# Prediction of pancreatic adenocarcinoma patient risk status using alternative splicing events

**DOI:** 10.1101/2021.06.02.446780

**Authors:** Rajesh Kumar, Anjali Lathwal, Gajendra P.S. Raghava

## Abstract

In literature, several mRNA, miRNA, lncRNA based biomarkers are identified by genomic analysis to stratify the patients into high and low risk groups of pancreatic adenocarcinoma (PAAD). The identified biomarkers are of limited use in terms of sensitivity and prediction ability. Thus, we aimed to identify the prognostic alternative splicing events and their related mutations in the PAAD. PAAD splicing data of 174 samples (17874 AS events in 6209 genes) and corresponding clinical information was obtained from the SpliceSeq and The Cancer Genome Atlas (TCGA), respectively. Prognostic-index based modeling was used to obtain the best predictive models for the seven AS types. However, model based on multiple spliced events genes (*APP*; *LATS1*; *MRPL4*; *LAS1L*; *STARD10*; *PHF21A*; *NMRAL1*) outperformed the single event models with a remarkable HR of 9.13 (p-value = 6.42e-10) as well as other existing models. Results from g:Profiler suggest that transcription factors *ZF5, ER81, E2F-1/2/3, ER81, Erg*, and *PEA3* are most related to the prognostic spliced genes. We also identified 565 mutations across 160 spliced genes that have a strong association with the prognostic AS events. The analysis revealed that around 560 of these mutations were not reported before in context to splice event/region. Overall, we conclude that altered AS events may serve as strong indicators for overall survival in pancreatic cancer patients, and novel linkage of the known mutations to the survival-related AS events may provide a new dimension to the advancement of diagnostic and therapeutic interventions in these patients.

## 1. Introduction

Despite the improvement in screening, diagnosis, curative resection, and preventive strategies, pancreatic cancer remains the leading cause of morbidity among the developed countries (Bray et al., 2018). The incidence of pancreatic cancer is more in men than women. Multiple risk factors have been identified that include obesity, diet, smoking, diabetes mellitus, cirrhosis, and bacterial infection (Rawla, Sunkara, & Gaduputi, 2019). Pancreatic cancer is considered as an intractable malignancy with an estimated 57,600 new cases and 47,050 deaths in 2019 (https://www.cancer.org/cancer/pancreatic-cancer/about/key-statistics.html). It is one of the fatal cancer types with an overall five-year survival rate of nearly 8% (Miller et al., 2018).

In the early stage, pancreatic cancer is almost asymptomatic and difficult to assess. Whereas, in advance stage, it gradually starts showing symptoms that include abnormal weight loss, fatigue, light-coloured stool, jaundice, pruritus, and abdominal pain (Saad, Turk, Al-Husseini, & Abdel-Rahman, 2018). The serum tumor antigen (CA 19-9) and computed tomography are the only available approved diagnostic standards for pancreatic cancer (Kim et al., 2004). However, they may miss the early stage of the cancer onset (Freelove & Walling, 2006). Surgical resection is the only available standard protocol for the treatment of pancreatic cancer. However, the use of drugs such as Gemcitabine, Leucovorin, Fluorouracil, and adjuvant chemotherapy can improve overall survival by several months (McGuigan et al., 2018; Neoptolemos et al., 2010). Even today, there is no definite cure for pancreatic cancer. The silent nature of the disease, poor prognosis, high mortality, and no available cure, opens an avenue to the researchers for the development of new therapies and markers.

Over the past decade, advancement in high-throughput technologies opened a new platform for genomic cancer studies. With the application of RNA-seq and gene expression studies, various molecular markers have been identified (Doherty et al., 2019; Kumar, Patiyal, Kumar, Nagpal, & Raghava, 2019). However, such studies, despite having promising results, were mainly confined to alteration at the transcriptome level, whereas systematic analysis of the post-transcriptional modification process is mostly neglected, especially in pancreatic cancer.

Alternative splicing (AS) is a post-transcriptional modification process that contributes to the generation of protein diversity in the eukaryotes (Pan, Shai, Lee, Frey, & Blencowe, 2008). Almost 95% of the human protein-coding gene undergoes splicing and produces protein isoforms in a tissue-specific manner (Nilsen & Graveley, 2010). Emerging data suggests the association of splice isoform with cancer progression, metastasis, and therapy resistance (Climente-González, Porta-Pardo, Godzik, & Eyras, 2017). One study identifies that the splice variant of the *MUC4* gene has been associated with the progress of pancreatic cancer (Jahan et al., 2018). Another study associates the *KLF6* gene splice variant with prognosis and tumor grade of pancreatic cancer (Hartel et al., 2008). An additional study showed the prognostic potential of the secretin receptor protein splice variant (Hayes, Carrigan, Dong, Reubi, & Miller, 2007). Thus, it is evident that cancer-specific splice isoforms can be used for diagnosis, prognosis, and therapeutic purposes.

Due to the technical limitations of experimental setups, the functional role of AS events in the progression of pancreatic cancer has been studied individually. Thus, analysis of all splicing isoforms with the pancreatic cancer progression needs to be considered to find out the reliable prognostic markers. Nowadays, machine learning techniques have been widely used for the identification of prognostic markers and the development of prediction classifiers (Chang & Chen, 2019; Lathwal, Arora, & Raghava, 2019). Therefore, in the current study, we have illuminated the role of AS events in the prognosis of pancreatic adenocarcinoma (PAAD). We have developed various machine learning and prognostic-index based models to gain insight into the prognostic potential of PAAD-specific AS patterns. We hope that the results obtained from this study will help in developing the novel and potent prognostic markers against pancreatic cancer.

## 2. Materials and Methods

### 2.1. Data Retrieval and Pre-processing

We have retrieved the clinical information of PAAD from the TCGA (https://tcga-data.nci.nih.gov/) using TCGA Assembler (Zhu, Qiu, & Ji, 2014). The downloaded dataset consists of a total of 178 samples. All the data on alternative splicing events in PAAD was obtained using a java based application, TCGA SpliceSeq (https://bioinformatics.mdanderson.org/TCGASpliceSeq). It covers seven types of AS events which include i) Alternate Acceptor Site (AA), ii) Alternate Promoter (AP), iii) Alternate Donor Site (AD) (iv) Alternate Terminator (AT), v) Exon Skip (ES), vi) Mutually Exclusive Exons (ME), and vii) Retained Intron (RI). SpliceSeq measures the percentage splice index (PSI) value corresponding to each sample, which is indicative of the ratio of transcripts mapped on the parent gene to all AS events. We have used stringent criteria for creating reliable data by considering only those samples which have PSI value greater than equal to 75% and the average PSI value of 50% or more.

### 2.2. Prognosis-associated AS Events, UpSet Plot and Functional Enrichment Analysis

We have identified the AS events having a significant relation with the overall survival (OS) (p-value < 0.05) in PAAD patients using Univariate Cox regression analysis implemented via ‘survival’ package in R (V.4.0.0, The R Foundation). The patients were scrutinized into high and low-risk groups on the basis of median PSI value of the AS events under consideration. For each AS event, the samples with PSI value higher than the median (PSI) were grouped in high-risk and samples with PSI value less than equal to median (PSI) were termed as low-risk samples. UpSet plot is used for the visualization of different intersections among all the types of AS using Intervene web application (https://asntech.shinyapps.io/intervene) (Khan & Mathelier, 2017). Using ggforest function in ‘survminer’ package available in R version 4.0.0, we have generated the forest plot for the top 10 events (Concordance) in each AS type. Functional enrichment of all the genes with survival-related AS events was done using the g:Profiler (https://biit.cs.ut.ee/gprofiler) (Reimand, Arak, & Vilo, 2011). It provides comprehensive data on functional evidence of genes under consideration which includes the related Gene Ontology (GO) terms, biological pathways, regulatory motifs of transcription factors and miRNAs, human diseases annotations and protein-protein interactions from multiple resources such as Reactome, KEGG, WikiPathways, Transfac, miTarBase, CORUM and Human Phenotype Ontology (HPO).

### 2.3. Prognostic-Index based Prediction Models using AS Events

Similar to studies found in literature, we have used the prognostic index to construct the different models for each AS type. The general idea of the formulation of the Prognostic Index (PI) is as follows:-

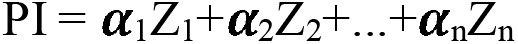

Where ***α*** is the regression coefficient for an event Z, which was retrieved via univariate cox model.

For each set of AS types, PI at median cut-offs of PSI values was used to classify patients in high- and low-risk groups. The patient with PI > median (PI) was grouped under high-risk set, and patients with PI ≤ median (PI) were termed as the low-risk group. For each AS type, we have selected the top 20 survival-associated events and iteratively searched for the minimum possible event-set that can be used to build the predictive models to further enhance its scalability in the clinical setup. Then, considering all the events in the top models within each AS type, we built our final highest performing prognostic model using the same iterative approach, as explained earlier. The calculation of all the statistical metrics such as Hazard ratio (HR), p-value, concordance, and standard error was done to screen the best performing final model.

### 2.4. Identification of Mutations in AS Events and Drug Targets

In order to identify the mutations linked with the survival-associated AS events, we first downloaded the mutation data from cBioPortal (https://www.cbioportal.org/datasets) (Cerami et al., 2012). Exon numbers comprising the AS events were obtained from SpliceSeq and their corresponding coordinate information was retrieved using the AceView by selecting human species option in the dropdown menu (https://www.ncbi.nlm.nih.gov/IEB/Research/Acembly/) (Thierry-Mieg & Thierry-Mieg, 2006). We identified 565 AS-event associated mutations in different exons among 160 genes. DrugBank (https://www.drugbank.ca/) was utilized to retrieve all the information on the drug targets against the genes having AS-events associated with OS.

### 2.5. Evaluation Metrics

All the statistical metrics of the survival models such as HR, log-rank test, Wald test, and concordance index were calculated to assess the performance of the models. HR depicts the risk of death linked to the high and low-risk groups. A Log-rank test was done to verify the statistical significance of the survival curves of the two risk sets. The Wald test was conducted to illustrate the importance of explanatory variables used to obtain HR. The prediction ability (PA) in terms of concordance index evaluates the model’s ability to categorize among the high- and low-risk sets. Lower log-rank, p-value (<0.05), and a higher concordance value (>0.5) (Chaudhary, Poirion, Lu, & Garmire, 2018; Han, Zhang, & Shao, 2017) account for a prognostic model with good predictive power. The complete workflow of our study is in Fig. 1.

**Fig. 1.**
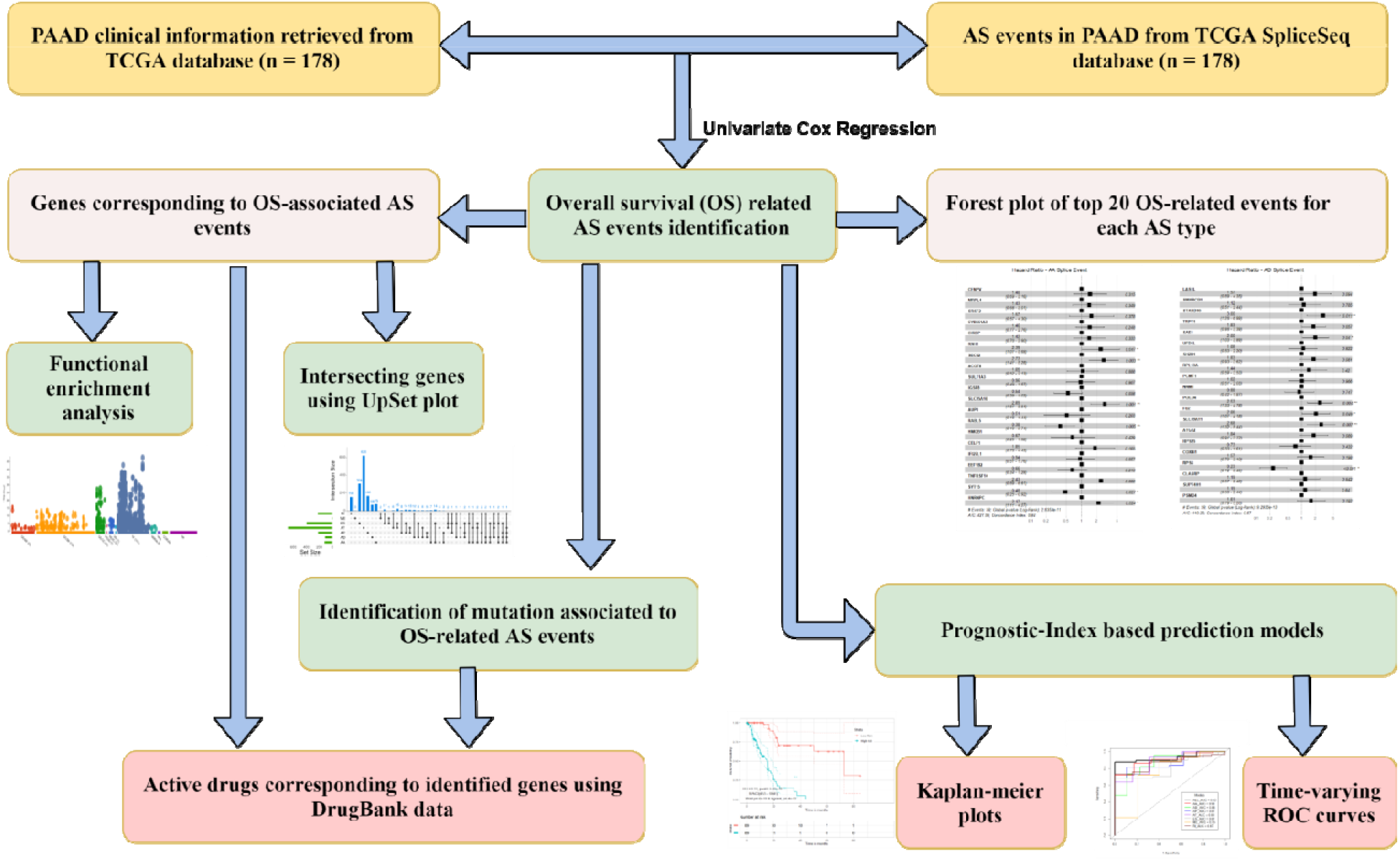
The overall flow of the study

## 3. Results

### 3.1. Overview of Alternating Splicing Events in TCGA PAAD Dataset

We retrieved a total of 17,874 AS events against 6,209 genes in 178 PAAD samples downloaded from TCGA SpliceSeq. We observed that there were many genes exhibiting more than one type of AS event. We detected 1099 AA events in 865 genes, 966 AD events in 707 genes, 2411 AP events in 967 genes, 7466 AT events in 3257 genes, 4566 ES events in 2582 genes, 58 ME events in 58 genes and 1318 RI events in 962 genes. Clearly, AT was the most common event found in 3257 genes (largest), whereas ME was found to be the rarest among all the subtypes (58 genes).

### 3.2. Survival-associated AS Events and Functional Enrichment Analysis

We have done cox univariate analysis to detect the events related to the overall OS of the PAAD patients. The patients were grouped among low and high-risk sets on the basis of median cut-off of the PSI value for the event under consideration. We have identified a total of 2523 AS events in 1559 genes that are significantly associated with the OS of the patients. In total, 122 AS events in 117 genes, 117 AD events in 109 genes, 372 AP events in 218 genes, 1189 AT events in 683 genes, 443 ES events in 395 genes, 6 ME events in 6 genes, and 274 RI events in 223 genes were found to be significant prognostic markers (p-value < 0.05) in PAAD patients. Statistical parameters such as HR, p-value, concordance, group size, and 95%CI obtained from the analysis of the AS events related to OS in PAAD patients can be found in the Supplementary S1 Table 1. Fig. 2 shows the intersections among the genes exhibiting alternative splicing events. Clearly, one gene may undergo more than one type of AS event.

For each AS subtype, further analysis was done using top 20 prognostic markers with the best distinguishability (Concordance Index) among the high- and low-risk groups for PAAD patients. We have implemented the coxph multivariate regression using these top events, and ‘ggforest’ function in ‘survminer’ package was used to generate the forest plot (top ten) for each event type, as shown in Fig. 3.

**Fig. 2.**
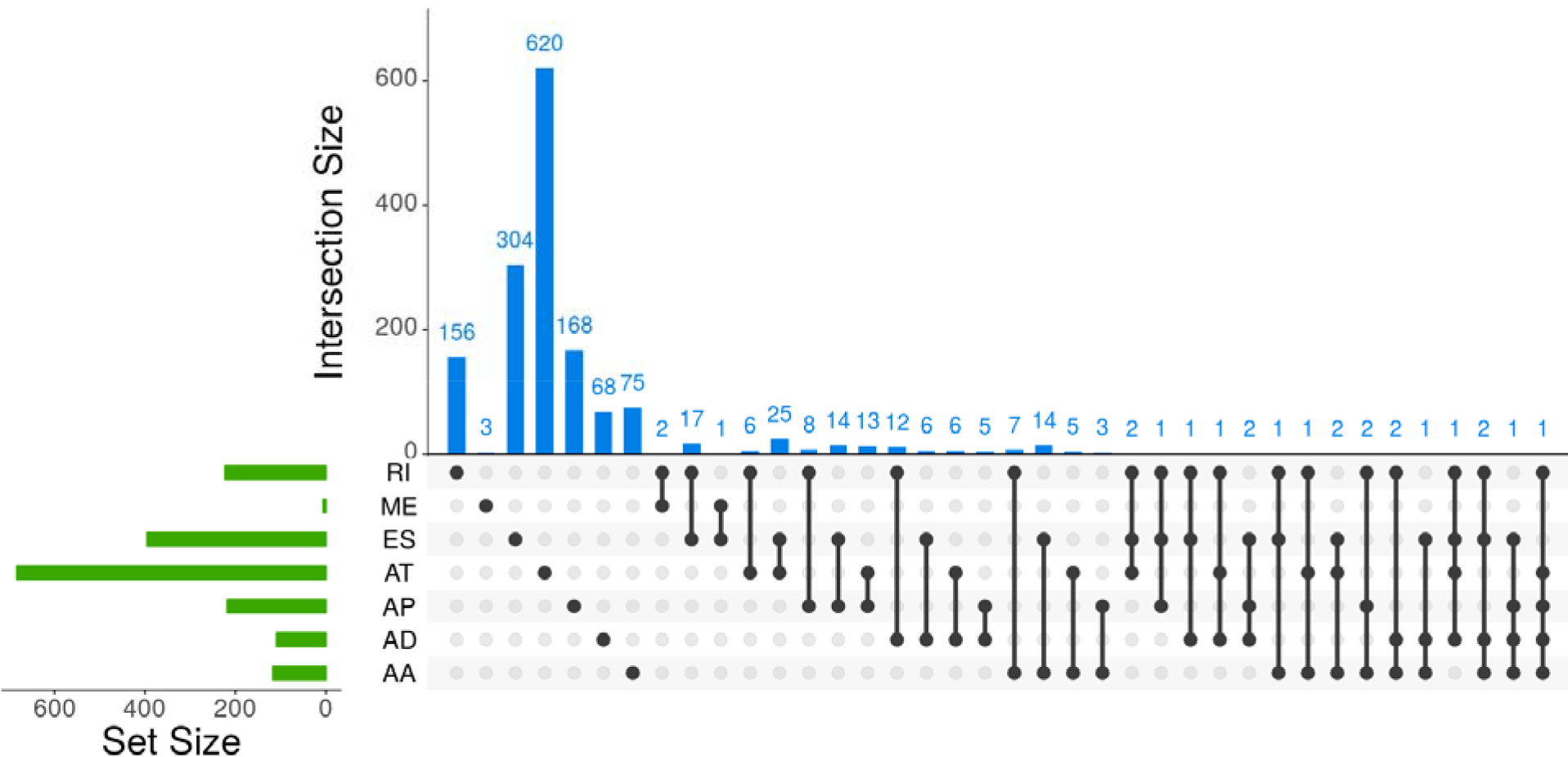
UpSet plot showing intersecting genes with prognosis-related AS events. Y-axis depicts the number of intersecting genes with a particular AS event. The X-axis describes the amount of each type of AS event (green bar) and the intersecting AS events (black line) in specific genes.

**Fig. 3.**
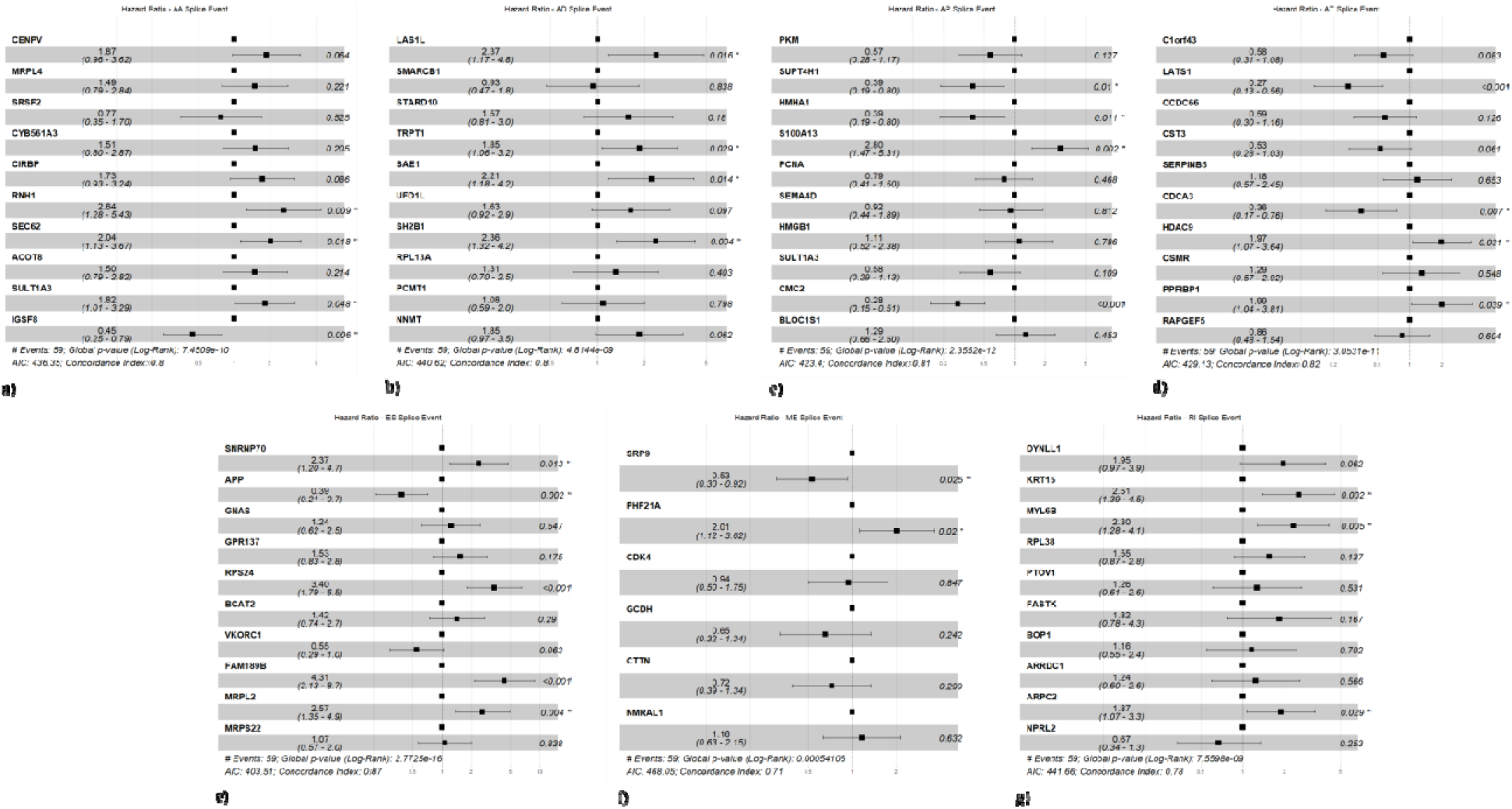
Forest plot of top 10 most significant AS events in PAAD a) Alternate Acceptor (AA), b) Alternate Donor (AD), c) Alternate Promoter (AP), d) Alternate Terminator (AT), e) Exon Skip (ES), f) Mutually Exclusive (ME), and g) Retained Intron (RI).

We have analyzed the functional importance of the parent genes of the prognosis-associated AS-events. g:Profiler was used to conduct the functional enrichment analysis of the genes and results are shown in a tabular form for all related Gene Ontology (GO) terms, biological pathways, regulatory motifs of transcription factors and miRNAs, human diseases annotations and protein-protein interactions obtained from several reliable resources like Reactome, KEGG, WikiPathways, Transfac, miTarBase, CORUM and Human Phenotype Ontology (HPO). We have given the details of the functional enrichment analysis for the parent genes against each process in Supplementary S1 Table 2. Functional enrichment analysis reveals that the majority of genes are enriched in mRNA metabolic pathways. It is well-observed in the literature that pancreatic cancer often exhibits reprogramming of metabolic pathways to sustain its growth. PAAD also utilizes the reprogrammed metabolic pathways to exhibit the chemoresistance (Qin et al., 2020). Molecular functional enrichment analysis revealed that the identified genes are the major components of the ribosome, which involved in protein synthesis machinery. Alteration within the protein synthesis machinery leads to the formation of faulty proteins and thus causing aggressive behaviour of cells which is the root cause of tumorigenesis (Halbrook & Lyssiotis, 2017). Somatic mutations in ribosomal proteins may impair ribosome biogenesis and proliferation, thus posing a role in the tumor progression (Sulima, Hofman, De Keersmaecker, & Dinman, 2017). Functional enrichment analysis suggests strong evidence against the role of identified prognostic spliced genes in the ribosomal machinery. Ribosomal proteins such as *RPL15*, *MRPL51* were also found to have mutations in PAAD patients, refer Supplementary S1 Table 3. Enrichment analysis depicts that majority of spliced genes are involved in the cellular processes related to the membrane-bound organelles such as the highly conserved cytoskeletal genes like the *ACT*, which is associated with the pancreatic tumorigenesis in past studies (Liu, Zhang, Go, & Hu, 2010). We observed that nearly 22 spliced genes are associated with microRNA, miR-652, which is responsible for inhibiting the transition from epithelial to mesenchymal via acidic microenvironment in pancreatic cancer cells (Deng et al., 2015). Results also suggest the strong association of transcription factors such as *ZF5, ER81, E2F-1/2/3, ER81, Erg*, and *PEA3* to the prognostic AS genes. Overall, the strong literature evidence against the identified prognostic spliced genes depicts their functional importance in the pancreatic cancer tumorigenesis.

### 3.3. Prognostic-index based Prediction Models

We screened for the top 20 prognostic markers within each subtype based on their prediction ability, i.e., concordance index, HR and p-values. For each AS type, using an iterative search process, we screened the minimum possible event signature that can be exploited to build the best predictive models. This way, we improved the scalability of the identified prognostic signature in the research and clinical setup.

A refined model was developed using the same iterative approach as explained above, on the combination of AS events obtained from best models in each subtype. We have generated the Kaplan-Meier plots for all the top models to analyse the survival curves in the two risk groups, which can be seen in Fig. 4. All the statistical and event-specific details such as HR, 95% confidence interval, p-value, and prediction ability (concordance index) of the these models against each splicing type can be found in Table 1. We observed that almost all the models are performing well with reasonably good prediction accuracy of nearly 70% except ES and ME. Combined model based on the seven spliced gene events (*APP*-ES; *LATS1*-AT; *MRPL4*-AA; *LAS1L*-AD; *STARD10*-AD; *PHF21A*-ME; *NMRAL1*-ME) outperformed the other subtype-specific models with an overall HR of 9.13, P-value of 6.42e-10 and prediction ability of 74% followed by the AP & AD splice type models with HR of 6.80, P-value of 1.63e-08 and prediction ability of 72.2% and HR of 6.17, P-value of 8.11e-08 and prediction ability of 72.00%, respectively.

**Table 1:**
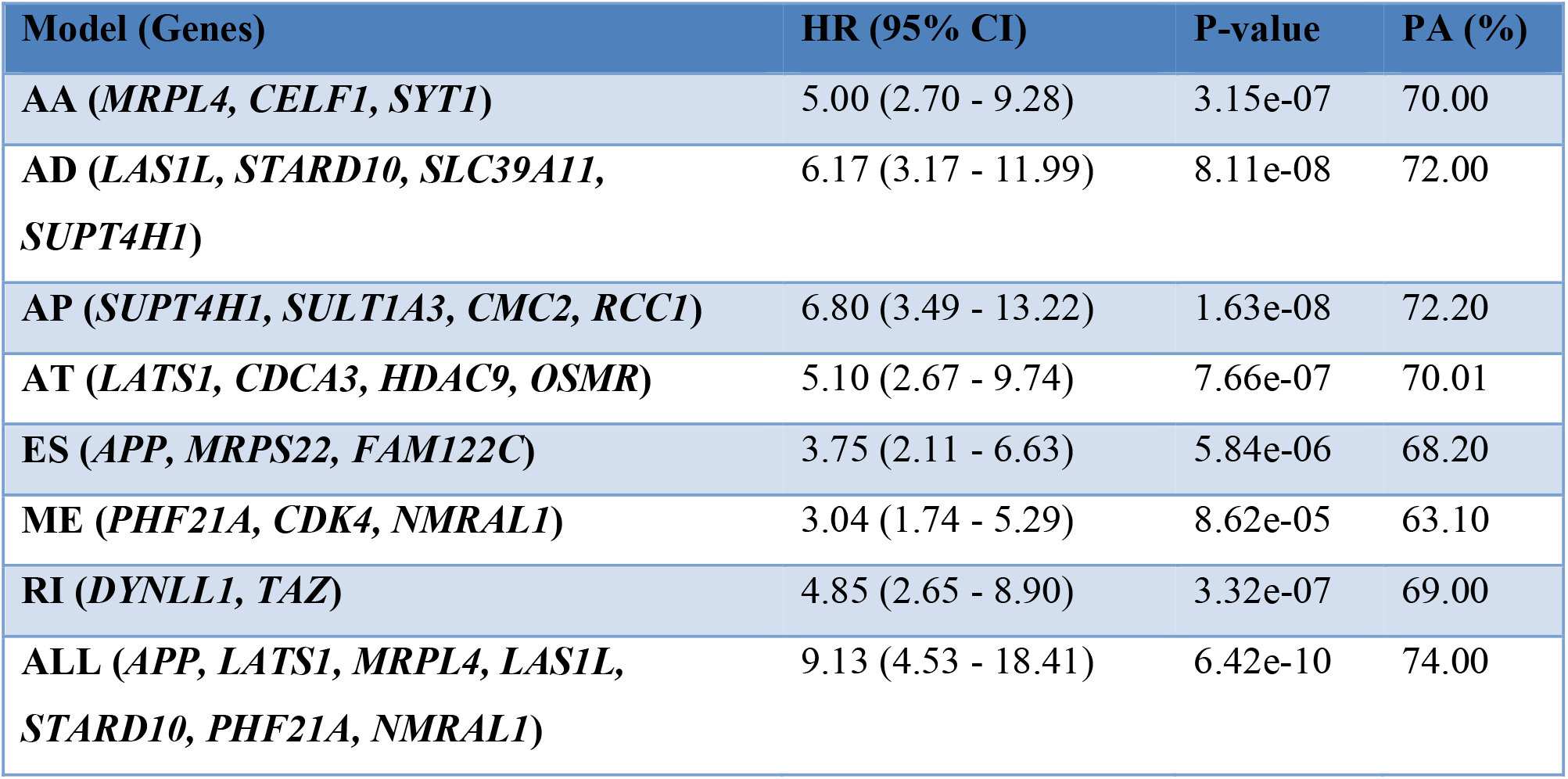
Statistical details of the top predictive models for each subtype and overall final model.

**Fig. 4.**
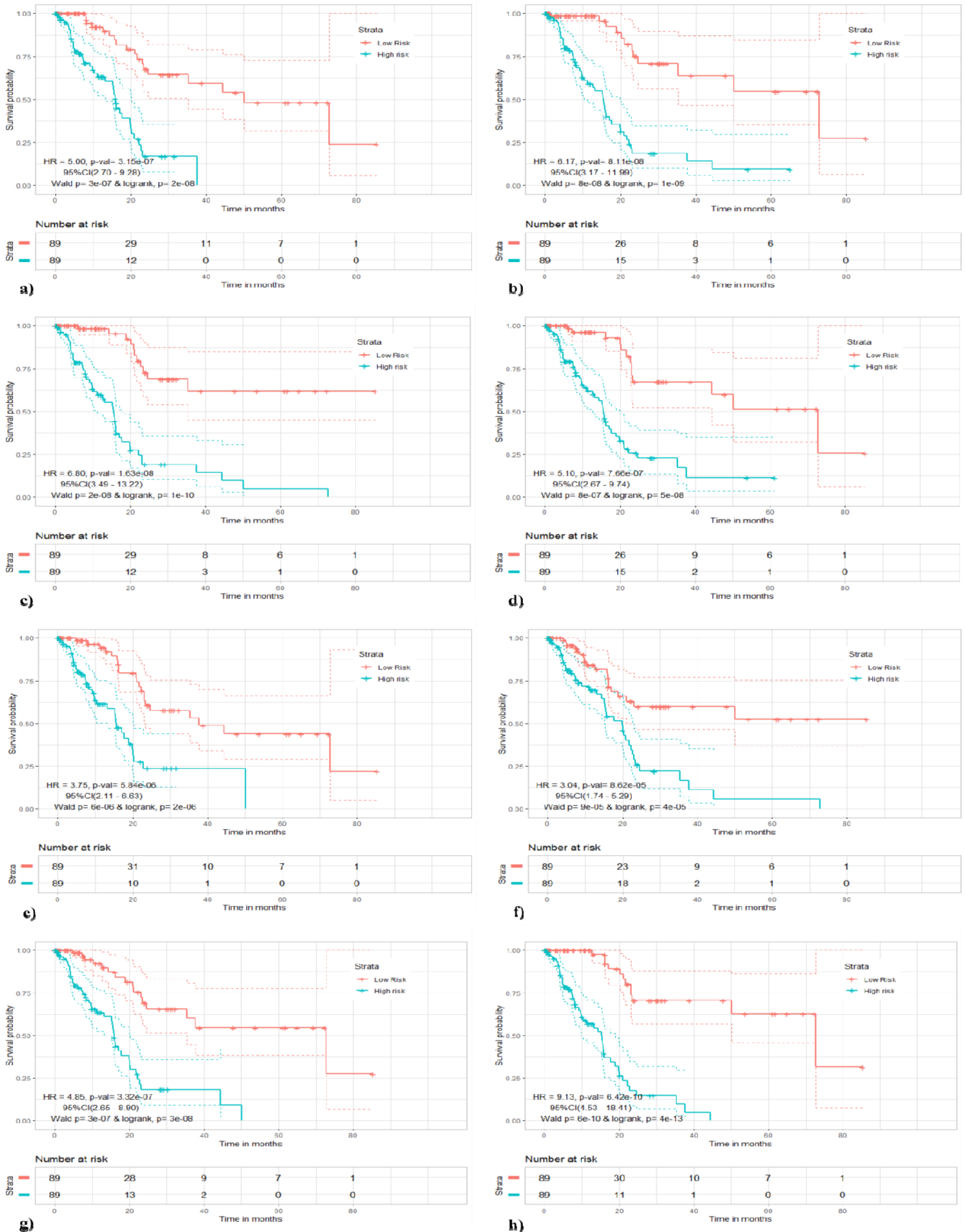
Kaplan-Meier plots against the best predictive models: a) Alternate Acceptor, HR = 5.0 & P-value = 3.15e-07; b) Alternate Donor, HR = 6.17 & P-value = 8.11e-08; c) Alternate Promoter, HR = 6.80 & P-value = 1.63e-08; d) Alternate Terminator, HR = 5.10 & P-value = 7.66e-07; e) Exon Skip, HR = 3.75 & P-value = 5.84e-06; f) Mutually Exclusive, HR = 3.04 & P-value = 8.62e-05; g) Retained Intron, HR = 4.85 & P-value = 3.32e-07; and h) Combined/ All, HR = 9.13 & P-value = 6.42e-10.

We have also assessed the stability and efficiency of our different predictive signatures using the time-varying ROC estimation at 4 and 5 years. The ROC Curve for 4 & 5 years, as shown in Fig. 5 indicated that our seven spliced gene-based prognostic model outperformed the other models with an overall AUC of 0.91 and 0.93, respectively. The study again confirms that the prognostic model based on the multiple splice events is better and reliable when compared to the individual AS type based models.

**Fig. 5.**
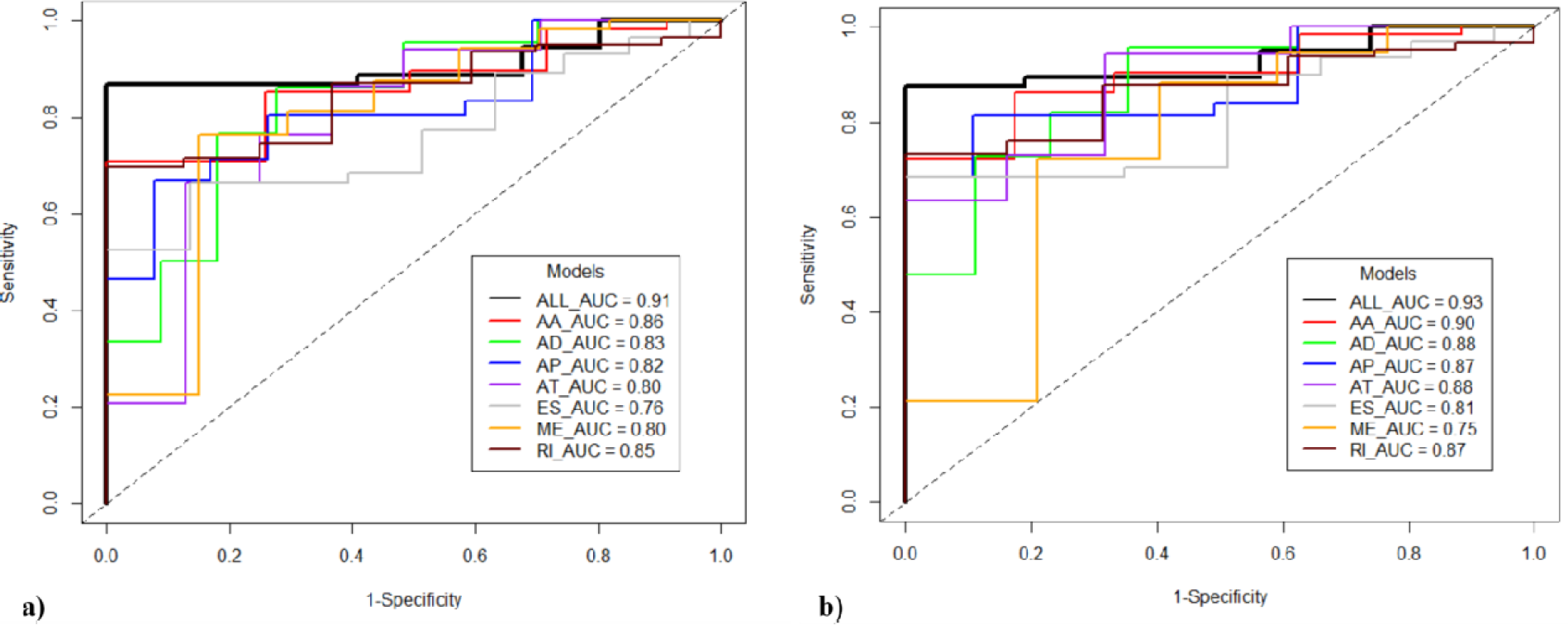
Time-varying ROC curves along with estimated AUCs of the prognostic models built using a single type of event or all events for risk stratification among the PAAD patients at a) 4 years, and b) 5 years.

### 3.4. Mutations in Overall Survival related Alternative Splicing Events

Aberrations and abnormalities in the alternative splicing are now widely accepted as one of the new indicators of initiation of carcinogenic process. In this regard, our goal is to identify the possible cancerous mutations that may be present in the spliced gene regions showing association with the OS of the PAAD patients. Firstly, we retrieved the respective exon numbers of the OS-related splicing events from the TCGA SpliceSeq, and to get the coordinates of these exons, we mapped them to the exon information obtained from the AceView. Further, the PAAD specific mutation data downloaded from the cBioPortal was utilized to identify the possible mutations within those exonic regions where the survival-related splice events have been observed. cBioPotal mutation file consisted of nearly 31,394 PAAD specific mutations across 12,829 unique genes.

We identified a total of 565 mutations (including 36 high impact, 126 moderate, 335 low, and 68 modifier mutations) in different exons involved in OS-related splicing events across 160 unique genes. We have summarized the results along with other details such as coordinate information of exons and mutations, type of variant, its biotype, etc. in the Supplementary S1 Table 3. We have shown the distribution of various mutation types available in literature within these AS events in the form of a pie-chart (Fig. 6). The majority of the mutations are missense mutations followed by silent mutations. The data analysis revealed that the known mutations associated with the splice sites or regions in 160 unique genes were only 5. However, based on our above study, it may not be wrong to say that all the 565 mutations are linked to splice sites or regions. In this way, we may conclude that ~ 560 mutations were identified to have novel associations with prognostic AS events in PAAD cancer patients. However, we emphasize that strong experimental evidence will be required to validate the results. We then searched for the therapeutic potential of the total 1559 OS-related spliced genes using the DrugBank data and found that 228 pharmacologically active drugs are available against 54 genes.

**Fig. 6.**
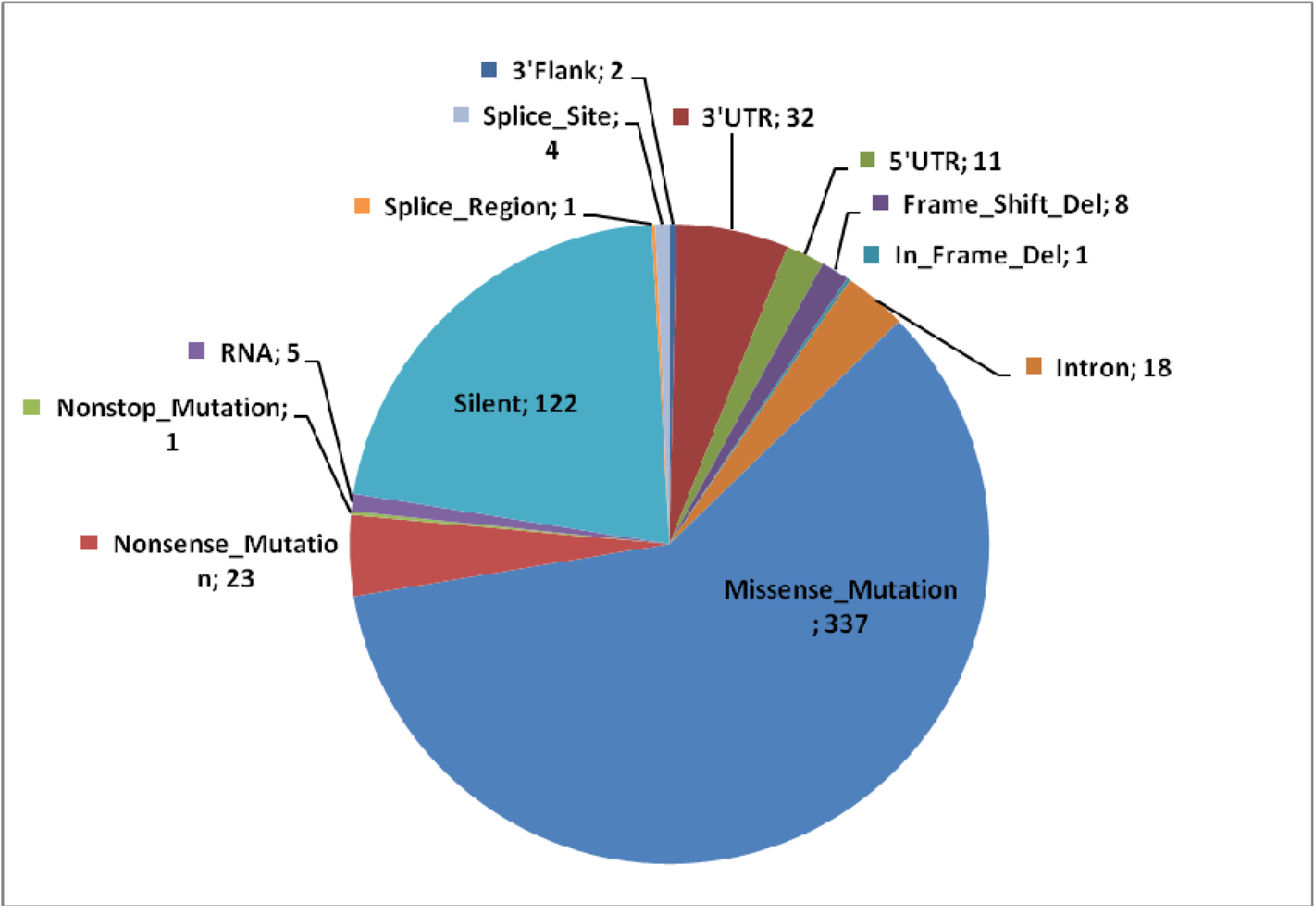
Categorical distribution of associated mutations (variant type, Freq) found in literature across the OS-related AS events in PAAD.

Further analysis revealed that only eight genes (out of 54) were having both carcinogenic mutations, and active drug targets designed against these genes. Table 2 showed the details of these genes and corresponding drug targets, however, the complete results against the 54 genes can be found in the Supplementary S1 Table 4.

**Table 2:** Drugs against OS-related spliced genes that also express carcinogenic mutations.

| Genes | UniProt ID | Drug Name (DrugBank IDs) |
| --- | --- | --- |
| **ACAA1** | P09110 | Trimetazidine (DB09069) |
| **APP** | P05067 | Florbetapir (DB09149), Flutemetamol (DB09151), Florbetaben (DB09148), |
| **CD44** | P16070 | Hyaluronic acid (DB08818) |
| **PPARD** | Q03181 | Icosapent (DB00159), Bezafibrate (DB01393), Glycerin (DB09462), Treprostinil (DB00374) |
| **PRLR** | P16471 | Fluoxymesterone (DB01185), Somatotropin (DB00052) |
| **SDHA** | P31040 | Ubidecarenone (DB09270) |
| **SHMT1** | P34896 | Mimosine (DB01055) |
| **TYMS** | P04818 | Pemetrexed (DB00642), Methotrexate (DB00563), Fluorouracil (DB00544), Trifluridine (DB00432), Tegafur-uracil (DB09327), Pralatrexate (DB06813), Tegafur (DB09256), Raltitrexed (DB00293), Floxuridine (DB00322), |

### 3.5. Improved Performance over Existing PAAD Prognostic Signatures

We have analyzed the performance of our final prognostic model in comparison with the previously identified prognostic markers obtained from several published studies. We observed that our splice event based seven gene prognostic model *(HR = 9.13; P-value = 6.42e-10, Concordance = 0.7; and 5yr AUC = 0.93)* outperformed the existing PAAD specific prognostic markers such as five miRNAs *(HR = 0.13; P < 0.001)* (Liang et al., 2018) and four miRNAs based prognostic models *(HR= 2.8; P < 0.001; 2yr AUC = 0.79)* (Wang, Deng, & Ma, 2019), 20 gene-based prognostic signature *(HR = 2.01; P = 0.007)* (Canlı et al., 2020), three lncRNA signature *(P < 0.001; AUC = 0.72)* (Shi et al., 2018), 29 gene-based ceRNA signature *(HR = 1.66; P < 1.0e-04)* (Zhao & Liu, 2017); 9 gene prognostic signature *(P < 0.0001; 1yr AUC = 0.70)* (Wu, Li, Zhang, Liu, & Zhao, 2019), and 7 gene-based AS event prognostic signature *(AUC = 0.89)* (Yu et al., 2019). Our splice-event based prognostic marker also outperformed an exisiting study in terms of HR and AUC that evaluates the risk assessment capability of PAAD related AS events (Xu, Pan, Ding, & Pan, 2020). All the above mentioned comparative analysis to existing studies highlights the reliability, better performance and strength of our identified prognostic signature in a more clear and robust way.

## 4. Discussion and conclusion

Alternative splicing is a crucial process in eukaryotic organisms for generating protein diversity. Emerging studies highlight the involvement of splicing events in the onset and progression of cancer. Isoform switching is commonly observed in pancreatic cancer, which contributes to the aggressiveness and development of chemoresistance (Calabretta et al., 2016). The functional contribution of AS events in tumorigenesis in the context of their prognosis related potential has mostly been unexplored. The present study focuses on the exploration of AS events for the identification of prognostic spliced gene biomarkers in PAAD. We have utilized univariate and multivariate cox proportional hazard methods to build predictive models that can stratify the patients into high and low-risk groups. Statistical metrics such as HR, P-value, and concordance revealed that the model developed on a combination of AS types outperformed the subtype-specific models. Functional enrichment analysis revealed the role of the majority of spliced genes in ribosomal machinery, maintaining membrane integrity, and any mutations in these genes may aid in carcinogenesis. Recent literature evidence also highlights the importance of somatic mutations in the progression and sustenance of cancer. In this study, we have made an effort to link the known mutations to the prognostic AS events. We identified 560 novel linkages of the known mutations to the survival-related splicing events. We have also reported genes such as *APP*, which is a remarkable OS-related spliced gene and have numerous mutational changes associated with it’s AS events. Mutations in *APP* gene may further explain the onset of dementia observed in the majority of PAAD patients and are widely targeted for therapeutic interventions (Woods & Padmanabhan, 2013). We have listed the active drug targets against the OS-associated spliced genes in our study. Results of linking known mutations to the AS events may further enhance our understanding of disease etiology. We may like to highlight the limitation of our study that includes the lack of independent datasets for the validation and experimental proof of the identified results. Despite having such restrictions in the study, we hope that our findings may open novel insights and aid in the advancement of diagnostic, prognostic and therapeutic approaches against the pancreatic cancer patients

## Author’s Contribution

**Rajesh Kumar:** Conceptualization, Methodology, Validation, Formal analysis, Investigation, Data curation, Writing original draft, Reviewing and editing original draft. **Anjali Lathwal:** Conceptualization, Methodology, Validation, Formal analysis, Investigation, Data curation, Writing original draft, Reviewing and editing original draft. **Gajendra Pal Singh Raghava:** Supervision, Project administration, Writing original draft, Reviewing and editing original draft, Investigation, Validation.

## Data Availability

All the data is freely available on TCGA, TCGA SpliceSeq, cBioPortal and AceView websites.

## Acknowledgment

Authors are thankful for the funding agencies, Council of scientific and industrial research (CSIR), University Grants Commission (UGC), and Indraprastha Institute of Information Technology (IIIT-D) Delhi for providing necessary facility and infrastructure to carry out this research work.

