## Supplementary files for "Prediction of pancreatic adenocarcinoma patient risk status using alternative splicing events"

| Gene | Splice | GP1 | GP2 | Beta | HR | se.beta. | z | P.value |
|---|---|---|---|---|---|---|---|---|
| PKM | AP | 89 | 89 | 1.389054 | 4.011055 | 0.295491 | 4.700831 | 2.59E-06 |
| SNRNP70 | ES | 89 | 89 | -1.37215 | 0.253561 | 0.301736 | -4.54753 | 5.43E-06 |
| APP | ES | 90 | 88 | 1.101446 | 3.008512 | 0.276431 | 3.984517 | 6.76E-05 |
| C1orf43 | AT | 90 | 88 | 1.179614 | 3.253118 | 0.284638 | 4.144258 | 3.41E-05 |
| SUPT4H1 | AP | 89 | 89 | 1.349446 | 3.855288 | 0.316017 | 4.27017 | 1.95E-05 |
| C1orf43 | AT | 90 | 88 | -1.16062 | 0.313291 | 0.284883 | -4.07404 | 4.62E-05 |
| LATS1 | AT | 89 | 89 | -1.14885 | 0.317001 | 0.278001 | -4.13254 | 3.59E-05 |
| LATS1 | AT | 89 | 89 | 1.14885 | 3.154564 | 0.278001 | 4.132538 | 3.59E-05 |
| CCDC66 | AT | 89 | 89 | 1.145656 | 3.144505 | 0.281661 | 4.067497 | 4.75E-05 |
| DYNLL1 | RI | 89 | 89 | -1.28908 | 0.275524 | 0.291324 | -4.4249 | 9.65E-06 |
| CST3 | AT | 91 | 87 | 1.127711 | 3.088579 | 0.272706 | 4.135269 | 3.55E-05 |
| GNAS | ES | 90 | 88 | -1.1274 | 0.323876 | 0.284478 | -3.96303 | 7.40E-05 |
| GPR137 | ES | 89 | 89 | -1.08713 | 0.337184 | 0.289323 | -3.75748 | 0.000172 |
| RPS24 | ES | 89 | 89 | -1.05048 | 0.349771 | 0.279124 | -3.76348 | 0.000168 |
| HMHA1 | AP | 89 | 89 | 1.218744 | 3.382935 | 0.294754 | 4.134778 | 3.55E-05 |
| SERPINB5 | AT | 90 | 88 | -1.12244 | 0.325483 | 0.28597 | -3.92505 | 8.67E-05 |
| S100A13 | AP | 89 | 89 | -1.15519 | 0.314999 | 0.280775 | -4.11428 | 3.88E-05 |
| BCAT2 | ES | 89 | 89 | -0.97645 | 0.376648 | 0.274764 | -3.55376 | 0.00038 |
| GNAS | ES | 89 | 89 | -1.08556 | 0.337712 | 0.280913 | -3.86441 | 0.000111 |
| CENPV | AA | 89 | 89 | -1.17757 | 0.308025 | 0.29076 | -4.04999 | 5.12E-05 |
| MRPL4 | AA | 90 | 88 | -1.16818 | 0.310932 | 0.289282 | -4.03821 | 5.39E-05 |
| CST3 | AT | 92 | 86 | -1.10977 | 0.329634 | 0.277789 | -3.99501 | 6.47E-05 |
| FAM189B | ES | 89 | 89 | -0.95181 | 0.386043 | 0.272298 | -3.49546 | 0.000473 |
| VKORC1 | ES | 89 | 89 | 0.835929 | 2.306956 | 0.277252 | 3.015051 | 0.002569 |
| MRPL2 | ES | 89 | 89 | -0.98602 | 0.37306 | 0.284816 | -3.46194 | 0.000536 |
| CDCA3 | AT | 89 | 89 | 1.222952 | 3.397201 | 0.302482 | 4.043051 | 5.28E-05 |
| CDCA3 | AT | 89 | 89 | -1.22295 | 0.29436 | 0.302482 | -4.04305 | 5.28E-05 |
| PCNA | AP | 89 | 89 | -0.98638 | 0.372923 | 0.275256 | -3.58351 | 0.000339 |
| PCNA | AP | 89 | 89 | 0.986384 | 2.681519 | 0.275256 | 3.583508 | 0.000339 |
| SRSF2 | AA | 89 | 89 | -1.15439 | 0.315249 | 0.29344 | -3.93399 | 8.35E-05 |
| MRPS22 | ES | 89 | 89 | -0.99961 | 0.368023 | 0.279658 | -3.5744 | 0.000351 |
| HDAC9 | AT | 89 | 89 | -0.84673 | 0.428814 | 0.276059 | -3.06722 | 0.002161 |
| KRT15 | RI | 89 | 89 | -1.0532 | 0.348821 | 0.278104 | -3.78706 | 0.000152 |
| SUPT4H1 | AP | 89 | 89 | -0.99875 | 0.36834 | 0.289342 | -3.45179 | 0.000557 |
| OSMR | AT | 89 | 89 | 0.984242 | 2.675782 | 0.283375 | 3.473283 | 0.000514 |
| OSMR | AT | 89 | 89 | -0.98424 | 0.373722 | 0.283375 | -3.47328 | 0.000514 |
| PPFIBP1 | AT | 89 | 89 | -0.9464 | 0.388135 | 0.273115 | -3.46522 | 0.00053 |
| SEMA4D | AP | 89 | 89 | -1.14794 | 0.317289 | 0.294541 | -3.89739 | 9.72E-05 |
| LAS1L | AD | 89 | 89 | -1.10562 | 0.331005 | 0.295405 | -3.74273 | 0.000182 |
| HMGB1 | AP | 91 | 87 | -1.10671 | 0.330646 | 0.297363 | -3.72174 | 0.000198 |
| GNAS | ES | 89 | 89 | -1.11908 | 0.326581 | 0.282927 | -3.95536 | 7.64E-05 |
| SULF2 | ES | 89 | 89 | -1.05704 | 0.347485 | 0.283457 | -3.72908 | 0.000192 |
| CYB561A3 | AA | 89 | 89 | -0.87427 | 0.417169 | 0.278941 | -3.13423 | 0.001723 |
| PKM | AP | 90 | 88 | -0.89104 | 0.410229 | 0.276342 | -3.22441 | 0.001262 |
| BAIAP2 | ES | 89 | 89 | 0.814875 | 2.258894 | 0.274224 | 2.971566 | 0.002963 |
| S100A13 | AP | 89 | 89 | 1.003001 | 2.726451 | 0.274773 | 3.650287 | 0.000262 |

| | | | | | | | | |
|---|---|---|---|---|---|---|---|---|
| **RAPGEF5** | AT | 89 | 89 | 0.717523 | 2.049351 | 0.265352 | 2.704039 | 0.00685 |
| **RAPGEF5** | AT | 89 | 89 | -0.71752 | 0.487959 | 0.265352 | -2.70404 | 0.00685 |
| **SSU72** | AT | 89 | 89 | -0.91421 | 0.400834 | 0.284291 | -3.21574 | 0.001301 |
| **SSU72** | AT | 89 | 89 | 0.914207 | 2.494796 | 0.284291 | 3.21574 | 0.001301 |
| **MYL6B** | RI | 90 | 88 | -1.14711 | 0.317552 | 0.279973 | -4.09723 | 4.18E-05 |
| **FAM122C** | ES | 89 | 89 | -1.01077 | 0.36394 | 0.275085 | -3.67437 | 0.000238 |
| **SRP9** | ME | 89 | 89 | 0.863851 | 2.372279 | 0.27487 | 3.142758 | 0.001674 |
| **IQCG** | AT | 89 | 89 | -1.05134 | 0.349468 | 0.274425 | -3.83108 | 0.000128 |
| **IQCG** | AT | 89 | 89 | 1.051345 | 2.861496 | 0.274425 | 3.831075 | 0.000128 |
| **TMEM67** | AT | 89 | 89 | -0.73981 | 0.477204 | 0.26995 | -2.74055 | 0.006134 |
| **TMEM67** | AT | 89 | 89 | 0.739812 | 2.095541 | 0.26995 | 2.740555 | 0.006134 |
| **SULT1A3** | AP | 89 | 89 | 1.038946 | 2.826236 | 0.282461 | 3.678187 | 0.000235 |
| **UAP1** | ES | 89 | 89 | 0.878533 | 2.407365 | 0.274266 | 3.203217 | 0.001359 |
| **PILRB** | ES | 89 | 89 | -0.78194 | 0.457519 | 0.272624 | -2.86819 | 0.004128 |
| **FAM72A** | AT | 89 | 89 | -0.6966 | 0.498278 | 0.270163 | -2.57843 | 0.009925 |
| **CMC2** | AP | 89 | 89 | 0.985219 | 2.678397 | 0.286444 | 3.439477 | 0.000583 |
| **HDAC9** | AT | 89 | 89 | 0.822253 | 2.275621 | 0.273596 | 3.005354 | 0.002653 |
| **YBX3** | ES | 89 | 89 | -0.93696 | 0.391819 | 0.287192 | -3.26248 | 0.001104 |
| **BLOC1S1** | AP | 89 | 89 | -0.77041 | 0.462821 | 0.270658 | -2.84645 | 0.004421 |
| **BLOC1S1** | AP | 89 | 89 | 0.770415 | 2.160663 | 0.270658 | 2.846448 | 0.004421 |
| **CACYBP** | AP | 89 | 89 | 0.93538 | 2.548181 | 0.277435 | 3.371526 | 0.000748 |
| **Sep-09** | AP | 89 | 89 | 0.950111 | 2.585997 | 0.279567 | 3.39851 | 0.000678 |
| **Sep-09** | AP | 89 | 89 | -0.89306 | 0.409401 | 0.282604 | -3.16011 | 0.001577 |
| **CIRBP** | AA | 89 | 89 | -0.82697 | 0.437373 | 0.274517 | -3.01246 | 0.002591 |
| **RHOF** | AT | 89 | 89 | -1.07847 | 0.340115 | 0.286977 | -3.75804 | 0.000171 |
| **RHOF** | AT | 89 | 89 | 1.078472 | 2.940183 | 0.286977 | 3.758036 | 0.000171 |
| **SMARCB1** | AD | 89 | 89 | -0.7757 | 0.46038 | 0.270757 | -2.86494 | 0.004171 |
| **RPL38** | RI | 91 | 87 | -0.92799 | 0.395348 | 0.271099 | -3.42306 | 0.000619 |
| **PRR13** | AT | 93 | 85 | 0.883715 | 2.419872 | 0.275535 | 3.207268 | 0.00134 |
| **STARD10** | AD | 89 | 89 | -0.88823 | 0.411385 | 0.273587 | -3.24659 | 0.001168 |
| **RNH1** | AA | 91 | 87 | -1.37196 | 0.25361 | 0.31192 | -4.39842 | 1.09E-05 |
| **DNASE1L1** | AP | 89 | 89 | -0.96284 | 0.381805 | 0.28546 | -3.37296 | 0.000744 |
| **KIF4A** | AT | 89 | 89 | -0.7812 | 0.457858 | 0.27739 | -2.81624 | 0.004859 |
| **KIF4A** | AT | 89 | 89 | 0.781197 | 2.184085 | 0.27739 | 2.816245 | 0.004859 |
| **RBCK1** | AT | 89 | 89 | 1.096399 | 2.993369 | 0.286225 | 3.830545 | 0.000128 |
| **RBCK1** | AT | 89 | 89 | -1.0964 | 0.334072 | 0.286225 | -3.83055 | 0.000128 |
| **PTOV1** | RI | 89 | 89 | -1.02925 | 0.357275 | 0.287007 | -3.58615 | 0.000336 |
| **FASTK** | RI | 89 | 89 | -1.25763 | 0.284328 | 0.297494 | -4.2274 | 2.36E-05 |
| **TMEM135** | AT | 89 | 89 | 0.764484 | 2.147887 | 0.277248 | 2.757399 | 0.005826 |
| **TMEM135** | AT | 89 | 89 | -0.76448 | 0.465574 | 0.277248 | -2.7574 | 0.005826 |
| **RNF32** | AT | 89 | 89 | 0.909821 | 2.483877 | 0.281873 | 3.227769 | 0.001248 |
| **RNF32** | AT | 89 | 89 | -0.90982 | 0.402596 | 0.281873 | -3.22777 | 0.001248 |
| **LY75** | AT | 89 | 89 | -0.78814 | 0.454691 | 0.281631 | -2.79847 | 0.005134 |
| **LY75** | AT | 89 | 89 | 0.788137 | 2.199295 | 0.281631 | 2.798474 | 0.005134 |
| **ATG16L2** | AT | 89 | 89 | -1.04586 | 0.351389 | 0.27616 | -3.78715 | 0.000152 |
| **ATG16L2** | AT | 89 | 89 | 1.04586 | 2.845845 | 0.27616 | 3.78715 | 0.000152 |
| **RCC1** | AP | 89 | 89 | -0.95487 | 0.384861 | 0.279772 | -3.41303 | 0.000642 |

| | | | | | | | | |
|---|---|---|---|---|---|---|---|---|
| **RCC1** | AP | 89 | 89 | 0.954872 | 2.598339 | 0.279772 | 3.413033 | 0.000642 |
| **SNRPB** | ES | 89 | 89 | -0.97912 | 0.375642 | 0.293089 | -3.34069 | 0.000836 |
| **PARD3** | AT | 89 | 89 | -0.93708 | 0.39177 | 0.281016 | -3.33462 | 0.000854 |
| **TCAIM** | AT | 89 | 89 | -1.27043 | 0.28071 | 0.311882 | -4.07344 | 4.63E-05 |
| **TCAIM** | AT | 89 | 89 | 1.270433 | 3.562394 | 0.311882 | 4.073443 | 4.63E-05 |
| **SLC3A2** | AP | 90 | 88 | 0.801264 | 2.228355 | 0.269779 | 2.970074 | 0.002977 |
| **DHPS** | AP | 89 | 89 | -0.78762 | 0.454924 | 0.272566 | -2.88967 | 0.003856 |
| **DHPS** | AP | 89 | 89 | 0.787625 | 2.198169 | 0.272566 | 2.88967 | 0.003856 |
| **ACTR6** | AT | 89 | 89 | -0.80727 | 0.446074 | 0.268512 | -3.00646 | 0.002643 |
| **ACTR6** | AT | 89 | 89 | 0.807271 | 2.241782 | 0.268512 | 3.006458 | 0.002643 |
| **SERPINB5** | AT | 90 | 88 | 0.973144 | 2.646252 | 0.279126 | 3.486403 | 0.00049 |
| **MAP3K7CL** | AP | 95 | 83 | 0.828439 | 2.289741 | 0.268663 | 3.083556 | 0.002045 |
| **SEC62** | AA | 89 | 89 | -0.7344 | 0.479793 | 0.27744 | -2.64706 | 0.00812 |
| **SET** | AP | 89 | 89 | -0.937 | 0.391801 | 0.277367 | -3.3782 | 0.00073 |
| **CTSC** | AT | 90 | 88 | 0.742364 | 2.100897 | 0.272331 | 2.725963 | 0.006411 |
| **CTSC** | AT | 90 | 88 | -0.74236 | 0.475987 | 0.272331 | -2.72596 | 0.006411 |
| **ZNF235** | AT | 89 | 89 | 0.664728 | 1.943961 | 0.267164 | 2.488092 | 0.012843 |
| **BOP1** | RI | 89 | 89 | -1.0874 | 0.337093 | 0.280293 | -3.87949 | 0.000105 |
| **ARRDC1** | RI | 89 | 89 | -1.12299 | 0.325305 | 0.279555 | -4.01706 | 5.89E-05 |
| **TRPT1** | AD | 89 | 89 | -0.62082 | 0.537506 | 0.267296 | -2.32258 | 0.020202 |
| **KLC1** | ES | 89 | 89 | -0.81787 | 0.441371 | 0.273163 | -2.99407 | 0.002753 |
| **RBP5** | AT | 89 | 89 | -0.84164 | 0.431002 | 0.267389 | -3.14763 | 0.001646 |
| **RBP5** | AT | 89 | 89 | 0.841642 | 2.320173 | 0.267389 | 3.14763 | 0.001646 |
| **ZC3HAV1** | AT | 89 | 89 | 0.805339 | 2.237454 | 0.272925 | 2.950765 | 0.00317 |
| **ZC3HAV1** | AT | 89 | 89 | -0.80534 | 0.446937 | 0.272925 | -2.95076 | 0.00317 |
| **UAP1** | ES | 89 | 89 | 0.876843 | 2.403299 | 0.277825 | 3.1561 | 0.001599 |
| **SCAF11** | AT | 89 | 89 | 0.898809 | 2.456676 | 0.276295 | 3.253073 | 0.001142 |
| **DBI** | AP | 90 | 88 | -0.79082 | 0.453474 | 0.284692 | -2.7778 | 0.005473 |
| **ARPC2** | RI | 91 | 87 | -0.67062 | 0.51139 | 0.267478 | -2.5072 | 0.012169 |
| **PMFBP1** | AT | 89 | 89 | 1.004611 | 2.730844 | 0.277341 | 3.622289 | 0.000292 |
| **PMFBP1** | AT | 89 | 89 | -1.00461 | 0.366187 | 0.277341 | -3.62229 | 0.000292 |
| **LIMK2** | AT | 89 | 89 | 0.963858 | 2.621792 | 0.288829 | 3.337128 | 0.000846 |
| **LIMK2** | AT | 89 | 89 | -0.96386 | 0.381418 | 0.288829 | -3.33713 | 0.000846 |
| **VDAC3** | ES | 89 | 89 | -0.87858 | 0.415373 | 0.274926 | -3.1957 | 0.001395 |
| **NOB1** | ES | 89 | 89 | 0.801002 | 2.227772 | 0.272053 | 2.944286 | 0.003237 |
| **WDR19** | AT | 90 | 88 | 0.77277 | 2.165757 | 0.266808 | 2.896351 | 0.003775 |
| **PRR13** | AT | 91 | 87 | -0.84476 | 0.42966 | 0.281744 | -2.99833 | 0.002715 |
| **SIRT6** | AP | 89 | 89 | -0.85506 | 0.425258 | 0.268674 | -3.18252 | 0.00146 |
| **SIRT6** | AP | 89 | 89 | 0.855059 | 2.351513 | 0.268674 | 3.18252 | 0.00146 |
| **NUMA1** | ES | 89 | 89 | -0.80716 | 0.446125 | 0.275284 | -2.93209 | 0.003367 |
| **GPATCH2L** | AT | 89 | 89 | -0.71199 | 0.490666 | 0.268845 | -2.64833 | 0.008089 |
| **SUMF2** | ES | 90 | 88 | -0.6083 | 0.544277 | 0.272257 | -2.23427 | 0.025465 |
| **BUD31** | ES | 89 | 89 | -1.12184 | 0.325681 | 0.287171 | -3.90651 | 9.36E-05 |
| **SAE1** | AD | 92 | 86 | -1.07009 | 0.342978 | 0.292457 | -3.65896 | 0.000253 |
| **NPRL2** | RI | 89 | 89 | -0.69552 | 0.498815 | 0.273508 | -2.54295 | 0.010992 |
| **OSGEP** | RI | 89 | 89 | -0.96348 | 0.381563 | 0.274055 | -3.51564 | 0.000439 |
| **TBL2** | AT | 89 | 89 | 0.92924 | 2.532585 | 0.280387 | 3.314131 | 0.000919 |

| | | | | | | | | |
|---|---|---|---|---|---|---|---|---|
| **TBL2** | AT | 89 | 89 | -0.92924 | 0.394854 | 0.280387 | -3.31413 | 0.000919 |
| **PPFIBP1** | AT | 89 | 89 | 0.835442 | 2.305832 | 0.274469 | 3.043844 | 0.002336 |
| **ARHGAP6** | AT | 94 | 84 | -0.73294 | 0.480493 | 0.274403 | -2.67104 | 0.007562 |
| **SCARB1** | ES | 89 | 89 | 0.827524 | 2.287648 | 0.272175 | 3.040417 | 0.002363 |
| **OSGEP** | AP | 89 | 89 | 1.037025 | 2.820812 | 0.280986 | 3.690657 | 0.000224 |
| **OSGEP** | AP | 89 | 89 | -1.03702 | 0.354508 | 0.280986 | -3.69066 | 0.000224 |
| **CRYGS** | AP | 89 | 89 | -0.69605 | 0.498552 | 0.271061 | -2.56787 | 0.010233 |
| **CRYGS** | AP | 89 | 89 | 0.696047 | 2.005809 | 0.271061 | 2.567867 | 0.010233 |
| **CTTN** | RI | 89 | 89 | 0.842353 | 2.321824 | 0.276992 | 3.041074 | 0.002357 |
| **TSPAN11** | AT | 89 | 89 | 0.755565 | 2.128813 | 0.271228 | 2.785717 | 0.005341 |
| **TSPAN11** | AT | 89 | 89 | -0.75556 | 0.469745 | 0.271228 | -2.78572 | 0.005341 |
| **ITSN1** | AT | 89 | 89 | 0.864779 | 2.374482 | 0.281752 | 3.069298 | 0.002146 |
| **EIF5A** | AP | 89 | 89 | 0.944571 | 2.571711 | 0.283293 | 3.334254 | 0.000855 |
| **TNFSF10** | ES | 90 | 88 | -0.73464 | 0.479677 | 0.268214 | -2.73902 | 0.006162 |
| **COMMD5** | AP | 89 | 89 | 0.756553 | 2.130919 | 0.288747 | 2.62012 | 0.00879 |
| **COMMD5** | AP | 89 | 89 | -0.75655 | 0.469281 | 0.288747 | -2.62012 | 0.00879 |
| **SEMA4D** | AP | 89 | 89 | 0.931508 | 2.538335 | 0.294416 | 3.163922 | 0.001557 |
| **RUNX1** | AP | 89 | 89 | 0.65507 | 1.925277 | 0.265354 | 2.468663 | 0.013562 |
| **CHID1** | ES | 89 | 89 | -0.89291 | 0.409464 | 0.27262 | -3.27527 | 0.001056 |
| **NUDT22** | ES | 90 | 88 | 0.921318 | 2.5126 | 0.277047 | 3.325495 | 0.000883 |
| **AMZ2** | ES | 89 | 89 | 0.692963 | 1.999632 | 0.276634 | 2.504978 | 0.012246 |
| **UFD1L** | AD | 91 | 87 | -0.56797 | 0.566673 | 0.262133 | -2.16674 | 0.030255 |
| **USP38** | AT | 90 | 88 | -0.85951 | 0.423368 | 0.270413 | -3.17853 | 0.00148 |
| **C11orf73** | RI | 89 | 89 | -1.09186 | 0.335593 | 0.284458 | -3.83837 | 0.000124 |
| **TAZ** | RI | 89 | 89 | -0.90217 | 0.405689 | 0.278795 | -3.23596 | 0.001212 |
| **THYN1** | RI | 89 | 89 | -0.94702 | 0.387897 | 0.289895 | -3.26675 | 0.001088 |
| **DBNDD1** | AP | 89 | 89 | -0.71175 | 0.490785 | 0.275886 | -2.57987 | 0.009884 |
| **SH2B1** | AD | 89 | 89 | -0.81192 | 0.444004 | 0.274743 | -2.9552 | 0.003125 |
| **SLC3A2** | AP | 89 | 89 | -0.82473 | 0.438353 | 0.267982 | -3.07755 | 0.002087 |
| **NOP16** | AT | 89 | 89 | 0.818381 | 2.266827 | 0.275853 | 2.966725 | 0.00301 |
| **NOP16** | AT | 89 | 89 | -0.81838 | 0.441145 | 0.275853 | -2.96672 | 0.00301 |
| **RPL13A** | AD | 90 | 88 | -0.89348 | 0.409229 | 0.280973 | -3.17995 | 0.001473 |
| **ARHGEF39** | AP | 89 | 89 | -0.83414 | 0.434247 | 0.272946 | -3.05607 | 0.002243 |
| **ARHGEF39** | AP | 89 | 89 | 0.834142 | 2.302837 | 0.272946 | 3.056067 | 0.002243 |
| **GSTK1** | RI | 89 | 89 | -0.96455 | 0.381156 | 0.287789 | -3.35158 | 0.000804 |
| **TTLL5** | AT | 89 | 89 | -0.94697 | 0.387914 | 0.283032 | -3.34581 | 0.00082 |
| **LDHA** | ES | 90 | 88 | -0.93602 | 0.392187 | 0.275638 | -3.39581 | 0.000684 |
| **PSMA6** | AP | 89 | 89 | 0.922317 | 2.51511 | 0.283133 | 3.25754 | 0.001124 |
| **PSMA6** | AP | 89 | 89 | -0.92232 | 0.397597 | 0.283133 | -3.25754 | 0.001124 |
| **ACOT8** | AA | 89 | 89 | -0.78399 | 0.456579 | 0.27146 | -2.88807 | 0.003876 |
| **AP3B2** | AT | 89 | 89 | 0.678917 | 1.97174 | 0.2686 | 2.527608 | 0.011484 |
| **AP3B2** | AT | 89 | 89 | -0.67892 | 0.507166 | 0.2686 | -2.52761 | 0.011484 |
| **RPS9** | RI | 90 | 88 | -0.55219 | 0.575688 | 0.274782 | -2.00955 | 0.044478 |
| **SLC35G1** | AT | 89 | 89 | 0.854141 | 2.349354 | 0.272937 | 3.129447 | 0.001751 |
| **RCE1** | ES | 89 | 89 | 1.023633 | 2.783287 | 0.285849 | 3.581022 | 0.000342 |
| **MRPL33** | ES | 89 | 89 | 0.692657 | 1.999019 | 0.267318 | 2.591137 | 0.009566 |
| **FAM72A** | AT | 89 | 89 | 0.580915 | 1.787674 | 0.267747 | 2.169642 | 0.030034 |

| | | | | | | | | |
|---|---|---|---|---|---|---|---|---|
| **ZNF410** | ES | 89 | 89 | -0.86401 | 0.421471 | 0.270891 | -3.18949 | 0.001425 |
| **RUVBL2** | ES | 89 | 89 | 0.825061 | 2.28202 | 0.272859 | 3.023766 | 0.002496 |
| **PCMT1** | AD | 89 | 89 | -0.66095 | 0.516358 | 0.266207 | -2.48286 | 0.013033 |
| **EBPL** | AT | 89 | 89 | 0.767448 | 2.154261 | 0.27307 | 2.810441 | 0.004947 |
| **EBPL** | AT | 89 | 89 | -0.76745 | 0.464196 | 0.27307 | -2.81044 | 0.004947 |
| **TESC** | ES | 89 | 89 | -0.85358 | 0.425887 | 0.27235 | -3.13414 | 0.001724 |
| **SNRNP70** | ES | 89 | 89 | -0.83088 | 0.435664 | 0.276805 | -3.00169 | 0.002685 |
| **SULT1A3** | AA | 90 | 88 | -0.74422 | 0.475104 | 0.268619 | -2.77055 | 0.005596 |
| **IGSF8** | AA | 89 | 89 | 0.949648 | 2.584799 | 0.276687 | 3.432204 | 0.000599 |
| **NUDT5** | RI | 90 | 88 | -0.98118 | 0.37487 | 0.286007 | -3.4306 | 0.000602 |
| **RBPMS** | AT | 89 | 89 | -0.92585 | 0.396195 | 0.279408 | -3.3136 | 0.000921 |
| **NNMT** | AD | 90 | 88 | -0.92152 | 0.397916 | 0.285015 | -3.23322 | 0.001224 |
| **SNRPN** | ES | 89 | 89 | -0.71951 | 0.486992 | 0.268545 | -2.67928 | 0.007378 |
| **SPAG7** | AP | 89 | 89 | 1.027241 | 2.793349 | 0.291491 | 3.524097 | 0.000425 |
| **SPAG7** | AP | 89 | 89 | -1.02724 | 0.357993 | 0.291491 | -3.5241 | 0.000425 |
| **C17orf104** | AT | 89 | 89 | 0.588753 | 1.801741 | 0.266158 | 2.212043 | 0.026964 |
| **C17orf104** | AT | 89 | 89 | -0.58875 | 0.555019 | 0.266158 | -2.21204 | 0.026964 |
| **WDR62** | RI | 90 | 88 | -0.79172 | 0.453066 | 0.276456 | -2.86381 | 0.004186 |
| **RPS9** | RI | 89 | 89 | -0.64439 | 0.524984 | 0.284704 | -2.26336 | 0.023613 |
| **IL1RAP** | AT | 89 | 89 | -0.89474 | 0.408716 | 0.276191 | -3.23955 | 0.001197 |
| **STYXL1** | ES | 90 | 88 | 0.926811 | 2.52644 | 0.275213 | 3.367613 | 0.000758 |
| **RFX5** | AT | 89 | 89 | 0.826874 | 2.286162 | 0.280375 | 2.949174 | 0.003186 |
| **RFX5** | AT | 89 | 89 | -0.82687 | 0.437414 | 0.280375 | -2.94917 | 0.003186 |
| **POLD4** | AD | 89 | 89 | -0.69529 | 0.498931 | 0.268481 | -2.5897 | 0.009606 |
| **PLEKHA5** | AT | 89 | 89 | -0.70156 | 0.495813 | 0.26999 | -2.59845 | 0.009364 |
| **RUNX1** | AP | 89 | 89 | -0.64796 | 0.523111 | 0.265818 | -2.43761 | 0.014785 |
| **RANBP1** | RI | 89 | 89 | -0.91888 | 0.398966 | 0.275927 | -3.33015 | 0.000868 |
| **UBE2L6** | AP | 89 | 89 | 0.970576 | 2.639466 | 0.271991 | 3.568418 | 0.000359 |
| **UBE2L6** | AP | 89 | 89 | -0.97058 | 0.378865 | 0.271991 | -3.56842 | 0.000359 |
| **VRK3** | AT | 89 | 89 | 0.918982 | 2.506738 | 0.283077 | 3.246407 | 0.001169 |
| **WIBG** | AP | 89 | 89 | -0.82529 | 0.438109 | 0.270235 | -3.05397 | 0.002258 |
| **LZTFL1** | ES | 89 | 89 | -0.87958 | 0.414959 | 0.270539 | -3.25119 | 0.001149 |
| **FAM175A** | AT | 89 | 89 | -0.78806 | 0.454726 | 0.268759 | -2.93222 | 0.003365 |
| **FAM175A** | AT | 89 | 89 | 0.78806 | 2.199127 | 0.268759 | 2.932223 | 0.003365 |
| **ZC3H14** | ES | 89 | 89 | 0.719125 | 2.052635 | 0.273213 | 2.632106 | 0.008486 |
| **MRPL2** | ES | 89 | 89 | 0.751697 | 2.120595 | 0.273405 | 2.74939 | 0.005971 |
| **FIS1** | AP | 90 | 88 | -0.54769 | 0.578286 | 0.275437 | -1.98843 | 0.046765 |
| **SLC15A3** | ES | 89 | 89 | -0.57082 | 0.565061 | 0.270065 | -2.11365 | 0.034545 |
| **PPM1K** | AT | 89 | 89 | 0.804983 | 2.236659 | 0.268107 | 3.002473 | 0.002678 |
| **TUBA1C** | AP | 98 | 80 | -0.82312 | 0.43906 | 0.278469 | -2.95587 | 0.003118 |
| **IRF9** | AT | 89 | 89 | 0.756752 | 2.131343 | 0.28023 | 2.700471 | 0.006924 |
| **IRF9** | AT | 89 | 89 | -0.75675 | 0.469188 | 0.28023 | -2.70047 | 0.006924 |
| **DST** | AT | 89 | 89 | 0.934384 | 2.545645 | 0.273803 | 3.41262 | 0.000643 |
| **ANTXR2** | RI | 90 | 88 | 0.612258 | 1.844591 | 0.267963 | 2.284863 | 0.022321 |
| **STIP1** | AP | 89 | 89 | 0.94725 | 2.57861 | 0.27498 | 3.444795 | 0.000571 |
| **STIP1** | AP | 89 | 89 | -0.94725 | 0.387806 | 0.27498 | -3.44479 | 0.000571 |
| **LRRC1** | AT | 90 | 88 | -0.78676 | 0.455318 | 0.283574 | -2.77444 | 0.00553 |

| | | | | | | | | |
|---|---|---|---|---|---|---|---|---|
| **PHF19** | AT | 89 | 89 | 0.896931 | 2.452067 | 0.290382 | 3.088802 | 0.00201 |
| **PHF19** | AT | 89 | 89 | -0.89693 | 0.407819 | 0.290382 | -3.0888 | 0.00201 |
| **GRAMD1C** | AP | 89 | 89 | 0.843556 | 2.324619 | 0.280516 | 3.007156 | 0.002637 |
| **VCAN** | AT | 89 | 89 | -0.71416 | 0.489602 | 0.264121 | -2.70392 | 0.006853 |
| **VCAN** | AT | 89 | 89 | 0.714161 | 2.042473 | 0.264121 | 2.70392 | 0.006853 |
| **PCBP2** | ES | 89 | 89 | -0.98061 | 0.375083 | 0.286798 | -3.41916 | 0.000628 |
| **LPCAT3** | RI | 89 | 89 | -1.0641 | 0.345039 | 0.288749 | -3.68519 | 0.000229 |
| **CD44** | ES | 89 | 89 | 0.855604 | 2.352794 | 0.273661 | 3.126504 | 0.001769 |
| **TM9SF1** | AT | 89 | 89 | 0.901961 | 2.464431 | 0.282842 | 3.18892 | 0.001428 |
| **TM9SF1** | AT | 89 | 89 | -0.90196 | 0.405773 | 0.282842 | -3.18892 | 0.001428 |
| **MAGED2** | AP | 89 | 89 | -0.67169 | 0.510843 | 0.269029 | -2.49673 | 0.012534 |
| **SON** | AT | 89 | 89 | -0.85665 | 0.424584 | 0.278211 | -3.07912 | 0.002076 |
| **TET2** | AT | 89 | 89 | -0.65936 | 0.517182 | 0.268685 | -2.45403 | 0.014127 |
| **TET2** | AT | 89 | 89 | 0.65936 | 1.933554 | 0.268685 | 2.454025 | 0.014127 |
| **UQCRQ** | RI | 89 | 89 | -0.94595 | 0.38831 | 0.283663 | -3.33477 | 0.000854 |
| **SNF8** | ES | 89 | 89 | -0.83977 | 0.431812 | 0.272124 | -3.08597 | 0.002029 |
| **GRK6** | AT | 89 | 89 | -0.69137 | 0.500891 | 0.279109 | -2.47705 | 0.013247 |
| **HMGA2** | AT | 89 | 89 | 0.931543 | 2.538423 | 0.272122 | 3.423262 | 0.000619 |
| **CDC37** | ES | 89 | 89 | -0.626 | 0.534728 | 0.269719 | -2.32093 | 0.020291 |
| **RPS16** | RI | 91 | 87 | -0.72625 | 0.483719 | 0.270035 | -2.68947 | 0.007157 |
| **CPSF3L** | ES | 90 | 88 | 1.082744 | 2.95277 | 0.2894 | 3.741335 | 0.000183 |
| **TSG101** | AT | 89 | 89 | 0.962984 | 2.619502 | 0.281522 | 3.42064 | 0.000625 |
| **TSG101** | AT | 89 | 89 | -0.96298 | 0.381752 | 0.281522 | -3.42064 | 0.000625 |
| **OBSL1** | AT | 89 | 89 | 0.798437 | 2.222066 | 0.276972 | 2.882736 | 0.003942 |
| **TTC21B** | AT | 89 | 89 | 0.979549 | 2.663255 | 0.282982 | 3.461524 | 0.000537 |
| **TTC21B** | AT | 89 | 89 | -0.97955 | 0.37548 | 0.282982 | -3.46152 | 0.000537 |
| **PTPN13** | AT | 89 | 89 | -0.72919 | 0.482297 | 0.272842 | -2.67259 | 0.007527 |
| **PTPN13** | AT | 89 | 89 | 0.729194 | 2.07341 | 0.272842 | 2.672591 | 0.007527 |
| **UBR4** | AP | 89 | 89 | -0.78012 | 0.458349 | 0.280199 | -2.78418 | 0.005366 |
| **PSMA1** | AP | 89 | 89 | -0.71754 | 0.48795 | 0.266401 | -2.69346 | 0.007071 |
| **PSMA1** | AP | 89 | 89 | 0.717542 | 2.049391 | 0.266401 | 2.693465 | 0.007071 |
| **YLPM1** | AT | 89 | 89 | -0.77115 | 0.462482 | 0.264207 | -2.91873 | 0.003515 |
| **YLPM1** | AT | 89 | 89 | 0.771148 | 2.162247 | 0.264207 | 2.918726 | 0.003515 |
| **PHF21A** | ME | 89 | 89 | -0.83955 | 0.431905 | 0.27946 | -3.00418 | 0.002663 |
| **CHCHD1** | RI | 89 | 89 | -0.93331 | 0.393251 | 0.290273 | -3.21527 | 0.001303 |
| **SLC39A11** | AD | 90 | 88 | -0.90475 | 0.404645 | 0.275192 | -3.28769 | 0.00101 |
| **FUZ** | AD | 89 | 89 | -0.87874 | 0.415307 | 0.274793 | -3.19781 | 0.001385 |
| **RGS3** | AP | 89 | 89 | 0.810011 | 2.247933 | 0.274292 | 2.953097 | 0.003146 |
| **FAM3A** | AT | 89 | 89 | 0.828013 | 2.288765 | 0.279335 | 2.964223 | 0.003034 |
| **FAM3A** | AT | 89 | 89 | -0.82801 | 0.436917 | 0.279335 | -2.96422 | 0.003034 |
| **PSMA3** | ES | 89 | 89 | 0.575214 | 1.777512 | 0.265473 | 2.166752 | 0.030254 |
| **MUC1** | ES | 90 | 88 | -0.61296 | 0.541743 | 0.272999 | -2.2453 | 0.024749 |
| **METTL17** | RI | 89 | 89 | -1.08947 | 0.336396 | 0.294987 | -3.69327 | 0.000221 |
| **FAM173A** | RI | 89 | 89 | -0.95162 | 0.386115 | 0.288718 | -3.29602 | 0.000981 |
| **ECSIT** | RI | 90 | 88 | -0.82701 | 0.437356 | 0.281054 | -2.94253 | 0.003255 |
| **EPB41L5** | AT | 89 | 89 | 0.776318 | 2.173455 | 0.276247 | 2.810236 | 0.004951 |
| **EPB41L5** | AT | 89 | 89 | -0.77632 | 0.460097 | 0.276247 | -2.81024 | 0.004951 |

| | | | | | | | | |
|---|---|---|---|---|---|---|---|---|
| **ATG12** | AD | 90 | 88 | -0.7317 | 0.48109 | 0.272754 | -2.68264 | 0.007304 |
| **PPRC1** | ES | 89 | 89 | 0.751999 | 2.121235 | 0.269309 | 2.792332 | 0.005233 |
| **RPS25** | AD | 90 | 88 | -0.82793 | 0.436955 | 0.278571 | -2.97204 | 0.002958 |
| **NLRP1** | AT | 89 | 89 | -0.84477 | 0.429654 | 0.269624 | -3.13316 | 0.001729 |
| **ZBTB44** | RI | 89 | 89 | 0.752354 | 2.121989 | 0.27262 | 2.759712 | 0.005785 |
| **NIP7** | AT | 89 | 89 | -0.87865 | 0.415343 | 0.273629 | -3.2111 | 0.001322 |
| **NIP7** | AT | 89 | 89 | 0.878652 | 2.407651 | 0.273629 | 3.211102 | 0.001322 |
| **TTC9C** | AT | 89 | 89 | -0.83336 | 0.434585 | 0.270236 | -3.08384 | 0.002043 |
| **TTC9C** | AT | 89 | 89 | 0.833365 | 2.301048 | 0.270236 | 3.083839 | 0.002043 |
| **COX4I1** | AD | 89 | 89 | -0.6575 | 0.518146 | 0.266434 | -2.46777 | 0.013596 |
| **PLCB2** | AT | 89 | 89 | 0.766454 | 2.152121 | 0.268205 | 2.857718 | 0.004267 |
| **PLCB2** | AT | 89 | 89 | -0.76645 | 0.464658 | 0.268205 | -2.85772 | 0.004267 |
| **DUSP22** | RI | 89 | 89 | -0.81922 | 0.440775 | 0.280233 | -2.92336 | 0.003463 |
| **SLC25A10** | AA | 89 | 89 | -0.63262 | 0.531196 | 0.264442 | -2.3923 | 0.016743 |
| **UBR4** | AP | 89 | 89 | 0.765436 | 2.149931 | 0.271916 | 2.814969 | 0.004878 |
| **NUDT22** | AT | 89 | 89 | 0.754442 | 2.126424 | 0.271609 | 2.777678 | 0.005475 |
| **NUDT22** | AT | 89 | 89 | -0.75444 | 0.470273 | 0.271609 | -2.77768 | 0.005475 |
| **TMX2** | ES | 89 | 89 | -0.60961 | 0.543561 | 0.272763 | -2.23496 | 0.02542 |
| **PLEKHA5** | AT | 89 | 89 | 0.775565 | 2.171818 | 0.267585 | 2.89838 | 0.003751 |
| **SPOCK2** | AT | 89 | 89 | -0.78338 | 0.456858 | 0.275193 | -2.84666 | 0.004418 |
| **SPOCK2** | AT | 89 | 89 | 0.783382 | 2.188863 | 0.275193 | 2.84666 | 0.004418 |
| **CMC2** | AP | 89 | 89 | -0.714 | 0.48968 | 0.272685 | -2.61841 | 0.008834 |
| **THEM4** | AT | 89 | 89 | 0.768316 | 2.156132 | 0.272993 | 2.814415 | 0.004887 |
| **THEM4** | AT | 89 | 89 | -0.76832 | 0.463794 | 0.272993 | -2.81442 | 0.004887 |
| **RDH11** | ES | 89 | 89 | -0.7302 | 0.481814 | 0.266909 | -2.73575 | 0.006224 |
| **DBNDD2** | AP | 89 | 89 | 0.72137 | 2.05725 | 0.2766 | 2.60799 | 0.009108 |
| **FAM92A1** | AT | 89 | 89 | -1.02385 | 0.35921 | 0.284028 | -3.60474 | 0.000312 |
| **SDF4** | AT | 92 | 86 | 0.597825 | 1.818159 | 0.269454 | 2.218652 | 0.02651 |
| **NISCH** | AT | 89 | 89 | 1.004838 | 2.731465 | 0.285177 | 3.523562 | 0.000426 |
| **NISCH** | AT | 89 | 89 | -1.00484 | 0.366104 | 0.285177 | -3.52356 | 0.000426 |
| **TMED4** | AT | 89 | 89 | 0.826552 | 2.285425 | 0.300368 | 2.751794 | 0.005927 |
| **TMED4** | AT | 89 | 89 | -0.82655 | 0.437555 | 0.300368 | -2.75179 | 0.005927 |
| **CENPK** | AT | 89 | 89 | 0.680927 | 1.975708 | 0.266681 | 2.55334 | 0.01067 |
| **CENPK** | AT | 89 | 89 | -0.68093 | 0.506148 | 0.266681 | -2.55334 | 0.01067 |
| **RPS6** | AD | 110 | 68 | 0.696825 | 2.00737 | 0.263389 | 2.645613 | 0.008154 |
| **RPL29** | AT | 96 | 82 | 0.694387 | 2.002482 | 0.269832 | 2.573403 | 0.01007 |
| **SULT1A3** | AP | 89 | 89 | -0.88681 | 0.411967 | 0.28004 | -3.16674 | 0.001542 |
| **CD99L2** | AT | 89 | 89 | 0.810638 | 2.249344 | 0.272284 | 2.977178 | 0.002909 |
| **CD99L2** | AT | 89 | 89 | -0.81064 | 0.444574 | 0.272284 | -2.97718 | 0.002909 |
| **GGPS1** | AP | 89 | 89 | 0.807311 | 2.241872 | 0.276402 | 2.920786 | 0.003491 |
| **RTEL1** | AT | 89 | 89 | -0.58949 | 0.554608 | 0.267572 | -2.20312 | 0.027586 |
| **RTEL1** | AT | 89 | 89 | 0.589493 | 1.803074 | 0.267572 | 2.203118 | 0.027586 |
| **COASY** | RI | 89 | 89 | 0.854429 | 2.350031 | 0.277714 | 3.076648 | 0.002093 |
| **SRSF1** | RI | 89 | 89 | 0.916146 | 2.499638 | 0.277017 | 3.30718 | 0.000942 |
| **AUP1** | AA | 90 | 88 | -0.95687 | 0.384093 | 0.279337 | -3.42551 | 0.000614 |
| **SAP30BP** | ES | 89 | 89 | -0.79304 | 0.452465 | 0.273393 | -2.90075 | 0.003723 |
| **YIPF3** | RI | 90 | 88 | -0.96303 | 0.381736 | 0.28298 | -3.40316 | 0.000666 |

| | | | | | | | | |
|---|---|---|---|---|---|---|---|---|
| **FOXRED1** | ES | 89 | 89 | 0.711217 | 2.036468 | 0.273 | 2.605192 | 0.009182 |
| **RABL5** | AA | 89 | 89 | 0.695201 | 2.004113 | 0.268298 | 2.591157 | 0.009565 |
| **TRNT1** | ES | 89 | 89 | -0.81582 | 0.442277 | 0.268216 | -3.04165 | 0.002353 |
| **C11orf31** | AT | 89 | 89 | 0.849835 | 2.339261 | 0.276923 | 3.068845 | 0.002149 |
| **C11orf31** | AT | 89 | 89 | -0.84983 | 0.427485 | 0.276923 | -3.06885 | 0.002149 |
| **KXD1** | ES | 89 | 89 | 0.861571 | 2.366877 | 0.275339 | 3.12913 | 0.001753 |
| **ITGB1BP1** | RI | 89 | 89 | -0.88893 | 0.411095 | 0.273257 | -3.25309 | 0.001142 |
| **MED15** | RI | 89 | 89 | -0.76031 | 0.467521 | 0.27579 | -2.75685 | 0.005836 |
| **PARD6G** | AT | 89 | 89 | 0.939706 | 2.559229 | 0.283747 | 3.311777 | 0.000927 |
| **PARD6G** | AT | 89 | 89 | -0.93971 | 0.390743 | 0.283747 | -3.31178 | 0.000927 |
| **SH3YL1** | ES | 89 | 89 | -0.61928 | 0.538333 | 0.264683 | -2.3397 | 0.019299 |
| **IP6K2** | RI | 89 | 89 | -0.95361 | 0.385348 | 0.282105 | -3.38033 | 0.000724 |
| **CLASRP** | AD | 89 | 89 | -0.79314 | 0.45242 | 0.273879 | -2.89596 | 0.00378 |
| **JPH2** | AT | 89 | 89 | -0.71366 | 0.489846 | 0.268685 | -2.65613 | 0.007904 |
| **JPH2** | AT | 89 | 89 | 0.713664 | 2.041457 | 0.268685 | 2.656135 | 0.007904 |
| **C1orf198** | AP | 89 | 89 | 0.626304 | 1.870683 | 0.277882 | 2.253843 | 0.024206 |
| **UGGT2** | AT | 89 | 89 | 0.756448 | 2.130695 | 0.268377 | 2.818606 | 0.004823 |
| **MICAL1** | AP | 89 | 89 | -0.74796 | 0.473329 | 0.288878 | -2.58921 | 0.00962 |
| **MICAL1** | AP | 89 | 89 | 0.747964 | 2.112694 | 0.288878 | 2.589205 | 0.00962 |
| **HS2ST1** | AT | 89 | 89 | 0.710533 | 2.035075 | 0.272709 | 2.605459 | 0.009175 |
| **PQLC1** | ES | 89 | 89 | -0.74371 | 0.475349 | 0.271126 | -2.74303 | 0.006087 |
| **UBE2I** | AP | 89 | 89 | -0.84613 | 0.429073 | 0.279849 | -3.02351 | 0.002499 |
| **SUPT4H1** | AD | 89 | 89 | -0.85143 | 0.426804 | 0.286273 | -2.97419 | 0.002938 |
| **SRSF2** | ES | 89 | 89 | -0.83475 | 0.433981 | 0.274397 | -3.04214 | 0.002349 |
| **KCNMA1** | AT | 89 | 89 | -0.61855 | 0.538722 | 0.26841 | -2.30451 | 0.021194 |
| **SLC39A13** | RI | 89 | 89 | -0.72599 | 0.483847 | 0.272696 | -2.66226 | 0.007762 |
| **YIF1A** | RI | 89 | 89 | -0.83363 | 0.43447 | 0.278909 | -2.98889 | 0.0028 |
| **THYN1** | RI | 89 | 89 | -0.68448 | 0.504351 | 0.275177 | -2.48743 | 0.012867 |
| **PRMT2** | ES | 90 | 88 | -0.67511 | 0.509102 | 0.265736 | -2.54052 | 0.011069 |
| **EIF3M** | ES | 89 | 89 | 0.55577 | 1.743284 | 0.265163 | 2.095955 | 0.036086 |
| **MAGT1** | AT | 89 | 89 | 0.807053 | 2.241292 | 0.271285 | 2.974922 | 0.002931 |
| **MAGT1** | AT | 89 | 89 | -0.80705 | 0.446171 | 0.271285 | -2.97492 | 0.002931 |
| **JUP** | AT | 89 | 89 | -0.65604 | 0.5189 | 0.270245 | -2.42759 | 0.015199 |
| **JUP** | AT | 89 | 89 | 0.656044 | 1.927153 | 0.270245 | 2.427593 | 0.015199 |
| **RANBP1** | RI | 89 | 89 | -0.89241 | 0.409669 | 0.27573 | -3.23652 | 0.00121 |
| **CDC42** | AT | 93 | 85 | -1.03931 | 0.353698 | 0.297137 | -3.49776 | 0.000469 |
| **TBX6** | AT | 89 | 89 | -0.70731 | 0.492969 | 0.277498 | -2.54887 | 0.010807 |
| **TBX6** | AT | 89 | 89 | 0.707308 | 2.028523 | 0.277498 | 2.548875 | 0.010807 |
| **TMEM205** | RI | 89 | 89 | -1.01549 | 0.362224 | 0.290079 | -3.50074 | 0.000464 |
| **XBP1** | RI | 89 | 89 | -0.81746 | 0.441553 | 0.283615 | -2.88227 | 0.003948 |
| **DST** | AT | 89 | 89 | 0.718812 | 2.051994 | 0.270554 | 2.656819 | 0.007888 |
| **HMGB1** | AA | 89 | 89 | -0.86439 | 0.421309 | 0.294658 | -2.93353 | 0.003351 |
| **SRSF2** | RI | 89 | 89 | 0.823421 | 2.27828 | 0.269647 | 3.0537 | 0.00226 |
| **PTK2** | AT | 89 | 89 | 0.727713 | 2.070341 | 0.271341 | 2.681913 | 0.00732 |
| **PTK2** | AT | 89 | 89 | -0.72771 | 0.483012 | 0.271341 | -2.68191 | 0.00732 |
| **CELF1** | AA | 89 | 89 | -0.87927 | 0.415087 | 0.280202 | -3.13798 | 0.001701 |
| **EIF5A** | AP | 90 | 88 | -0.7979 | 0.450274 | 0.276482 | -2.88591 | 0.003903 |

| | | | | | | | | |
|---|---|---|---|---|---|---|---|---|
| **AUNIP** | AT | 90 | 88 | -0.78714 | 0.455146 | 0.27454 | -2.86711 | 0.004142 |
| **C14orf2** | ES | 89 | 89 | 0.880952 | 2.413195 | 0.274595 | 3.208189 | 0.001336 |
| **GUK1** | AP | 90 | 88 | -0.67878 | 0.507235 | 0.272674 | -2.48935 | 0.012798 |
| **CENPN** | AT | 90 | 88 | 0.737389 | 2.090471 | 0.267521 | 2.756376 | 0.005845 |
| **CARD14** | AT | 90 | 88 | 0.985228 | 2.678423 | 0.286468 | 3.439224 | 0.000583 |
| **ATF7** | AT | 89 | 89 | -0.94572 | 0.388401 | 0.280046 | -3.37701 | 0.000733 |
| **PCYT2** | ES | 89 | 89 | 0.862277 | 2.368549 | 0.275429 | 3.130674 | 0.001744 |
| **FBXW9** | AT | 89 | 89 | 0.895904 | 2.449548 | 0.289445 | 3.095244 | 0.001967 |
| **FBXW9** | AT | 89 | 89 | -0.8959 | 0.408239 | 0.289445 | -3.09524 | 0.001967 |
| **USE1** | AT | 89 | 89 | 0.806168 | 2.239311 | 0.284117 | 2.837456 | 0.004547 |
| **USE1** | AT | 89 | 89 | -0.80617 | 0.446566 | 0.284117 | -2.83746 | 0.004547 |
| **UQCRQ** | RI | 92 | 86 | -0.85201 | 0.426557 | 0.293518 | -2.90275 | 0.003699 |
| **HSBP1L1** | ES | 89 | 89 | -0.79329 | 0.452356 | 0.268845 | -2.95072 | 0.00317 |
| **AGPAT4** | AT | 89 | 89 | 0.529469 | 1.69803 | 0.264964 | 1.998271 | 0.045687 |
| **AGPAT4** | AT | 89 | 89 | -0.52947 | 0.588918 | 0.264964 | -1.99827 | 0.045687 |
| **IFI27L1** | AA | 89 | 89 | 0.630481 | 1.878514 | 0.265168 | 2.377671 | 0.017422 |
| **MAF** | RI | 89 | 89 | 0.637709 | 1.892141 | 0.266884 | 2.389463 | 0.016873 |
| **ASF1B** | AT | 89 | 89 | 0.748232 | 2.113261 | 0.269667 | 2.774649 | 0.005526 |
| **ASF1B** | AT | 89 | 89 | -0.74823 | 0.473202 | 0.269667 | -2.77465 | 0.005526 |
| **CCDC25** | ES | 89 | 89 | 0.77 | 2.159766 | 0.270283 | 2.848864 | 0.004388 |
| **DEF8** | ES | 89 | 89 | -0.84998 | 0.427425 | 0.273226 | -3.11089 | 0.001865 |
| **EEF1B2** | AA | 89 | 89 | 0.696117 | 2.005949 | 0.273016 | 2.549729 | 0.010781 |
| **PPP1CC** | RI | 89 | 89 | 0.8667 | 2.379048 | 0.27301 | 3.174608 | 0.0015 |
| **PDLIM5** | AT | 89 | 89 | 0.743906 | 2.104139 | 0.274365 | 2.711378 | 0.0067 |
| **EDF1** | RI | 89 | 89 | -0.92089 | 0.398163 | 0.277609 | -3.31723 | 0.000909 |
| **GADL1** | AT | 107 | 71 | -0.96382 | 0.381435 | 0.318239 | -3.02859 | 0.002457 |
| **RFX3** | AT | 89 | 89 | -0.89568 | 0.40833 | 0.277431 | -3.22847 | 0.001245 |
| **PPA1** | ES | 89 | 89 | -0.83534 | 0.433726 | 0.271863 | -3.07266 | 0.002122 |
| **SNRNP70** | ES | 89 | 89 | -0.90195 | 0.405777 | 0.282213 | -3.19599 | 0.001394 |
| **USP38** | AT | 90 | 88 | 0.789173 | 2.201576 | 0.268817 | 2.935732 | 0.003328 |
| **DGKE** | AT | 89 | 89 | 1.059143 | 2.883899 | 0.278991 | 3.796332 | 0.000147 |
| **DGKE** | AT | 89 | 89 | -1.05914 | 0.346753 | 0.278991 | -3.79633 | 0.000147 |
| **C21orf59** | RI | 89 | 89 | -0.8945 | 0.408813 | 0.275423 | -3.24773 | 0.001163 |
| **TNFRSF14** | AA | 89 | 89 | -0.78277 | 0.457137 | 0.276155 | -2.83454 | 0.004589 |
| **ENDOV** | AT | 89 | 89 | -0.65181 | 0.5211 | 0.267827 | -2.4337 | 0.014945 |
| **VAMP8** | ES | 89 | 89 | -0.77254 | 0.461838 | 0.2769 | -2.78996 | 0.005271 |
| **CAMTA1** | AT | 89 | 89 | 0.725195 | 2.065135 | 0.266887 | 2.717242 | 0.006583 |
| **SCAF11** | AT | 89 | 89 | -0.75569 | 0.469686 | 0.272868 | -2.76944 | 0.005615 |
| **CAPG** | AP | 89 | 89 | 0.540347 | 1.716603 | 0.265682 | 2.033812 | 0.041971 |
| **CAPG** | AP | 89 | 89 | -0.54035 | 0.582546 | 0.265682 | -2.03381 | 0.041971 |
| **SYT15** | AA | 89 | 89 | 0.552704 | 1.737945 | 0.264605 | 2.088787 | 0.036727 |
| **CRIP1** | AT | 89 | 89 | 0.836635 | 2.308585 | 0.276128 | 3.029883 | 0.002446 |
| **CRIP1** | AT | 89 | 89 | -0.83663 | 0.433166 | 0.276128 | -3.02988 | 0.002446 |
| **PPIL2** | RI | 89 | 89 | 0.743953 | 2.104237 | 0.266137 | 2.795374 | 0.005184 |
| **PSMD4** | AD | 89 | 89 | -0.92105 | 0.398103 | 0.27488 | -3.35072 | 0.000806 |
| **TROAP** | AT | 89 | 89 | -0.88179 | 0.41404 | 0.288014 | -3.06164 | 0.002201 |
| **RPL28** | RI | 89 | 89 | -0.71052 | 0.491387 | 0.269813 | -2.63339 | 0.008454 |

| | | | | | | | | |
|---|---|---|---|---|---|---|---|---|
| **CLDND1** | AT | 90 | 88 | -0.83616 | 0.433372 | 0.277038 | -3.01821 | 0.002543 |
| **CDC42** | AT | 90 | 88 | 0.891645 | 2.439139 | 0.282361 | 3.157819 | 0.00159 |
| **VIM** | RI | 89 | 89 | -0.89886 | 0.407035 | 0.302266 | -2.97373 | 0.002942 |
| **GSTK1** | AT | 89 | 89 | 0.768924 | 2.157443 | 0.280261 | 2.743599 | 0.006077 |
| **GSTK1** | AT | 89 | 89 | -0.76892 | 0.463512 | 0.280261 | -2.7436 | 0.006077 |
| **LIG1** | AD | 89 | 89 | -0.53128 | 0.58785 | 0.269638 | -1.97036 | 0.048797 |
| **PSMA7** | AT | 89 | 89 | 0.797419 | 2.219805 | 0.277626 | 2.872279 | 0.004075 |
| **PSMA7** | AT | 89 | 89 | -0.79742 | 0.45049 | 0.277626 | -2.87228 | 0.004075 |
| **CDC14B** | AT | 89 | 89 | -0.74291 | 0.475729 | 0.270763 | -2.74376 | 0.006074 |
| **CDC14B** | AT | 89 | 89 | 0.742908 | 2.102039 | 0.270763 | 2.743758 | 0.006074 |
| **SET** | AP | 89 | 89 | 0.778879 | 2.179029 | 0.278751 | 2.794179 | 0.005203 |
| **TIMM17B** | RI | 89 | 89 | -0.85546 | 0.425087 | 0.281892 | -3.03472 | 0.002408 |
| **TMEM126I** | ES | 89 | 89 | 0.675913 | 1.965827 | 0.270568 | 2.498123 | 0.012485 |
| **XAF1** | AT | 90 | 88 | 0.718375 | 2.051097 | 0.26608 | 2.699843 | 0.006937 |
| **APEH** | ES | 90 | 88 | 0.865577 | 2.376376 | 0.278185 | 3.111519 | 0.001861 |
| **WDR19** | AT | 91 | 87 | -0.5568 | 0.573038 | 0.265574 | -2.0966 | 0.036029 |
| **HYI** | AT | 89 | 89 | -0.84063 | 0.431438 | 0.282915 | -2.97132 | 0.002965 |
| **HYI** | AT | 89 | 89 | 0.840631 | 2.317829 | 0.282915 | 2.971319 | 0.002965 |
| **C10orf11** | AP | 120 | 58 | -0.79857 | 0.449973 | 0.335708 | -2.37875 | 0.017371 |
| **SLC35G1** | AT | 89 | 89 | -0.68406 | 0.504564 | 0.265251 | -2.57891 | 0.009911 |
| **RPL21** | AD | 89 | 89 | -0.6331 | 0.530945 | 0.26967 | -2.34767 | 0.018891 |
| **RPS9** | RI | 90 | 88 | -0.78945 | 0.454097 | 0.287997 | -2.74116 | 0.006122 |
| **DBNDD2** | AP | 89 | 89 | -0.77126 | 0.462431 | 0.276133 | -2.79307 | 0.005221 |
| **TXNL4A** | ES | 89 | 89 | 0.635247 | 1.887489 | 0.268985 | 2.361644 | 0.018194 |
| **CAPN10** | AT | 89 | 89 | 0.858138 | 2.358764 | 0.272023 | 3.154648 | 0.001607 |
| **CAPN10** | AT | 89 | 89 | -0.85814 | 0.423951 | 0.272023 | -3.15465 | 0.001607 |
| **RBM33** | AT | 89 | 89 | 0.785219 | 2.192886 | 0.279565 | 2.808719 | 0.004974 |
| **RBM33** | AT | 89 | 89 | -0.78522 | 0.45602 | 0.279565 | -2.80872 | 0.004974 |
| **LYPLA2** | RI | 89 | 89 | -0.76123 | 0.467092 | 0.273661 | -2.78165 | 0.005408 |
| **NUTF2** | AP | 89 | 89 | -0.7678 | 0.464035 | 0.270578 | -2.83762 | 0.004545 |
| **NUTF2** | AP | 89 | 89 | 0.767796 | 2.155012 | 0.270578 | 2.837618 | 0.004545 |
| **PODXL** | ES | 89 | 89 | -0.76055 | 0.467408 | 0.270874 | -2.80777 | 0.004989 |
| **CAMLG** | ES | 89 | 89 | 0.73493 | 2.085337 | 0.268154 | 2.740699 | 0.006131 |
| **EPB41L4B** | AT | 89 | 89 | -0.77068 | 0.462697 | 0.27184 | -2.83506 | 0.004582 |
| **EPB41L4B** | AT | 89 | 89 | 0.770683 | 2.161242 | 0.27184 | 2.83506 | 0.004582 |
| **CALM1** | ES | 89 | 89 | 0.801758 | 2.229458 | 0.273201 | 2.934685 | 0.003339 |
| **TCF4** | AP | 89 | 89 | 0.686933 | 1.98761 | 0.269416 | 2.549714 | 0.010781 |
| **UCK2** | ES | 90 | 88 | 0.605181 | 1.831583 | 0.267647 | 2.261113 | 0.023752 |
| **PPOX** | AT | 90 | 88 | 0.863525 | 2.371505 | 0.271485 | 3.180741 | 0.001469 |
| **TMPO** | AT | 89 | 89 | -0.76535 | 0.465169 | 0.274365 | -2.78955 | 0.005278 |
| **ATPAF2** | AT | 89 | 89 | 0.836729 | 2.308802 | 0.272563 | 3.069858 | 0.002142 |
| **LSM12** | ES | 89 | 89 | -0.83501 | 0.433872 | 0.268543 | -3.10939 | 0.001875 |
| **C17orf62** | ES | 89 | 89 | -0.68948 | 0.501836 | 0.271786 | -2.53685 | 0.011185 |
| **PRMT1** | ES | 89 | 89 | -0.71689 | 0.488269 | 0.2727 | -2.62886 | 0.008567 |
| **TTLL5** | AT | 89 | 89 | 0.83124 | 2.296164 | 0.273993 | 3.033796 | 0.002415 |
| **TMEM205** | RI | 89 | 89 | -0.81094 | 0.444441 | 0.282504 | -2.87053 | 0.004098 |
| **CASC4** | AT | 91 | 87 | 0.602693 | 1.827033 | 0.269735 | 2.23439 | 0.025457 |

| | | | | | | | | |
|---|---|---|---|---|---|---|---|---|
| **EXOSC9** | ES | 89 | 89 | 0.729652 | 2.074359 | 0.268239 | 2.720156 | 0.006525 |
| **STOML2** | RI | 89 | 89 | -0.94856 | 0.387298 | 0.283599 | -3.34472 | 0.000824 |
| **WNK1** | AT | 89 | 89 | 0.856344 | 2.354536 | 0.271259 | 3.156928 | 0.001594 |
| **C19orf54** | AP | 89 | 89 | -0.65852 | 0.517616 | 0.270118 | -2.43791 | 0.014773 |
| **ZNF7** | AT | 89 | 89 | -0.72845 | 0.482659 | 0.268782 | -2.71017 | 0.006725 |
| **TXNDC17** | AD | 91 | 87 | 0.654395 | 1.923978 | 0.265707 | 2.462846 | 0.013784 |
| **ARL6IP4** | RI | 89 | 89 | -0.81963 | 0.440594 | 0.273319 | -2.99881 | 0.00271 |
| **PIGL** | AT | 89 | 89 | -0.72242 | 0.485575 | 0.273162 | -2.64466 | 0.008177 |
| **BOP1** | AT | 89 | 89 | 0.816595 | 2.262781 | 0.275962 | 2.959086 | 0.003086 |
| **BOP1** | AT | 89 | 89 | -0.81659 | 0.441934 | 0.275962 | -2.95909 | 0.003086 |
| **C19orf48** | RI | 89 | 89 | -0.89525 | 0.408504 | 0.28108 | -3.18506 | 0.001447 |
| **STRADA** | RI | 89 | 89 | -0.90403 | 0.404935 | 0.283005 | -3.19439 | 0.001401 |
| **PPAN** | AT | 89 | 89 | 0.761609 | 2.141719 | 0.275044 | 2.769045 | 0.005622 |
| **PPAN** | AT | 89 | 89 | -0.76161 | 0.466915 | 0.275044 | -2.76905 | 0.005622 |
| **ZFAND2B** | RI | 89 | 89 | -0.78571 | 0.455794 | 0.279704 | -2.80909 | 0.004968 |
| **AP1S3** | AT | 89 | 89 | -0.83345 | 0.434547 | 0.268706 | -3.10173 | 0.001924 |
| **ACAD10** | AT | 111 | 67 | -0.64458 | 0.524885 | 0.289911 | -2.22335 | 0.026192 |
| **HNRNPC** | AA | 89 | 89 | -0.69151 | 0.500821 | 0.273916 | -2.52452 | 0.011586 |
| **CYB561** | AA | 90 | 88 | -0.77942 | 0.458673 | 0.276265 | -2.82127 | 0.004783 |
| **SHARPIN** | ES | 89 | 89 | 0.82082 | 2.272361 | 0.279558 | 2.936136 | 0.003323 |
| **ZNF644** | ES | 89 | 89 | -0.56982 | 0.565626 | 0.264334 | -2.1557 | 0.031107 |
| **TRPT1** | ES | 89 | 89 | 0.719431 | 2.053264 | 0.269773 | 2.666802 | 0.007658 |
| **CBFA2T2** | AP | 89 | 89 | -0.83668 | 0.433147 | 0.275441 | -3.03759 | 0.002385 |
| **STAU1** | ES | 89 | 89 | -0.80546 | 0.44688 | 0.268974 | -2.99458 | 0.002748 |
| **SLC9B2** | AT | 89 | 89 | 0.719485 | 2.053376 | 0.269534 | 2.669369 | 0.007599 |
| **SLC9B2** | AT | 89 | 89 | -0.71949 | 0.487003 | 0.269534 | -2.66937 | 0.007599 |
| **C15orf57** | AT | 89 | 89 | -0.72891 | 0.482432 | 0.270056 | -2.69912 | 0.006952 |
| **COX4I1** | RI | 90 | 88 | -1.11079 | 0.3293 | 0.297582 | -3.7327 | 0.000189 |
| **SUGT1** | ES | 89 | 89 | -0.60075 | 0.548399 | 0.267023 | -2.24982 | 0.024461 |
| **HYPK** | AP | 89 | 89 | -0.79102 | 0.453382 | 0.278 | -2.84539 | 0.004436 |
| **HYPK** | AP | 89 | 89 | 0.791019 | 2.205644 | 0.278 | 2.845388 | 0.004436 |
| **GRAMD1C** | AP | 89 | 89 | -0.69023 | 0.50146 | 0.277393 | -2.48828 | 0.012836 |
| **UPP1** | ES | 89 | 89 | -0.61043 | 0.543119 | 0.273089 | -2.23527 | 0.0254 |
| **SRP9** | ES | 89 | 89 | -0.61219 | 0.542162 | 0.267772 | -2.28624 | 0.02224 |
| **CARS** | AD | 89 | 89 | -0.99627 | 0.369254 | 0.292165 | -3.40996 | 0.00065 |
| **MEF2BNB** | ES | 89 | 89 | -0.82656 | 0.437552 | 0.271824 | -3.04079 | 0.00236 |
| **ZNF250** | AT | 89 | 89 | -0.77064 | 0.462716 | 0.271425 | -2.83925 | 0.004522 |
| **RPL36** | AD | 89 | 89 | -0.74429 | 0.475072 | 0.265853 | -2.79962 | 0.005116 |
| **MOB1A** | AT | 89 | 89 | 0.906896 | 2.476624 | 0.270944 | 3.347169 | 0.000816 |
| **MOB1A** | AT | 89 | 89 | -0.9069 | 0.403775 | 0.270944 | -3.34717 | 0.000816 |
| **NFAM1** | AT | 89 | 89 | -0.67519 | 0.509058 | 0.272498 | -2.47779 | 0.01322 |
| **NFAM1** | AT | 89 | 89 | 0.675193 | 1.964411 | 0.272498 | 2.477791 | 0.01322 |
| **CDK11A** | AD | 89 | 89 | 0.617294 | 1.853904 | 0.26532 | 2.326597 | 0.019987 |
| **NUMA1** | ES | 89 | 89 | 0.735174 | 2.085846 | 0.270302 | 2.719824 | 0.006532 |
| **RBM3** | RI | 89 | 89 | -0.78543 | 0.455923 | 0.269947 | -2.90958 | 0.003619 |
| **COX4I1** | AD | 91 | 87 | -0.91461 | 0.400671 | 0.283388 | -3.22743 | 0.001249 |
| **SEPW1** | AA | 90 | 88 | -1.03401 | 0.355578 | 0.305411 | -3.38564 | 0.00071 |

| | | | | | | | | |
|---|---|---|---|---|---|---|---|---|
| **RPL28** | RI | 93 | 85 | -0.56309 | 0.569444 | 0.275184 | -2.04625 | 0.040732 |
| **MDH1** | AP | 89 | 89 | 0.745098 | 2.106648 | 0.274371 | 2.715655 | 0.006614 |
| **MDH1** | AP | 89 | 89 | -0.7451 | 0.474688 | 0.274371 | -2.71565 | 0.006614 |
| **LAMP2** | AT | 89 | 89 | -0.74261 | 0.47587 | 0.270411 | -2.74624 | 0.006028 |
| **ZNF197** | AT | 89 | 89 | -0.90282 | 0.405424 | 0.272767 | -3.30987 | 0.000933 |
| **ZNF197** | AT | 89 | 89 | 0.902821 | 2.466552 | 0.272767 | 3.309867 | 0.000933 |
| **TMEM180** | ES | 89 | 89 | -0.62733 | 0.534017 | 0.269724 | -2.32581 | 0.020029 |
| **COPZ1** | AA | 90 | 88 | 0.705679 | 2.025221 | 0.270837 | 2.605547 | 0.009173 |
| **INO80E** | ES | 89 | 89 | -0.56708 | 0.567179 | 0.268275 | -2.11381 | 0.034532 |
| **CCNL2** | AT | 89 | 89 | -0.77472 | 0.460831 | 0.271272 | -2.85589 | 0.004292 |
| **CCNL2** | AT | 89 | 89 | 0.774723 | 2.169992 | 0.271272 | 2.855889 | 0.004292 |
| **MAPKAPK5** | AT | 89 | 89 | -0.61775 | 0.539155 | 0.264441 | -2.33607 | 0.019488 |
| **MAPKAPK5** | AT | 89 | 89 | 0.617752 | 1.854754 | 0.264441 | 2.33607 | 0.019488 |
| **DGUOK** | ES | 89 | 89 | -0.79366 | 0.452187 | 0.281711 | -2.81728 | 0.004843 |
| **HSP90AB1** | AP | 90 | 88 | 0.656873 | 1.928751 | 0.27215 | 2.413644 | 0.015794 |
| **DUT** | AP | 90 | 88 | -0.66503 | 0.514259 | 0.269846 | -2.46447 | 0.013721 |
| **XAF1** | AT | 90 | 88 | -0.69596 | 0.498598 | 0.266495 | -2.61151 | 0.009014 |
| **SYNGR2** | AA | 89 | 89 | -0.76901 | 0.463473 | 0.27593 | -2.78696 | 0.00532 |
| **CEP70** | AT | 89 | 89 | -0.81204 | 0.443954 | 0.27202 | -2.9852 | 0.002834 |
| **TMEM134** | ES | 89 | 89 | -0.66145 | 0.516102 | 0.269036 | -2.4586 | 0.013948 |
| **LINC00908** | AT | 127 | 51 | -0.68194 | 0.505635 | 0.326183 | -2.09067 | 0.036558 |
| **LRRFIP1** | AT | 90 | 88 | 0.825172 | 2.282273 | 0.281401 | 2.932368 | 0.003364 |
| **FAM219B** | RI | 89 | 89 | 0.675024 | 1.964081 | 0.273512 | 2.467985 | 0.013588 |
| **LRRFIP1** | AT | 90 | 88 | -0.82476 | 0.438341 | 0.281407 | -2.93084 | 0.00338 |
| **RAB2A** | ES | 90 | 88 | -0.75542 | 0.469814 | 0.275544 | -2.74156 | 0.006115 |
| **PSMA6** | ES | 89 | 89 | 0.538964 | 1.71423 | 0.269691 | 1.998451 | 0.045668 |
| **ZNF524** | AP | 90 | 88 | -0.89683 | 0.407861 | 0.278323 | -3.22225 | 0.001272 |
| **NLRP1** | AT | 89 | 89 | 0.764013 | 2.146875 | 0.267883 | 2.852038 | 0.004344 |
| **ATP5J** | AT | 91 | 87 | 0.608031 | 1.83681 | 0.265998 | 2.285849 | 0.022263 |
| **ELP6** | ES | 89 | 89 | 0.86955 | 2.385836 | 0.274385 | 3.169091 | 0.001529 |
| **RMDN1** | ES | 90 | 88 | -0.75279 | 0.471052 | 0.269739 | -2.7908 | 0.005258 |
| **TPM2** | AT | 89 | 89 | -1.02159 | 0.360022 | 0.281875 | -3.62426 | 0.00029 |
| **TPM2** | AT | 89 | 89 | 1.02159 | 2.777608 | 0.281875 | 3.624264 | 0.00029 |
| **Sep-09** | AP | 89 | 89 | -0.7937 | 0.45217 | 0.280594 | -2.82863 | 0.004675 |
| **TMEM205** | ES | 89 | 89 | 0.583752 | 1.792752 | 0.264959 | 2.203179 | 0.027582 |
| **GPBP1** | AA | 89 | 89 | -0.72485 | 0.484399 | 0.273322 | -2.65198 | 0.008002 |
| **GPALPP1** | AT | 89 | 89 | 0.740286 | 2.096536 | 0.268252 | 2.759671 | 0.005786 |
| **GPALPP1** | AT | 89 | 89 | -0.74029 | 0.476977 | 0.268252 | -2.75967 | 0.005786 |
| **PDLIM5** | AT | 89 | 89 | -0.79297 | 0.452501 | 0.274841 | -2.88518 | 0.003912 |
| **SIAH1** | AP | 89 | 89 | -0.84721 | 0.42861 | 0.280892 | -3.01613 | 0.00256 |
| **TSFM** | AT | 89 | 89 | 0.697579 | 2.008884 | 0.278833 | 2.501782 | 0.012357 |
| **PCNXL4** | AT | 89 | 89 | 0.921865 | 2.513974 | 0.289359 | 3.185888 | 0.001443 |
| **USE1** | RI | 89 | 89 | -0.68306 | 0.505069 | 0.274223 | -2.49089 | 0.012742 |
| **GGCX** | AT | 89 | 89 | -0.85979 | 0.42325 | 0.277584 | -3.09742 | 0.001952 |
| **GGCX** | AT | 89 | 89 | 0.859793 | 2.36267 | 0.277584 | 3.097419 | 0.001952 |
| **ZEB2** | AT | 89 | 89 | 0.799535 | 2.224507 | 0.276146 | 2.895332 | 0.003788 |
| **ZEB2** | AT | 89 | 89 | -0.79954 | 0.449538 | 0.276146 | -2.89533 | 0.003788 |

| | | | | | | | | |
|---|---|---|---|---|---|---|---|---|
| **ASPH** | AT | 90 | 88 | 0.717847 | 2.050014 | 0.277333 | 2.588393 | 0.009643 |
| **TPM3** | AT | 90 | 88 | -0.67561 | 0.508844 | 0.27174 | -2.48625 | 0.01291 |
| **TRMT112** | RI | 92 | 86 | -0.72318 | 0.485208 | 0.28034 | -2.57964 | 0.00989 |
| **PNPO** | ES | 89 | 89 | -0.71768 | 0.487885 | 0.272514 | -2.63353 | 0.00845 |
| **RPTOR** | AT | 89 | 89 | 0.812086 | 2.252601 | 0.271963 | 2.98602 | 0.002826 |
| **CHPF** | AT | 89 | 89 | -0.54532 | 0.579655 | 0.273718 | -1.99228 | 0.04634 |
| **CHPF** | AT | 89 | 89 | 0.545322 | 1.725164 | 0.273718 | 1.992279 | 0.04634 |
| **STRIP2** | AT | 89 | 89 | 0.610104 | 1.840623 | 0.269856 | 2.260852 | 0.023768 |
| **STRIP2** | AT | 89 | 89 | -0.6101 | 0.543294 | 0.269856 | -2.26085 | 0.023768 |
| **ZNF625** | AT | 89 | 89 | -0.68368 | 0.504757 | 0.275541 | -2.48122 | 0.013093 |
| **ZNF625** | AT | 89 | 89 | 0.683679 | 1.981152 | 0.275541 | 2.481223 | 0.013093 |
| **DST** | AT | 89 | 89 | -0.74153 | 0.476383 | 0.271547 | -2.73077 | 0.006319 |
| **RPS20** | RI | 92 | 86 | -0.86373 | 0.421588 | 0.276343 | -3.12556 | 0.001775 |
| **GABPB1** | AT | 89 | 89 | -0.61353 | 0.541438 | 0.26895 | -2.28119 | 0.022537 |
| **GABPB1** | AT | 89 | 89 | 0.613527 | 1.846933 | 0.26895 | 2.281194 | 0.022537 |
| **NIT2** | AD | 89 | 89 | -0.79947 | 0.449566 | 0.274352 | -2.91403 | 0.003568 |
| **LYNX1** | AT | 89 | 89 | -0.7875 | 0.454981 | 0.27744 | -2.83845 | 0.004533 |
| **LYNX1** | AT | 89 | 89 | 0.7875 | 2.197894 | 0.27744 | 2.838448 | 0.004533 |
| **MPDU1** | AT | 91 | 87 | -0.58251 | 0.558495 | 0.268054 | -2.1731 | 0.029773 |
| **TWSG1** | AT | 89 | 89 | -0.79661 | 0.450853 | 0.268892 | -2.96258 | 0.003051 |
| **TWSG1** | AT | 89 | 89 | 0.796613 | 2.218016 | 0.268892 | 2.96258 | 0.003051 |
| **CLN3** | ES | 89 | 89 | -0.75714 | 0.469006 | 0.275407 | -2.74917 | 0.005975 |
| **PSMA1** | AD | 89 | 89 | -0.58243 | 0.558537 | 0.271755 | -2.14324 | 0.032094 |
| **MAPK13** | ES | 89 | 89 | -0.68309 | 0.505056 | 0.266719 | -2.56107 | 0.010435 |
| **VPS28** | RI | 89 | 89 | -0.90192 | 0.405788 | 0.281013 | -3.20955 | 0.001329 |
| **ZSWIM7** | RI | 89 | 89 | -0.6232 | 0.536224 | 0.2681 | -2.32452 | 0.020098 |
| **RPS15** | RI | 105 | 73 | -0.80769 | 0.445887 | 0.283111 | -2.8529 | 0.004332 |
| **DCTD** | ES | 90 | 88 | -0.64505 | 0.524635 | 0.268469 | -2.40271 | 0.016274 |
| **UBXN1** | RI | 89 | 89 | -0.75795 | 0.468626 | 0.273368 | -2.77263 | 0.005561 |
| **ZNF814** | AT | 89 | 89 | 0.729651 | 2.074356 | 0.267324 | 2.729459 | 0.006344 |
| **ERLEC1** | ES | 89 | 89 | -0.59562 | 0.551221 | 0.266848 | -2.23206 | 0.025611 |
| **GMPPB** | RI | 89 | 89 | -0.88957 | 0.410832 | 0.277526 | -3.20536 | 0.001349 |
| **HSPB1** | RI | 89 | 89 | -0.70887 | 0.492199 | 0.268685 | -2.6383 | 0.008332 |
| **HHLA3** | AT | 90 | 88 | 0.574625 | 1.776464 | 0.276516 | 2.07809 | 0.037701 |
| **VWA2** | AT | 89 | 89 | 0.754253 | 2.126023 | 0.280716 | 2.686887 | 0.007212 |
| **VWA2** | AT | 89 | 89 | -0.75425 | 0.470362 | 0.280716 | -2.68689 | 0.007212 |
| **LRRC27** | AT | 89 | 89 | 0.532257 | 1.702772 | 0.262738 | 2.025814 | 0.042784 |
| **MVD** | ES | 90 | 88 | -0.9274 | 0.395582 | 0.28373 | -3.2686 | 0.001081 |
| **SAT1** | RI | 89 | 89 | -0.64103 | 0.526748 | 0.281294 | -2.27887 | 0.022675 |
| **ERCC5** | AP | 90 | 88 | -0.68911 | 0.502021 | 0.266763 | -2.58324 | 0.009788 |
| **EMC4** | AT | 89 | 89 | -0.94195 | 0.389867 | 0.28526 | -3.30207 | 0.00096 |
| **IL32** | RI | 89 | 89 | -0.80051 | 0.4491 | 0.274932 | -2.91166 | 0.003595 |
| **CAPN10** | ES | 89 | 89 | 0.614339 | 1.848435 | 0.267645 | 2.295353 | 0.021713 |
| **SEMA3C** | AT | 94 | 84 | 0.945723 | 2.574675 | 0.279297 | 3.386082 | 0.000709 |
| **LMBR1** | ES | 89 | 89 | -0.77088 | 0.462607 | 0.271707 | -2.83717 | 0.004552 |
| **MCTS1** | AP | 89 | 89 | 0.827299 | 2.287133 | 0.281376 | 2.940188 | 0.00328 |
| **MCTS1** | AP | 89 | 89 | -0.8273 | 0.437229 | 0.281376 | -2.94019 | 0.00328 |

| | | | | | | | | |
|---|---|---|---|---|---|---|---|---|
| **PPOX** | AT | 90 | 88 | -0.82665 | 0.437513 | 0.271577 | -3.04388 | 0.002335 |
| **NAP1L1** | ES | 91 | 87 | 0.618392 | 1.855942 | 0.264471 | 2.33822 | 0.019376 |
| **SERPINB1** | ES | 89 | 89 | -0.61764 | 0.539217 | 0.267993 | -2.30467 | 0.021185 |
| **PRKRIP1** | ES | 89 | 89 | -0.60735 | 0.544793 | 0.267777 | -2.26812 | 0.023322 |
| **DDOST** | ES | 90 | 88 | -0.62598 | 0.534737 | 0.268827 | -2.32856 | 0.019882 |
| **TPM3** | ES | 89 | 89 | 0.691883 | 1.997474 | 0.275587 | 2.510581 | 0.012053 |
| **SYNGR1** | AT | 90 | 88 | 0.631413 | 1.880266 | 0.271967 | 2.321654 | 0.020252 |
| **C12orf76** | AT | 89 | 89 | -0.57561 | 0.56236 | 0.266431 | -2.16046 | 0.030737 |
| **RANGRF** | RI | 89 | 89 | -0.83808 | 0.432541 | 0.279097 | -3.00283 | 0.002675 |
| **LSR** | ES | 89 | 89 | -0.57195 | 0.564426 | 0.267677 | -2.1367 | 0.032622 |
| **MTA3** | AT | 89 | 89 | -0.72483 | 0.484408 | 0.270981 | -2.67483 | 0.007477 |
| **MTA3** | AT | 89 | 89 | 0.724827 | 2.064374 | 0.270981 | 2.674828 | 0.007477 |
| **GGPS1** | AP | 89 | 89 | -0.73882 | 0.477675 | 0.273211 | -2.70422 | 0.006846 |
| **NUDT5** | AT | 89 | 89 | -0.56606 | 0.56776 | 0.272611 | -2.07642 | 0.037855 |
| **SRSF2** | AD | 89 | 89 | -0.7724 | 0.461903 | 0.284944 | -2.7107 | 0.006714 |
| **DOCK10** | AT | 89 | 89 | -0.81117 | 0.444338 | 0.276271 | -2.93613 | 0.003323 |
| **DOCK10** | AT | 89 | 89 | 0.81117 | 2.250539 | 0.276271 | 2.936133 | 0.003323 |
| **CHCHD7** | ES | 89 | 89 | -0.62854 | 0.533371 | 0.2696 | -2.33137 | 0.019734 |
| **HS2ST1** | AT | 89 | 89 | -0.70569 | 0.493767 | 0.268603 | -2.62727 | 0.008607 |
| **DCN** | ES | 89 | 89 | -0.83349 | 0.434528 | 0.273692 | -3.04538 | 0.002324 |
| **MEIS2** | AP | 89 | 89 | 0.613029 | 1.846014 | 0.275171 | 2.227813 | 0.025893 |
| **ZNF524** | AP | 90 | 88 | 0.864601 | 2.374059 | 0.274777 | 3.146561 | 0.001652 |
| **HMGA1** | AD | 89 | 89 | -0.88733 | 0.411755 | 0.278341 | -3.18791 | 0.001433 |
| **HNRNPC** | AP | 92 | 86 | -0.80801 | 0.445743 | 0.27817 | -2.90474 | 0.003676 |
| **PSME2** | AA | 89 | 89 | -0.83423 | 0.434208 | 0.284418 | -2.93312 | 0.003356 |
| **PHTF2** | AT | 89 | 89 | -0.67083 | 0.511285 | 0.268579 | -2.4977 | 0.0125 |
| **RBPMS** | AT | 89 | 89 | 0.89263 | 2.441542 | 0.277779 | 3.213458 | 0.001311 |
| **PABPC1** | AD | 90 | 88 | -0.69949 | 0.496839 | 0.27428 | -2.55028 | 0.010764 |
| **RRNAD1** | RI | 89 | 89 | -0.60602 | 0.545518 | 0.269025 | -2.25265 | 0.024281 |
| **PTPRB** | AT | 89 | 89 | 0.727648 | 2.070205 | 0.269161 | 2.703389 | 0.006864 |
| **PTPRB** | AT | 89 | 89 | -0.72765 | 0.483044 | 0.269161 | -2.70339 | 0.006864 |
| **EMC4** | AT | 89 | 89 | 0.825749 | 2.283591 | 0.284504 | 2.902412 | 0.003703 |
| **DBNL** | AT | 89 | 89 | 0.568944 | 1.766402 | 0.268766 | 2.116879 | 0.03427 |
| **NAT9** | AA | 89 | 89 | -0.53704 | 0.584475 | 0.264244 | -2.03237 | 0.042116 |
| **BCAM** | AT | 89 | 89 | -0.58264 | 0.55842 | 0.266227 | -2.18852 | 0.028632 |
| **BCAM** | AT | 89 | 89 | 0.582644 | 1.790767 | 0.266227 | 2.188521 | 0.028632 |
| **ZNF836** | AT | 89 | 89 | 0.705442 | 2.024741 | 0.268434 | 2.627991 | 0.008589 |
| **ZNF836** | AT | 89 | 89 | -0.70544 | 0.49389 | 0.268434 | -2.62799 | 0.008589 |
| **RBP1** | AT | 90 | 88 | -0.62617 | 0.534633 | 0.276838 | -2.26188 | 0.023705 |
| **RAD1** | ES | 89 | 89 | -0.64954 | 0.522285 | 0.277418 | -2.34138 | 0.019212 |
| **H2AFV** | AT | 90 | 88 | -0.7554 | 0.469822 | 0.272798 | -2.76909 | 0.005621 |
| **RBMX** | RI | 89 | 89 | -0.82579 | 0.437887 | 0.274662 | -3.00658 | 0.002642 |
| **BLVRB** | AD | 89 | 89 | -0.65912 | 0.517308 | 0.272981 | -2.41452 | 0.015756 |
| **TOMM40** | RI | 89 | 89 | -0.94307 | 0.389429 | 0.284276 | -3.31746 | 0.000908 |
| **PPP3CB** | AT | 89 | 89 | -0.57032 | 0.565347 | 0.274053 | -2.08105 | 0.03743 |
| **PPP3CB** | AT | 89 | 89 | 0.570316 | 1.768825 | 0.274053 | 2.081045 | 0.03743 |
| **BRIP1** | AT | 89 | 89 | -0.86291 | 0.421932 | 0.272835 | -3.16276 | 0.001563 |

| | | | | | | | | |
|---|---|---|---|---|---|---|---|---|
| **BRIP1** | AT | 89 | 89 | 0.86291 | 2.370048 | 0.272835 | 3.162758 | 0.001563 |
| **MEF2BNB** | AT | 89 | 89 | 0.764143 | 2.147154 | 0.271567 | 2.81383 | 0.004896 |
| **MEF2BNB** | AT | 89 | 89 | -0.76414 | 0.465733 | 0.271567 | -2.81383 | 0.004896 |
| **SERPINB6** | AP | 89 | 89 | -0.88388 | 0.413178 | 0.276005 | -3.2024 | 0.001363 |
| **HSP90AB1** | AP | 91 | 87 | -0.67709 | 0.508092 | 0.275346 | -2.45906 | 0.01393 |
| **RABEP2** | ES | 89 | 89 | 0.535388 | 1.708111 | 0.266186 | 2.011332 | 0.04429 |
| **TADA3** | AP | 89 | 89 | -0.63786 | 0.528424 | 0.27131 | -2.35102 | 0.018722 |
| **TADA3** | AP | 89 | 89 | 0.637856 | 1.892418 | 0.27131 | 2.35102 | 0.018722 |
| **TPM3** | ES | 94 | 84 | 0.631447 | 1.88033 | 0.266458 | 2.369781 | 0.017799 |
| **ZNF20** | AT | 89 | 89 | -0.66396 | 0.514808 | 0.266235 | -2.49389 | 0.012635 |
| **ZNF20** | AT | 89 | 89 | 0.663961 | 1.942472 | 0.266235 | 2.493894 | 0.012635 |
| **ATP6V1F** | ES | 89 | 89 | 0.639931 | 1.89635 | 0.267397 | 2.393184 | 0.016703 |
| **AUNIP** | AT | 91 | 87 | 0.732234 | 2.079722 | 0.272247 | 2.689595 | 0.007154 |
| **WAC** | AP | 89 | 89 | 0.706029 | 2.02593 | 0.266367 | 2.650585 | 0.008035 |
| **PSME2** | ES | 90 | 88 | -0.73827 | 0.477939 | 0.273386 | -2.70048 | 0.006924 |
| **CARD14** | AT | 90 | 88 | -0.86064 | 0.422893 | 0.283617 | -3.0345 | 0.002409 |
| **SLC33A1** | RI | 89 | 89 | -0.63452 | 0.530192 | 0.270114 | -2.34906 | 0.018821 |
| **RALGPS1** | AT | 90 | 88 | 0.67078 | 1.955762 | 0.270587 | 2.478977 | 0.013176 |
| **ZNF706** | AP | 89 | 89 | -0.57846 | 0.560761 | 0.267604 | -2.16163 | 0.030646 |
| **COX4I1** | ES | 89 | 89 | -0.75086 | 0.471958 | 0.268364 | -2.79793 | 0.005143 |
| **ZNF687** | ES | 89 | 89 | -0.66719 | 0.513148 | 0.268605 | -2.48391 | 0.012995 |
| **KLC1** | ES | 90 | 88 | -0.80747 | 0.445984 | 0.270519 | -2.98491 | 0.002837 |
| **PDCD5** | RI | 89 | 89 | -0.83282 | 0.434821 | 0.276149 | -3.01584 | 0.002563 |
| **GTF2IRD1** | RI | 89 | 89 | -0.65548 | 0.519191 | 0.267009 | -2.45491 | 0.014092 |
| **NXN** | AP | 107 | 71 | -0.83393 | 0.434339 | 0.300984 | -2.77068 | 0.005594 |
| **EIF5A** | RI | 90 | 88 | -0.68114 | 0.506039 | 0.2773 | -2.45634 | 0.014036 |
| **KLK10** | AP | 89 | 89 | -0.64163 | 0.526436 | 0.268619 | -2.38861 | 0.016912 |
| **TXNL4A** | ES | 90 | 88 | -0.71154 | 0.490885 | 0.267327 | -2.6617 | 0.007775 |
| **EPS15L1** | AT | 89 | 89 | 0.89189 | 2.439736 | 0.288508 | 3.091392 | 0.001992 |
| **NUDT1** | AP | 89 | 89 | 0.665799 | 1.946045 | 0.271485 | 2.452434 | 0.014189 |
| **POT1** | ES | 89 | 89 | 0.531337 | 1.701205 | 0.270062 | 1.967461 | 0.04913 |
| **TRIM16** | ES | 90 | 88 | 0.800612 | 2.226904 | 0.289078 | 2.769537 | 0.005614 |
| **ANAPC11** | AP | 90 | 88 | 0.993037 | 2.699421 | 0.290632 | 3.416825 | 0.000634 |
| **POLR3H** | RI | 89 | 89 | -0.53802 | 0.583903 | 0.263523 | -2.04165 | 0.041187 |
| **NOMO2** | RI | 89 | 89 | 0.715716 | 2.045651 | 0.271683 | 2.634377 | 0.008429 |
| **RAD23A** | AA | 89 | 89 | -0.61946 | 0.538233 | 0.265758 | -2.33094 | 0.019757 |
| **GNAS** | AT | 89 | 89 | -0.81059 | 0.444594 | 0.274994 | -2.94768 | 0.003202 |
| **GNAS** | AT | 89 | 89 | 0.810594 | 2.249245 | 0.274994 | 2.947679 | 0.003202 |
| **Sep-11** | AT | 89 | 89 | -0.57129 | 0.564794 | 0.265891 | -2.1486 | 0.031666 |
| **HSD17B10** | ES | 89 | 89 | 0.8576 | 2.357495 | 0.275698 | 3.110643 | 0.001867 |
| **B2M** | ES | 89 | 89 | 0.707264 | 2.028434 | 0.270293 | 2.616655 | 0.00888 |
| **ABHD17A** | AD | 89 | 89 | -0.70797 | 0.492641 | 0.271755 | -2.60519 | 0.009182 |
| **GGT5** | AA | 89 | 89 | -0.55287 | 0.575297 | 0.268778 | -2.05697 | 0.039689 |
| **DDX41** | AA | 90 | 88 | -0.72092 | 0.486306 | 0.275616 | -2.61566 | 0.008906 |
| **NT5C** | RI | 90 | 88 | -0.89624 | 0.408101 | 0.289226 | -3.09876 | 0.001943 |
| **USP45** | AT | 89 | 89 | 0.677593 | 1.969132 | 0.272492 | 2.486655 | 0.012895 |
| **CHCHD7** | ES | 89 | 89 | -0.53138 | 0.587793 | 0.265396 | -2.00222 | 0.045261 |

| | | | | | | | | |
|---|---|---|---|---|---|---|---|---|
| **DYNLL1** | RI | 89 | 89 | -0.70029 | 0.496441 | 0.271403 | -2.58026 | 0.009873 |
| **PSME1** | RI | 91 | 87 | -0.64079 | 0.526874 | 0.279022 | -2.29657 | 0.021643 |
| **SLC16A1** | AT | 89 | 89 | -0.95527 | 0.384708 | 0.281247 | -3.39656 | 0.000682 |
| **SLC16A1** | AT | 89 | 89 | 0.955271 | 2.599376 | 0.281247 | 3.396556 | 0.000682 |
| **HNRNPC** | AP | 90 | 88 | 0.761075 | 2.140575 | 0.274777 | 2.769787 | 0.005609 |
| **ILF3** | AA | 91 | 87 | 0.566377 | 1.761872 | 0.26514 | 2.13614 | 0.032668 |
| **PKP4** | ES | 89 | 89 | -0.66661 | 0.513444 | 0.275889 | -2.41625 | 0.015681 |
| **AGAP3** | AP | 91 | 87 | 0.752971 | 2.123299 | 0.269081 | 2.798301 | 0.005137 |
| **INADL** | AT | 89 | 89 | -0.66954 | 0.511945 | 0.270874 | -2.47177 | 0.013445 |
| **PDE4DIP** | AP | 89 | 89 | 0.72302 | 2.060647 | 0.273273 | 2.64578 | 0.00815 |
| **AP1S3** | AT | 89 | 89 | 0.784533 | 2.191382 | 0.269009 | 2.916376 | 0.003541 |
| **ZCCHC4** | AT | 89 | 89 | -0.62618 | 0.534628 | 0.267901 | -2.33737 | 0.01942 |
| **ZCCHC4** | AT | 89 | 89 | 0.626185 | 1.870461 | 0.267901 | 2.337372 | 0.01942 |
| **MDH2** | AT | 89 | 89 | 0.730339 | 2.075783 | 0.27309 | 2.674348 | 0.007487 |
| **MDH2** | AT | 89 | 89 | -0.73034 | 0.481746 | 0.27309 | -2.67435 | 0.007487 |
| **STXBP2** | AP | 89 | 89 | 0.634917 | 1.886865 | 0.2691 | 2.359406 | 0.018304 |
| **STXBP2** | AP | 89 | 89 | -0.63492 | 0.52998 | 0.2691 | -2.35941 | 0.018304 |
| **C21orf59** | RI | 89 | 89 | -0.83562 | 0.433604 | 0.281024 | -2.97349 | 0.002944 |
| **EEF1D** | ES | 89 | 89 | -0.97491 | 0.377227 | 0.291002 | -3.35018 | 0.000808 |
| **HSPH1** | ES | 89 | 89 | -0.6095 | 0.543624 | 0.266445 | -2.28752 | 0.022166 |
| **CHCHD5** | RI | 89 | 89 | -0.74676 | 0.473899 | 0.280544 | -2.66184 | 0.007772 |
| **NAA10** | AA | 90 | 88 | -0.5353 | 0.585492 | 0.267515 | -2.00102 | 0.04539 |
| **C11orf48** | RI | 90 | 88 | -0.57248 | 0.564125 | 0.266677 | -2.14672 | 0.031816 |
| **SNRPN** | AP | 100 | 78 | -0.67654 | 0.508372 | 0.284782 | -2.37565 | 0.017518 |
| **SGK1** | AT | 89 | 89 | -0.87067 | 0.418672 | 0.273991 | -3.17773 | 0.001484 |
| **SGK1** | AT | 89 | 89 | 0.870668 | 2.388506 | 0.273991 | 3.17773 | 0.001484 |
| **RRP8** | RI | 89 | 89 | -0.87417 | 0.417208 | 0.289652 | -3.018 | 0.002544 |
| **OAS3** | AT | 89 | 89 | -0.69902 | 0.497074 | 0.267736 | -2.61084 | 0.009032 |
| **MED6** | AT | 89 | 89 | -0.77863 | 0.459033 | 0.268149 | -2.90374 | 0.003687 |
| **MED6** | AT | 89 | 89 | 0.778634 | 2.178494 | 0.268149 | 2.903738 | 0.003687 |
| **SEC31A** | AD | 89 | 89 | -0.8191 | 0.44083 | 0.269419 | -3.04023 | 0.002364 |
| **CARHSP1** | AT | 89 | 89 | -0.62046 | 0.5377 | 0.272195 | -2.27945 | 0.02264 |
| **CARHSP1** | AT | 89 | 89 | 0.620455 | 1.859774 | 0.272195 | 2.279448 | 0.02264 |
| **MT1E** | RI | 89 | 89 | -1.02696 | 0.358094 | 0.301902 | -3.40163 | 0.00067 |
| **RPL18** | RI | 92 | 86 | -0.60199 | 0.547721 | 0.275341 | -2.18634 | 0.028791 |
| **ATP10D** | AT | 89 | 89 | 0.707698 | 2.029314 | 0.273541 | 2.587176 | 0.009677 |
| **ATP10D** | AT | 89 | 89 | -0.7077 | 0.492777 | 0.273541 | -2.58718 | 0.009677 |
| **CNST** | AT | 89 | 89 | -0.73913 | 0.477531 | 0.272326 | -2.71412 | 0.006645 |
| **CNST** | AT | 89 | 89 | 0.739125 | 2.094103 | 0.272326 | 2.714117 | 0.006645 |
| **LLGL2** | ES | 90 | 88 | 0.561633 | 1.753534 | 0.265841 | 2.112662 | 0.03463 |
| **PRLR** | AT | 141 | 37 | -1.55042 | 0.21216 | 0.473958 | -3.27121 | 0.001071 |
| **EGFR** | AT | 110 | 68 | 0.676386 | 1.966757 | 0.262497 | 2.576743 | 0.009974 |
| **CASZ1** | AT | 89 | 89 | 0.792279 | 2.208424 | 0.280874 | 2.820763 | 0.004791 |
| **CASZ1** | AT | 89 | 89 | -0.79228 | 0.452812 | 0.280874 | -2.82076 | 0.004791 |
| **SH3GLB1** | ES | 89 | 89 | -0.69245 | 0.500349 | 0.267427 | -2.5893 | 0.009617 |
| **CHEK1** | AT | 89 | 89 | 0.832401 | 2.298831 | 0.276191 | 3.013856 | 0.002579 |
| **CHEK1** | AT | 89 | 89 | -0.8324 | 0.435004 | 0.276191 | -3.01386 | 0.002579 |

| | | | | | | | | |
|---|---|---|---|---|---|---|---|---|
| **SSH2** | RI | 89 | 89 | -0.55042 | 0.576709 | 0.265869 | -2.07026 | 0.038428 |
| **COX17** | AT | 89 | 89 | 0.733463 | 2.082279 | 0.271367 | 2.702844 | 0.006875 |
| **CAMK2D** | AT | 89 | 89 | -0.79965 | 0.449487 | 0.266393 | -3.00176 | 0.002684 |
| **CAMK2D** | AT | 89 | 89 | 0.799648 | 2.224759 | 0.266393 | 3.001764 | 0.002684 |
| **COX5A** | ES | 89 | 89 | -0.59662 | 0.550667 | 0.266062 | -2.24242 | 0.024934 |
| **HNRNPLL** | AT | 89 | 89 | 0.672829 | 1.959774 | 0.272286 | 2.471039 | 0.013472 |
| **SUMO3** | AT | 89 | 89 | -0.84928 | 0.427722 | 0.273529 | -3.1049 | 0.001903 |
| **SUMO3** | AT | 89 | 89 | 0.849282 | 2.337968 | 0.273529 | 3.104904 | 0.001903 |
| **HINT1** | AT | 89 | 89 | -0.65943 | 0.517147 | 0.278994 | -2.3636 | 0.018099 |
| **HINT1** | AT | 89 | 89 | 0.659428 | 1.933687 | 0.278994 | 2.363596 | 0.018099 |
| **C5orf45** | AT | 90 | 88 | 0.761502 | 2.141491 | 0.269769 | 2.822794 | 0.004761 |
| **FASTK** | RI | 89 | 89 | -0.86547 | 0.420855 | 0.281156 | -3.07824 | 0.002082 |
| **PHF20L1** | AT | 89 | 89 | 0.718158 | 2.050652 | 0.270146 | 2.658403 | 0.007851 |
| **PHF20L1** | AT | 89 | 89 | -0.71816 | 0.48765 | 0.270146 | -2.6584 | 0.007851 |
| **SET** | AP | 89 | 89 | 0.744336 | 2.105044 | 0.271161 | 2.744993 | 0.006051 |
| **ZFAND4** | AT | 89 | 89 | 0.740423 | 2.096822 | 0.271628 | 2.725876 | 0.006413 |
| **HNRNPA1** | RI | 89 | 89 | -0.64367 | 0.525362 | 0.277246 | -2.32165 | 0.020252 |
| **LGMN** | ES | 89 | 89 | 0.554476 | 1.741029 | 0.263694 | 2.102728 | 0.03549 |
| **SIAH1** | AT | 89 | 89 | 0.790996 | 2.205592 | 0.289147 | 2.735621 | 0.006226 |
| **SIAH1** | AT | 89 | 89 | -0.791 | 0.453393 | 0.289147 | -2.73562 | 0.006226 |
| **TADA3** | AT | 89 | 89 | -0.83172 | 0.4353 | 0.288249 | -2.88542 | 0.003909 |
| **TADA3** | AT | 89 | 89 | 0.831719 | 2.297265 | 0.288249 | 2.885416 | 0.003909 |
| **CD151** | ES | 89 | 89 | 0.866574 | 2.378746 | 0.280319 | 3.091381 | 0.001992 |
| **CD59** | AP | 90 | 88 | 0.780324 | 2.182179 | 0.27119 | 2.877411 | 0.00401 |
| **HIGD1A** | AT | 89 | 89 | 0.726036 | 2.066871 | 0.270912 | 2.67997 | 0.007363 |
| **HIGD1A** | AT | 89 | 89 | -0.72604 | 0.483823 | 0.270912 | -2.67997 | 0.007363 |
| **WAC** | AP | 89 | 89 | -0.67606 | 0.508618 | 0.265599 | -2.54541 | 0.010915 |
| **POLR2C** | ES | 89 | 89 | -0.69011 | 0.501521 | 0.273761 | -2.52084 | 0.011707 |
| **C5orf45** | AT | 90 | 88 | -0.7938 | 0.452124 | 0.271912 | -2.91932 | 0.003508 |
| **FKBP3** | ES | 89 | 89 | -0.65323 | 0.52036 | 0.273126 | -2.3917 | 0.016771 |
| **NDUFV2** | AP | 91 | 87 | 0.542285 | 1.719932 | 0.264332 | 2.05153 | 0.040215 |
| **EGLN2** | AP | 89 | 89 | -0.74399 | 0.475216 | 0.277311 | -2.68286 | 0.0073 |
| **RPL15** | AD | 90 | 88 | 0.665723 | 1.945896 | 0.27195 | 2.447956 | 0.014367 |
| **SIRT5** | ES | 89 | 89 | -0.5394 | 0.583101 | 0.266818 | -2.02158 | 0.043219 |
| **CD44** | ES | 89 | 89 | 0.628116 | 1.874077 | 0.274052 | 2.291961 | 0.021908 |
| **TMEM134** | AA | 89 | 89 | -0.76504 | 0.465315 | 0.273648 | -2.79571 | 0.005179 |
| **INTS4** | AT | 89 | 89 | 0.622094 | 1.862824 | 0.268412 | 2.31768 | 0.020467 |
| **RPS15** | RI | 89 | 89 | -0.71925 | 0.487118 | 0.266515 | -2.69872 | 0.006961 |
| **KCTD7** | AT | 89 | 89 | 0.885533 | 2.424275 | 0.27452 | 3.225754 | 0.001256 |
| **KCTD7** | AT | 89 | 89 | -0.88553 | 0.412494 | 0.27452 | -3.22575 | 0.001256 |
| **B3GAT3** | AA | 89 | 89 | 0.687571 | 1.988878 | 0.268181 | 2.563833 | 0.010352 |
| **RPL38** | RI | 89 | 89 | -0.62394 | 0.535831 | 0.263912 | -2.36418 | 0.01807 |
| **RFXANK** | ES | 89 | 89 | -0.62557 | 0.534956 | 0.267712 | -2.33674 | 0.019453 |
| **ZNF506** | ES | 89 | 89 | 0.533186 | 1.704353 | 0.271553 | 1.96347 | 0.049592 |
| **SPINT2** | ES | 89 | 89 | -0.69815 | 0.497505 | 0.273007 | -2.55726 | 0.01055 |
| **CCDC66** | AT | 89 | 89 | -0.70568 | 0.493773 | 0.273633 | -2.57893 | 0.009911 |
| **REEP5** | ES | 90 | 88 | -0.61707 | 0.539524 | 0.270732 | -2.27926 | 0.022652 |

| | | | | | | | | |
|---|---|---|---|---|---|---|---|---|
| **ASH2L** | AP | 89 | 89 | 0.826926 | 2.28628 | 0.272956 | 3.029524 | 0.002449 |
| **ASH2L** | AP | 89 | 89 | -0.82693 | 0.437392 | 0.272956 | -3.02952 | 0.002449 |
| **FTH1** | AT | 117 | 61 | -1.01223 | 0.363408 | 0.349122 | -2.89936 | 0.003739 |
| **PFDN5** | ES | 89 | 89 | -0.65317 | 0.520394 | 0.270414 | -2.41544 | 0.015716 |
| **TCTN1** | AT | 89 | 89 | -0.65143 | 0.521302 | 0.27034 | -2.40966 | 0.015967 |
| **TCTN1** | AT | 89 | 89 | 0.651427 | 1.918276 | 0.27034 | 2.409659 | 0.015967 |
| **MTHFS** | AD | 89 | 89 | -0.76726 | 0.464284 | 0.274435 | -2.79578 | 0.005178 |
| **TMEM101** | AT | 91 | 87 | 0.642705 | 1.901619 | 0.276303 | 2.326089 | 0.020014 |
| **SMARCA4** | ES | 89 | 89 | -0.60601 | 0.545523 | 0.267764 | -2.26323 | 0.023622 |
| **C11orf49** | AA | 89 | 89 | 0.532282 | 1.702814 | 0.267436 | 1.990313 | 0.046556 |
| **PAFAH1B2** | AT | 90 | 88 | -0.65266 | 0.52066 | 0.26971 | -2.41985 | 0.015527 |
| **S100A4** | AP | 133 | 45 | 0.980757 | 2.666474 | 0.290703 | 3.373738 | 0.000742 |
| **PLAC9** | ES | 90 | 88 | 0.547085 | 1.728209 | 0.269517 | 2.029874 | 0.042369 |
| **COA3** | AT | 92 | 86 | -0.81609 | 0.442158 | 0.285152 | -2.86194 | 0.004211 |
| **ZNF343** | AT | 89 | 89 | 0.680519 | 1.974903 | 0.268628 | 2.533317 | 0.011299 |
| **ZNF343** | AT | 89 | 89 | -0.68052 | 0.506354 | 0.268628 | -2.53332 | 0.011299 |
| **RPL29** | AT | 90 | 88 | -0.54173 | 0.581739 | 0.272299 | -1.98948 | 0.046648 |
| **FAM65B** | AT | 89 | 89 | 0.949862 | 2.585354 | 0.278361 | 3.412336 | 0.000644 |
| **MPC2** | AP | 90 | 88 | 0.648633 | 1.912924 | 0.270644 | 2.396628 | 0.016547 |
| **SNRPF** | AT | 89 | 89 | -0.6574 | 0.518194 | 0.269536 | -2.43902 | 0.014727 |
| **TXNRD2** | AT | 89 | 89 | -0.65389 | 0.520018 | 0.264128 | -2.47566 | 0.013299 |
| **TXNRD2** | AT | 89 | 89 | 0.653891 | 1.923009 | 0.264128 | 2.475664 | 0.013299 |
| **TYMS** | ES | 89 | 89 | 0.804748 | 2.236132 | 0.2726 | 2.952117 | 0.003156 |
| **C8orf59** | AP | 89 | 89 | -0.79562 | 0.451303 | 0.277652 | -2.86552 | 0.004163 |
| **C8orf59** | AP | 89 | 89 | 0.795616 | 2.215806 | 0.277652 | 2.865516 | 0.004163 |
| **RBFOX2** | ES | 106 | 72 | -0.75705 | 0.469046 | 0.300128 | -2.52244 | 0.011654 |
| **PTK7** | AT | 89 | 89 | 0.576398 | 1.779617 | 0.270003 | 2.134784 | 0.032779 |
| **GTF2IRD2** | AT | 89 | 89 | 0.70638 | 2.026642 | 0.273303 | 2.584608 | 0.009749 |
| **GBA2** | RI | 89 | 89 | 0.550699 | 1.734465 | 0.267076 | 2.061956 | 0.039212 |
| **EIF3M** | RI | 89 | 89 | -0.77018 | 0.46293 | 0.280418 | -2.74655 | 0.006023 |
| **COPS4** | ES | 89 | 89 | -0.69269 | 0.500228 | 0.274729 | -2.52137 | 0.01169 |
| **TMEM39B** | AT | 89 | 89 | -0.74394 | 0.475238 | 0.276719 | -2.68843 | 0.007179 |
| **TMEM39B** | AT | 89 | 89 | 0.74394 | 2.104211 | 0.276719 | 2.68843 | 0.007179 |
| **PPIH** | AD | 91 | 87 | -0.6399 | 0.527343 | 0.270621 | -2.36457 | 0.018051 |
| **YIF1A** | ES | 90 | 88 | -0.67903 | 0.507111 | 0.272481 | -2.49201 | 0.012702 |
| **ARHGEF1** | ES | 89 | 89 | -0.79867 | 0.449926 | 0.283632 | -2.81588 | 0.004864 |
| **SYNGR1** | AT | 89 | 89 | -0.64006 | 0.527261 | 0.272301 | -2.35056 | 0.018745 |
| **DALRD3** | RI | 90 | 88 | -0.75177 | 0.47153 | 0.276158 | -2.72226 | 0.006484 |
| **NSUN5** | RI | 89 | 89 | -0.91425 | 0.400817 | 0.279613 | -3.2697 | 0.001077 |
| **DAZAP2** | AP | 94 | 84 | 0.618049 | 1.855305 | 0.265387 | 2.328856 | 0.019867 |
| **CLN3** | AT | 89 | 89 | -0.76645 | 0.46466 | 0.281913 | -2.71874 | 0.006553 |
| **CLN3** | AT | 89 | 89 | 0.766448 | 2.152109 | 0.281913 | 2.718744 | 0.006553 |
| **DRG2** | AD | 89 | 89 | 0.882096 | 2.415958 | 0.279811 | 3.152469 | 0.001619 |
| **SON** | AT | 89 | 89 | 0.678038 | 1.970008 | 0.271672 | 2.4958 | 0.012567 |
| **BRF1** | AT | 89 | 89 | 0.742471 | 2.101121 | 0.271569 | 2.734003 | 0.006257 |
| **BRF1** | AT | 89 | 89 | -0.74247 | 0.475936 | 0.271569 | -2.734 | 0.006257 |
| **ZFPL1** | AT | 89 | 89 | 0.683927 | 1.981645 | 0.273623 | 2.499524 | 0.012436 |

| | | | | | | | | |
|---|---|---|---|---|---|---|---|---|
| **ZFPL1** | AT | 89 | 89 | -0.68393 | 0.504631 | 0.273623 | -2.49952 | 0.012436 |
| **COPZ1** | ES | 90 | 88 | 0.773642 | 2.167647 | 0.265983 | 2.908618 | 0.00363 |
| **OARD1** | ES | 89 | 89 | -0.70345 | 0.494875 | 0.268033 | -2.62449 | 0.008678 |
| **SLC25A23** | AT | 89 | 89 | -0.76493 | 0.465366 | 0.275806 | -2.77344 | 0.005547 |
| **TPM4** | AP | 89 | 89 | 0.75695 | 2.131764 | 0.269545 | 2.808253 | 0.004981 |
| **TPM4** | AP | 89 | 89 | -0.75695 | 0.469095 | 0.269545 | -2.80825 | 0.004981 |
| **FKBP5** | AT | 90 | 88 | -0.56642 | 0.567553 | 0.266024 | -2.12921 | 0.033237 |
| **FAM102A** | AP | 89 | 89 | -0.80669 | 0.446332 | 0.27439 | -2.93995 | 0.003283 |
| **FAM102A** | AP | 89 | 89 | 0.806692 | 2.240483 | 0.27439 | 2.939949 | 0.003283 |
| **TRPM4** | ES | 89 | 89 | -0.68376 | 0.504715 | 0.271213 | -2.52112 | 0.011698 |
| **MFSD10** | ES | 89 | 89 | 0.742855 | 2.101927 | 0.279019 | 2.662379 | 0.007759 |
| **MRPS18C** | ES | 89 | 89 | -0.81799 | 0.441316 | 0.280167 | -2.91967 | 0.003504 |
| **CACYBP** | AP | 90 | 88 | -0.58339 | 0.558002 | 0.265094 | -2.2007 | 0.027757 |
| **GMPR2** | RI | 89 | 89 | -0.62646 | 0.534483 | 0.265915 | -2.35585 | 0.01848 |
| **MCFD2** | AP | 89 | 89 | -0.70426 | 0.494475 | 0.272781 | -2.58177 | 0.00983 |
| **ZSCAN9** | AT | 89 | 89 | -0.88316 | 0.413475 | 0.272602 | -3.23973 | 0.001196 |
| **ZSCAN9** | AT | 89 | 89 | 0.883158 | 2.418526 | 0.272602 | 3.239729 | 0.001196 |
| **PFKFB2** | AT | 89 | 89 | -0.82608 | 0.437761 | 0.277228 | -2.9798 | 0.002884 |
| **PFKFB2** | AT | 89 | 89 | 0.826082 | 2.284352 | 0.277228 | 2.979796 | 0.002884 |
| **RIC8A** | AA | 89 | 89 | 0.567577 | 1.763987 | 0.269619 | 2.10511 | 0.035282 |
| **TMPO** | AT | 89 | 89 | 0.67008 | 1.954394 | 0.271758 | 2.465722 | 0.013674 |
| **LGALS3** | AT | 89 | 89 | 0.643561 | 1.903247 | 0.270209 | 2.381714 | 0.017232 |
| **LGALS3** | AT | 89 | 89 | -0.64356 | 0.525418 | 0.270209 | -2.38171 | 0.017232 |
| **TCP1** | AA | 89 | 89 | -0.67018 | 0.511616 | 0.272366 | -2.46059 | 0.013871 |
| **SYTL1** | AA | 89 | 89 | -0.6153 | 0.540478 | 0.267964 | -2.29621 | 0.021664 |
| **SLC35B1** | AA | 89 | 89 | -0.75912 | 0.46808 | 0.272854 | -2.78214 | 0.0054 |
| **SCAP** | ES | 89 | 89 | -0.87021 | 0.418863 | 0.279926 | -3.10872 | 0.001879 |
| **CD74** | ES | 92 | 86 | 0.61676 | 1.852914 | 0.271781 | 2.269325 | 0.023249 |
| **PHF6** | AT | 89 | 89 | -0.76072 | 0.46733 | 0.270004 | -2.81744 | 0.004841 |
| **PARK7** | AP | 90 | 88 | 0.666506 | 1.947422 | 0.270545 | 2.463572 | 0.013756 |
| **ADAM19** | AT | 89 | 89 | -0.59404 | 0.552094 | 0.269738 | -2.20227 | 0.027646 |
| **MPDU1** | AT | 89 | 89 | 0.684065 | 1.981918 | 0.271146 | 2.522871 | 0.01164 |
| **SERF1A** | AP | 89 | 89 | 0.559717 | 1.750177 | 0.269614 | 2.075994 | 0.037895 |
| **SERF1A** | AP | 89 | 89 | -0.55972 | 0.571371 | 0.269614 | -2.07599 | 0.037895 |
| **SMARCC2** | AD | 89 | 89 | 0.567462 | 1.763784 | 0.275482 | 2.059886 | 0.039409 |
| **NFKBIA** | AA | 89 | 89 | 0.664989 | 1.944469 | 0.275503 | 2.413725 | 0.01579 |
| **CIRBP** | RI | 89 | 89 | -0.85454 | 0.42548 | 0.29337 | -2.91283 | 0.003582 |
| **SLC30A5** | AA | 89 | 89 | -0.72284 | 0.485372 | 0.267843 | -2.69875 | 0.00696 |
| **GNB5** | AT | 89 | 89 | -0.75501 | 0.470006 | 0.274887 | -2.74662 | 0.006021 |
| **GNB5** | AT | 89 | 89 | 0.755009 | 2.12763 | 0.274887 | 2.746619 | 0.006021 |
| **TRPV3** | AT | 89 | 89 | 0.698447 | 2.010629 | 0.268181 | 2.604391 | 0.009204 |
| **TRPV3** | AT | 89 | 89 | -0.69845 | 0.497357 | 0.268181 | -2.60439 | 0.009204 |
| **THADA** | AT | 90 | 88 | 0.589604 | 1.803273 | 0.267661 | 2.202796 | 0.027609 |
| **AP1S3** | AT | 89 | 89 | 0.629874 | 1.877374 | 0.271744 | 2.317893 | 0.020455 |
| **CBR1** | AT | 90 | 88 | 0.702157 | 2.018101 | 0.280308 | 2.504951 | 0.012247 |
| **PHC3** | AT | 89 | 89 | -0.83737 | 0.432849 | 0.26856 | -3.11799 | 0.001821 |
| **USP45** | AT | 89 | 89 | -0.64345 | 0.525477 | 0.272645 | -2.36002 | 0.018274 |

| | | | | | | | | |
|---|---|---|---|---|---|---|---|---|
| **COMMD4** | AT | 89 | 89 | -0.81962 | 0.440599 | 0.280942 | -2.9174 | 0.00353 |
| **COMMD4** | AT | 89 | 89 | 0.819619 | 2.269635 | 0.280942 | 2.917399 | 0.00353 |
| **CCDC40** | AT | 89 | 89 | -0.76488 | 0.465391 | 0.272248 | -2.80949 | 0.004962 |
| **PIGT** | ES | 89 | 89 | -0.67355 | 0.509896 | 0.266317 | -2.52912 | 0.011435 |
| **NHP2** | ES | 90 | 88 | 0.618925 | 1.856931 | 0.264689 | 2.338314 | 0.019371 |
| **MROH1** | AT | 89 | 89 | 0.637532 | 1.891806 | 0.270129 | 2.360099 | 0.01827 |
| **MROH1** | AT | 89 | 89 | -0.63753 | 0.528595 | 0.270129 | -2.3601 | 0.01827 |
| **CASC4** | AT | 92 | 86 | -0.57529 | 0.562542 | 0.274432 | -2.09629 | 0.036057 |
| **EIF2B3** | AT | 89 | 89 | 0.630744 | 1.879008 | 0.265694 | 2.373946 | 0.017599 |
| **EIF2B3** | AT | 89 | 89 | -0.63074 | 0.532196 | 0.265694 | -2.37395 | 0.017599 |
| **C16orf13** | ES | 90 | 88 | -0.67432 | 0.509502 | 0.271856 | -2.48044 | 0.013122 |
| **UBE2N** | RI | 90 | 88 | -0.61255 | 0.541967 | 0.272212 | -2.25027 | 0.024432 |
| **NFATC4** | AD | 89 | 89 | -0.72308 | 0.485253 | 0.273777 | -2.64114 | 0.008263 |
| **MFSD11** | RI | 89 | 89 | -0.71714 | 0.488148 | 0.272918 | -2.62766 | 0.008597 |
| **LFNG** | AT | 89 | 89 | 0.657617 | 1.930186 | 0.271647 | 2.420853 | 0.015484 |
| **LFNG** | AT | 89 | 89 | -0.65762 | 0.518085 | 0.271647 | -2.42085 | 0.015484 |
| **TTC12** | AT | 89 | 89 | -0.66017 | 0.516765 | 0.2734 | -2.41466 | 0.01575 |
| **PET112** | AT | 89 | 89 | -0.65097 | 0.521539 | 0.265694 | -2.45008 | 0.014282 |
| **ZNHIT3** | ES | 89 | 89 | 0.74922 | 2.115349 | 0.275639 | 2.71812 | 0.006565 |
| **ABHD17A** | ES | 90 | 88 | -0.56241 | 0.569834 | 0.284391 | -1.9776 | 0.047974 |
| **RPS11** | AT | 101 | 77 | 0.566902 | 1.762797 | 0.265717 | 2.133477 | 0.032886 |
| **PDCD6** | AT | 89 | 89 | -0.68773 | 0.502714 | 0.284183 | -2.42004 | 0.015519 |
| **NSUN5** | RI | 89 | 89 | -0.82722 | 0.437265 | 0.27704 | -2.9859 | 0.002827 |
| **MOK** | AP | 89 | 89 | -0.62742 | 0.533965 | 0.267291 | -2.34735 | 0.018908 |
| **XYLT2** | AT | 89 | 89 | -0.54507 | 0.5798 | 0.277004 | -1.96775 | 0.049097 |
| **XYLT2** | AT | 89 | 89 | 0.545073 | 1.724734 | 0.277004 | 1.967746 | 0.049097 |
| **FAM195B** | RI | 89 | 89 | -0.72615 | 0.483769 | 0.273346 | -2.65652 | 0.007895 |
| **MFF** | ES | 89 | 89 | -0.70289 | 0.495152 | 0.26963 | -2.60687 | 0.009138 |
| **SNX5** | AT | 89 | 89 | -0.77774 | 0.459442 | 0.269842 | -2.88221 | 0.003949 |
| **SNX5** | AT | 89 | 89 | 0.777743 | 2.176555 | 0.269842 | 2.882212 | 0.003949 |
| **ASPH** | AT | 91 | 87 | -0.59683 | 0.550553 | 0.274002 | -2.1782 | 0.029391 |
| **GPR56** | AP | 90 | 88 | -0.62239 | 0.536661 | 0.26862 | -2.31699 | 0.020504 |
| **ENY2** | AA | 90 | 88 | -0.63961 | 0.527499 | 0.274572 | -2.32947 | 0.019834 |
| **TMSB15B** | AD | 89 | 89 | -0.62199 | 0.536873 | 0.272171 | -2.28531 | 0.022295 |
| **ZNF81** | AT | 119 | 59 | -0.71579 | 0.488804 | 0.307257 | -2.32962 | 0.019826 |
| **NOP2** | ES | 89 | 89 | -0.71617 | 0.488623 | 0.26662 | -2.68609 | 0.007229 |
| **NUP133** | AT | 89 | 89 | -0.55655 | 0.573181 | 0.26781 | -2.07816 | 0.037694 |
| **NUP133** | AT | 89 | 89 | 0.556553 | 1.744649 | 0.26781 | 2.078163 | 0.037694 |
| **RAPGEF3** | AT | 89 | 89 | 0.609573 | 1.839646 | 0.270393 | 2.254401 | 0.024171 |
| **RAPGEF3** | AT | 89 | 89 | -0.60957 | 0.543583 | 0.270393 | -2.2544 | 0.024171 |
| **FAM32A** | ES | 89 | 89 | -0.65903 | 0.517352 | 0.270079 | -2.44014 | 0.014682 |
| **ZSCAN20** | AT | 89 | 89 | -0.70464 | 0.494286 | 0.268886 | -2.62059 | 0.008778 |
| **ZSCAN20** | AT | 89 | 89 | 0.704642 | 2.023122 | 0.268886 | 2.620592 | 0.008778 |
| **MAP3K8** | AT | 89 | 89 | -0.81095 | 0.444435 | 0.276262 | -2.93544 | 0.003331 |
| **MAP3K8** | AT | 89 | 89 | 0.810952 | 2.250049 | 0.276262 | 2.935444 | 0.003331 |
| **HAUS4** | ES | 89 | 89 | 0.519166 | 1.680625 | 0.264223 | 1.964874 | 0.049429 |
| **RPAIN** | ES | 90 | 88 | 0.636788 | 1.8904 | 0.270603 | 2.353219 | 0.018612 |

| | | | | | | | | |
|---|---|---|---|---|---|---|---|---|
| **COX20** | ES | 89 | 89 | 0.764533 | 2.147992 | 0.269643 | 2.835357 | 0.004577 |
| **MT2A** | RI | 89 | 89 | -0.66259 | 0.515512 | 0.277385 | -2.38872 | 0.016907 |
| **MRPS17** | AP | 89 | 89 | -0.55905 | 0.57175 | 0.265016 | -2.10951 | 0.0349 |
| **MRPS17** | AP | 89 | 89 | 0.559054 | 1.749018 | 0.265016 | 2.109512 | 0.0349 |
| **TMEM147** | RI | 89 | 89 | -0.60691 | 0.54503 | 0.26946 | -2.25234 | 0.024301 |
| **TMEM14B** | AT | 90 | 88 | -0.67917 | 0.507037 | 0.26948 | -2.5203 | 0.011726 |
| **COA1** | AT | 89 | 89 | -0.69804 | 0.49756 | 0.272365 | -2.56288 | 0.010381 |
| **KIAA1429** | AT | 89 | 89 | -0.95465 | 0.384947 | 0.278499 | -3.42784 | 0.000608 |
| **KIAA1429** | AT | 89 | 89 | 0.954649 | 2.597759 | 0.278499 | 3.427841 | 0.000608 |
| **SPAG7** | RI | 89 | 89 | -0.63695 | 0.528903 | 0.270729 | -2.35272 | 0.018637 |
| **SMN1** | AA | 89 | 89 | -1.0362 | 0.354802 | 0.285911 | -3.62419 | 0.00029 |
| **FAM72A** | AP | 89 | 89 | -0.78184 | 0.457565 | 0.272902 | -2.8649 | 0.004171 |
| **FAM72A** | AP | 89 | 89 | 0.781836 | 2.185482 | 0.272902 | 2.864897 | 0.004171 |
| **NEK2** | AT | 89 | 89 | 0.660137 | 1.935058 | 0.26812 | 2.462099 | 0.013813 |
| **NEK2** | AT | 89 | 89 | -0.66014 | 0.51678 | 0.26812 | -2.4621 | 0.013813 |
| **RPS24** | ES | 89 | 89 | -0.72908 | 0.482351 | 0.267656 | -2.72396 | 0.006451 |
| **ELP4** | ES | 89 | 89 | -0.73488 | 0.479565 | 0.274505 | -2.6771 | 0.007426 |
| **ACOT2** | AD | 89 | 89 | -0.85097 | 0.427002 | 0.278395 | -3.05669 | 0.002238 |
| **ZNF506** | AT | 89 | 89 | -0.72307 | 0.48526 | 0.270897 | -2.66917 | 0.007604 |
| **SH3YL1** | ES | 89 | 89 | 0.603121 | 1.827814 | 0.264633 | 2.279083 | 0.022662 |
| **USP40** | AT | 116 | 62 | 0.608854 | 1.838324 | 0.270821 | 2.248179 | 0.024565 |
| **PCID2** | RI | 89 | 89 | -0.58918 | 0.554782 | 0.272873 | -2.15918 | 0.030837 |
| **TTLL5** | AT | 89 | 89 | -0.6384 | 0.528136 | 0.269906 | -2.36527 | 0.018017 |
| **NDUFV2** | AP | 91 | 87 | -0.54872 | 0.577687 | 0.265369 | -2.06777 | 0.038661 |
| **TGOLN2** | RI | 89 | 89 | -0.68077 | 0.506227 | 0.272993 | -2.49372 | 0.012641 |
| **TSC22D3** | AP | 89 | 89 | -0.85895 | 0.423608 | 0.278193 | -3.08759 | 0.002018 |
| **NFYC** | AA | 89 | 89 | 0.646295 | 1.908456 | 0.267795 | 2.413392 | 0.015805 |
| **ZXDC** | AT | 89 | 89 | 0.691263 | 1.996234 | 0.267351 | 2.585603 | 0.009721 |
| **ZXDC** | AT | 89 | 89 | -0.69126 | 0.500943 | 0.267351 | -2.5856 | 0.009721 |
| **SGCD** | AT | 90 | 88 | -0.54988 | 0.577021 | 0.269629 | -2.03939 | 0.041411 |
| **COBL** | AT | 92 | 86 | 0.626471 | 1.870996 | 0.268701 | 2.33148 | 0.019728 |
| **PEX2** | ES | 89 | 89 | 0.650836 | 1.917142 | 0.273204 | 2.382234 | 0.017208 |
| **POLR1C** | AT | 89 | 89 | 0.60946 | 1.839438 | 0.272774 | 2.234305 | 0.025463 |
| **POLR1C** | AT | 89 | 89 | -0.60946 | 0.543644 | 0.272774 | -2.2343 | 0.025463 |
| **TNKS** | AP | 89 | 89 | -0.81281 | 0.44361 | 0.271911 | -2.98925 | 0.002797 |
| **TNKS** | AP | 89 | 89 | 0.81281 | 2.254233 | 0.271911 | 2.989253 | 0.002797 |
| **MDP1** | ES | 89 | 89 | 0.625114 | 1.86846 | 0.27095 | 2.307123 | 0.021048 |
| **GRAPL** | AT | 91 | 87 | 0.734444 | 2.084322 | 0.28169 | 2.607272 | 0.009127 |
| **SMIM4** | AD | 89 | 89 | 0.696718 | 2.007155 | 0.266467 | 2.614647 | 0.008932 |
| **RFX3** | AT | 89 | 89 | 0.831005 | 2.295624 | 0.27683 | 3.001859 | 0.002683 |
| **PRRC2B** | ES | 89 | 89 | -0.71418 | 0.489595 | 0.270668 | -2.63857 | 0.008326 |
| **RPL37A** | AT | 89 | 89 | 0.647329 | 1.91043 | 0.277379 | 2.333731 | 0.01961 |
| **PIGH** | AT | 90 | 88 | 0.747781 | 2.112308 | 0.274903 | 2.720168 | 0.006525 |
| **ELOF1** | AP | 91 | 87 | -0.64159 | 0.526455 | 0.270967 | -2.36778 | 0.017895 |
| **STX10** | RI | 89 | 89 | -0.7961 | 0.451083 | 0.270887 | -2.93887 | 0.003294 |
| **USP4** | AT | 90 | 88 | -0.76872 | 0.463606 | 0.276027 | -2.78495 | 0.005354 |
| **RNH1** | AP | 89 | 89 | -1.02624 | 0.35835 | 0.286143 | -3.58647 | 0.000335 |

| | | | | | | | | |
|---|---|---|---|---|---|---|---|---|
| **C12orf57** | AT | 90 | 88 | -0.79298 | 0.452493 | 0.280258 | -2.82948 | 0.004662 |
| **DCTN5** | AT | 89 | 89 | 0.691608 | 1.996925 | 0.270782 | 2.554112 | 0.010646 |
| **PEX26** | AT | 89 | 89 | -0.61795 | 0.53905 | 0.274677 | -2.24973 | 0.024466 |
| **PEX26** | AT | 89 | 89 | 0.617947 | 1.855116 | 0.274677 | 2.249727 | 0.024466 |
| **ADAM19** | AT | 89 | 89 | 0.731041 | 2.077242 | 0.274865 | 2.659642 | 0.007822 |
| **ZNF7** | AT | 89 | 89 | -0.80562 | 0.446813 | 0.270617 | -2.97696 | 0.002911 |
| **NDUFS3** | RI | 89 | 89 | -0.78157 | 0.457686 | 0.27458 | -2.84642 | 0.004421 |
| **RNASE1** | ES | 89 | 89 | 0.764635 | 2.148209 | 0.266547 | 2.868668 | 0.004122 |
| **DRG2** | AA | 89 | 89 | 0.722992 | 2.060588 | 0.276936 | 2.610684 | 0.009036 |
| **C17orf62** | AP | 89 | 89 | -0.62343 | 0.536102 | 0.277872 | -2.2436 | 0.024858 |
| **L3MBTL2** | AT | 89 | 89 | -0.7387 | 0.477734 | 0.274261 | -2.69342 | 0.007072 |
| **L3MBTL2** | AT | 89 | 89 | 0.738701 | 2.093214 | 0.274261 | 2.693424 | 0.007072 |
| **ERGIC1** | AT | 89 | 89 | 0.808779 | 2.245165 | 0.281989 | 2.868127 | 0.004129 |
| **C7orf10** | AT | 89 | 89 | 0.801469 | 2.228814 | 0.278827 | 2.87443 | 0.004048 |
| **C7orf10** | AT | 89 | 89 | -0.80147 | 0.448669 | 0.278827 | -2.87443 | 0.004048 |
| **NMNAT3** | AT | 132 | 46 | -1.17153 | 0.309891 | 0.383771 | -3.05269 | 0.002268 |
| **PHF6** | AT | 89 | 89 | 0.652598 | 1.920524 | 0.265461 | 2.458354 | 0.013958 |
| **ARL16** | ES | 89 | 89 | -0.57532 | 0.562527 | 0.267748 | -2.14872 | 0.031657 |
| **BTN3A2** | AT | 89 | 89 | 0.728823 | 2.072639 | 0.27597 | 2.640945 | 0.008268 |
| **BTN3A2** | AT | 89 | 89 | -0.72882 | 0.482477 | 0.27597 | -2.64094 | 0.008268 |
| **KPNA5** | AT | 90 | 88 | 0.555493 | 1.742799 | 0.265911 | 2.08902 | 0.036706 |
| **OXLD1** | AD | 89 | 89 | -0.56416 | 0.56884 | 0.265591 | -2.12415 | 0.033657 |
| **PFDN5** | ES | 89 | 89 | -0.56618 | 0.567688 | 0.266657 | -2.12326 | 0.033732 |
| **MPHOSPH** | AT | 89 | 89 | 0.660303 | 1.935379 | 0.269661 | 2.44864 | 0.01434 |
| **MPHOSPH** | AT | 89 | 89 | -0.6603 | 0.516695 | 0.269661 | -2.44864 | 0.01434 |
| **ATG12** | ES | 89 | 89 | -0.93714 | 0.391747 | 0.283577 | -3.30471 | 0.000951 |
| **LY6E** | AA | 89 | 89 | -0.59045 | 0.554079 | 0.267516 | -2.20715 | 0.027303 |
| **NUBP2** | RI | 89 | 89 | -0.61073 | 0.542955 | 0.269865 | -2.26309 | 0.02363 |
| **EIF4A1** | RI | 89 | 89 | -0.65458 | 0.519661 | 0.275866 | -2.37282 | 0.017653 |
| **RANGRF** | ES | 89 | 89 | 0.650816 | 1.917104 | 0.2759 | 2.358885 | 0.01833 |
| **ZNF223** | AT | 89 | 89 | -0.7534 | 0.470763 | 0.271508 | -2.77487 | 0.005522 |
| **MYL9** | ES | 89 | 89 | -0.87439 | 0.417116 | 0.275196 | -3.17734 | 0.001486 |
| **OARD1** | AD | 89 | 89 | 0.596485 | 1.815726 | 0.269132 | 2.21633 | 0.026669 |
| **GPATCH2L** | AT | 89 | 89 | 0.7383 | 2.092375 | 0.270496 | 2.729432 | 0.006344 |
| **STAU2** | AT | 89 | 89 | 0.576565 | 1.779915 | 0.264813 | 2.177257 | 0.029461 |
| **CD151** | ES | 89 | 89 | 0.688929 | 1.991582 | 0.278918 | 2.470003 | 0.013511 |
| **PFDN5** | AA | 93 | 85 | -0.6106 | 0.543028 | 0.270657 | -2.25597 | 0.024072 |
| **RPLP0** | RI | 92 | 86 | -0.75299 | 0.470956 | 0.269207 | -2.79707 | 0.005157 |
| **METTL5** | RI | 89 | 89 | -0.80834 | 0.445597 | 0.294096 | -2.74856 | 0.005986 |
| **C16orf13** | ES | 89 | 89 | 0.542797 | 1.720813 | 0.2697 | 2.012596 | 0.044157 |
| **PDCD5** | AP | 89 | 89 | -0.69987 | 0.496649 | 0.273168 | -2.56206 | 0.010405 |
| **PDCD5** | AP | 89 | 89 | 0.699873 | 2.013496 | 0.273168 | 2.562057 | 0.010405 |
| **C6orf132** | AT | 89 | 89 | -0.59546 | 0.551307 | 0.268768 | -2.21553 | 0.026724 |
| **C6orf132** | AT | 89 | 89 | 0.595463 | 1.813871 | 0.268768 | 2.215529 | 0.026724 |
| **FASTK** | RI | 89 | 89 | -0.75106 | 0.471864 | 0.278939 | -2.69258 | 0.00709 |
| **HAX1** | AA | 91 | 87 | -0.55037 | 0.576735 | 0.279063 | -1.97222 | 0.048585 |
| **NUDT5** | AT | 89 | 89 | 0.563035 | 1.755993 | 0.280292 | 2.008742 | 0.044564 |

| | | | | | | | | |
|---|---|---|---|---|---|---|---|---|
| **DNAJC12** | AT | 89 | 89 | 0.526895 | 1.693666 | 0.266372 | 1.978043 | 0.047924 |
| **DNAJC12** | AT | 89 | 89 | -0.5269 | 0.590435 | 0.266372 | -1.97804 | 0.047924 |
| **SLC35E1** | AT | 131 | 47 | -0.79307 | 0.452453 | 0.32762 | -2.4207 | 0.015491 |
| **TMEM108** | AT | 90 | 88 | 0.600146 | 1.822385 | 0.267459 | 2.243879 | 0.02484 |
| **FAM126A** | AT | 89 | 89 | -0.59046 | 0.55407 | 0.267403 | -2.20814 | 0.027234 |
| **FAM126A** | AT | 89 | 89 | 0.590464 | 1.804826 | 0.267403 | 2.208144 | 0.027234 |
| **RC3H2** | AT | 89 | 89 | 0.649722 | 1.915008 | 0.275719 | 2.356462 | 0.01845 |
| **ZMYM6** | AT | 89 | 89 | 0.701562 | 2.0169 | 0.269348 | 2.604666 | 0.009196 |
| **DGAT1** | RI | 89 | 89 | -0.67076 | 0.511319 | 0.273594 | -2.45167 | 0.01422 |
| **MPC2** | AP | 90 | 88 | -0.64545 | 0.524426 | 0.272681 | -2.36705 | 0.017931 |
| **KRAS** | AT | 89 | 89 | 0.757852 | 2.133688 | 0.27544 | 2.751424 | 0.005934 |
| **KRAS** | AT | 89 | 89 | -0.75785 | 0.468672 | 0.27544 | -2.75142 | 0.005934 |
| **OGFOD2** | ES | 89 | 89 | -0.702 | 0.495591 | 0.275996 | -2.54353 | 0.010974 |
| **MOK** | ES | 89 | 89 | -0.58169 | 0.558953 | 0.26458 | -2.19854 | 0.027911 |
| **GFOD1** | AT | 89 | 89 | 0.72603 | 2.066859 | 0.27658 | 2.625032 | 0.008664 |
| **GFOD1** | AT | 89 | 89 | -0.72603 | 0.483826 | 0.27658 | -2.62503 | 0.008664 |
| **KLHL7** | AT | 89 | 89 | -0.5837 | 0.557831 | 0.269778 | -2.16363 | 0.030493 |
| **ASH2L** | ES | 89 | 89 | -0.87254 | 0.417888 | 0.276726 | -3.15308 | 0.001616 |
| **CIZ1** | ES | 89 | 89 | -0.78191 | 0.457532 | 0.281992 | -2.77281 | 0.005558 |
| **METTL3** | RI | 89 | 89 | -0.72746 | 0.483136 | 0.272942 | -2.66525 | 0.007693 |
| **EIF3G** | ES | 89 | 89 | -0.68637 | 0.503401 | 0.270506 | -2.53735 | 0.011169 |
| **TSEN34** | AP | 89 | 89 | 0.696411 | 2.006539 | 0.263928 | 2.638636 | 0.008324 |
| **SSR3** | AP | 124 | 54 | 0.676111 | 1.966216 | 0.282397 | 2.394182 | 0.016657 |
| **MPV17** | ES | 89 | 89 | 0.559201 | 1.749274 | 0.263905 | 2.118949 | 0.034095 |
| **MRPL55** | ES | 89 | 89 | 0.726063 | 2.066928 | 0.276045 | 2.630234 | 0.008533 |
| **FTH1** | AA | 102 | 76 | -0.61841 | 0.538799 | 0.274031 | -2.25672 | 0.024025 |
| **SSB** | ES | 90 | 88 | -0.70568 | 0.493775 | 0.267546 | -2.63759 | 0.00835 |
| **ARIH2** | ES | 89 | 89 | -0.75066 | 0.472054 | 0.267814 | -2.80293 | 0.005064 |
| **H2AFV** | AT | 89 | 89 | 0.623155 | 1.864802 | 0.272202 | 2.289311 | 0.022061 |
| **RPL8** | RI | 94 | 84 | -0.52806 | 0.589746 | 0.265057 | -1.99227 | 0.046342 |
| **CREB3L4** | RI | 90 | 88 | -0.79873 | 0.449899 | 0.281551 | -2.8369 | 0.004555 |
| **LIMA1** | AP | 90 | 88 | -0.78031 | 0.458262 | 0.271742 | -2.87152 | 0.004085 |
| **ZNF280D** | AT | 89 | 89 | 0.702193 | 2.018173 | 0.271783 | 2.583654 | 0.009776 |
| **CALM2** | ES | 90 | 88 | -0.75131 | 0.471748 | 0.269452 | -2.78829 | 0.005299 |
| **PICK1** | ES | 89 | 89 | -0.61927 | 0.538335 | 0.266254 | -2.32587 | 0.020025 |
| **FBXL18** | AT | 89 | 89 | -0.72085 | 0.486339 | 0.268102 | -2.68871 | 0.007173 |
| **UBE2D4** | AT | 89 | 89 | 0.727385 | 2.069662 | 0.281056 | 2.588041 | 0.009652 |
| **UBE2D4** | AT | 89 | 89 | -0.72739 | 0.483171 | 0.281056 | -2.58804 | 0.009652 |
| **ECE1** | ES | 89 | 89 | -0.56341 | 0.569263 | 0.266477 | -2.1143 | 0.03449 |
| **ZNF512** | AT | 89 | 89 | -0.82697 | 0.437374 | 0.272096 | -3.03925 | 0.002372 |
| **ZNF512** | AT | 89 | 89 | 0.826966 | 2.286371 | 0.272096 | 3.039248 | 0.002372 |
| **OCIAD1** | AD | 89 | 89 | -0.6913 | 0.500923 | 0.293818 | -2.35283 | 0.018631 |
| **C1orf86** | ES | 89 | 89 | 0.676397 | 1.966779 | 0.289756 | 2.334371 | 0.019576 |
| **ASH1L** | ES | 89 | 89 | -0.64141 | 0.526551 | 0.265754 | -2.41354 | 0.015798 |
| **TCF25** | ES | 89 | 89 | -0.54633 | 0.579069 | 0.272921 | -2.0018 | 0.045306 |
| **RPS9** | AT | 93 | 85 | -0.54477 | 0.579975 | 0.272051 | -2.00246 | 0.045236 |
| **RNF181** | RI | 89 | 89 | -0.77398 | 0.461175 | 0.278717 | -2.77692 | 0.005488 |

| | | | | | | | | |
|---|---|---|---|---|---|---|---|---|
| **ARL6IP5** | ES | 89 | 89 | -0.6935 | 0.499823 | 0.268497 | -2.58289 | 0.009798 |
| **COA3** | AT | 90 | 88 | 0.731817 | 2.078855 | 0.27804 | 2.632053 | 0.008487 |
| **METTL5** | AD | 89 | 89 | -0.63471 | 0.53009 | 0.268847 | -2.36085 | 0.018233 |
| **SLMAP** | AT | 91 | 87 | -0.714 | 0.489679 | 0.268014 | -2.66406 | 0.00772 |
| **RBBP6** | AT | 89 | 89 | 0.750147 | 2.117312 | 0.288599 | 2.599271 | 0.009342 |
| **RBBP6** | AT | 89 | 89 | -0.75015 | 0.472297 | 0.288599 | -2.59927 | 0.009342 |
| **ADD1** | ES | 89 | 89 | -0.57827 | 0.560868 | 0.26402 | -2.19025 | 0.028506 |
| **OARD1** | AD | 89 | 89 | 0.708259 | 2.030453 | 0.268601 | 2.636841 | 0.008368 |
| **NIPAL3** | AT | 89 | 89 | -0.57357 | 0.56351 | 0.268362 | -2.1373 | 0.032573 |
| **KCNC4** | RI | 89 | 89 | -0.73661 | 0.478736 | 0.270083 | -2.72733 | 0.006385 |
| **ZNF248** | AT | 89 | 89 | -0.69094 | 0.501106 | 0.275423 | -2.50865 | 0.01212 |
| **ZNF248** | AT | 89 | 89 | 0.690938 | 1.995586 | 0.275423 | 2.508645 | 0.01212 |
| **TJP1** | AD | 89 | 89 | -0.55398 | 0.574659 | 0.267945 | -2.06751 | 0.038686 |
| **UGT8** | AP | 89 | 89 | -0.74079 | 0.476739 | 0.27697 | -2.67461 | 0.007482 |
| **UGT8** | AP | 89 | 89 | 0.740785 | 2.097582 | 0.27697 | 2.674605 | 0.007482 |
| **NREP** | AT | 91 | 87 | 0.627211 | 1.872382 | 0.266144 | 2.356662 | 0.01844 |
| **CHTOP** | AA | 89 | 89 | -0.63002 | 0.532582 | 0.277317 | -2.27184 | 0.023096 |
| **TSKU** | AP | 89 | 89 | -0.58841 | 0.55521 | 0.268908 | -2.18814 | 0.028659 |
| **TSKU** | AP | 89 | 89 | 0.588408 | 1.80112 | 0.268908 | 2.188144 | 0.028659 |
| **TPT1** | AD | 129 | 49 | 0.546685 | 1.727517 | 0.270924 | 2.017857 | 0.043606 |
| **POLR2M** | AT | 92 | 86 | 0.538696 | 1.713771 | 0.268012 | 2.009971 | 0.044434 |
| **BAX** | ES | 89 | 89 | 0.754728 | 2.127032 | 0.278882 | 2.70626 | 0.006805 |
| **KIAA0226** | AT | 89 | 89 | 0.604167 | 1.829728 | 0.266567 | 2.266474 | 0.023422 |
| **RPL35** | ES | 90 | 88 | -0.80017 | 0.449252 | 0.276445 | -2.8945 | 0.003798 |
| **UBA1** | AP | 89 | 89 | 0.980536 | 2.665886 | 0.282571 | 3.470051 | 0.00052 |
| **CDK4** | ME | 90 | 88 | 0.590512 | 1.804912 | 0.2702 | 2.185464 | 0.028855 |
| **DLG4** | AT | 89 | 89 | -0.94039 | 0.390477 | 0.278996 | -3.37061 | 0.00075 |
| **DLG4** | AT | 89 | 89 | 0.940385 | 2.560967 | 0.278996 | 3.370606 | 0.00075 |
| **NME1** | RI | 90 | 88 | -0.54525 | 0.579697 | 0.268626 | -2.02977 | 0.04238 |
| **SLC52A2** | ES | 91 | 87 | -0.6351 | 0.529884 | 0.26967 | -2.35509 | 0.018518 |
| **NTMT1** | AD | 89 | 89 | 0.653136 | 1.921558 | 0.266221 | 2.453357 | 0.014153 |
| **ERRFI1** | AT | 90 | 88 | -0.77574 | 0.460361 | 0.273006 | -2.84149 | 0.00449 |
| **ANAPC13** | AD | 89 | 89 | 0.620808 | 1.86043 | 0.267044 | 2.324741 | 0.020086 |
| **RNF8** | ES | 89 | 89 | -0.5788 | 0.560573 | 0.26763 | -2.16267 | 0.030567 |
| **TRPM6** | AT | 89 | 89 | 0.668523 | 1.951354 | 0.277666 | 2.407652 | 0.016055 |
| **TPM3** | AT | 89 | 89 | 0.551286 | 1.735483 | 0.268935 | 2.049887 | 0.040376 |
| **SMAGP** | ES | 90 | 88 | -0.82405 | 0.43865 | 0.273027 | -3.01822 | 0.002543 |
| **UBE2I** | AT | 90 | 88 | -0.62367 | 0.535972 | 0.276814 | -2.25305 | 0.024256 |
| **RPS20** | AT | 89 | 89 | -0.72826 | 0.48275 | 0.268461 | -2.71271 | 0.006673 |
| **RPS20** | AT | 89 | 89 | 0.728257 | 2.071467 | 0.268461 | 2.712713 | 0.006673 |
| **TSPAN31** | AT | 89 | 89 | -0.73985 | 0.477185 | 0.273643 | -2.70371 | 0.006857 |
| **TSPAN31** | AT | 89 | 89 | 0.739851 | 2.095623 | 0.273643 | 2.703707 | 0.006857 |
| **PPP2R5C** | AT | 89 | 89 | -0.59093 | 0.55381 | 0.266197 | -2.21991 | 0.026425 |
| **TMC7** | AT | 89 | 89 | -0.7774 | 0.4596 | 0.275052 | -2.82637 | 0.004708 |
| **TMC7** | AT | 89 | 89 | 0.7774 | 2.175807 | 0.275052 | 2.826374 | 0.004708 |
| **LRRC1** | AT | 89 | 89 | 0.608162 | 1.837052 | 0.269677 | 2.255148 | 0.024124 |
| **MUM1** | AT | 89 | 89 | 0.741903 | 2.099929 | 0.269431 | 2.753594 | 0.005894 |

| | | | | | | | | |
|---|---|---|---|---|---|---|---|---|
| **MUM1** | AT | 89 | 89 | -0.7419 | 0.476207 | 0.269431 | -2.75359 | 0.005894 |
| **SRSF7** | AA | 89 | 89 | -0.80337 | 0.447819 | 0.276301 | -2.90757 | 0.003642 |
| **MTIF2** | ES | 89 | 89 | -0.7564 | 0.469352 | 0.270307 | -2.79831 | 0.005137 |
| **TMEM9** | AP | 89 | 89 | -0.7094 | 0.491941 | 0.26776 | -2.64937 | 0.008064 |
| **NXF1** | RI | 89 | 89 | -0.58534 | 0.556918 | 0.28002 | -2.09035 | 0.036587 |
| **PIGH** | AT | 90 | 88 | -0.72731 | 0.483209 | 0.275098 | -2.6438 | 0.008198 |
| **EEF2K** | AT | 89 | 89 | -0.57995 | 0.559928 | 0.276189 | -2.09982 | 0.035744 |
| **EEF2K** | AT | 89 | 89 | 0.579948 | 1.785945 | 0.276189 | 2.099824 | 0.035744 |
| **NT5E** | AT | 89 | 89 | -0.84112 | 0.431227 | 0.26897 | -3.12719 | 0.001765 |
| **NT5E** | AT | 89 | 89 | 0.841121 | 2.318965 | 0.26897 | 3.127189 | 0.001765 |
| **FAM192A** | AP | 89 | 89 | -0.66214 | 0.515747 | 0.270319 | -2.44946 | 0.014307 |
| **SLC39A6** | AT | 89 | 89 | 0.59785 | 1.818205 | 0.264266 | 2.262305 | 0.023679 |
| **SLC39A6** | AT | 89 | 89 | -0.59785 | 0.549993 | 0.264266 | -2.26231 | 0.023679 |
| **ADRA1A** | AT | 89 | 89 | -0.56469 | 0.568537 | 0.280375 | -2.01405 | 0.044005 |
| **CDC42** | ES | 93 | 85 | -0.58862 | 0.555093 | 0.265309 | -2.21862 | 0.026513 |
| **APOLD1** | AT | 89 | 89 | -0.60592 | 0.545572 | 0.283091 | -2.14037 | 0.032325 |
| **APOLD1** | AT | 89 | 89 | 0.605921 | 1.832939 | 0.283091 | 2.140374 | 0.032325 |
| **GCDH** | ME | 89 | 89 | 0.685091 | 1.983952 | 0.268632 | 2.550299 | 0.010763 |
| **RALY** | ES | 89 | 89 | 0.561042 | 1.752498 | 0.270265 | 2.075896 | 0.037904 |
| **CBR1** | AT | 92 | 86 | -0.67991 | 0.506665 | 0.284497 | -2.38986 | 0.016855 |
| **VPS28** | RI | 90 | 88 | -0.78977 | 0.453951 | 0.276582 | -2.85545 | 0.004298 |
| **ZNF593** | RI | 89 | 89 | -0.57774 | 0.561166 | 0.270789 | -2.13354 | 0.032881 |
| **MARS** | RI | 89 | 89 | -0.61951 | 0.53821 | 0.283341 | -2.18644 | 0.028784 |
| **RPS16** | AD | 89 | 89 | -0.63005 | 0.532564 | 0.277773 | -2.26822 | 0.023316 |
| **GLYCTK** | RI | 89 | 89 | -0.55099 | 0.576377 | 0.267352 | -2.06093 | 0.03931 |
| **FAM131A** | AP | 89 | 89 | 0.670054 | 1.954343 | 0.276015 | 2.427599 | 0.015199 |
| **CFLAR** | AT | 89 | 89 | -0.55066 | 0.576571 | 0.266769 | -2.06417 | 0.039001 |
| **PCMT1** | AT | 89 | 89 | -0.54382 | 0.580526 | 0.269884 | -2.01502 | 0.043903 |
| **PCMT1** | AT | 89 | 89 | 0.54382 | 1.722575 | 0.269884 | 2.015016 | 0.043903 |
| **SYP** | AT | 89 | 89 | -0.56053 | 0.570904 | 0.267399 | -2.09625 | 0.03606 |
| **SYP** | AT | 89 | 89 | 0.560535 | 1.751609 | 0.267399 | 2.096246 | 0.03606 |
| **TCEAL3** | AP | 89 | 89 | 0.538312 | 1.713113 | 0.266127 | 2.022765 | 0.043097 |
| **TCEAL3** | AP | 89 | 89 | -0.53831 | 0.583733 | 0.266127 | -2.02277 | 0.043097 |
| **TRIM46** | AT | 89 | 89 | 0.807367 | 2.241998 | 0.269711 | 2.993456 | 0.002758 |
| **C11orf57** | RI | 89 | 89 | -0.8383 | 0.432444 | 0.279647 | -2.99772 | 0.00272 |
| **RPS9** | AD | 89 | 89 | -0.54207 | 0.581542 | 0.271097 | -1.99955 | 0.045548 |
| **LAMTOR5** | ES | 89 | 89 | 0.638269 | 1.893201 | 0.267392 | 2.387015 | 0.016986 |
| **ZNF365** | AT | 89 | 89 | 0.63941 | 1.895362 | 0.272971 | 2.342413 | 0.01916 |
| **GALT** | AA | 89 | 89 | -0.79616 | 0.451056 | 0.280346 | -2.83994 | 0.004512 |
| **NUDT22** | ES | 89 | 89 | 0.767192 | 2.153709 | 0.275428 | 2.785457 | 0.005345 |
| **RBM42** | ES | 89 | 89 | 0.651921 | 1.919225 | 0.270704 | 2.408241 | 0.01603 |
| **CLASP2** | AT | 91 | 87 | -0.60385 | 0.546704 | 0.275532 | -2.19157 | 0.02841 |
| **PCNP** | AA | 89 | 89 | -0.64867 | 0.522741 | 0.266605 | -2.43307 | 0.014971 |
| **VDAC1** | AP | 89 | 89 | 0.798423 | 2.222033 | 0.272638 | 2.928503 | 0.003406 |
| **VDAC1** | AP | 89 | 89 | -0.79842 | 0.450038 | 0.272638 | -2.9285 | 0.003406 |
| **CNTRL** | AT | 89 | 89 | -0.62903 | 0.533108 | 0.269203 | -2.33665 | 0.019458 |
| **FAM69B** | AP | 89 | 89 | -0.61613 | 0.540029 | 0.268938 | -2.29098 | 0.021964 |

| | | | | | | | | |
|---|---|---|---|---|---|---|---|---|
| **FAM69B** | AP | 89 | 89 | 0.616132 | 1.851751 | 0.268938 | 2.290982 | 0.021964 |
| **RER1** | AA | 89 | 89 | -1.02153 | 0.360042 | 0.291077 | -3.50949 | 0.000449 |
| **DOK4** | RI | 89 | 89 | -0.81533 | 0.442493 | 0.274254 | -2.9729 | 0.00295 |
| **NISCH** | RI | 90 | 88 | -0.62786 | 0.533733 | 0.283249 | -2.21664 | 0.026648 |
| **NEK11** | AT | 89 | 89 | 0.655231 | 1.925588 | 0.275856 | 2.37527 | 0.017536 |
| **RPL34** | ES | 89 | 89 | -0.61675 | 0.539694 | 0.269406 | -2.2893 | 0.022062 |
| **CREBZF** | RI | 89 | 89 | 0.756827 | 2.131502 | 0.273818 | 2.763975 | 0.00571 |
| **ZSWIM7** | RI | 90 | 88 | -0.6111 | 0.542754 | 0.272706 | -2.24087 | 0.025034 |
| **EIF2A** | ES | 89 | 89 | -0.84109 | 0.43124 | 0.269462 | -3.12137 | 0.0018 |
| **SOD2** | AT | 89 | 89 | 0.535719 | 1.708677 | 0.26898 | 1.991671 | 0.046407 |
| **SOD2** | AT | 89 | 89 | -0.53572 | 0.585248 | 0.26898 | -1.99167 | 0.046407 |
| **PIGL** | AT | 89 | 89 | -0.60278 | 0.547286 | 0.274308 | -2.19747 | 0.027987 |
| **GLI4** | AT | 89 | 89 | -0.68878 | 0.502188 | 0.267879 | -2.57124 | 0.010133 |
| **GLI4** | AT | 89 | 89 | 0.688781 | 1.991287 | 0.267879 | 2.571241 | 0.010133 |
| **TRPM6** | AT | 90 | 88 | -0.58418 | 0.557564 | 0.27211 | -2.14685 | 0.031805 |
| **NPIPB3** | AP | 89 | 89 | -0.72173 | 0.485912 | 0.270428 | -2.66883 | 0.007612 |
| **RPL37A** | AT | 89 | 89 | -0.55527 | 0.573919 | 0.274327 | -2.02411 | 0.042959 |
| **ZNF354A** | ES | 89 | 89 | -0.66885 | 0.512298 | 0.268388 | -2.49209 | 0.012699 |
| **IL32** | RI | 89 | 89 | -0.60262 | 0.547375 | 0.274151 | -2.19814 | 0.027939 |
| **NKIRAS2** | ES | 89 | 89 | -0.60678 | 0.545103 | 0.263049 | -2.30672 | 0.02107 |
| **NUP62** | AD | 89 | 89 | -0.52185 | 0.593422 | 0.264347 | -1.97411 | 0.048369 |
| **SCN11A** | AT | 103 | 75 | -0.62611 | 0.53467 | 0.29635 | -2.11272 | 0.034625 |
| **CPSF3L** | AA | 89 | 89 | -0.7151 | 0.489144 | 0.271937 | -2.62965 | 0.008547 |
| **RECQL5** | AT | 89 | 89 | -0.61568 | 0.540274 | 0.265423 | -2.31962 | 0.020362 |
| **RECQL5** | AT | 89 | 89 | 0.615679 | 1.850913 | 0.265423 | 2.319616 | 0.020362 |
| **ACAA1** | RI | 89 | 89 | -0.80753 | 0.445958 | 0.273379 | -2.95389 | 0.003138 |
| **RPL24** | AT | 90 | 88 | -0.80621 | 0.446549 | 0.287658 | -2.80265 | 0.005068 |
| **OLFM1** | AT | 111 | 67 | -0.58659 | 0.556218 | 0.276911 | -2.11835 | 0.034145 |
| **PSPC1** | AT | 89 | 89 | -0.63009 | 0.532545 | 0.269686 | -2.33637 | 0.019472 |
| **ZNF675** | AT | 89 | 89 | 0.647725 | 1.911188 | 0.266292 | 2.432386 | 0.015 |
| **OBSL1** | AT | 89 | 89 | -0.67433 | 0.5095 | 0.272865 | -2.47128 | 0.013463 |
| **PSMD7** | ES | 89 | 89 | -0.94371 | 0.389183 | 0.289312 | -3.2619 | 0.001107 |
| **TMEM205** | RI | 89 | 89 | -0.63142 | 0.531836 | 0.268197 | -2.35431 | 0.018557 |
| **NADSYN1** | AP | 89 | 89 | 0.539694 | 1.715482 | 0.272403 | 1.981232 | 0.047565 |
| **NADSYN1** | AP | 89 | 89 | -0.53969 | 0.582927 | 0.272403 | -1.98123 | 0.047565 |
| **PAM16** | AT | 89 | 89 | -0.57364 | 0.563471 | 0.265927 | -2.15713 | 0.030996 |
| **PAM16** | AT | 89 | 89 | 0.57364 | 1.774715 | 0.265927 | 2.15713 | 0.030996 |
| **DCAKD** | AT | 89 | 89 | 0.661862 | 1.938398 | 0.271582 | 2.437059 | 0.014807 |
| **DCAKD** | AT | 89 | 89 | -0.66186 | 0.51589 | 0.271582 | -2.43706 | 0.014807 |
| **SKA2** | ES | 89 | 89 | -0.62635 | 0.534541 | 0.267565 | -2.34092 | 0.019236 |
| **NEDD4L** | AP | 89 | 89 | 0.807413 | 2.242101 | 0.267289 | 3.020747 | 0.002522 |
| **CFLAR** | AP | 89 | 89 | 0.809742 | 2.247329 | 0.270509 | 2.993405 | 0.002759 |
| **FCHO2** | AT | 90 | 88 | 0.564997 | 1.759443 | 0.266183 | 2.122588 | 0.033788 |
| **ZNF511** | RI | 89 | 89 | -0.60438 | 0.546413 | 0.272759 | -2.2158 | 0.026705 |
| **TRMT112** | RI | 89 | 89 | -0.65971 | 0.517001 | 0.276873 | -2.38272 | 0.017185 |
| **TMEM101** | AT | 90 | 88 | -0.56742 | 0.566985 | 0.276427 | -2.0527 | 0.040102 |
| **ZNF585B** | AP | 90 | 88 | 0.544177 | 1.72319 | 0.26319 | 2.067625 | 0.038675 |

| | | | | | | | | |
|---|---|---|---|---|---|---|---|---|
| **ZNF283** | AT | 89 | 89 | 0.928302 | 2.53021 | 0.282145 | 3.290159 | 0.001001 |
| **IRF3** | RI | 89 | 89 | -0.73607 | 0.478995 | 0.276171 | -2.66525 | 0.007693 |
| **IFT172** | AT | 89 | 89 | -0.66375 | 0.514917 | 0.279043 | -2.37867 | 0.017375 |
| **IFT172** | AT | 89 | 89 | 0.66375 | 1.942062 | 0.279043 | 2.378669 | 0.017375 |
| **MRPL2** | AT | 89 | 89 | -0.76783 | 0.464021 | 0.272033 | -2.82254 | 0.004764 |
| **MRPL2** | AT | 89 | 89 | 0.767826 | 2.155077 | 0.272033 | 2.822545 | 0.004764 |
| **TRAPPC2** | RI | 89 | 89 | 0.516307 | 1.675827 | 0.263051 | 1.962766 | 0.049673 |
| **DYNLT3** | AT | 89 | 89 | 0.588455 | 1.801203 | 0.266321 | 2.209567 | 0.027135 |
| **DYNLT3** | AT | 89 | 89 | -0.58845 | 0.555185 | 0.266321 | -2.20957 | 0.027135 |
| **INADL** | AT | 89 | 89 | 0.577079 | 1.780829 | 0.269771 | 2.139143 | 0.032424 |
| **GTF2H1** | AP | 89 | 89 | -0.74957 | 0.472569 | 0.265269 | -2.8257 | 0.004718 |
| **TAF1D** | AT | 89 | 89 | -0.74083 | 0.476718 | 0.268145 | -2.7628 | 0.005731 |
| **TAF1D** | AT | 89 | 89 | 0.740831 | 2.097678 | 0.268145 | 2.762797 | 0.005731 |
| **TAF1D** | RI | 89 | 89 | -0.59196 | 0.553242 | 0.270091 | -2.1917 | 0.028401 |
| **CIRBP** | AA | 90 | 88 | -0.67423 | 0.50955 | 0.279973 | -2.40818 | 0.016032 |
| **NUP62** | AD | 89 | 89 | -0.65196 | 0.521023 | 0.275518 | -2.36631 | 0.017967 |
| **RNF130** | ES | 89 | 89 | 0.704229 | 2.022287 | 0.272962 | 2.579949 | 0.009882 |
| **PIGO** | RI | 89 | 89 | 0.719079 | 2.052541 | 0.270499 | 2.658342 | 0.007853 |
| **CYB561A3** | RI | 89 | 89 | -0.87643 | 0.416268 | 0.28432 | -3.08253 | 0.002052 |
| **C11orf73** | ES | 89 | 89 | -0.6039 | 0.546678 | 0.272651 | -2.21491 | 0.026767 |
| **PSME2** | ES | 91 | 87 | -0.61889 | 0.538544 | 0.270418 | -2.28863 | 0.022101 |
| **BFAR** | ES | 89 | 89 | -0.52704 | 0.590351 | 0.265311 | -1.98649 | 0.046979 |
| **D2HGDH** | ES | 89 | 89 | -0.69599 | 0.498582 | 0.270782 | -2.57028 | 0.010162 |
| **KDELR3** | AT | 89 | 89 | -0.69659 | 0.498282 | 0.269624 | -2.58355 | 0.009779 |
| **KDELR3** | AT | 89 | 89 | 0.696589 | 2.006896 | 0.269624 | 2.583553 | 0.009779 |
| **CLTA** | AT | 89 | 89 | -0.60889 | 0.543957 | 0.272812 | -2.23189 | 0.025622 |
| **CLTA** | AT | 89 | 89 | 0.608885 | 1.838381 | 0.272812 | 2.231887 | 0.025622 |
| **CC2D1B** | AT | 89 | 89 | 0.771662 | 2.163358 | 0.270508 | 2.852642 | 0.004336 |
| **CC2D1B** | AT | 89 | 89 | -0.77166 | 0.462244 | 0.270508 | -2.85264 | 0.004336 |
| **ARPC5** | AT | 89 | 89 | -0.58813 | 0.555365 | 0.270418 | -2.17489 | 0.029639 |
| **ARPC5** | AT | 89 | 89 | 0.588129 | 1.800616 | 0.270418 | 2.174886 | 0.029639 |
| **MYL6** | RI | 91 | 87 | -0.57059 | 0.565194 | 0.269902 | -2.11405 | 0.034511 |
| **SIN3A** | AT | 90 | 88 | 0.694372 | 2.002452 | 0.274274 | 2.531673 | 0.011352 |
| **KIAA1430** | AT | 90 | 88 | -0.74037 | 0.476938 | 0.270924 | -2.73276 | 0.006281 |
| **BRD8** | AT | 128 | 50 | -1.07218 | 0.342263 | 0.350843 | -3.056 | 0.002243 |
| **DBNL** | AD | 90 | 88 | -0.69922 | 0.496973 | 0.273025 | -2.56101 | 0.010437 |
| **NAA38** | RI | 89 | 89 | -0.65023 | 0.521926 | 0.283157 | -2.29636 | 0.021655 |
| **SLC35A2** | AT | 89 | 89 | -0.6692 | 0.512119 | 0.272809 | -2.45299 | 0.014167 |
| **UBR4** | AT | 89 | 89 | 0.607758 | 1.836309 | 0.276599 | 2.197254 | 0.028002 |
| **UBR4** | AT | 89 | 89 | -0.60776 | 0.544571 | 0.276599 | -2.19725 | 0.028002 |
| **RBM4** | AT | 89 | 89 | 0.807627 | 2.242579 | 0.276027 | 2.925895 | 0.003435 |
| **NPIPB3** | AP | 89 | 89 | 0.834647 | 2.304001 | 0.277158 | 3.011453 | 0.0026 |
| **TLCD1** | AP | 90 | 88 | 0.654622 | 1.924415 | 0.271088 | 2.414799 | 0.015744 |
| **ZNF69** | AT | 89 | 89 | 0.608008 | 1.836769 | 0.264843 | 2.295729 | 0.021691 |
| **ZNF69** | AT | 89 | 89 | -0.60801 | 0.544434 | 0.264843 | -2.29573 | 0.021691 |
| **FCHO2** | AT | 90 | 88 | -0.56282 | 0.569601 | 0.266208 | -2.11421 | 0.034497 |
| **POLR3H** | AD | 90 | 88 | -0.53086 | 0.588099 | 0.266648 | -1.99086 | 0.046496 |

| | | | | | | | | |
|---|---|---|---|---|---|---|---|---|
| **LAMP2** | AT | 89 | 89 | 0.678908 | 1.971723 | 0.2701 | 2.513546 | 0.011952 |
| **ARL6IP4** | AD | 89 | 89 | -0.63833 | 0.528176 | 0.26925 | -2.37076 | 0.017751 |
| **EP400NL** | AT | 89 | 89 | -0.55957 | 0.571455 | 0.27428 | -2.04014 | 0.041336 |
| **CLN3** | ES | 89 | 89 | -0.73879 | 0.477692 | 0.279828 | -2.64015 | 0.008287 |
| **ALDOA** | AD | 89 | 89 | -0.83887 | 0.432198 | 0.280286 | -2.99291 | 0.002763 |
| **SCRN2** | AA | 89 | 89 | -0.61288 | 0.541787 | 0.279145 | -2.19558 | 0.028122 |
| **SUPT4H1** | AP | 89 | 89 | -0.59787 | 0.549984 | 0.27682 | -2.15976 | 0.030791 |
| **TMEM108** | AT | 90 | 88 | -0.576 | 0.56214 | 0.267764 | -2.15116 | 0.031463 |
| **PLAC8** | AT | 89 | 89 | 0.746214 | 2.109001 | 0.277472 | 2.689328 | 0.00716 |
| **PLAC8** | AT | 89 | 89 | -0.74621 | 0.474158 | 0.277472 | -2.68933 | 0.00716 |
| **ST7** | AP | 89 | 89 | -0.54475 | 0.579985 | 0.264484 | -2.05968 | 0.039429 |
| **SLC35G1** | AT | 90 | 88 | -0.86212 | 0.422264 | 0.283203 | -3.04419 | 0.002333 |
| **GEMIN2** | ES | 90 | 88 | -0.60435 | 0.546428 | 0.270371 | -2.23527 | 0.0254 |
| **ALKBH6** | AD | 89 | 89 | -0.52307 | 0.592697 | 0.265488 | -1.97023 | 0.048812 |
| **ANAPC13** | AP | 104 | 74 | -0.69804 | 0.497561 | 0.299288 | -2.33233 | 0.019683 |
| **SAT1** | ES | 90 | 88 | 0.621867 | 1.862402 | 0.269249 | 2.309634 | 0.020908 |
| **LYPLAL1** | AA | 89 | 89 | -0.73935 | 0.477423 | 0.269744 | -2.74094 | 0.006126 |
| **ST8SIA1** | AT | 89 | 89 | 0.879852 | 2.410543 | 0.285678 | 3.079872 | 0.002071 |
| **ST8SIA1** | AT | 89 | 89 | -0.87985 | 0.414844 | 0.285678 | -3.07987 | 0.002071 |
| **SNX29** | AT | 124 | 54 | 0.550438 | 1.734013 | 0.269943 | 2.039091 | 0.041441 |
| **NDUFA11** | RI | 89 | 89 | -0.53831 | 0.583735 | 0.274092 | -1.96397 | 0.049534 |
| **KIAA1430** | AT | 90 | 88 | 0.667059 | 1.948499 | 0.267288 | 2.495655 | 0.012572 |
| **THOC3** | AT | 89 | 89 | -0.72353 | 0.485038 | 0.275024 | -2.63078 | 0.008519 |
| **THOC3** | AT | 89 | 89 | 0.723528 | 2.061694 | 0.275024 | 2.630784 | 0.008519 |
| **NRP1** | ES | 89 | 89 | -0.57325 | 0.563693 | 0.267966 | -2.13925 | 0.032416 |
| **AXIN1** | ES | 89 | 89 | -0.73958 | 0.477316 | 0.274421 | -2.69504 | 0.007038 |
| **HNRNPUL1** | AA | 89 | 89 | 0.665946 | 1.946331 | 0.272526 | 2.443604 | 0.014541 |
| **DBNDD2** | AP | 89 | 89 | -0.60729 | 0.544827 | 0.270107 | -2.24833 | 0.024555 |
| **PDE4DIP** | AT | 89 | 89 | 0.667799 | 1.94994 | 0.272479 | 2.450824 | 0.014253 |
| **IL32** | AD | 89 | 89 | -0.65777 | 0.518006 | 0.278295 | -2.36357 | 0.0181 |
| **TUBB3** | AP | 89 | 89 | 0.536425 | 1.709882 | 0.263783 | 2.03358 | 0.041994 |
| **TP53** | AT | 89 | 89 | -0.76327 | 0.466138 | 0.273773 | -2.78798 | 0.005304 |
| **KLK10** | AP | 89 | 89 | 0.827094 | 2.286665 | 0.283108 | 2.921481 | 0.003484 |
| **PSMD10** | RI | 89 | 89 | -0.54859 | 0.577761 | 0.266639 | -2.05745 | 0.039643 |
| **RPLP0** | AP | 89 | 89 | 0.567759 | 1.764308 | 0.263694 | 2.153101 | 0.031311 |
| **RPLP0** | AP | 89 | 89 | -0.56776 | 0.566794 | 0.263694 | -2.1531 | 0.031311 |
| **AP1G2** | RI | 89 | 89 | -0.6931 | 0.500022 | 0.268936 | -2.5772 | 0.00996 |
| **MOK** | AP | 89 | 89 | 0.648844 | 1.913328 | 0.270613 | 2.397685 | 0.016499 |
| **WASH4P** | AD | 89 | 89 | 0.735052 | 2.08559 | 0.27502 | 2.672722 | 0.007524 |
| **ATPAF2** | AT | 89 | 89 | -0.55753 | 0.572622 | 0.267525 | -2.08403 | 0.037157 |
| **NPRL2** | RI | 89 | 89 | -0.60336 | 0.546972 | 0.270359 | -2.23169 | 0.025636 |
| **RPS3A** | ES | 89 | 89 | 0.531526 | 1.701527 | 0.265179 | 2.004405 | 0.045027 |
| **MOSPD3** | AP | 141 | 37 | -1.61882 | 0.198133 | 0.595031 | -2.72056 | 0.006517 |
| **HDAC8** | AT | 89 | 89 | 0.643406 | 1.902952 | 0.272584 | 2.360397 | 0.018255 |
| **SERPINA1** | ES | 89 | 89 | -0.75939 | 0.467951 | 0.27335 | -2.7781 | 0.005468 |
| **SH3GL1** | ES | 89 | 89 | -0.63446 | 0.530221 | 0.270136 | -2.34868 | 0.01884 |
| **BTF3** | AD | 89 | 89 | -0.67208 | 0.510644 | 0.275021 | -2.44376 | 0.014535 |

| | | | | | | | | |
|---|---|---|---|---|---|---|---|---|
| **PML** | ES | 90 | 88 | 0.596801 | 1.816299 | 0.271752 | 2.196125 | 0.028083 |
| **TMEM199** | AT | 91 | 87 | -0.63598 | 0.529414 | 0.273267 | -2.32734 | 0.019947 |
| **CHCHD7** | AP | 90 | 88 | 0.533557 | 1.704986 | 0.269075 | 1.98293 | 0.047375 |
| **IFITM2** | AP | 89 | 89 | -0.70069 | 0.496243 | 0.27289 | -2.56767 | 0.010238 |
| **IFITM2** | AP | 89 | 89 | 0.700691 | 2.015144 | 0.27289 | 2.567669 | 0.010238 |
| **NAA38** | RI | 89 | 89 | -0.66273 | 0.515444 | 0.27538 | -2.40658 | 0.016103 |
| **SLC25A24** | AT | 89 | 89 | 0.587446 | 1.799387 | 0.265769 | 2.210367 | 0.02708 |
| **SLC25A24** | AT | 89 | 89 | -0.58745 | 0.555745 | 0.265769 | -2.21037 | 0.02708 |
| **LAMB4** | AT | 89 | 89 | 0.641224 | 1.898804 | 0.267557 | 2.396588 | 0.016549 |
| **LAMB4** | AT | 89 | 89 | -0.64122 | 0.526647 | 0.267557 | -2.39659 | 0.016549 |
| **OXR1** | AT | 89 | 89 | 0.666937 | 1.94826 | 0.272727 | 2.44544 | 0.014468 |
| **OXR1** | AT | 89 | 89 | -0.66694 | 0.513279 | 0.272727 | -2.44544 | 0.014468 |
| **TCP11L1** | AT | 89 | 89 | -0.55157 | 0.576042 | 0.266463 | -2.06999 | 0.038453 |
| **TCP11L1** | AT | 89 | 89 | 0.551575 | 1.735985 | 0.266463 | 2.069989 | 0.038453 |
| **NDUFA3** | ES | 89 | 89 | 0.588087 | 1.80054 | 0.27224 | 2.160174 | 0.030759 |
| **UBE2D3** | AP | 90 | 88 | -0.69578 | 0.498685 | 0.275348 | -2.52692 | 0.011507 |
| **C1orf63** | ES | 89 | 89 | -0.61977 | 0.538067 | 0.268413 | -2.30903 | 0.020942 |
| **EIF3C** | AP | 89 | 89 | 0.62523 | 1.868676 | 0.271132 | 2.305998 | 0.021111 |
| **EIF3C** | AP | 89 | 89 | -0.62523 | 0.535138 | 0.271132 | -2.306 | 0.021111 |
| **MGST2** | ES | 89 | 89 | -0.60759 | 0.544663 | 0.269132 | -2.25758 | 0.023972 |
| **C5orf24** | RI | 89 | 89 | -0.57288 | 0.563897 | 0.269764 | -2.12365 | 0.0337 |
| **SLC35A2** | AT | 89 | 89 | 0.669178 | 1.952632 | 0.27255 | 2.455247 | 0.014079 |
| **SLC37A4** | ES | 89 | 89 | -0.71107 | 0.49112 | 0.274458 | -2.5908 | 0.009575 |
| **INO80E** | AA | 89 | 89 | -0.56522 | 0.568236 | 0.273756 | -2.06468 | 0.038953 |
| **GATSL3** | ES | 89 | 89 | 0.596519 | 1.815786 | 0.268211 | 2.224065 | 0.026144 |
| **RBP1** | AT | 89 | 89 | 0.520671 | 1.683156 | 0.265548 | 1.960739 | 0.049909 |
| **PPP2R5C** | AT | 89 | 89 | 0.53149 | 1.701466 | 0.267566 | 1.986385 | 0.046991 |
| **ZNF396** | AT | 89 | 89 | 0.563879 | 1.757477 | 0.271561 | 2.076438 | 0.037853 |
| **FKBP5** | AT | 90 | 88 | 0.559469 | 1.749743 | 0.2649 | 2.111999 | 0.034687 |
| **COX19** | AT | 89 | 89 | 0.542484 | 1.720275 | 0.26516 | 2.045876 | 0.040769 |
| **COX19** | AT | 89 | 89 | -0.54248 | 0.581302 | 0.26516 | -2.04588 | 0.040769 |
| **AAMDC** | AT | 89 | 89 | 0.635006 | 1.887034 | 0.26973 | 2.354233 | 0.018561 |
| **SNAPC2** | AP | 90 | 88 | 0.538109 | 1.712765 | 0.265991 | 2.023038 | 0.043069 |
| **TKT** | ES | 89 | 89 | -0.70835 | 0.492458 | 0.270886 | -2.61492 | 0.008925 |
| **TMEM66** | ES | 90 | 88 | 0.632877 | 1.883021 | 0.272369 | 2.323604 | 0.020147 |
| **RPL24** | AT | 91 | 87 | 0.733886 | 2.083159 | 0.278921 | 2.631158 | 0.008509 |
| **C10orf128** | AT | 89 | 89 | -0.73894 | 0.477619 | 0.273761 | -2.69922 | 0.00695 |
| **RPS3** | ES | 95 | 83 | -0.55714 | 0.572845 | 0.269294 | -2.06889 | 0.038556 |
| **SNAPC2** | AP | 90 | 88 | -0.53723 | 0.584364 | 0.266005 | -2.01963 | 0.043422 |
| **SYTL4** | AT | 89 | 89 | -0.84925 | 0.427735 | 0.280407 | -3.02864 | 0.002457 |
| **SYTL4** | AT | 89 | 89 | 0.849251 | 2.337894 | 0.280407 | 3.028639 | 0.002457 |
| **LYRM5** | AT | 89 | 89 | 0.704711 | 2.023262 | 0.266445 | 2.644868 | 0.008172 |
| **LYRM5** | AT | 89 | 89 | -0.70471 | 0.494251 | 0.266445 | -2.64487 | 0.008172 |
| **FAM192A** | AP | 89 | 89 | 0.700469 | 2.014696 | 0.275007 | 2.547091 | 0.010863 |
| **DNAJC2** | AT | 89 | 89 | 0.624793 | 1.867859 | 0.272347 | 2.294104 | 0.021785 |
| **DNAJC2** | AT | 89 | 89 | -0.62479 | 0.535372 | 0.272347 | -2.2941 | 0.021785 |
| **LRRC27** | AT | 89 | 89 | -0.57674 | 0.561729 | 0.265171 | -2.17496 | 0.029633 |

| | | | | | | | | |
|---|---|---|---|---|---|---|---|---|
| **CD59** | AP | 91 | 87 | -0.71318 | 0.490082 | 0.268865 | -2.65257 | 0.007988 |
| **DENND1A** | AT | 89 | 89 | -0.6283 | 0.533496 | 0.271804 | -2.31161 | 0.020799 |
| **EPN1** | AP | 90 | 88 | 0.942042 | 2.565214 | 0.295352 | 3.189555 | 0.001425 |
| **MRPS24** | AT | 90 | 88 | 0.613132 | 1.846205 | 0.276623 | 2.21649 | 0.026658 |
| **RHEB** | ES | 89 | 89 | -0.52989 | 0.588669 | 0.265086 | -1.99894 | 0.045615 |
| **PAXIP1-AS** | AT | 89 | 89 | 0.735974 | 2.087513 | 0.27171 | 2.708675 | 0.006755 |
| **RDX** | AT | 89 | 89 | -0.75057 | 0.472099 | 0.269441 | -2.78564 | 0.005342 |
| **MAGOHB** | ES | 89 | 89 | -0.61912 | 0.538421 | 0.269642 | -2.29606 | 0.021672 |
| **HNRNPC** | AD | 89 | 89 | -0.64341 | 0.525497 | 0.271154 | -2.37286 | 0.017651 |
| **SOGA2** | AT | 89 | 89 | 0.72684 | 2.068533 | 0.271257 | 2.679524 | 0.007373 |
| **SOGA2** | AT | 89 | 89 | -0.72684 | 0.483434 | 0.271257 | -2.67952 | 0.007373 |
| **GSTK1** | ES | 89 | 89 | 0.808178 | 2.243815 | 0.275543 | 2.93304 | 0.003357 |
| **EEF1D** | ES | 89 | 89 | -0.72096 | 0.486286 | 0.276356 | -2.6088 | 0.009086 |
| **IDH3G** | RI | 89 | 89 | -0.67289 | 0.51023 | 0.273412 | -2.4611 | 0.013851 |
| **HPS1** | AT | 89 | 89 | 0.777347 | 2.175694 | 0.27824 | 2.793798 | 0.005209 |
| **HPS1** | AT | 89 | 89 | -0.77735 | 0.459624 | 0.27824 | -2.7938 | 0.005209 |
| **HMGB1** | AP | 89 | 89 | 0.718374 | 2.051095 | 0.280276 | 2.563096 | 0.010374 |
| **CREB3L2** | AT | 89 | 89 | -0.55366 | 0.574844 | 0.266027 | -2.08121 | 0.037415 |
| **STXBP2** | AA | 89 | 89 | -0.67437 | 0.50948 | 0.274281 | -2.45866 | 0.013946 |
| **RPS9** | AA | 89 | 89 | -0.65817 | 0.517796 | 0.272556 | -2.41483 | 0.015743 |
| **ITPA** | AD | 89 | 89 | -0.84203 | 0.430835 | 0.276666 | -3.04349 | 0.002339 |
| **METTL3** | RI | 89 | 89 | -0.66524 | 0.514151 | 0.272854 | -2.43808 | 0.014766 |
| **RGS12** | AT | 90 | 88 | 0.887959 | 2.430164 | 0.284806 | 3.11777 | 0.001822 |
| **NQO2** | ES | 89 | 89 | 0.648075 | 1.911858 | 0.263952 | 2.455273 | 0.014078 |
| **RNH1** | AA | 89 | 89 | -0.56918 | 0.565987 | 0.280689 | -2.02781 | 0.04258 |
| **EML3** | AA | 89 | 89 | -0.55836 | 0.572146 | 0.267133 | -2.0902 | 0.0366 |
| **DNAJA3** | ES | 89 | 89 | -0.66662 | 0.513439 | 0.268744 | -2.48052 | 0.013119 |
| **TLCD1** | AP | 90 | 88 | -0.66007 | 0.516814 | 0.272027 | -2.42649 | 0.015246 |
| **ZCCHC9** | RI | 89 | 89 | -0.62053 | 0.537658 | 0.272346 | -2.27848 | 0.022698 |
| **C7orf55-LU** | AP | 90 | 88 | -0.77404 | 0.461148 | 0.270986 | -2.85637 | 0.004285 |
| **ZMAT1** | ES | 89 | 89 | 0.598186 | 1.818816 | 0.271257 | 2.205237 | 0.027437 |
| **VPS25** | ES | 91 | 87 | -0.72807 | 0.48284 | 0.271875 | -2.67796 | 0.007407 |
| **SNRPD1** | ES | 89 | 89 | 0.53957 | 1.715269 | 0.265681 | 2.030895 | 0.042266 |
| **DCAF5** | AT | 89 | 89 | -0.55392 | 0.574692 | 0.272513 | -2.03264 | 0.042088 |
| **DCAF5** | AT | 89 | 89 | 0.553921 | 1.740063 | 0.272513 | 2.032644 | 0.042088 |
| **PCTP** | AA | 90 | 88 | -0.61962 | 0.538146 | 0.278545 | -2.22451 | 0.026114 |
| **ZNF585B** | AP | 90 | 88 | -0.52413 | 0.592072 | 0.263357 | -1.99018 | 0.046572 |
| **NDUFA3** | RI | 89 | 89 | -0.82212 | 0.439499 | 0.287064 | -2.86389 | 0.004185 |
| **UCHL5** | AT | 89 | 89 | 0.634758 | 1.886565 | 0.270747 | 2.344471 | 0.019054 |
| **CKLF** | ES | 89 | 89 | 0.712091 | 2.038248 | 0.27199 | 2.618072 | 0.008843 |
| **GRAMD1A** | AT | 89 | 89 | 0.618801 | 1.8567 | 0.271054 | 2.282944 | 0.022434 |
| **GRAMD1A** | AT | 89 | 89 | -0.6188 | 0.53859 | 0.271054 | -2.28294 | 0.022434 |
| **AKT1S1** | AD | 89 | 89 | -0.59172 | 0.553375 | 0.267404 | -2.21283 | 0.026909 |
| **NSFL1C** | ES | 90 | 88 | 0.772868 | 2.165969 | 0.268162 | 2.882096 | 0.00395 |
| **MKRN1** | ES | 89 | 89 | 0.659217 | 1.933279 | 0.266998 | 2.469001 | 0.013549 |
| **TUSC3** | AT | 108 | 70 | -0.6331 | 0.530943 | 0.28924 | -2.18884 | 0.028608 |
| **SMUG1** | RI | 90 | 88 | -0.68572 | 0.50373 | 0.275276 | -2.49101 | 0.012738 |

| | | | | | | | | |
|---|---|---|---|---|---|---|---|---|
| **INO80C** | ES | 89 | 89 | -0.84217 | 0.430774 | 0.269683 | -3.12282 | 0.001791 |
| **COMT** | AT | 91 | 87 | -0.68312 | 0.505037 | 0.27857 | -2.45225 | 0.014197 |
| **AIFM2** | AT | 89 | 89 | -0.62346 | 0.536088 | 0.275961 | -2.25922 | 0.023869 |
| **AIFM2** | AT | 89 | 89 | 0.623457 | 1.865366 | 0.275961 | 2.259225 | 0.023869 |
| **WIBG** | AP | 89 | 89 | 0.626172 | 1.870438 | 0.269697 | 2.321763 | 0.020246 |
| **GTF3A** | AD | 89 | 89 | -0.65952 | 0.517102 | 0.277342 | -2.37799 | 0.017408 |
| **HMOX2** | ES | 89 | 89 | -0.6979 | 0.497627 | 0.270681 | -2.57832 | 0.009928 |
| **FARS2** | AP | 89 | 89 | -0.70984 | 0.491723 | 0.273191 | -2.59833 | 0.009368 |
| **FARS2** | AP | 89 | 89 | 0.70984 | 2.033667 | 0.273191 | 2.598328 | 0.009368 |
| **COX4I1** | AA | 89 | 89 | -0.72766 | 0.483037 | 0.268908 | -2.70598 | 0.00681 |
| **EMC9** | AA | 89 | 89 | -0.64098 | 0.526774 | 0.273162 | -2.34654 | 0.018949 |
| **LSMD1** | RI | 89 | 89 | -0.67701 | 0.508132 | 0.284412 | -2.3804 | 0.017294 |
| **GRAPL** | AT | 90 | 88 | -0.62621 | 0.534616 | 0.280736 | -2.23059 | 0.025708 |
| **SPHK2** | AT | 89 | 89 | -0.65424 | 0.519836 | 0.267796 | -2.44306 | 0.014563 |
| **MAGED2** | AP | 89 | 89 | 0.561652 | 1.753567 | 0.270142 | 2.079096 | 0.037609 |
| **HMGA2** | AT | 89 | 89 | -0.63767 | 0.52852 | 0.267168 | -2.38679 | 0.016996 |
| **CD37** | AT | 89 | 89 | 0.558268 | 1.747644 | 0.273525 | 2.041011 | 0.04125 |
| **CD37** | AT | 89 | 89 | -0.55827 | 0.572199 | 0.273525 | -2.04101 | 0.04125 |
| **SLC6A9** | AT | 89 | 89 | 0.580023 | 1.78608 | 0.273526 | 2.120542 | 0.03396 |
| **SLC6A9** | AT | 89 | 89 | -0.58002 | 0.559885 | 0.273526 | -2.12054 | 0.03396 |
| **PCNXL4** | AT | 89 | 89 | -0.61351 | 0.541445 | 0.270153 | -2.27098 | 0.023148 |
| **CHURC1** | ES | 89 | 89 | -0.55642 | 0.57326 | 0.269457 | -2.06495 | 0.038928 |
| **CMC2** | AP | 89 | 89 | -0.76441 | 0.465607 | 0.283151 | -2.69967 | 0.006941 |
| **VRK3** | AT | 89 | 89 | -0.60535 | 0.545885 | 0.270977 | -2.23394 | 0.025487 |
| **DHRS4** | ES | 89 | 89 | 0.549324 | 1.732082 | 0.271194 | 2.025574 | 0.042808 |
| **IFI27** | ES | 91 | 87 | 0.823539 | 2.27855 | 0.275559 | 2.98861 | 0.002802 |
| **ARRB2** | AA | 91 | 87 | -0.5933 | 0.552501 | 0.277133 | -2.14085 | 0.032286 |
| **WDPCP** | AT | 92 | 86 | 0.678383 | 1.970689 | 0.270207 | 2.510611 | 0.012052 |
| **SMIM12** | AT | 92 | 86 | -0.65828 | 0.517739 | 0.275045 | -2.39337 | 0.016695 |
| **ACTR10** | ES | 89 | 89 | -0.63734 | 0.528699 | 0.265618 | -2.39945 | 0.01642 |
| **DIP2A** | AT | 90 | 88 | 0.638225 | 1.893118 | 0.270357 | 2.360672 | 0.018242 |
| **TRAK1** | AT | 89 | 89 | 0.542447 | 1.720211 | 0.271686 | 1.996593 | 0.045869 |
| **RBM6** | AT | 89 | 89 | 0.572456 | 1.772616 | 0.264994 | 2.160263 | 0.030752 |
| **RBM6** | AT | 89 | 89 | -0.57246 | 0.564138 | 0.264994 | -2.16026 | 0.030752 |
| **CNTRL** | AT | 89 | 89 | 0.563578 | 1.756947 | 0.268642 | 2.09788 | 0.035916 |
| **SNRPA1** | ES | 89 | 89 | 0.692273 | 1.998252 | 0.274011 | 2.526445 | 0.011522 |
| **NAA15** | AT | 89 | 89 | -0.69718 | 0.497989 | 0.27442 | -2.54055 | 0.011068 |
| **NAA15** | AT | 89 | 89 | 0.697177 | 2.008075 | 0.27442 | 2.540545 | 0.011068 |
| **PRTFDC1** | AT | 89 | 89 | 0.584244 | 1.793635 | 0.27905 | 2.093687 | 0.036288 |
| **PRTFDC1** | AT | 89 | 89 | -0.58424 | 0.557527 | 0.27905 | -2.09369 | 0.036288 |
| **MLST8** | AA | 91 | 87 | -0.61903 | 0.538467 | 0.275887 | -2.24378 | 0.024846 |
| **VKORC1** | AP | 91 | 87 | 0.594209 | 1.811597 | 0.266742 | 2.227656 | 0.025903 |
| **NDUFB3** | ES | 90 | 88 | 0.656177 | 1.927409 | 0.274789 | 2.387927 | 0.016944 |
| **HMGN1** | AP | 89 | 89 | 0.906993 | 2.476863 | 0.29119 | 3.114786 | 0.001841 |
| **NDUFB2** | AP | 89 | 89 | 0.599382 | 1.820994 | 0.271761 | 2.205548 | 0.027416 |
| **CDK2AP2** | AT | 89 | 89 | 0.626522 | 1.871091 | 0.271691 | 2.306012 | 0.02111 |
| **CDK2AP2** | AT | 89 | 89 | -0.62652 | 0.534448 | 0.271691 | -2.30601 | 0.02111 |

| | | | | | | | | |
|---|---|---|---|---|---|---|---|---|
| **HMBS** | RI | 89 | 89 | -0.68471 | 0.504237 | 0.270355 | -2.53263 | 0.011321 |
| **SRSF1** | RI | 89 | 89 | -0.71752 | 0.487962 | 0.271775 | -2.64012 | 0.008288 |
| **NDUFA10** | ES | 123 | 55 | -0.62779 | 0.533769 | 0.300261 | -2.09082 | 0.036544 |
| **C1orf213** | RI | 89 | 89 | 0.621232 | 1.861219 | 0.266791 | 2.328537 | 0.019884 |
| **RPS27L** | AT | 89 | 89 | 0.692932 | 1.99957 | 0.270006 | 2.56636 | 0.010277 |
| **RPS27L** | AT | 89 | 89 | -0.69293 | 0.500108 | 0.270006 | -2.56636 | 0.010277 |
| **MFGE8** | ES | 89 | 89 | -0.72358 | 0.485012 | 0.276214 | -2.61964 | 0.008802 |
| **B3GNTL1** | AT | 89 | 89 | 0.564764 | 1.759032 | 0.267451 | 2.11165 | 0.034716 |
| **GMFG** | ES | 90 | 88 | 0.589265 | 1.802663 | 0.269184 | 2.18908 | 0.028591 |
| **MFF** | ES | 89 | 89 | 0.574588 | 1.776398 | 0.267353 | 2.149169 | 0.031621 |
| **AZI2** | AT | 89 | 89 | 0.592207 | 1.807974 | 0.277247 | 2.136029 | 0.032677 |
| **AZI2** | AT | 89 | 89 | -0.59221 | 0.553105 | 0.277247 | -2.13603 | 0.032677 |
| **GPR56** | AP | 90 | 88 | -0.64521 | 0.524551 | 0.275357 | -2.34319 | 0.01912 |
| **DALRD3** | AA | 89 | 89 | -0.52696 | 0.5904 | 0.262675 | -2.00612 | 0.044844 |
| **RPL29** | RI | 89 | 89 | -0.596 | 0.55101 | 0.272448 | -2.18758 | 0.0287 |
| **REPS2** | ES | 89 | 89 | -0.66436 | 0.514605 | 0.271772 | -2.44453 | 0.014504 |
| **GOSR2** | AT | 89 | 89 | -0.67155 | 0.510916 | 0.26821 | -2.50381 | 0.012286 |
| **MYL12A** | AD | 89 | 89 | 0.55449 | 1.741052 | 0.268199 | 2.067459 | 0.038691 |
| **CHCHD7** | AD | 89 | 89 | -0.78398 | 0.456586 | 0.275303 | -2.8477 | 0.004404 |
| **TATDN1** | ES | 89 | 89 | -0.58962 | 0.554541 | 0.267428 | -2.20477 | 0.02747 |
| **DENND1A** | AT | 89 | 89 | 0.655579 | 1.926257 | 0.271933 | 2.41081 | 0.015917 |
| **PSMG2** | AP | 127 | 51 | -0.76128 | 0.467066 | 0.308528 | -2.46747 | 0.013607 |
| **MPV17** | AT | 90 | 88 | -0.61391 | 0.541231 | 0.276686 | -2.21879 | 0.026501 |
| **ITSN1** | AT | 89 | 89 | -0.58653 | 0.556253 | 0.270295 | -2.16997 | 0.030009 |
| **SGSM3** | RI | 90 | 88 | -0.69113 | 0.501011 | 0.278109 | -2.48509 | 0.012952 |
| **WBSCR16** | AT | 89 | 89 | -0.53987 | 0.582824 | 0.267698 | -2.01671 | 0.043726 |
| **ZBTB8OS** | AT | 89 | 89 | 0.619836 | 1.858622 | 0.265447 | 2.335063 | 0.01954 |
| **ZBTB8OS** | AT | 89 | 89 | -0.61984 | 0.538033 | 0.265447 | -2.33506 | 0.01954 |
| **RDX** | AT | 89 | 89 | 0.770639 | 2.161146 | 0.269473 | 2.859804 | 0.004239 |
| **SPG21** | AP | 89 | 89 | -0.70121 | 0.495984 | 0.274741 | -2.55226 | 0.010703 |
| **COX4I1** | RI | 89 | 89 | -0.75112 | 0.471839 | 0.272692 | -2.75445 | 0.005879 |
| **TBC1D22A** | AP | 90 | 88 | 0.599817 | 1.821785 | 0.267852 | 2.239357 | 0.025133 |
| **IP6K2** | ES | 89 | 89 | 0.555332 | 1.742519 | 0.265735 | 2.089797 | 0.036636 |
| **SLK** | ES | 89 | 89 | 0.543189 | 1.721488 | 0.266422 | 2.038831 | 0.041467 |
| **EBPL** | ES | 89 | 89 | -0.69283 | 0.500161 | 0.273178 | -2.53617 | 0.011207 |
| **UBE2I** | AT | 91 | 87 | 0.579397 | 1.784962 | 0.273697 | 2.116927 | 0.034266 |
| **RPS7** | AT | 91 | 87 | 0.569277 | 1.766989 | 0.266205 | 2.138489 | 0.032477 |
| **LAMTOR1** | AT | 90 | 88 | 0.750205 | 2.117433 | 0.281055 | 2.669244 | 0.007602 |
| **APAF1** | AT | 125 | 53 | 0.552301 | 1.737245 | 0.271307 | 2.035701 | 0.04178 |
| **KAT8** | RI | 90 | 88 | -0.67332 | 0.510011 | 0.27224 | -2.47327 | 0.013388 |
| **U2AF1L4** | AD | 89 | 89 | 0.612709 | 1.845424 | 0.270731 | 2.263168 | 0.023625 |
| **SPINT2** | AD | 90 | 88 | -0.82325 | 0.439002 | 0.279592 | -2.94448 | 0.003235 |
| **KIF13B** | AT | 112 | 66 | 0.723098 | 2.060807 | 0.271748 | 2.660915 | 0.007793 |
| **CARM1** | AA | 89 | 89 | -0.58933 | 0.554696 | 0.269509 | -2.18669 | 0.028765 |
| **C1QC** | AD | 89 | 89 | -0.57764 | 0.561221 | 0.269714 | -2.14167 | 0.03222 |
| **EPHA10** | AT | 89 | 89 | -0.79523 | 0.451478 | 0.272146 | -2.92207 | 0.003477 |
| **EPHA10** | AT | 89 | 89 | 0.795229 | 2.214947 | 0.272146 | 2.922068 | 0.003477 |

| | | | | | | | | |
|---|---|---|---|---|---|---|---|---|
| **TPT1** | RI | 92 | 86 | -0.64302 | 0.525703 | 0.267858 | -2.4006 | 0.016368 |
| **MAZ** | AP | 89 | 89 | 0.743048 | 2.102334 | 0.28072 | 2.646934 | 0.008123 |
| **C2ORF15** | AP | 89 | 89 | -0.59162 | 0.553432 | 0.263199 | -2.24779 | 0.024589 |
| **C2ORF15** | AP | 89 | 89 | 0.591616 | 1.806907 | 0.263199 | 2.247794 | 0.024589 |
| **FAM104B** | AD | 91 | 87 | 0.700883 | 2.015531 | 0.27219 | 2.574978 | 0.010025 |
| **CCDC24** | RI | 89 | 89 | -0.70644 | 0.493399 | 0.276553 | -2.55444 | 0.010636 |
| **TTC39C** | AT | 89 | 89 | 0.721263 | 2.057029 | 0.273281 | 2.639268 | 0.008309 |
| **TTC39C** | AT | 89 | 89 | -0.72126 | 0.486138 | 0.273281 | -2.63927 | 0.008309 |
| **ATL2** | ES | 89 | 89 | -0.56903 | 0.566077 | 0.265549 | -2.14283 | 0.032127 |
| **NCBP2** | ES | 90 | 88 | -0.71583 | 0.488789 | 0.26902 | -2.66086 | 0.007794 |
| **PSMG4** | RI | 89 | 89 | -0.6798 | 0.506718 | 0.27805 | -2.44489 | 0.01449 |
| **C14orf2** | RI | 89 | 89 | -0.79045 | 0.453641 | 0.274533 | -2.87925 | 0.003986 |
| **NAT9** | RI | 89 | 89 | -0.70871 | 0.492278 | 0.274852 | -2.57852 | 0.009922 |
| **TTLL3** | AT | 90 | 88 | -0.61965 | 0.538133 | 0.263425 | -2.35228 | 0.018659 |
| **TBCA** | RI | 89 | 89 | -0.64219 | 0.526137 | 0.271834 | -2.36245 | 0.018154 |
| **DDX10** | AT | 89 | 89 | -0.63585 | 0.529485 | 0.270466 | -2.35094 | 0.018726 |
| **DDX10** | AT | 89 | 89 | 0.635851 | 1.888628 | 0.270466 | 2.350943 | 0.018726 |
| **SKA2** | ES | 89 | 89 | -0.69334 | 0.499904 | 0.274875 | -2.52238 | 0.011656 |
| **NDUFA10** | AT | 89 | 89 | -0.63612 | 0.529343 | 0.272218 | -2.3368 | 0.01945 |
| **PLD1** | AT | 89 | 89 | -0.69635 | 0.498402 | 0.275637 | -2.52632 | 0.011527 |
| **PLD1** | AT | 89 | 89 | 0.696347 | 2.006411 | 0.275637 | 2.526318 | 0.011527 |
| **PAM16** | AP | 89 | 89 | 0.683229 | 1.980262 | 0.26882 | 2.541587 | 0.011035 |
| **POLR2E** | AT | 90 | 88 | 0.684464 | 1.98271 | 0.274359 | 2.494777 | 0.012604 |
| **MPV17** | AT | 89 | 89 | 0.627236 | 1.872427 | 0.273753 | 2.291247 | 0.021949 |
| **UBE2D3** | AP | 89 | 89 | 0.541453 | 1.718502 | 0.266712 | 2.030103 | 0.042346 |
| **PSMG4** | RI | 89 | 89 | -0.66769 | 0.512891 | 0.278464 | -2.39777 | 0.016495 |
| **SNX3** | AD | 89 | 89 | -0.53489 | 0.585732 | 0.268235 | -1.99412 | 0.046139 |
| **C17orf62** | AP | 89 | 89 | 0.66047 | 1.935702 | 0.279086 | 2.366542 | 0.017955 |
| **SYPL1** | ES | 90 | 88 | -0.64702 | 0.523601 | 0.272459 | -2.37476 | 0.01756 |
| **SLC37A3** | AT | 90 | 88 | 0.621904 | 1.86247 | 0.272383 | 2.283198 | 0.022419 |
| **USP15** | AT | 89 | 89 | 0.703944 | 2.021711 | 0.276827 | 2.542905 | 0.010994 |
| **USP15** | AT | 89 | 89 | -0.70394 | 0.49463 | 0.276827 | -2.5429 | 0.010994 |
| **BCL7B** | ES | 89 | 89 | -0.56258 | 0.569737 | 0.269732 | -2.0857 | 0.037006 |
| **RABL5** | ES | 147 | 31 | -0.95432 | 0.385075 | 0.432839 | -2.20479 | 0.027469 |
| **ENY2** | AA | 89 | 89 | -0.58348 | 0.557951 | 0.269144 | -2.16793 | 0.030164 |
| **TMEM41A** | ES | 89 | 89 | -0.59051 | 0.554045 | 0.269983 | -2.18721 | 0.028727 |
| **TAZ** | ES | 89 | 89 | -0.6752 | 0.509056 | 0.272653 | -2.4764 | 0.013272 |
| **EMC3** | AT | 89 | 89 | 0.689998 | 1.993712 | 0.274806 | 2.510851 | 0.012044 |
| **EMC3** | AT | 89 | 89 | -0.69 | 0.501577 | 0.274806 | -2.51085 | 0.012044 |
| **ZNF7** | AT | 89 | 89 | 0.625609 | 1.869384 | 0.265129 | 2.359639 | 0.018293 |
| **AKR1A1** | ES | 89 | 89 | 0.617775 | 1.854797 | 0.271743 | 2.273377 | 0.023003 |
| **PDDC1** | ES | 89 | 89 | 0.817985 | 2.26593 | 0.284896 | 2.871173 | 0.00409 |
| **ILK** | RI | 89 | 89 | -0.62516 | 0.535177 | 0.265909 | -2.35102 | 0.018722 |
| **MDK** | AD | 89 | 89 | 0.72623 | 2.067273 | 0.276842 | 2.623271 | 0.008709 |
| **PIWIL4** | AP | 89 | 89 | 0.898592 | 2.456141 | 0.27675 | 3.246939 | 0.001167 |
| **BLOC1S1** | RI | 89 | 89 | -0.56879 | 0.566208 | 0.278191 | -2.04462 | 0.040892 |
| **SIN3A** | AT | 90 | 88 | -0.60471 | 0.546233 | 0.27388 | -2.20793 | 0.027249 |

| | | | | | | | | |
|---|---|---|---|---|---|---|---|---|
| **THOP1** | AP | 89 | 89 | -0.61588 | 0.540164 | 0.268029 | -2.29782 | 0.021572 |
| **PRKCSH** | AD | 89 | 89 | 0.54453 | 1.723798 | 0.272385 | 1.99912 | 0.045595 |
| **ECT2** | ES | 89 | 89 | -0.65275 | 0.52061 | 0.266805 | -2.44657 | 0.014422 |
| **NFE2L1** | RI | 89 | 89 | -0.70825 | 0.492508 | 0.279603 | -2.53304 | 0.011308 |
| **GRK4** | AT | 89 | 89 | -0.69279 | 0.500181 | 0.276713 | -2.50363 | 0.012293 |
| **GRK4** | AT | 89 | 89 | 0.692785 | 1.999277 | 0.276713 | 2.503628 | 0.012293 |
| **TAF9** | AP | 89 | 89 | -0.90919 | 0.402849 | 0.286069 | -3.17822 | 0.001482 |
| **RNPS1** | ES | 89 | 89 | -0.74806 | 0.473286 | 0.272112 | -2.74907 | 0.005976 |
| **GPS2** | AT | 89 | 89 | 0.597337 | 1.817273 | 0.264726 | 2.256433 | 0.024044 |
| **GPS2** | AT | 89 | 89 | -0.59734 | 0.550275 | 0.264726 | -2.25643 | 0.024044 |
| **NAAA** | AT | 89 | 89 | 0.726366 | 2.067553 | 0.272826 | 2.662382 | 0.007759 |
| **FIG4** | AT | 89 | 89 | 0.611718 | 1.843595 | 0.26936 | 2.271006 | 0.023147 |
| **FIG4** | AT | 89 | 89 | -0.61172 | 0.542418 | 0.26936 | -2.27101 | 0.023147 |
| **C7orf55-LU** | AP | 89 | 89 | 0.702304 | 2.018397 | 0.27044 | 2.59689 | 0.009407 |
| **IAH1** | AT | 89 | 89 | -0.61005 | 0.543324 | 0.26663 | -2.28799 | 0.022138 |
| **IAH1** | AT | 89 | 89 | 0.610049 | 1.840521 | 0.26663 | 2.287994 | 0.022138 |
| **ACAA1** | AT | 90 | 88 | 0.654708 | 1.924581 | 0.272944 | 2.398692 | 0.016454 |
| **PNRC1** | AD | 89 | 89 | 0.560011 | 1.750692 | 0.272745 | 2.053238 | 0.040049 |
| **ARPC1A** | AT | 89 | 89 | -0.69183 | 0.500659 | 0.269236 | -2.56961 | 0.010181 |
| **ARPC1A** | AT | 89 | 89 | 0.69183 | 1.997368 | 0.269236 | 2.569606 | 0.010181 |
| **BTF3L4** | ES | 89 | 89 | -0.62559 | 0.534946 | 0.275433 | -2.27129 | 0.023129 |
| **UBA52** | AD | 90 | 88 | -0.65974 | 0.516988 | 0.270235 | -2.44134 | 0.014633 |
| **ZNF581** | AP | 89 | 89 | 0.587984 | 1.800355 | 0.267086 | 2.201475 | 0.027702 |
| **ACSM3** | AT | 89 | 89 | -0.59986 | 0.548886 | 0.268992 | -2.23004 | 0.025745 |
| **ACSM3** | AT | 89 | 89 | 0.599864 | 1.821871 | 0.268992 | 2.230043 | 0.025745 |
| **FHL2** | AT | 91 | 87 | -0.56979 | 0.565647 | 0.271248 | -2.10061 | 0.035676 |
| **ZFAND2B** | RI | 91 | 87 | -0.63031 | 0.532424 | 0.279051 | -2.25878 | 0.023897 |
| **KIF13A** | AT | 89 | 89 | -0.58967 | 0.554513 | 0.264741 | -2.22733 | 0.025925 |
| **C6orf141** | AT | 89 | 89 | 0.531029 | 1.700682 | 0.264771 | 2.005615 | 0.044897 |
| **RPS4X** | AT | 90 | 88 | 0.592247 | 1.808046 | 0.275545 | 2.149368 | 0.031605 |
| **AK2** | ES | 89 | 89 | -0.61808 | 0.538976 | 0.270848 | -2.28203 | 0.022487 |
| **CSF1** | AT | 89 | 89 | -0.61011 | 0.543292 | 0.266042 | -2.29328 | 0.021832 |
| **CSF1** | AT | 89 | 89 | 0.610108 | 1.84063 | 0.266042 | 2.293279 | 0.021832 |
| **VAPA** | ES | 90 | 88 | 0.522702 | 1.686578 | 0.266607 | 1.960572 | 0.049929 |
| **ACAA1** | AD | 89 | 89 | -0.62307 | 0.536296 | 0.268887 | -2.31721 | 0.020492 |
| **GUCY1A3** | AT | 89 | 89 | -0.60001 | 0.548804 | 0.270836 | -2.21541 | 0.026732 |
| **GUCY1A3** | AT | 89 | 89 | 0.600013 | 1.822143 | 0.270836 | 2.215408 | 0.026732 |
| **AIG1** | AT | 89 | 89 | 0.529377 | 1.697875 | 0.267763 | 1.977038 | 0.048037 |
| **BMP1** | AA | 89 | 89 | -0.56776 | 0.566794 | 0.268209 | -2.11686 | 0.034272 |
| **CAPN2** | AP | 89 | 89 | 0.534707 | 1.706948 | 0.266486 | 2.006512 | 0.044802 |
| **CAPN2** | AP | 89 | 89 | -0.53471 | 0.585841 | 0.266486 | -2.00651 | 0.044802 |
| **ACVR1B** | AT | 89 | 89 | 0.573137 | 1.773823 | 0.266035 | 2.154365 | 0.031212 |
| **MLH1** | AT | 89 | 89 | 0.701789 | 2.017359 | 0.2742 | 2.559407 | 0.010485 |
| **MLH1** | AT | 89 | 89 | -0.70179 | 0.495698 | 0.2742 | -2.55941 | 0.010485 |
| **PCMTD1** | AT | 89 | 89 | 0.611165 | 1.842576 | 0.266769 | 2.290989 | 0.021964 |
| **PCMTD1** | AT | 89 | 89 | -0.61116 | 0.542718 | 0.266769 | -2.29099 | 0.021964 |
| **ZNF280D** | AP | 89 | 89 | -0.68917 | 0.501992 | 0.269779 | -2.55457 | 0.010632 |

| | | | | | | | | |
|---|---|---|---|---|---|---|---|---|
| **ZNF280D** | AP | 89 | 89 | 0.689171 | 1.992063 | 0.269779 | 2.554572 | 0.010632 |
| **LRRC59** | AT | 90 | 88 | -0.57442 | 0.56303 | 0.271727 | -2.11397 | 0.034518 |
| **DTD2** | AD | 89 | 89 | 0.719538 | 2.053484 | 0.269501 | 2.669889 | 0.007588 |
| **VTI1B** | ES | 89 | 89 | -0.5977 | 0.550075 | 0.267278 | -2.23625 | 0.025335 |
| **CAMKK1** | AT | 89 | 89 | 0.54336 | 1.721782 | 0.264667 | 2.052993 | 0.040073 |
| **CAMKK1** | AT | 89 | 89 | -0.54336 | 0.580794 | 0.264667 | -2.05299 | 0.040073 |
| **SSC5D** | AT | 98 | 80 | -0.68734 | 0.502912 | 0.285523 | -2.40731 | 0.016071 |
| **SUMO1** | ES | 89 | 89 | -0.7798 | 0.458497 | 0.275593 | -2.82953 | 0.004662 |
| **SIRPB2** | AT | 89 | 89 | 0.537431 | 1.711604 | 0.264369 | 2.03288 | 0.042065 |
| **RGS12** | AP | 89 | 89 | -0.62374 | 0.535937 | 0.266391 | -2.34144 | 0.019209 |
| **TMSB15B** | AT | 89 | 89 | 0.524271 | 1.689227 | 0.265214 | 1.976785 | 0.048066 |
| **DHDDS** | AA | 89 | 89 | -0.60674 | 0.545127 | 0.268456 | -2.2601 | 0.023815 |
| **MRPS16** | AT | 90 | 88 | -0.5856 | 0.55677 | 0.26936 | -2.17406 | 0.029701 |
| **ARL2** | ES | 90 | 88 | -0.53991 | 0.5828 | 0.267643 | -2.01728 | 0.043666 |
| **ZNF276** | RI | 89 | 89 | 0.628001 | 1.873862 | 0.270336 | 2.323036 | 0.020177 |
| **RPTOR** | AT | 89 | 89 | -0.60677 | 0.545108 | 0.268168 | -2.26265 | 0.023657 |
| **SMIM12** | AT | 90 | 88 | 0.668997 | 1.952278 | 0.272317 | 2.456687 | 0.014022 |
| **NMRAL1** | RI | 89 | 89 | -0.74873 | 0.472967 | 0.290577 | -2.5767 | 0.009975 |
| **LRRC59** | AT | 96 | 82 | 0.550348 | 1.733857 | 0.267034 | 2.060965 | 0.039306 |
| **RBM42** | ES | 89 | 89 | 0.737412 | 2.090518 | 0.277577 | 2.656599 | 0.007893 |
| **PRKCE** | AT | 89 | 89 | -0.57625 | 0.562002 | 0.273266 | -2.10875 | 0.034966 |
| **PRKCE** | AT | 89 | 89 | 0.576249 | 1.779352 | 0.273266 | 2.108747 | 0.034966 |
| **COMMD1** | ES | 89 | 89 | -0.70301 | 0.495092 | 0.268011 | -2.62307 | 0.008714 |
| **FAM3A** | ES | 89 | 89 | 0.572713 | 1.773071 | 0.27118 | 2.111927 | 0.034693 |
| **XPNPEP1** | ES | 89 | 89 | -0.66958 | 0.511923 | 0.268239 | -2.49621 | 0.012553 |
| **CEP63** | AT | 89 | 89 | -0.68654 | 0.503313 | 0.280489 | -2.44767 | 0.014379 |
| **CEP63** | AT | 89 | 89 | 0.686542 | 1.986834 | 0.280489 | 2.447666 | 0.014379 |
| **TAF1D** | RI | 89 | 89 | -0.73395 | 0.48001 | 0.277989 | -2.64021 | 0.008286 |
| **TIAF1** | AP | 89 | 89 | 0.530535 | 1.699841 | 0.266904 | 1.98774 | 0.04684 |
| **TIAF1** | AP | 89 | 89 | -0.53053 | 0.58829 | 0.266904 | -1.98774 | 0.04684 |
| **RPS16** | AD | 91 | 87 | -0.55913 | 0.571704 | 0.26449 | -2.11401 | 0.034514 |
| **RPS23** | RI | 92 | 86 | -0.74016 | 0.477036 | 0.277695 | -2.66538 | 0.00769 |
| **RPL30** | ES | 89 | 89 | -0.60846 | 0.544186 | 0.283878 | -2.1434 | 0.032081 |
| **ERRFI1** | AT | 90 | 88 | 0.709869 | 2.033724 | 0.271275 | 2.616789 | 0.008876 |
| **MGRN1** | AA | 89 | 89 | -0.53915 | 0.583245 | 0.273181 | -1.97359 | 0.048428 |
| **RGS12** | AT | 90 | 88 | -0.84029 | 0.431586 | 0.285044 | -2.94792 | 0.003199 |
| **SDHA** | ES | 90 | 88 | -0.72156 | 0.485995 | 0.269341 | -2.67898 | 0.007385 |
| **ZNF483** | AT | 89 | 89 | -0.53926 | 0.58318 | 0.270716 | -1.99198 | 0.046374 |
| **FAM195B** | RI | 89 | 89 | -0.64687 | 0.523682 | 0.27253 | -2.37357 | 0.017617 |
| **REEP5** | ES | 89 | 89 | -0.53982 | 0.582852 | 0.267043 | -2.02147 | 0.043231 |
| **ZNF783** | AT | 89 | 89 | -0.69914 | 0.497013 | 0.267985 | -2.60887 | 0.009084 |
| **ZNF783** | AT | 89 | 89 | 0.699139 | 2.01202 | 0.267985 | 2.608874 | 0.009084 |
| **IDS** | AT | 89 | 89 | -0.68223 | 0.505488 | 0.280523 | -2.432 | 0.015016 |
| **SZT2** | AT | 89 | 89 | -0.67473 | 0.509294 | 0.272434 | -2.47667 | 0.013261 |
| **ZNF609** | AT | 91 | 87 | -0.56402 | 0.568915 | 0.270805 | -2.08276 | 0.037273 |
| **ABLIM2** | AT | 89 | 89 | 0.570937 | 1.769925 | 0.263653 | 2.165486 | 0.03035 |
| **ABLIM2** | AT | 89 | 89 | -0.57094 | 0.564996 | 0.263653 | -2.16549 | 0.03035 |

| | | | | | | | | |
|---|---|---|---|---|---|---|---|---|
| **SLC37A3** | AT | 91 | 87 | -0.61116 | 0.542723 | 0.272513 | -2.24266 | 0.024918 |
| **VPS28** | RI | 89 | 89 | -0.65735 | 0.518221 | 0.2727 | -2.41054 | 0.015929 |
| **SULT1A3** | AP | 89 | 89 | -0.56068 | 0.570823 | 0.266845 | -2.10113 | 0.035629 |
| **TAF1D** | AA | 89 | 89 | -0.591 | 0.553772 | 0.26871 | -2.1994 | 0.027849 |
| **C12orf57** | AA | 89 | 89 | -0.61103 | 0.54279 | 0.269625 | -2.26623 | 0.023437 |
| **SNF8** | AD | 90 | 88 | -0.62953 | 0.532841 | 0.277488 | -2.26868 | 0.023288 |
| **EPN1** | AP | 90 | 88 | -0.87838 | 0.415456 | 0.295199 | -2.97555 | 0.002925 |
| **STAG2** | AP | 89 | 89 | 0.578499 | 1.783359 | 0.271617 | 2.12983 | 0.033186 |
| **ZNF747** | AT | 90 | 88 | -0.79398 | 0.452041 | 0.273213 | -2.90609 | 0.00366 |
| **PSMD8** | AT | 90 | 88 | -0.53441 | 0.586014 | 0.272355 | -1.96219 | 0.049741 |
| **RQCD1** | ES | 89 | 89 | -0.67794 | 0.507661 | 0.264735 | -2.56083 | 0.010442 |
| **ZFAND2B** | RI | 89 | 89 | -0.68464 | 0.504271 | 0.276144 | -2.47929 | 0.013164 |
| **EIF4A1** | RI | 90 | 88 | -0.65986 | 0.516923 | 0.275829 | -2.39228 | 0.016744 |
| **NSRP1** | AT | 89 | 89 | 0.577828 | 1.782164 | 0.266848 | 2.165381 | 0.030359 |
| **NSRP1** | AT | 89 | 89 | -0.57783 | 0.561116 | 0.266848 | -2.16538 | 0.030359 |
| **NUP85** | AP | 89 | 89 | 0.613598 | 1.847065 | 0.272722 | 2.249901 | 0.024455 |
| **NUP85** | AP | 89 | 89 | -0.6136 | 0.5414 | 0.272722 | -2.2499 | 0.024455 |
| **IFRD2** | ES | 90 | 88 | 0.662218 | 1.939089 | 0.270004 | 2.452626 | 0.014182 |
| **LRRC23** | ES | 89 | 89 | -0.73276 | 0.480582 | 0.270868 | -2.70521 | 0.006826 |
| **ERCC5** | AP | 90 | 88 | 0.559289 | 1.749429 | 0.264332 | 2.115863 | 0.034356 |
| **RAB26** | AT | 89 | 89 | -0.63122 | 0.531943 | 0.267957 | -2.35567 | 0.018489 |
| **RAB26** | AT | 89 | 89 | 0.631219 | 1.879901 | 0.267957 | 2.355675 | 0.018489 |
| **GRN** | ES | 93 | 85 | -0.54204 | 0.581559 | 0.268023 | -2.02237 | 0.043138 |
| **ZNF606** | AT | 89 | 89 | -0.60619 | 0.545424 | 0.271597 | -2.23196 | 0.025618 |
| **GNAS** | AA | 89 | 89 | -0.65655 | 0.518636 | 0.270601 | -2.42628 | 0.015254 |
| **APP** | AP | 89 | 89 | 0.669672 | 1.953596 | 0.269712 | 2.482911 | 0.013031 |
| **ACHE** | ES | 89 | 89 | -0.59528 | 0.551407 | 0.269191 | -2.21137 | 0.02701 |
| **IDH3G** | RI | 89 | 89 | -0.58411 | 0.557604 | 0.270605 | -2.15852 | 0.030888 |
| **VWA5B1** | AT | 91 | 87 | -0.81346 | 0.443322 | 0.275831 | -2.94912 | 0.003187 |
| **PEX16** | AD | 89 | 89 | -0.63931 | 0.527657 | 0.278302 | -2.29717 | 0.021609 |
| **BSCL2** | ES | 89 | 89 | -0.57921 | 0.560339 | 0.265575 | -2.18098 | 0.029185 |
| **SUMF1** | AT | 90 | 88 | 0.606482 | 1.833967 | 0.269787 | 2.247998 | 0.024576 |
| **SUFU** | AT | 90 | 88 | -0.67105 | 0.511172 | 0.270867 | -2.47742 | 0.013234 |
| **CD44** | ES | 89 | 89 | -0.68933 | 0.50191 | 0.272276 | -2.53174 | 0.01135 |
| **FBXO34** | AT | 90 | 88 | 0.525544 | 1.691379 | 0.266071 | 1.9752 | 0.048245 |
| **ZNHIT3** | AT | 89 | 89 | -0.56171 | 0.570236 | 0.27109 | -2.07203 | 0.038263 |
| **ZNHIT3** | AT | 89 | 89 | 0.561705 | 1.753661 | 0.27109 | 2.072028 | 0.038263 |
| **ECE2** | AT | 89 | 89 | 0.565911 | 1.761052 | 0.268043 | 2.111269 | 0.034749 |
| **ECE2** | AT | 89 | 89 | -0.56591 | 0.567842 | 0.268043 | -2.11127 | 0.034749 |
| **ZNF33A** | AT | 89 | 89 | 0.59751 | 1.817587 | 0.26893 | 2.221806 | 0.026296 |
| **ZNF33A** | AT | 89 | 89 | -0.59751 | 0.55018 | 0.26893 | -2.22181 | 0.026296 |
| **NDUFA7** | ES | 91 | 87 | 0.571815 | 1.771479 | 0.265997 | 2.149702 | 0.031579 |
| **CCNG2** | AT | 89 | 89 | -0.6637 | 0.514942 | 0.27199 | -2.44016 | 0.014681 |
| **CCNG2** | AT | 89 | 89 | 0.6637 | 1.941965 | 0.27199 | 2.440164 | 0.014681 |
| **TM4SF1** | ES | 90 | 88 | -0.57502 | 0.562691 | 0.270864 | -2.12293 | 0.03376 |
| **PAFAH1B2** | AT | 90 | 88 | 0.531091 | 1.700787 | 0.265488 | 2.000434 | 0.045453 |
| **CYB5D2** | AD | 89 | 89 | -0.54979 | 0.577073 | 0.265495 | -2.07079 | 0.038378 |

| | | | | | | | | |
|---|---|---|---|---|---|---|---|---|
| **DNAH2** | AT | 89 | 89 | 0.521401 | 1.684385 | 0.265229 | 1.96585 | 0.049316 |
| **DNAH2** | AT | 89 | 89 | -0.5214 | 0.593688 | 0.265229 | -1.96585 | 0.049316 |
| **HN1** | ES | 90 | 88 | -0.62382 | 0.535891 | 0.266727 | -2.33881 | 0.019345 |
| **FLT3LG** | AT | 89 | 89 | 0.553702 | 1.739681 | 0.265843 | 2.082815 | 0.037268 |
| **LPP** | AT | 89 | 89 | 0.80088 | 2.227499 | 0.270723 | 2.958294 | 0.003093 |
| **LPP** | AT | 89 | 89 | -0.80088 | 0.448934 | 0.270723 | -2.95829 | 0.003093 |
| **RABEPK** | AT | 89 | 89 | 0.568198 | 1.765084 | 0.271097 | 2.095924 | 0.036089 |
| **RABEPK** | AT | 89 | 89 | -0.5682 | 0.566545 | 0.271097 | -2.09592 | 0.036089 |
| **SELENBP1** | ES | 89 | 89 | -0.66203 | 0.515805 | 0.265783 | -2.49085 | 0.012744 |
| **NUB1** | AT | 89 | 89 | 0.574129 | 1.775584 | 0.268223 | 2.14049 | 0.032315 |
| **NUB1** | AT | 89 | 89 | -0.57413 | 0.563195 | 0.268223 | -2.14049 | 0.032315 |
| **FDPS** | ES | 89 | 89 | 0.556401 | 1.744383 | 0.269358 | 2.065658 | 0.038861 |
| **OS9** | RI | 90 | 88 | -0.60014 | 0.548736 | 0.275413 | -2.17905 | 0.029328 |
| **ZNF581** | AP | 89 | 89 | -0.60502 | 0.546063 | 0.267417 | -2.26246 | 0.023669 |
| **ADCK3** | AP | 89 | 89 | 0.571804 | 1.77146 | 0.268645 | 2.128473 | 0.033298 |
| **ADCK3** | AP | 89 | 89 | -0.5718 | 0.564506 | 0.268645 | -2.12847 | 0.033298 |
| **DNAJC24** | AT | 89 | 89 | -0.56286 | 0.569577 | 0.274256 | -2.05232 | 0.040139 |
| **DYNLL1** | AP | 90 | 88 | 0.733672 | 2.082714 | 0.276817 | 2.650381 | 0.00804 |
| **PHC3** | AT | 89 | 89 | 0.75932 | 2.136824 | 0.267227 | 2.84148 | 0.00449 |
| **C12orf57** | AT | 93 | 85 | 0.673503 | 1.961095 | 0.271748 | 2.478407 | 0.013197 |
| **DYNC1I2** | ES | 89 | 89 | -0.60577 | 0.545652 | 0.269126 | -2.25089 | 0.024392 |
| **HSPB1** | AD | 89 | 89 | 0.533098 | 1.704204 | 0.265884 | 2.005 | 0.044963 |
| **GRHPR** | AP | 89 | 89 | -0.67583 | 0.508736 | 0.272558 | -2.47957 | 0.013154 |
| **GRHPR** | AP | 89 | 89 | 0.675827 | 1.965657 | 0.272558 | 2.479567 | 0.013154 |
| **LYPLA2** | RI | 89 | 89 | -0.56976 | 0.565662 | 0.269971 | -2.11044 | 0.03482 |
| **NDUFS2** | RI | 89 | 89 | -0.59662 | 0.550668 | 0.270386 | -2.20656 | 0.027345 |
| **TROAP** | AT | 89 | 89 | 0.641426 | 1.899186 | 0.269932 | 2.376252 | 0.017489 |
| **CARM1** | ES | 89 | 89 | -0.64758 | 0.523308 | 0.269974 | -2.3987 | 0.016454 |
| **GUSB** | AT | 91 | 87 | 0.60809 | 1.83692 | 0.274989 | 2.211328 | 0.027013 |
| **SLC39A14** | AT | 89 | 89 | 0.648923 | 1.913478 | 0.273585 | 2.371928 | 0.017696 |
| **SLC39A14** | AT | 89 | 89 | -0.64892 | 0.522609 | 0.273585 | -2.37193 | 0.017696 |
| **ANAPC11** | AD | 89 | 89 | -0.54304 | 0.580978 | 0.26805 | -2.0259 | 0.042775 |
| **MRPS33** | AP | 89 | 89 | -0.58539 | 0.556887 | 0.266646 | -2.19539 | 0.028136 |
| **EXOSC2** | ES | 89 | 89 | -0.56772 | 0.566818 | 0.270842 | -2.09611 | 0.036072 |
| **RBM7** | RI | 89 | 89 | -0.70945 | 0.491914 | 0.270855 | -2.6193 | 0.008811 |
| **PSMC5** | AA | 89 | 89 | -0.65605 | 0.518896 | 0.271061 | -2.42031 | 0.015507 |
| **RPS9** | AD | 89 | 89 | -0.54325 | 0.580856 | 0.276141 | -1.9673 | 0.049149 |
| **MPRIP** | AT | 89 | 89 | -0.56805 | 0.566631 | 0.265785 | -2.13724 | 0.032578 |
| **MPRIP** | AT | 89 | 89 | 0.568048 | 1.764818 | 0.265785 | 2.137243 | 0.032578 |
| **HNRNPM** | ES | 89 | 89 | 0.635608 | 1.88817 | 0.266887 | 2.381566 | 0.017239 |
| **ZNF816** | AT | 89 | 89 | 0.560556 | 1.751645 | 0.268976 | 2.084039 | 0.037157 |
| **ALS2** | AT | 89 | 89 | 0.535285 | 1.707934 | 0.264309 | 2.025222 | 0.042845 |
| **ALS2** | AT | 89 | 89 | -0.53528 | 0.585503 | 0.264309 | -2.02522 | 0.042845 |
| **ACSS1** | ES | 89 | 89 | -0.5916 | 0.553443 | 0.268139 | -2.20631 | 0.027363 |
| **MCUR1** | ES | 90 | 88 | -0.53645 | 0.58482 | 0.264638 | -2.02712 | 0.042651 |
| **NGDN** | RI | 89 | 89 | -0.62548 | 0.535002 | 0.276023 | -2.26606 | 0.023448 |
| **NIPA1** | AP | 89 | 89 | -0.66472 | 0.514416 | 0.271509 | -2.44825 | 0.014355 |

| | | | | | | | | |
|---|---|---|---|---|---|---|---|---|
| **GTF3C5** | AT | 89 | 89 | 0.623527 | 1.865496 | 0.27772 | 2.245166 | 0.024757 |
| **GTF3C5** | AT | 89 | 89 | -0.62353 | 0.53605 | 0.27772 | -2.24517 | 0.024757 |
| **C11orf57** | RI | 89 | 89 | -0.73155 | 0.481165 | 0.267529 | -2.73445 | 0.006248 |
| **PCNXL4** | AT | 89 | 89 | -0.70146 | 0.495862 | 0.277456 | -2.52817 | 0.011466 |
| **TP53** | AT | 89 | 89 | 0.597983 | 1.818447 | 0.271648 | 2.201318 | 0.027714 |
| **NEDD4L** | AP | 89 | 89 | -0.69727 | 0.497943 | 0.266068 | -2.62065 | 0.008776 |
| **AP2S1** | AP | 91 | 87 | -0.60217 | 0.547622 | 0.267794 | -2.24863 | 0.024536 |
| **TRAPPC2P1** | RI | 89 | 89 | -0.62782 | 0.533755 | 0.276899 | -2.26732 | 0.023371 |
| **DNAJC21** | AA | 89 | 89 | 0.565197 | 1.759794 | 0.268505 | 2.104978 | 0.035293 |
| **KCNH2** | AT | 89 | 89 | -0.58733 | 0.55581 | 0.267815 | -2.19304 | 0.028304 |
| **KCNH2** | AT | 89 | 89 | 0.587329 | 1.799176 | 0.267815 | 2.193043 | 0.028304 |
| **ME3** | ES | 89 | 89 | -0.73205 | 0.480923 | 0.271666 | -2.69466 | 0.007046 |
| **ANXA5** | AD | 90 | 88 | -0.54771 | 0.578273 | 0.271394 | -2.01813 | 0.043577 |
| **IK** | ES | 89 | 89 | -0.754 | 0.470483 | 0.269609 | -2.79662 | 0.005164 |
| **POLH** | AT | 90 | 88 | -0.61984 | 0.538031 | 0.265169 | -2.33752 | 0.019412 |
| **LRP10** | RI | 91 | 87 | 0.600767 | 1.823517 | 0.267861 | 2.24283 | 0.024908 |
| **CIRBP** | AP | 89 | 89 | -0.52414 | 0.592064 | 0.265081 | -1.97728 | 0.04801 |
| **PDCD5** | RI | 89 | 89 | -0.69285 | 0.50015 | 0.276256 | -2.50799 | 0.012142 |
| **SRSF7** | RI | 89 | 89 | -0.65359 | 0.520176 | 0.275906 | -2.36888 | 0.017842 |
| **SRSF7** | AD | 89 | 89 | -0.62586 | 0.534801 | 0.276864 | -2.26054 | 0.023788 |
| **CCT4** | ES | 89 | 89 | -0.65453 | 0.519686 | 0.27505 | -2.37968 | 0.017328 |
| **REEP5** | ES | 89 | 89 | -0.57479 | 0.562824 | 0.267505 | -2.1487 | 0.031658 |
| **APRT** | ES | 90 | 88 | 0.635599 | 1.888152 | 0.270622 | 2.34866 | 0.018841 |
| **DIP2A** | AT | 89 | 89 | -0.52587 | 0.591043 | 0.267036 | -1.96927 | 0.048922 |
| **RHBDF1** | AT | 89 | 89 | -0.55124 | 0.576232 | 0.272837 | -2.02042 | 0.04334 |
| **RHBDF1** | AT | 89 | 89 | 0.551245 | 1.735412 | 0.272837 | 2.020416 | 0.04334 |
| **ZNF843** | AT | 89 | 89 | 0.565048 | 1.759533 | 0.267227 | 2.114488 | 0.034474 |
| **ZNF843** | AT | 89 | 89 | -0.56505 | 0.568333 | 0.267227 | -2.11449 | 0.034474 |
| **ACAA1** | AT | 89 | 89 | -0.57035 | 0.565329 | 0.268292 | -2.12585 | 0.033516 |
| **SERPINB6** | AP | 89 | 89 | -0.55146 | 0.576111 | 0.267124 | -2.06442 | 0.038978 |
| **CDK2AP1** | AP | 89 | 89 | -0.58359 | 0.557893 | 0.268991 | -2.16954 | 0.030041 |
| **HAUS1** | ES | 89 | 89 | -0.54832 | 0.577921 | 0.265359 | -2.06633 | 0.038798 |
| **RPS7** | AT | 91 | 87 | -0.53308 | 0.586796 | 0.267998 | -1.98911 | 0.046689 |
| **FHL2** | ES | 89 | 89 | -0.6156 | 0.540317 | 0.271428 | -2.268 | 0.023329 |
| **NDUFC1** | AP | 89 | 89 | -0.64415 | 0.525108 | 0.278686 | -2.31138 | 0.020812 |
| **NDUFC1** | AP | 89 | 89 | 0.64415 | 1.904368 | 0.278686 | 2.311381 | 0.020812 |
| **CAMKMT** | AT | 129 | 49 | 0.569559 | 1.767487 | 0.286217 | 1.989956 | 0.046596 |
| **PKIG** | AT | 89 | 89 | -0.53119 | 0.587905 | 0.266675 | -1.9919 | 0.046383 |
| **PKIG** | AT | 89 | 89 | 0.531189 | 1.700954 | 0.266675 | 1.991895 | 0.046383 |
| **ATP11B** | AT | 101 | 77 | 0.658717 | 1.932312 | 0.266314 | 2.473466 | 0.013381 |
| **TAF9** | AP | 89 | 89 | 0.788288 | 2.199627 | 0.273442 | 2.882836 | 0.003941 |
| **ZNF345** | AT | 89 | 89 | -0.69716 | 0.497999 | 0.278273 | -2.5053 | 0.012235 |
| **ZNF345** | AT | 89 | 89 | 0.697157 | 2.008035 | 0.278273 | 2.505296 | 0.012235 |
| **ITGB1BP1** | AA | 89 | 89 | -0.76366 | 0.465959 | 0.275766 | -2.76923 | 0.005619 |
| **KLHL7** | AT | 89 | 89 | 0.628367 | 1.874547 | 0.271208 | 2.316916 | 0.020508 |
| **MRPL55** | ES | 89 | 89 | 0.622236 | 1.86309 | 0.2708 | 2.297769 | 0.021575 |
| **ATP5L** | AT | 101 | 77 | 0.532918 | 1.703897 | 0.264675 | 2.013483 | 0.044064 |

| | | | | | | | | |
|---|---|---|---|---|---|---|---|---|
| **MYL6** | AT | 89 | 89 | -0.5764 | 0.561917 | 0.267504 | -2.15474 | 0.031182 |
| **ANXA2** | ES | 96 | 82 | 0.52526 | 1.690898 | 0.263788 | 1.991219 | 0.046457 |
| **TBC1D5** | AP | 89 | 89 | 0.673984 | 1.962038 | 0.270247 | 2.493955 | 0.012633 |
| **EXOSC8** | AA | 89 | 89 | 0.556877 | 1.745213 | 0.267047 | 2.085311 | 0.037041 |
| **DUT** | RI | 91 | 87 | -0.57473 | 0.562856 | 0.274261 | -2.09556 | 0.036121 |
| **CRYZL1** | AT | 89 | 89 | 0.617462 | 1.854217 | 0.265246 | 2.327883 | 0.019918 |
| **DDX17** | AP | 89 | 89 | -0.79396 | 0.452052 | 0.278661 | -2.84919 | 0.004383 |
| **DDX17** | AP | 89 | 89 | 0.793958 | 2.212134 | 0.278661 | 2.849192 | 0.004383 |
| **SNX13** | AT | 89 | 89 | 0.802971 | 2.232162 | 0.281307 | 2.854433 | 0.004311 |
| **SNX13** | AT | 89 | 89 | -0.80297 | 0.447996 | 0.281307 | -2.85443 | 0.004311 |
| **SLC31A1** | AT | 89 | 89 | -0.69357 | 0.499791 | 0.275769 | -2.51503 | 0.011902 |
| **SLC31A1** | AT | 89 | 89 | 0.693566 | 2.000838 | 0.275769 | 2.515029 | 0.011902 |
| **LRRC23** | AT | 89 | 89 | 0.622836 | 1.864207 | 0.269094 | 2.314569 | 0.020637 |
| **LRRC23** | AT | 89 | 89 | -0.62284 | 0.536421 | 0.269094 | -2.31457 | 0.020637 |
| **TCFL5** | AT | 89 | 89 | -0.67016 | 0.511626 | 0.274748 | -2.43919 | 0.01472 |
| **TCFL5** | AT | 89 | 89 | 0.670162 | 1.954554 | 0.274748 | 2.439186 | 0.01472 |
| **UBE2L3** | AP | 89 | 89 | -0.64967 | 0.522217 | 0.274094 | -2.37025 | 0.017776 |
| **UBE2L3** | AP | 89 | 89 | 0.649672 | 1.914913 | 0.274094 | 2.370252 | 0.017776 |
| **IP6K2** | RI | 89 | 89 | -0.57066 | 0.56515 | 0.273937 | -2.0832 | 0.037233 |
| **ACTB** | ES | 96 | 82 | -0.59769 | 0.550082 | 0.28094 | -2.12746 | 0.033382 |
| **PELI3** | AT | 89 | 89 | 0.539688 | 1.715471 | 0.264299 | 2.041957 | 0.041156 |
| **PELI3** | AT | 89 | 89 | -0.53969 | 0.58293 | 0.264299 | -2.04196 | 0.041156 |
| **ATP5G2** | AA | 89 | 89 | -0.52984 | 0.588696 | 0.268272 | -1.97502 | 0.048265 |
| **TPT1** | AD | 89 | 89 | -0.58456 | 0.557354 | 0.267247 | -2.18732 | 0.028719 |
| **PABPN1** | RI | 89 | 89 | -0.62008 | 0.5379 | 0.272571 | -2.27494 | 0.02291 |
| **RPS15A** | AD | 89 | 89 | -0.58022 | 0.559775 | 0.268025 | -2.1648 | 0.030403 |
| **POLR2E** | AT | 90 | 88 | -0.65161 | 0.521208 | 0.275038 | -2.36915 | 0.017829 |
| **NDUFA7** | ES | 89 | 89 | 0.586609 | 1.797882 | 0.269284 | 2.178401 | 0.029376 |
| **QTRT1** | RI | 89 | 89 | -0.69725 | 0.497955 | 0.269922 | -2.58314 | 0.009791 |
| **CCDC74B** | AT | 89 | 89 | -0.60849 | 0.54417 | 0.27313 | -2.22785 | 0.025891 |
| **CCDC74B** | AT | 89 | 89 | 0.608493 | 1.83766 | 0.27313 | 2.227849 | 0.025891 |
| **RUVBL1** | AP | 90 | 88 | -0.5349 | 0.585725 | 0.266377 | -2.00807 | 0.044635 |
| **LAMTOR4** | RI | 90 | 88 | -0.61196 | 0.542285 | 0.280836 | -2.17908 | 0.029326 |
| **EEF1D** | AP | 89 | 89 | -0.55502 | 0.574058 | 0.273008 | -2.03299 | 0.042053 |
| **ZNFX1** | AT | 89 | 89 | -0.61159 | 0.542489 | 0.268084 | -2.28132 | 0.022529 |
| **ZNFX1** | AT | 89 | 89 | 0.611587 | 1.843355 | 0.268084 | 2.281324 | 0.022529 |
| **DUSP18** | AT | 89 | 89 | 0.682854 | 1.979519 | 0.276407 | 2.470462 | 0.013494 |
| **PLEKHA8** | AT | 89 | 89 | -0.56068 | 0.57082 | 0.267072 | -2.09936 | 0.035785 |
| **GTF2IRD2** | AT | 89 | 89 | -0.55963 | 0.571418 | 0.272123 | -2.05655 | 0.039729 |
| **CCDC171** | AT | 116 | 62 | -0.72671 | 0.483498 | 0.310146 | -2.34312 | 0.019123 |
| **RPLP0** | RI | 89 | 89 | -0.57326 | 0.563687 | 0.264736 | -2.16539 | 0.030358 |
| **PPP1R13B** | ES | 89 | 89 | -0.57636 | 0.56194 | 0.263399 | -2.18817 | 0.028657 |
| **RPS17** | RI | 89 | 89 | -0.70132 | 0.495932 | 0.280674 | -2.49868 | 0.012466 |
| **C19orf54** | AP | 89 | 89 | 0.543376 | 1.721809 | 0.27656 | 1.964769 | 0.049441 |
| **WDR45** | AA | 89 | 89 | -0.75951 | 0.467895 | 0.273564 | -2.77636 | 0.005497 |
| **EIF2AK4** | AT | 89 | 89 | -0.5295 | 0.588897 | 0.263408 | -2.0102 | 0.04441 |
| **DEF8** | AT | 89 | 89 | 0.713507 | 2.041137 | 0.275599 | 2.588931 | 0.009627 |

| | | | | | | | | |
|---|---|---|---|---|---|---|---|---|
| **DEF8** | AT | 89 | 89 | -0.71351 | 0.489923 | 0.275599 | -2.58893 | 0.009627 |
| **RPL36** | RI | 89 | 89 | -0.56061 | 0.570858 | 0.263879 | -2.12451 | 0.033627 |
| **HDAC10** | ES | 89 | 89 | 0.576657 | 1.780077 | 0.27242 | 2.116794 | 0.034277 |
| **LARP1B** | AT | 89 | 89 | 0.585432 | 1.795766 | 0.269455 | 2.172652 | 0.029807 |
| **ATP6V0B** | ES | 89 | 89 | -0.70849 | 0.492385 | 0.27708 | -2.557 | 0.010558 |
| **TRPT1** | AA | 89 | 89 | -0.55349 | 0.574941 | 0.269172 | -2.05626 | 0.039757 |
| **METTL23** | RI | 89 | 89 | -0.53767 | 0.584105 | 0.269944 | -1.9918 | 0.046393 |
| **NPIPB4** | ES | 89 | 89 | -0.63489 | 0.529992 | 0.273252 | -2.32347 | 0.020154 |
| **RCAN1** | AP | 128 | 50 | -0.62212 | 0.536808 | 0.316296 | -1.96688 | 0.049197 |
| **SEC13** | ES | 89 | 89 | 0.591486 | 1.806671 | 0.268243 | 2.205042 | 0.027451 |
| **H2AFV** | AT | 89 | 89 | -0.54291 | 0.581054 | 0.270061 | -2.01033 | 0.044396 |
| **RMDN1** | AA | 90 | 88 | 0.569425 | 1.767251 | 0.270742 | 2.103204 | 0.035448 |
| **C12orf73** | RI | 89 | 89 | -0.60906 | 0.543863 | 0.272825 | -2.23241 | 0.025588 |
| **NDUFB10** | RI | 89 | 89 | -0.52857 | 0.589449 | 0.267162 | -1.97845 | 0.047878 |
| **MFSD11** | RI | 89 | 89 | -0.65356 | 0.52019 | 0.269218 | -2.42763 | 0.015198 |
| **CAMTA1** | ES | 90 | 88 | 0.571419 | 1.770778 | 0.266382 | 2.14511 | 0.031944 |
| **ERAP2** | AT | 90 | 88 | -0.53579 | 0.585205 | 0.268778 | -1.99345 | 0.046213 |
| **CLSPN** | AT | 89 | 89 | 0.531165 | 1.700913 | 0.269867 | 1.968246 | 0.04904 |
| **CLSPN** | AT | 89 | 89 | -0.53117 | 0.587919 | 0.269867 | -1.96825 | 0.04904 |
| **ATPAF1** | ES | 89 | 89 | -0.71051 | 0.491394 | 0.278387 | -2.55224 | 0.010703 |
| **MRPL51** | AP | 90 | 88 | -0.64644 | 0.523907 | 0.272377 | -2.37333 | 0.017628 |
| **SFSWAP** | ES | 89 | 89 | 0.55897 | 1.74887 | 0.270336 | 2.067688 | 0.038669 |
| **TMEM98** | AT | 89 | 89 | -0.54202 | 0.581572 | 0.267346 | -2.02741 | 0.04262 |
| **TMEM98** | AT | 89 | 89 | 0.542021 | 1.719478 | 0.267346 | 2.027414 | 0.04262 |
| **ZNF586** | AT | 89 | 89 | -0.7247 | 0.484468 | 0.270694 | -2.6772 | 0.007424 |
| **CGREF1** | AT | 89 | 89 | -0.62242 | 0.536642 | 0.267884 | -2.32349 | 0.020153 |
| **CGREF1** | AT | 89 | 89 | 0.622424 | 1.86344 | 0.267884 | 2.323487 | 0.020153 |
| **NUDT22** | RI | 89 | 89 | -0.59072 | 0.553928 | 0.268116 | -2.20323 | 0.027578 |
| **MPND** | AA | 90 | 88 | -0.56792 | 0.566705 | 0.275313 | -2.06281 | 0.039131 |
| **ATP5E** | RI | 90 | 88 | -0.61881 | 0.538583 | 0.274767 | -2.25214 | 0.024314 |
| **SUMF1** | AT | 90 | 88 | -0.58588 | 0.556618 | 0.269913 | -2.17061 | 0.029961 |
| **TBC1D5** | AP | 89 | 89 | -0.64262 | 0.525911 | 0.26947 | -2.38477 | 0.01709 |
| **C1orf85** | AT | 89 | 89 | 0.785459 | 2.193413 | 0.282642 | 2.778989 | 0.005453 |
| **C1orf85** | AT | 89 | 89 | -0.78546 | 0.455911 | 0.282642 | -2.77899 | 0.005453 |
| **MRPL43** | AT | 89 | 89 | -0.57338 | 0.563615 | 0.265362 | -2.16076 | 0.030714 |
| **FTH1** | AT | 93 | 85 | 0.764068 | 2.146993 | 0.280289 | 2.725998 | 0.006411 |
| **SRSF2** | RI | 89 | 89 | -0.6699 | 0.51176 | 0.281092 | -2.38321 | 0.017163 |
| **ISOC2** | ES | 89 | 89 | -0.59684 | 0.550551 | 0.267882 | -2.22798 | 0.025882 |
| **MANBAL** | ES | 94 | 84 | 0.56208 | 1.754317 | 0.267914 | 2.097984 | 0.035907 |
| **WDR20** | AT | 89 | 89 | -0.58369 | 0.557838 | 0.274092 | -2.12952 | 0.033211 |
| **COMMD4** | AA | 89 | 89 | -0.66462 | 0.51447 | 0.273231 | -2.43244 | 0.014997 |
| **TMEM53** | AT | 89 | 89 | 0.608992 | 1.838577 | 0.26867 | 2.266691 | 0.023409 |
| **TMEM53** | AT | 89 | 89 | -0.60899 | 0.543899 | 0.26867 | -2.26669 | 0.023409 |
| **ILK** | RI | 89 | 89 | -0.61003 | 0.543334 | 0.27266 | -2.23733 | 0.025265 |
| **MRPL51** | AP | 90 | 88 | 0.607585 | 1.835993 | 0.269716 | 2.252683 | 0.024279 |
| **MCM7** | AP | 89 | 89 | -0.5309 | 0.588078 | 0.268862 | -1.97461 | 0.048313 |
| **MCM7** | AP | 89 | 89 | 0.530896 | 1.700455 | 0.268862 | 1.974606 | 0.048313 |

| | | | | | | | | |
|---|---|---|---|---|---|---|---|---|
| **C9orf64** | AP | 103 | 75 | -0.81748 | 0.441544 | 0.29718 | -2.75079 | 0.005945 |
| **PPIE** | ES | 89 | 89 | -0.54029 | 0.582579 | 0.273064 | -1.97862 | 0.047859 |
| **RCN1** | AP | 108 | 70 | -0.9486 | 0.387282 | 0.308365 | -3.07623 | 0.002096 |
| **IQCK** | AT | 89 | 89 | -0.6209 | 0.537458 | 0.274353 | -2.26316 | 0.023626 |
| **IQCK** | AT | 89 | 89 | 0.620904 | 1.860609 | 0.274353 | 2.263155 | 0.023626 |
| **NSMF** | RI | 90 | 88 | -0.6465 | 0.523875 | 0.271542 | -2.38086 | 0.017272 |
| **SETD4** | AT | 89 | 89 | -0.62556 | 0.534963 | 0.267464 | -2.33885 | 0.019343 |
| **SETD4** | AT | 89 | 89 | 0.625557 | 1.869286 | 0.267464 | 2.338848 | 0.019343 |
| **PHPT1** | ES | 90 | 88 | 0.575617 | 1.778228 | 0.268301 | 2.145419 | 0.031919 |
| **MED8** | AA | 90 | 88 | -0.77848 | 0.459103 | 0.284957 | -2.73192 | 0.006297 |
| **PARP6** | ES | 89 | 89 | -0.55792 | 0.572396 | 0.263968 | -2.1136 | 0.034549 |
| **KIAA0513** | AT | 89 | 89 | 0.563572 | 1.756937 | 0.268116 | 2.10197 | 0.035556 |
| **KIAA0513** | AT | 89 | 89 | -0.56357 | 0.569172 | 0.268116 | -2.10197 | 0.035556 |
| **ANAPC11** | AP | 89 | 89 | -0.63608 | 0.529363 | 0.276197 | -2.303 | 0.021279 |
| **RBM43** | AT | 89 | 89 | 0.551922 | 1.736588 | 0.266594 | 2.070273 | 0.038427 |
| **RBM43** | AT | 89 | 89 | -0.55192 | 0.575842 | 0.266594 | -2.07027 | 0.038427 |
| **EIF6** | AP | 89 | 89 | 0.631352 | 1.880151 | 0.265623 | 2.376869 | 0.01746 |
| **EIF6** | AP | 89 | 89 | -0.63135 | 0.531872 | 0.265623 | -2.37687 | 0.01746 |
| **FRMD8** | AT | 89 | 89 | -0.58255 | 0.558474 | 0.269028 | -2.16538 | 0.030359 |
| **FRMD8** | AT | 89 | 89 | 0.582548 | 1.790594 | 0.269028 | 2.165378 | 0.030359 |
| **ALDOA** | RI | 89 | 89 | -0.5472 | 0.578569 | 0.279107 | -1.96053 | 0.049934 |
| **LMNA** | AT | 89 | 89 | 0.609256 | 1.839062 | 0.270686 | 2.250785 | 0.024399 |
| **LMNA** | AT | 89 | 89 | -0.60926 | 0.543755 | 0.270686 | -2.25079 | 0.024399 |
| **C1orf198** | AP | 89 | 89 | -0.54179 | 0.581708 | 0.273236 | -1.98285 | 0.047384 |
| **ATP5L** | AT | 96 | 82 | -0.70404 | 0.494583 | 0.284921 | -2.471 | 0.013474 |
| **CDK2AP1** | AP | 89 | 89 | 0.64722 | 1.910223 | 0.268019 | 2.414828 | 0.015743 |
| **DCTN5** | ES | 89 | 89 | -0.60099 | 0.548269 | 0.266969 | -2.25116 | 0.024375 |
| **NLRP1** | AT | 89 | 89 | 0.587864 | 1.80014 | 0.266911 | 2.202473 | 0.027632 |
| **SPAG16** | AT | 89 | 89 | -0.59677 | 0.550587 | 0.268806 | -2.22008 | 0.026413 |
| **TMX2** | RI | 89 | 89 | -0.58909 | 0.554831 | 0.276937 | -2.12717 | 0.033406 |
| **NARF** | AP | 89 | 89 | -0.55705 | 0.572895 | 0.270963 | -2.05583 | 0.039799 |
| **NARF** | AP | 89 | 89 | 0.557053 | 1.745521 | 0.270963 | 2.055831 | 0.039799 |
| **TMEM205** | ES | 89 | 89 | 0.563473 | 1.756763 | 0.266471 | 2.114576 | 0.034466 |
| **SLC35A2** | AT | 89 | 89 | 0.584195 | 1.793546 | 0.264702 | 2.206988 | 0.027315 |
| **ROGDI** | AD | 89 | 89 | -0.70147 | 0.495858 | 0.273336 | -2.56632 | 0.010278 |
| **RNMT** | AA | 90 | 88 | -0.59744 | 0.550219 | 0.269539 | -2.21652 | 0.026656 |
| **PUF60** | ES | 89 | 89 | -0.60491 | 0.546125 | 0.266731 | -2.26785 | 0.023338 |
| **ACRC** | AP | 113 | 65 | 0.607477 | 1.835794 | 0.270117 | 2.248943 | 0.024516 |
| **ACOT8** | ES | 89 | 89 | 0.571588 | 1.771077 | 0.266283 | 2.146543 | 0.03183 |
| **FBXO34** | AT | 92 | 86 | -0.57744 | 0.561333 | 0.268987 | -2.14672 | 0.031815 |
| **MT1F** | AD | 90 | 88 | -0.66734 | 0.513073 | 0.27292 | -2.44517 | 0.014478 |
| **ZSCAN30** | AT | 89 | 89 | 0.622421 | 1.863435 | 0.265501 | 2.344327 | 0.019061 |
| **NDUFA10** | AT | 90 | 88 | 0.538574 | 1.713562 | 0.267374 | 2.01431 | 0.043977 |
| **ACY1** | AP | 89 | 89 | -0.64644 | 0.523909 | 0.281005 | -2.30044 | 0.021423 |
| **ACY1** | AP | 89 | 89 | 0.646437 | 1.908727 | 0.281005 | 2.300443 | 0.021423 |
| **GPR110** | AT | 89 | 89 | 0.540841 | 1.717451 | 0.266016 | 2.033118 | 0.042041 |
| **GPR110** | AT | 89 | 89 | -0.54084 | 0.582258 | 0.266016 | -2.03312 | 0.042041 |

| | | | | | | | | |
|---|---|---|---|---|---|---|---|---|
| **DYNLL1** | AP | 90 | 88 | -0.77721 | 0.459688 | 0.28291 | -2.74719 | 0.006011 |
| **DHX30** | ES | 89 | 89 | -0.59253 | 0.552924 | 0.266443 | -2.22387 | 0.026157 |
| **AP4M1** | AT | 89 | 89 | 0.534735 | 1.706996 | 0.264095 | 2.024786 | 0.042889 |
| **AP4M1** | AT | 89 | 89 | -0.53474 | 0.585824 | 0.264095 | -2.02479 | 0.042889 |
| **CD300LG** | AT | 123 | 55 | -0.67438 | 0.50947 | 0.320071 | -2.10698 | 0.035119 |
| **ATP9B** | AT | 93 | 85 | -0.68048 | 0.506373 | 0.278471 | -2.44364 | 0.01454 |
| **XYLB** | AT | 89 | 89 | -0.66352 | 0.515036 | 0.268327 | -2.4728 | 0.013406 |
| **XYLB** | AT | 89 | 89 | 0.663519 | 1.941613 | 0.268327 | 2.4728 | 0.013406 |
| **ABCC5** | AT | 89 | 89 | 0.560739 | 1.751968 | 0.270329 | 2.074283 | 0.038053 |
| **FAM65B** | AT | 89 | 89 | -0.62217 | 0.536776 | 0.271467 | -2.2919 | 0.021911 |
| **ZNF250** | AT | 89 | 89 | 0.54574 | 1.725885 | 0.264576 | 2.062697 | 0.039141 |
| **IMMP1L** | ES | 89 | 89 | 0.629338 | 1.876367 | 0.273079 | 2.304601 | 0.021189 |
| **MRPL48** | ES | 90 | 88 | 0.553755 | 1.739774 | 0.267748 | 2.068198 | 0.038621 |
| **SHMT1** | ES | 89 | 89 | -0.52868 | 0.589382 | 0.263922 | -2.00318 | 0.045158 |
| **FLJ27365** | AT | 89 | 89 | -0.53017 | 0.588507 | 0.267461 | -1.98222 | 0.047455 |
| **FLJ27365** | AT | 89 | 89 | 0.530166 | 1.699215 | 0.267461 | 1.982216 | 0.047455 |
| **IFI44** | AT | 89 | 89 | -0.54655 | 0.578941 | 0.271802 | -2.01086 | 0.044341 |
| **IFI44** | AT | 89 | 89 | 0.546555 | 1.727292 | 0.271802 | 2.010857 | 0.044341 |
| **SCRN2** | RI | 89 | 89 | -0.65075 | 0.521655 | 0.273699 | -2.37761 | 0.017425 |
| **AK9** | AT | 89 | 89 | -0.87646 | 0.416256 | 0.278128 | -3.15127 | 0.001626 |
| **COQ4** | AT | 89 | 89 | 0.535776 | 1.708774 | 0.268216 | 1.997553 | 0.045765 |
| **ANKRD10** | ES | 89 | 89 | -0.87958 | 0.414956 | 0.288575 | -3.04802 | 0.002304 |
| **ERAL1** | AD | 89 | 89 | -0.74204 | 0.476143 | 0.273473 | -2.71338 | 0.00666 |
| **SMARCE1** | ES | 89 | 89 | 0.657027 | 1.929049 | 0.27057 | 2.428309 | 0.015169 |
| **ACOT9** | AT | 89 | 89 | -0.56396 | 0.56895 | 0.2804 | -2.01128 | 0.044296 |
| **ACOT9** | AT | 89 | 89 | 0.563963 | 1.757624 | 0.2804 | 2.011276 | 0.044296 |
| **ARMCX4** | AT | 89 | 89 | 0.770887 | 2.161684 | 0.273969 | 2.813772 | 0.004896 |
| **INO80C** | AT | 89 | 89 | -0.56513 | 0.568284 | 0.280275 | -2.01636 | 0.043762 |
| **INO80C** | AT | 89 | 89 | 0.565135 | 1.759685 | 0.280275 | 2.01636 | 0.043762 |
| **MTHFD2L** | AT | 89 | 89 | -0.54849 | 0.577819 | 0.272622 | -2.01193 | 0.044228 |
| **ACTG1** | RI | 90 | 88 | -0.62969 | 0.532756 | 0.270908 | -2.32438 | 0.020105 |
| **EHMT1** | AT | 89 | 89 | 0.569053 | 1.766593 | 0.277165 | 2.053117 | 0.040061 |
| **LAMTOR1** | AT | 89 | 89 | -0.63244 | 0.531296 | 0.275211 | -2.298 | 0.021562 |
| **NUMB** | AP | 89 | 89 | 0.538058 | 1.712677 | 0.265521 | 2.026425 | 0.042721 |
| **NUMB** | AP | 89 | 89 | -0.53806 | 0.583881 | 0.265521 | -2.02642 | 0.042721 |
| **MRPL27** | RI | 89 | 89 | -0.73501 | 0.479501 | 0.286824 | -2.56258 | 0.01039 |
| **POLH** | AT | 90 | 88 | 0.5415 | 1.718583 | 0.262818 | 2.060363 | 0.039364 |
| **RAP1B** | AT | 89 | 89 | -0.54858 | 0.577769 | 0.265362 | -2.0673 | 0.038706 |
| **ZNF747** | AT | 90 | 88 | 0.702973 | 2.019748 | 0.269462 | 2.608799 | 0.009086 |
| **ZNF616** | AT | 89 | 89 | 0.522944 | 1.686988 | 0.265708 | 1.968117 | 0.049055 |
| **ZNF616** | AT | 89 | 89 | -0.52294 | 0.592773 | 0.265708 | -1.96812 | 0.049055 |
| **PRKCZ** | ES | 89 | 89 | -0.54277 | 0.581136 | 0.27099 | -2.00291 | 0.045187 |
| **CERS2** | AT | 91 | 87 | -0.61665 | 0.539748 | 0.275505 | -2.23827 | 0.025204 |
| **ANO6** | AT | 89 | 89 | -0.73378 | 0.480092 | 0.269152 | -2.72626 | 0.006406 |
| **VIMP** | AT | 89 | 89 | -0.62754 | 0.533902 | 0.274272 | -2.28804 | 0.022135 |
| **VIMP** | AT | 89 | 89 | 0.627543 | 1.873003 | 0.274272 | 2.288035 | 0.022135 |
| **SLC25A35** | AT | 89 | 89 | 0.553977 | 1.740159 | 0.27211 | 2.035853 | 0.041765 |

| | | | | | | | | |
|---|---|---|---|---|---|---|---|---|
| **SLC25A35** | AT | 89 | 89 | -0.55398 | 0.57466 | 0.27211 | -2.03585 | 0.041765 |
| **CLIP4** | AT | 89 | 89 | 0.60522 | 1.831656 | 0.272106 | 2.224208 | 0.026134 |
| **COMT** | AT | 90 | 88 | 0.536877 | 1.710656 | 0.270107 | 1.987642 | 0.046851 |
| **MFSD1** | ES | 89 | 89 | -0.6046 | 0.54629 | 0.27776 | -2.17672 | 0.029502 |
| **GANC** | AT | 89 | 89 | -0.54635 | 0.57906 | 0.26913 | -2.03006 | 0.042351 |
| **FLAD1** | RI | 89 | 89 | -0.54929 | 0.577358 | 0.272084 | -2.01884 | 0.043504 |
| **IFT81** | AT | 89 | 89 | 0.874499 | 2.397674 | 0.285309 | 3.065099 | 0.002176 |
| **IFT81** | AT | 89 | 89 | -0.8745 | 0.417071 | 0.285309 | -3.0651 | 0.002176 |
| **KLHL7** | AT | 89 | 89 | -0.56233 | 0.56988 | 0.269224 | -2.08871 | 0.036734 |
| **DPAGT1** | RI | 89 | 89 | -0.69555 | 0.498798 | 0.273357 | -2.54449 | 0.010944 |
| **DUSP11** | AT | 89 | 89 | 0.530415 | 1.699638 | 0.267426 | 1.983406 | 0.047322 |
| **DUSP11** | AT | 89 | 89 | -0.53042 | 0.588361 | 0.267426 | -1.98341 | 0.047322 |
| **PON2** | AA | 89 | 89 | -0.59911 | 0.549301 | 0.269935 | -2.21945 | 0.026456 |
| **BCAP29** | AT | 89 | 89 | -0.54543 | 0.579592 | 0.268221 | -2.03352 | 0.042 |
| **CNTLN** | AT | 89 | 89 | 0.544409 | 1.723589 | 0.272632 | 1.996861 | 0.04584 |
| **ARMCX4** | AT | 89 | 89 | -0.76928 | 0.463346 | 0.274771 | -2.79971 | 0.005115 |
| **DNAJB12** | AP | 89 | 89 | 0.777288 | 2.175563 | 0.278938 | 2.786595 | 0.005326 |
| **DNAJB12** | AP | 89 | 89 | -0.77729 | 0.459651 | 0.278938 | -2.7866 | 0.005326 |
| **C1QTNF6** | RI | 89 | 89 | 0.595468 | 1.81388 | 0.262895 | 2.265041 | 0.02351 |
| **PPIE** | AD | 89 | 89 | 0.540524 | 1.716905 | 0.271373 | 1.991813 | 0.046392 |
| **HMGB1** | RI | 89 | 89 | -0.83387 | 0.434364 | 0.281602 | -2.96118 | 0.003065 |
| **CDK10** | RI | 89 | 89 | -0.60428 | 0.546469 | 0.272741 | -2.21558 | 0.026721 |
| **DDX56** | AD | 89 | 89 | -0.57032 | 0.565343 | 0.271041 | -2.1042 | 0.035361 |
| **PSMD9** | ES | 90 | 88 | 0.57188 | 1.771594 | 0.267106 | 2.141019 | 0.032273 |
| **MPHOSPH** | AT | 89 | 89 | -0.56274 | 0.569645 | 0.265337 | -2.12086 | 0.033934 |
| **MPHOSPH** | AT | 89 | 89 | 0.562742 | 1.75548 | 0.265337 | 2.120861 | 0.033934 |
| **DPY30** | AD | 89 | 89 | 0.688108 | 1.989947 | 0.270774 | 2.541267 | 0.011045 |
| **GPR125** | AT | 89 | 89 | 0.521152 | 1.683966 | 0.264107 | 1.973261 | 0.048466 |
| **COQ2** | AT | 90 | 88 | 0.530034 | 1.69899 | 0.269254 | 1.96853 | 0.049007 |
| **UBE2A** | ES | 89 | 89 | -0.54127 | 0.582009 | 0.269873 | -2.00564 | 0.044894 |
| **METTL17** | RI | 89 | 89 | -0.53177 | 0.587563 | 0.266448 | -1.99578 | 0.045958 |
| **ITFG3** | AT | 90 | 88 | 0.697638 | 2.009002 | 0.273514 | 2.550648 | 0.010752 |
| **FCGRT** | AP | 89 | 89 | 0.537044 | 1.710941 | 0.264433 | 2.030922 | 0.042263 |
| **FCGRT** | AP | 89 | 89 | -0.53704 | 0.584474 | 0.264433 | -2.03092 | 0.042263 |
| **Sep-08** | AT | 89 | 89 | 0.662552 | 1.939736 | 0.27208 | 2.435137 | 0.014886 |
| **GPR137** | AA | 90 | 88 | -0.55087 | 0.576449 | 0.264838 | -2.08002 | 0.037524 |
| **MBD1** | AA | 89 | 89 | -0.53667 | 0.584694 | 0.269668 | -1.9901 | 0.04658 |
| **VIL1** | AT | 89 | 89 | 0.557143 | 1.745678 | 0.265491 | 2.098542 | 0.035857 |
| **VIL1** | AT | 89 | 89 | -0.55714 | 0.572843 | 0.265491 | -2.09854 | 0.035857 |
| **PKN2** | AT | 89 | 89 | -0.60605 | 0.545499 | 0.273013 | -2.21988 | 0.026427 |
| **PKN2** | AT | 89 | 89 | 0.606054 | 1.833184 | 0.273013 | 2.219877 | 0.026427 |
| **CREM** | AT | 89 | 89 | -0.54947 | 0.577253 | 0.268183 | -2.04888 | 0.040474 |
| **MRPS7** | RI | 89 | 89 | -0.59298 | 0.552676 | 0.273239 | -2.1702 | 0.029991 |
| **CEP70** | AT | 89 | 89 | 0.533405 | 1.704727 | 0.26597 | 2.005509 | 0.044909 |
| **TSC22D3** | AP | 90 | 88 | 0.781039 | 2.18374 | 0.271415 | 2.877657 | 0.004006 |
| **MED1** | AT | 89 | 89 | 0.581683 | 1.789047 | 0.267133 | 2.177507 | 0.029443 |
| **MED1** | AT | 89 | 89 | -0.58168 | 0.558957 | 0.267133 | -2.17751 | 0.029443 |

| | | | | | | | | |
|---|---|---|---|---|---|---|---|---|
| **SKA2** | ES | 89 | 89 | -0.61194 | 0.542298 | 0.26843 | -2.2797 | 0.022626 |
| **NDUFS7** | RI | 89 | 89 | -0.57568 | 0.56232 | 0.269947 | -2.13258 | 0.032959 |
| **CHFR** | AT | 89 | 89 | -0.59554 | 0.551266 | 0.26583 | -2.2403 | 0.025072 |
| **CHFR** | AT | 89 | 89 | 0.595537 | 1.814005 | 0.26583 | 2.240296 | 0.025072 |
| **NIPA1** | AP | 89 | 89 | 0.626963 | 1.871916 | 0.273961 | 2.28851 | 0.022108 |
| **IL32** | RI | 89 | 89 | -0.67573 | 0.508783 | 0.27522 | -2.45525 | 0.014079 |
| **METTL23** | AD | 89 | 89 | -0.5733 | 0.56366 | 0.272184 | -2.10631 | 0.035178 |
| **ZNF714** | AT | 89 | 89 | 0.664951 | 1.944395 | 0.281899 | 2.358825 | 0.018333 |
| **ZNF714** | AT | 89 | 89 | -0.66495 | 0.514299 | 0.281899 | -2.35883 | 0.018333 |
| **NR5A2** | AT | 89 | 89 | -0.58627 | 0.556396 | 0.264419 | -2.21722 | 0.026608 |
| **NR5A2** | AT | 89 | 89 | 0.586275 | 1.79728 | 0.264419 | 2.217221 | 0.026608 |
| **MRPS16** | AT | 90 | 88 | 0.532516 | 1.703213 | 0.268074 | 1.986453 | 0.046983 |
| **TUBGCP2** | AP | 89 | 89 | -0.66401 | 0.514781 | 0.270844 | -2.45164 | 0.014221 |
| **SPAG16** | AT | 90 | 88 | -0.60271 | 0.547328 | 0.275776 | -2.1855 | 0.028852 |
| **PPARD** | AT | 89 | 89 | -0.64384 | 0.525272 | 0.271915 | -2.3678 | 0.017894 |
| **PPARD** | AT | 89 | 89 | 0.643839 | 1.903776 | 0.271915 | 2.367798 | 0.017894 |
| **SWI5** | AA | 89 | 89 | 0.62551 | 1.869198 | 0.267759 | 2.336092 | 0.019486 |
| **MAZ** | AP | 89 | 89 | -0.56551 | 0.568073 | 0.272233 | -2.07728 | 0.037776 |
| **RNASEK** | ES | 90 | 88 | 0.535809 | 1.708831 | 0.266844 | 2.007951 | 0.044648 |
| **ARMCX4** | AT | 89 | 89 | -0.57513 | 0.562634 | 0.267646 | -2.14883 | 0.031648 |
| **ZC3H14** | AA | 89 | 89 | -0.56163 | 0.570277 | 0.269692 | -2.0825 | 0.037297 |
| **CLIP4** | AT | 89 | 89 | -0.56074 | 0.570784 | 0.270017 | -2.0767 | 0.03783 |
| **SNRNP40** | ES | 89 | 89 | -0.60839 | 0.544225 | 0.266605 | -2.282 | 0.022489 |
| **SSH3** | AA | 89 | 89 | 0.526492 | 1.692983 | 0.267729 | 1.966513 | 0.049239 |
| **CMC2** | ES | 89 | 89 | -0.68126 | 0.505977 | 0.270158 | -2.52172 | 0.011678 |
| **TTLL3** | AT | 91 | 87 | 0.518257 | 1.679099 | 0.262561 | 1.973858 | 0.048398 |
| **SLC39A13** | RI | 89 | 89 | -0.58862 | 0.555091 | 0.278513 | -2.11345 | 0.034562 |
| **TPM1** | AT | 89 | 89 | -0.65668 | 0.518572 | 0.270244 | -2.42994 | 0.015101 |
| **RPL27** | AP | 96 | 82 | -0.57244 | 0.564147 | 0.270303 | -2.11777 | 0.034194 |
| **NCK2** | ES | 89 | 89 | -0.7155 | 0.488945 | 0.271952 | -2.631 | 0.008513 |
| **SNAPC3** | AT | 89 | 89 | -0.64018 | 0.527198 | 0.272003 | -2.35357 | 0.018594 |
| **TSC22D3** | AP | 90 | 88 | -0.57002 | 0.565514 | 0.264507 | -2.15503 | 0.03116 |
| **KIAA1328** | AT | 89 | 89 | 0.543311 | 1.721698 | 0.269106 | 2.018951 | 0.043492 |
| **DYNC2LI1** | AT | 89 | 89 | 0.581809 | 1.789273 | 0.271629 | 2.14193 | 0.032199 |
| **DYNC2LI1** | AT | 89 | 89 | -0.58181 | 0.558886 | 0.271629 | -2.14193 | 0.032199 |
| **GNB2L1** | AP | 89 | 89 | 0.534868 | 1.707223 | 0.267426 | 2.000059 | 0.045494 |
| **GNB2L1** | AP | 89 | 89 | -0.53487 | 0.585747 | 0.267426 | -2.00006 | 0.045494 |
| **ZNF655** | AT | 89 | 89 | 0.609029 | 1.838646 | 0.268621 | 2.267245 | 0.023375 |
| **ZNF655** | AT | 89 | 89 | -0.60903 | 0.543878 | 0.268621 | -2.26724 | 0.023375 |
| **IFI6** | AD | 91 | 87 | -0.68301 | 0.505092 | 0.280306 | -2.43667 | 0.014823 |
| **DGUOK** | ES | 89 | 89 | 0.556795 | 1.745071 | 0.267941 | 2.078054 | 0.037704 |
| **UBE2F** | ES | 90 | 88 | -0.52553 | 0.591241 | 0.267531 | -1.96438 | 0.049486 |
| **GUSB** | AT | 90 | 88 | -0.57855 | 0.560712 | 0.278376 | -2.07829 | 0.037682 |
| **ZMYM6** | AT | 89 | 89 | -0.56696 | 0.56725 | 0.270572 | -2.0954 | 0.036136 |
| **SSSCA1** | AT | 89 | 89 | -0.6833 | 0.504947 | 0.267776 | -2.55176 | 0.010718 |
| **SSSCA1** | AT | 89 | 89 | 0.683301 | 1.980404 | 0.267776 | 2.551762 | 0.010718 |
| **DAD1** | AD | 89 | 89 | -0.66665 | 0.513425 | 0.273764 | -2.43513 | 0.014887 |

| | | | | | | | | |
|---|---|---|---|---|---|---|---|---|
| **C15orf40** | AT | 89 | 89 | -0.70624 | 0.493494 | 0.276716 | -2.55223 | 0.010703 |
| **MAP4** | ES | 89 | 89 | 0.650082 | 1.915698 | 0.274518 | 2.368089 | 0.01788 |
| **CORO1B** | RI | 89 | 89 | -0.60678 | 0.545104 | 0.284545 | -2.13245 | 0.03297 |
| **MSRB3** | AT | 90 | 88 | -0.6012 | 0.548152 | 0.267034 | -2.25141 | 0.02436 |
| **DCTN1** | ES | 89 | 89 | -0.68816 | 0.502501 | 0.267774 | -2.56992 | 0.010172 |
| **TIMM9** | AA | 90 | 88 | -0.77967 | 0.458558 | 0.27706 | -2.81408 | 0.004892 |
| **SERHL2** | AD | 89 | 89 | -0.58463 | 0.557311 | 0.270192 | -2.16377 | 0.030482 |
| **HYKK** | AT | 89 | 89 | 0.558639 | 1.748291 | 0.265505 | 2.104063 | 0.035373 |
| **OAZ1** | ES | 89 | 89 | 0.568153 | 1.765004 | 0.266323 | 2.133323 | 0.032898 |
| **BIN1** | ES | 89 | 89 | -0.54751 | 0.578386 | 0.266421 | -2.05507 | 0.039872 |
| **UQCRB** | ES | 89 | 89 | 0.600615 | 1.82324 | 0.270043 | 2.224145 | 0.026139 |
| **MED17** | AT | 89 | 89 | -0.61401 | 0.541177 | 0.271311 | -2.26311 | 0.023629 |
| **MED17** | AT | 89 | 89 | 0.614009 | 1.847824 | 0.271311 | 2.263114 | 0.023629 |
| **DHRS4L2** | ES | 89 | 89 | 0.705202 | 2.024256 | 0.27989 | 2.519566 | 0.01175 |
| **TPM1** | AT | 90 | 88 | 0.570294 | 1.768788 | 0.265049 | 2.151654 | 0.031425 |
| **SEMA5A** | AT | 126 | 52 | -0.6929 | 0.500125 | 0.328329 | -2.11038 | 0.034826 |
| **DCTN6** | ES | 89 | 89 | -0.54545 | 0.579583 | 0.264736 | -2.06035 | 0.039365 |
| **ZSCAN30** | AT | 89 | 89 | -0.63417 | 0.530377 | 0.266764 | -2.37727 | 0.017442 |
| **PHF20** | ES | 90 | 88 | -0.65279 | 0.520592 | 0.265023 | -2.46314 | 0.013773 |
| **WDR6** | RI | 89 | 89 | -0.57012 | 0.56546 | 0.269587 | -2.11477 | 0.034449 |
| **MRPL36** | AD | 89 | 89 | 0.535036 | 1.70751 | 0.266224 | 2.009719 | 0.044461 |
| **REEP4** | ES | 89 | 89 | 0.546213 | 1.726702 | 0.267372 | 2.042896 | 0.041063 |
| **RBM4B** | AT | 89 | 89 | 0.592129 | 1.807834 | 0.272111 | 2.176057 | 0.029551 |
| **RBM4B** | AT | 89 | 89 | -0.59213 | 0.553148 | 0.272111 | -2.17606 | 0.029551 |
| **FAM195B** | AA | 90 | 88 | -0.55013 | 0.576872 | 0.2697 | -2.0398 | 0.04137 |
| **SHPK** | AT | 90 | 88 | 0.678701 | 1.971315 | 0.271592 | 2.498976 | 0.012455 |
| **ETHE1** | AT | 89 | 89 | -0.61907 | 0.538445 | 0.267816 | -2.31155 | 0.020803 |
| **ETHE1** | AT | 89 | 89 | 0.619069 | 1.857199 | 0.267816 | 2.311547 | 0.020803 |
| **RPS4X** | AT | 93 | 85 | -0.57486 | 0.562785 | 0.279968 | -2.0533 | 0.040043 |
| **DIDO1** | AT | 89 | 89 | -0.53276 | 0.586985 | 0.270199 | -1.97172 | 0.048642 |
| **CTTN** | ME | 89 | 89 | 0.532346 | 1.702922 | 0.27034 | 1.96917 | 0.048934 |
| **MPDU1** | AT | 89 | 89 | -0.66188 | 0.515879 | 0.273126 | -2.42337 | 0.015377 |
| **SERPINE3** | AT | 89 | 89 | -0.58989 | 0.554387 | 0.269358 | -2.19 | 0.028524 |
| **SERPINE3** | AT | 89 | 89 | 0.589893 | 1.803795 | 0.269358 | 2.189997 | 0.028524 |
| **ZNF585A** | AT | 89 | 89 | 0.535098 | 1.707616 | 0.26543 | 2.015967 | 0.043803 |
| **IP6K2** | RI | 90 | 88 | -0.57155 | 0.56465 | 0.276099 | -2.07009 | 0.038444 |
| **SAA2** | AT | 89 | 89 | 0.539124 | 1.714505 | 0.269986 | 1.996857 | 0.045841 |
| **ZNF223** | AT | 89 | 89 | 0.711714 | 2.037481 | 0.27228 | 2.613907 | 0.008951 |
| **MRPL33** | AT | 89 | 89 | 0.612153 | 1.844397 | 0.271037 | 2.258555 | 0.023911 |
| **MRPL33** | AT | 89 | 89 | -0.61215 | 0.542182 | 0.271037 | -2.25856 | 0.023911 |
| **MFF** | ES | 89 | 89 | -0.57201 | 0.56439 | 0.266127 | -2.14939 | 0.031604 |
| **Sep-02** | ES | 89 | 89 | -0.54452 | 0.580119 | 0.263109 | -2.06957 | 0.038493 |
| **GGCT** | AP | 89 | 89 | -0.5795 | 0.560177 | 0.272245 | -2.12861 | 0.033287 |
| **GGCT** | AP | 89 | 89 | 0.579503 | 1.78515 | 0.272245 | 2.12861 | 0.033287 |
| **LY6E** | AA | 90 | 88 | 0.524142 | 1.689009 | 0.266068 | 1.969957 | 0.048843 |
| **NUBP2** | ES | 89 | 89 | 0.527116 | 1.694039 | 0.266337 | 1.979131 | 0.047801 |
| **STRADA** | RI | 89 | 89 | -0.6403 | 0.527132 | 0.269893 | -2.37244 | 0.017671 |

| | | | | | | | | |
|---|---|---|---|---|---|---|---|---|
| **OPA3** | AT | 90 | 88 | -0.57632 | 0.561962 | 0.269489 | -2.13857 | 0.03247 |
| **QDPR** | ES | 160 | 18 | -1.34387 | 0.260834 | 0.594399 | -2.26089 | 0.023766 |
| **FASTK** | AD | 90 | 88 | -0.54359 | 0.580663 | 0.265101 | -2.05049 | 0.040317 |
| **C12orf10** | AP | 89 | 89 | -0.59074 | 0.553918 | 0.268594 | -2.19938 | 0.027851 |
| **C12orf10** | AP | 89 | 89 | 0.590739 | 1.805322 | 0.268594 | 2.199378 | 0.027851 |
| **PRKCSH** | AD | 89 | 89 | -0.63239 | 0.531319 | 0.268241 | -2.35755 | 0.018396 |
| **SNRNP200** | ES | 89 | 89 | -0.5555 | 0.573785 | 0.268122 | -2.07182 | 0.038282 |
| **RPS14** | ES | 89 | 89 | -0.56008 | 0.571164 | 0.266197 | -2.104 | 0.035378 |
| **SERPINB6** | AP | 89 | 89 | 0.625617 | 1.8694 | 0.272042 | 2.299706 | 0.021465 |
| **RSBN1L** | AP | 98 | 80 | -0.82259 | 0.439293 | 0.295454 | -2.78415 | 0.005367 |
| **EIF3G** | RI | 89 | 89 | -0.64185 | 0.52632 | 0.287999 | -2.22864 | 0.025838 |
| **TARDBP** | ES | 89 | 89 | -0.5275 | 0.590081 | 0.267441 | -1.97238 | 0.048566 |
| **LSM14A** | ES | 89 | 89 | -0.52123 | 0.593787 | 0.264206 | -1.97283 | 0.048515 |
| **COQ4** | AT | 89 | 89 | -0.53393 | 0.586295 | 0.268311 | -1.98997 | 0.046594 |
| **DDB2** | ES | 89 | 89 | -0.5193 | 0.594937 | 0.263938 | -1.96751 | 0.049124 |
| **EMC4** | AD | 90 | 88 | -0.58204 | 0.558757 | 0.268317 | -2.16923 | 0.030065 |
| **SYNJ2** | AT | 89 | 89 | -0.5716 | 0.56462 | 0.271098 | -2.10847 | 0.034991 |
| **CNOT7** | AT | 89 | 89 | 0.654029 | 1.923274 | 0.265649 | 2.462007 | 0.013816 |
| **CNOT7** | AT | 89 | 89 | -0.65403 | 0.519947 | 0.265649 | -2.46201 | 0.013816 |
| **AP4S1** | AT | 89 | 89 | -0.64232 | 0.52607 | 0.268534 | -2.39196 | 0.016759 |
| **AP4S1** | AT | 89 | 89 | 0.642321 | 1.900889 | 0.268534 | 2.391957 | 0.016759 |
| **ITFG3** | AT | 91 | 87 | -0.71359 | 0.489884 | 0.276911 | -2.57696 | 0.009967 |
| **LTA4H** | AA | 89 | 89 | -0.53791 | 0.583966 | 0.267306 | -2.01235 | 0.044183 |
| **CLIP1** | AD | 89 | 89 | -0.58236 | 0.558578 | 0.269595 | -2.16013 | 0.030763 |
| **HNRNPC** | ES | 89 | 89 | -0.5873 | 0.555828 | 0.269351 | -2.18041 | 0.029227 |
| **PXK** | AT | 89 | 89 | 0.721142 | 2.05678 | 0.278879 | 2.585857 | 0.009714 |
| **PXK** | AT | 89 | 89 | -0.72114 | 0.486197 | 0.278879 | -2.58586 | 0.009714 |
| **RNF4** | AP | 89 | 89 | -0.59943 | 0.549122 | 0.268782 | -2.23018 | 0.025735 |
| **GSTO2** | AT | 89 | 89 | -0.59479 | 0.551678 | 0.266577 | -2.23121 | 0.025667 |
| **GSTO2** | AT | 89 | 89 | 0.59479 | 1.812651 | 0.266577 | 2.231211 | 0.025667 |
| **CKLF-CMTM** | AT | 90 | 88 | -0.56816 | 0.566568 | 0.268574 | -2.11546 | 0.034391 |
| **PKM** | AD | 93 | 85 | -0.70885 | 0.492209 | 0.275627 | -2.57178 | 0.010118 |
| **LLGL2** | AT | 89 | 89 | -0.59824 | 0.549778 | 0.272186 | -2.19791 | 0.027955 |
| **LLGL2** | AT | 89 | 89 | 0.598241 | 1.818917 | 0.272186 | 2.197912 | 0.027955 |
| **ABHD17A** | AP | 89 | 89 | 0.563597 | 1.756981 | 0.266328 | 2.116178 | 0.03433 |
| **ABHD17A** | AP | 89 | 89 | -0.5636 | 0.569158 | 0.266328 | -2.11618 | 0.03433 |
| **FLT3LG** | AT | 89 | 89 | -0.54487 | 0.579916 | 0.265424 | -2.05284 | 0.040088 |
| **RNF149** | AT | 89 | 89 | -0.53619 | 0.584973 | 0.264634 | -2.02616 | 0.042749 |
| **RNF149** | AT | 89 | 89 | 0.53619 | 1.709481 | 0.264634 | 2.026159 | 0.042749 |
| **URM1** | RI | 89 | 89 | -0.52647 | 0.590689 | 0.268143 | -1.96338 | 0.049602 |
| **LDHA** | ES | 89 | 89 | -0.60538 | 0.54587 | 0.26801 | -2.25878 | 0.023897 |
| **ARHGEF7** | AT | 89 | 89 | 0.589924 | 1.803851 | 0.269833 | 2.186258 | 0.028797 |
| **ARHGEF7** | AT | 89 | 89 | -0.58992 | 0.55437 | 0.269833 | -2.18626 | 0.028797 |
| **DAPK2** | AT | 90 | 88 | 0.668122 | 1.95057 | 0.26768 | 2.495973 | 0.012561 |
| **CSF2RA** | ES | 89 | 89 | -0.64265 | 0.525899 | 0.269579 | -2.38388 | 0.017131 |
| **RHOC** | AA | 89 | 89 | -0.54674 | 0.578834 | 0.273484 | -1.99916 | 0.045591 |
| **ACVR1B** | AT | 89 | 89 | -0.51682 | 0.596414 | 0.263682 | -1.96001 | 0.049995 |

| | | | | | | | | |
|---|---|---|---|---|---|---|---|---|
| **LDLRAD3** | ES | 89 | 89 | -0.60427 | 0.546474 | 0.268385 | -2.2515 | 0.024354 |
| **TMEM19** | AT | 89 | 89 | -0.56912 | 0.566026 | 0.271468 | -2.09643 | 0.036044 |
| **TMEM19** | AT | 89 | 89 | 0.569115 | 1.766703 | 0.271468 | 2.096435 | 0.036044 |
| **MIEN1** | RI | 89 | 89 | -0.61955 | 0.538184 | 0.275804 | -2.24636 | 0.024681 |
| **COX7C** | RI | 89 | 89 | -0.5869 | 0.556046 | 0.269688 | -2.17623 | 0.029538 |
| **TMEM98** | ES | 89 | 89 | 0.606624 | 1.834228 | 0.268527 | 2.259078 | 0.023879 |
| **HBS1L** | AP | 89 | 89 | 0.609945 | 1.84033 | 0.268886 | 2.268414 | 0.023304 |
| **HBS1L** | AP | 89 | 89 | -0.60994 | 0.543381 | 0.268886 | -2.26841 | 0.023304 |
| **RELA** | RI | 89 | 89 | -0.5935 | 0.552389 | 0.268576 | -2.20981 | 0.027118 |
| **B9D1** | AT | 89 | 89 | -0.61722 | 0.539441 | 0.276429 | -2.23284 | 0.02556 |
| **ABHD14B** | RI | 89 | 89 | -0.59299 | 0.552671 | 0.268988 | -2.20453 | 0.027487 |
| **NDUFS4** | ES | 90 | 88 | -0.60913 | 0.543824 | 0.272006 | -2.2394 | 0.02513 |
| **AHI1** | AT | 89 | 89 | -0.77319 | 0.461541 | 0.281591 | -2.74577 | 0.006037 |
| **GALK2** | ES | 89 | 89 | -0.56397 | 0.568948 | 0.268345 | -2.10165 | 0.035584 |
| **DRG2** | RI | 89 | 89 | -0.55539 | 0.573851 | 0.269703 | -2.05925 | 0.03947 |
| **ZNF283** | AT | 90 | 88 | -0.6586 | 0.517576 | 0.270113 | -2.43823 | 0.014759 |
| **AAMDC** | ES | 89 | 89 | -0.61391 | 0.541229 | 0.270832 | -2.26677 | 0.023405 |
| **LIN37** | RI | 89 | 89 | -0.56623 | 0.56766 | 0.274371 | -2.06375 | 0.039042 |
| **KLK10** | AP | 89 | 89 | 0.54427 | 1.723349 | 0.268434 | 2.027576 | 0.042604 |
| **SLC25A45** | AP | 89 | 89 | -0.59296 | 0.552691 | 0.276702 | -2.14295 | 0.032118 |
| **RPS3** | AT | 89 | 89 | 0.534419 | 1.706457 | 0.265076 | 2.016101 | 0.043789 |
| **ZFAND4** | AT | 89 | 89 | -0.85464 | 0.425437 | 0.283256 | -3.01719 | 0.002551 |
| **HMGN1** | AP | 89 | 89 | -0.7084 | 0.492432 | 0.286682 | -2.47102 | 0.013473 |
| **OAZ2** | ES | 90 | 88 | -0.53875 | 0.583478 | 0.271971 | -1.98091 | 0.047602 |
| **NPIPB6** | AP | 89 | 89 | 0.65737 | 1.929711 | 0.269979 | 2.434898 | 0.014896 |
| **NPIPB6** | AP | 89 | 89 | -0.65737 | 0.518212 | 0.269979 | -2.4349 | 0.014896 |
| **LRRC37A** | AP | 89 | 89 | -0.58228 | 0.558622 | 0.267404 | -2.17754 | 0.02944 |
| **LRRC37A** | AP | 89 | 89 | 0.582283 | 1.79012 | 0.267404 | 2.177539 | 0.02944 |
| **ANAPC11** | AP | 90 | 88 | -0.78056 | 0.458149 | 0.279466 | -2.79304 | 0.005221 |
| **WHSC1** | AT | 89 | 89 | -0.5387 | 0.583507 | 0.271294 | -1.98567 | 0.04707 |
| **WHSC1** | AT | 89 | 89 | 0.538699 | 1.713776 | 0.271294 | 1.985665 | 0.04707 |
| **DBN1** | RI | 105 | 73 | -0.62203 | 0.536852 | 0.274919 | -2.26261 | 0.02366 |
| **ASPH** | AT | 92 | 86 | 0.601818 | 1.825434 | 0.269501 | 2.233083 | 0.025544 |
| **CPNE1** | AA | 89 | 89 | -0.65299 | 0.520486 | 0.265815 | -2.45656 | 0.014027 |
| **SYNJ2** | AT | 89 | 89 | 0.557565 | 1.746414 | 0.272868 | 2.043349 | 0.041018 |
| **APOC1** | ES | 89 | 89 | 0.587643 | 1.799742 | 0.264521 | 2.221536 | 0.026315 |
| **CDKL3** | AT | 147 | 31 | -0.86116 | 0.42267 | 0.433704 | -1.9856 | 0.047078 |
| **C12orf66** | AT | 89 | 89 | -0.60652 | 0.545245 | 0.275496 | -2.20156 | 0.027697 |
| **C12orf66** | AT | 89 | 89 | 0.60652 | 1.834038 | 0.275496 | 2.201558 | 0.027697 |
| **ATP5G3** | RI | 89 | 89 | -0.57373 | 0.563422 | 0.267848 | -2.14199 | 0.032195 |
| **INPP4B** | AT | 89 | 89 | -0.54207 | 0.581543 | 0.269001 | -2.01513 | 0.043891 |
| **AHI1** | AT | 90 | 88 | 0.620778 | 1.860374 | 0.272846 | 2.275191 | 0.022894 |
| **FASTK** | RI | 89 | 89 | -0.68703 | 0.503068 | 0.273296 | -2.51387 | 0.011941 |
| **TMEM229I** | AT | 89 | 89 | 0.539411 | 1.714996 | 0.270612 | 1.993301 | 0.046228 |
| **TMEM229I** | AT | 89 | 89 | -0.53941 | 0.583092 | 0.270612 | -1.9933 | 0.046228 |
| **HNRNPL** | ES | 89 | 89 | -0.6036 | 0.546841 | 0.270488 | -2.23151 | 0.025648 |
| **PDE1A** | AT | 154 | 24 | 1.154082 | 3.171111 | 0.397234 | 2.905298 | 0.003669 |

| | | | | | | | | |
|---|---|---|---|---|---|---|---|---|
| **ENTPD6** | AA | 89 | 89 | -0.53708 | 0.58445 | 0.265512 | -2.02282 | 0.043092 |
| **NUDT9** | ES | 89 | 89 | -0.56287 | 0.569569 | 0.268306 | -2.09788 | 0.035915 |
| **FDFT1** | AP | 89 | 89 | -0.61331 | 0.541557 | 0.267162 | -2.29563 | 0.021697 |
| **APOBEC3F** | AT | 89 | 89 | 0.571083 | 1.770182 | 0.265104 | 2.154186 | 0.031226 |
| **APOBEC3F** | AT | 89 | 89 | -0.57108 | 0.564914 | 0.265104 | -2.15419 | 0.031226 |
| **MARK2** | ES | 89 | 89 | 0.730416 | 2.075944 | 0.278172 | 2.62577 | 0.008645 |
| **CD33** | AT | 89 | 89 | 0.659821 | 1.934446 | 0.268086 | 2.461226 | 0.013846 |
| **CD33** | AT | 89 | 89 | -0.65982 | 0.516944 | 0.268086 | -2.46123 | 0.013846 |
| **ANKRD39** | AT | 89 | 89 | 0.574335 | 1.77595 | 0.270511 | 2.123152 | 0.033741 |
| **ANKRD39** | AT | 89 | 89 | -0.57434 | 0.563079 | 0.270511 | -2.12315 | 0.033741 |
| **MAGOH** | ES | 89 | 89 | -0.63979 | 0.527404 | 0.271692 | -2.35483 | 0.018531 |
| **TMEM223** | AT | 89 | 89 | 0.590703 | 1.805257 | 0.277148 | 2.131365 | 0.033059 |
| **TMEM223** | AT | 89 | 89 | -0.5907 | 0.553938 | 0.277148 | -2.13136 | 0.033059 |
| **CHEK1** | AP | 89 | 89 | -0.58974 | 0.554473 | 0.266411 | -2.21363 | 0.026854 |
| **CHEK1** | AP | 89 | 89 | 0.589737 | 1.803514 | 0.266411 | 2.213635 | 0.026854 |
| **ANO6** | AT | 90 | 88 | 0.625618 | 1.8694 | 0.267667 | 2.337298 | 0.019424 |
| **PHF12** | RI | 89 | 89 | -0.54511 | 0.57978 | 0.266496 | -2.04546 | 0.040809 |
| **TXNL4A** | ES | 89 | 89 | -0.54214 | 0.581501 | 0.265433 | -2.04249 | 0.041103 |
| **MRPL4** | RI | 89 | 89 | -0.54062 | 0.582386 | 0.269408 | -2.0067 | 0.044782 |
| **GLE1** | AT | 89 | 89 | 0.599819 | 1.821789 | 0.270262 | 2.219399 | 0.02646 |
| **GLE1** | AT | 89 | 89 | -0.59982 | 0.548911 | 0.270262 | -2.2194 | 0.02646 |
| **UNC119** | RI | 89 | 89 | -0.68783 | 0.502665 | 0.270826 | -2.53975 | 0.011093 |
| **ACACA** | AP | 89 | 89 | -0.65751 | 0.518139 | 0.268707 | -2.44694 | 0.014407 |
| **ST7** | AT | 89 | 89 | 0.595952 | 1.814757 | 0.269891 | 2.208123 | 0.027236 |
| **ST7** | AT | 89 | 89 | -0.59595 | 0.551038 | 0.269891 | -2.20812 | 0.027236 |
| **ZRANB2** | ES | 89 | 89 | 0.620141 | 1.85919 | 0.26988 | 2.297842 | 0.021571 |
| **DAPK2** | AT | 90 | 88 | -0.65578 | 0.519038 | 0.267756 | -2.44916 | 0.014319 |
| **PAK1** | ES | 89 | 89 | 0.668241 | 1.950802 | 0.271757 | 2.458961 | 0.013934 |
| **STAMBP** | AT | 89 | 89 | 0.724574 | 2.063852 | 0.277503 | 2.611052 | 0.009026 |
| **STAMBP** | AT | 89 | 89 | -0.72457 | 0.484531 | 0.277503 | -2.61105 | 0.009026 |
| **SPAG16** | AT | 89 | 89 | 0.544286 | 1.723378 | 0.267946 | 2.03133 | 0.042222 |
| **ATP13A2** | ES | 89 | 89 | 0.760736 | 2.139851 | 0.276813 | 2.748199 | 0.005992 |
| **PBXIP1** | ES | 89 | 89 | -0.71647 | 0.488474 | 0.271696 | -2.63703 | 0.008364 |
| **ESRRA** | AA | 90 | 88 | 0.638491 | 1.893622 | 0.267694 | 2.385151 | 0.017072 |
| **ZNF652** | AT | 89 | 89 | 0.642268 | 1.900786 | 0.269176 | 2.38605 | 0.01703 |
| **ZNF652** | AT | 89 | 89 | -0.64227 | 0.526098 | 0.269176 | -2.38605 | 0.01703 |
| **RGS3** | AP | 89 | 89 | 0.546634 | 1.727428 | 0.274164 | 1.993817 | 0.046172 |
| **TXNDC12** | AT | 89 | 89 | 0.561971 | 1.754126 | 0.276869 | 2.029734 | 0.042384 |
| **TXNDC12** | AT | 89 | 89 | -0.56197 | 0.570085 | 0.276869 | -2.02973 | 0.042384 |
| **SEC14L1** | AP | 166 | 12 | -1.32495 | 0.265817 | 0.601935 | -2.20115 | 0.027725 |
| **RNF4** | AP | 89 | 89 | 0.529131 | 1.697456 | 0.267693 | 1.976633 | 0.048083 |
| **BYSL** | ES | 90 | 88 | 0.580508 | 1.786946 | 0.270909 | 2.142815 | 0.032128 |
| **FANCI** | AT | 159 | 19 | 0.798368 | 2.221911 | 0.327679 | 2.436433 | 0.014833 |
| **GINS4** | AT | 89 | 89 | 0.552599 | 1.737763 | 0.266541 | 2.073226 | 0.038151 |
| **GINS4** | AT | 89 | 89 | -0.5526 | 0.575452 | 0.266541 | -2.07323 | 0.038151 |
| **DHPS** | ES | 89 | 89 | -0.52025 | 0.594373 | 0.265361 | -1.96053 | 0.049934 |
| **RNF130** | AT | 89 | 89 | -0.68337 | 0.504914 | 0.272183 | -2.51069 | 0.01205 |

| | | | | | | | | |
|---|---|---|---|---|---|---|---|---|
| **RNF130** | AT | 89 | 89 | 0.683367 | 1.980536 | 0.272183 | 2.510686 | 0.01205 |
| **CSRP2BP** | AP | 142 | 36 | 0.730814 | 2.076771 | 0.291011 | 2.511298 | 0.012029 |
| **UBA6** | AT | 89 | 89 | -0.66648 | 0.51351 | 0.279532 | -2.38429 | 0.017112 |
| **UBA6** | AT | 89 | 89 | 0.666485 | 1.94738 | 0.279532 | 2.384288 | 0.017112 |
| **HIVEP1** | AT | 89 | 89 | -0.52595 | 0.590995 | 0.267202 | -1.96835 | 0.049027 |
| **HIVEP1** | AT | 89 | 89 | 0.525948 | 1.692062 | 0.267202 | 1.968355 | 0.049027 |
| **LUC7L3** | RI | 89 | 89 | -0.59148 | 0.553509 | 0.272663 | -2.16926 | 0.030063 |
| **TMED1** | AP | 108 | 70 | -0.7106 | 0.491349 | 0.288974 | -2.45904 | 0.013931 |
| **MDH2** | ES | 92 | 86 | 0.532382 | 1.702984 | 0.266119 | 2.000539 | 0.045442 |
| **RHBDL2** | AP | 89 | 89 | 0.638412 | 1.893471 | 0.271008 | 2.35569 | 0.018488 |
| **RHBDL2** | AP | 89 | 89 | -0.63841 | 0.528131 | 0.271008 | -2.35569 | 0.018488 |
| **MEIS2** | AP | 90 | 88 | -0.54263 | 0.581217 | 0.264413 | -2.05221 | 0.040149 |
| **EHBP1L1** | AP | 89 | 89 | 0.561831 | 1.753881 | 0.267311 | 2.101791 | 0.035572 |
| **EHBP1L1** | AP | 89 | 89 | -0.56183 | 0.570164 | 0.267311 | -2.10179 | 0.035572 |
| **DPP9** | AT | 89 | 89 | 0.724402 | 2.063498 | 0.275631 | 2.628164 | 0.008585 |
| **DPP9** | AT | 89 | 89 | -0.7244 | 0.484614 | 0.275631 | -2.62816 | 0.008585 |
| **IFNAR2** | ES | 89 | 89 | -0.63076 | 0.532186 | 0.277617 | -2.27206 | 0.023083 |
| **PTGR1** | AT | 89 | 89 | 0.549762 | 1.73284 | 0.278122 | 1.976692 | 0.048077 |
| **PTGR1** | AT | 89 | 89 | -0.54976 | 0.577087 | 0.278122 | -1.97669 | 0.048077 |
| **PCNXL2** | RI | 89 | 89 | 0.555316 | 1.742491 | 0.268927 | 2.064929 | 0.03893 |
| **SNRPN** | AP | 89 | 89 | 0.593606 | 1.810505 | 0.275938 | 2.151229 | 0.031458 |
| **SEC24C** | RI | 89 | 89 | -0.57641 | 0.56191 | 0.268641 | -2.14566 | 0.0319 |
| **TMEM159** | ES | 89 | 89 | -0.62734 | 0.534012 | 0.272607 | -2.30125 | 0.021378 |
| **PSMC5** | ES | 90 | 88 | 0.544266 | 1.723344 | 0.269052 | 2.022904 | 0.043083 |
| **GALNT10** | AT | 89 | 89 | 0.564355 | 1.758313 | 0.268971 | 2.098197 | 0.035888 |
| **GALNT10** | AT | 89 | 89 | -0.56435 | 0.568727 | 0.268971 | -2.0982 | 0.035888 |
| **EIF4A1** | ES | 89 | 89 | -0.54612 | 0.579193 | 0.26427 | -2.06652 | 0.03878 |
| **MRPL34** | AP | 157 | 21 | -1.10228 | 0.332112 | 0.472614 | -2.33231 | 0.019684 |
| **MYL6** | AD | 89 | 89 | -0.71117 | 0.491071 | 0.276152 | -2.57527 | 0.010016 |
| **C12orf76** | AT | 89 | 89 | 0.581406 | 1.788552 | 0.272226 | 2.135749 | 0.0327 |
| **EPG5** | AT | 154 | 24 | 0.804642 | 2.235895 | 0.368812 | 2.181714 | 0.029131 |
| **EIF3G** | RI | 89 | 89 | -0.56837 | 0.566446 | 0.283232 | -2.00674 | 0.044777 |
| **MVB12B** | AT | 89 | 89 | -0.58075 | 0.559481 | 0.274188 | -2.11806 | 0.03417 |
| **MVB12B** | AT | 89 | 89 | 0.580746 | 1.787372 | 0.274188 | 2.118057 | 0.03417 |
| **THADA** | AT | 89 | 89 | 0.630965 | 1.879424 | 0.273414 | 2.307729 | 0.021014 |
| **ADAMTS9** | AT | 89 | 89 | -0.56259 | 0.569731 | 0.270347 | -2.081 | 0.037434 |
| **ADAMTS9** | AT | 89 | 89 | 0.562592 | 1.755215 | 0.270347 | 2.080996 | 0.037434 |
| **C8orf82** | RI | 89 | 89 | -0.67278 | 0.51029 | 0.28152 | -2.3898 | 0.016858 |
| **FOXP1** | ES | 89 | 89 | -0.53209 | 0.587374 | 0.267268 | -1.99086 | 0.046496 |
| **PRPF31** | AA | 89 | 89 | 0.610427 | 1.841218 | 0.268191 | 2.276089 | 0.022841 |
| **HGSNAT** | AT | 89 | 89 | -0.55866 | 0.571974 | 0.272887 | -2.04723 | 0.040636 |
| **HGSNAT** | AT | 89 | 89 | 0.558661 | 1.748331 | 0.272887 | 2.047227 | 0.040636 |
| **RPL31** | RI | 89 | 89 | -0.58823 | 0.555308 | 0.267776 | -2.19673 | 0.02804 |
| **TP53I3** | RI | 89 | 89 | -0.62711 | 0.534131 | 0.267537 | -2.34403 | 0.019077 |
| **POLR2L** | AT | 89 | 89 | -0.60133 | 0.548084 | 0.271528 | -2.2146 | 0.026787 |
| **POLR2L** | AT | 89 | 89 | 0.601326 | 1.824537 | 0.271528 | 2.214605 | 0.026787 |
| **ZNF850** | AT | 89 | 89 | -0.59571 | 0.551174 | 0.266685 | -2.23374 | 0.0255 |

| | | | | | | | | |
|---|---|---|---|---|---|---|---|---|
| **ZNF850** | AT | 89 | 89 | 0.595705 | 1.81431 | 0.266685 | 2.233742 | 0.0255 |
| **PPP1R7** | ES | 89 | 89 | -0.54294 | 0.581038 | 0.266324 | -2.03864 | 0.041486 |
| **SERINC5** | AT | 89 | 89 | -0.58674 | 0.556135 | 0.267272 | -2.19531 | 0.028142 |
| **SERINC5** | AT | 89 | 89 | 0.586745 | 1.798125 | 0.267272 | 2.195307 | 0.028142 |
| **C16orf46** | AT | 90 | 88 | 0.767962 | 2.15537 | 0.279785 | 2.744833 | 0.006054 |
| **IFNAR1** | AP | 89 | 89 | -0.72382 | 0.484896 | 0.273414 | -2.64735 | 0.008113 |
| **IFNAR1** | AP | 89 | 89 | 0.72382 | 2.062297 | 0.273414 | 2.647347 | 0.008113 |
| **NDUFA8** | AT | 150 | 28 | -1.30957 | 0.269936 | 0.484344 | -2.7038 | 0.006855 |
| **HNRNPC** | AA | 89 | 89 | -0.63498 | 0.529948 | 0.271706 | -2.337 | 0.019439 |
| **MSRB1** | ES | 95 | 83 | 0.55148 | 1.73582 | 0.265368 | 2.078169 | 0.037694 |
| **MTHFD2L** | AT | 89 | 89 | 0.523758 | 1.688361 | 0.267119 | 1.960768 | 0.049906 |
| **ANKHD1** | AT | 89 | 89 | 0.630578 | 1.878696 | 0.273027 | 2.309584 | 0.020911 |
| **SSBP4** | AP | 89 | 89 | 0.709717 | 2.033416 | 0.272225 | 2.6071 | 0.009131 |
| **SSBP4** | AP | 89 | 89 | -0.70972 | 0.491783 | 0.272225 | -2.6071 | 0.009131 |
| **GMFG** | RI | 89 | 89 | -0.67587 | 0.508712 | 0.269664 | -2.50635 | 0.012198 |
| **TMEM107** | ES | 89 | 89 | 0.550275 | 1.73373 | 0.269625 | 2.040892 | 0.041262 |
| **RPS9** | AP | 89 | 89 | -0.53019 | 0.588493 | 0.266245 | -1.99136 | 0.046441 |
| **RPS9** | AP | 89 | 89 | 0.530191 | 1.699256 | 0.266245 | 1.991363 | 0.046441 |
| **PDXK** | AT | 89 | 89 | 0.630216 | 1.878016 | 0.273257 | 2.30631 | 0.021093 |
| **PDXK** | AT | 89 | 89 | -0.63022 | 0.532477 | 0.273257 | -2.30631 | 0.021093 |
| **SMS** | ES | 89 | 89 | -0.52827 | 0.589624 | 0.267666 | -1.97362 | 0.048425 |
| **NPIPB5** | ES | 89 | 89 | -0.5492 | 0.57741 | 0.27047 | -2.03055 | 0.042301 |
| **ACACA** | AP | 89 | 89 | 0.62248 | 1.863544 | 0.265989 | 2.340248 | 0.019271 |
| **H2AFY** | ES | 89 | 89 | -0.60774 | 0.544581 | 0.267902 | -2.26851 | 0.023298 |
| **GSK3A** | RI | 89 | 89 | -0.56397 | 0.568946 | 0.269942 | -2.08923 | 0.036687 |
| **TOMM7** | ES | 100 | 78 | 0.524329 | 1.689326 | 0.265568 | 1.974369 | 0.04834 |
| **ZNF195** | AT | 89 | 89 | 0.531934 | 1.702222 | 0.265273 | 2.005232 | 0.044938 |
| **ALDOA** | AD | 90 | 88 | -0.58038 | 0.559685 | 0.268954 | -2.15792 | 0.030934 |
| **AK9** | AT | 89 | 89 | 0.687445 | 1.988629 | 0.27343 | 2.514151 | 0.011932 |
| **ITGB1BP1** | ES | 90 | 88 | -0.63018 | 0.532496 | 0.271126 | -2.32431 | 0.020109 |
| **CERS2** | AT | 90 | 88 | 0.552766 | 1.738053 | 0.272794 | 2.026309 | 0.042733 |
| **RPN1** | AP | 89 | 89 | 0.5503 | 1.733773 | 0.26921 | 2.044132 | 0.040941 |
| **RPN1** | AP | 89 | 89 | -0.5503 | 0.576777 | 0.26921 | -2.04413 | 0.040941 |
| **CSF2RA** | ES | 89 | 89 | -0.62136 | 0.537212 | 0.26807 | -2.31791 | 0.020454 |
| **ZNF410** | ES | 89 | 89 | -0.54955 | 0.577209 | 0.265925 | -2.06657 | 0.038775 |
| **ZNF66** | AT | 89 | 89 | -0.53747 | 0.584227 | 0.270427 | -1.98747 | 0.04687 |
| **TRIM9** | AT | 125 | 53 | -0.81848 | 0.4411 | 0.348766 | -2.34679 | 0.018936 |
| **YIF1A** | AP | 89 | 89 | -0.55004 | 0.576927 | 0.269331 | -2.04224 | 0.041128 |
| **YIF1A** | AP | 89 | 89 | 0.550039 | 1.733321 | 0.269331 | 2.04224 | 0.041128 |
| **HNRNPC** | ES | 93 | 85 | -0.54418 | 0.580319 | 0.270043 | -2.01515 | 0.043889 |
| **KIAA1958** | AT | 89 | 89 | 0.548503 | 1.73066 | 0.264606 | 2.072902 | 0.038181 |
| **KIAA1958** | AT | 89 | 89 | -0.5485 | 0.577814 | 0.264606 | -2.0729 | 0.038181 |
| **SURF4** | ES | 90 | 88 | -0.59376 | 0.552246 | 0.275084 | -2.15847 | 0.030891 |
| **RPE** | ES | 89 | 89 | -0.64655 | 0.52385 | 0.274263 | -2.35741 | 0.018403 |
| **EDEM2** | AT | 89 | 89 | 0.540009 | 1.716022 | 0.273746 | 1.972664 | 0.048534 |
| **EDEM2** | AT | 89 | 89 | -0.54001 | 0.582743 | 0.273746 | -1.97266 | 0.048534 |
| **TVP23B** | AP | 89 | 89 | 0.567627 | 1.764077 | 0.271035 | 2.094296 | 0.036234 |

| | | | | | | | | |
|---|---|---|---|---|---|---|---|---|
| **GBGT1** | AT | 89 | 89 | 0.553251 | 1.738898 | 0.2684 | 2.061296 | 0.039275 |
| **GBGT1** | AT | 89 | 89 | -0.55325 | 0.575077 | 0.2684 | -2.0613 | 0.039275 |
| **RPAIN** | ES | 89 | 89 | -0.54316 | 0.580909 | 0.267216 | -2.03267 | 0.042086 |
| **CDK5RAP3** | AP | 89 | 89 | -0.60855 | 0.544142 | 0.269909 | -2.25463 | 0.024157 |
| **CDK5RAP3** | AP | 89 | 89 | 0.608546 | 1.837757 | 0.269909 | 2.254628 | 0.024157 |
| **PRMT1** | ES | 89 | 89 | -0.54138 | 0.581945 | 0.26837 | -2.01728 | 0.043666 |
| **ZNF816** | AT | 89 | 89 | -0.57165 | 0.564593 | 0.265831 | -2.15043 | 0.031521 |
| **CCDC74A** | AT | 126 | 52 | -0.76252 | 0.466489 | 0.335142 | -2.27522 | 0.022893 |
| **CCDC53** | ES | 89 | 89 | 0.54738 | 1.728718 | 0.272179 | 2.011105 | 0.044314 |
| **FAM178A** | AT | 89 | 89 | 0.607268 | 1.83541 | 0.271713 | 2.234959 | 0.02542 |
| **WWP2** | AT | 89 | 89 | 0.528336 | 1.696108 | 0.264676 | 1.996166 | 0.045916 |
| **WWP2** | AT | 89 | 89 | -0.52834 | 0.589585 | 0.264676 | -1.99617 | 0.045916 |
| **ZNF141** | AT | 89 | 89 | -0.54852 | 0.577805 | 0.265233 | -2.06807 | 0.038634 |
| **ZNF141** | AT | 89 | 89 | 0.548519 | 1.730688 | 0.265233 | 2.068066 | 0.038634 |
| **ZNF599** | AT | 89 | 89 | 0.540649 | 1.717121 | 0.267798 | 2.018869 | 0.043501 |
| **LIN7B** | ES | 90 | 88 | 0.699354 | 2.012452 | 0.272945 | 2.562253 | 0.0104 |
| **NPRL3** | ES | 89 | 89 | -0.56017 | 0.571114 | 0.266885 | -2.09891 | 0.035825 |
| **CNTNAP3** | AT | 89 | 89 | 0.603639 | 1.828761 | 0.272541 | 2.214857 | 0.02677 |
| **ST7** | AP | 89 | 89 | 0.58197 | 1.789561 | 0.274458 | 2.120433 | 0.03397 |
| **IL15RA** | AA | 89 | 89 | -0.52642 | 0.590718 | 0.26472 | -1.98858 | 0.046747 |
| **SDR39U1** | AP | 145 | 33 | -0.91662 | 0.399869 | 0.435832 | -2.10314 | 0.035453 |
| **ITGB2** | AT | 91 | 87 | -0.60421 | 0.546504 | 0.275853 | -2.19034 | 0.028499 |
| **SRSF5** | AA | 89 | 89 | -0.59871 | 0.549521 | 0.275234 | -2.17527 | 0.02961 |
| **LGALS9** | AA | 89 | 89 | -0.58954 | 0.554584 | 0.273082 | -2.15883 | 0.030863 |
| **CTNNB1** | RI | 89 | 89 | -0.60683 | 0.545075 | 0.268382 | -2.26107 | 0.023755 |
| **ABCB8** | AT | 89 | 89 | 0.553074 | 1.738589 | 0.26766 | 2.066334 | 0.038797 |
| **ABCB8** | AT | 89 | 89 | -0.55307 | 0.575179 | 0.26766 | -2.06633 | 0.038797 |
| **CEP152** | AT | 89 | 89 | 0.534274 | 1.706208 | 0.265204 | 2.014572 | 0.043949 |
| **CEP152** | AT | 89 | 89 | -0.53427 | 0.586095 | 0.265204 | -2.01457 | 0.043949 |
| **C17orf97** | AT | 89 | 89 | -0.53874 | 0.583483 | 0.266461 | -2.02183 | 0.043194 |
| **C17orf97** | AT | 89 | 89 | 0.538739 | 1.713845 | 0.266461 | 2.021832 | 0.043194 |
| **DIABLO** | RI | 89 | 89 | -0.53257 | 0.587095 | 0.271663 | -1.9604 | 0.049949 |
| **CD63** | AP | 90 | 88 | 0.642365 | 1.900972 | 0.273557 | 2.348195 | 0.018865 |
| **C15orf40** | AT | 89 | 89 | 0.661465 | 1.937629 | 0.273661 | 2.417095 | 0.015645 |
| **CYB5D2** | AT | 143 | 35 | -1.1223 | 0.325529 | 0.470791 | -2.38387 | 0.017132 |
| **HSD17B12** | AT | 89 | 89 | -0.63517 | 0.529844 | 0.268241 | -2.36791 | 0.017889 |
| **HSD17B12** | AT | 89 | 89 | 0.635172 | 1.887347 | 0.268241 | 2.367915 | 0.017889 |
| **PTMS** | AA | 89 | 89 | 0.528259 | 1.695976 | 0.267773 | 1.972783 | 0.04852 |
| **TVP23B** | AP | 89 | 89 | -0.548 | 0.578105 | 0.270943 | -2.02257 | 0.043118 |
| **PSMD11** | RI | 89 | 89 | -0.58973 | 0.554474 | 0.268715 | -2.19465 | 0.028189 |
| **CRTAC1** | AT | 90 | 88 | -0.57118 | 0.564859 | 0.27187 | -2.10093 | 0.035647 |
| **SCN8A** | AT | 164 | 14 | -1.54711 | 0.212863 | 0.725237 | -2.13324 | 0.032905 |
| **PAPOLA** | ES | 89 | 89 | 0.55384 | 1.739922 | 0.271244 | 2.04185 | 0.041166 |
| **MAP2K5** | AP | 173 | 5 | 2.686747 | 14.68383 | 0.648673 | 4.141911 | 3.44E-05 |
| **ZCCHC9** | RI | 89 | 89 | -0.52946 | 0.588925 | 0.263909 | -2.0062 | 0.044835 |
| **ZBTB43** | AP | 89 | 89 | -0.59643 | 0.550777 | 0.269905 | -2.20976 | 0.027122 |
| **ZBTB43** | AP | 89 | 89 | 0.596426 | 1.815617 | 0.269905 | 2.209763 | 0.027122 |

| | | | | | | | | |
|---|---|---|---|---|---|---|---|---|
| **WASH4P** | RI | 89 | 89 | -0.67692 | 0.508178 | 0.276678 | -2.44661 | 0.014421 |
| **PROM2** | AA | 90 | 88 | -0.54171 | 0.581753 | 0.265654 | -2.03916 | 0.041434 |
| **RAPH1** | AT | 89 | 89 | -0.55215 | 0.575709 | 0.269409 | -2.04949 | 0.040414 |
| **SMOX** | AP | 89 | 89 | 0.524882 | 1.690259 | 0.265953 | 1.973587 | 0.048429 |
| **SMOX** | AP | 89 | 89 | -0.52488 | 0.591625 | 0.265953 | -1.97359 | 0.048429 |
| **ZFYVE20** | AT | 89 | 89 | 0.628701 | 1.875173 | 0.272068 | 2.310825 | 0.020843 |
| **ZFYVE20** | AT | 89 | 89 | -0.6287 | 0.533284 | 0.272068 | -2.31082 | 0.020843 |
| **TRIM4** | AT | 89 | 89 | 0.538104 | 1.712757 | 0.267144 | 2.014286 | 0.04398 |
| **TRIM4** | AT | 89 | 89 | -0.5381 | 0.583854 | 0.267144 | -2.01429 | 0.04398 |
| **EIF3E** | AA | 89 | 89 | -0.57451 | 0.562981 | 0.266978 | -2.1519 | 0.031405 |
| **KXD1** | AP | 89 | 89 | -0.58153 | 0.55904 | 0.269347 | -2.15906 | 0.030846 |
| **PSMG2** | AT | 90 | 88 | 0.566654 | 1.762361 | 0.2721 | 2.082523 | 0.037295 |
| **MOK** | AP | 141 | 37 | -0.7659 | 0.464916 | 0.382269 | -2.00356 | 0.045118 |
| **NME1** | ES | 89 | 89 | 0.604759 | 1.830811 | 0.268732 | 2.250413 | 0.024423 |
| **ELP5** | AT | 89 | 89 | 0.587047 | 1.798669 | 0.26646 | 2.203131 | 0.027586 |
| **ELP5** | AT | 89 | 89 | -0.58705 | 0.555967 | 0.26646 | -2.20313 | 0.027586 |
| **MRPL55** | ES | 89 | 89 | 0.616491 | 1.852417 | 0.266552 | 2.312833 | 0.020732 |
| **SLC39A1** | AP | 89 | 89 | -0.56882 | 0.566191 | 0.26876 | -2.11647 | 0.034305 |
| **GIPC1** | ES | 90 | 88 | 0.586747 | 1.79813 | 0.270448 | 2.169541 | 0.030042 |
| **EXOSC10** | RI | 89 | 89 | -0.55393 | 0.574686 | 0.268015 | -2.06679 | 0.038754 |
| **NUMA1** | ES | 89 | 89 | -0.53264 | 0.587055 | 0.264374 | -2.01471 | 0.043935 |
| **VPS13A** | AT | 89 | 89 | -0.62488 | 0.535325 | 0.268451 | -2.32773 | 0.019927 |
| **BSCL2** | AP | 89 | 89 | -0.56347 | 0.569231 | 0.271124 | -2.07827 | 0.037685 |
| **PABPN1** | RI | 89 | 89 | -0.53073 | 0.588175 | 0.270313 | -1.9634 | 0.0496 |
| **CCDC79** | AT | 127 | 51 | -0.68628 | 0.503446 | 0.335684 | -2.04442 | 0.040912 |
| **HN1** | AD | 89 | 89 | 0.614151 | 1.848087 | 0.273834 | 2.242786 | 0.024911 |
| **SDCCAG8** | ES | 89 | 89 | -0.54741 | 0.578449 | 0.271706 | -2.0147 | 0.043936 |
| **N4BP2L2** | AA | 89 | 89 | 0.551419 | 1.735715 | 0.267862 | 2.058591 | 0.039533 |
| **TECPR2** | AT | 89 | 89 | -0.53404 | 0.586233 | 0.26964 | -1.98056 | 0.047641 |
| **TECPR2** | AT | 89 | 89 | 0.534038 | 1.705806 | 0.26964 | 1.980555 | 0.047641 |
| **IFRD2** | RI | 91 | 87 | -0.54649 | 0.578978 | 0.27247 | -2.00569 | 0.044889 |
| **AAK1** | AT | 89 | 89 | -0.61201 | 0.54226 | 0.272289 | -2.24765 | 0.024599 |
| **GCA** | AP | 89 | 89 | 0.544689 | 1.724072 | 0.274014 | 1.987814 | 0.046832 |
| **ATRX** | AT | 169 | 9 | 1.003833 | 2.728721 | 0.433548 | 2.315392 | 0.020591 |
| **ACP6** | AT | 89 | 89 | -0.636 | 0.529405 | 0.268262 | -2.37083 | 0.017748 |
| **ACP6** | AT | 89 | 89 | 0.636002 | 1.888915 | 0.268262 | 2.370828 | 0.017748 |
| **C4orf29** | AT | 89 | 89 | 0.62432 | 1.866975 | 0.273381 | 2.283697 | 0.022389 |
| **C4orf29** | AT | 89 | 89 | -0.62432 | 0.535626 | 0.273381 | -2.2837 | 0.022389 |
| **GCHFR** | AT | 89 | 89 | 0.581718 | 1.789109 | 0.275752 | 2.10957 | 0.034895 |
| **GCHFR** | AT | 89 | 89 | -0.58172 | 0.558937 | 0.275752 | -2.10957 | 0.034895 |
| **ROR1** | AT | 89 | 89 | 0.58388 | 1.792982 | 0.280839 | 2.079056 | 0.037612 |
| **ROR1** | AT | 89 | 89 | -0.58388 | 0.55773 | 0.280839 | -2.07906 | 0.037612 |
| **KLC1** | ES | 89 | 89 | -0.51664 | 0.59652 | 0.263409 | -1.96137 | 0.049836 |
| **MIB2** | AA | 90 | 88 | 0.522424 | 1.686109 | 0.262525 | 1.99 | 0.046591 |
| **FAM195A** | ES | 89 | 89 | -0.6401 | 0.52724 | 0.273569 | -2.33981 | 0.019294 |
| **GAPDH** | AP | 89 | 89 | -0.55131 | 0.576195 | 0.275682 | -1.99981 | 0.045521 |
| **ALG13** | AT | 89 | 89 | 0.580206 | 1.786406 | 0.267403 | 2.16978 | 0.030023 |

| | | | | | | | | |
|---|---|---|---|---|---|---|---|---|
| **ALG13** | AT | 89 | 89 | -0.58021 | 0.559783 | 0.267403 | -2.16978 | 0.030023 |
| **C16orf46** | AT | 90 | 88 | -0.65544 | 0.519215 | 0.271419 | -2.41486 | 0.015741 |
| **SMIM7** | AT | 89 | 89 | -0.56789 | 0.566719 | 0.267763 | -2.12088 | 0.033932 |
| **TKT** | ES | 91 | 87 | 0.578453 | 1.783278 | 0.270445 | 2.138897 | 0.032444 |
| **CD164** | AP | 89 | 89 | 0.5159 | 1.675145 | 0.263199 | 1.960114 | 0.049982 |
| **CD164** | AP | 89 | 89 | -0.5159 | 0.596963 | 0.263199 | -1.96011 | 0.049982 |
| **FAHD2A** | ES | 91 | 87 | 0.603519 | 1.828542 | 0.272827 | 2.212095 | 0.02696 |
| **NMRAL1** | ME | 89 | 89 | -0.54299 | 0.581006 | 0.266555 | -2.03708 | 0.041642 |
| **SLC41A3** | RI | 89 | 89 | -0.56153 | 0.570338 | 0.275989 | -2.0346 | 0.041892 |
| **IL13RA1** | AT | 89 | 89 | 0.569187 | 1.766831 | 0.268157 | 2.122586 | 0.033789 |
| **IL13RA1** | AT | 89 | 89 | -0.56919 | 0.565985 | 0.268157 | -2.12259 | 0.033789 |
| **TANK** | AT | 90 | 88 | 0.698636 | 2.011008 | 0.279852 | 2.496453 | 0.012544 |
| **PTGR1** | AA | 89 | 89 | -0.57875 | 0.560599 | 0.277379 | -2.0865 | 0.036934 |
| **DNASE1L1** | AP | 89 | 89 | 0.528176 | 1.695836 | 0.268862 | 1.964483 | 0.049474 |
| **LTA4H** | ES | 89 | 89 | -0.5371 | 0.584439 | 0.270441 | -1.98602 | 0.047031 |
| **FAU** | AD | 155 | 23 | 0.823067 | 2.277475 | 0.372858 | 2.207453 | 0.027282 |
| **ACAA1** | ES | 89 | 89 | 0.637886 | 1.892476 | 0.27158 | 2.348793 | 0.018834 |
| **Sep-15** | ES | 90 | 88 | 0.561366 | 1.753066 | 0.269254 | 2.084897 | 0.037079 |
| **IRF3** | AA | 89 | 89 | -0.56736 | 0.567022 | 0.271727 | -2.08796 | 0.036801 |
| **RGS12** | AP | 155 | 23 | 1.021654 | 2.777785 | 0.379018 | 2.695531 | 0.007028 |
| **CXorf36** | AT | 89 | 89 | 0.627355 | 1.87265 | 0.275999 | 2.273034 | 0.023024 |
| **CXorf36** | AT | 89 | 89 | -0.62735 | 0.534003 | 0.275999 | -2.27303 | 0.023024 |
| **TSTA3** | AD | 90 | 88 | 0.546102 | 1.726511 | 0.273143 | 1.999331 | 0.045573 |
| **TPM2** | AA | 89 | 89 | -0.6097 | 0.543513 | 0.274477 | -2.22132 | 0.026329 |
| **BLZF1** | AT | 89 | 89 | 0.658436 | 1.93177 | 0.269496 | 2.443214 | 0.014557 |
| **AKAP8L** | RI | 89 | 89 | -0.53877 | 0.583464 | 0.270212 | -1.99389 | 0.046164 |
| **Sep-06** | AT | 89 | 89 | -0.53575 | 0.58523 | 0.267711 | -2.00123 | 0.045368 |
| **ATL2** | ES | 89 | 89 | -0.56689 | 0.567289 | 0.268755 | -2.10931 | 0.034918 |
| **TTLL3** | AD | 89 | 89 | -0.59749 | 0.55019 | 0.271127 | -2.20374 | 0.027543 |
| **WDR83** | AA | 91 | 87 | 0.547727 | 1.729319 | 0.266717 | 2.053592 | 0.040015 |
| **ADD2** | AT | 89 | 89 | -0.63839 | 0.528141 | 0.267585 | -2.38575 | 0.017044 |
| **SLC52A2** | RI | 89 | 89 | -0.55897 | 0.571796 | 0.269892 | -2.0711 | 0.03835 |
| **TMEM107** | ES | 89 | 89 | -0.57483 | 0.562803 | 0.275106 | -2.08947 | 0.036666 |
| **DLGAP4** | AD | 91 | 87 | -0.52431 | 0.591966 | 0.265162 | -1.97731 | 0.048007 |
| **CYP46A1** | AT | 90 | 88 | -0.76071 | 0.467333 | 0.27248 | -2.79182 | 0.005241 |
| **ZNF44** | AT | 90 | 88 | -0.52566 | 0.591162 | 0.266823 | -1.97009 | 0.048828 |
| **OGFOD2** | AP | 89 | 89 | 0.524329 | 1.689325 | 0.265486 | 1.974976 | 0.048271 |
| **OGFOD2** | AP | 89 | 89 | -0.52433 | 0.591952 | 0.265486 | -1.97498 | 0.048271 |
| **TKT** | ES | 98 | 80 | 0.57822 | 1.782862 | 0.262788 | 2.200326 | 0.027784 |
| **FBXO16** | AT | 89 | 89 | -0.55646 | 0.573234 | 0.276725 | -2.01088 | 0.044338 |
| **DNAJC10** | AT | 89 | 89 | -0.57128 | 0.5648 | 0.280483 | -2.03678 | 0.041672 |
| **DNAJC10** | AT | 89 | 89 | 0.571284 | 1.770539 | 0.280483 | 2.036783 | 0.041672 |
| **RANGAP1** | AP | 161 | 17 | 0.735125 | 2.085742 | 0.36524 | 2.01272 | 0.044144 |
| **CERCAM** | RI | 89 | 89 | -0.53373 | 0.586416 | 0.270157 | -1.97562 | 0.048198 |
| **CCDC24** | RI | 89 | 89 | -0.52784 | 0.589879 | 0.267847 | -1.97067 | 0.048762 |
| **ETFA** | ES | 89 | 89 | -0.63258 | 0.531221 | 0.272296 | -2.32312 | 0.020173 |
| **TRIP10** | ES | 91 | 87 | -0.5723 | 0.564224 | 0.271975 | -2.10426 | 0.035356 |

| | | | | | | | | |
|---|---|---|---|---|---|---|---|---|
| **TMEM205** | AD | 89 | 89 | 0.631575 | 1.88057 | 0.274289 | 2.302585 | 0.021302 |
| **ZNF519** | AT | 89 | 89 | 0.596831 | 1.816354 | 0.275969 | 2.16267 | 0.030567 |
| **CRYAB** | AP | 89 | 89 | -0.53918 | 0.583226 | 0.27236 | -1.97966 | 0.047742 |
| **HEATR1** | AT | 89 | 89 | 0.541212 | 1.718088 | 0.27383 | 1.97645 | 0.048104 |
| **HEATR1** | AT | 89 | 89 | -0.54121 | 0.582042 | 0.27383 | -1.97645 | 0.048104 |
| **MFF** | ES | 90 | 88 | 0.645757 | 1.90743 | 0.269661 | 2.394699 | 0.016634 |
| **ZNF318** | AT | 117 | 61 | -0.79863 | 0.449947 | 0.349792 | -2.28314 | 0.022422 |
| **PAPOLA** | RI | 89 | 89 | -0.6025 | 0.547442 | 0.268513 | -2.24383 | 0.024843 |
| **CYP46A1** | AT | 90 | 88 | 0.709736 | 2.033454 | 0.270105 | 2.627634 | 0.008598 |
| **TANK** | AT | 90 | 88 | -0.62088 | 0.537469 | 0.274557 | -2.26141 | 0.023734 |
| **TMEM25** | AA | 89 | 89 | 0.603029 | 1.827646 | 0.271823 | 2.218463 | 0.026523 |
| **CYB561A3** | RI | 89 | 89 | -0.53185 | 0.587519 | 0.267955 | -1.98484 | 0.047163 |
| **FDFT1** | AP | 89 | 89 | 0.593513 | 1.810337 | 0.272049 | 2.18164 | 0.029136 |
| **C16orf52** | AT | 89 | 89 | -0.59139 | 0.553557 | 0.275053 | -2.1501 | 0.031548 |
| **DERA** | ES | 89 | 89 | -0.55429 | 0.574478 | 0.277709 | -1.99595 | 0.045939 |
| **EMR2** | AT | 89 | 89 | -0.55395 | 0.574676 | 0.266901 | -2.07548 | 0.037942 |
| **EMR2** | AT | 89 | 89 | 0.553949 | 1.74011 | 0.266901 | 2.075479 | 0.037942 |
| **TXNL4A** | AP | 89 | 89 | -0.58298 | 0.558233 | 0.272247 | -2.14136 | 0.032245 |
| **SFI1** | AA | 91 | 87 | 0.534727 | 1.706983 | 0.270629 | 1.97587 | 0.048169 |
| **ANAPC1** | AT | 174 | 4 | 1.361239 | 3.901025 | 0.597993 | 2.276346 | 0.022825 |
| **ME3** | AT | 89 | 89 | 0.572475 | 1.772649 | 0.272076 | 2.104102 | 0.03537 |
| **ME3** | AT | 89 | 89 | -0.57247 | 0.564127 | 0.272076 | -2.1041 | 0.03537 |
| **ADD2** | AT | 90 | 88 | 0.577961 | 1.7824 | 0.265863 | 2.173901 | 0.029713 |
| **COX15** | AT | 89 | 89 | 0.528271 | 1.695998 | 0.266688 | 1.980858 | 0.047607 |
| **COX15** | AT | 89 | 89 | -0.52827 | 0.589623 | 0.266688 | -1.98086 | 0.047607 |
| **RCAN1** | AP | 174 | 4 | 1.698868 | 5.467756 | 0.605682 | 2.804886 | 0.005033 |
| **FAM3A** | AA | 89 | 89 | -0.53758 | 0.584158 | 0.268107 | -2.00511 | 0.044951 |
| **FXR1** | AT | 168 | 10 | 1.082446 | 2.951891 | 0.535807 | 2.020217 | 0.043361 |
| **BABAM1** | AT | 177 | 1 | 3.344531 | 28.34727 | 1.095711 | 3.052384 | 0.00227 |
| **C20orf96** | ES | 90 | 88 | -0.58133 | 0.559156 | 0.272554 | -2.13289 | 0.032934 |
| **SYNE2** | AT | 177 | 1 | 2.656514 | 14.24654 | 1.049888 | 2.530284 | 0.011397 |
| **CNTN4** | AP | 177 | 1 | 2.163487 | 8.70143 | 1.032963 | 2.094447 | 0.03622 |
| **MOSPD2** | AA | 89 | 89 | -0.57877 | 0.560587 | 0.270835 | -2.13699 | 0.032599 |

| CI95..Lowe | CI95..Uppe | Concordan | se.concord | P.value_lik | P.value_wa | P.value_logrank |
|---|---|---|---|---|---|---|
| 2.247688 | 7.157827 | 0.675296 | 0.029801 | 6.26E-07 | 2.59E-06 | 4.74E-07 |
| 0.14036 | 0.458057 | 0.673671 | 0.030047 | 8.69E-07 | 5.43E-06 | 1.01E-06 |
| 1.75006 | 5.171906 | 0.671813 | 0.027403 | 4.12E-05 | 6.76E-05 | 3.03E-05 |
| 1.862153 | 5.683086 | 0.667054 | 0.028943 | 1.32E-05 | 3.41E-05 | 1.20E-05 |
| 2.075213 | 7.162272 | 0.665893 | 0.028668 | 2.53E-06 | 1.95E-05 | 4.49E-06 |
| 0.179248 | 0.547571 | 0.663455 | 0.028913 | 1.86E-05 | 4.62E-05 | 1.74E-05 |
| 0.183834 | 0.546633 | 0.66299 | 0.029212 | 2.15E-05 | 3.59E-05 | 1.47E-05 |
| 1.829382 | 5.439692 | 0.66299 | 0.029212 | 2.15E-05 | 3.59E-05 | 1.47E-05 |
| 1.810513 | 5.461384 | 0.662178 | 0.027065 | 2.90E-05 | 4.75E-05 | 2.03E-05 |
| 0.155663 | 0.48768 | 0.660436 | 0.030876 | 3.28E-06 | 9.65E-06 | 2.64E-06 |
| 1.809803 | 5.270916 | 0.657302 | 0.030825 | 2.65E-05 | 3.55E-05 | 1.48E-05 |
| 0.185452 | 0.565622 | 0.656257 | 0.027832 | 4.62E-05 | 7.40E-05 | 3.54E-05 |
| 0.191247 | 0.594483 | 0.651265 | 0.032001 | 7.18E-05 | 0.000172 | 8.23E-05 |
| 0.202392 | 0.60447 | 0.650104 | 0.03141 | 9.89E-05 | 0.000168 | 8.74E-05 |
| 1.898446 | 6.028219 | 0.648944 | 0.032756 | 1.16E-05 | 3.55E-05 | 1.28E-05 |
| 0.185828 | 0.570094 | 0.648827 | 0.030531 | 4.49E-05 | 8.67E-05 | 4.00E-05 |
| 0.181683 | 0.546141 | 0.648595 | 0.033728 | 2.43E-05 | 3.88E-05 | 1.63E-05 |
| 0.219814 | 0.645378 | 0.648247 | 0.030292 | 0.000258 | 0.00038 | 0.000226 |
| 0.19473 | 0.585679 | 0.64755 | 0.030046 | 7.86E-05 | 0.000111 | 5.74E-05 |
| 0.174217 | 0.544605 | 0.647202 | 0.029797 | 2.47E-05 | 5.12E-05 | 2.23E-05 |
| 0.176371 | 0.548154 | 0.646738 | 0.030658 | 1.91E-05 | 5.39E-05 | 2.03E-05 |
| 0.19124 | 0.568182 | 0.646738 | 0.031365 | 3.94E-05 | 6.47E-05 | 2.93E-05 |
| 0.226389 | 0.658288 | 0.646506 | 0.02951 | 0.000354 | 0.000473 | 0.000295 |
| 1.339806 | 3.972251 | 0.646506 | 0.031306 | 0.001889 | 0.002569 | 0.001938 |
| 0.213473 | 0.65195 | 0.64639 | 0.027773 | 0.000325 | 0.000536 | 0.00033 |
| 1.877793 | 6.146032 | 0.645229 | 0.031783 | 1.40E-05 | 5.28E-05 | 1.83E-05 |
| 0.162707 | 0.53254 | 0.645229 | 0.031783 | 1.40E-05 | 5.28E-05 | 1.83E-05 |
| 0.21743 | 0.639614 | 0.644416 | 0.03238 | 0.000231 | 0.000339 | 0.0002 |
| 1.563444 | 4.599171 | 0.644416 | 0.03238 | 0.000231 | 0.000339 | 0.0002 |
| 0.177369 | 0.560313 | 0.643952 | 0.03131 | 3.65E-05 | 8.35E-05 | 3.76E-05 |
| 0.21273 | 0.636678 | 0.642791 | 0.031715 | 0.000199 | 0.000351 | 0.000198 |
| 0.249624 | 0.736632 | 0.642675 | 0.03083 | 0.001674 | 0.002161 | 0.001611 |
| 0.202246 | 0.601624 | 0.641978 | 0.033407 | 0.000111 | 0.000152 | 8.18E-05 |
| 0.208911 | 0.649438 | 0.641514 | 0.031096 | 0.000277 | 0.000557 | 0.00033 |
| 1.53547 | 4.662943 | 0.640817 | 0.031276 | 0.000314 | 0.000514 | 0.000314 |
| 0.214457 | 0.651266 | 0.640817 | 0.031276 | 0.000314 | 0.000514 | 0.000314 |
| 0.227252 | 0.662915 | 0.640237 | 0.030171 | 0.000429 | 0.00053 | 0.000345 |
| 0.178132 | 0.565157 | 0.638612 | 0.032335 | 3.91E-05 | 9.72E-05 | 4.32E-05 |
| 0.185518 | 0.590587 | 0.638031 | 0.033141 | 6.99E-05 | 0.000182 | 8.52E-05 |
| 0.184606 | 0.592214 | 0.637799 | 0.032301 | 7.79E-05 | 0.000198 | 9.63E-05 |
| 0.18757 | 0.568616 | 0.637567 | 0.031758 | 4.76E-05 | 7.64E-05 | 3.61E-05 |
| 0.199368 | 0.60564 | 0.637335 | 0.032338 | 0.000107 | 0.000192 | 0.0001 |
| 0.241478 | 0.720686 | 0.63687 | 0.030476 | 0.001238 | 0.001723 | 0.00124 |
| 0.238673 | 0.705097 | 0.63687 | 0.032565 | 0.000869 | 0.001262 | 0.000872 |
| 1.319702 | 3.866482 | 0.635709 | 0.031186 | 0.002295 | 0.002963 | 0.002287 |
| 1.591147 | 4.671809 | 0.635013 | 0.033483 | 0.000193 | 0.000262 | 0.000153 |

| | | | | | | |
|---|---|---|---|---|---|---|
| 1.218282 | 3.447344 | 0.634781 | 0.03179 | 0.006658 | 0.00685 | 0.00581 |
| 0.290078 | 0.820828 | 0.634781 | 0.03179 | 0.006658 | 0.00685 | 0.00581 |
| 0.229602 | 0.699768 | 0.634665 | 0.031314 | 0.000783 | 0.001301 | 0.000884 |
| 1.429045 | 4.35536 | 0.634665 | 0.031314 | 0.000783 | 0.001301 | 0.000884 |
| 0.183443 | 0.549703 | 0.634665 | 0.034231 | 2.21E-05 | 4.18E-05 | 1.69E-05 |
| 0.212264 | 0.623997 | 0.634665 | 0.034398 | 0.000191 | 0.000238 | 0.000137 |
| 1.38419 | 4.065705 | 0.6342 | 0.031862 | 0.001273 | 0.001674 | 0.001203 |
| 0.204087 | 0.598409 | 0.633271 | 0.033681 | 9.27E-05 | 0.000128 | 6.43E-05 |
| 1.671097 | 4.899869 | 0.633271 | 0.033681 | 9.27E-05 | 0.000128 | 6.43E-05 |
| 0.28114 | 0.810001 | 0.632691 | 0.031965 | 0.005429 | 0.006134 | 0.005086 |
| 1.234567 | 3.55695 | 0.632691 | 0.031965 | 0.005429 | 0.006134 | 0.005086 |
| 1.624714 | 4.916319 | 0.632575 | 0.033916 | 0.000132 | 0.000235 | 0.000125 |
| 1.406327 | 4.120949 | 0.632343 | 0.03292 | 0.001 | 0.001359 | 0.000959 |
| 0.268133 | 0.78067 | 0.632343 | 0.032165 | 0.00326 | 0.004128 | 0.003276 |
| 0.293433 | 0.846125 | 0.631878 | 0.033275 | 0.008544 | 0.009925 | 0.008545 |
| 1.527753 | 4.695662 | 0.631878 | 0.033279 | 0.000328 | 0.000583 | 0.000354 |
| 1.331112 | 3.890321 | 0.631878 | 0.033151 | 0.002188 | 0.002653 | 0.002035 |
| 0.223166 | 0.687928 | 0.631646 | 0.031259 | 0.000663 | 0.001104 | 0.000738 |
| 0.272288 | 0.78668 | 0.630834 | 0.031895 | 0.003667 | 0.004421 | 0.003557 |
| 1.271166 | 3.672585 | 0.630834 | 0.031895 | 0.003667 | 0.004421 | 0.003557 |
| 1.479371 | 4.38918 | 0.630601 | 0.030941 | 0.000628 | 0.000748 | 0.000514 |
| 1.495066 | 4.47297 | 0.630485 | 0.033355 | 0.000454 | 0.000678 | 0.000433 |
| 0.235286 | 0.712364 | 0.630369 | 0.032782 | 0.001055 | 0.001577 | 0.001114 |
| 0.255377 | 0.749067 | 0.630253 | 0.030401 | 0.002 | 0.002591 | 0.001976 |
| 0.193798 | 0.5969 | 0.628628 | 0.033739 | 8.40E-05 | 0.000171 | 8.49E-05 |
| 1.675323 | 5.160004 | 0.628628 | 0.033739 | 8.40E-05 | 0.000171 | 8.49E-05 |
| 0.2708 | 0.782682 | 0.628512 | 0.030673 | 0.003448 | 0.004171 | 0.003337 |
| 0.232391 | 0.672574 | 0.628047 | 0.031454 | 0.000535 | 0.000619 | 0.000413 |
| 1.410122 | 4.152679 | 0.627815 | 0.034365 | 0.001007 | 0.00134 | 0.000941 |
| 0.240642 | 0.703278 | 0.627003 | 0.031572 | 0.000956 | 0.001168 | 0.000834 |
| 0.137613 | 0.467384 | 0.62677 | 0.035492 | 2.06E-06 | 1.09E-05 | 2.82E-06 |
| 0.218202 | 0.668076 | 0.62619 | 0.032428 | 0.000422 | 0.000744 | 0.000471 |
| 0.265837 | 0.788578 | 0.625842 | 0.033001 | 0.003758 | 0.004859 | 0.003893 |
| 1.268105 | 3.7617 | 0.625842 | 0.033001 | 0.003758 | 0.004859 | 0.003893 |
| 1.708145 | 5.245607 | 0.625726 | 0.034183 | 5.82E-05 | 0.000128 | 6.04E-05 |
| 0.190636 | 0.58543 | 0.625726 | 0.034183 | 5.82E-05 | 0.000128 | 6.04E-05 |
| 0.203564 | 0.627052 | 0.625261 | 0.034509 | 0.000189 | 0.000336 | 0.000191 |
| 0.158706 | 0.509386 | 0.625029 | 0.035658 | 7.55E-06 | 2.36E-05 | 7.82E-06 |
| 1.247433 | 3.698329 | 0.624681 | 0.033087 | 0.004576 | 0.005826 | 0.00475 |
| 0.270392 | 0.801647 | 0.624681 | 0.033087 | 0.004576 | 0.005826 | 0.00475 |
| 1.42955 | 4.315795 | 0.624565 | 0.034939 | 0.00082 | 0.001248 | 0.000853 |
| 0.231707 | 0.699521 | 0.624565 | 0.034939 | 0.00082 | 0.001248 | 0.000853 |
| 0.261813 | 0.789662 | 0.624449 | 0.031207 | 0.003809 | 0.005134 | 0.004122 |
| 1.266365 | 3.819516 | 0.624449 | 0.031207 | 0.003809 | 0.005134 | 0.004122 |
| 0.204513 | 0.603749 | 0.624216 | 0.034533 | 9.78E-05 | 0.000152 | 7.92E-05 |
| 1.656316 | 4.889667 | 0.624216 | 0.034533 | 9.78E-05 | 0.000152 | 7.92E-05 |
| 0.222414 | 0.665958 | 0.623984 | 0.033607 | 0.000509 | 0.000642 | 0.000425 |

| | | | | | | |
|---|---|---|---|---|---|---|
| 1.501596 | 4.496125 | 0.623984 | 0.033607 | 0.000509 | 0.000642 | 0.000425 |
| 0.211494 | 0.667194 | 0.623984 | 0.032897 | 0.000446 | 0.000836 | 0.000531 |
| 0.225855 | 0.679567 | 0.62352 | 0.033642 | 0.000583 | 0.000854 | 0.000568 |
| 0.152329 | 0.517288 | 0.62352 | 0.033819 | 1.13E-05 | 4.63E-05 | 1.65E-05 |
| 1.93316 | 6.56472 | 0.62352 | 0.033819 | 1.13E-05 | 4.63E-05 | 1.65E-05 |
| 1.313252 | 3.781121 | 0.623172 | 0.033955 | 0.002581 | 0.002977 | 0.002305 |
| 0.266643 | 0.776153 | 0.623055 | 0.032687 | 0.003228 | 0.003856 | 0.00306 |
| 1.288406 | 3.75033 | 0.623055 | 0.032687 | 0.003228 | 0.003856 | 0.00306 |
| 0.263541 | 0.755031 | 0.622823 | 0.033203 | 0.002246 | 0.002643 | 0.00203 |
| 1.324449 | 3.794474 | 0.622823 | 0.033203 | 0.002246 | 0.002643 | 0.00203 |
| 1.531225 | 4.573232 | 0.622475 | 0.03373 | 0.000349 | 0.00049 | 0.000308 |
| 1.352383 | 3.876798 | 0.622475 | 0.03413 | 0.001848 | 0.002045 | 0.001532 |
| 0.278546 | 0.82644 | 0.622475 | 0.031834 | 0.006526 | 0.00812 | 0.006823 |
| 0.227494 | 0.674777 | 0.622359 | 0.032809 | 0.000481 | 0.00073 | 0.000471 |
| 1.231959 | 3.582723 | 0.622243 | 0.03222 | 0.005478 | 0.006411 | 0.005325 |
| 0.279117 | 0.811715 | 0.622243 | 0.03222 | 0.005478 | 0.006411 | 0.005325 |
| 1.151536 | 3.28169 | 0.622243 | 0.032983 | 0.012291 | 0.012843 | 0.011335 |
| 0.19461 | 0.583897 | 0.622127 | 0.035961 | 6.15E-05 | 0.000105 | 4.98E-05 |
| 0.188076 | 0.562664 | 0.622011 | 0.034877 | 3.21E-05 | 5.89E-05 | 2.55E-05 |
| 0.318317 | 0.907624 | 0.621662 | 0.031965 | 0.018813 | 0.020202 | 0.018363 |
| 0.258397 | 0.753912 | 0.621546 | 0.032593 | 0.002124 | 0.002753 | 0.0021 |
| 0.255198 | 0.727916 | 0.62143 | 0.034598 | 0.00166 | 0.001646 | 0.001232 |
| 1.373784 | 3.918523 | 0.62143 | 0.034598 | 0.00166 | 0.001646 | 0.001232 |
| 1.310508 | 3.820046 | 0.62143 | 0.032461 | 0.002754 | 0.00317 | 0.002496 |
| 0.261777 | 0.763063 | 0.62143 | 0.032461 | 0.002754 | 0.00317 | 0.002496 |
| 1.394193 | 4.142788 | 0.621314 | 0.034174 | 0.001118 | 0.001599 | 0.001144 |
| 1.429436 | 4.222124 | 0.621198 | 0.033702 | 0.000779 | 0.001142 | 0.000778 |
| 0.259551 | 0.792288 | 0.621082 | 0.031991 | 0.003894 | 0.005473 | 0.004381 |
| 0.302743 | 0.863834 | 0.620966 | 0.032266 | 0.011214 | 0.012169 | 0.010716 |
| 1.585709 | 4.70295 | 0.620502 | 0.034978 | 0.000198 | 0.000292 | 0.000166 |
| 0.212632 | 0.630633 | 0.620502 | 0.034978 | 0.000198 | 0.000292 | 0.000166 |
| 1.488493 | 4.617955 | 0.620269 | 0.033591 | 0.000493 | 0.000846 | 0.000549 |
| 0.216546 | 0.67182 | 0.620269 | 0.033591 | 0.000493 | 0.000846 | 0.000549 |
| 0.242338 | 0.711959 | 0.620269 | 0.032391 | 0.001179 | 0.001395 | 0.001015 |
| 1.30707 | 3.79702 | 0.620153 | 0.033929 | 0.002688 | 0.003237 | 0.002519 |
| 1.283815 | 3.653569 | 0.619921 | 0.033404 | 0.003385 | 0.003775 | 0.003016 |
| 0.247346 | 0.746355 | 0.619689 | 0.033936 | 0.001904 | 0.002715 | 0.002037 |
| 0.251164 | 0.720026 | 0.619689 | 0.033491 | 0.001448 | 0.00146 | 0.001068 |
| 1.38884 | 3.981464 | 0.619689 | 0.033491 | 0.001448 | 0.00146 | 0.001068 |
| 0.260097 | 0.765206 | 0.619573 | 0.03362 | 0.002622 | 0.003367 | 0.002639 |
| 0.289697 | 0.831051 | 0.619573 | 0.032594 | 0.007734 | 0.008089 | 0.006935 |
| 0.319208 | 0.928038 | 0.619573 | 0.032311 | 0.022374 | 0.025465 | 0.023292 |
| 0.185503 | 0.571785 | 0.619341 | 0.035781 | 4.29E-05 | 9.36E-05 | 4.15E-05 |
| 0.193342 | 0.608423 | 0.619225 | 0.03416 | 0.000113 | 0.000253 | 0.000134 |
| 0.291829 | 0.85261 | 0.619108 | 0.031795 | 0.009227 | 0.010992 | 0.009523 |
| 0.222992 | 0.652893 | 0.618992 | 0.035071 | 0.000302 | 0.000439 | 0.000266 |
| 1.461833 | 4.387631 | 0.618992 | 0.035483 | 0.000631 | 0.000919 | 0.000616 |

| | | | | | | |
|---|---|---|---|---|---|---|
| 0.227913 | 0.684073 | 0.618992 | 0.035483 | 0.000631 | 0.000919 | 0.000616 |
| 1.346477 | 3.948721 | 0.61876 | 0.032957 | 0.001784 | 0.002336 | 0.001759 |
| 0.280617 | 0.822735 | 0.61876 | 0.032384 | 0.006215 | 0.007562 | 0.006375 |
| 1.34188 | 3.899999 | 0.618644 | 0.034323 | 0.001935 | 0.002363 | 0.001779 |
| 1.62629 | 4.892718 | 0.618528 | 0.035037 | 0.000122 | 0.000224 | 0.000118 |
| 0.204385 | 0.614897 | 0.618528 | 0.035037 | 0.000122 | 0.000224 | 0.000118 |
| 0.293078 | 0.848082 | 0.618528 | 0.033609 | 0.009793 | 0.010233 | 0.008862 |
| 1.179132 | 3.41206 | 0.618528 | 0.033609 | 0.009793 | 0.010233 | 0.008862 |
| 1.349128 | 3.995815 | 0.618296 | 0.034098 | 0.001731 | 0.002357 | 0.001746 |
| 1.25103 | 3.622491 | 0.61818 | 0.034184 | 0.004476 | 0.005341 | 0.00438 |
| 0.276053 | 0.799341 | 0.61818 | 0.034184 | 0.004476 | 0.005341 | 0.00438 |
| 1.366915 | 4.124737 | 0.61818 | 0.032271 | 0.001477 | 0.002146 | 0.001568 |
| 1.475987 | 4.480864 | 0.617948 | 0.03378 | 0.000528 | 0.000855 | 0.000564 |
| 0.28356 | 0.811433 | 0.617948 | 0.033908 | 0.005388 | 0.006162 | 0.005123 |
| 1.209998 | 3.752746 | 0.617948 | 0.032775 | 0.006275 | 0.00879 | 0.007304 |
| 0.266472 | 0.826448 | 0.617948 | 0.032775 | 0.006275 | 0.00879 | 0.007304 |
| 1.425417 | 4.520182 | 0.617948 | 0.033134 | 0.000832 | 0.001557 | 0.001046 |
| 1.14452 | 3.238642 | 0.617831 | 0.031726 | 0.013187 | 0.013562 | 0.012076 |
| 0.239972 | 0.698668 | 0.617715 | 0.034816 | 0.000837 | 0.001056 | 0.000724 |
| 1.459824 | 4.324601 | 0.617715 | 0.033803 | 0.000754 | 0.000883 | 0.000608 |
| 1.162729 | 3.438918 | 0.617599 | 0.033018 | 0.010097 | 0.012246 | 0.01062 |
| 0.339003 | 0.94724 | 0.617599 | 0.032727 | 0.030039 | 0.030255 | 0.028133 |
| 0.249197 | 0.719272 | 0.617483 | 0.033982 | 0.001314 | 0.00148 | 0.001094 |
| 0.192168 | 0.586062 | 0.617367 | 0.035767 | 6.58E-05 | 0.000124 | 5.99E-05 |
| 0.234899 | 0.700654 | 0.617251 | 0.034263 | 0.000852 | 0.001212 | 0.000836 |
| 0.219764 | 0.684661 | 0.617019 | 0.032802 | 0.00065 | 0.001088 | 0.000734 |
| 0.285797 | 0.842803 | 0.617019 | 0.03535 | 0.007962 | 0.009884 | 0.008448 |
| 0.259134 | 0.760763 | 0.616787 | 0.034221 | 0.002401 | 0.003125 | 0.002414 |
| 0.259249 | 0.741193 | 0.616671 | 0.034191 | 0.002071 | 0.002087 | 0.001587 |
| 1.320114 | 3.892469 | 0.616671 | 0.033698 | 0.002214 | 0.00301 | 0.002299 |
| 0.256906 | 0.75751 | 0.616671 | 0.033698 | 0.002214 | 0.00301 | 0.002299 |
| 0.23594 | 0.709791 | 0.616438 | 0.033192 | 0.000961 | 0.001473 | 0.001033 |
| 0.254334 | 0.741428 | 0.616322 | 0.033438 | 0.001789 | 0.002243 | 0.001694 |
| 1.348748 | 3.931835 | 0.616322 | 0.033438 | 0.001789 | 0.002243 | 0.001694 |
| 0.216838 | 0.66999 | 0.61609 | 0.034426 | 0.000464 | 0.000804 | 0.000514 |
| 0.22275 | 0.675542 | 0.615974 | 0.033717 | 0.000518 | 0.00082 | 0.000533 |
| 0.228491 | 0.673158 | 0.615626 | 0.034406 | 0.000487 | 0.000684 | 0.000447 |
| 1.443956 | 4.380867 | 0.615626 | 0.033679 | 0.000735 | 0.001124 | 0.000764 |
| 0.228265 | 0.692542 | 0.615626 | 0.033679 | 0.000735 | 0.001124 | 0.000764 |
| 0.268194 | 0.777289 | 0.615626 | 0.034115 | 0.003394 | 0.003876 | 0.003101 |
| 1.164707 | 3.337973 | 0.615394 | 0.032612 | 0.010777 | 0.011484 | 0.010108 |
| 0.299583 | 0.858585 | 0.615394 | 0.032612 | 0.010777 | 0.011484 | 0.010108 |
| 0.335964 | 0.986466 | 0.615394 | 0.033703 | 0.040611 | 0.044478 | 0.041907 |
| 1.376019 | 4.011184 | 0.615161 | 0.034276 | 0.001384 | 0.001751 | 0.00129 |
| 1.589435 | 4.873864 | 0.615161 | 0.03457 | 0.000175 | 0.000342 | 0.000192 |
| 1.183793 | 3.375656 | 0.615161 | 0.03523 | 0.009136 | 0.009566 | 0.008271 |
| 1.057746 | 3.021308 | 0.615045 | 0.033925 | 0.027717 | 0.030034 | 0.027827 |

| | | | | | | |
|---|---|---|---|---|---|---|
| 0.247847 | 0.716721 | 0.615045 | 0.034256 | 0.001121 | 0.001425 | 0.001017 |
| 1.336785 | 3.895626 | 0.614929 | 0.035243 | 0.002051 | 0.002496 | 0.001896 |
| 0.306447 | 0.870055 | 0.614929 | 0.03583 | 0.012482 | 0.013033 | 0.011497 |
| 1.261422 | 3.679052 | 0.614813 | 0.033806 | 0.003953 | 0.004947 | 0.004001 |
| 0.271809 | 0.792756 | 0.614813 | 0.033806 | 0.003953 | 0.004947 | 0.004001 |
| 0.249729 | 0.726304 | 0.614349 | 0.036293 | 0.001603 | 0.001724 | 0.001285 |
| 0.253241 | 0.749495 | 0.614117 | 0.033748 | 0.001964 | 0.002685 | 0.002034 |
| 0.280634 | 0.804336 | 0.614117 | 0.033858 | 0.004869 | 0.005596 | 0.004616 |
| 1.50283 | 4.445734 | 0.613884 | 0.035271 | 0.000456 | 0.000599 | 0.000386 |
| 0.214009 | 0.656644 | 0.613768 | 0.035579 | 0.000339 | 0.000602 | 0.000366 |
| 0.229127 | 0.685082 | 0.613768 | 0.034361 | 0.000679 | 0.000921 | 0.00063 |
| 0.227607 | 0.695658 | 0.613536 | 0.034822 | 0.000848 | 0.001224 | 0.000864 |
| 0.287697 | 0.824343 | 0.61342 | 0.033497 | 0.006488 | 0.007378 | 0.006231 |
| 1.57764 | 4.945868 | 0.613304 | 0.035485 | 0.000208 | 0.000425 | 0.000241 |
| 0.202189 | 0.633858 | 0.613304 | 0.035485 | 0.000208 | 0.000425 | 0.000241 |
| 1.069395 | 3.035615 | 0.613304 | 0.033571 | 0.025296 | 0.026964 | 0.024877 |
| 0.329423 | 0.935108 | 0.613304 | 0.033571 | 0.025296 | 0.026964 | 0.024877 |
| 0.263537 | 0.7789 | 0.613188 | 0.033678 | 0.003348 | 0.004186 | 0.003339 |
| 0.300473 | 0.917246 | 0.613072 | 0.033888 | 0.019334 | 0.023613 | 0.021333 |
| 0.237863 | 0.702289 | 0.613072 | 0.035748 | 0.000897 | 0.001197 | 0.000833 |
| 1.473151 | 4.332821 | 0.612956 | 0.034977 | 0.000593 | 0.000758 | 0.000506 |
| 1.319628 | 3.960614 | 0.61284 | 0.035507 | 0.00227 | 0.003186 | 0.002423 |
| 0.252486 | 0.757789 | 0.61284 | 0.035507 | 0.00227 | 0.003186 | 0.002423 |
| 0.294787 | 0.844448 | 0.61284 | 0.032459 | 0.009186 | 0.009606 | 0.008332 |
| 0.29208 | 0.841654 | 0.61284 | 0.035688 | 0.008038 | 0.009364 | 0.008022 |
| 0.310691 | 0.880763 | 0.61284 | 0.031946 | 0.014034 | 0.014785 | 0.013206 |
| 0.232309 | 0.685182 | 0.61284 | 0.035599 | 0.000638 | 0.000868 | 0.000578 |
| 1.548806 | 4.49816 | 0.612723 | 0.035785 | 0.000337 | 0.000359 | 0.000222 |
| 0.222313 | 0.645659 | 0.612723 | 0.035785 | 0.000337 | 0.000359 | 0.000222 |
| 1.439307 | 4.365804 | 0.612723 | 0.033826 | 0.000823 | 0.001169 | 0.00082 |
| 0.257963 | 0.744057 | 0.612375 | 0.033323 | 0.001908 | 0.002258 | 0.001717 |
| 0.244187 | 0.705162 | 0.612259 | 0.035578 | 0.000893 | 0.001149 | 0.000797 |
| 0.268523 | 0.770048 | 0.612259 | 0.034501 | 0.002877 | 0.003365 | 0.002649 |
| 1.298621 | 3.724071 | 0.612259 | 0.034501 | 0.002877 | 0.003365 | 0.002649 |
| 1.20158 | 3.506476 | 0.612143 | 0.033336 | 0.007309 | 0.008486 | 0.007218 |
| 1.240895 | 3.623936 | 0.612143 | 0.032507 | 0.005263 | 0.005971 | 0.004979 |
| 0.337047 | 0.992191 | 0.612143 | 0.03345 | 0.042886 | 0.046765 | 0.044171 |
| 0.332825 | 0.959345 | 0.612027 | 0.031234 | 0.031447 | 0.034545 | 0.032199 |
| 1.322473 | 3.782793 | 0.611795 | 0.034537 | 0.002603 | 0.002678 | 0.002113 |
| 0.254385 | 0.757805 | 0.611679 | 0.032498 | 0.002306 | 0.003118 | 0.002414 |
| 1.230613 | 3.691351 | 0.611563 | 0.034262 | 0.005268 | 0.006924 | 0.005698 |
| 0.270904 | 0.812603 | 0.611563 | 0.034262 | 0.005268 | 0.006924 | 0.005698 |
| 1.488459 | 4.353706 | 0.611563 | 0.034982 | 0.000577 | 0.000643 | 0.000434 |
| 1.090963 | 3.11882 | 0.61133 | 0.033387 | 0.020394 | 0.022321 | 0.020347 |
| 1.504257 | 4.420274 | 0.611214 | 0.03507 | 0.000442 | 0.000571 | 0.000367 |
| 0.22623 | 0.66478 | 0.611214 | 0.03507 | 0.000442 | 0.000571 | 0.000367 |
| 0.261178 | 0.793768 | 0.611098 | 0.032701 | 0.003889 | 0.00553 | 0.004438 |

| | | | | | | |
|---|---|---|---|---|---|---|
| 1.387903 | 4.332171 | 0.611098 | 0.034618 | 0.001204 | 0.00201 | 0.001426 |
| 0.230831 | 0.720512 | 0.611098 | 0.034618 | 0.001204 | 0.00201 | 0.001426 |
| 1.341454 | 4.028355 | 0.610982 | 0.034428 | 0.002052 | 0.002637 | 0.001999 |
| 0.291759 | 0.821606 | 0.610982 | 0.035038 | 0.006648 | 0.006853 | 0.005791 |
| 1.217128 | 3.427492 | 0.610982 | 0.035038 | 0.006648 | 0.006853 | 0.005791 |
| 0.213799 | 0.658037 | 0.610866 | 0.035265 | 0.000377 | 0.000628 | 0.000395 |
| 0.195923 | 0.607649 | 0.61075 | 0.035298 | 0.000115 | 0.000229 | 0.000121 |
| 1.376078 | 4.022768 | 0.610518 | 0.033816 | 0.001402 | 0.001769 | 0.001308 |
| 1.415667 | 4.290149 | 0.610518 | 0.034699 | 0.000995 | 0.001428 | 0.001013 |
| 0.233092 | 0.706381 | 0.610518 | 0.034699 | 0.000995 | 0.001428 | 0.001013 |
| 0.301502 | 0.865536 | 0.610402 | 0.033421 | 0.011658 | 0.012534 | 0.011035 |
| 0.246122 | 0.732448 | 0.610286 | 0.032034 | 0.001561 | 0.002076 | 0.001557 |
| 0.305449 | 0.875687 | 0.610286 | 0.034154 | 0.013555 | 0.014127 | 0.012589 |
| 1.141961 | 3.273869 | 0.610286 | 0.034154 | 0.013555 | 0.014127 | 0.012589 |
| 0.222702 | 0.677069 | 0.610286 | 0.034663 | 0.000487 | 0.000854 | 0.000544 |
| 0.253316 | 0.736083 | 0.610053 | 0.034589 | 0.001653 | 0.002029 | 0.001502 |
| 0.289845 | 0.865608 | 0.610053 | 0.033664 | 0.010533 | 0.013247 | 0.011532 |
| 1.489134 | 4.327074 | 0.609937 | 0.036275 | 0.000549 | 0.000619 | 0.000409 |
| 0.315172 | 0.907231 | 0.609937 | 0.03331 | 0.018411 | 0.020291 | 0.018416 |
| 0.284931 | 0.821197 | 0.609705 | 0.034698 | 0.006066 | 0.007157 | 0.005999 |
| 1.674525 | 5.206761 | 0.609589 | 0.035605 | 7.74E-05 | 0.000183 | 8.90E-05 |
| 1.508645 | 4.548314 | 0.609589 | 0.036224 | 0.000376 | 0.000625 | 0.000391 |
| 0.219862 | 0.662846 | 0.609589 | 0.036224 | 0.000376 | 0.000625 | 0.000391 |
| 1.291213 | 3.823984 | 0.609589 | 0.033184 | 0.00314 | 0.003942 | 0.003131 |
| 1.52946 | 4.637538 | 0.609473 | 0.036093 | 0.000328 | 0.000537 | 0.000329 |
| 0.215632 | 0.653826 | 0.609473 | 0.036093 | 0.000328 | 0.000537 | 0.000329 |
| 0.282535 | 0.8233 | 0.609473 | 0.033008 | 0.007088 | 0.007527 | 0.006434 |
| 1.214624 | 3.539391 | 0.609473 | 0.033008 | 0.007088 | 0.007527 | 0.006434 |
| 0.264661 | 0.793783 | 0.609357 | 0.034662 | 0.003995 | 0.005366 | 0.004309 |
| 0.289477 | 0.822501 | 0.609357 | 0.035028 | 0.006427 | 0.007071 | 0.005963 |
| 1.215804 | 3.454506 | 0.609357 | 0.035028 | 0.006427 | 0.007071 | 0.005963 |
| 0.275551 | 0.776226 | 0.609357 | 0.035252 | 0.003554 | 0.003515 | 0.002818 |
| 1.288285 | 3.629099 | 0.609357 | 0.035252 | 0.003554 | 0.003515 | 0.002818 |
| 0.249753 | 0.746905 | 0.609357 | 0.03445 | 0.001835 | 0.002663 | 0.001991 |
| 0.222633 | 0.694626 | 0.609241 | 0.034613 | 0.00078 | 0.001303 | 0.000892 |
| 0.235955 | 0.693933 | 0.609241 | 0.034843 | 0.000772 | 0.00101 | 0.000703 |
| 0.242362 | 0.711662 | 0.609241 | 0.035308 | 0.00116 | 0.001385 | 0.001001 |
| 1.313123 | 3.848231 | 0.609241 | 0.035099 | 0.003132 | 0.003146 | 0.002513 |
| 1.323825 | 3.957054 | 0.609241 | 0.033705 | 0.002116 | 0.003034 | 0.002302 |
| 0.252713 | 0.755387 | 0.609241 | 0.033705 | 0.002116 | 0.003034 | 0.002302 |
| 1.056431 | 2.990774 | 0.609125 | 0.033286 | 0.029749 | 0.030254 | 0.028108 |
| 0.317261 | 0.925061 | 0.609009 | 0.032384 | 0.021722 | 0.024749 | 0.022622 |
| 0.188694 | 0.599713 | 0.609009 | 0.035844 | 8.55E-05 | 0.000221 | 0.000109 |
| 0.21926 | 0.679946 | 0.609009 | 0.033543 | 0.000558 | 0.000981 | 0.000648 |
| 0.252116 | 0.758697 | 0.608893 | 0.0337 | 0.002296 | 0.003255 | 0.002507 |
| 1.264763 | 3.735014 | 0.608893 | 0.033924 | 0.004013 | 0.004951 | 0.004018 |
| 0.267737 | 0.790662 | 0.608893 | 0.033924 | 0.004013 | 0.004951 | 0.004018 |

| | | | | | | |
|---|---|---|---|---|---|---|
| 0.281875 | 0.821098 | 0.608893 | 0.033696 | 0.006279 | 0.007304 | 0.006142 |
| 1.251276 | 3.596041 | 0.608776 | 0.033502 | 0.00516 | 0.005233 | 0.004357 |
| 0.253114 | 0.754322 | 0.608776 | 0.034315 | 0.002214 | 0.002958 | 0.002263 |
| 0.253288 | 0.728825 | 0.608776 | 0.035067 | 0.001722 | 0.001729 | 0.0013 |
| 1.243621 | 3.620745 | 0.60866 | 0.034362 | 0.004924 | 0.005785 | 0.004767 |
| 0.242937 | 0.710101 | 0.608428 | 0.035579 | 0.001058 | 0.001322 | 0.000937 |
| 1.40825 | 4.116301 | 0.608428 | 0.035579 | 0.001058 | 0.001322 | 0.000937 |
| 0.255887 | 0.738074 | 0.608312 | 0.035212 | 0.001632 | 0.002043 | 0.001515 |
| 1.354878 | 3.907969 | 0.608312 | 0.035212 | 0.001632 | 0.002043 | 0.001515 |
| 0.307371 | 0.873456 | 0.608312 | 0.035207 | 0.012564 | 0.013596 | 0.012038 |
| 1.272244 | 3.640518 | 0.60808 | 0.035071 | 0.003682 | 0.004267 | 0.003432 |
| 0.274686 | 0.786013 | 0.60808 | 0.035071 | 0.003682 | 0.004267 | 0.003432 |
| 0.254497 | 0.763398 | 0.607964 | 0.034574 | 0.002614 | 0.003463 | 0.002707 |
| 0.316345 | 0.891966 | 0.607848 | 0.033087 | 0.015905 | 0.016743 | 0.015036 |
| 1.261737 | 3.663363 | 0.607732 | 0.035036 | 0.004124 | 0.004878 | 0.00395 |
| 1.248694 | 3.621125 | 0.607732 | 0.035095 | 0.004831 | 0.005475 | 0.004517 |
| 0.276157 | 0.800837 | 0.607732 | 0.035095 | 0.004831 | 0.005475 | 0.004517 |
| 0.318473 | 0.927736 | 0.607616 | 0.032875 | 0.022337 | 0.02542 | 0.023264 |
| 1.285447 | 3.669379 | 0.607616 | 0.036225 | 0.003563 | 0.003751 | 0.003 |
| 0.266401 | 0.783477 | 0.607499 | 0.034021 | 0.00373 | 0.004418 | 0.003583 |
| 1.276361 | 3.753736 | 0.607499 | 0.034021 | 0.00373 | 0.004418 | 0.003583 |
| 0.286947 | 0.835646 | 0.607499 | 0.034993 | 0.007302 | 0.008834 | 0.007508 |
| 1.262709 | 3.681692 | 0.607383 | 0.034543 | 0.004127 | 0.004887 | 0.003968 |
| 0.271614 | 0.791948 | 0.607383 | 0.034543 | 0.004127 | 0.004887 | 0.003968 |
| 0.285553 | 0.812967 | 0.607383 | 0.034999 | 0.005935 | 0.006224 | 0.005199 |
| 1.196313 | 3.537768 | 0.607383 | 0.03432 | 0.007279 | 0.009108 | 0.007744 |
| 0.205865 | 0.626778 | 0.607383 | 0.035154 | 0.000181 | 0.000312 | 0.000177 |
| 1.072191 | 3.083128 | 0.607267 | 0.034163 | 0.02385 | 0.02651 | 0.024343 |
| 1.561898 | 4.776817 | 0.607267 | 0.035678 | 0.000224 | 0.000426 | 0.000246 |
| 0.209344 | 0.640247 | 0.607267 | 0.035678 | 0.000224 | 0.000426 | 0.000246 |
| 1.268507 | 4.117571 | 0.607267 | 0.033257 | 0.003624 | 0.005927 | 0.004655 |
| 0.242862 | 0.788328 | 0.607267 | 0.033257 | 0.003624 | 0.005927 | 0.004655 |
| 1.17145 | 3.33213 | 0.607151 | 0.034184 | 0.009794 | 0.01067 | 0.009293 |
| 0.300108 | 0.853643 | 0.607151 | 0.034184 | 0.009794 | 0.01067 | 0.009293 |
| 1.197927 | 3.363756 | 0.607151 | 0.036689 | 0.008785 | 0.008154 | 0.00696 |
| 1.180013 | 3.398211 | 0.606803 | 0.034737 | 0.00867 | 0.01007 | 0.008672 |
| 0.237953 | 0.713235 | 0.606803 | 0.034599 | 0.001153 | 0.001542 | 0.001116 |
| 1.319128 | 3.835523 | 0.606687 | 0.035221 | 0.002616 | 0.002909 | 0.002281 |
| 0.260721 | 0.758076 | 0.606687 | 0.035221 | 0.002616 | 0.002909 | 0.002281 |
| 1.304178 | 3.85376 | 0.606571 | 0.034034 | 0.002601 | 0.003491 | 0.002715 |
| 0.328268 | 0.93701 | 0.606571 | 0.034179 | 0.026571 | 0.027586 | 0.025507 |
| 1.067224 | 3.04629 | 0.606571 | 0.034179 | 0.026571 | 0.027586 | 0.025507 |
| 1.363587 | 4.050086 | 0.606455 | 0.034847 | 0.001608 | 0.002093 | 0.001561 |
| 1.452378 | 4.302042 | 0.606339 | 0.035419 | 0.000633 | 0.000942 | 0.000629 |
| 0.222159 | 0.664061 | 0.606339 | 0.035722 | 0.0004 | 0.000614 | 0.000391 |
| 0.264772 | 0.773211 | 0.606222 | 0.035567 | 0.003249 | 0.003723 | 0.002981 |
| 0.219225 | 0.664716 | 0.606106 | 0.03521 | 0.00044 | 0.000666 | 0.00043 |

| | | | | | | |
|---|---|---|---|---|---|---|
| 1.192614 | 3.477405 | 0.60599 | 0.034939 | 0.00762 | 0.009182 | 0.007835 |
| 1.184532 | 3.390763 | 0.605874 | 0.035347 | 0.008488 | 0.009565 | 0.008241 |
| 0.26145 | 0.748169 | 0.605758 | 0.034559 | 0.002081 | 0.002353 | 0.001801 |
| 1.359443 | 4.025281 | 0.605642 | 0.035326 | 0.001542 | 0.002149 | 0.001588 |
| 0.24843 | 0.735595 | 0.605642 | 0.035326 | 0.001542 | 0.002149 | 0.001588 |
| 1.37977 | 4.060173 | 0.605642 | 0.035165 | 0.00142 | 0.001753 | 0.001295 |
| 0.240627 | 0.702326 | 0.605642 | 0.03591 | 0.000967 | 0.001142 | 0.000808 |
| 0.2723 | 0.8027 | 0.605642 | 0.034285 | 0.004514 | 0.005836 | 0.004764 |
| 1.467518 | 4.463081 | 0.60541 | 0.035817 | 0.000574 | 0.000927 | 0.000618 |
| 0.22406 | 0.681423 | 0.60541 | 0.035817 | 0.000574 | 0.000927 | 0.000618 |
| 0.320444 | 0.904376 | 0.605294 | 0.035247 | 0.019402 | 0.019299 | 0.017499 |
| 0.221679 | 0.669857 | 0.605294 | 0.035474 | 0.000528 | 0.000724 | 0.000475 |
| 0.264494 | 0.773871 | 0.605178 | 0.03434 | 0.002974 | 0.00378 | 0.002987 |
| 0.289304 | 0.829402 | 0.605178 | 0.034993 | 0.007303 | 0.007904 | 0.006724 |
| 1.205688 | 3.456571 | 0.605178 | 0.034993 | 0.007303 | 0.007904 | 0.006724 |
| 1.085091 | 3.225033 | 0.605062 | 0.032502 | 0.020879 | 0.024206 | 0.022047 |
| 1.259153 | 3.605488 | 0.605062 | 0.034444 | 0.004652 | 0.004823 | 0.004001 |
| 0.268702 | 0.83379 | 0.605062 | 0.034784 | 0.006941 | 0.00962 | 0.008062 |
| 1.199343 | 3.721601 | 0.605062 | 0.034784 | 0.006941 | 0.00962 | 0.008062 |
| 1.192477 | 3.47305 | 0.604945 | 0.034761 | 0.00797 | 0.009175 | 0.007859 |
| 0.279402 | 0.808714 | 0.604945 | 0.034108 | 0.005111 | 0.006087 | 0.005043 |
| 0.247927 | 0.742574 | 0.604829 | 0.03418 | 0.001808 | 0.002499 | 0.001896 |
| 0.24353 | 0.748005 | 0.604713 | 0.03549 | 0.00196 | 0.002938 | 0.00221 |
| 0.253457 | 0.743085 | 0.604713 | 0.034671 | 0.002088 | 0.002349 | 0.00181 |
| 0.318342 | 0.911667 | 0.604597 | 0.033967 | 0.019939 | 0.021194 | 0.019284 |
| 0.283523 | 0.82571 | 0.604597 | 0.035017 | 0.006684 | 0.007762 | 0.006557 |
| 0.251508 | 0.750529 | 0.604597 | 0.033934 | 0.00214 | 0.0028 | 0.002171 |
| 0.294104 | 0.864895 | 0.604597 | 0.0334 | 0.010964 | 0.012867 | 0.011261 |
| 0.30242 | 0.857038 | 0.604597 | 0.036352 | 0.010167 | 0.011069 | 0.009647 |
| 1.036718 | 2.931403 | 0.604365 | 0.034992 | 0.036058 | 0.036086 | 0.033845 |
| 1.316982 | 3.814319 | 0.604365 | 0.035238 | 0.002383 | 0.002931 | 0.002269 |
| 0.26217 | 0.759312 | 0.604365 | 0.035238 | 0.002383 | 0.002931 | 0.002269 |
| 0.305528 | 0.881285 | 0.604249 | 0.034811 | 0.013822 | 0.015199 | 0.013519 |
| 1.134707 | 3.273021 | 0.604249 | 0.034811 | 0.013822 | 0.015199 | 0.013519 |
| 0.238633 | 0.70329 | 0.604249 | 0.035891 | 0.000906 | 0.00121 | 0.000841 |
| 0.197564 | 0.633221 | 0.604133 | 0.03588 | 0.000211 | 0.000469 | 0.000266 |
| 0.286163 | 0.849233 | 0.604133 | 0.033724 | 0.008838 | 0.010807 | 0.009297 |
| 1.177533 | 3.494515 | 0.604133 | 0.033724 | 0.008838 | 0.010807 | 0.009297 |
| 0.205145 | 0.639577 | 0.604133 | 0.036351 | 0.000224 | 0.000464 | 0.000268 |
| 0.253262 | 0.769834 | 0.604133 | 0.035726 | 0.002683 | 0.003948 | 0.00306 |
| 1.207481 | 3.487157 | 0.604133 | 0.033974 | 0.007526 | 0.007888 | 0.006772 |
| 0.236476 | 0.75061 | 0.604017 | 0.034574 | 0.001981 | 0.003351 | 0.002509 |
| 1.343022 | 3.864834 | 0.604017 | 0.034879 | 0.002088 | 0.00226 | 0.001748 |
| 1.216399 | 3.523773 | 0.603901 | 0.035059 | 0.006532 | 0.00732 | 0.006189 |
| 0.283787 | 0.822099 | 0.603901 | 0.035059 | 0.006532 | 0.00732 | 0.006189 |
| 0.23968 | 0.718865 | 0.603668 | 0.034956 | 0.001223 | 0.001701 | 0.001231 |
| 0.2619 | 0.774137 | 0.603552 | 0.033907 | 0.00307 | 0.003903 | 0.003117 |

| | | | | | | |
|---|---|---|---|---|---|---|
| 0.265743 | 0.779542 | 0.603436 | 0.035874 | 0.003594 | 0.004142 | 0.00335 |
| 1.408825 | 4.133594 | 0.603436 | 0.035927 | 0.001238 | 0.001336 | 0.000979 |
| 0.297241 | 0.865585 | 0.60332 | 0.034264 | 0.011245 | 0.012798 | 0.011229 |
| 1.237455 | 3.531497 | 0.60332 | 0.036111 | 0.005555 | 0.005845 | 0.00486 |
| 1.527696 | 4.695926 | 0.60332 | 0.035675 | 0.000351 | 0.000583 | 0.000364 |
| 0.224339 | 0.672443 | 0.603088 | 0.036395 | 0.000496 | 0.000733 | 0.000476 |
| 1.380502 | 4.063757 | 0.602972 | 0.03488 | 0.001299 | 0.001744 | 0.001272 |
| 1.389024 | 4.319786 | 0.602856 | 0.034441 | 0.001173 | 0.001967 | 0.001386 |
| 0.231493 | 0.71993 | 0.602856 | 0.034441 | 0.001173 | 0.001967 | 0.001386 |
| 1.28314 | 3.908003 | 0.60274 | 0.035542 | 0.003171 | 0.004547 | 0.003564 |
| 0.255885 | 0.779338 | 0.60274 | 0.035542 | 0.003171 | 0.004547 | 0.003564 |
| 0.239958 | 0.758263 | 0.60274 | 0.033659 | 0.002213 | 0.003699 | 0.002789 |
| 0.267078 | 0.766164 | 0.602624 | 0.034563 | 0.002939 | 0.00317 | 0.002525 |
| 1.010202 | 2.85419 | 0.602624 | 0.034161 | 0.04533 | 0.045687 | 0.043406 |
| 0.350362 | 0.989901 | 0.602624 | 0.034161 | 0.04533 | 0.045687 | 0.043406 |
| 1.117129 | 3.158825 | 0.602508 | 0.035013 | 0.017311 | 0.017422 | 0.015779 |
| 1.121454 | 3.192459 | 0.602508 | 0.033704 | 0.016934 | 0.016873 | 0.015216 |
| 1.245695 | 3.585043 | 0.602508 | 0.035555 | 0.005064 | 0.005526 | 0.004573 |
| 0.278937 | 0.802765 | 0.602508 | 0.035555 | 0.005064 | 0.005526 | 0.004573 |
| 1.271573 | 3.668361 | 0.602391 | 0.034806 | 0.004017 | 0.004388 | 0.003568 |
| 0.250201 | 0.730181 | 0.602275 | 0.035573 | 0.001403 | 0.001865 | 0.001362 |
| 1.174703 | 3.425402 | 0.602275 | 0.033296 | 0.009315 | 0.010781 | 0.009393 |
| 1.39321 | 4.062467 | 0.602159 | 0.036185 | 0.001181 | 0.0015 | 0.001088 |
| 1.228952 | 3.602583 | 0.602159 | 0.034833 | 0.005524 | 0.0067 | 0.005559 |
| 0.231078 | 0.68606 | 0.602159 | 0.0364 | 0.000611 | 0.000909 | 0.000606 |
| 0.204425 | 0.711714 | 0.602043 | 0.031327 | 0.001103 | 0.002457 | 0.001687 |
| 0.237062 | 0.703334 | 0.602043 | 0.036237 | 0.001097 | 0.001245 | 0.00091 |
| 0.254569 | 0.738968 | 0.601927 | 0.036084 | 0.001699 | 0.002122 | 0.001591 |
| 0.233382 | 0.705518 | 0.601927 | 0.035253 | 0.000969 | 0.001394 | 0.001002 |
| 1.29992 | 3.728642 | 0.601927 | 0.035809 | 0.00312 | 0.003328 | 0.002671 |
| 1.669177 | 4.982621 | 0.601811 | 0.038069 | 9.42E-05 | 0.000147 | 7.79E-05 |
| 0.200698 | 0.599098 | 0.601811 | 0.038069 | 9.42E-05 | 0.000147 | 7.79E-05 |
| 0.238278 | 0.701399 | 0.601811 | 0.036166 | 0.000965 | 0.001163 | 0.000824 |
| 0.266062 | 0.785434 | 0.601695 | 0.035442 | 0.003489 | 0.004589 | 0.003665 |
| 0.308281 | 0.880839 | 0.601695 | 0.034351 | 0.013469 | 0.014945 | 0.013281 |
| 0.268406 | 0.794672 | 0.601695 | 0.035818 | 0.004102 | 0.005271 | 0.004252 |
| 1.223979 | 3.484357 | 0.601579 | 0.036217 | 0.006192 | 0.006583 | 0.005561 |
| 0.275133 | 0.801813 | 0.601579 | 0.035045 | 0.004516 | 0.005615 | 0.004591 |
| 1.019814 | 2.889475 | 0.601579 | 0.034113 | 0.039791 | 0.041971 | 0.039564 |
| 0.346084 | 0.980571 | 0.601579 | 0.034113 | 0.039791 | 0.041971 | 0.039564 |
| 1.034675 | 2.91923 | 0.601463 | 0.035578 | 0.035267 | 0.036727 | 0.034443 |
| 1.34371 | 3.966306 | 0.601463 | 0.035662 | 0.001772 | 0.002446 | 0.001828 |
| 0.252124 | 0.744208 | 0.601463 | 0.035662 | 0.001772 | 0.002446 | 0.001828 |
| 1.248988 | 3.545119 | 0.601463 | 0.034616 | 0.00468 | 0.005184 | 0.004255 |
| 0.232283 | 0.682297 | 0.601347 | 0.036081 | 0.00067 | 0.000806 | 0.000549 |
| 0.235442 | 0.728114 | 0.601347 | 0.03563 | 0.001309 | 0.002201 | 0.001575 |
| 0.289573 | 0.833853 | 0.601231 | 0.033229 | 0.00748 | 0.008454 | 0.007241 |

| | | | | | | |
|---|---|---|---|---|---|---|
| 0.251794 | 0.745892 | 0.601231 | 0.035821 | 0.001989 | 0.002543 | 0.001938 |
| 1.40246 | 4.242118 | 0.601114 | 0.035801 | 0.001066 | 0.00159 | 0.001123 |
| 0.225083 | 0.736072 | 0.601114 | 0.034057 | 0.001608 | 0.002942 | 0.002136 |
| 1.245606 | 3.736782 | 0.600998 | 0.03556 | 0.004572 | 0.006077 | 0.00494 |
| 0.26761 | 0.802822 | 0.600998 | 0.03556 | 0.004572 | 0.006077 | 0.00494 |
| 0.346538 | 0.997201 | 0.600882 | 0.033018 | 0.045074 | 0.048797 | 0.046218 |
| 1.288247 | 3.824991 | 0.600882 | 0.036048 | 0.003118 | 0.004075 | 0.003213 |
| 0.261439 | 0.776249 | 0.600882 | 0.036048 | 0.003118 | 0.004075 | 0.003213 |
| 0.279824 | 0.808785 | 0.600882 | 0.033973 | 0.005112 | 0.006074 | 0.00503 |
| 1.236423 | 3.57367 | 0.600882 | 0.033973 | 0.005112 | 0.006074 | 0.00503 |
| 1.261798 | 3.763015 | 0.600882 | 0.035485 | 0.004038 | 0.005203 | 0.00421 |
| 0.244642 | 0.738626 | 0.600766 | 0.036029 | 0.0018 | 0.002408 | 0.001805 |
| 1.156744 | 3.340823 | 0.60065 | 0.035371 | 0.012999 | 0.012485 | 0.011054 |
| 1.217582 | 3.455207 | 0.60065 | 0.03482 | 0.007375 | 0.006937 | 0.005936 |
| 1.377603 | 4.099269 | 0.60065 | 0.035897 | 0.001493 | 0.001861 | 0.001392 |
| 0.340507 | 0.964363 | 0.60065 | 0.034059 | 0.034036 | 0.036029 | 0.033717 |
| 0.2478 | 0.751167 | 0.600534 | 0.035861 | 0.001945 | 0.002965 | 0.002219 |
| 1.331262 | 4.035518 | 0.600534 | 0.035861 | 0.001945 | 0.002965 | 0.002219 |
| 0.233041 | 0.868845 | 0.600534 | 0.026652 | 0.010325 | 0.017371 | 0.014623 |
| 0.300009 | 0.848591 | 0.600418 | 0.034854 | 0.009343 | 0.009911 | 0.0086 |
| 0.312972 | 0.900727 | 0.600418 | 0.035132 | 0.016747 | 0.018891 | 0.017002 |
| 0.258229 | 0.798531 | 0.600418 | 0.03551 | 0.004148 | 0.006122 | 0.004922 |
| 0.269155 | 0.794495 | 0.600418 | 0.035452 | 0.004008 | 0.005221 | 0.004222 |
| 1.114099 | 3.197755 | 0.600302 | 0.034539 | 0.016481 | 0.018194 | 0.016398 |
| 1.384006 | 4.020048 | 0.600302 | 0.035526 | 0.00127 | 0.001607 | 0.001169 |
| 0.248753 | 0.72254 | 0.600302 | 0.035526 | 0.00127 | 0.001607 | 0.001169 |
| 1.267799 | 3.792992 | 0.600302 | 0.034996 | 0.003856 | 0.004974 | 0.00402 |
| 0.263644 | 0.788769 | 0.600302 | 0.034996 | 0.003856 | 0.004974 | 0.00402 |
| 0.273188 | 0.798625 | 0.60007 | 0.03546 | 0.004342 | 0.005408 | 0.004417 |
| 0.273045 | 0.788618 | 0.60007 | 0.036629 | 0.003772 | 0.004545 | 0.003665 |
| 1.268042 | 3.662401 | 0.60007 | 0.036629 | 0.003772 | 0.004545 | 0.003665 |
| 0.27487 | 0.794812 | 0.60007 | 0.034484 | 0.004154 | 0.004989 | 0.004059 |
| 1.232886 | 3.527196 | 0.599837 | 0.035706 | 0.005554 | 0.006131 | 0.005133 |
| 0.271585 | 0.788292 | 0.599837 | 0.035167 | 0.003807 | 0.004582 | 0.003712 |
| 1.268565 | 3.682086 | 0.599837 | 0.035167 | 0.003807 | 0.004582 | 0.003712 |
| 1.305119 | 3.808449 | 0.599721 | 0.035769 | 0.0026 | 0.003339 | 0.002597 |
| 1.172207 | 3.370221 | 0.599721 | 0.034636 | 0.010017 | 0.010781 | 0.009403 |
| 1.083939 | 3.094913 | 0.599605 | 0.035358 | 0.022394 | 0.023752 | 0.021733 |
| 1.392949 | 4.037503 | 0.599489 | 0.035519 | 0.001156 | 0.001469 | 0.001055 |
| 0.271688 | 0.796436 | 0.599489 | 0.034954 | 0.004391 | 0.005278 | 0.004347 |
| 1.353259 | 3.939059 | 0.599489 | 0.036554 | 0.00172 | 0.002142 | 0.001614 |
| 0.256317 | 0.734422 | 0.599489 | 0.035837 | 0.001656 | 0.001875 | 0.001405 |
| 0.294589 | 0.854884 | 0.599373 | 0.033968 | 0.010338 | 0.011185 | 0.009804 |
| 0.286112 | 0.833262 | 0.599373 | 0.034152 | 0.007676 | 0.008567 | 0.007373 |
| 1.342083 | 3.928498 | 0.599257 | 0.036156 | 0.001979 | 0.002415 | 0.001835 |
| 0.255474 | 0.773184 | 0.599257 | 0.034979 | 0.002992 | 0.004098 | 0.003228 |
| 1.076831 | 3.099883 | 0.599141 | 0.033774 | 0.023555 | 0.025457 | 0.023368 |

| | | | | | | |
|---|---|---|---|---|---|---|
| 1.226192 | 3.50921 | 0.599141 | 0.036385 | 0.006626 | 0.006525 | 0.005527 |
| 0.222149 | 0.675221 | 0.599141 | 0.035796 | 0.000534 | 0.000824 | 0.000551 |
| 1.383597 | 4.006831 | 0.599025 | 0.036267 | 0.001622 | 0.001594 | 0.001218 |
| 0.304848 | 0.878885 | 0.599025 | 0.035999 | 0.012952 | 0.014773 | 0.01309 |
| 0.285005 | 0.817388 | 0.599025 | 0.035559 | 0.005898 | 0.006725 | 0.00564 |
| 1.142957 | 3.238697 | 0.598909 | 0.035984 | 0.013716 | 0.013784 | 0.012318 |
| 0.257863 | 0.752815 | 0.598793 | 0.036763 | 0.002085 | 0.00271 | 0.002064 |
| 0.284276 | 0.829416 | 0.598793 | 0.034157 | 0.007442 | 0.008177 | 0.006992 |
| 1.317478 | 3.886347 | 0.598793 | 0.03558 | 0.002428 | 0.003086 | 0.002385 |
| 0.257311 | 0.759026 | 0.598793 | 0.03558 | 0.002428 | 0.003086 | 0.002385 |
| 0.235473 | 0.708682 | 0.598677 | 0.035515 | 0.000945 | 0.001447 | 0.001014 |
| 0.232536 | 0.705147 | 0.59856 | 0.03525 | 0.000927 | 0.001401 | 0.000977 |
| 1.249237 | 3.671809 | 0.59856 | 0.035994 | 0.004597 | 0.005622 | 0.004602 |
| 0.272345 | 0.800489 | 0.59856 | 0.035994 | 0.004597 | 0.005622 | 0.004602 |
| 0.263441 | 0.788595 | 0.59856 | 0.035522 | 0.003627 | 0.004968 | 0.003975 |
| 0.256634 | 0.735799 | 0.59856 | 0.034408 | 0.001698 | 0.001924 | 0.001446 |
| 0.297366 | 0.926483 | 0.598444 | 0.029645 | 0.021079 | 0.026192 | 0.023768 |
| 0.292769 | 0.856722 | 0.598444 | 0.03439 | 0.010163 | 0.011586 | 0.010124 |
| 0.266898 | 0.788243 | 0.598444 | 0.034755 | 0.003646 | 0.004783 | 0.003837 |
| 1.313764 | 3.930405 | 0.598444 | 0.03564 | 0.002338 | 0.003323 | 0.002548 |
| 0.336921 | 0.949577 | 0.598328 | 0.035071 | 0.029823 | 0.031107 | 0.028934 |
| 1.210079 | 3.48398 | 0.598328 | 0.034764 | 0.007517 | 0.007658 | 0.00657 |
| 0.252452 | 0.743175 | 0.598328 | 0.035548 | 0.001945 | 0.002385 | 0.001816 |
| 0.263779 | 0.757081 | 0.598328 | 0.035031 | 0.002439 | 0.002748 | 0.002143 |
| 1.210712 | 3.482538 | 0.598328 | 0.034257 | 0.006698 | 0.007599 | 0.00645 |
| 0.287147 | 0.82596 | 0.598328 | 0.034257 | 0.006698 | 0.007599 | 0.00645 |
| 0.284161 | 0.819047 | 0.598212 | 0.03515 | 0.006395 | 0.006952 | 0.00587 |
| 0.183776 | 0.590057 | 0.598212 | 0.036472 | 8.15E-05 | 0.000189 | 9.64E-05 |
| 0.324942 | 0.925522 | 0.598096 | 0.034489 | 0.022914 | 0.024461 | 0.022472 |
| 0.262924 | 0.781806 | 0.59798 | 0.036092 | 0.003408 | 0.004436 | 0.003528 |
| 1.279089 | 3.80338 | 0.59798 | 0.036092 | 0.003408 | 0.004436 | 0.003528 |
| 0.291151 | 0.863682 | 0.59798 | 0.034144 | 0.011248 | 0.012836 | 0.011244 |
| 0.318011 | 0.927574 | 0.59798 | 0.033801 | 0.022336 | 0.0254 | 0.023256 |
| 0.320776 | 0.916341 | 0.597864 | 0.035116 | 0.020933 | 0.02224 | 0.020272 |
| 0.208273 | 0.65466 | 0.597864 | 0.036396 | 0.000375 | 0.00065 | 0.000396 |
| 0.256834 | 0.745429 | 0.597748 | 0.036247 | 0.002125 | 0.00236 | 0.001813 |
| 0.271817 | 0.787683 | 0.597632 | 0.034267 | 0.003952 | 0.004522 | 0.003659 |
| 0.28214 | 0.799934 | 0.597516 | 0.036268 | 0.004616 | 0.005116 | 0.00419 |
| 1.456236 | 4.212 | 0.597516 | 0.036402 | 0.000666 | 0.000816 | 0.000554 |
| 0.237417 | 0.686702 | 0.597516 | 0.036402 | 0.000666 | 0.000816 | 0.000554 |
| 0.298413 | 0.868397 | 0.597516 | 0.03454 | 0.011192 | 0.01322 | 0.011586 |
| 1.151548 | 3.351065 | 0.597516 | 0.03454 | 0.011192 | 0.01322 | 0.011586 |
| 1.102164 | 3.118375 | 0.5974 | 0.035061 | 0.019035 | 0.019987 | 0.018168 |
| 1.228006 | 3.54294 | 0.5974 | 0.036203 | 0.006099 | 0.006532 | 0.005546 |
| 0.268604 | 0.773875 | 0.5974 | 0.035622 | 0.003093 | 0.003619 | 0.002878 |
| 0.229915 | 0.698246 | 0.597283 | 0.036102 | 0.000821 | 0.001249 | 0.000861 |
| 0.19542 | 0.646995 | 0.597283 | 0.036309 | 0.000324 | 0.00071 | 0.000427 |

| | | | | | | |
|---|---|---|---|---|---|---|
| 0.332058 | 0.976536 | 0.597283 | 0.032411 | 0.036035 | 0.040732 | 0.038145 |
| 1.230401 | 3.606928 | 0.597283 | 0.035084 | 0.005642 | 0.006614 | 0.005529 |
| 0.277244 | 0.812743 | 0.597283 | 0.035084 | 0.005642 | 0.006614 | 0.005529 |
| 0.280101 | 0.808466 | 0.597283 | 0.036061 | 0.005545 | 0.006028 | 0.005039 |
| 0.237537 | 0.691973 | 0.597167 | 0.035962 | 0.000775 | 0.000933 | 0.000637 |
| 1.445143 | 4.209878 | 0.597167 | 0.035962 | 0.000775 | 0.000933 | 0.000637 |
| 0.31475 | 0.906035 | 0.597051 | 0.034184 | 0.01819 | 0.020029 | 0.018171 |
| 1.191065 | 3.443575 | 0.597051 | 0.035977 | 0.007857 | 0.009173 | 0.007857 |
| 0.335246 | 0.959568 | 0.597051 | 0.035179 | 0.032026 | 0.034532 | 0.032243 |
| 0.270791 | 0.784241 | 0.596935 | 0.035497 | 0.003891 | 0.004292 | 0.003478 |
| 1.275119 | 3.692882 | 0.596935 | 0.035497 | 0.003891 | 0.004292 | 0.003478 |
| 0.321086 | 0.905328 | 0.596935 | 0.035709 | 0.018556 | 0.019488 | 0.017666 |
| 1.104572 | 3.11443 | 0.596935 | 0.035709 | 0.018556 | 0.019488 | 0.017666 |
| 0.260331 | 0.785436 | 0.596935 | 0.034981 | 0.003733 | 0.004843 | 0.003921 |
| 1.131415 | 3.287991 | 0.596935 | 0.036391 | 0.013506 | 0.015794 | 0.014009 |
| 0.303032 | 0.872721 | 0.596819 | 0.034873 | 0.012333 | 0.013721 | 0.012168 |
| 0.295739 | 0.840604 | 0.596819 | 0.034766 | 0.009506 | 0.009014 | 0.007866 |
| 0.269869 | 0.79597 | 0.596819 | 0.035003 | 0.004522 | 0.00532 | 0.00441 |
| 0.260492 | 0.756626 | 0.596819 | 0.035005 | 0.002642 | 0.002834 | 0.002221 |
| 0.304601 | 0.874458 | 0.596703 | 0.036453 | 0.013327 | 0.013948 | 0.01241 |
| 0.266802 | 0.958262 | 0.596703 | 0.025137 | 0.026402 | 0.036558 | 0.033183 |
| 1.314736 | 3.961838 | 0.596703 | 0.035269 | 0.002507 | 0.003364 | 0.002615 |
| 1.149067 | 3.35717 | 0.596587 | 0.033724 | 0.011972 | 0.013588 | 0.011992 |
| 0.25251 | 0.760932 | 0.596587 | 0.035251 | 0.002521 | 0.00338 | 0.002629 |
| 0.273768 | 0.806249 | 0.596587 | 0.035086 | 0.004746 | 0.006115 | 0.005009 |
| 1.010434 | 2.908241 | 0.596471 | 0.034304 | 0.046023 | 0.045668 | 0.043423 |
| 0.236376 | 0.703756 | 0.596471 | 0.03486 | 0.000881 | 0.001272 | 0.00089 |
| 1.269943 | 3.629355 | 0.596355 | 0.036107 | 0.004445 | 0.004344 | 0.003555 |
| 1.090553 | 3.093726 | 0.596355 | 0.036286 | 0.020793 | 0.022263 | 0.020317 |
| 1.393426 | 4.085048 | 0.596355 | 0.037982 | 0.00121 | 0.001529 | 0.00112 |
| 0.27763 | 0.799228 | 0.596355 | 0.03478 | 0.005218 | 0.005258 | 0.004413 |
| 0.207203 | 0.625549 | 0.596355 | 0.036661 | 0.000162 | 0.00029 | 0.000161 |
| 1.598595 | 4.82618 | 0.596355 | 0.036661 | 0.000162 | 0.00029 | 0.000161 |
| 0.260892 | 0.783689 | 0.596239 | 0.036593 | 0.003442 | 0.004675 | 0.003707 |
| 1.066564 | 3.013379 | 0.596239 | 0.035627 | 0.026404 | 0.027582 | 0.025515 |
| 0.283499 | 0.827667 | 0.596239 | 0.035375 | 0.006573 | 0.008002 | 0.006755 |
| 1.23927 | 3.546815 | 0.596123 | 0.035594 | 0.006025 | 0.005786 | 0.004835 |
| 0.281943 | 0.806926 | 0.596123 | 0.035594 | 0.006025 | 0.005786 | 0.004835 |
| 0.264043 | 0.775469 | 0.596123 | 0.036343 | 0.003132 | 0.003912 | 0.003091 |
| 0.247154 | 0.74329 | 0.596007 | 0.035032 | 0.001958 | 0.00256 | 0.001954 |
| 1.163086 | 3.469748 | 0.59589 | 0.035594 | 0.00976 | 0.012357 | 0.010693 |
| 1.425798 | 4.432651 | 0.59589 | 0.035566 | 0.000874 | 0.001443 | 0.000997 |
| 0.295075 | 0.86451 | 0.59589 | 0.035326 | 0.010891 | 0.012742 | 0.011133 |
| 0.24565 | 0.72925 | 0.59589 | 0.036639 | 0.001391 | 0.001952 | 0.001431 |
| 1.371272 | 4.070827 | 0.59589 | 0.036639 | 0.001391 | 0.001952 | 0.001431 |
| 1.294725 | 3.821994 | 0.59589 | 0.03659 | 0.003149 | 0.003788 | 0.003024 |
| 0.261644 | 0.772365 | 0.59589 | 0.03659 | 0.003149 | 0.003788 | 0.003024 |

| | | | | | | |
|---|---|---|---|---|---|---|
| 1.190394 | 3.530394 | 0.59589 | 0.034393 | 0.008071 | 0.009643 | 0.008285 |
| 0.29873 | 0.866743 | 0.595774 | 0.035368 | 0.011372 | 0.01291 | 0.011323 |
| 0.280093 | 0.840531 | 0.595774 | 0.034627 | 0.007752 | 0.00989 | 0.008401 |
| 0.285991 | 0.832305 | 0.595774 | 0.035777 | 0.006964 | 0.00845 | 0.007152 |
| 1.321872 | 3.838657 | 0.595774 | 0.036176 | 0.002457 | 0.002826 | 0.002197 |
| 0.338985 | 0.991194 | 0.595774 | 0.03454 | 0.042271 | 0.04634 | 0.043718 |
| 1.008884 | 2.949981 | 0.595774 | 0.03454 | 0.042271 | 0.04634 | 0.043718 |
| 1.084584 | 3.12368 | 0.595774 | 0.03537 | 0.021297 | 0.023768 | 0.021699 |
| 0.320135 | 0.922013 | 0.595774 | 0.03537 | 0.021297 | 0.023768 | 0.021699 |
| 0.294131 | 0.86621 | 0.595658 | 0.03458 | 0.010742 | 0.013093 | 0.01143 |
| 1.154455 | 3.399842 | 0.595658 | 0.03458 | 0.010742 | 0.013093 | 0.01143 |
| 0.279779 | 0.811143 | 0.595658 | 0.035586 | 0.006024 | 0.006319 | 0.005348 |
| 0.245282 | 0.724622 | 0.595658 | 0.036213 | 0.001356 | 0.001775 | 0.001294 |
| 0.319608 | 0.917232 | 0.595542 | 0.035279 | 0.021253 | 0.022537 | 0.020598 |
| 1.090237 | 3.128829 | 0.595542 | 0.035279 | 0.021253 | 0.022537 | 0.020598 |
| 0.262582 | 0.769702 | 0.595542 | 0.03616 | 0.002782 | 0.003568 | 0.002802 |
| 0.264141 | 0.783702 | 0.595542 | 0.036497 | 0.003434 | 0.004533 | 0.003627 |
| 1.275996 | 3.785859 | 0.595542 | 0.036497 | 0.003434 | 0.004533 | 0.003627 |
| 0.330256 | 0.944469 | 0.595426 | 0.035093 | 0.028204 | 0.029773 | 0.027596 |
| 0.266167 | 0.763689 | 0.595426 | 0.036694 | 0.002604 | 0.003051 | 0.002381 |
| 1.309434 | 3.75704 | 0.595426 | 0.036694 | 0.002604 | 0.003051 | 0.002381 |
| 0.27337 | 0.804647 | 0.59531 | 0.035621 | 0.004628 | 0.005975 | 0.004882 |
| 0.327895 | 0.951415 | 0.595194 | 0.034203 | 0.029351 | 0.032094 | 0.029789 |
| 0.299438 | 0.851866 | 0.595194 | 0.036043 | 0.009564 | 0.010435 | 0.009071 |
| 0.233938 | 0.703879 | 0.595194 | 0.036421 | 0.000856 | 0.001329 | 0.000919 |
| 0.317058 | 0.906887 | 0.595078 | 0.036609 | 0.019253 | 0.020098 | 0.018258 |
| 0.256 | 0.776623 | 0.595078 | 0.034393 | 0.003168 | 0.004332 | 0.003434 |
| 0.309982 | 0.887929 | 0.595078 | 0.035786 | 0.015235 | 0.016274 | 0.01457 |
| 0.274243 | 0.800789 | 0.594962 | 0.035615 | 0.004475 | 0.005561 | 0.00455 |
| 1.22839 | 3.502919 | 0.594962 | 0.034815 | 0.00612 | 0.006344 | 0.005358 |
| 0.326727 | 0.929966 | 0.594962 | 0.033864 | 0.024685 | 0.025611 | 0.023576 |
| 0.23847 | 0.707774 | 0.594962 | 0.03569 | 0.001056 | 0.001349 | 0.000964 |
| 0.290693 | 0.833385 | 0.594962 | 0.036197 | 0.007701 | 0.008332 | 0.007116 |
| 1.033203 | 3.054409 | 0.594846 | 0.034388 | 0.033394 | 0.037701 | 0.035149 |
| 1.226371 | 3.685652 | 0.594846 | 0.034901 | 0.005716 | 0.007212 | 0.006037 |
| 0.271322 | 0.815414 | 0.594846 | 0.034901 | 0.005716 | 0.007212 | 0.006037 |
| 1.017452 | 2.8497 | 0.594846 | 0.035153 | 0.043631 | 0.042784 | 0.0405 |
| 0.226843 | 0.689838 | 0.594846 | 0.035718 | 0.000682 | 0.001081 | 0.000737 |
| 0.303504 | 0.914199 | 0.594846 | 0.034555 | 0.018992 | 0.022675 | 0.020514 |
| 0.297613 | 0.84682 | 0.59473 | 0.035997 | 0.009209 | 0.009788 | 0.008505 |
| 0.222896 | 0.681916 | 0.59473 | 0.036996 | 0.000568 | 0.00096 | 0.000623 |
| 0.262011 | 0.769779 | 0.59473 | 0.035491 | 0.002815 | 0.003595 | 0.002835 |
| 1.093917 | 3.123372 | 0.59473 | 0.03583 | 0.020441 | 0.021713 | 0.019765 |
| 1.489307 | 4.451031 | 0.59473 | 0.036858 | 0.00054 | 0.000709 | 0.000472 |
| 0.271603 | 0.787933 | 0.59473 | 0.03589 | 0.003999 | 0.004552 | 0.003696 |
| 1.3176 | 3.97008 | 0.59473 | 0.035634 | 0.002317 | 0.00328 | 0.002531 |
| 0.251884 | 0.758956 | 0.59473 | 0.035634 | 0.002317 | 0.00328 | 0.002531 |

| | | | | | | |
|---|---|---|---|---|---|---|
| 0.256936 | 0.745003 | 0.594613 | 0.03556 | 0.001879 | 0.002335 | 0.001767 |
| 1.105213 | 3.116612 | 0.594613 | 0.035875 | 0.019453 | 0.019376 | 0.017563 |
| 0.318895 | 0.911759 | 0.594613 | 0.035957 | 0.019297 | 0.021185 | 0.019245 |
| 0.322329 | 0.920797 | 0.594613 | 0.035293 | 0.021769 | 0.023322 | 0.021381 |
| 0.315729 | 0.905663 | 0.594497 | 0.035258 | 0.018105 | 0.019882 | 0.018023 |
| 1.163861 | 3.42816 | 0.594497 | 0.034772 | 0.012495 | 0.012053 | 0.010536 |
| 1.103368 | 3.204189 | 0.594497 | 0.03474 | 0.01815 | 0.020252 | 0.018307 |
| 0.333602 | 0.947983 | 0.594381 | 0.035889 | 0.028932 | 0.030737 | 0.028563 |
| 0.250299 | 0.747471 | 0.594381 | 0.035088 | 0.001984 | 0.002675 | 0.002026 |
| 0.33401 | 0.953793 | 0.594381 | 0.034209 | 0.030724 | 0.032622 | 0.030438 |
| 0.284808 | 0.823893 | 0.594381 | 0.035897 | 0.006677 | 0.007477 | 0.006326 |
| 1.21375 | 3.511135 | 0.594381 | 0.035897 | 0.006677 | 0.007477 | 0.006326 |
| 0.279624 | 0.816 | 0.594265 | 0.035102 | 0.005579 | 0.006846 | 0.005704 |
| 0.33275 | 0.96875 | 0.594265 | 0.034517 | 0.034811 | 0.037855 | 0.035458 |
| 0.264244 | 0.807414 | 0.594265 | 0.035504 | 0.004878 | 0.006714 | 0.005487 |
| 0.258554 | 0.763618 | 0.594265 | 0.035823 | 0.002754 | 0.003323 | 0.002633 |
| 1.309555 | 3.867668 | 0.594265 | 0.035823 | 0.002754 | 0.003323 | 0.002633 |
| 0.314445 | 0.904718 | 0.594265 | 0.035453 | 0.018549 | 0.019734 | 0.017913 |
| 0.291667 | 0.835906 | 0.594149 | 0.036523 | 0.007967 | 0.008607 | 0.00737 |
| 0.254127 | 0.742993 | 0.594149 | 0.036179 | 0.001902 | 0.002324 | 0.001757 |
| 1.076488 | 3.165634 | 0.594149 | 0.035184 | 0.022917 | 0.025893 | 0.023727 |
| 1.385483 | 4.068007 | 0.594149 | 0.035022 | 0.001238 | 0.001652 | 0.001199 |
| 0.238624 | 0.710498 | 0.594149 | 0.036335 | 0.001013 | 0.001433 | 0.001005 |
| 0.258408 | 0.768888 | 0.594033 | 0.036111 | 0.002794 | 0.003676 | 0.002873 |
| 0.248657 | 0.758219 | 0.594033 | 0.035451 | 0.002239 | 0.003356 | 0.002563 |
| 0.302029 | 0.865521 | 0.594033 | 0.034669 | 0.011855 | 0.0125 | 0.011103 |
| 1.416507 | 4.208329 | 0.594033 | 0.036751 | 0.001046 | 0.001311 | 0.000947 |
| 0.290234 | 0.850516 | 0.594033 | 0.03586 | 0.009101 | 0.010764 | 0.009285 |
| 0.32197 | 0.924281 | 0.593917 | 0.035019 | 0.023361 | 0.024281 | 0.022341 |
| 1.221526 | 3.50852 | 0.593917 | 0.036127 | 0.006322 | 0.006864 | 0.005778 |
| 0.28502 | 0.818648 | 0.593917 | 0.036127 | 0.006322 | 0.006864 | 0.005778 |
| 1.307518 | 3.988311 | 0.593917 | 0.036514 | 0.002532 | 0.003703 | 0.002842 |
| 1.043075 | 2.991322 | 0.593917 | 0.036067 | 0.031981 | 0.03427 | 0.031927 |
| 0.34821 | 0.98105 | 0.593801 | 0.034583 | 0.04117 | 0.042116 | 0.039804 |
| 0.331396 | 0.940966 | 0.593801 | 0.034982 | 0.02977 | 0.028632 | 0.026671 |
| 1.062737 | 3.017534 | 0.593801 | 0.034982 | 0.02977 | 0.028632 | 0.026671 |
| 1.196405 | 3.426579 | 0.593801 | 0.035588 | 0.008175 | 0.008589 | 0.007379 |
| 0.291836 | 0.835837 | 0.593801 | 0.035588 | 0.008175 | 0.008589 | 0.007379 |
| 0.31075 | 0.919815 | 0.593801 | 0.035042 | 0.020412 | 0.023705 | 0.021532 |
| 0.303228 | 0.899593 | 0.593801 | 0.034613 | 0.016337 | 0.019212 | 0.017233 |
| 0.27525 | 0.801936 | 0.593801 | 0.036199 | 0.004783 | 0.005621 | 0.004623 |
| 0.255605 | 0.750162 | 0.593801 | 0.036761 | 0.002182 | 0.002642 | 0.002035 |
| 0.302961 | 0.883305 | 0.593685 | 0.035017 | 0.013498 | 0.015756 | 0.013994 |
| 0.223076 | 0.679836 | 0.593685 | 0.037118 | 0.000523 | 0.000908 | 0.000587 |
| 0.330401 | 0.967362 | 0.593569 | 0.033776 | 0.033787 | 0.03743 | 0.034959 |
| 1.033739 | 3.026627 | 0.593569 | 0.033776 | 0.033787 | 0.03743 | 0.034959 |
| 0.247176 | 0.720245 | 0.593569 | 0.037237 | 0.001228 | 0.001563 | 0.001135 |

| | | | | | | |
|---|---|---|---|---|---|---|
| 1.388417 | 4.045708 | 0.593569 | 0.037237 | 0.001228 | 0.001563 | 0.001135 |
| 1.260971 | 3.656127 | 0.593569 | 0.03661 | 0.004072 | 0.004896 | 0.003985 |
| 0.273513 | 0.79304 | 0.593569 | 0.03661 | 0.004072 | 0.004896 | 0.003985 |
| 0.240548 | 0.709696 | 0.593569 | 0.036845 | 0.001135 | 0.001363 | 0.000986 |
| 0.296188 | 0.8716 | 0.593569 | 0.036437 | 0.011475 | 0.01393 | 0.012212 |
| 1.013767 | 2.878019 | 0.593453 | 0.037015 | 0.042061 | 0.04429 | 0.041871 |
| 0.310487 | 0.899337 | 0.593453 | 0.034046 | 0.017166 | 0.018722 | 0.016914 |
| 1.11193 | 3.220748 | 0.593453 | 0.034046 | 0.017166 | 0.018722 | 0.016914 |
| 1.115384 | 3.169885 | 0.593336 | 0.035004 | 0.017217 | 0.017799 | 0.016106 |
| 0.30551 | 0.86749 | 0.593336 | 0.034707 | 0.011951 | 0.012635 | 0.011175 |
| 1.15275 | 3.273213 | 0.593336 | 0.034707 | 0.011951 | 0.012635 | 0.011175 |
| 1.122818 | 3.202783 | 0.593336 | 0.035651 | 0.015454 | 0.016703 | 0.015001 |
| 1.219743 | 3.54603 | 0.593104 | 0.036623 | 0.006536 | 0.007154 | 0.006067 |
| 1.201966 | 3.414732 | 0.593104 | 0.035396 | 0.008025 | 0.008035 | 0.006933 |
| 0.279683 | 0.816731 | 0.593104 | 0.034618 | 0.005653 | 0.006924 | 0.005778 |
| 0.242558 | 0.737302 | 0.593104 | 0.035793 | 0.001677 | 0.002409 | 0.001798 |
| 0.312256 | 0.900233 | 0.593104 | 0.036377 | 0.017238 | 0.018821 | 0.016974 |
| 1.150778 | 3.323843 | 0.593104 | 0.035256 | 0.011935 | 0.013176 | 0.011614 |
| 0.331889 | 0.947463 | 0.592988 | 0.036094 | 0.028296 | 0.030646 | 0.028419 |
| 0.278915 | 0.798612 | 0.592988 | 0.036063 | 0.00471 | 0.005143 | 0.004215 |
| 0.303113 | 0.86872 | 0.592872 | 0.034646 | 0.011651 | 0.012995 | 0.011453 |
| 0.262454 | 0.757853 | 0.592872 | 0.036947 | 0.002302 | 0.002837 | 0.002182 |
| 0.253077 | 0.747083 | 0.592872 | 0.037016 | 0.001862 | 0.002563 | 0.001925 |
| 0.307645 | 0.876205 | 0.592872 | 0.035697 | 0.013012 | 0.014092 | 0.012518 |
| 0.240786 | 0.783478 | 0.592756 | 0.032626 | 0.003369 | 0.005594 | 0.004395 |
| 0.293864 | 0.871408 | 0.592756 | 0.035271 | 0.011685 | 0.014036 | 0.012314 |
| 0.310954 | 0.891239 | 0.592756 | 0.0352 | 0.015672 | 0.016912 | 0.015237 |
| 0.290691 | 0.828951 | 0.59264 | 0.034999 | 0.007078 | 0.007775 | 0.006619 |
| 1.386005 | 4.294582 | 0.59264 | 0.036006 | 0.001284 | 0.001992 | 0.001449 |
| 1.143048 | 3.313152 | 0.59264 | 0.035099 | 0.012869 | 0.014189 | 0.012587 |
| 1.002027 | 2.888244 | 0.59264 | 0.034704 | 0.045398 | 0.04913 | 0.046563 |
| 1.263682 | 3.924327 | 0.592524 | 0.036143 | 0.003806 | 0.005614 | 0.004468 |
| 1.52716 | 4.771518 | 0.592524 | 0.036932 | 0.000347 | 0.000634 | 0.000391 |
| 0.348361 | 0.978705 | 0.592524 | 0.034876 | 0.04083 | 0.041187 | 0.038905 |
| 1.201087 | 3.484086 | 0.592408 | 0.036523 | 0.00729 | 0.008429 | 0.007154 |
| 0.31971 | 0.906115 | 0.592408 | 0.036239 | 0.019843 | 0.019757 | 0.017967 |
| 0.259351 | 0.762147 | 0.592408 | 0.03624 | 0.002852 | 0.003202 | 0.00254 |
| 1.312083 | 3.855779 | 0.592408 | 0.03624 | 0.002852 | 0.003202 | 0.00254 |
| 0.3354 | 0.951079 | 0.592408 | 0.036123 | 0.029819 | 0.031666 | 0.029453 |
| 1.373333 | 4.04693 | 0.592408 | 0.037169 | 0.001432 | 0.001867 | 0.001366 |
| 1.194227 | 3.445362 | 0.592292 | 0.034812 | 0.009376 | 0.00888 | 0.007694 |
| 0.289209 | 0.839169 | 0.592292 | 0.034747 | 0.008102 | 0.009182 | 0.007932 |
| 0.339709 | 0.974263 | 0.592292 | 0.035111 | 0.037775 | 0.039689 | 0.037328 |
| 0.283338 | 0.834669 | 0.592292 | 0.035729 | 0.007469 | 0.008906 | 0.007593 |
| 0.231515 | 0.719377 | 0.592059 | 0.03584 | 0.001233 | 0.001943 | 0.001404 |
| 1.154329 | 3.359077 | 0.592059 | 0.035636 | 0.011696 | 0.012895 | 0.011396 |
| 0.349397 | 0.988848 | 0.592059 | 0.034704 | 0.045187 | 0.045261 | 0.042895 |

| | | | | | | |
|---|---|---|---|---|---|---|
| 0.291641 | 0.845059 | 0.591943 | 0.036557 | 0.008857 | 0.009873 | 0.008535 |
| 0.304932 | 0.910354 | 0.591943 | 0.03488 | 0.017906 | 0.021643 | 0.019506 |
| 0.221683 | 0.667619 | 0.591827 | 0.037075 | 0.000415 | 0.000682 | 0.000432 |
| 1.49786 | 4.510939 | 0.591827 | 0.037075 | 0.000415 | 0.000682 | 0.000432 |
| 1.249222 | 3.667933 | 0.591827 | 0.036214 | 0.004579 | 0.005609 | 0.004586 |
| 1.04782 | 2.962525 | 0.591827 | 0.034866 | 0.033373 | 0.032668 | 0.030596 |
| 0.29899 | 0.881718 | 0.591827 | 0.034417 | 0.013031 | 0.015681 | 0.013879 |
| 1.253051 | 3.597938 | 0.591827 | 0.036331 | 0.004635 | 0.005137 | 0.004248 |
| 0.301061 | 0.870547 | 0.591711 | 0.035794 | 0.013856 | 0.013445 | 0.012058 |
| 1.206128 | 3.520578 | 0.591711 | 0.035727 | 0.006932 | 0.00815 | 0.00695 |
| 1.293412 | 3.712781 | 0.591711 | 0.03487 | 0.003037 | 0.003541 | 0.002805 |
| 0.316238 | 0.903835 | 0.591595 | 0.033998 | 0.01909 | 0.01942 | 0.017669 |
| 1.106396 | 3.16218 | 0.591595 | 0.033998 | 0.01909 | 0.01942 | 0.017669 |
| 1.215422 | 3.545168 | 0.591595 | 0.035515 | 0.006443 | 0.007487 | 0.006314 |
| 0.282074 | 0.82276 | 0.591595 | 0.035515 | 0.006443 | 0.007487 | 0.006314 |
| 1.11348 | 3.197416 | 0.591479 | 0.034653 | 0.016601 | 0.018304 | 0.016512 |
| 0.312753 | 0.898086 | 0.591479 | 0.034653 | 0.016601 | 0.018304 | 0.016512 |
| 0.249969 | 0.752145 | 0.591479 | 0.036117 | 0.002084 | 0.002944 | 0.002226 |
| 0.213256 | 0.667273 | 0.591479 | 0.036631 | 0.000455 | 0.000808 | 0.000518 |
| 0.322478 | 0.916425 | 0.591363 | 0.035683 | 0.020706 | 0.022166 | 0.020237 |
| 0.273455 | 0.821268 | 0.591363 | 0.035946 | 0.005974 | 0.007772 | 0.006468 |
| 0.346587 | 0.989077 | 0.591363 | 0.035705 | 0.042478 | 0.04539 | 0.0429 |
| 0.334487 | 0.951417 | 0.591247 | 0.034813 | 0.030722 | 0.031816 | 0.029613 |
| 0.29092 | 0.888359 | 0.591015 | 0.033822 | 0.013948 | 0.017518 | 0.015509 |
| 0.24471 | 0.7163 | 0.591015 | 0.036287 | 0.001518 | 0.001484 | 0.001112 |
| 1.396063 | 4.086463 | 0.591015 | 0.036287 | 0.001518 | 0.001484 | 0.001112 |
| 0.236484 | 0.736046 | 0.590899 | 0.036986 | 0.001569 | 0.002544 | 0.001855 |
| 0.294119 | 0.840077 | 0.590899 | 0.035709 | 0.008621 | 0.009032 | 0.007783 |
| 0.271391 | 0.776412 | 0.590899 | 0.036839 | 0.003165 | 0.003687 | 0.002924 |
| 1.287976 | 3.684724 | 0.590899 | 0.036839 | 0.003165 | 0.003687 | 0.002924 |
| 0.259981 | 0.747482 | 0.590899 | 0.03581 | 0.002094 | 0.002364 | 0.00182 |
| 0.315389 | 0.916712 | 0.590782 | 0.034838 | 0.020773 | 0.02264 | 0.020668 |
| 1.090855 | 3.170687 | 0.590782 | 0.034838 | 0.020773 | 0.02264 | 0.020668 |
| 0.198161 | 0.647109 | 0.590782 | 0.037079 | 0.00028 | 0.00067 | 0.000394 |
| 0.319292 | 0.939572 | 0.590782 | 0.034578 | 0.024916 | 0.028791 | 0.026468 |
| 1.187165 | 3.468867 | 0.590782 | 0.036007 | 0.008419 | 0.009677 | 0.008337 |
| 0.288279 | 0.842343 | 0.590782 | 0.036007 | 0.008419 | 0.009677 | 0.008337 |
| 0.280025 | 0.814341 | 0.590666 | 0.036018 | 0.005923 | 0.006645 | 0.00559 |
| 1.227986 | 3.571104 | 0.590666 | 0.036018 | 0.005923 | 0.006645 | 0.00559 |
| 1.041429 | 2.952561 | 0.590666 | 0.0347 | 0.032687 | 0.03463 | 0.032353 |
| 0.083797 | 0.537151 | 0.590666 | 0.025158 | 6.61E-05 | 0.001071 | 0.000349 |
| 1.175746 | 3.289942 | 0.590666 | 0.036737 | 0.011091 | 0.009974 | 0.008682 |
| 1.273509 | 3.829683 | 0.59055 | 0.036314 | 0.003536 | 0.004791 | 0.003812 |
| 0.261118 | 0.785232 | 0.59055 | 0.036314 | 0.003536 | 0.004791 | 0.003812 |
| 0.296237 | 0.8451 | 0.59055 | 0.035697 | 0.008802 | 0.009617 | 0.008325 |
| 1.337866 | 3.950039 | 0.59055 | 0.037195 | 0.001877 | 0.002579 | 0.00194 |
| 0.253162 | 0.747459 | 0.59055 | 0.037195 | 0.001877 | 0.002579 | 0.00194 |

| | | | | | | |
|---|---|---|---|---|---|---|
| 0.342491 | 0.971102 | 0.59055 | 0.036358 | 0.036379 | 0.038428 | 0.036082 |
| 1.22335 | 3.544271 | 0.59055 | 0.034165 | 0.006682 | 0.006875 | 0.00584 |
| 0.266663 | 0.757654 | 0.59055 | 0.03642 | 0.00261 | 0.002684 | 0.002105 |
| 1.319863 | 3.750049 | 0.59055 | 0.03642 | 0.00261 | 0.002684 | 0.002105 |
| 0.326901 | 0.927603 | 0.590434 | 0.03613 | 0.023348 | 0.024934 | 0.022904 |
| 1.149306 | 3.341766 | 0.590434 | 0.035497 | 0.012376 | 0.013472 | 0.011912 |
| 0.250226 | 0.731122 | 0.590434 | 0.036621 | 0.001435 | 0.001903 | 0.001395 |
| 1.367761 | 3.996384 | 0.590434 | 0.036621 | 0.001435 | 0.001903 | 0.001395 |
| 0.299319 | 0.893498 | 0.590434 | 0.035969 | 0.014764 | 0.018099 | 0.016115 |
| 1.119196 | 3.340919 | 0.590434 | 0.035969 | 0.014764 | 0.018099 | 0.016115 |
| 1.262084 | 3.633658 | 0.590434 | 0.036265 | 0.004343 | 0.004761 | 0.003887 |
| 0.242556 | 0.73022 | 0.590434 | 0.03667 | 0.001494 | 0.002082 | 0.001533 |
| 1.207655 | 3.482097 | 0.590434 | 0.035304 | 0.007957 | 0.007851 | 0.006777 |
| 0.287183 | 0.828051 | 0.590434 | 0.035304 | 0.007957 | 0.007851 | 0.006777 |
| 1.237223 | 3.581575 | 0.590434 | 0.035497 | 0.005086 | 0.006051 | 0.005013 |
| 1.231266 | 3.570848 | 0.590318 | 0.036053 | 0.005412 | 0.006413 | 0.005348 |
| 0.305117 | 0.904588 | 0.590318 | 0.035485 | 0.017293 | 0.020252 | 0.018222 |
| 1.038364 | 2.919192 | 0.590318 | 0.035676 | 0.035155 | 0.03549 | 0.033295 |
| 1.25142 | 3.887293 | 0.590202 | 0.0357 | 0.004276 | 0.006226 | 0.005009 |
| 0.257248 | 0.799093 | 0.590202 | 0.0357 | 0.004276 | 0.006226 | 0.005009 |
| 0.247418 | 0.765856 | 0.590202 | 0.036548 | 0.002504 | 0.003909 | 0.002999 |
| 1.305728 | 4.04175 | 0.590202 | 0.036548 | 0.002504 | 0.003909 | 0.002999 |
| 1.373219 | 4.12056 | 0.590086 | 0.036689 | 0.001402 | 0.001992 | 0.00147 |
| 1.282488 | 3.713022 | 0.590086 | 0.036113 | 0.003309 | 0.00401 | 0.003201 |
| 1.215382 | 3.514908 | 0.590086 | 0.035378 | 0.00648 | 0.007363 | 0.006255 |
| 0.284502 | 0.822787 | 0.590086 | 0.035378 | 0.00648 | 0.007363 | 0.006255 |
| 0.302214 | 0.855993 | 0.58997 | 0.035625 | 0.010867 | 0.010915 | 0.009604 |
| 0.293267 | 0.85766 | 0.58997 | 0.035508 | 0.009843 | 0.011707 | 0.010194 |
| 0.265342 | 0.770389 | 0.58997 | 0.036359 | 0.003047 | 0.003508 | 0.002779 |
| 0.304663 | 0.88877 | 0.589854 | 0.035242 | 0.015221 | 0.016771 | 0.015048 |
| 1.024499 | 2.887426 | 0.589854 | 0.03542 | 0.040187 | 0.040215 | 0.037882 |
| 0.275958 | 0.818349 | 0.589854 | 0.036783 | 0.005812 | 0.0073 | 0.006074 |
| 1.141918 | 3.315921 | 0.589854 | 0.035572 | 0.013003 | 0.014367 | 0.012755 |
| 0.345643 | 0.983693 | 0.589854 | 0.035508 | 0.042521 | 0.043219 | 0.040885 |
| 1.095252 | 3.206718 | 0.589738 | 0.034906 | 0.019407 | 0.021908 | 0.019964 |
| 0.272156 | 0.795566 | 0.589738 | 0.037202 | 0.00437 | 0.005179 | 0.00423 |
| 1.100776 | 3.152424 | 0.589738 | 0.035067 | 0.020356 | 0.020467 | 0.01865 |
| 0.288919 | 0.821281 | 0.589738 | 0.03578 | 0.006321 | 0.006961 | 0.005863 |
| 1.415502 | 4.151962 | 0.589738 | 0.036588 | 0.001003 | 0.001256 | 0.000889 |
| 0.24085 | 0.706463 | 0.589738 | 0.036588 | 0.001003 | 0.001256 | 0.000889 |
| 1.175797 | 3.364217 | 0.589622 | 0.036696 | 0.010145 | 0.010352 | 0.009042 |
| 0.319437 | 0.898815 | 0.589622 | 0.035847 | 0.017569 | 0.01807 | 0.016336 |
| 0.316549 | 0.904054 | 0.589622 | 0.035314 | 0.018704 | 0.019453 | 0.017652 |
| 1.000953 | 2.902056 | 0.589622 | 0.035224 | 0.046071 | 0.049592 | 0.046994 |
| 0.291349 | 0.849536 | 0.589622 | 0.034884 | 0.009222 | 0.01055 | 0.00914 |
| 0.288809 | 0.844197 | 0.589622 | 0.034216 | 0.009052 | 0.009911 | 0.008604 |
| 0.317368 | 0.917187 | 0.589622 | 0.035295 | 0.020967 | 0.022652 | 0.020749 |

| | | | | | | |
|---|---|---|---|---|---|---|
| 1.339026 | 3.903639 | 0.589622 | 0.036806 | 0.002097 | 0.002449 | 0.001874 |
| 0.256171 | 0.746811 | 0.589622 | 0.036806 | 0.002097 | 0.002449 | 0.001874 |
| 0.183325 | 0.720389 | 0.589505 | 0.029997 | 0.001365 | 0.003739 | 0.002528 |
| 0.306306 | 0.884117 | 0.589505 | 0.03581 | 0.013834 | 0.015716 | 0.013993 |
| 0.306885 | 0.885529 | 0.589389 | 0.03552 | 0.014387 | 0.015967 | 0.014296 |
| 1.129269 | 3.258552 | 0.589389 | 0.03552 | 0.014387 | 0.015967 | 0.014296 |
| 0.271134 | 0.795029 | 0.589389 | 0.036222 | 0.004381 | 0.005178 | 0.004244 |
| 1.106455 | 3.268233 | 0.589389 | 0.034835 | 0.016926 | 0.020014 | 0.018021 |
| 0.322769 | 0.922007 | 0.589389 | 0.035127 | 0.021622 | 0.023622 | 0.021598 |
| 1.008149 | 2.876135 | 0.589273 | 0.034735 | 0.043588 | 0.046556 | 0.044049 |
| 0.306886 | 0.883348 | 0.589273 | 0.036568 | 0.013641 | 0.015527 | 0.013795 |
| 1.508309 | 4.713945 | 0.589157 | 0.033821 | 0.001472 | 0.000742 | 0.000473 |
| 1.01902 | 2.930957 | 0.589157 | 0.036053 | 0.038906 | 0.042369 | 0.039868 |
| 0.252846 | 0.773214 | 0.589157 | 0.03539 | 0.003053 | 0.004211 | 0.00334 |
| 1.166513 | 3.343507 | 0.589157 | 0.035993 | 0.01049 | 0.011299 | 0.009868 |
| 0.299087 | 0.857256 | 0.589157 | 0.035993 | 0.01049 | 0.011299 | 0.009868 |
| 0.341151 | 0.991995 | 0.589157 | 0.034528 | 0.042328 | 0.046648 | 0.044043 |
| 1.49823 | 4.461302 | 0.589157 | 0.037475 | 0.0005 | 0.000644 | 0.000429 |
| 1.125447 | 3.251399 | 0.589041 | 0.035645 | 0.015082 | 0.016547 | 0.014809 |
| 0.305537 | 0.878865 | 0.589041 | 0.036127 | 0.012905 | 0.014727 | 0.013033 |
| 0.30988 | 0.872659 | 0.589041 | 0.036047 | 0.012913 | 0.013299 | 0.011795 |
| 1.145923 | 3.227061 | 0.589041 | 0.036047 | 0.012913 | 0.013299 | 0.011795 |
| 1.310569 | 3.815357 | 0.588925 | 0.036816 | 0.002985 | 0.003156 | 0.002523 |
| 0.261897 | 0.777689 | 0.588925 | 0.035929 | 0.003133 | 0.004163 | 0.003305 |
| 1.285861 | 3.818297 | 0.588925 | 0.035929 | 0.003133 | 0.004163 | 0.003305 |
| 0.260463 | 0.844666 | 0.588809 | 0.033207 | 0.007884 | 0.011654 | 0.009801 |
| 1.048334 | 3.02102 | 0.588809 | 0.035207 | 0.029779 | 0.032779 | 0.030469 |
| 1.186155 | 3.462683 | 0.588809 | 0.036139 | 0.008475 | 0.009749 | 0.008395 |
| 1.027614 | 2.927529 | 0.588809 | 0.035287 | 0.037959 | 0.039212 | 0.036886 |
| 0.267192 | 0.80206 | 0.588693 | 0.035868 | 0.004479 | 0.006023 | 0.00492 |
| 0.291956 | 0.857072 | 0.588577 | 0.036088 | 0.009793 | 0.01169 | 0.010185 |
| 0.276291 | 0.817438 | 0.588461 | 0.036037 | 0.005914 | 0.007179 | 0.006017 |
| 1.223335 | 3.619371 | 0.588461 | 0.036037 | 0.005914 | 0.007179 | 0.006017 |
| 0.31027 | 0.896286 | 0.588461 | 0.035572 | 0.016528 | 0.018051 | 0.016253 |
| 0.297281 | 0.865045 | 0.588461 | 0.036212 | 0.010732 | 0.012702 | 0.0111 |
| 0.258056 | 0.784456 | 0.588461 | 0.036406 | 0.003374 | 0.004864 | 0.003854 |
| 0.309203 | 0.899102 | 0.588461 | 0.035183 | 0.016755 | 0.018745 | 0.016873 |
| 0.274437 | 0.810168 | 0.588461 | 0.036037 | 0.005342 | 0.006484 | 0.005391 |
| 0.231707 | 0.693351 | 0.588461 | 0.03733 | 0.000751 | 0.001077 | 0.000735 |
| 1.102852 | 3.121143 | 0.588345 | 0.036687 | 0.019511 | 0.019867 | 0.018027 |
| 0.267406 | 0.807421 | 0.588345 | 0.035501 | 0.004976 | 0.006553 | 0.005392 |
| 1.238511 | 3.739632 | 0.588345 | 0.035501 | 0.004976 | 0.006553 | 0.005392 |
| 1.396091 | 4.180854 | 0.588345 | 0.036333 | 0.001173 | 0.001619 | 0.001176 |
| 1.1567 | 3.355175 | 0.588345 | 0.034537 | 0.010924 | 0.012567 | 0.011037 |
| 1.233931 | 3.57776 | 0.588228 | 0.037433 | 0.005745 | 0.006257 | 0.005256 |
| 0.279504 | 0.810418 | 0.588228 | 0.037433 | 0.005745 | 0.006257 | 0.005256 |
| 1.159091 | 3.387928 | 0.588112 | 0.036155 | 0.010479 | 0.012436 | 0.01087 |

| | | | | | | |
|---|---|---|---|---|---|---|
| 0.295166 | 0.862745 | 0.588112 | 0.036155 | 0.010479 | 0.012436 | 0.01087 |
| 1.287015 | 3.650847 | 0.588112 | 0.036141 | 0.0035 | 0.00363 | 0.002913 |
| 0.292648 | 0.836846 | 0.588112 | 0.036846 | 0.007933 | 0.008678 | 0.007467 |
| 0.271037 | 0.799027 | 0.587996 | 0.035566 | 0.004517 | 0.005547 | 0.00454 |
| 1.256904 | 3.615562 | 0.587996 | 0.035619 | 0.004322 | 0.004981 | 0.00408 |
| 0.276582 | 0.795605 | 0.587996 | 0.035619 | 0.004322 | 0.004981 | 0.00408 |
| 0.336951 | 0.955975 | 0.587996 | 0.035395 | 0.031337 | 0.033237 | 0.030994 |
| 0.260674 | 0.764221 | 0.587996 | 0.036644 | 0.002552 | 0.003283 | 0.002559 |
| 1.308521 | 3.836212 | 0.587996 | 0.036644 | 0.002552 | 0.003283 | 0.002559 |
| 0.296612 | 0.858822 | 0.58788 | 0.036874 | 0.010152 | 0.011698 | 0.010219 |
| 1.216511 | 3.631777 | 0.58788 | 0.035586 | 0.006197 | 0.007759 | 0.006514 |
| 0.254842 | 0.764236 | 0.58788 | 0.03602 | 0.002795 | 0.003504 | 0.002787 |
| 0.331885 | 0.938176 | 0.587764 | 0.035906 | 0.02702 | 0.027757 | 0.025734 |
| 0.317385 | 0.90008 | 0.587764 | 0.035811 | 0.018547 | 0.01848 | 0.016738 |
| 0.289703 | 0.843989 | 0.587764 | 0.035846 | 0.008549 | 0.00983 | 0.008462 |
| 0.242331 | 0.705487 | 0.587648 | 0.036341 | 0.001036 | 0.001196 | 0.000865 |
| 1.417461 | 4.126581 | 0.587648 | 0.036341 | 0.001036 | 0.001196 | 0.000865 |
| 0.25425 | 0.753726 | 0.587532 | 0.036396 | 0.002122 | 0.002884 | 0.002206 |
| 1.326741 | 3.933144 | 0.587532 | 0.036396 | 0.002122 | 0.002884 | 0.002206 |
| 1.03991 | 2.992231 | 0.587532 | 0.034614 | 0.032957 | 0.035282 | 0.032947 |
| 1.147337 | 3.329148 | 0.587532 | 0.034952 | 0.011944 | 0.013674 | 0.012085 |
| 1.120708 | 3.232197 | 0.587532 | 0.034875 | 0.015216 | 0.017232 | 0.015431 |
| 0.309387 | 0.892293 | 0.587532 | 0.034875 | 0.015216 | 0.017232 | 0.015431 |
| 0.299989 | 0.872534 | 0.587532 | 0.035565 | 0.01225 | 0.013871 | 0.012241 |
| 0.319659 | 0.913838 | 0.587416 | 0.035994 | 0.019762 | 0.021664 | 0.01971 |
| 0.2742 | 0.79905 | 0.587416 | 0.036335 | 0.004641 | 0.0054 | 0.004471 |
| 0.24199 | 0.725012 | 0.587416 | 0.036437 | 0.001447 | 0.001879 | 0.001375 |
| 1.087714 | 3.156428 | 0.587416 | 0.035645 | 0.020972 | 0.023249 | 0.021193 |
| 0.275293 | 0.793326 | 0.587416 | 0.036913 | 0.004418 | 0.004841 | 0.003961 |
| 1.145967 | 3.309391 | 0.5873 | 0.03568 | 0.012024 | 0.013756 | 0.012139 |
| 0.325396 | 0.93673 | 0.5873 | 0.035251 | 0.026093 | 0.027646 | 0.025557 |
| 1.164893 | 3.371982 | 0.587184 | 0.036367 | 0.010087 | 0.01164 | 0.010158 |
| 1.031778 | 2.968779 | 0.587184 | 0.0348 | 0.035278 | 0.037895 | 0.035591 |
| 0.336839 | 0.969201 | 0.587184 | 0.0348 | 0.035278 | 0.037895 | 0.035591 |
| 1.027909 | 3.026469 | 0.587068 | 0.035723 | 0.03481 | 0.039409 | 0.036856 |
| 1.133163 | 3.336643 | 0.587068 | 0.035745 | 0.013126 | 0.01579 | 0.013973 |
| 0.239421 | 0.75613 | 0.587068 | 0.036164 | 0.002134 | 0.003582 | 0.002689 |
| 0.287135 | 0.82047 | 0.587068 | 0.03524 | 0.006839 | 0.00696 | 0.005899 |
| 0.274233 | 0.805541 | 0.586951 | 0.035826 | 0.005885 | 0.006021 | 0.005123 |
| 1.241401 | 3.646532 | 0.586951 | 0.035826 | 0.005885 | 0.006021 | 0.005123 |
| 1.188656 | 3.401008 | 0.586951 | 0.035423 | 0.00853 | 0.009204 | 0.007913 |
| 0.294031 | 0.841287 | 0.586951 | 0.035423 | 0.00853 | 0.009204 | 0.007913 |
| 1.067156 | 3.047161 | 0.586951 | 0.0366 | 0.025414 | 0.027609 | 0.025463 |
| 1.102153 | 3.197864 | 0.586951 | 0.035414 | 0.019708 | 0.020455 | 0.01859 |
| 1.16505 | 3.495757 | 0.586951 | 0.035885 | 0.009765 | 0.012247 | 0.010585 |
| 0.255704 | 0.732715 | 0.586835 | 0.037297 | 0.001687 | 0.001821 | 0.001374 |
| 0.307948 | 0.896664 | 0.586835 | 0.035888 | 0.016754 | 0.018274 | 0.01652 |

| | | | | | | |
|---|---|---|---|---|---|---|
| 0.254042 | 0.764156 | 0.586719 | 0.036948 | 0.002537 | 0.00353 | 0.002721 |
| 1.308633 | 3.936354 | 0.586719 | 0.036948 | 0.002537 | 0.00353 | 0.002721 |
| 0.272948 | 0.793515 | 0.586719 | 0.036733 | 0.004198 | 0.004962 | 0.004028 |
| 0.302547 | 0.859352 | 0.586719 | 0.036166 | 0.0105 | 0.011435 | 0.009998 |
| 1.105331 | 3.119602 | 0.586719 | 0.036381 | 0.01844 | 0.019371 | 0.01756 |
| 1.114146 | 3.212263 | 0.586719 | 0.035856 | 0.017151 | 0.01827 | 0.016523 |
| 0.311307 | 0.897548 | 0.586719 | 0.035856 | 0.017151 | 0.01827 | 0.016523 |
| 0.328517 | 0.963279 | 0.586603 | 0.033866 | 0.032494 | 0.036057 | 0.033625 |
| 1.11627 | 3.16292 | 0.586487 | 0.035747 | 0.017875 | 0.017599 | 0.015986 |
| 0.316164 | 0.895841 | 0.586487 | 0.035747 | 0.017875 | 0.017599 | 0.015986 |
| 0.299048 | 0.868059 | 0.586487 | 0.036281 | 0.011564 | 0.013122 | 0.011524 |
| 0.317882 | 0.924017 | 0.586371 | 0.035793 | 0.022057 | 0.024432 | 0.022341 |
| 0.283746 | 0.829866 | 0.586371 | 0.035531 | 0.007376 | 0.008263 | 0.007084 |
| 0.285919 | 0.833412 | 0.586371 | 0.035389 | 0.007094 | 0.008597 | 0.007294 |
| 1.133374 | 3.287194 | 0.586371 | 0.035658 | 0.013735 | 0.015484 | 0.013751 |
| 0.304211 | 0.882321 | 0.586371 | 0.035658 | 0.013735 | 0.015484 | 0.013751 |
| 0.302395 | 0.883103 | 0.586255 | 0.035985 | 0.013592 | 0.01575 | 0.01395 |
| 0.309833 | 0.877901 | 0.586255 | 0.036303 | 0.013542 | 0.014282 | 0.012724 |
| 1.232417 | 3.630833 | 0.586139 | 0.036749 | 0.005123 | 0.006565 | 0.005415 |
| 0.326343 | 0.994998 | 0.586139 | 0.0349 | 0.041602 | 0.047974 | 0.04513 |
| 1.047185 | 2.967435 | 0.586139 | 0.036878 | 0.030992 | 0.032886 | 0.03064 |
| 0.288021 | 0.877441 | 0.586139 | 0.036306 | 0.012209 | 0.015519 | 0.013601 |
| 0.254055 | 0.752596 | 0.586139 | 0.036828 | 0.002072 | 0.002827 | 0.002155 |
| 0.316224 | 0.901636 | 0.586023 | 0.035037 | 0.017598 | 0.018908 | 0.017122 |
| 0.336893 | 0.997847 | 0.586023 | 0.034644 | 0.044163 | 0.049097 | 0.046393 |
| 1.002158 | 2.968302 | 0.586023 | 0.034644 | 0.044163 | 0.049097 | 0.046393 |
| 0.283117 | 0.826629 | 0.586023 | 0.036472 | 0.00649 | 0.007895 | 0.00666 |
| 0.291897 | 0.839941 | 0.586023 | 0.036073 | 0.008105 | 0.009138 | 0.00787 |
| 0.270732 | 0.779688 | 0.586023 | 0.037865 | 0.003559 | 0.003949 | 0.003204 |
| 1.282565 | 3.693686 | 0.586023 | 0.037865 | 0.003559 | 0.003949 | 0.003204 |
| 0.321787 | 0.941956 | 0.586023 | 0.035083 | 0.026211 | 0.029391 | 0.027098 |
| 0.316994 | 0.908552 | 0.585907 | 0.035526 | 0.019284 | 0.020504 | 0.018627 |
| 0.307968 | 0.903521 | 0.585907 | 0.034747 | 0.017198 | 0.019834 | 0.017928 |
| 0.31492 | 0.915259 | 0.585907 | 0.035647 | 0.020846 | 0.022295 | 0.020395 |
| 0.267668 | 0.892632 | 0.585791 | 0.03017 | 0.013916 | 0.019826 | 0.017347 |
| 0.289752 | 0.823987 | 0.585674 | 0.036062 | 0.006786 | 0.007229 | 0.006144 |
| 0.339103 | 0.968841 | 0.585558 | 0.03535 | 0.037012 | 0.037694 | 0.035462 |
| 1.032161 | 2.948958 | 0.585558 | 0.03535 | 0.037012 | 0.037694 | 0.035462 |
| 1.082869 | 3.125308 | 0.585558 | 0.035972 | 0.023711 | 0.024171 | 0.022316 |
| 0.319968 | 0.923473 | 0.585558 | 0.035972 | 0.023711 | 0.024171 | 0.022316 |
| 0.304716 | 0.878371 | 0.585558 | 0.036887 | 0.013364 | 0.014682 | 0.013033 |
| 0.291811 | 0.837249 | 0.585442 | 0.036194 | 0.008095 | 0.008778 | 0.007521 |
| 1.194388 | 3.426877 | 0.585442 | 0.036194 | 0.008095 | 0.008778 | 0.007521 |
| 0.258615 | 0.76377 | 0.585442 | 0.036384 | 0.002587 | 0.003331 | 0.002614 |
| 1.309294 | 3.866756 | 0.585442 | 0.036384 | 0.002587 | 0.003331 | 0.002614 |
| 1.001298 | 2.820839 | 0.585442 | 0.035077 | 0.049531 | 0.049429 | 0.047073 |
| 1.112284 | 3.212857 | 0.585442 | 0.036502 | 0.017985 | 0.018612 | 0.016834 |

| | | | | | | |
|---|---|---|---|---|---|---|
| 1.266229 | 3.643786 | 0.585326 | 0.036942 | 0.003969 | 0.004577 | 0.003724 |
| 0.299315 | 0.88787 | 0.585326 | 0.035489 | 0.014098 | 0.016907 | 0.015068 |
| 0.340113 | 0.961143 | 0.585326 | 0.035907 | 0.034151 | 0.0349 | 0.032729 |
| 1.040428 | 2.940196 | 0.585326 | 0.035907 | 0.034151 | 0.0349 | 0.032729 |
| 0.321408 | 0.92424 | 0.58521 | 0.035644 | 0.022249 | 0.024301 | 0.022297 |
| 0.298991 | 0.859849 | 0.58521 | 0.036935 | 0.010595 | 0.011726 | 0.010233 |
| 0.291748 | 0.848561 | 0.58521 | 0.035033 | 0.00932 | 0.010381 | 0.00902 |
| 0.223019 | 0.664446 | 0.58521 | 0.038159 | 0.000529 | 0.000608 | 0.000416 |
| 1.505013 | 4.483915 | 0.58521 | 0.038159 | 0.000529 | 0.000608 | 0.000416 |
| 0.311123 | 0.899127 | 0.585094 | 0.034816 | 0.017811 | 0.018637 | 0.016907 |
| 0.20259 | 0.621375 | 0.585094 | 0.038337 | 0.000182 | 0.00029 | 0.00017 |
| 0.268014 | 0.781173 | 0.584978 | 0.037342 | 0.003297 | 0.004171 | 0.003316 |
| 1.280125 | 3.731144 | 0.584978 | 0.037342 | 0.003297 | 0.004171 | 0.003316 |
| 1.144116 | 3.272788 | 0.584978 | 0.037055 | 0.012728 | 0.013813 | 0.012272 |
| 0.30555 | 0.874037 | 0.584978 | 0.037055 | 0.012728 | 0.013813 | 0.012272 |
| 0.285452 | 0.815065 | 0.584978 | 0.036796 | 0.006775 | 0.006451 | 0.005462 |
| 0.28002 | 0.821309 | 0.584978 | 0.037258 | 0.006156 | 0.007426 | 0.006219 |
| 0.247434 | 0.736885 | 0.584978 | 0.037239 | 0.001636 | 0.002238 | 0.001656 |
| 0.285356 | 0.825207 | 0.584978 | 0.036989 | 0.007386 | 0.007604 | 0.006502 |
| 1.088118 | 3.070353 | 0.584978 | 0.035434 | 0.021622 | 0.022662 | 0.02073 |
| 1.081182 | 3.125686 | 0.584978 | 0.036777 | 0.027849 | 0.024565 | 0.022523 |
| 0.324977 | 0.947092 | 0.584746 | 0.034812 | 0.027382 | 0.030837 | 0.028533 |
| 0.311173 | 0.896376 | 0.584746 | 0.035951 | 0.016506 | 0.018017 | 0.016204 |
| 0.343408 | 0.971796 | 0.584746 | 0.035254 | 0.038047 | 0.038661 | 0.036347 |
| 0.296465 | 0.864406 | 0.584746 | 0.036508 | 0.010674 | 0.012641 | 0.011053 |
| 0.245565 | 0.730739 | 0.584746 | 0.03632 | 0.001445 | 0.002018 | 0.001489 |
| 1.129106 | 3.225743 | 0.58463 | 0.036387 | 0.014273 | 0.015805 | 0.014097 |
| 1.182067 | 3.371171 | 0.58463 | 0.035973 | 0.009949 | 0.009721 | 0.008534 |
| 0.296633 | 0.845975 | 0.58463 | 0.035973 | 0.009949 | 0.009721 | 0.008534 |
| 0.34016 | 0.978813 | 0.58463 | 0.036238 | 0.038884 | 0.041411 | 0.038973 |
| 1.10498 | 3.168048 | 0.58463 | 0.036237 | 0.018289 | 0.019728 | 0.017919 |
| 1.122284 | 3.274959 | 0.58463 | 0.035118 | 0.015606 | 0.017208 | 0.01546 |
| 1.077704 | 3.139575 | 0.584514 | 0.036324 | 0.022394 | 0.025463 | 0.023306 |
| 0.318514 | 0.927898 | 0.584514 | 0.036324 | 0.022394 | 0.025463 | 0.023306 |
| 0.260346 | 0.755879 | 0.584514 | 0.036338 | 0.002263 | 0.002797 | 0.002157 |
| 1.322964 | 3.841047 | 0.584514 | 0.036338 | 0.002263 | 0.002797 | 0.002157 |
| 1.098629 | 3.177726 | 0.584397 | 0.035487 | 0.019136 | 0.021048 | 0.019177 |
| 1.200023 | 3.620263 | 0.584397 | 0.036361 | 0.007109 | 0.009127 | 0.007723 |
| 1.190593 | 3.383752 | 0.584397 | 0.036438 | 0.008404 | 0.008932 | 0.007711 |
| 1.334328 | 3.949471 | 0.584397 | 0.03751 | 0.002392 | 0.002683 | 0.002112 |
| 0.288035 | 0.832205 | 0.584397 | 0.03575 | 0.007663 | 0.008326 | 0.007132 |
| 1.10924 | 3.290311 | 0.584281 | 0.034608 | 0.016687 | 0.01961 | 0.017611 |
| 1.232423 | 3.620385 | 0.584165 | 0.036798 | 0.00537 | 0.006525 | 0.005408 |
| 0.309538 | 0.895382 | 0.584165 | 0.035856 | 0.016385 | 0.017895 | 0.016141 |
| 0.265263 | 0.767073 | 0.584165 | 0.036385 | 0.002815 | 0.003294 | 0.002607 |
| 0.269895 | 0.796349 | 0.584165 | 0.037323 | 0.004361 | 0.005354 | 0.004374 |
| 0.204523 | 0.627875 | 0.584049 | 0.037933 | 0.000191 | 0.000335 | 0.000192 |

| | | | | | | |
|---|---|---|---|---|---|---|
| 0.26125 | 0.783732 | 0.584049 | 0.03615 | 0.003592 | 0.004662 | 0.003751 |
| 1.17455 | 3.395095 | 0.584049 | 0.036766 | 0.009585 | 0.010646 | 0.009247 |
| 0.314647 | 0.923494 | 0.584049 | 0.035653 | 0.021632 | 0.024466 | 0.022341 |
| 1.082844 | 3.178164 | 0.584049 | 0.035653 | 0.021632 | 0.024466 | 0.022341 |
| 1.212054 | 3.560019 | 0.583933 | 0.036028 | 0.007126 | 0.007822 | 0.006705 |
| 0.262891 | 0.759409 | 0.583933 | 0.037289 | 0.002476 | 0.002911 | 0.002275 |
| 0.267205 | 0.783955 | 0.583817 | 0.036338 | 0.003511 | 0.004421 | 0.003554 |
| 1.274064 | 3.62211 | 0.583817 | 0.036932 | 0.003854 | 0.004122 | 0.003344 |
| 1.197466 | 3.545842 | 0.583817 | 0.036758 | 0.0073 | 0.009036 | 0.007652 |
| 0.310973 | 0.924213 | 0.583817 | 0.03437 | 0.021921 | 0.024858 | 0.022787 |
| 0.279084 | 0.817782 | 0.583817 | 0.035735 | 0.006219 | 0.007072 | 0.005968 |
| 1.22282 | 3.583149 | 0.583817 | 0.035735 | 0.006219 | 0.007072 | 0.005968 |
| 1.291871 | 3.901912 | 0.583817 | 0.037018 | 0.00301 | 0.004129 | 0.003248 |
| 1.290433 | 3.849568 | 0.583817 | 0.03693 | 0.003098 | 0.004048 | 0.003202 |
| 0.259769 | 0.774933 | 0.583817 | 0.03693 | 0.003098 | 0.004048 | 0.003202 |
| 0.146064 | 0.65747 | 0.583701 | 0.02964 | 0.000519 | 0.002268 | 0.001283 |
| 1.141455 | 3.231326 | 0.583701 | 0.035935 | 0.013231 | 0.013958 | 0.012411 |
| 0.33284 | 0.950717 | 0.583585 | 0.036979 | 0.029265 | 0.031657 | 0.029411 |
| 1.20675 | 3.559837 | 0.583585 | 0.035312 | 0.006569 | 0.008268 | 0.006965 |
| 0.280912 | 0.828672 | 0.583585 | 0.035312 | 0.006569 | 0.008268 | 0.006965 |
| 1.034913 | 2.934884 | 0.583585 | 0.035493 | 0.038621 | 0.036706 | 0.034596 |
| 0.338001 | 0.95733 | 0.583469 | 0.035864 | 0.033243 | 0.033657 | 0.031522 |
| 0.336613 | 0.957389 | 0.583353 | 0.034813 | 0.032317 | 0.033732 | 0.031561 |
| 1.140854 | 3.283235 | 0.583353 | 0.037289 | 0.013039 | 0.01434 | 0.012698 |
| 0.304578 | 0.876536 | 0.583353 | 0.037289 | 0.013039 | 0.01434 | 0.012698 |
| 0.224712 | 0.682947 | 0.583353 | 0.037886 | 0.000671 | 0.000951 | 0.000644 |
| 0.327991 | 0.936012 | 0.583353 | 0.035267 | 0.026315 | 0.027303 | 0.025236 |
| 0.319929 | 0.921453 | 0.583237 | 0.035938 | 0.021609 | 0.02363 | 0.021658 |
| 0.302623 | 0.892354 | 0.583237 | 0.03608 | 0.014797 | 0.017653 | 0.015752 |
| 1.116348 | 3.292244 | 0.583237 | 0.035052 | 0.015398 | 0.01833 | 0.016398 |
| 0.2765 | 0.801514 | 0.583237 | 0.036399 | 0.005188 | 0.005522 | 0.004594 |
| 0.243226 | 0.715326 | 0.583237 | 0.037463 | 0.00119 | 0.001486 | 0.001074 |
| 1.071432 | 3.07706 | 0.583237 | 0.036224 | 0.024511 | 0.026669 | 0.024591 |
| 1.231383 | 3.555381 | 0.583121 | 0.036843 | 0.006266 | 0.006344 | 0.005386 |
| 1.05923 | 2.990943 | 0.583121 | 0.035806 | 0.029238 | 0.029461 | 0.02744 |
| 1.152876 | 3.44044 | 0.583004 | 0.035565 | 0.010754 | 0.013511 | 0.011775 |
| 0.319476 | 0.923009 | 0.583004 | 0.035926 | 0.022229 | 0.024072 | 0.022052 |
| 0.277863 | 0.798233 | 0.583004 | 0.0373 | 0.004477 | 0.005157 | 0.004232 |
| 0.250385 | 0.793007 | 0.583004 | 0.037 | 0.003877 | 0.005986 | 0.00475 |
| 1.014296 | 2.91946 | 0.582888 | 0.035309 | 0.04124 | 0.044157 | 0.041747 |
| 0.290756 | 0.848341 | 0.582888 | 0.036019 | 0.008988 | 0.010405 | 0.00905 |
| 1.178771 | 3.439315 | 0.582888 | 0.036019 | 0.008988 | 0.010405 | 0.00905 |
| 0.32555 | 0.933618 | 0.582888 | 0.036228 | 0.025238 | 0.026724 | 0.024641 |
| 1.071101 | 3.071723 | 0.582888 | 0.036228 | 0.025238 | 0.026724 | 0.024641 |
| 0.273139 | 0.815173 | 0.582772 | 0.036633 | 0.005338 | 0.00709 | 0.005854 |
| 0.333762 | 0.996587 | 0.582656 | 0.034749 | 0.042742 | 0.048585 | 0.045828 |
| 1.013766 | 3.041641 | 0.582656 | 0.034906 | 0.039153 | 0.044564 | 0.041843 |

| | | | | | | |
|---|---|---|---|---|---|---|
| 1.004827 | 2.854722 | 0.582656 | 0.035261 | 0.046249 | 0.047924 | 0.045551 |
| 0.350297 | 0.995196 | 0.582656 | 0.035261 | 0.046249 | 0.047924 | 0.045551 |
| 0.238069 | 0.859894 | 0.582656 | 0.028946 | 0.009579 | 0.015491 | 0.013121 |
| 1.078893 | 3.078235 | 0.582656 | 0.03581 | 0.02388 | 0.02484 | 0.02283 |
| 0.328058 | 0.93579 | 0.582656 | 0.03799 | 0.026285 | 0.027234 | 0.025174 |
| 1.068616 | 3.04824 | 0.582656 | 0.03799 | 0.026285 | 0.027234 | 0.025174 |
| 1.115522 | 3.28748 | 0.582656 | 0.035129 | 0.015508 | 0.01845 | 0.016508 |
| 1.189638 | 3.419432 | 0.58254 | 0.036657 | 0.008496 | 0.009196 | 0.00792 |
| 0.299095 | 0.874129 | 0.58254 | 0.036438 | 0.012087 | 0.01422 | 0.012551 |
| 0.307311 | 0.894934 | 0.582424 | 0.035748 | 0.015974 | 0.017931 | 0.016097 |
| 1.243587 | 3.660882 | 0.582424 | 0.036958 | 0.00486 | 0.005934 | 0.004885 |
| 0.273158 | 0.804126 | 0.582424 | 0.036958 | 0.00486 | 0.005934 | 0.004885 |
| 0.288533 | 0.851238 | 0.582424 | 0.036228 | 0.009201 | 0.010974 | 0.009552 |
| 0.332786 | 0.938829 | 0.582424 | 0.035691 | 0.027211 | 0.027911 | 0.025878 |
| 1.201949 | 3.55415 | 0.582424 | 0.037294 | 0.0072 | 0.008664 | 0.007367 |
| 0.281361 | 0.831982 | 0.582424 | 0.037294 | 0.0072 | 0.008664 | 0.007367 |
| 0.328751 | 0.946537 | 0.582424 | 0.036787 | 0.02761 | 0.030493 | 0.028229 |
| 0.242946 | 0.718803 | 0.582424 | 0.037967 | 0.001226 | 0.001616 | 0.001167 |
| 0.263263 | 0.795158 | 0.582424 | 0.036862 | 0.0043 | 0.005558 | 0.004548 |
| 0.28297 | 0.824893 | 0.582308 | 0.036791 | 0.006636 | 0.007693 | 0.006502 |
| 0.29625 | 0.855399 | 0.582308 | 0.03684 | 0.009666 | 0.011169 | 0.009709 |
| 1.196165 | 3.365921 | 0.582308 | 0.03688 | 0.008313 | 0.008324 | 0.007159 |
| 1.130457 | 3.419859 | 0.582308 | 0.035715 | 0.021306 | 0.016657 | 0.014887 |
| 1.04285 | 2.934229 | 0.582192 | 0.03655 | 0.033275 | 0.034095 | 0.031902 |
| 1.203248 | 3.550547 | 0.582076 | 0.036609 | 0.006792 | 0.008533 | 0.007208 |
| 0.314899 | 0.921897 | 0.582076 | 0.034573 | 0.021733 | 0.024025 | 0.022011 |
| 0.292276 | 0.834189 | 0.582076 | 0.037356 | 0.007615 | 0.00835 | 0.007153 |
| 0.279273 | 0.797912 | 0.582076 | 0.03633 | 0.004536 | 0.005064 | 0.004153 |
| 1.09379 | 3.1793 | 0.582076 | 0.03618 | 0.019846 | 0.022061 | 0.020056 |
| 0.35079 | 0.991475 | 0.582076 | 0.034895 | 0.044706 | 0.046342 | 0.043942 |
| 0.259095 | 0.781217 | 0.58196 | 0.036212 | 0.003504 | 0.004555 | 0.003671 |
| 0.269033 | 0.780587 | 0.58196 | 0.036559 | 0.003368 | 0.004085 | 0.00327 |
| 1.184722 | 3.437956 | 0.58196 | 0.035907 | 0.008666 | 0.009776 | 0.00849 |
| 0.278196 | 0.79996 | 0.58196 | 0.03654 | 0.004608 | 0.005299 | 0.004362 |
| 0.31946 | 0.907171 | 0.58196 | 0.03658 | 0.018659 | 0.020025 | 0.018169 |
| 0.287561 | 0.822523 | 0.58196 | 0.036713 | 0.007026 | 0.007173 | 0.006088 |
| 1.193064 | 3.590334 | 0.58196 | 0.037048 | 0.007559 | 0.009652 | 0.008197 |
| 0.278526 | 0.838178 | 0.58196 | 0.037048 | 0.007559 | 0.009652 | 0.008197 |
| 0.337667 | 0.959707 | 0.581844 | 0.036007 | 0.033366 | 0.03449 | 0.032232 |
| 0.256593 | 0.745524 | 0.581844 | 0.036395 | 0.00242 | 0.002372 | 0.001853 |
| 1.341339 | 3.89722 | 0.581844 | 0.036395 | 0.00242 | 0.002372 | 0.001853 |
| 0.281626 | 0.890983 | 0.581844 | 0.035147 | 0.013887 | 0.018631 | 0.01644 |
| 1.11459 | 3.470532 | 0.581727 | 0.036581 | 0.015293 | 0.019576 | 0.017411 |
| 0.312774 | 0.886442 | 0.581727 | 0.036137 | 0.014978 | 0.015798 | 0.014161 |
| 0.339172 | 0.988647 | 0.581727 | 0.035452 | 0.041907 | 0.045306 | 0.042795 |
| 0.340282 | 0.988507 | 0.581727 | 0.035161 | 0.040972 | 0.045236 | 0.042639 |
| 0.267068 | 0.796363 | 0.581727 | 0.036969 | 0.004273 | 0.005488 | 0.004462 |

| | | | | | | |
|---|---|---|---|---|---|---|
| 0.295305 | 0.845984 | 0.581727 | 0.036572 | 0.008947 | 0.009798 | 0.008503 |
| 1.205468 | 3.585029 | 0.581611 | 0.036186 | 0.007049 | 0.008487 | 0.007229 |
| 0.312973 | 0.897827 | 0.581611 | 0.036699 | 0.016513 | 0.018233 | 0.016431 |
| 0.289586 | 0.828029 | 0.581611 | 0.036052 | 0.007031 | 0.00772 | 0.006582 |
| 1.202621 | 3.727699 | 0.581495 | 0.036956 | 0.007001 | 0.009342 | 0.007923 |
| 0.268262 | 0.831517 | 0.581495 | 0.036956 | 0.007001 | 0.009342 | 0.007923 |
| 0.334292 | 0.941011 | 0.581495 | 0.036203 | 0.029265 | 0.028506 | 0.026517 |
| 1.199386 | 3.437373 | 0.581495 | 0.035877 | 0.007632 | 0.008368 | 0.007187 |
| 0.333021 | 0.953524 | 0.581379 | 0.036299 | 0.03149 | 0.032573 | 0.030416 |
| 0.281969 | 0.812815 | 0.581379 | 0.036756 | 0.006038 | 0.006385 | 0.005369 |
| 0.292072 | 0.859745 | 0.581379 | 0.035782 | 0.010311 | 0.01212 | 0.01057 |
| 1.163135 | 3.423818 | 0.581379 | 0.035782 | 0.010311 | 0.01212 | 0.01057 |
| 0.339887 | 0.971595 | 0.581379 | 0.035603 | 0.037954 | 0.038686 | 0.03643 |
| 0.277028 | 0.820424 | 0.581379 | 0.036217 | 0.006153 | 0.007482 | 0.006281 |
| 1.218882 | 3.609742 | 0.581379 | 0.036217 | 0.006153 | 0.007482 | 0.006281 |
| 1.111354 | 3.154544 | 0.581379 | 0.036032 | 0.017141 | 0.01844 | 0.016643 |
| 0.309267 | 0.917147 | 0.581263 | 0.034804 | 0.019857 | 0.023096 | 0.020959 |
| 0.327765 | 0.940485 | 0.581263 | 0.03522 | 0.028185 | 0.028659 | 0.026683 |
| 1.063281 | 3.050964 | 0.581263 | 0.03522 | 0.028185 | 0.028659 | 0.026683 |
| 1.015808 | 2.937872 | 0.581263 | 0.036187 | 0.049535 | 0.043606 | 0.041058 |
| 1.013493 | 2.89791 | 0.581263 | 0.03536 | 0.044192 | 0.044434 | 0.042115 |
| 1.231372 | 3.674167 | 0.581263 | 0.036673 | 0.005104 | 0.006805 | 0.005596 |
| 1.085136 | 3.085239 | 0.581263 | 0.03634 | 0.023025 | 0.023422 | 0.021496 |
| 0.261324 | 0.772326 | 0.581263 | 0.037055 | 0.002847 | 0.003798 | 0.002979 |
| 1.532204 | 4.638383 | 0.581263 | 0.037475 | 0.000308 | 0.00052 | 0.000319 |
| 1.062824 | 3.06514 | 0.581147 | 0.036046 | 0.026094 | 0.028855 | 0.026646 |
| 0.226003 | 0.674649 | 0.581147 | 0.037349 | 0.000637 | 0.00075 | 0.000513 |
| 1.482253 | 4.424719 | 0.581147 | 0.037349 | 0.000637 | 0.00075 | 0.000513 |
| 0.34241 | 0.981423 | 0.581147 | 0.035955 | 0.04046 | 0.04238 | 0.039989 |
| 0.312347 | 0.898926 | 0.581147 | 0.035875 | 0.016807 | 0.018518 | 0.016731 |
| 1.140369 | 3.237886 | 0.581147 | 0.037436 | 0.013901 | 0.014153 | 0.012589 |
| 0.269597 | 0.786107 | 0.581031 | 0.036468 | 0.004092 | 0.00449 | 0.003684 |
| 1.102314 | 3.13994 | 0.581031 | 0.036103 | 0.019727 | 0.020086 | 0.018283 |
| 0.331761 | 0.947196 | 0.581031 | 0.03624 | 0.028225 | 0.030567 | 0.028343 |
| 1.132364 | 3.362683 | 0.581031 | 0.036139 | 0.01336 | 0.016055 | 0.014272 |
| 1.024478 | 2.939939 | 0.580799 | 0.035929 | 0.037863 | 0.040376 | 0.037927 |
| 0.256872 | 0.749063 | 0.580799 | 0.036897 | 0.002053 | 0.002543 | 0.001953 |
| 0.311543 | 0.922075 | 0.580799 | 0.036455 | 0.020909 | 0.024256 | 0.022062 |
| 0.285238 | 0.817027 | 0.580799 | 0.03668 | 0.005847 | 0.006673 | 0.005588 |
| 1.22395 | 3.50584 | 0.580799 | 0.03668 | 0.005847 | 0.006673 | 0.005588 |
| 0.279101 | 0.815854 | 0.580683 | 0.037219 | 0.005955 | 0.006857 | 0.005805 |
| 1.22571 | 3.582934 | 0.580683 | 0.037219 | 0.005955 | 0.006857 | 0.005805 |
| 0.32868 | 0.933143 | 0.580683 | 0.03796 | 0.026937 | 0.026425 | 0.024448 |
| 0.268074 | 0.78796 | 0.580683 | 0.036819 | 0.003807 | 0.004708 | 0.003789 |
| 1.2691 | 3.730312 | 0.580683 | 0.036819 | 0.003807 | 0.004708 | 0.003789 |
| 1.082859 | 3.116529 | 0.580683 | 0.035563 | 0.021616 | 0.024124 | 0.022038 |
| 1.23841 | 3.560775 | 0.580567 | 0.036245 | 0.005986 | 0.005894 | 0.004966 |

| | | | | | | |
|---|---|---|---|---|---|---|
| 0.280838 | 0.807487 | 0.580567 | 0.036245 | 0.005986 | 0.005894 | 0.004966 |
| 0.260564 | 0.769646 | 0.580567 | 0.036559 | 0.002901 | 0.003642 | 0.00287 |
| 0.27632 | 0.797231 | 0.580567 | 0.036336 | 0.004861 | 0.005137 | 0.00425 |
| 0.291069 | 0.83144 | 0.58045 | 0.037634 | 0.007936 | 0.008064 | 0.006917 |
| 0.32169 | 0.964149 | 0.58045 | 0.035794 | 0.0317 | 0.036587 | 0.033993 |
| 0.28182 | 0.828513 | 0.58045 | 0.036861 | 0.006834 | 0.008198 | 0.006933 |
| 0.325866 | 0.962109 | 0.58045 | 0.035739 | 0.031436 | 0.035744 | 0.033281 |
| 1.039383 | 3.068741 | 0.58045 | 0.035739 | 0.031436 | 0.035744 | 0.033281 |
| 0.254541 | 0.730556 | 0.58045 | 0.037348 | 0.001635 | 0.001765 | 0.00133 |
| 1.36882 | 3.92864 | 0.58045 | 0.037348 | 0.001635 | 0.001765 | 0.00133 |
| 0.303627 | 0.876059 | 0.580334 | 0.037037 | 0.013334 | 0.014307 | 0.012733 |
| 1.083177 | 3.052012 | 0.580334 | 0.036158 | 0.023943 | 0.023679 | 0.021811 |
| 0.327653 | 0.92321 | 0.580334 | 0.036158 | 0.023943 | 0.023679 | 0.021811 |
| 0.328173 | 0.984951 | 0.580334 | 0.035634 | 0.038637 | 0.044005 | 0.041295 |
| 0.330015 | 0.933679 | 0.580218 | 0.036667 | 0.025346 | 0.026513 | 0.024482 |
| 0.313245 | 0.95021 | 0.580218 | 0.034949 | 0.026971 | 0.032325 | 0.029779 |
| 1.052399 | 3.192386 | 0.580218 | 0.034949 | 0.026971 | 0.032325 | 0.029779 |
| 1.171849 | 3.358852 | 0.580218 | 0.036304 | 0.009583 | 0.010763 | 0.009359 |
| 1.031828 | 2.976512 | 0.580218 | 0.035779 | 0.034646 | 0.037904 | 0.035501 |
| 0.290106 | 0.88488 | 0.580218 | 0.035765 | 0.013364 | 0.016855 | 0.014872 |
| 0.263987 | 0.780613 | 0.580218 | 0.037405 | 0.003252 | 0.004298 | 0.003415 |
| 0.330062 | 0.954086 | 0.580102 | 0.034907 | 0.030337 | 0.032881 | 0.030673 |
| 0.308867 | 0.937846 | 0.580102 | 0.03611 | 0.023787 | 0.028784 | 0.026343 |
| 0.30898 | 0.917937 | 0.580102 | 0.035378 | 0.020061 | 0.023316 | 0.021184 |
| 0.3413 | 0.973368 | 0.580102 | 0.036491 | 0.038065 | 0.03931 | 0.036993 |
| 1.137775 | 3.356952 | 0.580102 | 0.035954 | 0.013843 | 0.015199 | 0.013554 |
| 0.341806 | 0.972583 | 0.579986 | 0.037642 | 0.038384 | 0.039001 | 0.03673 |
| 0.342055 | 0.985252 | 0.579986 | 0.034891 | 0.041251 | 0.043903 | 0.041431 |
| 1.014969 | 2.923504 | 0.579986 | 0.034891 | 0.041251 | 0.043903 | 0.041431 |
| 0.338028 | 0.964214 | 0.579986 | 0.035678 | 0.037061 | 0.03606 | 0.033889 |
| 1.037114 | 2.958338 | 0.579986 | 0.035678 | 0.037061 | 0.03606 | 0.033889 |
| 1.016854 | 2.886114 | 0.579986 | 0.036253 | 0.042387 | 0.043097 | 0.040738 |
| 0.346487 | 0.983426 | 0.579986 | 0.036253 | 0.042387 | 0.043097 | 0.040738 |
| 1.321469 | 3.803762 | 0.57987 | 0.036846 | 0.002646 | 0.002758 | 0.002183 |
| 0.249973 | 0.748111 | 0.57987 | 0.038075 | 0.00201 | 0.00272 | 0.002065 |
| 0.34184 | 0.989325 | 0.57987 | 0.036544 | 0.042183 | 0.045548 | 0.042984 |
| 1.120965 | 3.197433 | 0.579754 | 0.036964 | 0.01631 | 0.016986 | 0.015279 |
| 1.110041 | 3.236273 | 0.579754 | 0.035794 | 0.017158 | 0.01916 | 0.017291 |
| 0.260375 | 0.781377 | 0.579754 | 0.037007 | 0.003679 | 0.004512 | 0.003652 |
| 1.255286 | 3.695144 | 0.579638 | 0.037493 | 0.004942 | 0.005345 | 0.004442 |
| 1.129021 | 3.262495 | 0.579638 | 0.036981 | 0.014232 | 0.01603 | 0.014245 |
| 0.31858 | 0.938178 | 0.579638 | 0.036011 | 0.024579 | 0.02841 | 0.026105 |
| 0.309993 | 0.881498 | 0.579638 | 0.035894 | 0.014164 | 0.014971 | 0.013387 |
| 1.302208 | 3.791586 | 0.579638 | 0.036738 | 0.002786 | 0.003406 | 0.002688 |
| 0.263742 | 0.767927 | 0.579638 | 0.036738 | 0.002786 | 0.003406 | 0.002688 |
| 0.314535 | 0.903568 | 0.579638 | 0.03609 | 0.018292 | 0.019458 | 0.017638 |
| 0.318784 | 0.914824 | 0.579638 | 0.037142 | 0.020222 | 0.021964 | 0.019967 |

| | | | | | | |
|---|---|---|---|---|---|---|
| 1.093106 | 3.136915 | 0.579638 | 0.037142 | 0.020222 | 0.021964 | 0.019967 |
| 0.203511 | 0.636969 | 0.579522 | 0.03826 | 0.000217 | 0.000449 | 0.000259 |
| 0.2585 | 0.757446 | 0.579522 | 0.037454 | 0.002557 | 0.00295 | 0.002319 |
| 0.306353 | 0.929878 | 0.579522 | 0.035444 | 0.021864 | 0.026648 | 0.02427 |
| 1.121385 | 3.306527 | 0.579522 | 0.036839 | 0.015202 | 0.017536 | 0.0157 |
| 0.318294 | 0.915096 | 0.579522 | 0.03653 | 0.021121 | 0.022062 | 0.02017 |
| 1.246267 | 3.645528 | 0.579406 | 0.03706 | 0.004855 | 0.00571 | 0.004714 |
| 0.318036 | 0.926255 | 0.579406 | 0.036395 | 0.023389 | 0.025034 | 0.023024 |
| 0.254304 | 0.731283 | 0.579406 | 0.03734 | 0.001584 | 0.0018 | 0.001347 |
| 1.008565 | 2.894782 | 0.579406 | 0.035608 | 0.048239 | 0.046407 | 0.044179 |
| 0.345449 | 0.991508 | 0.579406 | 0.035608 | 0.048239 | 0.046407 | 0.044179 |
| 0.319686 | 0.936927 | 0.57929 | 0.03649 | 0.024927 | 0.027987 | 0.025741 |
| 0.297062 | 0.848956 | 0.57929 | 0.03726 | 0.009275 | 0.010133 | 0.00881 |
| 1.177918 | 3.3663 | 0.57929 | 0.03726 | 0.009275 | 0.010133 | 0.00881 |
| 0.327095 | 0.950418 | 0.57929 | 0.035931 | 0.028855 | 0.031805 | 0.029581 |
| 0.286002 | 0.825556 | 0.579173 | 0.036383 | 0.007161 | 0.007612 | 0.006476 |
| 0.33523 | 0.982557 | 0.579173 | 0.03466 | 0.039038 | 0.042959 | 0.040405 |
| 0.30274 | 0.866913 | 0.579173 | 0.036119 | 0.013141 | 0.012699 | 0.01133 |
| 0.319836 | 0.936791 | 0.579057 | 0.035291 | 0.024664 | 0.027939 | 0.025742 |
| 0.325515 | 0.912822 | 0.579057 | 0.036895 | 0.020937 | 0.02107 | 0.019237 |
| 0.353469 | 0.996267 | 0.579057 | 0.035507 | 0.046678 | 0.048369 | 0.045919 |
| 0.29911 | 0.955739 | 0.579057 | 0.033094 | 0.027797 | 0.034625 | 0.031955 |
| 0.287054 | 0.833507 | 0.578941 | 0.037128 | 0.007309 | 0.008547 | 0.007303 |
| 0.321134 | 0.908955 | 0.578941 | 0.03707 | 0.019414 | 0.020362 | 0.018534 |
| 1.100165 | 3.113968 | 0.578941 | 0.03707 | 0.019414 | 0.020362 | 0.018534 |
| 0.260971 | 0.762068 | 0.578941 | 0.037437 | 0.00255 | 0.003138 | 0.002461 |
| 0.254106 | 0.784738 | 0.578941 | 0.037119 | 0.003348 | 0.005068 | 0.003992 |
| 0.32325 | 0.957089 | 0.578941 | 0.03355 | 0.031366 | 0.034145 | 0.031964 |
| 0.313906 | 0.90347 | 0.578825 | 0.03617 | 0.017705 | 0.019472 | 0.017649 |
| 1.134058 | 3.220858 | 0.578825 | 0.036129 | 0.014727 | 0.015 | 0.013394 |
| 0.298456 | 0.869776 | 0.578825 | 0.035698 | 0.012188 | 0.013463 | 0.011926 |
| 0.220745 | 0.686145 | 0.578709 | 0.037935 | 0.00085 | 0.001107 | 0.000762 |
| 0.314404 | 0.899637 | 0.578709 | 0.036537 | 0.017422 | 0.018557 | 0.016747 |
| 1.00581 | 2.925878 | 0.578593 | 0.035551 | 0.04319 | 0.047565 | 0.044951 |
| 0.341778 | 0.994223 | 0.578593 | 0.035551 | 0.04319 | 0.047565 | 0.044951 |
| 0.334591 | 0.948919 | 0.578593 | 0.035971 | 0.031508 | 0.030996 | 0.028895 |
| 1.053831 | 2.988728 | 0.578593 | 0.035971 | 0.031508 | 0.030996 | 0.028895 |
| 1.138339 | 3.300762 | 0.578593 | 0.036056 | 0.012977 | 0.014807 | 0.013155 |
| 0.30296 | 0.878473 | 0.578593 | 0.036056 | 0.012977 | 0.014807 | 0.013155 |
| 0.316395 | 0.903093 | 0.578593 | 0.036269 | 0.017918 | 0.019236 | 0.017445 |
| 1.327817 | 3.785926 | 0.578593 | 0.03777 | 0.002338 | 0.002522 | 0.001955 |
| 1.322541 | 3.818775 | 0.578593 | 0.036699 | 0.002479 | 0.002759 | 0.002137 |
| 1.044238 | 2.964494 | 0.578593 | 0.036657 | 0.031905 | 0.033788 | 0.031539 |
| 0.320146 | 0.932597 | 0.578477 | 0.03625 | 0.023535 | 0.026705 | 0.024511 |
| 0.30048 | 0.889541 | 0.578477 | 0.036836 | 0.014494 | 0.017185 | 0.015285 |
| 0.329819 | 0.974691 | 0.578477 | 0.034784 | 0.035481 | 0.040102 | 0.037572 |
| 1.02874 | 2.886426 | 0.578477 | 0.036432 | 0.037759 | 0.038675 | 0.036372 |

| | | | | | | |
|---|---|---|---|---|---|---|
| 1.45544 | 4.398645 | 0.578477 | 0.037564 | 0.000642 | 0.001001 | 0.000664 |
| 0.278775 | 0.823016 | 0.578477 | 0.036474 | 0.006383 | 0.007693 | 0.006484 |
| 0.297999 | 0.889731 | 0.578477 | 0.035999 | 0.014125 | 0.017375 | 0.015428 |
| 1.123936 | 3.355713 | 0.578477 | 0.035999 | 0.014125 | 0.017375 | 0.015428 |
| 0.272259 | 0.790847 | 0.578477 | 0.038067 | 0.004131 | 0.004764 | 0.003916 |
| 1.264467 | 3.672974 | 0.578477 | 0.038067 | 0.004131 | 0.004764 | 0.003916 |
| 1.000737 | 2.806328 | 0.578477 | 0.036034 | 0.048614 | 0.049673 | 0.04723 |
| 1.068734 | 3.035678 | 0.578477 | 0.036689 | 0.026171 | 0.027135 | 0.025043 |
| 0.329416 | 0.935687 | 0.578477 | 0.036689 | 0.026171 | 0.027135 | 0.025043 |
| 1.049525 | 3.021705 | 0.578361 | 0.036117 | 0.031292 | 0.032424 | 0.030295 |
| 0.280975 | 0.794809 | 0.578361 | 0.036364 | 0.004576 | 0.004718 | 0.003877 |
| 0.281848 | 0.806319 | 0.578361 | 0.036126 | 0.005172 | 0.005731 | 0.004768 |
| 1.240204 | 3.548008 | 0.578361 | 0.036126 | 0.005172 | 0.005731 | 0.004768 |
| 0.325847 | 0.939328 | 0.578361 | 0.035951 | 0.025639 | 0.028401 | 0.026201 |
| 0.294357 | 0.882065 | 0.578361 | 0.035622 | 0.012999 | 0.016032 | 0.014174 |
| 0.303624 | 0.894085 | 0.578361 | 0.036294 | 0.015061 | 0.017967 | 0.01604 |
| 1.184395 | 3.452938 | 0.578361 | 0.036702 | 0.008848 | 0.009882 | 0.008563 |
| 1.207933 | 3.487716 | 0.578361 | 0.036734 | 0.007117 | 0.007853 | 0.00673 |
| 0.23843 | 0.726753 | 0.578245 | 0.038244 | 0.001412 | 0.002052 | 0.00151 |
| 0.320369 | 0.932851 | 0.578245 | 0.036758 | 0.023586 | 0.026767 | 0.024566 |
| 0.316986 | 0.914958 | 0.578245 | 0.035337 | 0.020338 | 0.022101 | 0.020133 |
| 0.350976 | 0.992987 | 0.578245 | 0.036082 | 0.046911 | 0.046979 | 0.044592 |
| 0.293256 | 0.84767 | 0.578245 | 0.03663 | 0.009422 | 0.010162 | 0.008848 |
| 0.293745 | 0.84524 | 0.578245 | 0.037018 | 0.009065 | 0.009779 | 0.008472 |
| 1.183096 | 3.404314 | 0.578245 | 0.037018 | 0.009065 | 0.009779 | 0.008472 |
| 0.318674 | 0.928501 | 0.578245 | 0.036552 | 0.022547 | 0.025622 | 0.023461 |
| 1.077005 | 3.138003 | 0.578245 | 0.036552 | 0.022547 | 0.025622 | 0.023461 |
| 1.273127 | 3.676079 | 0.578129 | 0.037602 | 0.003975 | 0.004336 | 0.003525 |
| 0.272029 | 0.785467 | 0.578129 | 0.037602 | 0.003975 | 0.004336 | 0.003525 |
| 0.326888 | 0.943538 | 0.578129 | 0.036026 | 0.027548 | 0.029639 | 0.027456 |
| 1.059841 | 3.059156 | 0.578129 | 0.036026 | 0.027548 | 0.029639 | 0.027456 |
| 0.333009 | 0.959264 | 0.578129 | 0.036291 | 0.031422 | 0.034511 | 0.032161 |
| 1.169768 | 3.427871 | 0.578129 | 0.036882 | 0.009633 | 0.011352 | 0.009834 |
| 0.280447 | 0.811097 | 0.578129 | 0.037376 | 0.005568 | 0.006281 | 0.005235 |
| 0.172077 | 0.680767 | 0.578129 | 0.03338 | 0.000718 | 0.002243 | 0.001394 |
| 0.291027 | 0.848656 | 0.578129 | 0.03653 | 0.009112 | 0.010437 | 0.009035 |
| 0.29963 | 0.909144 | 0.578129 | 0.036255 | 0.017431 | 0.021655 | 0.019457 |
| 0.300023 | 0.87415 | 0.578129 | 0.036487 | 0.01281 | 0.014167 | 0.012582 |
| 1.067836 | 3.157817 | 0.578013 | 0.035739 | 0.024358 | 0.028002 | 0.025684 |
| 0.316674 | 0.936473 | 0.578013 | 0.035739 | 0.024358 | 0.028002 | 0.025684 |
| 1.305548 | 3.852144 | 0.578013 | 0.037175 | 0.002552 | 0.003435 | 0.002662 |
| 1.338337 | 3.966428 | 0.578013 | 0.037361 | 0.001889 | 0.0026 | 0.001962 |
| 1.131224 | 3.273775 | 0.578013 | 0.035825 | 0.015038 | 0.015744 | 0.01416 |
| 1.092999 | 3.086663 | 0.578013 | 0.035895 | 0.021077 | 0.021691 | 0.019836 |
| 0.323974 | 0.914914 | 0.578013 | 0.035895 | 0.021077 | 0.021691 | 0.019836 |
| 0.338045 | 0.95977 | 0.578013 | 0.03661 | 0.032591 | 0.034497 | 0.032234 |
| 0.348722 | 0.991795 | 0.577896 | 0.036942 | 0.045895 | 0.046496 | 0.044202 |

| | | | | | | |
|---|---|---|---|---|---|---|
| 1.161279 | 3.347766 | 0.577896 | 0.036971 | 0.011144 | 0.011952 | 0.010518 |
| 0.311597 | 0.89529 | 0.57778 | 0.036113 | 0.0166 | 0.017751 | 0.015987 |
| 0.333822 | 0.978249 | 0.57778 | 0.034979 | 0.037489 | 0.041336 | 0.038805 |
| 0.276031 | 0.826681 | 0.57778 | 0.035554 | 0.006334 | 0.008287 | 0.006955 |
| 0.249519 | 0.748622 | 0.57778 | 0.037459 | 0.002052 | 0.002763 | 0.00211 |
| 0.313487 | 0.936347 | 0.57778 | 0.035635 | 0.023755 | 0.028122 | 0.025767 |
| 0.319684 | 0.946194 | 0.57778 | 0.035384 | 0.026937 | 0.030791 | 0.028402 |
| 0.332601 | 0.950092 | 0.57778 | 0.035728 | 0.030331 | 0.031463 | 0.029289 |
| 1.224311 | 3.63297 | 0.57778 | 0.036387 | 0.005625 | 0.00716 | 0.005979 |
| 0.275257 | 0.816786 | 0.57778 | 0.036387 | 0.005625 | 0.00716 | 0.005979 |
| 0.345372 | 0.973972 | 0.57778 | 0.036187 | 0.038516 | 0.039429 | 0.037163 |
| 0.242394 | 0.735609 | 0.577664 | 0.037532 | 0.001616 | 0.002333 | 0.001731 |
| 0.321657 | 0.928269 | 0.577664 | 0.03517 | 0.024003 | 0.0254 | 0.023413 |
| 0.352248 | 0.997279 | 0.577664 | 0.035941 | 0.048716 | 0.048812 | 0.046409 |
| 0.276753 | 0.894541 | 0.577664 | 0.033916 | 0.014232 | 0.019683 | 0.017328 |
| 1.098723 | 3.156886 | 0.577664 | 0.036956 | 0.019043 | 0.020908 | 0.019013 |
| 0.281382 | 0.810045 | 0.577548 | 0.037213 | 0.00572 | 0.006126 | 0.005174 |
| 1.377036 | 4.21973 | 0.577548 | 0.037225 | 0.001308 | 0.002071 | 0.001501 |
| 0.236982 | 0.726197 | 0.577548 | 0.037225 | 0.001308 | 0.002071 | 0.001501 |
| 1.02159 | 2.943257 | 0.5772 | 0.036698 | 0.046465 | 0.041441 | 0.038988 |
| 0.341121 | 0.998902 | 0.5772 | 0.036055 | 0.045304 | 0.049534 | 0.046886 |
| 1.153942 | 3.290156 | 0.5772 | 0.037089 | 0.012031 | 0.012572 | 0.011085 |
| 0.282928 | 0.831527 | 0.5772 | 0.036639 | 0.007108 | 0.008519 | 0.007221 |
| 1.202607 | 3.534474 | 0.5772 | 0.036639 | 0.007108 | 0.008519 | 0.007221 |
| 0.333388 | 0.953094 | 0.577084 | 0.035981 | 0.031319 | 0.032416 | 0.030246 |
| 0.278752 | 0.817323 | 0.577084 | 0.037647 | 0.005818 | 0.007038 | 0.005865 |
| 1.140885 | 3.320408 | 0.577084 | 0.036711 | 0.013959 | 0.014541 | 0.012947 |
| 0.32088 | 0.925067 | 0.577084 | 0.036123 | 0.022027 | 0.024555 | 0.022469 |
| 1.143106 | 3.326259 | 0.576968 | 0.03669 | 0.012123 | 0.014253 | 0.01256 |
| 0.300227 | 0.893757 | 0.576968 | 0.036353 | 0.015169 | 0.0181 | 0.016228 |
| 1.019608 | 2.867471 | 0.576968 | 0.036309 | 0.041043 | 0.041994 | 0.039667 |
| 0.27257 | 0.79717 | 0.576968 | 0.037292 | 0.004616 | 0.005304 | 0.004362 |
| 1.312867 | 3.982761 | 0.576968 | 0.037296 | 0.002328 | 0.003484 | 0.002658 |
| 0.342598 | 0.974342 | 0.576968 | 0.035755 | 0.037497 | 0.039643 | 0.037293 |
| 1.052248 | 2.958222 | 0.576852 | 0.036402 | 0.030539 | 0.031311 | 0.029169 |
| 0.338041 | 0.950346 | 0.576852 | 0.036402 | 0.030539 | 0.031311 | 0.029169 |
| 0.295169 | 0.847048 | 0.576852 | 0.036675 | 0.008838 | 0.00996 | 0.008614 |
| 1.125754 | 3.251887 | 0.576852 | 0.035743 | 0.014547 | 0.016499 | 0.014743 |
| 1.216554 | 3.575414 | 0.576852 | 0.037468 | 0.006464 | 0.007524 | 0.006374 |
| 0.338962 | 0.967354 | 0.576852 | 0.03529 | 0.035153 | 0.037157 | 0.034892 |
| 0.321985 | 0.929171 | 0.576852 | 0.036258 | 0.023036 | 0.025636 | 0.023523 |
| 1.011855 | 2.861273 | 0.576852 | 0.036532 | 0.044019 | 0.045027 | 0.042705 |
| 0.061726 | 0.635987 | 0.576852 | 0.023024 | 0.000497 | 0.006517 | 0.002528 |
| 1.115331 | 3.246771 | 0.576852 | 0.035794 | 0.016134 | 0.018255 | 0.016459 |
| 0.273857 | 0.799606 | 0.576736 | 0.036878 | 0.004789 | 0.005468 | 0.004514 |
| 0.31226 | 0.90032 | 0.576736 | 0.036665 | 0.017939 | 0.01884 | 0.017064 |
| 0.297865 | 0.875418 | 0.576736 | 0.036582 | 0.012503 | 0.014535 | 0.012839 |

| | | | | | | |
|---|---|---|---|---|---|---|
| 1.066281 | 3.093875 | 0.576619 | 0.036027 | 0.025525 | 0.028083 | 0.025878 |
| 0.309878 | 0.904483 | 0.576619 | 0.037291 | 0.017312 | 0.019947 | 0.018002 |
| 1.006199 | 2.889069 | 0.576619 | 0.03654 | 0.044637 | 0.047375 | 0.044837 |
| 0.290676 | 0.847185 | 0.576503 | 0.037363 | 0.008546 | 0.010238 | 0.00881 |
| 1.18038 | 3.440252 | 0.576503 | 0.037363 | 0.008546 | 0.010238 | 0.00881 |
| 0.300454 | 0.884272 | 0.576503 | 0.036781 | 0.013401 | 0.016103 | 0.014265 |
| 1.068813 | 3.029335 | 0.576387 | 0.036305 | 0.027603 | 0.02708 | 0.025075 |
| 0.330105 | 0.935617 | 0.576387 | 0.036305 | 0.027603 | 0.02708 | 0.025075 |
| 1.123919 | 3.207932 | 0.576387 | 0.037023 | 0.016224 | 0.016549 | 0.014893 |
| 0.311727 | 0.889744 | 0.576387 | 0.037023 | 0.016224 | 0.016549 | 0.014893 |
| 1.141567 | 3.325005 | 0.576387 | 0.037545 | 0.013131 | 0.014468 | 0.012873 |
| 0.300751 | 0.875989 | 0.576387 | 0.037545 | 0.013131 | 0.014468 | 0.012873 |
| 0.341697 | 0.971108 | 0.576271 | 0.036381 | 0.037206 | 0.038453 | 0.036115 |
| 1.029751 | 2.926573 | 0.576271 | 0.036381 | 0.037206 | 0.038453 | 0.036115 |
| 1.056018 | 3.06997 | 0.576271 | 0.035504 | 0.027303 | 0.030759 | 0.028437 |
| 0.290703 | 0.855466 | 0.576271 | 0.036549 | 0.009769 | 0.011507 | 0.009998 |
| 0.317953 | 0.910563 | 0.576155 | 0.0361 | 0.019101 | 0.020942 | 0.019032 |
| 1.098363 | 3.179231 | 0.576155 | 0.036258 | 0.018965 | 0.021111 | 0.019114 |
| 0.314541 | 0.910446 | 0.576155 | 0.036258 | 0.018965 | 0.021111 | 0.019114 |
| 0.321397 | 0.923026 | 0.576039 | 0.036383 | 0.021931 | 0.023972 | 0.021967 |
| 0.332336 | 0.956805 | 0.576039 | 0.035974 | 0.031418 | 0.0337 | 0.031398 |
| 1.144525 | 3.331314 | 0.576039 | 0.037076 | 0.01276 | 0.014079 | 0.012503 |
| 0.286793 | 0.841021 | 0.575923 | 0.037149 | 0.007959 | 0.009575 | 0.008223 |
| 0.332282 | 0.97174 | 0.575923 | 0.03679 | 0.035218 | 0.038953 | 0.036442 |
| 1.073404 | 3.071611 | 0.575923 | 0.035678 | 0.025591 | 0.026144 | 0.024151 |
| 1.000206 | 2.832431 | 0.575923 | 0.036882 | 0.04913 | 0.049909 | 0.047453 |
| 1.007094 | 2.874592 | 0.575807 | 0.036835 | 0.044946 | 0.046991 | 0.044502 |
| 1.032135 | 2.992557 | 0.575807 | 0.036352 | 0.034818 | 0.037853 | 0.035424 |
| 1.041096 | 2.940748 | 0.575807 | 0.036744 | 0.033299 | 0.034687 | 0.032455 |
| 1.023042 | 2.892694 | 0.575807 | 0.035658 | 0.039222 | 0.040769 | 0.03844 |
| 0.345698 | 0.977477 | 0.575807 | 0.035658 | 0.039222 | 0.040769 | 0.03844 |
| 1.112207 | 3.20165 | 0.575691 | 0.036874 | 0.017007 | 0.018561 | 0.016719 |
| 1.016919 | 2.884758 | 0.575691 | 0.036684 | 0.042371 | 0.043069 | 0.040708 |
| 0.289595 | 0.837429 | 0.575691 | 0.036434 | 0.007878 | 0.008925 | 0.007684 |
| 1.104115 | 3.211412 | 0.575691 | 0.036714 | 0.017473 | 0.020147 | 0.018168 |
| 1.205881 | 3.598658 | 0.575575 | 0.037459 | 0.00651 | 0.008509 | 0.007141 |
| 0.27929 | 0.816784 | 0.575459 | 0.037123 | 0.005655 | 0.00695 | 0.005801 |
| 0.33792 | 0.971093 | 0.575459 | 0.037441 | 0.036774 | 0.038556 | 0.036249 |
| 0.346944 | 0.984254 | 0.575459 | 0.036642 | 0.04272 | 0.043422 | 0.041057 |
| 0.246884 | 0.741067 | 0.575459 | 0.037684 | 0.001804 | 0.002457 | 0.00185 |
| 1.349405 | 4.05049 | 0.575459 | 0.037684 | 0.001804 | 0.002457 | 0.00185 |
| 1.200201 | 3.410752 | 0.575342 | 0.036352 | 0.007452 | 0.008172 | 0.006972 |
| 0.29319 | 0.833194 | 0.575342 | 0.036352 | 0.007452 | 0.008172 | 0.006972 |
| 1.17523 | 3.453793 | 0.575342 | 0.037547 | 0.009413 | 0.010863 | 0.00943 |
| 1.095271 | 3.185417 | 0.575342 | 0.036385 | 0.018976 | 0.021785 | 0.019741 |
| 0.313931 | 0.913016 | 0.575342 | 0.036385 | 0.018976 | 0.021785 | 0.019741 |
| 0.334051 | 0.944584 | 0.575226 | 0.037027 | 0.030883 | 0.029633 | 0.027646 |

| | | | | | | |
|---|---|---|---|---|---|---|
| 0.289341 | 0.830093 | 0.575226 | 0.037066 | 0.007046 | 0.007988 | 0.006795 |
| 0.313163 | 0.908846 | 0.575226 | 0.036373 | 0.018683 | 0.020799 | 0.018834 |
| 1.437869 | 4.576441 | 0.57511 | 0.038276 | 0.000756 | 0.001425 | 0.000951 |
| 1.073539 | 3.174986 | 0.57511 | 0.036358 | 0.023118 | 0.026658 | 0.02438 |
| 0.35013 | 0.989721 | 0.57511 | 0.035967 | 0.04461 | 0.045615 | 0.043287 |
| 1.225602 | 3.555569 | 0.57511 | 0.036891 | 0.005979 | 0.006755 | 0.005667 |
| 0.27841 | 0.800538 | 0.574994 | 0.036839 | 0.004646 | 0.005342 | 0.004401 |
| 0.317396 | 0.913359 | 0.574994 | 0.036957 | 0.019732 | 0.021672 | 0.019752 |
| 0.308861 | 0.894081 | 0.574994 | 0.037581 | 0.016557 | 0.017651 | 0.015954 |
| 1.215537 | 3.520114 | 0.574994 | 0.036648 | 0.006737 | 0.007373 | 0.006248 |
| 0.284082 | 0.822682 | 0.574994 | 0.036648 | 0.006737 | 0.007373 | 0.006248 |
| 1.307509 | 3.850609 | 0.574994 | 0.037126 | 0.002773 | 0.003357 | 0.002652 |
| 0.282916 | 0.835846 | 0.574994 | 0.036945 | 0.00726 | 0.009086 | 0.00772 |
| 0.298564 | 0.871955 | 0.574994 | 0.037349 | 0.012221 | 0.013851 | 0.012241 |
| 1.261128 | 3.7535 | 0.574878 | 0.037468 | 0.004061 | 0.005209 | 0.004216 |
| 0.266418 | 0.792941 | 0.574878 | 0.037468 | 0.004061 | 0.005209 | 0.004216 |
| 1.184172 | 3.552686 | 0.574878 | 0.03703 | 0.008163 | 0.010374 | 0.008846 |
| 0.341277 | 0.968261 | 0.574878 | 0.037412 | 0.036212 | 0.037415 | 0.035086 |
| 0.297617 | 0.872159 | 0.574762 | 0.037342 | 0.011964 | 0.013946 | 0.012265 |
| 0.3035 | 0.883402 | 0.574762 | 0.036753 | 0.013462 | 0.015743 | 0.01397 |
| 0.250503 | 0.740986 | 0.574762 | 0.036738 | 0.001819 | 0.002339 | 0.001762 |
| 0.301187 | 0.877696 | 0.574646 | 0.036976 | 0.013068 | 0.014766 | 0.013094 |
| 1.39062 | 4.246809 | 0.574646 | 0.037397 | 0.001225 | 0.001822 | 0.001315 |
| 1.139669 | 3.207246 | 0.574646 | 0.03737 | 0.014337 | 0.014078 | 0.012575 |
| 0.3265 | 0.981137 | 0.57453 | 0.036685 | 0.037313 | 0.04258 | 0.039904 |
| 0.33894 | 0.965808 | 0.57453 | 0.036373 | 0.033993 | 0.0366 | 0.034231 |
| 0.303203 | 0.869448 | 0.57453 | 0.036631 | 0.011762 | 0.013119 | 0.011571 |
| 0.303239 | 0.880815 | 0.57453 | 0.036002 | 0.014201 | 0.015246 | 0.013656 |
| 0.315272 | 0.91691 | 0.57453 | 0.036671 | 0.020464 | 0.022698 | 0.020679 |
| 0.271129 | 0.784339 | 0.57453 | 0.036905 | 0.003916 | 0.004285 | 0.003484 |
| 1.068795 | 3.095159 | 0.57453 | 0.037613 | 0.024918 | 0.027437 | 0.025235 |
| 0.283388 | 0.822666 | 0.574414 | 0.037597 | 0.006608 | 0.007407 | 0.006274 |
| 1.019024 | 2.887223 | 0.574414 | 0.037413 | 0.040962 | 0.042266 | 0.039848 |
| 0.336877 | 0.980389 | 0.574298 | 0.035761 | 0.037999 | 0.042088 | 0.039554 |
| 1.020004 | 2.968439 | 0.574298 | 0.035761 | 0.037999 | 0.042088 | 0.039554 |
| 0.311747 | 0.928962 | 0.574298 | 0.036596 | 0.022468 | 0.026114 | 0.023955 |
| 0.35335 | 0.992075 | 0.574298 | 0.036365 | 0.04553 | 0.046572 | 0.044165 |
| 0.250385 | 0.771448 | 0.574298 | 0.03628 | 0.002893 | 0.004185 | 0.00328 |
| 1.109716 | 3.207244 | 0.574182 | 0.036726 | 0.018203 | 0.019054 | 0.017303 |
| 1.19602 | 3.473567 | 0.574182 | 0.038236 | 0.007829 | 0.008843 | 0.007636 |
| 1.091491 | 3.158372 | 0.574182 | 0.03629 | 0.021079 | 0.022434 | 0.020536 |
| 0.316619 | 0.916178 | 0.574182 | 0.03629 | 0.021079 | 0.022434 | 0.020536 |
| 0.327646 | 0.934618 | 0.574182 | 0.036613 | 0.025249 | 0.026909 | 0.024861 |
| 1.280538 | 3.663631 | 0.574182 | 0.037443 | 0.003539 | 0.00395 | 0.003182 |
| 1.145581 | 3.262596 | 0.574182 | 0.036797 | 0.012984 | 0.013549 | 0.012002 |
| 0.301194 | 0.935943 | 0.574182 | 0.034742 | 0.023187 | 0.028608 | 0.026083 |
| 0.293686 | 0.863998 | 0.574065 | 0.036107 | 0.01089 | 0.012738 | 0.011153 |

| | | | | | | |
|---|---|---|---|---|---|---|
| 0.253919 | 0.73081 | 0.574065 | 0.037843 | 0.0015 | 0.001791 | 0.001326 |
| 0.292553 | 0.871852 | 0.574065 | 0.037133 | 0.011831 | 0.014197 | 0.012494 |
| 0.312131 | 0.920734 | 0.573949 | 0.035922 | 0.020413 | 0.023869 | 0.02172 |
| 1.08609 | 3.203778 | 0.573949 | 0.035922 | 0.020413 | 0.023869 | 0.02172 |
| 1.102495 | 3.17329 | 0.573949 | 0.036039 | 0.018427 | 0.020246 | 0.018391 |
| 0.300263 | 0.890533 | 0.573949 | 0.036797 | 0.014703 | 0.017408 | 0.015509 |
| 0.292752 | 0.845879 | 0.573949 | 0.037409 | 0.008945 | 0.009928 | 0.008583 |
| 0.287859 | 0.839965 | 0.573949 | 0.037306 | 0.008586 | 0.009368 | 0.008119 |
| 1.190526 | 3.473927 | 0.573949 | 0.037306 | 0.008586 | 0.009368 | 0.008119 |
| 0.285158 | 0.818231 | 0.573949 | 0.037023 | 0.005978 | 0.00681 | 0.005721 |
| 0.308396 | 0.899787 | 0.573833 | 0.036472 | 0.017357 | 0.018949 | 0.017172 |
| 0.290994 | 0.887296 | 0.573833 | 0.036373 | 0.013748 | 0.017294 | 0.015286 |
| 0.308375 | 0.926839 | 0.573833 | 0.036307 | 0.021721 | 0.025708 | 0.023424 |
| 0.307552 | 0.878649 | 0.573833 | 0.036326 | 0.01345 | 0.014563 | 0.012981 |
| 1.032706 | 2.97761 | 0.573833 | 0.036057 | 0.035187 | 0.037609 | 0.035246 |
| 0.313074 | 0.892227 | 0.573717 | 0.036921 | 0.016513 | 0.016996 | 0.015386 |
| 1.022416 | 2.987295 | 0.573717 | 0.036153 | 0.037404 | 0.04125 | 0.038693 |
| 0.334751 | 0.978075 | 0.573717 | 0.036153 | 0.037404 | 0.04125 | 0.038693 |
| 1.044901 | 3.052998 | 0.573601 | 0.037104 | 0.030496 | 0.03396 | 0.031541 |
| 0.327547 | 0.957028 | 0.573601 | 0.037104 | 0.030496 | 0.03396 | 0.031541 |
| 0.31886 | 0.91941 | 0.573601 | 0.036871 | 0.021334 | 0.023148 | 0.021138 |
| 0.338057 | 0.972108 | 0.573601 | 0.036454 | 0.036476 | 0.038928 | 0.036522 |
| 0.267301 | 0.811032 | 0.573601 | 0.036404 | 0.005287 | 0.006941 | 0.005754 |
| 0.320956 | 0.928447 | 0.573601 | 0.036176 | 0.024018 | 0.025487 | 0.023493 |
| 1.017952 | 2.947198 | 0.573485 | 0.035922 | 0.042454 | 0.042808 | 0.040601 |
| 1.327707 | 3.910346 | 0.573485 | 0.038061 | 0.00229 | 0.002802 | 0.002166 |
| 0.32095 | 0.951106 | 0.573485 | 0.036721 | 0.028357 | 0.032286 | 0.029872 |
| 1.160427 | 3.346713 | 0.573485 | 0.03691 | 0.011329 | 0.012052 | 0.01055 |
| 0.301989 | 0.887626 | 0.573369 | 0.036637 | 0.014443 | 0.016695 | 0.014881 |
| 0.314133 | 0.889821 | 0.573369 | 0.03696 | 0.015581 | 0.01642 | 0.014752 |
| 1.11442 | 3.215928 | 0.573369 | 0.036539 | 0.016719 | 0.018242 | 0.016431 |
| 1.010001 | 2.929824 | 0.573369 | 0.035278 | 0.042241 | 0.045869 | 0.04339 |
| 1.054512 | 2.979736 | 0.573369 | 0.035858 | 0.029451 | 0.030752 | 0.028601 |
| 0.3356 | 0.948306 | 0.573369 | 0.035858 | 0.029451 | 0.030752 | 0.028601 |
| 1.037745 | 2.974588 | 0.573369 | 0.036182 | 0.034178 | 0.035916 | 0.03363 |
| 1.167917 | 3.418917 | 0.573253 | 0.037246 | 0.010061 | 0.011522 | 0.010053 |
| 0.290826 | 0.852721 | 0.573253 | 0.037295 | 0.009381 | 0.011068 | 0.009572 |
| 1.172717 | 3.438482 | 0.573253 | 0.037295 | 0.009381 | 0.011068 | 0.009572 |
| 1.03802 | 3.09929 | 0.573137 | 0.035796 | 0.031245 | 0.036288 | 0.033728 |
| 0.322655 | 0.963372 | 0.573137 | 0.035796 | 0.031245 | 0.036288 | 0.033728 |
| 0.313562 | 0.924686 | 0.573137 | 0.035937 | 0.021942 | 0.024846 | 0.02273 |
| 1.074016 | 3.055714 | 0.573137 | 0.036682 | 0.025887 | 0.025903 | 0.023927 |
| 1.124794 | 3.302745 | 0.573137 | 0.037104 | 0.014683 | 0.016944 | 0.015117 |
| 1.39972 | 4.382915 | 0.573137 | 0.038482 | 0.001149 | 0.001841 | 0.001324 |
| 1.069018 | 3.10193 | 0.573137 | 0.036404 | 0.025357 | 0.027416 | 0.025313 |
| 1.098579 | 3.186825 | 0.573021 | 0.035626 | 0.018819 | 0.02111 | 0.019187 |
| 0.313792 | 0.910267 | 0.573021 | 0.035626 | 0.018819 | 0.02111 | 0.019187 |

| | | | | | | |
|---|---|---|---|---|---|---|
| 0.29683 | 0.856566 | 0.573021 | 0.036906 | 0.009805 | 0.011321 | 0.009848 |
| 0.286451 | 0.83123 | 0.573021 | 0.03703 | 0.007387 | 0.008288 | 0.007072 |
| 0.296327 | 0.96147 | 0.573021 | 0.031741 | 0.028752 | 0.036544 | 0.033632 |
| 1.103329 | 3.139713 | 0.572905 | 0.036577 | 0.019089 | 0.019884 | 0.018033 |
| 1.177897 | 3.394422 | 0.572905 | 0.037325 | 0.009247 | 0.010277 | 0.00889 |
| 0.294601 | 0.848971 | 0.572905 | 0.037325 | 0.009247 | 0.010277 | 0.00889 |
| 0.282253 | 0.833425 | 0.572905 | 0.037834 | 0.007591 | 0.008802 | 0.007559 |
| 1.041403 | 2.971178 | 0.572905 | 0.035923 | 0.035752 | 0.034716 | 0.032589 |
| 1.063616 | 3.055232 | 0.572905 | 0.038425 | 0.02707 | 0.028591 | 0.026479 |
| 1.051886 | 2.999936 | 0.572905 | 0.037401 | 0.030507 | 0.031621 | 0.029439 |
| 1.050025 | 3.113043 | 0.572905 | 0.036109 | 0.028717 | 0.032677 | 0.030259 |
| 0.321229 | 0.952359 | 0.572905 | 0.036109 | 0.028717 | 0.032677 | 0.030259 |
| 0.305776 | 0.899854 | 0.572788 | 0.036472 | 0.017204 | 0.01912 | 0.017371 |
| 0.352823 | 0.98795 | 0.572788 | 0.036422 | 0.045069 | 0.044844 | 0.042513 |
| 0.323036 | 0.939869 | 0.572788 | 0.036691 | 0.025389 | 0.0287 | 0.026439 |
| 0.302093 | 0.87661 | 0.572788 | 0.037287 | 0.013159 | 0.014504 | 0.012891 |
| 0.302029 | 0.864273 | 0.572672 | 0.036585 | 0.011752 | 0.012286 | 0.010833 |
| 1.02925 | 2.945119 | 0.572672 | 0.037498 | 0.036638 | 0.038691 | 0.03642 |
| 0.266185 | 0.783178 | 0.572672 | 0.037716 | 0.003664 | 0.004404 | 0.003547 |
| 0.328321 | 0.93663 | 0.572672 | 0.036076 | 0.025766 | 0.02747 | 0.0254 |
| 1.130432 | 3.282342 | 0.572672 | 0.036538 | 0.014142 | 0.015917 | 0.01417 |
| 0.255128 | 0.855064 | 0.572556 | 0.033543 | 0.009002 | 0.013607 | 0.011587 |
| 0.314679 | 0.93089 | 0.572556 | 0.036366 | 0.022975 | 0.026501 | 0.02423 |
| 0.327489 | 0.944817 | 0.572556 | 0.035755 | 0.027167 | 0.030009 | 0.027775 |
| 0.290482 | 0.86412 | 0.572556 | 0.037293 | 0.010712 | 0.012952 | 0.011306 |
| 0.344883 | 0.984924 | 0.572556 | 0.036782 | 0.041761 | 0.043726 | 0.041283 |
| 1.104695 | 3.127088 | 0.57244 | 0.036761 | 0.018998 | 0.01954 | 0.017787 |
| 0.319786 | 0.905228 | 0.57244 | 0.036761 | 0.018998 | 0.01954 | 0.017787 |
| 1.274408 | 3.664878 | 0.57244 | 0.0378 | 0.003658 | 0.004239 | 0.003421 |
| 0.289473 | 0.849823 | 0.57244 | 0.036045 | 0.010233 | 0.010703 | 0.009369 |
| 0.276489 | 0.805212 | 0.57244 | 0.037592 | 0.004938 | 0.005879 | 0.00488 |
| 1.077707 | 3.079594 | 0.57244 | 0.036741 | 0.023057 | 0.025133 | 0.023063 |
| 1.035103 | 2.933402 | 0.57244 | 0.037897 | 0.035188 | 0.036636 | 0.034394 |
| 1.021234 | 2.9019 | 0.572324 | 0.036409 | 0.039911 | 0.041467 | 0.039172 |
| 0.292806 | 0.854356 | 0.572324 | 0.037784 | 0.010584 | 0.011207 | 0.00986 |
| 1.043896 | 3.052111 | 0.572324 | 0.037124 | 0.030784 | 0.034266 | 0.031845 |
| 1.048672 | 2.977337 | 0.572324 | 0.037601 | 0.030599 | 0.032477 | 0.030256 |
| 1.220605 | 3.673198 | 0.572208 | 0.036841 | 0.005835 | 0.007602 | 0.006324 |
| 1.020761 | 2.95664 | 0.572208 | 0.035695 | 0.047733 | 0.04178 | 0.039332 |
| 0.299122 | 0.869582 | 0.572208 | 0.03721 | 0.011783 | 0.013388 | 0.011775 |
| 1.08555 | 3.137201 | 0.572208 | 0.037773 | 0.021803 | 0.023625 | 0.021621 |
| 0.253792 | 0.759373 | 0.572208 | 0.037847 | 0.002272 | 0.003235 | 0.002474 |
| 1.209832 | 3.510343 | 0.572208 | 0.036205 | 0.009301 | 0.007793 | 0.006653 |
| 0.327076 | 0.940724 | 0.572092 | 0.036709 | 0.026707 | 0.028765 | 0.026575 |
| 0.33079 | 0.952172 | 0.571976 | 0.036658 | 0.029997 | 0.03222 | 0.029949 |
| 0.264841 | 0.76964 | 0.571976 | 0.037772 | 0.003128 | 0.003477 | 0.002766 |
| 1.299309 | 3.775847 | 0.571976 | 0.037772 | 0.003128 | 0.003477 | 0.002766 |

| | | | | | | |
|---|---|---|---|---|---|---|
| 0.310985 | 0.888672 | 0.571976 | 0.03719 | 0.014791 | 0.016368 | 0.014633 |
| 1.212696 | 3.644613 | 0.571976 | 0.036784 | 0.006677 | 0.008123 | 0.006917 |
| 0.330392 | 0.927042 | 0.571976 | 0.037233 | 0.024899 | 0.024589 | 0.022675 |
| 1.0787 | 3.02671 | 0.571976 | 0.037233 | 0.024899 | 0.024589 | 0.022675 |
| 1.182228 | 3.436194 | 0.571976 | 0.037694 | 0.009514 | 0.010025 | 0.008768 |
| 0.286943 | 0.848399 | 0.57186 | 0.036794 | 0.008606 | 0.010636 | 0.009158 |
| 1.20399 | 3.514455 | 0.57186 | 0.036642 | 0.007104 | 0.008309 | 0.007105 |
| 0.284539 | 0.830572 | 0.57186 | 0.036642 | 0.007104 | 0.008309 | 0.007105 |
| 0.336388 | 0.9526 | 0.57186 | 0.036266 | 0.030771 | 0.032127 | 0.029958 |
| 0.28849 | 0.828155 | 0.57186 | 0.037768 | 0.006864 | 0.007794 | 0.006618 |
| 0.293825 | 0.873862 | 0.57186 | 0.037058 | 0.012092 | 0.01449 | 0.012759 |
| 0.264868 | 0.776955 | 0.571744 | 0.037991 | 0.003147 | 0.003986 | 0.003173 |
| 0.287247 | 0.843655 | 0.571744 | 0.036644 | 0.008253 | 0.009922 | 0.008548 |
| 0.321116 | 0.901816 | 0.571744 | 0.03709 | 0.018926 | 0.018659 | 0.016941 |
| 0.308826 | 0.896363 | 0.571744 | 0.03701 | 0.016211 | 0.018154 | 0.016299 |
| 0.311625 | 0.899653 | 0.571628 | 0.036856 | 0.016603 | 0.018726 | 0.016866 |
| 1.11154 | 3.208986 | 0.571628 | 0.036856 | 0.016603 | 0.018726 | 0.016866 |
| 0.291684 | 0.856763 | 0.571628 | 0.037715 | 0.009911 | 0.011656 | 0.010131 |
| 0.310474 | 0.902505 | 0.571628 | 0.036965 | 0.017397 | 0.01945 | 0.017543 |
| 0.290374 | 0.855467 | 0.571628 | 0.036349 | 0.009372 | 0.011527 | 0.009967 |
| 1.168953 | 3.443838 | 0.571628 | 0.036349 | 0.009372 | 0.011527 | 0.009967 |
| 1.169238 | 3.35384 | 0.571511 | 0.036901 | 0.011046 | 0.011035 | 0.009688 |
| 1.158042 | 3.394641 | 0.571511 | 0.03695 | 0.011092 | 0.012604 | 0.011087 |
| 1.094929 | 3.202019 | 0.571511 | 0.036782 | 0.019281 | 0.021949 | 0.019884 |
| 1.018883 | 2.898516 | 0.571511 | 0.037892 | 0.043455 | 0.042346 | 0.040078 |
| 0.297164 | 0.885226 | 0.571511 | 0.037119 | 0.013892 | 0.016495 | 0.01467 |
| 0.34624 | 0.990879 | 0.571511 | 0.036096 | 0.045292 | 0.046139 | 0.043794 |
| 1.120159 | 3.345008 | 0.571395 | 0.03539 | 0.01582 | 0.017955 | 0.016199 |
| 0.306961 | 0.893137 | 0.571395 | 0.036666 | 0.015109 | 0.01756 | 0.015696 |
| 1.092035 | 3.176449 | 0.571395 | 0.03685 | 0.020185 | 0.022419 | 0.020407 |
| 1.175124 | 3.4782 | 0.571279 | 0.036081 | 0.009599 | 0.010994 | 0.009619 |
| 0.287505 | 0.850974 | 0.571279 | 0.036081 | 0.009599 | 0.010994 | 0.009619 |
| 0.335798 | 0.966653 | 0.571279 | 0.036141 | 0.033789 | 0.037006 | 0.034601 |
| 0.164859 | 0.899453 | 0.571279 | 0.022625 | 0.012667 | 0.027469 | 0.022197 |
| 0.32923 | 0.945566 | 0.571279 | 0.036347 | 0.02831 | 0.030164 | 0.028071 |
| 0.326389 | 0.940492 | 0.571163 | 0.037203 | 0.027742 | 0.028727 | 0.026708 |
| 0.29832 | 0.868657 | 0.571163 | 0.036796 | 0.011604 | 0.013272 | 0.011732 |
| 1.163447 | 3.416474 | 0.571047 | 0.038153 | 0.010549 | 0.012044 | 0.010561 |
| 0.292699 | 0.859515 | 0.571047 | 0.038153 | 0.010549 | 0.012044 | 0.010561 |
| 1.111784 | 3.143235 | 0.571047 | 0.037084 | 0.01778 | 0.018293 | 0.01658 |
| 1.0889 | 3.1594 | 0.570931 | 0.037541 | 0.020765 | 0.023003 | 0.020954 |
| 1.296411 | 3.960503 | 0.570931 | 0.037785 | 0.003046 | 0.00409 | 0.00325 |
| 0.317801 | 0.901238 | 0.570931 | 0.037686 | 0.01781 | 0.018722 | 0.016967 |
| 1.201572 | 3.556688 | 0.570931 | 0.036985 | 0.006948 | 0.008709 | 0.007384 |
| 1.427852 | 4.224971 | 0.570931 | 0.038007 | 0.000973 | 0.001167 | 0.000836 |
| 0.328231 | 0.976724 | 0.570931 | 0.036104 | 0.036419 | 0.040892 | 0.038345 |
| 0.319338 | 0.934342 | 0.570931 | 0.036967 | 0.02421 | 0.027249 | 0.025012 |

| | | | | | | |
|---|---|---|---|---|---|---|
| 0.319432 | 0.913424 | 0.570931 | 0.036937 | 0.019686 | 0.021572 | 0.019626 |
| 1.010723 | 2.939954 | 0.570931 | 0.036566 | 0.041326 | 0.045595 | 0.043007 |
| 0.308608 | 0.878247 | 0.570931 | 0.037169 | 0.014172 | 0.014422 | 0.01286 |
| 0.284717 | 0.851945 | 0.570815 | 0.037357 | 0.008884 | 0.011308 | 0.009733 |
| 0.290796 | 0.86033 | 0.570815 | 0.036319 | 0.010696 | 0.012293 | 0.010797 |
| 1.162344 | 3.438833 | 0.570815 | 0.036319 | 0.010696 | 0.012293 | 0.010797 |
| 0.229953 | 0.705741 | 0.570815 | 0.037977 | 0.000968 | 0.001482 | 0.001044 |
| 0.277652 | 0.806763 | 0.570699 | 0.037803 | 0.005291 | 0.005976 | 0.004976 |
| 1.081645 | 3.053202 | 0.570699 | 0.036274 | 0.023394 | 0.024044 | 0.022115 |
| 0.327525 | 0.924518 | 0.570699 | 0.036274 | 0.023394 | 0.024044 | 0.022115 |
| 1.211231 | 3.529281 | 0.570699 | 0.037453 | 0.006895 | 0.007759 | 0.006599 |
| 1.087392 | 3.125684 | 0.570699 | 0.036308 | 0.021814 | 0.023147 | 0.021194 |
| 0.31993 | 0.919632 | 0.570699 | 0.036308 | 0.021814 | 0.023147 | 0.021194 |
| 1.187975 | 3.429302 | 0.570699 | 0.037203 | 0.008714 | 0.009407 | 0.008142 |
| 0.322183 | 0.916253 | 0.570583 | 0.037537 | 0.020669 | 0.022138 | 0.020213 |
| 1.091402 | 3.103824 | 0.570583 | 0.037537 | 0.020669 | 0.022138 | 0.020213 |
| 1.127213 | 3.285992 | 0.570583 | 0.036666 | 0.014618 | 0.016454 | 0.014697 |
| 1.025766 | 2.987935 | 0.570583 | 0.036384 | 0.03605 | 0.040049 | 0.037554 |
| 0.295371 | 0.848625 | 0.570583 | 0.036534 | 0.010168 | 0.010181 | 0.008885 |
| 1.178376 | 3.385573 | 0.570583 | 0.036534 | 0.010168 | 0.010181 | 0.008885 |
| 0.311789 | 0.917824 | 0.570467 | 0.037279 | 0.019733 | 0.023129 | 0.02099 |
| 0.304408 | 0.878022 | 0.570467 | 0.037129 | 0.012826 | 0.014633 | 0.012961 |
| 1.06663 | 3.038803 | 0.570467 | 0.037389 | 0.027009 | 0.027702 | 0.025751 |
| 0.323978 | 0.929927 | 0.570351 | 0.035912 | 0.023614 | 0.025745 | 0.023682 |
| 1.075353 | 3.086627 | 0.570351 | 0.035912 | 0.023614 | 0.025745 | 0.023682 |
| 0.332398 | 0.96257 | 0.570351 | 0.036721 | 0.032735 | 0.035676 | 0.033277 |
| 0.308127 | 0.919996 | 0.570351 | 0.036505 | 0.02059 | 0.023897 | 0.02178 |
| 0.330038 | 0.931665 | 0.570351 | 0.036098 | 0.0262 | 0.025925 | 0.024013 |
| 1.012161 | 2.857569 | 0.570351 | 0.037097 | 0.045441 | 0.044897 | 0.042532 |
| 1.053575 | 3.102797 | 0.570351 | 0.036526 | 0.028254 | 0.031605 | 0.029292 |
| 0.316974 | 0.916465 | 0.570235 | 0.037094 | 0.020746 | 0.022487 | 0.020529 |
| 0.322536 | 0.915142 | 0.570235 | 0.03762 | 0.022054 | 0.021832 | 0.020052 |
| 1.092726 | 3.100428 | 0.570235 | 0.03762 | 0.022054 | 0.021832 | 0.020052 |
| 1.000162 | 2.844084 | 0.570235 | 0.03649 | 0.049672 | 0.049929 | 0.047512 |
| 0.316612 | 0.908409 | 0.570235 | 0.036602 | 0.018682 | 0.020492 | 0.018611 |
| 0.322761 | 0.933155 | 0.570235 | 0.037156 | 0.025215 | 0.026732 | 0.024701 |
| 1.071633 | 3.098266 | 0.570235 | 0.037156 | 0.025215 | 0.026732 | 0.024701 |
| 1.004582 | 2.869629 | 0.570235 | 0.036011 | 0.047797 | 0.048037 | 0.045677 |
| 0.335062 | 0.958793 | 0.570235 | 0.037148 | 0.032557 | 0.034272 | 0.032 |
| 1.012482 | 2.877753 | 0.570002 | 0.036047 | 0.042541 | 0.044802 | 0.042384 |
| 0.347493 | 0.987672 | 0.570002 | 0.036047 | 0.042541 | 0.044802 | 0.042384 |
| 1.053078 | 2.987857 | 0.570002 | 0.037276 | 0.029928 | 0.031212 | 0.029089 |
| 1.178647 | 3.452887 | 0.570002 | 0.037047 | 0.008761 | 0.010485 | 0.009062 |
| 0.289613 | 0.84843 | 0.570002 | 0.037047 | 0.008761 | 0.010485 | 0.009062 |
| 1.092324 | 3.108133 | 0.570002 | 0.037094 | 0.020481 | 0.021964 | 0.020043 |
| 0.321737 | 0.915479 | 0.570002 | 0.037094 | 0.020481 | 0.021964 | 0.020043 |
| 0.295842 | 0.851792 | 0.56977 | 0.036915 | 0.010266 | 0.010632 | 0.009377 |

| | | | | | | |
|---|---|---|---|---|---|---|
| 1.173996 | 3.380178 | 0.56977 | 0.036915 | 0.010266 | 0.010632 | 0.009377 |
| 0.33055 | 0.959016 | 0.56977 | 0.036793 | 0.031639 | 0.034518 | 0.032159 |
| 1.210854 | 3.4825 | 0.569654 | 0.036806 | 0.006882 | 0.007588 | 0.006473 |
| 0.325773 | 0.928815 | 0.569654 | 0.03737 | 0.024665 | 0.025335 | 0.023443 |
| 1.024927 | 2.892432 | 0.569654 | 0.03751 | 0.038551 | 0.040073 | 0.03774 |
| 0.34573 | 0.975679 | 0.569654 | 0.03751 | 0.038551 | 0.040073 | 0.03774 |
| 0.287379 | 0.880094 | 0.569654 | 0.036299 | 0.012703 | 0.016071 | 0.014157 |
| 0.267148 | 0.786905 | 0.569654 | 0.037816 | 0.003693 | 0.004662 | 0.00377 |
| 1.019464 | 2.873655 | 0.569654 | 0.035955 | 0.041097 | 0.042065 | 0.039751 |
| 0.317952 | 0.903371 | 0.569654 | 0.036646 | 0.018258 | 0.019209 | 0.017442 |
| 1.004471 | 2.840787 | 0.569654 | 0.036416 | 0.046325 | 0.048066 | 0.045638 |
| 0.322098 | 0.922587 | 0.569538 | 0.036106 | 0.022636 | 0.023815 | 0.021918 |
| 0.328395 | 0.943964 | 0.569538 | 0.035977 | 0.029106 | 0.029701 | 0.027664 |
| 0.344906 | 0.984777 | 0.569538 | 0.036629 | 0.042281 | 0.043666 | 0.041283 |
| 1.10313 | 3.183086 | 0.569538 | 0.036031 | 0.01996 | 0.020177 | 0.018376 |
| 0.322269 | 0.922036 | 0.569538 | 0.037701 | 0.022321 | 0.023657 | 0.021659 |
| 1.144841 | 3.329185 | 0.569422 | 0.037371 | 0.012374 | 0.014022 | 0.012378 |
| 0.267603 | 0.83593 | 0.569422 | 0.037098 | 0.007704 | 0.009975 | 0.008506 |
| 1.027338 | 2.926261 | 0.569422 | 0.037448 | 0.037448 | 0.039306 | 0.036904 |
| 1.213332 | 3.601873 | 0.569422 | 0.036975 | 0.006498 | 0.007893 | 0.006659 |
| 0.328953 | 0.960158 | 0.569422 | 0.036641 | 0.031278 | 0.034966 | 0.032586 |
| 1.041495 | 3.039951 | 0.569422 | 0.036641 | 0.031278 | 0.034966 | 0.032586 |
| 0.292789 | 0.837177 | 0.569422 | 0.037024 | 0.008525 | 0.008714 | 0.007502 |
| 1.04207 | 3.016861 | 0.569422 | 0.037198 | 0.032764 | 0.034693 | 0.032437 |
| 0.302607 | 0.866025 | 0.569306 | 0.037559 | 0.011223 | 0.012553 | 0.011024 |
| 0.29046 | 0.872149 | 0.569306 | 0.037737 | 0.011621 | 0.014379 | 0.012587 |
| 1.146593 | 3.442816 | 0.569306 | 0.037737 | 0.011621 | 0.014379 | 0.012587 |
| 0.278373 | 0.827704 | 0.56919 | 0.037606 | 0.006592 | 0.008286 | 0.007016 |
| 1.007441 | 2.868119 | 0.56919 | 0.037076 | 0.046051 | 0.04684 | 0.044454 |
| 0.348661 | 0.992614 | 0.56919 | 0.037076 | 0.046051 | 0.04684 | 0.044454 |
| 0.340437 | 0.960075 | 0.56919 | 0.036689 | 0.035267 | 0.034514 | 0.032405 |
| 0.276806 | 0.822101 | 0.56919 | 0.037804 | 0.006063 | 0.00769 | 0.00646 |
| 0.311968 | 0.949259 | 0.56919 | 0.036188 | 0.026777 | 0.032081 | 0.029569 |
| 1.19504 | 3.461 | 0.569074 | 0.037095 | 0.008441 | 0.008876 | 0.007692 |
| 0.341444 | 0.996284 | 0.569074 | 0.035673 | 0.046991 | 0.048428 | 0.046022 |
| 0.246853 | 0.754566 | 0.569074 | 0.03743 | 0.002263 | 0.003199 | 0.002462 |
| 0.286661 | 0.823938 | 0.569074 | 0.038572 | 0.006808 | 0.007385 | 0.006256 |
| 0.343059 | 0.991371 | 0.569074 | 0.036658 | 0.043672 | 0.046374 | 0.043905 |
| 0.306966 | 0.893401 | 0.568958 | 0.0366 | 0.015162 | 0.017617 | 0.015753 |
| 0.345343 | 0.983708 | 0.568958 | 0.036919 | 0.044384 | 0.043231 | 0.040975 |
| 0.29394 | 0.840382 | 0.568958 | 0.03762 | 0.008279 | 0.009084 | 0.007832 |
| 1.189934 | 3.402057 | 0.568958 | 0.03762 | 0.008279 | 0.009084 | 0.007832 |
| 0.291695 | 0.875977 | 0.568958 | 0.037336 | 0.012169 | 0.015016 | 0.013187 |
| 0.298588 | 0.86869 | 0.568841 | 0.036562 | 0.012276 | 0.013261 | 0.011762 |
| 0.334609 | 0.967292 | 0.568841 | 0.036935 | 0.034816 | 0.037273 | 0.034926 |
| 1.055681 | 2.967406 | 0.568841 | 0.037027 | 0.030094 | 0.03035 | 0.028267 |
| 0.336995 | 0.947255 | 0.568841 | 0.037027 | 0.030094 | 0.03035 | 0.028267 |

| | | | | | | |
|---|---|---|---|---|---|---|
| 0.318137 | 0.925853 | 0.568841 | 0.036879 | 0.022535 | 0.024918 | 0.022827 |
| 0.303663 | 0.884377 | 0.568841 | 0.037451 | 0.013634 | 0.015929 | 0.014151 |
| 0.338348 | 0.963031 | 0.568841 | 0.036965 | 0.034439 | 0.035629 | 0.033354 |
| 0.327043 | 0.937687 | 0.568609 | 0.037114 | 0.025637 | 0.027849 | 0.025728 |
| 0.319982 | 0.92074 | 0.568609 | 0.037053 | 0.021624 | 0.023437 | 0.021408 |
| 0.309314 | 0.917901 | 0.568609 | 0.036079 | 0.020361 | 0.023288 | 0.021209 |
| 0.232944 | 0.740967 | 0.568609 | 0.038233 | 0.001722 | 0.002925 | 0.002147 |
| 1.047219 | 3.036965 | 0.568609 | 0.035936 | 0.030131 | 0.033186 | 0.030907 |
| 0.264618 | 0.772214 | 0.568493 | 0.03744 | 0.003174 | 0.00366 | 0.002917 |
| 0.343621 | 0.999395 | 0.568493 | 0.036315 | 0.046231 | 0.049741 | 0.047173 |
| 0.302156 | 0.852936 | 0.568493 | 0.037104 | 0.009857 | 0.010442 | 0.009087 |
| 0.293502 | 0.866399 | 0.568493 | 0.037974 | 0.011553 | 0.013164 | 0.011636 |
| 0.301051 | 0.88759 | 0.568377 | 0.036827 | 0.013974 | 0.016744 | 0.014885 |
| 1.056345 | 3.006696 | 0.568377 | 0.036834 | 0.028561 | 0.030359 | 0.028206 |
| 0.332591 | 0.94666 | 0.568377 | 0.036834 | 0.028561 | 0.030359 | 0.028206 |
| 1.082282 | 3.152272 | 0.568377 | 0.036698 | 0.022072 | 0.024455 | 0.022376 |
| 0.317232 | 0.923973 | 0.568377 | 0.036698 | 0.022072 | 0.024455 | 0.022376 |
| 1.142274 | 3.291738 | 0.568377 | 0.037846 | 0.013228 | 0.014182 | 0.012614 |
| 0.282621 | 0.817206 | 0.568261 | 0.037177 | 0.005771 | 0.006826 | 0.00571 |
| 1.04207 | 2.936945 | 0.568261 | 0.037293 | 0.034022 | 0.034356 | 0.032199 |
| 0.314615 | 0.899395 | 0.568261 | 0.036634 | 0.016786 | 0.018489 | 0.016663 |
| 1.111859 | 3.178487 | 0.568261 | 0.036634 | 0.016786 | 0.018489 | 0.016663 |
| 0.343915 | 0.983412 | 0.568261 | 0.036639 | 0.041736 | 0.043138 | 0.040767 |
| 0.320295 | 0.928791 | 0.568261 | 0.037182 | 0.023205 | 0.025618 | 0.023481 |
| 0.305159 | 0.881451 | 0.568261 | 0.037328 | 0.014925 | 0.015254 | 0.01372 |
| 1.151477 | 3.314471 | 0.568261 | 0.037718 | 0.013355 | 0.013031 | 0.011604 |
| 0.325339 | 0.934563 | 0.568261 | 0.036906 | 0.026375 | 0.02701 | 0.024995 |
| 0.328085 | 0.947688 | 0.568261 | 0.036801 | 0.028734 | 0.030888 | 0.028678 |
| 0.258185 | 0.761214 | 0.568145 | 0.038061 | 0.00265 | 0.003187 | 0.002511 |
| 0.305816 | 0.910423 | 0.568145 | 0.037297 | 0.018461 | 0.021609 | 0.019541 |
| 0.33296 | 0.942993 | 0.568145 | 0.037337 | 0.030939 | 0.029185 | 0.027207 |
| 1.080807 | 3.111967 | 0.568145 | 0.037244 | 0.02204 | 0.024576 | 0.022479 |
| 0.300611 | 0.869219 | 0.568029 | 0.03728 | 0.012309 | 0.013234 | 0.011732 |
| 0.29435 | 0.855831 | 0.568029 | 0.037199 | 0.009945 | 0.01135 | 0.009872 |
| 1.004062 | 2.849188 | 0.568029 | 0.036788 | 0.04805 | 0.048245 | 0.045843 |
| 0.335199 | 0.970077 | 0.568029 | 0.037377 | 0.035196 | 0.038263 | 0.035809 |
| 1.030846 | 2.983304 | 0.568029 | 0.037377 | 0.035196 | 0.038263 | 0.035809 |
| 1.04139 | 2.978043 | 0.568029 | 0.03692 | 0.033606 | 0.034749 | 0.032535 |
| 0.335791 | 0.960255 | 0.568029 | 0.03692 | 0.033606 | 0.034749 | 0.032535 |
| 1.072956 | 3.078994 | 0.567913 | 0.036922 | 0.024827 | 0.026296 | 0.024231 |
| 0.324781 | 0.932005 | 0.567913 | 0.036922 | 0.024827 | 0.026296 | 0.024231 |
| 1.051765 | 2.983687 | 0.567913 | 0.037892 | 0.029739 | 0.031579 | 0.029372 |
| 0.302162 | 0.87756 | 0.567913 | 0.036356 | 0.014527 | 0.014681 | 0.013149 |
| 1.139523 | 3.30948 | 0.567913 | 0.036356 | 0.014527 | 0.014681 | 0.013149 |
| 0.33091 | 0.95682 | 0.567797 | 0.037263 | 0.030692 | 0.03376 | 0.031453 |
| 1.010802 | 2.861763 | 0.567681 | 0.037738 | 0.04318 | 0.045453 | 0.042993 |
| 0.342958 | 0.971004 | 0.567681 | 0.036256 | 0.036306 | 0.038378 | 0.036021 |

| | | | | | | |
|---|---|---|---|---|---|---|
| 1.001562 | 2.832728 | 0.567681 | 0.036104 | 0.047598 | 0.049316 | 0.046885 |
| 0.353017 | 0.99844 | 0.567681 | 0.036104 | 0.047598 | 0.049316 | 0.046885 |
| 0.317715 | 0.903889 | 0.567681 | 0.037569 | 0.018008 | 0.019345 | 0.017527 |
| 1.033198 | 2.929244 | 0.567681 | 0.037265 | 0.03524 | 0.037268 | 0.034943 |
| 1.31032 | 3.786673 | 0.567681 | 0.038067 | 0.002633 | 0.003093 | 0.00243 |
| 0.264084 | 0.763172 | 0.567681 | 0.038067 | 0.002633 | 0.003093 | 0.00243 |
| 1.037546 | 3.00278 | 0.567681 | 0.037218 | 0.033122 | 0.036089 | 0.033676 |
| 0.333025 | 0.963813 | 0.567681 | 0.037218 | 0.033122 | 0.036089 | 0.033676 |
| 0.306374 | 0.868401 | 0.567564 | 0.037197 | 0.012047 | 0.012744 | 0.011265 |
| 1.049613 | 3.003678 | 0.567564 | 0.037695 | 0.031236 | 0.032315 | 0.030158 |
| 0.332925 | 0.952732 | 0.567564 | 0.037695 | 0.031236 | 0.032315 | 0.030158 |
| 1.028879 | 2.957464 | 0.567448 | 0.036527 | 0.036168 | 0.038861 | 0.036526 |
| 0.319839 | 0.941446 | 0.567448 | 0.03713 | 0.026137 | 0.029328 | 0.027072 |
| 0.323309 | 0.922293 | 0.567448 | 0.037715 | 0.022615 | 0.023669 | 0.021786 |
| 1.046309 | 2.999179 | 0.567332 | 0.036707 | 0.030835 | 0.033298 | 0.031045 |
| 0.333425 | 0.95574 | 0.567332 | 0.036707 | 0.030835 | 0.033298 | 0.031045 |
| 0.33274 | 0.97499 | 0.567332 | 0.036847 | 0.036351 | 0.040139 | 0.037626 |
| 1.210604 | 3.583085 | 0.567332 | 0.037699 | 0.006705 | 0.00804 | 0.006818 |
| 1.265624 | 3.60772 | 0.567332 | 0.038118 | 0.004191 | 0.00449 | 0.003674 |
| 1.151293 | 3.340499 | 0.567216 | 0.037223 | 0.011937 | 0.013197 | 0.011654 |
| 0.321985 | 0.924691 | 0.567216 | 0.037972 | 0.024974 | 0.024392 | 0.02262 |
| 1.012046 | 2.86974 | 0.567216 | 0.036896 | 0.04425 | 0.044963 | 0.042575 |
| 0.298188 | 0.867949 | 0.567216 | 0.036559 | 0.011474 | 0.013154 | 0.011612 |
| 1.152141 | 3.35359 | 0.567216 | 0.036559 | 0.011474 | 0.013154 | 0.011612 |
| 0.33324 | 0.960189 | 0.5671 | 0.036806 | 0.031714 | 0.03482 | 0.032466 |
| 0.324143 | 0.935498 | 0.5671 | 0.037465 | 0.024654 | 0.027345 | 0.025184 |
| 1.118926 | 3.223545 | 0.5671 | 0.037443 | 0.01545 | 0.017489 | 0.015669 |
| 0.308287 | 0.8883 | 0.5671 | 0.038047 | 0.015523 | 0.016454 | 0.014853 |
| 1.071567 | 3.148917 | 0.5671 | 0.03695 | 0.023965 | 0.027013 | 0.024807 |
| 1.119304 | 3.27114 | 0.5671 | 0.036547 | 0.01525 | 0.017696 | 0.015856 |
| 0.305704 | 0.893413 | 0.5671 | 0.036547 | 0.01525 | 0.017696 | 0.015856 |
| 0.343554 | 0.982481 | 0.566984 | 0.037317 | 0.040836 | 0.042775 | 0.040357 |
| 0.330215 | 0.939154 | 0.566984 | 0.036834 | 0.027415 | 0.028136 | 0.026157 |
| 0.333351 | 0.963797 | 0.566984 | 0.037566 | 0.033105 | 0.036072 | 0.033651 |
| 0.289292 | 0.836454 | 0.566868 | 0.037452 | 0.008096 | 0.008811 | 0.00757 |
| 0.305037 | 0.88269 | 0.566868 | 0.037515 | 0.013631 | 0.015507 | 0.01381 |
| 0.338078 | 0.997977 | 0.566868 | 0.036355 | 0.045345 | 0.049149 | 0.046617 |
| 0.336561 | 0.953975 | 0.566752 | 0.036845 | 0.031232 | 0.032578 | 0.030414 |
| 1.048246 | 2.971233 | 0.566752 | 0.036845 | 0.031232 | 0.032578 | 0.030414 |
| 1.119094 | 3.185778 | 0.566752 | 0.037822 | 0.017318 | 0.017239 | 0.015575 |
| 1.033936 | 2.967554 | 0.566752 | 0.036722 | 0.037847 | 0.037157 | 0.035138 |
| 1.017398 | 2.867157 | 0.566752 | 0.036338 | 0.042471 | 0.042845 | 0.040566 |
| 0.348778 | 0.9829 | 0.566752 | 0.036338 | 0.042471 | 0.042845 | 0.040566 |
| 0.327215 | 0.936081 | 0.566752 | 0.036963 | 0.025862 | 0.027363 | 0.025248 |
| 0.348147 | 0.982386 | 0.566752 | 0.036597 | 0.041667 | 0.042651 | 0.040338 |
| 0.311461 | 0.918982 | 0.566636 | 0.037383 | 0.020039 | 0.023448 | 0.021315 |
| 0.302139 | 0.875837 | 0.566636 | 0.037603 | 0.012573 | 0.014355 | 0.012727 |

| | | | | | | |
|---|---|---|---|---|---|---|
| 1.082428 | 3.215066 | 0.566636 | 0.036457 | 0.021368 | 0.024757 | 0.022572 |
| 0.311036 | 0.923849 | 0.566636 | 0.036457 | 0.021368 | 0.024757 | 0.022572 |
| 0.284821 | 0.812859 | 0.56652 | 0.037537 | 0.005854 | 0.006248 | 0.005262 |
| 0.287866 | 0.854146 | 0.56652 | 0.036907 | 0.009298 | 0.011466 | 0.009938 |
| 1.06776 | 3.096903 | 0.56652 | 0.037338 | 0.025626 | 0.027714 | 0.025593 |
| 0.295598 | 0.838796 | 0.56652 | 0.037979 | 0.008507 | 0.008776 | 0.007592 |
| 0.323992 | 0.925608 | 0.56652 | 0.036913 | 0.023163 | 0.024536 | 0.022499 |
| 0.310202 | 0.918415 | 0.56652 | 0.038085 | 0.020111 | 0.023371 | 0.021212 |
| 1.039705 | 2.978608 | 0.56652 | 0.037068 | 0.036135 | 0.035293 | 0.033116 |
| 0.328823 | 0.939487 | 0.56652 | 0.036757 | 0.026069 | 0.028304 | 0.026143 |
| 1.064411 | 3.041149 | 0.56652 | 0.036757 | 0.026069 | 0.028304 | 0.026143 |
| 0.282379 | 0.819064 | 0.566404 | 0.038008 | 0.00597 | 0.007046 | 0.005923 |
| 0.33972 | 0.984337 | 0.566404 | 0.036813 | 0.040292 | 0.043577 | 0.04105 |
| 0.277365 | 0.79806 | 0.566404 | 0.037951 | 0.004483 | 0.005164 | 0.004243 |
| 0.319959 | 0.904732 | 0.566404 | 0.037572 | 0.018471 | 0.019412 | 0.01761 |
| 1.078713 | 3.082574 | 0.566287 | 0.037934 | 0.023968 | 0.024908 | 0.022916 |
| 0.352153 | 0.99542 | 0.566287 | 0.037693 | 0.04632 | 0.04801 | 0.045589 |
| 0.291039 | 0.859507 | 0.566287 | 0.037851 | 0.009914 | 0.012142 | 0.010555 |
| 0.3029 | 0.89331 | 0.566287 | 0.037205 | 0.015475 | 0.017842 | 0.015991 |
| 0.310832 | 0.920151 | 0.566287 | 0.037491 | 0.020487 | 0.023788 | 0.021613 |
| 0.303122 | 0.890972 | 0.566287 | 0.037092 | 0.015022 | 0.017328 | 0.015484 |
| 0.333175 | 0.950764 | 0.566287 | 0.037452 | 0.029265 | 0.031658 | 0.029404 |
| 1.110921 | 3.209155 | 0.566171 | 0.037202 | 0.017261 | 0.018841 | 0.017007 |
| 0.350202 | 0.997518 | 0.566171 | 0.036842 | 0.046581 | 0.048922 | 0.04648 |
| 0.337565 | 0.983642 | 0.566055 | 0.037467 | 0.03919 | 0.04334 | 0.040799 |
| 1.01663 | 2.96239 | 0.566055 | 0.037467 | 0.03919 | 0.04334 | 0.040799 |
| 1.042158 | 2.970717 | 0.566055 | 0.037048 | 0.032526 | 0.034474 | 0.032243 |
| 0.336619 | 0.959548 | 0.566055 | 0.037048 | 0.032526 | 0.034474 | 0.032243 |
| 0.334141 | 0.956471 | 0.566055 | 0.036644 | 0.03105 | 0.033516 | 0.031248 |
| 0.341294 | 0.972484 | 0.566055 | 0.037296 | 0.038263 | 0.038978 | 0.036694 |
| 0.329295 | 0.945184 | 0.565939 | 0.037342 | 0.028908 | 0.030041 | 0.027932 |
| 0.343554 | 0.97217 | 0.565939 | 0.037272 | 0.038707 | 0.038798 | 0.036504 |
| 0.34703 | 0.992218 | 0.565939 | 0.037702 | 0.043724 | 0.046689 | 0.0442 |
| 0.317401 | 0.91979 | 0.565939 | 0.037228 | 0.021044 | 0.023329 | 0.02126 |
| 0.30411 | 0.906708 | 0.565939 | 0.037292 | 0.017767 | 0.020812 | 0.018796 |
| 1.102891 | 3.288283 | 0.565939 | 0.037292 | 0.017767 | 0.020812 | 0.018796 |
| 1.008621 | 3.097309 | 0.565823 | 0.034616 | 0.055678 | 0.046596 | 0.043786 |
| 0.348588 | 0.991521 | 0.565823 | 0.036908 | 0.045016 | 0.046383 | 0.043948 |
| 1.008552 | 2.868712 | 0.565823 | 0.036908 | 0.045016 | 0.046383 | 0.043948 |
| 1.146544 | 3.256595 | 0.565823 | 0.036903 | 0.014186 | 0.013381 | 0.011935 |
| 1.287049 | 3.759267 | 0.565823 | 0.038589 | 0.003438 | 0.003941 | 0.003173 |
| 0.288644 | 0.859202 | 0.565707 | 0.03719 | 0.011354 | 0.012235 | 0.010683 |
| 1.163871 | 3.464478 | 0.565707 | 0.03719 | 0.011354 | 0.012235 | 0.010683 |
| 0.271404 | 0.799982 | 0.565707 | 0.03728 | 0.004577 | 0.005619 | 0.004603 |
| 1.101649 | 3.189697 | 0.565707 | 0.036665 | 0.018631 | 0.020508 | 0.018664 |
| 1.095792 | 3.167666 | 0.565591 | 0.037518 | 0.01925 | 0.021575 | 0.019611 |
| 1.014266 | 2.86243 | 0.565591 | 0.037282 | 0.04336 | 0.044064 | 0.041644 |

| | | | | | | |
|---|---|---|---|---|---|---|
| 0.332639 | 0.949231 | 0.565591 | 0.0374 | 0.02957 | 0.031182 | 0.028956 |
| 1.008279 | 2.835659 | 0.565591 | 0.037248 | 0.044795 | 0.046457 | 0.044007 |
| 1.155242 | 3.332284 | 0.565591 | 0.037201 | 0.012017 | 0.012633 | 0.011172 |
| 1.03404 | 2.945503 | 0.565475 | 0.036904 | 0.035518 | 0.037041 | 0.034817 |
| 0.328811 | 0.963494 | 0.565475 | 0.037066 | 0.032561 | 0.036121 | 0.033683 |
| 1.10251 | 3.118449 | 0.565475 | 0.036697 | 0.020169 | 0.019918 | 0.018199 |
| 0.261814 | 0.780522 | 0.565475 | 0.037846 | 0.003522 | 0.004383 | 0.003539 |
| 1.281195 | 3.819511 | 0.565475 | 0.037846 | 0.003522 | 0.004383 | 0.003539 |
| 1.286107 | 3.874131 | 0.565475 | 0.038196 | 0.003478 | 0.004311 | 0.003457 |
| 0.258122 | 0.77754 | 0.565475 | 0.038196 | 0.003478 | 0.004311 | 0.003457 |
| 0.291108 | 0.85807 | 0.565475 | 0.03735 | 0.010385 | 0.011902 | 0.010429 |
| 1.165406 | 3.435156 | 0.565475 | 0.03735 | 0.010385 | 0.011902 | 0.010429 |
| 1.100123 | 3.158983 | 0.565359 | 0.036766 | 0.020144 | 0.020637 | 0.018825 |
| 0.316558 | 0.908989 | 0.565359 | 0.036766 | 0.020144 | 0.020637 | 0.018825 |
| 0.298597 | 0.876634 | 0.565359 | 0.037236 | 0.01267 | 0.01472 | 0.013009 |
| 1.140726 | 3.348991 | 0.565359 | 0.037236 | 0.01267 | 0.01472 | 0.013009 |
| 0.30517 | 0.893635 | 0.565359 | 0.0375 | 0.01545 | 0.017776 | 0.015895 |
| 1.119025 | 3.276864 | 0.565359 | 0.0375 | 0.01545 | 0.017776 | 0.015895 |
| 0.33036 | 0.966805 | 0.565359 | 0.037421 | 0.033594 | 0.037233 | 0.034761 |
| 0.317169 | 0.954033 | 0.565359 | 0.035998 | 0.028741 | 0.033382 | 0.03089 |
| 1.021907 | 2.879753 | 0.565243 | 0.037121 | 0.039615 | 0.041156 | 0.038795 |
| 0.347252 | 0.978562 | 0.565243 | 0.037121 | 0.039615 | 0.041156 | 0.038795 |
| 0.347967 | 0.995968 | 0.565243 | 0.036864 | 0.045251 | 0.048265 | 0.045769 |
| 0.330103 | 0.941048 | 0.565127 | 0.037409 | 0.027668 | 0.028719 | 0.0266 |
| 0.315274 | 0.917729 | 0.565127 | 0.037443 | 0.020017 | 0.02291 | 0.020832 |
| 0.331032 | 0.946578 | 0.565127 | 0.03782 | 0.028078 | 0.030403 | 0.028196 |
| 0.304017 | 0.893561 | 0.565127 | 0.03699 | 0.015929 | 0.017829 | 0.016067 |
| 1.060586 | 3.047731 | 0.565127 | 0.038006 | 0.027832 | 0.029376 | 0.027249 |
| 0.293381 | 0.845178 | 0.565127 | 0.037926 | 0.008684 | 0.009791 | 0.008476 |
| 0.3186 | 0.929445 | 0.565127 | 0.036814 | 0.023414 | 0.025891 | 0.023776 |
| 1.075911 | 3.13873 | 0.565127 | 0.036814 | 0.023414 | 0.025891 | 0.023776 |
| 0.347499 | 0.987267 | 0.565127 | 0.03737 | 0.042993 | 0.044635 | 0.042295 |
| 0.312738 | 0.94032 | 0.565127 | 0.037476 | 0.025021 | 0.029326 | 0.026934 |
| 0.336179 | 0.98026 | 0.565127 | 0.036826 | 0.03796 | 0.042053 | 0.039536 |
| 0.320773 | 0.917455 | 0.56501 | 0.037237 | 0.020588 | 0.022529 | 0.020551 |
| 1.089972 | 3.117474 | 0.56501 | 0.037237 | 0.020588 | 0.022529 | 0.020551 |
| 1.151546 | 3.402812 | 0.56501 | 0.036817 | 0.011789 | 0.013494 | 0.011922 |
| 0.338195 | 0.963455 | 0.56501 | 0.036605 | 0.035608 | 0.035785 | 0.033562 |
| 0.335214 | 0.974059 | 0.56501 | 0.037132 | 0.03662 | 0.039729 | 0.037288 |
| 0.263268 | 0.887956 | 0.56501 | 0.033584 | 0.013405 | 0.019123 | 0.016771 |
| 0.335502 | 0.947069 | 0.564894 | 0.037821 | 0.029079 | 0.030358 | 0.028212 |
| 0.335339 | 0.941662 | 0.564894 | 0.03667 | 0.02794 | 0.028657 | 0.026569 |
| 0.286096 | 0.859672 | 0.564894 | 0.037408 | 0.009952 | 0.012466 | 0.010796 |
| 1.00133 | 2.960691 | 0.564894 | 0.036511 | 0.044489 | 0.049441 | 0.046716 |
| 0.27371 | 0.799846 | 0.564894 | 0.038218 | 0.004668 | 0.005497 | 0.00452 |
| 0.35142 | 0.986854 | 0.564778 | 0.03731 | 0.04343 | 0.04441 | 0.042039 |
| 1.189274 | 3.503181 | 0.564778 | 0.038099 | 0.008084 | 0.009627 | 0.008251 |

| | | | | | | |
|---|---|---|---|---|---|---|
| 0.285455 | 0.840849 | 0.564778 | 0.038099 | 0.008084 | 0.009627 | 0.008251 |
| 0.340341 | 0.957509 | 0.564778 | 0.036823 | 0.032799 | 0.033627 | 0.03144 |
| 1.043649 | 3.036148 | 0.564778 | 0.036423 | 0.030608 | 0.034277 | 0.03188 |
| 1.058984 | 3.04516 | 0.564778 | 0.037975 | 0.029212 | 0.029807 | 0.027771 |
| 0.286058 | 0.847533 | 0.564662 | 0.037919 | 0.008532 | 0.010558 | 0.009094 |
| 0.339237 | 0.974412 | 0.564662 | 0.036376 | 0.036992 | 0.039757 | 0.037391 |
| 0.344123 | 0.991443 | 0.564662 | 0.03662 | 0.042775 | 0.046393 | 0.043854 |
| 0.310225 | 0.905444 | 0.564546 | 0.036938 | 0.017477 | 0.020154 | 0.018198 |
| 0.288793 | 0.997816 | 0.564546 | 0.032275 | 0.038257 | 0.049197 | 0.04578 |
| 1.067949 | 3.056381 | 0.564546 | 0.037428 | 0.027917 | 0.027451 | 0.0255 |
| 0.342247 | 0.98649 | 0.564546 | 0.037079 | 0.041733 | 0.044396 | 0.041925 |
| 1.039543 | 3.004373 | 0.564546 | 0.037078 | 0.032508 | 0.035448 | 0.033036 |
| 0.318611 | 0.928364 | 0.56443 | 0.037598 | 0.022488 | 0.025588 | 0.023428 |
| 0.349171 | 0.995073 | 0.56443 | 0.036904 | 0.044865 | 0.047878 | 0.045345 |
| 0.306905 | 0.8817 | 0.56443 | 0.037326 | 0.014715 | 0.015198 | 0.0137 |
| 1.050556 | 2.984757 | 0.564314 | 0.037781 | 0.031103 | 0.031944 | 0.029859 |
| 0.34556 | 0.991041 | 0.564314 | 0.037407 | 0.043508 | 0.046213 | 0.043677 |
| 1.002238 | 2.886647 | 0.564198 | 0.036708 | 0.045308 | 0.04904 | 0.046466 |
| 0.346423 | 0.997767 | 0.564198 | 0.036708 | 0.045308 | 0.04904 | 0.046466 |
| 0.284752 | 0.847993 | 0.564198 | 0.037797 | 0.008739 | 0.010703 | 0.009215 |
| 0.30719 | 0.893516 | 0.564198 | 0.036826 | 0.015173 | 0.017628 | 0.01576 |
| 1.02955 | 2.970762 | 0.564198 | 0.037521 | 0.036707 | 0.038669 | 0.036352 |
| 0.34438 | 0.982129 | 0.564198 | 0.037322 | 0.040673 | 0.04262 | 0.040178 |
| 1.018196 | 2.903768 | 0.564198 | 0.037322 | 0.040673 | 0.04262 | 0.040178 |
| 0.285003 | 0.823532 | 0.564198 | 0.038284 | 0.006538 | 0.007424 | 0.006308 |
| 0.31744 | 0.907209 | 0.564198 | 0.037122 | 0.018764 | 0.020153 | 0.018329 |
| 1.102281 | 3.150203 | 0.564198 | 0.037122 | 0.018764 | 0.020153 | 0.018329 |
| 0.327516 | 0.936858 | 0.564082 | 0.037388 | 0.02537 | 0.027578 | 0.025444 |
| 0.330377 | 0.972083 | 0.564082 | 0.03718 | 0.034538 | 0.039131 | 0.036576 |
| 0.314319 | 0.922859 | 0.564082 | 0.03669 | 0.021483 | 0.024314 | 0.022195 |
| 0.327949 | 0.94473 | 0.564082 | 0.037262 | 0.027107 | 0.029961 | 0.027714 |
| 0.310126 | 0.891837 | 0.564082 | 0.03724 | 0.016297 | 0.01709 | 0.015385 |
| 1.260478 | 3.816853 | 0.563966 | 0.037987 | 0.00406 | 0.005453 | 0.004413 |
| 0.261996 | 0.79335 | 0.563966 | 0.037987 | 0.00406 | 0.005453 | 0.004413 |
| 0.335047 | 0.948111 | 0.563966 | 0.037038 | 0.0289 | 0.030714 | 0.028506 |
| 1.239504 | 3.71889 | 0.563966 | 0.038065 | 0.004799 | 0.006411 | 0.005266 |
| 0.294985 | 0.887834 | 0.563966 | 0.037814 | 0.014064 | 0.017163 | 0.015235 |
| 0.325669 | 0.93072 | 0.563966 | 0.037399 | 0.02468 | 0.025882 | 0.023928 |
| 1.03767 | 2.965904 | 0.563966 | 0.037086 | 0.033871 | 0.035907 | 0.033662 |
| 0.325987 | 0.954588 | 0.56385 | 0.037187 | 0.029808 | 0.033211 | 0.030828 |
| 0.301152 | 0.87889 | 0.56385 | 0.037066 | 0.013613 | 0.014997 | 0.013389 |
| 1.085899 | 3.112964 | 0.563733 | 0.036946 | 0.023355 | 0.023409 | 0.021537 |
| 0.321237 | 0.920896 | 0.563733 | 0.036946 | 0.023355 | 0.023409 | 0.021537 |
| 0.318404 | 0.927161 | 0.563733 | 0.037471 | 0.022192 | 0.025265 | 0.02311 |
| 1.082151 | 3.114971 | 0.563733 | 0.036774 | 0.021764 | 0.024279 | 0.02219 |
| 0.3472 | 0.996071 | 0.563733 | 0.037448 | 0.04622 | 0.048313 | 0.045851 |
| 1.003944 | 2.880187 | 0.563733 | 0.037448 | 0.04622 | 0.048313 | 0.045851 |

| | | | | | | |
|---|---|---|---|---|---|---|
| 0.246612 | 0.790557 | 0.563733 | 0.037354 | 0.003878 | 0.005945 | 0.004764 |
| 0.341132 | 0.994919 | 0.563617 | 0.03671 | 0.044357 | 0.047859 | 0.045321 |
| 0.211615 | 0.708774 | 0.563617 | 0.036874 | 0.000989 | 0.002096 | 0.001437 |
| 0.313917 | 0.920184 | 0.563617 | 0.037488 | 0.02154 | 0.023626 | 0.02162 |
| 1.086739 | 3.185553 | 0.563617 | 0.037488 | 0.02154 | 0.023626 | 0.02162 |
| 0.307674 | 0.892 | 0.563617 | 0.037655 | 0.016405 | 0.017272 | 0.015587 |
| 0.316708 | 0.903628 | 0.563501 | 0.036914 | 0.018999 | 0.019343 | 0.01758 |
| 1.10665 | 3.157485 | 0.563501 | 0.036914 | 0.018999 | 0.019343 | 0.01758 |
| 1.051016 | 3.008606 | 0.563501 | 0.037007 | 0.030278 | 0.031919 | 0.029699 |
| 0.262636 | 0.80254 | 0.563385 | 0.03745 | 0.00484 | 0.006297 | 0.005206 |
| 0.341198 | 0.960255 | 0.563385 | 0.036835 | 0.034803 | 0.034549 | 0.032419 |
| 1.038808 | 2.97151 | 0.563385 | 0.036608 | 0.034972 | 0.035556 | 0.033386 |
| 0.336529 | 0.962642 | 0.563385 | 0.036608 | 0.034972 | 0.035556 | 0.033386 |
| 0.308073 | 0.909605 | 0.563385 | 0.037131 | 0.018606 | 0.021279 | 0.019286 |
| 1.029844 | 2.928342 | 0.563385 | 0.037227 | 0.038363 | 0.038427 | 0.03619 |
| 0.34149 | 0.971021 | 0.563385 | 0.037227 | 0.038363 | 0.038427 | 0.03619 |
| 1.117104 | 3.164402 | 0.563385 | 0.037312 | 0.016591 | 0.01746 | 0.01575 |
| 0.316015 | 0.895172 | 0.563385 | 0.037312 | 0.016591 | 0.01746 | 0.01575 |
| 0.329614 | 0.946237 | 0.563269 | 0.037802 | 0.028029 | 0.030359 | 0.028184 |
| 1.056818 | 3.033852 | 0.563269 | 0.037802 | 0.028029 | 0.030359 | 0.028184 |
| 0.334795 | 0.999842 | 0.563269 | 0.036178 | 0.044002 | 0.049934 | 0.047164 |
| 1.081903 | 3.126113 | 0.563153 | 0.037234 | 0.021877 | 0.024399 | 0.022333 |
| 0.319886 | 0.924298 | 0.563153 | 0.037234 | 0.021877 | 0.024399 | 0.022333 |
| 0.340507 | 0.993765 | 0.563153 | 0.036411 | 0.043049 | 0.047384 | 0.044805 |
| 0.282953 | 0.864499 | 0.563153 | 0.037431 | 0.010474 | 0.013474 | 0.011698 |
| 1.129655 | 3.230146 | 0.563153 | 0.037441 | 0.01421 | 0.015743 | 0.014042 |
| 0.3249 | 0.925204 | 0.563153 | 0.037071 | 0.023225 | 0.024375 | 0.02244 |
| 1.066869 | 3.037395 | 0.563153 | 0.037311 | 0.027163 | 0.027632 | 0.025595 |
| 0.325101 | 0.932468 | 0.563153 | 0.037724 | 0.025583 | 0.026413 | 0.024495 |
| 0.322427 | 0.954751 | 0.563037 | 0.037453 | 0.029373 | 0.033406 | 0.030958 |
| 0.336846 | 0.974358 | 0.563037 | 0.037523 | 0.036672 | 0.039799 | 0.037315 |
| 1.026317 | 2.968718 | 0.563037 | 0.037523 | 0.036672 | 0.039799 | 0.037315 |
| 1.04206 | 2.961648 | 0.563037 | 0.037554 | 0.033323 | 0.034466 | 0.032206 |
| 1.067573 | 3.013197 | 0.563037 | 0.03769 | 0.027532 | 0.027315 | 0.025349 |
| 0.290197 | 0.847268 | 0.562921 | 0.038251 | 0.009451 | 0.010278 | 0.008973 |
| 0.324417 | 0.933184 | 0.562921 | 0.037426 | 0.025566 | 0.026656 | 0.024624 |
| 0.32378 | 0.921158 | 0.562921 | 0.03726 | 0.021834 | 0.023338 | 0.021379 |
| 1.081185 | 3.117079 | 0.562921 | 0.036271 | 0.027122 | 0.024516 | 0.022446 |
| 1.050938 | 2.984682 | 0.562805 | 0.037554 | 0.029976 | 0.03183 | 0.029625 |
| 0.331328 | 0.951005 | 0.562689 | 0.036883 | 0.030145 | 0.031815 | 0.029611 |
| 0.300517 | 0.87597 | 0.562689 | 0.037125 | 0.012328 | 0.014478 | 0.012783 |
| 1.107437 | 3.135518 | 0.562689 | 0.037324 | 0.018473 | 0.019061 | 0.017309 |
| 1.014637 | 2.893935 | 0.562689 | 0.03742 | 0.042 | 0.043977 | 0.041518 |
| 0.30204 | 0.908758 | 0.562689 | 0.038076 | 0.017879 | 0.021423 | 0.019304 |
| 1.100403 | 3.310824 | 0.562689 | 0.038076 | 0.017879 | 0.021423 | 0.019304 |
| 1.019651 | 2.892794 | 0.562689 | 0.03712 | 0.039849 | 0.042041 | 0.039642 |
| 0.345687 | 0.980728 | 0.562689 | 0.03712 | 0.039849 | 0.042041 | 0.039642 |

| | | | | | | |
|---|---|---|---|---|---|---|
| 0.264028 | 0.800344 | 0.562573 | 0.037965 | 0.004541 | 0.006011 | 0.004922 |
| 0.327996 | 0.932099 | 0.562573 | 0.037435 | 0.024519 | 0.026157 | 0.024101 |
| 1.017266 | 2.864378 | 0.562573 | 0.036395 | 0.041905 | 0.042889 | 0.040557 |
| 0.349116 | 0.983027 | 0.562573 | 0.036395 | 0.041905 | 0.042889 | 0.040557 |
| 0.272066 | 0.954034 | 0.562456 | 0.031567 | 0.026274 | 0.035119 | 0.03209 |
| 0.293384 | 0.873988 | 0.562456 | 0.037997 | 0.012136 | 0.01454 | 0.012816 |
| 0.304395 | 0.87144 | 0.562456 | 0.036836 | 0.012648 | 0.013406 | 0.011934 |
| 1.147525 | 3.28521 | 0.562456 | 0.036836 | 0.012648 | 0.013406 | 0.011934 |
| 1.031386 | 2.975985 | 0.562456 | 0.037507 | 0.034787 | 0.038053 | 0.035649 |
| 0.315297 | 0.913831 | 0.562456 | 0.037545 | 0.020651 | 0.021911 | 0.020064 |
| 1.027553 | 2.898809 | 0.562456 | 0.037075 | 0.038788 | 0.039141 | 0.036922 |
| 1.098684 | 3.204519 | 0.562224 | 0.037404 | 0.01905 | 0.021189 | 0.019244 |
| 1.029403 | 2.940356 | 0.562224 | 0.03817 | 0.035949 | 0.038621 | 0.036239 |
| 0.351355 | 0.98866 | 0.562224 | 0.037507 | 0.043525 | 0.045158 | 0.04273 |
| 0.348408 | 0.994066 | 0.562224 | 0.037325 | 0.044457 | 0.047455 | 0.044938 |
| 1.005969 | 2.870198 | 0.562224 | 0.037325 | 0.044457 | 0.047455 | 0.044938 |
| 0.339841 | 0.986262 | 0.562108 | 0.037465 | 0.041036 | 0.044341 | 0.041819 |
| 1.013929 | 2.942549 | 0.562108 | 0.037465 | 0.041036 | 0.044341 | 0.041819 |
| 0.305077 | 0.891982 | 0.562108 | 0.037227 | 0.014993 | 0.017425 | 0.0156 |
| 0.241333 | 0.717965 | 0.562108 | 0.038761 | 0.001302 | 0.001626 | 0.001222 |
| 1.010133 | 2.890619 | 0.562108 | 0.037414 | 0.043756 | 0.045765 | 0.043315 |
| 0.235704 | 0.730529 | 0.561992 | 0.03844 | 0.001381 | 0.002304 | 0.001662 |
| 0.278584 | 0.813801 | 0.561992 | 0.038128 | 0.005414 | 0.00666 | 0.005538 |
| 1.135099 | 3.278331 | 0.561992 | 0.037063 | 0.013868 | 0.015169 | 0.013551 |
| 0.328395 | 0.985715 | 0.561992 | 0.037783 | 0.038912 | 0.044296 | 0.041583 |
| 1.014492 | 3.045111 | 0.561992 | 0.037783 | 0.038912 | 0.044296 | 0.041583 |
| 1.26354 | 3.698242 | 0.561992 | 0.038018 | 0.003907 | 0.004896 | 0.003964 |
| 0.328091 | 0.984318 | 0.561876 | 0.037308 | 0.038406 | 0.043762 | 0.041052 |
| 1.015932 | 3.047932 | 0.561876 | 0.037308 | 0.038406 | 0.043762 | 0.041052 |
| 0.338638 | 0.985934 | 0.561876 | 0.036222 | 0.040591 | 0.044228 | 0.041776 |
| 0.313279 | 0.905993 | 0.56176 | 0.037899 | 0.018472 | 0.020105 | 0.018232 |
| 1.026155 | 3.041306 | 0.56176 | 0.037028 | 0.03611 | 0.040061 | 0.037581 |
| 0.309796 | 0.911164 | 0.561644 | 0.036829 | 0.018945 | 0.021562 | 0.019554 |
| 1.017803 | 2.881955 | 0.561644 | 0.036722 | 0.041117 | 0.042721 | 0.040372 |
| 0.346987 | 0.982508 | 0.561644 | 0.036722 | 0.041117 | 0.042721 | 0.040372 |
| 0.273303 | 0.841268 | 0.561644 | 0.037672 | 0.007853 | 0.01039 | 0.008857 |
| 1.026738 | 2.876612 | 0.561644 | 0.03749 | 0.039033 | 0.039364 | 0.037087 |
| 0.343461 | 0.97192 | 0.561528 | 0.037681 | 0.038657 | 0.038706 | 0.036427 |
| 1.191051 | 3.425025 | 0.561528 | 0.037362 | 0.008632 | 0.009086 | 0.007846 |
| 1.002169 | 2.839769 | 0.561528 | 0.037672 | 0.046678 | 0.049055 | 0.04656 |
| 0.352141 | 0.997836 | 0.561528 | 0.037672 | 0.046678 | 0.049055 | 0.04656 |
| 0.341673 | 0.988429 | 0.561412 | 0.037664 | 0.04184 | 0.045187 | 0.042624 |
| 0.314544 | 0.926192 | 0.561412 | 0.036956 | 0.022291 | 0.025204 | 0.023069 |
| 0.283284 | 0.813631 | 0.561412 | 0.037988 | 0.005799 | 0.006406 | 0.005394 |
| 0.31189 | 0.913949 | 0.561412 | 0.037328 | 0.019467 | 0.022135 | 0.020079 |
| 1.094153 | 3.206262 | 0.561412 | 0.037328 | 0.019467 | 0.022135 | 0.020079 |
| 1.020865 | 2.966263 | 0.561412 | 0.037149 | 0.038564 | 0.041765 | 0.03929 |

| | | | | | | |
|---|---|---|---|---|---|---|
| 0.337125 | 0.979562 | 0.561412 | 0.037149 | 0.038564 | 0.041765 | 0.03929 |
| 1.07455 | 3.122201 | 0.561412 | 0.037002 | 0.024105 | 0.026134 | 0.024066 |
| 1.007504 | 2.904549 | 0.561412 | 0.037352 | 0.044116 | 0.046851 | 0.044352 |
| 0.316953 | 0.941571 | 0.561296 | 0.037757 | 0.025778 | 0.029502 | 0.027179 |
| 0.341695 | 0.981313 | 0.561179 | 0.036284 | 0.039938 | 0.042351 | 0.039982 |
| 0.338724 | 0.98411 | 0.561063 | 0.0369 | 0.040239 | 0.043504 | 0.041004 |
| 1.370677 | 4.194163 | 0.561063 | 0.038694 | 0.001618 | 0.002176 | 0.001603 |
| 0.238427 | 0.729567 | 0.561063 | 0.038694 | 0.001618 | 0.002176 | 0.001603 |
| 0.336217 | 0.965934 | 0.561063 | 0.0366 | 0.03464 | 0.036734 | 0.034502 |
| 0.291906 | 0.852327 | 0.560947 | 0.037622 | 0.009775 | 0.010944 | 0.009537 |
| 1.006289 | 2.870715 | 0.560947 | 0.037173 | 0.045259 | 0.047322 | 0.044823 |
| 0.348345 | 0.99375 | 0.560947 | 0.037173 | 0.045259 | 0.047322 | 0.044823 |
| 0.323624 | 0.932352 | 0.560947 | 0.037589 | 0.02381 | 0.026456 | 0.024306 |
| 0.34262 | 0.980465 | 0.560947 | 0.037444 | 0.039202 | 0.042 | 0.039582 |
| 1.01011 | 2.941024 | 0.560947 | 0.036334 | 0.045408 | 0.04584 | 0.043618 |
| 0.270408 | 0.793947 | 0.560947 | 0.037896 | 0.004301 | 0.005115 | 0.004178 |
| 1.259329 | 3.75841 | 0.560831 | 0.038805 | 0.004322 | 0.005326 | 0.004377 |
| 0.26607 | 0.794074 | 0.560831 | 0.038805 | 0.004322 | 0.005326 | 0.004377 |
| 1.083507 | 3.036583 | 0.560831 | 0.037216 | 0.02334 | 0.02351 | 0.021588 |
| 1.00868 | 2.922397 | 0.560715 | 0.03715 | 0.045507 | 0.046392 | 0.044143 |
| 0.250123 | 0.754316 | 0.560715 | 0.03865 | 0.002368 | 0.003065 | 0.002391 |
| 0.32019 | 0.932659 | 0.560715 | 0.037196 | 0.024205 | 0.026721 | 0.024573 |
| 0.332355 | 0.961662 | 0.560715 | 0.038255 | 0.033055 | 0.035361 | 0.033075 |
| 1.049549 | 2.990373 | 0.560599 | 0.037647 | 0.031342 | 0.032273 | 0.030177 |
| 0.338648 | 0.958207 | 0.560599 | 0.037233 | 0.032561 | 0.033934 | 0.031729 |
| 1.043616 | 2.952916 | 0.560599 | 0.037233 | 0.032561 | 0.033934 | 0.031729 |
| 1.170465 | 3.383173 | 0.560599 | 0.037662 | 0.010256 | 0.011045 | 0.009673 |
| 1.003518 | 2.8258 | 0.560599 | 0.037453 | 0.048016 | 0.048466 | 0.04611 |
| 1.002309 | 2.879917 | 0.560599 | 0.037432 | 0.04682 | 0.049007 | 0.046535 |
| 0.342937 | 0.987748 | 0.560599 | 0.037004 | 0.042772 | 0.044894 | 0.042477 |
| 0.34854 | 0.990502 | 0.560483 | 0.037732 | 0.043674 | 0.045958 | 0.043526 |
| 1.175343 | 3.433966 | 0.560483 | 0.037468 | 0.008989 | 0.010752 | 0.009296 |
| 1.018941 | 2.872905 | 0.560483 | 0.037499 | 0.040694 | 0.042263 | 0.03989 |
| 0.34808 | 0.981411 | 0.560483 | 0.037499 | 0.040694 | 0.042263 | 0.03989 |
| 1.138014 | 3.306263 | 0.560483 | 0.037579 | 0.013803 | 0.014886 | 0.01329 |
| 0.343029 | 0.968705 | 0.560367 | 0.037268 | 0.036052 | 0.037524 | 0.035235 |
| 0.344657 | 0.991906 | 0.560367 | 0.037298 | 0.045523 | 0.04658 | 0.044205 |
| 1.037476 | 2.937312 | 0.560367 | 0.037651 | 0.033866 | 0.035857 | 0.033545 |
| 0.340447 | 0.963878 | 0.560367 | 0.037651 | 0.033866 | 0.035857 | 0.033545 |
| 0.319452 | 0.9315 | 0.560251 | 0.037266 | 0.02393 | 0.026427 | 0.024296 |
| 1.073538 | 3.130363 | 0.560251 | 0.037266 | 0.02393 | 0.026427 | 0.024296 |
| 0.341262 | 0.976437 | 0.560251 | 0.03751 | 0.038596 | 0.040474 | 0.038094 |
| 0.323511 | 0.944173 | 0.560251 | 0.036637 | 0.026595 | 0.029991 | 0.027721 |
| 1.012187 | 2.871104 | 0.560251 | 0.037258 | 0.043242 | 0.044909 | 0.042545 |
| 1.282839 | 3.717318 | 0.560251 | 0.037782 | 0.00345 | 0.004006 | 0.003236 |
| 1.059835 | 3.01999 | 0.560135 | 0.036573 | 0.02933 | 0.029443 | 0.027357 |
| 0.331127 | 0.943543 | 0.560135 | 0.036573 | 0.02933 | 0.029443 | 0.027357 |

| | | | | | | |
|---|---|---|---|---|---|---|
| 0.320442 | 0.917754 | 0.560019 | 0.038084 | 0.020683 | 0.022626 | 0.020654 |
| 0.331287 | 0.954471 | 0.560019 | 0.03754 | 0.030708 | 0.032959 | 0.030679 |
| 0.327406 | 0.928188 | 0.559902 | 0.036927 | 0.023926 | 0.025072 | 0.02309 |
| 1.077367 | 3.054309 | 0.559902 | 0.036927 | 0.023926 | 0.025072 | 0.02309 |
| 1.094184 | 3.202451 | 0.559902 | 0.037061 | 0.019285 | 0.022108 | 0.020099 |
| 0.296664 | 0.87257 | 0.559902 | 0.038314 | 0.012263 | 0.014079 | 0.012525 |
| 0.330623 | 0.96095 | 0.559902 | 0.037073 | 0.032259 | 0.035178 | 0.032819 |
| 1.119004 | 3.378607 | 0.559902 | 0.037611 | 0.015368 | 0.018333 | 0.016431 |
| 0.29598 | 0.893652 | 0.559902 | 0.037611 | 0.015368 | 0.018333 | 0.016431 |
| 0.331368 | 0.934238 | 0.559786 | 0.037042 | 0.027152 | 0.026608 | 0.024579 |
| 1.070391 | 3.017792 | 0.559786 | 0.037042 | 0.027152 | 0.026608 | 0.024579 |
| 1.007126 | 2.880407 | 0.559786 | 0.036664 | 0.046766 | 0.046983 | 0.044662 |
| 0.302747 | 0.875318 | 0.559786 | 0.03788 | 0.012442 | 0.014221 | 0.012583 |
| 0.318792 | 0.939698 | 0.559786 | 0.037465 | 0.024988 | 0.028852 | 0.026543 |
| 0.308269 | 0.895032 | 0.559786 | 0.037215 | 0.015844 | 0.017894 | 0.016107 |
| 1.117279 | 3.243921 | 0.559786 | 0.037215 | 0.015844 | 0.017894 | 0.016107 |
| 1.105958 | 3.159164 | 0.559786 | 0.037212 | 0.019133 | 0.019486 | 0.017724 |
| 0.33318 | 0.968567 | 0.55967 | 0.036668 | 0.035805 | 0.037776 | 0.035541 |
| 1.012887 | 2.882949 | 0.55967 | 0.036544 | 0.043917 | 0.044648 | 0.042292 |
| 0.33297 | 0.950707 | 0.55967 | 0.037896 | 0.029253 | 0.031648 | 0.029399 |
| 0.336143 | 0.967493 | 0.559554 | 0.037258 | 0.034067 | 0.037297 | 0.034885 |
| 0.336227 | 0.968972 | 0.559554 | 0.036888 | 0.035909 | 0.03783 | 0.035522 |
| 0.322734 | 0.917725 | 0.559438 | 0.037713 | 0.020986 | 0.022489 | 0.020546 |
| 1.001755 | 2.86117 | 0.559438 | 0.037458 | 0.046196 | 0.049239 | 0.046713 |
| 0.297969 | 0.859192 | 0.559438 | 0.037792 | 0.011419 | 0.011678 | 0.010317 |
| 1.003655 | 2.809107 | 0.559438 | 0.037198 | 0.04988 | 0.048398 | 0.046035 |
| 0.321583 | 0.958153 | 0.559322 | 0.037561 | 0.030436 | 0.034562 | 0.03213 |
| 0.305336 | 0.880727 | 0.559322 | 0.038031 | 0.013253 | 0.015101 | 0.013405 |
| 0.332132 | 0.958241 | 0.559322 | 0.037458 | 0.032494 | 0.034194 | 0.031994 |
| 0.286929 | 0.833194 | 0.559322 | 0.038115 | 0.007559 | 0.008513 | 0.007265 |
| 0.309346 | 0.89847 | 0.55909 | 0.037839 | 0.017028 | 0.018594 | 0.016805 |
| 0.33674 | 0.949713 | 0.55909 | 0.037275 | 0.030386 | 0.03116 | 0.029046 |
| 1.016 | 2.917563 | 0.558974 | 0.037123 | 0.042481 | 0.043492 | 0.041134 |
| 1.050669 | 3.047103 | 0.558974 | 0.037475 | 0.030454 | 0.032199 | 0.03005 |
| 0.328181 | 0.951775 | 0.558974 | 0.037475 | 0.030454 | 0.032199 | 0.03005 |
| 1.01078 | 2.883526 | 0.558974 | 0.037769 | 0.042565 | 0.045494 | 0.042999 |
| 0.346798 | 0.989335 | 0.558974 | 0.037769 | 0.042565 | 0.045494 | 0.042999 |
| 1.086044 | 3.112782 | 0.558974 | 0.037208 | 0.021389 | 0.023375 | 0.021383 |
| 0.321256 | 0.920773 | 0.558974 | 0.037208 | 0.021389 | 0.023375 | 0.021383 |
| 0.29159 | 0.874919 | 0.558858 | 0.037818 | 0.012376 | 0.014823 | 0.013118 |
| 1.032147 | 2.950424 | 0.558858 | 0.037631 | 0.036428 | 0.037704 | 0.035418 |
| 0.349979 | 0.99882 | 0.558858 | 0.037092 | 0.048056 | 0.049486 | 0.047048 |
| 0.324927 | 0.967597 | 0.558858 | 0.036994 | 0.033355 | 0.037682 | 0.035183 |
| 0.333782 | 0.964019 | 0.558742 | 0.03711 | 0.034242 | 0.036136 | 0.03386 |
| 0.298755 | 0.853449 | 0.558742 | 0.038201 | 0.010239 | 0.010718 | 0.009353 |
| 1.171716 | 3.347228 | 0.558742 | 0.038201 | 0.010239 | 0.010718 | 0.009353 |
| 0.300226 | 0.878023 | 0.558742 | 0.037698 | 0.013441 | 0.014887 | 0.013264 |

| | | | | | | |
|---|---|---|---|---|---|---|
| 0.286907 | 0.848835 | 0.558742 | 0.037261 | 0.008916 | 0.010703 | 0.009297 |
| 1.118555 | 3.280929 | 0.558742 | 0.037857 | 0.015537 | 0.01788 | 0.016003 |
| 0.312086 | 0.952105 | 0.558625 | 0.036634 | 0.027608 | 0.03297 | 0.030458 |
| 0.324789 | 0.925126 | 0.558509 | 0.037235 | 0.024144 | 0.02436 | 0.022525 |
| 0.297309 | 0.849311 | 0.558509 | 0.037667 | 0.009591 | 0.010172 | 0.008891 |
| 0.266416 | 0.789275 | 0.558393 | 0.03855 | 0.003962 | 0.004892 | 0.003974 |
| 0.328178 | 0.946423 | 0.558393 | 0.037062 | 0.030804 | 0.030482 | 0.028465 |
| 1.039 | 2.94179 | 0.558277 | 0.037053 | 0.034471 | 0.035373 | 0.033192 |
| 1.047252 | 2.97468 | 0.558277 | 0.037349 | 0.031502 | 0.032898 | 0.030733 |
| 0.343115 | 0.974979 | 0.558277 | 0.037668 | 0.037763 | 0.039872 | 0.037522 |
| 1.073947 | 3.095316 | 0.558277 | 0.03743 | 0.025076 | 0.026139 | 0.024143 |
| 0.317979 | 0.921043 | 0.558161 | 0.038172 | 0.021325 | 0.023629 | 0.021546 |
| 1.085725 | 3.144861 | 0.558161 | 0.038172 | 0.021325 | 0.023629 | 0.021546 |
| 1.169559 | 3.503552 | 0.558161 | 0.037879 | 0.009251 | 0.01175 | 0.01015 |
| 1.05212 | 2.973624 | 0.558161 | 0.037861 | 0.030114 | 0.031425 | 0.029263 |
| 0.262787 | 0.951814 | 0.558161 | 0.030276 | 0.024983 | 0.034826 | 0.031581 |
| 0.344963 | 0.973775 | 0.558161 | 0.038003 | 0.037871 | 0.039365 | 0.037047 |
| 0.314423 | 0.894652 | 0.558045 | 0.037596 | 0.016168 | 0.017442 | 0.015696 |
| 0.309678 | 0.875157 | 0.558045 | 0.036806 | 0.013377 | 0.013773 | 0.012264 |
| 0.333372 | 0.959124 | 0.558045 | 0.037632 | 0.032717 | 0.034449 | 0.032214 |
| 1.013334 | 2.877226 | 0.558045 | 0.037046 | 0.043122 | 0.044461 | 0.042033 |
| 1.022421 | 2.916118 | 0.558045 | 0.037067 | 0.039154 | 0.041063 | 0.038644 |
| 1.060564 | 3.081627 | 0.557929 | 0.037462 | 0.026883 | 0.029551 | 0.027308 |
| 0.324504 | 0.942894 | 0.557929 | 0.037462 | 0.026883 | 0.029551 | 0.027308 |
| 0.340025 | 0.978698 | 0.557929 | 0.037886 | 0.03883 | 0.04137 | 0.038934 |
| 1.157649 | 3.356875 | 0.557813 | 0.037912 | 0.010819 | 0.012455 | 0.01093 |
| 0.318549 | 0.910137 | 0.557813 | 0.037825 | 0.01937 | 0.020803 | 0.01895 |
| 1.098735 | 3.139234 | 0.557813 | 0.037825 | 0.01937 | 0.020803 | 0.01895 |
| 0.325112 | 0.974207 | 0.557813 | 0.036579 | 0.034741 | 0.040043 | 0.037444 |
| 0.345647 | 0.996829 | 0.557697 | 0.037892 | 0.045886 | 0.048642 | 0.046131 |
| 1.002492 | 2.892737 | 0.557581 | 0.037728 | 0.046121 | 0.048934 | 0.046414 |
| 0.302039 | 0.881115 | 0.557581 | 0.037868 | 0.013619 | 0.015377 | 0.013677 |
| 0.32699 | 0.939919 | 0.557465 | 0.037087 | 0.02865 | 0.028524 | 0.026423 |
| 1.063921 | 3.058193 | 0.557465 | 0.037087 | 0.02865 | 0.028524 | 0.026423 |
| 1.014976 | 2.872928 | 0.557465 | 0.037789 | 0.043097 | 0.043803 | 0.041413 |
| 0.328673 | 0.970052 | 0.557465 | 0.037361 | 0.034681 | 0.038444 | 0.036003 |
| 1.01001 | 2.910393 | 0.557349 | 0.036695 | 0.043768 | 0.045841 | 0.043439 |
| 1.194892 | 3.47423 | 0.557349 | 0.037704 | 0.008642 | 0.008951 | 0.007743 |
| 1.084294 | 3.137342 | 0.557349 | 0.036767 | 0.022851 | 0.023911 | 0.021987 |
| 0.318741 | 0.922259 | 0.557349 | 0.036767 | 0.022851 | 0.023911 | 0.021987 |
| 0.335005 | 0.950839 | 0.557349 | 0.037513 | 0.02977 | 0.031604 | 0.029401 |
| 0.346384 | 0.971574 | 0.557349 | 0.037341 | 0.038724 | 0.038493 | 0.036263 |
| 0.328541 | 0.955125 | 0.557349 | 0.037344 | 0.029675 | 0.033287 | 0.030905 |
| 1.046983 | 3.043756 | 0.557349 | 0.037344 | 0.029675 | 0.033287 | 0.030905 |
| 1.002662 | 2.845178 | 0.557349 | 0.037092 | 0.04647 | 0.048843 | 0.046364 |
| 1.005118 | 2.855156 | 0.557232 | 0.037102 | 0.046072 | 0.047801 | 0.045417 |
| 0.310589 | 0.89465 | 0.557116 | 0.037791 | 0.01598 | 0.017671 | 0.015915 |

| | | | | | | |
|---|---|---|---|---|---|---|
| 0.331373 | 0.953007 | 0.557116 | 0.037253 | 0.030041 | 0.03247 | 0.030261 |
| 0.08136 | 0.836215 | 0.557116 | 0.017654 | 0.005469 | 0.023766 | 0.014982 |
| 0.345358 | 0.976288 | 0.557116 | 0.0374 | 0.038779 | 0.040317 | 0.037993 |
| 0.327203 | 0.937719 | 0.557 | 0.037907 | 0.026149 | 0.027851 | 0.02581 |
| 1.066418 | 3.056203 | 0.557 | 0.037907 | 0.026149 | 0.027851 | 0.02581 |
| 0.314071 | 0.89884 | 0.557 | 0.036973 | 0.016698 | 0.018396 | 0.01658 |
| 0.339253 | 0.970455 | 0.557 | 0.038229 | 0.035626 | 0.038282 | 0.035919 |
| 0.338979 | 0.962383 | 0.556884 | 0.037528 | 0.033402 | 0.035378 | 0.033098 |
| 1.09683 | 3.186142 | 0.556884 | 0.037607 | 0.019309 | 0.021465 | 0.019481 |
| 0.246186 | 0.783872 | 0.556768 | 0.038324 | 0.003618 | 0.005367 | 0.004322 |
| 0.299299 | 0.92554 | 0.556652 | 0.036923 | 0.020589 | 0.025838 | 0.023411 |
| 0.349354 | 0.996684 | 0.556536 | 0.037659 | 0.048275 | 0.048566 | 0.046177 |
| 0.353784 | 0.996606 | 0.556536 | 0.036828 | 0.048042 | 0.048515 | 0.046156 |
| 0.346521 | 0.991981 | 0.556536 | 0.037336 | 0.044527 | 0.046594 | 0.044128 |
| 0.354655 | 0.99801 | 0.55642 | 0.036761 | 0.048703 | 0.049124 | 0.046765 |
| 0.330241 | 0.945398 | 0.556304 | 0.03849 | 0.029551 | 0.030065 | 0.02801 |
| 0.331892 | 0.960541 | 0.556304 | 0.038201 | 0.032694 | 0.034991 | 0.032711 |
| 1.142669 | 3.237142 | 0.556304 | 0.037287 | 0.013074 | 0.013816 | 0.012275 |
| 0.308914 | 0.875144 | 0.556304 | 0.037287 | 0.013074 | 0.013816 | 0.012275 |
| 0.31079 | 0.890471 | 0.556188 | 0.037806 | 0.015172 | 0.016759 | 0.015025 |
| 1.123001 | 3.217607 | 0.556188 | 0.037806 | 0.015172 | 0.016759 | 0.015025 |
| 0.2847 | 0.842947 | 0.556188 | 0.037592 | 0.008016 | 0.009967 | 0.008542 |
| 0.345825 | 0.986095 | 0.556072 | 0.037485 | 0.041895 | 0.044183 | 0.041791 |
| 0.329309 | 0.947466 | 0.556072 | 0.037846 | 0.029161 | 0.030763 | 0.02861 |
| 0.327845 | 0.942351 | 0.556072 | 0.037917 | 0.028073 | 0.029227 | 0.02713 |
| 1.190709 | 3.552797 | 0.556072 | 0.038011 | 0.007913 | 0.009714 | 0.008316 |
| 0.281468 | 0.839836 | 0.556072 | 0.038011 | 0.007913 | 0.009714 | 0.008316 |
| 0.324251 | 0.929945 | 0.556072 | 0.037342 | 0.023607 | 0.025735 | 0.023668 |
| 0.327171 | 0.930244 | 0.555955 | 0.03724 | 0.024912 | 0.025667 | 0.023725 |
| 1.074987 | 3.056506 | 0.555955 | 0.03724 | 0.024912 | 0.025667 | 0.023725 |
| 0.334689 | 0.959097 | 0.555955 | 0.037435 | 0.034138 | 0.034391 | 0.03221 |
| 0.286771 | 0.844819 | 0.555839 | 0.038438 | 0.00815 | 0.010118 | 0.008659 |
| 0.322479 | 0.937287 | 0.555839 | 0.037501 | 0.025805 | 0.027955 | 0.025819 |
| 1.066909 | 3.100973 | 0.555839 | 0.037501 | 0.025805 | 0.027955 | 0.025819 |
| 1.042482 | 2.961188 | 0.555839 | 0.03782 | 0.033708 | 0.03433 | 0.03211 |
| 0.337702 | 0.95925 | 0.555839 | 0.03782 | 0.033708 | 0.03433 | 0.03211 |
| 0.344696 | 0.975651 | 0.555839 | 0.037512 | 0.037964 | 0.040088 | 0.037701 |
| 0.34824 | 0.982635 | 0.555839 | 0.03782 | 0.041168 | 0.042749 | 0.040376 |
| 1.017672 | 2.87158 | 0.555839 | 0.03782 | 0.041168 | 0.042749 | 0.040376 |
| 0.349233 | 0.999085 | 0.555839 | 0.037947 | 0.047455 | 0.049602 | 0.047096 |
| 0.322818 | 0.923038 | 0.555723 | 0.037819 | 0.022322 | 0.023897 | 0.021945 |
| 1.062964 | 3.061135 | 0.555723 | 0.037479 | 0.028205 | 0.028797 | 0.026796 |
| 0.326676 | 0.940765 | 0.555723 | 0.037479 | 0.028205 | 0.028797 | 0.026796 |
| 1.154282 | 3.296181 | 0.555723 | 0.038016 | 0.011248 | 0.012561 | 0.011026 |
| 0.310053 | 0.892008 | 0.555723 | 0.037805 | 0.016729 | 0.017131 | 0.015479 |
| 0.33866 | 0.989337 | 0.555607 | 0.037539 | 0.04217 | 0.045591 | 0.043094 |
| 0.355714 | 0.999988 | 0.555607 | 0.037883 | 0.049586 | 0.049995 | 0.04762 |

| | | | | | | |
|---|---|---|---|---|---|---|
| 0.322939 | 0.924738 | 0.555491 | 0.037735 | 0.023814 | 0.024354 | 0.022408 |
| 0.332478 | 0.96363 | 0.555491 | 0.037643 | 0.034155 | 0.036044 | 0.033801 |
| 1.037742 | 3.007722 | 0.555491 | 0.037643 | 0.034155 | 0.036044 | 0.033801 |
| 0.313448 | 0.92405 | 0.555491 | 0.037584 | 0.021157 | 0.024681 | 0.022499 |
| 0.327757 | 0.943343 | 0.555491 | 0.037883 | 0.02745 | 0.029538 | 0.027334 |
| 1.083634 | 3.104733 | 0.555375 | 0.037451 | 0.023315 | 0.023879 | 0.021941 |
| 1.086474 | 3.117252 | 0.555375 | 0.037271 | 0.021302 | 0.023304 | 0.021315 |
| 0.320795 | 0.920408 | 0.555375 | 0.037271 | 0.021302 | 0.023304 | 0.021315 |
| 0.326312 | 0.935099 | 0.555143 | 0.037612 | 0.024942 | 0.027118 | 0.025013 |
| 0.313796 | 0.927344 | 0.555143 | 0.037537 | 0.022951 | 0.02556 | 0.023487 |
| 0.326215 | 0.936332 | 0.555143 | 0.037628 | 0.026868 | 0.027487 | 0.025465 |
| 0.3191 | 0.926809 | 0.555143 | 0.037704 | 0.022539 | 0.02513 | 0.023071 |
| 0.265778 | 0.801495 | 0.555143 | 0.037893 | 0.005228 | 0.006037 | 0.005013 |
| 0.336246 | 0.962694 | 0.555027 | 0.037656 | 0.033836 | 0.035584 | 0.033292 |
| 0.338242 | 0.973577 | 0.555027 | 0.037979 | 0.038144 | 0.03947 | 0.037212 |
| 0.304827 | 0.878809 | 0.555027 | 0.037822 | 0.012937 | 0.014759 | 0.013077 |
| 0.318309 | 0.920267 | 0.554911 | 0.038012 | 0.021592 | 0.023405 | 0.021409 |
| 0.331545 | 0.971926 | 0.554911 | 0.037186 | 0.035108 | 0.039042 | 0.036617 |
| 1.018315 | 2.916517 | 0.554911 | 0.037412 | 0.041473 | 0.042604 | 0.040386 |
| 0.321331 | 0.950629 | 0.554795 | 0.037545 | 0.028177 | 0.032118 | 0.029693 |
| 1.014992 | 2.868984 | 0.554795 | 0.037675 | 0.042182 | 0.043789 | 0.041418 |
| 0.24419 | 0.741213 | 0.554678 | 0.038906 | 0.001908 | 0.002551 | 0.001976 |
| 0.280751 | 0.863716 | 0.554678 | 0.038208 | 0.010685 | 0.013473 | 0.011797 |
| 0.342391 | 0.99432 | 0.554562 | 0.037903 | 0.04849 | 0.047602 | 0.045411 |
| 1.136805 | 3.275657 | 0.554562 | 0.038066 | 0.014221 | 0.014896 | 0.013321 |
| 0.305282 | 0.879658 | 0.554562 | 0.038066 | 0.014221 | 0.014896 | 0.013321 |
| 0.330752 | 0.94348 | 0.554562 | 0.037071 | 0.028492 | 0.02944 | 0.027385 |
| 1.059906 | 3.02341 | 0.554562 | 0.037071 | 0.028492 | 0.02944 | 0.027385 |
| 0.264926 | 0.792298 | 0.554446 | 0.038129 | 0.005417 | 0.005221 | 0.004386 |
| 0.342862 | 0.993052 | 0.554446 | 0.037538 | 0.04481 | 0.04707 | 0.044651 |
| 1.006997 | 2.916621 | 0.554446 | 0.037538 | 0.04481 | 0.04707 | 0.044651 |
| 0.313215 | 0.920164 | 0.55433 | 0.038296 | 0.020777 | 0.02366 | 0.021631 |
| 1.076382 | 3.095749 | 0.55433 | 0.037645 | 0.025515 | 0.025544 | 0.023656 |
| 0.309134 | 0.876338 | 0.554214 | 0.038343 | 0.013287 | 0.014027 | 0.012483 |
| 1.023014 | 2.981349 | 0.554214 | 0.037999 | 0.037841 | 0.041018 | 0.038577 |
| 1.071641 | 3.022532 | 0.554098 | 0.037908 | 0.026078 | 0.026315 | 0.02435 |
| 0.180647 | 0.988942 | 0.554098 | 0.021076 | 0.026594 | 0.047078 | 0.040845 |
| 0.317753 | 0.935608 | 0.553982 | 0.037499 | 0.025008 | 0.027697 | 0.025548 |
| 1.068823 | 3.147101 | 0.553982 | 0.037499 | 0.025008 | 0.027697 | 0.025548 |
| 0.333305 | 0.952415 | 0.553982 | 0.037675 | 0.030549 | 0.032195 | 0.029956 |
| 0.343248 | 0.985271 | 0.553982 | 0.037659 | 0.040999 | 0.043891 | 0.041464 |
| 1.089816 | 3.175759 | 0.553982 | 0.037078 | 0.021797 | 0.022894 | 0.021017 |
| 0.29444 | 0.859521 | 0.553982 | 0.038448 | 0.010692 | 0.011941 | 0.010469 |
| 1.009062 | 2.914798 | 0.553866 | 0.036929 | 0.042613 | 0.046228 | 0.043714 |
| 0.343077 | 0.991019 | 0.553866 | 0.036929 | 0.042613 | 0.046228 | 0.043714 |
| 0.321826 | 0.929184 | 0.553866 | 0.0379 | 0.023063 | 0.025648 | 0.02354 |
| 1.455746 | 6.907762 | 0.553866 | 0.02745 | 0.010002 | 0.003669 | 0.002188 |

| | | | | | | |
|---|---|---|---|---|---|---|
| 0.347331 | 0.98345 | 0.553866 | 0.037193 | 0.042108 | 0.043092 | 0.040808 |
| 0.336639 | 0.963672 | 0.553866 | 0.037365 | 0.033353 | 0.035915 | 0.0336 |
| 0.320801 | 0.914226 | 0.553866 | 0.037538 | 0.02025 | 0.021697 | 0.019803 |
| 1.052838 | 2.976286 | 0.55375 | 0.037089 | 0.031749 | 0.031226 | 0.029097 |
| 0.335989 | 0.949814 | 0.55375 | 0.037089 | 0.031749 | 0.031226 | 0.029097 |
| 1.20347 | 3.580933 | 0.553518 | 0.038601 | 0.006885 | 0.008645 | 0.007344 |
| 1.143829 | 3.27154 | 0.553518 | 0.037576 | 0.013898 | 0.013846 | 0.012354 |
| 0.305666 | 0.874257 | 0.553518 | 0.037576 | 0.013898 | 0.013846 | 0.012354 |
| 1.045133 | 3.017796 | 0.553518 | 0.03703 | 0.030689 | 0.033741 | 0.031427 |
| 0.331368 | 0.956816 | 0.553518 | 0.03703 | 0.030689 | 0.033741 | 0.031427 |
| 0.309656 | 0.898273 | 0.553401 | 0.039017 | 0.017824 | 0.018531 | 0.016872 |
| 1.04865 | 3.107761 | 0.553401 | 0.037567 | 0.029069 | 0.033059 | 0.030628 |
| 0.321775 | 0.953607 | 0.553401 | 0.037567 | 0.029069 | 0.033059 | 0.030628 |
| 0.328936 | 0.934652 | 0.553401 | 0.037555 | 0.026161 | 0.026854 | 0.024899 |
| 1.069917 | 3.040109 | 0.553401 | 0.037555 | 0.026161 | 0.026854 | 0.024899 |
| 1.106276 | 3.158936 | 0.553401 | 0.038061 | 0.018069 | 0.019424 | 0.017624 |
| 0.343891 | 0.977472 | 0.553401 | 0.037709 | 0.039269 | 0.040809 | 0.038528 |
| 0.345632 | 0.978334 | 0.553401 | 0.0367 | 0.03956 | 0.041103 | 0.038781 |
| 0.343471 | 0.987488 | 0.553401 | 0.037552 | 0.042325 | 0.044782 | 0.042405 |
| 1.072632 | 3.09418 | 0.553401 | 0.037757 | 0.023799 | 0.02646 | 0.024317 |
| 0.323187 | 0.932286 | 0.553401 | 0.037757 | 0.023799 | 0.02646 | 0.024317 |
| 0.295632 | 0.854686 | 0.553285 | 0.037801 | 0.009861 | 0.011093 | 0.009701 |
| 0.306001 | 0.877346 | 0.553285 | 0.037558 | 0.013308 | 0.014407 | 0.012854 |
| 1.06927 | 3.079994 | 0.553285 | 0.037988 | 0.026279 | 0.027236 | 0.025246 |
| 0.324676 | 0.935218 | 0.553285 | 0.037988 | 0.026279 | 0.027236 | 0.025246 |
| 1.095473 | 3.155338 | 0.553169 | 0.037798 | 0.019653 | 0.021571 | 0.019666 |
| 0.307103 | 0.877231 | 0.553169 | 0.038016 | 0.012882 | 0.014319 | 0.012686 |
| 1.145231 | 3.323023 | 0.553053 | 0.038361 | 0.012143 | 0.013934 | 0.01232 |
| 1.19803 | 3.555407 | 0.553053 | 0.038013 | 0.008357 | 0.009026 | 0.007821 |
| 0.281262 | 0.834703 | 0.553053 | 0.038013 | 0.008357 | 0.009026 | 0.007821 |
| 1.019306 | 2.913778 | 0.553053 | 0.037621 | 0.041055 | 0.042222 | 0.039971 |
| 1.243828 | 3.681347 | 0.552937 | 0.038034 | 0.005305 | 0.005992 | 0.004983 |
| 0.286796 | 0.831973 | 0.552937 | 0.038608 | 0.007482 | 0.008364 | 0.00715 |
| 1.12055 | 3.200037 | 0.552937 | 0.037779 | 0.015452 | 0.017072 | 0.015301 |
| 1.121528 | 3.221487 | 0.552937 | 0.037409 | 0.016638 | 0.01703 | 0.015369 |
| 0.310416 | 0.891641 | 0.552937 | 0.037409 | 0.016638 | 0.01703 | 0.015369 |
| 1.009324 | 2.956441 | 0.552937 | 0.037321 | 0.041836 | 0.046172 | 0.043626 |
| 1.019505 | 3.01809 | 0.552821 | 0.037916 | 0.037817 | 0.042384 | 0.039765 |
| 0.331335 | 0.980868 | 0.552821 | 0.037916 | 0.037817 | 0.042384 | 0.039765 |
| 0.081699 | 0.864869 | 0.552821 | 0.01582 | 0.007653 | 0.027725 | 0.018771 |
| 1.004472 | 2.868529 | 0.552821 | 0.037356 | 0.045067 | 0.048083 | 0.045567 |
| 1.050784 | 3.038852 | 0.552821 | 0.038342 | 0.029675 | 0.032128 | 0.02996 |
| 1.168977 | 4.223255 | 0.552705 | 0.028143 | 0.024265 | 0.014833 | 0.012368 |
| 1.030649 | 2.930018 | 0.552705 | 0.03812 | 0.036079 | 0.038151 | 0.035831 |
| 0.341295 | 0.970262 | 0.552705 | 0.03812 | 0.036079 | 0.038151 | 0.035831 |
| 0.353333 | 0.99985 | 0.552589 | 0.036715 | 0.049329 | 0.049934 | 0.047563 |
| 0.296165 | 0.860796 | 0.552473 | 0.038127 | 0.010577 | 0.01205 | 0.010523 |

| | | | | | | |
|---|---|---|---|---|---|---|
| 1.161715 | 3.376491 | 0.552473 | 0.038127 | 0.010577 | 0.01205 | 0.010523 |
| 1.174032 | 3.673647 | 0.552357 | 0.029951 | 0.017591 | 0.012029 | 0.010333 |
| 0.296901 | 0.888152 | 0.552357 | 0.038401 | 0.015869 | 0.017112 | 0.015252 |
| 1.125933 | 3.36813 | 0.552357 | 0.038401 | 0.015869 | 0.017112 | 0.015252 |
| 0.350059 | 0.99776 | 0.552241 | 0.037623 | 0.047608 | 0.049027 | 0.046581 |
| 1.002245 | 2.85666 | 0.552241 | 0.037623 | 0.047608 | 0.049027 | 0.046581 |
| 0.324365 | 0.944531 | 0.552124 | 0.037973 | 0.026661 | 0.030063 | 0.027773 |
| 0.278878 | 0.865696 | 0.552124 | 0.036742 | 0.010406 | 0.013931 | 0.012094 |
| 1.010856 | 2.869007 | 0.552124 | 0.036928 | 0.04408 | 0.045442 | 0.042998 |
| 1.113207 | 3.220635 | 0.552008 | 0.037689 | 0.016388 | 0.018488 | 0.016653 |
| 0.310498 | 0.898306 | 0.552008 | 0.037689 | 0.016388 | 0.018488 | 0.016653 |
| 0.346154 | 0.975904 | 0.552008 | 0.037333 | 0.039196 | 0.040149 | 0.037862 |
| 1.03864 | 2.961661 | 0.551892 | 0.037496 | 0.034098 | 0.035572 | 0.033384 |
| 0.337648 | 0.962798 | 0.551892 | 0.037496 | 0.034098 | 0.035572 | 0.033384 |
| 1.202228 | 3.541776 | 0.551892 | 0.037993 | 0.007362 | 0.008585 | 0.007329 |
| 0.282344 | 0.831789 | 0.551892 | 0.037993 | 0.007362 | 0.008585 | 0.007329 |
| 0.308856 | 0.917005 | 0.55166 | 0.037823 | 0.019844 | 0.023083 | 0.020954 |
| 1.004663 | 2.988798 | 0.55166 | 0.037545 | 0.04323 | 0.048077 | 0.045428 |
| 0.334583 | 0.995358 | 0.55166 | 0.037545 | 0.04323 | 0.048077 | 0.045428 |
| 1.02863 | 2.951764 | 0.551544 | 0.037753 | 0.037576 | 0.03893 | 0.03664 |
| 1.054195 | 3.109414 | 0.551544 | 0.037428 | 0.027383 | 0.031458 | 0.029084 |
| 0.331893 | 0.951337 | 0.551312 | 0.037733 | 0.029489 | 0.0319 | 0.029674 |
| 0.312973 | 0.91116 | 0.551312 | 0.038329 | 0.019018 | 0.021378 | 0.019454 |
| 1.017078 | 2.920044 | 0.551312 | 0.036937 | 0.04021 | 0.043083 | 0.04067 |
| 1.037881 | 2.978825 | 0.551196 | 0.037473 | 0.03561 | 0.035888 | 0.033697 |
| 0.335703 | 0.963502 | 0.551196 | 0.037473 | 0.03561 | 0.035888 | 0.033697 |
| 0.345045 | 0.972234 | 0.55108 | 0.037876 | 0.037875 | 0.03878 | 0.036516 |
| 0.131521 | 0.838639 | 0.55108 | 0.022235 | 0.006995 | 0.019684 | 0.014486 |
| 0.285814 | 0.843734 | 0.550964 | 0.038206 | 0.008432 | 0.010016 | 0.008622 |
| 1.049017 | 3.049445 | 0.550964 | 0.037697 | 0.029711 | 0.0327 | 0.030462 |
| 1.085221 | 4.606642 | 0.550964 | 0.028572 | 0.045741 | 0.029131 | 0.025077 |
| 0.32514 | 0.986839 | 0.550964 | 0.036967 | 0.038429 | 0.044777 | 0.041994 |
| 0.326885 | 0.957579 | 0.550964 | 0.037902 | 0.033428 | 0.03417 | 0.031948 |
| 1.044301 | 3.059175 | 0.550964 | 0.037902 | 0.033428 | 0.03417 | 0.031948 |
| 1.099751 | 3.211849 | 0.550847 | 0.037501 | 0.018281 | 0.021014 | 0.019031 |
| 0.33539 | 0.967809 | 0.550847 | 0.037782 | 0.034211 | 0.037434 | 0.035044 |
| 1.033262 | 2.981607 | 0.550847 | 0.037782 | 0.034211 | 0.037434 | 0.035044 |
| 0.293891 | 0.886027 | 0.550847 | 0.038177 | 0.013805 | 0.016858 | 0.014957 |
| 0.347869 | 0.991777 | 0.550731 | 0.036488 | 0.045917 | 0.046496 | 0.044227 |
| 1.08848 | 3.114511 | 0.550615 | 0.037365 | 0.021903 | 0.022841 | 0.020897 |
| 0.335038 | 0.976468 | 0.550615 | 0.037859 | 0.036621 | 0.040636 | 0.038137 |
| 1.024099 | 2.984732 | 0.550615 | 0.037859 | 0.036621 | 0.040636 | 0.038137 |
| 0.32855 | 0.938567 | 0.550383 | 0.037768 | 0.027017 | 0.02804 | 0.025958 |
| 0.31617 | 0.902351 | 0.550267 | 0.037883 | 0.018317 | 0.019077 | 0.01728 |
| 0.321901 | 0.933194 | 0.550035 | 0.038304 | 0.025047 | 0.026787 | 0.024836 |
| 1.071588 | 3.106544 | 0.550035 | 0.038304 | 0.025047 | 0.026787 | 0.024836 |
| 0.326803 | 0.929589 | 0.550035 | 0.037284 | 0.024352 | 0.0255 | 0.023533 |

| | | | | | | |
|---|---|---|---|---|---|---|
| 1.075744 | 3.059948 | 0.550035 | 0.037284 | 0.024352 | 0.0255 | 0.023533 |
| 0.344754 | 0.979265 | 0.549919 | 0.037604 | 0.040186 | 0.041486 | 0.039099 |
| 0.329365 | 0.939037 | 0.549919 | 0.037741 | 0.02639 | 0.028142 | 0.026046 |
| 1.064921 | 3.036146 | 0.549919 | 0.037741 | 0.02639 | 0.028142 | 0.026046 |
| 1.245571 | 3.72971 | 0.549803 | 0.038569 | 0.006004 | 0.006054 | 0.0051 |
| 0.283739 | 0.828665 | 0.54957 | 0.03788 | 0.007838 | 0.008113 | 0.006997 |
| 1.206761 | 3.524368 | 0.54957 | 0.03788 | 0.007838 | 0.008113 | 0.006997 |
| 0.104469 | 0.697486 | 0.549454 | 0.027751 | 0.001502 | 0.006855 | 0.004251 |
| 0.31114 | 0.90263 | 0.549338 | 0.038141 | 0.018791 | 0.019439 | 0.01766 |
| 1.031865 | 2.920024 | 0.549338 | 0.037497 | 0.036789 | 0.037694 | 0.03548 |
| 1.000215 | 2.849951 | 0.549338 | 0.037446 | 0.048141 | 0.049906 | 0.047529 |
| 1.10016 | 3.20817 | 0.549338 | 0.038062 | 0.020351 | 0.020911 | 0.019167 |
| 1.192636 | 3.466923 | 0.549222 | 0.038317 | 0.007834 | 0.009131 | 0.007847 |
| 0.28844 | 0.838478 | 0.549222 | 0.038317 | 0.007834 | 0.009131 | 0.007847 |
| 0.29987 | 0.863 | 0.549222 | 0.037922 | 0.010889 | 0.012198 | 0.010717 |
| 1.02206 | 2.940942 | 0.549106 | 0.037712 | 0.039882 | 0.041262 | 0.03897 |
| 0.349231 | 0.991675 | 0.549106 | 0.037457 | 0.044135 | 0.046441 | 0.043995 |
| 1.008395 | 2.863434 | 0.549106 | 0.037457 | 0.044135 | 0.046441 | 0.043995 |
| 1.099265 | 3.208456 | 0.549106 | 0.037815 | 0.019621 | 0.021093 | 0.019234 |
| 0.311676 | 0.909699 | 0.549106 | 0.037815 | 0.019621 | 0.021093 | 0.019234 |
| 0.348929 | 0.996352 | 0.549106 | 0.037346 | 0.046341 | 0.048425 | 0.045923 |
| 0.339828 | 0.981089 | 0.54899 | 0.037792 | 0.03884 | 0.042301 | 0.039833 |
| 1.106444 | 3.138701 | 0.54899 | 0.037195 | 0.018734 | 0.019271 | 0.017541 |
| 0.322125 | 0.920664 | 0.548874 | 0.037906 | 0.022426 | 0.023298 | 0.021362 |
| 0.335194 | 0.965707 | 0.548758 | 0.038021 | 0.034804 | 0.036687 | 0.034397 |
| 1.003833 | 2.842925 | 0.548758 | 0.038096 | 0.048875 | 0.04834 | 0.045965 |
| 1.012081 | 2.862972 | 0.548642 | 0.037421 | 0.043854 | 0.044938 | 0.0426 |
| 0.330377 | 0.948152 | 0.548642 | 0.037772 | 0.028602 | 0.030934 | 0.028747 |
| 1.163615 | 3.398585 | 0.548642 | 0.03821 | 0.010722 | 0.011932 | 0.010563 |
| 0.312992 | 0.905939 | 0.548526 | 0.037777 | 0.017866 | 0.020109 | 0.018205 |
| 1.018263 | 2.966648 | 0.548293 | 0.037613 | 0.039468 | 0.042733 | 0.040259 |
| 1.022917 | 2.938622 | 0.548293 | 0.038336 | 0.040496 | 0.040941 | 0.038836 |
| 0.340296 | 0.977596 | 0.548293 | 0.038336 | 0.040496 | 0.040941 | 0.038836 |
| 0.317661 | 0.908506 | 0.548293 | 0.037588 | 0.020381 | 0.020454 | 0.018648 |
| 0.34275 | 0.972049 | 0.548177 | 0.03786 | 0.036692 | 0.038775 | 0.036426 |
| 0.34387 | 0.992588 | 0.548177 | 0.037767 | 0.0443 | 0.04687 | 0.044483 |
| 0.222673 | 0.873792 | 0.547945 | 0.032796 | 0.010794 | 0.018936 | 0.015916 |
| 0.340303 | 0.978084 | 0.547713 | 0.037785 | 0.040663 | 0.041128 | 0.039022 |
| 1.022407 | 2.938557 | 0.547713 | 0.037785 | 0.040663 | 0.041128 | 0.039022 |
| 0.341826 | 0.985209 | 0.547713 | 0.038092 | 0.042385 | 0.043889 | 0.041551 |
| 1.030335 | 2.906999 | 0.547713 | 0.037326 | 0.038972 | 0.038181 | 0.036021 |
| 0.343997 | 0.970558 | 0.547713 | 0.037326 | 0.038972 | 0.038181 | 0.036021 |
| 0.322092 | 0.946858 | 0.547597 | 0.03816 | 0.028123 | 0.030891 | 0.028695 |
| 0.306023 | 0.896726 | 0.547481 | 0.037859 | 0.017069 | 0.018403 | 0.016674 |
| 1.003483 | 2.934513 | 0.547481 | 0.038299 | 0.044367 | 0.048534 | 0.045887 |
| 0.340772 | 0.996529 | 0.547481 | 0.038299 | 0.044367 | 0.048534 | 0.045887 |
| 1.037079 | 3.000702 | 0.547365 | 0.037647 | 0.033025 | 0.036234 | 0.033877 |

| | | | | | | |
|---|---|---|---|---|---|---|
| 1.027571 | 2.942635 | 0.547365 | 0.038351 | 0.038579 | 0.039275 | 0.03704 |
| 0.339831 | 0.973169 | 0.547365 | 0.038351 | 0.038579 | 0.039275 | 0.03704 |
| 0.344076 | 0.980759 | 0.547249 | 0.037447 | 0.039888 | 0.042086 | 0.039726 |
| 0.320601 | 0.923548 | 0.547249 | 0.037588 | 0.024052 | 0.024157 | 0.022292 |
| 1.082781 | 3.119143 | 0.547249 | 0.037588 | 0.024052 | 0.024157 | 0.022292 |
| 0.34391 | 0.984735 | 0.547249 | 0.038704 | 0.041714 | 0.043666 | 0.041248 |
| 0.33532 | 0.950629 | 0.547133 | 0.038322 | 0.030726 | 0.031521 | 0.029438 |
| 0.241862 | 0.899735 | 0.547133 | 0.033785 | 0.014362 | 0.022893 | 0.019814 |
| 1.014017 | 2.947158 | 0.547016 | 0.038267 | 0.043201 | 0.044314 | 0.042021 |
| 1.077582 | 3.126194 | 0.546784 | 0.038021 | 0.024259 | 0.02542 | 0.023449 |
| 1.009628 | 2.84935 | 0.546784 | 0.038099 | 0.045511 | 0.045916 | 0.043612 |
| 0.350957 | 0.990464 | 0.546784 | 0.038099 | 0.045511 | 0.045916 | 0.043612 |
| 0.343569 | 0.971735 | 0.546784 | 0.03757 | 0.03942 | 0.038634 | 0.036486 |
| 1.029087 | 2.91062 | 0.546784 | 0.03757 | 0.03942 | 0.038634 | 0.036486 |
| 1.0159 | 2.902359 | 0.546668 | 0.037338 | 0.044568 | 0.043501 | 0.041249 |
| 1.178676 | 3.436029 | 0.546552 | 0.038278 | 0.009061 | 0.0104 | 0.008993 |
| 0.338493 | 0.963598 | 0.54632 | 0.037678 | 0.033826 | 0.035825 | 0.033557 |
| 1.071939 | 3.119926 | 0.54632 | 0.038238 | 0.024296 | 0.02677 | 0.024626 |
| 1.045026 | 3.064544 | 0.546204 | 0.038023 | 0.030527 | 0.03397 | 0.031582 |
| 0.351601 | 0.992453 | 0.546088 | 0.037772 | 0.045687 | 0.046747 | 0.044387 |
| 0.170191 | 0.939505 | 0.546088 | 0.02745 | 0.018287 | 0.035453 | 0.029833 |
| 0.318263 | 0.938427 | 0.545972 | 0.038529 | 0.024675 | 0.028499 | 0.026202 |
| 0.320409 | 0.942463 | 0.545856 | 0.038107 | 0.026892 | 0.02961 | 0.027447 |
| 0.324728 | 0.947141 | 0.545856 | 0.03861 | 0.02853 | 0.030863 | 0.028663 |
| 0.322114 | 0.922367 | 0.545856 | 0.038222 | 0.022393 | 0.023755 | 0.021756 |
| 1.02888 | 2.937848 | 0.545856 | 0.038167 | 0.036952 | 0.038797 | 0.036425 |
| 0.340385 | 0.971931 | 0.545856 | 0.038167 | 0.036952 | 0.038797 | 0.036425 |
| 1.014588 | 2.86929 | 0.545739 | 0.038301 | 0.042891 | 0.043949 | 0.041626 |
| 0.348518 | 0.985622 | 0.545739 | 0.038301 | 0.042891 | 0.043949 | 0.041626 |
| 0.346112 | 0.98365 | 0.545739 | 0.037612 | 0.04098 | 0.043194 | 0.040797 |
| 1.016622 | 2.88924 | 0.545739 | 0.037612 | 0.04098 | 0.043194 | 0.040797 |
| 0.344721 | 0.999881 | 0.545623 | 0.037343 | 0.046974 | 0.049949 | 0.047416 |
| 1.112048 | 3.249585 | 0.545507 | 0.03803 | 0.01685 | 0.018865 | 0.017014 |
| 1.133261 | 3.312923 | 0.545507 | 0.03783 | 0.013699 | 0.015645 | 0.013979 |
| 0.129375 | 0.819082 | 0.545391 | 0.028463 | 0.005576 | 0.017132 | 0.012226 |
| 0.313199 | 0.896346 | 0.545275 | 0.037867 | 0.017154 | 0.017889 | 0.016156 |
| 1.115641 | 3.192853 | 0.545275 | 0.037867 | 0.017154 | 0.017889 | 0.016156 |
| 1.003439 | 2.866479 | 0.545275 | 0.037451 | 0.04642 | 0.04852 | 0.046019 |
| 0.339922 | 0.983182 | 0.545159 | 0.037598 | 0.03959 | 0.043118 | 0.040645 |
| 0.327454 | 0.938884 | 0.545159 | 0.039145 | 0.028197 | 0.028189 | 0.026224 |
| 0.33153 | 0.962402 | 0.544811 | 0.037873 | 0.034247 | 0.035647 | 0.033464 |
| 0.051378 | 0.881908 | 0.544695 | 0.017194 | 0.006297 | 0.032905 | 0.019203 |
| 1.02246 | 2.96083 | 0.544695 | 0.037961 | 0.040174 | 0.041166 | 0.038914 |
| 4.118013 | 52.35896 | 0.544695 | 0.025247 | 0.001655 | 3.44E-05 | 3.93E-08 |
| 0.351091 | 0.987871 | 0.544695 | 0.03741 | 0.045007 | 0.044835 | 0.042537 |
| 0.324513 | 0.934801 | 0.544579 | 0.037744 | 0.026525 | 0.027122 | 0.025142 |
| 1.069747 | 3.081539 | 0.544579 | 0.037744 | 0.026525 | 0.027122 | 0.025142 |

| | | | | | | |
|---|---|---|---|---|---|---|
| 0.295466 | 0.874027 | 0.544463 | 0.038394 | 0.012639 | 0.014421 | 0.012805 |
| 0.345632 | 0.979181 | 0.544463 | 0.037525 | 0.041384 | 0.041434 | 0.039126 |
| 0.339533 | 0.976168 | 0.544463 | 0.038481 | 0.038518 | 0.040414 | 0.038074 |
| 1.00363 | 2.846642 | 0.544463 | 0.037727 | 0.04732 | 0.048429 | 0.046084 |
| 0.351291 | 0.996383 | 0.544463 | 0.037727 | 0.04732 | 0.048429 | 0.046084 |
| 1.100162 | 3.196141 | 0.544463 | 0.038138 | 0.020693 | 0.020843 | 0.019107 |
| 0.312877 | 0.908957 | 0.544463 | 0.038138 | 0.020693 | 0.020843 | 0.019107 |
| 1.014618 | 2.891273 | 0.544463 | 0.037873 | 0.041744 | 0.04398 | 0.041595 |
| 0.345868 | 0.985593 | 0.544463 | 0.037873 | 0.041744 | 0.04398 | 0.041595 |
| 0.333612 | 0.950048 | 0.544463 | 0.03784 | 0.030031 | 0.031405 | 0.029287 |
| 0.329742 | 0.947788 | 0.544346 | 0.038108 | 0.028497 | 0.030846 | 0.028669 |
| 1.033911 | 3.004047 | 0.54423 | 0.038548 | 0.034289 | 0.037295 | 0.034894 |
| 0.219779 | 0.983474 | 0.544114 | 0.029544 | 0.028625 | 0.045118 | 0.040226 |
| 1.08118 | 3.100195 | 0.544114 | 0.038063 | 0.02281 | 0.024423 | 0.022467 |
| 1.06694 | 3.032235 | 0.543766 | 0.03826 | 0.025852 | 0.027586 | 0.02548 |
| 0.32979 | 0.93726 | 0.543766 | 0.03826 | 0.025852 | 0.027586 | 0.02548 |
| 1.098624 | 3.123407 | 0.54365 | 0.03853 | 0.02079 | 0.020732 | 0.018921 |
| 0.334344 | 0.958809 | 0.543534 | 0.037684 | 0.031811 | 0.034305 | 0.032036 |
| 1.058317 | 3.055106 | 0.543418 | 0.038051 | 0.02719 | 0.030042 | 0.02781 |
| 0.339856 | 0.971774 | 0.543302 | 0.038015 | 0.036065 | 0.038754 | 0.036377 |
| 0.349658 | 0.98563 | 0.543186 | 0.037765 | 0.042929 | 0.043935 | 0.041596 |
| 0.316309 | 0.90599 | 0.543186 | 0.03801 | 0.018522 | 0.019927 | 0.018118 |
| 0.334585 | 0.968434 | 0.543069 | 0.038129 | 0.035273 | 0.037685 | 0.035351 |
| 0.34627 | 0.999073 | 0.543069 | 0.037891 | 0.046806 | 0.0496 | 0.047083 |
| 0.260746 | 0.972049 | 0.543069 | 0.033218 | 0.028953 | 0.040912 | 0.037179 |
| 1.080524 | 3.160896 | 0.542837 | 0.038211 | 0.022887 | 0.024911 | 0.022907 |
| 0.339616 | 0.985238 | 0.542721 | 0.038432 | 0.041235 | 0.043936 | 0.041512 |
| 1.026771 | 2.934156 | 0.542257 | 0.038147 | 0.036812 | 0.039533 | 0.037138 |
| 0.345583 | 0.994463 | 0.542141 | 0.038354 | 0.045549 | 0.047641 | 0.04521 |
| 1.005568 | 2.893663 | 0.542141 | 0.038354 | 0.045549 | 0.047641 | 0.04521 |
| 0.339418 | 0.987618 | 0.542141 | 0.038279 | 0.04207 | 0.044889 | 0.042442 |
| 0.318006 | 0.924656 | 0.542025 | 0.038122 | 0.023397 | 0.024599 | 0.022634 |
| 1.007661 | 2.949826 | 0.541909 | 0.037863 | 0.042964 | 0.046832 | 0.044347 |
| 1.166602 | 6.382572 | 0.541909 | 0.023464 | 0.04147 | 0.020591 | 0.015791 |
| 0.312927 | 0.895638 | 0.541792 | 0.038632 | 0.016485 | 0.017748 | 0.016034 |
| 1.116523 | 3.195634 | 0.541792 | 0.038632 | 0.016485 | 0.017748 | 0.016034 |
| 1.092537 | 3.190369 | 0.541676 | 0.03877 | 0.020168 | 0.022389 | 0.020408 |
| 0.313443 | 0.915301 | 0.541676 | 0.03877 | 0.020168 | 0.022389 | 0.020408 |
| 1.042117 | 3.071548 | 0.54156 | 0.038113 | 0.031389 | 0.034895 | 0.032527 |
| 0.325569 | 0.959585 | 0.54156 | 0.038113 | 0.031389 | 0.034895 | 0.032527 |
| 1.034011 | 3.109042 | 0.541444 | 0.038365 | 0.034225 | 0.037612 | 0.035153 |
| 0.321642 | 0.967107 | 0.541444 | 0.038365 | 0.034225 | 0.037612 | 0.035153 |
| 0.355968 | 0.99963 | 0.541444 | 0.037718 | 0.048754 | 0.049836 | 0.0474 |
| 1.007916 | 2.820636 | 0.541328 | 0.037422 | 0.046805 | 0.046591 | 0.044234 |
| 0.308423 | 0.901304 | 0.541328 | 0.038573 | 0.017689 | 0.019294 | 0.017517 |
| 0.335667 | 0.989076 | 0.541096 | 0.037898 | 0.040571 | 0.045521 | 0.042882 |
| 1.057709 | 3.017131 | 0.541096 | 0.038247 | 0.029479 | 0.030023 | 0.027934 |

| | | | | | | |
|---|---|---|---|---|---|---|
| 0.331441 | 0.945439 | 0.541096 | 0.038247 | 0.029479 | 0.030023 | 0.027934 |
| 0.305011 | 0.883852 | 0.540632 | 0.03871 | 0.015778 | 0.015741 | 0.014263 |
| 0.335311 | 0.957829 | 0.540632 | 0.038282 | 0.031918 | 0.033932 | 0.031706 |
| 1.049582 | 3.029856 | 0.540632 | 0.038881 | 0.030778 | 0.032444 | 0.030278 |
| 1.00004 | 2.806 | 0.540515 | 0.037505 | 0.048908 | 0.049982 | 0.04754 |
| 0.356379 | 0.99996 | 0.540515 | 0.037505 | 0.048908 | 0.049982 | 0.04754 |
| 1.071209 | 3.121299 | 0.540283 | 0.038335 | 0.026531 | 0.02696 | 0.024979 |
| 0.344579 | 0.979654 | 0.539587 | 0.038153 | 0.039466 | 0.041642 | 0.039269 |
| 0.332055 | 0.979613 | 0.539587 | 0.038009 | 0.037966 | 0.041892 | 0.039395 |
| 1.044573 | 2.988484 | 0.539587 | 0.038179 | 0.031304 | 0.033789 | 0.031509 |
| 0.334618 | 0.957329 | 0.539587 | 0.038179 | 0.031304 | 0.033789 | 0.031509 |
| 1.161994 | 3.480357 | 0.539122 | 0.03939 | 0.010861 | 0.012544 | 0.01106 |
| 0.325497 | 0.965512 | 0.539122 | 0.038001 | 0.033264 | 0.036934 | 0.034567 |
| 1.001216 | 2.872367 | 0.539122 | 0.037917 | 0.046437 | 0.049474 | 0.046988 |
| 0.343985 | 0.992977 | 0.53889 | 0.038245 | 0.044009 | 0.047031 | 0.044613 |
| 1.09667 | 4.729676 | 0.538658 | 0.025753 | 0.042891 | 0.027282 | 0.023268 |
| 1.111375 | 3.222552 | 0.538426 | 0.038644 | 0.016703 | 0.018834 | 0.016998 |
| 1.034211 | 2.97158 | 0.53831 | 0.038345 | 0.03468 | 0.037079 | 0.034697 |
| 0.332894 | 0.965817 | 0.538194 | 0.038326 | 0.03438 | 0.036801 | 0.034497 |
| 1.321534 | 5.83874 | 0.537845 | 0.023999 | 0.014852 | 0.007028 | 0.004944 |
| 1.09025 | 3.216527 | 0.537845 | 0.0386 | 0.020664 | 0.023024 | 0.021049 |
| 0.310894 | 0.917221 | 0.537845 | 0.0386 | 0.020664 | 0.023024 | 0.021049 |
| 1.010811 | 2.948958 | 0.537497 | 0.038035 | 0.042643 | 0.045573 | 0.043115 |
| 0.317376 | 0.930776 | 0.537265 | 0.038242 | 0.023139 | 0.026329 | 0.024182 |
| 1.139095 | 3.276052 | 0.537033 | 0.038583 | 0.013911 | 0.014557 | 0.013 |
| 0.343565 | 0.990876 | 0.537033 | 0.038102 | 0.042551 | 0.046164 | 0.043637 |
| 0.346299 | 0.989014 | 0.536568 | 0.037997 | 0.045096 | 0.045368 | 0.043026 |
| 0.334996 | 0.960657 | 0.53622 | 0.038052 | 0.032945 | 0.034918 | 0.032723 |
| 0.323392 | 0.936043 | 0.53622 | 0.037948 | 0.025955 | 0.027543 | 0.025488 |
| 1.025287 | 2.916787 | 0.536104 | 0.038657 | 0.037895 | 0.040015 | 0.037673 |
| 0.312594 | 0.892317 | 0.53564 | 0.038781 | 0.015808 | 0.017044 | 0.015342 |
| 0.336906 | 0.970452 | 0.535524 | 0.03809 | 0.035619 | 0.03835 | 0.036026 |
| 0.328236 | 0.965 | 0.535407 | 0.038708 | 0.033624 | 0.036666 | 0.034357 |
| 0.352039 | 0.995412 | 0.535291 | 0.038721 | 0.046954 | 0.048007 | 0.045654 |
| 0.273963 | 0.797189 | 0.535059 | 0.039065 | 0.004539 | 0.005241 | 0.004344 |
| 0.350418 | 0.997302 | 0.534943 | 0.038278 | 0.049507 | 0.048828 | 0.046588 |
| 1.003994 | 2.842468 | 0.534711 | 0.037919 | 0.047484 | 0.048271 | 0.045822 |
| 0.351807 | 0.996022 | 0.534711 | 0.037919 | 0.047484 | 0.048271 | 0.045822 |
| 1.065202 | 2.984032 | 0.534711 | 0.037648 | 0.028557 | 0.027784 | 0.025778 |
| 0.33326 | 0.986008 | 0.534595 | 0.037839 | 0.045042 | 0.044338 | 0.042006 |
| 0.325947 | 0.978684 | 0.534247 | 0.038463 | 0.036454 | 0.041672 | 0.039003 |
| 1.02178 | 3.067986 | 0.534247 | 0.038463 | 0.036454 | 0.041672 | 0.039003 |
| 1.019455 | 4.267299 | 0.534247 | 0.025352 | 0.064061 | 0.044144 | 0.039573 |
| 0.345341 | 0.995781 | 0.534247 | 0.038145 | 0.045962 | 0.048198 | 0.045734 |
| 0.348957 | 0.997137 | 0.533898 | 0.038223 | 0.046347 | 0.048762 | 0.046335 |
| 0.311528 | 0.905846 | 0.533202 | 0.038973 | 0.018096 | 0.020173 | 0.018245 |
| 0.33109 | 0.961516 | 0.533086 | 0.038795 | 0.032187 | 0.035356 | 0.033044 |

| | | | | | | |
|---|---|---|---|---|---|---|
| 1.098535 | 3.219326 | 0.53297 | 0.038962 | 0.019451 | 0.021302 | 0.019415 |
| 1.057535 | 3.119651 | 0.532853 | 0.038045 | 0.028815 | 0.030567 | 0.02839 |
| 0.341982 | 0.99465 | 0.532621 | 0.038902 | 0.044866 | 0.047742 | 0.045262 |
| 1.004525 | 2.938532 | 0.532505 | 0.038209 | 0.043745 | 0.048104 | 0.045539 |
| 0.340306 | 0.995496 | 0.532505 | 0.038209 | 0.043745 | 0.048104 | 0.045539 |
| 1.124379 | 3.23582 | 0.532505 | 0.037988 | 0.016176 | 0.016634 | 0.014975 |
| 0.226682 | 0.89311 | 0.532505 | 0.03585 | 0.013426 | 0.022422 | 0.019179 |
| 0.323429 | 0.926611 | 0.532389 | 0.038275 | 0.024322 | 0.024843 | 0.022896 |
| 1.197625 | 3.452611 | 0.532389 | 0.038749 | 0.00782 | 0.008598 | 0.007418 |
| 0.313798 | 0.920569 | 0.532389 | 0.039303 | 0.021207 | 0.023734 | 0.021775 |
| 1.072794 | 3.113639 | 0.532273 | 0.038407 | 0.025673 | 0.026523 | 0.024484 |
| 0.347487 | 0.993357 | 0.531809 | 0.038256 | 0.04698 | 0.047163 | 0.044846 |
| 1.062162 | 3.085516 | 0.531693 | 0.038677 | 0.026333 | 0.029136 | 0.026974 |
| 0.322877 | 0.949047 | 0.531577 | 0.038344 | 0.028978 | 0.031548 | 0.029272 |
| 0.33334 | 0.990056 | 0.531228 | 0.038304 | 0.041188 | 0.045939 | 0.043302 |
| 0.340593 | 0.969639 | 0.529835 | 0.038281 | 0.037853 | 0.037942 | 0.035715 |
| 1.031311 | 2.936052 | 0.529835 | 0.038281 | 0.037853 | 0.037942 | 0.035715 |
| 0.327399 | 0.951815 | 0.528558 | 0.03911 | 0.029294 | 0.032245 | 0.030021 |
| 1.004314 | 2.901273 | 0.528326 | 0.038213 | 0.046338 | 0.048169 | 0.045713 |
| 1.208276 | 12.5948 | 0.528094 | 0.020376 | 0.058177 | 0.022825 | 0.014053 |
| 1.039995 | 3.02144 | 0.527049 | 0.038512 | 0.032629 | 0.03537 | 0.033113 |
| 0.330968 | 0.961543 | 0.527049 | 0.038512 | 0.032629 | 0.03537 | 0.033113 |
| 1.058527 | 3.001293 | 0.526701 | 0.03845 | 0.028446 | 0.029713 | 0.027615 |
| 1.005588 | 2.860426 | 0.525656 | 0.037623 | 0.048106 | 0.047607 | 0.045272 |
| 0.349598 | 0.994443 | 0.525656 | 0.037623 | 0.048106 | 0.047607 | 0.045272 |
| 1.668214 | 17.92117 | 0.524843 | 0.016202 | 0.02396 | 0.005033 | 0.001609 |
| 0.345396 | 0.987969 | 0.521128 | 0.038929 | 0.042651 | 0.044951 | 0.042578 |
| 1.032811 | 8.43684 | 0.514976 | 0.013179 | 0.076895 | 0.043361 | 0.034014 |
| 3.310094 | 242.7628 | 0.51486 | 0.014558 | 0.032252 | 0.00227 | 2.77E-06 |
| 0.327744 | 0.953963 | 0.513815 | 0.038953 | 0.032516 | 0.032934 | 0.030849 |
| 1.819882 | 111.5259 | 0.513583 | 0.013344 | 0.066466 | 0.011397 | 0.00083 |
| 1.149028 | 65.89473 | 0.512073 | 0.011903 | 0.113434 | 0.03622 | 0.011488 |
| 0.329691 | 0.953187 | 0.508707 | 0.038818 | 0.0301 | 0.032599 | 0.030412 |

| # sources: | GO:BP | GO:CC | KEGG | REAC | WP |
|---|---|---|---|---|---|
| source | term_name | term_id | adjusted_p_value | negative_l | intersectio |
| GO:MF | structural constituent of | GO:0003735 | 2.03E-16 | 15.69205 | 52 |
| GO:MF | RNA binding | GO:0003723 | 2.68E-14 | 13.57145 | 229 |
| GO:MF | oxidoreductase activity, a | GO:0016655 | 3.51E-05 | 4.454154 | 19 |
| GO:MF | structural molecule activi | GO:0005198 | 5.52E-05 | 4.257874 | 85 |
| GO:MF | nucleic acid binding | GO:0003676 | 0.000254565 | 3.594201 | 380 |
| GO:MF | oxidoreductase activity, a | GO:0016651 | 0.000449659 | 3.347117 | 25 |
| GO:MF | protein binding | GO:0005515 | 0.000776712 | 3.10974 | 841 |
| GO:MF | cadherin binding | GO:0045296 | 0.00176236 | 2.753905 | 51 |
| GO:MF | translation regulator acti | GO:0090079 | 0.005039901 | 2.297578 | 19 |
| GO:MF | ubiquitin binding | GO:0043130 | 0.009011925 | 2.045182 | 14 |
| GO:MF | translation regulator acti | GO:0045182 | 0.010453021 | 1.980758 | 26 |
| GO:MF | NADH dehydrogenase (qu | GO:0050136 | 0.014120564 | 1.850148 | 13 |
| GO:MF | NADH dehydrogenase (ub | GO:0008137 | 0.014120564 | 1.850148 | 13 |
| GO:MF | NADH dehydrogenase act | GO:0003954 | 0.018047796 | 1.743576 | 13 |
| GO:MF | small protein activating e | GO:0044388 | 0.022062821 | 1.656339 | 4 |
| GO:MF | translation factor activity | GO:0008135 | 0.023059186 | 1.637156 | 15 |
| GO:MF | heterocyclic compound b | GO:1901363 | 0.026268886 | 1.580558 | 506 |
| GO:MF | ubiquitin-like protein bin | GO:0032182 | 0.02936259 | 1.532206 | 15 |
| GO:MF | cell adhesion molecule bi | GO:0050839 | 0.032930045 | 1.482408 | 31 |
| GO:MF | organic cyclic compound | GO:0097159 | 0.035970829 | 1.44405 | 511 |
| GO:MF | sulfate binding | GO:0043199 | 0.039383691 | 1.404684 | 2 |
| GO:MF | rRNA binding | GO:0019843 | 0.043781413 | 1.35871 | 14 |
| GO:MF | double-stranded RNA bin | GO:0003725 | 0.043910694 | 1.35743 | 11 |
| GO:BP | cotranslational protein ta | GO:0006613 | 2.36E-14 | 13.6271 | 37 |
| GO:BP | SRP-dependent cotransla | GO:0006614 | 2.67E-14 | 13.57274 | 38 |
| GO:BP | mRNA metabolic process | GO:0016071 | 3.05E-14 | 13.51573 | 144 |
| GO:BP | nuclear-transcribed mRN | GO:0000184 | 2.11E-13 | 12.6759 | 42 |
| GO:BP | protein localization to en | GO:0070972 | 4.79E-13 | 12.31988 | 42 |
| GO:BP | protein targeting to ER | GO:0045047 | 5.31E-13 | 12.2745 | 35 |
| GO:BP | viral process | GO:0016032 | 5.32E-13 | 12.27416 | 116 |
| GO:BP | translational initiation | GO:0006413 | 6.24E-13 | 12.20457 | 54 |
| GO:BP | establishment of protein | GO:0072599 | 1.86E-12 | 11.73158 | 35 |
| GO:BP | symbiotic process | GO:0044403 | 3.14E-12 | 11.50353 | 119 |
| GO:BP | intracellular protein trans | GO:0006886 | 3.22E-12 | 11.49208 | 174 |
| GO:BP | viral gene expression | GO:0019080 | 5.94E-12 | 11.22621 | 46 |
| GO:BP | cellular macromolecule c | GO:0044265 | 9.56E-12 | 11.01963 | 145 |
| GO:BP | translation | GO:0006412 | 1.19E-11 | 10.92507 | 116 |
| GO:BP | peptide biosynthetic proc | GO:0043043 | 2.19E-11 | 10.66017 | 118 |
| GO:BP | intracellular transport | GO:0046907 | 4.40E-11 | 10.35652 | 233 |
| GO:BP | mRNA catabolic process | GO:0006402 | 5.97E-11 | 10.22431 | 65 |
| GO:BP | aromatic compound cata | GO:0019439 | 8.21E-11 | 10.08564 | 107 |
| GO:BP | interspecies interaction b | GO:0044419 | 9.61E-11 | 10.01731 | 120 |
| GO:BP | cellular macromolecule lo | GO:0070727 | 1.93E-10 | 9.715416 | 250 |
| GO:BP | establishment of protein | GO:0072594 | 2.09E-10 | 9.679183 | 91 |
| GO:BP | nucleobase-containing co | GO:0034655 | 2.98E-10 | 9.525914 | 98 |

| | | | | | |
|---|---|---|---|---|---|
| GO:BP | cellular protein localizatio | GO:0034613 | 3.30E-10 | 9.481359 | 248 |
| GO:BP | macromolecule catabolic | GO:0009057 | 4.28E-10 | 9.369002 | 159 |
| GO:BP | viral transcription | GO:0019083 | 4.64E-10 | 9.333305 | 41 |
| GO:BP | peptide metabolic proces | GO:0006518 | 9.95E-10 | 9.002049 | 128 |
| GO:BP | organic cyclic compound | GO:1901361 | 1.11E-09 | 8.955745 | 108 |
| GO:BP | protein targeting | GO:0006605 | 1.15E-09 | 8.938333 | 81 |
| GO:BP | nuclear-transcribed mRN | GO:0000956 | 1.21E-09 | 8.917614 | 51 |
| GO:BP | heterocycle catabolic pro | GO:0046700 | 1.34E-09 | 8.871315 | 102 |
| GO:BP | cellular nitrogen compou | GO:0044270 | 1.49E-09 | 8.825461 | 102 |
| GO:BP | amide biosynthetic proce | GO:0043604 | 2.84E-09 | 8.545971 | 127 |
| GO:BP | RNA catabolic process | GO:0006401 | 3.47E-09 | 8.460077 | 66 |
| GO:BP | cellular localization | GO:0051641 | 7.50E-09 | 8.12501 | 328 |
| GO:BP | protein targeting to mem | GO:0006612 | 1.22E-08 | 7.913595 | 41 |
| GO:BP | protein localization to org | GO:0033365 | 1.66E-08 | 7.779757 | 123 |
| GO:BP | organic substance catabo | GO:1901575 | 2.40E-08 | 7.619716 | 268 |
| GO:BP | cellular amide metabolic | GO:0043603 | 6.56E-08 | 7.182908 | 149 |
| GO:BP | cellular protein-containin | GO:0034622 | 2.11E-07 | 6.676145 | 148 |
| GO:BP | protein-containing compl | GO:0043933 | 2.80E-07 | 6.553292 | 233 |
| GO:BP | cellular catabolic process | GO:0044248 | 3.15E-07 | 6.501475 | 270 |
| GO:BP | RNA processing | GO:0006396 | 6.49E-07 | 6.187767 | 132 |
| GO:BP | catabolic process | GO:0009056 | 1.06E-06 | 5.973747 | 303 |
| GO:BP | protein transport | GO:0015031 | 1.13E-06 | 5.945804 | 246 |
| GO:BP | establishment of protein | GO:0045184 | 1.59E-06 | 5.797756 | 256 |
| GO:BP | cellular component bioge | GO:0044085 | 2.02E-06 | 5.695666 | 319 |
| GO:BP | ribonucleoprotein comple | GO:0022613 | 2.62E-06 | 5.582008 | 72 |
| GO:BP | organonitrogen compour | GO:1901566 | 3.28E-06 | 5.484667 | 223 |
| GO:BP | establishment of protein | GO:0090150 | 3.62E-06 | 5.441458 | 51 |
| GO:BP | peptide transport | GO:0015833 | 4.23E-06 | 5.373854 | 247 |
| GO:BP | amide transport | GO:0042886 | 1.20E-05 | 4.919875 | 248 |
| GO:BP | protein localization | GO:0008104 | 1.53E-05 | 4.815637 | 314 |
| GO:BP | RNA splicing | GO:0008380 | 2.54E-05 | 4.594799 | 72 |
| GO:BP | nucleobase-containing co | GO:0006139 | 3.71E-05 | 4.430902 | 342 |
| GO:BP | protein-containing compl | GO:0065003 | 3.77E-05 | 4.42339 | 204 |
| GO:BP | cellular nitrogen compou | GO:0034641 | 4.07E-05 | 4.390239 | 554 |
| GO:BP | organelle organization | GO:0006996 | 5.00E-05 | 4.301135 | 367 |
| GO:BP | ribonucleoprotein comple | GO:0071826 | 5.43E-05 | 4.265277 | 40 |
| GO:BP | ribonucleoprotein comple | GO:0022618 | 5.95E-05 | 4.225786 | 39 |
| GO:BP | regulation of mRNA meta | GO:1903311 | 7.45E-05 | 4.127853 | 49 |
| GO:BP | RNA splicing, via transest | GO:0000375 | 0.00012764 | 3.894013 | 60 |
| GO:BP | protein localization to me | GO:0072657 | 0.000150858 | 3.821432 | 83 |
| GO:BP | mitochondrial respiratory | GO:0033108 | 0.000152305 | 3.817287 | 25 |
| GO:BP | mitochondrion organizati | GO:0007005 | 0.000153793 | 3.813065 | 77 |
| GO:BP | heterocycle metabolic pr | GO:0046483 | 0.000159868 | 3.796239 | 347 |
| GO:BP | modification-dependent | GO:0043632 | 0.000198107 | 3.703101 | 93 |
| GO:BP | organonitrogen compour | GO:1901565 | 0.000217657 | 3.662227 | 158 |
| GO:BP | RNA splicing, via transest | GO:0000377 | 0.00022287 | 3.651947 | 59 |
| GO:BP | mRNA splicing, via splicec | GO:0000398 | 0.00022287 | 3.651947 | 59 |

| | | | | | |
|---|---|---|---|---|---|
| GO:BP | cytoplasmic translation | GO:0002181 | 0.000250407 | 3.601354 | 22 |
| GO:BP | macromolecule localizati | GO:0033036 | 0.000315725 | 3.500691 | 339 |
| GO:BP | cellular protein metaboli | GO:0044267 | 0.000317066 | 3.498851 | 423 |
| GO:BP | cellular component asser | GO:0022607 | 0.000320778 | 3.493796 | 287 |
| GO:BP | cellular component orgar | GO:0071840 | 0.000328834 | 3.483024 | 622 |
| GO:BP | cellular nitrogen compou | GO:0044271 | 0.000332832 | 3.477775 | 308 |
| GO:BP | cellular aromatic compou | GO:0006725 | 0.000387929 | 3.411248 | 347 |
| GO:BP | cellular protein catabolic | GO:0044257 | 0.000477298 | 3.32121 | 86 |
| GO:BP | NADH dehydrogenase co | GO:0010257 | 0.000487173 | 3.312317 | 19 |
| GO:BP | mitochondrial respiratory | GO:0032981 | 0.000487173 | 3.312317 | 19 |
| GO:BP | ubiquitin-dependent prot | GO:0006511 | 0.000526289 | 3.278776 | 71 |
| GO:BP | mRNA processing | GO:0006397 | 0.000572832 | 3.241973 | 77 |
| GO:BP | cellular metabolic proces | GO:0044237 | 0.000732846 | 3.134987 | 972 |
| GO:BP | modification-dependent | GO:0019941 | 0.000752698 | 3.123379 | 71 |
| GO:BP | nucleic acid metabolic pr | GO:0090304 | 0.001233443 | 2.908881 | 307 |
| GO:BP | organic cyclic compound | GO:1901360 | 0.001235673 | 2.908096 | 383 |
| GO:BP | translational elongation | GO:0006414 | 0.001301397 | 2.88559 | 29 |
| GO:BP | nitrogen compound trans | GO:0071705 | 0.001328567 | 2.876616 | 265 |
| GO:BP | cellular macromolecule b | GO:0034645 | 0.001861393 | 2.730162 | 296 |
| GO:BP | protein catabolic process | GO:0030163 | 0.001891999 | 2.723079 | 97 |
| GO:BP | proteolysis involved in ce | GO:0051603 | 0.002001436 | 2.698658 | 77 |
| GO:BP | macromolecule biosynthe | GO:0009059 | 0.002629447 | 2.580136 | 303 |
| GO:BP | organonitrogen compour | GO:1901564 | 0.002673959 | 2.572845 | 590 |
| GO:BP | ribosome biogenesis | GO:0042254 | 0.003273501 | 2.484988 | 45 |
| GO:BP | organic substance biosyn | GO:1901576 | 0.003316251 | 2.479353 | 283 |
| GO:BP | cellular component orgar | GO:0016043 | 0.004049356 | 2.392614 | 598 |
| GO:BP | translational termination | GO:0006415 | 0.005198877 | 2.28409 | 23 |
| GO:BP | cellular component disas | GO:0022411 | 0.006152235 | 2.210967 | 76 |
| GO:BP | mitochondrial ATP synthe | GO:0042775 | 0.006813991 | 2.166598 | 23 |
| GO:BP | regulation of catabolic pr | GO:0009894 | 0.006965388 | 2.157055 | 100 |
| GO:BP | cellular biosynthetic proc | GO:0044249 | 0.007242037 | 2.140139 | 335 |
| GO:BP | ATP synthesis coupled ele | GO:0042773 | 0.008201807 | 2.08609 | 23 |
| GO:BP | protein metabolic proces | GO:0019538 | 0.010222878 | 1.990427 | 513 |
| GO:BP | ribonucleoprotein comple | GO:0071166 | 0.010692479 | 1.970922 | 27 |
| GO:BP | mitochondrial translation | GO:0070125 | 0.012682712 | 1.896788 | 20 |
| GO:BP | RNA metabolic process | GO:0016070 | 0.013090646 | 1.883039 | 276 |
| GO:BP | regulation of mRNA splici | GO:0048024 | 0.013921641 | 1.85631 | 24 |
| GO:BP | biosynthetic process | GO:0009058 | 0.014098415 | 1.85083 | 342 |
| GO:BP | alternative mRNA splicing | GO:0000380 | 0.016267811 | 1.788671 | 20 |
| GO:BP | metabolic process | GO:0008152 | 0.016832952 | 1.77384 | 1005 |
| GO:BP | regulation of organelle or | GO:0033043 | 0.017477897 | 1.757511 | 152 |
| GO:BP | nuclear transport | GO:0051169 | 0.017817477 | 1.749154 | 46 |
| GO:BP | regulation of mRNA proc | GO:0050684 | 0.018261071 | 1.738474 | 29 |
| GO:BP | respiratory electron trans | GO:0022904 | 0.018273081 | 1.738188 | 25 |
| GO:BP | protein export from nucle | GO:0006611 | 0.020869879 | 1.68048 | 33 |
| GO:BP | regulation of cellular cata | GO:0031329 | 0.022311067 | 1.65148 | 85 |
| GO:BP | regulation of alternative | GO:0000381 | 0.02619405 | 1.581797 | 18 |

| | | | | | |
|---|---|---|---|---|---|
| GO:BP | regulation of mRNA stabi | GO:0043488 | 0.027044347 | 1.567923 | 17 |
| GO:BP | ribonucleoprotein comple | GO:0071426 | 0.028270102 | 1.548673 | 26 |
| GO:BP | 2'-deoxyribonucleotide m | GO:0009394 | 0.030230948 | 1.519548 | 10 |
| GO:BP | mitochondrial electron tr | GO:0006120 | 0.030507445 | 1.515594 | 15 |
| GO:BP | nucleocytoplasmic transp | GO:0006913 | 0.031768565 | 1.498002 | 45 |
| GO:BP | regulation of mRNA catab | GO:0061013 | 0.033198474 | 1.478882 | 18 |
| GO:BP | cellular response to stres | GO:0033554 | 0.034570262 | 1.461297 | 165 |
| GO:BP | cytoplasmic translational | GO:0002183 | 0.0381589 | 1.418404 | 9 |
| GO:BP | cellular macromolecule m | GO:0044260 | 0.047296932 | 1.325167 | 434 |
| GO:BP | ribosomal large subunit b | GO:0042273 | 0.047772958 | 1.320818 | 14 |
| GO:CC | intracellular membrane-b | GO:0043231 | 2.31E-28 | 27.63634 | 1058 |
| GO:CC | cytosol | GO:0005829 | 1.09E-27 | 26.96382 | 591 |
| GO:CC | membrane-bounded orga | GO:0043227 | 7.67E-26 | 25.11521 | 1155 |
| GO:CC | intracellular organelle | GO:0043229 | 3.77E-25 | 24.42382 | 1170 |
| GO:CC | cytoplasm | GO:0005737 | 2.65E-24 | 23.57685 | 1084 |
| GO:CC | intracellular | GO:0005622 | 4.72E-24 | 23.3261 | 1258 |
| GO:CC | organelle | GO:0043226 | 3.16E-23 | 22.50063 | 1222 |
| GO:CC | ribonucleoprotein comple | GO:1990904 | 5.00E-21 | 20.30136 | 134 |
| GO:CC | membrane-enclosed lum | GO:0031974 | 4.77E-19 | 18.32156 | 576 |
| GO:CC | intracellular organelle lu | GO:0070013 | 4.77E-19 | 18.32156 | 576 |
| GO:CC | organelle lumen | GO:0043233 | 4.77E-19 | 18.32156 | 576 |
| GO:CC | nucleus | GO:0005634 | 1.16E-17 | 16.93425 | 745 |
| GO:CC | ribosomal subunit | GO:0044391 | 4.93E-17 | 16.30739 | 55 |
| GO:CC | nucleoplasm | GO:0005654 | 8.68E-16 | 15.06149 | 446 |
| GO:CC | ribosome | GO:0005840 | 8.91E-16 | 15.05023 | 59 |
| GO:CC | cytosolic ribosome | GO:0022626 | 8.30E-15 | 14.08105 | 37 |
| GO:CC | nuclear lumen | GO:0031981 | 9.02E-13 | 12.04495 | 462 |
| GO:CC | protein-containing compl | GO:0032991 | 4.47E-12 | 11.34994 | 499 |
| GO:CC | organelle envelope | GO:0031967 | 1.31E-11 | 10.88278 | 157 |
| GO:CC | envelope | GO:0031975 | 1.31E-11 | 10.88278 | 157 |
| GO:CC | small ribosomal subunit | GO:0015935 | 6.25E-11 | 10.20386 | 27 |
| GO:CC | cytosolic small ribosomal | GO:0022627 | 1.05E-09 | 8.978784 | 20 |
| GO:CC | mitochondrial inner mem | GO:0005743 | 5.94E-09 | 8.225884 | 75 |
| GO:CC | mitochondrial envelope | GO:0005740 | 8.90E-09 | 8.050643 | 103 |
| GO:CC | organelle inner membran | GO:0019866 | 2.02E-08 | 7.693908 | 80 |
| GO:CC | intracellular non-membra | GO:0043232 | 2.91E-08 | 7.536148 | 451 |
| GO:CC | non-membrane-bounded | GO:0043228 | 3.95E-08 | 7.402959 | 451 |
| GO:CC | catalytic complex | GO:1902494 | 5.06E-08 | 7.29545 | 155 |
| GO:CC | nucleolus | GO:0005730 | 1.36E-07 | 6.867553 | 121 |
| GO:CC | mitochondrion | GO:0005739 | 4.90E-07 | 6.309432 | 182 |
| GO:CC | mitochondrial protein co | GO:0098798 | 6.28E-07 | 6.202235 | 49 |
| GO:CC | mitochondrial membrane | GO:0031966 | 8.81E-07 | 6.054832 | 93 |
| GO:CC | large ribosomal subunit | GO:0015934 | 1.29E-06 | 5.889882 | 29 |
| GO:CC | extracellular vesicle | GO:1903561 | 1.70E-06 | 5.770237 | 119 |
| GO:CC | extracellular exosome | GO:0070062 | 1.76E-06 | 5.753981 | 118 |
| GO:CC | extracellular organelle | GO:0043230 | 1.94E-06 | 5.711779 | 119 |
| GO:CC | mitochondrial matrix | GO:0005759 | 3.40E-06 | 5.467901 | 66 |

| | | | | | |
|---|---|---|---|---|---|
| **GO:CC** | organelle membrane | GO:0031090 | 3.77E-06 | 5.423342 | 353 |
| **GO:CC** | focal adhesion | GO:0005925 | 7.29E-06 | 5.137515 | 63 |
| **GO:CC** | cell-substrate junction | GO:0030055 | 1.39E-05 | 4.856564 | 63 |
| **GO:CC** | cytosolic large ribosomal | GO:0022625 | 1.84E-05 | 4.736245 | 16 |
| **GO:CC** | vesicle | GO:0031982 | 0.000186693 | 3.728872 | 378 |
| **GO:CC** | spliceosomal complex | GO:0005681 | 0.000209505 | 3.678806 | 34 |
| **GO:CC** | respiratory chain comple | GO:0098803 | 0.000236252 | 3.626624 | 21 |
| **GO:CC** | mitochondrial respirason | GO:0005746 | 0.000267541 | 3.572609 | 22 |
| **GO:CC** | respirasome | GO:0070469 | 0.000707025 | 3.150565 | 23 |
| **GO:CC** | peptidase complex | GO:1905368 | 0.000762615 | 3.117695 | 18 |
| **GO:CC** | respiratory chain comple | GO:0045271 | 0.000848597 | 3.071299 | 15 |
| **GO:CC** | mitochondrial respiratory | GO:0005747 | 0.000848597 | 3.071299 | 15 |
| **GO:CC** | NADH dehydrogenase co | GO:0030964 | 0.000848597 | 3.071299 | 15 |
| **GO:CC** | proteasome complex | GO:0000502 | 0.001226581 | 2.911304 | 15 |
| **GO:CC** | catalytic step 2 spliceoso | GO:0071013 | 0.001287976 | 2.890092 | 20 |
| **GO:CC** | endopeptidase complex | GO:1905369 | 0.00149291 | 2.825966 | 15 |
| **GO:CC** | organelle envelope lumer | GO:0031970 | 0.001861145 | 2.73022 | 17 |
| **GO:CC** | inner mitochondrial mem | GO:0098800 | 0.001866483 | 2.728976 | 26 |
| **GO:CC** | neuron projection cytopl | GO:0120111 | 0.003038008 | 2.517411 | 7 |
| **GO:CC** | axon cytoplasm | GO:1904115 | 0.003411421 | 2.467065 | 6 |
| **GO:CC** | mitochondrial ribosome | GO:0005761 | 0.003905963 | 2.408272 | 19 |
| **GO:CC** | organellar ribosome | GO:0000313 | 0.003905963 | 2.408272 | 19 |
| **GO:CC** | proteasome core comple | GO:0019773 | 0.004087724 | 2.388518 | 4 |
| **GO:CC** | mitochondrial intermemb | GO:0005758 | 0.006039351 | 2.21901 | 18 |
| **GO:CC** | muscle thin filament trop | GO:0005862 | 0.006250408 | 2.204092 | 4 |
| **GO:CC** | polysome | GO:0005844 | 0.006708149 | 2.173397 | 15 |
| **GO:CC** | cell leading edge | GO:0031252 | 0.006715582 | 2.172916 | 20 |
| **GO:CC** | nuclear outer membrane | GO:0042175 | 0.008271472 | 2.082417 | 90 |
| **GO:CC** | endoplasmic reticulum m | GO:0005789 | 0.010729813 | 1.969408 | 88 |
| **GO:CC** | spindle midzone | GO:0051233 | 0.01273414 | 1.89503 | 4 |
| **GO:CC** | nuclear envelope | GO:0005635 | 0.013353824 | 1.874394 | 62 |
| **GO:CC** | intrinsic component of or | GO:0031300 | 0.013794329 | 1.860299 | 43 |
| **GO:CC** | anchoring junction | GO:0070161 | 0.018060173 | 1.743278 | 88 |
| **GO:CC** | pyruvate kinase complex | GO:1902912 | 0.018940542 | 1.722608 | 1 |
| **GO:CC** | lamellipodium | GO:0030027 | 0.019278855 | 1.714919 | 32 |
| **GO:CC** | mitotic spindle | GO:0072686 | 0.021244668 | 1.67275 | 21 |
| **GO:CC** | lysosomal membrane | GO:0005765 | 0.022624079 | 1.645429 | 49 |
| **GO:CC** | lytic vacuole membrane | GO:0098852 | 0.022624079 | 1.645429 | 49 |
| **GO:CC** | endoplasmic reticulum | GO:0005783 | 0.027521805 | 1.560323 | 136 |
| **GO:CC** | preribosome, large subur | GO:0030687 | 0.027656123 | 1.558209 | 5 |
| **GO:CC** | cilium | GO:0005929 | 0.029836935 | 1.525246 | 4 |
| **GO:CC** | nuclear periphery | GO:0034399 | 0.033719604 | 1.472118 | 21 |
| **GO:CC** | whole membrane | GO:0098805 | 0.034195703 | 1.466028 | 137 |
| **GO:CC** | vacuolar membrane | GO:0005774 | 0.035903818 | 1.444859 | 54 |
| **GO:CC** | integral component of or | GO:0031301 | 0.044647221 | 1.350206 | 39 |
| **KEGG** | Ribosome | KEGG:03010 | 0.000391839 | 3.406892 | 45 |
| **REAC** | Eukaryotic Translation Te | REAC:R-HSA-72764 | 1.34E-08 | 7.873057 | 34 |

| | | | | | |
|---|---|---|---|---|---|
| REAC | Eukaryotic Translation Elo | REAC:R-HSA-156842 | 1.34E-08 | 7.873057 | 34 |
| REAC | Nonsense Mediated Deca | REAC:R-HSA-975956 | 2.62E-08 | 7.581128 | 34 |
| REAC | Formation of a pool of fre | REAC:R-HSA-72689 | 3.79E-08 | 7.421359 | 35 |
| REAC | L13a-mediated translatio | REAC:R-HSA-156827 | 3.86E-08 | 7.413825 | 37 |
| REAC | Peptide chain elongation | REAC:R-HSA-156902 | 8.64E-08 | 7.063335 | 32 |
| REAC | Viral mRNA Translation | REAC:R-HSA-192823 | 8.64E-08 | 7.063335 | 32 |
| REAC | Eukaryotic Translation Ini | REAC:R-HSA-72613 | 8.79E-08 | 7.056022 | 38 |
| REAC | Cap-dependent Translatio | REAC:R-HSA-72737 | 8.79E-08 | 7.056022 | 38 |
| REAC | Regulation of expression | REAC:R-HSA-9010553 | 1.47E-07 | 6.83337 | 47 |
| REAC | GTP hydrolysis and joinin | REAC:R-HSA-72706 | 2.20E-07 | 6.656913 | 36 |
| REAC | Selenocysteine synthesis | REAC:R-HSA-2408557 | 3.12E-07 | 6.506122 | 32 |
| REAC | Nonsense Mediated Deca | REAC:R-HSA-975957 | 4.98E-07 | 6.303198 | 36 |
| REAC | Nonsense-Mediated Deca | REAC:R-HSA-927802 | 4.98E-07 | 6.303198 | 36 |
| REAC | SRP-dependent cotransla | REAC:R-HSA-1799339 | 9.02E-07 | 6.044672 | 35 |
| REAC | Translation | REAC:R-HSA-72766 | 9.57E-07 | 6.019015 | 71 |
| REAC | Influenza Viral RNA Trans | REAC:R-HSA-168273 | 2.91E-06 | 5.535837 | 38 |
| REAC | Influenza Infection | REAC:R-HSA-168254 | 4.68E-06 | 5.32976 | 41 |
| REAC | Influenza Life Cycle | REAC:R-HSA-168255 | 6.23E-06 | 5.205392 | 39 |
| REAC | Signaling by ROBO recept | REAC:R-HSA-376176 | 9.38E-06 | 5.027715 | 51 |
| REAC | Selenoamino acid metabo | REAC:R-HSA-2408522 | 9.44E-06 | 5.024979 | 34 |
| REAC | Translation initiation com | REAC:R-HSA-72649 | 0.000111429 | 3.953003 | 24 |
| REAC | Formation of the ternary | REAC:R-HSA-72695 | 0.000156812 | 3.804621 | 22 |
| REAC | Activation of the mRNA u | REAC:R-HSA-72662 | 0.000162128 | 3.790141 | 24 |
| REAC | Ribosomal scanning and s | REAC:R-HSA-72702 | 0.000490507 | 3.309355 | 23 |
| REAC | rRNA processing in the nu | REAC:R-HSA-8868773 | 0.000584005 | 3.233583 | 43 |
| REAC | rRNA processing | REAC:R-HSA-72312 | 0.000967596 | 3.014306 | 44 |
| REAC | Infectious disease | REAC:R-HSA-5663205 | 0.001537087 | 2.813302 | 68 |
| REAC | Major pathway of rRNA p | REAC:R-HSA-6791226 | 0.00226892 | 2.644181 | 40 |
| REAC | Metabolism of RNA | REAC:R-HSA-8953854 | 0.030171861 | 1.520398 | 104 |
| WP | Cytoplasmic Ribosomal P | WP:WP477 | 0.002762601 | 2.558682 | 31 |
| TF | Factor: ZF5; motif: GGSG( | TF:M10438_1 | 2.36E-48 | 47.62619 | 1317 |
| TF | Factor: ZF5; motif: GSGC( | TF:M00716_1 | 5.29E-44 | 43.27675 | 1249 |
| TF | Factor: ER81; motif: RCC( | TF:M02065_1 | 1.69E-43 | 42.77102 | 753 |
| TF | Factor: ZF5; motif: GGSG( | TF:M10438 | 1.94E-42 | 41.71231 | 1359 |
| TF | Factor: ZF5; motif: GGSG( | TF:M10438_0 | 1.94E-42 | 41.71231 | 1359 |
| TF | Factor: ZF5; motif: GSGC( | TF:M00716 | 4.35E-41 | 40.36169 | 1364 |
| TF | Factor: ZF5; motif: GSGC( | TF:M00716_0 | 4.35E-41 | 40.36169 | 1364 |
| TF | Factor: E2F-4; motif: SNG | TF:M09894_1 | 8.10E-39 | 38.09125 | 1226 |
| TF | Factor: Fli-1; motif: NACC | TF:M11400_0 | 1.63E-38 | 37.78874 | 970 |
| TF | Factor: Fli-1; motif: NACC | TF:M11400 | 1.63E-38 | 37.78874 | 970 |
| TF | Factor: E2F-3:HES-7; mot | TF:M08526_1 | 2.28E-38 | 37.64259 | 1287 |
| TF | Factor: Erm; motif: ACCG | TF:M02069_0 | 9.20E-38 | 37.03613 | 740 |
| TF | Factor: Erm; motif: ACCG | TF:M02069 | 9.20E-38 | 37.03613 | 740 |
| TF | Factor: Erg; motif: NACC( | TF:M11396_1 | 1.41E-37 | 36.85064 | 582 |
| TF | Factor: Fli-1; motif: NACC | TF:M11404 | 1.61E-37 | 36.79329 | 892 |
| TF | Factor: Fli-1; motif: NACC | TF:M11404_0 | 1.61E-37 | 36.79329 | 892 |
| TF | Factor: E2F-1:Elk-1; motif | TF:M08205_0 | 2.00E-37 | 36.69805 | 1248 |

| | | | | | |
|---|---|---|---|---|---|
| **TF** | Factor: E2F-1:Elk-1; motif | TF:M08205 | 2.00E-37 | 36.69805 | 1248 |
| **TF** | Factor: SAP-1; motif: NN( | TF:M09905_0 | 4.48E-37 | 36.34835 | 853 |
| **TF** | Factor: SAP-1; motif: NN( | TF:M09905 | 4.48E-37 | 36.34835 | 853 |
| **TF** | Factor: ETV4; motif: ACC( | TF:M03982 | 5.71E-37 | 36.24373 | 734 |
| **TF** | Factor: ETV4; motif: ACC( | TF:M03982_0 | 5.71E-37 | 36.24373 | 734 |
| **TF** | Factor: Erg; motif: NACC( | TF:M11392_0 | 2.29E-36 | 35.63935 | 845 |
| **TF** | Factor: Erg; motif: NACC( | TF:M11392 | 2.29E-36 | 35.63935 | 845 |
| **TF** | Factor: E2F-4; motif: SNG | TF:M09894 | 4.32E-36 | 35.36407 | 1346 |
| **TF** | Factor: E2F-4; motif: SNG | TF:M09894_0 | 4.32E-36 | 35.36407 | 1346 |
| **TF** | Factor: FLI-1; motif: NAYT | TF:M08913 | 6.39E-36 | 35.19461 | 853 |
| **TF** | Factor: FLI-1; motif: NAYT | TF:M08913_0 | 6.39E-36 | 35.19461 | 853 |
| **TF** | Factor: ETS1; motif: ACC( | TF:M03977 | 7.76E-36 | 35.11008 | 1001 |
| **TF** | Factor: ETS1; motif: ACC( | TF:M03977_0 | 7.76E-36 | 35.11008 | 1001 |
| **TF** | Factor: nerf; motif: NNN( | TF:M09901 | 1.79E-35 | 34.74689 | 610 |
| **TF** | Factor: nerf; motif: NNN( | TF:M09901_0 | 1.79E-35 | 34.74689 | 610 |
| **TF** | Factor: ETV7; motif: NCC( | TF:M02071 | 1.86E-35 | 34.73031 | 973 |
| **TF** | Factor: ETV7; motif: NCC( | TF:M02071_0 | 1.86E-35 | 34.73031 | 973 |
| **TF** | Factor: Elk-1; motif: NNN | TF:M00025_0 | 2.23E-35 | 34.65156 | 985 |
| **TF** | Factor: Elk-1; motif: NNN | TF:M00025 | 2.23E-35 | 34.65156 | 985 |
| **TF** | Factor: E2F-2; motif: GCG | TF:M11529_1 | 4.34E-35 | 34.36224 | 1265 |
| **TF** | Factor: ERG; motif: ACCG | TF:M03975_0 | 1.50E-34 | 33.82301 | 974 |
| **TF** | Factor: ERG; motif: ACCG | TF:M03975 | 1.50E-34 | 33.82301 | 974 |
| **TF** | Factor: Elk-1; motif: NNC( | TF:M11435_1 | 1.70E-34 | 33.76957 | 637 |
| **TF** | Factor: E2F; motif: GGCG | TF:M00803 | 1.81E-34 | 33.74286 | 1180 |
| **TF** | Factor: E2F; motif: GGCG | TF:M00803_0 | 1.81E-34 | 33.74286 | 1180 |
| **TF** | Factor: EHF; motif: CSCG( | TF:M02052_1 | 1.96E-34 | 33.70717 | 495 |
| **TF** | Factor: E2F-1:HES-7; mot | TF:M08525_1 | 2.41E-34 | 33.61818 | 1311 |
| **TF** | Factor: E2F-2; motif: GCG | TF:M11529_0 | 2.45E-34 | 33.61073 | 1346 |
| **TF** | Factor: E2F-2; motif: GCG | TF:M11529 | 2.45E-34 | 33.61073 | 1346 |
| **TF** | Factor: Elf-1; motif: NAM | TF:M11447 | 4.01E-34 | 33.39721 | 783 |
| **TF** | Factor: Elf-1; motif: NAM | TF:M11447_0 | 4.01E-34 | 33.39721 | 783 |
| **TF** | Factor: FLI1; motif: ACCG | TF:M03988 | 8.14E-34 | 33.08925 | 848 |
| **TF** | Factor: FLI1; motif: ACCG | TF:M03988_0 | 8.14E-34 | 33.08925 | 848 |
| **TF** | Factor: ER81; motif: RCC( | TF:M02065 | 1.44E-33 | 32.84212 | 1143 |
| **TF** | Factor: ER81; motif: RCC( | TF:M02065_0 | 1.44E-33 | 32.84212 | 1143 |
| **TF** | Factor: Erg; motif: ACCGG | TF:M02062_0 | 2.42E-33 | 32.61629 | 816 |
| **TF** | Factor: Erg; motif: ACCGG | TF:M02062 | 2.42E-33 | 32.61629 | 816 |
| **TF** | Factor: Erg; motif: NACC( | TF:M11396_0 | 4.12E-33 | 32.38554 | 1012 |
| **TF** | Factor: Erg; motif: NACC( | TF:M11396 | 4.12E-33 | 32.38554 | 1012 |
| **TF** | Factor: ER81; motif: NNC | TF:M11419_0 | 5.83E-33 | 32.23443 | 660 |
| **TF** | Factor: ER81; motif: NNC | TF:M11419 | 5.83E-33 | 32.23443 | 660 |
| **TF** | Factor: PEA3; motif: NAC | TF:M11428 | 8.31E-33 | 32.08039 | 747 |
| **TF** | Factor: PEA3; motif: NAC | TF:M11428_0 | 8.31E-33 | 32.08039 | 747 |
| **TF** | Factor: p300; motif: ACN | TF:M04826_1 | 9.57E-33 | 32.01897 | 1295 |
| **TF** | Factor: Elk-1; motif: NNN | TF:M00025_1 | 1.21E-32 | 31.9169 | 533 |
| **TF** | Factor: ELK-1; motif: ACC | TF:M02059_1 | 1.74E-32 | 31.75892 | 348 |
| **TF** | Factor: PEA3; motif: RCC( | TF:M02066 | 2.44E-32 | 31.61326 | 737 |

| | | | | | |
|---|---|---|---|---|---|
| **TF** | Factor: PEA3; motif: RCC( | TF:M02066_0 | 2.44E-32 | 31.61326 | 737 |
| **TF** | Factor: c-Ets-2; motif: NC | TF:M08759_0 | 4.51E-32 | 31.3463 | 831 |
| **TF** | Factor: c-Ets-2; motif: NC | TF:M08759 | 4.51E-32 | 31.3463 | 831 |
| **TF** | Factor: PEA3; motif: NAC( | TF:M11425_0 | 5.85E-32 | 31.23317 | 763 |
| **TF** | Factor: PEA3; motif: NAC( | TF:M11425 | 5.85E-32 | 31.23317 | 763 |
| **TF** | Factor: ER71; motif: ACC( | TF:M02067_0 | 6.73E-32 | 31.17201 | 656 |
| **TF** | Factor: ER71; motif: ACC( | TF:M02067 | 6.73E-32 | 31.17201 | 656 |
| **TF** | Factor: GABPA; motif: AC | TF:M03990_0 | 7.46E-32 | 31.12747 | 590 |
| **TF** | Factor: GABPA; motif: AC | TF:M03990 | 7.46E-32 | 31.12747 | 590 |
| **TF** | Factor: Erm; motif: NNSC | TF:M11424_0 | 1.44E-31 | 30.84174 | 790 |
| **TF** | Factor: Erm; motif: NNSC | TF:M11424 | 1.44E-31 | 30.84174 | 790 |
| **TF** | Factor: PEA3; motif: NAC( | TF:M11430 | 1.49E-31 | 30.82592 | 1025 |
| **TF** | Factor: PEA3; motif: NAC( | TF:M11430_0 | 1.49E-31 | 30.82592 | 1025 |
| **TF** | Factor: Elf-1; motif: ACTT | TF:M04714_0 | 1.50E-31 | 30.82515 | 790 |
| **TF** | Factor: Elf-1; motif: ACTT | TF:M04714 | 1.50E-31 | 30.82515 | 790 |
| **TF** | Factor: Elf-1; motif: NAM | TF:M11447_1 | 2.10E-31 | 30.67731 | 346 |
| **TF** | Factor: ERG; motif: ACCG | TF:M01752 | 3.61E-31 | 30.44217 | 915 |
| **TF** | Factor: ERG; motif: ACCG | TF:M01752_0 | 3.61E-31 | 30.44217 | 915 |
| **TF** | Factor: Elf-1; motif: NAN( | TF:M11451_0 | 5.98E-31 | 30.2233 | 719 |
| **TF** | Factor: Elf-1; motif: NAN( | TF:M11451 | 5.98E-31 | 30.2233 | 719 |
| **TF** | Factor: c-ets-1; motif: AC | TF:M02063 | 1.60E-30 | 29.79714 | 772 |
| **TF** | Factor: c-ets-1; motif: AC | TF:M02063_0 | 1.60E-30 | 29.79714 | 772 |
| **TF** | Factor: Elk-1; motif: RCC( | TF:M09904_0 | 2.95E-30 | 29.52988 | 814 |
| **TF** | Factor: Elk-1; motif: RCC( | TF:M09904 | 2.95E-30 | 29.52988 | 814 |
| **TF** | Factor: c-Ets-1; motif: NN | TF:M01078_1 | 4.77E-30 | 29.32139 | 336 |
| **TF** | Factor: c-Ets-1; motif: NN | TF:M01078_0 | 5.97E-30 | 29.22387 | 765 |
| **TF** | Factor: c-Ets-1; motif: NN | TF:M01078 | 5.97E-30 | 29.22387 | 765 |
| **TF** | Factor: Sp1; motif: NGGG | TF:M00196_0 | 5.99E-30 | 29.22249 | 959 |
| **TF** | Factor: Sp1; motif: NGGG | TF:M00196 | 5.99E-30 | 29.22249 | 959 |
| **TF** | Factor: FLI-1; motif: NAYT | TF:M08913_1 | 6.87E-30 | 29.16305 | 392 |
| **TF** | Factor: PEA3; motif: NAC( | TF:M11425_1 | 9.96E-30 | 29.00159 | 248 |
| **TF** | Factor: PEA3; motif: NAC( | TF:M11427_0 | 1.03E-29 | 28.98752 | 999 |
| **TF** | Factor: PEA3; motif: NAC( | TF:M11427 | 1.03E-29 | 28.98752 | 999 |
| **TF** | Factor: E2F-2; motif: GCG | TF:M11531_0 | 1.09E-29 | 28.96087 | 1162 |
| **TF** | Factor: E2F-2; motif: GCG | TF:M11531 | 1.09E-29 | 28.96087 | 1162 |
| **TF** | Factor: E2F-3:HES-7; mot | TF:M08526 | 1.20E-29 | 28.92075 | 1373 |
| **TF** | Factor: E2F-3:HES-7; mot | TF:M08526_0 | 1.20E-29 | 28.92075 | 1373 |
| **TF** | Factor: c-Ets-2; motif: NC | TF:M08759_1 | 1.37E-29 | 28.86343 | 376 |
| **TF** | Factor: Elk-1; motif: NNC( | TF:M11435 | 1.79E-29 | 28.746 | 1060 |
| **TF** | Factor: Elk-1; motif: NNC( | TF:M11435_0 | 1.79E-29 | 28.746 | 1060 |
| **TF** | Factor: E2F-3; motif: GGC | TF:M02089_0 | 1.92E-29 | 28.71736 | 1153 |
| **TF** | Factor: E2F-3; motif: GGC | TF:M02089 | 1.92E-29 | 28.71736 | 1153 |
| **TF** | Factor: ERG; motif: ACCG | TF:M01752_1 | 2.15E-29 | 28.66792 | 337 |
| **TF** | Factor: Fli-1; motif: ACCG | TF:M02073_0 | 2.41E-29 | 28.61761 | 630 |
| **TF** | Factor: Fli-1; motif: ACCG | TF:M02073 | 2.41E-29 | 28.61761 | 630 |
| **TF** | Factor: Elf-1; motif: NAN( | TF:M11451_1 | 2.59E-29 | 28.58744 | 295 |
| **TF** | Factor: Erm; motif: NNSC | TF:M11424_1 | 3.04E-29 | 28.5166 | 319 |

| | | | | | | |
|---|---|---|---|---|---|---|
| **TF** | Factor: GABP-alpha; moti | TF:M01660_1 | | 5.28E-29 | 28.27768 | 501 |
| **TF** | Factor: Elk-1; motif: NRS( | TF:M11434 | | 5.66E-29 | 28.24693 | 786 |
| **TF** | Factor: Elk-1; motif: NRS( | TF:M11434_0 | | 5.66E-29 | 28.24693 | 786 |
| **TF** | Factor: ELK-1; motif: ACC | TF:M02059_0 | | 5.95E-29 | 28.22534 | 905 |
| **TF** | Factor: ELK-1; motif: ACC | TF:M02059 | | 5.95E-29 | 28.22534 | 905 |
| **TF** | Factor: PEA3; motif: NAC | TF:M11427_1 | | 6.66E-29 | 28.17652 | 529 |
| **TF** | Factor: TEL1; motif: CNC( | TF:M02070_0 | | 1.29E-28 | 27.88884 | 996 |
| **TF** | Factor: TEL1; motif: CNC( | TF:M02070 | | 1.29E-28 | 27.88884 | 996 |
| **TF** | Factor: Elk-1; motif: RCC( | TF:M09904_1 | | 1.55E-28 | 27.80893 | 359 |
| **TF** | Factor: E2F-2; motif: NW | TF:M11530 | | 2.03E-28 | 27.69226 | 1255 |
| **TF** | Factor: E2F-2; motif: NW | TF:M11530_0 | | 2.03E-28 | 27.69226 | 1255 |
| **TF** | Factor: PEA3; motif: NAC | TF:M11428_1 | | 2.07E-28 | 27.68466 | 225 |
| **TF** | Factor: Sp1; motif: NGGG | TF:M07395 | | 2.13E-28 | 27.67235 | 995 |
| **TF** | Factor: Sp1; motif: NGGG | TF:M07395_0 | | 2.13E-28 | 27.67235 | 995 |
| **TF** | Factor: SAP-1; motif: NR( | TF:M11437_0 | | 2.33E-28 | 27.63311 | 311 |
| **TF** | Factor: SAP-1; motif: NR( | TF:M11437 | | 2.33E-28 | 27.63311 | 311 |
| **TF** | Factor: ETS1; motif: ACC( | TF:M03977_1 | | 3.26E-28 | 27.48615 | 536 |
| **TF** | Factor: E2F-1:Elk-1; motif | TF:M08205_1 | | 6.73E-28 | 27.17226 | 979 |
| **TF** | Factor: SAP-1; motif: NN( | TF:M09905_1 | | 6.78E-28 | 27.16891 | 383 |
| **TF** | Factor: nerf; motif: NNN( | TF:M09901_1 | | 7.96E-28 | 27.09898 | 224 |
| **TF** | Factor: PEA3; motif: NAC | TF:M11430_1 | | 1.12E-27 | 26.9503 | 549 |
| **TF** | Factor: ELK1; motif: ACC( | TF:M03970_0 | | 1.75E-27 | 26.75671 | 538 |
| **TF** | Factor: ELK1; motif: ACC( | TF:M03970 | | 1.75E-27 | 26.75671 | 538 |
| **TF** | Factor: Net; motif: NRCC( | TF:M11439_0 | | 2.34E-27 | 26.6313 | 291 |
| **TF** | Factor: Net; motif: NRCC( | TF:M11439 | | 2.34E-27 | 26.6313 | 291 |
| **TF** | Factor: ELF1; motif: NCC( | TF:M02053_0 | | 2.60E-27 | 26.58509 | 646 |
| **TF** | Factor: ELF1; motif: NCC( | TF:M02053 | | 2.60E-27 | 26.58509 | 646 |
| **TF** | Factor: E2F-4; motif: NTT | TF:M07380_0 | | 2.85E-27 | 26.54578 | 1114 |
| **TF** | Factor: E2F-4; motif: NTT | TF:M07380 | | 2.85E-27 | 26.54578 | 1114 |
| **TF** | Factor: Elf-1; motif: ACTT | TF:M04714_1 | | 3.40E-27 | 26.46828 | 256 |
| **TF** | Factor: Fli-1; motif: NACC | TF:M11400_1 | | 5.35E-27 | 26.27165 | 491 |
| **TF** | Factor: E2F-1; motif: NG( | TF:M07206 | | 5.35E-27 | 26.27155 | 1231 |
| **TF** | Factor: E2F-1; motif: NG( | TF:M07206_0 | | 5.35E-27 | 26.27155 | 1231 |
| **TF** | Factor: Elk-1; motif: NRS( | TF:M11434_1 | | 5.42E-27 | 26.26589 | 373 |
| **TF** | Factor: ETV5; motif: NCC( | TF:M03983_0 | | 6.24E-27 | 26.20462 | 335 |
| **TF** | Factor: ETV5; motif: NCC( | TF:M03983 | | 6.24E-27 | 26.20462 | 335 |
| **TF** | Factor: nerf; motif: NRRN | TF:M09606 | | 8.81E-27 | 26.05504 | 649 |
| **TF** | Factor: nerf; motif: NRRN | TF:M09606_0 | | 8.81E-27 | 26.05504 | 649 |
| **TF** | Factor: GABP-alpha; moti | TF:M11415 | | 1.12E-26 | 25.95103 | 343 |
| **TF** | Factor: GABP-alpha; moti | TF:M11415_0 | | 1.12E-26 | 25.95103 | 343 |
| **TF** | Factor: Erm; motif: NRCM | TF:M11422 | | 1.69E-26 | 25.77086 | 443 |
| **TF** | Factor: Erm; motif: NRCM | TF:M11422_0 | | 1.69E-26 | 25.77086 | 443 |
| **TF** | Factor: Elf-1; motif: NAAI | TF:M09605 | | 1.74E-26 | 25.75986 | 625 |
| **TF** | Factor: Elf-1; motif: NAAI | TF:M09605_0 | | 1.74E-26 | 25.75986 | 625 |
| **TF** | Factor: E2F-1; motif: NNN | TF:M09892 | | 2.90E-26 | 25.53686 | 1165 |
| **TF** | Factor: E2F-1; motif: NNN | TF:M09892_0 | | 2.90E-26 | 25.53686 | 1165 |
| **TF** | Factor: Erg; motif: NACC( | TF:M11392_1 | | 2.97E-26 | 25.52744 | 305 |

| | | | | | | |
|---|---|---|---|---|---|---|
| **TF** | Factor: ERG; motif: ACCG | TF:M03975_1 | | 4.40E-26 | 25.35653 | 348 |
| **TF** | Factor: GABP-alpha; moti | TF:M09935_0 | | 6.13E-26 | 25.21275 | 502 |
| **TF** | Factor: GABP-alpha; moti | TF:M09935 | | 6.13E-26 | 25.21275 | 502 |
| **TF** | Factor: ETV7; motif: NCC( | TF:M02071_1 | | 6.53E-26 | 25.18515 | 510 |
| **TF** | Factor: GABPalpha; motif | TF:M02074_0 | | 6.54E-26 | 25.18454 | 435 |
| **TF** | Factor: GABPalpha; motif | TF:M02074 | | 6.54E-26 | 25.18454 | 435 |
| **TF** | Factor: PEA3; motif: RCC( | TF:M02066_1 | | 6.86E-26 | 25.16351 | 310 |
| **TF** | Factor: E2F-2; motif: NWT | TF:M11530_1 | | 8.21E-26 | 25.08582 | 1171 |
| **TF** | Factor: TEL1; motif: CNC( | TF:M02070_1 | | 2.71E-25 | 24.56656 | 457 |
| **TF** | Factor: E2F-3:TBR2; moti | TF:M08207_1 | | 2.73E-25 | 24.56418 | 1301 |
| **TF** | Factor: GABP-ALPHA; mo | TF:M08917_0 | | 4.30E-25 | 24.36618 | 296 |
| **TF** | Factor: GABP-ALPHA; mo | TF:M08917 | | 4.30E-25 | 24.36618 | 296 |
| **TF** | Factor: c-ets-2; motif: AC | TF:M02064 | | 6.61E-25 | 24.18 | 993 |
| **TF** | Factor: c-ets-2; motif: AC | TF:M02064_0 | | 6.61E-25 | 24.18 | 993 |
| **TF** | Factor: ESE-1; motif: NSM | TF:M02055 | | 7.46E-25 | 24.12744 | 529 |
| **TF** | Factor: ESE-1; motif: NSM | TF:M02055_0 | | 7.46E-25 | 24.12744 | 529 |
| **TF** | Factor: ESE-1; motif: NSM | TF:M01214 | | 7.46E-25 | 24.12744 | 529 |
| **TF** | Factor: ESE-1; motif: NSM | TF:M01214_0 | | 7.46E-25 | 24.12744 | 529 |
| **TF** | Factor: GABP-alpha; moti | TF:M11414_0 | | 8.38E-25 | 24.07685 | 312 |
| **TF** | Factor: GABP-alpha; moti | TF:M11414 | | 8.38E-25 | 24.07685 | 312 |
| **TF** | Factor: Elf-1; motif: NNAI | TF:M09900 | | 1.52E-24 | 23.8183 | 571 |
| **TF** | Factor: Elf-1; motif: NNAI | TF:M09900_0 | | 1.52E-24 | 23.8183 | 571 |
| **TF** | Factor: Elk-1:HOXA1; mot | TF:M08451_1 | | 1.97E-24 | 23.70464 | 245 |
| **TF** | Factor: GABP-alpha; moti | TF:M04698 | | 2.22E-24 | 23.65398 | 331 |
| **TF** | Factor: GABP-alpha; moti | TF:M04698_0 | | 2.22E-24 | 23.65398 | 331 |
| **TF** | Factor: EHF; motif: CSCG( | TF:M02052 | | 2.41E-24 | 23.61824 | 919 |
| **TF** | Factor: EHF; motif: CSCG( | TF:M02052_0 | | 2.41E-24 | 23.61824 | 919 |
| **TF** | Factor: ETV3; motif: ACC( | TF:M03981_0 | | 2.53E-24 | 23.59658 | 265 |
| **TF** | Factor: ETV3; motif: ACC( | TF:M03981 | | 2.53E-24 | 23.59658 | 265 |
| **TF** | Factor: Elf-1; motif: AWC( | TF:M07415_0 | | 2.67E-24 | 23.57366 | 663 |
| **TF** | Factor: Elf-1; motif: AWC( | TF:M07415 | | 2.67E-24 | 23.57366 | 663 |
| **TF** | Factor: GABP-alpha; moti | TF:M01660 | | 2.79E-24 | 23.55475 | 1015 |
| **TF** | Factor: GABP-alpha; moti | TF:M01660_0 | | 2.79E-24 | 23.55475 | 1015 |
| **TF** | Factor: ELF2; motif: CCCG | TF:M02054 | | 3.00E-24 | 23.52295 | 377 |
| **TF** | Factor: ELF2; motif: CCCG | TF:M02054_0 | | 3.00E-24 | 23.52295 | 377 |
| **TF** | Factor: Elk-1; motif: NRC( | TF:M11433_0 | | 3.12E-24 | 23.50601 | 546 |
| **TF** | Factor: Elk-1; motif: NRC( | TF:M11433 | | 3.12E-24 | 23.50601 | 546 |
| **TF** | Factor: IRX-1; motif: NAC | TF:M11018_1 | | 3.26E-24 | 23.48638 | 1298 |
| **TF** | Factor: meis1:Elf-1; motif | TF:M08377 | | 3.82E-24 | 23.4179 | 1005 |
| **TF** | Factor: meis1:Elf-1; motif | TF:M08377_0 | | 3.82E-24 | 23.4179 | 1005 |
| **TF** | Factor: Sp1; motif: GGGG | TF:M00931_0 | | 3.88E-24 | 23.41124 | 953 |
| **TF** | Factor: Sp1; motif: GGGG | TF:M00931 | | 3.88E-24 | 23.41124 | 953 |
| **TF** | Factor: c-ets-1; motif: AC | TF:M02063_1 | | 8.31E-24 | 23.08051 | 231 |
| **TF** | Factor: Sp1; motif: NNGG | TF:M00932 | | 9.03E-24 | 23.04447 | 962 |
| **TF** | Factor: Sp1; motif: NNGG | TF:M00932_0 | | 9.03E-24 | 23.04447 | 962 |
| **TF** | Factor: ER81; motif: NNC | TF:M09913_0 | | 1.21E-23 | 22.91771 | 287 |
| **TF** | Factor: ER81; motif: NNC | TF:M09913 | | 1.21E-23 | 22.91771 | 287 |

| | | | | | | |
|---|---|---|---|---|---|---|
| **TF** | Factor: GABP; motif: CAC | TF:M01258_0 | | 1.79E-23 | 22.7474 | 354 |
| **TF** | Factor: GABP; motif: CAC | TF:M01258 | | 1.79E-23 | 22.7474 | 354 |
| **TF** | Factor: E2F-4; motif: NNT | TF:M04823_0 | | 2.51E-23 | 22.59953 | 1111 |
| **TF** | Factor: E2F-4; motif: NNT | TF:M04823 | | 2.51E-23 | 22.59953 | 1111 |
| **TF** | Factor: ETV4; motif: ACC | TF:M03982_1 | | 2.64E-23 | 22.5788 | 210 |
| **TF** | Factor: Elf-1; motif: NAN | TF:M11452 | | 2.65E-23 | 22.57645 | 623 |
| **TF** | Factor: Elf-1; motif: NAN | TF:M11452_0 | | 2.65E-23 | 22.57645 | 623 |
| **TF** | Factor: ZF5; motif: NRNG | TF:M00333_0 | | 2.79E-23 | 22.55476 | 1270 |
| **TF** | Factor: ZF5; motif: NRNG | TF:M00333 | | 2.79E-23 | 22.55476 | 1270 |
| **TF** | Factor: ELF1; motif: AAC | TF:M03966 | | 1.03E-22 | 21.98818 | 666 |
| **TF** | Factor: ELF1; motif: AAC | TF:M03966_0 | | 1.03E-22 | 21.98818 | 666 |
| **TF** | Factor: ETF; motif: CCCC | TF:M07039 | | 2.08E-22 | 21.68136 | 1344 |
| **TF** | Factor: ETF; motif: CCCC | TF:M07039_0 | | 2.08E-22 | 21.68136 | 1344 |
| **TF** | Factor: Erg; motif: ACCG | TF:M02062_1 | | 2.33E-22 | 21.632 | 255 |
| **TF** | Factor: GKLF; motif: NNC | TF:M12173_0 | | 2.35E-22 | 21.62944 | 923 |
| **TF** | Factor: GKLF; motif: NNC | TF:M12173 | | 2.35E-22 | 21.62944 | 923 |
| **TF** | Factor: FLI1; motif: ACCG | TF:M03988_1 | | 2.37E-22 | 21.62571 | 361 |
| **TF** | Factor: ELF4; motif: NCC | TF:M02056 | | 3.34E-22 | 21.47566 | 569 |
| **TF** | Factor: ELF4; motif: NCC | TF:M02056_0 | | 3.34E-22 | 21.47566 | 569 |
| **TF** | Factor: Elk-1; motif: NAC | TF:M11432_0 | | 3.89E-22 | 21.4098 | 636 |
| **TF** | Factor: Elk-1; motif: NAC | TF:M11432 | | 3.89E-22 | 21.4098 | 636 |
| **TF** | Factor: c-ets-2; motif: AC | TF:M02064_1 | | 4.12E-22 | 21.38547 | 356 |
| **TF** | Factor: Elf-1; motif: NAN | TF:M11448 | | 5.97E-22 | 21.22376 | 729 |
| **TF** | Factor: Elf-1; motif: NAN | TF:M11448_0 | | 5.97E-22 | 21.22376 | 729 |
| **TF** | Factor: Pet-1; motif: ACC | TF:M02072 | | 7.10E-22 | 21.14889 | 263 |
| **TF** | Factor: Pet-1; motif: ACC | TF:M02072_0 | | 7.10E-22 | 21.14889 | 263 |
| **TF** | Factor: Elf-1; motif: NAN | TF:M11448_1 | | 1.12E-21 | 20.94976 | 304 |
| **TF** | Factor: ER71; motif: NAC | TF:M11413 | | 1.30E-21 | 20.8856 | 240 |
| **TF** | Factor: ER71; motif: NAC | TF:M11413_0 | | 1.30E-21 | 20.8856 | 240 |
| **TF** | Factor: ERF; motif: ACCG | TF:M02061_0 | | 1.53E-21 | 20.81391 | 298 |
| **TF** | Factor: ERF; motif: ACCG | TF:M02061 | | 1.53E-21 | 20.81391 | 298 |
| **TF** | Factor: ER81; motif: NAC | TF:M11421_0 | | 2.20E-21 | 20.65845 | 271 |
| **TF** | Factor: ER81; motif: NAC | TF:M11421 | | 2.20E-21 | 20.65845 | 271 |
| **TF** | Factor: ER71; motif: ACC | TF:M02067_1 | | 3.35E-21 | 20.47481 | 172 |
| **TF** | Factor: ER81; motif: NNC | TF:M11419_1 | | 3.49E-21 | 20.45773 | 229 |
| **TF** | Factor: Fli-1; motif: NACC | TF:M11404_1 | | 2.20E-20 | 19.6569 | 374 |
| **TF** | Factor: Sp1; motif: GGGG | TF:M00008 | | 2.35E-20 | 19.62851 | 898 |
| **TF** | Factor: Sp1; motif: GGGG | TF:M00008_0 | | 2.35E-20 | 19.62851 | 898 |
| **TF** | Factor: Elf-1; motif: NAA | TF:M09605_1 | | 2.36E-20 | 19.62787 | 197 |
| **TF** | Factor: TCF-1; motif: ACA | TF:M11603_1 | | 3.30E-20 | 19.48193 | 1140 |
| **TF** | Factor: ER81; motif: NAC | TF:M11420 | | 4.93E-20 | 19.30697 | 320 |
| **TF** | Factor: ER81; motif: NAC | TF:M11420_0 | | 4.93E-20 | 19.30697 | 320 |
| **TF** | Factor: GKLF; motif: GCC | TF:M01588_0 | | 5.16E-20 | 19.28694 | 843 |
| **TF** | Factor: GKLF; motif: GCC | TF:M01588 | | 5.16E-20 | 19.28694 | 843 |
| **TF** | Factor: E2F-3:TBR2; moti | TF:M08207_0 | | 8.06E-20 | 19.09382 | 1358 |
| **TF** | Factor: E2F-3:TBR2; moti | TF:M08207 | | 8.06E-20 | 19.09382 | 1358 |
| **TF** | Factor: E2F; motif: GGCG | TF:M00803_1 | | 9.45E-20 | 19.02477 | 929 |

| | | | | | |
|---|---|---|---|---|---|
| **TF** | Factor: MAZ; motif: GGGI | TF:M09636 | 9.73E-20 | 19.01202 | 1314 |
| **TF** | Factor: MAZ; motif: GGGI | TF:M09636_0 | 9.73E-20 | 19.01202 | 1314 |
| **TF** | Factor: ESE-1; motif: NSM | TF:M02055_1 | 1.03E-19 | 18.98666 | 281 |
| **TF** | Factor: ESE-1; motif: NSM | TF:M01214_1 | 1.03E-19 | 18.98666 | 281 |
| **TF** | Factor: Fli-1; motif: ACCG | TF:M02073_1 | 2.20E-19 | 18.6581 | 158 |
| **TF** | Factor: Elf-1; motif: NNAI | TF:M09900_1 | 2.65E-19 | 18.57604 | 179 |
| **TF** | Factor: Elf-1; motif: NANS | TF:M11450 | 2.70E-19 | 18.56787 | 556 |
| **TF** | Factor: Elf-1; motif: NANS | TF:M11450_0 | 2.70E-19 | 18.56787 | 556 |
| **TF** | Factor: E2F-1:HES-7; mot | TF:M08525 | 3.16E-19 | 18.50065 | 1375 |
| **TF** | Factor: E2F-1:HES-7; mot | TF:M08525_0 | 3.16E-19 | 18.50065 | 1375 |
| **TF** | Factor: ELK1; motif: ACCC | TF:M03970_1 | 3.19E-19 | 18.49647 | 130 |
| **TF** | Factor: TCF-1; motif: ACA | TF:M11601_0 | 3.41E-19 | 18.46766 | 1243 |
| **TF** | Factor: TCF-1; motif: ACA | TF:M11601 | 3.41E-19 | 18.46766 | 1243 |
| **TF** | Factor: sp4; motif: NNGN | TF:M10072_0 | 4.42E-19 | 18.35469 | 938 |
| **TF** | Factor: sp4; motif: NNGN | TF:M10072 | 4.42E-19 | 18.35469 | 938 |
| **TF** | Factor: Elf-1; motif: NANC | TF:M11452_1 | 4.47E-19 | 18.34998 | 228 |
| **TF** | Factor: SP1; motif: RCCM | TF:M03920_0 | 4.76E-19 | 18.32225 | 679 |
| **TF** | Factor: SP1; motif: RCCM | TF:M03920 | 4.76E-19 | 18.32225 | 679 |
| **TF** | Factor: E2F-2; motif: GCG | TF:M11531_1 | 5.11E-19 | 18.2913 | 1075 |
| **TF** | Factor: Elf-1; motif: NANC | TF:M11449 | 5.13E-19 | 18.28973 | 472 |
| **TF** | Factor: Elf-1; motif: NANC | TF:M11449_0 | 5.13E-19 | 18.28973 | 472 |
| **TF** | Factor: etv3; motif: ACCG | TF:M02068_0 | 5.87E-19 | 18.23107 | 334 |
| **TF** | Factor: etv3; motif: ACCG | TF:M02068 | 5.87E-19 | 18.23107 | 334 |
| **TF** | Factor: Net; motif: NACC | TF:M11442 | 6.13E-19 | 18.21224 | 225 |
| **TF** | Factor: Net; motif: NACC | TF:M11442_0 | 6.13E-19 | 18.21224 | 225 |
| **TF** | Factor: Elk-1; motif: NACI | TF:M11432_1 | 6.52E-19 | 18.1857 | 170 |
| **TF** | Factor: ER71:Pax-5; motif | TF:M08266_0 | 7.43E-19 | 18.12886 | 1214 |
| **TF** | Factor: ER71:Pax-5; motif | TF:M08266 | 7.43E-19 | 18.12886 | 1214 |
| **TF** | Factor: SAP-1; motif: ACC | TF:M09607 | 8.37E-19 | 18.07724 | 354 |
| **TF** | Factor: SAP-1; motif: ACC | TF:M09607_0 | 8.37E-19 | 18.07724 | 354 |
| **TF** | Factor: meis1:Elf-1; motif | TF:M08377_1 | 1.11E-18 | 17.95484 | 532 |
| **TF** | Factor: E2F-4; motif: GCG | TF:M02090_0 | 1.14E-18 | 17.94477 | 1054 |
| **TF** | Factor: E2F-4; motif: GCG | TF:M02090 | 1.14E-18 | 17.94477 | 1054 |
| **TF** | Factor: Elk-1; motif: NRC( | TF:M11433_1 | 1.84E-18 | 17.73606 | 207 |
| **TF** | Factor: c-Ets-1(p54); mot | TF:M00032_1 | 1.90E-18 | 17.72238 | 207 |
| **TF** | Factor: E2F-1; motif: NNN | TF:M07250_0 | 1.91E-18 | 17.71937 | 963 |
| **TF** | Factor: E2F-1; motif: NNN | TF:M07250 | 1.91E-18 | 17.71937 | 963 |
| **TF** | Factor: E2F-7; motif: GRG | TF:M09896 | 2.27E-18 | 17.64466 | 909 |
| **TF** | Factor: E2F-7; motif: GRG | TF:M09896_0 | 2.27E-18 | 17.64466 | 909 |
| **TF** | Factor: ER71:Pax-5; motif | TF:M08266_1 | 2.37E-18 | 17.62471 | 832 |
| **TF** | Factor: Sp1; motif: CCCCC | TF:M00933 | 3.18E-18 | 17.49735 | 888 |
| **TF** | Factor: Sp1; motif: CCCCC | TF:M00933_0 | 3.18E-18 | 17.49735 | 888 |
| **TF** | Factor: ELF3; motif: WAC | TF:M03967 | 4.18E-18 | 17.37847 | 678 |
| **TF** | Factor: ELF3; motif: WAC | TF:M03967_0 | 4.18E-18 | 17.37847 | 678 |
| **TF** | Factor: E2F-3; motif: GGC | TF:M02089_1 | 5.33E-18 | 17.27355 | 862 |
| **TF** | Factor: GABP-alpha; moti | TF:M09935_1 | 7.55E-18 | 17.12223 | 161 |
| **TF** | Factor: E2F-1; motif: WW | TF:M04515_0 | 8.07E-18 | 17.09338 | 1136 |

| | | | | | |
|---|---|---|---|---|---|
| **TF** | Factor: E2F-1; motif: WW | TF:M04515 | 8.07E-18 | 17.09338 | 1136 |
| **TF** | Factor: GABPalpha_GABP | TF:M03791 | 8.32E-18 | 17.08 | 348 |
| **TF** | Factor: GABPalpha_GABP | TF:M03791_0 | 8.32E-18 | 17.08 | 348 |
| **TF** | Factor: BTEB4; motif: NC( | TF:M12186_0 | 1.18E-17 | 16.92974 | 846 |
| **TF** | Factor: BTEB4; motif: NC( | TF:M12186 | 1.18E-17 | 16.92974 | 846 |
| **TF** | Factor: MAZ; motif: GGGI | TF:M09636_1 | 1.21E-17 | 16.91735 | 1174 |
| **TF** | Factor: Elf-1; motif: AWC | TF:M07415_1 | 1.26E-17 | 16.89991 | 216 |
| **TF** | Factor: Erm; motif: ACCG | TF:M02069_1 | 1.87E-17 | 16.72931 | 192 |
| **TF** | Factor: Elf-1; motif: NANS | TF:M11450_1 | 2.09E-17 | 16.67953 | 192 |
| **TF** | Factor: ETF; motif: GVGG | TF:M00695 | 2.14E-17 | 16.67057 | 937 |
| **TF** | Factor: ETF; motif: GVGG | TF:M00695_0 | 2.14E-17 | 16.67057 | 937 |
| **TF** | Factor: E2F-4; motif: TTT | TF:M11535_0 | 2.43E-17 | 16.61481 | 966 |
| **TF** | Factor: E2F-4; motif: TTT | TF:M11535 | 2.43E-17 | 16.61481 | 966 |
| **TF** | Factor: BEN; motif: CAGC | TF:M01240_1 | 2.55E-17 | 16.59429 | 1131 |
| **TF** | Factor: SP2; motif: GNNG | TF:M03807 | 2.56E-17 | 16.59251 | 779 |
| **TF** | Factor: SP2; motif: GNNG | TF:M03807_0 | 2.56E-17 | 16.59251 | 779 |
| **TF** | Factor: BEN; motif: CAGC | TF:M01240_0 | 2.77E-17 | 16.5581 | 1191 |
| **TF** | Factor: BEN; motif: CAGC | TF:M01240 | 2.77E-17 | 16.5581 | 1191 |
| **TF** | Factor: TCF-1; motif: ACA | TF:M11601_1 | 3.60E-17 | 16.44325 | 983 |
| **TF** | Factor: E2F-1; motif: NGC | TF:M07206_1 | 5.00E-17 | 16.30131 | 967 |
| **TF** | Factor: c-Ets-1(p54); mot | TF:M00032 | 7.38E-17 | 16.13169 | 516 |
| **TF** | Factor: c-Ets-1(p54); mot | TF:M00032_0 | 7.38E-17 | 16.13169 | 516 |
| **TF** | Factor: ELF1; motif: NCC( | TF:M02053_1 | 8.19E-17 | 16.08673 | 152 |
| **TF** | Factor: nerf; motif: NRRN | TF:M09606_1 | 1.17E-16 | 15.93303 | 199 |
| **TF** | Factor: Elk-1:Pax-5; motif | TF:M08225_1 | 1.29E-16 | 15.88901 | 754 |
| **TF** | Factor: Elk-1:HOXA1; mot | TF:M08451 | 2.03E-16 | 15.69167 | 584 |
| **TF** | Factor: Elk-1:HOXA1; mot | TF:M08451_0 | 2.03E-16 | 15.69167 | 584 |
| **TF** | Factor: GABPA; motif: AC | TF:M03990_1 | 2.32E-16 | 15.63505 | 132 |
| **TF** | Factor: Sp2; motif: GGGG | TF:M10435_0 | 2.45E-16 | 15.6117 | 695 |
| **TF** | Factor: Sp1; motif: GGGG | TF:M07063_0 | 2.45E-16 | 15.6117 | 695 |
| **TF** | Factor: Sp2; motif: GGGG | TF:M10435 | 2.45E-16 | 15.6117 | 695 |
| **TF** | Factor: Sp1; motif: GGGG | TF:M07063 | 2.45E-16 | 15.6117 | 695 |
| **TF** | Factor: E2F; motif: TTTCG | TF:M00425_0 | 3.25E-16 | 15.48853 | 980 |
| **TF** | Factor: E2F; motif: TTTCG | TF:M00425 | 3.25E-16 | 15.48853 | 980 |
| **TF** | Factor: ZF5; motif: NRNG | TF:M00333_1 | 3.35E-16 | 15.47468 | 1072 |
| **TF** | Factor: IRX-1; motif: NAC | TF:M11018_0 | 5.13E-16 | 15.29019 | 1353 |
| **TF** | Factor: IRX-1; motif: NAC | TF:M11018 | 5.13E-16 | 15.29019 | 1353 |
| **TF** | Factor: ETF; motif: CCCC( | TF:M07039_1 | 7.40E-16 | 15.13073 | 1152 |
| **TF** | Factor: Kaiso; motif: TCT( | TF:M04691_1 | 7.71E-16 | 15.11291 | 939 |
| **TF** | Factor: CHD2; motif: TCT( | TF:M04710_0 | 9.64E-16 | 15.01578 | 1166 |
| **TF** | Factor: CHD2; motif: TCT( | TF:M04710 | 9.64E-16 | 15.01578 | 1166 |
| **TF** | Factor: ESE-1; motif: NWT | TF:M11382 | 1.35E-15 | 14.86904 | 883 |
| **TF** | Factor: ESE-1; motif: NWT | TF:M11382_0 | 1.35E-15 | 14.86904 | 883 |
| **TF** | Factor: Sp1; motif: NWRG | TF:M12151 | 1.49E-15 | 14.82607 | 661 |
| **TF** | Factor: Sp1; motif: NWRG | TF:M12151_0 | 1.49E-15 | 14.82607 | 661 |
| **TF** | Factor: CHD2; motif: TCT( | TF:M04710_1 | 1.55E-15 | 14.80997 | 994 |
| **TF** | Factor: Sp1; motif: NWRG | TF:M12152 | 1.66E-15 | 14.78032 | 700 |

| | | | | | |
|---|---|---|---|---|---|
| **TF** | Factor: Sp1; motif: NWRG | TF:M12152_0 | 1.66E-15 | 14.78032 | 700 |
| **TF** | Factor: ELF1; motif: AACC | TF:M03966_1 | 2.42E-15 | 14.61625 | 200 |
| **TF** | Factor: E2F-1; motif: WW | TF:M04515_1 | 2.86E-15 | 14.54432 | 1051 |
| **TF** | Factor: p300; motif: ACN | TF:M04826 | 4.49E-15 | 14.34806 | 1391 |
| **TF** | Factor: p300; motif: ACN | TF:M04826_0 | 4.49E-15 | 14.34806 | 1391 |
| **TF** | Factor: E2F-4; motif: NGG | TF:M07084_0 | 4.79E-15 | 14.31927 | 959 |
| **TF** | Factor: E2F-4; motif: NGG | TF:M07084 | 4.79E-15 | 14.31927 | 959 |
| **TF** | Factor: ELF3; motif: WAC | TF:M03967_1 | 5.15E-15 | 14.28859 | 246 |
| **TF** | Factor: BTEB2; motif: RG | TF:M07277_0 | 5.61E-15 | 14.25122 | 784 |
| **TF** | Factor: BTEB2; motif: RG | TF:M07277 | 5.61E-15 | 14.25122 | 784 |
| **TF** | Factor: E2F-1:TBR2; moti | TF:M08523 | 7.27E-15 | 14.13874 | 1036 |
| **TF** | Factor: E2F-1:TBR2; moti | TF:M08523_0 | 7.27E-15 | 14.13874 | 1036 |
| **TF** | Factor: TCF-1; motif: ACA | TF:M11603 | 1.05E-14 | 13.97905 | 1322 |
| **TF** | Factor: TCF-1; motif: ACA | TF:M11603_0 | 1.05E-14 | 13.97905 | 1322 |
| **TF** | Factor: Egr-1; motif: GCG | TF:M04869_1 | 2.55E-14 | 13.59383 | 911 |
| **TF** | Factor: Sp2; motif: WGG | TF:M05332 | 3.16E-14 | 13.49966 | 788 |
| **TF** | Factor: Sp6; motif: WGG | TF:M05361 | 3.16E-14 | 13.49966 | 788 |
| **TF** | Factor: Sp2; motif: WGG | TF:M05332_0 | 3.16E-14 | 13.49966 | 788 |
| **TF** | Factor: Sp6; motif: WGG | TF:M05361_0 | 3.16E-14 | 13.49966 | 788 |
| **TF** | Factor: CPBP; motif: NGG | TF:M05444_0 | 3.16E-14 | 13.49966 | 788 |
| **TF** | Factor: CPBP; motif: NGG | TF:M05444 | 3.16E-14 | 13.49966 | 788 |
| **TF** | Factor: Sp2; motif: NYSG | TF:M03567 | 3.21E-14 | 13.49396 | 712 |
| **TF** | Factor: Sp2; motif: NYSG | TF:M03567_0 | 3.21E-14 | 13.49396 | 712 |
| **TF** | Factor: Erg; motif: NNAC | TF:M09608_0 | 4.35E-14 | 13.36178 | 749 |
| **TF** | Factor: Erg; motif: NNAC | TF:M09608 | 4.35E-14 | 13.36178 | 749 |
| **TF** | Factor: Elk-1; motif: AAC | TF:M01163_0 | 5.76E-14 | 13.23991 | 439 |
| **TF** | Factor: Elk-1; motif: AAC | TF:M01163 | 5.76E-14 | 13.23991 | 439 |
| **TF** | Factor: SP2; motif: GGGC | TF:M01783 | 6.03E-14 | 13.21956 | 793 |
| **TF** | Factor: SP2; motif: GGGC | TF:M01783_0 | 6.03E-14 | 13.21956 | 793 |
| **TF** | Factor: SAP-1; motif: NRN | TF:M11436 | 6.70E-14 | 13.17405 | 152 |
| **TF** | Factor: SAP-1; motif: NRN | TF:M11436_0 | 6.70E-14 | 13.17405 | 152 |
| **TF** | Factor: Sp1; motif: NGGG | TF:M10071 | 6.92E-14 | 13.1596 | 770 |
| **TF** | Factor: Sp1; motif: NGGG | TF:M10071_0 | 6.92E-14 | 13.1596 | 770 |
| **TF** | Factor: ETV7; motif: NNS | TF:M08760_0 | 9.07E-14 | 13.04237 | 269 |
| **TF** | Factor: ETV7; motif: NNS | TF:M08760 | 9.07E-14 | 13.04237 | 269 |
| **TF** | Factor: E2F-1; motif: TTT | TF:M04516 | 1.00E-13 | 13.00007 | 988 |
| **TF** | Factor: E2F-1; motif: TTT | TF:M04516_0 | 1.00E-13 | 13.00007 | 988 |
| **TF** | Factor: ESE-1; motif: NW | TF:M11382_1 | 1.02E-13 | 12.99173 | 416 |
| **TF** | Factor: E2F-4:DP-1; motif | TF:M00738 | 1.14E-13 | 12.94175 | 879 |
| **TF** | Factor: E2F-4:DP-1; motif | TF:M00738_0 | 1.14E-13 | 12.94175 | 879 |
| **TF** | Factor: sp4; motif: NNNG | TF:M10530_0 | 1.56E-13 | 12.80774 | 759 |
| **TF** | Factor: sp4; motif: NNNG | TF:M10530 | 1.56E-13 | 12.80774 | 759 |
| **TF** | Factor: E2F-4; motif: NTT | TF:M07380_1 | 1.57E-13 | 12.80333 | 739 |
| **TF** | Factor: E2F-1; motif: NTT | TF:M00940 | 1.90E-13 | 12.72064 | 726 |
| **TF** | Factor: E2F-1; motif: NTT | TF:M00940_0 | 1.90E-13 | 12.72064 | 726 |
| **TF** | Factor: Elk-1:Pax-5; motif | TF:M08225_0 | 2.16E-13 | 12.66597 | 1115 |
| **TF** | Factor: Elk-1:Pax-5; motif | TF:M08225 | 2.16E-13 | 12.66597 | 1115 |

| | | | | | |
|---|---|---|---|---|---|
| **TF** | Factor: FEV; motif: ACCG( | TF:M03987 | 2.19E-13 | 12.65973 | 154 |
| **TF** | Factor: FEV; motif: ACCG( | TF:M03987_0 | 2.19E-13 | 12.65973 | 154 |
| **TF** | Factor: E2F-3:FOXI1; mot | TF:M08208_0 | 2.48E-13 | 12.60524 | 1067 |
| **TF** | Factor: E2F-3:FOXI1; mot | TF:M08208 | 2.48E-13 | 12.60524 | 1067 |
| **TF** | Factor: TIEG1; motif: NC( | TF:M12351 | 2.57E-13 | 12.59039 | 1036 |
| **TF** | Factor: TIEG1; motif: NC( | TF:M12351_0 | 2.57E-13 | 12.59039 | 1036 |
| **TF** | Factor: E2F-1; motif: NTT | TF:M11533 | 2.73E-13 | 12.56305 | 1102 |
| **TF** | Factor: E2F-1; motif: NTT | TF:M11533_0 | 2.73E-13 | 12.56305 | 1102 |
| **TF** | Factor: SAP-1a; motif: NA | TF:M01167_0 | 3.10E-13 | 12.50863 | 205 |
| **TF** | Factor: SAP-1a; motif: NA | TF:M01167 | 3.10E-13 | 12.50863 | 205 |
| **TF** | Factor: E2F-1:DP-1; motif | TF:M00736 | 4.52E-13 | 12.34514 | 919 |
| **TF** | Factor: E2F-1:DP-1; motif | TF:M00736_0 | 4.52E-13 | 12.34514 | 919 |
| **TF** | Factor: KLF3; motif: NNN | TF:M09970 | 5.23E-13 | 12.28189 | 717 |
| **TF** | Factor: KLF3; motif: NNN | TF:M09970_0 | 5.23E-13 | 12.28189 | 717 |
| **TF** | Factor: SP4; motif: NWRG | TF:M03922_0 | 5.45E-13 | 12.26335 | 581 |
| **TF** | Factor: SP4; motif: NWRG | TF:M03922 | 5.45E-13 | 12.26335 | 581 |
| **TF** | Factor: TEF-3:ER81; motif | TF:M08423_0 | 6.75E-13 | 12.17094 | 409 |
| **TF** | Factor: TEF-3:ER81; motif | TF:M08423 | 6.75E-13 | 12.17094 | 409 |
| **TF** | Factor: SP1; motif: NRGK( | TF:M09765_0 | 9.39E-13 | 12.02718 | 688 |
| **TF** | Factor: SP1; motif: NRGK( | TF:M09765 | 9.39E-13 | 12.02718 | 688 |
| **TF** | Factor: E2F; motif: NNTTT | TF:M08875 | 1.11E-12 | 11.95388 | 598 |
| **TF** | Factor: E2F; motif: NNTTT | TF:M08875_0 | 1.11E-12 | 11.95388 | 598 |
| **TF** | Factor: BTEB1; motif: GG( | TF:M09723 | 1.23E-12 | 11.91094 | 884 |
| **TF** | Factor: BTEB1; motif: GG( | TF:M09723_0 | 1.23E-12 | 11.91094 | 884 |
| **TF** | Factor: ELK3; motif: ACC( | TF:M03972_0 | 1.39E-12 | 11.85771 | 143 |
| **TF** | Factor: ELK3; motif: ACC( | TF:M03972 | 1.39E-12 | 11.85771 | 143 |
| **TF** | Factor: Elk-1:Pax-9; motif | TF:M08226_1 | 1.42E-12 | 11.84909 | 549 |
| **TF** | Factor: Erg; motif: NNAC( | TF:M09608_1 | 1.42E-12 | 11.84806 | 198 |
| **TF** | Factor: Sp1; motif: NGGG | TF:M00196_1 | 1.53E-12 | 11.81516 | 566 |
| **TF** | Factor: ER81; motif: NNN | TF:M09611_0 | 1.64E-12 | 11.78642 | 236 |
| **TF** | Factor: ER81; motif: NNN | TF:M09611 | 1.64E-12 | 11.78642 | 236 |
| **TF** | Factor: Hey1; motif: NGC | TF:M11058_0 | 1.74E-12 | 11.75873 | 1077 |
| **TF** | Factor: Hey1; motif: NGC | TF:M11058 | 1.74E-12 | 11.75873 | 1077 |
| **TF** | Factor: Churchill; motif: ( | TF:M00986_0 | 3.35E-12 | 11.47511 | 1159 |
| **TF** | Factor: Churchill; motif: ( | TF:M00986 | 3.35E-12 | 11.47511 | 1159 |
| **TF** | Factor: Rb:E2F-1:DP-1; m | TF:M00740_0 | 3.87E-12 | 11.41184 | 916 |
| **TF** | Factor: Rb:E2F-1:DP-1; m | TF:M00740 | 3.87E-12 | 11.41184 | 916 |
| **TF** | Factor: Elk-1; motif: RAC( | TF:M01165_0 | 4.47E-12 | 11.34977 | 505 |
| **TF** | Factor: Elk-1; motif: RAC( | TF:M01165 | 4.47E-12 | 11.34977 | 505 |
| **TF** | Factor: EKLF; motif: NGG( | TF:M09969_0 | 4.70E-12 | 11.32807 | 465 |
| **TF** | Factor: EKLF; motif: NGG( | TF:M09969 | 4.70E-12 | 11.32807 | 465 |
| **TF** | Factor: Pet-1; motif: NAC | TF:M11406 | 5.68E-12 | 11.24582 | 147 |
| **TF** | Factor: Pet-1; motif: NAC | TF:M11406_0 | 5.68E-12 | 11.24582 | 147 |
| **TF** | Factor: Fli-1; motif: NAC( | TF:M11399_0 | 5.76E-12 | 11.23962 | 1095 |
| **TF** | Factor: Fli-1; motif: NAC( | TF:M11399 | 5.76E-12 | 11.23962 | 1095 |
| **TF** | Factor: ELF5; motif: ANSN | TF:M03969_0 | 5.99E-12 | 11.22266 | 349 |
| **TF** | Factor: ELF5; motif: ANSN | TF:M03969 | 5.99E-12 | 11.22266 | 349 |

| | | | | | |
|---|---|---|---|---|---|
| **TF** | Factor: Sp2; motif: NTAA( | TF:M12156 | 6.58E-12 | 11.18145 | 424 |
| **TF** | Factor: Sp2; motif: NTAA( | TF:M12156_0 | 6.58E-12 | 11.18145 | 424 |
| **TF** | Factor: BTEB2; motif: GC( | TF:M07409_0 | 6.83E-12 | 11.16586 | 493 |
| **TF** | Factor: BTEB2; motif: GC( | TF:M07409 | 6.83E-12 | 11.16586 | 493 |
| **TF** | Factor: Elk-1; motif: RAC( | TF:M01165_1 | 6.89E-12 | 11.16192 | 184 |
| **TF** | Factor: Tel-2; motif: YTA( | TF:M00678 | 7.00E-12 | 11.15474 | 364 |
| **TF** | Factor: Tel-2; motif: YTA( | TF:M00678_0 | 7.00E-12 | 11.15474 | 364 |
| **TF** | Factor: Sp2; motif: NTWA | TF:M12155 | 7.26E-12 | 11.13896 | 454 |
| **TF** | Factor: Sp2; motif: NTWA | TF:M12155_0 | 7.26E-12 | 11.13896 | 454 |
| **TF** | Factor: pax-6; motif: NYA | TF:M11882_1 | 7.35E-12 | 11.13394 | 965 |
| **TF** | Factor: Egr-1; motif: GCG | TF:M04869_0 | 9.07E-12 | 11.04234 | 970 |
| **TF** | Factor: Egr-1; motif: GCG | TF:M04869 | 9.07E-12 | 11.04234 | 970 |
| **TF** | Factor: Fli-1; motif: NAC( | TF:M11402_0 | 9.78E-12 | 11.00955 | 295 |
| **TF** | Factor: Fli-1; motif: NAC( | TF:M11402 | 9.78E-12 | 11.00955 | 295 |
| **TF** | Factor: E2F-1; motif: NKT | TF:M00428 | 1.03E-11 | 10.98864 | 717 |
| **TF** | Factor: E2F-1; motif: NKT | TF:M00428_0 | 1.03E-11 | 10.98864 | 717 |
| **TF** | Factor: Pax-5; motif: RRM | TF:M00144_0 | 1.08E-11 | 10.96727 | 1091 |
| **TF** | Factor: Pax-5; motif: RRM | TF:M00144 | 1.08E-11 | 10.96727 | 1091 |
| **TF** | Factor: Hey1; motif: NGC | TF:M11058_1 | 1.16E-11 | 10.93667 | 664 |
| **TF** | Factor: FLI-1; motif: CMG | TF:M01208 | 1.59E-11 | 10.79895 | 419 |
| **TF** | Factor: FLI-1; motif: CMG | TF:M01208_0 | 1.59E-11 | 10.79895 | 419 |
| **TF** | Factor: Fli-1; motif: RCM( | TF:M09613 | 1.61E-11 | 10.79337 | 364 |
| **TF** | Factor: Fli-1; motif: RCM( | TF:M09613_0 | 1.61E-11 | 10.79337 | 364 |
| **TF** | Factor: AP-2gamma:Elk-1 | TF:M08441_1 | 1.75E-11 | 10.75651 | 1006 |
| **TF** | Factor: Sp1; motif: NNGG | TF:M00932_1 | 2.09E-11 | 10.67898 | 577 |
| **TF** | Factor: GCMa:PEA3; mot | TF:M08313_1 | 2.36E-11 | 10.62735 | 859 |
| **TF** | Factor: Erm; motif: NRCM | TF:M11422_1 | 2.58E-11 | 10.58801 | 101 |
| **TF** | Factor: Sp2; motif: GYCC( | TF:M07129_0 | 2.77E-11 | 10.55763 | 569 |
| **TF** | Factor: Sp2; motif: GYCC( | TF:M07129 | 2.77E-11 | 10.55763 | 569 |
| **TF** | Factor: pax-6; motif: NYA | TF:M11882_0 | 2.99E-11 | 10.52463 | 1257 |
| **TF** | Factor: pax-6; motif: NYA | TF:M11882 | 2.99E-11 | 10.52463 | 1257 |
| **TF** | Factor: Kaiso; motif: TCT( | TF:M04691 | 3.18E-11 | 10.49777 | 1163 |
| **TF** | Factor: Kaiso; motif: TCT( | TF:M04691_0 | 3.18E-11 | 10.49777 | 1163 |
| **TF** | Factor: TR4; motif: ACCC( | TF:M04934_1 | 3.60E-11 | 10.44404 | 1058 |
| **TF** | Factor: E2F; motif: TTTCG | TF:M00425_1 | 3.62E-11 | 10.4408 | 524 |
| **TF** | Factor: E2F-1; motif: NNN | TF:M09892_1 | 4.42E-11 | 10.35425 | 817 |
| **TF** | Factor: Sp1; motif: NGGG | TF:M07395_1 | 4.97E-11 | 10.30369 | 591 |
| **TF** | Factor: GKLF; motif: GCCI | TF:M01588_1 | 5.77E-11 | 10.2388 | 405 |
| **TF** | Factor: WT1; motif: GNG( | TF:M03893 | 6.08E-11 | 10.21639 | 739 |
| **TF** | Factor: WT1; motif: GNG( | TF:M03893_0 | 6.08E-11 | 10.21639 | 739 |
| **TF** | Factor: Elk-1:HOXB13; m( | TF:M08222_1 | 7.28E-11 | 10.13786 | 180 |
| **TF** | Factor: AP-2gamma:Elk-1 | TF:M08441_0 | 7.48E-11 | 10.12624 | 1308 |
| **TF** | Factor: AP-2gamma:Elk-1 | TF:M08441 | 7.48E-11 | 10.12624 | 1308 |
| **TF** | Factor: BTEB3; motif: CCI | TF:M09826 | 8.08E-11 | 10.09249 | 977 |
| **TF** | Factor: BTEB3; motif: CCI | TF:M09826_0 | 8.08E-11 | 10.09249 | 977 |
| **TF** | Factor: E2F-1; motif: TTG | TF:M00938 | 8.43E-11 | 10.07432 | 664 |
| **TF** | Factor: E2F-1; motif: TTG | TF:M00938_0 | 8.43E-11 | 10.07432 | 664 |

| | | | | | |
|---|---|---|---|---|---|
| **TF** | Factor: Elk-1:TBR2; motif | TF:M08213_1 | 1.02E-10 | 9.991187 | 181 |
| **TF** | Factor: GKLF; motif: WGG | TF:M09971 | 1.18E-10 | 9.928466 | 534 |
| **TF** | Factor: GKLF; motif: WGG | TF:M09971_0 | 1.18E-10 | 9.928466 | 534 |
| **TF** | Factor: Elf-1; motif: NAN | TF:M11449_1 | 1.51E-10 | 9.82139 | 105 |
| **TF** | Factor: Elk-1:HOXB13; mo | TF:M08222 | 1.90E-10 | 9.720797 | 479 |
| **TF** | Factor: Elk-1:HOXB13; mo | TF:M08222_0 | 1.90E-10 | 9.720797 | 479 |
| **TF** | Factor: etv3; motif: ACCG | TF:M02068_1 | 2.01E-10 | 9.695752 | 75 |
| **TF** | Factor: Elk-1:Pax-1; motif | TF:M08224_0 | 2.02E-10 | 9.694369 | 783 |
| **TF** | Factor: Elk-1:Pax-1; motif | TF:M08224 | 2.02E-10 | 9.694369 | 783 |
| **TF** | Factor: ER71:E2A; motif: | TF:M08568_1 | 2.07E-10 | 9.684431 | 1142 |
| **TF** | Factor: E2F; motif: TTTSG | TF:M00426 | 2.21E-10 | 9.654707 | 1010 |
| **TF** | Factor: E2F; motif: TTTSG | TF:M00426_0 | 2.21E-10 | 9.654707 | 1010 |
| **TF** | Factor: GKLF; motif: NNC | TF:M12173_1 | 2.43E-10 | 9.615282 | 488 |
| **TF** | Factor: E2F1; motif: NNN | TF:M08874 | 2.69E-10 | 9.569752 | 662 |
| **TF** | Factor: E2F1; motif: NNN | TF:M08874_0 | 2.69E-10 | 9.569752 | 662 |
| **TF** | Factor: Sp3; motif: NGCC | TF:M12153_0 | 2.98E-10 | 9.525818 | 580 |
| **TF** | Factor: Sp3; motif: NGCC | TF:M12153 | 2.98E-10 | 9.525818 | 580 |
| **TF** | Factor: AP-2rep; motif: N | TF:M09967 | 3.24E-10 | 9.489403 | 393 |
| **TF** | Factor: AP-2rep; motif: N | TF:M09967_0 | 3.24E-10 | 9.489403 | 393 |
| **TF** | Factor: GCMa:PEA3; mot | TF:M08313 | 3.76E-10 | 9.424449 | 1247 |
| **TF** | Factor: GCMa:PEA3; mot | TF:M08313_0 | 3.76E-10 | 9.424449 | 1247 |
| **TF** | Factor: GABPalpha; motif | TF:M02074_1 | 4.18E-10 | 9.378905 | 102 |
| **TF** | Factor: ELF2; motif: CCCG | TF:M02054_1 | 4.34E-10 | 9.362939 | 78 |
| **TF** | Factor: Elk-1:TBR2; motif | TF:M08213 | 5.14E-10 | 9.288928 | 583 |
| **TF** | Factor: Elk-1:TBR2; motif | TF:M08213_0 | 5.14E-10 | 9.288928 | 583 |
| **TF** | Factor: Erg; motif: NACC | TF:M11393 | 5.20E-10 | 9.283983 | 1163 |
| **TF** | Factor: Erg; motif: NACC | TF:M11393_0 | 5.20E-10 | 9.283983 | 1163 |
| **TF** | Factor: YY1; motif: NNCG | TF:M03924_0 | 6.79E-10 | 9.167976 | 855 |
| **TF** | Factor: YY1; motif: NNCG | TF:M03924 | 6.79E-10 | 9.167976 | 855 |
| **TF** | Factor: Fli-1; motif: NACC | TF:M11401_0 | 7.76E-10 | 9.110103 | 1014 |
| **TF** | Factor: Fli-1; motif: NACC | TF:M11401 | 7.76E-10 | 9.110103 | 1014 |
| **TF** | Factor: KLF17; motif: NG | TF:M05386 | 8.62E-10 | 9.064612 | 714 |
| **TF** | Factor: KLF17; motif: NG | TF:M05386_0 | 8.62E-10 | 9.064612 | 714 |
| **TF** | Factor: LKLF; motif: NGG | TF:M05499 | 8.62E-10 | 9.064612 | 714 |
| **TF** | Factor: LKLF; motif: NGG | TF:M05499_0 | 8.62E-10 | 9.064612 | 714 |
| **TF** | Factor: Sp3; motif: GGGG | TF:M07615_0 | 9.26E-10 | 9.033213 | 599 |
| **TF** | Factor: Sp3; motif: GGGG | TF:M07615 | 9.26E-10 | 9.033213 | 599 |
| **TF** | Factor: Erg; motif: NACC | TF:M11393_1 | 9.99E-10 | 9.000602 | 1131 |
| **TF** | Factor: SP1; motif: NCCC | TF:M07226 | 1.13E-09 | 8.948062 | 741 |
| **TF** | Factor: SP1; motif: NCCC | TF:M07226_0 | 1.13E-09 | 8.948062 | 741 |
| **TF** | Factor: Fli-1; motif: NACC | TF:M11401_1 | 1.26E-09 | 8.901292 | 1002 |
| **TF** | Factor: Sp1; motif: GGGG | TF:M00931_1 | 1.33E-09 | 8.874774 | 556 |
| **TF** | Factor: HOXB2:Elk-1; mot | TF:M08341 | 1.34E-09 | 8.872749 | 420 |
| **TF** | Factor: HOXB2:Elk-1; mot | TF:M08341_0 | 1.34E-09 | 8.872749 | 420 |
| **TF** | Factor: WT1; motif: CGC | TF:M02036 | 1.85E-09 | 8.731732 | 838 |
| **TF** | Factor: WT1; motif: CGC | TF:M02036_0 | 1.85E-09 | 8.731732 | 838 |
| **TF** | Factor: Elk-1:T-bet; motif | TF:M08231_1 | 1.93E-09 | 8.714532 | 192 |

| | | | | | |
|---|---|---|---|---|---|
| TF | Factor: SAP-1; motif: ACC | TF:M09607_1 | 2.71E-09 | 8.566928 | 128 |
| TF | Factor: FLI1; motif: ACCG | TF:M03989_0 | 3.65E-09 | 8.438286 | 902 |
| TF | Factor: FLI1; motif: ACCG | TF:M03989 | 3.65E-09 | 8.438286 | 902 |
| TF | Factor: DP-1; motif: NRNI | TF:M10098_0 | 4.29E-09 | 8.367906 | 645 |
| TF | Factor: DP-1; motif: NRNI | TF:M10098 | 4.29E-09 | 8.367906 | 645 |
| TF | Factor: Erg; motif: NACC( | TF:M11397 | 6.06E-09 | 8.217411 | 1013 |
| TF | Factor: Erg; motif: NACC( | TF:M11397_0 | 6.06E-09 | 8.217411 | 1013 |
| TF | Factor: GABP; motif: VCC | TF:M00341_0 | 6.18E-09 | 8.209291 | 726 |
| TF | Factor: GABP; motif: VCC | TF:M00341 | 6.18E-09 | 8.209291 | 726 |
| TF | Factor: Sp1; motif: GGGG | TF:M00008_1 | 7.27E-09 | 8.138643 | 453 |
| TF | Factor: Hey2; motif: NNC | TF:M11052_0 | 8.48E-09 | 8.071822 | 1003 |
| TF | Factor: Hey2; motif: NNC | TF:M11052 | 8.48E-09 | 8.071822 | 1003 |
| TF | Factor: ELK4; motif: NCR( | TF:M07209 | 8.97E-09 | 8.047161 | 110 |
| TF | Factor: ELK4; motif: NCR( | TF:M07209_0 | 8.97E-09 | 8.047161 | 110 |
| TF | Factor: HES-7; motif: GN( | TF:M11066 | 8.99E-09 | 8.046291 | 935 |
| TF | Factor: HES-7; motif: GN( | TF:M11066_0 | 8.99E-09 | 8.046291 | 935 |
| TF | Factor: Hey2; motif: NNC | TF:M11052_1 | 9.60E-09 | 8.017803 | 796 |
| TF | Factor: Fli-1; motif: NACC | TF:M11399_1 | 9.72E-09 | 8.012489 | 976 |
| TF | Factor: HOXB2:Elk-1; mot | TF:M08341_1 | 9.99E-09 | 8.000441 | 166 |
| TF | Factor: NRF-1; motif: SYG | TF:M09641 | 1.01E-08 | 7.99673 | 329 |
| TF | Factor: NRF-1; motif: SYG | TF:M09641_0 | 1.01E-08 | 7.99673 | 329 |
| TF | Factor: IRX2a; motif: ACR | TF:M11022 | 1.14E-08 | 7.943459 | 826 |
| TF | Factor: IRX2a; motif: ACR | TF:M11022_0 | 1.14E-08 | 7.943459 | 826 |
| TF | Factor: Erg; motif: NACC( | TF:M11397_1 | 1.19E-08 | 7.925148 | 992 |
| TF | Factor: YY1; motif: NNCG | TF:M03924_1 | 1.64E-08 | 7.784695 | 362 |
| TF | Factor: BTEB2; motif: NC( | TF:M12165_0 | 1.80E-08 | 7.745144 | 449 |
| TF | Factor: BTEB2; motif: NC( | TF:M12165 | 1.80E-08 | 7.745144 | 449 |
| TF | Factor: GCMa:Erg; motif: | TF:M08487_0 | 1.84E-08 | 7.735338 | 1140 |
| TF | Factor: GCMa:Erg; motif: | TF:M08487 | 1.84E-08 | 7.735338 | 1140 |
| TF | Factor: E2F; motif: TTTSG | TF:M00427 | 2.02E-08 | 7.693583 | 827 |
| TF | Factor: E2F; motif: TTTSG | TF:M00427_0 | 2.02E-08 | 7.693583 | 827 |
| TF | Factor: E2F; motif: TTTSG | TF:M00050_0 | 2.81E-08 | 7.551671 | 637 |
| TF | Factor: E2F; motif: TTTSG | TF:M00050 | 2.81E-08 | 7.551671 | 637 |
| TF | Factor: CREB,; motif: NTG | TF:M00981_0 | 2.93E-08 | 7.532966 | 533 |
| TF | Factor: CREB,; motif: NTG | TF:M00981 | 2.93E-08 | 7.532966 | 533 |
| TF | Factor: Elk-1:Pax-9; motif | TF:M08226_0 | 3.19E-08 | 7.496453 | 955 |
| TF | Factor: Elk-1:Pax-9; motif | TF:M08226 | 3.19E-08 | 7.496453 | 955 |
| TF | Factor: SP1; motif: GGGG | TF:M01303 | 3.25E-08 | 7.488471 | 704 |
| TF | Factor: SP1; motif: GGGG | TF:M01303_0 | 3.25E-08 | 7.488471 | 704 |
| TF | Factor: TR4; motif: ACCC( | TF:M04934 | 3.25E-08 | 7.487773 | 1402 |
| TF | Factor: TR4; motif: ACCC( | TF:M04934_0 | 3.25E-08 | 7.487773 | 1402 |
| TF | Factor: sp4; motif: NNGN | TF:M10072_1 | 3.28E-08 | 7.484557 | 576 |
| TF | Factor: Pet-1; motif: GCN | TF:M09918_1 | 3.37E-08 | 7.472775 | 625 |
| TF | Factor: E2F-3:FOXI1; mot | TF:M08208_1 | 3.51E-08 | 7.45483 | 609 |
| TF | Factor: ER71:SREBP-2; m( | TF:M08457 | 3.66E-08 | 7.436204 | 1009 |
| TF | Factor: ER71:SREBP-2; m( | TF:M08457_0 | 3.66E-08 | 7.436204 | 1009 |
| TF | Factor: KLF15; motif: RCC | TF:M12160 | 3.87E-08 | 7.412428 | 1066 |

| | | | | | |
|---|---|---|---|---|---|
| **TF** | Factor: KLF15; motif: RCC | TF:M12160_0 | 3.87E-08 | 7.412428 | 1066 |
| **TF** | Factor: ERG; motif: ACCG | TF:M03976_1 | 4.41E-08 | 7.355598 | 698 |
| **TF** | Factor: TEF-3:Elk-1; motif | TF:M08418_0 | 4.68E-08 | 7.329475 | 975 |
| **TF** | Factor: TEF-3:Elk-1; motif | TF:M08418 | 4.68E-08 | 7.329475 | 975 |
| **TF** | Factor: ER81; motif: NRC | TF:M11418_0 | 5.12E-08 | 7.290605 | 278 |
| **TF** | Factor: ER81; motif: NRC | TF:M11418 | 5.12E-08 | 7.290605 | 278 |
| **TF** | Factor: GABP-beta; motif | TF:M01876 | 5.50E-08 | 7.259404 | 169 |
| **TF** | Factor: GABP-beta; motif | TF:M01876_0 | 5.50E-08 | 7.259404 | 169 |
| **TF** | Factor: Elk-1; motif: AAC | TF:M01163_1 | 5.60E-08 | 7.251468 | 102 |
| **TF** | Factor: Pax-3; motif: NNN | TF:M00327_1 | 5.76E-08 | 7.239846 | 525 |
| **TF** | Factor: SP4; motif: SCCC | TF:M01273_0 | 6.07E-08 | 7.216518 | 336 |
| **TF** | Factor: SP4; motif: SCCC | TF:M01273 | 6.07E-08 | 7.216518 | 336 |
| **TF** | Factor: BTEB4; motif: NC | TF:M12186_1 | 6.27E-08 | 7.202736 | 378 |
| **TF** | Factor: Elk-1:Pax-1; motif | TF:M08224_1 | 6.61E-08 | 7.179773 | 289 |
| **TF** | Factor: CPBP; motif: SNC | TF:M01822_1 | 6.61E-08 | 7.179498 | 1430 |
| **TF** | Factor: Pax-5; motif: RRM | TF:M00144_1 | 6.95E-08 | 7.158191 | 594 |
| **TF** | Factor: GABP-alpha; moti | TF:M11414_1 | 7.76E-08 | 7.110104 | 57 |
| **TF** | Factor: Pet-1; motif: GCN | TF:M09918 | 8.28E-08 | 7.082145 | 1106 |
| **TF** | Factor: Pet-1; motif: GCN | TF:M09918_0 | 8.28E-08 | 7.082145 | 1106 |
| **TF** | Factor: E2F-1; motif: NTT | TF:M00430_1 | 9.64E-08 | 7.015759 | 266 |
| **TF** | Factor: ETV1; motif: ACC | TF:M03979 | 1.03E-07 | 6.988967 | 121 |
| **TF** | Factor: ETV1; motif: ACC | TF:M03979_0 | 1.03E-07 | 6.988967 | 121 |
| **TF** | Factor: E2F-1; motif: TTT | TF:M04516_1 | 1.10E-07 | 6.958006 | 872 |
| **TF** | Factor: Elk-1:T-bet; motif | TF:M08231 | 1.12E-07 | 6.951963 | 596 |
| **TF** | Factor: Elk-1:T-bet; motif | TF:M08231_0 | 1.12E-07 | 6.951963 | 596 |
| **TF** | Factor: CREB1; motif: NR | TF:M09887_0 | 1.22E-07 | 6.912839 | 383 |
| **TF** | Factor: CREB1; motif: NR | TF:M09887 | 1.22E-07 | 6.912839 | 383 |
| **TF** | Factor: RREB-1; motif: GG | TF:M05439_0 | 1.47E-07 | 6.834011 | 665 |
| **TF** | Factor: RREB-1; motif: GG | TF:M05439 | 1.47E-07 | 6.834011 | 665 |
| **TF** | Factor: SP1; motif: NRGK | TF:M09765_1 | 1.55E-07 | 6.810212 | 322 |
| **TF** | Factor: E2F-1:DP-2; motif | TF:M00737_0 | 1.64E-07 | 6.784263 | 761 |
| **TF** | Factor: E2F-1:DP-2; motif | TF:M00737 | 1.64E-07 | 6.784263 | 761 |
| **TF** | Factor: CREM; motif: TGA | TF:M01820 | 1.95E-07 | 6.710666 | 576 |
| **TF** | Factor: CREM; motif: TGA | TF:M01820_0 | 1.95E-07 | 6.710666 | 576 |
| **TF** | Factor: GKLF; motif: NNN | TF:M07289 | 1.96E-07 | 6.70769 | 1184 |
| **TF** | Factor: GKLF; motif: NNN | TF:M07289_0 | 1.96E-07 | 6.70769 | 1184 |
| **TF** | Factor: SP2; motif: GGGC | TF:M01783_1 | 2.04E-07 | 6.689752 | 402 |
| **TF** | Factor: ER81; motif: NAC | TF:M11420_1 | 2.05E-07 | 6.688063 | 59 |
| **TF** | Factor: E2F; motif: TTTSG | TF:M00426_1 | 2.13E-07 | 6.671383 | 540 |
| **TF** | Factor: E2F-6; motif: NGG | TF:M09895 | 2.15E-07 | 6.668213 | 692 |
| **TF** | Factor: E2F-6; motif: NGG | TF:M09895_0 | 2.15E-07 | 6.668213 | 692 |
| **TF** | Factor: Rb:E2F-1:DP-1; m | TF:M00740_1 | 2.18E-07 | 6.662136 | 409 |
| **TF** | Factor: E2F-4:DP-1; motif | TF:M00738_1 | 2.19E-07 | 6.658601 | 392 |
| **TF** | Factor: KLF15; motif: NCC | TF:M12158_0 | 2.21E-07 | 6.655797 | 907 |
| **TF** | Factor: KLF15; motif: NCC | TF:M12158 | 2.21E-07 | 6.655797 | 907 |
| **TF** | Factor: ER81; motif: NNC | TF:M09913_1 | 2.36E-07 | 6.627829 | 54 |
| **TF** | Factor: elf5; motif: GAMC | TF:M11389 | 2.36E-07 | 6.626312 | 154 |

| | | | | | |
|---|---|---|---|---|---|
| **TF** | Factor: elf5; motif: GAM( | TF:M11389_0 | 2.36E-07 | 6.626312 | 154 |
| **TF** | Factor: AP-2; motif: MKC( | TF:M00189_0 | 2.49E-07 | 6.604407 | 896 |
| **TF** | Factor: AP-2; motif: MKC( | TF:M00189 | 2.49E-07 | 6.604407 | 896 |
| **TF** | Factor: Elf5; motif: CSMG | TF:M02057_1 | 2.51E-07 | 6.601049 | 346 |
| **TF** | Factor: FLI-1; motif: CMG | TF:M01208_1 | 2.52E-07 | 6.598882 | 110 |
| **TF** | Factor: Fli-1; motif: RCM( | TF:M09613_1 | 3.02E-07 | 6.519778 | 83 |
| **TF** | Factor: Erg; motif: NACC( | TF:M11395 | 3.02E-07 | 6.519694 | 763 |
| **TF** | Factor: Erg; motif: NACC( | TF:M11395_0 | 3.02E-07 | 6.519694 | 763 |
| **TF** | Factor: E2F-1:TBR2; moti | TF:M08523_1 | 3.04E-07 | 6.517423 | 647 |
| **TF** | Factor: ETV5; motif: NCC( | TF:M03983_1 | 3.11E-07 | 6.507084 | 62 |
| **TF** | Factor: E2F-1; motif: TTTS | TF:M00431 | 3.46E-07 | 6.460944 | 650 |
| **TF** | Factor: E2F-1; motif: TTTS | TF:M00431_0 | 3.46E-07 | 6.460944 | 650 |
| **TF** | Factor: Elf-1; motif: RANS | TF:M11446 | 3.55E-07 | 6.449732 | 228 |
| **TF** | Factor: Elf-1; motif: RANS | TF:M11446_0 | 3.55E-07 | 6.449732 | 228 |
| **TF** | Factor: Erg; motif: NACC( | TF:M11395_1 | 3.79E-07 | 6.421151 | 705 |
| **TF** | Factor: Cdx-2; motif: NRT | TF:M10849_1 | 4.31E-07 | 6.365634 | 306 |
| **TF** | Factor: Sp1; motif: CCCC( | TF:M00933_1 | 4.49E-07 | 6.348223 | 485 |
| **TF** | Factor: CPBP; motif: GNN | TF:M09973_0 | 4.51E-07 | 6.346262 | 915 |
| **TF** | Factor: CPBP; motif: GNN | TF:M09973 | 4.51E-07 | 6.346262 | 915 |
| **TF** | Factor: ETS1; motif: GCC( | TF:M03978 | 4.66E-07 | 6.331614 | 704 |
| **TF** | Factor: ETS1; motif: GCC( | TF:M03978_0 | 4.66E-07 | 6.331614 | 704 |
| **TF** | Factor: ZIC4; motif: NNC( | TF:M12227_0 | 4.81E-07 | 6.317752 | 750 |
| **TF** | Factor: ZIC4; motif: NNC( | TF:M12227 | 4.81E-07 | 6.317752 | 750 |
| **TF** | Factor: Sp1; motif: NWR( | TF:M12152_1 | 4.85E-07 | 6.313836 | 294 |
| **TF** | Factor: ESE-1; motif: SATI | TF:M11381_1 | 4.99E-07 | 6.301539 | 837 |
| **TF** | Factor: ATF-3; motif: GG( | TF:M04744_1 | 5.59E-07 | 6.252371 | 458 |
| **TF** | Factor: pax-6; motif: NYA | TF:M11883_0 | 6.00E-07 | 6.221971 | 966 |
| **TF** | Factor: pax-6; motif: NYA | TF:M11883 | 6.00E-07 | 6.221971 | 966 |
| **TF** | Factor: ETV6; motif: NSC( | TF:M03985_0 | 6.03E-07 | 6.219821 | 138 |
| **TF** | Factor: ETV6; motif: NSC( | TF:M03985 | 6.03E-07 | 6.219821 | 138 |
| **TF** | Factor: GABP-alpha; moti | TF:M11415_1 | 6.65E-07 | 6.177082 | 62 |
| **TF** | Factor: PSF; motif: KWAT | TF:M07612 | 6.95E-07 | 6.157874 | 511 |
| **TF** | Factor: PSF; motif: KWAT | TF:M07612_0 | 6.95E-07 | 6.157874 | 511 |
| **TF** | Factor: Sp2; motif: NTAA( | TF:M12156_1 | 7.56E-07 | 6.121644 | 128 |
| **TF** | Factor: TAFII250; motif: R | TF:M10086_0 | 8.15E-07 | 6.088986 | 772 |
| **TF** | Factor: TAFII250; motif: R | TF:M10086 | 8.15E-07 | 6.088986 | 772 |
| **TF** | Factor: Egr-1; motif: ACC( | TF:M04918 | 8.17E-07 | 6.087585 | 459 |
| **TF** | Factor: Egr-1; motif: ACC( | TF:M04918_0 | 8.17E-07 | 6.087585 | 459 |
| **TF** | Factor: Churchill; motif: ( | TF:M00986_1 | 8.52E-07 | 6.069747 | 865 |
| **TF** | Factor: GABP-alpha; moti | TF:M04698_1 | 9.75E-07 | 6.010793 | 86 |
| **TF** | Factor: E2F-1; motif: TTTS | TF:M00516 | 1.13E-06 | 5.945193 | 620 |
| **TF** | Factor: E2F-1; motif: TTTS | TF:M00516_0 | 1.13E-06 | 5.945193 | 620 |
| **TF** | Factor: ATF-3; motif: GG( | TF:M04744 | 1.14E-06 | 5.943671 | 854 |
| **TF** | Factor: ATF-3; motif: GG( | TF:M04744_0 | 1.14E-06 | 5.943671 | 854 |
| **TF** | Factor: LBP-1; motif: NR( | TF:M11490 | 1.22E-06 | 5.915279 | 586 |
| **TF** | Factor: LBP-1; motif: NR( | TF:M11490_0 | 1.22E-06 | 5.915279 | 586 |
| **TF** | Factor: E2F-1; motif: NTT | TF:M00430 | 1.26E-06 | 5.898396 | 686 |

| | | | | | |
|---|---|---|---|---|---|
| **TF** | Factor: E2F-1; motif: NTT | TF:M00430_0 | 1.26E-06 | 5.898396 | 686 |
| **TF** | Factor: GABP; motif: CAC | TF:M01258_1 | 1.28E-06 | 5.893589 | 69 |
| **TF** | Factor: Egr-3; motif: NTG | TF:M00245 | 1.29E-06 | 5.887818 | 642 |
| **TF** | Factor: Egr-3; motif: NTG | TF:M00245_0 | 1.29E-06 | 5.887818 | 642 |
| **TF** | Factor: BEN; motif: CWG( | TF:M01241 | 1.35E-06 | 5.868625 | 577 |
| **TF** | Factor: BEN; motif: CWG( | TF:M01241_0 | 1.35E-06 | 5.868625 | 577 |
| **TF** | Factor: E2F-4; motif: NNT | TF:M04823_1 | 1.37E-06 | 5.864283 | 692 |
| **TF** | Factor: TIEG1; motif: NC( | TF:M12351_1 | 1.40E-06 | 5.853385 | 590 |
| **TF** | Factor: XBP-1; motif: WN | TF:M01770 | 1.57E-06 | 5.802871 | 916 |
| **TF** | Factor: XBP-1; motif: WN | TF:M01770_0 | 1.57E-06 | 5.802871 | 916 |
| **TF** | Factor: CREB1; motif: TK/ | TF:M09600 | 1.58E-06 | 5.802261 | 344 |
| **TF** | Factor: CREB1; motif: TK/ | TF:M09600_0 | 1.58E-06 | 5.802261 | 344 |
| **TF** | Factor: GABP-ALPHA; mo | TF:M08917_1 | 1.70E-06 | 5.770348 | 64 |
| **TF** | Factor: E2F-4; motif: GCG | TF:M02090_1 | 1.92E-06 | 5.717513 | 577 |
| **TF** | Factor: sp4; motif: NNNG | TF:M10530_1 | 2.11E-06 | 5.675604 | 365 |
| **TF** | Factor: E2F; motif: TTTSG | TF:M00427_1 | 2.39E-06 | 5.621572 | 376 |
| **TF** | Factor: E2F-1; motif: NTT | TF:M11533_1 | 2.45E-06 | 5.609982 | 753 |
| **TF** | Factor: ERG; motif: ACCG | TF:M03976_0 | 2.82E-06 | 5.549892 | 1053 |
| **TF** | Factor: ERG; motif: ACCG | TF:M03976 | 2.82E-06 | 5.549892 | 1053 |
| **TF** | Factor: SAP1A; motif: NC( | TF:M02058_0 | 2.85E-06 | 5.544613 | 87 |
| **TF** | Factor: SAP1A; motif: NC( | TF:M02058 | 2.85E-06 | 5.544613 | 87 |
| **TF** | Factor: ELK1; motif: NACT | TF:M03971 | 2.86E-06 | 5.543866 | 928 |
| **TF** | Factor: ELK1; motif: NACT | TF:M03971_0 | 2.86E-06 | 5.543866 | 928 |
| **TF** | Factor: GCMa:Erg; motif: | TF:M08487_1 | 2.96E-06 | 5.528023 | 748 |
| **TF** | Factor: SP1; motif: NCCC( | TF:M07226_1 | 3.16E-06 | 5.50007 | 380 |
| **TF** | Factor: NRF-1; motif: CN( | TF:M04694 | 3.30E-06 | 5.48155 | 175 |
| **TF** | Factor: NRF-1; motif: CN( | TF:M04694_0 | 3.30E-06 | 5.48155 | 175 |
| **TF** | Factor: BTEB2; motif: RG( | TF:M07277_1 | 3.40E-06 | 5.468657 | 399 |
| **TF** | Factor: ELF4; motif: NCC( | TF:M02056_1 | 3.63E-06 | 5.439787 | 106 |
| **TF** | Factor: E2F-4; motif: TTT | TF:M11535_1 | 3.77E-06 | 5.423678 | 586 |
| **TF** | Factor: Egr-1; motif: GCG | TF:M07354_0 | 3.78E-06 | 5.422856 | 652 |
| **TF** | Factor: Egr-1; motif: GCG | TF:M07354 | 3.78E-06 | 5.422856 | 652 |
| **TF** | Factor: Kaiso; motif: GCM | TF:M03876 | 3.91E-06 | 5.40767 | 1102 |
| **TF** | Factor: Kaiso; motif: GCM | TF:M03876_0 | 3.91E-06 | 5.40767 | 1102 |
| **TF** | Factor: NRF-1; motif: GC( | TF:M07052_0 | 4.15E-06 | 5.382281 | 224 |
| **TF** | Factor: NRF-1; motif: GC( | TF:M07052 | 4.15E-06 | 5.382281 | 224 |
| **TF** | Factor: Net; motif: CACTT | TF:M03832_0 | 4.21E-06 | 5.3754 | 89 |
| **TF** | Factor: Net; motif: CACTT | TF:M03832 | 4.21E-06 | 5.3754 | 89 |
| **TF** | Factor: ELF5; motif: ANSN | TF:M03969_1 | 4.61E-06 | 5.336377 | 68 |
| **TF** | Factor: ELK4; motif: ACC( | TF:M03973 | 4.61E-06 | 5.336051 | 69 |
| **TF** | Factor: ELK4; motif: ACC( | TF:M03973_0 | 4.61E-06 | 5.336051 | 69 |
| **TF** | Factor: c-Myc:Max; motif | TF:M00322_0 | 4.79E-06 | 5.319251 | 470 |
| **TF** | Factor: c-Myc:Max; motif | TF:M00322 | 4.79E-06 | 5.319251 | 470 |
| **TF** | Factor: HES-1; motif: GG( | TF:M11061 | 4.96E-06 | 5.304398 | 545 |
| **TF** | Factor: HES-1; motif: GG( | TF:M11061_0 | 4.96E-06 | 5.304398 | 545 |
| **TF** | Factor: ZFX; motif: SNAG( | TF:M10121_0 | 5.09E-06 | 5.293622 | 336 |
| **TF** | Factor: ZFX; motif: SNAG( | TF:M10121 | 5.09E-06 | 5.293622 | 336 |

| | | | | | | |
|---|---|---|---|---|---|---|
| **TF** | Factor: HOXB2:PEA3; mo | TF:M08345_0 | | 5.35E-06 | 5.271558 | 490 |
| **TF** | Factor: HOXB2:PEA3; mo | TF:M08345 | | 5.35E-06 | 5.271558 | 490 |
| **TF** | Factor: Erg; motif: NACC | TF:M11391 | | 5.41E-06 | 5.266509 | 855 |
| **TF** | Factor: Erg; motif: NACC | TF:M11391_0 | | 5.41E-06 | 5.266509 | 855 |
| **TF** | Factor: GABP; motif: VCC | TF:M00341_1 | | 5.82E-06 | 5.235291 | 173 |
| **TF** | Factor: HOXB2:PEA3; mo | TF:M08345_1 | | 5.92E-06 | 5.227491 | 169 |
| **TF** | Factor: ER71:SREBP-2; m | TF:M08457_1 | | 6.47E-06 | 5.189405 | 513 |
| **TF** | Factor: E2F; motif: TTTSG | TF:M00918_0 | | 6.47E-06 | 5.189405 | 513 |
| **TF** | Factor: E2F; motif: TTTSG | TF:M00918 | | 6.47E-06 | 5.189405 | 513 |
| **TF** | Factor: Cdx-2; motif: NRT | TF:M10849 | | 7.71E-06 | 5.113042 | 848 |
| **TF** | Factor: Cdx-2; motif: NRT | TF:M10849_0 | | 7.71E-06 | 5.113042 | 848 |
| **TF** | Factor: E2F-1:DP-1; motif | TF:M00736_1 | | 7.92E-06 | 5.101303 | 437 |
| **TF** | Factor: Sp1; motif: GGGG | TF:M07063_1 | | 8.50E-06 | 5.070831 | 339 |
| **TF** | Factor: Sp2; motif: GGGG | TF:M10435_1 | | 8.50E-06 | 5.070831 | 339 |
| **TF** | Factor: LBP-1; motif: NR( | TF:M11490_1 | | 9.31E-06 | 5.030865 | 541 |
| **TF** | Factor: c-Myc; motif: RA( | TF:M01145_0 | | 9.54E-06 | 5.020268 | 496 |
| **TF** | Factor: c-Myc; motif: RA( | TF:M01145 | | 9.54E-06 | 5.020268 | 496 |
| **TF** | Factor: SP2; motif: GNNG | TF:M03807_1 | | 9.62E-06 | 5.0167 | 395 |
| **TF** | Factor: Pet-1; motif: NAC | TF:M11407_0 | | 1.14E-05 | 4.942483 | 74 |
| **TF** | Factor: Pet-1; motif: NAC | TF:M11407 | | 1.14E-05 | 4.942483 | 74 |
| **TF** | Factor: KLF14; motif: NR( | TF:M12183_0 | | 1.19E-05 | 4.924825 | 602 |
| **TF** | Factor: KLF14; motif: NR( | TF:M12183 | | 1.19E-05 | 4.924825 | 602 |
| **TF** | Factor: E2F-2; motif: NTT | TF:M11528 | | 1.21E-05 | 4.918388 | 864 |
| **TF** | Factor: E2F-2; motif: NTT | TF:M11528_0 | | 1.21E-05 | 4.918388 | 864 |
| **TF** | Factor: YY1; motif: CAAN | TF:M10436_0 | | 1.22E-05 | 4.915094 | 631 |
| **TF** | Factor: YY1; motif: CAAN | TF:M10436 | | 1.22E-05 | 4.915094 | 631 |
| **TF** | Factor: Elf5; motif: CSMG | TF:M02057_0 | | 1.25E-05 | 4.90152 | 848 |
| **TF** | Factor: Elf5; motif: CSMG | TF:M02057 | | 1.25E-05 | 4.90152 | 848 |
| **TF** | Factor: E2F; motif: TTTSG | TF:M00939_0 | | 1.29E-05 | 4.890976 | 576 |
| **TF** | Factor: E2F; motif: TTTSG | TF:M00939 | | 1.29E-05 | 4.890976 | 576 |
| **TF** | Factor: Elk-1:OC-2; motif | TF:M08223_0 | | 1.37E-05 | 4.862391 | 1026 |
| **TF** | Factor: Elk-1:OC-2; motif | TF:M08223 | | 1.37E-05 | 4.862391 | 1026 |
| **TF** | Factor: NRF-1; motif: CNS | TF:M10018_0 | | 1.48E-05 | 4.829092 | 196 |
| **TF** | Factor: NRF-1; motif: CNS | TF:M10018 | | 1.48E-05 | 4.829092 | 196 |
| **TF** | Factor: FLI1; motif: ACCG | TF:M03989_1 | | 1.51E-05 | 4.821028 | 512 |
| **TF** | Factor: E2F-1; motif: NTT | TF:M00940_1 | | 1.58E-05 | 4.801154 | 269 |
| **TF** | Factor: E2F-7; motif: GRG | TF:M09896_1 | | 1.86E-05 | 4.730532 | 425 |
| **TF** | Factor: MOVO-B; motif: ( | TF:M01104_0 | | 1.99E-05 | 4.700207 | 679 |
| **TF** | Factor: MOVO-B; motif: ( | TF:M01104 | | 1.99E-05 | 4.700207 | 679 |
| **TF** | Factor: BTEB1; motif: GG( | TF:M09723_1 | | 2.03E-05 | 4.69328 | 533 |
| **TF** | Factor: ATF-1; motif: TNA | TF:M01861 | | 2.07E-05 | 4.684348 | 383 |
| **TF** | Factor: ATF-1; motif: TNA | TF:M01861_0 | | 2.07E-05 | 4.684348 | 383 |
| **TF** | Factor: RUNX3; motif: NR | TF:M04110_0 | | 2.16E-05 | 4.665528 | 790 |
| **TF** | Factor: RUNX3; motif: NR | TF:M04110 | | 2.16E-05 | 4.665528 | 790 |
| **TF** | Factor: SP1; motif: RCCM | TF:M03920_1 | | 2.23E-05 | 4.652178 | 288 |
| **TF** | Factor: Fli-1; motif: NACC | TF:M11402_1 | | 2.44E-05 | 4.613021 | 62 |
| **TF** | Factor: HES-7; motif: GN( | TF:M11066_1 | | 2.48E-05 | 4.605136 | 680 |

| | | | | | | |
|---|---|---|---|---|---|---|
| **TF** | Factor: pax-2; motif: NCG | TF:M11877_0 | | 2.73E-05 | 4.564188 | 855 |
| **TF** | Factor: pax-2; motif: NCG | TF:M11877 | | 2.73E-05 | 4.564188 | 855 |
| **TF** | Factor: KLF3; motif: NNN | TF:M09970_1 | | 2.90E-05 | 4.537937 | 359 |
| **TF** | Factor: ERF; motif: ACCG( | TF:M03974 | | 2.92E-05 | 4.534825 | 84 |
| **TF** | Factor: ERF; motif: ACCG( | TF:M03974_0 | | 2.92E-05 | 4.534825 | 84 |
| **TF** | Factor: ETV3; motif: ACC( | TF:M03981_1 | | 3.02E-05 | 4.520245 | 39 |
| **TF** | Factor: ETS1; motif: GCC( | TF:M03978_1 | | 3.08E-05 | 4.511487 | 568 |
| **TF** | Factor: BTEB2; motif: NM | TF:M12164 | | 3.12E-05 | 4.506087 | 252 |
| **TF** | Factor: BTEB2; motif: NM | TF:M12164_0 | | 3.12E-05 | 4.506087 | 252 |
| **TF** | Factor: ATF; motif: CNST( | TF:M00017_1 | | 3.36E-05 | 4.47313 | 220 |
| **TF** | Factor: KLF15; motif: RCC | TF:M12157 | | 3.59E-05 | 4.445205 | 347 |
| **TF** | Factor: KLF15; motif: RCC | TF:M12157_0 | | 3.59E-05 | 4.445205 | 347 |
| **TF** | Factor: hoxa9; motif: RTC | TF:M10785_0 | | 3.74E-05 | 4.427279 | 433 |
| **TF** | Factor: hoxa9; motif: RTC | TF:M10785 | | 3.74E-05 | 4.427279 | 433 |
| **TF** | Factor: E2F-3:FOXO6; mo | TF:M08209 | | 3.92E-05 | 4.406978 | 1036 |
| **TF** | Factor: E2F-3:FOXO6; mo | TF:M08209_0 | | 3.92E-05 | 4.406978 | 1036 |
| **TF** | Factor: Erg; motif: NACC( | TF:M11391_1 | | 3.92E-05 | 4.406231 | 775 |
| **TF** | Factor: Sp1; motif: NWR( | TF:M12151_1 | | 4.46E-05 | 4.350337 | 248 |
| **TF** | Factor: Elk-1:OC-2; motif: | TF:M08223_1 | | 4.52E-05 | 4.345111 | 521 |
| **TF** | Factor: E2F-1; motif: NKT | TF:M00428_1 | | 4.58E-05 | 4.339351 | 345 |
| **TF** | Factor: ATF-3; motif: GGT | TF:M09868_1 | | 4.58E-05 | 4.338856 | 222 |
| **TF** | Factor: RUNX2; motif: WI | TF:M04107 | | 4.79E-05 | 4.319236 | 1013 |
| **TF** | Factor: RUNX2; motif: WI | TF:M04107_0 | | 4.79E-05 | 4.319236 | 1013 |
| **TF** | Factor: pax-6; motif: NYA | TF:M11883_1 | | 5.07E-05 | 4.295284 | 661 |
| **TF** | Factor: CREB; motif: NST( | TF:M00178_0 | | 5.16E-05 | 4.287004 | 359 |
| **TF** | Factor: CREB; motif: NST( | TF:M00178 | | 5.16E-05 | 4.287004 | 359 |
| **TF** | Factor: CP2; motif: AACC( | TF:M11485_0 | | 5.27E-05 | 4.278586 | 611 |
| **TF** | Factor: CP2; motif: AACC( | TF:M11485 | | 5.27E-05 | 4.278586 | 611 |
| **TF** | Factor: YY2; motif: NCCG( | TF:M03925_0 | | 5.35E-05 | 4.272042 | 425 |
| **TF** | Factor: YY2; motif: NCCG( | TF:M03925 | | 5.35E-05 | 4.272042 | 425 |
| **TF** | Factor: XBP-1; motif: WN | TF:M01770_1 | | 5.44E-05 | 4.264346 | 419 |
| **TF** | Factor: Sp2; motif: NYSG( | TF:M03567_1 | | 5.49E-05 | 4.26076 | 332 |
| **TF** | Factor: Zfx; motif: SNSCA | TF:M01593_0 | | 5.81E-05 | 4.235795 | 331 |
| **TF** | Factor: Zfx; motif: SNSCA | TF:M01593 | | 5.81E-05 | 4.235795 | 331 |
| **TF** | Factor: Sp1; motif: NGGG | TF:M10071_1 | | 5.89E-05 | 4.229904 | 409 |
| **TF** | Factor: MECP2; motif: CC | TF:M01299 | | 6.74E-05 | 4.17166 | 444 |
| **TF** | Factor: MECP2; motif: CC | TF:M01299_0 | | 6.74E-05 | 4.17166 | 444 |
| **TF** | Factor: A-Myb:Elf-1; moti | TF:M08390 | | 7.19E-05 | 4.143031 | 867 |
| **TF** | Factor: A-Myb:Elf-1; moti | TF:M08390_0 | | 7.19E-05 | 4.143031 | 867 |
| **TF** | Factor: pax-6; motif: NYA | TF:M11881 | | 7.77E-05 | 4.109844 | 1057 |
| **TF** | Factor: pax-6; motif: NYA | TF:M11881_0 | | 7.77E-05 | 4.109844 | 1057 |
| **TF** | Factor: ZNF76; motif: RNI | TF:M10169_0 | | 7.85E-05 | 4.10535 | 156 |
| **TF** | Factor: ZNF76; motif: RNI | TF:M10169 | | 7.85E-05 | 4.10535 | 156 |
| **TF** | Factor: TEF-3:Elk-1; motif | TF:M08418_1 | | 8.60E-05 | 4.065711 | 477 |
| **TF** | Factor: CREB,; motif: NTG | TF:M00981_1 | | 9.77E-05 | 4.010035 | 219 |
| **TF** | Factor: c-Ets-2; motif: NA | TF:M11417_0 | | 9.98E-05 | 4.000905 | 1065 |
| **TF** | Factor: c-Ets-2; motif: NA | TF:M11417 | | 9.98E-05 | 4.000905 | 1065 |

| | | | | | |
|---|---|---|---|---|---|
| **TF** | Factor: KLF17; motif: NGI | TF:M12188 | 0.00010111 | 3.995207 | 350 |
| **TF** | Factor: KLF17; motif: NGI | TF:M12188_0 | 0.00010111 | 3.995207 | 350 |
| **TF** | Factor: ER81; motif: NAC | TF:M11421_1 | 0.000106307 | 3.973437 | 41 |
| **TF** | Factor: E2F-1; motif: NNN | TF:M07250_1 | 0.000111271 | 3.953617 | 421 |
| **TF** | Factor: NRF-1; motif: YGC | TF:M04059 | 0.000111498 | 3.952733 | 161 |
| **TF** | Factor: NRF-1; motif: YGC | TF:M04059_0 | 0.000111498 | 3.952733 | 161 |
| **TF** | Factor: Egr-1; motif: NGC | TF:M04950_0 | 0.000112391 | 3.949269 | 895 |
| **TF** | Factor: Egr-1; motif: NGC | TF:M04950 | 0.000112391 | 3.949269 | 895 |
| **TF** | Factor: NRF-1; motif: YGC | TF:M02102_0 | 0.000121327 | 3.916042 | 209 |
| **TF** | Factor: NRF-1; motif: YGC | TF:M02102 | 0.000121327 | 3.916042 | 209 |
| **TF** | Factor: Erg; motif: NACM | TF:M11394 | 0.00012495 | 3.903263 | 267 |
| **TF** | Factor: Erg; motif: NACM | TF:M11394_0 | 0.00012495 | 3.903263 | 267 |
| **TF** | Factor: etv3; motif: NNC | TF:M11410 | 0.000136483 | 3.864921 | 113 |
| **TF** | Factor: etv3; motif: NNC | TF:M11410_0 | 0.000136483 | 3.864921 | 113 |
| **TF** | Factor: CREB; motif: NNG | TF:M00113 | 0.000137551 | 3.861537 | 429 |
| **TF** | Factor: CREB; motif: NNG | TF:M00113_0 | 0.000137551 | 3.861537 | 429 |
| **TF** | Factor: CREB; motif: NNT | TF:M00916_0 | 0.000146686 | 3.833612 | 357 |
| **TF** | Factor: CREB; motif: NNT | TF:M00916 | 0.000146686 | 3.833612 | 357 |
| **TF** | Factor: DEAF1; motif: NC | TF:M01001 | 0.000156949 | 3.804242 | 195 |
| **TF** | Factor: DEAF1; motif: NC | TF:M01001_0 | 0.000156949 | 3.804242 | 195 |
| **TF** | Factor: ELK1; motif: NACT | TF:M03971_1 | 0.000171806 | 3.764961 | 618 |
| **TF** | Factor: Pax-3; motif: NNN | TF:M00327 | 0.000177073 | 3.751849 | 946 |
| **TF** | Factor: Pax-3; motif: NNN | TF:M00327_0 | 0.000177073 | 3.751849 | 946 |
| **TF** | Factor: GABP-alpha; moti | TF:M04748_1 | 0.000185291 | 3.732145 | 749 |
| **TF** | Factor: Tax/CREB; motif: | TF:M00114_1 | 0.000195432 | 3.709003 | 143 |
| **TF** | Factor: KLF15; motif: RCC | TF:M12160_1 | 0.000218161 | 3.661223 | 698 |
| **TF** | Factor: ER71; motif: NAC | TF:M11413_1 | 0.000224484 | 3.648814 | 43 |
| **TF** | Factor: SP1:SP3; motif: C | TF:M01219_0 | 0.000232919 | 3.632795 | 323 |
| **TF** | Factor: SP1:SP3; motif: C | TF:M01219 | 0.000232919 | 3.632795 | 323 |
| **TF** | Factor: SAP-1a; motif: NA | TF:M01167_1 | 0.000237272 | 3.624753 | 28 |
| **TF** | Factor: Sp2; motif: GGSN | TF:M09658_0 | 0.000250626 | 3.600973 | 476 |
| **TF** | Factor: Sp2; motif: GGSN | TF:M09658 | 0.000250626 | 3.600973 | 476 |
| **TF** | Factor: SAP-1; motif: NRC | TF:M11437_1 | 0.000268065 | 3.57176 | 63 |
| **TF** | Factor: Net; motif: ACCG | TF:M02060 | 0.000276305 | 3.558612 | 54 |
| **TF** | Factor: Net; motif: ACCG | TF:M02060_0 | 0.000276305 | 3.558612 | 54 |
| **TF** | Factor: CP2; motif: AACC | TF:M11485_1 | 0.00028756 | 3.541271 | 546 |
| **TF** | Factor: LKLF; motif: NRC | TF:M12172 | 0.000301066 | 3.521339 | 210 |
| **TF** | Factor: LKLF; motif: NRC | TF:M12172_0 | 0.000301066 | 3.521339 | 210 |
| **TF** | Factor: GABPalpha_GABP | TF:M03791_1 | 0.000305989 | 3.514295 | 56 |
| **TF** | Factor: PAX5; motif: RNG | TF:M09817_1 | 0.000311013 | 3.507221 | 520 |
| **TF** | Factor: KROX; motif: CCC | TF:M00982 | 0.000323644 | 3.489933 | 662 |
| **TF** | Factor: KROX; motif: CCC | TF:M00982_0 | 0.000323644 | 3.489933 | 662 |
| **TF** | Factor: CPBP; motif: GNN | TF:M09973_1 | 0.000360008 | 3.443688 | 578 |
| **TF** | Factor: E2F-4:DP-2; motif | TF:M00739 | 0.000383095 | 3.416694 | 504 |
| **TF** | Factor: E2F-4:DP-2; motif | TF:M00739_0 | 0.000383095 | 3.416694 | 504 |
| **TF** | Factor: nerf; motif: NANS | TF:M11460_0 | 0.000385236 | 3.414274 | 91 |
| **TF** | Factor: nerf; motif: NANS | TF:M11460 | 0.000385236 | 3.414274 | 91 |

| | | | | | |
|---|---|---|---|---|---|
| **TF** | Factor: KLF16; motif: GM | TF:M03914_0 | 0.000421835 | 3.374858 | 319 |
| **TF** | Factor: KLF16; motif: GM | TF:M03914 | 0.000421835 | 3.374858 | 319 |
| **TF** | Factor: E2F; motif: NCSC( | TF:M00919 | 0.000422812 | 3.373853 | 543 |
| **TF** | Factor: E2F; motif: NCSC( | TF:M00919_0 | 0.000422812 | 3.373853 | 543 |
| **TF** | Factor: E2F-1:DP-2; motif | TF:M00737_1 | 0.000426632 | 3.369947 | 285 |
| **TF** | Factor: Pet-1; motif: ACC( | TF:M02072_1 | 0.000427376 | 3.36919 | 49 |
| **TF** | Factor: Elk-1:SREBP-2; mo | TF:M08229_1 | 0.000446566 | 3.350114 | 232 |
| **TF** | Factor: Net; motif: NRCC( | TF:M11439_1 | 0.000457713 | 3.339407 | 55 |
| **TF** | Factor: BTEB2; motif: WG | TF:M09972 | 0.000461698 | 3.335642 | 777 |
| **TF** | Factor: BTEB2; motif: WG | TF:M09972_0 | 0.000461698 | 3.335642 | 777 |
| **TF** | Factor: MAZ; motif: GGGI | TF:M10432_0 | 0.000482323 | 3.316662 | 343 |
| **TF** | Factor: MAZ; motif: GGGI | TF:M10432 | 0.000482323 | 3.316662 | 343 |
| **TF** | Factor: CTCF; motif: NCCI | TF:M08911_0 | 0.000491045 | 3.308879 | 756 |
| **TF** | Factor: CTCF; motif: NCCI | TF:M08911 | 0.000491045 | 3.308879 | 756 |
| **TF** | Factor: SP1; motif: CCCCK | TF:M02281 | 0.000504569 | 3.297079 | 567 |
| **TF** | Factor: SP1; motif: CCCCK | TF:M02281_0 | 0.000504569 | 3.297079 | 567 |
| **TF** | Factor: USF; motif: GYCA( | TF:M00187_0 | 0.000512153 | 3.290601 | 349 |
| **TF** | Factor: USF; motif: GYCA( | TF:M00187 | 0.000512153 | 3.290601 | 349 |
| **TF** | Factor: KLF15; motif: GC( | TF:M12159_1 | 0.0005136 | 3.289375 | 33 |
| **TF** | Factor: PDEF; motif: MC( | TF:M02075_0 | 0.000520393 | 3.283669 | 262 |
| **TF** | Factor: PDEF; motif: MC( | TF:M02075 | 0.000520393 | 3.283669 | 262 |
| **TF** | Factor: TEF-3:ER81; motif | TF:M08423_1 | 0.000542895 | 3.265284 | 38 |
| **TF** | Factor: E2F; motif: TTTSG | TF:M00050_1 | 0.000554283 | 3.256268 | 196 |
| **TF** | Factor: Sp2; motif: NTWA | TF:M12155_1 | 0.000557194 | 3.253994 | 114 |
| **TF** | Factor: Sp1; motif: GGND | TF:M04953 | 0.000571402 | 3.243058 | 325 |
| **TF** | Factor: Sp1; motif: GGND | TF:M04953_0 | 0.000571402 | 3.243058 | 325 |
| **TF** | Factor: BTEB3; motif: CCN | TF:M09826_1 | 0.000603432 | 3.219372 | 655 |
| **TF** | Factor: EGR; motif: CGCC | TF:M08878_0 | 0.000606807 | 3.21695 | 615 |
| **TF** | Factor: EGR; motif: CGCC | TF:M08878 | 0.000606807 | 3.21695 | 615 |
| **TF** | Factor: Sp2; motif: GYCC( | TF:M07129_1 | 0.000615297 | 3.210915 | 113 |
| **TF** | Factor: E2F-3:T-bet; moti | TF:M08529 | 0.000623967 | 3.204839 | 920 |
| **TF** | Factor: E2F-3:T-bet; moti | TF:M08529_0 | 0.000623967 | 3.204839 | 920 |
| **TF** | Factor: KLF15; motif: GC( | TF:M12159 | 0.000648324 | 3.188208 | 150 |
| **TF** | Factor: KLF15; motif: GC( | TF:M12159_0 | 0.000648324 | 3.188208 | 150 |
| **TF** | Factor: ATF2; motif: VGT( | TF:M00179 | 0.000660782 | 3.179941 | 363 |
| **TF** | Factor: ATF2; motif: VGT( | TF:M00179_0 | 0.000660782 | 3.179941 | 363 |
| **TF** | Factor: c-Myc; motif: KA( | TF:M01154_1 | 0.000664119 | 3.177754 | 340 |
| **TF** | Factor: ATF-1; motif: NNI | TF:M09866_0 | 0.000666343 | 3.176302 | 187 |
| **TF** | Factor: ATF-1; motif: NNI | TF:M09866 | 0.000666343 | 3.176302 | 187 |
| **TF** | Factor: ZIC4; motif: NNC( | TF:M12227_1 | 0.000677966 | 3.168792 | 231 |
| **TF** | Factor: Sp4; motif: SCCC( | TF:M07617 | 0.000679188 | 3.16801 | 340 |
| **TF** | Factor: Sp4; motif: SCCC( | TF:M07617_0 | 0.000679188 | 3.16801 | 340 |
| **TF** | Factor: Egr-1; motif: GCG | TF:M01873 | 0.000684014 | 3.164935 | 560 |
| **TF** | Factor: Egr-1; motif: GCG | TF:M01873_0 | 0.000684014 | 3.164935 | 560 |
| **TF** | Factor: E2F-3:Prrxl1; mot | TF:M08206_1 | 0.000705478 | 3.151516 | 428 |
| **TF** | Factor: YY1; motif: GCCG( | TF:M02044 | 0.000763577 | 3.117147 | 525 |
| **TF** | Factor: YY1; motif: GCCG( | TF:M02044_0 | 0.000763577 | 3.117147 | 525 |

| | | | | | |
|---|---|---|---|---|---|
| **TF** | Factor: CREB; motif: CNN | TF:M00917 | 0.000766217 | 3.115648 | 338 |
| **TF** | Factor: CREB; motif: CNN | TF:M00917_0 | 0.000766217 | 3.115648 | 338 |
| **TF** | Factor: REX-1; motif: GGC | TF:M10119 | 0.00082942 | 3.081226 | 195 |
| **TF** | Factor: REX-1; motif: GGC | TF:M10119_0 | 0.00082942 | 3.081226 | 195 |
| **TF** | Factor: ER81; motif: NRCI | TF:M11418_1 | 0.000892756 | 3.049267 | 58 |
| **TF** | Factor: CREM; motif: NNT | TF:M11225_1 | 0.000967864 | 3.014186 | 246 |
| **TF** | Factor: Kaiso; motif: GNT | TF:M09632_0 | 0.000989207 | 3.004713 | 267 |
| **TF** | Factor: Kaiso; motif: GNT | TF:M09632 | 0.000989207 | 3.004713 | 267 |
| **TF** | Factor: RREB-1; motif: GC | TF:M05439_1 | 0.0010037 | 2.998396 | 291 |
| **TF** | Factor: SREBP-1; motif: R | TF:M11081_0 | 0.001018218 | 2.992159 | 898 |
| **TF** | Factor: SREBP-1; motif: R | TF:M11081 | 0.001018218 | 2.992159 | 898 |
| **TF** | Factor: ER71:E2A; motif: | TF:M08568 | 0.001020403 | 2.991228 | 1364 |
| **TF** | Factor: ER71:E2A; motif: | TF:M08568_0 | 0.001020403 | 2.991228 | 1364 |
| **TF** | Factor: YY1; motif: CAAN/ | TF:M10436_1 | 0.001064959 | 2.972667 | 199 |
| **TF** | Factor: USF1; motif: NNN | TF:M07067 | 0.001075796 | 2.96827 | 301 |
| **TF** | Factor: USF1; motif: NNN | TF:M07067_0 | 0.001075796 | 2.96827 | 301 |
| **TF** | Factor: ELFR; motif: NAN( | TF:M11454 | 0.001089462 | 2.962788 | 120 |
| **TF** | Factor: ELFR; motif: NAN( | TF:M11454_0 | 0.001089462 | 2.962788 | 120 |
| **TF** | Factor: CP2; motif: AACC( | TF:M11486_1 | 0.001097265 | 2.959688 | 283 |
| **TF** | Factor: BTEB2; motif: GC( | TF:M07409_1 | 0.00118044 | 2.927956 | 190 |
| **TF** | Factor: Tax/CREB; motif: | TF:M00114_0 | 0.001197026 | 2.921897 | 334 |
| **TF** | Factor: Tax/CREB; motif: | TF:M00114 | 0.001197026 | 2.921897 | 334 |
| **TF** | Factor: TFDP1; motif: NS( | TF:M09789 | 0.001290246 | 2.889328 | 199 |
| **TF** | Factor: TFDP1; motif: NS( | TF:M09789_0 | 0.001290246 | 2.889328 | 199 |
| **TF** | Factor: foxl2; motif: NAC( | TF:M11578_0 | 0.001355086 | 2.868033 | 183 |
| **TF** | Factor: foxl2; motif: NAC( | TF:M11578 | 0.001355086 | 2.868033 | 183 |
| **TF** | Factor: Net; motif: NRCC( | TF:M11440 | 0.001367992 | 2.863916 | 44 |
| **TF** | Factor: Net; motif: NRCC( | TF:M11440_0 | 0.001367992 | 2.863916 | 44 |
| **TF** | Factor: SP4; motif: SCCCC | TF:M01273_1 | 0.001387924 | 2.857634 | 105 |
| **TF** | Factor: ZBP89; motif: CC( | TF:M07397 | 0.001391699 | 2.856455 | 615 |
| **TF** | Factor: ZBP89; motif: CC( | TF:M07397_0 | 0.001391699 | 2.856455 | 615 |
| **TF** | Factor: CREB; motif: TGA( | TF:M00039 | 0.001420565 | 2.847539 | 318 |
| **TF** | Factor: CREB; motif: TGA( | TF:M00039_0 | 0.001420565 | 2.847539 | 318 |
| **TF** | Factor: ELFR; motif: NANS | TF:M11455_0 | 0.001438581 | 2.842066 | 79 |
| **TF** | Factor: ELFR; motif: NANS | TF:M11455 | 0.001438581 | 2.842066 | 79 |
| **TF** | Factor: ZSCAN18; motif: T | TF:M05432 | 0.001438862 | 2.841981 | 186 |
| **TF** | Factor: ZSCAN18; motif: T | TF:M05432_0 | 0.001438862 | 2.841981 | 186 |
| **TF** | Factor: c-Ets-2; motif: NA | TF:M11417_1 | 0.00148634 | 2.827882 | 1039 |
| **TF** | Factor: TFAP2A; motif: N( | TF:M04148_0 | 0.001510792 | 2.820795 | 828 |
| **TF** | Factor: TFAP2A; motif: N( | TF:M04148 | 0.001510792 | 2.820795 | 828 |
| **TF** | Factor: ATF-2; motif: NNT | TF:M11307_1 | 0.001583251 | 2.80045 | 180 |
| **TF** | Factor: CP2; motif: AACC( | TF:M11486_0 | 0.001609263 | 2.793373 | 340 |
| **TF** | Factor: CP2; motif: AACC( | TF:M11486 | 0.001609263 | 2.793373 | 340 |
| **TF** | Factor: ESE-1; motif: ACC | TF:M11384 | 0.001649957 | 2.782527 | 217 |
| **TF** | Factor: ESE-1; motif: ACC | TF:M11384_0 | 0.001649957 | 2.782527 | 217 |
| **TF** | Factor: ESE-1; motif: SATI | TF:M11381 | 0.001825112 | 2.73871 | 1208 |
| **TF** | Factor: ESE-1; motif: SATI | TF:M11381_0 | 0.001825112 | 2.73871 | 1208 |

| | | | | | |
|---|---|---|---|---|---|
| **TF** | Factor: YY1; motif: NAAN | TF:M08982_1 | 0.001864334 | 2.729476 | 161 |
| **TF** | Factor: GKLF; motif: NNC | TF:M12174_0 | 0.001905484 | 2.719995 | 242 |
| **TF** | Factor: GKLF; motif: NNC | TF:M12174 | 0.001905484 | 2.719995 | 242 |
| **TF** | Factor: E2F-4; motif: NGG | TF:M07084_1 | 0.001968314 | 2.705906 | 482 |
| **TF** | Factor: CREB1; motif: NR | TF:M11223_0 | 0.00205422 | 2.687353 | 272 |
| **TF** | Factor: CREB1; motif: NR | TF:M11223 | 0.00205422 | 2.687353 | 272 |
| **TF** | Factor: CREB1; motif: NR | TF:M11223_1 | 0.002103114 | 2.677137 | 261 |
| **TF** | Factor: ETV7; motif: NNS | TF:M08760_1 | 0.002106521 | 2.676434 | 48 |
| **TF** | Factor: SP4; motif: NWRG | TF:M03922_1 | 0.002186311 | 2.660288 | 207 |
| **TF** | Factor: nerf; motif: NANG | TF:M11457 | 0.002257575 | 2.646358 | 118 |
| **TF** | Factor: nerf; motif: NANG | TF:M11457_0 | 0.002257575 | 2.646358 | 118 |
| **TF** | Factor: YY1; motif: GCCA | TF:M00793 | 0.002335222 | 2.631672 | 453 |
| **TF** | Factor: YY1; motif: GCCA | TF:M00793_0 | 0.002335222 | 2.631672 | 453 |
| **TF** | Factor: BTEB2; motif: GN | TF:M03814_0 | 0.002360189 | 2.627053 | 199 |
| **TF** | Factor: BTEB2; motif: GN | TF:M03814 | 0.002360189 | 2.627053 | 199 |
| **TF** | Factor: E2F; motif: NKCG | TF:M00920_0 | 0.002378843 | 2.623634 | 500 |
| **TF** | Factor: E2F; motif: NKCG | TF:M00920 | 0.002378843 | 2.623634 | 500 |
| **TF** | Factor: AP-2rep; motif: N | TF:M09967_1 | 0.002402226 | 2.619386 | 144 |
| **TF** | Factor: CREB1; motif: RT | TF:M11221_0 | 0.002906099 | 2.53669 | 795 |
| **TF** | Factor: CREB1; motif: RT | TF:M11221 | 0.002906099 | 2.53669 | 795 |
| **TF** | Factor: ZFX; motif: SNAG | TF:M10121_1 | 0.002946756 | 2.530656 | 94 |
| **TF** | Factor: ctcf; motif: CCRS | TF:M10426 | 0.003089844 | 2.510063 | 776 |
| **TF** | Factor: ctcf; motif: CCRS | TF:M10426_0 | 0.003089844 | 2.510063 | 776 |
| **TF** | Factor: ERF; motif: ACCG | TF:M02061_1 | 0.00314536 | 2.50233 | 40 |
| **TF** | Factor: KLF14; motif: NR | TF:M12183_1 | 0.003154389 | 2.501085 | 232 |
| **TF** | Factor: AML3; motif: NN | TF:M12000 | 0.003205559 | 2.494096 | 418 |
| **TF** | Factor: AML3; motif: NN | TF:M12000_0 | 0.003205559 | 2.494096 | 418 |
| **TF** | Factor: PATZ; motif: GGG | TF:M10026_0 | 0.003466668 | 2.460088 | 706 |
| **TF** | Factor: PATZ; motif: GGG | TF:M10026 | 0.003466668 | 2.460088 | 706 |
| **TF** | Factor: IRX2a; motif: ACR | TF:M11022_1 | 0.003558352 | 2.448751 | 533 |
| **TF** | Factor: CREM; motif: TGA | TF:M01820_1 | 0.003575262 | 2.446692 | 194 |
| **TF** | Factor: Tel-2; motif: YTA | TF:M00678_1 | 0.00386277 | 2.413101 | 92 |
| **TF** | Factor: CTCF; motif: WG | TF:M03895 | 0.003865796 | 2.412761 | 771 |
| **TF** | Factor: CTCF; motif: WG | TF:M03895_0 | 0.003865796 | 2.412761 | 771 |
| **TF** | Factor: HES-1; motif: GN | TF:M08767_0 | 0.004045232 | 2.393057 | 640 |
| **TF** | Factor: HES-1; motif: GN | TF:M08767 | 0.004045232 | 2.393057 | 640 |
| **TF** | Factor: AHR; motif: CACG | TF:M01855 | 0.00405395 | 2.392122 | 707 |
| **TF** | Factor: AHR; motif: CACG | TF:M01855_0 | 0.00405395 | 2.392122 | 707 |
| **TF** | Factor: ESE-1; motif: ACC | TF:M11379 | 0.004129082 | 2.384147 | 293 |
| **TF** | Factor: ESE-1; motif: ACC | TF:M11379_0 | 0.004129082 | 2.384147 | 293 |
| **TF** | Factor: AhR,; motif: NRC | TF:M00976 | 0.004150727 | 2.381876 | 388 |
| **TF** | Factor: AhR,; motif: NRC | TF:M00976_0 | 0.004150727 | 2.381876 | 388 |
| **TF** | Factor: CREB1; motif: NN | TF:M03544 | 0.004270998 | 2.369471 | 282 |
| **TF** | Factor: CREB1; motif: NN | TF:M03544_0 | 0.004270998 | 2.369471 | 282 |
| **TF** | Factor: hoxa9; motif: RT | TF:M10785_1 | 0.004289761 | 2.367567 | 112 |
| **TF** | Factor: pax-6; motif: NYA | TF:M11881_1 | 0.004306142 | 2.365912 | 601 |
| **TF** | Factor: CREB1; motif: NN | TF:M11220_0 | 0.0044938 | 2.347386 | 161 |

| | | | | | |
|---|---|---|---|---|---|
| **TF** | Factor: CREB1; motif: NN | TF:M11220 | 0.0044938 | 2.347386 | 161 |
| **TF** | Factor: CREM; motif: NNT | TF:M11225_0 | 0.004498825 | 2.346901 | 261 |
| **TF** | Factor: CREM; motif: NNT | TF:M11225 | 0.004498825 | 2.346901 | 261 |
| **TF** | Factor: RUNX2; motif: WI | TF:M04107_1 | 0.004947262 | 2.305635 | 555 |
| **TF** | Factor: HIF-1alpha; motif | TF:M02012_0 | 0.00501613 | 2.299631 | 406 |
| **TF** | Factor: HIF-1alpha; motif | TF:M02012 | 0.00501613 | 2.299631 | 406 |
| **TF** | Factor: HES-1; motif: GN( | TF:M08767_1 | 0.005040068 | 2.297564 | 550 |
| **TF** | Factor: Arnt; motif: NNNI | TF:M00539_1 | 0.00516259 | 2.287132 | 523 |
| **TF** | Factor: Arnt; motif: NNNI | TF:M00539_0 | 0.00516259 | 2.287132 | 523 |
| **TF** | Factor: Arnt; motif: NNNI | TF:M00539 | 0.00516259 | 2.287132 | 523 |
| **TF** | Factor: ZNF777; motif: G | TF:M08993_0 | 0.00529266 | 2.276326 | 311 |
| **TF** | Factor: ZNF777; motif: G | TF:M08993 | 0.00529266 | 2.276326 | 311 |
| **TF** | Factor: USF1; motif: NNC | TF:M11094 | 0.005375977 | 2.269543 | 193 |
| **TF** | Factor: USF1; motif: NNC | TF:M11094_0 | 0.005375977 | 2.269543 | 193 |
| **TF** | Factor: NRF-1; motif: CN( | TF:M04694_1 | 0.005376872 | 2.26947 | 62 |
| **TF** | Factor: Elk-1; motif: NAA | TF:M00007 | 0.005794576 | 2.236978 | 355 |
| **TF** | Factor: Elk-1; motif: NAA | TF:M00007_0 | 0.005794576 | 2.236978 | 355 |
| **TF** | Factor: c-Myc; motif: NS( | TF:M04743_1 | 0.005820931 | 2.235008 | 197 |
| **TF** | Factor: RUNX2; motif: NR | TF:M04106 | 0.005975645 | 2.223615 | 1092 |
| **TF** | Factor: RUNX2; motif: NR | TF:M04106_0 | 0.005975645 | 2.223615 | 1092 |
| **TF** | Factor: YY1; motif: NAAN | TF:M08982_0 | 0.006117195 | 2.213448 | 506 |
| **TF** | Factor: YY1; motif: NAAN | TF:M08982 | 0.006117195 | 2.213448 | 506 |
| **TF** | Factor: E2F-1; motif: TWS | TF:M00024_0 | 0.006147839 | 2.211278 | 747 |
| **TF** | Factor: E2F-1; motif: TWS | TF:M00024 | 0.006147839 | 2.211278 | 747 |
| **TF** | Factor: CREBPA; motif: N | TF:M11313_1 | 0.006364093 | 2.196264 | 183 |
| **TF** | Factor: Egr-1; motif: NGC | TF:M04950_1 | 0.006375635 | 2.195477 | 487 |
| **TF** | Factor: CREB; motif: CGT( | TF:M00801 | 0.006497603 | 2.187247 | 374 |
| **TF** | Factor: CREB; motif: CGT( | TF:M00801_0 | 0.006497603 | 2.187247 | 374 |
| **TF** | Factor: nerf; motif: AAM( | TF:M08757 | 0.006535654 | 2.184711 | 68 |
| **TF** | Factor: nerf; motif: AAM( | TF:M08757_0 | 0.006535654 | 2.184711 | 68 |
| **TF** | Factor: VDR; motif: GGGK | TF:M00444_1 | 0.006643758 | 2.177586 | 186 |
| **TF** | Factor: SREBP-1; motif: R | TF:M11081_1 | 0.00666736 | 2.176046 | 667 |
| **TF** | Factor: YY1; motif: NNNN | TF:M07370 | 0.006928141 | 2.159383 | 499 |
| **TF** | Factor: YY1; motif: NNNN | TF:M07370_0 | 0.006928141 | 2.159383 | 499 |
| **TF** | Factor: CREB1; motif: NN | TF:M07248_1 | 0.006958765 | 2.157468 | 126 |
| **TF** | Factor: Kaiso; motif: GNT | TF:M09632_1 | 0.007216559 | 2.14167 | 49 |
| **TF** | Factor: CREB1; motif: NN | TF:M11220_1 | 0.007448503 | 2.127931 | 159 |
| **TF** | Factor: TFAP2A; motif: N( | TF:M04148_1 | 0.007837001 | 2.10585 | 790 |
| **TF** | Factor: CREB1; motif: NN | TF:M03544_1 | 0.008284812 | 2.081717 | 124 |
| **TF** | Factor: USF2; motif: NNN | TF:M03860_1 | 0.008406088 | 2.075406 | 100 |
| **TF** | Factor: TEF-3:Fli-1; motif: | TF:M08510_0 | 0.008440923 | 2.07361 | 565 |
| **TF** | Factor: TEF-3:Fli-1; motif: | TF:M08510 | 0.008440923 | 2.07361 | 565 |
| **TF** | Factor: ATF; motif: NTGA | TF:M00338 | 0.008691709 | 2.060895 | 239 |
| **TF** | Factor: ATF; motif: NTGA | TF:M00338_0 | 0.008691709 | 2.060895 | 239 |
| **TF** | Factor: USF; motif: NNRY | TF:M00121 | 0.009134175 | 2.039331 | 238 |
| **TF** | Factor: USF; motif: NNRY | TF:M00121_0 | 0.009134175 | 2.039331 | 238 |
| **TF** | Factor: USF; motif: NNRY | TF:M00121_1 | 0.009134175 | 2.039331 | 238 |

| | | | | | |
|---|---|---|---|---|---|
| **TF** | Factor: Elk-1; motif: GGA/ | TF:M03819_1 | 0.009135046 | 2.039289 | 461 |
| **TF** | Factor: ATF-2; motif: NNT | TF:M11307_0 | 0.009558256 | 2.019621 | 208 |
| **TF** | Factor: ATF-2; motif: NNT | TF:M11307 | 0.009558256 | 2.019621 | 208 |
| **TF** | Factor: Elk-1; motif: NAA/ | TF:M00007_1 | 0.009789047 | 2.00926 | 90 |
| **TF** | Factor: HOXB2:Elf-1; mot | TF:M08339_0 | 0.009909739 | 2.003938 | 122 |
| **TF** | Factor: HOXB2:Elf-1; mot | TF:M08339 | 0.009909739 | 2.003938 | 122 |
| **TF** | Factor: Elk-1:HOXA3; mot | TF:M08220 | 0.01053764 | 1.977257 | 982 |
| **TF** | Factor: Elk-1:HOXA3; mot | TF:M08220_0 | 0.01053764 | 1.977257 | 982 |
| **TF** | Factor: ZNF614; motif: N( | TF:M09862 | 0.011084737 | 1.955275 | 766 |
| **TF** | Factor: ZNF614; motif: N( | TF:M09862_0 | 0.011084737 | 1.955275 | 766 |
| **TF** | Factor: A-Myb:Elf-1; moti | TF:M08390_1 | 0.01136175 | 1.944555 | 423 |
| **TF** | Factor: ZNF76; motif: RNI | TF:M10169_1 | 0.011394101 | 1.94332 | 45 |
| **TF** | Factor: YY1; motif: GGCS( | TF:M12273_0 | 0.011408561 | 1.942769 | 529 |
| **TF** | Factor: YY1; motif: GGCS( | TF:M12273 | 0.011408561 | 1.942769 | 529 |
| **TF** | Factor: Sp3; motif: GGGG | TF:M07615_1 | 0.011412044 | 1.942637 | 275 |
| **TF** | Factor: YY1; motif: CAAR/ | TF:M10104_0 | 0.011537876 | 1.937874 | 517 |
| **TF** | Factor: YY1; motif: CAAR/ | TF:M10104 | 0.011537876 | 1.937874 | 517 |
| **TF** | Factor: ATF-3; motif: RTK | TF:M01863 | 0.011710822 | 1.931413 | 188 |
| **TF** | Factor: ATF-3; motif: RTK | TF:M01863_0 | 0.011710822 | 1.931413 | 188 |
| **TF** | Factor: Pax-4; motif: NNN | TF:M00378_1 | 0.012514318 | 1.902593 | 1154 |
| **TF** | Factor: Elk-1:TEF; motif: I | TF:M08232 | 0.012539095 | 1.901734 | 461 |
| **TF** | Factor: Elk-1:TEF; motif: I | TF:M08232_0 | 0.012539095 | 1.901734 | 461 |
| **TF** | Factor: Elk-1:TEF; motif: I | TF:M08232_1 | 0.012733311 | 1.895059 | 131 |
| **TF** | Factor: ZNF37A; motif: C( | TF:M12354 | 0.012863621 | 1.890637 | 879 |
| **TF** | Factor: ZNF37A; motif: C( | TF:M12354_0 | 0.012863621 | 1.890637 | 879 |
| **TF** | Factor: LBP-1; motif: AAC | TF:M11489_1 | 0.013738037 | 1.862075 | 164 |
| **TF** | Factor: Kaiso; motif: SARI | TF:M09966_0 | 0.013935385 | 1.855881 | 202 |
| **TF** | Factor: Kaiso; motif: SARI | TF:M09966 | 0.013935385 | 1.855881 | 202 |
| **TF** | Factor: GKLF; motif: WG( | TF:M09971_1 | 0.014253919 | 1.846066 | 142 |
| **TF** | Factor: TEF-3:Max; motif: | TF:M08647 | 0.014420013 | 1.841034 | 4 |
| **TF** | Factor: TEF-3:Max; motif: | TF:M08647_0 | 0.014420013 | 1.841034 | 4 |
| **TF** | Factor: CREB1; motif: TG/ | TF:M02279_0 | 0.014534387 | 1.837603 | 159 |
| **TF** | Factor: CREB1; motif: TG/ | TF:M02279 | 0.014534387 | 1.837603 | 159 |
| **TF** | Factor: ATF-1; motif: NNN | TF:M07034_1 | 0.014611464 | 1.835306 | 121 |
| **TF** | Factor: ATF-2; motif: NRT | TF:M11309_1 | 0.01735583 | 1.760555 | 224 |
| **TF** | Factor: c-Myc:Max; motif | TF:M00615_0 | 0.017380558 | 1.759936 | 551 |
| **TF** | Factor: c-Myc:Max; motif | TF:M00615 | 0.017380558 | 1.759936 | 551 |
| **TF** | Factor: c-Myc:Max; motif | TF:M00615_1 | 0.017380558 | 1.759936 | 551 |
| **TF** | Factor: CPBP; motif: NGC | TF:M12166 | 0.017563869 | 1.75538 | 123 |
| **TF** | Factor: CPBP; motif: NGC | TF:M12166_0 | 0.017563869 | 1.75538 | 123 |
| **TF** | Factor: CREB; motif: NNT | TF:M00916_1 | 0.019136713 | 1.718133 | 172 |
| **TF** | Factor: ATF-2; motif: NNT | TF:M07312 | 0.019383284 | 1.712573 | 278 |
| **TF** | Factor: ATF-2; motif: NNT | TF:M07312_0 | 0.019383284 | 1.712573 | 278 |
| **TF** | Factor: CREB; motif: TGA( | TF:M00039_1 | 0.019815267 | 1.703 | 146 |
| **TF** | Factor: ER81; motif: NNN | TF:M09611_1 | 0.020902676 | 1.679798 | 33 |
| **TF** | Factor: Erm:HOXB13; mo | TF:M08467_0 | 0.021374079 | 1.670113 | 69 |
| **TF** | Factor: Erm:HOXB13; mo | TF:M08467 | 0.021374079 | 1.670113 | 69 |

| | | | | | |
|---|---|---|---|---|---|
| **TF** | Factor: rax; motif: NNCRT | TF:M10948 | 0.022156623 | 1.654496 | 85 |
| **TF** | Factor: rax; motif: NNCRT | TF:M10948_0 | 0.022156623 | 1.654496 | 85 |
| **TF** | Factor: RNF96; motif: BC( | TF:M01199 | 0.023450273 | 1.629852 | 580 |
| **TF** | Factor: RNF96; motif: BC( | TF:M01199_0 | 0.023450273 | 1.629852 | 580 |
| **TF** | Factor: Net; motif: NACC( | TF:M11442_1 | 0.023517737 | 1.628604 | 35 |
| **TF** | Factor: USF; motif: NCAC( | TF:M00217_0 | 0.023944332 | 1.620797 | 344 |
| **TF** | Factor: USF; motif: NCAC( | TF:M00217 | 0.023944332 | 1.620797 | 344 |
| **TF** | Factor: Kaiso; motif: GCM | TF:M03876_1 | 0.024056276 | 1.618772 | 514 |
| **TF** | Factor: Pax-3; motif: TCG | TF:M00360_1 | 0.024351268 | 1.613478 | 422 |
| **TF** | Factor: AP-2gamma:Max; | TF:M08664 | 0.024446722 | 1.611779 | 207 |
| **TF** | Factor: AP-2gamma:Max; | TF:M08664_0 | 0.024446722 | 1.611779 | 207 |
| **TF** | Factor: Egr-1; motif: WTG | TF:M00243_0 | 0.025722526 | 1.589686 | 467 |
| **TF** | Factor: Egr-1; motif: WTG | TF:M00243 | 0.025722526 | 1.589686 | 467 |
| **TF** | Factor: Net; motif: CACTT | TF:M03832_1 | 0.027820927 | 1.555628 | 10 |
| **TF** | Factor: ATF-3; motif: SNT | TF:M07313 | 0.028258835 | 1.548846 | 275 |
| **TF** | Factor: ATF-3; motif: SNT | TF:M07313_0 | 0.028258835 | 1.548846 | 275 |
| **TF** | Factor: PPARalpha:RXRal | TF:M00518_0 | 0.029199499 | 1.534625 | 670 |
| **TF** | Factor: PPARalpha:RXRal | TF:M00518 | 0.029199499 | 1.534625 | 670 |
| **TF** | Factor: LHX4; motif: NNC | TF:M11005_0 | 0.029465448 | 1.530687 | 731 |
| **TF** | Factor: LHX4; motif: NNC | TF:M11005 | 0.029465448 | 1.530687 | 731 |
| **TF** | Factor: ATF-3; motif: GGT | TF:M09868_0 | 0.029598621 | 1.528729 | 547 |
| **TF** | Factor: ATF-3; motif: GGT | TF:M09868 | 0.029598621 | 1.528729 | 547 |
| **TF** | Factor: E2F-3; motif: NNG | TF:M09893 | 0.030177867 | 1.520311 | 498 |
| **TF** | Factor: E2F-3; motif: NNG | TF:M09893_0 | 0.030177867 | 1.520311 | 498 |
| **TF** | Factor: Fli-1:Prrxl1; motif | TF:M08282 | 0.030220986 | 1.519691 | 174 |
| **TF** | Factor: Fli-1:Prrxl1; motif | TF:M08282_0 | 0.030220986 | 1.519691 | 174 |
| **TF** | Factor: pax-6; motif: NYA | TF:M11880_0 | 0.030638619 | 1.513731 | 912 |
| **TF** | Factor: pax-6; motif: NYA | TF:M11880 | 0.030638619 | 1.513731 | 912 |
| **TF** | Factor: Elk-1:HOXA3; mot | TF:M08220_1 | 0.030841084 | 1.51087 | 466 |
| **TF** | Factor: YY1; motif: CAAR | TF:M07234 | 0.031913431 | 1.496027 | 512 |
| **TF** | Factor: YY1; motif: CAAR | TF:M07234_0 | 0.031913431 | 1.496027 | 512 |
| **TF** | Factor: ATF; motif: CNSTC | TF:M00017 | 0.033722985 | 1.472074 | 612 |
| **TF** | Factor: ATF; motif: CNSTC | TF:M00017_0 | 0.033722985 | 1.472074 | 612 |
| **TF** | Factor: PAX5; motif: RNG | TF:M09817_0 | 0.033911993 | 1.469647 | 983 |
| **TF** | Factor: PAX5; motif: RNG | TF:M09817 | 0.033911993 | 1.469647 | 983 |
| **TF** | Factor: YY1; motif: GCCG( | TF:M04696 | 0.033936441 | 1.469334 | 646 |
| **TF** | Factor: YY1; motif: GCCG( | TF:M04696_0 | 0.033936441 | 1.469334 | 646 |
| **TF** | Factor: ZNF501; motif: NI | TF:M12085 | 0.034044253 | 1.467956 | 693 |
| **TF** | Factor: ZNF501; motif: NI | TF:M12085_0 | 0.034044253 | 1.467956 | 693 |
| **TF** | Factor: Foxn2; motif: NN( | TF:M11541 | 0.034193309 | 1.466059 | 494 |
| **TF** | Factor: Foxn2; motif: NN( | TF:M11541_0 | 0.034193309 | 1.466059 | 494 |
| **TF** | Factor: ETF; motif: GVGG | TF:M00695_1 | 0.037851332 | 1.421919 | 592 |
| **TF** | Factor: ATF-2; motif: NRT | TF:M11309_0 | 0.038193651 | 1.418009 | 369 |
| **TF** | Factor: ATF-2; motif: NRT | TF:M11309 | 0.038193651 | 1.418009 | 369 |
| **TF** | Factor: c-Myc; motif: KAC | TF:M01154_0 | 0.039049399 | 1.408386 | 390 |
| **TF** | Factor: c-Myc; motif: KAC | TF:M01154 | 0.039049399 | 1.408386 | 390 |
| **TF** | Factor: C-Myc; motif: NG | TF:M07601 | 0.041072774 | 1.386446 | 426 |

| | | | | | |
|---|---|---|---|---|---|
| **TF** | Factor: C-Myc; motif: NG( | TF:M07601_0 | 0.041072774 | 1.386446 | 426 |
| **TF** | Factor: SAP-1; motif: NTC | TF:M11438_0 | 0.041736063 | 1.379489 | 510 |
| **TF** | Factor: SAP-1; motif: NTC | TF:M11438 | 0.041736063 | 1.379489 | 510 |
| **TF** | Factor: ATF-1; motif: TNA | TF:M01861_1 | 0.042169372 | 1.375003 | 140 |
| **TF** | Factor: USF; motif: NNRN | TF:M00122 | 0.04266516 | 1.369927 | 211 |
| **TF** | Factor: USF; motif: NNRN | TF:M00122_1 | 0.04266516 | 1.369927 | 211 |
| **TF** | Factor: USF; motif: NNRN | TF:M00122_0 | 0.04266516 | 1.369927 | 211 |
| **TF** | Factor: GKLF; motif: NNR | TF:M07040 | 0.042841799 | 1.368132 | 1013 |
| **TF** | Factor: GKLF; motif: NNR | TF:M07040_0 | 0.042841799 | 1.368132 | 1013 |
| **TF** | Factor: PDEF; motif: NAM | TF:M11470_0 | 0.043375697 | 1.362754 | 63 |
| **TF** | Factor: PDEF; motif: NAM | TF:M11470 | 0.043375697 | 1.362754 | 63 |
| **TF** | Factor: CREB1; motif: NN | TF:M07248 | 0.043420069 | 1.362309 | 285 |
| **TF** | Factor: CREB1; motif: NN | TF:M07248_0 | 0.043420069 | 1.362309 | 285 |
| **TF** | Factor: pax-6; motif: NYA | TF:M11880_1 | 0.043591927 | 1.360594 | 518 |
| **TF** | Factor: Osx; motif: CCNC( | TF:M07329 | 0.044291634 | 1.353678 | 565 |
| **TF** | Factor: Osx; motif: CCNC( | TF:M07329_0 | 0.044291634 | 1.353678 | 565 |
| **TF** | Factor: BMAL1; motif: NS | TF:M09877 | 0.044908516 | 1.347671 | 206 |
| **TF** | Factor: BMAL1; motif: NS | TF:M09877_0 | 0.044908516 | 1.347671 | 206 |
| **TF** | Factor: TEF-3:Elf-1; motif | TF:M08415_0 | 0.045122255 | 1.345609 | 541 |
| **TF** | Factor: TEF-3:Elf-1; motif | TF:M08415 | 0.045122255 | 1.345609 | 541 |
| **TF** | Factor: CREB; motif: CNN | TF:M00917_1 | 0.045319906 | 1.343711 | 153 |
| **TF** | Factor: HIF1; motif: NGTA | TF:M00466_0 | 0.046983645 | 1.328053 | 146 |
| **TF** | Factor: HIF1; motif: NGTA | TF:M00466 | 0.046983645 | 1.328053 | 146 |
| **TF** | Factor: USF2; motif: NNC | TF:M11096_0 | 0.047618859 | 1.322221 | 154 |
| **TF** | Factor: USF2; motif: NNC | TF:M11096 | 0.047618859 | 1.322221 | 154 |
| **TF** | Factor: Fli-1:ETV7; motif: | TF:M08283 | 0.047739726 | 1.32112 | 251 |
| **TF** | Factor: Fli-1:ETV7; motif: | TF:M08283_0 | 0.047739726 | 1.32112 | 251 |
| **TF** | Factor: LBP9; motif: ACYF | TF:M11492_1 | 0.048870313 | 1.310955 | 499 |
| **TF** | Factor: YY1; motif: NGCC( | TF:M12275_0 | 0.049780541 | 1.30294 | 585 |
| **TF** | Factor: YY1; motif: NGCC( | TF:M12275 | 0.049780541 | 1.30294 | 585 |
| **MIRNA** | hsa-miR-652-3p | MIRNA:hsa-miR-652-3p | 0.000977972 | 3.009674 | 22 |
| **MIRNA** | hsa-miR-615-3p | MIRNA:hsa-miR-615-3p | 0.001136625 | 2.944383 | 131 |
| **MIRNA** | hsa-miR-320a | MIRNA:hsa-miR-320a | 0.005328762 | 2.273374 | 69 |
| **MIRNA** | hsa-miR-1301-3p | MIRNA:hsa-miR-1301-3 | 0.02425461 | 1.615206 | 25 |
| **MIRNA** | hsa-miR-328-3p | MIRNA:hsa-miR-328-3p | 0.03535343 | 1.451568 | 3 |
| **MIRNA** | hsa-miR-296-3p | MIRNA:hsa-miR-296-3p | 0.036557566 | 1.437023 | 29 |
| **MIRNA** | hsa-miR-1208 | MIRNA:hsa-miR-1208 | 0.047481554 | 1.323475 | 5 |
| **MIRNA** | hsa-miR-92a-3p | MIRNA:hsa-miR-92a-3p | 0.047680095 | 1.321663 | 182 |

| TF | MIRNA | HPA | CORUM |
|---|---|---|---|
| intersections | | | |
| RPS24,MRPL4,MRPL2,MRPS22,RPL38,RPL13A,RPS9,MRPL | | | |
| PKM,SNRNP70,RPS24,MRPL4,MRPL2,SRSF2,LAS1L,HMGB1 | | | |
| NDUFV2,CBR1,NDUFS3,NDUFA3,NQO2,AIFM2,DHRS4,NDU | | | |
| RPS24,MRPL4,MRPL2,MRPS22,KRT15,MYL6B,RPL38,ARPC | | | |
| PKM,SNRNP70,APP,RPS24,MRPL4,MRPL2,PCNA,SRSF2,LA | | | |
| MICAL1,TXNDC17,NXN,NDUFV2,TXNRD2,GMPR2,CBR1,ND | | | |
| PKM,SNRNP70,APP,SUPT4H1,LATS1,CCDC66,DYNLL1,CST3 | | | |
| PKM,PPFIBP1,BAIAP2,SLC3A2,ZC3HAV1,CTTN,LDHA,PCMT | | | |
| EIF5A,EIF3M,CELF1,EEF1B2,TSFM,MCTS1,PABPC1,EEF1D,T | | | |
| RBCK1,UBE2L6,TSG101,AUP1,ZFAND2B,SHARPIN,VPS28,U | | | |
| CIRBP,EIF5A,RPS9,EIF3M,CELF1,EEF1B2,TSFM,MCTS1,PAB | | | |
| NDUFV2,NDUFS3,NDUFA3,NDUFB3,NDUFB2,NDUFA10,ND | | | |
| NDUFV2,NDUFS3,NDUFA3,NDUFB3,NDUFB2,NDUFA10,ND | | | |
| NDUFV2,NDUFS3,NDUFA3,NDUFB3,NDUFB2,NDUFA10,ND | | | |
| SAE1,UBE2I,PARK7,SUMO1 | | | |
| EIF5A,EIF3M,EEF1B2,TSFM,MCTS1,EEF1D,BRF1,EIF2B3,EL | | | |
| PKM,SNRNP70,APP,LATS1,GNAS,RPS24,MRPL4,MRPL2,PC | | | |
| RBCK1,UBE2L6,TSG101,AUP1,ZFAND2B,SHARPIN,VPS28,U | | | |
| PKM,PPFIBP1,HMGB1,BAIAP2,SLC3A2,ZC3HAV1,CTTN,LD | | | |
| PKM,SNRNP70,APP,LATS1,GNAS,RPS24,MRPL4,MRPL2,PC | | | |
| SULT1A3,RCC1 | | | |
| CIRBP,RPS9,PPAN,MRPS18C,RPS11,MRPS17,RPLP0,RPL8,R | | | |
| SLC3A2,VIM,STAU1,HSP90AB1,LRRFIP1,PRKRIP1,ILF3,OAS | | | |
| RPS24,SRP9,RPL38,SEC62,RPL13A,RPS9,RPS16,RPS25,RPS | | | |
| RPS24,SRP9,RPL38,RPL13A,RPS9,RPS16,RPS25,RPS6,RPL2 | | | |
| SNRNP70,APP,SUPT4H1,RPS24,SRSF2,SSU72,YBX3,CIRBP,F | | | |
| RPS24,RPL38,RPL13A,RPS9,RPS16,RPS25,RPS6,RPL29,RPL | | | |
| RPS24,SRP9,RPL38,SEC62,RPL13A,RPS9,RPS16,RPS25,RPS | | | |
| RPS24,SRP9,RPL38,SEC62,RPL13A,RPS9,RPS16,RPS25,RPS | | | |
| SUPT4H1,DYNLL1,RPS24,PCNA,SMARCB1,RPL38,RBCK1,AT | | | |
| RPS24,RPL38,RPL13A,RPS9,RPS16,RPS25,RPS6,RPL29,EIF3 | | | |
| RPS24,SRP9,RPL38,SEC62,RPL13A,RPS9,RPS16,RPS25,RPS | | | |
| SUPT4H1,DYNLL1,RPS24,PCNA,SMARCB1,RPL38,RBCK1,AT | | | |
| DYNLL1,RPS24,SRSF2,SRP9,RPL38,PARD3,SEC62,WDR19,S | | | |
| SUPT4H1,RPS24,SMARCB1,RPL38,RPL13A,RPS9,PCBP2,H | | | |
| LATS1,CST3,RPS24,HMGB1,TMEM67,YBX3,CIRBP,RPL38,R | | | |
| APP,RPS24,MRPL4,MRPL2,MRPS22,SRP9,YBX3,CIRBP,RPL | | | |
| APP,RPS24,MRPL4,MRPL2,MRPS22,SRP9,YBX3,CIRBP,RPL | | | |
| APP,DYNLL1,GNAS,RPS24,S100A13,SRSF2,SRP9,BLOC1S1,F | | | |
| RPS24,YBX3,CIRBP,RPL38,RNH1,SET,ZC3HAV1,RPL13A,PS | | | |
| RPS24,HMGB1,SULT1A3,YBX3,CIRBP,RPL38,RNH1,DNASE1 | | | |
| SUPT4H1,DYNLL1,RPS24,PCNA,SMARCB1,RPL38,RBCK1,AT | | | |
| LATS1,CCDC66,DYNLL1,RPS24,SRSF2,SRP9,RPL38,PARD3,S | | | |
| RPS24,SRP9,RPL38,SEC62,SAE1,RPL13A,GSTK1,ACOT8,RPS | | | |
| RPS24,HMGB1,YBX3,CIRBP,RPL38,RNH1,DNASE1L1,SET,ZC | | | |

| |
|---|
| LATS1,CCDC66,DYNLL1,RPS24,SRSF2,SRP9,RPL38,PARD3,S |
| SNRNP70,LATS1,CST3,RPS24,HMGB1,TMEM67,YBX3,CIRB |
| SUPT4H1,RPS24,SMARCB1,RPL38,RPL13A,RPS9,HMGA2,R |
| APP,RPS24,MRPL4,MRPL2,MRPS22,SRP9,YBX3,CIRBP,RPL3 |
| RPS24,HMGB1,SULT1A3,YBX3,CIRBP,RPL38,RNH1,DNASE1 |
| RPS24,SRP9,RPL38,PARD3,SEC62,SAE1,RPL13A,GSTK1,ACC |
| RPS24,RPL38,RPL13A,RPS9,RPS16,RPS25,RPS6,RPL29,RPL2 |
| RPS24,HMGB1,YBX3,CIRBP,RPL38,RNH1,DNASE1L1,SET,ZC |
| RPS24,HMGB1,YBX3,CIRBP,RPL38,RNH1,DNASE1L1,SET,ZC |
| APP,RPS24,MRPL4,MRPL2,MRPS22,SRP9,YBX3,CIRBP,RPL3 |
| RPS24,YBX3,CIRBP,RPL38,RNH1,SET,ZC3HAV1,RPL13A,PSM |
| APP,LATS1,CCDC66,DYNLL1,GNAS,RPS24,S100A13,SRSF2,S |
| RPS24,SRP9,RPL38,PARD3,SEC62,RPL13A,RPS9,FIS1,RPS16 |
| CCDC66,RPS24,SRP9,RPL38,SEC62,WDR19,NUMA1,SAE1,R |
| PKM,SNRNP70,APP,LATS1,CST3,RPS24,BCAT2,HMGB1,TM |
| APP,RPS24,MRPL4,MRPL2,MRPS22,SRP9,YBX3,CIRBP,RPL3 |
| LATS1,CENPV,HMGB1,BAIAP2,PILRB,SMARCB1,RPL38,SNR |
| PKM,LATS1,CENPV,MRPL4,MRPL2,MRPS22,HMGB1,BAIAP |
| SNRNP70,APP,LATS1,DYNLL1,CST3,RPS24,BCAT2,HMGB1, |
| SNRNP70,APP,SUPT4H1,RPS24,SRSF2,LAS1L,SSU72,CIRBP, |
| PKM,SNRNP70,APP,LATS1,DYNLL1,CST3,RPS24,BCAT2,HM |
| DYNLL1,GNAS,RPS24,S100A13,SRSF2,HDAC9,HMGB1,SRP9 |
| DYNLL1,GNAS,RPS24,S100A13,SRSF2,HDAC9,HMGB1,SRP9 |
| PKM,APP,LATS1,CCDC66,DYNLL1,RPS24,CENPV,SEMA4D,L |
| RPS24,LAS1L,RPL38,SNRPB,BOP1,SCAF11,NOB1,NOP16,RP |
| PKM,APP,RPS24,BCAT2,MRPL4,MRPL2,MRPS22,SRP9,YBX3 |
| RPS24,SRP9,RPL38,PARD3,SEC62,RPL13A,RPS9,FIS1,LRRC1 |
| DYNLL1,GNAS,RPS24,S100A13,SRSF2,HDAC9,HMGB1,SRP9 |
| DYNLL1,GNAS,RPS24,S100A13,SRSF2,HDAC9,HMGB1,SRP9 |
| LATS1,CCDC66,DYNLL1,GNAS,RPS24,S100A13,SRSF2,HDAC |
| SNRNP70,SRSF2,CIRBP,FASTK,SNRPB,TRPT1,SCAF11,BUD3 |
| PKM,SNRNP70,APP,SUPT4H1,GNAS,RPS24,PCNA,SRSF2,HD |
| PKM,LATS1,CENPV,HMGB1,BAIAP2,PILRB,SMARCB1,RPL38 |
| PKM,SNRNP70,APP,SUPT4H1,GNAS,RPS24,MRPL4,MRPL2, |
| SUPT4H1,LATS1,CCDC66,DYNLL1,CENPV,PCNA,HDAC9,KRT |
| RPL38,SNRPB,BOP1,KLC1,SCAF11,RPL13A,RUVBL2,SRSF1,E |
| RPL38,SNRPB,BOP1,SCAF11,RPL13A,RUVBL2,SRSF1,EIF3M |
| SNRNP70,SRSF2,YBX3,CIRBP,SET,ZC3HAV1,PSMA6,ZC3H14 |
| SNRNP70,SRSF2,CIRBP,SNRPB,SCAF11,BUD31,SNRPN,PCB |
| RPS24,SRP9,RPL38,PARD3,SEC62,WDR19,RPL13A,RPS9,TE |
| TAZ,ECSIT,FOXRED1,TMEM126B,COX17,NDUFV2,COA3,CC |
| TMEM135,SAE1,CTTN,TNFSF10,TAZ,FIS1,CDC37,ECSIT,ATG |
| PKM,SNRNP70,APP,SUPT4H1,GNAS,RPS24,PCNA,SRSF2,HD |
| LATS1,TMEM67,RBCK1,ARRDC1,SIRT6,UFD1L,USP38,PSMA |
| SNRNP70,LATS1,CST3,BCAT2,TMEM67,SULT1A3,RBCK1,CT |
| SNRNP70,SRSF2,CIRBP,SNRPB,SCAF11,BUD31,SNRPN,PCB |
| SNRNP70,SRSF2,CIRBP,SNRPB,SCAF11,BUD31,SNRPN,PCB |

| |
|---|
| YBX3,RPL38,RPL13A,RPL29,EIF3M,RPL36,MCTS1,RPL18,RP |
| LATS1,CCDC66,DYNLL1,GNAS,RPS24,S100A13,SRSF2,HDAC |
| APP,LATS1,CST3,RPS24,SERPINB5,MRPL4,VKORC1,MRPL2, |
| PKM,APP,LATS1,CCDC66,DYNLL1,CENPV,SEMA4D,HMGB1, |
| PKM,APP,SUPT4H1,LATS1,CCDC66,DYNLL1,CST3,RPS24,SE |
| PKM,APP,SUPT4H1,RPS24,MRPL4,MRPL2,PCNA,SRSF2,MR |
| PKM,SNRNP70,APP,SUPT4H1,GNAS,RPS24,PCNA,SRSF2,HD |
| LATS1,TMEM67,RBCK1,CTSC,ARRDC1,SIRT6,UFD1L,USP38, |
| TAZ,ECSIT,FOXRED1,TMEM126B,NDUFV2,COA1,NDUFS3,N |
| TAZ,ECSIT,FOXRED1,TMEM126B,NDUFV2,COA1,NDUFS3,N |
| LATS1,TMEM67,RBCK1,ARRDC1,SIRT6,UFD1L,USP38,PSMA |
| SNRNP70,APP,SUPT4H1,SRSF2,SSU72,CIRBP,SNRPB,SCAF1 |
| PKM,SNRNP70,APP,SUPT4H1,LATS1,DYNLL1,CST3,GNAS,R |
| LATS1,TMEM67,RBCK1,ARRDC1,SIRT6,UFD1L,USP38,PSMA |
| SNRNP70,APP,SUPT4H1,GNAS,RPS24,PCNA,SRSF2,HDAC9, |
| PKM,SNRNP70,APP,SUPT4H1,GNAS,RPS24,PCNA,SRSF2,HD |
| MRPL4,MRPL2,MRPS22,SRP9,EIF5A,MRPL33,CHCHD1,EEF |
| DYNLL1,GNAS,RPS24,S100A13,SRSF2,HDAC9,HMGB1,SRP9 |
| APP,SUPT4H1,CST3,RPS24,MRPL4,MRPL2,PCNA,MRPS22,H |
| SNRNP70,LATS1,CST3,TMEM67,RBCK1,CTSC,ARRDC1,SIRT |
| LATS1,TMEM67,RBCK1,CTSC,ARRDC1,SIRT6,UFD1L,USP38, |
| APP,SUPT4H1,CST3,RPS24,MRPL4,MRPL2,PCNA,SRSF2,MR |
| PKM,SNRNP70,APP,LATS1,CST3,RPS24,SERPINB5,BCAT2,M |
| RPS24,LAS1L,RPL38,BOP1,NOB1,NOP16,RPS9,RPS16,NIP7, |
| PKM,APP,SUPT4H1,CST3,RPS24,BCAT2,MRPL4,MRPL2,PCN |
| PKM,APP,SUPT4H1,LATS1,CCDC66,DYNLL1,CST3,SERPINB5 |
| MRPL4,MRPL2,MRPS22,EIF5A,MRPL33,CHCHD1,APEH,TRM |
| CST3,MRPL4,MRPL2,MRPS22,HMGB1,SMARCB1,SET,KLC1, |
| TAZ,UQCRQ,COX4I1,COX5A,NDUFV2,PARK7,NDUFS3,NDU |
| SNRNP70,APP,LATS1,CST3,HMGB1,YBX3,CIRBP,SET,CTSC,Z |
| PKM,APP,SUPT4H1,CST3,RPS24,BCAT2,MRPL4,MRPL2,PCN |
| TAZ,UQCRQ,COX4I1,COX5A,NDUFV2,PARK7,NDUFS3,NDU |
| SNRNP70,APP,LATS1,CST3,RPS24,SERPINB5,MRPL4,VKORC |
| SRSF2,EIF5A,SRSF1,RPS15,ENY2,NUP133,PCID2,SSB,CHTOP |
| MRPL4,MRPL2,MRPS22,MRPL33,CHCHD1,TSFM,MRPS18C |
| SNRNP70,APP,SUPT4H1,RPS24,PCNA,SRSF2,HDAC9,SEMA4 |
| SNRNP70,SRSF2,CIRBP,SON,SRSF1,CELF1,RBM3,RBMX,HN |
| PKM,APP,SUPT4H1,CST3,RPS24,BCAT2,MRPL4,MRPL2,PCN |
| SRSF2,SRSF1,CELF1,RBMX,HNRNPA1,RBFOX2,SRSF7,RBM4 |
| PKM,SNRNP70,APP,SUPT4H1,LATS1,DYNLL1,CST3,GNAS,R |
| LATS1,CENPV,BAIAP2,TMEM67,RHOF,SMARCB1,TMEM135 |
| SRSF2,SNRPB,EIF5A,SRSF1,JUP,PPP1CC,NUTF2,STRADA,HS |
| SNRNP70,SRSF2,CIRBP,ZC3H14,SON,SRSF1,CELF1,CDK11A, |
| TAZ,UQCRQ,COX4I1,COX5A,NDUFV2,PARK7,NDUFS3,NDU |
| SRSF2,EIF5A,SRSF1,NUTF2,STRADA,RPS15,PARK7,ENY2,NU |
| SNRNP70,APP,LATS1,CST3,HMGB1,YBX3,CIRBP,SET,CTSC,Z |
| SRSF2,SRSF1,CELF1,RBMX,HNRNPA1,RBFOX2,SRSF7,RBM4 |

| |
|---|
| YBX3,CIRBP,SET,PSMA6,ZC3H14,PSMA1,PSMA3,PSMD4,V |
| SRSF2,EIF5A,SRSF1,RPS15,ENY2,NUP133,PCID2,SSB,CHTO |
| GUK1,DGUOK,DUT,DCTD,NUDT1,NT5C,TYMS,SMUG1,SH |
| NDUFV2,PARK7,NDUFS3,NDUFA11,NDUFA3,NDUFB3,NDU |
| SRSF2,SNRPB,EIF5A,SRSF1,JUP,PPP1CC,NUTF2,STRADA,HS |
| YBX3,CIRBP,SET,ZC3HAV1,PSMA6,ZC3H14,PSMA1,PSMA3, |
| APP,CST3,PCNA,HMGB1,TMEM67,SULT1A3,YBX3,SMARC |
| RPL13A,EIF3M,MCTS1,EIF2B3,EIF4A1,METTL3,EIF3G,RBM |
| APP,SUPT4H1,LATS1,CST3,GNAS,RPS24,SERPINB5,MRPL4, |
| LAS1L,RPL38,BOP1,NOP16,NIP7,FBXW9,PPAN,NHP2,ZNH |
| PKM,SNRNP70,APP,SUPT4H1,DYNLL1,CST3,GNAS,GPR137 |
| PKM,APP,LATS1,CCDC66,DYNLL1,GNAS,RPS24,S100A13,C |
| PKM,SNRNP70,APP,SUPT4H1,DYNLL1,CST3,GNAS,GPR137 |
| PKM,SNRNP70,APP,SUPT4H1,LATS1,CCDC66,DYNLL1,CST3 |
| PKM,APP,LATS1,CCDC66,DYNLL1,CST3,GNAS,GPR137,RPS |
| PKM,SNRNP70,APP,SUPT4H1,LATS1,CCDC66,DYNLL1,CST3 |
| PKM,SNRNP70,APP,SUPT4H1,LATS1,CCDC66,DYNLL1,CST3 |
| SNRNP70,RPS24,MRPL4,MRPL2,SRSF2,MRPS22,LAS1L,SRP |
| PKM,SNRNP70,APP,SUPT4H1,CST3,RPS24,S100A13,BCAT2 |
| PKM,SNRNP70,APP,SUPT4H1,CST3,RPS24,S100A13,BCAT2 |
| PKM,SNRNP70,APP,SUPT4H1,CST3,RPS24,S100A13,BCAT2 |
| PKM,SNRNP70,APP,SUPT4H1,DYNLL1,CST3,GNAS,RPS24,S |
| RPS24,MRPL4,MRPL2,MRPS22,RPL38,RPL13A,RPS9,MRPL |
| SNRNP70,SUPT4H1,RPS24,S100A13,CENPV,PCNA,SRSF2,H |
| RPS24,MRPL4,MRPL2,MRPS22,RPL38,RPL13A,RPS9,MRPL |
| RPS24,RPL38,RPL13A,RPS9,RPS16,RPS25,RPS6,RPL29,RPL |
| SNRNP70,SUPT4H1,RPS24,S100A13,CENPV,PCNA,SRSF2,H |
| PKM,SNRNP70,APP,SUPT4H1,DYNLL1,GNAS,RPS24,MRPL4 |
| APP,CST3,MRPL4,MRPL2,MRPS22,BLOC1S1,CACYBP,TME |
| APP,CST3,MRPL4,MRPL2,MRPS22,BLOC1S1,CACYBP,TME |
| RPS24,MRPS22,RPS9,RPS16,RPS25,RPS6,RPS20,RPS15,MC |
| RPS24,RPS9,RPS16,RPS25,RPS6,RPS20,RPS15,MCTS1,RPS1 |
| MRPL4,MRPL2,MRPS22,TAZ,MRPL33,UQCRQ,CHCHD1,EC |
| MRPL4,MRPL2,MRPS22,BLOC1S1,TMEM135,VDAC3,TAZ, |
| MRPL4,MRPL2,MRPS22,TAZ,MRPL33,UQCRQ,CHCHD1,EC |
| APP,LATS1,CCDC66,DYNLL1,CST3,RPS24,S100A13,CENPV, |
| APP,LATS1,CCDC66,DYNLL1,CST3,RPS24,S100A13,CENPV, |
| PKM,DYNLL1,GNAS,PCNA,HDAC9,LAS1L,CACYBP,SMARCB |
| S100A13,LAS1L,CYB561A3,SMARCB1,BOP1,SCAF11,NOP16 |
| PKM,DYNLL1,BCAT2,MRPL4,MRPL2,MRPS22,FAM72A,CM |
| MRPL4,MRPL2,MRPS22,MRPL33,UQCRQ,COX4I1,FOXRED |
| MRPL4,MRPL2,MRPS22,TMEM135,VDAC3,TAZ,MRPL33,F |
| MRPL4,MRPL2,RPL38,RPL13A,MRPL33,RPL29,RPL28,RPL |
| PKM,APP,CST3,GNAS,PCNA,KRT15,BAIAP2,MYL6B,IQCG,C |
| PKM,APP,CST3,GNAS,PCNA,KRT15,BAIAP2,MYL6B,IQCG,C |
| PKM,APP,CST3,GNAS,PCNA,KRT15,BAIAP2,MYL6B,IQCG,C |
| BCAT2,MRPL4,MRPL2,MRPS22,BLOC1S1,FASTK,GSTK1,MR |

| |
|---|
| APP,DYNLL1,CST3,GNAS,GPR137,MRPL4,VKORC1,MRPL2,[illegible] |
| PPFIBP1,RPL38,ARPC2,CTTN,RPL13A,RPS9,DST,PCBP2,CD4[illegible] |
| PPFIBP1,RPL38,ARPC2,CTTN,RPL13A,RPS9,DST,PCBP2,CD4[illegible] |
| RPL38,RPL13A,RPL29,RPL28,RPL21,RPL36,RPL18,RPL15,R[illegible] |
| PKM,APP,DYNLL1,CST3,GNAS,PCNA,KRT15,HMGB1,CYB56[illegible] |
| SNRNP70,SRSF2,CIRBP,SNRPB,BUD31,SNRPN,SRSF1,TXNL[illegible] |
| UQCRQ,COX4I1,FOXRED1,COX5A,NDUFV2,COA3,NDUFS3,[illegible] |
| UQCRQ,COX4I1,FOXRED1,COX5A,NDUFV2,COA3,NDUFS3,[illegible] |
| UQCRQ,COX4I1,FOXRED1,COX5A,HIGD1A,NDUFV2,COA3,[illegible] |
| PSMA6,PSMA1,PSMA3,PSMD4,PSMA7,ZFAND2B,HSPB1,P[illegible] |
| FOXRED1,NDUFV2,NDUFS3,NDUFA11,NDUFA3,NDUFB3,N[illegible] |
| FOXRED1,NDUFV2,NDUFS3,NDUFA11,NDUFA3,NDUFB3,N[illegible] |
| FOXRED1,NDUFV2,NDUFS3,NDUFA11,NDUFA3,NDUFB3,N[illegible] |
| PSMA6,PSMA1,PSMA3,PSMD4,PSMA7,ZFAND2B,HSPB1,P[illegible] |
| SNRPB,BUD31,SNRPN,SRSF1,HNRNPC,PABPC1,RBMX,DDX[illegible] |
| PSMA6,PSMA1,PSMA3,PSMD4,PSMA7,ZFAND2B,HSPB1,P[illegible] |
| APP,BLOC1S1,CACYBP,THEM4,PPOX,STOML2,CHCHD7,CH[illegible] |
| UQCRQ,COX4I1,FOXRED1,C14ORF2,TIMM17B,ATP5J,COX[illegible] |
| BAIAP2,BLOC1S1,KIF4A,AP3B2,RANBP1,ZC3H14,DST |
| BLOC1S1,KIF4A,AP3B2,RANBP1,ZC3H14,DST |
| MRPL4,MRPL2,MRPS22,MRPL33,CHCHD1,MRPS18C,MRP[illegible] |
| MRPL4,MRPL2,MRPS22,MRPL33,CHCHD1,MRPS18C,MRP[illegible] |
| PSMA6,PSMA1,PSMA3,PSMA7 |
| BLOC1S1,THEM4,PPOX,STOML2,CHCHD7,CHCHD5,COX17,[illegible] |
| TPM2,TPM3,TPM4,TPM1 |
| RPL38,PSMA6,PSMA1,VIM,RPL36,RPL18,RPL8,RPL24,NAA[illegible] |
| APP,GNAS,BAIAP2,ARPC2,CTTN,ITSN1,TESC,DST,CD44,PTP[illegible] |
| VKORC1,SRP9,TMEM67,SEC62,TBL2,EIF5A,UFD1L,SLC35G[illegible] |
| VKORC1,SRP9,TMEM67,SEC62,TBL2,EIF5A,UFD1L,SLC35G[illegible] |
| APP,CENPV,NUMA1,CTTN |
| APP,CST3,CACYBP,RCC1,ITSN1,EIF5A,RANBP1,DST,UBE2I,[illegible] |
| SEC62,TBL2,RCE1,FIS1,AUP1,SLC39A13,YIF1A,XBP1,TIMM[illegible] |
| APP,PPFIBP1,YBX3,RPL38,STARD10,PARD3,ARPC2,CTTN,R[illegible] |
| PKM |
| APP,ARPC2,CTTN,ITSN1,TESC,CD44,PTPN13,DUSP22,ITGB[illegible] |
| DYNLL1,LIMK2,NUMA1,CTTN,CDC42,CAPG,CDC14B,TADA[illegible] |
| GPR137,CYB561A3,BLOC1S1,SLC3A2,NPRL2,SCARB1,SLC1[illegible] |
| GPR137,CYB561A3,BLOC1S1,SLC3A2,NPRL2,SCARB1,SLC1[illegible] |
| APP,CST3,VKORC1,SULF2,BAIAP2,SRP9,TMEM67,DNASE1L[illegible] |
| LAS1L,BOP1,NIP7,FBXW9,PPAN |
| PKM,APP,CCDC66,DYNLL1 |
| PCNA,KIF4A,NUMA1,PSMA6,RUVBL2,ATF7,VIM,PSPC1,TP[illegible] |
| APP,DYNLL1,GPR137,CYB561A3,BLOC1S1,RHOF,TMEM13[illegible] |
| GPR137,CYB561A3,BLOC1S1,ATG16L2,SLC3A2,NPRL2,SCA[illegible] |
| SEC62,TBL2,RCE1,FIS1,AUP1,SLC39A13,YIF1A,XBP1,TIMM[illegible] |
| RPS24,MRPL4,MRPL2,RPL38,RPL13A,RPS9,MRPL33,RPS16[illegible] |
| RPS24,RPL38,RPL13A,RPS9,RPS16,RPS25,RPS6,RPL29,RPL[illegible] |

| |
|---|
| RPS24,RPL38,RPL13A,RPS9,RPS16,RPS25,RPS6,RPL29,EEF1 |
| RPS24,RPL38,RPL13A,RPS9,RPS16,RPS25,RPS6,RPL29,RPL |
| RPS24,RPL38,RPL13A,RPS9,RPS16,RPS25,RPS6,RPL29,EIF3 |
| RPS24,RPL38,RPL13A,RPS9,RPS16,RPS25,RPS6,RPL29,EIF3 |
| RPS24,RPL38,RPL13A,RPS9,RPS16,RPS25,RPS6,RPL29,RPL |
| RPS24,RPL38,RPL13A,RPS9,RPS16,RPS25,RPS6,RPL29,RPL |
| RPS24,RPL38,RPL13A,RPS9,RPS16,RPS25,RPS6,RPL29,EIF3 |
| RPS24,RPL38,RPL13A,RPS9,RPS16,RPS25,RPS6,RPL29,EIF3 |
| RPS24,RPL38,RPL13A,PSMA6,RPS9,RPS16,PSMA1,PSMA3, |
| RPS24,RPL38,RPL13A,RPS9,RPS16,RPS25,RPS6,RPL29,EIF3 |
| RPS24,RPL38,RPL13A,RPS9,RPS16,RPS25,RPS6,RPL29,RPL |
| RPS24,RPL38,RPL13A,RPS9,RPS16,RPS25,RPS6,RPL29,RPL |
| RPS24,RPL38,RPL13A,RPS9,RPS16,RPS25,RPS6,RPL29,RPL |
| RPS24,SRP9,RPL38,RPL13A,RPS9,RPS16,RPS25,RPS6,RPL2 |
| RPS24,MRPL4,MRPL2,MRPS22,SRP9,RPL38,RPL13A,RPS9, |
| RPS24,RPL38,RPL13A,RPS9,RPS16,RPS25,RPS6,RPL29,RPL |
| RPS24,RPL38,RPL13A,RPS9,RPS16,RPS25,RPS6,RPL29,RPL |
| RPS24,RPL38,RPL13A,RPS9,RPS16,RPS25,RPS6,RPL29,RPL |
| RPS24,RPL38,RPL13A,PSMA6,RPS9,RPS16,PSMA1,PSMA3, |
| RPS24,RPL38,RPL13A,RPS9,NNMT,RPS16,RPS25,RPS6,RPL |
| RPS24,RPS9,RPS16,RPS25,RPS6,EIF3M,RPS20,RPS15,PABP |
| RPS24,RPS9,RPS16,RPS25,RPS6,EIF3M,RPS20,RPS15,RPS1 |
| RPS24,RPS9,RPS16,RPS25,RPS6,EIF3M,RPS20,RPS15,PABP |
| RPS24,RPS9,RPS16,RPS25,RPS6,EIF3M,RPS20,RPS15,RPS1 |
| RPS24,LAS1L,RPL38,BOP1,NOB1,RPL13A,RPS9,RPS16,RPS2 |
| RPS24,LAS1L,RPL38,BOP1,NOB1,RPL13A,RPS9,RPS16,RPS2 |
| SUPT4H1,RPS24,RPL38,RCC1,RPL13A,PSMA6,RPS9,RANBP |
| RPS24,LAS1L,RPL38,BOP1,NOB1,RPL13A,RPS9,RPS16,RPS2 |
| SNRNP70,RPS24,SRSF2,LAS1L,RPL38,SNRPB,SET,BOP1,NO |
| RPS24,RPL38,RPL13A,RPS9,RPS16,RPS25,RPS6,RPL29,RPL |
| PKM,SNRNP70,APP,C1ORF43,SUPT4H1,LATS1,DYNLL1,CST |
| PKM,SNRNP70,APP,C1ORF43,SUPT4H1,LATS1,DYNLL1,CST |
| C1ORF43,SUPT4H1,CST3,GNAS,GPR137,SERPINB5,MRPL4, |
| PKM,SNRNP70,APP,C1ORF43,SUPT4H1,LATS1,CCDC66,DY |
| PKM,SNRNP70,APP,C1ORF43,SUPT4H1,LATS1,CCDC66,DY |
| PKM,SNRNP70,APP,C1ORF43,SUPT4H1,LATS1,CCDC66,DY |
| PKM,SNRNP70,APP,C1ORF43,SUPT4H1,LATS1,CCDC66,DY |
| PKM,SNRNP70,APP,C1ORF43,SUPT4H1,LATS1,CCDC66,DY |
| SNRNP70,C1ORF43,SUPT4H1,LATS1,CST3,GNAS,RPS24,SE |
| SNRNP70,C1ORF43,SUPT4H1,LATS1,CST3,GNAS,RPS24,SE |
| PKM,SNRNP70,APP,C1ORF43,SUPT4H1,LATS1,CCDC66,DY |
| SUPT4H1,LATS1,CST3,GNAS,RPS24,SERPINB5,S100A13,BC |
| SUPT4H1,LATS1,CST3,GNAS,RPS24,SERPINB5,S100A13,BC |
| C1ORF43,SUPT4H1,CST3,GPR137,SERPINB5,MRPS22,LAS1 |
| SNRNP70,SUPT4H1,CST3,GNAS,RPS24,SERPINB5,S100A13 |
| SNRNP70,SUPT4H1,CST3,GNAS,RPS24,SERPINB5,S100A13 |
| PKM,SNRNP70,APP,C1ORF43,SUPT4H1,LATS1,CCDC66,DY |

| |
|---|
| PKM,SNRNP70,APP,C1ORF43,SUPT4H1,LATS1,CCDC66,DYI |
| SUPT4H1,CST3,GNAS,RPS24,SERPINB5,S100A13,BCAT2,FA |
| SUPT4H1,CST3,GNAS,RPS24,SERPINB5,S100A13,BCAT2,FA |
| SNRNP70,SUPT4H1,CST3,GNAS,RPS24,SERPINB5,S100A13, |
| SNRNP70,SUPT4H1,CST3,GNAS,RPS24,SERPINB5,S100A13, |
| C1ORF43,SUPT4H1,CST3,GNAS,RPS24,SERPINB5,S100A13, |
| C1ORF43,SUPT4H1,CST3,GNAS,RPS24,SERPINB5,S100A13, |
| PKM,SNRNP70,APP,C1ORF43,SUPT4H1,LATS1,CCDC66,DYI |
| PKM,SNRNP70,APP,C1ORF43,SUPT4H1,LATS1,CCDC66,DYI |
| SNRNP70,C1ORF43,SUPT4H1,CST3,GNAS,GPR137,RPS24,S |
| SNRNP70,C1ORF43,SUPT4H1,CST3,GNAS,GPR137,RPS24,S |
| C1ORF43,SUPT4H1,LATS1,CST3,GNAS,GPR137,RPS24,SERF |
| C1ORF43,SUPT4H1,LATS1,CST3,GNAS,GPR137,RPS24,SERF |
| SUPT4H1,CST3,GNAS,SERPINB5,BCAT2,FAM189B,CDCA3,N |
| SUPT4H1,CST3,GNAS,SERPINB5,BCAT2,FAM189B,CDCA3,N |
| C1ORF43,SUPT4H1,LATS1,DYNLL1,CST3,GNAS,RPS24,SERP |
| C1ORF43,SUPT4H1,LATS1,DYNLL1,CST3,GNAS,RPS24,SERP |
| SNRNP70,C1ORF43,SUPT4H1,DYNLL1,CST3,GNAS,GPR137, |
| SNRNP70,C1ORF43,SUPT4H1,DYNLL1,CST3,GNAS,GPR137, |
| PKM,SNRNP70,APP,C1ORF43,SUPT4H1,LATS1,DYNLL1,CST |
| SNRNP70,C1ORF43,SUPT4H1,CST3,GNAS,RPS24,SERPINB5 |
| SNRNP70,C1ORF43,SUPT4H1,CST3,GNAS,RPS24,SERPINB5 |
| SUPT4H1,CST3,GNAS,GPR137,RPS24,MRPL4,FAM189B,SRS |
| PKM,SNRNP70,APP,C1ORF43,SUPT4H1,LATS1,CCDC66,DYI |
| PKM,SNRNP70,APP,C1ORF43,SUPT4H1,LATS1,CCDC66,DYI |
| SUPT4H1,GNAS,GPR137,S100A13,CDCA3,SRSF2,MRPS22,L |
| PKM,SNRNP70,APP,C1ORF43,SUPT4H1,LATS1,CCDC66,DYI |
| PKM,SNRNP70,APP,C1ORF43,SUPT4H1,LATS1,CCDC66,DYI |
| PKM,SNRNP70,APP,C1ORF43,SUPT4H1,LATS1,CCDC66,DYI |
| SUPT4H1,CST3,GNAS,GPR137,SERPINB5,S100A13,BCAT2,F |
| SUPT4H1,CST3,GNAS,GPR137,SERPINB5,S100A13,BCAT2,F |
| SNRNP70,SUPT4H1,CST3,GNAS,RPS24,SERPINB5,S100A13, |
| SNRNP70,SUPT4H1,CST3,GNAS,RPS24,SERPINB5,S100A13, |
| SNRNP70,C1ORF43,SUPT4H1,CCDC66,CST3,GNAS,GPR137 |
| SNRNP70,C1ORF43,SUPT4H1,CCDC66,CST3,GNAS,GPR137 |
| SNRNP70,SUPT4H1,CST3,GNAS,RPS24,SERPINB5,S100A13, |
| SNRNP70,SUPT4H1,CST3,GNAS,RPS24,SERPINB5,S100A13, |
| SNRNP70,C1ORF43,SUPT4H1,LATS1,CST3,GNAS,GPR137,R |
| SNRNP70,C1ORF43,SUPT4H1,LATS1,CST3,GNAS,GPR137,R |
| SUPT4H1,LATS1,CST3,GNAS,SERPINB5,S100A13,BCAT2,VK |
| SUPT4H1,LATS1,CST3,GNAS,SERPINB5,S100A13,BCAT2,VK |
| SNRNP70,SUPT4H1,LATS1,CST3,GNAS,RPS24,SERPINB5,S1 |
| SNRNP70,SUPT4H1,LATS1,CST3,GNAS,RPS24,SERPINB5,S1 |
| PKM,SNRNP70,APP,C1ORF43,SUPT4H1,LATS1,CCDC66,DYI |
| C1ORF43,SUPT4H1,CST3,BCAT2,MRPS22,LAS1L,HMGB1,B |
| SUPT4H1,CST3,GPR137,SERPINB5,MRPS22,LAS1L,HMGB1, |
| SNRNP70,SUPT4H1,CST3,GNAS,GPR137,RPS24,SERPINB5,S |

| |
|---|
| SNRNP70,SUPT4H1,CST3,GNAS,GPR137,RPS24,SERPINB5,S |
| SNRNP70,SUPT4H1,LATS1,CST3,GNAS,GPR137,SERPINB5,S |
| SNRNP70,SUPT4H1,LATS1,CST3,GNAS,GPR137,SERPINB5,S |
| SNRNP70,SUPT4H1,CST3,GNAS,GPR137,SERPINB5,S100A1 |
| SNRNP70,SUPT4H1,CST3,GNAS,GPR137,SERPINB5,S100A1 |
| SNRNP70,SUPT4H1,SERPINB5,S100A13,BCAT2,CDCA3,SRS |
| SNRNP70,SUPT4H1,SERPINB5,S100A13,BCAT2,CDCA3,SRS |
| SUPT4H1,LATS1,CST3,GNAS,RPS24,SERPINB5,S100A13,BC |
| SUPT4H1,LATS1,CST3,GNAS,RPS24,SERPINB5,S100A13,BC |
| SUPT4H1,LATS1,CST3,GNAS,GPR137,SERPINB5,S100A13,B |
| SUPT4H1,LATS1,CST3,GNAS,GPR137,SERPINB5,S100A13,B |
| SNRNP70,C1ORF43,SUPT4H1,LATS1,CST3,GNAS,GPR137,R |
| SNRNP70,C1ORF43,SUPT4H1,LATS1,CST3,GNAS,GPR137,R |
| SNRNP70,SUPT4H1,CST3,GNAS,GPR137,RPS24,SERPINB5,S |
| SNRNP70,SUPT4H1,CST3,GNAS,GPR137,RPS24,SERPINB5,S |
| SUPT4H1,CST3,GNAS,GPR137,SRSF2,MRPS22,LAS1L,HMGI |
| SNRNP70,SUPT4H1,CST3,GNAS,GPR137,RPS24,SERPINB5,S |
| SNRNP70,SUPT4H1,CST3,GNAS,GPR137,RPS24,SERPINB5,S |
| SUPT4H1,GPR137,SERPINB5,BCAT2,FAM189B,CDCA3,SRSI |
| SUPT4H1,GPR137,SERPINB5,BCAT2,FAM189B,CDCA3,SRSI |
| SNRNP70,SUPT4H1,LATS1,CST3,GNAS,GPR137,RPS24,SERI |
| SNRNP70,SUPT4H1,LATS1,CST3,GNAS,GPR137,RPS24,SERI |
| SNRNP70,SUPT4H1,CST3,GNAS,GPR137,RPS24,SERPINB5,S |
| SNRNP70,SUPT4H1,CST3,GNAS,GPR137,RPS24,SERPINB5,S |
| SUPT4H1,CST3,MRPS22,LAS1L,HMGB1,CMC2,BLOC1S1,RB |
| SNRNP70,SUPT4H1,LATS1,CST3,GNAS,GPR137,SERPINB5,S |
| SNRNP70,SUPT4H1,LATS1,CST3,GNAS,GPR137,SERPINB5,S |
| APP,C1ORF43,SUPT4H1,DYNLL1,CST3,GNAS,GPR137,SERP |
| APP,C1ORF43,SUPT4H1,DYNLL1,CST3,GNAS,GPR137,SERP |
| SUPT4H1,CST3,SERPINB5,MRPS22,LAS1L,HMGB1,MYL6B,C |
| SUPT4H1,CST3,MRPS22,LAS1L,HMGB1,BLOC1S1,RBCK1,LY |
| SNRNP70,C1ORF43,SUPT4H1,LATS1,CST3,GNAS,GPR137,R |
| SNRNP70,C1ORF43,SUPT4H1,LATS1,CST3,GNAS,GPR137,R |
| PKM,SNRNP70,APP,C1ORF43,LATS1,DYNLL1,CST3,GNAS,G |
| PKM,SNRNP70,APP,C1ORF43,LATS1,DYNLL1,CST3,GNAS,G |
| PKM,SNRNP70,APP,C1ORF43,SUPT4H1,LATS1,CCDC66,DYI |
| PKM,SNRNP70,APP,C1ORF43,SUPT4H1,LATS1,CCDC66,DYI |
| SUPT4H1,CST3,GPR137,S100A13,MRPS22,LAS1L,HMGB1,N |
| SNRNP70,SUPT4H1,LATS1,DYNLL1,CST3,GNAS,GPR137,RP |
| SNRNP70,SUPT4H1,LATS1,DYNLL1,CST3,GNAS,GPR137,RP |
| SNRNP70,APP,C1ORF43,SUPT4H1,LATS1,CCDC66,DYNLL1, |
| SNRNP70,APP,C1ORF43,SUPT4H1,LATS1,CCDC66,DYNLL1, |
| SUPT4H1,CST3,SERPINB5,MRPS22,LAS1L,HMGB1,MYL6B,C |
| SUPT4H1,RPS24,SERPINB5,S100A13,BCAT2,CDCA3,MRPS2 |
| SUPT4H1,RPS24,SERPINB5,S100A13,BCAT2,CDCA3,MRPS2 |
| SUPT4H1,GPR137,SRSF2,MRPS22,LAS1L,HMGB1,CMC2,BL |
| SUPT4H1,CST3,GNAS,MRPS22,LAS1L,HMGB1,SSU72,BLOC |

| |
|---|
| SUPT4H1,CST3,GNAS,GPR137,RPS24,MRPL4,FAM189B,SR |
| SNRNP70,SUPT4H1,CST3,GNAS,GPR137,SERPINB5,S100A1 |
| SNRNP70,SUPT4H1,CST3,GNAS,GPR137,SERPINB5,S100A1 |
| SNRNP70,SUPT4H1,CST3,GNAS,GPR137,RPS24,SERPINB5, |
| SNRNP70,SUPT4H1,CST3,GNAS,GPR137,RPS24,SERPINB5, |
| SUPT4H1,CST3,GPR137,S100A13,VKORC1,MRPS22,LAS1L, |
| PKM,SNRNP70,APP,C1ORF43,SUPT4H1,GNAS,GPR137,RPS |
| PKM,SNRNP70,APP,C1ORF43,SUPT4H1,GNAS,GPR137,RPS |
| SUPT4H1,MRPS22,LAS1L,HMGB1,MYL6B,BLOC1S1,CIRBP, |
| PKM,SNRNP70,APP,C1ORF43,LATS1,CCDC66,DYNLL1,CST3 |
| PKM,SNRNP70,APP,C1ORF43,LATS1,CCDC66,DYNLL1,CST3 |
| SUPT4H1,CST3,MRPS22,LAS1L,HMGB1,MYL6B,BLOC1S1,R |
| APP,C1ORF43,SUPT4H1,DYNLL1,CST3,GNAS,GPR137,RPS2 |
| APP,C1ORF43,SUPT4H1,DYNLL1,CST3,GNAS,GPR137,RPS2 |
| SUPT4H1,BCAT2,MRPS22,SEMA4D,LAS1L,HMGB1,SSU72, |
| SUPT4H1,BCAT2,MRPS22,SEMA4D,LAS1L,HMGB1,SSU72, |
| C1ORF43,SUPT4H1,CST3,SRSF2,MRPS22,LAS1L,HMGB1,B |
| PKM,SNRNP70,APP,C1ORF43,SUPT4H1,CST3,GNAS,GPR13 |
| SUPT4H1,CST3,MRPS22,LAS1L,HMGB1,MYL6B,CMC2,BLO |
| SUPT4H1,MRPS22,LAS1L,HMGB1,BLOC1S1,RBCK1,SNRPB, |
| SUPT4H1,CST3,GPR137,S100A13,VKORC1,MRPS22,LAS1L, |
| SNRNP70,SUPT4H1,RPS24,SERPINB5,BCAT2,MRPS22,SEM |
| SNRNP70,SUPT4H1,RPS24,SERPINB5,BCAT2,MRPS22,SEM |
| SUPT4H1,BCAT2,MRPS22,LAS1L,HMGB1,SSU72,MYL6B,T |
| SUPT4H1,BCAT2,MRPS22,LAS1L,HMGB1,SSU72,MYL6B,T |
| SUPT4H1,LATS1,RPS24,SERPINB5,BCAT2,CDCA3,SRSF2,M |
| SUPT4H1,LATS1,RPS24,SERPINB5,BCAT2,CDCA3,SRSF2,M |
| PKM,SNRNP70,APP,C1ORF43,SUPT4H1,LATS1,CCDC66,DY |
| PKM,SNRNP70,APP,C1ORF43,SUPT4H1,LATS1,CCDC66,DY |
| SUPT4H1,CST3,GPR137,MRPS22,LAS1L,HMGB1,SSU72,MY |
| C1ORF43,SUPT4H1,CST3,SERPINB5,MRPS22,LAS1L,HMGB |
| PKM,SNRNP70,APP,C1ORF43,SUPT4H1,LATS1,CCDC66,DY |
| PKM,SNRNP70,APP,C1ORF43,SUPT4H1,LATS1,CCDC66,DY |
| SUPT4H1,CST3,GNAS,MRPS22,LAS1L,HMGB1,BLOC1S1,CA |
| SUPT4H1,SERPINB5,BCAT2,MRPS22,SEMA4D,LAS1L,HMG |
| SUPT4H1,SERPINB5,BCAT2,MRPS22,SEMA4D,LAS1L,HMG |
| SUPT4H1,CST3,GNAS,SERPINB5,S100A13,BCAT2,MRPL4,C |
| SUPT4H1,CST3,GNAS,SERPINB5,S100A13,BCAT2,MRPL4,C |
| SUPT4H1,CST3,GNAS,SERPINB5,S100A13,BCAT2,MRPS22, |
| SUPT4H1,CST3,GNAS,SERPINB5,S100A13,BCAT2,MRPS22, |
| SUPT4H1,CST3,GNAS,SERPINB5,S100A13,BCAT2,MRPS22, |
| SUPT4H1,CST3,GNAS,SERPINB5,S100A13,BCAT2,MRPS22, |
| SUPT4H1,CST3,GNAS,BCAT2,FAM189B,CDCA3,MRPS22,LA |
| SUPT4H1,CST3,GNAS,BCAT2,FAM189B,CDCA3,MRPS22,LA |
| PKM,SNRNP70,SUPT4H1,LATS1,CCDC66,DYNLL1,CST3,GN |
| PKM,SNRNP70,SUPT4H1,LATS1,CCDC66,DYNLL1,CST3,GN |
| SUPT4H1,CST3,SERPINB5,MRPS22,LAS1L,HMGB1,MYL6B, |

| |
|---|
| SUPT4H1,CST3,SERPINB5,SRSF2,MRPS22,SEMA4D,LAS1L, |
| SUPT4H1,CST3,GNAS,GPR137,BCAT2,MRPS22,LAS1L,HMG |
| SUPT4H1,CST3,GNAS,GPR137,BCAT2,MRPS22,LAS1L,HMG |
| C1ORF43,SUPT4H1,CST3,BCAT2,SRSF2,MRPS22,LAS1L,HM |
| SUPT4H1,RPS24,SERPINB5,BCAT2,SRSF2,MRPS22,HDAC9, |
| SUPT4H1,RPS24,SERPINB5,BCAT2,SRSF2,MRPS22,HDAC9, |
| SUPT4H1,CST3,MRPS22,LAS1L,HMGB1,MYL6B,CMC2,BLO |
| PKM,SNRNP70,APP,C1ORF43,LATS1,CCDC66,DYNLL1,CST3 |
| SNRNP70,APP,SUPT4H1,GNAS,RPS24,BCAT2,FAM189B,SR |
| PKM,SNRNP70,APP,C1ORF43,SUPT4H1,LATS1,CCDC66,DY |
| SUPT4H1,BCAT2,SRSF2,MRPS22,LAS1L,HMGB1,TMEM67, |
| SUPT4H1,BCAT2,SRSF2,MRPS22,LAS1L,HMGB1,TMEM67, |
| SNRNP70,SUPT4H1,LATS1,CST3,GNAS,GPR137,RPS24,SER |
| SNRNP70,SUPT4H1,LATS1,CST3,GNAS,GPR137,RPS24,SER |
| SUPT4H1,GPR137,S100A13,BCAT2,CDCA3,SRSF2,MRPS22, |
| SUPT4H1,GPR137,S100A13,BCAT2,CDCA3,SRSF2,MRPS22, |
| SUPT4H1,GPR137,S100A13,BCAT2,CDCA3,SRSF2,MRPS22, |
| SUPT4H1,GPR137,S100A13,BCAT2,CDCA3,SRSF2,MRPS22, |
| SUPT4H1,SERPINB5,BCAT2,MRPS22,LAS1L,HMGB1,SSU72, |
| SUPT4H1,SERPINB5,BCAT2,MRPS22,LAS1L,HMGB1,SSU72, |
| SUPT4H1,GNAS,BCAT2,MRPS22,LAS1L,HMGB1,SSU72,TM |
| SUPT4H1,GNAS,BCAT2,MRPS22,LAS1L,HMGB1,SSU72,TM |
| SNRNP70,C1ORF43,SUPT4H1,LATS1,GPR137,MRPS22,LAS |
| SUPT4H1,GPR137,BCAT2,MRPS22,LAS1L,HMGB1,TMEM6 |
| SUPT4H1,GPR137,BCAT2,MRPS22,LAS1L,HMGB1,TMEM6 |
| SNRNP70,SUPT4H1,GNAS,GPR137,RPS24,SERPINB5,S100A |
| SNRNP70,SUPT4H1,GNAS,GPR137,RPS24,SERPINB5,S100A |
| SNRNP70,SUPT4H1,BCAT2,MRPS22,SEMA4D,LAS1L,HMGB |
| SNRNP70,SUPT4H1,BCAT2,MRPS22,SEMA4D,LAS1L,HMGB |
| SNRNP70,SUPT4H1,CST3,GNAS,SERPINB5,S100A13,BCAT2 |
| SNRNP70,SUPT4H1,CST3,GNAS,SERPINB5,S100A13,BCAT2 |
| SNRNP70,APP,SUPT4H1,DYNLL1,CST3,GNAS,GPR137,RPS2 |
| SNRNP70,APP,SUPT4H1,DYNLL1,CST3,GNAS,GPR137,RPS2 |
| SUPT4H1,BCAT2,CDCA3,SRSF2,MRPS22,LAS1L,HMGB1,BA |
| SUPT4H1,BCAT2,CDCA3,SRSF2,MRPS22,LAS1L,HMGB1,BA |
| SNRNP70,SUPT4H1,LATS1,CST3,GNAS,RPS24,SERPINB5,S1 |
| SNRNP70,SUPT4H1,LATS1,CST3,GNAS,RPS24,SERPINB5,S1 |
| PKM,SNRNP70,APP,C1ORF43,SUPT4H1,LATS1,CCDC66,DY |
| C1ORF43,SUPT4H1,LATS1,DYNLL1,CST3,GNAS,GPR137,RP |
| C1ORF43,SUPT4H1,LATS1,DYNLL1,CST3,GNAS,GPR137,RP |
| APP,C1ORF43,SUPT4H1,DYNLL1,GNAS,GPR137,SERPINB5, |
| APP,C1ORF43,SUPT4H1,DYNLL1,GNAS,GPR137,SERPINB5, |
| SUPT4H1,MRPS22,LAS1L,HMGB1,MYL6B,CMC2,BLOC1S1, |
| PKM,APP,C1ORF43,SUPT4H1,DYNLL1,GNAS,GPR137,SERP |
| PKM,APP,C1ORF43,SUPT4H1,DYNLL1,GNAS,GPR137,SERP |
| SUPT4H1,CST3,GNAS,S100A13,BCAT2,MRPS22,LAS1L,HM |
| SUPT4H1,CST3,GNAS,S100A13,BCAT2,MRPS22,LAS1L,HM |

| |
|---|
| SUPT4H1,CST3,GNAS,BCAT2,MRPS22,LAS1L,HMGB1,SSU7 |
| SUPT4H1,CST3,GNAS,BCAT2,MRPS22,LAS1L,HMGB1,SSU7 |
| PKM,SNRNP70,SUPT4H1,LATS1,CCDC66,DYNLL1,CST3,GN |
| PKM,SNRNP70,SUPT4H1,LATS1,CCDC66,DYNLL1,CST3,GN |
| SUPT4H1,CST3,MRPS22,LAS1L,HMGB1,MYL6B,CMC2,BLO |
| SNRNP70,C1ORF43,SUPT4H1,GPR137,SERPINB5,BCAT2,SR |
| SNRNP70,C1ORF43,SUPT4H1,GPR137,SERPINB5,BCAT2,SR |
| PKM,SNRNP70,APP,C1ORF43,SUPT4H1,LATS1,CCDC66,DY |
| PKM,SNRNP70,APP,C1ORF43,SUPT4H1,LATS1,CCDC66,DY |
| SNRNP70,SUPT4H1,LATS1,GPR137,RPS24,S100A13,BCAT2 |
| SNRNP70,SUPT4H1,LATS1,GPR137,RPS24,S100A13,BCAT2 |
| PKM,SNRNP70,APP,C1ORF43,SUPT4H1,LATS1,DYNLL1,CST |
| PKM,SNRNP70,APP,C1ORF43,SUPT4H1,LATS1,DYNLL1,CST |
| SUPT4H1,CST3,SERPINB5,MRPS22,LAS1L,HMGB1,MYL6B, |
| SNRNP70,APP,C1ORF43,SUPT4H1,LATS1,DYNLL1,GNAS,G |
| SNRNP70,APP,C1ORF43,SUPT4H1,LATS1,DYNLL1,GNAS,G |
| SUPT4H1,CST3,SERPINB5,MRPS22,LAS1L,HMGB1,MYL6B, |
| SUPT4H1,GPR137,RPS24,SERPINB5,S100A13,BCAT2,SRSF2 |
| SUPT4H1,GPR137,RPS24,SERPINB5,S100A13,BCAT2,SRSF2 |
| PKM,SUPT4H1,CST3,GNAS,SERPINB5,S100A13,BCAT2,MR |
| PKM,SUPT4H1,CST3,GNAS,SERPINB5,S100A13,BCAT2,MR |
| SUPT4H1,SERPINB5,VKORC1,MRPS22,SEMA4D,LAS1L,HM |
| SNRNP70,C1ORF43,SUPT4H1,GNAS,GPR137,SERPINB5,BC |
| SNRNP70,C1ORF43,SUPT4H1,GNAS,GPR137,SERPINB5,BC |
| SUPT4H1,BCAT2,MRPS22,SEMA4D,LAS1L,HMGB1,TMEM6 |
| SUPT4H1,BCAT2,MRPS22,SEMA4D,LAS1L,HMGB1,TMEM6 |
| GNAS,GPR137,SRSF2,MRPS22,HMGB1,SSU72,TMEM67,CA |
| SUPT4H1,BCAT2,MRPS22,LAS1L,HMGB1,TMEM67,CMC2, |
| SUPT4H1,BCAT2,MRPS22,LAS1L,HMGB1,TMEM67,CMC2, |
| SNRNP70,SUPT4H1,MRPS22,LAS1L,HMGB1,TMEM67,CM |
| SNRNP70,SUPT4H1,MRPS22,LAS1L,HMGB1,TMEM67,CM |
| SUPT4H1,SERPINB5,BCAT2,MRPS22,LAS1L,HMGB1,TMEM |
| SUPT4H1,SERPINB5,BCAT2,MRPS22,LAS1L,HMGB1,TMEM |
| SUPT4H1,MRPS22,LAS1L,HMGB1,CMC2,BLOC1S1,CACYBP, |
| SUPT4H1,MRPS22,LAS1L,HMGB1,BLOC1S1,RBCK1,SNRPB, |
| SUPT4H1,CST3,SERPINB5,MRPS22,LAS1L,HMGB1,MYL6B, |
| SNRNP70,C1ORF43,CCDC66,DYNLL1,CST3,GNAS,GPR137, |
| SNRNP70,C1ORF43,CCDC66,DYNLL1,CST3,GNAS,GPR137, |
| SUPT4H1,CST3,GNAS,MRPS22,LAS1L,HMGB1,BLOC1S1,CI |
| PKM,SNRNP70,APP,C1ORF43,SUPT4H1,CCDC66,DYNLL1,C |
| SUPT4H1,SERPINB5,BCAT2,MRPS22,SEMA4D,LAS1L,HMG |
| SUPT4H1,SERPINB5,BCAT2,MRPS22,SEMA4D,LAS1L,HMG |
| PKM,SNRNP70,C1ORF43,SUPT4H1,CCDC66,DYNLL1,CST3, |
| PKM,SNRNP70,C1ORF43,SUPT4H1,CCDC66,DYNLL1,CST3, |
| PKM,SNRNP70,APP,C1ORF43,SUPT4H1,LATS1,CCDC66,DY |
| PKM,SNRNP70,APP,C1ORF43,SUPT4H1,LATS1,CCDC66,DY |
| PKM,SNRNP70,APP,C1ORF43,LATS1,DYNLL1,CST3,GNAS,G |

| |
|---|
| PKM,SNRNP70,APP,C1ORF43,SUPT4H1,LATS1,CCDC66,DY |
| PKM,SNRNP70,APP,C1ORF43,SUPT4H1,LATS1,CCDC66,DY |
| SUPT4H1,GPR137,CDCA3,MRPS22,LAS1L,HMGB1,SSU72,C |
| SUPT4H1,GPR137,CDCA3,MRPS22,LAS1L,HMGB1,SSU72,C |
| SUPT4H1,MRPS22,LAS1L,HMGB1,MYL6B,CMC2,BLOC1S1, |
| SUPT4H1,GNAS,MRPS22,LAS1L,HMGB1,CIRBP,RPL38,RBC |
| SNRNP70,SUPT4H1,GPR137,BCAT2,SRSF2,MRPS22,OSMR, |
| SNRNP70,SUPT4H1,GPR137,BCAT2,SRSF2,MRPS22,OSMR, |
| PKM,SNRNP70,APP,C1ORF43,SUPT4H1,LATS1,CCDC66,DY |
| PKM,SNRNP70,APP,C1ORF43,SUPT4H1,LATS1,CCDC66,DY |
| SUPT4H1,MRPS22,LAS1L,HMGB1,MYL6B,BLOC1S1,RBCK1, |
| PKM,SNRNP70,APP,C1ORF43,SUPT4H1,CCDC66,DYNLL1,C |
| PKM,SNRNP70,APP,C1ORF43,SUPT4H1,CCDC66,DYNLL1,C |
| PKM,SNRNP70,APP,C1ORF43,SUPT4H1,DYNLL1,CST3,GNA |
| PKM,SNRNP70,APP,C1ORF43,SUPT4H1,DYNLL1,CST3,GNA |
| GPR137,SRSF2,MRPS22,HMGB1,TMEM67,PRR13,RBCK1,L |
| C1ORF43,LATS1,DYNLL1,GPR137,RPS24,SERPINB5,BCAT2, |
| C1ORF43,LATS1,DYNLL1,GPR137,RPS24,SERPINB5,BCAT2, |
| PKM,SNRNP70,APP,C1ORF43,LATS1,DYNLL1,CST3,GNAS,G |
| SUPT4H1,GNAS,BCAT2,SRSF2,MRPS22,LAS1L,HMGB1,RAP |
| SUPT4H1,GNAS,BCAT2,SRSF2,MRPS22,LAS1L,HMGB1,RAP |
| SNRNP70,SUPT4H1,SERPINB5,BCAT2,MRPS22,LAS1L,HMG |
| SNRNP70,SUPT4H1,SERPINB5,BCAT2,MRPS22,LAS1L,HMG |
| SUPT4H1,MRPS22,LAS1L,HMGB1,TMEM67,CMC2,CIRBP,R |
| SUPT4H1,MRPS22,LAS1L,HMGB1,TMEM67,CMC2,CIRBP,R |
| SUPT4H1,GNAS,SERPINB5,MRPS22,LAS1L,HMGB1,BLOC1S |
| PKM,SNRNP70,APP,C1ORF43,SUPT4H1,LATS1,CCDC66,DY |
| PKM,SNRNP70,APP,C1ORF43,SUPT4H1,LATS1,CCDC66,DY |
| SNRNP70,SUPT4H1,GPR137,BCAT2,SRSF2,MRPS22,LAS1L, |
| SNRNP70,SUPT4H1,GPR137,BCAT2,SRSF2,MRPS22,LAS1L, |
| C1ORF43,SUPT4H1,LATS1,CST3,GNAS,GPR137,BCAT2,MR |
| PKM,SNRNP70,APP,C1ORF43,SUPT4H1,LATS1,CCDC66,CS |
| PKM,SNRNP70,APP,C1ORF43,SUPT4H1,LATS1,CCDC66,CS |
| SUPT4H1,LATS1,SRSF2,MRPS22,LAS1L,HMGB1,SSU72,MYL |
| SUPT4H1,CDCA3,MRPS22,LAS1L,HMGB1,SRP9,BLOC1S1,C |
| SNRNP70,APP,C1ORF43,SUPT4H1,LATS1,CST3,GNAS,GPR1 |
| SNRNP70,APP,C1ORF43,SUPT4H1,LATS1,CST3,GNAS,GPR1 |
| PKM,SNRNP70,SUPT4H1,LATS1,DYNLL1,CST3,GNAS,GPR1 |
| PKM,SNRNP70,SUPT4H1,LATS1,DYNLL1,CST3,GNAS,GPR1 |
| PKM,SNRNP70,APP,C1ORF43,SUPT4H1,LATS1,DYNLL1,CS |
| PKM,APP,C1ORF43,SUPT4H1,DYNLL1,CST3,GNAS,GPR137, |
| PKM,APP,C1ORF43,SUPT4H1,DYNLL1,CST3,GNAS,GPR137, |
| SNRNP70,SUPT4H1,LATS1,GNAS,GPR137,RPS24,S100A13, |
| SNRNP70,SUPT4H1,LATS1,GNAS,GPR137,RPS24,S100A13, |
| SNRNP70,APP,C1ORF43,SUPT4H1,DYNLL1,CST3,GNAS,GP |
| SUPT4H1,MRPS22,LAS1L,HMGB1,RBCK1,SNRPB,DHPS,DBI, |
| PKM,SNRNP70,APP,C1ORF43,LATS1,CCDC66,DYNLL1,CST3 |

| |
|---|
| PKM,SNRNP70,APP,C1ORF43,LATS1,CCDC66,DYNLL1,CST3 |
| SUPT4H1,CST3,GNAS,SERPINB5,S100A13,MRPS22,LAS1L, |
| SUPT4H1,CST3,GNAS,SERPINB5,S100A13,MRPS22,LAS1L, |
| SNRNP70,C1ORF43,LATS1,DYNLL1,GPR137,RPS24,SERPIN |
| SNRNP70,C1ORF43,LATS1,DYNLL1,GPR137,RPS24,SERPIN |
| PKM,SNRNP70,APP,C1ORF43,SUPT4H1,LATS1,DYNLL1,CST |
| SUPT4H1,CST3,CDCA3,MRPS22,LAS1L,HMGB1,MYL6B,CM |
| SUPT4H1,MRPS22,LAS1L,HMGB1,MYL6B,CMC2,BLOC1S1, |
| GPR137,MRPS22,HMGB1,RBCK1,LY75,SNRPB,DHPS,ZNF23 |
| PKM,SNRNP70,APP,C1ORF43,SUPT4H1,LATS1,CCDC66,DY |
| PKM,SNRNP70,APP,C1ORF43,SUPT4H1,LATS1,CCDC66,DY |
| APP,C1ORF43,LATS1,CCDC66,DYNLL1,CST3,GNAS,GPR137, |
| APP,C1ORF43,LATS1,CCDC66,DYNLL1,CST3,GNAS,GPR137, |
| PKM,SNRNP70,APP,C1ORF43,SUPT4H1,LATS1,CCDC66,DY |
| SNRNP70,APP,C1ORF43,SUPT4H1,DYNLL1,GNAS,GPR137, |
| SNRNP70,APP,C1ORF43,SUPT4H1,DYNLL1,GNAS,GPR137, |
| PKM,SNRNP70,APP,C1ORF43,SUPT4H1,LATS1,CCDC66,DY |
| PKM,SNRNP70,APP,C1ORF43,SUPT4H1,LATS1,CCDC66,DY |
| PKM,APP,C1ORF43,SUPT4H1,CCDC66,CST3,GNAS,RPS24,B |
| PKM,SNRNP70,APP,C1ORF43,SUPT4H1,LATS1,CST3,GNAS, |
| PKM,SUPT4H1,LATS1,GNAS,SERPINB5,BCAT2,VKORC1,CD |
| PKM,SUPT4H1,LATS1,GNAS,SERPINB5,BCAT2,VKORC1,CD |
| SUPT4H1,MRPS22,LAS1L,MYL6B,CMC2,RNH1,RBCK1,RCC1 |
| SUPT4H1,GNAS,HMGB1,SSU72,SNRPB,ZNF235,BOP1,ARR |
| PKM,APP,C1ORF43,SUPT4H1,CCDC66,DYNLL1,CST3,GPR1 |
| SNRNP70,C1ORF43,SUPT4H1,LATS1,DYNLL1,CST3,GPR137 |
| SNRNP70,C1ORF43,SUPT4H1,LATS1,DYNLL1,CST3,GPR137 |
| SUPT4H1,MRPS22,LAS1L,HMGB1,MYL6B,CMC2,RBCK1,RC |
| APP,C1ORF43,SUPT4H1,DYNLL1,GNAS,GPR137,SERPINB5, |
| APP,C1ORF43,SUPT4H1,DYNLL1,GNAS,GPR137,SERPINB5, |
| APP,C1ORF43,SUPT4H1,DYNLL1,GNAS,GPR137,SERPINB5, |
| APP,C1ORF43,SUPT4H1,DYNLL1,GNAS,GPR137,SERPINB5, |
| SNRNP70,APP,C1ORF43,SUPT4H1,LATS1,DYNLL1,CST3,GN |
| SNRNP70,APP,C1ORF43,SUPT4H1,LATS1,DYNLL1,CST3,GN |
| PKM,SNRNP70,APP,C1ORF43,SUPT4H1,LATS1,DYNLL1,CST |
| PKM,SNRNP70,APP,C1ORF43,SUPT4H1,LATS1,CCDC66,DY |
| PKM,SNRNP70,APP,C1ORF43,SUPT4H1,LATS1,CCDC66,DY |
| PKM,SNRNP70,APP,C1ORF43,SUPT4H1,LATS1,DYNLL1,CST |
| PKM,SNRNP70,APP,C1ORF43,SUPT4H1,CCDC66,CST3,GNA |
| PKM,SNRNP70,APP,C1ORF43,SUPT4H1,LATS1,CCDC66,CS |
| PKM,SNRNP70,APP,C1ORF43,SUPT4H1,LATS1,CCDC66,CS |
| LATS1,DYNLL1,CST3,GNAS,GPR137,RPS24,BCAT2,CENPV, |
| LATS1,DYNLL1,CST3,GNAS,GPR137,RPS24,BCAT2,CENPV, |
| PKM,SNRNP70,C1ORF43,DYNLL1,GNAS,GPR137,RPS24,SE |
| PKM,SNRNP70,C1ORF43,DYNLL1,GNAS,GPR137,RPS24,SE |
| PKM,SNRNP70,APP,C1ORF43,SUPT4H1,LATS1,CCDC66,CS |
| SNRNP70,C1ORF43,LATS1,DYNLL1,GNAS,GPR137,RPS24,S |

| |
|---|
| SNRNP70,C1ORF43,LATS1,DYNLL1,GNAS,GPR137,RPS24,S |
| SRSF2,MRPS22,HMGB1,CIRBP,RBCK1,SNRPB,ACTR6,KLC1, |
| SNRNP70,APP,C1ORF43,CCDC66,DYNLL1,CST3,GNAS,GPR1 |
| PKM,SNRNP70,APP,C1ORF43,SUPT4H1,LATS1,CCDC66,DY |
| PKM,SNRNP70,APP,C1ORF43,SUPT4H1,LATS1,CCDC66,DY |
| PKM,SNRNP70,SUPT4H1,LATS1,CCDC66,DYNLL1,CST3,GN |
| PKM,SNRNP70,SUPT4H1,LATS1,CCDC66,DYNLL1,CST3,GN |
| GNAS,SRSF2,MRPS22,HMGB1,PILRB,BLOC1S1,CIRBP,RPL3 |
| PKM,APP,C1ORF43,SUPT4H1,DYNLL1,GNAS,GPR137,SERP |
| PKM,APP,C1ORF43,SUPT4H1,DYNLL1,GNAS,GPR137,SERP |
| PKM,SNRNP70,APP,C1ORF43,SUPT4H1,LATS1,GNAS,RPS2 |
| PKM,SNRNP70,APP,C1ORF43,SUPT4H1,LATS1,GNAS,RPS2 |
| PKM,SNRNP70,APP,C1ORF43,SUPT4H1,CCDC66,DYNLL1,C |
| PKM,SNRNP70,APP,C1ORF43,SUPT4H1,CCDC66,DYNLL1,C |
| SNRNP70,APP,C1ORF43,SUPT4H1,LATS1,CST3,GNAS,GPR1 |
| PKM,C1ORF43,LATS1,DYNLL1,GNAS,GPR137,BCAT2,CENP |
| PKM,C1ORF43,LATS1,DYNLL1,GNAS,GPR137,BCAT2,CENP |
| PKM,C1ORF43,LATS1,DYNLL1,GNAS,GPR137,BCAT2,CENP |
| PKM,C1ORF43,LATS1,DYNLL1,GNAS,GPR137,BCAT2,CENP |
| PKM,C1ORF43,LATS1,DYNLL1,GNAS,GPR137,BCAT2,CENP |
| PKM,C1ORF43,LATS1,DYNLL1,GNAS,GPR137,BCAT2,CENP |
| APP,DYNLL1,GNAS,GPR137,SERPINB5,CENPV,FAM189B,M |
| APP,DYNLL1,GNAS,GPR137,SERPINB5,CENPV,FAM189B,M |
| SNRNP70,SUPT4H1,LATS1,RPS24,SERPINB5,BCAT2,SRSF2, |
| SNRNP70,SUPT4H1,LATS1,RPS24,SERPINB5,BCAT2,SRSF2, |
| SNRNP70,SUPT4H1,LATS1,RPS24,SERPINB5,BCAT2,SRSF2, |
| SNRNP70,SUPT4H1,LATS1,RPS24,SERPINB5,BCAT2,SRSF2, |
| C1ORF43,DYNLL1,CST3,GPR137,RPS24,SERPINB5,BCAT2,C |
| C1ORF43,DYNLL1,CST3,GPR137,RPS24,SERPINB5,BCAT2,C |
| SUPT4H1,MRPS22,LAS1L,HMGB1,TMEM67,CMC2,CIRBP,K |
| SUPT4H1,MRPS22,LAS1L,HMGB1,TMEM67,CMC2,CIRBP,K |
| PKM,C1ORF43,DYNLL1,CST3,GNAS,GPR137,SERPINB5,CEN |
| PKM,C1ORF43,DYNLL1,CST3,GNAS,GPR137,SERPINB5,CEN |
| SUPT4H1,GPR137,BCAT2,MRPS22,LAS1L,HMGB1,CMC2,R |
| SUPT4H1,GPR137,BCAT2,MRPS22,LAS1L,HMGB1,CMC2,R |
| SNRNP70,APP,C1ORF43,LATS1,DYNLL1,CST3,GNAS,GPR13 |
| SNRNP70,APP,C1ORF43,LATS1,DYNLL1,CST3,GNAS,GPR13 |
| GPR137,RPS24,BCAT2,MRPL4,VKORC1,SRSF2,LAS1L,HMG |
| SNRNP70,APP,C1ORF43,SUPT4H1,LATS1,DYNLL1,GNAS,G |
| SNRNP70,APP,C1ORF43,SUPT4H1,LATS1,DYNLL1,GNAS,G |
| PKM,SNRNP70,C1ORF43,SUPT4H1,DYNLL1,GPR137,SERPI |
| PKM,SNRNP70,C1ORF43,SUPT4H1,DYNLL1,GPR137,SERPI |
| PKM,SNRNP70,APP,SUPT4H1,LATS1,CST3,GNAS,GPR137,R |
| PKM,SNRNP70,APP,C1ORF43,LATS1,GPR137,BCAT2,CENP |
| PKM,SNRNP70,APP,C1ORF43,LATS1,GPR137,BCAT2,CENP |
| PKM,SNRNP70,APP,C1ORF43,SUPT4H1,CCDC66,DYNLL1,C |
| PKM,SNRNP70,APP,C1ORF43,SUPT4H1,CCDC66,DYNLL1,C |

| |
|---|
| SUPT4H1,MRPS22,LAS1L,HMGB1,TMEM67,CMC2,CIRBP,R |
| SUPT4H1,MRPS22,LAS1L,HMGB1,TMEM67,CMC2,CIRBP,R |
| PKM,SNRNP70,APP,C1ORF43,SUPT4H1,LATS1,DYNLL1,CST |
| PKM,SNRNP70,APP,C1ORF43,SUPT4H1,LATS1,DYNLL1,CST |
| PKM,APP,C1ORF43,SUPT4H1,LATS1,DYNLL1,CST3,GNAS,G |
| PKM,APP,C1ORF43,SUPT4H1,LATS1,DYNLL1,CST3,GNAS,G |
| PKM,SNRNP70,APP,LATS1,CCDC66,DYNLL1,CST3,GNAS,GP |
| PKM,SNRNP70,APP,LATS1,CCDC66,DYNLL1,CST3,GNAS,GP |
| SUPT4H1,SERPINB5,MRPS22,LAS1L,HMGB1,TMEM67,CM( |
| SUPT4H1,SERPINB5,MRPS22,LAS1L,HMGB1,TMEM67,CM( |
| PKM,SNRNP70,APP,C1ORF43,SUPT4H1,LATS1,CCDC66,CST |
| PKM,SNRNP70,APP,C1ORF43,SUPT4H1,LATS1,CCDC66,CST |
| C1ORF43,DYNLL1,GPR137,SERPINB5,CENPV,FAM189B,MR |
| C1ORF43,DYNLL1,GPR137,SERPINB5,CENPV,FAM189B,MR |
| APP,C1ORF43,DYNLL1,GNAS,GPR137,SERPINB5,BCAT2,VK( |
| APP,C1ORF43,DYNLL1,GNAS,GPR137,SERPINB5,BCAT2,VK( |
| SUPT4H1,SERPINB5,S100A13,BCAT2,SRSF2,MRPS22,LAS1L |
| SUPT4H1,SERPINB5,S100A13,BCAT2,SRSF2,MRPS22,LAS1L |
| PKM,C1ORF43,DYNLL1,GPR137,SERPINB5,CENPV,FAM189 |
| PKM,C1ORF43,DYNLL1,GPR137,SERPINB5,CENPV,FAM189 |
| PKM,SNRNP70,APP,C1ORF43,LATS1,GPR137,BCAT2,CENP' |
| PKM,SNRNP70,APP,C1ORF43,LATS1,GPR137,BCAT2,CENP' |
| PKM,APP,C1ORF43,SUPT4H1,DYNLL1,CST3,GNAS,GPR137, |
| PKM,APP,C1ORF43,SUPT4H1,DYNLL1,CST3,GNAS,GPR137, |
| SUPT4H1,MRPS22,LAS1L,HMGB1,TMEM67,CMC2,CIRBP,R |
| SUPT4H1,MRPS22,LAS1L,HMGB1,TMEM67,CMC2,CIRBP,R |
| PKM,APP,DYNLL1,GPR137,RPS24,BCAT2,VKORC1,SRSF2,M |
| SUPT4H1,SERPINB5,SRSF2,MRPS22,OSMR,SEMA4D,LAS1L, |
| C1ORF43,DYNLL1,GNAS,GPR137,SERPINB5,MRPL2,CDCA3, |
| GNAS,MRPS22,LAS1L,HMGB1,SSU72,CIRBP,KIF4A,RBCK1,A |
| GNAS,MRPS22,LAS1L,HMGB1,SSU72,CIRBP,KIF4A,RBCK1,A |
| PKM,SNRNP70,APP,C1ORF43,LATS1,GNAS,RPS24,SERPINB |
| PKM,SNRNP70,APP,C1ORF43,LATS1,GNAS,RPS24,SERPINB |
| PKM,SNRNP70,APP,C1ORF43,SUPT4H1,LATS1,DYNLL1,CST |
| PKM,SNRNP70,APP,C1ORF43,SUPT4H1,LATS1,DYNLL1,CST |
| SNRNP70,APP,C1ORF43,SUPT4H1,LATS1,DYNLL1,GNAS,GF |
| SNRNP70,APP,C1ORF43,SUPT4H1,LATS1,DYNLL1,GNAS,GF |
| SNRNP70,SUPT4H1,LATS1,RPS24,SERPINB5,BCAT2,SRSF2,I |
| SNRNP70,SUPT4H1,LATS1,RPS24,SERPINB5,BCAT2,SRSF2,I |
| C1ORF43,DYNLL1,GPR137,SERPINB5,CENPV,FAM189B,PCI |
| C1ORF43,DYNLL1,GPR137,SERPINB5,CENPV,FAM189B,PCI |
| SUPT4H1,MRPS22,LAS1L,HMGB1,TMEM67,CMC2,CIRBP,R |
| SUPT4H1,MRPS22,LAS1L,HMGB1,TMEM67,CMC2,CIRBP,R |
| PKM,SNRNP70,APP,C1ORF43,LATS1,DYNLL1,CST3,GNAS,G |
| PKM,SNRNP70,APP,C1ORF43,LATS1,DYNLL1,CST3,GNAS,G |
| GNAS,BCAT2,HMGB1,RAPGEF5,TMEM67,CMC2,CACYBP,C |
| GNAS,BCAT2,HMGB1,RAPGEF5,TMEM67,CMC2,CACYBP,C |

| |
|---|
| C1ORF43,MRPL2,MRPS22,OSMR,HMGB1,BAIAP2,MYL6B, |
| C1ORF43,MRPL2,MRPS22,OSMR,HMGB1,BAIAP2,MYL6B, |
| C1ORF43,SUPT4H1,DYNLL1,CST3,GPR137,SERPINB5,CENP |
| C1ORF43,SUPT4H1,DYNLL1,CST3,GPR137,SERPINB5,CENP |
| SUPT4H1,SRSF2,MRPS22,LAS1L,HMGB1,RAPGEF5,SSU72, |
| SNRNP70,SUPT4H1,GNAS,SERPINB5,BCAT2,CDCA3,MRPS2 |
| SNRNP70,SUPT4H1,GNAS,SERPINB5,BCAT2,CDCA3,MRPS2 |
| PKM,GPR137,SERPINB5,MRPL2,PCNA,MRPS22,OSMR,BAI |
| PKM,GPR137,SERPINB5,MRPL2,PCNA,MRPS22,OSMR,BAI |
| PKM,SNRNP70,C1ORF43,DYNLL1,CST3,GNAS,GPR137,RPS |
| SNRNP70,APP,C1ORF43,SUPT4H1,LATS1,CST3,GNAS,GPR1 |
| SNRNP70,APP,C1ORF43,SUPT4H1,LATS1,CST3,GNAS,GPR1 |
| SUPT4H1,RPS24,BCAT2,SRSF2,MRPS22,KRT15,SEMA4D,LA |
| SUPT4H1,RPS24,BCAT2,SRSF2,MRPS22,KRT15,SEMA4D,LA |
| PKM,SNRNP70,APP,LATS1,DYNLL1,GNAS,GPR137,RPS24,S |
| PKM,SNRNP70,APP,LATS1,DYNLL1,GNAS,GPR137,RPS24,S |
| PKM,SNRNP70,APP,C1ORF43,SUPT4H1,LATS1,CCDC66,DY |
| PKM,SNRNP70,APP,C1ORF43,SUPT4H1,LATS1,CCDC66,DY |
| PKM,SNRNP70,APP,C1ORF43,GNAS,RPS24,SERPINB5,S100 |
| APP,SUPT4H1,CST3,SERPINB5,BCAT2,FAM189B,CDCA3,M |
| APP,SUPT4H1,CST3,SERPINB5,BCAT2,FAM189B,CDCA3,M |
| SUPT4H1,DYNLL1,GNAS,SERPINB5,BCAT2,MRPS22,LAS1L, |
| SUPT4H1,DYNLL1,GNAS,SERPINB5,BCAT2,MRPS22,LAS1L, |
| PKM,SNRNP70,APP,C1ORF43,CCDC66,DYNLL1,CST3,GNAS, |
| C1ORF43,DYNLL1,GNAS,GPR137,SERPINB5,MRPL2,CDCA3, |
| PKM,SNRNP70,SUPT4H1,LATS1,GNAS,GPR137,RPS24,SER |
| SUPT4H1,GNAS,SERPINB5,MRPS22,LAS1L,HMGB1,RBCK1, |
| DYNLL1,CST3,GPR137,SERPINB5,CENPV,FAM189B,MRPL2, |
| DYNLL1,CST3,GPR137,SERPINB5,CENPV,FAM189B,MRPL2, |
| PKM,SNRNP70,APP,C1ORF43,SUPT4H1,LATS1,CCDC66,DY |
| PKM,SNRNP70,APP,C1ORF43,SUPT4H1,LATS1,CCDC66,DY |
| PKM,SNRNP70,APP,C1ORF43,SUPT4H1,CCDC66,CST3,GNA |
| PKM,SNRNP70,APP,C1ORF43,SUPT4H1,CCDC66,CST3,GNA |
| PKM,SNRNP70,APP,C1ORF43,SUPT4H1,LATS1,CCDC66,DY |
| SNRNP70,APP,C1ORF43,SUPT4H1,LATS1,GNAS,GPR137,CE |
| PKM,SNRNP70,SUPT4H1,LATS1,CST3,GNAS,GPR137,BCAT |
| C1ORF43,DYNLL1,GNAS,GPR137,SERPINB5,MRPL2,CDCA3, |
| DYNLL1,GPR137,SERPINB5,FAM189B,MRPL2,PCNA,SRSF2, |
| APP,SUPT4H1,DYNLL1,GNAS,GPR137,RPS24,CENPV,FAM1 |
| APP,SUPT4H1,DYNLL1,GNAS,GPR137,RPS24,CENPV,FAM1 |
| SUPT4H1,GPR137,FAM189B,MRPS22,LAS1L,SRP9,TMEM6 |
| PKM,SNRNP70,APP,C1ORF43,LATS1,CCDC66,DYNLL1,CST3 |
| PKM,SNRNP70,APP,C1ORF43,LATS1,CCDC66,DYNLL1,CST3 |
| PKM,SNRNP70,APP,C1ORF43,SUPT4H1,LATS1,DYNLL1,CST |
| PKM,SNRNP70,APP,C1ORF43,SUPT4H1,LATS1,DYNLL1,CST |
| PKM,SNRNP70,C1ORF43,LATS1,GNAS,BCAT2,CENPV,MRPL |
| PKM,SNRNP70,C1ORF43,LATS1,GNAS,BCAT2,CENPV,MRPL |

| |
|---|
| SUPT4H1,CDCA3,HMGB1,BLOC1S1,RHOF,DNASE1L1,RBCK |
| PKM,SNRNP70,SUPT4H1,CCDC66,DYNLL1,CST3,GPR137,R |
| PKM,SNRNP70,SUPT4H1,CCDC66,DYNLL1,CST3,GPR137,R |
| SRSF2,MRPS22,HMGB1,CIRBP,RBCK1,ACTR6,BUD31,NPRL |
| PKM,SNRNP70,C1ORF43,SUPT4H1,CST3,GPR137,RPS24,SE |
| PKM,SNRNP70,C1ORF43,SUPT4H1,CST3,GPR137,RPS24,SE |
| SUPT4H1,MRPS22,LAS1L,RBCK1,ZNF235,DBI,ARPC2,LIMK2 |
| PKM,SNRNP70,APP,C1ORF43,CCDC66,DYNLL1,CST3,GPR1 |
| PKM,SNRNP70,APP,C1ORF43,CCDC66,DYNLL1,CST3,GPR1 |
| PKM,SNRNP70,APP,C1ORF43,SUPT4H1,LATS1,CCDC66,DY |
| SNRNP70,APP,C1ORF43,LATS1,DYNLL1,CST3,GNAS,GPR13 |
| SNRNP70,APP,C1ORF43,LATS1,DYNLL1,CST3,GNAS,GPR13 |
| SNRNP70,DYNLL1,GNAS,RPS24,SERPINB5,BCAT2,MRPL2,C |
| PKM,SNRNP70,APP,C1ORF43,SUPT4H1,LATS1,DYNLL1,GP |
| PKM,SNRNP70,APP,C1ORF43,SUPT4H1,LATS1,DYNLL1,GP |
| SNRNP70,C1ORF43,LATS1,DYNLL1,GNAS,GPR137,RPS24,S |
| SNRNP70,C1ORF43,LATS1,DYNLL1,GNAS,GPR137,RPS24,S |
| C1ORF43,DYNLL1,GPR137,SERPINB5,CENPV,FAM189B,MR |
| C1ORF43,DYNLL1,GPR137,SERPINB5,CENPV,FAM189B,MR |
| PKM,SNRNP70,APP,C1ORF43,SUPT4H1,LATS1,CCDC66,DY |
| PKM,SNRNP70,APP,C1ORF43,SUPT4H1,LATS1,CCDC66,DY |
| MRPS22,LAS1L,HMGB1,MYL6B,CMC2,RBCK1,RCC1,SNRPB, |
| SUPT4H1,MRPS22,LAS1L,CMC2,RBCK1,ACTR6,BUD31,NPR |
| PKM,SUPT4H1,CST3,SERPINB5,CDCA3,SRSF2,MRPS22,LAS |
| PKM,SUPT4H1,CST3,SERPINB5,CDCA3,SRSF2,MRPS22,LAS |
| PKM,SNRNP70,APP,C1ORF43,LATS1,DYNLL1,CST3,GPR137 |
| PKM,SNRNP70,APP,C1ORF43,LATS1,DYNLL1,CST3,GPR137 |
| C1ORF43,LATS1,GNAS,RPS24,BCAT2,MRPL4,FAM189B,VK |
| C1ORF43,LATS1,GNAS,RPS24,BCAT2,MRPL4,FAM189B,VK |
| PKM,APP,C1ORF43,LATS1,DYNLL1,CST3,GPR137,RPS24,S1 |
| PKM,APP,C1ORF43,LATS1,DYNLL1,CST3,GPR137,RPS24,S1 |
| PKM,SNRNP70,C1ORF43,SUPT4H1,LATS1,GNAS,GPR137,S |
| PKM,SNRNP70,C1ORF43,SUPT4H1,LATS1,GNAS,GPR137,S |
| PKM,SNRNP70,C1ORF43,SUPT4H1,LATS1,GNAS,GPR137,S |
| PKM,SNRNP70,C1ORF43,SUPT4H1,LATS1,GNAS,GPR137,S |
| APP,C1ORF43,SUPT4H1,DYNLL1,GNAS,GPR137,SERPINB5, |
| APP,C1ORF43,SUPT4H1,DYNLL1,GNAS,GPR137,SERPINB5, |
| PKM,SNRNP70,APP,C1ORF43,LATS1,DYNLL1,CST3,GPR137 |
| PKM,C1ORF43,DYNLL1,CST3,GNAS,GPR137,SERPINB5,CEN |
| PKM,C1ORF43,DYNLL1,CST3,GNAS,GPR137,SERPINB5,CEN |
| PKM,APP,C1ORF43,LATS1,DYNLL1,CST3,GPR137,RPS24,S1 |
| C1ORF43,DYNLL1,GNAS,GPR137,SERPINB5,MRPL2,CDCA3, |
| SNRNP70,C1ORF43,SUPT4H1,LATS1,CST3,GPR137,RPS24, |
| SNRNP70,C1ORF43,SUPT4H1,LATS1,CST3,GPR137,RPS24, |
| SNRNP70,APP,SUPT4H1,DYNLL1,CST3,GNAS,GPR137,RPS2 |
| SNRNP70,APP,SUPT4H1,DYNLL1,CST3,GNAS,GPR137,RPS2 |
| SUPT4H1,HMGB1,CMC2,BLOC1S1,RHOF,PRR13,RBCK1,RN |

| |
|---|
| GPR137,MRPS22,RBCK1,RCC1,SNRPB,DHPS,DBI,ARPC2,LI |
| PKM,SNRNP70,APP,C1ORF43,DYNLL1,CST3,GNAS,GPR137, |
| PKM,SNRNP70,APP,C1ORF43,DYNLL1,CST3,GNAS,GPR137, |
| PKM,SUPT4H1,LATS1,CST3,GNAS,GPR137,RPS24,CENPV, |
| PKM,SUPT4H1,LATS1,CST3,GNAS,GPR137,RPS24,CENPV, |
| APP,C1ORF43,LATS1,DYNLL1,CST3,GNAS,GPR137,RPS24, |
| APP,C1ORF43,LATS1,DYNLL1,CST3,GNAS,GPR137,RPS24, |
| PKM,SNRNP70,SUPT4H1,LATS1,DYNLL1,CST3,GNAS,RPS24 |
| PKM,SNRNP70,SUPT4H1,LATS1,DYNLL1,CST3,GNAS,RPS24 |
| C1ORF43,CST3,GNAS,SERPINB5,MRPL2,PCNA,SRSF2,HMG |
| PKM,SNRNP70,APP,C1ORF43,GNAS,RPS24,SERPINB5,S100 |
| PKM,SNRNP70,APP,C1ORF43,GNAS,RPS24,SERPINB5,S100 |
| SUPT4H1,BCAT2,MRPS22,LAS1L,HMGB1,CMC2,RBCK1,SN |
| SUPT4H1,BCAT2,MRPS22,LAS1L,HMGB1,CMC2,RBCK1,SN |
| PKM,SNRNP70,APP,C1ORF43,GNAS,RPS24,SERPINB5,S100 |
| PKM,SNRNP70,APP,C1ORF43,GNAS,RPS24,SERPINB5,S100 |
| PKM,SNRNP70,APP,C1ORF43,RPS24,SERPINB5,S100A13,C |
| SNRNP70,APP,C1ORF43,LATS1,DYNLL1,CST3,GNAS,GPR13 |
| C1ORF43,SUPT4H1,MRPS22,CMC2,BLOC1S1,RCC1,SNRPB, |
| SNRNP70,APP,SUPT4H1,GPR137,BCAT2,PCNA,SRSF2,PPFI |
| SNRNP70,APP,SUPT4H1,GPR137,BCAT2,PCNA,SRSF2,PPFI |
| PKM,APP,C1ORF43,DYNLL1,CST3,GNAS,GPR137,RPS24,BC |
| PKM,APP,C1ORF43,DYNLL1,CST3,GNAS,GPR137,RPS24,BC |
| APP,C1ORF43,LATS1,DYNLL1,CST3,GNAS,GPR137,RPS24, |
| C1ORF43,GNAS,RPS24,VKORC1,CDCA3,SRSF2,SSU72,MYL |
| SNRNP70,LATS1,RPS24,SERPINB5,BCAT2,VKORC1,CDCA3, |
| SNRNP70,LATS1,RPS24,SERPINB5,BCAT2,VKORC1,CDCA3, |
| PKM,SNRNP70,APP,C1ORF43,SUPT4H1,LATS1,DYNLL1,CST |
| PKM,SNRNP70,APP,C1ORF43,SUPT4H1,LATS1,DYNLL1,CST |
| SNRNP70,APP,C1ORF43,LATS1,DYNLL1,CST3,GNAS,GPR13 |
| SNRNP70,APP,C1ORF43,LATS1,DYNLL1,CST3,GNAS,GPR13 |
| SNRNP70,APP,C1ORF43,SUPT4H1,LATS1,DYNLL1,GNAS,G |
| SNRNP70,APP,C1ORF43,SUPT4H1,LATS1,DYNLL1,GNAS,G |
| APP,C1ORF43,SUPT4H1,LATS1,CCDC66,DYNLL1,CST3,RPS2 |
| APP,C1ORF43,SUPT4H1,LATS1,CCDC66,DYNLL1,CST3,RPS2 |
| PKM,SNRNP70,APP,C1ORF43,SUPT4H1,LATS1,CCDC66,DY |
| PKM,SNRNP70,APP,C1ORF43,SUPT4H1,LATS1,CCDC66,DY |
| PKM,APP,SUPT4H1,LATS1,DYNLL1,CST3,GNAS,GPR137,SE |
| PKM,APP,SUPT4H1,LATS1,DYNLL1,CST3,GNAS,GPR137,SE |
| PKM,SNRNP70,APP,C1ORF43,SUPT4H1,LATS1,CCDC66,DY |
| PKM,SNRNP70,APP,C1ORF43,SUPT4H1,LATS1,CCDC66,DY |
| C1ORF43,DYNLL1,CST3,GNAS,GPR137,SERPINB5,CENPV,F |
| PKM,SNRNP70,SUPT4H1,DYNLL1,CST3,GNAS,SERPINB5,S1 |
| SNRNP70,APP,LATS1,GNAS,GPR137,RPS24,BCAT2,CENPV, |
| PKM,SNRNP70,APP,C1ORF43,SUPT4H1,LATS1,CCDC66,DY |
| PKM,SNRNP70,APP,C1ORF43,SUPT4H1,LATS1,CCDC66,DY |
| PKM,SNRNP70,C1ORF43,LATS1,DYNLL1,CST3,GNAS,GPR13 |

| |
|---|
| PKM,SNRNP70,C1ORF43,LATS1,DYNLL1,CST3,GNAS,GPR13 |
| APP,C1ORF43,DYNLL1,GPR137,RPS24,BCAT2,FAM189B,V |
| SNRNP70,C1ORF43,SUPT4H1,LATS1,DYNLL1,CST3,GPR137 |
| SNRNP70,C1ORF43,SUPT4H1,LATS1,DYNLL1,CST3,GPR137 |
| PKM,SUPT4H1,GNAS,SERPINB5,MRPS22,LAS1L,HMGB1,C |
| PKM,SUPT4H1,GNAS,SERPINB5,MRPS22,LAS1L,HMGB1,C |
| SUPT4H1,SERPINB5,LAS1L,HMGB1,TMEM67,DNASE1L1,K |
| SUPT4H1,SERPINB5,LAS1L,HMGB1,TMEM67,DNASE1L1,K |
| SUPT4H1,SRSF2,MRPS22,LAS1L,CMC2,RBCK1,RCC1,SNRPB |
| PKM,C1ORF43,SUPT4H1,CCDC66,SERPINB5,MRPL4,FAM1 |
| C1ORF43,GPR137,SERPINB5,CENPV,MRPL2,PCNA,SRSF2,B |
| C1ORF43,GPR137,SERPINB5,CENPV,MRPL2,PCNA,SRSF2,B |
| SNRNP70,RPS24,SERPINB5,BCAT2,MRPL2,CDCA3,PCNA,SR |
| PKM,C1ORF43,CCDC66,GPR137,RPS24,BCAT2,SRSF2,HDA |
| PKM,SNRNP70,APP,C1ORF43,SUPT4H1,LATS1,CCDC66,DY |
| SNRNP70,APP,C1ORF43,LATS1,CST3,RPS24,BCAT2,MRPL4, |
| SUPT4H1,MRPS22,LAS1L,HMGB1,RBCK1,RCC1,SNRPB,BU |
| PKM,SNRNP70,APP,SUPT4H1,LATS1,CCDC66,DYNLL1,CST3 |
| PKM,SNRNP70,APP,SUPT4H1,LATS1,CCDC66,DYNLL1,CST3 |
| SNRNP70,APP,C1ORF43,LATS1,SERPINB5,CENPV,SRSF2,LA |
| LAS1L,HMGB1,TMEM67,KIF4A,RBCK1,SNRPB,DHPS,GPAT |
| LAS1L,HMGB1,TMEM67,KIF4A,RBCK1,SNRPB,DHPS,GPAT |
| APP,C1ORF43,DYNLL1,CST3,GNAS,GPR137,BCAT2,CENPV, |
| PKM,SUPT4H1,DYNLL1,CST3,SERPINB5,CDCA3,MRPS22,LA |
| PKM,SUPT4H1,DYNLL1,CST3,SERPINB5,CDCA3,MRPS22,LA |
| C1ORF43,SUPT4H1,LATS1,CCDC66,DYNLL1,SERPINB5,CEN |
| C1ORF43,SUPT4H1,LATS1,CCDC66,DYNLL1,SERPINB5,CEN |
| SNRNP70,APP,LATS1,DYNLL1,CST3,GNAS,MRPL4,FAM189 |
| SNRNP70,APP,LATS1,DYNLL1,CST3,GNAS,MRPL4,FAM189 |
| C1ORF43,DYNLL1,SERPINB5,MRPL2,CDCA3,PCNA,SRSF2,B |
| SNRNP70,APP,C1ORF43,SUPT4H1,LATS1,DYNLL1,GNAS,G |
| SNRNP70,APP,C1ORF43,SUPT4H1,LATS1,DYNLL1,GNAS,G |
| SUPT4H1,LATS1,CCDC66,DYNLL1,RPS24,SERPINB5,CENPV, |
| SUPT4H1,LATS1,CCDC66,DYNLL1,RPS24,SERPINB5,CENPV, |
| PKM,SNRNP70,APP,C1ORF43,SUPT4H1,LATS1,DYNLL1,CST |
| PKM,SNRNP70,APP,C1ORF43,SUPT4H1,LATS1,DYNLL1,CST |
| DYNLL1,GPR137,SERPINB5,FAM189B,MRPL2,PCNA,SRSF2, |
| SUPT4H1,MRPS22,LAS1L,RBCK1,RCC1,SNRPB,BUD31,NPR |
| SNRNP70,APP,LATS1,GNAS,GPR137,SERPINB5,CENPV,PCN |
| PKM,SNRNP70,SUPT4H1,LATS1,CST3,GNAS,GPR137,RPS2 |
| PKM,SNRNP70,SUPT4H1,LATS1,CST3,GNAS,GPR137,RPS2 |
| SNRNP70,APP,SUPT4H1,GNAS,GPR137,VKORC1,PCNA,SRS |
| SNRNP70,APP,C1ORF43,SUPT4H1,GNAS,GPR137,VKORC1, |
| PKM,SNRNP70,C1ORF43,LATS1,DYNLL1,CST3,GNAS,GPR13 |
| PKM,SNRNP70,C1ORF43,LATS1,DYNLL1,CST3,GNAS,GPR13 |
| SUPT4H1,MRPS22,LAS1L,RBCK1,CTTN,COMMD5,NUDT5,S |
| HMGB1,CMC2,RPL38,KIF4A,TRPT1,NPRL2,CTTN,NUDT22, |

| |
|---|
| HMGB1,CMC2,RPL38,KIF4A,TRPT1,NPRL2,CTTN,NUDT22, |
| PKM,APP,LATS1,DYNLL1,CST3,GNAS,GPR137,BCAT2,CENP |
| PKM,APP,LATS1,DYNLL1,CST3,GNAS,GPR137,BCAT2,CENP |
| RPS24,S100A13,FAM189B,CDCA3,SRSF2,MRPS22,KRT15,B |
| SUPT4H1,MRPS22,CYB561A3,RAPGEF5,CIRBP,RPL38,RBCK |
| SUPT4H1,SERPINB5,MRPS22,LAS1L,TMEM67,CIRBP,RPL38 |
| APP,C1ORF43,DYNLL1,GPR137,RPS24,S100A13,BCAT2,MR |
| APP,C1ORF43,DYNLL1,GPR137,RPS24,S100A13,BCAT2,MR |
| PKM,SNRNP70,SUPT4H1,RPS24,BCAT2,CENPV,MRPL4,PC |
| SUPT4H1,MRPS22,LAS1L,RBCK1,ARPC2,LIMK2,BUD31,NP |
| SNRNP70,APP,C1ORF43,LATS1,GNAS,GPR137,SERPINB5,V |
| SNRNP70,APP,C1ORF43,LATS1,GNAS,GPR137,SERPINB5,V |
| SUPT4H1,OSMR,PILRB,CMC2,CIRBP,RPL38,LY75,SNRPB,ZN |
| SUPT4H1,OSMR,PILRB,CMC2,CIRBP,RPL38,LY75,SNRPB,ZN |
| APP,C1ORF43,DYNLL1,GPR137,RPS24,S100A13,BCAT2,MR |
| PKM,APP,LATS1,CCDC66,DYNLL1,GPR137,SRSF2,HDAC9,O |
| C1ORF43,DYNLL1,GNAS,GPR137,SERPINB5,MRPL2,CDCA3, |
| PKM,C1ORF43,SUPT4H1,DYNLL1,CST3,GNAS,GPR137,SER |
| PKM,C1ORF43,SUPT4H1,DYNLL1,CST3,GNAS,GPR137,SER |
| PKM,SNRNP70,C1ORF43,SUPT4H1,DYNLL1,GNAS,GPR137, |
| PKM,SNRNP70,C1ORF43,SUPT4H1,DYNLL1,GNAS,GPR137, |
| PKM,SNRNP70,APP,C1ORF43,DYNLL1,CST3,GNAS,GPR137, |
| PKM,SNRNP70,APP,C1ORF43,DYNLL1,CST3,GNAS,GPR137, |
| SNRNP70,DYNLL1,SERPINB5,MRPL2,CDCA3,PCNA,SRSF2,B |
| PKM,SNRNP70,APP,C1ORF43,LATS1,CCDC66,GPR137,RPS |
| PKM,APP,C1ORF43,LATS1,CST3,S100A13,CENPV,MRPL4,V |
| PKM,C1ORF43,SUPT4H1,LATS1,DYNLL1,GNAS,GPR137,RP |
| PKM,C1ORF43,SUPT4H1,LATS1,DYNLL1,GNAS,GPR137,RP |
| HMGB1,SNRPB,ZNF235,BOP1,TRPT1,ARPC2,PMFBP1,LIMK |
| HMGB1,SNRPB,ZNF235,BOP1,TRPT1,ARPC2,PMFBP1,LIMK |
| SUPT4H1,MRPS22,LAS1L,HMGB1,RBCK1,BUD31,NPRL2,CT |
| PKM,APP,LATS1,CST3,GNAS,GPR137,RPS24,S100A13,BCA |
| PKM,APP,LATS1,CST3,GNAS,GPR137,RPS24,S100A13,BCA |
| MRPL2,BAIAP2,MYL6B,PRR13,RNH1,ARRDC1,NUMA1,TBL |
| APP,C1ORF43,LATS1,DYNLL1,GNAS,RPS24,CENPV,FAM18 |
| APP,C1ORF43,LATS1,DYNLL1,GNAS,RPS24,CENPV,FAM18 |
| PKM,LATS1,CENPV,FAM189B,PCNA,SRSF2,SULF2,BAIAP2, |
| PKM,LATS1,CENPV,FAM189B,PCNA,SRSF2,SULF2,BAIAP2, |
| PKM,SNRNP70,APP,C1ORF43,SUPT4H1,LATS1,DYNLL1,CST |
| MRPS22,RBCK1,SNRPB,ARPC2,LIMK2,NPRL2,CTTN,STYXL1, |
| SNRNP70,APP,LATS1,DYNLL1,GNAS,GPR137,SERPINB5,CE |
| SNRNP70,APP,LATS1,DYNLL1,GNAS,GPR137,SERPINB5,CE |
| PKM,APP,C1ORF43,LATS1,CCDC66,DYNLL1,CST3,SERPINB |
| PKM,APP,C1ORF43,LATS1,CCDC66,DYNLL1,CST3,SERPINB |
| PKM,C1ORF43,LATS1,GPR137,RPS24,FAM189B,PCNA,SRS |
| PKM,C1ORF43,LATS1,GPR137,RPS24,FAM189B,PCNA,SRS |
| SNRNP70,APP,C1ORF43,SUPT4H1,LATS1,CCDC66,CST3,GN |

| |
|---|
| SNRNP70,APP,C1ORF43,SUPT4H1,LATS1,CCDC66,CST3,GN |
| MRPS22,LAS1L,RBCK1,NPRL2,CTTN,COMMD5,NUDT5,STY |
| SNRNP70,APP,C1ORF43,CCDC66,DYNLL1,GNAS,RPS24,SER |
| SNRNP70,APP,C1ORF43,CCDC66,DYNLL1,GNAS,RPS24,SER |
| PKM,SNRNP70,APP,SUPT4H1,LATS1,CST3,GNAS,MRPL4,V |
| PKM,SNRNP70,APP,SUPT4H1,LATS1,CST3,GNAS,MRPL4,V |
| PKM,SNRNP70,SUPT4H1,LATS1,CST3,GNAS,GPR137,CENP |
| APP,C1ORF43,DYNLL1,CST3,GNAS,GPR137,SERPINB5,CEN |
| PKM,SNRNP70,APP,LATS1,CCDC66,CST3,RPS24,SERPINB5, |
| PKM,SNRNP70,APP,LATS1,CCDC66,CST3,RPS24,SERPINB5, |
| C1ORF43,SUPT4H1,LATS1,CCDC66,DYNLL1,SERPINB5,CEN |
| C1ORF43,SUPT4H1,LATS1,CCDC66,DYNLL1,SERPINB5,CEN |
| MRPS22,LAS1L,CMC2,RBCK1,RCC1,SNRPB,ARPC2,LIMK2,B |
| PKM,SNRNP70,C1ORF43,LATS1,GNAS,S100A13,CENPV,FA |
| DYNLL1,GPR137,SERPINB5,MRPL2,CDCA3,PCNA,SRSF2,H |
| SNRNP70,APP,GNAS,GPR137,CENPV,PCNA,SRSF2,OSMR,R |
| SNRNP70,APP,DYNLL1,GNAS,GPR137,BCAT2,CENPV,VKOR |
| PKM,SNRNP70,APP,C1ORF43,DYNLL1,CST3,GNAS,GPR137, |
| PKM,SNRNP70,APP,C1ORF43,DYNLL1,CST3,GNAS,GPR137, |
| SUPT4H1,MRPS22,LAS1L,HMGB1,TMEM67,CMC2,RBCK1, |
| SUPT4H1,MRPS22,LAS1L,HMGB1,TMEM67,CMC2,RBCK1, |
| SNRNP70,C1ORF43,SUPT4H1,LATS1,DYNLL1,GNAS,GPR13 |
| SNRNP70,C1ORF43,SUPT4H1,LATS1,DYNLL1,GNAS,GPR13 |
| PKM,C1ORF43,SUPT4H1,LATS1,DYNLL1,CST3,GNAS,GPR13 |
| DYNLL1,GNAS,GPR137,SERPINB5,MRPL2,CDCA3,PCNA,SR |
| SNRNP70,SUPT4H1,PCNA,SRSF2,FAM122C,IQCG,RHOF,SM |
| SNRNP70,SUPT4H1,PCNA,SRSF2,FAM122C,IQCG,RHOF,SM |
| DYNLL1,GNAS,GPR137,SERPINB5,MRPL2,CDCA3,PCNA,SR |
| MRPS22,MYL6B,CMC2,RNH1,RBCK1,RCC1,SNRPB,ARPC2,L |
| C1ORF43,CCDC66,DYNLL1,GNAS,GPR137,BCAT2,CENPV,M |
| APP,SUPT4H1,DYNLL1,CST3,GNAS,GPR137,RPS24,S100A1 |
| APP,SUPT4H1,DYNLL1,CST3,GNAS,GPR137,RPS24,S100A1 |
| PKM,SNRNP70,APP,C1ORF43,LATS1,DYNLL1,CST3,GNAS,G |
| PKM,SNRNP70,APP,C1ORF43,LATS1,DYNLL1,CST3,GNAS,G |
| SNRNP70,GPR137,BCAT2,PCNA,SRSF2,PPFIBP1,FAM122C, |
| SNRNP70,GPR137,BCAT2,PCNA,SRSF2,PPFIBP1,FAM122C, |
| BCAT2,HMGB1,TMEM67,RBCK1,SNRPB,BUD31,CTTN,EIF5 |
| BCAT2,HMGB1,TMEM67,RBCK1,SNRPB,BUD31,CTTN,EIF5 |
| RPL38,NPRL2,CTTN,RPL13A,TSG101,RPS6,RPL29,TRNT1,M |
| LAS1L,HMGB1,TMEM67,KIF4A,RBCK1,RCC1,SNRPB,DHPS, |
| LAS1L,HMGB1,TMEM67,KIF4A,RBCK1,RCC1,SNRPB,DHPS, |
| APP,C1ORF43,LATS1,GNAS,SERPINB5,CENPV,MRPL4,CDCA |
| APP,C1ORF43,LATS1,GNAS,SERPINB5,CENPV,MRPL4,CDCA |
| PKM,SNRNP70,APP,C1ORF43,SERPINB5,MRPL4,MRPL2,PC |
| PKM,SNRNP70,APP,C1ORF43,SERPINB5,MRPL4,MRPL2,PC |
| C1ORF43,CST3,GNAS,CENPV,FAM189B,BAIAP2,YBX3,BLO |
| C1ORF43,CST3,GNAS,CENPV,FAM189B,BAIAP2,YBX3,BLO |

| |
|---|
| PKM,APP,SUPT4H1,LATS1,CST3,GPR137,RPS24,SERPINB5, |
| PKM,APP,SUPT4H1,LATS1,CST3,GPR137,RPS24,SERPINB5, |
| C1ORF43,LATS1,DYNLL1,GNAS,GPR137,RPS24,S100A13,B |
| C1ORF43,LATS1,DYNLL1,GNAS,GPR137,RPS24,S100A13,B |
| SUPT4H1,GNAS,MRPS22,PPFIBP1,LAS1L,SSU72,SULT1A3, |
| S100A13,BCAT2,MRPS22,LAS1L,MYL6B,FAM122C,CMC2,R |
| PKM,SNRNP70,C1ORF43,CENPV,FAM189B,CDCA3,PCNA,L |
| SNRNP70,APP,LATS1,DYNLL1,CST3,GNAS,GPR137,SERPIN |
| SNRNP70,APP,LATS1,DYNLL1,CST3,GNAS,GPR137,SERPIN |
| PKM,SNRNP70,APP,C1ORF43,SUPT4H1,LATS1,CCDC66,DY |
| PKM,SNRNP70,APP,C1ORF43,SUPT4H1,LATS1,CCDC66,DY |
| SNRNP70,APP,LATS1,GNAS,GPR137,FAM189B,PCNA,SRSF |
| DYNLL1,GNAS,GPR137,SERPINB5,MRPL2,CDCA3,PCNA,SR |
| DYNLL1,GNAS,GPR137,SERPINB5,MRPL2,CDCA3,PCNA,SR |
| PKM,C1ORF43,LATS1,GPR137,RPS24,FAM189B,PCNA,SRS |
| PKM,SNRNP70,APP,C1ORF43,LATS1,CST3,RPS24,SERPINB |
| PKM,SNRNP70,APP,C1ORF43,LATS1,CST3,RPS24,SERPINB |
| DYNLL1,GNAS,GPR137,SERPINB5,MRPL2,PCNA,SRSF2,BAI |
| SUPT4H1,MRPS22,LAS1L,HMGB1,TMEM67,CMC2,RBCK1, |
| SUPT4H1,MRPS22,LAS1L,HMGB1,TMEM67,CMC2,RBCK1, |
| DYNLL1,GNAS,GPR137,RPS24,SERPINB5,BCAT2,CENPV,M |
| DYNLL1,GNAS,GPR137,RPS24,SERPINB5,BCAT2,CENPV,M |
| SNRNP70,APP,LATS1,DYNLL1,CST3,GPR137,SERPINB5,BCA |
| SNRNP70,APP,LATS1,DYNLL1,CST3,GPR137,SERPINB5,BCA |
| C1ORF43,LATS1,GNAS,GPR137,RPS24,S100A13,BCAT2,FA |
| C1ORF43,LATS1,GNAS,GPR137,RPS24,S100A13,BCAT2,FA |
| SNRNP70,APP,SUPT4H1,GNAS,GPR137,RPS24,S100A13,B |
| SNRNP70,APP,SUPT4H1,GNAS,GPR137,RPS24,S100A13,B |
| SNRNP70,APP,LATS1,DYNLL1,CST3,GNAS,GPR137,SERPIN |
| SNRNP70,APP,LATS1,DYNLL1,CST3,GNAS,GPR137,SERPIN |
| C1ORF43,SUPT4H1,LATS1,DYNLL1,CST3,GPR137,RPS24,SE |
| C1ORF43,SUPT4H1,LATS1,DYNLL1,CST3,GPR137,RPS24,SE |
| SNRNP70,SUPT4H1,PCNA,SRSF2,FAM122C,IQCG,RHOF,SM |
| SNRNP70,SUPT4H1,PCNA,SRSF2,FAM122C,IQCG,RHOF,SM |
| C1ORF43,GPR137,S100A13,BCAT2,FAM189B,VKORC1,CD |
| SNRNP70,APP,C1ORF43,LATS1,CENPV,CDCA3,PCNA,SSU7 |
| PKM,LATS1,CST3,GNAS,GPR137,MRPL4,MRPL2,CDCA3,SR |
| PKM,SNRNP70,APP,C1ORF43,SUPT4H1,LATS1,DYNLL1,CST |
| PKM,SNRNP70,APP,C1ORF43,SUPT4H1,LATS1,DYNLL1,CST |
| C1ORF43,DYNLL1,GNAS,GPR137,SERPINB5,FAM189B,MR |
| APP,C1ORF43,SUPT4H1,LATS1,CCDC66,DYNLL1,CST3,RPS2 |
| APP,C1ORF43,SUPT4H1,LATS1,CCDC66,DYNLL1,CST3,RPS2 |
| PKM,SNRNP70,APP,LATS1,CST3,GNAS,GPR137,SERPINB5, |
| PKM,SNRNP70,APP,LATS1,CST3,GNAS,GPR137,SERPINB5, |
| SERPINB5,MRPL2,PCNA,SRSF2,HMGB1,BAIAP2,RAPGEF5, |
| MRPS22,SEMA4D,LAS1L,RBCK1,RCC1,SNRPB,DHPS,BUD31 |
| PKM,SNRNP70,APP,C1ORF43,RPS24,SERPINB5,CENPV,MR |

| |
|---|
| PKM,SNRNP70,C1ORF43,CCDC66,CST3,GNAS,GPR137,RPS |
| PKM,SNRNP70,C1ORF43,CCDC66,CST3,GNAS,GPR137,RPS |
| DYNLL1,GPR137,SERPINB5,MRPL2,PCNA,SRSF2,BAIAP2,RA |
| RPS24,LAS1L,HMGB1,MYL6B,TMEM67,KIF4A,RBCK1,RCC1, |
| RPS24,LAS1L,HMGB1,MYL6B,TMEM67,KIF4A,RBCK1,RCC1, |
| MRPS22,RBCK1,RCC1,SNRPB,DBI,ARPC2,LIMK2,BUD31,NP |
| PKM,SNRNP70,C1ORF43,SUPT4H1,DYNLL1,GNAS,CENPV,F |
| SNRNP70,LATS1,RPS24,SERPINB5,BCAT2,CDCA3,SMARCB1 |
| SNRNP70,LATS1,RPS24,SERPINB5,BCAT2,CDCA3,SMARCB1 |
| CCDC66,SERPINB5,MRPL4,PCNA,PPFIBP1,LAS1L,FAM122C, |
| GPR137,RPS24,SERPINB5,BCAT2,MRPL2,CDCA3,PCNA,BAI |
| GPR137,RPS24,SERPINB5,BCAT2,MRPL2,CDCA3,PCNA,BAI |
| PKM,SNRNP70,APP,SUPT4H1,LATS1,CCDC66,DYNLL1,GPR |
| PKM,SNRNP70,APP,SUPT4H1,LATS1,CCDC66,DYNLL1,GPR |
| PKM,APP,C1ORF43,SUPT4H1,LATS1,DYNLL1,CST3,GNAS,G |
| PKM,APP,C1ORF43,SUPT4H1,LATS1,DYNLL1,CST3,GNAS,G |
| C1ORF43,LATS1,GNAS,GPR137,RPS24,S100A13,BCAT2,MR |
| DYNLL1,SERPINB5,MRPL2,PCNA,SRSF2,BAIAP2,MYL6B,RH |
| C1ORF43,LATS1,DYNLL1,CST3,RPS24,MRPL4,FAM189B,VK |
| PKM,APP,DYNLL1,GPR137,PCNA,SRSF2,BAIAP2,RAPGEF5, |
| PKM,C1ORF43,SERPINB5,CENPV,MRPL4,MRPL2,FAM122C, |
| PKM,SNRNP70,APP,LATS1,CCDC66,CST3,GNAS,GPR137,SE |
| PKM,SNRNP70,APP,LATS1,CCDC66,CST3,GNAS,GPR137,SE |
| PKM,C1ORF43,DYNLL1,GPR137,RPS24,SERPINB5,BCAT2,C |
| C1ORF43,LATS1,CCDC66,DYNLL1,SERPINB5,CENPV,MRPL4 |
| C1ORF43,LATS1,CCDC66,DYNLL1,SERPINB5,CENPV,MRPL4 |
| PKM,C1ORF43,LATS1,DYNLL1,GPR137,S100A13,FAM189B, |
| PKM,C1ORF43,LATS1,DYNLL1,GPR137,S100A13,FAM189B, |
| C1ORF43,LATS1,GNAS,BCAT2,FAM189B,VKORC1,CDCA3,S |
| C1ORF43,LATS1,GNAS,BCAT2,FAM189B,VKORC1,CDCA3,S |
| PKM,SNRNP70,CST3,RPS24,SERPINB5,FAM189B,MRPL2,C |
| DYNLL1,GPR137,SERPINB5,MRPL2,CDCA3,PCNA,SRSF2,BA |
| PKM,C1ORF43,CST3,GNAS,CENPV,FAM189B,KRT15,HMGB |
| PKM,C1ORF43,CST3,GNAS,CENPV,FAM189B,KRT15,HMGB |
| DYNLL1,GNAS,SERPINB5,CENPV,MRPL2,CDCA3,PCNA,SRS |
| PKM,SNRNP70,C1ORF43,DYNLL1,GNAS,GPR137,CENPV,FA |
| PKM,SNRNP70,C1ORF43,DYNLL1,GNAS,GPR137,CENPV,FA |
| PKM,SNRNP70,APP,SUPT4H1,CCDC66,DYNLL1,RPS24,S10 |
| PKM,SNRNP70,APP,SUPT4H1,CCDC66,DYNLL1,RPS24,S10 |
| PKM,C1ORF43,SUPT4H1,LATS1,DYNLL1,GPR137,RPS24,SE |
| PKM,C1ORF43,SUPT4H1,LATS1,DYNLL1,GPR137,RPS24,SE |
| FAM189B,VKORC1,CDCA3,PCNA,SRSF2,MRPS22,MYL6B,F |
| FAM189B,VKORC1,CDCA3,PCNA,SRSF2,MRPS22,MYL6B,F |
| SNRNP70,SUPT4H1,LATS1,CST3,GPR137,RPS24,S100A13, |
| C1ORF43,SERPINB5,MRPL4,FAM189B,CDCA3,PPFIBP1,BAI |
| SNRNP70,SUPT4H1,LATS1,DYNLL1,CST3,GNAS,GPR137,RP |
| SNRNP70,SUPT4H1,LATS1,DYNLL1,CST3,GNAS,GPR137,RP |

| |
|---|
| PKM,SNRNP70,DYNLL1,RPS24,SERPINB5,BCAT2,CDCA3,SR |
| PKM,SNRNP70,DYNLL1,RPS24,SERPINB5,BCAT2,CDCA3,SR |
| MRPS22,LAS1L,RBCK1,RCC1,SNRPB,BUD31,NPRL2,CTTN,EI |
| SNRNP70,APP,C1ORF43,LATS1,GNAS,GPR137,CENPV,PCN |
| SNRNP70,PCNA,SRSF2,SMARCB1,STARD10,RCC1,TCAIM,S |
| SNRNP70,PCNA,SRSF2,SMARCB1,STARD10,RCC1,TCAIM,S |
| SNRNP70,APP,C1ORF43,LATS1,DYNLL1,CST3,GNAS,RPS24, |
| SNRNP70,APP,C1ORF43,LATS1,DYNLL1,CST3,GNAS,RPS24, |
| SNRNP70,SUPT4H1,GPR137,PCNA,FAM122C,RHOF,STARD |
| SNRNP70,SUPT4H1,GPR137,PCNA,FAM122C,RHOF,STARD |
| PKM,SUPT4H1,GNAS,SERPINB5,SRSF2,MRPS22,KRT15,SEN |
| PKM,SUPT4H1,GNAS,SERPINB5,SRSF2,MRPS22,KRT15,SEN |
| CST3,MRPS22,MYL6B,IQCG,KIF4A,TCAIM,SET,DBI,SIRT6,IT |
| CST3,MRPS22,MYL6B,IQCG,KIF4A,TCAIM,SET,DBI,SIRT6,IT |
| SUPT4H1,CCDC66,DYNLL1,SERPINB5,S100A13,MRPL4,FAN |
| SUPT4H1,CCDC66,DYNLL1,SERPINB5,S100A13,MRPL4,FAN |
| APP,SUPT4H1,LATS1,CCDC66,DYNLL1,CST3,RPS24,SERPIN |
| APP,SUPT4H1,LATS1,CCDC66,DYNLL1,CST3,RPS24,SERPIN |
| SNRNP70,SRSF2,BAIAP2,RNH1,RBCK1,SNRPB,TCAIM,SLC3 |
| SNRNP70,SRSF2,BAIAP2,RNH1,RBCK1,SNRPB,TCAIM,SLC3 |
| C1ORF43,SUPT4H1,DYNLL1,GNAS,GPR137,SERPINB5,BCAT |
| PKM,C1ORF43,SUPT4H1,LATS1,CCDC66,DYNLL1,GNAS,RPS |
| PKM,C1ORF43,SUPT4H1,LATS1,CCDC66,DYNLL1,GNAS,RPS |
| PKM,SNRNP70,APP,LATS1,DYNLL1,CST3,GNAS,GPR137,RP |
| DYNLL1,CST3,SERPINB5,MRPL4,FAM189B,HMGB1,IQCG,C |
| PKM,SNRNP70,C1ORF43,LATS1,DYNLL1,GNAS,GPR137,RPS |
| MRPS22,LAS1L,RBCK1,RCC1,SNRPB,BUD31,NPRL2,CTTN,C |
| CST3,GNAS,GPR137,RPS24,CENPV,FAM189B,MRPL2,PCNA |
| CST3,GNAS,GPR137,RPS24,CENPV,FAM189B,MRPL2,PCNA |
| SUPT4H1,MRPS22,LAS1L,SNRPB,BUD31,CTTN,EIF5A,RPL2 |
| DYNLL1,GNAS,GPR137,SERPINB5,MRPL2,CDCA3,PCNA,SRS |
| DYNLL1,GNAS,GPR137,SERPINB5,MRPL2,CDCA3,PCNA,SRS |
| MRPS22,LAS1L,HMGB1,RBCK1,BUD31,NPRL2,CTTN,EIF5A, |
| HMGB1,TMEM67,RBCK1,RCC1,SNRPB,DHPS,GPATCH2L,BU |
| HMGB1,TMEM67,RBCK1,RCC1,SNRPB,DHPS,GPATCH2L,BU |
| PKM,C1ORF43,LATS1,DYNLL1,GPR137,S100A13,FAM189B, |
| RPS24,SERPINB5,BCAT2,CDCA3,SMARCB1,RNH1,DNASE1L |
| RPS24,SERPINB5,BCAT2,CDCA3,SMARCB1,RNH1,DNASE1L |
| SUPT4H1,MRPS22,LAS1L,HMGB1,LY75,BUD31,EIF5A,ACOT |
| PKM,C1ORF43,SUPT4H1,CCDC66,CST3,RPS24,BCAT2,MRP |
| APP,SUPT4H1,DYNLL1,CST3,GNAS,GPR137,RPS24,CENPV, |
| APP,SUPT4H1,DYNLL1,CST3,GNAS,GPR137,RPS24,CENPV, |
| PKM,C1ORF43,DYNLL1,CST3,GNAS,GPR137,SERPINB5,FAN |
| SNRNP70,APP,C1ORF43,LATS1,GPR137,PCNA,OSMR,PPFIE |
| SNRNP70,APP,C1ORF43,LATS1,GPR137,PCNA,OSMR,PPFIE |
| CMC2,TRPT1,PMFBP1,NPRL2,CTTN,NUDT22,RPL13A,ARH |
| CMC2,TRPT1,PMFBP1,NPRL2,CTTN,NUDT22,RPL13A,ARH |

| |
|---|
| GPR137,RPS24,SERPINB5,MRPL2,CDCA3,PCNA,BAIAP2,MY |
| GPR137,RPS24,SERPINB5,MRPL2,CDCA3,PCNA,BAIAP2,MY |
| SNRNP70,APP,LATS1,DYNLL1,CST3,GNAS,GPR137,SERPIN |
| SNRNP70,APP,LATS1,DYNLL1,CST3,GNAS,GPR137,SERPIN |
| SNRNP70,SUPT4H1,LATS1,GNAS,PCNA,OSMR,SRP9,SULT1 |
| MRPS22,LAS1L,RBCK1,RCC1,SNRPB,DBI,BUD31,NPRL2,CTT |
| PKM,SNRNP70,SERPINB5,SRSF2,MRPS22,OSMR,SEMA4D, |
| MRPS22,LAS1L,HMGB1,RBCK1,BUD31,NPRL2,CTTN,EIF5A, |
| PKM,C1ORF43,SUPT4H1,LATS1,CCDC66,DYNLL1,CST3,GPR |
| PKM,C1ORF43,SUPT4H1,LATS1,CCDC66,DYNLL1,CST3,GPR |
| PKM,APP,C1ORF43,SUPT4H1,DYNLL1,CST3,GNAS,GPR137, |
| PKM,APP,C1ORF43,SUPT4H1,DYNLL1,CST3,GNAS,GPR137, |
| APP,CCDC66,DYNLL1,CST3,GNAS,GPR137,S100A13,BCAT2 |
| APP,CCDC66,DYNLL1,CST3,GNAS,GPR137,S100A13,BCAT2 |
| PKM,DYNLL1,CST3,GNAS,GPR137,MRPL2,CDCA3,PCNA,M |
| PKM,DYNLL1,CST3,GNAS,GPR137,MRPL2,CDCA3,PCNA,M |
| PKM,C1ORF43,LATS1,RPS24,SERPINB5,FAM189B,VKORC1, |
| PKM,C1ORF43,LATS1,RPS24,SERPINB5,FAM189B,VKORC1, |
| MRPL2,MYL6B,PTOV1,ARRDC1,PRMT2,PPP1CC,LIG1,TIM |
| SNRNP70,LATS1,FAM189B,VKORC1,PCNA,SEMA4D,PTOV1 |
| SNRNP70,LATS1,FAM189B,VKORC1,PCNA,SEMA4D,PTOV1 |
| MRPS22,LAS1L,CMC2,RNH1,RBCK1,RCC1,SNRPB,ZC3HAV1 |
| SNRNP70,APP,SUPT4H1,GNAS,PCNA,SRSF2,OSMR,RPL38, |
| MRPL2,BAIAP2,MYL6B,FAM72A,PRR13,ARRDC1,NOP16,T |
| PKM,APP,C1ORF43,DYNLL1,CST3,GNAS,GPR137,BCAT2,CE |
| PKM,APP,C1ORF43,DYNLL1,CST3,GNAS,GPR137,BCAT2,CE |
| C1ORF43,DYNLL1,CST3,GNAS,GPR137,SERPINB5,FAM189 |
| APP,SUPT4H1,LATS1,DYNLL1,CST3,GNAS,RPS24,S100A13, |
| APP,SUPT4H1,LATS1,DYNLL1,CST3,GNAS,RPS24,S100A13, |
| DYNLL1,GPR137,MRPL2,CDCA3,PCNA,BAIAP2,RAPGEF5,M |
| PKM,C1ORF43,SUPT4H1,LATS1,CST3,GNAS,GPR137,RPS24 |
| PKM,C1ORF43,SUPT4H1,LATS1,CST3,GNAS,GPR137,RPS24 |
| SERPINB5,MRPL2,CDCA3,MYL6B,PRR13,PTOV1,ARRDC1,LI |
| SERPINB5,MRPL2,CDCA3,MYL6B,PRR13,PTOV1,ARRDC1,LI |
| LATS1,CCDC66,DYNLL1,CST3,RPS24,SERPINB5,MRPL4,FA |
| LATS1,CCDC66,DYNLL1,CST3,RPS24,SERPINB5,MRPL4,FA |
| PKM,SNRNP70,APP,C1ORF43,SERPINB5,SRSF2,SEMA4D,R |
| CCDC66,DYNLL1,CENPV,MRPL4,PPFIBP1,SSU72,FAM122C, |
| CCDC66,DYNLL1,CENPV,MRPL4,PPFIBP1,SSU72,FAM122C, |
| SNRNP70,APP,C1ORF43,CST3,GNAS,S100A13,CENPV,MRP |
| C1ORF43,DYNLL1,GNAS,GPR137,MRPL2,CDCA3,PCNA,BAI |
| C1ORF43,DYNLL1,GNAS,GPR137,MRPL2,CDCA3,PCNA,BAI |
| APP,SUPT4H1,DYNLL1,GNAS,S100A13,CENPV,FAM189B,M |
| APP,SUPT4H1,DYNLL1,GNAS,S100A13,CENPV,FAM189B,M |
| CCDC66,GPR137,RPS24,VKORC1,CDCA3,PCNA,SRSF2,KRT1 |
| C1ORF43,LATS1,GPR137,RPS24,VKORC1,MRPL2,CDCA3,SR |
| C1ORF43,LATS1,GPR137,RPS24,VKORC1,MRPL2,CDCA3,SR |

| |
|---|
| APP,SUPT4H1,LATS1,CCDC66,DYNLL1,CST3,RPS24,SERPIN |
| APP,SUPT4H1,LATS1,CCDC66,DYNLL1,CST3,RPS24,SERPIN |
| LATS1,GPR137,SRSF2,SSU72,SRP9,IQCG,YBX3,CACYBP,DH |
| LATS1,GPR137,SRSF2,SSU72,SRP9,IQCG,YBX3,CACYBP,DH |
| SUPT4H1,SERPINB5,MRPS22,LAS1L,RPL38,DHPS,BUD31,E |
| PKM,C1ORF43,LATS1,SERPINB5,MRPL4,FAM122C,BLOC1S |
| C1ORF43,CENPV,MRPL4,CDCA3,SRSF2,SULF2,BAIAP2,RAP |
| C1ORF43,CENPV,MRPL4,CDCA3,SRSF2,SULF2,BAIAP2,RAP |
| APP,DYNLL1,CST3,GNAS,SRSF2,PPFIBP1,SULF2,SSU72,CAC |
| PKM,SNRNP70,APP,C1ORF43,LATS1,GNAS,RPS24,SERPINB |
| PKM,SNRNP70,APP,C1ORF43,LATS1,GNAS,RPS24,SERPINB |
| PKM,SNRNP70,APP,C1ORF43,SUPT4H1,LATS1,CCDC66,DY |
| PKM,SNRNP70,APP,C1ORF43,SUPT4H1,LATS1,CCDC66,DY |
| C1ORF43,GNAS,GPR137,BCAT2,VKORC1,SRSF2,SSU72,TM |
| PKM,C1ORF43,SERPINB5,SEMA4D,HMGB1,YBX3,BLOC1S1, |
| PKM,C1ORF43,SERPINB5,SEMA4D,HMGB1,YBX3,BLOC1S1, |
| CMC2,RPL38,TRPT1,PMFBP1,NPRL2,CTTN,NUDT22,RPL13 |
| CMC2,RPL38,TRPT1,PMFBP1,NPRL2,CTTN,NUDT22,RPL13 |
| PKM,C1ORF43,LATS1,GPR137,FAM189B,PCNA,SSU72,FAM |
| SERPINB5,MRPL2,PCNA,SRSF2,HMGB1,BAIAP2,RAPGEF5, |
| APP,DYNLL1,CST3,GPR137,RPS24,SERPINB5,MRPL4,FAM1 |
| APP,DYNLL1,CST3,GPR137,RPS24,SERPINB5,MRPL4,FAM1 |
| CST3,GNAS,PCNA,BAIAP2,SRP9,FAM72A,BLOC1S1,STARD1 |
| CST3,GNAS,PCNA,BAIAP2,SRP9,FAM72A,BLOC1S1,STARD1 |
| PKM,APP,SUPT4H1,CCDC66,S100A13,MRPL4,VKORC1,PC |
| PKM,APP,SUPT4H1,CCDC66,S100A13,MRPL4,VKORC1,PC |
| HMGB1,TMEM67,RBCK1,SNRPB,DHPS,BUD31,CTTN,EIF5A, |
| HMGB1,TMEM67,RBCK1,SNRPB,DHPS,BUD31,CTTN,EIF5A, |
| SERPINB5,MRPL2,PCNA,BAIAP2,RAPGEF5,MYL6B,RHOF,P |
| PKM,C1ORF43,DYNLL1,CST3,GNAS,GPR137,BCAT2,MRPL2 |
| PKM,C1ORF43,DYNLL1,CST3,GNAS,GPR137,BCAT2,MRPL2 |
| APP,SUPT4H1,LATS1,CCDC66,DYNLL1,CST3,RPS24,SERPIN |
| APP,SUPT4H1,LATS1,CCDC66,DYNLL1,CST3,RPS24,SERPIN |
| CMC2,TRPT1,PMFBP1,NPRL2,CTTN,NUDT22,RPL13A,ARH |
| CMC2,TRPT1,PMFBP1,NPRL2,CTTN,NUDT22,RPL13A,ARH |
| APP,C1ORF43,CCDC66,DYNLL1,GNAS,CDCA3,SULF2,BAIAP |
| APP,C1ORF43,CCDC66,DYNLL1,GNAS,CDCA3,SULF2,BAIAP |
| SNRNP70,LATS1,DYNLL1,CST3,GNAS,GPR137,RPS24,BCAT |
| PKM,C1ORF43,SUPT4H1,LATS1,CCDC66,DYNLL1,CST3,GN |
| PKM,C1ORF43,SUPT4H1,LATS1,CCDC66,DYNLL1,CST3,GN |
| PKM,LATS1,SERPINB5,MRPL4,FAM122C,DNASE1L1,SNRPB |
| PKM,C1ORF43,LATS1,GPR137,FAM189B,PCNA,SULF2,SSU |
| PKM,C1ORF43,LATS1,GPR137,FAM189B,PCNA,SULF2,SSU |
| C1ORF43,GNAS,RPS24,SRSF2,RAPGEF5,PILRB,CMC2,CACY |
| C1ORF43,GNAS,RPS24,SRSF2,RAPGEF5,PILRB,CMC2,CACY |
| PKM,SNRNP70,APP,C1ORF43,LATS1,CCDC66,DYNLL1,GNA |
| PKM,SNRNP70,APP,C1ORF43,LATS1,CCDC66,DYNLL1,GNA |

| |
|---|
| GPR137,RPS24,VKORC1,SRSF2,MYL6B,SRP9,IQCG,ATG16L |
| LATS1,RPS24,SERPINB5,BCAT2,CDCA3,SMARCB1,RNH1,DN |
| LATS1,RPS24,SERPINB5,BCAT2,CDCA3,SMARCB1,RNH1,DN |
| PKM,SNRNP70,SUPT4H1,LATS1,CST3,GNAS,CENPV,MRPL4 |
| PKM,LATS1,DYNLL1,SERPINB5,MRPL4,MRPL2,FAM122C,B |
| PKM,LATS1,DYNLL1,SERPINB5,MRPL4,MRPL2,FAM122C,B |
| PKM,LATS1,DYNLL1,SERPINB5,MRPL4,MRPL2,FAM122C,B |
| SNRPB,ARPC2,CTTN,TSG101,RPS6,RTEL1,SLC39A13,USE1, |
| MRPL2,PCNA,SRSF2,BAIAP2,MYL6B,RHOF,PRR13,RNH1,DN |
| CMC2,RPL38,TRPT1,PMFBP1,NPRL2,CTTN,NUDT22,RPL13 |
| CMC2,RPL38,TRPT1,PMFBP1,NPRL2,CTTN,NUDT22,RPL13 |
| C1ORF43,SUPT4H1,LATS1,CCDC66,GPR137,RPS24,CDCA3, |
| C1ORF43,SUPT4H1,LATS1,CCDC66,GPR137,RPS24,CDCA3, |
| PKM,SNRNP70,C1ORF43,CST3,GNAS,GPR137,RPS24,CENP |
| PKM,SNRNP70,C1ORF43,CST3,GNAS,GPR137,RPS24,CENP |
| SNRNP70,APP,LATS1,DYNLL1,CST3,GNAS,GPR137,SERPIN |
| SNRNP70,APP,LATS1,DYNLL1,CST3,GNAS,GPR137,SERPIN |
| SERPINB5,MRPL2,PCNA,SRSF2,BAIAP2,RAPGEF5,MYL6B,R |
| PKM,SNRNP70,APP,C1ORF43,CCDC66,DYNLL1,CST3,RPS24 |
| PKM,SNRNP70,APP,C1ORF43,CCDC66,DYNLL1,CST3,RPS24 |
| C1ORF43,BAIAP2,CIRBP,RNH1,RNF32,ATG16L2,BOP1,TBL2 |
| APP,SUPT4H1,CCDC66,DYNLL1,CST3,GNAS,GPR137,S100A |
| APP,SUPT4H1,CCDC66,DYNLL1,CST3,GNAS,GPR137,S100A |
| MRPS22,LAS1L,SNRPB,DBI,ARPC2,LIMK2,BUD31,EIF5A,VR |
| DYNLL1,MRPL2,CDCA3,PCNA,BAIAP2,MYL6B,RHOF,SMAR |
| APP,CCDC66,DYNLL1,CST3,GPR137,SERPINB5,BCAT2,CEN |
| APP,CCDC66,DYNLL1,CST3,GPR137,SERPINB5,BCAT2,CEN |
| PKM,SUPT4H1,DYNLL1,CST3,GNAS,GPR137,SERPINB5,CEN |
| PKM,SUPT4H1,DYNLL1,CST3,GNAS,GPR137,SERPINB5,CEN |
| PKM,C1ORF43,DYNLL1,CST3,GNAS,RPS24,CENPV,VKORC1, |
| LATS1,CCDC66,SERPINB5,MRPL4,CDCA3,PPFIBP1,BAIAP2, |
| SUPT4H1,MRPS22,CIRBP,RPL38,DNASE1L1,RBCK1,DHPS,A |
| APP,C1ORF43,SUPT4H1,CCDC66,DYNLL1,GNAS,GPR137,B |
| APP,C1ORF43,SUPT4H1,CCDC66,DYNLL1,GNAS,GPR137,B |
| PKM,SNRNP70,APP,C1ORF43,RPS24,SERPINB5,CENPV,VK |
| PKM,SNRNP70,APP,C1ORF43,RPS24,SERPINB5,CENPV,VK |
| PKM,C1ORF43,DYNLL1,CST3,GNAS,SERPINB5,MRPL4,VKO |
| PKM,C1ORF43,DYNLL1,CST3,GNAS,SERPINB5,MRPL4,VKO |
| C1ORF43,RPS24,SRSF2,PILRB,CMC2,CACYBP,CIRBP,RPL38, |
| C1ORF43,RPS24,SRSF2,PILRB,CMC2,CACYBP,CIRBP,RPL38, |
| SNRNP70,APP,C1ORF43,DYNLL1,MRPL4,PCNA,SSU72,FAM |
| SNRNP70,APP,C1ORF43,DYNLL1,MRPL4,PCNA,SSU72,FAM |
| LATS1,DYNLL1,RPS24,SERPINB5,MRPL4,PPFIBP1,BAIAP2,S |
| LATS1,DYNLL1,RPS24,SERPINB5,MRPL4,PPFIBP1,BAIAP2,S |
| PKM,LATS1,DYNLL1,SRSF2,HDAC9,IQCG,TMEM67,RNF32, |
| PKM,C1ORF43,DYNLL1,RPS24,SERPINB5,BCAT2,CENPV,M |
| LATS1,SERPINB5,MRPL4,FAM122C,DNASE1L1,SNRPB,TCA |

| |
|---|
| LATS1,SERPINB5,MRPL4,FAM122C,DNASE1L1,SNRPB,TCAI |
| PKM,C1ORF43,LATS1,SERPINB5,MRPL4,MRPL2,FAM122C, |
| PKM,C1ORF43,LATS1,SERPINB5,MRPL4,MRPL2,FAM122C, |
| SNRNP70,APP,LATS1,CST3,GNAS,GPR137,SERPINB5,CENP |
| PKM,SNRNP70,APP,VKORC1,CDCA3,SRSF2,MRPS22,HDAC |
| PKM,SNRNP70,APP,VKORC1,CDCA3,SRSF2,MRPS22,HDAC |
| PKM,SNRNP70,APP,C1ORF43,RPS24,SERPINB5,CENPV,VKC |
| PKM,SNRNP70,APP,C1ORF43,RPS24,SERPINB5,CENPV,VKC |
| PKM,SNRNP70,APP,C1ORF43,RPS24,SERPINB5,CENPV,VKC |
| PKM,SNRNP70,APP,C1ORF43,RPS24,SERPINB5,CENPV,VKC |
| PKM,APP,CCDC66,DYNLL1,CST3,GPR137,RPS24,SERPINB5, |
| PKM,APP,CCDC66,DYNLL1,CST3,GPR137,RPS24,SERPINB5, |
| C1ORF43,SERPINB5,BLOC1S1,DHPS,SET,BOP1,SAE1,OSGEF |
| C1ORF43,SERPINB5,BLOC1S1,DHPS,SET,BOP1,SAE1,OSGEF |
| SNRNP70,PCNA,TCAIM,SCAF11,VDAC3,NOB1,SAE1,EIF5A, |
| PKM,SUPT4H1,LATS1,CST3,SERPINB5,BCAT2,OSMR,SEMA |
| PKM,SUPT4H1,LATS1,CST3,SERPINB5,BCAT2,OSMR,SEMA |
| PKM,APP,SERPINB5,SRSF2,SEMA4D,RAPGEF5,YBX3,STARD |
| PKM,SNRNP70,APP,C1ORF43,LATS1,CCDC66,DYNLL1,CST3 |
| PKM,SNRNP70,APP,C1ORF43,LATS1,CCDC66,DYNLL1,CST3 |
| C1ORF43,SUPT4H1,LATS1,CCDC66,GPR137,RPS24,BCAT2, |
| C1ORF43,SUPT4H1,LATS1,CCDC66,GPR137,RPS24,BCAT2, |
| SNRNP70,APP,SUPT4H1,LATS1,RPS24,SERPINB5,CENPV,M |
| SNRNP70,APP,SUPT4H1,LATS1,RPS24,SERPINB5,CENPV,M |
| PKM,LATS1,SERPINB5,MRPL4,FAM122C,DNASE1L1,SNRPB |
| SNRNP70,C1ORF43,DYNLL1,GNAS,RPS24,MRPL4,VKORC1, |
| SUPT4H1,CCDC66,DYNLL1,GPR137,RPS24,CENPV,MRPL4,F |
| SUPT4H1,CCDC66,DYNLL1,GPR137,RPS24,CENPV,MRPL4,F |
| BCAT2,CMC2,NPRL2,CTTN,ARHGEF39,PSMA6,ACOT8,SNRF |
| BCAT2,CMC2,NPRL2,CTTN,ARHGEF39,PSMA6,ACOT8,SNRF |
| APP,C1ORF43,LATS1,CCDC66,DYNLL1,CST3,GNAS,GPR137, |
| PKM,SNRNP70,APP,C1ORF43,GNAS,RPS24,SERPINB5,CENF |
| C1ORF43,SUPT4H1,LATS1,CCDC66,GPR137,RPS24,MRPL4, |
| C1ORF43,SUPT4H1,LATS1,CCDC66,GPR137,RPS24,MRPL4, |
| SERPINB5,MRPL4,FAM122C,TCAIM,KLC1,OSGEP,LDHA,RCE |
| MRPL4,CACYBP,SEC62,KLC1,VDAC3,NPRL2,OSGEP,ITSN1,A |
| SERPINB5,MRPL4,FAM122C,DNASE1L1,SNRPB,TCAIM,KLC |
| PKM,C1ORF43,SUPT4H1,LATS1,CCDC66,DYNLL1,CST3,GNA |
| SERPINB5,MRPL4,FAM122C,TCAIM,KLC1,OSGEP,LDHA,RCE |
| C1ORF43,SERPINB5,BLOC1S1,RNH1,SAE1,OSGEP,EIF5A,RP |
| PKM,SNRNP70,LATS1,CCDC66,SERPINB5,BCAT2,VKORC1,S |
| PKM,SNRNP70,LATS1,CCDC66,SERPINB5,BCAT2,VKORC1,S |
| LATS1,CCDC66,DYNLL1,CST3,SERPINB5,MRPL4,PPFIBP1,BA |
| LATS1,CCDC66,DYNLL1,CST3,SERPINB5,MRPL4,PPFIBP1,BA |
| PKM,SNRNP70,C1ORF43,SERPINB5,SRSF2,BLOC1S1,CIRBP, |
| PKM,SNRNP70,C1ORF43,SERPINB5,SRSF2,BLOC1S1,CIRBP, |
| PKM,SNRNP70,C1ORF43,SERPINB5,SRSF2,BLOC1S1,CIRBP, |

| |
|---|
| SNRNP70,SUPT4H1,GPR137,SERPINB5,CDCA3,MRPS22,HD |
| PKM,LATS1,SERPINB5,MRPL4,FAM122C,DNASE1L1,SNRPB |
| PKM,LATS1,SERPINB5,MRPL4,FAM122C,DNASE1L1,SNRPB |
| RPL38,DNASE1L1,SLC3A2,DHPS,TRPT1,BUD31,NPRL2,CTT |
| SUPT4H1,RPS24,MRPS22,LAS1L,HMGB1,IQCG,PILRB,RPL3 |
| SUPT4H1,RPS24,MRPS22,LAS1L,HMGB1,IQCG,PILRB,RPL3 |
| PKM,SNRNP70,CCDC66,DYNLL1,GPR137,RPS24,S100A13, |
| PKM,SNRNP70,CCDC66,DYNLL1,GPR137,RPS24,S100A13, |
| PKM,APP,C1ORF43,SUPT4H1,DYNLL1,CST3,GNAS,GPR137, |
| PKM,APP,C1ORF43,SUPT4H1,DYNLL1,CST3,GNAS,GPR137, |
| SUPT4H1,CCDC66,DYNLL1,S100A13,CENPV,MRPL2,SRSF2, |
| STARD10,ZNF235,NOP16,FIS1,PPRC1,RTEL1,MED15,TBX6, |
| C1ORF43,SUPT4H1,LATS1,CCDC66,GPR137,RPS24,SERPIN |
| C1ORF43,SUPT4H1,LATS1,CCDC66,GPR137,RPS24,SERPIN |
| DYNLL1,GNAS,GPR137,SERPINB5,MRPL2,CDCA3,PCNA,SRS |
| C1ORF43,SUPT4H1,LATS1,CCDC66,GPR137,RPS24,VKORC1 |
| C1ORF43,SUPT4H1,LATS1,CCDC66,GPR137,RPS24,VKORC1 |
| SUPT4H1,SERPINB5,MRPL4,PPFIBP1,SSU72,FAM122C,RNH |
| SUPT4H1,SERPINB5,MRPL4,PPFIBP1,SSU72,FAM122C,RNH |
| PKM,SNRNP70,APP,C1ORF43,SUPT4H1,LATS1,CCDC66,DY |
| SNRNP70,APP,DYNLL1,VKORC1,CDCA3,HDAC9,OSMR,LAS1 |
| SNRNP70,APP,DYNLL1,VKORC1,CDCA3,HDAC9,OSMR,LAS1 |
| DYNLL1,FAM122C,IQCG,PRR13,RCC1,TCAIM,ZC3HAV1,DBI |
| PKM,SNRNP70,APP,DYNLL1,GNAS,GPR137,SERPINB5,S10 |
| PKM,SNRNP70,APP,DYNLL1,GNAS,GPR137,SERPINB5,S10 |
| PKM,C1ORF43,GPR137,PCNA,SSU72,FAM122C,TMEM67,F |
| SNRNP70,C1ORF43,MRPL4,CDCA3,SRSF2,MYL6B,FAM72A, |
| SNRNP70,C1ORF43,MRPL4,CDCA3,SRSF2,MYL6B,FAM72A, |
| SERPINB5,KRT15,CIRBP,SMARCB1,RNH1,PTOV1,FASTK,AR |
| PKM,SNRNP70,APP,C1ORF43 |
| PKM,SNRNP70,APP,C1ORF43 |
| LATS1,SERPINB5,MRPL4,FAM122C,DNASE1L1,SNRPB,TCAI |
| LATS1,SERPINB5,MRPL4,FAM122C,DNASE1L1,SNRPB,TCAI |
| MRPL4,FAM122C,TCAIM,KLC1,OSGEP,LDHA,RPS9,RCE1,ZN |
| SERPINB5,MRPL4,MRPL2,PPFIBP1,FAM122C,CACYBP,TCAI |
| PKM,SNRNP70,APP,C1ORF43,RPS24,SERPINB5,CENPV,FAN |
| PKM,SNRNP70,APP,C1ORF43,RPS24,SERPINB5,CENPV,FAN |
| PKM,SNRNP70,APP,C1ORF43,RPS24,SERPINB5,CENPV,FAN |
| SNRNP70,SERPINB5,BCAT2,CDCA3,SMARCB1,RNH1,DNASE |
| SNRNP70,SERPINB5,BCAT2,CDCA3,SMARCB1,RNH1,DNASE |
| SERPINB5,MRPL4,PPFIBP1,FAM122C,CACYBP,TCAIM,KLC1, |
| LATS1,DYNLL1,SERPINB5,MRPL4,SSU72,FAM122C,TMEM6 |
| LATS1,DYNLL1,SERPINB5,MRPL4,SSU72,FAM122C,TMEM6 |
| SERPINB5,MRPL4,PPFIBP1,FAM122C,TCAIM,KLC1,OSGEP,L |
| MRPS22,SAE1,CTTN,EIF5A,COMMD5,IGSF8,NUDT5,CPSF3 |
| CST3,HMGB1,NOB1,NUMA1,ARHGAP6,AMZ2,FIS1,PPM1K, |
| CST3,HMGB1,NOB1,NUMA1,ARHGAP6,AMZ2,FIS1,PPM1K, |

| |
|---|
| PKM,SNRNP70,C1ORF43,SUPT4H1,LATS1,CCDC66,DYNLL1 |
| PKM,SNRNP70,C1ORF43,SUPT4H1,LATS1,CCDC66,DYNLL1 |
| PKM,SNRNP70,APP,C1ORF43,CST3,GNAS,BCAT2,CENPV,F |
| PKM,SNRNP70,APP,C1ORF43,CST3,GNAS,BCAT2,CENPV,F |
| MRPS22,LAS1L,RBCK1,BUD31,CTTN,EIF5A,COMMD5,CPSF |
| PKM,SNRNP70,APP,C1ORF43,RPS24,SERPINB5,VKORC1,SR |
| PKM,SNRNP70,APP,C1ORF43,RPS24,SERPINB5,VKORC1,SR |
| PKM,SNRNP70,APP,C1ORF43,LATS1,DYNLL1,CST3,GNAS,G |
| C1ORF43,SUPT4H1,LATS1,CCDC66,CST3,GNAS,SERPINB5,S |
| C1ORF43,VKORC1,CDCA3,SRSF2,OSMR,BAIAP2,YBX3,BLO |
| C1ORF43,VKORC1,CDCA3,SRSF2,OSMR,BAIAP2,YBX3,BLO |
| PKM,APP,C1ORF43,SUPT4H1,LATS1,CCDC66,DYNLL1,GNA |
| PKM,APP,C1ORF43,SUPT4H1,LATS1,CCDC66,DYNLL1,GNA |
| STYXL1,CDK11A,TRMT112,DDOST,TOMM40,MDH2,SEC31 |
| SUPT4H1,CCDC66,DYNLL1,SERPINB5,MRPL4,PPFIBP1,BAIA |
| SUPT4H1,CCDC66,DYNLL1,SERPINB5,MRPL4,PPFIBP1,BAIA |
| PKM,SNRNP70,APP,C1ORF43,CCDC66,GNAS,RPS24,SERPIN |
| PKM,SNRNP70,APP,C1ORF43,CCDC66,GNAS,RPS24,SERPIN |
| PKM,SNRNP70,C1ORF43,SUPT4H1,LATS1,DYNLL1,RPS24,S |
| PKM,SNRNP70,C1ORF43,SUPT4H1,LATS1,DYNLL1,RPS24,S |
| PKM,SNRNP70,C1ORF43,SUPT4H1,CCDC66,DYNLL1,SERPI |
| PKM,SNRNP70,C1ORF43,SUPT4H1,CCDC66,DYNLL1,SERPI |
| PKM,SNRNP70,SUPT4H1,LATS1,CST3,GNAS,BCAT2,FAM18 |
| PKM,SNRNP70,SUPT4H1,LATS1,CST3,GNAS,BCAT2,FAM18 |
| C1ORF43,SUPT4H1,CST3,SERPINB5,CENPV,FAM189B,MRP |
| C1ORF43,SUPT4H1,CST3,SERPINB5,CENPV,FAM189B,MRP |
| PKM,SNRNP70,C1ORF43,SUPT4H1,LATS1,DYNLL1,GNAS,G |
| PKM,SNRNP70,C1ORF43,SUPT4H1,LATS1,DYNLL1,GNAS,G |
| SNRNP70,FAM189B,VKORC1,MRPL2,CDCA3,SRSF2,MRPS2 |
| C1ORF43,SUPT4H1,LATS1,CCDC66,GPR137,RPS24,VKORC1 |
| C1ORF43,SUPT4H1,LATS1,CCDC66,GPR137,RPS24,VKORC1 |
| APP,SUPT4H1,LATS1,CCDC66,DYNLL1,CST3,GNAS,RPS24,S |
| APP,SUPT4H1,LATS1,CCDC66,DYNLL1,CST3,GNAS,RPS24,S |
| PKM,SNRNP70,APP,C1ORF43,SUPT4H1,CCDC66,DYNLL1,C |
| PKM,SNRNP70,APP,C1ORF43,SUPT4H1,CCDC66,DYNLL1,C |
| C1ORF43,SUPT4H1,LATS1,GPR137,RPS24,BCAT2,MRPL2,C |
| C1ORF43,SUPT4H1,LATS1,GPR137,RPS24,BCAT2,MRPL2,C |
| APP,C1ORF43,LATS1,CCDC66,GNAS,RPS24,S100A13,CENP |
| APP,C1ORF43,LATS1,CCDC66,GNAS,RPS24,S100A13,CENP |
| PKM,APP,C1ORF43,LATS1,RPS24,FAM189B,VKORC1,CDCA |
| PKM,APP,C1ORF43,LATS1,RPS24,FAM189B,VKORC1,CDCA |
| PKM,SNRNP70,APP,C1ORF43,LATS1,CCDC66,DYNLL1,GNA |
| PKM,APP,SUPT4H1,LATS1,CCDC66,DYNLL1,CST3,RPS24,SE |
| PKM,APP,SUPT4H1,LATS1,CCDC66,DYNLL1,CST3,RPS24,SE |
| PKM,SNRNP70,APP,C1ORF43,RPS24,SERPINB5,CENPV,SRS |
| PKM,SNRNP70,APP,C1ORF43,RPS24,SERPINB5,CENPV,SRS |
| PKM,SNRNP70,APP,C1ORF43,RPS24,SERPINB5,CENPV,VK |

| |
|---|
| PKM,SNRNP70,APP,C1ORF43,RPS24,SERPINB5,CENPV,VKC |
| SNRNP70,APP,CCDC66,GNAS,RPS24,SERPINB5,S100A13,VI |
| SNRNP70,APP,CCDC66,GNAS,RPS24,SERPINB5,S100A13,VI |
| SERPINB5,MRPL4,PPFIBP1,SSU72,FAM122C,CACYBP,DNAS |
| PKM,SNRNP70,C1ORF43,SERPINB5,CENPV,SRSF2,BLOC1S1 |
| PKM,SNRNP70,C1ORF43,SERPINB5,CENPV,SRSF2,BLOC1S1 |
| PKM,SNRNP70,C1ORF43,SERPINB5,CENPV,SRSF2,BLOC1S1 |
| PKM,C1ORF43,SUPT4H1,DYNLL1,CST3,GNAS,GPR137,BCA |
| PKM,C1ORF43,SUPT4H1,DYNLL1,CST3,GNAS,GPR137,BCA |
| LATS1,SEMA4D,SNRPB,BUD31,CTTN,AMZ2,MRPL33,WDR6 |
| LATS1,SEMA4D,SNRPB,BUD31,CTTN,AMZ2,MRPL33,WDR6 |
| LATS1,DYNLL1,CST3,RPS24,SERPINB5,MRPL4,BAIAP2,SSU7 |
| LATS1,DYNLL1,CST3,RPS24,SERPINB5,MRPL4,BAIAP2,SSU7 |
| PKM,SNRNP70,C1ORF43,SUPT4H1,DYNLL1,GPR137,RPS24 |
| C1ORF43,SUPT4H1,DYNLL1,GNAS,GPR137,BCAT2,FAM189 |
| C1ORF43,SUPT4H1,DYNLL1,GNAS,GPR137,BCAT2,FAM189 |
| PKM,C1ORF43,SERPINB5,RAPGEF5,IQCG,YBX3,STARD10,R |
| PKM,C1ORF43,SERPINB5,RAPGEF5,IQCG,YBX3,STARD10,R |
| PKM,SNRNP70,APP,C1ORF43,LATS1,CCDC66,DYNLL1,CST3 |
| PKM,SNRNP70,APP,C1ORF43,LATS1,CCDC66,DYNLL1,CST3 |
| SERPINB5,MRPL4,PPFIBP1,FAM122C,CACYBP,TCAIM,KLC1, |
| PKM,VKORC1,MRPL2,OSMR,PPFIBP1,RAPGEF5,IQCG,SULT |
| PKM,VKORC1,MRPL2,OSMR,PPFIBP1,RAPGEF5,IQCG,SULT |
| C1ORF43,SERPINB5,BLOC1S1,DHPS,SET,SAE1,OSGEP,TESC, |
| C1ORF43,SERPINB5,BLOC1S1,DHPS,SET,SAE1,OSGEP,TESC, |
| SNRNP70,GPR137,S100A13,PCNA,OSMR,FAM72A,CIRBP,R |
| SNRNP70,GPR137,S100A13,PCNA,OSMR,FAM72A,CIRBP,R |
| PKM,C1ORF43,LATS1,GPR137,FAM189B,VKORC1,PCNA,OS |
| C1ORF43,SUPT4H1,LATS1,CCDC66,GPR137,RPS24,SERPINI |
| C1ORF43,SUPT4H1,LATS1,CCDC66,GPR137,RPS24,SERPINI |
| HMGB1,YBX3,TAZ,LRRC1,GRK6,RPS16,ZBTB44,RPS6,RPL29 |
| SNRNP70,GNAS,BCAT2,PCNA,ZNF235,ZC3HAV1,SAE1,SCAF |
| PKM,GNAS,VKORC1,CDCA3,HMGB1,SNRPB,RPL13A,PCBP2 |
| PKM,RPS24,ZNF410,RUVBL2,DST,PHF19,SON,CELF1,VIM,T |
| PKM,APP,DYNLL1 |
| PKM,C1ORF43,SET,BOP1,NOB1,ITSN1,EIF5A,TUBA1C,UBR4 |
| SRSF2,OSMR,TMEM67,SET,NUMA1 |
| PKM,SNRNP70,DYNLL1,RPS24,BCAT2,FAM189B,MRPL2,OS |

| HP |
| --- |
| 33,RPS16,RPS25,RPS6,RPL29,RPL28,RPL21,RPL36,RPS20,RPS15,RPL18,RPL15,MRPS18C,RPS11,MRPS17,RF |
| .,SRP9,YBX3,CIRBP,RPL38,FASTK,SNRPB,SLC3A2,BOP1,ZC3HAV1,SCAF11,TBL2,EIF5A,NOP16,RPL13A,PSMA |
| JFB3,NDUFB2,NDUFA10,NDUFA7,NDUFS2,NDUFC1,CRYZL1,NDUFB10,NDUFS7,NDUFS4,TP53I3,NDUFA8 |
| 2,NUMA1,CRYGS,RPL13A,RPS9,MRPL33,TUBA1C,DST,VCAN,RPS16,RPS25,RPS6,RPL29,JUP,RPL28,VIM,RPL |
| 31L,HMGB1,SRP9,YBX3,CIRBP,SMARCB1,RPL38,DNASE1L1,KIF4A,FASTK,RCC1,SNRPB,SLC3A2,SET,ZNF235, |
| )UFS3,NDUFA3,NQO2,AIFM2,DHRS4,NDUFB3,NDUFB2,NDUFA10,NDUFA7,NDUFS2,NDUFC1,CRYZL1,NDUI |
| ,GNAS,RPS24,SERPINB5,S100A13,BCAT2,MRPL4,FAM189B,CDCA3,PCNA,SRSF2,HDAC9,KRT15,OSMR,PPFI |
| 1,RANBP1,RPL29,JUP,PDLIM5,CAPG,LYPLA2,TMPO,HSP90AB1,LRRFIP1,PTPRB,DBNL,EPS15L1,PKP4,EEF1D |
| YMS,BRF1,EIF2B3,ELOF1,EIF4A1,EIF3G,MTIF2,EEF2K,EIF2A,EIF3C,RPS27L |
| BXN1,HSPB1,RAD23A,UBE2N,RNF8,NUP62,NSFL1C |
| PC1,EEF1D,TYMS,BRF1,EIF2B3,ELOF1,EIF4A1,EIF3G,MTIF2,EEF2K,EIF2A,EIF3C,RPS27L,EIF2AK4,EIF6,SHMT |
| )UFA7,NDUFS2,NDUFC1,NDUFB10,NDUFS7,NDUFS4,NDUFA8 |
| )UFA7,NDUFS2,NDUFC1,NDUFB10,NDUFS7,NDUFS4,NDUFA8 |
| )UFA7,NDUFS2,NDUFC1,NDUFB10,NDUFS7,NDUFS4,NDUFA8 |
| )F1,EIF4A1,EIF3G,MTIF2,EEF2K,EIF2A,EIF3C |
| NA,SRSF2,LAS1L,HMGB1,SRP9,YBX3,CIRBP,RHOF,SMARCB1,RPL38,DNASE1L1,KIF4A,FASTK,RCC1,SNRPB,SI |
| BXN1,HSPB1,RAD23A,UBE2N,RNF8,NUP62,PML,NSFL1C |
| IA,PCMT1,RANBP1,DST,NISCH,RPL29,ITGB1BP1,JUP,PDLIM5,CAPG,LYPLA2,CAMLG,TMPO,HSP90AB1,LRRF |
| NA,SRSF2,LAS1L,HMGB1,SRP9,YBX3,CIRBP,RHOF,SMARCB1,RPL38,DNASE1L1,KIF4A,FASTK,RCC1,SNRPB,SI |
| PS3,GTF3A,RPS4X,ERAL1,MRPS7,RPS14 |
| 3,EIF4A1,STAU2,RC3H2 |
| ,RPL29,RPL28,RPL21,ZFAND2B,RPL36,RPS20,RPS15,RPL18,RPL15,RPS11,RPL37A,RPLP0,SSR3,RPL8,RPL35 |
| ),RPL28,RPL21,ZFAND2B,RPL36,RPS20,RPS15,RPL18,RPL15,RPS11,RPL37A,RPLP0,SSR3,RPL8,RPL35,RPL34 |
| RPL38,RNH1,SNRPB,SET,ZC3HAV1,SCAF11,BUD31,RPL13A,PSMA6,RPS9,SNRPN,ZC3H14,PCBP2,SON,RPS16 |
| 28,RPL21,RPL36,RPS20,RPS15,PABPC1,RPL18,RPL15,RPS11,RPL37A,RPLP0,RPL8,RPL35,RPL34,RPL24,RPS3 |
| ,RPL29,RPL28,RPL21,ZFAND2B,RPL36,RPS20,RPS15,RPL18,RPL15,RPS11,RPL37A,RPLP0,SSR3,RPL8,RPL35 |
| ,RPL29,RPL28,RPL21,ZFAND2B,RPL36,RPS20,RPS15,RPL18,RPL15,RPS11,RPL37A,RPLP0,SSR3,RPL8,RPL35 |
| G16L2,RCC1,SET,KLC1,ZC3HAV1,SCARB1,ITSN1,RPL13A,ACOT8,RPS9,RANBP1,PCBP2,SNF8,HMGA2,RPS16 |
| M,RPL28,RPL21,RPL36,RPS20,RPS15,HSPB1,MCTS1,PABPC1,RPL18,RPL15,BRF1,EIF2B3,RPS11,RPL37A,EIF |
| ,RPL29,RPL28,RPL21,ZFAND2B,RPL36,RPS20,RPS15,RPL18,RPL15,RPS11,RPL37A,RPLP0,SSR3,RPL8,RPL35 |
| G16L2,RCC1,SET,KLC1,ZC3HAV1,SCARB1,ITSN1,RPL13A,ACOT8,RPS9,RANBP1,PCBP2,SNF8,HMGA2,RPS16 |
| AE1,CTTN,EIF5A,UFD1L,RPL13A,GSTK1,ACOT8,AP3B2,RPS9,FIS1,LRRC1,SNF8,CDC37,RPS16,TTC21B,RPS25 |
| IGA2,RPS16,RPS25,RPS6,RPL29,RPL28,RPL21,RPL36,RPS20,RPS15,MCTS1,RPL18,POLR2C,RPL15,SMARCA |
| NH1,DNASE1L1,RBCK1,SET,CTSC,ARRDC1,ZC3HAV1,SIRT6,UFD1L,USP38,RPL13A,PSMA6,RPS9,UBE2L6,ZC3 |
| 38,DHPS,EIF5A,RPL13A,RPS9,MRPL33,RPS16,CHCHD1,METTL17,RPS25,RPS6,RPL29,EIF3M,CELF1,EEF1B2,F |
| 38,DHPS,EIF5A,RPL13A,RPS9,MRPL33,RPS16,CHCHD1,METTL17,RPS25,RPS6,RPL29,EIF3M,CELF1,EEF1B2,F |
| RPL38,KIF4A,SNRPB,PARD3,SEC62,CTSC,WDR19,NUMA1,SAE1,CTTN,EIF5A,UFD1L,RPL13A,GSTK1,ACOT8,A |
| IA6,RPS9,ZC3H14,RPS16,PSMA1,PSMA3,RPS25,RPS6,RPL29,PSMD4,RPL28,VIM,PSMA7,RPL21,EXOSC9,HI |
| L1,SET,ZC3HAV1,RPL13A,PSMA6,RPS9,NUDT5,ZC3H14,TET2,RPS16,PSMA1,PSMA3,RPS25,RPS6,RPL29,PS |
| G16L2,RCC1,SET,KLC1,ZC3HAV1,SCARB1,ITSN1,RPL13A,ACOT8,RPS9,RANBP1,PCBP2,SNF8,HMGA2,RPS16 |
| EC62,WDR19,NUMA1,SAE1,CTTN,ITSN1,EIF5A,UFD1L,RPL13A,GSTK1,ACOT8,AP3B2,RPS9,RUVBL2,TESC,L |
| ),RUVBL2,FIS1,LRRC1,SNF8,RPS16,TSG101,RPS25,RPS6,RPL29,JUP,RPL28,TIMM17B,RPL21,NUTF2,ZFAND |
| 3HAV1,RPL13A,PSMA6,RPS9,NUDT5,ZC3H14,RPS16,PSMA1,PSMA3,RPS25,RPS6,RPL29,PSMD4,RPL28,VIN |

EC62,WDR19,NUMA1,SAE1,CTTN,ITSN1,EIF5A,UFD1L,RPL13A,GSTK1,ACOT8,AP3B2,RPS9,RUVBL2,TESC,LZ
P,RPL38,RNH1,DNASE1L1,RBCK1,SET,CTSC,ARRDC1,ZC3HAV1,SIRT6,UFD1L,USP38,RPL13A,PSMA6,RPS9,U
PS16,RPS25,RPS6,RPL29,RPL28,RPL21,RPL36,RPS20,RPS15,RPL18,POLR2C,RPL15,SMARCA4,RPS11,NUP13
38,DHPS,EIF5A,RPL13A,GSTK1,RPS9,MRPL33,RPS16,CHCHD1,METTL17,RPS25,RPS6,RPL29,EIF3M,CELF1,EE
L1,SET,ZC3HAV1,SCARB1,RPL13A,PSMA6,RPS9,NUDT5,ZC3H14,TET2,RPS16,PSMA1,PSMA3,RPS25,RPS6,R
OT8,RPS9,FIS1,CDC37,RPS16,RPS25,RPS6,RPL29,ITGB1BP1,RPL28,TIMM17B,RPL21,ZFAND2B,AP1S3,RPL36
28,RPL21,EXOSC9,RPL36,RPS20,RPS15,PABPC1,RPL18,RPL15,RPS11,PCID2,RPL37A,RPLP0,SSB,RPL8,RPL35
C3HAV1,RPL13A,PSMA6,RPS9,NUDT5,ZC3H14,TET2,RPS16,PSMA1,PSMA3,RPS25,RPS6,RPL29,PSMD4,RPL2
C3HAV1,RPL13A,PSMA6,RPS9,NUDT5,ZC3H14,TET2,RPS16,PSMA1,PSMA3,RPS25,RPS6,RPL29,PSMD4,RPL2
38,DHPS,EIF5A,RPL13A,RPS9,MRPL33,RPS16,CHCHD1,METTL17,RPS25,RPS6,RPL29,EIF3M,CELF1,EEF1B2,R
MA6,RPS9,ZC3H14,RPS16,PSMA1,PSMA3,RPS25,RPS6,RPL29,PSMD4,RPL28,VIM,PSMA7,RPL21,EXOSC9,HN
SRP9,TMEM67,BLOC1S1,RPL38,KIF4A,SNRPB,PARD3,SEC62,CTSC,WDR19,NUMA1,SAE1,CTTN,ITSN1,EIF5A
6,RPS25,RPS6,RPL29,ITGB1BP1,RPL28,RPL21,ZFAND2B,RPL36,RPS20,RPS15,RPL18,RPL15,RPS11,MFF,RPL3
RPL13A,GSTK1,ACOT8,RPS9,RUVBL2,LZTFL1,FIS1,LRRC1,SNF8,RPS16,TSG101,OBSL1,TTC21B,RPS25,RPS6,R
EM67,SULT1A3,YBX3,CIRBP,RPL38,RNH1,DNASE1L1,RBCK1,SET,CTSC,ARRDC1,ZC3HAV1,SIRT6,SCARB1,UF
38,DHPS,DBI,EIF5A,RPL13A,GSTK1,ACOT8,RPS9,MRPL33,RPS16,CHCHD1,METTL17,RPS25,THEM4,RPS6,RP
PB,SET,BOP1,SCAF11,ARPC2,NUMA1,ARHGAP6,CTTN,TAZ,RPL13A,RUVBL2,ECSIT,CENPK,SRSF1,FOXRED1,
2,PILRB,SMARCB1,RPL38,RCC1,SNRPB,PARD3,ACTR6,SET,CTSC,BOP1,KLC1,SCAF11,ARPC2,WDR19,NUMA
TMEM67,SULT1A3,YBX3,CIRBP,RPL38,RNH1,DNASE1L1,RBCK1,ATG16L2,SET,CTSC,ARRDC1,ZC3HAV1,SIRT
FASTK,SNRPB,BOP1,TRPT1,SCAF11,NOB1,BUD31,OSGEP,RPS9,RBPMS,SNRPN,ZC3H14,PCBP2,SON,RPS16,
GB1,TMEM67,SULT1A3,YBX3,CIRBP,RPL38,RNH1,DNASE1L1,RBCK1,ATG16L2,SET,CTSC,ARRDC1,ZC3HAV1
9,RPL38,ATG16L2,PARD3,SEC62,ARRDC1,WDR19,SAE1,CTTN,ITSN1,EIF5A,UFD1L,RPL13A,GSTK1,ACOT8,AP
9,RPL38,ATG16L2,PARD3,SEC62,ARRDC1,WDR19,SAE1,CTTN,ITSN1,EIF5A,UFD1L,RPL13A,GSTK1,ACOT8,AP
AS1L,HMGB1,BAIAP2,IQCG,TMEM67,PILRB,CIRBP,RHOF,SMARCB1,RPL38,KIF4A,ATG16L2,RCC1,SNRPB,PA
PL13A,RPS9,RUVBL2,RPS16,NIP7,RPS6,SRSF1,EIF3M,CELF1,FBXW9,TXNL4A,EXOSC9,PPAN,HSP90AB1,TRM
3,CIRBP,RPL38,STARD10,DHPS,EIF5A,TAZ,RPL13A,GSTK1,ACOT8,RPS9,MRPL33,NNMT,IL1RAP,VCAN,TET2,
,RPS16,RPS25,RPS6,RPL29,ITGB1BP1,RPL28,RPL21,ZFAND2B,RPL36,RPS20,RPS15,PDCD5,RPL18,CNST,EG
9,RPL38,ATG16L2,PARD3,SEC62,ARRDC1,WDR19,SAE1,CTTN,ITSN1,EIF5A,UFD1L,RPL13A,GSTK1,ACOT8,AP
9,RPL38,ATG16L2,PARD3,SEC62,ARRDC1,WDR19,SAE1,CTTN,ITSN1,EIF5A,UFD1L,RPL13A,GSTK1,ACOT8,AP
C9,HMGB1,BAIAP2,SRP9,RPL38,ATG16L2,PARD3,SEC62,ARRDC1,WDR19,NUMA1,SAE1,CTTN,ITSN1,EIF5A,
1,SNRPN,PCBP2,SON,SRSF1,CLASRP,CELF1,TXNL4A,ARL6IP4,HNRNPC,RBM3,PRKRIP1,PABPC1,RBMX,DDX4
DAC9,SEMA4D,LAS1L,HMGB1,SSU72,SULT1A3,UAP1,YBX3,CIRBP,SMARCB1,RPL38,RNH1,DNASE1L1,RBCK1
3,RCC1,SNRPB,PARD3,SET,CTSC,BOP1,SCAF11,ARPC2,WDR19,NUMA1,ARHGAP6,CTTN,TAZ,RPL13A,RUVBL
,PCNA,SRSF2,MRPS22,HDAC9,SEMA4D,LAS1L,HMGB1,SSU72,SRP9,SULT1A3,UAP1,YBX3,CIRBP,SMARCB1,
T15,HMGB1,BAIAP2,IQCG,TMEM67,BLOC1S1,CIRBP,RHOF,SMARCB1,RPL38,KIF4A,TMEM135,ATG16L2,RC
EIF3M,CELF1,TXNL4A,PPAN,HSP90AB1,RPS15,MCTS1,RBMX,SNRPF,ZNHIT3,NOP2,USP4,RPLP0,EIF3G,RPL2
,CELF1,TXNL4A,PPAN,HSP90AB1,RPS15,MCTS1,RBMX,SNRPF,ZNHIT3,NOP2,USP4,RPLP0,EIF3G,RPL24,GEM
4,SON,PSMA1,PSMA3,SRSF1,CELF1,PSMD4,VIM,PSMA7,EXOSC9,HNRNPC,CDK11A,RBM3,HSPB1,PSME2,PA
P2,SON,SRSF1,CELF1,TXNL4A,HNRNPC,RBM3,PABPC1,RBMX,DDX41,HNRNPA1,POLR2C,SNRPF,RBFOX2,PP
SC,LZTFL1,FIS1,LRRC1,TM9SF1,RPS16,RPS25,RPS6,RPL29,ITGB1BP1,JUP,VAMP8,PPIL2,RPL28,RPL21,ZFAN
OX20,COA1,NDUFS3,NDUFA11,NDUFA3,COX19,NDUFB3,NDUFB2,NDUFA10,NDUFA7,NDUFS2,NDUFC1,ND
G12,PPRC1,THEM4,FOXRED1,TIMM17B,TMEM126B,STOML2,SHARPIN,ATP5J,GABPB1,DCN,TOMM40,PDC
DAC9,SEMA4D,LAS1L,HMGB1,SSU72,SULT1A3,UAP1,YBX3,CIRBP,SMARCB1,RPL38,RNH1,DNASE1L1,RBCK1
A6,UBE2L6,PCBP2,SNF8,TSG101,UBR4,PSMA1,PSMA3,AUP1,UBE2I,XBP1,PTK2,PSMD4,PSMA7,CAMLG,EXO
TSC,ARRDC1,SIRT6,SCARB1,UFD1L,USP38,PSMA6,UBE2L6,PPM1K,VCAN,PCBP2,CD44,TET2,SNF8,TSG101,U
P2,SON,SRSF1,CELF1,TXNL4A,HNRNPC,RBM3,PABPC1,RBMX,DDX41,HNRNPA1,POLR2C,SNRPF,RBFOX2,PP
P2,SON,SRSF1,CELF1,TXNL4A,HNRNPC,RBM3,PABPC1,RBMX,DDX41,HNRNPA1,POLR2C,SNRPF,RBFOX2,PP

L15,DRG2,EIF2B3,EIF4A1,RPLP0,METTL3,EIF3G,RPL8,RPL24,RBM4,EIF3C,RPS23,RPL30,DNAJC24
9,HMGB1,BAIAP2,SRP9,RPL38,STARD10,ATG16L2,PARD3,SEC62,ARRDC1,WDR19,NUMA1,SAE1,SCARB1,
CDCA3,PCNA,MRPS22,HDAC9,SEMA4D,HMGB1,SSU72,SRP9,TMEM67,SULT1A3,PILRB,YBX3,BLOC1S1,CIR
,BAIAP2,IQCG,TMEM67,PILRB,CIRBP,RHOF,SMARCB1,RPL38,KIF4A,ATG16L2,RCC1,SNRPB,PARD3,SET,CTS
RPINB5,S100A13,CENPV,MRPL4,MRPL2,PCNA,MRPS22,HDAC9,KRT15,PPFIBP1,SEMA4D,LAS1L,HMGB1,S
PS22,HDAC9,SEMA4D,HMGB1,SSU72,SRP9,UAP1,YBX3,CIRBP,SMARCB1,RPL38,RBCK1,PTOV1,SNRPB,DH
AC9,SEMA4D,LAS1L,HMGB1,SSU72,SULT1A3,UAP1,YBX3,CIRBP,SMARCB1,RPL38,RNH1,DNASE1L1,RBCK
,PSMA6,UBE2L6,PCBP2,SNF8,TSG101,UBR4,PSMA1,PSMA3,AUP1,UGGT2,UBE2I,XBP1,PTK2,PSMD4,PSMA
NDUFA11,NDUFA3,NDUFB3,NDUFB2,NDUFA10,NDUFA7,NDUFS2,NDUFC1,NDUFB10,NDUFS7,NDUFS4,ND
NDUFA11,NDUFA3,NDUFB3,NDUFB2,NDUFA10,NDUFA7,NDUFS2,NDUFC1,NDUFB10,NDUFS7,NDUFS4,ND
A6,UBE2L6,PCBP2,SNF8,TSG101,UBR4,PSMA1,PSMA3,AUP1,UBE2I,XBP1,PTK2,PSMD4,PSMA7,CAMLG,ZF
1,BUD31,SNRPN,ZC3H14,PCBP2,SON,SRSF1,CLASRP,CELF1,TXNL4A,ARL6IP4,HNRNPC,CDK11A,RBM3,PRK
PS24,SERPINB5,BCAT2,MRPL4,VKORC1,MRPL2,CDCA3,PCNA,SRSF2,MRPS22,HDAC9,SEMA4D,LAS1L,HMG
A6,UBE2L6,PCBP2,SNF8,TSG101,UBR4,PSMA1,PSMA3,AUP1,UBE2I,XBP1,PTK2,PSMD4,PSMA7,CAMLG,ZF
SEMA4D,LAS1L,HMGB1,SSU72,YBX3,CIRBP,SMARCB1,RPL38,RNH1,DNASE1L1,RBCK1,PTOV1,FASTK,SNRP
AC9,SEMA4D,LAS1L,HMGB1,SSU72,SULT1A3,UAP1,YBX3,CIRBP,SMARCB1,RPL38,RNH1,DNASE1L1,RBCK
1B2,TSFM,EEF1D,MRPS18C,MRPS17,ELOF1,MRPL55,EEF2K,MRPS24,MRPS16,RPL30,DNAJC24,MRPS33,M
,RPL38,ATG16L2,PARD3,SLC3A2,SEC62,ARRDC1,VDAC3,WDR19,SAE1,CTTN,ITSN1,EIF5A,UFD1L,RPL13A,
HDAC9,SEMA4D,HMGB1,SSU72,SRP9,YBX3,CACYBP,CIRBP,SMARCB1,RPL38,RBCK1,PTOV1,SNRPB,DHPS,S
6,UFD1L,USP38,PSMA6,UBE2L6,PCBP2,SNF8,TSG101,UBR4,PSMA1,PSMA3,AUP1,UGGT2,UBE2I,XBP1,PTK
,PSMA6,UBE2L6,PCBP2,SNF8,TSG101,UBR4,PSMA1,PSMA3,AUP1,UGGT2,UBE2I,XBP1,PTK2,PSMD4,PSMA
PS22,HDAC9,SEMA4D,HMGB1,SSU72,SRP9,YBX3,CACYBP,CIRBP,SMARCB1,RPL38,RBCK1,PTOV1,SNRPB,
MRPL4,VKORC1,MRPL2,CDCA3,PCNA,MRPS22,HDAC9,SEMA4D,HMGB1,SULF2,SSU72,SRP9,TMEM67,SULT
RPS6,FBXW9,EXOSC9,PPAN,TRMT112,RPS15,RRNAD1,ZCCHC4,RRP8,NSUN5,NHP2,ZNHIT3,NOP2,MPHOS
NA,SRSF2,MRPS22,HDAC9,SEMA4D,HMGB1,SSU72,SRP9,UAP1,YBX3,CACYBP,CIRBP,SMARCB1,RPL38,STA
,S100A13,CENPV,MRPL4,MRPL2,PCNA,MRPS22,HDAC9,KRT15,PPFIBP1,SEMA4D,HMGB1,SULF2,BAIAP2,
MT112,MRPS18C,MRPS17,MRPL55,MRPS24,MRPS16,MRPS33,MRPL51,MRPL43,MRPL48,ERAL1,MRPL27,
,CTTN,EIF5A,MRPL33,FIS1,CD44,CDC37,TSG101,CHCHD1,ATG12,MICAL1,VAMP8,CAPG,APEH,CAPN10,SH
FA11,NDUFA3,NDUFB3,NDUFB2,NDUFA10,SDHA,NDUFA7,NDUFS2,NDUFC1,NDUFB10,NDUFS7,UQCRB,C
C3HAV1,SIRT6,NPRL2,SCARB1,CTTN,PSMA6,ZC3H14,SNF8,CDC37,PSMA1,PSMA3,XBP1,PTK2,PSMD4,VIM
NA,SRSF2,MRPS22,HDAC9,SEMA4D,HMGB1,SSU72,SRP9,UAP1,YBX3,CACYBP,CIRBP,SMARCB1,RPL38,STA
FA11,NDUFA3,NDUFB3,NDUFB2,NDUFA10,SDHA,NDUFA7,NDUFS2,NDUFC1,NDUFB10,NDUFS7,UQCRB,C
1,MRPL2,CDCA3,PCNA,MRPS22,HDAC9,SEMA4D,HMGB1,SULF2,SSU72,SRP9,TMEM67,SULT1A3,PILRB,Y
,SRSF7,NXF1,ZNF593,NUP62,THOC3,MAGOHB,U2AF1L4,NCBP2,RNPS1,NUP85,PABPN1,SEC13,EIF6,MAG
,MRPS17,MRPL55,MRPS24,MRPS16,MRPS33,MRPL51,MRPL43,MRPL48,ERAL1,MRPL27,MRPS7,MRPL36,
4D,LAS1L,HMGB1,SSU72,YBX3,CIRBP,SMARCB1,RPL38,RNH1,RBCK1,PTOV1,FASTK,SNRPB,SET,ZNF235,BO
RNPA1,RBFOX2,SRSF7,RBM42,RBM4,RNPS1,NSRP1,RBM7,IK,DDX17,SFSWAP,RBM4B,HNRNPL,MAGOH,SR
NA,SRSF2,MRPS22,HDAC9,SEMA4D,HMGB1,SSU72,SRP9,UAP1,YBX3,CACYBP,CIRBP,SMARCB1,RPL38,STA
,RNPS1,NSRP1,RBM7,HNRNPM,IK,DDX17,SFSWAP,RBM4B,HNRNPL,MAGOH,SRSF5,FXR1
PS24,SERPINB5,BCAT2,MRPL4,VKORC1,MRPL2,CDCA3,PCNA,SRSF2,MRPS22,HDAC9,SEMA4D,LAS1L,HMG
5,RCC1,SET,ARPC2,VDAC3,SIRT6,NUMA1,SAE1,NPRL2,ARHGAP6,CTTN,TNFSF10,SH2B1,RUVBL2,IL1RAP,ST
P90AB1,RPS15,HMGA1,HNRNPA1,SNRPF,PARK7,NFKBIA,ENY2,NUP133,RAPGEF3,RPAIN,PCID2,KPNA5,SS
,RBM3,RBMX,HNRNPA1,RBFOX2,SRSF7,RBM42,RBM4,NCBP2,RNPS1,NSRP1,RBM7,IK,DDX17,PABPN1,SFS
FA11,NDUFA3,AIFM2,NDUFB3,NDUFB2,NDUFA10,SDHA,NDUFA7,NDUFS2,NDUFC1,NDUFB10,NDUFS7,U
P133,RAPGEF3,PCID2,SSB,CHTOP,SRSF7,NXF1,ZNF593,NUP62,THOC3,TP53,MAGOHB,IFI27,U2AF1L4,NC
C3HAV1,SIRT6,NPRL2,SCARB1,CTTN,PSMA6,ZC3H14,CDC37,PSMA1,PSMA3,XBP1,PTK2,PSMD4,VIM,PSM
,RNPS1,NSRP1,RBM7,IK,DDX17,RBM4B,HNRNPL,MAGOH,SRSF5,FXR1

M,PSMA7,EXOSC9,HNRNPC,HSPB1,PSME2,PABPC1,PSME1,CARHSP1
P,SRSF7,NXF1,ZNF593,NUP62,THOC3,MAGOHB,U2AF1L4,NCBP2,RNPS1,NUP85,PABPN1,SEC13,EIF6,MAG
IT1,AK9
FB2,NDUFA10,NDUFA7,NDUFS2,NDUFC1,NDUFB10,NDUFS7,NDUFS4,NDUFA8
P90AB1,RPS15,HNRNPA1,SNRPF,PARK7,NFKBIA,ENY2,NUP133,RAPGEF3,RPAIN,PCID2,KPNA5,SSB,CHTOP,
PSMD4,VIM,PSMA7,EXOSC9,HNRNPC,HSPB1,PSME2,PABPC1,PSME1,CARHSP1
1,SIRT6,NPRL2,TBL2,UFD1L,PSMA6,RUVBL2,POLD4,UBE2L6,CD44,HMGA2,CDC37,PSMA1,PSMA3,MUC1,
I,EIF3C
VKORC1,MRPL2,CDCA3,PCNA,MRPS22,HDAC9,SEMA4D,HMGB1,SULF2,SSU72,SRP9,TMEM67,SULT1A3,PI
T3,NOP2,RPLP0,RPL35,RPL24,GTF3A
,RPS24,S100A13,BCAT2,CENPV,MRPL4,VKORC1,MRPL2,PCNA,SRSF2,MRPS22,HDAC9,KRT15,LAS1L,HMGB
NPV,CDCA3,SRSF2,KRT15,PPFIBP1,LAS1L,CYB561A3,BAIAP2,SSU72,MYL6B,SRP9,SULT1A3,UAP1,FAM72A
,RPS24,S100A13,BCAT2,CENPV,MRPL4,VKORC1,MRPL2,PCNA,SRSF2,MRPS22,HDAC9,KRT15,LAS1L,HMGB
,GNAS,GPR137,RPS24,S100A13,BCAT2,CENPV,MRPL4,VKORC1,MRPL2,PCNA,SRSF2,MRPS22,HDAC9,KRT1
24,SERPINB5,S100A13,BCAT2,CENPV,MRPL4,VKORC1,MRPL2,CDCA3,SRSF2,MRPS22,HDAC9,KRT15,PPFIB
,GNAS,GPR137,RPS24,SERPINB5,S100A13,BCAT2,CENPV,MRPL4,VKORC1,MRPL2,CDCA3,PCNA,SRSF2,MR
,GNAS,GPR137,RPS24,S100A13,BCAT2,CENPV,MRPL4,VKORC1,MRPL2,PCNA,SRSF2,MRPS22,HDAC9,KRT1
9,CIRBP,RPL38,SNRPB,BOP1,NOB1,BUD31,RPL13A,PSMA6,RPS9,MRPL33,RUVBL2,SNRPN,ZC3H14,RPS16,
,CENPV,MRPL4,MRPL2,PCNA,SRSF2,MRPS22,HDAC9,LAS1L,HMGB1,CYB561A3,BAIAP2,RAPGEF5,SSU72,U
,CENPV,MRPL4,MRPL2,PCNA,SRSF2,MRPS22,HDAC9,LAS1L,HMGB1,CYB561A3,BAIAP2,RAPGEF5,SSU72,U
,CENPV,MRPL4,MRPL2,PCNA,SRSF2,MRPS22,HDAC9,LAS1L,HMGB1,CYB561A3,BAIAP2,RAPGEF5,SSU72,U
100A13,CENPV,PCNA,SRSF2,HDAC9,KRT15,LAS1L,HMGB1,CYB561A3,BAIAP2,RAPGEF5,SSU72,UAP1,YBX3
33,RPS16,RPS25,RPS6,RPL29,RPL28,RPL21,RPL36,RBM3,RPS20,RPS15,MCTS1,RPL18,RPL15,MRPS18C,RPS
DAC9,LAS1L,HMGB1,CYB561A3,BAIAP2,RAPGEF5,SSU72,UAP1,CACYBP,CIRBP,SMARCB1,PRR13,RNH1,KIF
33,RPS16,CHCHD1,METTL17,RPS25,RPS6,RPL29,RPL28,RPL21,RPL36,RBM3,RPS20,RPS15,MCTS1,RPL18,R
28,RPL21,RPL36,RPS20,RPS15,MCTS1,RPL18,RPL15,RPS11,RPL37A,RPLP0,RPL8,RPL35,RPL34,EIF2A,RPL24,
DAC9,LAS1L,HMGB1,CYB561A3,BAIAP2,RAPGEF5,SSU72,UAP1,CACYBP,CIRBP,SMARCB1,PRR13,RNH1,KIF
,MRPL2,PCNA,SRSF2,MRPS22,HDAC9,OSMR,LAS1L,HMGB1,SSU72,MYL6B,SRP9,TMEM67,BLOC1S1,CACY
M135,RCC1,VDAC3,ITSN1,EIF5A,TAZ,MRPL33,RANBP1,FIS1,DST,UQCRQ,CHCHD1,ECSIT,COX4I1,SLC25A10,
M135,RCC1,VDAC3,ITSN1,EIF5A,TAZ,MRPL33,RANBP1,FIS1,DST,UQCRQ,CHCHD1,ECSIT,COX4I1,SLC25A10,
TS1,MRPS18C,RPS11,MRPS17,EIF2A,RPS3A,RPS3,MRPS24,RPS27L,RPS7,UBA52,RPS4X,MRPS16,RPS23,MF
1,EIF2A,RPS3A,RPS3,RPS27L,RPS7,UBA52,RPS4X,RPS23,RPS15A,RPS17,RPS14
IT,COX4I1,SLC25A10,THEM4,FOXRED1,C14ORF2,TIMM17B,TMEM126B,PPOX,STOML2,SLC9B2,ATP5J,TO
MRPL33,FIS1,UQCRQ,CHCHD1,ECSIT,COX4I1,SLC25A10,THEM4,COASY,FOXRED1,C14ORF2,AGPAT4,PPP1C
IT,COX4I1,SLC25A10,THEM4,FOXRED1,C14ORF2,TIMM17B,TMEM126B,NUTF2,PPOX,TMPO,STOML2,SLC
MRPL4,MRPL2,PCNA,MRPS22,KRT15,LAS1L,HMGB1,CYB561A3,BAIAP2,MYL6B,IQCG,TMEM67,CIRBP,RHO
MRPL4,MRPL2,PCNA,MRPS22,KRT15,LAS1L,HMGB1,CYB561A3,BAIAP2,MYL6B,IQCG,TMEM67,CIRBP,RHO
L,RBCK1,SNRPB,ACTR6,BUD31,SAE1,PSMA6,RUVBL2,SNRPN,POLD4,UBE2L6,PHF19,UQCRQ,PSMA1,PHF21
,RPL13A,RPS9,VRK3,ZC3H14,MAGED2,TSG101,PTPN13,CHCHD1,RPS25,NIP7,RPS6,IP6K2,PRMT2,FBXW9,
C2,BLOC1S1,FASTK,TMEM135,TCAIM,VDAC3,TAZ,GSTK1,MRPL33,STYXL1,FIS1,PPM1K,UQCRQ,CHCHD1,M
1,C14ORF2,TIMM17B,ATP5J,TOMM40,HSD17B10,COX5A,NDUFV2,COA3,MRPS18C,MRPS17,NDUFS3,MRF
S1,UQCRQ,CHCHD1,ECSIT,COX4I1,SLC25A10,THEM4,COASY,FOXRED1,C14ORF2,AGPAT4,PPP1CC,TIMM17
1,RPL36,RBM3,RPL18,RPL15,RPL37A,RPLP0,MRPL55,RPL8,RPL35,RPL34,RPL24,UBA52,RPL30,MRPL51,MR
ACYBP,RHOF,RNH1,LY75,SLC3A2,CTSC,ARRDC1,RBP5,DBI,ARPC2,VDAC3,NUMA1,SCARB1,TNFSF10,CHID1,
ACYBP,RHOF,RNH1,LY75,SLC3A2,CTSC,ARRDC1,RBP5,DBI,ARPC2,VDAC3,NUMA1,SCARB1,TNFSF10,CHID1,
ACYBP,RHOF,RNH1,LY75,SLC3A2,CTSC,ARRDC1,RBP5,DBI,ARPC2,VDAC3,NUMA1,SCARB1,TNFSF10,CHID1,
PL33,STYXL1,PPM1K,CHCHD1,THEM4,COASY,TRNT1,ACAD10,DGUOK,TSFM,CHPF,NUDT1,HSD17B10,MD

MRPS22,CYB561A3,SRP9,TMEM67,BLOC1S1,RHOF,TMEM135,ATG16L2,RCC1,SLC3A2,SEC62,CTSC,VDAC3,
4,RPS16,EPB41L5,CD99L2,JUP,CDC42,PTK2,PPP1CC,VIM,RPS15,HSPB1,HMGA1,PABPC1,B2M,RPL18,EGFR
4,RPS16,EPB41L5,CD99L2,JUP,CDC42,PTK2,PPP1CC,VIM,RPS15,HSPB1,HMGA1,PABPC1,B2M,RPL18,EGFR
PL37A,RPLP0,RPL8,RPL35,RPL34,RPL24,UBA52,RPL30
1A3,BAIAP2,MYL6B,IQCG,TMEM67,CACYBP,RHOF,RNH1,DNASE1L1,RNF32,LY75,SLC3A2,CTSC,ARRDC1,KL
4A,HNRNPC,RBM3,PRKRIP1,PABPC1,RBMX,WAC,DDX41,HNRNPA1,SNRPF,PPIH,LGALS3,RALY,GEMIN2,RHE
NDUFA11,NDUFA3,NDUFB3,NDUFB2,NDUFA10,SDHA,NDUFA7,NDUFS2,NDUFC1,NDUFB10,NDUFS7,UQCI
NDUFA11,NDUFA3,NDUFB3,NDUFB2,NDUFA10,SDHA,NDUFA7,NDUFS2,NDUFC1,NDUFB10,NDUFS7,UQCI
NDUFS3,NDUFA11,NDUFA3,NDUFB3,NDUFB2,NDUFA10,SDHA,NDUFA7,NDUFS2,NDUFC1,NDUFB10,NDUI
SME2,TADA3,RAD23A,PSME1,ENY2,PSMD7,PSMD10,UCHL5,TAF9,PSMD8,PSMC5
DUFB2,NDUFA10,NDUFA7,NDUFS2,NDUFC1,NDUFB10,NDUFS7,NDUFS4,NDUFA8
DUFB2,NDUFA10,NDUFA7,NDUFS2,NDUFC1,NDUFB10,NDUFS7,NDUFS4,NDUFA8
DUFB2,NDUFA10,NDUFA7,NDUFS2,NDUFC1,NDUFB10,NDUFS7,NDUFS4,NDUFA8
SME2,RAD23A,PSME1,PSMD7,PSMD10,UCHL5,PSMD8,PSMC5
41,HNRNPA1,SNRPF,RALY,MAGOHB,SNRPD1,SNRPA1,HNRNPM,PPIE,SNRNP40,SNRNP200,MAGOH,WDR8
SME2,RAD23A,PSME1,PSMD7,PSMD10,UCHL5,PSMD8,PSMC5
CHD5,COX17,SIRT5,PARK7,HAX1,COX19,THOP1,AK2,CAPN2,ARL2
5A,NDUFV2,COA3,NDUFS3,PAM16,NDUFA11,NDUFA3,NDUFB3,NDUFB2,NDUFA10,SDHA,NDUFA7,NDUFS

517,MRPL55,MRPS24,MRPS16,NDUFA7,MRPS33,MRPL51,MRPL43,MRPL48,MRPL27,MRPS7,MRPL36,MRI
517,MRPL55,MRPS24,MRPS16,NDUFA7,MRPS33,MRPL51,MRPL43,MRPL48,MRPL27,MRPS7,MRPL36,MRI

SIRT5,PARK7,HAX1,COX19,THOP1,AK2,CAPN2,ARL2,TIMM9,SEC14L1,NDUFA8

38,BTF3,RPS3,RPS4X,RPS23,RPL30,EIF2AK4
N13,EPB41L5,DUSP22,THEM4,ITGB1BP1,SH3YL1,CDC42,PTK2,CAPG,VIM,PODXL
L,RCE1,EBPL,DST,ANTXR2,GRAMD1C,LPCAT3,RDH11,TMED4,AUP1,JPH2,YIF1A,MAGT1,CDC42,XBP1,PCYT
L,RCE1,EBPL,DST,ANTXR2,GRAMD1C,LPCAT3,RDH11,TMED4,AUP1,JPH2,YIF1A,MAGT1,CDC42,XBP1,PCYT

PEH,TXNL4A,NUTF2,TMPO,BRIP1,NUDT1,EGFR,MFSD10,ENY2,NUP133,PCID2,TNKS,DCTN5,KPNA5,HAX1,
17B,PPOX,LAMP2,EMC4,SYNGR1,TOMM40,B2M,ABHD17A,COA3,MPC2,SLC35B1,CD74,PIGT,LFNG,MFF,C
PL13A,RPS9,DST,LRRC1,PCBP2,CD44,RPS16,OBSL1,EPB41L5,CD99L2,PARD6G,JUP,CDC42,PTK2,PPP1CC,VI

1BP1,PTK2,CAPG,PODXL,DBNL,RAPGEF3,HAX1,SLC39A6,ARPC5,TUBB3,RDX,ILK,SLC39A14,ALS2,ACTB,VIL1
3,PKP4,TNKS,CLASP2,NUP62,DYNLT3,TAF1D,RPS3,EML3,ECT2,STAG2,IK,MAP4,DCTN1
5A3,TM9SF1,KXD1,MAGT1,VAMP8,AP1S3,CYB561,NFAM1,LAMP2,HSP90AB1,RAB2A,RPTOR,DDOST,SYNG
5A3,TM9SF1,KXD1,MAGT1,VAMP8,AP1S3,CYB561,NFAM1,LAMP2,HSP90AB1,RAB2A,RPTOR,DDOST,SYNG
1,SEC62,SET,CTSC,DBI,SUMF2,TBL2,EIF5A,UFD1L,SLC35G1,RCE1,EBPL,FIS1,DST,ANTXR2,GRAMD1C,VCAN

53,PML,RGS12,KAT8,STAG2,HNRNPM,RUVBL1,NSMF,EIF6,LMNA,NARF,PRKCZ,MBD1
5,ATG16L2,SLC3A2,VDAC3,NPRL2,SCARB1,EIF5A,AP3B2,FIS1,SLC15A3,ANTXR2,CD44,TM9SF1,SNF8,TSG10
RB1,SLC15A3,TM9SF1,KXD1,MAGT1,VAMP8,AP1S3,CYB561,NFAM1,LAMP2,HSP90AB1,RAB2A,RPTOR,DD
17B,PPOX,LAMP2,EMC4,SYNGR1,TOMM40,B2M,COA3,MPC2,SLC35B1,CD74,PIGT,LFNG,MFF,COA1,PEX2,
,RPS25,RPS6,RPL29,RPL28,RPL21,RPL36,RPS20,RPS15,RPL18,RPL15,MRPS18C,RPS11,MRPS17,RPL37A,RP
28,APEH,RPL21,RPL36,TRMT112,RPS20,RPS15,RPL18,RPL15,RPS11,RPL37A,RPLP0,RPL8,RPL35,RPL34,RPL

1B2,RPL28,RPL21,RPL36,RPS20,RPS15,EEF1D,RPL18,RPL15,RPS11,RPL37A,RPLP0,RPL8,RPL35,RPL34,RPL2[illegible]
28,RPL21,RPL36,RPS20,RPS15,PABPC1,RPL18,RPL15,RPS11,RPL37A,RPLP0,RPL8,RPL35,RPL34,RPL24,RPS3[illegible]
M,RPL28,RPL21,RPL36,RPS20,RPS15,RPL18,RPL15,RPS11,RPL37A,RPLP0,EIF3G,RPL8,RPL35,RPL34,RPL24,[illegible]
M,RPL28,RPL21,RPL36,RPS20,RPS15,PABPC1,RPL18,RPL15,RPS11,RPL37A,EIF4A1,RPLP0,EIF3G,RPL8,RPL3[illegible]
28,RPL21,RPL36,RPS20,RPS15,RPL18,RPL15,RPS11,RPL37A,RPLP0,RPL8,RPL35,RPL34,RPL24,RPS3A,RPS3,[illegible]
28,RPL21,RPL36,RPS20,RPS15,RPL18,RPL15,RPS11,RPL37A,RPLP0,RPL8,RPL35,RPL34,RPL24,RPS3A,RPS3,[illegible]
M,RPL28,RPL21,RPL36,RPS20,RPS15,PABPC1,RPL18,RPL15,EIF2B3,RPS11,RPL37A,EIF4A1,RPLP0,EIF3G,RP[illegible]
M,RPL28,RPL21,RPL36,RPS20,RPS15,PABPC1,RPL18,RPL15,EIF2B3,RPS11,RPL37A,EIF4A1,RPLP0,EIF3G,RP[illegible]
RPS25,RPS6,RPL29,PSMD4,RPL28,PSMA7,RPL21,RPL36,RPS20,RPS15,PSME2,PABPC1,PSME1,RPL18,RPL1[illegible]
M,RPL28,RPL21,RPL36,RPS20,RPS15,RPL18,RPL15,RPS11,RPL37A,EIF4A1,RPLP0,EIF3G,RPL8,RPL35,RPL34,
28,RPL21,RPL36,RPS20,RPS15,RPL18,RPL15,RPS11,RPL37A,RPLP0,RPL8,RPL35,RPL34,RPL24,RPS3A,RPS3,[illegible]
28,RPL21,RPL36,RPS20,RPS15,PABPC1,RPL18,RPL15,RPS11,RPL37A,RPLP0,RPL8,RPL35,RPL34,RPL24,RPS3[illegible]
28,RPL21,RPL36,RPS20,RPS15,PABPC1,RPL18,RPL15,RPS11,RPL37A,RPLP0,RPL8,RPL35,RPL34,RPL24,RPS3[illegible]
9,RPL28,RPL21,RPL36,RPS20,RPS15,DDOST,RPL18,RPL15,RPS11,RPL37A,RPLP0,SSR3,RPL8,RPL35,RPL34,R[illegible]
MRPL33,RPS16,CHCHD1,RPS25,RPS6,RPL29,EIF3M,EEF1B2,PPA1,RPL28,APEH,RPL21,CARS,RPL36,TSFM,T[illegible]
28,RPL21,RPL36,RPS20,RPS15,RPL18,POLR2C,RPL15,RPS11,NUP133,RPL37A,RPLP0,RPL8,RPL35,RPL34,NU[illegible]
28,RPL21,RPL36,RPS20,RPS15,RPL18,POLR2C,RPL15,RPS11,NUP133,RPL37A,KPNA5,RPLP0,RPL8,RPL35,RP[illegible]
28,RPL21,RPL36,RPS20,RPS15,RPL18,POLR2C,RPL15,RPS11,NUP133,RPL37A,RPLP0,RPL8,RPL35,RPL34,NU[illegible]
RPS25,RPS6,RPL29,CDC42,PSMD4,RPL28,PSMA7,RPL21,RPL36,RPS20,RPS15,PSME2,PABPC1,PPP3CB,PSM[illegible]
29,RPL28,RPL21,RPL36,RPS20,RPS15,RPL18,RPL15,RPS11,RPL37A,RPLP0,RPL8,RPL35,MARS,RPL34,RPL24,
C1,RPS11,EIF4A1,EIF3G,RPS3A,EIF3C,RPS3,RPS27L,RPS7,RPS4X,RPS23,RPS15A,RPS17,RPS14,EIF3E,FAU
1,EIF3G,RPS3A,EIF3C,RPS3,RPS27L,RPS7,RPS4X,RPS23,RPS15A,RPS17,RPS14,EIF3E,FAU
C1,RPS11,EIF4A1,EIF3G,RPS3A,EIF3C,RPS3,RPS27L,RPS7,RPS4X,RPS23,RPS15A,RPS17,RPS14,EIF3E,FAU
1,EIF4A1,EIF3G,RPS3A,EIF3C,RPS3,RPS27L,RPS7,RPS4X,RPS23,RPS15A,RPS17,RPS14,EIF3E,FAU
25,NIP7,RPS6,RPL29,RPL28,RPL21,EXOSC9,RPL36,TRMT112,RPS20,RPS15,RPL18,RPL15,NHP2,RPS11,NOP2[illegible]
25,NIP7,RPS6,RPL29,RPL28,RPL21,EXOSC9,RPL36,TRMT112,RPS20,RPS15,HSD17B10,RPL18,RPL15,NHP2,[illegible]
1,RPS16,TSG101,PSMA1,PSMA3,RPS25,RPS6,RPL29,PSMD4,RPL28,LIG1,PSMA7,RPL21,AP1S3,RPL36,HSP[illegible]
25,NIP7,RPS6,RPL29,RPL28,RPL21,EXOSC9,RPL36,RPS20,RPS15,RPL18,RPL15,RPS11,RPL37A,MPHOSPH10,
31,BUD31,OSGEP,RPL13A,PSMA6,RPS9,SNRPN,PCBP2,RPS16,PSMA1,PSMA3,RPS25,NIP7,RPS6,RPL29,SRS[illegible]
28,RPL21,RPL36,RPS20,RPS15,RPL18,RPL15,RPS11,RPL37A,RPLP0,RPL8,RPL35,RPL34,RPL24,RPS3A,RPS3,[illegible]
T3,GNAS,GPR137,RPS24,SERPINB5,S100A13,BCAT2,CENPV,MRPL4,FAM189B,VKORC1,MRPL2,CDCA3,PCN[illegible]
T3,GNAS,GPR137,RPS24,BCAT2,CENPV,MRPL4,FAM189B,VKORC1,MRPL2,CDCA3,PCNA,SRSF2,MRPS22,OS[illegible]
SRSF2,MRPS22,LAS1L,HMGB1,BAIAP2,RAPGEF5,SSU72,MYL6B,FAM122C,TMEM67,FAM72A,CMC2,BLOC1[illegible]
NLL1,CST3,GNAS,GPR137,RPS24,SERPINB5,S100A13,BCAT2,CENPV,MRPL4,FAM189B,VKORC1,MRPL2,CD[illegible]
NLL1,CST3,GNAS,GPR137,RPS24,SERPINB5,S100A13,BCAT2,CENPV,MRPL4,FAM189B,VKORC1,MRPL2,CD[illegible]
NLL1,CST3,GNAS,GPR137,RPS24,BCAT2,CENPV,MRPL4,FAM189B,VKORC1,MRPL2,CDCA3,PCNA,SRSF2,MR[illegible]
NLL1,CST3,GNAS,GPR137,RPS24,BCAT2,CENPV,MRPL4,FAM189B,VKORC1,MRPL2,CDCA3,PCNA,SRSF2,MR[illegible]
NLL1,CST3,GNAS,GPR137,RPS24,SERPINB5,BCAT2,CENPV,MRPL4,FAM189B,MRPL2,CDCA3,PCNA,SRSF2,O[illegible]
RPINB5,S100A13,BCAT2,FAM189B,CDCA3,SRSF2,MRPS22,HDAC9,SEMA4D,LAS1L,HMGB1,BAIAP2,RAPGE[illegible]
RPINB5,S100A13,BCAT2,FAM189B,CDCA3,SRSF2,MRPS22,HDAC9,SEMA4D,LAS1L,HMGB1,BAIAP2,RAPGE[illegible]
NLL1,CST3,GNAS,GPR137,RPS24,BCAT2,CENPV,MRPL4,FAM189B,VKORC1,MRPL2,CDCA3,PCNA,SRSF2,MR[illegible]
AT2,FAM189B,CDCA3,SRSF2,MRPS22,SEMA4D,LAS1L,HMGB1,BAIAP2,SSU72,MYL6B,TMEM67,PILRB,CMC[illegible]
AT2,FAM189B,CDCA3,SRSF2,MRPS22,SEMA4D,LAS1L,HMGB1,BAIAP2,SSU72,MYL6B,TMEM67,PILRB,CMC[illegible]
L,HMGB1,BAIAP2,MYL6B,FAM122C,CMC2,BLOC1S1,CACYBP,RHOF,SMARCB1,RPL38,RNH1,DNASE1L1,RB[illegible]
,BCAT2,FAM189B,CDCA3,SRSF2,MRPS22,HDAC9,SEMA4D,LAS1L,HMGB1,BAIAP2,RAPGEF5,SSU72,MYL6B,
,BCAT2,FAM189B,CDCA3,SRSF2,MRPS22,HDAC9,SEMA4D,LAS1L,HMGB1,BAIAP2,RAPGEF5,SSU72,MYL6B,
NLL1,CST3,GNAS,GPR137,RPS24,SERPINB5,S100A13,BCAT2,MRPL4,VKORC1,MRPL2,CDCA3,PCNA,SRSF2,[illegible]

NLL1,CST3,GNAS,GPR137,RPS24,SERPINB5,S100A13,BCAT2,MRPL4,VKORC1,MRPL2,CDCA3,PCNA,SRSF2,N
M189B,CDCA3,SRSF2,MRPS22,LAS1L,HMGB1,BAIAP2,SSU72,MYL6B,FAM122C,TMEM67,PILRB,CMC2,BLC
M189B,CDCA3,SRSF2,MRPS22,LAS1L,HMGB1,BAIAP2,SSU72,MYL6B,FAM122C,TMEM67,PILRB,CMC2,BLC
,BCAT2,FAM189B,CDCA3,SRSF2,MRPS22,SEMA4D,LAS1L,HMGB1,BAIAP2,SSU72,MYL6B,TMEM67,CMC2,E
,BCAT2,FAM189B,CDCA3,SRSF2,MRPS22,SEMA4D,LAS1L,HMGB1,BAIAP2,SSU72,MYL6B,TMEM67,CMC2,E
BCAT2,FAM189B,CDCA3,SRSF2,MRPS22,HDAC9,SEMA4D,LAS1L,HMGB1,BAIAP2,RAPGEF5,SSU72,MYL6B,
BCAT2,FAM189B,CDCA3,SRSF2,MRPS22,HDAC9,SEMA4D,LAS1L,HMGB1,BAIAP2,RAPGEF5,SSU72,MYL6B,
NLL1,CST3,GNAS,GPR137,RPS24,SERPINB5,BCAT2,CENPV,MRPL4,FAM189B,VKORC1,MRPL2,CDCA3,PCNA
NLL1,CST3,GNAS,GPR137,RPS24,SERPINB5,BCAT2,CENPV,MRPL4,FAM189B,VKORC1,MRPL2,CDCA3,PCNA
ERPINB5,S100A13,BCAT2,CDCA3,SRSF2,MRPS22,HDAC9,SEMA4D,LAS1L,HMGB1,BAIAP2,SSU72,MYL6B,F
ERPINB5,S100A13,BCAT2,CDCA3,SRSF2,MRPS22,HDAC9,SEMA4D,LAS1L,HMGB1,BAIAP2,SSU72,MYL6B,F
INB5,S100A13,BCAT2,FAM189B,CDCA3,SRSF2,MRPS22,HDAC9,SEMA4D,LAS1L,HMGB1,BAIAP2,RAPGEF5
INB5,S100A13,BCAT2,FAM189B,CDCA3,SRSF2,MRPS22,HDAC9,SEMA4D,LAS1L,HMGB1,BAIAP2,RAPGEF5
RPS22,LAS1L,HMGB1,SSU72,MYL6B,TMEM67,FAM72A,CMC2,BLOC1S1,CIRBP,RHOF,SMARCB1,RNH1,KIF
RPS22,LAS1L,HMGB1,SSU72,MYL6B,TMEM67,FAM72A,CMC2,BLOC1S1,CIRBP,RHOF,SMARCB1,RNH1,KIF
INB5,S100A13,BCAT2,FAM189B,VKORC1,CDCA3,PCNA,SRSF2,MRPS22,HDAC9,LAS1L,HMGB1,BAIAP2,RAF
INB5,S100A13,BCAT2,FAM189B,VKORC1,CDCA3,PCNA,SRSF2,MRPS22,HDAC9,LAS1L,HMGB1,BAIAP2,RAF
,RPS24,SERPINB5,S100A13,BCAT2,FAM189B,CDCA3,SRSF2,MRPS22,HDAC9,LAS1L,HMGB1,BAIAP2,RAPGE
,RPS24,SERPINB5,S100A13,BCAT2,FAM189B,CDCA3,SRSF2,MRPS22,HDAC9,LAS1L,HMGB1,BAIAP2,RAPGE
3,GNAS,GPR137,RPS24,BCAT2,CENPV,MRPL4,FAM189B,VKORC1,MRPL2,CDCA3,PCNA,SRSF2,MRPS22,OS
,S100A13,BCAT2,FAM189B,CDCA3,SRSF2,MRPS22,HDAC9,KRT15,SEMA4D,LAS1L,HMGB1,BAIAP2,RAPGEI
,S100A13,BCAT2,FAM189B,CDCA3,SRSF2,MRPS22,HDAC9,KRT15,SEMA4D,LAS1L,HMGB1,BAIAP2,RAPGEI
SF2,MRPS22,LAS1L,HMGB1,RAPGEF5,SSU72,MYL6B,TMEM67,CMC2,BLOC1S1,CACYBP,SMARCB1,PRR13,F
NLL1,CST3,GNAS,GPR137,RPS24,SERPINB5,BCAT2,CENPV,MRPL4,FAM189B,VKORC1,PCNA,SRSF2,OSMR,P
NLL1,CST3,GNAS,GPR137,RPS24,SERPINB5,BCAT2,CENPV,MRPL4,FAM189B,VKORC1,PCNA,SRSF2,OSMR,P
AS1L,HMGB1,SSU72,CMC2,BLOC1S1,CACYBP,CIRBP,RPL38,PRR13,RNH1,RBCK1,LY75,RCC1,SNRPB,SLC3A2
NLL1,CST3,GNAS,GPR137,RPS24,SERPINB5,S100A13,BCAT2,CENPV,MRPL4,FAM189B,VKORC1,MRPL2,CDC
NLL1,CST3,GNAS,GPR137,RPS24,SERPINB5,S100A13,BCAT2,CENPV,MRPL4,FAM189B,VKORC1,MRPL2,CDC
NLL1,CST3,GNAS,GPR137,RPS24,SERPINB5,S100A13,BCAT2,CENPV,MRPL4,FAM189B,VKORC1,MRPL2,CDC
AM189B,CDCA3,SRSF2,MRPS22,LAS1L,HMGB1,CYB561A3,BAIAP2,SSU72,MYL6B,TMEM67,PILRB,FAM72A
AM189B,CDCA3,SRSF2,MRPS22,LAS1L,HMGB1,CYB561A3,BAIAP2,SSU72,MYL6B,TMEM67,PILRB,FAM72A
,BCAT2,CDCA3,SRSF2,MRPS22,HDAC9,SEMA4D,LAS1L,HMGB1,BAIAP2,RAPGEF5,SSU72,MYL6B,TMEM67,I
,BCAT2,CDCA3,SRSF2,MRPS22,HDAC9,SEMA4D,LAS1L,HMGB1,BAIAP2,RAPGEF5,SSU72,MYL6B,TMEM67,I
,RPS24,SERPINB5,S100A13,BCAT2,MRPL4,FAM189B,CDCA3,SRSF2,MRPS22,HDAC9,SEMA4D,LAS1L,HMGE
,RPS24,SERPINB5,S100A13,BCAT2,MRPL4,FAM189B,CDCA3,SRSF2,MRPS22,HDAC9,SEMA4D,LAS1L,HMGE
,BCAT2,FAM189B,CDCA3,SRSF2,MRPS22,SEMA4D,LAS1L,HMGB1,BAIAP2,RAPGEF5,SSU72,MYL6B,TMEM6
,BCAT2,FAM189B,CDCA3,SRSF2,MRPS22,SEMA4D,LAS1L,HMGB1,BAIAP2,RAPGEF5,SSU72,MYL6B,TMEM6
PS24,SERPINB5,S100A13,BCAT2,FAM189B,CDCA3,SRSF2,MRPS22,HDAC9,SEMA4D,LAS1L,HMGB1,BAIAP2
PS24,SERPINB5,S100A13,BCAT2,FAM189B,CDCA3,SRSF2,MRPS22,HDAC9,SEMA4D,LAS1L,HMGB1,BAIAP2
ORC1,CDCA3,PCNA,MRPS22,SEMA4D,LAS1L,HMGB1,BAIAP2,SSU72,MYL6B,TMEM67,CMC2,BLOC1S1,CAC
ORC1,CDCA3,PCNA,MRPS22,SEMA4D,LAS1L,HMGB1,BAIAP2,SSU72,MYL6B,TMEM67,CMC2,BLOC1S1,CAC
00A13,BCAT2,FAM189B,CDCA3,MRPS22,SEMA4D,LAS1L,HMGB1,BAIAP2,SSU72,MYL6B,TMEM67,CMC2,E
00A13,BCAT2,FAM189B,CDCA3,MRPS22,SEMA4D,LAS1L,HMGB1,BAIAP2,SSU72,MYL6B,TMEM67,CMC2,E
NLL1,CST3,GPR137,SERPINB5,BCAT2,CENPV,MRPL4,FAM189B,VKORC1,MRPL2,CDCA3,PCNA,SRSF2,MRPS
AIAP2,SSU72,MYL6B,FAM122C,CMC2,BLOC1S1,CACYBP,RHOF,SMARCB1,RPL38,PRR13,DNASE1L1,RBCK1,
MYL6B,TMEM67,CMC2,BLOC1S1,CACYBP,RHOF,SMARCB1,PRR13,RBCK1,RNF32,LY75,RCC1,SNRPB,DHPS,
S100A13,BCAT2,CDCA3,SRSF2,MRPS22,SEMA4D,LAS1L,HMGB1,BAIAP2,RAPGEF5,SSU72,MYL6B,TMEM67,

S100A13,BCAT2,CDCA3,SRSF2,MRPS22,SEMA4D,LAS1L,HMGB1,BAIAP2,RAPGEF5,SSU72,MYL6B,TMEM67,

S100A13,BCAT2,CDCA3,SRSF2,MRPS22,SEMA4D,LAS1L,HMGB1,BAIAP2,SSU72,MYL6B,TMEM67,PILRB,CM

S100A13,BCAT2,CDCA3,SRSF2,MRPS22,SEMA4D,LAS1L,HMGB1,BAIAP2,SSU72,MYL6B,TMEM67,PILRB,CM

3,BCAT2,FAM189B,CDCA3,MRPS22,SEMA4D,LAS1L,HMGB1,BAIAP2,SSU72,MYL6B,TMEM67,CMC2,BLOC

3,BCAT2,FAM189B,CDCA3,MRPS22,SEMA4D,LAS1L,HMGB1,BAIAP2,SSU72,MYL6B,TMEM67,CMC2,BLOC

F2,MRPS22,HDAC9,LAS1L,HMGB1,BAIAP2,SSU72,MYL6B,TMEM67,CMC2,BLOC1S1,CACYBP,CIRBP,RHOF,

F2,MRPS22,HDAC9,LAS1L,HMGB1,BAIAP2,SSU72,MYL6B,TMEM67,CMC2,BLOC1S1,CACYBP,CIRBP,RHOF,

AT2,CDCA3,SRSF2,MRPS22,SEMA4D,LAS1L,HMGB1,SULF2,BAIAP2,SSU72,MYL6B,TMEM67,CMC2,BLOC1S

AT2,CDCA3,SRSF2,MRPS22,SEMA4D,LAS1L,HMGB1,SULF2,BAIAP2,SSU72,MYL6B,TMEM67,CMC2,BLOC1S

CAT2,FAM189B,CDCA3,SRSF2,MRPS22,SEMA4D,LAS1L,HMGB1,BAIAP2,RAPGEF5,SSU72,MYL6B,TMEM67

CAT2,FAM189B,CDCA3,SRSF2,MRPS22,SEMA4D,LAS1L,HMGB1,BAIAP2,RAPGEF5,SSU72,MYL6B,TMEM67

PS24,SERPINB5,S100A13,BCAT2,FAM189B,VKORC1,CDCA3,PCNA,SRSF2,MRPS22,SEMA4D,LAS1L,HMGB1,

PS24,SERPINB5,S100A13,BCAT2,FAM189B,VKORC1,CDCA3,PCNA,SRSF2,MRPS22,SEMA4D,LAS1L,HMGB1,

S100A13,BCAT2,FAM189B,CDCA3,SRSF2,MRPS22,LAS1L,HMGB1,BAIAP2,SSU72,MYL6B,TMEM67,PILRB,C

S100A13,BCAT2,FAM189B,CDCA3,SRSF2,MRPS22,LAS1L,HMGB1,BAIAP2,SSU72,MYL6B,TMEM67,PILRB,C

B1,MYL6B,CMC2,BLOC1S1,CACYBP,RBCK1,RNF32,LY75,RCC1,SNRPB,ACTR6,ZNF235,BOP1,ARRDC1,TRPT1,

S100A13,BCAT2,FAM189B,CDCA3,SRSF2,MRPS22,SEMA4D,LAS1L,HMGB1,BAIAP2,RAPGEF5,SSU72,MYL6

S100A13,BCAT2,FAM189B,CDCA3,SRSF2,MRPS22,SEMA4D,LAS1L,HMGB1,BAIAP2,RAPGEF5,SSU72,MYL6

F2,MRPS22,LAS1L,HMGB1,BAIAP2,SSU72,MYL6B,TMEM67,PILRB,FAM72A,CMC2,BLOC1S1,CACYBP,CIRBP,

F2,MRPS22,LAS1L,HMGB1,BAIAP2,SSU72,MYL6B,TMEM67,PILRB,FAM72A,CMC2,BLOC1S1,CACYBP,CIRBP,

PINB5,S100A13,BCAT2,CDCA3,SRSF2,MRPS22,SEMA4D,LAS1L,HMGB1,BAIAP2,SSU72,MYL6B,TMEM67,CM

PINB5,S100A13,BCAT2,CDCA3,SRSF2,MRPS22,SEMA4D,LAS1L,HMGB1,BAIAP2,SSU72,MYL6B,TMEM67,CM

S100A13,BCAT2,FAM189B,CDCA3,SRSF2,MRPS22,PPFIBP1,LAS1L,HMGB1,BAIAP2,SSU72,MYL6B,TMEM67

S100A13,BCAT2,FAM189B,CDCA3,SRSF2,MRPS22,PPFIBP1,LAS1L,HMGB1,BAIAP2,SSU72,MYL6B,TMEM67

CK1,LY75,RCC1,SNRPB,DHPS,ACTR6,ZNF235,ARRDC1,TRPT1,DBI,ARPC2,LIMK2,GPATCH2L,BUD31,NPRL2,

S100A13,BCAT2,VKORC1,CDCA3,PCNA,MRPS22,SEMA4D,LAS1L,HMGB1,BAIAP2,SSU72,MYL6B,TMEM67,C

S100A13,BCAT2,VKORC1,CDCA3,PCNA,MRPS22,SEMA4D,LAS1L,HMGB1,BAIAP2,SSU72,MYL6B,TMEM67,C

INB5,CENPV,MRPL4,FAM189B,MRPL2,CDCA3,PCNA,SRSF2,MRPS22,KRT15,PPFIBP1,HMGB1,SULF2,BAIAP

INB5,CENPV,MRPL4,FAM189B,MRPL2,CDCA3,PCNA,SRSF2,MRPS22,KRT15,PPFIBP1,HMGB1,SULF2,BAIAP

CMC2,BLOC1S1,RHOF,SMARCB1,RPL38,RBCK1,LY75,RCC1,SNRPB,ACTR6,TRPT1,ZC3HAV1,DBI,ARPC2,LIMK

75,RCC1,SNRPB,DHPS,ACTR6,TRPT1,DBI,ARPC2,LIMK2,GPATCH2L,BUD31,NPRL2,CTTN,EIF5A,COMMD5,N

PS24,SERPINB5,S100A13,BCAT2,FAM189B,VKORC1,CDCA3,PCNA,SRSF2,MRPS22,SEMA4D,LAS1L,HMGB1,

PS24,SERPINB5,S100A13,BCAT2,FAM189B,VKORC1,CDCA3,PCNA,SRSF2,MRPS22,SEMA4D,LAS1L,HMGB1,

PR137,RPS24,S100A13,BCAT2,CENPV,MRPL4,VKORC1,MRPL2,CDCA3,PCNA,SRSF2,MRPS22,OSMR,PPFIBP

PR137,RPS24,S100A13,BCAT2,CENPV,MRPL4,VKORC1,MRPL2,CDCA3,PCNA,SRSF2,MRPS22,OSMR,PPFIBP

NLL1,CST3,GNAS,GPR137,RPS24,S100A13,BCAT2,CENPV,MRPL4,FAM189B,VKORC1,MRPL2,CDCA3,PCNA,

NLL1,CST3,GNAS,GPR137,RPS24,S100A13,BCAT2,CENPV,MRPL4,FAM189B,VKORC1,MRPL2,CDCA3,PCNA,

MYL6B,BLOC1S1,SMARCB1,RBCK1,LY75,RCC1,SNRPB,DHPS,ACTR6,BOP1,TRPT1,ZC3HAV1,DBI,ARPC2,LIMK

S24,SERPINB5,S100A13,BCAT2,MRPL4,FAM189B,CDCA3,SRSF2,MRPS22,SEMA4D,LAS1L,HMGB1,SULF2,C

S24,SERPINB5,S100A13,BCAT2,MRPL4,FAM189B,CDCA3,SRSF2,MRPS22,SEMA4D,LAS1L,HMGB1,SULF2,C

CST3,GNAS,GPR137,RPS24,SERPINB5,BCAT2,CENPV,MRPL4,FAM189B,VKORC1,MRPL2,CDCA3,PCNA,SRSF

CST3,GNAS,GPR137,RPS24,SERPINB5,BCAT2,CENPV,MRPL4,FAM189B,VKORC1,MRPL2,CDCA3,PCNA,SRSF

CMC2,BLOC1S1,CACYBP,RHOF,SMARCB1,PRR13,RBCK1,RNF32,LY75,RCC1,SNRPB,DHPS,ACTR6,ZNF235,BO

2,SEMA4D,LAS1L,HMGB1,BAIAP2,RAPGEF5,SSU72,MYL6B,TMEM67,CMC2,BLOC1S1,CACYBP,CIRBP,RHOF

2,SEMA4D,LAS1L,HMGB1,BAIAP2,RAPGEF5,SSU72,MYL6B,TMEM67,CMC2,BLOC1S1,CACYBP,CIRBP,RHOF

OC1S1,RBCK1,SNRPB,ACTR6,ZNF235,BOP1,ARRDC1,TRPT1,DBI,ARPC2,LIMK2,BUD31,NPRL2,CTTN,EIF5A,C

1S1,RBCK1,RNF32,LY75,SNRPB,ACTR6,ZNF235,BOP1,TRPT1,ARPC2,LIMK2,SIRT6,NUMA1,GPATCH2L,BUD3

SF2,MRPS22,LAS1L,HMGB1,RAPGEF5,SSU72,TMEM67,BLOC1S1,CACYBP,PRR13,RBCK1,RNF32,LY75,SNRPB
13,BCAT2,MRPL4,FAM189B,CDCA3,SRSF2,MRPS22,LAS1L,HMGB1,SULF2,BAIAP2,RAPGEF5,SSU72,MYL6B,T
13,BCAT2,MRPL4,FAM189B,CDCA3,SRSF2,MRPS22,LAS1L,HMGB1,SULF2,BAIAP2,RAPGEF5,SSU72,MYL6B,T
S100A13,BCAT2,FAM189B,CDCA3,SRSF2,MRPS22,SEMA4D,LAS1L,HMGB1,BAIAP2,RAPGEF5,SSU72,MYL6B
S100A13,BCAT2,FAM189B,CDCA3,SRSF2,MRPS22,SEMA4D,LAS1L,HMGB1,BAIAP2,RAPGEF5,SSU72,MYL6B
HMGB1,MYL6B,PILRB,CMC2,BLOC1S1,CACYBP,SMARCB1,PRR13,RNH1,RBCK1,RNF32,LY75,RCC1,SNRPB,D
S24,S100A13,BCAT2,FAM189B,VKORC1,CDCA3,SRSF2,MRPS22,HDAC9,LAS1L,HMGB1,SULF2,CYB561A3,BA
S24,S100A13,BCAT2,FAM189B,VKORC1,CDCA3,SRSF2,MRPS22,HDAC9,LAS1L,HMGB1,SULF2,CYB561A3,BA
SMARCB1,RPL38,RBCK1,LY75,RCC1,SNRPB,DHPS,ACTR6,TRPT1,DBI,ARPC2,LIMK2,SIRT6,GPATCH2L,BUD31
,GNAS,GPR137,RPS24,SERPINB5,BCAT2,CENPV,MRPL4,FAM189B,VKORC1,MRPL2,CDCA3,PCNA,SRSF2,MR
,GNAS,GPR137,RPS24,SERPINB5,BCAT2,CENPV,MRPL4,FAM189B,VKORC1,MRPL2,CDCA3,PCNA,SRSF2,MR
BCK1,LY75,RCC1,SNRPB,DHPS,ACTR6,TRPT1,DBI,ARPC2,LIMK2,GPATCH2L,BUD31,NPRL2,CTTN,EIF5A,COM
4,SERPINB5,CENPV,MRPL4,FAM189B,MRPL2,CDCA3,PCNA,SRSF2,MRPS22,KRT15,OSMR,PPFIBP1,HMGB1
4,SERPINB5,CENPV,MRPL4,FAM189B,MRPL2,CDCA3,PCNA,SRSF2,MRPS22,KRT15,OSMR,PPFIBP1,HMGB1
MYL6B,TMEM67,CMC2,CIRBP,RNH1,KIF4A,RBCK1,RCC1,SNRPB,DHPS,ACTR6,DBI,LIMK2,SIRT6,NUMA1,BUI
MYL6B,TMEM67,CMC2,CIRBP,RNH1,KIF4A,RBCK1,RCC1,SNRPB,DHPS,ACTR6,DBI,LIMK2,SIRT6,NUMA1,BUI
IAP2,SSU72,MYL6B,FAM122C,CMC2,BLOC1S1,RHOF,SMARCB1,RPL38,RNH1,DNASE1L1,RBCK1,RNF32,LY7
7,RPS24,SERPINB5,BCAT2,MRPL4,VKORC1,CDCA3,PCNA,SRSF2,MRPS22,OSMR,LAS1L,HMGB1,SULF2,BAIA
C1S1,RHOF,SMARCB1,RBCK1,LY75,RCC1,SNRPB,ACTR6,BOP1,TRPT1,ARPC2,LIMK2,NUMA1,GPATCH2L,BUI
ACTR6,TRPT1,ARPC2,LIMK2,BUD31,NPRL2,CTTN,COMMD5,NUDT22,THYN1,RPL13A,PSMA6,ACOT8,RCE1,
HMGB1,MYL6B,PILRB,CMC2,BLOC1S1,CACYBP,RHOF,SMARCB1,PRR13,RNH1,RBCK1,RNF32,LY75,RCC1,SN
A4D,LAS1L,HMGB1,SSU72,MYL6B,TMEM67,CMC2,BLOC1S1,CIRBP,RPL38,RNH1,KIF4A,RBCK1,RNF32,RCC1
A4D,LAS1L,HMGB1,SSU72,MYL6B,TMEM67,CMC2,BLOC1S1,CIRBP,RPL38,RNH1,KIF4A,RBCK1,RNF32,RCC1
MEM67,CMC2,CIRBP,RNH1,KIF4A,RBCK1,RCC1,SNRPB,DHPS,ACTR6,DBI,LIMK2,SIRT6,BUD31,NPRL2,CTTN,E
MEM67,CMC2,CIRBP,RNH1,KIF4A,RBCK1,RCC1,SNRPB,DHPS,ACTR6,DBI,LIMK2,SIRT6,BUD31,NPRL2,CTTN,E
RPS22,HDAC9,SEMA4D,LAS1L,HMGB1,BAIAP2,MYL6B,TMEM67,PILRB,CMC2,BLOC1S1,CACYBP,CIRBP,SMA
RPS22,HDAC9,SEMA4D,LAS1L,HMGB1,BAIAP2,MYL6B,TMEM67,PILRB,CMC2,BLOC1S1,CACYBP,CIRBP,SMA
NLL1,CST3,GNAS,GPR137,RPS24,SERPINB5,BCAT2,CENPV,MRPL4,FAM189B,VKORC1,MRPL2,CDCA3,PCNA
NLL1,CST3,GNAS,GPR137,RPS24,SERPINB5,BCAT2,CENPV,MRPL4,FAM189B,VKORC1,MRPL2,CDCA3,PCNA
L6B,CMC2,BLOC1S1,CACYBP,SMARCB1,PRR13,RBCK1,RNF32,LY75,RCC1,SNRPB,DHPS,ACTR6,BOP1,TRPT1
1,BAIAP2,MYL6B,FAM122C,CMC2,BLOC1S1,RHOF,SMARCB1,RPL38,DNASE1L1,RBCK1,RNF32,LY75,RCC1,S
NLL1,CST3,GNAS,GPR137,RPS24,SERPINB5,BCAT2,CENPV,MRPL4,FAM189B,MRPL2,CDCA3,PCNA,SRSF2,KI
NLL1,CST3,GNAS,GPR137,RPS24,SERPINB5,BCAT2,CENPV,MRPL4,FAM189B,MRPL2,CDCA3,PCNA,SRSF2,KI
CYBP,RBCK1,LY75,SNRPB,DHPS,ACTR6,ZNF235,BOP1,TRPT1,DBI,ARPC2,LIMK2,SIRT6,NUMA1,GPATCH2L,E
31,SSU72,TMEM67,CMC2,CIRBP,RNH1,KIF4A,RBCK1,RNF32,RCC1,SNRPB,PARD3,DHPS,ACTR6,ZNF235,TRP
31,SSU72,TMEM67,CMC2,CIRBP,RNH1,KIF4A,RBCK1,RNF32,RCC1,SNRPB,PARD3,DHPS,ACTR6,ZNF235,TRP
DCA3,MRPS22,LAS1L,HMGB1,SULF2,RAPGEF5,SSU72,MYL6B,TMEM67,FAM72A,CMC2,RHOF,SMARCB1,KI
DCA3,MRPS22,LAS1L,HMGB1,SULF2,RAPGEF5,SSU72,MYL6B,TMEM67,FAM72A,CMC2,RHOF,SMARCB1,KI
LAS1L,HMGB1,SSU72,MYL6B,TMEM67,CMC2,CIRBP,RHOF,RNH1,KIF4A,RBCK1,RNF32,RCC1,SNRPB,PARD3
LAS1L,HMGB1,SSU72,MYL6B,TMEM67,CMC2,CIRBP,RHOF,RNH1,KIF4A,RBCK1,RNF32,RCC1,SNRPB,PARD3
LAS1L,HMGB1,SSU72,MYL6B,TMEM67,PILRB,CMC2,CIRBP,RPL38,RNH1,DNASE1L1,KIF4A,RBCK1,RNF32,LY
LAS1L,HMGB1,SSU72,MYL6B,TMEM67,PILRB,CMC2,CIRBP,RPL38,RNH1,DNASE1L1,KIF4A,RBCK1,RNF32,LY
S1L,HMGB1,BAIAP2,SSU72,TMEM67,CMC2,BLOC1S1,CIRBP,SMARCB1,RPL38,RNH1,KIF4A,RBCK1,LY75,RC
S1L,HMGB1,BAIAP2,SSU72,TMEM67,CMC2,BLOC1S1,CIRBP,SMARCB1,RPL38,RNH1,KIF4A,RBCK1,LY75,RC
AS,GPR137,RPS24,SERPINB5,BCAT2,CENPV,MRPL4,FAM189B,MRPL2,CDCA3,PCNA,SRSF2,OSMR,PPFIBP1,I
AS,GPR137,RPS24,SERPINB5,BCAT2,CENPV,MRPL4,FAM189B,MRPL2,CDCA3,PCNA,SRSF2,OSMR,PPFIBP1,I
FAM122C,CMC2,BLOC1S1,RHOF,RPL38,DNASE1L1,RBCK1,RNF32,LY75,RCC1,SNRPB,DHPS,ACTR6,TRPT1,ZC

HMGB1,MYL6B,FAM122C,CMC2,BLOC1S1,CACYBP,RHOF,SMARCB1,RPL38,DNASE1L1,RBCK1,RNF32,LY75,
B1,SSU72,TMEM67,CMC2,CIRBP,RPL38,RNH1,KIF4A,RBCK1,LY75,RCC1,SNRPB,DHPS,ACTR6,ZNF235,BOP1
B1,SSU72,TMEM67,CMC2,CIRBP,RPL38,RNH1,KIF4A,RBCK1,LY75,RCC1,SNRPB,DHPS,ACTR6,ZNF235,BOP1
GB1,BAIAP2,SSU72,MYL6B,FAM122C,FAM72A,CMC2,BLOC1S1,RHOF,SMARCB1,RPL38,DNASE1L1,RBCK1,
LAS1L,HMGB1,SSU72,MYL6B,TMEM67,CMC2,CACYBP,CIRBP,RHOF,RPL38,PRR13,RNH1,DNASE1L1,KIF4A,
LAS1L,HMGB1,SSU72,MYL6B,TMEM67,CMC2,CACYBP,CIRBP,RHOF,RPL38,PRR13,RNH1,DNASE1L1,KIF4A,
C1S1,CACYBP,RBCK1,RNF32,LY75,RCC1,SNRPB,ACTR6,ZNF235,TRPT1,DBI,ARPC2,LIMK2,GPATCH2L,BUD31
,GNAS,GPR137,RPS24,BCAT2,CENPV,MRPL4,FAM189B,VKORC1,MRPL2,CDCA3,PCNA,SRSF2,MRPS22,OSM
SF2,MRPS22,LAS1L,HMGB1,CYB561A3,BAIAP2,SSU72,FAM72A,CMC2,CACYBP,SMARCB1,RPL38,RNH1,RB
NLL1,CST3,GNAS,GPR137,RPS24,SERPINB5,BCAT2,CENPV,MRPL4,FAM189B,VKORC1,MRPL2,CDCA3,PCNA
CMC2,CACYBP,CIRBP,PRR13,RNH1,KIF4A,RBCK1,RCC1,SNRPB,DHPS,ACTR6,DBI,ARPC2,PMFBP1,LIMK2,GP
CMC2,CACYBP,CIRBP,PRR13,RNH1,KIF4A,RBCK1,RCC1,SNRPB,DHPS,ACTR6,DBI,ARPC2,PMFBP1,LIMK2,GP
PINB5,S100A13,BCAT2,VKORC1,CDCA3,PCNA,SRSF2,MRPS22,KRT15,SEMA4D,LAS1L,HMGB1,SULF2,BAIAP
PINB5,S100A13,BCAT2,VKORC1,CDCA3,PCNA,SRSF2,MRPS22,KRT15,SEMA4D,LAS1L,HMGB1,SULF2,BAIAP
LAS1L,HMGB1,BAIAP2,SSU72,MYL6B,TMEM67,CMC2,BLOC1S1,CACYBP,CIRBP,RPL38,PRR13,RNH1,KIF4A,
LAS1L,HMGB1,BAIAP2,SSU72,MYL6B,TMEM67,CMC2,BLOC1S1,CACYBP,CIRBP,RPL38,PRR13,RNH1,KIF4A,
LAS1L,HMGB1,BAIAP2,SSU72,MYL6B,TMEM67,CMC2,BLOC1S1,CACYBP,CIRBP,RPL38,PRR13,RNH1,KIF4A,
LAS1L,HMGB1,BAIAP2,SSU72,MYL6B,TMEM67,CMC2,BLOC1S1,CACYBP,CIRBP,RPL38,PRR13,RNH1,KIF4A,
,MYL6B,TMEM67,CMC2,CIRBP,RNH1,KIF4A,RBCK1,RNF32,RCC1,SNRPB,PARD3,DHPS,ACTR6,DBI,ARPC2,LI
,MYL6B,TMEM67,CMC2,CIRBP,RNH1,KIF4A,RBCK1,RNF32,RCC1,SNRPB,PARD3,DHPS,ACTR6,DBI,ARPC2,LI
EM67,CMC2,CIRBP,SMARCB1,RPL38,RNH1,KIF4A,RBCK1,LY75,RCC1,SNRPB,DHPS,ACTR6,ZNF235,ARRDC1,
EM67,CMC2,CIRBP,SMARCB1,RPL38,RNH1,KIF4A,RBCK1,LY75,RCC1,SNRPB,DHPS,ACTR6,ZNF235,ARRDC1,
1L,HMGB1,MYL6B,TMEM67,CMC2,RBCK1,LY75,SNRPB,DHPS,ACTR6,ZNF235,ZC3HAV1,DBI,ARPC2,PMFBP
7,CMC2,CIRBP,RPL38,RNH1,RBCK1,RCC1,SNRPB,DHPS,ACTR6,ZNF235,BOP1,TRPT1,DBI,ARPC2,PMFBP1,LI
7,CMC2,CIRBP,RPL38,RNH1,RBCK1,RCC1,SNRPB,DHPS,ACTR6,ZNF235,BOP1,TRPT1,DBI,ARPC2,PMFBP1,LI
13,BCAT2,MRPL4,FAM189B,CDCA3,SRSF2,MRPS22,HDAC9,LAS1L,HMGB1,SULF2,CYB561A3,BAIAP2,SSU7
13,BCAT2,MRPL4,FAM189B,CDCA3,SRSF2,MRPS22,HDAC9,LAS1L,HMGB1,SULF2,CYB561A3,BAIAP2,SSU7
1,TMEM67,CMC2,CIRBP,RPL38,RNH1,RBCK1,RCC1,SNRPB,DHPS,ACTR6,ZNF235,TRPT1,DBI,ARPC2,PMFB
1,TMEM67,CMC2,CIRBP,RPL38,RNH1,RBCK1,RCC1,SNRPB,DHPS,ACTR6,ZNF235,TRPT1,DBI,ARPC2,PMFB
,FAM189B,CDCA3,SRSF2,MRPS22,HDAC9,LAS1L,HMGB1,BAIAP2,SSU72,MYL6B,TMEM67,PILRB,CMC2,BL
,FAM189B,CDCA3,SRSF2,MRPS22,HDAC9,LAS1L,HMGB1,BAIAP2,SSU72,MYL6B,TMEM67,PILRB,CMC2,BL
4,SERPINB5,S100A13,BCAT2,MRPL4,FAM189B,CDCA3,SRSF2,MRPS22,LAS1L,HMGB1,SULF2,CYB561A3,B
4,SERPINB5,S100A13,BCAT2,MRPL4,FAM189B,CDCA3,SRSF2,MRPS22,LAS1L,HMGB1,SULF2,CYB561A3,B
IAP2,TMEM67,CMC2,BLOC1S1,CACYBP,CIRBP,RNH1,KIF4A,RBCK1,RCC1,SNRPB,DHPS,ACTR6,ARRDC1,TRP
IAP2,TMEM67,CMC2,BLOC1S1,CACYBP,CIRBP,RNH1,KIF4A,RBCK1,RCC1,SNRPB,DHPS,ACTR6,ARRDC1,TRP
00A13,BCAT2,VKORC1,CDCA3,SRSF2,MRPS22,OSMR,SEMA4D,LAS1L,HMGB1,SULF2,BAIAP2,SSU72,MYL6
00A13,BCAT2,VKORC1,CDCA3,SRSF2,MRPS22,OSMR,SEMA4D,LAS1L,HMGB1,SULF2,BAIAP2,SSU72,MYL6
NLL1,CST3,GNAS,GPR137,RPS24,SERPINB5,S100A13,BCAT2,CENPV,MRPL4,FAM189B,VKORC1,MRPL2,CD
S24,S100A13,BCAT2,MRPL4,FAM189B,VKORC1,MRPL2,SRSF2,MRPS22,HDAC9,KRT15,LAS1L,HMGB1,SULF
S24,S100A13,BCAT2,MRPL4,FAM189B,VKORC1,MRPL2,SRSF2,MRPS22,HDAC9,KRT15,LAS1L,HMGB1,SULF
CENPV,MRPL4,FAM189B,MRPL2,CDCA3,PCNA,SRSF2,MRPS22,KRT15,PPFIBP1,HMGB1,SULF2,CYB561A3,
CENPV,MRPL4,FAM189B,MRPL2,CDCA3,PCNA,SRSF2,MRPS22,KRT15,PPFIBP1,HMGB1,SULF2,CYB561A3,
CACYBP,RBCK1,LY75,RCC1,SNRPB,DHPS,ACTR6,ZNF235,TRPT1,DBI,ARPC2,LIMK2,GPATCH2L,BUD31,NPRL
INB5,CENPV,MRPL4,FAM189B,MRPL2,CDCA3,PCNA,SRSF2,MRPS22,KRT15,PPFIBP1,HMGB1,SULF2,BAIAP
INB5,CENPV,MRPL4,FAM189B,MRPL2,CDCA3,PCNA,SRSF2,MRPS22,KRT15,PPFIBP1,HMGB1,SULF2,BAIAP
GB1,SSU72,MYL6B,CMC2,CIRBP,RNH1,KIF4A,RBCK1,LY75,RCC1,SNRPB,DHPS,ACTR6,ARRDC1,RBP5,DBI,LI
GB1,SSU72,MYL6B,CMC2,CIRBP,RNH1,KIF4A,RBCK1,LY75,RCC1,SNRPB,DHPS,ACTR6,ARRDC1,RBP5,DBI,LI

2,TMEM67,CMC2,CIRBP,RNH1,KIF4A,RBCK1,LY75,RCC1,SNRPB,DHPS,ACTR6,ARRDC1,DBI,LIMK2,SIRT6,NU
2,TMEM67,CMC2,CIRBP,RNH1,KIF4A,RBCK1,LY75,RCC1,SNRPB,DHPS,ACTR6,ARRDC1,DBI,LIMK2,SIRT6,NU
AS,GPR137,RPS24,SERPINB5,BCAT2,CENPV,MRPL4,FAM189B,MRPL2,CDCA3,PCNA,SRSF2,OSMR,PPFIBP1,
AS,GPR137,RPS24,SERPINB5,BCAT2,CENPV,MRPL4,FAM189B,MRPL2,CDCA3,PCNA,SRSF2,OSMR,PPFIBP1,
C1S1,RBCK1,LY75,RCC1,SNRPB,ACTR6,TRPT1,DBI,ARPC2,LIMK2,BUD31,NPRL2,CTTN,EIF5A,COMMD5,NUD
SF2,MRPS22,LAS1L,HMGB1,SULF2,SSU72,TMEM67,FAM72A,CMC2,CACYBP,CIRBP,SMARCB1,RPL38,PRR1
SF2,MRPS22,LAS1L,HMGB1,SULF2,SSU72,TMEM67,FAM72A,CMC2,CACYBP,CIRBP,SMARCB1,RPL38,PRR1
NLL1,CST3,GNAS,GPR137,RPS24,S100A13,BCAT2,CENPV,MRPL4,VKORC1,MRPL2,CDCA3,PCNA,SRSF2,MRF
NLL1,CST3,GNAS,GPR137,RPS24,S100A13,BCAT2,CENPV,MRPL4,VKORC1,MRPL2,CDCA3,PCNA,SRSF2,MRF
,SRSF2,MRPS22,LAS1L,HMGB1,RAPGEF5,TMEM67,PILRB,CMC2,BLOC1S1,CACYBP,CIRBP,SMARCB1,RPL38
,SRSF2,MRPS22,LAS1L,HMGB1,RAPGEF5,TMEM67,PILRB,CMC2,BLOC1S1,CACYBP,CIRBP,SMARCB1,RPL38
3,GNAS,GPR137,RPS24,SERPINB5,S100A13,BCAT2,CENPV,MRPL4,FAM189B,MRPL2,CDCA3,PCNA,SRSF2,M
3,GNAS,GPR137,RPS24,SERPINB5,S100A13,BCAT2,CENPV,MRPL4,FAM189B,MRPL2,CDCA3,PCNA,SRSF2,M
CMC2,BLOC1S1,CACYBP,RHOF,RBCK1,RNF32,LY75,RCC1,SNRPB,ACTR6,ZNF235,TRPT1,DBI,ARPC2,LIMK2,B
PR137,RPS24,SERPINB5,BCAT2,CENPV,FAM189B,VKORC1,MRPL2,CDCA3,PCNA,SRSF2,MRPS22,KRT15,PPF
PR137,RPS24,SERPINB5,BCAT2,CENPV,FAM189B,VKORC1,MRPL2,CDCA3,PCNA,SRSF2,MRPS22,KRT15,PPF
CMC2,BLOC1S1,RHOF,SMARCB1,RPL38,RBCK1,RNF32,LY75,RCC1,SNRPB,ACTR6,TRPT1,ZC3HAV1,DBI,ARPC
,MRPS22,HDAC9,LAS1L,HMGB1,MYL6B,TMEM67,PILRB,CMC2,CACYBP,CIRBP,SMARCB1,RPL38,PRR13,RN
,MRPS22,HDAC9,LAS1L,HMGB1,MYL6B,TMEM67,PILRB,CMC2,CACYBP,CIRBP,SMARCB1,RPL38,PRR13,RN
PS22,SEMA4D,LAS1L,HMGB1,SSU72,MYL6B,SRP9,TMEM67,PILRB,CMC2,BLOC1S1,CACYBP,CIRBP,RHOF,SN
PS22,SEMA4D,LAS1L,HMGB1,SSU72,MYL6B,SRP9,TMEM67,PILRB,CMC2,BLOC1S1,CACYBP,CIRBP,RHOF,SN
GB1,MYL6B,CMC2,BLOC1S1,CACYBP,RHOF,SMARCB1,RNH1,DNASE1L1,RBCK1,RNF32,LY75,RCC1,SNRPB,D
AT2,MRPL4,SRSF2,MRPS22,LAS1L,HMGB1,SULF2,CYB561A3,RAPGEF5,SSU72,TMEM67,FAM72A,CMC2,CA
AT2,MRPL4,SRSF2,MRPS22,LAS1L,HMGB1,SULF2,CYB561A3,RAPGEF5,SSU72,TMEM67,FAM72A,CMC2,CA
7,CMC2,CIRBP,RNH1,KIF4A,RBCK1,RCC1,SNRPB,DHPS,ACTR6,DBI,LIMK2,GPATCH2L,BUD31,NPRL2,CTTN,E
7,CMC2,CIRBP,RNH1,KIF4A,RBCK1,RCC1,SNRPB,DHPS,ACTR6,DBI,LIMK2,GPATCH2L,BUD31,NPRL2,CTTN,E
ACYBP,PRR13,RBCK1,LY75,SNRPB,DHPS,ZNF235,TRPT1,DBI,ARPC2,LIMK2,GPATCH2L,BUD31,NPRL2,CTTN,N
CIRBP,RNH1,KIF4A,RBCK1,RCC1,SNRPB,DHPS,ACTR6,DBI,LIMK2,GPATCH2L,BUD31,NPRL2,CTTN,EIF5A,CON
CIRBP,RNH1,KIF4A,RBCK1,RCC1,SNRPB,DHPS,ACTR6,DBI,LIMK2,GPATCH2L,BUD31,NPRL2,CTTN,EIF5A,CON
2,CIRBP,RPL38,RNH1,KIF4A,RBCK1,RCC1,SNRPB,DHPS,ACTR6,ZNF235,TRPT1,DBI,ARPC2,PMFBP1,LIMK2,G
2,CIRBP,RPL38,RNH1,KIF4A,RBCK1,RCC1,SNRPB,DHPS,ACTR6,ZNF235,TRPT1,DBI,ARPC2,PMFBP1,LIMK2,G
67,CMC2,CIRBP,RNH1,KIF4A,RBCK1,RCC1,SNRPB,DHPS,ACTR6,DBI,ARPC2,LIMK2,GPATCH2L,BUD31,NPRL
67,CMC2,CIRBP,RNH1,KIF4A,RBCK1,RCC1,SNRPB,DHPS,ACTR6,DBI,ARPC2,LIMK2,GPATCH2L,BUD31,NPRL
,RHOF,RPL38,RBCK1,RCC1,SNRPB,ACTR6,ZNF235,TRPT1,DBI,ARPC2,LIMK2,BUD31,NPRL2,CTTN,EIF5A,CON
ACTR6,ZNF235,TRPT1,DBI,ARPC2,LIMK2,BUD31,NPRL2,CTTN,EIF5A,COMMD5,CHID1,NUDT22,PSMA6,RPS
CMC2,BLOC1S1,RHOF,SMARCB1,RPL38,RBCK1,RNF32,LY75,RCC1,SNRPB,DHPS,ACTR6,TRPT1,DBI,ARPC2,LI
RPS24,SERPINB5,CENPV,FAM189B,MRPL2,CDCA3,PCNA,SRSF2,KRT15,PPFIBP1,HMGB1,SULF2,BAIAP2,RAP
RPS24,SERPINB5,CENPV,FAM189B,MRPL2,CDCA3,PCNA,SRSF2,KRT15,PPFIBP1,HMGB1,SULF2,BAIAP2,RAP
RBP,RBCK1,ACTR6,TRPT1,DBI,BUD31,NPRL2,CTTN,EIF5A,COMMD5,NUDT22,THYN1,PSMA6,NUDT5,STYXL1
ST3,GNAS,RPS24,SERPINB5,BCAT2,CENPV,MRPL4,FAM189B,MRPL2,CDCA3,PCNA,SRSF2,KRT15,OSMR,PPF
31,SSU72,TMEM67,CMC2,CIRBP,RNH1,KIF4A,RBCK1,RNF32,RCC1,SNRPB,PARD3,SLC3A2,DHPS,ACTR6,DBI,
31,SSU72,TMEM67,CMC2,CIRBP,RNH1,KIF4A,RBCK1,RNF32,RCC1,SNRPB,PARD3,SLC3A2,DHPS,ACTR6,DBI,
GPR137,RPS24,SERPINB5,S100A13,BCAT2,CENPV,FAM189B,MRPL2,CDCA3,PCNA,SRSF2,KRT15,PPFIBP1,SE
GPR137,RPS24,SERPINB5,S100A13,BCAT2,CENPV,FAM189B,MRPL2,CDCA3,PCNA,SRSF2,KRT15,PPFIBP1,SE
NLL1,CST3,GNAS,GPR137,RPS24,SERPINB5,BCAT2,CENPV,MRPL4,FAM189B,VKORC1,MRPL2,CDCA3,PCNA
NLL1,CST3,GNAS,GPR137,RPS24,SERPINB5,BCAT2,CENPV,MRPL4,FAM189B,VKORC1,MRPL2,CDCA3,PCNA
PR137,RPS24,BCAT2,CENPV,MRPL4,FAM189B,VKORC1,PCNA,SRSF2,OSMR,PPFIBP1,HMGB1,SULF2,BAIAP

NLL1,CST3,GNAS,GPR137,SERPINB5,S100A13,BCAT2,CENPV,MRPL4,FAM189B,VKORC1,MRPL2,CDCA3,PC
NLL1,CST3,GNAS,GPR137,SERPINB5,S100A13,BCAT2,CENPV,MRPL4,FAM189B,VKORC1,MRPL2,CDCA3,PC
MC2,BLOC1S1,CIRBP,RPL38,PRR13,RNH1,RBCK1,RCC1,SNRPB,SLC3A2,DHPS,ACTR6,TRPT1,DBI,ARPC2,LIM
MC2,BLOC1S1,CIRBP,RPL38,PRR13,RNH1,RBCK1,RCC1,SNRPB,SLC3A2,DHPS,ACTR6,TRPT1,DBI,ARPC2,LIM
RHOF,RBCK1,RNF32,RCC1,SNRPB,ACTR6,DBI,BUD31,NPRL2,CTTN,EIF5A,COMMD5,SH2B1,PSMA6,ACOT8,
K1,SNRPB,ARPC2,LIMK2,NPRL2,CTTN,COMMD5,THYN1,RPL13A,PSMA6,NUDT5,STYXL1,IRF9,CPSF3L,TSG1
LAS1L,HMGB1,SULF2,TMEM67,FAM72A,CMC2,CACYBP,CIRBP,SMARCB1,RPL38,PRR13,RNH1,KIF4A,RBCK
LAS1L,HMGB1,SULF2,TMEM67,FAM72A,CMC2,CACYBP,CIRBP,SMARCB1,RPL38,PRR13,RNH1,KIF4A,RBCK
NLL1,CST3,GNAS,GPR137,RPS24,SERPINB5,S100A13,BCAT2,CENPV,MRPL4,FAM189B,VKORC1,MRPL2,CD
NLL1,CST3,GNAS,GPR137,RPS24,SERPINB5,S100A13,BCAT2,CENPV,MRPL4,FAM189B,VKORC1,MRPL2,CD
RCC1,SNRPB,DHPS,TRPT1,DBI,ARPC2,LIMK2,BUD31,NPRL2,CTTN,EIF5A,COMMD5,NUDT22,PSMA6,NUDT
ST3,GNAS,GPR137,RPS24,SERPINB5,BCAT2,CENPV,MRPL4,FAM189B,VKORC1,MRPL2,CDCA3,PCNA,SRSF2
ST3,GNAS,GPR137,RPS24,SERPINB5,BCAT2,CENPV,MRPL4,FAM189B,VKORC1,MRPL2,CDCA3,PCNA,SRSF2
S,GPR137,SERPINB5,CENPV,MRPL4,FAM189B,MRPL2,CDCA3,PCNA,SRSF2,MRPS22,PPFIBP1,HMGB1,BAIA
S,GPR137,SERPINB5,CENPV,MRPL4,FAM189B,MRPL2,CDCA3,PCNA,SRSF2,MRPS22,PPFIBP1,HMGB1,BAIA
Y75,SNRPB,DHPS,ZNF235,TRPT1,DBI,ARPC2,LIMK2,NPRL2,CTTN,NUDT22,SH2B1,RPL13A,SLC35G1,RCE1,Z
CENPV,FAM189B,MRPL2,CDCA3,PCNA,SRSF2,KRT15,PPFIBP1,HMGB1,BAIAP2,RAPGEF5,MYL6B,IQCG,BLO
CENPV,FAM189B,MRPL2,CDCA3,PCNA,SRSF2,KRT15,PPFIBP1,HMGB1,BAIAP2,RAPGEF5,MYL6B,IQCG,BLO
PR137,RPS24,S100A13,BCAT2,CENPV,MRPL4,VKORC1,PCNA,SRSF2,MRPS22,OSMR,PPFIBP1,HMGB1,SULF
GEF5,TMEM67,CMC2,CIRBP,SMARCB1,RPL38,RNH1,KIF4A,RBCK1,LY75,RCC1,SNRPB,DHPS,ACTR6,TRPT1,
GEF5,TMEM67,CMC2,CIRBP,SMARCB1,RPL38,RNH1,KIF4A,RBCK1,LY75,RCC1,SNRPB,DHPS,ACTR6,TRPT1,
B1,SSU72,TMEM67,CMC2,CACYBP,CIRBP,RPL38,RNH1,KIF4A,RBCK1,RNF32,SNRPB,PARD3,DHPS,ACTR6,Z
B1,SSU72,TMEM67,CMC2,CACYBP,CIRBP,RPL38,RNH1,KIF4A,RBCK1,RNF32,SNRPB,PARD3,DHPS,ACTR6,Z
NH1,KIF4A,RBCK1,RCC1,SNRPB,DHPS,ACTR6,DBI,LIMK2,BUD31,NPRL2,CTTN,EIF5A,COMMD5,UFD1L,ARH
NH1,KIF4A,RBCK1,RCC1,SNRPB,DHPS,ACTR6,DBI,LIMK2,BUD31,NPRL2,CTTN,EIF5A,COMMD5,UFD1L,ARH
1,RPL38,RBCK1,LY75,RCC1,SNRPB,DHPS,DBI,ARPC2,LIMK2,GPATCH2L,BUD31,NPRL2,CTTN,EIF5A,COMM
NLL1,CST3,GPR137,RPS24,S100A13,BCAT2,CENPV,VKORC1,MRPL2,CDCA3,PCNA,SRSF2,MRPS22,KRT15,O
NLL1,CST3,GPR137,RPS24,S100A13,BCAT2,CENPV,VKORC1,MRPL2,CDCA3,PCNA,SRSF2,MRPS22,KRT15,O
HMGB1,TMEM67,CMC2,CACYBP,CIRBP,RPL38,PRR13,RNH1,DNASE1L1,RBCK1,RCC1,SNRPB,DHPS,ACTR6,
HMGB1,TMEM67,CMC2,CACYBP,CIRBP,RPL38,PRR13,RNH1,DNASE1L1,RBCK1,RCC1,SNRPB,DHPS,ACTR6,
L4,SRSF2,MRPS22,LAS1L,RAPGEF5,SSU72,MYL6B,FAM122C,CMC2,BLOC1S1,CACYBP,RHOF,PRR13,RNH1,
T3,GNAS,GPR137,RPS24,S100A13,CENPV,MRPL4,FAM189B,VKORC1,MRPL2,CDCA3,PCNA,SRSF2,OSMR,PP
T3,GNAS,GPR137,RPS24,S100A13,CENPV,MRPL4,FAM189B,VKORC1,MRPL2,CDCA3,PCNA,SRSF2,OSMR,PP
6B,CMC2,BLOC1S1,DNASE1L1,RBCK1,LY75,RCC1,SNRPB,DHPS,ACTR6,ZNF235,TRPT1,DBI,ARPC2,LIMK2,B
IRBP,RPL38,RNH1,RBCK1,RCC1,SNRPB,SLC3A2,DHPS,ACTR6,TRPT1,NUMA1,GPATCH2L,SUMF2,BUD31,NP
37,RPS24,SERPINB5,BCAT2,CENPV,VKORC1,CDCA3,PCNA,SRSF2,OSMR,HMGB1,SULF2,RAPGEF5,SSU72,F
37,RPS24,SERPINB5,BCAT2,CENPV,VKORC1,CDCA3,PCNA,SRSF2,OSMR,HMGB1,SULF2,RAPGEF5,SSU72,F
37,RPS24,SERPINB5,BCAT2,CENPV,MRPL4,MRPL2,CDCA3,PCNA,SRSF2,OSMR,HMGB1,SULF2,CYB561A3,B
37,RPS24,SERPINB5,BCAT2,CENPV,MRPL4,MRPL2,CDCA3,PCNA,SRSF2,OSMR,HMGB1,SULF2,CYB561A3,B
T3,GPR137,RPS24,BCAT2,CDCA3,SRSF2,MRPS22,KRT15,OSMR,LAS1L,BAIAP2,RAPGEF5,SSU72,MYL6B,IQC
SERPINB5,CENPV,FAM189B,MRPL2,CDCA3,PCNA,SRSF2,MRPS22,KRT15,PPFIBP1,HMGB1,SULF2,BAIAP2,
SERPINB5,CENPV,FAM189B,MRPL2,CDCA3,PCNA,SRSF2,MRPS22,KRT15,PPFIBP1,HMGB1,SULF2,BAIAP2,
BCAT2,VKORC1,CDCA3,SRSF2,MRPS22,LAS1L,HMGB1,SULF2,CYB561A3,RAPGEF5,SSU72,SRP9,IQCG,TME
BCAT2,VKORC1,CDCA3,SRSF2,MRPS22,LAS1L,HMGB1,SULF2,CYB561A3,RAPGEF5,SSU72,SRP9,IQCG,TME
137,SERPINB5,CENPV,MRPL2,CDCA3,PCNA,SRSF2,HMGB1,SULF2,BAIAP2,RAPGEF5,MYL6B,FAM72A,BLO
,ARPC2,LIMK2,NPRL2,CTTN,EIF5A,COMMD5,THYN1,ZNF410,NUDT5,STYXL1,VRK3,LRRC1,CPSF3L,TSG101,
,GNAS,GPR137,BCAT2,CENPV,MRPL4,FAM189B,VKORC1,MRPL2,CDCA3,PCNA,SRSF2,MRPS22,PPFIBP1,LA

,GNAS,GPR137,BCAT2,CENPV,MRPL4,FAM189B,VKORC1,MRPL2,CDCA3,PCNA,SRSF2,MRPS22,PPFIBP1,LA
HMGB1,SSU72,MYL6B,TMEM67,CMC2,CIRBP,RHOF,SMARCB1,RNH1,KIF4A,RBCK1,RNF32,LY75,SNRPB,PAR
HMGB1,SSU72,MYL6B,TMEM67,CMC2,CIRBP,RHOF,SMARCB1,RNH1,KIF4A,RBCK1,RNF32,LY75,SNRPB,PAR
35,BCAT2,CENPV,FAM189B,VKORC1,MRPL2,CDCA3,PCNA,SRSF2,KRT15,PPFIBP1,HMGB1,CYB561A3,BAIAP
35,BCAT2,CENPV,FAM189B,VKORC1,MRPL2,CDCA3,PCNA,SRSF2,KRT15,PPFIBP1,HMGB1,CYB561A3,BAIAP
T3,GNAS,GPR137,SERPINB5,S100A13,BCAT2,CENPV,MRPL4,FAM189B,VKORC1,MRPL2,CDCA3,PCNA,SRSF2
C2,BLOC1S1,CIRBP,SMARCB1,RBCK1,LY75,RCC1,SNRPB,DHPS,ACTR6,TRPT1,NUMA1,BUD31,NPRL2,CTTN,
RHOF,SMARCB1,RBCK1,LY75,RCC1,SNRPB,ACTR6,ARPC2,BUD31,NPRL2,CTTN,EIF5A,COMMD5,THYN1,PSM
5,DBI,ARPC2,LIMK2,NPRL2,CTTN,SH2B1,RCE1,STYXL1,VRK3,CPSF3L,TSG101,PSMA1,FAM3A,RPS25,TTC9C
NLL1,CST3,GNAS,S100A13,CENPV,MRPL4,FAM189B,MRPL2,CDCA3,BAIAP2,RAPGEF5,SSU72,FAM122C,TM
NLL1,CST3,GNAS,S100A13,CENPV,MRPL4,FAM189B,MRPL2,CDCA3,BAIAP2,RAPGEF5,SSU72,FAM122C,TM
,BCAT2,CENPV,MRPL4,FAM189B,VKORC1,MRPL2,CDCA3,PCNA,SRSF2,MRPS22,KRT15,OSMR,LAS1L,HMGB
,BCAT2,CENPV,MRPL4,FAM189B,VKORC1,MRPL2,CDCA3,PCNA,SRSF2,MRPS22,KRT15,OSMR,LAS1L,HMGB
NLL1,CST3,GNAS,GPR137,SERPINB5,S100A13,BCAT2,CENPV,MRPL4,FAM189B,VKORC1,PCNA,SRSF2,PPFIB
SERPINB5,CENPV,FAM189B,MRPL2,CDCA3,PCNA,SRSF2,MRPS22,PPFIBP1,HMGB1,BAIAP2,RAPGEF5,MYL6
SERPINB5,CENPV,FAM189B,MRPL2,CDCA3,PCNA,SRSF2,MRPS22,PPFIBP1,HMGB1,BAIAP2,RAPGEF5,MYL6
NLL1,CST3,GNAS,GPR137,SERPINB5,S100A13,BCAT2,CENPV,MRPL4,FAM189B,VKORC1,MRPL2,CDCA3,PCN
NLL1,CST3,GNAS,GPR137,SERPINB5,S100A13,BCAT2,CENPV,MRPL4,FAM189B,VKORC1,MRPL2,CDCA3,PCN
CAT2,CENPV,MRPL4,FAM189B,MRPL2,CDCA3,PCNA,SRSF2,KRT15,OSMR,LAS1L,HMGB1,SULF2,BAIAP2,RA
GPR137,RPS24,BCAT2,CENPV,MRPL4,FAM189B,MRPL2,CDCA3,PCNA,SRSF2,OSMR,PPFIBP1,SULF2,CYB56
CA3,PCNA,MRPS22,SEMA4D,LAS1L,HMGB1,BAIAP2,RAPGEF5,SSU72,MYL6B,SRP9,TMEM67,PILRB,CMC2,B
CA3,PCNA,MRPS22,SEMA4D,LAS1L,HMGB1,BAIAP2,RAPGEF5,SSU72,MYL6B,SRP9,TMEM67,PILRB,CMC2,B
,SNRPB,ACTR6,TRPT1,ARPC2,LIMK2,BUD31,NPRL2,CTTN,COMMD5,NUDT22,SH2B1,PSMA6,RPS9,MRPL33
OC1,TRPT1,ARPC2,LIMK2,GPATCH2L,BUD31,CTTN,COMMD5,NUDT22,THYN1,RPL13A,RCE1,NUDT5,STYXL1
37,BCAT2,VKORC1,PCNA,SRSF2,MRPS22,OSMR,BAIAP2,RAPGEF5,SSU72,MYL6B,IQCG,PILRB,CMC2,RHOF,S
,SERPINB5,S100A13,BCAT2,FAM189B,PCNA,MRPS22,LAS1L,HMGB1,CYB561A3,BAIAP2,RAPGEF5,MYL6B,S
,SERPINB5,S100A13,BCAT2,FAM189B,PCNA,MRPS22,LAS1L,HMGB1,CYB561A3,BAIAP2,RAPGEF5,MYL6B,S
C1,SNRPB,ACTR6,BUD31,NPRL2,CTTN,EIF5A,COMMD5,PSMA6,RPS9,MRPL33,NUDT5,STYXL1,PPM1K,SON
CENPV,FAM189B,MRPL2,CDCA3,PCNA,SRSF2,MRPS22,KRT15,PPFIBP1,HMGB1,SULF2,BAIAP2,RAPGEF5,M
CENPV,FAM189B,MRPL2,CDCA3,PCNA,SRSF2,MRPS22,KRT15,PPFIBP1,HMGB1,SULF2,BAIAP2,RAPGEF5,M
CENPV,FAM189B,MRPL2,CDCA3,PCNA,SRSF2,MRPS22,KRT15,PPFIBP1,HMGB1,SULF2,BAIAP2,RAPGEF5,M
CENPV,FAM189B,MRPL2,CDCA3,PCNA,SRSF2,MRPS22,KRT15,PPFIBP1,HMGB1,SULF2,BAIAP2,RAPGEF5,M
AS,GPR137,RPS24,SERPINB5,BCAT2,CENPV,MRPL4,FAM189B,VKORC1,MRPL2,CDCA3,PCNA,SRSF2,OSMR,
AS,GPR137,RPS24,SERPINB5,BCAT2,CENPV,MRPL4,FAM189B,VKORC1,MRPL2,CDCA3,PCNA,SRSF2,OSMR,
T3,GNAS,GPR137,S100A13,BCAT2,CENPV,MRPL4,VKORC1,CDCA3,PCNA,SRSF2,OSMR,PPFIBP1,LAS1L,HMG
NLL1,CST3,GNAS,GPR137,RPS24,SERPINB5,S100A13,BCAT2,CENPV,MRPL4,FAM189B,VKORC1,MRPL2,CDC
NLL1,CST3,GNAS,GPR137,RPS24,SERPINB5,S100A13,BCAT2,CENPV,MRPL4,FAM189B,VKORC1,MRPL2,CDC
T3,GNAS,GPR137,SERPINB5,BCAT2,CENPV,MRPL4,FAM189B,MRPL2,CDCA3,PCNA,SRSF2,KRT15,OSMR,PPF
S,GPR137,RPS24,S100A13,CENPV,MRPL4,MRPL2,CDCA3,PCNA,SRSF2,MRPS22,HDAC9,OSMR,SULF2,BAIA
T3,GNAS,GPR137,RPS24,S100A13,CENPV,MRPL4,VKORC1,MRPL2,CDCA3,PCNA,SRSF2,MRPS22,HDAC9,OS
T3,GNAS,GPR137,RPS24,S100A13,CENPV,MRPL4,VKORC1,MRPL2,CDCA3,PCNA,SRSF2,MRPS22,HDAC9,OS
MRPL4,FAM189B,VKORC1,PCNA,SRSF2,MRPS22,LAS1L,HMGB1,SULF2,CYB561A3,RAPGEF5,SSU72,TMEM6
MRPL4,FAM189B,VKORC1,PCNA,SRSF2,MRPS22,LAS1L,HMGB1,SULF2,CYB561A3,RAPGEF5,SSU72,TMEM6
RPINB5,BCAT2,MRPL2,CDCA3,PCNA,SRSF2,PPFIBP1,HMGB1,BAIAP2,MYL6B,IQCG,FAM72A,YBX3,BLOC1S1
RPINB5,BCAT2,MRPL2,CDCA3,PCNA,SRSF2,PPFIBP1,HMGB1,BAIAP2,MYL6B,IQCG,FAM72A,YBX3,BLOC1S1
T3,GNAS,GPR137,RPS24,S100A13,CENPV,MRPL4,VKORC1,MRPL2,CDCA3,PCNA,SRSF2,MRPS22,OSMR,HM
ERPINB5,BCAT2,VKORC1,MRPL2,CDCA3,PCNA,SRSF2,MRPS22,HMGB1,BAIAP2,MYL6B,IQCG,YBX3,BLOC1S

ERPINB5,BCAT2,VKORC1,MRPL2,CDCA3,PCNA,SRSF2,MRPS22,HMGB1,BAIAP2,MYL6B,IQCG,YBX3,BLOC1S
DBI,BUD31,NPRL2,TBL2,CTTN,COMMD5,SH2B1,NOP16,RPL13A,PSMA6,RPS9,STYXL1,FIS1,CPSF3L,TSG101,
137,BCAT2,CENPV,MRPL4,FAM189B,VKORC1,MRPL2,PCNA,SRSF2,MRPS22,PPFIBP1,LAS1L,HMGB1,SULF2,
NLL1,CST3,GPR137,SERPINB5,S100A13,BCAT2,CENPV,MRPL4,FAM189B,VKORC1,MRPL2,CDCA3,PCNA,SRS
NLL1,CST3,GPR137,SERPINB5,S100A13,BCAT2,CENPV,MRPL4,FAM189B,VKORC1,MRPL2,CDCA3,PCNA,SRS
AS,GPR137,RPS24,BCAT2,CENPV,MRPL4,FAM189B,MRPL2,CDCA3,PCNA,SRSF2,OSMR,PPFIBP1,BAIAP2,RA
AS,GPR137,RPS24,BCAT2,CENPV,MRPL4,FAM189B,MRPL2,CDCA3,PCNA,SRSF2,OSMR,PPFIBP1,BAIAP2,RA
3,RBCK1,SNRPB,ACTR6,TRPT1,BUD31,NPRL2,TBL2,CTTN,EIF5A,COMMD5,NUDT22,USP38,SH2B1,RPL13A,
INB5,CENPV,FAM189B,MRPL2,CDCA3,PCNA,SRSF2,MRPS22,PPFIBP1,HMGB1,SULF2,BAIAP2,RAPGEF5,MY
INB5,CENPV,FAM189B,MRPL2,CDCA3,PCNA,SRSF2,MRPS22,PPFIBP1,HMGB1,SULF2,BAIAP2,RAPGEF5,MY
4,BCAT2,CENPV,MRPL4,FAM189B,VKORC1,PCNA,SRSF2,MRPS22,KRT15,OSMR,HMGB1,SULF2,BAIAP2,RAI
4,BCAT2,CENPV,MRPL4,FAM189B,VKORC1,PCNA,SRSF2,MRPS22,KRT15,OSMR,HMGB1,SULF2,BAIAP2,RAI
ST3,GNAS,GPR137,RPS24,SERPINB5,BCAT2,CENPV,MRPL4,FAM189B,VKORC1,MRPL2,CDCA3,PCNA,SRSF2
ST3,GNAS,GPR137,RPS24,SERPINB5,BCAT2,CENPV,MRPL4,FAM189B,VKORC1,MRPL2,CDCA3,PCNA,SRSF2
137,BCAT2,CENPV,MRPL4,VKORC1,CDCA3,PCNA,SRSF2,OSMR,PPFIBP1,HMGB1,SULF2,BAIAP2,RAPGEF5,S
V,MRPL4,VKORC1,MRPL2,CDCA3,PCNA,SRSF2,OSMR,PPFIBP1,HMGB1,BAIAP2,RAPGEF5,MYL6B,IQCG,TME
V,MRPL4,VKORC1,MRPL2,CDCA3,PCNA,SRSF2,OSMR,PPFIBP1,HMGB1,BAIAP2,RAPGEF5,MYL6B,IQCG,TME
V,MRPL4,VKORC1,MRPL2,CDCA3,PCNA,SRSF2,OSMR,PPFIBP1,HMGB1,BAIAP2,RAPGEF5,MYL6B,IQCG,TME
V,MRPL4,VKORC1,MRPL2,CDCA3,PCNA,SRSF2,OSMR,PPFIBP1,HMGB1,BAIAP2,RAPGEF5,MYL6B,IQCG,TME
V,MRPL4,VKORC1,MRPL2,CDCA3,PCNA,SRSF2,OSMR,PPFIBP1,HMGB1,BAIAP2,RAPGEF5,MYL6B,IQCG,TME
V,MRPL4,VKORC1,MRPL2,CDCA3,PCNA,SRSF2,OSMR,PPFIBP1,HMGB1,BAIAP2,RAPGEF5,MYL6B,IQCG,TME
RPL2,CDCA3,PCNA,SRSF2,MRPS22,PPFIBP1,HMGB1,BAIAP2,RAPGEF5,MYL6B,IQCG,BLOC1S1,CACYBP,RHC
RPL2,CDCA3,PCNA,SRSF2,MRPS22,PPFIBP1,HMGB1,BAIAP2,RAPGEF5,MYL6B,IQCG,BLOC1S1,CACYBP,RHC
MRPS22,KRT15,OSMR,SEMA4D,LAS1L,HMGB1,RAPGEF5,SSU72,MYL6B,IQCG,TMEM67,PILRB,CMC2,BLOC
MRPS22,KRT15,OSMR,SEMA4D,LAS1L,HMGB1,RAPGEF5,SSU72,MYL6B,IQCG,TMEM67,PILRB,CMC2,BLOC
MRPS22,OSMR,SEMA4D,LAS1L,HMGB1,SSU72,MYL6B,IQCG,TMEM67,CMC2,CIRBP,RPL38,RNH1,DNASE1L
MRPS22,OSMR,SEMA4D,LAS1L,HMGB1,SSU72,MYL6B,IQCG,TMEM67,CMC2,CIRBP,RPL38,RNH1,DNASE1L
ENPV,MRPL4,FAM189B,VKORC1,MRPL2,PCNA,SRSF2,KRT15,PPFIBP1,HMGB1,BAIAP2,RAPGEF5,MYL6B,FA
ENPV,MRPL4,FAM189B,VKORC1,MRPL2,PCNA,SRSF2,KRT15,PPFIBP1,HMGB1,BAIAP2,RAPGEF5,MYL6B,FA
IF4A,RBCK1,SNRPB,DHPS,ZNF235,TRPT1,DBI,ARPC2,PMFBP1,LIMK2,BUD31,CTTN,EIF5A,COMMD5,NUDT2
IF4A,RBCK1,SNRPB,DHPS,ZNF235,TRPT1,DBI,ARPC2,PMFBP1,LIMK2,BUD31,CTTN,EIF5A,COMMD5,NUDT2
IPV,FAM189B,MRPL2,CDCA3,PCNA,SRSF2,MRPS22,KRT15,PPFIBP1,HMGB1,BAIAP2,RAPGEF5,MYL6B,IQCG
IPV,FAM189B,MRPL2,CDCA3,PCNA,SRSF2,MRPS22,KRT15,PPFIBP1,HMGB1,BAIAP2,RAPGEF5,MYL6B,IQCG
3CK1,LY75,SNRPB,ZNF235,BOP1,TRPT1,ARPC2,PMFBP1,LIMK2,GPATCH2L,BUD31,NPRL2,CTTN,EIF5A,COM
3CK1,LY75,SNRPB,ZNF235,BOP1,TRPT1,ARPC2,PMFBP1,LIMK2,GPATCH2L,BUD31,NPRL2,CTTN,EIF5A,COM
7,BCAT2,CENPV,MRPL4,FAM189B,VKORC1,MRPL2,PCNA,SRSF2,MRPS22,PPFIBP1,LAS1L,HMGB1,SULF2,B
7,BCAT2,CENPV,MRPL4,FAM189B,VKORC1,MRPL2,PCNA,SRSF2,MRPS22,PPFIBP1,LAS1L,HMGB1,SULF2,B
31,RAPGEF5,TMEM67,CMC2,BLOC1S1,CACYBP,CIRBP,PRR13,KIF4A,LY75,ATG16L2,SNRPB,PARD3,SLC3A2,
R137,SERPINB5,FAM189B,VKORC1,CDCA3,PCNA,SRSF2,OSMR,PPFIBP1,LAS1L,HMGB1,SULF2,RAPGEF5,S
R137,SERPINB5,FAM189B,VKORC1,CDCA3,PCNA,SRSF2,OSMR,PPFIBP1,LAS1L,HMGB1,SULF2,RAPGEF5,S
NB5,CENPV,FAM189B,MRPL2,CDCA3,PCNA,SRSF2,MRPS22,OSMR,PPFIBP1,HMGB1,BAIAP2,RAPGEF5,MYL
NB5,CENPV,FAM189B,MRPL2,CDCA3,PCNA,SRSF2,MRPS22,OSMR,PPFIBP1,HMGB1,BAIAP2,RAPGEF5,MYL
PS24,BCAT2,MRPL4,CDCA3,PCNA,SRSF2,OSMR,CYB561A3,BAIAP2,RAPGEF5,FAM122C,SRP9,IQCG,SULT1
V,VKORC1,CDCA3,PCNA,SRSF2,OSMR,PPFIBP1,RAPGEF5,SSU72,SRP9,IQCG,TMEM67,CACYBP,SMARCB1,R
V,VKORC1,CDCA3,PCNA,SRSF2,OSMR,PPFIBP1,RAPGEF5,SSU72,SRP9,IQCG,TMEM67,CACYBP,SMARCB1,R
ST3,GPR137,RPS24,S100A13,BCAT2,MRPL4,FAM189B,VKORC1,MRPL2,CDCA3,PCNA,SRSF2,MRPS22,KRT1
ST3,GPR137,RPS24,S100A13,BCAT2,MRPL4,FAM189B,VKORC1,MRPL2,CDCA3,PCNA,SRSF2,MRPS22,KRT1

NH1,RBCK1,SNRPB,DHPS,ACTR6,DBI,LIMK2,BUD31,CTTN,EIF5A,UFD1L,ARHGEF39,ACOT8,STYXL1,LZTFL1,
NH1,RBCK1,SNRPB,DHPS,ACTR6,DBI,LIMK2,BUD31,CTTN,EIF5A,UFD1L,ARHGEF39,ACOT8,STYXL1,LZTFL1,
T3,GNAS,GPR137,RPS24,SERPINB5,BCAT2,CENPV,MRPL4,FAM189B,VKORC1,MRPL2,PCNA,SRSF2,KRT15,O
T3,GNAS,GPR137,RPS24,SERPINB5,BCAT2,CENPV,MRPL4,FAM189B,VKORC1,MRPL2,PCNA,SRSF2,KRT15,O
PR137,RPS24,SERPINB5,CENPV,FAM189B,MRPL2,CDCA3,PCNA,SRSF2,MRPS22,KRT15,PPFIBP1,HMGB1,B
PR137,RPS24,SERPINB5,CENPV,FAM189B,MRPL2,CDCA3,PCNA,SRSF2,MRPS22,KRT15,PPFIBP1,HMGB1,B
R137,BCAT2,CENPV,FAM189B,VKORC1,MRPL2,PCNA,SRSF2,MRPS22,KRT15,OSMR,LAS1L,HMGB1,SULF2,
R137,BCAT2,CENPV,FAM189B,VKORC1,MRPL2,PCNA,SRSF2,MRPS22,KRT15,OSMR,LAS1L,HMGB1,SULF2,
C2,CIRBP,RNH1,KIF4A,RBCK1,RNF32,RCC1,SNRPB,PARD3,DHPS,ACTR6,ARPC2,LIMK2,GPATCH2L,BUD31,N
C2,CIRBP,RNH1,KIF4A,RBCK1,RNF32,RCC1,SNRPB,PARD3,DHPS,ACTR6,ARPC2,LIMK2,GPATCH2L,BUD31,N
T3,GNAS,GPR137,RPS24,CENPV,FAM189B,VKORC1,CDCA3,PCNA,SRSF2,OSMR,PPFIBP1,HMGB1,SULF2,BA
T3,GNAS,GPR137,RPS24,CENPV,FAM189B,VKORC1,CDCA3,PCNA,SRSF2,OSMR,PPFIBP1,HMGB1,SULF2,BA
PL2,CDCA3,PCNA,SRSF2,KRT15,PPFIBP1,HMGB1,BAIAP2,RAPGEF5,MYL6B,IQCG,BLOC1S1,CACYBP,CIRBP,
PL2,CDCA3,PCNA,SRSF2,KRT15,PPFIBP1,HMGB1,BAIAP2,RAPGEF5,MYL6B,IQCG,BLOC1S1,CACYBP,CIRBP,
ORC1,MRPL2,CDCA3,PCNA,SRSF2,HMGB1,BAIAP2,MYL6B,YBX3,BLOC1S1,RHOF,SMARCB1,PRR13,RNH1,D
ORC1,MRPL2,CDCA3,PCNA,SRSF2,HMGB1,BAIAP2,MYL6B,YBX3,BLOC1S1,RHOF,SMARCB1,PRR13,RNH1,D
,HMGB1,MYL6B,TMEM67,CMC2,CACYBP,CIRBP,PRR13,RNH1,RBCK1,RCC1,SNRPB,DHPS,ZNF235,TRPT1,Z
,HMGB1,MYL6B,TMEM67,CMC2,CACYBP,CIRBP,PRR13,RNH1,RBCK1,RCC1,SNRPB,DHPS,ZNF235,TRPT1,Z
B,MRPL2,CDCA3,PCNA,SRSF2,MRPS22,PPFIBP1,HMGB1,BAIAP2,RAPGEF5,SSU72,MYL6B,IQCG,BLOC1S1,R
B,MRPL2,CDCA3,PCNA,SRSF2,MRPS22,PPFIBP1,HMGB1,BAIAP2,RAPGEF5,SSU72,MYL6B,IQCG,BLOC1S1,R
V,VKORC1,CDCA3,PCNA,SRSF2,OSMR,PPFIBP1,SSU72,SRP9,IQCG,TMEM67,CACYBP,SMARCB1,RPL38,PTO
V,VKORC1,CDCA3,PCNA,SRSF2,OSMR,PPFIBP1,SSU72,SRP9,IQCG,TMEM67,CACYBP,SMARCB1,RPL38,PTO
RPS24,SERPINB5,S100A13,CENPV,FAM189B,MRPL2,CDCA3,PCNA,SRSF2,MRPS22,PPFIBP1,HMGB1,SULF2
RPS24,SERPINB5,S100A13,CENPV,FAM189B,MRPL2,CDCA3,PCNA,SRSF2,MRPS22,PPFIBP1,HMGB1,SULF2
NH1,RBCK1,SNRPB,DHPS,ACTR6,DBI,LIMK2,BUD31,CTTN,EIF5A,UFD1L,ARHGEF39,ACOT8,STYXL1,LZTFL1,
NH1,RBCK1,SNRPB,DHPS,ACTR6,DBI,LIMK2,BUD31,CTTN,EIF5A,UFD1L,ARHGEF39,ACOT8,STYXL1,LZTFL1,
RPS22,HDAC9,OSMR,IQCG,PILRB,CMC2,RHOF,PRR13,STARD10,RNH1,DNASE1L1,RNF32,SLC3A2,ACTR6,S
,HMGB1,MYL6B,CMC2,BLOC1S1,RBCK1,RNF32,RCC1,SNRPB,DHPS,ACTR6,DBI,ARPC2,LIMK2,BUD31,NPRL
,PCNA,SRSF2,SULF2,BAIAP2,RAPGEF5,MYL6B,RHOF,SMARCB1,PRR13,RNH1,RBCK1,PTOV1,FASTK,RNF32,L
CTR6,BOP1,TRPT1,RBP5,LIMK2,SIRT6,NUMA1,SAE1,CTTN,TSPAN11,EIF5A,COMMD5,CHID1,NUDT22,UFD
CTR6,BOP1,TRPT1,RBP5,LIMK2,SIRT6,NUMA1,SAE1,CTTN,TSPAN11,EIF5A,COMMD5,CHID1,NUDT22,UFD
5,S100A13,CENPV,MRPL4,VKORC1,MRPL2,CDCA3,PCNA,SRSF2,OSMR,PPFIBP1,SEMA4D,HMGB1,SULF2,B
5,S100A13,CENPV,MRPL4,VKORC1,MRPL2,CDCA3,PCNA,SRSF2,OSMR,PPFIBP1,SEMA4D,HMGB1,SULF2,B
T3,GNAS,GPR137,BCAT2,CENPV,MRPL4,FAM189B,VKORC1,MRPL2,CDCA3,PCNA,SRSF2,MRPS22,OSMR,PP
T3,GNAS,GPR137,BCAT2,CENPV,MRPL4,FAM189B,VKORC1,MRPL2,CDCA3,PCNA,SRSF2,MRPS22,OSMR,PP
R137,SERPINB5,CENPV,MRPL4,FAM189B,VKORC1,PCNA,SRSF2,OSMR,PPFIBP1,LAS1L,HMGB1,SULF2,RAP
R137,SERPINB5,CENPV,MRPL4,FAM189B,VKORC1,PCNA,SRSF2,OSMR,PPFIBP1,LAS1L,HMGB1,SULF2,RAP
MRPS22,OSMR,SEMA4D,LAS1L,HMGB1,RAPGEF5,SSU72,MYL6B,IQCG,TMEM67,CMC2,BLOC1S1,CIRBP,RP
MRPS22,OSMR,SEMA4D,LAS1L,HMGB1,RAPGEF5,SSU72,MYL6B,IQCG,TMEM67,CMC2,BLOC1S1,CIRBP,RP
NA,SRSF2,KRT15,PPFIBP1,HMGB1,BAIAP2,RAPGEF5,MYL6B,IQCG,BLOC1S1,CACYBP,CIRBP,RHOF,SMARCB
NA,SRSF2,KRT15,PPFIBP1,HMGB1,BAIAP2,RAPGEF5,MYL6B,IQCG,BLOC1S1,CACYBP,CIRBP,RHOF,SMARCB
NH1,KIF4A,RBCK1,SNRPB,DHPS,ACTR6,DBI,LIMK2,BUD31,CTTN,EIF5A,COMMD5,UFD1L,USP38,ARHGEF3
NH1,KIF4A,RBCK1,SNRPB,DHPS,ACTR6,DBI,LIMK2,BUD31,CTTN,EIF5A,COMMD5,UFD1L,USP38,ARHGEF3
PR137,RPS24,S100A13,BCAT2,CENPV,MRPL4,FAM189B,VKORC1,MRPL2,CDCA3,PCNA,MRPS22,OSMR,SE
PR137,RPS24,S100A13,BCAT2,CENPV,MRPL4,FAM189B,VKORC1,MRPL2,CDCA3,PCNA,MRPS22,OSMR,SE
IRBP,SMARCB1,RPL38,RNH1,KIF4A,RBCK1,LY75,RCC1,TRPT1,PMFBP1,NPRL2,CRYGS,CTTN,COMMD5,NUD
IRBP,SMARCB1,RPL38,RNH1,KIF4A,RBCK1,LY75,RCC1,TRPT1,PMFBP1,NPRL2,CRYGS,CTTN,COMMD5,NUD

FAM72A,BLOC1S1,RHOF,PRR13,RNH1,RBCK1,FASTK,TMEM135,SET,ARRDC1,TRPT1,RBP5,LIMK2,NUMA1,G
FAM72A,BLOC1S1,RHOF,PRR13,RNH1,RBCK1,FASTK,TMEM135,SET,ARRDC1,TRPT1,RBP5,LIMK2,NUMA1,G
V,FAM189B,MRPL2,PCNA,SRSF2,PPFIBP1,HMGB1,CYB561A3,BAIAP2,RAPGEF5,MYL6B,IQCG,BLOC1S1,CAC
V,FAM189B,MRPL2,PCNA,SRSF2,PPFIBP1,HMGB1,CYB561A3,BAIAP2,RAPGEF5,MYL6B,IQCG,BLOC1S1,CAC
TMEM67,CMC2,DNASE1L1,KIF4A,RBCK1,RCC1,SNRPB,DHPS,ZNF235,DBI,ARPC2,LIMK2,BUD31,NPRL2,CTTN
2,PPFIBP1,LAS1L,HMGB1,RAPGEF5,TMEM67,PILRB,FAM72A,CMC2,BLOC1S1,CIRBP,RPL38,RNH1,DNASE1
2,PPFIBP1,LAS1L,HMGB1,RAPGEF5,TMEM67,PILRB,FAM72A,CMC2,BLOC1S1,CIRBP,RPL38,RNH1,DNASE1
AP2,MYL6B,FAM72A,BLOC1S1,RHOF,SMARCB1,PRR13,STARD10,RNH1,RBCK1,PTOV1,TMEM135,RNF32,BC
AP2,MYL6B,FAM72A,BLOC1S1,RHOF,SMARCB1,PRR13,STARD10,RNH1,RBCK1,PTOV1,TMEM135,RNF32,BC
24,SERPINB5,S100A13,BCAT2,CENPV,MRPL4,FAM189B,VKORC1,MRPL2,CDCA3,PCNA,SRSF2,OSMR,PPFIBP
137,BCAT2,CENPV,MRPL4,VKORC1,MRPL2,CDCA3,PCNA,SRSF2,OSMR,PPFIBP1,HMGB1,SULF2,BAIAP2,RAP
137,BCAT2,CENPV,MRPL4,VKORC1,MRPL2,CDCA3,PCNA,SRSF2,OSMR,PPFIBP1,HMGB1,SULF2,BAIAP2,RAP
S1L,HMGB1,MYL6B,TMEM67,CMC2,BLOC1S1,CIRBP,PRR13,RNH1,DNASE1L1,KIF4A,RBCK1,RCC1,SNRPB,D
S1L,HMGB1,MYL6B,TMEM67,CMC2,BLOC1S1,CIRBP,PRR13,RNH1,DNASE1L1,KIF4A,RBCK1,RCC1,SNRPB,D
ERPINB5,CENPV,VKORC1,MRPL2,PCNA,SRSF2,LAS1L,HMGB1,SULF2,BAIAP2,RAPGEF5,SSU72,SRP9,IQCG,T
ERPINB5,CENPV,VKORC1,MRPL2,PCNA,SRSF2,LAS1L,HMGB1,SULF2,BAIAP2,RAPGEF5,SSU72,SRP9,IQCG,T
NLL1,CST3,GNAS,GPR137,RPS24,S100A13,BCAT2,CENPV,MRPL4,FAM189B,VKORC1,PCNA,SRSF2,LAS1L,H
NLL1,CST3,GNAS,GPR137,RPS24,S100A13,BCAT2,CENPV,MRPL4,FAM189B,VKORC1,PCNA,SRSF2,LAS1L,H
A13,CENPV,MRPL4,VKORC1,CDCA3,PCNA,SRSF2,PPFIBP1,SEMA4D,HMGB1,SULF2,BAIAP2,RAPGEF5,SSU7
RPS22,PPFIBP1,LAS1L,HMGB1,CYB561A3,RAPGEF5,MYL6B,IQCG,TMEM67,PILRB,CMC2,BLOC1S1,CIRBP,R
RPS22,PPFIBP1,LAS1L,HMGB1,CYB561A3,RAPGEF5,MYL6B,IQCG,TMEM67,PILRB,CMC2,BLOC1S1,CIRBP,R
HMGB1,SSU72,SRP9,TMEM67,PILRB,CMC2,CIRBP,RPL38,RNH1,DNASE1L1,KIF4A,RBCK1,RNF32,SNRPB,SLC
HMGB1,SSU72,SRP9,TMEM67,PILRB,CMC2,CIRBP,RPL38,RNH1,DNASE1L1,KIF4A,RBCK1,RNF32,SNRPB,SLC
,GPR137,RPS24,S100A13,BCAT2,CENPV,MRPL4,FAM189B,VKORC1,CDCA3,PCNA,SRSF2,MRPS22,KRT15,O
,PCNA,SRSF2,HMGB1,BAIAP2,RAPGEF5,MYL6B,RHOF,SMARCB1,PRR13,RNH1,RBCK1,PTOV1,FASTK,RNF32
PINB5,BCAT2,CENPV,MRPL4,FAM189B,VKORC1,PCNA,SRSF2,MRPS22,KRT15,OSMR,SEMA4D,LAS1L,HMGB
DHPS,ARPC2,LIMK2,BUD31,NPRL2,CTTN,EIF5A,COMMD5,ZNF410,NUDT5,STYXL1,VRK3,LZTFL1,IRF9,CPSF
CDCA3,PCNA,SRSF2,HMGB1,BAIAP2,RAPGEF5,MYL6B,IQCG,FAM72A,BLOC1S1,RHOF,SMARCB1,PRR13,R
CDCA3,PCNA,SRSF2,HMGB1,BAIAP2,RAPGEF5,MYL6B,IQCG,FAM72A,BLOC1S1,RHOF,SMARCB1,PRR13,R
NLL1,CST3,GNAS,GPR137,RPS24,SERPINB5,S100A13,BCAT2,CENPV,MRPL4,FAM189B,VKORC1,MRPL2,CDC
NLL1,CST3,GNAS,GPR137,RPS24,SERPINB5,S100A13,BCAT2,CENPV,MRPL4,FAM189B,VKORC1,MRPL2,CDC
S,GPR137,RPS24,S100A13,BCAT2,CENPV,MRPL4,FAM189B,MRPL2,CDCA3,PCNA,SRSF2,MRPS22,HDAC9,C
S,GPR137,RPS24,S100A13,BCAT2,CENPV,MRPL4,FAM189B,MRPL2,CDCA3,PCNA,SRSF2,MRPS22,HDAC9,C
NLL1,CST3,GNAS,GPR137,RPS24,S100A13,BCAT2,CENPV,MRPL4,FAM189B,VKORC1,CDCA3,PCNA,SRSF2,M
ENPV,VKORC1,PCNA,SRSF2,PPFIBP1,LAS1L,BAIAP2,RAPGEF5,SSU72,SRP9,YBX3,BLOC1S1,CACYBP,SMARCB
2,CENPV,MRPL4,FAM189B,MRPL2,CDCA3,PCNA,SRSF2,OSMR,BAIAP2,RAPGEF5,SSU72,SRP9,SULT1A3,PIL
,PCNA,SRSF2,BAIAP2,RAPGEF5,MYL6B,FAM72A,RHOF,SMARCB1,PRR13,RNH1,RBCK1,PTOV1,FASTK,RNF3
KRT15,HMGB1,BAIAP2,RAPGEF5,MYL6B,CIRBP,RHOF,SMARCB1,PRR13,RNH1,RBCK1,PTOV1,FASTK,RNF32
89B,MRPL2,CDCA3,PCNA,SRSF2,MRPS22,BAIAP2,RAPGEF5,SSU72,MYL6B,SRP9,SULT1A3,FAM72A,RHOF,S
89B,MRPL2,CDCA3,PCNA,SRSF2,MRPS22,BAIAP2,RAPGEF5,SSU72,MYL6B,SRP9,SULT1A3,FAM72A,RHOF,S
7,CMC2,BLOC1S1,RBCK1,RCC1,DHPS,ZC3HAV1,DBI,LIMK2,NOB1,NUMA1,BUD31,NPRL2,CTTN,EIF5A,COM
,GNAS,GPR137,RPS24,SERPINB5,S100A13,BCAT2,CENPV,MRPL4,FAM189B,VKORC1,MRPL2,CDCA3,PCNA,
,GNAS,GPR137,RPS24,SERPINB5,S100A13,BCAT2,CENPV,MRPL4,FAM189B,VKORC1,MRPL2,CDCA3,PCNA,
3,GNAS,GPR137,SERPINB5,CENPV,FAM189B,MRPL2,CDCA3,PCNA,SRSF2,MRPS22,KRT15,PPFIBP1,HMGB1
3,GNAS,GPR137,SERPINB5,CENPV,FAM189B,MRPL2,CDCA3,PCNA,SRSF2,MRPS22,KRT15,PPFIBP1,HMGB1
4,MRPL2,CDCA3,PCNA,SRSF2,OSMR,LAS1L,HMGB1,SULF2,RAPGEF5,SSU72,SRP9,IQCG,YBX3,BLOC1S1,CA
4,MRPL2,CDCA3,PCNA,SRSF2,OSMR,LAS1L,HMGB1,SULF2,RAPGEF5,SSU72,SRP9,IQCG,YBX3,BLOC1S1,CA

1,RNF32,RCC1,DHPS,ACTR6,DBI,GPATCH2L,BUD31,EIF5A,COMMD5,SH2B1,RPL13A,NUDT5,WDR62,VRK3,

PS24,SERPINB5,BCAT2,FAM189B,CDCA3,PCNA,SRSF2,KRT15,HMGB1,CYB561A3,RAPGEF5,CACYBP,CIRBP,

PS24,SERPINB5,BCAT2,FAM189B,CDCA3,PCNA,SRSF2,KRT15,HMGB1,CYB561A3,RAPGEF5,CACYBP,CIRBP,

2,CTTN,RPL13A,STYXL1,FIS1,CPSF3L,TSG101,TTC9C,RPS6,RPL29,TRNT1,MED15,SH3YL1,EIF3M,CDC42,ASF

RPINB5,S100A13,FAM189B,CDCA3,SRSF2,MRPS22,SEMA4D,LAS1L,HMGB1,SSU72,MYL6B,SRP9,IQCG,TM

RPINB5,S100A13,FAM189B,CDCA3,SRSF2,MRPS22,SEMA4D,LAS1L,HMGB1,SSU72,MYL6B,SRP9,IQCG,TM

,BUD31,CTTN,EIF5A,COMMD5,NUDT5,STYXL1,VRK3,TSG101,RPS6,RPL29,TRNT1,CLASRP,FBXW9,ASF1B,E

37,RPS24,S100A13,BCAT2,VKORC1,MRPL2,SRSF2,MRPS22,HDAC9,KRT15,OSMR,HMGB1,RAPGEF5,SSU72,

37,RPS24,S100A13,BCAT2,VKORC1,MRPL2,SRSF2,MRPS22,HDAC9,KRT15,OSMR,HMGB1,RAPGEF5,SSU72,

NLL1,CST3,GPR137,RPS24,S100A13,BCAT2,CENPV,MRPL4,FAM189B,VKORC1,MRPL2,CDCA3,PCNA,SRSF2,

7,RPS24,SERPINB5,CENPV,MRPL4,FAM189B,VKORC1,PCNA,SRSF2,HDAC9,KRT15,OSMR,PPFIBP1,SEMA4D

7,RPS24,SERPINB5,CENPV,MRPL4,FAM189B,VKORC1,PCNA,SRSF2,HDAC9,KRT15,OSMR,PPFIBP1,SEMA4D

DCA3,PCNA,SRSF2,BAIAP2,RAPGEF5,SSU72,MYL6B,IQCG,RHOF,SMARCB1,PRR13,RNH1,DNASE1L1,RBCK1,

R137,CDCA3,PCNA,OSMR,PPFIBP1,HMGB1,SSU72,SRP9,IQCG,CACYBP,SMARCB1,RPL38,RBCK1,PTOV1,FAS

R137,CDCA3,PCNA,OSMR,PPFIBP1,HMGB1,SSU72,SRP9,IQCG,CACYBP,SMARCB1,RPL38,RBCK1,PTOV1,FAS

ERPINB5,BCAT2,MRPL4,MRPL2,CDCA3,PCNA,SRSF2,KRT15,BAIAP2,SSU72,MYL6B,CIRBP,RHOF,SMARCB1,

ERPINB5,BCAT2,MRPL4,MRPL2,CDCA3,PCNA,SRSF2,KRT15,BAIAP2,SSU72,MYL6B,CIRBP,RHOF,SMARCB1,

PL2,PCNA,SRSF2,HMGB1,BAIAP2,RAPGEF5,MYL6B,IQCG,BLOC1S1,RHOF,PRR13,RNH1,RBCK1,PTOV1,RNF

PL2,PCNA,SRSF2,HMGB1,BAIAP2,RAPGEF5,MYL6B,IQCG,BLOC1S1,RHOF,PRR13,RNH1,RBCK1,PTOV1,RNF

NLL1,GNAS,GPR137,RPS24,SERPINB5,S100A13,BCAT2,CENPV,MRPL4,FAM189B,VKORC1,MRPL2,CDCA3,P

NLL1,GNAS,GPR137,RPS24,SERPINB5,S100A13,BCAT2,CENPV,MRPL4,FAM189B,VKORC1,MRPL2,CDCA3,P

,ACTR6,DBI,ARPC2,BUD31,NPRL2,CTTN,EIF5A,COMMD5,STYXL1,SON,UQCRQ,SNF8,CPSF3L,TSG101,METT

L2,CTTN,EIF5A,COMMD5,PSMA6,RPS9,STYXL1,CPSF3L,MED15,UBE2I,EIF3M,CDC42,FBXW9,PPIL2,APEH,L

1L,HMGB1,BAIAP2,RAPGEF5,MYL6B,TMEM67,CMC2,BLOC1S1,CACYBP,RHOF,PRR13,STARD10,DNASE1L1,

1L,HMGB1,BAIAP2,RAPGEF5,MYL6B,TMEM67,CMC2,BLOC1S1,CACYBP,RHOF,PRR13,STARD10,DNASE1L1,

,RPS24,S100A13,BCAT2,MRPL4,FAM189B,VKORC1,MRPL2,CDCA3,PCNA,SRSF2,MRPS22,OSMR,SEMA4D,L

,RPS24,S100A13,BCAT2,MRPL4,FAM189B,VKORC1,MRPL2,CDCA3,PCNA,SRSF2,MRPS22,OSMR,SEMA4D,L

ORC1,CDCA3,PCNA,SRSF2,MRPS22,KRT15,HMGB1,SSU72,MYL6B,SRP9,IQCG,TMEM67,UAP1,FAM72A,CA

ORC1,CDCA3,PCNA,SRSF2,MRPS22,KRT15,HMGB1,SSU72,MYL6B,SRP9,IQCG,TMEM67,UAP1,FAM72A,CA

00A13,BCAT2,MRPL4,FAM189B,VKORC1,MRPL2,CDCA3,PCNA,MRPS22,OSMR,SEMA4D,LAS1L,HMGB1,SU

00A13,BCAT2,MRPL4,FAM189B,VKORC1,MRPL2,CDCA3,PCNA,MRPS22,OSMR,SEMA4D,LAS1L,HMGB1,SU

100A13,BCAT2,MRPL4,FAM189B,MRPL2,SRSF2,SEMA4D,SULF2,RAPGEF5,SSU72,SRP9,IQCG,PILRB,FAM72

100A13,BCAT2,MRPL4,FAM189B,MRPL2,SRSF2,SEMA4D,SULF2,RAPGEF5,SSU72,SRP9,IQCG,PILRB,FAM72

100A13,BCAT2,MRPL4,FAM189B,MRPL2,SRSF2,SEMA4D,SULF2,RAPGEF5,SSU72,SRP9,IQCG,PILRB,FAM72

100A13,BCAT2,MRPL4,FAM189B,MRPL2,SRSF2,SEMA4D,SULF2,RAPGEF5,SSU72,SRP9,IQCG,PILRB,FAM72

CENPV,FAM189B,MRPL2,CDCA3,PCNA,SRSF2,MRPS22,PPFIBP1,HMGB1,BAIAP2,RAPGEF5,MYL6B,IQCG,BL

CENPV,FAM189B,MRPL2,CDCA3,PCNA,SRSF2,MRPS22,PPFIBP1,HMGB1,BAIAP2,RAPGEF5,MYL6B,IQCG,BL

,RPS24,S100A13,BCAT2,MRPL4,FAM189B,VKORC1,MRPL2,CDCA3,PCNA,SRSF2,MRPS22,OSMR,SEMA4D,L

PV,FAM189B,MRPL2,CDCA3,PCNA,SRSF2,MRPS22,PPFIBP1,HMGB1,BAIAP2,RAPGEF5,MYL6B,IQCG,FAM7

PV,FAM189B,MRPL2,CDCA3,PCNA,SRSF2,MRPS22,PPFIBP1,HMGB1,BAIAP2,RAPGEF5,MYL6B,IQCG,FAM7

00A13,BCAT2,MRPL4,FAM189B,VKORC1,MRPL2,CDCA3,PCNA,MRPS22,OSMR,SEMA4D,LAS1L,HMGB1,SU

,PCNA,SRSF2,SULF2,BAIAP2,RAPGEF5,MYL6B,RHOF,SMARCB1,PRR13,RNH1,RBCK1,PTOV1,FASTK,RNF32,L

SERPINB5,S100A13,BCAT2,FAM189B,MRPS22,LAS1L,HMGB1,SSU72,MYL6B,FAM122C,TMEM67,PILRB,CM

SERPINB5,S100A13,BCAT2,FAM189B,MRPS22,LAS1L,HMGB1,SSU72,MYL6B,FAM122C,TMEM67,PILRB,CM

4,S100A13,CENPV,FAM189B,MRPL2,CDCA3,PCNA,SRSF2,OSMR,PPFIBP1,SULF2,BAIAP2,RAPGEF5,SSU72,

4,S100A13,CENPV,FAM189B,MRPL2,CDCA3,PCNA,SRSF2,OSMR,PPFIBP1,SULF2,BAIAP2,RAPGEF5,SSU72,

F32,LY75,SNRPB,DHPS,ACTR6,DBI,GPATCH2L,BUD31,CTTN,EIF5A,COMMD5,SH2B1,RPL13A,NUDT5,VRK3,

/IK2,GPATCH2L,BUD31,NPRL2,CTTN,RPL13A,STYXL1,VRK3,LZTFL1,FIS1,CPSF3L,TSG101,RPS25,NIP7,DBNDI
,RPS24,S100A13,BCAT2,MRPL4,FAM189B,VKORC1,MRPL2,CDCA3,PCNA,MRPS22,OSMR,SEMA4D,HMGB1
,RPS24,S100A13,BCAT2,MRPL4,FAM189B,VKORC1,MRPL2,CDCA3,PCNA,MRPS22,OSMR,SEMA4D,HMGB1
/IRPL4,CDCA3,PCNA,SRSF2,OSMR,CYB561A3,BAIAP2,FAM122C,SRP9,SULT1A3,PILRB,FAM72A,BLOC1S1,CA
/IRPL4,CDCA3,PCNA,SRSF2,OSMR,CYB561A3,BAIAP2,FAM122C,SRP9,SULT1A3,PILRB,FAM72A,BLOC1S1,CA
100A13,BCAT2,MRPL4,FAM189B,VKORC1,MRPL2,CDCA3,PCNA,MRPS22,OSMR,SEMA4D,LAS1L,HMGB1,SU
100A13,BCAT2,MRPL4,FAM189B,VKORC1,MRPL2,CDCA3,PCNA,MRPS22,OSMR,SEMA4D,LAS1L,HMGB1,SU
-,SERPINB5,BCAT2,MRPS22,PPFIBP1,LAS1L,HMGB1,SSU72,TMEM67,SULT1A3,PILRB,FAM72A,CMC2,CIRBF
-,SERPINB5,BCAT2,MRPS22,PPFIBP1,LAS1L,HMGB1,SSU72,TMEM67,SULT1A3,PILRB,FAM72A,CMC2,CIRBF
31,SULF2,BAIAP2,RAPGEF5,MYL6B,BLOC1S1,CACYBP,RHOF,PRR13,RNH1,RBCK1,PTOV1,FASTK,RNF32,LY7
)A13,CENPV,MRPL4,VKORC1,MRPL2,CDCA3,PCNA,SRSF2,MRPS22,HDAC9,OSMR,PPFIBP1,SEMA4D,HMGB
)A13,CENPV,MRPL4,VKORC1,MRPL2,CDCA3,PCNA,SRSF2,MRPS22,HDAC9,OSMR,PPFIBP1,SEMA4D,HMGB
RPB,ACTR6,DBI,BUD31,CTTN,EIF5A,COMMD5,UFD1L,ARHGEF39,NUDT5,STYXL1,FIS1,MAGED2,SON,TSG10
RPB,ACTR6,DBI,BUD31,CTTN,EIF5A,COMMD5,UFD1L,ARHGEF39,NUDT5,STYXL1,FIS1,MAGED2,SON,TSG10
)A13,CENPV,MRPL4,VKORC1,MRPL2,CDCA3,PCNA,SRSF2,MRPS22,HDAC9,OSMR,PPFIBP1,SEMA4D,HMGB
)A13,CENPV,MRPL4,VKORC1,MRPL2,CDCA3,PCNA,SRSF2,MRPS22,HDAC9,OSMR,PPFIBP1,SEMA4D,HMGB
ENPV,MRPL4,VKORC1,PCNA,SRSF2,OSMR,PPFIBP1,SEMA4D,HMGB1,SULF2,RAPGEF5,FAM122C,IQCG,YBX
7,RPS24,S100A13,BCAT2,MRPL4,FAM189B,VKORC1,MRPL2,CDCA3,PCNA,MRPS22,OSMR,SEMA4D,LAS1L,
ACTR6,ZNF235,DBI,ARPC2,BUD31,NPRL2,COMMD5,THYN1,NUDT5,STYXL1,LZTFL1,IRF9,DST,VCAN,MAGEI
3P1,FAM122C,RHOF,SMARCB1,STARD10,TMEM135,RNF32,RCC1,TCAIM,SET,CTSC,BOP1,ARRDC1,SCAF11,
3P1,FAM122C,RHOF,SMARCB1,STARD10,TMEM135,RNF32,RCC1,TCAIM,SET,CTSC,BOP1,ARRDC1,SCAF11,
AT2,CENPV,MRPL4,VKORC1,PCNA,SRSF2,MRPS22,OSMR,LAS1L,HMGB1,SULF2,BAIAP2,RAPGEF5,SSU72,IC
AT2,CENPV,MRPL4,VKORC1,PCNA,SRSF2,MRPS22,OSMR,LAS1L,HMGB1,SULF2,BAIAP2,RAPGEF5,SSU72,IC
100A13,BCAT2,MRPL4,FAM189B,VKORC1,MRPL2,CDCA3,PCNA,MRPS22,OSMR,SEMA4D,LAS1L,HMGB1,SU
5B,SRP9,IQCG,TMEM67,FASTK,ATG16L2,RCC1,DHPS,SET,BOP1,RBP5,ZC3HAV1,ARPC2,NUMA1,SAE1,SCARI
PCNA,MRPS22,KRT15,SSU72,IQCG,SMARCB1,RNH1,DNASE1L1,PTOV1,RNF32,RCC1,TCAIM,DHPS,BOP1,VD
PCNA,MRPS22,KRT15,SSU72,IQCG,SMARCB1,RNH1,DNASE1L1,PTOV1,RNF32,RCC1,TCAIM,DHPS,BOP1,VD
T3,GNAS,GPR137,RPS24,S100A13,BCAT2,CENPV,MRPL4,MRPL2,CDCA3,SRSF2,MRPS22,KRT15,OSMR,PPFIE
T3,GNAS,GPR137,RPS24,S100A13,BCAT2,CENPV,MRPL4,MRPL2,CDCA3,SRSF2,MRPS22,KRT15,OSMR,PPFIE
7,SERPINB5,CENPV,FAM189B,VKORC1,PCNA,SRSF2,OSMR,PPFIBP1,HMGB1,RAPGEF5,SSU72,FAM122C,SR
7,SERPINB5,CENPV,FAM189B,VKORC1,PCNA,SRSF2,OSMR,PPFIBP1,HMGB1,RAPGEF5,SSU72,FAM122C,SR
PR137,SERPINB5,FAM189B,PCNA,SRSF2,OSMR,PPFIBP1,HMGB1,SULF2,RAPGEF5,SSU72,SRP9,CACYBP,SM
PR137,SERPINB5,FAM189B,PCNA,SRSF2,OSMR,PPFIBP1,HMGB1,SULF2,RAPGEF5,SSU72,SRP9,CACYBP,SM
24,SERPINB5,CENPV,MRPL4,FAM189B,VKORC1,MRPL2,CDCA3,PCNA,OSMR,PPFIBP1,LAS1L,HMGB1,BAIAP
24,SERPINB5,CENPV,MRPL4,FAM189B,VKORC1,MRPL2,CDCA3,PCNA,OSMR,PPFIBP1,LAS1L,HMGB1,BAIAP
NLL1,CST3,GPR137,RPS24,BCAT2,VKORC1,PCNA,SRSF2,MRPS22,HDAC9,KRT15,OSMR,LAS1L,HMGB1,BAIA
NLL1,CST3,GPR137,RPS24,BCAT2,VKORC1,PCNA,SRSF2,MRPS22,HDAC9,KRT15,OSMR,LAS1L,HMGB1,BAIA
RPINB5,BCAT2,CENPV,FAM189B,MRPL2,CDCA3,PCNA,SRSF2,KRT15,PPFIBP1,SEMA4D,HMGB1,SULF2,BAIA
RPINB5,BCAT2,CENPV,FAM189B,MRPL2,CDCA3,PCNA,SRSF2,KRT15,PPFIBP1,SEMA4D,HMGB1,SULF2,BAIA
NLL1,CST3,GNAS,GPR137,RPS24,S100A13,BCAT2,CENPV,MRPL4,FAM189B,VKORC1,MRPL2,CDCA3,PCNA,S
NLL1,CST3,GNAS,GPR137,RPS24,S100A13,BCAT2,CENPV,MRPL4,FAM189B,VKORC1,MRPL2,CDCA3,PCNA,S
AM189B,MRPL2,CDCA3,PCNA,SRSF2,BAIAP2,RAPGEF5,MYL6B,IQCG,RHOF,SMARCB1,RPL38,PRR13,RNH1,I
00A13,CDCA3,MRPS22,PPFIBP1,LAS1L,HMGB1,RAPGEF5,SSU72,SULT1A3,PILRB,FAM72A,CMC2,YBX3,CIRI
MRPL4,FAM189B,VKORC1,SRSF2,SULF2,BAIAP2,RAPGEF5,SSU72,SULT1A3,FAM72A,YBX3,CACYBP,SMARC
NLL1,CENPV,MRPL4,FAM189B,CDCA3,PCNA,PPFIBP1,LAS1L,HMGB1,SULF2,RAPGEF5,SSU72,MYL6B,FAM1
NLL1,CENPV,MRPL4,FAM189B,CDCA3,PCNA,PPFIBP1,LAS1L,HMGB1,SULF2,RAPGEF5,SSU72,MYL6B,FAM1
37,RPS24,SERPINB5,BCAT2,CENPV,MRPL4,FAM189B,MRPL2,CDCA3,PCNA,SRSF2,MRPS22,KRT15,OSMR,PI

37,RPS24,SERPINB5,BCAT2,CENPV,MRPL4,FAM189B,MRPL2,CDCA3,PCNA,SRSF2,MRPS22,KRT15,OSMR,P
ORC1,CDCA3,PCNA,MRPS22,OSMR,SEMA4D,LAS1L,HMGB1,SULF2,RAPGEF5,SSU72,FAM122C,IQCG,SULT
,RPS24,S100A13,BCAT2,VKORC1,MRPL2,CDCA3,PCNA,SRSF2,MRPS22,HDAC9,OSMR,PPFIBP1,LAS1L,HMG
,RPS24,S100A13,BCAT2,VKORC1,MRPL2,CDCA3,PCNA,SRSF2,MRPS22,HDAC9,OSMR,PPFIBP1,LAS1L,HMG
B561A3,SSU72,SRP9,TMEM67,PILRB,CMC2,CACYBP,CIRBP,RPL38,RNH1,DNASE1L1,KIF4A,RBCK1,SNRPB,S
B561A3,SSU72,SRP9,TMEM67,PILRB,CMC2,CACYBP,CIRBP,RPL38,RNH1,DNASE1L1,KIF4A,RBCK1,SNRPB,S
F4A,RBCK1,RNF32,SNRPB,PARD3,DHPS,MAP3K7CL,ZNF235,TRPT1,ARPC2,PMFBP1,LIMK2,NUMA1,BUD31
F4A,RBCK1,RNF32,SNRPB,PARD3,DHPS,MAP3K7CL,ZNF235,TRPT1,ARPC2,PMFBP1,LIMK2,NUMA1,BUD31
,DHPS,DBI,ARPC2,LIMK2,BUD31,NPRL2,CTTN,EIF5A,COMMD5,RPL13A,NUDT5,STYXL1,VRK3,LZTFL1,FIS1,
39B,MRPL2,PCNA,SRSF2,LAS1L,HMGB1,BAIAP2,RAPGEF5,SSU72,IQCG,TMEM67,FAM72A,BLOC1S1,RHOF,
AIAP2,RAPGEF5,MYL6B,RHOF,PRR13,RNH1,RBCK1,PTOV1,RNF32,LY75,ATG16L2,SLC3A2,ARRDC1,RBP5,LI
AIAP2,RAPGEF5,MYL6B,RHOF,PRR13,RNH1,RBCK1,PTOV1,RNF32,LY75,ATG16L2,SLC3A2,ARRDC1,RBP5,LI
SF2,BAIAP2,RAPGEF5,MYL6B,IQCG,RHOF,SMARCB1,PRR13,RNH1,RBCK1,PTOV1,RNF32,LY75,DHPS,ARRD
C9,KRT15,MYL6B,PILRB,CMC2,RHOF,STARD10,RNH1,CTSC,RBP5,BUD31,SAE1,CTTN,TSPAN11,EIF5A,UFD1L
NLL1,CST3,GNAS,GPR137,RPS24,SERPINB5,S100A13,BCAT2,CENPV,MRPL4,FAM189B,VKORC1,MRPL2,CD
,FAM189B,PCNA,SRSF2,LAS1L,HMGB1,BAIAP2,SSU72,IQCG,FAM72A,YBX3,CACYBP,CIRBP,RHOF,SMARCB1
D31,NPRL2,CTTN,EIF5A,COMMD5,NUDT5,STYXL1,CPSF3L,TSG101,RPL29,TRNT1,CDC42,FBXW9,ENDOV,LY
,GNAS,RPS24,SERPINB5,S100A13,BCAT2,MRPL4,FAM189B,MRPL2,CDCA3,SRSF2,MRPS22,HDAC9,PPFIBP
,GNAS,RPS24,SERPINB5,S100A13,BCAT2,MRPL4,FAM189B,MRPL2,CDCA3,SRSF2,MRPS22,HDAC9,PPFIBP
S1L,RAPGEF5,FAM72A,BLOC1S1,CACYBP,RNH1,FASTK,SNRPB,BOP1,SCAF11,SCARB1,UFD1L,NOP16,AP3B
H2L,BUD31,CTTN,EIF5A,COMMD5,UFD1L,ACOT8,AP3B2,STYXL1,SNF8,CPSF3L,NIP7,RTEL1,ITGB1BP1,CLA
H2L,BUD31,CTTN,EIF5A,COMMD5,UFD1L,ACOT8,AP3B2,STYXL1,SNF8,CPSF3L,NIP7,RTEL1,ITGB1BP1,CLA
FAM189B,VKORC1,MRPL2,PCNA,SRSF2,MRPS22,LAS1L,HMGB1,SULF2,BAIAP2,RAPGEF5,SSU72,FAM122C,
S1L,HMGB1,BAIAP2,RAPGEF5,MYL6B,TMEM67,CMC2,BLOC1S1,CIRBP,RHOF,PRR13,STARD10,DNASE1L1,
S1L,HMGB1,BAIAP2,RAPGEF5,MYL6B,TMEM67,CMC2,BLOC1S1,CIRBP,RHOF,PRR13,STARD10,DNASE1L1,
PV,MRPL4,FAM189B,VKORC1,MRPL2,CDCA3,PCNA,PPFIBP1,SSU72,FAM122C,IQCG,TMEM67,CACYBP,STA
PV,MRPL4,FAM189B,VKORC1,MRPL2,CDCA3,PCNA,PPFIBP1,SSU72,FAM122C,IQCG,TMEM67,CACYBP,STA
,CDCA3,SRSF2,OSMR,PPFIBP1,HMGB1,SULF2,BAIAP2,SSU72,FAM122C,SRP9,UAP1,CACYBP,CIRBP,RHOF,
,CDCA3,SRSF2,OSMR,PPFIBP1,HMGB1,SULF2,BAIAP2,SSU72,FAM122C,SRP9,UAP1,CACYBP,CIRBP,RHOF,
AIAP2,RAPGEF5,MYL6B,RHOF,SMARCB1,PRR13,RNH1,RBCK1,PTOV1,FASTK,RNF32,LY75,ARRDC1,LIMK2,N
R137,CENPV,PCNA,SRSF2,OSMR,PPFIBP1,HMGB1,SULF2,BAIAP2,SSU72,FAM122C,SRP9,SULT1A3,FAM72
R137,CENPV,PCNA,SRSF2,OSMR,PPFIBP1,HMGB1,SULF2,BAIAP2,SSU72,FAM122C,SRP9,SULT1A3,FAM72
MRPL4,FAM189B,MRPL2,CDCA3,PCNA,SRSF2,OSMR,PPFIBP1,HMGB1,BAIAP2,FAM122C,IQCG,TMEM67,C
MRPL4,FAM189B,MRPL2,CDCA3,PCNA,SRSF2,OSMR,PPFIBP1,HMGB1,BAIAP2,FAM122C,IQCG,TMEM67,C
3,GNAS,GPR137,RPS24,SERPINB5,S100A13,BCAT2,CENPV,MRPL4,FAM189B,VKORC1,MRPL2,CDCA3,PCN
3,GNAS,GPR137,RPS24,SERPINB5,S100A13,BCAT2,CENPV,MRPL4,FAM189B,VKORC1,MRPL2,CDCA3,PCN
KRT15,BAIAP2,RAPGEF5,MYL6B,FAM122C,PILRB,BLOC1S1,RHOF,PRR13,RNH1,RBCK1,PTOV1,FASTK,RNF3
2,CTTN,EIF5A,COMMD5,NUDT5,STYXL1,LZTFL1,CPSF3L,RPS25,RPL29,TRNT1,CDC42,FBXW9,ENDOV,LYPL
A,SRSF2,OSMR,PPFIBP1,RAPGEF5,SSU72,SRP9,BLOC1S1,CACYBP,SMARCB1,RPL38,RNH1,PTOV1,FASTK,AT
,BCAT2,CENPV,MRPL4,MRPL2,CDCA3,PCNA,SRSF2,OSMR,BAIAP2,SRP9,IQCG,TMEM67,SULT1A3,PILRB,F
,BCAT2,CENPV,MRPL4,MRPL2,CDCA3,PCNA,SRSF2,OSMR,BAIAP2,SRP9,IQCG,TMEM67,SULT1A3,PILRB,F
F2,OSMR,RAPGEF5,SSU72,SRP9,CACYBP,SMARCB1,RPL38,RNH1,PTOV1,SNRPB,PARD3,TCAIM,ACTR6,SCA
PCNA,SRSF2,OSMR,RAPGEF5,SSU72,SRP9,SMARCB1,RPL38,PTOV1,SNRPB,ACTR6,CTSC,ARRDC1,KLC1,SCA
37,RPS24,SERPINB5,BCAT2,MRPL4,MRPL2,CDCA3,PCNA,SRSF2,MRPS22,KRT15,PPFIBP1,HMGB1,BAIAP2,R
37,RPS24,SERPINB5,BCAT2,MRPL4,MRPL2,CDCA3,PCNA,SRSF2,MRPS22,KRT15,PPFIBP1,HMGB1,BAIAP2,R
TYXL1,CPSF3L,TSG101,FBXW9,ENDOV,LYPLA2,C17ORF62,PRMT1,PPAN,HYPK,ZNF197,LRRFIP1,TRMT112,
RPL13A,ARHGEF39,ACOT8,TSG101,FUZ,NIP7,DUSP22,RPL29,COASY,AUP1,FOXRED1,MED15,SH3YL1,EIF3

RPL13A,ARHGEF39,ACOT8,TSG101,FUZ,NIP7,DUSP22,RPL29,COASY,AUP1,FOXRED1,MED15,SH3YL1,EIF3M

V,FAM189B,MRPL2,PCNA,HDAC9,PPFIBP1,LAS1L,HMGB1,SULF2,CYB561A3,BAIAP2,RAPGEF5,SSU72,PILRB

V,FAM189B,MRPL2,PCNA,HDAC9,PPFIBP1,LAS1L,HMGB1,SULF2,CYB561A3,BAIAP2,RAPGEF5,SSU72,PILRB

AIAP2,RAPGEF5,SSU72,BLOC1S1,CACYBP,CIRBP,RPL38,DNASE1L1,RNF32,ATG16L2,RCC1,SNRPB,SEC62,CT

1,DHPS,ACTR6,TRPT1,DBI,BUD31,NPRL2,CTTN,EIF5A,COMMD5,NUDT22,RBPMS,STYXL1,VRK3,LZTFL1,PPM

,DNASE1L1,RBCK1,DHPS,TRPT1,CTTN,EIF5A,COMMD5,NUDT22,RPL13A,RBPMS,STYXL1,VRK3,LZTFL1,TUB

PL4,FAM189B,MRPL2,CDCA3,PCNA,MRPS22,OSMR,SEMA4D,LAS1L,HMGB1,SULF2,RAPGEF5,SSU72,FAM1

PL4,FAM189B,MRPL2,CDCA3,PCNA,MRPS22,OSMR,SEMA4D,LAS1L,HMGB1,SULF2,RAPGEF5,SSU72,FAM1

IA,MRPS22,OSMR,SULF2,BAIAP2,RAPGEF5,SSU72,IQCG,FAM72A,CACYBP,SMARCB1,PRR13,RNH1,DNASE1

L2,CTTN,EIF5A,COMMD5,NUDT5,STYXL1,VRK3,CPSF3L,TSG101,RPS6,RPL29,TRNT1,CDC42,FBXW9,ASF1B,

VKORC1,PCNA,SRSF2,OSMR,PPFIBP1,RAPGEF5,SSU72,FAM122C,SRP9,IQCG,TMEM67,PILRB,CACYBP,SMARC

VKORC1,PCNA,SRSF2,OSMR,PPFIBP1,RAPGEF5,SSU72,FAM122C,SRP9,IQCG,TMEM67,PILRB,CACYBP,SMARC

IF235,BOP1,TRPT1,ARPC2,PMFBP1,LIMK2,GPATCH2L,NPRL2,CRYGS,CTTN,COMMD5,NUDT22,USP38,SH2B

IF235,BOP1,TRPT1,ARPC2,PMFBP1,LIMK2,GPATCH2L,NPRL2,CRYGS,CTTN,COMMD5,NUDT22,USP38,SH2B

PL4,FAM189B,CDCA3,PCNA,MRPS22,OSMR,SEMA4D,LAS1L,HMGB1,SULF2,RAPGEF5,SSU72,FAM122C,SU

SMR,LAS1L,RAPGEF5,MYL6B,SRP9,IQCG,TMEM67,SULT1A3,RHOF,STARD10,DNASE1L1,KIF4A,PTOV1,RNF3

,PCNA,SRSF2,BAIAP2,RAPGEF5,MYL6B,RHOF,SMARCB1,PRR13,RNH1,RBCK1,PTOV1,FASTK,RNF32,LY75,PA

PINB5,S100A13,CENPV,FAM189B,MRPL2,CDCA3,PCNA,SRSF2,KRT15,PPFIBP1,HMGB1,SULF2,BAIAP2,RAPG

PINB5,S100A13,CENPV,FAM189B,MRPL2,CDCA3,PCNA,SRSF2,KRT15,PPFIBP1,HMGB1,SULF2,BAIAP2,RAPG

RPS24,S100A13,BCAT2,CENPV,MRPL4,FAM189B,MRPL2,PCNA,SRSF2,MRPS22,KRT15,PPFIBP1,LAS1L,HMG

RPS24,S100A13,BCAT2,CENPV,MRPL4,FAM189B,MRPL2,PCNA,SRSF2,MRPS22,KRT15,PPFIBP1,LAS1L,HMG

,S100A13,BCAT2,CENPV,MRPL4,FAM189B,VKORC1,MRPL2,SRSF2,KRT15,OSMR,LAS1L,SULF2,BAIAP2,RAP

,S100A13,BCAT2,CENPV,MRPL4,FAM189B,VKORC1,MRPL2,SRSF2,KRT15,OSMR,LAS1L,SULF2,BAIAP2,RAP

AIAP2,MYL6B,RHOF,PRR13,RNH1,DNASE1L1,RBCK1,PTOV1,FASTK,LY75,SET,ARRDC1,RBP5,LIMK2,NUMA1

24,SERPINB5,S100A13,BCAT2,CENPV,MRPL4,VKORC1,MRPL2,PCNA,SRSF2,HDAC9,OSMR,HMGB1,BAIAP2,

KORC1,MRPL2,HMGB1,SSU72,IQCG,YBX3,BLOC1S1,CACYBP,RHOF,PRR13,RNH1,DNASE1L1,PTOV1,RNF32,

24,SERPINB5,S100A13,BCAT2,CENPV,MRPL4,VKORC1,MRPL2,CDCA3,PCNA,SRSF2,MRPS22,OSMR,PPFIBP

24,SERPINB5,S100A13,BCAT2,CENPV,MRPL4,VKORC1,MRPL2,CDCA3,PCNA,SRSF2,MRPS22,OSMR,PPFIBP

2,GPATCH2L,BUD31,CTTN,EIF5A,NUDT22,UFD1L,SH2B1,RPL13A,PCMT1,VRK3,IRF9,TM9SF1,RPS16,TSG10

2,GPATCH2L,BUD31,CTTN,EIF5A,NUDT22,UFD1L,SH2B1,RPL13A,PCMT1,VRK3,IRF9,TM9SF1,RPS16,TSG10

TN,EIF5A,COMMD5,NUDT5,STYXL1,SNF8,CPSF3L,TSG101,RPL29,TRNT1,CDC42,FBXW9,RFX3,ENDOV,LYPL

T2,FAM189B,VKORC1,SRSF2,OSMR,PPFIBP1,LAS1L,BAIAP2,TMEM67,FAM72A,YBX3,CACYBP,CIRBP,KIF4A,

T2,FAM189B,VKORC1,SRSF2,OSMR,PPFIBP1,LAS1L,BAIAP2,TMEM67,FAM72A,YBX3,CACYBP,CIRBP,KIF4A,

2,NOP16,RUVBL2,EBPL,WDR62,UBE2L6,LPCAT3,CDC37,SLC39A11,ZBTB44,TTC9C,CD99L2,IP6K2,HS2ST1,U

B,PCNA,HMGB1,BAIAP2,RAPGEF5,SSU72,MYL6B,SRP9,IQCG,TMEM67,FAM72A,YBX3,RHOF,SMARCB1,RP

B,PCNA,HMGB1,BAIAP2,RAPGEF5,SSU72,MYL6B,SRP9,IQCG,TMEM67,FAM72A,YBX3,RHOF,SMARCB1,RP

MYL6B,SULT1A3,FAM72A,BLOC1S1,CACYBP,STARD10,RNH1,KIF4A,RBCK1,PTOV1,FASTK,RNF32,LY75,SNRP

MYL6B,SULT1A3,FAM72A,BLOC1S1,CACYBP,STARD10,RNH1,KIF4A,RBCK1,PTOV1,FASTK,RNF32,LY75,SNRP

3,GNAS,GPR137,BCAT2,CENPV,MRPL4,VKORC1,MRPL2,PCNA,SRSF2,MRPS22,OSMR,SULF2,BAIAP2,RAPG

,VRK3,CPSF3L,TSG101,DBNDD2,RPS6,RTEL1,SLC39A13,CDC42,USE1,LIG1,DGUOK,TRMT112,MPDU1,ERCC

NPV,VKORC1,PCNA,SRSF2,OSMR,SULF2,RAPGEF5,SSU72,SRP9,IQCG,TMEM67,PILRB,CACYBP,SMARCB1,RP

NPV,VKORC1,PCNA,SRSF2,OSMR,SULF2,RAPGEF5,SSU72,SRP9,IQCG,TMEM67,PILRB,CACYBP,SMARCB1,RP

,S100A13,CENPV,MRPL4,VKORC1,MRPL2,CDCA3,PCNA,HDAC9,OSMR,PPFIBP1,LAS1L,HMGB1,SSU72,FAM

,S100A13,CENPV,MRPL4,VKORC1,MRPL2,CDCA3,PCNA,HDAC9,OSMR,PPFIBP1,LAS1L,HMGB1,SSU72,FAM

2,OSMR,SULF2,BAIAP2,SSU72,FAM122C,TMEM67,PILRB,FAM72A,BLOC1S1,CIRBP,RHOF,RPL38,KIF4A,RB

2,OSMR,SULF2,BAIAP2,SSU72,FAM122C,TMEM67,PILRB,FAM72A,BLOC1S1,CIRBP,RHOF,RPL38,KIF4A,RB

AS,RPS24,SERPINB5,BCAT2,CENPV,PCNA,SRSF2,OSMR,LAS1L,RAPGEF5,SSU72,FAM122C,TMEM67,FAM72

IAS,RPS24,SERPINB5,BCAT2,CENPV,PCNA,SRSF2,OSMR,LAS1L,RAPGEF5,SSU72,FAM122C,TMEM67,FAM72

XL1,CPSF3L,CDC42,FBXW9,RFX3,ENDOV,LYPLA2,C17ORF62,PRMT1,PPAN,ZNF197,LRRFIP1,TRMT112,DDO

PINB5,S100A13,BCAT2,MRPL4,CDCA3,PCNA,SRSF2,HMGB1,SULF2,CYB561A3,BAIAP2,SSU72,IQCG,YBX3,C

PINB5,S100A13,BCAT2,MRPL4,CDCA3,PCNA,SRSF2,HMGB1,SULF2,CYB561A3,BAIAP2,SSU72,IQCG,YBX3,C

KORC1,PCNA,SRSF2,MRPS22,HDAC9,OSMR,HMGB1,CYB561A3,BAIAP2,RAPGEF5,SSU72,MYL6B,FAM122C,

KORC1,PCNA,SRSF2,MRPS22,HDAC9,OSMR,HMGB1,CYB561A3,BAIAP2,RAPGEF5,SSU72,MYL6B,FAM122C,

V,MRPL4,FAM189B,CDCA3,SRSF2,OSMR,BAIAP2,SSU72,SRP9,SULT1A3,PILRB,FAM72A,BLOC1S1,CACYBP,F

PV,FAM189B,MRPL2,CDCA3,PCNA,SRSF2,HMGB1,BAIAP2,RAPGEF5,MYL6B,BLOC1S1,CACYBP,CIRBP,RHOF

S100A13,BCAT2,FAM189B,VKORC1,MRPL2,CDCA3,PCNA,SRSF2,MRPS22,KRT15,PPFIBP1,SEMA4D,HMGB1

S100A13,BCAT2,FAM189B,VKORC1,MRPL2,CDCA3,PCNA,SRSF2,MRPS22,KRT15,PPFIBP1,SEMA4D,HMGB1

PV,MRPL4,FAM189B,VKORC1,MRPL2,CDCA3,PCNA,PPFIBP1,SSU72,FAM122C,IQCG,TMEM67,CACYBP,STA

PV,MRPL4,FAM189B,VKORC1,MRPL2,CDCA3,PCNA,PPFIBP1,SSU72,FAM122C,IQCG,TMEM67,CACYBP,STA

UD31,NPRL2,CTTN,EIF5A,COMMD5,STYXL1,UQCRQ,CPSF3L,TSG101,ATG12,NIP7,RPS6,CDC42,FBXW9,RBN

M189B,MRPL2,SRSF2,OSMR,BAIAP2,MYL6B,SRP9,SULT1A3,FAM72A,BLOC1S1,CIRBP,RHOF,RPL38,DNASE1

MGB1,BAIAP2,RAPGEF5,MYL6B,FAM72A,RHOF,SMARCB1,PRR13,RNH1,RBCK1,PTOV1,RNF32,ATG16L2,SET

APGEF5,SSU72,SRP9,BLOC1S1,CACYBP,SMARCB1,RPL38,RNH1,PTOV1,SNRPB,ACTR6,SET,SCAF11,LIMK2,N

C1,MRPL2,PCNA,SRSF2,MRPS22,LAS1L,HMGB1,SULF2,BAIAP2,RAPGEF5,SSU72,FAM122C,IQCG,PILRB,YBX

,RPS24,S100A13,BCAT2,MRPL4,FAM189B,VKORC1,MRPL2,CDCA3,PCNA,MRPS22,OSMR,SEMA4D,LAS1L,H

,RPS24,S100A13,BCAT2,MRPL4,FAM189B,VKORC1,MRPL2,CDCA3,PCNA,MRPS22,OSMR,SEMA4D,LAS1L,H

SNRPB,DHPS,BUD31,CTTN,EIF5A,UFD1L,ARHGEF39,ACOT8,STYXL1,FIS1,MAGED2,CPSF3L,RTEL1,RABL5,ITG

SNRPB,DHPS,BUD31,CTTN,EIF5A,UFD1L,ARHGEF39,ACOT8,STYXL1,FIS1,MAGED2,CPSF3L,RTEL1,RABL5,ITG

7,SERPINB5,BCAT2,MRPL4,VKORC1,SRSF2,MRPS22,KRT15,OSMR,PPFIBP1,LAS1L,HMGB1,SULF2,BAIAP2,RA

7,SERPINB5,BCAT2,MRPL4,VKORC1,SRSF2,MRPS22,KRT15,OSMR,PPFIBP1,LAS1L,HMGB1,SULF2,BAIAP2,RA

37,RPS24,S100A13,BCAT2,CENPV,MRPL4,MRPL2,CDCA3,SRSF2,MRPS22,KRT15,OSMR,PPFIBP1,SULF2,BAIA

SF2,HMGB1,BAIAP2,RAPGEF5,MYL6B,RHOF,SMARCB1,PRR13,RNH1,RBCK1,PTOV1,RNF32,ARRDC1,KLC1,LI

IARCB1,STARD10,RCC1,TCAIM,ARRDC1,SCAF11,VDAC3,NOB1,SUMF2,SAE1,NPRL2,EIF5A,AMZ2,RPL13A,TT

IARCB1,STARD10,RCC1,TCAIM,ARRDC1,SCAF11,VDAC3,NOB1,SUMF2,SAE1,NPRL2,EIF5A,AMZ2,RPL13A,TT

SF2,BAIAP2,RAPGEF5,MYL6B,RHOF,SMARCB1,PRR13,RNH1,RBCK1,PTOV1,FASTK,RNF32,LY75,TCAIM,ARRD

LIMK2,GPATCH2L,BUD31,NPRL2,CTTN,UFD1L,SH2B1,RUVBL2,NUDT5,STYXL1,VRK3,IRF9,DST,ANTXR2,SON,

IRPL2,PCNA,SRSF2,MRPS22,LAS1L,HMGB1,FAM122C,IQCG,PILRB,SMARCB1,RPL38,RNH1,FASTK,PARD3,TC

3,CENPV,FAM189B,MRPL2,SRSF2,BAIAP2,RAPGEF5,SSU72,MYL6B,SULT1A3,FAM72A,RHOF,SMARCB1,PRR

3,CENPV,FAM189B,MRPL2,SRSF2,BAIAP2,RAPGEF5,SSU72,MYL6B,SULT1A3,FAM72A,RHOF,SMARCB1,PRR

PR137,RPS24,S100A13,BCAT2,CENPV,VKORC1,MRPL2,CDCA3,PCNA,SRSF2,KRT15,OSMR,PPFIBP1,LAS1L,H

PR137,RPS24,S100A13,BCAT2,CENPV,VKORC1,MRPL2,CDCA3,PCNA,SRSF2,KRT15,OSMR,PPFIBP1,LAS1L,H

RHOF,SMARCB1,STARD10,TMEM135,RNF32,RCC1,TCAIM,SET,CTSC,BOP1,ARRDC1,SCAF11,ARPC2,VDAC3,

RHOF,SMARCB1,STARD10,TMEM135,RNF32,RCC1,TCAIM,SET,CTSC,BOP1,ARRDC1,SCAF11,ARPC2,VDAC3,

A,COMMD5,UFD1L,USP38,ACOT8,NUDT5,STYXL1,TSG101,RPS25,THEM4,RTEL1,TRNT1,ITGB1BP1,UGGT2,C

A,COMMD5,UFD1L,USP38,ACOT8,NUDT5,STYXL1,TSG101,RPS25,THEM4,RTEL1,TRNT1,ITGB1BP1,UGGT2,C

ED15,SH3YL1,EIF3M,CDC42,ASF1B,LYPLA2,TRMT112,SAT1,LSR,PSME2,TADA3,RAD23A,SEC31A,CNST,PAF

GPATCH2L,BUD31,CTTN,EIF5A,COMMD5,UFD1L,ACOT8,STYXL1,CPSF3L,NIP7,RTEL1,ITGB1BP1,CLASRP,C1C

GPATCH2L,BUD31,CTTN,EIF5A,COMMD5,UFD1L,ACOT8,STYXL1,CPSF3L,NIP7,RTEL1,ITGB1BP1,CLASRP,C1C

A3,PCNA,SRSF2,MRPS22,OSMR,PPFIBP1,SEMA4D,BAIAP2,RAPGEF5,YBX3,SMARCB1,RPL38,STARD10,RNH1

A3,PCNA,SRSF2,MRPS22,OSMR,PPFIBP1,SEMA4D,BAIAP2,RAPGEF5,YBX3,SMARCB1,RPL38,STARD10,RNH1

NA,SRSF2,OSMR,PPFIBP1,SEMA4D,HMGB1,YBX3,BLOC1S1,CACYBP,STARD10,RNH1,FASTK,TMEM135,RNF

NA,SRSF2,OSMR,PPFIBP1,SEMA4D,HMGB1,YBX3,BLOC1S1,CACYBP,STARD10,RNH1,FASTK,TMEM135,RNF

C1S1,CIRBP,RNH1,PTOV1,RNF32,LY75,ATG16L2,SNRPB,PARD3,SEC62,SET,BOP1,ARRDC1,ZC3HAV1,WDR19

C1S1,CIRBP,RNH1,PTOV1,RNF32,LY75,ATG16L2,SNRPB,PARD3,SEC62,SET,BOP1,ARRDC1,ZC3HAV1,WDR19

S100A13,BCAT2,MRPL2,SRSF2,MRPS22,HDAC9,KRT15,OSMR,SEMA4D,LAS1L,HMGB1,MYL6B,FAM122C,TI
S100A13,BCAT2,MRPL2,SRSF2,MRPS22,HDAC9,KRT15,OSMR,SEMA4D,LAS1L,HMGB1,MYL6B,FAM122C,TI
CAT2,MRPL4,FAM189B,VKORC1,MRPL2,CDCA3,PCNA,MRPS22,OSMR,SEMA4D,HMGB1,SULF2,RAPGEF5,S
CAT2,MRPL4,FAM189B,VKORC1,MRPL2,CDCA3,PCNA,MRPS22,OSMR,SEMA4D,HMGB1,SULF2,RAPGEF5,S
IRBP,RPL38,DNASE1L1,FASTK,RNF32,LY75,PARD3,ACTR6,ARRDC1,SIRT6,CTTN,COMMD5,TAZ,THYN1,RPS
BCK1,RCC1,SNRPB,TCAIM,ACTR6,ZC3HAV1,DBI,ARPC2,PMFBP1,LIMK2,BUD31,TSPAN11,UFD1L,ARHGEF3
AS1L,HMGB1,SULF2,RAPGEF5,SSU72,SRP9,PILRB,FAM72A,YBX3,BLOC1S1,CACYBP,RNH1,DNASE1L1,PTOV
35,SRSF2,OSMR,RAPGEF5,SSU72,SRP9,YBX3,BLOC1S1,CACYBP,SMARCB1,RPL38,RNH1,PTOV1,FASTK,LY75,
35,SRSF2,OSMR,RAPGEF5,SSU72,SRP9,YBX3,BLOC1S1,CACYBP,SMARCB1,RPL38,RNH1,PTOV1,FASTK,LY75,
NLL1,CST3,GNAS,GPR137,RPS24,SERPINB5,BCAT2,CENPV,VKORC1,MRPL2,SRSF2,HDAC9,OSMR,SEMA4D,L
NLL1,CST3,GNAS,GPR137,RPS24,SERPINB5,BCAT2,CENPV,VKORC1,MRPL2,SRSF2,HDAC9,OSMR,SEMA4D,L
2,SSU72,SRP9,IQCG,FAM72A,SMARCB1,RPL38,PTOV1,FASTK,SNRPB,PARD3,ACTR6,SEC62,ARRDC1,SCAF11
SF2,BAIAP2,RAPGEF5,MYL6B,RHOF,SMARCB1,PRR13,RNH1,RBCK1,PTOV1,RNF32,LY75,TCAIM,ARRDC1,LIN
SF2,BAIAP2,RAPGEF5,MYL6B,RHOF,SMARCB1,PRR13,RNH1,RBCK1,PTOV1,RNF32,LY75,TCAIM,ARRDC1,LIN
F2,OSMR,SULF2,BAIAP2,SSU72,TMEM67,PILRB,FAM72A,BLOC1S1,CIRBP,RHOF,RPL38,KIF4A,RBCK1,FASTK
5,CENPV,MRPL4,MRPL2,SRSF2,OSMR,SEMA4D,SULF2,RAPGEF5,SSU72,YBX3,BLOC1S1,CIRBP,SMARCB1,PR
5,CENPV,MRPL4,MRPL2,SRSF2,OSMR,SEMA4D,SULF2,RAPGEF5,SSU72,YBX3,BLOC1S1,CIRBP,SMARCB1,PR
AP2,RAPGEF5,MYL6B,RHOF,SMARCB1,PRR13,RNH1,RBCK1,PTOV1,FASTK,RNF32,LY75,PARD3,TCAIM,ARRD
SNRPB,DHPS,BUD31,CTTN,EIF5A,UFD1L,ARHGEF39,ACOT8,STYXL1,FIS1,MAGED2,CPSF3L,RTEL1,RABL5,ITG
SNRPB,DHPS,BUD31,CTTN,EIF5A,UFD1L,ARHGEF39,ACOT8,STYXL1,FIS1,MAGED2,CPSF3L,RTEL1,RABL5,ITG
RPL2,CDCA3,PCNA,MRPS22,BAIAP2,RAPGEF5,MYL6B,CIRBP,RHOF,SMARCB1,PRR13,RNH1,DNASE1L1,RBCK
RPL2,CDCA3,PCNA,MRPS22,BAIAP2,RAPGEF5,MYL6B,CIRBP,RHOF,SMARCB1,PRR13,RNH1,DNASE1L1,RBCK
T2,CENPV,FAM189B,MRPL2,PCNA,SRSF2,KRT15,OSMR,LAS1L,HMGB1,SULF2,RAPGEF5,SSU72,FAM122C,S
T2,CENPV,FAM189B,MRPL2,PCNA,SRSF2,KRT15,OSMR,LAS1L,HMGB1,SULF2,RAPGEF5,SSU72,FAM122C,S
M189B,VKORC1,MRPL2,CDCA3,SRSF2,MRPS22,PPFIBP1,SEMA4D,LAS1L,HMGB1,CYB561A3,SSU72,MYL6B,
M189B,VKORC1,MRPL2,CDCA3,SRSF2,MRPS22,PPFIBP1,SEMA4D,LAS1L,HMGB1,CYB561A3,SSU72,MYL6B,
CAT2,FAM189B,VKORC1,MRPL2,CDCA3,SRSF2,MRPS22,HDAC9,KRT15,SEMA4D,LAS1L,CYB561A3,BAIAP2,R
CAT2,FAM189B,VKORC1,MRPL2,CDCA3,SRSF2,MRPS22,HDAC9,KRT15,SEMA4D,LAS1L,CYB561A3,BAIAP2,R
35,BCAT2,VKORC1,PCNA,SRSF2,RAPGEF5,SSU72,SRP9,IQCG,BLOC1S1,CACYBP,SMARCB1,RPL38,RNH1,PTO
35,BCAT2,VKORC1,PCNA,SRSF2,RAPGEF5,SSU72,SRP9,IQCG,BLOC1S1,CACYBP,SMARCB1,RPL38,RNH1,PTO
RPINB5,S100A13,BCAT2,MRPL4,FAM189B,VKORC1,MRPL2,SRSF2,MRPS22,HDAC9,KRT15,OSMR,LAS1L,HN
RPINB5,S100A13,BCAT2,MRPL4,FAM189B,VKORC1,MRPL2,SRSF2,MRPS22,HDAC9,KRT15,OSMR,LAS1L,HN
IARCB1,STARD10,RNF32,RCC1,TCAIM,ARRDC1,SCAF11,VDAC3,NOB1,SIRT6,SUMF2,SAE1,NPRL2,EIF5A,AM
IARCB1,STARD10,RNF32,RCC1,TCAIM,ARRDC1,SCAF11,VDAC3,NOB1,SIRT6,SUMF2,SAE1,NPRL2,EIF5A,AM
CA3,MRPS22,OSMR,SEMA4D,HMGB1,SULF2,RAPGEF5,SSU72,FAM122C,SULT1A3,PILRB,FAM72A,YBX3,CA
2,SRP9,SMARCB1,RPL38,STARD10,PTOV1,FASTK,SNRPB,PARD3,ACTR6,SCAF11,LIMK2,VDAC3,SIRT6,CTTN,
SF2,BAIAP2,SRP9,SULT1A3,PILRB,FAM72A,BLOC1S1,CACYBP,STARD10,RBCK1,PTOV1,FASTK,TMEM135,AT
T3,GNAS,GPR137,CENPV,MRPL4,FAM189B,MRPL2,CDCA3,PCNA,SRSF2,MRPS22,OSMR,HMGB1,SULF2,CYB
T3,GNAS,GPR137,CENPV,MRPL4,FAM189B,MRPL2,CDCA3,PCNA,SRSF2,MRPS22,OSMR,HMGB1,SULF2,CYB
PL2,CDCA3,PCNA,SRSF2,BAIAP2,RAPGEF5,MYL6B,CACYBP,CIRBP,RHOF,SMARCB1,PRR13,RNH1,DNASE1L1,
24,SERPINB5,CENPV,MRPL4,FAM189B,MRPL2,CDCA3,PCNA,PPFIBP1,LAS1L,BAIAP2,SSU72,FAM122C,TMEI
24,SERPINB5,CENPV,MRPL4,FAM189B,MRPL2,CDCA3,PCNA,PPFIBP1,LAS1L,BAIAP2,SSU72,FAM122C,TMEI
BCAT2,CENPV,MRPL4,FAM189B,MRPS22,KRT15,OSMR,SULF2,BAIAP2,RAPGEF5,SSU72,FAM122C,TMEM67
BCAT2,CENPV,MRPL4,FAM189B,MRPS22,KRT15,OSMR,SULF2,BAIAP2,RAPGEF5,SSU72,FAM122C,TMEM67
MYL6B,RHOF,SMARCB1,PRR13,RNH1,RBCK1,PTOV1,FASTK,RNF32,SLC3A2,CTSC,ARRDC1,KLC1,LIMK2,NUM
,NPRL2,CTTN,EIF5A,COMMD5,UFD1L,STYXL1,LZTFL1,VCAN,UQCRQ,CPSF3L,TSG101,RPS25,CDC42,FBXW9,
PL4,VKORC1,PCNA,SRSF2,PPFIBP1,SEMA4D,HMGB1,SULF2,RAPGEF5,IQCG,YBX3,BLOC1S1,CACYBP,CIRBP,

24,SERPINB5,S100A13,BCAT2,MRPL4,FAM189B,VKORC1,MRPL2,CDCA3,PCNA,SRSF2,HDAC9,PPFIBP1,LAS
24,SERPINB5,S100A13,BCAT2,MRPL4,FAM189B,VKORC1,MRPL2,CDCA3,PCNA,SRSF2,HDAC9,PPFIBP1,LAS
PGEF5,MYL6B,CACYBP,RHOF,PRR13,RNH1,RBCK1,PTOV1,FASTK,RNF32,LY75,PARD3,SET,BOP1,ARRDC1,K
,SNRPB,DHPS,GPATCH2L,BUD31,CTTN,EIF5A,COMMD5,UFD1L,ACOT8,STYXL1,CPSF3L,NIP7,RTEL1,ITGB1B
,SNRPB,DHPS,GPATCH2L,BUD31,CTTN,EIF5A,COMMD5,UFD1L,ACOT8,STYXL1,CPSF3L,NIP7,RTEL1,ITGB1B
RL2,CTTN,STYXL1,CPSF3L,TSG101,NIP7,RPS6,CDC42,TRMT112,CHPF,DDOST,LSR,TOMM40,TADA3,MDH2,
AM189B,MRPS22,KRT15,HMGB1,RAPGEF5,FAM122C,SRP9,FAM72A,RHOF,RNH1,DNASE1L1,RBCK1,PTOV
L,RNH1,DNASE1L1,PTOV1,RNF32,TCAIM,DHPS,BOP1,VDAC3,NOB1,GPATCH2L,SUMF2,OSGEP,AMZ2,TAZ,S
L,RNH1,DNASE1L1,PTOV1,RNF32,TCAIM,DHPS,BOP1,VDAC3,NOB1,GPATCH2L,SUMF2,OSGEP,AMZ2,TAZ,S
,RNH1,DNASE1L1,RNF32,TCAIM,KLC1,SCAF11,NPRL2,OSGEP,TAZ,NOP16,LDHA,RPS9,RCE1,MRPL33,ZNF41
AP2,MYL6B,RHOF,SMARCB1,PRR13,RNH1,DNASE1L1,RBCK1,PTOV1,LY75,DHPS,ARRDC1,RBP5,LIMK2,NUM
AP2,MYL6B,RHOF,SMARCB1,PRR13,RNH1,DNASE1L1,RBCK1,PTOV1,LY75,DHPS,ARRDC1,RBP5,LIMK2,NUM
137,RPS24,SERPINB5,MRPL2,SRSF2,HDAC9,OSMR,SULF2,CYB561A3,RAPGEF5,MYL6B,FAM122C,SRP9,IQC
137,RPS24,SERPINB5,MRPL2,SRSF2,HDAC9,OSMR,SULF2,CYB561A3,RAPGEF5,MYL6B,FAM122C,SRP9,IQC
PR137,S100A13,BCAT2,CENPV,MRPL4,FAM189B,VKORC1,MRPL2,CDCA3,PCNA,SRSF2,MRPS22,KRT15,OS
PR137,S100A13,BCAT2,CENPV,MRPL4,FAM189B,VKORC1,MRPL2,CDCA3,PCNA,SRSF2,MRPS22,KRT15,OS
PL4,FAM189B,VKORC1,MRPL2,CDCA3,PCNA,MRPS22,SEMA4D,HMGB1,SULF2,RAPGEF5,SSU72,FAM122C
OF,PRR13,RNH1,RBCK1,PTOV1,FASTK,LY75,ARRDC1,RBP5,LIMK2,NUMA1,TBL2,SCARB1,ITSN1,AMZ2,RUV
ORC1,SRSF2,MRPS22,LAS1L,HMGB1,SULF2,BAIAP2,IQCG,CMC2,CACYBP,SMARCB1,RPL38,STARD10,KIF4A
SU72,IQCG,SMARCB1,RNH1,PTOV1,FASTK,PARD3,ACTR6,CTSC,SCAF11,VDAC3,SUMF2,OSGEP,CTTN,EIF5
,BLOC1S1,TCAIM,DHPS,KLC1,SCAF11,WDR19,SAE1,OSGEP,EIF5A,RPL13A,LDHA,IGSF8,WDR62,RANBP1,VR
RPINB5,BCAT2,CENPV,MRPL4,FAM189B,VKORC1,MRPL2,CDCA3,PCNA,SRSF2,MRPS22,KRT15,OSMR,LAS1
RPINB5,BCAT2,CENPV,MRPL4,FAM189B,VKORC1,MRPL2,CDCA3,PCNA,SRSF2,MRPS22,KRT15,OSMR,LAS1
ENPV,MRPL4,VKORC1,CDCA3,PCNA,SRSF2,OSMR,PPFIBP1,LAS1L,HMGB1,BAIAP2,RAPGEF5,SSU72,FAM12
,FAM189B,CDCA3,PCNA,PPFIBP1,HMGB1,SSU72,FAM122C,IQCG,TMEM67,CACYBP,STARD10,RNH1,DNAS
,FAM189B,CDCA3,PCNA,PPFIBP1,HMGB1,SSU72,FAM122C,IQCG,TMEM67,CACYBP,STARD10,RNH1,DNAS
,VKORC1,CDCA3,PCNA,HMGB1,SULF2,SSU72,MYL6B,FAM122C,TMEM67,SULT1A3,UAP1,PILRB,FAM72A,C
,VKORC1,CDCA3,PCNA,HMGB1,SULF2,SSU72,MYL6B,FAM122C,TMEM67,SULT1A3,UAP1,PILRB,FAM72A,C
RSF2,HMGB1,SSU72,MYL6B,SRP9,IQCG,TMEM67,CACYBP,PRR13,DNASE1L1,PTOV1,FASTK,TMEM135,ATG
RSF2,HMGB1,SSU72,MYL6B,SRP9,IQCG,TMEM67,CACYBP,PRR13,DNASE1L1,PTOV1,FASTK,TMEM135,ATG
CA3,SRSF2,PPFIBP1,SEMA4D,HMGB1,SULF2,BAIAP2,PILRB,YBX3,CACYBP,RPL38,STARD10,RNH1,PTOV1,F
IAP2,RAPGEF5,MYL6B,RHOF,SMARCB1,PRR13,RNH1,RBCK1,PTOV1,FASTK,RNF32,ARRDC1,LIMK2,TBL2,SC
1,BAIAP2,SSU72,FAM72A,YBX3,BLOC1S1,RNH1,DNASE1L1,PTOV1,FASTK,RNF32,ATG16L2,SNRPB,PARD3,
1,BAIAP2,SSU72,FAM72A,YBX3,BLOC1S1,RNH1,DNASE1L1,PTOV1,FASTK,RNF32,ATG16L2,SNRPB,PARD3,
2,BAIAP2,RAPGEF5,MYL6B,CIRBP,RHOF,PRR13,RNH1,RBCK1,PTOV1,FASTK,RNF32,LY75,SLC3A2,SET,ARRD
M189B,VKORC1,CDCA3,MRPS22,OSMR,SULF2,CYB561A3,BAIAP2,SSU72,MYL6B,IQCG,CMC2,YBX3,BLOC1
M189B,VKORC1,CDCA3,MRPS22,OSMR,SULF2,CYB561A3,BAIAP2,SSU72,MYL6B,IQCG,CMC2,YBX3,BLOC1
A13,BCAT2,CENPV,MRPL4,VKORC1,MRPL2,CDCA3,SRSF2,MRPS22,OSMR,PPFIBP1,LAS1L,HMGB1,SULF2,
A13,BCAT2,CENPV,MRPL4,VKORC1,MRPL2,CDCA3,SRSF2,MRPS22,OSMR,PPFIBP1,LAS1L,HMGB1,SULF2,
RPINB5,S100A13,BCAT2,CENPV,MRPL4,FAM189B,VKORC1,MRPL2,CDCA3,PCNA,SRSF2,OSMR,PPFIBP1,LA
RPINB5,S100A13,BCAT2,CENPV,MRPL4,FAM189B,VKORC1,MRPL2,CDCA3,PCNA,SRSF2,OSMR,PPFIBP1,LA
M72A,RPL38,STARD10,PTOV1,ZNF235,SCAF11,NPRL2,EIF5A,UFD1L,THYN1,NOP16,VRK3,FIS1,OBSL1,PPR
M72A,RPL38,STARD10,PTOV1,ZNF235,SCAF11,NPRL2,EIF5A,UFD1L,THYN1,NOP16,VRK3,FIS1,OBSL1,PPR
KORC1,PCNA,SRSF2,MRPS22,PPFIBP1,LAS1L,RAPGEF5,SSU72,MYL6B,TMEM67,FAM72A,CMC2,BLOC1S1,
AP2,SSU72,FAM122C,IQCG,CACYBP,TCAIM,KLC1,SIRT6,NUMA1,OSGEP,GSTK1,LDHA,RPS9,RCE1,ZNF410,P
S24,S100A13,BCAT2,CENPV,MRPL4,CDCA3,PCNA,SRSF2,MRPS22,KRT15,OSMR,PPFIBP1,HMGB1,SULF2,CY
S24,S100A13,BCAT2,CENPV,MRPL4,CDCA3,PCNA,SRSF2,MRPS22,KRT15,OSMR,PPFIBP1,HMGB1,SULF2,CY

SF2,RAPGEF5,CIRBP,SMARCB1,RNH1,DNASE1L1,PTOV1,FASTK,TMEM135,ATG16L2,DHPS,ZNF235,KLC1,ZC
SF2,RAPGEF5,CIRBP,SMARCB1,RNH1,DNASE1L1,PTOV1,FASTK,TMEM135,ATG16L2,DHPS,ZNF235,KLC1,ZC
IF5A,COMMD5,STYXL1,CPSF3L,CDC42,FBXW9,LYPLA2,ZNF197,TRMT112,DDOST,TOMM40,TADA3,MDH2,S
A,SRSF2,SULF2,RAPGEF5,SSU72,SRP9,BLOC1S1,CACYBP,SMARCB1,RPL38,RNH1,PTOV1,PARD3,TCAIM,ACT
CAF11,VDAC3,NOB1,SIRT6,SUMF2,SAE1,NPRL2,EIF5A,AMZ2,RPL13A,TTLL5,RPS9,SLC35G1,PCMT1,SNRPN,
CAF11,VDAC3,NOB1,SIRT6,SUMF2,SAE1,NPRL2,EIF5A,AMZ2,RPL13A,TTLL5,RPS9,SLC35G1,PCMT1,SNRPN,
SERPINB5,BCAT2,MRPL4,VKORC1,CDCA3,PCNA,MRPS22,OSMR,PPFIBP1,LAS1L,SULF2,CYB561A3,BAIAP2,
SERPINB5,BCAT2,MRPL4,VKORC1,CDCA3,PCNA,MRPS22,OSMR,PPFIBP1,LAS1L,SULF2,CYB561A3,BAIAP2,
10,RNH1,RCC1,TCAIM,SET,BOP1,ARRDC1,SCAF11,ARPC2,VDAC3,NOB1,WDR19,SIRT6,SUMF2,SAE1,EIF5A,
10,RNH1,RCC1,TCAIM,SET,BOP1,ARRDC1,SCAF11,ARPC2,VDAC3,NOB1,WDR19,SIRT6,SUMF2,SAE1,EIF5A,
MA4D,LAS1L,HMGB1,TMEM67,SULT1A3,PILRB,CMC2,BLOC1S1,CACYBP,CIRBP,RPL38,PRR13,RNH1,DNASE1
MA4D,LAS1L,HMGB1,TMEM67,SULT1A3,PILRB,CMC2,BLOC1S1,CACYBP,CIRBP,RPL38,PRR13,RNH1,DNASE1
SN1,UFD1L,USP38,ZNF410,TESC,SON,FAM3A,RPS25,MED15,MICAL1,HS2ST1,EIF3M,TBX6,PDLIM5,RFX3,A
SN1,UFD1L,USP38,ZNF410,TESC,SON,FAM3A,RPS25,MED15,MICAL1,HS2ST1,EIF3M,TBX6,PDLIM5,RFX3,A
M189B,MRPL2,CDCA3,OSMR,PPFIBP1,LAS1L,BAIAP2,MYL6B,IQCG,CACYBP,RHOF,SMARCB1,STARD10,RNH1
M189B,MRPL2,CDCA3,OSMR,PPFIBP1,LAS1L,BAIAP2,MYL6B,IQCG,CACYBP,RHOF,SMARCB1,STARD10,RNH1
35,CENPV,MRPL4,MRPL2,PPFIBP1,LAS1L,BAIAP2,SSU72,FAM122C,TMEM67,PILRB,FAM72A,CACYBP,RNH1
35,CENPV,MRPL4,MRPL2,PPFIBP1,LAS1L,BAIAP2,SSU72,FAM122C,TMEM67,PILRB,FAM72A,CACYBP,RNH1
A2,KLC1,SCAF11,DBI,ARPC2,NUMA1,EIF5A,DBNDD1,SH2B1,RUVBL2,TESC,NUDT5,RANBP1,DST,LRRC1,PHF
A2,KLC1,SCAF11,DBI,ARPC2,NUMA1,EIF5A,DBNDD1,SH2B1,RUVBL2,TESC,NUDT5,RANBP1,DST,LRRC1,PHF
T2,VKORC1,SRSF2,MRPS22,OSMR,HMGB1,SULF2,BAIAP2,RAPGEF5,MYL6B,IQCG,PILRB,YBX3,RHOF,SMARC
S24,SERPINB5,MRPL4,FAM189B,VKORC1,MRPL2,PCNA,SRSF2,PPFIBP1,SEMA4D,LAS1L,HMGB1,SULF2,BAI
S24,SERPINB5,MRPL4,FAM189B,VKORC1,MRPL2,PCNA,SRSF2,PPFIBP1,SEMA4D,LAS1L,HMGB1,SULF2,BAI
S24,S100A13,MRPL4,FAM189B,VKORC1,MRPL2,CDCA3,PCNA,MRPS22,OSMR,LAS1L,SULF2,BAIAP2,SSU72
ACYBP,RNH1,PTOV1,DHPS,BOP1,KLC1,ZC3HAV1,WDR19,NUMA1,OSGEP,TBL2,ARHGAP6,EIF5A,CHID1,AM
S24,SERPINB5,BCAT2,FAM189B,MRPL2,CDCA3,PCNA,SRSF2,KRT15,HMGB1,BAIAP2,RAPGEF5,SSU72,MYL6
OMMD5,STYXL1,CPSF3L,CDC42,FBXW9,TRMT112,DDOST,TOMM40,TADA3,MDH2,SEC31A,B3GAT3,PAFAH
A,SRSF2,OSMR,SULF2,BAIAP2,RAPGEF5,SSU72,MYL6B,FAM72A,SMARCB1,PRR13,RNH1,RBCK1,PTOV1,FAS
A,SRSF2,OSMR,SULF2,BAIAP2,RAPGEF5,SSU72,MYL6B,FAM72A,SMARCB1,PRR13,RNH1,RBCK1,PTOV1,FAS
9,FBXW9,LYPLA2,ZNF197,TRMT112,DDOST,RRNAD1,TOMM40,ANAPC11,COX17,B3GAT3,OGFOD2,PICK1,T
SF2,PPFIBP1,HMGB1,BAIAP2,RAPGEF5,MYL6B,IQCG,BLOC1S1,RHOF,SMARCB1,PRR13,RNH1,RBCK1,PTOV1
SF2,PPFIBP1,HMGB1,BAIAP2,RAPGEF5,MYL6B,IQCG,BLOC1S1,RHOF,SMARCB1,PRR13,RNH1,RBCK1,PTOV1
COMMD5,STYXL1,SNF8,CPSF3L,TSG101,CDC42,FBXW9,ENDOV,LYPLA2,PRMT1,PPAN,TRMT112,DDOST,TC
JD31,CTTN,EIF5A,UFD1L,ACOT8,STYXL1,CPSF3L,NIP7,RTEL1,ITGB1BP1,C1ORF198,UGGT2,CDC42,RPL28,LI
JD31,CTTN,EIF5A,UFD1L,ACOT8,STYXL1,CPSF3L,NIP7,RTEL1,ITGB1BP1,C1ORF198,UGGT2,CDC42,RPL28,LI
,VKORC1,CDCA3,PCNA,HMGB1,SULF2,SSU72,MYL6B,FAM122C,TMEM67,SULT1A3,UAP1,PILRB,FAM72A,C
1,PTOV1,RNF32,DHPS,NOB1,GPATCH2L,OSGEP,AMZ2,TAZ,MRPL33,RUVBL2,SNRPN,SNF8,UBR4,CHCHD1,E
1,PTOV1,RNF32,DHPS,NOB1,GPATCH2L,OSGEP,AMZ2,TAZ,MRPL33,RUVBL2,SNRPN,SNF8,UBR4,CHCHD1,E
T8,ZNF410,NUDT5,SNF8,RPL29,MED15,C1ORF198,FBXW9,LYPLA2,PPAN,ZNF197,TRMT112,DDOST,TOMM
L4,MRPL2,PCNA,SRSF2,RAPGEF5,TMEM67,PILRB,FAM72A,BLOC1S1,CACYBP,RHOF,DNASE1L1,PTOV1,FAS
FAM189B,MRPL2,CDCA3,PCNA,SRSF2,BAIAP2,RAPGEF5,MYL6B,SULT1A3,FAM72A,RHOF,SMARCB1,PRR13
FAM189B,MRPL2,CDCA3,PCNA,SRSF2,BAIAP2,RAPGEF5,MYL6B,SULT1A3,FAM72A,RHOF,SMARCB1,PRR13
M189B,MRPL2,CDCA3,PCNA,SRSF2,PPFIBP1,BAIAP2,RAPGEF5,MYL6B,PILRB,CACYBP,CIRBP,RHOF,PRR13,R
BP1,HMGB1,BAIAP2,SSU72,SULT1A3,CACYBP,SMARCB1,RPL38,KIF4A,PTOV1,FASTK,TMEM135,SNRPB,PAR
BP1,HMGB1,BAIAP2,SSU72,SULT1A3,CACYBP,SMARCB1,RPL38,KIF4A,PTOV1,FASTK,TMEM135,SNRPB,PAR
GEF39,ACOT8,VRK3,IRF9,RPS16,TSG101,FUZ,DBNDD2,RPS6,MED15,CDC42,ASF1B,CAPG,PPIL2,LIG1,ACAD1
GEF39,ACOT8,VRK3,IRF9,RPS16,TSG101,FUZ,DBNDD2,RPS6,MED15,CDC42,ASF1B,CAPG,PPIL2,LIG1,ACAD1

L6B,RHOF,PRR13,RNH1,RBCK1,PTOV1,LY75,ARRDC1,RBP5,LIMK2,NUMA1,SCARB1,CTTN,TSPAN11,COMM
L6B,RHOF,PRR13,RNH1,RBCK1,PTOV1,LY75,ARRDC1,RBP5,LIMK2,NUMA1,SCARB1,CTTN,TSPAN11,COMM
35,BCAT2,VKORC1,PCNA,SRSF2,RAPGEF5,SSU72,SRP9,IQCG,BLOC1S1,CACYBP,SMARCB1,RPL38,RNH1,PTO
35,BCAT2,VKORC1,PCNA,SRSF2,RAPGEF5,SSU72,SRP9,IQCG,BLOC1S1,CACYBP,SMARCB1,RPL38,RNH1,PTO
A3,RPL38,KIF4A,PTOV1,SNRPB,PARD3,ACTR6,SEC62,SCAF11,WDR19,NUMA1,OSGEP,SH2B1,LDHA,AP3B2,
N,EIF5A,COMMD5,STYXL1,CPSF3L,CDC42,FBXW9,TRMT112,DDOST,TOMM40,TADA3,MDH2,CHCHD5,SEC
LAS1L,HMGB1,SULF2,CYB561A3,SSU72,TMEM67,SULT1A3,BLOC1S1,CACYBP,RPL38,RNH1,DNASE1L1,RCC
COMMD5,STYXL1,CPSF3L,TSG101,CDC42,FBXW9,ENDOV,PPAN,TRMT112,DDOST,TOMM40,TADA3,MDH
137,SERPINB5,S100A13,BCAT2,CENPV,FAM189B,MRPL2,CDCA3,PCNA,SRSF2,KRT15,PPFIBP1,SEMA4D,HM
137,SERPINB5,S100A13,BCAT2,CENPV,FAM189B,MRPL2,CDCA3,PCNA,SRSF2,KRT15,PPFIBP1,SEMA4D,HM
MRPL4,MRPL2,CDCA3,SRSF2,MRPS22,KRT15,PPFIBP1,HMGB1,CYB561A3,BAIAP2,RAPGEF5,MYL6B,TMEM
MRPL4,MRPL2,CDCA3,SRSF2,MRPS22,KRT15,PPFIBP1,HMGB1,CYB561A3,BAIAP2,RAPGEF5,MYL6B,TMEM
,CENPV,MRPL2,PCNA,SRSF2,HMGB1,SULF2,BAIAP2,RAPGEF5,SSU72,SRP9,TMEM67,PILRB,BLOC1S1,RHOF
,CENPV,MRPL2,PCNA,SRSF2,HMGB1,SULF2,BAIAP2,RAPGEF5,SSU72,SRP9,TMEM67,PILRB,BLOC1S1,RHOF
RPS22,BAIAP2,RAPGEF5,MYL6B,FAM72A,RHOF,SMARCB1,RPL38,PRR13,RNH1,RBCK1,PTOV1,LY75,PARD3,
RPS22,BAIAP2,RAPGEF5,MYL6B,FAM72A,RHOF,SMARCB1,RPL38,PRR13,RNH1,RBCK1,PTOV1,LY75,PARD3,
MRPL2,HMGB1,SRP9,YBX3,BLOC1S1,CACYBP,PRR13,STARD10,RNH1,FASTK,RCC1,DHPS,SET,BOP1,ARRDC
MRPL2,HMGB1,SRP9,YBX3,BLOC1S1,CACYBP,PRR13,STARD10,RNH1,FASTK,RCC1,DHPS,SET,BOP1,ARRDC
17B,ZFAND2B,LRRFIP1,TWSG1,RABEP2,RALGPS1,NXN,OARD1,PFKFB2,ZXDC,TSEN34,SLC39A6,NDUFA11,
,LY75,SNRPB,DHPS,BUD31,NPRL2,SCARB1,CTTN,AMZ2,USP38,LDHA,MRPL33,PCMT1,WDR62,STYXL1,UBE
,LY75,SNRPB,DHPS,BUD31,NPRL2,SCARB1,CTTN,AMZ2,USP38,LDHA,MRPL33,PCMT1,WDR62,STYXL1,UBE
,ARPC2,LIMK2,BUD31,NPRL2,CTTN,UFD1L,STYXL1,UQCRQ,CPSF3L,METTL17,ATG12,NIP7,FOXRED1,IP6K2,
SNRPB,ACTR6,SCAF11,NUMA1,OSGEP,RPS9,WDR62,RANBP1,SLC15A3,PHF19,LPCAT3,CD44,TM9SF1,SON,
LL5,RUVBL2,WDR62,UBE2L6,ANTXR2,LPCAT3,GRK6,ZBTB44,TTC9C,HS2ST1,AUNIP,C14ORF2,ATF7,PCYT2,
NPV,MRPL2,CDCA3,PCNA,SRSF2,MRPS22,HMGB1,BAIAP2,RAPGEF5,MYL6B,BLOC1S1,RHOF,SMARCB1,PR
NPV,MRPL2,CDCA3,PCNA,SRSF2,MRPS22,HMGB1,BAIAP2,RAPGEF5,MYL6B,BLOC1S1,RHOF,SMARCB1,PR
,MRPL2,CDCA3,PCNA,SRSF2,HMGB1,BAIAP2,RAPGEF5,MYL6B,CACYBP,CIRBP,RHOF,SMARCB1,PRR13,RN
CENPV,FAM189B,MRPL2,SRSF2,BAIAP2,RAPGEF5,MYL6B,SRP9,SULT1A3,FAM72A,RHOF,SMARCB1,PRR13,
CENPV,FAM189B,MRPL2,SRSF2,BAIAP2,RAPGEF5,MYL6B,SRP9,SULT1A3,FAM72A,RHOF,SMARCB1,PRR13,
YL6B,RHOF,SMARCB1,PRR13,RNH1,RBCK1,PTOV1,ARRDC1,LIMK2,NUMA1,TBL2,SCARB1,CTTN,ITSN1,DBN
,S100A13,CENPV,MRPL4,FAM189B,VKORC1,MRPL2,CDCA3,PCNA,SRSF2,MRPS22,OSMR,HMGB1,SULF2,B
,S100A13,CENPV,MRPL4,FAM189B,VKORC1,MRPL2,CDCA3,PCNA,SRSF2,MRPS22,OSMR,HMGB1,SULF2,B
MK2,NUMA1,SCARB1,TSPAN11,AMZ2,NOP16,PSMA6,MRPL33,RUVBL2,EBPL,STIP1,SLC39A11,ZBTB44,SLC
MK2,NUMA1,SCARB1,TSPAN11,AMZ2,NOP16,PSMA6,MRPL33,RUVBL2,EBPL,STIP1,SLC39A11,ZBTB44,SLC
189B,MRPL2,CDCA3,PCNA,PPFIBP1,LAS1L,HMGB1,BAIAP2,SSU72,FAM122C,IQCG,TMEM67,PILRB,CACYB
189B,MRPL2,CDCA3,PCNA,PPFIBP1,LAS1L,HMGB1,BAIAP2,SSU72,FAM122C,IQCG,TMEM67,PILRB,CACYB
APGEF5,YBX3,CIRBP,STARD10,RNH1,FASTK,ATG16L2,RCC1,SET,BOP1,DBI,GPATCH2L,SAE1,OSGEP,SCARB1,
TMEM67,CACYBP,RNH1,PTOV1,ATG16L2,TCAIM,BOP1,ARRDC1,KLC1,SCAF11,GPATCH2L,NPRL2,OSGEP,A
TMEM67,CACYBP,RNH1,PTOV1,ATG16L2,TCAIM,BOP1,ARRDC1,KLC1,SCAF11,GPATCH2L,NPRL2,OSGEP,A
L4,MRPL2,SRSF2,LAS1L,BAIAP2,RAPGEF5,SULT1A3,CACYBP,RHOF,KIF4A,PTOV1,FASTK,TMEM135,LY75,PA
AP2,MYL6B,FAM72A,RHOF,SMARCB1,PRR13,RNH1,RBCK1,PTOV1,LY75,ATG16L2,SLC3A2,ARRDC1,RBP5,L
AP2,MYL6B,FAM72A,RHOF,SMARCB1,PRR13,RNH1,RBCK1,PTOV1,LY75,ATG16L2,SLC3A2,ARRDC1,RBP5,L
IRPL2,SRSF2,BAIAP2,RAPGEF5,MYL6B,SULT1A3,FAM72A,SMARCB1,PRR13,STARD10,RNH1,PTOV1,LY75,P
IRPL2,SRSF2,BAIAP2,RAPGEF5,MYL6B,SULT1A3,FAM72A,SMARCB1,PRR13,STARD10,RNH1,PTOV1,LY75,P
.5,SEMA4D,LAS1L,CYB561A3,PRR13,RBCK1,PTOV1,RNF32,TCAIM,ACTR6,SEC62,SET,SCAF11,ARPC2,PMFBP
SF2,MRPS22,SSU72,SRP9,IQCG,UAP1,CMC2,YBX3,CACYBP,RPL38,RBCK1,FASTK,ATG16L2,RCC1,SNRPB,PA
SF2,MRPS22,SSU72,SRP9,IQCG,UAP1,CMC2,YBX3,CACYBP,RPL38,RBCK1,FASTK,ATG16L2,RCC1,SNRPB,PA

35,MRPL4,MRPL2,PPFIBP1,LAS1L,BAIAP2,SSU72,FAM122C,TMEM67,PILRB,CACYBP,RNH1,DNASE1L1,PTO
35,MRPL4,MRPL2,PPFIBP1,LAS1L,BAIAP2,SSU72,FAM122C,TMEM67,PILRB,CACYBP,RNH1,DNASE1L1,PTO
PS,SET,BOP1,KLC1,PMFBP1,NOB1,NOP16,RPL13A,NNMT,STYXL1,PHF19,LPCAT3,TET2,YLPM1,FUZ,ECSIT,P
PS,SET,BOP1,KLC1,PMFBP1,NOB1,NOP16,RPL13A,NNMT,STYXL1,PHF19,LPCAT3,TET2,YLPM1,FUZ,ECSIT,P
F5A,USP38,LZTFL1,SLC15A3,CPSF3L,FBXW9,CCDC25,TROAP,LYPLA2,PPAN,CDK11A,ZNF197,ATP5J,TRMT1
1,DNASE1L1,SNRPB,TCAIM,DHPS,KLC1,SCAF11,WDR19,SAE1,OSGEP,TAZ,LDHA,RCE1,ZNF410,IGSF8,NNMT
GEF5,MYL6B,FAM72A,CACYBP,RHOF,LY75,ATG16L2,RCC1,SLC3A2,DHPS,SEC62,KLC1,SCAF11,LIMK2,VDAC
GEF5,MYL6B,FAM72A,CACYBP,RHOF,LY75,ATG16L2,RCC1,SLC3A2,DHPS,SEC62,KLC1,SCAF11,LIMK2,VDAC
YBP,RHOF,RNH1,KIF4A,RBCK1,ATG16L2,SNRPB,PARD3,TCAIM,ARRDC1,KLC1,RBP5,SCAF11,DBI,ARPC2,WD
5,S100A13,CENPV,MRPL4,VKORC1,MRPL2,CDCA3,PCNA,SRSF2,MRPS22,PPFIBP1,SEMA4D,HMGB1,SULF2,
5,S100A13,CENPV,MRPL4,VKORC1,MRPL2,CDCA3,PCNA,SRSF2,MRPS22,PPFIBP1,SEMA4D,HMGB1,SULF2,
NLL1,CST3,GNAS,GPR137,RPS24,SERPINB5,S100A13,BCAT2,CENPV,MRPL4,FAM189B,VKORC1,MRPL2,CDC
NLL1,CST3,GNAS,GPR137,RPS24,SERPINB5,S100A13,BCAT2,CENPV,MRPL4,FAM189B,VKORC1,MRPL2,CDC
EM67,RNF32,ATG16L2,SET,BOP1,ARPC2,OSGEP,SCARB1,USP38,SH2B1,NOP16,RPL13A,SPAG7,STYXL1,SON
,STARD10,RNH1,DNASE1L1,FASTK,DHPS,MAP3K7CL,SET,BOP1,RBP5,SAE1,OSGEP,CTTN,EIF5A,COMMD5,C
,STARD10,RNH1,DNASE1L1,FASTK,DHPS,MAP3K7CL,SET,BOP1,RBP5,SAE1,OSGEP,CTTN,EIF5A,COMMD5,C
A,ARHGEF39,ACOT8,VRK3,IRF9,RPS16,TSG101,FUZ,NIP7,TTC9C,DBNDD2,RPS6,MED15,SLC39A13,CDC42,A
A,ARHGEF39,ACOT8,VRK3,IRF9,RPS16,TSG101,FUZ,NIP7,TTC9C,DBNDD2,RPS6,MED15,SLC39A13,CDC42,A
1122C,TMEM67,UAP1,FAM72A,RNF32,SLC3A2,CTSC,RBP5,ARPC2,NUMA1,GPATCH2L,EIF5A,CHID1,AP3B2,
MYL6B,RHOF,PRR13,RNH1,RBCK1,PTOV1,RNF32,ARRDC1,KLC1,LIMK2,NUMA1,TBL2,SCARB1,CTTN,DBNDD
89B,CDCA3,PCNA,SRSF2,OSMR,LAS1L,HMGB1,BAIAP2,RAPGEF5,MYL6B,IQCG,BLOC1S1,CACYBP,SMARCB1
89B,CDCA3,PCNA,SRSF2,OSMR,LAS1L,HMGB1,BAIAP2,RAPGEF5,MYL6B,IQCG,BLOC1S1,CACYBP,SMARCB1
.0,DNASE1L1,PTOV1,FASTK,SNRPB,ACTR6,WDR19,SIRT6,NUMA1,NPRL2,USP38,TAZ,AP3B2,RANBP1,FIS1,S
.0,DNASE1L1,PTOV1,FASTK,SNRPB,ACTR6,WDR19,SIRT6,NUMA1,NPRL2,USP38,TAZ,AP3B2,RANBP1,FIS1,S
IA,MRPS22,HDAC9,LAS1L,HMGB1,CYB561A3,FAM122C,TMEM67,PILRB,CMC2,BLOC1S1,CACYBP,CIRBP,RH
IA,MRPS22,HDAC9,LAS1L,HMGB1,CYB561A3,FAM122C,TMEM67,PILRB,CMC2,BLOC1S1,CACYBP,CIRBP,RH
,UFD1L,ACOT8,STYXL1,CPSF3L,RTEL1,ITGB1BP1,C1ORF198,UGGT2,CDC42,LIG1,CYB561,ZNF197,TRMT112
,UFD1L,ACOT8,STYXL1,CPSF3L,RTEL1,ITGB1BP1,C1ORF198,UGGT2,CDC42,LIG1,CYB561,ZNF197,TRMT112
R13,RNH1,RBCK1,PTOV1,ARRDC1,LIMK2,SCARB1,DBNDD1,RCE1,EBPL,TESC,IGSF8,SPAG7,WDR62,UBE2L6
,CDCA3,PCNA,MRPS22,BAIAP2,RAPGEF5,MYL6B,FAM72A,CIRBP,RHOF,SMARCB1,PRR13,RNH1,RBCK1,PTC
,CDCA3,PCNA,MRPS22,BAIAP2,RAPGEF5,MYL6B,FAM72A,CIRBP,RHOF,SMARCB1,PRR13,RNH1,RBCK1,PTC
35,MRPL4,MRPL2,PPFIBP1,LAS1L,BAIAP2,SSU72,FAM122C,TMEM67,PILRB,CACYBP,RNH1,DNASE1L1,PTO
35,MRPL4,MRPL2,PPFIBP1,LAS1L,BAIAP2,SSU72,FAM122C,TMEM67,PILRB,CACYBP,RNH1,DNASE1L1,PTO
GEF39,ACOT8,VRK3,TSG101,FUZ,RPS6,MED15,CDC42,ASF1B,PPIL2,LIG1,ZNF197,RAB2A,TRMT112,UBXN1,
GEF39,ACOT8,VRK3,TSG101,FUZ,RPS6,MED15,CDC42,ASF1B,PPIL2,LIG1,ZNF197,RAB2A,TRMT112,UBXN1,
2,RAPGEF5,SSU72,MYL6B,PILRB,YBX3,CACYBP,SMARCB1,KIF4A,FASTK,RCC1,TCAIM,DHPS,ACTR6,SET,TRPT
2,RAPGEF5,SSU72,MYL6B,PILRB,YBX3,CACYBP,SMARCB1,KIF4A,FASTK,RCC1,TCAIM,DHPS,ACTR6,SET,TRPT
2,CENPV,MRPL4,CDCA3,SRSF2,MRPS22,KRT15,OSMR,HMGB1,SULF2,CYB561A3,RAPGEF5,SSU72,FAM122
AS,GPR137,CENPV,MRPL4,FAM189B,VKORC1,PCNA,SRSF2,KRT15,SEMA4D,HMGB1,SULF2,BAIAP2,RAPGEF
AS,GPR137,CENPV,MRPL4,FAM189B,VKORC1,PCNA,SRSF2,KRT15,SEMA4D,HMGB1,SULF2,BAIAP2,RAPGEF
,TCAIM,KLC1,SCAF11,WDR19,OSGEP,TAZ,LDHA,RCE1,ZNF410,IGSF8,NNMT,PLEKHA5,VRK3,PCBP2,TET2,G
72,FAM122C,TMEM67,UAP1,PILRB,FAM72A,CIRBP,RNF32,SLC3A2,CTSC,RBP5,ARPC2,NUMA1,GPATCH2L,
72,FAM122C,TMEM67,UAP1,PILRB,FAM72A,CIRBP,RNF32,SLC3A2,CTSC,RBP5,ARPC2,NUMA1,GPATCH2L,
BP,CIRBP,RPL38,RCC1,ACTR6,CTSC,TRPT1,ARPC2,PMFBP1,BUD31,NPRL2,CRYGS,CTTN,NUDT22,USP38,RPL
BP,CIRBP,RPL38,RCC1,ACTR6,CTSC,TRPT1,ARPC2,PMFBP1,BUD31,NPRL2,CRYGS,CTTN,NUDT22,USP38,RPL
S,GPR137,RPS24,SERPINB5,S100A13,BCAT2,CENPV,MRPL4,FAM189B,VKORC1,MRPL2,CDCA3,PCNA,SRSF
S,GPR137,RPS24,SERPINB5,S100A13,BCAT2,CENPV,MRPL4,FAM189B,VKORC1,MRPL2,CDCA3,PCNA,SRSF

2,SET,ARPC2,NOB1,SAE1,SCARB1,USP38,NOP16,RPL13A,TESC,NNMT,STYXL1,TET2,UBR4,FUZ,ECSIT,PPRC1

NASE1L1,PTOV1,RNF32,DHPS,BOP1,VDAC3,NOB1,GPATCH2L,OSGEP,AMZ2,TAZ,MRPL33,RUVBL2,SNRPN,S

NASE1L1,PTOV1,RNF32,DHPS,BOP1,VDAC3,NOB1,GPATCH2L,OSGEP,AMZ2,TAZ,MRPL33,RUVBL2,SNRPN,S

,FAM189B,CDCA3,SRSF2,OSMR,BAIAP2,SRP9,SULT1A3,PILRB,FAM72A,RPL38,STARD10,KIF4A,RBCK1,PTO

LOC1S1,CACYBP,DNASE1L1,SNRPB,TCAIM,KLC1,SCAF11,WDR19,SAE1,OSGEP,TAZ,LDHA,RCE1,ZNF410,IGS

LOC1S1,CACYBP,DNASE1L1,SNRPB,TCAIM,KLC1,SCAF11,WDR19,SAE1,OSGEP,TAZ,LDHA,RCE1,ZNF410,IGS

LOC1S1,CACYBP,DNASE1L1,SNRPB,TCAIM,KLC1,SCAF11,WDR19,SAE1,OSGEP,TAZ,LDHA,RCE1,ZNF410,IGS

LIG1,DGUOK,ATP5J,TRMT112,ERCC5,EMC4,DDOST,LSR,TOMM40,TADA3,ABHD17A,SEC31A,CNST,TCP1,UE

NASE1L1,RBCK1,PTOV1,FASTK,LY75,SET,ARRDC1,RBP5,NUMA1,TBL2,SCARB1,ITSN1,AMZ2,TAZ,RUVBL2,EB

A,ARHGEF39,ACOT8,VRK3,IRF9,RPS16,TSG101,FUZ,NIP7,TTC9C,DBNDD2,RPS6,MED15,SLC39A13,CDC42,A

A,ARHGEF39,ACOT8,VRK3,IRF9,RPS16,TSG101,FUZ,NIP7,TTC9C,DBNDD2,RPS6,MED15,SLC39A13,CDC42,A

SRSF2,OSMR,PPFIBP1,SSU72,MYL6B,SRP9,IQCG,UAP1,CMC2,YBX3,CACYBP,RBCK1,FASTK,SNRPB,PARD3,D

SRSF2,OSMR,PPFIBP1,SSU72,MYL6B,SRP9,IQCG,UAP1,CMC2,YBX3,CACYBP,RBCK1,FASTK,SNRPB,PARD3,D

V,FAM189B,MRPL2,CDCA3,HMGB1,BAIAP2,MYL6B,SRP9,PILRB,CIRBP,SMARCB1,PRR13,STARD10,TMEM1

V,FAM189B,MRPL2,CDCA3,HMGB1,BAIAP2,MYL6B,SRP9,PILRB,CIRBP,SMARCB1,PRR13,STARD10,TMEM1

35,CENPV,VKORC1,PCNA,SRSF2,RAPGEF5,SSU72,SRP9,IQCG,BLOC1S1,CACYBP,SMARCB1,PTOV1,FASTK,LY

35,CENPV,VKORC1,PCNA,SRSF2,RAPGEF5,SSU72,SRP9,IQCG,BLOC1S1,CACYBP,SMARCB1,PTOV1,FASTK,LY

HOF,PRR13,RNH1,RBCK1,PTOV1,ARRDC1,LIMK2,NUMA1,TBL2,SCARB1,DBNDD1,RCE1,EBPL,TESC,IGSF8,SP

4,S100A13,CENPV,FAM189B,VKORC1,MRPL2,CDCA3,HDAC9,SEMA4D,LAS1L,SULF2,SSU72,MYL6B,TMEM6

4,S100A13,CENPV,FAM189B,VKORC1,MRPL2,CDCA3,HDAC9,SEMA4D,LAS1L,SULF2,SSU72,MYL6B,TMEM6

2,COASY,RABL5,TRNT1,HSBP1L1,EDF1,DGKE,CAMLG,EPB41L4B,ZNF7,ARL6IP4,STAU1,SYNGR2,RPTOR,LYN

13,BCAT2,CENPV,MRPL4,FAM189B,MRPL2,PCNA,SRSF2,OSMR,HMGB1,BAIAP2,SSU72,SRP9,PILRB,RHOF,

13,BCAT2,CENPV,MRPL4,FAM189B,MRPL2,PCNA,SRSF2,OSMR,HMGB1,BAIAP2,SSU72,SRP9,PILRB,RHOF,

K3,NIP7,RPS6,FBXW9,LYPLA2,TRMT112,DDOST,TOMM40,CNST,B3GAT3,DCTN5,NDUFS3,OGFOD2,ARIH2,

CB1,PRR13,RNH1,DNASE1L1,RBCK1,PTOV1,SET,ARRDC1,CTTN,ITSN1,COMMD5,AMZ2,TAZ,RUVBL2,STIP1,

PV,FAM189B,SRSF2,HDAC9,KRT15,OSMR,CYB561A3,BAIAP2,RAPGEF5,FAM122C,TMEM67,FAM72A,YBX3,

PV,FAM189B,SRSF2,HDAC9,KRT15,OSMR,CYB561A3,BAIAP2,RAPGEF5,FAM122C,TMEM67,FAM72A,YBX3,

IPV,FAM189B,MRPL2,CDCA3,PCNA,SRSF2,SEMA4D,HMGB1,SULF2,BAIAP2,RAPGEF5,MYL6B,IQCG,CIRBP,R

IPV,FAM189B,MRPL2,CDCA3,PCNA,SRSF2,SEMA4D,HMGB1,SULF2,BAIAP2,RAPGEF5,MYL6B,IQCG,CIRBP,R

,PCNA,SRSF2,MRPS22,OSMR,HMGB1,SULF2,BAIAP2,RAPGEF5,SSU72,IQCG,YBX3,CIRBP,RHOF,RNH1,PTOV

FAM122C,IQCG,CACYBP,DNASE1L1,TCAIM,KLC1,SCAF11,NUMA1,OSGEP,TAZ,LDHA,RCE1,ZNF410,PCMT1,I

CTR6,TRPT1,NPRL2,CTTN,EIF5A,COMMD5,NUDT22,STYXL1,VRK3,LZTFL1,IRF9,CPSF3L,TSG101,RPS25,TTC

CAT2,CENPV,FAM189B,CDCA3,SRSF2,HMGB1,RAPGEF5,SSU72,SRP9,IQCG,TMEM67,YBX3,BLOC1S1,RHOF,

CAT2,CENPV,FAM189B,CDCA3,SRSF2,HMGB1,RAPGEF5,SSU72,SRP9,IQCG,TMEM67,YBX3,BLOC1S1,RHOF,

ORC1,MRPL2,PCNA,SRSF2,OSMR,SEMA4D,HMGB1,SULF2,RAPGEF5,IQCG,YBX3,BLOC1S1,CIRBP,SMARCB1,

ORC1,MRPL2,PCNA,SRSF2,OSMR,SEMA4D,HMGB1,SULF2,RAPGEF5,IQCG,YBX3,BLOC1S1,CIRBP,SMARCB1,

RC1,KRT15,LAS1L,SULF2,CYB561A3,RAPGEF5,SSU72,MYL6B,IQCG,PILRB,FAM72A,YBX3,CACYBP,RHOF,PRR

RC1,KRT15,LAS1L,SULF2,CYB561A3,RAPGEF5,SSU72,MYL6B,IQCG,PILRB,FAM72A,YBX3,CACYBP,RHOF,PRR

RBCK1,RCC1,ACTR6,CTSC,TRPT1,PMFBP1,BUD31,NPRL2,CRYGS,CTTN,NUDT22,USP38,RPL13A,ARHGEF39,

RBCK1,RCC1,ACTR6,CTSC,TRPT1,PMFBP1,BUD31,NPRL2,CRYGS,CTTN,NUDT22,USP38,RPL13A,ARHGEF39,

122C,CACYBP,CIRBP,STARD10,TMEM135,LY75,RCC1,CTSC,ZNF235,BOP1,ARRDC1,KLC1,RBP5,ARPC2,VDA

122C,CACYBP,CIRBP,STARD10,TMEM135,LY75,RCC1,CTSC,ZNF235,BOP1,ARRDC1,KLC1,RBP5,ARPC2,VDA

SU72,FAM122C,TMEM67,PILRB,CACYBP,RNH1,DNASE1L1,PTOV1,SNRPB,TCAIM,SEC62,BOP1,KLC1,SCAF1

SU72,FAM122C,TMEM67,PILRB,CACYBP,RNH1,DNASE1L1,PTOV1,SNRPB,TCAIM,SEC62,BOP1,KLC1,SCAF1

RCC1,TBL2,ARHGAP6,UFD1L,DST,TTC21B,PSMA1,FAM3A,RPS25,TMX2,CENPK,SRSF1,KXD1,C1ORF198,PPA

RPL4,VKORC1,CDCA3,PCNA,SRSF2,OSMR,LAS1L,BAIAP2,RAPGEF5,SSU72,FAM122C,SRP9,UAP1,CACYBP,CI

M,KLC1,SCAF11,OSGEP,TAZ,LDHA,RCE1,ZNF410,IGSF8,NNMT,VRK3,PCBP2,TET2,GRK6,CDC37,PSMA3,ME

M,KLC1,SCAF11,OSGEP,TAZ,LDHA,RCE1,ZNF410,IGSF8,NNMT,VRK3,PCBP2,TET2,GRK6,CDC37,PSMA3,MET
BLOC1S1,DNASE1L1,SNRPB,TCAIM,DHPS,KLC1,SCAF11,WDR19,SAE1,OSGEP,TBL2,TAZ,LDHA,RCE1,ZNF410
BLOC1S1,DNASE1L1,SNRPB,TCAIM,DHPS,KLC1,SCAF11,WDR19,SAE1,OSGEP,TBL2,TAZ,LDHA,RCE1,ZNF410
V,VKORC1,CDCA3,SRSF2,OSMR,LAS1L,SULF2,CYB561A3,BAIAP2,RAPGEF5,SSU72,FAM122C,FAM72A,YBX3,
9,OSMR,SEMA4D,HMGB1,SULF2,BAIAP2,RAPGEF5,SSU72,IQCG,TMEM67,CMC2,YBX3,SMARCB1,RPL38,ST
9,OSMR,SEMA4D,HMGB1,SULF2,BAIAP2,RAPGEF5,SSU72,IQCG,TMEM67,CMC2,YBX3,SMARCB1,RPL38,ST
ORC1,MRPL2,PCNA,SRSF2,OSMR,SEMA4D,HMGB1,SULF2,RAPGEF5,IQCG,YBX3,BLOC1S1,CIRBP,SMARCB1,
ORC1,MRPL2,SRSF2,SEMA4D,HMGB1,SULF2,RAPGEF5,IQCG,YBX3,BLOC1S1,CIRBP,SMARCB1,STARD10,RNI
ORC1,MRPL2,SRSF2,SEMA4D,HMGB1,SULF2,RAPGEF5,IQCG,YBX3,BLOC1S1,CIRBP,SMARCB1,STARD10,RNI
ORC1,MRPL2,SRSF2,SEMA4D,HMGB1,SULF2,RAPGEF5,IQCG,YBX3,BLOC1S1,CIRBP,SMARCB1,STARD10,RNI
S100A13,BCAT2,MRPL4,FAM189B,MRPL2,CDCA3,PCNA,SRSF2,MRPS22,KRT15,OSMR,PPFIBP1,SEMA4D,H
S100A13,BCAT2,MRPL4,FAM189B,MRPL2,CDCA3,PCNA,SRSF2,MRPS22,KRT15,OSMR,PPFIBP1,SEMA4D,H
P,CHID1,TESC,SPAG7,WDR62,STYXL1,RANBP1,LZTFL1,FIS1,STIP1,TET2,GRK6,CDC37,SLC39A11,FAM3A,PPR
P,CHID1,TESC,SPAG7,WDR62,STYXL1,RANBP1,LZTFL1,FIS1,STIP1,TET2,GRK6,CDC37,SLC39A11,FAM3A,PPR
RPL13A,TTLL5,RPS9,CPSF3L,TMED4,ITGB1BP1,SH3YL1,PRMT2,PPA1,FAM219B,ELP6,LRRC27,SSH2,COX17,I
4D,HMGB1,RAPGEF5,TMEM67,CMC2,CIRBP,RPL38,RNH1,DNASE1L1,KIF4A,RBCK1,FASTK,RNF32,LY75,RCC
4D,HMGB1,RAPGEF5,TMEM67,CMC2,CIRBP,RPL38,RNH1,DNASE1L1,KIF4A,RBCK1,FASTK,RNF32,LY75,RCC
010,RNH1,FASTK,ATG16L2,RCC1,BOP1,DBI,GPATCH2L,SAE1,OSGEP,SCARB1,CTTN,EIF5A,COMMD5,CHID1,I
,GNAS,RPS24,SERPINB5,BCAT2,CENPV,VKORC1,MRPL2,CDCA3,PCNA,SRSF2,KRT15,OSMR,PPFIBP1,LAS1L,
,GNAS,RPS24,SERPINB5,BCAT2,CENPV,VKORC1,MRPL2,CDCA3,PCNA,SRSF2,KRT15,OSMR,PPFIBP1,LAS1L,
MRPL4,VKORC1,CDCA3,SRSF2,MRPS22,OSMR,PPFIBP1,SSU72,MYL6B,SRP9,IQCG,TMEM67,UAP1,FAM72A
MRPL4,VKORC1,CDCA3,SRSF2,MRPS22,OSMR,PPFIBP1,SSU72,MYL6B,SRP9,IQCG,TMEM67,UAP1,FAM72A
RPL4,MRPL2,CDCA3,PCNA,SRSF2,HDAC9,PPFIBP1,HMGB1,SSU72,SRP9,IQCG,TMEM67,FAM72A,CMC2,CA
RPL4,MRPL2,CDCA3,PCNA,SRSF2,HDAC9,PPFIBP1,HMGB1,SSU72,SRP9,IQCG,TMEM67,FAM72A,CMC2,CA
,TCAIM,KLC1,SCAF11,WDR19,OSGEP,TAZ,LDHA,RCE1,ZNF410,IGSF8,NNMT,VRK3,PCBP2,TET2,GRK6,CDC3
CDCA3,PCNA,OSMR,PPFIBP1,LAS1L,SULF2,RAPGEF5,SSU72,CACYBP,CIRBP,STARD10,RNH1,DNASE1L1,RNF
FAM189B,VKORC1,MRPL2,SSU72,FAM122C,CACYBP,PRR13,RNH1,DNASE1L1,TMEM135,ACTR6,ZNF235,BC
FAM189B,VKORC1,MRPL2,SSU72,FAM122C,CACYBP,PRR13,RNH1,DNASE1L1,TMEM135,ACTR6,ZNF235,BC
PN,STYXL1,UQCRQ,FUZ,TRNT1,MED15,EIF3M,CDC42,PPIL2,LIG1,CDC14B,C17ORF62,ZNF197,RAB2A,TRMT
PN,STYXL1,UQCRQ,FUZ,TRNT1,MED15,EIF3M,CDC42,PPIL2,LIG1,CDC14B,C17ORF62,ZNF197,RAB2A,TRMT
,S100A13,FAM189B,PCNA,KRT15,BAIAP2,SSU72,MYL6B,PILRB,FAM72A,CIRBP,RPL38,RNH1,RBCK1,PTOV1
PV,VKORC1,MRPL2,SRSF2,MRPS22,PPFIBP1,SEMA4D,HMGB1,SULF2,RAPGEF5,IQCG,YBX3,BLOC1S1,CIRBP,
VKORC1,CDCA3,SRSF2,OSMR,PPFIBP1,HMGB1,SSU72,MYL6B,SRP9,IQCG,TMEM67,UAP1,CMC2,YBX3,CAC
VKORC1,CDCA3,SRSF2,OSMR,PPFIBP1,HMGB1,SSU72,MYL6B,SRP9,IQCG,TMEM67,UAP1,CMC2,YBX3,CAC
E1,ZNF410,IGSF8,NNMT,WDR62,VRK3,GRK6,CDC37,FAM3A,PSMA3,METTL17,NIP7,TTC9C,COX4I1,AUP1,F
AP3B2,RPS9,STYXL1,CDC37,ECSIT,RPS6,CD99L2,COASY,YIPF3,UBE2I,EIF3M,C14ORF2,ATF7,CRIP1,CASC4,CY
1,SCAF11,OSGEP,TAZ,LDHA,RCE1,ZNF410,IGSF8,NNMT,VRK3,PCBP2,TET2,GRK6,CDC37,PSMA3,METTL17,
AS,GPR137,CENPV,MRPL4,FAM189B,VKORC1,PCNA,SRSF2,KRT15,HMGB1,SULF2,BAIAP2,RAPGEF5,SSU72,
E1,ZNF410,IGSF8,NNMT,WDR62,VRK3,TET2,GRK6,CDC37,FAM3A,PSMA3,METTL17,NIP7,TTC9C,COX4I1,AU
L13A,SPAG7,WDR62,RANBP1,FIS1,TUBA1C,TET2,GRK6,CDC37,TTC21B,FAM3A,PPRC1,C1ORF198,EEF1B2,I
RSF2,MRPS22,KRT15,OSMR,SEMA4D,LAS1L,HMGB1,RAPGEF5,SSU72,MYL6B,FAM122C,TMEM67,CMC2,Y
RSF2,MRPS22,KRT15,OSMR,SEMA4D,LAS1L,HMGB1,RAPGEF5,SSU72,MYL6B,FAM122C,TMEM67,CMC2,Y
AIAP2,FAM122C,CACYBP,RNH1,DNASE1L1,SNRPB,TCAIM,BOP1,ARRDC1,KLC1,SCAF11,GPATCH2L,NPRL2,O
AIAP2,FAM122C,CACYBP,RNH1,DNASE1L1,SNRPB,TCAIM,BOP1,ARRDC1,KLC1,SCAF11,GPATCH2L,NPRL2,O
RNH1,FASTK,DHPS,SET,BOP1,SAE1,OSGEP,EIF5A,CHID1,RPL13A,MRPL33,SNRPN,WDR62,RANBP1,FIS1,TU
RNH1,FASTK,DHPS,SET,BOP1,SAE1,OSGEP,EIF5A,CHID1,RPL13A,MRPL33,SNRPN,WDR62,RANBP1,FIS1,TU
RNH1,FASTK,DHPS,SET,BOP1,SAE1,OSGEP,EIF5A,CHID1,RPL13A,MRPL33,SNRPN,WDR62,RANBP1,FIS1,TU

AC9,OSMR,PPFIBP1,LAS1L,RAPGEF5,SULT1A3,PILRB,CIRBP,RPL38,PRR13,STARD10,DNASE1L1,RBCK1,LY7

,TCAIM,KLC1,SCAF11,WDR19,OSGEP,TAZ,LDHA,RCE1,ZNF410,IGSF8,NNMT,PLEKHA5,VRK3,PCBP2,TET2,G

,TCAIM,KLC1,SCAF11,WDR19,OSGEP,TAZ,LDHA,RCE1,ZNF410,IGSF8,NNMT,PLEKHA5,VRK3,PCBP2,TET2,G

N,EIF5A,NUDT22,VRK3,LZTFL1,CPSF3L,OBSL1,MUC1,PPRC1,RPS25,PLCB2,RPS6,SLC39A13,TBX6,PCYT2,TRC

3,RNH1,RCC1,PMFBP1,LIMK2,ARHGAP6,TNFSF10,SH2B1,SNRPN,STYXL1,VRK3,FIS1,DST,LRRC1,TM9SF1,M

3,RNH1,RCC1,PMFBP1,LIMK2,ARHGAP6,TNFSF10,SH2B1,SNRPN,STYXL1,VRK3,FIS1,DST,LRRC1,TM9SF1,M

3CAT2,CENPV,FAM189B,VKORC1,MRPL2,CDCA3,PCNA,SRSF2,MRPS22,HDAC9,KRT15,OSMR,PPFIBP1,LAS1

3CAT2,CENPV,FAM189B,VKORC1,MRPL2,CDCA3,PCNA,SRSF2,MRPS22,HDAC9,KRT15,OSMR,PPFIBP1,LAS1

,SERPINB5,CENPV,VKORC1,MRPL2,CDCA3,SRSF2,OSMR,SEMA4D,HMGB1,BAIAP2,RAPGEF5,MYL6B,TMEM

,SERPINB5,CENPV,VKORC1,MRPL2,CDCA3,SRSF2,OSMR,SEMA4D,HMGB1,BAIAP2,RAPGEF5,MYL6B,TMEM

MRPS22,OSMR,LAS1L,HMGB1,RAPGEF5,MYL6B,IQCG,FAM72A,BLOC1S1,CACYBP,CIRBP,SMARCB1,PRR13,

CLDND1,CAPN10,NUTF2,WNK1,RPL36,PABPC1,HSPH1,ENY2,ZNF223,DGAT1,MARS,ZNF354A,DCAKD,NAA

35,CDCA3,SRSF2,MRPS22,OSMR,PPFIBP1,SSU72,MYL6B,SRP9,IQCG,UAP1,CMC2,YBX3,CACYBP,RBCK1,FAS

35,CDCA3,SRSF2,MRPS22,OSMR,PPFIBP1,SSU72,MYL6B,SRP9,IQCG,UAP1,CMC2,YBX3,CACYBP,RBCK1,FAS

SF2,BAIAP2,RAPGEF5,MYL6B,RHOF,SMARCB1,PRR13,RNH1,RBCK1,PTOV1,RNF32,LY75,ARRDC1,LIMK2,NU

L,CDCA3,SRSF2,OSMR,PPFIBP1,SSU72,MYL6B,SRP9,IQCG,UAP1,CMC2,YBX3,CACYBP,RBCK1,FASTK,ATG16L

L,CDCA3,SRSF2,OSMR,PPFIBP1,SSU72,MYL6B,SRP9,IQCG,UAP1,CMC2,YBX3,CACYBP,RBCK1,FASTK,ATG16L

I1,TCAIM,BOP1,ARRDC1,KLC1,BUD31,LDHA,RPS9,RCE1,ZNF410,IGSF8,VRK3,TET2,GRK6,CDC37,TSG101,UE

I1,TCAIM,BOP1,ARRDC1,KLC1,BUD31,LDHA,RPS9,RCE1,ZNF410,IGSF8,VRK3,TET2,GRK6,CDC37,TSG101,UE

NLL1,CST3,GNAS,GPR137,S100A13,BCAT2,CENPV,MRPL4,FAM189B,CDCA3,PCNA,SRSF2,KRT15,PPFIBP1,S

LL,HMGB1,RAPGEF5,SSU72,FAM122C,IQCG,CMC2,YBX3,BLOC1S1,CACYBP,CIRBP,PRR13,RNH1,KIF4A,RBCK

LL,HMGB1,RAPGEF5,SSU72,FAM122C,IQCG,CMC2,YBX3,BLOC1S1,CACYBP,CIRBP,PRR13,RNH1,KIF4A,RBCK

,BUD31,CTTN,AP3B2,RPS9,ZNF410,NNMT,WDR62,PPM1K,DST,TM9SF1,SON,UQCRQ,TSG101,PHF21A,FAM

0A13,BCAT2,CENPV,FAM189B,CDCA3,MRPS22,KRT15,OSMR,PPFIBP1,SULF2,BAIAP2,MYL6B,FAM122C,IQC

0A13,BCAT2,CENPV,FAM189B,CDCA3,MRPS22,KRT15,OSMR,PPFIBP1,SULF2,BAIAP2,MYL6B,FAM122C,IQC

AM72A,RNF32,SLC3A2,RBP5,NUMA1,GPATCH2L,EIF5A,CHID1,AP3B2,ZNF410,NUDT5,TUBA1C,MAGED2,S

,CACYBP,RHOF,TMEM135,LY75,RCC1,SEC62,KLC1,SCAF11,LIMK2,VDAC3,BUD31,OSGEP,ITSN1,COMMD5,N

,CACYBP,RHOF,TMEM135,LY75,RCC1,SEC62,KLC1,SCAF11,LIMK2,VDAC3,BUD31,OSGEP,ITSN1,COMMD5,N

RDC1,KLC1,GPATCH2L,TBL2,ITSN1,AMZ2,TAZ,RUVBL2,PPM1K,LPCAT3,MAGED2,GRK6,CPSF3L,SLC39A11,F

M,KLC1,SCAF11,OSGEP,TAZ,LDHA,RCE1,ZNF410,IGSF8,NNMT,VRK3,PCBP2,TET2,GRK6,CDC37,PSMA3,ME

M,KLC1,SCAF11,OSGEP,TAZ,LDHA,RCE1,ZNF410,IGSF8,NNMT,VRK3,PCBP2,TET2,GRK6,CDC37,PSMA3,ME

IF410,IGSF8,NNMT,WDR62,VRK3,TM9SF1,CDC37,FAM3A,PSMA3,NIP7,TTC9C,COX4I1,AUP1,FOXRED1,TR

M,KLC1,SCAF11,OSGEP,LDHA,RPS9,RCE1,ZNF410,IGSF8,NNMT,WDR62,VRK3,TM9SF1,TET2,GRK6,CDC37,

M189B,VKORC1,SRSF2,SEMA4D,HMGB1,SULF2,RAPGEF5,IQCG,YBX3,BLOC1S1,CIRBP,RHOF,SMARCB1,RPL

M189B,VKORC1,SRSF2,SEMA4D,HMGB1,SULF2,RAPGEF5,IQCG,YBX3,BLOC1S1,CIRBP,RHOF,SMARCB1,RPL

M189B,VKORC1,SRSF2,SEMA4D,HMGB1,SULF2,RAPGEF5,IQCG,YBX3,BLOC1S1,CIRBP,RHOF,SMARCB1,RPL

E1L1,PTOV1,DHPS,SUMF2,AMZ2,TAZ,MRPL33,RUVBL2,OBSL1,UBR4,NLRP1,PLCB2,SLC25A10,TMX2,CD99L

E1L1,PTOV1,DHPS,SUMF2,AMZ2,TAZ,MRPL33,RUVBL2,OBSL1,UBR4,NLRP1,PLCB2,SLC25A10,TMX2,CD99L

,NUMA1,OSGEP,LDHA,RPS9,RCE1,ZNF410,IGSF8,NNMT,WDR62,VRK3,TM9SF1,TET2,GRK6,CDC37,FAM3A,

7,CACYBP,RNH1,DNASE1L1,PTOV1,SNRPB,TCAIM,BOP1,KLC1,SCAF11,GPATCH2L,NPRL2,OSGEP,ARHGAP6

7,CACYBP,RNH1,DNASE1L1,PTOV1,SNRPB,TCAIM,BOP1,KLC1,SCAF11,GPATCH2L,NPRL2,OSGEP,ARHGAP6

LDHA,RPS9,RCE1,ZNF410,IGSF8,NNMT,WDR62,VRK3,TM9SF1,TET2,GRK6,CDC37,FAM3A,PSMA3,METTL17

L,TSG101,FBXW9,CCDC25,ENDOV,LYPLA2,C17ORF62,PPAN,ATP5J,PPP3CB,TADA3,NT5C,GBA2,NSUN5,UBE

,TM9SF1,SON,GRK6,TSG101,RPS25,RDH11,COASY,CARD14,USE1,PPIL2,LIG1,WNK1,TRMT112,SAT1,MCTS

,TM9SF1,SON,GRK6,TSG101,RPS25,RDH11,COASY,CARD14,USE1,PPIL2,LIG1,WNK1,TRMT112,SAT1,MCTS

,RPS24,SERPINB5,BCAT2,CENPV,MRPL4,FAM189B,VKORC1,MRPL2,CDCA3,PCNA,SRSF2,MRPS22,HDAC9,K
,RPS24,SERPINB5,BCAT2,CENPV,MRPL4,FAM189B,VKORC1,MRPL2,CDCA3,PCNA,SRSF2,MRPS22,HDAC9,K
AM189B,MRPL2,CDCA3,PCNA,SRSF2,KRT15,BAIAP2,RAPGEF5,SSU72,MYL6B,YBX3,BLOC1S1,RHOF,RNH1,D
AM189B,MRPL2,CDCA3,PCNA,SRSF2,KRT15,BAIAP2,RAPGEF5,SSU72,MYL6B,YBX3,BLOC1S1,RHOF,RNH1,D
3L,CDC42,FBXW9,TRMT112,DDOST,TOMM40,TADA3,PAFAH1B2,UBE2N,FAM32A,OGFOD2,ARIH2,PICK1,T
RSF2,SEMA4D,HMGB1,RAPGEF5,IQCG,YBX3,SMARCB1,STARD10,RNH1,DNASE1L1,FASTK,SET,BOP1,DBI,GP
RSF2,SEMA4D,HMGB1,RAPGEF5,IQCG,YBX3,SMARCB1,STARD10,RNH1,DNASE1L1,FASTK,SET,BOP1,DBI,GP
PR137,BCAT2,CENPV,MRPL2,CDCA3,SRSF2,KRT15,HMGB1,SULF2,BAIAP2,RAPGEF5,SSU72,MYL6B,YBX3,B
S100A13,MRPL4,CDCA3,SRSF2,OSMR,HMGB1,SSU72,IQCG,FAM72A,CMC2,RHOF,RNH1,DNASE1L1,PTOV1,
C1S1,STARD10,ATG16L2,SET,BOP1,ARRDC1,KLC1,CTTN,COMMD5,RUNX1,RPL13A,LDHA,SLC35G1,TESC,RA
C1S1,STARD10,ATG16L2,SET,BOP1,ARRDC1,KLC1,CTTN,COMMD5,RUNX1,RPL13A,LDHA,SLC35G1,TESC,RA
S,GPR137,RPS24,S100A13,BCAT2,MRPL4,LAS1L,BAIAP2,SMARCB1,STARD10,RNH1,SEC62,ZNF235,ARRDC1
S,GPR137,RPS24,S100A13,BCAT2,MRPL4,LAS1L,BAIAP2,SMARCB1,STARD10,RNH1,SEC62,ZNF235,ARRDC1
A,PDCD6,FAM32A,TSPAN31
AP2,MYL6B,IQCG,CACYBP,RHOF,STARD10,RNH1,PTOV1,RNF32,TCAIM,KLC1,SIRT6,BUD31,OSGEP,ARHGAP
AP2,MYL6B,IQCG,CACYBP,RHOF,STARD10,RNH1,PTOV1,RNF32,TCAIM,KLC1,SIRT6,BUD31,OSGEP,ARHGAP
NB5,S100A13,BCAT2,CENPV,MRPL4,FAM189B,VKORC1,PCNA,SRSF2,OSMR,PPFIBP1,SEMA4D,HMGB1,CYB
NB5,S100A13,BCAT2,CENPV,MRPL4,FAM189B,VKORC1,PCNA,SRSF2,OSMR,PPFIBP1,SEMA4D,HMGB1,CYB
ERPINB5,BCAT2,CENPV,MRPL4,VKORC1,MRPL2,CDCA3,PCNA,SRSF2,MRPS22,HDAC9,KRT15,PPFIBP1,SEM
ERPINB5,BCAT2,CENPV,MRPL4,VKORC1,MRPL2,CDCA3,PCNA,SRSF2,MRPS22,HDAC9,KRT15,PPFIBP1,SEM
NB5,CENPV,MRPL4,MRPL2,PPFIBP1,SSU72,FAM122C,TMEM67,BLOC1S1,CACYBP,RHOF,STARD10,RNH1,PT
NB5,CENPV,MRPL4,MRPL2,PPFIBP1,SSU72,FAM122C,TMEM67,BLOC1S1,CACYBP,RHOF,STARD10,RNH1,PT
9B,CDCA3,PCNA,OSMR,PPFIBP1,SULF2,BAIAP2,MYL6B,SRP9,IQCG,TMEM67,FAM72A,BLOC1S1,SMARCB1,
9B,CDCA3,PCNA,OSMR,PPFIBP1,SULF2,BAIAP2,MYL6B,SRP9,IQCG,TMEM67,FAM72A,BLOC1S1,SMARCB1,
S22,HMGB1,TMEM67,BLOC1S1,RNH1,SNRPB,TCAIM,ZNF235,LIMK2,ARHGAP6,TSPAN11,USP38,SH2B1,NC
S22,HMGB1,TMEM67,BLOC1S1,RNH1,SNRPB,TCAIM,ZNF235,LIMK2,ARHGAP6,TSPAN11,USP38,SH2B1,NC
PR137,RPS24,SERPINB5,BCAT2,CENPV,MRPL4,VKORC1,MRPL2,CDCA3,PCNA,SRSF2,MRPS22,OSMR,LAS1L,
PR137,RPS24,SERPINB5,BCAT2,CENPV,MRPL4,VKORC1,MRPL2,CDCA3,PCNA,SRSF2,MRPS22,OSMR,LAS1L,
2,OSMR,PPFIBP1,SSU72,FAM122C,IQCG,TMEM67,CMC2,SMARCB1,RPL38,PRR13,RNH1,RNF32,RCC1,SNR
L,CDCA3,SRSF2,OSMR,PPFIBP1,SSU72,MYL6B,SRP9,IQCG,UAP1,CMC2,YBX3,CACYBP,RBCK1,FASTK,SNRPB,
L,CDCA3,SRSF2,OSMR,PPFIBP1,SSU72,MYL6B,SRP9,IQCG,UAP1,CMC2,YBX3,CACYBP,RBCK1,FASTK,SNRPB,
ERPINB5,CENPV,MRPL4,FAM189B,MRPL2,PCNA,PPFIBP1,LAS1L,HMGB1,BAIAP2,RAPGEF5,SSU72,FAM122
ERPINB5,CENPV,MRPL4,FAM189B,MRPL2,PCNA,PPFIBP1,LAS1L,HMGB1,BAIAP2,RAPGEF5,SSU72,FAM122
ST3,GNAS,GPR137,RPS24,SERPINB5,BCAT2,CENPV,MRPL4,FAM189B,VKORC1,MRPL2,PCNA,SRSF2,MRPS2
ST3,GNAS,GPR137,RPS24,SERPINB5,BCAT2,CENPV,MRPL4,FAM189B,VKORC1,MRPL2,PCNA,SRSF2,MRPS2
DCA3,SRSF2,MRPS22,OSMR,SSU72,MYL6B,SRP9,IQCG,TMEM67,UAP1,CMC2,YBX3,CACYBP,CIRBP,RPL38,
DCA3,SRSF2,MRPS22,OSMR,SSU72,MYL6B,SRP9,IQCG,TMEM67,UAP1,CMC2,YBX3,CACYBP,CIRBP,RPL38,
V,MRPL4,VKORC1,SRSF2,OSMR,PPFIBP1,SEMA4D,LAS1L,CYB561A3,BAIAP2,SSU72,MYL6B,FAM122C,TMEI
V,MRPL4,VKORC1,SRSF2,OSMR,PPFIBP1,SEMA4D,LAS1L,CYB561A3,BAIAP2,SSU72,MYL6B,FAM122C,TMEI
3,PCNA,LAS1L,HMGB1,SSU72,CACYBP,CIRBP,RPL38,STARD10,RNH1,KIF4A,RBCK1,RNF32,ATG16L2,PARD3
3,PCNA,LAS1L,HMGB1,SSU72,CACYBP,CIRBP,RPL38,STARD10,RNH1,KIF4A,RBCK1,RNF32,ATG16L2,PARD3
S,CENPV,FAM189B,BAIAP2,RAPGEF5,SSU72,YBX3,RHOF,SMARCB1,DNASE1L1,RBCK1,FASTK,ATG16L2,SNR
RPINB5,MRPL4,MRPL2,PPFIBP1,LAS1L,BAIAP2,SSU72,FAM122C,TMEM67,PILRB,CACYBP,RNH1,DNASE1L1
RPINB5,MRPL4,MRPL2,PPFIBP1,LAS1L,BAIAP2,SSU72,FAM122C,TMEM67,PILRB,CACYBP,RNH1,DNASE1L1
F2,SEMA4D,HMGB1,RAPGEF5,FAM122C,IQCG,YBX3,CIRBP,STARD10,RNH1,FASTK,ATG16L2,RCC1,DHPS,SE
F2,SEMA4D,HMGB1,RAPGEF5,FAM122C,IQCG,YBX3,CIRBP,STARD10,RNH1,FASTK,ATG16L2,RCC1,DHPS,SE
ORC1,SRSF2,SEMA4D,HMGB1,SULF2,RAPGEF5,IQCG,YBX3,BLOC1S1,CIRBP,SMARCB1,STARD10,RNH1,DNA

ORC1,SRSF2,SEMA4D,HMGB1,SULF2,RAPGEF5,IQCG,YBX3,BLOC1S1,CIRBP,SMARCB1,STARD10,RNH1,DNA

KORC1,MRPL2,PCNA,SRSF2,MRPS22,HDAC9,OSMR,LAS1L,SSU72,SRP9,TMEM67,CMC2,CACYBP,SMARCB1

KORC1,MRPL2,PCNA,SRSF2,MRPS22,HDAC9,OSMR,LAS1L,SSU72,SRP9,TMEM67,CMC2,CACYBP,SMARCB1

E1L1,TCAIM,KLC1,OSGEP,TAZ,LDHA,RPS9,RCE1,ZNF410,IGSF8,NNMT,WDR62,VRK3,TM9SF1,TET2,GRK6,C

L,CIRBP,FASTK,RCC1,DHPS,SET,BOP1,SAE1,OSGEP,CHID1,RPL13A,MRPL33,SNRPN,WDR62,FIS1,TUBA1C,IR

L,CIRBP,FASTK,RCC1,DHPS,SET,BOP1,SAE1,OSGEP,CHID1,RPL13A,MRPL33,SNRPN,WDR62,FIS1,TUBA1C,IR

L,CIRBP,FASTK,RCC1,DHPS,SET,BOP1,SAE1,OSGEP,CHID1,RPL13A,MRPL33,SNRPN,WDR62,FIS1,TUBA1C,IR

T2,CENPV,MRPL4,FAM189B,VKORC1,MRPL2,CDCA3,SRSF2,HDAC9,KRT15,PPFIBP1,SEMA4D,HMGB1,SULF

T2,CENPV,MRPL4,FAM189B,VKORC1,MRPL2,CDCA3,SRSF2,HDAC9,KRT15,PPFIBP1,SEMA4D,HMGB1,SULF

62,ZC3H14,PPM1K,SON,UQCRQ,GRK6,PHF21A,TMX2,GGPS1,YIPF3,YIF1A,ASF1B,RFX3,TNFRSF14,PRMT1,S

62,ZC3H14,PPM1K,SON,UQCRQ,GRK6,PHF21A,TMX2,GGPS1,YIPF3,YIF1A,ASF1B,RFX3,TNFRSF14,PRMT1,S

72,FAM122C,TMEM67,PILRB,CACYBP,RNH1,DNASE1L1,PTOV1,SNRPB,TCAIM,SEC62,BOP1,KLC1,SCAF11,G

72,FAM122C,TMEM67,PILRB,CACYBP,RNH1,DNASE1L1,PTOV1,SNRPB,TCAIM,SEC62,BOP1,KLC1,SCAF11,G

,SERPINB5,BCAT2,CENPV,MRPL4,VKORC1,MRPL2,CDCA3,PCNA,SRSF2,OSMR,LAS1L,BAIAP2,SSU72,FAM12

9B,MRPL2,CDCA3,SRSF2,OSMR,PPFIBP1,HMGB1,BAIAP2,RAPGEF5,MYL6B,SULT1A3,PRR13,STARD10,RNH1

9B,MRPL2,CDCA3,SRSF2,OSMR,PPFIBP1,HMGB1,BAIAP2,RAPGEF5,MYL6B,SULT1A3,PRR13,STARD10,RNH1

NH1,FASTK,SET,BOP1,OSGEP,EIF5A,CHID1,RPL13A,LDHA,SPAG7,STYXL1,LZTFL1,FIS1,TUBA1C,IRF9,STIP1,T

NH1,FASTK,SET,BOP1,OSGEP,EIF5A,CHID1,RPL13A,LDHA,SPAG7,STYXL1,LZTFL1,FIS1,TUBA1C,IRF9,STIP1,T

,GPR137,RPS24,SERPINB5,FAM189B,PCNA,SRSF2,MRPS22,KRT15,LAS1L,SULF2,CYB561A3,SSU72,FAM122

,GPR137,RPS24,SERPINB5,FAM189B,PCNA,SRSF2,MRPS22,KRT15,LAS1L,SULF2,CYB561A3,SSU72,FAM122

,OSGEP,LDHA,RPS9,RCE1,ZNF410,IGSF8,NNMT,WDR62,VRK3,TM9SF1,TET2,GRK6,CDC37,FAM3A,PSMA3,

1A3,STARD10,RNH1,LY75,ATG16L2,SEC62,TRPT1,ARPC2,NOB1,BUD31,CTTN,NUDT22,AMZ2,LDHA,PSMA6

1A3,STARD10,RNH1,LY75,ATG16L2,SEC62,TRPT1,ARPC2,NOB1,BUD31,CTTN,NUDT22,AMZ2,LDHA,PSMA6

,SPAG7,WDR62,RANBP1,FIS1,STIP1,TET2,GRK6,CDC37,FAM3A,PPRC1,TRNT1,C1ORF198,EEF1B2,PPA1,TN

,SPAG7,WDR62,RANBP1,FIS1,STIP1,TET2,GRK6,CDC37,FAM3A,PPRC1,TRNT1,C1ORF198,EEF1B2,PPA1,TN

NH1,DNASE1L1,TMEM135,LY75,SNRPB,PARD3,DBI,NOB1,WDR19,SIRT6,ITSN1,COMMD5,TAZ,THYN1,DBN

NH1,DNASE1L1,TMEM135,LY75,SNRPB,PARD3,DBI,NOB1,WDR19,SIRT6,ITSN1,COMMD5,TAZ,THYN1,DBN

SMR,HMGB1,SULF2,SSU72,TMEM67,SULT1A3,PILRB,FAM72A,RHOF,RPL38,DNASE1L1,RBCK1,RNF32,LY75,

35,VKORC1,CDCA3,PCNA,SRSF2,OSMR,PPFIBP1,LAS1L,HMGB1,SSU72,MYL6B,SRP9,IQCG,UAP1,CMC2,YBX

35,VKORC1,CDCA3,PCNA,SRSF2,OSMR,PPFIBP1,LAS1L,HMGB1,SSU72,MYL6B,SRP9,IQCG,UAP1,CMC2,YBX

,UBE2I,RPL21,CARS,GGCX,SAT1,MCTS1,MTA3,NXN,AGAP3,C11ORF48,TCP1,SNX5,COX20

RB1,SPAG7,WDR62,DST,PCBP2,TM9SF1,CDC37,NIP7,RPL29,RTEL1,YIPF3,EIF3M,JUP,EDF1,RFX3,DGKE,TRO

,CD44,GRK6,RPS16,CPSF3L,METTL17,AUP1,XBP1,VIM,TMPO,ATPAF2,LSM12,STOML2,WNK1,RPL36,HSP9

XNL4A,NUTF2,TMPO,ATP5J,DCTD,SYNGR1,HMGA1,POLR3H,MDH2,HIGD1A,NOP2,NUP133,ARIH2,UBA1,R

4,PPRC1,TMX2,EDF1,LYPLA2,C19ORF54,HSP90AB1,TRMT112,MTA3,HMGA1,ZNF20,GBA2,FAM102A,PHC3

SMR,HMGB1,YBX3,RNH1,RCC1,SLC3A2,ZC3HAV1,SCAF11,ARPC2,WDR19,SCARB1,CTTN,RPL13A,RUVBL2,T

PL37A,RPLP0,MRPL55,RPL8,RPL35,RPL34,RPL24,RPS3A,RPS3,MRPS24,RPS27L,RPS7,UBA52,RPS4X,MRPS
A6,RPS9,NUDT5,RBPMS,SNRPN,ZC3H14,STIP1,PCBP2,SON,RPS16,PSMA1,YLPM1,CHCHD1,PPRC1,RPS25,N

L21,NUTF2,EPB41L4B,RPL36,TPM2,ASPH,RPS20,RPS15,DCN,HMGA1,SEC31A,RPL18,RPL15,TPM4,MRPS1
BOP1,ZC3HAV1,SCAF11,TBL2,EIF5A,RUNX1,NOP16,RPL13A,PSMA6,RPS9,ZNF410,RUVBL2,NUDT5,RBPM

BP1,SEMA4D,HMGB1,CYB561A3,BAIAP2,RAPGEF5,SSU72,MYL6B,SRP9,IQCG,TMEM67,SULT1A3,UAP1,P
,EGFR,SH3GLB1,RPL15,PARK7,SNX5,LIMA1,ADD1,TJP1,RPL34,EIF2A,RPL24,SH3GL1,RDX,SLK,VAPA,LRRC

LC3A2,SET,ZNF235,BOP1,ZC3HAV1,SCAF11,DBI,LIMK2,VDAC3,SIRT6,TBL2,EIF5A,RUNX1,NOP16,RPL13A,

LC3A2,SET,ZNF235,BOP1,ZC3HAV1,SCAF11,DBI,LIMK2,VDAC3,SIRT6,TBL2,EIF5A,RUNX1,NOP16,RPL13A,

,RPL24,RPS3A,RPS3,RPS7,UBA52,RPS4X,RPS23,RPL30,RPS15A,RPS17,RPL27,RPS14,RPL31,FAU
,PSMA1,PSMA3,RPS25,RPS6,RPL29,SRSF1,CLASRP,CELF1,PSMD4,RPL28,VIM,PSMA7,RPL21,TXNL4A,EXO
A,RPS3,MAGOHB,RPS7,NCBP2,RNPS1,UBA52,RPS4X,RPS23,RPL30,RPS15A,RPS17,RPL27,RPS14,MAGOH,
,RER1,RPL34,RPL24,KDELR3,RPS3A,RPS3,RPS7,UBA52,RPS4X,VAPA,RPS23,RPL30,OS9,RPS15A,RPS17,BC

,TSG101,UBR4,PSMA3,RPS25,NLRP1,RPS6,RPL29,UBE2I,CDC42,ATF7,TNFRSF14,VAMP8,RPL28,VIM,PSM
4A1,RPLP0,METTL3,EIF3G,RPL8,RPL35,MTIF2,RPL34,EIF2A,RPL24,RBM4,RPS3A,EIF3C,RPS3,RPS7,NCBP2,

,TSG101,UBR4,PSMA3,RPS25,NLRP1,RPS6,RPL29,UBE2I,CDC42,ATF7,TNFRSF14,VAMP8,RPL28,VIM,PSM
,TMED4,RPS6,RPL29,SRSF1,AUP1,RABL5,ITGB1BP1,JUP,CDC42,RPL28,TIMM17B,RPL21,NUTF2,STRADA,
,RPS11,NUP133,RPL37A,RPLP0,EIF3G,SSB,RPL8,RPL35,RPL34,NUP62,RPL24,RPS3A,RPS3,RPS7,POLR2E,
H14,PCBP2,SNF8,RPS16,TSG101,UBR4,PSMA1,PSMA3,RPS25,RPS6,RPL29,AUP1,UGGT2,UBE2I,XBP1,PT
PA1,RPL28,VIM,APEH,RPL21,CARS,RPL36,RBM3,MAPKAPK5,TSFM,TRMT112,RPS20,RPS15,HSPB1,MCTS
PA1,RPL28,VIM,APEH,RPL21,CARS,RPL36,RBM3,MAPKAPK5,TSFM,TRMT112,RPS20,RPS15,HSPB1,MCTS
P3B2,RPS9,RANBP1,FIS1,TUBA1C,DST,LRRC1,SNF8,CDC37,RPS16,TSG101,TTC21B,RPS25,TMED4,RPS6,R
NRNPC,RPL36,RPS20,RPS15,HSPB1,PSME2,PABPC1,PSME1,CARHSP1,RPL18,RPL15,RPS11,PCID2,RPL37A,
MD4,RPL28,VIM,PSMA7,RPL21,EXOSC9,HNRNPC,UPP1,RPL36,DUT,RPS20,RPS15,HSPB1,PSME2,PABPC1,
,TSG101,UBR4,PSMA3,RPS25,NLRP1,RPS6,RPL29,UBE2I,CDC42,ATF7,TNFRSF14,VAMP8,RPL28,VIM,PSM
TFL1,FIS1,LRRC1,TM9SF1,SNF8,CDC37,RPS16,TSG101,OBSL1,TTC21B,RPS25,TMED4,RPS6,RPL29,SRSF1,
2B,RPL36,LAMP2,HSP90AB1,RPS20,VPS28,RPS15,TOMM40,PDCD5,POT1,RPL18,SH3GLB1,RPL15,TCP1,N
M,PSMA7,RPL21,EXOSC9,HNRNPC,UPP1,RPL36,DUT,RPS20,RPS15,HSPB1,PSME2,PABPC1,NUDT1,NT5C,P

ZTFL1,FIS1,LRRC1,TM9SF1,SNF8,CDC37,RPS16,TSG101,OBSL1,TTC21B,RPS25,TMED4,RPS6,RPL29,SRSF1,
BE2L6,ZC3H14,VCAN,PCBP2,CD44,SNF8,RPS16,TSG101,UBR4,PSMA1,PSMA3,RPS25,RPS6,RPL29,AUP1,U
3,RPL37A,RPLP0,RPL8,RPL35,RPL34,NUP62,RPL24,RPS3A,RPS3,RPS7,POLR2E,UBA52,RPS4X,RPS23,RPL3
EF1B2,PPA1,RPL28,VIM,APEH,RPL21,CARS,RPL36,RBM3,MAPKAPK5,TSFM,TRMT112,RPS20,RPS15,HSPB1
RPL29,PSMD4,RPL28,VIM,PSMA7,RPL21,EXOSC9,HNRNPC,UPP1,RPL36,DUT,RPS20,RPS15,HSPB1,PSME2,
,LAMP2,RPS20,RPS15,SYNGR1,TOMM40,PDCD5,RPL18,SH3GLB1,RPL15,RPS11,MFF,ACOT2,PEX2,RPL37
,RPL34,RPL24,RPS3A,RPS3,MAGOHB,RPS7,NCBP2,RNPS1,UBA52,RPS4X,MLH1,RPS23,RPL30,RQCD1,EXO
28,VIM,PSMA7,RPL21,EXOSC9,HNRNPC,UPP1,RPL36,DUT,RPS20,RPS15,HSPB1,PSME2,PABPC1,BLVRB,N
28,VIM,PSMA7,RPL21,EXOSC9,HNRNPC,UPP1,RPL36,DUT,RPS20,RPS15,HSPB1,PSME2,PABPC1,BLVRB,N
PPA1,RPL28,VIM,APEH,RPL21,CARS,RPL36,RBM3,MAPKAPK5,TSFM,TRMT112,RPS20,RPS15,HSPB1,MCTS
NRNPC,RPL36,RPS20,RPS15,HSPB1,PSME2,PABPC1,PSME1,CARHSP1,RPL18,RPL15,RPS11,PCID2,RPL37A,
,UFD1L,RPL13A,GSTK1,ACOT8,AP3B2,RPS9,RUVBL2,TESC,RANBP1,LZTFL1,FIS1,TUBA1C,DST,LRRC1,TM9
7A,PEX26,RPLP0,SSR3,RPL8,RPL35,RPL34,RPL24,RPS3A,RPS3,RPS7,UBA52,RPS4X,RPS23,RPL30,PEX16,R
PL29,JUP,RPL28,TIMM17B,RPL21,NUTF2,ZFAND2B,RPL36,LAMP2,HSP90AB1,RPS20,VPS28,RPS15,TOMM
D1L,USP38,RPL13A,LDHA,PSMA6,ACOT8,RPS9,NUDT5,UBE2L6,ZC3H14,PPM1K,VCAN,PCBP2,CD44,TET2,
L29,EIF3M,CELF1,EEF1B2,PPA1,RPL28,VIM,APEH,RPL21,CARS,RPL36,RBM3,MAPKAPK5,TSFM,TRMT112,
EIF3M,PTK2,CELF1,CENPN,ASF1B,VAMP8,CAPG,PSMD4,TMEM126B,TXNL4A,TCF4,ATPAF2,PPAN,HSP90
1,ARHGAP6,SCARB1,CTTN,EIF5A,TAZ,RPL13A,MRPL33,RUVBL2,IL1RAP,FIS1,TSG101,CHCHD1,ECSIT,NLRP
6,NPRL2,SCARB1,CTTN,UFD1L,USP38,RPL13A,PSMA6,ACOT8,RPS9,NUDT5,UBE2L6,ZC3H14,FIS1,PPM1K,
CPSF3L,RPS6,SRSF1,TRNT1,CLASRP,CELF1,FBXW9,TXNL4A,EXOSC9,ARL6IP4,HNRNPC,CDK11A,RBM3,ELP
,SIRT6,NPRL2,SCARB1,CTTN,UFD1L,USP38,RPL13A,LDHA,PSMA6,ACOT8,RPS9,NUDT5,UBE2L6,ZC3H14,F
P3B2,RPS9,TESC,IL1RAP,FIS1,SLC15A3,LRRC1,SNF8,CDC37,RPS16,TSG101,TTC21B,FUZ,RPS25,NLRP1,TME
P3B2,RPS9,RUVBL2,TESC,IL1RAP,FIS1,SLC15A3,LRRC1,SNF8,CDC37,RPS16,TSG101,TTC21B,FUZ,EPB41L5,
ARD3,SET,CTSC,BOP1,ARRDC1,SCAF11,ARPC2,VDAC3,NOB1,WDR19,SIRT6,NUMA1,NPRL2,ARHGAP6,CTT
T112,RPS15,MCTS1,RRNAD1,RBMX,ZCCHC4,RRP8,SNRPF,NSUN5,NHP2,ZNHIT3,NOP2,USP4,MPHOSPH1
RPS16,CHCHD1,MUC1,METTL17,ATG12,RPS25,RPS6,RPL29,COASY,UGGT2,HS2ST1,EIF3M,MAGT1,CELF1
FR,SH3GLB1,RPL15,RPS11,MFF,RPL37A,PEX26,RPLP0,SSR3,RPL8,BAX,RPL35,RPL34,RPL24,TP53,RPS3A,R
P3B2,RPS9,TESC,IL1RAP,FIS1,SLC15A3,LRRC1,SNF8,CDC37,RPS16,TSG101,TTC21B,FUZ,RPS25,NLRP1,TME
P3B2,RPS9,TESC,IL1RAP,FIS1,SLC15A3,LRRC1,SNF8,CDC37,RPS16,TSG101,TTC21B,FUZ,RPS25,NLRP1,TME
UFD1L,RPL13A,GSTK1,ACOT8,AP3B2,RPS9,RUVBL2,TESC,IL1RAP,LZTFL1,FIS1,SLC15A3,LRRC1,TM9SF1,SN
41,HNRNPLL,HNRNPA1,POLR2C,SNRPF,RBFOX2,PPIH,LGALS3,USP4,MPHOSPH10,METTL3,TSEN34,SRSF7,
1,PTOV1,FASTK,SNRPB,SET,ZNF235,BOP1,TRPT1,ZC3HAV1,SCAF11,DBI,NOB1,SIRT6,NUMA1,BUD31,OSG
L2,IL1RAP,FIS1,ECSIT,NLRP1,CENPK,SRSF1,FOXRED1,ITGB1BP1,EIF3M,CDC42,PTK2,CELF1,CENPN,ASF1B,
RPL38,RNH1,DNASE1L1,RBCK1,PTOV1,FASTK,SNRPB,DHPS,SET,ZNF235,BOP1,TRPT1,ZC3HAV1,SCAF11,D
C1,PARD3,ACTR6,SET,CTSC,BOP1,KLC1,ARPC2,LIMK2,VDAC3,WDR19,SIRT6,NUMA1,SAE1,NPRL2,ARHGA
4,GEMIN2,EIF3C,SNRPD1,RPS27L,TAF9,RUVBL1,SFSWAP,EIF6,DHX30,ERAL1,MRPS7,SNRNP200,RPS14,L
MIN2,EIF3C,SNRPD1,RPS27L,TAF9,RUVBL1,SFSWAP,EIF6,DHX30,ERAL1,MRPS7,SNRNP200,RPS14,LUC7L3
ABPC1,RBMX,PSME1,CARHSP1,HNRNPA1,RBFOX2,PCID2,METTL3,SRSF7,RBM42,PSMD7,RBM4,PSMD10,
PIH,USP4,MPHOSPH10,METTL3,SRSF7,RALY,RBM42,PSPC1,RBM4,GEMIN2,HNRNPUL1,MAGOHB,SNRPD1
D2B,RPL36,RPS20,RPS15,RANGRF,PDCD5,ABHD17A,RPL18,CNST,EGFR,SH3GLB1,RPL15,LGALS3,RPS11,M

D5,HSD17B10,SH3GLB1,COX17,HIGD1A,NDUFV2,SIRT5,COA3,PARK7,MFF,COX20,COA1,NDUFS3,HAX1,
1,PTOV1,FASTK,SNRPB,SET,ZNF235,BOP1,TRPT1,ZC3HAV1,SCAF11,DBI,NOB1,SIRT6,NUMA1,BUD31,OSG
OSC9,ZFAND2B,SHARPIN,HSP90AB1,SIAH1,VPS28,UBXN1,ERLEC1,PSME2,WAC,ANAPC11,RAD23A,USP45
UBR4,PSMA1,PSMA3,AUP1,UGGT2,UBE2I,XBP1,PTK2,USE1,GADL1,PSMD4,PSMA7,LYPLA2,CAMLG,ZFAN
PIH,USP4,METTL3,SRSF7,RALY,RBM42,PSPC1,RBM4,GEMIN2,HNRNPUL1,MAGOHB,SNRPD1,SNRPA1,U2A
PIH,USP4,METTL3,SRSF7,RALY,RBM42,PSPC1,RBM4,GEMIN2,HNRNPUL1,MAGOHB,SNRPD1,SNRPA1,U2A

CTTN,ITSN1,EIF5A,UFD1L,RPL13A,GSTK1,ACOT8,AP3B2,RPS9,RUVBL2,TESC,IL1RAP,LZTFL1,FIS1,SLC15A3,
BP,SMARCB1,RPL38,RBCK1,FASTK,SNRPB,PARD3,DHPS,SET,CTSC,ARRDC1,TRPT1,ZC3HAV1,LIMK2,SIRT6,
C,BOP1,SCAF11,ARPC2,VDAC3,WDR19,SIRT6,NUMA1,NPRL2,ARHGAP6,CTTN,RUNX1,TAZ,SH2B1,RPL13A
ULF2,BAIAP2,IQCG,TMEM67,PILRB,BLOC1S1,CIRBP,RHOF,SMARCB1,RPL38,KIF4A,TMEM135,ATG16L2,RC
PS,SET,ZNF235,SIRT6,NUMA1,BUD31,EIF5A,RUNX1,UFD1L,TAZ,SH2B1,RPL13A,PSMA6,RPS9,MRPL33,RU
1,PTOV1,FASTK,SNRPB,SET,ZNF235,BOP1,TRPT1,ZC3HAV1,SCAF11,DBI,NOB1,SIRT6,NUMA1,BUD31,OSG
A7,CAMLG,ZFAND2B,SHARPIN,LAMP2,HSP90AB1,SIAH1,VPS28,UBXN1,ERLEC1,PSME2,WAC,ANAPC11,RA

AND2B,SHARPIN,HSP90AB1,SIAH1,VPS28,UBXN1,ERLEC1,PSME2,WAC,ANAPC11,RAD23A,USP45,PSME1,
RIP1,PABPC1,RBMX,DDX41,HNRNPLL,HNRNPA1,POLR2C,SNRPF,RBFOX2,PPIH,LGALS3,USP4,METTL3,TSE
B1,SULF2,SSU72,SRP9,TMEM67,SULT1A3,UAP1,PILRB,YBX3,BLOC1S1,CACYBP,CIRBP,SMARCB1,RPL38,ST
AND2B,SHARPIN,HSP90AB1,SIAH1,VPS28,UBXN1,ERLEC1,PSME2,WAC,ANAPC11,RAD23A,USP45,PSME1,
B,SET,ZNF235,BOP1,TRPT1,ZC3HAV1,SCAF11,NOB1,SIRT6,NUMA1,BUD31,OSGEP,RUNX1,UFD1L,SH2B1,
1,PTOV1,FASTK,SNRPB,SET,ZNF235,BOP1,TRPT1,ZC3HAV1,SCAF11,DBI,NOB1,SIRT6,NUMA1,BUD31,OSG

GSTK1,ACOT8,AP3B2,RPS9,TESC,IL1RAP,FIS1,SLC15A3,LRRC1,SNF8,CDC37,RPS16,TSG101,TTC21B,FUZ,RP
ET,ZNF235,SIRT6,NUMA1,BUD31,EIF5A,RUNX1,UFD1L,SH2B1,RPL13A,PSMA6,ACOT8,RPS9,MRPL33,RUV
2,USE1,PSMD4,PSMA7,LYPLA2,CAMLG,ZFAND2B,SHARPIN,LAMP2,HSP90AB1,SIAH1,VPS28,UBXN1,ERLE
A7,CAMLG,ZFAND2B,SHARPIN,HSP90AB1,SIAH1,VPS28,UBXN1,ERLEC1,PSME2,WAC,ANAPC11,RAD23A,U
DHPS,SET,ZNF235,SIRT6,NUMA1,BUD31,EIF5A,RUNX1,UFD1L,SH2B1,RPL13A,PSMA6,ACOT8,RPS9,MRPL3
1A3,PILRB,YBX3,BLOC1S1,CIRBP,SMARCB1,RPL38,STARD10,RBCK1,FASTK,SNRPB,PARD3,DHPS,SET,CTSC,
PH10,RPLP0,RPL35,ZNF593,EIF2A,RPL24,GTF3A,RPS27L,RPS7,DDX10,EXOSC2,NGDN,EXOSC8,DDX17,RPS
RD10,RBCK1,PTOV1,SNRPB,DHPS,SET,ZNF235,SIRT6,NUMA1,BUD31,SCARB1,EIF5A,RUNX1,UFD1L,TAZ,SH
QCG,TMEM67,PILRB,BLOC1S1,CIRBP,RHOF,SMARCB1,RPL38,KIF4A,TMEM135,ATG16L2,RCC1,SNRPB,PA

ARPIN,LAMP2,ASPH,TRMT112,MCTS1,HMGA1,SMARCA4,MRPS18C,PARK7,SMARCC2,MRPS17,HAX1,MR

,PSMA7,CAMLG,EXOSC9,HNRNPC,HSP90AB1,RPTOR,VPS28,UBXN1,ERLEC1,HSPB1,DCN,PSME2,PABPC1,
RD10,RBCK1,PTOV1,SNRPB,DHPS,SET,ZNF235,SIRT6,NUMA1,BUD31,SCARB1,EIF5A,RUNX1,UFD1L,TAZ,SH

BX3,BLOC1S1,CIRBP,SMARCB1,RPL38,RBCK1,FASTK,SNRPB,PARD3,DHPS,SET,CTSC,ARRDC1,TRPT1,ZC3HA

P1,TRPT1,ZC3HAV1,SCAF11,NOB1,SIRT6,NUMA1,BUD31,OSGEP,RUNX1,RPL13A,PSMA6,RPS9,RUVBL2,TI

RD10,RBCK1,PTOV1,SNRPB,DHPS,SET,ZNF235,SIRT6,NUMA1,BUD31,SCARB1,EIF5A,RUNX1,UFD1L,TAZ,SH

B1,SULF2,CYB561A3,SSU72,SRP9,TMEM67,SULT1A3,UAP1,PILRB,YBX3,BLOC1S1,CACYBP,CIRBP,SMARCB
YXL1,RANBP1,FIS1,PHF19,CDC37,TSG101,OBSL1,TTC21B,YLPM1,FUZ,MUC1,RTEL1,ITGB1BP1,CDC42,PTK
3,CHTOP,SRSF7,NXF1,ZNF593,NUP62,RBM4,GEMIN2,THOC3,TP53,PML,MAGOHB,SNRPD1,IFI27,U2AF1L

A7,CAMLG,EXOSC9,HNRNPC,HSP90AB1,RPTOR,UBXN1,ERLEC1,HSPB1,DCN,PSME2,PABPC1,WAC,RAD23

,SRSF7,NXF1,ZNF593,NUP62,RBM4,GEMIN2,THOC3,TP53,PML,MAGOHB,SNRPD1,IFI27,U2AF1L4,NCBP2,

DUSP22,RTEL1,AUP1,UGGT2,YIF1A,CDC42,XBP1,AUNIP,HSBP1L1,ENDOV,CRIP1,PSMD4,LIG1,PSMA7,CDC

LRB,YBX3,BLOC1S1,CACYBP,CIRBP,SMARCB1,RPL38,RNH1,DNASE1L1,RBCK1,PTOV1,FASTK,SNRPB,PARD3

1,SULF2,CYB561A3,BAIAP2,RAPGEF5,SSU72,SRP9,TMEM67,UAP1,FAM72A,CMC2,YBX3,BLOC1S1,CACYB
,YBX3,BLOC1S1,CACYBP,RHOF,RPL38,PRR13,STARD10,RNH1,KIF4A,RBCK1,SNRPB,PARD3,SLC3A2,DHPS,N
1,SULF2,CYB561A3,BAIAP2,RAPGEF5,SSU72,MYL6B,SRP9,IQCG,TMEM67,UAP1,FAM72A,CMC2,YBX3,BLO
.5,LAS1L,HMGB1,SULF2,CYB561A3,BAIAP2,RAPGEF5,SSU72,MYL6B,SRP9,IQCG,TMEM67,UAP1,FAM72A,
P1,LAS1L,HMGB1,SULF2,CYB561A3,BAIAP2,SSU72,MYL6B,SRP9,IQCG,TMEM67,SULT1A3,UAP1,FAM72A,
PS22,HDAC9,KRT15,PPFIBP1,LAS1L,HMGB1,SULF2,CYB561A3,BAIAP2,RAPGEF5,SSU72,MYL6B,SRP9,IQCG
.5,LAS1L,HMGB1,SULF2,CYB561A3,BAIAP2,RAPGEF5,SSU72,MYL6B,SRP9,IQCG,TMEM67,UAP1,FAM72A,
PSMA1,RPS25,NIP7,RPS6,RPL29,SRSF1,EIF3M,CELF1,FBXW9,RPL28,VIM,RPL21,TXNL4A,PPAN,HNRNPC,R
IAP1,BLOC1S1,CACYBP,CIRBP,SMARCB1,PRR13,RNH1,DNASE1L1,KIF4A,PTOV1,FASTK,ATG16L2,RCC1,SNF
IAP1,BLOC1S1,CACYBP,CIRBP,SMARCB1,PRR13,RNH1,DNASE1L1,KIF4A,PTOV1,FASTK,ATG16L2,RCC1,SNF
IAP1,BLOC1S1,CACYBP,CIRBP,SMARCB1,PRR13,RNH1,DNASE1L1,KIF4A,PTOV1,FASTK,ATG16L2,RCC1,SNF
,CACYBP,CIRBP,SMARCB1,PRR13,RNH1,DNASE1L1,KIF4A,PTOV1,ATG16L2,RCC1,SNRPB,SLC3A2,ACTR6,M
11,MRPS17,RPL37A,RPLP0,MRPL55,RPL8,RPL35,RPL34,EIF2A,RPL24,RPS3A,RPS3,MRPS24,RPS27L,RPS7,
4A,PTOV1,ATG16L2,RCC1,SNRPB,SLC3A2,ACTR6,SET,CTSC,BOP1,SCAF11,NOB1,SIRT6,NUMA1,BUD31,SA
PL15,MRPS18C,RPS11,MRPS17,RPL37A,RPLP0,MRPL55,RPL8,RPL35,RPL34,EIF2A,RPL24,RPS3A,BTF3,RPS

4A,PTOV1,ATG16L2,RCC1,SNRPB,SLC3A2,ACTR6,SET,CTSC,BOP1,SCAF11,NOB1,SIRT6,NUMA1,BUD31,SA
BP,CIRBP,SMARCB1,RPL38,RNH1,KIF4A,RBCK1,PTOV1,RCC1,SNRPB,PARD3,SLC3A2,ACTR6,SET,BOP1,KLC
THEM4,COASY,FOXRED1,UBE2I,C14ORF2,AGPAT4,PPP1CC,TIMM17B,TMEM126B,APEH,TXNL4A,NUTF2,I
THEM4,COASY,FOXRED1,UBE2I,C14ORF2,AGPAT4,PPP1CC,TIMM17B,TMEM126B,APEH,TXNL4A,NUTF2,I

MM40,COX5A,HIGD1A,NDUFV2,SIRT5,COA3,MPC2,TYMS,SLC25A23,MRPS18C,LGALS3,COX20,MRPS17,T
C,TIMM17B,TMEM126B,PPOX,STOML2,SLC9B2,ATP5J,CHCHD7,TOMM40,CHCHD5,SH3GLB1,COX17,COX
9B2,ATP5J,TOMM40,COX5A,HIGD1A,NDUFV2,SIRT5,COA3,MPC2,TYMS,SLC25A23,MFSD10,MRPS18C,LG
F,SMARCB1,RPL38,KIF4A,TMEM135,RCC1,PARD3,ACTR6,SET,CTSC,BOP1,KLC1,SCAF11,ARPC2,LIMK2,WD
F,SMARCB1,RPL38,KIF4A,TMEM135,RCC1,PARD3,ACTR6,SET,CTSC,BOP1,KLC1,SCAF11,ARPC2,LIMK2,WD
.A,PSMA3,ATG12,SRSF1,AUP1,FOXRED1,UBE2I,MAGT1,PPP1CC,PSMD4,PSMA7,PRMT1,EXOSC9,ZFAND2I
PPP1CC,EDF1,ENDOV,CAMTA1,CAPG,CDC14B,PODXL,EXOSC9,ARL6IP4,PPAN,RPL36,MOB1A,INO80E,ZEB
IETTL17,ECSIT,COX4I1,SLC25A10,THEM4,COASY,FOXRED1,TRNT1,C14ORF2,AGPAT4,IFI27L1,PPP1CC,LIG1
L55,BAX,VDAC1,PAM16,NDUFA11,NDUFA3,MRPS24,NDUFB3,NDUFB2,NDUFA10,MRPS16,SDHA,NDUFA
7B,TMEM126B,PPOX,STOML2,SLC9B2,ATP5J,TOMM40,SH3GLB1,COX5A,HIGD1A,NDUFV2,SIRT5,COA3,M

GSTK1,LDHA,PSMA6,RPS9,RUVBL2,PCMT1,IGSF8,NUDT5,PCBP2,CD44,SNF8,CDC37,RPS16,TSG101,PTPN
GSTK1,LDHA,PSMA6,RPS9,RUVBL2,PCMT1,IGSF8,NUDT5,PCBP2,CD44,SNF8,CDC37,RPS16,TSG101,PTPN
GSTK1,LDHA,PSMA6,RPS9,RUVBL2,PCMT1,IGSF8,NUDT5,PCBP2,CD44,SNF8,CDC37,RPS16,TSG101,PTPN
H2,SIRT5,MTHFS,TXNRD2,TYMS,MRPS18C,PARK7,MRPS17,ACOT2,NDUFS3,NMNAT3,MRPL55,GCDH,VDA

NUMA1,NPRL2,TBL2,SCARB1,EIF5A,UFD1L,TAZ,AP3B2,SLC35G1,RCE1,MRPL33,EBPL,FIS1,SLC15A3,DST,A

,CD151,PTK7,TPM4,RPS11,RPL37A,RPLP0,KRAS,RPL8,LIMA1,ADD1,CLASP2,ARPC5,ALKBH6,NRP1,RPS3A,

,CD151,PTK7,TPM4,RPS11,RPL37A,RPLP0,KRAS,RPL8,LIMA1,ADD1,CLASP2,ARPC5,ALKBH6,NRP1,RPS3A,

C1,RBP5,DBI,ARPC2,VDAC3,NUMA1,SCARB1,ITSN1,TNFSF10,CHID1,GSTK1,LDHA,PSMA6,AP3B2,RPS9,RL

B,MAGOHB,SNRPD1,SNRPA1,U2AF1L4,HNRNPM,IK,PPIE,SNRNP40,SNRNP200,MAGOH,LUC7L3,PRPF31

2,USE1,AGPAT4,NUTF2,C17ORF62,PIGL,ZFAND2B,COPZ1,RAB2A,GGCX,ASPH,MPDU1,UBXN1,EMC4,DDO

2,USE1,AGPAT4,C17ORF62,PIGL,ZFAND2B,COPZ1,RAB2A,GGCX,ASPH,MPDU1,UBXN1,EMC4,DDOST,SLC3

CREB3L4,POLR2M,BAX,CDK4,NXF1,ADRA1A,NUP62,TP53,MGST2,DNAJC2,IFI27,VAPA,LRRC59,SUMO1,N

DA1,PEX2,STX10,PEX26,ERGIC1,ECE1,SYP,RER1,BFAR,SLC35A2,GRAMD1A,EMC9,SLC6A9,SYPL1,SLC37A3,

M,PODXL,EPB41L4B,RPS15,HSPB1,RANGRF,LSR,HMGA1,PABPC1,B2M,PKP4,RPL18,EGFR,CD151,PTK7,TP

R1,CD74,ECE1,UBA1,TMEM9,LAMTOR5,CLTA,RHEB,SPHK2,LAMTOR1,PLD1,UBA52,VAPA,VTI1B,SZT2,GR

R1,CD74,ECE1,UBA1,TMEM9,LAMTOR5,CLTA,RHEB,SPHK2,LAMTOR1,PLD1,UBA52,VAPA,VTI1B,SZT2,GR

,LPCAT3,RDH11,SDF4,TMED4,RPS6,AUP1,JPH2,UGGT2,SLC39A13,YIF1A,MAGT1,XBP1,USE1,AGPAT4,EEF

1,UBR4,ATG12,COASY,KXD1,KCNMA1,MAGT1,AGPAT4,PPP1CC,VAMP8,NUTF2,PODXL,STOML2,AP1S3,C

OST,SYNGR1,ATP6V1F,SH3GLB1,CD74,ECE1,UBA1,TMEM9,LAMTOR5,CLTA,TMEM199,RHEB,SPHK2,LAM

STX10,PEX26,ERGIC1,SYP,RER1,BFAR,SLC35A2,EMC9,SLC6A9,SYPL1,SLC37A3,EMC3,PEX16,BSCL2,MCUR

LP0,RPL8,RPL35,RPL34,RPL24,RPS3A,RPS3,RPS27L,RPS7,UBA52,RPS4X,MRPS16,RPS23,RPL30,RPS15A,RF

L8,RPL35,RPL34,RPL24,RPS3A,EIF3C,RPS3,RPS27L,RPS7,UBA52,RPS4X,RPS23,RPL30,RPS15A,RPS17
L8,RPL35,RPL34,RPL24,RPS3A,EIF3C,RPS3,RPS27L,RPS7,UBA52,RPS4X,RPS23,RPL30,RPS15A,RPS17
5,RPS11,RPL37A,RPLP0,RPL8,RPL35,RPL34,RPL24,PSMD7,PSMD10,RPS3A,RPS3,MAGOHB,RPS27L,RPS7,N

RMT112,RPS20,RPS15,DDOST,PABPC1,EEF1D,RPL18,RPL15,MRPS18C,EIF2B3,RPS11,MRPS17,RPL37A,EIF4
P62,RPL24,RPS3A,RPS3,RPS27L,RPS7,POLR2E,UBA52,RPS4X,RPS23,RPL30,NUP85,RPS15A,RPS17,SEC13
L34,NUP62,RPL24,CLTA,RPS3A,RPS3,RPS27L,RPS7,POLR2E,UBA52,RPS4X,RPS23,RPL30,NUP85,PABPN1,I
P62,RPL24,CLTA,RPS3A,RPS3,RPS27L,RPS7,POLR2E,UBA52,RPS4X,RPS23,RPL30,NUP85,RPS15A,RPS17,SI
IE1,RPL18,RPL15,RPS11,RPL37A,RPLP0,RPL8,RPL35,CLASP2,RPL34,RPL24,PSMD7,NRP1,PSMD10,RPS3A,I

2,RPL37A,MPHOSPH10,RPLP0,RPL8,RPL35,RPL34,RPL24,RPS3A,RPS3,RPS27L,RPS7,UBA52,RPS4X,RPS23,I
RPS11,NOP2,RPL37A,MPHOSPH10,RPLP0,RPL8,RPL35,RPL34,RPL24,RPS3A,RPS3,RPS27L,RPS7,UBA52,RPS
0AB1,RPS20,VPS28,RPS15,HMGA1,PSME2,B2M,PSME1,RPL18,POLR2C,RPL15,RPS11,NUP133,RPL37A,KI
RPLP0,RPL8,RPL35,RPL34,RPL24,RPS3A,RPS3,RPS27L,RPS7,UBA52,RPS4X,RPS23,RPL30,EXOSC2,EXOSC8,
F1,TRNT1,PSMD4,RPL28,PSMA7,RPL21,TXNL4A,EXOSC9,HNRNPC,RPL36,TRMT112,RPS20,RPS15,HSPB1,

A,SRSF2,MRPS22,KRT15,OSMR,PPFIBP1,SEMA4D,LAS1L,HMGB1,SULF2,BAIAP2,RAPGEF5,SSU72,MYL6B,F
MR,LAS1L,HMGB1,SULF2,BAIAP2,RAPGEF5,SSU72,MYL6B,SRP9,IQCG,SULT1A3,FAM72A,CMC2,YBX3,BLC
1S1,CACYBP,RHOF,SMARCB1,RPL38,PRR13,RNH1,DNASE1L1,RBCK1,RNF32,LY75,RCC1,SNRPB,DHPS,ACTR
CA3,PCNA,SRSF2,MRPS22,KRT15,OSMR,PPFIBP1,SEMA4D,LAS1L,HMGB1,SULF2,CYB561A3,BAIAP2,RAPG
CA3,PCNA,SRSF2,MRPS22,KRT15,OSMR,PPFIBP1,SEMA4D,LAS1L,HMGB1,SULF2,CYB561A3,BAIAP2,RAPG
RPS22,OSMR,PPFIBP1,LAS1L,HMGB1,SULF2,CYB561A3,BAIAP2,RAPGEF5,SSU72,MYL6B,FAM122C,SRP9,IC
RPS22,OSMR,PPFIBP1,LAS1L,HMGB1,SULF2,CYB561A3,BAIAP2,RAPGEF5,SSU72,MYL6B,FAM122C,SRP9,IC
SMR,PPFIBP1,HMGB1,SULF2,CYB561A3,BAIAP2,RAPGEF5,SSU72,MYL6B,SRP9,IQCG,TMEM67,SULT1A3,F
F5,SSU72,MYL6B,FAM122C,TMEM67,PILRB,CMC2,BLOC1S1,CACYBP,CIRBP,RHOF,SMARCB1,RPL38,PRR13
F5,SSU72,MYL6B,FAM122C,TMEM67,PILRB,CMC2,BLOC1S1,CACYBP,CIRBP,RHOF,SMARCB1,RPL38,PRR13
RPS22,OSMR,PPFIBP1,LAS1L,HMGB1,SULF2,BAIAP2,RAPGEF5,SSU72,FAM122C,SRP9,IQCG,TMEM67,PILR
2,BLOC1S1,CACYBP,CIRBP,RHOF,SMARCB1,PRR13,RNH1,KIF4A,RBCK1,RNF32,LY75,RCC1,SNRPB,PARD3,I
2,BLOC1S1,CACYBP,CIRBP,RHOF,SMARCB1,PRR13,RNH1,KIF4A,RBCK1,RNF32,LY75,RCC1,SNRPB,PARD3,I
CK1,RNF32,LY75,RCC1,SNRPB,DHPS,ACTR6,ZNF235,BOP1,TRPT1,ZC3HAV1,DBI,ARPC2,LIMK2,NUMA1,GP
,TMEM67,PILRB,CMC2,BLOC1S1,CACYBP,CIRBP,RHOF,SMARCB1,RPL38,PRR13,STARD10,RNH1,DNASE1L1
,TMEM67,PILRB,CMC2,BLOC1S1,CACYBP,CIRBP,RHOF,SMARCB1,RPL38,PRR13,STARD10,RNH1,DNASE1L1
MRPS22,OSMR,LAS1L,HMGB1,SULF2,BAIAP2,RAPGEF5,SSU72,MYL6B,FAM122C,IQCG,TMEM67,FAM72A,

/IRPS22,OSMR,LAS1L,HMGB1,SULF2,BAIAP2,RAPGEF5,SSU72,MYL6B,FAM122C,IQCG,TMEM67,FAM72A,

)C1S1,CACYBP,CIRBP,RHOF,SMARCB1,RPL38,PRR13,RNH1,DNASE1L1,KIF4A,RBCK1,RNF32,LY75,RCC1,SN

)C1S1,CACYBP,CIRBP,RHOF,SMARCB1,RPL38,PRR13,RNH1,DNASE1L1,KIF4A,RBCK1,RNF32,LY75,RCC1,SN

3LOC1S1,CACYBP,CIRBP,RHOF,SMARCB1,RPL38,PRR13,RNH1,KIF4A,RBCK1,RNF32,LY75,RCC1,SNRPB,PAR

3LOC1S1,CACYBP,CIRBP,RHOF,SMARCB1,RPL38,PRR13,RNH1,KIF4A,RBCK1,RNF32,LY75,RCC1,SNRPB,PAR

FAM122C,TMEM67,CMC2,BLOC1S1,CACYBP,CIRBP,RHOF,SMARCB1,RPL38,PRR13,STARD10,RNH1,DNAS

FAM122C,TMEM67,CMC2,BLOC1S1,CACYBP,CIRBP,RHOF,SMARCB1,RPL38,PRR13,STARD10,RNH1,DNAS

,SRSF2,MRPS22,KRT15,OSMR,PPFIBP1,HMGB1,SULF2,CYB561A3,BAIAP2,RAPGEF5,SSU72,MYL6B,FAM12

,SRSF2,MRPS22,KRT15,OSMR,PPFIBP1,HMGB1,SULF2,CYB561A3,BAIAP2,RAPGEF5,SSU72,MYL6B,FAM12

AM122C,TMEM67,PILRB,CMC2,BLOC1S1,CACYBP,CIRBP,RHOF,SMARCB1,RPL38,PRR13,RNH1,DNASE1L1,

AM122C,TMEM67,PILRB,CMC2,BLOC1S1,CACYBP,CIRBP,RHOF,SMARCB1,RPL38,PRR13,RNH1,DNASE1L1,

,SSU72,MYL6B,FAM122C,TMEM67,PILRB,FAM72A,CMC2,YBX3,BLOC1S1,CACYBP,CIRBP,RHOF,SMARCB1

,SSU72,MYL6B,FAM122C,TMEM67,PILRB,FAM72A,CMC2,YBX3,BLOC1S1,CACYBP,CIRBP,RHOF,SMARCB1

F4A,RBCK1,LY75,RCC1,SNRPB,DHPS,ACTR6,ZNF235,ARRDC1,TRPT1,RBP5,ZC3HAV1,DBI,ARPC2,PMFBP1,L

F4A,RBCK1,LY75,RCC1,SNRPB,DHPS,ACTR6,ZNF235,ARRDC1,TRPT1,RBP5,ZC3HAV1,DBI,ARPC2,PMFBP1,L

PGEF5,SSU72,MYL6B,FAM122C,TMEM67,PILRB,FAM72A,CMC2,YBX3,BLOC1S1,CACYBP,CIRBP,RHOF,SMA

PGEF5,SSU72,MYL6B,FAM122C,TMEM67,PILRB,FAM72A,CMC2,YBX3,BLOC1S1,CACYBP,CIRBP,RHOF,SMA

F5,SSU72,MYL6B,FAM122C,TMEM67,PILRB,FAM72A,CMC2,YBX3,BLOC1S1,CACYBP,CIRBP,RHOF,SMARC

F5,SSU72,MYL6B,FAM122C,TMEM67,PILRB,FAM72A,CMC2,YBX3,BLOC1S1,CACYBP,CIRBP,RHOF,SMARC

MR,PPFIBP1,HMGB1,SULF2,BAIAP2,RAPGEF5,SSU72,MYL6B,FAM122C,SRP9,IQCG,SULT1A3,PILRB,FAM7

F5,SSU72,MYL6B,FAM122C,TMEM67,PILRB,CMC2,BLOC1S1,CACYBP,CIRBP,RHOF,SMARCB1,RPL38,PRR1

F5,SSU72,MYL6B,FAM122C,TMEM67,PILRB,CMC2,BLOC1S1,CACYBP,CIRBP,RHOF,SMARCB1,RPL38,PRR1

RBCK1,RNF32,LY75,RCC1,SNRPB,PARD3,SLC3A2,DHPS,ACTR6,ZNF235,BOP1,ARRDC1,TRPT1,RBP5,DBI,AR

PFIBP1,LAS1L,HMGB1,SULF2,CYB561A3,BAIAP2,RAPGEF5,SSU72,FAM122C,IQCG,TMEM67,FAM72A,YBX

PFIBP1,LAS1L,HMGB1,SULF2,CYB561A3,BAIAP2,RAPGEF5,SSU72,FAM122C,IQCG,TMEM67,FAM72A,YBX

2,DHPS,ACTR6,ZNF235,BOP1,ARRDC1,TRPT1,RBP5,ZC3HAV1,DBI,ARPC2,LIMK2,SIRT6,NUMA1,GPATCH2L

CA3,PCNA,SRSF2,MRPS22,OSMR,PPFIBP1,LAS1L,HMGB1,SULF2,BAIAP2,RAPGEF5,SSU72,MYL6B,FAM122

CA3,PCNA,SRSF2,MRPS22,OSMR,PPFIBP1,SEMA4D,HMGB1,SULF2,CYB561A3,BAIAP2,RAPGEF5,SSU72,M

CA3,PCNA,SRSF2,MRPS22,OSMR,PPFIBP1,SEMA4D,HMGB1,SULF2,CYB561A3,BAIAP2,RAPGEF5,SSU72,M

A,CMC2,BLOC1S1,CACYBP,CIRBP,RHOF,SMARCB1,RPL38,PRR13,RNH1,KIF4A,RBCK1,RNF32,LY75,RCC1,SN

A,CMC2,BLOC1S1,CACYBP,CIRBP,RHOF,SMARCB1,RPL38,PRR13,RNH1,KIF4A,RBCK1,RNF32,LY75,RCC1,SN

PILRB,CMC2,BLOC1S1,CACYBP,CIRBP,RHOF,SMARCB1,RPL38,PRR13,STARD10,RNH1,DNASE1L1,KIF4A,RB

PILRB,CMC2,BLOC1S1,CACYBP,CIRBP,RHOF,SMARCB1,RPL38,PRR13,STARD10,RNH1,DNASE1L1,KIF4A,RB

31,SULF2,CYB561A3,BAIAP2,RAPGEF5,SSU72,MYL6B,FAM122C,TMEM67,PILRB,FAM72A,CMC2,YBX3,BLC

31,SULF2,CYB561A3,BAIAP2,RAPGEF5,SSU72,MYL6B,FAM122C,TMEM67,PILRB,FAM72A,CMC2,YBX3,BLC

57,CMC2,BLOC1S1,CACYBP,CIRBP,RHOF,SMARCB1,RPL38,PRR13,STARD10,RNH1,KIF4A,RBCK1,RNF32,LY7

57,CMC2,BLOC1S1,CACYBP,CIRBP,RHOF,SMARCB1,RPL38,PRR13,STARD10,RNH1,KIF4A,RBCK1,RNF32,LY7

2,RAPGEF5,SSU72,MYL6B,FAM122C,TMEM67,PILRB,CMC2,BLOC1S1,CACYBP,CIRBP,RHOF,SMARCB1,RPL3

2,RAPGEF5,SSU72,MYL6B,FAM122C,TMEM67,PILRB,CMC2,BLOC1S1,CACYBP,CIRBP,RHOF,SMARCB1,RPL3

CYBP,CIRBP,RHOF,SMARCB1,RPL38,RNH1,KIF4A,RBCK1,RNF32,LY75,RCC1,SNRPB,PARD3,DHPS,ACTR6,ZN

CYBP,CIRBP,RHOF,SMARCB1,RPL38,RNH1,KIF4A,RBCK1,RNF32,LY75,RCC1,SNRPB,PARD3,DHPS,ACTR6,ZN

3LOC1S1,CIRBP,RHOF,SMARCB1,RPL38,PRR13,RNH1,DNASE1L1,KIF4A,RBCK1,RNF32,LY75,RCC1,SNRPB,P

3LOC1S1,CIRBP,RHOF,SMARCB1,RPL38,PRR13,RNH1,DNASE1L1,KIF4A,RBCK1,RNF32,LY75,RCC1,SNRPB,P

22,OSMR,LAS1L,HMGB1,SULF2,BAIAP2,RAPGEF5,SSU72,MYL6B,FAM122C,SRP9,IQCG,TMEM67,SULT1A3

RNF32,LY75,RCC1,SNRPB,DHPS,ACTR6,ARRDC1,TRPT1,ZC3HAV1,DBI,ARPC2,LIMK2,GPATCH2L,BUD31,NF

ACTR6,ZNF235,BOP1,TRPT1,ZC3HAV1,DBI,ARPC2,LIMK2,NUMA1,GPATCH2L,BUD31,NPRL2,ARHGAP6,CT

,CMC2,BLOC1S1,CACYBP,CIRBP,RHOF,SMARCB1,RPL38,PRR13,RNH1,KIF4A,RBCK1,RNF32,LY75,RCC1,SNI

,CMC2,BLOC1S1,CACYBP,CIRBP,RHOF,SMARCB1,RPL38,PRR13,RNH1,KIF4A,RBCK1,RNF32,LY75,RCC1,SNI
C2,BLOC1S1,CACYBP,CIRBP,RHOF,SMARCB1,RPL38,PRR13,RNH1,DNASE1L1,KIF4A,RBCK1,RNF32,LY75,R(
C2,BLOC1S1,CACYBP,CIRBP,RHOF,SMARCB1,RPL38,PRR13,RNH1,DNASE1L1,KIF4A,RBCK1,RNF32,LY75,R(
1S1,CIRBP,RHOF,SMARCB1,RPL38,RNH1,DNASE1L1,KIF4A,RBCK1,RNF32,LY75,RCC1,SNRPB,PARD3,SLC3A
1S1,CIRBP,RHOF,SMARCB1,RPL38,RNH1,DNASE1L1,KIF4A,RBCK1,RNF32,LY75,RCC1,SNRPB,PARD3,SLC3A
RPL38,PRR13,RNH1,DNASE1L1,KIF4A,RBCK1,RNF32,RCC1,SNRPB,PARD3,DHPS,ACTR6,CTSC,ZNF235,ARRD
RPL38,PRR13,RNH1,DNASE1L1,KIF4A,RBCK1,RNF32,RCC1,SNRPB,PARD3,DHPS,ACTR6,CTSC,ZNF235,ARRD
1,CACYBP,CIRBP,RHOF,PRR13,RNH1,DNASE1L1,KIF4A,RBCK1,RNF32,RCC1,SNRPB,PARD3,SLC3A2,DHPS,A
1,CACYBP,CIRBP,RHOF,PRR13,RNH1,DNASE1L1,KIF4A,RBCK1,RNF32,RCC1,SNRPB,PARD3,SLC3A2,DHPS,A
,CMC2,BLOC1S1,CACYBP,CIRBP,RHOF,SMARCB1,RNH1,KIF4A,RBCK1,RNF32,LY75,RCC1,SNRPB,PARD3,DH
,CMC2,BLOC1S1,CACYBP,CIRBP,RHOF,SMARCB1,RNH1,KIF4A,RBCK1,RNF32,LY75,RCC1,SNRPB,PARD3,DH
,SULF2,BAIAP2,RAPGEF5,SSU72,MYL6B,SRP9,TMEM67,PILRB,CMC2,BLOC1S1,CACYBP,CIRBP,RHOF,SMAI
,SULF2,BAIAP2,RAPGEF5,SSU72,MYL6B,SRP9,TMEM67,PILRB,CMC2,BLOC1S1,CACYBP,CIRBP,RHOF,SMAI
MC2,BLOC1S1,CACYBP,CIRBP,RHOF,SMARCB1,RPL38,PRR13,RNH1,KIF4A,RBCK1,RNF32,LY75,RCC1,SNRPI
MC2,BLOC1S1,CACYBP,CIRBP,RHOF,SMARCB1,RPL38,PRR13,RNH1,KIF4A,RBCK1,RNF32,LY75,RCC1,SNRPI
,DBI,ARPC2,LIMK2,GPATCH2L,BUD31,NPRL2,CTTN,EIF5A,COMMD5,NUDT22,USP38,THYN1,SH2B1,NOP1
3,TMEM67,PILRB,CMC2,BLOC1S1,CACYBP,CIRBP,RHOF,SMARCB1,RPL38,PRR13,RNH1,DNASE1L1,KIF4A,R
3,TMEM67,PILRB,CMC2,BLOC1S1,CACYBP,CIRBP,RHOF,SMARCB1,RPL38,PRR13,RNH1,DNASE1L1,KIF4A,R
,RHOF,SMARCB1,RPL38,PRR13,RNH1,KIF4A,RBCK1,RNF32,LY75,RCC1,SNRPB,PARD3,DHPS,ACTR6,ZNF23
,RHOF,SMARCB1,RPL38,PRR13,RNH1,KIF4A,RBCK1,RNF32,LY75,RCC1,SNRPB,PARD3,DHPS,ACTR6,ZNF23
MC2,BLOC1S1,CACYBP,CIRBP,RHOF,SMARCB1,RPL38,PRR13,RNH1,DNASE1L1,KIF4A,RBCK1,RNF32,LY75,R
MC2,BLOC1S1,CACYBP,CIRBP,RHOF,SMARCB1,RPL38,PRR13,RNH1,DNASE1L1,KIF4A,RBCK1,RNF32,LY75,R
,CMC2,BLOC1S1,CACYBP,CIRBP,RHOF,SMARCB1,RPL38,RNH1,DNASE1L1,KIF4A,RBCK1,FASTK,RNF32,LY7
,CMC2,BLOC1S1,CACYBP,CIRBP,RHOF,SMARCB1,RPL38,RNH1,DNASE1L1,KIF4A,RBCK1,FASTK,RNF32,LY7
CTTN,EIF5A,COMMD5,CHID1,NUDT22,SH2B1,NOP16,RPL13A,PSMA6,ACOT8,RPS9,MRPL33,ZNF410,IGSF
CMC2,BLOC1S1,CACYBP,CIRBP,RHOF,SMARCB1,RPL38,PRR13,RNH1,KIF4A,RBCK1,RNF32,LY75,RCC1,SNRF
CMC2,BLOC1S1,CACYBP,CIRBP,RHOF,SMARCB1,RPL38,PRR13,RNH1,KIF4A,RBCK1,RNF32,LY75,RCC1,SNRF
2,RAPGEF5,SSU72,MYL6B,FAM122C,IQCG,FAM72A,BLOC1S1,CACYBP,CIRBP,RHOF,SMARCB1,PRR13,RNH
2,RAPGEF5,SSU72,MYL6B,FAM122C,IQCG,FAM72A,BLOC1S1,CACYBP,CIRBP,RHOF,SMARCB1,PRR13,RNH
2,GPATCH2L,BUD31,NPRL2,CTTN,EIF5A,COMMD5,NUDT22,USP38,THYN1,SH2B1,NOP16,PSMA6,ACOT8,
IUDT22,USP38,THYN1,SH2B1,NOP16,PSMA6,ACOT8,RPS9,MRPL33,ZNF410,NUDT5,STYXL1,POLD4,VRK3,
,SULF2,BAIAP2,RAPGEF5,SSU72,MYL6B,SRP9,TMEM67,PILRB,FAM72A,CMC2,BLOC1S1,CACYBP,CIRBP,RH
,SULF2,BAIAP2,RAPGEF5,SSU72,MYL6B,SRP9,TMEM67,PILRB,FAM72A,CMC2,BLOC1S1,CACYBP,CIRBP,RH
1,HMGB1,SULF2,BAIAP2,RAPGEF5,SSU72,MYL6B,FAM122C,IQCG,FAM72A,YBX3,CACYBP,RHOF,SMARCB
1,HMGB1,SULF2,BAIAP2,RAPGEF5,SSU72,MYL6B,FAM122C,IQCG,FAM72A,YBX3,CACYBP,RHOF,SMARCB
SRSF2,MRPS22,OSMR,PPFIBP1,SEMA4D,LAS1L,HMGB1,SULF2,BAIAP2,RAPGEF5,SSU72,MYL6B,FAM122C
SRSF2,MRPS22,OSMR,PPFIBP1,SEMA4D,LAS1L,HMGB1,SULF2,BAIAP2,RAPGEF5,SSU72,MYL6B,FAM122C
2,SIRT6,GPATCH2L,BUD31,NPRL2,CTTN,EIF5A,COMMD5,NUDT22,USP38,THYN1,SH2B1,NOP16,RPL13A,
YB561A3,BAIAP2,RAPGEF5,SSU72,MYL6B,TMEM67,PILRB,FAM72A,CMC2,BLOC1S1,CACYBP,CIRBP,RHOF,
YB561A3,BAIAP2,RAPGEF5,SSU72,MYL6B,TMEM67,PILRB,FAM72A,CMC2,BLOC1S1,CACYBP,CIRBP,RHOF,
2,MRPS22,KRT15,PPFIBP1,HMGB1,SULF2,CYB561A3,BAIAP2,RAPGEF5,SSU72,MYL6B,SRP9,IQCG,SULT1A
2,MRPS22,KRT15,PPFIBP1,HMGB1,SULF2,CYB561A3,BAIAP2,RAPGEF5,SSU72,MYL6B,SRP9,IQCG,SULT1A
P1,TRPT1,ZC3HAV1,DBI,ARPC2,LIMK2,GPATCH2L,BUD31,NPRL2,CTTN,EIF5A,COMMD5,CHID1,NUDT22,U
,STARD10,RNH1,KIF4A,RBCK1,RNF32,LY75,RCC1,SNRPB,PARD3,SLC3A2,DHPS,ACTR6,ARRDC1,TRPT1,KL(
,STARD10,RNH1,KIF4A,RBCK1,RNF32,LY75,RCC1,SNRPB,PARD3,SLC3A2,DHPS,ACTR6,ARRDC1,TRPT1,KL(
COMMD5,NUDT22,SH2B1,NOP16,RPL13A,PSMA6,ACOT8,RPS9,RCE1,ZNF410,NUDT5,STYXL1,VRK3,SNF8,
1,NPRL2,CTTN,EIF5A,COMMD5,CHID1,NUDT22,AMZ2,THYN1,SH2B1,NOP16,RPL13A,PSMA6,ACOT8,RPS

3,PARD3,SLC3A2,DHPS,ACTR6,ZNF235,BOP1,ARRDC1,TRPT1,RBP5,DBI,ARPC2,LIMK2,SIRT6,NUMA1,GPA
TMEM67,FAM72A,CMC2,BLOC1S1,CACYBP,CIRBP,RHOF,SMARCB1,RPL38,RNH1,KIF4A,RBCK1,RNF32,LY7
TMEM67,FAM72A,CMC2,BLOC1S1,CACYBP,CIRBP,RHOF,SMARCB1,RPL38,RNH1,KIF4A,RBCK1,RNF32,LY7
3,TMEM67,PILRB,FAM72A,CMC2,BLOC1S1,CACYBP,CIRBP,RHOF,SMARCB1,RPL38,PRR13,RNH1,KIF4A,RB
3,TMEM67,PILRB,FAM72A,CMC2,BLOC1S1,CACYBP,CIRBP,RHOF,SMARCB1,RPL38,PRR13,RNH1,KIF4A,RB
HPS,ACTR6,ZNF235,BOP1,TRPT1,ZC3HAV1,DBI,ARPC2,LIMK2,SIRT6,NUMA1,GPATCH2L,BUD31,NPRL2,AI
IAP2,SSU72,MYL6B,TMEM67,PILRB,FAM72A,CMC2,YBX3,BLOC1S1,CACYBP,CIRBP,SMARCB1,RPL38,RNH
IAP2,SSU72,MYL6B,TMEM67,PILRB,FAM72A,CMC2,YBX3,BLOC1S1,CACYBP,CIRBP,SMARCB1,RPL38,RNH
,NPRL2,CTTN,EIF5A,COMMD5,NUDT22,THYN1,SH2B1,NOP16,RPL13A,PSMA6,ACOT8,RPS9,ZNF410,IGSF
RPS22,OSMR,PPFIBP1,LAS1L,HMGB1,SULF2,CYB561A3,BAIAP2,RAPGEF5,SSU72,MYL6B,FAM122C,SRP9,I
RPS22,OSMR,PPFIBP1,LAS1L,HMGB1,SULF2,CYB561A3,BAIAP2,RAPGEF5,SSU72,MYL6B,FAM122C,SRP9,I
MMD5,NUDT22,USP38,THYN1,NOP16,PSMA6,ACOT8,RPS9,MRPL33,ZNF410,NUDT5,STYXL1,VRK3,LZTFL1
,SULF2,BAIAP2,RAPGEF5,SSU72,MYL6B,FAM122C,IQCG,FAM72A,YBX3,BLOC1S1,CACYBP,CIRBP,RHOF,SM
,SULF2,BAIAP2,RAPGEF5,SSU72,MYL6B,FAM122C,IQCG,FAM72A,YBX3,BLOC1S1,CACYBP,CIRBP,RHOF,SM
D31,NPRL2,CTTN,TSPAN11,EIF5A,COMMD5,CHID1,UFD1L,ARHGEF39,PSMA6,ACOT8,AP3B2,ZNF410,NUD
D31,NPRL2,CTTN,TSPAN11,EIF5A,COMMD5,CHID1,UFD1L,ARHGEF39,PSMA6,ACOT8,AP3B2,ZNF410,NUD
75,RCC1,SNRPB,DHPS,ACTR6,ARRDC1,TRPT1,KLC1,ZC3HAV1,DBI,ARPC2,LIMK2,NUMA1,BUD31,NPRL2,CT
AP2,RAPGEF5,SSU72,IQCG,FAM72A,YBX3,CACYBP,CIRBP,RHOF,SMARCB1,RPL38,STARD10,RNH1,DNASE1
D31,NPRL2,CTTN,EIF5A,COMMD5,NUDT22,USP38,THYN1,SH2B1,NOP16,RPL13A,PSMA6,ACOT8,RPS9,ZN
ZNF410,NUDT5,STYXL1,SNF8,CDC37,CPSF3L,TSG101,UBR4,RPS6,RPL29,RTEL1,AUP1,YIPF3,TRNT1,KXD1,
IRPB,DHPS,ACTR6,ZNF235,BOP1,ARRDC1,TRPT1,ZC3HAV1,DBI,ARPC2,LIMK2,SIRT6,GPATCH2L,BUD31,NP
L,SNRPB,PARD3,SLC3A2,DHPS,ACTR6,ZNF235,ARRDC1,TRPT1,SCAF11,DBI,ARPC2,PMFBP1,LIMK2,SIRT6,N
L,SNRPB,PARD3,SLC3A2,DHPS,ACTR6,ZNF235,ARRDC1,TRPT1,SCAF11,DBI,ARPC2,PMFBP1,LIMK2,SIRT6,N
EIF5A,COMMD5,CHID1,UFD1L,ARHGEF39,PSMA6,ACOT8,AP3B2,ZNF410,NUDT5,SNRPN,STYXL1,POLD4,L
EIF5A,COMMD5,CHID1,UFD1L,ARHGEF39,PSMA6,ACOT8,AP3B2,ZNF410,NUDT5,SNRPN,STYXL1,POLD4,L
RCB1,RPL38,PRR13,RNH1,DNASE1L1,KIF4A,RBCK1,RCC1,SNRPB,DHPS,ACTR6,CTSC,ZNF235,ARRDC1,TRP
RCB1,RPL38,PRR13,RNH1,DNASE1L1,KIF4A,RBCK1,RCC1,SNRPB,DHPS,ACTR6,CTSC,ZNF235,ARRDC1,TRP
,SRSF2,MRPS22,OSMR,HMGB1,SULF2,CYB561A3,BAIAP2,RAPGEF5,SSU72,FAM122C,SRP9,IQCG,TMEM6
,SRSF2,MRPS22,OSMR,HMGB1,SULF2,CYB561A3,BAIAP2,RAPGEF5,SSU72,FAM122C,SRP9,IQCG,TMEM6
,DBI,ARPC2,LIMK2,GPATCH2L,BUD31,NPRL2,CTTN,EIF5A,COMMD5,NUDT22,USP38,THYN1,SH2B1,NOP1
NRPB,DHPS,ACTR6,TRPT1,ZC3HAV1,DBI,ARPC2,LIMK2,NUMA1,BUD31,NPRL2,CTTN,EIF5A,COMMD5,NU
RT15,OSMR,PPFIBP1,SULF2,CYB561A3,BAIAP2,RAPGEF5,SSU72,FAM122C,SRP9,IQCG,TMEM67,SULT1A3
RT15,OSMR,PPFIBP1,SULF2,CYB561A3,BAIAP2,RAPGEF5,SSU72,FAM122C,SRP9,IQCG,TMEM67,SULT1A3
BUD31,NPRL2,CTTN,EIF5A,COMMD5,CHID1,NUDT22,SH2B1,NOP16,RPL13A,PSMA6,ACOT8,RPS9,ZNF41
T1,DBI,ARPC2,PMFBP1,LIMK2,SIRT6,BUD31,NPRL2,CTTN,EIF5A,COMMD5,NUDT22,UFD1L,RPL13A,ARH
T1,DBI,ARPC2,PMFBP1,LIMK2,SIRT6,BUD31,NPRL2,CTTN,EIF5A,COMMD5,NUDT22,UFD1L,RPL13A,ARH
F4A,RBCK1,RNF32,LY75,RCC1,SNRPB,SLC3A2,ACTR6,MAP3K7CL,ZNF235,BOP1,ARRDC1,TRPT1,RBP5,ZC3
F4A,RBCK1,RNF32,LY75,RCC1,SNRPB,SLC3A2,ACTR6,MAP3K7CL,ZNF235,BOP1,ARRDC1,TRPT1,RBP5,ZC3
,DHPS,ACTR6,RBP5,ZC3HAV1,DBI,ARPC2,LIMK2,SIRT6,BUD31,NPRL2,CTTN,EIF5A,COMMD5,CHID1,UFD1
,DHPS,ACTR6,RBP5,ZC3HAV1,DBI,ARPC2,LIMK2,SIRT6,BUD31,NPRL2,CTTN,EIF5A,COMMD5,CHID1,UFD1
75,RCC1,SNRPB,PARD3,SLC3A2,DHPS,ACTR6,ZNF235,TRPT1,RBP5,DBI,ARPC2,PMFBP1,LIMK2,SIRT6,BUD
75,RCC1,SNRPB,PARD3,SLC3A2,DHPS,ACTR6,ZNF235,TRPT1,RBP5,DBI,ARPC2,PMFBP1,LIMK2,SIRT6,BUD
C1,SNRPB,DHPS,ACTR6,ARRDC1,TRPT1,RBP5,SCAF11,DBI,ARPC2,LIMK2,NOB1,SIRT6,NUMA1,SUMF2,BU
C1,SNRPB,DHPS,ACTR6,ARRDC1,TRPT1,RBP5,SCAF11,DBI,ARPC2,LIMK2,NOB1,SIRT6,NUMA1,SUMF2,BU
HMGB1,CYB561A3,BAIAP2,RAPGEF5,SSU72,FAM122C,SRP9,IQCG,TMEM67,SULT1A3,PILRB,FAM72A,YBX
HMGB1,CYB561A3,BAIAP2,RAPGEF5,SSU72,FAM122C,SRP9,IQCG,TMEM67,SULT1A3,PILRB,FAM72A,YBX
3HAV1,DBI,ARPC2,LIMK2,BUD31,NPRL2,CTTN,EIF5A,COMMD5,NUDT22,TAZ,THYN1,SH2B1,NOP16,PSM

RCC1,SNRPB,DHPS,ACTR6,ZNF235,TRPT1,ZC3HAV1,DBI,ARPC2,LIMK2,BUD31,NPRL2,CTTN,EIF5A,COMMI
,ARRDC1,TRPT1,DBI,ARPC2,PMFBP1,LIMK2,SIRT6,NUMA1,GPATCH2L,BUD31,NPRL2,CTTN,TSPAN11,EIF5
,ARRDC1,TRPT1,DBI,ARPC2,PMFBP1,LIMK2,SIRT6,NUMA1,GPATCH2L,BUD31,NPRL2,CTTN,TSPAN11,EIF5
RNF32,LY75,RCC1,SNRPB,DHPS,ACTR6,ARRDC1,KLC1,RBP5,ZC3HAV1,DBI,ARPC2,NUMA1,BUD31,NPRL2,
RBCK1,RCC1,SNRPB,DHPS,ACTR6,CTSC,DBI,ARPC2,LIMK2,SIRT6,GPATCH2L,BUD31,NPRL2,CTTN,EIF5A,CO
RBCK1,RCC1,SNRPB,DHPS,ACTR6,CTSC,DBI,ARPC2,LIMK2,SIRT6,GPATCH2L,BUD31,NPRL2,CTTN,EIF5A,CO
,NPRL2,CTTN,EIF5A,COMMD5,CHID1,NUDT22,SH2B1,NOP16,PSMA6,ACOT8,RPS9,ZNF410,IGSF8,NUDT5
MR,PPFIBP1,LAS1L,HMGB1,SULF2,CYB561A3,BAIAP2,RAPGEF5,SSU72,MYL6B,FAM122C,SRP9,IQCG,PILRB
K1,ATG16L2,RCC1,SNRPB,TCAIM,DHPS,ACTR6,SEC62,ZNF235,BOP1,ARRDC1,TRPT1,ZC3HAV1,ARPC2,LIN
,SRSF2,MRPS22,KRT15,OSMR,PPFIBP1,HMGB1,SULF2,BAIAP2,RAPGEF5,SSU72,MYL6B,FAM122C,SRP9,IC
ATCH2L,BUD31,NPRL2,CTTN,EIF5A,COMMD5,UFD1L,ARHGEF39,PSMA6,ACOT8,AP3B2,PCMT1,NUDT5,SN
ATCH2L,BUD31,NPRL2,CTTN,EIF5A,COMMD5,UFD1L,ARHGEF39,PSMA6,ACOT8,AP3B2,PCMT1,NUDT5,SN
2,RAPGEF5,SSU72,MYL6B,TMEM67,PILRB,CMC2,BLOC1S1,CACYBP,CIRBP,RHOF,SMARCB1,RPL38,PRR13,
2,RAPGEF5,SSU72,MYL6B,TMEM67,PILRB,CMC2,BLOC1S1,CACYBP,CIRBP,RHOF,SMARCB1,RPL38,PRR13,
RBCK1,RNF32,RCC1,SNRPB,SLC3A2,DHPS,ACTR6,ZNF235,BOP1,ARRDC1,TRPT1,ZC3HAV1,SCAF11,DBI,AR
RBCK1,RNF32,RCC1,SNRPB,SLC3A2,DHPS,ACTR6,ZNF235,BOP1,ARRDC1,TRPT1,ZC3HAV1,SCAF11,DBI,AR
RBCK1,RNF32,RCC1,SNRPB,SLC3A2,DHPS,ACTR6,ZNF235,BOP1,ARRDC1,TRPT1,ZC3HAV1,SCAF11,DBI,AR
RBCK1,RNF32,RCC1,SNRPB,SLC3A2,DHPS,ACTR6,ZNF235,BOP1,ARRDC1,TRPT1,ZC3HAV1,SCAF11,DBI,AR
MK2,GPATCH2L,BUD31,NPRL2,CTTN,EIF5A,COMMD5,CHID1,UFD1L,RPL13A,ARHGEF39,PSMA6,ACOT8,A
MK2,GPATCH2L,BUD31,NPRL2,CTTN,EIF5A,COMMD5,CHID1,UFD1L,RPL13A,ARHGEF39,PSMA6,ACOT8,A
TRPT1,DBI,ARPC2,PMFBP1,LIMK2,SIRT6,NUMA1,GPATCH2L,SUMF2,BUD31,NPRL2,CRYGS,CTTN,TSPAN1
TRPT1,DBI,ARPC2,PMFBP1,LIMK2,SIRT6,NUMA1,GPATCH2L,SUMF2,BUD31,NPRL2,CRYGS,CTTN,TSPAN1
1,LIMK2,NUMA1,GPATCH2L,BUD31,NPRL2,CTTN,EIF5A,COMMD5,USP38,SH2B1,NOP16,PSMA6,ZNF410,
MK2,GPATCH2L,BUD31,NPRL2,CTTN,EIF5A,COMMD5,NUDT22,UFD1L,SH2B1,RPL13A,ARHGEF39,PSMA6,
MK2,GPATCH2L,BUD31,NPRL2,CTTN,EIF5A,COMMD5,NUDT22,UFD1L,SH2B1,RPL13A,ARHGEF39,PSMA6,
2,MYL6B,TMEM67,FAM72A,CMC2,BLOC1S1,CACYBP,CIRBP,RHOF,RPL38,PRR13,RNH1,KIF4A,RBCK1,RNF
2,MYL6B,TMEM67,FAM72A,CMC2,BLOC1S1,CACYBP,CIRBP,RHOF,RPL38,PRR13,RNH1,KIF4A,RBCK1,RNF
1,LIMK2,GPATCH2L,BUD31,NPRL2,CTTN,EIF5A,COMMD5,NUDT22,UFD1L,ARHGEF39,PSMA6,ACOT8,PCI
1,LIMK2,GPATCH2L,BUD31,NPRL2,CTTN,EIF5A,COMMD5,NUDT22,UFD1L,ARHGEF39,PSMA6,ACOT8,PCI
OC1S1,CACYBP,CIRBP,RHOF,SMARCB1,RPL38,PRR13,RNH1,KIF4A,RBCK1,RNF32,LY75,RCC1,SNRPB,PARD
OC1S1,CACYBP,CIRBP,RHOF,SMARCB1,RPL38,PRR13,RNH1,KIF4A,RBCK1,RNF32,LY75,RCC1,SNRPB,PARD
IAP2,RAPGEF5,SSU72,MYL6B,TMEM67,FAM72A,CMC2,BLOC1S1,CACYBP,CIRBP,RHOF,SMARCB1,RPL38,
IAP2,RAPGEF5,SSU72,MYL6B,TMEM67,FAM72A,CMC2,BLOC1S1,CACYBP,CIRBP,RHOF,SMARCB1,RPL38,
T1,SCAF11,DBI,LIMK2,SUMF2,BUD31,NPRL2,CTTN,EIF5A,COMMD5,NUDT22,UFD1L,ARHGEF39,TTLL5,PS
T1,SCAF11,DBI,LIMK2,SUMF2,BUD31,NPRL2,CTTN,EIF5A,COMMD5,NUDT22,UFD1L,ARHGEF39,TTLL5,PS
3,IQCG,TMEM67,CMC2,BLOC1S1,CIRBP,RHOF,SMARCB1,RPL38,RNH1,DNASE1L1,KIF4A,RBCK1,RNF32,LY
3,IQCG,TMEM67,CMC2,BLOC1S1,CIRBP,RHOF,SMARCB1,RPL38,RNH1,DNASE1L1,KIF4A,RBCK1,RNF32,LY
A3,PCNA,SRSF2,MRPS22,HDAC9,OSMR,PPFIBP1,LAS1L,HMGB1,SULF2,BAIAP2,RAPGEF5,SSU72,MYL6B,F
2,CYB561A3,RAPGEF5,SSU72,MYL6B,FAM122C,TMEM67,CMC2,BLOC1S1,CACYBP,CIRBP,RHOF,SMARCB
2,CYB561A3,RAPGEF5,SSU72,MYL6B,FAM122C,TMEM67,CMC2,BLOC1S1,CACYBP,CIRBP,RHOF,SMARCB
BAIAP2,RAPGEF5,SSU72,MYL6B,IQCG,BLOC1S1,CACYBP,CIRBP,RHOF,SMARCB1,PRR13,RNH1,DNASE1L1,K
BAIAP2,RAPGEF5,SSU72,MYL6B,IQCG,BLOC1S1,CACYBP,CIRBP,RHOF,SMARCB1,PRR13,RNH1,DNASE1L1,K
2,CTTN,EIF5A,COMMD5,CHID1,NUDT22,SH2B1,RPL13A,PSMA6,ACOT8,RPS9,MRPL33,ZNF410,NUDT5,ST
2,RAPGEF5,SSU72,MYL6B,IQCG,FAM72A,BLOC1S1,CACYBP,CIRBP,RHOF,SMARCB1,PRR13,RNH1,DNASE1
2,RAPGEF5,SSU72,MYL6B,IQCG,FAM72A,BLOC1S1,CACYBP,CIRBP,RHOF,SMARCB1,PRR13,RNH1,DNASE1
MK2,SIRT6,NUMA1,BUD31,CTTN,TSPAN11,EIF5A,COMMD5,CHID1,UFD1L,THYN1,ARHGEF39,ACOT8,AP3
MK2,SIRT6,NUMA1,BUD31,CTTN,TSPAN11,EIF5A,COMMD5,CHID1,UFD1L,THYN1,ARHGEF39,ACOT8,AP3

MA1,BUD31,NPRL2,CTTN,TSPAN11,EIF5A,COMMD5,UFD1L,ARHGEF39,PSMA6,ACOT8,SLC35G1,IGSF8,N
MA1,BUD31,NPRL2,CTTN,TSPAN11,EIF5A,COMMD5,UFD1L,ARHGEF39,PSMA6,ACOT8,SLC35G1,IGSF8,N
YB561A3,BAIAP2,RAPGEF5,SSU72,FAM122C,SRP9,IQCG,TMEM67,SULT1A3,PILRB,FAM72A,BLOC1S1,CA
YB561A3,BAIAP2,RAPGEF5,SSU72,FAM122C,SRP9,IQCG,TMEM67,SULT1A3,PILRB,FAM72A,BLOC1S1,CA
T22,THYN1,SH2B1,NOP16,PSMA6,ACOT8,RPS9,MRPL33,ZNF410,NUDT5,STYXL1,VRK3,PPM1K,SON,UQC
3,RNH1,KIF4A,RBCK1,RNF32,LY75,RCC1,SNRPB,DHPS,ACTR6,MAP3K7CL,CTSC,ZNF235,BOP1,ARRDC1,TR
3,RNH1,KIF4A,RBCK1,RNF32,LY75,RCC1,SNRPB,DHPS,ACTR6,MAP3K7CL,CTSC,ZNF235,BOP1,ARRDC1,TR
S22,HDAC9,OSMR,PPFIBP1,LAS1L,HMGB1,SULF2,CYB561A3,BAIAP2,RAPGEF5,SSU72,MYL6B,FAM122C,
S22,HDAC9,OSMR,PPFIBP1,LAS1L,HMGB1,SULF2,CYB561A3,BAIAP2,RAPGEF5,SSU72,MYL6B,FAM122C,
,RNH1,KIF4A,RBCK1,LY75,RCC1,SNRPB,SLC3A2,DHPS,ACTR6,BOP1,TRPT1,KLC1,ZC3HAV1,DBI,ARPC2,PM
,RNH1,KIF4A,RBCK1,LY75,RCC1,SNRPB,SLC3A2,DHPS,ACTR6,BOP1,TRPT1,KLC1,ZC3HAV1,DBI,ARPC2,PM
RPS22,KRT15,OSMR,PPFIBP1,SEMA4D,HMGB1,SULF2,CYB561A3,BAIAP2,RAPGEF5,SSU72,MYL6B,FAM
RPS22,KRT15,OSMR,PPFIBP1,SEMA4D,HMGB1,SULF2,CYB561A3,BAIAP2,RAPGEF5,SSU72,MYL6B,FAM
UD31,NPRL2,CTTN,EIF5A,COMMD5,CHID1,NUDT22,THYN1,SH2B1,NOP16,PSMA6,ACOT8,RPS9,RCE1,ZN
IBP1,HMGB1,SULF2,CYB561A3,BAIAP2,RAPGEF5,SSU72,MYL6B,IQCG,BLOC1S1,CIRBP,RHOF,SMARCB1,P
IBP1,HMGB1,SULF2,CYB561A3,BAIAP2,RAPGEF5,SSU72,MYL6B,IQCG,BLOC1S1,CIRBP,RHOF,SMARCB1,P
2,LIMK2,BUD31,NPRL2,CTTN,EIF5A,COMMD5,NUDT22,USP38,THYN1,SH2B1,NOP16,PSMA6,ACOT8,RPS
H1,DNASE1L1,RBCK1,RCC1,SNRPB,DHPS,ACTR6,CTSC,ZNF235,BOP1,TRPT1,ZC3HAV1,DBI,ARPC2,PMFBP
H1,DNASE1L1,RBCK1,RCC1,SNRPB,DHPS,ACTR6,CTSC,ZNF235,BOP1,TRPT1,ZC3HAV1,DBI,ARPC2,PMFBP
ARCB1,RPL38,RNH1,DNASE1L1,KIF4A,RBCK1,RNF32,LY75,RCC1,SNRPB,PARD3,SLC3A2,DHPS,ACTR6,ZN
ARCB1,RPL38,RNH1,DNASE1L1,KIF4A,RBCK1,RNF32,LY75,RCC1,SNRPB,PARD3,SLC3A2,DHPS,ACTR6,ZN
HPS,ACTR6,ZNF235,TRPT1,RBP5,ZC3HAV1,DBI,ARPC2,LIMK2,SUMF2,BUD31,NPRL2,CTTN,EIF5A,COMM
CYBP,CIRBP,SMARCB1,RPL38,PRR13,RNH1,KIF4A,RBCK1,RNF32,LY75,RCC1,SNRPB,SLC3A2,DHPS,ACTR6,
CYBP,CIRBP,SMARCB1,RPL38,PRR13,RNH1,KIF4A,RBCK1,RNF32,LY75,RCC1,SNRPB,SLC3A2,DHPS,ACTR6,
IF5A,COMMD5,UFD1L,ARHGEF39,PSMA6,ACOT8,AP3B2,NUDT5,SNRPN,STYXL1,LZTFL1,FIS1,PPM1K,VC
IF5A,COMMD5,UFD1L,ARHGEF39,PSMA6,ACOT8,AP3B2,NUDT5,SNRPN,STYXL1,LZTFL1,FIS1,PPM1K,VC
UDT22,USP38,SH2B1,RPL13A,ARHGEF39,PSMA6,RCE1,ZNF410,STYXL1,VRK3,LRRC1,CDC37,RPS16,CPSF
MD5,UFD1L,ARHGEF39,PSMA6,ACOT8,NUDT5,SNRPN,STYXL1,LZTFL1,FIS1,MAGED2,SON,UQCRQ,CPSF
MD5,UFD1L,ARHGEF39,PSMA6,ACOT8,NUDT5,SNRPN,STYXL1,LZTFL1,FIS1,MAGED2,SON,UQCRQ,CPSF
PATCH2L,BUD31,CTTN,EIF5A,COMMD5,NUDT22,UFD1L,SH2B1,ARHGEF39,ACOT8,AP3B2,PCMT1,STYXL
PATCH2L,BUD31,CTTN,EIF5A,COMMD5,NUDT22,UFD1L,SH2B1,ARHGEF39,ACOT8,AP3B2,PCMT1,STYXL
2,CTTN,EIF5A,COMMD5,UFD1L,RPL13A,ARHGEF39,PSMA6,ACOT8,AP3B2,NUDT5,SNRPN,STYXL1,LZTFL1,
2,CTTN,EIF5A,COMMD5,UFD1L,RPL13A,ARHGEF39,PSMA6,ACOT8,AP3B2,NUDT5,SNRPN,STYXL1,LZTFL1,
MD5,CHID1,NUDT22,PSMA6,RPS9,NUDT5,STYXL1,VRK3,SON,UQCRQ,SNF8,CPSF3L,TSG101,ATG12,NIP
9,MRPL33,NUDT5,STYXL1,VRK3,PPM1K,SON,UQCRQ,SNF8,CPSF3L,TSG101,UBR4,RPS6,RPL29,RTEL1,AU
MK2,BUD31,NPRL2,CTTN,EIF5A,COMMD5,NUDT22,USP38,THYN1,SH2B1,NOP16,PSMA6,ACOT8,RPS9,R
GEF5,SSU72,MYL6B,IQCG,BLOC1S1,CACYBP,CIRBP,RHOF,SMARCB1,PRR13,RNH1,DNASE1L1,KIF4A,RBCK
GEF5,SSU72,MYL6B,IQCG,BLOC1S1,CACYBP,CIRBP,RHOF,SMARCB1,PRR13,RNH1,DNASE1L1,KIF4A,RBCK
L,TUBA1C,CPSF3L,TSG101,FUZ,RPS25,COX4I1,RPL29,AUP1,FOXRED1,TRNT1,KXD1,MED15,UBE2I,EIF3M,
IBP1,LAS1L,HMGB1,SULF2,BAIAP2,RAPGEF5,SSU72,FAM122C,SRP9,IQCG,SULT1A3,FAM72A,YBX3,BLOC
,ARPC2,LIMK2,GPATCH2L,BUD31,NPRL2,CTTN,EIF5A,COMMD5,UFD1L,USP38,DBNDD1,RPL13A,ARHGEF
,ARPC2,LIMK2,GPATCH2L,BUD31,NPRL2,CTTN,EIF5A,COMMD5,UFD1L,USP38,DBNDD1,RPL13A,ARHGEF
MA4D,HMGB1,CYB561A3,BAIAP2,RAPGEF5,MYL6B,IQCG,BLOC1S1,CACYBP,CIRBP,RHOF,SMARCB1,PRR
MA4D,HMGB1,CYB561A3,BAIAP2,RAPGEF5,MYL6B,IQCG,BLOC1S1,CACYBP,CIRBP,RHOF,SMARCB1,PRR
,SRSF2,MRPS22,KRT15,OSMR,PPFIBP1,SEMA4D,LAS1L,HMGB1,SULF2,CYB561A3,BAIAP2,RAPGEF5,SSU7
,SRSF2,MRPS22,KRT15,OSMR,PPFIBP1,SEMA4D,LAS1L,HMGB1,SULF2,CYB561A3,BAIAP2,RAPGEF5,SSU7
2,RAPGEF5,SSU72,IQCG,FAM72A,CACYBP,RHOF,SMARCB1,RPL38,PRR13,STARD10,RNH1,DNASE1L1,KIF

NA,SRSF2,MRPS22,KRT15,OSMR,PPFIBP1,SEMA4D,HMGB1,SULF2,CYB561A3,BAIAP2,RAPGEF5,SSU72,M

NA,SRSF2,MRPS22,KRT15,OSMR,PPFIBP1,SEMA4D,HMGB1,SULF2,CYB561A3,BAIAP2,RAPGEF5,SSU72,M

IK2,GPATCH2L,BUD31,NPRL2,CTTN,EIF5A,COMMD5,NUDT22,RPL13A,PSMA6,RPS9,RCE1,STYXL1,VRK3,L

IK2,GPATCH2L,BUD31,NPRL2,CTTN,EIF5A,COMMD5,NUDT22,RPL13A,PSMA6,RPS9,RCE1,STYXL1,VRK3,L

RPS9,RCE1,ZNF410,NUDT5,STYXL1,PPM1K,SON,UQCRQ,SNF8,CPSF3L,TSG101,NIP7,RPL29,COASY,YIPF3,T

)1,RPS25,RPS6,RPL29,RTEL1,AUP1,TRNT1,KXD1,MED15,CLASRP,UBE2I,EIF3M,CDC42,ASF1B,CCDC25,RF

1,LY75,RCC1,SNRPB,DHPS,ACTR6,MAP3K7CL,ZNF235,BOP1,ARRDC1,TRPT1,DBI,ARPC2,PMFBP1,LIMK2,G

1,LY75,RCC1,SNRPB,DHPS,ACTR6,MAP3K7CL,ZNF235,BOP1,ARRDC1,TRPT1,DBI,ARPC2,PMFBP1,LIMK2,G

CA3,PCNA,SRSF2,MRPS22,OSMR,PPFIBP1,SEMA4D,LAS1L,HMGB1,SULF2,BAIAP2,RAPGEF5,SSU72,MYL6B

CA3,PCNA,SRSF2,MRPS22,OSMR,PPFIBP1,SEMA4D,LAS1L,HMGB1,SULF2,BAIAP2,RAPGEF5,SSU72,MYL6B

5,STYXL1,VRK3,LZTFL1,PPM1K,SNF8,CPSF3L,TSG101,RPS25,NIP7,RPS6,RPL29,YIPF3,TRNT1,EIF3M,CDC42

,HDAC9,KRT15,OSMR,PPFIBP1,LAS1L,HMGB1,SULF2,BAIAP2,RAPGEF5,SSU72,FAM122C,SRP9,IQCG,SULT

,HDAC9,KRT15,OSMR,PPFIBP1,LAS1L,HMGB1,SULF2,BAIAP2,RAPGEF5,SSU72,FAM122C,SRP9,IQCG,SULT

P2,RAPGEF5,MYL6B,SRP9,IQCG,SULT1A3,FAM72A,BLOC1S1,CACYBP,CIRBP,RHOF,SMARCB1,RPL38,PRR1

P2,RAPGEF5,MYL6B,SRP9,IQCG,SULT1A3,FAM72A,BLOC1S1,CACYBP,CIRBP,RHOF,SMARCB1,RPL38,PRR1

NF410,STYXL1,VRK3,CPSF3L,TSG101,FAM3A,RPS25,TTC9C,TMX2,DBNDD2,RPS6,RPL29,RTEL1,SRSF1,SAP

C1S1,CIRBP,RHOF,SMARCB1,PRR13,RNH1,DNASE1L1,RBCK1,PTOV1,FASTK,RNF32,LY75,ATG16L2,SNRPB,

C1S1,CIRBP,RHOF,SMARCB1,PRR13,RNH1,DNASE1L1,RBCK1,PTOV1,FASTK,RNF32,LY75,ATG16L2,SNRPB,

F2,BAIAP2,RAPGEF5,SSU72,MYL6B,FAM122C,IQCG,FAM72A,YBX3,CACYBP,RHOF,SMARCB1,STARD10,RN

DBI,PMFBP1,LIMK2,BUD31,NPRL2,CRYGS,CTTN,EIF5A,COMMD5,NUDT22,UFD1L,USP38,SH2B1,RPL13A,A

DBI,PMFBP1,LIMK2,BUD31,NPRL2,CRYGS,CTTN,EIF5A,COMMD5,NUDT22,UFD1L,USP38,SH2B1,RPL13A,A

NF235,TRPT1,DBI,ARPC2,PMFBP1,LIMK2,SIRT6,BUD31,NPRL2,SCARB1,CTTN,EIF5A,COMMD5,CHID1,NUD

NF235,TRPT1,DBI,ARPC2,PMFBP1,LIMK2,SIRT6,BUD31,NPRL2,SCARB1,CTTN,EIF5A,COMMD5,CHID1,NUD

GEF39,ACOT8,AP3B2,NUDT5,STYXL1,LZTFL1,FIS1,MAGED2,SON,SNF8,CPSF3L,TSG101,FUZ,RPS25,THEM

GEF39,ACOT8,AP3B2,NUDT5,STYXL1,LZTFL1,FIS1,MAGED2,SON,SNF8,CPSF3L,TSG101,FUZ,RPS25,THEM

D5,RPL13A,PSMA6,ACOT8,ZNF410,NUDT5,STYXL1,VRK3,LZTFL1,PPM1K,SNF8,CPSF3L,TSG101,RPS25,NIP

SMR,SEMA4D,LAS1L,HMGB1,SULF2,BAIAP2,RAPGEF5,SSU72,MYL6B,IQCG,UAP1,PILRB,FAM72A,CMC2,BI

SMR,SEMA4D,LAS1L,HMGB1,SULF2,BAIAP2,RAPGEF5,SSU72,MYL6B,IQCG,UAP1,PILRB,FAM72A,CMC2,BI

ZNF235,BOP1,TRPT1,DBI,ARPC2,PMFBP1,LIMK2,GPATCH2L,BUD31,NPRL2,CRYGS,CTTN,EIF5A,COMMD5,

ZNF235,BOP1,TRPT1,DBI,ARPC2,PMFBP1,LIMK2,GPATCH2L,BUD31,NPRL2,CRYGS,CTTN,EIF5A,COMMD5,

RBCK1,FASTK,RNF32,LY75,SNRPB,PARD3,SLC3A2,ACTR6,BOP1,TRPT1,RBP5,ZC3HAV1,DBI,ARPC2,SIRT6,G

PFIBP1,HMGB1,CYB561A3,BAIAP2,SSU72,MYL6B,SRP9,SULT1A3,PILRB,FAM72A,BLOC1S1,CIRBP,RHOF,RP

PFIBP1,HMGB1,CYB561A3,BAIAP2,SSU72,MYL6B,SRP9,SULT1A3,PILRB,FAM72A,BLOC1S1,CIRBP,RHOF,RP

JD31,NPRL2,CTTN,EIF5A,COMMD5,NUDT22,THYN1,RPL13A,PSMA6,ACOT8,RPS9,MRPL33,ZNF410,NUDT

RL2,CTTN,EIF5A,COMMD5,NUDT22,RPL13A,PSMA6,ACOT8,RPS9,MRPL33,ZNF410,NUDT5,STYXL1,VRK3,I

AM122C,SRP9,IQCG,TMEM67,PILRB,FAM72A,YBX3,BLOC1S1,CACYBP,SMARCB1,RPL38,STARD10,RNH1,K

AM122C,SRP9,IQCG,TMEM67,PILRB,FAM72A,YBX3,BLOC1S1,CACYBP,SMARCB1,RPL38,STARD10,RNH1,K

AIAP2,FAM122C,SRP9,IQCG,TMEM67,SULT1A3,PILRB,FAM72A,BLOC1S1,CACYBP,CIRBP,STARD10,RNH1,D

AIAP2,FAM122C,SRP9,IQCG,TMEM67,SULT1A3,PILRB,FAM72A,BLOC1S1,CACYBP,CIRBP,STARD10,RNH1,D

G,PILRB,FAM72A,CMC2,BLOC1S1,CIRBP,RHOF,SMARCB1,PRR13,STARD10,RNH1,DNASE1L1,RBCK1,RNF32

RAPGEF5,SSU72,MYL6B,IQCG,FAM72A,BLOC1S1,CACYBP,CIRBP,RHOF,SMARCB1,PRR13,RNH1,KIF4A,RBC

RAPGEF5,SSU72,MYL6B,IQCG,FAM72A,BLOC1S1,CACYBP,CIRBP,RHOF,SMARCB1,PRR13,RNH1,KIF4A,RBC

M67,PILRB,FAM72A,CMC2,BLOC1S1,CACYBP,CIRBP,SMARCB1,RPL38,RNH1,KIF4A,RBCK1,TMEM135,LY75

M67,PILRB,FAM72A,CMC2,BLOC1S1,CACYBP,CIRBP,SMARCB1,RPL38,RNH1,KIF4A,RBCK1,TMEM135,LY75

C1S1,CACYBP,RHOF,SMARCB1,PRR13,RNH1,DNASE1L1,RBCK1,PTOV1,FASTK,RNF32,LY75,ATG16L2,SNRP

RPS6,RTEL1,ITGB1BP1,CLASRP,SLC39A13,CDC42,FBXW9,USE1,ASF1B,RFX3,ENDOV,LYPLA2,C17ORF62,PP

AS1L,HMGB1,SULF2,CYB561A3,BAIAP2,RAPGEF5,SSU72,MYL6B,FAM122C,SRP9,IQCG,PILRB,YBX3,CACYB

S1L,HMGB1,SULF2,CYB561A3,BAIAP2,RAPGEF5,SSU72,MYL6B,FAM122C,SRP9,IQCG,PILRB,YBX3,CACYB
D3,DHPS,ACTR6,RBP5,ZC3HAV1,DBI,ARPC2,LIMK2,SIRT6,BUD31,CTTN,EIF5A,COMMD5,CHID1,UFD1L,T
D3,DHPS,ACTR6,RBP5,ZC3HAV1,DBI,ARPC2,LIMK2,SIRT6,BUD31,CTTN,EIF5A,COMMD5,CHID1,UFD1L,T
2,RAPGEF5,SSU72,MYL6B,IQCG,BLOC1S1,RHOF,SMARCB1,PRR13,RNH1,DNASE1L1,KIF4A,RBCK1,PTOV1,
2,RAPGEF5,SSU72,MYL6B,IQCG,BLOC1S1,RHOF,SMARCB1,PRR13,RNH1,DNASE1L1,KIF4A,RBCK1,PTOV1,
,KRT15,OSMR,PPFIBP1,SEMA4D,HMGB1,SULF2,CYB561A3,BAIAP2,RAPGEF5,SSU72,MYL6B,SRP9,IQCG,S
EIF5A,COMMD5,NUDT22,USP38,THYN1,NOP16,RPL13A,PSMA6,ACOT8,RPS9,ZNF410,NUDT5,STYXL1,PO
A6,ACOT8,RPS9,MRPL33,ZNF410,IGSF8,NUDT5,WDR62,STYXL1,PPM1K,LRRC1,SON,UQCRQ,SNF8,CPSF3
,DBNDD2,RPS6,RTEL1,SRSF1,SAP30BP,TRNT1,ITGB1BP1,MED15,SLC39A13,EIF3M,CDC42,USE1,ASF1B,LI
EM67,YBX3,BLOC1S1,CACYBP,RHOF,SMARCB1,PRR13,RNH1,DNASE1L1,KIF4A,RBCK1,PTOV1,FASTK,TME
EM67,YBX3,BLOC1S1,CACYBP,RHOF,SMARCB1,PRR13,RNH1,DNASE1L1,KIF4A,RBCK1,PTOV1,FASTK,TME
1,SULF2,RAPGEF5,SSU72,FAM122C,SRP9,IQCG,PILRB,YBX3,CACYBP,SMARCB1,RPL38,RNH1,DNASE1L1,
1,SULF2,RAPGEF5,SSU72,FAM122C,SRP9,IQCG,PILRB,YBX3,CACYBP,SMARCB1,RPL38,RNH1,DNASE1L1,
P1,LAS1L,HMGB1,SULF2,BAIAP2,RAPGEF5,SSU72,MYL6B,SRP9,IQCG,TMEM67,SULT1A3,YBX3,BLOC1S1,
B,IQCG,BLOC1S1,CIRBP,RHOF,SMARCB1,PRR13,RNH1,RBCK1,PTOV1,FASTK,RNF32,LY75,ATG16L2,PARD
B,IQCG,BLOC1S1,CIRBP,RHOF,SMARCB1,PRR13,RNH1,RBCK1,PTOV1,FASTK,RNF32,LY75,ATG16L2,PARD
A,SRSF2,MRPS22,HDAC9,KRT15,OSMR,PPFIBP1,LAS1L,HMGB1,SULF2,CYB561A3,BAIAP2,RAPGEF5,SSU
A,SRSF2,MRPS22,HDAC9,KRT15,OSMR,PPFIBP1,LAS1L,HMGB1,SULF2,CYB561A3,BAIAP2,RAPGEF5,SSU
PGEF5,SSU72,FAM122C,SRP9,SULT1A3,FAM72A,YBX3,CACYBP,RHOF,SMARCB1,RPL38,RNH1,DNASE1L1
1A3,BAIAP2,RAPGEF5,SSU72,FAM122C,SRP9,IQCG,SULT1A3,FAM72A,BLOC1S1,CACYBP,RHOF,SMARCB1
LOC1S1,CACYBP,CIRBP,RPL38,RNH1,DNASE1L1,KIF4A,RBCK1,RCC1,SNRPB,SLC3A2,DHPS,ACTR6,ARRDC1
LOC1S1,CACYBP,CIRBP,RPL38,RNH1,DNASE1L1,KIF4A,RBCK1,RCC1,SNRPB,SLC3A2,DHPS,ACTR6,ARRDC1
,ZNF410,RUVBL2,NUDT5,WDR62,STYXL1,PPM1K,IRF9,DST,SON,UQCRQ,CPSF3L,TSG101,ATG12,NIP7,RP
,LRRC1,CDC37,RPS16,CPSF3L,TSG101,RPS6,RPL29,RTEL1,SRSF1,AUP1,TRNT1,KXD1,MED15,SH3YL1,C1O
MARCB1,PRR13,STARD10,RNH1,DNASE1L1,KIF4A,RNF32,SNRPB,SLC3A2,ACTR6,SEC62,SET,CTSC,ZNF23
RP9,IQCG,TMEM67,CMC2,BLOC1S1,CACYBP,CIRBP,RHOF,RPL38,PRR13,RNH1,KIF4A,RBCK1,RNF32,LY75
RP9,IQCG,TMEM67,CMC2,BLOC1S1,CACYBP,CIRBP,RHOF,RPL38,PRR13,RNH1,KIF4A,RBCK1,RNF32,LY75
,UQCRQ,SNF8,HMGA2,CPSF3L,TSG101,NIP7,RPL29,AUP1,YIPF3,TRNT1,MED15,UBE2I,EIF3M,CDC42,FBX
YL6B,IQCG,BLOC1S1,CACYBP,CIRBP,RHOF,SMARCB1,PRR13,RNH1,KIF4A,RBCK1,PTOV1,FASTK,RNF32,LY
YL6B,IQCG,BLOC1S1,CACYBP,CIRBP,RHOF,SMARCB1,PRR13,RNH1,KIF4A,RBCK1,PTOV1,FASTK,RNF32,LY
YL6B,IQCG,BLOC1S1,CACYBP,CIRBP,RHOF,SMARCB1,PRR13,RNH1,KIF4A,RBCK1,PTOV1,FASTK,RNF32,LY
YL6B,IQCG,BLOC1S1,CACYBP,CIRBP,RHOF,SMARCB1,PRR13,RNH1,KIF4A,RBCK1,PTOV1,FASTK,RNF32,LY
PPFIBP1,SEMA4D,LAS1L,BAIAP2,RAPGEF5,SSU72,MYL6B,FAM122C,SRP9,IQCG,TMEM67,PILRB,FAM72A,
PPFIBP1,SEMA4D,LAS1L,BAIAP2,RAPGEF5,SSU72,MYL6B,FAM122C,SRP9,IQCG,TMEM67,PILRB,FAM72A,
B1,SULF2,BAIAP2,RAPGEF5,SSU72,MYL6B,FAM122C,IQCG,SULT1A3,FAM72A,CMC2,YBX3,CACYBP,CIRBP
A3,PCNA,SRSF2,MRPS22,HDAC9,OSMR,PPFIBP1,LAS1L,HMGB1,SULF2,BAIAP2,RAPGEF5,SSU72,MYL6B,
A3,PCNA,SRSF2,MRPS22,HDAC9,OSMR,PPFIBP1,LAS1L,HMGB1,SULF2,BAIAP2,RAPGEF5,SSU72,MYL6B,
IBP1,SEMA4D,HMGB1,SULF2,CYB561A3,BAIAP2,RAPGEF5,SSU72,MYL6B,FAM122C,SRP9,IQCG,PILRB,FA
P2,SSU72,MYL6B,SRP9,IQCG,PILRB,FAM72A,CMC2,YBX3,CACYBP,RHOF,RPL38,PRR13,STARD10,RNH1,D
MR,HMGB1,SULF2,BAIAP2,RAPGEF5,SSU72,MYL6B,FAM122C,SRP9,IQCG,UAP1,PILRB,FAM72A,YBX3,CA
MR,HMGB1,SULF2,BAIAP2,RAPGEF5,SSU72,MYL6B,FAM122C,SRP9,IQCG,UAP1,PILRB,FAM72A,YBX3,CA
7,FAM72A,CMC2,BLOC1S1,CACYBP,CIRBP,RHOF,SMARCB1,RPL38,PRR13,RNH1,KIF4A,RNF32,LY75,ATG1
7,FAM72A,CMC2,BLOC1S1,CACYBP,CIRBP,RHOF,SMARCB1,RPL38,PRR13,RNH1,KIF4A,RNF32,LY75,ATG1
,CIRBP,RHOF,SMARCB1,PRR13,RNH1,DNASE1L1,RBCK1,PTOV1,FASTK,LY75,ATG16L2,TCAIM,SLC3A2,DH
,CIRBP,RHOF,SMARCB1,PRR13,RNH1,DNASE1L1,RBCK1,PTOV1,FASTK,LY75,ATG16L2,TCAIM,SLC3A2,DH
GB1,SULF2,BAIAP2,RAPGEF5,MYL6B,FAM122C,SRP9,IQCG,UAP1,PILRB,FAM72A,YBX3,CACYBP,RHOF,RP
1,RHOF,SMARCB1,PRR13,RNH1,DNASE1L1,RBCK1,PTOV1,FASTK,LY75,ATG16L2,TCAIM,SLC3A2,DHPS,SE

1,RHOF,SMARCB1,PRR13,RNH1,DNASE1L1,RBCK1,PTOV1,FASTK,LY75,ATG16L2,TCAIM,SLC3A2,DHPS,SE
,TTC9C,RPS6,RPL29,SAP30BP,YIPF3,TRNT1,MED15,SH3YL1,SLC39A13,YIF1A,EIF3M,CDC42,TMEM205,AS
BAIAP2,RAPGEF5,SSU72,FAM122C,SRP9,IQCG,PILRB,YBX3,RHOF,SMARCB1,RPL38,STARD10,DNASE1L1,
F2,MRPS22,KRT15,OSMR,PPFIBP1,SEMA4D,LAS1L,HMGB1,SULF2,CYB561A3,BAIAP2,RAPGEF5,SSU72,M
F2,MRPS22,KRT15,OSMR,PPFIBP1,SEMA4D,LAS1L,HMGB1,SULF2,CYB561A3,BAIAP2,RAPGEF5,SSU72,M
PGEF5,SSU72,SRP9,IQCG,TMEM67,SULT1A3,PILRB,FAM72A,BLOC1S1,CACYBP,CIRBP,SMARCB1,RPL38,S
PGEF5,SSU72,SRP9,IQCG,TMEM67,SULT1A3,PILRB,FAM72A,BLOC1S1,CACYBP,CIRBP,SMARCB1,RPL38,S
TTLL5,PSMA6,RCE1,MRPL33,STYXL1,FIS1,CPSF3L,TSG101,NIP7,TTC9C,TMX2,DBNDD2,TMED4,RPS6,RPL2
L6B,IQCG,PILRB,FAM72A,BLOC1S1,CACYBP,CIRBP,RHOF,SMARCB1,PRR13,RNH1,KIF4A,RBCK1,PTOV1,FA
L6B,IQCG,PILRB,FAM72A,BLOC1S1,CACYBP,CIRBP,RHOF,SMARCB1,PRR13,RNH1,KIF4A,RBCK1,PTOV1,FA
GEF5,SSU72,FAM122C,IQCG,SULT1A3,FAM72A,CACYBP,CIRBP,RHOF,SMARCB1,PRR13,RNH1,DNASE1L1
GEF5,SSU72,FAM122C,IQCG,SULT1A3,FAM72A,CACYBP,CIRBP,RHOF,SMARCB1,PRR13,RNH1,DNASE1L1
,HDAC9,KRT15,OSMR,PPFIBP1,LAS1L,HMGB1,SULF2,BAIAP2,RAPGEF5,SSU72,FAM122C,SRP9,IQCG,SULT
,HDAC9,KRT15,OSMR,PPFIBP1,LAS1L,HMGB1,SULF2,BAIAP2,RAPGEF5,SSU72,FAM122C,SRP9,IQCG,SULT
U72,FAM122C,SRP9,IQCG,YBX3,CACYBP,RHOF,SMARCB1,STARD10,RNH1,RBCK1,PTOV1,FASTK,TMEM1
M67,SULT1A3,PILRB,YBX3,BLOC1S1,CACYBP,CIRBP,RHOF,SMARCB1,RPL38,KIF4A,RBCK1,PTOV1,FASTK,
M67,SULT1A3,PILRB,YBX3,BLOC1S1,CACYBP,CIRBP,RHOF,SMARCB1,RPL38,KIF4A,RBCK1,PTOV1,FASTK,
M67,SULT1A3,PILRB,YBX3,BLOC1S1,CACYBP,CIRBP,RHOF,SMARCB1,RPL38,KIF4A,RBCK1,PTOV1,FASTK,
M67,SULT1A3,PILRB,YBX3,BLOC1S1,CACYBP,CIRBP,RHOF,SMARCB1,RPL38,KIF4A,RBCK1,PTOV1,FASTK,
M67,SULT1A3,PILRB,YBX3,BLOC1S1,CACYBP,CIRBP,RHOF,SMARCB1,RPL38,KIF4A,RBCK1,PTOV1,FASTK,
M67,SULT1A3,PILRB,YBX3,BLOC1S1,CACYBP,CIRBP,RHOF,SMARCB1,RPL38,KIF4A,RBCK1,PTOV1,FASTK,
F,SMARCB1,PRR13,RNH1,KIF4A,RBCK1,PTOV1,FASTK,RNF32,LY75,ATG16L2,PARD3,TCAIM,ACTR6,SET,B
F,SMARCB1,PRR13,RNH1,KIF4A,RBCK1,PTOV1,FASTK,RNF32,LY75,ATG16L2,PARD3,TCAIM,ACTR6,SET,B
1S1,CACYBP,CIRBP,RHOF,SMARCB1,RPL38,PRR13,STARD10,RNH1,DNASE1L1,KIF4A,RBCK1,RNF32,RCC1,
1S1,CACYBP,CIRBP,RHOF,SMARCB1,RPL38,PRR13,STARD10,RNH1,DNASE1L1,KIF4A,RBCK1,RNF32,RCC1,
1,KIF4A,RBCK1,RNF32,RCC1,SNRPB,PARD3,SLC3A2,DHPS,ACTR6,ZNF235,TRPT1,DBI,ARPC2,PMFBP1,LIM
1,KIF4A,RBCK1,RNF32,RCC1,SNRPB,PARD3,SLC3A2,DHPS,ACTR6,ZNF235,TRPT1,DBI,ARPC2,PMFBP1,LIM
M122C,IQCG,PILRB,FAM72A,BLOC1S1,CACYBP,CIRBP,RHOF,SMARCB1,PRR13,RNH1,KIF4A,RBCK1,PTOV
M122C,IQCG,PILRB,FAM72A,BLOC1S1,CACYBP,CIRBP,RHOF,SMARCB1,PRR13,RNH1,KIF4A,RBCK1,PTOV
2,UFD1L,ARHGEF39,ACOT8,AP3B2,PCMT1,STYXL1,LZTFL1,FIS1,MAGED2,SNF8,CPSF3L,TSG101,FUZ,RTEL
2,UFD1L,ARHGEF39,ACOT8,AP3B2,PCMT1,STYXL1,LZTFL1,FIS1,MAGED2,SNF8,CPSF3L,TSG101,FUZ,RTEL
,BLOC1S1,CACYBP,CIRBP,RHOF,SMARCB1,PRR13,RNH1,DNASE1L1,KIF4A,RBCK1,PTOV1,FASTK,TMEM13
,BLOC1S1,CACYBP,CIRBP,RHOF,SMARCB1,PRR13,RNH1,DNASE1L1,KIF4A,RBCK1,PTOV1,FASTK,TMEM13
MD5,NUDT22,UFD1L,SH2B1,RPL13A,ARHGEF39,PCMT1,NUDT5,STYXL1,RANBP1,VRK3,FIS1,IRF9,STIP1,L
MD5,NUDT22,UFD1L,SH2B1,RPL13A,ARHGEF39,PCMT1,NUDT5,STYXL1,RANBP1,VRK3,FIS1,IRF9,STIP1,L
IAP2,RAPGEF5,SSU72,FAM122C,SRP9,IQCG,PILRB,YBX3,CACYBP,RHOF,SMARCB1,RPL38,STARD10,DNA
IAP2,RAPGEF5,SSU72,FAM122C,SRP9,IQCG,PILRB,YBX3,CACYBP,RHOF,SMARCB1,RPL38,STARD10,DNA
ZNF235,BOP1,ARRDC1,TRPT1,RBP5,ARPC2,LIMK2,NOB1,GPATCH2L,BUD31,NPRL2,SCARB1,CTTN,NUDT2
U72,SRP9,IQCG,TMEM67,PILRB,CACYBP,SMARCB1,RPL38,RNH1,KIF4A,PTOV1,FASTK,TMEM135,LY75,A
U72,SRP9,IQCG,TMEM67,PILRB,CACYBP,SMARCB1,RPL38,RNH1,KIF4A,PTOV1,FASTK,TMEM135,LY75,A
6B,SRP9,IQCG,FAM72A,BLOC1S1,CIRBP,RHOF,SMARCB1,PRR13,RNH1,RBCK1,PTOV1,RNF32,LY75,ATG16
6B,SRP9,IQCG,FAM72A,BLOC1S1,CIRBP,RHOF,SMARCB1,PRR13,RNH1,RBCK1,PTOV1,RNF32,LY75,ATG16
3,PILRB,FAM72A,BLOC1S1,SMARCB1,STARD10,RNH1,DNASE1L1,RBCK1,PTOV1,FASTK,TMEM135,LY75,
L38,STARD10,RNH1,DNASE1L1,PTOV1,FASTK,TMEM135,SNRPB,PARD3,DHPS,ACTR6,MAP3K7CL,SET,AR
L38,STARD10,RNH1,DNASE1L1,PTOV1,FASTK,TMEM135,SNRPB,PARD3,DHPS,ACTR6,MAP3K7CL,SET,AR
5,OSMR,LAS1L,HMGB1,SULF2,BAIAP2,RAPGEF5,SSU72,MYL6B,FAM122C,IQCG,PILRB,FAM72A,CMC2,YB
5,OSMR,LAS1L,HMGB1,SULF2,BAIAP2,RAPGEF5,SSU72,MYL6B,FAM122C,IQCG,PILRB,FAM72A,CMC2,YB

FIS1,MAGED2,SON,CPSF3L,FUZ,RTEL1,YIPF3,RABL5,ITGB1BP1,MED15,C1ORF198,UGGT2,CDC42,CARD14

FIS1,MAGED2,SON,CPSF3L,FUZ,RTEL1,YIPF3,RABL5,ITGB1BP1,MED15,C1ORF198,UGGT2,CDC42,CARD14

SMR,SEMA4D,LAS1L,SULF2,CYB561A3,BAIAP2,RAPGEF5,SSU72,FAM122C,SRP9,IQCG,TMEM67,SULT1A3,

SMR,SEMA4D,LAS1L,SULF2,CYB561A3,BAIAP2,RAPGEF5,SSU72,FAM122C,SRP9,IQCG,TMEM67,SULT1A3,

AIAP2,RAPGEF5,MYL6B,IQCG,FAM72A,BLOC1S1,CACYBP,CIRBP,RHOF,SMARCB1,RPL38,PRR13,STARD10,

AIAP2,RAPGEF5,MYL6B,IQCG,FAM72A,BLOC1S1,CACYBP,CIRBP,RHOF,SMARCB1,RPL38,PRR13,STARD10,

CYB561A3,BAIAP2,RAPGEF5,SSU72,MYL6B,FAM122C,SRP9,IQCG,PILRB,YBX3,CACYBP,CIRBP,RHOF,SMAR

CYB561A3,BAIAP2,RAPGEF5,SSU72,MYL6B,FAM122C,SRP9,IQCG,PILRB,YBX3,CACYBP,CIRBP,RHOF,SMAR

PRL2,CTTN,EIF5A,COMMD5,UFD1L,RPL13A,ARHGEF39,ACOT8,AP3B2,NUDT5,STYXL1,LZTFL1,FIS1,MAGED

PRL2,CTTN,EIF5A,COMMD5,UFD1L,RPL13A,ARHGEF39,ACOT8,AP3B2,NUDT5,STYXL1,LZTFL1,FIS1,MAGED

IAP2,RAPGEF5,SSU72,SRP9,IQCG,TMEM67,SULT1A3,FAM72A,BLOC1S1,CACYBP,RHOF,SMARCB1,RPL38,

IAP2,RAPGEF5,SSU72,SRP9,IQCG,TMEM67,SULT1A3,FAM72A,BLOC1S1,CACYBP,RHOF,SMARCB1,RPL38,

RHOF,SMARCB1,PRR13,RNH1,RBCK1,PTOV1,FASTK,RNF32,LY75,ATG16L2,SNRPB,PARD3,SLC3A2,SET,ZNF

RHOF,SMARCB1,PRR13,RNH1,RBCK1,PTOV1,FASTK,RNF32,LY75,ATG16L2,SNRPB,PARD3,SLC3A2,SET,ZNF

NASE1L1,RBCK1,PTOV1,FASTK,LY75,ATG16L2,TCAIM,DHPS,SET,ARRDC1,RBP5,LIMK2,NUMA1,TBL2,SCAR

NASE1L1,RBCK1,PTOV1,FASTK,LY75,ATG16L2,TCAIM,DHPS,SET,ARRDC1,RBP5,LIMK2,NUMA1,TBL2,SCAR

C3HAV1,DBI,ARPC2,LIMK2,GPATCH2L,BUD31,NPRL2,ARHGAP6,CTTN,EIF5A,COMMD5,NUDT22,UFD1L,AF

C3HAV1,DBI,ARPC2,LIMK2,GPATCH2L,BUD31,NPRL2,ARHGAP6,CTTN,EIF5A,COMMD5,NUDT22,UFD1L,AF

RHOF,SMARCB1,PRR13,RNH1,RBCK1,PTOV1,FASTK,RNF32,LY75,ATG16L2,SLC3A2,SEC62,SET,CTSC,ARRDC

RHOF,SMARCB1,PRR13,RNH1,RBCK1,PTOV1,FASTK,RNF32,LY75,ATG16L2,SLC3A2,SEC62,SET,CTSC,ARRDC

V1,FASTK,TMEM135,SNRPB,PARD3,DHPS,ACTR6,MAP3K7CL,SET,BOP1,ARRDC1,KLC1,SCAF11,LIMK2,VDA

V1,FASTK,TMEM135,SNRPB,PARD3,DHPS,ACTR6,MAP3K7CL,SET,BOP1,ARRDC1,KLC1,SCAF11,LIMK2,VDA

,BAIAP2,RAPGEF5,SSU72,MYL6B,IQCG,BLOC1S1,CACYBP,CIRBP,RHOF,SMARCB1,PRR13,RNH1,DNASE1L1

,BAIAP2,RAPGEF5,SSU72,MYL6B,IQCG,BLOC1S1,CACYBP,CIRBP,RHOF,SMARCB1,PRR13,RNH1,DNASE1L1

FIS1,MAGED2,SON,CPSF3L,FUZ,RTEL1,RABL5,ITGB1BP1,MED15,C1ORF198,UGGT2,CDC42,CARD14,FBXW

FIS1,MAGED2,SON,CPSF3L,FUZ,RTEL1,RABL5,ITGB1BP1,MED15,C1ORF198,UGGT2,CDC42,CARD14,FBXW

EC62,SET,CTSC,ZNF235,BOP1,RBP5,SIRT6,BUD31,SAE1,TSPAN11,EIF5A,UFD1L,USP38,TAZ,SH2B1,RPL13A

2,CTTN,EIF5A,COMMD5,UFD1L,USP38,SH2B1,NOP16,RPL13A,ZNF410,RUVBL2,IGSF8,NUDT5,STYXL1,POL

LY75,PARD3,TCAIM,SET,BOP1,ARRDC1,TRPT1,KLC1,LIMK2,SIRT6,NUMA1,BUD31,TBL2,SCARB1,CTTN,ITSN

1L,THYN1,AP3B2,SLC35G1,RCE1,IGSF8,NUDT5,IL1RAP,POLD4,FIS1,IRF9,PHF19,MAGED2,CDC37,CPSF3L,

1L,THYN1,AP3B2,SLC35G1,RCE1,IGSF8,NUDT5,IL1RAP,POLD4,FIS1,IRF9,PHF19,MAGED2,CDC37,CPSF3L,

AIAP2,RAPGEF5,SSU72,FAM122C,IQCG,FAM72A,YBX3,BLOC1S1,CACYBP,CIRBP,RHOF,SMARCB1,RPL38,S

AIAP2,RAPGEF5,SSU72,FAM122C,IQCG,FAM72A,YBX3,BLOC1S1,CACYBP,CIRBP,RHOF,SMARCB1,RPL38,S

FIBP1,SEMA4D,HMGB1,SULF2,CYB561A3,BAIAP2,RAPGEF5,SSU72,SRP9,FAM72A,YBX3,CACYBP,CIRBP,RH

FIBP1,SEMA4D,HMGB1,SULF2,CYB561A3,BAIAP2,RAPGEF5,SSU72,SRP9,FAM72A,YBX3,CACYBP,CIRBP,RH

GEF5,SSU72,SRP9,IQCG,TMEM67,PILRB,YBX3,CACYBP,SMARCB1,RPL38,STARD10,RNH1,KIF4A,PTOV1,FA

GEF5,SSU72,SRP9,IQCG,TMEM67,PILRB,YBX3,CACYBP,SMARCB1,RPL38,STARD10,RNH1,KIF4A,PTOV1,FA

L38,RNH1,DNASE1L1,KIF4A,RBCK1,RNF32,RCC1,SNRPB,PARD3,SLC3A2,DHPS,ACTR6,CTSC,ZNF235,BOP1,

L38,RNH1,DNASE1L1,KIF4A,RBCK1,RNF32,RCC1,SNRPB,PARD3,SLC3A2,DHPS,ACTR6,CTSC,ZNF235,BOP1,

1,PRR13,RNH1,DNASE1L1,RBCK1,PTOV1,FASTK,RNF32,LY75,ATG16L2,SLC3A2,ARRDC1,TRPT1,KLC1,LIMK

1,PRR13,RNH1,DNASE1L1,RBCK1,PTOV1,FASTK,RNF32,LY75,ATG16L2,SLC3A2,ARRDC1,TRPT1,KLC1,LIMK

,ACOT8,STYXL1,LZTFL1,FIS1,MAGED2,SON,CPSF3L,FUZ,RPS25,RTEL1,RABL5,ITGB1BP1,MED15,C1ORF19

,ACOT8,STYXL1,LZTFL1,FIS1,MAGED2,SON,CPSF3L,FUZ,RPS25,RTEL1,RABL5,ITGB1BP1,MED15,C1ORF19

MA4D,LAS1L,HMGB1,SULF2,CYB561A3,BAIAP2,RAPGEF5,SSU72,MYL6B,FAM122C,SRP9,IQCG,TMEM67,S

MA4D,LAS1L,HMGB1,SULF2,CYB561A3,BAIAP2,RAPGEF5,SSU72,MYL6B,FAM122C,SRP9,IQCG,TMEM67,S

T22,USP38,SH2B1,RPL13A,ARHGEF39,PSMA6,ACOT8,ZNF410,NUDT5,SNRPN,STYXL1,VRK3,IRF9,UQCRQ,

T22,USP38,SH2B1,RPL13A,ARHGEF39,PSMA6,ACOT8,ZNF410,NUDT5,SNRPN,STYXL1,VRK3,IRF9,UQCRQ,

PATCH2L,BUD31,TBL2,SCARB1,TSPAN11,ITSN1,EIF5A,COMMD5,NUDT22,NOP16,TTLL5,PSMA6,RUVBL2,
PATCH2L,BUD31,TBL2,SCARB1,TSPAN11,ITSN1,EIF5A,COMMD5,NUDT22,NOP16,TTLL5,PSMA6,RUVBL2,
YBP,CIRBP,RHOF,SMARCB1,PRR13,RNH1,RBCK1,PTOV1,RNF32,LY75,ATG16L2,RCC1,SLC3A2,ARRDC1,KL
YBP,CIRBP,RHOF,SMARCB1,PRR13,RNH1,RBCK1,PTOV1,RNF32,LY75,ATG16L2,RCC1,SLC3A2,ARRDC1,KL
,EIF5A,COMMD5,DBNDD1,SH2B1,NOP16,RPL13A,NUDT5,STYXL1,POLD4,VRK3,LZTFL1,FIS1,SNF8,CPSF3
L1,RBCK1,FASTK,RCC1,SNRPB,SLC3A2,DHPS,ACTR6,TRPT1,DBI,LIMK2,SIRT6,NUMA1,BUD31,NPRL2,TBL2,
L1,RBCK1,FASTK,RCC1,SNRPB,SLC3A2,DHPS,ACTR6,TRPT1,DBI,LIMK2,SIRT6,NUMA1,BUD31,NPRL2,TBL2,
P1,ARRDC1,RBP5,NUMA1,TBL2,SCARB1,CTTN,TSPAN11,EIF5A,COMMD5,RUNX1,NOP16,TTLL5,PSMA6,
P1,ARRDC1,RBP5,NUMA1,TBL2,SCARB1,CTTN,TSPAN11,EIF5A,COMMD5,RUNX1,NOP16,TTLL5,PSMA6,
1,LAS1L,HMGB1,BAIAP2,RAPGEF5,SSU72,SRP9,IQCG,PILRB,FAM72A,YBX3,CACYBP,RHOF,SMARCB1,RPL
GEF5,SSU72,FAM122C,SRP9,IQCG,YBX3,CACYBP,CIRBP,RHOF,SMARCB1,STARD10,RNH1,RBCK1,PTOV1,
GEF5,SSU72,FAM122C,SRP9,IQCG,YBX3,CACYBP,CIRBP,RHOF,SMARCB1,STARD10,RNH1,RBCK1,PTOV1,
HPS,ACTR6,MAP3K7CL,DBI,LIMK2,GPATCH2L,BUD31,NPRL2,CTTN,EIF5A,COMMD5,UFD1L,USP38,ARHG
HPS,ACTR6,MAP3K7CL,DBI,LIMK2,GPATCH2L,BUD31,NPRL2,CTTN,EIF5A,COMMD5,UFD1L,USP38,ARHG
MEM67,PILRB,YBX3,CACYBP,RHOF,SMARCB1,RPL38,RNH1,PTOV1,FASTK,LY75,ATG16L2,PARD3,TCAIM,S
MEM67,PILRB,YBX3,CACYBP,RHOF,SMARCB1,RPL38,RNH1,PTOV1,FASTK,LY75,ATG16L2,PARD3,TCAIM,S
MGB1,BAIAP2,RAPGEF5,SSU72,MYL6B,SRP9,IQCG,SULT1A3,FAM72A,YBX3,CACYBP,CIRBP,RHOF,SMARC
MGB1,BAIAP2,RAPGEF5,SSU72,MYL6B,SRP9,IQCG,SULT1A3,FAM72A,YBX3,CACYBP,CIRBP,RHOF,SMARC
2,FAM122C,IQCG,FAM72A,YBX3,BLOC1S1,CACYBP,CIRBP,RHOF,SMARCB1,RPL38,STARD10,RNH1,DNASE
L38,RNH1,DNASE1L1,RBCK1,LY75,RCC1,SNRPB,SLC3A2,DHPS,ACTR6,TRPT1,DBI,LIMK2,SIRT6,NUMA1,G
L38,RNH1,DNASE1L1,RBCK1,LY75,RCC1,SNRPB,SLC3A2,DHPS,ACTR6,TRPT1,DBI,LIMK2,SIRT6,NUMA1,G
3A2,DHPS,ACTR6,TRPT1,ZC3HAV1,DBI,SIRT6,NUMA1,BUD31,CRYGS,CTTN,TSPAN11,EIF5A,COMMD5,NU
3A2,DHPS,ACTR6,TRPT1,ZC3HAV1,DBI,SIRT6,NUMA1,BUD31,CRYGS,CTTN,TSPAN11,EIF5A,COMMD5,NU
SMR,PPFIBP1,HMGB1,SULF2,CYB561A3,BAIAP2,RAPGEF5,SSU72,MYL6B,IQCG,TMEM67,FAM72A,YBX3,B
,LY75,PARD3,TCAIM,ACTR6,SET,BOP1,ARRDC1,KLC1,LIMK2,SIRT6,NUMA1,BUD31,TBL2,SCARB1,CTTN,IT
1,CYB561A3,BAIAP2,RAPGEF5,SSU72,FAM122C,SRP9,IQCG,TMEM67,SULT1A3,PILRB,BLOC1S1,CACYBP,
L,TSG101,RPS25,COX4I1,RPS6,RPL29,TRNT1,CDC42,FBXW9,ASF1B,CCDC25,ENDOV,LYPLA2,PPAN,RPL36
H1,RBCK1,PTOV1,TMEM135,RNF32,LY75,ATG16L2,SLC3A2,ARRDC1,RBP5,LIMK2,NUMA1,TBL2,SCARB1,
H1,RBCK1,PTOV1,TMEM135,RNF32,LY75,ATG16L2,SLC3A2,ARRDC1,RBP5,LIMK2,NUMA1,TBL2,SCARB1,
A3,PCNA,SRSF2,MRPS22,OSMR,PPFIBP1,SEMA4D,LAS1L,HMGB1,SULF2,BAIAP2,RAPGEF5,SSU72,FAM12
A3,PCNA,SRSF2,MRPS22,OSMR,PPFIBP1,SEMA4D,LAS1L,HMGB1,SULF2,BAIAP2,RAPGEF5,SSU72,FAM12
SMR,PPFIBP1,SEMA4D,HMGB1,SULF2,BAIAP2,SSU72,MYL6B,FAM122C,SRP9,IQCG,UAP1,PILRB,FAM72
SMR,PPFIBP1,SEMA4D,HMGB1,SULF2,BAIAP2,SSU72,MYL6B,FAM122C,SRP9,IQCG,UAP1,PILRB,FAM72
RPS22,HDAC9,KRT15,OSMR,PPFIBP1,SEMA4D,LAS1L,HMGB1,SULF2,CYB561A3,BAIAP2,RAPGEF5,SSU72
1,STARD10,RNH1,PTOV1,ATG16L2,SNRPB,PARD3,TCAIM,ACTR6,SET,CTSC,BOP1,SCAF11,DBI,LIMK2,NPR
B,FAM72A,BLOC1S1,CACYBP,RPL38,STARD10,DNASE1L1,KIF4A,RBCK1,PTOV1,FASTK,TMEM135,ATG16
2,LY75,ATG16L2,PARD3,TCAIM,SLC3A2,SET,ARRDC1,KLC1,LIMK2,SIRT6,NUMA1,BUD31,TBL2,SCARB1,CT
,ATG16L2,ARRDC1,KLC1,LIMK2,WDR19,SIRT6,NUMA1,GPATCH2L,TBL2,SCARB1,CTTN,ITSN1,AMZ2,TAZ,
MARCB1,PRR13,RNH1,RBCK1,PTOV1,LY75,ATG16L2,PARD3,DHPS,SEC62,SET,ARRDC1,TRPT1,RBP5,ZC3H
MARCB1,PRR13,RNH1,RBCK1,PTOV1,LY75,ATG16L2,PARD3,DHPS,SEC62,SET,ARRDC1,TRPT1,RBP5,ZC3H
MD5,NOP16,RPL13A,PSMA6,SNRPN,STYXL1,LZTFL1,FIS1,SON,UQCRQ,SNF8,RPS16,CPSF3L,TSG101,FAM
SRSF2,MRPS22,KRT15,OSMR,PPFIBP1,LAS1L,HMGB1,SULF2,CYB561A3,BAIAP2,RAPGEF5,SSU72,MYL6B,
SRSF2,MRPS22,KRT15,OSMR,PPFIBP1,LAS1L,HMGB1,SULF2,CYB561A3,BAIAP2,RAPGEF5,SSU72,MYL6B,
L,SULF2,BAIAP2,RAPGEF5,SSU72,MYL6B,IQCG,PILRB,FAM72A,YBX3,BLOC1S1,CACYBP,CIRBP,RHOF,SMAR
L,SULF2,BAIAP2,RAPGEF5,SSU72,MYL6B,IQCG,PILRB,FAM72A,YBX3,BLOC1S1,CACYBP,CIRBP,RHOF,SMAR
CYBP,SMARCB1,RPL38,PRR13,RNH1,DNASE1L1,KIF4A,PTOV1,FASTK,TMEM135,RCC1,PARD3,DHPS,SET,C
CYBP,SMARCB1,RPL38,PRR13,RNH1,DNASE1L1,KIF4A,PTOV1,FASTK,TMEM135,RCC1,PARD3,DHPS,SET,C

PPM1K,SON,SNF8,CPSF3L,METTL17,NIP7,COX4I1,THEM4,RPL29,FOXRED1,TRNT1,MICAL1,UBE2I,YIF1A,P
RHOF,SMARCB1,RNH1,DNASE1L1,RBCK1,PTOV1,FASTK,ATG16L2,RCC1,DHPS,SET,ZNF235,BOP1,ARRDC1,
RHOF,SMARCB1,RNH1,DNASE1L1,RBCK1,PTOV1,FASTK,ATG16L2,RCC1,DHPS,SET,ZNF235,BOP1,ARRDC1,
1B,LIG1,LYPLA2,CYB561,ATP5J,TRMT112,SAT1,DDOST,LSR,PSME2,TOMM40,RABEP2,TADA3,RAD23A,PD
EM67,PILRB,CMC2,BLOC1S1,CIRBP,SMARCB1,RPL38,PRR13,RNH1,RBCK1,RCC1,SNRPB,TCAIM,SLC3A2,DH
EM67,PILRB,CMC2,BLOC1S1,CIRBP,SMARCB1,RPL38,PRR13,RNH1,RBCK1,RCC1,SNRPB,TCAIM,SLC3A2,DH
NDOV,LYPLA2,PPAN,SUGT1,ZNF197,INO80E,TRMT112,CHPF,DDOST,LSR,TOMM40,TADA3,ANAPC11,MD
MYL6B,FAM122C,IQCG,TMEM67,PILRB,CMC2,CIRBP,RHOF,PRR13,STARD10,RNH1,KIF4A,RNF32,RCC1,SN
MYL6B,FAM122C,IQCG,TMEM67,PILRB,CMC2,CIRBP,RHOF,PRR13,STARD10,RNH1,KIF4A,RNF32,RCC1,SN
MRPS22,OSMR,PPFIBP1,SEMA4D,LAS1L,HMGB1,SULF2,BAIAP2,RAPGEF5,SSU72,FAM122C,IQCG,TMEM6
,HMGB1,BAIAP2,RAPGEF5,SSU72,MYL6B,FAM122C,SRP9,IQCG,TMEM67,PILRB,FAM72A,CMC2,YBX3,BLO
,HMGB1,BAIAP2,RAPGEF5,SSU72,MYL6B,FAM122C,SRP9,IQCG,TMEM67,PILRB,FAM72A,CMC2,YBX3,BLO
,PTOV1,FASTK,RNF32,LY75,PARD3,DHPS,SET,BOP1,ARRDC1,KLC1,LIMK2,VDAC3,NOB1,NUMA1,BUD31,T
TK,TMEM135,ATG16L2,SNRPB,PARD3,ACTR6,MAP3K7CL,SET,KLC1,SCAF11,ARPC2,LIMK2,VDAC3,WDR1
TK,TMEM135,ATG16L2,SNRPB,PARD3,ACTR6,MAP3K7CL,SET,KLC1,SCAF11,ARPC2,LIMK2,VDAC3,WDR1
PRR13,RNH1,DNASE1L1,RBCK1,PTOV1,FASTK,LY75,RCC1,DHPS,SET,CTSC,ARRDC1,RBP5,LIMK2,NOB1,NU
PRR13,RNH1,DNASE1L1,RBCK1,PTOV1,FASTK,LY75,RCC1,DHPS,SET,CTSC,ARRDC1,RBP5,LIMK2,NOB1,NU
32,LY75,ATG16L2,SLC3A2,ARRDC1,RBP5,LIMK2,SIRT6,NUMA1,TBL2,SCARB1,CTTN,TSPAN11,DBNDD1,NC
32,LY75,ATG16L2,SLC3A2,ARRDC1,RBP5,LIMK2,SIRT6,NUMA1,TBL2,SCARB1,CTTN,TSPAN11,DBNDD1,NC
CNA,SRSF2,MRPS22,HDAC9,KRT15,OSMR,PPFIBP1,SEMA4D,LAS1L,HMGB1,SULF2,CYB561A3,BAIAP2,RA
CNA,SRSF2,MRPS22,HDAC9,KRT15,OSMR,PPFIBP1,SEMA4D,LAS1L,HMGB1,SULF2,CYB561A3,BAIAP2,RA
L17,NIP7,TRNT1,MED15,CLASRP,EIF3M,CDC42,FBXW9,EEF1B2,ENDOV,PPIL2,RPL28,LYPLA2,C17ORF62,Z
YPLA2,C17ORF62,CARS,LRRFIP1,ATP5J,TRMT112,DDOST,TOMM40,TADA3,MDH2,SEC31A,PAFAH1B2,PPI
KIF4A,RBCK1,RNF32,LY75,RCC1,SNRPB,PARD3,DHPS,ACTR6,ZNF235,DBI,ARPC2,SIRT6,GPATCH2L,BUD31
KIF4A,RBCK1,RNF32,LY75,RCC1,SNRPB,PARD3,DHPS,ACTR6,ZNF235,DBI,ARPC2,SIRT6,GPATCH2L,BUD31
LAS1L,HMGB1,SULF2,BAIAP2,RAPGEF5,SSU72,FAM122C,SRP9,IQCG,TMEM67,SULT1A3,PILRB,FAM72A,C
LAS1L,HMGB1,SULF2,BAIAP2,RAPGEF5,SSU72,FAM122C,SRP9,IQCG,TMEM67,SULT1A3,PILRB,FAM72A,C
CYBP,SMARCB1,RPL38,PRR13,STARD10,DNASE1L1,PTOV1,FASTK,TMEM135,ATG16L2,RCC1,SNRPB,PARD
CYBP,SMARCB1,RPL38,PRR13,STARD10,DNASE1L1,PTOV1,FASTK,TMEM135,ATG16L2,RCC1,SNRPB,PARD
LF2,BAIAP2,RAPGEF5,SSU72,FAM122C,SRP9,IQCG,TMEM67,SULT1A3,PILRB,FAM72A,CMC2,YBX3,BLOC1
LF2,BAIAP2,RAPGEF5,SSU72,FAM122C,SRP9,IQCG,TMEM67,SULT1A3,PILRB,FAM72A,CMC2,YBX3,BLOC1
A,CMC2,YBX3,CACYBP,RHOF,STARD10,RNH1,DNASE1L1,RBCK1,PTOV1,RNF32,ATG16L2,SNRPB,PARD3,S
A,CMC2,YBX3,CACYBP,RHOF,STARD10,RNH1,DNASE1L1,RBCK1,PTOV1,RNF32,ATG16L2,SNRPB,PARD3,S
A,CMC2,YBX3,CACYBP,RHOF,STARD10,RNH1,DNASE1L1,RBCK1,PTOV1,RNF32,ATG16L2,SNRPB,PARD3,S
A,CMC2,YBX3,CACYBP,RHOF,STARD10,RNH1,DNASE1L1,RBCK1,PTOV1,RNF32,ATG16L2,SNRPB,PARD3,S
LOC1S1,CACYBP,CIRBP,RHOF,SMARCB1,PRR13,RNH1,KIF4A,RBCK1,PTOV1,FASTK,RNF32,LY75,ATG16L2,P
LOC1S1,CACYBP,CIRBP,RHOF,SMARCB1,PRR13,RNH1,KIF4A,RBCK1,PTOV1,FASTK,RNF32,LY75,ATG16L2,P
LAS1L,HMGB1,SULF2,BAIAP2,RAPGEF5,SSU72,FAM122C,SRP9,IQCG,TMEM67,SULT1A3,PILRB,FAM72A,C
2A,BLOC1S1,CIRBP,RHOF,SMARCB1,PRR13,RNH1,RBCK1,PTOV1,FASTK,RNF32,LY75,ATG16L2,PARD3,TC
2A,BLOC1S1,CIRBP,RHOF,SMARCB1,PRR13,RNH1,RBCK1,PTOV1,FASTK,RNF32,LY75,ATG16L2,PARD3,TC
LF2,BAIAP2,RAPGEF5,SSU72,FAM122C,SRP9,IQCG,TMEM67,SULT1A3,PILRB,FAM72A,CMC2,YBX3,BLOC1
Y75,PARD3,TCAIM,ACTR6,SET,BOP1,ARRDC1,KLC1,LIMK2,SIRT6,NUMA1,BUD31,TBL2,SCARB1,CTTN,ITS
C2,BLOC1S1,CIRBP,RNH1,KIF4A,RBCK1,RNF32,ATG16L2,RCC1,SNRPB,DHPS,ACTR6,ZNF235,DBI,ARPC2,LI
C2,BLOC1S1,CIRBP,RNH1,KIF4A,RBCK1,RNF32,ATG16L2,RCC1,SNRPB,DHPS,ACTR6,ZNF235,DBI,ARPC2,LI
MYL6B,SRP9,SULT1A3,FAM72A,RHOF,SMARCB1,PRR13,STARD10,RNH1,DNASE1L1,RBCK1,PTOV1,TMEM
MYL6B,SRP9,SULT1A3,FAM72A,RHOF,SMARCB1,PRR13,STARD10,RNH1,DNASE1L1,RBCK1,PTOV1,TMEM
PPM1K,SON,SNF8,METTL17,NIP7,THEM4,RPL29,TRNT1,MICAL1,UBE2I,YIF1A,EIF3M,PCYT2,FBXW9,IFI27L

D2,RPS6,RTEL1,ITGB1BP1,SLC39A13,CDC42,USE1,CCDC25,RPL28,LIG1,LYPLA2,NUTF2,CDK11A,DGUOK,TF
,SULF2,BAIAP2,RAPGEF5,SSU72,FAM122C,SRP9,IQCG,TMEM67,SULT1A3,PILRB,FAM72A,CMC2,YBX3,CA
,SULF2,BAIAP2,RAPGEF5,SSU72,FAM122C,SRP9,IQCG,TMEM67,SULT1A3,PILRB,FAM72A,CMC2,YBX3,CA
ACYBP,CIRBP,STARD10,DNASE1L1,KIF4A,RBCK1,PTOV1,FASTK,TMEM135,LY75,ATG16L2,SNRPB,PARD3,DI
ACYBP,CIRBP,STARD10,DNASE1L1,KIF4A,RBCK1,PTOV1,FASTK,TMEM135,LY75,ATG16L2,SNRPB,PARD3,DI
ULF2,BAIAP2,RAPGEF5,SSU72,FAM122C,SRP9,IQCG,TMEM67,SULT1A3,PILRB,FAM72A,CMC2,YBX3,BLOC
ULF2,BAIAP2,RAPGEF5,SSU72,FAM122C,SRP9,IQCG,TMEM67,SULT1A3,PILRB,FAM72A,CMC2,YBX3,BLOC
P,RHOF,RPL38,PRR13,STARD10,DNASE1L1,KIF4A,RBCK1,FASTK,TMEM135,RNF32,LY75,RCC1,SNRPB,PARD
P,RHOF,RPL38,PRR13,STARD10,DNASE1L1,KIF4A,RBCK1,FASTK,TMEM135,RNF32,LY75,RCC1,SNRPB,PARD
5,ATG16L2,SNRPB,PARD3,DHPS,ARRDC1,KLC1,ARPC2,LIMK2,NOB1,SIRT6,NUMA1,GPATCH2L,TBL2,SCAR
1,SULF2,BAIAP2,RAPGEF5,SSU72,FAM122C,IQCG,PILRB,YBX3,BLOC1S1,CACYBP,CIRBP,RHOF,SMARCB1,R
1,SULF2,BAIAP2,RAPGEF5,SSU72,FAM122C,IQCG,PILRB,YBX3,BLOC1S1,CACYBP,CIRBP,RHOF,SMARCB1,R
D1,THEM4,RTEL1,RABL5,TRNT1,ITGB1BP1,MED15,UGGT2,FBXW9,CCDC25,EEF1B2,ENDOV,PPIL2,LIG1,AP
D1,THEM4,RTEL1,RABL5,TRNT1,ITGB1BP1,MED15,UGGT2,FBXW9,CCDC25,EEF1B2,ENDOV,PPIL2,LIG1,AP
1,SULF2,BAIAP2,RAPGEF5,SSU72,FAM122C,IQCG,YBX3,BLOC1S1,CACYBP,CIRBP,RHOF,SMARCB1,RPL38,S
1,SULF2,BAIAP2,RAPGEF5,SSU72,FAM122C,IQCG,YBX3,BLOC1S1,CACYBP,CIRBP,RHOF,SMARCB1,RPL38,S
3,BLOC1S1,CACYBP,CIRBP,SMARCB1,RPL38,STARD10,RNH1,DNASE1L1,FASTK,TMEM135,RNF32,LY75,AT
HMGB1,SULF2,BAIAP2,RAPGEF5,SSU72,FAM122C,TMEM67,SULT1A3,PILRB,FAM72A,CMC2,YBX3,BLOC1
D2,SON,SNF8,CPSF3L,NIP7,THEM4,RPL29,SAP30BP,YIPF3,SH3YL1,CLASRP,C1ORF198,EIF3M,IFI27L1,END
ARPC2,VDAC3,NOB1,SIRT6,SUMF2,SAE1,NPRL2,EIF5A,AMZ2,USP38,DBNDD1,RPL13A,ARHGEF39,TTLL5,F
ARPC2,VDAC3,NOB1,SIRT6,SUMF2,SAE1,NPRL2,EIF5A,AMZ2,USP38,DBNDD1,RPL13A,ARHGEF39,TTLL5,F
QCG,PILRB,YBX3,CIRBP,RHOF,SMARCB1,RNH1,PTOV1,FASTK,RNF32,LY75,RCC1,SNRPB,TCAIM,SLC3A2,DH
QCG,PILRB,YBX3,CIRBP,RHOF,SMARCB1,RNH1,PTOV1,FASTK,RNF32,LY75,RCC1,SNRPB,TCAIM,SLC3A2,DH
ULF2,BAIAP2,RAPGEF5,SSU72,FAM122C,SRP9,IQCG,TMEM67,SULT1A3,PILRB,FAM72A,CMC2,YBX3,BLOC
B1,SH2B1,NOP16,RPL13A,PSMA6,SPAG7,WDR62,RANBP1,UBE2L6,SLC15A3,PHF19,SON,SNF8,PTPN13,UI
AC3,NOB1,GPATCH2L,SUMF2,OSGEP,CRYGS,ITSN1,AMZ2,TAZ,SH2B1,GSTK1,RCE1,MRPL33,ZNF410,RUVI
AC3,NOB1,GPATCH2L,SUMF2,OSGEP,CRYGS,ITSN1,AMZ2,TAZ,SH2B1,GSTK1,RCE1,MRPL33,ZNF410,RUVI
BP1,LAS1L,HMGB1,SULF2,CYB561A3,BAIAP2,RAPGEF5,SSU72,FAM122C,SRP9,IQCG,SULT1A3,PILRB,FAM7
BP1,LAS1L,HMGB1,SULF2,CYB561A3,BAIAP2,RAPGEF5,SSU72,FAM122C,SRP9,IQCG,SULT1A3,PILRB,FAM7
RP9,IQCG,TMEM67,PILRB,BLOC1S1,CACYBP,SMARCB1,RPL38,STARD10,RNH1,KIF4A,PTOV1,FASTK,RNF32
RP9,IQCG,TMEM67,PILRB,BLOC1S1,CACYBP,SMARCB1,RPL38,STARD10,RNH1,KIF4A,PTOV1,FASTK,RNF32
ARCB1,RPL38,KIF4A,PTOV1,FASTK,ATG16L2,SNRPB,PARD3,ACTR6,MAP3K7CL,SEC62,CTSC,ARRDC1,KLC1,
ARCB1,RPL38,KIF4A,PTOV1,FASTK,ATG16L2,SNRPB,PARD3,ACTR6,MAP3K7CL,SEC62,CTSC,ARRDC1,KLC1,
2,SSU72,MYL6B,FAM122C,IQCG,TMEM67,PILRB,FAM72A,CACYBP,RHOF,STARD10,RNH1,DNASE1L1,PTO
2,SSU72,MYL6B,FAM122C,IQCG,TMEM67,PILRB,FAM72A,CACYBP,RHOF,STARD10,RNH1,DNASE1L1,PTO
AP2,SSU72,MYL6B,IQCG,PILRB,FAM72A,CMC2,CIRBP,RHOF,PRR13,STARD10,RNH1,DNASE1L1,KIF4A,RBCK
AP2,SSU72,MYL6B,IQCG,PILRB,FAM72A,CMC2,CIRBP,RHOF,PRR13,STARD10,RNH1,DNASE1L1,KIF4A,RBCK
AP2,RAPGEF5,MYL6B,IQCG,CACYBP,CIRBP,RHOF,SMARCB1,PRR13,RNH1,KIF4A,RBCK1,PTOV1,FASTK,TME
AP2,RAPGEF5,MYL6B,IQCG,CACYBP,CIRBP,RHOF,SMARCB1,PRR13,RNH1,KIF4A,RBCK1,PTOV1,FASTK,TME
SRSF2,MRPS22,HDAC9,KRT15,OSMR,PPFIBP1,SEMA4D,LAS1L,HMGB1,SULF2,CYB561A3,BAIAP2,RAPGEF5
SRSF2,MRPS22,HDAC9,KRT15,OSMR,PPFIBP1,SEMA4D,LAS1L,HMGB1,SULF2,CYB561A3,BAIAP2,RAPGEF5
RBCK1,PTOV1,FASTK,RNF32,LY75,ATG16L2,SNRPB,PARD3,TCAIM,SEC62,SET,ARRDC1,TRPT1,KLC1,LIMK2
BP,RPL38,PRR13,STARD10,RNH1,DNASE1L1,KIF4A,RBCK1,FASTK,TMEM135,LY75,SNRPB,TCAIM,ACTR6,SI
B1,RPL38,PRR13,RNH1,DNASE1L1,KIF4A,PTOV1,FASTK,RNF32,LY75,RCC1,PARD3,DHPS,ACTR6,SEC62,SET
22C,SRP9,IQCG,SULT1A3,UAP1,PILRB,FAM72A,YBX3,BLOC1S1,CACYBP,RHOF,PRR13,RNH1,DNASE1L1,KI
22C,SRP9,IQCG,SULT1A3,UAP1,PILRB,FAM72A,YBX3,BLOC1S1,CACYBP,RHOF,PRR13,RNH1,DNASE1L1,KI
PFIBP1,HMGB1,CYB561A3,BAIAP2,RAPGEF5,SSU72,MYL6B,SRP9,IQCG,FAM72A,BLOC1S1,CIRBP,RHOF,SN

PFIBP1,HMGB1,CYB561A3,BAIAP2,RAPGEF5,SSU72,MYL6B,SRP9,IQCG,FAM72A,BLOC1S1,CIRBP,RHOF,SM
1A3,PILRB,FAM72A,YBX3,CACYBP,CIRBP,RHOF,RNH1,DNASE1L1,KIF4A,RBCK1,FASTK,RCC1,SNRPB,PARD3
B1,SULF2,BAIAP2,RAPGEF5,SSU72,MYL6B,SRP9,IQCG,TMEM67,UAP1,FAM72A,CMC2,BLOC1S1,CACYBP,
B1,SULF2,BAIAP2,RAPGEF5,SSU72,MYL6B,SRP9,IQCG,TMEM67,UAP1,FAM72A,CMC2,BLOC1S1,CACYBP,
LC3A2,DHPS,ACTR6,RBP5,SCAF11,DBI,LIMK2,SIRT6,GPATCH2L,BUD31,CRYGS,CTTN,EIF5A,COMMD5,RUN
LC3A2,DHPS,ACTR6,RBP5,SCAF11,DBI,LIMK2,SIRT6,GPATCH2L,BUD31,CRYGS,CTTN,EIF5A,COMMD5,RUN
,ARHGAP6,CTTN,EIF5A,COMMD5,NUDT22,UFD1L,USP38,ACOT8,PCMT1,NUDT5,STYXL1,VRK3,LZTFL1,LR
,ARHGAP6,CTTN,EIF5A,COMMD5,NUDT22,UFD1L,USP38,ACOT8,PCMT1,NUDT5,STYXL1,VRK3,LZTFL1,LR
SNF8,CPSF3L,TSG101,RPS25,NIP7,RPS6,RPL29,AUP1,TRNT1,YIF1A,CDC42,FBXW9,ASF1B,LYPLA2,SUGT1,C
RPL38,PRR13,RNH1,KIF4A,FASTK,RNF32,LY75,ATG16L2,SNRPB,DHPS,SEC62,SET,ARRDC1,TRPT1,SCAF11,
MK2,SIRT6,NUMA1,SCARB1,CTTN,TSPAN11,DBNDD1,NOP16,PSMA6,ACOT8,AP3B2,RCE1,EBPL,TESC,IGS
MK2,SIRT6,NUMA1,SCARB1,CTTN,TSPAN11,DBNDD1,NOP16,PSMA6,ACOT8,AP3B2,RCE1,EBPL,TESC,IGS
C1,LIMK2,VDAC3,NUMA1,TBL2,SCARB1,CTTN,ITSN1,EIF5A,AMZ2,TAZ,DBNDD1,GSTK1,RCE1,RUVBL2,EBP
L,USP38,SH2B1,RPL13A,TTLL5,LDHA,PSMA6,AP3B2,SLC35G1,MRPL33,ZNF410,IGSF8,RANBP1,LPCAT3,CD
CA3,PCNA,SRSF2,MRPS22,HDAC9,KRT15,OSMR,PPFIBP1,SEMA4D,LAS1L,HMGB1,SULF2,CYB561A3,BAIAP
,RNH1,PTOV1,RNF32,RCC1,DHPS,ACTR6,SEC62,SET,ZNF235,BOP1,ARRDC1,KLC1,SCAF11,LIMK2,WDR19,
PLA2,ZNF197,TRMT112,DDOST,TOMM40,TADA3,ANAPC11,MDH2,SEC31A,COX17,B3GAT3,PAFAH1B2,DA
1,LAS1L,HMGB1,SULF2,CYB561A3,RAPGEF5,SSU72,MYL6B,SRP9,TMEM67,SULT1A3,PILRB,FAM72A,CMC
1,LAS1L,HMGB1,SULF2,CYB561A3,RAPGEF5,SSU72,MYL6B,SRP9,TMEM67,SULT1A3,PILRB,FAM72A,CMC
2,IGSF8,RANBP1,ZC3H14,LRRC1,PHF19,TM9SF1,SON,RPS16,UBR4,CHCHD1,PSMA3,RPS25,SPOCK2,THEM
SRP,C1ORF198,UGGT2,UBE2I,CDC42,PCYT2,FBXW9,RPL28,LIG1,LYPLA2,STOML2,CYB561,ZNF197,COPZ1,
SRP,C1ORF198,UGGT2,UBE2I,CDC42,PCYT2,FBXW9,RPL28,LIG1,LYPLA2,STOML2,CYB561,ZNF197,COPZ1,
IQCG,PILRB,YBX3,RHOF,SMARCB1,RPL38,STARD10,DNASE1L1,RBCK1,PTOV1,FASTK,TMEM135,RNF32,LY
KIF4A,RBCK1,RNF32,LY75,RCC1,SNRPB,PARD3,DHPS,ACTR6,KLC1,DBI,ARPC2,NOB1,SIRT6,GPATCH2L,BUD
KIF4A,RBCK1,RNF32,LY75,RCC1,SNRPB,PARD3,DHPS,ACTR6,KLC1,DBI,ARPC2,NOB1,SIRT6,GPATCH2L,BUD
RD10,RNH1,DNASE1L1,PTOV1,ATG16L2,SNRPB,TCAIM,BOP1,ARRDC1,KLC1,ZC3HAV1,SCAF11,SIRT6,NUM
RD10,RNH1,DNASE1L1,PTOV1,ATG16L2,SNRPB,TCAIM,BOP1,ARRDC1,KLC1,ZC3HAV1,SCAF11,SIRT6,NUM
RPL38,RNH1,KIF4A,RBCK1,FASTK,TMEM135,LY75,ATG16L2,SNRPB,PARD3,TCAIM,ARRDC1,TRPT1,KLC1,R
RPL38,RNH1,KIF4A,RBCK1,FASTK,TMEM135,LY75,ATG16L2,SNRPB,PARD3,TCAIM,ARRDC1,TRPT1,KLC1,R
UMA1,TBL2,SCARB1,CTTN,ITSN1,DBNDD1,NOP16,ACOT8,AP3B2,RCE1,RUVBL2,EBPL,TESC,IGSF8,SPAG7,
A,CACYBP,SMARCB1,RPL38,RNH1,KIF4A,PTOV1,FASTK,TMEM135,SNRPB,PARD3,TCAIM,ACTR6,MAP3K7
A,CACYBP,SMARCB1,RPL38,RNH1,KIF4A,PTOV1,FASTK,TMEM135,SNRPB,PARD3,TCAIM,ACTR6,MAP3K7
ACYBP,RHOF,STARD10,RNH1,DNASE1L1,PTOV1,RNF32,ATG16L2,TCAIM,BOP1,KLC1,SCAF11,WDR19,SIR
ACYBP,RHOF,STARD10,RNH1,DNASE1L1,PTOV1,RNF32,ATG16L2,TCAIM,BOP1,KLC1,SCAF11,WDR19,SIR
A,SRSF2,MRPS22,KRT15,OSMR,PPFIBP1,SEMA4D,HMGB1,CYB561A3,BAIAP2,RAPGEF5,SSU72,MYL6B,FAM
A,SRSF2,MRPS22,KRT15,OSMR,PPFIBP1,SEMA4D,HMGB1,CYB561A3,BAIAP2,RAPGEF5,SSU72,MYL6B,FAM
2,LY75,ARRDC1,KLC1,RBP5,DBI,LIMK2,NUMA1,OSGEP,TBL2,ARHGAP6,SCARB1,CTTN,TSPAN11,ITSN1,CO
A2,TXNDC17,ZNF197,TRMT112,DDOST,TOMM40,TADA3,ANAPC11,MDH2,SEC31A,COX17,B3GAT3,PAFA
TG16L2,SNRPB,PARD3,SLC3A2,ACTR6,SET,SCAF11,LIMK2,NUMA1,GPATCH2L,SAE1,OSGEP,TBL2,SCARB1,
AM72A,BLOC1S1,CACYBP,CIRBP,STARD10,RNH1,DNASE1L1,KIF4A,RBCK1,PTOV1,FASTK,TMEM135,LY75,
AM72A,BLOC1S1,CACYBP,CIRBP,STARD10,RNH1,DNASE1L1,KIF4A,RBCK1,PTOV1,FASTK,TMEM135,LY75,
F11,DBI,LIMK2,SIRT6,NUMA1,NPRL2,OSGEP,EIF5A,UFD1L,DBNDD1,RPL13A,GSTK1,RPS9,SLC35G1,MRPL
F11,LIMK2,VDAC3,NUMA1,OSGEP,EIF5A,UFD1L,DBNDD1,RPS9,MRPL33,RUVBL2,TESC,WDR62,POLD4,PL
APGEF5,SSU72,MYL6B,BLOC1S1,CIRBP,RHOF,SMARCB1,PRR13,RNH1,DNASE1L1,RBCK1,PTOV1,RNF32,L
APGEF5,SSU72,MYL6B,BLOC1S1,CIRBP,RHOF,SMARCB1,PRR13,RNH1,DNASE1L1,RBCK1,PTOV1,RNF32,L
DDOST,LSR,TOMM40,RABEP2,TADA3,MDH2,SEC31A,GBA2,MFSD10,RIC8A,UBE2N,LFNG,FAM32A,C12OR
1,CDC42,ASF1B,PPIL2,LIG1,LYPLA2,SHARPIN,ZNF197,CCNL2,RAB2A,ATP5J,TRMT112,UBXN1,SAT1,DDOST

1,CDC42,ASF1B,PPIL2,LIG1,LYPLA2,SHARPIN,ZNF197,CCNL2,RAB2A,ATP5J,TRMT112,UBXN1,SAT1,DDOST
3,FAM72A,YBX3,CACYBP,CIRBP,RHOF,SMARCB1,RPL38,PRR13,STARD10,RNH1,DNASE1L1,KIF4A,RBCK1,P
3,FAM72A,YBX3,CACYBP,CIRBP,RHOF,SMARCB1,RPL38,PRR13,STARD10,RNH1,DNASE1L1,KIF4A,RBCK1,P
SC,ARRDC1,TRPT1,ZC3HAV1,DBI,NOB1,WDR19,NPRL2,ITSN1,COMMD5,NUDT22,USP38,TAZ,RPL13A,PSN
M1K,IRF9,CPSF3L,TSG101,MUC1,RPS25,COX4I1,MED15,SH3YL1,CDC42,PCYT2,FBXW9,CCDC25,ENDOV,TF
A1C,IRF9,CPSF3L,TSG101,COX4I1,MED15,UGGT2,FBXW9,CCDC25,ENDOV,PPAN,CYB561,CDK11A,ATP5J,
L22C,SULT1A3,PILRB,FAM72A,CMC2,YBX3,CACYBP,CIRBP,RHOF,RNH1,DNASE1L1,KIF4A,RBCK1,RCC1,SNF
L22C,SULT1A3,PILRB,FAM72A,CMC2,YBX3,CACYBP,CIRBP,RHOF,RNH1,DNASE1L1,KIF4A,RBCK1,RCC1,SNF
L1,KIF4A,PTOV1,TMEM135,RNF32,RCC1,PARD3,TCAIM,SLC3A2,DHPS,ACTR6,SEC62,SET,CTSC,ARRDC1,Z
ENDOV,LYPLA2,PPAN,ZNF197,TRMT112,DDOST,TOMM40,TADA3,ANAPC11,MDH2,SEC31A,CNST,COX17
CB1,RPL38,STARD10,RNH1,PTOV1,FASTK,RNF32,LY75,ATG16L2,SNRPB,PARD3,SLC3A2,ACTR6,MAP3K7CI
CB1,RPL38,STARD10,RNH1,PTOV1,FASTK,RNF32,LY75,ATG16L2,SNRPB,PARD3,SLC3A2,ACTR6,MAP3K7CI
31,RPL13A,ARHGEF39,ACOT8,VRK3,IRF9,ANTXR2,STIP1,CD44,TM9SF1,MAGED2,RPS16,TSG101,UBR4,PSN
31,RPL13A,ARHGEF39,ACOT8,VRK3,IRF9,ANTXR2,STIP1,CD44,TM9SF1,MAGED2,RPS16,TSG101,UBR4,PSN
LT1A3,PILRB,FAM72A,CMC2,YBX3,CACYBP,CIRBP,RHOF,RNH1,DNASE1L1,KIF4A,RBCK1,RCC1,SNRPB,PAR
32,ATG16L2,RCC1,SNRPB,DHPS,ACTR6,SET,BOP1,KLC1,PMFBP1,VDAC3,NUMA1,BUD31,OSGEP,ARHGAP
RD3,TCAIM,SET,ARRDC1,LIMK2,SIRT6,NUMA1,TBL2,SCARB1,CTTN,ITSN1,EIF5A,THYN1,DBNDD1,NOP16,
GEF5,MYL6B,IQCG,PILRB,BLOC1S1,CACYBP,CIRBP,RHOF,SMARCB1,PRR13,RNH1,DNASE1L1,KIF4A,RBCK1,
GEF5,MYL6B,IQCG,PILRB,BLOC1S1,CACYBP,CIRBP,RHOF,SMARCB1,PRR13,RNH1,DNASE1L1,KIF4A,RBCK1,
GB1,SULF2,BAIAP2,RAPGEF5,MYL6B,FAM122C,SRP9,IQCG,SULT1A3,FAM72A,CMC2,BLOC1S1,CIRBP,RHC
GB1,SULF2,BAIAP2,RAPGEF5,MYL6B,FAM122C,SRP9,IQCG,SULT1A3,FAM72A,CMC2,BLOC1S1,CIRBP,RHC
GEF5,SSU72,MYL6B,IQCG,TMEM67,SULT1A3,PILRB,FAM72A,CMC2,YBX3,CACYBP,CIRBP,RHOF,SMARCB1
GEF5,SSU72,MYL6B,IQCG,TMEM67,SULT1A3,PILRB,FAM72A,CMC2,YBX3,CACYBP,CIRBP,RHOF,SMARCB1
,TBL2,SCARB1,CTTN,ITSN1,AMZ2,TAZ,RUVBL2,EBPL,IGSF8,SPAG7,WDR62,RANBP1,STIP1,LPCAT3,FAM3A
RAPGEF5,SSU72,SRP9,PILRB,FAM72A,BLOC1S1,CACYBP,CIRBP,RHOF,SMARCB1,PRR13,STARD10,RNH1,R
SNRPB,TCAIM,DHPS,SET,KLC1,SCAF11,NUMA1,SUMF2,SAE1,OSGEP,EIF5A,AMZ2,TAZ,RPL13A,TTLL5,LDH
1,LAS1L,HMGB1,BAIAP2,RAPGEF5,SSU72,FAM122C,SRP9,IQCG,UAP1,FAM72A,YBX3,CACYBP,CIRBP,RHC
1,LAS1L,HMGB1,BAIAP2,RAPGEF5,SSU72,FAM122C,SRP9,IQCG,UAP1,FAM72A,YBX3,CACYBP,CIRBP,RHC
1,UBR4,ATG12,DBNDD2,RPS6,RTEL1,FOXRED1,CLASRP,UGGT2,USE1,ASF1B,CAPG,LIG1,PSMA7,ACAD10,
1,UBR4,ATG12,DBNDD2,RPS6,RTEL1,FOXRED1,CLASRP,UGGT2,USE1,ASF1B,CAPG,LIG1,PSMA7,ACAD10,
A2,PPAN,ZNF197,TRMT112,DDOST,LSR,TOMM40,TADA3,ANAPC11,MDH2,EEF1D,SEC31A,COX17,PAFAH
TOV1,TMEM135,RCC1,SNRPB,PARD3,TCAIM,DHPS,SEC62,SET,TRPT1,NOB1,SIRT6,GPATCH2L,SAE1,OSG
TOV1,TMEM135,RCC1,SNRPB,PARD3,TCAIM,DHPS,SEC62,SET,TRPT1,NOB1,SIRT6,GPATCH2L,SAE1,OSG
BE2I,AUNIP,C14ORF2,PCYT2,PPP1CC,DGKE,CAMTA1,LIG1,TIMM17B,PRMT1,ZFAND2B,RPL36,LAMP2,LR
L38,PRR13,KIF4A,RBCK1,PTOV1,FASTK,TMEM135,SNRPB,PARD3,TCAIM,SLC3A2,DHPS,ACTR6,SEC62,SET,
L38,PRR13,KIF4A,RBCK1,PTOV1,FASTK,TMEM135,SNRPB,PARD3,TCAIM,SLC3A2,DHPS,ACTR6,SEC62,SET,
B,PARD3,SLC3A2,DHPS,SEC62,BOP1,TRPT1,KLC1,RBP5,ZC3HAV1,SCAF11,WDR19,GPATCH2L,SUMF2,SAE
B,PARD3,SLC3A2,DHPS,SEC62,BOP1,TRPT1,KLC1,RBP5,ZC3HAV1,SCAF11,WDR19,GPATCH2L,SUMF2,SAE
EF5,SSU72,FAM72A,CACYBP,CIRBP,RHOF,RPL38,RNH1,RBCK1,PTOV1,FASTK,TMEM135,RNF32,LY75,PAR
5,EMC4,DDOST,LSR,NAT9,TOMM40,TADA3,ABHD17A,MDH2,SEC31A,CNST,PAFAH1B2,NSUN5,TCP1,PIG
L38,RNH1,PTOV1,FASTK,LY75,ATG16L2,PARD3,TCAIM,SLC3A2,ACTR6,SET,CTSC,BOP1,SCAF11,DBI,PMFB
L38,RNH1,PTOV1,FASTK,LY75,ATG16L2,PARD3,TCAIM,SLC3A2,ACTR6,SET,CTSC,BOP1,SCAF11,DBI,PMFB
1122C,IQCG,TMEM67,SULT1A3,PILRB,YBX3,BLOC1S1,CACYBP,RHOF,PRR13,RNH1,DNASE1L1,PTOV1,RNF
1122C,IQCG,TMEM67,SULT1A3,PILRB,YBX3,BLOC1S1,CACYBP,RHOF,PRR13,RNH1,DNASE1L1,PTOV1,RNF
CK1,FASTK,RNF32,ATG16L2,PARD3,SLC3A2,SET,CTSC,BOP1,ARRDC1,RBP5,SCAF11,DBI,ARPC2,VDAC3,NU
CK1,FASTK,RNF32,ATG16L2,PARD3,SLC3A2,SET,CTSC,BOP1,ARRDC1,RBP5,SCAF11,DBI,ARPC2,VDAC3,NU
A,BLOC1S1,CACYBP,SMARCB1,RPL38,STARD10,RNH1,KIF4A,PTOV1,FASTK,RNF32,ATG16L2,SNRPB,PARD

A,BLOC1S1,CACYBP,SMARCB1,RPL38,STARD10,RNH1,KIF4A,PTOV1,FASTK,RNF32,ATG16L2,SNRPB,PARD
ST,LSR,TOMM40,TADA3,MDH2,SEC31A,PFDN5,PAFAH1B2,GBA2,NSUN5,MFSD10,UBE2N,LFNG,FAM32A
IRBP,RNH1,PTOV1,LY75,ATG16L2,SLC3A2,DHPS,CTSC,ZNF235,BOP1,ARRDC1,KLC1,RBP5,ARPC2,LIMK2,N
IRBP,RNH1,PTOV1,LY75,ATG16L2,SLC3A2,DHPS,CTSC,ZNF235,BOP1,ARRDC1,KLC1,RBP5,ARPC2,LIMK2,N
SRP9,TMEM67,YBX3,BLOC1S1,CACYBP,RHOF,SMARCB1,RPL38,DNASE1L1,KIF4A,FASTK,TMEM135,LY75,
SRP9,TMEM67,YBX3,BLOC1S1,CACYBP,RHOF,SMARCB1,RPL38,DNASE1L1,KIF4A,FASTK,TMEM135,LY75,
RPL38,STARD10,DNASE1L1,KIF4A,RBCK1,PTOV1,FASTK,TMEM135,ATG16L2,SNRPB,PARD3,DHPS,ACTR6,S
,SMARCB1,RPL38,PRR13,RNH1,RBCK1,PTOV1,FASTK,TMEM135,RNF32,LY75,ATG16L2,PARD3,ACTR6,SEC
L,SULF2,CYB561A3,BAIAP2,RAPGEF5,SRP9,IQCG,TMEM67,PILRB,CMC2,YBX3,BLOC1S1,CACYBP,RHOF,SM
L,SULF2,CYB561A3,BAIAP2,RAPGEF5,SRP9,IQCG,TMEM67,PILRB,CMC2,YBX3,BLOC1S1,CACYBP,RHOF,SM
RD10,RNH1,DNASE1L1,PTOV1,ATG16L2,SNRPB,TCAIM,BOP1,ARRDC1,KLC1,ZC3HAV1,SCAF11,NUMA1,G
RD10,RNH1,DNASE1L1,PTOV1,ATG16L2,SNRPB,TCAIM,BOP1,ARRDC1,KLC1,ZC3HAV1,SCAF11,NUMA1,G
M3,TRMT112,DDOST,TOMM40,TADA3,MDH2,SEC31A,CNST,B3GAT3,PAFAH1B2,NSUN5,UBE2N,FAM32A,
L1,RBCK1,PTOV1,FASTK,ATG16L2,SNRPB,SLC3A2,ACTR6,SET,CTSC,ARRDC1,KLC1,RBP5,ZC3HAV1,SCAF11
,ARRDC1,TRPT1,LIMK2,NUMA1,TBL2,SCARB1,CTTN,ITSN1,COMMD5,NUDT22,DBNDD1,TTLL5,RCE1,EBPL
UMA1,OSGEP,SCARB1,EIF5A,UFD1L,DBNDD1,MRPL33,ZNF410,RUVBL2,WDR62,PLEKHA5,RANBP1,LZTFL
3,SMARCB1,RPL38,RBCK1,PTOV1,FASTK,RNF32,LY75,ATG16L2,PARD3,TCAIM,SLC3A2,DHPS,ACTR6,SEC6
IMGB1,SULF2,BAIAP2,RAPGEF5,SSU72,FAM122C,SRP9,IQCG,TMEM67,SULT1A3,PILRB,FAM72A,CMC2,YB
IMGB1,SULF2,BAIAP2,RAPGEF5,SSU72,FAM122C,SRP9,IQCG,TMEM67,SULT1A3,PILRB,FAM72A,CMC2,YB
B1BP1,MED15,C1ORF198,UGGT2,CDC42,FBXW9,CCDC25,EEF1B2,LIG1,LYPLA2,STOML2,PPAN,CYB561,Z
B1BP1,MED15,C1ORF198,UGGT2,CDC42,FBXW9,CCDC25,EEF1B2,LIG1,LYPLA2,STOML2,PPAN,CYB561,Z
APGEF5,SSU72,MYL6B,FAM122C,IQCG,SULT1A3,PILRB,CMC2,YBX3,CACYBP,RHOF,SMARCB1,RPL38,PRR1
APGEF5,SSU72,MYL6B,FAM122C,IQCG,SULT1A3,PILRB,CMC2,YBX3,CACYBP,RHOF,SMARCB1,RPL38,PRR1
P2,SSU72,FAM122C,IQCG,SULT1A3,FAM72A,YBX3,CACYBP,CIRBP,RHOF,SMARCB1,RPL38,PRR13,RNH1,
MK2,NUMA1,TBL2,SCARB1,CTTN,ITSN1,EIF5A,DBNDD1,NOP16,TTLL5,AP3B2,RCE1,EBPL,TESC,IGSF8,SPA
LL5,PSMA6,RPS9,SLC35G1,SNRPN,ZC3H14,RPS16,CPSF3L,ZBTB44,TMED4,SRSF1,ITGB1BP1,SH3YL1,PRM
LL5,PSMA6,RPS9,SLC35G1,SNRPN,ZC3H14,RPS16,CPSF3L,ZBTB44,TMED4,SRSF1,ITGB1BP1,SH3YL1,PRM
C1,KLC1,LIMK2,SIRT6,NUMA1,TBL2,SCARB1,CTTN,ITSN1,THYN1,DBNDD1,NOP16,RCE1,RUVBL2,EBPL,TE
UQCRQ,CPSF3L,TSG101,ATG12,NIP7,RPS6,RTEL1,SAP30BP,FOXRED1,TRNT1,MED15,CLASRP,CDC42,USE
AIM,SLC3A2,DHPS,SET,CTSC,BOP1,ARRDC1,TRPT1,ARPC2,PMFBP1,LIMK2,VDAC3,WDR19,SIRT6,NUMA1
13,STARD10,RNH1,RBCK1,PTOV1,LY75,PARD3,DHPS,SEC62,SET,ARRDC1,TRPT1,ZC3HAV1,WDR19,NUM
13,STARD10,RNH1,RBCK1,PTOV1,LY75,PARD3,DHPS,SEC62,SET,ARRDC1,TRPT1,ZC3HAV1,WDR19,NUM
IMGB1,SULF2,BAIAP2,RAPGEF5,SSU72,MYL6B,SRP9,IQCG,TMEM67,SULT1A3,PILRB,FAM72A,YBX3,BLOC
IMGB1,SULF2,BAIAP2,RAPGEF5,SSU72,MYL6B,SRP9,IQCG,TMEM67,SULT1A3,PILRB,FAM72A,YBX3,BLOC
NOB1,SIRT6,SUMF2,SAE1,NPRL2,EIF5A,AMZ2,RPL13A,TTLL5,PSMA6,RPS9,SLC35G1,VRK3,ZC3H14,RPS16
NOB1,SIRT6,SUMF2,SAE1,NPRL2,EIF5A,AMZ2,RPL13A,TTLL5,PSMA6,RPS9,SLC35G1,VRK3,ZC3H14,RPS16
DC42,PPIL2,LIG1,APEH,RBM33,C17ORF62,CYB561,ZNF250,CDK11A,ZNF197,DGUOK,TSFM,TRMT112,CH
DC42,PPIL2,LIG1,APEH,RBM33,C17ORF62,CYB561,ZNF250,CDK11A,ZNF197,DGUOK,TSFM,TRMT112,CH
AH1B2,ZFPL1,GMPR2,RIC8A,FAM32A,NDUFS3,OXLD1,PICK1,RNF181,RALY,GLYCTK,NEK11,EIF2A,PSMD7,
RF198,UGGT2,UBE2I,CDC42,FBXW9,RPL28,LIG1,TIMM17B,CYB561,ZNF197,TRMT112,DDOST,RRNAD1,
RF198,UGGT2,UBE2I,CDC42,FBXW9,RPL28,LIG1,TIMM17B,CYB561,ZNF197,TRMT112,DDOST,RRNAD1,
,FASTK,RNF32,ATG16L2,SET,BOP1,ARRDC1,TRPT1,KLC1,ZC3HAV1,SCAF11,DBI,WDR19,GPATCH2L,SUMF
,FASTK,RNF32,ATG16L2,SET,BOP1,ARRDC1,TRPT1,KLC1,ZC3HAV1,SCAF11,DBI,WDR19,GPATCH2L,SUMF
32,LY75,RCC1,DHPS,SET,CTSC,BOP1,SCAF11,WDR19,SUMF2,BUD31,SAE1,OSGEP,TBL2,ARHGAP6,SCARB
32,LY75,RCC1,DHPS,SET,CTSC,BOP1,SCAF11,WDR19,SUMF2,BUD31,SAE1,OSGEP,TBL2,ARHGAP6,SCARB
,NUMA1,TBL2,SCARB1,ITSN1,COMMD5,AMZ2,UFD1L,THYN1,DBNDD1,RCE1,ZNF410,RBPMS,SPAG7,RAN
,NUMA1,TBL2,SCARB1,ITSN1,COMMD5,AMZ2,UFD1L,THYN1,DBNDD1,RCE1,ZNF410,RBPMS,SPAG7,RAN

MEM67,SULT1A3,CMC2,BLOC1S1,CACYBP,CIRBP,SMARCB1,RPL38,PRR13,RNH1,RBCK1,PTOV1,RCC1,SNR

MEM67,SULT1A3,CMC2,BLOC1S1,CACYBP,CIRBP,SMARCB1,RPL38,PRR13,RNH1,RBCK1,PTOV1,RCC1,SNR

SU72,FAM122C,SULT1A3,PILRB,FAM72A,CMC2,YBX3,BLOC1S1,CACYBP,CIRBP,RHOF,RNH1,DNASE1L1,KIF

SU72,FAM122C,SULT1A3,PILRB,FAM72A,CMC2,YBX3,BLOC1S1,CACYBP,CIRBP,RHOF,RNH1,DNASE1L1,KIF

,IGSF8,NUDT5,STYXL1,POLD4,RANBP1,SLC15A3,TUBA1C,GRK6,CPSF3L,COX4I1,NISCH,RTEL1,TRNT1,MED

9,ACOT8,RUVBL2,NUDT5,STYXL1,LRRC1,VCAN,SON,UQCRQ,CPSF3L,TSG101,METTL17,NIP7,COX4I1,DUSP

1,LY75,ATG16L2,SNRPB,SLC3A2,SEC62,SET,ZC3HAV1,LIMK2,SAE1,OSGEP,ARHGAP6,CTTN,ITSN1,COMMD

ATG16L2,PARD3,SLC3A2,ACTR6,SEC62,CTSC,DBI,ARPC2,PMFBP1,VDAC3,NUMA1,GPATCH2L,OSGEP,SCA

ATG16L2,PARD3,SLC3A2,ACTR6,SEC62,CTSC,DBI,ARPC2,PMFBP1,VDAC3,NUMA1,GPATCH2L,OSGEP,SCA

AS1L,SULF2,CYB561A3,BAIAP2,RAPGEF5,MYL6B,FAM122C,SRP9,IQCG,TMEM67,SULT1A3,BLOC1S1,CIRB

AS1L,SULF2,CYB561A3,BAIAP2,RAPGEF5,MYL6B,FAM122C,SRP9,IQCG,TMEM67,SULT1A3,BLOC1S1,CIRB

,LIMK2,SIRT6,NUMA1,SAE1,OSGEP,CTTN,EIF5A,COMMD5,TTLL5,LDHA,AP3B2,MRPL33,EBPL,WDR62,RA

MK2,SIRT6,NUMA1,TBL2,SCARB1,CTTN,ITSN1,THYN1,DBNDD1,NOP16,RCE1,EBPL,TESC,IGSF8,SPAG7,WD

MK2,SIRT6,NUMA1,TBL2,SCARB1,CTTN,ITSN1,THYN1,DBNDD1,NOP16,RCE1,EBPL,TESC,IGSF8,SPAG7,WD

,RNF32,ATG16L2,PARD3,SLC3A2,SET,CTSC,BOP1,ARRDC1,RBP5,SCAF11,DBI,ARPC2,VDAC3,GPATCH2L,BU

R13,STARD10,FASTK,ATG16L2,RCC1,DHPS,SET,BOP1,DBI,NOB1,GPATCH2L,SUMF2,SAE1,OSGEP,SCARB1,

R13,STARD10,FASTK,ATG16L2,RCC1,DHPS,SET,BOP1,DBI,NOB1,GPATCH2L,SUMF2,SAE1,OSGEP,SCARB1,

C1,TRPT1,KLC1,LIMK2,SIRT6,NUMA1,TBL2,SCARB1,CTTN,ITSN1,NUDT22,UFD1L,THYN1,DBNDD1,NOP16

B1BP1,MED15,C1ORF198,UGGT2,CDC42,FBXW9,CCDC25,EEF1B2,LIG1,PPAN,CYB561,ZNF197,TRMT112,

B1BP1,MED15,C1ORF198,UGGT2,CDC42,FBXW9,CCDC25,EEF1B2,LIG1,PPAN,CYB561,ZNF197,TRMT112,

1,PTOV1,LY75,TCAIM,DHPS,SET,CTSC,ARRDC1,RBP5,LIMK2,NOB1,NUMA1,GPATCH2L,OSGEP,TBL2,SCAR

1,PTOV1,LY75,TCAIM,DHPS,SET,CTSC,ARRDC1,RBP5,LIMK2,NOB1,NUMA1,GPATCH2L,OSGEP,TBL2,SCAR

RP9,IQCG,PILRB,YBX3,CACYBP,CIRBP,SMARCB1,RPL38,STARD10,RNH1,KIF4A,RBCK1,PTOV1,FASTK,TMEM

RP9,IQCG,PILRB,YBX3,CACYBP,CIRBP,SMARCB1,RPL38,STARD10,RNH1,KIF4A,RBCK1,PTOV1,FASTK,TMEM

,SRP9,IQCG,TMEM67,UAP1,FAM72A,YBX3,CACYBP,RPL38,PRR13,PTOV1,FASTK,TMEM135,RNF32,ATG16

,SRP9,IQCG,TMEM67,UAP1,FAM72A,YBX3,CACYBP,RPL38,PRR13,PTOV1,FASTK,TMEM135,RNF32,ATG16

APGEF5,SSU72,MYL6B,SRP9,SULT1A3,PILRB,CMC2,YBX3,BLOC1S1,CACYBP,CIRBP,RPL38,DNASE1L1,RBC

APGEF5,SSU72,MYL6B,SRP9,SULT1A3,PILRB,CMC2,YBX3,BLOC1S1,CACYBP,CIRBP,RPL38,DNASE1L1,RBC

V1,FASTK,LY75,ATG16L2,PARD3,SLC3A2,ACTR6,SEC62,SET,BOP1,TRPT1,SCAF11,DBI,PMFBP1,LIMK2,VDA

V1,FASTK,LY75,ATG16L2,PARD3,SLC3A2,ACTR6,SEC62,SET,BOP1,TRPT1,SCAF11,DBI,PMFBP1,LIMK2,VDA

MGB1,SULF2,BAIAP2,RAPGEF5,SSU72,MYL6B,FAM122C,SRP9,IQCG,SULT1A3,CMC2,YBX3,CACYBP,SMARC

MGB1,SULF2,BAIAP2,RAPGEF5,SSU72,MYL6B,FAM122C,SRP9,IQCG,SULT1A3,CMC2,YBX3,CACYBP,SMARC

Z2,USP38,RPL13A,TTLL5,PSMA6,RPS9,SLC35G1,SNRPN,ZC3H14,RPS16,CPSF3L,ZBTB44,TMED4,SRSF1,ITG

Z2,USP38,RPL13A,TTLL5,PSMA6,RPS9,SLC35G1,SNRPN,ZC3H14,RPS16,CPSF3L,ZBTB44,TMED4,SRSF1,ITG

CYBP,CIRBP,RHOF,RNH1,DNASE1L1,RBCK1,PTOV1,FASTK,RCC1,SNRPB,PARD3,ACTR6,SET,KLC1,SIRT6,BUD

MRPL33,WDR62,IL1RAP,RANBP1,LZTFL1,ZC3H14,FIS1,STIP1,LRRC1,PHF19,TM9SF1,SON,TET2,RPS16,FUZ

G16L2,SNRPB,PARD3,DHPS,ACTR6,ARRDC1,KLC1,SCAF11,DBI,VDAC3,SIRT6,NUMA1,SAE1,OSGEP,TBL2,A

561A3,BAIAP2,RAPGEF5,FAM72A,YBX3,CACYBP,CIRBP,RHOF,SMARCB1,RPL38,RNH1,PTOV1,FASTK,TME

561A3,BAIAP2,RAPGEF5,FAM72A,YBX3,CACYBP,CIRBP,RHOF,SMARCB1,RPL38,RNH1,PTOV1,FASTK,TME

RBCK1,PTOV1,FASTK,RNF32,LY75,PARD3,SLC3A2,SET,ARRDC1,KLC1,LIMK2,SIRT6,NUMA1,TBL2,SCARB1,

M67,PILRB,FAM72A,CACYBP,RHOF,STARD10,RNH1,DNASE1L1,PTOV1,ATG16L2,SNRPB,TCAIM,SEC62,BO

M67,PILRB,FAM72A,CACYBP,RHOF,STARD10,RNH1,DNASE1L1,PTOV1,ATG16L2,SNRPB,TCAIM,SEC62,BO

7,FAM72A,CMC2,YBX3,CACYBP,RHOF,SMARCB1,RPL38,PRR13,DNASE1L1,KIF4A,RBCK1,PTOV1,FASTK,RN

7,FAM72A,CMC2,YBX3,CACYBP,RHOF,SMARCB1,RPL38,PRR13,DNASE1L1,KIF4A,RBCK1,PTOV1,FASTK,RN

A1,TBL2,SCARB1,CTTN,ITSN1,AMZ2,DBNDD1,AP3B2,RCE1,RUVBL2,EBPL,TESC,IGSF8,SPAG7,WDR62,RFX

,CDK11A,RBM3,TRMT112,RPS20,GABPB1,PRKRIP1,DDOST,LSR,PSME2,RRNAD1,TOMM40,TADA3,MDH2,

SMARCB1,STARD10,RNH1,DNASE1L1,FASTK,RNF32,LY75,ATG16L2,RCC1,DHPS,SET,BOP1,ARRDC1,SCAF1

1L,HMGB1,SULF2,RAPGEF5,SSU72,MYL6B,SRP9,IQCG,TMEM67,UAP1,PILRB,FAM72A,YBX3,BLOC1S1,CAC

1L,HMGB1,SULF2,RAPGEF5,SSU72,MYL6B,SRP9,IQCG,TMEM67,UAP1,PILRB,FAM72A,YBX3,BLOC1S1,CAC

LC1,LIMK2,NUMA1,TBL2,SCARB1,CTTN,ITSN1,DBNDD1,ACOT8,AP3B2,RCE1,PCMT1,EBPL,TESC,IGSF8,SPA

P1,CLASRP,C1ORF198,UGGT2,UBE2I,CDC42,FBXW9,RPL28,LIG1,TIMM17B,AP1S3,CYB561,ZNF197,COPZ1

P1,CLASRP,C1ORF198,UGGT2,UBE2I,CDC42,FBXW9,RPL28,LIG1,TIMM17B,AP1S3,CYB561,ZNF197,COPZ1

SEC31A,PAFAH1B2,NSUN5,TCP1,UBE2N,FAM32A,DCTN5,PEX26,ARIH2,FBXL18,TSPAN31,ZNF593,GLYCTK

1,FASTK,TMEM135,RNF32,ATG16L2,SLC3A2,SEC62,SET,ZNF235,ARRDC1,ZC3HAV1,SCAF11,ARPC2,LIMK2

H2B1,MRPL33,RUVBL2,SNRPN,SLC15A3,SNF8,OBSL1,UBR4,CHCHD1,ECSIT,NLRP1,NIP7,PLCB2,SLC25A10

H2B1,MRPL33,RUVBL2,SNRPN,SLC15A3,SNF8,OBSL1,UBR4,CHCHD1,ECSIT,NLRP1,NIP7,PLCB2,SLC25A10

.0,PCMT1,IGSF8,NNMT,WDR62,VRK3,TM9SF1,TET2,GRK6,CDC37,TSG101,OBSL1,FAM3A,PSMA3,METTL1

MA1,OSGEP,SCARB1,CTTN,TSPAN11,AMZ2,TAZ,SH2B1,NOP16,PSMA6,MRPL33,RUVBL2,EBPL,TESC,SNRPN

MA1,OSGEP,SCARB1,CTTN,TSPAN11,AMZ2,TAZ,SH2B1,NOP16,PSMA6,MRPL33,RUVBL2,EBPL,TESC,SNRPN

G,TMEM67,CIRBP,SMARCB1,PRR13,KIF4A,RNF32,ATG16L2,RCC1,TCAIM,ACTR6,SEC62,SET,ARRDC1,KLC1

G,TMEM67,CIRBP,SMARCB1,PRR13,KIF4A,RNF32,ATG16L2,RCC1,TCAIM,ACTR6,SEC62,SET,ARRDC1,KLC1

MR,PPFIBP1,SULF2,RAPGEF5,SSU72,MYL6B,IQCG,SULT1A3,FAM72A,YBX3,BLOC1S1,CIRBP,RHOF,SMARCI

MR,PPFIBP1,SULF2,RAPGEF5,SSU72,MYL6B,IQCG,SULT1A3,FAM72A,YBX3,BLOC1S1,CIRBP,RHOF,SMARCI

,SULT1A3,PILRB,FAM72A,CMC2,YBX3,BLOC1S1,CACYBP,CIRBP,RHOF,RNH1,DNASE1L1,KIF4A,RBCK1,FAST

3L2,EBPL,IGSF8,SNRPN,SPAG7,WDR62,STIP1,LPCAT3,SLC39A11,FAM3A,ZBTB44,NISCH,CD99L2,PARD6G,

,RNF32,RCC1,ACTR6,CTSC,ZNF235,TRPT1,KLC1,ZC3HAV1,DBI,ARPC2,VDAC3,NUMA1,GPATCH2L,BUD31,

A,AMZ2,DBNDD1,SH2B1,PSMA6,RPS9,MRPL33,RUVBL2,WDR62,STYXL1,PLEKHA5,SLC15A3,PHF19,LPCAT

K3,TET2,GRK6,CDC37,FAM3A,METTL17,NLRP1,NIP7,TRNT1,MED15,XBP1,GUK1,CENPN,PCYT2,IFI27L1,P

L,SULF2,CYB561A3,BAIAP2,RAPGEF5,SSU72,FAM122C,TMEM67,PILRB,FAM72A,CMC2,YBX3,BLOC1S1,CA

L,SULF2,CYB561A3,BAIAP2,RAPGEF5,SSU72,FAM122C,TMEM67,PILRB,FAM72A,CMC2,YBX3,BLOC1S1,CA

2C,SRP9,UAP1,RHOF,SMARCB1,PRR13,STARD10,RNH1,DNASE1L1,PTOV1,RNF32,LY75,RCC1,PARD3,TCAI

E1L1,PTOV1,ATG16L2,SNRPB,TCAIM,ARRDC1,KLC1,ZC3HAV1,SCAF11,NUMA1,BUD31,NPRL2,OSGEP,TBL

E1L1,PTOV1,ATG16L2,SNRPB,TCAIM,ARRDC1,KLC1,ZC3HAV1,SCAF11,NUMA1,BUD31,NPRL2,OSGEP,TBL

IRBP,RHOF,RPL38,DNASE1L1,RNF32,ATG16L2,SNRPB,PARD3,TCAIM,SLC3A2,CTSC,RBP5,ARPC2,PMFBP1,

IRBP,RHOF,RPL38,DNASE1L1,RNF32,ATG16L2,SNRPB,PARD3,TCAIM,SLC3A2,CTSC,RBP5,ARPC2,PMFBP1,

16L2,DHPS,SEC62,SET,BOP1,ZC3HAV1,DBI,ARPC2,OSGEP,SCARB1,TAZ,SH2B1,NOP16,RPL13A,RCE1,PCM

16L2,DHPS,SEC62,SET,BOP1,ZC3HAV1,DBI,ARPC2,OSGEP,SCARB1,TAZ,SH2B1,NOP16,RPL13A,RCE1,PCM

ASTK,LY75,SLC3A2,SEC62,CTSC,BOP1,ARRDC1,TRPT1,SCAF11,DBI,ARPC2,EIF5A,CHID1,NUDT22,UFD1L,U

ARB1,CTTN,ITSN1,DBNDD1,ACOT8,AP3B2,RCE1,EBPL,TESC,IGSF8,SPAG7,WDR62,POLD4,UBE2L6,PPM1K,

SET,BOP1,ARRDC1,TBL2,ARHGAP6,SCARB1,ITSN1,COMMD5,UFD1L,TAZ,DBNDD1,RCE1,ZNF410,EBPL,IGS

SET,BOP1,ARRDC1,TBL2,ARHGAP6,SCARB1,ITSN1,COMMD5,UFD1L,TAZ,DBNDD1,RCE1,ZNF410,EBPL,IGS

C1,KLC1,LIMK2,NUMA1,TBL2,SCARB1,CTTN,ITSN1,AMZ2,DBNDD1,ACOT8,AP3B2,RCE1,TESC,IGSF8,SPAG

S1,CACYBP,CIRBP,SMARCB1,RPL38,SNRPB,DHPS,SEC62,ZNF235,RBP5,ZC3HAV1,DBI,NUMA1,SUMF2,NPR

S1,CACYBP,CIRBP,SMARCB1,RPL38,SNRPB,DHPS,SEC62,ZNF235,RBP5,ZC3HAV1,DBI,NUMA1,SUMF2,NPR

BAIAP2,RAPGEF5,MYL6B,IQCG,PILRB,FAM72A,BLOC1S1,CACYBP,CIRBP,RHOF,SMARCB1,PRR13,STARD10,

BAIAP2,RAPGEF5,MYL6B,IQCG,PILRB,FAM72A,BLOC1S1,CACYBP,CIRBP,RHOF,SMARCB1,PRR13,STARD10,

S1L,HMGB1,BAIAP2,RAPGEF5,SSU72,FAM122C,SRP9,IQCG,UAP1,PILRB,YBX3,CACYBP,CIRBP,RHOF,SMAR

S1L,HMGB1,BAIAP2,RAPGEF5,SSU72,FAM122C,SRP9,IQCG,UAP1,PILRB,YBX3,CACYBP,CIRBP,RHOF,SMAR

C1,DUSP22,GGPS1,RTEL1,AUP1,KXD1,MED15,JPH2,TBX6,RFX3,PPIL2,PSMD4,CLDND1,LIG1,CDC14B,CAPN

C1,DUSP22,GGPS1,RTEL1,AUP1,KXD1,MED15,JPH2,TBX6,RFX3,PPIL2,PSMD4,CLDND1,LIG1,CDC14B,CAPN

CIRBP,RHOF,PRR13,RNH1,RNF32,SNRPB,SLC3A2,DHPS,ACTR6,ARRDC1,RBP5,DBI,ARPC2,NOB1,SIRT6,GPA

CMT1,IGSF8,NNMT,WDR62,VRK3,TM9SF1,SON,TET2,UQCRQ,GRK6,CDC37,OBSL1,UBR4,PHF21A,FAM3A

B561A3,RAPGEF5,SSU72,FAM122C,IQCG,TMEM67,SULT1A3,UAP1,FAM72A,CMC2,BLOC1S1,CIRBP,RHO

B561A3,RAPGEF5,SSU72,FAM122C,IQCG,TMEM67,SULT1A3,UAP1,FAM72A,CMC2,BLOC1S1,CIRBP,RHO

C3HAV1,DBI,GPATCH2L,SUMF2,BUD31,TBL2,ARHGAP6,CHID1,AMZ2,UFD1L,TAZ,AP3B2,MRPL33,RUVBL2,
C3HAV1,DBI,GPATCH2L,SUMF2,BUD31,TBL2,ARHGAP6,CHID1,AMZ2,UFD1L,TAZ,AP3B2,MRPL33,RUVBL2,
SEC31A,B3GAT3,PAFAH1B2,DALRD3,NSUN5,UBE2N,FAM32A,OGFOD2,ARIH2,PICK1,TSPAN31,GLYCTK,TP
R6,SET,BOP1,ARRDC1,KLC1,SCAF11,LIMK2,VDAC3,NUMA1,OSGEP,ARHGAP6,SCARB1,ITSN1,EIF5A,UFD1
ZC3H14,RPS16,CPSF3L,ZBTB44,TMED4,ITGB1BP1,SH3YL1,UGGT2,MAGT1,CENPN,PPA1,ACAD10,FAM21
ZC3H14,RPS16,CPSF3L,ZBTB44,TMED4,ITGB1BP1,SH3YL1,UGGT2,MAGT1,CENPN,PPA1,ACAD10,FAM21
RAPGEF5,SSU72,MYL6B,FAM122C,SRP9,SULT1A3,FAM72A,YBX3,CACYBP,CIRBP,STARD10,RNH1,DNASE1
RAPGEF5,SSU72,MYL6B,FAM122C,SRP9,SULT1A3,FAM72A,YBX3,CACYBP,CIRBP,STARD10,RNH1,DNASE1
AMZ2,USP38,NOP16,RPL13A,PSMA6,RPS9,SLC35G1,SNRPN,ZC3H14,PPM1K,HMGA2,CPSF3L,ZBTB44,TM
AMZ2,USP38,NOP16,RPL13A,PSMA6,RPS9,SLC35G1,SNRPN,ZC3H14,PPM1K,HMGA2,CPSF3L,ZBTB44,TM
1L1,RBCK1,RCC1,SNRPB,TCAIM,SLC3A2,DHPS,ACTR6,MAP3K7CL,DBI,PMFBP1,LIMK2,GPATCH2L,SUMF2,
1L1,RBCK1,RCC1,SNRPB,TCAIM,SLC3A2,DHPS,ACTR6,MAP3K7CL,DBI,PMFBP1,LIMK2,GPATCH2L,SUMF2,
PEH,LYPLA2,C17ORF62,PRMT1,EXOSC9,HNRNPC,STAU1,RPL36,CDK11A,RAB2A,STRIP2,UBXN1,HHLA3,ER
PEH,LYPLA2,C17ORF62,PRMT1,EXOSC9,HNRNPC,STAU1,RPL36,CDK11A,RAB2A,STRIP2,UBXN1,HHLA3,ER
1,DNASE1L1,PTOV1,RNF32,TCAIM,BOP1,KLC1,WDR19,SIRT6,NUMA1,BUD31,OSGEP,TBL2,ARHGAP6,EIF5
1,DNASE1L1,PTOV1,RNF32,TCAIM,BOP1,KLC1,WDR19,SIRT6,NUMA1,BUD31,OSGEP,TBL2,ARHGAP6,EIF5
L,DNASE1L1,PTOV1,ATG16L2,SNRPB,TCAIM,SEC62,BOP1,ARRDC1,KLC1,SCAF11,NUMA1,GPATCH2L,BUD
L,DNASE1L1,PTOV1,ATG16L2,SNRPB,TCAIM,SEC62,BOP1,ARRDC1,KLC1,SCAF11,NUMA1,GPATCH2L,BUD
19,LPCAT3,MAGED2,GRK6,CPSF3L,TSG101,PTPN13,PHF21A,FAM3A,EPB41L5,RPL29,SRSF1,RABL5,TRNT1
19,LPCAT3,MAGED2,GRK6,CPSF3L,TSG101,PTPN13,PHF21A,FAM3A,EPB41L5,RPL29,SRSF1,RABL5,TRNT1
CB1,RPL38,STARD10,DNASE1L1,RBCK1,FASTK,LY75,RCC1,PARD3,DHPS,SEC62,ZNF235,ARRDC1,TRPT1,RB
AP2,RAPGEF5,SSU72,SRP9,IQCG,TMEM67,SULT1A3,FAM72A,BLOC1S1,CIRBP,RHOF,RPL38,PRR13,RNH1,
AP2,RAPGEF5,SSU72,SRP9,IQCG,TMEM67,SULT1A3,FAM72A,BLOC1S1,CIRBP,RHOF,RPL38,PRR13,RNH1,
,MYL6B,SRP9,IQCG,TMEM67,PILRB,FAM72A,CMC2,YBX3,CACYBP,SMARCB1,STARD10,DNASE1L1,KIF4A,
Z2,THYN1,TTLL5,RCE1,MRPL33,IGSF8,VRK3,FIS1,SON,UQCRQ,PSMA1,RGS3,DBNDD2,COASY,SRSF1,UBE2
B,CIRBP,RHOF,SMARCB1,PRR13,RNH1,DNASE1L1,RBCK1,PTOV1,FASTK,LY75,PARD3,SLC3A2,SET,CTSC,Z
1B2,NSUN5,UBE2N,FAM32A,OGFOD2,ARIH2,PICK1,TSPAN31,GLYCTK,TP53,TMEM199,EPN1,ZCCHC9,TB
TK,LY75,PARD3,SEC62,SET,ARRDC1,TRPT1,RBP5,SCAF11,LIMK2,NUMA1,BUD31,SAE1,TBL2,SCARB1,CTT
TK,LY75,PARD3,SEC62,SET,ARRDC1,TRPT1,RBP5,SCAF11,LIMK2,NUMA1,BUD31,SAE1,TBL2,SCARB1,CTT

L,FASTK,LY75,ARRDC1,LIMK2,NUMA1,TBL2,SCARB1,CTTN,ITSN1,COMMD5,UFD1L,DBNDD1,NOP16,ARH
L,FASTK,LY75,ARRDC1,LIMK2,NUMA1,TBL2,SCARB1,CTTN,ITSN1,COMMD5,UFD1L,DBNDD1,NOP16,ARH
MM40,TADA3,MDH2,SEC31A,PAFAH1B2,NSUN5,MCFD2,RIC8A,UBE2N,LFNG,FAM32A,C12ORF57,SLC35
G1,TIMM17B,CYB561,ZNF197,TRMT112,DDOST,TOMM40,MDH2,POLR2C,B3GAT3,PAFAH1B2,NSUN5,T
G1,TIMM17B,CYB561,ZNF197,TRMT112,DDOST,TOMM40,MDH2,POLR2C,B3GAT3,PAFAH1B2,NSUN5,T
IRBP,RPL38,DNASE1L1,RNF32,ATG16L2,SLC3A2,CTSC,RBP5,ARPC2,PMFBP1,VDAC3,NUMA1,GPATCH2L,
ECSIT,NLRP1,NIP7,PLCB2,SLC25A10,TMX2,RPL29,CD99L2,PARD6G,CLASRP,C1ORF198,UGGT2,MICAL1,PR
ECSIT,NLRP1,NIP7,PLCB2,SLC25A10,TMX2,RPL29,CD99L2,PARD6G,CLASRP,C1ORF198,UGGT2,MICAL1,PR
40,ANAPC11,COX17,GBA2,MCFD2,RIC8A,COX20,PEX2,C12ORF57,SLC35E1,OGFOD2,PICK1,TSPAN31,RAL
TK,RNF32,LY75,RCC1,PARD3,DHPS,SCAF11,DBI,SIRT6,GPATCH2L,BUD31,OSGEP,ARHGAP6,SCARB1,CTTN,
,RNH1,RBCK1,PTOV1,LY75,PARD3,SLC3A2,DHPS,SEC62,SET,ARRDC1,TRPT1,KLC1,NOB1,NUMA1,TBL2,SC
,RNH1,RBCK1,PTOV1,LY75,PARD3,SLC3A2,DHPS,SEC62,SET,ARRDC1,TRPT1,KLC1,NOB1,NUMA1,TBL2,SC
NH1,RBCK1,PTOV1,FASTK,RNF32,PARD3,SLC3A2,SEC62,SET,ARRDC1,TRPT1,KLC1,SCAF11,ARPC2,LIMK2,
D3,ACTR6,MAP3K7CL,SEC62,KLC1,SCAF11,ARPC2,WDR19,SIRT6,NUMA1,SAE1,EIF5A,DBNDD1,LDHA,AP
D3,ACTR6,MAP3K7CL,SEC62,KLC1,SCAF11,ARPC2,WDR19,SIRT6,NUMA1,SAE1,EIF5A,DBNDD1,LDHA,AP
10,ZNF197,RAB2A,TRMT112,UBXN1,DDOST,MEIS2,TOMM40,TADA3,KLK10,HSPH1,CNST,PFDN5,PAFAH1
10,ZNF197,RAB2A,TRMT112,UBXN1,DDOST,MEIS2,TOMM40,TADA3,KLK10,HSPH1,CNST,PFDN5,PAFAH1

MD5,AMZ2,NOP16,PSMA6,MRPL33,RUVBL2,EBPL,TESC,SPAG7,STIP1,PHF19,MAGED2,TET2,CHCHD1,SLC3
MD5,AMZ2,NOP16,PSMA6,MRPL33,RUVBL2,EBPL,TESC,SPAG7,STIP1,PHF19,MAGED2,TET2,CHCHD1,SLC3
V1,FASTK,LY75,ATG16L2,PARD3,SLC3A2,ACTR6,SEC62,SET,BOP1,TRPT1,SCAF11,DBI,PMFBP1,LIMK2,VDA
V1,FASTK,LY75,ATG16L2,PARD3,SLC3A2,ACTR6,SEC62,SET,BOP1,TRPT1,SCAF11,DBI,PMFBP1,LIMK2,VDA
RPS9,SLC35G1,WDR62,RFX5,ZC3H14,SLC15A3,PPM1K,LRRC1,LPCAT3,CD44,TM9SF1,SON,CDC37,RPS16,
31A,B3GAT3,PAFAH1B2,NSUN5,UBE2N,FAM32A,OGFOD2,ARIH2,PICK1,TSPAN31,GLYCTK,TP53,TMEM19
1,SNRPB,PARD3,ARPC2,OSGEP,CTTN,EIF5A,COMMD5,UFD1L,USP38,ACOT8,RPS9,ZNF410,RUVBL2,TESC,
2,SEC31A,PAFAH1B2,NSUN5,RIC8A,UBE2N,FAM32A,C12ORF57,SLC35E1,OGFOD2,ARIH2,PICK1,TSPAN31,
MGB1,CYB561A3,RAPGEF5,IQCG,PILRB,CACYBP,CIRBP,RHOF,SMARCB1,STARD10,RNH1,DNASE1L1,RBCK1
MGB1,CYB561A3,RAPGEF5,IQCG,PILRB,CACYBP,CIRBP,RHOF,SMARCB1,STARD10,RNH1,DNASE1L1,RBCK1
M67,PILRB,FAM72A,CIRBP,RHOF,SMARCB1,RBCK1,PTOV1,FASTK,TMEM135,RNF32,LY75,PARD3,TCAIM,S
M67,PILRB,FAM72A,CIRBP,RHOF,SMARCB1,RBCK1,PTOV1,FASTK,TMEM135,RNF32,LY75,PARD3,TCAIM,S
,SMARCB1,RPL38,PRR13,STARD10,RNH1,DNASE1L1,RBCK1,PTOV1,FASTK,TMEM135,LY75,ATG16L2,PAR
,SMARCB1,RPL38,PRR13,STARD10,RNH1,DNASE1L1,RBCK1,PTOV1,FASTK,TMEM135,LY75,ATG16L2,PAR
SET,ARRDC1,RBP5,SCAF11,LIMK2,NUMA1,BUD31,ARHGAP6,SCARB1,CTTN,TSPAN11,ITSN1,EIF5A,DBND
SET,ARRDC1,RBP5,SCAF11,LIMK2,NUMA1,BUD31,ARHGAP6,SCARB1,CTTN,TSPAN11,ITSN1,EIF5A,DBND
1,SCAF11,SAE1,OSGEP,SCARB1,EIF5A,CHID1,RPS9,RCE1,MRPL33,TESC,SPAG7,WDR62,RANBP1,LZTFL1,FI
1,SCAF11,SAE1,OSGEP,SCARB1,EIF5A,CHID1,RPS9,RCE1,MRPL33,TESC,SPAG7,WDR62,RANBP1,LZTFL1,FI
SNAPC2,AKT1S1,TCFL5,ANKRD10,RAP1B,ZNF616,UBE2A,CREM,SWI5,OAZ1,PHF20,MARK2,TMED1
2L6,ZC3H14,PPM1K,LRRC1,CD44,MAGED2,SON,GRK6,CDC37,PHF21A,CHCHD1,RPS25,NIP7,TTC9C,TMX2
2L6,ZC3H14,PPM1K,LRRC1,CD44,MAGED2,SON,GRK6,CDC37,PHF21A,CHCHD1,RPS25,NIP7,TTC9C,TMX2
CDC42,EEF1B2,RPL28,LYPLA2,C17ORF62,RBM3,RAB2A,ATP5J,TRMT112,EMC4,DDOST,LSR,RRNAD1,RAD
UQCRQ,PTPN13,FAM3A,NIP7,COX4I1,GGPS1,RTEL1,SRSF1,FOXRED1,SH3YL1,PPP1CC,RFX3,PSMD4,VIM,
PPP1CC,DGKE,CAMTA1,RPL28,LIG1,CAMLG,PRMT1,ZFAND2B,RPL36,HSP90AB1,ZNF524,RMDN1,PSME2,
R13,RNH1,RBCK1,PTOV1,TMEM135,LY75,PARD3,SLC3A2,SET,CTSC,ARRDC1,TRPT1,KLC1,RBP5,SCAF11,LI
R13,RNH1,RBCK1,PTOV1,TMEM135,LY75,PARD3,SLC3A2,SET,CTSC,ARRDC1,TRPT1,KLC1,RBP5,SCAF11,LI
H1,RBCK1,PTOV1,FASTK,RNF32,LY75,ATG16L2,PARD3,TCAIM,SLC3A2,SEC62,SET,ARRDC1,TRPT1,KLC1,SC
STARD10,RNH1,PTOV1,LY75,PARD3,DHPS,SEC62,SET,BOP1,ARRDC1,TRPT1,ZC3HAV1,WDR19,NUMA1,SA
STARD10,RNH1,PTOV1,LY75,PARD3,DHPS,SEC62,SET,BOP1,ARRDC1,TRPT1,ZC3HAV1,WDR19,NUMA1,SA
IDD1,RCE1,TESC,IGSF8,SPAG7,UBE2L6,STIP1,PHF19,GRK6,METTL17,EPB41L5,ZBTB44,DUSP22,NISCH,CD9
AIAP2,RAPGEF5,SSU72,SRP9,TMEM67,SULT1A3,PILRB,CMC2,YBX3,BLOC1S1,CACYBP,SMARCB1,RPL38,P
AIAP2,RAPGEF5,SSU72,SRP9,TMEM67,SULT1A3,PILRB,CMC2,YBX3,BLOC1S1,CACYBP,SMARCB1,RPL38,P
25A10,TMED4,CD99L2,SH3YL1,IP6K2,CLASRP,JPH2,C1ORF198,MICAL1,PRMT2,TBX6,PCYT2,USE1,PPP1C
25A10,TMED4,CD99L2,SH3YL1,IP6K2,CLASRP,JPH2,C1ORF198,MICAL1,PRMT2,TBX6,PCYT2,USE1,PPP1C
P,RNH1,DNASE1L1,PTOV1,ATG16L2,SNRPB,TCAIM,BOP1,ARRDC1,KLC1,ZC3HAV1,SCAF11,NUMA1,GPAT
P,RNH1,DNASE1L1,PTOV1,ATG16L2,SNRPB,TCAIM,BOP1,ARRDC1,KLC1,ZC3HAV1,SCAF11,NUMA1,GPAT
,CTTN,EIF5A,COMMD5,CHID1,RPL13A,LDHA,RPS9,TESC,SPAG7,WDR62,STYXL1,RFX5,RANBP1,LZTFL1,FIS
RHGAP6,EIF5A,RPL13A,LDHA,RPS9,RCE1,ZNF410,PCMT1,IGSF8,NNMT,WDR62,RFX5,VRK3,PPM1K,TM9S
RHGAP6,EIF5A,RPL13A,LDHA,RPS9,RCE1,ZNF410,PCMT1,IGSF8,NNMT,WDR62,RFX5,VRK3,PPM1K,TM9S
RD3,TCAIM,ACTR6,SET,BOP1,ARRDC1,TRPT1,KLC1,SCAF11,LIMK2,SIRT6,GPATCH2L,BUD31,NPRL2,TBL2,
MK2,NUMA1,SCARB1,CTTN,TSPAN11,ITSN1,DBNDD1,NOP16,PSMA6,AP3B2,EBPL,TESC,IGSF8,SPAG7,PP
MK2,NUMA1,SCARB1,CTTN,TSPAN11,ITSN1,DBNDD1,NOP16,PSMA6,AP3B2,EBPL,TESC,IGSF8,SPAG7,PP
RD3,DHPS,SEC62,SET,ARRDC1,TRPT1,ZC3HAV1,WDR19,NUMA1,TBL2,CTTN,TSPAN11,ITSN1,EIF5A,COM
RD3,DHPS,SEC62,SET,ARRDC1,TRPT1,ZC3HAV1,WDR19,NUMA1,TBL2,CTTN,TSPAN11,ITSN1,EIF5A,COM
1,NOB1,WDR19,SIRT6,SAE1,TBL2,ARHGAP6,SCARB1,TNFSF10,COMMD5,USP38,THYN1,DBNDD1,GSTK1,
RD3,DHPS,SEC62,SET,BOP1,KLC1,ARPC2,PMFBP1,NOB1,SIRT6,NUMA1,GPATCH2L,SAE1,NPRL2,SCARB1,
RD3,DHPS,SEC62,SET,BOP1,KLC1,ARPC2,PMFBP1,NOB1,SIRT6,NUMA1,GPATCH2L,SAE1,NPRL2,SCARB1,

V1,ATG16L2,SNRPB,TCAIM,SEC62,BOP1,ARRDC1,KLC1,SCAF11,GPATCH2L,BUD31,NPRL2,OSGEP,ARHGAI

V1,ATG16L2,SNRPB,TCAIM,SEC62,BOP1,ARRDC1,KLC1,SCAF11,GPATCH2L,BUD31,NPRL2,OSGEP,ARHGAI

PRC1,NIP7,NISCH,CENPK,GGPS1,SRSF1,AUP1,YIPF3,IP6K2,MAGT1,CCDC25,EEF1B2,PPP1CC,RFX3,CAMTA

PRC1,NIP7,NISCH,CENPK,GGPS1,SRSF1,AUP1,YIPF3,IP6K2,MAGT1,CCDC25,EEF1B2,PPP1CC,RFX3,CAMTA

12,DDOST,TOMM40,ZNF687,CARHSP1,COX17,FTH1,GMPR2,RIC8A,PDCD6,FAM32A,USP40,PEX2,C12ORI

T,SNRPN,WDR62,PLEKHA5,VRK3,PCBP2,TET2,GRK6,CDC37,PTPN13,FAM3A,PSMA3,METTL17,TTC9C,COX

3,NUMA1,BUD31,NPRL2,OSGEP,ITSN1,EIF5A,COMMD5,DBNDD1,NOP16,AP3B2,RPS9,SLC35G1,ZNF410,

3,NUMA1,BUD31,NPRL2,OSGEP,ITSN1,EIF5A,COMMD5,DBNDD1,NOP16,AP3B2,RPS9,SLC35G1,ZNF410,

R19,EIF5A,DBNDD1,SH2B1,NOP16,RPL13A,TTLL5,ACOT8,ZNF410,PCMT1,IGSF8,RBPMS,WDR62,RANBP1

,RAPGEF5,SSU72,MYL6B,IQCG,TMEM67,YBX3,BLOC1S1,CACYBP,CIRBP,RHOF,SMARCB1,RPL38,STARD10,

,RAPGEF5,SSU72,MYL6B,IQCG,TMEM67,YBX3,BLOC1S1,CACYBP,CIRBP,RHOF,SMARCB1,RPL38,STARD10,

CA3,PCNA,SRSF2,MRPS22,OSMR,PPFIBP1,SEMA4D,LAS1L,HMGB1,SULF2,BAIAP2,RAPGEF5,SSU72,MYL6B

CA3,PCNA,SRSF2,MRPS22,OSMR,PPFIBP1,SEMA4D,LAS1L,HMGB1,SULF2,BAIAP2,RAPGEF5,SSU72,MYL6B

N,TET2,FUZ,PPRC1,RDH11,NISCH,SRSF1,AUP1,KXD1,IP6K2,YIF1A,MAGT1,GUK1,USE1,PPP1CC,RPL28,APEI

CHID1,TAZ,RPL13A,MRPL33,TESC,SPAG7,WDR62,STYXL1,RFX5,RANBP1,LZTFL1,FIS1,TUBA1C,IRF9,STIP1,T

CHID1,TAZ,RPL13A,MRPL33,TESC,SPAG7,WDR62,STYXL1,RFX5,RANBP1,LZTFL1,FIS1,TUBA1C,IRF9,STIP1,T

SF1B,CAPG,PPIL2,TROAP,LIG1,ACAD10,ZNF197,RAB2A,ZNF524,TRMT112,UBXN1,DDOST,MEIS2,TOMM4

SF1B,CAPG,PPIL2,TROAP,LIG1,ACAD10,ZNF197,RAB2A,ZNF524,TRMT112,UBXN1,DDOST,MEIS2,TOMM4

,ZNF410,EBPL,NUDT5,ZC3H14,TUBA1C,CD44,MAGED2,SNF8,CDC37,CHCHD1,RGS3,METTL17,ZBTB44,NIF

1,RCE1,EBPL,TESC,IGSF8,SPAG7,WDR62,POLD4,UBE2L6,STIP1,PHF19,METTL17,ZBTB44,CD99L2,PARD6G

L,PRR13,RNH1,DNASE1L1,KIF4A,PTOV1,TMEM135,RNF32,ATG16L2,TCAIM,DHPS,ACTR6,SET,BOP1,KLC1,I

L,PRR13,RNH1,DNASE1L1,KIF4A,PTOV1,TMEM135,RNF32,ATG16L2,TCAIM,DHPS,ACTR6,SET,BOP1,KLC1,I

STIP1,LPCAT3,TM9SF1,CDC37,RPS16,YLPM1,METTL17,DUSP22,SRSF1,AUP1,RABL5,PARD6G,UGGT2,TBX6

STIP1,LPCAT3,TM9SF1,CDC37,RPS16,YLPM1,METTL17,DUSP22,SRSF1,AUP1,RABL5,PARD6G,UGGT2,TBX6

HOF,SMARCB1,RPL38,RNH1,KIF4A,PTOV1,SNRPB,PARD3,TCAIM,SEC62,BOP1,ZC3HAV1,SCAF11,NOB1,WD

HOF,SMARCB1,RPL38,RNH1,KIF4A,PTOV1,SNRPB,PARD3,TCAIM,SEC62,BOP1,ZC3HAV1,SCAF11,NOB1,WD

,DDOST,TOMM40,MDH2,POLR2C,PAFAH1B2,NSUN5,FAM32A,C12ORF57,ZNF223,OGFOD2,EIF3G,SLC52

,DDOST,TOMM40,MDH2,POLR2C,PAFAH1B2,NSUN5,FAM32A,C12ORF57,ZNF223,OGFOD2,EIF3G,SLC52

5,PHF19,ZBTB44,CD99L2,PARD6G,SH3YL1,IP6K2,UBE2I,AUNIP,C14ORF2,PCYT2,PPP1CC,DGKE,CAMTA1,LI

OV1,TMEM135,LY75,PARD3,SLC3A2,SET,ARRDC1,TRPT1,RBP5,LIMK2,NUMA1,BUD31,ARHGAP6,SCARB1,

OV1,TMEM135,LY75,PARD3,SLC3A2,SET,ARRDC1,TRPT1,RBP5,LIMK2,NUMA1,BUD31,ARHGAP6,SCARB1,

V1,SNRPB,TCAIM,SEC62,BOP1,ARRDC1,KLC1,SCAF11,GPATCH2L,NPRL2,OSGEP,ARHGAP6,EIF5A,AMZ2,TA

V1,SNRPB,TCAIM,SEC62,BOP1,ARRDC1,KLC1,SCAF11,GPATCH2L,NPRL2,OSGEP,ARHGAP6,EIF5A,AMZ2,TA

DDOST,MEIS2,TOMM40,TADA3,KLK10,HSPH1,CNST,PFDN5,PAFAH1B2,ZFPL1,GMPR2,RIC8A,TCP1,FAM3

DDOST,MEIS2,TOMM40,TADA3,KLK10,HSPH1,CNST,PFDN5,PAFAH1B2,ZFPL1,GMPR2,RIC8A,TCP1,FAM3

T1,SCAF11,LIMK2,SCARB1,ITSN1,NUDT22,USP38,DBNDD1,NOP16,TTLL5,LDHA,EBPL,TESC,NUDT5,RBPMS

T1,SCAF11,LIMK2,SCARB1,ITSN1,NUDT22,USP38,DBNDD1,NOP16,TTLL5,LDHA,EBPL,TESC,NUDT5,RBPMS

C,IQCG,TMEM67,SULT1A3,FAM72A,CMC2,BLOC1S1,CIRBP,RHOF,SMARCB1,PRR13,RNH1,DNASE1L1,KIF4

F5,SSU72,SRP9,IQCG,SULT1A3,UAP1,PILRB,FAM72A,YBX3,CACYBP,CIRBP,RHOF,SMARCB1,PRR13,RNH1,D

F5,SSU72,SRP9,IQCG,SULT1A3,UAP1,PILRB,FAM72A,YBX3,CACYBP,CIRBP,RHOF,SMARCB1,PRR13,RNH1,D

RK6,CDC37,PTPN13,PSMA3,METTL17,TTC9C,COX4I1,THEM4,CENPK,AUP1,TRNT1,MED15,MAGT1,C14OF

ARHGAP6,EIF5A,CHID1,AP3B2,ZNF410,RUVBL2,PCMT1,EBPL,NUDT5,RANBP1,ZC3H14,TUBA1C,CD44,MA

ARHGAP6,EIF5A,CHID1,AP3B2,ZNF410,RUVBL2,PCMT1,EBPL,NUDT5,RANBP1,ZC3H14,TUBA1C,CD44,MA

L13A,ARHGEF39,PSMA6,ACOT8,RCE1,NUDT5,SNRPN,STYXL1,POLD4,VRK3,IRF9,DST,PCBP2,UQCRQ,RPS16

L13A,ARHGEF39,PSMA6,ACOT8,RCE1,NUDT5,SNRPN,STYXL1,POLD4,VRK3,IRF9,DST,PCBP2,UQCRQ,RPS16

2,MRPS22,HDAC9,OSMR,SEMA4D,LAS1L,HMGB1,BAIAP2,RAPGEF5,SSU72,MYL6B,FAM122C,SRP9,IQCG,T

2,MRPS22,HDAC9,OSMR,SEMA4D,LAS1L,HMGB1,BAIAP2,RAPGEF5,SSU72,MYL6B,FAM122C,SRP9,IQCG,T

,NISCH,RPS6,SRSF1,FOXRED1,IP6K2,MAGT1,RFX3,RPL21,CAMLG,C19ORF54,C19ORF48,CBFA2T2,HYPK,M
LC15A3,SNF8,UBR4,CHCHD1,ECSIT,NLRP1,NIP7,PLCB2,SLC25A10,TMX2,SDF4,RPL29,CD99L2,PARD6G,CL
LC15A3,SNF8,UBR4,CHCHD1,ECSIT,NLRP1,NIP7,PLCB2,SLC25A10,TMX2,SDF4,RPL29,CD99L2,PARD6G,CL
V1,FASTK,TMEM135,ATG16L2,SNRPB,ACTR6,SET,BOP1,ARRDC1,KLC1,SCAF11,DBI,LIMK2,VDAC3,SIRT6,N
F8,NNMT,WDR62,PLEKHA5,VRK3,PPM1K,PCBP2,TET2,GRK6,CDC37,PTPN13,FAM3A,PSMA3,METTL17,EP
F8,NNMT,WDR62,PLEKHA5,VRK3,PPM1K,PCBP2,TET2,GRK6,CDC37,PTPN13,FAM3A,PSMA3,METTL17,EP
F8,NNMT,WDR62,PLEKHA5,VRK3,PPM1K,PCBP2,TET2,GRK6,CDC37,PTPN13,FAM3A,PSMA3,METTL17,EP
3E2N,FAM32A,DCTN5,OXLD1,ARIH2,FBXL18,ZNF512,ZNF593,GLYCTK,EIF2A,GLI4,SLC35A2,EPN1,ZBTB8OS
PL,SPAG7,WDR62,RANBP1,STIP1,LPCAT3,CPSF3L,ZBTB44,NISCH,CD99L2,PARD6G,SH3YL1,IP6K2,MICAL1
SF1B,CAPG,PPIL2,LIG1,ACAD10,CDK11A,ZNF197,RAB2A,ZNF524,TRMT112,UBXN1,SAT1,DDOST,MEIS2,T
SF1B,CAPG,PPIL2,LIG1,ACAD10,CDK11A,ZNF197,RAB2A,ZNF524,TRMT112,UBXN1,SAT1,DDOST,MEIS2,T
HPS,SEC62,SET,BOP1,TRPT1,KLC1,ARPC2,PMFBP1,NOB1,SIRT6,NUMA1,GPATCH2L,SAE1,SCARB1,TSPAN
HPS,SEC62,SET,BOP1,TRPT1,KLC1,ARPC2,PMFBP1,NOB1,SIRT6,NUMA1,GPATCH2L,SAE1,SCARB1,TSPAN
35,ATG16L2,PARD3,TCAIM,SLC3A2,SET,CTSC,BOP1,ARRDC1,TRPT1,DBI,LIMK2,NUMA1,ARHGAP6,SCARB
35,ATG16L2,PARD3,TCAIM,SLC3A2,SET,CTSC,BOP1,ARRDC1,TRPT1,DBI,LIMK2,NUMA1,ARHGAP6,SCARB
75,ATG16L2,PARD3,SLC3A2,ACTR6,SET,SCAF11,DBI,PMFBP1,LIMK2,VDAC3,NUMA1,GPATCH2L,CTTN,TSP
75,ATG16L2,PARD3,SLC3A2,ACTR6,SET,SCAF11,DBI,PMFBP1,LIMK2,VDAC3,NUMA1,GPATCH2L,CTTN,TSP
AG7,WDR62,POLD4,UBE2L6,STIP1,PHF19,ZBTB44,CD99L2,PARD6G,SH3YL1,IP6K2,UBE2I,AUNIP,C14ORF
7,UAP1,FAM72A,BLOC1S1,CACYBP,CIRBP,RHOF,RPL38,STARD10,PTOV1,FASTK,TMEM135,ATG16L2,RCC
7,UAP1,FAM72A,BLOC1S1,CACYBP,CIRBP,RHOF,RPL38,STARD10,PTOV1,FASTK,TMEM135,ATG16L2,RCC
1,TWSG1,GMPPB,LMBR1,NAP1L1,CHCHD7,PDCD5,C11ORF48,LLGL2,B3GAT3,BRF1,OARD1,SMARCC2,N
SMARCB1,RPL38,PRR13,STARD10,RNH1,DNASE1L1,PTOV1,FASTK,TMEM135,LY75,ATG16L2,PARD3,TCAI
SMARCB1,RPL38,PRR13,STARD10,RNH1,DNASE1L1,PTOV1,FASTK,TMEM135,LY75,ATG16L2,PARD3,TCAI
ICK1,FBXL18,TSPAN31,GLYCTK,EIF2A,TMEM199,EPN1,ZCCHC9,AZI2,LAMTOR1,U2AF1L4,UBA52,SUMO1
ECSIT,ZBTB44,NIP7,NISCH,CD99L2,PARD6G,IP6K2,CLASRP,JPH2,MICAL1,HS2ST1,PRMT2,TMEM205,PTK2,
CACYBP,RPL38,PRR13,KIF4A,PTOV1,RNF32,ATG16L2,RCC1,PARD3,DHPS,ACTR6,MAP3K7CL,SEC62,ZNF23
CACYBP,RPL38,PRR13,KIF4A,PTOV1,RNF32,ATG16L2,RCC1,PARD3,DHPS,ACTR6,MAP3K7CL,SEC62,ZNF23
HOF,SMARCB1,RPL38,PRR13,RNH1,DNASE1L1,KIF4A,RBCK1,PTOV1,RNF32,LY75,ATG16L2,PARD3,SLC3A
HOF,SMARCB1,RPL38,PRR13,RNH1,DNASE1L1,KIF4A,RBCK1,PTOV1,RNF32,LY75,ATG16L2,PARD3,SLC3A
1,FASTK,RNF32,RCC1,ACTR6,SET,CTSC,SCAF11,DBI,ARPC2,NUMA1,BUD31,NPRL2,TBL2,CTTN,ITSN1,AMZ
GSF8,NNMT,WDR62,VRK3,TET2,CDC37,OBSL1,FAM3A,PSMA3,NLRP1,TTC9C,COX4I1,DBNDD2,COASY,SR
C,CDC42,TMEM205,CCDC25,TROAP,LIG1,CYB561,CDK11A,RAB2A,TRMT112,CHPF,ZNF814,ERCC5,DDOS
RPL38,STARD10,RNH1,DNASE1L1,RBCK1,PTOV1,FASTK,TMEM135,LY75,ATG16L2,TCAIM,SLC3A2,MAP3K
RPL38,STARD10,RNH1,DNASE1L1,RBCK1,PTOV1,FASTK,TMEM135,LY75,ATG16L2,TCAIM,SLC3A2,MAP3K
STARD10,RNH1,DNASE1L1,FASTK,TMEM135,LY75,ATG16L2,RCC1,DHPS,SET,CTSC,BOP1,SCAF11,DBI,WD
STARD10,RNH1,DNASE1L1,FASTK,TMEM135,LY75,ATG16L2,RCC1,DHPS,SET,CTSC,BOP1,SCAF11,DBI,WD
13,RNH1,RBCK1,RNF32,LY75,DHPS,ACTR6,SEC62,CTSC,ZNF235,BOP1,ARRDC1,TRPT1,KLC1,RBP5,ZC3HA
13,RNH1,RBCK1,RNF32,LY75,DHPS,ACTR6,SEC62,CTSC,ZNF235,BOP1,ARRDC1,TRPT1,KLC1,RBP5,ZC3HA
PSMA6,ACOT8,RCE1,NUDT5,SNRPN,STYXL1,POLD4,VRK3,IRF9,DST,UQCRQ,RPS16,TSG101,SLC39A11,FU
PSMA6,ACOT8,RCE1,NUDT5,SNRPN,STYXL1,POLD4,VRK3,IRF9,DST,UQCRQ,RPS16,TSG101,SLC39A11,FU
C3,WDR19,NUMA1,TBL2,SCARB1,TSPAN11,CHID1,AMZ2,DBNDD1,RPL13A,AP3B2,ZNF410,PCMT1,EBPL,T
C3,WDR19,NUMA1,TBL2,SCARB1,TSPAN11,CHID1,AMZ2,DBNDD1,RPL13A,AP3B2,ZNF410,PCMT1,EBPL,T
1,GPATCH2L,NPRL2,OSGEP,ARHGAP6,EIF5A,AMZ2,TAZ,NOP16,LDHA,RCE1,ZNF410,PCMT1,IGSF8,NNMT,
1,GPATCH2L,NPRL2,OSGEP,ARHGAP6,EIF5A,AMZ2,TAZ,NOP16,LDHA,RCE1,ZNF410,PCMT1,IGSF8,NNMT,
1,CAPG,PSMD4,TIMM17B,TMEM126B,PODXL,EXOSC9,ACAD10,DUT,ZNF524,PNPO,ZNF625,RPS20,MPDU
RBP,SMARCB1,PRR13,STARD10,RNH1,DNASE1L1,PTOV1,RNF32,LY75,SEC62,SET,ARRDC1,SCAF11,ARPC2,
TTL17,TTC9C,COX4I1,THEM4,CENPK,AUP1,TRNT1,MED15,MAGT1,C14ORF2,GUK1,CENPN,PCYT2,MAF,PF

TTL17,TTC9C,COX4I1,THEM4,CENPK,AUP1,TRNT1,MED15,MAGT1,C14ORF2,GUK1,CENPN,PCYT2,MAF,P
),IGSF8,NNMT,SNRPN,WDR62,PLEKHA5,VRK3,PCBP2,TET2,GRK6,CDC37,PTPN13,FAM3A,PSMA3,METTL1
),IGSF8,NNMT,SNRPN,WDR62,PLEKHA5,VRK3,PCBP2,TET2,GRK6,CDC37,PTPN13,FAM3A,PSMA3,METTL1
,CACYBP,RHOF,SMARCB1,RPL38,PRR13,KIF4A,RBCK1,PTOV1,RNF32,LY75,ATG16L2,RCC1,PARD3,MAP3K
ARD10,RNH1,DNASE1L1,KIF4A,LY75,ATG16L2,RCC1,SNRPB,DHPS,BOP1,TRPT1,DBI,NOB1,CTTN,TSPAN11
ARD10,RNH1,DNASE1L1,KIF4A,LY75,ATG16L2,RCC1,SNRPB,DHPS,BOP1,TRPT1,DBI,NOB1,CTTN,TSPAN11
STARD10,RNH1,DNASE1L1,FASTK,TMEM135,LY75,ATG16L2,RCC1,DHPS,SET,CTSC,BOP1,SCAF11,DBI,WD
H1,DNASE1L1,FASTK,TMEM135,ATG16L2,RCC1,SLC3A2,DHPS,SET,BOP1,SCAF11,DBI,WDR19,GPATCH2L,
H1,DNASE1L1,FASTK,TMEM135,ATG16L2,RCC1,SLC3A2,DHPS,SET,BOP1,SCAF11,DBI,WDR19,GPATCH2L,
H1,DNASE1L1,FASTK,TMEM135,ATG16L2,RCC1,SLC3A2,DHPS,SET,BOP1,SCAF11,DBI,WDR19,GPATCH2L,
MGB1,SULF2,BAIAP2,RAPGEF5,SSU72,MYL6B,FAM122C,SRP9,IQCG,PILRB,FAM72A,YBX3,CACYBP,CIRBP,
MGB1,SULF2,BAIAP2,RAPGEF5,SSU72,MYL6B,FAM122C,SRP9,IQCG,PILRB,FAM72A,YBX3,CACYBP,CIRBP,
C1,SLC25A10,SDF4,TRNT1,ITGB1BP1,C1ORF198,EEF1B2,PPA1,TNFRSF14,CAPN10,PRMT1,C19ORF54,C1
C1,SLC25A10,SDF4,TRNT1,ITGB1BP1,C1ORF198,EEF1B2,PPA1,TNFRSF14,CAPN10,PRMT1,C19ORF54,C1
HNRNPA1,RFXANK,SPINT2,MOK,ELOF1,DCTN5,TCF25,ZNF248,NXF1,NUP62,PSPC1,NDUFA3,C1ORF63,DN
1,SNRPB,PARD3,SLC3A2,DHPS,ACTR6,ZNF235,TRPT1,DBI,PMFBP1,BUD31,NPRL2,CRYGS,CTTN,EIF5A,CHI
1,SNRPB,PARD3,SLC3A2,DHPS,ACTR6,ZNF235,TRPT1,DBI,PMFBP1,BUD31,NPRL2,CRYGS,CTTN,EIF5A,CHI
RPL13A,LDHA,RPS9,TESC,SPAG7,STYXL1,RFX5,RANBP1,LZTFL1,FIS1,TUBA1C,STIP1,TET2,GRK6,HMGA2,CD
HMGB1,SULF2,CYB561A3,BAIAP2,RAPGEF5,SSU72,FAM122C,IQCG,TMEM67,SULT1A3,FAM72A,YBX3,BL
HMGB1,SULF2,CYB561A3,BAIAP2,RAPGEF5,SSU72,FAM122C,IQCG,TMEM67,SULT1A3,FAM72A,YBX3,BL
,CMC2,YBX3,CACYBP,SMARCB1,RPL38,RBCK1,PTOV1,FASTK,ATG16L2,RCC1,PARD3,DHPS,SEC62,SET,BO
,CMC2,YBX3,CACYBP,SMARCB1,RPL38,RBCK1,PTOV1,FASTK,ATG16L2,RCC1,PARD3,DHPS,SEC62,SET,BO
CYBP,RPL38,KIF4A,PTOV1,RNF32,ATG16L2,SNRPB,PARD3,ACTR6,MAP3K7CL,CTSC,ZNF235,TRPT1,ZC3HA
CYBP,RPL38,KIF4A,PTOV1,RNF32,ATG16L2,SNRPB,PARD3,ACTR6,MAP3K7CL,CTSC,ZNF235,TRPT1,ZC3HA
7,PTPN13,PSMA3,METTL17,TTC9C,COX4I1,THEM4,CENPK,AUP1,TRNT1,MED15,MAGT1,C14ORF2,GUK1,
32,LY75,TCAIM,SET,CTSC,BOP1,ARRDC1,KLC1,ZC3HAV1,SCAF11,ARPC2,VDAC3,NOB1,WDR19,BUD31,N
P1,KLC1,RBP5,GPATCH2L,SUMF2,BUD31,ITSN1,EIF5A,TAZ,NOP16,GSTK1,IGSF8,NUDT5,SNRPN,WDR62,
P1,KLC1,RBP5,GPATCH2L,SUMF2,BUD31,ITSN1,EIF5A,TAZ,NOP16,GSTK1,IGSF8,NUDT5,SNRPN,WDR62,
112,DDOST,MEIS2,TOMM40,TADA3,PDCD5,MDH2,SEC31A,PFDN5,PAFAH1B2,NSUN5,GMPR2,RIC8A,TC
112,DDOST,MEIS2,TOMM40,TADA3,PDCD5,MDH2,SEC31A,PFDN5,PAFAH1B2,NSUN5,GMPR2,RIC8A,TC
,FASTK,TMEM135,ATG16L2,TCAIM,SLC3A2,BOP1,TRPT1,KLC1,RBP5,DBI,ARPC2,SIRT6,GPATCH2L,CTTN,E
,SMARCB1,STARD10,RNH1,DNASE1L1,FASTK,ATG16L2,RCC1,PARD3,DHPS,SET,BOP1,SCAF11,DBI,WDR19
YBP,RBCK1,FASTK,SNRPB,DHPS,SEC62,SET,BOP1,TRPT1,KLC1,ARPC2,PMFBP1,NOB1,SIRT6,NUMA1,SAE1
YBP,RBCK1,FASTK,SNRPB,DHPS,SEC62,SET,BOP1,TRPT1,KLC1,ARPC2,PMFBP1,NOB1,SIRT6,NUMA1,SAE1
OXRED1,TRNT1,MAGT1,GUK1,CENPN,MAF,PPA1,PPIL2,ZFAND2B,MAPKAPK5,HSP90AB1,CEP70,RAB2A,
B561,MOB1A,SYNGR2,GPBP1,ASPH,RPS15,HHLA3,SYNGR1,MTA3,PSME1,MDH2,HSPH1,SIRT5,ASH2L,S
TTC9C,COX4I1,THEM4,CENPK,AUP1,TRNT1,MED15,MAGT1,C14ORF2,GUK1,CENPN,PCYT2,MAF,PPA1,PP
SRP9,IQCG,SULT1A3,UAP1,PILRB,YBX3,CACYBP,CIRBP,RHOF,SMARCB1,PRR13,RNH1,DNASE1L1,RBCK1,P
JP1,FOXRED1,TRNT1,MAGT1,GUK1,CENPN,MAF,PPA1,PPIL2,ZFAND2B,MAPKAPK5,HSP90AB1,CEP70,RA
PPA1,CAPN10,C19ORF54,C19ORF48,UPP1,RPL36,NFAM1,COPZ1,ASPH,PNPO,ZNF814,C12ORF76,SLC33A
BX3,BLOC1S1,CACYBP,CIRBP,RHOF,RPL38,PRR13,KIF4A,RBCK1,RNF32,RCC1,SNRPB,TCAIM,SLC3A2,DHPS
BX3,BLOC1S1,CACYBP,CIRBP,RHOF,RPL38,PRR13,KIF4A,RBCK1,RNF32,RCC1,SNRPB,TCAIM,SLC3A2,DHPS
SGEP,ARHGAP6,TAZ,LDHA,RPS9,RCE1,ZNF410,PCMT1,IGSF8,NNMT,WDR62,VRK3,PPM1K,PHF19,PCBP2,
SGEP,ARHGAP6,TAZ,LDHA,RPS9,RCE1,ZNF410,PCMT1,IGSF8,NNMT,WDR62,VRK3,PPM1K,PHF19,PCBP2,
BA1C,IRF9,STIP1,TET2,HMGA2,CDC37,UBR4,FAM3A,PPRC1,TRNT1,ITGB1BP1,C1ORF198,PCYT2,USE1,CC
BA1C,IRF9,STIP1,TET2,HMGA2,CDC37,UBR4,FAM3A,PPRC1,TRNT1,ITGB1BP1,C1ORF198,PCYT2,USE1,CC
BA1C,IRF9,STIP1,TET2,HMGA2,CDC37,UBR4,FAM3A,PPRC1,TRNT1,ITGB1BP1,C1ORF198,PCYT2,USE1,CC

5,SNRPB,DHPS,ACTR6,ZNF235,BOP1,TRPT1,ZC3HAV1,DBI,ARPC2,PMFBP1,LIMK2,VDAC3,SIRT6,NUMA1,G
RK6,CDC37,PTPN13,PSMA3,METTL17,TTC9C,COX4I1,THEM4,CENPK,AUP1,TRNT1,MED15,MAGT1,C14OF
RK6,CDC37,PTPN13,PSMA3,METTL17,TTC9C,COX4I1,THEM4,CENPK,AUP1,TRNT1,MED15,MAGT1,C14OF
DAP,PRMT1,STOML2,HNRNPC,CYB561,MOB1A,CDK11A,DGUOK,UBXN1,ERCC5,LMBR1,MCTS1,DDOST,LSI
AGED2,RPS16,TSG101,PHF21A,METTL17,RPS25,COX4I1,PLCB2,TMX2,CENPK,FOXRED1,CDC42,PTK2,GUK
AGED2,RPS16,TSG101,PHF21A,METTL17,RPS25,COX4I1,PLCB2,TMX2,CENPK,FOXRED1,CDC42,PTK2,GUK
L,SULF2,BAIAP2,SSU72,FAM122C,IQCG,TMEM67,PILRB,CMC2,BLOC1S1,CACYBP,RHOF,SMARCB1,RPL38,
L,SULF2,BAIAP2,SSU72,FAM122C,IQCG,TMEM67,PILRB,CMC2,BLOC1S1,CACYBP,RHOF,SMARCB1,RPL38,
67,SULT1A3,PILRB,FAM72A,YBX3,BLOC1S1,CACYBP,CIRBP,RHOF,SMARCB1,RPL38,PRR13,RNH1,RBCK1,P
67,SULT1A3,PILRB,FAM72A,YBX3,BLOC1S1,CACYBP,CIRBP,RHOF,SMARCB1,RPL38,PRR13,RNH1,RBCK1,P
,RNH1,ATG16L2,SNRPB,SEC62,SET,CTSC,ZNF235,ARRDC1,RBP5,ZC3HAV1,PMFBP1,NUMA1,BUD31,ARHG
38,NDUFA3,SMUG1,GRK4,VAPA,MGRN1,ABLIM2,IFRD2,DYNC1I2,AP2S1,ZNF345,NDUFB10,DNAJB12,CO
TK,SNRPB,PARD3,DHPS,SEC62,SET,BOP1,KLC1,SCAF11,ARPC2,PMFBP1,NOB1,SIRT6,NUMA1,GPATCH2L,
TK,SNRPB,PARD3,DHPS,SEC62,SET,BOP1,KLC1,SCAF11,ARPC2,PMFBP1,NOB1,SIRT6,NUMA1,GPATCH2L,
MA1,TBL2,SCARB1,CTTN,ITSN1,THYN1,DBNDD1,RCE1,EBPL,TESC,IGSF8,SPAG7,WDR62,POLD4,UBE2L6,S
2,SNRPB,PARD3,DHPS,SEC62,SET,BOP1,TRPT1,KLC1,ARPC2,PMFBP1,NOB1,SIRT6,NUMA1,GPATCH2L,SA
2,SNRPB,PARD3,DHPS,SEC62,SET,BOP1,TRPT1,KLC1,ARPC2,PMFBP1,NOB1,SIRT6,NUMA1,GPATCH2L,SA
3R4,FAM3A,PSMA3,METTL17,PPRC1,COX4I1,AUP1,TRNT1,ITGB1BP1,MAGT1,GUK1,CENPN,CARD14,PCY
3R4,FAM3A,PSMA3,METTL17,PPRC1,COX4I1,AUP1,TRNT1,ITGB1BP1,MAGT1,GUK1,CENPN,CARD14,PCY
EMA4D,LAS1L,HMGB1,CYB561A3,BAIAP2,RAPGEF5,SSU72,MYL6B,FAM122C,SRP9,IQCG,TMEM67,SULT1
1,FASTK,ATG16L2,RCC1,SNRPB,TCAIM,SLC3A2,DHPS,ZC3HAV1,DBI,GPATCH2L,BUD31,OSGEP,ARHGAP6,
1,FASTK,ATG16L2,RCC1,SNRPB,TCAIM,SLC3A2,DHPS,ZC3HAV1,DBI,GPATCH2L,BUD31,OSGEP,ARHGAP6,
M3A,TTC9C,MED15,UGGT2,TMEM205,EEF1B2,PPA1,ENDOV,RPL28,VIM,C19ORF54,HNRNPC,CARS,DGUO
G,SULT1A3,CACYBP,CIRBP,RHOF,SMARCB1,RPL38,RNH1,DNASE1L1,RBCK1,PTOV1,FASTK,TMEM135,RN
G,SULT1A3,CACYBP,CIRBP,RHOF,SMARCB1,RPL38,RNH1,DNASE1L1,RBCK1,PTOV1,FASTK,TMEM135,RN
NF8,CDC37,CHCHD1,ZBTB44,CD99L2,COASY,SRSF1,MED15,IP6K2,KCNMA1,CDC42,GUK1,EDF1,DGKE,CRI
NOP16,AP3B2,RPS9,SLC35G1,TESC,NUDT5,WDR62,IL1RAP,STYXL1,POLD4,PPM1K,LRRC1,HMGA2,CPSF3L
NOP16,AP3B2,RPS9,SLC35G1,TESC,NUDT5,WDR62,IL1RAP,STYXL1,POLD4,PPM1K,LRRC1,HMGA2,CPSF3L
AM3A,NLRP1,PLCB2,FOXRED1,KXD1,PRMT2,JUP,GUK1,ATF7,PCYT2,EDF1,LIG1,APEH,NUTF2,SHARPIN,ZN

TTL17,TTC9C,COX4I1,THEM4,CENPK,AUP1,TRNT1,MED15,MAGT1,C14ORF2,GUK1,CENPN,PCYT2,MAF,PF
TTL17,TTC9C,COX4I1,THEM4,CENPK,AUP1,TRNT1,MED15,MAGT1,C14ORF2,GUK1,CENPN,PCYT2,MAF,PF
NT1,MAGT1,GUK1,CENPN,MAF,PPA1,PPIL2,ZFAND2B,MAPKAPK5,HSP90AB1,RAB2A,GPBP1,CHPF,MVD,IL
FAM3A,PSMA3,METTL17,NIP7,TTC9C,COX4I1,THEM4,AUP1,FOXRED1,TRNT1,MED15,MAGT1,C14ORF2,G
38,STARD10,RNH1,DNASE1L1,FASTK,ATG16L2,RCC1,DHPS,SET,BOP1,DBI,GPATCH2L,SAE1,NPRL2,OSGEP,
38,STARD10,RNH1,DNASE1L1,FASTK,ATG16L2,RCC1,DHPS,SET,BOP1,DBI,GPATCH2L,SAE1,NPRL2,OSGEP,
38,STARD10,RNH1,DNASE1L1,FASTK,ATG16L2,RCC1,DHPS,SET,BOP1,DBI,GPATCH2L,SAE1,NPRL2,OSGEP,
2,SAP30BP,CLASRP,UGGT2,MICAL1,PRMT2,EIF3M,JUP,PTK2,ATF7,TIMM17B,LSM12,ZFAND2B,SHARPIN,
2,SAP30BP,CLASRP,UGGT2,MICAL1,PRMT2,EIF3M,JUP,PTK2,ATF7,TIMM17B,LSM12,ZFAND2B,SHARPIN,
,PSMA3,METTL17,NIP7,TTC9C,COX4I1,SRSF1,AUP1,FOXRED1,TRNT1,MAGT1,GUK1,CENPN,AGPAT4,MAF
,TAZ,LDHA,RCE1,ZNF410,PCMT1,IGSF8,NNMT,WDR62,RFX5,VRK3,PPM1K,PHF19,PCBP2,TM9SF1,TET2,G
,TAZ,LDHA,RCE1,ZNF410,PCMT1,IGSF8,NNMT,WDR62,RFX5,VRK3,PPM1K,PHF19,PCBP2,TM9SF1,TET2,G
,NIP7,TTC9C,COX4I1,AUP1,FOXRED1,TRNT1,MAGT1,GUK1,CENPN,AGPAT4,IFI27L1,MAF,PPA1,ENDOV,P

1,LSR,NAT9,ZNF687,NOMO2,C11ORF48,RRP8,MT1E,ZNF506,DALRD3,MROH1,PDCD6,MPHOSPH10,C6OF
1,LSR,NAT9,ZNF687,NOMO2,C11ORF48,RRP8,MT1E,ZNF506,DALRD3,MROH1,PDCD6,MPHOSPH10,C6OF

RT15,PPFIBP1,SEMA4D,LAS1L,HMGB1,SULF2,CYB561A3,BAIAP2,RAPGEF5,SSU72,MYL6B,FAM122C,SRP9
RT15,PPFIBP1,SEMA4D,LAS1L,HMGB1,SULF2,CYB561A3,BAIAP2,RAPGEF5,SSU72,MYL6B,FAM122C,SRP9
NASE1L1,KIF4A,RBCK1,PTOV1,FASTK,RNF32,LY75,ATG16L2,SNRPB,PARD3,ACTR6,SET,ZNF235,BOP1,ARR
NASE1L1,KIF4A,RBCK1,PTOV1,FASTK,RNF32,LY75,ATG16L2,SNRPB,PARD3,ACTR6,SET,ZNF235,BOP1,ARR
SPAN31,GLYCTK,TMEM199,EPN1,ZCCHC9,AZI2,TBC1D22A,SZT2,MRPL51,MED8,MRPL27,MED17,SEC14L
ATCH2L,SAE1,OSGEP,SCARB1,EIF5A,CHID1,TAZ,RPL13A,LDHA,RPS9,MRPL33,TESC,SPAG7,STYXL1,RANBP
ATCH2L,SAE1,OSGEP,SCARB1,EIF5A,CHID1,TAZ,RPL13A,LDHA,RPS9,MRPL33,TESC,SPAG7,STYXL1,RANBP
LOC1S1,RHOF,RPL38,PRR13,STARD10,RNH1,DNASE1L1,RBCK1,PTOV1,LY75,ATG16L2,PARD3,DHPS,ACTR
FASTK,TMEM135,SNRPB,PARD3,ZNF235,TRPT1,KLC1,RBP5,ZC3HAV1,SCAF11,DBI,NUMA1,TBL2,CTTN,TS
NBP1,TET2,UBR4,FUZ,ZBTB44,SLC25A10,RDH11,NISCH,TMED4,FOXRED1,ITGB1BP1,MED15,SH3YL1,CLA
NBP1,TET2,UBR4,FUZ,ZBTB44,SLC25A10,RDH11,NISCH,TMED4,FOXRED1,ITGB1BP1,MED15,SH3YL1,CLA
,KLC1,LIMK2,VDAC3,WDR19,NUMA1,BUD31,SAE1,OSGEP,SCARB1,TSPAN11,ITSN1,EIF5A,COMMD5,AM
,KLC1,LIMK2,VDAC3,WDR19,NUMA1,BUD31,SAE1,OSGEP,SCARB1,TSPAN11,ITSN1,EIF5A,COMMD5,AM

6,AMZ2,LDHA,RCE1,ZNF410,IGSF8,IL1RAP,RFX5,RANBP1,VRK3,FIS1,PPM1K,TET2,UQCRQ,GRK6,OBSL1,F
6,AMZ2,LDHA,RCE1,ZNF410,IGSF8,IL1RAP,RFX5,RANBP1,VRK3,FIS1,PPM1K,TET2,UQCRQ,GRK6,OBSL1,F
561A3,MYL6B,FAM122C,SRP9,SULT1A3,UAP1,CMC2,YBX3,BLOC1S1,CIRBP,RPL38,PRR13,RNH1,DNASE1L
561A3,MYL6B,FAM122C,SRP9,SULT1A3,UAP1,CMC2,YBX3,BLOC1S1,CIRBP,RPL38,PRR13,RNH1,DNASE1L
A4D,LAS1L,HMGB1,SULF2,CYB561A3,BAIAP2,RAPGEF5,SSU72,MYL6B,FAM122C,IQCG,TMEM67,UAP1,F
A4D,LAS1L,HMGB1,SULF2,CYB561A3,BAIAP2,RAPGEF5,SSU72,MYL6B,FAM122C,IQCG,TMEM67,UAP1,F
TOV1,TMEM135,TCAIM,DHPS,SET,CTSC,BOP1,ARRDC1,KLC1,SCAF11,VDAC3,WDR19,NUMA1,SAE1,NPRL
TOV1,TMEM135,TCAIM,DHPS,SET,CTSC,BOP1,ARRDC1,KLC1,SCAF11,VDAC3,WDR19,NUMA1,SAE1,NPRL
,RPL38,STARD10,DNASE1L1,RBCK1,PTOV1,FASTK,ATG16L2,SNRPB,ACTR6,MAP3K7CL,ARRDC1,SCAF11,LI
,RPL38,STARD10,DNASE1L1,RBCK1,PTOV1,FASTK,ATG16L2,SNRPB,ACTR6,MAP3K7CL,ARRDC1,SCAF11,LI
P16,LDHA,MRPL33,SNRPN,FIS1,CHCHD1,RPS25,TMX2,RDH11,CENPK,COASY,EIF3M,CDC42,XBP1,PTK2,C
P16,LDHA,MRPL33,SNRPN,FIS1,CHCHD1,RPS25,TMX2,RDH11,CENPK,COASY,EIF3M,CDC42,XBP1,PTK2,C
,HMGB1,BAIAP2,RAPGEF5,SSU72,FAM122C,SRP9,IQCG,UAP1,PILRB,YBX3,CACYBP,CIRBP,RHOF,SMARCB
,HMGB1,BAIAP2,RAPGEF5,SSU72,FAM122C,SRP9,IQCG,UAP1,PILRB,YBX3,CACYBP,CIRBP,RHOF,SMARCB
PB,TCAIM,DHPS,MAP3K7CL,CTSC,ZNF235,DBI,WDR19,NUMA1,OSGEP,TBL2,CRYGS,ITSN1,EIF5A,USP38,S
PARD3,DHPS,SEC62,SET,BOP1,TRPT1,KLC1,ARPC2,PMFBP1,NOB1,SIRT6,NUMA1,GPATCH2L,SAE1,SCARB
PARD3,DHPS,SEC62,SET,BOP1,TRPT1,KLC1,ARPC2,PMFBP1,NOB1,SIRT6,NUMA1,GPATCH2L,SAE1,SCARB
C,TMEM67,PILRB,CACYBP,RHOF,RNH1,DNASE1L1,PTOV1,RNF32,ATG16L2,SNRPB,TCAIM,SEC62,BOP1,A
C,TMEM67,PILRB,CACYBP,RHOF,RNH1,DNASE1L1,PTOV1,RNF32,ATG16L2,SNRPB,TCAIM,SEC62,BOP1,A
2,HDAC9,SEMA4D,LAS1L,HMGB1,SULF2,BAIAP2,RAPGEF5,FAM122C,IQCG,TMEM67,SULT1A3,PILRB,FA
2,HDAC9,SEMA4D,LAS1L,HMGB1,SULF2,BAIAP2,RAPGEF5,FAM122C,IQCG,TMEM67,SULT1A3,PILRB,FA
RBCK1,FASTK,ATG16L2,SNRPB,PARD3,SLC3A2,DHPS,ACTR6,MAP3K7CL,SEC62,SET,CTSC,BOP1,TRPT1,KLC
RBCK1,FASTK,ATG16L2,SNRPB,PARD3,SLC3A2,DHPS,ACTR6,MAP3K7CL,SEC62,SET,CTSC,BOP1,TRPT1,KLC
M67,CMC2,YBX3,CACYBP,CIRBP,RHOF,SMARCB1,RPL38,RNH1,PTOV1,RNF32,ATG16L2,SNRPB,PARD3,SL
M67,CMC2,YBX3,CACYBP,CIRBP,RHOF,SMARCB1,RPL38,RNH1,PTOV1,RNF32,ATG16L2,SNRPB,PARD3,SL
,DHPS,SET,ARPC2,NOB1,SIRT6,ARHGAP6,CTTN,EIF5A,THYN1,DBNDD1,GSTK1,RCE1,PCMT1,TESC,IGSF8,N
,DHPS,SET,ARPC2,NOB1,SIRT6,ARHGAP6,CTTN,EIF5A,THYN1,DBNDD1,GSTK1,RCE1,PCMT1,TESC,IGSF8,N
PB,PARD3,ACTR6,SEC62,SET,BOP1,ARRDC1,TRPT1,KLC1,SCAF11,DBI,ARPC2,LIMK2,BUD31,SAE1,OSGEP,
,PTOV1,ATG16L2,SNRPB,TCAIM,SEC62,BOP1,ARRDC1,KLC1,ZC3HAV1,SCAF11,WDR19,GPATCH2L,BUD31
,PTOV1,ATG16L2,SNRPB,TCAIM,SEC62,BOP1,ARRDC1,KLC1,ZC3HAV1,SCAF11,WDR19,GPATCH2L,BUD31
T,BOP1,DBI,GPATCH2L,SAE1,OSGEP,SCARB1,CTTN,EIF5A,COMMD5,CHID1,RPL13A,LDHA,AP3B2,RPS9,M
T,BOP1,DBI,GPATCH2L,SAE1,OSGEP,SCARB1,CTTN,EIF5A,COMMD5,CHID1,RPL13A,LDHA,AP3B2,RPS9,M
SE1L1,FASTK,ATG16L2,RCC1,DHPS,SET,BOP1,DBI,GPATCH2L,SAE1,OSGEP,SCARB1,CTTN,EIF5A,COMMD5

SE1L1,FASTK,ATG16L2,RCC1,DHPS,SET,BOP1,DBI,GPATCH2L,SAE1,OSGEP,SCARB1,CTTN,EIF5A,COMMD5
,PRR13,STARD10,RNH1,RBCK1,RNF32,ATG16L2,RCC1,SNRPB,TCAIM,ACTR6,CTSC,ZNF235,KLC1,RBP5,ZC3
,PRR13,STARD10,RNH1,RBCK1,RNF32,ATG16L2,RCC1,SNRPB,TCAIM,ACTR6,CTSC,ZNF235,KLC1,RBP5,ZC3
DC37,UBR4,FAM3A,PSMA3,METTL17,PPRC1,NLRP1,NIP7,TTC9C,COX4I1,AUP1,FOXRED1,TRNT1,ITGB1BP
F9,STIP1,HMGA2,UBR4,FAM3A,SDF4,RPL29,ITGB1BP1,C1ORF198,PCYT2,USE1,CCDC25,EEF1B2,PPA1,TIM
F9,STIP1,HMGA2,UBR4,FAM3A,SDF4,RPL29,ITGB1BP1,C1ORF198,PCYT2,USE1,CCDC25,EEF1B2,PPA1,TIM
F9,STIP1,HMGA2,UBR4,FAM3A,SDF4,RPL29,ITGB1BP1,C1ORF198,PCYT2,USE1,CCDC25,EEF1B2,PPA1,TIM
2,CYB561A3,BAIAP2,RAPGEF5,SSU72,MYL6B,IQCG,SULT1A3,PILRB,FAM72A,YBX3,BLOC1S1,CACYBP,CIRB
2,CYB561A3,BAIAP2,RAPGEF5,SSU72,MYL6B,IQCG,SULT1A3,PILRB,FAM72A,YBX3,BLOC1S1,CACYBP,CIRB
TOML2,C19ORF54,PPAN,HNRNPC,RPS20,GABPB1,DCTD,ERLEC1,PSME2,PHTF2,RRNAD1,WAC,PRLR,CAM
TOML2,C19ORF54,PPAN,HNRNPC,RPS20,GABPB1,DCTD,ERLEC1,PSME2,PHTF2,RRNAD1,WAC,PRLR,CAM
PATCH2L,NPRL2,OSGEP,ARHGAP6,EIF5A,AMZ2,TAZ,NOP16,LDHA,RPS9,RCE1,ZNF410,PCMT1,IGSF8,NNM
PATCH2L,NPRL2,OSGEP,ARHGAP6,EIF5A,AMZ2,TAZ,NOP16,LDHA,RPS9,RCE1,ZNF410,PCMT1,IGSF8,NNM
2C,PILRB,YBX3,CACYBP,SMARCB1,PRR13,STARD10,DNASE1L1,PTOV1,RNF32,LY75,ATG16L2,TCAIM,DHP
L,TMEM135,LY75,ATG16L2,PARD3,SEC62,SET,ARRDC1,TRPT1,SCAF11,DBI,LIMK2,NUMA1,BUD31,TBL2,S
L,TMEM135,LY75,ATG16L2,PARD3,SEC62,SET,ARRDC1,TRPT1,SCAF11,DBI,LIMK2,NUMA1,BUD31,TBL2,S
ET2,UQCRQ,GRK6,CDC37,SLC39A11,FUZ,PPRC1,SLC25A10,SDF4,ITGB1BP1,SH3YL1,C1ORF198,C14ORF2,
ET2,UQCRQ,GRK6,CDC37,SLC39A11,FUZ,PPRC1,SLC25A10,SDF4,ITGB1BP1,SH3YL1,C1ORF198,C14ORF2,
2C,SRP9,UAP1,FAM72A,CIRBP,SMARCB1,PRR13,STARD10,KIF4A,FASTK,LY75,RCC1,SNRPB,TCAIM,SLC3A2,
2C,SRP9,UAP1,FAM72A,CIRBP,SMARCB1,PRR13,STARD10,KIF4A,FASTK,LY75,RCC1,SNRPB,TCAIM,SLC3A2,
METTL17,NIP7,TTC9C,COX4I1,AUP1,FOXRED1,TRNT1,MAGT1,GUK1,CENPN,AGPAT4,MAF,PPA1,ENDOV,
,AP3B2,SLC35G1,RCE1,RUVBL2,EBPL,WDR62,IRF9,TET2,UQCRQ,SNF8,GRK6,OBSL1,FUZ,METTL17,ZBTB4
,AP3B2,SLC35G1,RCE1,RUVBL2,EBPL,WDR62,IRF9,TET2,UQCRQ,SNF8,GRK6,OBSL1,FUZ,METTL17,ZBTB4
FRSF14,CAPN10,PRMT1,C19ORF54,C19ORF48,RPL36,RBM3,SYNGR2,TPM2,PNPO,RPS15,UBXN1,ZNF814,
FRSF14,CAPN10,PRMT1,C19ORF54,C19ORF48,RPL36,RBM3,SYNGR2,TPM2,PNPO,RPS15,UBXN1,ZNF814,
NDD1,GSTK1,MRPL33,ZNF410,EBPL,RBPMS,RANBP1,UBE2L6,LZTFL1,ANTXR2,VCAN,MAGED2,SON,OBSL1,
NDD1,GSTK1,MRPL33,ZNF410,EBPL,RBPMS,RANBP1,UBE2L6,LZTFL1,ANTXR2,VCAN,MAGED2,SON,OBSL1,
,ATG16L2,PARD3,SLC3A2,CTSC,BOP1,RBP5,SCAF11,DBI,ARPC2,NUMA1,GPATCH2L,BUD31,SAE1,OSGEP,T
3,CACYBP,RBCK1,FASTK,SNRPB,PARD3,DHPS,SEC62,SET,BOP1,TRPT1,KLC1,ARPC2,PMFBP1,NOB1,SIRT6,
3,CACYBP,RBCK1,FASTK,SNRPB,PARD3,DHPS,SEC62,SET,BOP1,TRPT1,KLC1,ARPC2,PMFBP1,NOB1,SIRT6,

AP,VIM,LIG1,TMEM126B,APEH,RPL21,CAMLG,TMPO,LSM12,STOML2,WNK1,C19ORF54,HNRNPC,RPL36,
0AB1,SYNGR2,RAB2A,ZEB2,NIT2,RAD23A,HSD17B10,NT5C,SLC16A1,ILF3,MDH2,KCTD7,TMEM101,FAM1

UBA1C,PCBP2,TET2,SNF8,HMGA2,CDC37,PTPN13,FAM3A,RPS25,COX4I1,SPOCK2,AUP1,SAP30BP,RABL5,

16,RPS23,RPL30,NDUFA7,MRPS33,RPS15A,RPS17,MRPL51,MRPL43,MRPL27,MRPS7,RPL27,MRPL36,RPS
NIP7,RPS6,RPL29,SRSF1,TRNT1,UBE2I,EIF3M,CELF1,EEF1B2,PPP1CC,EDF1,ENDOV,PSMD4,RPL28,VIM,APE

8C,RPS11,NUP133,MRPS17,RPL37A,MYL9,RPLP0,MRPL55,RPL8,ADD1,RPL35,RPL34,NUP62,RPL24,CLTA,
5,SNRPN,SPAG7,RFX5,ZC3H14,IRF9,STIP1,PHF19,PCBP2,SON,TET2,HMGA2,RPS16,TSG101,PSMA1,YLPM1

ILRB,FAM72A,YBX3,BLOC1S1,CACYBP,CIRBP,RHOF,SMARCB1,PRR13,STARD10,RNH1,KIF4A,RBCK1,FASTK
59,MPRIP,ANXA2,RUVBL1,PUF60,NUMB,PKN2,CORO1B,DBN1,MARK2,CTNNB1,EIF3E,GIPC1,RANGAP1

TTLL5,LDHA,PSMA6,RPS9,ZNF410,RUVBL2,NUDT5,RBPMS,SNRPN,SPAG7,RFX5,UBE2L6,VRK3,ZC3H14,TU

TTLL5,LDHA,PSMA6,RPS9,ZNF410,RUVBL2,NUDT5,RBPMS,SNRPN,SPAG7,RFX5,UBE2L6,VRK3,ZC3H14,TU

SC9,ARL6IP4,HNRNPC,RPL36,CDK11A,RBM3,RPS20,RPS15,HSPB1,PRKRIP1,PSME2,PABPC1,RBMX,DDX41

A7,RPL21,CAMLG,AP1S3,STAU1,RPL36,HSP90AB1,RPTOR,RPS20,VPS28,RPS15,UBXN1,MCTS1,HMGA1,P
UBA52,RPS4X,RPS23,RPL30,RPS15A,RPS17,EIF2AK4,EIF6,RPL27,NCK2,RPS14,GLE1,RPL31,EIF3E,FAU

A7,RPL21,CAMLG,AP1S3,STAU1,SUGT1,RPL36,HSP90AB1,RPTOR,RPS20,VPS28,RPS15,UBXN1,MCTS1,HN
FAND2B,AP1S3,RPL36,LAMP2,COPZ1,HSP90AB1,RAB2A,RPS20,VPS28,RPS15,ERLEC1,HSPB1,SYNGR1,RA

2,PSMD4,RPL28,VIM,PSMA7,RPL21,CAMLG,EXOSC9,ZFAND2B,HNRNPC,SHARPIN,RPL36,LAMP2,HSP90A
1,PABPC1,ZNF706,GGT5,ILF3,ZCCHC4,EEF1D,RPL18,EGFR,HNRNPLL,RPL15,COA3,TYMS,DALRD3,DRG2,B
1,PABPC1,ZNF706,GGT5,ILF3,ZCCHC4,EEF1D,RPL18,EGFR,HNRNPLL,RPL15,COA3,TYMS,DALRD3,DRG2,B
PL29,SRSF1,AUP1,RABL5,ITGB1BP1,MICAL1,YIF1A,JUP,CDC42,PTK2,PPP1CC,VAMP8,RPL28,TIMM17B,RP
RPLP0,METTL3,SSB,RPL8,RPL35,RPL34,RPL24,PSMD7,PSMD10,RPS3A,RPS3,MAGOHB,RPS7,NCBP2,RNPS
BLVRB,NUDT1,NT5C,PSME1,CARHSP1,RPL18,HINT1,RPL15,MTHFS,RPS11,PCID2,RPL37A,RPLP0,METTL3,
A7,RPL21,CAMLG,AP1S3,STAU1,SUGT1,RPL36,HSP90AB1,RPTOR,RPS20,VPS28,RPS15,UBXN1,MCTS1,HN
AUP1,RABL5,ITGB1BP1,JUP,CDC42,VAMP8,PPIL2,RPL28,TIMM17B,RPL21,NUTF2,STRADA,ZFAND2B,AP1S
FKBIA,RPS11,MFF,NUP133,RPAIN,ACOT2,PEX2,RPL37A,PEX26,KPNA5,RPLP0,HAX1,SSR3,MPV17,ARIH2,
SME1,CARHSP1,RPL18,HINT1,RPL15,RPS11,PCID2,RPL37A,RPLP0,METTL3,SSB,RPL8,BAX,RPL35,NT5E,GA

AUP1,RABL5,ITGB1BP1,JUP,CDC42,VAMP8,PPIL2,RPL28,TIMM17B,RPL21,NUTF2,STRADA,ZFAND2B,AP1S
JGGT2,UBE2I,XBP1,PTK2,USE1,PSMD4,RPL28,VIM,PSMA7,RPL21,LYPLA2,CAMLG,EXOSC9,ZFAND2B,HNRI

1,MCTS1,PABPC1,ZNF706,GGT5,ILF3,ZCCHC4,EEF1D,RPL18,EGFR,HNRNPLL,RPL15,COA3,TYMS,DALRD3,D
.PABPC1,BLVRB,NUDT1,NT5C,PSME1,CARHSP1,RPL18,HINT1,RPL15,MTHFS,RPS11,PCID2,RPL37A,RPLP0,
A,PEX26,RPLP0,HAX1,SSR3,MPV17,ARIH2,RPL8,RPL35,RPL34,ACAA1,RPL24,PAM16,RPS3A,UBE2D3,RPS3
SC2,EXOSC8,RPS15A,RPS17,RPL27,RPS14,CNOT7,HBS1L,MAGOH,RPL31,EIF3E,EXOSC10,FAU
JDT1,NT5C,PSME1,CARHSP1,RPL18,HINT1,RPL15,MTHFS,RPS11,PCID2,RPL37A,RPLP0,METTL3,SSB,RPL8,
JDT1,NT5C,PSME1,CARHSP1,RPL18,HINT1,RPL15,MTHFS,RPS11,PCID2,RPL37A,RPLP0,METTL3,SSB,RPL8,
;1,PABPC1,ZNF706,GGT5,ILF3,ZCCHC4,EEF1D,RPL18,EGFR,HNRNPLL,RPL15,MTHFS,COA3,MPC2,TYMS,DA
RPLP0,METTL3,SSB,RPL8,RPL35,RPL34,RPL24,PSMD7,PSMD10,RPS3A,RPS3,MAGOHB,RPS7,NCBP2,RNPS
;F1,SNF8,CDC37,RPS16,TSG101,OBSL1,TTC21B,RPS25,PLCB2,TMED4,RPS6,RPL29,SRSF1,AUP1,RABL5,KXI

M40,PDCD5,POT1,ABHD17A,RPL18,SH3GLB1,RPL15,TCTN1,TCP1,PARK7,NFKBIA,RPS11,MFF,NUP133,RPA
,SNF8,RPS16,TSG101,UBR4,PSMA1,PSMA3,RPS25,PLCB2,RPS6,RPL29,AUP1,UGGT2,UBE2I,XBP1,PTK2,US
RPS20,NIT2,RPS15,HSPB1,MVD,MCTS1,PABPC1,ZNF706,GGT5,ILF3,ZCCHC4,EEF1D,RPL18,EGFR,HNRNPL
AB1,RPS15,ERCC5,MCTS1,NAP1L1,RANGRF,DBNL,RBMX,PDE4DIP,MED6,PRLR,SSH2,COX17,NDUFV2,SMA
'1,CENPK,SRSF1,FOXRED1,ITGB1BP1,MICAL1,EIF3M,CDC42,PTK2,CELF1,CENPN,ASF1B,VAMP8,CAPG,PSM
VCAN,PCBP2,CD44,TM9SF1,TET2,SNF8,CDC37,RPS16,TSG101,UBR4,PSMA1,PSMA3,ATG12,RPS25,RPS6,F
6,TRMT112,RPS15,PRKRIP1,PABPC1,RRNAD1,RBMX,HSD17B10,DDX41,ZCCHC4,RRP8,EGFR,HNRNPLL,HN
IS1,PPM1K,VCAN,PCBP2,CD44,TM9SF1,TET2,SNF8,CDC37,RPS16,TSG101,UBR4,PSMA1,PSMA3,ATG12,RI
ED4,RPS6,RPL29,SRSF1,AUP1,RABL5,ITGB1BP1,YIF1A,JUP,CDC42,XBP1,GUK1,USE1,RFX3,TNFRSF14,VAMI
RPS25,NLRP1,TMED4,RPS6,RPL29,SRSF1,AUP1,RABL5,ITGB1BP1,YIF1A,JUP,CDC42,XBP1,GUK1,USE1,RFX
N,RUNX1,TAZ,SH2B1,NOP16,RPL13A,RPS9,RUVBL2,WDR62,IL1RAP,STYXL1,FIS1,DST,SNF8,HMGA2,RPS16
),RPLP0,EIF3G,RPL35,ZNF593,EIF2A,RPL24,GEMIN2,EIF3C,SNRPD1,GTF3A,RPS27L,RPS7,DDX10,TAF9,EXC
,GUK1,AGPAT4,EEF1B2,GADL1,PPA1,RPL28,VIM,APEH,RPL21,UCK2,PPOX,PRMT1,STOML2,PIGL,UPP1,CA

ED4,RPS6,RPL29,SRSF1,AUP1,RABL5,ITGB1BP1,YIF1A,JUP,CDC42,XBP1,GUK1,USE1,RFX3,TNFRSF14,VAMI
ED4,RPS6,RPL29,SRSF1,AUP1,RABL5,ITGB1BP1,YIF1A,JUP,CDC42,XBP1,GUK1,USE1,RFX3,TNFRSF14,VAMI
IF8,CDC37,RPS16,TSG101,OBSL1,TTC21B,FUZ,EPB41L5,RPS25,NLRP1,TMED4,RPS6,RPL29,SRSF1,AUP1,RA
RALY,RBM42,PSPC1,RBM4,GEMIN2,THOC3,HNRNPUL1,MAGOHB,SNRPD1,SNRPA1,ZBTB8OS,U2AF1L4,N
EP,RUNX1,UFD1L,TAZ,SH2B1,RPL13A,LDHA,PSMA6,ACOT8,RPS9,RUVBL2,TESC,NUDT5,RBPMS,NNMT,SN
VAMP8,CAPG,PSMD4,TMEM126B,TXNL4A,TCF4,ATPAF2,PRMT1,STOML2,PPAN,HSP90AB1,RPS15,VWA2,
)BI,NOB1,SIRT6,NUMA1,BUD31,OSGEP,EIF5A,RUNX1,UFD1L,TAZ,SH2B1,RPL13A,GSTK1,LDHA,PSMA6,AC
P6,CTTN,TNFSF10,TAZ,SH2B1,GSTK1,ACOT8,RUVBL2,NUDT5,WDR62,IL1RAP,STYXL1,POLD4,RANBP1,FIS

NCBP2,RNPS1,UBA52,PSMD8,NSRP1,EXOSC2,RBM7,PSMC5,HNRNPM,IK,EXOSC8,DDX17,PABPN1,SFSWA
.,SNRPA1,U2AF1L4,NCBP2,POLR2E,RNPS1,NSRP1,RBM7,HNRNPM,IK,DDX17,PABPN1,SFSWAP,PPIE,PUF6
IFF,RPL37A,PEX26,RPLP0,FAM126A,SSR3,RPL8,PICK1,SLMAP,BAX,RPL35,DLG4,CLASP2,RER1,RPL34,RPL2

MPV17,ARIH2,BAX,VDAC1,PAM16,NDUFA11,TP53,PSMD10,NDUFA3,UBE2D3,COX19,DNAJA3,AIFM2,ARR
EP,RUNX1,UFD1L,TAZ,SH2B1,RPL13A,LDHA,PSMA6,ACOT8,RPS9,RUVBL2,TESC,NUDT5,RBPMS,NNMT,SN
,PSME1,AGAP3,PARK7,USP40,USP4,ARIH2,FBXL18,UBE2D4,RBBP6,UBA1,RNF8,PPP2R5C,PCNP,PSMD7,N
)2B,SHARPIN,UPP1,LAMP2,HSP90AB1,DUT,SIAH1,VPS28,UBXN1,ERLEC1,SAT1,DCN,PSME2,BLVRB,WAC,I
.F1L4,NCBP2,POLR2E,RNPS1,NSRP1,RBM7,HNRNPM,IK,DDX17,PABPN1,SFSWAP,PPIE,PUF60,SNRNP40,RI
.F1L4,NCBP2,POLR2E,RNPS1,NSRP1,RBM7,HNRNPM,IK,DDX17,PABPN1,SFSWAP,PPIE,PUF60,SNRNP40,RI

LRRC1,GRAMD1C,TM9SF1,SNF8,CDC37,RPS16,TSG101,OBSL1,TTC21B,FUZ,EPB41L5,RPS25,NLRP1,TMED
SUMF2,SAE1,NPRL2,EIF5A,TNFSF10,UFD1L,USP38,RPL13A,TTLL5,PSMA6,RPS9,RCE1,MRPL33,RUVBL2,PC
,RUVBL2,WDR62,IL1RAP,STYXL1,FIS1,DST,SNF8,HMGA2,TSG101,OBSL1,TTC21B,FUZ,ECSIT,EPB41L5,ATG
C1,SNRPB,PARD3,ACTR6,SET,CTSC,BOP1,ARRDC1,KLC1,SCAF11,ARPC2,LIMK2,VDAC3,NOB1,WDR19,SIRT
VBL2,TESC,NUDT5,RBPMS,NNMT,IL1RAP,RFX5,POLD4,UBE2L6,IRF9,PHF19,TET2,SNF8,HMGA2,RPS16,CPS
EP,RUNX1,UFD1L,TAZ,SH2B1,RPL13A,LDHA,PSMA6,ACOT8,RPS9,RUVBL2,TESC,NUDT5,RBPMS,NNMT,SN
D23A,USP45,PSME1,AGAP3,LGMN,PARK7,SNX5,USP40,USP4,ARIH2,FBXL18,UBE2D4,RBBP6,UBA1,RNF8,

AGAP3,PARK7,USP40,USP4,ARIH2,FBXL18,UBE2D4,RBBP6,UBA1,RNF8,PPP2R5C,PCNP,PSMD7,NEDD4L,R
N34,RBBP6,CHTOP,SRSF7,RALY,RBM42,PSPC1,GTF2H1,RBM4,GEMIN2,THOC3,HNRNPUL1,MAGOHB,SNF
TARD10,RNH1,DNASE1L1,RBCK1,PTOV1,FASTK,ATG16L2,SNRPB,PARD3,DHPS,SET,CTSC,ZNF235,BOP1,AR
AGAP3,PARK7,USP40,USP4,ARIH2,FBXL18,UBE2D4,RBBP6,UBA1,RNF8,PPP2R5C,PCNP,PSMD7,NEDD4L,R
RPL13A,PSMA6,RPS9,RUVBL2,TESC,RBPMS,SNRPN,IL1RAP,RFX5,POLD4,UBE2L6,ZC3H14,IRF9,PHF19,PCB
EP,SCARB1,RUNX1,UFD1L,TAZ,SH2B1,RPL13A,LDHA,PSMA6,ACOT8,RPS9,RUVBL2,EBPL,TESC,NUDT5,RBF

S25,NLRP1,NISCH,TMED4,RPS6,RPL29,SRSF1,AUP1,RABL5,ITGB1BP1,YIF1A,JUP,CDC42,XBP1,GUK1,USE1
BL2,TESC,IL1RAP,RFX5,POLD4,UBE2L6,IRF9,PHF19,VCAN,TET2,SNF8,HMGA2,RPS16,CPSF3L,TSG101,TTC
C1,PSME2,WAC,ANAPC11,RAD23A,ABHD17A,USP45,PSME1,AGAP3,EGFR,LGMN,EGLN2,PARK7,SNX5,US
SP45,PSME1,AGAP3,LGMN,PARK7,USP40,USP4,ARIH2,FBXL18,UBE2D4,RBBP6,UBA1,RNF8,PPP2R5C,FAN
3,RUVBL2,TESC,RBPMS,IL1RAP,RFX5,POLD4,UBE2L6,IRF9,PHF19,VCAN,TET2,SNF8,HMGA2,RPS16,CPSF3
ARRDC1,TRPT1,ZC3HAV1,DBI,LIMK2,SIRT6,SUMF2,SAE1,NPRL2,SCARB1,EIF5A,TNFSF10,AMZ2,UFD1L,US

H2B1,RPL13A,GSTK1,PSMA6,ACOT8,RPS9,MRPL33,RUVBL2,TESC,NUDT5,RBPMS,NNMT,IL1RAP,RFX5,POL
RD3,ACTR6,SET,CTSC,BOP1,KLC1,SCAF11,ARPC2,LIMK2,VDAC3,WDR19,SIRT6,NUMA1,SAE1,NPRL2,ARHC

PL55,LIMA1,ADD1,BAX,MTIF2,CLASP2,VDAC1,PAM16,AXIN1,TP53,SH3GL1,MRPS24,RDX,APAF1,C1QC,BN

WAC,RAD23A,PSME1,CARHSP1,EGFR,SH3GLB1,EGLN2,PAFAH1B2,PFKFB2,PARK7,SNX5,NUP133,PCID2,H
H2B1,RPL13A,GSTK1,PSMA6,ACOT8,RPS9,MRPL33,RUVBL2,TESC,NUDT5,RBPMS,NNMT,IL1RAP,RFX5,POL

V1,LIMK2,SIRT6,SUMF2,SAE1,NPRL2,EIF5A,TNFSF10,AMZ2,UFD1L,USP38,RPL13A,TTLL5,PSMA6,ACOT8,

ESC,RBPMS,SNRPN,IL1RAP,RFX5,ZC3H14,IRF9,PHF19,PCBP2,SON,TET2,SNF8,HMGA2,RPS16,CPSF3L,TSG1

H2B1,RPL13A,GSTK1,PSMA6,ACOT8,RPS9,MRPL33,RUVBL2,TESC,NUDT5,RBPMS,NNMT,IL1RAP,RFX5,POL

1,RPL38,STARD10,RNH1,DNASE1L1,RBCK1,PTOV1,FASTK,ATG16L2,SNRPB,PARD3,SLC3A2,DHPS,SET,CTS
2,CAPG,CDC14B,HNRNPC,MAPKAPK5,CEP70,ZEB2,RANGRF,DCN,HMGA1,DBNL,RAD1,RABEP2,TADA3,PD

A,PSME1,CARHSP1,SH3GLB1,PAFAH1B2,PFKFB2,PARK7,SNX5,PCID2,HAX1,METTL3,ARIH2,BAX,FAM192A

14B,PRMT1,STOML2,STRADA,STAU1,UPP1,LAMP2,INO80E,MAPKAPK5,HSP90AB1,ZEB2,RPTOR,MAPK13

3,DHPS,SET,CTSC,ZNF235,ARRDC1,TRPT1,ZC3HAV1,LIMK2,SIRT6,NUMA1,SUMF2,BUD31,SAE1,NPRL2,EI

P,CIRBP,RHOF,SMARCB1,PRR13,RNH1,DNASE1L1,KIF4A,PTOV1,FASTK,TMEM135,ATG16L2,RCC1,SNRPB,
AP3K7CL,SEC62,SET,KLC1,ZC3HAV1,ARPC2,NOB1,NUMA1,ARHGAP6,CTTN,ITSN1,EIF5A,COMMD5,UFD
C1S1,CACYBP,CIRBP,RHOF,SMARCB1,PRR13,RNH1,DNASE1L1,KIF4A,PTOV1,FASTK,TMEM135,RNF32,LY
CMC2,YBX3,BLOC1S1,CACYBP,CIRBP,RHOF,SMARCB1,RPL38,PRR13,RNH1,DNASE1L1,KIF4A,PTOV1,FAST
CMC2,YBX3,BLOC1S1,CACYBP,CIRBP,RHOF,RPL38,PRR13,STARD10,RNH1,DNASE1L1,KIF4A,RBCK1,PTOV
,TMEM67,SULT1A3,UAP1,FAM72A,CMC2,YBX3,BLOC1S1,CACYBP,CIRBP,RHOF,SMARCB1,RPL38,PRR13,
CMC2,YBX3,BLOC1S1,CACYBP,CIRBP,RHOF,SMARCB1,RPL38,PRR13,STARD10,RNH1,DNASE1L1,KIF4A,PT
PL36,RBM3,RPS20,RPS15,MCTS1,PRKRIP1,PABPC1,RBMX,WAC,HSD17B10,DDX41,ILF3,RPL18,HNRNPA1,
RPB,SLC3A2,ACTR6,SET,CTSC,BOP1,SCAF11,DBI,NOB1,SIRT6,NUMA1,SUMF2,BUD31,SAE1,OSGEP,COMM
RPB,SLC3A2,ACTR6,SET,CTSC,BOP1,SCAF11,DBI,NOB1,SIRT6,NUMA1,SUMF2,BUD31,SAE1,OSGEP,COMM
RPB,SLC3A2,ACTR6,SET,CTSC,BOP1,SCAF11,DBI,NOB1,SIRT6,NUMA1,SUMF2,BUD31,SAE1,OSGEP,COMM
AP3K7CL,SET,CTSC,ZNF235,BOP1,ZC3HAV1,SCAF11,ARPC2,LIMK2,VDAC3,NOB1,SIRT6,NUMA1,BUD31,S
UBA52,RPS4X,MRPS16,RPS23,RPL30,MRPS33,RPS15A,RPS17,MRPL51,MRPL43,MRPL48,MRPL27,MRPS7
E1,OSGEP,COMMD5,RUNX1,NUDT22,UFD1L,NOP16,PSMA6,RPS9,RUVBL2,TESC,RBPMS,SNRPN,POLD4,P
3,MRPS24,RPS27L,RPS7,UBA52,RPS4X,MRPS16,RPS23,RPL30,NDUFA7,MRPS33,DNAJC21,RPS15A,RPS17

E1,OSGEP,COMMD5,RUNX1,NUDT22,UFD1L,NOP16,RPL13A,PSMA6,RPS9,RUVBL2,TESC,RBPMS,SNRPN,
1,ARPC2,VDAC3,NOB1,WDR19,NUMA1,BUD31,SAE1,NPRL2,OSGEP,CTTN,EIF5A,RUNX1,UFD1L,RPL13A,P
PPOX,TMPO,STOML2,SLC9B2,ATP5J,CHCHD7,TOMM40,BRIP1,NUDT1,CHCHD5,EGFR,SH3GLB1,COX17,C
PPOX,TMPO,STOML2,SLC9B2,ATP5J,CHCHD7,TOMM40,BRIP1,NUDT1,CHCHD5,EGFR,SH3GLB1,COX17,C

MEM14B,COA1,NDUFS3,MPV17,MRPL55,PAM16,NDUFA11,SLC25A24,NDUFA3,RPS3,MRPS24,SPHK2,N
5A,HIGD1A,NDUFV2,SIRT5,COA3,MPC2,TYMS,SLC25A23,MRPS18C,LGALS3,PARK7,MFF,COX20,MRPS17,
ALS3,COX20,MRPS17,TMEM14B,COA1,NDUFS3,MPV17,MRPL55,PAM16,NDUFA11,SLC25A24,NDUFA3,
R19,SIRT6,NUMA1,BUD31,ARHGAP6,CTTN,ITSN1,RUNX1,NOP16,RPL13A,TTLL5,PSMA6,RPS9,MRPL33,R
R19,SIRT6,NUMA1,BUD31,ARHGAP6,CTTN,ITSN1,RUNX1,NOP16,RPL13A,TTLL5,PSMA6,RPS9,MRPL33,R
B,HNRNPC,SHARPIN,SUGT1,INO80E,HSPB1,DDOST,MTA3,PSME2,PABPC1,RBMX,PPP3CB,TADA3,ANAPC1
2,PRKRIP1,ANAPC11,ILF3,ZCCHC4,EEF1D,NAA10,RRP8,RPL18,INTS4,ZNF506,SMARCA4,PAFAH1B2,TYMS
,TIMM17B,TMEM126B,XAF1,CAPN10,PPOX,ATPAF2,STOML2,ACAD10,SLC9B2,DGUOK,HSP90AB1,DUT,
7,NDUFS2,MRPS33,NDUFC1,NDUFB10,MRPL51,MRPL43,IMMP1L,MRPL48,MRPL27,MRPS7,NDUFS7,TI
PC2,TYMS,SLC25A23,MRPS18C,LGALS3,MFF,COX20,MRPS17,TMEM14B,COA1,NDUFS3,HAX1,MPV17,M

13,PSMA1,FUZ,PSMA3,MUC1,RPS25,SDF4,COASY,AUP1,JUP,CDC42,PPA1,VAMP8,CAPG,RPL28,VIM,PSM
13,PSMA1,FUZ,PSMA3,MUC1,RPS25,SDF4,COASY,AUP1,JUP,CDC42,PPA1,VAMP8,CAPG,RPL28,VIM,PSM
13,PSMA1,FUZ,PSMA3,MUC1,RPS25,SDF4,COASY,AUP1,JUP,CDC42,PPA1,VAMP8,CAPG,RPL28,VIM,PSM
C1,PAM16,D2HGDH,TP53,RPS3,MRPS24,IDH3G,DNAJA3,FARS2,NDUFA10,ACSM3,LRRC59,MRPS16,ARL

ANTXR2,GRAMD1C,LPCAT3,CD44,TM9SF1,UQCRQ,SNF8,TSG101,UBR4,CHCHD1,ECSIT,ATG12,COX4I1,SLC
RPS3,RDX,RPS7,ILK,FHL2,RPS4X,CAPN2,ARL2,RPL30,LPP,MPRIP,ANXA5,ACTB,RPS17,ACTG1,NUMB,RPL2
RPS3,RDX,RPS7,ILK,FHL2,RPS4X,CAPN2,ARL2,RPL30,LPP,MPRIP,ANXA5,ACTB,RPS17,ACTG1,NUMB,RPL2

JVBL2,PCMT1,IGSF8,NUDT5,TUBA1C,DST,ANTXR2,PCBP2,CD44,TM9SF1,MAGED2,SNF8,CDC37,RPS16,TS

ST,SLC33A1,NOMO2,B2M,SGK1,SEC31A,ATP10D,EGFR,CAMK2D,GBA2,MCFD2,SLC35B1,SCAP,CD74,THA
3A1,NOMO2,B2M,SGK1,SEC31A,ATP10D,EGFR,CAMK2D,GBA2,MCFD2,SLC35B1,SCAP,CD74,THADA,PIG

UP85,ATP11B,SEC13,NSMF,PARP6,LMNA,ROGDI,AK9,PRKCZ,DNAJB12,DCTN1,BIN1,PHF20,CLIP1,CPNE1,

M4,RPS11,RPL37A,RPLP0,KRAS,RPL8,LIMA1,ASH1L,ADD1,TJP1,UBA1,CDK4,CLASP2,ARPC5,ALKBH6,NRP1

N,AP2S1,ATP11B,ANXA2,LAMTOR4,SEC13,RAP1B,CPNE1,C12ORF66,ATP13A2,HGSNAT,SURF4,NPRL3,CD
N,AP2S1,ATP11B,ANXA2,LAMTOR4,SEC13,RAP1B,CPNE1,C12ORF66,ATP13A2,HGSNAT,SURF4,NPRL3,CD
1B2,RPL21,CAMLG,C17ORF62,PIGL,ZFAND2B,STAU1,RAB2A,GGCX,ASPH,LYNX1,UBXN1,ERLEC1,DDOST,R

YB561,SLC9B2,NFAM1,LAMP2,COPZ1,HSP90AB1,SYNGR2,RAB2A,RPTOR,VPS28,DDOST,SYNGR1,RANGRF
TOR1,PLD1,UBA52,VAPA,VTI1B,SZT2,GRN,AP2S1,ATP11B,ANXA2,LAMTOR4,ATP6V0B,SEC13,RAP1B,CPN

4A1,RPLP0,EIF3G,SSR3,MRPL55,RPL8,RPL35,MTIF2,MARS,RPL34,RPL24,RPS3A,EIF3C,RPS3,MRPS24,FARS

RPS3,MAGOHB,RPS27L,RPS7,NCBP2,RNPS1,UBA52,RPS4X,RPS23,RPL30,PSMD8,PSMC5,RPS15A,RPS17

PNA5,RPLP0,RPL8,RPL35,RPL34,NUP62,RPL24,PSMD7,NEDD4L,GTF2H1,CLTA,PSMD10,RPS3A,SH3GL1,RF

PSME2,PABPC1,RBMX,HSD17B10,PSME1,RPL18,HNRNPA1,POLR2C,RPL15,SNRPF,PPIH,THADA,NHP2,RPS

FAM122C,SRP9,IQCG,PILRB,FAM72A,YBX3,BLOC1S1,CACYBP,CIRBP,RHOF,SMARCB1,RPL38,PRR13,STARD
OC1S1,CACYBP,RHOF,SMARCB1,RPL38,PRR13,STARD10,RNH1,DNASE1L1,KIF4A,RBCK1,PTOV1,FASTK,TM
R6,CTSC,ZNF235,BOP1,ARRDC1,TRPT1,KLC1,ZC3HAV1,DBI,ARPC2,LIMK2,SIRT6,NUMA1,GPATCH2L,BUD3
EF5,SSU72,MYL6B,FAM122C,SRP9,IQCG,TMEM67,SULT1A3,PILRB,FAM72A,CMC2,YBX3,BLOC1S1,CACYB
EF5,SSU72,MYL6B,FAM122C,SRP9,IQCG,TMEM67,SULT1A3,PILRB,FAM72A,CMC2,YBX3,BLOC1S1,CACYB
QCG,SULT1A3,PILRB,FAM72A,CMC2,YBX3,BLOC1S1,CACYBP,RHOF,SMARCB1,RPL38,PRR13,STARD10,RNH
QCG,SULT1A3,PILRB,FAM72A,CMC2,YBX3,BLOC1S1,CACYBP,RHOF,SMARCB1,RPL38,PRR13,STARD10,RNH
PILRB,FAM72A,YBX3,BLOC1S1,CACYBP,CIRBP,RHOF,SMARCB1,RPL38,PRR13,STARD10,RNH1,DNASE1L1,K
3,STARD10,RNH1,DNASE1L1,KIF4A,RBCK1,RNF32,LY75,ATG16L2,RCC1,SNRPB,PARD3,TCAIM,SLC3A2,DHF
3,STARD10,RNH1,DNASE1L1,KIF4A,RBCK1,RNF32,LY75,ATG16L2,RCC1,SNRPB,PARD3,TCAIM,SLC3A2,DHF
B,FAM72A,YBX3,BLOC1S1,CACYBP,RHOF,SMARCB1,RPL38,PRR13,STARD10,RNH1,DNASE1L1,KIF4A,RBCK
DHPS,ACTR6,BOP1,ARRDC1,TRPT1,RBP5,ZC3HAV1,SCAF11,DBI,ARPC2,LIMK2,SIRT6,NUMA1,GPATCH2L,S
DHPS,ACTR6,BOP1,ARRDC1,TRPT1,RBP5,ZC3HAV1,SCAF11,DBI,ARPC2,LIMK2,SIRT6,NUMA1,GPATCH2L,S
ATCH2L,BUD31,NPRL2,SCARB1,CTTN,EIF5A,COMMD5,CHID1,NUDT22,USP38,TAZ,THYN1,SH2B1,NOP16,
1,KIF4A,RBCK1,RNF32,LY75,RCC1,SNRPB,PARD3,SLC3A2,DHPS,ACTR6,CTSC,ZNF235,BOP1,ARRDC1,TRPT
1,KIF4A,RBCK1,RNF32,LY75,RCC1,SNRPB,PARD3,SLC3A2,DHPS,ACTR6,CTSC,ZNF235,BOP1,ARRDC1,TRPT
YBX3,CACYBP,CIRBP,RHOF,SMARCB1,RPL38,PRR13,STARD10,RNH1,DNASE1L1,KIF4A,RBCK1,PTOV1,FAST

YBX3,CACYBP,CIRBP,RHOF,SMARCB1,RPL38,PRR13,STARD10,RNH1,DNASE1L1,KIF4A,RBCK1,PTOV1,FAST

RPB,PARD3,TCAIM,DHPS,ACTR6,CTSC,ZNF235,BOP1,ARRDC1,TRPT1,RBP5,ZC3HAV1,SCAF11,DBI,ARPC2,

RPB,PARD3,TCAIM,DHPS,ACTR6,CTSC,ZNF235,BOP1,ARRDC1,TRPT1,RBP5,ZC3HAV1,SCAF11,DBI,ARPC2,

D3,DHPS,ACTR6,ZNF235,BOP1,ARRDC1,TRPT1,RBP5,ZC3HAV1,SCAF11,DBI,ARPC2,PMFBP1,LIMK2,SIRT6

D3,DHPS,ACTR6,ZNF235,BOP1,ARRDC1,TRPT1,RBP5,ZC3HAV1,SCAF11,DBI,ARPC2,PMFBP1,LIMK2,SIRT6

E1L1,KIF4A,RBCK1,RNF32,LY75,ATG16L2,RCC1,SNRPB,PARD3,TCAIM,DHPS,ACTR6,SEC62,CTSC,ZNF235,B

E1L1,KIF4A,RBCK1,RNF32,LY75,ATG16L2,RCC1,SNRPB,PARD3,TCAIM,DHPS,ACTR6,SEC62,CTSC,ZNF235,B

2C,SRP9,IQCG,TMEM67,SULT1A3,PILRB,FAM72A,CMC2,YBX3,BLOC1S1,CACYBP,CIRBP,RHOF,SMARCB1,

2C,SRP9,IQCG,TMEM67,SULT1A3,PILRB,FAM72A,CMC2,YBX3,BLOC1S1,CACYBP,CIRBP,RHOF,SMARCB1,

,KIF4A,RBCK1,RNF32,LY75,RCC1,SNRPB,PARD3,TCAIM,DHPS,ACTR6,CTSC,ZNF235,BOP1,ARRDC1,TRPT1,

,KIF4A,RBCK1,RNF32,LY75,RCC1,SNRPB,PARD3,TCAIM,DHPS,ACTR6,CTSC,ZNF235,BOP1,ARRDC1,TRPT1,

,RPL38,PRR13,STARD10,RNH1,DNASE1L1,KIF4A,RBCK1,FASTK,RNF32,LY75,ATG16L2,RCC1,SNRPB,PARD3

,RPL38,PRR13,STARD10,RNH1,DNASE1L1,KIF4A,RBCK1,FASTK,RNF32,LY75,ATG16L2,RCC1,SNRPB,PARD3

IMK2,SIRT6,NUMA1,GPATCH2L,SUMF2,BUD31,SAE1,NPRL2,CTTN,TSPAN11,EIF5A,COMMD5,CHID1,NUD

IMK2,SIRT6,NUMA1,GPATCH2L,SUMF2,BUD31,SAE1,NPRL2,CTTN,TSPAN11,EIF5A,COMMD5,CHID1,NUD

RCB1,RPL38,PRR13,STARD10,RNH1,DNASE1L1,KIF4A,RBCK1,FASTK,RNF32,LY75,ATG16L2,RCC1,SNRPB,

RCB1,RPL38,PRR13,STARD10,RNH1,DNASE1L1,KIF4A,RBCK1,FASTK,RNF32,LY75,ATG16L2,RCC1,SNRPB,

B1,RPL38,PRR13,STARD10,RNH1,DNASE1L1,KIF4A,RBCK1,FASTK,RNF32,LY75,ATG16L2,RCC1,SNRPB,PAR

B1,RPL38,PRR13,STARD10,RNH1,DNASE1L1,KIF4A,RBCK1,FASTK,RNF32,LY75,ATG16L2,RCC1,SNRPB,PAR

2A,CMC2,YBX3,CACYBP,CIRBP,RHOF,SMARCB1,RPL38,PRR13,STARD10,RNH1,DNASE1L1,KIF4A,RBCK1,P

3,STARD10,RNH1,DNASE1L1,KIF4A,RBCK1,RNF32,LY75,RCC1,SNRPB,PARD3,TCAIM,SLC3A2,DHPS,ACTR6,

3,STARD10,RNH1,DNASE1L1,KIF4A,RBCK1,RNF32,LY75,RCC1,SNRPB,PARD3,TCAIM,SLC3A2,DHPS,ACTR6,

PC2,LIMK2,SIRT6,NUMA1,GPATCH2L,BUD31,SAE1,NPRL2,SCARB1,CTTN,EIF5A,COMMD5,CHID1,NUDT22

3,CACYBP,RHOF,SMARCB1,RPL38,PRR13,STARD10,RNH1,DNASE1L1,KIF4A,RBCK1,PTOV1,FASTK,TMEM1

3,CACYBP,RHOF,SMARCB1,RPL38,PRR13,STARD10,RNH1,DNASE1L1,KIF4A,RBCK1,PTOV1,FASTK,TMEM1

,BUD31,NPRL2,CTTN,EIF5A,COMMD5,CHID1,NUDT22,UFD1L,USP38,NOP16,RPL13A,PSMA6,ACOT8,RPS

C,SRP9,IQCG,TMEM67,PILRB,FAM72A,YBX3,BLOC1S1,CACYBP,RHOF,SMARCB1,RPL38,PRR13,STARD10,

YL6B,FAM122C,SRP9,IQCG,TMEM67,SULT1A3,PILRB,FAM72A,CMC2,YBX3,BLOC1S1,CACYBP,CIRBP,RHO

YL6B,FAM122C,SRP9,IQCG,TMEM67,SULT1A3,PILRB,FAM72A,CMC2,YBX3,BLOC1S1,CACYBP,CIRBP,RHO

IRPB,PARD3,DHPS,ACTR6,MAP3K7CL,ZNF235,BOP1,ARRDC1,TRPT1,RBP5,ZC3HAV1,SCAF11,DBI,ARPC2,P

IRPB,PARD3,DHPS,ACTR6,MAP3K7CL,ZNF235,BOP1,ARRDC1,TRPT1,RBP5,ZC3HAV1,SCAF11,DBI,ARPC2,P

CK1,RNF32,LY75,RCC1,SNRPB,PARD3,SLC3A2,DHPS,ACTR6,CTSC,ZNF235,ARRDC1,TRPT1,KLC1,RBP5,ZC3

CK1,RNF32,LY75,RCC1,SNRPB,PARD3,SLC3A2,DHPS,ACTR6,CTSC,ZNF235,ARRDC1,TRPT1,KLC1,RBP5,ZC3

C1S1,CACYBP,CIRBP,RHOF,SMARCB1,RPL38,PRR13,STARD10,RNH1,DNASE1L1,KIF4A,RBCK1,FASTK,RNF

C1S1,CACYBP,CIRBP,RHOF,SMARCB1,RPL38,PRR13,STARD10,RNH1,DNASE1L1,KIF4A,RBCK1,FASTK,RNF

75,RCC1,SNRPB,PARD3,SLC3A2,DHPS,ACTR6,ZNF235,BOP1,ARRDC1,TRPT1,KLC1,RBP5,ZC3HAV1,SCAF11,

75,RCC1,SNRPB,PARD3,SLC3A2,DHPS,ACTR6,ZNF235,BOP1,ARRDC1,TRPT1,KLC1,RBP5,ZC3HAV1,SCAF11,

38,PRR13,STARD10,RNH1,DNASE1L1,KIF4A,RBCK1,RNF32,LY75,ATG16L2,RCC1,SNRPB,PARD3,TCAIM,SLC

38,PRR13,STARD10,RNH1,DNASE1L1,KIF4A,RBCK1,RNF32,LY75,ATG16L2,RCC1,SNRPB,PARD3,TCAIM,SLC

IF235,ARRDC1,TRPT1,RBP5,ZC3HAV1,SCAF11,DBI,ARPC2,PMFBP1,LIMK2,SIRT6,NUMA1,GPATCH2L,SUM

IF235,ARRDC1,TRPT1,RBP5,ZC3HAV1,SCAF11,DBI,ARPC2,PMFBP1,LIMK2,SIRT6,NUMA1,GPATCH2L,SUM

ARD3,SLC3A2,DHPS,ACTR6,ZNF235,BOP1,ARRDC1,TRPT1,RBP5,ZC3HAV1,SCAF11,DBI,ARPC2,PMFBP1,LI

ARD3,SLC3A2,DHPS,ACTR6,ZNF235,BOP1,ARRDC1,TRPT1,RBP5,ZC3HAV1,SCAF11,DBI,ARPC2,PMFBP1,LI

,FAM72A,CMC2,YBX3,BLOC1S1,CACYBP,CIRBP,RHOF,SMARCB1,RPL38,PRR13,RNH1,DNASE1L1,KIF4A,RB

RL2,SCARB1,CTTN,EIF5A,COMMD5,CHID1,NUDT22,USP38,TAZ,THYN1,SH2B1,NOP16,RPL13A,GSTK1,PS

TN,EIF5A,COMMD5,CHID1,NUDT22,USP38,THYN1,SH2B1,NOP16,RPL13A,TTLL5,PSMA6,ACOT8,RPS9,RC

RPB,PARD3,DHPS,ACTR6,ZNF235,BOP1,ARRDC1,TRPT1,RBP5,ZC3HAV1,SCAF11,DBI,ARPC2,PMFBP1,LIMK

RPB,PARD3,DHPS,ACTR6,ZNF235,BOP1,ARRDC1,TRPT1,RBP5,ZC3HAV1,SCAF11,DBI,ARPC2,PMFBP1,LIMK

CC1,SNRPB,PARD3,DHPS,ACTR6,ZNF235,BOP1,ARRDC1,TRPT1,RBP5,ZC3HAV1,SCAF11,DBI,ARPC2,PMFB

CC1,SNRPB,PARD3,DHPS,ACTR6,ZNF235,BOP1,ARRDC1,TRPT1,RBP5,ZC3HAV1,SCAF11,DBI,ARPC2,PMFB

A2,DHPS,ACTR6,ZNF235,BOP1,ARRDC1,TRPT1,RBP5,ZC3HAV1,SCAF11,DBI,ARPC2,PMFBP1,LIMK2,SIRT6,N

A2,DHPS,ACTR6,ZNF235,BOP1,ARRDC1,TRPT1,RBP5,ZC3HAV1,SCAF11,DBI,ARPC2,PMFBP1,LIMK2,SIRT6,N

DC1,TRPT1,RBP5,ZC3HAV1,SCAF11,DBI,ARPC2,PMFBP1,LIMK2,SIRT6,NUMA1,GPATCH2L,SUMF2,BUD31,

DC1,TRPT1,RBP5,ZC3HAV1,SCAF11,DBI,ARPC2,PMFBP1,LIMK2,SIRT6,NUMA1,GPATCH2L,SUMF2,BUD31,

ACTR6,BOP1,ARRDC1,TRPT1,RBP5,ZC3HAV1,SCAF11,DBI,ARPC2,LIMK2,SIRT6,NUMA1,GPATCH2L,SUMF2,

ACTR6,BOP1,ARRDC1,TRPT1,RBP5,ZC3HAV1,SCAF11,DBI,ARPC2,LIMK2,SIRT6,NUMA1,GPATCH2L,SUMF2,

HPS,ACTR6,MAP3K7CL,ZNF235,BOP1,ARRDC1,TRPT1,RBP5,ZC3HAV1,SCAF11,DBI,ARPC2,PMFBP1,LIMK2,

HPS,ACTR6,MAP3K7CL,ZNF235,BOP1,ARRDC1,TRPT1,RBP5,ZC3HAV1,SCAF11,DBI,ARPC2,PMFBP1,LIMK2,

RCB1,RPL38,PRR13,RNH1,DNASE1L1,KIF4A,RBCK1,RNF32,LY75,ATG16L2,RCC1,SNRPB,PARD3,SLC3A2,DH

RCB1,RPL38,PRR13,RNH1,DNASE1L1,KIF4A,RBCK1,RNF32,LY75,ATG16L2,RCC1,SNRPB,PARD3,SLC3A2,DH

B,PARD3,DHPS,ACTR6,ZNF235,BOP1,ARRDC1,TRPT1,RBP5,ZC3HAV1,SCAF11,DBI,ARPC2,PMFBP1,LIMK2,

B,PARD3,DHPS,ACTR6,ZNF235,BOP1,ARRDC1,TRPT1,RBP5,ZC3HAV1,SCAF11,DBI,ARPC2,PMFBP1,LIMK2,

6,RPL13A,PSMA6,ACOT8,RPS9,RCE1,ZNF410,NUDT5,STYXL1,VRK3,SNF8,CDC37,CPSF3L,TSG101,UBR4,YL

RBCK1,RNF32,LY75,RCC1,SNRPB,PARD3,SLC3A2,DHPS,ACTR6,ZNF235,BOP1,ARRDC1,TRPT1,KLC1,RBP5,ZC

RBCK1,RNF32,LY75,RCC1,SNRPB,PARD3,SLC3A2,DHPS,ACTR6,ZNF235,BOP1,ARRDC1,TRPT1,KLC1,RBP5,ZC

5,BOP1,ARRDC1,TRPT1,ZC3HAV1,SCAF11,DBI,ARPC2,PMFBP1,LIMK2,SIRT6,NUMA1,GPATCH2L,SUMF2,B

5,BOP1,ARRDC1,TRPT1,ZC3HAV1,SCAF11,DBI,ARPC2,PMFBP1,LIMK2,SIRT6,NUMA1,GPATCH2L,SUMF2,B

CC1,SNRPB,PARD3,SLC3A2,DHPS,ACTR6,ZNF235,BOP1,ARRDC1,TRPT1,RBP5,ZC3HAV1,SCAF11,DBI,ARPC

CC1,SNRPB,PARD3,SLC3A2,DHPS,ACTR6,ZNF235,BOP1,ARRDC1,TRPT1,RBP5,ZC3HAV1,SCAF11,DBI,ARPC

5,RCC1,SNRPB,PARD3,DHPS,ACTR6,ZNF235,BOP1,ARRDC1,TRPT1,RBP5,ZC3HAV1,SCAF11,DBI,ARPC2,PM

5,RCC1,SNRPB,PARD3,DHPS,ACTR6,ZNF235,BOP1,ARRDC1,TRPT1,RBP5,ZC3HAV1,SCAF11,DBI,ARPC2,PM

8,NUDT5,STYXL1,VRK3,PPM1K,SON,UQCRQ,SNF8,CPSF3L,TSG101,UBR4,NIP7,TTC9C,TMX2,DBNDD2,RPS

PB,PARD3,DHPS,ACTR6,ZNF235,BOP1,ARRDC1,TRPT1,RBP5,ZC3HAV1,SCAF11,DBI,ARPC2,PMFBP1,LIMK2

PB,PARD3,DHPS,ACTR6,ZNF235,BOP1,ARRDC1,TRPT1,RBP5,ZC3HAV1,SCAF11,DBI,ARPC2,PMFBP1,LIMK2

H1,DNASE1L1,KIF4A,RBCK1,PTOV1,FASTK,TMEM135,RNF32,LY75,ATG16L2,SNRPB,PARD3,TCAIM,SLC3A2

H1,DNASE1L1,KIF4A,RBCK1,PTOV1,FASTK,TMEM135,RNF32,LY75,ATG16L2,SNRPB,PARD3,TCAIM,SLC3A2

,RPS9,ZNF410,RUVBL2,IGSF8,NUDT5,STYXL1,VRK3,PPM1K,IRF9,ANTXR2,SON,UQCRQ,SNF8,RPS16,CPSF3

LZTFL1,PPM1K,IRF9,SON,SNF8,CPSF3L,TSG101,UBR4,RPS25,NIP7,COX4I1,THEM4,RPS6,RPL29,RTEL1,AU

HOF,SMARCB1,RPL38,PRR13,RNH1,DNASE1L1,KIF4A,RBCK1,RNF32,LY75,RCC1,SNRPB,PARD3,SLC3A2,DH

HOF,SMARCB1,RPL38,PRR13,RNH1,DNASE1L1,KIF4A,RBCK1,RNF32,LY75,RCC1,SNRPB,PARD3,SLC3A2,DH

1,STARD10,RNH1,DNASE1L1,KIF4A,RBCK1,PTOV1,FASTK,TMEM135,RNF32,LY75,ATG16L2,RCC1,PARD3,

1,STARD10,RNH1,DNASE1L1,KIF4A,RBCK1,PTOV1,FASTK,TMEM135,RNF32,LY75,ATG16L2,RCC1,PARD3,

,SRP9,IQCG,TMEM67,SULT1A3,PILRB,FAM72A,CMC2,YBX3,BLOC1S1,CACYBP,CIRBP,RHOF,SMARCB1,RP

,SRP9,IQCG,TMEM67,SULT1A3,PILRB,FAM72A,CMC2,YBX3,BLOC1S1,CACYBP,CIRBP,RHOF,SMARCB1,RP

GSTK1,PSMA6,ACOT8,RPS9,MRPL33,ZNF410,IGSF8,NUDT5,STYXL1,VRK3,PPM1K,IRF9,STIP1,LRRC1,SON,

,SMARCB1,RPL38,PRR13,RNH1,KIF4A,RBCK1,PTOV1,RNF32,LY75,ATG16L2,RCC1,SNRPB,PARD3,TCAIM,SL

,SMARCB1,RPL38,PRR13,RNH1,KIF4A,RBCK1,PTOV1,RNF32,LY75,ATG16L2,RCC1,SNRPB,PARD3,TCAIM,SL

A3,FAM72A,BLOC1S1,CACYBP,CIRBP,RHOF,SMARCB1,RPL38,PRR13,RNH1,DNASE1L1,KIF4A,RBCK1,PTOV1

A3,FAM72A,BLOC1S1,CACYBP,CIRBP,RHOF,SMARCB1,RPL38,PRR13,RNH1,DNASE1L1,KIF4A,RBCK1,PTOV1

USP38,THYN1,SH2B1,NOP16,RPL13A,PSMA6,ACOT8,RPS9,RCE1,MRPL33,ZNF410,RUVBL2,IGSF8,NUDT5,S

C1,RBP5,ZC3HAV1,SCAF11,DBI,ARPC2,LIMK2,NOB1,SIRT6,NUMA1,GPATCH2L,SUMF2,BUD31,NPRL2,CTTN

C1,RBP5,ZC3HAV1,SCAF11,DBI,ARPC2,LIMK2,NOB1,SIRT6,NUMA1,GPATCH2L,SUMF2,BUD31,NPRL2,CTTN

CPSF3L,TSG101,UBR4,FAM3A,TTC9C,THEM4,DBNDD2,RPS6,RPL29,RTEL1,SAP30BP,YIPF3,FOXRED1,TRNT

S9,RCE1,MRPL33,ZNF410,IGSF8,NUDT5,STYXL1,VRK3,PPM1K,LRRC1,SON,UQCRQ,SNF8,CDC37,RPS16,CPS

TCH2L,BUD31,SAE1,NPRL2,SCARB1,CTTN,EIF5A,COMMD5,CHID1,NUDT22,USP38,THYN1,SH2B1,NOP16,[illegible]
5,RCC1,SNRPB,PARD3,SLC3A2,DHPS,ACTR6,MAP3K7CL,CTSC,ZNF235,BOP1,ARRDC1,TRPT1,RBP5,ZC3HA[illegible]
5,RCC1,SNRPB,PARD3,SLC3A2,DHPS,ACTR6,MAP3K7CL,CTSC,ZNF235,BOP1,ARRDC1,TRPT1,RBP5,ZC3HA[illegible]
CK1,RNF32,LY75,RCC1,SNRPB,PARD3,DHPS,ACTR6,MAP3K7CL,CTSC,ZNF235,BOP1,ARRDC1,TRPT1,KLC1,[illegible]
CK1,RNF32,LY75,RCC1,SNRPB,PARD3,DHPS,ACTR6,MAP3K7CL,CTSC,ZNF235,BOP1,ARRDC1,TRPT1,KLC1,[illegible]
RHGAP6,CTTN,EIF5A,COMMD5,CHID1,NUDT22,USP38,THYN1,SH2B1,NOP16,RPL13A,GSTK1,PSMA6,ACO
I1,DNASE1L1,KIF4A,RBCK1,PTOV1,FASTK,RNF32,ATG16L2,RCC1,SNRPB,TCAIM,SLC3A2,DHPS,ACTR6,SEC[illegible]
I1,DNASE1L1,KIF4A,RBCK1,PTOV1,FASTK,RNF32,ATG16L2,RCC1,SNRPB,TCAIM,SLC3A2,DHPS,ACTR6,SEC[illegible]
8,NUDT5,STYXL1,POLD4,VRK3,IRF9,STIP1,LRRC1,UQCRQ,SNF8,CPSF3L,TSG101,UBR4,RPS25,NIP7,COX4I1
QCG,PILRB,FAM72A,YBX3,CACYBP,RHOF,SMARCB1,RPL38,PRR13,STARD10,RNH1,DNASE1L1,KIF4A,RBCK
QCG,PILRB,FAM72A,YBX3,CACYBP,RHOF,SMARCB1,RPL38,PRR13,STARD10,RNH1,DNASE1L1,KIF4A,RBCK
,PPM1K,SON,UQCRQ,SNF8,CPSF3L,TSG101,UBR4,RPS25,NIP7,THEM4,RPS6,RPL29,RTEL1,AUP1,YIPF3,FO
MARCB1,PRR13,STARD10,RNH1,KIF4A,RBCK1,PTOV1,FASTK,TMEM135,RNF32,LY75,ATG16L2,SNRPB,PAR
MARCB1,PRR13,STARD10,RNH1,KIF4A,RBCK1,PTOV1,FASTK,TMEM135,RNF32,LY75,ATG16L2,SNRPB,PAR
DT5,SNRPN,STYXL1,POLD4,LZTFL1,FIS1,PPM1K,MAGED2,SON,UQCRQ,SNF8,CDC37,CPSF3L,TSG101,FUZ,[illegible]
DT5,SNRPN,STYXL1,POLD4,LZTFL1,FIS1,PPM1K,MAGED2,SON,UQCRQ,SNF8,CDC37,CPSF3L,TSG101,FUZ,[illegible]
TN,EIF5A,COMMD5,NUDT22,USP38,TAZ,THYN1,DBNDD1,SH2B1,NOP16,GSTK1,PSMA6,ACOT8,RPS9,RC[illegible]
L1,RBCK1,PTOV1,FASTK,TMEM135,RNF32,ATG16L2,RCC1,SNRPB,PARD3,TCAIM,DHPS,ACTR6,SEC62,SET[illegible]
NF410,RUVBL2,IGSF8,NUDT5,STYXL1,ANTXR2,LRRC1,SON,UQCRQ,SNF8,RPS16,CPSF3L,TSG101,UBR4,ATG
MED15,CLASRP,C1ORF198,UBE2I,EIF3M,CDC42,CARD14,PCYT2,FBXW9,ASF1B,RFX3,ENDOV,CAMTA1,PP[illegible]
RL2,ARHGAP6,CTTN,EIF5A,COMMD5,CHID1,NUDT22,USP38,THYN1,SH2B1,NOP16,RPL13A,GSTK1,PSMA
NUMA1,GPATCH2L,BUD31,NPRL2,CTTN,TSPAN11,EIF5A,COMMD5,CHID1,NUDT22,UFD1L,USP38,DBNDD
NUMA1,GPATCH2L,BUD31,NPRL2,CTTN,TSPAN11,EIF5A,COMMD5,CHID1,NUDT22,UFD1L,USP38,DBNDD
ZTFL1,FIS1,MAGED2,SON,UQCRQ,SNF8,CDC37,CPSF3L,TSG101,FUZ,FAM3A,RPS25,THEM4,RPS6,RTEL1,S[illegible]
ZTFL1,FIS1,MAGED2,SON,UQCRQ,SNF8,CDC37,CPSF3L,TSG101,FUZ,FAM3A,RPS25,THEM4,RPS6,RTEL1,S[illegible]
T1,ZC3HAV1,SCAF11,DBI,ARPC2,PMFBP1,LIMK2,NUMA1,GPATCH2L,SUMF2,BUD31,NPRL2,ARHGAP6,CT
T1,ZC3HAV1,SCAF11,DBI,ARPC2,PMFBP1,LIMK2,NUMA1,GPATCH2L,SUMF2,BUD31,NPRL2,ARHGAP6,CT
7,SULT1A3,PILRB,FAM72A,YBX3,BLOC1S1,CACYBP,CIRBP,RHOF,SMARCB1,RPL38,STARD10,RNH1,DNASE[illegible]
7,SULT1A3,PILRB,FAM72A,YBX3,BLOC1S1,CACYBP,CIRBP,RHOF,SMARCB1,RPL38,STARD10,RNH1,DNASE[illegible]
16,RPL13A,PSMA6,ACOT8,RPS9,RCE1,ZNF410,NUDT5,STYXL1,VRK3,STIP1,LRRC1,SON,UQCRQ,SNF8,CPSF[illegible]
DT22,USP38,TAZ,THYN1,SH2B1,NOP16,PSMA6,ACOT8,RPS9,RCE1,MRPL33,ZNF410,RUVBL2,IGSF8,NUDT
,PILRB,FAM72A,YBX3,BLOC1S1,CACYBP,CIRBP,RHOF,SMARCB1,RPL38,STARD10,RNH1,DNASE1L1,KIF4A,[illegible]
,PILRB,FAM72A,YBX3,BLOC1S1,CACYBP,CIRBP,RHOF,SMARCB1,RPL38,STARD10,RNH1,DNASE1L1,KIF4A,[illegible]
),IGSF8,NUDT5,STYXL1,VRK3,LRRC1,SNF8,CDC37,RPS16,CPSF3L,TSG101,UBR4,FUZ,FAM3A,ATG12,RPS25
GEF39,PSMA6,ACOT8,AP3B2,ZNF410,PCMT1,NUDT5,SNRPN,STYXL1,POLD4,VRK3,LZTFL1,FIS1,PPM1K,M[illegible]
GEF39,PSMA6,ACOT8,AP3B2,ZNF410,PCMT1,NUDT5,SNRPN,STYXL1,POLD4,VRK3,LZTFL1,FIS1,PPM1K,M[illegible]
HAV1,SCAF11,DBI,ARPC2,PMFBP1,LIMK2,SIRT6,NUMA1,GPATCH2L,SUMF2,BUD31,SAE1,NPRL2,OSGEP,[illegible]
HAV1,SCAF11,DBI,ARPC2,PMFBP1,LIMK2,SIRT6,NUMA1,GPATCH2L,SUMF2,BUD31,SAE1,NPRL2,OSGEP,[illegible]
L,THYN1,RPL13A,ARHGEF39,PSMA6,ACOT8,AP3B2,ZNF410,IGSF8,NUDT5,SNRPN,STYXL1,POLD4,LZTFL1,
L,THYN1,RPL13A,ARHGEF39,PSMA6,ACOT8,AP3B2,ZNF410,IGSF8,NUDT5,SNRPN,STYXL1,POLD4,LZTFL1,
)31,NPRL2,CRYGS,CTTN,EIF5A,COMMD5,CHID1,NUDT22,UFD1L,USP38,THYN1,RPL13A,ARHGEF39,PSMA
)31,NPRL2,CRYGS,CTTN,EIF5A,COMMD5,CHID1,NUDT22,UFD1L,USP38,THYN1,RPL13A,ARHGEF39,PSMA
D31,SAE1,NPRL2,CRYGS,CTTN,TSPAN11,EIF5A,COMMD5,CHID1,NUDT22,UFD1L,USP38,THYN1,SH2B1,R[illegible]
D31,SAE1,NPRL2,CRYGS,CTTN,TSPAN11,EIF5A,COMMD5,CHID1,NUDT22,UFD1L,USP38,THYN1,SH2B1,R[illegible]
3,BLOC1S1,CACYBP,CIRBP,RHOF,SMARCB1,RPL38,STARD10,RNH1,DNASE1L1,KIF4A,RBCK1,PTOV1,FAST[illegible]
3,BLOC1S1,CACYBP,CIRBP,RHOF,SMARCB1,RPL38,STARD10,RNH1,DNASE1L1,KIF4A,RBCK1,PTOV1,FAST[illegible]
A6,ACOT8,RPS9,RCE1,ZNF410,IGSF8,NUDT5,STYXL1,VRK3,PPM1K,TUBA1C,LRRC1,VCAN,SON,UQCRQ,SN

D5,CHID1,NUDT22,UFD1L,USP38,TAZ,THYN1,SH2B1,NOP16,RPL13A,PSMA6,ACOT8,RPS9,RCE1,ZNF410,R

5A,COMMD5,CHID1,NUDT22,UFD1L,THYN1,SH2B1,RPL13A,ARHGEF39,GSTK1,PSMA6,ACOT8,SLC35G1,ZI

5A,COMMD5,CHID1,NUDT22,UFD1L,THYN1,SH2B1,RPL13A,ARHGEF39,GSTK1,PSMA6,ACOT8,SLC35G1,ZI

CTTN,EIF5A,COMMD5,USP38,TAZ,THYN1,SH2B1,NOP16,RPL13A,GSTK1,PSMA6,ACOT8,RPS9,RCE1,MRPL

MMD5,CHID1,UFD1L,TAZ,SH2B1,ARHGEF39,PSMA6,ACOT8,AP3B2,ZNF410,TESC,IGSF8,NUDT5,SNRPN,SI

MMD5,CHID1,UFD1L,TAZ,SH2B1,ARHGEF39,PSMA6,ACOT8,AP3B2,ZNF410,TESC,IGSF8,NUDT5,SNRPN,SI

,STYXL1,VRK3,PPM1K,IRF9,ANTXR2,UQCRQ,SNF8,CPSF3L,TSG101,UBR4,ATG12,NIP7,RPS6,RPL29,RTEL1,

,YBX3,RHOF,SMARCB1,RPL38,PRR13,STARD10,RNH1,DNASE1L1,KIF4A,RBCK1,PTOV1,FASTK,TMEM135,F

MK2,GPATCH2L,BUD31,SAE1,NPRL2,OSGEP,CTTN,ITSN1,EIF5A,COMMD5,NUDT22,USP38,TAZ,SH2B1,NOI

QCG,TMEM67,SULT1A3,FAM72A,YBX3,BLOC1S1,CACYBP,RHOF,SMARCB1,RPL38,PRR13,STARD10,RNH1,I

NRPN,STYXL1,VRK3,LZTFL1,FIS1,STIP1,VCAN,MAGED2,SON,UQCRQ,SNF8,CPSF3L,TSG101,FUZ,FAM3A,AT

NRPN,STYXL1,VRK3,LZTFL1,FIS1,STIP1,VCAN,MAGED2,SON,UQCRQ,SNF8,CPSF3L,TSG101,FUZ,FAM3A,AT

,STARD10,RNH1,DNASE1L1,KIF4A,RBCK1,RNF32,LY75,RCC1,SNRPB,PARD3,SLC3A2,DHPS,ACTR6,MAP3K7

,STARD10,RNH1,DNASE1L1,KIF4A,RBCK1,RNF32,LY75,RCC1,SNRPB,PARD3,SLC3A2,DHPS,ACTR6,MAP3K7

PC2,PMFBP1,LIMK2,SIRT6,NUMA1,GPATCH2L,SUMF2,BUD31,SAE1,NPRL2,SCARB1,CRYGS,CTTN,EIF5A,C

PC2,PMFBP1,LIMK2,SIRT6,NUMA1,GPATCH2L,SUMF2,BUD31,SAE1,NPRL2,SCARB1,CRYGS,CTTN,EIF5A,C

PC2,PMFBP1,LIMK2,SIRT6,NUMA1,GPATCH2L,SUMF2,BUD31,SAE1,NPRL2,SCARB1,CRYGS,CTTN,EIF5A,C

PC2,PMFBP1,LIMK2,SIRT6,NUMA1,GPATCH2L,SUMF2,BUD31,SAE1,NPRL2,SCARB1,CRYGS,CTTN,EIF5A,C

P3B2,NUDT5,SNRPN,STYXL1,POLD4,LZTFL1,FIS1,MAGED2,SON,UQCRQ,SNF8,CDC37,CPSF3L,TSG101,FUZ

P3B2,NUDT5,SNRPN,STYXL1,POLD4,LZTFL1,FIS1,MAGED2,SON,UQCRQ,SNF8,CDC37,CPSF3L,TSG101,FUZ

1,EIF5A,COMMD5,NUDT22,UFD1L,USP38,THYN1,SH2B1,RPL13A,ARHGEF39,GSTK1,PSMA6,ACOT8,RPS9,

1,EIF5A,COMMD5,NUDT22,UFD1L,USP38,THYN1,SH2B1,RPL13A,ARHGEF39,GSTK1,PSMA6,ACOT8,RPS9,

NUDT5,STYXL1,VRK3,LZTFL1,FIS1,STIP1,LRRC1,CDC37,CPSF3L,TSG101,UBR4,PHF21A,FUZ,METTL17,ATG1

,ACOT8,PCMT1,NUDT5,SNRPN,STYXL1,RANBP1,VRK3,LZTFL1,FIS1,IRF9,ANTXR2,STIP1,LRRC1,TM9SF1,MA

,ACOT8,PCMT1,NUDT5,SNRPN,STYXL1,RANBP1,VRK3,LZTFL1,FIS1,IRF9,ANTXR2,STIP1,LRRC1,TM9SF1,MA

32,LY75,RCC1,SNRPB,TCAIM,SLC3A2,DHPS,ACTR6,SEC62,ZNF235,BOP1,ARRDC1,TRPT1,RBP5,ZC3HAV1,S

32,LY75,RCC1,SNRPB,TCAIM,SLC3A2,DHPS,ACTR6,SEC62,ZNF235,BOP1,ARRDC1,TRPT1,RBP5,ZC3HAV1,S

MT1,NUDT5,SNRPN,STYXL1,LZTFL1,FIS1,PPM1K,MAGED2,SON,UQCRQ,CPSF3L,TSG101,FUZ,FAM3A,RPS2

MT1,NUDT5,SNRPN,STYXL1,LZTFL1,FIS1,PPM1K,MAGED2,SON,UQCRQ,CPSF3L,TSG101,FUZ,FAM3A,RPS2

3,SLC3A2,DHPS,ACTR6,BOP1,ARRDC1,TRPT1,RBP5,ZC3HAV1,SCAF11,DBI,ARPC2,LIMK2,SIRT6,NUMA1,GF

3,SLC3A2,DHPS,ACTR6,BOP1,ARRDC1,TRPT1,RBP5,ZC3HAV1,SCAF11,DBI,ARPC2,LIMK2,SIRT6,NUMA1,GF

PRR13,RNH1,KIF4A,RBCK1,PTOV1,RNF32,LY75,ATG16L2,RCC1,SNRPB,PARD3,TCAIM,SLC3A2,DHPS,ACTR

PRR13,RNH1,KIF4A,RBCK1,PTOV1,RNF32,LY75,ATG16L2,RCC1,SNRPB,PARD3,TCAIM,SLC3A2,DHPS,ACTR

MA6,ACOT8,AP3B2,RPS9,RCE1,MRPL33,NUDT5,SNRPN,STYXL1,LZTFL1,FIS1,PPM1K,MAGED2,SON,UQCR

MA6,ACOT8,AP3B2,RPS9,RCE1,MRPL33,NUDT5,SNRPN,STYXL1,LZTFL1,FIS1,PPM1K,MAGED2,SON,UQCR

75,RCC1,SNRPB,PARD3,SLC3A2,DHPS,ACTR6,ZNF235,BOP1,ARRDC1,TRPT1,RBP5,ZC3HAV1,SCAF11,DBI,A

75,RCC1,SNRPB,PARD3,SLC3A2,DHPS,ACTR6,ZNF235,BOP1,ARRDC1,TRPT1,RBP5,ZC3HAV1,SCAF11,DBI,A

AM122C,SRP9,IQCG,TMEM67,FAM72A,CMC2,YBX3,CACYBP,RHOF,SMARCB1,RPL38,PRR13,STARD10,RN

1,RPL38,PRR13,RNH1,DNASE1L1,RBCK1,FASTK,TMEM135,RNF32,LY75,ATG16L2,RCC1,SNRPB,PARD3,TC

1,RPL38,PRR13,RNH1,DNASE1L1,RBCK1,FASTK,TMEM135,RNF32,LY75,ATG16L2,RCC1,SNRPB,PARD3,TC

IF4A,RBCK1,PTOV1,FASTK,RNF32,LY75,ATG16L2,SNRPB,PARD3,TCAIM,SLC3A2,ACTR6,SEC62,SET,CTSC,Z

IF4A,RBCK1,PTOV1,FASTK,RNF32,LY75,ATG16L2,SNRPB,PARD3,TCAIM,SLC3A2,ACTR6,SEC62,SET,CTSC,Z

YXL1,VRK3,LZTFL1,PPM1K,SON,UQCRQ,SNF8,CPSF3L,TSG101,UBR4,METTL17,ATG12,RPS25,NIP7,TMX2,I

L1,KIF4A,RBCK1,PTOV1,FASTK,RNF32,LY75,ATG16L2,SNRPB,PARD3,TCAIM,SLC3A2,ACTR6,SEC62,SET,CT

L1,KIF4A,RBCK1,PTOV1,FASTK,RNF32,LY75,ATG16L2,SNRPB,PARD3,TCAIM,SLC3A2,ACTR6,SEC62,SET,CT

32,SLC35G1,ZNF410,IGSF8,NUDT5,STYXL1,RFX5,POLD4,LZTFL1,FIS1,PPM1K,MAGED2,SON,CDC37,CPSF3L

32,SLC35G1,ZNF410,IGSF8,NUDT5,STYXL1,RFX5,POLD4,LZTFL1,FIS1,PPM1K,MAGED2,SON,CDC37,CPSF3L

IUDT5,SNRPN,STYXL1,POLD4,LZTFL1,FIS1,PPM1K,MAGED2,SON,UQCRQ,CPSF3L,TSG101,FUZ,FAM3A,RPS
IUDT5,SNRPN,STYXL1,POLD4,LZTFL1,FIS1,PPM1K,MAGED2,SON,UQCRQ,CPSF3L,TSG101,FUZ,FAM3A,RPS
CYBP,CIRBP,RHOF,SMARCB1,RPL38,STARD10,RNH1,DNASE1L1,KIF4A,RBCK1,PTOV1,FASTK,TMEM135,LY
CYBP,CIRBP,RHOF,SMARCB1,RPL38,STARD10,RNH1,DNASE1L1,KIF4A,RBCK1,PTOV1,FASTK,TMEM135,LY
RQ,SNF8,CPSF3L,TSG101,ATG12,NIP7,THEM4,RPS6,RPL29,AUP1,YIPF3,FOXRED1,TRNT1,KXD1,MED15,C1
PT1,DBI,ARPC2,PMFBP1,LIMK2,GPATCH2L,BUD31,NPRL2,SCARB1,CTTN,EIF5A,COMMD5,NUDT22,UFD1L
PT1,DBI,ARPC2,PMFBP1,LIMK2,GPATCH2L,BUD31,NPRL2,SCARB1,CTTN,EIF5A,COMMD5,NUDT22,UFD1L
SRP9,IQCG,SULT1A3,FAM72A,CMC2,YBX3,BLOC1S1,CACYBP,CIRBP,RHOF,SMARCB1,RPL38,PRR13,STARD
SRP9,IQCG,SULT1A3,FAM72A,CMC2,YBX3,BLOC1S1,CACYBP,CIRBP,RHOF,SMARCB1,RPL38,PRR13,STARD
FBP1,LIMK2,NOB1,WDR19,SUMF2,BUD31,NPRL2,OSGEP,TBL2,CRYGS,CTTN,EIF5A,COMMD5,NUDT22,AN
FBP1,LIMK2,NOB1,WDR19,SUMF2,BUD31,NPRL2,OSGEP,TBL2,CRYGS,CTTN,EIF5A,COMMD5,NUDT22,AN
122C,SRP9,IQCG,TMEM67,SULT1A3,UAP1,PILRB,FAM72A,YBX3,BLOC1S1,CACYBP,CIRBP,RHOF,SMARCB1
122C,SRP9,IQCG,TMEM67,SULT1A3,UAP1,PILRB,FAM72A,YBX3,BLOC1S1,CACYBP,CIRBP,RHOF,SMARCB1
F410,IGSF8,NUDT5,STYXL1,VRK3,PPM1K,LRRC1,SON,UQCRQ,SNF8,CPSF3L,TSG101,METTL17,ATG12,NIP
RR13,RNH1,DNASE1L1,RBCK1,PTOV1,FASTK,RNF32,LY75,ATG16L2,RCC1,SNRPB,PARD3,TCAIM,SLC3A2,D
RR13,RNH1,DNASE1L1,RBCK1,PTOV1,FASTK,RNF32,LY75,ATG16L2,RCC1,SNRPB,PARD3,TCAIM,SLC3A2,D
9,RCE1,ZNF410,RUVBL2,IGSF8,NUDT5,STYXL1,VRK3,PPM1K,LRRC1,SON,UQCRQ,SNF8,CPSF3L,TSG101,M
1,LIMK2,GPATCH2L,BUD31,NPRL2,ARHGAP6,CTTN,EIF5A,COMMD5,NUDT22,UFD1L,USP38,TAZ,SH2B1,R
1,LIMK2,GPATCH2L,BUD31,NPRL2,ARHGAP6,CTTN,EIF5A,COMMD5,NUDT22,UFD1L,USP38,TAZ,SH2B1,R
F235,ARRDC1,TRPT1,RBP5,ZC3HAV1,DBI,ARPC2,PMFBP1,LIMK2,SIRT6,NUMA1,GPATCH2L,BUD31,NPRL2
F235,ARRDC1,TRPT1,RBP5,ZC3HAV1,DBI,ARPC2,PMFBP1,LIMK2,SIRT6,NUMA1,GPATCH2L,BUD31,NPRL2
)5,CHID1,NUDT22,UFD1L,USP38,THYN1,SH2B1,RPL13A,PSMA6,ACOT8,AP3B2,RPS9,RCE1,MRPL33,ZNF4
MAP3K7CL,CTSC,ZNF235,BOP1,ARRDC1,TRPT1,RBP5,DBI,ARPC2,PMFBP1,LIMK2,SIRT6,NUMA1,GPATCH
MAP3K7CL,CTSC,ZNF235,BOP1,ARRDC1,TRPT1,RBP5,DBI,ARPC2,PMFBP1,LIMK2,SIRT6,NUMA1,GPATCH
N,MAGED2,SON,UQCRQ,SNF8,CPSF3L,TSG101,FUZ,FAM3A,RPS25,NIP7,THEM4,RPS6,RTEL1,SAP30BP,YI
N,MAGED2,SON,UQCRQ,SNF8,CPSF3L,TSG101,FUZ,FAM3A,RPS25,NIP7,THEM4,RPS6,RTEL1,SAP30BP,YI
3L,TSG101,CHCHD1,FUZ,FAM3A,RPS25,TTC9C,TMX2,DBNDD2,RPS6,RPL29,RTEL1,SRSF1,SAP30BP,FOXRE
3L,TSG101,FUZ,FAM3A,RPS25,NIP7,THEM4,RPS6,RTEL1,SAP30BP,YIPF3,RABL5,TRNT1,ITGB1BP1,MED15,
3L,TSG101,FUZ,FAM3A,RPS25,NIP7,THEM4,RPS6,RTEL1,SAP30BP,YIPF3,RABL5,TRNT1,ITGB1BP1,MED15,
1,VRK3,LZTFL1,FIS1,MAGED2,SON,SNF8,CPSF3L,TSG101,FUZ,METTL17,ATG12,NIP7,RPS6,RTEL1,FOXRED
1,VRK3,LZTFL1,FIS1,MAGED2,SON,SNF8,CPSF3L,TSG101,FUZ,METTL17,ATG12,NIP7,RPS6,RTEL1,FOXRED
FIS1,MAGED2,SON,UQCRQ,SNF8,CPSF3L,TSG101,FUZ,FAM3A,RPS25,NIP7,THEM4,RPS6,RTEL1,SAP30BP,
FIS1,MAGED2,SON,UQCRQ,SNF8,CPSF3L,TSG101,FUZ,FAM3A,RPS25,NIP7,THEM4,RPS6,RTEL1,SAP30BP,
7,RPS6,RPL29,YIPF3,FOXRED1,TRNT1,MED15,CLASRP,UBE2I,EIF3M,CDC42,FBXW9,USE1,ASF1B,ENDOV,P
P1,YIPF3,TRNT1,MED15,CLASRP,UBE2I,EIF3M,CDC42,FBXW9,USE1,ASF1B,RFX3,ENDOV,PPIL2,APEH,LYPL
CE1,MRPL33,ZNF410,RUVBL2,IGSF8,NUDT5,STYXL1,VRK3,PPM1K,LRRC1,SON,UQCRQ,SNF8,CPSF3L,TSG1
1,PTOV1,FASTK,TMEM135,RNF32,LY75,ATG16L2,SNRPB,PARD3,TCAIM,SLC3A2,DHPS,ACTR6,SEC62,CTSC
1,PTOV1,FASTK,TMEM135,RNF32,LY75,ATG16L2,SNRPB,PARD3,TCAIM,SLC3A2,DHPS,ACTR6,SEC62,CTSC
CDC42,FBXW9,CCDC25,RFX3,ENDOV,TROAP,APEH,LYPLA2,C17ORF62,PRMT1,PPAN,ACAD10,CYB561,SUG
1S1,CACYBP,CIRBP,RHOF,SMARCB1,RPL38,PRR13,STARD10,RNH1,DNASE1L1,KIF4A,RBCK1,PTOV1,FASTK
39,PSMA6,ACOT8,AP3B2,NUDT5,SNRPN,STYXL1,POLD4,LZTFL1,FIS1,PPM1K,MAGED2,SON,UQCRQ,SNF8,
39,PSMA6,ACOT8,AP3B2,NUDT5,SNRPN,STYXL1,POLD4,LZTFL1,FIS1,PPM1K,MAGED2,SON,UQCRQ,SNF8,
13,RNH1,DNASE1L1,RBCK1,PTOV1,FASTK,RNF32,LY75,ATG16L2,RCC1,SLC3A2,DHPS,SET,ZNF235,BOP1,A
13,RNH1,DNASE1L1,RBCK1,PTOV1,FASTK,RNF32,LY75,ATG16L2,RCC1,SLC3A2,DHPS,SET,ZNF235,BOP1,A
2,MYL6B,FAM122C,SRP9,IQCG,TMEM67,SULT1A3,UAP1,PILRB,FAM72A,YBX3,BLOC1S1,CACYBP,RHOF,SN
2,MYL6B,FAM122C,SRP9,IQCG,TMEM67,SULT1A3,UAP1,PILRB,FAM72A,YBX3,BLOC1S1,CACYBP,RHOF,SN
4A,RBCK1,PTOV1,FASTK,TMEM135,RNF32,ATG16L2,RCC1,PARD3,TCAIM,SLC3A2,DHPS,ACTR6,SEC62,SET

YL6B,FAM122C,SRP9,IQCG,TMEM67,SULT1A3,PILRB,FAM72A,YBX3,BLOC1S1,CACYBP,CIRBP,RHOF,SMAF

YL6B,FAM122C,SRP9,IQCG,TMEM67,SULT1A3,PILRB,FAM72A,YBX3,BLOC1S1,CACYBP,CIRBP,RHOF,SMAF

ZTFL1,IRF9,UQCRQ,CPSF3L,TSG101,UBR4,ATG12,RPS25,NIP7,DBNDD2,RPS6,RTEL1,FOXRED1,KXD1,ITGB1

ZTFL1,IRF9,UQCRQ,CPSF3L,TSG101,UBR4,ATG12,RPS25,NIP7,DBNDD2,RPS6,RTEL1,FOXRED1,KXD1,ITGB1

RNT1,KXD1,MED15,C1ORF198,UBE2I,EIF3M,CDC42,FBXW9,EEF1B2,ENDOV,PPIL2,APEH,LYPLA2,C17ORF

3,ENDOV,PPIL2,LYPLA2,C17ORF62,PRMT1,PPAN,ACAD10,CYB561,SUGT1,CARS,CDK11A,ZNF197,DGUOK

PATCH2L,BUD31,NPRL2,CRYGS,CTTN,EIF5A,COMMD5,CHID1,NUDT22,UFD1L,SH2B1,RPL13A,ARHGEF39,

PATCH2L,BUD31,NPRL2,CRYGS,CTTN,EIF5A,COMMD5,CHID1,NUDT22,UFD1L,SH2B1,RPL13A,ARHGEF39,

,FAM122C,SRP9,IQCG,TMEM67,PILRB,FAM72A,YBX3,BLOC1S1,CACYBP,CIRBP,RHOF,SMARCB1,RPL38,PF

,FAM122C,SRP9,IQCG,TMEM67,PILRB,FAM72A,YBX3,BLOC1S1,CACYBP,CIRBP,RHOF,SMARCB1,RPL38,PF

,FBXW9,ASF1B,ENDOV,PPIL2,LYPLA2,C17ORF62,PRMT1,TXNDC17,PPAN,CARS,CDK11A,ZNF197,LRRFIP1,

1A3,PILRB,FAM72A,YBX3,BLOC1S1,CACYBP,CIRBP,RHOF,SMARCB1,RPL38,PRR13,STARD10,RNH1,DNASE

1A3,PILRB,FAM72A,YBX3,BLOC1S1,CACYBP,CIRBP,RHOF,SMARCB1,RPL38,PRR13,STARD10,RNH1,DNASE

.3,RNH1,RBCK1,PTOV1,FASTK,TMEM135,RNF32,LY75,ATG16L2,SNRPB,PARD3,TCAIM,SLC3A2,DHPS,SEC6

.3,RNH1,RBCK1,PTOV1,FASTK,TMEM135,RNF32,LY75,ATG16L2,SNRPB,PARD3,TCAIM,SLC3A2,DHPS,SEC6

30BP,FOXRED1,TRNT1,ITGB1BP1,MED15,CLASRP,SLC39A13,EIF3M,CDC42,USE1,ASF1B,DGKE,ENDOV,PPI

SLC3A2,DHPS,SET,CTSC,ZNF235,ARRDC1,TRPT1,KLC1,RBP5,LIMK2,NOB1,SIRT6,NUMA1,GPATCH2L,OSGE

SLC3A2,DHPS,SET,CTSC,ZNF235,ARRDC1,TRPT1,KLC1,RBP5,LIMK2,NOB1,SIRT6,NUMA1,GPATCH2L,OSGE

H1,DNASE1L1,KIF4A,RBCK1,PTOV1,FASTK,TMEM135,RNF32,LY75,ATG16L2,RCC1,PARD3,TCAIM,SLC3A2,

RHGEF39,PSMA6,ACOT8,ZNF410,NUDT5,SNRPN,STYXL1,POLD4,VRK3,LZTFL1,FIS1,IRF9,DST,MAGED2,SC

RHGEF39,PSMA6,ACOT8,ZNF410,NUDT5,SNRPN,STYXL1,POLD4,VRK3,LZTFL1,FIS1,IRF9,DST,MAGED2,SC

DT22,UFD1L,RPL13A,ARHGEF39,ACOT8,AP3B2,ZNF410,PCMT1,NUDT5,STYXL1,POLD4,VRK3,LZTFL1,FIS1,I

DT22,UFD1L,RPL13A,ARHGEF39,ACOT8,AP3B2,ZNF410,PCMT1,NUDT5,STYXL1,POLD4,VRK3,LZTFL1,FIS1,I

,RPS6,RTEL1,SAP30BP,YIPF3,RABL5,TRNT1,ITGB1BP1,MED15,CLASRP,C1ORF198,UGGT2,UBE2I,EIF3M,C

,RPS6,RTEL1,SAP30BP,YIPF3,RABL5,TRNT1,ITGB1BP1,MED15,CLASRP,C1ORF198,UGGT2,UBE2I,EIF3M,C

,COX4I1,RPS6,RPL29,YIPF3,TRNT1,MED15,C1ORF198,MICAL1,EIF3M,CDC42,FBXW9,ASF1B,CCDC25,RFX

LOC1S1,CIRBP,RHOF,SMARCB1,PRR13,STARD10,RNH1,DNASE1L1,KIF4A,RBCK1,PTOV1,FASTK,RNF32,LY7

LOC1S1,CIRBP,RHOF,SMARCB1,PRR13,STARD10,RNH1,DNASE1L1,KIF4A,RBCK1,PTOV1,FASTK,RNF32,LY7

NUDT22,UFD1L,USP38,SH2B1,RPL13A,ARHGEF39,PSMA6,ACOT8,ZNF410,PCMT1,NUDT5,SNRPN,STYXL1

NUDT22,UFD1L,USP38,SH2B1,RPL13A,ARHGEF39,PSMA6,ACOT8,ZNF410,PCMT1,NUDT5,SNRPN,STYXL1

PATCH2L,BUD31,NPRL2,NUDT22,NOP16,RPL13A,TTLL5,LDHA,ACOT8,RPS9,SLC35G1,MRPL33,ZNF410,EE

L38,STARD10,RNH1,DNASE1L1,RBCK1,PTOV1,FASTK,RNF32,ATG16L2,RCC1,SNRPB,PARD3,SLC3A2,DHPS,

L38,STARD10,RNH1,DNASE1L1,RBCK1,PTOV1,FASTK,RNF32,ATG16L2,RCC1,SNRPB,PARD3,SLC3A2,DHPS,

T5,STYXL1,POLD4,VRK3,LZTFL1,FIS1,PPM1K,SON,SNF8,GRK6,HMGA2,CPSF3L,TSG101,RPS25,NIP7,SLC25A

LZTFL1,PPM1K,TUBA1C,IRF9,CD44,SON,UQCRQ,SNF8,GRK6,CPSF3L,TSG101,MUC1,RPS25,COX4I1,SLC25

IF4A,PTOV1,FASTK,TMEM135,RNF32,LY75,ATG16L2,SNRPB,PARD3,TCAIM,SLC3A2,ACTR6,SET,CTSC,BOP1

IF4A,PTOV1,FASTK,TMEM135,RNF32,LY75,ATG16L2,SNRPB,PARD3,TCAIM,SLC3A2,ACTR6,SET,CTSC,BOP1

NASE1L1,KIF4A,RBCK1,PTOV1,FASTK,TMEM135,RNF32,LY75,ATG16L2,SNRPB,PARD3,TCAIM,SLC3A2,DH

NASE1L1,KIF4A,RBCK1,PTOV1,FASTK,TMEM135,RNF32,LY75,ATG16L2,SNRPB,PARD3,TCAIM,SLC3A2,DH

,RCC1,SNRPB,SLC3A2,ACTR6,SEC62,SET,CTSC,ZNF235,BOP1,ARRDC1,KLC1,RBP5,SCAF11,DBI,ARPC2,LIMI

K1,PTOV1,FASTK,RNF32,LY75,ATG16L2,SNRPB,PARD3,TCAIM,SLC3A2,SEC62,SET,CTSC,ZNF235,ARRDC1,

K1,PTOV1,FASTK,RNF32,LY75,ATG16L2,SNRPB,PARD3,TCAIM,SLC3A2,SEC62,SET,CTSC,ZNF235,ARRDC1,

,RCC1,SNRPB,SLC3A2,DHPS,ACTR6,BOP1,TRPT1,ZC3HAV1,DBI,PMFBP1,LIMK2,SIRT6,NUMA1,BUD31,NPI

,RCC1,SNRPB,SLC3A2,DHPS,ACTR6,BOP1,TRPT1,ZC3HAV1,DBI,PMFBP1,LIMK2,SIRT6,NUMA1,BUD31,NPI

B,PARD3,TCAIM,ACTR6,SEC62,SET,CTSC,BOP1,ARRDC1,TRPT1,KLC1,DBI,ARPC2,LIMK2,SIRT6,NUMA1,BUI

AN,SUGT1,ZNF197,DGUOK,LRRFIP1,TRMT112,CHPF,ERCC5,EMC4,DDOST,LSR,TOMM40,RABEP2,TADA3,

,RHOF,SMARCB1,RPL38,STARD10,RNH1,DNASE1L1,KIF4A,RBCK1,PTOV1,FASTK,TMEM135,RNF32,LY75,

P,RHOF,SMARCB1,RPL38,STARD10,RNH1,DNASE1L1,KIF4A,RBCK1,PTOV1,FASTK,TMEM135,RNF32,LY75,A
HYN1,RPL13A,ARHGEF39,ACOT8,AP3B2,SLC35G1,ZNF410,IGSF8,NUDT5,STYXL1,RFX5,POLD4,LZTFL1,FIS1,
HYN1,RPL13A,ARHGEF39,ACOT8,AP3B2,SLC35G1,ZNF410,IGSF8,NUDT5,STYXL1,RFX5,POLD4,LZTFL1,FIS1,
,FASTK,RNF32,LY75,ATG16L2,RCC1,SNRPB,TCAIM,SLC3A2,DHPS,ZNF235,BOP1,ARRDC1,KLC1,RBP5,ARPC
,FASTK,RNF32,LY75,ATG16L2,RCC1,SNRPB,TCAIM,SLC3A2,DHPS,ZNF235,BOP1,ARRDC1,KLC1,RBP5,ARPC
SULT1A3,PILRB,FAM72A,YBX3,BLOC1S1,CACYBP,CIRBP,RHOF,SMARCB1,RPL38,PRR13,RNH1,DNASE1L1,K
LD4,IRF9,SON,SNF8,CDC37,CPSF3L,TSG101,RPS25,NIP7,TTC9C,SLC25A10,THEM4,RPL29,AUP1,YIPF3,FOX
3L,TSG101,NIP7,THEM4,RPL29,AUP1,SAP30BP,YIPF3,FOXRED1,TRNT1,KXD1,MED15,C1ORF198,UBE2I,EIF
G1,APEH,LYPLA2,C17ORF62,INO80E,CCNL2,DGUOK,ATP5J,TRMT112,MPDU1,SAT1,ERCC5,EMC4,SERPINE
EM135,LY75,ATG16L2,SNRPB,PARD3,TCAIM,ACTR6,SEC62,SET,BOP1,ARRDC1,TRPT1,KLC1,SCAF11,DBI,AF
EM135,LY75,ATG16L2,SNRPB,PARD3,TCAIM,ACTR6,SEC62,SET,BOP1,ARRDC1,TRPT1,KLC1,SCAF11,DBI,AF
KIF4A,RBCK1,PTOV1,FASTK,TMEM135,ATG16L2,PARD3,TCAIM,SLC3A2,DHPS,ACTR6,SEC62,SET,CTSC,BOP
KIF4A,RBCK1,PTOV1,FASTK,TMEM135,ATG16L2,PARD3,TCAIM,SLC3A2,DHPS,ACTR6,SEC62,SET,CTSC,BOP
CACYBP,RHOF,SMARCB1,RPL38,STARD10,RNH1,DNASE1L1,KIF4A,RBCK1,PTOV1,FASTK,TMEM135,RNF32
3,TCAIM,SLC3A2,SEC62,SET,BOP1,ARRDC1,TRPT1,KLC1,RBP5,DBI,LIMK2,SIRT6,NUMA1,GPATCH2L,BUD3
3,TCAIM,SLC3A2,SEC62,SET,BOP1,ARRDC1,TRPT1,KLC1,RBP5,DBI,LIMK2,SIRT6,NUMA1,GPATCH2L,BUD3
72,MYL6B,FAM122C,SRP9,IQCG,TMEM67,SULT1A3,PILRB,FAM72A,YBX3,BLOC1S1,CACYBP,RHOF,SMARC
72,MYL6B,FAM122C,SRP9,IQCG,TMEM67,SULT1A3,PILRB,FAM72A,YBX3,BLOC1S1,CACYBP,RHOF,SMARC
,PTOV1,FASTK,TMEM135,RNF32,LY75,ATG16L2,RCC1,SNRPB,PARD3,SLC3A2,DHPS,ACTR6,SEC62,SET,BO
,RPL38,STARD10,RNH1,DNASE1L1,KIF4A,RBCK1,PTOV1,FASTK,TMEM135,ATG16L2,RCC1,SNRPB,PARD3,
,TRPT1,RBP5,SCAF11,DBI,ARPC2,LIMK2,SIRT6,NUMA1,GPATCH2L,SUMF2,BUD31,NPRL2,CRYGS,CTTN,TS
,TRPT1,RBP5,SCAF11,DBI,ARPC2,LIMK2,SIRT6,NUMA1,GPATCH2L,SUMF2,BUD31,NPRL2,CRYGS,CTTN,TS
S6,SAP30BP,YIPF3,FOXRED1,TRNT1,MED15,CLASRP,UBE2I,EIF3M,CDC42,FBXW9,CCDC25,RFX3,ENDOV,P
RF198,UBE2I,USE1,ASF1B,ENDOV,CAMTA1,APEH,LYPLA2,C17ORF62,PRMT1,PPAN,ACAD10,CDK11A,ZNF
5,BOP1,ARRDC1,KLC1,RBP5,SCAF11,ARPC2,SIRT6,BUD31,SAE1,NPRL2,CTTN,TSPAN11,EIF5A,COMMD5,UI
,RCC1,SNRPB,PARD3,DHPS,ACTR6,CTSC,ZNF235,BOP1,ARRDC1,TRPT1,RBP5,ZC3HAV1,SCAF11,DBI,ARPC
,RCC1,SNRPB,PARD3,DHPS,ACTR6,CTSC,ZNF235,BOP1,ARRDC1,TRPT1,RBP5,ZC3HAV1,SCAF11,DBI,ARPC
W9,RFX3,ENDOV,PPIL2,APEH,LYPLA2,C17ORF62,PRMT1,STOML2,C19ORF54,TXNDC17,PPAN,HNRNPC,CA
75,ATG16L2,SNRPB,PARD3,TCAIM,SLC3A2,SEC62,ARRDC1,KLC1,RBP5,LIMK2,SIRT6,NUMA1,GPATCH2L,N
75,ATG16L2,SNRPB,PARD3,TCAIM,SLC3A2,SEC62,ARRDC1,KLC1,RBP5,LIMK2,SIRT6,NUMA1,GPATCH2L,N
75,ATG16L2,SNRPB,PARD3,TCAIM,SLC3A2,SEC62,ARRDC1,KLC1,RBP5,LIMK2,SIRT6,NUMA1,GPATCH2L,N
75,ATG16L2,SNRPB,PARD3,TCAIM,SLC3A2,SEC62,ARRDC1,KLC1,RBP5,LIMK2,SIRT6,NUMA1,GPATCH2L,N
,CMC2,YBX3,BLOC1S1,CACYBP,SMARCB1,RPL38,STARD10,RNH1,KIF4A,PTOV1,FASTK,RNF32,LY75,ATG16
,CMC2,YBX3,BLOC1S1,CACYBP,SMARCB1,RPL38,STARD10,RNH1,KIF4A,PTOV1,FASTK,RNF32,LY75,ATG16
P,RHOF,SMARCB1,RPL38,STARD10,RNH1,DNASE1L1,KIF4A,RBCK1,PTOV1,FASTK,TMEM135,RNF32,ATG16
FAM122C,SRP9,IQCG,TMEM67,PILRB,FAM72A,CMC2,YBX3,CACYBP,RHOF,SMARCB1,RPL38,PRR13,STARD
FAM122C,SRP9,IQCG,TMEM67,PILRB,FAM72A,CMC2,YBX3,CACYBP,RHOF,SMARCB1,RPL38,PRR13,STARD
M72A,BLOC1S1,CACYBP,CIRBP,RHOF,SMARCB1,RPL38,PRR13,RNH1,DNASE1L1,KIF4A,RBCK1,PTOV1,FAS
NASE1L1,PTOV1,TMEM135,RNF32,RCC1,SNRPB,PARD3,TCAIM,SLC3A2,DHPS,ACTR6,SEC62,SET,CTSC,ARR
CYBP,RHOF,RPL38,PRR13,STARD10,RNH1,DNASE1L1,PTOV1,FASTK,TMEM135,RNF32,LY75,ATG16L2,RCC
CYBP,RHOF,RPL38,PRR13,STARD10,RNH1,DNASE1L1,PTOV1,FASTK,TMEM135,RNF32,LY75,ATG16L2,RCC
6L2,SNRPB,PARD3,SLC3A2,DHPS,MAP3K7CL,SET,CTSC,ZNF235,BOP1,ARRDC1,TRPT1,KLC1,RBP5,ZC3HAV
6L2,SNRPB,PARD3,SLC3A2,DHPS,MAP3K7CL,SET,CTSC,ZNF235,BOP1,ARRDC1,TRPT1,KLC1,RBP5,ZC3HAV
PS,SET,CTSC,ARRDC1,TRPT1,RBP5,LIMK2,NOB1,NUMA1,GPATCH2L,OSGEP,TBL2,SCARB1,CTTN,TSPAN11,
PS,SET,CTSC,ARRDC1,TRPT1,RBP5,LIMK2,NOB1,NUMA1,GPATCH2L,OSGEP,TBL2,SCARB1,CTTN,TSPAN11,
L38,PRR13,RNH1,DNASE1L1,PTOV1,TMEM135,RNF32,LY75,ATG16L2,RCC1,SNRPB,PARD3,TCAIM,SLC3A2
T,CTSC,ARRDC1,TRPT1,RBP5,LIMK2,VDAC3,NOB1,NUMA1,GPATCH2L,SUMF2,OSGEP,TBL2,SCARB1,CTTN,

T,CTSC,ARRDC1,TRPT1,RBP5,LIMK2,VDAC3,NOB1,NUMA1,GPATCH2L,SUMF2,OSGEP,TBL2,SCARB1,CTTN,

F1B,LIG1,LYPLA2,C17ORF62,PRMT1,C19ORF54,ARL6IP4,HNRNPC,ZNF197,MAPKAPK5,ATP5J,TRMT112,C

RBCK1,PTOV1,FASTK,TMEM135,RNF32,LY75,ATG16L2,PARD3,TCAIM,SLC3A2,DHPS,ACTR6,SEC62,SET,CT

YL6B,FAM122C,SRP9,IQCG,TMEM67,SULT1A3,PILRB,FAM72A,CMC2,YBX3,BLOC1S1,CACYBP,CIRBP,RHO

YL6B,FAM122C,SRP9,IQCG,TMEM67,SULT1A3,PILRB,FAM72A,CMC2,YBX3,BLOC1S1,CACYBP,CIRBP,RHO

TARD10,RNH1,DNASE1L1,KIF4A,RBCK1,PTOV1,FASTK,TMEM135,LY75,ATG16L2,RCC1,SNRPB,PARD3,SLC

TARD10,RNH1,DNASE1L1,KIF4A,RBCK1,PTOV1,FASTK,TMEM135,LY75,ATG16L2,RCC1,SNRPB,PARD3,SLC

9,SAP30BP,TRNT1,MED15,SH3YL1,C1ORF198,SLC39A13,YIF1A,EIF3M,CDC42,TMEM205,CARD14,ASF1B,L

STK,RNF32,LY75,ATG16L2,PARD3,TCAIM,SLC3A2,SEC62,BOP1,ARRDC1,KLC1,RBP5,LIMK2,VDAC3,SIRT6,N

STK,RNF32,LY75,ATG16L2,PARD3,TCAIM,SLC3A2,SEC62,BOP1,ARRDC1,KLC1,RBP5,LIMK2,VDAC3,SIRT6,N

,KIF4A,RBCK1,PTOV1,FASTK,TMEM135,RNF32,RCC1,SNRPB,PARD3,TCAIM,SLC3A2,DHPS,ACTR6,SEC62,S

,KIF4A,RBCK1,PTOV1,FASTK,TMEM135,RNF32,RCC1,SNRPB,PARD3,TCAIM,SLC3A2,DHPS,ACTR6,SEC62,S

1A3,PILRB,FAM72A,YBX3,BLOC1S1,CACYBP,CIRBP,RHOF,SMARCB1,RPL38,PRR13,STARD10,RNH1,DNASE

1A3,PILRB,FAM72A,YBX3,BLOC1S1,CACYBP,CIRBP,RHOF,SMARCB1,RPL38,PRR13,STARD10,RNH1,DNASE

35,RNF32,LY75,ATG16L2,RCC1,PARD3,TCAIM,DHPS,SET,CTSC,BOP1,ARRDC1,KLC1,ZC3HAV1,SCAF11,DBI,

Y75,SNRPB,PARD3,TCAIM,ACTR6,CTSC,ZNF235,BOP1,ARRDC1,KLC1,SCAF11,LIMK2,WDR19,SUMF2,BUD

Y75,SNRPB,PARD3,TCAIM,ACTR6,CTSC,ZNF235,BOP1,ARRDC1,KLC1,SCAF11,LIMK2,WDR19,SUMF2,BUD

Y75,SNRPB,PARD3,TCAIM,ACTR6,CTSC,ZNF235,BOP1,ARRDC1,KLC1,SCAF11,LIMK2,WDR19,SUMF2,BUD

Y75,SNRPB,PARD3,TCAIM,ACTR6,CTSC,ZNF235,BOP1,ARRDC1,KLC1,SCAF11,LIMK2,WDR19,SUMF2,BUD

Y75,SNRPB,PARD3,TCAIM,ACTR6,CTSC,ZNF235,BOP1,ARRDC1,KLC1,SCAF11,LIMK2,WDR19,SUMF2,BUD

Y75,SNRPB,PARD3,TCAIM,ACTR6,CTSC,ZNF235,BOP1,ARRDC1,KLC1,SCAF11,LIMK2,WDR19,SUMF2,BUD

OP1,ARRDC1,TRPT1,KLC1,RBP5,LIMK2,SIRT6,NUMA1,TBL2,SCARB1,CTTN,TSPAN11,ITSN1,EIF5A,COMMD

OP1,ARRDC1,TRPT1,KLC1,RBP5,LIMK2,SIRT6,NUMA1,TBL2,SCARB1,CTTN,TSPAN11,ITSN1,EIF5A,COMMD

SNRPB,PARD3,SLC3A2,DHPS,ACTR6,MAP3K7CL,ZNF235,TRPT1,KLC1,ZC3HAV1,DBI,ARPC2,PMFBP1,LIMK

SNRPB,PARD3,SLC3A2,DHPS,ACTR6,MAP3K7CL,ZNF235,TRPT1,KLC1,ZC3HAV1,DBI,ARPC2,PMFBP1,LIMK

K2,GPATCH2L,BUD31,NPRL2,TBL2,CTTN,EIF5A,COMMD5,NUDT22,UFD1L,USP38,THYN1,DBNDD1,NOP1

K2,GPATCH2L,BUD31,NPRL2,TBL2,CTTN,EIF5A,COMMD5,NUDT22,UFD1L,USP38,THYN1,DBNDD1,NOP1

1,FASTK,TMEM135,RNF32,LY75,ATG16L2,SLC3A2,DHPS,ACTR6,SET,BOP1,ARRDC1,TRPT1,KLC1,RBP5,DBI,

1,FASTK,TMEM135,RNF32,LY75,ATG16L2,SLC3A2,DHPS,ACTR6,SET,BOP1,ARRDC1,TRPT1,KLC1,RBP5,DBI,

1,RABL5,ITGB1BP1,MED15,CLASRP,C1ORF198,UGGT2,UBE2I,CDC42,PCYT2,FBXW9,CCDC25,EEF1B2,LIG1

1,RABL5,ITGB1BP1,MED15,CLASRP,C1ORF198,UGGT2,UBE2I,CDC42,PCYT2,FBXW9,CCDC25,EEF1B2,LIG1

5,RNF32,LY75,ATG16L2,SNRPB,PARD3,SLC3A2,SET,ARRDC1,TRPT1,KLC1,ZC3HAV1,DBI,ARPC2,LIMK2,SIR

5,RNF32,LY75,ATG16L2,SNRPB,PARD3,SLC3A2,SET,ARRDC1,TRPT1,KLC1,ZC3HAV1,DBI,ARPC2,LIMK2,SIR

RRC1,TM9SF1,MAGED2,RPS16,TSG101,UBR4,PSMA1,ATG12,THEM4,DBNDD2,SDF4,NISCH,RPS6,RTEL1,F

RRC1,TM9SF1,MAGED2,RPS16,TSG101,UBR4,PSMA1,ATG12,THEM4,DBNDD2,SDF4,NISCH,RPS6,RTEL1,F

E1L1,RBCK1,PTOV1,FASTK,TMEM135,RNF32,LY75,ATG16L2,PARD3,TCAIM,SLC3A2,DHPS,ACTR6,SEC62,S

E1L1,RBCK1,PTOV1,FASTK,TMEM135,RNF32,LY75,ATG16L2,PARD3,TCAIM,SLC3A2,DHPS,ACTR6,SEC62,S

2,USP38,SH2B1,NOP16,RPL13A,ARHGEF39,PSMA6,ACOT8,RPS9,RCE1,ZNF410,NUDT5,SPAG7,STYXL1,RF

TG16L2,SNRPB,PARD3,SLC3A2,ACTR6,MAP3K7CL,SEC62,SET,CTSC,ARRDC1,KLC1,SCAF11,DBI,ARPC2,PMF

TG16L2,SNRPB,PARD3,SLC3A2,ACTR6,MAP3K7CL,SEC62,SET,CTSC,ARRDC1,KLC1,SCAF11,DBI,ARPC2,PMF

L2,PARD3,SLC3A2,SET,ARRDC1,TRPT1,KLC1,RBP5,LIMK2,SIRT6,NUMA1,GPATCH2L,TBL2,SCARB1,CTTN,T

L2,PARD3,SLC3A2,SET,ARRDC1,TRPT1,KLC1,RBP5,LIMK2,SIRT6,NUMA1,GPATCH2L,TBL2,SCARB1,CTTN,T

ATG16L2,SNRPB,PARD3,TCAIM,SLC3A2,DHPS,ACTR6,SET,BOP1,ARRDC1,KLC1,SCAF11,DBI,LIMK2,VDAC3,

RDC1,KLC1,SCAF11,LIMK2,VDAC3,WDR19,SIRT6,NUMA1,GPATCH2L,BUD31,NPRL2,ARHGAP6,CTTN,ITSN

RDC1,KLC1,SCAF11,LIMK2,VDAC3,WDR19,SIRT6,NUMA1,GPATCH2L,BUD31,NPRL2,ARHGAP6,CTTN,ITSN

X3,BLOC1S1,CIRBP,RHOF,SMARCB1,PRR13,STARD10,RNH1,DNASE1L1,KIF4A,RNF32,RCC1,SNRPB,PARD3,

X3,BLOC1S1,CIRBP,RHOF,SMARCB1,PRR13,STARD10,RNH1,DNASE1L1,KIF4A,RNF32,RCC1,SNRPB,PARD3,

,FBXW9,CCDC25,EEF1B2,ENDOV,LIG1,LYPLA2,STOML2,ZNF7,PPAN,CYB561,SUGT1,HYPK,ZNF197,INO80E
,FBXW9,CCDC25,EEF1B2,ENDOV,LIG1,LYPLA2,STOML2,ZNF7,PPAN,CYB561,SUGT1,HYPK,ZNF197,INO80E
,UAP1,FAM72A,YBX3,BLOC1S1,CACYBP,SMARCB1,RPL38,PRR13,RNH1,DNASE1L1,KIF4A,PTOV1,FASTK,RI
,UAP1,FAM72A,YBX3,BLOC1S1,CACYBP,SMARCB1,RPL38,PRR13,RNH1,DNASE1L1,KIF4A,PTOV1,FASTK,RI
RNH1,KIF4A,RBCK1,PTOV1,FASTK,TMEM135,RNF32,LY75,ATG16L2,SNRPB,PARD3,SLC3A2,DHPS,ACTR6,S
RNH1,KIF4A,RBCK1,PTOV1,FASTK,TMEM135,RNF32,LY75,ATG16L2,SNRPB,PARD3,SLC3A2,DHPS,ACTR6,S
CB1,RPL38,STARD10,RNH1,DNASE1L1,KIF4A,RBCK1,PTOV1,FASTK,RNF32,LY75,ATG16L2,RCC1,PARD3,TC
CB1,RPL38,STARD10,RNH1,DNASE1L1,KIF4A,RBCK1,PTOV1,FASTK,RNF32,LY75,ATG16L2,RCC1,PARD3,TC
D2,SON,SNF8,CPSF3L,FUZ,RPS25,NIP7,COX4I1,RPS6,RPL29,RTEL1,RABL5,TRNT1,ITGB1BP1,MED15,CLASR
D2,SON,SNF8,CPSF3L,FUZ,RPS25,NIP7,COX4I1,RPS6,RPL29,RTEL1,RABL5,TRNT1,ITGB1BP1,MED15,CLASR
STARD10,DNASE1L1,KIF4A,RBCK1,PTOV1,FASTK,TMEM135,RCC1,SNRPB,PARD3,DHPS,ACTR6,MAP3K7CL
STARD10,DNASE1L1,KIF4A,RBCK1,PTOV1,FASTK,TMEM135,RCC1,SNRPB,PARD3,DHPS,ACTR6,MAP3K7CL
F235,BOP1,ARRDC1,TRPT1,KLC1,DBI,ARPC2,LIMK2,SIRT6,NUMA1,GPATCH2L,BUD31,TBL2,SCARB1,CTTN,
F235,BOP1,ARRDC1,TRPT1,KLC1,DBI,ARPC2,LIMK2,SIRT6,NUMA1,GPATCH2L,BUD31,TBL2,SCARB1,CTTN,
B1,CTTN,TSPAN11,ITSN1,COMMD5,AMZ2,TAZ,DBNDD1,NOP16,PSMA6,ACOT8,AP3B2,MRPL33,RUVBL2,
B1,CTTN,TSPAN11,ITSN1,COMMD5,AMZ2,TAZ,DBNDD1,NOP16,PSMA6,ACOT8,AP3B2,MRPL33,RUVBL2,
RHGEF39,PSMA6,ACOT8,RUVBL2,PCMT1,NUDT5,WDR62,STYXL1,LZTFL1,FIS1,DST,STIP1,VCAN,MAGED2,
RHGEF39,PSMA6,ACOT8,RUVBL2,PCMT1,NUDT5,WDR62,STYXL1,LZTFL1,FIS1,DST,STIP1,VCAN,MAGED2,
1,TRPT1,RBP5,LIMK2,NOB1,SIRT6,NUMA1,TBL2,SCARB1,CTTN,TSPAN11,ITSN1,COMMD5,NUDT22,DBNI
1,TRPT1,RBP5,LIMK2,NOB1,SIRT6,NUMA1,TBL2,SCARB1,CTTN,TSPAN11,ITSN1,COMMD5,NUDT22,DBNI
C3,WDR19,SIRT6,NUMA1,BUD31,OSGEP,ARHGAP6,CTTN,ITSN1,EIF5A,USP38,DBNDD1,SH2B1,AP3B2,RP
C3,WDR19,SIRT6,NUMA1,BUD31,OSGEP,ARHGAP6,CTTN,ITSN1,EIF5A,USP38,DBNDD1,SH2B1,AP3B2,RP
,KIF4A,RBCK1,PTOV1,FASTK,TMEM135,RNF32,LY75,ATG16L2,PARD3,SLC3A2,SEC62,SET,ZNF235,ARRDC
,KIF4A,RBCK1,PTOV1,FASTK,TMEM135,RNF32,LY75,ATG16L2,PARD3,SLC3A2,SEC62,SET,ZNF235,ARRDC
/9,CCDC25,EEF1B2,ENDOV,LIG1,LYPLA2,STOML2,ZNF7,PPAN,CYB561,SUGT1,HYPK,ZNF197,INO80E,HSP9
/9,CCDC25,EEF1B2,ENDOV,LIG1,LYPLA2,STOML2,ZNF7,PPAN,CYB561,SUGT1,HYPK,ZNF197,INO80E,HSP9
,GSTK1,TTLL5,PSMA6,AP3B2,SLC35G1,MRPL33,ZNF410,IGSF8,NNMT,WDR62,IL1RAP,RANBP1,UBE2L6,ZC
D4,VRK3,LZTFL1,FIS1,STIP1,VCAN,LPCAT3,SON,UQCRQ,SNF8,CPSF3L,TSG101,METTL17,ATG12,RPS25,NII
1,COMMD5,NUDT22,UFD1L,THYN1,DBNDD1,NOP16,ACOT8,AP3B2,RCE1,ZNF410,RUVBL2,PCMT1,EBPL,
TSG101,SLC39A11,NIP7,COX4I1,RTEL1,RABL5,TRNT1,ITGB1BP1,MED15,CLASRP,UGGT2,UBE2I,PRMT2,CD
TSG101,SLC39A11,NIP7,COX4I1,RTEL1,RABL5,TRNT1,ITGB1BP1,MED15,CLASRP,UGGT2,UBE2I,PRMT2,CD
TARD10,RNH1,DNASE1L1,KIF4A,PTOV1,FASTK,TMEM135,RNF32,LY75,ATG16L2,RCC1,PARD3,TCAIM,DHF
TARD10,RNH1,DNASE1L1,KIF4A,PTOV1,FASTK,TMEM135,RNF32,LY75,ATG16L2,RCC1,PARD3,TCAIM,DHF
HOF,SMARCB1,RPL38,RNH1,DNASE1L1,KIF4A,RBCK1,PTOV1,FASTK,TMEM135,RNF32,LY75,ATG16L2,PAR
HOF,SMARCB1,RPL38,RNH1,DNASE1L1,KIF4A,RBCK1,PTOV1,FASTK,TMEM135,RNF32,LY75,ATG16L2,PAR
STK,LY75,ATG16L2,SNRPB,PARD3,TCAIM,SLC3A2,ACTR6,MAP3K7CL,SEC62,SET,CTSC,BOP1,ARRDC1,KLC
STK,LY75,ATG16L2,SNRPB,PARD3,TCAIM,SLC3A2,ACTR6,MAP3K7CL,SEC62,SET,CTSC,BOP1,ARRDC1,KLC
ARRDC1,TRPT1,DBI,ARPC2,PMFBP1,LIMK2,NUMA1,GPATCH2L,BUD31,NPRL2,TBL2,SCARB1,CTTN,EIF5A,
ARRDC1,TRPT1,DBI,ARPC2,PMFBP1,LIMK2,NUMA1,GPATCH2L,BUD31,NPRL2,TBL2,SCARB1,CTTN,EIF5A,
2,SIRT6,BUD31,TBL2,SCARB1,CTTN,ITSN1,NUDT22,AMZ2,TAZ,DBNDD1,NOP16,ACOT8,AP3B2,RCE1,MRP
2,SIRT6,BUD31,TBL2,SCARB1,CTTN,ITSN1,NUDT22,AMZ2,TAZ,DBNDD1,NOP16,ACOT8,AP3B2,RCE1,MRP
8,UGGT2,UBE2I,CDC42,FBXW9,CCDC25,EEF1B2,LIG1,LYPLA2,STOML2,ZNF7,PPAN,CYB561,SUGT1,HYPK,Z
8,UGGT2,UBE2I,CDC42,FBXW9,CCDC25,EEF1B2,LIG1,LYPLA2,STOML2,ZNF7,PPAN,CYB561,SUGT1,HYPK,Z
ULT1A3,PILRB,FAM72A,CMC2,YBX3,BLOC1S1,CACYBP,CIRBP,RHOF,RPL38,PRR13,RNH1,DNASE1L1,KIF4A
ULT1A3,PILRB,FAM72A,CMC2,YBX3,BLOC1S1,CACYBP,CIRBP,RHOF,RPL38,PRR13,RNH1,DNASE1L1,KIF4A
RPS16,TSG101,SLC39A11,FUZ,FAM3A,RPS25,NIP7,TTC9C,PLCB2,DUSP22,THEM4,DBNDD2,RPS6,RPL29,C
RPS16,TSG101,SLC39A11,FUZ,FAM3A,RPS25,NIP7,TTC9C,PLCB2,DUSP22,THEM4,DBNDD2,RPS6,RPL29,C

,EBPL,WDR62,UBE2L6,VRK3,ANTXR2,STIP1,LPCAT3,GRK6,CDC37,RPS16,SLC39A11,MUC1,ECSIT,EPB41L5,
,EBPL,WDR62,UBE2L6,VRK3,ANTXR2,STIP1,LPCAT3,GRK6,CDC37,RPS16,SLC39A11,MUC1,ECSIT,EPB41L5,
C1,RBP5,LIMK2,WDR19,SIRT6,NUMA1,GPATCH2L,TBL2,SCARB1,CTTN,TSPAN11,ITSN1,DBNDD1,NOP16,T
C1,RBP5,LIMK2,WDR19,SIRT6,NUMA1,GPATCH2L,TBL2,SCARB1,CTTN,TSPAN11,ITSN1,DBNDD1,NOP16,T
L,TSG101,FAM3A,METTL17,RPS25,NIP7,CENPK,RPS6,RPL29,AUP1,SAP30BP,TRNT1,MED15,SLC39A13,YIF
,CRYGS,CTTN,EIF5A,COMMD5,NUDT22,UFD1L,USP38,TAZ,RPL13A,ARHGEF39,PSMA6,ACOT8,RPS9,ZNF4
,CRYGS,CTTN,EIF5A,COMMD5,NUDT22,UFD1L,USP38,TAZ,RPL13A,ARHGEF39,PSMA6,ACOT8,RPS9,ZNF4
RCE1,RUVBL2,EBPL,WDR62,UBE2L6,VRK3,IRF9,ANTXR2,STIP1,LPCAT3,MAGED2,GRK6,CDC37,SLC39A11,
RCE1,RUVBL2,EBPL,WDR62,UBE2L6,VRK3,IRF9,ANTXR2,STIP1,LPCAT3,MAGED2,GRK6,CDC37,SLC39A11,
38,PRR13,STARD10,RNH1,DNASE1L1,KIF4A,PTOV1,FASTK,TMEM135,RNF32,LY75,ATG16L2,RCC1,PARD3
ASTK,TMEM135,RNF32,LY75,ATG16L2,RCC1,PARD3,TCAIM,DHPS,SET,CTSC,BOP1,ARRDC1,KLC1,ZC3HAV
ASTK,TMEM135,RNF32,LY75,ATG16L2,RCC1,PARD3,TCAIM,DHPS,SET,CTSC,BOP1,ARRDC1,KLC1,ZC3HAV
EF39,PSMA6,ACOT8,AP3B2,ZNF410,NUDT5,SNRPN,STYXL1,POLD4,VRK3,LZTFL1,FIS1,PPM1K,STIP1,VCAN
EF39,PSMA6,ACOT8,AP3B2,ZNF410,NUDT5,SNRPN,STYXL1,POLD4,VRK3,LZTFL1,FIS1,PPM1K,STIP1,VCAN
LC3A2,DHPS,ACTR6,SEC62,SET,CTSC,ARRDC1,SCAF11,PMFBP1,LIMK2,VDAC3,NOB1,NUMA1,GPATCH2L,
LC3A2,DHPS,ACTR6,SEC62,SET,CTSC,ARRDC1,SCAF11,PMFBP1,LIMK2,VDAC3,NOB1,NUMA1,GPATCH2L,
31,PRR13,STARD10,RNH1,DNASE1L1,KIF4A,RBCK1,PTOV1,FASTK,RNF32,LY75,ATG16L2,RCC1,PARD3,TCA
31,PRR13,STARD10,RNH1,DNASE1L1,KIF4A,RBCK1,PTOV1,FASTK,RNF32,LY75,ATG16L2,RCC1,PARD3,TCA
1L1,KIF4A,FASTK,TMEM135,RNF32,LY75,ATG16L2,RCC1,PARD3,TCAIM,DHPS,SET,CTSC,BOP1,ARRDC1,T
PATCH2L,BUD31,NPRL2,CRYGS,CTTN,EIF5A,COMMD5,CHID1,NUDT22,UFD1L,USP38,SH2B1,RPL13A,ARH
PATCH2L,BUD31,NPRL2,CRYGS,CTTN,EIF5A,COMMD5,CHID1,NUDT22,UFD1L,USP38,SH2B1,RPL13A,ARH
DT22,UFD1L,USP38,TAZ,RPL13A,ARHGEF39,ACOT8,AP3B2,ZNF410,RUVBL2,EBPL,IGSF8,NUDT5,RBPMS,
DT22,UFD1L,USP38,TAZ,RPL13A,ARHGEF39,ACOT8,AP3B2,ZNF410,RUVBL2,EBPL,IGSF8,NUDT5,RBPMS,
LOC1S1,CACYBP,CIRBP,SMARCB1,STARD10,RNH1,DNASE1L1,KIF4A,RBCK1,PTOV1,FASTK,TMEM135,RNF
SN1,EIF5A,COMMD5,UFD1L,THYN1,DBNDD1,NOP16,ACOT8,AP3B2,RCE1,ZNF410,RUVBL2,PCMT1,EBPL,T
RHOF,SMARCB1,PRR13,STARD10,RNH1,PTOV1,FASTK,RNF32,LY75,RCC1,SNRPB,PARD3,TCAIM,DHPS,SEC
,CDK11A,ZNF197,ATP5J,TRMT112,MCTS1,DDOST,LSR,TOMM40,TADA3,ZNF687,ANAPC11,MDH2,SEC31A
CTTN,TSPAN11,ITSN1,COMMD5,DBNDD1,NOP16,TTLL5,PSMA6,ACOT8,AP3B2,RCE1,RUVBL2,EBPL,TESC,
CTTN,TSPAN11,ITSN1,COMMD5,DBNDD1,NOP16,TTLL5,PSMA6,ACOT8,AP3B2,RCE1,RUVBL2,EBPL,TESC,
2C,SRP9,IQCG,UAP1,PILRB,FAM72A,YBX3,CACYBP,CIRBP,RHOF,SMARCB1,RPL38,PRR13,STARD10,RNH1,
2C,SRP9,IQCG,UAP1,PILRB,FAM72A,YBX3,CACYBP,CIRBP,RHOF,SMARCB1,RPL38,PRR13,STARD10,RNH1,
A,CMC2,YBX3,BLOC1S1,CACYBP,CIRBP,RHOF,RPL38,PRR13,STARD10,RNH1,DNASE1L1,PTOV1,FASTK,TME
A,CMC2,YBX3,BLOC1S1,CACYBP,CIRBP,RHOF,RPL38,PRR13,STARD10,RNH1,DNASE1L1,PTOV1,FASTK,TME
,MYL6B,FAM122C,SULT1A3,PILRB,FAM72A,YBX3,CACYBP,CIRBP,RHOF,RPL38,PRR13,STARD10,RNH1,DN
L2,OSGEP,TBL2,SCARB1,ITSN1,EIF5A,UFD1L,DBNDD1,NOP16,RPL13A,GSTK1,LDHA,RPS9,SLC35G1,MRPL3
L2,SNRPB,PARD3,DHPS,ACTR6,SEC62,SET,CTSC,BOP1,ARRDC1,KLC1,ZC3HAV1,SCAF11,DBI,LIMK2,VDAC3,
TN,ITSN1,EIF5A,UFD1L,THYN1,DBNDD1,NOP16,ACOT8,AP3B2,RCE1,ZNF410,RUVBL2,PCMT1,EBPL,TESC,I
DBNDD1,ACOT8,AP3B2,RCE1,RUVBL2,EBPL,TESC,IGSF8,SNRPN,SPAG7,WDR62,POLD4,UBE2L6,FIS1,PPM1
AV1,LIMK2,NOB1,NUMA1,TBL2,SCARB1,CTTN,TSPAN11,ITSN1,EIF5A,COMMD5,NUDT22,DBNDD1,NOP1
AV1,LIMK2,NOB1,NUMA1,TBL2,SCARB1,CTTN,TSPAN11,ITSN1,EIF5A,COMMD5,NUDT22,DBNDD1,NOP1
3A,PSMA3,RPS25,COX4I1,TMX2,THEM4,AUP1,RABL5,ITGB1BP1,MED15,UGGT2,EIF3M,CDC42,CARD14,FE
FAM122C,IQCG,TMEM67,SULT1A3,UAP1,PILRB,FAM72A,CMC2,YBX3,BLOC1S1,CACYBP,CIRBP,RHOF,SMA
FAM122C,IQCG,TMEM67,SULT1A3,UAP1,PILRB,FAM72A,CMC2,YBX3,BLOC1S1,CACYBP,CIRBP,RHOF,SMA
RCB1,RPL38,PRR13,RNH1,KIF4A,RBCK1,PTOV1,FASTK,TMEM135,RNF32,LY75,ATG16L2,PARD3,TCAIM,SLC
RCB1,RPL38,PRR13,RNH1,KIF4A,RBCK1,PTOV1,FASTK,TMEM135,RNF32,LY75,ATG16L2,PARD3,TCAIM,SLC
TSC,BOP1,KLC1,SCAF11,ARPC2,LIMK2,VDAC3,SIRT6,NUMA1,SUMF2,BUD31,SAE1,NPRL2,TBL2,ARHGAP6
TSC,BOP1,KLC1,SCAF11,ARPC2,LIMK2,VDAC3,SIRT6,NUMA1,SUMF2,BUD31,SAE1,NPRL2,TBL2,ARHGAP6

CYT2,FBXW9,RFX3,DGKE,RPL28,APEH,CAPN10,LYPLA2,C19ORF54,ZNF7,PIGL,PPAN,CDK11A,ZNF197,COF

TRPT1,KLC1,WDR19,GPATCH2L,SUMF2,BUD31,TBL2,ITSN1,EIF5A,TNFSF10,CHID1,NUDT22,AMZ2,UFD1L,

TRPT1,KLC1,WDR19,GPATCH2L,SUMF2,BUD31,TBL2,ITSN1,EIF5A,TNFSF10,CHID1,NUDT22,AMZ2,UFD1L,

E4DIP,MDH2,SEC31A,CNST,PAFAH1B2,NSUN5,ZFPL1,GMPR2,RIC8A,UBE2N,MFSD11,FAM32A,TMEM147

HPS,ACTR6,ZNF235,RBP5,ZC3HAV1,SCAF11,DBI,ARPC2,PMFBP1,LIMK2,NOB1,WDR19,NUMA1,SUMF2,BU

HPS,ACTR6,ZNF235,RBP5,ZC3HAV1,SCAF11,DBI,ARPC2,PMFBP1,LIMK2,NOB1,WDR19,NUMA1,SUMF2,BU

H2,SEC31A,CNST,COX17,HNRNPLL,HIGD1A,OARD1,TCP1,UBE2N,FAM32A,COA1,C12ORF57,DCTN5,NDUF

NRPB,ACTR6,SEC62,SET,CTSC,ARRDC1,KLC1,RBP5,SCAF11,DBI,ARPC2,SUMF2,BUD31,SAE1,NPRL2,CRYGS,

NRPB,ACTR6,SEC62,SET,CTSC,ARRDC1,KLC1,RBP5,SCAF11,DBI,ARPC2,SUMF2,BUD31,SAE1,NPRL2,CRYGS,

57,PILRB,FAM72A,CMC2,YBX3,BLOC1S1,CACYBP,CIRBP,RHOF,SMARCB1,PRR13,STARD10,RNH1,DNASE1L

OC1S1,CACYBP,RHOF,SMARCB1,RPL38,STARD10,RNH1,KIF4A,RBCK1,PTOV1,FASTK,LY75,ATG16L2,SNRPB

OC1S1,CACYBP,RHOF,SMARCB1,RPL38,STARD10,RNH1,KIF4A,RBCK1,PTOV1,FASTK,LY75,ATG16L2,SNRPB

BL2,SCARB1,CTTN,ITSN1,AMZ2,TAZ,DBNDD1,GSTK1,ACOT8,AP3B2,RCE1,RUVBL2,EBPL,TESC,IGSF8,SNRP

9,SIRT6,NUMA1,BUD31,OSGEP,ARHGAP6,ITSN1,EIF5A,USP38,SH2B1,GSTK1,AP3B2,RPS9,RCE1,MRPL33,E

9,SIRT6,NUMA1,BUD31,OSGEP,ARHGAP6,ITSN1,EIF5A,USP38,SH2B1,GSTK1,AP3B2,RPS9,RCE1,MRPL33,E

MA1,GPATCH2L,OSGEP,SCARB1,CTTN,TSPAN11,ITSN1,AMZ2,TAZ,SH2B1,NOP16,PSMA6,ACOT8,MRPL33,

MA1,GPATCH2L,OSGEP,SCARB1,CTTN,TSPAN11,ITSN1,AMZ2,TAZ,SH2B1,NOP16,PSMA6,ACOT8,MRPL33,

OP16,PSMA6,ACOT8,AP3B2,RCE1,EBPL,TESC,IGSF8,SPAG7,WDR62,POLD4,UBE2L6,FIS1,STIP1,PHF19,CDC

OP16,PSMA6,ACOT8,AP3B2,RCE1,EBPL,TESC,IGSF8,SPAG7,WDR62,POLD4,UBE2L6,FIS1,STIP1,PHF19,CDC

GEF5,SSU72,FAM122C,SRP9,IQCG,TMEM67,SULT1A3,UAP1,PILRB,FAM72A,CMC2,YBX3,BLOC1S1,CACYE

GEF5,SSU72,FAM122C,SRP9,IQCG,TMEM67,SULT1A3,UAP1,PILRB,FAM72A,CMC2,YBX3,BLOC1S1,CACYE

NF7,TXNDC17,PPAN,ZFAND2B,RBM3,ATP5J,TRMT112,DDOST,LSR,RRNAD1,DBNL,TOMM40,RABEP2,TAD

H,NSUN5,UBE2N,FAM32A,PEX26,OGFOD2,SSR3,ARIH2,PICK1,SLC52A2,TSPAN31,GLYCTK,ACAA1,PIGO,A

,ARHGAP6,SCARB1,CTTN,TSPAN11,EIF5A,COMMD5,CHID1,UFD1L,USP38,TAZ,THYN1,DBNDD1,SH2B1,RP

,ARHGAP6,SCARB1,CTTN,TSPAN11,EIF5A,COMMD5,CHID1,UFD1L,USP38,TAZ,THYN1,DBNDD1,SH2B1,RP

MC2,YBX3,BLOC1S1,CACYBP,CIRBP,RHOF,RPL38,PRR13,STARD10,RNH1,DNASE1L1,KIF4A,RBCK1,PTOV1,

MC2,YBX3,BLOC1S1,CACYBP,CIRBP,RHOF,RPL38,PRR13,STARD10,RNH1,DNASE1L1,KIF4A,RBCK1,PTOV1,

3,DHPS,SEC62,SET,CTSC,BOP1,TRPT1,KLC1,RBP5,ZC3HAV1,DBI,ARPC2,PMFBP1,LIMK2,NUMA1,SAE1,NPF

3,DHPS,SEC62,SET,CTSC,BOP1,TRPT1,KLC1,RBP5,ZC3HAV1,DBI,ARPC2,PMFBP1,LIMK2,NUMA1,SAE1,NPF

LS1,CACYBP,CIRBP,RHOF,PRR13,RNH1,DNASE1L1,KIF4A,RBCK1,PTOV1,FASTK,RNF32,ATG16L2,RCC1,SNR

LS1,CACYBP,CIRBP,RHOF,PRR13,RNH1,DNASE1L1,KIF4A,RBCK1,PTOV1,FASTK,RNF32,ATG16L2,RCC1,SNR

LC3A2,DHPS,ACTR6,SET,CTSC,BOP1,ARRDC1,TRPT1,KLC1,RBP5,ARPC2,NOB1,NUMA1,GPATCH2L,SUMF2,

LC3A2,DHPS,ACTR6,SET,CTSC,BOP1,ARRDC1,TRPT1,KLC1,RBP5,ARPC2,NOB1,NUMA1,GPATCH2L,SUMF2,

LC3A2,DHPS,ACTR6,SET,CTSC,BOP1,ARRDC1,TRPT1,KLC1,RBP5,ARPC2,NOB1,NUMA1,GPATCH2L,SUMF2,

LC3A2,DHPS,ACTR6,SET,CTSC,BOP1,ARRDC1,TRPT1,KLC1,RBP5,ARPC2,NOB1,NUMA1,GPATCH2L,SUMF2,

ARD3,TCAIM,SLC3A2,ARRDC1,KLC1,RBP5,LIMK2,SIRT6,NUMA1,NPRL2,TBL2,SCARB1,CTTN,TSPAN11,ITSN

ARD3,TCAIM,SLC3A2,ARRDC1,KLC1,RBP5,LIMK2,SIRT6,NUMA1,NPRL2,TBL2,SCARB1,CTTN,TSPAN11,ITSN

MC2,YBX3,CACYBP,CIRBP,RHOF,RPL38,PRR13,STARD10,RNH1,DNASE1L1,KIF4A,RBCK1,PTOV1,FASTK,TM

AIM,SLC3A2,SEC62,SET,ARRDC1,KLC1,RBP5,LIMK2,SIRT6,NUMA1,GPATCH2L,TBL2,ARHGAP6,SCARB1,CTT

AIM,SLC3A2,SEC62,SET,ARRDC1,KLC1,RBP5,LIMK2,SIRT6,NUMA1,GPATCH2L,TBL2,ARHGAP6,SCARB1,CTT

LS1,CACYBP,CIRBP,RHOF,PRR13,RNH1,DNASE1L1,KIF4A,RBCK1,PTOV1,FASTK,RNF32,ATG16L2,RCC1,SNR

N1,EIF5A,COMMD5,UFD1L,THYN1,DBNDD1,NOP16,ACOT8,AP3B2,RCE1,ZNF410,RUVBL2,PCMT1,EBPL,TE

MK2,WDR19,BUD31,SAE1,NPRL2,CTTN,TSPAN11,COMMD5,UFD1L,USP38,THYN1,RPL13A,ARHGEF39,GS

MK2,WDR19,BUD31,SAE1,NPRL2,CTTN,TSPAN11,COMMD5,UFD1L,USP38,THYN1,RPL13A,ARHGEF39,GS

135,RNF32,LY75,ATG16L2,PARD3,TCAIM,SLC3A2,DHPS,SEC62,SET,ARRDC1,TRPT1,RBP5,LIMK2,WDR19,N

135,RNF32,LY75,ATG16L2,PARD3,TCAIM,SLC3A2,DHPS,SEC62,SET,ARRDC1,TRPT1,RBP5,LIMK2,WDR19,N

1,EEF1B2,RFX3,DGKE,RPL28,APEH,CAPN10,LYPLA2,C19ORF54,ZNF7,PIGL,PPAN,CDK11A,ZNF197,COPZ1,

RMT112,CHPF,MPDU1,ERCC5,EMC4,DDOST,LSR,PSME2,NAT9,TOMM40,TADA3,ABHD17A,MDH2,CHCHD

CYBP,CIRBP,RHOF,PRR13,STARD10,RNH1,DNASE1L1,KIF4A,RBCK1,PTOV1,FASTK,ATG16L2,RCC1,SNRPB,

CYBP,CIRBP,RHOF,PRR13,STARD10,RNH1,DNASE1L1,KIF4A,RBCK1,PTOV1,FASTK,ATG16L2,RCC1,SNRPB,

HPS,ACTR6,ARRDC1,SCAF11,DBI,VDAC3,WDR19,SIRT6,NUMA1,SAE1,NPRL2,OSGEP,TBL2,ITSN1,EIF5A,CC

HPS,ACTR6,ARRDC1,SCAF11,DBI,VDAC3,WDR19,SIRT6,NUMA1,SAE1,NPRL2,OSGEP,TBL2,ITSN1,EIF5A,CC

C1S1,CACYBP,CIRBP,RHOF,PRR13,RNH1,DNASE1L1,KIF4A,RBCK1,PTOV1,FASTK,RNF32,ATG16L2,RCC1,SN

C1S1,CACYBP,CIRBP,RHOF,PRR13,RNH1,DNASE1L1,KIF4A,RBCK1,PTOV1,FASTK,RNF32,ATG16L2,RCC1,SN

D3,ACTR6,ZNF235,ARRDC1,TRPT1,ZC3HAV1,ARPC2,NOB1,SIRT6,NUMA1,GPATCH2L,BUD31,SAE1,NPRL2,

D3,ACTR6,ZNF235,ARRDC1,TRPT1,ZC3HAV1,ARPC2,NOB1,SIRT6,NUMA1,GPATCH2L,BUD31,SAE1,NPRL2,

B1,CTTN,ITSN1,TNFSF10,COMMD5,UFD1L,DBNDD1,NOP16,PSMA6,ACOT8,RCE1,RUVBL2,EBPL,TESC,IGS

PL38,STARD10,RNH1,DNASE1L1,KIF4A,FASTK,TMEM135,RNF32,LY75,ATG16L2,RCC1,SNRPB,PARD3,TCA

PL38,STARD10,RNH1,DNASE1L1,KIF4A,FASTK,TMEM135,RNF32,LY75,ATG16L2,RCC1,SNRPB,PARD3,TCA

EH,LYPLA2,C17ORF62,PPAN,CYB561,HYPK,ZNF250,ZNF197,INO80E,HSP90AB1,ATP5J,TRMT112,CHPF,ER

EH,LYPLA2,C17ORF62,PPAN,CYB561,HYPK,ZNF250,ZNF197,INO80E,HSP90AB1,ATP5J,TRMT112,CHPF,ER

STARD10,RNH1,DNASE1L1,FASTK,TMEM135,RNF32,LY75,ATG16L2,RCC1,PARD3,TCAIM,DHPS,SET,CTSC,E

STARD10,RNH1,DNASE1L1,FASTK,TMEM135,RNF32,LY75,ATG16L2,RCC1,PARD3,TCAIM,DHPS,SET,CTSC,E

TG16L2,RCC1,TCAIM,DHPS,SET,CTSC,BOP1,ARRDC1,TRPT1,SCAF11,DBI,ARPC2,VDAC3,WDR19,GPATCH2L

S1,CACYBP,CIRBP,RHOF,RPL38,RNH1,DNASE1L1,KIF4A,RBCK1,PTOV1,FASTK,RNF32,ATG16L2,RCC1,SNR

OV,PPIL2,LIG1,CDC14B,TIMM17B,CAPN10,C17ORF62,C19ORF48,PPAN,SUGT1,COPZ1,INO80E,DGUOK,TR

PSMA6,RPS9,SLC35G1,EBPL,IGSF8,SNRPN,SPAG7,WDR62,RANBP1,VRK3,ZC3H14,FIS1,PPM1K,LRRC1,PHF

PSMA6,RPS9,SLC35G1,EBPL,IGSF8,SNRPN,SPAG7,WDR62,RANBP1,VRK3,ZC3H14,FIS1,PPM1K,LRRC1,PHF

IPS,ACTR6,SET,CTSC,ARRDC1,TRPT1,ZC3HAV1,SCAF11,DBI,ARPC2,VDAC3,NUMA1,BUD31,SAE1,NPRL2,O

IPS,ACTR6,SET,CTSC,ARRDC1,TRPT1,ZC3HAV1,SCAF11,DBI,ARPC2,VDAC3,NUMA1,BUD31,SAE1,NPRL2,O

C1S1,CACYBP,CIRBP,RHOF,RNH1,DNASE1L1,KIF4A,RBCK1,PTOV1,FASTK,RNF32,ATG16L2,RCC1,SNRPB,PA

BR4,FUZ,ECSIT,PPRC1,COX4I1,TMX2,SDF4,NISCH,RPS6,GGPS1,SRSF1,AUP1,SAP30BP,FOXRED1,RABL5,KX

BL2,EBPL,TESC,SNRPN,SLC15A3,GRAMD1C,SNF8,RPS16,OBSL1,PTPN13,UBR4,PSMA1,CHCHD1,ECSIT,NLR

BL2,EBPL,TESC,SNRPN,SLC15A3,GRAMD1C,SNF8,RPS16,OBSL1,PTPN13,UBR4,PSMA1,CHCHD1,ECSIT,NLR

72A,CMC2,YBX3,BLOC1S1,CACYBP,CIRBP,RHOF,SMARCB1,RPL38,PRR13,STARD10,RNH1,RBCK1,PTOV1,F

72A,CMC2,YBX3,BLOC1S1,CACYBP,CIRBP,RHOF,SMARCB1,RPL38,PRR13,STARD10,RNH1,RBCK1,PTOV1,F

,LY75,ATG16L2,SNRPB,PARD3,SLC3A2,ACTR6,MAP3K7CL,SET,CTSC,BOP1,SCAF11,DBI,PMFBP1,LIMK2,VD

,LY75,ATG16L2,SNRPB,PARD3,SLC3A2,ACTR6,MAP3K7CL,SET,CTSC,BOP1,SCAF11,DBI,PMFBP1,LIMK2,VD

,SCAF11,DBI,ARPC2,LIMK2,VDAC3,WDR19,SIRT6,NUMA1,OSGEP,SCARB1,CTTN,COMMD5,UFD1L,RPL13A

,SCAF11,DBI,ARPC2,LIMK2,VDAC3,WDR19,SIRT6,NUMA1,OSGEP,SCARB1,CTTN,COMMD5,UFD1L,RPL13A

V1,RNF32,ATG16L2,SNRPB,TCAIM,SEC62,SET,BOP1,ARRDC1,KLC1,ZC3HAV1,SCAF11,SIRT6,NUMA1,GPAT

V1,RNF32,ATG16L2,SNRPB,TCAIM,SEC62,SET,BOP1,ARRDC1,KLC1,ZC3HAV1,SCAF11,SIRT6,NUMA1,GPAT

K1,RNF32,RCC1,SNRPB,PARD3,SLC3A2,ACTR6,SEC62,SET,CTSC,ZNF235,BOP1,ARRDC1,KLC1,RBP5,SCAF11

K1,RNF32,RCC1,SNRPB,PARD3,SLC3A2,ACTR6,SEC62,SET,CTSC,ZNF235,BOP1,ARRDC1,KLC1,RBP5,SCAF11

EM135,RNF32,LY75,ATG16L2,PARD3,TCAIM,SEC62,SET,CTSC,ARRDC1,TRPT1,KLC1,LIMK2,WDR19,SIRT6,N

EM135,RNF32,LY75,ATG16L2,PARD3,TCAIM,SEC62,SET,CTSC,ARRDC1,TRPT1,KLC1,LIMK2,WDR19,SIRT6,N

5,SSU72,MYL6B,FAM122C,SRP9,IQCG,TMEM67,SULT1A3,PILRB,FAM72A,CMC2,YBX3,BLOC1S1,CACYBP,C

5,SSU72,MYL6B,FAM122C,SRP9,IQCG,TMEM67,SULT1A3,PILRB,FAM72A,CMC2,YBX3,BLOC1S1,CACYBP,C

,WDR19,SIRT6,NUMA1,GPATCH2L,TBL2,SCARB1,CTTN,ITSN1,COMMD5,NUDT22,UFD1L,THYN1,DBNDD1

EC62,BOP1,ARRDC1,TRPT1,KLC1,ZC3HAV1,SCAF11,DBI,LIMK2,SIRT6,GPATCH2L,ARHGAP6,SCARB1,CTTN,

T,BOP1,TRPT1,ZC3HAV1,VDAC3,NOB1,SIRT6,NUMA1,GPATCH2L,SUMF2,OSGEP,CRYGS,CTTN,EIF5A,COM

F4A,RBCK1,PTOV1,FASTK,TMEM135,RNF32,LY75,ATG16L2,RCC1,SNRPB,SLC3A2,ACTR6,SEC62,SET,BOP1,

F4A,RBCK1,PTOV1,FASTK,TMEM135,RNF32,LY75,ATG16L2,RCC1,SNRPB,SLC3A2,ACTR6,SEC62,SET,BOP1,

MARCB1,RPL38,PRR13,RNH1,DNASE1L1,KIF4A,RBCK1,PTOV1,FASTK,TMEM135,RNF32,LY75,RCC1,PARD3,

MARCB1,RPL38,PRR13,RNH1,DNASE1L1,KIF4A,RBCK1,PTOV1,FASTK,TMEM135,RNF32,LY75,RCC1,PARD3,
3,TCAIM,DHPS,ACTR6,SET,ZNF235,ARRDC1,KLC1,DBI,LIMK2,NOB1,BUD31,SAE1,NPRL2,OSGEP,TBL2,EIF5
CIRBP,RHOF,PRR13,STARD10,RNH1,TMEM135,RNF32,LY75,SNRPB,PARD3,TCAIM,SLC3A2,DHPS,ACTR6,N
CIRBP,RHOF,PRR13,STARD10,RNH1,TMEM135,RNF32,LY75,SNRPB,PARD3,TCAIM,SLC3A2,DHPS,ACTR6,N
NX1,UFD1L,USP38,RPL13A,ARHGEF39,ACOT8,AP3B2,MRPL33,ZNF410,EBPL,IGSF8,STYXL1,POLD4,VRK3,L
NX1,UFD1L,USP38,RPL13A,ARHGEF39,ACOT8,AP3B2,MRPL33,ZNF410,EBPL,IGSF8,STYXL1,POLD4,VRK3,L
RC1,CDC37,CPSF3L,TSG101,FAM3A,METTL17,ATG12,RPS25,COX4I1,RPS6,RPL29,RTEL1,SRSF1,FOXRED1,
RC1,CDC37,CPSF3L,TSG101,FAM3A,METTL17,ATG12,RPS25,COX4I1,RPS6,RPL29,RTEL1,SRSF1,FOXRED1,
DK11A,ZNF197,TRMT112,CHPF,GABPB1,EMC4,DDOST,LSR,PSME2,RRNAD1,TOMM40,TADA3,ZNF687,A
VDAC3,WDR19,NUMA1,SUMF2,SAE1,NPRL2,OSGEP,ARHGAP6,SCARB1,CRYGS,EIF5A,TNFSF10,CHID1,NU
F8,SPAG7,WDR62,POLD4,UBE2L6,STIP1,PHF19,SLC39A11,FAM3A,EPB41L5,ZBTB44,NISCH,TMED4,CD99L
F8,SPAG7,WDR62,POLD4,UBE2L6,STIP1,PHF19,SLC39A11,FAM3A,EPB41L5,ZBTB44,NISCH,TMED4,CD99L
L,TESC,IGSF8,SPAG7,WDR62,POLD4,UBE2L6,STIP1,PHF19,RPS16,OBSL1,CHCHD1,SLC39A11,FAM3A,NLR
44,MAGED2,GRK6,HMGA2,PSMA1,ECSIT,PPRC1,NIP7,TTC9C,TMED4,RPS6,COASY,SRSF1,SAP30BP,YIPF3,
2,RAPGEF5,SSU72,MYL6B,FAM122C,SRP9,IQCG,TMEM67,SULT1A3,PILRB,FAM72A,CMC2,YBX3,BLOC1S1
NUMA1,NPRL2,OSGEP,TBL2,SCARB1,CRYGS,TSPAN11,TNFSF10,RUNX1,AMZ2,UFD1L,USP38,THYN1,RPL1
LRD3,NSUN5,RIC8A,UBE2N,FAM32A,C12ORF57,OGFOD2,ARIH2,PICK1,TSPAN31,GLYCTK,TRAPPC2,AXIN
2,YBX3,CIRBP,RHOF,RPL38,PRR13,STARD10,RNH1,DNASE1L1,KIF4A,RBCK1,FASTK,TMEM135,RNF32,LY7
2,YBX3,CIRBP,RHOF,RPL38,PRR13,STARD10,RNH1,DNASE1L1,KIF4A,RBCK1,FASTK,TMEM135,RNF32,LY7
4,GGPS1,SRSF1,TRNT1,SH3YL1,IP6K2,UGGT2,PRMT2,MAGT1,GUK1,CENPN,ATF7,ASF1B,ENDOV,RPL28,V
ATP5J,TRMT112,VWA2,LMBR1,DDOST,RRNAD1,RAD1,TOMM40,ZNF687,MDH2,HNRNPLL,POLR2C,B3GA
ATP5J,TRMT112,VWA2,LMBR1,DDOST,RRNAD1,RAD1,TOMM40,ZNF687,MDH2,HNRNPLL,POLR2C,B3GA
75,ATG16L2,PARD3,TCAIM,SLC3A2,DHPS,ACTR6,SEC62,SET,CTSC,BOP1,ARRDC1,TRPT1,KLC1,RBP5,ARPC
D31,ARHGAP6,SCARB1,CTTN,TSPAN11,EIF5A,COMMD5,CHID1,UFD1L,USP38,THYN1,DBNDD1,SH2B1,RPL
D31,ARHGAP6,SCARB1,CTTN,TSPAN11,EIF5A,COMMD5,CHID1,UFD1L,USP38,THYN1,DBNDD1,SH2B1,RPL
MA1,GPATCH2L,SUMF2,BUD31,NPRL2,OSGEP,TBL2,ARHGAP6,EIF5A,UFD1L,TAZ,RPL13A,GSTK1,TTLL5,LD
MA1,GPATCH2L,SUMF2,BUD31,NPRL2,OSGEP,TBL2,ARHGAP6,EIF5A,UFD1L,TAZ,RPL13A,GSTK1,TTLL5,LD
BP5,SCAF11,DBI,ARPC2,NOB1,WDR19,SIRT6,GPATCH2L,SUMF2,NPRL2,TBL2,ARHGAP6,EIF5A,COMMD5,
BP5,SCAF11,DBI,ARPC2,NOB1,WDR19,SIRT6,GPATCH2L,SUMF2,NPRL2,TBL2,ARHGAP6,EIF5A,COMMD5,
WDR62,POLD4,UBE2L6,PPM1K,STIP1,PHF19,LPCAT3,GRK6,CPSF3L,FAM3A,ZBTB44,NISCH,CD99L2,GGPS
L,SEC62,ZNF235,ARRDC1,KLC1,SCAF11,ARPC2,LIMK2,WDR19,SIRT6,NUMA1,GPATCH2L,SAE1,NPRL2,OS
L,SEC62,ZNF235,ARRDC1,KLC1,SCAF11,ARPC2,LIMK2,WDR19,SIRT6,NUMA1,GPATCH2L,SAE1,NPRL2,OS
T6,NUMA1,GPATCH2L,OSGEP,TBL2,ARHGAP6,EIF5A,AMZ2,UFD1L,TAZ,THYN1,NOP16,GSTK1,TTLL5,LDHA
T6,NUMA1,GPATCH2L,OSGEP,TBL2,ARHGAP6,EIF5A,AMZ2,UFD1L,TAZ,THYN1,NOP16,GSTK1,TTLL5,LDHA
M122C,SRP9,IQCG,TMEM67,SULT1A3,UAP1,PILRB,FAM72A,YBX3,BLOC1S1,CACYBP,CIRBP,RHOF,SMARC
M122C,SRP9,IQCG,TMEM67,SULT1A3,UAP1,PILRB,FAM72A,YBX3,BLOC1S1,CACYBP,CIRBP,RHOF,SMARC
MMD5,CHID1,UFD1L,DBNDD1,ACOT8,AP3B2,RCE1,RUVBL2,EBPL,TESC,IGSF8,SPAG7,WDR62,POLD4,PLE
H1B2,DALRD3,NSUN5,GMPR2,UBE2N,PDCD6,FAM32A,C12ORF57,OGFOD2,ARIH2,PICK1,TSPAN31,GLYCT
CTTN,EIF5A,UFD1L,THYN1,DBNDD1,NOP16,RPL13A,RPS9,MRPL33,ZNF410,RUVBL2,EBPL,IGSF8,RBPMS,W
TG16L2,SNRPB,PARD3,SLC3A2,DHPS,ACTR6,ARRDC1,KLC1,SCAF11,DBI,VDAC3,WDR19,SIRT6,NUMA1,SU
TG16L2,SNRPB,PARD3,SLC3A2,DHPS,ACTR6,ARRDC1,KLC1,SCAF11,DBI,VDAC3,WDR19,SIRT6,NUMA1,SU
33,RUVBL2,WDR62,PLEKHA5,RANBP1,FIS1,SLC15A3,STIP1,PHF19,LPCAT3,CD44,TM9SF1,SON,TET2,UQC
EKHA5,RANBP1,LZTFL1,FIS1,SLC15A3,LRRC1,PHF19,LPCAT3,CD44,TM9SF1,SON,TET2,UQCRQ,PTPN13,P
75,RCC1,PARD3,SLC3A2,DHPS,SET,CTSC,ZNF235,ARRDC1,TRPT1,KLC1,RBP5,SCAF11,LIMK2,NOB1,NUM
75,RCC1,PARD3,SLC3A2,DHPS,SET,CTSC,ZNF235,ARRDC1,TRPT1,KLC1,RBP5,SCAF11,LIMK2,NOB1,NUM
F57,METTL5,SLC35E1,OGFOD2,TSEN34,ARIH2,PICK1,TSPAN31,GLYCTK,PIGO,AXIN1,TMEM199,EPN1,HP
,CHCHD7,MEIS2,PABPC1,ZNF836,TOMM40,RABEP2,TADA3,KLK10,TRIM16,RAD23A,STXBP2,HSPH1,CNS

,CHCHD7,MEIS2,PABPC1,ZNF836,TOMM40,RABEP2,TADA3,KLK10,TRIM16,RAD23A,STXBP2,HSPH1,CNST
TOV1,FASTK,RNF32,LY75,ATG16L2,PARD3,DHPS,ACTR6,SEC62,SET,BOP1,ARRDC1,TRPT1,KLC1,SCAF11,A
TOV1,FASTK,RNF32,LY75,ATG16L2,PARD3,DHPS,ACTR6,SEC62,SET,BOP1,ARRDC1,TRPT1,KLC1,SCAF11,A
MA6,AP3B2,RPS9,RCE1,NUDT5,RBPMS,STYXL1,TUBA1C,IRF9,SON,GRK6,CDC37,TSG101,TTC21B,UBR4,PS
ROAP,C17ORF62,PRMT1,C19ORF54,CYB561,SUGT1,HYPK,CDK11A,RAB2A,TRMT112,CHPF,RPS20,DCTD,Z
TPM2,TSFM,TRMT112,DDOST,LSR,TOMM40,ZNF20,SLC33A1,ZNF687,NXN,PDE4DIP,MDH2,STXBP2,SEC3
RPB,PARD3,TCAIM,ACTR6,SEC62,SET,CTSC,ARRDC1,KLC1,RBP5,DBI,LIMK2,NOB1,NUMA1,GPATCH2L,BUD
RPB,PARD3,TCAIM,ACTR6,SEC62,SET,CTSC,ARRDC1,KLC1,RBP5,DBI,LIMK2,NOB1,NUMA1,GPATCH2L,BUD
C3HAV1,VDAC3,NOB1,WDR19,SIRT6,NUMA1,SUMF2,TBL2,CRYGS,ITSN1,EIF5A,CHID1,USP38,TAZ,DBNDD
,PAFAH1B2,DALRD3,NSUN5,UBE2N,FAM32A,C12ORF57,DCTN5,OXLD1,SLC35E1,OGFOD2,ARIH2,PICK1,T
L,SET,BOP1,SCAF11,DBI,PMFBP1,LIMK2,VDAC3,WDR19,SIRT6,NUMA1,GPATCH2L,OSGEP,ARHGAP6,SCA
L,SET,BOP1,SCAF11,DBI,PMFBP1,LIMK2,VDAC3,WDR19,SIRT6,NUMA1,GPATCH2L,OSGEP,ARHGAP6,SCA
MA1,SLC39A11,FUZ,ATG12,NIP7,TTC9C,PLCB2,DBNDD2,SDF4,NISCH,RPS6,RTEL1,TRNT1,MED15,SLC39A1
MA1,SLC39A11,FUZ,ATG12,NIP7,TTC9C,PLCB2,DBNDD2,SDF4,NISCH,RPS6,RTEL1,TRNT1,MED15,SLC39A1
D3,TCAIM,ACTR6,SEC62,SET,CTSC,ARRDC1,KLC1,LIMK2,NOB1,NUMA1,BUD31,SAE1,NPRL2,OSGEP,ARH
,UFD1L,USP38,TAZ,NOP16,TTLL5,NUDT5,NNMT,SPAG7,WDR62,STYXL1,RFX5,DST,MAGED2,TET2,CDC37
TTLL5,AP3B2,RCE1,ZNF410,RUVBL2,PCMT1,EBPL,TESC,IGSF8,SPAG7,WDR62,POLD4,UBE2L6,PPM1K,STI
PTOV1,FASTK,TMEM135,RNF32,LY75,ATG16L2,SNRPB,PARD3,SLC3A2,DHPS,ACTR6,SEC62,SET,ZNF235,B
PTOV1,FASTK,TMEM135,RNF32,LY75,ATG16L2,SNRPB,PARD3,SLC3A2,DHPS,ACTR6,SEC62,SET,ZNF235,B
F,SMARCB1,RPL38,PRR13,RNH1,DNASE1L1,RBCK1,PTOV1,FASTK,TMEM135,RNF32,ATG16L2,RCC1,SNR
F,SMARCB1,RPL38,PRR13,RNH1,DNASE1L1,RBCK1,PTOV1,FASTK,TMEM135,RNF32,ATG16L2,RCC1,SNR
,RPL38,RNH1,KIF4A,PTOV1,FASTK,TMEM135,RNF32,LY75,PARD3,TCAIM,ACTR6,SEC62,SET,BOP1,ARRDC
,RPL38,RNH1,KIF4A,PTOV1,FASTK,TMEM135,RNF32,LY75,PARD3,TCAIM,ACTR6,SEC62,SET,BOP1,ARRDC
,ZBTB44,NISCH,CD99L2,PARD6G,SH3YL1,IP6K2,CLASRP,JPH2,MICAL1,UBE2I,PRMT2,EIF3M,TMEM205,P
BCK1,PTOV1,FASTK,TMEM135,RNF32,ATG16L2,SNRPB,TCAIM,SLC3A2,DHPS,MAP3K7CL,SEC62,SET,ZNF2
A,RPS9,RCE1,ZNF410,PCMT1,IGSF8,WDR62,STYXL1,PLEKHA5,RANBP1,VRK3,LZTFL1,FIS1,PPM1K,PHF19,
F,SMARCB1,RPL38,PRR13,STARD10,RNH1,DNASE1L1,KIF4A,PTOV1,RNF32,LY75,ATG16L2,RCC1,PARD3,T
F,SMARCB1,RPL38,PRR13,STARD10,RNH1,DNASE1L1,KIF4A,PTOV1,RNF32,LY75,ATG16L2,RCC1,PARD3,T
CYB561,DGUOK,TRMT112,MPDU1,UBXN1,EMC4,MCTS1,DDOST,NAT9,TOMM40,TADA3,KLK10,HSPH1,C
CYB561,DGUOK,TRMT112,MPDU1,UBXN1,EMC4,MCTS1,DDOST,NAT9,TOMM40,TADA3,KLK10,HSPH1,C
1B2,GBA2,DALRD3,NSUN5,MCFD2,RIC8A,UBE2N,FAM32A,C12ORF57,SLC35E1,OGFOD2,ARIH2,PICK1,TS
EP,SCARB1,CTTN,NUDT22,UFD1L,SH2B1,NOP16,GSTK1,LDHA,PSMA6,ACOT8,AP3B2,RCE1,MRPL33,IGSF8
EP,SCARB1,CTTN,NUDT22,UFD1L,SH2B1,NOP16,GSTK1,LDHA,PSMA6,ACOT8,AP3B2,RCE1,MRPL33,IGSF8
RFIP1,RMDN1,TWSG1,LRRC27,RANGRF,CHCHD7,PSME2,PABPC1,BCAM,ZNF836,BLVRB,TOMM40,RABEP
BOP1,ARRDC1,SCAF11,ARPC2,LIMK2,VDAC3,NOB1,WDR19,SIRT6,GPATCH2L,SUMF2,BUD31,SAE1,ARHG
BOP1,ARRDC1,SCAF11,ARPC2,LIMK2,VDAC3,NOB1,WDR19,SIRT6,GPATCH2L,SUMF2,BUD31,SAE1,ARHG
1,SCARB1,TSPAN11,COMMD5,NUDT22,AMZ2,USP38,THYN1,NOP16,RPL13A,ARHGEF39,GSTK1,TTLL5,L
1,SCARB1,TSPAN11,COMMD5,NUDT22,AMZ2,USP38,THYN1,NOP16,RPL13A,ARHGEF39,GSTK1,TTLL5,L
D3,DHPS,ACTR6,SET,BOP1,ARRDC1,TRPT1,KLC1,SCAF11,DBI,ARPC2,LIMK2,WDR19,NUMA1,BUD31,NPRL
T,UBE2N,FAM32A,TNKS,DCTN5,PEX26,OXLD1,ARIH2,PICK1,FBXL18,ZNF512,TSPAN31,ZNF593,GLYCTK,EI
P1,LIMK2,VDAC3,NUMA1,GPATCH2L,OSGEP,ARHGAP6,SCARB1,CTTN,EIF5A,UFD1L,DBNDD1,GSTK1,PSM
P1,LIMK2,VDAC3,NUMA1,GPATCH2L,OSGEP,ARHGAP6,SCARB1,CTTN,EIF5A,UFD1L,DBNDD1,GSTK1,PSM
32,LY75,SNRPB,TCAIM,DHPS,SEC62,SET,CTSC,BOP1,ARRDC1,KLC1,SCAF11,WDR19,NUMA1,SUMF2,BUD
32,LY75,SNRPB,TCAIM,DHPS,SEC62,SET,CTSC,BOP1,ARRDC1,KLC1,SCAF11,WDR19,NUMA1,SUMF2,BUD
MA1,GPATCH2L,BUD31,TBL2,ARHGAP6,CTTN,EIF5A,COMMD5,CHID1,UFD1L,USP38,THYN1,DBNDD1,PS
MA1,GPATCH2L,BUD31,TBL2,ARHGAP6,CTTN,EIF5A,COMMD5,CHID1,UFD1L,USP38,THYN1,DBNDD1,PS
3,SEC62,SET,BOP1,TRPT1,SCAF11,DBI,VDAC3,NOB1,SIRT6,SAE1,OSGEP,TBL2,SCARB1,CTTN,TSPAN11,IT

3,SEC62,SET,BOP1,TRPT1,SCAF11,DBI,VDAC3,NOB1,SIRT6,SAE1,OSGEP,TBL2,SCARB1,CTTN,TSPAN11,IT
,C12ORF57,METTL5,SLC35E1,OGFOD2,ARIH2,PICK1,TSPAN31,GLYCTK,PIGO,AXIN1,TMEM199,EPN1,HPS
IOB1,NUMA1,BUD31,SAE1,OSGEP,TBL2,ARHGAP6,SCARB1,TSPAN11,ITSN1,COMMD5,USP38,THYN1,DB
IOB1,NUMA1,BUD31,SAE1,OSGEP,TBL2,ARHGAP6,SCARB1,TSPAN11,ITSN1,COMMD5,USP38,THYN1,DB
RCC1,SNRPB,SLC3A2,ACTR6,SET,CTSC,ZNF235,ARRDC1,TRPT1,KLC1,RBP5,ZC3HAV1,SCAF11,ARPC2,LIMK
RCC1,SNRPB,SLC3A2,ACTR6,SET,CTSC,ZNF235,ARRDC1,TRPT1,KLC1,RBP5,ZC3HAV1,SCAF11,ARPC2,LIMK
SET,BOP1,ARRDC1,KLC1,SCAF11,DBI,LIMK2,VDAC3,SIRT6,NUMA1,BUD31,SAE1,OSGEP,TBL2,TSPAN11,IT
C62,SET,BOP1,ARRDC1,TRPT1,KLC1,RBP5,SCAF11,ARPC2,LIMK2,VDAC3,NUMA1,TBL2,SCARB1,CTTN,TSP
IARCB1,RPL38,PRR13,STARD10,RNH1,KIF4A,RBCK1,PTOV1,FASTK,RNF32,LY75,ATG16L2,RCC1,SNRPB,SLC
IARCB1,RPL38,PRR13,STARD10,RNH1,KIF4A,RBCK1,PTOV1,FASTK,RNF32,LY75,ATG16L2,RCC1,SNRPB,SLC
PATCH2L,SUMF2,BUD31,NPRL2,OSGEP,TBL2,ARHGAP6,EIF5A,TAZ,RPL13A,GSTK1,TTLL5,LDHA,RPS9,RCE
PATCH2L,SUMF2,BUD31,NPRL2,OSGEP,TBL2,ARHGAP6,EIF5A,TAZ,RPL13A,GSTK1,TTLL5,LDHA,RPS9,RCE
,DCTN5,OGFOD2,ARIH2,PICK1,TSPAN31,GLYCTK,TP53,TMEM199,EPN1,ZCCHC9,AZI2,TBC1D22A,SZT2,M
L,DBI,ARPC2,LIMK2,VDAC3,WDR19,SIRT6,SAE1,NPRL2,OSGEP,ARHGAP6,TSPAN11,ITSN1,COMMD5,USP3
,TESC,IGSF8,SPAG7,WDR62,POLD4,RANBP1,UBE2L6,STIP1,PHF19,SON,OBSL1,FAM3A,MUC1,EPB41L5,P
1,FIS1,STIP1,LRRC1,PHF19,LPCAT3,CD44,TM9SF1,SON,TET2,UQCRQ,CDC37,PTPN13,YLPM1,FUZ,FAM3A,
2,SET,CTSC,BOP1,TRPT1,RBP5,ARPC2,PMFBP1,LIMK2,VDAC3,NUMA1,GPATCH2L,SUMF2,BUD31,TBL2,A
3X3,CACYBP,CIRBP,RHOF,SMARCB1,RPL38,PRR13,STARD10,RNH1,DNASE1L1,KIF4A,RBCK1,PTOV1,FASTK,
3X3,CACYBP,CIRBP,RHOF,SMARCB1,RPL38,PRR13,STARD10,RNH1,DNASE1L1,KIF4A,RBCK1,PTOV1,FASTK,
NF197,ATP5J,TRMT112,EMC4,DDOST,TOMM40,ANAPC11,MDH2,NAA10,CARHSP1,COX17,POLR2C,PFDN
NF197,ATP5J,TRMT112,EMC4,DDOST,TOMM40,ANAPC11,MDH2,NAA10,CARHSP1,COX17,POLR2C,PFDN
13,STARD10,DNASE1L1,RBCK1,FASTK,LY75,RCC1,SNRPB,PARD3,TCAIM,SLC3A2,DHPS,ACTR6,SEC62,CTSC,
13,STARD10,DNASE1L1,RBCK1,FASTK,LY75,RCC1,SNRPB,PARD3,TCAIM,SLC3A2,DHPS,ACTR6,SEC62,CTSC,
RBCK1,PTOV1,FASTK,LY75,ATG16L2,SNRPB,PARD3,DHPS,SET,BOP1,ARRDC1,TRPT1,KLC1,SCAF11,DBI,VD
G7,WDR62,POLD4,UBE2L6,STIP1,PHF19,MAGED2,SON,GRK6,PHF21A,MUC1,ATG12,PPRC1,ZBTB44,DBN
T2,PPA1,ENDOV,CDK11A,INO80E,FAM219B,ZNF524,ELP6,SIAH1,TRMT112,MPDU1,RPS15,LRRC27,MVD,
T2,PPA1,ENDOV,CDK11A,INO80E,FAM219B,ZNF524,ELP6,SIAH1,TRMT112,MPDU1,RPS15,LRRC27,MVD,
SC,IGSF8,SPAG7,WDR62,POLD4,UBE2L6,PPM1K,STIP1,PHF19,MAGED2,SON,GRK6,CDC37,RPS16,CPSF3L
1,CCDC25,ENDOV,PPIL2,RPL28,LYPLA2,C17ORF62,ZNF7,ZFAND2B,RBM3,DGUOK,RAB2A,TRMT112,UBXN
,SUMF2,BUD31,SAE1,NPRL2,TBL2,ARHGAP6,SCARB1,ITSN1,COMMD5,NUDT22,AMZ2,USP38,DBNDD1,S
1,SAE1,TBL2,SCARB1,CTTN,TSPAN11,ITSN1,EIF5A,COMMD5,NUDT22,DBNDD1,SH2B1,NOP16,TTLL5,AC
1,SAE1,TBL2,SCARB1,CTTN,TSPAN11,ITSN1,EIF5A,COMMD5,NUDT22,DBNDD1,SH2B1,NOP16,TTLL5,AC
1S1,CACYBP,CIRBP,RHOF,SMARCB1,RPL38,PRR13,STARD10,RNH1,DNASE1L1,RBCK1,PTOV1,FASTK,TME
1S1,CACYBP,CIRBP,RHOF,SMARCB1,RPL38,PRR13,STARD10,RNH1,DNASE1L1,RBCK1,PTOV1,FASTK,TME
,CPSF3L,DUSP22,SLC25A10,TMED4,YIPF3,ITGB1BP1,SH3YL1,YIF1A,PRMT2,PTK2,CENPN,PPA1,ATPAF2,Z
,CPSF3L,DUSP22,SLC25A10,TMED4,YIPF3,ITGB1BP1,SH3YL1,YIF1A,PRMT2,PTK2,CENPN,PPA1,ATPAF2,Z
IPF,DDOST,LSR,DOCK10,TOMM40,TADA3,RAD23A,ABHD17A,MDH2,SEC31A,POLR2C,PAFAH1B2,NSUN5,
IPF,DDOST,LSR,DOCK10,TOMM40,TADA3,RAD23A,ABHD17A,MDH2,SEC31A,POLR2C,PAFAH1B2,NSUN5,
TRAPPC2,CLTA,SLC35A2,SLC25A24,HPS1,ITPA,DNAJA3,SNRPA1,AZI2,TATDN1,TBC1D22A,ZNF581,ZNF48
TOMM40,ZNF687,MDH2,POLR2C,B3GAT3,PAFAH1B2,TXNRD2,NSUN5,THADA,ZNF81,FAM32A,C12ORF57
TOMM40,ZNF687,MDH2,POLR2C,B3GAT3,PAFAH1B2,TXNRD2,NSUN5,THADA,ZNF81,FAM32A,C12ORF57
2,BUD31,SAE1,OSGEP,TBL2,SCARB1,CTTN,EIF5A,COMMD5,CHID1,NUDT22,AMZ2,DBNDD1,SH2B1,LDHA,
2,BUD31,SAE1,OSGEP,TBL2,SCARB1,CTTN,EIF5A,COMMD5,CHID1,NUDT22,AMZ2,DBNDD1,SH2B1,LDHA,
1,CTTN,ITSN1,EIF5A,COMMD5,CHID1,AMZ2,LDHA,SLC35G1,MRPL33,ZNF410,TESC,NUDT5,SPAG7,WDR6
1,CTTN,ITSN1,EIF5A,COMMD5,CHID1,AMZ2,LDHA,SLC35G1,MRPL33,ZNF410,TESC,NUDT5,SPAG7,WDR6
IBP1,ZC3H14,PPM1K,STIP1,GRAMD1C,LPCAT3,SON,RPS16,OBSL1,FAM3A,ECSIT,ATG12,COX4I1,DUSP22,
IBP1,ZC3H14,PPM1K,STIP1,GRAMD1C,LPCAT3,SON,RPS16,OBSL1,FAM3A,ECSIT,ATG12,COX4I1,DUSP22,

PB,TCAIM,DHPS,ACTR6,MAP3K7CL,RBP5,ZC3HAV1,DBI,ARPC2,PMFBP1,LIMK2,WDR19,SIRT6,GPATCH2L,

PB,TCAIM,DHPS,ACTR6,MAP3K7CL,RBP5,ZC3HAV1,DBI,ARPC2,PMFBP1,LIMK2,WDR19,SIRT6,GPATCH2L,

F4A,RBCK1,PTOV1,FASTK,RNF32,ATG16L2,RCC1,SNRPB,PARD3,ACTR6,SEC62,SET,CTSC,BOP1,ARRDC1,KL

F4A,RBCK1,PTOV1,FASTK,RNF32,ATG16L2,RCC1,SNRPB,PARD3,ACTR6,SEC62,SET,CTSC,BOP1,ARRDC1,KL

D15,SH3YL1,CLASRP,UGGT2,MICAL1,SLC39A13,PRMT2,TBX6,FBXW9,ASF1B,CCDC25,DGKE,ENDOV,LIG1,C

P22,RPL29,TRNT1,MED15,IP6K2,C1ORF198,EIF3M,CDC42,TMEM205,USE1,IFI27L1,ASF1B,CCDC25,EEF1B

D5,CHID1,AMZ2,UFD1L,TAZ,THYN1,SH2B1,NOP16,RPL13A,LDHA,SLC35G1,MRPL33,RUVBL2,IGSF8,WDR6

RB1,CTTN,TSPAN11,EIF5A,UFD1L,USP38,DBNDD1,PSMA6,RUVBL2,TESC,IGSF8,NNMT,PLEKHA5,RANBP1,

RB1,CTTN,TSPAN11,EIF5A,UFD1L,USP38,DBNDD1,PSMA6,RUVBL2,TESC,IGSF8,NNMT,PLEKHA5,RANBP1,

P,RHOF,SMARCB1,RPL38,PRR13,STARD10,DNASE1L1,KIF4A,PTOV1,RNF32,ATG16L2,RCC1,SNRPB,TCAIM

P,RHOF,SMARCB1,RPL38,PRR13,STARD10,DNASE1L1,KIF4A,PTOV1,RNF32,ATG16L2,RCC1,SNRPB,TCAIM

NBP1,ZC3H14,FIS1,PPM1K,STIP1,LRRC1,LPCAT3,TM9SF1,SON,TET2,CDC37,RPS16,OBSL1,TTC21B,UBR4,F

R62,POLD4,UBE2L6,STIP1,PHF19,GRK6,RPS16,SLC39A11,MUC1,ATG12,ZBTB44,NISCH,CD99L2,GGPS1,AU

R62,POLD4,UBE2L6,STIP1,PHF19,GRK6,RPS16,SLC39A11,MUC1,ATG12,ZBTB44,NISCH,CD99L2,GGPS1,AU

JD31,TBL2,ARHGAP6,EIF5A,COMMD5,CHID1,UFD1L,THYN1,DBNDD1,PSMA6,AP3B2,SLC35G1,ZNF410,PC

CRYGS,CTTN,COMMD5,CHID1,NOP16,RPL13A,LDHA,PSMA6,RPS9,MRPL33,ZNF410,EBPL,TESC,NNMT,SN

CRYGS,CTTN,COMMD5,CHID1,NOP16,RPL13A,LDHA,PSMA6,RPS9,MRPL33,ZNF410,EBPL,TESC,NNMT,SN

,ACOT8,AP3B2,RCE1,ZNF410,EBPL,TESC,IGSF8,SPAG7,WDR62,POLD4,UBE2L6,PPM1K,STIP1,PHF19,LPCA

EMC4,DDOST,TOMM40,ANAPC11,MDH2,NAA10,CARHSP1,COX17,POLR2C,PAFAH1B2,NSUN5,FAM32A,

EMC4,DDOST,TOMM40,ANAPC11,MDH2,NAA10,CARHSP1,COX17,POLR2C,PAFAH1B2,NSUN5,FAM32A,

RB1,CTTN,TSPAN11,ITSN1,COMMD5,AMZ2,TAZ,SH2B1,NOP16,TTLL5,PSMA6,MRPL33,RUVBL2,EBPL,TESC

RB1,CTTN,TSPAN11,ITSN1,COMMD5,AMZ2,TAZ,SH2B1,NOP16,TTLL5,PSMA6,MRPL33,RUVBL2,EBPL,TESC

M135,RNF32,LY75,PARD3,TCAIM,SLC3A2,DHPS,ACTR6,SEC62,SET,CTSC,BOP1,TRPT1,RBP5,ZC3HAV1,PMF

M135,RNF32,LY75,PARD3,TCAIM,SLC3A2,DHPS,ACTR6,SEC62,SET,CTSC,BOP1,TRPT1,RBP5,ZC3HAV1,PMF

L2,SLC3A2,DHPS,ACTR6,SEC62,SET,BOP1,TRPT1,KLC1,ZC3HAV1,ARPC2,PMFBP1,LIMK2,NOB1,SIRT6,GPA

L2,SLC3A2,DHPS,ACTR6,SEC62,SET,BOP1,TRPT1,KLC1,ZC3HAV1,ARPC2,PMFBP1,LIMK2,NOB1,SIRT6,GPA

K1,RNF32,LY75,ATG16L2,RCC1,SNRPB,PARD3,TCAIM,SLC3A2,DHPS,ACTR6,SEC62,CTSC,BOP1,ARRDC1,TF

K1,RNF32,LY75,ATG16L2,RCC1,SNRPB,PARD3,TCAIM,SLC3A2,DHPS,ACTR6,SEC62,CTSC,BOP1,ARRDC1,TF

AC3,NUMA1,GPATCH2L,OSGEP,ARHGAP6,SCARB1,CTTN,TSPAN11,ITSN1,EIF5A,NUDT22,UFD1L,DBNDD1,

AC3,NUMA1,GPATCH2L,OSGEP,ARHGAP6,SCARB1,CTTN,TSPAN11,ITSN1,EIF5A,NUDT22,UFD1L,DBNDD1,

CB1,RPL38,STARD10,DNASE1L1,KIF4A,FASTK,TMEM135,RNF32,RCC1,SNRPB,TCAIM,ACTR6,SET,CTSC,ZNF

CB1,RPL38,STARD10,DNASE1L1,KIF4A,FASTK,TMEM135,RNF32,RCC1,SNRPB,TCAIM,ACTR6,SET,CTSC,ZNF

GB1BP1,SH3YL1,PRMT2,MAGT1,CENPN,PPA1,ENDOV,CDK11A,INO80E,FAM219B,ZNF524,ELP6,SIAH1,TRN

GB1BP1,SH3YL1,PRMT2,MAGT1,CENPN,PPA1,ENDOV,CDK11A,INO80E,FAM219B,ZNF524,ELP6,SIAH1,TRN

D31,SAE1,NPRL2,OSGEP,TBL2,EIF5A,UFD1L,DBNDD1,SH2B1,NOP16,TTLL5,ACOT8,SLC35G1,MRPL33,ZNF

,COX4I1,DUSP22,TMED4,CENPK,GGPS1,RTEL1,SRSF1,AUP1,FOXRED1,SH3YL1,UGGT2,YIF1A,TBX6,AUNIP,

RHGAP6,SCARB1,ITSN1,COMMD5,UFD1L,USP38,SH2B1,NOP16,AP3B2,RUVBL2,TESC,RBPMS,WDR62,PLE

M135,RNF32,LY75,ATG16L2,PARD3,TCAIM,DHPS,ACTR6,SET,BOP1,ARRDC1,TRPT1,KLC1,SCAF11,DBI,ARF

M135,RNF32,LY75,ATG16L2,PARD3,TCAIM,DHPS,ACTR6,SET,BOP1,ARRDC1,TRPT1,KLC1,SCAF11,DBI,ARF

CTTN,TSPAN11,ITSN1,COMMD5,AMZ2,TAZ,DBNDD1,ACOT8,AP3B2,RCE1,EBPL,TESC,IGSF8,RBPMS,SPAG

P1,ARRDC1,KLC1,ZC3HAV1,SCAF11,WDR19,NUMA1,GPATCH2L,NPRL2,OSGEP,TBL2,ARHGAP6,EIF5A,AM

P1,ARRDC1,KLC1,ZC3HAV1,SCAF11,WDR19,NUMA1,GPATCH2L,NPRL2,OSGEP,TBL2,ARHGAP6,EIF5A,AM

F32,LY75,ATG16L2,RCC1,PARD3,DHPS,MAP3K7CL,SEC62,ZNF235,ARRDC1,TRPT1,KLC1,ARPC2,LIMK2,VD

F32,LY75,ATG16L2,RCC1,PARD3,DHPS,MAP3K7CL,SEC62,ZNF235,ARRDC1,TRPT1,KLC1,ARPC2,LIMK2,VD

5,POLD4,UBE2L6,STIP1,PHF19,GRK6,RPS16,CPSF3L,SLC39A11,ZBTB44,NISCH,CD99L2,AUP1,PARD6G,SH3

SEC31A,COX17,B3GAT3,PAFAH1B2,NSUN5,GMPR2,UBE2N,PDCD6,FAM32A,TNKS,PEX26,OGFOD2,ARIH2

1,DBI,WDR19,GPATCH2L,SUMF2,SAE1,OSGEP,TBL2,SCARB1,CTTN,TSPAN11,EIF5A,COMMD5,CHID1,AMZ

CYBP,RHOF,SMARCB1,PRR13,STARD10,RNH1,DNASE1L1,KIF4A,RBCK1,PTOV1,RNF32,ATG16L2,RCC1,TCA
CYBP,RHOF,SMARCB1,PRR13,STARD10,RNH1,DNASE1L1,KIF4A,RBCK1,PTOV1,RNF32,ATG16L2,RCC1,TCA
AG7,WDR62,RFX5,POLD4,UBE2L6,PHF19,LPCAT3,GRK6,CPSF3L,FAM3A,METTL17,EPB41L5,ATG12,ZBTB4
L,TRMT112,DDOST,PABPC1,RRNAD1,TOMM40,TADA3,ZNF687,MDH2,POLR2C,B3GAT3,PAFAH1B2,TXNR
L,TRMT112,DDOST,PABPC1,RRNAD1,TOMM40,TADA3,ZNF687,MDH2,POLR2C,B3GAT3,PAFAH1B2,TXNR

2,SUMF2,BUD31,SAE1,NPRL2,OSGEP,SCARB1,CTTN,ITSN1,EIF5A,TNFSF10,COMMD5,USP38,TAZ,THYN1,D
,TMX2,SDF4,RPL29,CD99L2,SAP30BP,PARD6G,CLASRP,JPH2,C1ORF198,UGGT2,MICAL1,PRMT2,EIF3M,JL
,TMX2,SDF4,RPL29,CD99L2,SAP30BP,PARD6G,CLASRP,JPH2,C1ORF198,UGGT2,MICAL1,PRMT2,EIF3M,JL
17,NIP7,TTC9C,COX4I1,DBNDD2,COASY,SRSF1,AUP1,FOXRED1,TRNT1,MAGT1,GUK1,CENPN,AGPAT4,IFI2
N,SPAG7,STIP1,PHF19,OBSL1,UBR4,SLC39A11,FAM3A,EPB41L5,NLRP1,ZBTB44,SLC25A10,DBNDD2,NISCH
N,SPAG7,STIP1,PHF19,OBSL1,UBR4,SLC39A11,FAM3A,EPB41L5,NLRP1,ZBTB44,SLC25A10,DBNDD2,NISCH
,PMFBP1,WDR19,NUMA1,GPATCH2L,BUD31,SAE1,OSGEP,TBL2,ARHGAP6,SCARB1,CHID1,UFD1L,SH2B1,
,PMFBP1,WDR19,NUMA1,GPATCH2L,BUD31,SAE1,OSGEP,TBL2,ARHGAP6,SCARB1,CHID1,UFD1L,SH2B1,
B1,RPL38,PRR13,STARD10,RNH1,DNASE1L1,KIF4A,RBCK1,PTOV1,FASTK,TMEM135,RNF32,ATG16L2,SNRI
B1,RPL38,PRR13,STARD10,RNH1,DNASE1L1,KIF4A,RBCK1,PTOV1,FASTK,TMEM135,RNF32,ATG16L2,SNRI
TK,RNF32,ATG16L2,RCC1,SNRPB,PARD3,ACTR6,SEC62,SET,CTSC,BOP1,ARRDC1,KLC1,RBP5,DBI,ARPC2,LIN
SH3YL1,IP6K2,CLASRP,MICAL1,UBE2I,PRMT2,PTK2,AUNIP,C14ORF2,GUK1,PCYT2,PPP1CC,PPA1,DGKE,CA
CTTN,ITSN1,EIF5A,RUNX1,NUDT22,UFD1L,SH2B1,NOP16,TTLL5,LDHA,MRPL33,RUVBL2,NUDT5,WDR62,V
3,TM9SF1,SON,TET2,TTC21B,PPRC1,RPS25,NIP7,TTC9C,COX4I1,SPOCK2,TMED4,CENPK,RTEL1,AUP1,SAP
PIL2,TXNL4A,CAPN10,EXOSC9,C19ORF54,ZFAND2B,RPL36,CDK11A,MAPKAPK5,TMEM134,GPBP1,TRMT1
ACYBP,CIRBP,RHOF,SMARCB1,RPL38,PRR13,DNASE1L1,KIF4A,RBCK1,PTOV1,FASTK,RNF32,LY75,ATG16L2
ACYBP,CIRBP,RHOF,SMARCB1,RPL38,PRR13,DNASE1L1,KIF4A,RBCK1,PTOV1,FASTK,RNF32,LY75,ATG16L2
M,SEC62,SET,BOP1,ARRDC1,SCAF11,ARPC2,LIMK2,VDAC3,SIRT6,NUMA1,SUMF2,OSGEP,TBL2,ARHGAP6
L2,ARHGAP6,EIF5A,AMZ2,TAZ,NOP16,RPL13A,TTLL5,LDHA,RPS9,RCE1,ZNF410,IGSF8,NNMT,WDR62,STYX
L2,ARHGAP6,EIF5A,AMZ2,TAZ,NOP16,RPL13A,TTLL5,LDHA,RPS9,RCE1,ZNF410,IGSF8,NNMT,WDR62,STYX
,VDAC3,NUMA1,GPATCH2L,BUD31,NPRL2,OSGEP,ARHGAP6,TSPAN11,EIF5A,CHID1,UFD1L,TAZ,THYN1,RI
,VDAC3,NUMA1,GPATCH2L,BUD31,NPRL2,OSGEP,ARHGAP6,TSPAN11,EIF5A,CHID1,UFD1L,TAZ,THYN1,RI
T1,TESC,SPAG7,WDR62,POLD4,PLEKHA5,RANBP1,LZTFL1,CD44,SON,CDC37,TSG101,PTPN13,PSMA1,YLP
T1,TESC,SPAG7,WDR62,POLD4,PLEKHA5,RANBP1,LZTFL1,CD44,SON,CDC37,TSG101,PTPN13,PSMA1,YLP
SP38,PSMA6,AP3B2,RPS9,SLC35G1,RUVBL2,PCMT1,SPAG7,WDR62,STYXL1,RANBP1,FIS1,PPM1K,TUBA1
,STIP1,PHF19,CD44,GRK6,CPSF3L,FAM3A,EPB41L5,ATG12,ZBTB44,NISCH,CD99L2,GGPS1,FOXRED1,PARD
F8,RBPMS,SPAG7,ZC3H14,FIS1,TUBA1C,STIP1,PHF19,TET2,GRK6,OBSL1,ECSIT,ATG12,NLRP1,COX4I1,PLC
F8,RBPMS,SPAG7,ZC3H14,FIS1,TUBA1C,STIP1,PHF19,TET2,GRK6,OBSL1,ECSIT,ATG12,NLRP1,COX4I1,PLC
G7,WDR62,POLD4,RANBP1,UBE2L6,PPM1K,TUBA1C,STIP1,PHF19,LPCAT3,CD44,CPSF3L,OBSL1,YLPM1,SL
RL2,ARHGAP6,EIF5A,USP38,THYN1,DBNDD1,SH2B1,NOP16,RPL13A,ARHGEF39,LDHA,PSMA6,RPS9,IGSF8,
RL2,ARHGAP6,EIF5A,USP38,THYN1,DBNDD1,SH2B1,NOP16,RPL13A,ARHGEF39,LDHA,PSMA6,RPS9,IGSF8,
RNH1,DNASE1L1,RBCK1,PTOV1,FASTK,RNF32,ATG16L2,SNRPB,TCAIM,SLC3A2,ACTR6,SEC62,SET,CTSC,ZN
RNH1,DNASE1L1,RBCK1,PTOV1,FASTK,RNF32,ATG16L2,SNRPB,TCAIM,SLC3A2,ACTR6,SEC62,SET,CTSC,ZN
CB1,PRR13,STARD10,RNH1,DNASE1L1,PTOV1,RNF32,LY75,ATG16L2,RCC1,PARD3,TCAIM,DHPS,MAP3K7
CB1,PRR13,STARD10,RNH1,DNASE1L1,PTOV1,RNF32,LY75,ATG16L2,RCC1,PARD3,TCAIM,DHPS,MAP3K7
N10,RBM33,NUTF2,ATPAF2,PRMT1,WNK1,STAU1,CARS,RPL36,MDH1,INO80E,ELP6,PABPC1,ATP6V1F,TR
N10,RBM33,NUTF2,ATPAF2,PRMT1,WNK1,STAU1,CARS,RPL36,MDH1,INO80E,ELP6,PABPC1,ATP6V1F,TR
ATCH2L,BUD31,SAE1,NPRL2,OSGEP,UFD1L,USP38,SH2B1,NOP16,ACOT8,RPS9,MRPL33,ZNF410,RUVBL2,N
,PSMA3,METTL17,PPRC1,RPS25,NIP7,TTC9C,COX4I1,DBNDD2,CENPK,RTEL1,COASY,SRSF1,AUP1,FOXRED
F,SMARCB1,PRR13,RNH1,DNASE1L1,KIF4A,RBCK1,PTOV1,FASTK,TMEM135,RNF32,LY75,ATG16L2,SNRPB
F,SMARCB1,PRR13,RNH1,DNASE1L1,KIF4A,RBCK1,PTOV1,FASTK,TMEM135,RNF32,LY75,ATG16L2,SNRPB

,SNRPN,SPAG7,PPM1K,ANTXR2,LPCAT3,MAGED2,SON,GRK6,CPSF3L,OBSL1,UBR4,FAM3A,NLRP1,PLCB2,S
,SNRPN,SPAG7,PPM1K,ANTXR2,LPCAT3,MAGED2,SON,GRK6,CPSF3L,OBSL1,UBR4,FAM3A,NLRP1,PLCB2,S

L,DBNDD1,MRPL33,RUVBL2,TESC,RANBP1,VRK3,LZTFL1,FIS1,PPM1K,LRRC1,PHF19,LPCAT3,TM9SF1,SON
9B,ZNF524,ATP5J,ELP6,TRMT112,RPS15,LRRC27,MVD,PABPC1,NAT9,PDCD5,ANAPC11,DDX41,CNST,CASZ
9B,ZNF524,ATP5J,ELP6,TRMT112,RPS15,LRRC27,MVD,PABPC1,NAT9,PDCD5,ANAPC11,DDX41,CNST,CASZ
L1,KIF4A,TMEM135,RNF32,LY75,RCC1,PARD3,TCAIM,SLC3A2,DHPS,ACTR6,SET,CTSC,BOP1,ARRDC1,KLC1
L1,KIF4A,TMEM135,RNF32,LY75,RCC1,PARD3,TCAIM,SLC3A2,DHPS,ACTR6,SET,CTSC,BOP1,ARRDC1,KLC1
IED4,GGPS1,SRSF1,ITGB1BP1,SH3YL1,YIF1A,PRMT2,MAGT1,PTK2,CENPN,PPA1,ENDOV,ATPAF2,ZNF644,C
IED4,GGPS1,SRSF1,ITGB1BP1,SH3YL1,YIF1A,PRMT2,MAGT1,PTK2,CENPN,PPA1,ENDOV,ATPAF2,ZNF644,C
BUD31,SAE1,CRYGS,CTTN,EIF5A,UFD1L,USP38,ARHGEF39,ACOT8,ZNF410,EBPL,STYXL1,VRK3,LZTFL1,FIS1
BUD31,SAE1,CRYGS,CTTN,EIF5A,UFD1L,USP38,ARHGEF39,ACOT8,ZNF410,EBPL,STYXL1,VRK3,LZTFL1,FIS1
CC5,LMBR1,PPP3CB,CHCHD5,CNST,SH3GLB1,HNRNPA1,INTS4,SPINT2,ARHGEF1,DRG2,MRPS18C,GNB5,U
CC5,LMBR1,PPP3CB,CHCHD5,CNST,SH3GLB1,HNRNPA1,INTS4,SPINT2,ARHGEF1,DRG2,MRPS18C,GNB5,U
5A,AMZ2,UFD1L,TAZ,THYN1,GSTK1,TTLL5,LDHA,RPS9,RCE1,ZNF410,EBPL,IGSF8,WDR62,RFX5,RANBP1,VR
5A,AMZ2,UFD1L,TAZ,THYN1,GSTK1,TTLL5,LDHA,RPS9,RCE1,ZNF410,EBPL,IGSF8,WDR62,RFX5,RANBP1,VR
31,NPRL2,OSGEP,ARHGAP6,EIF5A,AMZ2,TAZ,NOP16,RPL13A,GSTK1,TTLL5,LDHA,RPS9,RCE1,ZNF410,PCN
31,NPRL2,OSGEP,ARHGAP6,EIF5A,AMZ2,TAZ,NOP16,RPL13A,GSTK1,TTLL5,LDHA,RPS9,RCE1,ZNF410,PCN
L,PRMT2,JUP,TBX6,C14ORF2,IFI27L1,CCDC25,EEF1B2,DGKE,CRIP1,PSMD4,RPL28,CLDND1,VIM,CDC14B,T
L,PRMT2,JUP,TBX6,C14ORF2,IFI27L1,CCDC25,EEF1B2,DGKE,CRIP1,PSMD4,RPL28,CLDND1,VIM,CDC14B,T
P5,ZC3HAV1,LIMK2,VDAC3,SIRT6,BUD31,SAE1,ARHGAP6,SCARB1,CTTN,TSPAN11,TNFSF10,COMMD5,NU
DNASE1L1,KIF4A,PTOV1,FASTK,TMEM135,RNF32,LY75,ATG16L2,SNRPB,PARD3,TCAIM,SLC3A2,DHPS,AC
DNASE1L1,KIF4A,PTOV1,FASTK,TMEM135,RNF32,LY75,ATG16L2,SNRPB,PARD3,TCAIM,SLC3A2,DHPS,AC
PTOV1,RNF32,LY75,ATG16L2,SNRPB,TCAIM,SLC3A2,DHPS,ACTR6,MAP3K7CL,SEC62,SET,CTSC,ZNF235,BC
I,JUP,C14ORF2,RFX3,DGKE,TXNL4A,PRMT1,ARL6IP4,ZFAND2B,ACAD10,SUGT1,INO80E,CEP70,ELP6,GPB
NF235,ARRDC1,TRPT1,KLC1,RBP5,SCAF11,NUMA1,GPATCH2L,BUD31,TBL2,ARHGAP6,SCARB1,CTTN,TSPA

N,ITSN1,EIF5A,COMMD5,NUDT22,DBNDD1,NOP16,TTLL5,AP3B2,RCE1,RUVBL2,EBPL,TESC,NNMT,SPAG7,
N,ITSN1,EIF5A,COMMD5,NUDT22,DBNDD1,NOP16,TTLL5,AP3B2,RCE1,RUVBL2,EBPL,TESC,NNMT,SPAG7,

GEF39,PSMA6,AP3B2,RCE1,TESC,IGSF8,SPAG7,WDR62,PLEKHA5,RANBP1,UBE2L6,TUBA1C,ANTXR2,STIP1
GEF39,PSMA6,AP3B2,RCE1,TESC,IGSF8,SPAG7,WDR62,PLEKHA5,RANBP1,UBE2L6,TUBA1C,ANTXR2,STIP1
SE1,OGFOD2,ARIH2,PICK1,TPT1,TSPAN31,GLYCTK,TRAPPC2,AXIN1,TMEM199,EPN1,ZCCHC9,SNRPD1,AZI
HADA,FAM32A,C12ORF57,ZNF223,OGFOD2,METTL3,EIF3G,SLC52A2,NTMT1,TSPAN31,GLYCTK,CFLAR,TRI
HADA,FAM32A,C12ORF57,ZNF223,OGFOD2,METTL3,EIF3G,SLC52A2,NTMT1,TSPAN31,GLYCTK,CFLAR,TRI
BUD31,OSGEP,ARHGAP6,TSPAN11,EIF5A,CHID1,UFD1L,TAZ,THYN1,LDHA,ACOT8,AP3B2,ZNF410,RUVBL2,
MT2,EIF3M,JUP,GUK1,ATF7,CAPG,PPIL2,TIMM17B,LSM12,PIGL,C19ORF48,ZFAND2B,SHARPIN,CDK11A,L
MT2,EIF3M,JUP,GUK1,ATF7,CAPG,PPIL2,TIMM17B,LSM12,PIGL,C19ORF48,ZFAND2B,SHARPIN,CDK11A,L
LY,GLYCTK,TRAPPC2,AXIN1,TMEM199,EPN1,ITPA,ZCCHC9,SNRPD1,AZI2,GOSR2,MYL12A,LAMTOR1,BCL7
,TSPAN11,EIF5A,COMMD5,AMZ2,TAZ,THYN1,SH2B1,RPL13A,PSMA6,ACOT8,AP3B2,SLC35G1,MRPL33,TE
ARB1,CTTN,TSPAN11,ITSN1,EIF5A,COMMD5,NUDT22,DBNDD1,NOP16,TTLL5,PSMA6,ACOT8,AP3B2,RCE
ARB1,CTTN,TSPAN11,ITSN1,EIF5A,COMMD5,NUDT22,DBNDD1,NOP16,TTLL5,PSMA6,ACOT8,AP3B2,RCE
SIRT6,NUMA1,GPATCH2L,BUD31,TBL2,SCARB1,CTTN,ITSN1,EIF5A,COMMD5,NUDT22,UFD1L,DBNDD1,SH
3B2,RPS9,RCE1,MRPL33,TESC,RBPMS,WDR62,POLD4,RANBP1,ZC3H14,FIS1,SLC15A3,PPM1K,STIP1,PHF1
3B2,RPS9,RCE1,MRPL33,TESC,RBPMS,WDR62,POLD4,RANBP1,ZC3H14,FIS1,SLC15A3,PPM1K,STIP1,PHF1
B2,ZFPL1,GMPR2,RIC8A,TCP1,PIGT,FAM32A,PEX2,ARL16,OXLD1,SSR3,ARIH2,PICK1,ZNF512,RNF181,TSP
B2,ZFPL1,GMPR2,RIC8A,TCP1,PIGT,FAM32A,PEX2,ARL16,OXLD1,SSR3,ARIH2,PICK1,ZNF512,RNF181,TSP

9A11,FAM3A,EPB41L5,NLRP1,ZBTB44,SLC25A10,DBNDD2,NISCH,TMED4,CD99L2,TRNT1,PARD6G,SH3YL

9A11,FAM3A,EPB41L5,NLRP1,ZBTB44,SLC25A10,DBNDD2,NISCH,TMED4,CD99L2,TRNT1,PARD6G,SH3YL

AC3,NUMA1,GPATCH2L,OSGEP,SCARB1,CTTN,TSPAN11,ITSN1,EIF5A,NUDT22,UFD1L,DBNDD1,GSTK1,PSM

AC3,NUMA1,GPATCH2L,OSGEP,SCARB1,CTTN,TSPAN11,ITSN1,EIF5A,NUDT22,UFD1L,DBNDD1,GSTK1,PSM

TTC21B,RPS25,COX4I1,DUSP22,GGPS1,SRSF1,FOXRED1,ITGB1BP1,SH3YL1,TBX6,PTK2,CENPN,ATF7,HSBP

99,EPN1,ZCCHC9,AZI2,TBC1D22A,SZT2,MRPL51,C9ORF64,MED8,MRPL27,DDX56,MED17,URM1,SEC14L1

GSF8,NUDT5,IL1RAP,STYXL1,RFX5,VRK3,STIP1,LRRC1,LPCAT3,TM9SF1,SON,UBR4,PSMA1,CHCHD1,RPS25

,GLYCTK,TRAPPC2,AXIN1,TMEM199,EPN1,ZCCHC9,SNRPD1,AZI2,TBC1D22A,BCL7B,SZT2,MRPL51,C9ORF

,PTOV1,FASTK,TMEM135,RNF32,LY75,ATG16L2,RCC1,SLC3A2,DHPS,SEC62,SET,CTSC,ZNF235,ARRDC1,TR

,PTOV1,FASTK,TMEM135,RNF32,LY75,ATG16L2,RCC1,SLC3A2,DHPS,SEC62,SET,CTSC,ZNF235,ARRDC1,TR

LC3A2,ACTR6,SET,BOP1,ARRDC1,TRPT1,RBP5,ZC3HAV1,DBI,VDAC3,NOB1,NUMA1,GPATCH2L,BUD31,NP

LC3A2,ACTR6,SET,BOP1,ARRDC1,TRPT1,RBP5,ZC3HAV1,DBI,VDAC3,NOB1,NUMA1,GPATCH2L,BUD31,NP

D3,TCAIM,SLC3A2,DHPS,SEC62,SET,ARRDC1,TRPT1,RBP5,ZC3HAV1,SCAF11,DBI,LIMK2,NOB1,WDR19,NU

D3,TCAIM,SLC3A2,DHPS,SEC62,SET,ARRDC1,TRPT1,RBP5,ZC3HAV1,SCAF11,DBI,LIMK2,NOB1,WDR19,NU

D1,SH2B1,NOP16,TTLL5,PSMA6,AP3B2,RUVBL2,EBPL,IGSF8,SPAG7,WDR62,IL1RAP,STYXL1,RANBP1,UBE2

D1,SH2B1,NOP16,TTLL5,PSMA6,AP3B2,RUVBL2,EBPL,IGSF8,SPAG7,WDR62,IL1RAP,STYXL1,RANBP1,UBE2

S1,TUBA1C,STIP1,TET2,GRK6,CDC37,RPS16,OBSL1,PTPN13,SLC39A11,FAM3A,PPRC1,RPS25,SLC25A10,SD

S1,TUBA1C,STIP1,TET2,GRK6,CDC37,RPS16,OBSL1,PTPN13,SLC39A11,FAM3A,PPRC1,RPS25,SLC25A10,SD

,THEM4,GGPS1,YIPF3,IP6K2,UBE2I,KCNMA1,YIF1A,TMEM205,XBP1,GUK1,CARD14,FBXW9,ASF1B,PPP1C

,THEM4,GGPS1,YIPF3,IP6K2,UBE2I,KCNMA1,YIF1A,TMEM205,XBP1,GUK1,CARD14,FBXW9,ASF1B,PPP1C

TMPO,PRMT1,CASC4,EXOSC9,WNK1,ZNF7,HNRNPC,ZNF644,STAU1,CCNL2,LRRFIP1,ZNF524,GPBP1,SIAH1

NAT9,TOMM40,RABEP2,ZNF20,RALGPS1,RAD23A,ABHD17A,ILF3,COX5A,PHF20L1,HNRNPA1,FKBP3,RFX

MK2,NUMA1,BUD31,OSGEP,ARHGAP6,SCARB1,CTTN,TSPAN11,ITSN1,EIF5A,COMMD5,NUDT22,UFD1L,U

MK2,NUMA1,BUD31,OSGEP,ARHGAP6,SCARB1,CTTN,TSPAN11,ITSN1,EIF5A,COMMD5,NUDT22,UFD1L,U

AF11,LIMK2,SIRT6,GPATCH2L,TBL2,ARHGAP6,SCARB1,CTTN,TSPAN11,ITSN1,EIF5A,COMMD5,NUDT22,T

AE1,OSGEP,TBL2,CTTN,TSPAN11,ITSN1,EIF5A,COMMD5,NUDT22,DBNDD1,SH2B1,NOP16,TTLL5,ACOT8,A

AE1,OSGEP,TBL2,CTTN,TSPAN11,ITSN1,EIF5A,COMMD5,NUDT22,DBNDD1,SH2B1,NOP16,TTLL5,ACOT8,A

99L2,GGPS1,PARD6G,SH3YL1,IP6K2,MICAL1,HS2ST1,UBE2I,TBX6,TMEM205,C14ORF2,PCYT2,PPP1CC,PPA

RR13,STARD10,RNH1,DNASE1L1,KIF4A,PTOV1,FASTK,TMEM135,RNF32,ATG16L2,RCC1,TCAIM,SLC3A2,D

RR13,STARD10,RNH1,DNASE1L1,KIF4A,PTOV1,FASTK,TMEM135,RNF32,ATG16L2,RCC1,TCAIM,SLC3A2,D

C,DGKE,CAMTA1,LIG1,TIMM17B,APEH,CAPN10,ATPAF2,CASC4,ZFAND2B,SHARPIN,SYNGR2,CEP70,LRRFI

C,DGKE,CAMTA1,LIG1,TIMM17B,APEH,CAPN10,ATPAF2,CASC4,ZFAND2B,SHARPIN,SYNGR2,CEP70,LRRFI

CH2L,NPRL2,OSGEP,TBL2,ARHGAP6,EIF5A,AMZ2,TAZ,NOP16,RPL13A,TTLL5,LDHA,RPS9,RCE1,ZNF410,PC

CH2L,NPRL2,OSGEP,TBL2,ARHGAP6,EIF5A,AMZ2,TAZ,NOP16,RPL13A,TTLL5,LDHA,RPS9,RCE1,ZNF410,PC

1,TUBA1C,IRF9,STIP1,TET2,GRK6,HMGA2,CDC37,TTC21B,UBR4,SLC39A11,PSMA3,PPRC1,SLC25A10,SDF4

F1,GRK6,CDC37,CPSF3L,TSG101,UBR4,FAM3A,METTL17,NLRP1,NIP7,TTC9C,COX4I1,THEM4,DBNDD2,CE

F1,GRK6,CDC37,CPSF3L,TSG101,UBR4,FAM3A,METTL17,NLRP1,NIP7,TTC9C,COX4I1,THEM4,DBNDD2,CE

SCARB1,CTTN,TSPAN11,ITSN1,COMMD5,NUDT22,UFD1L,USP38,THYN1,DBNDD1,SH2B1,LDHA,AP3B2,RP

M1K,STIP1,PHF19,MAGED2,GRK6,YLPM1,PHF21A,SLC39A11,FAM3A,MUC1,EPB41L5,ZBTB44,NISCH,TME

M1K,STIP1,PHF19,MAGED2,GRK6,YLPM1,PHF21A,SLC39A11,FAM3A,MUC1,EPB41L5,ZBTB44,NISCH,TME

MD5,NUDT22,SH2B1,NOP16,TTLL5,ACOT8,AP3B2,RUVBL2,EBPL,RBPMS,SPAG7,IL1RAP,STYXL1,RFX5,PLE

MD5,NUDT22,SH2B1,NOP16,TTLL5,ACOT8,AP3B2,RUVBL2,EBPL,RBPMS,SPAG7,IL1RAP,STYXL1,RFX5,PLE

TTLL5,ACOT8,AP3B2,MRPL33,PCMT1,EBPL,NNMT,WDR62,DST,STIP1,LRRC1,PHF19,LPCAT3,CD44,SON,T

TSPAN11,USP38,SH2B1,NOP16,RPL13A,PSMA6,RPS9,SLC35G1,RCE1,RUVBL2,NNMT,SPAG7,WDR62,STYX

TSPAN11,USP38,SH2B1,NOP16,RPL13A,PSMA6,RPS9,SLC35G1,RCE1,RUVBL2,NNMT,SPAG7,WDR62,STYX

P6,EIF5A,AMZ2,TAZ,NOP16,GSTK1,TTLL5,LDHA,RPS9,RCE1,ZNF410,PCMT1,IGSF8,NNMT,WDR62,STYXL1,

P6,EIF5A,AMZ2,TAZ,NOP16,GSTK1,TTLL5,LDHA,RPS9,RCE1,ZNF410,PCMT1,IGSF8,NNMT,WDR62,STYXL1,

1,PSMA7,XAF1,LYPLA2,CAMLG,ATPAF2,LSM12,PRMT1,C19ORF54,HNRNPC,ZNF644,RBM3,CCNL2,HSP90

1,PSMA7,XAF1,LYPLA2,CAMLG,ATPAF2,LSM12,PRMT1,C19ORF54,HNRNPC,ZNF644,RBM3,CCNL2,HSP90

F57,ZNF223,SLC35E1,OGFOD2,MPV17,PICK1,TSPAN31,FAM192A,GLYCTK,RBM42,GTF2H1,AXIN1,TMEM1

4I1,THEM4,CENPK,AUP1,TRNT1,MED15,MAGT1,C14ORF2,GUK1,CENPN,PCYT2,MAF,PPA1,PPIL2,TXNL4A

TESC,STYXL1,PPM1K,TET2,SNF8,CDC37,CPSF3L,CHCHD1,MUC1,ECSIT,PPRC1,DBNDD2,RPS6,CD99L2,COA

TESC,STYXL1,PPM1K,TET2,SNF8,CDC37,CPSF3L,CHCHD1,MUC1,ECSIT,PPRC1,DBNDD2,RPS6,CD99L2,COA

,VRK3,STIP1,LRRC1,LPCAT3,TET2,PHF21A,COX4I1,NISCH,TMED4,RPS6,SRSF1,YIPF3,TRNT1,ITGB1BP1,MI

,RNH1,DNASE1L1,PTOV1,FASTK,TMEM135,RNF32,ATG16L2,RCC1,SNRPB,PARD3,SLC3A2,DHPS,SET,BOP1

,RNH1,DNASE1L1,PTOV1,FASTK,TMEM135,RNF32,ATG16L2,RCC1,SNRPB,PARD3,SLC3A2,DHPS,SET,BOP1

,FAM122C,SRP9,IQCG,TMEM67,PILRB,FAM72A,CMC2,YBX3,BLOC1S1,CACYBP,CIRBP,RHOF,SMARCB1,PR

,FAM122C,SRP9,IQCG,TMEM67,PILRB,FAM72A,CMC2,YBX3,BLOC1S1,CACYBP,CIRBP,RHOF,SMARCB1,PR

H,LYPLA2,NUTF2,C19ORF48,SUGT1,HYPK,UPP1,MOB1A,INO80E,CCNL2,TMEM134,RAB2A,ZNF524,ATP5J,

ET2,GRK6,CDC37,UBR4,SLC39A11,FAM3A,PPRC1,SLC25A10,THEM4,SDF4,TRNT1,ITGB1BP1,C1ORF198,C

ET2,GRK6,CDC37,UBR4,SLC39A11,FAM3A,PPRC1,SLC25A10,THEM4,SDF4,TRNT1,ITGB1BP1,C1ORF198,C

40,TADA3,KLK10,TRIM16,HSPH1,CNST,COX5A,PFDN5,PAFAH1B2,COA3,DAZAP2,ZFPL1,GMPR2,RIC8A,TC

40,TADA3,KLK10,TRIM16,HSPH1,CNST,COX5A,PFDN5,PAFAH1B2,COA3,DAZAP2,ZFPL1,GMPR2,RIC8A,TC

P7,THEM4,CENPK,CD99L2,COASY,SRSF1,KXD1,MED15,IP6K2,KCNMA1,CDC42,XBP1,GUK1,CENPN,DGKE,T

,SH3YL1,IP6K2,UBE2I,AUNIP,C14ORF2,PCYT2,PPP1CC,DGKE,CAMTA1,LIG1,TIMM17B,APEH,NUTF2,C17O

RBP5,ZC3HAV1,NOB1,WDR19,NUMA1,GPATCH2L,SUMF2,NPRL2,OSGEP,TBL2,ARHGAP6,CRYGS,EIF5A,CH

RBP5,ZC3HAV1,NOB1,WDR19,NUMA1,GPATCH2L,SUMF2,NPRL2,OSGEP,TBL2,ARHGAP6,CRYGS,EIF5A,CH

,TMEM205,C14ORF2,CENPN,HSBP1L1,ASF1B,EEF1B2,PPP1CC,PPIL2,LIG1,TMEM126B,UCK2,C19ORF54,T

,TMEM205,C14ORF2,CENPN,HSBP1L1,ASF1B,EEF1B2,PPP1CC,PPIL2,LIG1,TMEM126B,UCK2,C19ORF54,T

DR19,SIRT6,NUMA1,SUMF2,TSPAN11,EIF5A,COMMD5,UFD1L,TTLL5,LDHA,ZNF410,NUDT5,NNMT,SNRPN

DR19,SIRT6,NUMA1,SUMF2,TSPAN11,EIF5A,COMMD5,UFD1L,TTLL5,LDHA,ZNF410,NUDT5,NNMT,SNRPN

A2,TSPAN31,GLYCTK,CFLAR,TRIM46,EIF2A,NKIRAS2,ACAA1,GTF2H1,BFAR,TMEM199,ZNF396

A2,TSPAN31,GLYCTK,CFLAR,TRIM46,EIF2A,NKIRAS2,ACAA1,GTF2H1,BFAR,TMEM199,ZNF396

IG1,C17ORF62,ZNF7,ZFAND2B,LRRFIP1,RAB2A,LYNX1,LMBR1,BCAM,RABEP2,RALGPS1,EEF1D,NAA10,LLG

CTTN,TSPAN11,ITSN1,EIF5A,NUDT22,DBNDD1,SH2B1,NOP16,TTLL5,PSMA6,AP3B2,RUVBL2,EBPL,IGSF8,S

CTTN,TSPAN11,ITSN1,EIF5A,NUDT22,DBNDD1,SH2B1,NOP16,TTLL5,PSMA6,AP3B2,RUVBL2,EBPL,IGSF8,S

AZ,NOP16,GSTK1,LDHA,RPS9,RCE1,ZNF410,PCMT1,IGSF8,NNMT,WDR62,STYXL1,RFX5,RANBP1,VRK3,PPM

AZ,NOP16,GSTK1,LDHA,RPS9,RCE1,ZNF410,PCMT1,IGSF8,NNMT,WDR62,STYXL1,RFX5,RANBP1,VRK3,PPM

2A,PEX2,ARL16,OXLD1,SSR3,ARIH2,PICK1,RNF181,TSPAN31,GLYCTK,PCNP,CREBZF,EIF2A,NKIRAS2,PIGO,S

2A,PEX2,ARL16,OXLD1,SSR3,ARIH2,PICK1,RNF181,TSPAN31,GLYCTK,PCNP,CREBZF,EIF2A,NKIRAS2,PIGO,S

,POLD4,PLEKHA5,VRK3,LZTFL1,FIS1,PPM1K,STIP1,LPCAT3,UQCRQ,GRK6,CPSF3L,TSG101,PTPN13,UBR4,P

,POLD4,PLEKHA5,VRK3,LZTFL1,FIS1,PPM1K,STIP1,LPCAT3,UQCRQ,GRK6,CPSF3L,TSG101,PTPN13,UBR4,P

A,RBCK1,PTOV1,FASTK,TMEM135,RNF32,LY75,ATG16L2,SNRPB,PARD3,TCAIM,SLC3A2,DHPS,ACTR6,MA

DNASE1L1,RBCK1,PTOV1,FASTK,RNF32,LY75,ATG16L2,RCC1,PARD3,TCAIM,ACTR6,SET,BOP1,ARRDC1,TRP

DNASE1L1,RBCK1,PTOV1,FASTK,RNF32,LY75,ATG16L2,RCC1,PARD3,TCAIM,ACTR6,SET,BOP1,ARRDC1,TRP

RF2,GUK1,CENPN,PCYT2,MAF,PPA1,PPIL2,TXNL4A,EXOSC9,ZFAND2B,SUGT1,ZNF250,NFAM1,CDK11A,MA

GED2,SNF8,CDC37,CPSF3L,CHCHD1,RGS3,MUC1,METTL17,ZBTB44,NIP7,SLC25A10,THEM4,CENPK,CD99L

GED2,SNF8,CDC37,CPSF3L,CHCHD1,RGS3,MUC1,METTL17,ZBTB44,NIP7,SLC25A10,THEM4,CENPK,CD99L

6,TSG101,YLPM1,PHF21A,SLC39A11,FUZ,RPS25,NIP7,TTC9C,PLCB2,THEM4,DBNDD2,RPS6,RTEL1,COASY,

6,TSG101,YLPM1,PHF21A,SLC39A11,FUZ,RPS25,NIP7,TTC9C,PLCB2,THEM4,DBNDD2,RPS6,RTEL1,COASY,

TMEM67,UAP1,PILRB,FAM72A,CMC2,YBX3,BLOC1S1,CACYBP,CIRBP,RHOF,SMARCB1,PRR13,STARD10,RN

TMEM67,UAP1,PILRB,FAM72A,CMC2,YBX3,BLOC1S1,CACYBP,CIRBP,RHOF,SMARCB1,PRR13,STARD10,RN

MOB1A,INO80E,HSP90AB1,RAB2A,ZNF524,MPDU1,ZSWIM7,ERLEC1,C12ORF76,HMGA1,NAT9,RBMX,PPP
ASRP,C1ORF198,UGGT2,MICAL1,SLC39A13,PRMT2,EIF3M,JUP,GUK1,ATF7,CAPG,PPIL2,LIG1,TIMM17B,L
ASRP,C1ORF198,UGGT2,MICAL1,SLC39A13,PRMT2,EIF3M,JUP,GUK1,ATF7,CAPG,PPIL2,LIG1,TIMM17B,L
UMA1,SAE1,TSPAN11,ITSN1,EIF5A,UFD1L,USP38,SH2B1,LDHA,AP3B2,RPS9,RCE1,RBPMS,WDR62,PLEKH
B41L5,TTC9C,COX4I1,THEM4,CENPK,SRSF1,AUP1,TRNT1,MED15,MAGT1,C14ORF2,GUK1,CENPN,PCYT2,
B41L5,TTC9C,COX4I1,THEM4,CENPK,SRSF1,AUP1,TRNT1,MED15,MAGT1,C14ORF2,GUK1,CENPN,PCYT2,
B41L5,TTC9C,COX4I1,THEM4,CENPK,SRSF1,AUP1,TRNT1,MED15,MAGT1,C14ORF2,GUK1,CENPN,PCYT2,
,U2AF1L4,RNPS1,ZNF483,AP4M1,MRPL27,SSH3,RPS14,RNF4,MAGOH,ZRANB2,SEC14L1,FANCI
,UBE2I,PRMT2,PTK2,AUNIP,C14ORF2,GUK1,PCYT2,PPP1CC,PPA1,DGKE,CAMTA1,CRIP1,LIG1,TIMM17B,A
OMM40,TADA3,KLK10,TRIM16,HSPH1,CNST,COX5A,PFDN5,PAFAH1B2,COA3,DAZAP2,ZFPL1,GMPR2,RIC
OMM40,TADA3,KLK10,TRIM16,HSPH1,CNST,COX5A,PFDN5,PAFAH1B2,COA3,DAZAP2,ZFPL1,GMPR2,RIC
11,NUDT22,USP38,NOP16,RPL13A,PSMA6,RPS9,RUVBL2,TESC,NNMT,SPAG7,STYXL1,LZTFL1,PPM1K,STIP
11,NUDT22,USP38,NOP16,RPL13A,PSMA6,RPS9,RUVBL2,TESC,NNMT,SPAG7,STYXL1,LZTFL1,PPM1K,STIP
1,CTTN,ITSN1,COMMD5,NUDT22,SH2B1,NOP16,TTLL5,PSMA6,RUVBL2,EBPL,IGSF8,PLEKHA5,UBE2L6,IRF
1,CTTN,ITSN1,COMMD5,NUDT22,SH2B1,NOP16,TTLL5,PSMA6,RUVBL2,EBPL,IGSF8,PLEKHA5,UBE2L6,IRF
AN11,ITSN1,EIF5A,UFD1L,DBNDD1,GSTK1,PSMA6,RUVBL2,EBPL,TESC,IGSF8,NNMT,PLEKHA5,RANBP1,V
AN11,ITSN1,EIF5A,UFD1L,DBNDD1,GSTK1,PSMA6,RUVBL2,EBPL,TESC,IGSF8,NNMT,PLEKHA5,RANBP1,V
2,PCYT2,PPP1CC,DGKE,CAMTA1,LIG1,TIMM17B,APEH,C17ORF62,ZNF7,ZFAND2B,TMEM134,LRRFIP1,RA
1,DHPS,SET,BOP1,ARRDC1,SCAF11,VDAC3,WDR19,NUMA1,BUD31,SAE1,OSGEP,ARHGAP6,CTTN,TSPAN1
1,DHPS,SET,BOP1,ARRDC1,SCAF11,VDAC3,WDR19,NUMA1,BUD31,SAE1,OSGEP,ARHGAP6,CTTN,TSPAN1
KBIA,TRPV3,LFNG,HAUS4,PCID2,PRRC2B,EIF4A1,TSEN34,ARIH2,RBBP6,LAMTOR5,ACAA1,MYL6,AXIN1,R
M,SLC3A2,SEC62,SET,BOP1,ARRDC1,TRPT1,KLC1,RBP5,ZC3HAV1,SCAF11,DBI,LIMK2,NOB1,WDR19,NUM
M,SLC3A2,SEC62,SET,BOP1,ARRDC1,TRPT1,KLC1,RBP5,ZC3HAV1,SCAF11,DBI,LIMK2,NOB1,WDR19,NUM

,AUNIP,PCYT2,PPP1CC,PPA1,DGKE,CAMTA1,LIG1,TIMM17B,APEH,PODXL,C17ORF62,PRMT1,CASC4,ZFAN
5,ARRDC1,ARPC2,LIMK2,VDAC3,NOB1,SIRT6,NUMA1,SUMF2,BUD31,SAE1,TBL2,ARHGAP6,SCARB1,CTTN
5,ARRDC1,ARPC2,LIMK2,VDAC3,NOB1,SIRT6,NUMA1,SUMF2,BUD31,SAE1,TBL2,ARHGAP6,SCARB1,CTTN
2,DHPS,MAP3K7CL,SEC62,SET,BOP1,ARRDC1,TRPT1,KLC1,ZC3HAV1,SCAF11,DBI,NUMA1,ARHGAP6,SCAR
2,DHPS,MAP3K7CL,SEC62,SET,BOP1,ARRDC1,TRPT1,KLC1,ZC3HAV1,SCAF11,DBI,NUMA1,ARHGAP6,SCAR
2,THYN1,DBNDD1,SH2B1,RPL13A,TTLL5,LDHA,ACOT8,RPS9,SLC35G1,RCE1,RUVBL2,EBPL,TESC,NUDT5,S
SF1,AUP1,TRNT1,PRMT2,MAGT1,GUK1,CENPN,AGPAT4,MAF,EDF1,RFX3,PPA1,PPIL2,TXNL4A,UCK2,PRM
T,LSR,TOMM40,TADA3,SLC33A1,PDE4DIP,MDH2,STXBP2,SEC31A,MT1E,COX17,PAFAH1B2,NSUN5,GMPF
7CL,SEC62,SET,CTSC,BOP1,TRPT1,ZC3HAV1,SCAF11,LIMK2,NOB1,WDR19,NUMA1,BUD31,SAE1,NPRL2,T
7CL,SEC62,SET,CTSC,BOP1,TRPT1,ZC3HAV1,SCAF11,LIMK2,NOB1,WDR19,NUMA1,BUD31,SAE1,NPRL2,T
R19,GPATCH2L,SUMF2,BUD31,SAE1,OSGEP,SCARB1,CTTN,ITSN1,EIF5A,COMMD5,CHID1,AMZ2,TAZ,RPL1
R19,GPATCH2L,SUMF2,BUD31,SAE1,OSGEP,SCARB1,CTTN,ITSN1,EIF5A,COMMD5,CHID1,AMZ2,TAZ,RPL1
/1,ARPC2,VDAC3,NOB1,WDR19,SIRT6,NUMA1,GPATCH2L,NPRL2,TBL2,ARHGAP6,SCARB1,CTTN,TSPAN1
/1,ARPC2,VDAC3,NOB1,WDR19,SIRT6,NUMA1,GPATCH2L,NPRL2,TBL2,ARHGAP6,SCARB1,CTTN,TSPAN1
Z,RPS25,NIP7,TTC9C,PLCB2,THEM4,DBNDD2,RPS6,COASY,AUP1,SAP30BP,FOXRED1,MED15,SH3YL1,SLC
Z,RPS25,NIP7,TTC9C,PLCB2,THEM4,DBNDD2,RPS6,COASY,AUP1,SAP30BP,FOXRED1,MED15,SH3YL1,SLC
TESC,SPAG7,RANBP1,ZC3H14,STIP1,VCAN,LPCAT3,RPS16,ATG12,ZBTB44,DUSP22,SPOCK2,THEM4,SDF4,
TESC,SPAG7,RANBP1,ZC3H14,STIP1,VCAN,LPCAT3,RPS16,ATG12,ZBTB44,DUSP22,SPOCK2,THEM4,SDF4,
WDR62,STYXL1,RFX5,VRK3,PPM1K,PHF19,PCBP2,TM9SF1,TET2,GRK6,CDC37,TSG101,OBSL1,FAM3A,PS
WDR62,STYXL1,RFX5,VRK3,PPM1K,PHF19,PCBP2,TM9SF1,TET2,GRK6,CDC37,TSG101,OBSL1,FAM3A,PS
J1,TWSG1,ZSWIM7,ERLEC1,MVD,MCTS1,DCN,TOMM40,ATP6V1F,SLC33A1,NT5C,MDH2,SH3GLB1,SSH2,
SIRT6,NUMA1,SUMF2,BUD31,OSGEP,TBL2,ARHGAP6,CTTN,TSPAN11,ITSN1,EIF5A,AMZ2,USP38,TAZ,DB
A1,PPIL2,TXNL4A,EXOSC9,ZFAND2B,ZNF250,NFAM1,MAPKAPK5,DGUOK,HSP90AB1,RAB2A,RMDN1,GP

PA1,PPIL2,TXNL4A,EXOSC9,ZFAND2B,ZNF250,NFAM1,MAPKAPK5,DGUOK,HSP90AB1,RAB2A,RMDN1,GPE

7,EPB41L5,TTC9C,COX4I1,THEM4,CENPK,AUP1,TRNT1,MED15,MAGT1,C14ORF2,GUK1,CENPN,PCYT2,HS

7,EPB41L5,TTC9C,COX4I1,THEM4,CENPK,AUP1,TRNT1,MED15,MAGT1,C14ORF2,GUK1,CENPN,PCYT2,HS

7CL,SEC62,ARRDC1,TRPT1,KLC1,ARPC2,VDAC3,SIRT6,SUMF2,BUD31,SAE1,CTTN,EIF5A,NUDT22,AMZ2,DE

,COMMD5,CHID1,NUDT22,TAZ,GSTK1,LDHA,PSMA6,RPS9,RCE1,MRPL33,PCMT1,EBPL,SPAG7,STYXL1,RF

,COMMD5,CHID1,NUDT22,TAZ,GSTK1,LDHA,PSMA6,RPS9,RCE1,MRPL33,PCMT1,EBPL,SPAG7,STYXL1,RF

R19,GPATCH2L,SUMF2,BUD31,SAE1,OSGEP,SCARB1,CTTN,EIF5A,COMMD5,CHID1,AMZ2,TAZ,RPL13A,LD

SAE1,OSGEP,SCARB1,CTTN,EIF5A,COMMD5,RUNX1,CHID1,TAZ,RPL13A,LDHA,AP3B2,RPS9,MRPL33,TESC,

SAE1,OSGEP,SCARB1,CTTN,EIF5A,COMMD5,RUNX1,CHID1,TAZ,RPL13A,LDHA,AP3B2,RPS9,MRPL33,TESC,

SAE1,OSGEP,SCARB1,CTTN,EIF5A,COMMD5,RUNX1,CHID1,TAZ,RPL13A,LDHA,AP3B2,RPS9,MRPL33,TESC,

,SMARCB1,RPL38,PRR13,RNH1,DNASE1L1,KIF4A,RBCK1,PTOV1,TMEM135,RNF32,PARD3,TCAIM,SLC3A2,

,SMARCB1,RPL38,PRR13,RNH1,DNASE1L1,KIF4A,RBCK1,PTOV1,TMEM135,RNF32,PARD3,TCAIM,SLC3A2,

ORF48,UPP1,RPL36,RBM3,SYNGR2,RAB2A,TPM2,SIAH1,ASPH,PNPO,NIT2,RPS15,UBXN1,ZNF814,VWA2,

ORF48,UPP1,RPL36,RBM3,SYNGR2,RAB2A,TPM2,SIAH1,ASPH,PNPO,NIT2,RPS15,UBXN1,ZNF814,VWA2,

IAJA3,AKT1S1,MLST8,C2ORF15,IAH1,SDHA,ZNF483,PEX16,HNRNPM,POLH,EHMT1,MBD1,NDUFS7,DAD1,

D1,NUDT22,UFD1L,USP38,DBNDD1,TTLL5,ACOT8,AP3B2,ZNF410,NUDT5,RBPMS,STYXL1,POLD4,VRK3,LZ

D1,NUDT22,UFD1L,USP38,DBNDD1,TTLL5,ACOT8,AP3B2,ZNF410,NUDT5,RBPMS,STYXL1,POLD4,VRK3,LZ

C37,TTC21B,UBR4,SLC39A11,METTL17,PPRC1,SLC25A10,SDF4,TMED4,RPL29,TRNT1,ITGB1BP1,SH3YL1,

OC1S1,CACYBP,CIRBP,RHOF,SMARCB1,RPL38,RNH1,DNASE1L1,PTOV1,FASTK,RNF32,LY75,RCC1,PARD3,T

OC1S1,CACYBP,CIRBP,RHOF,SMARCB1,RPL38,RNH1,DNASE1L1,PTOV1,FASTK,RNF32,LY75,RCC1,PARD3,T

P1,KLC1,ARPC2,PMFBP1,NOB1,SIRT6,NUMA1,GPATCH2L,SAE1,SCARB1,CRYGS,TSPAN11,COMMD5,CHID1

P1,KLC1,ARPC2,PMFBP1,NOB1,SIRT6,NUMA1,GPATCH2L,SAE1,SCARB1,CRYGS,TSPAN11,COMMD5,CHID1

V1,SCAF11,DBI,LIMK2,VDAC3,WDR19,SIRT6,NUMA1,SUMF2,BUD31,NPRL2,OSGEP,CRYGS,CTTN,TSPAN1

V1,SCAF11,DBI,LIMK2,VDAC3,WDR19,SIRT6,NUMA1,SUMF2,BUD31,NPRL2,OSGEP,CRYGS,CTTN,TSPAN1

,CENPN,PCYT2,MAF,PPA1,PPIL2,TXNL4A,EXOSC9,ZFAND2B,SUGT1,ZNF250,NFAM1,RBM3,MAPKAPK5,DG

RL2,TBL2,SCARB1,CTTN,CHID1,AMZ2,TAZ,LDHA,AP3B2,ZNF410,PCMT1,EBPL,TESC,IGSF8,NUDT5,RANBP

RFX5,RANBP1,LRRC1,TET2,UQCRQ,SNF8,GRK6,CDC37,TSG101,TTC21B,UBR4,PSMA1,SLC39A11,TTC9C,CO

RFX5,RANBP1,LRRC1,TET2,UQCRQ,SNF8,GRK6,CDC37,TSG101,TTC21B,UBR4,PSMA1,SLC39A11,TTC9C,CO

P1,FAM32A,PEX2,SSR3,ARIH2,TSPAN31,GLYCTK,CREBZF,EIF2A,GLI4,NKIRAS2,PIGO,BFAR,SLC25A24,EIF3C

P1,FAM32A,PEX2,SSR3,ARIH2,TSPAN31,GLYCTK,CREBZF,EIF2A,GLI4,NKIRAS2,PIGO,BFAR,SLC25A24,EIF3C

IF5A,COMMD5,NUDT22,USP38,DBNDD1,SH2B1,RCE1,TESC,IGSF8,STYXL1,RANBP1,SLC15A3,PPM1K,PHF1

,GPATCH2L,SAE1,OSGEP,SCARB1,CTTN,ITSN1,EIF5A,COMMD5,CHID1,AMZ2,TAZ,RPL13A,LDHA,RPS9,MR

,SCARB1,COMMD5,NUDT22,USP38,NOP16,RPL13A,TTLL5,PSMA6,MRPL33,RUVBL2,NNMT,SPAG7,WDR6

,SCARB1,COMMD5,NUDT22,USP38,NOP16,RPL13A,TTLL5,PSMA6,MRPL33,RUVBL2,NNMT,SPAG7,WDR6

GPBP1,CHPF,MVD,IL32,SYNGR1,RRNAD1,ZNF687,POT1,RAD23A,PKP4,HSPH1,RPL18,CHEK1,PHF20L1,FKB

IL2,TXNL4A,EXOSC9,ZFAND2B,ZNF250,NFAM1,MAPKAPK5,DGUOK,HSP90AB1,RAB2A,RMDN1,GPBP1,GP

TOV1,FASTK,RNF32,LY75,ATG16L2,RCC1,PARD3,TCAIM,ACTR6,SET,BOP1,ARRDC1,TRPT1,KLC1,RBP5,ZC3

B2A,GPBP1,CHPF,MVD,IL32,SYNGR1,RRNAD1,ZNF687,POT1,RAD23A,PKP4,HSPH1,RPL18,CHEK1,PHF20L1

1,ZNF706,RAD23A,SLC16A1,ILF3,AGAP3,EEF1D,SUMO3,PHF20L1,EGLN2,SIRT5,RFXANK,TMEM101,PPIH,

,ACTR6,MAP3K7CL,SEC62,ZNF235,BOP1,TRPT1,KLC1,RBP5,ZC3HAV1,SCAF11,DBI,ARPC2,PMFBP1,VDAC3

,ACTR6,MAP3K7CL,SEC62,ZNF235,BOP1,TRPT1,KLC1,RBP5,ZC3HAV1,SCAF11,DBI,ARPC2,PMFBP1,VDAC3

TM9SF1,SON,TET2,GRK6,CDC37,PHF21A,FAM3A,PSMA3,METTL17,NLRP1,NIP7,TTC9C,COX4I1,THEM4,D

TM9SF1,SON,TET2,GRK6,CDC37,PHF21A,FAM3A,PSMA3,METTL17,NLRP1,NIP7,TTC9C,COX4I1,THEM4,D

DC25,EEF1B2,PPA1,TIMM17B,TMEM126B,CAPN10,ATPAF2,C19ORF54,ARL6IP4,C19ORF48,CBFA2T2,UPF

DC25,EEF1B2,PPA1,TIMM17B,TMEM126B,CAPN10,ATPAF2,C19ORF54,ARL6IP4,C19ORF48,CBFA2T2,UPF

DC25,EEF1B2,PPA1,TIMM17B,TMEM126B,CAPN10,ATPAF2,C19ORF54,ARL6IP4,C19ORF48,CBFA2T2,UPF

PATCH2L,BUD31,NPRL2,CRYGS,CTTN,EIF5A,TNFSF10,COMMD5,NUDT22,SH2B1,NOP16,RPL13A,ZNF410
F2,GUK1,CENPN,PCYT2,IFI27L1,MAF,PPA1,PPIL2,TXNL4A,EXOSC9,ZFAND2B,SUGT1,ZNF250,NFAM1,CDK
F2,GUK1,CENPN,PCYT2,IFI27L1,MAF,PPA1,PPIL2,TXNL4A,EXOSC9,ZFAND2B,SUGT1,ZNF250,NFAM1,CDK
R,PSME2,TOMM40,BRIP1,TADA3,SLC33A1,CARHSP1,CNST,HINT1,KCTD7,RFXANK,NSUN5,OARD1,GMPR2
1,CARD14,USE1,PPA1,PPIL2,VIM,LIG1,WNK1,CBFA2T2,SUGT1,DGUOK,CHPF,STRIP2,DCTD,SAT1,ERCC5,LI
1,CARD14,USE1,PPA1,PPIL2,VIM,LIG1,WNK1,CBFA2T2,SUGT1,DGUOK,CHPF,STRIP2,DCTD,SAT1,ERCC5,LI
PRR13,RNH1,RBCK1,FASTK,RNF32,LY75,RCC1,SNRPB,TCAIM,DHPS,ACTR6,MAP3K7CL,SEC62,CTSC,ZNF23
PRR13,RNH1,RBCK1,FASTK,RNF32,LY75,RCC1,SNRPB,TCAIM,DHPS,ACTR6,MAP3K7CL,SEC62,CTSC,ZNF23
TOV1,FASTK,RNF32,LY75,ATG16L2,PARD3,TCAIM,DHPS,ACTR6,SEC62,BOP1,ARRDC1,TRPT1,KLC1,RBP5,
TOV1,FASTK,RNF32,LY75,ATG16L2,PARD3,TCAIM,DHPS,ACTR6,SEC62,BOP1,ARRDC1,TRPT1,KLC1,RBP5,
AP6,SCARB1,ITSN1,EIF5A,COMMD5,SH2B1,RUVBL2,WDR62,IL1RAP,PLEKHA5,LZTFL1,ANTXR2,STIP1,GRA

SAE1,SCARB1,TSPAN11,COMMD5,USP38,NOP16,RPL13A,PSMA6,RPS9,ZNF410,RUVBL2,NNMT,SPAG7,ST
SAE1,SCARB1,TSPAN11,COMMD5,USP38,NOP16,RPL13A,PSMA6,RPS9,ZNF410,RUVBL2,NNMT,SPAG7,ST
TIP1,PHF19,GRK6,RPS16,SLC39A11,ATG12,ZBTB44,CD99L2,GGPS1,PARD6G,SH3YL1,IP6K2,MICAL1,HS2S
E1,SCARB1,TSPAN11,COMMD5,NUDT22,USP38,NOP16,RPL13A,PSMA6,RPS9,RUVBL2,TESC,NNMT,SPAG
E1,SCARB1,TSPAN11,COMMD5,NUDT22,USP38,NOP16,RPL13A,PSMA6,RPS9,RUVBL2,TESC,NNMT,SPAG
T2,MAF,PPP1CC,PPA1,PPIL2,TXNL4A,EXOSC9,ZFAND2B,UPP1,MAPKAPK5,HSP90AB1,CEP70,RMDN1,GPB
T2,MAF,PPP1CC,PPA1,PPIL2,TXNL4A,EXOSC9,ZFAND2B,UPP1,MAPKAPK5,HSP90AB1,CEP70,RMDN1,GPB
A3,PILRB,FAM72A,YBX3,CACYBP,CIRBP,RHOF,RPL38,PRR13,DNASE1L1,RBCK1,PTOV1,FASTK,TMEM135,
,CRYGS,CTTN,ITSN1,RUNX1,AMZ2,UFD1L,USP38,DBNDD1,RPL13A,TTLL5,LDHA,AP3B2,RPS9,ZNF410,RUV
,CRYGS,CTTN,ITSN1,RUNX1,AMZ2,UFD1L,USP38,DBNDD1,RPL13A,TTLL5,LDHA,AP3B2,RPS9,ZNF410,RUV
K,TRMT112,RPS20,ZSWIM7,EMC4,MCTS1,DDOST,LSR,PABPC1,RAD1,ZNF687,SLC16A1,NAA10,SEC31A,F
32,LY75,ATG16L2,RCC1,PARD3,TCAIM,SLC3A2,DHPS,ACTR6,SET,CTSC,BOP1,ARRDC1,KLC1,DBI,VDAC3,B
32,LY75,ATG16L2,RCC1,PARD3,TCAIM,SLC3A2,DHPS,ACTR6,SET,CTSC,BOP1,ARRDC1,KLC1,DBI,VDAC3,B
P1,TROAP,VIM,ATPAF2,CASC4,PIGL,STAU1,SLC9B2,CDK11A,RBM3,ZNF197,ZNF524,TPM2,TPM3,ZNF625
,MUC1,ECSIT,TMED4,RPS6,CD99L2,COASY,YIPF3,ITGB1BP1,SH3YL1,EIF3M,AUNIP,C14ORF2,GUK1,ATF7,
,MUC1,ECSIT,TMED4,RPS6,CD99L2,COASY,YIPF3,ITGB1BP1,SH3YL1,EIF3M,AUNIP,C14ORF2,GUK1,ATF7,
F250,SYNGR2,NIT2,TWSG1,IL32,CHCHD7,ZNF836,BLVRB,RABEP2,RALGPS1,NXN,NUDT1,RAD23A,ABHD1

A1,PPIL2,TXNL4A,EXOSC9,ZFAND2B,ZNF250,NFAM1,MAPKAPK5,DGUOK,HSP90AB1,RAB2A,RMDN1,GP
A1,PPIL2,TXNL4A,EXOSC9,ZFAND2B,ZNF250,NFAM1,MAPKAPK5,DGUOK,HSP90AB1,RAB2A,RMDN1,GP
32,SYNGR1,RRNAD1,ZNF687,POT1,RAD23A,PKP4,HSPH1,RPL18,CHEK1,PHF20L1,FKBP3,B3GAT3,GTF2IR
UK1,CENPN,PCYT2,AGPAT4,IFI27L1,MAF,PPA1,ENDOV,PPIL2,TXNL4A,EXOSC9,ZFAND2B,RBM3,MAPKAP
SCARB1,CTTN,EIF5A,COMMD5,CHID1,UFD1L,TAZ,RPL13A,LDHA,RPS9,MRPL33,TESC,SNRPN,SPAG7,WDR
SCARB1,CTTN,EIF5A,COMMD5,CHID1,UFD1L,TAZ,RPL13A,LDHA,RPS9,MRPL33,TESC,SNRPN,SPAG7,WDR
SCARB1,CTTN,EIF5A,COMMD5,CHID1,UFD1L,TAZ,RPL13A,LDHA,RPS9,MRPL33,TESC,SNRPN,SPAG7,WDR
RBM3,SYNGR2,ZNF524,ASPH,TWSG1,MAPK13,VPS28,DBNL,TOMM40,SERPINB6,RABEP2,RALGPS1,ZNF6
RBM3,SYNGR2,ZNF524,ASPH,TWSG1,MAPK13,VPS28,DBNL,TOMM40,SERPINB6,RABEP2,RALGPS1,ZNF6
,PPA1,ENDOV,PPIL2,ZFAND2B,MAPKAPK5,HSP90AB1,CEP70,RAB2A,GPBP1,GPALPP1,CHPF,GMPPB,MVD
RK6,CDC37,TSG101,FAM3A,PSMA3,METTL17,NIP7,TTC9C,COX4I1,THEM4,DBNDD2,CENPK,COASY,SRSF1
RK6,CDC37,TSG101,FAM3A,PSMA3,METTL17,NIP7,TTC9C,COX4I1,THEM4,DBNDD2,CENPK,COASY,SRSF1
PIL2,ZFAND2B,MAPKAPK5,HSP90AB1,CEP70,RAB2A,GPBP1,GPALPP1,CHPF,MVD,IL32,SYNGR1,RRNAD1,

F132,FAM126A,RNF181,CHTOP,ZNF593,LAMTOR5,RBM42,ACAA1,NEDD4L,ZNF283,GTF2H1,NAA38,ZN
F132,FAM126A,RNF181,CHTOP,ZNF593,LAMTOR5,RBM42,ACAA1,NEDD4L,ZNF283,GTF2H1,NAA38,ZN

),IQCG,TMEM67,UAP1,FAM72A,CMC2,YBX3,CACYBP,CIRBP,RPL38,PRR13,STARD10,RNH1,DNASE1L1,KIF
),IQCG,TMEM67,UAP1,FAM72A,CMC2,YBX3,CACYBP,CIRBP,RPL38,PRR13,STARD10,RNH1,DNASE1L1,KIF
DC1,TRPT1,KLC1,ZC3HAV1,SCAF11,VDAC3,SIRT6,NUMA1,ARHGAP6,TSPAN11,ITSN1,EIF5A,COMMD5,CH
DC1,TRPT1,KLC1,ZC3HAV1,SCAF11,VDAC3,SIRT6,NUMA1,ARHGAP6,TSPAN11,ITSN1,EIF5A,COMMD5,CH

1,LZTFL1,FIS1,TUBA1C,IRF9,STIP1,TET2,UQCRQ,GRK6,HMGA2,CDC37,TTC21B,SLC39A11,FUZ,PSMA3,PPF
1,LZTFL1,FIS1,TUBA1C,IRF9,STIP1,TET2,UQCRQ,GRK6,HMGA2,CDC37,TTC21B,SLC39A11,FUZ,PSMA3,PPF
6,SET,BOP1,ARRDC1,TRPT1,KLC1,RBP5,SCAF11,ARPC2,VDAC3,SIRT6,NUMA1,SUMF2,SAE1,ARHGAP6,SC
PAN11,NUDT22,AMZ2,USP38,TAZ,THYN1,TTLL5,AP3B2,RPS9,RUVBL2,PCMT1,NUDT5,WDR62,RFX5,RAN
SRP,JUP,PCYT2,DEF8,CAMTA1,LIG1,TMEM126B,CAPN10,UCK2,TMPO,PRMT1,EXOSC9,ARL6IP4,STRADA,
SRP,JUP,PCYT2,DEF8,CAMTA1,LIG1,TMEM126B,CAPN10,UCK2,TMPO,PRMT1,EXOSC9,ARL6IP4,STRADA,
Z2,USP38,DBNDD1,AP3B2,ZNF410,IGSF8,RBPMS,SPAG7,STYXL1,RANBP1,UBE2L6,VRK3,TUBA1C,ANTXR2,
Z2,USP38,DBNDD1,AP3B2,ZNF410,IGSF8,RBPMS,SPAG7,STYXL1,RANBP1,UBE2L6,VRK3,TUBA1C,ANTXR2,

AM3A,PPRC1,NLRP1,TTC9C,COX4I1,RPS6,SRSF1,AUP1,SAP30BP,TRNT1,KXD1,ITGB1BP1,MED15,YIF1A,PR
AM3A,PPRC1,NLRP1,TTC9C,COX4I1,RPS6,SRSF1,AUP1,SAP30BP,TRNT1,KXD1,ITGB1BP1,MED15,YIF1A,PR
1,KIF4A,RBCK1,TMEM135,RNF32,LY75,RCC1,SLC3A2,DHPS,ACTR6,MAP3K7CL,SEC62,CTSC,BOP1,TRPT1,
1,KIF4A,RBCK1,TMEM135,RNF32,LY75,RCC1,SLC3A2,DHPS,ACTR6,MAP3K7CL,SEC62,CTSC,BOP1,TRPT1,
AM72A,CACYBP,CIRBP,RPL38,PRR13,STARD10,RNH1,DNASE1L1,KIF4A,TMEM135,ATG16L2,RCC1,SNRPB,
AM72A,CACYBP,CIRBP,RPL38,PRR13,STARD10,RNH1,DNASE1L1,KIF4A,TMEM135,ATG16L2,RCC1,SNRPB,
2,OSGEP,TBL2,ARHGAP6,EIF5A,CHID1,USP38,RPL13A,LDHA,RPS9,RCE1,ZNF410,PCMT1,IGSF8,SNRPN,WD
2,OSGEP,TBL2,ARHGAP6,EIF5A,CHID1,USP38,RPL13A,LDHA,RPS9,RCE1,ZNF410,PCMT1,IGSF8,SNRPN,WD
MK2,VDAC3,WDR19,SIRT6,NUMA1,GPATCH2L,BUD31,SAE1,NPRL2,EIF5A,USP38,TAZ,GSTK1,TTLL5,AP3B
MK2,VDAC3,WDR19,SIRT6,NUMA1,GPATCH2L,BUD31,SAE1,NPRL2,EIF5A,USP38,TAZ,GSTK1,TTLL5,AP3B
C14ORF2,CARD14,PCYT2,USE1,ENDOV,PPIL2,RPL28,VIM,LIG1,TMEM126B,STOML2,C19ORF54,PPAN,ACA
C14ORF2,CARD14,PCYT2,USE1,ENDOV,PPIL2,RPL28,VIM,LIG1,TMEM126B,STOML2,C19ORF54,PPAN,ACA
1,PRR13,STARD10,DNASE1L1,KIF4A,PTOV1,TMEM135,RNF32,LY75,ATG16L2,RCC1,PARD3,TCAIM,SLC3A
1,PRR13,STARD10,DNASE1L1,KIF4A,PTOV1,TMEM135,RNF32,LY75,ATG16L2,RCC1,PARD3,TCAIM,SLC3A
H2B1,RPL13A,ARHGEF39,LDHA,SLC35G1,ZNF410,UBE2L6,LZTFL1,PPM1K,TUBA1C,DST,LRRC1,GRAMD1C,
1,TSPAN11,COMMD5,NUDT22,USP38,NOP16,RPL13A,PSMA6,RPS9,RUVBL2,TESC,NNMT,SPAG7,STYXL1,
1,TSPAN11,COMMD5,NUDT22,USP38,NOP16,RPL13A,PSMA6,RPS9,RUVBL2,TESC,NNMT,SPAG7,STYXL1,
RRDC1,KLC1,SCAF11,GPATCH2L,NPRL2,OSGEP,TBL2,ARHGAP6,ITSN1,EIF5A,AMZ2,TAZ,NOP16,GSTK1,TTL
RRDC1,KLC1,SCAF11,GPATCH2L,NPRL2,OSGEP,TBL2,ARHGAP6,ITSN1,EIF5A,AMZ2,TAZ,NOP16,GSTK1,TTL
M72A,YBX3,BLOC1S1,CACYBP,CIRBP,RHOF,SMARCB1,RPL38,PRR13,RNH1,DNASE1L1,RBCK1,PTOV1,FAST
M72A,YBX3,BLOC1S1,CACYBP,CIRBP,RHOF,SMARCB1,RPL38,PRR13,RNH1,DNASE1L1,RBCK1,PTOV1,FAST
1,ZC3HAV1,SCAF11,ARPC2,PMFBP1,NOB1,SIRT6,NUMA1,GPATCH2L,SAE1,SCARB1,TSPAN11,COMMD5,
1,ZC3HAV1,SCAF11,ARPC2,PMFBP1,NOB1,SIRT6,NUMA1,GPATCH2L,SAE1,SCARB1,TSPAN11,COMMD5,
3A2,ACTR6,SEC62,SET,CTSC,ZNF235,BOP1,ZC3HAV1,ARPC2,NOB1,WDR19,SIRT6,NUMA1,GPATCH2L,SU
3A2,ACTR6,SEC62,SET,CTSC,ZNF235,BOP1,ZC3HAV1,ARPC2,NOB1,WDR19,SIRT6,NUMA1,GPATCH2L,SU
UDT5,STYXL1,POLD4,RANBP1,ZC3H14,FIS1,SLC15A3,ANTXR2,LRRC1,SNF8,GRK6,RPS16,TSG101,OBSL1,Z
UDT5,STYXL1,POLD4,RANBP1,ZC3H14,FIS1,SLC15A3,ANTXR2,LRRC1,SNF8,GRK6,RPS16,TSG101,OBSL1,Z
ARHGAP6,CTTN,ITSN1,EIF5A,NUDT22,AMZ2,UFD1L,USP38,TAZ,DBNDD1,SH2B1,AP3B2,PCMT1,EBPL,TES
,NPRL2,OSGEP,ARHGAP6,EIF5A,AMZ2,TAZ,NOP16,RPL13A,GSTK1,LDHA,RPS9,RCE1,ZNF410,PCMT1,IGS
,NPRL2,OSGEP,ARHGAP6,EIF5A,AMZ2,TAZ,NOP16,RPL13A,GSTK1,LDHA,RPS9,RCE1,ZNF410,PCMT1,IGS
RPL33,TESC,SNRPN,SPAG7,WDR62,STYXL1,RFX5,RANBP1,LZTFL1,FIS1,TUBA1C,IRF9,STIP1,TET2,UQCRQ,
RPL33,TESC,SNRPN,SPAG7,WDR62,STYXL1,RFX5,RANBP1,LZTFL1,FIS1,TUBA1C,IRF9,STIP1,TET2,UQCRQ,
,CHID1,TAZ,RPL13A,LDHA,RPS9,MRPL33,TESC,SNRPN,SPAG7,WDR62,STYXL1,RFX5,RANBP1,LZTFL1,FIS1,

,CHID1,TAZ,RPL13A,LDHA,RPS9,MRPL33,TESC,SNRPN,SPAG7,WDR62,STYXL1,RFX5,RANBP1,LZTFL1,FIS1,

3HAV1,ARPC2,LIMK2,WDR19,SIRT6,GPATCH2L,BUD31,SAE1,TBL2,ARHGAP6,SCARB1,CRYGS,TSPAN11,UF

3HAV1,ARPC2,LIMK2,WDR19,SIRT6,GPATCH2L,BUD31,SAE1,TBL2,ARHGAP6,SCARB1,CRYGS,TSPAN11,UF

P1,MAGT1,GUK1,CENPN,AGPAT4,IFI27L1,MAF,PPA1,ENDOV,PPIL2,ZFAND2B,MAPKAPK5,HSP90AB1,CEP7

MM17B,TMEM126B,CAPN10,EPB41L4B,ATPAF2,C19ORF54,ARL6IP4,CBFA2T2,UPP1,RPL36,ZNF197,COPZ

MM17B,TMEM126B,CAPN10,EPB41L4B,ATPAF2,C19ORF54,ARL6IP4,CBFA2T2,UPP1,RPL36,ZNF197,COPZ

MM17B,TMEM126B,CAPN10,EPB41L4B,ATPAF2,C19ORF54,ARL6IP4,CBFA2T2,UPP1,RPL36,ZNF197,COPZ

P,RHOF,SMARCB1,RPL38,PRR13,RNH1,DNASE1L1,KIF4A,RBCK1,PTOV1,FASTK,TMEM135,LY75,RCC1,PAR

P,RHOF,SMARCB1,RPL38,PRR13,RNH1,DNASE1L1,KIF4A,RBCK1,PTOV1,FASTK,TMEM135,LY75,RCC1,PAR

K2D,ZFAND4,BRF1,PARK7,CCDC40,XYLT2,POLR1C,EIF4A1,TMEM108,TSEN34,ZNF512,UBA1,ANAPC13,NK

K2D,ZFAND4,BRF1,PARK7,CCDC40,XYLT2,POLR1C,EIF4A1,TMEM108,TSEN34,ZNF512,UBA1,ANAPC13,NK

IT,WDR62,STYXL1,RFX5,VRK3,PPM1K,PHF19,PCBP2,TM9SF1,TET2,GRK6,CDC37,TSG101,OBSL1,FAM3A,P

IT,WDR62,STYXL1,RFX5,VRK3,PPM1K,PHF19,PCBP2,TM9SF1,TET2,GRK6,CDC37,TSG101,OBSL1,FAM3A,P

S,ACTR6,SEC62,SET,ARRDC1,SCAF11,ARPC2,LIMK2,NUMA1,SUMF2,OSGEP,TBL2,ARHGAP6,SCARB1,CTTN

CARB1,CTTN,TSPAN11,ITSN1,EIF5A,COMMD5,RUNX1,NUDT22,DBNDD1,NOP16,TTLL5,AP3B2,SLC35G1,R

CARB1,CTTN,TSPAN11,ITSN1,EIF5A,COMMD5,RUNX1,NUDT22,DBNDD1,NOP16,TTLL5,AP3B2,SLC35G1,R

CCDC25,EEF1B2,PPA1,TNFRSF14,CAPN10,C19ORF54,C19ORF48,UPP1,NFAM1,COPZ1,SYNGR2,RAB2A,EL

CCDC25,EEF1B2,PPA1,TNFRSF14,CAPN10,C19ORF54,C19ORF48,UPP1,NFAM1,COPZ1,SYNGR2,RAB2A,EL

,DHPS,MAP3K7CL,SEC62,SET,CTSC,ZNF235,BOP1,ARRDC1,TRPT1,KLC1,RBP5,SCAF11,ARPC2,PMFBP1,LIM

,DHPS,MAP3K7CL,SEC62,SET,CTSC,ZNF235,BOP1,ARRDC1,TRPT1,KLC1,RBP5,SCAF11,ARPC2,PMFBP1,LIM

PIL2,ZFAND2B,MAPKAPK5,HSP90AB1,CEP70,RAB2A,GPBP1,CHPF,MVD,IL32,SYNGR1,RRNAD1,ZNF687,P

4,COX4I1,SLC25A10,SPOCK2,SDF4,NISCH,TMED4,CENPK,RABL5,SH3YL1,UGGT2,UBE2I,PRMT2,XBP1,PTK

4,COX4I1,SLC25A10,SPOCK2,SDF4,NISCH,TMED4,CENPK,RABL5,SH3YL1,UGGT2,UBE2I,PRMT2,XBP1,PTK

VWA2,RANGRF,SLC33A1,GTF2IRD1,RAD23A,GGT5,SLC16A1,ILF3,AGAP3,EEF1D,CARHSP1,SUMO3,PHF20

VWA2,RANGRF,SLC33A1,GTF2IRD1,RAD23A,GGT5,SLC16A1,ILF3,AGAP3,EEF1D,CARHSP1,SUMO3,PHF20

,TTC21B,PHF21A,SLC39A11,MUC1,ATG12,NLRP1,PLCB2,THEM4,RABL5,PARD6G,IP6K2,UBE2I,YIF1A,PRM

,TTC21B,PHF21A,SLC39A11,MUC1,ATG12,NLRP1,PLCB2,THEM4,RABL5,PARD6G,IP6K2,UBE2I,YIF1A,PRM

TBL2,ARHGAP6,EIF5A,COMMD5,CHID1,UFD1L,THYN1,DBNDD1,AP3B2,SLC35G1,RCE1,ZNF410,PCMT1,NU

NUMA1,GPATCH2L,SAE1,SCARB1,TSPAN11,COMMD5,NUDT22,USP38,NOP16,RPL13A,TTLL5,PSMA6,RPS

NUMA1,GPATCH2L,SAE1,SCARB1,TSPAN11,COMMD5,NUDT22,USP38,NOP16,RPL13A,TTLL5,PSMA6,RPS

MAPKAPK5,HSP90AB1,TPM3,RPS15,UBXN1,H2AFV,TOMM40,ZNF687,PDCD5,POLR3H,RAD23A,SLC16A1,

02A,TCP1,SMARCC2,COMMD4,EIF2B3,DCTN5,RPL8,ADD1,CDK4,SRSF7,RALY,VDAC1,EIF2A,RNF130,ALKB

,PPP1CC,CLDND1,PSMA7,UCK2,PPAN,CBFA2T2,CARS,CDK11A,LAMP2,HSP90AB1,ATP5J,GGCX,TPM3,RPS

EH,RPL21,RBM33,PRMT1,EXOSC9,ARL6IP4,PPAN,HNRNPC,STAU1,CARS,RPL36,RBM3,HSP90AB1,DUT,LRF

ARPC5,MYL6,TUBB3,RPS3A,LAMB4,RPS3,MRPS24,VPS25,RPS27L,MFGE8,RPS7,UBA52,RPS4X,MRPS16,RF
L,PHF21A,CHCHD1,MUC1,PPRC1,RPS25,ZBTB44,NIP7,RPS6,RPL29,RTEL1,SRSF1,TRNT1,JPH2,UBE2I,EIF3M

,RNF32,ATG16L2,RCC1,SNRPB,PARD3,SLC3A2,DHPS,ACTR6,MAP3K7CL,SET,CTSC,BOP1,ARRDC1,KLC1,RB

BA1C,IRF9,STIP1,PHF19,PCBP2,SON,TET2,GRK6,HMGA2,RPS16,TSG101,PSMA1,YLPM1,PHF21A,CHCHD1,

BA1C,IRF9,STIP1,PHF19,GRAMD1C,PCBP2,SON,TET2,GRK6,HMGA2,RPS16,TSG101,PSMA1,YLPM1,PHF21

L,PSME1,CARHSP1,RPL18,HNRNPLL,HNRNPA1,POLR2C,RPL15,SNRPF,RBFOX2,PPIH,LGALS3,RPS11,PCID2,

ABPC1,RAD23A,B2M,ILF3,OAS3,RPL18,EGFR,HNRNPA1,POLR2C,RPL15,SMARCA4,TYMS,PPIH,CD74,NFKB

MGA1,PABPC1,RAD23A,B2M,ILF3,OAS3,RPL18,EGFR,HNRNPA1,POLR2C,RPL15,SMARCA4,TYMS,PPIH,TCP
NGRF,TOMM40,PDCD5,SEC31A,RPL18,SH3GLB1,RPL15,B3GAT3,SYTL1,CD74,PARK7,NFKBIA,RPS11,PDCD

B1,SIAH1,RPS20,VPS28,RPS15,UBXN1,ERLEC1,HSPB1,PSME2,PABPC1,WAC,ANAPC11,RAD23A,USP45,PS
RF1,MRPS18C,EIF2B3,RPS11,MRPS17,RPL37A,ELOF1,EIF4A1,RPLP0,METTL3,EIF3G,MRPL55,RPL8,RPL35,
RF1,MRPS18C,EIF2B3,RPS11,MRPS17,RPL37A,ELOF1,EIF4A1,RPLP0,METTL3,EIF3G,MRPL55,RPL8,RPL35,
L21,CAPN10,NUTF2,STOML2,STRADA,ZFAND2B,AP1S3,RPL36,LAMP2,COPZ1,HSP90AB1,RAB2A,ATP5J,RF
1,UBA52,RPS4X,MLH1,RPS23,RPL30,PSMD8,RQCD1,EXOSC2,PSMC5,HNRNPM,EXOSC8,RPS15A,RPS17
SSB,RPL8,BAX,RPL35,NT5E,GALT,RPL34,ACAA1,RPL24,PSMD7,PAM16,PSMD10,RPS3A,RPS3,MAGOHB,IT
MGA1,PABPC1,RAD23A,B2M,ILF3,OAS3,RPL18,EGFR,HNRNPA1,POLR2C,RPL15,SMARCA4,TYMS,PPIH,TCP
3,RPL36,LAMP2,COPZ1,HSP90AB1,RAB2A,RPS20,VPS28,RPS15,ERLEC1,HSPB1,SYNGR1,RANGRF,TOMM4
RPL8,BAX,RPL35,RPL34,NUP62,ACAA1,RPL24,PAM16,TP53,RPS3A,PML,UBE2D3,RPS3,VPS25,DHRS4,GOS
LT,RPL34,RPL24,PSMD7,PAM16,PSMD10,RPS3A,RPS3,MAGOHB,ITPA,SMUG1,RPS7,APAF1,NCBP2,RNPS

S3,RPL36,LAMP2,COPZ1,HSP90AB1,RAB2A,RPS20,VPS28,RPS15,ERLEC1,HSPB1,SYNGR1,RANGRF,TOMM4
NPC,SHARPIN,RPL36,LAMP2,HSP90AB1,SIAH1,RPS20,VPS28,RPS15,UBXN1,ERLEC1,HSPB1,DCN,PSME2,P

DRG2,BRF1,MRPS18C,PARK7,EIF2B3,RPS11,MRPS17,RPL37A,ELOF1,EIF4A1,RPLP0,METTL3,EIF3G,MRPL5
METTL3,SSB,RPL8,BAX,RPL35,NT5E,GALT,RPL34,ACAA1,RPL24,PSMD7,PAM16,PSMD10,RPS3A,RPS3,MA
,DHRS4,TRAK1,GOSR2,RPS7,KIF13B,UBA52,RPS4X,VTI1B,RPS23,RPL30,PEX16,OS9,UBE2L3,RPS15A,RPS1

,BAX,RPL35,NT5E,GALT,RPL34,RPL24,PSMD7,PAM16,PSMD10,RPS3A,RPS3,MAGOHB,ITPA,SMUG1,HMO
,BAX,RPL35,NT5E,GALT,RPL34,RPL24,PSMD7,PAM16,PSMD10,RPS3A,RPS3,MAGOHB,ITPA,SMUG1,HMO
ALRD3,DRG2,BRF1,MRPS18C,EIF2B3,RPS11,MRPS17,RPL37A,ELOF1,EIF4A1,RPLP0,METTL3,EIF3G,MRPL5
1,UBA52,RPS4X,MLH1,RPS23,RPL30,PSMD8,RQCD1,EXOSC2,RBM7,PSMC5,HNRNPM,EXOSC8,RPS15A,R
D1,ITGB1BP1,PARD6G,JPH2,MICAL1,YIF1A,JUP,CDC42,PTK2,DEF8,PPP1CC,VAMP8,PPIL2,RPL28,TIMM17

IN,ACOT2,PEX2,TNKS,RPL37A,PEX26,KPNA5,RPLP0,HAX1,CIZ1,SSR3,MPV17,SSB,ARIH2,RPL8,BAX,RPL35,
E1,GADL1,PSMD4,RPL28,VIM,PSMA7,RPL21,LYPLA2,CAMLG,EXOSC9,ZFAND2B,ACAD10,HNRNPC,SHARP
L,RPL15,MTHFS,COA3,MPC2,TYMS,GBA2,DALRD3,DRG2,BRF1,MRPS18C,PARK7,EIF2B3,RPS11,MRPS17,A
ARCA4,COA3,SNRPF,GBA2,BRF1,TCP1,CCDC40,ZNHIT3,NOP2,COX20,COA1,COBL,USP4,NDUFS3,RPLP0,HA
MD4,TMEM126B,APEH,TXNL4A,TCF4,ATPAF2,PRMT1,STOML2,PPAN,LAMP2,HSP90AB1,ASPH,TRMT112,
RPL29,AUP1,UGGT2,UBE2I,XBP1,PTK2,GADL1,VAMP8,PSMD4,RPL28,VIM,PSMA7,RPL21,CAMLG,EXOSC9
IRNPA1,POLR2C,INTS4,SNRPF,RBFOX2,PPIH,NSUN5,LGALS3,THADA,NHP2,ZNHIT3,NOP2,ELP4,USP4,MPH
PS25,PLCB2,RPS6,RPL29,AUP1,UGGT2,UBE2I,XBP1,PTK2,USE1,GADL1,VAMP8,PSMD4,RPL28,VIM,PSMA
P8,RPL28,TIMM17B,RPL21,CAPN10,NUTF2,STRADA,ZFAND2B,AP1S3,SLC9B2,RPL36,LAMP2,COPZ1,HSP9
3,TNFRSF14,VAMP8,RPL28,TIMM17B,RPL21,CAPN10,NUTF2,STRADA,ZFAND2B,AP1S3,SLC9B2,RPL36,LA
,TSG101,OBSL1,TTC21B,FUZ,ECSIT,EPB41L5,ATG12,NLRP1,NIP7,DUSP22,SPOCK2,CENPK,RPS6,RTEL1,SR
OSC2,NGDN,EXOSC8,DDX17,RUVBL1,RPS17,SFSWAP,EIF6,DHX30,ERAL1,DDX56,MRPS7,RPL27,MRPL36,S
ARS,RPL36,RBM3,MAPKAPK5,DGUOK,DUT,ATP5J,TSFM,TRMT112,PNPO,CHPF,RPS20,MPDU1,RPS15,DCT

P8,RPL28,TIMM17B,RPL21,CAPN10,NUTF2,STRADA,ZFAND2B,AP1S3,SLC9B2,RPL36,LAMP2,COPZ1,HSP9
P8,RPL28,TIMM17B,RPL21,CAPN10,NUTF2,STRADA,ZFAND2B,AP1S3,SLC9B2,RPL36,LAMP2,COPZ1,HSP9
ABL5,ITGB1BP1,YIF1A,JUP,CDC42,XBP1,GUK1,USE1,RFX3,TNFRSF14,VAMP8,PPIL2,RPL28,TIMM17B,RPL2
CBP2,POLR2E,RNPS1,NSRP1,RBM7,HNRNPM,IK,DDX17,PABPN1,SFSWAP,PPIE,PUF60,SNRNP40,RBM4B,
IRPN,IL1RAP,RFX5,POLD4,UBE2L6,ZC3H14,IRF9,PHF19,PCBP2,SON,TET2,SNF8,HMGA2,RPS16,CPSF3L,TS
,ERCC5,MCTS1,NAP1L1,RANGRF,HMGA1,DBNL,RBMX,HSD17B10,B2M,PDE4DIP,MED6,SEC31A,PRLR,SH3
OT8,RPS9,MRPL33,RUVBL2,TESC,NUDT5,RBPMS,NNMT,SNRPN,IL1RAP,RFX5,POLD4,UBE2L6,ZC3H14,IRF
1,TUBA1C,DST,PHF19,SON,TET2,SNF8,HMGA2,CDC37,TSG101,OBSL1,TTC21B,YLPM1,PHF21A,FUZ,MUC1

0,SNRNP40,RBM4B,SNRNP200,HNRNPL,MAGOH,LUC7L3,PRPF31,POLR2L,SRSF5,PAPOLA,WDR83,FXR1
4,FCHO2,TP53,RPS3A,RPS3,RDX,DNAJA3,MYL12A,RPS7,UBA52,KIF13A,RPS4X,VTI1B,PRKCE,RPS23,RPL30

B2,TRAK1,NDUFB3,NDUFB2,NDUFA10,NDUFA7,NDUFS2,NDUFC1,CAMKMT,UBE2L3,PPP1R13B,WDR45,
IRPN,IL1RAP,RFX5,POLD4,UBE2L6,ZC3H14,IRF9,PHF19,PCBP2,SON,TET2,SNF8,HMGA2,RPS16,CPSF3L,TS
IEDD4L,RNF130,BFAR,AXIN1,PSMD10,PML,UBE2D3,VPS25,UCHL5,NSFL1C,IFI27,ARRB2,RPS7,USP15,TAF
NUDT1,ANAPC11,RAD23A,HSD17B10,ABHD17A,GGT5,NT5C,USP45,PSME1,AGAP3,EGFR,HINT1,LGMN,E

)4,RPS6,RPL29,SRSF1,AUP1,RABL5,ITGB1BP1,YIF1A,JUP,CDC42,XBP1,GUK1,USE1,RFX3,TNFRSF14,VAMP8
CMT1,TESC,RBPMS,STYXL1,UBE2L6,VRK3,FIS1,PPM1K,PHF19,VCAN,PCBP2,CD44,TET2,SNF8,GRK6,HMGA
12,NLRP1,NIP7,DUSP22,SPOCK2,CENPK,RTEL1,SRSF1,FOXRED1,RABL5,ITGB1BP1,PARD6G,SH3YL1,MICAL
T6,NUMA1,SAE1,NPRL2,ARHGAP6,SCARB1,CTTN,ITSN1,EIF5A,TNFSF10,RUNX1,TAZ,SH2B1,NOP16,RPL13A
SF3L,TSG101,TTC21B,PSMA1,PHF21A,CHCHD1,PSMA3,MUC1,METTL17,PPRC1,RPS25,DUSP22,RPS6,RPL2
IRPN,IL1RAP,RFX5,POLD4,UBE2L6,ZC3H14,IRF9,PHF19,PCBP2,SON,TET2,SNF8,HMGA2,RPS16,CPSF3L,TSG
,PPP2R5C,FAM192A,PCNP,PSMD7,NEDD4L,RNF130,BFAR,CLTA,AXIN1,PSMD10,PML,TMEM199,UBE2D3,

RNF130,BFAR,AXIN1,PSMD10,PML,UBE2D3,VPS25,UCHL5,NSFL1C,IFI27,ARRB2,RPS7,USP15,TAF9,UBA52,
RPD1,SNRPA1,U2AF1L4,NCBP2,POLR2E,RNPS1,NSRP1,RBM7,HNRNPM,IK,DDX17,PABPN1,SFSWAP,PPIE,F
RDC1,TRPT1,ZC3HAV1,SCAF11,DBI,LIMK2,NOB1,SIRT6,NUMA1,SUMF2,BUD31,SAE1,NPRL2,OSGEP,SCAF
RNF130,BFAR,AXIN1,PSMD10,PML,UBE2D3,VPS25,UCHL5,NSFL1C,IFI27,ARRB2,RPS7,USP15,TAF9,UBA52,
3P2,SON,TET2,SNF8,HMGA2,RPS16,CPSF3L,TSG101,TTC21B,PSMA1,YLPM1,PHF21A,PSMA3,MUC1,PPRC1
PMS,NNMT,SNRPN,IL1RAP,RFX5,POLD4,UBE2L6,ZC3H14,IRF9,PHF19,PCBP2,SON,TET2,SNF8,HMGA2,RPS

,RFX3,TNFRSF14,VAMP8,RPL28,TIMM17B,RPL21,CAPN10,NUTF2,STRADA,ZFAND2B,AP1S3,SLC9B2,RPL3
21B,PSMA1,PHF21A,CHCHD1,PSMA3,MUC1,METTL17,ATG12,PPRC1,RPS25,DUSP22,RPS6,RPL29,RTEL1,S
P40,USP4,ARIH2,FBXL18,UBE2D4,RBBP6,UBA1,RNF8,PPP2R5C,FAM192A,PCNP,PSMD7,NEDD4L,RNF130
M192A,PCNP,PSMD7,NEDD4L,RNF130,BFAR,AXIN1,PSMD10,PML,UBE2D3,VPS25,UCHL5,NSFL1C,IFI27,AF
L,TSG101,TTC21B,PSMA1,PHF21A,CHCHD1,PSMA3,MUC1,METTL17,ATG12,PPRC1,RPS25,DUSP22,RPS6,
SP38,TAZ,RPL13A,GSTK1,TTLL5,LDHA,PSMA6,ACOT8,RPS9,RCE1,MRPL33,RUVBL2,PCMT1,TESC,RBPMS,N

LD4,UBE2L6,IRF9,PHF19,VCAN,LPCAT3,TET2,SNF8,HMGA2,RPS16,CPSF3L,TSG101,TTC21B,PTPN13,PSMA
GAP6,SCARB1,CTTN,ITSN1,EIF5A,TNFSF10,RUNX1,TAZ,SH2B1,RPL13A,GSTK1,ACOT8,MRPL33,RUVBL2,NU

MP1,CAPN2,MRPS16,MRPS33,WDR45,MRPL51,MRPL43,MRPL48,ERAL1,SMARCE1,MRPL27,VIL1,MRPS7,

IAX1,METTL3,ARIH2,BAX,CDK4,FAM192A,ADRA1A,LAMTOR5,VDAC1,NUP62,PSMD7,PAM16,NEDD4L,AXI
LD4,UBE2L6,IRF9,PHF19,VCAN,LPCAT3,TET2,SNF8,HMGA2,RPS16,CPSF3L,TSG101,TTC21B,PTPN13,PSMA

RPS9,RCE1,MRPL33,RUVBL2,PCMT1,TESC,RBPMS,IL1RAP,STYXL1,UBE2L6,VRK3,FIS1,PPM1K,PHF19,VCAN

L01,TTC21B,PSMA1,PHF21A,PSMA3,MUC1,PPRC1,RPS25,DUSP22,RPS6,RPL29,SRSF1,SAP30BP,TRNT1,ITG

LD4,UBE2L6,IRF9,PHF19,VCAN,LPCAT3,TET2,SNF8,HMGA2,RPS16,CPSF3L,TSG101,TTC21B,PTPN13,PSMA

C,ZNF235,BOP1,ARRDC1,TRPT1,ZC3HAV1,SCAF11,DBI,LIMK2,NOB1,SIRT6,NUMA1,SUMF2,BUD31,SAE1,I
DCD5,POT1,ANAPC11,PDE4DIP,NAA10,SH3GLB1,CHEK1,SSH2,CAMK2D,HNRNPA1,HIGD1A,GBA2,TCP1,PA

,ADRA1A,LAMTOR5,VDAC1,PSMD7,PAM16,AXIN1,TP53,PSMD10,PML,RHEB,RDX,UCHL5,MLST8,RPS7,LA

;,ZSWIM7,UBXN1,ERLEC1,HSPB1,ERCC5,MCTS1,HMGA1,PSME2,DBNL,RAD1,BRIP1,SERPINB6,WAC,NUDT

F5A,TNFSF10,RUNX1,UFD1L,USP38,SH2B1,RPL13A,TTLL5,PSMA6,ACOT8,RPS9,RCE1,MRPL33,RUVBL2,PCI

TCAIM,SLC3A2,ACTR6,MAP3K7CL,SEC62,SET,CTSC,ZNF235,BOP1,ZC3HAV1,SCAF11,DBI,ARPC2,LIMK2,VD
1L,SH2B1,RPL13A,GSTK1,TTLL5,LDHA,PSMA6,ACOT8,RPS9,RCE1,RUVBL2,PCMT1,TESC,NUDT5,RBPMS,NI
75,ATG16L2,RCC1,SNRPB,TCAIM,SLC3A2,ACTR6,MAP3K7CL,SEC62,SET,CTSC,ZNF235,BOP1,ARRDC1,KLC
K,TMEM135,RNF32,ATG16L2,RCC1,SNRPB,PARD3,TCAIM,SLC3A2,ACTR6,MAP3K7CL,SEC62,SET,CTSC,ZNI
1,FASTK,TMEM135,RNF32,ATG16L2,RCC1,SNRPB,PARD3,TCAIM,SLC3A2,DHPS,ACTR6,MAP3K7CL,SEC62,
STARD10,RNH1,DNASE1L1,KIF4A,RBCK1,PTOV1,FASTK,TMEM135,RNF32,ATG16L2,RCC1,SNRPB,PARD3,T
OV1,FASTK,TMEM135,RNF32,LY75,ATG16L2,RCC1,SNRPB,PARD3,TCAIM,SLC3A2,ACTR6,MAP3K7CL,SEC6
RPL15,SNRPF,PPIH,MRPS18C,LGALS3,NHP2,ZNHIT3,RPS11,MRPS17,RPL37A,MPHOSPH10,RPLP0,EIF3G,N
D5,RUNX1,CHID1,NUDT22,UFD1L,NOP16,RPL13A,GSTK1,PSMA6,ACOT8,RPS9,MRPL33,RUVBL2,TESC,RB
D5,RUNX1,CHID1,NUDT22,UFD1L,NOP16,RPL13A,GSTK1,PSMA6,ACOT8,RPS9,MRPL33,RUVBL2,TESC,RB
D5,RUNX1,CHID1,NUDT22,UFD1L,NOP16,RPL13A,GSTK1,PSMA6,ACOT8,RPS9,MRPL33,RUVBL2,TESC,RB
AE1,OSGEP,ITSN1,EIF5A,COMMD5,RUNX1,CHID1,NUDT22,UFD1L,THYN1,SH2B1,NOP16,RPL13A,TTLL5,L

PLEKHA5,UBE2L6,VRK3,ZC3H14,IRF9,DST,PHF19,PCBP2,MAGED2,SON,SNF8,HMGA2,RPS16,CPSF3L,PTPN

RFX5,POLD4,PLEKHA5,UBE2L6,VRK3,ZC3H14,IRF9,DST,PHF19,PCBP2,MAGED2,SON,SNF8,HMGA2,RPS16
SMA6,AP3B2,RPS9,MRPL33,RUVBL2,SNRPN,POLD4,RANBP1,UBE2L6,ZC3H14,FIS1,STIP1,LRRC1,PHF19,C
X5A,HIGD1A,NDUFV2,SIRT5,COA3,MPC2,TYMS,SLC25A23,MFSD10,MRPS18C,LGALS3,PARK7,MFF,ENY2
X5A,HIGD1A,NDUFV2,SIRT5,COA3,MPC2,TYMS,SLC25A23,MFSD10,MRPS18C,LGALS3,PARK7,MFF,ENY2

DUFB3,NDUFB2,NDUFA10,MRPS16,SDHA,NDUFA7,NDUFS2,MRPS33,MCUR1,NDUFC1,DUSP18,NDUFB10
TMEM14B,COA1,NDUFS3,HAX1,MPV17,MRPL55,BAX,NME1,VDAC1,PAM16,NDUFA11,NRP1,SLC25A24,N
RPS3,MRPS24,SPHK2,IFI27,NDUFB3,NDUFB2,NDUFA10,MRPS16,SDHA,NDUFA7,NDUFS2,MRPS33,MCUR
UVBL2,RBPMS,WDR62,RFX5,POLD4,RANBP1,VRK3,ZC3H14,TUBA1C,IRF9,DST,MAGED2,HMGA2,RPS16,T
UVBL2,RBPMS,WDR62,RFX5,POLD4,RANBP1,VRK3,ZC3H14,TUBA1C,IRF9,DST,MAGED2,HMGA2,RPS16,T
1,POLR3H,RAD23A,HSD17B10,DDX41,PSME1,NAA10,MED6,CARHSP1,HINT1,HNRNPA1,POLR2C,NDUFV
,NSUN5,OARD1,PHF6,SLC30A5,NHP2,UBE2N,RPS11,NOP2,FAM32A,RPAIN,NEK2,POLR1C,MPHOSPH10,S
ATP5J,TSFM,CHPF,CHCHD7,TOMM40,NUDT1,HSD17B10,ILF3,MDH2,CHCHD5,SGK1,SH3GLB1,COX17,COX

RPL55,BAX,NME1,VDAC1,PAM16,NDUFA11,NRP1,SLC25A24,NDUFA3,RPS3,MRPS24,AIFM2,SPHK2,IFI27,

A7,APEH,LYPLA2,NUTF2,PODXL,EXOSC9,TXNDC17,HNRNPC,STAU1,MOB1A,MDH1,LAMP2,HSP90AB1,DL
A7,APEH,LYPLA2,NUTF2,PODXL,EXOSC9,TXNDC17,HNRNPC,STAU1,MOB1A,MDH1,LAMP2,HSP90AB1,DL
A7,APEH,LYPLA2,NUTF2,PODXL,EXOSC9,TXNDC17,HNRNPC,STAU1,MOB1A,MDH1,LAMP2,HSP90AB1,DL
2,NDUFA7,NDUFS2,MRPS33,ACSS1,ME3,MRPL51,MRPL43,DHX30,MRPL48,ERAL1,ACOT9,MTHFD2L,MRF

25A10,THEM4,RDH11,TMED4,COASY,AUP1,FOXRED1,KXD1,JPH2,HS2ST1,SLC39A13,YIF1A,MAGT1,CDC4

G101,PTPN13,UBR4,PSMA1,FUZ,PSMA3,MUC1,ATG12,RPS25,SDF4,NISCH,TMED4,COASY,AUP1,YIPF3,YI

DA,PIGT,PDCD6,TMEM147,PIGH,ERGIC1,DGAT1,SSR3,CREB3L4,ARL6IP5,BAX,UBA1,FAM69B,NUP62,PIG
T,PDCD6,TMEM147,PIGH,ERGIC1,DGAT1,SSR3,CREB3L4,ARL6IP5,BAX,UBA1,FAM69B,PIGO,BFAR,KDELR3

,RPS3A,RPS3,RDX,SGSM3,RPS7,ILK,ECT2,FHL2,RPS4X,VAPA,CAPN2,ARL2,RPL30,LPP,MPRIP,ANXA5,ACTB

ANGRF,PHTF2,SLC33A1,NOMO2,B2M,DDX41,EEF1D,SGK1,SEC31A,RPL18,ATP10D,EGFR,CAMK2D,REEP5

,PTPRB,DBNL,TOMM40,SERPINB6,ATP6V1F,B2M,ABHD17A,STXBP2,SEC31A,EGFR,SH3GLB1,CAMK2D,M

;2,RPS27L,RPS7,UBA52,RPS4X,MRPS16,RPS23,RPL30,MRPS33,RPS15A,RPS17,MRPL51,MRPL43,MRPL48,

PS3,RPS27L,RPS7,NCBP2,POLR2E,TAF9,UBA52,RPS4X,RPS23,RPL30,PSMD8,NUP85,PSMC5,AP2S1,PABPN

;11,NOP2,NUP133,RPL37A,MPHOSPH10,EIF4A1,RPLP0,METTL3,TSEN34,RPL8,CHTOP,RPL35,SRSF7,NXF1

)10,RNH1,DNASE1L1,KIF4A,RBCK1,PTOV1,FASTK,TMEM135,RNF32,LY75,ATG16L2,RCC1,SNRPB,PARD3,T(
EM135,RNF32,LY75,ATG16L2,RCC1,PARD3,TCAIM,SLC3A2,DHPS,ACTR6,SEC62,SET,CTSC,BOP1,ARRDC1,T
1,NPRL2,ARHGAP6,SCARB1,CTTN,EIF5A,COMMD5,CHID1,NUDT22,AMZ2,UFD1L,USP38,TAZ,THYN1,SH2B
P,CIRBP,RHOF,SMARCB1,RPL38,PRR13,STARD10,RNH1,DNASE1L1,KIF4A,RBCK1,PTOV1,FASTK,TMEM135
P,CIRBP,RHOF,SMARCB1,RPL38,PRR13,STARD10,RNH1,DNASE1L1,KIF4A,RBCK1,PTOV1,FASTK,TMEM135
H1,DNASE1L1,KIF4A,RBCK1,PTOV1,FASTK,TMEM135,RNF32,LY75,ATG16L2,RCC1,PARD3,TCAIM,SLC3A2,I
H1,DNASE1L1,KIF4A,RBCK1,PTOV1,FASTK,TMEM135,RNF32,LY75,ATG16L2,RCC1,PARD3,TCAIM,SLC3A2,I
(IF4A,RBCK1,PTOV1,FASTK,TMEM135,RNF32,LY75,ATG16L2,RCC1,SNRPB,PARD3,TCAIM,SLC3A2,DHPS,A(
PS,ACTR6,SEC62,CTSC,ZNF235,BOP1,ARRDC1,TRPT1,KLC1,RBP5,ZC3HAV1,SCAF11,DBI,ARPC2,PMFBP1,LII
PS,ACTR6,SEC62,CTSC,ZNF235,BOP1,ARRDC1,TRPT1,KLC1,RBP5,ZC3HAV1,SCAF11,DBI,ARPC2,PMFBP1,LII
(1,PTOV1,FASTK,TMEM135,RNF32,LY75,ATG16L2,RCC1,SNRPB,PARD3,TCAIM,SLC3A2,DHPS,ACTR6,SEC6
;UMF2,BUD31,SAE1,NPRL2,CTTN,TSPAN11,EIF5A,COMMD5,CHID1,NUDT22,AMZ2,UFD1L,USP38,THYN1,
;UMF2,BUD31,SAE1,NPRL2,CTTN,TSPAN11,EIF5A,COMMD5,CHID1,NUDT22,AMZ2,UFD1L,USP38,THYN1,
GSTK1,PSMA6,ACOT8,RPS9,RCE1,MRPL33,ZNF410,RUVBL2,IGSF8,NUDT5,WDR62,STYXL1,VRK3,PPM1K,T
L,KLC1,RBP5,ZC3HAV1,SCAF11,DBI,ARPC2,PMFBP1,LIMK2,NOB1,SIRT6,NUMA1,GPATCH2L,SUMF2,BUD3
L,KLC1,RBP5,ZC3HAV1,SCAF11,DBI,ARPC2,PMFBP1,LIMK2,NOB1,SIRT6,NUMA1,GPATCH2L,SUMF2,BUD3
TK,TMEM135,RNF32,LY75,ATG16L2,RCC1,SNRPB,PARD3,TCAIM,SLC3A2,DHPS,ACTR6,SEC62,SET,CTSC,ZN

TK,TMEM135,RNF32,LY75,ATG16L2,RCC1,SNRPB,PARD3,TCAIM,SLC3A2,DHPS,ACTR6,SEC62,SET,CTSC,ZN

PMFBP1,LIMK2,SIRT6,NUMA1,GPATCH2L,SUMF2,BUD31,SAE1,NPRL2,ARHGAP6,CTTN,TSPAN11,EIF5A,C

PMFBP1,LIMK2,SIRT6,NUMA1,GPATCH2L,SUMF2,BUD31,SAE1,NPRL2,ARHGAP6,CTTN,TSPAN11,EIF5A,C

,NUMA1,GPATCH2L,SUMF2,BUD31,SAE1,NPRL2,CTTN,TSPAN11,EIF5A,COMMD5,CHID1,NUDT22,AMZ2,

,NUMA1,GPATCH2L,SUMF2,BUD31,SAE1,NPRL2,CTTN,TSPAN11,EIF5A,COMMD5,CHID1,NUDT22,AMZ2,

OP1,ARRDC1,TRPT1,KLC1,RBP5,ZC3HAV1,SCAF11,DBI,ARPC2,PMFBP1,LIMK2,NOB1,SIRT6,NUMA1,GPAT

OP1,ARRDC1,TRPT1,KLC1,RBP5,ZC3HAV1,SCAF11,DBI,ARPC2,PMFBP1,LIMK2,NOB1,SIRT6,NUMA1,GPAT

RPL38,PRR13,STARD10,RNH1,DNASE1L1,KIF4A,RBCK1,PTOV1,FASTK,TMEM135,RNF32,LY75,ATG16L2,R

RPL38,PRR13,STARD10,RNH1,DNASE1L1,KIF4A,RBCK1,PTOV1,FASTK,TMEM135,RNF32,LY75,ATG16L2,R

RBP5,ZC3HAV1,SCAF11,DBI,ARPC2,PMFBP1,LIMK2,SIRT6,NUMA1,GPATCH2L,SUMF2,BUD31,SAE1,NPRL2

RBP5,ZC3HAV1,SCAF11,DBI,ARPC2,PMFBP1,LIMK2,SIRT6,NUMA1,GPATCH2L,SUMF2,BUD31,SAE1,NPRL2

,TCAIM,SLC3A2,DHPS,ACTR6,SEC62,CTSC,ZNF235,BOP1,ARRDC1,TRPT1,KLC1,RBP5,ZC3HAV1,SCAF11,DE

,TCAIM,SLC3A2,DHPS,ACTR6,SEC62,CTSC,ZNF235,BOP1,ARRDC1,TRPT1,KLC1,RBP5,ZC3HAV1,SCAF11,DE

T22,UFD1L,THYN1,SH2B1,RPL13A,ARHGEF39,GSTK1,TTLL5,PSMA6,ACOT8,RPS9,SLC35G1,RCE1,MRPL33,

T22,UFD1L,THYN1,SH2B1,RPL13A,ARHGEF39,GSTK1,TTLL5,PSMA6,ACOT8,RPS9,SLC35G1,RCE1,MRPL33,

ARD3,TCAIM,SLC3A2,DHPS,ACTR6,SEC62,CTSC,BOP1,ARRDC1,TRPT1,KLC1,RBP5,ZC3HAV1,SCAF11,DBI,A

ARD3,TCAIM,SLC3A2,DHPS,ACTR6,SEC62,CTSC,BOP1,ARRDC1,TRPT1,KLC1,RBP5,ZC3HAV1,SCAF11,DBI,A

D3,TCAIM,SLC3A2,DHPS,ACTR6,SEC62,CTSC,ZNF235,BOP1,ARRDC1,TRPT1,KLC1,RBP5,ZC3HAV1,SCAF11,

D3,TCAIM,SLC3A2,DHPS,ACTR6,SEC62,CTSC,ZNF235,BOP1,ARRDC1,TRPT1,KLC1,RBP5,ZC3HAV1,SCAF11,

TOV1,FASTK,TMEM135,RNF32,LY75,ATG16L2,SNRPB,PARD3,TCAIM,SLC3A2,DHPS,ACTR6,SEC62,SET,CTS

MAP3K7CL,CTSC,ZNF235,BOP1,ARRDC1,TRPT1,KLC1,RBP5,ZC3HAV1,SCAF11,DBI,ARPC2,PMFBP1,LIMK2,

MAP3K7CL,CTSC,ZNF235,BOP1,ARRDC1,TRPT1,KLC1,RBP5,ZC3HAV1,SCAF11,DBI,ARPC2,PMFBP1,LIMK2,

,AMZ2,USP38,THYN1,SH2B1,NOP16,RPL13A,ARHGEF39,TTLL5,PSMA6,ACOT8,RPS9,SLC35G1,RCE1,MRP

35,RNF32,LY75,ATG16L2,RCC1,SNRPB,PARD3,TCAIM,SLC3A2,DHPS,ACTR6,SEC62,SET,CTSC,BOP1,ARRD

35,RNF32,LY75,ATG16L2,RCC1,SNRPB,PARD3,TCAIM,SLC3A2,DHPS,ACTR6,SEC62,SET,CTSC,BOP1,ARRD

9,RCE1,ZNF410,NUDT5,STYXL1,VRK3,TUBA1C,DST,STIP1,LRRC1,SON,UQCRQ,SNF8,CDC37,CPSF3L,TSG10

NH1,DNASE1L1,KIF4A,RBCK1,PTOV1,FASTK,TMEM135,RNF32,LY75,ATG16L2,RCC1,PARD3,TCAIM,SLC3A

,SMARCB1,RPL38,PRR13,STARD10,RNH1,DNASE1L1,KIF4A,RBCK1,PTOV1,FASTK,TMEM135,RNF32,LY75,

,SMARCB1,RPL38,PRR13,STARD10,RNH1,DNASE1L1,KIF4A,RBCK1,PTOV1,FASTK,TMEM135,RNF32,LY75,

MFBP1,LIMK2,SIRT6,NUMA1,GPATCH2L,SUMF2,BUD31,SAE1,NPRL2,SCARB1,CTTN,TSPAN11,EIF5A,COM

MFBP1,LIMK2,SIRT6,NUMA1,GPATCH2L,SUMF2,BUD31,SAE1,NPRL2,SCARB1,CTTN,TSPAN11,EIF5A,COM

HAV1,SCAF11,DBI,ARPC2,PMFBP1,LIMK2,NOB1,SIRT6,NUMA1,GPATCH2L,SUMF2,BUD31,SAE1,NPRL2,A

HAV1,SCAF11,DBI,ARPC2,PMFBP1,LIMK2,NOB1,SIRT6,NUMA1,GPATCH2L,SUMF2,BUD31,SAE1,NPRL2,A

32,LY75,ATG16L2,RCC1,SNRPB,PARD3,TCAIM,SLC3A2,DHPS,ACTR6,MAP3K7CL,SEC62,SET,CTSC,ZNF235,

32,LY75,ATG16L2,RCC1,SNRPB,PARD3,TCAIM,SLC3A2,DHPS,ACTR6,MAP3K7CL,SEC62,SET,CTSC,ZNF235,

DBI,ARPC2,PMFBP1,LIMK2,SIRT6,NUMA1,GPATCH2L,SUMF2,BUD31,SAE1,NPRL2,SCARB1,CTTN,TSPAN1

DBI,ARPC2,PMFBP1,LIMK2,SIRT6,NUMA1,GPATCH2L,SUMF2,BUD31,SAE1,NPRL2,SCARB1,CTTN,TSPAN1

3A2,DHPS,ACTR6,CTSC,ZNF235,BOP1,ARRDC1,TRPT1,KLC1,RBP5,ZC3HAV1,SCAF11,DBI,ARPC2,PMFBP1,

3A2,DHPS,ACTR6,CTSC,ZNF235,BOP1,ARRDC1,TRPT1,KLC1,RBP5,ZC3HAV1,SCAF11,DBI,ARPC2,PMFBP1,

F2,BUD31,NPRL2,CTTN,TSPAN11,EIF5A,COMMD5,CHID1,NUDT22,AMZ2,UFD1L,THYN1,RPL13A,ARHGEF

F2,BUD31,NPRL2,CTTN,TSPAN11,EIF5A,COMMD5,CHID1,NUDT22,AMZ2,UFD1L,THYN1,RPL13A,ARHGEF

MK2,SIRT6,NUMA1,GPATCH2L,SUMF2,BUD31,SAE1,NPRL2,CTTN,TSPAN11,EIF5A,COMMD5,CHID1,NUD

MK2,SIRT6,NUMA1,GPATCH2L,SUMF2,BUD31,SAE1,NPRL2,CTTN,TSPAN11,EIF5A,COMMD5,CHID1,NUD

CK1,PTOV1,FASTK,TMEM135,RNF32,LY75,ATG16L2,RCC1,SNRPB,PARD3,TCAIM,DHPS,ACTR6,SEC62,SET

MA6,ACOT8,RPS9,RCE1,ZNF410,RUVBL2,PCMT1,IGSF8,NUDT5,STYXL1,VRK3,TUBA1C,IRF9,ANTXR2,STIP1

E1,ZNF410,RUVBL2,IGSF8,NUDT5,SPAG7,STYXL1,VRK3,PPM1K,IRF9,ANTXR2,STIP1,LRRC1,SON,UQCRQ,S

2,SIRT6,NUMA1,GPATCH2L,SUMF2,BUD31,SAE1,NPRL2,CTTN,TSPAN11,EIF5A,COMMD5,CHID1,NUDT22

K2,SIRT6,NUMA1,GPATCH2L,SUMF2,BUD31,SAE1,NPRL2,CTTN,TSPAN11,EIF5A,COMMD5,CHID1,NUDT22
P1,LIMK2,SIRT6,NUMA1,GPATCH2L,SUMF2,BUD31,SAE1,NPRL2,CRYGS,CTTN,TSPAN11,EIF5A,COMMD5,
P1,LIMK2,SIRT6,NUMA1,GPATCH2L,SUMF2,BUD31,SAE1,NPRL2,CRYGS,CTTN,TSPAN11,EIF5A,COMMD5,
NUMA1,GPATCH2L,SUMF2,BUD31,SAE1,NPRL2,CRYGS,CTTN,TSPAN11,EIF5A,COMMD5,CHID1,NUDT22,A
NUMA1,GPATCH2L,SUMF2,BUD31,SAE1,NPRL2,CRYGS,CTTN,TSPAN11,EIF5A,COMMD5,CHID1,NUDT22,A
NPRL2,SCARB1,CTTN,TSPAN11,EIF5A,COMMD5,CHID1,NUDT22,UFD1L,TAZ,SH2B1,RPL13A,ARHGEF39,TT
NPRL2,SCARB1,CTTN,TSPAN11,EIF5A,COMMD5,CHID1,NUDT22,UFD1L,TAZ,SH2B1,RPL13A,ARHGEF39,TT
,BUD31,NPRL2,CTTN,TSPAN11,EIF5A,COMMD5,CHID1,NUDT22,AMZ2,UFD1L,TAZ,THYN1,DBNDD1,RPL13
,BUD31,NPRL2,CTTN,TSPAN11,EIF5A,COMMD5,CHID1,NUDT22,AMZ2,UFD1L,TAZ,THYN1,DBNDD1,RPL13
,SIRT6,NUMA1,GPATCH2L,SUMF2,BUD31,SAE1,NPRL2,ARHGAP6,CTTN,TSPAN11,EIF5A,COMMD5,CHID1,
,SIRT6,NUMA1,GPATCH2L,SUMF2,BUD31,SAE1,NPRL2,ARHGAP6,CTTN,TSPAN11,EIF5A,COMMD5,CHID1,
PS,ACTR6,MAP3K7CL,ZNF235,BOP1,ARRDC1,TRPT1,KLC1,RBP5,ZC3HAV1,SCAF11,DBI,ARPC2,PMFBP1,LI
PS,ACTR6,MAP3K7CL,ZNF235,BOP1,ARRDC1,TRPT1,KLC1,RBP5,ZC3HAV1,SCAF11,DBI,ARPC2,PMFBP1,LI
SIRT6,NUMA1,GPATCH2L,SUMF2,BUD31,SAE1,NPRL2,ARHGAP6,CTTN,TSPAN11,EIF5A,COMMD5,CHID1,
SIRT6,NUMA1,GPATCH2L,SUMF2,BUD31,SAE1,NPRL2,ARHGAP6,CTTN,TSPAN11,EIF5A,COMMD5,CHID1,
PM1,FAM3A,ATG12,NIP7,TTC9C,THEM4,DBNDD2,RPS6,RPL29,RTEL1,SRSF1,AUP1,SAP30BP,YIPF3,FOXRE
C3HAV1,SCAF11,DBI,ARPC2,PMFBP1,LIMK2,SIRT6,NUMA1,GPATCH2L,SUMF2,BUD31,SAE1,NPRL2,ARHG
C3HAV1,SCAF11,DBI,ARPC2,PMFBP1,LIMK2,SIRT6,NUMA1,GPATCH2L,SUMF2,BUD31,SAE1,NPRL2,ARHG
UD31,SAE1,NPRL2,CTTN,TSPAN11,EIF5A,COMMD5,CHID1,NUDT22,UFD1L,USP38,THYN1,SH2B1,NOP16,
UD31,SAE1,NPRL2,CTTN,TSPAN11,EIF5A,COMMD5,CHID1,NUDT22,UFD1L,USP38,THYN1,SH2B1,NOP16,
C2,PMFBP1,LIMK2,SIRT6,NUMA1,GPATCH2L,SUMF2,BUD31,NPRL2,SCARB1,CTTN,TSPAN11,EIF5A,COMM
C2,PMFBP1,LIMK2,SIRT6,NUMA1,GPATCH2L,SUMF2,BUD31,NPRL2,SCARB1,CTTN,TSPAN11,EIF5A,COMM
MFBP1,LIMK2,SIRT6,NUMA1,GPATCH2L,SUMF2,BUD31,SAE1,NPRL2,CRYGS,CTTN,TSPAN11,EIF5A,COMM
MFBP1,LIMK2,SIRT6,NUMA1,GPATCH2L,SUMF2,BUD31,SAE1,NPRL2,CRYGS,CTTN,TSPAN11,EIF5A,COMM
6,RPL29,RTEL1,AUP1,YIPF3,TRNT1,KXD1,ITGB1BP1,MED15,CLASRP,C1ORF198,UBE2I,SLC39A13,EIF3M,C
,SIRT6,NUMA1,GPATCH2L,SUMF2,BUD31,NPRL2,SCARB1,CTTN,TSPAN11,EIF5A,COMMD5,CHID1,NUDT2
,SIRT6,NUMA1,GPATCH2L,SUMF2,BUD31,NPRL2,SCARB1,CTTN,TSPAN11,EIF5A,COMMD5,CHID1,NUDT2
,ACTR6,SEC62,SET,CTSC,ZNF235,BOP1,ARRDC1,TRPT1,KLC1,RBP5,DBI,LIMK2,NOB1,SIRT6,NUMA1,GPAT
,ACTR6,SEC62,SET,CTSC,ZNF235,BOP1,ARRDC1,TRPT1,KLC1,RBP5,DBI,LIMK2,NOB1,SIRT6,NUMA1,GPAT
L,TSG101,UBR4,METTL17,ATG12,NIP7,DUSP22,RPS6,RPL29,RTEL1,AUP1,SAP30BP,YIPF3,FOXRED1,TRNT
P1,YIPF3,FOXRED1,TRNT1,KXD1,ITGB1BP1,MED15,CLASRP,C1ORF198,UBE2I,EIF3M,CDC42,CARD14,PCY
PS,ACTR6,MAP3K7CL,ZNF235,BOP1,ARRDC1,TRPT1,KLC1,RBP5,ZC3HAV1,SCAF11,DBI,ARPC2,PMFBP1,LIN
PS,ACTR6,MAP3K7CL,ZNF235,BOP1,ARRDC1,TRPT1,KLC1,RBP5,ZC3HAV1,SCAF11,DBI,ARPC2,PMFBP1,LIN
TCAIM,SLC3A2,DHPS,ACTR6,SET,CTSC,ZNF235,BOP1,ARRDC1,TRPT1,KLC1,RBP5,ZC3HAV1,SCAF11,DBI,AF
TCAIM,SLC3A2,DHPS,ACTR6,SET,CTSC,ZNF235,BOP1,ARRDC1,TRPT1,KLC1,RBP5,ZC3HAV1,SCAF11,DBI,AF
38,PRR13,STARD10,RNH1,DNASE1L1,KIF4A,RBCK1,PTOV1,FASTK,TMEM135,RNF32,LY75,ATG16L2,RCC1
38,PRR13,STARD10,RNH1,DNASE1L1,KIF4A,RBCK1,PTOV1,FASTK,TMEM135,RNF32,LY75,ATG16L2,RCC1
UQCRQ,SNF8,CPSF3L,TSG101,UBR4,RPS25,NIP7,COX4I1,SLC25A10,TMX2,DBNDD2,RPS6,RPL29,RTEL1,AU
LC3A2,DHPS,ACTR6,MAP3K7CL,CTSC,ZNF235,BOP1,ARRDC1,TRPT1,KLC1,RBP5,ZC3HAV1,SCAF11,DBI,ARF
LC3A2,DHPS,ACTR6,MAP3K7CL,CTSC,ZNF235,BOP1,ARRDC1,TRPT1,KLC1,RBP5,ZC3HAV1,SCAF11,DBI,ARF
L,FASTK,TMEM135,RNF32,LY75,ATG16L2,SNRPB,PARD3,TCAIM,SLC3A2,DHPS,ACTR6,SEC62,SET,CTSC,ZN
L,FASTK,TMEM135,RNF32,LY75,ATG16L2,SNRPB,PARD3,TCAIM,SLC3A2,DHPS,ACTR6,SEC62,SET,CTSC,ZN
STYXL1,VRK3,LZTFL1,PPM1K,IRF9,ANTXR2,STIP1,LRRC1,SON,UQCRQ,SNF8,RPS16,CPSF3L,TSG101,UBR4,N
N,TSPAN11,EIF5A,COMMD5,CHID1,NUDT22,UFD1L,DBNDD1,SH2B1,NOP16,RPL13A,ARHGEF39,TTLL5,PS
N,TSPAN11,EIF5A,COMMD5,CHID1,NUDT22,UFD1L,DBNDD1,SH2B1,NOP16,RPL13A,ARHGEF39,TTLL5,PS
T1,KXD1,MED15,CLASRP,UBE2I,SLC39A13,EIF3M,CDC42,CARD14,PCYT2,FBXW9,USE1,ASF1B,ENDOV,PPIL
SF3L,TSG101,UBR4,CHCHD1,FAM3A,THEM4,RPS6,RPL29,RTEL1,AUP1,YIPF3,FOXRED1,TRNT1,KXD1,MED1

RPL13A,ARHGEF39,TTLL5,PSMA6,ACOT8,RPS9,SLC35G1,RCE1,ZNF410,IGSF8,NUDT5,SPAG7,STYXL1,VRK3
V1,SCAF11,DBI,ARPC2,PMFBP1,LIMK2,SIRT6,NUMA1,GPATCH2L,SUMF2,BUD31,SAE1,NPRL2,OSGEP,ARH
V1,SCAF11,DBI,ARPC2,PMFBP1,LIMK2,SIRT6,NUMA1,GPATCH2L,SUMF2,BUD31,SAE1,NPRL2,OSGEP,ARH
RBP5,ZC3HAV1,SCAF11,DBI,ARPC2,PMFBP1,LIMK2,SIRT6,NUMA1,GPATCH2L,SUMF2,BUD31,SAE1,NPRL2
RBP5,ZC3HAV1,SCAF11,DBI,ARPC2,PMFBP1,LIMK2,SIRT6,NUMA1,GPATCH2L,SUMF2,BUD31,SAE1,NPRL2
T8,RPS9,MRPL33,ZNF410,RUVBL2,IGSF8,NUDT5,WDR62,STYXL1,VRK3,LZTFL1,PPM1K,IRF9,ANTXR2,STIP
2,CTSC,ZNF235,BOP1,ARRDC1,TRPT1,KLC1,ZC3HAV1,SCAF11,DBI,ARPC2,PMFBP1,LIMK2,WDR19,SIRT6,N
2,CTSC,ZNF235,BOP1,ARRDC1,TRPT1,KLC1,ZC3HAV1,SCAF11,DBI,ARPC2,PMFBP1,LIMK2,WDR19,SIRT6,N
,SLC25A10,TMX2,DBNDD2,RPS6,RPL29,RTEL1,AUP1,SAP30BP,YIPF3,FOXRED1,TRNT1,KXD1,ITGB1BP1,M
1,PTOV1,FASTK,TMEM135,RNF32,LY75,ATG16L2,RCC1,PARD3,TCAIM,SLC3A2,DHPS,ACTR6,SEC62,SET,C
1,PTOV1,FASTK,TMEM135,RNF32,LY75,ATG16L2,RCC1,PARD3,TCAIM,SLC3A2,DHPS,ACTR6,SEC62,SET,C
XRED1,TRNT1,KXD1,MED15,C1ORF198,UBE2I,EIF3M,CDC42,CARD14,PCYT2,FBXW9,USE1,ASF1B,RFX3,E
D3,TCAIM,SLC3A2,ACTR6,SEC62,SET,ZNF235,BOP1,ARRDC1,TRPT1,KLC1,RBP5,DBI,LIMK2,SIRT6,NUMA1,
D3,TCAIM,SLC3A2,ACTR6,SEC62,SET,ZNF235,BOP1,ARRDC1,TRPT1,KLC1,RBP5,DBI,LIMK2,SIRT6,NUMA1,
AM3A,RPS25,THEM4,RPS6,RTEL1,SAP30BP,YIPF3,RABL5,TRNT1,ITGB1BP1,MED15,CLASRP,C1ORF198,U
AM3A,RPS25,THEM4,RPS6,RTEL1,SAP30BP,YIPF3,RABL5,TRNT1,ITGB1BP1,MED15,CLASRP,C1ORF198,U
E1,MRPL33,ZNF410,RUVBL2,PCMT1,IGSF8,NUDT5,WDR62,STYXL1,RFX5,PLEKHA5,PPM1K,TUBA1C,ANTX
,CTSC,BOP1,ARRDC1,TRPT1,KLC1,RBP5,SCAF11,DBI,ARPC2,LIMK2,VDAC3,NOB1,SIRT6,NUMA1,SUMF2,B
12,RPS25,NIP7,THEM4,RPS6,RPL29,RTEL1,AUP1,SAP30BP,YIPF3,FOXRED1,TRNT1,KXD1,MED15,IP6K2,C
IL2,LYPLA2,C17ORF62,PRMT1,PPAN,ACAD10,SUGT1,HYPK,CARS,ZNF197,DGUOK,LRRFIP1,ZNF524,ATP5J,
6,ACOT8,RPS9,RCE1,MRPL33,ZNF410,RUVBL2,IGSF8,NUDT5,RBPMS,SPAG7,WDR62,STYXL1,VRK3,PPM1
1,RPL13A,ARHGEF39,PSMA6,ACOT8,AP3B2,MRPL33,ZNF410,PCMT1,NUDT5,SNRPN,STYXL1,POLD4,VRK3
1,RPL13A,ARHGEF39,PSMA6,ACOT8,AP3B2,MRPL33,ZNF410,PCMT1,NUDT5,SNRPN,STYXL1,POLD4,VRK3
AP30BP,YIPF3,RABL5,TRNT1,ITGB1BP1,MED15,CLASRP,C1ORF198,UGGT2,UBE2I,EIF3M,CDC42,TBX6,CA
AP30BP,YIPF3,RABL5,TRNT1,ITGB1BP1,MED15,CLASRP,C1ORF198,UGGT2,UBE2I,EIF3M,CDC42,TBX6,CA
TN,TSPAN11,EIF5A,COMMD5,NUDT22,AMZ2,UFD1L,USP38,TAZ,SH2B1,RPL13A,ARHGEF39,TTLL5,PSMA6
TN,TSPAN11,EIF5A,COMMD5,NUDT22,AMZ2,UFD1L,USP38,TAZ,SH2B1,RPL13A,ARHGEF39,TTLL5,PSMA6
1L1,KIF4A,RBCK1,PTOV1,FASTK,TMEM135,LY75,ATG16L2,RCC1,SNRPB,PARD3,TCAIM,SLC3A2,DHPS,ACT
1L1,KIF4A,RBCK1,PTOV1,FASTK,TMEM135,LY75,ATG16L2,RCC1,SNRPB,PARD3,TCAIM,SLC3A2,DHPS,ACT
3L,TSG101,UBR4,ATG12,NIP7,THEM4,DBNDD2,RPS6,RPL29,RTEL1,AUP1,YIPF3,FOXRED1,TRNT1,KXD1,IT
5,WDR62,STYXL1,VRK3,PPM1K,TUBA1C,ANTXR2,STIP1,LRRC1,VCAN,SON,UQCRQ,SNF8,GRK6,CDC37,CP
RBCK1,PTOV1,FASTK,TMEM135,LY75,ATG16L2,RCC1,SNRPB,PARD3,TCAIM,SLC3A2,DHPS,ACTR6,MAP3K
RBCK1,PTOV1,FASTK,TMEM135,LY75,ATG16L2,RCC1,SNRPB,PARD3,TCAIM,SLC3A2,DHPS,ACTR6,MAP3K
,NIP7,TTC9C,SLC25A10,RPS6,RPL29,RTEL1,AUP1,YIPF3,FOXRED1,TRNT1,ITGB1BP1,MED15,SH3YL1,CLAS
AGED2,SON,UQCRQ,SNF8,CPSF3L,TSG101,FUZ,FAM3A,ATG12,RPS25,COX4I1,THEM4,RPS6,RPL29,RTEL1,
AGED2,SON,UQCRQ,SNF8,CPSF3L,TSG101,FUZ,FAM3A,ATG12,RPS25,COX4I1,THEM4,RPS6,RPL29,RTEL1,
CTTN,TSPAN11,ITSN1,EIF5A,COMMD5,CHID1,NUDT22,AMZ2,UFD1L,THYN1,SH2B1,NOP16,RPL13A,ARHG
CTTN,TSPAN11,ITSN1,EIF5A,COMMD5,CHID1,NUDT22,AMZ2,UFD1L,THYN1,SH2B1,NOP16,RPL13A,ARHG
FIS1,MAGED2,SON,UQCRQ,SNF8,CDC37,CPSF3L,TSG101,FUZ,FAM3A,RPS25,COX4I1,THEM4,RPS6,RPL29
FIS1,MAGED2,SON,UQCRQ,SNF8,CDC37,CPSF3L,TSG101,FUZ,FAM3A,RPS25,COX4I1,THEM4,RPS6,RPL29
6,ACOT8,AP3B2,ZNF410,PCMT1,EBPL,IGSF8,NUDT5,SNRPN,STYXL1,POLD4,VRK3,LZTFL1,FIS1,TUBA1C,IR
6,ACOT8,AP3B2,ZNF410,PCMT1,EBPL,IGSF8,NUDT5,SNRPN,STYXL1,POLD4,VRK3,LZTFL1,FIS1,TUBA1C,IR
PL13A,ARHGEF39,GSTK1,TTLL5,PSMA6,ACOT8,RPS9,SLC35G1,RCE1,MRPL33,ZNF410,IGSF8,NUDT5,RBPM
PL13A,ARHGEF39,GSTK1,TTLL5,PSMA6,ACOT8,RPS9,SLC35G1,RCE1,MRPL33,ZNF410,IGSF8,NUDT5,RBPM
,TMEM135,RNF32,LY75,ATG16L2,RCC1,SNRPB,PARD3,TCAIM,SLC3A2,DHPS,ACTR6,MAP3K7CL,SEC62,SE
,TMEM135,RNF32,LY75,ATG16L2,RCC1,SNRPB,PARD3,TCAIM,SLC3A2,DHPS,ACTR6,MAP3K7CL,SEC62,SE
F8,CDC37,CPSF3L,TSG101,UBR4,METTL17,ATG12,RPS25,NIP7,COX4I1,THEM4,RPS6,RPL29,COASY,AUP1,

RUVBL2,IGSF8,NUDT5,STYXL1,POLD4,VRK3,LZTFL1,FIS1,PPM1K,TUBA1C,LRRC1,VCAN,SON,UQCRQ,SNF8,
NF410,PCMT1,IGSF8,NUDT5,SNRPN,STYXL1,RFX5,POLD4,RANBP1,VRK3,LZTFL1,FIS1,PPM1K,IRF9,ANTXR
NF410,PCMT1,IGSF8,NUDT5,SNRPN,STYXL1,RFX5,POLD4,RANBP1,VRK3,LZTFL1,FIS1,PPM1K,IRF9,ANTXR
33,ZNF410,RUVBL2,IGSF8,NUDT5,STYXL1,RFX5,LZTFL1,TUBA1C,IRF9,ANTXR2,STIP1,LRRC1,VCAN,SON,U
PAG7,WDR62,STYXL1,POLD4,LZTFL1,FIS1,TUBA1C,ANTXR2,STIP1,LRRC1,VCAN,MAGED2,SON,UQCRQ,SN
PAG7,WDR62,STYXL1,POLD4,LZTFL1,FIS1,TUBA1C,ANTXR2,STIP1,LRRC1,VCAN,MAGED2,SON,UQCRQ,SN
AUP1,SAP30BP,YIPF3,FOXRED1,TRNT1,ITGB1BP1,MED15,CLASRP,C1ORF198,UBE2I,EIF3M,CDC42,CARD
RNF32,LY75,ATG16L2,RCC1,PARD3,TCAIM,SLC3A2,DHPS,ACTR6,SEC62,SET,CTSC,BOP1,ARRDC1,TRPT1,KL
P16,ARHGEF39,PSMA6,ACOT8,RPS9,RCE1,MRPL33,ZNF410,RUVBL2,TESC,NUDT5,STYXL1,LZTFL1,PPM1K,
DNASE1L1,KIF4A,RBCK1,PTOV1,FASTK,TMEM135,RNF32,LY75,ATG16L2,RCC1,PARD3,TCAIM,SLC3A2,DHI
G12,RPS25,NIP7,THEM4,RPS6,RTEL1,SAP30BP,YIPF3,FOXRED1,RABL5,TRNT1,ITGB1BP1,MED15,CLASRP,
G12,RPS25,NIP7,THEM4,RPS6,RTEL1,SAP30BP,YIPF3,FOXRED1,RABL5,TRNT1,ITGB1BP1,MED15,CLASRP,
CL,ZNF235,BOP1,ARRDC1,TRPT1,KLC1,RBP5,ZC3HAV1,SCAF11,DBI,ARPC2,PMFBP1,LIMK2,NOB1,SIRT6,N
CL,ZNF235,BOP1,ARRDC1,TRPT1,KLC1,RBP5,ZC3HAV1,SCAF11,DBI,ARPC2,PMFBP1,LIMK2,NOB1,SIRT6,N
OMMD5,CHID1,NUDT22,UFD1L,USP38,SH2B1,RPL13A,ARHGEF39,TTLL5,PSMA6,ACOT8,RPS9,RCE1,MRP
OMMD5,CHID1,NUDT22,UFD1L,USP38,SH2B1,RPL13A,ARHGEF39,TTLL5,PSMA6,ACOT8,RPS9,RCE1,MRP
OMMD5,CHID1,NUDT22,UFD1L,USP38,SH2B1,RPL13A,ARHGEF39,TTLL5,PSMA6,ACOT8,RPS9,RCE1,MRP
OMMD5,CHID1,NUDT22,UFD1L,USP38,SH2B1,RPL13A,ARHGEF39,TTLL5,PSMA6,ACOT8,RPS9,RCE1,MRP
,FAM3A,RPS25,NIP7,COX4I1,THEM4,RPS6,RPL29,RTEL1,SAP30BP,YIPF3,RABL5,TRNT1,ITGB1BP1,MED15
,FAM3A,RPS25,NIP7,COX4I1,THEM4,RPS6,RPL29,RTEL1,SAP30BP,YIPF3,RABL5,TRNT1,ITGB1BP1,MED15
SLC35G1,ZNF410,PCMT1,NUDT5,RBPMS,SNRPN,STYXL1,RFX5,POLD4,LZTFL1,FIS1,PPM1K,TUBA1C,IRF9,I
SLC35G1,ZNF410,PCMT1,NUDT5,RBPMS,SNRPN,STYXL1,RFX5,POLD4,LZTFL1,FIS1,PPM1K,TUBA1C,IRF9,I
12,RPS25,NIP7,TTC9C,TMX2,RPS6,RPL29,YIPF3,TRNT1,ITGB1BP1,MED15,EIF3M,CDC42,CARD14,PCYT2,F
AGED2,SON,UQCRQ,RPS16,CPSF3L,TSG101,UBR4,FUZ,FAM3A,ATG12,RPS25,NIP7,THEM4,DBNDD2,RPS6,
AGED2,SON,UQCRQ,RPS16,CPSF3L,TSG101,UBR4,FUZ,FAM3A,ATG12,RPS25,NIP7,THEM4,DBNDD2,RPS6,
CAF11,DBI,ARPC2,PMFBP1,LIMK2,SIRT6,NUMA1,GPATCH2L,SUMF2,BUD31,SAE1,NPRL2,OSGEP,ARHGA
CAF11,DBI,ARPC2,PMFBP1,LIMK2,SIRT6,NUMA1,GPATCH2L,SUMF2,BUD31,SAE1,NPRL2,OSGEP,ARHGA
5,NIP7,THEM4,RPS6,RTEL1,SAP30BP,YIPF3,RABL5,TRNT1,ITGB1BP1,MED15,C1ORF198,UGGT2,SLC39A1
5,NIP7,THEM4,RPS6,RTEL1,SAP30BP,YIPF3,RABL5,TRNT1,ITGB1BP1,MED15,C1ORF198,UGGT2,SLC39A1
ATCH2L,SUMF2,BUD31,SAE1,NPRL2,TBL2,ARHGAP6,CRYGS,CTTN,TSPAN11,EIF5A,COMMD5,CHID1,NUD
ATCH2L,SUMF2,BUD31,SAE1,NPRL2,TBL2,ARHGAP6,CRYGS,CTTN,TSPAN11,EIF5A,COMMD5,CHID1,NUD
6,MAP3K7CL,CTSC,ZNF235,BOP1,ARRDC1,TRPT1,RBP5,ZC3HAV1,SCAF11,DBI,ARPC2,PMFBP1,LIMK2,SIR
6,MAP3K7CL,CTSC,ZNF235,BOP1,ARRDC1,TRPT1,RBP5,ZC3HAV1,SCAF11,DBI,ARPC2,PMFBP1,LIMK2,SIR
Q,SNF8,CPSF3L,TSG101,UBR4,FUZ,FAM3A,EPB41L5,ATG12,RPS25,NIP7,DUSP22,SLC25A10,THEM4,RPS6,
Q,SNF8,CPSF3L,TSG101,UBR4,FUZ,FAM3A,EPB41L5,ATG12,RPS25,NIP7,DUSP22,SLC25A10,THEM4,RPS6,
RPC2,PMFBP1,LIMK2,SIRT6,NUMA1,GPATCH2L,SUMF2,BUD31,NPRL2,TBL2,CTTN,TSPAN11,EIF5A,COM
RPC2,PMFBP1,LIMK2,SIRT6,NUMA1,GPATCH2L,SUMF2,BUD31,NPRL2,TBL2,CTTN,TSPAN11,EIF5A,COM
H1,DNASE1L1,KIF4A,RBCK1,PTOV1,TMEM135,RNF32,LY75,ATG16L2,RCC1,PARD3,TCAIM,SLC3A2,DHPS,
IM,SLC3A2,DHPS,ACTR6,SEC62,ZNF235,BOP1,ARRDC1,TRPT1,RBP5,ZC3HAV1,DBI,ARPC2,LIMK2,VDAC3,
IM,SLC3A2,DHPS,ACTR6,SEC62,ZNF235,BOP1,ARRDC1,TRPT1,RBP5,ZC3HAV1,DBI,ARPC2,LIMK2,VDAC3,
NF235,BOP1,ARRDC1,TRPT1,KLC1,RBP5,ZC3HAV1,DBI,ARPC2,LIMK2,NOB1,SIRT6,NUMA1,GPATCH2L,BU
NF235,BOP1,ARRDC1,TRPT1,KLC1,RBP5,ZC3HAV1,DBI,ARPC2,LIMK2,NOB1,SIRT6,NUMA1,GPATCH2L,BU
RPS6,RPL29,RTEL1,AUP1,YIPF3,FOXRED1,TRNT1,ITGB1BP1,MED15,CLASRP,C1ORF198,UBE2I,EIF3M,CDC
C,ZNF235,BOP1,ARRDC1,TRPT1,KLC1,RBP5,ZC3HAV1,DBI,ARPC2,LIMK2,NOB1,SIRT6,NUMA1,GPATCH2L
C,ZNF235,BOP1,ARRDC1,TRPT1,KLC1,RBP5,ZC3HAV1,DBI,ARPC2,LIMK2,NOB1,SIRT6,NUMA1,GPATCH2L
,TSG101,FUZ,NIP7,COX4I1,THEM4,RPS6,RTEL1,AUP1,SAP30BP,RABL5,TRNT1,KXD1,ITGB1BP1,MED15,CL
,TSG101,FUZ,NIP7,COX4I1,THEM4,RPS6,RTEL1,AUP1,SAP30BP,RABL5,TRNT1,KXD1,ITGB1BP1,MED15,CL

25,THEM4,RPS6,RPL29,RTEL1,AUP1,SAP30BP,YIPF3,RABL5,TRNT1,KXD1,ITGB1BP1,MED15,CLASRP,C1O
25,THEM4,RPS6,RPL29,RTEL1,AUP1,SAP30BP,YIPF3,RABL5,TRNT1,KXD1,ITGB1BP1,MED15,CLASRP,C1O
75,ATG16L2,RCC1,SNRPB,PARD3,SLC3A2,DHPS,ACTR6,MAP3K7CL,SEC62,SET,CTSC,BOP1,ARRDC1,KLC1,
75,ATG16L2,RCC1,SNRPB,PARD3,SLC3A2,DHPS,ACTR6,MAP3K7CL,SEC62,SET,CTSC,BOP1,ARRDC1,KLC1,
ORF198,UBE2I,EIF3M,CDC42,CARD14,PCYT2,FBXW9,USE1,ASF1B,RFX3,ENDOV,CAMTA1,PPIL2,APEH,CA
,SH2B1,RPL13A,ARHGEF39,LDHA,PSMA6,ACOT8,RPS9,SLC35G1,RCE1,ZNF410,RUVBL2,PCMT1,TESC,NU
,SH2B1,RPL13A,ARHGEF39,LDHA,PSMA6,ACOT8,RPS9,SLC35G1,RCE1,ZNF410,RUVBL2,PCMT1,TESC,NU
10,RNH1,DNASE1L1,KIF4A,RBCK1,PTOV1,FASTK,TMEM135,RNF32,LY75,ATG16L2,RCC1,PARD3,TCAIM,S
10,RNH1,DNASE1L1,KIF4A,RBCK1,PTOV1,FASTK,TMEM135,RNF32,LY75,ATG16L2,RCC1,PARD3,TCAIM,S
Z2,UFD1L,USP38,TAZ,THYN1,DBNDD1,SH2B1,NOP16,RPL13A,ARHGEF39,TTLL5,PSMA6,ACOT8,RPS9,M
Z2,UFD1L,USP38,TAZ,THYN1,DBNDD1,SH2B1,NOP16,RPL13A,ARHGEF39,TTLL5,PSMA6,ACOT8,RPS9,M
,RPL38,PRR13,STARD10,RNH1,DNASE1L1,KIF4A,RBCK1,PTOV1,FASTK,TMEM135,RNF32,LY75,ATG16L2,R
,RPL38,PRR13,STARD10,RNH1,DNASE1L1,KIF4A,RBCK1,PTOV1,FASTK,TMEM135,RNF32,LY75,ATG16L2,R
7,THEM4,RPS6,RPL29,AUP1,YIPF3,FOXRED1,TRNT1,KXD1,MED15,CLASRP,C1ORF198,UBE2I,EIF3M,CDC4
HPS,SET,CTSC,BOP1,ARRDC1,TRPT1,KLC1,RBP5,LIMK2,VDAC3,NOB1,SIRT6,NUMA1,GPATCH2L,SUMF2,B
HPS,SET,CTSC,BOP1,ARRDC1,TRPT1,KLC1,RBP5,LIMK2,VDAC3,NOB1,SIRT6,NUMA1,GPATCH2L,SUMF2,B
ETTL17,ATG12,RPS25,NIP7,DUSP22,RPS6,RPL29,COASY,AUP1,SAP30BP,YIPF3,FOXRED1,TRNT1,MED15,
PL13A,ARHGEF39,PSMA6,ACOT8,ZNF410,RUVBL2,PCMT1,TESC,IGSF8,NUDT5,SNRPN,SPAG7,WDR62,STY
PL13A,ARHGEF39,PSMA6,ACOT8,ZNF410,RUVBL2,PCMT1,TESC,IGSF8,NUDT5,SNRPN,SPAG7,WDR62,STY
,CRYGS,CTTN,TSPAN11,EIF5A,COMMD5,CHID1,NUDT22,UFD1L,USP38,THYN1,RPL13A,ARHGEF39,PSMA
,CRYGS,CTTN,TSPAN11,EIF5A,COMMD5,CHID1,NUDT22,UFD1L,USP38,THYN1,RPL13A,ARHGEF39,PSMA
10,RUVBL2,PCMT1,IGSF8,NUDT5,WDR62,STYXL1,POLD4,VRK3,LZTFL1,FIS1,PPM1K,TUBA1C,STIP1,VCAN,
L,BUD31,NPRL2,OSGEP,SCARB1,CTTN,ITSN1,EIF5A,COMMD5,CHID1,NUDT22,AMZ2,UFD1L,USP38,SH2
L,BUD31,NPRL2,OSGEP,SCARB1,CTTN,ITSN1,EIF5A,COMMD5,CHID1,NUDT22,AMZ2,UFD1L,USP38,SH2
PF3,RABL5,TRNT1,ITGB1BP1,MED15,CLASRP,C1ORF198,UGGT2,UBE2I,EIF3M,CDC42,CARD14,PCYT2,FBX
PF3,RABL5,TRNT1,ITGB1BP1,MED15,CLASRP,C1ORF198,UGGT2,UBE2I,EIF3M,CDC42,CARD14,PCYT2,FBX
D1,TRNT1,ITGB1BP1,MED15,CLASRP,SLC39A13,EIF3M,CDC42,C14ORF2,USE1,ASF1B,DGKE,ENDOV,PPIL
,CLASRP,C1ORF198,UGGT2,UBE2I,EIF3M,CDC42,CARD14,FBXW9,CCDC25,EEF1B2,RFX3,ENDOV,PPIL2,RP
,CLASRP,C1ORF198,UGGT2,UBE2I,EIF3M,CDC42,CARD14,FBXW9,CCDC25,EEF1B2,RFX3,ENDOV,PPIL2,RP
1,RABL5,ITGB1BP1,MED15,CLASRP,C1ORF198,UGGT2,UBE2I,SLC39A13,CDC42,CARD14,PCYT2,FBXW9,U
1,RABL5,ITGB1BP1,MED15,CLASRP,C1ORF198,UGGT2,UBE2I,SLC39A13,CDC42,CARD14,PCYT2,FBXW9,U
,YIPF3,RABL5,TRNT1,ITGB1BP1,MED15,CLASRP,C1ORF198,UGGT2,UBE2I,EIF3M,CDC42,CARD14,PCYT2,F
,YIPF3,RABL5,TRNT1,ITGB1BP1,MED15,CLASRP,C1ORF198,UGGT2,UBE2I,EIF3M,CDC42,CARD14,PCYT2,F
PIL2,RPL28,APEH,LYPLA2,C17ORF62,PRMT1,ZNF7,TXNDC17,C19ORF48,PPAN,ZFAND2B,SUGT1,CARS,RB
A2,C17ORF62,PRMT1,STOML2,TXNDC17,C19ORF48,PPAN,CARS,ZNF197,INO80E,CCNL2,DGUOK,HSP90A
01,METTL17,ATG12,RPS25,NIP7,THEM4,RPS6,RPL29,COASY,AUP1,SAP30BP,YIPF3,FOXRED1,TRNT1,KXD
,ZNF235,BOP1,ARRDC1,TRPT1,KLC1,RBP5,ZC3HAV1,ARPC2,LIMK2,NOB1,SIRT6,NUMA1,GPATCH2L,OSG
,ZNF235,BOP1,ARRDC1,TRPT1,KLC1,RBP5,ZC3HAV1,ARPC2,LIMK2,NOB1,SIRT6,NUMA1,GPATCH2L,OSG
T1,HYPK,CDK11A,ZNF197,LRRFIP1,RAB2A,ATP5J,TRMT112,CHPF,RPS20,SAT1,ERCC5,SERPINB1,DDOST,L
,TMEM135,RNF32,LY75,ATG16L2,RCC1,SNRPB,PARD3,TCAIM,SLC3A2,DHPS,ACTR6,SEC62,SET,CTSC,BOP
,HMGA2,CPSF3L,TSG101,FUZ,FAM3A,RPS25,NIP7,COX4I1,THEM4,RPS6,RPL29,RTEL1,SAP30BP,YIPF3,RA
,HMGA2,CPSF3L,TSG101,FUZ,FAM3A,RPS25,NIP7,COX4I1,THEM4,RPS6,RPL29,RTEL1,SAP30BP,YIPF3,RA
RRDC1,TRPT1,KLC1,RBP5,DBI,LIMK2,NOB1,WDR19,SIRT6,NUMA1,GPATCH2L,SUMF2,BUD31,TBL2,SCAR
RRDC1,TRPT1,KLC1,RBP5,DBI,LIMK2,NOB1,WDR19,SIRT6,NUMA1,GPATCH2L,SUMF2,BUD31,TBL2,SCAR
MARCB1,RPL38,PRR13,STARD10,RNH1,DNASE1L1,KIF4A,RBCK1,PTOV1,FASTK,TMEM135,RNF32,LY75,AT
MARCB1,RPL38,PRR13,STARD10,RNH1,DNASE1L1,KIF4A,RBCK1,PTOV1,FASTK,TMEM135,RNF32,LY75,AT
,CTSC,BOP1,ARRDC1,TRPT1,KLC1,ZC3HAV1,SCAF11,ARPC2,LIMK2,VDAC3,NOB1,WDR19,SIRT6,NUMA1,S

RCB1,RPL38,PRR13,RNH1,DNASE1L1,KIF4A,RBCK1,PTOV1,FASTK,TMEM135,RNF32,LY75,ATG16L2,SNRPB

RCB1,RPL38,PRR13,RNH1,DNASE1L1,KIF4A,RBCK1,PTOV1,FASTK,TMEM135,RNF32,LY75,ATG16L2,SNRPB

LBP1,MED15,CLASRP,MICAL1,UBE2I,SLC39A13,EIF3M,CDC42,TBX6,PTK2,CARD14,FBXW9,USE1,CCDC25,I

LBP1,MED15,CLASRP,MICAL1,UBE2I,SLC39A13,EIF3M,CDC42,TBX6,PTK2,CARD14,FBXW9,USE1,CCDC25,I

62,PRMT1,TXNDC17,PIGL,PPAN,CARS,CDK11A,ZNF197,COPZ1,INO80E,CCNL2,LRRFIP1,ATP5J,TRMT112,C

,LRRFIP1,RAB2A,ATP5J,TRMT112,SAT1,ERCC5,EMC4,DDOST,LSR,PSME2,TOMM40,RABEP2,TADA3,ZNF6

,PSMA6,ACOT8,RCE1,PCMT1,NUDT5,SNRPN,STYXL1,RANBP1,VRK3,LZTFL1,FIS1,IRF9,ANTXR2,STIP1,LRRC

,PSMA6,ACOT8,RCE1,PCMT1,NUDT5,SNRPN,STYXL1,RANBP1,VRK3,LZTFL1,FIS1,IRF9,ANTXR2,STIP1,LRRC

RR13,STARD10,RNH1,DNASE1L1,KIF4A,RBCK1,PTOV1,FASTK,TMEM135,RNF32,LY75,ATG16L2,RCC1,PARD

RR13,STARD10,RNH1,DNASE1L1,KIF4A,RBCK1,PTOV1,FASTK,TMEM135,RNF32,LY75,ATG16L2,RCC1,PARD

ATP5J,TRMT112,CHPF,RPS20,DDOST,LSR,RRNAD1,TOMM40,TADA3,ANAPC11,NT5C,MDH2,STXBP2,SEC3

E1L1,RBCK1,PTOV1,FASTK,TMEM135,RNF32,LY75,ATG16L2,RCC1,SNRPB,PARD3,TCAIM,SLC3A2,DHPS,AC

E1L1,RBCK1,PTOV1,FASTK,TMEM135,RNF32,LY75,ATG16L2,RCC1,SNRPB,PARD3,TCAIM,SLC3A2,DHPS,AC

62,SET,ARRDC1,TRPT1,KLC1,RBP5,SCAF11,LIMK2,VDAC3,WDR19,SIRT6,NUMA1,GPATCH2L,NPRL2,OSGEP

62,SET,ARRDC1,TRPT1,KLC1,RBP5,SCAF11,LIMK2,VDAC3,WDR19,SIRT6,NUMA1,GPATCH2L,NPRL2,OSGEP

IL2,RPL28,LIG1,APEH,LYPLA2,ATPAF2,C17ORF62,AP1S3,ZNF197,INO80E,CCNL2,DGUOK,LRRFIP1,ATP5J,T

EP,TBL2,SCARB1,CTTN,TSPAN11,ITSN1,EIF5A,NUDT22,AMZ2,TAZ,DBNDD1,NOP16,GSTK1,TTLL5,PSMA6,A

EP,TBL2,SCARB1,CTTN,TSPAN11,ITSN1,EIF5A,NUDT22,AMZ2,TAZ,DBNDD1,NOP16,GSTK1,TTLL5,PSMA6,A

DHPS,ACTR6,SET,CTSC,BOP1,ARRDC1,TRPT1,KLC1,RBP5,ZC3HAV1,SCAF11,DBI,ARPC2,VDAC3,NOB1,WDR

ON,UQCRQ,RPS16,CPSF3L,TSG101,SLC39A11,FUZ,FAM3A,RPS25,TTC9C,PLCB2,DUSP22,THEM4,DBNDD2,

ON,UQCRQ,RPS16,CPSF3L,TSG101,SLC39A11,FUZ,FAM3A,RPS25,TTC9C,PLCB2,DUSP22,THEM4,DBNDD2,

MAGED2,SON,SNF8,CPSF3L,TSG101,CHCHD1,FUZ,ATG12,RPS25,NIP7,COX4I1,THEM4,RPS6,RPL29,RTEL1,

MAGED2,SON,SNF8,CPSF3L,TSG101,CHCHD1,FUZ,ATG12,RPS25,NIP7,COX4I1,THEM4,RPS6,RPL29,RTEL1,

DC42,CARD14,PCYT2,FBXW9,CCDC25,EEF1B2,RFX3,ENDOV,PPIL2,LIG1,APEH,LYPLA2,STOML2,ZNF7,PPAI

DC42,CARD14,PCYT2,FBXW9,CCDC25,EEF1B2,RFX3,ENDOV,PPIL2,LIG1,APEH,LYPLA2,STOML2,ZNF7,PPAI

3,ENDOV,APEH,LYPLA2,C17ORF62,PRMT1,PPAN,RPL36,CDK11A,ZNF197,LRRFIP1,ATP5J,TRMT112,CHPF,

5,ATG16L2,RCC1,SNRPB,PARD3,TCAIM,SLC3A2,DHPS,ACTR6,MAP3K7CL,SEC62,SET,CTSC,ZNF235,BOP1,A

5,ATG16L2,RCC1,SNRPB,PARD3,TCAIM,SLC3A2,DHPS,ACTR6,MAP3K7CL,SEC62,SET,CTSC,ZNF235,BOP1,A

,POLD4,RANBP1,VRK3,LZTFL1,FIS1,IRF9,ANTXR2,STIP1,LRRC1,CD44,TM9SF1,MAGED2,SON,UQCRQ,RPS1

,POLD4,RANBP1,VRK3,LZTFL1,FIS1,IRF9,ANTXR2,STIP1,LRRC1,CD44,TM9SF1,MAGED2,SON,UQCRQ,RPS1

BPL,IGSF8,NUDT5,SPAG7,FIS1,TUBA1C,DST,ANTXR2,STIP1,LRRC1,VCAN,TM9SF1,SON,UQCRQ,SNF8,GRK6

,ACTR6,MAP3K7CL,SEC62,SET,CTSC,BOP1,ARRDC1,TRPT1,KLC1,RBP5,ZC3HAV1,SCAF11,DBI,ARPC2,LIMK

,ACTR6,MAP3K7CL,SEC62,SET,CTSC,BOP1,ARRDC1,TRPT1,KLC1,RBP5,ZC3HAV1,SCAF11,DBI,ARPC2,LIMK

A10,RPS6,RPL29,AUP1,YIPF3,TRNT1,MED15,IP6K2,C1ORF198,UBE2I,YIF1A,EIF3M,CDC42,FBXW9,ASF1B,E

A10,TMX2,YIPF3,MED15,MICAL1,UBE2I,EIF3M,CDC42,FBXW9,CCDC25,RFX3,ENDOV,PPIL2,TROAP,RPL28,

1,ARRDC1,TRPT1,KLC1,RBP5,SCAF11,DBI,ARPC2,PMFBP1,LIMK2,VDAC3,NOB1,WDR19,NUMA1,GPATCH2

1,ARRDC1,TRPT1,KLC1,RBP5,SCAF11,DBI,ARPC2,PMFBP1,LIMK2,VDAC3,NOB1,WDR19,NUMA1,GPATCH2

HPS,ACTR6,SEC62,SET,BOP1,ARRDC1,KLC1,SCAF11,DBI,LIMK2,VDAC3,WDR19,SIRT6,NUMA1,GPATCH2L,S

HPS,ACTR6,SEC62,SET,BOP1,ARRDC1,KLC1,SCAF11,DBI,LIMK2,VDAC3,WDR19,SIRT6,NUMA1,GPATCH2L,S

K2,WDR19,SIRT6,NUMA1,SUMF2,BUD31,SAE1,NPRL2,CTTN,TSPAN11,EIF5A,COMMD5,CHID1,UFD1L,USP

KLC1,RBP5,LIMK2,NOB1,SIRT6,NUMA1,GPATCH2L,BUD31,NPRL2,TBL2,ARHGAP6,SCARB1,CTTN,TSPAN11

KLC1,RBP5,LIMK2,NOB1,SIRT6,NUMA1,GPATCH2L,BUD31,NPRL2,TBL2,ARHGAP6,SCARB1,CTTN,TSPAN11

RL2,OSGEP,TBL2,CRYGS,CTTN,TSPAN11,EIF5A,COMMD5,NUDT22,AMZ2,UFD1L,USP38,TAZ,THYN1,SH2B

RL2,OSGEP,TBL2,CRYGS,CTTN,TSPAN11,EIF5A,COMMD5,NUDT22,AMZ2,UFD1L,USP38,TAZ,THYN1,SH2B

D31,OSGEP,TBL2,SCARB1,CTTN,TSPAN11,ITSN1,EIF5A,COMMD5,NUDT22,AMZ2,UFD1L,TAZ,THYN1,DBN

ABHD17A,MDH2,SEC31A,CNST,ZNF506,PAFAH1B2,GBA2,NSUN5,ZFPL1,MFSD10,RIC8A,TCP1,PIGT,UBE2N

ATG16L2,PARD3,TCAIM,SLC3A2,DHPS,ACTR6,SEC62,SET,CTSC,BOP1,ARRDC1,TRPT1,KLC1,RBP5,ARPC2,PI

ATG16L2,PARD3,TCAIM,SLC3A2,DHPS,ACTR6,SEC62,SET,CTSC,BOP1,ARRDC1,TRPT1,KLC1,RBP5,ARPC2,PI

,MAGED2,SON,SNF8,CDC37,CPSF3L,TSG101,FUZ,COX4I1,RPL29,RTEL1,AUP1,RABL5,TRNT1,ITGB1BP1,ME

,MAGED2,SON,SNF8,CDC37,CPSF3L,TSG101,FUZ,COX4I1,RPL29,RTEL1,AUP1,RABL5,TRNT1,ITGB1BP1,ME

2,LIMK2,VDAC3,NOB1,SIRT6,NUMA1,GPATCH2L,SUMF2,OSGEP,TBL2,SCARB1,CRYGS,CTTN,TSPAN11,ITS

2,LIMK2,VDAC3,NOB1,SIRT6,NUMA1,GPATCH2L,SUMF2,OSGEP,TBL2,SCARB1,CRYGS,CTTN,TSPAN11,ITS

IF4A,RBCK1,PTOV1,FASTK,TMEM135,RNF32,LY75,ATG16L2,SNRPB,PARD3,TCAIM,SLC3A2,DHPS,ACTR6,S

RED1,TRNT1,KXD1,MED15,C1ORF198,UBE2I,EIF3M,CDC42,CARD14,PCYT2,FBXW9,CCDC25,RFX3,ENDOV

3M,CDC42,CARD14,PCYT2,FBXW9,CCDC25,RFX3,ENDOV,CAMTA1,PPIL2,APEH,CAPN10,LYPLA2,C17ORF6

31,DDOST,LSR,PABPC1,RRNAD1,NAT9,TOMM40,RABEP2,TADA3,RAD23A,ABHD17A,MDH2,HSPH1,CHCHI

RPC2,LIMK2,SIRT6,GPATCH2L,SUMF2,BUD31,SAE1,NPRL2,OSGEP,TBL2,ARHGAP6,CTTN,TSPAN11,ITSN1,E

RPC2,LIMK2,SIRT6,GPATCH2L,SUMF2,BUD31,SAE1,NPRL2,OSGEP,TBL2,ARHGAP6,CTTN,TSPAN11,ITSN1,E

1,ARRDC1,TRPT1,KLC1,RBP5,SCAF11,ARPC2,PMFBP1,LIMK2,VDAC3,WDR19,SIRT6,NUMA1,SUMF2,BUD

1,ARRDC1,TRPT1,KLC1,RBP5,SCAF11,ARPC2,PMFBP1,LIMK2,VDAC3,WDR19,SIRT6,NUMA1,SUMF2,BUD

,LY75,ATG16L2,SNRPB,PARD3,TCAIM,SLC3A2,DHPS,ACTR6,MAP3K7CL,SEC62,SET,CTSC,ZNF235,BOP1,AI

1,NPRL2,TBL2,SCARB1,CTTN,TSPAN11,ITSN1,EIF5A,COMMD5,NUDT22,AMZ2,UFD1L,THYN1,DBNDD1,NC

1,NPRL2,TBL2,SCARB1,CTTN,TSPAN11,ITSN1,EIF5A,COMMD5,NUDT22,AMZ2,UFD1L,THYN1,DBNDD1,NC

B1,RPL38,PRR13,STARD10,RNH1,DNASE1L1,KIF4A,RBCK1,PTOV1,FASTK,TMEM135,RNF32,LY75,ATG16L

B1,RPL38,PRR13,STARD10,RNH1,DNASE1L1,KIF4A,RBCK1,PTOV1,FASTK,TMEM135,RNF32,LY75,ATG16L

P1,ARRDC1,TRPT1,KLC1,RBP5,SCAF11,DBI,ARPC2,LIMK2,VDAC3,NOB1,WDR19,SIRT6,SUMF2,BUD31,NPI

TCAIM,SLC3A2,DHPS,ACTR6,SET,BOP1,ARRDC1,KLC1,SCAF11,DBI,ARPC2,LIMK2,VDAC3,SIRT6,NUMA1,BU

PAN11,EIF5A,COMMD5,RUNX1,CHID1,NUDT22,AMZ2,UFD1L,USP38,DBNDD1,RPL13A,ARHGEF39,TTLL5,

PAN11,EIF5A,COMMD5,RUNX1,CHID1,NUDT22,AMZ2,UFD1L,USP38,DBNDD1,RPL13A,ARHGEF39,TTLL5,

PIL2,RPL28,APEH,LYPLA2,C17ORF62,PRMT1,STOML2,ZNF7,PPAN,ZFAND2B,HNRNPC,CARS,RBM3,COPZ1

197,INO80E,CCNL2,LRRFIP1,ZNF524,ATP5J,RMDN1,TRMT112,ERCC5,SERPINB1,DDOST,LSR,PABPC1,RRN

FD1L,USP38,TAZ,SH2B1,NOP16,RPL13A,ARHGEF39,GSTK1,TTLL5,LDHA,PSMA6,ACOT8,AP3B2,RPS9,SLC3

2,PMFBP1,LIMK2,VDAC3,NUMA1,GPATCH2L,BUD31,SAE1,NPRL2,ARHGAP6,SCARB1,CTTN,EIF5A,COMMI

2,PMFBP1,LIMK2,VDAC3,NUMA1,GPATCH2L,BUD31,SAE1,NPRL2,ARHGAP6,SCARB1,CTTN,EIF5A,COMMI

ARS,ZNF197,LRRFIP1,ATP5J,TRMT112,CHPF,RPS20,DCTD,DDOST,LSR,CHCHD7,PABPC1,RRNAD1,TOMM4C

IPRL2,TBL2,SCARB1,CTTN,TSPAN11,ITSN1,EIF5A,THYN1,DBNDD1,NOP16,GSTK1,TTLL5,PSMA6,ACOT8,AP

IPRL2,TBL2,SCARB1,CTTN,TSPAN11,ITSN1,EIF5A,THYN1,DBNDD1,NOP16,GSTK1,TTLL5,PSMA6,ACOT8,AP

IPRL2,TBL2,SCARB1,CTTN,TSPAN11,ITSN1,EIF5A,THYN1,DBNDD1,NOP16,GSTK1,TTLL5,PSMA6,ACOT8,AP

IPRL2,TBL2,SCARB1,CTTN,TSPAN11,ITSN1,EIF5A,THYN1,DBNDD1,NOP16,GSTK1,TTLL5,PSMA6,ACOT8,AP

L2,RCC1,SNRPB,PARD3,TCAIM,SLC3A2,ACTR6,SET,CTSC,BOP1,TRPT1,SCAF11,DBI,PMFBP1,LIMK2,VDAC3

L2,RCC1,SNRPB,PARD3,TCAIM,SLC3A2,ACTR6,SET,CTSC,BOP1,TRPT1,SCAF11,DBI,PMFBP1,LIMK2,VDAC3

L2,PARD3,TCAIM,SLC3A2,DHPS,ACTR6,SEC62,SET,CTSC,ZNF235,BOP1,ARRDC1,TRPT1,KLC1,RBP5,ZC3HA

010,RNH1,DNASE1L1,KIF4A,RBCK1,PTOV1,FASTK,TMEM135,RNF32,LY75,ATG16L2,RCC1,PARD3,TCAIM,S

010,RNH1,DNASE1L1,KIF4A,RBCK1,PTOV1,FASTK,TMEM135,RNF32,LY75,ATG16L2,RCC1,PARD3,TCAIM,S

TK,TMEM135,RNF32,LY75,ATG16L2,RCC1,SNRPB,PARD3,TCAIM,SLC3A2,DHPS,ACTR6,SEC62,SET,CTSC,B

RDC1,KLC1,RBP5,ZC3HAV1,SCAF11,DBI,LIMK2,VDAC3,WDR19,NUMA1,GPATCH2L,BUD31,SAE1,NPRL2,O

1,SNRPB,PARD3,TCAIM,SLC3A2,DHPS,ACTR6,SEC62,SET,CTSC,BOP1,ARRDC1,KLC1,RBP5,ZC3HAV1,SCAF1

1,SNRPB,PARD3,TCAIM,SLC3A2,DHPS,ACTR6,SEC62,SET,CTSC,BOP1,ARRDC1,KLC1,RBP5,ZC3HAV1,SCAF1

1,SCAF11,DBI,ARPC2,PMFBP1,LIMK2,NOB1,SIRT6,NUMA1,GPATCH2L,SUMF2,BUD31,SAE1,NPRL2,OSGEI

1,SCAF11,DBI,ARPC2,PMFBP1,LIMK2,NOB1,SIRT6,NUMA1,GPATCH2L,SUMF2,BUD31,SAE1,NPRL2,OSGEI

ITSN1,TNFSF10,COMMD5,NUDT22,AMZ2,TAZ,NOP16,PSMA6,ACOT8,AP3B2,RCE1,MRPL33,RUVBL2,EBPI

ITSN1,TNFSF10,COMMD5,NUDT22,AMZ2,TAZ,NOP16,PSMA6,ACOT8,AP3B2,RCE1,MRPL33,RUVBL2,EBPI

,DHPS,ACTR6,SEC62,SET,CTSC,BOP1,ARRDC1,KLC1,RBP5,ZC3HAV1,SCAF11,DBI,ARPC2,LIMK2,VDAC3,NO

,TSPAN11,ITSN1,COMMD5,NUDT22,AMZ2,TAZ,SH2B1,NOP16,TTLL5,PSMA6,ACOT8,AP3B2,RCE1,MRPL3

,TSPAN11,ITSN1,COMMD5,NUDT22,AMZ2,TAZ,SH2B1,NOP16,TTLL5,PSMA6,ACOT8,AP3B2,RCE1,MRPL3
HPF,RPS20,NIT2,ZNF814,SAT1,ERCC5,EMC4,DDOST,LSR,MEIS2,PSME2,RRNAD1,TOMM40,RABEP2,TADA
C,BOP1,ARRDC1,TRPT1,KLC1,RBP5,ARPC2,PMFBP1,LIMK2,WDR19,SIRT6,NUMA1,GPATCH2L,SUMF2,BU
F,SMARCB1,RPL38,PRR13,RNH1,DNASE1L1,KIF4A,RBCK1,PTOV1,FASTK,TMEM135,RNF32,LY75,ATG16L2,
F,SMARCB1,RPL38,PRR13,RNH1,DNASE1L1,KIF4A,RBCK1,PTOV1,FASTK,TMEM135,RNF32,LY75,ATG16L2,
3A2,DHPS,ACTR6,MAP3K7CL,SEC62,SET,BOP1,ARRDC1,KLC1,SCAF11,DBI,LIMK2,VDAC3,WDR19,SIRT6,NU
3A2,DHPS,ACTR6,MAP3K7CL,SEC62,SET,BOP1,ARRDC1,KLC1,SCAF11,DBI,LIMK2,VDAC3,WDR19,SIRT6,NU
IG1,TMEM126B,LYPLA2,CAMLG,PRMT1,STOML2,C19ORF54,ZNF7,ARL6IP4,HNRNPC,CYB561,CCNL2,MA
UMA1,GPATCH2L,BUD31,NPRL2,TBL2,SCARB1,CTTN,TSPAN11,ITSN1,EIF5A,UFD1L,THYN1,DBNDD1,NOP
UMA1,GPATCH2L,BUD31,NPRL2,TBL2,SCARB1,CTTN,TSPAN11,ITSN1,EIF5A,UFD1L,THYN1,DBNDD1,NOP
ET,CTSC,BOP1,ARRDC1,ZC3HAV1,SCAF11,DBI,ARPC2,PMFBP1,LIMK2,VDAC3,NOB1,WDR19,SIRT6,NUMA
ET,CTSC,BOP1,ARRDC1,ZC3HAV1,SCAF11,DBI,ARPC2,PMFBP1,LIMK2,VDAC3,NOB1,WDR19,SIRT6,NUMA
1L1,KIF4A,RBCK1,PTOV1,FASTK,TMEM135,RNF32,LY75,ATG16L2,RCC1,SNRPB,PARD3,TCAIM,SLC3A2,DH
1L1,KIF4A,RBCK1,PTOV1,FASTK,TMEM135,RNF32,LY75,ATG16L2,RCC1,SNRPB,PARD3,TCAIM,SLC3A2,DH
ARPC2,LIMK2,VDAC3,NOB1,WDR19,SIRT6,NUMA1,SUMF2,SAE1,NPRL2,OSGEP,TBL2,SCARB1,CTTN,TSPA
31,TBL2,SCARB1,CTTN,TSPAN11,ITSN1,EIF5A,AMZ2,UFD1L,DBNDD1,SH2B1,NOP16,GSTK1,PSMA6,ACOT
31,TBL2,SCARB1,CTTN,TSPAN11,ITSN1,EIF5A,AMZ2,UFD1L,DBNDD1,SH2B1,NOP16,GSTK1,PSMA6,ACOT
31,TBL2,SCARB1,CTTN,TSPAN11,ITSN1,EIF5A,AMZ2,UFD1L,DBNDD1,SH2B1,NOP16,GSTK1,PSMA6,ACOT
31,TBL2,SCARB1,CTTN,TSPAN11,ITSN1,EIF5A,AMZ2,UFD1L,DBNDD1,SH2B1,NOP16,GSTK1,PSMA6,ACOT
31,TBL2,SCARB1,CTTN,TSPAN11,ITSN1,EIF5A,AMZ2,UFD1L,DBNDD1,SH2B1,NOP16,GSTK1,PSMA6,ACOT
31,TBL2,SCARB1,CTTN,TSPAN11,ITSN1,EIF5A,AMZ2,UFD1L,DBNDD1,SH2B1,NOP16,GSTK1,PSMA6,ACOT
5,NUDT22,AMZ2,UFD1L,DBNDD1,NOP16,TTLL5,PSMA6,ACOT8,AP3B2,RCE1,ZNF410,RUVBL2,EBPL,TESC
5,NUDT22,AMZ2,UFD1L,DBNDD1,NOP16,TTLL5,PSMA6,ACOT8,AP3B2,RCE1,ZNF410,RUVBL2,EBPL,TESC
,NOB1,GPATCH2L,BUD31,SAE1,NPRL2,TBL2,ARHGAP6,CTTN,EIF5A,COMMD5,NUDT22,UFD1L,USP38,TH
,NOB1,GPATCH2L,BUD31,SAE1,NPRL2,TBL2,ARHGAP6,CTTN,EIF5A,COMMD5,NUDT22,UFD1L,USP38,TH
,RPL13A,ARHGEF39,PSMA6,ACOT8,AP3B2,ZNF410,RUVBL2,PCMT1,NUDT5,SNRPN,STYXL1,POLD4,VRK3,
,RPL13A,ARHGEF39,PSMA6,ACOT8,AP3B2,ZNF410,RUVBL2,PCMT1,NUDT5,SNRPN,STYXL1,POLD4,VRK3,
,LIMK2,NOB1,SIRT6,NUMA1,BUD31,OSGEP,TBL2,ARHGAP6,SCARB1,CTTN,TSPAN11,ITSN1,COMMD5,CHI
,LIMK2,NOB1,SIRT6,NUMA1,BUD31,OSGEP,TBL2,ARHGAP6,SCARB1,CTTN,TSPAN11,ITSN1,COMMD5,CHI
,LYPLA2,STOML2,ZNF7,PPAN,CYB561,ZNF197,COPZ1,ATP5J,TRMT112,UBXN1,EMC4,DDOST,RRNAD1,TC
,LYPLA2,STOML2,ZNF7,PPAN,CYB561,ZNF197,COPZ1,ATP5J,TRMT112,UBXN1,EMC4,DDOST,RRNAD1,TC
T6,NUMA1,BUD31,TBL2,SCARB1,CTTN,TSPAN11,ITSN1,COMMD5,NUDT22,AMZ2,UFD1L,TAZ,DBNDD1,N
T6,NUMA1,BUD31,TBL2,SCARB1,CTTN,TSPAN11,ITSN1,COMMD5,NUDT22,AMZ2,UFD1L,TAZ,DBNDD1,N
OXRED1,RABL5,TRNT1,MED15,CLASRP,UGGT2,SLC39A13,FBXW9,USE1,CCDC25,EEF1B2,ENDOV,CAPG,P
OXRED1,RABL5,TRNT1,MED15,CLASRP,UGGT2,SLC39A13,FBXW9,USE1,CCDC25,EEF1B2,ENDOV,CAPG,P
ET,CTSC,BOP1,ARRDC1,TRPT1,KLC1,RBP5,ARPC2,PMFBP1,LIMK2,WDR19,SIRT6,NUMA1,GPATCH2L,SUM
ET,CTSC,BOP1,ARRDC1,TRPT1,KLC1,RBP5,ARPC2,PMFBP1,LIMK2,WDR19,SIRT6,NUMA1,GPATCH2L,SUM
5,VRK3,TET2,UQCRQ,SNF8,CDC37,RPS16,TSG101,CHCHD1,ATG12,PPRC1,NIP7,TTC9C,RPS6,RPL29,SRSF
BP1,LIMK2,VDAC3,WDR19,SIRT6,NUMA1,GPATCH2L,BUD31,OSGEP,ARHGAP6,SCARB1,CTTN,EIF5A,COM
BP1,LIMK2,VDAC3,WDR19,SIRT6,NUMA1,GPATCH2L,BUD31,OSGEP,ARHGAP6,SCARB1,CTTN,EIF5A,COM
SPAN11,ITSN1,COMMD5,NUDT22,DBNDD1,SH2B1,NOP16,ARHGEF39,TTLL5,PSMA6,ACOT8,AP3B2,RCE1
SPAN11,ITSN1,COMMD5,NUDT22,DBNDD1,SH2B1,NOP16,ARHGEF39,TTLL5,PSMA6,ACOT8,AP3B2,RCE1
SIRT6,NUMA1,BUD31,SAE1,NPRL2,TBL2,ARHGAP6,ITSN1,EIF5A,COMMD5,UFD1L,USP38,TAZ,SH2B1,RPL
1,EIF5A,USP38,TAZ,DBNDD1,AP3B2,RPS9,MRPL33,EBPL,TESC,RBPMS,WDR62,IL1RAP,POLD4,PLEKHA5,R
1,EIF5A,USP38,TAZ,DBNDD1,AP3B2,RPS9,MRPL33,EBPL,TESC,RBPMS,WDR62,IL1RAP,POLD4,PLEKHA5,R
,SLC3A2,ACTR6,SEC62,SET,CTSC,ZNF235,BOP1,ARRDC1,TRPT1,KLC1,RBP5,ZC3HAV1,SCAF11,DBI,ARPC2,L
,SLC3A2,ACTR6,SEC62,SET,CTSC,ZNF235,BOP1,ARRDC1,TRPT1,KLC1,RBP5,ZC3HAV1,SCAF11,DBI,ARPC2,L

,HSP90AB1,ATP5J,TRMT112,ZNF814,HSPB1,ERCC5,EMC4,DDOST,TOMM40,TADA3,ANAPC11,MDH2,EEF
,HSP90AB1,ATP5J,TRMT112,ZNF814,HSPB1,ERCC5,EMC4,DDOST,TOMM40,TADA3,ANAPC11,MDH2,EEF
NF32,LY75,ATG16L2,RCC1,PARD3,SLC3A2,DHPS,ACTR6,SEC62,SET,CTSC,BOP1,ARRDC1,TRPT1,KLC1,RBP5
NF32,LY75,ATG16L2,RCC1,PARD3,SLC3A2,DHPS,ACTR6,SEC62,SET,CTSC,BOP1,ARRDC1,TRPT1,KLC1,RBP5
EC62,SET,ZNF235,BOP1,ARRDC1,TRPT1,KLC1,RBP5,ZC3HAV1,SCAF11,DBI,ARPC2,LIMK2,VDAC3,WDR19,
EC62,SET,ZNF235,BOP1,ARRDC1,TRPT1,KLC1,RBP5,ZC3HAV1,SCAF11,DBI,ARPC2,LIMK2,VDAC3,WDR19,
AIM,SLC3A2,DHPS,ACTR6,SEC62,SET,CTSC,BOP1,ARRDC1,TRPT1,KLC1,RBP5,SCAF11,ARPC2,PMFBP1,LIM
AIM,SLC3A2,DHPS,ACTR6,SEC62,SET,CTSC,BOP1,ARRDC1,TRPT1,KLC1,RBP5,SCAF11,ARPC2,PMFBP1,LIM
P,C1ORF198,UGGT2,UBE2I,EIF3M,CDC42,CARD14,PCYT2,FBXW9,ASF1B,CCDC25,EEF1B2,RFX3,RPL28,LIG
P,C1ORF198,UGGT2,UBE2I,EIF3M,CDC42,CARD14,PCYT2,FBXW9,ASF1B,CCDC25,EEF1B2,RFX3,RPL28,LIG
,SEC62,SET,ARRDC1,KLC1,SCAF11,ARPC2,PMFBP1,LIMK2,VDAC3,WDR19,SIRT6,NUMA1,GPATCH2L,SAE1
,SEC62,SET,ARRDC1,KLC1,SCAF11,ARPC2,PMFBP1,LIMK2,VDAC3,WDR19,SIRT6,NUMA1,GPATCH2L,SAE1
TSPAN11,ITSN1,EIF5A,NUDT22,AMZ2,UFD1L,DBNDD1,NOP16,ACOT8,AP3B2,RCE1,RUVBL2,PCMT1,EBPL
TSPAN11,ITSN1,EIF5A,NUDT22,AMZ2,UFD1L,DBNDD1,NOP16,ACOT8,AP3B2,RCE1,RUVBL2,PCMT1,EBPL
,EBPL,TESC,IGSF8,SNRPN,SPAG7,WDR62,RANBP1,UBE2L6,FIS1,SLC15A3,PPM1K,TUBA1C,ANTXR2,STIP1,
,EBPL,TESC,IGSF8,SNRPN,SPAG7,WDR62,RANBP1,UBE2L6,FIS1,SLC15A3,PPM1K,TUBA1C,ANTXR2,STIP1,
UQCRQ,CPSF3L,TSG101,FUZ,FAM3A,METTL17,EPB41L5,ATG12,RPS25,NIP7,DUSP22,THEM4,RPS6,RTEL1,
UQCRQ,CPSF3L,TSG101,FUZ,FAM3A,METTL17,EPB41L5,ATG12,RPS25,NIP7,DUSP22,THEM4,RPS6,RTEL1,
D1,NOP16,TTLL5,PSMA6,ACOT8,AP3B2,RCE1,ZNF410,RUVBL2,EBPL,TESC,IGSF8,SNRPN,SPAG7,WDR62,
D1,NOP16,TTLL5,PSMA6,ACOT8,AP3B2,RCE1,ZNF410,RUVBL2,EBPL,TESC,IGSF8,SNRPN,SPAG7,WDR62,
S9,MRPL33,EBPL,TESC,WDR62,IL1RAP,POLD4,PLEKHA5,RANBP1,UBE2L6,VRK3,LZTFL1,ZC3H14,FIS1,TUB
S9,MRPL33,EBPL,TESC,WDR62,IL1RAP,POLD4,PLEKHA5,RANBP1,UBE2L6,VRK3,LZTFL1,ZC3H14,FIS1,TUB
1,TRPT1,KLC1,ZC3HAV1,LIMK2,SIRT6,NUMA1,GPATCH2L,BUD31,OSGEP,TBL2,ARHGAP6,SCARB1,CTTN,T
1,TRPT1,KLC1,ZC3HAV1,LIMK2,SIRT6,NUMA1,GPATCH2L,BUD31,OSGEP,TBL2,ARHGAP6,SCARB1,CTTN,T
0AB1,ATP5J,TRMT112,ZNF814,ERCC5,EMC4,DDOST,TOMM40,ANAPC11,MDH2,EEF1D,NAA10,CARHSP1,
0AB1,ATP5J,TRMT112,ZNF814,ERCC5,EMC4,DDOST,TOMM40,ANAPC11,MDH2,EEF1D,NAA10,CARHSP1,
3H14,PHF19,VCAN,LPCAT3,CD44,MAGED2,GRK6,HMGA2,TTC21B,PSMA1,ECSIT,PPRC1,RPS25,NIP7,TTC
P7,CENPK,RPS6,RPL29,AUP1,SAP30BP,YIPF3,FOXRED1,TRNT1,CLASRP,YIF1A,EIF3M,CDC42,FBXW9,USE1,
,TESC,IGSF8,SPAG7,WDR62,POLD4,RANBP1,UBE2L6,PPM1K,STIP1,PHF19,LPCAT3,CD44,MAGED2,SON,G
C42,TBX6,CENPN,FBXW9,CCDC25,EEF1B2,RFX3,DGKE,TNFRSF14,ENDOV,LIG1,APEH,CAPN10,LYPLA2,C1
C42,TBX6,CENPN,FBXW9,CCDC25,EEF1B2,RFX3,DGKE,TNFRSF14,ENDOV,LIG1,APEH,CAPN10,LYPLA2,C1
S,SET,CTSC,BOP1,ARRDC1,TRPT1,KLC1,ZC3HAV1,SCAF11,DBI,ARPC2,VDAC3,NOB1,WDR19,NUMA1,GPA
S,SET,CTSC,BOP1,ARRDC1,TRPT1,KLC1,ZC3HAV1,SCAF11,DBI,ARPC2,VDAC3,NOB1,WDR19,NUMA1,GPA
D3,TCAIM,SLC3A2,DHPS,ACTR6,SET,BOP1,ARRDC1,TRPT1,KLC1,ZC3HAV1,SCAF11,DBI,ARPC2,LIMK2,VDA
D3,TCAIM,SLC3A2,DHPS,ACTR6,SET,BOP1,ARRDC1,TRPT1,KLC1,ZC3HAV1,SCAF11,DBI,ARPC2,LIMK2,VDA
1,SCAF11,DBI,ARPC2,PMFBP1,LIMK2,VDAC3,NOB1,WDR19,SIRT6,NUMA1,GPATCH2L,NPRL2,OSGEP,TBL
1,SCAF11,DBI,ARPC2,PMFBP1,LIMK2,VDAC3,NOB1,WDR19,SIRT6,NUMA1,GPATCH2L,NPRL2,OSGEP,TBL
COMMD5,NUDT22,AMZ2,UFD1L,USP38,THYN1,DBNDD1,SH2B1,NOP16,RPL13A,ARHGEF39,PSMA6,ACO
COMMD5,NUDT22,AMZ2,UFD1L,USP38,THYN1,DBNDD1,SH2B1,NOP16,RPL13A,ARHGEF39,PSMA6,ACO
L33,EBPL,TESC,IGSF8,SPAG7,RFX5,POLD4,RANBP1,UBE2L6,FIS1,PPM1K,PHF19,LPCAT3,CD44,MAGED2,S
L33,EBPL,TESC,IGSF8,SPAG7,RFX5,POLD4,RANBP1,UBE2L6,FIS1,PPM1K,PHF19,LPCAT3,CD44,MAGED2,S
ZNF197,DGUOK,ATP5J,TRMT112,ZNF814,EMC4,DDOST,RRNAD1,TOMM40,ZNF687,ANAPC11,MDH2,NAA
ZNF197,DGUOK,ATP5J,TRMT112,ZNF814,EMC4,DDOST,RRNAD1,TOMM40,ZNF687,ANAPC11,MDH2,NAA
,RBCK1,PTOV1,FASTK,RNF32,ATG16L2,RCC1,SNRPB,PARD3,TCAIM,DHPS,ACTR6,SEC62,SET,CTSC,BOP1,
,RBCK1,PTOV1,FASTK,RNF32,ATG16L2,RCC1,SNRPB,PARD3,TCAIM,DHPS,ACTR6,SEC62,SET,CTSC,BOP1,
OASY,AUP1,SAP30BP,YIPF3,FOXRED1,TRNT1,MED15,SH3YL1,SLC39A13,EIF3M,CDC42,PTK2,MAF,ASF1B,
OASY,AUP1,SAP30BP,YIPF3,FOXRED1,TRNT1,MED15,SH3YL1,SLC39A13,EIF3M,CDC42,PTK2,MAF,ASF1B,

,ZBTB44,TTC9C,THEM4,DBNDD2,NISCH,TMED4,CD99L2,KXD1,PARD6G,SH3YL1,IP6K2,CLASRP,JPH2,MICA
,ZBTB44,TTC9C,THEM4,DBNDD2,NISCH,TMED4,CD99L2,KXD1,PARD6G,SH3YL1,IP6K2,CLASRP,JPH2,MICA
TLL5,PSMA6,ACOT8,AP3B2,RCE1,EBPL,TESC,IGSF8,SPAG7,WDR62,RFX5,POLD4,UBE2L6,FIS1,STIP1,PHF1
TLL5,PSMA6,ACOT8,AP3B2,RCE1,EBPL,TESC,IGSF8,SPAG7,WDR62,RFX5,POLD4,UBE2L6,FIS1,STIP1,PHF1
1A,EIF3M,CDC42,FBXW9,ASF1B,ENDOV,CLDND1,LYPLA2,C17ORF62,PRMT1,CASC4,ZNF7,TXNDC17,SUGT
10,EBPL,NUDT5,RBPMS,SNRPN,STYXL1,POLD4,VRK3,LZTFL1,FIS1,TUBA1C,IRF9,DST,GRAMD1C,PCBP2,MA
10,EBPL,NUDT5,RBPMS,SNRPN,STYXL1,POLD4,VRK3,LZTFL1,FIS1,TUBA1C,IRF9,DST,GRAMD1C,PCBP2,MA
MUC1,ECSIT,ZBTB44,TTC9C,DUSP22,NISCH,TMED4,CD99L2,YIPF3,KXD1,PARD6G,SH3YL1,IP6K2,CLASRP,J
MUC1,ECSIT,ZBTB44,TTC9C,DUSP22,NISCH,TMED4,CD99L2,YIPF3,KXD1,PARD6G,SH3YL1,IP6K2,CLASRP,J
,TCAIM,SLC3A2,DHPS,SEC62,SET,BOP1,ARRDC1,TRPT1,KLC1,SCAF11,ARPC2,LIMK2,VDAC3,NOB1,SIRT6,N
1,SCAF11,DBI,ARPC2,LIMK2,VDAC3,NOB1,WDR19,SIRT6,NUMA1,SUMF2,SAE1,NPRL2,OSGEP,TBL2,SCAR
1,SCAF11,DBI,ARPC2,LIMK2,VDAC3,NOB1,WDR19,SIRT6,NUMA1,SUMF2,SAE1,NPRL2,OSGEP,TBL2,SCAR
,MAGED2,SON,UQCRQ,CPSF3L,TSG101,FUZ,FAM3A,METTL17,ATG12,RPS25,NIP7,THEM4,RPS6,RTEL1,SA
,MAGED2,SON,UQCRQ,CPSF3L,TSG101,FUZ,FAM3A,METTL17,ATG12,RPS25,NIP7,THEM4,RPS6,RTEL1,SA
SUMF2,BUD31,SAE1,NPRL2,OSGEP,TBL2,ARHGAP6,SCARB1,CTTN,EIF5A,COMMD5,AMZ2,UFD1L,USP38,T
SUMF2,BUD31,SAE1,NPRL2,OSGEP,TBL2,ARHGAP6,SCARB1,CTTN,EIF5A,COMMD5,AMZ2,UFD1L,USP38,T
IM,DHPS,ACTR6,MAP3K7CL,SEC62,SET,ZNF235,BOP1,ARRDC1,KLC1,RBP5,SCAF11,DBI,ARPC2,LIMK2,VDA
IM,DHPS,ACTR6,MAP3K7CL,SEC62,SET,ZNF235,BOP1,ARRDC1,KLC1,RBP5,SCAF11,DBI,ARPC2,LIMK2,VDA
RPT1,KLC1,SCAF11,DBI,ARPC2,VDAC3,WDR19,NUMA1,GPATCH2L,SUMF2,SAE1,OSGEP,TBL2,ARHGAP6,S
GEF39,TTLL5,LDHA,PSMA6,ACOT8,RCE1,ZNF410,EBPL,NUDT5,RBPMS,SNRPN,STYXL1,RFX5,POLD4,VRK3,
GEF39,TTLL5,LDHA,PSMA6,ACOT8,RCE1,ZNF410,EBPL,NUDT5,RBPMS,SNRPN,STYXL1,RFX5,POLD4,VRK3,
STYXL1,POLD4,PLEKHA5,VRK3,LZTFL1,FIS1,SLC15A3,TUBA1C,IRF9,PHF19,PCBP2,MAGED2,SON,UQCRQ,H
STYXL1,POLD4,PLEKHA5,VRK3,LZTFL1,FIS1,SLC15A3,TUBA1C,IRF9,PHF19,PCBP2,MAGED2,SON,UQCRQ,H
32,LY75,RCC1,SNRPB,PARD3,TCAIM,SLC3A2,ACTR6,MAP3K7CL,SEC62,SET,BOP1,ARRDC1,TRPT1,KLC1,RE
TESC,IGSF8,SPAG7,WDR62,RFX5,POLD4,RANBP1,UBE2L6,PPM1K,STIP1,PHF19,LPCAT3,CD44,MAGED2,SC
62,SET,BOP1,ARRDC1,TRPT1,ZC3HAV1,DBI,ARPC2,PMFBP1,LIMK2,VDAC3,NOB1,WDR19,SIRT6,NUMA1,
A,CNST,COX17,ZNF506,FTH1,PAFAH1B2,GBA2,DALRD3,NSUN5,GMPR2,RIC8A,TCP1,UBE2N,PDCD6,FAM3
IGSF8,SPAG7,WDR62,IL1RAP,POLD4,UBE2L6,FIS1,PPM1K,STIP1,PHF19,VCAN,MAGED2,SON,GRK6,CDC3
IGSF8,SPAG7,WDR62,IL1RAP,POLD4,UBE2L6,FIS1,PPM1K,STIP1,PHF19,VCAN,MAGED2,SON,GRK6,CDC3
,DNASE1L1,KIF4A,RBCK1,PTOV1,FASTK,TMEM135,RNF32,LY75,ATG16L2,RCC1,PARD3,TCAIM,SLC3A2,DH
,DNASE1L1,KIF4A,RBCK1,PTOV1,FASTK,TMEM135,RNF32,LY75,ATG16L2,RCC1,PARD3,TCAIM,SLC3A2,DH
EM135,RNF32,LY75,RCC1,SNRPB,PARD3,TCAIM,SLC3A2,DHPS,ACTR6,SEC62,SET,CTSC,ZNF235,BOP1,ARR
EM135,RNF32,LY75,RCC1,SNRPB,PARD3,TCAIM,SLC3A2,DHPS,ACTR6,SEC62,SET,CTSC,ZNF235,BOP1,ARR
ASE1L1,RBCK1,PTOV1,FASTK,TMEM135,RNF32,LY75,ATG16L2,RCC1,SNRPB,SLC3A2,DHPS,ACTR6,SET,CT
33,ZNF410,RUVBL2,EBPL,IGSF8,WDR62,STYXL1,PLEKHA5,RANBP1,LZTFL1,PPM1K,TUBA1C,STIP1,LRRC1,P
SIRT6,NUMA1,BUD31,SAE1,NPRL2,OSGEP,TBL2,TSPAN11,ITSN1,EIF5A,COMMD5,UFD1L,USP38,TAZ,SH2
GSF8,SPAG7,WDR62,POLD4,PLEKHA5,RANBP1,UBE2L6,ZC3H14,PPM1K,STIP1,PHF19,LPCAT3,CD44,MAG
1K,STIP1,PHF19,LPCAT3,MAGED2,GRK6,CPSF3L,SLC39A11,FAM3A,METTL17,ATG12,NLRP1,ZBTB44,PLCB
6,TTLL5,PSMA6,ACOT8,AP3B2,RCE1,ZNF410,RUVBL2,EBPL,TESC,IGSF8,NNMT,SPAG7,WDR62,IL1RAP,STY
6,TTLL5,PSMA6,ACOT8,AP3B2,RCE1,ZNF410,RUVBL2,EBPL,TESC,IGSF8,NNMT,SPAG7,WDR62,IL1RAP,STY
BXW9,RFX3,ENDOV,PPIL2,RPL28,LIG1,CDC14B,TMEM126B,CAPN10,LYPLA2,C19ORF48,PPAN,STAU1,CAR
ARCB1,RPL38,STARD10,RNH1,DNASE1L1,KIF4A,RBCK1,PTOV1,FASTK,TMEM135,RNF32,LY75,RCC1,SNRPE
ARCB1,RPL38,STARD10,RNH1,DNASE1L1,KIF4A,RBCK1,PTOV1,FASTK,TMEM135,RNF32,LY75,RCC1,SNRPE
C3A2,SEC62,SET,ZNF235,ARRDC1,TRPT1,KLC1,RBP5,ZC3HAV1,SCAF11,LIMK2,SIRT6,NUMA1,GPATCH2L,B
C3A2,SEC62,SET,ZNF235,ARRDC1,TRPT1,KLC1,RBP5,ZC3HAV1,SCAF11,LIMK2,SIRT6,NUMA1,GPATCH2L,B
,SCARB1,CRYGS,CTTN,TSPAN11,ITSN1,COMMD5,USP38,TAZ,THYN1,SH2B1,GSTK1,LDHA,PSMA6,AP3B2,I
,SCARB1,CRYGS,CTTN,TSPAN11,ITSN1,COMMD5,USP38,TAZ,THYN1,SH2B1,GSTK1,LDHA,PSMA6,AP3B2,I

PZ1,LRRFIP1,TRMT112,CHPF,ZNF625,GABPB1,VWA2,DDOST,LSR,RRNAD1,RAD1,TOMM40,TADA3,ATP6V
,TAZ,TTLL5,AP3B2,MRPL33,RUVBL2,IGSF8,SNRPN,WDR62,RFX5,UBE2L6,FIS1,SLC15A3,PPM1K,ANTXR2,L
,TAZ,TTLL5,AP3B2,MRPL33,RUVBL2,IGSF8,SNRPN,WDR62,RFX5,UBE2L6,FIS1,SLC15A3,PPM1K,ANTXR2,L
,C12ORF57,DCTN5,NDUFS3,OXLD1,ZNF223,ARIH2,PICK1,RNF181,TSPAN31,RALY,GLYCTK,RBM42,PSMD7
JD31,SAE1,NPRL2,ARHGAP6,CRYGS,CTTN,TSPAN11,EIF5A,COMMD5,AMZ2,UFD1L,USP38,DBNDD1,SH2B
JD31,SAE1,NPRL2,ARHGAP6,CRYGS,CTTN,TSPAN11,EIF5A,COMMD5,AMZ2,UFD1L,USP38,DBNDD1,SH2B
FS3,OXLD1,SLC35E1,OGFOD2,EIF3G,ARIH2,PICK1,FBXL18,TSPAN31,ZNF593,GLYCTK,CFLAR,RER1,EIF2A,K
CTTN,TSPAN11,EIF5A,RUNX1,UFD1L,USP38,DBNDD1,SH2B1,RPL13A,ARHGEF39,GSTK1,TTLL5,LDHA,PSM
CTTN,TSPAN11,EIF5A,RUNX1,UFD1L,USP38,DBNDD1,SH2B1,RPL13A,ARHGEF39,GSTK1,TTLL5,LDHA,PSM
1,KIF4A,RBCK1,PTOV1,RNF32,LY75,ATG16L2,RCC1,SNRPB,PARD3,TCAIM,DHPS,ACTR6,MAP3K7CL,SEC62
,PARD3,SLC3A2,DHPS,ACTR6,MAP3K7CL,SET,CTSC,TRPT1,SCAF11,DBI,PMFBP1,LIMK2,VDAC3,NOB1,WD
,PARD3,SLC3A2,DHPS,ACTR6,MAP3K7CL,SET,CTSC,TRPT1,SCAF11,DBI,PMFBP1,LIMK2,VDAC3,NOB1,WD
N,SPAG7,WDR62,POLD4,UBE2L6,STIP1,PHF19,LPCAT3,GRK6,RPS16,CPSF3L,OBSL1,SLC39A11,FAM3A,AT
EBPL,TESC,RBPMS,WDR62,IL1RAP,POLD4,PLEKHA5,RANBP1,UBE2L6,VRK3,LZTFL1,FIS1,SLC15A3,TUBA1C,
EBPL,TESC,RBPMS,WDR62,IL1RAP,POLD4,PLEKHA5,RANBP1,UBE2L6,VRK3,LZTFL1,FIS1,SLC15A3,TUBA1C,
,RUVBL2,EBPL,TESC,SNRPN,SPAG7,WDR62,RANBP1,UBE2L6,PPM1K,ANTXR2,STIP1,PHF19,VCAN,LPCAT3,
,RUVBL2,EBPL,TESC,SNRPN,SPAG7,WDR62,RANBP1,UBE2L6,PPM1K,ANTXR2,STIP1,PHF19,VCAN,LPCAT3,
37,RPS16,SLC39A11,FAM3A,EPB41L5,ATG12,ZBTB44,NISCH,TMED4,CD99L2,GGPS1,TRNT1,PARD6G,SH3
37,RPS16,SLC39A11,FAM3A,EPB41L5,ATG12,ZBTB44,NISCH,TMED4,CD99L2,GGPS1,TRNT1,PARD6G,SH3
3P,CIRBP,RHOF,SMARCB1,PRR13,STARD10,RNH1,PTOV1,FASTK,TMEM135,RNF32,LY75,RCC1,SNRPB,PAR
3P,CIRBP,RHOF,SMARCB1,PRR13,STARD10,RNH1,PTOV1,FASTK,TMEM135,RNF32,LY75,RCC1,SNRPB,PAR
A3,TRIM16,MDH2,CHCHD5,SEC31A,CARHSP1,RPL15,B3GAT3,ZNF506,PAFAH1B2,GBA2,DALRD3,NSUN5,
XIN1,TMEM199,EPN1,HPS1,ITPA,ZCCHC9,VPS25,CHURC1,AZI2,TBC1D22A,ZNF276,SZT2,PEX16,MRPL51,
L13A,ARHGEF39,GSTK1,PSMA6,ACOT8,AP3B2,SLC35G1,MRPL33,ZNF410,PCMT1,NUDT5,SNRPN,SPAG7,
L13A,ARHGEF39,GSTK1,PSMA6,ACOT8,AP3B2,SLC35G1,MRPL33,ZNF410,PCMT1,NUDT5,SNRPN,SPAG7,
FASTK,TMEM135,LY75,ATG16L2,RCC1,SNRPB,PARD3,TCAIM,SLC3A2,DHPS,ACTR6,MAP3K7CL,SEC62,SET,
FASTK,TMEM135,LY75,ATG16L2,RCC1,SNRPB,PARD3,TCAIM,SLC3A2,DHPS,ACTR6,MAP3K7CL,SEC62,SET,
RL2,OSGEP,TBL2,SCARB1,CRYGS,ITSN1,TNFSF10,COMMD5,CHID1,NUDT22,UFD1L,USP38,TAZ,THYN1,DB
RL2,OSGEP,TBL2,SCARB1,CRYGS,ITSN1,TNFSF10,COMMD5,CHID1,NUDT22,UFD1L,USP38,TAZ,THYN1,DB
PB,PARD3,TCAIM,DHPS,ACTR6,SEC62,SET,CTSC,BOP1,ARRDC1,KLC1,RBP5,ZC3HAV1,DBI,ARPC2,LIMK2,N
PB,PARD3,TCAIM,DHPS,ACTR6,SEC62,SET,CTSC,BOP1,ARRDC1,KLC1,RBP5,ZC3HAV1,DBI,ARPC2,LIMK2,N
,BUD31,NPRL2,ARHGAP6,CRYGS,TSPAN11,ITSN1,COMMD5,NUDT22,AMZ2,UFD1L,TAZ,THYN1,SH2B1,NC
,BUD31,NPRL2,ARHGAP6,CRYGS,TSPAN11,ITSN1,COMMD5,NUDT22,AMZ2,UFD1L,TAZ,THYN1,SH2B1,NC
,BUD31,NPRL2,ARHGAP6,CRYGS,TSPAN11,ITSN1,COMMD5,NUDT22,AMZ2,UFD1L,TAZ,THYN1,SH2B1,NC
,BUD31,NPRL2,ARHGAP6,CRYGS,TSPAN11,ITSN1,COMMD5,NUDT22,AMZ2,UFD1L,TAZ,THYN1,SH2B1,NC
N1,THYN1,DBNDD1,NOP16,GSTK1,PSMA6,ACOT8,AP3B2,RCE1,ZNF410,EBPL,TESC,IGSF8,SPAG7,WDR62,
N1,THYN1,DBNDD1,NOP16,GSTK1,PSMA6,ACOT8,AP3B2,RCE1,ZNF410,EBPL,TESC,IGSF8,SPAG7,WDR62,
EM135,ATG16L2,RCC1,SNRPB,PARD3,TCAIM,SLC3A2,DHPS,ACTR6,MAP3K7CL,SEC62,SET,CTSC,ZNF235,
TN,TSPAN11,ITSN1,EIF5A,DBNDD1,SH2B1,NOP16,GSTK1,TTLL5,PSMA6,ACOT8,AP3B2,RCE1,RUVBL2,EBP
TN,TSPAN11,ITSN1,EIF5A,DBNDD1,SH2B1,NOP16,GSTK1,TTLL5,PSMA6,ACOT8,AP3B2,RCE1,RUVBL2,EBP
PB,PARD3,TCAIM,DHPS,ACTR6,SEC62,SET,CTSC,BOP1,ARRDC1,KLC1,RBP5,ZC3HAV1,DBI,ARPC2,LIMK2,N
SC,IGSF8,SPAG7,WDR62,POLD4,UBE2L6,ZC3H14,PPM1K,STIP1,PHF19,LPCAT3,CD44,SON,GRK6,CDC37,R
TK1,LDHA,PSMA6,ACOT8,RPS9,SLC35G1,NUDT5,NNMT,SNRPN,SPAG7,WDR62,STYXL1,VRK3,LZTFL1,FIS1
TK1,LDHA,PSMA6,ACOT8,RPS9,SLC35G1,NUDT5,NNMT,SNRPN,SPAG7,WDR62,STYXL1,VRK3,LZTFL1,FIS1
NUMA1,SUMF2,SAE1,TBL2,SCARB1,CTTN,TSPAN11,ITSN1,EIF5A,COMMD5,NUDT22,AMZ2,TAZ,DBNDD1,
NUMA1,SUMF2,SAE1,TBL2,SCARB1,CTTN,TSPAN11,ITSN1,EIF5A,COMMD5,NUDT22,AMZ2,TAZ,DBNDD1,
INO80E,CCNL2,LRRFIP1,TRMT112,CHPF,ZNF625,GABPB1,ZNF814,VWA2,DDOST,LSR,RRNAD1,RAD1,TOM

5,SEC31A,CNST,COX17,PAFAH1B2,NSUN5,GMPR2,TCP1,PIGT,UBE2N,NFATC4,PDCD6,FAM32A,TGOLN2,

PARD3,TCAIM,ACTR6,SEC62,SET,CTSC,ZNF235,BOP1,KLC1,RBP5,ZC3HAV1,DBI,ARPC2,PMFBP1,LIMK2,NO

PARD3,TCAIM,ACTR6,SEC62,SET,CTSC,ZNF235,BOP1,KLC1,RBP5,ZC3HAV1,DBI,ARPC2,PMFBP1,LIMK2,NO

OMMD5,UFD1L,USP38,TAZ,SH2B1,NOP16,ARHGEF39,PSMA6,ACOT8,AP3B2,NUDT5,RBPMS,SPAG7,WDR6

OMMD5,UFD1L,USP38,TAZ,SH2B1,NOP16,ARHGEF39,PSMA6,ACOT8,AP3B2,NUDT5,RBPMS,SPAG7,WDR6

RPB,PARD3,TCAIM,DHPS,ACTR6,SEC62,SET,CTSC,BOP1,ARRDC1,TRPT1,KLC1,RBP5,DBI,ARPC2,LIMK2,VDA

RPB,PARD3,TCAIM,DHPS,ACTR6,SEC62,SET,CTSC,BOP1,ARRDC1,TRPT1,KLC1,RBP5,DBI,ARPC2,LIMK2,VDA

ARHGAP6,CRYGS,CTTN,TSPAN11,EIF5A,COMMD5,NUDT22,UFD1L,USP38,TAZ,THYN1,RPL13A,ARHGEF39

ARHGAP6,CRYGS,CTTN,TSPAN11,EIF5A,COMMD5,NUDT22,UFD1L,USP38,TAZ,THYN1,RPL13A,ARHGEF39

F8,SPAG7,WDR62,POLD4,UBE2L6,STIP1,PHF19,GRK6,CDC37,RPS16,TSG101,OBSL1,SLC39A11,FAM3A,MU

IM,DHPS,SET,CTSC,BOP1,ARRDC1,TRPT1,KLC1,SCAF11,DBI,ARPC2,VDAC3,WDR19,GPATCH2L,SUMF2,BU

IM,DHPS,SET,CTSC,BOP1,ARRDC1,TRPT1,KLC1,SCAF11,DBI,ARPC2,VDAC3,WDR19,GPATCH2L,SUMF2,BU

CC5,EMC4,DDOST,LSR,TOMM40,TADA3,ANAPC11,RAD23A,ABHD17A,MDH2,NAA10,SEC31A,CARHSP1,C

CC5,EMC4,DDOST,LSR,TOMM40,TADA3,ANAPC11,RAD23A,ABHD17A,MDH2,NAA10,SEC31A,CARHSP1,C

BOP1,ARRDC1,TRPT1,KLC1,SCAF11,DBI,VDAC3,WDR19,GPATCH2L,SUMF2,BUD31,SAE1,OSGEP,TBL2,ARH

BOP1,ARRDC1,TRPT1,KLC1,SCAF11,DBI,VDAC3,WDR19,GPATCH2L,SUMF2,BUD31,SAE1,OSGEP,TBL2,ARH

,SUMF2,SAE1,OSGEP,TBL2,SCARB1,CTTN,TSPAN11,EIF5A,COMMD5,CHID1,NUDT22,AMZ2,TAZ,RPL13A,L

PB,PARD3,TCAIM,DHPS,ACTR6,SEC62,SET,CTSC,BOP1,ARRDC1,KLC1,RBP5,DBI,ARPC2,LIMK2,VDAC3,NOB

RMT112,DDOST,LSR,TOMM40,TADA3,PDCD5,MDH2,CARHSP1,LLGL2,COX17,FKBP3,B3GAT3,PAFAH1B2,G

19,HMGA2,RPS16,CPSF3L,UBR4,ZBTB44,DUSP22,SLC25A10,TMED4,SRSF1,YIPF3,ITGB1BP1,SH3YL1,YIF1A

19,HMGA2,RPS16,CPSF3L,UBR4,ZBTB44,DUSP22,SLC25A10,TMED4,SRSF1,YIPF3,ITGB1BP1,SH3YL1,YIF1A

SGEP,TBL2,CTTN,ITSN1,COMMD5,CHID1,NUDT22,AMZ2,UFD1L,USP38,THYN1,DBNDD1,SH2B1,RPL13A,T

SGEP,TBL2,CTTN,ITSN1,COMMD5,CHID1,NUDT22,AMZ2,UFD1L,USP38,THYN1,DBNDD1,SH2B1,RPL13A,T

RD3,TCAIM,DHPS,ACTR6,SEC62,SET,CTSC,BOP1,ARRDC1,TRPT1,KLC1,RBP5,ARPC2,LIMK2,VDAC3,NOB1,V

D1,IP6K2,UGGT2,MAGT1,PCYT2,IFI27L1,CCDC25,PPP1CC,RFX3,RPL28,PSMA7,TIMM17B,APEH,RPL21,NU

P1,NIP7,PLCB2,DUSP22,SLC25A10,TMX2,DBNDD2,SDF4,RPL29,CD99L2,SAP30BP,RABL5,TRNT1,ITGB1BP

P1,NIP7,PLCB2,DUSP22,SLC25A10,TMX2,DBNDD2,SDF4,RPL29,CD99L2,SAP30BP,RABL5,TRNT1,ITGB1BP

ASTK,RNF32,LY75,ATG16L2,RCC1,SNRPB,PARD3,TCAIM,SLC3A2,DHPS,ACTR6,SET,CTSC,BOP1,ARRDC1,TR

ASTK,RNF32,LY75,ATG16L2,RCC1,SNRPB,PARD3,TCAIM,SLC3A2,DHPS,ACTR6,SET,CTSC,BOP1,ARRDC1,TR

DAC3,NOB1,WDR19,SIRT6,NUMA1,GPATCH2L,OSGEP,ARHGAP6,SCARB1,CTTN,TSPAN11,EIF5A,AMZ2,UFD

DAC3,NOB1,WDR19,SIRT6,NUMA1,GPATCH2L,OSGEP,ARHGAP6,SCARB1,CTTN,TSPAN11,EIF5A,AMZ2,UFD

A,TTLL5,AP3B2,RPS9,SLC35G1,ZNF410,TESC,RBPMS,SPAG7,WDR62,RFX5,POLD4,RANBP1,ZC3H14,FIS1,SL

A,TTLL5,AP3B2,RPS9,SLC35G1,ZNF410,TESC,RBPMS,SPAG7,WDR62,RFX5,POLD4,RANBP1,ZC3H14,FIS1,SL

TCH2L,SUMF2,BUD31,NPRL2,OSGEP,TBL2,ARHGAP6,EIF5A,CHID1,AMZ2,UFD1L,USP38,TAZ,THYN1,SH2B1

TCH2L,SUMF2,BUD31,NPRL2,OSGEP,TBL2,ARHGAP6,EIF5A,CHID1,AMZ2,UFD1L,USP38,TAZ,THYN1,SH2B1

,DBI,ARPC2,SIRT6,SUMF2,BUD31,SAE1,NPRL2,OSGEP,TBL2,ARHGAP6,TSPAN11,EIF5A,TNFSF10,RUNX1,U

,DBI,ARPC2,SIRT6,SUMF2,BUD31,SAE1,NPRL2,OSGEP,TBL2,ARHGAP6,TSPAN11,EIF5A,TNFSF10,RUNX1,U

NUMA1,GPATCH2L,TBL2,SCARB1,CTTN,TSPAN11,ITSN1,EIF5A,NUDT22,DBNDD1,NOP16,GSTK1,TTLL5,PSM

NUMA1,GPATCH2L,TBL2,SCARB1,CTTN,TSPAN11,ITSN1,EIF5A,NUDT22,DBNDD1,NOP16,GSTK1,TTLL5,PSM

CIRBP,RHOF,RPL38,PRR13,STARD10,RNH1,DNASE1L1,KIF4A,RBCK1,PTOV1,FASTK,TMEM135,RNF32,LY75,

CIRBP,RHOF,RPL38,PRR13,STARD10,RNH1,DNASE1L1,KIF4A,RBCK1,PTOV1,FASTK,TMEM135,RNF32,LY75,

,NOP16,ACOT8,AP3B2,RCE1,RUVBL2,EBPL,TESC,IGSF8,NNMT,SPAG7,WDR62,IL1RAP,RANBP1,UBE2L6,FI

,TSPAN11,TNFSF10,COMMD5,RUNX1,CHID1,NUDT22,USP38,TAZ,THYN1,DBNDD1,SH2B1,NOP16,ARHGE

MD5,NUDT22,UFD1L,TAZ,NOP16,SLC35G1,RCE1,EBPL,IGSF8,RBPMS,IL1RAP,PLEKHA5,RANBP1,UBE2L6,F

,ARRDC1,KLC1,RBP5,ZC3HAV1,DBI,ARPC2,LIMK2,NOB1,WDR19,SIRT6,NUMA1,GPATCH2L,BUD31,SAE1,N

,ARRDC1,KLC1,RBP5,ZC3HAV1,DBI,ARPC2,LIMK2,NOB1,WDR19,SIRT6,NUMA1,GPATCH2L,BUD31,SAE1,N

,TCAIM,SLC3A2,DHPS,SET,CTSC,ZNF235,ARRDC1,TRPT1,KLC1,RBP5,SCAF11,ARPC2,LIMK2,NOB1,NUMA1,

,TCAIM,SLC3A2,DHPS,SET,CTSC,ZNF235,ARRDC1,TRPT1,KLC1,RBP5,SCAF11,ARPC2,LIMK2,NOB1,NUMA1,

A,COMMD5,CHID1,UFD1L,DBNDD1,SH2B1,NOP16,RPL13A,GSTK1,TTLL5,ACOT8,SLC35G1,MRPL33,ZNF41

MAP3K7CL,CTSC,ZNF235,BOP1,ARRDC1,RBP5,ZC3HAV1,SCAF11,DBI,ARPC2,PMFBP1,LIMK2,NOB1,WDR1

MAP3K7CL,CTSC,ZNF235,BOP1,ARRDC1,RBP5,ZC3HAV1,SCAF11,DBI,ARPC2,PMFBP1,LIMK2,NOB1,WDR1

ZTFL1,FIS1,SLC15A3,PPM1K,IRF9,PHF19,PCBP2,MAGED2,SON,SNF8,CPSF3L,TSG101,SLC39A11,FUZ,METT

ZTFL1,FIS1,SLC15A3,PPM1K,IRF9,PHF19,PCBP2,MAGED2,SON,SNF8,CPSF3L,TSG101,SLC39A11,FUZ,METT

TRNT1,ITGB1BP1,CLASRP,C1ORF198,UGGT2,UBE2I,CDC42,FBXW9,USE1,ASF1B,RPL28,LIG1,RBM33,LYPL

TRNT1,ITGB1BP1,CLASRP,C1ORF198,UGGT2,UBE2I,CDC42,FBXW9,USE1,ASF1B,RPL28,LIG1,RBM33,LYPL

NAPC11,ABHD17A,MDH2,SEC31A,CNST,COX17,HNRNPLL,B3GAT3,PAFAH1B2,DALRD3,NSUN5,MRPS18C,

DT22,USP38,THYN1,NOP16,TTLL5,AP3B2,RPS9,MRPL33,PCMT1,WDR62,PLEKHA5,RANBP1,UBE2L6,VRK3

.2,TRNT1,PARD6G,SH3YL1,IP6K2,CLASRP,JPH2,C1ORF198,MICAL1,UBE2I,TBX6,XBP1,PTK2,AUNIP,C14ORF

.2,TRNT1,PARD6G,SH3YL1,IP6K2,CLASRP,JPH2,C1ORF198,MICAL1,UBE2I,TBX6,XBP1,PTK2,AUNIP,C14ORF

P1,ZBTB44,NISCH,RPL29,CD99L2,TRNT1,PARD6G,SH3YL1,IP6K2,CLASRP,JPH2,C1ORF198,MICAL1,UBE2I,P

FOXRED1,ITGB1BP1,EIF3M,CDC42,AUNIP,CENPN,CARD14,PCYT2,EEF1B2,DGKE,CAMTA1,PPIL2,PSMD4,R

.,CACYBP,CIRBP,RHOF,SMARCB1,RPL38,PRR13,STARD10,RNH1,DNASE1L1,KIF4A,RBCK1,PTOV1,FASTK,TM

.3A,GSTK1,LDHA,RPS9,RUVBL2,PCMT1,EBPL,IGSF8,WDR62,STYXL1,POLD4,RANBP1,ZC3H14,SLC15A3,STI

5,RCC1,SNRPB,PARD3,TCAIM,SLC3A2,ACTR6,MAP3K7CL,SEC62,CTSC,ZNF235,BOP1,ARRDC1,TRPT1,KLC1,

5,RCC1,SNRPB,PARD3,TCAIM,SLC3A2,ACTR6,MAP3K7CL,SEC62,CTSC,ZNF235,BOP1,ARRDC1,TRPT1,KLC1,

IM,TMPO,PRMT1,ZNF7,TXNDC17,ARL6IP4,COPZ1,CCNL2,CEP70,TPM2,SIAH1,GGCX,STRIP2,LYNX1,TWSG

.T3,RFXANK,FTH1,PFDN5,PAFAH1B2,TXNRD2,DALRD3,NSUN5,MRPS18C,SMARCC2,ZNF81,FAM32A,STX1

.T3,RFXANK,FTH1,PFDN5,PAFAH1B2,TXNRD2,DALRD3,NSUN5,MRPS18C,SMARCC2,ZNF81,FAM32A,STX1

C2,PMFBP1,LIMK2,WDR19,NUMA1,SUMF2,BUD31,SAE1,NPRL2,ARHGAP6,SCARB1,CTTN,TSPAN11,ITSN1,

.13A,GSTK1,PSMA6,ACOT8,AP3B2,SLC35G1,MRPL33,ZNF410,PCMT1,IGSF8,NUDT5,SNRPN,SPAG7,WDR6

.13A,GSTK1,PSMA6,ACOT8,AP3B2,SLC35G1,MRPL33,ZNF410,PCMT1,IGSF8,NUDT5,SNRPN,SPAG7,WDR6

HA,RPS9,RCE1,ZNF410,PCMT1,IGSF8,NUDT5,NNMT,SNRPN,WDR62,RFX5,RANBP1,VRK3,SLC15A3,PPM1

HA,RPS9,RCE1,ZNF410,PCMT1,IGSF8,NUDT5,NNMT,SNRPN,WDR62,RFX5,RANBP1,VRK3,SLC15A3,PPM1

NUDT22,DBNDD1,SH2B1,NOP16,RPL13A,GSTK1,TTLL5,LDHA,ACOT8,SLC35G1,ZNF410,PCMT1,TESC,IGSF

NUDT22,DBNDD1,SH2B1,NOP16,RPL13A,GSTK1,TTLL5,LDHA,ACOT8,SLC35G1,ZNF410,PCMT1,TESC,IGSF

1,PARD6G,SH3YL1,IP6K2,MICAL1,HS2ST1,UBE2I,TMEM205,AUNIP,C14ORF2,GUK1,PCYT2,PPP1CC,PPA1,

GEP,CTTN,ITSN1,EIF5A,COMMD5,AMZ2,SH2B1,RPL13A,GSTK1,TTLL5,LDHA,AP3B2,RPS9,SLC35G1,RCE1,

GEP,CTTN,ITSN1,EIF5A,COMMD5,AMZ2,SH2B1,RPL13A,GSTK1,TTLL5,LDHA,AP3B2,RPS9,SLC35G1,RCE1,

,RCE1,ZNF410,PCMT1,IGSF8,NNMT,WDR62,STYXL1,PLEKHA5,RANBP1,VRK3,FIS1,SLC15A3,PPM1K,ANTX

,RCE1,ZNF410,PCMT1,IGSF8,NNMT,WDR62,STYXL1,PLEKHA5,RANBP1,VRK3,FIS1,SLC15A3,PPM1K,ANTX

B1,RPL38,STARD10,DNASE1L1,KIF4A,RBCK1,PTOV1,FASTK,TMEM135,RNF32,LY75,ATG16L2,RCC1,SNRPB

B1,RPL38,STARD10,DNASE1L1,KIF4A,RBCK1,PTOV1,FASTK,TMEM135,RNF32,LY75,ATG16L2,RCC1,SNRPB

KHA5,RANBP1,UBE2L6,ANTXR2,STIP1,PHF19,LPCAT3,MAGED2,SON,GRK6,RPS16,CPSF3L,FAM3A,METTL1

K,GTF2H1,AXIN1,TP53,TMEM199,SLC25A24,EPN1,ITPA,ZCCHC9,SNRPD1,AZI2,GOSR2,TBC1D22A,LAMTC

VDR62,PLEKHA5,RANBP1,LZTFL1,ZC3H14,FIS1,STIP1,LRRC1,PHF19,LPCAT3,CD44,TM9SF1,SON,TET2,UQC

JMF2,BUD31,SAE1,NPRL2,SCARB1,ITSN1,EIF5A,UFD1L,USP38,TAZ,DBNDD1,SH2B1,ARHGEF39,GSTK1,PS

JMF2,BUD31,SAE1,NPRL2,SCARB1,ITSN1,EIF5A,UFD1L,USP38,TAZ,DBNDD1,SH2B1,ARHGEF39,GSTK1,PS

RQ,PTPN13,UBR4,PSMA1,FUZ,FAM3A,PPRC1,NIP7,COX4I1,DUSP22,SDF4,TMED4,GGPS1,RTEL1,SRSF1,AU

SMA1,FUZ,FAM3A,PPRC1,NIP7,COX4I1,DUSP22,TMED4,CENPK,GGPS1,RTEL1,SRSF1,AUP1,FOXRED1,SH3

A1,GPATCH2L,BUD31,OSGEP,ARHGAP6,SCARB1,CTTN,TSPAN11,ITSN1,TNFSF10,COMMD5,CHID1,NUDT2

A1,GPATCH2L,BUD31,OSGEP,ARHGAP6,SCARB1,CTTN,TSPAN11,ITSN1,TNFSF10,COMMD5,CHID1,NUDT2

T,ZNF506,PFDN5,PAFAH1B2,PPIH,DAZAP2,ZFPL1,GMPR2,RIC8A,TCP1,SMARCC2,FAM32A,COX20,PEX2,C1

T,ZNF506,PFDN5,PAFAH1B2,PPIH,DAZAP2,ZFPL1,GMPR2,RIC8A,TCP1,SMARCC2,FAM32A,COX20,PEX2,C1

RPC2,LIMK2,VDAC3,NUMA1,GPATCH2L,BUD31,TBL2,ARHGAP6,SCARB1,CTTN,TSPAN11,ITSN1,EIF5A,COM

RPC2,LIMK2,VDAC3,NUMA1,GPATCH2L,BUD31,TBL2,ARHGAP6,SCARB1,CTTN,TSPAN11,ITSN1,EIF5A,COM

MA1,YLPM1,CHCHD1,TTC9C,COX4I1,THEM4,DBNDD2,RPS6,RPL29,AUP1,SAP30BP,YIPF3,FOXRED1,TRNT

NF814,ERCC5,MCTS1,SERPINB1,DDOST,LSR,ZNF836,TOMM40,SERPINB6,TADA3,SLC33A1,MDH2,SEC31A

1A,COX17,B3GAT3,ASH2L,GMPR2,ZSCAN9,RIC8A,PARK7,UBE2N,PDCD6,FAM32A,USP40,C12ORF57,OXLD

D31,SAE1,NPRL2,OSGEP,ARHGAP6,EIF5A,COMMD5,RUNX1,CHID1,UFD1L,TAZ,DBNDD1,NOP16,GSTK1,TTL

D31,SAE1,NPRL2,OSGEP,ARHGAP6,EIF5A,COMMD5,RUNX1,CHID1,UFD1L,TAZ,DBNDD1,NOP16,GSTK1,TTL

D1,SH2B1,RPL13A,PSMA6,ACOT8,AP3B2,RPS9,SLC35G1,RCE1,PCMT1,EBPL,NUDT5,RBPMS,WDR62,PLEKH

TSPAN31,GLYCTK,RER1,AXIN1,TMEM199,EPN1,HPS1,ITPA,ZCCHC9,SNRPD1,SNRPA1,AZI2,GOSR2,TBC1D2

RB1,CTTN,EIF5A,AMZ2,UFD1L,DBNDD1,ARHGEF39,PSMA6,ACOT8,AP3B2,MRPL33,ZNF410,RUVBL2,EBPL,

RB1,CTTN,EIF5A,AMZ2,UFD1L,DBNDD1,ARHGEF39,PSMA6,ACOT8,AP3B2,MRPL33,ZNF410,RUVBL2,EBPL,

L3,JUP,CDC42,PTK2,USE1,ASF1B,PPP1CC,VAMP8,CAPG,PPIL2,TROAP,LIG1,PSMA7,C17ORF62,ACAD10,HN

L3,JUP,CDC42,PTK2,USE1,ASF1B,PPP1CC,VAMP8,CAPG,PPIL2,TROAP,LIG1,PSMA7,C17ORF62,ACAD10,HN

GAP6,EIF5A,RUNX1,CHID1,UFD1L,TAZ,DBNDD1,NOP16,GSTK1,TTLL5,ACOT8,AP3B2,SLC35G1,MRPL33,ZN

,TTC21B,UBR4,PSMA1,FAM3A,EPB41L5,RPS25,TTC9C,TMX2,THEM4,NISCH,TMED4,CENPK,RPS6,GGPS1,

P1,PHF19,LPCAT3,SON,GRK6,RPS16,CPSF3L,PHF21A,SLC39A11,MUC1,ATG12,PPRC1,ZBTB44,DBNDD2,NI

OP1,ARRDC1,TRPT1,KLC1,RBP5,ZC3HAV1,SCAF11,DBI,ARPC2,LIMK2,WDR19,SIRT6,NUMA1,GPATCH2L,B

OP1,ARRDC1,TRPT1,KLC1,RBP5,ZC3HAV1,SCAF11,DBI,ARPC2,LIMK2,WDR19,SIRT6,NUMA1,GPATCH2L,B

PB,PARD3,SLC3A2,ACTR6,MAP3K7CL,SEC62,SET,CTSC,ZNF235,BOP1,ARRDC1,TRPT1,KLC1,ZC3HAV1,SCAF

PB,PARD3,SLC3A2,ACTR6,MAP3K7CL,SEC62,SET,CTSC,ZNF235,BOP1,ARRDC1,TRPT1,KLC1,ZC3HAV1,SCAF

C1,TRPT1,KLC1,SCAF11,DBI,ARPC2,LIMK2,VDAC3,NOB1,WDR19,SIRT6,GPATCH2L,SUMF2,BUD31,SAE1,NP

C1,TRPT1,KLC1,SCAF11,DBI,ARPC2,LIMK2,VDAC3,NOB1,WDR19,SIRT6,GPATCH2L,SUMF2,BUD31,SAE1,NP

TK2,AUNIP,GUK1,PCYT2,PPP1CC,PPA1,DGKE,CAMTA1,CRIP1,PPIL2,LIG1,TIMM17B,APEH,NUTF2,C17ORF

35,BOP1,KLC1,ZC3HAV1,SCAF11,DBI,ARPC2,PMFBP1,LIMK2,NOB1,NUMA1,GPATCH2L,SUMF2,BUD31,O

SON,TET2,SNF8,GRK6,HMGA2,CDC37,CPSF3L,OBSL1,PSMA1,FAM3A,PSMA3,METTL17,NLRP1,ZBTB44,NI

TCAIM,SLC3A2,DHPS,ACTR6,SEC62,SET,ZNF235,BOP1,ARRDC1,KLC1,ZC3HAV1,SCAF11,ARPC2,PMFBP1,LI

TCAIM,SLC3A2,DHPS,ACTR6,SEC62,SET,ZNF235,BOP1,ARRDC1,KLC1,ZC3HAV1,SCAF11,ARPC2,PMFBP1,LI

HCHD5,CNST,RPL15,PAFAH1B2,NSUN5,ZFPL1,PIGT,UBE2N,NFATC4,FAM32A,ZSCAN20,TGOLN2,RPL37A,

HCHD5,CNST,RPL15,PAFAH1B2,NSUN5,ZFPL1,PIGT,UBE2N,NFATC4,FAM32A,ZSCAN20,TGOLN2,RPL37A,

PAN31,RALY,GLYCTK,TRAPPC2,AXIN1,TMEM199,EPN1,ITPA,ZCCHC9,SNRPD1,AZI2,GOSR2,MYL12A,TBC1

3,SPAG7,IL1RAP,ZC3H14,FIS1,SLC15A3,TUBA1C,IRF9,STIP1,GRAMD1C,VCAN,CD44,MAGED2,UQCRQ,SNF8

3,SPAG7,IL1RAP,ZC3H14,FIS1,SLC15A3,TUBA1C,IRF9,STIP1,GRAMD1C,VCAN,CD44,MAGED2,UQCRQ,SNF8

2,RALGPS1,ABHD17A,PHF20L1,KCTD7,RFXANK,PTK7,TMEM39B,BRF1,ZFPL1,PFKFB2,COMMD4,XYLT2,ZX

AP6,SCARB1,CTTN,ITSN1,EIF5A,COMMD5,RUNX1,CHID1,UFD1L,USP38,DBNDD1,NOP16,RPL13A,GSTK1,

AP6,SCARB1,CTTN,ITSN1,EIF5A,COMMD5,RUNX1,CHID1,UFD1L,USP38,DBNDD1,NOP16,RPL13A,GSTK1,

DHA,AP3B2,ZNF410,RUVBL2,PCMT1,IGSF8,SPAG7,WDR62,IL1RAP,POLD4,PLEKHA5,RANBP1,UBE2L6,ZC3H

DHA,AP3B2,ZNF410,RUVBL2,PCMT1,IGSF8,SPAG7,WDR62,IL1RAP,POLD4,PLEKHA5,RANBP1,UBE2L6,ZC3H

2,TBL2,CTTN,TSPAN11,ITSN1,EIF5A,COMMD5,NUDT22,AMZ2,UFD1L,THYN1,DBNDD1,SH2B1,RPL13A,LD

F2A,GLI4,SLC35A2,TMEM199,EPN1,HPS1,ITPA,ZBTB8OS,TBC1D22A,U2AF1L4,RNPS1,SUMO1,ZNF483,SZT

IA6,ACOT8,MRPL33,RUVBL2,EBPL,TESC,NNMT,WDR62,RFX5,PLEKHA5,RANBP1,LZTFL1,ZC3H14,TUBA1C,

IA6,ACOT8,MRPL33,RUVBL2,EBPL,TESC,NNMT,WDR62,RFX5,PLEKHA5,RANBP1,LZTFL1,ZC3H14,TUBA1C,

31,SAE1,OSGEP,TBL2,ARHGAP6,CRYGS,EIF5A,CHID1,AMZ2,TAZ,NOP16,RPL13A,GSTK1,TTLL5,LDHA,RPS9,

31,SAE1,OSGEP,TBL2,ARHGAP6,CRYGS,EIF5A,CHID1,AMZ2,TAZ,NOP16,RPL13A,GSTK1,TTLL5,LDHA,RPS9,

MA6,AP3B2,SLC35G1,ZNF410,PCMT1,EBPL,TESC,IGSF8,NUDT5,STYXL1,RFX5,PLEKHA5,RANBP1,FIS1,TUB

MA6,AP3B2,SLC35G1,ZNF410,PCMT1,EBPL,TESC,IGSF8,NUDT5,STYXL1,RFX5,PLEKHA5,RANBP1,FIS1,TUB

SN1,EIF5A,NUDT22,AMZ2,UFD1L,USP38,DBNDD1,NOP16,RPL13A,ARHGEF39,TTLL5,ACOT8,AP3B2,SLC35

N1,EIF5A,NUDT22,AMZ2,UFD1L,USP38,DBNDD1,NOP16,RPL13A,ARHGEF39,TTLL5,ACOT8,AP3B2,SLC35
1,ITPA,SNRPD1,CHURC1,SNRPA1,TBC1D22A,ZNF581,SZT2,LRP10,ACTB,C9ORF64,MED8,RNMT,MRPL27,
NDD1,SH2B1,AP3B2,RPS9,SLC35G1,ZNF410,EBPL,IGSF8,SPAG7,IL1RAP,RANBP1,UBE2L6,LZTFL1,ZC3H14,
NDD1,SH2B1,AP3B2,RPS9,SLC35G1,ZNF410,EBPL,IGSF8,SPAG7,IL1RAP,RANBP1,UBE2L6,LZTFL1,ZC3H14,
2,VDAC3,SIRT6,NPRL2,TBL2,ARHGAP6,SCARB1,CTTN,ITSN1,TNFSF10,COMMD5,RUNX1,NUDT22,TAZ,SH2
2,VDAC3,SIRT6,NPRL2,TBL2,ARHGAP6,SCARB1,CTTN,ITSN1,TNFSF10,COMMD5,RUNX1,NUDT22,TAZ,SH2
N1,EIF5A,COMMD5,UFD1L,USP38,TAZ,SH2B1,NOP16,LDHA,AP3B2,RPS9,RCE1,ZNF410,TESC,RBPMS,WD
N11,ITSN1,EIF5A,COMMD5,NUDT22,UFD1L,DBNDD1,SH2B1,NOP16,TTLL5,ACOT8,AP3B2,RCE1,RUVBL2,
3A2,DHPS,SEC62,SET,CTSC,BOP1,ARRDC1,TRPT1,KLC1,SCAF11,DBI,ARPC2,VDAC3,SIRT6,BUD31,NPRL2,C
3A2,DHPS,SEC62,SET,CTSC,BOP1,ARRDC1,TRPT1,KLC1,SCAF11,DBI,ARPC2,VDAC3,SIRT6,BUD31,NPRL2,C
1,ZNF410,PCMT1,IGSF8,NUDT5,NNMT,SNRPN,WDR62,RFX5,RANBP1,VRK3,PPM1K,PCBP2,TM9SF1,MAG
1,ZNF410,PCMT1,IGSF8,NUDT5,NNMT,SNRPN,WDR62,RFX5,RANBP1,VRK3,PPM1K,PCBP2,TM9SF1,MAG
RPL51,C9ORF64,MED8,AP4M1,MRPL27,DDX56,MED17,URM1,MARK2,SEC14L1,DPP9,SEC24C,TVP23B,SR
8,TAZ,NOP16,RPL13A,GSTK1,TTLL5,LDHA,AP3B2,RCE1,NUDT5,RBPMS,WDR62,POLD4,UBE2L6,FIS1,SLC1
PRC1,ZBTB44,TMX2,NISCH,CD99L2,GGPS1,SRSF1,PARD6G,SH3YL1,IP6K2,CLASRP,MICAL1,HS2ST1,UBE2I,
,PSMA3,NIP7,COX4I1,DUSP22,GGPS1,RTEL1,SRSF1,AUP1,FOXRED1,SH3YL1,UGGT2,HS2ST1,PRMT2,MAG
RHGAP6,SCARB1,CTTN,ITSN1,EIF5A,COMMD5,NUDT22,AMZ2,USP38,THYN1,DBNDD1,SH2B1,NOP16,AR
,RNF32,LY75,ATG16L2,RCC1,SNRPB,PARD3,TCAIM,SLC3A2,DHPS,ACTR6,SEC62,SET,CTSC,ZNF235,BOP1,A
,RNF32,LY75,ATG16L2,RCC1,SNRPB,PARD3,TCAIM,SLC3A2,DHPS,ACTR6,SEC62,SET,CTSC,ZNF235,BOP1,A
5,PAFAH1B2,NSUN5,FAM32A,PCID2,PEX2,C12ORF57,DCTN5,ZNF223,OGFOD2,EIF3G,PICK1,FBXL18,SLC5
5,PAFAH1B2,NSUN5,FAM32A,PCID2,PEX2,C12ORF57,DCTN5,ZNF223,OGFOD2,EIF3G,PICK1,FBXL18,SLC5
ZNF235,ARRDC1,TRPT1,RBP5,ZC3HAV1,DBI,ARPC2,LIMK2,VDAC3,NOB1,SIRT6,GPATCH2L,SUMF2,BUD31
ZNF235,ARRDC1,TRPT1,RBP5,ZC3HAV1,DBI,ARPC2,LIMK2,VDAC3,NOB1,SIRT6,GPATCH2L,SUMF2,BUD31
AC3,NOB1,NUMA1,GPATCH2L,SUMF2,SAE1,NPRL2,TBL2,ARHGAP6,SCARB1,CTTN,TSPAN11,ITSN1,EIF5A,
DD2,NISCH,CD99L2,GGPS1,PARD6G,SH3YL1,IP6K2,CLASRP,MICAL1,HS2ST1,UBE2I,AUNIP,C14ORF2,PCYT
SYNGR1,PABPC1,WAC,PDCD5,ANAPC11,USP45,ILF3,SSH2,COX17,HINT1,HNRNPA1,EGLN2,SIRT5,RFXANK
SYNGR1,PABPC1,WAC,PDCD5,ANAPC11,USP45,ILF3,SSH2,COX17,HINT1,HNRNPA1,EGLN2,SIRT5,RFXANK
,YLPM1,FAM3A,MUC1,ATG12,ZBTB44,DBNDD2,CD99L2,GGPS1,AUP1,KXD1,PARD6G,SH3YL1,IP6K2,C1O
1,ERLEC1,EMC4,DDOST,LSR,RRNAD1,DBNL,TOMM40,RABEP2,TADA3,TRIM16,MDH2,CHCHD5,SEC31A,C
H2B1,ARHGEF39,TTLL5,PSMA6,RPS9,SLC35G1,RCE1,RUVBL2,PCMT1,IGSF8,NUDT5,NNMT,WDR62,IL1RA
T8,AP3B2,RCE1,RUVBL2,EBPL,RBPMS,NNMT,SPAG7,IL1RAP,RFX5,PLEKHA5,UBE2L6,VRK3,ZC3H14,PPM
T8,AP3B2,RCE1,RUVBL2,EBPL,RBPMS,NNMT,SPAG7,IL1RAP,RFX5,PLEKHA5,UBE2L6,VRK3,ZC3H14,PPM
135,RNF32,LY75,ATG16L2,PARD3,TCAIM,SLC3A2,DHPS,ACTR6,MAP3K7CL,SEC62,SET,CTSC,ZNF235,BO
135,RNF32,LY75,ATG16L2,PARD3,TCAIM,SLC3A2,DHPS,ACTR6,MAP3K7CL,SEC62,SET,CTSC,ZNF235,BO
NF644,CDK11A,INO80E,FAM219B,ZNF524,ELP6,SIAH1,TRMT112,GABPB1,MPDU1,RPS15,LRRC27,MVD,S
NF644,CDK11A,INO80E,FAM219B,ZNF524,ELP6,SIAH1,TRMT112,GABPB1,MPDU1,RPS15,LRRC27,MVD,S
MFSD10,GMPR2,ZSCAN9,UBE2N,LFNG,PDCD6,FAM32A,NFYC,C12ORF57,OXLD1,ZNF223,OGFOD2,EIF3G,
MFSD10,GMPR2,ZSCAN9,UBE2N,LFNG,PDCD6,FAM32A,NFYC,C12ORF57,OXLD1,ZNF223,OGFOD2,EIF3G,
,SZT2,LAMTOR4,WDR45,C9ORF64,IQCK,SETD4,MED8,GANC,SSH3,C12ORF10,SNRNP200,RPS14,ZRANB2
,ZNF223,OGFOD2,METTL3,EIF3G,PICK1,SLC52A2,NTMT1,TSPAN31,GLYCTK,CFLAR,TRIM46,EIF2A,NKIRAS
,ZNF223,OGFOD2,METTL3,EIF3G,PICK1,SLC52A2,NTMT1,TSPAN31,GLYCTK,CFLAR,TRIM46,EIF2A,NKIRAS
RPS9,SLC35G1,RCE1,MRPL33,ZNF410,TESC,NUDT5,RBPMS,SPAG7,IL1RAP,STYXL1,RANBP1,UBE2L6,LZTF
RPS9,SLC35G1,RCE1,MRPL33,ZNF410,TESC,NUDT5,RBPMS,SPAG7,IL1RAP,STYXL1,RANBP1,UBE2L6,LZTF
2,STYXL1,RFX5,RANBP1,VRK3,LZTFL1,FIS1,TUBA1C,STIP1,TET2,SNF8,GRK6,CDC37,OBSL1,UBR4,SLC39A1
2,STYXL1,RFX5,RANBP1,VRK3,LZTFL1,FIS1,TUBA1C,STIP1,TET2,SNF8,GRK6,CDC37,OBSL1,UBR4,SLC39A1
LC25A10,DBNDD2,SDF4,CD99L2,GGPS1,COASY,SAP30BP,RABL5,TRNT1,ITGB1BP1,PARD6G,C1ORF198,
LC25A10,DBNDD2,SDF4,CD99L2,GGPS1,COASY,SAP30BP,RABL5,TRNT1,ITGB1BP1,PARD6G,C1ORF198,

,BUD31,NPRL2,CTTN,TSPAN11,ITSN1,EIF5A,COMMD5,UFD1L,SH2B1,NOP16,ARHGEF39,LDHA,PSMA6,AC

,BUD31,NPRL2,CTTN,TSPAN11,ITSN1,EIF5A,COMMD5,UFD1L,SH2B1,NOP16,ARHGEF39,LDHA,PSMA6,AC

C1,RBP5,DBI,ARPC2,LIMK2,VDAC3,NOB1,WDR19,SIRT6,NUMA1,GPATCH2L,BUD31,SAE1,NPRL2,ARHGAP

C1,RBP5,DBI,ARPC2,LIMK2,VDAC3,NOB1,WDR19,SIRT6,NUMA1,GPATCH2L,BUD31,SAE1,NPRL2,ARHGAP

C17ORF62,PRMT1,TXNDC17,PPAN,ACAD10,CYB561,RPL36,CDK11A,MDH1,ZNF197,DGUOK,LRRFIP1,ZNF5

2,PPA1,DGKE,ENDOV,PPIL2,RPL28,LIG1,TMEM126B,RPL21,LYPLA2,C17ORF62,CASC4,ZNF7,PPAN,CYB561

2,RFX5,PLEKHA5,RANBP1,VRK3,DST,ANTXR2,GRAMD1C,MAGED2,GRK6,HMGA2,TSG101,PTPN13,UBR4,

UBE2L6,VRK3,LZTFL1,ZC3H14,TUBA1C,ANTXR2,LRRC1,PHF19,LPCAT3,CD44,TM9SF1,MAGED2,SON,TET2

UBE2L6,VRK3,LZTFL1,ZC3H14,TUBA1C,ANTXR2,LRRC1,PHF19,LPCAT3,CD44,TM9SF1,MAGED2,SON,TET2

,DHPS,ACTR6,SEC62,SET,BOP1,KLC1,RBP5,ZC3HAV1,SCAF11,ARPC2,PMFBP1,VDAC3,NOB1,WDR19,NUM

,DHPS,ACTR6,SEC62,SET,BOP1,KLC1,RBP5,ZC3HAV1,SCAF11,ARPC2,PMFBP1,VDAC3,NOB1,WDR19,NUM

UZ,PPRC1,RPS25,COX4I1,DUSP22,SPOCK2,GGPS1,SRSF1,AUP1,FOXRED1,TRNT1,ITGB1BP1,PARD6G,SH3

JP1,PARD6G,SH3YL1,IP6K2,MICAL1,HS2ST1,UBE2I,AUNIP,C14ORF2,PCYT2,PPP1CC,RFX3,PPA1,DGKE,CAM

JP1,PARD6G,SH3YL1,IP6K2,MICAL1,HS2ST1,UBE2I,AUNIP,C14ORF2,PCYT2,PPP1CC,RFX3,PPA1,DGKE,CAM

CMT1,EBPL,TESC,IGSF8,NUDT5,STYXL1,PLEKHA5,RANBP1,FIS1,TUBA1C,STIP1,LRRC1,PHF19,LPCAT3,MAG

IRPN,SPAG7,WDR62,STYXL1,RFX5,RANBP1,LZTFL1,FIS1,SLC15A3,TUBA1C,IRF9,STIP1,TM9SF1,TET2,UQCR

IRPN,SPAG7,WDR62,STYXL1,RFX5,RANBP1,LZTFL1,FIS1,SLC15A3,TUBA1C,IRF9,STIP1,TM9SF1,TET2,UQCR

AT3,CD44,GRK6,RPS16,CPSF3L,FAM3A,MUC1,EPB41L5,ATG12,ZBTB44,NISCH,CD99L2,GGPS1,AUP1,FOXR

PCID2,PEX2,C12ORF57,ZNF223,OGFOD2,EIF3G,PICK1,FBXL18,SLC52A2,TSPAN31,FAM192A,GLYCTK,CFLA

PCID2,PEX2,C12ORF57,ZNF223,OGFOD2,EIF3G,PICK1,FBXL18,SLC52A2,TSPAN31,FAM192A,GLYCTK,CFLA

C,SNRPN,SPAG7,WDR62,IL1RAP,STYXL1,RANBP1,UBE2L6,PPM1K,TUBA1C,STIP1,PHF19,VCAN,SON,TET2,C

C,SNRPN,SPAG7,WDR62,IL1RAP,STYXL1,RANBP1,UBE2L6,PPM1K,TUBA1C,STIP1,PHF19,VCAN,SON,TET2,C

FBP1,LIMK2,WDR19,NUMA1,GPATCH2L,BUD31,SAE1,OSGEP,TBL2,SCARB1,CTTN,ITSN1,EIF5A,TNFSF10,N

FBP1,LIMK2,WDR19,NUMA1,GPATCH2L,BUD31,SAE1,OSGEP,TBL2,SCARB1,CTTN,ITSN1,EIF5A,TNFSF10,N

TCH2L,SAE1,OSGEP,SCARB1,TSPAN11,ITSN1,EIF5A,RUNX1,NUDT22,UFD1L,USP38,SH2B1,NOP16,RPL13A

TCH2L,SAE1,OSGEP,SCARB1,TSPAN11,ITSN1,EIF5A,RUNX1,NUDT22,UFD1L,USP38,SH2B1,NOP16,RPL13A

RPT1,KLC1,ZC3HAV1,SCAF11,DBI,LIMK2,NOB1,WDR19,SIRT6,NUMA1,SUMF2,SAE1,NPRL2,OSGEP,SCARB1

RPT1,KLC1,ZC3HAV1,SCAF11,DBI,LIMK2,NOB1,WDR19,SIRT6,NUMA1,SUMF2,SAE1,NPRL2,OSGEP,SCARB1

PSMA6,AP3B2,RUVBL2,EBPL,TESC,IGSF8,NNMT,RFX5,PLEKHA5,RANBP1,UBE2L6,VRK3,LZTFL1,ZC3H14,TU

PSMA6,AP3B2,RUVBL2,EBPL,TESC,IGSF8,NNMT,RFX5,PLEKHA5,RANBP1,UBE2L6,VRK3,LZTFL1,ZC3H14,TU

F235,BOP1,TRPT1,KLC1,ZC3HAV1,DBI,ARPC2,VDAC3,WDR19,NUMA1,GPATCH2L,SUMF2,BUD31,SAE1,AR

F235,BOP1,TRPT1,KLC1,ZC3HAV1,DBI,ARPC2,VDAC3,WDR19,NUMA1,GPATCH2L,SUMF2,BUD31,SAE1,AR

MT112,MPDU1,RPS15,LRRC27,MVD,SYNGR1,PABPC1,NAT9,WAC,PDCD5,ANAPC11,USP45,ILF3,EEF1D,CA

MT112,MPDU1,RPS15,LRRC27,MVD,SYNGR1,PABPC1,NAT9,WAC,PDCD5,ANAPC11,USP45,ILF3,EEF1D,CA

410,RUVBL2,PCMT1,NUDT5,WDR62,STYXL1,RFX5,RANBP1,UBE2L6,PPM1K,DST,STIP1,LRRC1,VCAN,LPCAT

,GUK1,PPP1CC,ENDOV,PSMD4,LIG1,PSMA7,TMEM126B,TXNL4A,NUTF2,TMPO,LSM12,PRMT1,CASC4,WI

EKHA5,RANBP1,FIS1,STIP1,LPCAT3,TM9SF1,MAGED2,HMGA2,CDC37,YLPM1,SLC39A11,FAM3A,MUC1,M

PC2,LIMK2,VDAC3,WDR19,BUD31,NPRL2,OSGEP,TBL2,CTTN,ITSN1,EIF5A,RUNX1,CHID1,NUDT22,AMZ2,U

PC2,LIMK2,VDAC3,WDR19,BUD31,NPRL2,OSGEP,TBL2,CTTN,ITSN1,EIF5A,RUNX1,CHID1,NUDT22,AMZ2,U

7,WDR62,STYXL1,RANBP1,UBE2L6,PPM1K,TUBA1C,STIP1,PHF19,VCAN,LPCAT3,CD44,MAGED2,SON,GRK

Z2,TAZ,NOP16,RPL13A,GSTK1,LDHA,RPS9,RCE1,ZNF410,PCMT1,IGSF8,NNMT,WDR62,STYXL1,RFX5,RANB

Z2,TAZ,NOP16,RPL13A,GSTK1,LDHA,RPS9,RCE1,ZNF410,PCMT1,IGSF8,NNMT,WDR62,STYXL1,RFX5,RANB

AC3,NOB1,GPATCH2L,SUMF2,BUD31,SAE1,NPRL2,OSGEP,TBL2,ARHGAP6,SCARB1,CTTN,TSPAN11,ITSN1,

AC3,NOB1,GPATCH2L,SUMF2,BUD31,SAE1,NPRL2,OSGEP,TBL2,ARHGAP6,SCARB1,CTTN,TSPAN11,ITSN1,

3YL1,IP6K2,MICAL1,UBE2I,PRMT2,JUP,AUNIP,C14ORF2,GUK1,PCYT2,PPP1CC,RFX3,DGKE,CAMTA1,CRIP1,

2,PICK1,TSPAN31,APOLD1,GLYCTK,GTF2H1,TP53,TMEM199,SLC25A24,EPN1,ZCCHC9,MKRN1,GTF3A,AZI2

Z2,TAZ,RPL13A,LDHA,RPS9,SLC35G1,MRPL33,EBPL,TESC,NUDT5,SNRPN,SPAG7,WDR62,STYXL1,RFX5,RAN

IM,DHPS,MAP3K7CL,SEC62,SET,BOP1,KLC1,SCAF11,WDR19,SIRT6,NUMA1,GPATCH2L,SUMF2,BUD31,SA
IM,DHPS,MAP3K7CL,SEC62,SET,BOP1,KLC1,SCAF11,WDR19,SIRT6,NUMA1,GPATCH2L,SUMF2,BUD31,SA
4,CD99L2,GGPS1,AUP1,FOXRED1,PARD6G,SH3YL1,IP6K2,MICAL1,HS2ST1,UBE2I,SLC39A13,AUNIP,C14OF
D2,NSUN5,DRG2,MRPS18C,SMARCC2,THADA,ZNF81,FAM32A,C12ORF57,PEX26,ZNF223,OGFOD2,METTL
D2,NSUN5,DRG2,MRPS18C,SMARCC2,THADA,ZNF81,FAM32A,C12ORF57,PEX26,ZNF223,OGFOD2,METTL

BNDD1,RPL13A,LDHA,PSMA6,AP3B2,RPS9,SLC35G1,RCE1,ZNF410,NUDT5,RBPMS,SPAG7,IL1RAP,POLD4,
JP,PTK2,GUK1,ATF7,CAPG,PPIL2,RPL28,LIG1,TIMM17B,APEH,NUTF2,LSM12,PIGL,C19ORF48,ZFAND2B,SH
JP,PTK2,GUK1,ATF7,CAPG,PPIL2,RPL28,LIG1,TIMM17B,APEH,NUTF2,LSM12,PIGL,C19ORF48,ZFAND2B,SH
7L1,MAF,PPA1,ENDOV,PPIL2,PSMA7,TMPO,PRMT1,TXNDC17,ARL6IP4,ZFAND2B,INO80E,MAPKAPK5,HSI
H,TMED4,CD99L2,TRNT1,PARD6G,SH3YL1,IP6K2,CLASRP,JPH2,C1ORF198,MICAL1,UBE2I,PRMT2,JUP,TBX
H,TMED4,CD99L2,TRNT1,PARD6G,SH3YL1,IP6K2,CLASRP,JPH2,C1ORF198,MICAL1,UBE2I,PRMT2,JUP,TBX
NOP16,GSTK1,AP3B2,MRPL33,ZNF410,NUDT5,WDR62,IL1RAP,STYXL1,DST,STIP1,CD44,TET2,CDC37,RPS
NOP16,GSTK1,AP3B2,MRPL33,ZNF410,NUDT5,WDR62,IL1RAP,STYXL1,DST,STIP1,CD44,TET2,CDC37,RPS
PB,PARD3,TCAIM,SLC3A2,DHPS,ACTR6,SET,CTSC,ZNF235,BOP1,ARRDC1,TRPT1,KLC1,ZC3HAV1,SCAF11,A
PB,PARD3,TCAIM,SLC3A2,DHPS,ACTR6,SET,CTSC,ZNF235,BOP1,ARRDC1,TRPT1,KLC1,ZC3HAV1,SCAF11,A
MK2,NOB1,SIRT6,NUMA1,GPATCH2L,BUD31,SAE1,NPRL2,ARHGAP6,ITSN1,EIF5A,COMMD5,RUNX1,UFD1
MTA1,CRIP1,LIG1,TIMM17B,APEH,NUTF2,C17ORF62,CASC4,ZFAND2B,CYB561,SHARPIN,RPL36,LAMP2,S
RK3,FIS1,IRF9,LRRC1,PHF19,VCAN,MAGED2,UQCRQ,TSG101,PTPN13,UBR4,PSMA1,FAM3A,PSMA3,RPS2
30BP,FOXRED1,TRNT1,ITGB1BP1,SH3YL1,MICAL1,JUP,TBX6,AUNIP,GUK1,CENPN,HSBP1L1,ASF1B,CCDC2
12,CHPF,GMPPB,MVD,IL32,SYNGR1,C12ORF76,RANGRF,RRNAD1,SLC33A1,ZNF687,POT1,POLR3H,RAD2
,RCC1,SNRPB,PARD3,TCAIM,SLC3A2,DHPS,ACTR6,MAP3K7CL,SEC62,CTSC,ZNF235,BOP1,ARRDC1,TRPT1,
,RCC1,SNRPB,PARD3,TCAIM,SLC3A2,DHPS,ACTR6,MAP3K7CL,SEC62,CTSC,ZNF235,BOP1,ARRDC1,TRPT1,
,SCARB1,CTTN,TSPAN11,ITSN1,EIF5A,AMZ2,USP38,TAZ,DBNDD1,RPL13A,GSTK1,ACOT8,SLC35G1,RCE1,N
L1,RFX5,VRK3,PPM1K,PCBP2,TM9SF1,MAGED2,TET2,UQCRQ,GRK6,CDC37,CPSF3L,TSG101,UBR4,FAM3
L1,RFX5,VRK3,PPM1K,PCBP2,TM9SF1,MAGED2,TET2,UQCRQ,GRK6,CDC37,CPSF3L,TSG101,UBR4,FAM3
PL13A,LDHA,ACOT8,AP3B2,ZNF410,RUVBL2,PCMT1,EBPL,NUDT5,WDR62,RANBP1,UBE2L6,VRK3,ZC3H14
PL13A,LDHA,ACOT8,AP3B2,ZNF410,RUVBL2,PCMT1,EBPL,NUDT5,WDR62,RANBP1,UBE2L6,VRK3,ZC3H14
M1,FUZ,FAM3A,METTL17,ATG12,PPRC1,ZBTB44,TMX2,RDH11,SDF4,NISCH,CENPK,RTEL1,SRSF1,AUP1,S
M1,FUZ,FAM3A,METTL17,ATG12,PPRC1,ZBTB44,TMX2,RDH11,SDF4,NISCH,CENPK,RTEL1,SRSF1,AUP1,S
C,TM9SF1,TET2,SNF8,CPSF3L,TSG101,OBSL1,UBR4,FAM3A,PSMA3,METTL17,ATG12,PPRC1,SLC25A10,SD
6G,SH3YL1,IP6K2,MICAL1,HS2ST1,UBE2I,TMEM205,AUNIP,C14ORF2,GUK1,PCYT2,AGPAT4,PPP1CC,EDF
B2,DUSP22,SPOCK2,DBNDD2,SDF4,NISCH,CD99L2,GGPS1,RABL5,TRNT1,PARD6G,IP6K2,CLASRP,C1ORF1
B2,DUSP22,SPOCK2,DBNDD2,SDF4,NISCH,CD99L2,GGPS1,RABL5,TRNT1,PARD6G,IP6K2,CLASRP,C1ORF1
C39A11,FAM3A,METTL17,EPB41L5,ATG12,ZBTB44,CD99L2,PARD6G,SH3YL1,IP6K2,CLASRP,MICAL1,HS2S
,NUDT5,RBPMS,NNMT,SPAG7,STYXL1,PLEKHA5,UBE2L6,PPM1K,TUBA1C,IRF9,DST,ANTXR2,STIP1,LRRC1,
,NUDT5,RBPMS,NNMT,SPAG7,STYXL1,PLEKHA5,UBE2L6,PPM1K,TUBA1C,IRF9,DST,ANTXR2,STIP1,LRRC1,
NF235,ARRDC1,TRPT1,KLC1,RBP5,ZC3HAV1,ARPC2,PMFBP1,NUMA1,SUMF2,BUD31,NPRL2,OSGEP,ARHG
NF235,ARRDC1,TRPT1,KLC1,RBP5,ZC3HAV1,ARPC2,PMFBP1,NUMA1,SUMF2,BUD31,NPRL2,OSGEP,ARHG
CL,SEC62,SET,ZNF235,ARRDC1,TRPT1,KLC1,ZC3HAV1,SCAF11,ARPC2,PMFBP1,LIMK2,VDAC3,NOB1,SIRT
CL,SEC62,SET,ZNF235,ARRDC1,TRPT1,KLC1,ZC3HAV1,SCAF11,ARPC2,PMFBP1,LIMK2,VDAC3,NOB1,SIRT
IM16,HSPH1,ZFAND4,LGMN,RPL15,KCTD7,COA3,ZNF343,OARD1,GMPR2,PHC3,COMMD4,UBE2N,MFSD
IM16,HSPH1,ZFAND4,LGMN,RPL15,KCTD7,COA3,ZNF343,OARD1,GMPR2,PHC3,COMMD4,UBE2N,MFSD
NUDT5,STYXL1,RFX5,PLEKHA5,FIS1,IRF9,DST,STIP1,VCAN,LPCAT3,MAGED2,SON,UQCRQ,HMGA2,CDC37,
1,TRNT1,ITGB1BP1,MAGT1,GUK1,CENPN,AGPAT4,IFI27L1,MAF,PPA1,ENDOV,PPIL2,TXNL4A,TMPO,ZFAN
,PARD3,TCAIM,SLC3A2,DHPS,ACTR6,MAP3K7CL,SEC62,SET,CTSC,ZNF235,BOP1,ARRDC1,TRPT1,KLC1,RBI
,PARD3,TCAIM,SLC3A2,DHPS,ACTR6,MAP3K7CL,SEC62,SET,CTSC,ZNF235,BOP1,ARRDC1,TRPT1,KLC1,RBI

SLC25A10,TMX2,THEM4,RDH11,SDF4,CD99L2,AUP1,SAP30BP,KXD1,CLASRP,C1ORF198,UGGT2,MICAL1,F
SLC25A10,TMX2,THEM4,RDH11,SDF4,CD99L2,AUP1,SAP30BP,KXD1,CLASRP,C1ORF198,UGGT2,MICAL1,F

,TET2,GRK6,CDC37,CPSF3L,OBSL1,YLPM1,FAM3A,PSMA3,ECSIT,EPB41L5,PPRC1,NIP7,COX4I1,SPOCK2,CE
Z1,SSH2,COX17,HINT1,HNRNPA1,EGLN2,SIRT5,RFXANK,SMARCA4,MPC2,OARD1,MRPS18C,MCFD2,SCAP,
Z1,SSH2,COX17,HINT1,HNRNPA1,EGLN2,SIRT5,RFXANK,SMARCA4,MPC2,OARD1,MRPS18C,MCFD2,SCAP,
L,RBP5,ZC3HAV1,SCAF11,DBI,ARPC2,VDAC3,NOB1,WDR19,NUMA1,GPATCH2L,BUD31,NPRL2,OSGEP,TBL
L,RBP5,ZC3HAV1,SCAF11,DBI,ARPC2,VDAC3,NOB1,WDR19,NUMA1,GPATCH2L,BUD31,NPRL2,OSGEP,TBL
CDK11A,INO80E,FAM219B,ZNF524,ELP6,SIAH1,TRMT112,MPDU1,RPS15,HSPB1,LRRC27,SYNGR1,NAT9,W
CDK11A,INO80E,FAM219B,ZNF524,ELP6,SIAH1,TRMT112,MPDU1,RPS15,HSPB1,LRRC27,SYNGR1,NAT9,W
,IRF9,STIP1,VCAN,PCBP2,MAGED2,SON,UQCRQ,SNF8,CPSF3L,TSG101,SLC39A11,FUZ,METTL17,ATG12,RI
,IRF9,STIP1,VCAN,PCBP2,MAGED2,SON,UQCRQ,SNF8,CPSF3L,TSG101,SLC39A11,FUZ,METTL17,ATG12,RI
UBE2N,NDUFS3,KPNA5,ZNF223,DGAT1,METTL3,EIF3G,ZNF512,RNF181,ARL6IP5,RPL35,NME1,FAM192A,
UBE2N,NDUFS3,KPNA5,ZNF223,DGAT1,METTL3,EIF3G,ZNF512,RNF181,ARL6IP5,RPL35,NME1,FAM192A,
RK3,FIS1,SLC15A3,PPM1K,GRAMD1C,SON,UQCRQ,SNF8,GRK6,CDC37,RPS16,OBSL1,PSMA1,PHF21A,FAM
RK3,FIS1,SLC15A3,PPM1K,GRAMD1C,SON,UQCRQ,SNF8,GRK6,CDC37,RPS16,OBSL1,PSMA1,PHF21A,FAM
MT1,IGSF8,NNMT,WDR62,STYXL1,RFX5,RANBP1,VRK3,PPM1K,PHF19,PCBP2,TM9SF1,SON,TET2,GRK6,CD
MT1,IGSF8,NNMT,WDR62,STYXL1,RFX5,RANBP1,VRK3,PPM1K,PHF19,PCBP2,TM9SF1,SON,TET2,GRK6,CD
TIMM17B,TMEM126B,UCK2,C17ORF62,ZNF7,PIGL,UPP1,RBM3,DUT,SYNGR2,FAM219B,RMDN1,STRIP2,R
TIMM17B,TMEM126B,UCK2,C17ORF62,ZNF7,PIGL,UPP1,RBM3,DUT,SYNGR2,FAM219B,RMDN1,STRIP2,R
UDT22,UFD1L,USP38,TAZ,SH2B1,ARHGEF39,TTLL5,LDHA,ACOT8,SLC35G1,ZNF410,PCMT1,EBPL,IGSF8,RBF
TR6,MAP3K7CL,SEC62,SET,CTSC,ARRDC1,TRPT1,KLC1,RBP5,ZC3HAV1,SCAF11,DBI,PMFBP1,VDAC3,WDR1
TR6,MAP3K7CL,SEC62,SET,CTSC,ARRDC1,TRPT1,KLC1,RBP5,ZC3HAV1,SCAF11,DBI,PMFBP1,VDAC3,WDR1
DP1,KLC1,RBP5,ZC3HAV1,SCAF11,ARPC2,PMFBP1,LIMK2,VDAC3,NOB1,WDR19,NUMA1,BUD31,SAE1,NPI
P1,ZSWIM7,ZNF814,NAP1L1,HMGA1,PABPC1,DBNL,ZNF836,ZNF706,ZNF687,NXN,NUDT1,TRIM16,ANAP
AN11,ITSN1,EIF5A,TNFSF10,COMMD5,NUDT22,AMZ2,TAZ,DBNDD1,SH2B1,TTLL5,AP3B2,RUVBL2,PCMT1,

WDR62,IL1RAP,STYXL1,PLEKHA5,RANBP1,UBE2L6,ZC3H14,ANTXR2,STIP1,LRRC1,PHF19,GRAMD1C,TET2,
WDR62,IL1RAP,STYXL1,PLEKHA5,RANBP1,UBE2L6,ZC3H14,ANTXR2,STIP1,LRRC1,PHF19,GRAMD1C,TET2,

,PHF19,VCAN,CD44,CDC37,RPS16,CPSF3L,YLPM1,PHF21A,SLC39A11,FAM3A,EPB41L5,ATG12,ZBTB44,PL
,PHF19,VCAN,CD44,CDC37,RPS16,CPSF3L,YLPM1,PHF21A,SLC39A11,FAM3A,EPB41L5,ATG12,ZBTB44,PL
2,TBC1D22A,BCL7B,ZNF581,SZT2,MRPL51,C9ORF64,SETD4,MED8,PUF60,MRPL27,DDX56,MED17,RPS14,

,PCMT1,EBPL,NUDT5,WDR62,RANBP1,VRK3,ZC3H14,TUBA1C,ANTXR2,STIP1,LRRC1,CD44,MAGED2,SON,
LAMP2,SYNGR2,ASPH,TRMT112,RPTOR,TWSG1,VPS28,ZSWIM7,HSPB1,MVD,EMC4,IL32,SEMA3C,C12ORI
LAMP2,SYNGR2,ASPH,TRMT112,RPTOR,TWSG1,VPS28,ZSWIM7,HSPB1,MVD,EMC4,IL32,SEMA3C,C12ORI

SC,RBPMS,NNMT,WDR62,STYXL1,RANBP1,FIS1,SLC15A3,IRF9,GRAMD1C,TM9SF1,TET2,GRK6,CPSF3L,TT
1,EBPL,RBPMS,SPAG7,WDR62,IL1RAP,RFX5,PLEKHA5,UBE2L6,ZC3H14,TUBA1C,ANTXR2,STIP1,LRRC1,PHF
1,EBPL,RBPMS,SPAG7,WDR62,IL1RAP,RFX5,PLEKHA5,UBE2L6,ZC3H14,TUBA1C,ANTXR2,STIP1,LRRC1,PHF
H2B1,ACOT8,AP3B2,RCE1,RUVBL2,PCMT1,IGSF8,RBPMS,SPAG7,WDR62,IL1RAP,STYXL1,RFX5,POLD4,PLEK
9,LPCAT3,CD44,TM9SF1,MAGED2,SON,HMGA2,CDC37,TSG101,TTC21B,UBR4,PSMA1,FUZ,METTL17,EPB4
9,LPCAT3,CD44,TM9SF1,MAGED2,SON,HMGA2,CDC37,TSG101,TTC21B,UBR4,PSMA1,FUZ,METTL17,EPB4
AN31,GLYCTK,PCNP,NEK11,CREBZF,EIF2A,NKIRAS2,PIGO,CLTA,SLC35A2,SLC25A24,ZNF396,HPS1,ITPA,SN
AN31,GLYCTK,PCNP,NEK11,CREBZF,EIF2A,NKIRAS2,PIGO,CLTA,SLC35A2,SLC25A24,ZNF396,HPS1,ITPA,SN

1,IP6K2,CLASRP,JPH2,C1ORF198,MICAL1,UBE2I,SLC39A13,PRMT2,TBX6,TMEM205,PTK2,AUNIP,C14ORF
1,IP6K2,CLASRP,JPH2,C1ORF198,MICAL1,UBE2I,SLC39A13,PRMT2,TBX6,TMEM205,PTK2,AUNIP,C14ORF
MA6,AP3B2,RUVBL2,TESC,IGSF8,NNMT,PLEKHA5,RANBP1,VRK3,LZTFL1,ZC3H14,TUBA1C,ANTXR2,LRRC1,
MA6,AP3B2,RUVBL2,TESC,IGSF8,NNMT,PLEKHA5,RANBP1,VRK3,LZTFL1,ZC3H14,TUBA1C,ANTXR2,LRRC1,
1L1,ASF1B,DEF8,PPP1CC,ENDOV,UCK2,TMPO,PRMT1,EXOSC9,WNK1,C19ORF54,ZNF7,ARL6IP4,STRADA,

,COX4I1,RTEL1,COASY,TRNT1,ITGB1BP1,CLASRP,UGGT2,UBE2I,JUP,TBX6,GUK1,CARD14,IFI27L1,EEF1B2

RPT1,KLC1,ZC3HAV1,DBI,ARPC2,LIMK2,VDAC3,WDR19,SIRT6,GPATCH2L,TBL2,ARHGAP6,CTTN,EIF5A,TNF
RPT1,KLC1,ZC3HAV1,DBI,ARPC2,LIMK2,VDAC3,WDR19,SIRT6,GPATCH2L,TBL2,ARHGAP6,CTTN,EIF5A,TNF
RL2,ARHGAP6,SCARB1,CTTN,TSPAN11,ITSN1,EIF5A,COMMD5,NUDT22,UFD1L,THYN1,DBNDD1,SH2B1,T
RL2,ARHGAP6,SCARB1,CTTN,TSPAN11,ITSN1,EIF5A,COMMD5,NUDT22,UFD1L,THYN1,DBNDD1,SH2B1,T
JMA1,BUD31,SAE1,TBL2,CTTN,TSPAN11,ITSN1,COMMD5,NUDT22,UFD1L,USP38,TAZ,THYN1,DBNDD1,SH
JMA1,BUD31,SAE1,TBL2,CTTN,TSPAN11,ITSN1,COMMD5,NUDT22,UFD1L,USP38,TAZ,THYN1,DBNDD1,SH
2L6,PPM1K,STIP1,PHF19,VCAN,MAGED2,SON,GRK6,OBSL1,UBR4,YLPM1,PHF21A,CHCHD1,SLC39A11,FAM
2L6,PPM1K,STIP1,PHF19,VCAN,MAGED2,SON,GRK6,OBSL1,UBR4,YLPM1,PHF21A,CHCHD1,SLC39A11,FAM
DF4,RPL29,AUP1,TRNT1,MED15,C1ORF198,UGGT2,HS2ST1,XBP1,C14ORF2,PCYT2,ASF1B,EEF1B2,PPA1,D
DF4,RPL29,AUP1,TRNT1,MED15,C1ORF198,UGGT2,HS2ST1,XBP1,C14ORF2,PCYT2,ASF1B,EEF1B2,PPA1,D

C,RFX3,TNFRSF14,PSMD4,RPL28,TXNL4A,NUTF2,PODXL,PRMT1,STOML2,C19ORF54,PPAN,HNRNPC,SHA
C,RFX3,TNFRSF14,PSMD4,RPL28,TXNL4A,NUTF2,PODXL,PRMT1,STOML2,C19ORF54,PPAN,HNRNPC,SHA

,ZEB2,STRIP2,RPS20,MPDU1,RPS15,ERLEC1,NAP1L1,MEIS2,PHTF2,NAT9,RBP1,RAD1,H2AFV,SERPINB6,T
ANK,PAFAH1B2,TMEM39B,DAZAP2,ZFPL1,TRPM4,MFSD10,PFKFB2,NDUFS3,NUBP2,STAU2,TSEN34,MRP
SP38,DBNDD1,SH2B1,NOP16,TTLL5,LDHA,PSMA6,AP3B2,RCE1,RUVBL2,EBPL,TESC,IGSF8,SPAG7,WDR62
SP38,DBNDD1,SH2B1,NOP16,TTLL5,LDHA,PSMA6,AP3B2,RCE1,RUVBL2,EBPL,TESC,IGSF8,SPAG7,WDR62
HYN1,DBNDD1,TTLL5,PSMA6,ACOT8,RCE1,RUVBL2,PCMT1,EBPL,TESC,IGSF8,RBPMS,WDR62,IL1RAP,STY
P3B2,RUVBL2,EBPL,RBPMS,SPAG7,IL1RAP,RFX5,PLEKHA5,RANBP1,UBE2L6,VRK3,PPM1K,TUBA1C,STIP1,
P3B2,RUVBL2,EBPL,RBPMS,SPAG7,IL1RAP,RFX5,PLEKHA5,RANBP1,UBE2L6,VRK3,PPM1K,TUBA1C,STIP1,
1,DGKE,CAMTA1,LIG1,TIMM17B,APEH,PODXL,CAMLG,ATPAF2,C17ORF62,PRMT1,CASC4,ZNF7,ZFAND2
HPS,ACTR6,SEC62,CTSC,BOP1,ARRDC1,TRPT1,KLC1,SCAF11,DBI,ARPC2,VDAC3,SIRT6,NUMA1,NPRL2,TBL
HPS,ACTR6,SEC62,CTSC,BOP1,ARRDC1,TRPT1,KLC1,SCAF11,DBI,ARPC2,VDAC3,SIRT6,NUMA1,NPRL2,TBL
P1,TPM2,SIAH1,STRIP2,TWSG1,LMBR1,NAP1L1,HMGA1,RABEP2,ZNF20,RALGPS1,ZNF687,NXN,HSD17B1
P1,TPM2,SIAH1,STRIP2,TWSG1,LMBR1,NAP1L1,HMGA1,RABEP2,ZNF20,RALGPS1,ZNF687,NXN,HSD17B1
MT1,IGSF8,NNMT,WDR62,STYXL1,RFX5,RANBP1,VRK3,PPM1K,PHF19,PCBP2,TM9SF1,SON,TET2,UQCRQ
MT1,IGSF8,NNMT,WDR62,STYXL1,RFX5,RANBP1,VRK3,PPM1K,PHF19,PCBP2,TM9SF1,SON,TET2,UQCRQ
,RPS6,RPL29,TRNT1,ITGB1BP1,SH3YL1,C1ORF198,C14ORF2,PCYT2,USE1,AGPAT4,CCDC25,EEF1B2,PPA1,
NPK,SRSF1,AUP1,SAP30BP,FOXRED1,TRNT1,KXD1,MED15,SLC39A13,MAGT1,C14ORF2,GUK1,CENPN,PCY
NPK,SRSF1,AUP1,SAP30BP,FOXRED1,TRNT1,KXD1,MED15,SLC39A13,MAGT1,C14ORF2,GUK1,CENPN,PCY
S9,RCE1,TESC,RBPMS,SPAG7,WDR62,IL1RAP,PLEKHA5,ZC3H14,SLC15A3,PPM1K,ANTXR2,STIP1,PHF19,V
D4,CD99L2,GGPS1,TRNT1,PARD6G,SH3YL1,IP6K2,CLASRP,JPH2,C1ORF198,MICAL1,HS2ST1,UBE2I,PRMT
D4,CD99L2,GGPS1,TRNT1,PARD6G,SH3YL1,IP6K2,CLASRP,JPH2,C1ORF198,MICAL1,HS2ST1,UBE2I,PRMT
KHA5,UBE2L6,TUBA1C,STIP1,LRRC1,PHF19,LPCAT3,SON,TET2,GRK6,CPSF3L,OBSL1,PTPN13,UBR4,YLPM
KHA5,UBE2L6,TUBA1C,STIP1,LRRC1,PHF19,LPCAT3,SON,TET2,GRK6,CPSF3L,OBSL1,PTPN13,UBR4,YLPM
ET2,GRK6,OBSL1,YLPM1,FUZ,PSMA3,EPB41L5,ATG12,PPRC1,RPS25,NIP7,TTC9C,COX4I1,PLCB2,RDH11,S
L1,UBE2L6,LZTFL1,STIP1,PHF19,PCBP2,LPCAT3,SON,TET2,HMGA2,RPS16,UBR4,FUZ,ECSIT,EPB41L5,ATG
L1,UBE2L6,LZTFL1,STIP1,PHF19,PCBP2,LPCAT3,SON,TET2,HMGA2,RPS16,UBR4,FUZ,ECSIT,EPB41L5,ATG

RFX5,RANBP1,VRK3,PPM1K,PHF19,PCBP2,TM9SF1,SON,TET2,GRK6,CDC37,TSG101,OBSL1,PHF21A,FAM3

RFX5,RANBP1,VRK3,PPM1K,PHF19,PCBP2,TM9SF1,SON,TET2,GRK6,CDC37,TSG101,OBSL1,PHF21A,FAM3

AB1,ZNF524,GPBP1,RPTOR,MPDU1,ZSWIM7,ZNF814,RAD1,RBMX,WAC,EPS15L1,RAD23A,HSD17B10,ILF3

AB1,ZNF524,GPBP1,RPTOR,MPDU1,ZSWIM7,ZNF814,RAD1,RBMX,WAC,EPS15L1,RAD23A,HSD17B10,ILF3

99,IFITM2,SLC25A24,EPN1,ZCCHC9,SNRPD1,AZI2,GOSR2,LAMTOR1,PRKCSH,ZNF276,TIAF1,SZT2

CAPN10,EXOSC9,C19ORF54,ZFAND2B,SUGT1,ZNF250,RPL36,NFAM1,CDK11A,MAPKAPK5,DGUOK,HSP9

SY,YIPF3,ITGB1BP1,MICAL1,HS2ST1,UBE2I,EIF3M,PTK2,AUNIP,C14ORF2,GUK1,ATF7,MAF,DGKE,ENDOV,

SY,YIPF3,ITGB1BP1,MICAL1,HS2ST1,UBE2I,EIF3M,PTK2,AUNIP,C14ORF2,GUK1,ATF7,MAF,DGKE,ENDOV,

CAL1,UBE2I,TBX6,TMEM205,PTK2,CENPN,FBXW9,ASF1B,CCDC25,CAMTA1,APEH,PPOX,TMPO,ATPAF2,ZN

,TRPT1,SCAF11,DBI,WDR19,GPATCH2L,SUMF2,BUD31,SAE1,OSGEP,TBL2,SCARB1,CTTN,TSPAN11,ITSN1,

,TRPT1,SCAF11,DBI,WDR19,GPATCH2L,SUMF2,BUD31,SAE1,OSGEP,TBL2,SCARB1,CTTN,TSPAN11,ITSN1,

R13,STARD10,RNH1,DNASE1L1,KIF4A,RBCK1,PTOV1,TMEM135,RNF32,LY75,ATG16L2,RCC1,SNRPB,PARD

R13,STARD10,RNH1,DNASE1L1,KIF4A,RBCK1,PTOV1,TMEM135,RNF32,LY75,ATG16L2,RCC1,SNRPB,PARD

,SIAH1,MPDU1,ZSWIM7,UBXN1,ERLEC1,NAT9,RAD1,RBMX,PPP3CB,ANAPC11,RAD23A,ABHD17A,ILF3,ZC

14ORF2,PCYT2,USE1,CCDC25,EEF1B2,PPA1,TNFRSF14,TIMM17B,APEH,TXNL4A,CAPN10,PRMT1,C19ORF

14ORF2,PCYT2,USE1,CCDC25,EEF1B2,PPA1,TNFRSF14,TIMM17B,APEH,TXNL4A,CAPN10,PRMT1,C19ORF

1,PIGT,FAM32A,PEX2,NDUFS3,ARL16,OXLD1,SSR3,ARIH2,PICK1,ZNF512,RNF181,CHTOP,TSPAN31,GLYC

1,PIGT,FAM32A,PEX2,NDUFS3,ARL16,OXLD1,SSR3,ARIH2,PICK1,ZNF512,RNF181,CHTOP,TSPAN31,GLYC

NFRSF14,CRIP1,PPIL2,TROAP,VIM,LIG1,CAPN10,NUTF2,ATPAF2,CASC4,PIGL,STAU1,SLC9B2,CARS,CDK11

RF62,ZNF7,ZFAND2B,HSP90AB1,TMEM134,LRRFIP1,RAB2A,ZNF524,NIT2,LYNX1,LMBR1,CHCHD7,PHTF2,

ID1,AMZ2,UFD1L,TAZ,THYN1,NOP16,RPL13A,TTLL5,RCE1,MRPL33,PCMT1,EBPL,IGSF8,NNMT,WDR62,R

ID1,AMZ2,UFD1L,TAZ,THYN1,NOP16,RPL13A,TTLL5,RCE1,MRPL33,PCMT1,EBPL,IGSF8,NNMT,WDR62,R

XNDC17,STRADA,AP1S3,ZNF644,CARS,ZNF250,NFAM1,SYNGR2,ZNF524,SIAH1,STRIP2,GABPB1,UBXN1,N

XNDC17,STRADA,AP1S3,ZNF644,CARS,ZNF250,NFAM1,SYNGR2,ZNF524,SIAH1,STRIP2,GABPB1,UBXN1,N

,SPAG7,POLD4,PLEKHA5,VRK3,TUBA1C,DST,PHF19,GRAMD1C,VCAN,TM9SF1,SON,UQCRQ,CDC37,OBSL1

,SPAG7,POLD4,PLEKHA5,VRK3,TUBA1C,DST,PHF19,GRAMD1C,VCAN,TM9SF1,SON,UQCRQ,CDC37,OBSL1

L2,COX17,FKBP3,EGLN2,KCTD7,SPINT2,BRF1,FAM102A,PFKFB2,RIC8A,LFNG,XYLT2,PCID2,ZXDC,COBL,M

PAG7,WDR62,IL1RAP,STYXL1,RFX5,RANBP1,UBE2L6,PPM1K,STIP1,PHF19,VCAN,MAGED2,SON,UQCRQ,

PAG7,WDR62,IL1RAP,STYXL1,RFX5,RANBP1,UBE2L6,PPM1K,STIP1,PHF19,VCAN,MAGED2,SON,UQCRQ,

M1K,PHF19,PCBP2,TM9SF1,SON,TET2,GRK6,CDC37,TSG101,OBSL1,PHF21A,FAM3A,PSMA3,METTL17,PP

M1K,PHF19,PCBP2,TM9SF1,SON,TET2,GRK6,CDC37,TSG101,OBSL1,PHF21A,FAM3A,PSMA3,METTL17,PP

LC35A2,SLC25A24,ZNF396,HPS1,ITPA,SNRPD1,AZI2,TATDN1,KAT8,NCBP2,SUMO1,ZNF483,SZT2,BSCL2,

LC35A2,SLC25A24,ZNF396,HPS1,ITPA,SNRPD1,AZI2,TATDN1,KAT8,NCBP2,SUMO1,ZNF483,SZT2,BSCL2,

HF21A,FUZ,FAM3A,METTL17,PPRC1,RPS25,ZBTB44,COX4I1,DUSP22,TMX2,CENPK,AUP1,RABL5,ITGB1B

HF21A,FUZ,FAM3A,METTL17,PPRC1,RPS25,ZBTB44,COX4I1,DUSP22,TMX2,CENPK,AUP1,RABL5,ITGB1B

P3K7CL,SEC62,SET,CTSC,ZNF235,BOP1,ARRDC1,TRPT1,KLC1,RBP5,ZC3HAV1,SCAF11,DBI,ARPC2,LIMK2,N

T1,KLC1,RBP5,ZC3HAV1,SCAF11,DBI,ARPC2,LIMK2,VDAC3,SIRT6,SAE1,NPRL2,TBL2,SCARB1,TSPAN11,ITS

T1,KLC1,RBP5,ZC3HAV1,SCAF11,DBI,ARPC2,LIMK2,VDAC3,SIRT6,SAE1,NPRL2,TBL2,SCARB1,TSPAN11,ITS

PKAPK5,DGUOK,HSP90AB1,RAB2A,RMDN1,GPBP1,GPALPP1,CHPF,ERLEC1,GMPPB,MVD,IL32,SYNGR1,R

2,COASY,SRSF1,KXD1,MED15,IP6K2,KCNMA1,CDC42,XBP1,GUK1,CENPN,EDF1,DGKE,TNFRSF14,CRIP1,P

2,COASY,SRSF1,KXD1,MED15,IP6K2,KCNMA1,CDC42,XBP1,GUK1,CENPN,EDF1,DGKE,TNFRSF14,CRIP1,P

AUP1,SAP30BP,FOXRED1,MED15,SH3YL1,SLC39A13,PRMT2,EIF3M,CDC42,PTK2,ASF1B,ENDOV,CAPG,PP

AUP1,SAP30BP,FOXRED1,MED15,SH3YL1,SLC39A13,PRMT2,EIF3M,CDC42,PTK2,ASF1B,ENDOV,CAPG,PP

IH1,DNASE1L1,KIF4A,RBCK1,PTOV1,FASTK,TMEM135,RNF32,LY75,ATG16L2,RCC1,SNRPB,PARD3,TCAIM,

IH1,DNASE1L1,KIF4A,RBCK1,PTOV1,FASTK,TMEM135,RNF32,LY75,ATG16L2,RCC1,SNRPB,PARD3,TCAIM,

3CB,RAD23A,ABHD17A,USP45,ILF3,MDH2,HNRNPA1,FKBP3,FTH1,PFDN5,MTHFS,MPC2,COPS4,DALRD3,I
SM12,PIGL,C19ORF48,ZFAND2B,SHARPIN,UPP1,CDK11A,LAMP2,SYNGR2,ZEB2,ASPH,TRMT112,RPTOR,TV
SM12,PIGL,C19ORF48,ZFAND2B,SHARPIN,UPP1,CDK11A,LAMP2,SYNGR2,ZEB2,ASPH,TRMT112,RPTOR,TV
A5,RANBP1,FIS1,STIP1,LPCAT3,TM9SF1,SON,HMGA2,CDC37,OBSL1,TTC21B,YLPM1,SLC39A11,FAM3A,M
IFI27L1,MAF,PPA1,PPIL2,TXNL4A,CAPN10,PRMT1,EXOSC9,ZFAND2B,SUGT1,ZNF250,RPL36,NFAM1,CDK1
IFI27L1,MAF,PPA1,PPIL2,TXNL4A,CAPN10,PRMT1,EXOSC9,ZFAND2B,SUGT1,ZNF250,RPL36,NFAM1,CDK1
IFI27L1,MAF,PPA1,PPIL2,TXNL4A,CAPN10,PRMT1,EXOSC9,ZFAND2B,SUGT1,ZNF250,RPL36,NFAM1,CDK1

APEH,C17ORF62,ZFAND2B,SHARPIN,RPL36,LAMP2,SYNGR2,TMEM134,LRRFIP1,ASPH,NIT2,TWSG1,HSPB1
C8A,TCP1,PIGT,FAM32A,PEX2,TNKS,NDUFS3,ARL16,OXLD1,SSR3,ARIH2,PICK1,ZNF512,RNF181,CHTOP,TS
C8A,TCP1,PIGT,FAM32A,PEX2,TNKS,NDUFS3,ARL16,OXLD1,SSR3,ARIH2,PICK1,ZNF512,RNF181,CHTOP,TS
P1,PHF19,PCBP2,LPCAT3,TET2,UQCRQ,RPS16,UBR4,YLPM1,CHCHD1,FUZ,ECSIT,ATG12,PPRC1,ZBTB44,NIF
P1,PHF19,PCBP2,LPCAT3,TET2,UQCRQ,RPS16,UBR4,YLPM1,CHCHD1,FUZ,ECSIT,ATG12,PPRC1,ZBTB44,NIF
F9,STIP1,PHF19,LPCAT3,MAGED2,SON,GRK6,CDC37,OBSL1,PTPN13,UBR4,PHF21A,SLC39A11,MUC1,ATG1
F9,STIP1,PHF19,LPCAT3,MAGED2,SON,GRK6,CDC37,OBSL1,PTPN13,UBR4,PHF21A,SLC39A11,MUC1,ATG1
RK3,LZTFL1,ZC3H14,TUBA1C,ANTXR2,LRRC1,PHF19,LPCAT3,CD44,TM9SF1,SON,TET2,UQCRQ,GRK6,CPSF
RK3,LZTFL1,ZC3H14,TUBA1C,ANTXR2,LRRC1,PHF19,LPCAT3,CD44,TM9SF1,SON,TET2,UQCRQ,GRK6,CPSF
B2A,ZNF524,LYNX1,LMBR1,PABPC1,BCAM,BLVRB,RABEP2,RALGPS1,EPS15L1,EEF1D,NAA10,LLGL2,COX1
1,ITSN1,TNFSF10,COMMD5,RUNX1,CHID1,AMZ2,UFD1L,USP38,THYN1,NOP16,RPL13A,GSTK1,LDHA,PSM
1,ITSN1,TNFSF10,COMMD5,RUNX1,CHID1,AMZ2,UFD1L,USP38,THYN1,NOP16,RPL13A,GSTK1,LDHA,PSM
PS3,RHEB,NQO2,EML3,PCTP,HMOX2,NAA15,CDK2AP2,CARM1,MAZ,UBA52,ACVR1B,VTI1B,NMRAL1,XPN
A1,SUMF2,BUD31,SAE1,TBL2,ARHGAP6,SCARB1,CTTN,ITSN1,COMMD5,NUDT22,AMZ2,TAZ,THYN1,DBND
A1,SUMF2,BUD31,SAE1,TBL2,ARHGAP6,SCARB1,CTTN,ITSN1,COMMD5,NUDT22,AMZ2,TAZ,THYN1,DBND

ND2B,HNRNPC,SHARPIN,SYNGR2,LRRFIP1,TPM2,ZEB2,ASPH,TWSG1,LRRC27,LMBR1,NAP1L1,RANGRF,LSF
N,TSPAN11,EIF5A,UFD1L,THYN1,DBNDD1,SH2B1,NOP16,ARHGEF39,GSTK1,ACOT8,SLC35G1,ZNF410,PCM
N,TSPAN11,EIF5A,UFD1L,THYN1,DBNDD1,SH2B1,NOP16,ARHGEF39,GSTK1,ACOT8,SLC35G1,ZNF410,PCM
RB1,CTTN,ITSN1,EIF5A,COMMD5,RUNX1,NUDT22,UFD1L,DBNDD1,SH2B1,ARHGEF39,LDHA,AP3B2,RCE1,I
RB1,CTTN,ITSN1,EIF5A,COMMD5,RUNX1,NUDT22,UFD1L,DBNDD1,SH2B1,ARHGEF39,LDHA,AP3B2,RCE1,I
PAG7,WDR62,RANBP1,LZTFL1,FIS1,STIP1,PHF19,TET2,SNF8,OBSL1,FAM3A,MUC1,EPB41L5,ATG12,RPS25
T1,ZFAND2B,ZNF250,INO80E,MAPKAPK5,HSP90AB1,RAB2A,GPBP1,CHPF,UBXN1,GMPPB,MVD,IL32,NAP
R2,UBE2N,PDCD6,FAM32A,DCTN5,OXLD1,ZNF223,ARIH2,TSPAN31,GLYCTK,TRIM46,RBM42,RER1,ZNF28
BL2,CTTN,TSPAN11,EIF5A,NUDT22,AMZ2,THYN1,DBNDD1,GSTK1,PSMA6,AP3B2,SLC35G1,RCE1,ZNF410,
BL2,CTTN,TSPAN11,EIF5A,NUDT22,AMZ2,THYN1,DBNDD1,GSTK1,PSMA6,AP3B2,SLC35G1,RCE1,ZNF410,
L3A,LDHA,RPS9,SLC35G1,MRPL33,ZNF410,TESC,NUDT5,SNRPN,SPAG7,WDR62,STYXL1,RFX5,RANBP1,VRI
L3A,LDHA,RPS9,SLC35G1,MRPL33,ZNF410,TESC,NUDT5,SNRPN,SPAG7,WDR62,STYXL1,RFX5,RANBP1,VRI
1,TNFSF10,RUNX1,CHID1,NUDT22,DBNDD1,SH2B1,ACOT8,RPS9,SLC35G1,RCE1,ZNF410,PCMT1,EBPL,IGS
1,TNFSF10,RUNX1,CHID1,NUDT22,DBNDD1,SH2B1,ACOT8,RPS9,SLC35G1,RCE1,ZNF410,PCMT1,EBPL,IGS
9A13,PRMT2,EIF3M,CDC42,TMEM205,PTK2,ASF1B,DGKE,CAPG,PPIL2,RPL28,LIG1,CDC14B,TMEM126B,I
9A13,PRMT2,EIF3M,CDC42,TMEM205,PTK2,ASF1B,DGKE,CAPG,PPIL2,RPL28,LIG1,CDC14B,TMEM126B,I
NISCH,SRSF1,YIPF3,KXD1,SH3YL1,IP6K2,UGGT2,MICAL1,HS2ST1,UBE2I,YIF1A,XBP1,PTK2,AUNIP,PCYT2,PF
NISCH,SRSF1,YIPF3,KXD1,SH3YL1,IP6K2,UGGT2,MICAL1,HS2ST1,UBE2I,YIF1A,XBP1,PTK2,AUNIP,PCYT2,PF
MA3,METTL17,NLRP1,NIP7,TTC9C,COX4I1,THEM4,DBNDD2,CENPK,COASY,SRSF1,AUP1,SAP30BP,FOXRED
MA3,METTL17,NLRP1,NIP7,TTC9C,COX4I1,THEM4,DBNDD2,CENPK,COASY,SRSF1,AUP1,SAP30BP,FOXRED
COX17,HNRNPA1,LGMN,NDUFV2,INTS4,ASH2L,PLAC9,SNRPF,NSUN5,GMPR2,PHC3,MFSD11,ELP4,DCTN5
NDD1,RPL13A,GSTK1,SLC35G1,RCE1,MRPL33,EBPL,NUDT5,RBPMS,SPAG7,WDR62,RANBP1,SLC15A3,TUB
3P1,GPALPP1,CHPF,ERLEC1,GMPPB,MVD,IL32,SYNGR1,RANGRF,ZNF687,POT1,RAD23A,PKP4,HSPH1,RPL

3P1,GPALPP1,CHPF,ERLEC1,GMPPB,MVD,IL32,SYNGR1,RANGRF,ZNF687,POT1,RAD23A,PKP4,HSPH1,RPL
5BP1L1,IFI27L1,MAF,PPA1,PPIL2,TXNL4A,CAPN10,EXOSC9,C19ORF54,ZFAND2B,SUGT1,ZNF250,RPL36,NF
5BP1L1,IFI27L1,MAF,PPA1,PPIL2,TXNL4A,CAPN10,EXOSC9,C19ORF54,ZFAND2B,SUGT1,ZNF250,RPL36,NF
3NDD1,SH2B1,NOP16,RPL13A,ACOT8,ZNF410,EBPL,STYXL1,POLD4,UBE2L6,TUBA1C,ANTXR2,STIP1,LRRC1
X5,POLD4,LZTFL1,ZC3H14,IRF9,PCBP2,SON,TET2,UQCRQ,SNF8,GRK6,HMGA2,OBSL1,TTC21B,UBR4,CHCH
X5,POLD4,LZTFL1,ZC3H14,IRF9,PCBP2,SON,TET2,UQCRQ,SNF8,GRK6,HMGA2,OBSL1,TTC21B,UBR4,CHCH
HA,RPS9,SLC35G1,MRPL33,ZNF410,TESC,NUDT5,SNRPN,SPAG7,WDR62,STYXL1,RFX5,RANBP1,VRK3,LZTI
,SNRPN,SPAG7,WDR62,STYXL1,RFX5,RANBP1,LZTFL1,FIS1,TUBA1C,IRF9,STIP1,LPCAT3,SON,TET2,UQCRQ
,SNRPN,SPAG7,WDR62,STYXL1,RFX5,RANBP1,LZTFL1,FIS1,TUBA1C,IRF9,STIP1,LPCAT3,SON,TET2,UQCRQ
,SNRPN,SPAG7,WDR62,STYXL1,RFX5,RANBP1,LZTFL1,FIS1,TUBA1C,IRF9,STIP1,LPCAT3,SON,TET2,UQCRQ
ACTR6,SEC62,SET,CTSC,ZNF235,BOP1,TRPT1,KLC1,SCAF11,DBI,ARPC2,WDR19,SIRT6,SUMF2,NPRL2,OSG
ACTR6,SEC62,SET,CTSC,ZNF235,BOP1,TRPT1,KLC1,SCAF11,DBI,ARPC2,WDR19,SIRT6,SUMF2,NPRL2,OSG
RANGRF,SLC33A1,GTF2IRD1,NUDT1,POLR3H,RAD23A,GGT5,SLC16A1,ILF3,AGAP3,MDH2,EEF1D,HSPH1,N
RANGRF,SLC33A1,GTF2IRD1,NUDT1,POLR3H,RAD23A,GGT5,SLC16A1,ILF3,AGAP3,MDH2,EEF1D,HSPH1,N
,C15ORF40,UQCRB,DCTN6,HBS1L,HNRNPL,ZNF652,NDUFA8,NPRL3,C17ORF97,PAPOLA,ELP5,SDCCAG8,M
ZTFL1,ZC3H14,PPM1K,TUBA1C,ANTXR2,SON,HMGA2,CDC37,CPSF3L,TSG101,OBSL1,CHCHD1,SLC39A11,F
ZTFL1,ZC3H14,PPM1K,TUBA1C,ANTXR2,SON,HMGA2,CDC37,CPSF3L,TSG101,OBSL1,CHCHD1,SLC39A11,F
C1ORF198,C14ORF2,USE1,AGPAT4,EEF1B2,PPA1,TMEM126B,ATPAF2,PRMT1,ARL6IP4,C19ORF48,CYB56
TCAIM,SLC3A2,DHPS,ACTR6,MAP3K7CL,SEC62,SET,BOP1,ARRDC1,KLC1,RBP5,SCAF11,LIMK2,VDAC3,NOB
TCAIM,SLC3A2,DHPS,ACTR6,MAP3K7CL,SEC62,SET,BOP1,ARRDC1,KLC1,RBP5,SCAF11,LIMK2,VDAC3,NOB
1,USP38,THYN1,SH2B1,NOP16,RPL13A,PSMA6,RPS9,RUVBL2,EBPL,TESC,NNMT,SPAG7,WDR62,STYXL1,PI
1,USP38,THYN1,SH2B1,NOP16,RPL13A,PSMA6,RPS9,RUVBL2,EBPL,TESC,NNMT,SPAG7,WDR62,STYXL1,PI
.1,TNFSF10,RUNX1,NUDT22,USP38,THYN1,SH2B1,RPL13A,GSTK1,TTLL5,PSMA6,AP3B2,RPS9,MRPL33,ZN
.1,TNFSF10,RUNX1,NUDT22,USP38,THYN1,SH2B1,RPL13A,GSTK1,TTLL5,PSMA6,AP3B2,RPS9,MRPL33,ZN
5UOK,HSP90AB1,RAB2A,RMDN1,GPBP1,GPALPP1,CHPF,ERLEC1,GMPPB,MVD,IL32,SYNGR1,RANGRF,H2A
1,UBE2L6,LZTFL1,ZC3H14,ANTXR2,STIP1,SON,UQCRQ,CDC37,CPSF3L,METTL17,ZBTB44,DUSP22,SLC25A1
DX4I1,DBNDD2,TMED4,CENPK,SAP30BP,FOXRED1,MED15,SH3YL1,MICAL1,HS2ST1,UBE2I,YIF1A,PRMT2,J
DX4I1,DBNDD2,TMED4,CENPK,SAP30BP,FOXRED1,MED15,SH3YL1,MICAL1,HS2ST1,UBE2I,YIF1A,PRMT2,J
C,ZNF396,DNAJC2,HPS1,ITPA,SNRPD1,MKRN1,GTF3A,IFI27,RPS27L,AZI2,KAT8,NCBP2,ARPC1A,ZNF581,SS
C,ZNF396,DNAJC2,HPS1,ITPA,SNRPD1,MKRN1,GTF3A,IFI27,RPS27L,AZI2,KAT8,NCBP2,ARPC1A,ZNF581,SS
19,CD44,MAGED2,CDC37,RPS16,OBSL1,PTPN13,YLPM1,PHF21A,FUZ,MUC1,ZBTB44,RPL29,CD99L2,GGPS
PL33,TESC,NUDT5,SNRPN,SPAG7,WDR62,IL1RAP,STYXL1,RFX5,RANBP1,LZTFL1,FIS1,TUBA1C,IRF9,STIP1,
52,STYXL1,LZTFL1,ZC3H14,PHF19,PCBP2,LPCAT3,MAGED2,SON,TET2,PTPN13,UBR4,YLPM1,CHCHD1,FUZ,
52,STYXL1,LZTFL1,ZC3H14,PHF19,PCBP2,LPCAT3,MAGED2,SON,TET2,PTPN13,UBR4,YLPM1,CHCHD1,FUZ,
P3,B3GAT3,GTF2IRD2,COPS4,TPM4,THADA,MROH1,SNX5,NUP133,TMEM147,ELP4,ZXDC,COBL,ARL16,SL

ALPP1,CHPF,ERLEC1,GMPPB,MVD,IL32,SYNGR1,RANGRF,ZNF687,POT1,RAD23A,PKP4,HSPH1,RPL18,CNS
HAV1,SCAF11,DBI,ARPC2,LIMK2,VDAC3,SIRT6,SAE1,NPRL2,TBL2,SCARB1,TSPAN11,ITSN1,COMMD5,RUN
1,FKBP3,B3GAT3,GTF2IRD2,COPS4,TPM4,THADA,SNX5,NUP133,TMEM147,ELP4,ZXDC,COBL,ARL16,SLC3
FKBP5,MFSD10,CBR1,SNX5,TMEM147,SMIM4,C6ORF132,SSR3,TCF25,BAX,CDK4,NME1,EEF2K,GTF2H1,TA
3,GPATCH2L,BUD31,NPRL2,TBL2,CRYGS,CTTN,TSPAN11,EIF5A,NUDT22,UFD1L,THYN1,NOP16,RPL13A,LDI
3,GPATCH2L,BUD31,NPRL2,TBL2,CRYGS,CTTN,TSPAN11,EIF5A,NUDT22,UFD1L,THYN1,NOP16,RPL13A,LDI
BNDD2,TMED4,CENPK,COASY,SRSF1,AUP1,FOXRED1,TRNT1,KXD1,ITGB1BP1,MED15,SLC39A13,PRMT2,N
BNDD2,TMED4,CENPK,COASY,SRSF1,AUP1,FOXRED1,TRNT1,KXD1,ITGB1BP1,MED15,SLC39A13,PRMT2,N
P1,RPL36,RBM3,COPZ1,SIAH1,ASPH,PNPO,NIT2,MPDU1,RPS15,UBXN1,ZNF814,GMPPB,IL32,C12ORF76,R
P1,RPL36,RBM3,COPZ1,SIAH1,ASPH,PNPO,NIT2,MPDU1,RPS15,UBXN1,ZNF814,GMPPB,IL32,C12ORF76,R
P1,RPL36,RBM3,COPZ1,SIAH1,ASPH,PNPO,NIT2,MPDU1,RPS15,UBXN1,ZNF814,GMPPB,IL32,C12ORF76,R

,TESC,NUDT5,RBPMS,STYXL1,RFX5,POLD4,RANBP1,VRK3,LZTFL1,FIS1,IRF9,DST,ANTXR2,STIP1,LRRC1,GR

K11A,MAPKAPK5,DGUOK,HSP90AB1,RAB2A,RMDN1,GPBP1,GPALPP1,CHPF,ERLEC1,GMPPB,HHLA3,MVD,

K11A,MAPKAPK5,DGUOK,HSP90AB1,RAB2A,RMDN1,GPBP1,GPALPP1,CHPF,ERLEC1,GMPPB,HHLA3,MVD,

2,ZSCAN9,RIC8A,PIGT,EIF2B3,PDCD6,FAM32A,TMEM147,USP40,PEX2,C12ORF57,OXLD1,ZNF223,OGFOD

MBR1,MCTS1,DDOST,DOCK10,MEIS2,RAD23A,MDH2,CHCHD5,PRLR,SSH2,HINT1,RPL15,ASH2L,GMPR2,P

MBR1,MCTS1,DDOST,DOCK10,MEIS2,RAD23A,MDH2,CHCHD5,PRLR,SSH2,HINT1,RPL15,ASH2L,GMPR2,P

5,BOP1,TRPT1,ZC3HAV1,DBI,ARPC2,PMFBP1,LIMK2,WDR19,SIRT6,NUMA1,GPATCH2L,BUD31,SAE1,OSG

5,BOP1,TRPT1,ZC3HAV1,DBI,ARPC2,PMFBP1,LIMK2,WDR19,SIRT6,NUMA1,GPATCH2L,BUD31,SAE1,OSG

DBI,ARPC2,LIMK2,VDAC3,WDR19,NUMA1,GPATCH2L,SAE1,TBL2,SCARB1,CTTN,TSPAN11,ITSN1,EIF5A,CH

DBI,ARPC2,LIMK2,VDAC3,WDR19,NUMA1,GPATCH2L,SAE1,TBL2,SCARB1,CTTN,TSPAN11,ITSN1,EIF5A,CH

MD1C,CD44,UQCRQ,CDC37,CPSF3L,TSG101,TTC21B,UBR4,YLPM1,ATG12,ZBTB44,SLC25A10,TMX2,SPOC

YXL1,LZTFL1,STIP1,PHF19,PCBP2,LPCAT3,CD44,MAGED2,SON,TET2,UQCRQ,CDC37,RPS16,UBR4,YLPM1,

YXL1,LZTFL1,STIP1,PHF19,PCBP2,LPCAT3,CD44,MAGED2,SON,TET2,UQCRQ,CDC37,RPS16,UBR4,YLPM1,

T1,UBE2I,AUNIP,C14ORF2,PCYT2,PPP1CC,PPA1,DGKE,CAMTA1,LIG1,TIMM17B,APEH,CAPN10,NUTF2,PO

7,WDR62,STYXL1,LZTFL1,PPM1K,STIP1,PHF19,PCBP2,LPCAT3,SON,TET2,UQCRQ,RPS16,UBR4,YLPM1,CH

7,WDR62,STYXL1,LZTFL1,PPM1K,STIP1,PHF19,PCBP2,LPCAT3,SON,TET2,UQCRQ,RPS16,UBR4,YLPM1,CH

P1,TRMT112,CHPF,MAPK13,ERLEC1,GMPPB,EMC4,IL32,SYNGR1,H2AFV,RALGPS1,ZNF687,POT1,RAD23A

P1,TRMT112,CHPF,MAPK13,ERLEC1,GMPPB,EMC4,IL32,SYNGR1,H2AFV,RALGPS1,ZNF687,POT1,RAD23A

RNF32,LY75,ATG16L2,RCC1,SNRPB,PARD3,SLC3A2,DHPS,ACTR6,SEC62,SET,CTSC,BOP1,ARRDC1,TRPT1,KL

BL2,TESC,IGSF8,NNMT,WDR62,IL1RAP,STYXL1,PPM1K,DST,ANTXR2,PHF19,VCAN,TM9SF1,SON,UQCRQ,C

BL2,TESC,IGSF8,NNMT,WDR62,IL1RAP,STYXL1,PPM1K,DST,ANTXR2,PHF19,VCAN,TM9SF1,SON,UQCRQ,C

KBP3,INTS4,B3GAT3,TCTN1,SNRPF,GBA2,RIC8A,COMMD4,UBE2N,NOP2,USP40,PCID2,TGOLN2,PEX2,MD

UD31,NPRL2,OSGEP,TBL2,ARHGAP6,SCARB1,CTTN,TSPAN11,ITSN1,EIF5A,COMMD5,AMZ2,UFD1L,USP38

UD31,NPRL2,OSGEP,TBL2,ARHGAP6,SCARB1,CTTN,TSPAN11,ITSN1,EIF5A,COMMD5,AMZ2,UFD1L,USP38

,RPS15,MVD,ERCC5,NAP1L1,MEIS2,PSME2,RRNAD1,DBNL,BCAM,ATP6V1F,NXN,KLK10,ANAPC11,USP45,

FBXW9,DEF8,CRIP1,TMEM126B,HYI,EPB41L4B,TCF4,PPOX,CASC4,C19ORF54,SHARPIN,SLC9B2,ZNF250,M

FBXW9,DEF8,CRIP1,TMEM126B,HYI,EPB41L4B,TCF4,PPOX,CASC4,C19ORF54,SHARPIN,SLC9B2,ZNF250,M

7A,STXBP2,ATP10D,CD151,RFXANK,FTH1,PAFAH1B2,TYMS,TMEM39B,DALRD3,BRF1,OARD1,MFSD10,U

BP1,GPALPP1,CHPF,ERLEC1,GMPPB,MVD,IL32,SYNGR1,RANGRF,ZNF687,POT1,RAD23A,PKP4,HSPH1,RPL

BP1,GPALPP1,CHPF,ERLEC1,GMPPB,MVD,IL32,SYNGR1,RANGRF,ZNF687,POT1,RAD23A,PKP4,HSPH1,RPL

D2,COPS4,TPM4,THADA,MROH1,SNX5,NUP133,TMEM147,ELP4,ZXDC,ARL16,SLC35E1,MRPL55,ECE1,OC

K5,HSP90AB1,CEP70,RAB2A,RMDN1,GPBP1,GPALPP1,CHPF,ERLEC1,GMPPB,HHLA3,MVD,IL32,SYNGR1,R

62,IL1RAP,STYXL1,RFX5,POLD4,RANBP1,LZTFL1,FIS1,TUBA1C,IRF9,STIP1,LPCAT3,SON,TET2,UQCRQ,GRK

62,IL1RAP,STYXL1,RFX5,POLD4,RANBP1,LZTFL1,FIS1,TUBA1C,IRF9,STIP1,LPCAT3,SON,TET2,UQCRQ,GRK

62,IL1RAP,STYXL1,RFX5,POLD4,RANBP1,LZTFL1,FIS1,TUBA1C,IRF9,STIP1,LPCAT3,SON,TET2,UQCRQ,GRK

87,PDCD5,NXN,HSD17B10,ABHD17A,GGT5,DDX41,ZCCHC4,STXBP2,SEC31A,SUMO3,LGMN,SMARCA4,TM

87,PDCD5,NXN,HSD17B10,ABHD17A,GGT5,DDX41,ZCCHC4,STXBP2,SEC31A,SUMO3,LGMN,SMARCA4,TM

,IL32,SYNGR1,RRNAD1,ZNF687,POT1,RAD23A,PKP4,HSPH1,RPL18,CHEK1,PHF20L1,FKBP3,B3GAT3,ZNF5

L,AUP1,SAP30BP,FOXRED1,TRNT1,ITGB1BP1,MED15,SLC39A13,MAGT1,C14ORF2,GUK1,CENPN,CARD14,

L,AUP1,SAP30BP,FOXRED1,TRNT1,ITGB1BP1,MED15,SLC39A13,MAGT1,C14ORF2,GUK1,CENPN,CARD14,

ZNF687,POT1,RAD23A,PKP4,HSPH1,MT1E,RPL18,CHEK1,PHF20L1,FKBP3,B3GAT3,GTF2IRD2,COPS4,TPM

396,DNAJC2,DNAJA3,VPS25,NDUFB3,RPS27L,GOSR2,ZBTB8OS,ILK,PRKCSH,BTF3L4,ZNF581,RPS23,MGR

396,DNAJC2,DNAJA3,VPS25,NDUFB3,RPS27L,GOSR2,ZBTB8OS,ILK,PRKCSH,BTF3L4,ZNF581,RPS23,MGR

4A,FASTK,TMEM135,RNF32,LY75,ATG16L2,RCC1,SNRPB,PARD3,TCAIM,DHPS,ACTR6,MAP3K7CL,SEC62,S
4A,FASTK,TMEM135,RNF32,LY75,ATG16L2,RCC1,SNRPB,PARD3,TCAIM,DHPS,ACTR6,MAP3K7CL,SEC62,S
ID1,NUDT22,AMZ2,UFD1L,TAZ,DBNDD1,RPL13A,TTLL5,RCE1,ZNF410,EBPL,TESC,RBPMS,IL1RAP,POLD4,F
ID1,NUDT22,AMZ2,UFD1L,TAZ,DBNDD1,RPL13A,TTLL5,RCE1,ZNF410,EBPL,TESC,RBPMS,IL1RAP,POLD4,F

C1,SLC25A10,SDF4,TMED4,RPS6,RTEL1,TRNT1,ITGB1BP1,SH3YL1,C1ORF198,C14ORF2,PCYT2,USE1,AGP
C1,SLC25A10,SDF4,TMED4,RPS6,RTEL1,TRNT1,ITGB1BP1,SH3YL1,C1ORF198,C14ORF2,PCYT2,USE1,AGP
RB1,CTTN,ITSN1,EIF5A,NUDT22,USP38,TAZ,THYN1,DBNDD1,SH2B1,TTLL5,LDHA,AP3B2,RCE1,PCMT1,EB
P1,PPM1K,ANTXR2,STIP1,CD44,TM9SF1,MAGED2,UQCRQ,GRK6,CDC37,TSG101,PTPN13,FAM3A,ZBTB4
HNRNPC,CYB561,SLC9B2,NFAM1,CDK11A,HSP90AB1,ASPH,PNPO,MPDU1,SAT1,PABPC1,NUDT1,GGT5,NT
HNRNPC,CYB561,SLC9B2,NFAM1,CDK11A,HSP90AB1,ASPH,PNPO,MPDU1,SAT1,PABPC1,NUDT1,GGT5,NT
,STIP1,LRRC1,TET2,OBSL1,TTC21B,UBR4,YLPM1,PHF21A,ATG12,RPS25,ZBTB44,DUSP22,THEM4,RDH11,D
,STIP1,LRRC1,TET2,OBSL1,TTC21B,UBR4,YLPM1,PHF21A,ATG12,RPS25,ZBTB44,DUSP22,THEM4,RDH11,D

MT2,MAGT1,CENPN,AGPAT4,MAF,PPP1CC,RFX3,PPA1,PPIL2,TIMM17B,RPL21,EXOSC9,ARL6IP4,C19ORF
MT2,MAGT1,CENPN,AGPAT4,MAF,PPP1CC,RFX3,PPA1,PPIL2,TIMM17B,RPL21,EXOSC9,ARL6IP4,C19ORF
KLC1,ZC3HAV1,ARPC2,LIMK2,NOB1,WDR19,SIRT6,NUMA1,GPATCH2L,SUMF2,NPRL2,ARHGAP6,SCARB1,
KLC1,ZC3HAV1,ARPC2,LIMK2,NOB1,WDR19,SIRT6,NUMA1,GPATCH2L,SUMF2,NPRL2,ARHGAP6,SCARB1,
PARD3,TCAIM,DHPS,ACTR6,SEC62,SET,CTSC,ZNF235,BOP1,ARRDC1,TRPT1,KLC1,ZC3HAV1,DBI,VDAC3,NC
PARD3,TCAIM,DHPS,ACTR6,SEC62,SET,CTSC,ZNF235,BOP1,ARRDC1,TRPT1,KLC1,ZC3HAV1,DBI,VDAC3,NC
R62,RFX5,POLD4,PLEKHA5,RANBP1,VRK3,FIS1,PPM1K,IRF9,STIP1,TM9SF1,TET2,GRK6,HMGA2,CDC37,C
R62,RFX5,POLD4,PLEKHA5,RANBP1,VRK3,FIS1,PPM1K,IRF9,STIP1,TM9SF1,TET2,GRK6,HMGA2,CDC37,C
2,RBPMS,WDR62,RANBP1,UBE2L6,FIS1,PPM1K,STIP1,PHF19,GRAMD1C,LPCAT3,CD44,TM9SF1,MAGED2,
2,RBPMS,WDR62,RANBP1,UBE2L6,FIS1,PPM1K,STIP1,PHF19,GRAMD1C,LPCAT3,CD44,TM9SF1,MAGED2,
D10,CBFA2T2,SUGT1,INO80E,MAPKAPK5,DGUOK,HSP90AB1,ATP5J,GGCX,TRMT112,PNPO,CHPF,RPS20,
D10,CBFA2T2,SUGT1,INO80E,MAPKAPK5,DGUOK,HSP90AB1,ATP5J,GGCX,TRMT112,PNPO,CHPF,RPS20,
,DHPS,ACTR6,SEC62,SET,ZNF235,BOP1,ARRDC1,TRPT1,KLC1,ZC3HAV1,SCAF11,ARPC2,PMFBP1,LIMK2,V
,DHPS,ACTR6,SEC62,SET,ZNF235,BOP1,ARRDC1,TRPT1,KLC1,ZC3HAV1,SCAF11,ARPC2,PMFBP1,LIMK2,V
,PCBP2,CD44,TM9SF1,MAGED2,SON,SNF8,CDC37,TSG101,UBR4,PSMA1,METTL17,ECSIT,ATG12,NIP7,TT
LZTFL1,PPM1K,STIP1,PHF19,PCBP2,LPCAT3,CD44,SON,TET2,CDC37,RPS16,UBR4,YLPM1,CHCHD1,FUZ,E
LZTFL1,PPM1K,STIP1,PHF19,PCBP2,LPCAT3,CD44,SON,TET2,CDC37,RPS16,UBR4,YLPM1,CHCHD1,FUZ,E
L5,LDHA,RPS9,RCE1,MRPL33,ZNF410,PCMT1,IGSF8,NNMT,SNRPN,WDR62,STYXL1,RFX5,PLEKHA5,RANB
L5,LDHA,RPS9,RCE1,MRPL33,ZNF410,PCMT1,IGSF8,NNMT,SNRPN,WDR62,STYXL1,RFX5,PLEKHA5,RANB
K,TMEM135,RNF32,LY75,RCC1,PARD3,DHPS,ACTR6,MAP3K7CL,SEC62,TRPT1,RBP5,SCAF11,DBI,PMFBP1,
K,TMEM135,RNF32,LY75,RCC1,PARD3,DHPS,ACTR6,MAP3K7CL,SEC62,TRPT1,RBP5,SCAF11,DBI,PMFBP1,
RUNX1,NUDT22,USP38,SH2B1,NOP16,RPL13A,PSMA6,RPS9,SLC35G1,MRPL33,RUVBL2,TESC,NNMT,SPA
RUNX1,NUDT22,USP38,SH2B1,NOP16,RPL13A,PSMA6,RPS9,SLC35G1,MRPL33,RUVBL2,TESC,NNMT,SPA
MF2,OSGEP,TBL2,CTTN,ITSN1,EIF5A,COMMD5,CHID1,UFD1L,USP38,DBNDD1,NOP16,ARHGEF39,LDHA,
MF2,OSGEP,TBL2,CTTN,ITSN1,EIF5A,COMMD5,CHID1,UFD1L,USP38,DBNDD1,NOP16,ARHGEF39,LDHA,
BTB44,NIP7,TTC9C,PLCB2,DUSP22,SLC25A10,DBNDD2,RTEL1,SAP30BP,FOXRED1,ITGB1BP1,PARD6G,SH
BTB44,NIP7,TTC9C,PLCB2,DUSP22,SLC25A10,DBNDD2,RTEL1,SAP30BP,FOXRED1,ITGB1BP1,PARD6G,SH
C,RBPMS,WDR62,IL1RAP,PLEKHA5,RANBP1,LZTFL1,FIS1,SLC15A3,PPM1K,STIP1,LRRC1,GRAMD1C,TM9SF
8,NNMT,WDR62,STYXL1,RFX5,PLEKHA5,RANBP1,VRK3,PPM1K,PHF19,PCBP2,TM9SF1,SON,TET2,GRK6,C
8,NNMT,WDR62,STYXL1,RFX5,PLEKHA5,RANBP1,VRK3,PPM1K,PHF19,PCBP2,TM9SF1,SON,TET2,GRK6,C
GRK6,HMGA2,CDC37,OBSL1,TTC21B,UBR4,SLC39A11,FUZ,PSMA3,PPRC1,SLC25A10,SDF4,TMED4,RPS6,R
GRK6,HMGA2,CDC37,OBSL1,TTC21B,UBR4,SLC39A11,FUZ,PSMA3,PPRC1,SLC25A10,SDF4,TMED4,RPS6,R
TUBA1C,IRF9,STIP1,TET2,UQCRQ,GRK6,HMGA2,CDC37,TTC21B,UBR4,SLC39A11,FUZ,FAM3A,PSMA3,ME

TUBA1C,IRF9,STIP1,TET2,UQCRQ,GRK6,HMGA2,CDC37,TTC21B,UBR4,SLC39A11,FUZ,FAM3A,PSMA3,ME
D1L,USP38,NOP16,AP3B2,RUVBL2,STYXL1,ZC3H14,PPM1K,DST,STIP1,VCAN,CD44,SON,UQCRQ,CDC37,RI
D1L,USP38,NOP16,AP3B2,RUVBL2,STYXL1,ZC3H14,PPM1K,DST,STIP1,VCAN,CD44,SON,UQCRQ,CDC37,RI
70,RAB2A,GPBP1,GPALPP1,CHPF,GMPPB,MVD,ERCC5,IL32,SYNGR1,RRNAD1,ZNF687,POT1,RAD23A,PKP
1,SIAH1,ASPH,PNPO,NIT2,LYNX1,MPDU1,RPS15,ZNF814,GMPPB,IL32,NAP1L1,SERPINB1,C12ORF76,RAN
1,SIAH1,ASPH,PNPO,NIT2,LYNX1,MPDU1,RPS15,ZNF814,GMPPB,IL32,NAP1L1,SERPINB1,C12ORF76,RAN
1,SIAH1,ASPH,PNPO,NIT2,LYNX1,MPDU1,RPS15,ZNF814,GMPPB,IL32,NAP1L1,SERPINB1,C12ORF76,RAN
RD3,TCAIM,SLC3A2,ACTR6,MAP3K7CL,SEC62,SET,CTSC,ZNF235,BOP1,ARRDC1,TRPT1,KLC1,RBP5,ZC3HAV
RD3,TCAIM,SLC3A2,ACTR6,MAP3K7CL,SEC62,SET,CTSC,ZNF235,BOP1,ARRDC1,TRPT1,KLC1,RBP5,ZC3HAV
KIRAS2,SKA2,AXIN1,AP1G2,EML3,UCHL5,TUSC3,FARS2,ARRB2,SNRPA1,CARM1,ECT2,DTD2,XPNPEP1,CEP
KIRAS2,SKA2,AXIN1,AP1G2,EML3,UCHL5,TUSC3,FARS2,ARRB2,SNRPA1,CARM1,ECT2,DTD2,XPNPEP1,CEP
SMA3,METTL17,NIP7,TTC9C,COX4I1,THEM4,DBNDD2,CENPK,COASY,SRSF1,AUP1,SAP30BP,FOXRED1,TR
SMA3,METTL17,NIP7,TTC9C,COX4I1,THEM4,DBNDD2,CENPK,COASY,SRSF1,AUP1,SAP30BP,FOXRED1,TR
N,TSPAN11,ITSN1,EIF5A,AMZ2,USP38,TAZ,RPL13A,GSTK1,ACOT8,SLC35G1,RCE1,MRPL33,IGSF8,NUDT5,R
UVBL2,PCMT1,NNMT,IL1RAP,STYXL1,PLEKHA5,SLC15A3,STIP1,LRRC1,PHF19,MAGED2,TET2,GRK6,RPS16
UVBL2,PCMT1,NNMT,IL1RAP,STYXL1,PLEKHA5,SLC15A3,STIP1,LRRC1,PHF19,MAGED2,TET2,GRK6,RPS16
P6,TPM2,GPALPP1,SIAH1,ASPH,PNPO,NIT2,RPS15,UBXN1,ZNF814,GMPPB,IL32,C12ORF76,RANGRF,PAB
P6,TPM2,GPALPP1,SIAH1,ASPH,PNPO,NIT2,RPS15,UBXN1,ZNF814,GMPPB,IL32,C12ORF76,RANGRF,PAB
1K2,VDAC3,WDR19,SIRT6,GPATCH2L,OSGEP,CTTN,ITSN1,EIF5A,COMMD5,NUDT22,USP38,THYN1,SH2B1,
1K2,VDAC3,WDR19,SIRT6,GPATCH2L,OSGEP,CTTN,ITSN1,EIF5A,COMMD5,NUDT22,USP38,THYN1,SH2B1,
OT1,RAD23A,PKP4,HSPH1,RPL18,CHEK1,PHF20L1,FKBP3,B3GAT3,ZNF506,TCTN1,GTF2IRD2,COPS4,ZFPL
2,FBXW9,USE1,HSBP1L1,IFI27L1,CCDC25,EEF1B2,EDF1,PPIL2,LIG1,PSMA7,TXNL4A,NUTF2,TCF4,C17ORF
2,FBXW9,USE1,HSBP1L1,IFI27L1,CCDC25,EEF1B2,EDF1,PPIL2,LIG1,PSMA7,TXNL4A,NUTF2,TCF4,C17ORF
L1,EGLN2,RFXANK,TMEM101,FKBP5,MRPS18C,TRPV3,CBR1,PHC3,COMMD4,SNX5,SMIM4,NMNAT3,BT
L1,EGLN2,RFXANK,TMEM101,FKBP5,MRPS18C,TRPV3,CBR1,PHC3,COMMD4,SNX5,SMIM4,NMNAT3,BT
T2,PTK2,C14ORF2,CARD14,HSBP1L1,IFI27L1,CCDC25,PPP1CC,ENDOV,PPIL2,TROAP,RPL28,APEH,TMPO,C
T2,PTK2,C14ORF2,CARD14,HSBP1L1,IFI27L1,CCDC25,PPP1CC,ENDOV,PPIL2,TROAP,RPL28,APEH,TMPO,C
JDT5,SNRPN,WDR62,IL1RAP,PLEKHA5,FIS1,TUBA1C,IRF9,DST,STIP1,LRRC1,PHF19,MAGED2,SNF8,CDC37,
9,RUVBL2,TESC,NNMT,SPAG7,STYXL1,LZTFL1,PPM1K,STIP1,PHF19,PCBP2,LPCAT3,SON,TET2,UQCRQ,CD
9,RUVBL2,TESC,NNMT,SPAG7,STYXL1,LZTFL1,PPM1K,STIP1,PHF19,PCBP2,LPCAT3,SON,TET2,UQCRQ,CD

,ILF3,MDH2,HNRNPA1,RPL15,SMARCA4,PAFAH1B2,TYMS,TCP1,SMARCC2,EIF2B3,XYLT2,NOP2,PRRC2B,R
H6,NRP1,AXIN1,HNRNPUL1,PSMD10,SH3GL1,NDUFA3,EIF3C,IDH3G,NQO2,NSFL1C,SMUG1,CDK2AP2,BC

15,DCTD,LRRC27,PHTF2,SLC33A1,ZNF687,NXN,HSD17B10,PDE4DIP,HSPH1,CHCHD5,CHEK1,HNRNPLL,EG

RFIP1,TSFM,RPS20,RPS15,HSPB1,MCTS1,NAP1L1,PRKRIP1,DCN,PABPC1,RBMX,NUDT1,HSD17B10,DDX41

PS23,RPL30,NUP85,NDUFA7,MRPS33,ACTB,RPS15A,RPS17,SEC13,MRPL51,MRPL43,ACTG1,MRPL27,MRP

1,TBX6,XBP1,CELF1,AUNIP,ATF7,MAF,EEF1B2,PPP1CC,EDF1,RFX3,ENDOV,CAMTA1,CRIP1,PSMD4,RPL28,

P5,ZC3HAV1,SCAF11,DBI,ARPC2,LIMK2,VDAC3,WDR19,SIRT6,NUMA1,GPATCH2L,BUD31,SAE1,NPRL2,O

,MUC1,PPRC1,RPS25,NLRP1,ZBTB44,NIP7,RPS6,RPL29,RTEL1,COASY,SRSF1,RABL5,TRNT1,IP6K2,JPH2,MI

A,CHCHD1,MUC1,PPRC1,RPS25,NLRP1,ZBTB44,NIP7,RPS6,RPL29,RTEL1,COASY,SRSF1,RABL5,TRNT1,IP6

RPL37A,USP4,RPLP0,METTL3,TSEN34,SSB,RPL8,RBBP6,CHTOP,RPL35,SRSF7,RALY,RBM42,RPL34,RPL24,

IA,RPS11,NUP133,RPL37A,KPNA5,EIF4A1,RPLP0,EIF3G,SSB,RPL8,BAX,RPL35,SLC52A2,NXF1,CFLAR,LAMT

1,CD74,NFKBIA,RPS11,NUP133,RPL37A,KPNA5,EIF4A1,RPLP0,EIF3G,SSB,RPL8,BAX,RPL35,SLC52A2,NXF1

06,MFF,SNX5,ENY2,NUP133,RAPGEF3,RPAIN,ACOT2,PCID2,PEX2,RPL37A,STX10,PEX26,KPNA5,RPLP0,HA

ME1,AGAP3,CARHSP1,RPL18,LGMN,RPL15,PARK7,RPS11,SNX5,USP40,PCID2,RPL37A,USP4,RPLP0,METT

CDK4,MTIF2,EEF2K,MARS,RPL34,EIF2A,RPL24,RBM4,RPS3A,EIF3C,RPS3,MRPS24,FARS2,RPS27L,RPS7,NC

CDK4,MTIF2,EEF2K,MARS,RPL34,EIF2A,RPL24,RBM4,RPS3A,EIF3C,MGST2,RPS3,MRPS24,FARS2,RPS27L,R

PS20,VPS28,RPS15,ERLEC1,HSPB1,SYNGR1,RANGRF,HMGA1,TOMM40,PDCD5,STXBP2,SEC31A,RPL18,SH

TPA,SMUG1,COMT,HMOX2,RPS7,APAF1,NCBP2,RNPS1,UBA52,RPS4X,MLH1,RPS23,RPL30,PSMD8,RQCD1

1,CD74,NFKBIA,RPS11,NUP133,RPL37A,KPNA5,EIF4A1,RPLP0,EIF3G,SSB,RPL8,BAX,RPL35,SLC52A2,NXF1

40,PDCD5,POT1,ABHD17A,SEC31A,RPL18,CNST,EGFR,SH3GLB1,RPL15,B3GAT3,TCTN1,LGALS3,TCP1,SYTL

R2,RPS7,ECT2,UBA52,RPS4X,VTI1B,RPS23,RPL30,NUP85,PEX16,SUFU,CCT4,PKIG,UBE2L3,RPS15A,PPP1R

1,UBA52,RPS4X,MLH1,RPS23,RPL30,PSMD8,RQCD1,EXOSC2,RBM7,PSMC5,HNRNPM,EXOSC8,RPS15A,RF

40,PDCD5,POT1,ABHD17A,SEC31A,RPL18,CNST,EGFR,SH3GLB1,RPL15,B3GAT3,TCTN1,LGALS3,TCP1,SYTL
ABPC1,WAC,ANAPC11,RAD23A,ABHD17A,USP45,PSME1,AGAP3,CARHSP1,RPL18,EGFR,SH3GLB1,LGMN,

5,RPL8,ECE1,RPL35,CDK4,MTIF2,EEF2K,MARS,RPL34,EIF2A,RPL24,RBM4,RPS3A,EIF3C,MGST2,RPS3,MRP
GOHB,ITPA,SMUG1,COMT,HMOX2,RPS7,APAF1,NCBP2,RNPS1,UBA52,RPS4X,MLH1,RPS23,RPL30,PSMD
7,AP4M1,IMMP1L,RPL27,TIMM9,RPS14,AP4S1,PAK1,RPL31,GSK3A,TOMM7,ITGB2,GIPC1,VPS13A,FAU

X2,RPS7,APAF1,NCBP2,RNPS1,UBA52,RPS4X,MLH1,RPS23,RPL30,PSMD8,RQCD1,EXOSC2,RBM7,PSMC5,
X2,RPS7,APAF1,NCBP2,RNPS1,UBA52,RPS4X,MLH1,RPS23,RPL30,PSMD8,RQCD1,EXOSC2,RBM7,PSMC5,
5,RPL8,UGT8,RPL35,CDK4,MTIF2,EEF2K,GCDH,MARS,RPL34,EIF2A,RPL24,RBM4,RPS3A,EIF3C,MGST2,RPS

3,RPL21,CAPN10,NUTF2,STOML2,STRADA,ZFAND2B,AP1S3,SLC9B2,RPL36,LAMP2,COPZ1,HSP90AB1,CEP

DLG4,LAMTOR5,RER1,RPL34,NUP62,ACAA1,RPL24,PAM16,KDELR3,SIN3A,NRP1,TP53,RPS3A,HDAC8,PM
IN,UPP1,RPL36,LAMP2,HSP90AB1,DUT,SIAH1,RPS20,VPS28,RPS15,UBXN1,ERLEC1,HSPB1,SAT1,DCN,PSN
ACOT2,RPL37A,ELOF1,EIF4A1,RPLP0,DGAT1,METTL3,EIF3G,MRPL55,RPL8,ECE1,UGT8,RPL35,CDK4,MTIF2
X1,EIF3G,PICK1,ADD1,CLASP2,RPL24,FCHO2,GTF2H1,CLTA,ARPC5,GEMIN2,NDUFA11,AXIN1,TP53,PSMD
RPS15,VWA2,ERCC5,MCTS1,NAP1L1,RANGRF,HMGA1,DBNL,RBMX,HSD17B10,B2M,PDE4DIP,MED6,SEC3
,ZFAND2B,ACAD10,HNRNPC,SHARPIN,UPP1,RPL36,LAMP2,HSP90AB1,DUT,SIAH1,RPTOR,RPS20,VPS28,R
OSPH10,METTL3,TSEN34,SSB,RBBP6,CHTOP,RPL35,SRSF7,RALY,RBM42,PSPC1,GTF2H1,RBM4,GEMIN2,T
7,RPL21,LYPLA2,CAMLG,EXOSC9,ZFAND2B,ACAD10,HNRNPC,SHARPIN,UPP1,RPL36,LAMP2,HSP90AB1,D
0AB1,RAB2A,RPS20,VPS28,RPS15,ERLEC1,HSPB1,SERPINB1,SYNGR1,RANGRF,TOMM40,PPP3CB,RABEP2,
MP2,COPZ1,HSP90AB1,RAB2A,RPS20,VPS28,RPS15,ERLEC1,HSPB1,SERPINB1,SYNGR1,RANGRF,TOMM4
SF1,FOXRED1,RABL5,ITGB1BP1,PARD6G,SH3YL1,MICAL1,EIF3M,JUP,CDC42,PTK2,CELF1,CENPN,FBXW9,

D,HSPB1,MVD,SAT1,MCTS1,DDOST,DCN,PABPC1,ZNF706,ABHD17A,GGT5,ILF3,ZCCHC4,EEF1D,RPL18,EG

0AB1,RAB2A,RPS20,VPS28,RPS15,ERLEC1,HSPB1,SERPINB1,SYNGR1,RANGRF,TOMM40,PPP3CB,RABEP2,
0AB1,RAB2A,RPS20,VPS28,RPS15,ERLEC1,HSPB1,SERPINB1,SYNGR1,RANGRF,TOMM40,PPP3CB,RABEP2,
1,CAPN10,NUTF2,STRADA,ZFAND2B,AP1S3,SLC9B2,RPL36,LAMP2,COPZ1,HSP90AB1,RAB2A,RPS20,VPS2
SNRNP200,HNRNPL,MAGOH,ZRANB2,LUC7L3,PRPF31,POLR2L,SRSF5,PAPOLA,AKAP8L,WDR83,FXR1
G101,TTC21B,PSMA1,YLPM1,PHF21A,PSMA3,MUC1,PPRC1,RPS25,DUSP22,THEM4,RPS6,RPL29,RTEL1,C
GLB1,SSH2,COX17,NDUFV2,KCTD7,ASH2L,SMARCA4,COA3,SNRPF,GBA2,PPIH,BRF1,TRPM4,MCFD2,LGA
9,PHF19,PCBP2,SON,TET2,SNF8,HMGA2,RPS16,CPSF3L,TSG101,TTC21B,PSMA1,YLPM1,PHF21A,CHCHD1
,ECSIT,EPB41L5,ATG12,PPRC1,NIP7,THEM4,NISCH,TMED4,CENPK,RTEL1,FOXRED1,RABL5,ITGB1BP1,PAR

,RAB26,PEX16,ANXA2,ACTB,RPS15A,PPP1R13B,RPS17,AP4M1,NUMB,PRKCZ,VIL1,RPL27,TIMM9,RPS14,

NDUFB10,ATPAF1,LMNA,IMMP1L,NDUFS7,IFI6,TIMM9,UQCRB,GGCT,OPA3,NDUFS4,MARK2,ATP13A2,ES
G101,TTC21B,PSMA1,YLPM1,PHF21A,PSMA3,MUC1,PPRC1,RPS25,DUSP22,THEM4,RPS6,RPL29,RTEL1,C
9,UBA52,SUMO1,COMMD1,PSMD8,SUFU,NUB1,OS9,EXOSC2,PSMC5,EXOSC8,UBE2L3,DNAJB12,UBE2A,
GLN2,MTHFS,GBA2,PARK7,SNX5,USP40,USP4,ARIH2,FBXL18,UBE2D4,ECE1,RBBP6,UBA1,RNF8,PPP2R5C,

3,PPIL2,RPL28,TIMM17B,RPL21,CAPN10,NUTF2,STOML2,STRADA,ZFAND2B,AP1S3,SLC9B2,RPL36,LAMP2
.2,CDC37,RPS16,TSG101,OBSL1,PTPN13,UBR4,PSMA1,CHCHD1,RGS3,PSMA3,MUC1,METTL17,ATG12,RPS
L1,EIF3M,JUP,CDC42,PTK2,CELF1,CENPN,ASF1B,DEF8,PDLIM5,RFX3,VAMP8,CAPG,PSMD4,CDC14B,TMEN
A,GSTK1,ACOT8,RPS9,MRPL33,RUVBL2,NUDT5,WDR62,IL1RAP,STYXL1,POLD4,RANBP1,FIS1,TUBA1C,DST
29,COASY,SAP30BP,ITGB1BP1,MED15,UBE2I,PRMT2,EIF3M,JUP,TBX6,XBP1,CELF1,GUK1,CARD14,ATF7,H
G101,TTC21B,PSMA1,YLPM1,PHF21A,PSMA3,MUC1,PPRC1,RPS25,DUSP22,THEM4,RPS6,RPL29,RTEL1,CC
,RDX,VPS25,UCHL5,NSFL1C,IFI27,ARRB2,RPS7,USP15,NFE2L1,TAF9,UBA52,CAPN2,SUMO1,COMMD1,PSN

RNMT,PUF60,SNRNP40,RBM4B,SNRNP200,HNRNPL,MAGOH,ZRANB2,LUC7L3,PRPF31,POLR2L,SRSF5,PAF
RB1,CTTN,ITSN1,EIF5A,TNFSF10,RUNX1,UFD1L,USP38,TAZ,SH2B1,RPL13A,GSTK1,TTLL5,LDHA,PSMA6,ACO

L,RPS25,DUSP22,RPS6,RPL29,RTEL1,SRSF1,SAP30BP,TRNT1,ITGB1BP1,MED15,CLASRP,UBE2I,PRMT2,JUP
16,CPSF3L,TSG101,TTC21B,PSMA1,YLPM1,PHF21A,PSMA3,MUC1,PPRC1,RPS25,DUSP22,THEM4,RPS6,RI

6,LAMP2,COPZ1,HSP90AB1,RAB2A,RPS20,VPS28,RPS15,ERLEC1,HSPB1,SERPINB1,SYNGR1,RANGRF,TOM
SAP30BP,ITGB1BP1,MED15,UGGT2,HS2ST1,UBE2I,PRMT2,EIF3M,MAGT1,JUP,CDC42,TBX6,XBP1,CELF1,CA
),BFAR,CLTA,LYPLAL1,AXIN1,PSMD10,PML,TMEM199,UBE2D3,RDX,VPS25,UCHL5,NSFL1C,IFI27,ARRB2,SC
RRB2,RPS7,USP15,NFE2L1,TAF9,UBA52,CAPN2,SUMO1,COMMD1,PSMD8,SUFU,NUB1,OS9,PSMC5
RPL29,RTEL1,SAP30BP,ITGB1BP1,MED15,UGGT2,HS2ST1,UBE2I,PRMT2,EIF3M,MAGT1,JUP,CDC42,TBX6,
NMT,IL1RAP,STYXL1,UBE2L6,VRK3,FIS1,PPM1K,PHF19,VCAN,PCBP2,LPCAT3,CD44,TET2,SNF8,GRK6,HMG

.1,PHF21A,CHCHD1,FAM3A,PSMA3,MUC1,METTL17,ATG12,PPRC1,RPS25,PLCB2,DUSP22,SLC25A10,RPS6
JDT5,WDR62,IL1RAP,STYXL1,POLD4,RANBP1,FIS1,TUBA1C,DST,PHF19,VCAN,CD44,SON,TET2,SNF8,HMGA

DCTN1,SEMA5A,MRPL36,MARK2,GLE1,ATP13A2,MRPL34,EPG5,GSK3A,TOMM7,CDK5RAP3,CTNNB1,AKA

IN1,TP53,PSMD10,PML,RHEB,RDX,UCHL5,MLST8,SGSM3,RPS7,LAMTOR1,APAF1,KAT8,SNX3,NFE2L1,TAF
.1,PHF21A,CHCHD1,FAM3A,PSMA3,MUC1,METTL17,ATG12,PPRC1,RPS25,DUSP22,RPS6,RPL29,GGPS1,RT

N,PCBP2,CD44,TET2,SNF8,GRK6,HMGA2,CDC37,RPS16,TSG101,OBSL1,PTPN13,UBR4,PSMA1,CHCHD1,RG

GB1BP1,MED15,CLASRP,UBE2I,PRMT2,JUP,TBX6,XBP1,CELF1,CARD14,ATF7,FBXW9,HSBP1L1,MAF,EDF1,

.1,PHF21A,CHCHD1,FAM3A,PSMA3,MUC1,METTL17,ATG12,PPRC1,RPS25,PLCB2,DUSP22,SLC25A10,RPS6

NPRL2,OSGEP,SCARB1,CTTN,ITSN1,EIF5A,TNFSF10,RUNX1,CHID1,AMZ2,UFD1L,USP38,TAZ,SH2B1,RPL13A
RK7,UBE2N,MFF,RAPGEF3,NEK2,PCID2,TNKS,HAX1,MPV17,ARIH2,LIMA1,PICK1,ADD1,TJP1,CHTOP,BAX,C

AMTOR1,APAF1,KAT8,NFE2L1,TAF9,UBA52,SUMO1,COMMD1,PSMD8,SUFU,NUB1,OS9,EXOSC2,PSMC5,H

1,POT1,RAD23A,B2M,USP45,PSME1,AGAP3,HSPH1,SGK1,RRP8,SEC31A,EGFR,SH3GLB1,CHEK1,CAMK2D

MT1,TESC,RBPMS,IL1RAP,STYXL1,RFX5,POLD4,UBE2L6,VRK3,ZC3H14,FIS1,PPM1K,IRF9,PHF19,VCAN,PCB

AC3,NOB1,SIRT6,NUMA1,SUMF2,BUD31,SAE1,NPRL2,OSGEP,TBL2,SCARB1,CTTN,ITSN1,EIF5A,COMMD5
NMT,WDR62,PLEKHA5,RANBP1,UBE2L6,LZTFL1,IRF9,DST,STIP1,LRRC1,PCBP2,CD44,MAGED2,SNF8,CDC3
1,RBP5,ZC3HAV1,SCAF11,DBI,ARPC2,LIMK2,VDAC3,NOB1,SIRT6,NUMA1,SUMF2,BUD31,SAE1,NPRL2,OS
F235,BOP1,ARRDC1,KLC1,ZC3HAV1,SCAF11,DBI,ARPC2,LIMK2,VDAC3,NOB1,WDR19,SIRT6,NUMA1,SUM
SET,CTSC,ARRDC1,KLC1,RBP5,ZC3HAV1,DBI,ARPC2,PMFBP1,LIMK2,VDAC3,NOB1,WDR19,SIRT6,NUMA1,
TCAIM,SLC3A2,DHPS,ACTR6,MAP3K7CL,SEC62,SET,CTSC,ZNF235,BOP1,ARRDC1,KLC1,RBP5,ZC3HAV1,SC
2,SET,CTSC,ZNF235,BOP1,ARRDC1,KLC1,RBP5,ZC3HAV1,SCAF11,DBI,ARPC2,PMFBP1,LIMK2,VDAC3,NOB
MRPL55,SSB,RPL8,RPL35,RALY,ZNF593,RBM42,RPL34,EIF2A,NUP62,RPL24,NAA38,GEMIN2,RPS3A,BTF3,
PMS,SNRPN,STYXL1,RFX5,POLD4,PLEKHA5,UBE2L6,VRK3,ZC3H14,PPM1K,IRF9,DST,PHF19,VCAN,PCBP2,
PMS,SNRPN,STYXL1,RFX5,POLD4,PLEKHA5,UBE2L6,VRK3,ZC3H14,PPM1K,IRF9,DST,PHF19,VCAN,PCBP2,
PMS,SNRPN,STYXL1,RFX5,POLD4,PLEKHA5,UBE2L6,VRK3,ZC3H14,PPM1K,IRF9,DST,PHF19,VCAN,PCBP2,
DHA,PSMA6,RPS9,ZNF410,RUVBL2,TESC,NUDT5,RBPMS,SNRPN,SPAG7,WDR62,RFX5,POLD4,PLEKHA5,R

13,UBR4,PSMA1,YLPM1,PHF21A,CHCHD1,PSMA3,METTL17,ECSIT,EPB41L5,PPRC1,RPS25,NLRP1,NIP7,C

,CPSF3L,TSG101,TTC21B,PTPN13,UBR4,PSMA1,YLPM1,PHF21A,CHCHD1,PSMA3,MUC1,METTL17,ECSIT,
D44,UQCRQ,SNF8,HMGA2,CDC37,RPS16,CPSF3L,TSG101,OBSL1,TTC21B,PSMA1,PHF21A,PSMA3,ATG12,
,NUP133,COX20,MRPS17,TMEM14B,COA1,PCID2,TNKS,DCTN5,NDUFS3,KPNA5,HAX1,MPV17,MRPL55,C
,NUP133,COX20,MRPS17,TMEM14B,COA1,PCID2,TNKS,DCTN5,NDUFS3,KPNA5,HAX1,MPV17,MRPL55,C

,MRPL51,MRPL43,IMMP1L,MRPL48,COQ4,ERAL1,MTHFD2L,MRPL27,SLC25A35,COQ2,MRPS7,NDUFS7,
NDUFA3,COX19,RPS3,MRPS24,AIFM2,SPHK2,IFI27,TRAK1,NDUFB3,NDUFB2,NDUFA10,THOP1,UBA52,AK
1,NDUFC1,ATP11B,DUSP18,NDUFB10,MRPL51,MRPL43,IMMP1L,MRPL48,COQ4,ERAL1,MTHFD2L,MRPL
SG101,OBSL1,TTC21B,PTPN13,UBR4,PSMA1,CHCHD1,FUZ,MUC1,METTL17,EPB41L5,RPS25,NIP7,DUSP2
SG101,OBSL1,TTC21B,PTPN13,UBR4,PSMA1,CHCHD1,FUZ,MUC1,METTL17,EPB41L5,RPS25,NIP7,DUSP2
2,ASH2L,SMARCA4,SNRPF,GMPR2,SMARCC2,PHC3,PIGT,NHP2,UBE2N,PDCD6,ENY2,PEX2,POLR1C,PIGH,
TAU2,KLHL7,TSEN34,RBBP6,CHTOP,RPL35,CDK4,ZNF593,MARS,NEK11,ZNF354A,PSPC1,TAF1D,SIN3A,RB
5A,HIGD1A,NDUFV2,SIRT5,MTHFS,COA3,MPC2,TXNRD2,TYMS,SLC25A23,MRPS18C,LGALS3,PARK7,MFF,

,TRAK1,NDUFB3,NDUFB2,NDUFA10,UBA52,MRPS16,SDHA,NDUFA7,NDUFS2,MRPS33,MCUR1,NDUFC1,

T,SYNGR2,RAB2A,TPM3,RPS20,NIT2,VPS28,HSPB1,VWA2,SEMA3C,SERPINB1,PRKRIP1,LSR,DOCK10,PSM
T,SYNGR2,RAB2A,TPM3,RPS20,NIT2,VPS28,HSPB1,VWA2,SEMA3C,SERPINB1,PRKRIP1,LSR,DOCK10,PSM
T,SYNGR2,RAB2A,TPM3,RPS20,NIT2,VPS28,HSPB1,VWA2,SEMA3C,SERPINB1,PRKRIP1,LSR,DOCK10,PSM

-2,XBP1,C14ORF2,PCYT2,USE1,AGPAT4,PPP1CC,VAMP8,TIMM17B,TMEM126B,APEH,TXNL4A,NUTF2,PP[illegible]

F1A,MAGT1,JUP,CDC42,USE1,PPA1,VAMP8,CAPG,RPL28,VIM,PSMA7,APEH,LYPLA2,NUTF2,PODXL,EXOS[illegible]

[illegible],BFAR,KDELR3,MOSPD3,PML,TMEM199,MGST2,RHEB,CREB3L2,GRAMD1A,TUSC3,HMOX2,EMC9,DHRS[illegible]
[illegible],MOSPD3,PML,TMEM199,MGST2,RHEB,CREB3L2,GRAMD1A,TUSC3,HMOX2,EMC9,DHRS4,IFI27,GOSR2,[illegible]

[illegible],RPS17,ACTG1,NUMB,RAP1B,PRKCZ,PKN2,RPL27,CORO1B,RPS14,ARHGEF7,AHI1,DBN1,PAK1,RPL31,LIN[illegible]

,GBA2,TRPM4,MCFD2,SLC35B1,SCAP,CD74,PARK7,PIGT,PDCD6,TMEM147,TGOLN2,SGCD,PIGH,ERGIC1,[illegible]

CFD2,LGALS3,SCAP,CD74,PARK7,SLC30A5,PDCD6,MFF,SNX5,GPR56,TGOLN2,PEX2,STX10,PEX26,HAX1,T[illegible]

,RPL34,NUP62,RPL24,PSMD7,GTF2H1,GEMIN2,THOC3,HNRNPUL1,PSMD10,RPS3A,RPS3,MAGOHB,SNRP

CAIM,SLC3A2,DHPS,ACTR6,SEC62,SET,CTSC,BOP1,ARRDC1,TRPT1,KLC1,RBP5,ZC3HAV1,SCAF11,DBI,ARPC
TRPT1,KLC1,RBP5,ZC3HAV1,SCAF11,DBI,ARPC2,LIMK2,VDAC3,NOB1,WDR19,SIRT6,NUMA1,GPATCH2L,SU
31,NOP16,RPL13A,ARHGEF39,GSTK1,TTLL5,PSMA6,ACOT8,RPS9,SLC35G1,RCE1,MRPL33,ZNF410,RUVBL2
5,RNF32,LY75,ATG16L2,RCC1,SNRPB,PARD3,TCAIM,SLC3A2,DHPS,ACTR6,SEC62,SET,CTSC,ZNF235,BOP1,A
5,RNF32,LY75,ATG16L2,RCC1,SNRPB,PARD3,TCAIM,SLC3A2,DHPS,ACTR6,SEC62,SET,CTSC,ZNF235,BOP1,A
DHPS,ACTR6,SEC62,SET,CTSC,ZNF235,BOP1,ARRDC1,TRPT1,KLC1,RBP5,ZC3HAV1,SCAF11,DBI,ARPC2,LIM
DHPS,ACTR6,SEC62,SET,CTSC,ZNF235,BOP1,ARRDC1,TRPT1,KLC1,RBP5,ZC3HAV1,SCAF11,DBI,ARPC2,LIM
CTR6,SEC62,SET,CTSC,ZNF235,BOP1,ARRDC1,TRPT1,KLC1,ZC3HAV1,SCAF11,DBI,ARPC2,LIMK2,VDAC3,SIF
MK2,NOB1,SIRT6,NUMA1,GPATCH2L,SUMF2,BUD31,SAE1,NPRL2,OSGEP,ARHGAP6,SCARB1,CTTN,TSPAN
MK2,NOB1,SIRT6,NUMA1,GPATCH2L,SUMF2,BUD31,SAE1,NPRL2,OSGEP,ARHGAP6,SCARB1,CTTN,TSPAN
2,SET,CTSC,ZNF235,BOP1,ARRDC1,TRPT1,KLC1,ZC3HAV1,SCAF11,DBI,ARPC2,LIMK2,VDAC3,NOB1,WDR1
SH2B1,RPL13A,ARHGEF39,TTLL5,PSMA6,ACOT8,AP3B2,RPS9,SLC35G1,RCE1,MRPL33,ZNF410,RUVBL2,IG
SH2B1,RPL13A,ARHGEF39,TTLL5,PSMA6,ACOT8,AP3B2,RPS9,SLC35G1,RCE1,MRPL33,ZNF410,RUVBL2,IG
TUBA1C,IRF9,ANTXR2,STIP1,LRRC1,VCAN,TM9SF1,SON,UQCRQ,SNF8,GRK6,HMGA2,CDC37,RPS16,CPSF3I
1,SAE1,NPRL2,ARHGAP6,CTTN,TSPAN11,EIF5A,COMMD5,CHID1,NUDT22,AMZ2,UFD1L,USP38,TAZ,THYN
1,SAE1,NPRL2,ARHGAP6,CTTN,TSPAN11,EIF5A,COMMD5,CHID1,NUDT22,AMZ2,UFD1L,USP38,TAZ,THYN
IF235,BOP1,ARRDC1,TRPT1,KLC1,RBP5,ZC3HAV1,SCAF11,DBI,ARPC2,LIMK2,VDAC3,NOB1,SIRT6,NUMA1,

IF235,BOP1,ARRDC1,TRPT1,KLC1,RBP5,ZC3HAV1,SCAF11,DBI,ARPC2,LIMK2,VDAC3,NOB1,SIRT6,NUMA1,

OMMD5,CHID1,NUDT22,UFD1L,USP38,TAZ,THYN1,SH2B1,NOP16,RPL13A,ARHGEF39,GSTK1,TTLL5,LDHA

OMMD5,CHID1,NUDT22,UFD1L,USP38,TAZ,THYN1,SH2B1,NOP16,RPL13A,ARHGEF39,GSTK1,TTLL5,LDHA

UFD1L,USP38,THYN1,SH2B1,NOP16,RPL13A,ARHGEF39,GSTK1,TTLL5,PSMA6,ACOT8,AP3B2,RPS9,SLC35

UFD1L,USP38,THYN1,SH2B1,NOP16,RPL13A,ARHGEF39,GSTK1,TTLL5,PSMA6,ACOT8,AP3B2,RPS9,SLC35

TCH2L,SUMF2,BUD31,SAE1,NPRL2,OSGEP,CTTN,TSPAN11,EIF5A,COMMD5,CHID1,NUDT22,UFD1L,USP38

TCH2L,SUMF2,BUD31,SAE1,NPRL2,OSGEP,CTTN,TSPAN11,EIF5A,COMMD5,CHID1,NUDT22,UFD1L,USP38

CC1,SNRPB,PARD3,TCAIM,SLC3A2,DHPS,ACTR6,MAP3K7CL,SEC62,SET,CTSC,ZNF235,BOP1,ARRDC1,TRPT

CC1,SNRPB,PARD3,TCAIM,SLC3A2,DHPS,ACTR6,MAP3K7CL,SEC62,SET,CTSC,ZNF235,BOP1,ARRDC1,TRPT

2,ARHGAP6,CTTN,TSPAN11,EIF5A,COMMD5,CHID1,NUDT22,UFD1L,USP38,TAZ,THYN1,SH2B1,NOP16,RPL

2,ARHGAP6,CTTN,TSPAN11,EIF5A,COMMD5,CHID1,NUDT22,UFD1L,USP38,TAZ,THYN1,SH2B1,NOP16,RPL

BI,ARPC2,PMFBP1,LIMK2,NOB1,SIRT6,NUMA1,GPATCH2L,SUMF2,BUD31,SAE1,NPRL2,OSGEP,ARHGAP6,

BI,ARPC2,PMFBP1,LIMK2,NOB1,SIRT6,NUMA1,GPATCH2L,SUMF2,BUD31,SAE1,NPRL2,OSGEP,ARHGAP6,

,ZNF410,PCMT1,IGSF8,NUDT5,SNRPN,STYXL1,RFX5,POLD4,VRK3,LZTFL1,FIS1,PPM1K,IRF9,LRRC1,TM9SF

,ZNF410,PCMT1,IGSF8,NUDT5,SNRPN,STYXL1,RFX5,POLD4,VRK3,LZTFL1,FIS1,PPM1K,IRF9,LRRC1,TM9SF

RPC2,LIMK2,NOB1,SIRT6,NUMA1,GPATCH2L,SUMF2,BUD31,SAE1,NPRL2,OSGEP,ARHGAP6,SCARB1,CTT

RPC2,LIMK2,NOB1,SIRT6,NUMA1,GPATCH2L,SUMF2,BUD31,SAE1,NPRL2,OSGEP,ARHGAP6,SCARB1,CTT

DBI,ARPC2,PMFBP1,LIMK2,SIRT6,NUMA1,GPATCH2L,SUMF2,BUD31,SAE1,NPRL2,OSGEP,ARHGAP6,SCA

DBI,ARPC2,PMFBP1,LIMK2,SIRT6,NUMA1,GPATCH2L,SUMF2,BUD31,SAE1,NPRL2,OSGEP,ARHGAP6,SCA

C,ZNF235,BOP1,ARRDC1,TRPT1,KLC1,RBP5,ZC3HAV1,SCAF11,DBI,ARPC2,LIMK2,VDAC3,NOB1,WDR19,SI

,NOB1,SIRT6,NUMA1,GPATCH2L,SUMF2,BUD31,SAE1,NPRL2,OSGEP,ARHGAP6,SCARB1,CTTN,TSPAN11,E

,NOB1,SIRT6,NUMA1,GPATCH2L,SUMF2,BUD31,SAE1,NPRL2,OSGEP,ARHGAP6,SCARB1,CTTN,TSPAN11,E

L33,ZNF410,RUVBL2,IGSF8,NUDT5,SPAG7,WDR62,STYXL1,VRK3,PPM1K,IRF9,DST,ANTXR2,STIP1,LRRC1,S

C1,TRPT1,KLC1,ZC3HAV1,SCAF11,DBI,ARPC2,LIMK2,VDAC3,NOB1,WDR19,SIRT6,NUMA1,SUMF2,BUD31,

C1,TRPT1,KLC1,ZC3HAV1,SCAF11,DBI,ARPC2,LIMK2,VDAC3,NOB1,WDR19,SIRT6,NUMA1,SUMF2,BUD31,

1,UBR4,CHCHD1,FUZ,FAM3A,ATG12,RPS25,NIP7,TTC9C,SLC25A10,THEM4,DBNDD2,RPS6,RPL29,RTEL1,S

2,DHPS,ACTR6,SEC62,SET,CTSC,BOP1,ARRDC1,TRPT1,KLC1,ZC3HAV1,SCAF11,DBI,ARPC2,LIMK2,VDAC3,

ATG16L2,RCC1,SNRPB,PARD3,TCAIM,SLC3A2,DHPS,ACTR6,SEC62,SET,CTSC,ZNF235,BOP1,ARRDC1,TRPT

ATG16L2,RCC1,SNRPB,PARD3,TCAIM,SLC3A2,DHPS,ACTR6,SEC62,SET,CTSC,ZNF235,BOP1,ARRDC1,TRPT

MD5,CHID1,NUDT22,UFD1L,USP38,THYN1,SH2B1,NOP16,RPL13A,ARHGEF39,GSTK1,TTLL5,PSMA6,ACO

MD5,CHID1,NUDT22,UFD1L,USP38,THYN1,SH2B1,NOP16,RPL13A,ARHGEF39,GSTK1,TTLL5,PSMA6,ACO

RHGAP6,CTTN,TSPAN11,EIF5A,COMMD5,CHID1,NUDT22,UFD1L,USP38,TAZ,THYN1,DBNDD1,SH2B1,NOP

RHGAP6,CTTN,TSPAN11,EIF5A,COMMD5,CHID1,NUDT22,UFD1L,USP38,TAZ,THYN1,DBNDD1,SH2B1,NOP

BOP1,ARRDC1,TRPT1,KLC1,RBP5,ZC3HAV1,SCAF11,DBI,ARPC2,PMFBP1,LIMK2,SIRT6,NUMA1,GPATCH2L,

BOP1,ARRDC1,TRPT1,KLC1,RBP5,ZC3HAV1,SCAF11,DBI,ARPC2,PMFBP1,LIMK2,SIRT6,NUMA1,GPATCH2L,

1,EIF5A,COMMD5,CHID1,NUDT22,AMZ2,UFD1L,USP38,THYN1,DBNDD1,SH2B1,NOP16,RPL13A,ARHGEF

1,EIF5A,COMMD5,CHID1,NUDT22,AMZ2,UFD1L,USP38,THYN1,DBNDD1,SH2B1,NOP16,RPL13A,ARHGEF

LIMK2,NOB1,SIRT6,NUMA1,GPATCH2L,SUMF2,BUD31,SAE1,NPRL2,OSGEP,ARHGAP6,SCARB1,CTTN,TSP

LIMK2,NOB1,SIRT6,NUMA1,GPATCH2L,SUMF2,BUD31,SAE1,NPRL2,OSGEP,ARHGAP6,SCARB1,CTTN,TSP

39,TTLL5,PSMA6,ACOT8,AP3B2,RPS9,RCE1,MRPL33,ZNF410,PCMT1,IGSF8,NUDT5,SNRPN,WDR62,STYXL

39,TTLL5,PSMA6,ACOT8,AP3B2,RPS9,RCE1,MRPL33,ZNF410,PCMT1,IGSF8,NUDT5,SNRPN,WDR62,STYXL

T22,AMZ2,UFD1L,USP38,THYN1,SH2B1,NOP16,RPL13A,ARHGEF39,GSTK1,TTLL5,PSMA6,ACOT8,AP3B2,R

T22,AMZ2,UFD1L,USP38,THYN1,SH2B1,NOP16,RPL13A,ARHGEF39,GSTK1,TTLL5,PSMA6,ACOT8,AP3B2,R

,CTSC,ZNF235,BOP1,ARRDC1,TRPT1,KLC1,ZC3HAV1,SCAF11,DBI,ARPC2,PMFBP1,LIMK2,VDAC3,NOB1,SI

,LRRC1,VCAN,TM9SF1,SON,UQCRQ,SNF8,CDC37,RPS16,CPSF3L,TSG101,UBR4,ATG12,RPS25,NIP7,TTC9C

NF8,CDC37,RPS16,CPSF3L,TSG101,UBR4,FAM3A,METTL17,ATG12,NIP7,DUSP22,THEM4,DBNDD2,RPS6,

2,UFD1L,USP38,THYN1,SH2B1,NOP16,RPL13A,ARHGEF39,TTLL5,PSMA6,ACOT8,AP3B2,RPS9,SLC35G1,RC

2,UFD1L,USP38,THYN1,SH2B1,NOP16,RPL13A,ARHGEF39,TTLL5,PSMA6,ACOT8,AP3B2,RPS9,SLC35G1,RC
CHID1,NUDT22,AMZ2,UFD1L,USP38,TAZ,THYN1,DBNDD1,SH2B1,NOP16,RPL13A,ARHGEF39,GSTK1,TTLL5
CHID1,NUDT22,AMZ2,UFD1L,USP38,TAZ,THYN1,DBNDD1,SH2B1,NOP16,RPL13A,ARHGEF39,GSTK1,TTLL5
MZ2,UFD1L,USP38,THYN1,SH2B1,NOP16,RPL13A,ARHGEF39,GSTK1,TTLL5,PSMA6,ACOT8,AP3B2,RPS9,S
MZ2,UFD1L,USP38,THYN1,SH2B1,NOP16,RPL13A,ARHGEF39,GSTK1,TTLL5,PSMA6,ACOT8,AP3B2,RPS9,S
LL5,PSMA6,ACOT8,AP3B2,RPS9,RCE1,MRPL33,ZNF410,PCMT1,TESC,NUDT5,SNRPN,SPAG7,STYXL1,POLD
LL5,PSMA6,ACOT8,AP3B2,RPS9,RCE1,MRPL33,ZNF410,PCMT1,TESC,NUDT5,SNRPN,SPAG7,STYXL1,POLD
3A,ARHGEF39,GSTK1,TTLL5,PSMA6,ACOT8,AP3B2,RPS9,RCE1,MRPL33,ZNF410,IGSF8,NUDT5,SNRPN,WD
3A,ARHGEF39,GSTK1,TTLL5,PSMA6,ACOT8,AP3B2,RPS9,RCE1,MRPL33,ZNF410,IGSF8,NUDT5,SNRPN,WD
,NUDT22,AMZ2,UFD1L,USP38,THYN1,SH2B1,NOP16,RPL13A,ARHGEF39,GSTK1,TTLL5,PSMA6,ACOT8,AP3
,NUDT22,AMZ2,UFD1L,USP38,THYN1,SH2B1,NOP16,RPL13A,ARHGEF39,GSTK1,TTLL5,PSMA6,ACOT8,AP3
MK2,SIRT6,NUMA1,GPATCH2L,SUMF2,BUD31,SAE1,NPRL2,TBL2,ARHGAP6,SCARB1,CTTN,TSPAN11,EIF5A
MK2,SIRT6,NUMA1,GPATCH2L,SUMF2,BUD31,SAE1,NPRL2,TBL2,ARHGAP6,SCARB1,CTTN,TSPAN11,EIF5A
NUDT22,UFD1L,USP38,THYN1,SH2B1,NOP16,RPL13A,ARHGEF39,GSTK1,TTLL5,PSMA6,ACOT8,AP3B2,RPS
NUDT22,UFD1L,USP38,THYN1,SH2B1,NOP16,RPL13A,ARHGEF39,GSTK1,TTLL5,PSMA6,ACOT8,AP3B2,RPS
D1,TRNT1,KXD1,ITGB1BP1,MED15,CLASRP,C1ORF198,MICAL1,UBE2I,SLC39A13,EIF3M,CDC42,CARD14,F
AP6,SCARB1,CTTN,TSPAN11,EIF5A,COMMD5,CHID1,NUDT22,AMZ2,UFD1L,USP38,THYN1,DBNDD1,SH2B
AP6,SCARB1,CTTN,TSPAN11,EIF5A,COMMD5,CHID1,NUDT22,AMZ2,UFD1L,USP38,THYN1,DBNDD1,SH2B
,RPL13A,ARHGEF39,GSTK1,TTLL5,PSMA6,ACOT8,AP3B2,RPS9,SLC35G1,RCE1,MRPL33,ZNF410,PCMT1,NU
,RPL13A,ARHGEF39,GSTK1,TTLL5,PSMA6,ACOT8,AP3B2,RPS9,SLC35G1,RCE1,MRPL33,ZNF410,PCMT1,NU
1D5,CHID1,NUDT22,AMZ2,UFD1L,USP38,THYN1,DBNDD1,SH2B1,RPL13A,ARHGEF39,TTLL5,PSMA6,ACOT
1D5,CHID1,NUDT22,AMZ2,UFD1L,USP38,THYN1,DBNDD1,SH2B1,RPL13A,ARHGEF39,TTLL5,PSMA6,ACOT
D5,CHID1,NUDT22,UFD1L,USP38,TAZ,THYN1,SH2B1,NOP16,RPL13A,ARHGEF39,GSTK1,TTLL5,PSMA6,AC
D5,CHID1,NUDT22,UFD1L,USP38,TAZ,THYN1,SH2B1,NOP16,RPL13A,ARHGEF39,GSTK1,TTLL5,PSMA6,AC
CDC42,CARD14,FBXW9,USE1,ASF1B,RFX3,ENDOV,CAMTA1,PPIL2,RPL28,TIMM17B,APEH,LYPLA2,C17ORF
22,AMZ2,UFD1L,THYN1,SH2B1,NOP16,RPL13A,ARHGEF39,TTLL5,PSMA6,ACOT8,AP3B2,RPS9,SLC35G1,RC
22,AMZ2,UFD1L,THYN1,SH2B1,NOP16,RPL13A,ARHGEF39,TTLL5,PSMA6,ACOT8,AP3B2,RPS9,SLC35G1,RC
CH2L,BUD31,NPRL2,TBL2,SCARB1,CTTN,TSPAN11,ITSN1,EIF5A,COMMD5,NUDT22,AMZ2,UFD1L,TAZ,THY
CH2L,BUD31,NPRL2,TBL2,SCARB1,CTTN,TSPAN11,ITSN1,EIF5A,COMMD5,NUDT22,AMZ2,UFD1L,TAZ,THY
1,KXD1,ITGB1BP1,MED15,CLASRP,C1ORF198,UBE2I,EIF3M,CDC42,CARD14,PCYT2,FBXW9,USE1,IFI27L1,
T2,FBXW9,USE1,ASF1B,CCDC25,RFX3,ENDOV,CAMTA1,PPIL2,APEH,CAPN10,LYPLA2,C17ORF62,PRMT1,ZI
MK2,SIRT6,NUMA1,GPATCH2L,SUMF2,BUD31,SAE1,NPRL2,TBL2,ARHGAP6,SCARB1,CTTN,TSPAN11,EIF5A
MK2,SIRT6,NUMA1,GPATCH2L,SUMF2,BUD31,SAE1,NPRL2,TBL2,ARHGAP6,SCARB1,CTTN,TSPAN11,EIF5A
RPC2,LIMK2,VDAC3,NOB1,WDR19,SIRT6,NUMA1,SUMF2,BUD31,SAE1,NPRL2,OSGEP,TBL2,ARHGAP6,SCA
RPC2,LIMK2,VDAC3,NOB1,WDR19,SIRT6,NUMA1,SUMF2,BUD31,SAE1,NPRL2,OSGEP,TBL2,ARHGAP6,SCA
,SNRPB,PARD3,TCAIM,SLC3A2,DHPS,ACTR6,SEC62,SET,CTSC,ZNF235,BOP1,ARRDC1,TRPT1,KLC1,RBP5,ZC
,SNRPB,PARD3,TCAIM,SLC3A2,DHPS,ACTR6,SEC62,SET,CTSC,ZNF235,BOP1,ARRDC1,TRPT1,KLC1,RBP5,ZC
UP1,SAP30BP,YIPF3,TRNT1,KXD1,ITGB1BP1,MED15,CLASRP,C1ORF198,UBE2I,SLC39A13,EIF3M,CDC42,CA
PC2,PMFBP1,LIMK2,SIRT6,NUMA1,GPATCH2L,SUMF2,BUD31,SAE1,NPRL2,OSGEP,ARHGAP6,SCARB1,CTT
PC2,PMFBP1,LIMK2,SIRT6,NUMA1,GPATCH2L,SUMF2,BUD31,SAE1,NPRL2,OSGEP,ARHGAP6,SCARB1,CTT
F235,BOP1,ARRDC1,TRPT1,KLC1,RBP5,ZC3HAV1,SCAF11,DBI,ARPC2,LIMK2,VDAC3,NOB1,SIRT6,NUMA1,
F235,BOP1,ARRDC1,TRPT1,KLC1,RBP5,ZC3HAV1,SCAF11,DBI,ARPC2,LIMK2,VDAC3,NOB1,SIRT6,NUMA1,
METTL17,ATG12,RPS25,NIP7,DUSP22,THEM4,DBNDD2,RPS6,RPL29,RTEL1,AUP1,SAP30BP,YIPF3,FOXRED
MA6,ACOT8,AP3B2,RPS9,SLC35G1,RCE1,MRPL33,ZNF410,IGSF8,NUDT5,SNRPN,STYXL1,RFX5,POLD4,LZT
MA6,ACOT8,AP3B2,RPS9,SLC35G1,RCE1,MRPL33,ZNF410,IGSF8,NUDT5,SNRPN,STYXL1,RFX5,POLD4,LZT
L2,LIG1,APEH,CAPN10,LYPLA2,C17ORF62,PRMT1,ZNF7,ARL6IP4,PPAN,ACAD10,SUGT1,HYPK,CARS,CDK11
15,CLASRP,C1ORF198,MICAL1,UBE2I,SLC39A13,EIF3M,CDC42,C14ORF2,CARD14,PCYT2,FBXW9,USE1,ASF

3,LRRC1,SNF8,CDC37,RPS16,CPSF3L,TSG101,UBR4,CHCHD1,FUZ,FAM3A,ATG12,RPS25,NIP7,TTC9C,SLC25

IGAP6,SCARB1,CTTN,TSPAN11,EIF5A,COMMD5,CHID1,NUDT22,AMZ2,UFD1L,USP38,THYN1,SH2B1,NOP1

IGAP6,SCARB1,CTTN,TSPAN11,EIF5A,COMMD5,CHID1,NUDT22,AMZ2,UFD1L,USP38,THYN1,SH2B1,NOP1

2,ARHGAP6,SCARB1,CTTN,TSPAN11,EIF5A,COMMD5,CHID1,NUDT22,UFD1L,USP38,THYN1,SH2B1,NOP16

2,ARHGAP6,SCARB1,CTTN,TSPAN11,EIF5A,COMMD5,CHID1,NUDT22,UFD1L,USP38,THYN1,SH2B1,NOP16

1,LRRC1,LPCAT3,CD44,SON,UQCRQ,SNF8,GRK6,HMGA2,CDC37,CPSF3L,TSG101,UBR4,FAM3A,ATG12,RP

NUMA1,GPATCH2L,SUMF2,BUD31,SAE1,NPRL2,OSGEP,TBL2,ARHGAP6,SCARB1,CTTN,TSPAN11,ITSN1,EIF

NUMA1,GPATCH2L,SUMF2,BUD31,SAE1,NPRL2,OSGEP,TBL2,ARHGAP6,SCARB1,CTTN,TSPAN11,ITSN1,EIF

IED15,CLASRP,C1ORF198,UBE2I,SLC39A13,EIF3M,CDC42,CARD14,PCYT2,FBXW9,USE1,ASF1B,CCDC25,RF

TSC,BOP1,ARRDC1,TRPT1,KLC1,RBP5,ZC3HAV1,SCAF11,ARPC2,PMFBP1,LIMK2,VDAC3,NOB1,WDR19,SIR

TSC,BOP1,ARRDC1,TRPT1,KLC1,RBP5,ZC3HAV1,SCAF11,ARPC2,PMFBP1,LIMK2,VDAC3,NOB1,WDR19,SIR

NDOV,CAMTA1,PPIL2,APEH,CAPN10,LYPLA2,C17ORF62,PRMT1,STOML2,C19ORF54,ZNF7,TXNDC17,ARL6

,GPATCH2L,BUD31,NPRL2,TBL2,SCARB1,CTTN,TSPAN11,ITSN1,EIF5A,COMMD5,NUDT22,AMZ2,UFD1L,US

,GPATCH2L,BUD31,NPRL2,TBL2,SCARB1,CTTN,TSPAN11,ITSN1,EIF5A,COMMD5,NUDT22,AMZ2,UFD1L,US

GGT2,UBE2I,EIF3M,CDC42,TBX6,CARD14,PCYT2,FBXW9,CCDC25,EEF1B2,RFX3,ENDOV,PPIL2,LIG1,CDC14

GGT2,UBE2I,EIF3M,CDC42,TBX6,CARD14,PCYT2,FBXW9,CCDC25,EEF1B2,RFX3,ENDOV,PPIL2,LIG1,CDC14

R2,STIP1,LRRC1,VCAN,MAGED2,SON,UQCRQ,SNF8,GRK6,CDC37,CPSF3L,TSG101,UBR4,PHF21A,ATG12,R

UD31,SAE1,NPRL2,TBL2,ARHGAP6,SCARB1,CTTN,ITSN1,EIF5A,COMMD5,CHID1,NUDT22,UFD1L,USP38,T

LASRP,C1ORF198,UBE2I,SLC39A13,EIF3M,CDC42,CARD14,PCYT2,FBXW9,USE1,IFI27L1,ASF1B,CCDC25,RF

,TRMT112,RPS20,SAT1,ERCC5,EMC4,SERPINB1,DDOST,LSR,TOMM40,RABEP2,TADA3,ZNF20,WAC,RAD23

K,IRF9,ANTXR2,STIP1,LRRC1,LPCAT3,CD44,MAGED2,SON,UQCRQ,SNF8,GRK6,HMGA2,CDC37,RPS16,CPS

3,LZTFL1,FIS1,PPM1K,MAGED2,SON,UQCRQ,SNF8,HMGA2,CDC37,CPSF3L,TSG101,FUZ,FAM3A,ATG12,RP

3,LZTFL1,FIS1,PPM1K,MAGED2,SON,UQCRQ,SNF8,HMGA2,CDC37,CPSF3L,TSG101,FUZ,FAM3A,ATG12,RP

RD14,PCYT2,FBXW9,CCDC25,EEF1B2,RFX3,ENDOV,PPIL2,LIG1,CDC14B,APEH,CAPN10,LYPLA2,C17ORF62,

RD14,PCYT2,FBXW9,CCDC25,EEF1B2,RFX3,ENDOV,PPIL2,LIG1,CDC14B,APEH,CAPN10,LYPLA2,C17ORF62,

5,ACOT8,RPS9,RCE1,MRPL33,ZNF410,RUVBL2,PCMT1,TESC,IGSF8,NUDT5,SNRPN,SPAG7,WDR62,STYXL1,

5,ACOT8,RPS9,RCE1,MRPL33,ZNF410,RUVBL2,PCMT1,TESC,IGSF8,NUDT5,SNRPN,SPAG7,WDR62,STYXL1,

R6,SET,BOP1,ARRDC1,KLC1,SCAF11,DBI,PMFBP1,LIMK2,VDAC3,WDR19,SIRT6,NUMA1,SUMF2,BUD31,SA

R6,SET,BOP1,ARRDC1,KLC1,SCAF11,DBI,PMFBP1,LIMK2,VDAC3,WDR19,SIRT6,NUMA1,SUMF2,BUD31,SA

GB1BP1,MED15,CLASRP,C1ORF198,MICAL1,UBE2I,SLC39A13,EIF3M,CDC42,CARD14,PCYT2,FBXW9,USE1

SF3L,TSG101,UBR4,PHF21A,METTL17,ATG12,RPS25,NIP7,COX4I1,DUSP22,THEM4,RPS6,RPL29,COASY,AU

7CL,SEC62,SET,CTSC,BOP1,ARRDC1,KLC1,RBP5,SCAF11,DBI,ARPC2,LIMK2,VDAC3,NOB1,WDR19,SIRT6,NU

7CL,SEC62,SET,CTSC,BOP1,ARRDC1,KLC1,RBP5,SCAF11,DBI,ARPC2,LIMK2,VDAC3,NOB1,WDR19,SIRT6,NU

RP,C1ORF198,MICAL1,UBE2I,SLC39A13,EIF3M,CDC42,CARD14,PCYT2,FBXW9,USE1,ASF1B,RFX3,ENDOV,

SAP30BP,YIPF3,FOXRED1,RABL5,TRNT1,ITGB1BP1,MED15,CLASRP,C1ORF198,UGGT2,UBE2I,EIF3M,CDC4

SAP30BP,YIPF3,FOXRED1,RABL5,TRNT1,ITGB1BP1,MED15,CLASRP,C1ORF198,UGGT2,UBE2I,EIF3M,CDC4

GEF39,TTLL5,LDHA,ACOT8,AP3B2,RPS9,SLC35G1,RCE1,ZNF410,PCMT1,TESC,IGSF8,NUDT5,RBPMS,STYXL1

GEF39,TTLL5,LDHA,ACOT8,AP3B2,RPS9,SLC35G1,RCE1,ZNF410,PCMT1,TESC,IGSF8,NUDT5,RBPMS,STYXL1

,RTEL1,AUP1,SAP30BP,YIPF3,RABL5,TRNT1,ITGB1BP1,MED15,CLASRP,C1ORF198,UGGT2,UBE2I,EIF3M,C

,RTEL1,AUP1,SAP30BP,YIPF3,RABL5,TRNT1,ITGB1BP1,MED15,CLASRP,C1ORF198,UGGT2,UBE2I,EIF3M,C

F9,PCBP2,MAGED2,SON,UQCRQ,SNF8,CDC37,CPSF3L,TSG101,SLC39A11,FUZ,FAM3A,METTL17,ATG12,R

F9,PCBP2,MAGED2,SON,UQCRQ,SNF8,CDC37,CPSF3L,TSG101,SLC39A11,FUZ,FAM3A,METTL17,ATG12,R

MS,SNRPN,STYXL1,RFX5,POLD4,LZTFL1,FIS1,PPM1K,TUBA1C,IRF9,LRRC1,TM9SF1,MAGED2,SON,UQCRQ,G

MS,SNRPN,STYXL1,RFX5,POLD4,LZTFL1,FIS1,PPM1K,TUBA1C,IRF9,LRRC1,TM9SF1,MAGED2,SON,UQCRQ,G

ET,CTSC,ZNF235,BOP1,ARRDC1,KLC1,ZC3HAV1,SCAF11,DBI,ARPC2,LIMK2,VDAC3,WDR19,SIRT6,NUMA1,S

ET,CTSC,ZNF235,BOP1,ARRDC1,KLC1,ZC3HAV1,SCAF11,DBI,ARPC2,LIMK2,VDAC3,WDR19,SIRT6,NUMA1,S

YIPF3,FOXRED1,TRNT1,KXD1,MED15,IP6K2,CLASRP,C1ORF198,UBE2I,EIF3M,CDC42,CARD14,PCYT2,FBXV

CPSF3L,TSG101,METTL17,ATG12,RPS25,NIP7,COX4I1,DUSP22,THEM4,RPS6,RPL29,COASY,AUP1,SAP30BI
2,LRRC1,TM9SF1,MAGED2,SON,UQCRQ,SNF8,RPS16,CPSF3L,TSG101,UBR4,FUZ,FAM3A,ATG12,RPS25,NI
2,LRRC1,TM9SF1,MAGED2,SON,UQCRQ,SNF8,RPS16,CPSF3L,TSG101,UBR4,FUZ,FAM3A,ATG12,RPS25,NI
QCRQ,SNF8,CDC37,CPSF3L,TSG101,UBR4,PHF21A,RPS25,NIP7,COX4I1,THEM4,RPL29,COASY,AUP1,SAP3(
F8,CDC37,RPS16,CPSF3L,TSG101,YLPM1,FUZ,FAM3A,METTL17,EPB41L5,ATG12,RPS25,NIP7,THEM4,CEN
F8,CDC37,RPS16,CPSF3L,TSG101,YLPM1,FUZ,FAM3A,METTL17,EPB41L5,ATG12,RPS25,NIP7,THEM4,CEN
14,PCYT2,FBXW9,USE1,ASF1B,RFX3,ENDOV,CAMTA1,PPIL2,APEH,CAPN10,LYPLA2,C17ORF62,PRMT1,ZNI
C1,RBP5,ZC3HAV1,SCAF11,ARPC2,PMFBP1,LIMK2,VDAC3,NOB1,WDR19,SIRT6,NUMA1,GPATCH2L,SUMI
IRF9,DST,VCAN,SON,UQCRQ,RPS16,CPSF3L,TSG101,UBR4,CHCHD1,FAM3A,PSMA3,MUC1,ATG12,RPS25,
S,ACTR6,SEC62,SET,CTSC,ZNF235,BOP1,ARRDC1,TRPT1,KLC1,ZC3HAV1,SCAF11,DBI,ARPC2,PMFBP1,VD/
C1ORF198,UGGT2,UBE2I,EIF3M,CDC42,CARD14,PCYT2,FBXW9,USE1,CCDC25,EEF1B2,RFX3,ENDOV,PPIL2
C1ORF198,UGGT2,UBE2I,EIF3M,CDC42,CARD14,PCYT2,FBXW9,USE1,CCDC25,EEF1B2,RFX3,ENDOV,PPIL2
UMA1,GPATCH2L,SUMF2,BUD31,SAE1,NPRL2,ARHGAP6,SCARB1,CTTN,TSPAN11,EIF5A,COMMD5,RUNX
UMA1,GPATCH2L,SUMF2,BUD31,SAE1,NPRL2,ARHGAP6,SCARB1,CTTN,TSPAN11,EIF5A,COMMD5,RUNX
L33,ZNF410,RUVBL2,PCMT1,NUDT5,SNRPN,STYXL1,POLD4,RANBP1,VRK3,LZTFL1,FIS1,PPM1K,TUBA1C,II
L33,ZNF410,RUVBL2,PCMT1,NUDT5,SNRPN,STYXL1,POLD4,RANBP1,VRK3,LZTFL1,FIS1,PPM1K,TUBA1C,II
L33,ZNF410,RUVBL2,PCMT1,NUDT5,SNRPN,STYXL1,POLD4,RANBP1,VRK3,LZTFL1,FIS1,PPM1K,TUBA1C,II
L33,ZNF410,RUVBL2,PCMT1,NUDT5,SNRPN,STYXL1,POLD4,RANBP1,VRK3,LZTFL1,FIS1,PPM1K,TUBA1C,II
,CLASRP,C1ORF198,UGGT2,UBE2I,EIF3M,CDC42,TBX6,CARD14,PCYT2,FBXW9,ASF1B,CCDC25,EEF1B2,RF
,CLASRP,C1ORF198,UGGT2,UBE2I,EIF3M,CDC42,TBX6,CARD14,PCYT2,FBXW9,ASF1B,CCDC25,EEF1B2,RF
LRRC1,TM9SF1,MAGED2,SON,UQCRQ,GRK6,RPS16,CPSF3L,TSG101,SLC39A11,FUZ,FAM3A,RPS25,NIP7,C
LRRC1,TM9SF1,MAGED2,SON,UQCRQ,GRK6,RPS16,CPSF3L,TSG101,SLC39A11,FUZ,FAM3A,RPS25,NIP7,C
3XW9,USE1,ASF1B,EEF1B2,PPP1CC,ENDOV,PPIL2,PSMD4,RPL28,CAPN10,LYPLA2,C17ORF62,PRMT1,STOI
RTEL1,SAP30BP,YIPF3,FOXRED1,RABL5,TRNT1,ITGB1BP1,MED15,CLASRP,C1ORF198,UGGT2,SLC39A13,E
RTEL1,SAP30BP,YIPF3,FOXRED1,RABL5,TRNT1,ITGB1BP1,MED15,CLASRP,C1ORF198,UGGT2,SLC39A13,E
P6,SCARB1,CTTN,TSPAN11,ITSN1,EIF5A,COMMD5,CHID1,NUDT22,AMZ2,UFD1L,USP38,THYN1,SH2B1,NC
P6,SCARB1,CTTN,TSPAN11,ITSN1,EIF5A,COMMD5,CHID1,NUDT22,AMZ2,UFD1L,USP38,THYN1,SH2B1,NC
3,EIF3M,CDC42,CARD14,FBXW9,CCDC25,EEF1B2,RFX3,ENDOV,PPIL2,RPL28,LIG1,CDC14B,TIMM17B,APE
3,EIF3M,CDC42,CARD14,FBXW9,CCDC25,EEF1B2,RFX3,ENDOV,PPIL2,RPL28,LIG1,CDC14B,TIMM17B,APE
T22,UFD1L,USP38,THYN1,SH2B1,NOP16,RPL13A,ARHGEF39,GSTK1,TTLL5,PSMA6,ACOT8,AP3B2,RPS9,SI
T22,UFD1L,USP38,THYN1,SH2B1,NOP16,RPL13A,ARHGEF39,GSTK1,TTLL5,PSMA6,ACOT8,AP3B2,RPS9,SI
T6,NUMA1,GPATCH2L,SUMF2,BUD31,SAE1,NPRL2,OSGEP,ARHGAP6,SCARB1,CTTN,TSPAN11,ITSN1,EIF5/
T6,NUMA1,GPATCH2L,SUMF2,BUD31,SAE1,NPRL2,OSGEP,ARHGAP6,SCARB1,CTTN,TSPAN11,ITSN1,EIF5/
,RTEL1,COASY,AUP1,SAP30BP,YIPF3,FOXRED1,RABL5,TRNT1,KXD1,ITGB1BP1,MED15,CLASRP,C1ORF198,
,RTEL1,COASY,AUP1,SAP30BP,YIPF3,FOXRED1,RABL5,TRNT1,KXD1,ITGB1BP1,MED15,CLASRP,C1ORF198,
MD5,CHID1,NUDT22,AMZ2,UFD1L,USP38,TAZ,THYN1,DBNDD1,NOP16,RPL13A,ARHGEF39,GSTK1,TTLL5,I
MD5,CHID1,NUDT22,AMZ2,UFD1L,USP38,TAZ,THYN1,DBNDD1,NOP16,RPL13A,ARHGEF39,GSTK1,TTLL5,I
ACTR6,SEC62,SET,CTSC,ZNF235,BOP1,ARRDC1,TRPT1,KLC1,ZC3HAV1,SCAF11,DBI,ARPC2,LIMK2,VDAC3,N
NOB1,SIRT6,GPATCH2L,SUMF2,BUD31,NPRL2,OSGEP,ARHGAP6,SCARB1,CTTN,EIF5A,CHID1,NUDT22,AN
NOB1,SIRT6,GPATCH2L,SUMF2,BUD31,NPRL2,OSGEP,ARHGAP6,SCARB1,CTTN,EIF5A,CHID1,NUDT22,AN
D31,NPRL2,TBL2,SCARB1,CTTN,TSPAN11,ITSN1,EIF5A,COMMD5,NUDT22,AMZ2,UFD1L,TAZ,THYN1,DBN
D31,NPRL2,TBL2,SCARB1,CTTN,TSPAN11,ITSN1,EIF5A,COMMD5,NUDT22,AMZ2,UFD1L,TAZ,THYN1,DBN
42,FBXW9,USE1,ASF1B,RFX3,ENDOV,PPIL2,APEH,LYPLA2,C17ORF62,PRMT1,STOML2,C19ORF54,TXNDC1
,BUD31,NPRL2,TBL2,SCARB1,CTTN,TSPAN11,ITSN1,EIF5A,COMMD5,NUDT22,AMZ2,UFD1L,TAZ,THYN1,D
,BUD31,NPRL2,TBL2,SCARB1,CTTN,TSPAN11,ITSN1,EIF5A,COMMD5,NUDT22,AMZ2,UFD1L,TAZ,THYN1,D
ASRP,C1ORF198,UGGT2,MICAL1,UBE2I,PRMT2,CDC42,TBX6,FBXW9,ASF1B,CCDC25,EEF1B2,RFX3,ENDO
ASRP,C1ORF198,UGGT2,MICAL1,UBE2I,PRMT2,CDC42,TBX6,FBXW9,ASF1B,CCDC25,EEF1B2,RFX3,ENDO

RF198,UGGT2,MICAL1,UBE2I,EIF3M,CDC42,TBX6,CARD14,FBXW9,ASF1B,CCDC25,EEF1B2,RFX3,ENDOV,F

RF198,UGGT2,MICAL1,UBE2I,EIF3M,CDC42,TBX6,CARD14,FBXW9,ASF1B,CCDC25,EEF1B2,RFX3,ENDOV,F

ZC3HAV1,SCAF11,DBI,ARPC2,LIMK2,VDAC3,WDR19,SIRT6,NUMA1,SUMF2,BUD31,SAE1,NPRL2,OSGEP,TI

ZC3HAV1,SCAF11,DBI,ARPC2,LIMK2,VDAC3,WDR19,SIRT6,NUMA1,SUMF2,BUD31,SAE1,NPRL2,OSGEP,TI

PN10,LYPLA2,C17ORF62,PRMT1,ZNF7,ARL6IP4,PPAN,SUGT1,HYPK,CARS,RBM3,ZNF197,LRRFIP1,ATP5J,R

DT5,SNRPN,STYXL1,RANBP1,VRK3,LZTFL1,FIS1,IRF9,ANTXR2,STIP1,LRRC1,CD44,TM9SF1,MAGED2,SON,U

DT5,SNRPN,STYXL1,RANBP1,VRK3,LZTFL1,FIS1,IRF9,ANTXR2,STIP1,LRRC1,CD44,TM9SF1,MAGED2,SON,U

LC3A2,DHPS,ACTR6,SEC62,SET,CTSC,ZNF235,BOP1,ARRDC1,TRPT1,KLC1,RBP5,ZC3HAV1,SCAF11,DBI,ARP

LC3A2,DHPS,ACTR6,SEC62,SET,CTSC,ZNF235,BOP1,ARRDC1,TRPT1,KLC1,RBP5,ZC3HAV1,SCAF11,DBI,ARP

RPL33,RUVBL2,NUDT5,SNRPN,STYXL1,POLD4,VRK3,LZTFL1,FIS1,TUBA1C,IRF9,DST,LRRC1,TM9SF1,MAGEI

RPL33,RUVBL2,NUDT5,SNRPN,STYXL1,POLD4,VRK3,LZTFL1,FIS1,TUBA1C,IRF9,DST,LRRC1,TM9SF1,MAGEI

CC1,SNRPB,PARD3,TCAIM,SLC3A2,DHPS,ACTR6,MAP3K7CL,SEC62,SET,CTSC,BOP1,ARRDC1,TRPT1,KLC1,I

CC1,SNRPB,PARD3,TCAIM,SLC3A2,DHPS,ACTR6,MAP3K7CL,SEC62,SET,CTSC,BOP1,ARRDC1,TRPT1,KLC1,I

2,CARD14,PCYT2,FBXW9,USE1,ASF1B,EEF1B2,RFX3,ENDOV,CAMTA1,PPIL2,RPL28,APEH,CAPN10,LYPLA2,

UD31,OSGEP,TBL2,SCARB1,CRYGS,CTTN,TSPAN11,ITSN1,NUDT22,AMZ2,UFD1L,TAZ,DBNDD1,SH2B1,NOI

UD31,OSGEP,TBL2,SCARB1,CRYGS,CTTN,TSPAN11,ITSN1,NUDT22,AMZ2,UFD1L,TAZ,DBNDD1,SH2B1,NOI

CLASRP,C1ORF198,UBE2I,EIF3M,CDC42,CARD14,PCYT2,FBXW9,USE1,IFI27L1,ASF1B,CCDC25,EEF1B2,RFX

YXL1,VRK3,LZTFL1,FIS1,TUBA1C,IRF9,DST,ANTXR2,STIP1,VCAN,LPCAT3,TM9SF1,MAGED2,SON,UQCRQ,RF

YXL1,VRK3,LZTFL1,FIS1,TUBA1C,IRF9,DST,ANTXR2,STIP1,VCAN,LPCAT3,TM9SF1,MAGED2,SON,UQCRQ,RF

5,ACOT8,AP3B2,SLC35G1,MRPL33,ZNF410,PCMT1,EBPL,IGSF8,NUDT5,SNRPN,STYXL1,RFX5,POLD4,VRK3,

5,ACOT8,AP3B2,SLC35G1,MRPL33,ZNF410,PCMT1,EBPL,IGSF8,NUDT5,SNRPN,STYXL1,RFX5,POLD4,VRK3,

CD44,MAGED2,SON,UQCRQ,SNF8,GRK6,HMGA2,CDC37,CPSF3L,TSG101,MUC1,METTL17,ATG12,RPS25,N

31,RPL13A,ARHGEF39,LDHA,PSMA6,ACOT8,RPS9,SLC35G1,RCE1,ZNF410,RUVBL2,PCMT1,TESC,IGSF8,NUI

31,RPL13A,ARHGEF39,LDHA,PSMA6,ACOT8,RPS9,SLC35G1,RCE1,ZNF410,RUVBL2,PCMT1,TESC,IGSF8,NUI

W9,CCDC25,EEF1B2,RFX3,ENDOV,PPIL2,RPL28,LIG1,CDC14B,TIMM17B,APEH,LYPLA2,C17ORF62,STOML

W9,CCDC25,EEF1B2,RFX3,ENDOV,PPIL2,RPL28,LIG1,CDC14B,TIMM17B,APEH,LYPLA2,C17ORF62,STOML

2,RPL28,LIG1,APEH,LYPLA2,NUTF2,ATPAF2,C17ORF62,AP1S3,HYPK,MOB1A,CDK11A,INO80E,CCNL2,DGU

L28,LIG1,CDC14B,TIMM17B,APEH,LYPLA2,C17ORF62,STOML2,ZNF7,PPAN,CYB561,SUGT1,HYPK,CARS,ZN

L28,LIG1,CDC14B,TIMM17B,APEH,LYPLA2,C17ORF62,STOML2,ZNF7,PPAN,CYB561,SUGT1,HYPK,CARS,ZN

SE1,ASF1B,CCDC25,EEF1B2,ENDOV,RPL28,CLDND1,LIG1,TIMM17B,LYPLA2,STOML2,ZNF7,PPAN,AP1S3,C

SE1,ASF1B,CCDC25,EEF1B2,ENDOV,RPL28,CLDND1,LIG1,TIMM17B,LYPLA2,STOML2,ZNF7,PPAN,AP1S3,C

BXW9,ASF1B,CCDC25,EEF1B2,RFX3,ENDOV,PPIL2,RPL28,LIG1,CDC14B,TIMM17B,APEH,LYPLA2,C17ORF6

BXW9,ASF1B,CCDC25,EEF1B2,RFX3,ENDOV,PPIL2,RPL28,LIG1,CDC14B,TIMM17B,APEH,LYPLA2,C17ORF6

M3,ZNF197,INO80E,CCNL2,LRRFIP1,ATP5J,TRMT112,CHPF,RPS20,DDOST,LSR,DBNL,TOMM40,RABEP2,TA

B1,LRRFIP1,ATP5J,ELP6,TRMT112,CHPF,RPS20,DCTD,DDOST,LSR,RRNAD1,TOMM40,TADA3,ANAPC11,N

1,MED15,CLASRP,C1ORF198,UBE2I,EIF3M,CDC42,CARD14,PCYT2,FBXW9,USE1,IFI27L1,ASF1B,CCDC25,EI

EP,TBL2,ARHGAP6,SCARB1,CTTN,TSPAN11,ITSN1,EIF5A,TNFSF10,COMMD5,CHID1,NUDT22,UFD1L,TAZ,D

EP,TBL2,ARHGAP6,SCARB1,CTTN,TSPAN11,ITSN1,EIF5A,TNFSF10,COMMD5,CHID1,NUDT22,UFD1L,TAZ,D

SR,TOMM40,SERPINB6,RABEP2,TADA3,WAC,SLC33A1,ZNF687,RAD23A,NT5C,MDH2,STXBP2,SEC31A,SIR

1,ARRDC1,TRPT1,KLC1,RBP5,ZC3HAV1,SCAF11,DBI,ARPC2,LIMK2,VDAC3,NOB1,WDR19,SIRT6,NUMA1,SI

BL5,TRNT1,ITGB1BP1,MED15,CLASRP,C1ORF198,UGGT2,UBE2I,EIF3M,CDC42,TBX6,CARD14,PCYT2,FBXW

BL5,TRNT1,ITGB1BP1,MED15,CLASRP,C1ORF198,UGGT2,UBE2I,EIF3M,CDC42,TBX6,CARD14,PCYT2,FBXW

31,CTTN,TSPAN11,ITSN1,EIF5A,TNFSF10,CHID1,NUDT22,AMZ2,UFD1L,TAZ,DBNDD1,NOP16,TTLL5,PSMA

31,CTTN,TSPAN11,ITSN1,EIF5A,TNFSF10,CHID1,NUDT22,AMZ2,UFD1L,TAZ,DBNDD1,NOP16,TTLL5,PSMA

G16L2,RCC1,SNRPB,PARD3,TCAIM,SLC3A2,DHPS,ACTR6,SEC62,SET,CTSC,ZNF235,BOP1,ARRDC1,TRPT1,K

G16L2,RCC1,SNRPB,PARD3,TCAIM,SLC3A2,DHPS,ACTR6,SEC62,SET,CTSC,ZNF235,BOP1,ARRDC1,TRPT1,K

UMF2,BUD31,SAE1,TBL2,ARHGAP6,SCARB1,CTTN,TSPAN11,ITSN1,EIF5A,COMMD5,NUDT22,AMZ2,TAZ,

,PARD3,TCAIM,SLC3A2,DHPS,ACTR6,SEC62,SET,CTSC,ZNF235,BOP1,ARRDC1,TRPT1,KLC1,RBP5,ZC3HAV1

,PARD3,TCAIM,SLC3A2,DHPS,ACTR6,SEC62,SET,CTSC,ZNF235,BOP1,ARRDC1,TRPT1,KLC1,RBP5,ZC3HAV1

ENDOV,PPIL2,TROAP,RPL28,LIG1,TMEM126B,APEH,CAPN10,LYPLA2,C17ORF62,TXNDC17,PPAN,CYB561,

ENDOV,PPIL2,TROAP,RPL28,LIG1,TMEM126B,APEH,CAPN10,LYPLA2,C17ORF62,TXNDC17,PPAN,CYB561,

CHPF,RPS20,DCTD,ERCC5,DDOST,PABPC1,RRNAD1,DBNL,TOMM40,TADA3,ATP6V1F,ANAPC11,NT5C,MD

87,RAD23A,NT5C,MDH2,STXBP2,SEC31A,PFDN5,MTHFS,PAFAH1B2,NSUN5,ZFPL1,GMPR2,RIC8A,UBE2N,

1,TM9SF1,MAGED2,SON,TET2,UQCRQ,SNF8,CDC37,RPS16,CPSF3L,TSG101,UBR4,PSMA1,SLC39A11,FUZ,

1,TM9SF1,MAGED2,SON,TET2,UQCRQ,SNF8,CDC37,RPS16,CPSF3L,TSG101,UBR4,PSMA1,SLC39A11,FUZ,

D3,TCAIM,SLC3A2,DHPS,ACTR6,SEC62,SET,CTSC,ZNF235,BOP1,ARRDC1,TRPT1,KLC1,ZC3HAV1,SCAF11,DE

D3,TCAIM,SLC3A2,DHPS,ACTR6,SEC62,SET,CTSC,ZNF235,BOP1,ARRDC1,TRPT1,KLC1,ZC3HAV1,SCAF11,DE

31A,CNST,COX17,HIGD1A,B3GAT3,ZNF506,MTHFS,PAFAH1B2,ZNF343,PPIH,DALRD3,NSUN5,GMPR2,MC

TR6,SEC62,SET,CTSC,BOP1,ARRDC1,TRPT1,KLC1,RBP5,ZC3HAV1,SCAF11,DBI,ARPC2,LIMK2,VDAC3,NOB1

TR6,SEC62,SET,CTSC,BOP1,ARRDC1,TRPT1,KLC1,RBP5,ZC3HAV1,SCAF11,DBI,ARPC2,LIMK2,VDAC3,NOB1

P,TBL2,SCARB1,CTTN,TSPAN11,ITSN1,EIF5A,COMMD5,RUNX1,NUDT22,UFD1L,THYN1,DBNDD1,SH2B1,N

P,TBL2,SCARB1,CTTN,TSPAN11,ITSN1,EIF5A,COMMD5,RUNX1,NUDT22,UFD1L,THYN1,DBNDD1,SH2B1,N

RMT112,MPDU1,SAT1,ERCC5,EMC4,SERPINB1,DDOST,LSR,RRNAD1,DBNL,NAT9,TOMM40,RABEP2,TADA

ACOT8,AP3B2,RCE1,MRPL33,RUVBL2,EBPL,TESC,IGSF8,SNRPN,SPAG7,WDR62,RFX5,POLD4,UBE2L6,FIS1,

ACOT8,AP3B2,RCE1,MRPL33,RUVBL2,EBPL,TESC,IGSF8,SNRPN,SPAG7,WDR62,RFX5,POLD4,UBE2L6,FIS1,

R19,SIRT6,NUMA1,BUD31,SAE1,NPRL2,TBL2,ARHGAP6,SCARB1,CRYGS,CTTN,TSPAN11,ITSN1,EIF5A,COM

RPS6,RPL29,RTEL1,COASY,AUP1,SAP30BP,YIPF3,FOXRED1,RABL5,TRNT1,ITGB1BP1,MED15,SH3YL1,C1OR

RPS6,RPL29,RTEL1,COASY,AUP1,SAP30BP,YIPF3,FOXRED1,RABL5,TRNT1,ITGB1BP1,MED15,SH3YL1,C1OR

,FOXRED1,RABL5,TRNT1,ITGB1BP1,MED15,CLASRP,C1ORF198,UGGT2,UBE2I,SLC39A13,EIF3M,CDC42,TB

,FOXRED1,RABL5,TRNT1,ITGB1BP1,MED15,CLASRP,C1ORF198,UGGT2,UBE2I,SLC39A13,EIF3M,CDC42,TB

N,CYB561,SUGT1,HYPK,ZNF250,ZNF197,COPZ1,INO80E,HSP90AB1,RAB2A,ATP5J,TRMT112,CHPF,ZNF814

N,CYB561,SUGT1,HYPK,ZNF250,ZNF197,COPZ1,INO80E,HSP90AB1,RAB2A,ATP5J,TRMT112,CHPF,ZNF814

RPS20,DCTD,MCTS1,DDOST,LSR,TOMM40,TADA3,ZNF687,ANAPC11,NT5C,MDH2,EEF1D,CHCHD5,SEC31

ARRDC1,TRPT1,KLC1,RBP5,ZC3HAV1,SCAF11,DBI,ARPC2,LIMK2,VDAC3,NOB1,WDR19,SIRT6,NUMA1,GPA

ARRDC1,TRPT1,KLC1,RBP5,ZC3HAV1,SCAF11,DBI,ARPC2,LIMK2,VDAC3,NOB1,WDR19,SIRT6,NUMA1,GPA

16,CPSF3L,TSG101,UBR4,SLC39A11,FUZ,FAM3A,PSMA3,METTL17,ATG12,RPS25,NIP7,TTC9C,PLCB2,TMX2

16,CPSF3L,TSG101,UBR4,SLC39A11,FUZ,FAM3A,PSMA3,METTL17,ATG12,RPS25,NIP7,TTC9C,PLCB2,TMX2

,HMGA2,CDC37,RPS16,CPSF3L,TSG101,UBR4,YLPM1,CHCHD1,FUZ,PSMA3,RPS25,NIP7,TTC9C,COX4I1,T

2,VDAC3,WDR19,SIRT6,NUMA1,BUD31,SAE1,NPRL2,OSGEP,ARHGAP6,CRYGS,TSPAN11,ITSN1,EIF5A,CO

2,VDAC3,WDR19,SIRT6,NUMA1,BUD31,SAE1,NPRL2,OSGEP,ARHGAP6,CRYGS,TSPAN11,ITSN1,EIF5A,CO

DF1,RFX3,ENDOV,PPIL2,APEH,LYPLA2,C17ORF62,PRMT1,C19ORF54,TXNDC17,PPAN,SUGT1,CARS,CDK1

,TIMM17B,APEH,LYPLA2,LSM12,C17ORF62,PRMT1,STOML2,C19ORF54,TXNDC17,PIGL,PPAN,CYB561,HY

2L,BUD31,OSGEP,ARHGAP6,SCARB1,CTTN,TSPAN11,ITSN1,EIF5A,NUDT22,AMZ2,UFD1L,DBNDD1,NOP16,

2L,BUD31,OSGEP,ARHGAP6,SCARB1,CTTN,TSPAN11,ITSN1,EIF5A,NUDT22,AMZ2,UFD1L,DBNDD1,NOP16,

SUMF2,BUD31,SAE1,NPRL2,OSGEP,TBL2,ARHGAP6,SCARB1,TSPAN11,ITSN1,EIF5A,COMMD5,UFD1L,USP

SUMF2,BUD31,SAE1,NPRL2,OSGEP,TBL2,ARHGAP6,SCARB1,TSPAN11,ITSN1,EIF5A,COMMD5,UFD1L,USP

P38,TAZ,DBNDD1,SH2B1,NOP16,RPL13A,GSTK1,TTLL5,LDHA,PSMA6,ACOT8,AP3B2,SLC35G1,MRPL33,ZN

,ITSN1,EIF5A,THYN1,DBNDD1,SH2B1,NOP16,GSTK1,TTLL5,PSMA6,ACOT8,AP3B2,RCE1,ZNF410,RUVBL2,

,ITSN1,EIF5A,THYN1,DBNDD1,SH2B1,NOP16,GSTK1,TTLL5,PSMA6,ACOT8,AP3B2,RCE1,ZNF410,RUVBL2,

1,RPL13A,ARHGEF39,TTLL5,PSMA6,ACOT8,RPS9,RCE1,MRPL33,ZNF410,RUVBL2,NUDT5,SNRPN,WDR62,

1,RPL13A,ARHGEF39,TTLL5,PSMA6,ACOT8,RPS9,RCE1,MRPL33,ZNF410,RUVBL2,NUDT5,SNRPN,WDR62,

DD1,NOP16,PSMA6,ACOT8,AP3B2,RCE1,ZNF410,RUVBL2,PCMT1,EBPL,TESC,IGSF8,RBPMS,SPAG7,WDR6

N,ZNF81,FAM32A,TNKS,ELOF1,STX10,C12ORF57,DCTN5,PEX26,NDUFS3,OXLD1,STAU2,METTL5,SLC35E1,

MFBP1,LIMK2,VDAC3,WDR19,SIRT6,NUMA1,GPATCH2L,SUMF2,BUD31,SAE1,NPRL2,TBL2,ARHGAP6,SCA

MFBP1,LIMK2,VDAC3,WDR19,SIRT6,NUMA1,GPATCH2L,SUMF2,BUD31,SAE1,NPRL2,TBL2,ARHGAP6,SCA
D15,CLASRP,C1ORF198,UGGT2,UBE2I,CDC42,TBX6,PCYT2,FBXW9,MAF,ASF1B,CCDC25,EEF1B2,RFX3,EN
D15,CLASRP,C1ORF198,UGGT2,UBE2I,CDC42,TBX6,PCYT2,FBXW9,MAF,ASF1B,CCDC25,EEF1B2,RFX3,EN
N1,EIF5A,COMMD5,AMZ2,TAZ,DBNDD1,SH2B1,NOP16,GSTK1,PSMA6,ACOT8,AP3B2,RCE1,MRPL33,ZNF
N1,EIF5A,COMMD5,AMZ2,TAZ,DBNDD1,SH2B1,NOP16,GSTK1,PSMA6,ACOT8,AP3B2,RCE1,MRPL33,ZNF
EC62,SET,CTSC,BOP1,ARRDC1,TRPT1,KLC1,RBP5,ZC3HAV1,SCAF11,DBI,ARPC2,LIMK2,VDAC3,NOB1,SIRT
,CAMTA1,PPIL2,LIG1,APEH,RPL21,CAPN10,LYPLA2,LSM12,C17ORF62,PRMT1,C19ORF54,ZNF7,ARL6IP4,
2,PRMT1,STOML2,ZNF7,TXNDC17,PPAN,HNRNPC,SUGT1,CARS,RBM3,ZNF197,COPZ1,LRRFIP1,ATP5J,TR
5,SEC31A,CNST,ZNF506,PAFAH1B2,DALRD3,NSUN5,ZFPL1,RIC8A,TCP1,PIGT,UBE2N,NFATC4,MFSD11,P
IF5A,COMMD5,CHID1,NUDT22,AMZ2,UFD1L,USP38,TAZ,THYN1,DBNDD1,SH2B1,TTLL5,LDHA,ACOT8,AP
IF5A,COMMD5,CHID1,NUDT22,AMZ2,UFD1L,USP38,TAZ,THYN1,DBNDD1,SH2B1,TTLL5,LDHA,ACOT8,AP
31,SAE1,NPRL2,OSGEP,TBL2,ARHGAP6,SCARB1,CRYGS,CTTN,ITSN1,EIF5A,COMMD5,CHID1,NUDT22,AM
31,SAE1,NPRL2,OSGEP,TBL2,ARHGAP6,SCARB1,CRYGS,CTTN,ITSN1,EIF5A,COMMD5,CHID1,NUDT22,AM
RDC1,TRPT1,KLC1,ZC3HAV1,SCAF11,DBI,ARPC2,LIMK2,VDAC3,NOB1,WDR19,NUMA1,SUMF2,BUD31,N
P16,GSTK1,TTLL5,PSMA6,ACOT8,AP3B2,RCE1,MRPL33,ZNF410,EBPL,TESC,IGSF8,NNMT,SPAG7,WDR62,
P16,GSTK1,TTLL5,PSMA6,ACOT8,AP3B2,RCE1,MRPL33,ZNF410,EBPL,TESC,IGSF8,NNMT,SPAG7,WDR62,
2,RCC1,SNRPB,PARD3,TCAIM,SLC3A2,DHPS,ACTR6,MAP3K7CL,SEC62,SET,CTSC,ZNF235,BOP1,ARRDC1,T
2,RCC1,SNRPB,PARD3,TCAIM,SLC3A2,DHPS,ACTR6,MAP3K7CL,SEC62,SET,CTSC,ZNF235,BOP1,ARRDC1,T
L2,OSGEP,TBL2,ARHGAP6,SCARB1,ITSN1,EIF5A,TNFSF10,COMMD5,RUNX1,CHID1,NUDT22,AMZ2,UFD1
D31,SAE1,NPRL2,OSGEP,TBL2,ARHGAP6,SCARB1,TSPAN11,ITSN1,EIF5A,COMMD5,AMZ2,UFD1L,USP38,
PSMA6,ACOT8,AP3B2,RPS9,RCE1,MRPL33,ZNF410,EBPL,NUDT5,RBPMS,SNRPN,WDR62,STYXL1,POLD4,
PSMA6,ACOT8,AP3B2,RPS9,RCE1,MRPL33,ZNF410,EBPL,NUDT5,RBPMS,SNRPN,WDR62,STYXL1,POLD4,
,LRRFIP1,RAB2A,ATP5J,TRMT112,CHPF,RPS20,GABPB1,UBXN1,ERLEC1,EMC4,DDOST,LSR,PSME2,PABPC
AD1,DBNL,NAT9,TOMM40,RABEP2,TADA3,ZNF20,WAC,RAD23A,NT5C,MDH2,STXBP2,HSPH1,CHCHD5,N
5G1,MRPL33,RUVBL2,EBPL,TESC,IGSF8,NUDT5,NNMT,SPAG7,WDR62,STYXL1,RANBP1,ZC3H14,IRF9,STIP
D5,NUDT22,AMZ2,UFD1L,USP38,THYN1,DBNDD1,SH2B1,NOP16,RPL13A,ARHGEF39,TTLL5,PSMA6,ACOT
D5,NUDT22,AMZ2,UFD1L,USP38,THYN1,DBNDD1,SH2B1,NOP16,RPL13A,ARHGEF39,TTLL5,PSMA6,ACOT
,TADA3,ANAPC11,NT5C,MDH2,STXBP2,EEF1D,CHCHD5,SEC31A,COX17,HIGD1A,B3GAT3,PAFAH1B2,GB
3B2,RCE1,ZNF410,RUVBL2,EBPL,TESC,IGSF8,SPAG7,WDR62,IL1RAP,RFX5,POLD4,UBE2L6,ZC3H14,FIS1,P
3B2,RCE1,ZNF410,RUVBL2,EBPL,TESC,IGSF8,SPAG7,WDR62,IL1RAP,RFX5,POLD4,UBE2L6,ZC3H14,FIS1,P
3B2,RCE1,ZNF410,RUVBL2,EBPL,TESC,IGSF8,SPAG7,WDR62,IL1RAP,RFX5,POLD4,UBE2L6,ZC3H14,FIS1,P
3B2,RCE1,ZNF410,RUVBL2,EBPL,TESC,IGSF8,SPAG7,WDR62,IL1RAP,RFX5,POLD4,UBE2L6,ZC3H14,FIS1,P
,NOB1,WDR19,SIRT6,NUMA1,GPATCH2L,NPRL2,OSGEP,TBL2,ARHGAP6,SCARB1,CRYGS,CTTN,TSPAN11,I
,NOB1,WDR19,SIRT6,NUMA1,GPATCH2L,NPRL2,OSGEP,TBL2,ARHGAP6,SCARB1,CRYGS,CTTN,TSPAN11,I
V1,SCAF11,DBI,ARPC2,LIMK2,VDAC3,NOB1,SIRT6,NUMA1,BUD31,OSGEP,TBL2,SCARB1,CTTN,TSPAN11,I
LC3A2,DHPS,ACTR6,SEC62,SET,CTSC,ZNF235,BOP1,ARRDC1,TRPT1,KLC1,ZC3HAV1,SCAF11,DBI,ARPC2,LI
LC3A2,DHPS,ACTR6,SEC62,SET,CTSC,ZNF235,BOP1,ARRDC1,TRPT1,KLC1,ZC3HAV1,SCAF11,DBI,ARPC2,LI
OP1,ARRDC1,TRPT1,KLC1,RBP5,ZC3HAV1,SCAF11,DBI,ARPC2,LIMK2,VDAC3,WDR19,SIRT6,NUMA1,GPAT
GEP,TBL2,SCARB1,CRYGS,CTTN,ITSN1,EIF5A,COMMD5,AMZ2,UFD1L,USP38,TAZ,THYN1,DBNDD1,SH2B1
1,DBI,ARPC2,LIMK2,VDAC3,NOB1,WDR19,SIRT6,NUMA1,GPATCH2L,SUMF2,BUD31,SAE1,NPRL2,OSGEP
1,DBI,ARPC2,LIMK2,VDAC3,NOB1,WDR19,SIRT6,NUMA1,GPATCH2L,SUMF2,BUD31,SAE1,NPRL2,OSGEP
,TBL2,SCARB1,CTTN,TSPAN11,EIF5A,COMMD5,CHID1,NUDT22,AMZ2,USP38,SH2B1,NOP16,RPL13A,AR
,TBL2,SCARB1,CTTN,TSPAN11,EIF5A,COMMD5,CHID1,NUDT22,AMZ2,USP38,SH2B1,NOP16,RPL13A,AR
L,TESC,IGSF8,SNRPN,SPAG7,WDR62,RANBP1,UBE2L6,FIS1,PPM1K,TUBA1C,ANTXR2,STIP1,PHF19,LPCAT3
L,TESC,IGSF8,SNRPN,SPAG7,WDR62,RANBP1,UBE2L6,FIS1,PPM1K,TUBA1C,ANTXR2,STIP1,PHF19,LPCAT3
B1,WDR19,SIRT6,NUMA1,GPATCH2L,SUMF2,BUD31,SAE1,NPRL2,OSGEP,TBL2,TSPAN11,ITSN1,EIF5A,CO
3,RUVBL2,EBPL,TESC,IGSF8,SNRPN,SPAG7,WDR62,RANBP1,UBE2L6,FIS1,PPM1K,TUBA1C,ANTXR2,STIP1,

3,RUVBL2,EBPL,TESC,IGSF8,SNRPN,SPAG7,WDR62,RANBP1,UBE2L6,FIS1,PPM1K,TUBA1C,ANTXR2,STIP1,

3,KLK10,RAD23A,ZCCHC4,MDH2,STXBP2,CHCHD5,NAA10,SEC31A,CNST,COX5A,HINT1,SIRT5,PAFAH1B2,D

D31,SAE1,NPRL2,TBL2,ARHGAP6,SCARB1,CRYGS,CTTN,TSPAN11,ITSN1,EIF5A,COMMD5,CHID1,NUDT22,

,RCC1,SNRPB,PARD3,TCAIM,SLC3A2,DHPS,ACTR6,SEC62,SET,CTSC,ZNF235,BOP1,ARRDC1,TRPT1,KLC1,RE

,RCC1,SNRPB,PARD3,TCAIM,SLC3A2,DHPS,ACTR6,SEC62,SET,CTSC,ZNF235,BOP1,ARRDC1,TRPT1,KLC1,RE

JMA1,SUMF2,BUD31,SAE1,NPRL2,ARHGAP6,SCARB1,CTTN,TSPAN11,ITSN1,EIF5A,COMMD5,UFD1L,USP3

JMA1,SUMF2,BUD31,SAE1,NPRL2,ARHGAP6,SCARB1,CTTN,TSPAN11,ITSN1,EIF5A,COMMD5,UFD1L,USP3

PKAPK5,ATP5J,TRMT112,CHPF,RPS20,DCTD,ZNF814,SAT1,ERCC5,EMC4,LMBR1,DDOST,LSR,MEIS2,PSME

16,TTLL5,PSMA6,ACOT8,AP3B2,RCE1,ZNF410,RUVBL2,EBPL,TESC,IGSF8,RBPMS,SNRPN,SPAG7,WDR62,I

16,TTLL5,PSMA6,ACOT8,AP3B2,RCE1,ZNF410,RUVBL2,EBPL,TESC,IGSF8,RBPMS,SNRPN,SPAG7,WDR62,I

1,SUMF2,BUD31,SAE1,TBL2,CRYGS,CTTN,TSPAN11,ITSN1,EIF5A,COMMD5,CHID1,AMZ2,USP38,TAZ,DBN

1,SUMF2,BUD31,SAE1,TBL2,CRYGS,CTTN,TSPAN11,ITSN1,EIF5A,COMMD5,CHID1,AMZ2,USP38,TAZ,DBN

HPS,ACTR6,SEC62,SET,CTSC,BOP1,ARRDC1,TRPT1,KLC1,RBP5,ZC3HAV1,SCAF11,DBI,ARPC2,LIMK2,VDAC3

HPS,ACTR6,SEC62,SET,CTSC,BOP1,ARRDC1,TRPT1,KLC1,RBP5,ZC3HAV1,SCAF11,DBI,ARPC2,LIMK2,VDAC3

N11,ITSN1,EIF5A,COMMD5,AMZ2,UFD1L,USP38,DBNDD1,SH2B1,NOP16,RPL13A,ARHGEF39,TTLL5,PSM

8,AP3B2,ZNF410,RUVBL2,PCMT1,TESC,IGSF8,RBPMS,SNRPN,SPAG7,WDR62,STYXL1,UBE2L6,VRK3,ZC3H

8,AP3B2,ZNF410,RUVBL2,PCMT1,TESC,IGSF8,RBPMS,SNRPN,SPAG7,WDR62,STYXL1,UBE2L6,VRK3,ZC3H

8,AP3B2,ZNF410,RUVBL2,PCMT1,TESC,IGSF8,RBPMS,SNRPN,SPAG7,WDR62,STYXL1,UBE2L6,VRK3,ZC3H

8,AP3B2,ZNF410,RUVBL2,PCMT1,TESC,IGSF8,RBPMS,SNRPN,SPAG7,WDR62,STYXL1,UBE2L6,VRK3,ZC3H

8,AP3B2,ZNF410,RUVBL2,PCMT1,TESC,IGSF8,RBPMS,SNRPN,SPAG7,WDR62,STYXL1,UBE2L6,VRK3,ZC3H

8,AP3B2,ZNF410,RUVBL2,PCMT1,TESC,IGSF8,RBPMS,SNRPN,SPAG7,WDR62,STYXL1,UBE2L6,VRK3,ZC3H

,IGSF8,SNRPN,SPAG7,WDR62,IL1RAP,POLD4,RANBP1,UBE2L6,FIS1,PPM1K,ANTXR2,STIP1,PHF19,VCAN,L

,IGSF8,SNRPN,SPAG7,WDR62,IL1RAP,POLD4,RANBP1,UBE2L6,FIS1,PPM1K,ANTXR2,STIP1,PHF19,VCAN,L

YN1,DBNDD1,SH2B1,NOP16,RPL13A,ARHGEF39,PSMA6,ACOT8,AP3B2,RCE1,ZNF410,RUVBL2,PCMT1,IG

YN1,DBNDD1,SH2B1,NOP16,RPL13A,ARHGEF39,PSMA6,ACOT8,AP3B2,RCE1,ZNF410,RUVBL2,PCMT1,IG

,LZTFL1,FIS1,PPM1K,LPCAT3,MAGED2,SON,UQCRQ,SNF8,HMGA2,CPSF3L,TSG101,FUZ,FAM3A,PSMA3,M

,LZTFL1,FIS1,PPM1K,LPCAT3,MAGED2,SON,UQCRQ,SNF8,HMGA2,CPSF3L,TSG101,FUZ,FAM3A,PSMA3,M

D1,NUDT22,AMZ2,UFD1L,THYN1,DBNDD1,NOP16,ARHGEF39,PSMA6,ACOT8,AP3B2,RCE1,MRPL33,RUV

D1,NUDT22,AMZ2,UFD1L,THYN1,DBNDD1,NOP16,ARHGEF39,PSMA6,ACOT8,AP3B2,RCE1,MRPL33,RUV

MM40,ZNF687,ANAPC11,MDH2,HSPH1,NAA10,CARHSP1,CNST,COX17,HNRNPLL,POLR2C,B3GAT3,RFXA

MM40,ZNF687,ANAPC11,MDH2,HSPH1,NAA10,CARHSP1,CNST,COX17,HNRNPLL,POLR2C,B3GAT3,RFXA

OP16,ARHGEF39,PSMA6,ACOT8,AP3B2,RCE1,MRPL33,RUVBL2,EBPL,TESC,IGSF8,RBPMS,NNMT,SPAG7,W

OP16,ARHGEF39,PSMA6,ACOT8,AP3B2,RCE1,MRPL33,RUVBL2,EBPL,TESC,IGSF8,RBPMS,NNMT,SPAG7,W

PIL2,LIG1,PSMA7,APEH,LYPLA2,NUTF2,C17ORF62,STOML2,PPAN,ACAD10,CYB561,ZNF250,RPL36,CDK11

PIL2,LIG1,PSMA7,APEH,LYPLA2,NUTF2,C17ORF62,STOML2,PPAN,ACAD10,CYB561,ZNF250,RPL36,CDK11

IF2,BUD31,SAE1,NPRL2,ARHGAP6,SCARB1,CRYGS,CTTN,TSPAN11,ITSN1,EIF5A,COMMD5,CHID1,NUDT22

IF2,BUD31,SAE1,NPRL2,ARHGAP6,SCARB1,CRYGS,CTTN,TSPAN11,ITSN1,EIF5A,COMMD5,CHID1,NUDT22

1,YIPF3,FOXRED1,TRNT1,MED15,SH3YL1,CLASRP,MICAL1,SLC39A13,EIF3M,C14ORF2,FBXW9,USE1,ASF1

IMD5,UFD1L,DBNDD1,NOP16,RPL13A,TTLL5,PSMA6,AP3B2,RPS9,SLC35G1,MRPL33,ZNF410,RUVBL2,EB

IMD5,UFD1L,DBNDD1,NOP16,RPL13A,TTLL5,PSMA6,AP3B2,RPS9,SLC35G1,MRPL33,ZNF410,RUVBL2,EB

,RUVBL2,PCMT1,EBPL,TESC,IGSF8,NNMT,SPAG7,WDR62,IL1RAP,POLD4,RANBP1,UBE2L6,ZC3H14,FIS1,P

,RUVBL2,PCMT1,EBPL,TESC,IGSF8,NNMT,SPAG7,WDR62,IL1RAP,POLD4,RANBP1,UBE2L6,ZC3H14,FIS1,P

13A,ARHGEF39,GSTK1,PSMA6,AP3B2,SLC35G1,RUVBL2,EBPL,TESC,NUDT5,RBPMS,SPAG7,WDR62,IL1RA

ANBP1,UBE2L6,LZTFL1,ZC3H14,FIS1,SLC15A3,TUBA1C,STIP1,LRRC1,PHF19,VCAN,LPCAT3,CD44,TM9SF1,

ANBP1,UBE2L6,LZTFL1,ZC3H14,FIS1,SLC15A3,TUBA1C,STIP1,LRRC1,PHF19,VCAN,LPCAT3,CD44,TM9SF1,

IMK2,VDAC3,WDR19,SIRT6,NUMA1,SUMF2,BUD31,SAE1,NPRL2,CRYGS,CTTN,TSPAN11,ITSN1,EIF5A,CO

IMK2,VDAC3,WDR19,SIRT6,NUMA1,SUMF2,BUD31,SAE1,NPRL2,CRYGS,CTTN,TSPAN11,ITSN1,EIF5A,CO

1D,NAA10,CARHSP1,COX17,HIGD1A,POLR2C,PFDN5,PAFAH1B2,DALRD3,NSUN5,OARD1,RIC8A,UBE2N,F
1D,NAA10,CARHSP1,COX17,HIGD1A,POLR2C,PFDN5,PAFAH1B2,DALRD3,NSUN5,OARD1,RIC8A,UBE2N,F
,ZC3HAV1,SCAF11,DBI,LIMK2,VDAC3,NOB1,WDR19,SIRT6,NUMA1,GPATCH2L,SUMF2,BUD31,NPRL2,OS
,ZC3HAV1,SCAF11,DBI,LIMK2,VDAC3,NOB1,WDR19,SIRT6,NUMA1,GPATCH2L,SUMF2,BUD31,NPRL2,OS
SIRT6,NUMA1,GPATCH2L,BUD31,NPRL2,OSGEP,TBL2,ARHGAP6,SCARB1,CTTN,TSPAN11,ITSN1,EIF5A,CO
SIRT6,NUMA1,GPATCH2L,BUD31,NPRL2,OSGEP,TBL2,ARHGAP6,SCARB1,CTTN,TSPAN11,ITSN1,EIF5A,CO
IK2,VDAC3,NOB1,WDR19,NUMA1,GPATCH2L,SUMF2,BUD31,NPRL2,OSGEP,TBL2,ARHGAP6,SCARB1,CTT
IK2,VDAC3,NOB1,WDR19,NUMA1,GPATCH2L,SUMF2,BUD31,NPRL2,OSGEP,TBL2,ARHGAP6,SCARB1,CTT
1,TIMM17B,LYPLA2,STOML2,PPAN,CYB561,SUGT1,HYPK,ZNF197,COPZ1,ATP5J,TRMT112,ZNF814,VWA
1,TIMM17B,LYPLA2,STOML2,PPAN,CYB561,SUGT1,HYPK,ZNF197,COPZ1,ATP5J,TRMT112,ZNF814,VWA
,NPRL2,OSGEP,ARHGAP6,CTTN,ITSN1,EIF5A,COMMD5,USP38,TAZ,DBNDD1,SH2B1,RPL13A,GSTK1,TTLL5
,NPRL2,OSGEP,ARHGAP6,CTTN,ITSN1,EIF5A,COMMD5,USP38,TAZ,DBNDD1,SH2B1,RPL13A,GSTK1,TTLL5
,TESC,IGSF8,SNRPN,SPAG7,WDR62,RFX5,POLD4,RANBP1,UBE2L6,FIS1,PPM1K,TUBA1C,ANTXR2,STIP1,P
,TESC,IGSF8,SNRPN,SPAG7,WDR62,RFX5,POLD4,RANBP1,UBE2L6,FIS1,PPM1K,TUBA1C,ANTXR2,STIP1,P
HF19,LPCAT3,GRK6,CDC37,RPS16,CPSF3L,OBSL1,UBR4,PHF21A,SLC39A11,FAM3A,MUC1,EPB41L5,ATG1
HF19,LPCAT3,GRK6,CDC37,RPS16,CPSF3L,OBSL1,UBR4,PHF21A,SLC39A11,FAM3A,MUC1,EPB41L5,ATG1
YIPF3,FOXRED1,RABL5,TRNT1,ITGB1BP1,MED15,IP6K2,CLASRP,C1ORF198,UGGT2,EIF3M,CDC42,TBX6,C
YIPF3,FOXRED1,RABL5,TRNT1,ITGB1BP1,MED15,IP6K2,CLASRP,C1ORF198,UGGT2,EIF3M,CDC42,TBX6,C
IL1RAP,POLD4,RANBP1,UBE2L6,FIS1,PPM1K,ANTXR2,STIP1,PHF19,VCAN,LPCAT3,MAGED2,SON,GRK6,C
IL1RAP,POLD4,RANBP1,UBE2L6,FIS1,PPM1K,ANTXR2,STIP1,PHF19,VCAN,LPCAT3,MAGED2,SON,GRK6,C
A1C,STIP1,LRRC1,PHF19,VCAN,LPCAT3,CD44,TM9SF1,SON,TET2,CDC37,CPSF3L,TSG101,OBSL1,TTC21B,P
A1C,STIP1,LRRC1,PHF19,VCAN,LPCAT3,CD44,TM9SF1,SON,TET2,CDC37,CPSF3L,TSG101,OBSL1,TTC21B,P
PAN11,ITSN1,EIF5A,COMMD5,NUDT22,AMZ2,UFD1L,TAZ,DBNDD1,SH2B1,NOP16,ARHGEF39,GSTK1,PS
PAN11,ITSN1,EIF5A,COMMD5,NUDT22,AMZ2,UFD1L,TAZ,DBNDD1,SH2B1,NOP16,ARHGEF39,GSTK1,PS
,COX17,HIGD1A,POLR2C,PFDN5,PAFAH1B2,NSUN5,OARD1,RIC8A,UBE2N,FAM32A,TMEM147,COA1,USP
,COX17,HIGD1A,POLR2C,PFDN5,PAFAH1B2,NSUN5,OARD1,RIC8A,UBE2N,FAM32A,TMEM147,COA1,USP
9C,COX4I1,SLC25A10,SPOCK2,NISCH,TMED4,RPS6,CD99L2,COASY,SRSF1,YIPF3,FOXRED1,KXD1,ITGB1BP1
IFI27L1,ASF1B,CCDC25,EEF1B2,RPL28,LYPLA2,C17ORF62,C19ORF54,SHARPIN,SUGT1,CDK11A,RBM3,ZN
RK6,CDC37,RPS16,CPSF3L,PHF21A,SLC39A11,FAM3A,MUC1,METTL17,EPB41L5,ATG12,ZBTB44,DUSP22,
7ORF62,PRMT1,STOML2,ZNF7,TXNDC17,ARL6IP4,PPAN,CYB561,STAU1,ZNF250,RPL36,INO80E,HSP90AB
7ORF62,PRMT1,STOML2,ZNF7,TXNDC17,ARL6IP4,PPAN,CYB561,STAU1,ZNF250,RPL36,INO80E,HSP90AB
TCH2L,SUMF2,BUD31,SAE1,OSGEP,TBL2,ARHGAP6,SCARB1,CTTN,TSPAN11,ITSN1,EIF5A,COMMD5,CHID
TCH2L,SUMF2,BUD31,SAE1,OSGEP,TBL2,ARHGAP6,SCARB1,CTTN,TSPAN11,ITSN1,EIF5A,COMMD5,CHID
C3,WDR19,SIRT6,NUMA1,GPATCH2L,BUD31,NPRL2,OSGEP,TBL2,SCARB1,CTTN,TSPAN11,ITSN1,EIF5A,C
C3,WDR19,SIRT6,NUMA1,GPATCH2L,BUD31,NPRL2,OSGEP,TBL2,SCARB1,CTTN,TSPAN11,ITSN1,EIF5A,C
2,ARHGAP6,SCARB1,CTTN,EIF5A,COMMD5,UFD1L,USP38,DBNDD1,SH2B1,NOP16,RPL13A,GSTK1,TTLL5,L
2,ARHGAP6,SCARB1,CTTN,EIF5A,COMMD5,UFD1L,USP38,DBNDD1,SH2B1,NOP16,RPL13A,GSTK1,TTLL5,L
T8,AP3B2,ZNF410,RUVBL2,PCMT1,EBPL,NUDT5,SNRPN,STYXL1,POLD4,VRK3,LZTFL1,FIS1,PPM1K,LPCAT3
T8,AP3B2,ZNF410,RUVBL2,PCMT1,EBPL,NUDT5,SNRPN,STYXL1,POLD4,VRK3,LZTFL1,FIS1,PPM1K,LPCAT3
ON,GRK6,CDC37,RPS16,CPSF3L,SLC39A11,FAM3A,METTL17,ATG12,NLRP1,ZBTB44,PLCB2,SLC25A10,THE
ON,GRK6,CDC37,RPS16,CPSF3L,SLC39A11,FAM3A,METTL17,ATG12,NLRP1,ZBTB44,PLCB2,SLC25A10,THE
10,CARHSP1,COX17,POLR2C,B3GAT3,PFDN5,PAFAH1B2,NSUN5,OARD1,GMPR2,RIC8A,UBE2N,PDCD6,Z
10,CARHSP1,COX17,POLR2C,B3GAT3,PFDN5,PAFAH1B2,NSUN5,OARD1,GMPR2,RIC8A,UBE2N,PDCD6,Z
RRDC1,KLC1,RBP5,DBI,ARPC2,LIMK2,VDAC3,NOB1,WDR19,SIRT6,NUMA1,GPATCH2L,BUD31,SAE1,NPRL
RRDC1,KLC1,RBP5,DBI,ARPC2,LIMK2,VDAC3,NOB1,WDR19,SIRT6,NUMA1,GPATCH2L,BUD31,SAE1,NPRL
RFX3,ENDOV,VAMP8,CAMTA1,CAPG,PPIL2,LIG1,CDC14B,APEH,LYPLA2,LSM12,C17ORF62,PRMT1,STOM
RFX3,ENDOV,VAMP8,CAMTA1,CAPG,PPIL2,LIG1,CDC14B,APEH,LYPLA2,LSM12,C17ORF62,PRMT1,STOM

L1,HS2ST1,UBE2I,EIF3M,TBX6,AUNIP,C14ORF2,ATF7,PCYT2,USE1,IFI27L1,PPP1CC,PPA1,DGKE,CAMTA1,
L1,HS2ST1,UBE2I,EIF3M,TBX6,AUNIP,C14ORF2,ATF7,PCYT2,USE1,IFI27L1,PPP1CC,PPA1,DGKE,CAMTA1,
9,MAGED2,CDC37,RPS16,SLC39A11,FAM3A,METTL17,EPB41L5,ATG12,RPS25,ZBTB44,DBNDD2,NISCH,T
9,MAGED2,CDC37,RPS16,SLC39A11,FAM3A,METTL17,EPB41L5,ATG12,RPS25,ZBTB44,DBNDD2,NISCH,T
1,RPL36,CDK11A,ZNF197,COPZ1,INO80E,CCNL2,DGUOK,ATP5J,TRMT112,CHPF,GABPB1,MPDU1,VWA2,
GED2,SON,UQCRQ,CPSF3L,TSG101,SLC39A11,FUZ,FAM3A,MUC1,METTL17,RPS25,ZBTB44,TTC9C,COX4
GED2,SON,UQCRQ,CPSF3L,TSG101,SLC39A11,FUZ,FAM3A,MUC1,METTL17,RPS25,ZBTB44,TTC9C,COX4
PH2,MICAL1,HS2ST1,UBE2I,EIF3M,TBX6,TMEM205,AUNIP,C14ORF2,ATF7,PCYT2,USE1,IFI27L1,PPP1CC,
PH2,MICAL1,HS2ST1,UBE2I,EIF3M,TBX6,TMEM205,AUNIP,C14ORF2,ATF7,PCYT2,USE1,IFI27L1,PPP1CC,
UMA1,SUMF2,BUD31,SAE1,NPRL2,OSGEP,TBL2,ARHGAP6,SCARB1,TSPAN11,ITSN1,EIF5A,NUDT22,AMZ
B1,CTTN,TSPAN11,ITSN1,EIF5A,COMMD5,AMZ2,UFD1L,USP38,DBNDD1,SH2B1,NOP16,RPL13A,ARHGEF
B1,CTTN,TSPAN11,ITSN1,EIF5A,COMMD5,AMZ2,UFD1L,USP38,DBNDD1,SH2B1,NOP16,RPL13A,ARHGEF
P30BP,YIPF3,FOXRED1,RABL5,TRNT1,ITGB1BP1,MED15,CLASRP,C1ORF198,UGGT2,UBE2I,EIF3M,MAGT
P30BP,YIPF3,FOXRED1,RABL5,TRNT1,ITGB1BP1,MED15,CLASRP,C1ORF198,UGGT2,UBE2I,EIF3M,MAGT
HYN1,DBNDD1,SH2B1,RPL13A,LDHA,PSMA6,ACOT8,RPS9,RCE1,MRPL33,RUVBL2,PCMT1,EBPL,TESC,WD
HYN1,DBNDD1,SH2B1,RPL13A,LDHA,PSMA6,ACOT8,RPS9,RCE1,MRPL33,RUVBL2,PCMT1,EBPL,TESC,WD
C3,WDR19,NUMA1,SAE1,NPRL2,OSGEP,TBL2,ARHGAP6,SCARB1,CRYGS,CTTN,TSPAN11,ITSN1,TNFSF10,
C3,WDR19,NUMA1,SAE1,NPRL2,OSGEP,TBL2,ARHGAP6,SCARB1,CRYGS,CTTN,TSPAN11,ITSN1,TNFSF10,
CARB1,CTTN,TSPAN11,ITSN1,EIF5A,COMMD5,CHID1,NUDT22,AMZ2,TAZ,DBNDD1,SH2B1,RPL13A,LDHA,
LZTFL1,FIS1,PPM1K,TUBA1C,IRF9,DST,LRRC1,PHF19,PCBP2,MAGED2,SON,UQCRQ,SNF8,CDC37,CPSF3L,
LZTFL1,FIS1,PPM1K,TUBA1C,IRF9,DST,LRRC1,PHF19,PCBP2,MAGED2,SON,UQCRQ,SNF8,CDC37,CPSF3L,
MGA2,CPSF3L,TSG101,OBSL1,YLPM1,SLC39A11,FUZ,METTL17,RPS25,ZBTB44,COX4I1,PLCB2,TMX2,THE
MGA2,CPSF3L,TSG101,OBSL1,YLPM1,SLC39A11,FUZ,METTL17,RPS25,ZBTB44,COX4I1,PLCB2,TMX2,THE
P5,ZC3HAV1,SCAF11,DBI,ARPC2,LIMK2,SIRT6,NUMA1,GPATCH2L,BUD31,SAE1,NPRL2,TBL2,SCARB1,CTT
N,GRK6,CDC37,RPS16,CPSF3L,PHF21A,SLC39A11,FAM3A,MUC1,METTL17,EPB41L5,ATG12,ZBTB44,DUS
GPATCH2L,SUMF2,OSGEP,TBL2,SCARB1,CTTN,ITSN1,TNFSF10,COMMD5,NUDT22,AMZ2,UFD1L,USP38,T
2A,USP40,C12ORF57,DCTN5,NDUFS3,OXLD1,ZNF223,SLC35E1,OGFOD2,MPV17,ARIH2,PICK1,FBXL18,TS
7,RPS16,YLPM1,PHF21A,SLC39A11,FAM3A,MUC1,METTL17,EPB41L5,ATG12,PPRC1,ZBTB44,TTC9C,PLCB
7,RPS16,YLPM1,PHF21A,SLC39A11,FAM3A,MUC1,METTL17,EPB41L5,ATG12,PPRC1,ZBTB44,TTC9C,PLCB
PS,SEC62,SET,BOP1,ARRDC1,TRPT1,KLC1,ZC3HAV1,SCAF11,ARPC2,PMFBP1,LIMK2,VDAC3,NOB1,WDR1
PS,SEC62,SET,BOP1,ARRDC1,TRPT1,KLC1,ZC3HAV1,SCAF11,ARPC2,PMFBP1,LIMK2,VDAC3,NOB1,WDR1
DC1,TRPT1,KLC1,RBP5,ZC3HAV1,SCAF11,DBI,ARPC2,LIMK2,VDAC3,NOB1,WDR19,SIRT6,NUMA1,GPATC
DC1,TRPT1,KLC1,RBP5,ZC3HAV1,SCAF11,DBI,ARPC2,LIMK2,VDAC3,NOB1,WDR19,SIRT6,NUMA1,GPATC
SC,BOP1,ARRDC1,TRPT1,KLC1,RBP5,ZC3HAV1,SCAF11,DBI,ARPC2,LIMK2,VDAC3,NOB1,WDR19,SIRT6,NU
HF19,LPCAT3,CD44,TM9SF1,SON,TET2,UQCRQ,CDC37,PTPN13,UBR4,YLPM1,FAM3A,PSMA3,PPRC1,NIP
B1,NOP16,TTLL5,LDHA,PSMA6,AP3B2,RPS9,RCE1,ZNF410,PCMT1,TESC,IGSF8,RBPMS,WDR62,PLEKHA5,
ED2,SON,GRK6,RPS16,CPSF3L,PHF21A,SLC39A11,FAM3A,MUC1,METTL17,EPB41L5,ATG12,ZBTB44,TMX
2,CD99L2,FOXRED1,KXD1,PARD6G,SH3YL1,IP6K2,CLASRP,C1ORF198,MICAL1,UBE2I,SLC39A13,PRMT2,EI
XL1,PLEKHA5,RANBP1,UBE2L6,PPM1K,ANTXR2,STIP1,LRRC1,PHF19,GRAMD1C,VCAN,MAGED2,SON,TET
XL1,PLEKHA5,RANBP1,UBE2L6,PPM1K,ANTXR2,STIP1,LRRC1,PHF19,GRAMD1C,VCAN,MAGED2,SON,TET
S,INO80E,DGUOK,ATP5J,RMDN1,GGCX,TRMT112,STRIP2,RPS20,LYNX1,DCTD,ERCC5,DDOST,LSR,CHCHD
,PARD3,TCAIM,SLC3A2,DHPS,ACTR6,MAP3K7CL,SEC62,SET,CTSC,ZNF235,BOP1,ARRDC1,TRPT1,KLC1,RB
,PARD3,TCAIM,SLC3A2,DHPS,ACTR6,MAP3K7CL,SEC62,SET,CTSC,ZNF235,BOP1,ARRDC1,TRPT1,KLC1,RB
UD31,NPRL2,TBL2,ARHGAP6,SCARB1,CTTN,TSPAN11,ITSN1,EIF5A,COMMD5,NUDT22,THYN1,DBNDD1,S
UD31,NPRL2,TBL2,ARHGAP6,SCARB1,CTTN,TSPAN11,ITSN1,EIF5A,COMMD5,NUDT22,THYN1,DBNDD1,S
RPS9,SLC35G1,ZNF410,RUVBL2,PCMT1,EBPL,TESC,RBPMS,WDR62,STYXL1,POLD4,PLEKHA5,RANBP1,LZT
RPS9,SLC35G1,ZNF410,RUVBL2,PCMT1,EBPL,TESC,RBPMS,WDR62,STYXL1,POLD4,PLEKHA5,RANBP1,LZT

1F,WAC,ANAPC11,NT5C,STXBP2,EEF1D,MED6,CHEK1,POLR2C,FKBP3,INTS4,B3GAT3,RFXANK,SPINT2,GBA
PCAT3,CD44,MAGED2,SON,GRK6,CPSF3L,OBSL1,UBR4,SLC39A11,FAM3A,MUC1,METTL17,ATG12,RPS25,
PCAT3,CD44,MAGED2,SON,GRK6,CPSF3L,OBSL1,UBR4,SLC39A11,FAM3A,MUC1,METTL17,ATG12,RPS25,
7,ZNF283,TRAPPC2,CLTA,SLC35A2,AXIN1,TP53,TMEM199,SLC25A24,HPS1,ITPA,VPS25,SNRPA1,AZI2,TATI
1,NOP16,RPL13A,ARHGEF39,PSMA6,ACOT8,RPS9,MRPL33,ZNF410,SNRPN,STYXL1,RFX5,VRK3,LZTFL1,FIS
1,NOP16,RPL13A,ARHGEF39,PSMA6,ACOT8,RPS9,MRPL33,ZNF410,SNRPN,STYXL1,RFX5,VRK3,LZTFL1,FIS
DELR3,AXIN1,TMEM199,EPN1,HPS1,ITPA,ZCCHC9,SNRPD1,AZI2,GOSR2,TATDN1,LAMTOR1,U2AF1L4
IA6,AP3B2,RPS9,SLC35G1,MRPL33,ZNF410,RUVBL2,TESC,IGSF8,NUDT5,RBPMS,NNMT,SPAG7,WDR62,ST
IA6,AP3B2,RPS9,SLC35G1,MRPL33,ZNF410,RUVBL2,TESC,IGSF8,NUDT5,RBPMS,NNMT,SPAG7,WDR62,ST
,SET,ZNF235,BOP1,ARRDC1,TRPT1,KLC1,SCAF11,ARPC2,PMFBP1,LIMK2,VDAC3,NOB1,SIRT6,NUMA1,GPA
R19,SIRT6,NUMA1,GPATCH2L,SAE1,OSGEP,TBL2,ARHGAP6,SCARB1,CRYGS,CTTN,TSPAN11,ITSN1,EIF5A,
R19,SIRT6,NUMA1,GPATCH2L,SAE1,OSGEP,TBL2,ARHGAP6,SCARB1,CRYGS,CTTN,TSPAN11,ITSN1,EIF5A,
G12,ZBTB44,TMX2,RPL29,CD99L2,GGPS1,AUP1,FOXRED1,TRNT1,ITGB1BP1,PARD6G,SH3YL1,IP6K2,CLAS
,STIP1,LRRC1,PHF19,LPCAT3,CD44,TM9SF1,SON,TET2,CDC37,CPSF3L,TSG101,OBSL1,TTC21B,UBR4,PSMA
,STIP1,LRRC1,PHF19,LPCAT3,CD44,TM9SF1,SON,TET2,CDC37,CPSF3L,TSG101,OBSL1,TTC21B,UBR4,PSMA
,MAGED2,GRK6,CPSF3L,OBSL1,PTPN13,UBR4,CHCHD1,SLC39A11,FAM3A,MUC1,EPB41L5,NLRP1,ZBTB44
,MAGED2,GRK6,CPSF3L,OBSL1,PTPN13,UBR4,CHCHD1,SLC39A11,FAM3A,MUC1,EPB41L5,NLRP1,ZBTB44
YL1,IP6K2,CLASRP,JPH2,C1ORF198,MICAL1,HS2ST1,UBE2I,TBX6,XBP1,PTK2,AUNIP,C14ORF2,PCYT2,USE1
YL1,IP6K2,CLASRP,JPH2,C1ORF198,MICAL1,HS2ST1,UBE2I,TBX6,XBP1,PTK2,AUNIP,C14ORF2,PCYT2,USE1
D3,TCAIM,SLC3A2,DHPS,ACTR6,SEC62,SET,CTSC,BOP1,ARRDC1,TRPT1,RBP5,ZC3HAV1,DBI,ARPC2,PMFB
D3,TCAIM,SLC3A2,DHPS,ACTR6,SEC62,SET,CTSC,BOP1,ARRDC1,TRPT1,RBP5,ZC3HAV1,DBI,ARPC2,PMFB
GMPR2,MCFD2,RIC8A,COMMD4,UBE2N,FAM32A,TMEM14B,C12ORF57,PEX26,NDUFS3,ZNF223,SLC35E
C9ORF64,MED8,RNMT,ANKRD10,MRPL27,DDX56,ZNF655,MED17,RNF4,URM1,SEC14L1,DPP9,SEC24C,RP
WDR62,STYXL1,VRK3,LZTFL1,FIS1,PPM1K,IRF9,ANTXR2,LRRC1,SON,SNF8,HMGA2,CDC37,CPSF3L,OBSL1,
WDR62,STYXL1,VRK3,LZTFL1,FIS1,PPM1K,IRF9,ANTXR2,LRRC1,SON,SNF8,HMGA2,CDC37,CPSF3L,OBSL1,
,CTSC,ZNF235,BOP1,ARRDC1,TRPT1,KLC1,RBP5,ZC3HAV1,DBI,ARPC2,PMFBP1,LIMK2,NOB1,WDR19,SIRT
,CTSC,ZNF235,BOP1,ARRDC1,TRPT1,KLC1,RBP5,ZC3HAV1,DBI,ARPC2,PMFBP1,LIMK2,NOB1,WDR19,SIRT
NDD1,SH2B1,NOP16,RPL13A,PSMA6,ACOT8,RPS9,SLC35G1,RCE1,RUVBL2,PCMT1,EBPL,TESC,IGSF8,NUD
NDD1,SH2B1,NOP16,RPL13A,PSMA6,ACOT8,RPS9,SLC35G1,RCE1,RUVBL2,PCMT1,EBPL,TESC,IGSF8,NUD
OB1,WDR19,SIRT6,NUMA1,GPATCH2L,BUD31,SAE1,NPRL2,OSGEP,TBL2,ARHGAP6,SCARB1,CRYGS,CTTN,
OB1,WDR19,SIRT6,NUMA1,GPATCH2L,BUD31,SAE1,NPRL2,OSGEP,TBL2,ARHGAP6,SCARB1,CRYGS,CTTN,
OP16,GSTK1,LDHA,AP3B2,RCE1,ZNF410,EBPL,RBPMS,STYXL1,RANBP1,UBE2L6,VRK3,LZTFL1,SLC15A3,IRF
OP16,GSTK1,LDHA,AP3B2,RCE1,ZNF410,EBPL,RBPMS,STYXL1,RANBP1,UBE2L6,VRK3,LZTFL1,SLC15A3,IRF
OP16,GSTK1,LDHA,AP3B2,RCE1,ZNF410,EBPL,RBPMS,STYXL1,RANBP1,UBE2L6,VRK3,LZTFL1,SLC15A3,IRF
OP16,GSTK1,LDHA,AP3B2,RCE1,ZNF410,EBPL,RBPMS,STYXL1,RANBP1,UBE2L6,VRK3,LZTFL1,SLC15A3,IRF
L1RAP,POLD4,UBE2L6,FIS1,PPM1K,STIP1,PHF19,CD44,SON,GRK6,CDC37,RPS16,TTC21B,YLPM1,PHF21A,
L1RAP,POLD4,UBE2L6,FIS1,PPM1K,STIP1,PHF19,CD44,SON,GRK6,CDC37,RPS16,TTC21B,YLPM1,PHF21A,
BOP1,ARRDC1,KLC1,RBP5,ZC3HAV1,DBI,ARPC2,PMFBP1,LIMK2,NOB1,WDR19,SIRT6,NUMA1,GPATCH2L,
L,TESC,IGSF8,SPAG7,WDR62,IL1RAP,RFX5,POLD4,RANBP1,UBE2L6,FIS1,PPM1K,STIP1,PHF19,VCAN,MAG
L,TESC,IGSF8,SPAG7,WDR62,IL1RAP,RFX5,POLD4,RANBP1,UBE2L6,FIS1,PPM1K,STIP1,PHF19,VCAN,MAG
OB1,WDR19,SIRT6,NUMA1,GPATCH2L,BUD31,SAE1,NPRL2,OSGEP,TBL2,ARHGAP6,CRYGS,CTTN,ITSN1,EI
PS16,CPSF3L,PHF21A,SLC39A11,FAM3A,MUC1,METTL17,EPB41L5,ATG12,ZBTB44,DUSP22,TMX2,NISCH,
,PPM1K,IRF9,DST,GRAMD1C,VCAN,PCBP2,LPCAT3,MAGED2,SON,UQCRQ,SNF8,HMGA2,CPSF3L,TSG101,
,PPM1K,IRF9,DST,GRAMD1C,VCAN,PCBP2,LPCAT3,MAGED2,SON,UQCRQ,SNF8,HMGA2,CPSF3L,TSG101,
SH2B1,NOP16,ARHGEF39,TTLL5,LDHA,ACOT8,AP3B2,RCE1,MRPL33,RUVBL2,EBPL,TESC,RBPMS,NNMT,SP
SH2B1,NOP16,ARHGEF39,TTLL5,LDHA,ACOT8,AP3B2,RCE1,MRPL33,RUVBL2,EBPL,TESC,RBPMS,NNMT,SP
MM40,TADA3,ATP6V1F,WAC,ANAPC11,NT5C,STXBP2,EEF1D,MED6,CARHSP1,POLR2C,INTS4,B3GAT3,RFX

PEX2,TNKS,ELOF1,DCTN5,PEX26,OXLD1,ZNF223,MPV17,ARIH2,PICK1,FBXL18,ZNF512,NTMT1,TSPAN31,Z
B1,WDR19,SIRT6,NUMA1,GPATCH2L,BUD31,SAE1,NPRL2,OSGEP,TBL2,CRYGS,ITSN1,EIF5A,COMMD5,UF
B1,WDR19,SIRT6,NUMA1,GPATCH2L,BUD31,SAE1,NPRL2,OSGEP,TBL2,CRYGS,ITSN1,EIF5A,COMMD5,UF
52,IL1RAP,PLEKHA5,RANBP1,FIS1,ANTXR2,STIP1,LPCAT3,TM9SF1,MAGED2,GRK6,HMGA2,CDC37,RPS16,
52,IL1RAP,PLEKHA5,RANBP1,FIS1,ANTXR2,STIP1,LPCAT3,TM9SF1,MAGED2,GRK6,HMGA2,CDC37,RPS16,
AC3,NOB1,WDR19,SIRT6,NUMA1,GPATCH2L,BUD31,SAE1,NPRL2,OSGEP,TBL2,ARHGAP6,SCARB1,CRYGS,
AC3,NOB1,WDR19,SIRT6,NUMA1,GPATCH2L,BUD31,SAE1,NPRL2,OSGEP,TBL2,ARHGAP6,SCARB1,CRYGS,
,GSTK1,LDHA,ACOT8,AP3B2,RPS9,SLC35G1,RCE1,MRPL33,RUVBL2,PCMT1,EBPL,TESC,IGSF8,NUDT5,RBP
,GSTK1,LDHA,ACOT8,AP3B2,RPS9,SLC35G1,RCE1,MRPL33,RUVBL2,PCMT1,EBPL,TESC,IGSF8,NUDT5,RBP
JC1,METTL17,ECSIT,ZBTB44,DUSP22,NISCH,CD99L2,GGPS1,PARD6G,SH3YL1,IP6K2,CLASRP,C1ORF198,M
D31,SAE1,OSGEP,TBL2,ARHGAP6,SCARB1,CTTN,TSPAN11,ITSN1,EIF5A,COMMD5,CHID1,NUDT22,AMZ2,T
D31,SAE1,OSGEP,TBL2,ARHGAP6,SCARB1,CTTN,TSPAN11,ITSN1,EIF5A,COMMD5,CHID1,NUDT22,AMZ2,T
OX17,PFDN5,PAFAH1B2,NSUN5,MFSD10,ZSCAN9,UBE2N,LFNG,PDCD6,FAM32A,PCID2,PEX2,C12ORF57,
OX17,PFDN5,PAFAH1B2,NSUN5,MFSD10,ZSCAN9,UBE2N,LFNG,PDCD6,FAM32A,PCID2,PEX2,C12ORF57,
GAP6,SCARB1,CTTN,TSPAN11,ITSN1,EIF5A,COMMD5,CHID1,NUDT22,AMZ2,TAZ,DBNDD1,SH2B1,RPL13
GAP6,SCARB1,CTTN,TSPAN11,ITSN1,EIF5A,COMMD5,CHID1,NUDT22,AMZ2,TAZ,DBNDD1,SH2B1,RPL13
LDHA,PSMA6,AP3B2,RPS9,SLC35G1,MRPL33,ZNF410,RUVBL2,EBPL,TESC,NUDT5,RBPMS,SNRPN,SPAG7,W
1,WDR19,SIRT6,NUMA1,GPATCH2L,BUD31,SAE1,NPRL2,OSGEP,TBL2,ARHGAP6,SCARB1,CRYGS,CTTN,ITS
BA2,PPIH,DALRD3,THADA,PDCD6,ZSCAN20,MAP3K8,USP40,PEX2,POLR1C,MDP1,C12ORF57,BTN3A2,FA
,PRMT2,PTK2,AUNIP,CENPN,PCYT2,PPA1,ENDOV,CAMTA1,PSMA7,ATPAF2,ZNF7,ARL6IP4,PIGL,PPAN,ZN
,PRMT2,PTK2,AUNIP,CENPN,PCYT2,PPA1,ENDOV,CAMTA1,PSMA7,ATPAF2,ZNF7,ARL6IP4,PIGL,PPAN,ZN
TLL5,LDHA,ACOT8,RPS9,SLC35G1,RCE1,ZNF410,RUVBL2,EBPL,TESC,NUDT5,RBPMS,SPAG7,WDR62,RANB
TLL5,LDHA,ACOT8,RPS9,SLC35G1,RCE1,ZNF410,RUVBL2,EBPL,TESC,NUDT5,RBPMS,SPAG7,WDR62,RANB
VDR19,SIRT6,NUMA1,GPATCH2L,BUD31,SAE1,NPRL2,OSGEP,TBL2,ARHGAP6,SCARB1,CRYGS,CTTN,ITSN1
TF2,C19ORF54,C19ORF48,ZNF644,HYPK,UPP1,MOB1A,RBM3,INO80E,CCNL2,CEP70,TMEM134,FAM219
1,PARD6G,CLASRP,JPH2,C1ORF198,UGGT2,MICAL1,SLC39A13,PRMT2,EIF3M,JUP,TMEM205,PTK2,GUK1
1,PARD6G,CLASRP,JPH2,C1ORF198,UGGT2,MICAL1,SLC39A13,PRMT2,EIF3M,JUP,TMEM205,PTK2,GUK1
PT1,KLC1,RBP5,ZC3HAV1,SCAF11,DBI,VDAC3,NOB1,SIRT6,NUMA1,GPATCH2L,SUMF2,BUD31,SAE1,NPRL
PT1,KLC1,RBP5,ZC3HAV1,SCAF11,DBI,VDAC3,NOB1,SIRT6,NUMA1,GPATCH2L,SUMF2,BUD31,SAE1,NPRL
1L,DBNDD1,RPL13A,ARHGEF39,PSMA6,ACOT8,AP3B2,RPS9,SLC35G1,MRPL33,ZNF410,RUVBL2,EBPL,TE
1L,DBNDD1,RPL13A,ARHGEF39,PSMA6,ACOT8,AP3B2,RPS9,SLC35G1,MRPL33,ZNF410,RUVBL2,EBPL,TE
C15A3,TUBA1C,ANTXR2,STIP1,LRRC1,PHF19,LPCAT3,CD44,TM9SF1,MAGED2,SON,TET2,UQCRQ,CDC37,
C15A3,TUBA1C,ANTXR2,STIP1,LRRC1,PHF19,LPCAT3,CD44,TM9SF1,MAGED2,SON,TET2,UQCRQ,CDC37,
,NOP16,RPL13A,GSTK1,TTLL5,LDHA,RPS9,RCE1,ZNF410,RUVBL2,PCMT1,IGSF8,NNMT,SNRPN,WDR62,ST
,NOP16,RPL13A,GSTK1,TTLL5,LDHA,RPS9,RCE1,ZNF410,RUVBL2,PCMT1,IGSF8,NNMT,SNRPN,WDR62,ST
FD1L,USP38,TAZ,DBNDD1,SH2B1,RPL13A,ARHGEF39,GSTK1,TTLL5,LDHA,PSMA6,AP3B2,RPS9,SLC35G1,
FD1L,USP38,TAZ,DBNDD1,SH2B1,RPL13A,ARHGEF39,GSTK1,TTLL5,LDHA,PSMA6,AP3B2,RPS9,SLC35G1,
MA6,ACOT8,AP3B2,RCE1,ZNF410,TESC,IGSF8,SPAG7,WDR62,IL1RAP,RFX5,POLD4,UBE2L6,ZC3H14,FIS1,S
MA6,ACOT8,AP3B2,RCE1,ZNF410,TESC,IGSF8,SPAG7,WDR62,IL1RAP,RFX5,POLD4,UBE2L6,ZC3H14,FIS1,S
,ATG16L2,RCC1,SNRPB,PARD3,TCAIM,SLC3A2,DHPS,ACTR6,MAP3K7CL,SEC62,SET,CTSC,ZNF235,BOP1,A
,ATG16L2,RCC1,SNRPB,PARD3,TCAIM,SLC3A2,DHPS,ACTR6,MAP3K7CL,SEC62,SET,CTSC,ZNF235,BOP1,A
S1,PPM1K,ANTXR2,STIP1,PHF19,LPCAT3,SON,TET2,GRK6,CPSF3L,YLPM1,PHF21A,FAM3A,MUC1,EPB41L
F39,TTLL5,LDHA,AP3B2,EBPL,IGSF8,NUDT5,RBPMS,IL1RAP,STYXL1,RFX5,POLD4,RANBP1,SLC15A3,TUBA1
IS1,TUBA1C,GRAMD1C,CD44,SON,SNF8,GRK6,RPS16,CPSF3L,TTC21B,PSMA1,PHF21A,CHCHD1,SLC39A1
PRL2,OSGEP,TBL2,ARHGAP6,CTTN,ITSN1,EIF5A,COMMD5,RUNX1,CHID1,AMZ2,UFD1L,USP38,TAZ,THYN
PRL2,OSGEP,TBL2,ARHGAP6,CTTN,ITSN1,EIF5A,COMMD5,RUNX1,CHID1,AMZ2,UFD1L,USP38,TAZ,THYN
,GPATCH2L,BUD31,NPRL2,OSGEP,TBL2,ARHGAP6,SCARB1,CTTN,TSPAN11,ITSN1,EIF5A,TNFSF10,COMM

,GPATCH2L,BUD31,NPRL2,OSGEP,TBL2,ARHGAP6,SCARB1,CTTN,TSPAN11,ITSN1,EIF5A,TNFSF10,COMMD

10,RUVBL2,PCMT1,TESC,NUDT5,RBPMS,WDR62,STYXL1,RFX5,RANBP1,UBE2L6,LZTFL1,PPM1K,DST,STIP1

9,SIRT6,NUMA1,GPATCH2L,SUMF2,BUD31,SAE1,NPRL2,OSGEP,TBL2,SCARB1,RUNX1,UFD1L,USP38,SH2B

9,SIRT6,NUMA1,GPATCH2L,SUMF2,BUD31,SAE1,NPRL2,OSGEP,TBL2,SCARB1,RUNX1,UFD1L,USP38,SH2B

L17,RPS25,COX4I1,PLCB2,RTEL1,RABL5,ITGB1BP1,MED15,CLASRP,JPH2,C1ORF198,UGGT2,MICAL1,HS2S

L17,RPS25,COX4I1,PLCB2,RTEL1,RABL5,ITGB1BP1,MED15,CLASRP,JPH2,C1ORF198,UGGT2,MICAL1,HS2S

A2,C17ORF62,ACAD10,CYB561,CBFA2T2,NFAM1,CDK11A,MDH1,ZNF197,DGUOK,TSFM,TRMT112,UBXN1

A2,C17ORF62,ACAD10,CYB561,CBFA2T2,NFAM1,CDK11A,MDH1,ZNF197,DGUOK,TSFM,TRMT112,UBXN1

GMPR2,TCP1,PIGT,UBE2N,MFSD11,PDCD6,FAM32A,ELOF1,DCTN5,PEX26,OXLD1,ZNF223,OGFOD2,ARIH

,FIS1,STIP1,PHF19,TM9SF1,SON,TET2,HMGA2,PSMA1,CHCHD1,SLC39A11,FUZ,FAM3A,PSMA3,ECSIT,EPB

F2,PCYT2,USE1,PPP1CC,RFX3,DGKE,CAMTA1,LIG1,TIMM17B,APEH,HYI,CAPN10,NUTF2,ATPAF2,C17ORF6

F2,PCYT2,USE1,PPP1CC,RFX3,DGKE,CAMTA1,LIG1,TIMM17B,APEH,HYI,CAPN10,NUTF2,ATPAF2,C17ORF6

RMT2,EIF3M,TMEM205,PTK2,AUNIP,C14ORF2,PCYT2,PPP1CC,RFX3,DGKE,CAMTA1,CAPG,CRIP1,PPIL2,C

PL28,TIMM17B,TXNL4A,LYPLA2,CAMLG,EPB41L4B,UCK2,TMPO,ATPAF2,C17ORF62,WNK1,ARL6IP4,C19

MEM135,RNF32,LY75,ATG16L2,RCC1,SNRPB,PARD3,TCAIM,SLC3A2,DHPS,ACTR6,SEC62,SET,CTSC,ZNF235

P1,PHF19,VCAN,TM9SF1,SON,SNF8,GRK6,RPS16,UBR4,YLPM1,SLC39A11,FAM3A,PSMA3,METTL17,ECSIT

RBP5,ZC3HAV1,SCAF11,DBI,ARPC2,LIMK2,NOB1,SIRT6,NUMA1,GPATCH2L,BUD31,SAE1,NPRL2,ARHGAP

RBP5,ZC3HAV1,SCAF11,DBI,ARPC2,LIMK2,NOB1,SIRT6,NUMA1,GPATCH2L,BUD31,SAE1,NPRL2,ARHGAP

1,RPS15,SAT1,ERCC5,MCTS1,PRKRIP1,CHCHD7,MEIS2,DBNL,H2AFV,RBMX,RABEP2,TADA3,WAC,NXN,B2

0,C12ORF57,DCTN5,ZNF223,OGFOD2,METTL3,EIF3G,PICK1,SLC52A2,NTMT1,TSPAN31,MTIF2,GLYCTK,CF

0,C12ORF57,DCTN5,ZNF223,OGFOD2,METTL3,EIF3G,PICK1,SLC52A2,NTMT1,TSPAN31,MTIF2,GLYCTK,CF

,EIF5A,COMMD5,NUDT22,AMZ2,USP38,TAZ,THYN1,DBNDD1,SH2B1,ARHGEF39,GSTK1,TTLL5,LDHA,AP3B

2,STYXL1,RFX5,VRK3,LZTFL1,FIS1,PPM1K,ANTXR2,LRRC1,SON,SNF8,HMGA2,RPS16,CPSF3L,OBSL1,FAM3

2,STYXL1,RFX5,VRK3,LZTFL1,FIS1,PPM1K,ANTXR2,LRRC1,SON,SNF8,HMGA2,RPS16,CPSF3L,OBSL1,FAM3

K,PCBP2,TM9SF1,TET2,UQCRQ,GRK6,CDC37,CPSF3L,TSG101,UBR4,SLC39A11,FAM3A,PSMA3,METTL17,N

K,PCBP2,TM9SF1,TET2,UQCRQ,GRK6,CDC37,CPSF3L,TSG101,UBR4,SLC39A11,FAM3A,PSMA3,METTL17,N

8,RBPMS,SPAG7,WDR62,STYXL1,PLEKHA5,RANBP1,VRK3,SLC15A3,TUBA1C,ANTXR2,STIP1,LRRC1,PHF19,

8,RBPMS,SPAG7,WDR62,STYXL1,PLEKHA5,RANBP1,VRK3,SLC15A3,TUBA1C,ANTXR2,STIP1,LRRC1,PHF19,

DGKE,CAMTA1,CRIP1,LIG1,TIMM17B,APEH,CAPN10,NUTF2,PODXL,C17ORF62,PRMT1,CASC4,ZNF7,ZFAN

TESC,RBPMS,SPAG7,WDR62,RFX5,POLD4,PLEKHA5,RANBP1,ZC3H14,FIS1,SLC15A3,PPM1K,TUBA1C,IRF9,

TESC,RBPMS,SPAG7,WDR62,RFX5,POLD4,PLEKHA5,RANBP1,ZC3H14,FIS1,SLC15A3,PPM1K,TUBA1C,IRF9,

R2,GRAMD1C,PCBP2,TM9SF1,MAGED2,SON,TET2,GRK6,CDC37,RPS16,CPSF3L,TSG101,OBSL1,FAM3A,PS

R2,GRAMD1C,PCBP2,TM9SF1,MAGED2,SON,TET2,GRK6,CDC37,RPS16,CPSF3L,TSG101,OBSL1,FAM3A,PS

,PARD3,TCAIM,SLC3A2,DHPS,ACTR6,SET,CTSC,ZNF235,BOP1,ARRDC1,TRPT1,KLC1,RBP5,SCAF11,DBI,ARP

,PARD3,TCAIM,SLC3A2,DHPS,ACTR6,SET,CTSC,ZNF235,BOP1,ARRDC1,TRPT1,KLC1,RBP5,SCAF11,DBI,ARP

L7,EPB41L5,ATG12,ZBTB44,CD99L2,FOXRED1,TRNT1,PARD6G,SH3YL1,IP6K2,UBE2I,SLC39A13,PRMT2,CD

RQ,CDC37,RPS16,PTPN13,FUZ,FAM3A,PPRC1,NIP7,COX4I1,DUSP22,DBNDD2,RPL29,GGPS1,RTEL1,SRSF1

MA6,AP3B2,RPS9,RCE1,NUDT5,RBPMS,PLEKHA5,RANBP1,FIS1,ANTXR2,STIP1,LPCAT3,TM9SF1,MAGED2,

MA6,AP3B2,RPS9,RCE1,NUDT5,RBPMS,PLEKHA5,RANBP1,FIS1,ANTXR2,STIP1,LPCAT3,TM9SF1,MAGED2,

P1,FOXRED1,SH3YL1,HS2ST1,YIF1A,PRMT2,EIF3M,MAGT1,TBX6,TMEM205,AUNIP,GUK1,ATF7,EEF1B2,P

YL1,HS2ST1,YIF1A,EIF3M,TBX6,TMEM205,AUNIP,GUK1,FBXW9,PPP1CC,RFX3,ENDOV,PSMD4,VIM,LIG1,P

2,AMZ2,UFD1L,TAZ,DBNDD1,SH2B1,NOP16,TTLL5,PSMA6,ACOT8,AP3B2,RCE1,MRPL33,RUVBL2,PCMT1,

2,AMZ2,UFD1L,TAZ,DBNDD1,SH2B1,NOP16,TTLL5,PSMA6,ACOT8,AP3B2,RCE1,MRPL33,RUVBL2,PCMT1,

12ORF57,NDUFS3,ARL16,OXLD1,METTL5,SSR3,ARIH2,PICK1,TPT1,TSPAN31,MTIF2,RALY,GLYCTK,PCNP,CF

12ORF57,NDUFS3,ARL16,OXLD1,METTL5,SSR3,ARIH2,PICK1,TPT1,TSPAN31,MTIF2,RALY,GLYCTK,PCNP,C
MMD5,RUNX1,CHID1,NUDT22,AMZ2,UFD1L,USP38,TAZ,THYN1,DBNDD1,GSTK1,LDHA,PSMA6,AP3B2,SLC
MMD5,RUNX1,CHID1,NUDT22,AMZ2,UFD1L,USP38,TAZ,THYN1,DBNDD1,GSTK1,LDHA,PSMA6,AP3B2,SLC
1,KXD1,MED15,SH3YL1,IP6K2,MICAL1,UBE2I,EIF3M,PTK2,CENPN,PCYT2,USE1,ASF1B,RFX3,DGKE,PPIL2,R
,COX17,HIGD1A,B3GAT3,PFDN5,PAFAH1B2,GBA2,NSUN5,MFSD10,GMPR2,ZSCAN9,RIC8A,UBE2N,ZNHIT
D1,ZNF223,SLC35E1,OGFOD2,TSEN34,ARIH2,PICK1,SLC52A2,TSPAN31,ZNF593,GLYCTK,RER1,GTF2H1,KD
LL5,ACOT8,AP3B2,SLC35G1,MRPL33,ZNF410,RUVBL2,PCMT1,EBPL,TESC,IGSF8,NUDT5,RBPMS,NNMT,SN
LL5,ACOT8,AP3B2,SLC35G1,MRPL33,ZNF410,RUVBL2,PCMT1,EBPL,TESC,IGSF8,NUDT5,RBPMS,NNMT,SN
HA5,RANBP1,VRK3,FIS1,SLC15A3,TUBA1C,GRAMD1C,VCAN,LPCAT3,UQCRQ,SNF8,GRK6,HMGA2,CDC37,

,TESC,WDR62,STYXL1,PLEKHA5,RANBP1,LZTFL1,ZC3H14,FIS1,STIP1,LRRC1,PHF19,LPCAT3,CD44,TM9SF1,
,TESC,WDR62,STYXL1,PLEKHA5,RANBP1,LZTFL1,ZC3H14,FIS1,STIP1,LRRC1,PHF19,LPCAT3,CD44,TM9SF1,
IRNPC,CYB561,STAU1,CDK11A,ZNF197,DGUOK,RAB2A,ZNF524,ATP5J,TRMT112,MPDU1,UBXN1,SAT1,EN
IRNPC,CYB561,STAU1,CDK11A,ZNF197,DGUOK,RAB2A,ZNF524,ATP5J,TRMT112,MPDU1,UBXN1,SAT1,EN
F410,RUVBL2,PCMT1,EBPL,TESC,IGSF8,NUDT5,RBPMS,NNMT,SNRPN,WDR62,STYXL1,RFX5,RANBP1,UBE
COASY,TRNT1,KXD1,ITGB1BP1,C1ORF198,UGGT2,HS2ST1,YIF1A,TMEM205,C14ORF2,GUK1,ATF7,MAF,EE
SCH,CD99L2,GGPS1,AUP1,PARD6G,SH3YL1,IP6K2,CLASRP,MICAL1,HS2ST1,UBE2I,PRMT2,AUNIP,C14ORF
UD31,NPRL2,TBL2,ARHGAP6,SCARB1,CTTN,TSPAN11,ITSN1,EIF5A,COMMD5,NUDT22,UFD1L,DBNDD1,S
UD31,NPRL2,TBL2,ARHGAP6,SCARB1,CTTN,TSPAN11,ITSN1,EIF5A,COMMD5,NUDT22,UFD1L,DBNDD1,S
11,DBI,ARPC2,LIMK2,SIRT6,NUMA1,GPATCH2L,SUMF2,BUD31,SAE1,NPRL2,OSGEP,ARHGAP6,SCARB1,CT
11,DBI,ARPC2,LIMK2,SIRT6,NUMA1,GPATCH2L,SUMF2,BUD31,SAE1,NPRL2,OSGEP,ARHGAP6,SCARB1,CT
PRL2,TBL2,ARHGAP6,SCARB1,CTTN,TSPAN11,ITSN1,EIF5A,COMMD5,CHID1,NUDT22,AMZ2,UFD1L,USP38
PRL2,TBL2,ARHGAP6,SCARB1,CTTN,TSPAN11,ITSN1,EIF5A,COMMD5,CHID1,NUDT22,AMZ2,UFD1L,USP38
62,ZFAND2B,SHARPIN,UPP1,RPL36,LAMP2,SYNGR2,TMEM134,LRRFIP1,RAB2A,ZNF524,ASPH,LYNX1,TW
SGEP,TBL2,SCARB1,EIF5A,THYN1,DBNDD1,SH2B1,NOP16,RPL13A,ARHGEF39,GSTK1,LDHA,PSMA6,ACOT
P7,DUSP22,SPOCK2,DBNDD2,CENPK,RPS6,COASY,AUP1,SAP30BP,FOXRED1,TRNT1,KXD1,MED15,UGGT2,
MK2,VDAC3,NOB1,SIRT6,NUMA1,SUMF2,BUD31,SAE1,OSGEP,TBL2,ARHGAP6,SCARB1,CRYGS,CTTN,TSP
MK2,VDAC3,NOB1,SIRT6,NUMA1,SUMF2,BUD31,SAE1,OSGEP,TBL2,ARHGAP6,SCARB1,CRYGS,CTTN,TSP
STX10,C12ORF57,DCTN5,ARL16,OXLD1,ZNF223,STAU2,OGFOD2,PICK1,ZNF512,RNF181,TSPAN31,SLC39
STX10,C12ORF57,DCTN5,ARL16,OXLD1,ZNF223,STAU2,OGFOD2,PICK1,ZNF512,RNF181,TSPAN31,SLC39

,HMGA2,OBSL1,TTC21B,PTPN13,PSMA1,YLPM1,PSMA3,ECSIT,RPS25,TTC9C,SLC25A10,TMX2,THEM4,RD
,HMGA2,OBSL1,TTC21B,PTPN13,PSMA1,YLPM1,PSMA3,ECSIT,RPS25,TTC9C,SLC25A10,TMX2,THEM4,RD
DC,NDUFS3,NUBP2,TSEN34,MRPL55,CREB3L4,TCF25,TJP1,CDK4,RALY,FAM69B,DOK4,NUP62,PSMD7,ZN
LDHA,AP3B2,RPS9,RCE1,ZNF410,RUVBL2,EBPL,TESC,NUDT5,RBPMS,WDR62,IL1RAP,PLEKHA5,RANBP1,Z
LDHA,AP3B2,RPS9,RCE1,ZNF410,RUVBL2,EBPL,TESC,NUDT5,RBPMS,WDR62,IL1RAP,PLEKHA5,RANBP1,Z
H14,TUBA1C,IRF9,LRRC1,GRAMD1C,MAGED2,SON,UQCRQ,CPSF3L,PHF21A,CHCHD1,SLC39A11,MUC1,EP
H14,TUBA1C,IRF9,LRRC1,GRAMD1C,MAGED2,SON,UQCRQ,CPSF3L,PHF21A,CHCHD1,SLC39A11,MUC1,EP
HA,PSMA6,AP3B2,SLC35G1,RCE1,MRPL33,ZNF410,PCMT1,TESC,IGSF8,RBPMS,NNMT,SPAG7,WDR62,IL1
2,PSMD8,SETD4,MED8,EIF6,DHX30,AP4M1,MRPL27,CHFR,SSH3,DHRS4L2,RPS14,RNF4,MAGOH,ZRANB2
ANTXR2,LRRC1,PHF19,LPCAT3,CD44,TM9SF1,MAGED2,SON,TET2,UQCRQ,GRK6,CPSF3L,OBSL1,YLPM1,C
ANTXR2,LRRC1,PHF19,LPCAT3,CD44,TM9SF1,MAGED2,SON,TET2,UQCRQ,GRK6,CPSF3L,OBSL1,YLPM1,C
RCE1,MRPL33,ZNF410,RUVBL2,PCMT1,EBPL,TESC,IGSF8,NNMT,SNRPN,WDR62,IL1RAP,STYXL1,PLEKHA5
RCE1,MRPL33,ZNF410,RUVBL2,PCMT1,EBPL,TESC,IGSF8,NNMT,SNRPN,WDR62,IL1RAP,STYXL1,PLEKHA5
A1C,STIP1,LRRC1,PHF19,LPCAT3,MAGED2,SNF8,CDC37,OBSL1,TTC21B,CHCHD1,MUC1,METTL17,EPB41L
A1C,STIP1,LRRC1,PHF19,LPCAT3,MAGED2,SNF8,CDC37,OBSL1,TTC21B,CHCHD1,MUC1,METTL17,EPB41L
G1,ZNF410,IGSF8,NUDT5,RBPMS,NNMT,SPAG7,WDR62,STYXL1,RANBP1,VRK3,LZTFL1,ZC3H14,FIS1,SLC1

G1,ZNF410,IGSF8,NUDT5,RBPMS,NNMT,SPAG7,WDR62,STYXL1,RANBP1,VRK3,LZTFL1,ZC3H14,FIS1,SLC1

LC15A3,TUBA1C,ANTXR2,STIP1,LPCAT3,TET2,HMGA2,RPS16,OBSL1,TTC21B,UBR4,PHF21A,ATG12,DUSP
LC15A3,TUBA1C,ANTXR2,STIP1,LPCAT3,TET2,HMGA2,RPS16,OBSL1,TTC21B,UBR4,PHF21A,ATG12,DUSP
B1,NOP16,RPL13A,TTLL5,ACOT8,SLC35G1,RCE1,EBPL,NUDT5,WDR62,IL1RAP,RANBP1,UBE2L6,VRK3,ZC3
B1,NOP16,RPL13A,TTLL5,ACOT8,SLC35G1,RCE1,EBPL,NUDT5,WDR62,IL1RAP,RANBP1,UBE2L6,VRK3,ZC3
R62,PLEKHA5,RANBP1,FIS1,PPM1K,DST,ANTXR2,STIP1,PHF19,LPCAT3,CD44,TM9SF1,MAGED2,SON,TET
,PCMT1,EBPL,TESC,IGSF8,RBPMS,SPAG7,WDR62,IL1RAP,STYXL1,RFX5,POLD4,PLEKHA5,RANBP1,UBE2L6,
SGEP,SCARB1,CTTN,TSPAN11,ITSN1,EIF5A,CHID1,NUDT22,AMZ2,UFD1L,USP38,GSTK1,LDHA,PSMA6,AP
SGEP,SCARB1,CTTN,TSPAN11,ITSN1,EIF5A,CHID1,NUDT22,AMZ2,UFD1L,USP38,GSTK1,LDHA,PSMA6,AP
ED2,TET2,GRK6,CDC37,CPSF3L,TSG101,UBR4,SLC39A11,FAM3A,PSMA3,METTL17,RPS25,NLRP1,NIP7,TT
ED2,TET2,GRK6,CDC37,CPSF3L,TSG101,UBR4,SLC39A11,FAM3A,PSMA3,METTL17,RPS25,NLRP1,NIP7,TT

5A3,ANTXR2,STIP1,PHF19,VCAN,LPCAT3,CD44,TM9SF1,SON,TET2,HMGA2,CDC37,TSG101,UBR4,PSMA1,
JUP,TBX6,TMEM205,PTK2,AUNIP,C14ORF2,GUK1,PCYT2,USE1,PPP1CC,PPA1,DGKE,CAMTA1,LIG1,TIMM
T1,TBX6,TMEM205,AUNIP,GUK1,CENPN,PPP1CC,RFX3,ENDOV,PSMD4,VIM,LIG1,PSMA7,TMPO,LSM12,P
HGEF39,LDHA,ACOT8,RPS9,SLC35G1,RCE1,RUVBL2,IGSF8,NUDT5,RBPMS,NNMT,WDR62,IL1RAP,POLD4,
RRDC1,KLC1,RBP5,ZC3HAV1,SCAF11,DBI,ARPC2,PMFBP1,LIMK2,VDAC3,NOB1,WDR19,NUMA1,GPATCH
RRDC1,KLC1,RBP5,ZC3HAV1,SCAF11,DBI,ARPC2,PMFBP1,LIMK2,VDAC3,NOB1,WDR19,NUMA1,GPATCH
2A2,TSPAN31,FAM192A,GLYCTK,CFLAR,TRIM46,EIF2A,NKIRAS2,ACAA1,GTF2H1,BFAR,BTF3,TMEM199,S
2A2,TSPAN31,FAM192A,GLYCTK,CFLAR,TRIM46,EIF2A,NKIRAS2,ACAA1,GTF2H1,BFAR,BTF3,TMEM199,S
L,SAE1,TBL2,ARHGAP6,SCARB1,CTTN,TSPAN11,ITSN1,EIF5A,TNFSF10,COMMD5,RUNX1,CHID1,NUDT22,
L,SAE1,TBL2,ARHGAP6,SCARB1,CTTN,TSPAN11,ITSN1,EIF5A,TNFSF10,COMMD5,RUNX1,CHID1,NUDT22,
COMMD5,NUDT22,THYN1,SH2B1,GSTK1,TTLL5,LDHA,PSMA6,ACOT8,SLC35G1,RCE1,ZNF410,PCMT1,RBP
2,USE1,AGPAT4,PPP1CC,RFX3,PPA1,DGKE,CAMTA1,CRIP1,VIM,LIG1,TIMM17B,APEH,NUTF2,PODXL,CAM
,SPINT2,SMARCA4,MPC2,MCFD2,CBR1,PHC3,MFSD11,MOK,ZNF81,TMEM14B,RPL37A,ELOF1,DCTN5,DG
,SPINT2,SMARCA4,MPC2,MCFD2,CBR1,PHC3,MFSD11,MOK,ZNF81,TMEM14B,RPL37A,ELOF1,DCTN5,DG
RF198,MICAL1,HS2ST1,UBE2I,TBX6,AUNIP,C14ORF2,GUK1,PCYT2,USE1,PPP1CC,EDF1,PPA1,DGKE,CAMT
ARHSP1,CNST,FKBP3,RPL15,PAFAH1B2,NSUN5,ZFPL1,GMPR2,PIGT,UBE2N,FAM32A,TMEM14B,PEX2,DC
P,STYXL1,FIS1,SLC15A3,TUBA1C,LRRC1,PHF19,TM9SF1,SON,SNF8,GRK6,RPS16,OBSL1,PTPN13,YLPM1,EP
1K,TUBA1C,STIP1,LRRC1,PHF19,GRAMD1C,LPCAT3,SON,TET2,GRK6,OBSL1,PTPN13,UBR4,YLPM1,PHF21
1K,TUBA1C,STIP1,LRRC1,PHF19,GRAMD1C,LPCAT3,SON,TET2,GRK6,OBSL1,PTPN13,UBR4,YLPM1,PHF21
1,ARRDC1,TRPT1,KLC1,RBP5,ZC3HAV1,SCAF11,DBI,ARPC2,LIMK2,VDAC3,WDR19,SIRT6,NUMA1,SUMF2,
1,ARRDC1,TRPT1,KLC1,RBP5,ZC3HAV1,SCAF11,DBI,ARPC2,LIMK2,VDAC3,WDR19,SIRT6,NUMA1,SUMF2,
YNGR1,PABPC1,NAT9,TOMM40,WAC,PDCD5,ANAPC11,NOMO2,USP45,ILF3,EEF1D,CASZ1,SSH2,COX17,
YNGR1,PABPC1,NAT9,TOMM40,WAC,PDCD5,ANAPC11,NOMO2,USP45,ILF3,EEF1D,CASZ1,SSH2,COX17,
MPV17,SLC52A2,TSPAN31,APOLD1,ZNF593,GLYCTK,CFLAR,TRIM46,EIF2A,GLI4,NKIRAS2,ACAA1,GTF2H1,
MPV17,SLC52A2,TSPAN31,APOLD1,ZNF593,GLYCTK,CFLAR,TRIM46,EIF2A,GLI4,NKIRAS2,ACAA1,GTF2H1,

L1,FIS1,SLC15A3,TUBA1C,STIP1,LPCAT3,CD44,TET2,UQCRQ,SNF8,GRK6,CDC37,OBSL1,SLC39A11,RGS3,M
L1,FIS1,SLC15A3,TUBA1C,STIP1,LPCAT3,CD44,TET2,UQCRQ,SNF8,GRK6,CDC37,OBSL1,SLC39A11,RGS3,M
1,RGS3,FAM3A,MUC1,ATG12,PPRC1,SLC25A10,SPOCK2,SDF4,NISCH,RPL29,AUP1,FOXRED1,TRNT1,KXD1
1,RGS3,FAM3A,MUC1,ATG12,PPRC1,SLC25A10,SPOCK2,SDF4,NISCH,RPL29,AUP1,FOXRED1,TRNT1,KXD1
UBE2I,EIF3M,TBX6,PTK2,C14ORF2,CENPN,HSBP1L1,EDF1,RFX3,DGKE,CAMTA1,CLDND1,VIM,LIG1,PSMA7,
BE2I,EIF3M,TBX6,PTK2,C14ORF2,CENPN,HSBP1L1,EDF1,RFX3,DGKE,CAMTA1,CLDND1,VIM,LIG1,PSMA7,

OT8,AP3B2,RPS9,ZNF410,RUVBL2,IGSF8,NUDT5,RBPMS,SNRPN,STYXL1,ZC3H14,FIS1,TUBA1C,IRF9,DST,S

OT8,AP3B2,RPS9,ZNF410,RUVBL2,IGSF8,NUDT5,RBPMS,SNRPN,STYXL1,ZC3H14,FIS1,TUBA1C,IRF9,DST,S

6,ITSN1,EIF5A,COMMD5,RUNX1,UFD1L,TAZ,DBNDD1,NOP16,RPL13A,GSTK1,TTLL5,ACOT8,AP3B2,SLC35

6,ITSN1,EIF5A,COMMD5,RUNX1,UFD1L,TAZ,DBNDD1,NOP16,RPL13A,GSTK1,TTLL5,ACOT8,AP3B2,SLC35

24,ATP5J,GPBP1,SIAH1,TSFM,TRMT112,CHPF,VPS28,DCTD,UBXN1,SAT1,ERCC5,SERPINB1,DDOST,LSR,PS

L,SLC9B2,ZNF250,RBM3,HSP90AB1,LRRFIP1,ATP5J,RMDN1,ZEB2,TRMT112,ZNF625,RPS20,ERLEC1,EMC4,

SMA1,PHF21A,CHCHD1,METTL17,PPRC1,RPS25,NIP7,COX4I1,TMX2,SPOCK2,RDH11,DBNDD2,SRSF1,AUI

,UQCRQ,YLPM1,CHCHD1,FUZ,FAM3A,PPRC1,RPS25,NIP7,COX4I1,SPOCK2,CENPK,GGPS1,RTEL1,COASY,S

,UQCRQ,YLPM1,CHCHD1,FUZ,FAM3A,PPRC1,RPS25,NIP7,COX4I1,SPOCK2,CENPK,GGPS1,RTEL1,COASY,S

A1,BUD31,SAE1,OSGEP,TBL2,ARHGAP6,SCARB1,CTTN,ITSN1,TNFSF10,RUNX1,CHID1,AMZ2,UFD1L,USP3

A1,BUD31,SAE1,OSGEP,TBL2,ARHGAP6,SCARB1,CTTN,ITSN1,TNFSF10,RUNX1,CHID1,AMZ2,UFD1L,USP3

YL1,CLASRP,JPH2,UGGT2,HS2ST1,CDC42,TBX6,TMEM205,XBP1,AUNIP,C14ORF2,CENPN,ATF7,HSBP1L1,A

MTA1,CRIP1,LIG1,TIMM17B,APEH,CAPN10,NUTF2,PODXL,ATPAF2,C17ORF62,PRMT1,CASC4,ZNF7,ZFAND

MTA1,CRIP1,LIG1,TIMM17B,APEH,CAPN10,NUTF2,PODXL,ATPAF2,C17ORF62,PRMT1,CASC4,ZNF7,ZFAND

ED2,SNF8,CDC37,TTC21B,CHCHD1,MUC1,EPB41L5,ZBTB44,NIP7,COX4I1,DUSP22,SLC25A10,SPOCK2,THE

RQ,SNF8,GRK6,HMGA2,CDC37,TTC21B,UBR4,PSMA1,SLC39A11,PSMA3,PPRC1,PLCB2,SLC25A10,SDF4,TM

RQ,SNF8,GRK6,HMGA2,CDC37,TTC21B,UBR4,PSMA1,SLC39A11,PSMA3,PPRC1,PLCB2,SLC25A10,SDF4,TM

ED1,PARD6G,SH3YL1,IP6K2,CLASRP,MICAL1,HS2ST1,UBE2I,TMEM205,AUNIP,C14ORF2,GUK1,PCYT2,PPF

R,TRIM46,EIF2A,NKIRAS2,ACAA1,GTF2H1,BFAR,BTF3,TMEM199,SLC25A24,ZNF396,EPN1,ZCCHC9,CHUR

R,TRIM46,EIF2A,NKIRAS2,ACAA1,GTF2H1,BFAR,BTF3,TMEM199,SLC25A24,ZNF396,EPN1,ZCCHC9,CHUR

GRK6,OBSL1,UBR4,YLPM1,PHF21A,SLC39A11,FAM3A,MUC1,ECSIT,EPB41L5,PPRC1,RPS25,NLRP1,ZBTB44

GRK6,OBSL1,UBR4,YLPM1,PHF21A,SLC39A11,FAM3A,MUC1,ECSIT,EPB41L5,PPRC1,RPS25,NLRP1,ZBTB44

UDT22,AMZ2,USP38,THYN1,DBNDD1,SH2B1,NOP16,TTLL5,LDHA,PSMA6,ACOT8,RPS9,SLC35G1,RCE1,ZN

UDT22,AMZ2,USP38,THYN1,DBNDD1,SH2B1,NOP16,TTLL5,LDHA,PSMA6,ACOT8,RPS9,SLC35G1,RCE1,ZN

A,PSMA6,ACOT8,RPS9,SLC35G1,RCE1,PCMT1,TESC,NNMT,SPAG7,WDR62,STYXL1,POLD4,PLEKHA5,RANBI

A,PSMA6,ACOT8,RPS9,SLC35G1,RCE1,PCMT1,TESC,NNMT,SPAG7,WDR62,STYXL1,POLD4,PLEKHA5,RANBI

1,CTTN,TSPAN11,ITSN1,EIF5A,COMMD5,CHID1,NUDT22,USP38,TAZ,THYN1,SH2B1,RPL13A,TTLL5,PSMA6

1,CTTN,TSPAN11,ITSN1,EIF5A,COMMD5,CHID1,NUDT22,USP38,TAZ,THYN1,SH2B1,RPL13A,TTLL5,PSMA6

UBA1C,ANTXR2,LRRC1,PHF19,LPCAT3,CD44,TM9SF1,SON,TET2,UQCRQ,CPSF3L,OBSL1,YLPM1,CHCHD1,F

UBA1C,ANTXR2,LRRC1,PHF19,LPCAT3,CD44,TM9SF1,SON,TET2,UQCRQ,CPSF3L,OBSL1,YLPM1,CHCHD1,F

HGAP6,CRYGS,CTTN,ITSN1,EIF5A,COMMD5,RUNX1,NUDT22,AMZ2,UFD1L,USP38,TAZ,DBNDD1,SH2B1,N

HGAP6,CRYGS,CTTN,ITSN1,EIF5A,COMMD5,RUNX1,NUDT22,AMZ2,UFD1L,USP38,TAZ,DBNDD1,SH2B1,N

SZ1,SSH2,COX17,HINT1,HNRNPA1,EGLN2,SIRT5,RFXANK,SPINT2,SMARCA4,MPC2,OARD1,MRPS18C,MCI

SZ1,SSH2,COX17,HINT1,HNRNPA1,EGLN2,SIRT5,RFXANK,SPINT2,SMARCA4,MPC2,OARD1,MRPS18C,MCI

T3,CD44,SON,TET2,HMGA2,CDC37,CPSF3L,TTC21B,PSMA1,FUZ,PSMA3,METTL17,ZBTB44,DUSP22,SLC25

NK1,C19ORF54,ZNF7,PPAN,HNRNPC,ZNF644,ZNF250,CCNL2,LRRFIP1,ZNF524,SIAH1,CHPF,STRIP2,RPS20,

ETTL17,EPB41L5,PPRC1,ZBTB44,DUSP22,RTEL1,SRSF1,YIPF3,UGGT2,HS2ST1,UBE2I,PRMT2,TBX6,TMEM2

FD1L,THYN1,DBNDD1,SH2B1,NOP16,RPL13A,LDHA,AP3B2,RPS9,RCE1,MRPL33,ZNF410,PCMT1,EBPL,TE

FD1L,THYN1,DBNDD1,SH2B1,NOP16,RPL13A,LDHA,AP3B2,RPS9,RCE1,MRPL33,ZNF410,PCMT1,EBPL,TE

6,CPSF3L,YLPM1,PHF21A,SLC39A11,MUC1,METTL17,EPB41L5,ATG12,ZBTB44,PLCB2,DUSP22,DBNDD2,C

P1,VRK3,PPM1K,PHF19,PCBP2,TM9SF1,MAGED2,SON,TET2,GRK6,CDC37,CPSF3L,TSG101,OBSL1,UBR4,F

P1,VRK3,PPM1K,PHF19,PCBP2,TM9SF1,MAGED2,SON,TET2,GRK6,CDC37,CPSF3L,TSG101,OBSL1,UBR4,F

,EIF5A,CHID1,NUDT22,UFD1L,TAZ,DBNDD1,SH2B1,RPL13A,ARHGEF39,GSTK1,ACOT8,SLC35G1,ZNF410,P

,EIF5A,CHID1,NUDT22,UFD1L,TAZ,DBNDD1,SH2B1,RPL13A,ARHGEF39,GSTK1,ACOT8,SLC35G1,ZNF410,P

,PPIL2,LIG1,CDC14B,TIMM17B,APEH,HYI,CAPN10,NUTF2,ATPAF2,C17ORF62,ZNF7,PPAN,ZFAND2B,RPL3

NBP1,LZTFL1,FIS1,TUBA1C,IRF9,STIP1,LPCAT3,TET2,UQCRQ,GRK6,HMGA2,CDC37,TTC21B,UBR4,SLC39A1

E1,NPRL2,OSGEP,TBL2,ARHGAP6,SCARB1,CTTN,TSPAN11,ITSN1,EIF5A,COMMD5,AMZ2,UFD1L,USP38,TA
E1,NPRL2,OSGEP,TBL2,ARHGAP6,SCARB1,CTTN,TSPAN11,ITSN1,EIF5A,COMMD5,AMZ2,UFD1L,USP38,TA
RF2,GUK1,PCYT2,PPP1CC,EDF1,RFX3,PPA1,DGKE,VAMP8,CAMTA1,CLDND1,LIG1,TIMM17B,APEH,HYI,CAI
L3,EIF3G,PICK1,SLC52A2,NTMT1,TSPAN31,ZNF593,GLYCTK,CFLAR,TRIM46,EIF2A,NKIRAS2,ACAA1,GTF2H
L3,EIF3G,PICK1,SLC52A2,NTMT1,TSPAN31,ZNF593,GLYCTK,CFLAR,TRIM46,EIF2A,NKIRAS2,ACAA1,GTF2H

,LZTFL1,FIS1,SLC15A3,DST,STIP1,LRRC1,PHF19,GRAMD1C,VCAN,CD44,TM9SF1,MAGED2,SON,GRK6,HM
HARPIN,UPP1,NFAM1,CDK11A,RBM3,LAMP2,SYNGR2,ZNF524,ASPH,TRMT112,RPTOR,TWSG1,MAPK13,V
HARPIN,UPP1,NFAM1,CDK11A,RBM3,LAMP2,SYNGR2,ZNF524,ASPH,TRMT112,RPTOR,TWSG1,MAPK13,V
P90AB1,CEP70,RAB2A,GPBP1,GPALPP1,PNPO,CHPF,NIT2,ZSWIM7,GMPPB,MVD,IL32,SYNGR1,RANGRF,R
6,TMEM205,PTK2,AUNIP,C14ORF2,PCYT2,USE1,PPP1CC,DGKE,CAMTA1,RPL28,LIG1,TIMM17B,APEH,HYI,
6,TMEM205,PTK2,AUNIP,C14ORF2,PCYT2,USE1,PPP1CC,DGKE,CAMTA1,RPL28,LIG1,TIMM17B,APEH,HYI,
16,TTC21B,PSMA1,FAM3A,ECSIT,ATG12,PPRC1,RPS25,NIP7,TTC9C,COX4I1,TMX2,SPOCK2,RDH11,TMED4
16,TTC21B,PSMA1,FAM3A,ECSIT,ATG12,PPRC1,RPS25,NIP7,TTC9C,COX4I1,TMX2,SPOCK2,RDH11,TMED4
RPC2,LIMK2,VDAC3,WDR19,SIRT6,NUMA1,GPATCH2L,SUMF2,BUD31,SAE1,OSGEP,TBL2,ARHGAP6,SCAR
RPC2,LIMK2,VDAC3,WDR19,SIRT6,NUMA1,GPATCH2L,SUMF2,BUD31,SAE1,OSGEP,TBL2,ARHGAP6,SCAR
L,TAZ,DBNDD1,NOP16,RPL13A,GSTK1,TTLL5,ACOT8,AP3B2,SLC35G1,MRPL33,ZNF410,RUVBL2,PCMT1,E
YNGR2,TMEM134,LRRFIP1,RAB2A,ZNF524,ASPH,LYNX1,TWSG1,HSPB1,LRRC27,LMBR1,CHCHD7,PSME2,
25,COX4I1,THEM4,RDH11,NISCH,TMED4,AUP1,SAP30BP,FOXRED1,RABL5,TRNT1,MED15,SH3YL1,IP6K2,J
5,DGKE,PSMD4,LIG1,PSMA7,APEH,CAPN10,NUTF2,PODXL,LSM12,C19ORF54,PIGL,PPAN,CYB561,SLC9B2
3A,SLC16A1,ILF3,HSPH1,RPL18,CNST,SUMO3,PHF20L1,HNRNPA1,NDUFV2,EGLN2,B3GAT3,SPINT2,TYMS,
KLC1,ZC3HAV1,SCAF11,ARPC2,LIMK2,VDAC3,NOB1,WDR19,SIRT6,NUMA1,GPATCH2L,SUMF2,BUD31,SA
KLC1,ZC3HAV1,SCAF11,ARPC2,LIMK2,VDAC3,NOB1,WDR19,SIRT6,NUMA1,GPATCH2L,SUMF2,BUD31,SA
MRPL33,EBPL,NUDT5,RBPMS,WDR62,RANBP1,SLC15A3,TUBA1C,DST,ANTXR2,LRRC1,VCAN,TM9SF1,SON,
A,PSMA3,METTL17,NLRP1,NIP7,TTC9C,COX4I1,THEM4,CENPK,COASY,SRSF1,AUP1,SAP30BP,FOXRED1,TR
A,PSMA3,METTL17,NLRP1,NIP7,TTC9C,COX4I1,THEM4,CENPK,COASY,SRSF1,AUP1,SAP30BP,FOXRED1,TR
4,TUBA1C,ANTXR2,STIP1,LRRC1,CD44,MAGED2,SON,SNF8,CDC37,CPSF3L,OBSL1,CHCHD1,RGS3,MUC1,M
4,TUBA1C,ANTXR2,STIP1,LRRC1,CD44,MAGED2,SON,SNF8,CDC37,CPSF3L,OBSL1,CHCHD1,RGS3,MUC1,M
AP30BP,YIPF3,FOXRED1,KXD1,SH3YL1,CLASRP,C1ORF198,MICAL1,HS2ST1,YIF1A,MAGT1,CARD14,USE1,E
AP30BP,YIPF3,FOXRED1,KXD1,SH3YL1,CLASRP,C1ORF198,MICAL1,HS2ST1,YIF1A,MAGT1,CARD14,USE1,E
DF4,COASY,SRSF1,FOXRED1,TRNT1,MED15,SH3YL1,C1ORF198,UGGT2,MICAL1,SLC39A13,YIF1A,PRMT2,M
1,PPA1,DGKE,CAMTA1,LIG1,TIMM17B,APEH,HYI,CAPN10,NUTF2,PODXL,C17ORF62,PRMT1,CASC4,ZNF7,
98,MICAL1,JUP,TBX6,XBP1,PTK2,C14ORF2,CENPN,PCYT2,HSBP1L1,DEF8,PPP1CC,EDF1,RFX3,DGKE,CAMT
98,MICAL1,JUP,TBX6,XBP1,PTK2,C14ORF2,CENPN,PCYT2,HSBP1L1,DEF8,PPP1CC,EDF1,RFX3,DGKE,CAMT
ST1,UBE2I,SLC39A13,PTK2,AUNIP,C14ORF2,PCYT2,USE1,PPP1CC,EDF1,RFX3,PPA1,DGKE,CAMTA1,CRIP1,
VCAN,LPCAT3,SON,TET2,UQCRQ,PSMA1,PHF21A,FAM3A,MUC1,EPB41L5,ATG12,NLRP1,NIP7,TTC9C,CO
VCAN,LPCAT3,SON,TET2,UQCRQ,PSMA1,PHF21A,FAM3A,MUC1,EPB41L5,ATG12,NLRP1,NIP7,TTC9C,CO
GAP6,SCARB1,CTTN,TSPAN11,ITSN1,EIF5A,COMMD5,RUNX1,CHID1,NUDT22,AMZ2,UFD1L,USP38,SH2B1,
GAP6,SCARB1,CTTN,TSPAN11,ITSN1,EIF5A,COMMD5,RUNX1,CHID1,NUDT22,AMZ2,UFD1L,USP38,SH2B1,
5,NUMA1,SUMF2,BUD31,SAE1,NPRL2,OSGEP,TBL2,ARHGAP6,SCARB1,CRYGS,CTTN,TSPAN11,ITSN1,EIF5
5,NUMA1,SUMF2,BUD31,SAE1,NPRL2,OSGEP,TBL2,ARHGAP6,SCARB1,CRYGS,CTTN,TSPAN11,ITSN1,EIF5
11,ENY2,ZSCAN20,RPAIN,TSC22D3,PEX2,PEX26,ZNF223,DNAJC12,DGAT1,OGFOD2,SRSF7,APOLD1,MARS,
11,ENY2,ZSCAN20,RPAIN,TSC22D3,PEX2,PEX26,ZNF223,DNAJC12,DGAT1,OGFOD2,SRSF7,APOLD1,MARS,
CPSF3L,TSG101,PSMA1,YLPM1,PHF21A,NIP7,NISCH,TMED4,CENPK,AUP1,SAP30BP,YIPF3,TRNT1,IP6K2,Y
ND2B,ZNF250,CDK11A,INO80E,MAPKAPK5,HSP90AB1,SYNGR2,CEP70,RAB2A,GPBP1,GPALPP1,PNPO,CHP
P5,ZC3HAV1,SCAF11,DBI,ARPC2,LIMK2,NOB1,WDR19,SIRT6,NUMA1,GPATCH2L,SUMF2,BUD31,SAE1,NP
P5,ZC3HAV1,SCAF11,DBI,ARPC2,LIMK2,NOB1,WDR19,SIRT6,NUMA1,GPATCH2L,SUMF2,BUD31,SAE1,NP

RMT2,EIF3M,JUP,TMEM205,XBP1,PTK2,GUK1,ATF7,AGPAT4,EDF1,RFX3,CRIP1,CLDND1,LIG1,TIMM17B,
RMT2,EIF3M,JUP,TMEM205,XBP1,PTK2,GUK1,ATF7,AGPAT4,EDF1,RFX3,CRIP1,CLDND1,LIG1,TIMM17B,

ENPK,RTEL1,SRSF1,AUP1,YIPF3,FOXRED1,KXD1,ITGB1BP1,SH3YL1,IP6K2,UGGT2,HS2ST1,UBE2I,EIF3M,TB
PHC3,MFSD11,MOK,ZNF81,TMEM14B,RPL37A,ELOF1,DCTN5,METTL3,TCF25,ZNF248,ANAPC13,SRSF7,N
PHC3,MFSD11,MOK,ZNF81,TMEM14B,RPL37A,ELOF1,DCTN5,METTL3,TCF25,ZNF248,ANAPC13,SRSF7,N
2,ARHGAP6,SCARB1,CTTN,TSPAN11,ITSN1,EIF5A,COMMD5,CHID1,AMZ2,USP38,TAZ,DBNDD1,RPL13A,LI
2,ARHGAP6,SCARB1,CTTN,TSPAN11,ITSN1,EIF5A,COMMD5,CHID1,AMZ2,USP38,TAZ,DBNDD1,RPL13A,LI
VAC,POLR3H,NOMO2,USP45,ILF3,EEF1D,CNST,CASZ1,COX17,HINT1,HNRNPA1,SIRT5,RFXANK,SPINT2,SM
VAC,POLR3H,NOMO2,USP45,ILF3,EEF1D,CNST,CASZ1,COX17,HINT1,HNRNPA1,SIRT5,RFXANK,SPINT2,SM
PS25,NIP7,TTC9C,COX4I1,PLCB2,DUSP22,RPS6,RTEL1,FOXRED1,RABL5,ITGB1BP1,MED15,C1ORF198,UG
PS25,NIP7,TTC9C,COX4I1,PLCB2,DUSP22,RPS6,RTEL1,FOXRED1,RABL5,ITGB1BP1,MED15,C1ORF198,UG
ZNF593,MARS,C11ORF57,PCNP,EIF2A,GTF2H1,KDELR3,SLC35A2,SCRN2,RPS3A,EIF3C,DENND1A,ITPA,ZC
ZNF593,MARS,C11ORF57,PCNP,EIF2A,GTF2H1,KDELR3,SLC35A2,SCRN2,RPS3A,EIF3C,DENND1A,ITPA,ZC
3A,PPRC1,NLRP1,TTC9C,RDH11,DBNDD2,RPS6,COASY,SRSF1,AUP1,SAP30BP,TRNT1,KXD1,MED15,UBE2I
3A,PPRC1,NLRP1,TTC9C,RDH11,DBNDD2,RPS6,COASY,SRSF1,AUP1,SAP30BP,TRNT1,KXD1,MED15,UBE2I
C37,TSG101,OBSL1,UBR4,PHF21A,FAM3A,PSMA3,METTL17,NLRP1,NIP7,TTC9C,COX4I1,DUSP22,THEM4,
C37,TSG101,OBSL1,UBR4,PHF21A,FAM3A,PSMA3,METTL17,NLRP1,NIP7,TTC9C,COX4I1,DUSP22,THEM4,
PS20,HHLA3,LMBR1,DDOST,RRNAD1,RABEP2,WAC,TRIM16,NT5C,STXBP2,EEF1D,NAA10,LLGL2,PRLR,SS
PS20,HHLA3,LMBR1,DDOST,RRNAD1,RABEP2,WAC,TRIM16,NT5C,STXBP2,EEF1D,NAA10,LLGL2,PRLR,SS
MS,IL1RAP,STYXL1,PLEKHA5,RANBP1,UBE2L6,ZC3H14,FIS1,DST,ANTXR2,STIP1,LRRC1,GRAMD1C,CD44,
9,NUMA1,GPATCH2L,SUMF2,SAE1,NPRL2,OSGEP,TBL2,ARHGAP6,SCARB1,CRYGS,TSPAN11,ITSN1,EIF5A,
9,NUMA1,GPATCH2L,SUMF2,SAE1,NPRL2,OSGEP,TBL2,ARHGAP6,SCARB1,CRYGS,TSPAN11,ITSN1,EIF5A,
RL2,TBL2,ARHGAP6,CTTN,ITSN1,EIF5A,TNFSF10,COMMD5,AMZ2,UFD1L,TAZ,THYN1,DBNDD1,SH2B1,NO
C11,AGAP3,C11ORF48,CARHSP1,SUMO3,PHF20L1,ZFAND4,INTS4,REEP5,FTH1,DAZAP2,BRF1,TPM4,SLC3
,EBPL,IGSF8,SPAG7,WDR62,STYXL1,POLD4,RANBP1,UBE2L6,PPM1K,STIP1,PHF19,VCAN,MAGED2,SON,T

GRK6,HMGA2,OBSL1,UBR4,YLPM1,PHF21A,CHCHD1,FAM3A,MUC1,METTL17,EPB41L5,ATG12,ZBTB44,T
GRK6,HMGA2,OBSL1,UBR4,YLPM1,PHF21A,CHCHD1,FAM3A,MUC1,METTL17,EPB41L5,ATG12,ZBTB44,T

CB2,SLC25A10,NISCH,RPL29,CD99L2,GGPS1,SRSF1,TRNT1,PARD6G,SH3YL1,IP6K2,CLASRP,JPH2,MICAL1,
CB2,SLC25A10,NISCH,RPL29,CD99L2,GGPS1,SRSF1,TRNT1,PARD6G,SH3YL1,IP6K2,CLASRP,JPH2,MICAL1,

SNF8,CDC37,CPSF3L,OBSL1,CHCHD1,RGS3,MUC1,METTL17,NLRP1,ZBTB44,NIP7,COX4I1,SLC25A10,THE
F76,SERPINB6,RABEP2,RALGPS1,ZNF687,PDCD5,NXN,HSD17B10,ABHD17A,GGT5,SLC16A1,ZCCHC4,STXB
F76,SERPINB6,RABEP2,RALGPS1,ZNF687,PDCD5,NXN,HSD17B10,ABHD17A,GGT5,SLC16A1,ZCCHC4,STXB

C21B,PTPN13,PSMA1,CHCHD1,FAM3A,ECSIT,EPB41L5,PPRC1,ZBTB44,TTC9C,SLC25A10,SPOCK2,SDF4,T
19,SON,TET2,GRK6,CDC37,CPSF3L,TSG101,UBR4,YLPM1,PHF21A,SLC39A11,FUZ,MUC1,METTL17,ECSIT,
19,SON,TET2,GRK6,CDC37,CPSF3L,TSG101,UBR4,YLPM1,PHF21A,SLC39A11,FUZ,MUC1,METTL17,ECSIT,
HA5,RANBP1,UBE2L6,ZC3H14,FIS1,SLC15A3,PPM1K,STIP1,PHF19,VCAN,LPCAT3,MAGED2,SON,GRK6,CD
1L5,RPS25,DUSP22,SPOCK2,RDH11,TMED4,GGPS1,SRSF1,AUP1,FOXRED1,RABL5,ITGB1BP1,CLASRP,JPH
1L5,RPS25,DUSP22,SPOCK2,RDH11,TMED4,GGPS1,SRSF1,AUP1,FOXRED1,RABL5,ITGB1BP1,CLASRP,JPH
RPD1,AZI2,TATDN1,KAT8,NCBP2,SUMO1,ZNF483,SZT2,BSCL2,PSMC5,C12ORF73,IQCK,SETD4,MED8,MB
RPD1,AZI2,TATDN1,KAT8,NCBP2,SUMO1,ZNF483,SZT2,BSCL2,PSMC5,C12ORF73,IQCK,SETD4,MED8,MB

2,PCYT2,USE1,DEF8,PPP1CC,RFX3,DGKE,CAMTA1,LIG1,CDC14B,TIMM17B,APEH,HYI,CAPN10,LYPLA2,PO
2,PCYT2,USE1,DEF8,PPP1CC,RFX3,DGKE,CAMTA1,LIG1,CDC14B,TIMM17B,APEH,HYI,CAPN10,LYPLA2,PO
PHF19,LPCAT3,CD44,TM9SF1,SON,TET2,UQCRQ,CPSF3L,OBSL1,YLPM1,CHCHD1,FUZ,RGS3,FAM3A,PPRC
PHF19,LPCAT3,CD44,TM9SF1,SON,TET2,UQCRQ,CPSF3L,OBSL1,YLPM1,CHCHD1,FUZ,RGS3,FAM3A,PPRC
HNRNPC,CYB561,ZNF644,STAU1,UPP1,ZNF250,LRRFIP1,ZNF524,RPTOR,ZNF625,RPS20,GABPB1,NIT2,RPS

,EDF1,CAMTA1,LIG1,TXNL4A,CAPN10,NUTF2,C19ORF54,ZNF7,ZFAND2B,CYB561,ZNF250,RPL36,NFAM1,

SF10,CHID1,NUDT22,UFD1L,TAZ,DBNDD1,SH2B1,NOP16,ARHGEF39,TTLL5,ACOT8,AP3B2,RPS9,SLC35G1,
SF10,CHID1,NUDT22,UFD1L,TAZ,DBNDD1,SH2B1,NOP16,ARHGEF39,TTLL5,ACOT8,AP3B2,RPS9,SLC35G1,
TLL5,LDHA,ZNF410,RUVBL2,PCMT1,EBPL,IGSF8,NUDT5,RBPMS,WDR62,IL1RAP,STYXL1,RFX5,POLD4,RAN
TLL5,LDHA,ZNF410,RUVBL2,PCMT1,EBPL,IGSF8,NUDT5,RBPMS,WDR62,IL1RAP,STYXL1,RFX5,POLD4,RAN
2B1,GSTK1,PSMA6,ACOT8,AP3B2,ZNF410,RUVBL2,EBPL,TESC,WDR62,IL1RAP,PLEKHA5,UBE2L6,VRK3,Z
2B1,GSTK1,PSMA6,ACOT8,AP3B2,ZNF410,RUVBL2,EBPL,TESC,WDR62,IL1RAP,PLEKHA5,UBE2L6,VRK3,Z
M3A,MUC1,ECSIT,EPB41L5,ATG12,PPRC1,ZBTB44,PLCB2,DUSP22,DBNDD2,NISCH,TMED4,CD99L2,GGPS1
M3A,MUC1,ECSIT,EPB41L5,ATG12,PPRC1,ZBTB44,PLCB2,DUSP22,DBNDD2,NISCH,TMED4,CD99L2,GGPS1
GKE,TNFRSF14,RPL28,TIMM17B,RPL21,CAPN10,ATPAF2,PRMT1,C19ORF54,C19ORF48,HNRNPC,CBFA2T
GKE,TNFRSF14,RPL28,TIMM17B,RPL21,CAPN10,ATPAF2,PRMT1,C19ORF54,C19ORF48,HNRNPC,CBFA2T

RPIN,ZNF644,CARS,RPL36,ZNF197,HSP90AB1,SYNGR2,ELP6,RPS20,GABPB1,NIT2,TWSG1,ZSWIM7,DCTD,
RPIN,ZNF644,CARS,RPL36,ZNF197,HSP90AB1,SYNGR2,ELP6,RPS20,GABPB1,NIT2,TWSG1,ZSWIM7,DCTD,

ADA3,WAC,ZNF687,ILF3,HSPH1,OAS3,LLGL2,PRLR,CHEK1,COX17,HNRNPLL,HNRNPA1,INTS4,SMARCA4,O
L55,CREB3L4,TCF25,TJP1,CDK4,RALY,TRIM46,DOK4,NUP62,PSMD7,ZNF511,IFT172,CC2D1B,ZNF69,AXIN
,IL1RAP,STYXL1,RFX5,RANBP1,UBE2L6,ZC3H14,SLC15A3,PPM1K,IRF9,DST,STIP1,PHF19,VCAN,MAGED2,S
,IL1RAP,STYXL1,RFX5,RANBP1,UBE2L6,ZC3H14,SLC15A3,PPM1K,IRF9,DST,STIP1,PHF19,VCAN,MAGED2,S
XL1,POLD4,RANBP1,UBE2L6,PPM1K,TUBA1C,STIP1,PHF19,VCAN,LPCAT3,MAGED2,SON,TET2,GRK6,HMG
LRRC1,PHF19,CD44,SON,TET2,GRK6,CDC37,OBSL1,PTPN13,UBR4,YLPM1,PHF21A,FUZ,MUC1,ECSIT,EPB4
LRRC1,PHF19,CD44,SON,TET2,GRK6,CDC37,OBSL1,PTPN13,UBR4,YLPM1,PHF21A,FUZ,MUC1,ECSIT,EPB4
3,SYNGR2,TMEM134,LRRFIP1,RAB2A,ZNF524,TPM2,ZEB2,CHPF,LYNX1,LRRC27,LMBR1,NAP1L1,CHCHD7,
2,CTTN,ITSN1,EIF5A,COMMD5,CHID1,NUDT22,UFD1L,USP38,TAZ,DBNDD1,RPL13A,GSTK1,PSMA6,ACOT
2,CTTN,ITSN1,EIF5A,COMMD5,CHID1,NUDT22,UFD1L,USP38,TAZ,DBNDD1,RPL13A,GSTK1,PSMA6,ACOT
0,ILF3,STXBP2,MED6,FKBP3,EGLN2,SIRT5,ZNF506,SPINT2,SMARCA4,PTK7,TMEM39B,BRF1,OARD1,MCF
0,ILF3,STXBP2,MED6,FKBP3,EGLN2,SIRT5,ZNF506,SPINT2,SMARCA4,PTK7,TMEM39B,BRF1,OARD1,MCF
,GRK6,CDC37,TSG101,OBSL1,UBR4,PHF21A,SLC39A11,FAM3A,PSMA3,METTL17,NLRP1,NIP7,TTC9C,COX
,GRK6,CDC37,TSG101,OBSL1,UBR4,PHF21A,SLC39A11,FAM3A,PSMA3,METTL17,NLRP1,NIP7,TTC9C,COX
,TIMM17B,TMEM126B,TXNL4A,CAPN10,ATPAF2,PRMT1,C19ORF54,ARL6IP4,C19ORF48,HNRNPC,CYB561
T2,IFI27L1,MAF,PPA1,ENDOV,PPIL2,RPL21,TXNL4A,PRMT1,EXOSC9,C19ORF48,ZFAND2B,HYPK,ZNF250,
T2,IFI27L1,MAF,PPA1,ENDOV,PPIL2,RPL21,TXNL4A,PRMT1,EXOSC9,C19ORF48,ZFAND2B,HYPK,ZNF250,
CAN,LPCAT3,TM9SF1,SNF8,RPS16,CPSF3L,TSG101,OBSL1,TTC21B,UBR4,PHF21A,RGS3,EPB41L5,ZBTB44,
2,TBX6,PTK2,AUNIP,C14ORF2,PCYT2,USE1,AGPAT4,PPP1CC,RFX3,PPA1,DGKE,CAMTA1,LIG1,TIMM17B,A
2,TBX6,PTK2,AUNIP,C14ORF2,PCYT2,USE1,AGPAT4,PPP1CC,RFX3,PPA1,DGKE,CAMTA1,LIG1,TIMM17B,A
1,PHF21A,FUZ,MUC1,ECSIT,EPB41L5,ATG12,PPRC1,ZBTB44,NIP7,TTC9C,PLCB2,DUSP22,SLC25A10,DBND
1,PHF21A,FUZ,MUC1,ECSIT,EPB41L5,ATG12,PPRC1,ZBTB44,NIP7,TTC9C,PLCB2,DUSP22,SLC25A10,DBND
F4,TMED4,CD99L2,GGPS1,FOXRED1,MED15,JPH2,C1ORF198,HS2ST1,YIF1A,JUP,XBP1,GUK1,CENPN,AT
12,PPRC1,NIP7,COX4I1,PLCB2,TMX2,RDH11,NISCH,CENPK,RPS6,GGPS1,SRSF1,AUP1,SAP30BP,YIPF3,TRN
12,PPRC1,NIP7,COX4I1,PLCB2,TMX2,RDH11,NISCH,CENPK,RPS6,GGPS1,SRSF1,AUP1,SAP30BP,YIPF3,TRN

3A,PSMA3,METTL17,PPRC1,NLRP1,NIP7,TTC9C,COX4I1,DUSP22,THEM4,DBNDD2,TMED4,CENPK,COASY,S
3A,PSMA3,METTL17,PPRC1,NLRP1,NIP7,TTC9C,COX4I1,DUSP22,THEM4,DBNDD2,TMED4,CENPK,COASY,S
3,ZCCHC4,MDH2,STXBP2,HNRNPA1,FKBP3,ZNF506,FTH1,COPS4,DAZAP2,OARD1,GNB5,MFSD11,RPS11,N
3,ZCCHC4,MDH2,STXBP2,HNRNPA1,FKBP3,ZNF506,FTH1,COPS4,DAZAP2,OARD1,GNB5,MFSD11,RPS11,N

0AB1,RAB2A,RMDN1,GPBP1,GPALPP1,CHPF,ERLEC1,GMPPB,MVD,IL32,SYNGR1,RANGRF,H2AFV,SLC33A
CRIP1,VIM,LIG1,HYI,EPB41L4B,TCF4,PPOX,TMPO,CASC4,ARL6IP4,PPAN,CYB561,SHARPIN,MOB1A,MAPKA
CRIP1,VIM,LIG1,HYI,EPB41L4B,TCF4,PPOX,TMPO,CASC4,ARL6IP4,PPAN,CYB561,SHARPIN,MOB1A,MAPKA
NF7,TXNDC17,C19ORF48,PPAN,AP1S3,SHARPIN,STAU1,MDH1,INO80E,MAPKAPK5,CEP70,LRRFIP1,ZNF52
EIF5A,COMMD5,RUNX1,CHID1,NUDT22,AMZ2,TAZ,THYN1,RPL13A,LDHA,RPS9,MRPL33,ZNF410,RUVBL2
EIF5A,COMMD5,RUNX1,CHID1,NUDT22,AMZ2,TAZ,THYN1,RPL13A,LDHA,RPS9,MRPL33,ZNF410,RUVBL2
D3,TCAIM,SLC3A2,DHPS,ACTR6,MAP3K7CL,SEC62,SET,CTSC,ZNF235,BOP1,ARRDC1,TRPT1,KLC1,RBP5,SCA
D3,TCAIM,SLC3A2,DHPS,ACTR6,MAP3K7CL,SEC62,SET,CTSC,ZNF235,BOP1,ARRDC1,TRPT1,KLC1,RBP5,SCA
CHC4,MDH2,STXBP2,HSPH1,CHCHD5,COX5A,HNRNPLL,SUMO3,HNRNPA1,FKBP3,RPL15,INTS4,REEP5,FT
54,C19ORF48,CBFA2T2,UPP1,RPL36,NFAM1,CDK11A,RBM3,COPZ1,SYNGR2,ELP6,TPM2,SIAH1,ASPH,PNF
54,C19ORF48,CBFA2T2,UPP1,RPL36,NFAM1,CDK11A,RBM3,COPZ1,SYNGR2,ELP6,TPM2,SIAH1,ASPH,PNF
TK,PCNP,NEK11,CREBZF,EIF2A,NKIRAS2,GTF2H1,PIGO,CLTA,SLC35A2,SLC25A24,ZNF396,HPS1,ITPA,SNRP
TK,PCNP,NEK11,CREBZF,EIF2A,NKIRAS2,GTF2H1,PIGO,CLTA,SLC35A2,SLC25A24,ZNF396,HPS1,ITPA,SNRP
A,RBM3,MDH1,ZNF197,LRRFIP1,ZNF524,ATP5J,TPM2,GPALPP1,ZNF625,GABPB1,VPS28,RPS15,ERLEC1,H
,PABPC1,NAT9,BCAM,BLVRB,RABEP2,RALGPS1,PDCD5,EPS15L1,EEF1D,NAA10,LLGL2,COX17,FKBP3,EGLN
FX5,PLEKHA5,RANBP1,UBE2L6,VRK3,ZC3H14,FIS1,SLC15A3,PPM1K,ANTXR2,GRAMD1C,SON,UQCRQ,SNF8
FX5,PLEKHA5,RANBP1,UBE2L6,VRK3,ZC3H14,FIS1,SLC15A3,PPM1K,ANTXR2,GRAMD1C,SON,UQCRQ,SNF8
IAP1L1,SYNGR1,LSR,MTA3,PABPC1,RRNAD1,ZNF836,H2AFV,WAC,SLC33A1,ANAPC11,PKP4,ZCCHC4,OAS3
IAP1L1,SYNGR1,LSR,MTA3,PABPC1,RRNAD1,ZNF836,H2AFV,WAC,SLC33A1,ANAPC11,PKP4,ZCCHC4,OAS3
L,UBR4,PSMA1,PHF21A,FUZ,RGS3,FAM3A,PSMA3,ECSIT,ATG12,PPRC1,NIP7,TTC9C,COX4I1,SPOCK2,RDH1
L,UBR4,PSMA1,PHF21A,FUZ,RGS3,FAM3A,PSMA3,ECSIT,ATG12,PPRC1,NIP7,TTC9C,COX4I1,SPOCK2,RDH1

YL9,TSEN34,TJP1,NADSYN1,ZNF511,D2HGDH,SNX29,AXIN1,AAMDC,SNAPC2,TKT,DENND1A,MRPS24,ZM
GRK6,OBSL1,PTPN13,UBR4,YLPM1,PHF21A,SLC39A11,FAM3A,MUC1,ECSIT,EPB41L5,ATG12,PPRC1,ZBTB4
GRK6,OBSL1,PTPN13,UBR4,YLPM1,PHF21A,SLC39A11,FAM3A,MUC1,ECSIT,EPB41L5,ATG12,PPRC1,ZBTB4
RC1,NLRP1,NIP7,TTC9C,COX4I1,DUSP22,THEM4,DBNDD2,TMED4,CENPK,COASY,SRSF1,AUP1,SAP30BP,F
RC1,NLRP1,NIP7,TTC9C,COX4I1,DUSP22,THEM4,DBNDD2,TMED4,CENPK,COASY,SRSF1,AUP1,SAP30BP,F
PSMC5,IQCK,SETD4,MED8,MBD1,CHFR,C12ORF10,RNF4,C12ORF66,ZRANB2,DPP9,SEC24C,RPL31,TVP23B
PSMC5,IQCK,SETD4,MED8,MBD1,CHFR,C12ORF10,RNF4,C12ORF66,ZRANB2,DPP9,SEC24C,RPL31,TVP23B
P1,MED15,PARD6G,SH3YL1,MICAL1,YIF1A,TBX6,PTK2,C14ORF2,GUK1,PCYT2,IFI27L1,EDF1,DGKE,CAPG,C
P1,MED15,PARD6G,SH3YL1,MICAL1,YIF1A,TBX6,PTK2,C14ORF2,GUK1,PCYT2,IFI27L1,EDF1,DGKE,CAPG,C
IOB1,WDR19,SIRT6,NUMA1,GPATCH2L,SUMF2,BUD31,SAE1,NPRL2,OSGEP,TBL2,ARHGAP6,SCARB1,CTTN
SN1,COMMD5,RUNX1,CHID1,NUDT22,AMZ2,UFD1L,USP38,TAZ,THYN1,DBNDD1,LDHA,AP3B2,RCE1,PCM
SN1,COMMD5,RUNX1,CHID1,NUDT22,AMZ2,UFD1L,USP38,TAZ,THYN1,DBNDD1,LDHA,AP3B2,RCE1,PCM
ANGRF,H2AFV,ZNF687,POT1,RAD23A,PKP4,HSPH1,RPL18,CNST,HNRNPA1,FKBP3,NDUFV2,B3GAT3,GTF2
PIL2,TROAP,VIM,LIG1,CAPN10,NUTF2,ATPAF2,CASC4,PIGL,CYB561,STAU1,SLC9B2,CARS,CDK11A,RBM3,N
PIL2,TROAP,VIM,LIG1,CAPN10,NUTF2,ATPAF2,CASC4,PIGL,CYB561,STAU1,SLC9B2,CARS,CDK11A,RBM3,N
IL2,LIG1,LYPLA2,LSM12,C19ORF54,ARL6IP4,ACAD10,SUGT1,ZNF250,RBM3,MDH1,ZNF197,CCNL2,HSP90A
IL2,LIG1,LYPLA2,LSM12,C19ORF54,ARL6IP4,ACAD10,SUGT1,ZNF250,RBM3,MDH1,ZNF197,CCNL2,HSP90A
SLC3A2,DHPS,ACTR6,MAP3K7CL,SEC62,SET,ZNF235,BOP1,ARRDC1,TRPT1,KLC1,ZC3HAV1,SCAF11,DBI,AR
SLC3A2,DHPS,ACTR6,MAP3K7CL,SEC62,SET,ZNF235,BOP1,ARRDC1,TRPT1,KLC1,ZC3HAV1,SCAF11,DBI,AR

DAZAP2,OARD1,RIC8A,SMARCC2,GNB5,CCDC40,RPS11,NUP133,RPAIN,COX20,ACOT2,NFYC,ZXDC,SGCD,
WSG1,VPS28,ZSWIM7,HSPB1,MVD,SAT1,EMC4,IL32,SEMA3C,C12ORF76,SERPINB6,RABEP2,RALGPS1,ZN
WSG1,VPS28,ZSWIM7,HSPB1,MVD,SAT1,EMC4,IL32,SEMA3C,C12ORF76,SERPINB6,RABEP2,RALGPS1,ZN
ETTL17,EPB41L5,RPS25,DUSP22,RDH11,DBNDD2,SRSF1,ITGB1BP1,PARD6G,SH3YL1,JPH2,UGGT2,UBE2I,
L1A,MAPKAPK5,DGUOK,HSP90AB1,RAB2A,RMDN1,GPBP1,GPALPP1,CHPF,RPS20,ZSWIM7,ERLEC1,GMPP
L1A,MAPKAPK5,DGUOK,HSP90AB1,RAB2A,RMDN1,GPBP1,GPALPP1,CHPF,RPS20,ZSWIM7,ERLEC1,GMPP
L1A,MAPKAPK5,DGUOK,HSP90AB1,RAB2A,RMDN1,GPBP1,GPALPP1,CHPF,ZSWIM7,ERLEC1,GMPPB,MVD

L,LRRC27,LMBR1,LSR,CHCHD7,PSME2,PHTF2,PABPC1,BCAM,ZNF836,BLVRB,RABEP2,RALGPS1,ZNF687,PD
PAN31,GLYCTK,PCNP,NEK11,CREBZF,EIF2A,NKIRAS2,PIGO,CLTA,SLC35A2,SLC25A24,ZNF396,HPS1,ITPA,S
PAN31,GLYCTK,PCNP,NEK11,CREBZF,EIF2A,NKIRAS2,PIGO,CLTA,SLC35A2,SLC25A24,ZNF396,HPS1,ITPA,S
P7,TMX2,RDH11,DBNDD2,NISCH,CENPK,RPS6,RPL29,GGPS1,SRSF1,AUP1,SAP30BP,YIPF3,FOXRED1,TRNT
P7,TMX2,RDH11,DBNDD2,NISCH,CENPK,RPS6,RPL29,GGPS1,SRSF1,AUP1,SAP30BP,YIPF3,FOXRED1,TRNT
12,PPRC1,ZBTB44,DBNDD2,SDF4,NISCH,TMED4,RPL29,SH3YL1,IP6K2,CLASRP,JPH2,MICAL1,TMEM205,XE
12,PPRC1,ZBTB44,DBNDD2,SDF4,NISCH,TMED4,RPL29,SH3YL1,IP6K2,CLASRP,JPH2,MICAL1,TMEM205,XE
3L,OBSL1,CHCHD1,FAM3A,PPRC1,RPS25,NIP7,COX4I1,SPOCK2,SDF4,CENPK,CD99L2,GGPS1,RTEL1,SRSF1
3L,OBSL1,CHCHD1,FAM3A,PPRC1,RPS25,NIP7,COX4I1,SPOCK2,SDF4,CENPK,CD99L2,GGPS1,RTEL1,SRSF1
7,FKBP3,EGLN2,KCTD7,B3GAT3,SPINT2,BRF1,FAM102A,PFKFB2,RIC8A,MFSD11,LFNG,XYLT2,PCID2,ZXDC,
MA6,SLC35G1,PCMT1,NUDT5,RBPMS,SPAG7,WDR62,PLEKHA5,VRK3,LZTFL1,FIS1,PPM1K,DST,STIP1,LRRC
MA6,SLC35G1,PCMT1,NUDT5,RBPMS,SPAG7,WDR62,PLEKHA5,VRK3,LZTFL1,FIS1,PPM1K,DST,STIP1,LRRC
NPEP1,RQCD1,EXOSC2,DNAJC21,CDK2AP1,NDUFC1,ACTB,PPP1R13B,HDAC10,LARP1B,KIAA0513,NARF,AF
DD1,GSTK1,PSMA6,AP3B2,MRPL33,RUVBL2,EBPL,TESC,RBPMS,SPAG7,WDR62,IL1RAP,POLD4,PLEKHA5,U
DD1,GSTK1,PSMA6,AP3B2,MRPL33,RUVBL2,EBPL,TESC,RBPMS,SPAG7,WDR62,IL1RAP,POLD4,PLEKHA5,U

R,CHCHD7,HMGA1,PSME2,PABPC1,NAT9,ZNF836,PPP3CB,RABEP2,TADA3,RALGPS1,ZNF687,GTF2IRD1,N
IT1,TESC,STYXL1,POLD4,UBE2L6,DST,ANTXR2,STIP1,LRRC1,GRAMD1C,VCAN,LPCAT3,HMGA2,PTPN13,UB
IT1,TESC,STYXL1,POLD4,UBE2L6,DST,ANTXR2,STIP1,LRRC1,GRAMD1C,VCAN,LPCAT3,HMGA2,PTPN13,UB
RUVBL2,TESC,IGSF8,RBPMS,IL1RAP,RFX5,PLEKHA5,RANBP1,UBE2L6,ZC3H14,SLC15A3,TUBA1C,DST,ANT
RUVBL2,TESC,IGSF8,RBPMS,IL1RAP,RFX5,PLEKHA5,RANBP1,UBE2L6,ZC3H14,SLC15A3,TUBA1C,DST,ANT
,NIP7,SLC25A10,TMX2,SDF4,CD99L2,RTEL1,SRSF1,AUP1,FOXRED1,TRNT1,ITGB1BP1,SH3YL1,CLASRP,C1
1L1,SYNGR1,RRNAD1,ZNF687,POT1,RAD23A,SLC16A1,PKP4,HSPH1,SEC31A,RPL18,PHF20L1,FKBP3,NDU
3,GTF2H1,SNX29,TMEM199,SLC25A24,OXR1,HPS1,VPS25,SMUG1,AIFM2,GTF3A,SNRPA1,TBC1D22A,PRK
RUVBL2,TESC,NNMT,SPAG7,WDR62,IL1RAP,POLD4,PLEKHA5,UBE2L6,VRK3,LZTFL1,FIS1,SLC15A3,PPM1K
RUVBL2,TESC,NNMT,SPAG7,WDR62,IL1RAP,POLD4,PLEKHA5,UBE2L6,VRK3,LZTFL1,FIS1,SLC15A3,PPM1K
K3,LZTFL1,FIS1,SLC15A3,TUBA1C,IRF9,STIP1,LPCAT3,TET2,UQCRQ,SNF8,GRK6,HMGA2,CDC37,TTC21B,UE
K3,LZTFL1,FIS1,SLC15A3,TUBA1C,IRF9,STIP1,LPCAT3,TET2,UQCRQ,SNF8,GRK6,HMGA2,CDC37,TTC21B,UE
F8,NUDT5,STYXL1,POLD4,RANBP1,UBE2L6,LZTFL1,ANTXR2,PHF19,PCBP2,CD44,SON,UQCRQ,GRK6,HMG
F8,NUDT5,STYXL1,POLD4,RANBP1,UBE2L6,LZTFL1,ANTXR2,PHF19,PCBP2,CD44,SON,UQCRQ,GRK6,HMG
LYPLA2,LSM12,CASC4,WNK1,ACAD10,STAU1,ZNF250,CDK11A,RBM3,MDH1,ZNF197,CCNL2,HSP90AB1,R
LYPLA2,LSM12,CASC4,WNK1,ACAD10,STAU1,ZNF250,CDK11A,RBM3,MDH1,ZNF197,CCNL2,HSP90AB1,R
A1,DGKE,TNFRSF14,ENDOV,PSMD4,PSMA7,APEH,RPL21,TXNL4A,TCF4,ATPAF2,C17ORF62,C19ORF54,AF
A1,DGKE,TNFRSF14,ENDOV,PSMD4,PSMA7,APEH,RPL21,TXNL4A,TCF4,ATPAF2,C17ORF62,C19ORF54,AF
1,TRNT1,KXD1,ITGB1BP1,MED15,SLC39A13,MAGT1,C14ORF2,GUK1,CENPN,CARD14,PCYT2,HSBP1L1,AG
1,TRNT1,KXD1,ITGB1BP1,MED15,SLC39A13,MAGT1,C14ORF2,GUK1,CENPN,CARD14,PCYT2,HSBP1L1,AG
5,PEX26,NDUFS3,SSR3,UBA1,SLC39A6,LAMTOR5,DCAKD,SKA2,IFT172,DYNLT3,PIGO,BFAR,CLTA,LYPLAL1,
A1C,ANTXR2,VCAN,SON,TET2,SNF8,HMGA2,CDC37,RPS16,OBSL1,UBR4,RGS3,ATG12,RPS25,ZBTB44,NIP
18,CNST,HNRNPA1,FKBP3,NDUFV2,B3GAT3,GTF2IRD2,TMEM39B,THADA,RPS11,SNX5,NUP133,TMEM1

18,CNST,HNRNPA1,FKBP3,NDUFV2,B3GAT3,GTF2IRD2,TMEM39B,THADA,RPS11,SNX5,NUP133,TMEM14

AM1,CDK11A,MAPKAPK5,DGUOK,HSP90AB1,RAB2A,ZNF524,RMDN1,GPBP1,GPALPP1,TRMT112,CHPF,Z

AM1,CDK11A,MAPKAPK5,DGUOK,HSP90AB1,RAB2A,ZNF524,RMDN1,GPBP1,GPALPP1,TRMT112,CHPF,Z

1,LPCAT3,TM9SF1,HMGA2,CPSF3L,OBSL1,PTPN13,UBR4,FAM3A,MUC1,RPS25,ZBTB44,PLCB2,DUSP22,TM

D1,FUZ,METTL17,ZBTB44,NIP7,COX4I1,DUSP22,SLC25A10,SPOCK2,SDF4,TMED4,RPS6,RPL29,CD99L2,G

D1,FUZ,METTL17,ZBTB44,NIP7,COX4I1,DUSP22,SLC25A10,SPOCK2,SDF4,TMED4,RPS6,RPL29,CD99L2,G

FL1,FIS1,SLC15A3,TUBA1C,IRF9,STIP1,LPCAT3,TET2,UQCRQ,SNF8,GRK6,HMGA2,CDC37,TTC21B,UBR4,SL

,GRK6,HMGA2,CDC37,TTC21B,UBR4,CHCHD1,SLC39A11,FUZ,FAM3A,PSMA3,METTL17,PPRC1,SLC25A10

,GRK6,HMGA2,CDC37,TTC21B,UBR4,CHCHD1,SLC39A11,FUZ,FAM3A,PSMA3,METTL17,PPRC1,SLC25A10

,GRK6,HMGA2,CDC37,TTC21B,UBR4,CHCHD1,SLC39A11,FUZ,FAM3A,PSMA3,METTL17,PPRC1,SLC25A10

EP,TBL2,ARHGAP6,SCARB1,CTTN,ITSN1,EIF5A,RUNX1,CHID1,NUDT22,USP38,TAZ,DBNDD1,SH2B1,NOP1

EP,TBL2,ARHGAP6,SCARB1,CTTN,ITSN1,EIF5A,RUNX1,CHID1,NUDT22,USP38,TAZ,DBNDD1,SH2B1,NOP1

NAA10,CARHSP1,SUMO3,PHF20L1,EGLN2,RFXANK,TMEM101,FKBP5,MRPS18C,TRPV3,CBR1,PHC3,COMM

NAA10,CARHSP1,SUMO3,PHF20L1,EGLN2,RFXANK,TMEM101,FKBP5,MRPS18C,TRPV3,CBR1,PHC3,COMM

UZ,MUC1,METTL17,PPRC1,RPS25,ZBTB44,PLCB2,TMX2,SDF4,CENPK,RPS6,RTEL1,FOXRED1,TRNT1,ITGB1

UZ,MUC1,METTL17,PPRC1,RPS25,ZBTB44,PLCB2,TMX2,SDF4,CENPK,RPS6,RTEL1,FOXRED1,TRNT1,ITGB1

1,UPP1,NFAM1,RBM3,ZNF197,COPZ1,HSP90AB1,SYNGR2,RAB2A,ATP5J,ELP6,TPM2,SIAH1,ASPH,PNPO,N

1,SIRT6,NUMA1,GPATCH2L,SUMF2,BUD31,SAE1,NPRL2,OSGEP,TBL2,ARHGAP6,SCARB1,CTTN,TSPAN11,I

1,SIRT6,NUMA1,GPATCH2L,SUMF2,BUD31,SAE1,NPRL2,OSGEP,TBL2,ARHGAP6,SCARB1,CTTN,TSPAN11,I

LEKHA5,UBE2L6,LZTFL1,STIP1,PHF19,PCBP2,LPCAT3,CD44,SON,TET2,UQCRQ,RPS16,PTPN13,UBR4,YLPM

LEKHA5,UBE2L6,LZTFL1,STIP1,PHF19,PCBP2,LPCAT3,CD44,SON,TET2,UQCRQ,RPS16,PTPN13,UBR4,YLPM

F410,PCMT1,TESC,IGSF8,NUDT5,RBPMS,NNMT,IL1RAP,RFX5,POLD4,RANBP1,VRK3,ZC3H14,FIS1,PPM1K,

F410,PCMT1,TESC,IGSF8,NUDT5,RBPMS,NNMT,IL1RAP,RFX5,POLD4,RANBP1,VRK3,ZC3H14,FIS1,PPM1K,

FV,ZNF687,POT1,RAD23A,PKP4,HSPH1,RPL18,CNST,HNRNPA1,FKBP3,NDUFV2,B3GAT3,GTF2IRD2,TMEM

0,SPOCK2,THEM4,DBNDD2,SDF4,NISCH,TMED4,RTEL1,SRSF1,FOXRED1,TRNT1,KXD1,MED15,SH3YL1,JP

UP,TBX6,TMEM205,AUNIP,GUK1,FBXW9,CCDC25,DEF8,EEF1B2,GADL1,PPA1,CAMTA1,CRIP1,PPIL2,XAF

UP,TBX6,TMEM205,AUNIP,GUK1,FBXW9,CCDC25,DEF8,EEF1B2,GADL1,PPA1,CAMTA1,CRIP1,PPIL2,XAF

1,SRSF1,AUP1,MED15,SH3YL1,JPH2,PRMT2,JUP,TBX6,C14ORF2,ATF7,HSBP1L1,AGPAT4,ASF1B,RFX3,PP

LPCAT3,TET2,UQCRQ,GRK6,HMGA2,CDC37,TTC21B,UBR4,CHCHD1,SLC39A11,FUZ,FAM3A,PSMA3,METT

ATG12,PPRC1,ZBTB44,NIP7,TMX2,THEM4,RDH11,DBNDD2,NISCH,CENPK,RPS6,RPL29,CD99L2,GGPS1,S

ATG12,PPRC1,ZBTB44,NIP7,TMX2,THEM4,RDH11,DBNDD2,NISCH,CENPK,RPS6,RPL29,CD99L2,GGPS1,S

C35E1,ECE1,OCIAD1,RBBP6,TPT1,TMEM9,ZNF593,GLYCTK,RBM42,CLASP2,CREBZF,ZNF511,GTF2H1,MY

T,HNRNPA1,FKBP3,NDUFV2,B3GAT3,GTF2IRD2,TMEM39B,THADA,RPS11,SNX5,NUP133,TMEM147,NFY

X1,CHID1,NUDT22,AMZ2,UFD1L,USP38,TAZ,THYN1,DBNDD1,LDHA,AP3B2,RCE1,PCMT1,TESC,RBPMS,NN

5E1,ECE1,OCIAD1,RBBP6,TPT1,TMEM9,ZNF593,GLYCTK,RBM42,CLASP2,CREBZF,ZNF511,GTF2H1,MYL6,

AF1D,D2HGDH,ST7,RGS12,PCTP,CKLF,GRAMD1A,AKT1S1,COMT,GTF3A,DHRS4,NDUFB2,USP15,AKR1A1,

HA,ACOT8,RUVBL2,IGSF8,NUDT5,NNMT,WDR62,IL1RAP,STYXL1,UBE2L6,LZTFL1,ZC3H14,FIS1,PPM1K,TU

HA,ACOT8,RUVBL2,IGSF8,NUDT5,NNMT,WDR62,IL1RAP,STYXL1,UBE2L6,LZTFL1,ZC3H14,FIS1,PPM1K,TU

MAGT1,C14ORF2,GUK1,CENPN,CARD14,PCYT2,FBXW9,HSBP1L1,AGPAT4,IFI27L1,MAF,PPA1,ENDOV,PPIL

MAGT1,C14ORF2,GUK1,CENPN,CARD14,PCYT2,FBXW9,HSBP1L1,AGPAT4,IFI27L1,MAF,PPA1,ENDOV,PPIL

ANGRF,RAD1,SLC33A1,ZNF706,GTF2IRD1,POLR3H,GGT5,SLC16A1,ILF3,AGAP3,EEF1D,SUMO3,PHF20L1,

ANGRF,RAD1,SLC33A1,ZNF706,GTF2IRD1,POLR3H,GGT5,SLC16A1,ILF3,AGAP3,EEF1D,SUMO3,PHF20L1,

ANGRF,RAD1,SLC33A1,ZNF706,GTF2IRD1,POLR3H,GGT5,SLC16A1,ILF3,AGAP3,EEF1D,SUMO3,PHF20L1,

AMD1C,VCAN,PCBP2,CD44,TM9SF1,CDC37,RPS16,CPSF3L,TSG101,TTC21B,PSMA1,PHF21A,SLC39A11,FU
,IL32,SYNGR1,RANGRF,H2AFV,ZNF687,POT1,RAD23A,PKP4,HSPH1,RPL18,CNST,HNRNPA1,FKBP3,NDUFV
,IL32,SYNGR1,RANGRF,H2AFV,ZNF687,POT1,RAD23A,PKP4,HSPH1,RPL18,CNST,HNRNPA1,FKBP3,NDUFV
2,MPV17,ARIH2,TSPAN31,PPP2R5C,ZNF593,GLYCTK,TRIM46,RPL34,FCHO2,GTF2H1,CC2D1B,ARPC5,ALKE
FKFB2,LGALS3,PARK7,NFKBIA,PDCD6,NOP2,FAM32A,ZSCAN20,TNKS,ELOF1,C12ORF57,EIF4A1,RPLP0,ZM
FKFB2,LGALS3,PARK7,NFKBIA,PDCD6,NOP2,FAM32A,ZSCAN20,TNKS,ELOF1,C12ORF57,EIF4A1,RPLP0,ZM
EP,TBL2,ARHGAP6,CRYGS,CTTN,ITSN1,EIF5A,TNFSF10,COMMD5,RUNX1,NUDT22,AMZ2,UFD1L,USP38,T
EP,TBL2,ARHGAP6,CRYGS,CTTN,ITSN1,EIF5A,TNFSF10,COMMD5,RUNX1,NUDT22,AMZ2,UFD1L,USP38,T
ID1,NUDT22,UFD1L,USP38,DBNDD1,SH2B1,NOP16,GSTK1,TTLL5,PSMA6,ACOT8,AP3B2,RCE1,RUVBL2,P
ID1,NUDT22,UFD1L,USP38,DBNDD1,SH2B1,NOP16,GSTK1,TTLL5,PSMA6,ACOT8,AP3B2,RCE1,RUVBL2,P
K2,TMED4,RPS6,RTEL1,SRSF1,SAP30BP,KXD1,MED15,IP6K2,TBX6,PTK2,AUNIP,CENPN,USE1,IFI27L1,MA

CHCHD1,SLC39A11,FUZ,ECSIT,ATG12,PPRC1,NLRP1,ZBTB44,NIP7,TMX2,RDH11,DBNDD2,NISCH,CENPK,R
CHCHD1,SLC39A11,FUZ,ECSIT,ATG12,PPRC1,NLRP1,ZBTB44,NIP7,TMX2,RDH11,DBNDD2,NISCH,CENPK,R
DXL,C17ORF62,PRMT1,CASC4,ZNF7,ZFAND2B,CYB561,HSP90AB1,SYNGR2,TMEM134,LRRFIP1,RAB2A,ZN
CHD1,FUZ,ECSIT,ATG12,PPRC1,ZBTB44,NIP7,TMX2,RDH11,DBNDD2,NISCH,CENPK,RPS6,RPL29,GGPS1,S
CHD1,FUZ,ECSIT,ATG12,PPRC1,ZBTB44,NIP7,TMX2,RDH11,DBNDD2,NISCH,CENPK,RPS6,RPL29,GGPS1,S
,PKP4,AGAP3,HSPH1,MT1E,RPL18,SH3GLB1,FKBP3,B3GAT3,ZNF506,PAFAH1B2,GTF2IRD2,ZFPL1,GMPR2
,PKP4,AGAP3,HSPH1,MT1E,RPL18,SH3GLB1,FKBP3,B3GAT3,ZNF506,PAFAH1B2,GTF2IRD2,ZFPL1,GMPR2
C1,RBP5,ZC3HAV1,SCAF11,DBI,ARPC2,LIMK2,VDAC3,WDR19,NUMA1,GPATCH2L,SUMF2,NPRL2,TBL2,A
DC37,CPSF3L,TSG101,OBSL1,UBR4,PSMA1,YLPM1,PHF21A,FUZ,FAM3A,PSMA3,EPB41L5,TTC9C,TMX2,T
DC37,CPSF3L,TSG101,OBSL1,UBR4,PSMA1,YLPM1,PHF21A,FUZ,FAM3A,PSMA3,EPB41L5,TTC9C,TMX2,T
P1,PEX26,ARL16,EIF4A1,FBXL18,OCIAD1,POLR2M,NME1,ANAPC13,TRPM6,PPP2R5C,GLYCTK,NUP62,ZN
,TAZ,THYN1,DBNDD1,RPL13A,ARHGEF39,GSTK1,TTLL5,PSMA6,ACOT8,AP3B2,RPS9,SLC35G1,RCE1,ZNF4
,TAZ,THYN1,DBNDD1,RPL13A,ARHGEF39,GSTK1,TTLL5,PSMA6,ACOT8,AP3B2,RPS9,SLC35G1,RCE1,ZNF4
STXBP2,CHCHD5,SH3GLB1,SSH2,EGLN2,B3GAT3,REEP5,PTK7,TRPM4,SCAP,MROH1,TTC12,SNX5,ELP4,AC
OB1A,INO80E,TMEM134,GPBP1,SIAH1,ASPH,CHPF,NIT2,RPS15,HHLA3,SEMA3C,LMBR1,MEIS2,HMGA1,T
OB1A,INO80E,TMEM134,GPBP1,SIAH1,ASPH,CHPF,NIT2,RPS15,HHLA3,SEMA3C,LMBR1,MEIS2,HMGA1,T
E2N,ZXDC,NMNAT3,MPHOSPH10,MYL9,GFOD1,TSEN34,MRPL55,TCF25,CHTOP,NXF1,RALY,RER1,NUP6

18,HNRNPA1,FKBP3,NDUFV2,B3GAT3,GTF2IRD2,TMEM39B,THADA,RPS11,SNX5,NUP133,TMEM147,NFY
18,HNRNPA1,FKBP3,NDUFV2,B3GAT3,GTF2IRD2,TMEM39B,THADA,RPS11,SNX5,NUP133,TMEM147,NFY
IAD1,RBBP6,ZNF248,TPT1,TMEM9,ZNF593,GLYCTK,RBM42,CLASP2,CREBZF,ZNF511,GTF2H1,MYL6,RPS3
RNAD1,H2AFV,SLC33A1,ZNF687,POT1,RAD23A,PKP4,HSPH1,MT1E,RPL18,CNST,CHEK1,PHF20L1,HNRNP
6,HMGA2,CDC37,TTC21B,UBR4,SLC39A11,FUZ,FAM3A,PSMA3,METTL17,ATG12,PPRC1,NIP7,SLC25A10,T
6,HMGA2,CDC37,TTC21B,UBR4,SLC39A11,FUZ,FAM3A,PSMA3,METTL17,ATG12,PPRC1,NIP7,SLC25A10,T
6,HMGA2,CDC37,TTC21B,UBR4,SLC39A11,FUZ,FAM3A,PSMA3,METTL17,ATG12,PPRC1,NIP7,SLC25A10,T
MEM39B,DALRD3,NSUN5,OARD1,MFSD10,TRPV3,UBE2N,FAM32A,KPNA5,MPHOSPH10,MYL9,SLC35E1,G
MEM39B,DALRD3,NSUN5,OARD1,MFSD10,TRPV3,UBE2N,FAM32A,KPNA5,MPHOSPH10,MYL9,SLC35E1,G
06,GTF2IRD2,ZFPL1,TPM4,SLC30A5,THADA,MROH1,SNX5,NUP133,TMEM147,ELP4,ZXDC,COBL,ELOF1,A
PCYT2,HSBP1L1,AGPAT4,IFI27L1,MAF,PPA1,ENDOV,PPIL2,TXNL4A,PRMT1,EXOSC9,ZFAND2B,HYPK,ZNF2
PCYT2,HSBP1L1,AGPAT4,IFI27L1,MAF,PPA1,ENDOV,PPIL2,TXNL4A,PRMT1,EXOSC9,ZFAND2B,HYPK,ZNF2
4,SLC30A5,THADA,MROH1,SNX5,NUP133,TMEM147,ELP4,ZXDC,COBL,ARL16,SLC35E1,MRPL55,ECE1,OC

ET,CTSC,ZNF235,BOP1,ARRDC1,TRPT1,KLC1,ZC3HAV1,DBI,LIMK2,VDAC3,NOB1,WDR19,SIRT6,NUMA1,GI
ET,CTSC,ZNF235,BOP1,ARRDC1,TRPT1,KLC1,ZC3HAV1,DBI,LIMK2,VDAC3,NOB1,WDR19,SIRT6,NUMA1,GI
PLEKHA5,RANBP1,ZC3H14,FIS1,SLC15A3,TUBA1C,ANTXR2,STIP1,LRRC1,PHF19,TM9SF1,GRK6,HMGA2,RP
PLEKHA5,RANBP1,ZC3H14,FIS1,SLC15A3,TUBA1C,ANTXR2,STIP1,LRRC1,PHF19,TM9SF1,GRK6,HMGA2,RP

AT4,CCDC25,EEF1B2,PPA1,TNFRSF14,TIMM17B,TMEM126B,CAPN10,ATPAF2,PRMT1,C19ORF54,ARL6IP
AT4,CCDC25,EEF1B2,PPA1,TNFRSF14,TIMM17B,TMEM126B,CAPN10,ATPAF2,PRMT1,C19ORF54,ARL6IP
3PL,TESC,IGSF8,NUDT5,RBPMS,SPAG7,WDR62,STYXL1,POLD4,PLEKHA5,UBE2L6,LZTFL1,SLC15A3,TUBA1C
4,GGPS1,COASY,SRSF1,FOXRED1,RABL5,TRNT1,KXD1,MED15,SH3YL1,HS2ST1,UBE2I,YIF1A,CDC42,TMEM
T5C,ILF3,STXBP2,EEF1D,C11ORF48,CNST,LLGL2,EGFR,HINT1,NDUFV2,SIRT5,TMEM101,DRG2,SLC25A23,T
T5C,ILF3,STXBP2,EEF1D,C11ORF48,CNST,LLGL2,EGFR,HINT1,NDUFV2,SIRT5,TMEM101,DRG2,SLC25A23,T
DBNDD2,SDF4,TMED4,CENPK,RPS6,GGPS1,TRNT1,ITGB1BP1,SH3YL1,UGGT2,MICAL1,UBE2I,SLC39A13,PR
DBNDD2,SDF4,TMED4,CENPK,RPS6,GGPS1,TRNT1,ITGB1BP1,SH3YL1,UGGT2,MICAL1,UBE2I,SLC39A13,PR

48,ZFAND2B,UPP1,NFAM1,INO80E,MAPKAPK5,DGUOK,HSP90AB1,GPBP1,CHPF,MAPK13,ERLEC1,MVD,IL
48,ZFAND2B,UPP1,NFAM1,INO80E,MAPKAPK5,DGUOK,HSP90AB1,GPBP1,CHPF,MAPK13,ERLEC1,MVD,IL
CTTN,TSPAN11,ITSN1,EIF5A,RUNX1,CHID1,NUDT22,AMZ2,UFD1L,TAZ,DBNDD1,SH2B1,ACOT8,RPS9,SLC3
CTTN,TSPAN11,ITSN1,EIF5A,RUNX1,CHID1,NUDT22,AMZ2,UFD1L,TAZ,DBNDD1,SH2B1,ACOT8,RPS9,SLC3
OB1,WDR19,SIRT6,NUMA1,GPATCH2L,BUD31,SAE1,OSGEP,TBL2,ARHGAP6,SCARB1,CTTN,TSPAN11,RUN
OB1,WDR19,SIRT6,NUMA1,GPATCH2L,BUD31,SAE1,OSGEP,TBL2,ARHGAP6,SCARB1,CTTN,TSPAN11,RUN
PSF3L,TSG101,PTPN13,UBR4,PSMA1,CHCHD1,FAM3A,PSMA3,METTL17,EPB41L5,RPS25,NLRP1,ZBTB44,I
PSF3L,TSG101,PTPN13,UBR4,PSMA1,CHCHD1,FAM3A,PSMA3,METTL17,EPB41L5,RPS25,NLRP1,ZBTB44,I
,CDC37,RPS16,TTC21B,UBR4,PSMA1,YLPM1,FUZ,METTL17,ZBTB44,TTC9C,COX4I1,DUSP22,NISCH,CENPK
,CDC37,RPS16,TTC21B,UBR4,PSMA1,YLPM1,FUZ,METTL17,ZBTB44,TTC9C,COX4I1,DUSP22,NISCH,CENPK
SAT1,ERCC5,LMBR1,MCTS1,DDOST,NAT9,ZNF836,RBMX,WAC,POLR3H,DDX41,EEF1D,CHCHD5,NAA10,CN
SAT1,ERCC5,LMBR1,MCTS1,DDOST,NAT9,ZNF836,RBMX,WAC,POLR3H,DDX41,EEF1D,CHCHD5,NAA10,CN
DAC3,NOB1,SIRT6,NUMA1,SUMF2,BUD31,NPRL2,OSGEP,TBL2,ARHGAP6,SCARB1,CTTN,TSPAN11,ITSN1,I
DAC3,NOB1,SIRT6,NUMA1,SUMF2,BUD31,NPRL2,OSGEP,TBL2,ARHGAP6,SCARB1,CTTN,TSPAN11,ITSN1,I
C9C,DUSP22,THEM4,RDH11,CENPK,RPS6,RPL29,RTEL1,SAP30BP,FOXRED1,TRNT1,ITGB1BP1,SH3YL1,CLA
CSIT,ATG12,PPRC1,ZBTB44,NIP7,TMX2,RDH11,DBNDD2,NISCH,CENPK,RPS6,RPL29,GGPS1,SRSF1,AUP1,SA
CSIT,ATG12,PPRC1,ZBTB44,NIP7,TMX2,RDH11,DBNDD2,NISCH,CENPK,RPS6,RPL29,GGPS1,SRSF1,AUP1,SA
P1,VRK3,PPM1K,PHF19,PCBP2,TM9SF1,SON,TET2,GRK6,CDC37,TSG101,OBSL1,PHF21A,FAM3A,PSMA3,N
P1,VRK3,PPM1K,PHF19,PCBP2,TM9SF1,SON,TET2,GRK6,CDC37,TSG101,OBSL1,PHF21A,FAM3A,PSMA3,N
,VDAC3,NOB1,SIRT6,GPATCH2L,BUD31,SAE1,NPRL2,OSGEP,ARHGAP6,SCARB1,CRYGS,CTTN,TSPAN11,ITS
,VDAC3,NOB1,SIRT6,GPATCH2L,BUD31,SAE1,NPRL2,OSGEP,ARHGAP6,SCARB1,CRYGS,CTTN,TSPAN11,ITS
G7,WDR62,STYXL1,PLEKHA5,LZTFL1,STIP1,PHF19,PCBP2,LPCAT3,CD44,MAGED2,SON,TET2,UQCRQ,CDC3
G7,WDR62,STYXL1,PLEKHA5,LZTFL1,STIP1,PHF19,PCBP2,LPCAT3,CD44,MAGED2,SON,TET2,UQCRQ,CDC3
RPS9,SLC35G1,MRPL33,RUVBL2,PCMT1,EBPL,RBPMS,SPAG7,STYXL1,RANBP1,UBE2L6,ZC3H14,TUBA1C,PH
RPS9,SLC35G1,MRPL33,RUVBL2,PCMT1,EBPL,RBPMS,SPAG7,STYXL1,RANBP1,UBE2L6,ZC3H14,TUBA1C,PH
3YL1,UGGT2,HS2ST1,PRMT2,EIF3M,MAGT1,TBX6,AUNIP,C14ORF2,GUK1,FBXW9,IFI27L1,MAF,EEF1B2,PP
3YL1,UGGT2,HS2ST1,PRMT2,EIF3M,MAGT1,TBX6,AUNIP,C14ORF2,GUK1,FBXW9,IFI27L1,MAF,EEF1B2,PP
F1,SON,TET2,SNF8,GRK6,HMGA2,CPSF3L,OBSL1,PTPN13,UBR4,PSMA1,YLPM1,PHF21A,EPB41L5,ZBTB44,
DC37,TSG101,OBSL1,PTPN13,UBR4,PHF21A,FAM3A,PSMA3,METTL17,EPB41L5,PPRC1,NLRP1,NIP7,TTC9
DC37,TSG101,OBSL1,PTPN13,UBR4,PHF21A,FAM3A,PSMA3,METTL17,EPB41L5,PPRC1,NLRP1,NIP7,TTC9
RPL29,RTEL1,SRSF1,TRNT1,ITGB1BP1,SH3YL1,C1ORF198,C14ORF2,ATF7,PCYT2,USE1,AGPAT4,CCDC25,EE
RPL29,RTEL1,SRSF1,TRNT1,ITGB1BP1,SH3YL1,C1ORF198,C14ORF2,ATF7,PCYT2,USE1,AGPAT4,CCDC25,EE
TTL17,PPRC1,SLC25A10,TMX2,SDF4,TMED4,RPS6,RPL29,RTEL1,TRNT1,ITGB1BP1,SH3YL1,C1ORF198,CEL

TTL17,PPRC1,SLC25A10,TMX2,SDF4,TMED4,RPS6,RPL29,RTEL1,TRNT1,ITGB1BP1,SH3YL1,C1ORF198,CEL

PS16,CPSF3L,TSG101,PTPN13,FAM3A,PSMA3,METTL17,EPB41L5,ATG12,PPRC1,RPS25,NIP7,DUSP22,TM

PS16,CPSF3L,TSG101,PTPN13,FAM3A,PSMA3,METTL17,EPB41L5,ATG12,PPRC1,RPS25,NIP7,DUSP22,TM

4,HSPH1,MT1E,RPL18,CHEK1,PHF20L1,FKBP3,B3GAT3,ZNF506,GTF2IRD2,COPS4,TMEM39B,ZFPL1,TPM4,

GRF,RRNAD1,RAD1,SLC33A1,ZNF706,GTF2IRD1,NUDT1,ANAPC11,POLR3H,RAD23A,SLC16A1,ILF3,EEF1D,

GRF,RRNAD1,RAD1,SLC33A1,ZNF706,GTF2IRD1,NUDT1,ANAPC11,POLR3H,RAD23A,SLC16A1,ILF3,EEF1D,

GRF,RRNAD1,RAD1,SLC33A1,ZNF706,GTF2IRD1,NUDT1,ANAPC11,POLR3H,RAD23A,SLC16A1,ILF3,EEF1D,

1,SCAF11,DBI,ARPC2,VDAC3,NOB1,WDR19,SIRT6,NUMA1,GPATCH2L,BUD31,SAE1,NPRL2,OSGEP,TBL2,A

1,SCAF11,DBI,ARPC2,VDAC3,NOB1,WDR19,SIRT6,NUMA1,GPATCH2L,BUD31,SAE1,NPRL2,OSGEP,TBL2,A

NT1,KXD1,ITGB1BP1,MED15,SLC39A13,MAGT1,C14ORF2,GUK1,CENPN,CARD14,PCYT2,HSBP1L1,AGPAT

NT1,KXD1,ITGB1BP1,MED15,SLC39A13,MAGT1,C14ORF2,GUK1,CENPN,CARD14,PCYT2,HSBP1L1,AGPAT

BPMS,RANBP1,TUBA1C,DST,VCAN,TM9SF1,SON,TET2,SNF8,GRK6,HMGA2,CDC37,RPS16,OBSL1,UBR4,R

,PTPN13,UBR4,YLPM1,PHF21A,SLC39A11,MUC1,EPB41L5,ATG12,PPRC1,ZBTB44,COX4I1,DUSP22,SLC25

,PTPN13,UBR4,YLPM1,PHF21A,SLC39A11,MUC1,EPB41L5,ATG12,PPRC1,ZBTB44,COX4I1,DUSP22,SLC25

PC1,SLC33A1,ZNF706,PDCD5,GTF2IRD1,NUDT1,POLR3H,RAD23A,SLC16A1,ILF3,AGAP3,MDH2,EEF1D,HS

PC1,SLC33A1,ZNF706,PDCD5,GTF2IRD1,NUDT1,POLR3H,RAD23A,SLC16A1,ILF3,AGAP3,MDH2,EEF1D,HS

RPL13A,ARHGEF39,GSTK1,LDHA,PSMA6,ACOT8,SLC35G1,RCE1,MRPL33,ZNF410,RUVBL2,PCMT1,EBPL,T

RPL13A,ARHGEF39,GSTK1,LDHA,PSMA6,ACOT8,SLC35G1,RCE1,MRPL33,ZNF410,RUVBL2,PCMT1,EBPL,T

1,TPM4,SLC30A5,THADA,MROH1,SNX5,NUP133,TMEM147,ELP4,ZXDC,COBL,ELOF1,ARL16,SLC35E1,MRP

52,PRMT1,CASC4,EXOSC9,TXNDC17,PPAN,CYB561,SUGT1,CARS,COPZ1,DUT,TPM2,GPBP1,PNPO,CHPF,L

52,PRMT1,CASC4,EXOSC9,TXNDC17,PPAN,CYB561,SUGT1,CARS,COPZ1,DUT,TPM2,GPBP1,PNPO,CHPF,L

N3A2,C6ORF132,TMEM108,ZMYM6,SSR3,MPV17,CREB3L4,ZNF512,OCIAD1,TCF25,SLMAP,RBBP6,BAX,C

N3A2,C6ORF132,TMEM108,ZMYM6,SSR3,MPV17,CREB3L4,ZNF512,OCIAD1,TCF25,SLMAP,RBBP6,BAX,C

C17ORF62,STOML2,PIGL,ACAD10,HNRNPC,SLC9B2,HYPK,MOB1A,INO80E,DGUOK,LRRFIP1,ZNF524,GPBP

C17ORF62,STOML2,PIGL,ACAD10,HNRNPC,SLC9B2,HYPK,MOB1A,INO80E,DGUOK,LRRFIP1,ZNF524,GPBP

OBSL1,PSMA1,CHCHD1,MUC1,METTL17,NLRP1,ZBTB44,NIP7,TTC9C,COX4I1,DUSP22,SLC25A10,THEM4,

C37,RPS16,UBR4,YLPM1,CHCHD1,FUZ,ECSIT,ATG12,PPRC1,RPS25,ZBTB44,NIP7,COX4I1,TMX2,THEM4,R

C37,RPS16,UBR4,YLPM1,CHCHD1,FUZ,ECSIT,ATG12,PPRC1,RPS25,ZBTB44,NIP7,COX4I1,TMX2,THEM4,R

RPL37A,C12ORF57,BTN3A2,EIF4A1,CIZ1,SSR3,TJP1,TPT1,UBA1,NXF1,MARS,C11ORF57,VDAC1,NUP62,EIF

LN2,RPL15,KCTD7,RBFOX2,PTK7,BRF1,RIC8A,PARK7,CBR1,XYLT2,MFF,RPAIN,COX20,NEK2,PRRC2B,ELOF

,ILF3,MDH2,EEF1D,RRP8,OAS3,CARHSP1,RPL18,HNRNPLL,HNRNPA1,FKBP3,RPL15,SMARCA4,S100A4,SN

VIM,LIG1,APEH,RPL21,RBM33,TCF4,TMPO,PRMT1,EXOSC9,ZNF7,ARL6IP4,PPAN,HNRNPC,ZNF644,STAU1

SGEP,TBL2,ARHGAP6,SCARB1,CRYGS,CTTN,ITSN1,EIF5A,TNFSF10,COMMD5,RUNX1,CHID1,NUDT22,UFD1

CAL1,UBE2I,EIF3M,CDC42,TBX6,XBP1,PTK2,CELF1,AUNIP,GUK1,ATF7,MAF,EEF1B2,PPP1CC,EDF1,GADL1,

K2,JPH2,MICAL1,UBE2I,EIF3M,CDC42,TBX6,XBP1,PTK2,CELF1,AUNIP,GUK1,ATF7,MAF,EEF1B2,PPP1CC,ED

PSPC1,PSMD7,GTF2H1,RBM4,GEMIN2,THOC3,HNRNPUL1,TP53,PSMD10,RPS3A,RPS3,MAGOHB,SNRPD1,

TOR5,VDAC1,RPL34,NUP62,RPL24,NEDD4L,IRF3,TP53,AP1G2,RPS3A,PML,IFITM2,RPS3,IFI27,AZI2,RPS7,CA

,CFLAR,LAMTOR5,VDAC1,RPL34,NUP62,RPL24,NEDD4L,IRF3,THOC3,TP53,AP1G2,RPS3A,PML,IFITM2,RPS

X1,SSR3,MPV17,SSB,ARIH2,RPL8,PICK1,CHTOP,RPL35,SRSF7,NXF1,ZNF593,RPL34,NUP62,ACAA1,RPL24,F

L3,SSB,ARIH2,RPL8,FBXL18,UBE2D4,RBBP6,BAX,RPL35,UBA1,RNF8,PPP2R5C,FAM192A,PCNP,RPL34,RPL

BP2,PLD1,PIWIL4,UBA52,RPS4X,MRPS16,RPS23,RPL30,RQCD1,NDUFA7,DNAJC24,MRPS33,NGDN,RPS15A

RPS7,NCBP2,PLD1,PIWIL4,UBA52,RPS4X,MRPS16,RPS23,RPL30,RQCD1,NDUFA7,DNAJC24,MRPS33,NGDN

3GLB1,HNRNPA1,RPL15,B3GAT3,SNRPF,MCFD2,SYTL1,CD74,PARK7,NFKBIA,RPS11,PDCD6,MFF,SNX5,EN

.,EXOSC2,RBM7,PSMC5,HNRNPM,EXOSC8,RPS15A,RPS17,PON2,RPL27,QDPR,RPS14,CNOT7,HBS1L,PDE1

,CFLAR,LAMTOR5,VDAC1,RPL34,NUP62,RPL24,NEDD4L,IRF3,THOC3,TP53,AP1G2,RPS3A,PML,IFITM2,RPS

.1,CD74,PARK7,NFKBIA,RPS11,PDCD6,MFF,SNX5,ENY2,NUP133,RAPGEF3,RPAIN,ACOT2,PCID2,SGCD,PEX

PS17,RPL27,RPS14,CNOT7,HBS1L,PDE1A,ENTPD6,NUDT9,APOBEC3F,MAGOH,RPL31,GSK3A,PSMD11,EIF3

1,CD74,PARK7,NFKBIA,RPS11,PDCD6,MFF,SNX5,ENY2,NUP133,RAPGEF3,RPAIN,ACOT2,PCID2,SGCD,PEX
EGLN2,RPL15,PARK7,RPS11,SNX5,USP40,PCID2,RPL37A,USP4,RPLP0,METTL3,SSB,ARIH2,RPL8,FBXL18,UE

S24,FARS2,RPS27L,RPS7,NCBP2,PLD1,PIWIL4,THOP1,NFE2L1,UBA52,RPS4X,MRPS16,XPNPEP1,RPS23,RP
8,RQCD1,EXOSC2,RBM7,PSMC5,HNRNPM,EXOSC8,RPS15A,RPS17,RPL27,QDPR,RPS14,CNOT7,HBS1L,PDI

HNRNPM,EXOSC8,RPS15A,RPS17,RPL27,RPS14,CNOT7,HBS1L,PDE1A,ENTPD6,NUDT9,APOBEC3F,MAGOI
HNRNPM,EXOSC8,RPS15A,RPS17,RPL27,RPS14,CNOT7,HBS1L,PDE1A,ENTPD6,NUDT9,APOBEC3F,MAGOI
53,MRPS24,FARS2,SPHK2,RPS27L,RPS7,NCBP2,PLD1,PIWIL4,UBA52,RPS4X,MRPS16,NMRAL1,RPS23,RPL3

70,RAB2A,ATP5J,RPS20,VPS28,RPS15,ERLEC1,HSPB1,SYNGR1,RANGRF,HMGA1,TOMM40,PPP3CB,PDCD5

L,UBE2D3,RPS3,RDX,VPS25,NSFL1C,DHRS4,GOSR2,RPS7,CARM1,ECT2,UBA52,RPS4X,VAPA,VTI1B,SUMO
ME2,PABPC1,BLVRB,WAC,NUDT1,ANAPC11,RAD23A,HSD17B10,ABHD17A,GGT5,NT5C,USP45,PSME1,AG
,EEF2K,GCDH,MARS,RPL34,EIF2A,RPL24,RBM4,TP53,RPS3A,EIF3C,MGST2,RPS3,MRPS24,FARS2,SPHK2,R
D10,TMEM199,NDUFA3,EIF3C,COX19,RPS3,RDX,SNRPD1,MLST8,NDUFB3,NDUFB2,NDUFA10,RPS27L,GM
1A,PRLR,SH3GLB1,SSH2,COX17,NDUFV2,KCTD7,ASH2L,SMARCA4,COA3,SNRPF,GBA2,PPIH,BRF1,TRPM4,
PS15,UBXN1,ERLEC1,HSPB1,DCN,PSME2,PABPC1,BLVRB,TOMM40,WAC,NUDT1,ANAPC11,RAD23A,HSD
THOC3,HNRNPUL1,TP53,MAGOHB,SNRPD1,FARS2,RBM6,SNRPA1,ZBTB8OS,RPS7,U2AF1L4,NCBP2,DDX10
UT,SIAH1,RPTOR,RPS20,VPS28,RPS15,UBXN1,ERLEC1,HSPB1,SAT1,DCN,PSME2,PABPC1,BLVRB,TOMM40
ATP6V1F,PDCD5,TRIM16,SLC16A1,STXBP2,SEC31A,RPL18,CNST,LLGL2,EGFR,SH3GLB1,RPL15,B3GAT3,M
),PPP3CB,RABEP2,ATP6V1F,PDCD5,POT1,TRIM16,SLC16A1,STXBP2,SEC31A,RPL18,CNST,LLGL2,EGFR,SH3
ASF1B,DEF8,PDLIM5,RFX3,VAMP8,CAPG,PSMD4,CDC14B,TMEM126B,TXNL4A,PODXL,TCF4,ATPAF2,PRMT

FR,HNRNPLL,RPL15,B3GAT3,MTHFS,COA3,MPC2,TYMS,DALRD3,DRG2,BRF1,MRPS18C,GMPR2,MCFD2,S

ATP6V1F,PDCD5,TRIM16,SLC16A1,STXBP2,SEC31A,RPL18,CNST,LLGL2,EGFR,SH3GLB1,RPL15,B3GAT3,M
ATP6V1F,SLC33A1,PDCD5,TRIM16,SLC16A1,STXBP2,SEC31A,RPL18,CNST,LLGL2,EGFR,SH3GLB1,RPL15,B3
8,RPS15,ERLEC1,HSPB1,SERPINB1,SYNGR1,RANGRF,LSR,TOMM40,PPP3CB,RABEP2,ATP6V1F,PDCD5,POT

OASY,SRSF1,SAP30BP,TRNT1,ITGB1BP1,MED15,CLASRP,UGGT2,UBE2I,PRMT2,JUP,TBX6,XBP1,CELF1,AUN
LS3,TCP1,CD74,PARK7,CCDC40,ZNHIT3,PDCD6,MFF,NOP2,COX20,COA1,COBL,USP4,NDUFS3,RPLP0,HAX1
,PSMA3,MUC1,METTL17,PPRC1,RPS25,DUSP22,THEM4,RPS6,RPL29,RTEL1,COASY,SRSF1,SAP30BP,TRNT
D6G,MICAL1,PRMT2,JUP,CDC42,PTK2,AUNIP,CENPN,ASF1B,PDLIM5,RFX3,VAMP8,CAPG,VIM,CDC14B,TI

OASY,SRSF1,SAP30BP,TRNT1,ITGB1BP1,MED15,CLASRP,UGGT2,UBE2I,PRMT2,JUP,TBX6,XBP1,CELF1,AUN
CHFR,RNF4,RNF149,UBA6,MVB12B,GSK3A,TRIM9,EDEM2,WWP2,CTNNB1,C17ORF97,PSMD11,GIPC1,EXC
NT5E,FAM192A,GCDH,PCNP,PSMD7,NEDD4L,RNF130,BFAR,CLTA,LYPLAL1,AXIN1,PSMD10,PML,TMEM19

,COPZ1,HSP90AB1,RAB2A,ATP5J,RPS20,VPS28,RPS15,ERLEC1,HSPB1,SERPINB1,SYNGR1,RANGRF,LSR,TO
S25,NLRP1,DUSP22,SPOCK2,DBNDD2,RPS6,RPL29,AUP1,ITGB1BP1,UGGT2,MICAL1,UBE2I,PRMT2,EIF3M,
M126B,TXNL4A,PODXL,TCF4,ATPAF2,PRMT1,STOML2,PPAN,HSP90AB1,CEP70,VPS28,RPS15,VWA2,ERCC
,PHF19,VCAN,CD44,SON,TET2,SNF8,HMGA2,CDC37,RPS16,TSG101,OBSL1,TTC21B,YLPM1,PHF21A,CHCH
SBP1L1,AGPAT4,MAF,EEF1B2,EDF1,RFX3,PPA1,CAMTA1,PSMD4,RPL28,VIM,LIG1,PSMA7,APEH,RPL21,TC
OASY,SRSF1,SAP30BP,TRNT1,ITGB1BP1,MED15,CLASRP,UGGT2,UBE2I,PRMT2,JUP,TBX6,XBP1,CELF1,AUN

OT8,RPS9,RCE1,MRPL33,RUVBL2,PCMT1,TESC,NUDT5,RBPMS,NNMT,SNRPN,IL1RAP,STYXL1,RFX5,POLD

,TBX6,XBP1,CELF1,AUNIP,CARD14,ATF7,FBXW9,HSBP1L1,MAF,EDF1,RFX3,PPA1,ENDOV,CAMTA1,PSMD
PL29,GGPS1,RTEL1,COASY,SRSF1,SAP30BP,TRNT1,ITGB1BP1,MED15,CLASRP,UGGT2,UBE2I,PRMT2,JUP,T

IM40,PPP3CB,RABEP2,ATP6V1F,SLC33A1,PDCD5,TRIM16,SLC16A1,STXBP2,SEC31A,RPL18,CNST,LLGL2,E
ARD14,ATF7,MAF,EEF1B2,EDF1,RFX3,PPA1,CAMTA1,PSMD4,RPL28,VIM,LIG1,PSMA7,APEH,RPL21,TCF4,
GSM3,RPS7,SNX3,USP15,NFE2L1,TAF9,UBA52,CAPN2,SUMO1,COMMD1,PSMD8,RAB26,SUFU,NUB1,OS9

,XBP1,CELF1,CARD14,ATF7,HSBP1L1,MAF,EEF1B2,EDF1,RFX3,PPA1,CAMTA1,PSMD4,RPL28,VIM,LIG1,PS
GA2,CDC37,RPS16,TSG101,OBSL1,PTPN13,UBR4,PSMA1,CHCHD1,RGS3,PSMA3,MUC1,METTL17,ATG12,

5,RPL29,GGPS1,RTEL1,COASY,SAP30BP,ITGB1BP1,MED15,SH3YL1,IP6K2,UGGT2,HS2ST1,UBE2I,PRMT2,E
A2,CDC37,TSG101,OBSL1,TTC21B,YLPM1,PHF21A,CHCHD1,FUZ,MUC1,ECSIT,EPB41L5,ATG12,PPRC1,NLR

9,UBA52,SUMO1,PRKCE,COMMD1,PSMD8,NUP85,RAB26,BSCL2,SUFU,NUB1,OS9,EXOSC2,PSMC5,HNRN
TEL1,COASY,SAP30BP,ITGB1BP1,MED15,SH3YL1,UGGT2,HS2ST1,UBE2I,PRMT2,EIF3M,MAGT1,JUP,CDC4

S3,PSMA3,MUC1,METTL17,ATG12,RPS25,NLRP1,DUSP22,SPOCK2,DBNDD2,RPS6,RPL29,AUP1,ITGB1BP

RFX3,PPA1,ENDOV,CAMTA1,PSMD4,RPL28,VIM,PSMA7,RPL21,TXNL4A,TCF4,TMPO,EXOSC9,ZNF7,ARL6I

5,RPL29,GGPS1,RTEL1,COASY,SAP30BP,ITGB1BP1,MED15,SH3YL1,IP6K2,UGGT2,HS2ST1,UBE2I,PRMT2,E

A,GSTK1,TTLL5,LDHA,PSMA6,ACOT8,RPS9,RCE1,MRPL33,RUVBL2,PCMT1,EBPL,TESC,NUDT5,RBPMS,NN
CLASP2,VDAC1,CNTRL,NUP62,PAM16,SKA2,ARPC5,SIN3A,NRP1,TP53,PSMD10,HDAC8,PML,UBE2D3,RPS

,HNRNPA1,HIGD1A,POLR2C,EGLN2,ASH2L,TXNRD2,COPS4,OARD1,SLC25A23,SCAP,CD74,PARK7,NFKBIA,

P2,CD44,TET2,SNF8,GRK6,HMGA2,CDC37,RPS16,CPSF3L,TSG101,OBSL1,TTC21B,PTPN13,UBR4,PSMA1,Y

,RUNX1,CHID1,NUDT22,UFD1L,TAZ,THYN1,SH2B1,NOP16,RPL13A,GSTK1,TTLL5,LDHA,PSMA6,ACOT8,AP
7,RPS16,CPSF3L,TSG101,OBSL1,PTPN13,UBR4,PSMA1,YLPM1,CHCHD1,RGS3,PSMA3,ECSIT,EPB41L5,ATG
GEP,TBL2,SCARB1,CTTN,ITSN1,EIF5A,TNFSF10,COMMD5,RUNX1,CHID1,NUDT22,UFD1L,TAZ,THYN1,SH2E
F2,BUD31,SAE1,NPRL2,OSGEP,TBL2,ARHGAP6,SCARB1,CTTN,ITSN1,EIF5A,COMMD5,RUNX1,CHID1,NUD
SUMF2,SAE1,NPRL2,OSGEP,TBL2,ARHGAP6,SCARB1,CTTN,ITSN1,EIF5A,COMMD5,CHID1,UFD1L,TAZ,DBN
AF11,DBI,ARPC2,PMFBP1,LIMK2,VDAC3,NOB1,WDR19,SIRT6,NUMA1,SUMF2,BUD31,SAE1,NPRL2,OSGEP
1,WDR19,SIRT6,NUMA1,SUMF2,BUD31,SAE1,NPRL2,OSGEP,TBL2,ARHGAP6,SCARB1,CTTN,ITSN1,EIF5A,
EIF3C,RPS3,MRPS24,RHEB,MAGOHB,SNRPD1,SNRPA1,RPS27L,RPS7,U2AF1L4,TAF9,UBA52,RPS4X,MRPS1
MAGED2,SON,SNF8,HMGA2,RPS16,CPSF3L,TSG101,TTC21B,PTPN13,UBR4,PSMA1,YLPM1,PHF21A,CHCH
MAGED2,SON,SNF8,HMGA2,RPS16,CPSF3L,TSG101,TTC21B,PTPN13,UBR4,PSMA1,YLPM1,PHF21A,CHCH
MAGED2,SON,SNF8,HMGA2,RPS16,CPSF3L,TSG101,TTC21B,PTPN13,UBR4,PSMA1,YLPM1,PHF21A,CHCH
ANBP1,UBE2L6,VRK3,ZC3H14,TUBA1C,IRF9,DST,STIP1,PHF19,PCBP2,MAGED2,SON,TET2,SNF8,HMGA2,R

X4I1,SLC25A10,CENPK,RPS6,RTEL1,SRSF1,SAP30BP,TRNT1,ITGB1BP1,MED15,IP6K2,CLASRP,UBE2I,PRM

EPB41L5,PPRC1,RPS25,NLRP1,NIP7,COX4I1,SLC25A10,CENPK,RPS6,RTEL1,SRSF1,SAP30BP,TRNT1,ITGB1B
RPS25,NLRP1,NIP7,COX4I1,DUSP22,CENPK,RPS6,RPL29,SRSF1,AUP1,FOXRED1,RABL5,KXD1,MED15,PAR
REB3L4,POLR2M,BAX,CDK4,NME1,NXF1,ADRA1A,VDAC1,NUP62,PAM16,NDUFA11,NRP1,TP53,SLC25A2
REB3L4,POLR2M,BAX,CDK4,NME1,NXF1,ADRA1A,VDAC1,NUP62,PAM16,NDUFA11,NRP1,TP53,SLC25A2

2,CAPN2,MRPS16,ARL2,SDHA,NDUFA7,NDUFS2,MRPS33,MCUR1,NDUFC1,QTRT1,DUSP18,NDUFB10,MR
27,SLC25A35,COQ2,MRPS7,NDUFS7,IFI6,TIMM9,UQCRB,MRPL36,COX7C,NDUFS4,SLC25A45
2,CENPK,RPS6,RPL29,RTEL1,SAP30BP,RABL5,ITGB1BP1,IP6K2,JPH2,MICAL1,UBE2I,PRMT2,JUP,CDC42,TB
2,CENPK,RPS6,RPL29,RTEL1,SAP30BP,RABL5,ITGB1BP1,IP6K2,JPH2,MICAL1,UBE2I,PRMT2,JUP,CDC42,TB
NDUFS3,KLHL7,METTL3,TSEN34,ARIH2,FBXL18,UBE2D4,POLR2M,CDK4,ANAPC13,RNF8,PPP2R5C,RALY,C
M4,GEMIN2,TP53,RPS3A,PML,OXR1,RPS3,DNAJC2,IDH3G,RGS12,ZCCHC9,SMUG1,RPS7,TBCA,POLR2E,SL
ENY2,COX20,MRPS17,TMEM14B,COA1,ACOT2,NDUFS3,NMNAT3,HAX1,KRAS,MPV17,MRPL55,CREB3L4,

QTRT1,DUSP18,NDUFB10,MRPL51,MRPL43,IMMP1L,MRPL48,COQ4,ERAL1,MTHFD2L,MRPL27,SLC25A35

IE2,PABPC1,DBNL,BCAM,H2AFV,RBMX,BLVRB,SERPINB6,ATP6V1F,PDCD5,B2M,NT5C,PSME1,SLC16A1,M
IE2,PABPC1,DBNL,BCAM,H2AFV,RBMX,BLVRB,SERPINB6,ATP6V1F,PDCD5,B2M,NT5C,PSME1,SLC16A1,M
IE2,PABPC1,DBNL,BCAM,H2AFV,RBMX,BLVRB,SERPINB6,ATP6V1F,PDCD5,B2M,NT5C,PSME1,SLC16A1,M

X,TMPO,C17ORF62,STOML2,PIGL,ZFAND2B,AP1S3,CYB561,SLC9B2,NFAM1,LAMP2,COPZ1,HSP90AB1,S

9,TXNDC17,AP1S3,HNRNPC,CYB561,STAU1,SLC9B2,MOB1A,NFAM1,MDH1,LAMP2,COPZ1,HSP90AB1,D

TAU2,RPLP0,HAX1,DGAT1,SSR3,CREB3L4,ARL6IP5,SLMAP,BAX,UBA1,DLG4,SLC39A6,FAM69B,RER1,RPL

MEM108,DGAT1,KRAS,MPV17,PICK1,ECE1,ADD1,BAX,UBA1,DLG4,NME1,TMEM9,ADRA1A,CFLAR,SYP,LA

'D1,SNRPA1,RPS27L,ZBTB8OS,RPS7,U2AF1L4,NCBP2,POLR2E,RNPS1,UBA52,RPS4X,RPS23,RPL30,PSMD8,

C2,LIMK2,VDAC3,NOB1,WDR19,SIRT6,NUMA1,GPATCH2L,SUMF2,BUD31,SAE1,NPRL2,OSGEP,TBL2,ARHG
JMF2,BUD31,SAE1,NPRL2,OSGEP,TBL2,ARHGAP6,SCARB1,CRYGS,CTTN,TSPAN11,ITSN1,EIF5A,COMMD5,
,IGSF8,NUDT5,SPAG7,STYXL1,RFX5,VRK3,LZTFL1,PPM1K,TUBA1C,IRF9,DST,ANTXR2,STIP1,LRRC1,VCAN,C
ARRDC1,TRPT1,KLC1,RBP5,ZC3HAV1,SCAF11,DBI,ARPC2,LIMK2,VDAC3,NOB1,WDR19,SIRT6,NUMA1,GPA
ARRDC1,TRPT1,KLC1,RBP5,ZC3HAV1,SCAF11,DBI,ARPC2,LIMK2,VDAC3,NOB1,WDR19,SIRT6,NUMA1,GPA
K2,VDAC3,NOB1,WDR19,SIRT6,NUMA1,GPATCH2L,SUMF2,BUD31,SAE1,NPRL2,OSGEP,TBL2,ARHGAP6,S
K2,VDAC3,NOB1,WDR19,SIRT6,NUMA1,GPATCH2L,SUMF2,BUD31,SAE1,NPRL2,OSGEP,TBL2,ARHGAP6,S
RT6,NUMA1,SUMF2,BUD31,SAE1,NPRL2,OSGEP,TBL2,ARHGAP6,SCARB1,CTTN,TSPAN11,ITSN1,EIF5A,CO
I11,EIF5A,COMMD5,CHID1,NUDT22,AMZ2,UFD1L,USP38,TAZ,THYN1,DBNDD1,SH2B1,NOP16,RPL13A,AR
I11,EIF5A,COMMD5,CHID1,NUDT22,AMZ2,UFD1L,USP38,TAZ,THYN1,DBNDD1,SH2B1,NOP16,RPL13A,AR
.9,SIRT6,NUMA1,GPATCH2L,SUMF2,BUD31,SAE1,NPRL2,OSGEP,TBL2,ARHGAP6,SCARB1,CTTN,TSPAN11,
GSF8,NUDT5,SNRPN,WDR62,STYXL1,RFX5,POLD4,LZTFL1,FIS1,PPM1K,IRF9,DST,ANTXR2,STIP1,LRRC1,VCA
GSF8,NUDT5,SNRPN,WDR62,STYXL1,RFX5,POLD4,LZTFL1,FIS1,PPM1K,IRF9,DST,ANTXR2,STIP1,LRRC1,VCA
L,TSG101,UBR4,PHF21A,METTL17,ATG12,RPS25,NIP7,TTC9C,COX4I1,DUSP22,SLC25A10,TMX2,THEM4,R
N1,DBNDD1,SH2B1,NOP16,RPL13A,ARHGEF39,GSTK1,TTLL5,PSMA6,ACOT8,AP3B2,RPS9,SLC35G1,RCE1,N
N1,DBNDD1,SH2B1,NOP16,RPL13A,ARHGEF39,GSTK1,TTLL5,PSMA6,ACOT8,AP3B2,RPS9,SLC35G1,RCE1,N
,SUMF2,BUD31,SAE1,NPRL2,OSGEP,TBL2,ARHGAP6,SCARB1,CRYGS,CTTN,TSPAN11,ITSN1,EIF5A,COMMD

,SUMF2,BUD31,SAE1,NPRL2,OSGEP,TBL2,ARHGAP6,SCARB1,CRYGS,CTTN,TSPAN11,ITSN1,EIF5A,COMMD
,PSMA6,ACOT8,AP3B2,RPS9,SLC35G1,RCE1,MRPL33,ZNF410,RUVBL2,PCMT1,TESC,IGSF8,NUDT5,SNRPN
,PSMA6,ACOT8,AP3B2,RPS9,SLC35G1,RCE1,MRPL33,ZNF410,RUVBL2,PCMT1,TESC,IGSF8,NUDT5,SNRPN
G1,RCE1,MRPL33,ZNF410,PCMT1,IGSF8,NUDT5,SNRPN,STYXL1,RFX5,POLD4,VRK3,LZTFL1,FIS1,PPM1K,ST
G1,RCE1,MRPL33,ZNF410,PCMT1,IGSF8,NUDT5,SNRPN,STYXL1,RFX5,POLD4,VRK3,LZTFL1,FIS1,PPM1K,ST
,TAZ,THYN1,SH2B1,NOP16,RPL13A,ARHGEF39,GSTK1,TTLL5,PSMA6,ACOT8,AP3B2,RPS9,SLC35G1,RCE1,
,TAZ,THYN1,SH2B1,NOP16,RPL13A,ARHGEF39,GSTK1,TTLL5,PSMA6,ACOT8,AP3B2,RPS9,SLC35G1,RCE1,
1,KLC1,RBP5,ZC3HAV1,SCAF11,DBI,ARPC2,LIMK2,VDAC3,NOB1,WDR19,SIRT6,NUMA1,GPATCH2L,SUMF
1,KLC1,RBP5,ZC3HAV1,SCAF11,DBI,ARPC2,LIMK2,VDAC3,NOB1,WDR19,SIRT6,NUMA1,GPATCH2L,SUMF
13A,ARHGEF39,GSTK1,TTLL5,PSMA6,ACOT8,AP3B2,RPS9,SLC35G1,RCE1,MRPL33,ZNF410,RUVBL2,PCM
13A,ARHGEF39,GSTK1,TTLL5,PSMA6,ACOT8,AP3B2,RPS9,SLC35G1,RCE1,MRPL33,ZNF410,RUVBL2,PCM
SCARB1,CTTN,TSPAN11,EIF5A,COMMD5,CHID1,NUDT22,AMZ2,UFD1L,USP38,TAZ,THYN1,DBNDD1,SH2B
SCARB1,CTTN,TSPAN11,EIF5A,COMMD5,CHID1,NUDT22,AMZ2,UFD1L,USP38,TAZ,THYN1,DBNDD1,SH2B
1,MAGED2,SON,UQCRQ,SNF8,CDC37,RPS16,CPSF3L,TSG101,UBR4,FUZ,FAM3A,ATG12,RPS25,NIP7,COX4
1,MAGED2,SON,UQCRQ,SNF8,CDC37,RPS16,CPSF3L,TSG101,UBR4,FUZ,FAM3A,ATG12,RPS25,NIP7,COX4
N,TSPAN11,EIF5A,COMMD5,RUNX1,CHID1,NUDT22,UFD1L,USP38,TAZ,THYN1,DBNDD1,SH2B1,NOP16,R
N,TSPAN11,EIF5A,COMMD5,RUNX1,CHID1,NUDT22,UFD1L,USP38,TAZ,THYN1,DBNDD1,SH2B1,NOP16,R
RB1,CTTN,TSPAN11,EIF5A,COMMD5,CHID1,NUDT22,UFD1L,USP38,TAZ,THYN1,DBNDD1,SH2B1,NOP16,R
RB1,CTTN,TSPAN11,EIF5A,COMMD5,CHID1,NUDT22,UFD1L,USP38,TAZ,THYN1,DBNDD1,SH2B1,NOP16,R
RT6,NUMA1,SUMF2,BUD31,SAE1,NPRL2,TBL2,ARHGAP6,SCARB1,CTTN,TSPAN11,ITSN1,EIF5A,COMMD5,
IF5A,COMMD5,CHID1,NUDT22,UFD1L,USP38,TAZ,THYN1,DBNDD1,SH2B1,NOP16,RPL13A,ARHGEF39,GS
IF5A,COMMD5,CHID1,NUDT22,UFD1L,USP38,TAZ,THYN1,DBNDD1,SH2B1,NOP16,RPL13A,ARHGEF39,GS
ON,UQCRQ,SNF8,CDC37,RPS16,CPSF3L,TSG101,UBR4,YLPM1,PHF21A,CHCHD1,FUZ,FAM3A,ATG12,RPS2
SAE1,NPRL2,OSGEP,TBL2,ARHGAP6,SCARB1,CRYGS,CTTN,TSPAN11,ITSN1,EIF5A,COMMD5,NUDT22,AMZ
SAE1,NPRL2,OSGEP,TBL2,ARHGAP6,SCARB1,CRYGS,CTTN,TSPAN11,ITSN1,EIF5A,COMMD5,NUDT22,AMZ
RSF1,SAP30BP,YIPF3,FOXRED1,TRNT1,KXD1,ITGB1BP1,MED15,SH3YL1,IP6K2,CLASRP,C1ORF198,MICAL1
NOB1,WDR19,SIRT6,NUMA1,GPATCH2L,SUMF2,BUD31,SAE1,NPRL2,OSGEP,TBL2,ARHGAP6,SCARB1,CRY
1,KLC1,RBP5,ZC3HAV1,SCAF11,DBI,ARPC2,LIMK2,VDAC3,NOB1,WDR19,SIRT6,NUMA1,GPATCH2L,SUMF
1,KLC1,RBP5,ZC3HAV1,SCAF11,DBI,ARPC2,LIMK2,VDAC3,NOB1,WDR19,SIRT6,NUMA1,GPATCH2L,SUMF
T8,AP3B2,RPS9,SLC35G1,RCE1,MRPL33,ZNF410,PCMT1,IGSF8,NUDT5,SNRPN,STYXL1,RFX5,POLD4,VRK3,
T8,AP3B2,RPS9,SLC35G1,RCE1,MRPL33,ZNF410,PCMT1,IGSF8,NUDT5,SNRPN,STYXL1,RFX5,POLD4,VRK3,
16,RPL13A,ARHGEF39,GSTK1,TTLL5,PSMA6,ACOT8,AP3B2,RPS9,SLC35G1,RCE1,MRPL33,ZNF410,RUVBL
16,RPL13A,ARHGEF39,GSTK1,TTLL5,PSMA6,ACOT8,AP3B2,RPS9,SLC35G1,RCE1,MRPL33,ZNF410,RUVBL
,SUMF2,BUD31,SAE1,NPRL2,OSGEP,ARHGAP6,SCARB1,CTTN,TSPAN11,EIF5A,COMMD5,CHID1,NUDT22,
,SUMF2,BUD31,SAE1,NPRL2,OSGEP,ARHGAP6,SCARB1,CTTN,TSPAN11,EIF5A,COMMD5,CHID1,NUDT22,
39,GSTK1,TTLL5,PSMA6,ACOT8,AP3B2,RPS9,SLC35G1,RCE1,MRPL33,ZNF410,PCMT1,IGSF8,NUDT5,SNRP
39,GSTK1,TTLL5,PSMA6,ACOT8,AP3B2,RPS9,SLC35G1,RCE1,MRPL33,ZNF410,PCMT1,IGSF8,NUDT5,SNRP
AN11,EIF5A,COMMD5,CHID1,NUDT22,AMZ2,UFD1L,USP38,TAZ,THYN1,DBNDD1,SH2B1,NOP16,RPL13A,
AN11,EIF5A,COMMD5,CHID1,NUDT22,AMZ2,UFD1L,USP38,TAZ,THYN1,DBNDD1,SH2B1,NOP16,RPL13A,
1,POLD4,VRK3,LZTFL1,FIS1,PPM1K,CD44,TM9SF1,MAGED2,SON,UQCRQ,SNF8,GRK6,HMGA2,CDC37,RPS
1,POLD4,VRK3,LZTFL1,FIS1,PPM1K,CD44,TM9SF1,MAGED2,SON,UQCRQ,SNF8,GRK6,HMGA2,CDC37,RPS
PS9,SLC35G1,RCE1,MRPL33,ZNF410,PCMT1,IGSF8,NUDT5,SNRPN,WDR62,STYXL1,RFX5,POLD4,VRK3,LZT
PS9,SLC35G1,RCE1,MRPL33,ZNF410,PCMT1,IGSF8,NUDT5,SNRPN,WDR62,STYXL1,RFX5,POLD4,VRK3,LZT
RT6,NUMA1,GPATCH2L,SUMF2,BUD31,SAE1,NPRL2,OSGEP,TBL2,ARHGAP6,CTTN,TSPAN11,ITSN1,EIF5A,
,COX4I1,THEM4,DBNDD2,RPS6,RPL29,RTEL1,AUP1,SAP30BP,YIPF3,FOXRED1,TRNT1,KXD1,ITGB1BP1,ME
RPL29,RTEL1,AUP1,SAP30BP,YIPF3,FOXRED1,TRNT1,KXD1,ITGB1BP1,MED15,SH3YL1,CLASRP,C1ORF198,
E1,MRPL33,ZNF410,PCMT1,IGSF8,NUDT5,SNRPN,SPAG7,WDR62,STYXL1,RFX5,POLD4,RANBP1,VRK3,LZT

E1,MRPL33,ZNF410,PCMT1,IGSF8,NUDT5,SNRPN,SPAG7,WDR62,STYXL1,RFX5,POLD4,RANBP1,VRK3,LZT

5,PSMA6,ACOT8,AP3B2,RPS9,SLC35G1,RCE1,MRPL33,ZNF410,RUVBL2,PCMT1,IGSF8,NUDT5,RBPMS,SNR

5,PSMA6,ACOT8,AP3B2,RPS9,SLC35G1,RCE1,MRPL33,ZNF410,RUVBL2,PCMT1,IGSF8,NUDT5,RBPMS,SNR

LC35G1,RCE1,MRPL33,ZNF410,PCMT1,IGSF8,NUDT5,SNRPN,STYXL1,RFX5,POLD4,RANBP1,VRK3,LZTFL1,

LC35G1,RCE1,MRPL33,ZNF410,PCMT1,IGSF8,NUDT5,SNRPN,STYXL1,RFX5,POLD4,RANBP1,VRK3,LZTFL1,

D4,VRK3,LZTFL1,FIS1,PPM1K,TUBA1C,STIP1,VCAN,MAGED2,SON,UQCRQ,SNF8,CDC37,CPSF3L,TSG101,UB

D4,VRK3,LZTFL1,FIS1,PPM1K,TUBA1C,STIP1,VCAN,MAGED2,SON,UQCRQ,SNF8,CDC37,CPSF3L,TSG101,UB

R62,STYXL1,POLD4,LZTFL1,FIS1,PPM1K,VCAN,MAGED2,SON,UQCRQ,SNF8,GRK6,HMGA2,CDC37,CPSF3L,

R62,STYXL1,POLD4,LZTFL1,FIS1,PPM1K,VCAN,MAGED2,SON,UQCRQ,SNF8,GRK6,HMGA2,CDC37,CPSF3L,

3B2,RPS9,SLC35G1,RCE1,MRPL33,ZNF410,PCMT1,IGSF8,NUDT5,RBPMS,SNRPN,WDR62,STYXL1,RFX5,PO

3B2,RPS9,SLC35G1,RCE1,MRPL33,ZNF410,PCMT1,IGSF8,NUDT5,RBPMS,SNRPN,WDR62,STYXL1,RFX5,PO

A,COMMD5,CHID1,NUDT22,AMZ2,UFD1L,USP38,TAZ,THYN1,DBNDD1,SH2B1,NOP16,RPL13A,ARHGEF39,

A,COMMD5,CHID1,NUDT22,AMZ2,UFD1L,USP38,TAZ,THYN1,DBNDD1,SH2B1,NOP16,RPL13A,ARHGEF39,

S9,SLC35G1,RCE1,MRPL33,ZNF410,RUVBL2,PCMT1,IGSF8,NUDT5,SNRPN,STYXL1,RFX5,POLD4,RANBP1,V

S9,SLC35G1,RCE1,MRPL33,ZNF410,RUVBL2,PCMT1,IGSF8,NUDT5,SNRPN,STYXL1,RFX5,POLD4,RANBP1,V

PCYT2,FBXW9,USE1,ASF1B,RFX3,ENDOV,CAMTA1,PPIL2,LIG1,APEH,CAPN10,LYPLA2,C17ORF62,PRMT1,Z

1,NOP16,RPL13A,ARHGEF39,GSTK1,TTLL5,PSMA6,ACOT8,AP3B2,RPS9,SLC35G1,RCE1,MRPL33,ZNF410,

1,NOP16,RPL13A,ARHGEF39,GSTK1,TTLL5,PSMA6,ACOT8,AP3B2,RPS9,SLC35G1,RCE1,MRPL33,ZNF410,

UDT5,SNRPN,STYXL1,RFX5,POLD4,VRK3,LZTFL1,FIS1,PPM1K,IRF9,ANTXR2,LRRC1,TM9SF1,MAGED2,SON,

UDT5,SNRPN,STYXL1,RFX5,POLD4,VRK3,LZTFL1,FIS1,PPM1K,IRF9,ANTXR2,LRRC1,TM9SF1,MAGED2,SON,

8,AP3B2,RPS9,SLC35G1,RCE1,MRPL33,ZNF410,PCMT1,IGSF8,NUDT5,SNRPN,WDR62,STYXL1,RFX5,POLD

8,AP3B2,RPS9,SLC35G1,RCE1,MRPL33,ZNF410,PCMT1,IGSF8,NUDT5,SNRPN,WDR62,STYXL1,RFX5,POLD

OT8,AP3B2,RPS9,SLC35G1,RCE1,MRPL33,ZNF410,PCMT1,IGSF8,NUDT5,RBPMS,SNRPN,STYXL1,RFX5,POL

OT8,AP3B2,RPS9,SLC35G1,RCE1,MRPL33,ZNF410,PCMT1,IGSF8,NUDT5,RBPMS,SNRPN,STYXL1,RFX5,POL

62,PRMT1,STOML2,ARL6IP4,C19ORF48,PPAN,HNRNPC,SUGT1,CARS,RBM3,ZNF197,INO80E,CCNL2,DGU

CE1,MRPL33,ZNF410,PCMT1,IGSF8,NUDT5,SNRPN,WDR62,STYXL1,RFX5,POLD4,RANBP1,VRK3,LZTFL1,FIS

CE1,MRPL33,ZNF410,PCMT1,IGSF8,NUDT5,SNRPN,WDR62,STYXL1,RFX5,POLD4,RANBP1,VRK3,LZTFL1,FIS

N1,DBNDD1,NOP16,GSTK1,TTLL5,PSMA6,ACOT8,AP3B2,RCE1,ZNF410,RUVBL2,PCMT1,EBPL,TESC,IGSF8,

N1,DBNDD1,NOP16,GSTK1,TTLL5,PSMA6,ACOT8,AP3B2,RCE1,ZNF410,RUVBL2,PCMT1,EBPL,TESC,IGSF8,

ASF1B,CCDC25,EEF1B2,RFX3,ENDOV,CAMTA1,PPIL2,RPL28,APEH,CAPN10,LYPLA2,C17ORF62,PRMT1,ZN

NF7,TXNDC17,ARL6IP4,PPAN,ACAD10,SUGT1,HYPK,CARS,CDK11A,ZNF197,DGUOK,LRRFIP1,ZNF524,ATP5

A,COMMD5,CHID1,NUDT22,AMZ2,UFD1L,USP38,TAZ,THYN1,DBNDD1,SH2B1,NOP16,RPL13A,ARHGEF39,

A,COMMD5,CHID1,NUDT22,AMZ2,UFD1L,USP38,TAZ,THYN1,DBNDD1,SH2B1,NOP16,RPL13A,ARHGEF39,

RB1,CRYGS,CTTN,TSPAN11,ITSN1,EIF5A,COMMD5,CHID1,NUDT22,AMZ2,UFD1L,TAZ,THYN1,DBNDD1,SH

RB1,CRYGS,CTTN,TSPAN11,ITSN1,EIF5A,COMMD5,CHID1,NUDT22,AMZ2,UFD1L,TAZ,THYN1,DBNDD1,SH

C3HAV1,SCAF11,DBI,ARPC2,LIMK2,VDAC3,NOB1,WDR19,SIRT6,NUMA1,GPATCH2L,SUMF2,BUD31,SAE1,

C3HAV1,SCAF11,DBI,ARPC2,LIMK2,VDAC3,NOB1,WDR19,SIRT6,NUMA1,GPATCH2L,SUMF2,BUD31,SAE1,

ARD14,PCYT2,FBXW9,USE1,ASF1B,CCDC25,RFX3,ENDOV,CAMTA1,PPIL2,RPL28,LIG1,APEH,CAPN10,LYPL

N,TSPAN11,ITSN1,EIF5A,COMMD5,CHID1,NUDT22,AMZ2,UFD1L,USP38,THYN1,SH2B1,NOP16,RPL13A,A

N,TSPAN11,ITSN1,EIF5A,COMMD5,CHID1,NUDT22,AMZ2,UFD1L,USP38,THYN1,SH2B1,NOP16,RPL13A,A

GPATCH2L,BUD31,NPRL2,OSGEP,TBL2,ARHGAP6,SCARB1,CTTN,TSPAN11,ITSN1,EIF5A,COMMD5,RUNX1,

GPATCH2L,BUD31,NPRL2,OSGEP,TBL2,ARHGAP6,SCARB1,CTTN,TSPAN11,ITSN1,EIF5A,COMMD5,RUNX1,

1,TRNT1,KXD1,ITGB1BP1,MED15,CLASRP,C1ORF198,UBE2I,SLC39A13,EIF3M,CDC42,CARD14,PCYT2,FBX

FL1,FIS1,PPM1K,LRRC1,MAGED2,SON,UQCRQ,SNF8,HMGA2,CDC37,RPS16,CPSF3L,TSG101,UBR4,FUZ,FA

FL1,FIS1,PPM1K,LRRC1,MAGED2,SON,UQCRQ,SNF8,HMGA2,CDC37,RPS16,CPSF3L,TSG101,UBR4,FUZ,FA

LA,ZNF197,COPZ1,INO80E,CCNL2,DGUOK,LRRFIP1,ZNF524,ATP5J,TRMT112,CHPF,RPS20,SAT1,ERCC5,EM

F1B,RFX3,ENDOV,CAMTA1,PPIL2,RPL28,APEH,CAPN10,LYPLA2,C17ORF62,PRMT1,STOML2,C19ORF54,ZN

A10,TMX2,THEM4,DBNDD2,RPS6,RPL29,RTEL1,SRSF1,AUP1,SAP30BP,YIPF3,FOXRED1,TRNT1,KXD1,ITGB

16,RPL13A,ARHGEF39,TTLL5,PSMA6,ACOT8,AP3B2,RPS9,SLC35G1,RCE1,MRPL33,ZNF410,PCMT1,IGSF8,N

16,RPL13A,ARHGEF39,TTLL5,PSMA6,ACOT8,AP3B2,RPS9,SLC35G1,RCE1,MRPL33,ZNF410,PCMT1,IGSF8,N

,RPL13A,ARHGEF39,GSTK1,TTLL5,PSMA6,ACOT8,AP3B2,RPS9,SLC35G1,RCE1,MRPL33,ZNF410,RUVBL2,P

,RPL13A,ARHGEF39,GSTK1,TTLL5,PSMA6,ACOT8,AP3B2,RPS9,SLC35G1,RCE1,MRPL33,ZNF410,RUVBL2,P

S25,NIP7,SLC25A10,TMX2,THEM4,DBNDD2,RPS6,RPL29,RTEL1,AUP1,SAP30BP,YIPF3,FOXRED1,TRNT1,K

F5A,COMMD5,RUNX1,NUDT22,UFD1L,USP38,TAZ,THYN1,DBNDD1,SH2B1,NOP16,RPL13A,ARHGEF39,GS

F5A,COMMD5,RUNX1,NUDT22,UFD1L,USP38,TAZ,THYN1,DBNDD1,SH2B1,NOP16,RPL13A,ARHGEF39,GS

FX3,ENDOV,CAMTA1,PPIL2,LIG1,TMEM126B,APEH,RPL21,CAPN10,LYPLA2,C17ORF62,PRMT1,ARL6IP4,PP

T6,NUMA1,GPATCH2L,SUMF2,BUD31,SAE1,NPRL2,OSGEP,TBL2,ARHGAP6,SCARB1,CRYGS,CTTN,TSPAN1

T6,NUMA1,GPATCH2L,SUMF2,BUD31,SAE1,NPRL2,OSGEP,TBL2,ARHGAP6,SCARB1,CRYGS,CTTN,TSPAN1

IP4,PPAN,ACAD10,HNRNPC,SUGT1,HYPK,CARS,CDK11A,ZNF197,DGUOK,LRRFIP1,ATP5J,TRMT112,CHPF,

SP38,THYN1,DBNDD1,NOP16,GSTK1,TTLL5,PSMA6,ACOT8,AP3B2,RCE1,ZNF410,RUVBL2,PCMT1,EBPL,TE

SP38,THYN1,DBNDD1,NOP16,GSTK1,TTLL5,PSMA6,ACOT8,AP3B2,RCE1,ZNF410,RUVBL2,PCMT1,EBPL,TE

B,APEH,RPL21,CAPN10,LYPLA2,C17ORF62,PRMT1,STOML2,ZNF7,TXNDC17,ARL6IP4,PPAN,CYB561,SHAR

B,APEH,RPL21,CAPN10,LYPLA2,C17ORF62,PRMT1,STOML2,ZNF7,TXNDC17,ARL6IP4,PPAN,CYB561,SHAR

PS25,NIP7,COX4I1,THEM4,RPS6,RPL29,COASY,AUP1,SAP30BP,YIPF3,FOXRED1,TRNT1,KXD1,ITGB1BP1,M

AZ,THYN1,SH2B1,NOP16,RPL13A,TTLL5,ACOT8,AP3B2,RPS9,SLC35G1,RCE1,ZNF410,RUVBL2,PCMT1,EBP

FX3,ENDOV,CAMTA1,PPIL2,RPL28,APEH,RPL21,CAPN10,LYPLA2,C17ORF62,PRMT1,C19ORF54,ZNF7,TXND

A,NT5C,MDH2,STXBP2,EEF1D,CHCHD5,SEC31A,CNST,HIGD1A,PFDN5,PAFAH1B2,ZNF343,GBA2,PPIH,DA

F3L,TSG101,UBR4,FAM3A,ATG12,NIP7,SLC25A10,TMX2,THEM4,DBNDD2,RPS6,RPL29,RTEL1,AUP1,SAP3

S25,NIP7,COX4I1,SLC25A10,THEM4,RPS6,RPL29,RTEL1,COASY,SAP30BP,YIPF3,FOXRED1,RABL5,TRNT1,K

S25,NIP7,COX4I1,SLC25A10,THEM4,RPS6,RPL29,RTEL1,COASY,SAP30BP,YIPF3,FOXRED1,RABL5,TRNT1,K

,PRMT1,STOML2,ZNF7,TXNDC17,ARL6IP4,PPAN,CYB561,SHARPIN,SUGT1,HYPK,CARS,ZNF250,ZNF197,CC

,PRMT1,STOML2,ZNF7,TXNDC17,ARL6IP4,PPAN,CYB561,SHARPIN,SUGT1,HYPK,CARS,ZNF250,ZNF197,CC

VRK3,LZTFL1,FIS1,PPM1K,TUBA1C,IRF9,DST,ANTXR2,STIP1,LRRC1,VCAN,LPCAT3,MAGED2,SON,UQCRQ,S

VRK3,LZTFL1,FIS1,PPM1K,TUBA1C,IRF9,DST,ANTXR2,STIP1,LRRC1,VCAN,LPCAT3,MAGED2,SON,UQCRQ,S

E1,NPRL2,OSGEP,TBL2,ARHGAP6,SCARB1,CRYGS,TSPAN11,ITSN1,EIF5A,COMMD5,UFD1L,USP38,TAZ,TH

E1,NPRL2,OSGEP,TBL2,ARHGAP6,SCARB1,CRYGS,TSPAN11,ITSN1,EIF5A,COMMD5,UFD1L,USP38,TAZ,TH

,ASF1B,CCDC25,RFX3,ENDOV,CAMTA1,PPIL2,RPL28,LIG1,APEH,CAPN10,LYPLA2,C17ORF62,PRMT1,ZNF7

P1,SAP30BP,YIPF3,FOXRED1,TRNT1,KXD1,MED15,IP6K2,CLASRP,C1ORF198,UBE2I,SLC39A13,EIF3M,CD

MA1,SUMF2,BUD31,SAE1,NPRL2,OSGEP,TBL2,ARHGAP6,SCARB1,CTTN,TSPAN11,ITSN1,EIF5A,COMMD

MA1,SUMF2,BUD31,SAE1,NPRL2,OSGEP,TBL2,ARHGAP6,SCARB1,CTTN,TSPAN11,ITSN1,EIF5A,COMMD

CAMTA1,PPIL2,RPL28,PSMA7,APEH,CAPN10,LYPLA2,C17ORF62,PRMT1,ZNF7,ARL6IP4,C19ORF48,PPAN,

2,TBX6,CARD14,PCYT2,FBXW9,USE1,ASF1B,CCDC25,EEF1B2,RFX3,ENDOV,PPIL2,LIG1,CDC14B,APEH,CAP

2,TBX6,CARD14,PCYT2,FBXW9,USE1,ASF1B,CCDC25,EEF1B2,RFX3,ENDOV,PPIL2,LIG1,CDC14B,APEH,CAP

L,RFX5,POLD4,VRK3,LZTFL1,FIS1,PPM1K,IRF9,ANTXR2,LRRC1,TM9SF1,MAGED2,SNF8,CDC37,RPS16,CPSF

L,RFX5,POLD4,VRK3,LZTFL1,FIS1,PPM1K,IRF9,ANTXR2,LRRC1,TM9SF1,MAGED2,SNF8,CDC37,RPS16,CPSF

DC42,TBX6,CARD14,PCYT2,FBXW9,ASF1B,CCDC25,EEF1B2,RFX3,ENDOV,PPIL2,LIG1,CDC14B,APEH,RPL21

DC42,TBX6,CARD14,PCYT2,FBXW9,ASF1B,CCDC25,EEF1B2,RFX3,ENDOV,PPIL2,LIG1,CDC14B,APEH,RPL21

PS25,ZBTB44,COX4I1,PLCB2,THEM4,RPS6,RPL29,RTEL1,SAP30BP,YIPF3,FOXRED1,RABL5,TRNT1,ITGB1BP

PS25,ZBTB44,COX4I1,PLCB2,THEM4,RPS6,RPL29,RTEL1,SAP30BP,YIPF3,FOXRED1,RABL5,TRNT1,ITGB1BP

RK6,CDC37,CPSF3L,TSG101,UBR4,SLC39A11,FUZ,FAM3A,RPS25,NIP7,COX4I1,PLCB2,DUSP22,TMX2,THE

RK6,CDC37,CPSF3L,TSG101,UBR4,SLC39A11,FUZ,FAM3A,RPS25,NIP7,COX4I1,PLCB2,DUSP22,TMX2,THE

SUMF2,BUD31,SAE1,NPRL2,OSGEP,TBL2,ARHGAP6,SCARB1,CTTN,TSPAN11,ITSN1,EIF5A,COMMD5,AMZ

SUMF2,BUD31,SAE1,NPRL2,OSGEP,TBL2,ARHGAP6,SCARB1,CTTN,TSPAN11,ITSN1,EIF5A,COMMD5,AMZ

W9,USE1,ASF1B,EEF1B2,RFX3,ENDOV,CAMTA1,PPIL2,RPL28,APEH,RPL21,CAPN10,LYPLA2,C17ORF62,PR

P,YIPF3,FOXRED1,TRNT1,KXD1,MED15,IP6K2,CLASRP,C1ORF198,UGGT2,UBE2I,EIF3M,CDC42,CARD14,PC
P7,TTC9C,COX4I1,TMX2,THEM4,DBNDD2,RPS6,RPL29,RTEL1,AUP1,SAP30BP,YIPF3,FOXRED1,RABL5,TRN
P7,TTC9C,COX4I1,TMX2,THEM4,DBNDD2,RPS6,RPL29,RTEL1,AUP1,SAP30BP,YIPF3,FOXRED1,RABL5,TRN
0BP,YIPF3,FOXRED1,TRNT1,KXD1,ITGB1BP1,MED15,IP6K2,C1ORF198,MICAL1,UBE2I,EIF3M,CDC42,CARD
PK,RPS6,RTEL1,SRSF1,SAP30BP,YIPF3,FOXRED1,RABL5,TRNT1,ITGB1BP1,MED15,CLASRP,C1ORF198,UGG
PK,RPS6,RTEL1,SRSF1,SAP30BP,YIPF3,FOXRED1,RABL5,TRNT1,ITGB1BP1,MED15,CLASRP,C1ORF198,UGG
F7,ARL6IP4,C19ORF48,PPAN,CARS,RBM3,ZNF197,INO80E,CCNL2,DGUOK,LRRFIP1,ZNF524,ATP5J,RMDN1
F2,BUD31,SAE1,NPRL2,OSGEP,TBL2,ARHGAP6,SCARB1,CRYGS,CTTN,TSPAN11,ITSN1,EIF5A,COMMD5,CH
,NIP7,COX4I1,SPOCK2,THEM4,DBNDD2,RPS6,RPL29,RTEL1,COASY,AUP1,SAP30BP,FOXRED1,RABL5,TRNT
AC3,NOB1,WDR19,SIRT6,NUMA1,GPATCH2L,SUMF2,SAE1,NPRL2,OSGEP,TBL2,SCARB1,CRYGS,CTTN,TSPA
,RPL28,LIG1,CDC14B,TIMM17B,APEH,LYPLA2,C17ORF62,STOML2,ZNF7,PPAN,HNRNPC,CYB561,SUGT1,H
,RPL28,LIG1,CDC14B,TIMM17B,APEH,LYPLA2,C17ORF62,STOML2,ZNF7,PPAN,HNRNPC,CYB561,SUGT1,H
1,CHID1,NUDT22,AMZ2,UFD1L,USP38,TAZ,THYN1,DBNDD1,SH2B1,NOP16,RPL13A,ARHGEF39,GSTK1,TT
1,CHID1,NUDT22,AMZ2,UFD1L,USP38,TAZ,THYN1,DBNDD1,SH2B1,NOP16,RPL13A,ARHGEF39,GSTK1,TT
RF9,ANTXR2,STIP1,LRRC1,VCAN,CD44,TM9SF1,MAGED2,SON,UQCRQ,SNF8,CDC37,RPS16,CPSF3L,TSG10
RF9,ANTXR2,STIP1,LRRC1,VCAN,CD44,TM9SF1,MAGED2,SON,UQCRQ,SNF8,CDC37,RPS16,CPSF3L,TSG10
RF9,ANTXR2,STIP1,LRRC1,VCAN,CD44,TM9SF1,MAGED2,SON,UQCRQ,SNF8,CDC37,RPS16,CPSF3L,TSG10
RF9,ANTXR2,STIP1,LRRC1,VCAN,CD44,TM9SF1,MAGED2,SON,UQCRQ,SNF8,CDC37,RPS16,CPSF3L,TSG10
X3,ENDOV,PPIL2,RPL28,LIG1,CDC14B,TIMM17B,APEH,LYPLA2,C17ORF62,PRMT1,STOML2,ZNF7,TXNDC1
X3,ENDOV,PPIL2,RPL28,LIG1,CDC14B,TIMM17B,APEH,LYPLA2,C17ORF62,PRMT1,STOML2,ZNF7,TXNDC1
OX4I1,PLCB2,DUSP22,TMX2,THEM4,DBNDD2,RPS6,RPL29,RTEL1,COASY,AUP1,SAP30BP,YIPF3,FOXRED1,
OX4I1,PLCB2,DUSP22,TMX2,THEM4,DBNDD2,RPS6,RPL29,RTEL1,COASY,AUP1,SAP30BP,YIPF3,FOXRED1,
ML2,C19ORF54,ZNF7,C19ORF48,PPAN,AP1S3,ACAD10,CYB561,SUGT1,HYPK,ZNF197,INO80E,CCNL2,MA
IF3M,CDC42,CARD14,FBXW9,USE1,ASF1B,CCDC25,EEF1B2,RFX3,ENDOV,CAPG,PPIL2,LIG1,PSMA7,CDC1
IF3M,CDC42,CARD14,FBXW9,USE1,ASF1B,CCDC25,EEF1B2,RFX3,ENDOV,CAPG,PPIL2,LIG1,PSMA7,CDC1
P16,RPL13A,ARHGEF39,GSTK1,TTLL5,PSMA6,ACOT8,AP3B2,RPS9,SLC35G1,RCE1,MRPL33,ZNF410,RUVB
P16,RPL13A,ARHGEF39,GSTK1,TTLL5,PSMA6,ACOT8,AP3B2,RPS9,SLC35G1,RCE1,MRPL33,ZNF410,RUVB
H,LYPLA2,C17ORF62,STOML2,ZNF7,PPAN,AP1S3,CYB561,SUGT1,HYPK,CARS,ZNF250,RPL36,ZNF197,INO
H,LYPLA2,C17ORF62,STOML2,ZNF7,PPAN,AP1S3,CYB561,SUGT1,HYPK,CARS,ZNF250,RPL36,ZNF197,INO
C35G1,RCE1,MRPL33,ZNF410,RUVBL2,IGSF8,NUDT5,RBPMS,SNRPN,STYXL1,RFX5,POLD4,PLEKHA5,LZTF
C35G1,RCE1,MRPL33,ZNF410,RUVBL2,IGSF8,NUDT5,RBPMS,SNRPN,STYXL1,RFX5,POLD4,PLEKHA5,LZTF
,COMMD5,CHID1,NUDT22,AMZ2,UFD1L,USP38,THYN1,SH2B1,NOP16,RPL13A,ARHGEF39,GSTK1,TTLL5,
,COMMD5,CHID1,NUDT22,AMZ2,UFD1L,USP38,THYN1,SH2B1,NOP16,RPL13A,ARHGEF39,GSTK1,TTLL5,
,UGGT2,MICAL1,UBE2I,EIF3M,CDC42,CARD14,PCYT2,FBXW9,CCDC25,EEF1B2,RFX3,ENDOV,PPIL2,LIG1,C
,UGGT2,MICAL1,UBE2I,EIF3M,CDC42,CARD14,PCYT2,FBXW9,CCDC25,EEF1B2,RFX3,ENDOV,PPIL2,LIG1,C
PSMA6,ACOT8,AP3B2,RPS9,SLC35G1,RCE1,MRPL33,ZNF410,RUVBL2,PCMT1,IGSF8,NUDT5,SNRPN,STYXL
PSMA6,ACOT8,AP3B2,RPS9,SLC35G1,RCE1,MRPL33,ZNF410,RUVBL2,PCMT1,IGSF8,NUDT5,SNRPN,STYXL
NOB1,WDR19,SIRT6,NUMA1,SUMF2,BUD31,SAE1,NPRL2,OSGEP,TBL2,ARHGAP6,SCARB1,CRYGS,CTTN,TS
Z2,UFD1L,TAZ,THYN1,SH2B1,NOP16,RPL13A,ARHGEF39,GSTK1,TTLL5,LDHA,PSMA6,ACOT8,AP3B2,RPS9
Z2,UFD1L,TAZ,THYN1,SH2B1,NOP16,RPL13A,ARHGEF39,GSTK1,TTLL5,LDHA,PSMA6,ACOT8,AP3B2,RPS9
DD1,NOP16,GSTK1,TTLL5,PSMA6,ACOT8,AP3B2,RCE1,ZNF410,RUVBL2,PCMT1,EBPL,TESC,IGSF8,SNRPN,
DD1,NOP16,GSTK1,TTLL5,PSMA6,ACOT8,AP3B2,RCE1,ZNF410,RUVBL2,PCMT1,EBPL,TESC,IGSF8,SNRPN,
7,C19ORF48,PPAN,HNRNPC,SUGT1,CARS,CDK11A,RBM3,ZNF197,INO80E,CCNL2,DGUOK,LRRFIP1,ZNF52
DBNDD1,NOP16,GSTK1,TTLL5,PSMA6,ACOT8,AP3B2,RCE1,ZNF410,RUVBL2,PCMT1,EBPL,TESC,IGSF8,SNR
DBNDD1,NOP16,GSTK1,TTLL5,PSMA6,ACOT8,AP3B2,RCE1,ZNF410,RUVBL2,PCMT1,EBPL,TESC,IGSF8,SNR
V,PPIL2,LIG1,APEH,CAPN10,LYPLA2,C17ORF62,PRMT1,STOML2,ZNF7,TXNDC17,ARL6IP4,C19ORF48,PPA
V,PPIL2,LIG1,APEH,CAPN10,LYPLA2,C17ORF62,PRMT1,STOML2,ZNF7,TXNDC17,ARL6IP4,C19ORF48,PPA

PIL2,LIG1,CDC14B,APEH,CAPN10,LYPLA2,C17ORF62,PRMT1,STOML2,ZNF7,TXNDC17,PPAN,CYB561,SHA

PIL2,LIG1,CDC14B,APEH,CAPN10,LYPLA2,C17ORF62,PRMT1,STOML2,ZNF7,TXNDC17,PPAN,CYB561,SHA

BL2,ARHGAP6,SCARB1,CTTN,TSPAN11,ITSN1,EIF5A,COMMD5,AMZ2,UFD1L,USP38,TAZ,THYN1,DBNDD1,

BL2,ARHGAP6,SCARB1,CTTN,TSPAN11,ITSN1,EIF5A,COMMD5,AMZ2,UFD1L,USP38,TAZ,THYN1,DBNDD1,

MDN1,TRMT112,CHPF,RPS20,DCTD,SAT1,ERCC5,SERPINB1,DDOST,LSR,PABPC1,RRNAD1,DBNL,TOMM4

UQCRQ,SNF8,CDC37,RPS16,CPSF3L,TSG101,UBR4,YLPM1,FUZ,FAM3A,PSMA3,ECSIT,ATG12,RPS25,NIP7,T

UQCRQ,SNF8,CDC37,RPS16,CPSF3L,TSG101,UBR4,YLPM1,FUZ,FAM3A,PSMA3,ECSIT,ATG12,RPS25,NIP7,T

C2,LIMK2,VDAC3,NOB1,WDR19,SIRT6,NUMA1,GPATCH2L,SUMF2,BUD31,SAE1,OSGEP,TBL2,ARHGAP6,S

C2,LIMK2,VDAC3,NOB1,WDR19,SIRT6,NUMA1,GPATCH2L,SUMF2,BUD31,SAE1,OSGEP,TBL2,ARHGAP6,S

D2,SON,UQCRQ,SNF8,GRK6,RPS16,CPSF3L,TSG101,TTC21B,PHF21A,SLC39A11,FUZ,FAM3A,PSMA3,MUC

D2,SON,UQCRQ,SNF8,GRK6,RPS16,CPSF3L,TSG101,TTC21B,PHF21A,SLC39A11,FUZ,FAM3A,PSMA3,MUC

RBP5,ZC3HAV1,SCAF11,DBI,ARPC2,LIMK2,VDAC3,NOB1,WDR19,SIRT6,NUMA1,GPATCH2L,BUD31,NPRL2

RBP5,ZC3HAV1,SCAF11,DBI,ARPC2,LIMK2,VDAC3,NOB1,WDR19,SIRT6,NUMA1,GPATCH2L,BUD31,NPRL2

,C17ORF62,PRMT1,C19ORF54,ZNF7,TXNDC17,ARL6IP4,PIGL,C19ORF48,PPAN,SUGT1,HYPK,CARS,CDK11

P16,GSTK1,PSMA6,ACOT8,AP3B2,RCE1,MRPL33,ZNF410,RUVBL2,EBPL,TESC,IGSF8,SNRPN,SPAG7,WDR6

P16,GSTK1,PSMA6,ACOT8,AP3B2,RCE1,MRPL33,ZNF410,RUVBL2,EBPL,TESC,IGSF8,SNRPN,SPAG7,WDR6

3,ENDOV,CAMTA1,PPIL2,RPL28,APEH,RPL21,CAPN10,LYPLA2,C17ORF62,PRMT1,C19ORF54,ZNF7,TXNDC

S16,CPSF3L,TSG101,UBR4,YLPM1,FUZ,FAM3A,METTL17,EPB41L5,ATG12,RPS25,NIP7,THEM4,DBNDD2,

S16,CPSF3L,TSG101,UBR4,YLPM1,FUZ,FAM3A,METTL17,EPB41L5,ATG12,RPS25,NIP7,THEM4,DBNDD2,

,LZTFL1,FIS1,PPM1K,IRF9,PCBP2,MAGED2,SON,UQCRQ,SNF8,CDC37,CPSF3L,TSG101,SLC39A11,FUZ,FAM

,LZTFL1,FIS1,PPM1K,IRF9,PCBP2,MAGED2,SON,UQCRQ,SNF8,CDC37,CPSF3L,TSG101,SLC39A11,FUZ,FAM

NIP7,DUSP22,SLC25A10,RPS6,RPL29,COASY,AUP1,SAP30BP,YIPF3,FOXRED1,TRNT1,ITGB1BP1,MED15,IP

DT5,RBPMS,SNRPN,SPAG7,IL1RAP,STYXL1,RANBP1,UBE2L6,VRK3,LZTFL1,FIS1,IRF9,ANTXR2,STIP1,LRRC1,

DT5,RBPMS,SNRPN,SPAG7,IL1RAP,STYXL1,RANBP1,UBE2L6,VRK3,LZTFL1,FIS1,IRF9,ANTXR2,STIP1,LRRC1,

2,ZNF7,PPAN,CYB561,SUGT1,HYPK,CARS,ZNF250,ZNF197,COPZ1,INO80E,HSP90AB1,RAB2A,ATP5J,TRMT

2,ZNF7,PPAN,CYB561,SUGT1,HYPK,CARS,ZNF250,ZNF197,COPZ1,INO80E,HSP90AB1,RAB2A,ATP5J,TRMT

OK,LRRFIP1,ATP5J,RMDN1,TRMT112,CHPF,MPDU1,RPS15,UBXN1,HSPB1,SAT1,ERCC5,EMC4,SERPINB1,

F250,ZNF197,INO80E,HSP90AB1,RAB2A,ATP5J,TRMT112,CHPF,ZNF814,HSPB1,ERCC5,EMC4,DDOST,LSR

F250,ZNF197,INO80E,HSP90AB1,RAB2A,ATP5J,TRMT112,CHPF,ZNF814,HSPB1,ERCC5,EMC4,DDOST,LSR

YB561,SUGT1,HYPK,RPL36,ZNF197,COPZ1,INO80E,HSP90AB1,ATP5J,RMDN1,TRMT112,CHPF,UBXN1,ZN

YB561,SUGT1,HYPK,RPL36,ZNF197,COPZ1,INO80E,HSP90AB1,ATP5J,RMDN1,TRMT112,CHPF,UBXN1,ZN

2,STOML2,ZNF7,PPAN,CYB561,SUGT1,HYPK,CARS,ZNF250,ZNF197,COPZ1,INO80E,HSP90AB1,RAB2A,AT

2,STOML2,ZNF7,PPAN,CYB561,SUGT1,HYPK,CARS,ZNF250,ZNF197,COPZ1,INO80E,HSP90AB1,RAB2A,AT

ADA3,TRIM16,ANAPC11,NT5C,MDH2,EEF1D,CHCHD5,SEC31A,CARHSP1,CNST,COX17,HNRNPLL,HIGD1A,

T5C,MDH2,EEF1D,CHCHD5,SEC31A,CNST,PRLR,COX17,HNRNPLL,HIGD1A,B3GAT3,ZNF506,SPINT2,MTHF

EF1B2,RFX3,ENDOV,CAMTA1,PPIL2,RPL28,APEH,RPL21,CAPN10,LYPLA2,C17ORF62,PRMT1,C19ORF54,Z

BNDD1,SH2B1,NOP16,GSTK1,TTLL5,LDHA,PSMA6,ACOT8,AP3B2,SLC35G1,RCE1,RUVBL2,PCMT1,EBPL,T

BNDD1,SH2B1,NOP16,GSTK1,TTLL5,LDHA,PSMA6,ACOT8,AP3B2,SLC35G1,RCE1,RUVBL2,PCMT1,EBPL,T

T5,B3GAT3,PFDN5,PAFAH1B2,ZNF343,GBA2,PPIH,NSUN5,ZFPL1,MFSD10,GMPR2,RIC8A,UBE2N,LFNG,Z

UMF2,BUD31,NPRL2,OSGEP,TBL2,ARHGAP6,SCARB1,ITSN1,EIF5A,TNFSF10,COMMD5,RUNX1,CHID1,NU

/9,ASF1B,CCDC25,EEF1B2,RFX3,ENDOV,PPIL2,RPL28,LIG1,CDC14B,TIMM17B,APEH,LYPLA2,C17ORF62,P

/9,ASF1B,CCDC25,EEF1B2,RFX3,ENDOV,PPIL2,RPL28,LIG1,CDC14B,TIMM17B,APEH,LYPLA2,C17ORF62,P

6,ACOT8,AP3B2,RCE1,MRPL33,RUVBL2,PCMT1,EBPL,TESC,IGSF8,SNRPN,SPAG7,WDR62,RFX5,POLD4,RA

6,ACOT8,AP3B2,RCE1,MRPL33,RUVBL2,PCMT1,EBPL,TESC,IGSF8,SNRPN,SPAG7,WDR62,RFX5,POLD4,RA

LC1,RBP5,ZC3HAV1,SCAF11,DBI,ARPC2,PMFBP1,LIMK2,VDAC3,NOB1,WDR19,SIRT6,NUMA1,GPATCH2L,

LC1,RBP5,ZC3HAV1,SCAF11,DBI,ARPC2,PMFBP1,LIMK2,VDAC3,NOB1,WDR19,SIRT6,NUMA1,GPATCH2L,

THYN1,DBNDD1,SH2B1,RPL13A,TTLL5,LDHA,PSMA6,ACOT8,AP3B2,RPS9,SLC35G1,RCE1,MRPL33,PCMT1,

,SCAF11,DBI,ARPC2,LIMK2,VDAC3,NOB1,WDR19,SIRT6,NUMA1,GPATCH2L,BUD31,NPRL2,TBL2,ARHGAP

,SCAF11,DBI,ARPC2,LIMK2,VDAC3,NOB1,WDR19,SIRT6,NUMA1,GPATCH2L,BUD31,NPRL2,TBL2,ARHGAP

CARS,CDK11A,RBM3,CCNL2,DGUOK,LRRFIP1,ZNF524,ATP5J,TRMT112,CHPF,RPS20,MPDU1,UBXN1,ERCC

CARS,CDK11A,RBM3,CCNL2,DGUOK,LRRFIP1,ZNF524,ATP5J,TRMT112,CHPF,RPS20,MPDU1,UBXN1,ERCC

H2,STXBP2,EEF1D,CHCHD5,SEC31A,COX17,HIGD1A,B3GAT3,RFXANK,PAFAH1B2,ZNF343,GBA2,PPIH,ARH

ZNF81,FAM32A,TGOLN2,C12ORF57,DCTN5,PEX26,NDUFS3,OXLD1,ZNF223,METTL5,SLC35E1,OGFOD2,T

FAM3A,PSMA3,ECSIT,ATG12,RPS25,NIP7,TTC9C,SLC25A10,TMX2,THEM4,DBNDD2,SDF4,NISCH,TMED4,

FAM3A,PSMA3,ECSIT,ATG12,RPS25,NIP7,TTC9C,SLC25A10,TMX2,THEM4,DBNDD2,SDF4,NISCH,TMED4,

I,ARPC2,LIMK2,VDAC3,NOB1,WDR19,SIRT6,NUMA1,GPATCH2L,SUMF2,BUD31,SAE1,NPRL2,OSGEP,TBL2

I,ARPC2,LIMK2,VDAC3,NOB1,WDR19,SIRT6,NUMA1,GPATCH2L,SUMF2,BUD31,SAE1,NPRL2,OSGEP,TBL2

FD2,RIC8A,TCP1,THADA,UBE2N,LFNG,ZNHIT3,PDCD6,FAM32A,PEX2,POLR1C,C12ORF57,DCTN5,PEX26,N

,WDR19,SIRT6,NUMA1,GPATCH2L,SUMF2,BUD31,SAE1,NPRL2,OSGEP,TBL2,ARHGAP6,SCARB1,CTTN,TSP

,WDR19,SIRT6,NUMA1,GPATCH2L,SUMF2,BUD31,SAE1,NPRL2,OSGEP,TBL2,ARHGAP6,SCARB1,CTTN,TSP

OP16,ARHGEF39,GSTK1,TTLL5,PSMA6,ACOT8,AP3B2,RPS9,RCE1,RUVBL2,PCMT1,EBPL,TESC,IGSF8,NNMT

OP16,ARHGEF39,GSTK1,TTLL5,PSMA6,ACOT8,AP3B2,RPS9,RCE1,RUVBL2,PCMT1,EBPL,TESC,IGSF8,NNMT

3,RAD23A,ABHD17A,MDH2,HSPH1,CHCHD5,SEC31A,CNST,SSH2,COX17,POLR2C,ZNF506,PAFAH1B2,DAL

PPM1K,TUBA1C,ANTXR2,STIP1,PHF19,LPCAT3,MAGED2,GRK6,CDC37,RPS16,CPSF3L,TSG101,TTC21B,PTP

PPM1K,TUBA1C,ANTXR2,STIP1,PHF19,LPCAT3,MAGED2,GRK6,CDC37,RPS16,CPSF3L,TSG101,TTC21B,PTP

MD5,CHID1,NUDT22,AMZ2,UFD1L,TAZ,THYN1,DBNDD1,SH2B1,RPL13A,GSTK1,TTLL5,LDHA,AP3B2,SLC35

F198,UGGT2,EIF3M,CDC42,TMEM205,PTK2,AUNIP,CARD14,FBXW9,ASF1B,CCDC25,EEF1B2,RFX3,PPA1,

F198,UGGT2,EIF3M,CDC42,TMEM205,PTK2,AUNIP,CARD14,FBXW9,ASF1B,CCDC25,EEF1B2,RFX3,PPA1,

X6,CARD14,PCYT2,FBXW9,USE1,ASF1B,CCDC25,EEF1B2,RFX3,ENDOV,PPIL2,LIG1,APEH,CAPN10,LYPLA2,

X6,CARD14,PCYT2,FBXW9,USE1,ASF1B,CCDC25,EEF1B2,RFX3,ENDOV,PPIL2,LIG1,APEH,CAPN10,LYPLA2,

,HSPB1,ERCC5,EMC4,DDOST,LSR,MEIS2,RRNAD1,TOMM40,TADA3,ZNF687,PDCD5,ANAPC11,RAD23A,A

,HSPB1,ERCC5,EMC4,DDOST,LSR,MEIS2,RRNAD1,TOMM40,TADA3,ZNF687,PDCD5,ANAPC11,RAD23A,A

A,CARHSP1,CNST,COX17,HIGD1A,INTS4,B3GAT3,ZNF506,FTH1,MTHFS,PAFAH1B2,ZNF343,GBA2,PPIH,DA

TCH2L,SUMF2,BUD31,SAE1,NPRL2,OSGEP,TBL2,CRYGS,CTTN,TSPAN11,ITSN1,EIF5A,TNFSF10,COMMD5,

TCH2L,SUMF2,BUD31,SAE1,NPRL2,OSGEP,TBL2,CRYGS,CTTN,TSPAN11,ITSN1,EIF5A,TNFSF10,COMMD5,

,THEM4,DBNDD2,RPS6,RTEL1,SAP30BP,YIPF3,FOXRED1,RABL5,TRNT1,ITGB1BP1,MED15,CLASRP,C1ORF

,THEM4,DBNDD2,RPS6,RTEL1,SAP30BP,YIPF3,FOXRED1,RABL5,TRNT1,ITGB1BP1,MED15,CLASRP,C1ORF

MX2,SPOCK2,THEM4,DBNDD2,RTEL1,AUP1,SAP30BP,YIPF3,TRNT1,ITGB1BP1,MED15,IP6K2,CLASRP,UGG

MMD5,NUDT22,AMZ2,USP38,TAZ,THYN1,DBNDD1,SH2B1,NOP16,RPL13A,GSTK1,TTLL5,LDHA,PSMA6,AC

MMD5,NUDT22,AMZ2,USP38,TAZ,THYN1,DBNDD1,SH2B1,NOP16,RPL13A,GSTK1,TTLL5,LDHA,PSMA6,AC

LA,ZNF197,LRRFIP1,ATP5J,ELP6,TRMT112,CHPF,RPS20,GABPB1,DCTD,EMC4,DDOST,LSR,CHCHD7,PSME2

PK,CARS,CDK11A,HSP90AB1,LRRFIP1,RAB2A,ATP5J,TRMT112,CHPF,RPS20,VPS28,ERCC5,MCTS1,DDOST,

,ARHGEF39,LDHA,PSMA6,ACOT8,AP3B2,SLC35G1,RCE1,MRPL33,ZNF410,RUVBL2,EBPL,TESC,IGSF8,WDR

,ARHGEF39,LDHA,PSMA6,ACOT8,AP3B2,SLC35G1,RCE1,MRPL33,ZNF410,RUVBL2,EBPL,TESC,IGSF8,WDR

38,TAZ,THYN1,DBNDD1,SH2B1,NOP16,ARHGEF39,GSTK1,TTLL5,PSMA6,ACOT8,AP3B2,RCE1,ZNF410,RUV

38,TAZ,THYN1,DBNDD1,SH2B1,NOP16,ARHGEF39,GSTK1,TTLL5,PSMA6,ACOT8,AP3B2,RCE1,ZNF410,RUV

F410,RUVBL2,EBPL,TESC,IGSF8,NUDT5,RBPMS,SPAG7,WDR62,STYXL1,RANBP1,VRK3,ZC3H14,FIS1,PPM1

PCMT1,EBPL,TESC,IGSF8,SPAG7,WDR62,IL1RAP,RFX5,POLD4,RANBP1,UBE2L6,ZC3H14,FIS1,PPM1K,TUB

PCMT1,EBPL,TESC,IGSF8,SPAG7,WDR62,IL1RAP,RFX5,POLD4,RANBP1,UBE2L6,ZC3H14,FIS1,PPM1K,TUB

STYXL1,POLD4,VRK3,LZTFL1,FIS1,TUBA1C,IRF9,DST,ANTXR2,CD44,MAGED2,SON,UQCRQ,SNF8,GRK6,RPS

STYXL1,POLD4,VRK3,LZTFL1,FIS1,TUBA1C,IRF9,DST,ANTXR2,CD44,MAGED2,SON,UQCRQ,SNF8,GRK6,RPS

2,POLD4,PLEKHA5,RANBP1,UBE2L6,ZC3H14,FIS1,PPM1K,ANTXR2,STIP1,LRRC1,PHF19,VCAN,LPCAT3,CD

OGFOD2,EIF3G,ARIH2,PICK1,FBXL18,ZNF512,RNF181,TSPAN31,RALY,ZNF593,GLYCTK,RER1,NEK11,EIF2A

RB1,CRYGS,CTTN,TSPAN11,ITSN1,EIF5A,COMMD5,CHID1,NUDT22,AMZ2,USP38,TAZ,THYN1,DBNDD1,SH

RB1,CRYGS,CTTN,TSPAN11,ITSN1,EIF5A,COMMD5,CHID1,NUDT22,AMZ2,USP38,TAZ,THYN1,DBNDD1,SH

DOV,CAMTA1,LIG1,RPL21,CAPN10,LYPLA2,PRMT1,STOML2,ZNF7,TXNDC17,ARL6IP4,C19ORF48,PPAN,CY

DOV,CAMTA1,LIG1,RPL21,CAPN10,LYPLA2,PRMT1,STOML2,ZNF7,TXNDC17,ARL6IP4,C19ORF48,PPAN,CY

410,RUVBL2,EBPL,TESC,IGSF8,SNRPN,SPAG7,WDR62,IL1RAP,RFX5,POLD4,UBE2L6,FIS1,SLC15A3,TUBA1C

410,RUVBL2,EBPL,TESC,IGSF8,SNRPN,SPAG7,WDR62,IL1RAP,RFX5,POLD4,UBE2L6,FIS1,SLC15A3,TUBA1C

,NUMA1,GPATCH2L,BUD31,NPRL2,TBL2,ARHGAP6,SCARB1,CTTN,TSPAN11,ITSN1,EIF5A,COMMD5,CHID

PAN,CYB561,SUGT1,HYPK,CARS,CDK11A,RBM3,ZNF197,COPZ1,LRRFIP1,RAB2A,ATP5J,TRMT112,CHPF,R

MT112,CHPF,RPS20,DCTD,SAT1,ERCC5,SERPINB1,DDOST,LSR,PSME2,PABPC1,RRNAD1,DBNL,TOMM40,

DCD6,FAM32A,ZSCAN20,TGOLN2,PEX2,TNKS,C12ORF57,DCTN5,PEX26,NDUFS3,ARL16,OXLD1,C6ORF132

3B2,SLC35G1,PCMT1,EBPL,TESC,NUDT5,RBPMS,SPAG7,WDR62,IL1RAP,STYXL1,POLD4,PLEKHA5,RANBP1

3B2,SLC35G1,PCMT1,EBPL,TESC,NUDT5,RBPMS,SPAG7,WDR62,IL1RAP,STYXL1,POLD4,PLEKHA5,RANBP1

2,UFD1L,USP38,TAZ,THYN1,DBNDD1,SH2B1,ARHGEF39,GSTK1,TTLL5,LDHA,PSMA6,ACOT8,AP3B2,RPS9,

2,UFD1L,USP38,TAZ,THYN1,DBNDD1,SH2B1,ARHGEF39,GSTK1,TTLL5,LDHA,PSMA6,ACOT8,AP3B2,RPS9,

RL2,OSGEP,TBL2,ARHGAP6,SCARB1,CTTN,TSPAN11,ITSN1,EIF5A,COMMD5,CHID1,NUDT22,AMZ2,UFD1

IL1RAP,RFX5,POLD4,RANBP1,UBE2L6,FIS1,PPM1K,TUBA1C,ANTXR2,STIP1,PHF19,LPCAT3,CD44,SON,TET

IL1RAP,RFX5,POLD4,RANBP1,UBE2L6,FIS1,PPM1K,TUBA1C,ANTXR2,STIP1,PHF19,LPCAT3,CD44,SON,TET

RPT1,KLC1,RBP5,ZC3HAV1,SCAF11,DBI,ARPC2,LIMK2,VDAC3,NOB1,WDR19,SIRT6,NUMA1,SUMF2,BUD3

RPT1,KLC1,RBP5,ZC3HAV1,SCAF11,DBI,ARPC2,LIMK2,VDAC3,NOB1,WDR19,SIRT6,NUMA1,SUMF2,BUD3

L,USP38,TAZ,THYN1,DBNDD1,SH2B1,NOP16,RPL13A,TTLL5,LDHA,ACOT8,SLC35G1,RCE1,RUVBL2,PCMT1

TAZ,DBNDD1,SH2B1,NOP16,RPL13A,ARHGEF39,GSTK1,LDHA,PSMA6,AP3B2,RPS9,SLC35G1,RCE1,ZNF41

RK3,LZTFL1,FIS1,PPM1K,TUBA1C,IRF9,PCBP2,CD44,MAGED2,SON,UQCRQ,SNF8,GRK6,HMGA2,CDC37,C

RK3,LZTFL1,FIS1,PPM1K,TUBA1C,IRF9,PCBP2,CD44,MAGED2,SON,UQCRQ,SNF8,GRK6,HMGA2,CDC37,C

1,RRNAD1,DBNL,TOMM40,RABEP2,TADA3,TRIM16,NT5C,MDH2,STXBP2,SEC31A,CNST,ZFAND4,HIGD1A,

AA10,MED6,SEC31A,CARHSP1,CNST,COX17,HNRNPLL,SIRT5,FTH1,PFDN5,MTHFS,SMARCA4,C11ORF49,Z

1,LRRC1,VCAN,LPCAT3,CD44,MAGED2,SON,SNF8,GRK6,HMGA2,CDC37,CPSF3L,PTPN13,PSMA1,ECSIT,EP

8,AP3B2,RPS9,SLC35G1,RCE1,MRPL33,ZNF410,PCMT1,NUDT5,SNRPN,IL1RAP,STYXL1,RFX5,POLD4,VRK3

8,AP3B2,RPS9,SLC35G1,RCE1,MRPL33,ZNF410,PCMT1,NUDT5,SNRPN,IL1RAP,STYXL1,RFX5,POLD4,VRK3

2,PPIH,DALRD3,NSUN5,DRG2,MCFD2,RIC8A,PARK7,THADA,UBE2N,LFNG,ZNHIT3,FAM32A,PEX2,POLR1C

M1K,TUBA1C,STIP1,PHF19,VCAN,CD44,SON,TET2,GRK6,CDC37,RPS16,TSG101,TTC21B,YLPM1,PHF21A,

M1K,TUBA1C,STIP1,PHF19,VCAN,CD44,SON,TET2,GRK6,CDC37,RPS16,TSG101,TTC21B,YLPM1,PHF21A,

M1K,TUBA1C,STIP1,PHF19,VCAN,CD44,SON,TET2,GRK6,CDC37,RPS16,TSG101,TTC21B,YLPM1,PHF21A,

M1K,TUBA1C,STIP1,PHF19,VCAN,CD44,SON,TET2,GRK6,CDC37,RPS16,TSG101,TTC21B,YLPM1,PHF21A,

TSN1,EIF5A,NUDT22,AMZ2,UFD1L,USP38,THYN1,DBNDD1,SH2B1,NOP16,RPL13A,ARHGEF39,GSTK1,LDH

TSN1,EIF5A,NUDT22,AMZ2,UFD1L,USP38,THYN1,DBNDD1,SH2B1,NOP16,RPL13A,ARHGEF39,GSTK1,LDH

TSN1,EIF5A,COMMD5,NUDT22,AMZ2,UFD1L,USP38,TAZ,THYN1,DBNDD1,SH2B1,NOP16,RPL13A,ARHGE

MK2,VDAC3,NOB1,WDR19,SIRT6,NUMA1,SUMF2,BUD31,SAE1,NPRL2,OSGEP,TBL2,ARHGAP6,SCARB1,CR

MK2,VDAC3,NOB1,WDR19,SIRT6,NUMA1,SUMF2,BUD31,SAE1,NPRL2,OSGEP,TBL2,ARHGAP6,SCARB1,CR

CH2L,BUD31,NPRL2,TBL2,ARHGAP6,SCARB1,CTTN,TSPAN11,ITSN1,EIF5A,TNFSF10,COMMD5,NUDT22,A

,NOP16,TTLL5,LDHA,PSMA6,ACOT8,AP3B2,RPS9,SLC35G1,RCE1,MRPL33,ZNF410,RUVBL2,PCMT1,EBPL,

,TBL2,SCARB1,CTTN,TSPAN11,ITSN1,EIF5A,COMMD5,CHID1,AMZ2,UFD1L,USP38,TAZ,THYN1,DBNDD1,S

,TBL2,SCARB1,CTTN,TSPAN11,ITSN1,EIF5A,COMMD5,CHID1,AMZ2,UFD1L,USP38,TAZ,THYN1,DBNDD1,S

HGEF39,GSTK1,TTLL5,LDHA,PSMA6,ACOT8,AP3B2,RPS9,SLC35G1,RCE1,ZNF410,PCMT1,TESC,IGSF8,NUD

HGEF39,GSTK1,TTLL5,LDHA,PSMA6,ACOT8,AP3B2,RPS9,SLC35G1,RCE1,ZNF410,PCMT1,TESC,IGSF8,NUD

,MAGED2,SON,GRK6,CDC37,RPS16,CPSF3L,OBSL1,PTPN13,UBR4,CHCHD1,SLC39A11,FAM3A,MUC1,EPB

,MAGED2,SON,GRK6,CDC37,RPS16,CPSF3L,OBSL1,PTPN13,UBR4,CHCHD1,SLC39A11,FAM3A,MUC1,EPB

MMD5,CHID1,AMZ2,UFD1L,USP38,TAZ,DBNDD1,NOP16,RPL13A,ARHGEF39,GSTK1,TTLL5,LDHA,PSMA6,

PHF19,LPCAT3,MAGED2,SON,GRK6,CDC37,RPS16,CPSF3L,OBSL1,PTPN13,UBR4,PHF21A,CHCHD1,SLC39

PHF19,LPCAT3,MAGED2,SON,GRK6,CDC37,RPS16,CPSF3L,OBSL1,PTPN13,UBR4,PHF21A,CHCHD1,SLC39

DALRD3,NSUN5,ZFPL1,GMPR2,RIC8A,PIGT,UBE2N,MFSD11,FAM32A,ZSCAN20,TMEM147,TGOLN2,PEX2,I

AMZ2,USP38,TAZ,THYN1,DBNDD1,SH2B1,ARHGEF39,GSTK1,TTLL5,LDHA,PSMA6,AP3B2,RPS9,SLC35G1,F

3P5,ZC3HAV1,SCAF11,DBI,ARPC2,PMFBP1,LIMK2,VDAC3,NOB1,SIRT6,NUMA1,GPATCH2L,SUMF2,BUD31

3P5,ZC3HAV1,SCAF11,DBI,ARPC2,PMFBP1,LIMK2,VDAC3,NOB1,SIRT6,NUMA1,GPATCH2L,SUMF2,BUD31

38,TAZ,DBNDD1,SH2B1,ARHGEF39,GSTK1,TTLL5,LDHA,PSMA6,ACOT8,AP3B2,RPS9,RCE1,ZNF410,PCMT1,

38,TAZ,DBNDD1,SH2B1,ARHGEF39,GSTK1,TTLL5,LDHA,PSMA6,ACOT8,AP3B2,RPS9,RCE1,ZNF410,PCMT1,

2,TOMM40,RABEP2,TADA3,KLK10,RAD23A,PDE4DIP,ZCCHC4,MDH2,STXBP2,HSPH1,CHCHD5,NAA10,SEC

L1RAP,POLD4,RANBP1,UBE2L6,ZC3H14,FIS1,PPM1K,ANTXR2,STIP1,PHF19,VCAN,PCBP2,LPCAT3,CD44,M

L1RAP,POLD4,RANBP1,UBE2L6,ZC3H14,FIS1,PPM1K,ANTXR2,STIP1,PHF19,VCAN,PCBP2,LPCAT3,CD44,M

IDD1,SH2B1,NOP16,RPL13A,ARHGEF39,GSTK1,PSMA6,ACOT8,AP3B2,RPS9,SLC35G1,RCE1,PCMT1,EBPL,T

IDD1,SH2B1,NOP16,RPL13A,ARHGEF39,GSTK1,PSMA6,ACOT8,AP3B2,RPS9,SLC35G1,RCE1,PCMT1,EBPL,T

,NOB1,WDR19,SIRT6,NUMA1,GPATCH2L,SUMF2,BUD31,SAE1,NPRL2,OSGEP,TBL2,ARHGAP6,SCARB1,CR

,NOB1,WDR19,SIRT6,NUMA1,GPATCH2L,SUMF2,BUD31,SAE1,NPRL2,OSGEP,TBL2,ARHGAP6,SCARB1,CR

A6,AP3B2,RPS9,SLC35G1,RCE1,PCMT1,TESC,IGSF8,RBPMS,SNRPN,SPAG7,IL1RAP,STYXL1,POLD4,PLEKHA

14,FIS1,PPM1K,TUBA1C,DST,ANTXR2,PHF19,LPCAT3,TM9SF1,MAGED2,SON,TET2,UQCRQ,GRK6,CDC37,C

14,FIS1,PPM1K,TUBA1C,DST,ANTXR2,PHF19,LPCAT3,TM9SF1,MAGED2,SON,TET2,UQCRQ,GRK6,CDC37,C

14,FIS1,PPM1K,TUBA1C,DST,ANTXR2,PHF19,LPCAT3,TM9SF1,MAGED2,SON,TET2,UQCRQ,GRK6,CDC37,C

14,FIS1,PPM1K,TUBA1C,DST,ANTXR2,PHF19,LPCAT3,TM9SF1,MAGED2,SON,TET2,UQCRQ,GRK6,CDC37,C

14,FIS1,PPM1K,TUBA1C,DST,ANTXR2,PHF19,LPCAT3,TM9SF1,MAGED2,SON,TET2,UQCRQ,GRK6,CDC37,C

14,FIS1,PPM1K,TUBA1C,DST,ANTXR2,PHF19,LPCAT3,TM9SF1,MAGED2,SON,TET2,UQCRQ,GRK6,CDC37,C

LPCAT3,CD44,TM9SF1,MAGED2,SON,GRK6,CDC37,RPS16,CPSF3L,TTC21B,PTPN13,YLPM1,PHF21A,SLC39

LPCAT3,CD44,TM9SF1,MAGED2,SON,GRK6,CDC37,RPS16,CPSF3L,TTC21B,PTPN13,YLPM1,PHF21A,SLC39

SF8,NUDT5,SNRPN,WDR62,STYXL1,POLD4,VRK3,LZTFL1,FIS1,PPM1K,IRF9,DST,ANTXR2,STIP1,LRRC1,VCA

SF8,NUDT5,SNRPN,WDR62,STYXL1,POLD4,VRK3,LZTFL1,FIS1,PPM1K,IRF9,DST,ANTXR2,STIP1,LRRC1,VCA

IETTL17,ATG12,RPS25,NIP7,COX4I1,THEM4,NISCH,CENPK,RPS6,RPL29,RTEL1,AUP1,SAP30BP,YIPF3,FOXR

IETTL17,ATG12,RPS25,NIP7,COX4I1,THEM4,NISCH,CENPK,RPS6,RPL29,RTEL1,AUP1,SAP30BP,YIPF3,FOXR

BL2,EBPL,TESC,IGSF8,RBPMS,SNRPN,SPAG7,WDR62,IL1RAP,POLD4,PLEKHA5,RANBP1,UBE2L6,FIS1,PPM1

BL2,EBPL,TESC,IGSF8,RBPMS,SNRPN,SPAG7,WDR62,IL1RAP,POLD4,PLEKHA5,RANBP1,UBE2L6,FIS1,PPM1

NK,PFDN5,PAFAH1B2,TXNRD2,NSUN5,ZFPL1,MRPS18C,RIC8A,SMARCC2,UBE2N,ZNF81,FAM32A,TMEM1

NK,PFDN5,PAFAH1B2,TXNRD2,NSUN5,ZFPL1,MRPS18C,RIC8A,SMARCC2,UBE2N,ZNF81,FAM32A,TMEM1

VDR62,IL1RAP,RFX5,POLD4,RANBP1,UBE2L6,FIS1,PPM1K,TUBA1C,STIP1,PHF19,VCAN,LPCAT3,CD44,SON

VDR62,IL1RAP,RFX5,POLD4,RANBP1,UBE2L6,FIS1,PPM1K,TUBA1C,STIP1,PHF19,VCAN,LPCAT3,CD44,SON

A,INO80E,DGUOK,HSP90AB1,ATP5J,TRMT112,CHPF,MPDU1,UBXN1,SAT1,ERCC5,EMC4,MCTS1,DDOST,L

A,INO80E,DGUOK,HSP90AB1,ATP5J,TRMT112,CHPF,MPDU1,UBXN1,SAT1,ERCC5,EMC4,MCTS1,DDOST,L

2,AMZ2,USP38,TAZ,THYN1,DBNDD1,SH2B1,ARHGEF39,GSTK1,TTLL5,LDHA,AP3B2,RPS9,SLC35G1,RCE1,RU

2,AMZ2,USP38,TAZ,THYN1,DBNDD1,SH2B1,ARHGEF39,GSTK1,TTLL5,LDHA,AP3B2,RPS9,SLC35G1,RCE1,RU

3,PPP1CC,DGKE,ENDOV,CAMTA1,PPIL2,RPL28,LIG1,TXNL4A,LYPLA2,ATPAF2,C17ORF62,EXOSC9,WNK1,C

PL,TESC,RBPMS,SNRPN,SPAG7,WDR62,IL1RAP,STYXL1,RFX5,POLD4,PLEKHA5,RANBP1,VRK3,LZTFL1,ZC3H

PL,TESC,RBPMS,SNRPN,SPAG7,WDR62,IL1RAP,STYXL1,RFX5,POLD4,PLEKHA5,RANBP1,VRK3,LZTFL1,ZC3H

PM1K,STIP1,PHF19,VCAN,LPCAT3,MAGED2,SON,GRK6,CDC37,RPS16,OBSL1,PTPN13,YLPM1,PHF21A,SLC

PM1K,STIP1,PHF19,VCAN,LPCAT3,MAGED2,SON,GRK6,CDC37,RPS16,OBSL1,PTPN13,YLPM1,PHF21A,SLC

P,STYXL1,PLEKHA5,RANBP1,LZTFL1,FIS1,PPM1K,TUBA1C,STIP1,LPCAT3,TM9SF1,MAGED2,SON,SNF8,GRK

SON,TET2,CDC37,RPS16,CPSF3L,TSG101,OBSL1,TTC21B,UBR4,PSMA1,FUZ,FAM3A,PSMA3,METTL17,PPR

SON,TET2,CDC37,RPS16,CPSF3L,TSG101,OBSL1,TTC21B,UBR4,PSMA1,FUZ,FAM3A,PSMA3,METTL17,PPR

MMD5,NUDT22,UFD1L,USP38,TAZ,THYN1,DBNDD1,SH2B1,NOP16,RPL13A,ARHGEF39,GSTK1,TTLL5,LDHA

MMD5,NUDT22,UFD1L,USP38,TAZ,THYN1,DBNDD1,SH2B1,NOP16,RPL13A,ARHGEF39,GSTK1,TTLL5,LDHA

AM32A,TMEM147,COA1,USP40,PCID2,PEX2,POLR1C,TNKS,C12ORF57,DCTN5,ZNF223,OGFOD2,EIF3G,SS
AM32A,TMEM147,COA1,USP40,PCID2,PEX2,POLR1C,TNKS,C12ORF57,DCTN5,ZNF223,OGFOD2,EIF3G,SS
GEP,TBL2,ARHGAP6,CRYGS,CTTN,TSPAN11,ITSN1,EIF5A,COMMD5,CHID1,NUDT22,AMZ2,UFD1L,TAZ,DBN
GEP,TBL2,ARHGAP6,CRYGS,CTTN,TSPAN11,ITSN1,EIF5A,COMMD5,CHID1,NUDT22,AMZ2,UFD1L,TAZ,DBN
MMD5,NUDT22,UFD1L,DBNDD1,SH2B1,NOP16,ARHGEF39,GSTK1,TTLL5,PSMA6,ACOT8,AP3B2,RPS9,RCE
MMD5,NUDT22,UFD1L,DBNDD1,SH2B1,NOP16,ARHGEF39,GSTK1,TTLL5,PSMA6,ACOT8,AP3B2,RPS9,RCE
N,ITSN1,EIF5A,TNFSF10,COMMD5,NUDT22,AMZ2,UFD1L,USP38,TAZ,THYN1,DBNDD1,SH2B1,NOP16,ARH
N,ITSN1,EIF5A,TNFSF10,COMMD5,NUDT22,AMZ2,UFD1L,USP38,TAZ,THYN1,DBNDD1,SH2B1,NOP16,ARH
2,EMC4,LMBR1,DDOST,RRNAD1,RAD1,TOMM40,TADA3,ZNF687,ANAPC11,MDH2,EEF1D,NAA10,CARHSP
2,EMC4,LMBR1,DDOST,RRNAD1,RAD1,TOMM40,TADA3,ZNF687,ANAPC11,MDH2,EEF1D,NAA10,CARHSP
,LDHA,AP3B2,RPS9,RCE1,MRPL33,RUVBL2,EBPL,TESC,NUDT5,RBPMS,SPAG7,WDR62,IL1RAP,POLD4,PLE
,LDHA,AP3B2,RPS9,RCE1,MRPL33,RUVBL2,EBPL,TESC,NUDT5,RBPMS,SPAG7,WDR62,IL1RAP,POLD4,PLE
HF19,LPCAT3,CD44,TM9SF1,MAGED2,SON,GRK6,CDC37,RPS16,CPSF3L,TTC21B,YLPM1,PHF21A,CHCHD1,
HF19,LPCAT3,CD44,TM9SF1,MAGED2,SON,GRK6,CDC37,RPS16,CPSF3L,TTC21B,YLPM1,PHF21A,CHCHD1,
L2,NLRP1,ZBTB44,PLCB2,SLC25A10,TMX2,NISCH,TMED4,CD99L2,GGPS1,AUP1,TRNT1,KXD1,MED15,PAR
L2,NLRP1,ZBTB44,PLCB2,SLC25A10,TMX2,NISCH,TMED4,CD99L2,GGPS1,AUP1,TRNT1,KXD1,MED15,PAR
ENPN,CARD14,FBXW9,USE1,IFI27L1,CCDC25,EEF1B2,RFX3,ENDOV,PPIL2,RPL28,LIG1,CDC14B,TIMM17B,
ENPN,CARD14,FBXW9,USE1,IFI27L1,CCDC25,EEF1B2,RFX3,ENDOV,PPIL2,RPL28,LIG1,CDC14B,TIMM17B,
C37,RPS16,CPSF3L,PTPN13,YLPM1,PHF21A,SLC39A11,FAM3A,MUC1,METTL17,ECSIT,EPB41L5,ATG12,P
C37,RPS16,CPSF3L,PTPN13,YLPM1,PHF21A,SLC39A11,FAM3A,MUC1,METTL17,ECSIT,EPB41L5,ATG12,P
SMA1,FAM3A,PSMA3,METTL17,ECSIT,PPRC1,RPS25,COX4I1,DUSP22,SPOCK2,TMED4,CENPK,CD99L2,GG
SMA1,FAM3A,PSMA3,METTL17,ECSIT,PPRC1,RPS25,COX4I1,DUSP22,SPOCK2,TMED4,CENPK,CD99L2,GG
MA6,ACOT8,AP3B2,RCE1,MRPL33,RUVBL2,PCMT1,EBPL,TESC,IGSF8,RBPMS,SPAG7,WDR62,IL1RAP,STYX
MA6,ACOT8,AP3B2,RCE1,MRPL33,RUVBL2,PCMT1,EBPL,TESC,IGSF8,RBPMS,SPAG7,WDR62,IL1RAP,STYX
40,PCID2,PEX2,TNKS,C12ORF57,DCTN5,ZNF223,OGFOD2,EIF3G,ARIH2,PICK1,FBXL18,NME1,SLC52A2,TS
40,PCID2,PEX2,TNKS,C12ORF57,DCTN5,ZNF223,OGFOD2,EIF3G,ARIH2,PICK1,FBXL18,NME1,SLC52A2,TS
L,MED15,SH3YL1,UGGT2,MICAL1,YIF1A,PRMT2,EIF3M,CDC42,AUNIP,CENPN,CARD14,PCYT2,USE1,ASF1B
F197,COPZ1,INO80E,ATP5J,TRMT112,CHPF,RPS20,GABPB1,UBXN1,EMC4,PRKRIP1,DDOST,LSR,PSME2,RR
TMX2,NISCH,CD99L2,GGPS1,AUP1,FOXRED1,PARD6G,SH3YL1,IP6K2,CLASRP,C1ORF198,MICAL1,HS2ST1,
1,LRRFIP1,ZNF524,ATP5J,ELP6,TRMT112,RPS20,DCTD,SAT1,SERPINB1,DDOST,LSR,HMGA1,RRNAD1,DBN
1,LRRFIP1,ZNF524,ATP5J,ELP6,TRMT112,RPS20,DCTD,SAT1,SERPINB1,DDOST,LSR,HMGA1,RRNAD1,DBN
1,NUDT22,AMZ2,TAZ,DBNDD1,SH2B1,RPL13A,LDHA,PSMA6,AP3B2,RPS9,SLC35G1,RCE1,MRPL33,ZNF41
1,NUDT22,AMZ2,TAZ,DBNDD1,SH2B1,RPL13A,LDHA,PSMA6,AP3B2,RPS9,SLC35G1,RCE1,MRPL33,ZNF41
OMMD5,RUNX1,CHID1,NUDT22,AMZ2,UFD1L,USP38,TAZ,THYN1,DBNDD1,SH2B1,NOP16,RPL13A,ARHG
OMMD5,RUNX1,CHID1,NUDT22,AMZ2,UFD1L,USP38,TAZ,THYN1,DBNDD1,SH2B1,NOP16,RPL13A,ARHG
DHA,PSMA6,AP3B2,RPS9,SLC35G1,MRPL33,ZNF410,RUVBL2,EBPL,TESC,RBPMS,NNMT,SNRPN,SPAG7,W
DHA,PSMA6,AP3B2,RPS9,SLC35G1,MRPL33,ZNF410,RUVBL2,EBPL,TESC,RBPMS,NNMT,SNRPN,SPAG7,W
,MAGED2,SON,UQCRQ,SNF8,HMGA2,CPSF3L,TSG101,OBSL1,FUZ,FAM3A,PSMA3,MUC1,METTL17,ATG1
,MAGED2,SON,UQCRQ,SNF8,HMGA2,CPSF3L,TSG101,OBSL1,FUZ,FAM3A,PSMA3,MUC1,METTL17,ATG1
M4,DBNDD2,NISCH,TMED4,CD99L2,AUP1,FOXRED1,TRNT1,KXD1,ITGB1BP1,PARD6G,SH3YL1,IP6K2,JPH
M4,DBNDD2,NISCH,TMED4,CD99L2,AUP1,FOXRED1,TRNT1,KXD1,ITGB1BP1,PARD6G,SH3YL1,IP6K2,JPH
NF81,FAM32A,TMEM147,COA1,USP40,PCID2,NFYC,PEX2,TNKS,C12ORF57,DCTN5,ZNF223,RC3H2,OGFO
NF81,FAM32A,TMEM147,COA1,USP40,PCID2,NFYC,PEX2,TNKS,C12ORF57,DCTN5,ZNF223,RC3H2,OGFO
2,OSGEP,TBL2,ARHGAP6,SCARB1,CRYGS,CTTN,TSPAN11,ITSN1,EIF5A,COMMD5,RUNX1,CHID1,UFD1L,U
2,OSGEP,TBL2,ARHGAP6,SCARB1,CRYGS,CTTN,TSPAN11,ITSN1,EIF5A,COMMD5,RUNX1,CHID1,UFD1L,U
2,ARL6IP4,ACAD10,SHARPIN,SUGT1,HYPK,CARS,ZNF250,CDK11A,ZNF197,CCNL2,MAPKAPK5,RAB2A,ZN
2,ARL6IP4,ACAD10,SHARPIN,SUGT1,HYPK,CARS,ZNF250,CDK11A,ZNF197,CCNL2,MAPKAPK5,RAB2A,ZN

RPL28,LIG1,TIMM17B,APEH,HYI,CAPN10,CAMLG,ATPAF2,PRMT1,CASC4,PIGL,STRADA,ZFAND2B,CYB561,
RPL28,LIG1,TIMM17B,APEH,HYI,CAPN10,CAMLG,ATPAF2,PRMT1,CASC4,PIGL,STRADA,ZFAND2B,CYB561,
MED4,CD99L2,GGPS1,COASY,TRNT1,KXD1,PARD6G,SH3YL1,IP6K2,CLASRP,JPH2,C1ORF198,MICAL1,HS2S
MED4,CD99L2,GGPS1,COASY,TRNT1,KXD1,PARD6G,SH3YL1,IP6K2,CLASRP,JPH2,C1ORF198,MICAL1,HS2S
EMC4,MCTS1,DDOST,LSR,PSME2,PABPC1,RRNAD1,TOMM40,RABEP2,TADA3,ZNF687,ANAPC11,POLR3H,
I1,PLCB2,SLC25A10,TMX2,THEM4,SDF4,RPS6,RTEL1,SAP30BP,YIPF3,RABL5,TRNT1,ITGB1BP1,MED15,SH3
I1,PLCB2,SLC25A10,TMX2,THEM4,SDF4,RPS6,RTEL1,SAP30BP,YIPF3,RABL5,TRNT1,ITGB1BP1,MED15,SH3
GKE,CAMTA1,RPL28,LIG1,PSMA7,TIMM17B,APEH,HYI,CAPN10,CAMLG,ATPAF2,PRMT1,CASC4,PIGL,STR
GKE,CAMTA1,RPL28,LIG1,PSMA7,TIMM17B,APEH,HYI,CAPN10,CAMLG,ATPAF2,PRMT1,CASC4,PIGL,STR
2,USP38,TAZ,THYN1,DBNDD1,NOP16,RPL13A,GSTK1,ACOT8,RPS9,SLC35G1,RCE1,MRPL33,PCMT1,EBPL,
39,TTLL5,PSMA6,AP3B2,RPS9,SLC35G1,RCE1,PCMT1,TESC,IGSF8,NUDT5,RBPMS,SNRPN,SPAG7,IL1RAP,S
39,TTLL5,PSMA6,AP3B2,RPS9,SLC35G1,RCE1,PCMT1,TESC,IGSF8,NUDT5,RBPMS,SNRPN,SPAG7,IL1RAP,S
1,CDC42,CARD14,PCYT2,FBXW9,CCDC25,EEF1B2,RFX3,PPA1,ENDOV,PPIL2,RPL28,LIG1,CDC14B,TIMM17
1,CDC42,CARD14,PCYT2,FBXW9,CCDC25,EEF1B2,RFX3,PPA1,ENDOV,PPIL2,RPL28,LIG1,CDC14B,TIMM17
R62,IL1RAP,STYXL1,PLEKHA5,RANBP1,LZTFL1,ZC3H14,SLC15A3,PHF19,GRAMD1C,VCAN,LPCAT3,CD44,T
R62,IL1RAP,STYXL1,PLEKHA5,RANBP1,LZTFL1,ZC3H14,SLC15A3,PHF19,GRAMD1C,VCAN,LPCAT3,CD44,T
COMMD5,RUNX1,AMZ2,UFD1L,USP38,TAZ,THYN1,DBNDD1,RPL13A,GSTK1,TTLL5,LDHA,PSMA6,ACOT8,
COMMD5,RUNX1,AMZ2,UFD1L,USP38,TAZ,THYN1,DBNDD1,RPL13A,GSTK1,TTLL5,LDHA,PSMA6,ACOT8,
PSMA6,AP3B2,RPS9,SLC35G1,MRPL33,ZNF410,RUVBL2,PCMT1,EBPL,TESC,NUDT5,RBPMS,SNRPN,SPAG
SG101,OBSL1,UBR4,SLC39A11,FUZ,FAM3A,MUC1,METTL17,RPS25,ZBTB44,NIP7,COX4I1,PLCB2,TMX2,S
SG101,OBSL1,UBR4,SLC39A11,FUZ,FAM3A,MUC1,METTL17,RPS25,ZBTB44,NIP7,COX4I1,PLCB2,TMX2,S
4,CENPK,RTEL1,FOXRED1,RABL5,TRNT1,ITGB1BP1,MED15,CLASRP,JPH2,UGGT2,MICAL1,UBE2I,PRMT2,
4,CENPK,RTEL1,FOXRED1,RABL5,TRNT1,ITGB1BP1,MED15,CLASRP,JPH2,UGGT2,MICAL1,UBE2I,PRMT2,
N,TSPAN11,ITSN1,EIF5A,COMMD5,CHID1,NUDT22,AMZ2,USP38,TAZ,DBNDD1,SH2B1,NOP16,RPL13A,A
P22,TMX2,DBNDD2,NISCH,CD99L2,GGPS1,AUP1,FOXRED1,PARD6G,SH3YL1,IP6K2,CLASRP,C1ORF198,M
HYN1,NOP16,RPL13A,ARHGEF39,GSTK1,TTLL5,LDHA,PSMA6,ACOT8,AP3B2,RPS9,RCE1,ZNF410,RUVBL2,
AN31,PPP2R5C,GLYCTK,RBM42,CNTRL,RER1,TRAPPC2,GTF2H1,AXIN1,TMEM199,SLC25A24,MGST2,EP
2,DUSP22,SLC25A10,NISCH,TMED4,CD99L2,GGPS1,YIPF3,TRNT1,PARD6G,SH3YL1,IP6K2,CLASRP,JPH2,C1
2,DUSP22,SLC25A10,NISCH,TMED4,CD99L2,GGPS1,YIPF3,TRNT1,PARD6G,SH3YL1,IP6K2,CLASRP,JPH2,C1
,SIRT6,NUMA1,SUMF2,BUD31,SAE1,NPRL2,OSGEP,TBL2,ARHGAP6,SCARB1,CRYGS,CTTN,TSPAN11,ITSN1
,SIRT6,NUMA1,SUMF2,BUD31,SAE1,NPRL2,OSGEP,TBL2,ARHGAP6,SCARB1,CRYGS,CTTN,TSPAN11,ITSN1
2L,BUD31,SAE1,NPRL2,OSGEP,TBL2,ARHGAP6,SCARB1,CRYGS,CTTN,TSPAN11,ITSN1,EIF5A,COMMD5,N
2L,BUD31,SAE1,NPRL2,OSGEP,TBL2,ARHGAP6,SCARB1,CRYGS,CTTN,TSPAN11,ITSN1,EIF5A,COMMD5,N
MA1,GPATCH2L,BUD31,NPRL2,OSGEP,TBL2,ARHGAP6,SCARB1,CTTN,TSPAN11,ITSN1,EIF5A,COMMD5,C
,COX4I1,SDF4,GGPS1,RTEL1,SRSF1,AUP1,FOXRED1,SH3YL1,IP6K2,UGGT2,HS2ST1,PRMT2,MAGT1,JUP,T
ANBP1,UBE2L6,ZC3H14,FIS1,PPM1K,DST,ANTXR2,STIP1,PHF19,VCAN,LPCAT3,CD44,TM9SF1,MAGED2,S
2,NISCH,CD99L2,GGPS1,AUP1,FOXRED1,PARD6G,SH3YL1,IP6K2,MICAL1,HS2ST1,UBE2I,SLC39A13,YIF1A,
F3M,JUP,PTK2,AUNIP,C14ORF2,GUK1,ATF7,PCYT2,PPP1CC,EDF1,PPA1,DGKE,CAMTA1,CAPG,CLDND1,LI
2,GRK6,CPSF3L,OBSL1,PTPN13,UBR4,YLPM1,PHF21A,SLC39A11,FAM3A,MUC1,ECSIT,EPB41L5,ATG12,PP
2,GRK6,CPSF3L,OBSL1,PTPN13,UBR4,YLPM1,PHF21A,SLC39A11,FAM3A,MUC1,ECSIT,EPB41L5,ATG12,PP
7,MEIS2,PSME2,DBNL,NAT9,TOMM40,RABEP2,TADA3,ZNF687,ANAPC11,POLR3H,NOMO2,RAD23A,B2M
P5,ZC3HAV1,SCAF11,DBI,ARPC2,LIMK2,SIRT6,NUMA1,GPATCH2L,BUD31,SAE1,NPRL2,OSGEP,TBL2,ARH
P5,ZC3HAV1,SCAF11,DBI,ARPC2,LIMK2,SIRT6,NUMA1,GPATCH2L,BUD31,SAE1,NPRL2,OSGEP,TBL2,ARH
H2B1,NOP16,ARHGEF39,GSTK1,TTLL5,LDHA,PSMA6,ACOT8,AP3B2,SLC35G1,RCE1,RUVBL2,PCMT1,EBPL,
H2B1,NOP16,ARHGEF39,GSTK1,TTLL5,LDHA,PSMA6,ACOT8,AP3B2,SLC35G1,RCE1,RUVBL2,PCMT1,EBPL,
L1,ZC3H14,SLC15A3,PPM1K,TUBA1C,ANTXR2,PHF19,GRAMD1C,LPCAT3,CD44,TM9SF1,SON,SNF8,GRK6,
L1,ZC3H14,SLC15A3,PPM1K,TUBA1C,ANTXR2,PHF19,GRAMD1C,LPCAT3,CD44,TM9SF1,SON,SNF8,GRK6,

A2,PPIH,ARHGEF1,NSUN5,ZFPL1,OARD1,MFSD10,MRPS18C,RIC8A,LGALS3,SCAP,PIGT,PDCD6,COA1,C12C
,NLRP1,COX4I1,PLCB2,SLC25A10,TMX2,SPOCK2,THEM4,DBNDD2,CD99L2,COASY,AUP1,SAP30BP,FOXRED
,NLRP1,COX4I1,PLCB2,SLC25A10,TMX2,SPOCK2,THEM4,DBNDD2,CD99L2,COASY,AUP1,SAP30BP,FOXRED
DN1,TBC1D22A,ZNF581,ZNF483,SZT2,PSMC5,LRP10,WDR45,C9ORF64,IQCK,SETD4,MED8,RNMT,MRPL2
1,PPM1K,IRF9,DST,ANTXR2,STIP1,LRRC1,LPCAT3,TM9SF1,MAGED2,SON,UQCRQ,SNF8,GRK6,CDC37,RPS
1,PPM1K,IRF9,DST,ANTXR2,STIP1,LRRC1,LPCAT3,TM9SF1,MAGED2,SON,UQCRQ,SNF8,GRK6,CDC37,RPS

YXL1,RANBP1,UBE2L6,VRK3,ZC3H14,FIS1,SLC15A3,TUBA1C,IRF9,STIP1,PHF19,LPCAT3,CD44,MAGED2,U
YXL1,RANBP1,UBE2L6,VRK3,ZC3H14,FIS1,SLC15A3,TUBA1C,IRF9,STIP1,PHF19,LPCAT3,CD44,MAGED2,U
ATCH2L,SUMF2,BUD31,SAE1,OSGEP,TBL2,ARHGAP6,SCARB1,CRYGS,CTTN,TSPAN11,ITSN1,EIF5A,COMM
TNFSF10,CHID1,NUDT22,AMZ2,UFD1L,USP38,THYN1,DBNDD1,SH2B1,NOP16,RPL13A,GSTK1,PSMA6,AC
TNFSF10,CHID1,NUDT22,AMZ2,UFD1L,USP38,THYN1,DBNDD1,SH2B1,NOP16,RPL13A,GSTK1,PSMA6,AC
RP,JPH2,C1ORF198,MICAL1,UBE2I,SLC39A13,PRMT2,EIF3M,PTK2,AUNIP,C14ORF2,GUK1,PCYT2,AGPAT4
1,FUZ,PSMA3,METTL17,ECSIT,PPRC1,RPS25,ZBTB44,NIP7,COX4I1,DUSP22,SPOCK2,TMED4,CENPK,CD99
1,FUZ,PSMA3,METTL17,ECSIT,PPRC1,RPS25,ZBTB44,NIP7,COX4I1,DUSP22,SPOCK2,TMED4,CENPK,CD99
,PLCB2,SLC25A10,TMX2,DBNDD2,NISCH,TMED4,CD99L2,AUP1,FOXRED1,TRNT1,PARD6G,SH3YL1,IP6K2,
,PLCB2,SLC25A10,TMX2,DBNDD2,NISCH,TMED4,CD99L2,AUP1,FOXRED1,TRNT1,PARD6G,SH3YL1,IP6K2,
,PPP1CC,RFX3,PPA1,DGKE,CAMTA1,LIG1,TIMM17B,APEH,HYI,CAPN10,NUTF2,ATPAF2,C17ORF62,PRMT
,PPP1CC,RFX3,PPA1,DGKE,CAMTA1,LIG1,TIMM17B,APEH,HYI,CAPN10,NUTF2,ATPAF2,C17ORF62,PRMT
P1,LIMK2,VDAC3,NOB1,WDR19,SIRT6,NUMA1,GPATCH2L,SUMF2,BUD31,OSGEP,TBL2,ARHGAP6,SCARB
P1,LIMK2,VDAC3,NOB1,WDR19,SIRT6,NUMA1,GPATCH2L,SUMF2,BUD31,OSGEP,TBL2,ARHGAP6,SCARB
1,OGFOD2,METTL3,ARIH2,PICK1,NME1,TSPAN31,GLYCTK,RER1,EIF2A,ZNF283,TRAPPC2,PIGO,AXIN1,HN

FAM3A,METTL17,EPB41L5,ATG12,RPS25,NIP7,COX4I1,SPOCK2,THEM4,RPL29,RTEL1,SRSF1,AUP1,YIPF3,
FAM3A,METTL17,EPB41L5,ATG12,RPS25,NIP7,COX4I1,SPOCK2,THEM4,RPL29,RTEL1,SRSF1,AUP1,YIPF3,
6,NUMA1,GPATCH2L,BUD31,SAE1,NPRL2,OSGEP,TBL2,ARHGAP6,SCARB1,CRYGS,CTTN,TSPAN11,ITSN1,E
6,NUMA1,GPATCH2L,BUD31,SAE1,NPRL2,OSGEP,TBL2,ARHGAP6,SCARB1,CRYGS,CTTN,TSPAN11,ITSN1,E
T5,RBPMS,NNMT,SPAG7,WDR62,STYXL1,POLD4,PLEKHA5,RANBP1,UBE2L6,LZTFL1,SLC15A3,PHF19,LPCA
T5,RBPMS,NNMT,SPAG7,WDR62,STYXL1,POLD4,PLEKHA5,RANBP1,UBE2L6,LZTFL1,SLC15A3,PHF19,LPCA
,TSPAN11,ITSN1,EIF5A,COMMD5,RUNX1,CHID1,UFD1L,USP38,TAZ,THYN1,DBNDD1,NOP16,GSTK1,TTLL5
,TSPAN11,ITSN1,EIF5A,COMMD5,RUNX1,CHID1,UFD1L,USP38,TAZ,THYN1,DBNDD1,NOP16,GSTK1,TTLL5
9,ANTXR2,STIP1,PHF19,LPCAT3,CD44,TET2,GRK6,HMGA2,RPS16,CPSF3L,TSG101,TTC21B,PTPN13,PHF21
9,ANTXR2,STIP1,PHF19,LPCAT3,CD44,TET2,GRK6,HMGA2,RPS16,CPSF3L,TSG101,TTC21B,PTPN13,PHF21
9,ANTXR2,STIP1,PHF19,LPCAT3,CD44,TET2,GRK6,HMGA2,RPS16,CPSF3L,TSG101,TTC21B,PTPN13,PHF21
9,ANTXR2,STIP1,PHF19,LPCAT3,CD44,TET2,GRK6,HMGA2,RPS16,CPSF3L,TSG101,TTC21B,PTPN13,PHF21
SLC39A11,FAM3A,METTL17,ECSIT,EPB41L5,ATG12,ZBTB44,PLCB2,DBNDD2,NISCH,TMED4,CD99L2,GGPS
SLC39A11,FAM3A,METTL17,ECSIT,EPB41L5,ATG12,ZBTB44,PLCB2,DBNDD2,NISCH,TMED4,CD99L2,GGPS
BUD31,SAE1,NPRL2,OSGEP,TBL2,ARHGAP6,SCARB1,CTTN,TSPAN11,ITSN1,EIF5A,COMMD5,RUNX1,CHID
ED2,SON,GRK6,CDC37,RPS16,OBSL1,TTC21B,UBR4,YLPM1,PHF21A,SLC39A11,FAM3A,MUC1,METTL17,E
ED2,SON,GRK6,CDC37,RPS16,OBSL1,TTC21B,UBR4,YLPM1,PHF21A,SLC39A11,FAM3A,MUC1,METTL17,E
IF5A,COMMD5,RUNX1,CHID1,UFD1L,USP38,TAZ,THYN1,DBNDD1,NOP16,GSTK1,TTLL5,LDHA,ACOT8,AP3
CD99L2,GGPS1,AUP1,FOXRED1,PARD6G,SH3YL1,IP6K2,C1ORF198,MICAL1,HS2ST1,UBE2I,SLC39A13,YIF
OBSL1,TTC21B,YLPM1,PHF21A,FUZ,PSMA3,EPB41L5,RPS25,NIP7,DUSP22,SLC25A10,TMX2,THEM4,CENP
OBSL1,TTC21B,YLPM1,PHF21A,FUZ,PSMA3,EPB41L5,RPS25,NIP7,DUSP22,SLC25A10,TMX2,THEM4,CENP
AG7,WDR62,IL1RAP,RFX5,PLEKHA5,UBE2L6,VRK3,ZC3H14,PPM1K,TUBA1C,IRF9,STIP1,LRRC1,PHF19,GR
AG7,WDR62,IL1RAP,RFX5,PLEKHA5,UBE2L6,VRK3,ZC3H14,PPM1K,TUBA1C,IRF9,STIP1,LRRC1,PHF19,GR
ANK,SPINT2,GBA2,ARHGEF1,DALRD3,NSUN5,MFSD10,MRPS18C,RIC8A,LGALS3,COMMD4,PIGT,PDCD6,C

ZNF593,GLYCTK,NEK11,EIF2A,GLI4,ACAA1,ZNF511,GTF2H1,SLC35A2,TP53,TMEM199,SLC25A24,EPN1,HP
D1L,DBNDD1,SH2B1,NOP16,GSTK1,TTLL5,LDHA,ACOT8,AP3B2,SLC35G1,MRPL33,ZNF410,RUVBL2,PCMT
D1L,DBNDD1,SH2B1,NOP16,GSTK1,TTLL5,LDHA,ACOT8,AP3B2,SLC35G1,MRPL33,ZNF410,RUVBL2,PCMT
UBR4,PSMA1,YLPM1,CHCHD1,SLC39A11,FUZ,FAM3A,MUC1,METTL17,ECSIT,EPB41L5,ATG12,PPRC1,TTC
UBR4,PSMA1,YLPM1,CHCHD1,SLC39A11,FUZ,FAM3A,MUC1,METTL17,ECSIT,EPB41L5,ATG12,PPRC1,TTC
CTTN,ITSN1,EIF5A,COMMD5,RUNX1,CHID1,NUDT22,UFD1L,USP38,TAZ,DBNDD1,SH2B1,NOP16,RPL13A,
CTTN,ITSN1,EIF5A,COMMD5,RUNX1,CHID1,NUDT22,UFD1L,USP38,TAZ,DBNDD1,SH2B1,NOP16,RPL13A,
MS,SPAG7,IL1RAP,STYXL1,RFX5,POLD4,PLEKHA5,RANBP1,FIS1,SLC15A3,PPM1K,TUBA1C,IRF9,STIP1,LRR
MS,SPAG7,IL1RAP,STYXL1,RFX5,POLD4,PLEKHA5,RANBP1,FIS1,SLC15A3,PPM1K,TUBA1C,IRF9,STIP1,LRR
ICAL1,HS2ST1,UBE2I,PRMT2,TMEM205,AUNIP,C14ORF2,PCYT2,FBXW9,USE1,DEF8,PPP1CC,RFX3,PPA1,
TAZ,DBNDD1,SH2B1,RPL13A,LDHA,PSMA6,AP3B2,RPS9,SLC35G1,RCE1,MRPL33,ZNF410,RUVBL2,PCMT1,
TAZ,DBNDD1,SH2B1,RPL13A,LDHA,PSMA6,AP3B2,RPS9,SLC35G1,RCE1,MRPL33,ZNF410,RUVBL2,PCMT1,
DCTN5,ZNF223,ARIH2,PICK1,FBXL18,FAM192A,ZNF593,GLYCTK,C11ORF57,EIF2A,GLI4,NKIRAS2,SCN11A,
DCTN5,ZNF223,ARIH2,PICK1,FBXL18,FAM192A,ZNF593,GLYCTK,C11ORF57,EIF2A,GLI4,NKIRAS2,SCN11A,
A,LDHA,PSMA6,AP3B2,RPS9,SLC35G1,MRPL33,ZNF410,RUVBL2,EBPL,TESC,NUDT5,RBPMS,SNRPN,SPAG7
A,LDHA,PSMA6,AP3B2,RPS9,SLC35G1,MRPL33,ZNF410,RUVBL2,EBPL,TESC,NUDT5,RBPMS,SNRPN,SPAG7
WDR62,STYXL1,RFX5,RANBP1,VRK3,LZTFL1,ZC3H14,FIS1,TUBA1C,IRF9,STIP1,LPCAT3,TET2,UQCRQ,SNF8,
SN1,EIF5A,COMMD5,RUNX1,CHID1,UFD1L,TAZ,DBNDD1,SH2B1,NOP16,GSTK1,TTLL5,LDHA,ACOT8,AP3B
M126A,SSB,PICK1,ZNF512,CHTOP,POLR2M,SLC52A2,ANAPC13,TSPAN31,MTIF2,EEF2K,FAM192A,GLYCT
IF644,SLC9B2,CDK11A,RBM3,INO80E,FAM219B,ZNF524,ELP6,GPBP1,SIAH1,TRMT112,GABPB1,MPDU1,T
IF644,SLC9B2,CDK11A,RBM3,INO80E,FAM219B,ZNF524,ELP6,GPBP1,SIAH1,TRMT112,GABPB1,MPDU1,T
P1,UBE2L6,LZTFL1,ZC3H14,FIS1,SLC15A3,STIP1,PHF19,GRAMD1C,PCBP2,LPCAT3,TET2,UQCRQ,SNF8,GR
P1,UBE2L6,LZTFL1,ZC3H14,FIS1,SLC15A3,STIP1,PHF19,GRAMD1C,PCBP2,LPCAT3,TET2,UQCRQ,SNF8,GR
,EIF5A,COMMD5,RUNX1,CHID1,NUDT22,UFD1L,USP38,TAZ,DBNDD1,NOP16,RPL13A,GSTK1,TTLL5,LDH
B,RAB2A,ZNF524,ATP5J,ELP6,TPM2,SIAH1,GGCX,RPTOR,CHPF,ZNF625,MPDU1,ZSWIM7,UBXN1,ERLEC1,
,ATF7,EDF1,VAMP8,CAPG,PPIL2,RPL28,CLDND1,VIM,LIG1,TIMM17B,APEH,CAPN10,LYPLA2,NUTF2,POD
,ATF7,EDF1,VAMP8,CAPG,PPIL2,RPL28,CLDND1,VIM,LIG1,TIMM17B,APEH,CAPN10,LYPLA2,NUTF2,POD
L2,TBL2,ARHGAP6,SCARB1,CTTN,TSPAN11,ITSN1,EIF5A,TNFSF10,COMMD5,CHID1,NUDT22,AMZ2,UFD1L
L2,TBL2,ARHGAP6,SCARB1,CTTN,TSPAN11,ITSN1,EIF5A,TNFSF10,COMMD5,CHID1,NUDT22,AMZ2,UFD1L
SC,IGSF8,RBPMS,NNMT,WDR62,STYXL1,RFX5,PLEKHA5,RANBP1,VRK3,LZTFL1,ZC3H14,FIS1,TUBA1C,ANT
SC,IGSF8,RBPMS,NNMT,WDR62,STYXL1,RFX5,PLEKHA5,RANBP1,VRK3,LZTFL1,ZC3H14,FIS1,TUBA1C,ANT
TTC21B,PTPN13,UBR4,PSMA1,CHCHD1,FUZ,FAM3A,METTL17,RPS25,NIP7,COX4I1,DUSP22,SPOCK2,TME
TTC21B,PTPN13,UBR4,PSMA1,CHCHD1,FUZ,FAM3A,METTL17,RPS25,NIP7,COX4I1,DUSP22,SPOCK2,TME
TYXL1,RFX5,RANBP1,VRK3,SLC15A3,PPM1K,PHF19,GRAMD1C,PCBP2,CD44,TM9SF1,MAGED2,SON,TET2,
TYXL1,RFX5,RANBP1,VRK3,SLC15A3,PPM1K,PHF19,GRAMD1C,PCBP2,CD44,TM9SF1,MAGED2,SON,TET2,
MRPL33,ZNF410,RUVBL2,EBPL,TESC,IGSF8,NUDT5,RBPMS,NNMT,SPAG7,WDR62,IL1RAP,STYXL1,RANBP1
MRPL33,ZNF410,RUVBL2,EBPL,TESC,IGSF8,NUDT5,RBPMS,NNMT,SPAG7,WDR62,IL1RAP,STYXL1,RANBP1
LC15A3,TUBA1C,ANTXR2,STIP1,PHF19,VCAN,CD44,SON,TET2,UQCRQ,CDC37,RPS16,TTC21B,PTPN13,YLP
LC15A3,TUBA1C,ANTXR2,STIP1,PHF19,VCAN,CD44,SON,TET2,UQCRQ,CDC37,RPS16,TTC21B,PTPN13,YLP
RRDC1,TRPT1,KLC1,RBP5,ZC3HAV1,SCAF11,DBI,ARPC2,PMFBP1,LIMK2,VDAC3,NOB1,WDR19,SIRT6,NUM
RRDC1,TRPT1,KLC1,RBP5,ZC3HAV1,SCAF11,DBI,ARPC2,PMFBP1,LIMK2,VDAC3,NOB1,WDR19,SIRT6,NUM
5,ATG12,ZBTB44,PLCB2,DUSP22,TMX2,DBNDD2,NISCH,RPL29,CD99L2,GGPS1,AUP1,FOXRED1,PARD6G,S
LC,DST,STIP1,LRRC1,VCAN,LPCAT3,TM9SF1,UQCRQ,SNF8,GRK6,HMGA2,CDC37,RPS16,CPSF3L,TSG101,O
1,FUZ,MUC1,ECSIT,PPRC1,RPS25,ZBTB44,NIP7,COX4I1,PLCB2,DUSP22,SLC25A10,TMX2,SDF4,NISCH,RPS
1,DBNDD1,SH2B1,NOP16,RPL13A,TTLL5,LDHA,ACOT8,AP3B2,RPS9,SLC35G1,MRPL33,ZNF410,RUVBL2,E
1,DBNDD1,SH2B1,NOP16,RPL13A,TTLL5,LDHA,ACOT8,AP3B2,RPS9,SLC35G1,MRPL33,ZNF410,RUVBL2,E
D5,CHID1,NUDT22,AMZ2,TAZ,DBNDD1,SH2B1,NOP16,RPL13A,TTLL5,PSMA6,ACOT8,AP3B2,RCE1,MRPL3

05,CHID1,NUDT22,AMZ2,TAZ,DBNDD1,SH2B1,NOP16,RPL13A,TTLL5,PSMA6,ACOT8,AP3B2,RCE1,MRPL33
,LRRC1,VCAN,LPCAT3,CD44,SON,TET2,SNF8,HMGA2,CDC37,CPSF3L,TTC21B,PSMA1,FUZ,PSMA3,METTL1
1,NOP16,RPL13A,GSTK1,LDHA,PSMA6,ACOT8,AP3B2,RPS9,SLC35G1,RCE1,MRPL33,ZNF410,RUVBL2,PCM
1,NOP16,RPL13A,GSTK1,LDHA,PSMA6,ACOT8,AP3B2,RPS9,SLC35G1,RCE1,MRPL33,ZNF410,RUVBL2,PCM
ST1,UBE2I,PRMT2,CDC42,TBX6,PTK2,GUK1,PCYT2,FBXW9,ASF1B,CCDC25,EEF1B2,ENDOV,VAMP8,CAPG,
ST1,UBE2I,PRMT2,CDC42,TBX6,PTK2,GUK1,PCYT2,FBXW9,ASF1B,CCDC25,EEF1B2,ENDOV,VAMP8,CAPG,
,VWA2,LMBR1,MCTS1,DDOST,DOCK10,PSME2,RAD1,TOMM40,TADA3,ZNF687,ANAPC11,MDH2,HSPH1,
,VWA2,LMBR1,MCTS1,DDOST,DOCK10,PSME2,RAD1,TOMM40,TADA3,ZNF687,ANAPC11,MDH2,HSPH1,
2,PICK1,FBXL18,CDK4,TSPAN31,ZNF593,GLYCTK,EIF2A,ZNF283,GTF2H1,TP53,TMEM199,SLC25A24,C1OR
41L5,RPS25,ZBTB44,NIP7,TTC9C,COX4I1,RDH11,NISCH,TMED4,CENPK,CD99L2,RTEL1,SRSF1,RABL5,TRNT
2,PRMT1,CASC4,ZNF7,PPAN,ZFAND2B,ZNF250,CCNL2,SYNGR2,CEP70,TMEM134,LRRFIP1,RAB2A,TPM2,
2,PRMT1,CASC4,ZNF7,PPAN,ZFAND2B,ZNF250,CCNL2,SYNGR2,CEP70,TMEM134,LRRFIP1,RAB2A,TPM2,
LDND1,LIG1,CDC14B,TIMM17B,APEH,CAPN10,NUTF2,ATPAF2,C17ORF62,PRMT1,ZNF7,PIGL,PPAN,ZFAN
RF48,PPAN,ZFAND2B,SHARPIN,SLC9B2,HYPK,RPL36,CDK11A,RBM3,INO80E,CCNL2,SYNGR2,FAM219B,A
,BOP1,ARRDC1,TRPT1,KLC1,RBP5,ZC3HAV1,SCAF11,DBI,ARPC2,LIMK2,VDAC3,NOB1,WDR19,SIRT6,NUM
,RPS25,NIP7,TTC9C,COX4I1,SLC25A10,TMX2,THEM4,RDH11,SDF4,TMED4,GGPS1,RTEL1,COASY,SRSF1,FC

6,SCARB1,CRYGS,CTTN,TSPAN11,EIF5A,TNFSF10,COMMD5,RUNX1,CHID1,NUDT22,UFD1L,USP38,TAZ,TH
6,SCARB1,CRYGS,CTTN,TSPAN11,EIF5A,TNFSF10,COMMD5,RUNX1,CHID1,NUDT22,UFD1L,USP38,TAZ,TH
M,ILF3,PDE4DIP,EEF1D,RRP8,CARHSP1,CNST,SH3GLB1,CHEK1,HNRNPLL,PHF20L1,HNRNPA1,HIGD1A,PO
LAR,TRIM46,EIF2A,NKIRAS2,ACAA1,GTF2H1,BFAR,TP53,SH3GL1,TMEM199,UBE2D3,ZNF396,EPN1,ZCCH
LAR,TRIM46,EIF2A,NKIRAS2,ACAA1,GTF2H1,BFAR,TP53,SH3GL1,TMEM199,UBE2D3,ZNF396,EPN1,ZCCH
32,RPS9,SLC35G1,RCE1,RUVBL2,PCMT1,EBPL,IGSF8,NUDT5,NNMT,SPAG7,WDR62,IL1RAP,STYXL1,POLD4
A,METTL17,ATG12,RPS25,NLRP1,NIP7,COX4I1,SPOCK2,THEM4,CENPK,RPL29,RTEL1,COASY,SRSF1,AUP1,
A,METTL17,ATG12,RPS25,NLRP1,NIP7,COX4I1,SPOCK2,THEM4,CENPK,RPL29,RTEL1,COASY,SRSF1,AUP1,
NLRP1,NIP7,TTC9C,COX4I1,THEM4,DBNDD2,TMED4,CENPK,SRSF1,AUP1,SAP30BP,FOXRED1,TRNT1,KXD1
NLRP1,NIP7,TTC9C,COX4I1,THEM4,DBNDD2,TMED4,CENPK,SRSF1,AUP1,SAP30BP,FOXRED1,TRNT1,KXD1
LPCAT3,CD44,TM9SF1,MAGED2,TET2,CDC37,RPS16,CPSF3L,TSG101,PTPN13,UBR4,PSMA1,PHF21A,CHC
LPCAT3,CD44,TM9SF1,MAGED2,TET2,CDC37,RPS16,CPSF3L,TSG101,PTPN13,UBR4,PSMA1,PHF21A,CHC
D2B,CYB561,RPL36,LAMP2,SYNGR2,TMEM134,LRRFIP1,RAB2A,ZNF524,TPM2,ZEB2,CHPF,LYNX1,LRRC27
STIP1,LRRC1,PHF19,LPCAT3,CD44,TM9SF1,MAGED2,SON,TET2,HMGA2,CDC37,RPS16,TSG101,TTC21B,U
STIP1,LRRC1,PHF19,LPCAT3,CD44,TM9SF1,MAGED2,SON,TET2,HMGA2,CDC37,RPS16,TSG101,TTC21B,U
MA3,METTL17,PPRC1,RPS25,NLRP1,NIP7,TTC9C,COX4I1,THEM4,DBNDD2,SDF4,CENPK,RTEL1,COASY,SR
MA3,METTL17,PPRC1,RPS25,NLRP1,NIP7,TTC9C,COX4I1,THEM4,DBNDD2,SDF4,CENPK,RTEL1,COASY,SR
C2,VDAC3,WDR19,NUMA1,GPATCH2L,BUD31,SAE1,NPRL2,OSGEP,TBL2,ARHGAP6,SCARB1,CTTN,TSPAN
C2,VDAC3,WDR19,NUMA1,GPATCH2L,BUD31,SAE1,NPRL2,OSGEP,TBL2,ARHGAP6,SCARB1,CTTN,TSPAN
C42,TBX6,AUNIP,C14ORF2,GUK1,PCYT2,USE1,IFI27L1,PPP1CC,EDF1,RFX3,PPA1,DGKE,TNFRSF14,ENDOV

,AUP1,YIPF3,FOXRED1,SH3YL1,CLASRP,UGGT2,HS2ST1,PRMT2,MAGT1,JUP,TBX6,TMEM205,AUNIP,GUK
SON,HMGA2,CDC37,RPS16,OBSL1,UBR4,PSMA1,YLPM1,CHCHD1,SLC39A11,FUZ,FAM3A,MUC1,METTL17
SON,HMGA2,CDC37,RPS16,OBSL1,UBR4,PSMA1,YLPM1,CHCHD1,SLC39A11,FUZ,FAM3A,MUC1,METTL17
PP1CC,RFX3,ENDOV,PSMD4,CLDND1,VIM,LIG1,PSMA7,RPL21,TXNL4A,TMPO,LSM12,PRMT1,CASC4,EXO
SMA7,RPL21,TXNL4A,NUTF2,TCF4,TMPO,LSM12,PRMT1,CASC4,EXOSC9,WNK1,C19ORF54,ZNF7,HNRNP
EBPL,TESC,IGSF8,SNRPN,SPAG7,WDR62,IL1RAP,STYXL1,POLD4,RANBP1,UBE2L6,PPM1K,STIP1,LRRC1,PH
EBPL,TESC,IGSF8,SNRPN,SPAG7,WDR62,IL1RAP,STYXL1,POLD4,RANBP1,UBE2L6,PPM1K,STIP1,LRRC1,PH

REBZF,EIF2A,NKIRAS2,PSMD7,NADSYN1,DCAKD,PIGO,D2HGDH,KDELR3,SLC35A2,AXIN1,SLC25A24,UBE2D

REBZF,EIF2A,NKIRAS2,PSMD7,NADSYN1,DCAKD,PIGO,D2HGDH,KDELR3,SLC35A2,AXIN1,SLC25A24,UBE2D

C35G1,RCE1,MRPL33,ZNF410,RUVBL2,TESC,IGSF8,RBPMS,SPAG7,IL1RAP,STYXL1,POLD4,PLEKHA5,RANBP

C35G1,RCE1,MRPL33,ZNF410,RUVBL2,TESC,IGSF8,RBPMS,SPAG7,IL1RAP,STYXL1,POLD4,PLEKHA5,RANBP

PL28,LIG1,APEH,HYI,C17ORF62,C19ORF54,CYB561,SHARPIN,SLC9B2,CARS,ZNF250,CDK11A,CCNL2,MAPI

3,PDCD6,FAM32A,TMEM147,PEX2,C12ORF57,L3MBTL2,OXLD1,ZNF223,DGAT1,OGFOD2,TSEN34,ARIH2,

RPN,WDR62,STYXL1,RFX5,RANBP1,UBE2L6,LZTFL1,PPM1K,TUBA1C,DST,ANTXR2,STIP1,LRRC1,GRAMD1C

RPN,WDR62,STYXL1,RFX5,RANBP1,UBE2L6,LZTFL1,PPM1K,TUBA1C,DST,ANTXR2,STIP1,LRRC1,GRAMD1C

CPSF3L,TTC21B,PTPN13,UBR4,YLPM1,PHF21A,SLC39A11,RGS3,METTL17,ECSIT,EPB41L5,ATG12,NIP7,TTC

SON,TET2,CDC37,CPSF3L,OBSL1,TTC21B,PSMA1,YLPM1,FUZ,PSMA3,METTL17,PPRC1,RPS25,COX4I1,DUS

SON,TET2,CDC37,CPSF3L,OBSL1,TTC21B,PSMA1,YLPM1,FUZ,PSMA3,METTL17,PPRC1,RPS25,COX4I1,DUS

MC4,LMBR1,MCTS1,DDOST,SYNGR1,LSR,MEIS2,PSME2,PABPC1,PTPRB,NAT9,TOMM40,SERPINB6,TADA3,

MC4,LMBR1,MCTS1,DDOST,SYNGR1,LSR,MEIS2,PSME2,PABPC1,PTPRB,NAT9,TOMM40,SERPINB6,TADA3,

2L6,LZTFL1,PPM1K,TUBA1C,ANTXR2,STIP1,LRRC1,VCAN,PCBP2,LPCAT3,CD44,SON,TET2,SNF8,HMGA2,C

EF1B2,GADL1,PPA1,DGKE,PSMD4,LIG1,PSMA7,TIMM17B,TMEM126B,RPL21,TXNL4A,CAPN10,TCF4,ATPA

2,GUK1,PCYT2,USE1,AGPAT4,PPP1CC,RFX3,PPA1,DGKE,CAMTA1,CRIP1,VIM,LIG1,CDC14B,TIMM17B,APE

H2B1,NOP16,GSTK1,TTLL5,LDHA,ACOT8,AP3B2,SLC35G1,RCE1,ZNF410,RUVBL2,PCMT1,EBPL,TESC,IGSF8

H2B1,NOP16,GSTK1,TTLL5,LDHA,ACOT8,AP3B2,SLC35G1,RCE1,ZNF410,RUVBL2,PCMT1,EBPL,TESC,IGSF8

TN,ITSN1,EIF5A,TNFSF10,COMMD5,RUNX1,NUDT22,USP38,TAZ,THYN1,DBNDD1,SH2B1,NOP16,RPL13A

TN,ITSN1,EIF5A,TNFSF10,COMMD5,RUNX1,NUDT22,USP38,TAZ,THYN1,DBNDD1,SH2B1,NOP16,RPL13A

3,THYN1,DBNDD1,SH2B1,NOP16,RPL13A,ARHGEF39,GSTK1,LDHA,PSMA6,AP3B2,RPS9,RCE1,RUVBL2,EBP

3,THYN1,DBNDD1,SH2B1,NOP16,RPL13A,ARHGEF39,GSTK1,LDHA,PSMA6,AP3B2,RPS9,RCE1,RUVBL2,EBP

SG1,HSPB1,LRRC27,MVD,LMBR1,LSR,CHCHD7,PSME2,PHTF2,PABPC1,BCAM,ZNF836,BLVRB,RABEP2,RAL

3,RCE1,MRPL33,PCMT1,NUDT5,RBPMS,SPAG7,WDR62,UBE2L6,VRK3,LZTFL1,ZC3H14,FIS1,SLC15A3,PPM

,UBE2I,PRMT2,MAGT1,PTK2,C14ORF2,GUK1,CENPN,CARD14,PCYT2,IFI27L1,CCDC25,EDF1,PPA1,ENDOV,

AN11,ITSN1,EIF5A,COMMD5,AMZ2,UFD1L,USP38,TAZ,DBNDD1,RPL13A,GSTK1,TTLL5,PSMA6,ACOT8,AP3

AN11,ITSN1,EIF5A,COMMD5,AMZ2,UFD1L,USP38,TAZ,DBNDD1,RPL13A,GSTK1,TTLL5,PSMA6,ACOT8,AP3

A6,CFLAR,PCNP,VDAC1,RER1,NEK11,EIF2A,NKIRAS2,ZNF511,GTF2H1,BFAR,CLTA,SLC35A2,RPS3A,MOSPD

A6,CFLAR,PCNP,VDAC1,RER1,NEK11,EIF2A,NKIRAS2,ZNF511,GTF2H1,BFAR,CLTA,SLC35A2,RPS3A,MOSPD

DH11,TMED4,SRSF1,AUP1,TRNT1,PARD6G,C1ORF198,UGGT2,MICAL1,HS2ST1,YIF1A,JUP,TBX6,TMEM205

DH11,TMED4,SRSF1,AUP1,TRNT1,PARD6G,C1ORF198,UGGT2,MICAL1,HS2ST1,YIF1A,JUP,TBX6,TMEM205

F511,IRF3,IFT172,D2HGDH,CC2D1B,ALKBH6,AXIN1,MOSPD3,AAMDC,SNAPC2,NQO2,AKT1S1,EMC9,B3GN

C3H14,SLC15A3,STIP1,LRRC1,PHF19,VCAN,LPCAT3,TET2,SNF8,GRK6,HMGA2,CPSF3L,PTPN13,UBR4,YLPM

C3H14,SLC15A3,STIP1,LRRC1,PHF19,VCAN,LPCAT3,TET2,SNF8,GRK6,HMGA2,CPSF3L,PTPN13,UBR4,YLPM

B41L5,ATG12,PPRC1,ZBTB44,NIP7,COX4I1,DUSP22,SLC25A10,TMX2,RDH11,DBNDD2,NISCH,RPS6,RTEL1

B41L5,ATG12,PPRC1,ZBTB44,NIP7,COX4I1,DUSP22,SLC25A10,TMX2,RDH11,DBNDD2,NISCH,RPS6,RTEL1

RAP,STYXL1,RFX5,POLD4,PLEKHA5,RANBP1,UBE2L6,VRK3,ZC3H14,FIS1,SLC15A3,PPM1K,TUBA1C,ANTXR

HCHD1,FUZ,FAM3A,PSMA3,PPRC1,RPS25,NIP7,COX4I1,DUSP22,SPOCK2,SDF4,CENPK,CD99L2,GGPS1,RTE

HCHD1,FUZ,FAM3A,PSMA3,PPRC1,RPS25,NIP7,COX4I1,DUSP22,SPOCK2,SDF4,CENPK,CD99L2,GGPS1,RTE

,RANBP1,VRK3,LZTFL1,FIS1,PPM1K,TUBA1C,IRF9,STIP1,PHF19,PCBP2,TM9SF1,SON,TET2,UQCRQ,SNF8,G

,RANBP1,VRK3,LZTFL1,FIS1,PPM1K,TUBA1C,IRF9,STIP1,PHF19,PCBP2,TM9SF1,SON,TET2,UQCRQ,SNF8,G

5,ZBTB44,NIP7,COX4I1,DUSP22,SLC25A10,SPOCK2,THEM4,CENPK,CD99L2,GGPS1,RTEL1,COASY,SRSF1,S

5,ZBTB44,NIP7,COX4I1,DUSP22,SLC25A10,SPOCK2,THEM4,CENPK,CD99L2,GGPS1,RTEL1,COASY,SRSF1,S

5A3,PPM1K,TUBA1C,DST,STIP1,LRRC1,PHF19,CD44,TM9SF1,SON,TET2,CDC37,RPS16,TTC21B,UBR4,YLPM

5A3,PPM1K,TUBA1C,DST,STIP1,LRRC1,PHF19,CD44,TM9SF1,SON,TET2,CDC37,RPS16,TTC21B,UBR4,YLPM

22,SLC25A10,RDH11,DBNDD2,SDF4,TMED4,CENPK,RPS6,GGPS1,COASY,SRSF1,RABL5,TRNT1,ITGB1BP1,
22,SLC25A10,RDH11,DBNDD2,SDF4,TMED4,CENPK,RPS6,GGPS1,COASY,SRSF1,RABL5,TRNT1,ITGB1BP1,
H14,PPM1K,IRF9,DST,ANTXR2,STIP1,LRRC1,VCAN,LPCAT3,CD44,SON,TET2,HMGA2,RPS16,TSG101,TTC2
H14,PPM1K,IRF9,DST,ANTXR2,STIP1,LRRC1,VCAN,LPCAT3,CD44,SON,TET2,HMGA2,RPS16,TSG101,TTC2
2,UQCRQ,HMGA2,CDC37,CPSF3L,OBSL1,TTC21B,PSMA1,YLPM1,SLC39A11,FAM3A,METTL17,EPB41L5,PP
ZC3H14,PPM1K,TUBA1C,ANTXR2,STIP1,LRRC1,PHF19,VCAN,LPCAT3,CD44,SON,GRK6,CPSF3L,OBSL1,TTC
3B2,RPS9,SLC35G1,RCE1,RUVBL2,PCMT1,EBPL,TESC,IGSF8,RBPMS,SPAG7,WDR62,STYXL1,RFX5,POLD4,P
3B2,RPS9,SLC35G1,RCE1,RUVBL2,PCMT1,EBPL,TESC,IGSF8,RBPMS,SPAG7,WDR62,STYXL1,RFX5,POLD4,P
C9C,COX4I1,THEM4,DBNDD2,TMED4,CENPK,SRSF1,AUP1,SAP30BP,FOXRED1,TRNT1,KXD1,ITGB1BP1,ME
C9C,COX4I1,THEM4,DBNDD2,TMED4,CENPK,SRSF1,AUP1,SAP30BP,FOXRED1,TRNT1,KXD1,ITGB1BP1,ME

,SLC39A11,FAM3A,EPB41L5,ZBTB44,DUSP22,GGPS1,SRSF1,AUP1,SAP30BP,YIPF3,FOXRED1,MED15,SH3Y
17B,APEH,HYI,NUTF2,PODXL,C17ORF62,PRMT1,CASC4,ZNF7,STRADA,ZFAND2B,STAU1,RPL36,LAMP2,CC
RMT1,CASC4,WNK1,ZNF7,ARL6IP4,PPAN,HNRNPC,ZNF644,SUGT1,COPZ1,CCNL2,MAPKAPK5,CEP70,LRR
PLEKHA5,RANBP1,LZTFL1,SLC15A3,TUBA1C,STIP1,PHF19,VCAN,TM9SF1,SON,UQCRQ,SNF8,GRK6,RPS16,
2L,BUD31,SAE1,NPRL2,OSGEP,TBL2,CRYGS,CTTN,ITSN1,EIF5A,COMMD5,RUNX1,CHID1,UFD1L,USP38,TH
2L,BUD31,SAE1,NPRL2,OSGEP,TBL2,CRYGS,CTTN,ITSN1,EIF5A,COMMD5,RUNX1,CHID1,UFD1L,USP38,TH
LC25A24,ZNF396,EPN1,ZCCHC9,CHURC1,HMBS,AZI2,U2AF1L4,EMC3,PRKCSH,ZNF276,ZNF483,SZT2,PSM
LC25A24,ZNF396,EPN1,ZCCHC9,CHURC1,HMBS,AZI2,U2AF1L4,EMC3,PRKCSH,ZNF276,ZNF483,SZT2,PSM
JFD1L,USP38,TAZ,THYN1,DBNDD1,SH2B1,NOP16,ARHGEF39,GSTK1,TTLL5,LDHA,ACOT8,SLC35G1,RCE1,
JFD1L,USP38,TAZ,THYN1,DBNDD1,SH2B1,NOP16,ARHGEF39,GSTK1,TTLL5,LDHA,ACOT8,SLC35G1,RCE1,
PMS,IL1RAP,STYXL1,POLD4,PLEKHA5,RANBP1,ZC3H14,PPM1K,IRF9,ANTXR2,STIP1,LRRC1,PHF19,VCAN,T
1LG,EPB41L4B,TCF4,ATPAF2,C17ORF62,PRMT1,CASC4,ZNF7,PPAN,ZFAND2B,HNRNPC,STAU1,ZNF250,NF
AT1,RPL8,UBE2D4,ASH1L,TCF25,ZNF248,ANAPC13,NXF1,SYP,NUP62,PSPC1,NADSYN1,FCHO2,IRF3,D2H
AT1,RPL8,UBE2D4,ASH1L,TCF25,ZNF248,ANAPC13,NXF1,SYP,NUP62,PSPC1,NADSYN1,FCHO2,IRF3,D2H
A1,CLDND1,LIG1,TIMM17B,APEH,LYPLA2,NUTF2,PODXL,TCF4,C17ORF62,PRMT1,CASC4,ZNF7,PPAN,ZFA
TN5,PEX26,OXLD1,ZNF223,ARIH2,PICK1,ZNF512,POLR2M,TSPAN31,GLYCTK,C11ORF57,LAMTOR5,RER1,
B41L5,ZBTB44,NIP7,COX4I1,PLCB2,SLC25A10,DBNDD2,TMED4,CENPK,RPS6,CD99L2,GGPS1,COASY,SRSF
A,FUZ,FAM3A,MUC1,ECSIT,EPB41L5,ATG12,PPRC1,ZBTB44,NIP7,TTC9C,PLCB2,DUSP22,SLC25A10,SPOCK
A,FUZ,FAM3A,MUC1,ECSIT,EPB41L5,ATG12,PPRC1,ZBTB44,NIP7,TTC9C,PLCB2,DUSP22,SLC25A10,SPOCK
,SAE1,TBL2,ARHGAP6,SCARB1,CTTN,TSPAN11,ITSN1,EIF5A,COMMD5,RUNX1,NUDT22,AMZ2,USP38,TAZ,
,SAE1,TBL2,ARHGAP6,SCARB1,CTTN,TSPAN11,ITSN1,EIF5A,COMMD5,RUNX1,NUDT22,AMZ2,USP38,TAZ,
IINT1,HNRNPA1,EGLN2,SIRT5,RFXANK,SPINT2,SMARCA4,OARD1,MCFD2,ADAM19,CBR1,PHC3,MROH1,E
IINT1,HNRNPA1,EGLN2,SIRT5,RFXANK,SPINT2,SMARCA4,OARD1,MCFD2,ADAM19,CBR1,PHC3,MROH1,E
,PIGO,BFAR,SLC25A24,EIF3C,DNAJC2,HPS1,GTF3A,CHURC1,IFI27,ARRB2,AZI2,NCBP2,POLR2E,PRKCSH,R
,PIGO,BFAR,SLC25A24,EIF3C,DNAJC2,HPS1,GTF3A,CHURC1,IFI27,ARRB2,AZI2,NCBP2,POLR2E,PRKCSH,R

IUC1,PPRC1,RPS25,NLRP1,ZBTB44,NIP7,SLC25A10,SDF4,NISCH,TMED4,CD99L2,RTEL1,COASY,AUP1,YIPF
IUC1,PPRC1,RPS25,NLRP1,ZBTB44,NIP7,SLC25A10,SDF4,NISCH,TMED4,CD99L2,RTEL1,COASY,AUP1,YIPF
.,ITGB1BP1,SH3YL1,C1ORF198,SLC39A13,YIF1A,EIF3M,JUP,XBP1,C14ORF2,GUK1,PCYT2,FBXW9,USE1,AG
.,ITGB1BP1,SH3YL1,C1ORF198,SLC39A13,YIF1A,EIF3M,JUP,XBP1,C14ORF2,GUK1,PCYT2,FBXW9,USE1,AG
,CAMLG,EPB41L4B,UCK2,LSM12,PRMT1,CASC4,ZNF7,TXNDC17,ARL6IP4,ACAD10,STAU1,SLC9B2,ZNF250,
,CAMLG,EPB41L4B,UCK2,LSM12,PRMT1,CASC4,ZNF7,TXNDC17,ARL6IP4,ACAD10,STAU1,SLC9B2,ZNF250,

STIP1,LRRC1,GRAMD1C,VCAN,PCBP2,LPCAT3,TM9SF1,MAGED2,SON,UQCRQ,SNF8,GRK6,CDC37,RPS16,C
STIP1,LRRC1,GRAMD1C,VCAN,PCBP2,LPCAT3,TM9SF1,MAGED2,SON,UQCRQ,SNF8,GRK6,CDC37,RPS16,C
G1,MRPL33,ZNF410,RUVBL2,PCMT1,EBPL,TESC,NUDT5,RBPMS,NNMT,SNRPN,SPAG7,WDR62,STYXL1,PL
G1,MRPL33,ZNF410,RUVBL2,PCMT1,EBPL,TESC,NUDT5,RBPMS,NNMT,SNRPN,SPAG7,WDR62,STYXL1,PL
SME2,PHTF2,RRNAD1,BCAM,TOMM40,RABEP2,TADA3,ZNF20,ATP6V1F,WAC,SLC33A1,ZNF687,NXN,ANA
,DDOST,LSR,RRNAD1,DBNL,RAD1,TOMM40,SERPINB6,TADA3,ATP6V1F,TRIM16,ILF3,MDH2,EEF1D,CHCH
P1,RABL5,TRNT1,ITGB1BP1,MED15,CLASRP,MICAL1,HS2ST1,SLC39A13,MAGT1,TBX6,XBP1,PTK2,AUNIP,C
RSF1,AUP1,FOXRED1,ITGB1BP1,SH3YL1,IP6K2,JPH2,EIF3M,MAGT1,CDC42,TBX6,TMEM205,XBP1,AUNIP,
RSF1,AUP1,FOXRED1,ITGB1BP1,SH3YL1,IP6K2,JPH2,EIF3M,MAGT1,CDC42,TBX6,TMEM205,XBP1,AUNIP,
8,TAZ,THYN1,SH2B1,NOP16,RPL13A,GSTK1,TTLL5,ACOT8,AP3B2,SLC35G1,MRPL33,ZNF410,PCMT1,EBPL
8,TAZ,THYN1,SH2B1,NOP16,RPL13A,GSTK1,TTLL5,ACOT8,AP3B2,SLC35G1,MRPL33,ZNF410,PCMT1,EBPL
SF1B,PPP1CC,PPIL2,VIM,LIG1,TIMM17B,TMEM126B,APEH,RPL21,TXNL4A,UCK2,TMPO,C17ORF62,EXOS
2B,HNRNPC,CYB561,ZNF250,HSP90AB1,SYNGR2,TMEM134,LRRFIP1,RAB2A,ZNF524,TPM2,SIAH1,ZEB2,
2B,HNRNPC,CYB561,ZNF250,HSP90AB1,SYNGR2,TMEM134,LRRFIP1,RAB2A,ZNF524,TPM2,SIAH1,ZEB2,
M4,CENPK,CD99L2,GGPS1,RTEL1,COASY,SRSF1,SAP30BP,YIPF3,TRNT1,MED15,IP6K2,CLASRP,C1ORF198
ED4,RPS6,RPL29,COASY,SRSF1,SAP30BP,TRNT1,ITGB1BP1,SH3YL1,IP6K2,C1ORF198,UGGT2,YIF1A,EIF3M
ED4,RPS6,RPL29,COASY,SRSF1,SAP30BP,TRNT1,ITGB1BP1,SH3YL1,IP6K2,C1ORF198,UGGT2,YIF1A,EIF3M
1CC,EDF1,PPA1,DGKE,CAMTA1,LIG1,TIMM17B,APEH,HYI,CAPN10,NUTF2,PODXL,ATPAF2,C17ORF62,PRI

,NIP7,PLCB2,DUSP22,SLC25A10,TMX2,DBNDD2,NISCH,TMED4,CD99L2,GGPS1,YIPF3,TRNT1,PARD6G,SH
,NIP7,PLCB2,DUSP22,SLC25A10,TMX2,DBNDD2,NISCH,TMED4,CD99L2,GGPS1,YIPF3,TRNT1,PARD6G,SH
IF410,RUVBL2,PCMT1,IGSF8,NUDT5,RBPMS,NNMT,WDR62,STYXL1,PLEKHA5,RANBP1,FIS1,SLC15A3,TUB
IF410,RUVBL2,PCMT1,IGSF8,NUDT5,RBPMS,NNMT,WDR62,STYXL1,PLEKHA5,RANBP1,FIS1,SLC15A3,TUB
P1,LZTFL1,TUBA1C,DST,PHF19,LPCAT3,CD44,SON,TET2,SNF8,CPSF3L,TSG101,OBSL1,TTC21B,PTPN13,UB
P1,LZTFL1,TUBA1C,DST,PHF19,LPCAT3,CD44,SON,TET2,SNF8,CPSF3L,TSG101,OBSL1,TTC21B,PTPN13,UB
,ACOT8,AP3B2,RPS9,SLC35G1,RCE1,MRPL33,RUVBL2,EBPL,TESC,NUDT5,RBPMS,SNRPN,WDR62,IL1RAP,
,ACOT8,AP3B2,RPS9,SLC35G1,RCE1,MRPL33,RUVBL2,EBPL,TESC,NUDT5,RBPMS,SNRPN,WDR62,IL1RAP,
UZ,RGS3,FAM3A,PSMA3,PPRC1,RPS25,NIP7,COX4I1,SPOCK2,NISCH,CENPK,GGPS1,RTEL1,SRSF1,AUP1,FC
UZ,RGS3,FAM3A,PSMA3,PPRC1,RPS25,NIP7,COX4I1,SPOCK2,NISCH,CENPK,GGPS1,RTEL1,SRSF1,AUP1,FC
IOP16,TTLL5,LDHA,PSMA6,ACOT8,AP3B2,RPS9,RCE1,MRPL33,ZNF410,RUVBL2,NUDT5,NNMT,SPAG7,WD
IOP16,TTLL5,LDHA,PSMA6,ACOT8,AP3B2,RPS9,RCE1,MRPL33,ZNF410,RUVBL2,NUDT5,NNMT,SPAG7,WD
FD2,SCAP,CBR1,PHC3,MFSD11,MOK,ZNF81,TMEM14B,RPL37A,ELOF1,DCTN5,ARL16,DGAT1,RPL8,UBE2D
FD2,SCAP,CBR1,PHC3,MFSD11,MOK,ZNF81,TMEM14B,RPL37A,ELOF1,DCTN5,ARL16,DGAT1,RPL8,UBE2D
A10,NISCH,GGPS1,COASY,SRSF1,AUP1,YIPF3,FOXRED1,SH3YL1,IP6K2,JUP,TBX6,TMEM205,XBP1,PTK2,C1
MPDU1,RPS15,ERLEC1,NAP1L1,MEIS2,HMGA1,PHTF2,NAT9,H2AFV,SERPINB6,WAC,ANAPC11,ILF3,AGAP
205,ATF7,PCYT2,IFI27L1,ASF1B,EEF1B2,PPP1CC,EDF1,DGKE,ENDOV,CAMTA1,PPIL2,RPL28,LIG1,TXNL4A,N
SC,IGSF8,NUDT5,RBPMS,SPAG7,WDR62,IL1RAP,STYXL1,RFX5,POLD4,PLEKHA5,RANBP1,VRK3,LZTFL1,ZC3
SC,IGSF8,NUDT5,RBPMS,SPAG7,WDR62,IL1RAP,STYXL1,RFX5,POLD4,PLEKHA5,RANBP1,VRK3,LZTFL1,ZC3
CD99L2,AUP1,PARD6G,SH3YL1,IP6K2,CLASRP,MICAL1,HS2ST1,UBE2I,PRMT2,PTK2,AUNIP,C14ORF2,PCYT
PHF21A,FAM3A,PSMA3,METTL17,PPRC1,RPS25,NLRP1,NIP7,TTC9C,COX4I1,DUSP22,THEM4,DBNDD2,TM
PHF21A,FAM3A,PSMA3,METTL17,PPRC1,RPS25,NLRP1,NIP7,TTC9C,COX4I1,DUSP22,THEM4,DBNDD2,TM
CMT1,EBPL,RBPMS,WDR62,STYXL1,UBE2L6,VRK3,PPM1K,TUBA1C,DST,ANTXR2,STIP1,LRRC1,PHF19,GRA
CMT1,EBPL,RBPMS,WDR62,STYXL1,UBE2L6,VRK3,PPM1K,TUBA1C,DST,ANTXR2,STIP1,LRRC1,PHF19,GRA
,LAMP2,HSP90AB1,SYNGR2,TMEM134,LRRFIP1,RAB2A,ZNF524,ASPH,TRMT112,STRIP2,LYNX1,TWSG1,L

1,FUZ,FAM3A,PSMA3,METTL17,ATG12,PPRC1,ZBTB44,SLC25A10,TMX2,SDF4,NISCH,TMED4,RPS6,RPL29

AZ,THYN1,DBNDD1,SH2B1,LDHA,PSMA6,AP3B2,RPS9,SLC35G1,RCE1,MRPL33,ZNF410,RUVBL2,PCMT1,NU
AZ,THYN1,DBNDD1,SH2B1,LDHA,PSMA6,AP3B2,RPS9,SLC35G1,RCE1,MRPL33,ZNF410,RUVBL2,PCMT1,NU
PN10,NUTF2,PODXL,C17ORF62,PRMT1,CASC4,ZNF7,TXNDC17,ZFAND2B,CYB561,STAU1,UPP1,ZNF250,RF
1,BFAR,TP53,SH3GL1,TMEM199,ZNF396,SNAPC2,EPN1,ZCCHC9,MKRN1,COMT,CHURC1,RBM6
1,BFAR,TP53,SH3GL1,TMEM199,ZNF396,SNAPC2,EPN1,ZCCHC9,MKRN1,COMT,CHURC1,RBM6

GA2,CDC37,RPS16,PTPN13,UBR4,PHF21A,CHCHD1,SLC39A11,FUZ,FAM3A,MUC1,METTL17,RPS25,TTC9C,
PS28,ZSWIM7,HSPB1,MVD,EMC4,IL32,SEMA3C,SYNGR1,C12ORF76,DBNL,TOMM40,SERPINB6,RABEP2,F
PS28,ZSWIM7,HSPB1,MVD,EMC4,IL32,SEMA3C,SYNGR1,C12ORF76,DBNL,TOMM40,SERPINB6,RABEP2,F
RNAD1,SLC33A1,RALGPS1,ZNF687,POT1,RAD23A,ILF3,PKP4,HSPH1,MT1E,RPL18,CHEK1,SUMO3,PHF20L
,CAPN10,LYPLA2,PODXL,ATPAF2,C17ORF62,PRMT1,CASC4,ARL6IP4,ZFAND2B,SHARPIN,UPP1,NFAM1,DU
,CAPN10,LYPLA2,PODXL,ATPAF2,C17ORF62,PRMT1,CASC4,ARL6IP4,ZFAND2B,SHARPIN,UPP1,NFAM1,DU
,CENPK,RPS6,SRSF1,SAP30BP,KXD1,SH3YL1,C1ORF198,UGGT2,HS2ST1,YIF1A,TMEM205,CELF1,C14ORF2
,CENPK,RPS6,SRSF1,SAP30BP,KXD1,SH3YL1,C1ORF198,UGGT2,HS2ST1,YIF1A,TMEM205,CELF1,C14ORF2
B1,CRYGS,CTTN,TSPAN11,ITSN1,EIF5A,COMMD5,RUNX1,CHID1,NUDT22,AMZ2,TAZ,THYN1,DBNDD1,SH
B1,CRYGS,CTTN,TSPAN11,ITSN1,EIF5A,COMMD5,RUNX1,CHID1,NUDT22,AMZ2,TAZ,THYN1,DBNDD1,SH
BPL,TESC,NUDT5,RBPMS,NNMT,SNRPN,WDR62,STYXL1,PLEKHA5,RANBP1,UBE2L6,FIS1,PPM1K,TUBA1C,
PHTF2,PABPC1,NAT9,BCAM,ZNF836,BLVRB,RABEP2,RALGPS1,ZNF687,PDCD5,NXN,EPS15L1,POLR3H,RAD
PH2,UGGT2,MICAL1,SLC39A13,CDC42,TBX6,XBP1,PTK2,C14ORF2,GUK1,CENPN,ATF7,FBXW9,USE1,HSBP
,CARS,MOB1A,CDK11A,MDH1,SYNGR2,CEP70,ELP6,SIAH1,TSFM,GABPB1,TWSG1,ZSWIM7,RPS15,DCTD,I
,TMEM39B,FKBP5,MRPS18C,SLC30A5,THADA,COMMD4,SNX5,NUP133,ELP4,ZXDC,ELOF1,ARL16,RC3H2,I
E1,NPRL2,OSGEP,TBL2,ARHGAP6,SCARB1,CRYGS,CTTN,TSPAN11,EIF5A,COMMD5,CHID1,NUDT22,AMZ2,
E1,NPRL2,OSGEP,TBL2,ARHGAP6,SCARB1,CRYGS,CTTN,TSPAN11,EIF5A,COMMD5,CHID1,NUDT22,AMZ2,
TET2,SNF8,GRK6,HMGA2,CDC37,RPS16,OBSL1,UBR4,PHF21A,RGS3,ATG12,RPS25,ZBTB44,NIP7,TTC9C,C
NT1,KXD1,ITGB1BP1,MED15,SLC39A13,YIF1A,MAGT1,C14ORF2,GUK1,CENPN,CARD14,PCYT2,HSBP1L1,A
NT1,KXD1,ITGB1BP1,MED15,SLC39A13,YIF1A,MAGT1,C14ORF2,GUK1,CENPN,CARD14,PCYT2,HSBP1L1,A
ETTL17,ATG12,NLRP1,ZBTB44,NIP7,COX4I1,SLC25A10,THEM4,SDF4,CENPK,RPL29,CD99L2,COASY,SRSF1,
ETTL17,ATG12,NLRP1,ZBTB44,NIP7,COX4I1,SLC25A10,THEM4,SDF4,CENPK,RPL29,CD99L2,COASY,SRSF1,
EF1B2,PPP1CC,EDF1,RFX3,PPIL2,RPL28,TMEM126B,APEH,RPL21,TXNL4A,NUTF2,CAMLG,TMPO,ATPAF2,
EF1B2,PPP1CC,EDF1,RFX3,PPIL2,RPL28,TMEM126B,APEH,RPL21,TXNL4A,NUTF2,CAMLG,TMPO,ATPAF2,
MAGT1,PTK2,AUNIP,C14ORF2,GUK1,FBXW9,USE1,AGPAT4,IFI27L1,DEF8,EEF1B2,PPP1CC,EDF1,RFX3,PPA1
ZFAND2B,CYB561,UPP1,RPL36,HSP90AB1,SYNGR2,TMEM134,LRRFIP1,RAB2A,ZNF524,ATP5J,TPM2,SIAH
A1,CLDND1,VIM,LIG1,PSMA7,CDC14B,CAPN10,NUTF2,CAMLG,EPB41L4B,UCK2,ATPAF2,LSM12,CASC4,W
A1,CLDND1,VIM,LIG1,PSMA7,CDC14B,CAPN10,NUTF2,CAMLG,EPB41L4B,UCK2,ATPAF2,LSM12,CASC4,W
CLDND1,LIG1,PSMA7,TIMM17B,APEH,HYI,CAPN10,NUTF2,PODXL,C17ORF62,PRMT1,CASC4,ZNF7,ZFAND
K4I1,TMX2,THEM4,RDH11,SDF4,TMED4,CENPK,RTEL1,SRSF1,RABL5,TRNT1,ITGB1BP1,PARD6G,C1ORF19
K4I1,TMX2,THEM4,RDH11,SDF4,TMED4,CENPK,RTEL1,SRSF1,RABL5,TRNT1,ITGB1BP1,PARD6G,C1ORF19
NOP16,RPL13A,GSTK1,TTLL5,LDHA,PSMA6,RPS9,RCE1,MRPL33,ZNF410,RUVBL2,PCMT1,EBPL,WDR62,IL
NOP16,RPL13A,GSTK1,TTLL5,LDHA,PSMA6,RPS9,RCE1,MRPL33,ZNF410,RUVBL2,PCMT1,EBPL,WDR62,IL
A,COMMD5,NUDT22,AMZ2,UFD1L,USP38,TAZ,DBNDD1,RPL13A,GSTK1,TTLL5,PSMA6,ACOT8,AP3B2,SLC
A,COMMD5,NUDT22,AMZ2,UFD1L,USP38,TAZ,DBNDD1,RPL13A,GSTK1,TTLL5,PSMA6,ACOT8,AP3B2,SLC
,FAM131A,ZNF354A,PSMD7,DCAKD,ZNF283,BRD8,NAA38,SCRN2,THOC3,TP53,PSMD10,HDAC8,NDUFA3
,FAM131A,ZNF354A,PSMD7,DCAKD,ZNF283,BRD8,NAA38,SCRN2,THOC3,TP53,PSMD10,HDAC8,NDUFA3
IF1A,MAGT1,TMEM205,PCYT2,FBXW9,EEF1B2,DGKE,ENDOV,VAMP8,CAMTA1,PPIL2,RPL28,TIMM17B,TN
PF,MPDU1,MAPK13,GMPPB,MVD,ERCC5,IL32,NAP1L1,SYNGR1,RRNAD1,NAT9,H2AFV,BLVRB,SLC33A1,ZN
RL2,OSGEP,TBL2,ARHGAP6,SCARB1,CTTN,ITSN1,EIF5A,COMMD5,RUNX1,NUDT22,UFD1L,USP38,TAZ,THY
RL2,OSGEP,TBL2,ARHGAP6,SCARB1,CTTN,ITSN1,EIF5A,COMMD5,RUNX1,NUDT22,UFD1L,USP38,TAZ,THY

APEH,PPOX,LSM12,ZFAND2B,CYB561,SHARPIN,UPP1,ZNF250,RBM3,DGUOK,SYNGR2,FAM219B,RAB2A,
APEH,PPOX,LSM12,ZFAND2B,CYB561,SHARPIN,UPP1,ZNF250,RBM3,DGUOK,SYNGR2,FAM219B,RAB2A,

X6,AUNIP,GUK1,CENPN,ATF7,FBXW9,MAF,PPP1CC,RFX3,DGKE,ENDOV,CAMTA1,CRIP1,PSMD4,VIM,LIG1
XF1,SYP,TRIM46,NUP62,PSPC1,NADSYN1,FCHO2,IRF3,IFT172,RNF130,BFAR,D2HGDH,GEMIN2,LYPLAL1,T
XF1,SYP,TRIM46,NUP62,PSPC1,NADSYN1,FCHO2,IRF3,IFT172,RNF130,BFAR,D2HGDH,GEMIN2,LYPLAL1,T
DHA,ACOT8,AP3B2,RPS9,ZNF410,RUVBL2,PCMT1,EBPL,TESC,IGSF8,NUDT5,RBPMS,SPAG7,STYXL1,RFX5,
DHA,ACOT8,AP3B2,RPS9,ZNF410,RUVBL2,PCMT1,EBPL,TESC,IGSF8,NUDT5,RBPMS,SPAG7,STYXL1,RFX5,
ARCA4,MPC2,MRPS18C,MCFD2,SCAP,CBR1,PHC3,MROH1,EIF2B3,MOK,ZNF81,NOP2,ZXDC,RPL37A,ELOF
ARCA4,MPC2,MRPS18C,MCFD2,SCAP,CBR1,PHC3,MROH1,EIF2B3,MOK,ZNF81,NOP2,ZXDC,RPL37A,ELOF
GT2,MAGT1,CDC42,PTK2,FBXW9,CCDC25,EEF1B2,PPA1,VAMP8,PSMD4,TROAP,RPL28,LIG1,CDC14B,TIM
GT2,MAGT1,CDC42,PTK2,FBXW9,CCDC25,EEF1B2,PPA1,VAMP8,PSMD4,TROAP,RPL28,LIG1,CDC14B,TIM
CHC9,SNRPA1,AZI2,ZBTB8OS,TBC1D22A,NCBP2,TMEM41A,ZNF581,AK2,MRPS16,RAB26,ZNF33A,DNAJC2
CHC9,SNRPA1,AZI2,ZBTB8OS,TBC1D22A,NCBP2,TMEM41A,ZNF581,AK2,MRPS16,RAB26,ZNF33A,DNAJC2
,YIF1A,PRMT2,MAGT1,JUP,C14ORF2,GUK1,CENPN,FBXW9,AGPAT4,IFI27L1,PPP1CC,RFX3,PPA1,ENDOV,
,YIF1A,PRMT2,MAGT1,JUP,C14ORF2,GUK1,CENPN,FBXW9,AGPAT4,IFI27L1,PPP1CC,RFX3,PPA1,ENDOV,
,DBNDD2,TMED4,CENPK,COASY,SRSF1,AUP1,SAP30BP,FOXRED1,TRNT1,KXD1,ITGB1BP1,MED15,SLC39A
,DBNDD2,TMED4,CENPK,COASY,SRSF1,AUP1,SAP30BP,FOXRED1,TRNT1,KXD1,ITGB1BP1,MED15,SLC39A
H2,CAMK2D,ZFAND4,SIRT5,B3GAT3,ZNF343,GBA2,COPS4,PPIH,DALRD3,TPM4,MFSD10,MRPS18C,GMPR
H2,CAMK2D,ZFAND4,SIRT5,B3GAT3,ZNF343,GBA2,COPS4,PPIH,DALRD3,TPM4,MFSD10,MRPS18C,GMPR
TM9SF1,TET2,SNF8,HMGA2,CDC37,RPS16,CPSF3L,OBSL1,TTC21B,PSMA1,YLPM1,SLC39A11,FAM3A,EPB4
TNFSF10,COMMD5,CHID1,NUDT22,AMZ2,UFD1L,USP38,TAZ,THYN1,DBNDD1,NOP16,RPL13A,GSTK1,TT
TNFSF10,COMMD5,CHID1,NUDT22,AMZ2,UFD1L,USP38,TAZ,THYN1,DBNDD1,NOP16,RPL13A,GSTK1,TT
P16,RPL13A,ARHGEF39,TTLL5,AP3B2,MRPL33,ZNF410,RUVBL2,EBPL,TESC,NUDT5,RBPMS,SPAG7,WDR62
0A5,GNB5,TRPV3,THADA,EIF2B3,PIGH,NUBP2,RPLP0,HAX1,TSEN34,OCIAD1,TCF25,RBBP6,TPT1,CDK4,AN
ET2,GRK6,HMGA2,CPSF3L,OBSL1,PTPN13,UBR4,PHF21A,CHCHD1,SLC39A11,FAM3A,MUC1,ECSIT,ATG12,

TC9C,PLCB2,DUSP22,SLC25A10,TMX2,THEM4,NISCH,TMED4,RPL29,CD99L2,RTEL1,SRSF1,YIPF3,RABL5,P
TC9C,PLCB2,DUSP22,SLC25A10,TMX2,THEM4,NISCH,TMED4,RPL29,CD99L2,RTEL1,SRSF1,YIPF3,RABL5,P

HS2ST1,UBE2I,YIF1A,TBX6,PTK2,C14ORF2,ATF7,PCYT2,USE1,HSBP1L1,PPP1CC,RFX3,PPA1,DGKE,ENDOV,
HS2ST1,UBE2I,YIF1A,TBX6,PTK2,C14ORF2,ATF7,PCYT2,USE1,HSBP1L1,PPP1CC,RFX3,PPA1,DGKE,ENDOV,

M4,CENPK,RPL29,CD99L2,COASY,SRSF1,SAP30BP,KXD1,MED15,SH3YL1,IP6K2,CLASRP,HS2ST1,KCNMA1,
P2,C11ORF48,OAS3,SEC31A,CNST,SUMO3,LGMN,CD151,SMARCA4,TYMS,TMEM39B,DALRD3,DAZAP2,O
P2,C11ORF48,OAS3,SEC31A,CNST,SUMO3,LGMN,CD151,SMARCA4,TYMS,TMEM39B,DALRD3,DAZAP2,O

MED4,CD99L2,RTEL1,SRSF1,AUP1,MED15,SH3YL1,UGGT2,HS2ST1,UBE2I,SLC39A13,YIF1A,PRMT2,JUP,CD
EPB41L5,ATG12,PPRC1,ZBTB44,NIP7,TTC9C,PLCB2,DUSP22,SLC25A10,SPOCK2,DBNDD2,NISCH,TMED4,C
EPB41L5,ATG12,PPRC1,ZBTB44,NIP7,TTC9C,PLCB2,DUSP22,SLC25A10,SPOCK2,DBNDD2,NISCH,TMED4,C
C37,RPS16,CPSF3L,OBSL1,YLPM1,PHF21A,SLC39A11,MUC1,METTL17,EPB41L5,ATG12,ZBTB44,PLCB2,DU
2,YIF1A,CDC42,TBX6,TMEM205,PTK2,CELF1,C14ORF2,CENPN,ATF7,FBXW9,HSBP1L1,ASF1B,PPP1CC,EDF
2,YIF1A,CDC42,TBX6,TMEM205,PTK2,CELF1,C14ORF2,CENPN,ATF7,FBXW9,HSBP1L1,ASF1B,PPP1CC,EDF
D1,CHFR,SSH3,C12ORF10,RNF4,C12ORF66,ZRANB2,DPP9,SEC24C,RPL31,TVP23B,CDK5RAP3,WWP2
D1,CHFR,SSH3,C12ORF10,RNF4,C12ORF66,ZRANB2,DPP9,SEC24C,RPL31,TVP23B,CDK5RAP3,WWP2

DXL,ATPAF2,C17ORF62,PRMT1,CASC4,ARL6IP4,ZFAND2B,HNRNPC,SHARPIN,SYNGR2,CEP70,TMEM134,L

DXL,ATPAF2,C17ORF62,PRMT1,CASC4,ARL6IP4,ZFAND2B,HNRNPC,SHARPIN,SYNGR2,CEP70,TMEM134,L

1,RPS25,NIP7,COX4I1,SPOCK2,NISCH,CENPK,GGPS1,RTEL1,SRSF1,AUP1,FOXRED1,ITGB1BP1,SH3YL1,IP6K

1,RPS25,NIP7,COX4I1,SPOCK2,NISCH,CENPK,GGPS1,RTEL1,SRSF1,AUP1,FOXRED1,ITGB1BP1,SH3YL1,IP6K

15,ERLEC1,IL32,NAP1L1,MEIS2,PHTF2,PABPC1,NAT9,ZNF836,RBP1,H2AFV,SERPINB6,RABEP2,WAC,ZNF7

DGUOK,SYNGR2,ATP5J,GGCX,TRMT112,PNPO,CHPF,ZNF625,RPS20,DCTD,SAT1,LMBR1,MCTS1,LSR,CHCH

,RCE1,PCMT1,EBPL,TESC,IGSF8,RBPMS,SPAG7,WDR62,RFX5,POLD4,RANBP1,UBE2L6,FIS1,SLC15A3,PPM

,RCE1,PCMT1,EBPL,TESC,IGSF8,RBPMS,SPAG7,WDR62,RFX5,POLD4,RANBP1,UBE2L6,FIS1,SLC15A3,PPM

BP1,UBE2L6,VRK3,ZC3H14,PPM1K,ANTXR2,STIP1,LRRC1,PHF19,PCBP2,LPCAT3,CD44,TM9SF1,MAGED2,

BP1,UBE2L6,VRK3,ZC3H14,PPM1K,ANTXR2,STIP1,LRRC1,PHF19,PCBP2,LPCAT3,CD44,TM9SF1,MAGED2,

C3H14,FIS1,IRF9,STIP1,PHF19,GRAMD1C,LPCAT3,CD44,MAGED2,SON,TET2,GRK6,HMGA2,CDC37,OBSL1,

C3H14,FIS1,IRF9,STIP1,PHF19,GRAMD1C,LPCAT3,CD44,MAGED2,SON,TET2,GRK6,HMGA2,CDC37,OBSL1,

,COASY,SRSF1,YIPF3,TRNT1,PARD6G,SH3YL1,IP6K2,CLASRP,JPH2,C1ORF198,MICAL1,HS2ST1,UBE2I,PRM

,COASY,SRSF1,YIPF3,TRNT1,PARD6G,SH3YL1,IP6K2,CLASRP,JPH2,C1ORF198,MICAL1,HS2ST1,UBE2I,PRM

2,UPP1,CARS,RPL36,NFAM1,RBM3,ZNF197,COPZ1,MAPKAPK5,SYNGR2,TMEM134,RAB2A,ELP6,TPM2,SI

2,UPP1,CARS,RPL36,NFAM1,RBM3,ZNF197,COPZ1,MAPKAPK5,SYNGR2,TMEM134,RAB2A,ELP6,TPM2,SI

,UBXN1,ERLEC1,HSPB1,HHLA3,EMC4,RANGRF,PSME2,PHTF2,RAD1,RBMX,SERPINB6,WAC,RALGPS1,KLK1

,UBXN1,ERLEC1,HSPB1,HHLA3,EMC4,RANGRF,PSME2,PHTF2,RAD1,RBMX,SERPINB6,WAC,RALGPS1,KLK1

ARD1,TPM4,MRPS18C,TCP1,NFATC4,MFSD11,PDCD6,MFF,FAM32A,COX20,COBL,PEX2,ELOF1,C12ORF57

1,SLC37A4,AAMDC,SNAPC2,NQO2,AKT1S1,B3GNTL1,KAT8,CARM1,MDK,ZNF276,LRRC23,BSCL2,CCNG2,

ON,TET2,GRK6,HMGA2,CPSF3L,OBSL1,PTPN13,UBR4,YLPM1,PHF21A,CHCHD1,SLC39A11,FAM3A,MUC1,

ON,TET2,GRK6,HMGA2,CPSF3L,OBSL1,PTPN13,UBR4,YLPM1,PHF21A,CHCHD1,SLC39A11,FAM3A,MUC1,

A2,RPS16,CPSF3L,OBSL1,TTC21B,YLPM1,PHF21A,SLC39A11,MUC1,METTL17,EPB41L5,ATG12,ZBTB44,PL

1L5,ATG12,PPRC1,ZBTB44,NIP7,TTC9C,PLCB2,DUSP22,DBNDD2,NISCH,TMED4,GGPS1,RTEL1,RABL5,TRN

1L5,ATG12,PPRC1,ZBTB44,NIP7,TTC9C,PLCB2,DUSP22,DBNDD2,NISCH,TMED4,GGPS1,RTEL1,RABL5,TRN

HMGA1,PSME2,PABPC1,RRNAD1,NAT9,BCAM,BLVRB,PPP3CB,RABEP2,TADA3,RALGPS1,ZNF687,PDCD5,

8,AP3B2,RPS9,SLC35G1,RCE1,RUVBL2,EBPL,TESC,IGSF8,RBPMS,SNRPN,WDR62,IL1RAP,POLD4,PLEKHA5,

8,AP3B2,RPS9,SLC35G1,RCE1,RUVBL2,EBPL,TESC,IGSF8,RBPMS,SNRPN,WDR62,IL1RAP,POLD4,PLEKHA5,

D2,PFKFB2,TRPV3,EIF2B3,ZXDC,SMIM4,KPNA5,TSEN34,KCNC4,TJP1,ANAPC13,SLC39A6,DOK4,SCN11A,N

D2,PFKFB2,TRPV3,EIF2B3,ZXDC,SMIM4,KPNA5,TSEN34,KCNC4,TJP1,ANAPC13,SLC39A6,DOK4,SCN11A,N

4I1,DUSP22,THEM4,DBNDD2,CENPK,RTEL1,COASY,SRSF1,AUP1,SAP30BP,FOXRED1,TRNT1,KXD1,ITGB1B

4I1,DUSP22,THEM4,DBNDD2,CENPK,RTEL1,COASY,SRSF1,AUP1,SAP30BP,FOXRED1,TRNT1,KXD1,ITGB1B

,CBFA2T2,UPP1,NFAM1,RBM3,ZNF197,COPZ1,HSP90AB1,SYNGR2,RAB2A,ELP6,TPM2,SIAH1,ASPH,PNPC

DK11A,INO80E,MAPKAPK5,HSP90AB1,SYNGR2,TMEM134,RAB2A,GPBP1,GPALPP1,TRMT112,CHPF,ZSW

DK11A,INO80E,MAPKAPK5,HSP90AB1,SYNGR2,TMEM134,RAB2A,GPBP1,GPALPP1,TRMT112,CHPF,ZSW

COX4I1,DUSP22,SLC25A10,TMX2,SPOCK2,NISCH,RPS6,CD99L2,GGPS1,AUP1,TRNT1,PARD6G,SH3YL1,MI

PEH,HYI,CAPN10,PODXL,ATPAF2,C17ORF62,PRMT1,CASC4,ZFAND2B,CYB561,STAU1,DUT,SYNGR2,CEP7

PEH,HYI,CAPN10,PODXL,ATPAF2,C17ORF62,PRMT1,CASC4,ZFAND2B,CYB561,STAU1,DUT,SYNGR2,CEP7

D2,NISCH,TMED4,GGPS1,RTEL1,RABL5,TRNT1,PARD6G,SH3YL1,CLASRP,C1ORF198,UGGT2,MICAL1,HS2S

D2,NISCH,TMED4,GGPS1,RTEL1,RABL5,TRNT1,PARD6G,SH3YL1,CLASRP,C1ORF198,UGGT2,MICAL1,HS2S

7,ASF1B,CCDC25,EEF1B2,TNFRSF14,CRIP1,PSMD4,RPL28,LIG1,PSMA7,CDC14B,TMEM126B,TMPO,C17O

T1,KXD1,SH3YL1,IP6K2,CLASRP,YIF1A,PRMT2,EIF3M,MAGT1,CELF1,USE1,CCDC25,EEF1B2,PPP1CC,RFX3,

T1,KXD1,SH3YL1,IP6K2,CLASRP,YIF1A,PRMT2,EIF3M,MAGT1,CELF1,USE1,CCDC25,EEF1B2,PPP1CC,RFX3,

SRSF1,AUP1,SAP30BP,FOXRED1,TRNT1,KXD1,ITGB1BP1,MED15,SLC39A13,YIF1A,PRMT2,MAGT1,C14OR
SRSF1,AUP1,SAP30BP,FOXRED1,TRNT1,KXD1,ITGB1BP1,MED15,SLC39A13,YIF1A,PRMT2,MAGT1,C14OR
MOK,SNX5,ZNF81,RPAIN,COX20,TMEM14B,ELP4,ACOT2,NFYC,ZXDC,POLR1C,TNKS,NDUFS3,L3MBTL2,KP
MOK,SNX5,ZNF81,RPAIN,COX20,TMEM14B,ELP4,ACOT2,NFYC,ZXDC,POLR1C,TNKS,NDUFS3,L3MBTL2,KP

1,ZNF687,POT1,RAD23A,ILF3,PKP4,HSPH1,RPL18,CNST,PHF20L1,HNRNPA1,FKBP3,NDUFV2,B3GAT3,SPI
APK5,HSP90AB1,SYNGR2,ZNF524,ATP5J,GPBP1,SIAH1,ASPH,STRIP2,TWSG1,MAPK13,RPS15,HHLA3,LRRC
APK5,HSP90AB1,SYNGR2,ZNF524,ATP5J,GPBP1,SIAH1,ASPH,STRIP2,TWSG1,MAPK13,RPS15,HHLA3,LRRC
4,ATP5J,ELP6,GPBP1,SIAH1,ASPH,RPS15,DCTD,HSPB1,LRRC27,RANGRF,CHCHD7,HMGA1,PABPC1,DBNL,
,TESC,NUDT5,SNRPN,SPAG7,WDR62,IL1RAP,STYXL1,RFX5,RANBP1,UBE2L6,VRK3,LZTFL1,FIS1,SLC15A3,T
,TESC,NUDT5,SNRPN,SPAG7,WDR62,IL1RAP,STYXL1,RFX5,RANBP1,UBE2L6,VRK3,LZTFL1,FIS1,SLC15A3,T
AF11,DBI,ARPC2,PMFBP1,LIMK2,VDAC3,NOB1,WDR19,SIRT6,NUMA1,GPATCH2L,SUMF2,BUD31,SAE1,N
AF11,DBI,ARPC2,PMFBP1,LIMK2,VDAC3,NOB1,WDR19,SIRT6,NUMA1,GPATCH2L,SUMF2,BUD31,SAE1,N
H1,MTHFS,MPC2,TXNRD2,COPS4,DALRD3,ZFPL1,OARD1,RIC8A,SMARCC2,UBE2N,ZNHIT3,RPS11,PDCD6,
PO,NIT2,TWSG1,RPS15,UBXN1,ZNF814,GMPPB,VWA2,IL32,C12ORF76,RANGRF,RAD1,SLC33A1,ZNF706,G
PO,NIT2,TWSG1,RPS15,UBXN1,ZNF814,GMPPB,VWA2,IL32,C12ORF76,RANGRF,RAD1,SLC33A1,ZNF706,G
D1,INO80C,HMOX2,AZI2,TATDN1,LAMTOR1,APAF1,KAT8,NCBP2,LRRC59,SUMO1,DHDDS,NMRAL1,ZNF4
D1,INO80C,HMOX2,AZI2,TATDN1,LAMTOR1,APAF1,KAT8,NCBP2,LRRC59,SUMO1,DHDDS,NMRAL1,ZNF4
SPB1,MVD,ERCC5,NAP1L1,RANGRF,MEIS2,PSME2,RRNAD1,DBNL,BCAM,TOMM40,ZNF20,ATP6V1F,SLC3
2,KCTD7,B3GAT3,SPINT2,PTK7,BRF1,ZFPL1,FAM102A,MFSD10,PFKFB2,RIC8A,MFSD11,LFNG,XYLT2,PCID
8,GRK6,RPS16,OBSL1,PSMA1,CHCHD1,RGS3,FAM3A,PPRC1,RPS25,NLRP1,THEM4,RDH11,DBNDD2,RPS6,
8,GRK6,RPS16,OBSL1,PSMA1,CHCHD1,RGS3,FAM3A,PPRC1,RPS25,NLRP1,THEM4,RDH11,DBNDD2,RPS6,
3,HNRNPLL,HIGD1A,B3GAT3,COA3,TYMS,SLC25A23,MRPS18C,TCP1,TRPV3,PHC3,MROH1,NFATC4,RPS11
3,HNRNPLL,HIGD1A,B3GAT3,COA3,TYMS,SLC25A23,MRPS18C,TCP1,TRPV3,PHC3,MROH1,NFATC4,RPS11
11,DBNDD2,SDF4,RPS6,COASY,AUP1,SAP30BP,YIPF3,RABL5,TRNT1,KXD1,ITGB1BP1,SH3YL1,HS2ST1,UBE
11,DBNDD2,SDF4,RPS6,COASY,AUP1,SAP30BP,YIPF3,RABL5,TRNT1,KXD1,ITGB1BP1,SH3YL1,HS2ST1,UBE

AT1,AKT1S1,B3GNTL1,CARM1,MDK,IAH1,AIG1,CAMKK1,ZNF276,ANXA2,TCFL5,HDAC10,METTL23,PHPT1
44,PLCB2,DUSP22,DBNDD2,NISCH,TMED4,CD99L2,GGPS1,COASY,SRSF1,YIPF3,TRNT1,PARD6G,SH3YL1,IP
44,PLCB2,DUSP22,DBNDD2,NISCH,TMED4,CD99L2,GGPS1,COASY,SRSF1,YIPF3,TRNT1,PARD6G,SH3YL1,IP
OXRED1,TRNT1,KXD1,ITGB1BP1,MED15,SLC39A13,PRMT2,MAGT1,C14ORF2,GUK1,CENPN,CARD14,PCYT
OXRED1,TRNT1,KXD1,ITGB1BP1,MED15,SLC39A13,PRMT2,MAGT1,C14ORF2,GUK1,CENPN,CARD14,PCYT

RIP1,CAPN10,EPB41L4B,UCK2,TMPO,ATPAF2,PRMT1,WNK1,ARL6IP4,PIGL,C19ORF48,STRADA,AP1S3,SH
RIP1,CAPN10,EPB41L4B,UCK2,TMPO,ATPAF2,PRMT1,WNK1,ARL6IP4,PIGL,C19ORF48,STRADA,AP1S3,SH
,ITSN1,EIF5A,COMMD5,RUNX1,NUDT22,UFD1L,USP38,TAZ,THYN1,DBNDD1,SH2B1,RPL13A,ARHGEF39,
T1,TESC,IGSF8,RBPMS,NNMT,SPAG7,IL1RAP,RFX5,POLD4,PLEKHA5,RANBP1,VRK3,ZC3H14,FIS1,PPM1K,T
T1,TESC,IGSF8,RBPMS,NNMT,SPAG7,IL1RAP,RFX5,POLD4,PLEKHA5,RANBP1,VRK3,ZC3H14,FIS1,PPM1K,T
IRD2,TMEM39B,THADA,RPS11,SNX5,NUP133,TMEM147,ELP4,NFYC,ZXDC,L3MBTL2,EIF4A1,RC3H2,KLHL
MDH1,ZNF197,LRRFIP1,ZNF524,ATP5J,TPM2,GPALPP1,SIAH1,TPM3,ZNF625,GABPB1,VPS28,RPS15,ERLE
MDH1,ZNF197,LRRFIP1,ZNF524,ATP5J,TPM2,GPALPP1,SIAH1,TPM3,ZNF625,GABPB1,VPS28,RPS15,ERLE
AB1,RAB2A,ZNF524,ATP5J,ELP6,TRMT112,CHPF,UBXN1,SAT1,DDOST,LSR,MEIS2,PSME2,PABPC1,RBP1,T
AB1,RAB2A,ZNF524,ATP5J,ELP6,TRMT112,CHPF,UBXN1,SAT1,DDOST,LSR,MEIS2,PSME2,PABPC1,RBP1,T
RPC2,PMFBP1,LIMK2,NOB1,SIRT6,NUMA1,GPATCH2L,SUMF2,BUD31,SAE1,NPRL2,OSGEP,TBL2,SCARB1,C
RPC2,PMFBP1,LIMK2,NOB1,SIRT6,NUMA1,GPATCH2L,SUMF2,BUD31,SAE1,NPRL2,OSGEP,TBL2,SCARB1,C

USP4,L3MBTL2,OXLD1,RPLP0,OGFOD2,METTL3,ASH1L,SLMAP,ADD1,CHTOP,TSKU,TPT1,BAX,TSPAN31,S
F687,PDCD5,NXN,KLK10,HSD17B10,ABHD17A,GGT5,SLC16A1,ZCCHC4,STXBP2,C11ORF48,OAS3,MED6,S
F687,PDCD5,NXN,KLK10,HSD17B10,ABHD17A,GGT5,SLC16A1,ZCCHC4,STXBP2,C11ORF48,OAS3,MED6,S
PRMT2,TMEM205,PTK2,C14ORF2,CENPN,ATF7,ASF1B,DEF8,EEF1B2,PPP1CC,RFX3,ENDOV,CAMTA1,PPIL
B,MVD,EMC4,IL32,SYNGR1,RANGRF,H2AFV,SLC33A1,ZNF687,POT1,RAD23A,ILF3,PKP4,HSPH1,RPL18,C
B,MVD,EMC4,IL32,SYNGR1,RANGRF,H2AFV,SLC33A1,ZNF687,POT1,RAD23A,ILF3,PKP4,HSPH1,RPL18,C
,EMC4,IL32,SYNGR1,RANGRF,H2AFV,SLC33A1,ZNF687,POT1,RAD23A,ILF3,PKP4,HSPH1,RPL18,CNST,SH

CD5,NXN,EPS15L1,ABHD17A,CHCHD5,NAA10,C11ORF48,LLGL2,SUMO3,FKBP3,SIRT5,KCTD7,RFXANK,SP
NRPD1,INO80C,HMOX2,AZI2,TATDN1,LAMTOR1,KAT8,NCBP2,LRRC59,SUMO1,NMRAL1,ZNF483,SZT2,BS
NRPD1,INO80C,HMOX2,AZI2,TATDN1,LAMTOR1,KAT8,NCBP2,LRRC59,SUMO1,NMRAL1,ZNF483,SZT2,BS
1,SH3YL1,IP6K2,PRMT2,EIF3M,MAGT1,CCDC25,EEF1B2,PPP1CC,PDLIM5,RFX3,VAMP8,PSMA7,TIMM17B
1,SH3YL1,IP6K2,PRMT2,EIF3M,MAGT1,CCDC25,EEF1B2,PPP1CC,PDLIM5,RFX3,VAMP8,PSMA7,TIMM17B
P1,PTK2,ATF7,PCYT2,MAF,ASF1B,PPP1CC,EDF1,RFX3,PPA1,DGKE,TNFRSF14,CAMTA1,CAPG,CLDND1,LI
P1,PTK2,ATF7,PCYT2,MAF,ASF1B,PPP1CC,EDF1,RFX3,PPA1,DGKE,TNFRSF14,CAMTA1,CAPG,CLDND1,LI
,AUP1,FOXRED1,ITGB1BP1,SH3YL1,IP6K2,JPH2,HS2ST1,MAGT1,CDC42,TBX6,AUNIP,GUK1,CARD14,FBX
,AUP1,FOXRED1,ITGB1BP1,SH3YL1,IP6K2,JPH2,HS2ST1,MAGT1,CDC42,TBX6,AUNIP,GUK1,CARD14,FBX
,COBL,ERGIC1,MYL9,TSEN34,TJP1,NUP62,NADSYN1,ZNF511,D2HGDH,SNX29,AXIN1,AAMDC,SNAPC2,TK
1,PHF19,PCBP2,LPCAT3,TM9SF1,MAGED2,SON,TET2,UQCRQ,SNF8,CDC37,RPS16,CPSF3L,TSG101,TTC21
1,PHF19,PCBP2,LPCAT3,TM9SF1,MAGED2,SON,TET2,UQCRQ,SNF8,CDC37,RPS16,CPSF3L,TSG101,TTC21
4M1,ATP9B,ANKRD10,ACTG1,CERS2,BCAP29,CNTLN,MRPS7,SERHL2,OAZ1,BIN1,RNF4,GSTO2,LDLRAD3,
BE2L6,ZC3H14,FIS1,SLC15A3,PPM1K,IRF9,STIP1,LRRC1,PHF19,GRAMD1C,LPCAT3,CD44,SON,TET2,GRK6
BE2L6,ZC3H14,FIS1,SLC15A3,PPM1K,IRF9,STIP1,LRRC1,PHF19,GRAMD1C,LPCAT3,CD44,SON,TET2,GRK6

XN,KLK10,RAD23A,ABHD17A,STXBP2,EEF1D,NAA10,C11ORF48,MED6,LGMN,CD151,EGLN2,SIRT5,KCTD7
R4,PSMA1,YLPM1,SLC39A11,PSMA3,MUC1,METTL17,RPS25,ZBTB44,SPOCK2,SDF4,NISCH,CENPK,RPL29,
R4,PSMA1,YLPM1,SLC39A11,PSMA3,MUC1,METTL17,RPS25,ZBTB44,SPOCK2,SDF4,NISCH,CENPK,RPL29,
R2,STIP1,LRRC1,PHF19,VCAN,CD44,MAGED2,TET2,GRK6,HMGA2,CDC37,RPS16,CPSF3L,OBSL1,PTPN13,
R2,STIP1,LRRC1,PHF19,VCAN,CD44,MAGED2,TET2,GRK6,HMGA2,CDC37,RPS16,CPSF3L,OBSL1,PTPN13,
ORF198,MICAL1,UBE2I,SLC39A13,YIF1A,PRMT2,EIF3M,JUP,XBP1,C14ORF2,GUK1,FBXW9,IFI27L1,MAF,DE
FV2,B3GAT3,ZNF506,PAFAH1B2,GTF2IRD2,DAZAP2,ZFPL1,TPM4,THADA,MROH1,SNX5,NUP133,RPAIN,T
CSH,SSC5D,ZNF276,COMMD1,SZT2,PSMC5,AP2S1,LRP10,SLC31A1,ACTB,IQCK,SETD4,MED8,MRPL27,RA
,TUBA1C,IRF9,STIP1,LRRC1,PHF19,GRAMD1C,LPCAT3,TM9SF1,MAGED2,SON,TET2,GRK6,HMGA2,RPS16
,TUBA1C,IRF9,STIP1,LRRC1,PHF19,GRAMD1C,LPCAT3,TM9SF1,MAGED2,SON,TET2,GRK6,HMGA2,RPS16
3R4,SLC39A11,FUZ,RGS3,FAM3A,PSMA3,METTL17,PPRC1,SLC25A10,TMX2,SPOCK2,SDF4,NISCH,TMED4,
3R4,SLC39A11,FUZ,RGS3,FAM3A,PSMA3,METTL17,PPRC1,SLC25A10,TMX2,SPOCK2,SDF4,NISCH,TMED4,
A2,CDC37,OBSL1,PTPN13,YLPM1,PHF21A,CHCHD1,FAM3A,MUC1,ZBTB44,COX4I1,SLC25A10,DBNDD2,S
A2,CDC37,OBSL1,PTPN13,YLPM1,PHF21A,CHCHD1,FAM3A,MUC1,ZBTB44,COX4I1,SLC25A10,DBNDD2,S
AB2A,ZNF524,ATP5J,ELP6,TRMT112,CHPF,UBXN1,SAT1,EMC4,LMBR1,DDOST,LSR,MEIS2,PSME2,RBP1,T
AB2A,ZNF524,ATP5J,ELP6,TRMT112,CHPF,UBXN1,SAT1,EMC4,LMBR1,DDOST,LSR,MEIS2,PSME2,RBP1,T
L6IP4,C19ORF48,STRADA,HNRNPC,SHARPIN,SLC9B2,CARS,ZNF250,RBM3,MDH1,INO80E,MAPKAPK5,SY
L6IP4,C19ORF48,STRADA,HNRNPC,SHARPIN,SLC9B2,CARS,ZNF250,RBM3,MDH1,INO80E,MAPKAPK5,SY
PAT4,IFI27L1,MAF,PPA1,ENDOV,PPIL2,TXNL4A,PRMT1,EXOSC9,ARL6IP4,ZFAND2B,HYPK,ZNF250,NFAM
PAT4,IFI27L1,MAF,PPA1,ENDOV,PPIL2,TXNL4A,PRMT1,EXOSC9,ARL6IP4,ZFAND2B,HYPK,ZNF250,NFAM
PSMD10,UCHL5,MYL12A,TATDN1,ZBTB8OS,SLK,CARM1,POLR2E,EMC3,PIWIL4,THOP1,TAF9,DTD2,PSMD
7,TTC9C,SLC25A10,NISCH,TMED4,CENPK,RPS6,RPL29,CD99L2,GGPS1,COASY,SRSF1,AUP1,FOXRED1,RAB
7,NFYC,ZXDC,L3MBTL2,EIF4A1,RC3H2,PICK1,ECE1,OCIAD1,RBBP6,NREP,CHTOP,TSPAN31,TMEM9,ADRA

17,NFYC,ZXDC,L3MBTL2,EIF4A1,RC3H2,PICK1,ECE1,OCIAD1,RBBP6,NREP,CHTOP,TSPAN31,TMEM9,ADRA
NF814,ERLEC1,GMPPB,MVD,IL32,SYNGR1,RANGRF,H2AFV,SLC33A1,ZNF687,POT1,POLR3H,RAD23A,ILF3
NF814,ERLEC1,GMPPB,MVD,IL32,SYNGR1,RANGRF,H2AFV,SLC33A1,ZNF687,POT1,POLR3H,RAD23A,ILF3
MX2,THEM4,DBNDD2,NISCH,RPL29,GGPS1,RTEL1,COASY,SRSF1,FOXRED1,TRNT1,MED15,SH3YL1,IP6K2,C
GPS1,ITGB1BP1,SH3YL1,C1ORF198,UBE2I,PRMT2,XBP1,PTK2,CENPN,PCYT2,USE1,HSBP1L1,AGPAT4,IFI27
GPS1,ITGB1BP1,SH3YL1,C1ORF198,UBE2I,PRMT2,XBP1,PTK2,CENPN,PCYT2,USE1,HSBP1L1,AGPAT4,IFI27
C39A11,FUZ,RGS3,FAM3A,PSMA3,METTL17,PPRC1,SLC25A10,TMX2,SPOCK2,SDF4,TMED4,RPS6,RPL29,R
,TMX2,SDF4,TMED4,RPS6,RPL29,RTEL1,TRNT1,ITGB1BP1,SH3YL1,C1ORF198,KCNMA1,XBP1,CELF1,C14O
,TMX2,SDF4,TMED4,RPS6,RPL29,RTEL1,TRNT1,ITGB1BP1,SH3YL1,C1ORF198,KCNMA1,XBP1,CELF1,C14O
,TMX2,SDF4,TMED4,RPS6,RPL29,RTEL1,TRNT1,ITGB1BP1,SH3YL1,C1ORF198,KCNMA1,XBP1,CELF1,C14O
5,RPL13A,GSTK1,TTLL5,ACOT8,AP3B2,RPS9,RCE1,MRPL33,ZNF410,RUVBL2,PCMT1,TESC,IGSF8,NUDT5,RI
5,RPL13A,GSTK1,TTLL5,ACOT8,AP3B2,RPS9,RCE1,MRPL33,ZNF410,RUVBL2,PCMT1,TESC,IGSF8,NUDT5,RI
MD4,SNX5,SMIM4,NMNAT3,BTN3A2,C6ORF132,TMEM108,ZMYM6,SSR3,MPV17,MRPL55,ARIH2,CREB3L
MD4,SNX5,SMIM4,NMNAT3,BTN3A2,C6ORF132,TMEM108,ZMYM6,SSR3,MPV17,MRPL55,ARIH2,CREB3L

BP1,MED15,CLASRP,C1ORF198,UGGT2,HS2ST1,UBE2I,SLC39A13,YIF1A,CDC42,TBX6,TMEM205,XBP1,PT
BP1,MED15,CLASRP,C1ORF198,UGGT2,HS2ST1,UBE2I,SLC39A13,YIF1A,CDC42,TBX6,TMEM205,XBP1,PT
NIT2,MPDU1,ZNF814,VWA2,IL32,LMBR1,SERPINB1,DDOST,C12ORF76,RANGRF,PABPC1,RAD1,BLVRB,ZNF
TSN1,EIF5A,COMMD5,RUNX1,CHID1,AMZ2,UFD1L,USP38,TAZ,THYN1,DBNDD1,SH2B1,NOP16,RPL13A,A
TSN1,EIF5A,COMMD5,RUNX1,CHID1,AMZ2,UFD1L,USP38,TAZ,THYN1,DBNDD1,SH2B1,NOP16,RPL13A,A
I1,CHCHD1,FUZ,RGS3,ECSIT,EPB41L5,ATG12,PPRC1,NIP7,COX4I1,TMX2,RDH11,DBNDD2,NISCH,CENPK,RI
I1,CHCHD1,FUZ,RGS3,ECSIT,EPB41L5,ATG12,PPRC1,NIP7,COX4I1,TMX2,RDH11,DBNDD2,NISCH,CENPK,RI
IRF9,LRRC1,PHF19,GRAMD1C,VCAN,PCBP2,LPCAT3,CD44,TM9SF1,SON,TET2,HMGA2,RPS16,TTC21B,PTF
IRF9,LRRC1,PHF19,GRAMD1C,VCAN,PCBP2,LPCAT3,CD44,TM9SF1,SON,TET2,HMGA2,RPS16,TTC21B,PTF
M39B,PPIH,THADA,RPS11,SNX5,NUP133,TMEM147,ELP4,NFYC,ZXDC,L3MBTL2,EIF4A1,RC3H2,KLHL7,PICK
H2,UGGT2,HS2ST1,UBE2I,SLC39A13,YIF1A,CDC42,XBP1,PTK2,AUNIP,PCYT2,HSBP1L1,IFI27L1,EDF1,RFX3,I
L,APEH,TXNL4A,RBM33,EXOSC9,WNK1,C19ORF54,ZFAND2B,HNRNPC,SLC9B2,SUGT1,HYPK,ZNF250,CDK1
L,APEH,TXNL4A,RBM33,EXOSC9,WNK1,C19ORF54,ZFAND2B,HNRNPC,SLC9B2,SUGT1,HYPK,ZNF250,CDK1

A1,TNFRSF14,CRIP1,TIMM17B,XAF1,APEH,HYI,CAMLG,UCK2,PPOX,TMPO,ATPAF2,PRMT1,WNK1,C19ORF
L17,ECSIT,PPRC1,ZBTB44,SLC25A10,TMX2,SDF4,NISCH,TMED4,RPS6,RPL29,RTEL1,FOXRED1,TRNT1,ITGB
RSF1,AUP1,SAP30BP,YIPF3,FOXRED1,TRNT1,SH3YL1,IP6K2,PRMT2,MAGT1,TMEM205,CCDC25,PPP1CC,PI
RSF1,AUP1,SAP30BP,YIPF3,FOXRED1,TRNT1,SH3YL1,IP6K2,PRMT2,MAGT1,TMEM205,CCDC25,PPP1CC,PI
L6,RPS3A,MAGOHB,IDH3G,CREB3L2,AKT1S1,SPHK2,HMGN1,NDUFB2,CDK2AP2,NDUFA10,AZI2,U2AF1L4,

C,ZXDC,L3MBTL2,EIF4A1,RC3H2,PICK1,ECE1,OCIAD1,RBBP6,NREP,CHTOP,TSPAN31,TMEM9,ADRA1A,ZNF
NMT,SPAG7,IL1RAP,RFX5,POLD4,PLEKHA5,RANBP1,ZC3H14,FIS1,PPM1K,TUBA1C,IRF9,DST,ANTXR2,STIP1
MAGOHB,IDH3G,CREB3L2,AKT1S1,SPHK2,HMGN1,NDUFB2,CDK2AP2,NDUFA10,AZI2,U2AF1L4,ATL2,DDX
GPS2,SUMF1,CCNG2,RABEPK,GRHPR,QTRT1,RUVBL1,ATP6V0B,METTL23,SETD4,PARP6,KIAA0513,SHMT1
BA1C,STIP1,LRRC1,VCAN,CD44,UQCRQ,SNF8,HMGA2,CDC37,RPS16,CPSF3L,TSG101,UBR4,PSMA1,PHF21
BA1C,STIP1,LRRC1,VCAN,CD44,UQCRQ,SNF8,HMGA2,CDC37,RPS16,CPSF3L,TSG101,UBR4,PSMA1,PHF21
2,TIMM17B,TXNL4A,TMPO,PRMT1,EXOSC9,ARL6IP4,ZFAND2B,HYPK,ZNF250,NFAM1,RBM3,MAPKAPK5,
2,TIMM17B,TXNL4A,TMPO,PRMT1,EXOSC9,ARL6IP4,ZFAND2B,HYPK,ZNF250,NFAM1,RBM3,MAPKAPK5,
HNRNPA1,EGLN2,SIRT5,RFXANK,TMEM101,S100A4,TXNRD2,FKBP5,TRPM4,MFSD10,MRPS18C,SYTL1,SLC
HNRNPA1,EGLN2,SIRT5,RFXANK,TMEM101,S100A4,TXNRD2,FKBP5,TRPM4,MFSD10,MRPS18C,SYTL1,SLC
HNRNPA1,EGLN2,SIRT5,RFXANK,TMEM101,S100A4,TXNRD2,FKBP5,TRPM4,MFSD10,MRPS18C,SYTL1,SLC

Z,FAM3A,MUC1,ATG12,RPS25,ZBTB44,NIP7,TTC9C,SLC25A10,TMX2,DBNDD2,SDF4,TMED4,CENPK,RPS6

2,B3GAT3,GTF2IRD2,TMEM39B,BRF1,THADA,RPS11,SNX5,NUP133,TMEM147,ELP4,NFYC,ZXDC,L3MBTL

2,B3GAT3,GTF2IRD2,TMEM39B,BRF1,THADA,RPS11,SNX5,NUP133,TMEM147,ELP4,NFYC,ZXDC,L3MBTL

3H6,BTF3,PML,IFITM2,SLC25A24,ZCCHC9,VPS25,SNRPD1,CD37,AZI2,ZBTB8OS,LAMTOR1,NCBP2,PLD1,BC

IYM6,OGFOD2,EIF3G,ARIH2,RNF181,CHTOP,PPP2R5C,ZNF593,GLYCTK,FAM131A,TCEAL3,C11ORF57,CNT

IYM6,OGFOD2,EIF3G,ARIH2,RNF181,CHTOP,PPP2R5C,ZNF593,GLYCTK,FAM131A,TCEAL3,C11ORF57,CNT

AZ,THYN1,SH2B1,RPL13A,ARHGEF39,GSTK1,TTLL5,LDHA,PSMA6,SLC35G1,RCE1,MRPL33,ZNF410,PCMT1

AZ,THYN1,SH2B1,RPL13A,ARHGEF39,GSTK1,TTLL5,LDHA,PSMA6,SLC35G1,RCE1,MRPL33,ZNF410,PCMT1

CMT1,EBPL,IGSF8,RBPMS,NNMT,SPAG7,WDR62,POLD4,RANBP1,VRK3,ZC3H14,PPM1K,TUBA1C,ANTXR2,

CMT1,EBPL,IGSF8,RBPMS,NNMT,SPAG7,WDR62,POLD4,RANBP1,VRK3,ZC3H14,PPM1K,TUBA1C,ANTXR2,

F,EEF1B2,PPP1CC,RFX3,DGKE,PSMD4,RPL28,VIM,TMEM126B,CAPN10,EPB41L4B,C17ORF62,PRMT1,STO

PS6,RPL29,CD99L2,GGPS1,SRSF1,AUP1,SAP30BP,YIPF3,TRNT1,SH3YL1,IP6K2,PRMT2,EIF3M,MAGT1,TM

PS6,RPL29,CD99L2,GGPS1,SRSF1,AUP1,SAP30BP,YIPF3,TRNT1,SH3YL1,IP6K2,PRMT2,EIF3M,MAGT1,TM

F524,SIAH1,ZEB2,STRIP2,LYNX1,LMBR1,NAP1L1,SERPINB1,RANGRF,CHCHD7,PSME2,PABPC1,NAT9,BCA

RSF1,AUP1,SAP30BP,YIPF3,FOXRED1,TRNT1,SH3YL1,IP6K2,YIF1A,PRMT2,EIF3M,MAGT1,CCDC25,EEF1B2,

RSF1,AUP1,SAP30BP,YIPF3,FOXRED1,TRNT1,SH3YL1,IP6K2,YIF1A,PRMT2,EIF3M,MAGT1,CCDC25,EEF1B2,

,THADA,MFF,SNX5,NUP133,MAP3K8,MRPS17,TMEM147,TSC22D3,ELOF1,NDUFS3,RC3H2,MRPL55,ECE1

,THADA,MFF,SNX5,NUP133,MAP3K8,MRPS17,TMEM147,TSC22D3,ELOF1,NDUFS3,RC3H2,MRPL55,ECE1

RHGAP6,SCARB1,TSPAN11,ITSN1,EIF5A,TNFSF10,COMMD5,CHID1,NUDT22,AMZ2,USP38,TAZ,DBNDD1,S

MED4,RPS6,FOXRED1,RABL5,MED15,C1ORF198,UGGT2,YIF1A,TMEM205,XBP1,PTK2,CELF1,AUNIP,C14O

MED4,RPS6,FOXRED1,RABL5,MED15,C1ORF198,UGGT2,YIF1A,TMEM205,XBP1,PTK2,CELF1,AUNIP,C14O

511,BFAR,SIN3A,RBM4,THOC3,TP53,TMEM199,OXR1,EPN1,CREB3L2,ZCCHC9,PCTP,AKT1S1,MKRN1,SM

10,RUVBL2,EBPL,TESC,IGSF8,RBPMS,WDR62,IL1RAP,STYXL1,POLD4,RANBP1,LZTFL1,FIS1,SLC15A3,PPM1

10,RUVBL2,EBPL,TESC,IGSF8,RBPMS,WDR62,IL1RAP,STYXL1,POLD4,RANBP1,LZTFL1,FIS1,SLC15A3,PPM1

OT2,TGOLN2,PEX2,DCTN5,L3MBTL2,OXLD1,EIF4A1,ARIH2,RPL8,ECE1,ZNF512,ARL6IP5,ZNF248,UBA1,N

ADA3,ZNF706,POLR3H,RAD23A,MDH2,HSPH1,CHCHD5,RPL18,CAMK2D,HINT1,SIRT5,SPINT2,ASH2L,TCT

ADA3,ZNF706,POLR3H,RAD23A,MDH2,HSPH1,CHCHD5,RPL18,CAMK2D,HINT1,SIRT5,SPINT2,ASH2L,TCT

2,SCN11A,PSMD7,ZNF511,IRF3,SLC35A2,NDUFA11,TUBB3,SLC25A24,COX19,SNAPC2,NQO2,ZMAT1,B3G

C,ZXDC,L3MBTL2,EIF4A1,RC3H2,PICK1,ECE1,OCIAD1,RBBP6,NREP,CHTOP,TSPAN31,TMEM9,ADRA1A,Z

C,ZXDC,L3MBTL2,EIF4A1,RC3H2,PICK1,ECE1,OCIAD1,RBBP6,NREP,CHTOP,TSPAN31,TMEM9,ADRA1A,Z

A,MAGOHB,IDH3G,CREB3L2,AKT1S1,SPHK2,HMGN1,NDUFB2,NDUFA10,AZI2,U2AF1L4,DDX10,EMC3,RN

A1,FKBP3,B3GAT3,GTF2IRD2,COPS4,TMEM39B,ZFPL1,TPM4,SLC30A5,THADA,MROH1,SNX5,NUP133,T

MX2,SDF4,TMED4,RPS6,RPL29,RTEL1,FOXRED1,TRNT1,ITGB1BP1,SH3YL1,C1ORF198,KCNMA1,XBP1,CEL

MX2,SDF4,TMED4,RPS6,RPL29,RTEL1,FOXRED1,TRNT1,ITGB1BP1,SH3YL1,C1ORF198,KCNMA1,XBP1,CEL

MX2,SDF4,TMED4,RPS6,RPL29,RTEL1,FOXRED1,TRNT1,ITGB1BP1,SH3YL1,C1ORF198,KCNMA1,XBP1,CEL

FOD1,TCF25,CHTOP,ANAPC13,SLC39A6,CNTRL,NUP62,SCN11A,RECQL5,IRF3,NDUFA11,TUBB3,MOSPD3,

FOD1,TCF25,CHTOP,ANAPC13,SLC39A6,CNTRL,NUP62,SCN11A,RECQL5,IRF3,NDUFA11,TUBB3,MOSPD3,

RL16,SLC35E1,MRPL55,ARIH2,ECE1,OCIAD1,TCF25,RBBP6,ZNF248,TPT1,SMAGP,TSPAN31,TMEM9,SLC3

50,NFAM1,MAPKAPK5,DGUOK,HSP90AB1,CEP70,RAB2A,RMDN1,GPBP1,GPALPP1,CHPF,MAPK13,ZSWI

50,NFAM1,MAPKAPK5,DGUOK,HSP90AB1,CEP70,RAB2A,RMDN1,GPBP1,GPALPP1,CHPF,MAPK13,ZSWI

AD1,TCF25,RBBP6,ZNF248,TPT1,TMEM9,ZNF593,GLYCTK,RBM42,CLASP2,CREBZF,ZNF511,GTF2H1,MYL

S16,CPSF3L,OBSL1,TTC21B,PTPN13,PSMA1,YLPM1,PHF21A,SLC39A11,PPRC1,RPS25,COX4I1,SPOCK2,THE
S16,CPSF3L,OBSL1,TTC21B,PTPN13,PSMA1,YLPM1,PHF21A,SLC39A11,PPRC1,RPS25,COX4I1,SPOCK2,THE

4,C19ORF48,CYB561,CBFA2T2,UPP1,NFAM1,RBM3,COPZ1,HSP90AB1,SYNGR2,RAB2A,ATP5J,ELP6,TPM2,
4,C19ORF48,CYB561,CBFA2T2,UPP1,NFAM1,RBM3,COPZ1,HSP90AB1,SYNGR2,RAB2A,ATP5J,ELP6,TPM2,
C,STIP1,LRRC1,PHF19,GRAMD1C,VCAN,LPCAT3,CD44,MAGED2,SNF8,HMGA2,CPSF3L,OBSL1,PSMA1,YLPM
I205,XBP1,C14ORF2,FBXW9,IFI27L1,ASF1B,EDF1,RFX3,TNFRSF14,ENDOV,CAPG,CRIP1,LIG1,PSMA7,CDC1
RPM4,GMPR2,TCP1,ADAM19,CBR1,EIF2B3,MFSD11,SNX5,NOP2,RAPGEF3,TMEM147,TMEM14B,ZXDC,C
RPM4,GMPR2,TCP1,ADAM19,CBR1,EIF2B3,MFSD11,SNX5,NOP2,RAPGEF3,TMEM147,TMEM14B,ZXDC,C
MT2,MAGT1,CDC42,TBX6,TMEM205,PTK2,AUNIP,CENPN,PCYT2,IFI27L1,MAF,ASF1B,CCDC25,DEF8,PPP1
MT2,MAGT1,CDC42,TBX6,TMEM205,PTK2,AUNIP,CENPN,PCYT2,IFI27L1,MAF,ASF1B,CCDC25,DEF8,PPP1

32,NAP1L1,SYNGR1,MTA3,H2AFV,BLVRB,ZNF706,ZNF687,POT1,RAD23A,PKP4,AGAP3,HSPH1,NAA10,CA
32,NAP1L1,SYNGR1,MTA3,H2AFV,BLVRB,ZNF706,ZNF687,POT1,RAD23A,PKP4,AGAP3,HSPH1,NAA10,CA
5G1,RCE1,ZNF410,PCMT1,TESC,IGSF8,NUDT5,RBPMS,WDR62,IL1RAP,STYXL1,RFX5,POLD4,PLEKHA5,RAN
5G1,RCE1,ZNF410,PCMT1,TESC,IGSF8,NUDT5,RBPMS,WDR62,IL1RAP,STYXL1,RFX5,POLD4,PLEKHA5,RAN
X1,NUDT22,UFD1L,USP38,TAZ,THYN1,DBNDD1,ARHGEF39,TTLL5,PSMA6,ACOT8,AP3B2,RPS9,SLC35G1,R
X1,NUDT22,UFD1L,USP38,TAZ,THYN1,DBNDD1,ARHGEF39,TTLL5,PSMA6,ACOT8,AP3B2,RPS9,SLC35G1,R
NIP7,COX4I1,DUSP22,THEM4,DBNDD2,TMED4,CENPK,RTEL1,SRSF1,AUP1,SAP30BP,FOXRED1,TRNT1,KXD
NIP7,COX4I1,DUSP22,THEM4,DBNDD2,TMED4,CENPK,RTEL1,SRSF1,AUP1,SAP30BP,FOXRED1,TRNT1,KXD
,CD99L2,SRSF1,AUP1,RABL5,TRNT1,ITGB1BP1,PARD6G,SH3YL1,CLASRP,JPH2,UGGT2,TBX6,TMEM205,PT
,CD99L2,SRSF1,AUP1,RABL5,TRNT1,ITGB1BP1,PARD6G,SH3YL1,CLASRP,JPH2,UGGT2,TBX6,TMEM205,PT
NST,SSH2,COX17,HINT1,POLR2C,SIRT5,RFXANK,SPINT2,ASH2L,PAFAH1B2,S100A4,COPS4,DALRD3,GMPR
NST,SSH2,COX17,HINT1,POLR2C,SIRT5,RFXANK,SPINT2,ASH2L,PAFAH1B2,S100A4,COPS4,DALRD3,GMPR
EIF5A,COMMD5,NUDT22,AMZ2,USP38,TAZ,THYN1,DBNDD1,RPL13A,GSTK1,TTLL5,ACOT8,AP3B2,RPS9,S
EIF5A,COMMD5,NUDT22,AMZ2,USP38,TAZ,THYN1,DBNDD1,RPL13A,GSTK1,TTLL5,ACOT8,AP3B2,RPS9,S
SRP,YIF1A,EIF3M,JUP,CDC42,AUNIP,CENPN,ATF7,USE1,MAF,CCDC25,EEF1B2,ENDOV,PSMD4,RPL28,CLD
AP30BP,YIPF3,FOXRED1,TRNT1,SH3YL1,IP6K2,PRMT2,EIF3M,MAGT1,TMEM205,CCDC25,EEF1B2,PPP1CC
AP30BP,YIPF3,FOXRED1,TRNT1,SH3YL1,IP6K2,PRMT2,EIF3M,MAGT1,TMEM205,CCDC25,EEF1B2,PPP1CC
METTL17,PPRC1,RPS25,NLRP1,NIP7,TTC9C,COX4I1,DUSP22,THEM4,DBNDD2,TMED4,CENPK,RTEL1,COAS
METTL17,PPRC1,RPS25,NLRP1,NIP7,TTC9C,COX4I1,DUSP22,THEM4,DBNDD2,TMED4,CENPK,RTEL1,COAS
N1,EIF5A,COMMD5,CHID1,NUDT22,AMZ2,UFD1L,TAZ,THYN1,SH2B1,NOP16,RPL13A,ARHGEF39,TTLL5,L
N1,EIF5A,COMMD5,CHID1,NUDT22,AMZ2,UFD1L,TAZ,THYN1,SH2B1,NOP16,RPL13A,ARHGEF39,TTLL5,L
7,RPS16,CPSF3L,UBR4,YLPM1,FUZ,METTL17,ECSIT,ATG12,PPRC1,RPS25,ZBTB44,NIP7,PLCB2,TMX2,RDH1
7,RPS16,CPSF3L,UBR4,YLPM1,FUZ,METTL17,ECSIT,ATG12,PPRC1,RPS25,ZBTB44,NIP7,PLCB2,TMX2,RDH1
HF19,MAGED2,UQCRQ,HMGA2,CDC37,CPSF3L,OBSL1,TTC21B,UBR4,PSMA1,PHF21A,PSMA3,MUC1,ECSI
HF19,MAGED2,UQCRQ,HMGA2,CDC37,CPSF3L,OBSL1,TTC21B,UBR4,PSMA1,PHF21A,PSMA3,MUC1,ECSI
P1CC,DGKE,ENDOV,CRIP1,LIG1,PSMA7,APEH,RPL21,TXNL4A,CAPN10,LYPLA2,PODXL,EPB41L4B,UCK2,TM
P1CC,DGKE,ENDOV,CRIP1,LIG1,PSMA7,APEH,RPL21,TXNL4A,CAPN10,LYPLA2,PODXL,EPB41L4B,UCK2,TM
COX4I1,DUSP22,SLC25A10,DBNDD2,NISCH,CD99L2,GGPS1,SRSF1,AUP1,TRNT1,MED15,C1ORF198,MICA
C,COX4I1,DUSP22,THEM4,DBNDD2,TMED4,CENPK,COASY,SRSF1,AUP1,SAP30BP,FOXRED1,TRNT1,KXD1,
C,COX4I1,DUSP22,THEM4,DBNDD2,TMED4,CENPK,COASY,SRSF1,AUP1,SAP30BP,FOXRED1,TRNT1,KXD1,
F1B2,PPA1,TNFRSF14,TIMM17B,TMEM126B,TXNL4A,CAPN10,EPB41L4B,ATPAF2,PRMT1,C19ORF54,ARL
F1B2,PPA1,TNFRSF14,TIMM17B,TMEM126B,TXNL4A,CAPN10,EPB41L4B,ATPAF2,PRMT1,C19ORF54,ARL
F1,C14ORF2,ATF7,PCYT2,USE1,AGPAT4,CCDC25,EEF1B2,PPA1,TNFRSF14,VAMP8,TIMM17B,TMEM126B,

F1,C14ORF2,ATF7,PCYT2,USE1,AGPAT4,CCDC25,EEF1B2,PPA1,TNFRSF14,VAMP8,TIMM17B,TMEM126B,

ED4,CENPK,RPS6,RPL29,CD99L2,GGPS1,RTEL1,SRSF1,AUP1,SAP30BP,FOXRED1,TRNT1,KXD1,IP6K2,CLASF

ED4,CENPK,RPS6,RPL29,CD99L2,GGPS1,RTEL1,SRSF1,AUP1,SAP30BP,FOXRED1,TRNT1,KXD1,IP6K2,CLASF

,SLC30A5,THADA,MROH1,SNX5,NUP133,TMEM147,ELP4,ZXDC,COBL,ELOF1,ARL16,SLC35E1,TSEN34,MR

,SUMO3,PHF20L1,HNRNPA1,EGLN2,RFXANK,TMEM101,S100A4,TXNRD2,NSUN5,FKBP5,TRPM4,SYTL1,SL

,SUMO3,PHF20L1,HNRNPA1,EGLN2,RFXANK,TMEM101,S100A4,TXNRD2,NSUN5,FKBP5,TRPM4,SYTL1,SL

,SUMO3,PHF20L1,HNRNPA1,EGLN2,RFXANK,TMEM101,S100A4,TXNRD2,NSUN5,FKBP5,TRPM4,SYTL1,SL

RHGAP6,SCARB1,CTTN,TSPAN11,ITSN1,EIF5A,TNFSF10,COMMD5,NUDT22,UFD1L,USP38,DBNDD1,SH2B

RHGAP6,SCARB1,CTTN,TSPAN11,ITSN1,EIF5A,TNFSF10,COMMD5,NUDT22,UFD1L,USP38,DBNDD1,SH2B

4,IFI27L1,MAF,PPA1,ENDOV,PPIL2,TXNL4A,PRMT1,EXOSC9,ARL6IP4,ZFAND2B,HYPK,ZNF250,NFAM1,MA

4,IFI27L1,MAF,PPA1,ENDOV,PPIL2,TXNL4A,PRMT1,EXOSC9,ARL6IP4,ZFAND2B,HYPK,ZNF250,NFAM1,MA

GS3,ECSIT,ATG12,PPRC1,NIP7,TTC9C,SLC25A10,SDF4,NISCH,TMED4,CENPK,RPL29,CD99L2,GGPS1,COASY

A10,SPOCK2,NISCH,CD99L2,GGPS1,RABL5,PARD6G,SH3YL1,CLASRP,C1ORF198,MICAL1,HS2ST1,UBE2I,JU

A10,SPOCK2,NISCH,CD99L2,GGPS1,RABL5,PARD6G,SH3YL1,CLASRP,C1ORF198,MICAL1,HS2ST1,UBE2I,JU

PH1,NAA10,CARHSP1,SUMO3,EGLN2,SIRT5,RFXANK,TMEM101,PPIH,FKBP5,MFSD10,PHC3,EIF2B3,XYLT2,

PH1,NAA10,CARHSP1,SUMO3,EGLN2,SIRT5,RFXANK,TMEM101,PPIH,FKBP5,MFSD10,PHC3,EIF2B3,XYLT2,

ESC,NNMT,SNRPN,SPAG7,STYXL1,PLEKHA5,UBE2L6,VRK3,LZTFL1,IRF9,DST,ANTXR2,LRRC1,GRAMD1C,VC

ESC,NNMT,SNRPN,SPAG7,STYXL1,PLEKHA5,UBE2L6,VRK3,LZTFL1,IRF9,DST,ANTXR2,LRRC1,GRAMD1C,VC

L55,ARIH2,ECE1,OCIAD1,TCF25,RBBP6,ZNF248,TPT1,TSPAN31,TMEM9,SLC39A6,ZNF593,GLYCTK,RBM42

MBR1,MCTS1,NAP1L1,DDOST,RANGRF,HMGA1,PABPC1,DBNL,SERPINB6,RABEP2,TADA3,ZNF20,ATP6V1F,

MBR1,MCTS1,NAP1L1,DDOST,RANGRF,HMGA1,PABPC1,DBNL,SERPINB6,RABEP2,TADA3,ZNF20,ATP6V1F,

K4,ANAPC13,FAM192A,TCEAL3,GALT,VDAC1,GTF2H1,TAF1D,D2HGDH,CC2D1B,ARPC5,SH3GL1,TKT,RHE

K4,ANAPC13,FAM192A,TCEAL3,GALT,VDAC1,GTF2H1,TAF1D,D2HGDH,CC2D1B,ARPC5,SH3GL1,TKT,RHE

1,SIAH1,ASPH,PNPO,CHPF,LYNX1,MPDU1,ZSWIM7,DCTD,LMBR1,HMGA1,PHTF2,SERPINB6,TADA3,ZNF20

1,SIAH1,ASPH,PNPO,CHPF,LYNX1,MPDU1,ZSWIM7,DCTD,LMBR1,HMGA1,PHTF2,SERPINB6,TADA3,ZNF20

RPL29,CD99L2,GGPS1,COASY,SRSF1,SAP30BP,TRNT1,KXD1,MED15,IP6K2,UGGT2,KCNMA1,JUP,CDC42,TN

H11,DBNDD2,NISCH,CENPK,RPS6,RPL29,GGPS1,SRSF1,AUP1,SAP30BP,YIPF3,FOXRED1,TRNT1,KXD1,SH3

H11,DBNDD2,NISCH,CENPK,RPS6,RPL29,GGPS1,SRSF1,AUP1,SAP30BP,YIPF3,FOXRED1,TRNT1,KXD1,SH3

3C,RPS3,SNRPD1,ARRB2,NDUFA10,SLK,MAZ,SYPL1,THOP1,RNPS1,UBA52,ZNF609,PSMD8,IFRD2,ZNF33A,

1,C12ORF57,ERGIC1,EIF3G,RPL8,KCNC4,CHTOP,TPT1,TSPAN31,SLC39A6,APOLD1,MARS,C11ORF57,FAM6

RPF,TYMS,RBFOX2,ARHGEF1,NSUN5,DRG2,BRF1,MRPS18C,LGALS3,TCP1,PHF6,PARK7,NHP2,EIF2B3,UBE

,CARS,ZNF250,RPL36,RBM3,ZNF197,HSP90AB1,DUT,LRRFIP1,ZNF524,GPBP1,TSFM,ZEB2,RPTOR,ZNF625,

LL,SH2B1,ARHGEF39,GSTK1,LDHA,PSMA6,ACOT8,AP3B2,RPS9,SLC35G1,ZNF410,RUVBL2,PCMT1,TESC,IG

RFX3,DGKE,ENDOV,CAMTA1,CRIP1,PSMD4,RPL28,VIM,LIG1,APEH,RPL21,RBM33,TCF4,UCK2,PPOX,TMP

DF1,GADL1,RFX3,DGKE,ENDOV,CAMTA1,CRIP1,PSMD4,RPL28,VIM,LIG1,APEH,RPL21,RBM33,TCF4,UCK2,

,SNRPA1,RPS7,U2AF1L4,NCBP2,POLR2E,RNPS1,UBA52,RPS4X,MLH1,RPS23,RPL30,PSMD8,RQCD1,NSRP1

ARM1,POLR2E,SNX3,GPS2,UBA52,RPS4X,VAPA,SUMO1,RPS23,RPL30,NUP85,FDPS,DYNC1I2,AP2S1,IK,AN

S3,IFI27,AZI2,RPS7,CARM1,POLR2E,SNX3,GPS2,UBA52,RPS4X,VAPA,SUMO1,RPS23,RPL30,NUP85,FDPS,D

PAM16,IFT172,CLTA,THOC3,TP53,AP1G2,RPS3A,PML,UBE2D3,RPS3,SYTL4,MAGOHB,VPS25,DHRS4,IFI27,

24,PSMD7,PAM16,NEDD4L,RNF130,BFAR,CLTA,AXIN1,PSMD10,RPS3A,PML,TMEM199,UBE2D3,RPS3,RD

A,RPS17,EIF2AK4,MRPL51,MRPL43,EIF6,MRPL48,SHMT1,ERAL1,MRPL27,MRPS7,RPL27,NCK2,MRPL36,R

N,RPS15A,RPS17,EIF2AK4,MRPL51,MRPL43,EIF6,MRPL48,SHMT1,ERAL1,MRPL27,MRPS7,RPL27,NCK2,MR

Y2,NUP133,RAPGEF3,RPAIN,ACOT2,PCID2,PEX2,RPL37A,STX10,DCTN5,PEX26,ERGIC1,KPNA5,RPLP0,HAX

A,ENTPD6,NUDT9,APOBEC3F,MAGOH,RPL31,GSK3A,PSMD11,EIF3E,EXOSC10,FAU,DERA,FXR1

S3,IFI27,AZI2,RPS7,CARM1,POLR2E,SNX3,GPS2,UBA52,RPS4X,VAPA,SSC5D,SUMO1,RPS23,RPL30,NUP85,

2,TNKS,RPL37A,STX10,USP4,PEX26,KPNA5,RPLP0,HAX1,FAM126A,CIZ1,SSR3,MPV17,SSB,ARIH2,RPL8,PI

2,TNKS,RPL37A,STX10,USP4,PEX26,KPNA5,RPLP0,HAX1,FAM126A,CIZ1,SSR3,MPV17,SSB,ARIH2,RPL8,PI(
3E2D4,RBBP6,BAX,RPL35,UBA1,RNF8,PPP2R5C,FAM192A,PCNP,RPL34,RPL24,PSMD7,PAM16,NEDD4L,RI

L30,RQCD1,NDUFA7,DNAJC24,MRPS33,NGDN,RPS15A,RPS17,EIF2AK4,ERAP2,MRPL51,MRPL43,EIF6,MR
E1A,ENTPD6,NUDT9,APOBEC3F,MAGOH,RPL31,GSK3A,PSMD11,EIF3E,EXOSC10,FAU,CYP46A1,DERA,FXR

30,RQCD1,NDUFA7,DNAJC24,MRPS33,ACSS1,NGDN,RPS15A,RPS17,EIF2AK4,MRPL51,MRPL43,EIF6,MRPL

5,POT1,ABHD17A,STXBP2,SEC31A,RPL18,CNST,LLGL2,EGFR,SH3GLB1,CAMK2D,HNRNPA1,RPL15,B3GAT3

1,ARL2,RPS23,RPL30,SZT2,NUP85,PEX16,SUFU,OS9,IK,CCT4,PKIG,UBE2L3,RPS15A,LAMTOR4,PPP1R13B,I
AP3,CARHSP1,RPL18,EGFR,SH3GLB1,HINT1,LGMN,EGLN2,RPL15,MTHFS,PAFAH1B2,GBA2,PFKFB2,PARK7
PS27L,RPS7,NCBP2,PLD1,PIWIL4,THOP1,NFE2L1,UBA52,ACSM3,RPS4X,MRPS16,NMRAL1,XPNPEP1,RPS2
FG,PSMG2,PSMG4,TBCA,TAF9,ARPC1A,UBA52,ARL2,PRKCE,NDUFA7,NDUFS2,PSMC5,GTF3C5,AP2S1,ND
,MRPS18C,MCFD2,LGALS3,TCP1,CD74,PARK7,SMARCC2,CCDC40,ZNHIT3,PDCD6,MFF,NOP2,NUP133,CO
17B10,GGT5,NT5C,USP45,PSME1,AGAP3,CARHSP1,RPL18,SH3GLB1,HINT1,LGMN,RPL15,MTHFS,PAFAH1
),POLR2E,RNPS1,NSRP1,EXOSC2,RBM7,HNRNPM,NGDN,IK,EXOSC8,DDX17,PABPN1,QTRT1,RPS17,SFSW
,WAC,NUDT1,ANAPC11,RAD23A,HSD17B10,ABHD17A,GGT5,NT5C,USP45,PSME1,AGAP3,CARHSP1,RPL1
PC2,TRPM4,MCFD2,PFKFB2,TCP1,SYTL1,CD74,PARK7,NFKBIA,RPS11,PDCD6,MFF,SNX5,ENY2,NUP133,R
3GLB1,RPL15,B3GAT3,MPC2,TRPM4,MCFD2,PFKFB2,TCP1,SYTL1,CD74,PARK7,NFKBIA,RPS11,PDCD6,MFF
T1,EXOSC9,STOML2,PPAN,HSP90AB1,CEP70,TRMT112,VPS28,RPS15,VWA2,ERCC5,MCTS1,NAP1L1,RAN(

CAP,CD74,PARK7,PIGT,EIF2B3,LFNG,RPS11,XYLT2,MRPS17,RPL37A,PIGH,ELOF1,NMNAT3,EIF4A1,RPLP0,

PC2,TRPM4,MCFD2,PFKFB2,TCP1,SYTL1,CD74,PARK7,NFKBIA,RPS11,PDCD6,MFF,SNX5,ENY2,NUP133,R
3GAT3,MPC2,TRPM4,MCFD2,PFKFB2,TCP1,SYTL1,CD74,PARK7,NFKBIA,RPS11,PDCD6,MFF,SNX5,ENY2,N
T1,TRIM16,ABHD17A,SLC16A1,STXBP2,SEC31A,RPL18,CNST,LLGL2,EGFR,SH3GLB1,RPL15,B3GAT3,TCTN1

NIP,GUK1,CARD14,ATF7,FBXW9,HSBP1L1,AGPAT4,MAF,EDF1,RFX3,PPA1,ENDOV,CAMTA1,PSMD4,RPL28,
1,DGAT1,EIF3G,PICK1,ADD1,KCNC4,BAX,DLG4,TRPM6,CLASP2,RPL24,FCHO2,TRAPPC2,GTF2H1,CLTA,ARF
1,ITGB1BP1,MED15,CLASRP,UGGT2,UBE2I,PRMT2,EIF3M,JUP,TBX6,XBP1,CELF1,AUNIP,GUK1,CARD14,A
MM17B,TMEM126B,CAPN10,EPB41L4B,PRMT1,STOML2,PPAN,HNRNPC,SHARPIN,INO80E,MAPKAPK5,H

NIP,GUK1,CARD14,ATF7,FBXW9,HSBP1L1,AGPAT4,MAF,EDF1,RFX3,PPA1,ENDOV,CAMTA1,PSMD4,RPL28,

)9,UBE2D3,RDX,ITPA,VPS25,UCHL5,NSFL1C,SMUG1,COMT,HMOX2,IFI27,ARRB2,SGSM3,RPS7,SNX3,USP1

MM40,PPP3CB,RABEP2,ATP6V1F,PDCD5,POT1,TRIM16,ABHD17A,SLC16A1,STXBP2,SEC31A,RPL18,ATP1
,MAGT1,CDC42,XBP1,PTK2,CELF1,CARD14,FBXW9,EEF1B2,PPP1CC,PPA1,TNFRSF14,CAMTA1,PPIL2,PSM
5,MCTS1,NAP1L1,RANGRF,LSR,HMGA1,DBNL,RBMX,BRIP1,RABEP2,POT1,HSD17B10,B2M,PKP4,PDE4DIP
D1,FUZ,MUC1,ECSIT,EPB41L5,ATG12,PPRC1,NLRP1,NIP7,DUSP22,SPOCK2,THEM4,NISCH,TMED4,CENPK
F4,UCK2,PPOX,TMPO,EXOSC9,STOML2,ZNF7,HNRNPC,ZNF644,CBFA2T2,UPP1,CARS,ZNF250,RPL36,NFA
IP,GUK1,CARD14,ATF7,FBXW9,HSBP1L1,AGPAT4,MAF,EDF1,RFX3,PPA1,ENDOV,CAMTA1,PSMD4,RPL28

,UBE2L6,VRK3,ZC3H14,FIS1,PPM1K,IRF9,PHF19,VCAN,PCBP2,LPCAT3,CD44,TM9SF1,SON,TET2,UQCRQ,

,RPL28,VIM,LIG1,PSMA7,CDC14B,RPL21,TXNL4A,TCF4,TMPO,EXOSC9,ZNF7,ARL6IP4,HNRNPC,ZNF644,C
BX6,XBP1,CELF1,AUNIP,GUK1,CARD14,ATF7,FBXW9,HSBP1L1,AGPAT4,MAF,EDF1,RFX3,PPA1,ENDOV,C

GFR,SH3GLB1,HNRNPA1,RPL15,B3GAT3,MPC2,SLC25A23,TRPM4,MCFD2,PFKFB2,TCP1,SYTL1,SLC35B1,C
TMPO,PRMT1,EXOSC9,STOML2,ZNF7,PIGL,HNRNPC,ZNF644,CBFA2T2,CARS,ZNF250,RPL36,NFAM1,CDK1

MA7,APEH,RPL21,TCF4,TMPO,PRMT1,EXOSC9,STOML2,ZNF7,PIGL,HNRNPC,ZNF644,CBFA2T2,CARS,ZNF
PS25,NLRP1,DUSP22,SPOCK2,THEM4,DBNDD2,RPS6,RPL29,COASY,AUP1,ITGB1BP1,UGGT2,MICAL1,HS2

IF3M,MAGT1,JUP,CDC42,TBX6,XBP1,CELF1,GUK1,CARD14,ATF7,PCYT2,HSBP1L1,AGPAT4,MAF,EEF1B2,E
P1,NIP7,DUSP22,SPOCK2,THEM4,NISCH,TMED4,CENPK,RTEL1,SRSF1,FOXRED1,RABL5,ITGB1BP1,PARD6

2,TBX6,XBP1,CELF1,GUK1,CARD14,ATF7,PCYT2,HSBP1L1,AGPAT4,MAF,EEF1B2,EDF1,GADL1,RFX3,PPA1,

,UGGT2,MICAL1,HS2ST1,UBE2I,PRMT2,EIF3M,MAGT1,CDC42,XBP1,PTK2,CELF1,CARD14,FBXW9,USE1,E

P4,HNRNPC,ZNF644,CBFA2T2,CARS,ZNF250,RPL36,NFAM1,CDK11A,RBM3,ZNF197,CCNL2,LRRFIP1,ZNF5

IF3M,MAGT1,JUP,CDC42,TBX6,XBP1,CELF1,GUK1,CARD14,ATF7,PCYT2,HSBP1L1,AGPAT4,MAF,EEF1B2,E

MT,SNRPN,IL1RAP,STYXL1,RFX5,POLD4,UBE2L6,VRK3,ZC3H14,FIS1,PPM1K,IRF9,PHF19,VCAN,PCBP2,LPC
3,RDX,NSFL1C,SPHK2,ARRB2,MLST8,CDK2AP2,GMFG,PSMG2,ILK,MDK,ECT2,NFE2L1,ARPC1A,CAPN2,AR

UBE2N,NFATC4,PDCD6,NUP133,MAP3K8,RPAIN,KRAS,METTL3,MPV17,CREB3L4,PICK1,ARL6IP5,RBBP6,A

'LPM1,PHF21A,CHCHD1,RGS3,PSMA3,MUC1,METTL17,ATG12,PPRC1,RPS25,NLRP1,DUSP22,SPOCK2,DBI

'3B2,RPS9,SLC35G1,RCE1,MRPL33,ZNF410,RUVBL2,EBPL,TESC,IGSF8,NUDT5,RBPMS,SNRPN,SPAG7,WDR
i12,RPS25,NLRP1,NIP7,COX4I1,PLCB2,DUSP22,THEM4,NISCH,CENPK,RPS6,RPL29,GGPS1,KXD1,ITGB1BP1
31,NOP16,RPL13A,GSTK1,TTLL5,LDHA,PSMA6,ACOT8,AP3B2,RPS9,SLC35G1,RCE1,MRPL33,ZNF410,RUVB
T22,UFD1L,TAZ,THYN1,SH2B1,NOP16,RPL13A,GSTK1,TTLL5,LDHA,PSMA6,ACOT8,AP3B2,RPS9,SLC35G1,F
NDD1,SH2B1,RPL13A,GSTK1,TTLL5,LDHA,PSMA6,ACOT8,AP3B2,RPS9,SLC35G1,RCE1,MRPL33,RUVBL2,PC
',TBL2,ARHGAP6,SCARB1,CTTN,ITSN1,EIF5A,COMMD5,RUNX1,CHID1,NUDT22,UFD1L,TAZ,THYN1,DBNDD
TNFSF10,COMMD5,RUNX1,CHID1,NUDT22,UFD1L,TAZ,THYN1,SH2B1,NOP16,RPL13A,GSTK1,TTLL5,LDHA
6,RPS23,RPL30,NSRP1,MRPS33,HNRNPM,NGDN,IK,DDX17,ACTB,PABPN1,RPS15A,RUVBL1,RPS17,EIF2AK
D1,PSMA3,MUC1,METTL17,ECSIT,EPB41L5,PPRC1,RPS25,NLRP1,NIP7,COX4I1,SLC25A10,THEM4,SDF4,CI
D1,PSMA3,MUC1,METTL17,ECSIT,EPB41L5,PPRC1,RPS25,NLRP1,NIP7,COX4I1,SLC25A10,THEM4,SDF4,CI
D1,PSMA3,MUC1,METTL17,ECSIT,EPB41L5,PPRC1,RPS25,NLRP1,NIP7,COX4I1,SLC25A10,THEM4,SDF4,CI
PS16,CPSF3L,TSG101,TTC21B,PTPN13,UBR4,PSMA1,YLPM1,PHF21A,CHCHD1,SLC39A11,RGS3,PSMA3,M

T2,XBP1,CELF1,CENPN,ATF7,ASF1B,PPP1CC,EDF1,DGKE,CAPG,PPIL2,PSMD4,LIG1,PSMA7,CDC14B,XAF1,

iP1,MED15,IP6K2,CLASRP,UBE2I,PRMT2,TBX6,XBP1,CELF1,CENPN,ATF7,FBXW9,MAF,ASF1B,PPP1CC,EDF
D6G,JPH2,UGGT2,UBE2I,KCNMA1,EIF3M,MAGT1,JUP,CDC42,CELF1,C14ORF2,FBXW9,USE1,ASF1B,EEF1B
I,NDUFA3,MGST2,COX19,RPS3,DNAJC2,MRPS24,AIFM2,SPHK2,IFI27,TRAK1,NDUFB3,NDUFB2,NDUFA10,
I,NDUFA3,MGST2,COX19,RPS3,DNAJC2,MRPS24,AIFM2,SPHK2,IFI27,TRAK1,NDUFB3,NDUFB2,NDUFA10,

PL51,MRPL43,IMMP1L,MRPL48,COQ4,ERAL1,MTHFD2L,MRPL27,SLC25A35,COQ2,MRPS7,NDUFS7,IFI6,T

X6,PTK2,AUNIP,CENPN,ATF7,FBXW9,MAF,ASF1B,PPP1CC,PDLIM5,EDF1,RFX3,ENDOV,CAMTA1,CAPG,RPL
X6,PTK2,AUNIP,CENPN,ATF7,FBXW9,MAF,ASF1B,PPP1CC,PDLIM5,EDF1,RFX3,ENDOV,CAMTA1,CAPG,RPL
FLAR,RECQL5,PSMD7,DYNLT3,GTF2H1,SIN3A,BRD8,NAA38,NDUFA11,TP53,PSMD10,HDAC8,NDUFA3,UB
JMO1,ARL2,SDHA,SELENBP1,NUB1,EXOSC2,HNRNPM,NGDN,DNAJC21,EXOSC8,DDX17,EIF6,NARF,RNMT,
PICK1,BAX,UBA1,NME1,MTIF2,GCDH,GLYCTK,VDAC1,CREBZF,PAM16,D2HGDH,BRD8,NDUFA11,NRP1,TP

DH2,STXBP2,HSPH1,HINT1,HNRNPA1,LGMN,B3GAT3,FTH1,PAFAH1B2,S100A4,TPM4,FKBP5,LGALS3,TCP
DH2,STXBP2,HSPH1,HINT1,HNRNPA1,LGMN,B3GAT3,FTH1,PAFAH1B2,S100A4,TPM4,FKBP5,LGALS3,TCP
DH2,STXBP2,HSPH1,HINT1,HNRNPA1,LGMN,B3GAT3,FTH1,PAFAH1B2,S100A4,TPM4,FKBP5,LGALS3,TCP

YNGR2,RAB2A,ATP5J,GGCX,ASPH,RPTOR,CHPF,MPDU1,VPS28,UBXN1,EMC4,DDOST,SYNGR1,PTPRB,DBN

JT,SYNGR2,RAB2A,SIAH1,TPM3,RPS20,NIT2,VPS28,HSPB1,VWA2,SEMA3C,NAP1L1,SERPINB1,PRKRIP1,D

34,RPL24,OLFM1,TRAPPC2,PIGO,BFAR,KDELR3,SLC35A2,TP53,RPS3A,SERPINA1,PML,TMEM199,MGST2,F

MTOR5,VDAC1,NUP62,KDELR3,CLTA,AP1G2,SH3GL1,PML,TMEM199,UBE2D3,SYTL4,DENND1A,RHEB,VPS

,RQCD1,NUP85,EXOSC2,PSMC5,HNRNPM,EXOSC8,PABPN1,RPS15A,QTRT1,RPS17,SEC13,PPIE,RNMT,PUF

GAP6,SCARB1,CRYGS,CTTN,TSPAN11,ITSN1,EIF5A,COMMD5,CHID1,NUDT22,AMZ2,UFD1L,TAZ,THYN1,DB
,NUDT22,AMZ2,UFD1L,USP38,TAZ,DBNDD1,SH2B1,NOP16,RPL13A,ARHGEF39,GSTK1,TTLL5,LDHA,PSMA
CD44,TM9SF1,SON,UQCRQ,SNF8,GRK6,CDC37,RPS16,CPSF3L,TSG101,UBR4,YLPM1,PHF21A,CHCHD1,FUZ
ATCH2L,SUMF2,BUD31,SAE1,NPRL2,OSGEP,TBL2,ARHGAP6,SCARB1,CRYGS,CTTN,TSPAN11,ITSN1,EIF5A,C
ATCH2L,SUMF2,BUD31,SAE1,NPRL2,OSGEP,TBL2,ARHGAP6,SCARB1,CRYGS,CTTN,TSPAN11,ITSN1,EIF5A,C
SCARB1,CRYGS,CTTN,TSPAN11,ITSN1,EIF5A,COMMD5,NUDT22,AMZ2,UFD1L,USP38,TAZ,THYN1,DBNDD1
SCARB1,CRYGS,CTTN,TSPAN11,ITSN1,EIF5A,COMMD5,NUDT22,AMZ2,UFD1L,USP38,TAZ,THYN1,DBNDD1
MMD5,NUDT22,AMZ2,UFD1L,USP38,TAZ,THYN1,DBNDD1,SH2B1,NOP16,RPL13A,ARHGEF39,GSTK1,TTLL
RHGEF39,GSTK1,TTLL5,LDHA,PSMA6,ACOT8,AP3B2,RPS9,SLC35G1,RCE1,MRPL33,ZNF410,RUVBL2,PCMT1
RHGEF39,GSTK1,TTLL5,LDHA,PSMA6,ACOT8,AP3B2,RPS9,SLC35G1,RCE1,MRPL33,ZNF410,RUVBL2,PCMT1
ITSN1,EIF5A,COMMD5,CHID1,NUDT22,AMZ2,UFD1L,USP38,TAZ,THYN1,DBNDD1,SH2B1,NOP16,RPL13A,
AN,LPCAT3,MAGED2,SON,UQCRQ,SNF8,GRK6,HMGA2,CDC37,CPSF3L,TSG101,UBR4,YLPM1,PHF21A,FUZ,
AN,LPCAT3,MAGED2,SON,UQCRQ,SNF8,GRK6,HMGA2,CDC37,CPSF3L,TSG101,UBR4,YLPM1,PHF21A,FUZ,
PS6,RPL29,RTEL1,COASY,AUP1,SAP30BP,YIPF3,FOXRED1,TRNT1,KXD1,ITGB1BP1,MED15,IP6K2,CLASRP,C
MRPL33,ZNF410,RUVBL2,PCMT1,TESC,IGSF8,NUDT5,SNRPN,SPAG7,WDR62,STYXL1,RFX5,POLD4,VRK3,LZ
MRPL33,ZNF410,RUVBL2,PCMT1,TESC,IGSF8,NUDT5,SNRPN,SPAG7,WDR62,STYXL1,RFX5,POLD4,VRK3,LZ
D5,CHID1,NUDT22,UFD1L,USP38,TAZ,THYN1,DBNDD1,SH2B1,NOP16,RPL13A,ARHGEF39,GSTK1,TTLL5,LD

D5,CHID1,NUDT22,UFD1L,USP38,TAZ,THYN1,DBNDD1,SH2B1,NOP16,RPL13A,ARHGEF39,GSTK1,TTLL5,LD

I,SPAG7,WDR62,STYXL1,RFX5,POLD4,RANBP1,LZTFL1,FIS1,PPM1K,TUBA1C,IRF9,DST,ANTXR2,STIP1,LRRC

I,SPAG7,WDR62,STYXL1,RFX5,POLD4,RANBP1,LZTFL1,FIS1,PPM1K,TUBA1C,IRF9,DST,ANTXR2,STIP1,LRRC

IP1,LRRC1,VCAN,MAGED2,SON,UQCRQ,SNF8,GRK6,CDC37,CPSF3L,TSG101,UBR4,FUZ,FAM3A,METTL17,

IP1,LRRC1,VCAN,MAGED2,SON,UQCRQ,SNF8,GRK6,CDC37,CPSF3L,TSG101,UBR4,FUZ,FAM3A,METTL17,

MRPL33,ZNF410,PCMT1,TESC,IGSF8,NUDT5,SNRPN,SPAG7,WDR62,STYXL1,RFX5,POLD4,VRK3,LZTFL1,FIS

MRPL33,ZNF410,PCMT1,TESC,IGSF8,NUDT5,SNRPN,SPAG7,WDR62,STYXL1,RFX5,POLD4,VRK3,LZTFL1,FIS

2,BUD31,SAE1,NPRL2,OSGEP,TBL2,ARHGAP6,SCARB1,CTTN,TSPAN11,ITSN1,EIF5A,COMMD5,RUNX1,NU

2,BUD31,SAE1,NPRL2,OSGEP,TBL2,ARHGAP6,SCARB1,CTTN,TSPAN11,ITSN1,EIF5A,COMMD5,RUNX1,NU

T1,TESC,IGSF8,NUDT5,SNRPN,SPAG7,WDR62,STYXL1,RFX5,POLD4,RANBP1,VRK3,LZTFL1,FIS1,PPM1K,TU

T1,TESC,IGSF8,NUDT5,SNRPN,SPAG7,WDR62,STYXL1,RFX5,POLD4,RANBP1,VRK3,LZTFL1,FIS1,PPM1K,TU

1,NOP16,RPL13A,ARHGEF39,GSTK1,TTLL5,LDHA,PSMA6,ACOT8,AP3B2,RPS9,SLC35G1,RCE1,MRPL33,ZN

1,NOP16,RPL13A,ARHGEF39,GSTK1,TTLL5,LDHA,PSMA6,ACOT8,AP3B2,RPS9,SLC35G1,RCE1,MRPL33,ZN

I1,DUSP22,THEM4,DBNDD2,RPS6,RPL29,RTEL1,COASY,AUP1,SAP30BP,YIPF3,FOXRED1,RABL5,TRNT1,KX

I1,DUSP22,THEM4,DBNDD2,RPS6,RPL29,RTEL1,COASY,AUP1,SAP30BP,YIPF3,FOXRED1,RABL5,TRNT1,KX

PL13A,ARHGEF39,GSTK1,TTLL5,LDHA,PSMA6,ACOT8,AP3B2,RPS9,SLC35G1,RCE1,MRPL33,ZNF410,RUVB

PL13A,ARHGEF39,GSTK1,TTLL5,LDHA,PSMA6,ACOT8,AP3B2,RPS9,SLC35G1,RCE1,MRPL33,ZNF410,RUVB

PL13A,ARHGEF39,GSTK1,TTLL5,PSMA6,ACOT8,AP3B2,RPS9,SLC35G1,RCE1,MRPL33,ZNF410,RUVBL2,PC

PL13A,ARHGEF39,GSTK1,TTLL5,PSMA6,ACOT8,AP3B2,RPS9,SLC35G1,RCE1,MRPL33,ZNF410,RUVBL2,PC

NUDT22,AMZ2,UFD1L,USP38,TAZ,THYN1,DBNDD1,SH2B1,NOP16,RPL13A,ARHGEF39,GSTK1,TTLL5,LDH

TK1,TTLL5,PSMA6,ACOT8,AP3B2,RPS9,SLC35G1,RCE1,MRPL33,ZNF410,RUVBL2,PCMT1,TESC,IGSF8,NUD

TK1,TTLL5,PSMA6,ACOT8,AP3B2,RPS9,SLC35G1,RCE1,MRPL33,ZNF410,RUVBL2,PCMT1,TESC,IGSF8,NUD

5,NIP7,TTC9C,SLC25A10,TMX2,THEM4,DBNDD2,RPS6,RPL29,RTEL1,SRSF1,AUP1,SAP30BP,YIPF3,FOXRE

2,UFD1L,USP38,TAZ,THYN1,DBNDD1,SH2B1,RPL13A,ARHGEF39,TTLL5,LDHA,PSMA6,ACOT8,AP3B2,RPS

2,UFD1L,USP38,TAZ,THYN1,DBNDD1,SH2B1,RPL13A,ARHGEF39,TTLL5,LDHA,PSMA6,ACOT8,AP3B2,RPS

UBE2I,SLC39A13,EIF3M,CDC42,TBX6,C14ORF2,CARD14,PCYT2,FBXW9,USE1,ASF1B,DGKE,ENDOV,PPIL2,

GS,CTTN,TSPAN11,ITSN1,EIF5A,COMMD5,CHID1,NUDT22,AMZ2,UFD1L,USP38,TAZ,THYN1,DBNDD1,SH2

2,BUD31,SAE1,NPRL2,OSGEP,TBL2,ARHGAP6,SCARB1,CTTN,TSPAN11,ITSN1,EIF5A,COMMD5,RUNX1,NU

2,BUD31,SAE1,NPRL2,OSGEP,TBL2,ARHGAP6,SCARB1,CTTN,TSPAN11,ITSN1,EIF5A,COMMD5,RUNX1,NU

LZTFL1,FIS1,PPM1K,IRF9,ANTXR2,STIP1,LRRC1,VCAN,CD44,TM9SF1,MAGED2,SON,UQCRQ,SNF8,CDC37,

LZTFL1,FIS1,PPM1K,IRF9,ANTXR2,STIP1,LRRC1,VCAN,CD44,TM9SF1,MAGED2,SON,UQCRQ,SNF8,CDC37,

2,PCMT1,TESC,IGSF8,NUDT5,SNRPN,SPAG7,WDR62,STYXL1,RFX5,POLD4,VRK3,LZTFL1,FIS1,PPM1K,TUBA

2,PCMT1,TESC,IGSF8,NUDT5,SNRPN,SPAG7,WDR62,STYXL1,RFX5,POLD4,VRK3,LZTFL1,FIS1,PPM1K,TUBA

AMZ2,UFD1L,USP38,TAZ,THYN1,DBNDD1,SH2B1,NOP16,RPL13A,ARHGEF39,GSTK1,TTLL5,LDHA,PSMA6,

AMZ2,UFD1L,USP38,TAZ,THYN1,DBNDD1,SH2B1,NOP16,RPL13A,ARHGEF39,GSTK1,TTLL5,LDHA,PSMA6,

N,SPAG7,STYXL1,RFX5,POLD4,VRK3,LZTFL1,FIS1,PPM1K,STIP1,LRRC1,VCAN,MAGED2,SON,UQCRQ,SNF8,

N,SPAG7,STYXL1,RFX5,POLD4,VRK3,LZTFL1,FIS1,PPM1K,STIP1,LRRC1,VCAN,MAGED2,SON,UQCRQ,SNF8,

ARHGEF39,GSTK1,TTLL5,LDHA,PSMA6,ACOT8,AP3B2,RPS9,SLC35G1,RCE1,MRPL33,ZNF410,RUVBL2,PCM

ARHGEF39,GSTK1,TTLL5,LDHA,PSMA6,ACOT8,AP3B2,RPS9,SLC35G1,RCE1,MRPL33,ZNF410,RUVBL2,PCM

16,CPSF3L,TSG101,UBR4,PHF21A,CHCHD1,FUZ,FAM3A,ATG12,RPS25,NIP7,COX4I1,DUSP22,SLC25A10,TI

16,CPSF3L,TSG101,UBR4,PHF21A,CHCHD1,FUZ,FAM3A,ATG12,RPS25,NIP7,COX4I1,DUSP22,SLC25A10,TI

FL1,FIS1,PPM1K,IRF9,LRRC1,TM9SF1,MAGED2,SON,UQCRQ,SNF8,GRK6,HMGA2,CDC37,RPS16,CPSF3L,T

FL1,FIS1,PPM1K,IRF9,LRRC1,TM9SF1,MAGED2,SON,UQCRQ,SNF8,GRK6,HMGA2,CDC37,RPS16,CPSF3L,T

TNFSF10,COMMD5,CHID1,NUDT22,AMZ2,UFD1L,USP38,TAZ,THYN1,DBNDD1,SH2B1,NOP16,RPL13A,GST

D15,IP6K2,CLASRP,C1ORF198,MICAL1,UBE2I,SLC39A13,EIF3M,CDC42,CARD14,PCYT2,FBXW9,USE1,IFI27

UBE2I,SLC39A13,EIF3M,CDC42,CARD14,PCYT2,FBXW9,USE1,IFI27L1,ASF1B,CCDC25,EEF1B2,RFX3,ENDO

FL1,FIS1,PPM1K,IRF9,ANTXR2,STIP1,VCAN,LPCAT3,TM9SF1,MAGED2,SON,UQCRQ,SNF8,CDC37,RPS16,C

FL1,FIS1,PPM1K,IRF9,ANTXR2,STIP1,VCAN,LPCAT3,TM9SF1,MAGED2,SON,UQCRQ,SNF8,CDC37,RPS16,C

RPN,WDR62,STYXL1,RFX5,POLD4,RANBP1,VRK3,LZTFL1,FIS1,PPM1K,IRF9,ANTXR2,STIP1,LRRC1,VCAN,LPC

RPN,WDR62,STYXL1,RFX5,POLD4,RANBP1,VRK3,LZTFL1,FIS1,PPM1K,IRF9,ANTXR2,STIP1,LRRC1,VCAN,LPC

FIS1,PPM1K,IRF9,ANTXR2,LRRC1,TM9SF1,MAGED2,SON,UQCRQ,SNF8,GRK6,CDC37,RPS16,CPSF3L,TSG1

FIS1,PPM1K,IRF9,ANTXR2,LRRC1,TM9SF1,MAGED2,SON,UQCRQ,SNF8,GRK6,CDC37,RPS16,CPSF3L,TSG1

BR4,CHCHD1,FUZ,FAM3A,METTL17,EPB41L5,ATG12,RPS25,NIP7,COX4I1,DUSP22,SLC25A10,THEM4,CEN

BR4,CHCHD1,FUZ,FAM3A,METTL17,EPB41L5,ATG12,RPS25,NIP7,COX4I1,DUSP22,SLC25A10,THEM4,CEN

,TSG101,UBR4,PHF21A,FUZ,FAM3A,EPB41L5,ATG12,RPS25,NIP7,COX4I1,DUSP22,SLC25A10,TMX2,THEM

,TSG101,UBR4,PHF21A,FUZ,FAM3A,EPB41L5,ATG12,RPS25,NIP7,COX4I1,DUSP22,SLC25A10,TMX2,THEM

LD4,VRK3,LZTFL1,FIS1,PPM1K,IRF9,ANTXR2,LRRC1,TM9SF1,MAGED2,SON,UQCRQ,SNF8,GRK6,CDC37,RP

LD4,VRK3,LZTFL1,FIS1,PPM1K,IRF9,ANTXR2,LRRC1,TM9SF1,MAGED2,SON,UQCRQ,SNF8,GRK6,CDC37,RP

,GSTK1,TTLL5,PSMA6,ACOT8,AP3B2,RPS9,SLC35G1,RCE1,MRPL33,ZNF410,RUVBL2,PCMT1,EBPL,IGSF8,N

,GSTK1,TTLL5,PSMA6,ACOT8,AP3B2,RPS9,SLC35G1,RCE1,MRPL33,ZNF410,RUVBL2,PCMT1,EBPL,IGSF8,N

RK3,LZTFL1,FIS1,PPM1K,IRF9,DST,ANTXR2,STIP1,LRRC1,VCAN,CD44,TM9SF1,MAGED2,SON,UQCRQ,SNF

RK3,LZTFL1,FIS1,PPM1K,IRF9,DST,ANTXR2,STIP1,LRRC1,VCAN,CD44,TM9SF1,MAGED2,SON,UQCRQ,SNF

NF7,TXNDC17,ARL6IP4,PPAN,ACAD10,SUGT1,HYPK,CARS,CDK11A,RBM3,ZNF197,COPZ1,INO80E,CCNL2,

RUVBL2,PCMT1,IGSF8,NUDT5,SNRPN,SPAG7,WDR62,STYXL1,RFX5,POLD4,RANBP1,VRK3,LZTFL1,FIS1,PPN

RUVBL2,PCMT1,IGSF8,NUDT5,SNRPN,SPAG7,WDR62,STYXL1,RFX5,POLD4,RANBP1,VRK3,LZTFL1,FIS1,PPN

UQCRQ,SNF8,CDC37,RPS16,CPSF3L,TSG101,UBR4,FUZ,FAM3A,PSMA3,ATG12,RPS25,NIP7,TTC9C,COX4I1

UQCRQ,SNF8,CDC37,RPS16,CPSF3L,TSG101,UBR4,FUZ,FAM3A,PSMA3,ATG12,RPS25,NIP7,TTC9C,COX4I1

4,RANBP1,VRK3,LZTFL1,FIS1,PPM1K,IRF9,ANTXR2,STIP1,LRRC1,VCAN,TM9SF1,MAGED2,SON,UQCRQ,SN

4,RANBP1,VRK3,LZTFL1,FIS1,PPM1K,IRF9,ANTXR2,STIP1,LRRC1,VCAN,TM9SF1,MAGED2,SON,UQCRQ,SN

LD4,RANBP1,VRK3,LZTFL1,FIS1,PPM1K,TUBA1C,IRF9,DST,ANTXR2,STIP1,LRRC1,LPCAT3,TM9SF1,MAGED2

LD4,RANBP1,VRK3,LZTFL1,FIS1,PPM1K,TUBA1C,IRF9,DST,ANTXR2,STIP1,LRRC1,LPCAT3,TM9SF1,MAGED2

OK,HSP90AB1,LRRFIP1,ZNF524,ATP5J,ELP6,TRMT112,CHPF,RPS20,MPDU1,DCTD,UBXN1,ERCC5,EMC4,D

1,PPM1K,IRF9,ANTXR2,STIP1,LRRC1,VCAN,TM9SF1,MAGED2,SON,UQCRQ,SNF8,GRK6,CDC37,RPS16,CPS

1,PPM1K,IRF9,ANTXR2,STIP1,LRRC1,VCAN,TM9SF1,MAGED2,SON,UQCRQ,SNF8,GRK6,CDC37,RPS16,CPS

,NNMT,SNRPN,SPAG7,WDR62,IL1RAP,RFX5,POLD4,RANBP1,UBE2L6,ZC3H14,FIS1,PPM1K,TUBA1C,ANTX

,NNMT,SNRPN,SPAG7,WDR62,IL1RAP,RFX5,POLD4,RANBP1,UBE2L6,ZC3H14,FIS1,PPM1K,TUBA1C,ANTX

F7,TXNDC17,ARL6IP4,PIGL,C19ORF48,PPAN,ZFAND2B,ACAD10,SUGT1,HYPK,CARS,CDK11A,RBM3,ZNF19

J,TRMT112,CHPF,RPS20,MPDU1,DCTD,SAT1,ERCC5,SERPINB1,DDOST,LSR,PSME2,PABPC1,RRNAD1,NAT

GSTK1,TTLL5,PSMA6,ACOT8,AP3B2,RPS9,SLC35G1,RCE1,MRPL33,ZNF410,RUVBL2,PCMT1,IGSF8,NUDT5

GSTK1,TTLL5,PSMA6,ACOT8,AP3B2,RPS9,SLC35G1,RCE1,MRPL33,ZNF410,RUVBL2,PCMT1,IGSF8,NUDT5

H2B1,RPL13A,ARHGEF39,GSTK1,TTLL5,LDHA,ACOT8,AP3B2,SLC35G1,RCE1,ZNF410,RUVBL2,PCMT1,EBPL,

H2B1,RPL13A,ARHGEF39,GSTK1,TTLL5,LDHA,ACOT8,AP3B2,SLC35G1,RCE1,ZNF410,RUVBL2,PCMT1,EBPL,

NPRL2,OSGEP,TBL2,ARHGAP6,SCARB1,CTTN,TSPAN11,ITSN1,EIF5A,COMMD5,CHID1,NUDT22,AMZ2,UFD

NPRL2,OSGEP,TBL2,ARHGAP6,SCARB1,CTTN,TSPAN11,ITSN1,EIF5A,COMMD5,CHID1,NUDT22,AMZ2,UFD

A2,C17ORF62,PRMT1,STOML2,C19ORF54,TXNDC17,ARL6IP4,PPAN,ACAD10,SUGT1,HYPK,CARS,CDK11A,

RHGEF39,GSTK1,TTLL5,LDHA,PSMA6,ACOT8,AP3B2,RPS9,SLC35G1,RCE1,MRPL33,ZNF410,RUVBL2,PCM

RHGEF39,GSTK1,TTLL5,LDHA,PSMA6,ACOT8,AP3B2,RPS9,SLC35G1,RCE1,MRPL33,ZNF410,RUVBL2,PCM

,NUDT22,AMZ2,UFD1L,USP38,TAZ,THYN1,DBNDD1,NOP16,ARHGEF39,GSTK1,TTLL5,LDHA,PSMA6,ACOT

,NUDT22,AMZ2,UFD1L,USP38,TAZ,THYN1,DBNDD1,NOP16,ARHGEF39,GSTK1,TTLL5,LDHA,PSMA6,ACOT

W9,USE1,IFI27L1,ASF1B,CCDC25,EEF1B2,RFX3,ENDOV,CAMTA1,PPIL2,RPL28,APEH,CAPN10,LYPLA2,C17

M3A,METTL17,ATG12,RPS25,NIP7,COX4I1,DUSP22,SLC25A10,SPOCK2,THEM4,CENPK,RPS6,RPL29,RTEL1

M3A,METTL17,ATG12,RPS25,NIP7,COX4I1,DUSP22,SLC25A10,SPOCK2,THEM4,CENPK,RPS6,RPL29,RTEL1

C4,SERPINB1,DDOST,LSR,RRNAD1,DBNL,TOMM40,RABEP2,TADA3,WAC,KLK10,ANAPC11,RAD23A,NT5C,

F7,TXNDC17,ARL6IP4,C19ORF48,PPAN,ACAD10,SUGT1,HYPK,CARS,CDK11A,ZNF197,INO80E,CCNL2,MA

1BP1,MED15,SH3YL1,IP6K2,CLASRP,C1ORF198,MICAL1,UBE2I,SLC39A13,EIF3M,CDC42,AUNIP,C14ORF2,
NUDT5,RBPMS,SNRPN,SPAG7,STYXL1,RFX5,POLD4,RANBP1,VRK3,LZTFL1,FIS1,PPM1K,IRF9,ANTXR2,LRRC
NUDT5,RBPMS,SNRPN,SPAG7,STYXL1,RFX5,POLD4,RANBP1,VRK3,LZTFL1,FIS1,PPM1K,IRF9,ANTXR2,LRRC
PCMT1,IGSF8,NUDT5,SNRPN,SPAG7,WDR62,STYXL1,RFX5,POLD4,RANBP1,VRK3,LZTFL1,FIS1,PPM1K,IRF9
PCMT1,IGSF8,NUDT5,SNRPN,SPAG7,WDR62,STYXL1,RFX5,POLD4,RANBP1,VRK3,LZTFL1,FIS1,PPM1K,IRF9
XD1,ITGB1BP1,MED15,IP6K2,CLASRP,C1ORF198,UBE2I,SLC39A13,YIF1A,EIF3M,CDC42,CARD14,PCYT2,FB
TK1,TTLL5,PSMA6,ACOT8,RPS9,SLC35G1,RCE1,MRPL33,ZNF410,RUVBL2,PCMT1,EBPL,TESC,IGSF8,NUDT5
TK1,TTLL5,PSMA6,ACOT8,RPS9,SLC35G1,RCE1,MRPL33,ZNF410,RUVBL2,PCMT1,EBPL,TESC,IGSF8,NUDT5
PAN,ACAD10,CYB561,SUGT1,HYPK,CARS,CDK11A,RBM3,ZNF197,DGUOK,LRRFIP1,RAB2A,ZNF524,ATP5J,T
1,ITSN1,EIF5A,COMMD5,CHID1,NUDT22,AMZ2,UFD1L,USP38,TAZ,THYN1,DBNDD1,SH2B1,NOP16,RPL13
1,ITSN1,EIF5A,COMMD5,CHID1,NUDT22,AMZ2,UFD1L,USP38,TAZ,THYN1,DBNDD1,SH2B1,NOP16,RPL13
RPS20,DCTD,SAT1,ERCC5,SERPINB1,DDOST,LSR,PSME2,PABPC1,RRNAD1,TOMM40,RABEP2,TADA3,ZNF
SC,IGSF8,SNRPN,SPAG7,WDR62,IL1RAP,RFX5,POLD4,PLEKHA5,RANBP1,UBE2L6,ZC3H14,FIS1,PPM1K,TU
SC,IGSF8,SNRPN,SPAG7,WDR62,IL1RAP,RFX5,POLD4,PLEKHA5,RANBP1,UBE2L6,ZC3H14,FIS1,PPM1K,TU
PIN,SUGT1,HYPK,CARS,ZNF250,ZNF197,COPZ1,INO80E,HSP90AB1,RAB2A,ATP5J,TRMT112,CHPF,GABPB
PIN,SUGT1,HYPK,CARS,ZNF250,ZNF197,COPZ1,INO80E,HSP90AB1,RAB2A,ATP5J,TRMT112,CHPF,GABPB
ED15,IP6K2,C1ORF198,UBE2I,SLC39A13,EIF3M,CDC42,CARD14,PCYT2,FBXW9,USE1,IFI27L1,ASF1B,CCDC
L,TESC,IGSF8,NUDT5,RBPMS,WDR62,IL1RAP,STYXL1,RFX5,POLD4,PLEKHA5,RANBP1,VRK3,LZTFL1,ZC3H1
C17,ARL6IP4,PPAN,ZFAND2B,ACAD10,SUGT1,HYPK,CARS,ZNF250,CDK11A,RBM3,ZNF197,COPZ1,DGUO
LRD3,NSUN5,ZFPL1,MFSD10,RIC8A,UBE2N,LFNG,ZNHIT3,PDCD6,FAM32A,TGOLN2,POLR1C,PIGH,STX10,
0BP,YIPF3,FOXRED1,TRNT1,KXD1,ITGB1BP1,MED15,IP6K2,CLASRP,C1ORF198,UBE2I,SLC39A13,YIF1A,EIF
XD1,ITGB1BP1,MED15,IP6K2,CLASRP,C1ORF198,UGGT2,MICAL1,UBE2I,SLC39A13,EIF3M,CDC42,TBX6,C
XD1,ITGB1BP1,MED15,IP6K2,CLASRP,C1ORF198,UGGT2,MICAL1,UBE2I,SLC39A13,EIF3M,CDC42,TBX6,C
PZ1,INO80E,HSP90AB1,RAB2A,ATP5J,TRMT112,CHPF,ZNF814,HSPB1,ERCC5,EMC4,DDOST,LSR,MEIS2,R
PZ1,INO80E,HSP90AB1,RAB2A,ATP5J,TRMT112,CHPF,ZNF814,HSPB1,ERCC5,EMC4,DDOST,LSR,MEIS2,R
SNF8,GRK6,CPSF3L,TSG101,UBR4,YLPM1,PHF21A,FUZ,FAM3A,METTL17,EPB41L5,ATG12,RPS25,NIP7,DU
SNF8,GRK6,CPSF3L,TSG101,UBR4,YLPM1,PHF21A,FUZ,FAM3A,METTL17,EPB41L5,ATG12,RPS25,NIP7,DU
YN1,DBNDD1,SH2B1,NOP16,RPL13A,ARHGEF39,GSTK1,LDHA,PSMA6,ACOT8,AP3B2,RPS9,SLC35G1,RCE1
YN1,DBNDD1,SH2B1,NOP16,RPL13A,ARHGEF39,GSTK1,LDHA,PSMA6,ACOT8,AP3B2,RPS9,SLC35G1,RCE1
,ARL6IP4,PPAN,ACAD10,SUGT1,HYPK,CARS,CDK11A,RBM3,ZNF197,COPZ1,CCNL2,DGUOK,LRRFIP1,ZNF5
C42,CARD14,PCYT2,FBXW9,USE1,IFI27L1,ASF1B,CCDC25,EEF1B2,RFX3,DGKE,ENDOV,CAMTA1,PPIL2,RPL
5,AMZ2,UFD1L,USP38,TAZ,THYN1,DBNDD1,SH2B1,NOP16,RPL13A,ARHGEF39,GSTK1,TTLL5,LDHA,PSMA
5,AMZ2,UFD1L,USP38,TAZ,THYN1,DBNDD1,SH2B1,NOP16,RPL13A,ARHGEF39,GSTK1,TTLL5,LDHA,PSMA
ACAD10,SUGT1,HYPK,CARS,CDK11A,ZNF197,INO80E,CCNL2,MAPKAPK5,DGUOK,LRRFIP1,ZNF524,ATP5J,
PN10,LYPLA2,C17ORF62,PRMT1,STOML2,ZNF7,TXNDC17,PPAN,CYB561,SHARPIN,SUGT1,HYPK,CARS,ZNF
PN10,LYPLA2,C17ORF62,PRMT1,STOML2,ZNF7,TXNDC17,PPAN,CYB561,SHARPIN,SUGT1,HYPK,CARS,ZNF
3L,TSG101,UBR4,CHCHD1,FUZ,FAM3A,ECSIT,ATG12,RPS25,TTC9C,COX4I1,DUSP22,SLC25A10,THEM4,DE
3L,TSG101,UBR4,CHCHD1,FUZ,FAM3A,ECSIT,ATG12,RPS25,TTC9C,COX4I1,DUSP22,SLC25A10,THEM4,DE
,CAPN10,LYPLA2,C17ORF62,PRMT1,STOML2,ZNF7,TXNDC17,ARL6IP4,C19ORF48,PPAN,CYB561,SHARPI
,CAPN10,LYPLA2,C17ORF62,PRMT1,STOML2,ZNF7,TXNDC17,ARL6IP4,C19ORF48,PPAN,CYB561,SHARPI
1,MED15,CLASRP,C1ORF198,UGGT2,MICAL1,UBE2I,SLC39A13,EIF3M,CDC42,TBX6,PTK2,CARD14,PCYT2,
1,MED15,CLASRP,C1ORF198,UGGT2,MICAL1,UBE2I,SLC39A13,EIF3M,CDC42,TBX6,PTK2,CARD14,PCYT2,
M4,DBNDD2,RPS6,RPL29,RTEL1,COASY,AUP1,SAP30BP,YIPF3,FOXRED1,RABL5,TRNT1,KXD1,ITGB1BP1,
M4,DBNDD2,RPS6,RPL29,RTEL1,COASY,AUP1,SAP30BP,YIPF3,FOXRED1,RABL5,TRNT1,KXD1,ITGB1BP1,
2,UFD1L,USP38,TAZ,THYN1,DBNDD1,SH2B1,NOP16,ARHGEF39,GSTK1,TTLL5,LDHA,PSMA6,ACOT8,AP3B2
2,UFD1L,USP38,TAZ,THYN1,DBNDD1,SH2B1,NOP16,ARHGEF39,GSTK1,TTLL5,LDHA,PSMA6,ACOT8,AP3B2
MT1,C19ORF54,ZNF7,TXNDC17,ARL6IP4,PIGL,C19ORF48,PPAN,ZFAND2B,SLC9B2,SUGT1,HYPK,CARS,ZNF

YT2,FBXW9,USE1,IFI27L1,ASF1B,CCDC25,EEF1B2,RFX3,ENDOV,CAMTA1,PPIL2,RPL28,TMEM126B,APEH,

T1,KXD1,ITGB1BP1,MED15,CLASRP,C1ORF198,UGGT2,UBE2I,SLC39A13,EIF3M,CDC42,TBX6,CARD14,PCY

T1,KXD1,ITGB1BP1,MED15,CLASRP,C1ORF198,UGGT2,UBE2I,SLC39A13,EIF3M,CDC42,TBX6,CARD14,PCY

14,PCYT2,FBXW9,USE1,IFI27L1,ASF1B,CCDC25,EEF1B2,RFX3,DGKE,ENDOV,CAMTA1,PPIL2,PSMD4,RPL2

T2,UBE2I,EIF3M,CDC42,TBX6,TMEM205,CARD14,PCYT2,FBXW9,USE1,IFI27L1,CCDC25,EEF1B2,PDLIM5,

T2,UBE2I,EIF3M,CDC42,TBX6,TMEM205,CARD14,PCYT2,FBXW9,USE1,IFI27L1,CCDC25,EEF1B2,PDLIM5,

,TRMT112,CHPF,RPS20,MPDU1,DCTD,SERPINB1,DDOST,LSR,PABPC1,RRNAD1,DBNL,NAT9,TOMM40,TA

ID1,NUDT22,AMZ2,USP38,TAZ,THYN1,DBNDD1,SH2B1,RPL13A,ARHGEF39,GSTK1,TTLL5,LDHA,PSMA6,A

1,KXD1,ITGB1BP1,MED15,SH3YL1,IP6K2,CLASRP,MICAL1,UBE2I,EIF3M,CDC42,TBX6,TMEM205,CARD14,

N11,ITSN1,EIF5A,COMMD5,CHID1,NUDT22,AMZ2,UFD1L,USP38,TAZ,DBNDD1,SH2B1,NOP16,RPL13A,A

YPK,CARS,ZNF250,RBM3,ZNF197,COPZ1,INO80E,HSP90AB1,RAB2A,ATP5J,TRMT112,CHPF,ZNF814,HSPB

YPK,CARS,ZNF250,RBM3,ZNF197,COPZ1,INO80E,HSP90AB1,RAB2A,ATP5J,TRMT112,CHPF,ZNF814,HSPB

LL5,PSMA6,ACOT8,AP3B2,RPS9,SLC35G1,RCE1,MRPL33,ZNF410,RUVBL2,PCMT1,IGSF8,NUDT5,SNRPN,S

LL5,PSMA6,ACOT8,AP3B2,RPS9,SLC35G1,RCE1,MRPL33,ZNF410,RUVBL2,PCMT1,IGSF8,NUDT5,SNRPN,S

1,UBR4,YLPM1,CHCHD1,SLC39A11,FUZ,FAM3A,PSMA3,METTL17,EPB41L5,ATG12,RPS25,ZBTB44,NIP7,T

1,UBR4,YLPM1,CHCHD1,SLC39A11,FUZ,FAM3A,PSMA3,METTL17,EPB41L5,ATG12,RPS25,ZBTB44,NIP7,T

1,UBR4,YLPM1,CHCHD1,SLC39A11,FUZ,FAM3A,PSMA3,METTL17,EPB41L5,ATG12,RPS25,ZBTB44,NIP7,T

1,UBR4,YLPM1,CHCHD1,SLC39A11,FUZ,FAM3A,PSMA3,METTL17,EPB41L5,ATG12,RPS25,ZBTB44,NIP7,T

7,ARL6IP4,PPAN,CYB561,SHARPIN,SUGT1,HYPK,CARS,ZNF250,ZNF197,COPZ1,INO80E,HSP90AB1,RAB2A

7,ARL6IP4,PPAN,CYB561,SHARPIN,SUGT1,HYPK,CARS,ZNF250,ZNF197,COPZ1,INO80E,HSP90AB1,RAB2A

RABL5,TRNT1,KXD1,ITGB1BP1,MED15,SH3YL1,CLASRP,C1ORF198,UGGT2,MICAL1,UBE2I,SLC39A13,EIF3

RABL5,TRNT1,KXD1,ITGB1BP1,MED15,SH3YL1,CLASRP,C1ORF198,UGGT2,MICAL1,UBE2I,SLC39A13,EIF3

KAPK5,DGUOK,LRRFIP1,ATP5J,RMDN1,TRMT112,CHPF,MPDU1,SAT1,ERCC5,EMC4,LMBR1,MCTS1,SERP

4B,APEH,LYPLA2,NUTF2,TMPO,C17ORF62,STOML2,WNK1,ZNF7,PPAN,ACAD10,CYB561,SUGT1,HYPK,CA

4B,APEH,LYPLA2,NUTF2,TMPO,C17ORF62,STOML2,WNK1,ZNF7,PPAN,ACAD10,CYB561,SUGT1,HYPK,CA

L2,PCMT1,IGSF8,NUDT5,RBPMS,SNRPN,SPAG7,IL1RAP,STYXL1,RFX5,POLD4,RANBP1,UBE2L6,VRK3,LZTF

L2,PCMT1,IGSF8,NUDT5,RBPMS,SNRPN,SPAG7,IL1RAP,STYXL1,RFX5,POLD4,RANBP1,UBE2L6,VRK3,LZTF

80E,HSP90AB1,RAB2A,ATP5J,TRMT112,CHPF,GABPB1,DCTD,UBXN1,ZNF814,HSPB1,ERCC5,EMC4,DDOST

80E,HSP90AB1,RAB2A,ATP5J,TRMT112,CHPF,GABPB1,DCTD,UBXN1,ZNF814,HSPB1,ERCC5,EMC4,DDOST

L1,FIS1,PPM1K,TUBA1C,IRF9,DST,STIP1,LRRC1,VCAN,MAGED2,SON,UQCRQ,SNF8,CDC37,CPSF3L,TSG101

L1,FIS1,PPM1K,TUBA1C,IRF9,DST,STIP1,LRRC1,VCAN,MAGED2,SON,UQCRQ,SNF8,CDC37,CPSF3L,TSG101

LDHA,PSMA6,ACOT8,AP3B2,RPS9,SLC35G1,RCE1,MRPL33,ZNF410,PCMT1,EBPL,TESC,IGSF8,NUDT5,RBP

LDHA,PSMA6,ACOT8,AP3B2,RPS9,SLC35G1,RCE1,MRPL33,ZNF410,PCMT1,EBPL,TESC,IGSF8,NUDT5,RBP

DC14B,APEH,LYPLA2,C17ORF62,STOML2,ZNF7,PPAN,ZFAND2B,CYB561,SUGT1,HYPK,CARS,ZNF250,ZNF1

DC14B,APEH,LYPLA2,C17ORF62,STOML2,ZNF7,PPAN,ZFAND2B,CYB561,SUGT1,HYPK,CARS,ZNF250,ZNF1

1,RFX5,POLD4,VRK3,LZTFL1,FIS1,PPM1K,TUBA1C,LPCAT3,MAGED2,SON,UQCRQ,SNF8,GRK6,HMGA2,CD

1,RFX5,POLD4,VRK3,LZTFL1,FIS1,PPM1K,TUBA1C,LPCAT3,MAGED2,SON,UQCRQ,SNF8,GRK6,HMGA2,CD

PAN11,ITSN1,EIF5A,COMMD5,CHID1,NUDT22,AMZ2,UFD1L,USP38,TAZ,THYN1,DBNDD1,SH2B1,RPL13A,

,SLC35G1,RCE1,MRPL33,ZNF410,PCMT1,EBPL,IGSF8,NUDT5,RBPMS,SNRPN,SPAG7,WDR62,IL1RAP,STYX

,SLC35G1,RCE1,MRPL33,ZNF410,PCMT1,EBPL,IGSF8,NUDT5,RBPMS,SNRPN,SPAG7,WDR62,IL1RAP,STYX

SPAG7,WDR62,IL1RAP,RFX5,POLD4,RANBP1,UBE2L6,ZC3H14,FIS1,PPM1K,TUBA1C,ANTXR2,STIP1,LRRC1

SPAG7,WDR62,IL1RAP,RFX5,POLD4,RANBP1,UBE2L6,ZC3H14,FIS1,PPM1K,TUBA1C,ANTXR2,STIP1,LRRC1

4,ATP5J,TRMT112,CHPF,RPS20,MPDU1,DCTD,EMC4,DDOST,LSR,PSME2,PABPC1,RRNAD1,DBNL,NAT9,T

PN,SPAG7,WDR62,IL1RAP,RFX5,POLD4,RANBP1,UBE2L6,ZC3H14,FIS1,PPM1K,TUBA1C,ANTXR2,STIP1,LR

PN,SPAG7,WDR62,IL1RAP,RFX5,POLD4,RANBP1,UBE2L6,ZC3H14,FIS1,PPM1K,TUBA1C,ANTXR2,STIP1,LR

N,ACAD10,CYB561,SHARPIN,SUGT1,HYPK,ZNF250,ZNF197,INO80E,HSP90AB1,LRRFIP1,ATP5J,ELP6,TRMT

N,ACAD10,CYB561,SHARPIN,SUGT1,HYPK,ZNF250,ZNF197,INO80E,HSP90AB1,LRRFIP1,ATP5J,ELP6,TRMT

RPIN,SUGT1,HYPK,CARS,ZNF250,ZNF197,INO80E,HSP90AB1,LRRFIP1,RAB2A,ATP5J,TRMT112,CHPF,DCTD
RPIN,SUGT1,HYPK,CARS,ZNF250,ZNF197,INO80E,HSP90AB1,LRRFIP1,RAB2A,ATP5J,TRMT112,CHPF,DCTD
SH2B1,NOP16,ARHGEF39,GSTK1,TTLL5,LDHA,PSMA6,ACOT8,AP3B2,RPS9,RCE1,MRPL33,ZNF410,RUVBL
SH2B1,NOP16,ARHGEF39,GSTK1,TTLL5,LDHA,PSMA6,ACOT8,AP3B2,RPS9,RCE1,MRPL33,ZNF410,RUVBL
0,RABEP2,TADA3,ZNF20,WAC,ANAPC11,RAD23A,NT5C,MDH2,STXBP2,EEF1D,CHCHD5,SEC31A,CNST,CO
TC9C,DUSP22,SLC25A10,TMX2,THEM4,DBNDD2,RPS6,RPL29,RTEL1,COASY,SRSF1,AUP1,SAP30BP,YIPF3,
TC9C,DUSP22,SLC25A10,TMX2,THEM4,DBNDD2,RPS6,RPL29,RTEL1,COASY,SRSF1,AUP1,SAP30BP,YIPF3,
CARB1,CTTN,TSPAN11,ITSN1,EIF5A,COMMD5,RUNX1,NUDT22,AMZ2,UFD1L,USP38,TAZ,THYN1,DBNDD1
CARB1,CTTN,TSPAN11,ITSN1,EIF5A,COMMD5,RUNX1,NUDT22,AMZ2,UFD1L,USP38,TAZ,THYN1,DBNDD1
1,ATG12,RPS25,NIP7,TTC9C,PLCB2,DUSP22,SLC25A10,TMX2,THEM4,DBNDD2,RPS6,RPL29,GGPS1,RTEL1,
1,ATG12,RPS25,NIP7,TTC9C,PLCB2,DUSP22,SLC25A10,TMX2,THEM4,DBNDD2,RPS6,RPL29,GGPS1,RTEL1,
,OSGEP,TBL2,ARHGAP6,SCARB1,CTTN,TSPAN11,ITSN1,EIF5A,TNFSF10,COMMD5,RUNX1,NUDT22,AMZ2,
,OSGEP,TBL2,ARHGAP6,SCARB1,CTTN,TSPAN11,ITSN1,EIF5A,TNFSF10,COMMD5,RUNX1,NUDT22,AMZ2,
,RBM3,ZNF197,COPZ1,INO80E,CCNL2,LRRFIP1,ATP5J,RMDN1,TRMT112,CHPF,RPS20,DCTD,SAT1,ERCC5,
2,IL1RAP,RFX5,POLD4,RANBP1,UBE2L6,FIS1,SLC15A3,PPM1K,ANTXR2,STIP1,PHF19,GRAMD1C,LPCAT3,C
2,IL1RAP,RFX5,POLD4,RANBP1,UBE2L6,FIS1,SLC15A3,PPM1K,ANTXR2,STIP1,PHF19,GRAMD1C,LPCAT3,C
17,ARL6IP4,PIGL,PPAN,ZFAND2B,SHARPIN,SUGT1,HYPK,CARS,CDK11A,RBM3,ZNF197,COPZ1,INO80E,CC
RPS6,RTEL1,SRSF1,SAP30BP,YIPF3,FOXRED1,RABL5,TRNT1,ITGB1BP1,MED15,IP6K2,CLASRP,C1ORF198,U
RPS6,RTEL1,SRSF1,SAP30BP,YIPF3,FOXRED1,RABL5,TRNT1,ITGB1BP1,MED15,IP6K2,CLASRP,C1ORF198,U
3A,METTL17,ATG12,RPS25,NIP7,COX4I1,PLCB2,THEM4,RPS6,RPL29,RTEL1,COASY,AUP1,SAP30BP,YIPF3,
3A,METTL17,ATG12,RPS25,NIP7,COX4I1,PLCB2,THEM4,RPS6,RPL29,RTEL1,COASY,AUP1,SAP30BP,YIPF3,
6K2,CLASRP,C1ORF198,MICAL1,UBE2I,SLC39A13,YIF1A,EIF3M,CDC42,GUK1,FBXW9,USE1,IFI27L1,ASF1B,
,CD44,TM9SF1,MAGED2,SON,UQCRQ,SNF8,CDC37,RPS16,CPSF3L,TSG101,PTPN13,UBR4,YLPM1,CHCHD
,CD44,TM9SF1,MAGED2,SON,UQCRQ,SNF8,CDC37,RPS16,CPSF3L,TSG101,PTPN13,UBR4,YLPM1,CHCHD
112,CHPF,GABPB1,DCTD,ZNF814,HSPB1,ERCC5,EMC4,DDOST,LSR,MEIS2,RRNAD1,TOMM40,TADA3,ZNF
112,CHPF,GABPB1,DCTD,ZNF814,HSPB1,ERCC5,EMC4,DDOST,LSR,MEIS2,RRNAD1,TOMM40,TADA3,ZNF
DOST,LSR,MEIS2,RRNAD1,DBNL,NAT9,TOMM40,RABEP2,TADA3,POLR3H,RAD23A,B2M,ABHD17A,MDH
,MEIS2,RRNAD1,TOMM40,TADA3,ZNF687,PDCD5,ANAPC11,RAD23A,ABHD17A,MDH2,EEF1D,NAA10,SE
,MEIS2,RRNAD1,TOMM40,TADA3,ZNF687,PDCD5,ANAPC11,RAD23A,ABHD17A,MDH2,EEF1D,NAA10,SE
F814,ERCC5,EMC4,MCTS1,DDOST,LSR,RRNAD1,DBNL,TOMM40,TADA3,ZNF687,TRIM16,ANAPC11,MDH
F814,ERCC5,EMC4,MCTS1,DDOST,LSR,RRNAD1,DBNL,TOMM40,TADA3,ZNF687,TRIM16,ANAPC11,MDH
P5J,TRMT112,CHPF,ZNF814,HSPB1,VWA2,ERCC5,EMC4,LMBR1,DDOST,LSR,MEIS2,RRNAD1,RAD1,TOMM
P5J,TRMT112,CHPF,ZNF814,HSPB1,VWA2,ERCC5,EMC4,LMBR1,DDOST,LSR,MEIS2,RRNAD1,RAD1,TOMM
3GAT3,ZNF506,SPINT2,MTHFS,PAFAH1B2,ZNF343,GBA2,PPIH,DALRD3,NSUN5,OARD1,GMPR2,MCFD2,
S,PAFAH1B2,ZNF343,GBA2,PPIH,DALRD3,NSUN5,OARD1,MCFD2,RIC8A,TCP1,PARK7,UBE2N,LFNG,ZNHIT
F7,TXNDC17,ARL6IP4,PIGL,PPAN,ZFAND2B,SHARPIN,SUGT1,HYPK,CARS,CDK11A,RBM3,ZNF197,COPZ1,
SC,IGSF8,RBPMS,SPAG7,WDR62,IL1RAP,STYXL1,RFX5,POLD4,RANBP1,UBE2L6,ZC3H14,FIS1,SLC15A3,PP
SC,IGSF8,RBPMS,SPAG7,WDR62,IL1RAP,STYXL1,RFX5,POLD4,RANBP1,UBE2L6,ZC3H14,FIS1,SLC15A3,PP
NHIT3,PDCD6,FAM32A,COX20,C12ORF57,PEX26,OXLD1,ZNF223,METTL5,SLC35E1,OGFOD2,TSEN34,ARIH
DT22,AMZ2,UFD1L,USP38,TAZ,THYN1,DBNDD1,SH2B1,NOP16,RPL13A,GSTK1,TTLL5,LDHA,ACOT8,RPS9,S
MT1,STOML2,C19ORF54,ZNF7,TXNDC17,PPAN,CYB561,SHARPIN,SUGT1,HYPK,CARS,ZNF250,ZNF197,CO
MT1,STOML2,C19ORF54,ZNF7,TXNDC17,PPAN,CYB561,SHARPIN,SUGT1,HYPK,CARS,ZNF250,ZNF197,CO
NBP1,UBE2L6,FIS1,SLC15A3,PPM1K,ANTXR2,STIP1,PHF19,LPCAT3,CD44,TM9SF1,MAGED2,SON,GRK6,CD
NBP1,UBE2L6,FIS1,SLC15A3,PPM1K,ANTXR2,STIP1,PHF19,LPCAT3,CD44,TM9SF1,MAGED2,SON,GRK6,CD
SUMF2,BUD31,SAE1,NPRL2,OSGEP,TBL2,SCARB1,CRYGS,CTTN,TSPAN11,ITSN1,EIF5A,COMMD5,RUNX1,
SUMF2,BUD31,SAE1,NPRL2,OSGEP,TBL2,SCARB1,CRYGS,CTTN,TSPAN11,ITSN1,EIF5A,COMMD5,RUNX1,
,EBPL,TESC,RBPMS,SPAG7,WDR62,IL1RAP,RFX5,PLEKHA5,RANBP1,UBE2L6,LZTFL1,ZC3H14,FIS1,SLC15A3

P6,SCARB1,CTTN,TSPAN11,ITSN1,EIF5A,TNFSF10,COMMD5,RUNX1,CHID1,NUDT22,AMZ2,UFD1L,USP38,T
P6,SCARB1,CTTN,TSPAN11,ITSN1,EIF5A,TNFSF10,COMMD5,RUNX1,CHID1,NUDT22,AMZ2,UFD1L,USP38,T
C5,EMC4,MCTS1,DDOST,LSR,PABPC1,RRNAD1,DBNL,NAT9,TOMM40,RABEP2,TADA3,ZNF687,ABHD17A,N
C5,EMC4,MCTS1,DDOST,LSR,PABPC1,RRNAD1,DBNL,NAT9,TOMM40,RABEP2,TADA3,ZNF687,ABHD17A,N
HGEF1,DALRD3,NSUN5,DRG2,MFSD10,MCFD2,RIC8A,THADA,UBE2N,LFNG,ZNHIT3,FAM32A,COX20,COA1
SEN34,ARIH2,RPL8,PICK1,ZNF512,RNF181,TSPAN31,RALY,GLYCTK,RBM42,VDAC1,RER1,DOK4,EIF2A,ACA
RPS6,RPL29,RTEL1,COASY,SRSF1,SAP30BP,YIPF3,FOXRED1,RABL5,TRNT1,ITGB1BP1,MED15,SH3YL1,CLAS
RPS6,RPL29,RTEL1,COASY,SRSF1,SAP30BP,YIPF3,FOXRED1,RABL5,TRNT1,ITGB1BP1,MED15,SH3YL1,CLAS
2,ARHGAP6,SCARB1,CRYGS,CTTN,TSPAN11,ITSN1,EIF5A,COMMD5,CHID1,NUDT22,AMZ2,UFD1L,USP38,T
2,ARHGAP6,SCARB1,CRYGS,CTTN,TSPAN11,ITSN1,EIF5A,COMMD5,CHID1,NUDT22,AMZ2,UFD1L,USP38,T
DUFS3,OXLD1,METTL5,SLC35E1,OGFOD2,METTL3,ARIH2,PICK1,FBXL18,RNF181,TPT1,SLC52A2,TSPAN31,
PAN11,ITSN1,EIF5A,TNFSF10,COMMD5,RUNX1,CHID1,NUDT22,AMZ2,UFD1L,USP38,TAZ,THYN1,DBNDD1
PAN11,ITSN1,EIF5A,TNFSF10,COMMD5,RUNX1,CHID1,NUDT22,AMZ2,UFD1L,USP38,TAZ,THYN1,DBNDD1
T,SPAG7,WDR62,IL1RAP,RFX5,POLD4,RANBP1,UBE2L6,VRK3,ZC3H14,FIS1,PPM1K,TUBA1C,IRF9,ANTXR2,S
T,SPAG7,WDR62,IL1RAP,RFX5,POLD4,RANBP1,UBE2L6,VRK3,ZC3H14,FIS1,PPM1K,TUBA1C,IRF9,ANTXR2,S
LRD3,NSUN5,ZFPL1,RIC8A,TCP1,SMARCC2,PIGT,UBE2N,NFATC4,MFSD11,PDCD6,FAM32A,ZSCAN20,COX2
N13,UBR4,SLC39A11,FAM3A,MUC1,METTL17,ECSIT,EPB41L5,ATG12,NLRP1,ZBTB44,NIP7,SLC25A10,TM
N13,UBR4,SLC39A11,FAM3A,MUC1,METTL17,ECSIT,EPB41L5,ATG12,NLRP1,ZBTB44,NIP7,SLC25A10,TM
5G1,RCE1,ZNF410,PCMT1,EBPL,TESC,NUDT5,RBPMS,SPAG7,WDR62,IL1RAP,STYXL1,RFX5,PLEKHA5,RANB
ENDOV,VAMP8,CAMTA1,CAPG,PPIL2,LIG1,CDC14B,APEH,LYPLA2,LSM12,C17ORF62,PRMT1,STOML2,ZNF
ENDOV,VAMP8,CAMTA1,CAPG,PPIL2,LIG1,CDC14B,APEH,LYPLA2,LSM12,C17ORF62,PRMT1,STOML2,ZNF
NUTF2,C17ORF62,PRMT1,STOML2,ZNF7,TXNDC17,C19ORF48,PPAN,AP1S3,CYB561,SHARPIN,SUGT1,HYP
NUTF2,C17ORF62,PRMT1,STOML2,ZNF7,TXNDC17,C19ORF48,PPAN,AP1S3,CYB561,SHARPIN,SUGT1,HYP
BHD17A,MDH2,EEF1D,NAA10,CARHSP1,COX17,HNRNPLL,HIGD1A,POLR2C,B3GAT3,RFXANK,PFDN5,PAFA
BHD17A,MDH2,EEF1D,NAA10,CARHSP1,COX17,HNRNPLL,HIGD1A,POLR2C,B3GAT3,RFXANK,PFDN5,PAFA
ALRD3,NSUN5,MFSD10,GMPR2,MCFD2,RIC8A,TCP1,THADA,UBE2N,LFNG,ZNHIT3,PDCD6,FAM32A,COX20
CHID1,NUDT22,UFD1L,USP38,TAZ,DBNDD1,SH2B1,NOP16,RPL13A,ARHGEF39,GSTK1,TTLL5,LDHA,PSMA6
CHID1,NUDT22,UFD1L,USP38,TAZ,DBNDD1,SH2B1,NOP16,RPL13A,ARHGEF39,GSTK1,TTLL5,LDHA,PSMA6
F198,UGGT2,SLC39A13,YIF1A,EIF3M,CDC42,PTK2,CARD14,FBXW9,USE1,ASF1B,CCDC25,EEF1B2,EDF1,RFX
F198,UGGT2,SLC39A13,YIF1A,EIF3M,CDC42,PTK2,CARD14,FBXW9,USE1,ASF1B,CCDC25,EEF1B2,EDF1,RFX
T2,MICAL1,SLC39A13,EIF3M,TBX6,AUNIP,C14ORF2,GUK1,PCYT2,USE1,ASF1B,EEF1B2,PPP1CC,RFX3,END
OT8,AP3B2,RPS9,SLC35G1,RCE1,MRPL33,ZNF410,RUVBL2,TESC,NUDT5,RBPMS,WDR62,IL1RAP,STYXL1,P
OT8,AP3B2,RPS9,SLC35G1,RCE1,MRPL33,ZNF410,RUVBL2,TESC,NUDT5,RBPMS,WDR62,IL1RAP,STYXL1,P
,PABPC1,RRNAD1,TOMM40,RABEP2,TADA3,ZNF687,ANAPC11,POLR3H,ABHD17A,NT5C,MDH2,STXBP2,E
LSR,PSME2,RRNAD1,TOMM40,TADA3,WAC,SLC33A1,ZNF687,ANAPC11,NT5C,PDE4DIP,ZCCHC4,MDH2,S
62,IL1RAP,STYXL1,POLD4,PLEKHA5,RANBP1,VRK3,LZTFL1,ZC3H14,FIS1,SLC15A3,PPM1K,TUBA1C,ANTXR2
62,IL1RAP,STYXL1,POLD4,PLEKHA5,RANBP1,VRK3,LZTFL1,ZC3H14,FIS1,SLC15A3,PPM1K,TUBA1C,ANTXR2
VBL2,TESC,IGSF8,NUDT5,RBPMS,SPAG7,WDR62,IL1RAP,PLEKHA5,RANBP1,FIS1,SLC15A3,ANTXR2,STIP1,LR
VBL2,TESC,IGSF8,NUDT5,RBPMS,SPAG7,WDR62,IL1RAP,PLEKHA5,RANBP1,FIS1,SLC15A3,ANTXR2,STIP1,LR
K,IRF9,DST,ANTXR2,STIP1,LRRC1,VCAN,LPCAT3,MAGED2,SON,SNF8,GRK6,HMGA2,CDC37,OBSL1,TTC21B
A1C,ANTXR2,STIP1,PHF19,VCAN,LPCAT3,CD44,MAGED2,SON,TET2,GRK6,CDC37,RPS16,CPSF3L,TSG101,C
A1C,ANTXR2,STIP1,PHF19,VCAN,LPCAT3,CD44,MAGED2,SON,TET2,GRK6,CDC37,RPS16,CPSF3L,TSG101,C
S16,CPSF3L,TSG101,TTC21B,PHF21A,SLC39A11,FUZ,FAM3A,MUC1,RPS25,ZBTB44,NIP7,TTC9C,PLCB2,DUS
S16,CPSF3L,TSG101,TTC21B,PHF21A,SLC39A11,FUZ,FAM3A,MUC1,RPS25,ZBTB44,NIP7,TTC9C,PLCB2,DUS
44,TM9SF1,SON,TET2,GRK6,HMGA2,CDC37,RPS16,CPSF3L,OBSL1,TTC21B,PTPN13,YLPM1,PHF21A,SLC39
A,GLI4,PIGO,CLTA,SLC35A2,AXIN1,TMEM199,EPN1,HPS1,ITPA,ZCCHC9,SNRPD1,MKRN1,HMOX2,SNRPA1,
H2B1,NOP16,ARHGEF39,GSTK1,TTLL5,LDHA,PSMA6,AP3B2,RPS9,SLC35G1,RCE1,MRPL33,RUVBL2,PCMT1

2B1,NOP16,ARHGEF39,GSTK1,TTLL5,LDHA,PSMA6,AP3B2,RPS9,SLC35G1,RCE1,MRPL33,RUVBL2,PCMT1
B561,SHARPIN,SUGT1,HYPK,RBM3,ZNF197,COPZ1,ATP5J,ELP6,TRMT112,DCTD,ZNF814,VWA2,EMC4,LM
B561,SHARPIN,SUGT1,HYPK,RBM3,ZNF197,COPZ1,ATP5J,ELP6,TRMT112,DCTD,ZNF814,VWA2,EMC4,LM
,STIP1,PHF19,GRAMD1C,MAGED2,SNF8,CDC37,RPS16,TSG101,OBSL1,TTC21B,PTPN13,UBR4,PSMA1,CH
,STIP1,PHF19,GRAMD1C,MAGED2,SNF8,CDC37,RPS16,TSG101,OBSL1,TTC21B,PTPN13,UBR4,PSMA1,CH
1,NUDT22,AMZ2,UFD1L,USP38,TAZ,THYN1,DBNDD1,SH2B1,NOP16,ARHGEF39,GSTK1,TTLL5,LDHA,PSM
PS20,DCTD,ZNF814,SAT1,ERCC5,SERPINB1,DDOST,LSR,PSME2,PABPC1,RRNAD1,RBP1,TOMM40,SERPINE
ABEP2,TADA3,ZNF20,ANAPC11,RAD23A,NT5C,MDH2,STXBP2,EEF1D,CHCHD5,SEC31A,COX17,HIGD1A,S
,ARIH2,PICK1,FBXL18,ZNF512,RNF181,RPL35,TSPAN31,EEF2K,RALY,ZNF593,GLYCTK,RBM42,NEK11,EIF2
,VRK3,LZTFL1,ZC3H14,FIS1,SLC15A3,PPM1K,TUBA1C,DST,ANTXR2,STIP1,LRRC1,PHF19,GRAMD1C,CD44,
,VRK3,LZTFL1,ZC3H14,FIS1,SLC15A3,PPM1K,TUBA1C,DST,ANTXR2,STIP1,LRRC1,PHF19,GRAMD1C,CD44,
SLC35G1,RCE1,RUVBL2,PCMT1,EBPL,IGSF8,NUDT5,NNMT,SPAG7,WDR62,IL1RAP,STYXL1,POLD4,PLEKHA
SLC35G1,RCE1,RUVBL2,PCMT1,EBPL,IGSF8,NUDT5,NNMT,SPAG7,WDR62,IL1RAP,STYXL1,POLD4,PLEKHA
L,USP38,TAZ,DBNDD1,SH2B1,NOP16,RPL13A,ARHGEF39,GSTK1,TTLL5,LDHA,PSMA6,ACOT8,AP3B2,SLC3
2,GRK6,CDC37,RPS16,CPSF3L,TTC21B,PTPN13,YLPM1,PHF21A,SLC39A11,FAM3A,MUC1,METTL17,EPB41
2,GRK6,CDC37,RPS16,CPSF3L,TTC21B,PTPN13,YLPM1,PHF21A,SLC39A11,FAM3A,MUC1,METTL17,EPB41
1,SAE1,NPRL2,OSGEP,TBL2,ARHGAP6,SCARB1,CTTN,TSPAN11,ITSN1,EIF5A,TNFSF10,COMMD5,RUNX1,C
1,SAE1,NPRL2,OSGEP,TBL2,ARHGAP6,SCARB1,CTTN,TSPAN11,ITSN1,EIF5A,TNFSF10,COMMD5,RUNX1,C
,EBPL,TESC,IGSF8,NUDT5,RBPMS,WDR62,IL1RAP,POLD4,PLEKHA5,RANBP1,UBE2L6,LZTFL1,ZC3H14,SLC1
,RUVBL2,PCMT1,TESC,IGSF8,NUDT5,RBPMS,WDR62,IL1RAP,STYXL1,POLD4,PLEKHA5,RANBP1,UBE2L6,
PSF3L,TSG101,UBR4,PHF21A,SLC39A11,FUZ,FAM3A,MUC1,METTL17,RPS25,ZBTB44,NIP7,COX4I1,PLCB2
PSF3L,TSG101,UBR4,PHF21A,SLC39A11,FUZ,FAM3A,MUC1,METTL17,RPS25,ZBTB44,NIP7,COX4I1,PLCB2
FKBP3,B3GAT3,PAFAH1B2,PPIH,NSUN5,DRG2,ZFPL1,GMPR2,THADA,PIGT,UBE2N,LFNG,ZNHIT3,NOP2,FA
NF343,GBA2,DALRD3,NSUN5,ZFPL1,MFSD10,MCFD2,RIC8A,SMARCC2,PHC3,UBE2N,NFATC4,PDCD6,FAM
B41L5,PPRC1,RPS25,NLRP1,NIP7,TTC9C,COX4I1,SLC25A10,THEM4,DBNDD2,NISCH,TMED4,RPS6,RPL29,
,LZTFL1,FIS1,PPM1K,IRF9,ANTXR2,STIP1,LRRC1,VCAN,LPCAT3,TM9SF1,MAGED2,SON,UQCRQ,SNF8,CDC
,LZTFL1,FIS1,PPM1K,IRF9,ANTXR2,STIP1,LRRC1,VCAN,LPCAT3,TM9SF1,MAGED2,SON,UQCRQ,SNF8,CDC
,C12ORF57,PEX26,EIF4A1,METTL5,SLC35E1,OGFOD2,METTL3,TSEN34,SSR3,ARIH2,PICK1,TPT1,SLC52A2,
LC39A11,FAM3A,MUC1,METTL17,ECSIT,EPB41L5,ATG12,PPRC1,ZBTB44,PLCB2,DUSP22,DBNDD2,NISCH
LC39A11,FAM3A,MUC1,METTL17,ECSIT,EPB41L5,ATG12,PPRC1,ZBTB44,PLCB2,DUSP22,DBNDD2,NISCH
LC39A11,FAM3A,MUC1,METTL17,ECSIT,EPB41L5,ATG12,PPRC1,ZBTB44,PLCB2,DUSP22,DBNDD2,NISCH
LC39A11,FAM3A,MUC1,METTL17,ECSIT,EPB41L5,ATG12,PPRC1,ZBTB44,PLCB2,DUSP22,DBNDD2,NISCH
A,PSMA6,ACOT8,AP3B2,RPS9,SLC35G1,MRPL33,ZNF410,RUVBL2,EBPL,TESC,IGSF8,NUDT5,NNMT,SNRP
A,PSMA6,ACOT8,AP3B2,RPS9,SLC35G1,MRPL33,ZNF410,RUVBL2,EBPL,TESC,IGSF8,NUDT5,NNMT,SNRP
F39,GSTK1,TTLL5,LDHA,AP3B2,RPS9,SLC35G1,RCE1,MRPL33,RUVBL2,PCMT1,EBPL,TESC,IGSF8,NUDT5,R
YGS,CTTN,TSPAN11,ITSN1,EIF5A,COMMD5,CHID1,NUDT22,AMZ2,UFD1L,USP38,TAZ,THYN1,DBNDD1,S
YGS,CTTN,TSPAN11,ITSN1,EIF5A,COMMD5,CHID1,NUDT22,AMZ2,UFD1L,USP38,TAZ,THYN1,DBNDD1,S
MZ2,UFD1L,USP38,TAZ,THYN1,DBNDD1,SH2B1,NOP16,TTLL5,PSMA6,ACOT8,AP3B2,RPS9,RCE1,ZNF410,
TESC,NUDT5,SNRPN,SPAG7,IL1RAP,STYXL1,RFX5,POLD4,PLEKHA5,UBE2L6,LZTFL1,ZC3H14,FIS1,PPM1K,T
H2B1,NOP16,RPL13A,ARHGEF39,GSTK1,TTLL5,LDHA,PSMA6,AP3B2,RPS9,SLC35G1,RCE1,MRPL33,ZNF41
H2B1,NOP16,RPL13A,ARHGEF39,GSTK1,TTLL5,LDHA,PSMA6,AP3B2,RPS9,SLC35G1,RCE1,MRPL33,ZNF41
T5,RBPMS,SPAG7,STYXL1,RFX5,RANBP1,UBE2L6,VRK3,LZTFL1,IRF9,STIP1,LRRC1,GRAMD1C,CD44,MAGE
T5,RBPMS,SPAG7,STYXL1,RFX5,RANBP1,UBE2L6,VRK3,LZTFL1,IRF9,STIP1,LRRC1,GRAMD1C,CD44,MAGE
41L5,ATG12,NLRP1,ZBTB44,PLCB2,SLC25A10,TMX2,THEM4,NISCH,TMED4,CD99L2,GGPS1,SAP30BP,TRN
41L5,ATG12,NLRP1,ZBTB44,PLCB2,SLC25A10,TMX2,THEM4,NISCH,TMED4,CD99L2,GGPS1,SAP30BP,TRN
AP3B2,RPS9,SLC35G1,MRPL33,ZNF410,RUVBL2,TESC,NUDT5,RBPMS,SNRPN,SPAG7,IL1RAP,STYXL1,RFX
11,FAM3A,MUC1,EPB41L5,ATG12,NLRP1,ZBTB44,PLCB2,SLC25A10,TMX2,THEM4,DBNDD2,NISCH,TME

A11,FAM3A,MUC1,EPB41L5,ATG12,NLRP1,ZBTB44,PLCB2,SLC25A10,TMX2,THEM4,DBNDD2,NISCH,TMED
POLR1C,TNKS,C12ORF57,DCTN5,PEX26,NDUFS3,OXLD1,EIF4A1,ZNF223,MPV17,ARIH2,PICK1,ZNF512,RN
RCE1,RUVBL2,PCMT1,EBPL,TESC,IGSF8,NUDT5,RBPMS,NNMT,SPAG7,WDR62,IL1RAP,STYXL1,RFX5,POLD4
,SAE1,NPRL2,OSGEP,TBL2,ARHGAP6,SCARB1,CRYGS,CTTN,TSPAN11,ITSN1,EIF5A,TNFSF10,COMMD5,RUI
,SAE1,NPRL2,OSGEP,TBL2,ARHGAP6,SCARB1,CRYGS,CTTN,TSPAN11,ITSN1,EIF5A,TNFSF10,COMMD5,RUI
,NUDT5,RBPMS,WDR62,POLD4,PLEKHA5,RANBP1,UBE2L6,ZC3H14,FIS1,PPM1K,DST,ANTXR2,STIP1,PHF1
,NUDT5,RBPMS,WDR62,POLD4,PLEKHA5,RANBP1,UBE2L6,ZC3H14,FIS1,PPM1K,DST,ANTXR2,STIP1,PHF1
31A,CNST,COX5A,HINT1,SPINT2,PAFAH1B2,PPIH,DALRD3,NSUN5,ZFPL1,GMPR2,RIC8A,THADA,UBE2N,M
AGED2,SON,TET2,GRK6,CDC37,RPS16,CPSF3L,OBSL1,YLPM1,PHF21A,SLC39A11,FAM3A,MUC1,METTL17
AGED2,SON,TET2,GRK6,CDC37,RPS16,CPSF3L,OBSL1,YLPM1,PHF21A,SLC39A11,FAM3A,MUC1,METTL17
TESC,IGSF8,NUDT5,RBPMS,NNMT,WDR62,IL1RAP,STYXL1,RFX5,PLEKHA5,RANBP1,VRK3,ZC3H14,FIS1,SLC
TESC,IGSF8,NUDT5,RBPMS,NNMT,WDR62,IL1RAP,STYXL1,RFX5,PLEKHA5,RANBP1,VRK3,ZC3H14,FIS1,SLC
YGS,CTTN,TSPAN11,ITSN1,EIF5A,TNFSF10,COMMD5,RUNX1,CHID1,NUDT22,AMZ2,UFD1L,USP38,TAZ,TH
YGS,CTTN,TSPAN11,ITSN1,EIF5A,TNFSF10,COMMD5,RUNX1,CHID1,NUDT22,AMZ2,UFD1L,USP38,TAZ,TH
5,RANBP1,UBE2L6,VRK3,LZTFL1,ZC3H14,FIS1,SLC15A3,PPM1K,TUBA1C,IRF9,ANTXR2,STIP1,LRRC1,PHF19
CPSF3L,TSG101,OBSL1,PTPN13,PSMA1,CHCHD1,FUZ,FAM3A,MUC1,METTL17,EPB41L5,ATG12,PPRC1,RPS
CPSF3L,TSG101,OBSL1,PTPN13,PSMA1,CHCHD1,FUZ,FAM3A,MUC1,METTL17,EPB41L5,ATG12,PPRC1,RPS
CPSF3L,TSG101,OBSL1,PTPN13,PSMA1,CHCHD1,FUZ,FAM3A,MUC1,METTL17,EPB41L5,ATG12,PPRC1,RPS
CPSF3L,TSG101,OBSL1,PTPN13,PSMA1,CHCHD1,FUZ,FAM3A,MUC1,METTL17,EPB41L5,ATG12,PPRC1,RPS
CPSF3L,TSG101,OBSL1,PTPN13,PSMA1,CHCHD1,FUZ,FAM3A,MUC1,METTL17,EPB41L5,ATG12,PPRC1,RPS
A11,FAM3A,METTL17,ECSIT,EPB41L5,ATG12,ZBTB44,PLCB2,DUSP22,DBNDD2,NISCH,TMED4,CD99L2,GG
A11,FAM3A,METTL17,ECSIT,EPB41L5,ATG12,ZBTB44,PLCB2,DUSP22,DBNDD2,NISCH,TMED4,CD99L2,GG
N,LPCAT3,MAGED2,SON,UQCRQ,SNF8,HMGA2,RPS16,CPSF3L,TSG101,YLPM1,FUZ,FAM3A,PSMA3,METT
N,LPCAT3,MAGED2,SON,UQCRQ,SNF8,HMGA2,RPS16,CPSF3L,TSG101,YLPM1,FUZ,FAM3A,PSMA3,METT
ED1,RABL5,TRNT1,ITGB1BP1,MED15,CLASRP,C1ORF198,UGGT2,UBE2I,SLC39A13,YIF1A,EIF3M,CDC42,C
ED1,RABL5,TRNT1,ITGB1BP1,MED15,CLASRP,C1ORF198,UGGT2,UBE2I,SLC39A13,YIF1A,EIF3M,CDC42,C
1K,ANTXR2,STIP1,LRRC1,PHF19,LPCAT3,CD44,TM9SF1,MAGED2,SON,GRK6,CDC37,RPS16,CPSF3L,CHCHD
1K,ANTXR2,STIP1,LRRC1,PHF19,LPCAT3,CD44,TM9SF1,MAGED2,SON,GRK6,CDC37,RPS16,CPSF3L,CHCHD
47,USP40,PCID2,PEX2,TNKS,RPL37A,STX10,C12ORF57,DCTN5,OXLD1,ZNF223,OGFOD2,EIF3G,PICK1,FBX
47,USP40,PCID2,PEX2,TNKS,RPL37A,STX10,C12ORF57,DCTN5,OXLD1,ZNF223,OGFOD2,EIF3G,PICK1,FBX
,TET2,GRK6,CDC37,RPS16,CPSF3L,TSG101,OBSL1,TTC21B,PTPN13,YLPM1,CHCHD1,SLC39A11,FAM3A,MI
,TET2,GRK6,CDC37,RPS16,CPSF3L,TSG101,OBSL1,TTC21B,PTPN13,YLPM1,CHCHD1,SLC39A11,FAM3A,MI
SR,RRNAD1,PTPRB,NAT9,TOMM40,TADA3,TRIM16,ANAPC11,RAD23A,ABHD17A,MDH2,HSPH1,CHCHD5,
SR,RRNAD1,PTPRB,NAT9,TOMM40,TADA3,TRIM16,ANAPC11,RAD23A,ABHD17A,MDH2,HSPH1,CHCHD5,
UVBL2,PCMT1,EBPL,IGSF8,NUDT5,NNMT,SPAG7,WDR62,IL1RAP,STYXL1,RFX5,POLD4,PLEKHA5,RANBP1,
UVBL2,PCMT1,EBPL,IGSF8,NUDT5,NNMT,SPAG7,WDR62,IL1RAP,STYXL1,RFX5,POLD4,PLEKHA5,RANBP1,
19ORF54,TXNDC17,ARL6IP4,STRADA,AP1S3,ACAD10,SHARPIN,SUGT1,HYPK,RPL36,MOB1A,CDK11A,MDH
14,FIS1,SLC15A3,TUBA1C,ANTXR2,STIP1,LRRC1,PHF19,LPCAT3,CD44,TM9SF1,MAGED2,SON,TET2,UQCR
14,FIS1,SLC15A3,TUBA1C,ANTXR2,STIP1,LRRC1,PHF19,LPCAT3,CD44,TM9SF1,MAGED2,SON,TET2,UQCR
39A11,FAM3A,MUC1,METTL17,ECSIT,EPB41L5,ATG12,PPRC1,ZBTB44,PLCB2,DUSP22,TMX2,DBNDD2,NI
39A11,FAM3A,MUC1,METTL17,ECSIT,EPB41L5,ATG12,PPRC1,ZBTB44,PLCB2,DUSP22,TMX2,DBNDD2,NI
6,HMGA2,CDC37,RPS16,OBSL1,TTC21B,PSMA1,YLPM1,SLC39A11,FUZ,FAM3A,METTL17,EPB41L5,PPRC1
C1,RPS25,NIP7,COX4I1,DUSP22,SLC25A10,SPOCK2,TMED4,CENPK,GGPS1,RTEL1,SRSF1,AUP1,YIPF3,FOX
C1,RPS25,NIP7,COX4I1,DUSP22,SLC25A10,SPOCK2,TMED4,CENPK,GGPS1,RTEL1,SRSF1,AUP1,YIPF3,FOX
A,PSMA6,ACOT8,AP3B2,RPS9,SLC35G1,MRPL33,ZNF410,RUVBL2,EBPL,TESC,IGSF8,NUDT5,RBPMS,NNMT
A,PSMA6,ACOT8,AP3B2,RPS9,SLC35G1,MRPL33,ZNF410,RUVBL2,EBPL,TESC,IGSF8,NUDT5,RBPMS,NNMT

R3,ARIH2,PICK1,FBXL18,NME1,SLC52A2,TSPAN31,FAM192A,GLYCTK,CFLAR,TRIM46,C11ORF57,RBM42,C
R3,ARIH2,PICK1,FBXL18,NME1,SLC52A2,TSPAN31,FAM192A,GLYCTK,CFLAR,TRIM46,C11ORF57,RBM42,C
NDD1,SH2B1,NOP16,RPL13A,GSTK1,PSMA6,ACOT8,AP3B2,RPS9,SLC35G1,RCE1,MRPL33,ZNF410,RUVBL2
NDD1,SH2B1,NOP16,RPL13A,GSTK1,PSMA6,ACOT8,AP3B2,RPS9,SLC35G1,RCE1,MRPL33,ZNF410,RUVBL2
E1,MRPL33,ZNF410,RUVBL2,PCMT1,EBPL,TESC,IGSF8,RBPMS,NNMT,SPAG7,WDR62,IL1RAP,STYXL1,RFX5
E1,MRPL33,ZNF410,RUVBL2,PCMT1,EBPL,TESC,IGSF8,RBPMS,NNMT,SPAG7,WDR62,IL1RAP,STYXL1,RFX5
HGEF39,GSTK1,TTLL5,LDHA,ACOT8,RPS9,SLC35G1,RCE1,ZNF410,RUVBL2,PCMT1,EBPL,IGSF8,NUDT5,RBP
HGEF39,GSTK1,TTLL5,LDHA,ACOT8,RPS9,SLC35G1,RCE1,ZNF410,RUVBL2,PCMT1,EBPL,IGSF8,NUDT5,RBP
1,COX17,HNRNPLL,POLR2C,B3GAT3,RFXANK,FTH1,PFDN5,PAFAH1B2,TXNRD2,DALRD3,NSUN5,MRPS18C
1,COX17,HNRNPLL,POLR2C,B3GAT3,RFXANK,FTH1,PFDN5,PAFAH1B2,TXNRD2,DALRD3,NSUN5,MRPS18C
KHA5,RANBP1,LZTFL1,ZC3H14,FIS1,SLC15A3,PPM1K,IRF9,DST,STIP1,LRRC1,PHF19,LPCAT3,CD44,TM9SF1
KHA5,RANBP1,LZTFL1,ZC3H14,FIS1,SLC15A3,PPM1K,IRF9,DST,STIP1,LRRC1,PHF19,LPCAT3,CD44,TM9SF1
SLC39A11,FAM3A,MUC1,METTL17,EPB41L5,ATG12,ZBTB44,PLCB2,THEM4,NISCH,TMED4,CD99L2,GGPS1
SLC39A11,FAM3A,MUC1,METTL17,EPB41L5,ATG12,ZBTB44,PLCB2,THEM4,NISCH,TMED4,CD99L2,GGPS1
D6G,SH3YL1,IP6K2,CLASRP,JPH2,UGGT2,MICAL1,HS2ST1,UBE2I,YIF1A,PRMT2,EIF3M,JUP,TBX6,TMEM20
D6G,SH3YL1,IP6K2,CLASRP,JPH2,UGGT2,MICAL1,HS2ST1,UBE2I,YIF1A,PRMT2,EIF3M,JUP,TBX6,TMEM20
TMEM126B,APEH,LYPLA2,C17ORF62,CASC4,STOML2,PIGL,PPAN,ZFAND2B,HNRNPC,CYB561,CARS,ZNF25
TMEM126B,APEH,LYPLA2,C17ORF62,CASC4,STOML2,PIGL,PPAN,ZFAND2B,HNRNPC,CYB561,CARS,ZNF25
PRC1,ZBTB44,PLCB2,DUSP22,NISCH,TMED4,CD99L2,GGPS1,SRSF1,AUP1,YIPF3,FOXRED1,TRNT1,KXD1,PA
PRC1,ZBTB44,PLCB2,DUSP22,NISCH,TMED4,CD99L2,GGPS1,SRSF1,AUP1,YIPF3,FOXRED1,TRNT1,KXD1,PA
PS1,RTEL1,SRSF1,AUP1,YIPF3,FOXRED1,KXD1,ITGB1BP1,MED15,SH3YL1,CLASRP,HS2ST1,YIF1A,TBX6,XB
PS1,RTEL1,SRSF1,AUP1,YIPF3,FOXRED1,KXD1,ITGB1BP1,MED15,SH3YL1,CLASRP,HS2ST1,YIF1A,TBX6,XB
L1,RFX5,RANBP1,UBE2L6,FIS1,PPM1K,TUBA1C,ANTXR2,STIP1,PHF19,GRAMD1C,VCAN,LPCAT3,CD44,MA
L1,RFX5,RANBP1,UBE2L6,FIS1,PPM1K,TUBA1C,ANTXR2,STIP1,PHF19,GRAMD1C,VCAN,LPCAT3,CD44,MA
AN31,FAM192A,GLYCTK,CFLAR,TRIM46,C11ORF57,RBM42,CREBZF,EIF2A,NKIRAS2,SCN11A,ACAA1,GTF2
AN31,FAM192A,GLYCTK,CFLAR,TRIM46,C11ORF57,RBM42,CREBZF,EIF2A,NKIRAS2,SCN11A,ACAA1,GTF2
,DEF8,EEF1B2,PPA1,DGKE,CAMTA1,CRIP1,PPIL2,PSMD4,RPL28,CLDND1,TIMM17B,TMEM126B,RPL21,TX
NAD1,DBNL,RBP1,RAD1,TOMM40,TADA3,ZNF687,TRIM16,ANAPC11,ABHD17A,MDH2,STXBP2,EEF1D,CH
UBE2I,SLC39A13,PRMT2,CDC42,TBX6,TMEM205,AUNIP,C14ORF2,GUK1,PCYT2,AGPAT4,IFI27L1,PPP1CC,
L,RAD1,TOMM40,PPP3CB,TADA3,ZNF20,WAC,ZNF687,PDCD5,NXN,ANAPC11,ABHD17A,NT5C,STXBP2,CH
L,RAD1,TOMM40,PPP3CB,TADA3,ZNF20,WAC,ZNF687,PDCD5,NXN,ANAPC11,ABHD17A,NT5C,STXBP2,CH
0,RUVBL2,PCMT1,EBPL,TESC,NUDT5,RBPMS,SNRPN,SPAG7,WDR62,STYXL1,RFX5,POLD4,PLEKHA5,RANB
0,RUVBL2,PCMT1,EBPL,TESC,NUDT5,RBPMS,SNRPN,SPAG7,WDR62,STYXL1,RFX5,POLD4,PLEKHA5,RANB
EF39,GSTK1,TTLL5,LDHA,PSMA6,ACOT8,AP3B2,RPS9,SLC35G1,RCE1,MRPL33,ZNF410,RUVBL2,PCMT1,EB
EF39,GSTK1,TTLL5,LDHA,PSMA6,ACOT8,AP3B2,RPS9,SLC35G1,RCE1,MRPL33,ZNF410,RUVBL2,PCMT1,EB
DR62,STYXL1,RFX5,POLD4,PLEKHA5,RANBP1,LZTFL1,ZC3H14,FIS1,SLC15A3,PPM1K,TUBA1C,ANTXR2,STI
DR62,STYXL1,RFX5,POLD4,PLEKHA5,RANBP1,LZTFL1,ZC3H14,FIS1,SLC15A3,PPM1K,TUBA1C,ANTXR2,STI
2,RPS25,NIP7,COX4I1,DUSP22,THEM4,RDH11,NISCH,CENPK,RPS6,RPL29,RTEL1,COASY,SRSF1,AUP1,SAP3
2,RPS25,NIP7,COX4I1,DUSP22,THEM4,RDH11,NISCH,CENPK,RPS6,RPL29,RTEL1,COASY,SRSF1,AUP1,SAP3
2,UGGT2,MICAL1,UBE2I,SLC39A13,YIF1A,PRMT2,EIF3M,JUP,TBX6,AUNIP,C14ORF2,GUK1,ATF7,PCYT2,PP
2,UGGT2,MICAL1,UBE2I,SLC39A13,YIF1A,PRMT2,EIF3M,JUP,TBX6,AUNIP,C14ORF2,GUK1,ATF7,PCYT2,PP
2,EIF3G,MPV17,PICK1,FBXL18,NME1,SLC52A2,TSPAN31,FAM192A,GLYCTK,CFLAR,TRIM46,C11ORF57,RI
2,EIF3G,MPV17,PICK1,FBXL18,NME1,SLC52A2,TSPAN31,FAM192A,GLYCTK,CFLAR,TRIM46,C11ORF57,RI
P38,TAZ,DBNDD1,SH2B1,NOP16,RPL13A,GSTK1,TTLL5,LDHA,ACOT8,AP3B2,SLC35G1,MRPL33,ZNF410,R
P38,TAZ,DBNDD1,SH2B1,NOP16,RPL13A,GSTK1,TTLL5,LDHA,ACOT8,AP3B2,SLC35G1,MRPL33,ZNF410,R
524,ATP5J,TRMT112,CHPF,RPS20,UBXN1,HSPB1,SAT1,LMBR1,DDOST,LSR,CHCHD7,MEIS2,PSME2,PABPC
524,ATP5J,TRMT112,CHPF,RPS20,UBXN1,HSPB1,SAT1,LMBR1,DDOST,LSR,CHCHD7,MEIS2,PSME2,PABPC

,ZNF644,UPP1,ZNF250,RPL36,LAMP2,HSP90AB1,SYNGR2,CEP70,LRRFIP1,ZNF524,ELP6,RMDN1,TPM2,GF

,ZNF644,UPP1,ZNF250,RPL36,LAMP2,HSP90AB1,SYNGR2,CEP70,LRRFIP1,ZNF524,ELP6,RMDN1,TPM2,GF

T1,UBE2I,EIF3M,TBX6,XBP1,PTK2,AUNIP,C14ORF2,PCYT2,USE1,PPP1CC,RFX3,PPA1,DGKE,CAMTA1,CLDN

T1,UBE2I,EIF3M,TBX6,XBP1,PTK2,AUNIP,C14ORF2,PCYT2,USE1,PPP1CC,RFX3,PPA1,DGKE,CAMTA1,CLDN

,ABHD17A,NT5C,MDH2,STXBP2,HSPH1,SEC31A,CARHSP1,CNST,COX17,HNRNPLL,HIGD1A,B3GAT3,RFXAN

3YL1,C1ORF198,UGGT2,YIF1A,EIF3M,CDC42,TBX6,TMEM205,PTK2,AUNIP,CARD14,FBXW9,AGPAT4,CCDC

3YL1,C1ORF198,UGGT2,YIF1A,EIF3M,CDC42,TBX6,TMEM205,PTK2,AUNIP,CARD14,FBXW9,AGPAT4,CCDC

ADA,ZFAND2B,ZNF644,UPP1,ZNF250,RPL36,CDK11A,DGUOK,HSP90AB1,SYNGR2,CEP70,TMEM134,LRRFI

ADA,ZFAND2B,ZNF644,UPP1,ZNF250,RPL36,CDK11A,DGUOK,HSP90AB1,SYNGR2,CEP70,TMEM134,LRRFI

TESC,IGSF8,RBPMS,WDR62,STYXL1,PLEKHA5,RANBP1,VRK3,LZTFL1,ZC3H14,FIS1,SLC15A3,PPM1K,TUBA1

STYXL1,POLD4,PLEKHA5,RANBP1,UBE2L6,VRK3,LZTFL1,ZC3H14,FIS1,SLC15A3,PPM1K,TUBA1C,IRF9,ANTX

STYXL1,POLD4,PLEKHA5,RANBP1,UBE2L6,VRK3,LZTFL1,ZC3H14,FIS1,SLC15A3,PPM1K,TUBA1C,IRF9,ANTX

B,TMEM126B,APEH,RBM33,LYPLA2,UCK2,C17ORF62,STOML2,ZNF7,PPAN,ZFAND2B,HNRNPC,CYB561,CB

B,TMEM126B,APEH,RBM33,LYPLA2,UCK2,C17ORF62,STOML2,ZNF7,PPAN,ZFAND2B,HNRNPC,CYB561,CB

M9SF1,SON,TET2,GRK6,RPS16,CPSF3L,OBSL1,TTC21B,PSMA1,YLPM1,FUZ,RGS3,PSMA3,ECSIT,EPB41L5,P

M9SF1,SON,TET2,GRK6,RPS16,CPSF3L,OBSL1,TTC21B,PSMA1,YLPM1,FUZ,RGS3,PSMA3,ECSIT,EPB41L5,P

AP3B2,RPS9,SLC35G1,RCE1,MRPL33,ZNF410,RUVBL2,PCMT1,EBPL,TESC,IGSF8,NUDT5,SNRPN,WDR62,ST

AP3B2,RPS9,SLC35G1,RCE1,MRPL33,ZNF410,RUVBL2,PCMT1,EBPL,TESC,IGSF8,NUDT5,SNRPN,WDR62,ST

7,WDR62,STYXL1,RFX5,PLEKHA5,RANBP1,VRK3,LZTFL1,ZC3H14,FIS1,SLC15A3,TUBA1C,IRF9,STIP1,GRAMI

POCK2,THEM4,DBNDD2,RPS6,RPL29,RTEL1,SAP30BP,YIPF3,RABL5,TRNT1,KXD1,ITGB1BP1,MED15,SH3YL

POCK2,THEM4,DBNDD2,RPS6,RPL29,RTEL1,SAP30BP,YIPF3,RABL5,TRNT1,KXD1,ITGB1BP1,MED15,SH3YL

,CDC42,TBX6,PTK2,GUK1,CENPN,FBXW9,ASF1B,CCDC25,EEF1B2,DGKE,TNFRSF14,ENDOV,VAMP8,PPIL2,T

,CDC42,TBX6,PTK2,GUK1,CENPN,FBXW9,ASF1B,CCDC25,EEF1B2,DGKE,TNFRSF14,ENDOV,VAMP8,PPIL2,T

RHGEF39,GSTK1,LDHA,ACOT8,AP3B2,SLC35G1,RCE1,ZNF410,RUVBL2,PCMT1,EBPL,TESC,IGSF8,NUDT5,SN

ICAL1,HS2ST1,UBE2I,SLC39A13,YIF1A,PRMT2,CDC42,AUNIP,C14ORF2,GUK1,PCYT2,USE1,AGPAT4,PPP1C

PCMT1,EBPL,TESC,IGSF8,NUDT5,NNMT,SPAG7,IL1RAP,STYXL1,PLEKHA5,RANBP1,VRK3,LZTFL1,ZC3H14,PF

I1,HPS1,ITPA,ZCCHC9,VPS25,SNRPD1,GTF3A,AZI2,GOSR2,TATDN1,TBC1D22A,LAMTOR1,U2AF1L4,C2ORF

ORF198,MICAL1,HS2ST1,UBE2I,TBX6,TMEM205,XBP1,PTK2,AUNIP,C14ORF2,PCYT2,USE1,ASF1B,PPP1CC

ORF198,MICAL1,HS2ST1,UBE2I,TBX6,TMEM205,XBP1,PTK2,AUNIP,C14ORF2,PCYT2,USE1,ASF1B,PPP1CC

L,EIF5A,NUDT22,AMZ2,UFD1L,USP38,TAZ,THYN1,DBNDD1,SH2B1,NOP16,RPL13A,ARHGEF39,GSTK1,TTLL

L,EIF5A,NUDT22,AMZ2,UFD1L,USP38,TAZ,THYN1,DBNDD1,SH2B1,NOP16,RPL13A,ARHGEF39,GSTK1,TTLL

UDT22,AMZ2,UFD1L,USP38,TAZ,THYN1,DBNDD1,SH2B1,NOP16,ARHGEF39,GSTK1,TTLL5,LDHA,PSMA6,A

UDT22,AMZ2,UFD1L,USP38,TAZ,THYN1,DBNDD1,SH2B1,NOP16,ARHGEF39,GSTK1,TTLL5,LDHA,PSMA6,A

HID1,NUDT22,AMZ2,UFD1L,TAZ,THYN1,DBNDD1,SH2B1,RPL13A,GSTK1,TTLL5,LDHA,PSMA6,ACOT8,AP3B

BX6,TMEM205,AUNIP,GUK1,CENPN,ATF7,ASF1B,EEF1B2,ENDOV,PPIL2,PSMD4,VIM,LIG1,PSMA7,APEH,H

ON,TET2,UQCRQ,HMGA2,CDC37,RPS16,CPSF3L,OBSL1,TTC21B,UBR4,PSMA1,YLPM1,PHF21A,SLC39A11,

PRMT2,CDC42,TBX6,AUNIP,C14ORF2,GUK1,PCYT2,AGPAT4,IFI27L1,PPP1CC,EDF1,RFX3,PPA1,DGKE,CAM

G1,TIMM17B,APEH,NUTF2,C17ORF62,CASC4,ZNF7,ZFAND2B,SHARPIN,STAU1,UPP1,ZNF250,RPL36,LAMP

PRC1,ZBTB44,NIP7,TTC9C,PLCB2,DUSP22,SLC25A10,DBNDD2,NISCH,TMED4,CD99L2,GGPS1,RTEL1,YIPF3,

PRC1,ZBTB44,NIP7,TTC9C,PLCB2,DUSP22,SLC25A10,DBNDD2,NISCH,TMED4,CD99L2,GGPS1,RTEL1,YIPF3,

,MDH2,EEF1D,CHCHD5,NAA10,C11ORF48,OAS3,CNST,HIGD1A,POLR2C,INTS4,ZNF506,ASH2L,PAFAH1B2,

GAP6,SCARB1,CTTN,TSPAN11,ITSN1,EIF5A,TNFSF10,COMMD5,CHID1,NUDT22,AMZ2,UFD1L,USP38,TAZ,D

GAP6,SCARB1,CTTN,TSPAN11,ITSN1,EIF5A,TNFSF10,COMMD5,CHID1,NUDT22,AMZ2,UFD1L,USP38,TAZ,D

TESC,IGSF8,RBPMS,NNMT,SPAG7,WDR62,IL1RAP,STYXL1,RFX5,POLD4,PLEKHA5,RANBP1,UBE2L6,ZC3H1

TESC,IGSF8,RBPMS,NNMT,SPAG7,WDR62,IL1RAP,STYXL1,RFX5,POLD4,PLEKHA5,RANBP1,UBE2L6,ZC3H1

,HMGA2,RPS16,CPSF3L,TSG101,OBSL1,TTC21B,PSMA1,YLPM1,ECSIT,EPB41L5,ATG12,PPRC1,ZBTB44,NIP

,HMGA2,RPS16,CPSF3L,TSG101,OBSL1,TTC21B,PSMA1,YLPM1,ECSIT,EPB41L5,ATG12,PPRC1,ZBTB44,NIP

RF57,NDUFS3,NUBP2,OGFOD2,TSEN34,PICK1,ZNF512,CDK4,SLC52A2,TSPAN31,FAM192A,TRIM46,C11C
D1,KXD1,ITGB1BP1,CLASRP,UGGT2,MICAL1,SLC39A13,PRMT2,EIF3M,MAGT1,JUP,TMEM205,PTK2,AUNIF
D1,KXD1,ITGB1BP1,CLASRP,UGGT2,MICAL1,SLC39A13,PRMT2,EIF3M,MAGT1,JUP,TMEM205,PTK2,AUNIF
7,GANC,DYNC2LI1,ZNF655,C12ORF10,SNRNP200,RPS14,ARHGEF7,ZRANB2,SEC14L1,UBA6,DPP9,SEC24C,
16,CPSF3L,TSG101,PSMA1,FUZ,FAM3A,PSMA3,METTL17,ECSIT,ATG12,RPS25,NLRP1,ZBTB44,NIP7,TTC9C
16,CPSF3L,TSG101,PSMA1,FUZ,FAM3A,PSMA3,METTL17,ECSIT,ATG12,RPS25,NLRP1,ZBTB44,NIP7,TTC9C

QCRQ,GRK6,HMGA2,CDC37,CPSF3L,TSG101,OBSL1,PTPN13,PSMA1,PHF21A,CHCHD1,FUZ,RGS3,METTL17
QCRQ,GRK6,HMGA2,CDC37,CPSF3L,TSG101,OBSL1,PTPN13,PSMA1,PHF21A,CHCHD1,FUZ,RGS3,METTL17
D5,CHID1,NUDT22,AMZ2,UFD1L,USP38,TAZ,THYN1,DBNDD1,SH2B1,NOP16,RPL13A,ARHGEF39,GSTK1,T
OT8,AP3B2,RPS9,SLC35G1,MRPL33,ZNF410,RUVBL2,EBPL,TESC,IGSF8,NUDT5,RBPMS,NNMT,WDR62,IL1F
OT8,AP3B2,RPS9,SLC35G1,MRPL33,ZNF410,RUVBL2,EBPL,TESC,IGSF8,NUDT5,RBPMS,NNMT,WDR62,IL1F
,PPP1CC,EDF1,RFX3,PPA1,DGKE,CAMTA1,CAPG,PPIL2,CLDND1,LIG1,TIMM17B,APEH,CAPN10,NUTF2,PO
L2,GGPS1,RTEL1,SRSF1,AUP1,YIPF3,FOXRED1,KXD1,ITGB1BP1,MED15,SH3YL1,CLASRP,HS2ST1,YIF1A,TB
L2,GGPS1,RTEL1,SRSF1,AUP1,YIPF3,FOXRED1,KXD1,ITGB1BP1,MED15,SH3YL1,CLASRP,HS2ST1,YIF1A,TB
CLASRP,JPH2,C1ORF198,UGGT2,MICAL1,HS2ST1,UBE2I,SLC39A13,PRMT2,JUP,TBX6,TMEM205,PTK2,AU
CLASRP,JPH2,C1ORF198,UGGT2,MICAL1,HS2ST1,UBE2I,SLC39A13,PRMT2,JUP,TBX6,TMEM205,PTK2,AU
1,CASC4,ZNF7,PPAN,ZFAND2B,ZNF250,RPL36,NFAM1,LAMP2,CCNL2,SYNGR2,CEP70,TMEM134,LRRFIP1,
1,CASC4,ZNF7,PPAN,ZFAND2B,ZNF250,RPL36,NFAM1,LAMP2,CCNL2,SYNGR2,CEP70,TMEM134,LRRFIP1,
1,CTTN,TSPAN11,ITSN1,TNFSF10,COMMD5,RUNX1,CHID1,NUDT22,AMZ2,UFD1L,USP38,TAZ,THYN1,SH2
1,CTTN,TSPAN11,ITSN1,TNFSF10,COMMD5,RUNX1,CHID1,NUDT22,AMZ2,UFD1L,USP38,TAZ,THYN1,SH2
RNPUL1,TP53,TMEM199,SNAPC2,EPN1,ITPA,ZCCHC9,SNRPD1,MKRN1,SMUG1,DHRS4,NDUFB3,AZI2,GOS

FOXRED1,RABL5,TRNT1,ITGB1BP1,PARD6G,CLASRP,C1ORF198,UGGT2,MICAL1,UBE2I,YIF1A,EIF3M,CDC4
FOXRED1,RABL5,TRNT1,ITGB1BP1,PARD6G,CLASRP,C1ORF198,UGGT2,MICAL1,UBE2I,YIF1A,EIF3M,CDC4
IF5A,COMMD5,RUNX1,CHID1,NUDT22,UFD1L,USP38,TAZ,THYN1,DBNDD1,SH2B1,NOP16,RPL13A,GSTK1
IF5A,COMMD5,RUNX1,CHID1,NUDT22,UFD1L,USP38,TAZ,THYN1,DBNDD1,SH2B1,NOP16,RPL13A,GSTK1
T3,CD44,SON,SNF8,GRK6,RPS16,TSG101,PTPN13,UBR4,PSMA1,YLPM1,CHCHD1,FUZ,RGS3,FAM3A,METT
T3,CD44,SON,SNF8,GRK6,RPS16,TSG101,PTPN13,UBR4,PSMA1,YLPM1,CHCHD1,FUZ,RGS3,FAM3A,METT
,LDHA,ACOT8,AP3B2,SLC35G1,MRPL33,ZNF410,RUVBL2,PCMT1,EBPL,TESC,IGSF8,NUDT5,RBPMS,NNMT
,LDHA,ACOT8,AP3B2,SLC35G1,MRPL33,ZNF410,RUVBL2,PCMT1,EBPL,TESC,IGSF8,NUDT5,RBPMS,NNMT
A,SLC39A11,FUZ,ECSIT,ATG12,ZBTB44,TTC9C,COX4I1,DUSP22,RDH11,DBNDD2,RPS6,RPL29,CD99L2,GGP
A,SLC39A11,FUZ,ECSIT,ATG12,ZBTB44,TTC9C,COX4I1,DUSP22,RDH11,DBNDD2,RPS6,RPL29,CD99L2,GGP
A,SLC39A11,FUZ,ECSIT,ATG12,ZBTB44,TTC9C,COX4I1,DUSP22,RDH11,DBNDD2,RPS6,RPL29,CD99L2,GGP
A,SLC39A11,FUZ,ECSIT,ATG12,ZBTB44,TTC9C,COX4I1,DUSP22,RDH11,DBNDD2,RPS6,RPL29,CD99L2,GGP
1,AUP1,TRNT1,PARD6G,SH3YL1,IP6K2,CLASRP,JPH2,MICAL1,HS2ST1,UBE2I,PRMT2,EIF3M,TBX6,XBP1,P
1,AUP1,TRNT1,PARD6G,SH3YL1,IP6K2,CLASRP,JPH2,MICAL1,HS2ST1,UBE2I,PRMT2,EIF3M,TBX6,XBP1,P
1,UFD1L,USP38,TAZ,THYN1,DBNDD1,SH2B1,NOP16,RPL13A,GSTK1,TTLL5,LDHA,ACOT8,AP3B2,SLC35G1,
CSIT,EPB41L5,ATG12,PPRC1,ZBTB44,PLCB2,DUSP22,DBNDD2,NISCH,TMED4,CD99L2,GGPS1,COASY,SRSF
CSIT,EPB41L5,ATG12,PPRC1,ZBTB44,PLCB2,DUSP22,DBNDD2,NISCH,TMED4,CD99L2,GGPS1,COASY,SRSF
B2,SLC35G1,MRPL33,ZNF410,RUVBL2,PCMT1,EBPL,TESC,IGSF8,NUDT5,RBPMS,NNMT,SNRPN,SPAG7,WD
LA,PRMT2,CDC42,AUNIP,C14ORF2,GUK1,PCYT2,AGPAT4,PPP1CC,EDF1,RFX3,PPA1,DGKE,CAMTA1,CRIP1,
K,RPS6,RPL29,RTEL1,COASY,SAP30BP,YIPF3,FOXRED1,RABL5,TRNT1,KXD1,MED15,SH3YL1,CLASRP,C1OR
K,RPS6,RPL29,RTEL1,COASY,SAP30BP,YIPF3,FOXRED1,RABL5,TRNT1,KXD1,MED15,SH3YL1,CLASRP,C1OR
AMD1C,MAGED2,SON,TET2,GRK6,CDC37,OBSL1,TTC21B,PTPN13,UBR4,YLPM1,PHF21A,SLC39A11,FUZ,F
AMD1C,MAGED2,SON,TET2,GRK6,CDC37,OBSL1,TTC21B,PTPN13,UBR4,YLPM1,PHF21A,SLC39A11,FUZ,F
12ORF57,NDUFS3,OXLD1,NUBP2,SLC35E1,OGFOD2,METTL3,TSEN34,PICK1,ZNF512,CDK4,NME1,SLC52A

S1,ITPA,GTF3A,ZBTB8OS,TBC1D22A,U2AF1L4,RNPS1,BTF3L4,SUMO1,ZNF483,SZT2,PSMD8,PSMC5,LRP1(
1,EBPL,TESC,NUDT5,RBPMS,NNMT,SNRPN,SPAG7,WDR62,IL1RAP,STYXL1,RFX5,RANBP1,UBE2L6,LZTFL1,
1,EBPL,TESC,NUDT5,RBPMS,NNMT,SNRPN,SPAG7,WDR62,IL1RAP,STYXL1,RFX5,RANBP1,UBE2L6,LZTFL1,
9C,DUSP22,SLC25A10,RDH11,DBNDD2,RTEL1,SRSF1,AUP1,RABL5,ITGB1BP1,MED15,PARD6G,C1ORF198,
9C,DUSP22,SLC25A10,RDH11,DBNDD2,RTEL1,SRSF1,AUP1,RABL5,ITGB1BP1,MED15,PARD6G,C1ORF198,
GSTK1,TTLL5,LDHA,ACOT8,AP3B2,SLC35G1,MRPL33,ZNF410,RUVBL2,PCMT1,EBPL,TESC,IGSF8,NUDT5,R
GSTK1,TTLL5,LDHA,ACOT8,AP3B2,SLC35G1,MRPL33,ZNF410,RUVBL2,PCMT1,EBPL,TESC,IGSF8,NUDT5,R
C1,PCBP2,TM9SF1,MAGED2,GRK6,CPSF3L,TSG101,OBSL1,PSMA1,YLPM1,PSMA3,MUC1,RPS25,NLRP1,NI
C1,PCBP2,TM9SF1,MAGED2,GRK6,CPSF3L,TSG101,OBSL1,PSMA1,YLPM1,PSMA3,MUC1,RPS25,NLRP1,NI
OGKE,TNFRSF14,CAMTA1,CRIP1,CLDND1,VIM,LIG1,CDC14B,TIMM17B,APEH,CAPN10,NUTF2,UCK2,ATPAI
EBPL,TESC,IGSF8,NUDT5,RBPMS,SNRPN,SPAG7,WDR62,IL1RAP,STYXL1,RFX5,POLD4,RANBP1,UBE2L6,VR
EBPL,TESC,IGSF8,NUDT5,RBPMS,SNRPN,SPAG7,WDR62,IL1RAP,STYXL1,RFX5,POLD4,RANBP1,UBE2L6,VR
GTF2H1,PIGO,AXIN1,BTF3,SLC25A24,EIF3C,DNAJC2,EPN1,HPS1,CREB3L2,ZCCHC9,AKT1S1,AIFM2,GTF3A,
GTF2H1,PIGO,AXIN1,BTF3,SLC25A24,EIF3C,DNAJC2,EPN1,HPS1,CREB3L2,ZCCHC9,AKT1S1,AIFM2,GTF3A,
,WDR62,IL1RAP,STYXL1,RFX5,POLD4,RANBP1,UBE2L6,VRK3,LZTFL1,ZC3H14,FIS1,SLC15A3,TUBA1C,IRF9,
,WDR62,IL1RAP,STYXL1,RFX5,POLD4,RANBP1,UBE2L6,VRK3,LZTFL1,ZC3H14,FIS1,SLC15A3,TUBA1C,IRF9,
GRK6,HMGA2,CDC37,TTC21B,UBR4,CHCHD1,SLC39A11,FUZ,RGS3,FAM3A,PSMA3,METTL17,ATG12,PPRC
2,SLC35G1,MRPL33,ZNF410,RUVBL2,PCMT1,EBPL,TESC,IGSF8,NUDT5,RBPMS,NNMT,SNRPN,WDR62,STY
K,RER1,NEK11,CREBZF,EIF2A,GLI4,SCN11A,RECQL5,PIGO,BFAR,THOC3,RPS3A,MOSPD3,HDAC8,EIF3C,TKT
WSG1,RPS15,HSPB1,LRRC27,MVD,SYNGR1,PABPC1,NAT9,TOMM40,ATP6V1F,WAC,PDCD5,ANAPC11,PO
WSG1,RPS15,HSPB1,LRRC27,MVD,SYNGR1,PABPC1,NAT9,TOMM40,ATP6V1F,WAC,PDCD5,ANAPC11,PO
K6,RPS16,CPSF3L,OBSL1,YLPM1,SLC39A11,FAM3A,PSMA3,MUC1,ECSIT,EPB41L5,ATG12,PPRC1,RPS25,N
K6,RPS16,CPSF3L,OBSL1,YLPM1,SLC39A11,FAM3A,PSMA3,MUC1,ECSIT,EPB41L5,ATG12,PPRC1,RPS25,N
,ACOT8,AP3B2,SLC35G1,MRPL33,ZNF410,RUVBL2,PCMT1,EBPL,TESC,IGSF8,NUDT5,RBPMS,NNMT,SNRP
HSPB1,ERCC5,C12ORF76,MEIS2,HMGA1,NAT9,RAD1,RBMX,PPP3CB,RABEP2,ATP6V1F,ZNF706,ZNF687,P
KL,TMPO,LSM12,ARL6IP4,PIGL,C19ORF48,STRADA,PPAN,ZFAND2B,SHARPIN,STAU1,UPP1,MOB1A,NFAM
KL,TMPO,LSM12,ARL6IP4,PIGL,C19ORF48,STRADA,PPAN,ZFAND2B,SHARPIN,STAU1,UPP1,MOB1A,NFAM
,THYN1,SH2B1,NOP16,RPL13A,ARHGEF39,GSTK1,TTLL5,LDHA,PSMA6,ACOT8,AP3B2,SLC35G1,RCE1,ZNF
,THYN1,SH2B1,NOP16,RPL13A,ARHGEF39,GSTK1,TTLL5,LDHA,PSMA6,ACOT8,AP3B2,SLC35G1,RCE1,ZNF
XR2,STIP1,LRRC1,PHF19,LPCAT3,CD44,TM9SF1,MAGED2,SON,TET2,UQCRQ,CDC37,CPSF3L,OBSL1,TTC21
XR2,STIP1,LRRC1,PHF19,LPCAT3,CD44,TM9SF1,MAGED2,SON,TET2,UQCRQ,CDC37,CPSF3L,OBSL1,TTC21
D4,GGPS1,RTEL1,SRSF1,AUP1,FOXRED1,TRNT1,ITGB1BP1,SH3YL1,IP6K2,CLASRP,MICAL1,HS2ST1,YIF1A,E
D4,GGPS1,RTEL1,SRSF1,AUP1,FOXRED1,TRNT1,ITGB1BP1,SH3YL1,IP6K2,CLASRP,MICAL1,HS2ST1,YIF1A,E
UQCRQ,GRK6,CDC37,RPS16,CPSF3L,TSG101,OBSL1,UBR4,PHF21A,SLC39A11,FAM3A,PSMA3,METTL17,P
UQCRQ,GRK6,CDC37,RPS16,CPSF3L,TSG101,OBSL1,UBR4,PHF21A,SLC39A11,FAM3A,PSMA3,METTL17,P
,UBE2L6,VRK3,ZC3H14,FIS1,PPM1K,TUBA1C,DST,ANTXR2,STIP1,PHF19,GRAMD1C,VCAN,LPCAT3,CD44,N
,UBE2L6,VRK3,ZC3H14,FIS1,PPM1K,TUBA1C,DST,ANTXR2,STIP1,PHF19,GRAMD1C,VCAN,LPCAT3,CD44,N
M1,PHF21A,SLC39A11,FAM3A,MUC1,METTL17,ECSIT,EPB41L5,ATG12,ZBTB44,PLCB2,DUSP22,SPOCK2,D
M1,PHF21A,SLC39A11,FAM3A,MUC1,METTL17,ECSIT,EPB41L5,ATG12,ZBTB44,PLCB2,DUSP22,SPOCK2,D
A1,GPATCH2L,BUD31,SAE1,NPRL2,OSGEP,TBL2,ARHGAP6,SCARB1,CRYGS,CTTN,TSPAN11,ITSN1,EIF5A,C
A1,GPATCH2L,BUD31,SAE1,NPRL2,OSGEP,TBL2,ARHGAP6,SCARB1,CRYGS,CTTN,TSPAN11,ITSN1,EIF5A,C
H3YL1,IP6K2,CLASRP,MICAL1,HS2ST1,UBE2I,JUP,TBX6,TMEM205,AUNIP,C14ORF2,GUK1,PCYT2,USE1,AS
BSL1,PSMA1,SLC39A11,FUZ,MUC1,RPS25,TTC9C,COX4I1,PLCB2,SLC25A10,TMX2,THEM4,SDF4,NISCH,CE
,RPL29,COASY,RABL5,TRNT1,PARD6G,SH3YL1,C1ORF198,UGGT2,MICAL1,SLC39A13,YIF1A,EIF3M,JUP,C
BPL,IGSF8,NUDT5,RBPMS,NNMT,WDR62,RFX5,POLD4,PLEKHA5,RANBP1,UBE2L6,VRK3,LZTFL1,ZC3H14,F
BPL,IGSF8,NUDT5,RBPMS,NNMT,WDR62,RFX5,POLD4,PLEKHA5,RANBP1,UBE2L6,VRK3,LZTFL1,ZC3H14,F
,RUVBL2,PCMT1,EBPL,TESC,IGSF8,SNRPN,SPAG7,WDR62,IL1RAP,STYXL1,RFX5,POLD4,RANBP1,UBE2L6,

3,RUVBL2,PCMT1,EBPL,TESC,IGSF8,SNRPN,SPAG7,WDR62,IL1RAP,STYXL1,RFX5,POLD4,RANBP1,UBE2L6,
.7,ZBTB44,NIP7,TTC9C,COX4I1,SLC25A10,NISCH,CENPK,RPL29,GGPS1,COASY,SRSF1,AUP1,YIPF3,FOXRED
MT1,TESC,IGSF8,NUDT5,RBPMS,SNRPN,SPAG7,IL1RAP,STYXL1,RFX5,PLEKHA5,UBE2L6,VRK3,LZTFL1,FIS1,
MT1,TESC,IGSF8,NUDT5,RBPMS,SNRPN,SPAG7,IL1RAP,STYXL1,RFX5,PLEKHA5,UBE2L6,VRK3,LZTFL1,FIS1,
TROAP,LIG1,APEH,RBM33,LYPLA2,PRMT1,STOML2,ZNF7,TXNDC17,PPAN,CYB561,SHARPIN,CBFA2T2,STA
TROAP,LIG1,APEH,RBM33,LYPLA2,PRMT1,STOML2,ZNF7,TXNDC17,PPAN,CYB561,SHARPIN,CBFA2T2,STA
CNST,COX17,HINT1,POLR2C,FTH1,MTHFS,SMARCA4,PAFAH1B2,TXNRD2,DALRD3,NSUN5,ZFPL1,MRPS18
CNST,COX17,HINT1,POLR2C,FTH1,MTHFS,SMARCA4,PAFAH1B2,TXNRD2,DALRD3,NSUN5,ZFPL1,MRPS18
F63,EPN1,HPS1,ITPA,ZCCHC9,MKRN1,GTF3A,CHURC1,AZI2,GOSR2,TBC1D22A,LAMTOR1,U2AF1L4
T1,KXD1,ITGB1BP1,MED15,PARD6G,UGGT2,HS2ST1,UBE2I,EIF3M,CDC42,TBX6,TMEM205,XBP1,C14ORF
GPBP1,SIAH1,ASPH,CHPF,STRIP2,LYNX1,LMBR1,NAP1L1,CHCHD7,HMGA1,PABPC1,NAT9,BCAM,RABEP2,
GPBP1,SIAH1,ASPH,CHPF,STRIP2,LYNX1,LMBR1,NAP1L1,CHCHD7,HMGA1,PABPC1,NAT9,BCAM,RABEP2,
D2B,MOB1A,NFAM1,CDK11A,LAMP2,COPZ1,SYNGR2,TMEM134,LRRFIP1,RAB2A,ZNF524,ELP6,SIAH1,AS
TP5J,ELP6,RPS15,ERLEC1,LRRC27,LMBR1,NAP1L1,DDOST,PSME2,RAD1,TOMM40,TADA3,ZNF20,WAC,SL
A1,GPATCH2L,SUMF2,BUD31,SAE1,NPRL2,OSGEP,TBL2,ARHGAP6,SCARB1,CTTN,TSPAN11,ITSN1,EIF5A,
XRED1,TRNT1,ITGB1BP1,PARD6G,SH3YL1,UGGT2,MICAL1,HS2ST1,UBE2I,EIF3M,CDC42,TBX6,TMEM205

YN1,DBNDD1,SH2B1,NOP16,RPL13A,ARHGEF39,GSTK1,TTLL5,LDHA,PSMA6,ACOT8,AP3B2,RPS9,SLC35G
YN1,DBNDD1,SH2B1,NOP16,RPL13A,ARHGEF39,GSTK1,TTLL5,LDHA,PSMA6,ACOT8,AP3B2,RPS9,SLC35G
LR2C,RPL15,INTS4,RFXANK,ZNF506,ASH2L,SMARCA4,COA3,MPC2,SNRPF,PTK7,PPIH,OARD1,MFSD10,MR
C9,UCHL5,COMT,CHURC1,AZI2,GOSR2,LAMTOR1,U2AF1L4,BCL7B,PRKCSH,UBA52,RPL30,ZNF483,SZT2,P
C9,UCHL5,COMT,CHURC1,AZI2,GOSR2,LAMTOR1,U2AF1L4,BCL7B,PRKCSH,UBA52,RPL30,ZNF483,SZT2,P
,PLEKHA5,RANBP1,VRK3,LZTFL1,ZC3H14,SLC15A3,TUBA1C,STIP1,LRRC1,PHF19,GRAMD1C,VCAN,LPCAT3
YIPF3,FOXRED1,RABL5,TRNT1,ITGB1BP1,PARD6G,CLASRP,C1ORF198,UGGT2,MICAL1,UBE2I,YIF1A,EIF3
YIPF3,FOXRED1,RABL5,TRNT1,ITGB1BP1,PARD6G,CLASRP,C1ORF198,UGGT2,MICAL1,UBE2I,YIF1A,EIF3
,MED15,SLC39A13,YIF1A,PRMT2,MAGT1,AUNIP,C14ORF2,GUK1,CENPN,PCYT2,FBXW9,HSBP1L1,AGPAT
,MED15,SLC39A13,YIF1A,PRMT2,MAGT1,AUNIP,C14ORF2,GUK1,CENPN,PCYT2,FBXW9,HSBP1L1,AGPAT
HD1,SLC39A11,FUZ,FAM3A,ATG12,PPRC1,COX4I1,PLCB2,DUSP22,TMX2,THEM4,NISCH,TMED4,CENPK,R
HD1,SLC39A11,FUZ,FAM3A,ATG12,PPRC1,COX4I1,PLCB2,DUSP22,TMX2,THEM4,NISCH,TMED4,CENPK,R
,LMBR1,NAP1L1,SERPINB1,RANGRF,CHCHD7,HMGA1,PSME2,PABPC1,DBNL,NAT9,BCAM,ZNF836,BLVRB
BR4,PSMA1,FUZ,FAM3A,METTL17,EPB41L5,PPRC1,RPS25,NIP7,COX4I1,DUSP22,SPOCK2,RDH11,DBNDD
BR4,PSMA1,FUZ,FAM3A,METTL17,EPB41L5,PPRC1,RPS25,NIP7,COX4I1,DUSP22,SPOCK2,RDH11,DBNDD
SF1,AUP1,SAP30BP,FOXRED1,TRNT1,KXD1,ITGB1BP1,MED15,UBE2I,SLC39A13,YIF1A,PRMT2,MAGT1,C1
SF1,AUP1,SAP30BP,FOXRED1,TRNT1,KXD1,ITGB1BP1,MED15,UBE2I,SLC39A13,YIF1A,PRMT2,MAGT1,C1
11,ITSN1,EIF5A,COMMD5,RUNX1,NUDT22,AMZ2,UFD1L,TAZ,THYN1,DBNDD1,SH2B1,RPL13A,ARHGEF39
11,ITSN1,EIF5A,COMMD5,RUNX1,NUDT22,AMZ2,UFD1L,TAZ,THYN1,DBNDD1,SH2B1,RPL13A,ARHGEF39
,CAMTA1,CRIP1,RPL28,CLDND1,LIG1,TIMM17B,APEH,CAPN10,NUTF2,PPOX,C17ORF62,PRMT1,ZNF7,AR

1,CARD14,ATF7,HSBP1L1,ASF1B,PPP1CC,EDF1,RFX3,ENDOV,PPIL2,PSMD4,VIM,LIG1,PSMA7,CAPN10,TM
,EPB41L5,ATG12,ZBTB44,DUSP22,THEM4,RDH11,DBNDD2,SRSF1,AUP1,SAP30BP,YIPF3,RABL5,ITGB1BP
,EPB41L5,ATG12,ZBTB44,DUSP22,THEM4,RDH11,DBNDD2,SRSF1,AUP1,SAP30BP,YIPF3,RABL5,ITGB1BP
SC9,WNK1,ZNF7,HNRNPC,ZNF644,STAU1,SUGT1,CDK11A,MDH1,COPZ1,INO80E,CCNL2,MAPKAPK5,LRR
C,ZNF644,STAU1,MOB1A,MDH1,CCNL2,MAPKAPK5,LRRFIP1,ZNF524,GPBP1,SIAH1,ZEB2,CHPF,STRIP2,RP
F19,VCAN,PCBP2,MAGED2,SON,TET2,GRK6,HMGA2,CPSF3L,OBSL1,PTPN13,UBR4,YLPM1,PHF21A,CHCH
F19,VCAN,PCBP2,MAGED2,SON,TET2,GRK6,HMGA2,CPSF3L,OBSL1,PTPN13,UBR4,YLPM1,PHF21A,CHCH

3,C1ORF63,ZNF396,COX19,HPS1,ITPA,VPS25,SNRPD1,NDUFB2,AZI2,TATDN1,SPG21,KAT8,NCBP2,UBA5

)3,C1ORF63,ZNF396,COX19,HPS1,ITPA,VPS25,SNRPD1,NDUFB2,AZI2,TATDN1,SPG21,KAT8,NCBP2,UBA5
1,UBE2L6,LZTFL1,ZC3H14,FIS1,SLC15A3,PPM1K,TUBA1C,IRF9,STIP1,LRRC1,PHF19,GRAMD1C,VCAN,LPCA
1,UBE2L6,LZTFL1,ZC3H14,FIS1,SLC15A3,PPM1K,TUBA1C,IRF9,STIP1,LRRC1,PHF19,GRAMD1C,VCAN,LPCA
KAPK5,LRRFIP1,RAB2A,ZNF524,ATP5J,GPBP1,TRMT112,CHPF,STRIP2,GABPB1,RPS15,ERLEC1,LRRC27,SAT
TSPAN31,ZNF593,GLYCTK,FAM131A,TRIM46,C11ORF57,RBM42,ZNF511,GTF2H1,PIGO,SNX29,THOC3,AX

C,VCAN,PCBP2,LPCAT3,CD44,SON,TET2,SNF8,HMGA2,CDC37,CPSF3L,TTC21B,UBR4,PSMA1,SLC39A11,FU
C,VCAN,PCBP2,LPCAT3,CD44,SON,TET2,SNF8,HMGA2,CDC37,CPSF3L,TTC21B,UBR4,PSMA1,SLC39A11,FU
9C,COX4I1,PLCB2,DUSP22,SLC25A10,DBNDD2,TMED4,CD99L2,GGPS1,RTEL1,COASY,SRSF1,TRNT1,PARD

SP22,SPOCK2,CENPK,RPL29,GGPS1,RTEL1,SRSF1,AUP1,FOXRED1,ITGB1BP1,SH3YL1,IP6K2,CLASRP,JPH2,U
SP22,SPOCK2,CENPK,RPL29,GGPS1,RTEL1,SRSF1,AUP1,FOXRED1,ITGB1BP1,SH3YL1,IP6K2,CLASRP,JPH2,U
,ZNF687,KLK10,TRIM16,PDE4DIP,HSPH1,CHCHD5,OAS3,SEC31A,CARHSP1,CNST,COX5A,HINT1,NDUFV2,F
,ZNF687,KLK10,TRIM16,PDE4DIP,HSPH1,CHCHD5,OAS3,SEC31A,CARHSP1,CNST,COX5A,HINT1,NDUFV2,F
DC37,CPSF3L,TTC21B,UBR4,PSMA1,FUZ,PSMA3,NLRP1,ZBTB44,DUSP22,SLC25A10,RDH11,NISCH,RPL29,
AF2,LSM12,EXOSC9,WNK1,ZNF7,TXNDC17,ACAD10,HNRNPC,CYB561,SHARPIN,SUGT1,CARS,ZNF250,MDH
EH,CAPN10,NUTF2,PODXL,CAMLG,EPB41L4B,TCF4,ATPAF2,C17ORF62,PRMT1,CASC4,ZNF7,PPAN,ZFAND
,RBPMS,SPAG7,WDR62,IL1RAP,STYXL1,RFX5,POLD4,PLEKHA5,RANBP1,UBE2L6,VRK3,ZC3H14,FIS1,SLC15
,RBPMS,SPAG7,WDR62,IL1RAP,STYXL1,RFX5,POLD4,PLEKHA5,RANBP1,UBE2L6,VRK3,ZC3H14,FIS1,SLC15
,GSTK1,TTLL5,LDHA,PSMA6,ACOT8,AP3B2,RPS9,SLC35G1,RCE1,MRPL33,ZNF410,RUVBL2,PCMT1,EBPL,T
,GSTK1,TTLL5,LDHA,PSMA6,ACOT8,AP3B2,RPS9,SLC35G1,RCE1,MRPL33,ZNF410,RUVBL2,PCMT1,EBPL,T
L,TESC,IGSF8,RBPMS,SPAG7,WDR62,IL1RAP,STYXL1,POLD4,PLEKHA5,UBE2L6,VRK3,ZC3H14,FIS1,SLC15A
L,TESC,IGSF8,RBPMS,SPAG7,WDR62,IL1RAP,STYXL1,POLD4,PLEKHA5,UBE2L6,VRK3,ZC3H14,FIS1,SLC15A
GPS1,ZNF687,PDCD5,NXN,EPS15L1,POLR3H,ABHD17A,ZCCHC4,STXBP2,CHCHD5,NAA10,C11ORF48,OAS
1K,TUBA1C,DST,ANTXR2,LRRC1,PHF19,GRAMD1C,CD44,MAGED2,SON,TET2,UQCRQ,GRK6,HMGA2,RPS1
PPIL2,PSMA7,TIMM17B,RPL21,TXNL4A,CAPN10,CAMLG,TMPO,EXOSC9,C19ORF54,ZNF7,TXNDC17,ZFAN
B2,SLC35G1,RCE1,MRPL33,ZNF410,RUVBL2,PCMT1,EBPL,IGSF8,NUDT5,RBPMS,SPAG7,WDR62,IL1RAP,S
B2,SLC35G1,RCE1,MRPL33,ZNF410,RUVBL2,PCMT1,EBPL,IGSF8,NUDT5,RBPMS,SPAG7,WDR62,IL1RAP,S
3,C1ORF63,RPS3,EPN1,HPS1,ITPA,SMUG1,EMC9,CHURC1,TATDN1,ZBTB8OS,PRKCSH,RNPS1,BTF3L4,SUM
3,C1ORF63,RPS3,EPN1,HPS1,ITPA,SMUG1,EMC9,CHURC1,TATDN1,ZBTB8OS,PRKCSH,RNPS1,BTF3L4,SUM

,XBP1,PTK2,AUNIP,GUK1,CENPN,ATF7,USE1,HSBP1L1,ASF1B,DEF8,EEF1B2,RFX3,TNFRSF14,ENDOV,CAM
,XBP1,PTK2,AUNIP,GUK1,CENPN,ATF7,USE1,HSBP1L1,ASF1B,DEF8,EEF1B2,RFX3,TNFRSF14,ENDOV,CAM
NTL1,KAT8,CARM1,CCDC24,MDK,CAMKK1,ZNF276,CCNG2,RABEPK,NUB1,MCUR1,TCFL5,PPP1R13B,TMEN
I1,PHF21A,CHCHD1,ECSIT,ATG12,PPRC1,RPS25,ZBTB44,NIP7,COX4I1,PLCB2,TMX2,RDH11,NISCH,CENPK,
I1,PHF21A,CHCHD1,ECSIT,ATG12,PPRC1,RPS25,ZBTB44,NIP7,COX4I1,PLCB2,TMX2,RDH11,NISCH,CENPK,
,COASY,SRSF1,AUP1,MED15,SH3YL1,IP6K2,UBE2I,YIF1A,PRMT2,TBX6,AUNIP,C14ORF2,CENPN,PCYT2,FB
,COASY,SRSF1,AUP1,MED15,SH3YL1,IP6K2,UBE2I,YIF1A,PRMT2,TBX6,AUNIP,C14ORF2,CENPN,PCYT2,FB
2,STIP1,LRRC1,PHF19,VCAN,LPCAT3,TM9SF1,MAGED2,SON,TET2,UQCRQ,GRK6,RPS16,OBSL1,PTPN13,U

EL1,SRSF1,AUP1,SAP30BP,FOXRED1,ITGB1BP1,PARD6G,SH3YL1,IP6K2,JPH2,HS2ST1,YIF1A,EIF3M,MAGT1
EL1,SRSF1,AUP1,SAP30BP,FOXRED1,ITGB1BP1,PARD6G,SH3YL1,IP6K2,JPH2,HS2ST1,YIF1A,EIF3M,MAGT1
RK6,HMGA2,CDC37,RPS16,CPSF3L,TSG101,OBSL1,TTC21B,PTPN13,PSMA1,PHF21A,CHCHD1,SLC39A11,F
RK6,HMGA2,CDC37,RPS16,CPSF3L,TSG101,OBSL1,TTC21B,PTPN13,PSMA1,PHF21A,CHCHD1,SLC39A11,F
AP30BP,YIPF3,RABL5,TRNT1,MED15,IP6K2,CLASRP,C1ORF198,UGGT2,UBE2I,YIF1A,JUP,CDC42,PTK2,GUK
AP30BP,YIPF3,RABL5,TRNT1,MED15,IP6K2,CLASRP,C1ORF198,UGGT2,UBE2I,YIF1A,JUP,CDC42,PTK2,GUK
M1,CHCHD1,SLC39A11,RGS3,PSMA3,RPS25,NIP7,COX4I1,SPOCK2,THEM4,RDH11,NISCH,RPS6,RPL29,CD9

M1,CHCHD1,SLC39A11,RGS3,PSMA3,RPS25,NIP7,COX4I1,SPOCK2,THEM4,RDH11,NISCH,RPS6,RPL29,CD9

PARD6G,SH3YL1,JPH2,UGGT2,MICAL1,HS2ST1,UBE2I,SLC39A13,PRMT2,MAGT1,JUP,CDC42,TBX6,TMEM2
PARD6G,SH3YL1,JPH2,UGGT2,MICAL1,HS2ST1,UBE2I,SLC39A13,PRMT2,MAGT1,JUP,CDC42,TBX6,TMEM2
1B,YLPM1,PHF21A,SLC39A11,PSMA3,MUC1,ATG12,PPRC1,ZBTB44,NIP7,TTC9C,COX4I1,DUSP22,SPOCK2,
1B,YLPM1,PHF21A,SLC39A11,PSMA3,MUC1,ATG12,PPRC1,ZBTB44,NIP7,TTC9C,COX4I1,DUSP22,SPOCK2,
PRC1,RPS25,DUSP22,THEM4,RDH11,DBNDD2,CD99L2,RTEL1,SRSF1,ITGB1BP1,PARD6G,SH3YL1,CLASRP,JF
21B,YLPM1,PHF21A,SLC39A11,FAM3A,MUC1,EPB41L5,ATG12,PPRC1,ZBTB44,PLCB2,DUSP22,SLC25A10,
LEKHA5,RANBP1,VRK3,LZTFL1,FIS1,SLC15A3,PPM1K,TUBA1C,STIP1,GRAMD1C,LPCAT3,CD44,TM9SF1,TE
LEKHA5,RANBP1,VRK3,LZTFL1,FIS1,SLC15A3,PPM1K,TUBA1C,STIP1,GRAMD1C,LPCAT3,CD44,TM9SF1,TE
D15,MICAL1,SLC39A13,YIF1A,PRMT2,MAGT1,AUNIP,C14ORF2,GUK1,CENPN,PCYT2,FBXW9,AGPAT4,IFI2
D15,MICAL1,SLC39A13,YIF1A,PRMT2,MAGT1,AUNIP,C14ORF2,GUK1,CENPN,PCYT2,FBXW9,AGPAT4,IFI2

L1,UGGT2,MICAL1,HS2ST1,UBE2I,SLC39A13,PRMT2,TMEM205,GUK1,ATF7,FBXW9,USE1,AGPAT4,MAF,A
NL2,HSP90AB1,DUT,SYNGR2,CEP70,TMEM134,LRRFIP1,RAB2A,ZNF524,TPM2,SIAH1,ZEB2,TRMT112,CHF
FIP1,RAB2A,ZNF524,TPM2,GPBP1,SIAH1,ZEB2,STRIP2,ZNF625,RPS20,LYNX1,MPDU1,TWSG1,RPS15,ERLE
CPSF3L,TSG101,OBSL1,PTPN13,YLPM1,FUZ,PSMA3,ECSIT,EPB41L5,NLRP1,ZBTB44,NIP7,COX4I1,PLCB2,SL
YN1,DBNDD1,SH2B1,NOP16,RPL13A,GSTK1,TTLL5,LDHA,ACOT8,AP3B2,SLC35G1,MRPL33,ZNF410,RUVBI
YN1,DBNDD1,SH2B1,NOP16,RPL13A,GSTK1,TTLL5,LDHA,ACOT8,AP3B2,SLC35G1,MRPL33,ZNF410,RUVBI

MRPL33,ZNF410,PCMT1,EBPL,IGSF8,RBPMS,NNMT,SNRPN,SPAG7,WDR62,IL1RAP,STYXL1,RFX5,PLEKHA5
MRPL33,ZNF410,PCMT1,EBPL,IGSF8,RBPMS,NNMT,SNRPN,SPAG7,WDR62,IL1RAP,STYXL1,RFX5,PLEKHA5
M9SF1,TET2,SNF8,CPSF3L,OBSL1,TTC21B,UBR4,PSMA1,YLPM1,PHF21A,SLC39A11,MUC1,ECSIT,EPB41L5,
AM1,MDH1,LAMP2,HSP90AB1,DUT,SYNGR2,TMEM134,LRRFIP1,RAB2A,ZNF524,TPM2,GPBP1,SIAH1,ZEB
GDH,LYPLAL1,THOC3,SH3GL1,BTF3,NDUFA3,C1ORF63,C5ORF24,SLC37A4,IDH3G,DNAJA3,CKLF,AKT1S1,IN
GDH,LYPLAL1,THOC3,SH3GL1,BTF3,NDUFA3,C1ORF63,C5ORF24,SLC37A4,IDH3G,DNAJA3,CKLF,AKT1S1,IN
ND2B,HNRNPC,CYB561,ZNF250,RPL36,HSP90AB1,DUT,SYNGR2,TMEM134,LRRFIP1,RAB2A,ZNF524,ATP5J
EIF2A,RECQL5,ZNF511,TRAPPC2,TAF1D,PIGO,SLC35A2,HNRNPUL1,TP53,RPS3A,TMEM199,SNAPC2,EPN1,
1,AUP1,FOXRED1,RABL5,TRNT1,ITGB1BP1,PARD6G,SH3YL1,JPH2,UGGT2,MICAL1,PRMT2,EIF3M,MAGT1,
2,DBNDD2,NISCH,TMED4,GGPS1,RTEL1,RABL5,TRNT1,PARD6G,SH3YL1,CLASRP,C1ORF198,UGGT2,MICA
2,DBNDD2,NISCH,TMED4,GGPS1,RTEL1,RABL5,TRNT1,PARD6G,SH3YL1,CLASRP,C1ORF198,UGGT2,MICA
THYN1,DBNDD1,SH2B1,NOP16,ARHGEF39,GSTK1,TTLL5,LDHA,PSMA6,ACOT8,AP3B2,SLC35G1,RCE1,ZNF
THYN1,DBNDD1,SH2B1,NOP16,ARHGEF39,GSTK1,TTLL5,LDHA,PSMA6,ACOT8,AP3B2,SLC35G1,RCE1,ZNF
IF2B3,MFSD11,MOK,ZNF81,TMEM14B,ZXDC,SMIM4,RPL37A,ELOF1,DCTN5,ARL16,LY6E,TMEM108,RPL8,
IF2B3,MFSD11,MOK,ZNF81,TMEM14B,ZXDC,SMIM4,RPL37A,ELOF1,DCTN5,ARL16,LY6E,TMEM108,RPL8,

3,FOXRED1,TRNT1,KXD1,ITGB1BP1,PARD6G,SH3YL1,CLASRP,C1ORF198,UGGT2,HS2ST1,KCNMA1,SLC39A
3,FOXRED1,TRNT1,KXD1,ITGB1BP1,PARD6G,SH3YL1,CLASRP,C1ORF198,UGGT2,HS2ST1,KCNMA1,SLC39A
PAT4,MAF,EEF1B2,PPA1,TNFRSF14,VAMP8,CAMTA1,PSMD4,RPL28,PSMA7,TIMM17B,RPL21,CAPN10,CA
PAT4,MAF,EEF1B2,PPA1,TNFRSF14,VAMP8,CAMTA1,PSMD4,RPL28,PSMA7,TIMM17B,RPL21,CAPN10,CA
,CDK11A,SYNGR2,LRRFIP1,RAB2A,ZNF524,ATP5J,SIAH1,RPTOR,STRIP2,LYNX1,MPDU1,TWSG1,ERLEC1,GN
,CDK11A,SYNGR2,LRRFIP1,RAB2A,ZNF524,ATP5J,SIAH1,RPTOR,STRIP2,LYNX1,MPDU1,TWSG1,ERLEC1,GN

PSF3L,TSG101,TTC21B,UBR4,PSMA1,YLPM1,SLC39A11,FUZ,FAM3A,PSMA3,METTL17,EPB41L5,ATG12,NI

PSF3L,TSG101,TTC21B,UBR4,PSMA1,YLPM1,SLC39A11,FUZ,FAM3A,PSMA3,METTL17,EPB41L5,ATG12,NI

EKHA5,RANBP1,UBE2L6,FIS1,PPM1K,TUBA1C,DST,ANTXR2,STIP1,LRRC1,PCBP2,LPCAT3,CD44,TM9SF1,SC

EKHA5,RANBP1,UBE2L6,FIS1,PPM1K,TUBA1C,DST,ANTXR2,STIP1,LRRC1,PCBP2,LPCAT3,CD44,TM9SF1,SC

PC11,NT5C,MDH2,STXBP2,SEC31A,CARHSP1,SPINT2,MTHFS,S100A4,TXNRD2,TYMS,GBA2,TMEM39B,DA

D5,C11ORF48,SEC31A,RPL15,B3GAT3,RFXANK,GTF2IRD2,DALRD3,NSUN5,ZFPL1,MRPS18C,GMPR2,ZSCA

C14ORF2,PCYT2,USE1,HSBP1L1,AGPAT4,MAF,CCDC25,EEF1B2,EDF1,PPA1,DGKE,CAMTA1,PPIL2,RPL28,LI

GUK1,CENPN,CARD14,FBXW9,CCDC25,PPP1CC,GADL1,RFX3,ENDOV,PSMD4,VIM,PSMA7,CDC14B,TMEM

GUK1,CENPN,CARD14,FBXW9,CCDC25,PPP1CC,GADL1,RFX3,ENDOV,PSMD4,VIM,PSMA7,CDC14B,TMEM

,NUDT5,NNMT,SNRPN,SPAG7,WDR62,IL1RAP,STYXL1,RFX5,POLD4,RANBP1,TUBA1C,DST,STIP1,GRAMD1

,NUDT5,NNMT,SNRPN,SPAG7,WDR62,IL1RAP,STYXL1,RFX5,POLD4,RANBP1,TUBA1C,DST,STIP1,GRAMD1

C9,WNK1,C19ORF54,ZNF7,ARL6IP4,STRADA,AP1S3,ACAD10,ZNF644,SLC9B2,HYPK,UPP1,ZNF250,MOB1A

ASPH,TRMT112,STRIP2,NIT2,LYNX1,TWSG1,LRRC27,MVD,LMBR1,NAP1L1,SERPINB1,RANGRF,CHCHD7,HN

ASPH,TRMT112,STRIP2,NIT2,LYNX1,TWSG1,LRRC27,MVD,LMBR1,NAP1L1,SERPINB1,RANGRF,CHCHD7,HN

,UGGT2,UBE2I,YIF1A,JUP,CDC42,PTK2,GUK1,CENPN,PCYT2,USE1,HSBP1L1,EEF1B2,EDF1,DGKE,TNFRSF14

M,TBX6,TMEM205,C14ORF2,ATF7,PCYT2,USE1,AGPAT4,CCDC25,EEF1B2,PPA1,ENDOV,PSMD4,PSMA7,TIM

M,TBX6,TMEM205,C14ORF2,ATF7,PCYT2,USE1,AGPAT4,CCDC25,EEF1B2,PPA1,ENDOV,PSMD4,PSMA7,TIM

MT1,CASC4,ZNF7,ZFAND2B,CYB561,UPP1,RPL36,LAMP2,SYNGR2,TMEM134,LRRFIP1,RAB2A,ZNF524,TPM

3YL1,IP6K2,CLASRP,JPH2,C1ORF198,MICAL1,HS2ST1,UBE2I,PRMT2,JUP,TBX6,TMEM205,PTK2,AUNIP,C14

3YL1,IP6K2,CLASRP,JPH2,C1ORF198,MICAL1,HS2ST1,UBE2I,PRMT2,JUP,TBX6,TMEM205,PTK2,AUNIP,C14

A1C,IRF9,DST,STIP1,LRRC1,PHF19,VCAN,PCBP2,LPCAT3,CD44,TM9SF1,MAGED2,SON,UQCRQ,SNF8,GRK6

A1C,IRF9,DST,STIP1,LRRC1,PHF19,VCAN,PCBP2,LPCAT3,CD44,TM9SF1,MAGED2,SON,UQCRQ,SNF8,GRK6

R4,YLPM1,CHCHD1,FUZ,FAM3A,METTL17,ECSIT,EPB41L5,ATG12,PPRC1,ZBTB44,NIP7,PLCB2,SLC25A10,T

R4,YLPM1,CHCHD1,FUZ,FAM3A,METTL17,ECSIT,EPB41L5,ATG12,PPRC1,ZBTB44,NIP7,PLCB2,SLC25A10,T

STYXL1,UBE2L6,PPM1K,TUBA1C,IRF9,DST,LRRC1,PHF19,GRAMD1C,VCAN,PCBP2,LPCAT3,SON,UQCRQ,SN

STYXL1,UBE2L6,PPM1K,TUBA1C,IRF9,DST,LRRC1,PHF19,GRAMD1C,VCAN,PCBP2,LPCAT3,SON,UQCRQ,SN

OXRED1,ITGB1BP1,SH3YL1,IP6K2,JPH2,HS2ST1,MAGT1,CDC42,TBX6,TMEM205,XBP1,AUNIP,GUK1,CENPN

OXRED1,ITGB1BP1,SH3YL1,IP6K2,JPH2,HS2ST1,MAGT1,CDC42,TBX6,TMEM205,XBP1,AUNIP,GUK1,CENPN

OR62,IL1RAP,STYXL1,RFX5,POLD4,PLEKHA5,RANBP1,UBE2L6,VRK3,LZTFL1,ZC3H14,FIS1,SLC15A3,PPM1K,

OR62,IL1RAP,STYXL1,RFX5,POLD4,PLEKHA5,RANBP1,UBE2L6,VRK3,LZTFL1,ZC3H14,FIS1,SLC15A3,PPM1K,

04,TCF25,ZNF248,ANAPC13,SRSF7,NXF1,SYP,TRIM46,NUP62,PSPC1,NADSYN1,FCHO2,IRF3,IFT172,D2HGD

04,TCF25,ZNF248,ANAPC13,SRSF7,NXF1,SYP,TRIM46,NUP62,PSPC1,NADSYN1,FCHO2,IRF3,IFT172,D2HGD

14ORF2,GUK1,CARD14,PCYT2,HSBP1L1,ASF1B,CCDC25,EEF1B2,PPA1,CAMTA1,PSMD4,RPL28,LIG1,TIMM

P3,EEF1D,NAA10,OAS3,CHEK1,COX17,HNRNPLL,SUMO3,HINT1,HIGD1A,INTS4,B3GAT3,ASH2L,SMARCA4,

NUTF2,CAMLG,EPB41L4B,UCK2,ATPAF2,LSM12,C17ORF62,PRMT1,CASC4,C19ORF54,ZNF7,STRADA,AP1S3

H14,FIS1,SLC15A3,PPM1K,TUBA1C,IRF9,ANTXR2,STIP1,LRRC1,PHF19,LPCAT3,CD44,TM9SF1,MAGED2,SC

H14,FIS1,SLC15A3,PPM1K,TUBA1C,IRF9,ANTXR2,STIP1,LRRC1,PHF19,LPCAT3,CD44,TM9SF1,MAGED2,SC

2,PPP1CC,EDF1,RFX3,PPA1,DGKE,VAMP8,CAMTA1,CRIP1,LIG1,TIMM17B,APEH,HYI,CAPN10,NUTF2,POD

IED4,CENPK,COASY,SRSF1,AUP1,SAP30BP,FOXRED1,TRNT1,KXD1,ITGB1BP1,MED15,SLC39A13,YIF1A,PRM

IED4,CENPK,COASY,SRSF1,AUP1,SAP30BP,FOXRED1,TRNT1,KXD1,ITGB1BP1,MED15,SLC39A13,YIF1A,PRM

MD1C,VCAN,LPCAT3,TM9SF1,UQCRQ,HMGA2,CPSF3L,OBSL1,UBR4,PHF21A,SLC39A11,FAM3A,MUC1,EP

MD1C,VCAN,LPCAT3,TM9SF1,UQCRQ,HMGA2,CPSF3L,OBSL1,UBR4,PHF21A,SLC39A11,FAM3A,MUC1,EP

RRC27,MVD,LMBR1,RANGRF,CHCHD7,PSME2,PHTF2,PABPC1,NAT9,BCAM,ZNF836,BLVRB,RABEP2,RALG

,RTEL1,SRSF1,AUP1,YIPF3,FOXRED1,TRNT1,ITGB1BP1,SH3YL1,IP6K2,C1ORF198,MICAL1,PRMT2,EIF3M,T

JDT5,RBPMS,SPAG7,WDR62,IL1RAP,STYXL1,PLEKHA5,RANBP1,UBE2L6,ZC3H14,FIS1,SLC15A3,TUBA1C,A
JDT5,RBPMS,SPAG7,WDR62,IL1RAP,STYXL1,PLEKHA5,RANBP1,UBE2L6,ZC3H14,FIS1,SLC15A3,TUBA1C,A
PL36,HSP90AB1,SYNGR2,TMEM134,LRRFIP1,RAB2A,ZNF524,TPM2,SIAH1,ZEB2,NIT2,LYNX1,TWSG1,ZSWI

,COX4I1,DUSP22,SPOCK2,DBNDD2,NISCH,GGPS1,COASY,AUP1,SAP30BP,TRNT1,PARD6G,IP6K2,CLASRP,L
RALGPS1,ZNF687,PDCD5,NXN,KLK10,HSD17B10,ABHD17A,GGT5,DDX41,SLC16A1,ZCCHC4,STXBP2,C11OF
RALGPS1,ZNF687,PDCD5,NXN,KLK10,HSD17B10,ABHD17A,GGT5,DDX41,SLC16A1,ZCCHC4,STXBP2,C11OF
.1,FKBP3,B3GAT3,TCTN1,GTF2IRD2,COPS4,TPM4,SLC35B1,SLC30A5,THADA,MROH1,SNX5,NUP133,TMEN
JT,SYNGR2,CEP70,TMEM134,LRRFIP1,TPM2,GPBP1,SIAH1,ASPH,CHPF,STRIP2,TWSG1,LMBR1,NAP1L1,CH
JT,SYNGR2,CEP70,TMEM134,LRRFIP1,TPM2,GPBP1,SIAH1,ASPH,CHPF,STRIP2,TWSG1,LMBR1,NAP1L1,CH
!,ATF7,FBXW9,IFI27L1,MAF,CCDC25,GADL1,PPA1,DGKE,CAPG,CRIP1,PSMD4,RPL28,VIM,LIG1,PSMA7,TIM
!,ATF7,FBXW9,IFI27L1,MAF,CCDC25,GADL1,PPA1,DGKE,CAPG,CRIP1,PSMD4,RPL28,VIM,LIG1,PSMA7,TIM
2B1,NOP16,RPL13A,ARHGEF39,GSTK1,TTLL5,LDHA,ACOT8,AP3B2,SLC35G1,RCE1,MRPL33,ZNF410,RUVBI
2B1,NOP16,RPL13A,ARHGEF39,GSTK1,TTLL5,LDHA,ACOT8,AP3B2,SLC35G1,RCE1,MRPL33,ZNF410,RUVBI
,DST,ANTXR2,STIP1,LRRC1,PCBP2,LPCAT3,CD44,TM9SF1,SON,TET2,SNF8,GRK6,HMGA2,CDC37,CPSF3L,T
D23A,ABHD17A,ZCCHC4,STXBP2,EEF1D,CHCHD5,NAA10,C11ORF48,ATP10D,LLGL2,SUMO3,FKBP3,EGLN2
1L1,MAF,CCDC25,DEF8,DGKE,PPIL2,PSMD4,RPL28,LIG1,TIMM17B,CAPN10,NUTF2,CAMLG,C17ORF62,CA
MVD,SERPINB1,SYNGR1,CHCHD7,MEIS2,PHTF2,RAD1,TOMM40,SERPINB6,WAC,ZNF706,GTF2IRD1,NXN,I
KLHL7,SSR3,ARIH2,PICK1,UBE2D4,ZNF512,OCIAD1,TCF25,SLMAP,RBBP6,ZNF248,TPT1,BAX,CDK4,TMEM
,UFD1L,USP38,TAZ,THYN1,DBNDD1,SH2B1,NOP16,RPL13A,ARHGEF39,GSTK1,LDHA,PSMA6,ACOT8,SLC3!
,UFD1L,USP38,TAZ,THYN1,DBNDD1,SH2B1,NOP16,RPL13A,ARHGEF39,GSTK1,LDHA,PSMA6,ACOT8,SLC3!
OX4I1,SLC25A10,RDH11,NISCH,TMED4,CENPK,RPS6,RPL29,CD99L2,GGPS1,COASY,SRSF1,AUP1,FOXRED1
AGPAT4,MAF,PPP1CC,RFX3,PPA1,PPIL2,RPL21,TXNL4A,LSM12,PRMT1,EXOSC9,ARL6IP4,C19ORF48,ZFANI
AGPAT4,MAF,PPP1CC,RFX3,PPA1,PPIL2,RPL21,TXNL4A,LSM12,PRMT1,EXOSC9,ARL6IP4,C19ORF48,ZFANI
,SAP30BP,TRNT1,KXD1,MED15,SH3YL1,IP6K2,CLASRP,HS2ST1,KCNMA1,CDC42,TMEM205,XBP1,CELF1,A
,SAP30BP,TRNT1,KXD1,MED15,SH3YL1,IP6K2,CLASRP,HS2ST1,KCNMA1,CDC42,TMEM205,XBP1,CELF1,A
PRMT1,C19ORF54,C19ORF48,ZNF644,SUGT1,HYPK,MOB1A,RBM3,MDH1,LAMP2,INO80E,CCNL2,HSP90A
PRMT1,C19ORF54,C19ORF48,ZNF644,SUGT1,HYPK,MOB1A,RBM3,MDH1,LAMP2,INO80E,CCNL2,HSP90A
1,DGKE,TNFRSF14,ENDOV,PPIL2,LIG1,PSMA7,CDC14B,RPL21,TCF4,TMPO,ATPAF2,C17ORF62,PRMT1,CAS
1,ZEB2,NIT2,LYNX1,TWSG1,LRRC27,MVD,LMBR1,NAP1L1,SERPINB1,RANGRF,CHCHD7,HMGA1,PSME2,P
VNK1,ZNF7,ARL6IP4,ACAD10,SHARPIN,STAU1,CDK11A,DUT,SYNGR2,TMEM134,LRRFIP1,RAB2A,ZNF524,(
VNK1,ZNF7,ARL6IP4,ACAD10,SHARPIN,STAU1,CDK11A,DUT,SYNGR2,TMEM134,LRRFIP1,RAB2A,ZNF524,(
2B,CYB561,STAU1,UPP1,ZNF250,RPL36,CCNL2,SYNGR2,CEP70,TMEM134,LRRFIP1,RAB2A,ZNF524,ATP5J
8,MICAL1,HS2ST1,EIF3M,MAGT1,CDC42,TMEM205,XBP1,PTK2,AUNIP,GUK1,ATF7,FBXW9,IFI27L1,MAF,C
8,MICAL1,HS2ST1,EIF3M,MAGT1,CDC42,TMEM205,XBP1,PTK2,AUNIP,GUK1,ATF7,FBXW9,IFI27L1,MAF,C
1RAP,PLEKHA5,RANBP1,UBE2L6,VRK3,LZTFL1,IRF9,ANTXR2,STIP1,PHF19,GRAMD1C,CD44,TM9SF1,SON,L
1RAP,PLEKHA5,RANBP1,UBE2L6,VRK3,LZTFL1,IRF9,ANTXR2,STIP1,PHF19,GRAMD1C,CD44,TM9SF1,SON,L
35G1,RCE1,MRPL33,ZNF410,RUVBL2,PCMT1,EBPL,IGSF8,NUDT5,RBPMS,SPAG7,WDR62,IL1RAP,STYXL1,P
35G1,RCE1,MRPL33,ZNF410,RUVBL2,PCMT1,EBPL,IGSF8,NUDT5,RBPMS,SPAG7,WDR62,IL1RAP,STYXL1,P
,EIF3C,DENND1A,EML3,DNAJA3,NSFL1C,SMUG1,TRAK1,RBM6,NDUFB2,MFGE8,AZI2,U2AF1L4,KIF13B,PC
,EIF3C,DENND1A,EML3,DNAJA3,NSFL1C,SMUG1,TRAK1,RBM6,NDUFB2,MFGE8,AZI2,U2AF1L4,KIF13B,PC
MEM126B,HYI,C17ORF62,CASC4,STOML2,WNK1,C19ORF54,ZFAND2B,STAU1,SLC9B2,SUGT1,HYPK,CARS,
IF687,POT1,TRIM16,RAD23A,PKP4,EEF1D,HSPH1,MT1E,RPL18,CHEK1,SUMO3,PHF20L1,ZFAND4,HNRNPA
(N1,DBNDD1,SH2B1,RPL13A,ARHGEF39,GSTK1,LDHA,PSMA6,ACOT8,AP3B2,RPS9,SLC35G1,RCE1,MRPL3
(N1,DBNDD1,SH2B1,RPL13A,ARHGEF39,GSTK1,LDHA,PSMA6,ACOT8,AP3B2,RPS9,SLC35G1,RCE1,MRPL3

ZNF524,ASPH,PNPO,NIT2,TWSG1,MAPK13,VPS28,ZSWIM7,DCN,DBNL,NAT9,BCAM,TOMM40,SERPINB6,
ZNF524,ASPH,PNPO,NIT2,TWSG1,MAPK13,VPS28,ZSWIM7,DCN,DBNL,NAT9,BCAM,TOMM40,SERPINB6,

,PSMA7,CDC14B,APEH,TXNL4A,CAPN10,NUTF2,EPB41L4B,UCK2,TMPO,ATPAF2,LSM12,PRMT1,CASC4,W
HOC3,SH3GL1,BTF3,NDUFA3,C1ORF63,SLC37A4,IDH3G,DNAJA3,CKLF,AKT1S1,GTF3A,ARRB2,DIP2A,MLS
HOC3,SH3GL1,BTF3,NDUFA3,C1ORF63,SLC37A4,IDH3G,DNAJA3,CKLF,AKT1S1,GTF3A,ARRB2,DIP2A,MLS
OLD4,RANBP1,UBE2L6,LZTFL1,ZC3H14,FIS1,PPM1K,TUBA1C,ANTXR2,STIP1,VCAN,LPCAT3,MAGED2,SON
OLD4,RANBP1,UBE2L6,LZTFL1,ZC3H14,FIS1,PPM1K,TUBA1C,ANTXR2,STIP1,VCAN,LPCAT3,MAGED2,SON
1,DCTN5,ARL16,RPL8,UBE2D4,ASH1L,TCF25,ZNF248,ANAPC13,SRSF7,NXF1,SYP,TRIM46,EIF2A,NUP62,P
1,DCTN5,ARL16,RPL8,UBE2D4,ASH1L,TCF25,ZNF248,ANAPC13,SRSF7,NXF1,SYP,TRIM46,EIF2A,NUP62,P
M17B,TMEM126B,RBM33,LYPLA2,UCK2,C17ORF62,STOML2,ZNF7,PPAN,ZFAND2B,HNRNPC,CYB561,CBF
M17B,TMEM126B,RBM33,LYPLA2,UCK2,C17ORF62,STOML2,ZNF7,PPAN,ZFAND2B,HNRNPC,CYB561,CBF
4,DYNC1I2,CAMKMT,CRYZL1,RPS15A,ATP6V0B,MPND,TMEM53,IQCK,RNMT,DHX30,MTHFD2L,MRPL27,
4,DYNC1I2,CAMKMT,CRYZL1,RPS15A,ATP6V0B,MPND,TMEM53,IQCK,RNMT,DHX30,MTHFD2L,MRPL27,
PIL2,TIMM17B,RPL21,TXNL4A,UCK2,PRMT1,EXOSC9,WNK1,TXNDC17,ARL6IP4,C19ORF48,ZFAND2B,SH
PIL2,TIMM17B,RPL21,TXNL4A,UCK2,PRMT1,EXOSC9,WNK1,TXNDC17,ARL6IP4,C19ORF48,ZFAND2B,SH
13,YIF1A,PRMT2,MAGT1,C14ORF2,GUK1,CENPN,CARD14,PCYT2,FBXW9,HSBP1L1,AGPAT4,IFI27L1,MAF,
13,YIF1A,PRMT2,MAGT1,C14ORF2,GUK1,CENPN,CARD14,PCYT2,FBXW9,HSBP1L1,AGPAT4,IFI27L1,MAF,
2,MCFD2,RIC8A,CBR1,PHC3,CCDC40,MROH1,MFSD11,SNX5,ENY2,FAM32A,HAUS4,PCID2,ZXDC,PEX2,TN
2,MCFD2,RIC8A,CBR1,PHC3,CCDC40,MROH1,MFSD11,SNX5,ENY2,FAM32A,HAUS4,PCID2,ZXDC,PEX2,TN
1L5,RPS25,ZBTB44,TTC9C,COX4I1,DUSP22,SLC25A10,THEM4,DBNDD2,SDF4,NISCH,RTEL1,AUP1,YIPF3,
L5,PSMA6,ACOT8,AP3B2,RPS9,SLC35G1,MRPL33,RUVBL2,PCMT1,EBPL,RBPMS,NNMT,SNRPN,SPAG7,W
L5,PSMA6,ACOT8,AP3B2,RPS9,SLC35G1,MRPL33,RUVBL2,PCMT1,EBPL,RBPMS,NNMT,SNRPN,SPAG7,W
,STYXL1,RFX5,POLD4,PLEKHA5,RANBP1,UBE2L6,VRK3,LZTFL1,FIS1,TUBA1C,DST,STIP1,LRRC1,PHF19,VC
APC13,SMAGP,TMEM9,NT5E,GLYCTK,TCEAL3,CLASP2,GLI4,ZNF675,TAF1D,BFAR,CC2D1B,PLAC8,NDUFA
,PPRC1,NLRP1,ZBTB44,NIP7,TTC9C,COX4I1,PLCB2,DUSP22,SLC25A10,TMX2,DBNDD2,NISCH,CD99L2,GG

ARD6G,SH3YL1,IP6K2,CLASRP,JPH2,C1ORF198,MICAL1,HS2ST1,UBE2I,SLC39A13,YIF1A,PRMT2,TMEM20
ARD6G,SH3YL1,IP6K2,CLASRP,JPH2,C1ORF198,MICAL1,HS2ST1,UBE2I,SLC39A13,YIF1A,PRMT2,TMEM20

CAMTA1,VIM,LIG1,PSMA7,XAF1,APEH,HYI,CAPN10,PODXL,EPB41L4B,UCK2,ATPAF2,C17ORF62,PRMT1,C
CAMTA1,VIM,LIG1,PSMA7,XAF1,APEH,HYI,CAPN10,PODXL,EPB41L4B,UCK2,ATPAF2,C17ORF62,PRMT1,C

DC42,TMEM205,XBP1,CELF1,AUNIP,C14ORF2,GUK1,CENPN,USE1,HSBP1L1,EEF1B2,EDF1,DGKE,TNFRSF
ARD1,MFSD10,PFKFB2,RIC8A,TRPV3,NHP2,FAM32A,TNKS,MDP1,PIGH,ELOF1,KPNA5,MPHOSPH10,MYL
ARD1,MFSD10,PFKFB2,RIC8A,TRPV3,NHP2,FAM32A,TNKS,MDP1,PIGH,ELOF1,KPNA5,MPHOSPH10,MYL

42,TBX6,TMEM205,XBP1,C14ORF2,PCYT2,HSBP1L1,IFI27L1,ASF1B,EDF1,DGKE,TNFRSF14,ENDOV,CAMT
D99L2,GGPS1,RTEL1,TRNT1,MED15,PARD6G,SH3YL1,IP6K2,CLASRP,JPH2,C1ORF198,MICAL1,HS2ST1,UB
D99L2,GGPS1,RTEL1,TRNT1,MED15,PARD6G,SH3YL1,IP6K2,CLASRP,JPH2,C1ORF198,MICAL1,HS2ST1,UB
SP22,DBNDD2,CD99L2,GGPS1,AUP1,KXD1,PARD6G,SH3YL1,JPH2,MICAL1,HS2ST1,UBE2I,SLC39A13,TBX
1,RFX3,CAMTA1,CRIP1,PSMD4,LIG1,TMEM126B,RPL21,TXNL4A,TCF4,UCK2,TMPO,C17ORF62,PRMT1,EX
1,RFX3,CAMTA1,CRIP1,PSMD4,LIG1,TMEM126B,RPL21,TXNL4A,TCF4,UCK2,TMPO,C17ORF62,PRMT1,EX

LRRFIP1,ZNF524,TPM2,GPBP1,SIAH1,ASPH,CHPF,STRIP2,TWSG1,LMBR1,NAP1L1,CHCHD7,HMGA1,PABPC
LRRFIP1,ZNF524,TPM2,GPBP1,SIAH1,ASPH,CHPF,STRIP2,TWSG1,LMBR1,NAP1L1,CHCHD7,HMGA1,PABPC
K2,JPH2,MAGT1,CDC42,TBX6,TMEM205,AUNIP,GUK1,CENPN,CARD14,FBXW9,CCDC25,PPP1CC,RFX3,END
K2,JPH2,MAGT1,CDC42,TBX6,TMEM205,AUNIP,GUK1,CENPN,CARD14,FBXW9,CCDC25,PPP1CC,RFX3,END
706,ZNF687,ILF3,PKP4,HSPH1,C11ORF48,OAS3,LLGL2,CHEK1,COX17,HNRNPLL,LGMN,B3GAT3,REEP5,SM

ID7,PHTF2,RAD1,PPP3CB,WAC,SLC33A1,GTF2IRD1,ANAPC11,NT5C,SLC16A1,ILF3,MDH2,MT1E,LLGL2,SH3

1K,IRF9,ANTXR2,STIP1,PHF19,VCAN,CD44,TM9SF1,MAGED2,SON,SNF8,GRK6,CPSF3L,TSG101,OBSL1,TTC
1K,IRF9,ANTXR2,STIP1,PHF19,VCAN,CD44,TM9SF1,MAGED2,SON,SNF8,GRK6,CPSF3L,TSG101,OBSL1,TTC
SON,GRK6,HMGA2,CDC37,RPS16,CPSF3L,OBSL1,TTC21B,PTPN13,YLPM1,PHF21A,SLC39A11,FUZ,FAM3A,
SON,GRK6,HMGA2,CDC37,RPS16,CPSF3L,OBSL1,TTC21B,PTPN13,YLPM1,PHF21A,SLC39A11,FUZ,FAM3A,
TTC21B,PTPN13,UBR4,PSMA1,YLPM1,PHF21A,FUZ,FAM3A,MUC1,EPB41L5,ATG12,PPRC1,ZBTB44,TTC9C
TTC21B,PTPN13,UBR4,PSMA1,YLPM1,PHF21A,FUZ,FAM3A,MUC1,EPB41L5,ATG12,PPRC1,ZBTB44,TTC9C
IT2,TBX6,TMEM205,PTK2,AUNIP,C14ORF2,PCYT2,USE1,AGPAT4,MAF,CCDC25,DEF8,PPP1CC,RFX3,PPA1,
IT2,TBX6,TMEM205,PTK2,AUNIP,C14ORF2,PCYT2,USE1,AGPAT4,MAF,CCDC25,DEF8,PPP1CC,RFX3,PPA1,
AH1,ASPH,TRMT112,PNPO,CHPF,NIT2,MPDU1,TWSG1,RPS15,DCTD,UBXN1,GMPPB,HSPB1,VWA2,MVD,I
AH1,ASPH,TRMT112,PNPO,CHPF,NIT2,MPDU1,TWSG1,RPS15,DCTD,UBXN1,GMPPB,HSPB1,VWA2,MVD,I

10,GGT5,AGAP3,MDH2,CNST,LLGL2,PRLR,CASZ1,CHEK1,CAMK2D,ZFAND4,RPL15,B3GAT3,C11ORF49,GTF
10,GGT5,AGAP3,MDH2,CNST,LLGL2,PRLR,CASZ1,CHEK1,CAMK2D,ZFAND4,RPL15,B3GAT3,C11ORF49,GTF

7,MPHOSPH10,EIF4A1,ZNF248,TPT1,POLR2M,SMAGP,SRSF7,NXF1,CFLAR,VDAC1,CNTRL,RPL34,CREBZF,E
RABEPK,NUB1,ZNF816,SLC31A1,ZNFX1,PPP1R13B,SETD4,PHPT1,LMNA,ANKRD10,RAP1B,PKN2,CREM,TU
,ECSIT,EPB41L5,ATG12,PPRC1,ZBTB44,COX4I1,PLCB2,DUSP22,SLC25A10,TMX2,DBNDD2,NISCH,TMED4,F
,ECSIT,EPB41L5,ATG12,PPRC1,ZBTB44,COX4I1,PLCB2,DUSP22,SLC25A10,TMX2,DBNDD2,NISCH,TMED4,F
LCB2,DUSP22,DBNDD2,CD99L2,GGPS1,AUP1,PARD6G,SH3YL1,IP6K2,CLASRP,C1ORF198,MICAL1,HS2ST1,
IT1,MED15,PARD6G,SH3YL1,CLASRP,C1ORF198,UGGT2,MICAL1,HS2ST1,UBE2I,YIF1A,TBX6,TMEM205,PT
IT1,MED15,PARD6G,SH3YL1,CLASRP,C1ORF198,UGGT2,MICAL1,HS2ST1,UBE2I,YIF1A,TBX6,TMEM205,PT
GTF2IRD1,NXN,KLK10,EPS15L1,NOMO2,ILF3,EEF1D,NAA10,LLGL2,COX17,SUMO3,FKBP3,EGLN2,KCTD7,B
RANBP1,ZC3H14,FIS1,SLC15A3,TUBA1C,DST,ANTXR2,STIP1,LPCAT3,CD44,TM9SF1,MAGED2,SON,SNF8,G
RANBP1,ZC3H14,FIS1,SLC15A3,TUBA1C,DST,ANTXR2,STIP1,LPCAT3,CD44,TM9SF1,MAGED2,SON,SNF8,G
ADSYN1,RNF130,ZNF69,NDUFA11,MOSPD3,AAMDC,SNAPC2,RHEB,AKT1S1,NSFL1C,TUSC3,NDUFA10,SPC
ADSYN1,RNF130,ZNF69,NDUFA11,MOSPD3,AAMDC,SNAPC2,RHEB,AKT1S1,NSFL1C,TUSC3,NDUFA10,SPC
P1,MED15,SLC39A13,YIF1A,PRMT2,MAGT1,C14ORF2,GUK1,CENPN,CARD14,PCYT2,FBXW9,HSBP1L1,AGI
P1,MED15,SLC39A13,YIF1A,PRMT2,MAGT1,C14ORF2,GUK1,CENPN,CARD14,PCYT2,FBXW9,HSBP1L1,AGI
),NIT2,MPDU1,ZNF814,GMPPB,VWA2,IL32,LMBR1,SERPINB1,DDOST,C12ORF76,RANGRF,PABPC1,RRNAD
IM7,GMPPB,MVD,SAT1,EMC4,IL32,SYNGR1,RRNAD1,TOMM40,ZNF687,POT1,ANAPC11,RAD23A,SLC16A
IM7,GMPPB,MVD,SAT1,EMC4,IL32,SYNGR1,RRNAD1,TOMM40,ZNF687,POT1,ANAPC11,RAD23A,SLC16A
CAL1,HS2ST1,UBE2I,SLC39A13,YIF1A,EIF3M,JUP,TBX6,XBP1,PTK2,C14ORF2,CENPN,ASF1B,DEF8,EEF1B2,F
0,TMEM134,LRRFIP1,TPM2,GPBP1,SIAH1,ZEB2,ASPH,TRMT112,CHPF,STRIP2,ZNF625,MAPK13,GMPPB,LI
0,TMEM134,LRRFIP1,TPM2,GPBP1,SIAH1,ZEB2,ASPH,TRMT112,CHPF,STRIP2,ZNF625,MAPK13,GMPPB,LI
T1,UBE2I,YIF1A,TBX6,TMEM205,PTK2,AUNIP,C14ORF2,GUK1,CENPN,PCYT2,USE1,PPP1CC,RFX3,PPA1,DC
T1,UBE2I,YIF1A,TBX6,TMEM205,PTK2,AUNIP,C14ORF2,GUK1,CENPN,PCYT2,USE1,PPP1CC,RFX3,PPA1,DC
RF62,PRMT1,WNK1,TXNDC17,ARL6IP4,PIGL,C19ORF48,HNRNPC,CBFA2T2,STAU1,SLC9B2,HYPK,CARS,MC
VAMP8,LIG1,PSMA7,TIMM17B,XAF1,RPL21,TXNL4A,LYPLA2,NUTF2,CAMLG,ATPAF2,LSM12,PRMT1,EXOS
VAMP8,LIG1,PSMA7,TIMM17B,XAF1,RPL21,TXNL4A,LYPLA2,NUTF2,CAMLG,ATPAF2,LSM12,PRMT1,EXOS

F2,GUK1,CENPN,CARD14,PCYT2,FBXW9,HSBP1L1,AGPAT4,IFI27L1,MAF,PPA1,ENDOV,PPIL2,TIMM17B,A
F2,GUK1,CENPN,CARD14,PCYT2,FBXW9,HSBP1L1,AGPAT4,IFI27L1,MAF,PPA1,ENDOV,PPIL2,TIMM17B,A
NA5,OXLD1,EIF4A1,GFOD1,METTL3,PICK1,FBXL18,ASH1L,SLMAP,CHTOP,POLR2M,BAX,SRSF7,GLYCTK,CN
NA5,OXLD1,EIF4A1,GFOD1,METTL3,PICK1,FBXL18,ASH1L,SLMAP,CHTOP,POLR2M,BAX,SRSF7,GLYCTK,CN

NT2,GTF2IRD2,TMEM39B,BRF1,FKBP5,MRPS18C,THADA,COMMD4,RPS11,SNX5,NUP133,TMEM147,ELP4
27,SEMA3C,SYNGR1,LSR,MTA3,MEIS2,HMGA1,TADA3,EPS15L1,RAD23A,NT5C,PSME1,MDH2,HSPH1,CH
27,SEMA3C,SYNGR1,LSR,MTA3,MEIS2,HMGA1,TADA3,EPS15L1,RAD23A,NT5C,PSME1,MDH2,HSPH1,CH
NAT9,BCAM,ZNF836,RBMX,TOMM40,TADA3,ATP6V1F,ZNF687,ILF3,EEF1D,LLGL2,CAMK2D,EGLN2,RPL15
UBA1C,IRF9,STIP1,VCAN,LPCAT3,SON,TET2,UQCRQ,GRK6,HMGA2,CDC37,CPSF3L,TTC21B,PTPN13,UBR4,
UBA1C,IRF9,STIP1,VCAN,LPCAT3,SON,TET2,UQCRQ,GRK6,HMGA2,CDC37,CPSF3L,TTC21B,PTPN13,UBR4,
PRL2,OSGEP,TBL2,ARHGAP6,SCARB1,CRYGS,CTTN,TSPAN11,ITSN1,EIF5A,COMMD5,CHID1,NUDT22,AMZ
PRL2,OSGEP,TBL2,ARHGAP6,SCARB1,CRYGS,CTTN,TSPAN11,ITSN1,EIF5A,COMMD5,CHID1,NUDT22,AMZ
SNX5,RPAIN,COX20,ACOT2,ZXDC,NDUFS3,L3MBTL2,OXLD1,MPHOSPH10,GFOD1,KLHL7,METTL3,MRPL5
TF2IRD1,POLR3H,RAD23A,GGT5,SLC16A1,ILF3,AGAP3,MDH2,EEF1D,HSPH1,NAA10,CARHSP1,COX5A,SU
TF2IRD1,POLR3H,RAD23A,GGT5,SLC16A1,ILF3,AGAP3,MDH2,EEF1D,HSPH1,NAA10,CARHSP1,COX5A,SU
83,SZT2,BSCL2,PSMC5,ACSS1,RUVBL1,LAMTOR4,RPS17,EIF2AK4,C12ORF73,IQCK,SETD4,MED8,MBD1,C
83,SZT2,BSCL2,PSMC5,ACSS1,RUVBL1,LAMTOR4,RPS17,EIF2AK4,C12ORF73,IQCK,SETD4,MED8,MBD1,C
3A1,NXN,KLK10,ANAPC11,USP45,PDE4DIP,STXBP2,CHCHD5,SGK1,SH3GLB1,SSH2,EGLN2,B3GAT3,RFXA
2,ZXDC,COBL,ERGIC1,MYL9,TSEN34,TCF25,TJP1,CHTOP,UBA1,GLI4,NUP62,SCN11A,NADSYN1,ZNF511,D
COASY,SRSF1,SAP30BP,FOXRED1,TRNT1,KXD1,ITGB1BP1,MED15,UGGT2,MICAL1,UBE2I,YIF1A,PRMT2,M
COASY,SRSF1,SAP30BP,FOXRED1,TRNT1,KXD1,ITGB1BP1,MED15,UGGT2,MICAL1,UBE2I,YIF1A,PRMT2,M
,GPR56,HAUS4,PEX2,PRRC2B,PIGH,NUBP2,RPLP0,C6ORF132,TSEN34,OCIAD1,TJP1,CHTOP,CDK4,SLC52A
,GPR56,HAUS4,PEX2,PRRC2B,PIGH,NUBP2,RPLP0,C6ORF132,TSEN34,OCIAD1,TJP1,CHTOP,CDK4,SLC52A
2I,SLC39A13,YIF1A,PRMT2,MAGT1,JUP,CDC42,TBX6,CARD14,FBXW9,USE1,ASF1B,EEF1B2,GADL1,DGKE,
2I,SLC39A13,YIF1A,PRMT2,MAGT1,JUP,CDC42,TBX6,CARD14,FBXW9,USE1,ASF1B,EEF1B2,GADL1,DGKE,

,PUF60,MT1F,ATP9B,ANKRD10,ACTG1,RAP1B,IFT81,UBE2A,PKN2,CREM,TUBGCP2,CORO1B,OAZ1,SEMA
6K2,CLASRP,JPH2,C1ORF198,MICAL1,HS2ST1,UBE2I,PRMT2,TBX6,TMEM205,PTK2,AUNIP,C14ORF2,PCY
6K2,CLASRP,JPH2,C1ORF198,MICAL1,HS2ST1,UBE2I,PRMT2,TBX6,TMEM205,PTK2,AUNIP,C14ORF2,PCY
2,FBXW9,HSBP1L1,AGPAT4,IFI27L1,MAF,PPA1,ENDOV,PPIL2,TIMM17B,APEH,TXNL4A,TMPO,PRMT1,EX
2,FBXW9,HSBP1L1,AGPAT4,IFI27L1,MAF,PPA1,ENDOV,PPIL2,TIMM17B,APEH,TXNL4A,TMPO,PRMT1,EX

ARPIN,STAU1,UPP1,CARS,ZNF250,RPL36,MOB1A,MDH1,ZNF197,INO80E,MAPKAPK5,LRRFIP1,RMDN1,G
ARPIN,STAU1,UPP1,CARS,ZNF250,RPL36,MOB1A,MDH1,ZNF197,INO80E,MAPKAPK5,LRRFIP1,RMDN1,G
GSTK1,LDHA,PSMA6,ACOT8,AP3B2,RPS9,SLC35G1,RCE1,MRPL33,ZNF410,PCMT1,TESC,IGSF8,NUDT5,RB
UBA1C,IRF9,DST,ANTXR2,STIP1,PHF19,LPCAT3,CD44,TM9SF1,TET2,GRK6,CDC37,CPSF3L,OBSL1,PTPN13,
UBA1C,IRF9,DST,ANTXR2,STIP1,PHF19,LPCAT3,CD44,TM9SF1,TET2,GRK6,CDC37,CPSF3L,OBSL1,PTPN13,
7,PICK1,ECE1,OCIAD1,RBBP6,NREP,CHTOP,TSPAN31,TMEM9,ADRA1A,ZNF593,GLYCTK,RBM42,CLASP2,
1,HSPB1,LRRC27,MVD,ERCC5,NAP1L1,PRKRIP1,RANGRF,LSR,MEIS2,PSME2,RRNAD1,DBNL,BCAM,TOM
1,HSPB1,LRRC27,MVD,ERCC5,NAP1L1,PRKRIP1,RANGRF,LSR,MEIS2,PSME2,RRNAD1,DBNL,BCAM,TOM
OMM40,SERPINB6,RABEP2,TADA3,PDCD5,KLK10,TRIM16,RAD23A,PDE4DIP,MDH2,HSPH1,CHCHD5,OAS
OMM40,SERPINB6,RABEP2,TADA3,PDCD5,KLK10,TRIM16,RAD23A,PDE4DIP,MDH2,HSPH1,CHCHD5,OAS
RYGS,CTTN,TSPAN11,ITSN1,EIF5A,COMMD5,CHID1,NUDT22,AMZ2,UFD1L,USP38,TAZ,THYN1,DBNDD1,S
RYGS,CTTN,TSPAN11,ITSN1,EIF5A,COMMD5,CHID1,NUDT22,AMZ2,UFD1L,USP38,TAZ,THYN1,DBNDD1,S

RSF7,RER1,RPL24,ZNF675,BRD8,SLC35A2,RPS3A,BTF3,TCP11L1,NDUFA3,MGST2,RPS3,CKLF,AKT1S1,FARS
EC31A,CNST,SUMO3,LGMN,CD151,KCTD7,SMARCA4,TYMS,TMEM39B,DALRD3,DAZAP2,OARD1,SLC25A2
EC31A,CNST,SUMO3,LGMN,CD151,KCTD7,SMARCA4,TYMS,TMEM39B,DALRD3,DAZAP2,OARD1,SLC25A2
2,VIM,LIG1,TMEM126B,NUTF2,CAMLG,EPB41L4B,TCF4,UCK2,TMPO,ATPAF2,C17ORF62,CASC4,C19ORF5
IST,SH3GLB1,PHF20L1,HNRNPA1,FKBP3,NDUFV2,B3GAT3,SPINT2,GTF2IRD2,TMEM39B,BRF1,TPM4,FKBF
IST,SH3GLB1,PHF20L1,HNRNPA1,FKBP3,NDUFV2,B3GAT3,SPINT2,GTF2IRD2,TMEM39B,BRF1,TPM4,FKBF
3GLB1,PHF20L1,HNRNPA1,FKBP3,NDUFV2,B3GAT3,SPINT2,GTF2IRD2,TMEM39B,BRF1,TPM4,FKBP5,GMF

PINT2,PTK7,TMEM39B,DALRD3,BRF1,OARD1,PFKFB2,NFKBIA,GNB5,LFNG,XYLT2,ZXDC,COBL,TNKS,SMIM
CL2,PSMC5,ACSS1,RUVBL1,LAMTOR4,EIF2AK4,C12ORF73,IQCK,SETD4,MED8,MBD1,CHFR,SSH3,OPA3,C
CL2,PSMC5,ACSS1,RUVBL1,LAMTOR4,EIF2AK4,C12ORF73,IQCK,SETD4,MED8,MBD1,CHFR,SSH3,OPA3,C
,XAF1,RPL21,LYPLA2,CAMLG,ATPAF2,LSM12,PRMT1,EXOSC9,C19ORF54,PIGL,C19ORF48,ZFAND2B,HNRN
,XAF1,RPL21,LYPLA2,CAMLG,ATPAF2,LSM12,PRMT1,EXOSC9,C19ORF54,PIGL,C19ORF48,ZFAND2B,HNRN
1,PSMA7,APEH,CAPN10,PODXL,EPB41L4B,TCF4,TMPO,ATPAF2,PRMT1,CASC4,ZNF7,C19ORF48,STRADA,
1,PSMA7,APEH,CAPN10,PODXL,EPB41L4B,TCF4,TMPO,ATPAF2,PRMT1,CASC4,ZNF7,C19ORF48,STRADA,
V9,CCDC25,PPP1CC,RFX3,ENDOV,PPIL2,PSMD4,VIM,LIG1,PSMA7,APEH,TXNL4A,NUTF2,PPOX,TMPO,ATP
V9,CCDC25,PPP1CC,RFX3,ENDOV,PPIL2,PSMD4,VIM,LIG1,PSMA7,APEH,TXNL4A,NUTF2,PPOX,TMPO,ATP
,DENND1A,MRPS24,ZMAT1,AKT1S1,FARS2,B3GNTL1,KAT8,KIF13B,CARM1,TTLL3,MDK,IAH1,AIG1,CAMK
3,PTPN13,UBR4,PSMA1,PHF21A,FUZ,RGS3,PSMA3,MUC1,METTL17,ECSIT,PPRC1,RPS25,NIP7,TTC9C,COX
3,PTPN13,UBR4,PSMA1,PHF21A,FUZ,RGS3,PSMA3,MUC1,METTL17,ECSIT,PPRC1,RPS25,NIP7,TTC9C,COX

,HMGA2,OBSL1,TTC21B,UBR4,PSMA1,PHF21A,FUZ,RGS3,FAM3A,MUC1,EPB41L5,ATG12,PPRC1,ZBTB44,
,HMGA2,OBSL1,TTC21B,UBR4,PSMA1,PHF21A,FUZ,RGS3,FAM3A,MUC1,EPB41L5,ATG12,PPRC1,ZBTB44,

,RFXANK,SPINT2,GTF2IRD2,TMEM39B,BRF1,OARD1,SLC25A23,FAM102A,GMPR2,PFKFB2,RIC8A,MROH1
,GGPS1,COASY,SRSF1,SAP30BP,FOXRED1,ITGB1BP1,MED15,PARD6G,SH3YL1,IP6K2,C1ORF198,HS2ST1,SI
,GGPS1,COASY,SRSF1,SAP30BP,FOXRED1,ITGB1BP1,MED15,PARD6G,SH3YL1,IP6K2,C1ORF198,HS2ST1,SI
YLPM1,PHF21A,CHCHD1,SLC39A11,FUZ,MUC1,METTL17,EPB41L5,ATG12,RPS25,ZBTB44,TTC9C,PLCB2,DI
YLPM1,PHF21A,CHCHD1,SLC39A11,FUZ,MUC1,METTL17,EPB41L5,ATG12,RPS25,ZBTB44,TTC9C,PLCB2,DI
F8,EDF1,GADL1,DGKE,TNFRSF14,ENDOV,CAMTA1,LIG1,PSMA7,TMEM126B,HYI,NUTF2,PODXL,CAMLG,E
MEM147,ELP4,ZXDC,COBL,ELOF1,ARL16,SLC35E1,TSEN34,ARIH2,RPL8,ECE1,OCIAD1,RBBP6,TPT1,TSPAN

,OBSL1,TTC21B,UBR4,YLPM1,FUZ,MUC1,METTL17,ATG12,PPRC1,COX4I1,SLC25A10,TMX2,THEM4,NISCH
,OBSL1,TTC21B,UBR4,YLPM1,FUZ,MUC1,METTL17,ATG12,PPRC1,COX4I1,SLC25A10,TMX2,THEM4,NISCH
RPS6,RPL29,RTEL1,AUP1,FOXRED1,TRNT1,KXD1,ITGB1BP1,SH3YL1,C1ORF198,SLC39A13,EIF3M,XBP1,CE
RPS6,RPL29,RTEL1,AUP1,FOXRED1,TRNT1,KXD1,ITGB1BP1,SH3YL1,C1ORF198,SLC39A13,EIF3M,XBP1,CE
DF4,NISCH,TMED4,GGPS1,RTEL1,COASY,SRSF1,SAP30BP,YIPF3,KXD1,ITGB1BP1,MED15,PARD6G,SH3YL1,
DF4,NISCH,TMED4,GGPS1,RTEL1,COASY,SRSF1,SAP30BP,YIPF3,KXD1,ITGB1BP1,MED15,PARD6G,SH3YL1,
OMM40,SERPINB6,RABEP2,TADA3,PDCD5,KLK10,TRIM16,RAD23A,PDE4DIP,MDH2,HSPH1,CHCHD5,OAS3
OMM40,SERPINB6,RABEP2,TADA3,PDCD5,KLK10,TRIM16,RAD23A,PDE4DIP,MDH2,HSPH1,CHCHD5,OAS3
NGR2,TMEM134,RAB2A,ELP6,GPBP1,TSFM,GGCX,TRMT112,RPS20,MPDU1,TWSG1,ZSWIM7,DCTD,GMPF
NGR2,TMEM134,RAB2A,ELP6,GPBP1,TSFM,GGCX,TRMT112,RPS20,MPDU1,TWSG1,ZSWIM7,DCTD,GMPF
L,MAPKAPK5,DGUOK,HSP90AB1,CEP70,RAB2A,RMDN1,GPBP1,GPALPP1,CHPF,MAPK13,ZSWIM7,ERLEC1
L,MAPKAPK5,DGUOK,HSP90AB1,CEP70,RAB2A,RMDN1,GPBP1,GPALPP1,CHPF,MAPK13,ZSWIM7,ERLEC1
08,SUFU,NDUFA7,NUB1,FDPS,OS9,RBM7,NGDN,DDX17,SLC31A1,DUSP18,PLEKHA8,EIF2AK4,MRPL51,WD
L5,ITGB1BP1,MED15,PARD6G,MICAL1,HS2ST1,UBE2I,KCNMA1,PRMT2,EIF3M,MAGT1,JUP,CDC42,TBX6,T
1A,ZNF593,GLYCTK,RBM42,CLASP2,CREBZF,RPL24,NADSYN1,ZNF511,GTF2H1,RNF130,MYL6,SIN3A,NAA

1A,ZNF593,GLYCTK,RBM42,CLASP2,CREBZF,RPL24,NADSYN1,ZNF511,GTF2H1,RNF130,MYL6,SIN3A,NAA
,PKP4,HSPH1,RPL18,CNST,PHF20L1,HNRNPA1,FKBP3,NDUFV2,B3GAT3,SPINT2,GTF2IRD2,TMEM39B,BRI
,PKP4,HSPH1,RPL18,CNST,PHF20L1,HNRNPA1,FKBP3,NDUFV2,B3GAT3,SPINT2,GTF2IRD2,TMEM39B,BRI
1ORF198,HS2ST1,UBE2I,SLC39A13,YIF1A,JUP,CDC42,PTK2,AUNIP,C14ORF2,GUK1,CARD14,PCYT2,HSBP1
L1,CCDC25,ENDOV,VAMP8,PPIL2,PSMD4,TIMM17B,TMEM126B,XAF1,TXNL4A,RBM33,NUTF2,CAMLG,U
L1,CCDC25,ENDOV,VAMP8,PPIL2,PSMD4,TIMM17B,TMEM126B,XAF1,TXNL4A,RBM33,NUTF2,CAMLG,U
TEL1,AUP1,FOXRED1,TRNT1,KXD1,ITGB1BP1,SH3YL1,C1ORF198,SLC39A13,EIF3M,XBP1,CELF1,C14ORF2
RF2,ATF7,PCYT2,USE1,AGPAT4,CCDC25,DEF8,EEF1B2,PPA1,TNFRSF14,VAMP8,TIMM17B,TMEM126B,AP
RF2,ATF7,PCYT2,USE1,AGPAT4,CCDC25,DEF8,EEF1B2,PPA1,TNFRSF14,VAMP8,TIMM17B,TMEM126B,AP
RF2,ATF7,PCYT2,USE1,AGPAT4,CCDC25,DEF8,EEF1B2,PPA1,TNFRSF14,VAMP8,TIMM17B,TMEM126B,AP
BPMS,SPAG7,WDR62,POLD4,PLEKHA5,RANBP1,UBE2L6,VRK3,ZC3H14,PPM1K,TUBA1C,IRF9,STIP1,LRRC1
BPMS,SPAG7,WDR62,POLD4,PLEKHA5,RANBP1,UBE2L6,VRK3,ZC3H14,PPM1K,TUBA1C,IRF9,STIP1,LRRC1
4,ZNF512,OCIAD1,TCF25,SLMAP,RBBP6,BAX,CDK4,NME1,ANAPC13,FAM192A,TCEAL3,GALT,VDAC1,ACA
4,ZNF512,OCIAD1,TCF25,SLMAP,RBBP6,BAX,CDK4,NME1,ANAPC13,FAM192A,TCEAL3,GALT,VDAC1,ACA

K2,AUNIP,C14ORF2,CARD14,PCYT2,FBXW9,USE1,CCDC25,EDF1,GADL1,DGKE,VAMP8,TROAP,LIG1,CDC1
K2,AUNIP,C14ORF2,CARD14,PCYT2,FBXW9,USE1,CCDC25,EDF1,GADL1,DGKE,VAMP8,TROAP,LIG1,CDC1
20,ZNF706,PDCD5,GTF2IRD1,NUDT1,ANAPC11,POLR3H,RAD23A,GGT5,NT5C,SLC16A1,ILF3,AGAP3,MDH
RHGEF39,GSTK1,ACOT8,AP3B2,RPS9,SLC35G1,RCE1,MRPL33,ZNF410,RUVBL2,EBPL,TESC,IGSF8,RBPMS,
RHGEF39,GSTK1,ACOT8,AP3B2,RPS9,SLC35G1,RCE1,MRPL33,ZNF410,RUVBL2,EBPL,TESC,IGSF8,RBPMS,
PS6,RPL29,GGPS1,RTEL1,SRSF1,AUP1,SAP30BP,YIPF3,FOXRED1,TRNT1,KXD1,SH3YL1,IP6K2,CLASRP,UGG
PS6,RPL29,GGPS1,RTEL1,SRSF1,AUP1,SAP30BP,YIPF3,FOXRED1,TRNT1,KXD1,SH3YL1,IP6K2,CLASRP,UGG
N13,UBR4,PSMA1,CHCHD1,FUZ,METTL17,EPB41L5,ATG12,PPRC1,ZBTB44,NIP7,TTC9C,COX4I1,THEM4,
N13,UBR4,PSMA1,CHCHD1,FUZ,METTL17,EPB41L5,ATG12,PPRC1,ZBTB44,NIP7,TTC9C,COX4I1,THEM4,
1,ECE1,OCIAD1,RBBP6,TJP1,NREP,CHTOP,TSPAN31,PPP2R5C,TMEM9,ADRA1A,ZNF593,GLYCTK,RBM42,
PPA1,DGKE,ENDOV,CAMTA1,RPL28,PSMA7,HYI,TXNL4A,NUTF2,PODXL,CAMLG,TCF4,UCK2,ATPAF2,LSM1
1A,RBM3,MDH1,LAMP2,MAPKAPK5,DGUOK,HSP90AB1,LRRFIP1,GPALPP1,TPM3,PNPO,ZNF625,GABPB1
1A,RBM3,MDH1,LAMP2,MAPKAPK5,DGUOK,HSP90AB1,LRRFIP1,GPALPP1,TPM3,PNPO,ZNF625,GABPB1

54,TXNDC17,AP1S3,CYB561,SHARPIN,UPP1,CARS,RPL36,MOB1A,NFAM1,RBM3,LAMP2,INO80E,CCNL2,
1BP1,MED15,SH3YL1,C1ORF198,MICAL1,HS2ST1,KCNMA1,EIF3M,MAGT1,TBX6,XBP1,CELF1,C14ORF2,A
LIM5,RFX3,VAMP8,PSMD4,PSMA7,TMEM126B,XAF1,RPL21,CAMLG,ATPAF2,LSM12,PRMT1,EXOSC9,ST
LIM5,RFX3,VAMP8,PSMD4,PSMA7,TMEM126B,XAF1,RPL21,CAMLG,ATPAF2,LSM12,PRMT1,EXOSC9,ST
,ATL2,DDX10,EMC3,RNPS1,MLH1,DTD2,XPNPEP1,ZNF483,ZNF783,FBXO34,CYB5D2,ALS2,EXOSC8,SLC31

593,GLYCTK,RBM42,CLASP2,CREBZF,RPL24,NADSYN1,ZNF511,GTF2H1,RNF130,MYL6,SIN3A,NAA38,TLC
1,PHF19,LPCAT3,CD44,TM9SF1,TET2,GRK6,CDC37,CPSF3L,OBSL1,PTPN13,YLPM1,PHF21A,CHCHD1,SLC3
10,EMC3,RNPS1,DTD2,XPNPEP1,ZNF483,ZNF783,FBXO34,CYB5D2,ALS2,EXOSC8,SLC31A1,RCAN1,C12OR
,MRPL27,MFSD1,FLAD1,FCGRT,RNASEK,SSH3,TIMM9,DHRS4L2,MRPL36,SYNJ2,URM1,MIEN1
A,PSMA3,METTL17,EPB41L5,ATG12,RPS25,NLRP1,NIP7,DUSP22,TMX2,RPS6,RPL29,RTEL1,COASY,FOXRE
A,PSMA3,METTL17,EPB41L5,ATG12,RPS25,NLRP1,NIP7,DUSP22,TMX2,RPS6,RPL29,RTEL1,COASY,FOXRE
DGUOK,HSP90AB1,CEP70,RAB2A,RMDN1,GPBP1,GPALPP1,CHPF,MAPK13,ZSWIM7,ERLEC1,GMPPB,MV
DGUOK,HSP90AB1,CEP70,RAB2A,RMDN1,GPBP1,GPALPP1,CHPF,MAPK13,ZSWIM7,ERLEC1,GMPPB,MV
30A5,TRPV3,THADA,PHC3,COMMD4,MFSD11,SNX5,COX20,TMEM147,SMIM4,PRRC2B,DCTN5,NMNAT3
30A5,TRPV3,THADA,PHC3,COMMD4,MFSD11,SNX5,COX20,TMEM147,SMIM4,PRRC2B,DCTN5,NMNAT3
30A5,TRPV3,THADA,PHC3,COMMD4,MFSD11,SNX5,COX20,TMEM147,SMIM4,PRRC2B,DCTN5,NMNAT3

,RPL29,RTEL1,SAP30BP,YIPF3,FOXRED1,RABL5,TRNT1,ITGB1BP1,SH3YL1,CLASRP,C1ORF198,MICAL1,HS2
2,EIF4A1,RC3H2,KLHL7,PICK1,ECE1,OCIAD1,RBBP6,NREP,CHTOP,TSPAN31,TMEM9,ADRA1A,ZNF593,GLY
2,EIF4A1,RC3H2,KLHL7,PICK1,ECE1,OCIAD1,RBBP6,NREP,CHTOP,TSPAN31,TMEM9,ADRA1A,ZNF593,GLY

RL,RER1,NEK11,RPL34,NKIRAS2,ACAA1,PIGO,MYL6,SLC35A2,RBM4,TP53,C1ORF63,C5ORF24,ZNF396,SN
RL,RER1,NEK11,RPL34,NKIRAS2,ACAA1,PIGO,MYL6,SLC35A2,RBM4,TP53,C1ORF63,C5ORF24,ZNF396,SN
,TESC,IGSF8,RBPMS,NNMT,SNRPN,WDR62,RFX5,UBE2L6,LZTFL1,FIS1,PPM1K,TUBA1C,DST,LRRC1,PHF19
,TESC,IGSF8,RBPMS,NNMT,SNRPN,WDR62,RFX5,UBE2L6,LZTFL1,FIS1,PPM1K,TUBA1C,DST,LRRC1,PHF19
STIP1,LRRC1,PHF19,VCAN,LPCAT3,CD44,TM9SF1,MAGED2,SON,TET2,UQCRQ,GRK6,CPSF3L,OBSL1,PTPN
STIP1,LRRC1,PHF19,VCAN,LPCAT3,CD44,TM9SF1,MAGED2,SON,TET2,UQCRQ,GRK6,CPSF3L,OBSL1,PTPN
ML2,C19ORF54,PIGL,STRADA,PPAN,AP1S3,SLC9B2,INO80E,CCNL2,MAPKAPK5,HSP90AB1,DUT,CEP70,LR

EM205,CELF1,CARD14,CCDC25,EEF1B2,PPP1CC,PDLIM5,RFX3,VAMP8,CRIP1,PSMA7,TMEM126B,XAF1,RI
EM205,CELF1,CARD14,CCDC25,EEF1B2,PPP1CC,PDLIM5,RFX3,VAMP8,CRIP1,PSMA7,TMEM126B,XAF1,RI
M,BLVRB,PPP3CB,RABEP2,TADA3,RALGPS1,ZNF706,ZNF687,PDCD5,GTF2IRD1,EPS15L1,POLR3H,ABHD17
,PPP1CC,PDLIM5,RFX3,PPA1,VAMP8,CRIP1,PSMA7,TIMM17B,XAF1,RPL21,TXNL4A,LYPLA2,NUTF2,CAML
,PPP1CC,PDLIM5,RFX3,PPA1,VAMP8,CRIP1,PSMA7,TIMM17B,XAF1,RPL21,TXNL4A,LYPLA2,NUTF2,CAML
,OCIAD1,TCF25,RBBP6,CHTOP,SMAGP,TSPAN31,TMEM9,ADRA1A,ZNF593,GLYCTK,FAM131A,RBM42,CL
,OCIAD1,TCF25,RBBP6,CHTOP,SMAGP,TSPAN31,TMEM9,ADRA1A,ZNF593,GLYCTK,FAM131A,RBM42,CL
H2B1,NOP16,RPL13A,ARHGEF39,GSTK1,TTLL5,LDHA,PSMA6,AP3B2,RPS9,SLC35G1,RCE1,ZNF410,RUVBL
RF2,CENPN,PCYT2,USE1,AGPAT4,MAF,EEF1B2,RFX3,PPA1,ENDOV,VAMP8,PPIL2,TROAP,RPL28,VIM,LIG1
RF2,CENPN,PCYT2,USE1,AGPAT4,MAF,EEF1B2,RFX3,PPA1,ENDOV,VAMP8,PPIL2,TROAP,RPL28,VIM,LIG1
UG1,NDUFB2,RPS7,APAF1,PLD1,ECT2,GPS2,PNRC1,MLH1,DTD2,SUMO1,COMMD1,CEP63,ZNF483,LRRC
K,TUBA1C,IRF9,ANTXR2,STIP1,LRRC1,PHF19,VCAN,PCBP2,LPCAT3,CD44,TM9SF1,MAGED2,SON,TET2,UQ
K,TUBA1C,IRF9,ANTXR2,STIP1,LRRC1,PHF19,VCAN,PCBP2,LPCAT3,CD44,TM9SF1,MAGED2,SON,TET2,UQ
ME1,PPP2R5C,MTIF2,GLYCTK,VDAC1,CREBZF,NKIRAS2,ACAA1,OLFM1,FCHO2,IRF3,KDELR3,CLTA,MYL6,PL
N1,PAFAH1B2,MPC2,TYMS,BRF1,TPM4,FKBP5,TRPM4,MRPS18C,PFKFB2,TRPV3,THADA,PIGT,NHP2,LFNG
N1,PAFAH1B2,MPC2,TYMS,BRF1,TPM4,FKBP5,TRPM4,MRPS18C,PFKFB2,TRPV3,THADA,PIGT,NHP2,LFNG
NTL1,REPS2,MYL12A,KAT8,CARM1,FAM104B,CCDC24,MDK,PNRC1,C6ORF141,ACVR1B,DTD2,CAMKK1,X

IF593,GLYCTK,RBM42,CLASP2,CREBZF,RPL24,NADSYN1,ZNF511,GTF2H1,RNF130,MYL6,SIN3A,NAA38,TL
IF593,GLYCTK,RBM42,CLASP2,CREBZF,RPL24,NADSYN1,ZNF511,GTF2H1,RNF130,MYL6,SIN3A,NAA38,TL
PS1,MLH1,DHDDS,XPNPEP1,ZNF483,ZNF783,FBXO34,CYB5D2,ALS2,EXOSC8,SLC31A1,NSMF,ATP9B,IMM
MEM147,ELP4,NFYC,ZXDC,COBL,ELOF1,L3MBTL2,ARL16,EIF4A1,SLC35E1,RC3H2,KLHL7,MRPL55,ARIH2,PI
F1,C14ORF2,ATF7,PCYT2,FBXW9,USE1,HSBP1L1,AGPAT4,CCDC25,DEF8,EEF1B2,PPA1,TNFRSF14,VAMP8,
F1,C14ORF2,ATF7,PCYT2,FBXW9,USE1,HSBP1L1,AGPAT4,CCDC25,DEF8,EEF1B2,PPA1,TNFRSF14,VAMP8,
F1,C14ORF2,ATF7,PCYT2,FBXW9,USE1,HSBP1L1,AGPAT4,CCDC25,DEF8,EEF1B2,PPA1,TNFRSF14,VAMP8,
,SNAPC2,EPN1,RDX,NSFL1C,TUSC3,B3GNTL1,SGSM3,APAF1,CARM1,CCDC24,TBCA,SNX3,IAH1,PNRC1,CA
,SNAPC2,EPN1,RDX,NSFL1C,TUSC3,B3GNTL1,SGSM3,APAF1,CARM1,CCDC24,TBCA,SNX3,IAH1,PNRC1,CA
A6,ZNF593,GLYCTK,TRIM46,RBM42,CLASP2,CREBZF,ZNF511,GTF2H1,MYL6,RPS3A,IFITM2,TCP11L1,MA
M7,ERLEC1,GMPPB,MVD,SAT1,EMC4,IL32,SYNGR1,RANGRF,RRNAD1,RBP1,H2AFV,ZNF687,POT1,ANAPC1
M7,ERLEC1,GMPPB,MVD,SAT1,EMC4,IL32,SYNGR1,RANGRF,RRNAD1,RBP1,H2AFV,ZNF687,POT1,ANAPC1
,RPS3A,MAGOHB,IDH3G,CREB3L2,AKT1S1,SPHK2,HMGN1,NDUFB2,CDK2AP2,NDUFA10,RPS27L,AZI2,SP

EM4,DBNDD2,NISCH,RPL29,CD99L2,GGPS1,RTEL1,SRSF1,AUP1,KXD1,ITGB1BP1,MED15,SH3YL1,IP6K2,JP
EM4,DBNDD2,NISCH,RPL29,CD99L2,GGPS1,RTEL1,SRSF1,AUP1,KXD1,ITGB1BP1,MED15,SH3YL1,IP6K2,JP

,GPALPP1,SIAH1,ASPH,PNPO,RPTOR,NIT2,MPDU1,TWSG1,RPS15,UBXN1,ZNF814,GMPPB,VWA2,IL32,LM
,GPALPP1,SIAH1,ASPH,PNPO,RPTOR,NIT2,MPDU1,TWSG1,RPS15,UBXN1,ZNF814,GMPPB,VWA2,IL32,LM
M1,PHF21A,CHCHD1,FAM3A,METTL17,EPB41L5,ATG12,ZBTB44,PLCB2,DUSP22,SLC25A10,SPOCK2,THEM
4B,TIMM17B,APEH,RPL21,RBM33,TMPO,LSM12,PRMT1,CASC4,TXNDC17,PIGL,HNRNPC,ZNF644,STAU1,
COBL,STX10,NMNAT3,KPNA5,MPHOSPH10,MYL9,TMEM108,OGFOD2,ZNF512,TCF25,RNF181,BAX,SLC52
COBL,STX10,NMNAT3,KPNA5,MPHOSPH10,MYL9,TMEM108,OGFOD2,ZNF512,TCF25,RNF181,BAX,SLC52
LCC,RFX3,PPA1,VIM,LIG1,PSMA7,PODXL,CAMLG,TCF4,TMPO,EXOSC9,WNK1,TXNDC17,ARL6IP4,PIGL,STR
LCC,RFX3,PPA1,VIM,LIG1,PSMA7,PODXL,CAMLG,TCF4,TMPO,EXOSC9,WNK1,TXNDC17,ARL6IP4,PIGL,STR

RHSP1,MT1E,RPL18,SUMO3,PHF20L1,FKBP3,B3GAT3,ZNF506,REEP5,GTF2IRD2,DAZAP2,ZFPL1,TPM4,SL
RHSP1,MT1E,RPL18,SUMO3,PHF20L1,FKBP3,B3GAT3,ZNF506,REEP5,GTF2IRD2,DAZAP2,ZFPL1,TPM4,SL
NBP1,UBE2L6,VRK3,SLC15A3,PPM1K,TUBA1C,IRF9,ANTXR2,STIP1,GRAMD1C,VCAN,PCBP2,MAGED2,TET2
NBP1,UBE2L6,VRK3,SLC15A3,PPM1K,TUBA1C,IRF9,ANTXR2,STIP1,GRAMD1C,VCAN,PCBP2,MAGED2,TET2
CE1,MRPL33,ZNF410,TESC,IGSF8,NUDT5,WDR62,IL1RAP,RFX5,PLEKHA5,RANBP1,LZTFL1,ZC3H14,PPM1K
CE1,MRPL33,ZNF410,TESC,IGSF8,NUDT5,WDR62,IL1RAP,RFX5,PLEKHA5,RANBP1,LZTFL1,ZC3H14,PPM1K
D1,MED15,SH3YL1,SLC39A13,MAGT1,XBP1,C14ORF2,GUK1,CENPN,PCYT2,IFI27L1,MAF,CCDC25,GADL1,P
D1,MED15,SH3YL1,SLC39A13,MAGT1,XBP1,C14ORF2,GUK1,CENPN,PCYT2,IFI27L1,MAF,CCDC25,GADL1,P
TK2,C14ORF2,CENPN,FBXW9,HSBP1L1,ASF1B,DEF8,EEF1B2,PPP1CC,EDF1,ENDOV,CAMTA1,CAPG,PPIL2,P
TK2,C14ORF2,CENPN,FBXW9,HSBP1L1,ASF1B,DEF8,EEF1B2,PPP1CC,EDF1,ENDOV,CAMTA1,CAPG,PPIL2,P
2,LGALS3,SCAP,NHP2,PDCD6,MOK,SNX5,TMEM147,USP40,TGOLN2,ELOF1,C12ORF57,HAX1,ZMYM6,OG
2,LGALS3,SCAP,NHP2,PDCD6,MOK,SNX5,TMEM147,USP40,TGOLN2,ELOF1,C12ORF57,HAX1,ZMYM6,OG
LC35G1,RCE1,MRPL33,ZNF410,EBPL,IGSF8,NUDT5,RBPMS,WDR62,IL1RAP,STYXL1,POLD4,RANBP1,VRK3,
LC35G1,RCE1,MRPL33,ZNF410,EBPL,IGSF8,NUDT5,RBPMS,WDR62,IL1RAP,STYXL1,POLD4,RANBP1,VRK3,
ND1,TIMM17B,TMEM126B,RBM33,LYPLA2,NUTF2,EPB41L4B,LSM12,PRMT1,ZNF7,ARL6IP4,C19ORF48,S
,PDLIM5,RFX3,PPA1,DGKE,VAMP8,CRIP1,PSMA7,TIMM17B,XAF1,RPL21,LYPLA2,CAMLG,ATPAF2,LSM12,
,PDLIM5,RFX3,PPA1,DGKE,VAMP8,CRIP1,PSMA7,TIMM17B,XAF1,RPL21,LYPLA2,CAMLG,ATPAF2,LSM12,
Y,SRSF1,AUP1,SAP30BP,FOXRED1,TRNT1,KXD1,ITGB1BP1,MED15,UGGT2,SLC39A13,PRMT2,MAGT1,C14
Y,SRSF1,AUP1,SAP30BP,FOXRED1,TRNT1,KXD1,ITGB1BP1,MED15,UGGT2,SLC39A13,PRMT2,MAGT1,C14
DHA,PSMA6,ACOT8,AP3B2,RPS9,SLC35G1,RCE1,MRPL33,RUVBL2,PCMT1,TESC,NUDT5,RBPMS,NNMT,SN
DHA,PSMA6,ACOT8,AP3B2,RPS9,SLC35G1,RCE1,MRPL33,RUVBL2,PCMT1,TESC,NUDT5,RBPMS,NNMT,SN
L1,DBNDD2,SDF4,NISCH,CENPK,RPS6,RPL29,GGPS1,RTEL1,SRSF1,AUP1,SAP30BP,YIPF3,FOXRED1,TRNT1,
L1,DBNDD2,SDF4,NISCH,CENPK,RPS6,RPL29,GGPS1,RTEL1,SRSF1,AUP1,SAP30BP,YIPF3,FOXRED1,TRNT1,
T,ATG12,PPRC1,ZBTB44,COX4I1,DUSP22,SLC25A10,TMX2,THEM4,RDH11,DBNDD2,NISCH,CENPK,RTEL1,
T,ATG12,PPRC1,ZBTB44,COX4I1,DUSP22,SLC25A10,TMX2,THEM4,RDH11,DBNDD2,NISCH,CENPK,RTEL1,
MPO,LSM12,C17ORF62,CASC4,C19ORF54,ZNF7,TXNDC17,C19ORF48,PPAN,UPP1,CDK11A,RBM3,MDH1,L
MPO,LSM12,C17ORF62,CASC4,C19ORF54,ZNF7,TXNDC17,C19ORF48,PPAN,UPP1,CDK11A,RBM3,MDH1,L
L1,UBE2I,SLC39A13,PTK2,C14ORF2,CENPN,PCYT2,MAF,CCDC25,DEF8,PPP1CC,RFX3,PPA1,DGKE,ENDOV,
,ITGB1BP1,MED15,UGGT2,SLC39A13,YIF1A,PRMT2,MAGT1,C14ORF2,GUK1,CENPN,CARD14,PCYT2,FBXW
,ITGB1BP1,MED15,UGGT2,SLC39A13,YIF1A,PRMT2,MAGT1,C14ORF2,GUK1,CENPN,CARD14,PCYT2,FBXW
6IP4,C19ORF48,ZFAND2B,HNRNPC,CYB561,CBFA2T2,SLC9B2,UPP1,RPL36,NFAM1,RBM3,ZNF197,COPZ1,
6IP4,C19ORF48,ZFAND2B,HNRNPC,CYB561,CBFA2T2,SLC9B2,UPP1,RPL36,NFAM1,RBM3,ZNF197,COPZ1,
CAPN10,EPB41L4B,ATPAF2,PRMT1,C19ORF54,ARL6IP4,C19ORF48,CYB561,CBFA2T2,SLC9B2,UPP1,RPL3

,CAPN10,EPB41L4B,ATPAF2,PRMT1,C19ORF54,ARL6IP4,C19ORF48,CYB561,CBFA2T2,SLC9B2,UPP1,RPL3
RP,C1ORF198,UGGT2,YIF1A,CDC42,TBX6,XBP1,C14ORF2,CENPN,ATF7,USE1,HSBP1L1,EEF1B2,GADL1,PPA
RP,C1ORF198,UGGT2,YIF1A,CDC42,TBX6,XBP1,C14ORF2,CENPN,ATF7,USE1,HSBP1L1,EEF1B2,GADL1,PPA
PL55,ARIH2,FBXL18,ECE1,OCIAD1,TCF25,RBBP6,ZNF248,TPT1,SMAGP,TSPAN31,TMEM9,NXF1,SLC39A6,
C30A5,PHC3,COMMD4,MFSD11,SNX5,COX20,TMEM147,GRAPL,SMIM4,PRRC2B,DCTN5,TMEM108,ZMY
C30A5,PHC3,COMMD4,MFSD11,SNX5,COX20,TMEM147,GRAPL,SMIM4,PRRC2B,DCTN5,TMEM108,ZMY
C30A5,PHC3,COMMD4,MFSD11,SNX5,COX20,TMEM147,GRAPL,SMIM4,PRRC2B,DCTN5,TMEM108,ZMY
1,NOP16,ARHGEF39,GSTK1,TTLL5,LDHA,PSMA6,AP3B2,SLC35G1,RCE1,MRPL33,RUVBL2,PCMT1,EBPL,TE
1,NOP16,ARHGEF39,GSTK1,TTLL5,LDHA,PSMA6,AP3B2,SLC35G1,RCE1,MRPL33,RUVBL2,PCMT1,EBPL,TE

PKAPK5,DGUOK,HSP90AB1,CEP70,RAB2A,RMDN1,GPBP1,GPALPP1,CHPF,MAPK13,ZSWIM7,ERLEC1,GM
PKAPK5,DGUOK,HSP90AB1,CEP70,RAB2A,RMDN1,GPBP1,GPALPP1,CHPF,MAPK13,ZSWIM7,ERLEC1,GM
,SRSF1,FOXRED1,MED15,PARD6G,SH3YL1,MICAL1,HS2ST1,UBE2I,KCNMA1,YIF1A,PRMT2,EIF3M,MAGT1
P,TBX6,TMEM205,PTK2,CENPN,PCYT2,USE1,HSBP1L1,MAF,ASF1B,PPP1CC,RFX3,PPA1,CAMTA1,CAPG,C
P,TBX6,TMEM205,PTK2,CENPN,PCYT2,USE1,HSBP1L1,MAF,ASF1B,PPP1CC,RFX3,PPA1,CAMTA1,CAPG,C
,GRAPL,DCTN5,NMNAT3,BTN3A2,MPHOSPH10,DNAJC12,TMEM108,ZMYM6,METTL3,SSR3,MPV17,MRP
,GRAPL,DCTN5,NMNAT3,BTN3A2,MPHOSPH10,DNAJC12,TMEM108,ZMYM6,METTL3,SSR3,MPV17,MRP
AN,TM9SF1,MAGED2,TET2,UQCRQ,SNF8,CDC37,RPS16,TSG101,TTC21B,PTPN13,PHF21A,FAM3A,METTL
AN,TM9SF1,MAGED2,TET2,UQCRQ,SNF8,CDC37,RPS16,TSG101,TTC21B,PTPN13,PHF21A,FAM3A,METTL
,CLASP2,CREBZF,ZNF511,GTF2H1,MYL6,RPS3A,MAGOHB,IDH3G,CREB3L2,AKT1S1,SPHK2,ACTR10,HMG
,PDCD5,NOMO2,NT5C,CHCHD5,NAA10,CARHSP1,MT1E,RPL18,ATP10D,CNST,LLGL2,EGFR,CASZ1,HNRNP
,PDCD5,NOMO2,NT5C,CHCHD5,NAA10,CARHSP1,MT1E,RPL18,ATP10D,CNST,LLGL2,EGFR,CASZ1,HNRNP
B,HPS1,RGS12,AKT1S1,NDUFB2,MYL12A,ZBTB8OS,SLC37A3,AKR1A1,PRKCSH,GPS2,VAPA,CAMKK1,DHD
B,HPS1,RGS12,AKT1S1,NDUFB2,MYL12A,ZBTB8OS,SLC37A3,AKR1A1,PRKCSH,GPS2,VAPA,CAMKK1,DHD
,ATP6V1F,NXN,POLR3H,STXBP2,CHCHD5,RRP8,OAS3,SH3GLB1,CHEK1,ZNF506,SPINT2,REEP5,ASH2L,TCT
,ATP6V1F,NXN,POLR3H,STXBP2,CHCHD5,RRP8,OAS3,SH3GLB1,CHEK1,ZNF506,SPINT2,REEP5,ASH2L,TCT
MEM205,PTK2,CELF1,GUK1,CENPN,PCYT2,USE1,HSBP1L1,EEF1B2,EDF1,DGKE,TNFRSF14,CAPG,CRIP1,PP
YL1,IP6K2,CLASRP,YIF1A,PRMT2,EIF3M,MAGT1,TMEM205,CARD14,CCDC25,EEF1B2,PPP1CC,PDLIM5,RF
YL1,IP6K2,CLASRP,YIF1A,PRMT2,EIF3M,MAGT1,TMEM205,CARD14,CCDC25,EEF1B2,PPP1CC,PDLIM5,RF

,HNRNPM,PABPN1,RPS17,CLSPN,MANBAL,MED8,LMNA,EHMT1,ANO6,MBD1,UBE2F,WDR6,REEP4,SHPK,

9B,ACAA1,RPL24,FCHO2,IFT172,TRAPPC2,DYNLT3,BFAR,CLTA,MYL6,SIN3A,GEMIN2,AXIN1,TUBB3,RPS3

2N,RPS11,NOP2,FAM32A,MRPS17,PCID2,PRRC2B,RPL37A,ELOF1,MPHOSPH10,EIF4A1,STAU2,RPLP0,RC[illegible]

,RPS20,GABPB1,RPS15,ZNF814,HSPB1,ERCC5,MCTS1,NAP1L1,PRKRIP1,MTA3,DCN,MEIS2,HMGA1,PHTF2

SF8,NUDT5,RBPMS,SNRPN,WDR62,IL1RAP,STYXL1,RFX5,POLD4,PLEKHA5,RANBP1,UBE2L6,VRK3,LZTFL1,

[illegible],PRMT1,EXOSC9,WNK1,ZNF7,ARL6IP4,STRADA,PPAN,ACAD10,HNRNPC,ZNF644,STAU1,CARS,ZNF250,R[illegible]

PPOX,TMPO,PRMT1,EXOSC9,WNK1,ZNF7,ARL6IP4,STRADA,PPAN,ACAD10,HNRNPC,ZNF644,STAU1,CAR[illegible]

[illegible],EXOSC2,RBM7,PSMC5,HNRNPM,IK,EXOSC8,DDX17,PABPN1,RPS15A,RPS17,SFSWAP,PPIE,RNMT,PUF60,

TRAK1,GOSR2,SGSM3,TBC1D22A,RPS7,U2AF1L4,KIF13B,NCBP2,SNX3,ECT2,RNPS1,UBA52,KIF13A,RPS4X[illegible]

X,MAGOHB,VPS25,UCHL5,NSFL1C,IFI27,ARRB2,RPS7,APAF1,NCBP2,USP15,NFE2L1,TAF9,RNPS1,UBA52,[illegible]

[illegible]1,TMEM108,SSR3,MPV17,SSB,ARIH2,RPL8,PICK1,CHTOP,RPL35,SRSF7,NXF1,ZNF593,TRIM46,RPL34,NU[illegible]

[illegible]K1,SLMAP,UGT8,CHTOP,BAX,RPL35,DLG4,SRSF7,NXF1,ZNF593,TRIM46,LAMTOR5,CLASP2,RER1,RPL34,[illegible]

CK1,SLMAP,UGT8,CHTOP,BAX,RPL35,DLG4,SRSF7,NXF1,ZNF593,TRIM46,LAMTOR5,CLASP2,RER1,RPL34,N
NF130,BFAR,CLTA,LYPLAL1,AXIN1,PSMD10,RPS3A,PML,TMEM199,UBE2D3,RPS3,RDX,MAGOHB,VPS25,U

PL48,SHMT1,ERAL1,MRPL27,MRPS7,RPL27,NCK2,BIN1,MRPL36,RBM4B,ETHE1,GGCT,RPS14,LSM14A,CN

.48,SHMT1,ERAL1,MRPL27,CERS2,MRPS7,RPL27,NCK2,MRPL36,RBM4B,GGCT,RPS14,LSM14A,CNOT7,HB

,FTH1,TCTN1,MPC2,SNRPF,TRPM4,MCFD2,LGALS3,TCP1,SYTL1,CD74,PARK7,NFKBIA,RPS11,PDCD6,MFF,

7,RPS11,SNX5,NUP133,USP40,PCID2,PEX2,RPL37A,USP4,RPLP0,METTL3,SSB,ARIH2,RPL8,FBXL18,UBE2D4
23,RPL30,RQCD1,NDUFA7,DNAJC24,MRPS33,ACSS1,NGDN,RPS15A,RPS17,EIF2AK4,ERAP2,MRPL51,MRPL
UFC1,RUVBL1,NDUFB10,ATPAF1,SFSWAP,MCM7,EIF6,DHX30,ERAL1,DNAJB12,VIL1,MRPS7,NDUFS7,TUB
X20,MRPS17,COA1,COBL,USP4,NDUFS3,RPLP0,HAX1,DGAT1,EIF3G,MRPL55,LIMA1,PICK1,ADD1,KCNC4,B
B2,GBA2,PFKFB2,PARK7,RPS11,SNX5,USP40,PCID2,PEX2,RPL37A,USP4,RPLP0,HAX1,METTL3,SSB,ARIH2,
AP,C9ORF64,PPIE,EIF6,RNMT,PUF60,DUSP11,DDX56,SNRNP40,RPL27,RBM4B,SNRNP200,RPS14,URM1,H
8,EGFR,SH3GLB1,HINT1,LGMN,EGLN2,RPL15,MTHFS,PAFAH1B2,GBA2,PFKFB2,PARK7,RPS11,SNX5,NUP1
APGEF3,RPAIN,ACOT2,PCID2,PEX2,TNKS,RPL37A,STX10,PEX26,BTN3A2,KPNA5,RPLP0,HAX1,SSR3,MPV17,
F,SNX5,ENY2,NUP133,RAPGEF3,RPAIN,ACOT2,PCID2,PEX2,TNKS,RPL37A,STX10,PEX26,BTN3A2,KPNA5,RF
GRF,LSR,HMGA1,RRNAD1,DBNL,RBMX,BRIP1,RABEP2,POT1,HSD17B10,B2M,PKP4,PDE4DIP,ZCCHC4,RRP8

METTL3,EIF3G,MRPL55,RPL8,UGT8,RPL35,CDK4,NME1,MTIF2,EEF2K,NT5E,GCDH,MARS,LAMTOR5,RPL34

APGEF3,RPAIN,ACOT2,PCID2,PEX2,TNKS,RPL37A,STX10,PEX26,BTN3A2,KPNA5,RPLP0,HAX1,SSR3,MPV17,
UP133,RAPGEF3,RPAIN,ACOT2,PCID2,PEX2,TNKS,RPL37A,STX10,PEX26,BTN3A2,KPNA5,RPLP0,HAX1,SSR
,MPC2,TRPM4,MCFD2,PFKFB2,LGALS3,TCP1,SYTL1,CD74,PARK7,NFKBIA,RPS11,PDCD6,MFF,SNX5,ENY2,

,VIM,LIG1,PSMA7,CDC14B,RPL21,TXNL4A,TCF4,UCK2,TMPO,EXOSC9,STOML2,ZNF7,ARL6IP4,HNRNPC,ZN
PC5,GEMIN2,NDUFA11,AXIN1,TP53,PSMD10,SERPINA1,PML,TMEM199,NDUFA3,EIF3C,COX19,RPS3,RDX,
TF7,FBXW9,HSBP1L1,AGPAT4,MAF,EEF1B2,EDF1,RFX3,PPA1,ENDOV,CAMTA1,PSMD4,RPL28,VIM,LIG1,P
SP90AB1,CEP70,RAB2A,ATP5J,TPM2,ZEB2,TPM3,STRIP2,GABPB1,VPS28,RPS15,MCTS1,NAP1L1,RANGRF,

,VIM,LIG1,PSMA7,CDC14B,RPL21,TXNL4A,TCF4,UCK2,PPOX,TMPO,EXOSC9,STOML2,ZNF7,ARL6IP4,HNRN

L5,NFE2L1,TAF9,UBA52,CAPN2,SUMO1,COMMD1,XPNPEP1,PSMD8,RAB26,SUFU,NUB1,OS9,GUSB,PSMC

0D,CNST,LLGL2,EGFR,SH3GLB1,HNRNPA1,RPL15,B3GAT3,TCTN1,MPC2,TRPM4,MCFD2,PFKFB2,LGALS3,T
D4,RPL28,VIM,PSMA7,CDC14B,APEH,RPL21,LYPLA2,CAMLG,PRMT1,STOML2,WNK1,PIGL,STRADA,ZFAND
,MED6,SEC31A,PRLR,SH3GLB1,SSH2,COX17,CD151,NDUFV2,KCTD7,ASH2L,TCTN1,SMARCA4,COA3,SNRP
,RPS6,RTEL1,SRSF1,FOXRED1,RABL5,ITGB1BP1,PARD6G,SH3YL1,IP6K2,MICAL1,KCNMA1,PRMT2,EIF3M,J
M1,CDK11A,RBM3,ZNF197,CCNL2,MAPKAPK5,DGUOK,HSP90AB1,DUT,LRRFIP1,ZNF524,ATP5J,GPBP1,TS
,VIM,LIG1,PSMA7,CDC14B,RPL21,TXNL4A,TCF4,UCK2,PPOX,TMPO,EXOSC9,STOML2,ZNF7,ARL6IP4,HNRN

SNF8,GRK6,HMGA2,CDC37,RPS16,CPSF3L,TSG101,OBSL1,TTC21B,PTPN13,UBR4,PSMA1,YLPM1,PHF21A,

BFA2T2,CARS,ZNF250,RPL36,NFAM1,CDK11A,RBM3,ZNF197,INO80E,CCNL2,MAPKAPK5,HSP90AB1,LRRF
MTA1,PSMD4,RPL28,VIM,LIG1,PSMA7,CDC14B,RPL21,TXNL4A,TCF4,UCK2,PPOX,TMPO,EXOSC9,STOML

D74,PARK7,NFKBIA,RPS11,PDCD6,MFF,SNX5,ENY2,NUP133,RAPGEF3,RPAIN,ACOT2,PCID2,PEX2,TNKS,RI
1A,RBM3,ZNF197,CCNL2,MAPKAPK5,HSP90AB1,DUT,LRRFIP1,ZNF524,GPBP1,TSFM,ZEB2,TRMT112,RPT

250,RPL36,NFAM1,CDK11A,RBM3,ZNF197,CCNL2,MAPKAPK5,HSP90AB1,DUT,LRRFIP1,ZNF524,GPBP1,TS
ST1,UBE2I,PRMT2,EIF3M,MAGT1,CDC42,XBP1,PTK2,CELF1,GUK1,CARD14,FBXW9,USE1,AGPAT4,EEF1B2

DF1,GADL1,RFX3,PPA1,DGKE,CAMTA1,PSMD4,RPL28,VIM,LIG1,PSMA7,APEH,RPL21,TCF4,UCK2,PPOX,TN
G,SH3YL1,IP6K2,MICAL1,KCNMA1,PRMT2,EIF3M,JUP,CDC42,TBX6,PTK2,CELF1,AUNIP,CENPN,ASF1B,DEF

DGKE,CAMTA1,PSMD4,RPL28,VIM,LIG1,PSMA7,APEH,RPL21,TCF4,UCK2,PPOX,TMPO,PRMT1,EXOSC9,ST

EF1B2,PPP1CC,PPA1,TNFRSF14,CAMTA1,PPIL2,PSMD4,RPL28,VIM,PSMA7,CDC14B,APEH,RPL21,CAPN1C

24,ELP6,GPBP1,TSFM,ZEB2,TRMT112,RPTOR,ZNF625,RPS20,GABPB1,TWSG1,RPS15,ZNF814,HSPB1,PRKI

DF1,GADL1,RFX3,PPA1,DGKE,CAMTA1,PSMD4,RPL28,VIM,LIG1,PSMA7,APEH,RPL21,TCF4,UCK2,PPOX,TN

AT3,CD44,TM9SF1,SON,TET2,UQCRQ,SNF8,GRK6,HMGA2,CDC37,RPS16,CPSF3L,TSG101,OBSL1,TTC21B,F
2,PRKCE,STAG2,GRN,IK,CCT4,ANXA2,UBE2L3,PPP1R13B,PHPT1,LMNA,ACTG1,CDK10,VIL1,SSH3,TPM1,N

ADD1,NREP,TPT1,BAX,UBA1,ERRFI1,RNF8,EEF2K,MARS,CFLAR,NEK11,EIF2A,NUP62,RECQL5,ZNF675,PSM

NDD2,RPS6,RPL29,RTEL1,AUP1,SAP30BP,ITGB1BP1,MED15,UGGT2,MICAL1,HS2ST1,UBE2I,PRMT2,EIF3M

R62,STYXL1,RFX5,POLD4,PLEKHA5,RANBP1,UBE2L6,VRK3,ZC3H14,FIS1,SLC15A3,PPM1K,TUBA1C,IRF9,DS
,PARD6G,C1ORF198,MICAL1,UBE2I,PRMT2,EIF3M,JUP,CDC42,PTK2,GUK1,CENPN,FBXW9,HSBP1L1,EEF1
L2,PCMT1,EBPL,TESC,IGSF8,NUDT5,RBPMS,SNRPN,SPAG7,WDR62,STYXL1,RFX5,POLD4,PLEKHA5,RANBP
RCE1,MRPL33,ZNF410,RUVBL2,EBPL,TESC,IGSF8,NUDT5,RBPMS,SNRPN,SPAG7,WDR62,STYXL1,RFX5,POL
MT1,EBPL,TESC,NUDT5,RBPMS,NNMT,SNRPN,WDR62,STYXL1,PLEKHA5,RANBP1,UBE2L6,VRK3,LZTFL1,Z
D1,SH2B1,NOP16,RPL13A,GSTK1,TTLL5,LDHA,PSMA6,ACOT8,AP3B2,RPS9,SLC35G1,RCE1,MRPL33,ZNF41
,PSMA6,ACOT8,AP3B2,RPS9,SLC35G1,RCE1,MRPL33,ZNF410,RUVBL2,PCMT1,EBPL,TESC,IGSF8,NUDT5,R
K4,MRPL51,MRPL43,PPIE,EIF6,MRPL48,MRPL27,MRPS7,SNRNP40,RPL27,MRPL36,SNRNP200,RPS14,HNR
ENPK,RPS6,RTEL1,COASY,SRSF1,SAP30BP,TRNT1,ITGB1BP1,MED15,IP6K2,CLASRP,UGGT2,UBE2I,PRMT2,
ENPK,RPS6,RTEL1,COASY,SRSF1,SAP30BP,TRNT1,ITGB1BP1,MED15,IP6K2,CLASRP,UGGT2,UBE2I,PRMT2,
ENPK,RPS6,RTEL1,COASY,SRSF1,SAP30BP,TRNT1,ITGB1BP1,MED15,IP6K2,CLASRP,UGGT2,UBE2I,PRMT2,
UC1,METTL17,ECSIT,EPB41L5,PPRC1,RPS25,NLRP1,ZBTB44,NIP7,COX4I1,DUSP22,SLC25A10,CENPK,RPS6

TXNL4A,NUTF2,ATPAF2,PRMT1,EXOSC9,ARL6IP4,STRADA,HNRNPC,HYPK,UPP1,MOB1A,RBM3,INO80E,CC

1,RFX3,DGKE,ENDOV,CAMTA1,CAPG,PPIL2,PSMD4,VIM,LIG1,PSMA7,CDC14B,XAF1,TXNL4A,NUTF2,POD
2,PPP1CC,PDLIM5,RFX3,VAMP8,CAPG,PSMD4,RPL28,VIM,PSMA7,TIMM17B,RPL21,TXNL4A,NUTF2,TCF4
,THOP1,UBA52,AK2,VAPA,CAPN2,LRRC59,SUMO1,MRPS16,ARL2,SDHA,NUP85,NDUFA7,NDUFS2,MRPS3
,THOP1,UBA52,AK2,VAPA,CAPN2,LRRC59,SUMO1,MRPS16,ARL2,SDHA,NUP85,NDUFA7,NDUFS2,MRPS3

L28,VIM,CDC14B,RPL21,PODXL,EPB41L4B,TCF4,TMPO,EXOSC9,STOML2,ARL6IP4,PPAN,HNRNPC,STAU1,S
L28,VIM,CDC14B,RPL21,PODXL,EPB41L4B,TCF4,TMPO,EXOSC9,STOML2,ARL6IP4,PPAN,HNRNPC,STAU1,S
E2D3,MAGOHB,SNRPD1,DCAF5,UCHL5,TUSC3,INO80C,SNRPA1,NAA15,NDUFB3,NDUFB2,NDUFA10,KAT
,DUSP11,DDX56,MED1,SNAPC3,ZNF655,RBM4B,RPS14,ABHD14B,GLE1,ACACA,ESRRA,BYSL,POLR2L,H2AF
P53,SLC25A24,OXR1,NDUFA3,COX19,RPS3,MRPS24,IDH3G,DNAJA3,UCHL5,TUSC3,AIFM2,FARS2,SPHK2,D

L,TOMM40,BRIP1,SERPINB6,ATP6V1F,SLC33A1,NUDT1,NOMO2,B2M,ABHD17A,STXBP2,SGK1,SEC31A,A

DOST,SYNGR1,LSR,DOCK10,PSME2,PABPC1,PTPRB,DBNL,BCAM,H2AFV,RBMX,BLVRB,SERPINB6,RABEP2,

RPS3,RHEB,CREB3L2,GRAMD1A,TUSC3,HMOX2,SPHK2,IFI27,MFGE8,GOSR2,ATL2,PLD1,SLC37A3,PRKCSH,

S25,AIFM2,HMOX2,SPHK2,SLC6A9,DHRS4,IFI27,GOSR2,SPG21,LAMTOR1,PLD1,SNX3,SYPL1,FIG4,UBA52,

NDD1,SH2B1,NOP16,RPL13A,ARHGEF39,GSTK1,TTLL5,LDHA,PSMA6,ACOT8,AP3B2,RPS9,SLC35G1,RCE1,

6,ACOT8,AP3B2,RPS9,SLC35G1,RCE1,MRPL33,ZNF410,RUVBL2,PCMT1,EBPL,TESC,IGSF8,NUDT5,RBPMS,

,FAM3A,PSMA3,METTL17,ATG12,RPS25,NIP7,TTC9C,COX4I1,DUSP22,TMX2,THEM4,DBNDD2,RPS6,RPL2

OMMD5,CHID1,NUDT22,AMZ2,UFD1L,TAZ,THYN1,DBNDD1,SH2B1,NOP16,RPL13A,ARHGEF39,GSTK1,TT

OMMD5,CHID1,NUDT22,AMZ2,UFD1L,TAZ,THYN1,DBNDD1,SH2B1,NOP16,RPL13A,ARHGEF39,GSTK1,TT

,SH2B1,NOP16,RPL13A,ARHGEF39,GSTK1,TTLL5,LDHA,PSMA6,ACOT8,AP3B2,RPS9,SLC35G1,RCE1,MRPL

,SH2B1,NOP16,RPL13A,ARHGEF39,GSTK1,TTLL5,LDHA,PSMA6,ACOT8,AP3B2,RPS9,SLC35G1,RCE1,MRPL

5,LDHA,PSMA6,ACOT8,AP3B2,RPS9,SLC35G1,RCE1,MRPL33,ZNF410,RUVBL2,PCMT1,EBPL,TESC,IGSF8,N

1,TESC,IGSF8,NUDT5,SNRPN,SPAG7,WDR62,STYXL1,RFX5,POLD4,VRK3,LZTFL1,FIS1,PPM1K,TUBA1C,IRF9,

1,TESC,IGSF8,NUDT5,SNRPN,SPAG7,WDR62,STYXL1,RFX5,POLD4,VRK3,LZTFL1,FIS1,PPM1K,TUBA1C,IRF9,

ARHGEF39,GSTK1,TTLL5,LDHA,PSMA6,ACOT8,AP3B2,RPS9,SLC35G1,RCE1,MRPL33,ZNF410,RUVBL2,PC

FAM3A,METTL17,EPB41L5,ATG12,RPS25,NIP7,COX4I1,DUSP22,SLC25A10,TMX2,THEM4,CENPK,RPS6,RP

FAM3A,METTL17,EPB41L5,ATG12,RPS25,NIP7,COX4I1,DUSP22,SLC25A10,TMX2,THEM4,CENPK,RPS6,RP

C1ORF198,UBE2I,SLC39A13,EIF3M,CDC42,CARD14,PCYT2,FBXW9,USE1,IFI27L1,ASF1B,CCDC25,EEF1B2,R

TFL1,FIS1,PPM1K,TUBA1C,IRF9,DST,ANTXR2,STIP1,LRRC1,VCAN,LPCAT3,MAGED2,SON,UQCRQ,SNF8,GR

TFL1,FIS1,PPM1K,TUBA1C,IRF9,DST,ANTXR2,STIP1,LRRC1,VCAN,LPCAT3,MAGED2,SON,UQCRQ,SNF8,GR

HA,PSMA6,ACOT8,AP3B2,RPS9,SLC35G1,RCE1,MRPL33,ZNF410,RUVBL2,PCMT1,EBPL,TESC,IGSF8,NUDT

HA,PSMA6,ACOT8,AP3B2,RPS9,SLC35G1,RCE1,MRPL33,ZNF410,RUVBL2,PCMT1,EBPL,TESC,IGSF8,NUDT

1,VCAN,LPCAT3,TM9SF1,MAGED2,SON,UQCRQ,SNF8,GRK6,CDC37,RPS16,CPSF3L,TSG101,UBR4,YLPM1,

1,VCAN,LPCAT3,TM9SF1,MAGED2,SON,UQCRQ,SNF8,GRK6,CDC37,RPS16,CPSF3L,TSG101,UBR4,YLPM1,

EPB41L5,ATG12,RPS25,NIP7,COX4I1,DUSP22,SLC25A10,THEM4,RPS6,RPL29,RTEL1,COASY,AUP1,SAP30E

EPB41L5,ATG12,RPS25,NIP7,COX4I1,DUSP22,SLC25A10,THEM4,RPS6,RPL29,RTEL1,COASY,AUP1,SAP30E

1,PPM1K,TUBA1C,ANTXR2,STIP1,LRRC1,VCAN,LPCAT3,CD44,TM9SF1,MAGED2,SON,UQCRQ,SNF8,GRK6,

1,PPM1K,TUBA1C,ANTXR2,STIP1,LRRC1,VCAN,LPCAT3,CD44,TM9SF1,MAGED2,SON,UQCRQ,SNF8,GRK6,

DT22,AMZ2,UFD1L,USP38,TAZ,THYN1,DBNDD1,SH2B1,NOP16,RPL13A,ARHGEF39,GSTK1,TTLL5,LDHA,PS

DT22,AMZ2,UFD1L,USP38,TAZ,THYN1,DBNDD1,SH2B1,NOP16,RPL13A,ARHGEF39,GSTK1,TTLL5,LDHA,PS

BA1C,IRF9,DST,ANTXR2,STIP1,LRRC1,VCAN,LPCAT3,TM9SF1,MAGED2,SON,UQCRQ,SNF8,GRK6,CDC37,R

BA1C,IRF9,DST,ANTXR2,STIP1,LRRC1,VCAN,LPCAT3,TM9SF1,MAGED2,SON,UQCRQ,SNF8,GRK6,CDC37,R

F410,RUVBL2,PCMT1,TESC,IGSF8,NUDT5,SNRPN,SPAG7,WDR62,STYXL1,RFX5,POLD4,PLEKHA5,VRK3,LZT

F410,RUVBL2,PCMT1,TESC,IGSF8,NUDT5,SNRPN,SPAG7,WDR62,STYXL1,RFX5,POLD4,PLEKHA5,VRK3,LZT

D1,ITGB1BP1,MED15,SH3YL1,IP6K2,CLASRP,C1ORF198,UGGT2,MICAL1,UBE2I,PRMT2,EIF3M,CDC42,TBX

D1,ITGB1BP1,MED15,SH3YL1,IP6K2,CLASRP,C1ORF198,UGGT2,MICAL1,UBE2I,PRMT2,EIF3M,CDC42,TBX

L2,PCMT1,TESC,IGSF8,NUDT5,SNRPN,SPAG7,WDR62,STYXL1,RFX5,POLD4,PLEKHA5,LZTFL1,FIS1,PPM1K,

L2,PCMT1,TESC,IGSF8,NUDT5,SNRPN,SPAG7,WDR62,STYXL1,RFX5,POLD4,PLEKHA5,LZTFL1,FIS1,PPM1K,

MT1,TESC,IGSF8,NUDT5,SNRPN,SPAG7,WDR62,STYXL1,RFX5,POLD4,RANBP1,VRK3,LZTFL1,FIS1,PPM1K,T

MT1,TESC,IGSF8,NUDT5,SNRPN,SPAG7,WDR62,STYXL1,RFX5,POLD4,RANBP1,VRK3,LZTFL1,FIS1,PPM1K,T

A,PSMA6,ACOT8,AP3B2,SLC35G1,RCE1,MRPL33,ZNF410,RUVBL2,PCMT1,EBPL,TESC,IGSF8,NUDT5,RBPM

DT5,SNRPN,SPAG7,WDR62,STYXL1,RFX5,POLD4,VRK3,LZTFL1,FIS1,PPM1K,TUBA1C,IRF9,DST,ANTXR2,STI

DT5,SNRPN,SPAG7,WDR62,STYXL1,RFX5,POLD4,VRK3,LZTFL1,FIS1,PPM1K,TUBA1C,IRF9,DST,ANTXR2,STI

D1,TRNT1,KXD1,ITGB1BP1,MED15,SH3YL1,IP6K2,CLASRP,C1ORF198,MICAL1,UBE2I,SLC39A13,YIF1A,EIF3

,SLC35G1,RCE1,MRPL33,ZNF410,RUVBL2,PCMT1,EBPL,TESC,IGSF8,NUDT5,RBPMS,SPAG7,WDR62,IL1RA

,SLC35G1,RCE1,MRPL33,ZNF410,RUVBL2,PCMT1,EBPL,TESC,IGSF8,NUDT5,RBPMS,SPAG7,WDR62,IL1RA

,RPL28,LIG1,PSMA7,TMEM126B,APEH,CAPN10,LYPLA2,NUTF2,ATPAF2,LSM12,C17ORF62,PRMT1,C19OR

B1,RPL13A,GSTK1,TTLL5,LDHA,PSMA6,ACOT8,AP3B2,RPS9,SLC35G1,RCE1,MRPL33,ZNF410,RUVBL2,PC

DT22,AMZ2,UFD1L,USP38,TAZ,THYN1,DBNDD1,SH2B1,NOP16,RPL13A,ARHGEF39,GSTK1,TTLL5,LDHA,PS

DT22,AMZ2,UFD1L,USP38,TAZ,THYN1,DBNDD1,SH2B1,NOP16,RPL13A,ARHGEF39,GSTK1,TTLL5,LDHA,PS

RPS16,CPSF3L,TSG101,UBR4,YLPM1,FUZ,FAM3A,PSMA3,EPB41L5,ATG12,RPS25,NIP7,TTC9C,COX4I1,DU

RPS16,CPSF3L,TSG101,UBR4,YLPM1,FUZ,FAM3A,PSMA3,EPB41L5,ATG12,RPS25,NIP7,TTC9C,COX4I1,DU

A1C,IRF9,DST,ANTXR2,STIP1,LRRC1,VCAN,LPCAT3,MAGED2,SON,UQCRQ,SNF8,HMGA2,CDC37,RPS16,CPS

A1C,IRF9,DST,ANTXR2,STIP1,LRRC1,VCAN,LPCAT3,MAGED2,SON,UQCRQ,SNF8,HMGA2,CDC37,RPS16,CPS

ACOT8,AP3B2,RPS9,SLC35G1,RCE1,MRPL33,ZNF410,RUVBL2,PCMT1,EBPL,TESC,IGSF8,NUDT5,RBPMS,SN

ACOT8,AP3B2,RPS9,SLC35G1,RCE1,MRPL33,ZNF410,RUVBL2,PCMT1,EBPL,TESC,IGSF8,NUDT5,RBPMS,SN

,HMGA2,CDC37,RPS16,CPSF3L,TSG101,UBR4,CHCHD1,FUZ,FAM3A,METTL17,EPB41L5,ATG12,RPS25,NIP

,HMGA2,CDC37,RPS16,CPSF3L,TSG101,UBR4,CHCHD1,FUZ,FAM3A,METTL17,EPB41L5,ATG12,RPS25,NIP

T1,TESC,IGSF8,NUDT5,SNRPN,SPAG7,WDR62,STYXL1,RFX5,POLD4,RANBP1,UBE2L6,VRK3,LZTFL1,FIS1,P

T1,TESC,IGSF8,NUDT5,SNRPN,SPAG7,WDR62,STYXL1,RFX5,POLD4,RANBP1,UBE2L6,VRK3,LZTFL1,FIS1,P

MX2,THEM4,DBNDD2,RPS6,RPL29,GGPS1,RTEL1,COASY,AUP1,SAP30BP,YIPF3,FOXRED1,RABL5,TRNT1,K

MX2,THEM4,DBNDD2,RPS6,RPL29,GGPS1,RTEL1,COASY,AUP1,SAP30BP,YIPF3,FOXRED1,RABL5,TRNT1,K

SG101,UBR4,PHF21A,FUZ,FAM3A,METTL17,ATG12,RPS25,NIP7,COX4I1,DUSP22,SLC25A10,TMX2,THEM

SG101,UBR4,PHF21A,FUZ,FAM3A,METTL17,ATG12,RPS25,NIP7,COX4I1,DUSP22,SLC25A10,TMX2,THEM

K1,TTLL5,LDHA,PSMA6,ACOT8,RPS9,SLC35G1,MRPL33,ZNF410,RUVBL2,PCMT1,EBPL,TESC,IGSF8,NUDT

L1,ASF1B,CCDC25,EEF1B2,RFX3,DGKE,ENDOV,CAMTA1,PPIL2,PSMD4,RPL28,APEH,RPL21,CAPN10,LYPLA

V,CAMTA1,PPIL2,RPL28,LIG1,APEH,CAPN10,LYPLA2,NUTF2,C17ORF62,PRMT1,ZNF7,ARL6IP4,PIGL,C19O

PSF3L,TSG101,UBR4,YLPM1,CHCHD1,FUZ,FAM3A,METTL17,EPB41L5,ATG12,RPS25,NIP7,TTC9C,COX4I1,

PSF3L,TSG101,UBR4,YLPM1,CHCHD1,FUZ,FAM3A,METTL17,EPB41L5,ATG12,RPS25,NIP7,TTC9C,COX4I1,I
CAT3,CD44,TM9SF1,MAGED2,SON,UQCRQ,SNF8,GRK6,HMGA2,CDC37,RPS16,CPSF3L,TSG101,TTC21B,UB
CAT3,CD44,TM9SF1,MAGED2,SON,UQCRQ,SNF8,GRK6,HMGA2,CDC37,RPS16,CPSF3L,TSG101,TTC21B,UB
01,UBR4,SLC39A11,FUZ,FAM3A,METTL17,ATG12,RPS25,NIP7,TTC9C,COX4I1,PLCB2,DUSP22,SLC25A10,T
01,UBR4,SLC39A11,FUZ,FAM3A,METTL17,ATG12,RPS25,NIP7,TTC9C,COX4I1,PLCB2,DUSP22,SLC25A10,T
PK,RPS6,RPL29,RTEL1,COASY,SRSF1,AUP1,SAP30BP,YIPF3,FOXRED1,RABL5,TRNT1,KXD1,ITGB1BP1,MED1
PK,RPS6,RPL29,RTEL1,COASY,SRSF1,AUP1,SAP30BP,YIPF3,FOXRED1,RABL5,TRNT1,KXD1,ITGB1BP1,MED1
I4,TMED4,RPS6,RPL29,RTEL1,COASY,AUP1,SAP30BP,YIPF3,FOXRED1,RABL5,TRNT1,KXD1,ITGB1BP1,MED
I4,TMED4,RPS6,RPL29,RTEL1,COASY,AUP1,SAP30BP,YIPF3,FOXRED1,RABL5,TRNT1,KXD1,ITGB1BP1,MED
S16,CPSF3L,TSG101,UBR4,YLPM1,PHF21A,CHCHD1,FUZ,FAM3A,ATG12,RPS25,NIP7,TTC9C,COX4I1,DUSP
S16,CPSF3L,TSG101,UBR4,YLPM1,PHF21A,CHCHD1,FUZ,FAM3A,ATG12,RPS25,NIP7,TTC9C,COX4I1,DUSP
IUDT5,RBPMS,SNRPN,SPAG7,WDR62,STYXL1,RFX5,POLD4,RANBP1,VRK3,LZTFL1,FIS1,PPM1K,TUBA1C,IR
IUDT5,RBPMS,SNRPN,SPAG7,WDR62,STYXL1,RFX5,POLD4,RANBP1,VRK3,LZTFL1,FIS1,PPM1K,TUBA1C,IR
8,CDC37,RPS16,CPSF3L,TSG101,UBR4,YLPM1,CHCHD1,FUZ,FAM3A,PSMA3,EPB41L5,ATG12,RPS25,NIP7,
8,CDC37,RPS16,CPSF3L,TSG101,UBR4,YLPM1,CHCHD1,FUZ,FAM3A,PSMA3,EPB41L5,ATG12,RPS25,NIP7,
DGUOK,LRRFIP1,ZNF524,ATP5J,TRMT112,CHPF,RPS20,SAT1,ERCC5,EMC4,SERPINB1,DDOST,LSR,PABPC1,
M1K,IRF9,DST,ANTXR2,STIP1,LRRC1,VCAN,LPCAT3,TM9SF1,MAGED2,SON,UQCRQ,SNF8,GRK6,HMGA2,CI
M1K,IRF9,DST,ANTXR2,STIP1,LRRC1,VCAN,LPCAT3,TM9SF1,MAGED2,SON,UQCRQ,SNF8,GRK6,HMGA2,CI
,DUSP22,SLC25A10,THEM4,DBNDD2,RPS6,RPL29,RTEL1,COASY,SRSF1,AUP1,SAP30BP,YIPF3,FOXRED1,R
,DUSP22,SLC25A10,THEM4,DBNDD2,RPS6,RPL29,RTEL1,COASY,SRSF1,AUP1,SAP30BP,YIPF3,FOXRED1,R
F8,GRK6,HMGA2,CDC37,RPS16,CPSF3L,TSG101,UBR4,PHF21A,CHCHD1,FUZ,FAM3A,METTL17,EPB41L5,A
F8,GRK6,HMGA2,CDC37,RPS16,CPSF3L,TSG101,UBR4,PHF21A,CHCHD1,FUZ,FAM3A,METTL17,EPB41L5,A
2,SON,UQCRQ,SNF8,CDC37,RPS16,CPSF3L,TSG101,UBR4,SLC39A11,FUZ,FAM3A,PSMA3,ATG12,RPS25,NI
2,SON,UQCRQ,SNF8,CDC37,RPS16,CPSF3L,TSG101,UBR4,SLC39A11,FUZ,FAM3A,PSMA3,ATG12,RPS25,NI
DOST,LSR,RRNAD1,NAT9,TOMM40,RABEP2,TADA3,ZNF20,WAC,KLK10,ANAPC11,RAD23A,ABHD17A,NT5
SF3L,TSG101,UBR4,PHF21A,CHCHD1,FUZ,FAM3A,PSMA3,ATG12,RPS25,NIP7,TTC9C,COX4I1,DUSP22,SLC
SF3L,TSG101,UBR4,PHF21A,CHCHD1,FUZ,FAM3A,PSMA3,ATG12,RPS25,NIP7,TTC9C,COX4I1,DUSP22,SLC
R2,STIP1,LRRC1,PHF19,VCAN,LPCAT3,CD44,TM9SF1,MAGED2,SON,TET2,GRK6,CDC37,RPS16,CPSF3L,TSG
R2,STIP1,LRRC1,PHF19,VCAN,LPCAT3,CD44,TM9SF1,MAGED2,SON,TET2,GRK6,CDC37,RPS16,CPSF3L,TSG
7,COPZ1,DGUOK,LRRFIP1,ZNF524,ATP5J,RMDN1,TRMT112,CHPF,RPS20,GABPB1,MPDU1,DCTD,UBXN1,E
9,TOMM40,RABEP2,TADA3,ZNF20,WAC,SLC33A1,ZNF687,ANAPC11,RAD23A,ABHD17A,NT5C,MDH2,STX
,SNRPN,SPAG7,WDR62,STYXL1,RFX5,POLD4,RANBP1,VRK3,LZTFL1,FIS1,PPM1K,TUBA1C,IRF9,DST,ANTXR
,SNRPN,SPAG7,WDR62,STYXL1,RFX5,POLD4,RANBP1,VRK3,LZTFL1,FIS1,PPM1K,TUBA1C,IRF9,DST,ANTXR
,TESC,NUDT5,RBPMS,SPAG7,WDR62,IL1RAP,STYXL1,RFX5,POLD4,PLEKHA5,RANBP1,UBE2L6,VRK3,LZTFL
,TESC,NUDT5,RBPMS,SPAG7,WDR62,IL1RAP,STYXL1,RFX5,POLD4,PLEKHA5,RANBP1,UBE2L6,VRK3,LZTFL
D1L,USP38,TAZ,THYN1,DBNDD1,SH2B1,NOP16,RPL13A,ARHGEF39,GSTK1,TTLL5,LDHA,PSMA6,ACOT8,AP
D1L,USP38,TAZ,THYN1,DBNDD1,SH2B1,NOP16,RPL13A,ARHGEF39,GSTK1,TTLL5,LDHA,PSMA6,ACOT8,AP
RBM3,ZNF197,DGUOK,DUT,LRRFIP1,RAB2A,ZNF524,ATP5J,TRMT112,CHPF,RPS20,MPDU1,DCTD,SAT1,ER
T1,EBPL,TESC,IGSF8,NUDT5,RBPMS,SNRPN,SPAG7,WDR62,IL1RAP,STYXL1,RFX5,POLD4,RANBP1,UBE2L6,
T1,EBPL,TESC,IGSF8,NUDT5,RBPMS,SNRPN,SPAG7,WDR62,IL1RAP,STYXL1,RFX5,POLD4,RANBP1,UBE2L6,
3,AP3B2,RPS9,RCE1,MRPL33,ZNF410,RUVBL2,PCMT1,EBPL,TESC,IGSF8,NUDT5,RBPMS,SNRPN,SPAG7,WI
3,AP3B2,RPS9,RCE1,MRPL33,ZNF410,RUVBL2,PCMT1,EBPL,TESC,IGSF8,NUDT5,RBPMS,SNRPN,SPAG7,WI
ORF62,PRMT1,C19ORF54,ZNF7,TXNDC17,ARL6IP4,PIGL,C19ORF48,PPAN,ZFAND2B,ACAD10,SUGT1,HYPK
L,COASY,AUP1,SAP30BP,YIPF3,RABL5,TRNT1,KXD1,ITGB1BP1,MED15,IP6K2,CLASRP,C1ORF198,UGGT2,M
L,COASY,AUP1,SAP30BP,YIPF3,RABL5,TRNT1,KXD1,ITGB1BP1,MED15,IP6K2,CLASRP,C1ORF198,UGGT2,M
,MDH2,STXBP2,CHCHD5,SEC31A,CNST,COX17,COX5A,HIGD1A,SIRT5,ZNF506,PFDN5,PAFAH1B2,ZNF343,I
PKAPK5,DGUOK,LRRFIP1,ZNF524,ATP5J,TRMT112,CHPF,RPS20,MPDU1,DCTD,UBXN1,ERLEC1,SAT1,ERCC

,CARD14,PCYT2,FBXW9,USE1,ASF1B,EEF1B2,PPP1CC,RFX3,DGKE,ENDOV,CAMTA1,PPIL2,RPL28,LIG1,PSM
1,TM9SF1,MAGED2,SON,UQCRQ,SNF8,CDC37,RPS16,CPSF3L,TSG101,UBR4,CHCHD1,FUZ,FAM3A,ECSIT,
1,TM9SF1,MAGED2,SON,UQCRQ,SNF8,CDC37,RPS16,CPSF3L,TSG101,UBR4,CHCHD1,FUZ,FAM3A,ECSIT,
,DST,ANTXR2,STIP1,LRRC1,VCAN,LPCAT3,CD44,TM9SF1,MAGED2,SON,UQCRQ,SNF8,CDC37,RPS16,CPSF
,DST,ANTXR2,STIP1,LRRC1,VCAN,LPCAT3,CD44,TM9SF1,MAGED2,SON,UQCRQ,SNF8,CDC37,RPS16,CPSF
XW9,USE1,IFI27L1,ASF1B,CCDC25,RFX3,TNFRSF14,ENDOV,CAMTA1,PPIL2,RPL28,LIG1,TIMM17B,TMEM
,SNRPN,SPAG7,WDR62,IL1RAP,STYXL1,RFX5,POLD4,PLEKHA5,UBE2L6,VRK3,LZTFL1,ZC3H14,FIS1,PPM1K
,SNRPN,SPAG7,WDR62,IL1RAP,STYXL1,RFX5,POLD4,PLEKHA5,UBE2L6,VRK3,LZTFL1,ZC3H14,FIS1,PPM1K
RMT112,CHPF,RPS20,MPDU1,ZNF814,SAT1,ERCC5,EMC4,SERPINB1,DDOST,LSR,PABPC1,RRNAD1,NAT9,
A,ARHGEF39,GSTK1,TTLL5,LDHA,PSMA6,ACOT8,AP3B2,RPS9,SLC35G1,RCE1,MRPL33,RUVBL2,PCMT1,EB
A,ARHGEF39,GSTK1,TTLL5,LDHA,PSMA6,ACOT8,AP3B2,RPS9,SLC35G1,RCE1,MRPL33,RUVBL2,PCMT1,EB
20,WAC,ANAPC11,RAD23A,NT5C,MDH2,STXBP2,EEF1D,CHCHD5,SEC31A,CNST,COX17,HIGD1A,SIRT5,INT
BA1C,ANTXR2,STIP1,LRRC1,PHF19,VCAN,LPCAT3,CD44,TM9SF1,MAGED2,SON,TET2,GRK6,CDC37,RPS16,
BA1C,ANTXR2,STIP1,LRRC1,PHF19,VCAN,LPCAT3,CD44,TM9SF1,MAGED2,SON,TET2,GRK6,CDC37,RPS16,
1,DCTD,ZNF814,HSPB1,ERCC5,EMC4,DDOST,LSR,MEIS2,RRNAD1,TOMM40,TADA3,ZNF20,ZNF687,PDCD5
1,DCTD,ZNF814,HSPB1,ERCC5,EMC4,DDOST,LSR,MEIS2,RRNAD1,TOMM40,TADA3,ZNF20,ZNF687,PDCD5
C25,EEF1B2,RFX3,DGKE,ENDOV,CAMTA1,PPIL2,PSMD4,RPL28,APEH,RPL21,CAPN10,LYPLA2,UCK2,C17OR
4,FIS1,SLC15A3,PPM1K,TUBA1C,DST,LRRC1,PHF19,GRAMD1C,LPCAT3,CD44,TM9SF1,MAGED2,SON,TET
K,LRRFIP1,ATP5J,RMDN1,TRMT112,CHPF,RPS20,UBXN1,ERLEC1,SAT1,ERCC5,EMC4,MCTS1,SERPINB1,DD
C12ORF57,DCTN5,PEX26,NDUFS3,ARL16,OXLD1,METTL5,SLC35E1,OGFOD2,EIF3G,TSEN34,ARIH2,PICK1,
3M,CDC42,CARD14,PCYT2,FBXW9,USE1,IFI27L1,MAF,ASF1B,CCDC25,EEF1B2,RFX3,TNFRSF14,ENDOV,C
ARD14,PCYT2,FBXW9,USE1,ASF1B,CCDC25,EEF1B2,RFX3,ENDOV,PPIL2,RPL28,LIG1,CDC14B,TIMM17B,A
ARD14,PCYT2,FBXW9,USE1,ASF1B,CCDC25,EEF1B2,RFX3,ENDOV,PPIL2,RPL28,LIG1,CDC14B,TIMM17B,A
RNAD1,TOMM40,TADA3,ZNF20,ZNF687,PDCD5,ANAPC11,RAD23A,ABHD17A,NT5C,MDH2,EEF1D,NAA10
RNAD1,TOMM40,TADA3,ZNF20,ZNF687,PDCD5,ANAPC11,RAD23A,ABHD17A,NT5C,MDH2,EEF1D,NAA10
SP22,SLC25A10,TMX2,THEM4,RPS6,RTEL1,COASY,SRSF1,AUP1,SAP30BP,YIPF3,FOXRED1,RABL5,TRNT1,K
SP22,SLC25A10,TMX2,THEM4,RPS6,RTEL1,COASY,SRSF1,AUP1,SAP30BP,YIPF3,FOXRED1,RABL5,TRNT1,K
,MRPL33,ZNF410,RUVBL2,PCMT1,EBPL,TESC,NUDT5,RBPMS,SPAG7,WDR62,IL1RAP,STYXL1,POLD4,PLEK
,MRPL33,ZNF410,RUVBL2,PCMT1,EBPL,TESC,NUDT5,RBPMS,SPAG7,WDR62,IL1RAP,STYXL1,POLD4,PLEK
24,ATP5J,TRMT112,CHPF,RPS20,MPDU1,UBXN1,SAT1,ERCC5,EMC4,SERPINB1,DDOST,LSR,PSME2,PABP
28,APEH,RPL21,CAPN10,LYPLA2,C17ORF62,PRMT1,STOML2,C19ORF54,ZNF7,TXNDC17,ARL6IP4,PIGL,C1
,ACOT8,AP3B2,RPS9,SLC35G1,RCE1,MRPL33,ZNF410,RUVBL2,PCMT1,EBPL,TESC,IGSF8,NUDT5,RBPMS,
,ACOT8,AP3B2,RPS9,SLC35G1,RCE1,MRPL33,ZNF410,RUVBL2,PCMT1,EBPL,TESC,IGSF8,NUDT5,RBPMS,
RMDN1,TRMT112,CHPF,RPS20,MPDU1,ERCC5,EMC4,SERPINB1,DDOST,LSR,PABPC1,RRNAD1,DBNL,NAT9
250,ZNF197,COPZ1,INO80E,HSP90AB1,RAB2A,ATP5J,TRMT112,CHPF,GABPB1,DCTD,UBXN1,ZNF814,HSP
250,ZNF197,COPZ1,INO80E,HSP90AB1,RAB2A,ATP5J,TRMT112,CHPF,GABPB1,DCTD,UBXN1,ZNF814,HSP
NDD2,RPS6,RPL29,GGPS1,RTEL1,COASY,SRSF1,AUP1,SAP30BP,FOXRED1,RABL5,TRNT1,KXD1,ITGB1BP1,
NDD2,RPS6,RPL29,GGPS1,RTEL1,COASY,SRSF1,AUP1,SAP30BP,FOXRED1,RABL5,TRNT1,KXD1,ITGB1BP1,
,SUGT1,HYPK,CARS,ZNF250,ZNF197,COPZ1,INO80E,HSP90AB1,RAB2A,ATP5J,ELP6,TRMT112,CHPF,DCT
,SUGT1,HYPK,CARS,ZNF250,ZNF197,COPZ1,INO80E,HSP90AB1,RAB2A,ATP5J,ELP6,TRMT112,CHPF,DCT
FBXW9,USE1,ASF1B,CCDC25,EEF1B2,RFX3,ENDOV,VAMP8,PPIL2,TROAP,LIG1,CDC14B,APEH,CAPN10,RB
FBXW9,USE1,ASF1B,CCDC25,EEF1B2,RFX3,ENDOV,VAMP8,PPIL2,TROAP,LIG1,CDC14B,APEH,CAPN10,RB
ED15,SH3YL1,CLASRP,C1ORF198,UGGT2,MICAL1,UBE2I,PRMT2,EIF3M,CDC42,TBX6,TMEM205,PTK2,CA
ED15,SH3YL1,CLASRP,C1ORF198,UGGT2,MICAL1,UBE2I,PRMT2,EIF3M,CDC42,TBX6,TMEM205,PTK2,CA
,RPS9,SLC35G1,RCE1,MRPL33,ZNF410,RUVBL2,PCMT1,TESC,IGSF8,NUDT5,RBPMS,SPAG7,WDR62,IL1RA
,RPS9,SLC35G1,RCE1,MRPL33,ZNF410,RUVBL2,PCMT1,TESC,IGSF8,NUDT5,RBPMS,SPAG7,WDR62,IL1RA
250,CDK11A,RBM3,ZNF197,COPZ1,INO80E,CCNL2,LRRFIP1,ATP5J,RMDN1,TRMT112,CHPF,STRIP2,RPS20,

,RPL21,CAPN10,LYPLA2,NUTF2,UCK2,C17ORF62,PRMT1,STOML2,C19ORF54,ZNF7,TXNDC17,ARL6IP4,PIG
T2,FBXW9,USE1,ASF1B,CCDC25,EEF1B2,RFX3,ENDOV,CAPG,PPIL2,LIG1,PSMA7,CDC14B,APEH,CAPN10,L
T2,FBXW9,USE1,ASF1B,CCDC25,EEF1B2,RFX3,ENDOV,CAPG,PPIL2,LIG1,PSMA7,CDC14B,APEH,CAPN10,L
,TIMM17B,APEH,RPL21,CAPN10,LYPLA2,UCK2,C17ORF62,PRMT1,CASC4,STOML2,C19ORF54,ZNF7,TXN
RFX3,ENDOV,PPIL2,RPL28,CLDND1,LIG1,CDC14B,TIMM17B,APEH,RPL21,CAPN10,LYPLA2,EPB41L4B,UCK
RFX3,ENDOV,PPIL2,RPL28,CLDND1,LIG1,CDC14B,TIMM17B,APEH,RPL21,CAPN10,LYPLA2,EPB41L4B,UCK
DA3,ZNF20,WAC,TRIM16,ANAPC11,ABHD17A,NT5C,MDH2,STXBP2,EEF1D,CHCHD5,SEC31A,CARHSP1,CN
COT8,AP3B2,RPS9,SLC35G1,RCE1,MRPL33,RUVBL2,PCMT1,EBPL,TESC,IGSF8,NUDT5,RBPMS,NNMT,SPAG
PCYT2,FBXW9,USE1,IFI27L1,ASF1B,CCDC25,EEF1B2,PPP1CC,RFX3,DGKE,ENDOV,CAMTA1,CRIP1,PPIL2,R
RHGEF39,GSTK1,TTLL5,LDHA,PSMA6,ACOT8,AP3B2,RPS9,SLC35G1,RCE1,MRPL33,ZNF410,RUVBL2,PCMT
1,ERCC5,EMC4,MCTS1,DDOST,LSR,MEIS2,RRNAD1,RBP1,TOMM40,TADA3,ZNF687,PDCD5,ANAPC11,RA
1,ERCC5,EMC4,MCTS1,DDOST,LSR,MEIS2,RRNAD1,RBP1,TOMM40,TADA3,ZNF687,PDCD5,ANAPC11,RA
PAG7,WDR62,STYXL1,RFX5,POLD4,PLEKHA5,VRK3,LZTFL1,FIS1,PPM1K,TUBA1C,IRF9,DST,ANTXR2,STIP1,
PAG7,WDR62,STYXL1,RFX5,POLD4,PLEKHA5,VRK3,LZTFL1,FIS1,PPM1K,TUBA1C,IRF9,DST,ANTXR2,STIP1,
TC9C,COX4I1,PLCB2,DUSP22,SLC25A10,TMX2,THEM4,RDH11,DBNDD2,RPS6,RPL29,RTEL1,COASY,AUP1,
TC9C,COX4I1,PLCB2,DUSP22,SLC25A10,TMX2,THEM4,RDH11,DBNDD2,RPS6,RPL29,RTEL1,COASY,AUP1,
TC9C,COX4I1,PLCB2,DUSP22,SLC25A10,TMX2,THEM4,RDH11,DBNDD2,RPS6,RPL29,RTEL1,COASY,AUP1,
TC9C,COX4I1,PLCB2,DUSP22,SLC25A10,TMX2,THEM4,RDH11,DBNDD2,RPS6,RPL29,RTEL1,COASY,AUP1,
,ATP5J,TRMT112,CHPF,ZNF814,HSPB1,VWA2,ERCC5,EMC4,LMBR1,DDOST,LSR,MEIS2,RRNAD1,RAD1,TO
,ATP5J,TRMT112,CHPF,ZNF814,HSPB1,VWA2,ERCC5,EMC4,LMBR1,DDOST,LSR,MEIS2,RRNAD1,RAD1,TO
M,CDC42,TBX6,TMEM205,PTK2,CARD14,FBXW9,USE1,HSBP1L1,ASF1B,CCDC25,EEF1B2,RFX3,ENDOV,VA
M,CDC42,TBX6,TMEM205,PTK2,CARD14,FBXW9,USE1,HSBP1L1,ASF1B,CCDC25,EEF1B2,RFX3,ENDOV,VA
INB1,DDOST,LSR,MEIS2,PSME2,NAT9,ZNF836,RAD1,TOMM40,RABEP2,TADA3,ZNF20,ZNF687,KLK10,AN
S,ZNF250,RPL36,CDK11A,ZNF197,INO80E,DGUOK,HSP90AB1,RAB2A,ZNF524,ATP5J,TRMT112,CHPF,MP
S,ZNF250,RPL36,CDK11A,ZNF197,INO80E,DGUOK,HSP90AB1,RAB2A,ZNF524,ATP5J,TRMT112,CHPF,MP
L1,FIS1,PPM1K,TUBA1C,IRF9,DST,ANTXR2,STIP1,LRRC1,VCAN,CD44,TM9SF1,MAGED2,SON,UQCRQ,SNF
L1,FIS1,PPM1K,TUBA1C,IRF9,DST,ANTXR2,STIP1,LRRC1,VCAN,CD44,TM9SF1,MAGED2,SON,UQCRQ,SNF
,LSR,MEIS2,TOMM40,TADA3,PDCD5,TRIM16,ANAPC11,RAD23A,ABHD17A,MDH2,EEF1D,HSPH1,CHCHD
,LSR,MEIS2,TOMM40,TADA3,PDCD5,TRIM16,ANAPC11,RAD23A,ABHD17A,MDH2,EEF1D,HSPH1,CHCHD
,UBR4,SLC39A11,FUZ,FAM3A,METTL17,EPB41L5,ATG12,RPS25,ZBTB44,NIP7,TTC9C,COX4I1,DUSP22,SLC
,UBR4,SLC39A11,FUZ,FAM3A,METTL17,EPB41L5,ATG12,RPS25,ZBTB44,NIP7,TTC9C,COX4I1,DUSP22,SLC
MS,SNRPN,SPAG7,IL1RAP,STYXL1,RFX5,POLD4,RANBP1,UBE2L6,VRK3,LZTFL1,FIS1,PPM1K,TUBA1C,IRF9,
MS,SNRPN,SPAG7,IL1RAP,STYXL1,RFX5,POLD4,RANBP1,UBE2L6,VRK3,LZTFL1,FIS1,PPM1K,TUBA1C,IRF9,
97,COPZ1,INO80E,CCNL2,HSP90AB1,LRRFIP1,RAB2A,ZNF524,ATP5J,TRMT112,CHPF,NIT2,ZNF814,HSPB1
97,COPZ1,INO80E,CCNL2,HSP90AB1,LRRFIP1,RAB2A,ZNF524,ATP5J,TRMT112,CHPF,NIT2,ZNF814,HSPB1
C37,CPSF3L,TSG101,UBR4,PSMA1,FUZ,FAM3A,PSMA3,METTL17,ATG12,RPS25,NIP7,COX4I1,DUSP22,SL
C37,CPSF3L,TSG101,UBR4,PSMA1,FUZ,FAM3A,PSMA3,METTL17,ATG12,RPS25,NIP7,COX4I1,DUSP22,SL
,ARHGEF39,TTLL5,LDHA,PSMA6,ACOT8,AP3B2,RPS9,SLC35G1,RCE1,MRPL33,ZNF410,RUVBL2,PCMT1,EB
L1,RFX5,POLD4,PLEKHA5,RANBP1,VRK3,LZTFL1,FIS1,PPM1K,TUBA1C,IRF9,DST,ANTXR2,STIP1,LRRC1,PH
L1,RFX5,POLD4,PLEKHA5,RANBP1,VRK3,LZTFL1,FIS1,PPM1K,TUBA1C,IRF9,DST,ANTXR2,STIP1,LRRC1,PH
,PHF19,VCAN,LPCAT3,CD44,TM9SF1,MAGED2,SON,TET2,GRK6,CDC37,RPS16,CPSF3L,TSG101,OBSL1,TT
,PHF19,VCAN,LPCAT3,CD44,TM9SF1,MAGED2,SON,TET2,GRK6,CDC37,RPS16,CPSF3L,TSG101,OBSL1,TT
MM40,RABEP2,TADA3,KLK10,ANAPC11,ABHD17A,NT5C,MDH2,STXBP2,EEF1D,CHCHD5,SEC31A,CNST,
RC1,PHF19,VCAN,LPCAT3,CD44,TM9SF1,MAGED2,SON,TET2,GRK6,CDC37,RPS16,CPSF3L,TSG101,OBSL1,
RC1,PHF19,VCAN,LPCAT3,CD44,TM9SF1,MAGED2,SON,TET2,GRK6,CDC37,RPS16,CPSF3L,TSG101,OBSL1,
112,CHPF,DCTD,ZNF814,SAT1,ERCC5,EMC4,SERPINB1,DDOST,LSR,RRNAD1,TOMM40,PPP3CB,RABEP2,T
112,CHPF,DCTD,ZNF814,SAT1,ERCC5,EMC4,SERPINB1,DDOST,LSR,RRNAD1,TOMM40,PPP3CB,RABEP2,T

,ZNF814,SAT1,ERCC5,EMC4,DDOST,LSR,MEIS2,RRNAD1,TOMM40,PPP3CB,RABEP2,TADA3,ZNF20,ZNF6

,ZNF814,SAT1,ERCC5,EMC4,DDOST,LSR,MEIS2,RRNAD1,TOMM40,PPP3CB,RABEP2,TADA3,ZNF20,ZNF6

2,PCMT1,TESC,IGSF8,NUDT5,RBPMS,SPAG7,WDR62,IL1RAP,STYXL1,POLD4,PLEKHA5,RANBP1,UBE2L6,V

2,PCMT1,TESC,IGSF8,NUDT5,RBPMS,SPAG7,WDR62,IL1RAP,STYXL1,POLD4,PLEKHA5,RANBP1,UBE2L6,V

X17,HIGD1A,SIRT5,INTS4,B3GAT3,ZNF506,PFDN5,PAFAH1B2,ZNF343,GBA2,PPIH,DALRD3,NSUN5,DRG2,

FOXRED1,RABL5,TRNT1,ITGB1BP1,MED15,SH3YL1,CLASRP,C1ORF198,UGGT2,SLC39A13,PRMT2,EIF3M,C

FOXRED1,RABL5,TRNT1,ITGB1BP1,MED15,SH3YL1,CLASRP,C1ORF198,UGGT2,SLC39A13,PRMT2,EIF3M,C

L,SH2B1,NOP16,RPL13A,ARHGEF39,GSTK1,TTLL5,LDHA,ACOT8,AP3B2,RPS9,SLC35G1,RCE1,MRPL33,ZNF

L,SH2B1,NOP16,RPL13A,ARHGEF39,GSTK1,TTLL5,LDHA,ACOT8,AP3B2,RPS9,SLC35G1,RCE1,MRPL33,ZNF

,COASY,AUP1,SAP30BP,YIPF3,FOXRED1,RABL5,TRNT1,KXD1,ITGB1BP1,MED15,SH3YL1,IP6K2,C1ORF198,

,COASY,AUP1,SAP30BP,YIPF3,FOXRED1,RABL5,TRNT1,KXD1,ITGB1BP1,MED15,SH3YL1,IP6K2,C1ORF198,

,UFD1L,USP38,TAZ,THYN1,DBNDD1,SH2B1,NOP16,RPL13A,ARHGEF39,GSTK1,TTLL5,LDHA,PSMA6,ACOT

,UFD1L,USP38,TAZ,THYN1,DBNDD1,SH2B1,NOP16,RPL13A,ARHGEF39,GSTK1,TTLL5,LDHA,PSMA6,ACOT

,SERPINB1,DDOST,LSR,PABPC1,RRNAD1,DBNL,TOMM40,RABEP2,TADA3,ZNF20,ATP6V1F,WAC,TRIM16,

D44,TM9SF1,MAGED2,SON,SNF8,GRK6,CDC37,RPS16,CPSF3L,OBSL1,PTPN13,UBR4,PSMA1,YLPM1,PHF2

D44,TM9SF1,MAGED2,SON,SNF8,GRK6,CDC37,RPS16,CPSF3L,OBSL1,PTPN13,UBR4,PSMA1,YLPM1,PHF2

NL2,LRRFIP1,ATP5J,RMDN1,TRMT112,CHPF,RPS20,GABPB1,DCTD,UBXN1,ERLEC1,SAT1,SERPINB1,DDOS

GGT2,EIF3M,CDC42,TBX6,CARD14,FBXW9,USE1,IFI27L1,ASF1B,CCDC25,EEF1B2,PDLIM5,RFX3,ENDOV,CA

GGT2,EIF3M,CDC42,TBX6,CARD14,FBXW9,USE1,IFI27L1,ASF1B,CCDC25,EEF1B2,PDLIM5,RFX3,ENDOV,CA

,FOXRED1,RABL5,TRNT1,KXD1,ITGB1BP1,MED15,IP6K2,CLASRP,C1ORF198,UGGT2,MICAL1,HS2ST1,UBE2

,FOXRED1,RABL5,TRNT1,KXD1,ITGB1BP1,MED15,IP6K2,CLASRP,C1ORF198,UGGT2,MICAL1,HS2ST1,UBE2

CCDC25,EEF1B2,EDF1,RFX3,ENDOV,CAMTA1,PPIL2,RPL28,TIMM17B,APEH,RPL21,LYPLA2,NUTF2,C17OR

1,FUZ,FAM3A,PSMA3,ECSIT,ATG12,RPS25,NIP7,TTC9C,DUSP22,SLC25A10,TMX2,THEM4,DBNDD2,RPS6,

1,FUZ,FAM3A,PSMA3,ECSIT,ATG12,RPS25,NIP7,TTC9C,DUSP22,SLC25A10,TMX2,THEM4,DBNDD2,RPS6,

687,PDCD5,ANAPC11,RAD23A,ABHD17A,MDH2,EEF1D,CHCHD5,NAA10,SEC31A,CARHSP1,COX17,HNRN

687,PDCD5,ANAPC11,RAD23A,ABHD17A,MDH2,EEF1D,CHCHD5,NAA10,SEC31A,CARHSP1,COX17,HNRN

2,STXBP2,EEF1D,HSPH1,CHCHD5,SEC31A,CARHSP1,CNST,COX17,POLR2C,ZNF506,SPINT2,SMARCA4,PAF

C31A,CARHSP1,COX17,HIGD1A,POLR2C,B3GAT3,SPINT2,PFDN5,PAFAH1B2,TXNRD2,DALRD3,NSUN5,OA

C31A,CARHSP1,COX17,HIGD1A,POLR2C,B3GAT3,SPINT2,PFDN5,PAFAH1B2,TXNRD2,DALRD3,NSUN5,OA

2,EEF1D,HSPH1,CHCHD5,NAA10,CARHSP1,CNST,COX17,HNRNPLL,HIGD1A,POLR2C,B3GAT3,RFXANK,ZNF

2,EEF1D,HSPH1,CHCHD5,NAA10,CARHSP1,CNST,COX17,HNRNPLL,HIGD1A,POLR2C,B3GAT3,RFXANK,ZNF

40,TADA3,ZNF687,PDCD5,ANAPC11,RAD23A,ABHD17A,MDH2,EEF1D,NAA10,SEC31A,CARHSP1,COX17,

40,TADA3,ZNF687,PDCD5,ANAPC11,RAD23A,ABHD17A,MDH2,EEF1D,NAA10,SEC31A,CARHSP1,COX17,

RIC8A,TCP1,THADA,COMMD4,PIGT,UBE2N,LFNG,ZNHIT3,FAM32A,TMEM14B,COA1,PEX2,POLR1C,C12OR

3,XYLT2,FAM32A,ZSCAN20,COA1,TGOLN2,PEX2,POLR1C,STX10,C12ORF57,DCTN5,PEX26,NDUFS3,OXLD

INO80E,CCNL2,LRRFIP1,ATP5J,RMDN1,TRMT112,CHPF,RPS20,GABPB1,DCTD,UBXN1,ERLEC1,SAT1,SERPI

M1K,TUBA1C,ANTXR2,STIP1,PHF19,GRAMD1C,VCAN,TM9SF1,MAGED2,TET2,SNF8,GRK6,HMGA2,CDC37

M1K,TUBA1C,ANTXR2,STIP1,PHF19,GRAMD1C,VCAN,TM9SF1,MAGED2,TET2,SNF8,GRK6,HMGA2,CDC37

I2,RPL8,PICK1,NIPAL3,TPT1,SLC52A2,TSPAN31,MTIF2,RALY,ZNF593,GLYCTK,RER1,DOK4,CREBZF,ACAA1,

LC35G1,RCE1,ZNF410,RUVBL2,PCMT1,EBPL,TESC,IGSF8,NUDT5,RBPMS,SPAG7,WDR62,IL1RAP,POLD4,P

PZ1,INO80E,DGUOK,HSP90AB1,RAB2A,ATP5J,TSFM,TRMT112,CHPF,GABPB1,DCTD,ZNF814,HSPB1,VWA

PZ1,INO80E,DGUOK,HSP90AB1,RAB2A,ATP5J,TSFM,TRMT112,CHPF,GABPB1,DCTD,ZNF814,HSPB1,VWA

C37,RPS16,CPSF3L,OBSL1,UBR4,CHCHD1,SLC39A11,FAM3A,MUC1,METTL17,EPB41L5,ATG12,RPS25,NL

C37,RPS16,CPSF3L,OBSL1,UBR4,CHCHD1,SLC39A11,FAM3A,MUC1,METTL17,EPB41L5,ATG12,RPS25,NL

HID1,NUDT22,AMZ2,UFD1L,USP38,TAZ,DBNDD1,SH2B1,NOP16,RPL13A,ARHGEF39,GSTK1,TTLL5,LDHA,

HID1,NUDT22,AMZ2,UFD1L,USP38,TAZ,DBNDD1,SH2B1,NOP16,RPL13A,ARHGEF39,GSTK1,TTLL5,LDHA,

,PPM1K,TUBA1C,STIP1,LRRC1,PHF19,GRAMD1C,VCAN,LPCAT3,CD44,TM9SF1,TET2,SNF8,GRK6,HMGA2,

TAZ,THYN1,DBNDD1,SH2B1,NOP16,RPL13A,ARHGEF39,GSTK1,TTLL5,LDHA,PSMA6,ACOT8,AP3B2,RPS9,S
TAZ,THYN1,DBNDD1,SH2B1,NOP16,RPL13A,ARHGEF39,GSTK1,TTLL5,LDHA,PSMA6,ACOT8,AP3B2,RPS9,S
IT5C,PDE4DIP,MDH2,STXBP2,CHCHD5,SEC31A,CNST,SSH2,COX5A,HIGD1A,INTS4,B3GAT3,ZNF506,MTHFS
IT5C,PDE4DIP,MDH2,STXBP2,CHCHD5,SEC31A,CNST,SSH2,COX5A,HIGD1A,INTS4,B3GAT3,ZNF506,MTHFS
,PEX2,POLR1C,PIGH,C12ORF57,PEX26,NDUFS3,METTL5,SLC35E1,OGFOD2,METTL3,SSR3,ARIH2,PICK1,TF
A1,PSMD7,TRAPPC2,PIGO,CLTA,SLC35A2,PLAC8,AXIN1,TMEM199,SLC25A24,EPN1,HPS1,ITPA,ZCCHC9,V
RP,C1ORF198,UGGT2,SLC39A13,EIF3M,JUP,CDC42,PTK2,CARD14,FBXW9,USE1,ASF1B,CCDC25,EEF1B2,P
RP,C1ORF198,UGGT2,SLC39A13,EIF3M,JUP,CDC42,PTK2,CARD14,FBXW9,USE1,ASF1B,CCDC25,EEF1B2,P
TAZ,THYN1,DBNDD1,SH2B1,RPL13A,ARHGEF39,GSTK1,TTLL5,LDHA,PSMA6,ACOT8,AP3B2,RPS9,SLC35G1,
TAZ,THYN1,DBNDD1,SH2B1,RPL13A,ARHGEF39,GSTK1,TTLL5,LDHA,PSMA6,ACOT8,AP3B2,RPS9,SLC35G1,
,RALY,ZNF593,GLYCTK,TRIM46,CNTRL,RER1,EIF2A,TRAPPC2,GTF2H1,PIGO,AXIN1,TP53,TMEM199,SLC25
,SH2B1,NOP16,RPL13A,TTLL5,LDHA,PSMA6,ACOT8,RPS9,SLC35G1,RCE1,ZNF410,RUVBL2,PCMT1,EBPL,T
,SH2B1,NOP16,RPL13A,TTLL5,LDHA,PSMA6,ACOT8,RPS9,SLC35G1,RCE1,ZNF410,RUVBL2,PCMT1,EBPL,T
STIP1,LRRC1,PHF19,GRAMD1C,VCAN,LPCAT3,CD44,MAGED2,SON,TET2,UQCRQ,GRK6,CDC37,RPS16,CPS
STIP1,LRRC1,PHF19,GRAMD1C,VCAN,LPCAT3,CD44,MAGED2,SON,TET2,UQCRQ,GRK6,CDC37,RPS16,CPS
20,TGOLN2,PEX2,TNKS,C12ORF57,DCTN5,PEX26,NDUFS3,ARL16,OXLD1,EIF3G,ARIH2,PICK1,FBXL18,ZNF5
X2,NISCH,TMED4,CENPK,RPL29,CD99L2,GGPS1,AUP1,TRNT1,PARD6G,SH3YL1,IP6K2,CLASRP,JPH2,C1ORF
X2,NISCH,TMED4,CENPK,RPL29,CD99L2,GGPS1,AUP1,TRNT1,PARD6G,SH3YL1,IP6K2,CLASRP,JPH2,C1ORF
P1,UBE2L6,VRK3,LZTFL1,ZC3H14,SLC15A3,PPM1K,TUBA1C,ANTXR2,STIP1,LRRC1,PHF19,GRAMD1C,VCAN
7,ARL6IP4,STRADA,PPAN,ACAD10,CYB561,SHARPIN,SUGT1,HYPK,CARS,ZNF250,MDH1,ZNF197,INO80E,C
7,ARL6IP4,STRADA,PPAN,ACAD10,CYB561,SHARPIN,SUGT1,HYPK,CARS,ZNF250,MDH1,ZNF197,INO80E,C
K,ZNF250,RPL36,MOB1A,ZNF197,COPZ1,INO80E,CCNL2,HSP90AB1,ATP5J,TRMT112,CHPF,ZNF625,RPS1
K,ZNF250,RPL36,MOB1A,ZNF197,COPZ1,INO80E,CCNL2,HSP90AB1,ATP5J,TRMT112,CHPF,ZNF625,RPS1
H1B2,TXNRD2,DALRD3,NSUN5,OARD1,MFSD10,MRPS18C,GMPR2,ZSCAN9,RIC8A,TCP1,SMARCC2,UBE2
H1B2,TXNRD2,DALRD3,NSUN5,OARD1,MFSD10,MRPS18C,GMPR2,ZSCAN9,RIC8A,TCP1,SMARCC2,UBE2
0,USP40,PEX2,POLR1C,C12ORF57,DCTN5,NDUFS3,ARL16,OXLD1,ZNF223,METTL5,SLC35E1,OGFOD2,MET
6,ACOT8,AP3B2,RPS9,SLC35G1,MRPL33,ZNF410,RUVBL2,PCMT1,EBPL,TESC,IGSF8,NUDT5,RBPMS,NNMT
6,ACOT8,AP3B2,RPS9,SLC35G1,MRPL33,ZNF410,RUVBL2,PCMT1,EBPL,TESC,IGSF8,NUDT5,RBPMS,NNMT
K3,ENDOV,VAMP8,CAPG,PPIL2,TROAP,RPL28,LIG1,PSMA7,CDC14B,TIMM17B,APEH,RBM33,LYPLA2,NUTF
K3,ENDOV,VAMP8,CAPG,PPIL2,TROAP,RPL28,LIG1,PSMA7,CDC14B,TIMM17B,APEH,RBM33,LYPLA2,NUTF
OV,CAMTA1,PPIL2,PSMD4,RPL28,LIG1,PSMA7,HYI,RPL21,CAPN10,LYPLA2,NUTF2,EPB41L4B,C17ORF62,C
POLD4,RANBP1,UBE2L6,VRK3,LZTFL1,ZC3H14,FIS1,SLC15A3,PPM1K,TUBA1C,IRF9,ANTXR2,STIP1,PHF19,V
POLD4,RANBP1,UBE2L6,VRK3,LZTFL1,ZC3H14,FIS1,SLC15A3,PPM1K,TUBA1C,IRF9,ANTXR2,STIP1,PHF19,V
EEF1D,CHCHD5,NAA10,SEC31A,CNST,COX17,HNRNPLL,HINT1,HIGD1A,B3GAT3,ZNF506,MTHFS,PAFAH1B
TXBP2,NAA10,SEC31A,CARHSP1,PRLR,COX17,ZFAND4,HIGD1A,POLR2C,SIRT5,B3GAT3,RFXANK,MTHFS,PA
2,LRRC1,PHF19,LPCAT3,TM9SF1,SON,TET2,UQCRQ,GRK6,HMGA2,CDC37,CPSF3L,OBSL1,PTPN13,YLPM1,C
2,LRRC1,PHF19,LPCAT3,TM9SF1,SON,TET2,UQCRQ,GRK6,HMGA2,CDC37,CPSF3L,OBSL1,PTPN13,YLPM1,C
PCAT3,TM9SF1,MAGED2,SON,TET2,GRK6,HMGA2,CDC37,RPS16,OBSL1,UBR4,PSMA1,YLPM1,CHCHD1,SL
PCAT3,TM9SF1,MAGED2,SON,TET2,GRK6,HMGA2,CDC37,RPS16,OBSL1,UBR4,PSMA1,YLPM1,CHCHD1,SL
3,UBR4,PHF21A,CHCHD1,RGS3,FAM3A,PSMA3,MUC1,ECSIT,EPB41L5,PPRC1,RPS25,NLRP1,ZBTB44,NIP7,
OBSL1,TTC21B,UBR4,YLPM1,PHF21A,SLC39A11,FAM3A,MUC1,METTL17,ECSIT,EPB41L5,ATG12,PPRC1,ZB
OBSL1,TTC21B,UBR4,YLPM1,PHF21A,SLC39A11,FAM3A,MUC1,METTL17,ECSIT,EPB41L5,ATG12,PPRC1,ZB
SP22,SLC25A10,TMX2,THEM4,DBNDD2,NISCH,TMED4,RPS6,RPL29,RTEL1,COASY,AUP1,SAP30BP,YIPF3,F
SP22,SLC25A10,TMX2,THEM4,DBNDD2,NISCH,TMED4,RPS6,RPL29,RTEL1,COASY,AUP1,SAP30BP,YIPF3,F
A11,FUZ,FAM3A,MUC1,METTL17,EPB41L5,ATG12,PPRC1,ZBTB44,PLCB2,DUSP22,SLC25A10,TMX2,THEM
AZI2,GOSR2,TATDN1,ZBTB8OS,TBC1D22A,LAMTOR1,U2AF1L4,RNPS1,BTF3L4,ZNF581,SUMO1,NMRAL1,
,EBPL,TESC,IGSF8,NUDT5,RBPMS,NNMT,SPAG7,WDR62,IL1RAP,STYXL1,RFX5,POLD4,PLEKHA5,RANBP1,V

,EBPL,TESC,IGSF8,NUDT5,RBPMS,NNMT,SPAG7,WDR62,IL1RAP,STYXL1,RFX5,POLD4,PLEKHA5,RANBP1,V

IBR1,DDOST,LSR,RRNAD1,RAD1,TOMM40,TADA3,ZNF20,ZNF687,ANAPC11,NT5C,MDH2,EEF1D,CHCHD5,

IBR1,DDOST,LSR,RRNAD1,RAD1,TOMM40,TADA3,ZNF20,ZNF687,ANAPC11,NT5C,MDH2,EEF1D,CHCHD5,

CHD1,SLC39A11,FAM3A,MUC1,METTL17,ECSIT,EPB41L5,ATG12,NLRP1,ZBTB44,NIP7,PLCB2,DUSP22,SLC

CHD1,SLC39A11,FAM3A,MUC1,METTL17,ECSIT,EPB41L5,ATG12,NLRP1,ZBTB44,NIP7,PLCB2,DUSP22,SLC

A6,ACOT8,AP3B2,SLC35G1,RCE1,MRPL33,ZNF410,RUVBL2,PCMT1,EBPL,TESC,IGSF8,NUDT5,RBPMS,SPA

36,RABEP2,TADA3,ZNF20,WAC,ZNF687,ANAPC11,RAD23A,NT5C,MDH2,STXBP2,EEF1D,CHCHD5,SEC31A,

IRT5,B3GAT3,PFDN5,C11ORF49,PAFAH1B2,ZNF343,GBA2,PPIH,DALRD3,NSUN5,DRG2,MFSD10,MCFD2,R

A,GLI4,NKIRAS2,RECQL5,ACAA1,IRF3,GTF2H1,CLTA,SLC35A2,AXIN1,MOSPD3,TMEM199,ZNF396,EPN1,H

TM9SF1,SON,TET2,UQCRQ,SNF8,GRK6,HMGA2,CDC37,RPS16,CPSF3L,OBSL1,TTC21B,PTPN13,UBR4,PSM

TM9SF1,SON,TET2,UQCRQ,SNF8,GRK6,HMGA2,CDC37,RPS16,CPSF3L,OBSL1,TTC21B,PTPN13,UBR4,PSM

A5,RANBP1,VRK3,LZTFL1,FIS1,SLC15A3,TUBA1C,STIP1,LRRC1,PHF19,VCAN,LPCAT3,TM9SF1,SON,SNF8,GR

A5,RANBP1,VRK3,LZTFL1,FIS1,SLC15A3,TUBA1C,STIP1,LRRC1,PHF19,VCAN,LPCAT3,TM9SF1,SON,SNF8,GR

5G1,RCE1,ZNF410,RUVBL2,PCMT1,EBPL,TESC,IGSF8,NUDT5,RBPMS,SPAG7,WDR62,IL1RAP,RFX5,POLD4,

L5,ATG12,ZBTB44,PLCB2,DUSP22,SLC25A10,DBNDD2,NISCH,TMED4,RPL29,CD99L2,GGPS1,AUP1,YIPF3,

L5,ATG12,ZBTB44,PLCB2,DUSP22,SLC25A10,DBNDD2,NISCH,TMED4,RPL29,CD99L2,GGPS1,AUP1,YIPF3,

HID1,NUDT22,AMZ2,UFD1L,USP38,TAZ,THYN1,DBNDD1,SH2B1,NOP16,RPL13A,ARHGEF39,GSTK1,TTLL5,

HID1,NUDT22,AMZ2,UFD1L,USP38,TAZ,THYN1,DBNDD1,SH2B1,NOP16,RPL13A,ARHGEF39,GSTK1,TTLL5,

5A3,PPM1K,TUBA1C,ANTXR2,STIP1,LRRC1,PHF19,LPCAT3,CD44,TM9SF1,TET2,UQCRQ,SNF8,GRK6,HMG

RK3,LZTFL1,ZC3H14,FIS1,PPM1K,DST,ANTXR2,STIP1,LRRC1,PHF19,VCAN,LPCAT3,CD44,TM9SF1,MAGED

,DUSP22,SLC25A10,TMX2,THEM4,RPS6,RPL29,GGPS1,RTEL1,COASY,AUP1,SAP30BP,YIPF3,RABL5,TRNT1,

,DUSP22,SLC25A10,TMX2,THEM4,RPS6,RPL29,GGPS1,RTEL1,COASY,AUP1,SAP30BP,YIPF3,RABL5,TRNT1,

M32A,PEX2,POLR1C,DCTN5,PEX26,EIF4A1,ZNF223,OGFOD2,METTL3,TSEN34,SSR3,ARIH2,PICK1,ZNF512

M32A,COX20,TGOLN2,PIGH,C12ORF57,DCTN5,PEX26,ARL16,OXLD1,STAU2,METTL5,SLC35E1,METTL3,ZN

CD99L2,COASY,SRSF1,SAP30BP,YIPF3,FOXRED1,RABL5,TRNT1,KXD1,ITGB1BP1,MED15,SH3YL1,UGGT2,M

37,RPS16,CPSF3L,TSG101,PTPN13,UBR4,PSMA1,YLPM1,PHF21A,SLC39A11,FUZ,FAM3A,PSMA3,METTL17

37,RPS16,CPSF3L,TSG101,PTPN13,UBR4,PSMA1,YLPM1,PHF21A,SLC39A11,FUZ,FAM3A,PSMA3,METTL17

NTMT1,ANAPC13,TSPAN31,RALY,GLYCTK,NKIRAS2,ACAA1,TRAPPC2,PIGO,AXIN1,TP53,BTF3,TMEM199,E

,TMED4,CD99L2,GGPS1,AUP1,YIPF3,TRNT1,PARD6G,SH3YL1,IP6K2,CLASRP,JPH2,C1ORF198,MICAL1,HS2

,TMED4,CD99L2,GGPS1,AUP1,YIPF3,TRNT1,PARD6G,SH3YL1,IP6K2,CLASRP,JPH2,C1ORF198,MICAL1,HS2

,TMED4,CD99L2,GGPS1,AUP1,YIPF3,TRNT1,PARD6G,SH3YL1,IP6K2,CLASRP,JPH2,C1ORF198,MICAL1,HS2

,TMED4,CD99L2,GGPS1,AUP1,YIPF3,TRNT1,PARD6G,SH3YL1,IP6K2,CLASRP,JPH2,C1ORF198,MICAL1,HS2

N,WDR62,IL1RAP,STYXL1,RFX5,PLEKHA5,RANBP1,UBE2L6,VRK3,LZTFL1,ZC3H14,FIS1,SLC15A3,PPM1K,TU

N,WDR62,IL1RAP,STYXL1,RFX5,PLEKHA5,RANBP1,UBE2L6,VRK3,LZTFL1,ZC3H14,FIS1,SLC15A3,PPM1K,TU

3PMS,WDR62,IL1RAP,STYXL1,RFX5,PLEKHA5,RANBP1,UBE2L6,VRK3,LZTFL1,ZC3H14,FIS1,SLC15A3,TUBA

H2B1,NOP16,RPL13A,ARHGEF39,GSTK1,TTLL5,LDHA,PSMA6,ACOT8,AP3B2,RPS9,SLC35G1,RCE1,MRPL33,

H2B1,NOP16,RPL13A,ARHGEF39,GSTK1,TTLL5,LDHA,PSMA6,ACOT8,AP3B2,RPS9,SLC35G1,RCE1,MRPL33,

RUVBL2,PCMT1,EBPL,TESC,IGSF8,NUDT5,RBPMS,NNMT,SPAG7,WDR62,IL1RAP,STYXL1,RFX5,POLD4,PLE

JBA1C,IRF9,ANTXR2,STIP1,LRRC1,PHF19,VCAN,LPCAT3,CD44,TM9SF1,SON,TET2,UQCRQ,SNF8,CDC37,R

0,RUVBL2,PCMT1,TESC,NUDT5,RBPMS,NNMT,SNRPN,SPAG7,WDR62,IL1RAP,STYXL1,RFX5,POLD4,PLEKH

0,RUVBL2,PCMT1,TESC,NUDT5,RBPMS,NNMT,SNRPN,SPAG7,WDR62,IL1RAP,STYXL1,RFX5,POLD4,PLEKH

D2,SON,TET2,UQCRQ,SNF8,HMGA2,CDC37,RPS16,CPSF3L,TSG101,TTC21B,PTPN13,PSMA1,YLPM1,CHCH

D2,SON,TET2,UQCRQ,SNF8,HMGA2,CDC37,RPS16,CPSF3L,TSG101,TTC21B,PTPN13,PSMA1,YLPM1,CHCH

T1,KXD1,MED15,PARD6G,SH3YL1,IP6K2,CLASRP,JPH2,UGGT2,MICAL1,HS2ST1,UBE2I,PRMT2,EIF3M,JUP,

T1,KXD1,MED15,PARD6G,SH3YL1,IP6K2,CLASRP,JPH2,UGGT2,MICAL1,HS2ST1,UBE2I,PRMT2,EIF3M,JUP,

5,POLD4,PLEKHA5,UBE2L6,LZTFL1,ZC3H14,FIS1,SLC15A3,PPM1K,TUBA1C,ANTXR2,STIP1,LRRC1,GRAMD1

D4,CD99L2,GGPS1,SAP30BP,TRNT1,KXD1,MED15,PARD6G,SH3YL1,IP6K2,CLASRP,JPH2,UGGT2,MICAL1,H

4,CD99L2,GGPS1,SAP30BP,TRNT1,KXD1,MED15,PARD6G,SH3YL1,IP6K2,CLASRP,JPH2,UGGT2,MICAL1,H

F181,NIPAL3,TSPAN31,RALY,ZNF593,GLYCTK,TRIM46,C11ORF57,LAMTOR5,RBM42,CLASP2,PCNP,NEK11

4,PLEKHA5,RANBP1,VRK3,LZTFL1,ZC3H14,FIS1,SLC15A3,PPM1K,TUBA1C,ANTXR2,STIP1,LRRC1,PHF19,GR

NX1,CHID1,NUDT22,AMZ2,UFD1L,USP38,TAZ,THYN1,DBNDD1,SH2B1,NOP16,RPL13A,ARHGEF39,GSTK1,

NX1,CHID1,NUDT22,AMZ2,UFD1L,USP38,TAZ,THYN1,DBNDD1,SH2B1,NOP16,RPL13A,ARHGEF39,GSTK1,

9,LPCAT3,CD44,TM9SF1,MAGED2,SON,UQCRQ,HMGA2,CDC37,RPS16,CPSF3L,TSG101,OBSL1,TTC21B,UE

9,LPCAT3,CD44,TM9SF1,MAGED2,SON,UQCRQ,HMGA2,CDC37,RPS16,CPSF3L,TSG101,OBSL1,TTC21B,UE

IFSD11,FAM32A,TMEM147,TGOLN2,PEX2,TNKS,C12ORF57,DCTN5,NDUFS3,ARL16,OXLD1,ZNF223,EIF3G,

,ECSIT,EPB41L5,ATG12,PPRC1,RPS25,ZBTB44,TTC9C,PLCB2,DUSP22,SPOCK2,THEM4,DBNDD2,NISCH,TM

,ECSIT,EPB41L5,ATG12,PPRC1,RPS25,ZBTB44,TTC9C,PLCB2,DUSP22,SPOCK2,THEM4,DBNDD2,NISCH,TM

15A3,PPM1K,TUBA1C,DST,GRAMD1C,VCAN,LPCAT3,SON,TET2,UQCRQ,SNF8,GRK6,HMGA2,CDC37,RPS1

15A3,PPM1K,TUBA1C,DST,GRAMD1C,VCAN,LPCAT3,SON,TET2,UQCRQ,SNF8,GRK6,HMGA2,CDC37,RPS1

YN1,DBNDD1,SH2B1,NOP16,RPL13A,GSTK1,TTLL5,LDHA,PSMA6,ACOT8,RPS9,SLC35G1,RCE1,ZNF410,RL

YN1,DBNDD1,SH2B1,NOP16,RPL13A,GSTK1,TTLL5,LDHA,PSMA6,ACOT8,RPS9,SLC35G1,RCE1,ZNF410,RL

9,GRAMD1C,VCAN,LPCAT3,CD44,TM9SF1,SON,TET2,UQCRQ,GRK6,HMGA2,CDC37,RPS16,CPSF3L,OBSL1,

25,ZBTB44,TTC9C,COX4I1,PLCB2,DUSP22,TMX2,THEM4,RDH11,TMED4,GGPS1,RTEL1,SRSF1,AUP1,SAP3

25,ZBTB44,TTC9C,COX4I1,PLCB2,DUSP22,TMX2,THEM4,RDH11,TMED4,GGPS1,RTEL1,SRSF1,AUP1,SAP3

25,ZBTB44,TTC9C,COX4I1,PLCB2,DUSP22,TMX2,THEM4,RDH11,TMED4,GGPS1,RTEL1,SRSF1,AUP1,SAP3

25,ZBTB44,TTC9C,COX4I1,PLCB2,DUSP22,TMX2,THEM4,RDH11,TMED4,GGPS1,RTEL1,SRSF1,AUP1,SAP3

25,ZBTB44,TTC9C,COX4I1,PLCB2,DUSP22,TMX2,THEM4,RDH11,TMED4,GGPS1,RTEL1,SRSF1,AUP1,SAP3

25,ZBTB44,TTC9C,COX4I1,PLCB2,DUSP22,TMX2,THEM4,RDH11,TMED4,GGPS1,RTEL1,SRSF1,AUP1,SAP3

PS1,AUP1,YIPF3,FOXRED1,TRNT1,PARD6G,SH3YL1,IP6K2,CLASRP,JPH2,MICAL1,HS2ST1,UBE2I,SLC39A13

PS1,AUP1,YIPF3,FOXRED1,TRNT1,PARD6G,SH3YL1,IP6K2,CLASRP,JPH2,MICAL1,HS2ST1,UBE2I,SLC39A13

L17,EPB41L5,ATG12,RPS25,NIP7,COX4I1,DUSP22,THEM4,NISCH,CENPK,RPS6,RPL29,RTEL1,COASY,AUP1,

L17,EPB41L5,ATG12,RPS25,NIP7,COX4I1,DUSP22,THEM4,NISCH,CENPK,RPS6,RPL29,RTEL1,COASY,AUP1,

ARD14,PCYT2,FBXW9,USE1,ASF1B,CCDC25,EEF1B2,EDF1,RFX3,ENDOV,VAMP8,PPIL2,RPL28,LIG1,CDC14

ARD14,PCYT2,FBXW9,USE1,ASF1B,CCDC25,EEF1B2,EDF1,RFX3,ENDOV,VAMP8,PPIL2,RPL28,LIG1,CDC14

1,SLC39A11,FAM3A,METTL17,ECSIT,EPB41L5,ATG12,PPRC1,ZBTB44,TTC9C,DUSP22,SLC25A10,THEM4,N

1,SLC39A11,FAM3A,METTL17,ECSIT,EPB41L5,ATG12,PPRC1,ZBTB44,TTC9C,DUSP22,SLC25A10,THEM4,N

L18,ZNF512,SLC52A2,TSPAN31,MTIF2,FAM192A,GLYCTK,CFLAR,TRIM46,C11ORF57,PCNP,RER1,CREBZF,

L18,ZNF512,SLC52A2,TSPAN31,MTIF2,FAM192A,GLYCTK,CFLAR,TRIM46,C11ORF57,PCNP,RER1,CREBZF,

ETTL17,ECSIT,EPB41L5,ATG12,NLRP1,ZBTB44,DUSP22,TMX2,THEM4,DBNDD2,NISCH,TMED4,RPL29,CD9

ETTL17,ECSIT,EPB41L5,ATG12,NLRP1,ZBTB44,DUSP22,TMX2,THEM4,DBNDD2,NISCH,TMED4,RPL29,CD9

,NAA10,SEC31A,CARHSP1,CNST,COX17,HIGD1A,RPL15,PFDN5,PAFAH1B2,COA3,NSUN5,ZFPL1,MFSD10,

,NAA10,SEC31A,CARHSP1,CNST,COX17,HIGD1A,RPL15,PFDN5,PAFAH1B2,COA3,NSUN5,ZFPL1,MFSD10,

VRK3,LZTFL1,ZC3H14,FIS1,SLC15A3,TUBA1C,STIP1,LRRC1,PHF19,GRAMD1C,VCAN,LPCAT3,TM9SF1,MAG

VRK3,LZTFL1,ZC3H14,FIS1,SLC15A3,TUBA1C,STIP1,LRRC1,PHF19,GRAMD1C,VCAN,LPCAT3,TM9SF1,MAG

1,ZNF197,INO80E,CCNL2,MAPKAPK5,LRRFIP1,RMDN1,TRMT112,CHPF,RPS20,MPDU1,MAPK13,RPS15,

Q,CDC37,CPSF3L,TSG101,OBSL1,TTC21B,PTPN13,UBR4,PSMA1,YLPM1,CHCHD1,FUZ,FAM3A,PSMA3,ME

Q,CDC37,CPSF3L,TSG101,OBSL1,TTC21B,PTPN13,UBR4,PSMA1,YLPM1,CHCHD1,FUZ,FAM3A,PSMA3,ME

SCH,TMED4,RPL29,CD99L2,GGPS1,SRSF1,TRNT1,PARD6G,SH3YL1,IP6K2,CLASRP,JPH2,MICAL1,HS2ST1,U

SCH,TMED4,RPL29,CD99L2,GGPS1,SRSF1,TRNT1,PARD6G,SH3YL1,IP6K2,CLASRP,JPH2,MICAL1,HS2ST1,U

,RPS25,NIP7,COX4I1,DUSP22,SLC25A10,DBNDD2,CENPK,GGPS1,RTEL1,SRSF1,AUP1,SAP30BP,RABL5,TR

RED1,RABL5,KXD1,ITGB1BP1,SH3YL1,CLASRP,JPH2,UGGT2,HS2ST1,YIF1A,EIF3M,TBX6,TMEM205,PTK2,A

RED1,RABL5,KXD1,ITGB1BP1,SH3YL1,CLASRP,JPH2,UGGT2,HS2ST1,YIF1A,EIF3M,TBX6,TMEM205,PTK2,A

,SPAG7,WDR62,STYXL1,RANBP1,UBE2L6,VRK3,LZTFL1,ZC3H14,FIS1,SLC15A3,PPM1K,TUBA1C,IRF9,DST,

,SPAG7,WDR62,STYXL1,RANBP1,UBE2L6,VRK3,LZTFL1,ZC3H14,FIS1,SLC15A3,PPM1K,TUBA1C,IRF9,DST,

REBZF,EIF2A,NKIRAS2,SCN11A,ACAA1,ZNF283,TRAPPC2,GTF2H1,BFAR,AXIN1,TP53,BTF3,TMEM199,SLC

REBZF,EIF2A,NKIRAS2,SCN11A,ACAA1,ZNF283,TRAPPC2,GTF2H1,BFAR,AXIN1,TP53,BTF3,TMEM199,SLC

,PCMT1,EBPL,TESC,IGSF8,NUDT5,RBPMS,SNRPN,SPAG7,WDR62,IL1RAP,POLD4,PLEKHA5,RANBP1,UBE2

,PCMT1,EBPL,TESC,IGSF8,NUDT5,RBPMS,SNRPN,SPAG7,WDR62,IL1RAP,POLD4,PLEKHA5,RANBP1,UBE2

,POLD4,PLEKHA5,RANBP1,UBE2L6,ZC3H14,FIS1,PPM1K,TUBA1C,IRF9,DST,ANTXR2,STIP1,LRRC1,PHF19,

,POLD4,PLEKHA5,RANBP1,UBE2L6,ZC3H14,FIS1,PPM1K,TUBA1C,IRF9,DST,ANTXR2,STIP1,LRRC1,PHF19,

MS,NNMT,SPAG7,WDR62,IL1RAP,STYXL1,POLD4,PLEKHA5,RANBP1,UBE2L6,VRK3,LZTFL1,ZC3H14,FIS1,S

MS,NNMT,SPAG7,WDR62,IL1RAP,STYXL1,POLD4,PLEKHA5,RANBP1,UBE2L6,VRK3,LZTFL1,ZC3H14,FIS1,S

,GMPR2,RIC8A,TCP1,SMARCC2,THADA,UBE2N,ZNF81,FAM32A,TMEM147,COA1,USP40,PCID2,PEX2,TN

,GMPR2,RIC8A,TCP1,SMARCC2,THADA,UBE2N,ZNF81,FAM32A,TMEM147,COA1,USP40,PCID2,PEX2,TN

,MAGED2,SON,TET2,HMGA2,CDC37,RPS16,CPSF3L,TSG101,OBSL1,TTC21B,UBR4,PSMA1,YLPM1,FUZ,PS

,MAGED2,SON,TET2,HMGA2,CDC37,RPS16,CPSF3L,TSG101,OBSL1,TTC21B,UBR4,PSMA1,YLPM1,FUZ,PS

1,AUP1,FOXRED1,TRNT1,KXD1,ITGB1BP1,PARD6G,SH3YL1,IP6K2,CLASRP,JPH2,MICAL1,HS2ST1,UBE2I,SL

1,AUP1,FOXRED1,TRNT1,KXD1,ITGB1BP1,PARD6G,SH3YL1,IP6K2,CLASRP,JPH2,MICAL1,HS2ST1,UBE2I,SL

5,PTK2,AUNIP,C14ORF2,GUK1,ATF7,PCYT2,USE1,PPP1CC,PPA1,DGKE,CAMTA1,CRIP1,LIG1,TIMM17B,TM

5,PTK2,AUNIP,C14ORF2,GUK1,ATF7,PCYT2,USE1,PPP1CC,PPA1,DGKE,CAMTA1,CRIP1,LIG1,TIMM17B,TM

0,RBM3,LAMP2,ZNF197,INO80E,HSP90AB1,RAB2A,ATP5J,RMDN1,TRMT112,CHPF,LYNX1,UBXN1,ZNF81

0,RBM3,LAMP2,ZNF197,INO80E,HSP90AB1,RAB2A,ATP5J,RMDN1,TRMT112,CHPF,LYNX1,UBXN1,ZNF81

ARD6G,SH3YL1,IP6K2,CLASRP,JPH2,C1ORF198,MICAL1,HS2ST1,UBE2I,YIF1A,EIF3M,CDC42,TBX6,TMEM2

ARD6G,SH3YL1,IP6K2,CLASRP,JPH2,C1ORF198,MICAL1,HS2ST1,UBE2I,YIF1A,EIF3M,CDC42,TBX6,TMEM2

P1,AUNIP,GUK1,FBXW9,MAF,ASF1B,EEF1B2,PPP1CC,GADL1,RFX3,DGKE,ENDOV,CAMTA1,CRIP1,PPIL2,PS

P1,AUNIP,GUK1,FBXW9,MAF,ASF1B,EEF1B2,PPP1CC,GADL1,RFX3,DGKE,ENDOV,CAMTA1,CRIP1,PPIL2,PS

GED2,SON,TET2,GRK6,HMGA2,CDC37,RPS16,CPSF3L,OBSL1,TTC21B,PTPN13,UBR4,YLPM1,PHF21A,CHC

GED2,SON,TET2,GRK6,HMGA2,CDC37,RPS16,CPSF3L,OBSL1,TTC21B,PTPN13,UBR4,YLPM1,PHF21A,CHC

2H1,BFAR,AXIN1,BTF3,TMEM199,SLC25A24,ZNF396,EPN1,CREB3L2,ITPA,ZCCHC9,VPS25,SNRPD1,AKT1S1

2H1,BFAR,AXIN1,BTF3,TMEM199,SLC25A24,ZNF396,EPN1,CREB3L2,ITPA,ZCCHC9,VPS25,SNRPD1,AKT1S1

NL4A,CAMLG,EPB41L4B,UCK2,TMPO,ATPAF2,LSM12,C17ORF62,WNK1,ZNF7,ARL6IP4,C19ORF48,PPAN,

HCHD5,NAA10,C11ORF48,SEC31A,CNST,COX17,HNRNPLL,FKBP3,RPL15,B3GAT3,RFXANK,ZNF506,C11ORF

EDF1,RFX3,PPA1,DGKE,CAMTA1,CRIP1,CLDND1,LIG1,PSMA7,CDC14B,TIMM17B,TMEM126B,APEH,HYI,C

HCHD5,NAA10,C11ORF48,MED6,SEC31A,CARHSP1,COX17,HNRNPLL,PHF20L1,SIRT5,B3GAT3,ASH2L,PFD

HCHD5,NAA10,C11ORF48,MED6,SEC31A,CARHSP1,COX17,HNRNPLL,PHF20L1,SIRT5,B3GAT3,ASH2L,PFD

P1,VRK3,LZTFL1,ZC3H14,FIS1,SLC15A3,TUBA1C,IRF9,STIP1,GRAMD1C,LPCAT3,TET2,UQCRQ,SNF8,GRK6,

P1,VRK3,LZTFL1,ZC3H14,FIS1,SLC15A3,TUBA1C,IRF9,STIP1,GRAMD1C,LPCAT3,TET2,UQCRQ,SNF8,GRK6,

PL,TESC,IGSF8,NUDT5,RBPMS,NNMT,SPAG7,WDR62,IL1RAP,STYXL1,RFX5,POLD4,PLEKHA5,RANBP1,UBE

PL,TESC,IGSF8,NUDT5,RBPMS,NNMT,SPAG7,WDR62,IL1RAP,STYXL1,RFX5,POLD4,PLEKHA5,RANBP1,UBE

P1,LRRC1,PHF19,LPCAT3,CD44,TM9SF1,MAGED2,SON,TET2,UQCRQ,GRK6,CDC37,CPSF3L,TSG101,OBSL1

P1,LRRC1,PHF19,LPCAT3,CD44,TM9SF1,MAGED2,SON,TET2,UQCRQ,GRK6,CDC37,CPSF3L,TSG101,OBSL1

30BP,YIPF3,FOXRED1,RABL5,TRNT1,KXD1,ITGB1BP1,MED15,CLASRP,C1ORF198,UGGT2,MICAL1,HS2ST1,

30BP,YIPF3,FOXRED1,RABL5,TRNT1,KXD1,ITGB1BP1,MED15,CLASRP,C1ORF198,UGGT2,MICAL1,HS2ST1,

P1CC,EDF1,PPA1,DGKE,VAMP8,CAMTA1,CAPG,CLDND1,LIG1,PSMA7,TIMM17B,XAF1,APEH,NUTF2,POD

P1CC,EDF1,PPA1,DGKE,VAMP8,CAMTA1,CAPG,CLDND1,LIG1,PSMA7,TIMM17B,XAF1,APEH,NUTF2,POD

BM42,CREBZF,EIF2A,NKIRAS2,ACAA1,GTF2H1,BFAR,BTF3,TMEM199,SLC25A24,ZNF396,EPN1,ITPA,ZCCH

BM42,CREBZF,EIF2A,NKIRAS2,ACAA1,GTF2H1,BFAR,BTF3,TMEM199,SLC25A24,ZNF396,EPN1,ITPA,ZCCH

UVBL2,PCMT1,EBPL,TESC,IGSF8,NUDT5,RBPMS,NNMT,SNRPN,SPAG7,WDR62,STYXL1,RFX5,RANBP1,UB

UVBL2,PCMT1,EBPL,TESC,IGSF8,NUDT5,RBPMS,NNMT,SNRPN,SPAG7,WDR62,STYXL1,RFX5,RANBP1,UB

1,ZNF836,TOMM40,SERPINB6,RABEP2,TADA3,ZNF687,PDCD5,KLK10,TRIM16,RAD23A,ABHD17A,PDE4D

1,ZNF836,TOMM40,SERPINB6,RABEP2,TADA3,ZNF687,PDCD5,KLK10,TRIM16,RAD23A,ABHD17A,PDE4D

BP1,SIAH1,ASPH,CHPF,STRIP2,LYNX1,TWSG1,ZSWIM7,RPS15,HSPB1,LRRC27,LMBR1,MCTS1,NAP1L1,RA

BP1,SIAH1,ASPH,CHPF,STRIP2,LYNX1,TWSG1,ZSWIM7,RPS15,HSPB1,LRRC27,LMBR1,MCTS1,NAP1L1,RA

D1,LIG1,TIMM17B,XAF1,APEH,HYI,CAPN10,NUTF2,ATPAF2,C17ORF62,PRMT1,CASC4,WNK1,ZNF7,STRA

D1,LIG1,TIMM17B,XAF1,APEH,HYI,CAPN10,NUTF2,ATPAF2,C17ORF62,PRMT1,CASC4,WNK1,ZNF7,STRA

NK,ZNF506,MTHFS,PAFAH1B2,DALRD3,NSUN5,MRPS18C,GMPR2,RIC8A,TCP1,THADA,PIGT,UBE2N,MFSD

25,EEF1B2,EDF1,RFX3,ENDOV,VAMP8,PPIL2,TROAP,LIG1,CDC14B,TMEM126B,APEH,RPL21,RBM33,LYPL

25,EEF1B2,EDF1,RFX3,ENDOV,VAMP8,PPIL2,TROAP,LIG1,CDC14B,TMEM126B,APEH,RPL21,RBM33,LYPL

P1,ZNF524,ELP6,RMDN1,TPM2,GPBP1,SIAH1,CHPF,STRIP2,RPS20,GABPB1,LYNX1,TWSG1,RPS15,DCTD,

P1,ZNF524,ELP6,RMDN1,TPM2,GPBP1,SIAH1,CHPF,STRIP2,RPS20,GABPB1,LYNX1,TWSG1,RPS15,DCTD,

C,DST,ANTXR2,STIP1,LRRC1,VCAN,TM9SF1,SON,TET2,UQCRQ,SNF8,GRK6,HMGA2,CDC37,RPS16,CPSF3L,

R2,STIP1,LRRC1,PHF19,GRAMD1C,VCAN,LPCAT3,CD44,TM9SF1,SON,TET2,UQCRQ,GRK6,HMGA2,CDC37,

R2,STIP1,LRRC1,PHF19,GRAMD1C,VCAN,LPCAT3,CD44,TM9SF1,SON,TET2,UQCRQ,GRK6,HMGA2,CDC37,

FA2T2,SUGT1,HYPK,CARS,ZNF250,CDK11A,RBM3,ZNF197,INO80E,DGUOK,HSP90AB1,RAB2A,ATP5J,TSF

FA2T2,SUGT1,HYPK,CARS,ZNF250,CDK11A,RBM3,ZNF197,INO80E,DGUOK,HSP90AB1,RAB2A,ATP5J,TSF

PRC1,RPS25,ZBTB44,NIP7,TTC9C,COX4I1,DUSP22,SLC25A10,SPOCK2,DBNDD2,NISCH,TMED4,CENPK,RPS

PRC1,RPS25,ZBTB44,NIP7,TTC9C,COX4I1,DUSP22,SLC25A10,SPOCK2,DBNDD2,NISCH,TMED4,CENPK,RPS

YXL1,POLD4,PLEKHA5,RANBP1,UBE2L6,VRK3,LZTFL1,ZC3H14,FIS1,SLC15A3,TUBA1C,STIP1,LRRC1,PHF19,

YXL1,POLD4,PLEKHA5,RANBP1,UBE2L6,VRK3,LZTFL1,ZC3H14,FIS1,SLC15A3,TUBA1C,STIP1,LRRC1,PHF19,

D1C,LPCAT3,TET2,UQCRQ,SNF8,GRK6,HMGA2,CDC37,TTC21B,UBR4,SLC39A11,FUZ,RGS3,FAM3A,PSMA3

1,C1ORF198,UGGT2,UBE2I,YIF1A,PRMT2,EIF3M,CDC42,TBX6,TMEM205,PTK2,CARD14,PCYT2,FBXW9,C

1,C1ORF198,UGGT2,UBE2I,YIF1A,PRMT2,EIF3M,CDC42,TBX6,TMEM205,PTK2,CARD14,PCYT2,FBXW9,C

TROAP,LIG1,APEH,RBM33,LYPLA2,C17ORF62,PRMT1,STOML2,C19ORF54,TXNDC17,PPAN,CYB561,SHARP

TROAP,LIG1,APEH,RBM33,LYPLA2,C17ORF62,PRMT1,STOML2,C19ORF54,TXNDC17,PPAN,CYB561,SHARP

NRPN,SPAG7,WDR62,IL1RAP,POLD4,PLEKHA5,VRK3,LZTFL1,ZC3H14,FIS1,PPM1K,TUBA1C,IRF9,ANTXR2,S

C,EDF1,RFX3,PPA1,DGKE,CAMTA1,CRIP1,CLDND1,LIG1,CDC14B,TIMM17B,APEH,HYI,CAPN10,NUTF2,PO

M1K,TUBA1C,IRF9,DST,ANTXR2,LRRC1,PHF19,GRAMD1C,VCAN,CD44,TM9SF1,MAGED2,TET2,SNF8,GR

,PPA1,DGKE,CAMTA1,CAPG,CLDND1,VIM,LIG1,PSMA7,TIMM17B,TMEM126B,APEH,HYI,CAPN10,PODXL,

,PPA1,DGKE,CAMTA1,CAPG,CLDND1,VIM,LIG1,PSMA7,TIMM17B,TMEM126B,APEH,HYI,CAPN10,PODXL,

5,PSMA6,ACOT8,AP3B2,RPS9,SLC35G1,RCE1,MRPL33,ZNF410,RUVBL2,PCMT1,EBPL,TESC,IGSF8,NUDT5,

5,PSMA6,ACOT8,AP3B2,RPS9,SLC35G1,RCE1,MRPL33,ZNF410,RUVBL2,PCMT1,EBPL,TESC,IGSF8,NUDT5,

COT8,AP3B2,RPS9,SLC35G1,RCE1,MRPL33,ZNF410,RUVBL2,PCMT1,EBPL,TESC,IGSF8,NUDT5,RBPMS,NN

COT8,AP3B2,RPS9,SLC35G1,RCE1,MRPL33,ZNF410,RUVBL2,PCMT1,EBPL,TESC,IGSF8,NUDT5,RBPMS,NN

2,RPS9,SLC35G1,RCE1,ZNF410,RUVBL2,PCMT1,TESC,IGSF8,NUDT5,RBPMS,NNMT,SNRPN,WDR62,IL1RA

YI,RPL21,LYPLA2,PPOX,TMPO,ATPAF2,LSM12,C17ORF62,PRMT1,CASC4,WNK1,PPAN,HNRNPC,STAU1,SU

FAM3A,MUC1,METTL17,ECSIT,EPB41L5,ATG12,PPRC1,RPS25,ZBTB44,DUSP22,SPOCK2,THEM4,RDH11,D

TA1,CRIP1,CLDND1,LIG1,PSMA7,CDC14B,TIMM17B,TMEM126B,APEH,HYI,CAPN10,NUTF2,PODXL,CAML

2,CCNL2,HSP90AB1,SYNGR2,TMEM134,LRRFIP1,RAB2A,ZNF524,ASPH,NIT2,LYNX1,TWSG1,ZSWIM7,HSP

RABL5,TRNT1,PARD6G,SH3YL1,IP6K2,CLASRP,JPH2,C1ORF198,MICAL1,HS2ST1,UBE2I,YIF1A,PRMT2,TBX

RABL5,TRNT1,PARD6G,SH3YL1,IP6K2,CLASRP,JPH2,C1ORF198,MICAL1,HS2ST1,UBE2I,YIF1A,PRMT2,TBX

,DALRD3,GMPR2,MCFD2,RIC8A,TCP1,SLC30A5,NFATC4,ZNHIT3,PDCD6,FAM32A,TMEM147,COA1,ELP4,Z

BNDD1,SH2B1,NOP16,RPL13A,ARHGEF39,GSTK1,TTLL5,LDHA,PSMA6,ACOT8,AP3B2,SLC35G1,RCE1,ZNF

BNDD1,SH2B1,NOP16,RPL13A,ARHGEF39,GSTK1,TTLL5,LDHA,PSMA6,ACOT8,AP3B2,SLC35G1,RCE1,ZNF

4,FIS1,SLC15A3,PPM1K,TUBA1C,STIP1,LRRC1,PHF19,GRAMD1C,VCAN,PCBP2,LPCAT3,CD44,MAGED2,SO

4,FIS1,SLC15A3,PPM1K,TUBA1C,STIP1,LRRC1,PHF19,GRAMD1C,VCAN,PCBP2,LPCAT3,CD44,MAGED2,SO

7,COX4I1,DUSP22,TMX2,TMED4,RPS6,CD99L2,GGPS1,SRSF1,AUP1,YIPF3,FOXRED1,RABL5,TRNT1,ITGB1

7,COX4I1,DUSP22,TMX2,TMED4,RPS6,CD99L2,GGPS1,SRSF1,AUP1,YIPF3,FOXRED1,RABL5,TRNT1,ITGB1

RF57,NEK11,EIF2A,PSPC1,ZNF283,TRAPPC2,ZNF69,ST7,THOC3,HNRNPUL1,TMEM199,UBE2D3,C1ORF63
,GUK1,ATF7,PCYT2,AGPAT4,PPP1CC,EDF1,TNFRSF14,VAMP8,CAPG,CRIP1,CLDND1,LIG1,PSMA7,TIMM1
,GUK1,ATF7,PCYT2,AGPAT4,PPP1CC,EDF1,TNFRSF14,VAMP8,CAPG,CRIP1,CLDND1,LIG1,PSMA7,TIMM1

,COX4I1,PLCB2,DUSP22,TMX2,SPOCK2,THEM4,RDH11,DBNDD2,CENPK,RPS6,RPL29,RTEL1,COASY,SRSF1
,COX4I1,PLCB2,DUSP22,TMX2,SPOCK2,THEM4,RDH11,DBNDD2,CENPK,RPS6,RPL29,RTEL1,COASY,SRSF1

7,ECSIT,PPRC1,RPS25,NIP7,TTC9C,COX4I1,DUSP22,SLC25A10,SPOCK2,THEM4,RDH11,TMED4,RPS6,RPL2
7,ECSIT,PPRC1,RPS25,NIP7,TTC9C,COX4I1,DUSP22,SLC25A10,SPOCK2,THEM4,RDH11,TMED4,RPS6,RPL2
TLL5,LDHA,PSMA6,ACOT8,AP3B2,RPS9,SLC35G1,RCE1,MRPL33,ZNF410,RUVBL2,PCMT1,EBPL,TESC,IGSF
AP,RFX5,PLEKHA5,RANBP1,UBE2L6,VRK3,LZTFL1,ZC3H14,FIS1,PPM1K,TUBA1C,IRF9,ANTXR2,STIP1,LRRC
AP,RFX5,PLEKHA5,RANBP1,UBE2L6,VRK3,LZTFL1,ZC3H14,FIS1,PPM1K,TUBA1C,IRF9,ANTXR2,STIP1,LRRC
DXL,ATPAF2,C17ORF62,CASC4,ZNF7,TXNDC17,PIGL,STRADA,PPAN,ZFAND2B,SHARPIN,UPP1,ZNF250,RPL
X6,XBP1,AUNIP,GUK1,FBXW9,IFI27L1,MAF,ASF1B,DEF8,PPP1CC,EDF1,GADL1,RFX3,ENDOV,CAMTA1,CA
X6,XBP1,AUNIP,GUK1,FBXW9,IFI27L1,MAF,ASF1B,DEF8,PPP1CC,EDF1,GADL1,RFX3,ENDOV,CAMTA1,CA
NIP,C14ORF2,GUK1,PCYT2,USE1,PPP1CC,PPA1,DGKE,VAMP8,CAMTA1,CRIP1,PPIL2,RPL28,LIG1,TIMM17
NIP,C14ORF2,GUK1,PCYT2,USE1,PPP1CC,PPA1,DGKE,VAMP8,CAMTA1,CRIP1,PPIL2,RPL28,LIG1,TIMM17
RAB2A,ZNF524,TPM2,GPBP1,SIAH1,ASPH,CHPF,STRIP2,LYNX1,MVD,LMBR1,NAP1L1,CHCHD7,HMGA1,P
RAB2A,ZNF524,TPM2,GPBP1,SIAH1,ASPH,CHPF,STRIP2,LYNX1,MVD,LMBR1,NAP1L1,CHCHD7,HMGA1,P
B1,NOP16,RPL13A,ARHGEF39,GSTK1,TTLL5,LDHA,PSMA6,ACOT8,AP3B2,RPS9,RCE1,ZNF410,RUVBL2,PC
B1,NOP16,RPL13A,ARHGEF39,GSTK1,TTLL5,LDHA,PSMA6,ACOT8,AP3B2,RPS9,RCE1,ZNF410,RUVBL2,PC

2,XBP1,PTK2,GUK1,CARD14,PCYT2,FBXW9,IFI27L1,ASF1B,CCDC25,EEF1B2,RFX3,DGKE,ENDOV,PSMD4,R
2,XBP1,PTK2,GUK1,CARD14,PCYT2,FBXW9,IFI27L1,ASF1B,CCDC25,EEF1B2,RFX3,DGKE,ENDOV,PSMD4,R
L,TTLL5,LDHA,ACOT8,AP3B2,SLC35G1,MRPL33,ZNF410,RUVBL2,PCMT1,EBPL,TESC,IGSF8,NUDT5,RBPMS,
L,TTLL5,LDHA,ACOT8,AP3B2,SLC35G1,MRPL33,ZNF410,RUVBL2,PCMT1,EBPL,TESC,IGSF8,NUDT5,RBPMS,
L17,ECSIT,EPB41L5,ATG12,PPRC1,RPS25,ZBTB44,NIP7,COX4I1,PLCB2,TMX2,SPOCK2,RDH11,SDF4,NISCH
L17,ECSIT,EPB41L5,ATG12,PPRC1,RPS25,ZBTB44,NIP7,COX4I1,PLCB2,TMX2,SPOCK2,RDH11,SDF4,NISCH
,SNRPN,SPAG7,WDR62,IL1RAP,STYXL1,RFX5,RANBP1,UBE2L6,VRK3,LZTFL1,FIS1,PPM1K,TUBA1C,DST,AN
,SNRPN,SPAG7,WDR62,IL1RAP,STYXL1,RFX5,RANBP1,UBE2L6,VRK3,LZTFL1,FIS1,PPM1K,TUBA1C,DST,AN
S1,COASY,AUP1,SAP30BP,FOXRED1,MED15,PARD6G,SH3YL1,MICAL1,HS2ST1,UBE2I,KCNMA1,SLC39A13
S1,COASY,AUP1,SAP30BP,FOXRED1,MED15,PARD6G,SH3YL1,MICAL1,HS2ST1,UBE2I,KCNMA1,SLC39A13
S1,COASY,AUP1,SAP30BP,FOXRED1,MED15,PARD6G,SH3YL1,MICAL1,HS2ST1,UBE2I,KCNMA1,SLC39A13
S1,COASY,AUP1,SAP30BP,FOXRED1,MED15,PARD6G,SH3YL1,MICAL1,HS2ST1,UBE2I,KCNMA1,SLC39A13
TK2,AUNIP,C14ORF2,PCYT2,USE1,AGPAT4,PPP1CC,EDF1,RFX3,PPA1,DGKE,CAMTA1,CAPG,CLDND1,LIG1,
TK2,AUNIP,C14ORF2,PCYT2,USE1,AGPAT4,PPP1CC,EDF1,RFX3,PPA1,DGKE,CAMTA1,CAPG,CLDND1,LIG1,
MRPL33,ZNF410,RUVBL2,PCMT1,EBPL,TESC,IGSF8,NUDT5,RBPMS,NNMT,SNRPN,WDR62,STYXL1,RFX5,R
1,YIPF3,TRNT1,PARD6G,SH3YL1,IP6K2,CLASRP,JPH2,C1ORF198,MICAL1,HS2ST1,UBE2I,PRMT2,EIF3M,TB
1,YIPF3,TRNT1,PARD6G,SH3YL1,IP6K2,CLASRP,JPH2,C1ORF198,MICAL1,HS2ST1,UBE2I,PRMT2,EIF3M,TB
R62,IL1RAP,STYXL1,RFX5,RANBP1,UBE2L6,VRK3,LZTFL1,FIS1,PPM1K,TUBA1C,DST,ANTXR2,STIP1,LRRC1
CLDND1,LIG1,CDC14B,TIMM17B,APEH,HYI,CAPN10,NUTF2,PODXL,ATPAF2,C17ORF62,PRMT1,CASC4,W
F198,HS2ST1,SLC39A13,PRMT2,EIF3M,TBX6,TMEM205,PTK2,AUNIP,CENPN,CARD14,FBXW9,USE1,HSB
F198,HS2ST1,SLC39A13,PRMT2,EIF3M,TBX6,TMEM205,PTK2,AUNIP,CENPN,CARD14,FBXW9,USE1,HSB
AM3A,MUC1,METTL17,ECSIT,EPB41L5,ATG12,PPRC1,ZBTB44,NIP7,TTC9C,PLCB2,DUSP22,SLC25A10,SPO
AM3A,MUC1,METTL17,ECSIT,EPB41L5,ATG12,PPRC1,ZBTB44,NIP7,TTC9C,PLCB2,DUSP22,SLC25A10,SPO
2,TSPAN31,FAM192A,GCDH,TRIM46,C11ORF57,GALT,EIF2A,PSPC1,ZNF283,TRAPPC2,ZNF69,ST7,THOC3,

,CAMKMT,ACTB,SETD4,MED8,EIF6,DHX30,AP4M1,MRPL27,CHFR,TUBGCP2,SSH3,RPL27,DHRS4L2,MRPL
,PPM1K,TUBA1C,DST,ANTXR2,STIP1,LRRC1,GRAMD1C,VCAN,PCBP2,LPCAT3,CD44,TM9SF1,SON,TET2,SN
,PPM1K,TUBA1C,DST,ANTXR2,STIP1,LRRC1,GRAMD1C,VCAN,PCBP2,LPCAT3,CD44,TM9SF1,SON,TET2,SN
UGGT2,UBE2I,SLC39A13,PRMT2,TBX6,TMEM205,PTK2,C14ORF2,CENPN,ATF7,PCYT2,HSBP1L1,IFI27L1,A
UGGT2,UBE2I,SLC39A13,PRMT2,TBX6,TMEM205,PTK2,C14ORF2,CENPN,ATF7,PCYT2,HSBP1L1,IFI27L1,A
BPMS,NNMT,SNRPN,SPAG7,WDR62,IL1RAP,STYXL1,RFX5,RANBP1,UBE2L6,LZTFL1,FIS1,PPM1K,TUBA1C,
BPMS,NNMT,SNRPN,SPAG7,WDR62,IL1RAP,STYXL1,RFX5,RANBP1,UBE2L6,LZTFL1,FIS1,PPM1K,TUBA1C,
P7,COX4I1,PLCB2,SLC25A10,TMX2,THEM4,DBNDD2,SDF4,NISCH,CENPK,RPL29,GGPS1,RTEL1,AUP1,YIPF
P7,COX4I1,PLCB2,SLC25A10,TMX2,THEM4,DBNDD2,SDF4,NISCH,CENPK,RPL29,GGPS1,RTEL1,AUP1,YIPF
F2,C17ORF62,PRMT1,CASC4,WNK1,ZNF7,TXNDC17,ARL6IP4,ZFAND2B,HNRNPC,UPP1,ZNF250,MOB1A,C
K3,LZTFL1,ZC3H14,FIS1,SLC15A3,TUBA1C,IRF9,STIP1,PCBP2,LPCAT3,CD44,TET2,UQCRQ,SNF8,GRK6,HM
K3,LZTFL1,ZC3H14,FIS1,SLC15A3,TUBA1C,IRF9,STIP1,PCBP2,LPCAT3,CD44,TET2,UQCRQ,SNF8,GRK6,HM

STIP1,LPCAT3,TET2,UQCRQ,SNF8,GRK6,HMGA2,CDC37,OBSL1,TTC21B,UBR4,CHCHD1,SLC39A11,FUZ,RG
STIP1,LPCAT3,TET2,UQCRQ,SNF8,GRK6,HMGA2,CDC37,OBSL1,TTC21B,UBR4,CHCHD1,SLC39A11,FUZ,RG
1,ZBTB44,DUSP22,SLC25A10,TMX2,THEM4,SDF4,NISCH,TMED4,RPS6,RPL29,RTEL1,SRSF1,AUP1,YIPF3,F
XL1,RFX5,RANBP1,UBE2L6,LZTFL1,FIS1,PPM1K,TUBA1C,DST,ANTXR2,STIP1,LRRC1,GRAMD1C,VCAN,PCB
,DENND1A,MRPS24,IDH3G,HPS1,ITPA,VPS25,SNRPD1,CD37,HMBS,RPS27L,TATDN1,ZBTB8OS,TBC1D22A
LR3H,NOMO2,RAD23A,USP45,SLC16A1,ILF3,EEF1D,NAA10,CNST,CASZ1,SSH2,COX17,HINT1,HNRNPA1,E
LR3H,NOMO2,RAD23A,USP45,SLC16A1,ILF3,EEF1D,NAA10,CNST,CASZ1,SSH2,COX17,HINT1,HNRNPA1,E
IP7,SLC25A10,TMX2,SDF4,NISCH,CENPK,RPL29,CD99L2,RTEL1,COASY,SRSF1,AUP1,FOXRED1,TRNT1,ITGB
IP7,SLC25A10,TMX2,SDF4,NISCH,CENPK,RPL29,CD99L2,RTEL1,COASY,SRSF1,AUP1,FOXRED1,TRNT1,ITGB
N,SPAG7,WDR62,IL1RAP,STYXL1,RFX5,RANBP1,UBE2L6,LZTFL1,FIS1,PPM1K,TUBA1C,DST,ANTXR2,STIP1,
DCD5,POT1,ANAPC11,RAD23A,ABHD17A,NT5C,USP45,ILF3,AGAP3,ZCCHC4,STXBP2,CHCHD5,MED6,HNR
1,CDK11A,RBM3,LAMP2,COPZ1,DUT,SYNGR2,TMEM134,RAB2A,ZNF524,ELP6,GPBP1,SIAH1,ZEB2,ASPH,
1,CDK11A,RBM3,LAMP2,COPZ1,DUT,SYNGR2,TMEM134,RAB2A,ZNF524,ELP6,GPBP1,SIAH1,ZEB2,ASPH,
410,RUVBL2,PCMT1,EBPL,TESC,NUDT5,RBPMS,SPAG7,WDR62,IL1RAP,STYXL1,POLD4,PLEKHA5,RANBP1,
410,RUVBL2,PCMT1,EBPL,TESC,NUDT5,RBPMS,SPAG7,WDR62,IL1RAP,STYXL1,POLD4,PLEKHA5,RANBP1,
B,PTPN13,UBR4,PSMA1,YLPM1,CHCHD1,FUZ,FAM3A,PSMA3,METTL17,PPRC1,RPS25,NIP7,COX4I1,DUS
B,PTPN13,UBR4,PSMA1,YLPM1,CHCHD1,FUZ,FAM3A,PSMA3,METTL17,PPRC1,RPS25,NIP7,COX4I1,DUS
IF3M,TBX6,TMEM205,XBP1,AUNIP,GUK1,FBXW9,ASF1B,DEF8,PPP1CC,EDF1,RFX3,ENDOV,CAMTA1,PSM
IF3M,TBX6,TMEM205,XBP1,AUNIP,GUK1,FBXW9,ASF1B,DEF8,PPP1CC,EDF1,RFX3,ENDOV,CAMTA1,PSM
RC1,RPS25,NLRP1,NIP7,TTC9C,COX4I1,DUSP22,THEM4,DBNDD2,TMED4,CENPK,RTEL1,COASY,SRSF1,AU
RC1,RPS25,NLRP1,NIP7,TTC9C,COX4I1,DUSP22,THEM4,DBNDD2,TMED4,CENPK,RTEL1,COASY,SRSF1,AU
MAGED2,SON,UQCRQ,GRK6,HMGA2,CDC37,CPSF3L,TSG101,OBSL1,TTC21B,PTPN13,PSMA1,YLPM1,PHF2
MAGED2,SON,UQCRQ,GRK6,HMGA2,CDC37,CPSF3L,TSG101,OBSL1,TTC21B,PTPN13,PSMA1,YLPM1,PHF2
BNDD2,NISCH,TMED4,CENPK,CD99L2,GGPS1,AUP1,YIPF3,TRNT1,KXD1,PARD6G,IP6K2,CLASRP,JPH2,MI
BNDD2,NISCH,TMED4,CENPK,CD99L2,GGPS1,AUP1,YIPF3,TRNT1,KXD1,PARD6G,IP6K2,CLASRP,JPH2,MI
OMMD5,CHID1,NUDT22,AMZ2,UFD1L,TAZ,THYN1,DBNDD1,SH2B1,NOP16,RPL13A,ARHGEF39,GSTK1,TT
OMMD5,CHID1,NUDT22,AMZ2,UFD1L,TAZ,THYN1,DBNDD1,SH2B1,NOP16,RPL13A,ARHGEF39,GSTK1,TT
F1B,DEF8,EEF1B2,PPP1CC,PPA1,DGKE,CAMTA1,CRIP1,LIG1,PSMA7,TIMM17B,APEH,HYI,CAPN10,NUTF2,
NPK,RPL29,GGPS1,RTEL1,AUP1,TRNT1,KXD1,MED15,SH3YL1,CLASRP,UGGT2,MICAL1,HS2ST1,UBE2I,SLC
C42,TMEM205,PTK2,GUK1,CENPN,PCYT2,USE1,CCDC25,DEF8,EEF1B2,EDF1,PPA1,DGKE,TNFRSF14,END
IS1,TUBA1C,DST,ANTXR2,STIP1,LRRC1,PHF19,GRAMD1C,LPCAT3,TM9SF1,MAGED2,SON,SNF8,GRK6,HM
IS1,TUBA1C,DST,ANTXR2,STIP1,LRRC1,PHF19,GRAMD1C,LPCAT3,TM9SF1,MAGED2,SON,SNF8,GRK6,HM
PPM1K,TUBA1C,IRF9,ANTXR2,STIP1,LRRC1,PHF19,VCAN,PCBP2,LPCAT3,MAGED2,SON,TET2,GRK6,HMG

PPM1K,TUBA1C,IRF9,ANTXR2,STIP1,LRRC1,PHF19,VCAN,PCBP2,LPCAT3,MAGED2,SON,TET2,GRK6,HMGA
1,ITGB1BP1,SH3YL1,IP6K2,SLC39A13,YIF1A,JUP,CDC42,TBX6,TMEM205,XBP1,PTK2,C14ORF2,GUK1,CARI
PPM1K,TUBA1C,IRF9,DST,ANTXR2,STIP1,LRRC1,VCAN,LPCAT3,CD44,MAGED2,SON,TET2,UQCRQ,SNF8,GI
PPM1K,TUBA1C,IRF9,DST,ANTXR2,STIP1,LRRC1,VCAN,LPCAT3,CD44,MAGED2,SON,TET2,UQCRQ,SNF8,GI
U1,SUGT1,HYPK,RPL36,CDK11A,ZNF197,COPZ1,DGUOK,ATP5J,TSFM,TRMT112,PNPO,RPS20,GABPB1,VP
U1,SUGT1,HYPK,RPL36,CDK11A,ZNF197,COPZ1,DGUOK,ATP5J,TSFM,TRMT112,PNPO,RPS20,GABPB1,VP
C,GMPR2,RIC8A,PIGT,RPS11,PDCD6,SNX5,ZNF81,FAM32A,NFYC,RPL37A,C12ORF57,DCTN5,ARL16,OXLD
C,GMPR2,RIC8A,PIGT,RPS11,PDCD6,SNX5,ZNF81,FAM32A,NFYC,RPL37A,C12ORF57,DCTN5,ARL16,OXLD

2,GUK1,PCYT2,HSBP1L1,AGPAT4,IFI27L1,ASF1B,DEF8,EEF1B2,EDF1,RFX3,TNFRSF14,ENDOV,CAMTA1,PPI
ZNF20,SLC33A1,RALGPS1,EPS15L1,POLR3H,ABHD17A,ILF3,EEF1D,NAA10,RRP8,MED6,LLGL2,CHEK1,COX
ZNF20,SLC33A1,RALGPS1,EPS15L1,POLR3H,ABHD17A,ILF3,EEF1D,NAA10,RRP8,MED6,LLGL2,CHEK1,COX
PH,TRMT112,STRIP2,ZNF625,LYNX1,TWSG1,LRRC27,MVD,IL32,LMBR1,C12ORF76,RANGRF,CHCHD7,HMG
C33A1,ABHD17A,GGT5,DDX41,NT5C,PSME1,HSPH1,CHCHD5,SGK1,SSH2,ZFAND4,RPL15,B3GAT3,RFXAN
TNFSF10,COMMD5,RUNX1,CHID1,NUDT22,AMZ2,UFD1L,USP38,TAZ,THYN1,DBNDD1,SH2B1,NOP16,RPL
,PTK2,C14ORF2,GUK1,PCYT2,IFI27L1,DEF8,PPP1CC,PPA1,DGKE,TNFRSF14,ENDOV,CAMTA1,PPIL2,RPL28

1,RCE1,MRPL33,RUVBL2,EBPL,TESC,IGSF8,NUDT5,RBPMS,SPAG7,WDR62,IL1RAP,STYXL1,RFX5,POLD4,PL
1,RCE1,MRPL33,RUVBL2,EBPL,TESC,IGSF8,NUDT5,RBPMS,SPAG7,WDR62,IL1RAP,STYXL1,RFX5,POLD4,PL
PS18C,MROH1,MFSD11,SNX5,FAM32A,RPAIN,ACOT2,PEX2,TNKS,SMIM4,PRRC2B,STX10,NMNAT3,MPH
SMD8,RQCD1,PEX16,ALS2,AP2S1,CAMKMT,RPS15A,METTL23,MRPL51,MRPL43,MCM7,C9ORF64,MED8,
SMD8,RQCD1,PEX16,ALS2,AP2S1,CAMKMT,RPS15A,METTL23,MRPL51,MRPL43,MCM7,C9ORF64,MED8,
,TM9SF1,SON,UQCRQ,SNF8,GRK6,HMGA2,CDC37,RPS16,CPSF3L,TSG101,OBSL1,PTPN13,YLPM1,PHF21A
,CDC42,XBP1,PTK2,GUK1,CARD14,PCYT2,FBXW9,USE1,IFI27L1,ASF1B,CCDC25,EEF1B2,RFX3,DGKE,END
,CDC42,XBP1,PTK2,GUK1,CARD14,PCYT2,FBXW9,USE1,IFI27L1,ASF1B,CCDC25,EEF1B2,RFX3,DGKE,END
4,IFI27L1,MAF,CCDC25,DEF8,PPA1,ENDOV,CAMTA1,PPIL2,RPL21,TXNL4A,LSM12,PRMT1,EXOSC9,C19OR
4,IFI27L1,MAF,CCDC25,DEF8,PPA1,ENDOV,CAMTA1,PPIL2,RPL21,TXNL4A,LSM12,PRMT1,EXOSC9,C19OR
PS6,CD99L2,GGPS1,SRSF1,AUP1,YIPF3,FOXRED1,TRNT1,ITGB1BP1,MED15,PARD6G,SH3YL1,CLASRP,C1O
PS6,CD99L2,GGPS1,SRSF1,AUP1,YIPF3,FOXRED1,TRNT1,ITGB1BP1,MED15,PARD6G,SH3YL1,CLASRP,C1O
,PPP3CB,RABEP2,TADA3,RALGPS1,ZNF687,PDCD5,GTF2IRD1,NXN,EPS15L1,POT1,POLR3H,RAD23A,ABH
2,TMED4,RPL29,GGPS1,RTEL1,SRSF1,AUP1,YIPF3,FOXRED1,RABL5,TRNT1,ITGB1BP1,SH3YL1,CLASRP,JPH
2,TMED4,RPL29,GGPS1,RTEL1,SRSF1,AUP1,YIPF3,FOXRED1,RABL5,TRNT1,ITGB1BP1,SH3YL1,CLASRP,JPH
4ORF2,GUK1,CENPN,CARD14,PCYT2,FBXW9,HSBP1L1,AGPAT4,IFI27L1,MAF,CCDC25,EEF1B2,PPP1CC,ED
4ORF2,GUK1,CENPN,CARD14,PCYT2,FBXW9,HSBP1L1,AGPAT4,IFI27L1,MAF,CCDC25,EEF1B2,PPP1CC,ED
,TTLL5,LDHA,ACOT8,AP3B2,RPS9,RCE1,MRPL33,ZNF410,RUVBL2,PCMT1,EBPL,TESC,IGSF8,NUDT5,RBPM
,TTLL5,LDHA,ACOT8,AP3B2,RPS9,RCE1,MRPL33,ZNF410,RUVBL2,PCMT1,EBPL,TESC,IGSF8,NUDT5,RBPM
L6IP4,ZFAND2B,ZNF644,UPP1,ZNF250,RPL36,LAMP2,CCNL2,HSP90AB1,SYNGR2,TMEM134,LRRFIP1,RAB

IPO,LSM12,C17ORF62,PRMT1,CASC4,STOML2,WNK1,C19ORF54,ZNF7,ARL6IP4,STRADA,HNRNPC,ZNF64
1,PARD6G,UGGT2,MICAL1,UBE2I,SLC39A13,YIF1A,PRMT2,TBX6,TMEM205,PTK2,C14ORF2,CENPN,ATF7,
1,PARD6G,UGGT2,MICAL1,UBE2I,SLC39A13,YIF1A,PRMT2,TBX6,TMEM205,PTK2,C14ORF2,CENPN,ATF7,
FIP1,ZNF524,GPBP1,SIAH1,ZEB2,TRMT112,STRIP2,ZNF625,RPS20,GABPB1,NIT2,LYNX1,MPDU1,RPS15,ER
S20,GABPB1,MPDU1,RPS15,ERLEC1,NAP1L1,DDOST,RANGRF,MEIS2,PHTF2,NAT9,ZNF836,RBP1,RAD1,H
D1,SLC39A11,FAM3A,MUC1,ECSIT,EPB41L5,ATG12,PPRC1,NLRP1,ZBTB44,NIP7,TTC9C,COX4I1,PLCB2,DU
D1,SLC39A11,FAM3A,MUC1,ECSIT,EPB41L5,ATG12,PPRC1,NLRP1,ZBTB44,NIP7,TTC9C,COX4I1,PLCB2,DU

2,LRRC59,SUMO1,ZNF483,SZT2,BSCL2,CYB5D2,RABEPK,PSMC5,ZNF816,HAUS1,LAMTOR4,WDR45,HDAC

2,LRRC59,SUMO1,ZNF483,SZT2,BSCL2,CYB5D2,RABEPK,PSMC5,ZNF816,HAUS1,LAMTOR4,WDR45,HDAC
AT3,CD44,SON,TET2,UQCRQ,SNF8,GRK6,HMGA2,CDC37,CPSF3L,TSG101,TTC21B,PTPN13,PSMA1,YLPM1,
AT3,CD44,SON,TET2,UQCRQ,SNF8,GRK6,HMGA2,CDC37,CPSF3L,TSG101,TTC21B,PTPN13,PSMA1,YLPM1,
1,ERCC5,IL32,LMBR1,MCTS1,SERPINB1,LSR,CHCHD7,MEIS2,PSME2,PHTF2,BCAM,ZNF836,PPP3CB,RABE

IZ,PSMA3,NLRP1,ZBTB44,DUSP22,SLC25A10,RDH11,NISCH,CENPK,RPL29,GGPS1,SRSF1,AUP1,SAP30BP,Y
IZ,PSMA3,NLRP1,ZBTB44,DUSP22,SLC25A10,RDH11,NISCH,CENPK,RPL29,GGPS1,SRSF1,AUP1,SAP30BP,Y
6G,SH3YL1,JPH2,UGGT2,MICAL1,SLC39A13,MAGT1,CDC42,XBP1,PTK2,AUNIP,GUK1,CENPN,PCYT2,FBXW

UGGT2,HS2ST1,PRMT2,JUP,CDC42,TBX6,TMEM205,AUNIP,GUK1,CENPN,FBXW9,ASF1B,CCDC25,PPP1CC,
UGGT2,HS2ST1,PRMT2,JUP,CDC42,TBX6,TMEM205,AUNIP,GUK1,CENPN,FBXW9,ASF1B,CCDC25,PPP1CC,
RPL15,B3GAT3,PFDN5,MTHFS,PAFAH1B2,COA3,NSUN5,DAZAP2,ZFPL1,MRPS18C,GMPR2,ZSCAN9,RIC8A,
RPL15,B3GAT3,PFDN5,MTHFS,PAFAH1B2,COA3,NSUN5,DAZAP2,ZFPL1,MRPS18C,GMPR2,ZSCAN9,RIC8A,
GGPS1,SRSF1,AUP1,YIPF3,FOXRED1,TRNT1,ITGB1BP1,SH3YL1,IP6K2,SLC39A13,YIF1A,JUP,CDC42,TBX6,X
1,ZNF197,DGUOK,DUT,CEP70,LRRFIP1,ZNF524,RMDN1,GPALPP1,TSFM,ZEB2,ASPH,ZNF625,RPS20,GAB
2B,HNRNPC,CYB561,ZNF250,RPL36,MOB1A,NFAM1,MDH1,LAMP2,HSP90AB1,DUT,SYNGR2,TMEM134,L
A3,PPM1K,TUBA1C,IRF9,ANTXR2,STIP1,PHF19,VCAN,PCBP2,LPCAT3,CD44,MAGED2,SON,GRK6,HMGA2,
A3,PPM1K,TUBA1C,IRF9,ANTXR2,STIP1,PHF19,VCAN,PCBP2,LPCAT3,CD44,MAGED2,SON,GRK6,HMGA2,
ESC,IGSF8,NUDT5,RBPMS,NNMT,SPAG7,IL1RAP,STYXL1,POLD4,PLEKHA5,UBE2L6,VRK3,LZTFL1,FIS1,SLC1
ESC,IGSF8,NUDT5,RBPMS,NNMT,SPAG7,IL1RAP,STYXL1,POLD4,PLEKHA5,UBE2L6,VRK3,LZTFL1,FIS1,SLC1
3,PPM1K,IRF9,ANTXR2,STIP1,PHF19,GRAMD1C,VCAN,LPCAT3,CD44,TM9SF1,SON,TET2,UQCRQ,SNF8,G
3,PPM1K,IRF9,ANTXR2,STIP1,PHF19,GRAMD1C,VCAN,LPCAT3,CD44,TM9SF1,SON,TET2,UQCRQ,SNF8,G
3,MED6,LLGL2,CHEK1,SUMO3,CD151,FKBP3,EGLN2,SIRT5,KCTD7,RFXANK,SPINT2,PTK7,TMEM39B,DALR
6,UBR4,PSMA1,YLPM1,FUZ,FAM3A,PSMA3,ECSIT,ATG12,PPRC1,RPS25,NIP7,TTC9C,COX4I1,SLC25A10,T
ID2B,SHARPIN,SUGT1,ZNF250,RPL36,CDK11A,COPZ1,INO80E,MAPKAPK5,HSP90AB1,DUT,TMEM134,RA
TYXL1,RFX5,POLD4,PLEKHA5,RANBP1,LZTFL1,FIS1,SLC15A3,PPM1K,TUBA1C,DST,ANTXR2,STIP1,LRRC1,V
TYXL1,RFX5,POLD4,PLEKHA5,RANBP1,LZTFL1,FIS1,SLC15A3,PPM1K,TUBA1C,DST,ANTXR2,STIP1,LRRC1,V
O1,SZT2,PSMD8,PEX16,PSMC5,NGDN,POLH,METTL23,C12ORF73,SETD4,MED8,AP4M1,MRPL27,GANC,
O1,SZT2,PSMD8,PEX16,PSMC5,NGDN,POLH,METTL23,C12ORF73,SETD4,MED8,AP4M1,MRPL27,GANC,

TA1,PSMD4,RPL28,PSMA7,RPL21,TXNL4A,PODXL,UCK2,LSM12,C17ORF62,PRMT1,CASC4,EXOSC9,WNK1
TA1,PSMD4,RPL28,PSMA7,RPL21,TXNL4A,PODXL,UCK2,LSM12,C17ORF62,PRMT1,CASC4,EXOSC9,WNK1
M53,PPIE,SETD4,PHPT1,ATP9B,ANKRD10,RAP1B,UBE2A,CREM,TUBGCP2,CORO1B,PHF20,AP4S1,C12ORF
RPS6,RPL29,CD99L2,SRSF1,AUP1,SAP30BP,FOXRED1,MED15,SH3YL1,IP6K2,CLASRP,HS2ST1,UBE2I,KCN
RPS6,RPL29,CD99L2,SRSF1,AUP1,SAP30BP,FOXRED1,MED15,SH3YL1,IP6K2,CLASRP,HS2ST1,UBE2I,KCN
KW9,MAF,ASF1B,CCDC25,PPP1CC,PDLIM5,RFX3,DGKE,TNFRSF14,ENDOV,CRIP1,CLDND1,PSMA7,TIMM1
KW9,MAF,ASF1B,CCDC25,PPP1CC,PDLIM5,RFX3,DGKE,TNFRSF14,ENDOV,CRIP1,CLDND1,PSMA7,TIMM1
BR4,YLPM1,PHF21A,FAM3A,METTL17,ATG12,PPRC1,RPS25,ZBTB44,TTC9C,COX4I1,PLCB2,DUSP22,SLC25

,JUP,CDC42,TBX6,TMEM205,AUNIP,GUK1,CENPN,ATF7,FBXW9,CCDC25,EEF1B2,PPP1CC,GADL1,RFX3,E
,JUP,CDC42,TBX6,TMEM205,AUNIP,GUK1,CENPN,ATF7,FBXW9,CCDC25,EEF1B2,PPP1CC,GADL1,RFX3,E
UZ,RGS3,FAM3A,PSMA3,MUC1,METTL17,ATG12,PPRC1,NLRP1,ZBTB44,NIP7,TTC9C,COX4I1,PLCB2,DUS
UZ,RGS3,FAM3A,PSMA3,MUC1,METTL17,ATG12,PPRC1,NLRP1,ZBTB44,NIP7,TTC9C,COX4I1,PLCB2,DUS
K1,CENPN,PCYT2,USE1,HSBP1L1,DEF8,EEF1B2,EDF1,DGKE,TNFRSF14,CAPG,CRIP1,PPIL2,TROAP,VIM,LIG1
K1,CENPN,PCYT2,USE1,HSBP1L1,DEF8,EEF1B2,EDF1,DGKE,TNFRSF14,CAPG,CRIP1,PPIL2,TROAP,VIM,LIG1
9L2,GGPS1,RTEL1,COASY,SRSF1,FOXRED1,TRNT1,ITGB1BP1,SH3YL1,IP6K2,UGGT2,HS2ST1,PRMT2,MAG

9L2,GGPS1,RTEL1,COASY,SRSF1,FOXRED1,TRNT1,ITGB1BP1,SH3YL1,IP6K2,UGGT2,HS2ST1,PRMT2,MAGT

205,PTK2,AUNIP,C14ORF2,GUK1,CENPN,ASF1B,CCDC25,DEF8,PPP1CC,RFX3,PPA1,DGKE,ENDOV,CAMTA1
205,PTK2,AUNIP,C14ORF2,GUK1,CENPN,ASF1B,CCDC25,DEF8,PPP1CC,RFX3,PPA1,DGKE,ENDOV,CAMTA1
,THEM4,DBNDD2,SDF4,TMED4,CENPK,GGPS1,RTEL1,SRSF1,SAP30BP,FOXRED1,RABL5,TRNT1,KXD1,ITGB
,THEM4,DBNDD2,SDF4,TMED4,CENPK,GGPS1,RTEL1,SRSF1,SAP30BP,FOXRED1,RABL5,TRNT1,KXD1,ITGB
PH2,UGGT2,MICAL1,UBE2I,SLC39A13,PRMT2,CDC42,TBX6,TMEM205,PTK2,C14ORF2,GUK1,CENPN,ATF7,
TMX2,SPOCK2,DBNDD2,NISCH,TMED4,RPL29,CD99L2,GGPS1,SRSF1,AUP1,PARD6G,SH3YL1,IP6K2,CLASR
T2,UQCRQ,SNF8,HMGA2,CDC37,RPS16,CPSF3L,TSG101,OBSL1,PTPN13,UBR4,PHF21A,SLC39A11,FUZ,FAI
T2,UQCRQ,SNF8,HMGA2,CDC37,RPS16,CPSF3L,TSG101,OBSL1,PTPN13,UBR4,PHF21A,SLC39A11,FUZ,FAI
7L1,MAF,PPA1,ENDOV,PPIL2,RPL21,TXNL4A,LSM12,PRMT1,EXOSC9,C19ORF48,ZFAND2B,HNRNPC,SLC9
7L1,MAF,PPA1,ENDOV,PPIL2,RPL21,TXNL4A,LSM12,PRMT1,EXOSC9,C19ORF48,ZFAND2B,HNRNPC,SLC9

ASF1B,EEF1B2,PPP1CC,RFX3,DGKE,ENDOV,CAMTA1,PPIL2,VIM,LIG1,CDC14B,TIMM17B,TMEM126B,APEH
PF,LYNX1,LRRC27,LMBR1,NAP1L1,SYNGR1,RANGRF,CHCHD7,HMGA1,PSME2,PHTF2,PABPC1,NAT9,BCAM
C1,SAT1,NAP1L1,DDOST,CHCHD7,MEIS2,PHTF2,DBNL,NAT9,ZNF836,RBP1,H2AFV,SERPINB6,RABEP2,TAD
C25A10,DBNDD2,CENPK,RPS6,RPL29,CD99L2,GGPS1,RTEL1,COASY,SRSF1,AUP1,FOXRED1,ITGB1BP1,PAI
L2,PCMT1,EBPL,TESC,IGSF8,NUDT5,RBPMS,NNMT,SNRPN,SPAG7,WDR62,STYXL1,RFX5,RANBP1,UBE2L6,
L2,PCMT1,EBPL,TESC,IGSF8,NUDT5,RBPMS,NNMT,SNRPN,SPAG7,WDR62,STYXL1,RFX5,RANBP1,UBE2L6,

,RANBP1,UBE2L6,VRK3,LZTFL1,ZC3H14,FIS1,IRF9,DST,ANTXR2,STIP1,LRRC1,GRAMD1C,CD44,TM9SF1,MA
,RANBP1,UBE2L6,VRK3,LZTFL1,ZC3H14,FIS1,IRF9,DST,ANTXR2,STIP1,LRRC1,GRAMD1C,CD44,TM9SF1,MA
ATG12,TTC9C,PLCB2,DUSP22,SLC25A10,TMX2,SPOCK2,DBNDD2,TMED4,CD99L2,GGPS1,SRSF1,AUP1,SAI
B2,ASPH,TRMT112,PNPO,LYNX1,TWSG1,GMPPB,LRRC27,LMBR1,MCTS1,NAP1L1,SERPINB1,RANGRF,MTA
NO80C,ARRB2,DIP2A,MLST8,HMGN1,CDK2AP2,MYL12A,MAZ,C2ORF15,TBCA,POLR2E,IAH1,PNRC1,ARPC1
NO80C,ARRB2,DIP2A,MLST8,HMGN1,CDK2AP2,MYL12A,MAZ,C2ORF15,TBCA,POLR2E,IAH1,PNRC1,ARPC1
I,TPM2,SIAH1,ZEB2,ASPH,TRMT112,NIT2,LYNX1,TWSG1,HSPB1,MVD,LMBR1,NAP1L1,SERPINB1,CHCHD7

,CDC42,TBX6,TMEM205,AUNIP,CENPN,FBXW9,USE1,HSBP1L1,IFI27L1,CCDC25,EEF1B2,EDF1,RFX3,DGKE,
L1,HS2ST1,UBE2I,SLC39A13,YIF1A,TBX6,TMEM205,PTK2,AUNIP,C14ORF2,GUK1,CENPN,PCYT2,USE1,AGF
L1,HS2ST1,UBE2I,SLC39A13,YIF1A,TBX6,TMEM205,PTK2,AUNIP,C14ORF2,GUK1,CENPN,PCYT2,USE1,AGF
F410,PCMT1,EBPL,TESC,IGSF8,NUDT5,RBPMS,NNMT,SPAG7,WDR62,IL1RAP,STYXL1,POLD4,PLEKHA5,RAN
F410,PCMT1,EBPL,TESC,IGSF8,NUDT5,RBPMS,NNMT,SPAG7,WDR62,IL1RAP,STYXL1,POLD4,PLEKHA5,RAN
,ASH1L,TCF25,ZNF248,ERRFI1,ANAPC13,NXF1,SYP,EIF2A,NUP62,PSPC1,NADSYN1,PAM16,FCHO2,IRF3,D2
,ASH1L,TCF25,ZNF248,ERRFI1,ANAPC13,NXF1,SYP,EIF2A,NUP62,PSPC1,NADSYN1,PAM16,FCHO2,IRF3,D2

A13,YIF1A,PRMT2,EIF3M,JUP,TBX6,C14ORF2,GUK1,FBXW9,USE1,AGPAT4,IFI27L1,MAF,ASF1B,DEF8,EEF1
A13,YIF1A,PRMT2,EIF3M,JUP,TBX6,C14ORF2,GUK1,FBXW9,USE1,AGPAT4,IFI27L1,MAF,ASF1B,DEF8,EEF1
AMLG,EPB41L4B,TCF4,PPOX,LSM12,PRMT1,WNK1,C19ORF54,C19ORF48,AP1S3,HNRNPC,CYB561,CBFA2T
AMLG,EPB41L4B,TCF4,PPOX,LSM12,PRMT1,WNK1,C19ORF54,C19ORF48,AP1S3,HNRNPC,CYB561,CBFA2T
MPPB,LRRC27,LMBR1,NAP1L1,SERPINB1,DDOST,SYNGR1,CHCHD7,HMGA1,RBP1,RAD1,H2AFV,SERPINB6,
MPPB,LRRC27,LMBR1,NAP1L1,SERPINB1,DDOST,SYNGR1,CHCHD7,HMGA1,RBP1,RAD1,H2AFV,SERPINB6,

LRP1,NIP7,TTC9C,COX4I1,DUSP22,THEM4,CENPK,RPS6,RPL29,RTEL1,COASY,AUP1,SAP30BP,YIPF3,FOXRE

LRP1,NIP7,TTC9C,COX4I1,DUSP22,THEM4,CENPK,RPS6,RPL29,RTEL1,COASY,AUP1,SAP30BP,YIPF3,FOXRE

ON,TET2,SNF8,GRK6,HMGA2,CDC37,CPSF3L,TTC21B,PTPN13,UBR4,PSMA1,PHF21A,SLC39A11,FUZ,PSMA

ON,TET2,SNF8,GRK6,HMGA2,CDC37,CPSF3L,TTC21B,PTPN13,UBR4,PSMA1,PHF21A,SLC39A11,FUZ,PSMA

ZAP2,ZFPL1,MFSD10,PARK7,PIGT,UBE2N,LFNG,PDCD6,FAM32A,COX20,PCID2,TGOLN2,C12ORF57,DCTN

N9,LGALS3,COMMD4,CCDC40,PIGT,UBE2N,PDCD6,FAM32A,ZSCAN20,MT2A,TMEM147,PEX2,TNKS,C12O

G1,PSMA7,CDC14B,TXNL4A,CAPN10,EPB41L4B,STOML2,WNK1,C19ORF54,ARL6IP4,C19ORF48,STRADA,

126B,NUTF2,PPOX,TMPO,LSM12,PRMT1,CASC4,STOML2,WNK1,PPAN,HNRNPC,CBFA2T2,STAU1,ZNF25

126B,NUTF2,PPOX,TMPO,LSM12,PRMT1,CASC4,STOML2,WNK1,PPAN,HNRNPC,CBFA2T2,STAU1,ZNF25

LC,VCAN,LPCAT3,CD44,TM9SF1,MAGED2,SON,TET2,SNF8,GRK6,CDC37,RPS16,TSG101,TTC21B,UBR4,PSM

LC,VCAN,LPCAT3,CD44,TM9SF1,MAGED2,SON,TET2,SNF8,GRK6,CDC37,RPS16,TSG101,TTC21B,UBR4,PSM

,MDH1,INO80E,LRRFIP1,ZNF524,GPALPP1,SIAH1,ASPH,RPTOR,STRIP2,ZNF625,RPS20,GABPB1,ZSWIM7,

MGA1,PSME2,PABPC1,DBNL,NAT9,BCAM,BLVRB,PPP3CB,RABEP2,TADA3,RALGPS1,ZNF706,ZNF687,PDCD

MGA1,PSME2,PABPC1,DBNL,NAT9,BCAM,BLVRB,PPP3CB,RABEP2,TADA3,RALGPS1,ZNF706,ZNF687,PDCD

4,CAPG,CRIP1,PPIL2,TROAP,VIM,LIG1,PSMA7,PODXL,EPB41L4B,PPOX,ATPAF2,CASC4,ZNF7,PIGL,AP1S3,A

MM17B,TMEM126B,TXNL4A,CAPN10,CAMLG,EPB41L4B,ATPAF2,PRMT1,C19ORF54,ARL6IP4,C19ORF48,

MM17B,TMEM126B,TXNL4A,CAPN10,CAMLG,EPB41L4B,ATPAF2,PRMT1,C19ORF54,ARL6IP4,C19ORF48,

M2,SIAH1,ZEB2,NIT2,LYNX1,TWSG1,ZSWIM7,HSPB1,LRRC27,LMBR1,NAP1L1,SERPINB1,RANGRF,CHCHD7

4ORF2,GUK1,PCYT2,USE1,AGPAT4,MAF,CCDC25,DEF8,PPP1CC,EDF1,RFX3,PPA1,DGKE,ENDOV,CAMTA1,

4ORF2,GUK1,PCYT2,USE1,AGPAT4,MAF,CCDC25,DEF8,PPP1CC,EDF1,RFX3,PPA1,DGKE,ENDOV,CAMTA1,

5,RPS16,TSG101,OBSL1,TTC21B,PTPN13,UBR4,PSMA1,YLPM1,FUZ,FAM3A,PSMA3,MUC1,ECSIT,EPB41L5,

5,RPS16,TSG101,OBSL1,TTC21B,PTPN13,UBR4,PSMA1,YLPM1,FUZ,FAM3A,PSMA3,MUC1,ECSIT,EPB41L5,

MX2,RDH11,SDF4,NISCH,TMED4,CENPK,RPS6,GGPS1,RTEL1,SRSF1,AUP1,SAP30BP,YIPF3,FOXRED1,TRNT

MX2,RDH11,SDF4,NISCH,TMED4,CENPK,RPS6,GGPS1,RTEL1,SRSF1,AUP1,SAP30BP,YIPF3,FOXRED1,TRNT

NF8,GRK6,CDC37,CPSF3L,TSG101,TTC21B,UBR4,PSMA1,YLPM1,CHCHD1,FAM3A,METTL17,EPB41L5,ATG1

NF8,GRK6,CDC37,CPSF3L,TSG101,TTC21B,UBR4,PSMA1,YLPM1,CHCHD1,FAM3A,METTL17,EPB41L5,ATG1

N,CARD14,FBXW9,CCDC25,PPP1CC,GADL1,RFX3,ENDOV,PPIL2,PSMD4,VIM,LIG1,PSMA7,CDC14B,APEH,T

N,CARD14,FBXW9,CCDC25,PPP1CC,GADL1,RFX3,ENDOV,PPIL2,PSMD4,VIM,LIG1,PSMA7,CDC14B,APEH,T

IRF9,ANTXR2,STIP1,LRRC1,PHF19,GRAMD1C,VCAN,PCBP2,CD44,MAGED2,SON,TET2,UQCRQ,GRK6,HMG

IRF9,ANTXR2,STIP1,LRRC1,PHF19,GRAMD1C,VCAN,PCBP2,CD44,MAGED2,SON,TET2,UQCRQ,GRK6,HMG

DH,ARPC5,GEMIN2,LYPLAL1,THOC3,SH3GL1,BTF3,NDUFA3,C1ORF63,C5ORF24,SLC37A4,IDH3G,DNAJA3,

DH,ARPC5,GEMIN2,LYPLAL1,THOC3,SH3GL1,BTF3,NDUFA3,C1ORF63,C5ORF24,SLC37A4,IDH3G,DNAJA3,

17B,TMEM126B,CAPN10,LYPLA2,TMPO,C17ORF62,WNK1,C19ORF54,ZNF7,PIGL,ZFAND2B,HNRNPC,ZNF

COA3,SNRPF,TYMS,ZFPL1,SLC25A23,TPM4,MRPS18C,RIC8A,TCP1,GNB5,COMMD4,NFATC4,PDCD6,FAM3

3,ZNF644,UPP1,ZNF250,NFAM1,MDH1,INO80E,CCNL2,MAPKAPK5,HSP90AB1,SYNGR2,LRRFIP1,ZNF524,

ON,TET2,UQCRQ,GRK6,RPS16,OBSL1,PTPN13,YLPM1,PHF21A,SLC39A11,FUZ,METTL17,EPB41L5,ATG12,P

ON,TET2,UQCRQ,GRK6,RPS16,OBSL1,PTPN13,YLPM1,PHF21A,SLC39A11,FUZ,METTL17,EPB41L5,ATG12,P

XL,EPB41L4B,TCF4,ATPAF2,C17ORF62,PRMT1,CASC4,WNK1,ZNF7,TXNDC17,ZFAND2B,HNRNPC,CYB561,

MT2,MAGT1,C14ORF2,GUK1,CENPN,CARD14,PCYT2,FBXW9,HSBP1L1,AGPAT4,IFI27L1,MAF,CCDC25,PPA1

MT2,MAGT1,C14ORF2,GUK1,CENPN,CARD14,PCYT2,FBXW9,HSBP1L1,AGPAT4,IFI27L1,MAF,CCDC25,PPA1

B41L5,ATG12,NLRP1,ZBTB44,COX4I1,DUSP22,TMX2,SPOCK2,DBNDD2,SDF4,NISCH,CENPK,GGPS1,RTEL1

B41L5,ATG12,NLRP1,ZBTB44,COX4I1,DUSP22,TMX2,SPOCK2,DBNDD2,SDF4,NISCH,CENPK,GGPS1,RTEL1

PS1,ZNF687,PDCD5,NXN,EPS15L1,POLR3H,RAD23A,ABHD17A,STXBP2,EEF1D,NAA10,LLGL2,CASZ1,COX1

BX6,PTK2,CELF1,C14ORF2,ATF7,PCYT2,USE1,AGPAT4,IFI27L1,CCDC25,DEF8,EEF1B2,PPP1CC,EDF1,RFX3,

NTXR2,STIP1,LRRC1,PHF19,GRAMD1C,PCBP2,CD44,TM9SF1,SON,GRK6,CDC37,RPS16,CPSF3L,TSG101,OE
NTXR2,STIP1,LRRC1,PHF19,GRAMD1C,PCBP2,CD44,TM9SF1,SON,GRK6,CDC37,RPS16,CPSF3L,TSG101,OE
M7,HSPB1,LRRC27,MVD,LMBR1,NAP1L1,RANGRF,CHCHD7,HMGA1,PABPC1,NAT9,BCAM,ZNF836,BLVRB

UGGT2,MICAL1,UBE2I,KCNMA1,YIF1A,EIF3M,XBP1,CELF1,AUNIP,HSBP1L1,AGPAT4,MAF,ASF1B,EEF1B2,P
RF48,OAS3,MED6,SEC31A,CNST,SUMO3,LGMN,CD151,SIRT5,RFXANK,SMARCA4,TYMS,TMEM39B,DALRD
RF48,OAS3,MED6,SEC31A,CNST,SUMO3,LGMN,CD151,SIRT5,RFXANK,SMARCA4,TYMS,TMEM39B,DALRD
M147,ELP4,TSC22D3,ZXDC,COBL,ARL16,SLC35E1,DGAT1,MRPL55,RPL8,FBXL18,ECE1,OCIAD1,TCF25,RBBI
HCHD7,HMGA1,PABPC1,BCAM,H2AFV,TOMM40,RABEP2,ZNF20,RALGPS1,ZNF687,PDCD5,NXN,KLK10,EPS
HCHD7,HMGA1,PABPC1,BCAM,H2AFV,TOMM40,RABEP2,ZNF20,RALGPS1,ZNF687,PDCD5,NXN,KLK10,EPS
MM17B,TMEM126B,XAF1,HYI,PODXL,TCF4,LSM12,CASC4,EXOSC9,PIGL,STRADA,ACAD10,ZNF644,CBFA2T
MM17B,TMEM126B,XAF1,HYI,PODXL,TCF4,LSM12,CASC4,EXOSC9,PIGL,STRADA,ACAD10,ZNF644,CBFA2T
L2,PCMT1,EBPL,TESC,NUDT5,RBPMS,NNMT,SNRPN,SPAG7,WDR62,STYXL1,RFX5,PLEKHA5,RANBP1,UBE2
L2,PCMT1,EBPL,TESC,NUDT5,RBPMS,NNMT,SNRPN,SPAG7,WDR62,STYXL1,RFX5,PLEKHA5,RANBP1,UBE2
TC21B,PTPN13,UBR4,PSMA1,SLC39A11,FUZ,PSMA3,MUC1,ATG12,NLRP1,ZBTB44,DUSP22,SLC25A10,TM
,SIRT5,KCTD7,RFXANK,SPINT2,PTK7,TMEM39B,DALRD3,BRF1,OARD1,SLC25A23,FAM102A,MFSD10,PFKI
ASC4,STOML2,C19ORF54,TXNDC17,PIGL,C19ORF48,PPAN,SLC9B2,SUGT1,ZNF250,RPL36,RBM3,MDH1,LA
POLR3H,NOMO2,B2M,ABHD17A,SLC16A1,ILF3,AGAP3,MDH2,EEF1D,SEC31A,CNST,SH3GLB1,CHEK1,COX
9,FAM192A,ADRA1A,ZNF593,GLYCTK,TCEAL3,RBM42,CLASP2,PCNP,VDAC1,ZNF511,GTF2H1,RNF130,D2H
5G1,ZNF410,PCMT1,EBPL,TESC,IGSF8,RBPMS,WDR62,STYXL1,POLD4,PLEKHA5,UBE2L6,VRK3,PPM1K,TUE
5G1,ZNF410,PCMT1,EBPL,TESC,IGSF8,RBPMS,WDR62,STYXL1,POLD4,PLEKHA5,UBE2L6,VRK3,PPM1K,TUE
L,RABL5,ITGB1BP1,MED15,PARD6G,CLASRP,UGGT2,MICAL1,HS2ST1,UBE2I,KCNMA1,PRMT2,EIF3M,MAG
D2B,UPP1,ZNF250,NFAM1,CDK11A,INO80E,CCNL2,MAPKAPK5,DGUOK,HSP90AB1,SYNGR2,CEP70,TMEM
D2B,UPP1,ZNF250,NFAM1,CDK11A,INO80E,CCNL2,MAPKAPK5,DGUOK,HSP90AB1,SYNGR2,CEP70,TMEM
UNIP,C14ORF2,GUK1,CENPN,USE1,HSBP1L1,EEF1B2,EDF1,DGKE,TNFRSF14,VAMP8,CAPG,CRIP1,PPIL2,TF
UNIP,C14ORF2,GUK1,CENPN,USE1,HSBP1L1,EEF1B2,EDF1,DGKE,TNFRSF14,VAMP8,CAPG,CRIP1,PPIL2,TF
AB1,TMEM134,FAM219B,ZNF524,ATP5J,SIAH1,GGCX,ZEB2,ZNF625,MPDU1,ZSWIM7,UBXN1,ERLEC1,HSP
AB1,TMEM134,FAM219B,ZNF524,ATP5J,SIAH1,GGCX,ZEB2,ZNF625,MPDU1,ZSWIM7,UBXN1,ERLEC1,HSP
C4,TXNDC17,C19ORF48,ZFAND2B,SHARPIN,HYPK,UPP1,CARS,RBM3,MDH1,COPZ1,CCNL2,CEP70,RAB2A
ABPC1,NAT9,BCAM,ZNF836,BLVRB,PPP3CB,RABEP2,RALGPS1,ZNF706,ZNF687,PDCD5,GTF2IRD1,NXN,EP
GPBP1,SIAH1,RPTOR,STRIP2,LYNX1,MPDU1,TWSG1,ZSWIM7,DCTD,GMPPB,MVD,LMBR1,NAP1L1,SERPIN
GPBP1,SIAH1,RPTOR,STRIP2,LYNX1,MPDU1,TWSG1,ZSWIM7,DCTD,GMPPB,MVD,LMBR1,NAP1L1,SERPIN
,TPM2,SIAH1,ZEB2,ASPH,CHPF,STRIP2,ZNF625,NIT2,LYNX1,TWSG1,GMPPB,HSPB1,LRRC27,LMBR1,NAP1
CDC25,EEF1B2,RFX3,DGKE,VAMP8,CAMTA1,PSMD4,TROAP,RPL28,TMEM126B,APEH,HYI,RPL21,TXNL4A
CDC25,EEF1B2,RFX3,DGKE,VAMP8,CAMTA1,PSMD4,TROAP,RPL28,TMEM126B,APEH,HYI,RPL21,TXNL4A
UQCRQ,GRK6,CDC37,CPSF3L,TSG101,OBSL1,TTC21B,PTPN13,UBR4,YLPM1,PHF21A,CHCHD1,SLC39A11,F
UQCRQ,GRK6,CDC37,CPSF3L,TSG101,OBSL1,TTC21B,PTPN13,UBR4,YLPM1,PHF21A,CHCHD1,SLC39A11,F
OLD4,PLEKHA5,RANBP1,FIS1,SLC15A3,PPM1K,TUBA1C,DST,ANTXR2,STIP1,LRRC1,VCAN,TM9SF1,SON,TE
OLD4,PLEKHA5,RANBP1,FIS1,SLC15A3,PPM1K,TUBA1C,DST,ANTXR2,STIP1,LRRC1,VCAN,TM9SF1,SON,TE
LR2E,SNX3,SLC37A3,ECT2,GRK4,BTF3L4,VAPA,MRPS16,RPS23,MGRN1,ABLIM2,IFRD2,GRN,ZNF606,ZNF
LR2E,SNX3,SLC37A3,ECT2,GRK4,BTF3L4,VAPA,MRPS16,RPS23,MGRN1,ABLIM2,IFRD2,GRN,ZNF606,ZNF
CDK11A,RBM3,MDH1,ZNF197,COPZ1,INO80E,CCNL2,DGUOK,HSP90AB1,CEP70,FAM219B,RMDN1,TSFM
A1,FKBP3,B3GAT3,ZNF506,TCTN1,TYMS,GTF2IRD2,COPS4,DAZAP2,ZFPL1,TPM4,RIC8A,SLC30A5,GNB5,TF
3,ZNF410,PCMT1,TESC,IGSF8,NUDT5,RBPMS,NNMT,SPAG7,IL1RAP,STYXL1,RFX5,POLD4,PLEKHA5,RANBI
3,ZNF410,PCMT1,TESC,IGSF8,NUDT5,RBPMS,NNMT,SPAG7,IL1RAP,STYXL1,RFX5,POLD4,PLEKHA5,RANBI

RABEP2,RALGPS1,ZNF687,PDCD5,GTF2IRD1,NXN,EPS15L1,NUDT1,POLR3H,HSD17B10,ABHD17A,GGT5,D
RABEP2,RALGPS1,ZNF687,PDCD5,GTF2IRD1,NXN,EPS15L1,NUDT1,POLR3H,HSD17B10,ABHD17A,GGT5,D

NK1,ARL6IP4,PPAN,AP1S3,HNRNPC,STAU1,ZNF250,MDH1,CCNL2,CEP70,ZNF524,TPM2,GPBP1,SIAH1,G
T8,HMGN1,CDK2AP2,U2AF1L4,C2ORF15,TBCA,DDX10,IAH1,PNRC1,ARPC1A,UBA52,CEP63,SDHA,ZNF483
T8,HMGN1,CDK2AP2,U2AF1L4,C2ORF15,TBCA,DDX10,IAH1,PNRC1,ARPC1A,UBA52,CEP63,SDHA,ZNF483
,TET2,UQCRQ,SNF8,GRK6,HMGA2,CDC37,RPS16,CPSF3L,OBSL1,PTPN13,PHF21A,CHCHD1,METTL17,ECS
,TET2,UQCRQ,SNF8,GRK6,HMGA2,CDC37,RPS16,CPSF3L,OBSL1,PTPN13,PHF21A,CHCHD1,METTL17,ECS
PC1,NADSYN1,FCHO2,IRF3,IFT172,D2HGDH,CC2D1B,ARPC5,GEMIN2,LYPLAL1,THOC3,SH3GL1,BTF3,NDU
PC1,NADSYN1,FCHO2,IRF3,IFT172,D2HGDH,CC2D1B,ARPC5,GEMIN2,LYPLAL1,THOC3,SH3GL1,BTF3,NDU
A2T2,STAU1,SUGT1,HYPK,CDK11A,RBM3,ZNF197,MAPKAPK5,DGUOK,ATP5J,TSFM,TRMT112,PNPO,ZNF
A2T2,STAU1,SUGT1,HYPK,CDK11A,RBM3,ZNF197,MAPKAPK5,DGUOK,ATP5J,TSFM,TRMT112,PNPO,ZNF

ARPIN,SUGT1,ZNF250,NFAM1,CDK11A,RBM3,INO80E,DGUOK,CEP70,LRRFIP1,FAM219B,RAB2A,GPBP1,G
ARPIN,SUGT1,ZNF250,NFAM1,CDK11A,RBM3,INO80E,DGUOK,CEP70,LRRFIP1,FAM219B,RAB2A,GPBP1,G
RFX3,PPA1,ENDOV,PPIL2,TIMM17B,APEH,RPL21,TXNL4A,TMPO,LSM12,PRMT1,EXOSC9,TXNDC17,ARL6I
RFX3,PPA1,ENDOV,PPIL2,TIMM17B,APEH,RPL21,TXNL4A,TMPO,LSM12,PRMT1,EXOSC9,TXNDC17,ARL6I
KS,RPL37A,PIGH,C12ORF57,PEX26,RNASE1,MYL9,STAU2,DGAT1,OGFOD2,KLHL7,EIF3G,TSEN34,SSB,CRE
KS,RPL37A,PIGH,C12ORF57,PEX26,RNASE1,MYL9,STAU2,DGAT1,OGFOD2,KLHL7,EIF3G,TSEN34,SSB,CRE
ED15,JPH2,UGGT2,MICAL1,HS2ST1,UBE2I,JUP,TBX6,TMEM205,PTK2,C14ORF2,PCYT2,FBXW9,USE1,IFI2
DR62,IL1RAP,STYXL1,POLD4,PLEKHA5,RANBP1,UBE2L6,VRK3,ZC3H14,FIS1,PPM1K,TUBA1C,ANTXR2,STIP
DR62,IL1RAP,STYXL1,POLD4,PLEKHA5,RANBP1,UBE2L6,VRK3,ZC3H14,FIS1,PPM1K,TUBA1C,ANTXR2,STIP
N,CD44,TM9SF1,MAGED2,SON,TET2,UQCRQ,HMGA2,CPSF3L,OBSL1,PSMA1,YLPM1,CHCHD1,RGS3,PSM
11,THOC3,IFITM2,TCP11L1,NDUFA3,RHEB,MAGOHB,ZMAT1,UCHL5,AKT1S1,GOSR2,TBCA,SLC37A3,EMC
PS1,SRSF1,PARD6G,SH3YL1,IP6K2,CLASRP,JPH2,C1ORF198,UGGT2,MICAL1,HS2ST1,UBE2I,SLC39A13,PRM

5,PTK2,GUK1,CENPN,PCYT2,USE1,AGPAT4,PPP1CC,EDF1,RFX3,PPA1,DGKE,TNFRSF14,ENDOV,CAMTA1,C
5,PTK2,GUK1,CENPN,PCYT2,USE1,AGPAT4,PPP1CC,EDF1,RFX3,PPA1,DGKE,TNFRSF14,ENDOV,CAMTA1,C

ASC4,WNK1,ZNF7,C19ORF48,ZFAND2B,AP1S3,HYPK,UPP1,MOB1A,NFAM1,INO80E,DUT,SYNGR2,CEP70,
ASC4,WNK1,ZNF7,C19ORF48,ZFAND2B,AP1S3,HYPK,UPP1,MOB1A,NFAM1,INO80E,DUT,SYNGR2,CEP70,

14,CAPG,CRIP1,PPIL2,TROAP,VIM,LIG1,PSMA7,RPL21,TXNL4A,CAPN10,NUTF2,PODXL,EPB41L4B,ATPAF2
,METTL5,FAM126A,RC3H2,OGFOD2,CIZ1,EIF3G,FBXL18,TCF25,ADD1,NIPAL3,CHTOP,ANAPC13,SLC39A6,
,METTL5,FAM126A,RC3H2,OGFOD2,CIZ1,EIF3G,FBXL18,TCF25,ADD1,NIPAL3,CHTOP,ANAPC13,SLC39A6,

A1,PPIL2,RPL28,CLDND1,VIM,LIG1,PSMA7,TIMM17B,APEH,TXNL4A,CAPN10,LYPLA2,EPB41L4B,ATPAF2,
E2I,SLC39A13,YIF1A,PRMT2,TBX6,TMEM205,PTK2,C14ORF2,CENPN,PCYT2,USE1,AGPAT4,DEF8,PPP1CC,
E2I,SLC39A13,YIF1A,PRMT2,TBX6,TMEM205,PTK2,C14ORF2,CENPN,PCYT2,USE1,AGPAT4,DEF8,PPP1CC,
,AUNIP,PCYT2,AGPAT4,PPP1CC,EDF1,RFX3,PPA1,DGKE,TNFRSF14,CAMTA1,CAPG,CRIP1,LIG1,PSMA7,TI
OSC9,C19ORF54,ZNF7,ARL6IP4,STRADA,AP1S3,HNRNPC,CYB561,ZNF644,STAU1,SLC9B2,SUGT1,UPP1,ZN
OSC9,C19ORF54,ZNF7,ARL6IP4,STRADA,AP1S3,HNRNPC,CYB561,ZNF644,STAU1,SLC9B2,SUGT1,UPP1,ZN

1,BCAM,RABEP2,ZNF20,RALGPS1,ZNF687,NXN,EPS15L1,HSD17B10,ABHD17A,USP45,ILF3,PKP4,STXBP2,

1,BCAM,RABEP2,ZNF20,RALGPS1,ZNF687,NXN,EPS15L1,HSD17B10,ABHD17A,USP45,ILF3,PKP4,STXBP2,

DOV,PPIL2,PSMD4,VIM,LIG1,PSMA7,CDC14B,APEH,TXNL4A,NUTF2,PPOX,TMPO,ATPAF2,LSM12,PRMT1,C

DOV,PPIL2,PSMD4,VIM,LIG1,PSMA7,CDC14B,APEH,TXNL4A,NUTF2,PPOX,TMPO,ATPAF2,LSM12,PRMT1,C

ARCA4,TYMS,GTF2IRD2,OARD1,TPM4,MRPS18C,RIC8A,TCP1,GNB5,CCDC40,UBE2N,NFATC4,PDCD6,MFF

3GLB1,COX5A,ZFAND4,HNRNPA1,HIGD1A,B3GAT3,RFXANK,SPINT2,RBFOX2,TMEM39B,DALRD3,DRG2,TF

21B,UBR4,YLPM1,PHF21A,SLC39A11,FAM3A,MUC1,METTL17,EPB41L5,ATG12,RPS25,NLRP1,ZBTB44,CO

21B,UBR4,YLPM1,PHF21A,SLC39A11,FAM3A,MUC1,METTL17,EPB41L5,ATG12,RPS25,NLRP1,ZBTB44,CO

MUC1,METTL17,EPB41L5,ATG12,RPS25,ZBTB44,TTC9C,PLCB2,DUSP22,TMX2,SPOCK2,DBNDD2,SDF4,NIS

MUC1,METTL17,EPB41L5,ATG12,RPS25,ZBTB44,TTC9C,PLCB2,DUSP22,TMX2,SPOCK2,DBNDD2,SDF4,NIS

,PLCB2,DUSP22,SLC25A10,TMX2,SDF4,NISCH,RPL29,YIPF3,FOXRED1,RABL5,PARD6G,SH3YL1,CLASRP,JPH

,PLCB2,DUSP22,SLC25A10,TMX2,SDF4,NISCH,RPL29,YIPF3,FOXRED1,RABL5,PARD6G,SH3YL1,CLASRP,JPH

DGKE,CAMTA1,CAPG,VIM,LIG1,TIMM17B,APEH,HYI,CAPN10,NUTF2,PODXL,CAMLG,EPB41L4B,TCF4,PPO

DGKE,CAMTA1,CAPG,VIM,LIG1,TIMM17B,APEH,HYI,CAPN10,NUTF2,PODXL,CAMLG,EPB41L4B,TCF4,PPO

L32,C12ORF76,RANGRF,RAD1,BLVRB,SLC33A1,ZNF706,PDCD5,GTF2IRD1,NUDT1,ANAPC11,POLR3H,RAD

L32,C12ORF76,RANGRF,RAD1,BLVRB,SLC33A1,ZNF706,PDCD5,GTF2IRD1,NUDT1,ANAPC11,POLR3H,RAD

2IRD2,DALRD3,TRPM4,TCP1,PHF6,PARK7,PHC3,CCDC40,XYLT2,NEK2,TGOLN2,POLR1C,PRRC2B,USP4,C1

2IRD2,DALRD3,TRPM4,TCP1,PHF6,PARK7,PHC3,CCDC40,XYLT2,NEK2,TGOLN2,POLR1C,PRRC2B,USP4,C1

IF2A,DCAKD,IRF3,TRAPPC2,RNF130,CLTA,SIN3A,NAA38,NDUFA11,TP53,BTF3,SYTL4,RDX,MAGOHB,INO8

3GCP2,CORO1B,PHF20,SHPK,C12ORF66,UNC119,ESRRA,GINS4,SEC24C,PPP1R7,C16ORF46,ANKHD1,KIAA

RPL29,CD99L2,GGPS1,SRSF1,YIPF3,TRNT1,PARD6G,SH3YL1,IP6K2,CLASRP,JPH2,C1ORF198,MICAL1,HS2ST

RPL29,CD99L2,GGPS1,SRSF1,YIPF3,TRNT1,PARD6G,SH3YL1,IP6K2,CLASRP,JPH2,C1ORF198,MICAL1,HS2ST

UBE2I,YIF1A,TBX6,PTK2,AUNIP,C14ORF2,GUK1,CENPN,PCYT2,USE1,AGPAT4,MAF,PPP1CC,EDF1,RFX3,PP

K2,AUNIP,C14ORF2,CENPN,PCYT2,USE1,DEF8,PPP1CC,EDF1,RFX3,PPA1,DGKE,ENDOV,CAMTA1,CLDND1

K2,AUNIP,C14ORF2,CENPN,PCYT2,USE1,DEF8,PPP1CC,EDF1,RFX3,PPA1,DGKE,ENDOV,CAMTA1,CLDND1

3GAT3,SPINT2,PAFAH1B2,GTF2IRD2,TMEM39B,BRF1,ZFPL1,FAM102A,MFSD10,PFKFB2,RIC8A

RK6,HMGA2,CDC37,CPSF3L,TSG101,OBSL1,TTC21B,PTPN13,PSMA1,YLPM1,PHF21A,SLC39A11,FUZ,RGS

RK6,HMGA2,CDC37,CPSF3L,TSG101,OBSL1,TTC21B,PTPN13,PSMA1,YLPM1,PHF21A,SLC39A11,FUZ,RGS

21,CCDC24,MDK,PNRC1,PCMTD1,CAMKK1,CEP63,ABLIM2,LRRC23,SUFU,CCNG2,HNRNPM,ME3,TCFL5,

21,CCDC24,MDK,PNRC1,PCMTD1,CAMKK1,CEP63,ABLIM2,LRRC23,SUFU,CCNG2,HNRNPM,ME3,TCFL5,

PAT4,IFI27L1,MAF,EEF1B2,PPA1,ENDOV,PPIL2,TIMM17B,APEH,RPL21,TXNL4A,CAMLG,LSM12,PRMT1,EX

PAT4,IFI27L1,MAF,EEF1B2,PPA1,ENDOV,PPIL2,TIMM17B,APEH,RPL21,TXNL4A,CAMLG,LSM12,PRMT1,EX

D1,RAD1,SLC33A1,ZNF706,PDCD5,GTF2IRD1,NUDT1,ANAPC11,POLR3H,RAD23A,GGT5,NT5C,SLC16A1,ILF

1,PKP4,HSPH1,NAA10,OAS3,RPL18,ATP10D,CHEK1,COX5A,SUMO3,PHF20L1,ZFAND4,HNRNPA1,CD151,

1,PKP4,HSPH1,NAA10,OAS3,RPL18,ATP10D,CHEK1,COX5A,SUMO3,PHF20L1,ZFAND4,HNRNPA1,CD151,

PP1CC,EDF1,RFX3,PPA1,DGKE,TNFRSF14,CAMTA1,CRIP1,VIM,LIG1,PSMA7,HYI,TXNL4A,CAPN10,NUTF2,

MBR1,NAP1L1,SERPINB1,SYNGR1,CHCHD7,HMGA1,PABPC1,BCAM,PPP3CB,RABEP2,TADA3,ZNF20,SLC33

MBR1,NAP1L1,SERPINB1,SYNGR1,CHCHD7,HMGA1,PABPC1,BCAM,PPP3CB,RABEP2,TADA3,ZNF20,SLC33

KE,CAMTA1,CLDND1,VIM,LIG1,PSMA7,TMEM126B,APEH,HYI,LYPLA2,NUTF2,PODXL,EPB41L4B,TCF4,UC

KE,CAMTA1,CLDND1,VIM,LIG1,PSMA7,TMEM126B,APEH,HYI,LYPLA2,NUTF2,PODXL,EPB41L4B,TCF4,UC

OB1A,CDK11A,MDH1,COPZ1,CEP70,FAM219B,ZNF524,ATP5J,GPBP1,TSFM,ZEB2,TRMT112,PNPO,GABPB

C9,STOML2,WNK1,C19ORF54,PIGL,C19ORF48,STRADA,ZFAND2B,HNRNPC,ZNF644,HYPK,MOB1A,CDK11

C9,STOML2,WNK1,C19ORF54,PIGL,C19ORF48,STRADA,ZFAND2B,HNRNPC,ZNF644,HYPK,MOB1A,CDK11

PEH,RPL21,TXNL4A,TMPO,PRMT1,EXOSC9,TXNDC17,ARL6IP4,C19ORF48,ZFAND2B,HYPK,ZNF250,NFAM1
PEH,RPL21,TXNL4A,TMPO,PRMT1,EXOSC9,TXNDC17,ARL6IP4,C19ORF48,ZFAND2B,HYPK,ZNF250,NFAM1
TRL,RER1,NKIRAS2,SCN11A,ZNF675,TAF1D,SLC35A2,RBM4,HNRNPUL1,BTF3,NDUFA3,AAMDC,MRPS24,R
TRL,RER1,NKIRAS2,SCN11A,ZNF675,TAF1D,SLC35A2,RBM4,HNRNPUL1,BTF3,NDUFA3,AAMDC,MRPS24,R

4,NFYC,ZXDC,L3MBTL2,MPHOSPH10,EIF4A1,RC3H2,KLHL7,SSR3,PICK1,UBE2D4,ECE1,ZNF512,OCIAD1,AR
CHD5,RRP8,RPL18,CAMK2D,ZFAND4,SIRT5,ASH2L,SMARCA4,PAFAH1B2,MPC2,GTF2IRD2,BRF1,FKBP5,MI
CHD5,RRP8,RPL18,CAMK2D,ZFAND4,SIRT5,ASH2L,SMARCA4,PAFAH1B2,MPC2,GTF2IRD2,BRF1,FKBP5,MI
5,ASH2L,PFDN5,TCTN1,PAFAH1B2,COA3,ZNF343,ARHGEF1,OARD1,TPM4,TRPM4,SMARCC2,PHC3,COMM
,CHCHD1,SLC39A11,FUZ,RGS3,FAM3A,PSMA3,MUC1,METTL17,ECSIT,EPB41L5,ATG12,PPRC1,ZBTB44,SLC
,CHCHD1,SLC39A11,FUZ,RGS3,FAM3A,PSMA3,MUC1,METTL17,ECSIT,EPB41L5,ATG12,PPRC1,ZBTB44,SLC
2,UFD1L,USP38,TAZ,THYN1,DBNDD1,SH2B1,NOP16,RPL13A,ARHGEF39,GSTK1,TTLL5,LDHA,PSMA6,ACOT
2,UFD1L,USP38,TAZ,THYN1,DBNDD1,SH2B1,NOP16,RPL13A,ARHGEF39,GSTK1,TTLL5,LDHA,PSMA6,ACOT
5,SSB,CREB3L4,OCIAD1,ASH1L,ARL6IP5,SLMAP,KCNC4,CHTOP,TSKU,TPT1,CDK4,TSPAN31,SRSF7,GLYCTK,
MO3,PHF20L1,EGLN2,RPL15,SIRT5,RFXANK,TMEM101,ZNF343,PPIH,FKBP5,TRPM4,MFSD10,MRPS18C,SI
MO3,PHF20L1,EGLN2,RPL15,SIRT5,RFXANK,TMEM101,ZNF343,PPIH,FKBP5,TRPM4,MFSD10,MRPS18C,SI
HFR,SSH3,OPA3,C12ORF10,RPS14,RNF4,C12ORF66,ZRANB2,UBA6,DPP9,SEC24C,RPL31,TVP23B,CDK5RAP
HFR,SSH3,OPA3,C12ORF10,RPS14,RNF4,C12ORF66,ZRANB2,UBA6,DPP9,SEC24C,RPL31,TVP23B,CDK5RAP
NK,REEP5,FTH1,PFDN5,TCTN1,TMEM101,C11ORF49,PTK7,TRPM4,SCAP,MROH1,TTC12,SNX5,MT2A,TMEM
2HGDH,KDELR3,CC2D1B,SNX29,AXIN1,COX19,AAMDC,SNAPC2,TKT,DENND1A,MRPS24,ZMAT1,GRAMD1
MAGT1,JUP,TBX6,XBP1,C14ORF2,GUK1,CENPN,PCYT2,IFI27L1,DEF8,EEF1B2,PPP1CC,EDF1,RFX3,DGKE,END
MAGT1,JUP,TBX6,XBP1,C14ORF2,GUK1,CENPN,PCYT2,IFI27L1,DEF8,EEF1B2,PPP1CC,EDF1,RFX3,DGKE,END
2,ANAPC13,NT5E,RALY,CLASP2,CNTRL,NUP62,PAM16,IRF3,GTF2H1,BFAR,MYL6,TLCD1,HNRNPUL1,BTF3,
2,ANAPC13,NT5E,RALY,CLASP2,CNTRL,NUP62,PAM16,IRF3,GTF2H1,BFAR,MYL6,TLCD1,HNRNPUL1,BTF3,
ENDOV,CAMTA1,CLDND1,PSMA7,TIMM17B,RPL21,PODXL,CAMLG,EPB41L4B,TMPO,PRMT1,EXOSC9,STO
ENDOV,CAMTA1,CLDND1,PSMA7,TIMM17B,RPL21,PODXL,CAMLG,EPB41L4B,TMPO,PRMT1,EXOSC9,STO

T2,USE1,AGPAT4,MAF,CCDC25,DEF8,PPP1CC,RFX3,PPA1,DGKE,CAMTA1,CAPG,VIM,LIG1,TIMM17B,APEH
T2,USE1,AGPAT4,MAF,CCDC25,DEF8,PPP1CC,RFX3,PPA1,DGKE,CAMTA1,CAPG,VIM,LIG1,TIMM17B,APEH
OSC9,TXNDC17,ARL6IP4,ZFAND2B,HYPK,ZNF250,NFAM1,RBM3,MAPKAPK5,DGUOK,HSP90AB1,CEP70,RA
OSC9,TXNDC17,ARL6IP4,ZFAND2B,HYPK,ZNF250,NFAM1,RBM3,MAPKAPK5,DGUOK,HSP90AB1,CEP70,RA

PBP1,SIAH1,ASPH,TRMT112,NIT2,ZSWIM7,ERLEC1,VWA2,LRRC27,SEMA3C,LMBR1,SERPINB1,LSR,MTA3,
PBP1,SIAH1,ASPH,TRMT112,NIT2,ZSWIM7,ERLEC1,VWA2,LRRC27,SEMA3C,LMBR1,SERPINB1,LSR,MTA3,
PMS,NNMT,SPAG7,IL1RAP,STYXL1,RFX5,POLD4,PLEKHA5,RANBP1,UBE2L6,VRK3,LZTFL1,ZC3H14,FIS1,SLC
,YLPM1,PHF21A,CHCHD1,SLC39A11,FAM3A,ECSIT,EPB41L5,ATG12,PPRC1,ZBTB44,TTC9C,COX4I1,PLCB2,
,YLPM1,PHF21A,CHCHD1,SLC39A11,FAM3A,ECSIT,EPB41L5,ATG12,PPRC1,ZBTB44,TTC9C,COX4I1,PLCB2,
CREBZF,EIF2A,RPL24,NADSYN1,ZNF511,GTF2H1,TAF1D,RNF130,MYL6,SIN3A,NAA38,TLCD1,GEMIN2,TUB
M40,ZNF20,ATP6V1F,SLC33A1,NXN,KLK10,ANAPC11,DDX41,USP45,PDE4DIP,STXBP2,HSPH1,CHCHD5,SGK
M40,ZNF20,ATP6V1F,SLC33A1,NXN,KLK10,ANAPC11,DDX41,USP45,PDE4DIP,STXBP2,HSPH1,CHCHD5,SGK
3,SEC31A,CARHSP1,CNST,LLGL2,COX17,COX5A,HINT1,HIGD1A,NDUFV2,B3GAT3,PFDN5,PAFAH1B2,COA3
3,SEC31A,CARHSP1,CNST,LLGL2,COX17,COX5A,HINT1,HIGD1A,NDUFV2,B3GAT3,PFDN5,PAFAH1B2,COA3
SH2B1,NOP16,RPL13A,ARHGEF39,GSTK1,TTLL5,LDHA,PSMA6,ACOT8,AP3B2,RPS9,RCE1,MRPL33,ZNF410,
SH2B1,NOP16,RPL13A,ARHGEF39,GSTK1,TTLL5,LDHA,PSMA6,ACOT8,AP3B2,RPS9,RCE1,MRPL33,ZNF410,

2,EMC9,CHURC1,NDUFA10,RPS7,MAZ,EMC3,PRKCSH,ACVR1B,PCMTD1,PSMD8,NSRP1,ZNF606,SUMF1,
3,MFSD10,GMPR2,PFKFB2,RIC8A,GNB5,TRPV3,NHP2,FAM32A,COX20,TNKS,MDP1,PRRC2B,PIGH,ELOF1,
3,MFSD10,GMPR2,PFKFB2,RIC8A,GNB5,TRPV3,NHP2,FAM32A,COX20,TNKS,MDP1,PRRC2B,PIGH,ELOF1,
4,ZNF7,ARL6IP4,PIGL,ZNF644,UPP1,ZNF250,NFAM1,MDH1,COPZ1,INO80E,CCNL2,MAPKAPK5,LRRFIP1,Z
5,GMPR2,THADA,RPS11,SNX5,NUP133,TMEM147,ELP4,NFYC,ZXDC,ELOF1,L3MBTL2,MPHOSPH10,EIF4A
5,GMPR2,THADA,RPS11,SNX5,NUP133,TMEM147,ELP4,NFYC,ZXDC,ELOF1,L3MBTL2,MPHOSPH10,EIF4A
R2,THADA,RPS11,SNX5,NUP133,TMEM147,ELP4,NFYC,ZXDC,ELOF1,L3MBTL2,MPHOSPH10,EIF4A1,RC3

,ERGIC1,KPNA5,MPHOSPH10,MYL9,TSEN34,UBE2D4,ECE1,TCF25,TJP1,SLC39A6,RALY,FAM69B,RER1,NU
12ORF10,RPS14,RNF4,C12ORF66,ZRANB2,UBA6,DPP9,SEC24C,RPL31,TVP23B,CDK5RAP3,WWP2,SRSF5
12ORF10,RPS14,RNF4,C12ORF66,ZRANB2,UBA6,DPP9,SEC24C,RPL31,TVP23B,CDK5RAP3,WWP2,SRSF5
NPC,ZNF644,CBFA2T2,HYPK,MOB1A,CDK11A,RBM3,INO80E,CCNL2,DGUOK,HSP90AB1,FAM219B,RAB2A,
NPC,ZNF644,CBFA2T2,HYPK,MOB1A,CDK11A,RBM3,INO80E,CCNL2,DGUOK,HSP90AB1,FAM219B,RAB2A,
,PPAN,MDH1,LAMP2,CCNL2,HSP90AB1,SYNGR2,RAB2A,ZNF524,ATP5J,ELP6,GPBP1,SIAH1,GGCX,ASPH,R
,PPAN,MDH1,LAMP2,CCNL2,HSP90AB1,SYNGR2,RAB2A,ZNF524,ATP5J,ELP6,GPBP1,SIAH1,GGCX,ASPH,R
AF2,LSM12,PRMT1,CASC4,STOML2,WNK1,PPAN,HNRNPC,ZNF644,STAU1,UPP1,ZNF250,MDH1,ZNF197,
AF2,LSM12,PRMT1,CASC4,STOML2,WNK1,PPAN,HNRNPC,ZNF644,STAU1,UPP1,ZNF250,MDH1,ZNF197,
K1,ZNF276,RQCD1,ANXA2,TCFL5,PPP1R13B,HDAC10,METTL23,TMEM53,PHPT1,PUF60,MT1F,ATP9B,AN
4I1,DUSP22,SDF4,TMED4,CENPK,RPS6,RPL29,CD99L2,GGPS1,AUP1,SAP30BP,FOXRED1,RABL5,KXD1,ITG
4I1,DUSP22,SDF4,TMED4,CENPK,RPS6,RPL29,CD99L2,GGPS1,AUP1,SAP30BP,FOXRED1,RABL5,KXD1,ITG

PLCB2,DUSP22,SLC25A10,TMX2,THEM4,DBNDD2,SDF4,NISCH,RPL29,GGPS1,AUP1,YIPF3,FOXRED1,RABL
PLCB2,DUSP22,SLC25A10,TMX2,THEM4,DBNDD2,SDF4,NISCH,RPL29,GGPS1,AUP1,YIPF3,FOXRED1,RABL

,LFNG,XYLT2,TMEM14B,ZXDC,COBL,TNKS,SMIM4,STX10,ERGIC1,MPHOSPH10,MYL9,STAU2,GFOD1,TSEN
LC39A13,YIF1A,JUP,TMEM205,XBP1,PTK2,AUNIP,C14ORF2,GUK1,CARD14,FBXW9,HSBP1L1,IFI27L1,MAF,
LC39A13,YIF1A,JUP,TMEM205,XBP1,PTK2,AUNIP,C14ORF2,GUK1,CARD14,FBXW9,HSBP1L1,IFI27L1,MAF,
USP22,TMX2,THEM4,DBNDD2,SDF4,NISCH,RPL29,CD99L2,GGPS1,COASY,SRSF1,PARD6G,SH3YL1,IP6K2,C
USP22,TMX2,THEM4,DBNDD2,SDF4,NISCH,RPL29,CD99L2,GGPS1,COASY,SRSF1,PARD6G,SH3YL1,IP6K2,C
PB41L4B,TCF4,UCK2,TMPO,STOML2,WNK1,ZNF7,ARL6IP4,PIGL,ZFAND2B,ACAD10,CYB561,ZNF644,STAU
31,TMEM9,NXF1,SLC39A6,ZNF593,GLYCTK,RBM42,CLASP2,CREBZF,GLI4,ZNF675,ZNF511,GTF2H1,MYL6,

,TMED4,RPL29,GGPS1,COASY,AUP1,FOXRED1,RABL5,PARD6G,SH3YL1,CLASRP,C1ORF198,MICAL1,UBE2
,TMED4,RPL29,GGPS1,COASY,AUP1,FOXRED1,RABL5,PARD6G,SH3YL1,CLASRP,C1ORF198,MICAL1,UBE2
LF1,C14ORF2,ATF7,PCYT2,USE1,AGPAT4,MAF,CCDC25,DEF8,EEF1B2,PPA1,TNFRSF14,VAMP8,CAMTA1,P
LF1,C14ORF2,ATF7,PCYT2,USE1,AGPAT4,MAF,CCDC25,DEF8,EEF1B2,PPA1,TNFRSF14,VAMP8,CAMTA1,P
CLASRP,JPH2,UGGT2,HS2ST1,UBE2I,SLC39A13,YIF1A,PRMT2,MAGT1,TMEM205,XBP1,PTK2,AUNIP,C14O
CLASRP,JPH2,UGGT2,HS2ST1,UBE2I,SLC39A13,YIF1A,PRMT2,MAGT1,TMEM205,XBP1,PTK2,AUNIP,C14O
,SEC31A,CNST,LLGL2,SSH2,COX5A,HINT1,HIGD1A,B3GAT3,PFDN5,PAFAH1B2,COA3,TMEM39B,PPIH,NSU
,SEC31A,CNST,LLGL2,SSH2,COX5A,HINT1,HIGD1A,B3GAT3,PFDN5,PAFAH1B2,COA3,TMEM39B,PPIH,NSU
B,HSPB1,HHLA3,LRRC27,MVD,DDOST,SYNGR1,MEIS2,RBP1,PPP3CB,WAC,ZNF706,ZNF687,GTF2IRD1,NX
B,HSPB1,HHLA3,LRRC27,MVD,DDOST,SYNGR1,MEIS2,RBP1,PPP3CB,WAC,ZNF706,ZNF687,GTF2IRD1,NX
,GMPPB,MVD,SAT1,EMC4,IL32,LMBR1,SYNGR1,RANGRF,RRNAD1,RBP1,H2AFV,ZNF687,POT1,ANAPC11,
,GMPPB,MVD,SAT1,EMC4,IL32,LMBR1,SYNGR1,RANGRF,RRNAD1,RBP1,H2AFV,ZNF687,POT1,ANAPC11,

MEM205,PTK2,AUNIP,ATF7,PCYT2,USE1,MAF,ASF1B,DEF8,EEF1B2,PPP1CC,RFX3,DGKE,ENDOV,CRIP1,PS
38,TLCD1,GEMIN2,TUBB3,MOSPD3,MAGOHB,CREB3L2,AIFM2,HMOX2,SPHK2,RBM6,HMGN1,NDUFB2,C

38,TLCD1,GEMIN2,TUBB3,MOSPD3,MAGOHB,CREB3L2,AIFM2,HMOX2,SPHK2,RBM6,HMGN1,NDUFB2,C
F1,FKBP5,MRPS18C,THADA,COMMD4,RPS11,SNX5,NUP133,TMEM147,ELP4,NFYC,ZXDC,L3MBTL2,MPHC
F1,FKBP5,MRPS18C,THADA,COMMD4,RPS11,SNX5,NUP133,TMEM147,ELP4,NFYC,ZXDC,L3MBTL2,MPHC
L1,IFI27L1,MAF,ASF1B,DEF8,EEF1B2,PPP1CC,RFX3,ENDOV,CRIP1,PPIL2,PSMD4,RPL28,PSMA7,TMEM12
CK2,C17ORF62,PRMT1,EXOSC9,TXNDC17,ARL6IP4,C19ORF48,STRADA,PPAN,CYB561,CBFA2T2,SUGT1,H
CK2,C17ORF62,PRMT1,EXOSC9,TXNDC17,ARL6IP4,C19ORF48,STRADA,PPAN,CYB561,CBFA2T2,SUGT1,H
,ATF7,PCYT2,USE1,AGPAT4,MAF,CCDC25,DEF8,EEF1B2,PPA1,TNFRSF14,VAMP8,CAMTA1,PSMA7,TIMM1
EH,TXNL4A,CAPN10,EPB41L4B,ATPAF2,LSM12,PRMT1,C19ORF54,ARL6IP4,C19ORF48,CYB561,CBFA2T2,
EH,TXNL4A,CAPN10,EPB41L4B,ATPAF2,LSM12,PRMT1,C19ORF54,ARL6IP4,C19ORF48,CYB561,CBFA2T2,
EH,TXNL4A,CAPN10,EPB41L4B,ATPAF2,LSM12,PRMT1,C19ORF54,ARL6IP4,C19ORF48,CYB561,CBFA2T2,
,GRAMD1C,VCAN,LPCAT3,CD44,MAGED2,SNF8,GRK6,RPS16,TSG101,OBSL1,PTPN13,UBR4,PSMA1,CHC
,GRAMD1C,VCAN,LPCAT3,CD44,MAGED2,SNF8,GRK6,RPS16,TSG101,OBSL1,PTPN13,UBR4,PSMA1,CHC
A1,ZNF675,GTF2H1,TAF1D,D2HGDH,CC2D1B,ARPC5,SH3GL1,TKT,DNAJC2,RHEB,HPS1,RGS12,AKT1S1,GT
A1,ZNF675,GTF2H1,TAF1D,D2HGDH,CC2D1B,ARPC5,SH3GL1,TKT,DNAJC2,RHEB,HPS1,RGS12,AKT1S1,GT

4B,APEH,RBM33,LYPLA2,C17ORF62,PRMT1,EXOSC9,STOML2,TXNDC17,PPAN,ACAD10,HNRNPC,CYB561,
4B,APEH,RBM33,LYPLA2,C17ORF62,PRMT1,EXOSC9,STOML2,TXNDC17,PPAN,ACAD10,HNRNPC,CYB561,
I2,STXBP2,EEF1D,HSPH1,NAA10,RRP8,CARHSP1,MT1E,CNST,LLGL2,EGFR,SUMO3,SIRT5,B3GAT3,PFDN5,
NRPN,WDR62,STYXL1,PLEKHA5,RANBP1,UBE2L6,VRK3,LZTFL1,ZC3H14,FIS1,SLC15A3,PPM1K,TUBA1C,D
NRPN,WDR62,STYXL1,PLEKHA5,RANBP1,UBE2L6,VRK3,LZTFL1,ZC3H14,FIS1,SLC15A3,PPM1K,TUBA1C,D
T2,YIF1A,PRMT2,MAGT1,GUK1,USE1,IFI27L1,CCDC25,EEF1B2,PPP1CC,PDLIM5,RFX3,PPA1,VAMP8,CRIP1
T2,YIF1A,PRMT2,MAGT1,GUK1,USE1,IFI27L1,CCDC25,EEF1B2,PPP1CC,PDLIM5,RFX3,PPA1,VAMP8,CRIP1
DH11,TMED4,CENPK,RPS6,RPL29,CD99L2,GGPS1,RTEL1,SRSF1,AUP1,SAP30BP,FOXRED1,TRNT1,SH3YL1,
DH11,TMED4,CENPK,RPS6,RPL29,CD99L2,GGPS1,RTEL1,SRSF1,AUP1,SAP30BP,FOXRED1,TRNT1,SH3YL1,
CLASP2,CREBZF,EIF2A,RPL24,NADSYN1,ZNF511,GTF2H1,RNF130,MYL6,SIN3A,NAA38,TLCD1,GEMIN2,TU
L2,C17ORF62,PRMT1,EXOSC9,C19ORF54,TXNDC17,ARL6IP4,C19ORF48,CYB561,SHARPIN,SLC9B2,RBM3,
,MPDU1,MAPK13,ZSWIM7,RPS15,HHLA3,MVD,SAT1,ERCC5,EMC4,NAP1L1,PRKRIP1,SYNGR1,C12ORF76,
,MPDU1,MAPK13,ZSWIM7,RPS15,HHLA3,MVD,SAT1,ERCC5,EMC4,NAP1L1,PRKRIP1,SYNGR1,C12ORF76,

MAPKAPK5,SYNGR2,TMEM134,LRRFIP1,FAM219B,ZNF524,TPM2,GPBP1,TRMT112,PNPO,CHPF,GABPB1,
TF7,PCYT2,USE1,AGPAT4,IFI27L1,CCDC25,DEF8,EEF1B2,PPP1CC,EDF1,PPA1,TNFRSF14,VAMP8,CAMTA1,
OML2,C19ORF54,C19ORF48,ZFAND2B,HNRNPC,ZNF644,CBFA2T2,HYPK,CARS,ZNF250,MOB1A,RBM3,CO
OML2,C19ORF54,C19ORF48,ZFAND2B,HNRNPC,ZNF644,CBFA2T2,HYPK,CARS,ZNF250,MOB1A,RBM3,CO
1,RCAN1,C12ORF73,NSMF,ATP9B,IMMP1L,RAP1B,IFT81,COQ2,TUBGCP2,SWI5,SNRNP40,DCTN6,WDR6,

D1,GEMIN2,TUBB3,MOSPD3,MAGOHB,CREB3L2,AIFM2,HMOX2,SPHK2,RBM6,HMGN1,NDUFB2,CDK2AP
A11,FAM3A,ECSIT,EPB41L5,ATG12,ZBTB44,TTC9C,COX4I1,PLCB2,DUSP22,SLC25A10,DBNDD2,SDF4,TM
F73,NSMF,ATP9B,IMMP1L,RAP1B,IFT81,COQ2,TUBGCP2,SWI5,SNRNP40,DCTN6,WDR6,OPA3,SNRNP20

D1,TRNT1,ITGB1BP1,SH3YL1,IP6K2,CLASRP,JPH2,C1ORF198,UGGT2,MICAL1,KCNMA1,SLC39A13,EIF3M,
D1,TRNT1,ITGB1BP1,SH3YL1,IP6K2,CLASRP,JPH2,C1ORF198,UGGT2,MICAL1,KCNMA1,SLC39A13,EIF3M,
,SAT1,ERCC5,IL32,LMBR1,SYNGR1,RANGRF,CHCHD7,RRNAD1,RBP1,RAD1,H2AFV,SLC33A1,ZNF687,NUD
,SAT1,ERCC5,IL32,LMBR1,SYNGR1,RANGRF,CHCHD7,RRNAD1,RBP1,RAD1,H2AFV,SLC33A1,ZNF687,NUD
,C6ORF132,TMEM108,ZMYM6,SSR3,MPV17,CREB3L4,UBE2D4,ZNF512,OCIAD1,TCF25,SLMAP,RBBP6,AD
,C6ORF132,TMEM108,ZMYM6,SSR3,MPV17,CREB3L4,UBE2D4,ZNF512,OCIAD1,TCF25,SLMAP,RBBP6,AD
,C6ORF132,TMEM108,ZMYM6,SSR3,MPV17,CREB3L4,UBE2D4,ZNF512,OCIAD1,TCF25,SLMAP,RBBP6,AD

ST1,UBE2I,SLC39A13,PRMT2,CDC42,TBX6,TMEM205,CELF1,AUNIP,CENPN,USE1,AGPAT4,CCDC25,EEF1B

CTK,RBM42,CLASP2,CREBZF,EIF2A,RPL24,NADSYN1,ZNF511,GTF2H1,TAF1D,RNF130,MYL6,SIN3A,NAA38

CTK,RBM42,CLASP2,CREBZF,EIF2A,RPL24,NADSYN1,ZNF511,GTF2H1,TAF1D,RNF130,MYL6,SIN3A,NAA38

APC2,RPS3,DNAJC2,EPN1,CREB3L2,ZMAT1,VPS25,SNRPD1,CKLF,TUSC3,SMUG1,AIFM2,CD37,SNRPA1,ZB

APC2,RPS3,DNAJC2,EPN1,CREB3L2,ZMAT1,VPS25,SNRPD1,CKLF,TUSC3,SMUG1,AIFM2,CD37,SNRPA1,ZB

,GRAMD1C,VCAN,PCBP2,CD44,TM9SF1,MAGED2,SON,UQCRQ,SNF8,CDC37,RPS16,CPSF3L,TSG101,UBR4

,GRAMD1C,VCAN,PCBP2,CD44,TM9SF1,MAGED2,SON,UQCRQ,SNF8,CDC37,RPS16,CPSF3L,TSG101,UBR4

13,UBR4,YLPM1,PHF21A,SLC39A11,FUZ,FAM3A,MUC1,EPB41L5,RPS25,NLRP1,ZBTB44,PLCB2,SLC25A10,

13,UBR4,YLPM1,PHF21A,SLC39A11,FUZ,FAM3A,MUC1,EPB41L5,RPS25,NLRP1,ZBTB44,PLCB2,SLC25A10,

RFIP1,TPM2,SIAH1,ZEB2,ASPH,TRMT112,RPTOR,STRIP2,GABPB1,RPS15,ERLEC1,LRRC27,MCTS1,NAP1L1,

PL21,LYPLA2,CAMLG,ATPAF2,LSM12,PRMT1,EXOSC9,STOML2,C19ORF54,ARL6IP4,PIGL,C19ORF48,ZFAN

PL21,LYPLA2,CAMLG,ATPAF2,LSM12,PRMT1,EXOSC9,STOML2,C19ORF54,ARL6IP4,PIGL,C19ORF48,ZFAN

A,AGAP3,EEF1D,NAA10,LLGL2,CHEK1,COX17,FKBP3,EGLN2,KCTD7,B3GAT3,SPINT2,TYMS,PTK7,BRF1,FAN

G,ATPAF2,LSM12,PRMT1,EXOSC9,STOML2,C19ORF54,PIGL,C19ORF48,ZFAND2B,HNRNPC,CYB561,ZNF64

G,ATPAF2,LSM12,PRMT1,EXOSC9,STOML2,C19ORF54,PIGL,C19ORF48,ZFAND2B,HNRNPC,CYB561,ZNF64

ASP2,PCNP,CREBZF,RPL24,ZNF675,NADSYN1,ZNF511,GTF2H1,BFAR,ARPC5,MYL6,TLCD1,MOSPD3,UBE2D

ASP2,PCNP,CREBZF,RPL24,ZNF675,NADSYN1,ZNF511,GTF2H1,BFAR,ARPC5,MYL6,TLCD1,MOSPD3,UBE2D

2,PCMT1,EBPL,TESC,IGSF8,NUDT5,RBPMS,SPAG7,WDR62,IL1RAP,STYXL1,RFX5,POLD4,RANBP1,UBE2L6,V

,PSMA7,TIMM17B,TMEM126B,LYPLA2,PODXL,CAMLG,UCK2,PRMT1,C19ORF54,ZNF7,ARL6IP4,C19ORF4

,PSMA7,TIMM17B,TMEM126B,LYPLA2,PODXL,CAMLG,UCK2,PRMT1,C19ORF54,ZNF7,ARL6IP4,C19ORF4

23,CCNG2,DNAJC24,MRPS33,PKIG,SLC31A1,SFSWAP,MPND,ROGDI,RNMT,AP4M1,SLC25A35,MFSD1,TUB

CRQ,GRK6,HMGA2,CPSF3L,OBSL1,PTPN13,YLPM1,PHF21A,CHCHD1,FUZ,RGS3,PSMA3,MUC1,EPB41L5,A

CRQ,GRK6,HMGA2,CPSF3L,OBSL1,PTPN13,YLPM1,PHF21A,CHCHD1,FUZ,RGS3,PSMA3,MUC1,EPB41L5,A

LAC8,SNX29,HNRNPUL1,LAMB4,UBE2D3,EIF3C,RPS3,DNAJC2,IDH3G,VPS25,PCTP,EMC9,NDUFB3,HMGN1

G,RPS11,SNX5,TGOLN2,POLR1C,USP4,EIF4A1,TMEM108,MPV17,SSB,RBBP6,NIPAL3,CFLAR,CNTRL,CREBZ

G,RPS11,SNX5,TGOLN2,POLR1C,USP4,EIF4A1,TMEM108,MPV17,SSB,RBBP6,NIPAL3,CFLAR,CNTRL,CREBZ

PNPEP1,SDHA,ABLIM2,DNAH2,ALS2,NIPA1,ANXA2,CRYZL1,TCFL5,NSMF,LMNA,ANKRD10,PRKCZ,FCGRT,T

CD1,GEMIN2,TUBB3,MOSPD3,MAGOHB,CREB3L2,HMOX2,SPHK2,RBM6,HMGN1,NDUFB2,CDK2AP2,NDU

CD1,GEMIN2,TUBB3,MOSPD3,MAGOHB,CREB3L2,HMOX2,SPHK2,RBM6,HMGN1,NDUFB2,CDK2AP2,NDU

P1L,RAP1B,IFT81,COQ2,TUBGCP2,SWI5,SNRNP40,DCTN6,WDR6,OPA3,SNRNP200,ARHGEF7,AHI1,LIN37,

CK1,ECE1,OCIAD1,TCF25,RBBP6,ZNF248,CHTOP,TPT1,SMAGP,TSPAN31,TMEM9,SLC39A6,ADRA1A,ZNF5

,CAPG,PPIL2,TIMM17B,TMEM126B,CAPN10,NUTF2,EPB41L4B,ATPAF2,PRMT1,C19ORF54,TXNDC17,ARL

,CAPG,PPIL2,TIMM17B,TMEM126B,CAPN10,NUTF2,EPB41L4B,ATPAF2,PRMT1,C19ORF54,TXNDC17,ARL

,CAPG,PPIL2,TIMM17B,TMEM126B,CAPN10,NUTF2,EPB41L4B,ATPAF2,PRMT1,C19ORF54,TXNDC17,ARL

MKK1,ZNF276,XPNPEP1,CEP63,SDHA,ABLIM2,FBXO34,HNRNPM,NIPA1,ANXA2,SLC31A1,TCFL5,ISOC2,RC

MKK1,ZNF276,XPNPEP1,CEP63,SDHA,ABLIM2,FBXO34,HNRNPM,NIPA1,ANXA2,SLC31A1,TCFL5,ISOC2,RC

GOHB,IDH3G,CREB3L2,AKT1S1,SPHK2,ACTR10,HMGN1,NDUFB2,CDK2AP2,NDUFA10,RPS27L,AZI2,U2AF1

1,RAD23A,SLC16A1,PKP4,EEF1D,HSPH1,NAA10,OAS3,SEC31A,RPL18,ATP10D,SH3GLB1,CHEK1,COX5A,PH

1,RAD23A,SLC16A1,PKP4,EEF1D,HSPH1,NAA10,OAS3,SEC31A,RPL18,ATP10D,SH3GLB1,CHEK1,COX5A,PH

G21,U2AF1L4,ATL2,DDX10,EMC3,RNPS1,MLH1,DTD2,DHDDS,XPNPEP1,ZNF483,ZNF783,FBXO34,CYB5D2

H2,C1ORF198,MICAL1,UBE2I,SLC39A13,PRMT2,JUP,TBX6,PTK2,C14ORF2,GUK1,CENPN,PCYT2,HSBP1L1,

H2,C1ORF198,MICAL1,UBE2I,SLC39A13,PRMT2,JUP,TBX6,PTK2,C14ORF2,GUK1,CENPN,PCYT2,HSBP1L1,

BR1,NAP1L1,C12ORF76,RANGRF,PABPC1,RAD1,BLVRB,ZNF20,SLC33A1,ZNF706,PDCD5,GTF2IRD1,NUDT

BR1,NAP1L1,C12ORF76,RANGRF,PABPC1,RAD1,BLVRB,ZNF20,SLC33A1,ZNF706,PDCD5,GTF2IRD1,NUDT

4,DBNDD2,SDF4,NISCH,TMED4,RPS6,CD99L2,RTEL1,COASY,AUP1,FOXRED1,TRNT1,PARD6G,SH3YL1,IP6K

UPP1,MOB1A,RBM3,LAMP2,CCNL2,MAPKAPK5,DUT,LRRFIP1,ZNF524,GPBP1,GABPB1,MPDU1,UBXN1,HI

A2,ANAPC13,EEF2K,TCEAL3,LAMTOR5,VDAC1,ZNF354A,NADSYN1,ZNF511,ZNF283,PIGO,BFAR,CC2D1B,A

A2,ANAPC13,EEF2K,TCEAL3,LAMTOR5,VDAC1,ZNF354A,NADSYN1,ZNF511,ZNF283,PIGO,BFAR,CC2D1B,A

ADA,AP1S3,ACAD10,CYB561,STAU1,SUGT1,HYPK,CARS,ZNF250,RPL36,MOB1A,NFAM1,MDH1,CCNL2,HS

ADA,AP1S3,ACAD10,CYB561,STAU1,SUGT1,HYPK,CARS,ZNF250,RPL36,MOB1A,NFAM1,MDH1,CCNL2,HS

C30A5,GNB5,THADA,CCDC40,MFF,SNX5,NUP133,MT2A,MRPS17,ELP4,TSC22D3,NFYC,ZXDC,MDP1,NDUF

C30A5,GNB5,THADA,CCDC40,MFF,SNX5,NUP133,MT2A,MRPS17,ELP4,TSC22D3,NFYC,ZXDC,MDP1,NDUF

2,UQCRQ,CDC37,RPS16,CPSF3L,OBSL1,PTPN13,CHCHD1,FAM3A,MUC1,EPB41L5,ATG12,PPRC1,RPS25,NL

2,UQCRQ,CDC37,RPS16,CPSF3L,OBSL1,PTPN13,CHCHD1,FAM3A,MUC1,EPB41L5,ATG12,PPRC1,RPS25,NL

,DST,ANTXR2,STIP1,GRAMD1C,VCAN,PCBP2,LPCAT3,TM9SF1,MAGED2,SON,UQCRQ,SNF8,RPS16,TSG10

,DST,ANTXR2,STIP1,GRAMD1C,VCAN,PCBP2,LPCAT3,TM9SF1,MAGED2,SON,UQCRQ,SNF8,RPS16,TSG10

PA1,TNFRSF14,ENDOV,CAMTA1,PPIL2,TIMM17B,RPL21,TXNL4A,CAPN10,CAMLG,PRMT1,EXOSC9,C19OF

PA1,TNFRSF14,ENDOV,CAMTA1,PPIL2,TIMM17B,RPL21,TXNL4A,CAPN10,CAMLG,PRMT1,EXOSC9,C19OF

SMD4,VIM,LIG1,TMEM126B,RPL21,CAMLG,UCK2,TMPO,LSM12,C17ORF62,EXOSC9,C19ORF54,ZNF7,TXN

SMD4,VIM,LIG1,TMEM126B,RPL21,CAMLG,UCK2,TMPO,LSM12,C17ORF62,EXOSC9,C19ORF54,ZNF7,TXN

OD2,ARIH2,PICK1,RNF181,NT5E,RALY,ZNF593,FAM131A,CFLAR,C11ORF57,CNTRL,RPL34,CREBZF,EIF2A,

OD2,ARIH2,PICK1,RNF181,NT5E,RALY,ZNF593,FAM131A,CFLAR,C11ORF57,CNTRL,RPL34,CREBZF,EIF2A,

LZTFL1,ZC3H14,FIS1,SLC15A3,TUBA1C,DST,ANTXR2,STIP1,LRRC1,PHF19,VCAN,LPCAT3,TM9SF1,SON,TET

LZTFL1,ZC3H14,FIS1,SLC15A3,TUBA1C,DST,ANTXR2,STIP1,LRRC1,PHF19,VCAN,LPCAT3,TM9SF1,SON,TET

TRADA,ZFAND2B,SUGT1,CDK11A,RBM3,MDH1,ZNF197,COPZ1,INO80E,CCNL2,TMEM134,RMDN1,SIAH1,

PRMT1,EXOSC9,STOML2,C19ORF54,ARL6IP4,PIGL,C19ORF48,ZFAND2B,HNRNPC,ZNF644,CBFA2T2,HYPK

PRMT1,EXOSC9,STOML2,C19ORF54,ARL6IP4,PIGL,C19ORF48,ZFAND2B,HNRNPC,ZNF644,CBFA2T2,HYPK

ORF2,GUK1,CENPN,CARD14,PCYT2,FBXW9,HSBP1L1,AGPAT4,IFI27L1,MAF,PPA1,ENDOV,PPIL2,CLDND1,

ORF2,GUK1,CENPN,CARD14,PCYT2,FBXW9,HSBP1L1,AGPAT4,IFI27L1,MAF,PPA1,ENDOV,PPIL2,CLDND1,

IRPN,SPAG7,WDR62,IL1RAP,STYXL1,POLD4,RANBP1,VRK3,ZC3H14,FIS1,SLC15A3,IRF9,ANTXR2,STIP1,LRR

IRPN,SPAG7,WDR62,IL1RAP,STYXL1,POLD4,RANBP1,VRK3,ZC3H14,FIS1,SLC15A3,IRF9,ANTXR2,STIP1,LRR

SH3YL1,IP6K2,HS2ST1,YIF1A,PRMT2,EIF3M,MAGT1,CELF1,USE1,CCDC25,EEF1B2,PPP1CC,PDLIM5,RFX3,

SH3YL1,IP6K2,HS2ST1,YIF1A,PRMT2,EIF3M,MAGT1,CELF1,USE1,CCDC25,EEF1B2,PPP1CC,PDLIM5,RFX3,

COASY,SRSF1,AUP1,YIPF3,TRNT1,ITGB1BP1,SH3YL1,CLASRP,JPH2,MICAL1,HS2ST1,YIF1A,PRMT2,EIF3M,JU

COASY,SRSF1,AUP1,YIPF3,TRNT1,ITGB1BP1,SH3YL1,CLASRP,JPH2,MICAL1,HS2ST1,YIF1A,PRMT2,EIF3M,JU

AMP2,ZNF197,INO80E,CCNL2,DGUOK,HSP90AB1,DUT,LRRFIP1,FAM219B,RAB2A,ZNF524,ATP5J,TPM2,GI

AMP2,ZNF197,INO80E,CCNL2,DGUOK,HSP90AB1,DUT,LRRFIP1,FAM219B,RAB2A,ZNF524,ATP5J,TPM2,GI

CAMTA1,PPIL2,RPL28,LIG1,PSMA7,HYI,TXNL4A,CAPN10,PODXL,EPB41L4B,TCF4,UCK2,TMPO,ATPAF2,EXC

V9,HSBP1L1,AGPAT4,IFI27L1,MAF,CCDC25,PPA1,ENDOV,PPIL2,TIMM17B,APEH,RPL21,TXNL4A,TMPO,LSI

V9,HSBP1L1,AGPAT4,IFI27L1,MAF,CCDC25,PPA1,ENDOV,PPIL2,TIMM17B,APEH,RPL21,TXNL4A,TMPO,LSI

,HSP90AB1,SYNGR2,RAB2A,ATP5J,ELP6,TPM2,GPALPP1,SIAH1,ASPH,PNPO,RPS20,NIT2,LYNX1,MPDU1,R

,HSP90AB1,SYNGR2,RAB2A,ATP5J,ELP6,TPM2,GPALPP1,SIAH1,ASPH,PNPO,RPS20,NIT2,LYNX1,MPDU1,R

5,NFAM1,RBM3,ZNF197,COPZ1,HSP90AB1,SYNGR2,RAB2A,ATP5J,ELP6,TPM2,GPALPP1,SIAH1,ASPH,PNP

6,NFAM1,RBM3,ZNF197,COPZ1,HSP90AB1,SYNGR2,RAB2A,ATP5J,ELP6,TPM2,GPALPP1,SIAH1,ASPH,PNP
.1,CAPG,PPIL2,RPL28,LIG1,TIMM17B,TMEM126B,XAF1,RPL21,TXNL4A,CAPN10,LYPLA2,PODXL,C17ORF62
.1,CAPG,PPIL2,RPL28,LIG1,TIMM17B,TMEM126B,XAF1,RPL21,TXNL4A,CAPN10,LYPLA2,PODXL,C17ORF62
ZNF593,GLYCTK,RBM42,CLASP2,CREBZF,ZNF511,GTF2H1,MYL6,RPS3A,TCP11L1,MAGOHB,IDH3G,CREB3L
M6,SSR3,CREB3L4,UBE2D4,ZNF512,TCF25,ADD1,NIPAL3,UGT8,BAX,RPL35,CDK4,NME1,ANAPC13,SRSF7,
M6,SSR3,CREB3L4,UBE2D4,ZNF512,TCF25,ADD1,NIPAL3,UGT8,BAX,RPL35,CDK4,NME1,ANAPC13,SRSF7,
M6,SSR3,CREB3L4,UBE2D4,ZNF512,TCF25,ADD1,NIPAL3,UGT8,BAX,RPL35,CDK4,NME1,ANAPC13,SRSF7,
SC,IGSF8,NUDT5,RBPMS,SPAG7,WDR62,IL1RAP,STYXL1,RFX5,POLD4,PLEKHA5,UBE2L6,VRK3,LZTFL1,ZC3
SC,IGSF8,NUDT5,RBPMS,SPAG7,WDR62,IL1RAP,STYXL1,RFX5,POLD4,PLEKHA5,UBE2L6,VRK3,LZTFL1,ZC3

PPB,MVD,SAT1,EMC4,IL32,LMBR1,SYNGR1,RANGRF,RRNAD1,RBP1,H2AFV,ZNF687,POT1,ANAPC11,RAD2
PPB,MVD,SAT1,EMC4,IL32,LMBR1,SYNGR1,RANGRF,RRNAD1,RBP1,H2AFV,ZNF687,POT1,ANAPC11,RAD2
,JUP,CDC42,TBX6,TMEM205,PTK2,AUNIP,CENPN,ATF7,PCYT2,MAF,ASF1B,DEF8,EEF1B2,PPP1CC,RFX3,PF
RIP1,CLDND1,LIG1,HYI,TXNL4A,CAPN10,LYPLA2,NUTF2,PODXL,CAMLG,TCF4,ATPAF2,C17ORF62,ZNF7,STF
RIP1,CLDND1,LIG1,HYI,TXNL4A,CAPN10,LYPLA2,NUTF2,PODXL,CAMLG,TCF4,ATPAF2,C17ORF62,ZNF7,STF
L55,SSB,ARIH2,CREB3L4,PICK1,TCF25,RBBP6,ADD1,CDK4,EEF2K,GALT,ACAA1,ZNF675,NADSYN1,DCAKD,C
L55,SSB,ARIH2,CREB3L4,PICK1,TCF25,RBBP6,ADD1,CDK4,EEF2K,GALT,ACAA1,ZNF675,NADSYN1,DCAKD,C
.17,ECSIT,EPB41L5,ATG12,RPS25,NLRP1,TTC9C,SLC25A10,TMX2,RDH11,DBNDD2,NISCH,TMED4,RPS6,RP
.17,ECSIT,EPB41L5,ATG12,RPS25,NLRP1,TTC9C,SLC25A10,TMX2,RDH11,DBNDD2,NISCH,TMED4,RPS6,RP
N1,NDUFB2,CDK2AP2,NDUFA10,RPS27L,AZI2,U2AF1L4,KIF13B,ATL2,DDX10,EMC3,RNPS1,MLH1,DTD2,DH
A1,LGMN,FKBP3,PFDN5,GTF2IRD2,PPIH,ARHGEF1,NSUN5,DRG2,MFSD10,RIC8A,TCP1,SCAP,CD74,PARK7
A1,LGMN,FKBP3,PFDN5,GTF2IRD2,PPIH,ARHGEF1,NSUN5,DRG2,MFSD10,RIC8A,TCP1,SCAP,CD74,PARK7
DS,ARL2,CEP63,ACHE,SUMF1,NDUFA7,CCNG2,RABEPK,GRHPR,SLC39A14,HNRNPM,MCUR1,QTRT1,LAMT
DS,ARL2,CEP63,ACHE,SUMF1,NDUFA7,CCNG2,RABEPK,GRHPR,SLC39A14,HNRNPM,MCUR1,QTRT1,LAMT
TN1,SNRPF,TYMS,PTK7,FAM102A,CD74,TRPV3,COMMD4,NFATC4,MFF,TMEM147,PCID2,TGOLN2,TSC22D
TN1,SNRPF,TYMS,PTK7,FAM102A,CD74,TRPV3,COMMD4,NFATC4,MFF,TMEM147,PCID2,TGOLN2,TSC22D
L2,TROAP,RPL28,VIM,LIG1,PSMA7,CAPN10,NUTF2,EPB41L4B,PPOX,ATPAF2,CASC4,PIGL,CYB561,STAU1,
X3,PPA1,VAMP8,CRIP1,PSMD4,PSMA7,TIMM17B,TMEM126B,XAF1,RPL21,TXNL4A,LYPLA2,NUTF2,CAML
X3,PPA1,VAMP8,CRIP1,PSMD4,PSMA7,TIMM17B,TMEM126B,XAF1,RPL21,TXNL4A,LYPLA2,NUTF2,CAML

,DIDO1,LSM14A,RNF149,DBN1,HNRNPL,ENTPD6,NUDT9,MAGOH,ATP13A2,ESRRA,UBA6,SEC24C,HGSNA

A,BTF3,EIF3C,C5ORF24,TKT,RHEB,CREB3L2,DNAJA3,SNRPD1,NSFL1C,CARM1,SLC37A3,TMEM41A,NFE2L1

3H2,METTL3,EIF3G,SSB,RPL8,RBBP6,ADD1,CHTOP,TPT1,RPL35,UBA1,NME1,SRSF7,MTIF2,NXF1,EEF2K,RA

2,PABPC1,ZNF836,RAD1,H2AFV,RBMX,BRIP1,ZNF20,ZNF687,PDCD5,GTF2IRD1,NUDT1,POT1,TRIM16,POL

ZC3H14,FIS1,SLC15A3,PPM1K,TUBA1C,IRF9,DST,ANTXR2,STIP1,LRRC1,PHF19,GRAMD1C,VCAN,PCBP2,CI

PL36,CDK11A,RBM3,MDH1,ZNF197,MAPKAPK5,DGUOK,HSP90AB1,DUT,LRRFIP1,RAB2A,ZNF524,GPBP1,

5,ZNF250,RPL36,CDK11A,RBM3,MDH1,ZNF197,MAPKAPK5,DGUOK,HSP90AB1,DUT,LRRFIP1,RAB2A,ZNF5

,MED1,PPARD,SNRNP40,RPL27,RBM4B,SNRNP200,RPS14,CNOT7,HBS1L,HNRNPL,MAGOH,ZRANB2,LUC7

,VTI1B,RPS23,RPL30,NUP85,RAB26,PEX16,SUFU,OS9,AP2S1,PKIG,TBC1D5,SNX13,UBE2L3,PABPN1,RPS15

RPS4X,CAPN2,MLH1,SUMO1,COMMD1,RPS23,RPL30,PSMD8,RQCD1,SUFU,NUB1,OS9,EXOSC2,RBM7,PSM

P62,ACAA1,RPL24,PAM16,IFT172,TRAPPC2,KDELR3,CLTA,RBM4,GEMIN2,THOC3,TP53,AP1G2,RPS3A,SER

NUP62,ACAA1,RPL24,PAM16,NEDD4L,FCHO2,IFT172,KDELR3,CLTA,SIN3A,THOC3,NRP1,TP53,AP1G2,RPS

NUP62,ACAA1,RPL24,PAM16,NEDD4L,FCHO2,IFT172,KDELR3,CLTA,SIN3A,THOC3,NRP1,TP53,AP1G2,RPS

CHL5,NSFL1C,IFI27,ARRB2,SGSM3,RPS7,APAF1,NCBP2,SNX3,USP15,NFE2L1,TAF9,RNPS1,UBA52,RPS4X,

SNX5,ENY2,NUP133,RAPGEF3,HAUS4,RPAIN,NEK2,ACOT2,PCID2,SGCD,PEX2,TNKS,RPL37A,STX10,USP4,

4,ECE1,RBBP6,BAX,RPL35,UBA1,CDK4,RNF8,PPP2R5C,NT5E,FAM192A,GCDH,GLYCTK,GALT,PCNP,RPL34,

43,EIF6,MRPL48,SHMT1,ERAL1,ACOT9,MTHFD2L,MRPL27,CERS2,MRPS7,RPL27,NCK2,BIN1,MRPL36,RB

GCP2,SWI5,SSH3,NCK2,CORO1B,DCTN1,BIN1,UQCRB,SNRNP200,RPS14,DDB2,CLIP1,RNF4,NDUFS4,DBN

AX,DLG4,RNF8,TRPM6,CLASP2,RPL24,FCHO2,TRAPPC2,GTF2H1,CLTA,ARPC5,GEMIN2,NDUFA11,AXIN1,

RPL8,FBXL18,UBE2D4,ECE1,RBBP6,BAX,RPL35,UBA1,RNF8,PPP2R5C,NT5E,FAM192A,ADRA1A,GCDH,LA

NRNPL,MAGOH,ZRANB2,BYSL,LUC7L3,PRPF31,POLR2L,SRSF5,PAPOLA,ELP5,EXOSC10,AKAP8L,WDR83,H

133,USP40,PCID2,PEX2,RPL37A,USP4,RPLP0,HAX1,METTL3,SSB,ARIH2,RPL8,FBXL18,UBE2D4,ECE1,RBBP

,SSB,ARIH2,RPL8,PICK1,CHTOP,RPL35,SRSF7,NXF1,ZNF593,RPL34,NUP62,ACAA1,RPL24,PAM16,IRF3,IFT1

PLP0,HAX1,SSR3,MPV17,SSB,ARIH2,RPL8,PICK1,CHTOP,BAX,RPL35,DLG4,SRSF7,NXF1,ZNF593,RPL34,NU

3,MED6,SEC31A,PRLR,SH3GLB1,SSH2,COX17,CD151,NDUFV2,KCTD7,ASH2L,TCTN1,SMARCA4,COA3,SNR

4,EIF2A,RPL24,NADSYN1,DCAKD,PIGO,RBM4,ST8SIA1,RPS3A,EIF3C,MGST2,RPS3,MRPS24,PCTP,TUSC3,F

,SSB,ARIH2,RPL8,PICK1,CHTOP,RPL35,SRSF7,NXF1,ZNF593,RPL34,NUP62,ACAA1,RPL24,PAM16,IRF3,IFT1

3,MPV17,SSB,ARIH2,RPL8,PICK1,CHTOP,RPL35,SRSF7,NXF1,ZNF593,RPL34,NUP62,ACAA1,RPL24,PAM16

NUP133,RAPGEF3,RPAIN,ACOT2,PCID2,SGCD,PEX2,TNKS,RPL37A,STX10,USP4,PEX26,BTN3A2,KPNA5,RP

NF644,CBFA2T2,UPP1,CARS,ZNF250,RPL36,NFAM1,CDK11A,RBM3,ZNF197,INO80E,CCNL2,MAPKAPK5,D

,SNRPD1,DHRS4,MLST8,NDUFB3,NDUFB2,NDUFA10,RPS27L,GMFG,REPS2,GOSR2,PSMG2,ATL2,PSMG4,T

SMA7,CDC14B,APEH,RPL21,TXNL4A,TCF4,UCK2,PPOX,TMPO,EXOSC9,STOML2,ZNF7,ARL6IP4,HNRNPC,Z

,MTA3,DCN,HMGA1,DBNL,RAD1,H2AFV,TOMM40,BRIP1,RABEP2,TADA3,ATP6V1F,WAC,PDCD5,POT1,TR

NPC,ZNF644,CBFA2T2,UPP1,CARS,ZNF250,RPL36,NFAM1,CDK11A,RBM3,ZNF197,INO80E,CCNL2,MAPKA

C5,AP2S1,RHBDF1,ANXA2,UBE2L3,ERAP2,SHMT1,DNAJB12,UBE2A,CHFR,HYKK,OAZ1,QDPR,LTA4H,RNF4,

TCP1,SYTL1,CD74,PARK7,NFKBIA,NHP2,ZNHIT3,RPS11,PDCD6,MFF,SNX5,ENY2,NUP133,RAPGEF3,RPAIN,
)2B,SHARPIN,CARS,RPL36,CDK11A,RBM3,LAMP2,INO80E,CCNL2,MAPKAPK5,DGUOK,HSP90AB1,RAB2A,S
'F,GBA2,PPIH,BRF1,TRPM4,MCFD2,LGALS3,TCP1,CD74,PARK7,CCDC40,ZNHIT3,PDCD6,MFF,NOP2,RAPGE
IUP,CDC42,TBX6,PTK2,CELF1,AUNIP,CENPN,FBXW9,ASF1B,DEF8,PDLIM5,RFX3,VAMP8,CAPG,PSMD4,VIN
;FM,ZEB2,TRMT112,PNPO,RPTOR,ZNF625,RPS20,GABPB1,TWSG1,RPS15,DCTD,ZNF814,GMPPB,HSPB1,S
IPC,ZNF644,CBFA2T2,UPP1,CARS,ZNF250,RPL36,NFAM1,CDK11A,RBM3,ZNF197,INO80E,CCNL2,MAPKAI

CHCHD1,RGS3,FAM3A,PSMA3,MUC1,METTL17,ATG12,PPRC1,RPS25,NLRP1,COX4I1,PLCB2,DUSP22,SLC2

FIP1,ZNF524,ELP6,GPBP1,TSFM,ZEB2,TRMT112,RPTOR,ZNF625,RPS20,GABPB1,TWSG1,ZSWIM7,RPS15,Z
2,ZNF7,ARL6IP4,HNRNPC,ZNF644,CBFA2T2,UPP1,CARS,ZNF250,RPL36,NFAM1,CDK11A,RBM3,ZNF197,IN

PL37A,STX10,PEX26,BTN3A2,KPNA5,RPLP0,HAX1,SSR3,MPV17,SSB,ARIH2,RPL8,PICK1,ARL6IP5,CHTOP,RI
OR,CHPF,ZNF625,RPS20,GABPB1,MPDU1,TWSG1,RPS15,ZNF814,HSPB1,MVD,MCTS1,NAP1L1,DDOST,M

;FM,ZEB2,TRMT112,RPTOR,CHPF,ZNF625,RPS20,GABPB1,MPDU1,TWSG1,RPS15,ZNF814,HSPB1,MVD,M
2,PPP1CC,GADL1,PPA1,TNFRSF14,CAMTA1,PPIL2,PSMD4,RPL28,VIM,PSMA7,CDC14B,APEH,RPL21,CAPN

MPO,PRMT1,EXOSC9,STOML2,ZNF7,PIGL,HNRNPC,ZNF644,CBFA2T2,UPP1,CARS,ZNF250,RPL36,NFAM1,C
8,PDLIM5,RFX3,VAMP8,CAPG,PSMD4,VIM,CDC14B,TIMM17B,TMEM126B,APEH,TXNL4A,CAPN10,LYPLA

OML2,ZNF7,PIGL,HNRNPC,ZNF644,CBFA2T2,UPP1,CARS,ZNF250,RPL36,NFAM1,CDK11A,RBM3,ZNF197,C

),LYPLA2,CAMLG,PRMT1,STOML2,WNK1,PIGL,STRADA,ZFAND2B,SHARPIN,CARS,RPL36,CDK11A,RBM3,L

RIP1,MTA3,DCN,MEIS2,HMGA1,PSME2,PABPC1,RRNAD1,ZNF836,H2AFV,RBMX,PPP3CB,BRIP1,TADA3,ZN

MPO,PRMT1,EXOSC9,STOML2,ZNF7,PIGL,HNRNPC,ZNF644,CBFA2T2,UPP1,CARS,ZNF250,RPL36,NFAM1,C

PTPN13,UBR4,PSMA1,YLPM1,PHF21A,CHCHD1,RGS3,FAM3A,PSMA3,MUC1,METTL17,ECSIT,EPB41L5,AT
CK2,DYNC2LI1,MAP4,CORO1B,DCTN1,BIN1,SEMA5A,CLIP1,RNF4,RHOC,DBN1,MARK2,PAK1,ATP13A2,H2

D7,IRF3,GTF2H1,BFAR,KDELR3,SIN3A,RBM4,ST8SIA1,AXIN1,TP53,PSMD10,PML,SLC25A24,OXR1,UBE2D3

,MAGT1,JUP,CDC42,TBX6,XBP1,PTK2,CELF1,AUNIP,CARD14,ATF7,FBXW9,MAF,EEF1B2,PPP1CC,EDF1,RF

T,ANTXR2,STIP1,PHF19,GRAMD1C,VCAN,PCBP2,LPCAT3,CD44,TM9SF1,MAGED2,SON,TET2,UQCRQ,SNF8
B2,PPP1CC,PDLIM5,EDF1,GADL1,PPA1,DGKE,VAMP8,CAMTA1,PSMD4,RPL28,VIM,PSMA7,XAF1,APEH,RP
1,UBE2L6,VRK3,ZC3H14,FIS1,SLC15A3,PPM1K,TUBA1C,IRF9,DST,ANTXR2,STIP1,PHF19,GRAMD1C,VCAN,
D4,PLEKHA5,RANBP1,UBE2L6,VRK3,ZC3H14,FIS1,SLC15A3,PPM1K,TUBA1C,IRF9,DST,ANTXR2,STIP1,PHF1
C3H14,FIS1,SLC15A3,PPM1K,TUBA1C,IRF9,DST,ANTXR2,STIP1,LRRC1,GRAMD1C,VCAN,PCBP2,LPCAT3,CD
0,RUVBL2,PCMT1,EBPL,TESC,IGSF8,NUDT5,RBPMS,NNMT,SNRPN,SPAG7,WDR62,STYXL1,RFX5,POLD4,PL
RBPMS,SNRPN,SPAG7,WDR62,STYXL1,RFX5,POLD4,PLEKHA5,RANBP1,UBE2L6,VRK3,ZC3H14,FIS1,SLC15A
NPL,APOBEC3F,MAGOH,BYSL,LUC7L3,MRPL34,PRPF31,RPL31,EIF3E,GAPDH,FAU,AKAP8L,WDR83,HEATR
JUP,TBX6,XBP1,CELF1,CENPN,ATF7,FBXW9,MAF,ASF1B,PPP1CC,EDF1,RFX3,DGKE,ENDOV,CAMTA1,CAPG
JUP,TBX6,XBP1,CELF1,CENPN,ATF7,FBXW9,MAF,ASF1B,PPP1CC,EDF1,RFX3,DGKE,ENDOV,CAMTA1,CAPG
JUP,TBX6,XBP1,CELF1,CENPN,ATF7,FBXW9,MAF,ASF1B,PPP1CC,EDF1,RFX3,DGKE,ENDOV,CAMTA1,CAPG
,RTEL1,SRSF1,SAP30BP,TRNT1,ITGB1BP1,MED15,PARD6G,IP6K2,CLASRP,JPH2,UBE2I,PRMT2,JUP,TBX6,X

CNL2,MAPKAPK5,HSP90AB1,DUT,TSFM,ZEB2,TRMT112,PNPO,RPTOR,RPS20,GABPB1,RPS15,ERCC5,RANG

XL,TCF4,ATPAF2,PRMT1,EXOSC9,ARL6IP4,STRADA,PPAN,HNRNPC,HYPK,UPP1,RPL36,MOB1A,RBM3,INO8
,PRMT1,EXOSC9,STOML2,PPAN,ZFAND2B,AP1S3,HNRNPC,SHARPIN,STAU1,SUGT1,HYPK,RPL36,RBM3,CC
3,MCUR1,NDUFC1,ATP11B,QTRT1,DUSP18,SEC13,NDUFB10,MRPL51,MRPL43,NSMF,PARP6,LMNA,ROGI
3,MCUR1,NDUFC1,ATP11B,QTRT1,DUSP18,SEC13,NDUFB10,MRPL51,MRPL43,NSMF,PARP6,LMNA,ROGI

SUGT1,HYPK,RPL36,MOB1A,RBM3,INO80E,SYNGR2,CEP70,LRRFIP1,RMDN1,TPM2,ZEB2,TPM3,RPTOR,RP
SUGT1,HYPK,RPL36,MOB1A,RBM3,INO80E,SYNGR2,CEP70,LRRFIP1,RMDN1,TPM2,ZEB2,TPM3,RPTOR,RP
3,POLR2E,TAF9,ACVR1B,SUMO1,DHDDS,COMMD1,SDHA,PSMD8,NDUFA7,CCNG2,DNAH2,OS9,DYNC1I2,

DHRS4,IFI27,TRAK1,NDUFB3,NDUFB2,NDUFA10,USP15,THOP1,GPS2,UBA52,ACSM3,AK2,CAPN2,LRRC59,

TP10D,EGFR,SH3GLB1,CAMK2D,COX5A,HIGD1A,NDUFV2,SIRT5,B3GAT3,COA3,MPC2,TYMS,GBA2,SLC25/

ATP6V1F,PDCD5,KLK10,NUDT1,B2M,ABHD17A,NT5C,PSME1,SLC16A1,MDH2,STXBP2,HSPH1,SEC31A,CN

,NFE2L1,FIG4,UBA52,CSF1,VAPA,CAPN2,LRRC59,DHDDS,PRKCE,RPS23,GRN,PEX16,BSCL2,SUMF1

KIF13A,VAPA,CAPN2,VTI1B,COMMD1,SZT2,RAB26,GRN,PEX16,RABEPK,AP2S1,ATP11B,ANXA2,TBC1D5,S

MRPL33,ZNF410,RUVBL2,PCMT1,EBPL,TESC,IGSF8,NUDT5,RBPMS,SPAG7,WDR62,IL1RAP,STYXL1,RFX5,P
NNMT,SPAG7,WDR62,IL1RAP,STYXL1,PLEKHA5,RANBP1,UBE2L6,VRK3,LZTFL1,ZC3H14,FIS1,SLC15A3,PPN
29,RTEL1,SRSF1,AUP1,SAP30BP,YIPF3,FOXRED1,TRNT1,KXD1,ITGB1BP1,MED15,SH3YL1,IP6K2,CLASRP,C1
TLL5,LDHA,PSMA6,ACOT8,AP3B2,RPS9,SLC35G1,RCE1,MRPL33,ZNF410,RUVBL2,PCMT1,EBPL,TESC,IGSF8
TLL5,LDHA,PSMA6,ACOT8,AP3B2,RPS9,SLC35G1,RCE1,MRPL33,ZNF410,RUVBL2,PCMT1,EBPL,TESC,IGSF8
33,ZNF410,RUVBL2,PCMT1,EBPL,TESC,IGSF8,NUDT5,RBPMS,NNMT,SNRPN,SPAG7,WDR62,IL1RAP,STYXL
33,ZNF410,RUVBL2,PCMT1,EBPL,TESC,IGSF8,NUDT5,RBPMS,NNMT,SNRPN,SPAG7,WDR62,IL1RAP,STYXL
IUDT5,RBPMS,SNRPN,SPAG7,WDR62,IL1RAP,STYXL1,POLD4,PLEKHA5,RANBP1,UBE2L6,VRK3,LZTFL1,ZC3
,DST,ANTXR2,STIP1,LRRC1,VCAN,LPCAT3,CD44,TM9SF1,MAGED2,SON,UQCRQ,SNF8,GRK6,HMGA2,CDC3
,DST,ANTXR2,STIP1,LRRC1,VCAN,LPCAT3,CD44,TM9SF1,MAGED2,SON,UQCRQ,SNF8,GRK6,HMGA2,CDC3
MT1,EBPL,TESC,IGSF8,NUDT5,RBPMS,SNRPN,SPAG7,WDR62,IL1RAP,STYXL1,RFX5,POLD4,PLEKHA5,RANBI
PL29,RTEL1,COASY,AUP1,SAP30BP,YIPF3,FOXRED1,RABL5,TRNT1,KXD1,ITGB1BP1,MED15,SH3YL1,IP6K2,C
PL29,RTEL1,COASY,AUP1,SAP30BP,YIPF3,FOXRED1,RABL5,TRNT1,KXD1,ITGB1BP1,MED15,SH3YL1,IP6K2,C
FX3,DGKE,ENDOV,CAMTA1,PPIL2,PSMD4,RPL28,APEH,RPL21,CAPN10,LYPLA2,NUTF2,C17ORF62,PRMT1,
K6,HMGA2,CDC37,RPS16,CPSF3L,TSG101,UBR4,YLPM1,FUZ,FAM3A,METTL17,EPB41L5,ATG12,RPS25,NI
K6,HMGA2,CDC37,RPS16,CPSF3L,TSG101,UBR4,YLPM1,FUZ,FAM3A,METTL17,EPB41L5,ATG12,RPS25,NI
5,RBPMS,SPAG7,WDR62,IL1RAP,STYXL1,RFX5,POLD4,PLEKHA5,RANBP1,VRK3,LZTFL1,ZC3H14,FIS1,SLC15

5,RBPMS,SPAG7,WDR62,IL1RAP,STYXL1,RFX5,POLD4,PLEKHA5,RANBP1,VRK3,LZTFL1,ZC3H14,FIS1,SLC15
PHF21A,FUZ,FAM3A,METTL17,EPB41L5,ATG12,RPS25,NIP7,COX4I1,DUSP22,SLC25A10,THEM4,DBNDD2,
PHF21A,FUZ,FAM3A,METTL17,EPB41L5,ATG12,RPS25,NIP7,COX4I1,DUSP22,SLC25A10,THEM4,DBNDD2,
3P,YIPF3,FOXRED1,RABL5,TRNT1,KXD1,ITGB1BP1,MED15,SH3YL1,IP6K2,CLASRP,C1ORF198,UGGT2,MICA
3P,YIPF3,FOXRED1,RABL5,TRNT1,KXD1,ITGB1BP1,MED15,SH3YL1,IP6K2,CLASRP,C1ORF198,UGGT2,MICA
,CDC37,RPS16,CPSF3L,TSG101,UBR4,YLPM1,FUZ,FAM3A,METTL17,EPB41L5,ATG12,RPS25,NIP7,TTC9C,C
,CDC37,RPS16,CPSF3L,TSG101,UBR4,YLPM1,FUZ,FAM3A,METTL17,EPB41L5,ATG12,RPS25,NIP7,TTC9C,C
MA6,ACOT8,AP3B2,RPS9,SLC35G1,RCE1,MRPL33,ZNF410,RUVBL2,PCMT1,EBPL,TESC,IGSF8,NUDT5,RBPI
MA6,ACOT8,AP3B2,RPS9,SLC35G1,RCE1,MRPL33,ZNF410,RUVBL2,PCMT1,EBPL,TESC,IGSF8,NUDT5,RBPI
PS16,CPSF3L,TSG101,UBR4,YLPM1,FUZ,FAM3A,PSMA3,METTL17,EPB41L5,ATG12,RPS25,NIP7,TTC9C,CO
PS16,CPSF3L,TSG101,UBR4,YLPM1,FUZ,FAM3A,PSMA3,METTL17,EPB41L5,ATG12,RPS25,NIP7,TTC9C,CO
FL1,FIS1,PPM1K,TUBA1C,IRF9,DST,ANTXR2,STIP1,LRRC1,VCAN,LPCAT3,CD44,TM9SF1,MAGED2,SON,UQ
FL1,FIS1,PPM1K,TUBA1C,IRF9,DST,ANTXR2,STIP1,LRRC1,VCAN,LPCAT3,CD44,TM9SF1,MAGED2,SON,UQ
6,CARD14,PCYT2,FBXW9,USE1,HSBP1L1,MAF,ASF1B,CCDC25,EEF1B2,RFX3,ENDOV,CAMTA1,CAPG,PPIL2
6,CARD14,PCYT2,FBXW9,USE1,HSBP1L1,MAF,ASF1B,CCDC25,EEF1B2,RFX3,ENDOV,CAMTA1,CAPG,PPIL2
TUBA1C,IRF9,DST,ANTXR2,STIP1,LRRC1,VCAN,LPCAT3,CD44,TM9SF1,MAGED2,SON,UQCRQ,SNF8,GRK6,
TUBA1C,IRF9,DST,ANTXR2,STIP1,LRRC1,VCAN,LPCAT3,CD44,TM9SF1,MAGED2,SON,UQCRQ,SNF8,GRK6,
UBA1C,IRF9,DST,ANTXR2,STIP1,LRRC1,VCAN,LPCAT3,CD44,TM9SF1,MAGED2,SON,UQCRQ,SNF8,GRK6,C
UBA1C,IRF9,DST,ANTXR2,STIP1,LRRC1,VCAN,LPCAT3,CD44,TM9SF1,MAGED2,SON,UQCRQ,SNF8,GRK6,C
S,SPAG7,WDR62,IL1RAP,STYXL1,RFX5,POLD4,PLEKHA5,RANBP1,UBE2L6,VRK3,LZTFL1,ZC3H14,SLC15A3,F
P1,LRRC1,VCAN,LPCAT3,CD44,TM9SF1,MAGED2,SON,UQCRQ,SNF8,GRK6,HMGA2,CDC37,RPS16,CPSF3L,
P1,LRRC1,VCAN,LPCAT3,CD44,TM9SF1,MAGED2,SON,UQCRQ,SNF8,GRK6,HMGA2,CDC37,RPS16,CPSF3L,
M,CDC42,AUNIP,C14ORF2,CARD14,PCYT2,FBXW9,USE1,ASF1B,CCDC25,EEF1B2,RFX3,DGKE,TNFRSF14,EI
P,STYXL1,RFX5,POLD4,PLEKHA5,RANBP1,UBE2L6,VRK3,LZTFL1,ZC3H14,FIS1,SLC15A3,PPM1K,TUBA1C,AN
P,STYXL1,RFX5,POLD4,PLEKHA5,RANBP1,UBE2L6,VRK3,LZTFL1,ZC3H14,FIS1,SLC15A3,PPM1K,TUBA1C,AN
F54,ZNF7,TXNDC17,ARL6IP4,C19ORF48,PPAN,ACAD10,CYB561,SHARPIN,SUGT1,HYPK,CARS,MOB1A,CDI
MT1,EBPL,TESC,IGSF8,NUDT5,RBPMS,SPAG7,WDR62,IL1RAP,STYXL1,RFX5,PLEKHA5,RANBP1,UBE2L6,VRK
MA6,ACOT8,AP3B2,RPS9,SLC35G1,RCE1,MRPL33,ZNF410,RUVBL2,PCMT1,EBPL,TESC,IGSF8,NUDT5,RBPI
MA6,ACOT8,AP3B2,RPS9,SLC35G1,RCE1,MRPL33,ZNF410,RUVBL2,PCMT1,EBPL,TESC,IGSF8,NUDT5,RBPI
SP22,SLC25A10,THEM4,DBNDD2,RPS6,RPL29,RTEL1,COASY,SRSF1,AUP1,SAP30BP,YIPF3,FOXRED1,RABL
SP22,SLC25A10,THEM4,DBNDD2,RPS6,RPL29,RTEL1,COASY,SRSF1,AUP1,SAP30BP,YIPF3,FOXRED1,RABL
SF3L,TSG101,UBR4,YLPM1,FUZ,FAM3A,METTL17,EPB41L5,ATG12,RPS25,NIP7,COX4I1,DUSP22,SLC25A1
SF3L,TSG101,UBR4,YLPM1,FUZ,FAM3A,METTL17,EPB41L5,ATG12,RPS25,NIP7,COX4I1,DUSP22,SLC25A1
RPN,SPAG7,WDR62,STYXL1,RFX5,POLD4,PLEKHA5,RANBP1,VRK3,LZTFL1,FIS1,PPM1K,TUBA1C,IRF9,DST,
RPN,SPAG7,WDR62,STYXL1,RFX5,POLD4,PLEKHA5,RANBP1,VRK3,LZTFL1,FIS1,PPM1K,TUBA1C,IRF9,DST,
7,COX4I1,DUSP22,SLC25A10,THEM4,CENPK,RPS6,RPL29,RTEL1,COASY,AUP1,SAP30BP,YIPF3,FOXRED1,R
7,COX4I1,DUSP22,SLC25A10,THEM4,CENPK,RPS6,RPL29,RTEL1,COASY,AUP1,SAP30BP,YIPF3,FOXRED1,R
M1K,TUBA1C,IRF9,DST,ANTXR2,STIP1,LRRC1,VCAN,LPCAT3,CD44,TM9SF1,MAGED2,SON,UQCRQ,SNF8,
M1K,TUBA1C,IRF9,DST,ANTXR2,STIP1,LRRC1,VCAN,LPCAT3,CD44,TM9SF1,MAGED2,SON,UQCRQ,SNF8,
XD1,ITGB1BP1,MED15,IP6K2,CLASRP,C1ORF198,UGGT2,MICAL1,UBE2I,SLC39A13,YIF1A,EIF3M,CDC42,TE
XD1,ITGB1BP1,MED15,IP6K2,CLASRP,C1ORF198,UGGT2,MICAL1,UBE2I,SLC39A13,YIF1A,EIF3M,CDC42,TE
4,DBNDD2,RPS6,RPL29,RTEL1,COASY,AUP1,SAP30BP,YIPF3,FOXRED1,RABL5,TRNT1,KXD1,ITGB1BP1,MED
4,DBNDD2,RPS6,RPL29,RTEL1,COASY,AUP1,SAP30BP,YIPF3,FOXRED1,RABL5,TRNT1,KXD1,ITGB1BP1,MED
,RBPMS,NNMT,SNRPN,SPAG7,WDR62,IL1RAP,STYXL1,RFX5,POLD4,PLEKHA5,RANBP1,UBE2L6,VRK3,LZTI
2,NUTF2,UCK2,C17ORF62,PRMT1,CASC4,STOML2,C19ORF54,ZNF7,TXNDC17,ARL6IP4,PIGL,C19ORF48,P
RF48,PPAN,ACAD10,SUGT1,HYPK,CARS,CDK11A,RBM3,ZNF197,COPZ1,INO80E,CCNL2,MAPKAPK5,DGUOI
DUSP22,SLC25A10,TMX2,THEM4,DBNDD2,RPS6,RPL29,RTEL1,COASY,AUP1,SAP30BP,YIPF3,FOXRED1,RA

DUSP22,SLC25A10,TMX2,THEM4,DBNDD2,RPS6,RPL29,RTEL1,COASY,AUP1,SAP30BP,YIPF3,FOXRED1,RA
3R4,PHF21A,SLC39A11,FUZ,FAM3A,PSMA3,ATG12,RPS25,NIP7,TTC9C,COX4I1,PLCB2,DUSP22,SLC25A10,
3R4,PHF21A,SLC39A11,FUZ,FAM3A,PSMA3,ATG12,RPS25,NIP7,TTC9C,COX4I1,PLCB2,DUSP22,SLC25A10,
MX2,THEM4,DBNDD2,RPS6,RPL29,RTEL1,COASY,AUP1,SAP30BP,YIPF3,FOXRED1,RABL5,TRNT1,KXD1,ITG
MX2,THEM4,DBNDD2,RPS6,RPL29,RTEL1,COASY,AUP1,SAP30BP,YIPF3,FOXRED1,RABL5,TRNT1,KXD1,ITG
L5,IP6K2,CLASRP,C1ORF198,UGGT2,MICAL1,UBE2I,SLC39A13,EIF3M,CDC42,TBX6,CARD14,PCYT2,FBXW9
L5,IP6K2,CLASRP,C1ORF198,UGGT2,MICAL1,UBE2I,SLC39A13,EIF3M,CDC42,TBX6,CARD14,PCYT2,FBXW9
15,CLASRP,C1ORF198,UGGT2,MICAL1,UBE2I,YIF1A,EIF3M,CDC42,TBX6,CARD14,PCYT2,FBXW9,ASF1B,C
15,CLASRP,C1ORF198,UGGT2,MICAL1,UBE2I,YIF1A,EIF3M,CDC42,TBX6,CARD14,PCYT2,FBXW9,ASF1B,C
22,SLC25A10,TMX2,THEM4,DBNDD2,RPS6,RPL29,GGPS1,RTEL1,COASY,SRSF1,AUP1,SAP30BP,YIPF3,FO
22,SLC25A10,TMX2,THEM4,DBNDD2,RPS6,RPL29,GGPS1,RTEL1,COASY,SRSF1,AUP1,SAP30BP,YIPF3,FO
F9,DST,ANTXR2,STIP1,LRRC1,VCAN,PCBP2,LPCAT3,CD44,TM9SF1,MAGED2,SON,UQCRQ,SNF8,GRK6,HM
F9,DST,ANTXR2,STIP1,LRRC1,VCAN,PCBP2,LPCAT3,CD44,TM9SF1,MAGED2,SON,UQCRQ,SNF8,GRK6,HM
TTC9C,COX4I1,DUSP22,SLC25A10,TMX2,THEM4,DBNDD2,RPS6,RPL29,RTEL1,COASY,AUP1,SAP30BP,YIP
TTC9C,COX4I1,DUSP22,SLC25A10,TMX2,THEM4,DBNDD2,RPS6,RPL29,RTEL1,COASY,AUP1,SAP30BP,YIP
,RRNAD1,DBNL,NAT9,TOMM40,RABEP2,TADA3,ZNF20,WAC,KLK10,ANAPC11,RAD23A,NT5C,MDH2,STXB
DC37,RPS16,CPSF3L,TSG101,UBR4,YLPM1,CHCHD1,FUZ,FAM3A,PSMA3,METTL17,EPB41L5,ATG12,RPS25
DC37,RPS16,CPSF3L,TSG101,UBR4,YLPM1,CHCHD1,FUZ,FAM3A,PSMA3,METTL17,EPB41L5,ATG12,RPS25
ABL5,TRNT1,KXD1,ITGB1BP1,MED15,SH3YL1,IP6K2,CLASRP,C1ORF198,UGGT2,MICAL1,UBE2I,SLC39A13,
ABL5,TRNT1,KXD1,ITGB1BP1,MED15,SH3YL1,IP6K2,CLASRP,C1ORF198,UGGT2,MICAL1,UBE2I,SLC39A13,
ATG12,RPS25,NIP7,TTC9C,COX4I1,DUSP22,SLC25A10,TMX2,THEM4,DBNDD2,RPS6,RPL29,RTEL1,COASY,
ATG12,RPS25,NIP7,TTC9C,COX4I1,DUSP22,SLC25A10,TMX2,THEM4,DBNDD2,RPS6,RPL29,RTEL1,COASY,
P7,TTC9C,COX4I1,PLCB2,DUSP22,SLC25A10,TMX2,THEM4,DBNDD2,SDF4,RPS6,RPL29,RTEL1,COASY,AUP
P7,TTC9C,COX4I1,PLCB2,DUSP22,SLC25A10,TMX2,THEM4,DBNDD2,SDF4,RPS6,RPL29,RTEL1,COASY,AUP
C,MDH2,STXBP2,EEF1D,CHCHD5,SEC31A,MT1E,CNST,PRLR,SH3GLB1,COX17,COX5A,HNRNPLL,HIGD1A,B
25A10,TMX2,THEM4,RDH11,DBNDD2,RPS6,RPL29,RTEL1,COASY,AUP1,SAP30BP,YIPF3,FOXRED1,RABL5,
25A10,TMX2,THEM4,RDH11,DBNDD2,RPS6,RPL29,RTEL1,COASY,AUP1,SAP30BP,YIPF3,FOXRED1,RABL5,
101,TTC21B,PTPN13,YLPM1,PHF21A,SLC39A11,FUZ,FAM3A,MUC1,METTL17,ECSIT,EPB41L5,ATG12,PPR
101,TTC21B,PTPN13,YLPM1,PHF21A,SLC39A11,FUZ,FAM3A,MUC1,METTL17,ECSIT,EPB41L5,ATG12,PPR
ERLEC1,SAT1,ERCC5,SERPINB1,DDOST,LSR,PSME2,PABPC1,RRNAD1,DBNL,NAT9,RAD1,TOMM40,RABEP2,
XBP2,EEF1D,CHCHD5,SEC31A,CNST,COX17,HIGD1A,SIRT5,INTS4,B3GAT3,ZNF506,PFDN5,MTHFS,PAFAH1
R2,STIP1,LRRC1,VCAN,LPCAT3,CD44,TM9SF1,MAGED2,SON,UQCRQ,SNF8,GRK6,HMGA2,CDC37,RPS16,C
R2,STIP1,LRRC1,VCAN,LPCAT3,CD44,TM9SF1,MAGED2,SON,UQCRQ,SNF8,GRK6,HMGA2,CDC37,RPS16,C
1,ZC3H14,FIS1,SLC15A3,PPM1K,TUBA1C,ANTXR2,STIP1,LRRC1,PHF19,GRAMD1C,VCAN,LPCAT3,CD44,T
1,ZC3H14,FIS1,SLC15A3,PPM1K,TUBA1C,ANTXR2,STIP1,LRRC1,PHF19,GRAMD1C,VCAN,LPCAT3,CD44,T
3B2,RPS9,SLC35G1,RCE1,MRPL33,ZNF410,RUVBL2,PCMT1,EBPL,TESC,IGSF8,NUDT5,RBPMS,NNMT,SNR
3B2,RPS9,SLC35G1,RCE1,MRPL33,ZNF410,RUVBL2,PCMT1,EBPL,TESC,IGSF8,NUDT5,RBPMS,NNMT,SNR
RCC5,EMC4,LMBR1,SERPINB1,DDOST,LSR,CHCHD7,PABPC1,RRNAD1,NAT9,TOMM40,RABEP2,TADA3,ZN
VRK3,LZTFL1,FIS1,PPM1K,TUBA1C,IRF9,DST,ANTXR2,STIP1,LRRC1,VCAN,CD44,TM9SF1,MAGED2,SON,U
VRK3,LZTFL1,FIS1,PPM1K,TUBA1C,IRF9,DST,ANTXR2,STIP1,LRRC1,VCAN,CD44,TM9SF1,MAGED2,SON,U
DR62,IL1RAP,STYXL1,RFX5,POLD4,PLEKHA5,RANBP1,UBE2L6,VRK3,LZTFL1,ZC3H14,FIS1,SLC15A3,PPM1K
DR62,IL1RAP,STYXL1,RFX5,POLD4,PLEKHA5,RANBP1,UBE2L6,VRK3,LZTFL1,ZC3H14,FIS1,SLC15A3,PPM1K
,CARS,CDK11A,RBM3,ZNF197,COPZ1,INO80E,CCNL2,DGUOK,LRRFIP1,ZNF524,ATP5J,RMDN1,TRMT112,
MICAL1,UBE2I,EIF3M,CDC42,TBX6,TMEM205,PTK2,CARD14,PCYT2,FBXW9,USE1,IFI27L1,ASF1B,CCDC25,E
MICAL1,UBE2I,EIF3M,CDC42,TBX6,TMEM205,PTK2,CARD14,PCYT2,FBXW9,USE1,IFI27L1,ASF1B,CCDC25,E
PPIH,DALRD3,NSUN5,ZFPL1,MFSD10,MCFD2,RIC8A,TCP1,PIGT,UBE2N,MFSD11,LFNG,ZNHIT3,PDCD6,ZN
5,EMC4,SERPINB1,DDOST,LSR,PSME2,RRNAD1,DBNL,NAT9,TOMM40,RABEP2,TADA3,ZNF20,WAC,KLK10

MA7,APEH,CAPN10,LYPLA2,NUTF2,ATPAF2,C17ORF62,PRMT1,C19ORF54,ZNF7,TXNDC17,ARL6IP4,C19OR
ATG12,RPS25,ZBTB44,NIP7,TTC9C,COX4I1,DUSP22,SLC25A10,TMX2,THEM4,DBNDD2,RPS6,RPL29,GGPS1
ATG12,RPS25,ZBTB44,NIP7,TTC9C,COX4I1,DUSP22,SLC25A10,TMX2,THEM4,DBNDD2,RPS6,RPL29,GGPS1
3L,TSG101,UBR4,YLPM1,CHCHD1,FUZ,FAM3A,PSMA3,METTL17,EPB41L5,ATG12,RPS25,ZBTB44,NIP7,TT
3L,TSG101,UBR4,YLPM1,CHCHD1,FUZ,FAM3A,PSMA3,METTL17,EPB41L5,ATG12,RPS25,ZBTB44,NIP7,TT
126B,APEH,CAPN10,LYPLA2,NUTF2,C17ORF62,PRMT1,STOML2,C19ORF54,ZNF7,TXNDC17,ARL6IP4,PIGL,
,TUBA1C,IRF9,DST,STIP1,LRRC1,GRAMD1C,VCAN,CD44,TM9SF1,MAGED2,SON,UQCRQ,CDC37,RPS16,CP
,TUBA1C,IRF9,DST,STIP1,LRRC1,GRAMD1C,VCAN,CD44,TM9SF1,MAGED2,SON,UQCRQ,CDC37,RPS16,CP
TOMM40,RABEP2,TADA3,ZNF20,WAC,SLC33A1,KLK10,ANAPC11,POLR3H,RAD23A,ABHD17A,NT5C,MDH
PL,TESC,IGSF8,NUDT5,RBPMS,NNMT,SPAG7,WDR62,IL1RAP,STYXL1,RFX5,POLD4,PLEKHA5,RANBP1,VRK
PL,TESC,IGSF8,NUDT5,RBPMS,NNMT,SPAG7,WDR62,IL1RAP,STYXL1,RFX5,POLD4,PLEKHA5,RANBP1,VRK
S4,B3GAT3,ZNF506,PFDN5,MTHFS,PAFAH1B2,ZNF343,GBA2,PPIH,DALRD3,NSUN5,DRG2,ZFPL1,MFSD10
CPSF3L,TSG101,OBSL1,TTC21B,PTPN13,YLPM1,PHF21A,SLC39A11,FUZ,FAM3A,MUC1,METTL17,ECSIT,EF
CPSF3L,TSG101,OBSL1,TTC21B,PTPN13,YLPM1,PHF21A,SLC39A11,FUZ,FAM3A,MUC1,METTL17,ECSIT,EF
,ANAPC11,POLR3H,RAD23A,ABHD17A,NT5C,MDH2,EEF1D,NAA10,MED6,SEC31A,CARHSP1,SSH2,COX17
,ANAPC11,POLR3H,RAD23A,ABHD17A,NT5C,MDH2,EEF1D,NAA10,MED6,SEC31A,CARHSP1,SSH2,COX17
F62,PRMT1,STOML2,C19ORF54,ZNF7,TXNDC17,ARL6IP4,PIGL,C19ORF48,PPAN,ZFAND2B,HNRNPC,SHAR
2,UQCRQ,GRK6,CDC37,RPS16,TSG101,TTC21B,PTPN13,UBR4,PSMA1,PHF21A,CHCHD1,SLC39A11,FAM3A
OST,LSR,PSME2,PABPC1,RRNAD1,DBNL,ZNF836,RAD1,TOMM40,RABEP2,TADA3,ZNF20,WAC,TRIM16,AI
NF512,RNF181,CHTOP,TPT1,SLC52A2,TSPAN31,MTIF2,RALY,GLYCTK,VDAC1,RER1,EIF2A,NKIRAS2,ACAA
AMTA1,PPIL2,PSMD4,RPL28,LIG1,TIMM17B,TMEM126B,APEH,CAPN10,LYPLA2,NUTF2,C17ORF62,PRMT1
PEH,RPL21,CAPN10,RBM33,LYPLA2,LSM12,C17ORF62,PRMT1,STOML2,C19ORF54,ZNF7,TXNDC17,ARL6II
PEH,RPL21,CAPN10,RBM33,LYPLA2,LSM12,C17ORF62,PRMT1,STOML2,C19ORF54,ZNF7,TXNDC17,ARL6II
,MED6,SEC31A,CARHSP1,COX17,HNRNPLL,PHF20L1,HIGD1A,POLR2C,B3GAT3,RFXANK,ZNF506,SPINT2,P
,MED6,SEC31A,CARHSP1,COX17,HNRNPLL,PHF20L1,HIGD1A,POLR2C,B3GAT3,RFXANK,ZNF506,SPINT2,P
XD1,ITGB1BP1,MED15,CLASRP,C1ORF198,UGGT2,MICAL1,UBE2I,YIF1A,EIF3M,CDC42,TMEM205,CARD1
XD1,ITGB1BP1,MED15,CLASRP,C1ORF198,UGGT2,MICAL1,UBE2I,YIF1A,EIF3M,CDC42,TMEM205,CARD1
HA5,RANBP1,LZTFL1,FIS1,SLC15A3,PPM1K,TUBA1C,ANTXR2,STIP1,LRRC1,PHF19,VCAN,LPCAT3,TM9SF1,
HA5,RANBP1,LZTFL1,FIS1,SLC15A3,PPM1K,TUBA1C,ANTXR2,STIP1,LRRC1,PHF19,VCAN,LPCAT3,TM9SF1,
1,RRNAD1,NAT9,TOMM40,RABEP2,TADA3,ZNF20,WAC,KLK10,ANAPC11,RAD23A,ABHD17A,NT5C,MDH
9ORF48,PPAN,ZFAND2B,HNRNPC,SHARPIN,SLC9B2,SUGT1,HYPK,CARS,ZNF250,CDK11A,RBM3,ZNF197,C
SPAG7,WDR62,IL1RAP,STYXL1,POLD4,PLEKHA5,RANBP1,UBE2L6,VRK3,LZTFL1,ZC3H14,FIS1,PPM1K,TUBA
SPAG7,WDR62,IL1RAP,STYXL1,POLD4,PLEKHA5,RANBP1,UBE2L6,VRK3,LZTFL1,ZC3H14,FIS1,PPM1K,TUBA
,TOMM40,RABEP2,TADA3,ZNF20,ATP6V1F,WAC,KLK10,ANAPC11,POLR3H,RAD23A,B2M,ABHD17A,DDX
B1,VWA2,ERCC5,EMC4,LMBR1,MCTS1,DDOST,LSR,MEIS2,RRNAD1,RAD1,TOMM40,TADA3,ZNF20,ZNF6
B1,VWA2,ERCC5,EMC4,LMBR1,MCTS1,DDOST,LSR,MEIS2,RRNAD1,RAD1,TOMM40,TADA3,ZNF20,ZNF6
MED15,SH3YL1,CLASRP,C1ORF198,UGGT2,UBE2I,SLC39A13,PRMT2,EIF3M,CDC42,TBX6,C14ORF2,GUK1,
MED15,SH3YL1,CLASRP,C1ORF198,UGGT2,UBE2I,SLC39A13,PRMT2,EIF3M,CDC42,TBX6,C14ORF2,GUK1,
,ZNF814,HSPB1,VWA2,ERCC5,EMC4,LMBR1,DDOST,LSR,MEIS2,RRNAD1,RAD1,TOMM40,TADA3,ZNF20,
,ZNF814,HSPB1,VWA2,ERCC5,EMC4,LMBR1,DDOST,LSR,MEIS2,RRNAD1,RAD1,TOMM40,TADA3,ZNF20,
M33,LYPLA2,C17ORF62,PRMT1,STOML2,C19ORF54,ZNF7,TXNDC17,ARL6IP4,C19ORF48,PPAN,CYB561,SH
M33,LYPLA2,C17ORF62,PRMT1,STOML2,C19ORF54,ZNF7,TXNDC17,ARL6IP4,C19ORF48,PPAN,CYB561,SH
RD14,PCYT2,FBXW9,USE1,HSBP1L1,MAF,ASF1B,CCDC25,EEF1B2,RFX3,ENDOV,VAMP8,CAMTA1,CAPG,PF
RD14,PCYT2,FBXW9,USE1,HSBP1L1,MAF,ASF1B,CCDC25,EEF1B2,RFX3,ENDOV,VAMP8,CAMTA1,CAPG,PF
P,STYXL1,POLD4,PLEKHA5,RANBP1,UBE2L6,VRK3,LZTFL1,ZC3H14,FIS1,PPM1K,TUBA1C,DST,ANTXR2,STIF
P,STYXL1,POLD4,PLEKHA5,RANBP1,UBE2L6,VRK3,LZTFL1,ZC3H14,FIS1,PPM1K,TUBA1C,DST,ANTXR2,STIF
,GABPB1,NIT2,DCTD,ERLEC1,HSPB1,SAT1,ERCC5,MCTS1,NAP1L1,SERPINB1,DDOST,LSR,PABPC1,RRNAD1

L,C19ORF48,PPAN,ZFAND2B,SHARPIN,SUGT1,HYPK,CARS,CDK11A,RBM3,ZNF197,COPZ1,INO80E,CCNL2,
YPLA2,C17ORF62,PRMT1,STOML2,ZNF7,TXNDC17,PPAN,ACAD10,CYB561,SHARPIN,SUGT1,HYPK,CARS,Z
YPLA2,C17ORF62,PRMT1,STOML2,ZNF7,TXNDC17,PPAN,ACAD10,CYB561,SHARPIN,SUGT1,HYPK,CARS,Z
C17,ARL6IP4,C19ORF48,PPAN,ZFAND2B,SHARPIN,SLC9B2,SUGT1,HYPK,CARS,ZNF250,CDK11A,RBM3,M
2,C17ORF62,PRMT1,CASC4,STOML2,ZNF7,TXNDC17,ARL6IP4,PIGL,PPAN,ZFAND2B,HNRNPC,CYB561,SHA
2,C17ORF62,PRMT1,CASC4,STOML2,ZNF7,TXNDC17,ARL6IP4,PIGL,PPAN,ZFAND2B,HNRNPC,CYB561,SHA
IST,COX17,HNRNPLL,HIGD1A,FKBP3,RPL15,INTS4,B3GAT3,ZNF506,SPINT2,PFDN5,MTHFS,PAFAH1B2,ZN
7,WDR62,IL1RAP,STYXL1,RFX5,POLD4,PLEKHA5,RANBP1,VRK3,LZTFL1,ZC3H14,FIS1,SLC15A3,PPM1K,TU
L28,LIG1,APEH,HYI,CAPN10,LYPLA2,NUTF2,UCK2,ATPAF2,C17ORF62,PRMT1,CASC4,STOML2,C19ORF54,
1,EBPL,TESC,IGSF8,NUDT5,RBPMS,NNMT,SNRPN,SPAG7,WDR62,IL1RAP,STYXL1,RFX5,POLD4,PLEKHA5,
D23A,ABHD17A,MDH2,EEF1D,HSPH1,CHCHD5,NAA10,SEC31A,CARHSP1,CNST,COX17,HNRNPLL,HIGD1A,
D23A,ABHD17A,MDH2,EEF1D,HSPH1,CHCHD5,NAA10,SEC31A,CARHSP1,CNST,COX17,HNRNPLL,HIGD1A,
RRC1,VCAN,LPCAT3,CD44,MAGED2,SON,TET2,UQCRQ,SNF8,GRK6,HMGA2,CDC37,RPS16,CPSF3L,TSG10
RRC1,VCAN,LPCAT3,CD44,MAGED2,SON,TET2,UQCRQ,SNF8,GRK6,HMGA2,CDC37,RPS16,CPSF3L,TSG10
AP30BP,YIPF3,FOXRED1,RABL5,TRNT1,KXD1,ITGB1BP1,MED15,IP6K2,CLASRP,C1ORF198,UGGT2,MICAL
AP30BP,YIPF3,FOXRED1,RABL5,TRNT1,KXD1,ITGB1BP1,MED15,IP6K2,CLASRP,C1ORF198,UGGT2,MICAL
AP30BP,YIPF3,FOXRED1,RABL5,TRNT1,KXD1,ITGB1BP1,MED15,IP6K2,CLASRP,C1ORF198,UGGT2,MICAL
AP30BP,YIPF3,FOXRED1,RABL5,TRNT1,KXD1,ITGB1BP1,MED15,IP6K2,CLASRP,C1ORF198,UGGT2,MICAL
MM40,TADA3,ZNF20,ZNF687,PDCD5,ANAPC11,RAD23A,ABHD17A,NT5C,MDH2,EEF1D,NAA10,SEC31A,C
MM40,TADA3,ZNF20,ZNF687,PDCD5,ANAPC11,RAD23A,ABHD17A,NT5C,MDH2,EEF1D,NAA10,SEC31A,C
MP8,CAPG,PPIL2,TROAP,LIG1,PSMA7,CDC14B,TMEM126B,APEH,RBM33,LYPLA2,C17ORF62,PRMT1,STO
MP8,CAPG,PPIL2,TROAP,LIG1,PSMA7,CDC14B,TMEM126B,APEH,RBM33,LYPLA2,C17ORF62,PRMT1,STO
PC11,RAD23A,ABHD17A,ZCCHC4,MDH2,STXBP2,EEF1D,HSPH1,CHCHD5,NAA10,OAS3,SEC31A,CNST,LLG
DU1,UBXN1,ZNF814,HSPB1,SAT1,ERCC5,EMC4,MCTS1,DDOST,LSR,MEIS2,RRNAD1,NAT9,TOMM40,RABE
DU1,UBXN1,ZNF814,HSPB1,SAT1,ERCC5,EMC4,MCTS1,DDOST,LSR,MEIS2,RRNAD1,NAT9,TOMM40,RABE
,CDC37,RPS16,CPSF3L,TSG101,PTPN13,UBR4,YLPM1,CHCHD1,FUZ,FAM3A,PSMA3,METTL17,ECSIT,EPB4
,CDC37,RPS16,CPSF3L,TSG101,PTPN13,UBR4,YLPM1,CHCHD1,FUZ,FAM3A,PSMA3,METTL17,ECSIT,EPB4
5,NAA10,SEC31A,CARHSP1,CNST,COX17,HIGD1A,POLR2C,B3GAT3,ZNF506,SPINT2,PFDN5,PAFAH1B2,DA
5,NAA10,SEC31A,CARHSP1,CNST,COX17,HIGD1A,POLR2C,B3GAT3,ZNF506,SPINT2,PFDN5,PAFAH1B2,DA
25A10,THEM4,RPS6,RPL29,RTEL1,COASY,AUP1,SAP30BP,YIPF3,FOXRED1,RABL5,TRNT1,KXD1,ITGB1BP1,
25A10,THEM4,RPS6,RPL29,RTEL1,COASY,AUP1,SAP30BP,YIPF3,FOXRED1,RABL5,TRNT1,KXD1,ITGB1BP1,
DST,ANTXR2,STIP1,LRRC1,CD44,TM9SF1,MAGED2,SON,UQCRQ,SNF8,HMGA2,CDC37,RPS16,CPSF3L,TSG
DST,ANTXR2,STIP1,LRRC1,CD44,TM9SF1,MAGED2,SON,UQCRQ,SNF8,HMGA2,CDC37,RPS16,CPSF3L,TSG
,ERCC5,EMC4,SERPINB1,DDOST,LSR,CHCHD7,MEIS2,PABPC1,RRNAD1,DBNL,ZNF836,RBP1,TOMM40,RA
,ERCC5,EMC4,SERPINB1,DDOST,LSR,CHCHD7,MEIS2,PABPC1,RRNAD1,DBNL,ZNF836,RBP1,TOMM40,RA
25A10,THEM4,NISCH,TMED4,CENPK,RPS6,RPL29,RTEL1,COASY,AUP1,SAP30BP,YIPF3,FOXRED1,RABL5,
25A10,THEM4,NISCH,TMED4,CENPK,RPS6,RPL29,RTEL1,COASY,AUP1,SAP30BP,YIPF3,FOXRED1,RABL5,
PL,TESC,NUDT5,RBPMS,SNRPN,SPAG7,WDR62,IL1RAP,STYXL1,RFX5,PLEKHA5,RANBP1,UBE2L6,VRK3,LZT
19,VCAN,PCBP2,LPCAT3,TM9SF1,MAGED2,SON,TET2,UQCRQ,SNF8,GRK6,HMGA2,CDC37,RPS16,CPSF3L,
19,VCAN,PCBP2,LPCAT3,TM9SF1,MAGED2,SON,TET2,UQCRQ,SNF8,GRK6,HMGA2,CDC37,RPS16,CPSF3L,
21B,YLPM1,PHF21A,CHCHD1,SLC39A11,FUZ,FAM3A,MUC1,METTL17,ECSIT,EPB41L5,ATG12,PPRC1,ZBT
21B,YLPM1,PHF21A,CHCHD1,SLC39A11,FUZ,FAM3A,MUC1,METTL17,ECSIT,EPB41L5,ATG12,PPRC1,ZBT
OX17,HNRNPLL,HIGD1A,B3GAT3,ZNF506,SPINT2,MTHFS,PAFAH1B2,ZNF343,GBA2,PPIH,DALRD3,NSUN
TTC21B,YLPM1,PHF21A,CHCHD1,SLC39A11,FUZ,FAM3A,MUC1,METTL17,ECSIT,EPB41L5,ATG12,PPRC1,
TTC21B,YLPM1,PHF21A,CHCHD1,SLC39A11,FUZ,FAM3A,MUC1,METTL17,ECSIT,EPB41L5,ATG12,PPRC1,
ADA3,ZNF20,ZNF687,PDCD5,ANAPC11,POLR3H,RAD23A,ABHD17A,NT5C,MDH2,STXBP2,NAA10,MED6,S
ADA3,ZNF20,ZNF687,PDCD5,ANAPC11,POLR3H,RAD23A,ABHD17A,NT5C,MDH2,STXBP2,NAA10,MED6,S

37,PDCD5,ANAPC11,POLR3H,RAD23A,ABHD17A,NT5C,MDH2,STXBP2,EEF1D,NAA10,SEC31A,CARHSP1,C
37,PDCD5,ANAPC11,POLR3H,RAD23A,ABHD17A,NT5C,MDH2,STXBP2,EEF1D,NAA10,SEC31A,CARHSP1,C
K3,LZTFL1,ZC3H14,FIS1,PPM1K,DST,ANTXR2,STIP1,LRRC1,PHF19,LPCAT3,CD44,TM9SF1,MAGED2,SON,T
K3,LZTFL1,ZC3H14,FIS1,PPM1K,DST,ANTXR2,STIP1,LRRC1,PHF19,LPCAT3,CD44,TM9SF1,MAGED2,SON,T
FPL1,MFSD10,MCFD2,RIC8A,TCP1,THADA,CCDC40,PIGT,UBE2N,LFNG,ZNHIT3,FAM32A,COX20,PEX2,POL
DC42,TMEM205,AUNIP,CARD14,FBXW9,USE1,MAF,ASF1B,CCDC25,EEF1B2,RFX3,DGKE,ENDOV,CAMTA1
DC42,TMEM205,AUNIP,CARD14,FBXW9,USE1,MAF,ASF1B,CCDC25,EEF1B2,RFX3,DGKE,ENDOV,CAMTA1
410,RUVBL2,PCMT1,EBPL,TESC,IGSF8,NUDT5,RBPMS,SPAG7,WDR62,IL1RAP,STYXL1,RFX5,POLD4,PLEKH
410,RUVBL2,PCMT1,EBPL,TESC,IGSF8,NUDT5,RBPMS,SPAG7,WDR62,IL1RAP,STYXL1,RFX5,POLD4,PLEKH
UGGT2,MICAL1,UBE2I,SLC39A13,YIF1A,EIF3M,CDC42,TBX6,TMEM205,XBP1,PTK2,AUNIP,C14ORF2,CARD
UGGT2,MICAL1,UBE2I,SLC39A13,YIF1A,EIF3M,CDC42,TBX6,TMEM205,XBP1,PTK2,AUNIP,C14ORF2,CARD
,AP3B2,RPS9,RCE1,ZNF410,RUVBL2,PCMT1,EBPL,TESC,IGSF8,NUDT5,RBPMS,NNMT,SPAG7,WDR62,IL1R
,AP3B2,RPS9,RCE1,ZNF410,RUVBL2,PCMT1,EBPL,TESC,IGSF8,NUDT5,RBPMS,NNMT,SPAG7,WDR62,IL1R
ANAPC11,RAD23A,NT5C,MDH2,STXBP2,EEF1D,CHCHD5,SEC31A,CARHSP1,CNST,COX17,HNRNPLL,HIGD1
1A,CHCHD1,SLC39A11,FAM3A,METTL17,ECSIT,EPB41L5,ATG12,NLRP1,ZBTB44,NIP7,PLCB2,DUSP22,SLC2
1A,CHCHD1,SLC39A11,FAM3A,METTL17,ECSIT,EPB41L5,ATG12,NLRP1,ZBTB44,NIP7,PLCB2,DUSP22,SLC2
T,LSR,PSME2,PABPC1,RRNAD1,DBNL,RAD1,TOMM40,RABEP2,TADA3,ZNF20,ATP6V1F,WAC,TRIM16,ANA
MTA1,CAPG,PPIL2,RPL28,CLDND1,LIG1,PSMA7,CDC14B,TIMM17B,TMEM126B,APEH,LYPLA2,EPB41L4B,
MTA1,CAPG,PPIL2,RPL28,CLDND1,LIG1,PSMA7,CDC14B,TIMM17B,TMEM126B,APEH,LYPLA2,EPB41L4B,
I,SLC39A13,EIF3M,CDC42,TBX6,PTK2,CARD14,PCYT2,FBXW9,USE1,MAF,ASF1B,CCDC25,EEF1B2,RFX3,EN
I,SLC39A13,EIF3M,CDC42,TBX6,PTK2,CARD14,PCYT2,FBXW9,USE1,MAF,ASF1B,CCDC25,EEF1B2,RFX3,EN
F62,PRMT1,STOML2,C19ORF54,ZNF7,TXNDC17,PIGL,C19ORF48,PPAN,HNRNPC,SHARPIN,SUGT1,CARS,C
PL29,GGPS1,RTEL1,COASY,SRSF1,AUP1,SAP30BP,YIPF3,FOXRED1,RABL5,TRNT1,ITGB1BP1,MED15,SH3Y
PL29,GGPS1,RTEL1,COASY,SRSF1,AUP1,SAP30BP,YIPF3,FOXRED1,RABL5,TRNT1,ITGB1BP1,MED15,SH3Y
LL,HIGD1A,POLR2C,B3GAT3,RFXANK,ZNF506,SPINT2,PFDN5,PAFAH1B2,TXNRD2,DALRD3,NSUN5,OARD
LL,HIGD1A,POLR2C,B3GAT3,RFXANK,ZNF506,SPINT2,PFDN5,PAFAH1B2,TXNRD2,DALRD3,NSUN5,OARD
AH1B2,ZNF343,DALRD3,NSUN5,ZFPL1,OARD1,MFSD10,RIC8A,TCP1,SMARCC2,PHC3,COMMD4,PIGT,UBE
D1,MFSD10,MRPS18C,GMPR2,ZSCAN9,RIC8A,TCP1,SMARCC2,THADA,UBE2N,LFNG,PDCD6,ZNF81,FAM
D1,MFSD10,MRPS18C,GMPR2,ZSCAN9,RIC8A,TCP1,SMARCC2,THADA,UBE2N,LFNG,PDCD6,ZNF81,FAM
506,PFDN5,PAFAH1B2,TXNRD2,GTF2IRD2,NSUN5,ZFPL1,OARD1,MRPS18C,RIC8A,TCP1,SMARCC2,GNB5,
506,PFDN5,PAFAH1B2,TXNRD2,GTF2IRD2,NSUN5,ZFPL1,OARD1,MRPS18C,RIC8A,TCP1,SMARCC2,GNB5,
HNRNPLL,HIGD1A,POLR2C,B3GAT3,RFXANK,SPINT2,FTH1,PFDN5,PAFAH1B2,TXNRD2,DALRD3,NSUN5,OA
HNRNPLL,HIGD1A,POLR2C,B3GAT3,RFXANK,SPINT2,FTH1,PFDN5,PAFAH1B2,TXNRD2,DALRD3,NSUN5,OA
F57,DCTN5,PEX26,NDUFS3,OXLD1,ZNF223,METTL5,SLC35E1,OGFOD2,METTL3,EIF3G,SSR3,ARIH2,PICK1,
1,EIF4A1,METTL5,SLC35E1,OGFOD2,EIF3G,TSEN34,SSR3,ARIH2,PICK1,FBXL18,ZNF512,RNF181,TPT1,UBA
NB1,DDOST,LSR,PSME2,PABPC1,RRNAD1,DBNL,TOMM40,RABEP2,TADA3,ZNF20,ATP6V1F,WAC,TRIM16,
,RPS16,CPSF3L,TSG101,OBSL1,TTC21B,PTPN13,CHCHD1,SLC39A11,FAM3A,PSMA3,MUC1,METTL17,ECS
,RPS16,CPSF3L,TSG101,OBSL1,TTC21B,PTPN13,CHCHD1,SLC39A11,FAM3A,PSMA3,MUC1,METTL17,ECS
PSMD7,TRAPPC2,PIGO,KDELR3,CLTA,SLC35A2,PLAC8,THOC3,AXIN1,TMEM199,SLC25A24,MGST2,EPN1,
LEKHA5,RANBP1,UBE2L6,LZTFL1,ZC3H14,FIS1,SLC15A3,PPM1K,TUBA1C,ANTXR2,STIP1,LRRC1,PHF19,GR
2,ERCC5,EMC4,LMBR1,DDOST,LSR,MEIS2,PSME2,RRNAD1,RAD1,TOMM40,TADA3,ZNF20,ZNF687,PDCD
2,ERCC5,EMC4,LMBR1,DDOST,LSR,MEIS2,PSME2,RRNAD1,RAD1,TOMM40,TADA3,ZNF20,ZNF687,PDCD
P1,ZBTB44,NIP7,PLCB2,SLC25A10,TMX2,SPOCK2,THEM4,DBNDD2,NISCH,TMED4,CD99L2,GGPS1,COASY
P1,ZBTB44,NIP7,PLCB2,SLC25A10,TMX2,SPOCK2,THEM4,DBNDD2,NISCH,TMED4,CD99L2,GGPS1,COASY
PSMA6,ACOT8,AP3B2,RPS9,SLC35G1,RCE1,MRPL33,ZNF410,RUVBL2,PCMT1,EBPL,TESC,IGSF8,NUDT5,RE
PSMA6,ACOT8,AP3B2,RPS9,SLC35G1,RCE1,MRPL33,ZNF410,RUVBL2,PCMT1,EBPL,TESC,IGSF8,NUDT5,RE
CDC37,CPSF3L,TSG101,OBSL1,TTC21B,PTPN13,PSMA1,PHF21A,CHCHD1,PSMA3,EPB41L5,ATG12,PPRC1,

LC35G1,RCE1,MRPL33,ZNF410,RUVBL2,PCMT1,EBPL,TESC,IGSF8,NUDT5,RBPMS,NNMT,SPAG7,WDR62,I

LC35G1,RCE1,MRPL33,ZNF410,RUVBL2,PCMT1,EBPL,TESC,IGSF8,NUDT5,RBPMS,NNMT,SPAG7,WDR62,I

S,PAFAH1B2,PPIH,NSUN5,OARD1,GMPR2,MCFD2,RIC8A,TCP1,THADA,PIGT,UBE2N,LFNG,ZNHIT3,PDCD6,

S,PAFAH1B2,PPIH,NSUN5,OARD1,GMPR2,MCFD2,RIC8A,TCP1,THADA,PIGT,UBE2N,LFNG,ZNHIT3,PDCD6,

T1,NME1,SLC52A2,TSPAN31,MTIF2,RALY,MARS,GLYCTK,TRIM46,C11ORF57,RER1,ACAA1,PSMD7,ZNF28

PS25,SNRPD1,MKRN1,SMUG1,CHURC1,SNRPA1,AZI2,MYL12A,TATDN1,TBC1D22A,LAMTOR1,ZNF581,RP

PP1CC,RFX3,DGKE,ENDOV,VAMP8,CAMTA1,CAPG,PPIL2,PSMD4,TROAP,RPL28,LIG1,PSMA7,CDC14B,API

PP1CC,RFX3,DGKE,ENDOV,VAMP8,CAMTA1,CAPG,PPIL2,PSMD4,TROAP,RPL28,LIG1,PSMA7,CDC14B,API

RCE1,MRPL33,ZNF410,RUVBL2,PCMT1,EBPL,TESC,IGSF8,NUDT5,RBPMS,NNMT,SPAG7,WDR62,IL1RAP,S

RCE1,MRPL33,ZNF410,RUVBL2,PCMT1,EBPL,TESC,IGSF8,NUDT5,RBPMS,NNMT,SPAG7,WDR62,IL1RAP,S

A24,EPN1,HPS1,ITPA,ZCCHC9,VPS25,SNRPD1,MKRN1,CHURC1,AZI2,GOSR2,TBC1D22A,LAMTOR1,U2AF1

ESC,IGSF8,NUDT5,RBPMS,NNMT,WDR62,IL1RAP,POLD4,PLEKHA5,RANBP1,UBE2L6,VRK3,LZTFL1,ZC3H1

ESC,IGSF8,NUDT5,RBPMS,NNMT,WDR62,IL1RAP,POLD4,PLEKHA5,RANBP1,UBE2L6,VRK3,LZTFL1,ZC3H1

F3L,OBSL1,TTC21B,PTPN13,PSMA1,YLPM1,PHF21A,SLC39A11,FAM3A,PSMA3,MUC1,METTL17,ECSIT,EPE

F3L,OBSL1,TTC21B,PTPN13,PSMA1,YLPM1,PHF21A,SLC39A11,FAM3A,PSMA3,MUC1,METTL17,ECSIT,EPE

12,RNF181,NTMT1,TSPAN31,RALY,ZNF593,GLYCTK,CFLAR,RER1,NEK11,EIF2A,GLI4,NKIRAS2,RECQL5,ACA

F198,UGGT2,MICAL1,HS2ST1,UBE2I,PRMT2,EIF3M,JUP,TBX6,TMEM205,XBP1,PTK2,AUNIP,C14ORF2,GUK

F198,UGGT2,MICAL1,HS2ST1,UBE2I,PRMT2,EIF3M,JUP,TBX6,TMEM205,XBP1,PTK2,AUNIP,C14ORF2,GUK

N,LPCAT3,CD44,TM9SF1,TET2,SNF8,GRK6,HMGA2,CDC37,RPS16,CPSF3L,TSG101,OBSL1,TTC21B,PTPN13

CNL2,MAPKAPK5,HSP90AB1,RAB2A,ATP5J,TRMT112,RPTOR,CHPF,RPS20,UBXN1,ZNF814,HSPB1,SAT1,E

CNL2,MAPKAPK5,HSP90AB1,RAB2A,ATP5J,TRMT112,RPTOR,CHPF,RPS20,UBXN1,ZNF814,HSPB1,SAT1,E

5,UBXN1,ZNF814,VWA2,ERCC5,EMC4,LMBR1,MCTS1,DDOST,LSR,RRNAD1,RAD1,TOMM40,TADA3,ZNF2

5,UBXN1,ZNF814,VWA2,ERCC5,EMC4,LMBR1,MCTS1,DDOST,LSR,RRNAD1,RAD1,TOMM40,TADA3,ZNF2

N,LFNG,PDCD6,ZNF81,FAM32A,TMEM147,COA1,USP40,PCID2,PEX2,POLR1C,TNKS,STX10,C12ORF57,DC

N,LFNG,PDCD6,ZNF81,FAM32A,TMEM147,COA1,USP40,PCID2,PEX2,POLR1C,TNKS,STX10,C12ORF57,DC

TL3,TSEN34,MPV17,ARIH2,PICK1,FBXL18,RNF181,TPT1,NTMT1,TSPAN31,PPP2R5C,FAM192A,RALY,GLY

,SPAG7,WDR62,STYXL1,PLEKHA5,RANBP1,UBE2L6,VRK3,LZTFL1,ZC3H14,FIS1,PPM1K,TUBA1C,IRF9,DST,

,SPAG7,WDR62,STYXL1,PLEKHA5,RANBP1,UBE2L6,VRK3,LZTFL1,ZC3H14,FIS1,PPM1K,TUBA1C,IRF9,DST,

F2,TMPO,C17ORF62,STOML2,WNK1,ZNF7,PPAN,AP1S3,ACAD10,CYB561,CBFA2T2,STAU1,SUGT1,HYPK,C

F2,TMPO,C17ORF62,STOML2,WNK1,ZNF7,PPAN,AP1S3,ACAD10,CYB561,CBFA2T2,STAU1,SUGT1,HYPK,C

C19ORF54,ZNF7,C19ORF48,PPAN,AP1S3,HNRNPC,STAU1,SUGT1,HYPK,CARS,CDK11A,RBM3,ZNF197,INO8

VCAN,LPCAT3,CD44,TM9SF1,SON,TET2,GRK6,HMGA2,CDC37,RPS16,CPSF3L,TSG101,TTC21B,PTPN13,UBR

VCAN,LPCAT3,CD44,TM9SF1,SON,TET2,GRK6,HMGA2,CDC37,RPS16,CPSF3L,TSG101,TTC21B,PTPN13,UBR

2,ZNF343,GBA2,PPIH,DALRD3,NSUN5,DRG2,MFSD10,MRPS18C,GMPR2,MCFD2,RIC8A,TCP1,PARK7,THA

AFAH1B2,ZNF343,PPIH,NSUN5,MFSD10,GMPR2,MCFD2,RIC8A,PARK7,THADA,UBE2N,LFNG,ZNHIT3,PDC

CHCHD1,FUZ,RGS3,FAM3A,PSMA3,MUC1,ECSIT,EPB41L5,ATG12,PPRC1,RPS25,NIP7,TTC9C,COX4I1,DUSP

CHCHD1,FUZ,RGS3,FAM3A,PSMA3,MUC1,ECSIT,EPB41L5,ATG12,PPRC1,RPS25,NIP7,TTC9C,COX4I1,DUSP

C39A11,FUZ,FAM3A,MUC1,METTL17,ECSIT,EPB41L5,ATG12,PPRC1,RPS25,ZBTB44,TTC9C,DUSP22,SLC25

C39A11,FUZ,FAM3A,MUC1,METTL17,ECSIT,EPB41L5,ATG12,PPRC1,RPS25,ZBTB44,TTC9C,DUSP22,SLC25

TTC9C,COX4I1,DUSP22,SLC25A10,SPOCK2,DBNDD2,NISCH,TMED4,RPS6,RPL29,CD99L2,RTEL1,COASY,SR

TB44,PLCB2,DUSP22,DBNDD2,NISCH,TMED4,CD99L2,GGPS1,COASY,SRSF1,AUP1,YIPF3,FOXRED1,TRNT1

TB44,PLCB2,DUSP22,DBNDD2,NISCH,TMED4,CD99L2,GGPS1,COASY,SRSF1,AUP1,YIPF3,FOXRED1,TRNT1

OXRED1,RABL5,TRNT1,ITGB1BP1,MED15,SH3YL1,IP6K2,C1ORF198,UGGT2,SLC39A13,YIF1A,EIF3M,CDC4

OXRED1,RABL5,TRNT1,ITGB1BP1,MED15,SH3YL1,IP6K2,C1ORF198,UGGT2,SLC39A13,YIF1A,EIF3M,CDC4

14,DBNDD2,NISCH,CD99L2,GGPS1,SRSF1,AUP1,FOXRED1,TRNT1,KXD1,PARD6G,SH3YL1,IP6K2,CLASRP,JP

ZNF483,SZT2,PSMD8,PEX16,CYB5D2,PSMC5,NGDN,LRP10,ACTB,LAMTOR4,MRPL51,C9ORF64,SETD4,ME

RK3,LZTFL1,ZC3H14,FIS1,SLC15A3,PPM1K,TUBA1C,ANTXR2,STIP1,LRRC1,PHF19,GRAMD1C,VCAN,PCBP2

RK3,LZTFL1,ZC3H14,FIS1,SLC15A3,PPM1K,TUBA1C,ANTXR2,STIP1,LRRC1,PHF19,GRAMD1C,VCAN,PCBP2

,NAA10,MED6,CARHSP1,SSH2,COX17,HNRNPLL,PHF20L1,POLR2C,INTS4,B3GAT3,RFXANK,ZNF506,FTH1,F

,NAA10,MED6,CARHSP1,SSH2,COX17,HNRNPLL,PHF20L1,POLR2C,INTS4,B3GAT3,RFXANK,ZNF506,FTH1,F

25A10,TMX2,RDH11,DBNDD2,SDF4,NISCH,TMED4,RPL29,CD99L2,GGPS1,AUP1,SAP30BP,RABL5,TRNT1,I

25A10,TMX2,RDH11,DBNDD2,SDF4,NISCH,TMED4,RPL29,CD99L2,GGPS1,AUP1,SAP30BP,RABL5,TRNT1,I

G7,WDR62,IL1RAP,STYXL1,RFX5,POLD4,PLEKHA5,RANBP1,UBE2L6,VRK3,ZC3H14,FIS1,SLC15A3,PPM1K,TU

RPL18,COX17,COX5A,HIGD1A,SIRT5,INTS4,B3GAT3,PFDN5,C11ORF49,PAFAH1B2,ZNF343,GBA2,PPIH,DA

IC8A,PARK7,THADA,UBE2N,LFNG,ZNHIT3,NOP2,FAM32A,COX20,PEX2,POLR1C,PIGH,C12ORF57,PEX26,N

PS1,ITPA,VPS25,SNRPD1,DHRS4,IFI27,SMIM12,TATDN1,ZBTB8OS,TBC1D22A,APAF1,U2AF1L4,TAF9,RNPS

A1,YLPM1,PHF21A,CHCHD1,FUZ,FAM3A,METTL17,EPB41L5,ATG12,PPRC1,ZBTB44,COX4I1,DUSP22,SLC2

A1,YLPM1,PHF21A,CHCHD1,FUZ,FAM3A,METTL17,EPB41L5,ATG12,PPRC1,ZBTB44,COX4I1,DUSP22,SLC2

K6,CDC37,RPS16,CPSF3L,OBSL1,PTPN13,UBR4,YLPM1,CHCHD1,FUZ,RGS3,MUC1,METTL17,ECSIT,EPB41L

K6,CDC37,RPS16,CPSF3L,OBSL1,PTPN13,UBR4,YLPM1,CHCHD1,FUZ,RGS3,MUC1,METTL17,ECSIT,EPB41L

PLEKHA5,RANBP1,UBE2L6,VRK3,LZTFL1,ZC3H14,FIS1,SLC15A3,PPM1K,TUBA1C,IRF9,DST,ANTXR2,STIP1,L

FOXRED1,TRNT1,KXD1,ITGB1BP1,PARD6G,SH3YL1,IP6K2,CLASRP,JPH2,C1ORF198,MICAL1,HS2ST1,UBE2I

FOXRED1,TRNT1,KXD1,ITGB1BP1,PARD6G,SH3YL1,IP6K2,CLASRP,JPH2,C1ORF198,MICAL1,HS2ST1,UBE2I

LDHA,PSMA6,ACOT8,AP3B2,SLC35G1,RCE1,MRPL33,ZNF410,RUVBL2,PCMT1,EBPL,TESC,IGSF8,NUDT5,R

LDHA,PSMA6,ACOT8,AP3B2,SLC35G1,RCE1,MRPL33,ZNF410,RUVBL2,PCMT1,EBPL,TESC,IGSF8,NUDT5,R

A2,RPS16,CPSF3L,TSG101,OBSL1,TTC21B,PTPN13,UBR4,PSMA1,YLPM1,PHF21A,CHCHD1,SLC39A11,FAM

2,SON,TET2,UQCRQ,GRK6,HMGA2,CDC37,RPS16,CPSF3L,TSG101,OBSL1,TTC21B,UBR4,PSMA1,YLPM1,PH

KXD1,ITGB1BP1,MED15,SH3YL1,IP6K2,CLASRP,C1ORF198,UGGT2,MICAL1,HS2ST1,UBE2I,SLC39A13,YIF1

KXD1,ITGB1BP1,MED15,SH3YL1,IP6K2,CLASRP,C1ORF198,UGGT2,MICAL1,HS2ST1,UBE2I,SLC39A13,YIF1

,RNF181,TPT1,POLR2M,SLC52A2,ANAPC13,TSPAN31,RALY,GLYCTK,C11ORF57,RER1,EIF2A,NKIRAS2,REC

F512,RNF181,TJP1,CHTOP,TPT1,NTMT1,MTIF2,RALY,GLYCTK,CFLAR,VDAC1,RER1,EIF2A,NKIRAS2,PSMD7,

ICAL1,UBE2I,SLC39A13,YIF1A,PRMT2,EIF3M,JUP,CDC42,XBP1,PTK2,AUNIP,GUK1,CENPN,CARD14,PCYT2,

,ECSIT,ATG12,RPS25,ZBTB44,NIP7,TTC9C,COX4I1,SLC25A10,TMX2,SPOCK2,THEM4,TMED4,RPS6,RPL29,

,ECSIT,ATG12,RPS25,ZBTB44,NIP7,TTC9C,COX4I1,SLC25A10,TMX2,SPOCK2,THEM4,TMED4,RPS6,RPL29,

PN1,HPS1,ITPA,EML3,ZCCHC9,VPS25,SNRPD1,UCHL5,MKRN1,CHURC1,AZI2,GOSR2,MYL12A,TBC1D22A,

ST1,UBE2I,PRMT2,EIF3M,TBX6,TMEM205,XBP1,PTK2,AUNIP,C14ORF2,GUK1,PCYT2,FBXW9,USE1,AGPA

ST1,UBE2I,PRMT2,EIF3M,TBX6,TMEM205,XBP1,PTK2,AUNIP,C14ORF2,GUK1,PCYT2,FBXW9,USE1,AGPA

ST1,UBE2I,PRMT2,EIF3M,TBX6,TMEM205,XBP1,PTK2,AUNIP,C14ORF2,GUK1,PCYT2,FBXW9,USE1,AGPA

ST1,UBE2I,PRMT2,EIF3M,TBX6,TMEM205,XBP1,PTK2,AUNIP,C14ORF2,GUK1,PCYT2,FBXW9,USE1,AGPA

BA1C,ANTXR2,STIP1,LRRC1,PHF19,LPCAT3,CD44,TM9SF1,MAGED2,SON,TET2,UQCRQ,GRK6,CDC37,RPS1

BA1C,ANTXR2,STIP1,LRRC1,PHF19,LPCAT3,CD44,TM9SF1,MAGED2,SON,TET2,UQCRQ,GRK6,CDC37,RPS1

1C,ANTXR2,STIP1,LRRC1,PHF19,GRAMD1C,VCAN,LPCAT3,CD44,TM9SF1,SON,TET2,UQCRQ,GRK6,HMGA2

ZNF410,RUVBL2,PCMT1,EBPL,TESC,IGSF8,NUDT5,RBPMS,NNMT,SNRPN,SPAG7,WDR62,IL1RAP,STYXL1,F

ZNF410,RUVBL2,PCMT1,EBPL,TESC,IGSF8,NUDT5,RBPMS,NNMT,SNRPN,SPAG7,WDR62,IL1RAP,STYXL1,F

KHA5,RANBP1,UBE2L6,VRK3,LZTFL1,ZC3H14,FIS1,SLC15A3,PPM1K,TUBA1C,IRF9,ANTXR2,STIP1,LRRC1,PH

S16,CPSF3L,PTPN13,UBR4,PSMA1,YLPM1,CHCHD1,FAM3A,PSMA3,METTL17,ECSIT,ATG12,PPRC1,NIP7,S

A5,UBE2L6,VRK3,LZTFL1,ZC3H14,FIS1,SLC15A3,PPM1K,TUBA1C,ANTXR2,STIP1,LRRC1,PHF19,GRAMD1C,

A5,UBE2L6,VRK3,LZTFL1,ZC3H14,FIS1,SLC15A3,PPM1K,TUBA1C,ANTXR2,STIP1,LRRC1,PHF19,GRAMD1C,

D1,FUZ,RGS3,FAM3A,PSMA3,ECSIT,ATG12,PPRC1,RPS25,ZBTB44,NIP7,TTC9C,DUSP22,SLC25A10,TMX2,S

D1,FUZ,RGS3,FAM3A,PSMA3,ECSIT,ATG12,PPRC1,RPS25,ZBTB44,NIP7,TTC9C,DUSP22,SLC25A10,TMX2,S

TBX6,TMEM205,PTK2,AUNIP,C14ORF2,GUK1,CENPN,ATF7,PCYT2,USE1,PPP1CC,PPA1,DGKE,TNFRSF14,C

TBX6,TMEM205,PTK2,AUNIP,C14ORF2,GUK1,CENPN,ATF7,PCYT2,USE1,PPP1CC,PPA1,DGKE,TNFRSF14,C

C,VCAN,LPCAT3,CD44,TM9SF1,SON,TET2,UQCRQ,SNF8,GRK6,HMGA2,CDC37,RPS16,CPSF3L,PTPN13,UB

S2ST1,UBE2I,PRMT2,EIF3M,JUP,TBX6,TMEM205,PTK2,AUNIP,C14ORF2,GUK1,ATF7,PCYT2,USE1,PPP1CC

S2ST1,UBE2I,PRMT2,EIF3M,JUP,TBX6,TMEM205,PTK2,AUNIP,C14ORF2,GUK1,ATF7,PCYT2,USE1,PPP1CC
,EIF2A,ACAA1,PSMD7,ZNF511,ZNF283,GTF2H1,PIGO,CLTA,SLC35A2,AXIN1,TP53,TMEM199,SLC25A24,C
AMD1C,VCAN,LPCAT3,TM9SF1,SON,UQCRQ,SNF8,GRK6,HMGA2,CDC37,RPS16,CPSF3L,TSG101,OBSL1,P
TTLL5,LDHA,PSMA6,ACOT8,AP3B2,RPS9,SLC35G1,RCE1,MRPL33,ZNF410,RUVBL2,PCMT1,EBPL,TESC,IGS
TTLL5,LDHA,PSMA6,ACOT8,AP3B2,RPS9,SLC35G1,RCE1,MRPL33,ZNF410,RUVBL2,PCMT1,EBPL,TESC,IGS
R4,PSMA1,YLPM1,PHF21A,CHCHD1,SLC39A11,FUZ,FAM3A,MUC1,METTL17,EPB41L5,ATG12,PPRC1,RPS
R4,PSMA1,YLPM1,PHF21A,CHCHD1,SLC39A11,FUZ,FAM3A,MUC1,METTL17,EPB41L5,ATG12,PPRC1,RPS
,MPV17,ARIH2,RPL8,PICK1,ZNF512,RNF181,TSPAN31,FAM192A,RALY,ZNF593,GLYCTK,TRIM46,C11ORF5
ED4,CD99L2,GGPS1,SRSF1,AUP1,YIPF3,TRNT1,KXD1,PARD6G,SH3YL1,IP6K2,CLASRP,JPH2,C1ORF198,MI
ED4,CD99L2,GGPS1,SRSF1,AUP1,YIPF3,TRNT1,KXD1,PARD6G,SH3YL1,IP6K2,CLASRP,JPH2,C1ORF198,MI
6,CPSF3L,OBSL1,TTC21B,PTPN13,UBR4,PSMA1,YLPM1,PHF21A,SLC39A11,FUZ,RGS3,FAM3A,MUC1,MET
6,CPSF3L,OBSL1,TTC21B,PTPN13,UBR4,PSMA1,YLPM1,PHF21A,SLC39A11,FUZ,RGS3,FAM3A,MUC1,MET
VBL2,PCMT1,EBPL,TESC,IGSF8,NUDT5,RBPMS,NNMT,SPAG7,WDR62,IL1RAP,STYXL1,POLD4,PLEKHA5,R
VBL2,PCMT1,EBPL,TESC,IGSF8,NUDT5,RBPMS,NNMT,SPAG7,WDR62,IL1RAP,STYXL1,POLD4,PLEKHA5,R
PTPN13,PSMA1,YLPM1,PHF21A,FAM3A,ECSIT,EPB41L5,ATG12,PPRC1,RPS25,ZBTB44,NIP7,COX4I1,DUSP
0BP,YIPF3,FOXRED1,KXD1,PARD6G,SH3YL1,IP6K2,CLASRP,C1ORF198,MICAL1,HS2ST1,YIF1A,PRMT2,CDC
0BP,YIPF3,FOXRED1,KXD1,PARD6G,SH3YL1,IP6K2,CLASRP,C1ORF198,MICAL1,HS2ST1,YIF1A,PRMT2,CDC
0BP,YIPF3,FOXRED1,KXD1,PARD6G,SH3YL1,IP6K2,CLASRP,C1ORF198,MICAL1,HS2ST1,YIF1A,PRMT2,CDC
0BP,YIPF3,FOXRED1,KXD1,PARD6G,SH3YL1,IP6K2,CLASRP,C1ORF198,MICAL1,HS2ST1,YIF1A,PRMT2,CDC
0BP,YIPF3,FOXRED1,KXD1,PARD6G,SH3YL1,IP6K2,CLASRP,C1ORF198,MICAL1,HS2ST1,YIF1A,PRMT2,CDC
0BP,YIPF3,FOXRED1,KXD1,PARD6G,SH3YL1,IP6K2,CLASRP,C1ORF198,MICAL1,HS2ST1,YIF1A,PRMT2,CDC
,EIF3M,TBX6,TMEM205,XBP1,PTK2,AUNIP,C14ORF2,GUK1,PCYT2,USE1,AGPAT4,PPP1CC,EDF1,RFX3,PP
,EIF3M,TBX6,TMEM205,XBP1,PTK2,AUNIP,C14ORF2,GUK1,PCYT2,USE1,AGPAT4,PPP1CC,EDF1,RFX3,PP
,SAP30BP,YIPF3,FOXRED1,RABL5,TRNT1,ITGB1BP1,MED15,CLASRP,C1ORF198,UGGT2,UBE2I,SLC39A13,
,SAP30BP,YIPF3,FOXRED1,RABL5,TRNT1,ITGB1BP1,MED15,CLASRP,C1ORF198,UGGT2,UBE2I,SLC39A13,
,TIMM17B,APEH,RBM33,LYPLA2,NUTF2,CAMLG,C17ORF62,STOML2,C19ORF54,ZNF7,TXNDC17,PPAN,A
,TIMM17B,APEH,RBM33,LYPLA2,NUTF2,CAMLG,C17ORF62,STOML2,C19ORF54,ZNF7,TXNDC17,PPAN,A
IISCH,TMED4,RPS6,CD99L2,GGPS1,RTEL1,COASY,AUP1,FOXRED1,TRNT1,KXD1,ITGB1BP1,MED15,PARD6
IISCH,TMED4,RPS6,CD99L2,GGPS1,RTEL1,COASY,AUP1,FOXRED1,TRNT1,KXD1,ITGB1BP1,MED15,PARD6
EIF2A,NKIRAS2,ACAA1,GTF2H1,BFAR,SLC35A2,SH3GL1,BTF3,TMEM199,SLC25A24,ZNF396,EPN1,ITPA,ZC
EIF2A,NKIRAS2,ACAA1,GTF2H1,BFAR,SLC35A2,SH3GL1,BTF3,TMEM199,SLC25A24,ZNF396,EPN1,ITPA,ZC
9L2,GGPS1,SRSF1,AUP1,FOXRED1,TRNT1,KXD1,PARD6G,SH3YL1,IP6K2,CLASRP,JPH2,C1ORF198,MICAL1,
9L2,GGPS1,SRSF1,AUP1,FOXRED1,TRNT1,KXD1,PARD6G,SH3YL1,IP6K2,CLASRP,JPH2,C1ORF198,MICAL1,
MPR2,ZSCAN9,TCP1,PIGT,UBE2N,NFATC4,LFNG,PDCD6,ZNF81,FAM32A,ZSCAN20,TMEM14B,PCID2,TGC
MPR2,ZSCAN9,TCP1,PIGT,UBE2N,NFATC4,LFNG,PDCD6,ZNF81,FAM32A,ZSCAN20,TMEM14B,PCID2,TGC
ED2,SON,UQCRQ,SNF8,GRK6,HMGA2,CDC37,RPS16,CPSF3L,TSG101,OBSL1,PTPN13,UBR4,YLPM1,PHF21
ED2,SON,UQCRQ,SNF8,GRK6,HMGA2,CDC37,RPS16,CPSF3L,TSG101,OBSL1,PTPN13,UBR4,YLPM1,PHF21
CTD,UBXN1,ERLEC1,HSPB1,HHLA3,SAT1,ERCC5,MCTS1,SERPINB1,DDOST,LSR,MEIS2,RRNAD1,NAT9,TO
TTL17,ECSIT,PPRC1,RPS25,NIP7,TTC9C,COX4I1,DUSP22,SLC25A10,SPOCK2,TMED4,CENPK,CD99L2,GGPS
TTL17,ECSIT,PPRC1,RPS25,NIP7,TTC9C,COX4I1,DUSP22,SLC25A10,SPOCK2,TMED4,CENPK,CD99L2,GGPS
BE2I,PRMT2,EIF3M,JUP,TBX6,TMEM205,XBP1,PTK2,AUNIP,C14ORF2,GUK1,CENPN,PCYT2,USE1,HSBP1L
BE2I,PRMT2,EIF3M,JUP,TBX6,TMEM205,XBP1,PTK2,AUNIP,C14ORF2,GUK1,CENPN,PCYT2,USE1,HSBP1L
T1,ITGB1BP1,PARD6G,CLASRP,UGGT2,MICAL1,UBE2I,PRMT2,EIF3M,JUP,CDC42,TBX6,TMEM205,PTK2,
UNIP,C14ORF2,GUK1,CENPN,FBXW9,HSBP1L1,MAF,ASF1B,PPP1CC,EDF1,GADL1,RFX3,DGKE,ENDOV,CA
UNIP,C14ORF2,GUK1,CENPN,FBXW9,HSBP1L1,MAF,ASF1B,PPP1CC,EDF1,GADL1,RFX3,DGKE,ENDOV,CA
ANTXR2,STIP1,LRRC1,GRAMD1C,VCAN,PCBP2,LPCAT3,CD44,TM9SF1,MAGED2,SON,TET2,UQCRQ,SNF8,G
ANTXR2,STIP1,LRRC1,GRAMD1C,VCAN,PCBP2,LPCAT3,CD44,TM9SF1,MAGED2,SON,TET2,UQCRQ,SNF8,G

25A24,ZNF396,EPN1,CREB3L2,ITPA,ZCCHC9,VPS25,SNRPD1,AKT1S1,AIFM2,CHURC1,SNRPA1,HMBS,AZI2

25A24,ZNF396,EPN1,CREB3L2,ITPA,ZCCHC9,VPS25,SNRPD1,AKT1S1,AIFM2,CHURC1,SNRPA1,HMBS,AZI2

L6,LZTFL1,ZC3H14,FIS1,TUBA1C,ANTXR2,STIP1,GRAMD1C,VCAN,LPCAT3,CD44,TM9SF1,MAGED2,SON,TE

L6,LZTFL1,ZC3H14,FIS1,TUBA1C,ANTXR2,STIP1,GRAMD1C,VCAN,LPCAT3,CD44,TM9SF1,MAGED2,SON,TE

GRAMD1C,VCAN,LPCAT3,CD44,MAGED2,SON,TET2,UQCRQ,GRK6,CDC37,RPS16,CPSF3L,OBSL1,TTC21B,P

GRAMD1C,VCAN,LPCAT3,CD44,MAGED2,SON,TET2,UQCRQ,GRK6,CDC37,RPS16,CPSF3L,OBSL1,TTC21B,P

LC15A3,TUBA1C,DST,STIP1,LRRC1,PHF19,VCAN,PCBP2,LPCAT3,CD44,TM9SF1,MAGED2,SON,UQCRQ,SN

LC15A3,TUBA1C,DST,STIP1,LRRC1,PHF19,VCAN,PCBP2,LPCAT3,CD44,TM9SF1,MAGED2,SON,UQCRQ,SN

KS,STX10,C12ORF57,DCTN5,OXLD1,ZNF223,OGFOD2,METTL3,EIF3G,PICK1,FBXL18,SLC52A2,NTMT1,TSP

KS,STX10,C12ORF57,DCTN5,OXLD1,ZNF223,OGFOD2,METTL3,EIF3G,PICK1,FBXL18,SLC52A2,NTMT1,TSP

MA3,METTL17,EPB41L5,PPRC1,RPS25,ZBTB44,COX4I1,DUSP22,SPOCK2,RDH11,DBNDD2,NISCH,TMED4,

MA3,METTL17,EPB41L5,PPRC1,RPS25,ZBTB44,COX4I1,DUSP22,SPOCK2,RDH11,DBNDD2,NISCH,TMED4,

C39A13,YIF1A,PRMT2,TBX6,XBP1,PTK2,AUNIP,C14ORF2,GUK1,PCYT2,USE1,AGPAT4,MAF,PPP1CC,EDF1,

C39A13,YIF1A,PRMT2,TBX6,XBP1,PTK2,AUNIP,C14ORF2,GUK1,PCYT2,USE1,AGPAT4,MAF,PPP1CC,EDF1,

EM126B,APEH,HYI,CAPN10,PODXL,ATPAF2,LSM12,C17ORF62,PRMT1,CASC4,PPAN,ZFAND2B,CYB561,SH

EM126B,APEH,HYI,CAPN10,PODXL,ATPAF2,LSM12,C17ORF62,PRMT1,CASC4,PPAN,ZFAND2B,CYB561,SH

4,ERLEC1,HSPB1,MVD,ERCC5,EMC4,DDOST,LSR,PABPC1,RRNAD1,DBNL,RBP1,RAD1,TOMM40,TADA3,PD

4,ERLEC1,HSPB1,MVD,ERCC5,EMC4,DDOST,LSR,PABPC1,RRNAD1,DBNL,RBP1,RAD1,TOMM40,TADA3,PD

05,XBP1,PTK2,AUNIP,C14ORF2,GUK1,PCYT2,USE1,AGPAT4,DEF8,PPP1CC,EDF1,PPA1,DGKE,VAMP8,CAM

05,XBP1,PTK2,AUNIP,C14ORF2,GUK1,PCYT2,USE1,AGPAT4,DEF8,PPP1CC,EDF1,PPA1,DGKE,VAMP8,CAM

SMD4,LIG1,PSMA7,TMEM126B,XAF1,APEH,HYI,TXNL4A,NUTF2,TCF4,UCK2,TMPO,ATPAF2,LSM12,C17OR

SMD4,LIG1,PSMA7,TMEM126B,XAF1,APEH,HYI,TXNL4A,NUTF2,TCF4,UCK2,TMPO,ATPAF2,LSM12,C17OR

HD1,SLC39A11,FAM3A,MUC1,METTL17,ECSIT,EPB41L5,ATG12,PPRC1,RPS25,NLRP1,ZBTB44,COX4I1,PLC

HD1,SLC39A11,FAM3A,MUC1,METTL17,ECSIT,EPB41L5,ATG12,PPRC1,RPS25,NLRP1,ZBTB44,COX4I1,PLC

L,AIFM2,CHURC1,SNRPA1,HMBS,AZI2,KAT8,U2AF1L4,EMC3,PRKCSH,ZNF276,COMMD1,ZNF483,SZT2,PS

L,AIFM2,CHURC1,SNRPA1,HMBS,AZI2,KAT8,U2AF1L4,EMC3,PRKCSH,ZNF276,COMMD1,ZNF483,SZT2,PS

ZFAND2B,AP1S3,ACAD10,HNRNPC,SHARPIN,SLC9B2,SUGT1,HYPK,CARS,ZNF250,RPL36,RBM3,MDH1,ZN

F49,PAFAH1B2,GTF2IRD2,ARHGEF1,DALRD3,NSUN5,ZFPL1,MRPS18C,GMPR2,TCP1,COMMD4,PIGT,UBE2

CAPN10,NUTF2,PODXL,CAMLG,ATPAF2,C17ORF62,PRMT1,CASC4,WNK1,ZNF7,TXNDC17,ZFAND2B,HNRN

N5,MTHFS,PAFAH1B2,SNRPF,GBA2,DALRD3,NSUN5,MFSD10,ZSCAN9,RIC8A,PARK7,SMARCC2,MROH1,U

N5,MTHFS,PAFAH1B2,SNRPF,GBA2,DALRD3,NSUN5,MFSD10,ZSCAN9,RIC8A,PARK7,SMARCC2,MROH1,U

HMGA2,CDC37,OBSL1,TTC21B,UBR4,CHCHD1,SLC39A11,FUZ,RGS3,FAM3A,PSMA3,MUC1,METTL17,ECSI

HMGA2,CDC37,OBSL1,TTC21B,UBR4,CHCHD1,SLC39A11,FUZ,RGS3,FAM3A,PSMA3,MUC1,METTL17,ECSI

2L6,VRK3,LZTFL1,ZC3H14,FIS1,SLC15A3,PPM1K,TUBA1C,IRF9,ANTXR2,STIP1,LRRC1,PHF19,GRAMD1C,V

2L6,VRK3,LZTFL1,ZC3H14,FIS1,SLC15A3,PPM1K,TUBA1C,IRF9,ANTXR2,STIP1,LRRC1,PHF19,GRAMD1C,V

,TTC21B,PTPN13,UBR4,PSMA1,YLPM1,CHCHD1,FUZ,FAM3A,PSMA3,METTL17,PPRC1,RPS25,NIP7,TTC9C

,TTC21B,PTPN13,UBR4,PSMA1,YLPM1,CHCHD1,FUZ,FAM3A,PSMA3,METTL17,PPRC1,RPS25,NIP7,TTC9C

UBE2I,SLC39A13,YIF1A,EIF3M,CDC42,TBX6,CELF1,CARD14,PCYT2,FBXW9,USE1,AGPAT4,ASF1B,CCDC25,

UBE2I,SLC39A13,YIF1A,EIF3M,CDC42,TBX6,CELF1,CARD14,PCYT2,FBXW9,USE1,AGPAT4,ASF1B,CCDC25,

XL,ATPAF2,LSM12,C17ORF62,CASC4,C19ORF54,ZNF7,PPAN,ZFAND2B,CYB561,SHARPIN,STAU1,ZNF250,R

XL,ATPAF2,LSM12,C17ORF62,CASC4,C19ORF54,ZNF7,PPAN,ZFAND2B,CYB561,SHARPIN,STAU1,ZNF250,R

C9,VPS25,SNRPD1,AIFM2,CHURC1,ARRB2,SNRPA1,HMBS,AZI2,APAF1,KAT8,U2AF1L4,EMC3,PRKCSH,UBA

C9,VPS25,SNRPD1,AIFM2,CHURC1,ARRB2,SNRPA1,HMBS,AZI2,APAF1,KAT8,U2AF1L4,EMC3,PRKCSH,UBA

E2L6,LZTFL1,FIS1,PPM1K,TUBA1C,DST,ANTXR2,STIP1,LRRC1,GRAMD1C,VCAN,PCBP2,LPCAT3,CD44,TM9

E2L6,LZTFL1,FIS1,PPM1K,TUBA1C,DST,ANTXR2,STIP1,LRRC1,GRAMD1C,VCAN,PCBP2,LPCAT3,CD44,TM9

DIP,MDH2,STXBP2,HSPH1,OAS3,SEC31A,CNST,COX17,COX5A,HINT1,HIGD1A,B3GAT3,ZNF506,SPINT2,PFD

DIP,MDH2,STXBP2,HSPH1,OAS3,SEC31A,CNST,COX17,COX5A,HINT1,HIGD1A,B3GAT3,ZNF506,SPINT2,PFD

NGRF,MTA3,CHCHD7,HMGA1,PSME2,PABPC1,NAT9,BCAM,ZNF836,BLVRB,TOMM40,PPP3CB,RABEP2,T
NGRF,MTA3,CHCHD7,HMGA1,PSME2,PABPC1,NAT9,BCAM,ZNF836,BLVRB,TOMM40,PPP3CB,RABEP2,T
A,PPAN,ZFAND2B,STAU1,ZNF250,RPL36,NFAM1,LAMP2,CCNL2,HSP90AB1,SYNGR2,CEP70,TMEM134,L
A,PPAN,ZFAND2B,STAU1,ZNF250,RPL36,NFAM1,LAMP2,CCNL2,HSP90AB1,SYNGR2,CEP70,TMEM134,L
11,ZNHIT3,PDCD6,NOP2,FAM32A,COA1,USP40,ELOF1,C12ORF57,DCTN5,PEX26,NDUFS3,OXLD1,ZNF223,
A2,NUTF2,PODXL,LSM12,C17ORF62,STOML2,C19ORF54,ZNF7,ARL6IP4,PPAN,CYB561,CBFA2T2,STAU1,S
A2,NUTF2,PODXL,LSM12,C17ORF62,STOML2,C19ORF54,ZNF7,ARL6IP4,PPAN,CYB561,CBFA2T2,STAU1,S
RLEC1,HSPB1,LMBR1,MCTS1,NAP1L1,PRKRIP1,RANGRF,HMGA1,PSME2,PABPC1,NAT9,ZNF836,RBP1,BL
RLEC1,HSPB1,LMBR1,MCTS1,NAP1L1,PRKRIP1,RANGRF,HMGA1,PSME2,PABPC1,NAT9,ZNF836,RBP1,BL
,OBSL1,TTC21B,PTPN13,UBR4,YLPM1,PHF21A,CHCHD1,RGS3,METTL17,ECSIT,EPB41L5,ATG12,PPRC1,RP
,RPS16,CPSF3L,OBSL1,PTPN13,UBR4,PSMA1,YLPM1,PHF21A,FAM3A,ECSIT,EPB41L5,ATG12,PPRC1,RPS2
,RPS16,CPSF3L,OBSL1,PTPN13,UBR4,PSMA1,YLPM1,PHF21A,FAM3A,ECSIT,EPB41L5,ATG12,PPRC1,RPS2
M,TRMT112,CHPF,ZNF625,RPS20,GABPB1,DCTD,ZNF814,HSPB1,ERCC5,EMC4,PRKRIP1,DDOST,LSR,DOCK
M,TRMT112,CHPF,ZNF625,RPS20,GABPB1,DCTD,ZNF814,HSPB1,ERCC5,EMC4,PRKRIP1,DDOST,LSR,DOCK
6,RTEL1,COASY,SRSF1,AUP1,SAP30BP,FOXRED1,TRNT1,ITGB1BP1,PARD6G,SH3YL1,CLASRP,JPH2,C1ORF
6,RTEL1,COASY,SRSF1,AUP1,SAP30BP,FOXRED1,TRNT1,ITGB1BP1,PARD6G,SH3YL1,CLASRP,JPH2,C1ORF
,VCAN,LPCAT3,TM9SF1,MAGED2,SON,TET2,UQCRQ,SNF8,GRK6,HMGA2,CDC37,RPS16,CPSF3L,OBSL1,TT
,VCAN,LPCAT3,TM9SF1,MAGED2,SON,TET2,UQCRQ,SNF8,GRK6,HMGA2,CDC37,RPS16,CPSF3L,OBSL1,TT
,METTL17,ECSIT,ATG12,PPRC1,ZBTB44,NIP7,DUSP22,SLC25A10,TMX2,SPOCK2,THEM4,SDF4,NISCH,TME
DC25,EEF1B2,RFX3,ENDOV,VAMP8,CAPG,CRIP1,PPIL2,TROAP,LIG1,CDC14B,APEH,RBM33,LYPLA2,PODX
DC25,EEF1B2,RFX3,ENDOV,VAMP8,CAPG,CRIP1,PPIL2,TROAP,LIG1,CDC14B,APEH,RBM33,LYPLA2,PODX
IN,CBFA2T2,STAU1,SUGT1,HYPK,CARS,ZNF250,RPL36,CDK11A,ZNF197,INO80E,DGUOK,HSP90AB1,SYNG
IN,CBFA2T2,STAU1,SUGT1,HYPK,CARS,ZNF250,RPL36,CDK11A,ZNF197,INO80E,DGUOK,HSP90AB1,SYNG
TIP1,LRRC1,PHF19,LPCAT3,CD44,MAGED2,SON,TET2,UQCRQ,SNF8,GRK6,HMGA2,CDC37,RPS16,CPSF3L,
DXL,ATPAF2,C17ORF62,PRMT1,CASC4,WNK1,ZNF7,TXNDC17,ZFAND2B,HNRNPC,CYB561,SHARPIN,STAU
6,CDC37,RPS16,TSG101,OBSL1,TTC21B,PTPN13,UBR4,PSMA1,CHCHD1,SLC39A11,FUZ,METTL17,EPB41L

,CAMLG,ATPAF2,C17ORF62,PRMT1,CASC4,WNK1,ZNF7,ZFAND2B,AP1S3,HNRNPC,CYB561,ZNF250,RPL3
,CAMLG,ATPAF2,C17ORF62,PRMT1,CASC4,WNK1,ZNF7,ZFAND2B,AP1S3,HNRNPC,CYB561,ZNF250,RPL3
,RBPMS,WDR62,STYXL1,POLD4,PLEKHA5,RANBP1,UBE2L6,VRK3,LZTFL1,ZC3H14,FIS1,SLC15A3,PPM1K,T
,RBPMS,WDR62,STYXL1,POLD4,PLEKHA5,RANBP1,UBE2L6,VRK3,LZTFL1,ZC3H14,FIS1,SLC15A3,PPM1K,T
MT,SNRPN,SPAG7,WDR62,IL1RAP,STYXL1,RFX5,POLD4,PLEKHA5,RANBP1,UBE2L6,VRK3,LZTFL1,ZC3H14,
MT,SNRPN,SPAG7,WDR62,IL1RAP,STYXL1,RFX5,POLD4,PLEKHA5,RANBP1,UBE2L6,VRK3,LZTFL1,ZC3H14,
P,STYXL1,RFX5,POLD4,PLEKHA5,RANBP1,UBE2L6,VRK3,LZTFL1,ZC3H14,FIS1,SLC15A3,PPM1K,TUBA1C,IR
GT1,ZNF250,RPL36,CDK11A,MDH1,COPZ1,INO80E,CCNL2,MAPKAPK5,CEP70,RAB2A,ZNF524,ELP6,TPM2
BNDD2,CD99L2,RTEL1,SRSF1,AUP1,SAP30BP,YIPF3,FOXRED1,ITGB1BP1,PARD6G,SH3YL1,CLASRP,JPH2,U
G,ATPAF2,C17ORF62,PRMT1,CASC4,ZNF7,TXNDC17,ZFAND2B,HNRNPC,CYB561,SHARPIN,ZNF644,STAU1
B1,MVD,SEMA3C,LMBR1,SERPINB1,CHCHD7,PHTF2,PABPC1,NAT9,BCAM,ZNF836,BLVRB,TOMM40,RABE
,TMEM205,PTK2,AUNIP,C14ORF2,GUK1,CENPN,PCYT2,USE1,AGPAT4,DEF8,PPP1CC,EDF1,PPA1,DGKE,C
,TMEM205,PTK2,AUNIP,C14ORF2,GUK1,CENPN,PCYT2,USE1,AGPAT4,DEF8,PPP1CC,EDF1,PPA1,DGKE,C
XDC,PEX2,ELOF1,DCTN5,PEX26,BTN3A2,OXLD1,EIF4A1,C6ORF132,EIF3G,ARIH2,PICK1,ADD1,CHTOP,TSP
410,RUVBL2,PCMT1,EBPL,TESC,IGSF8,NUDT5,NNMT,SNRPN,SPAG7,WDR62,IL1RAP,STYXL1,RFX5,POLD4,
410,RUVBL2,PCMT1,EBPL,TESC,IGSF8,NUDT5,NNMT,SNRPN,SPAG7,WDR62,IL1RAP,STYXL1,RFX5,POLD4,
N,TET2,GRK6,HMGA2,CDC37,RPS16,CPSF3L,OBSL1,TTC21B,PTPN13,UBR4,YLPM1,PHF21A,CHCHD1,SLC3
N,TET2,GRK6,HMGA2,CDC37,RPS16,CPSF3L,OBSL1,TTC21B,PTPN13,UBR4,YLPM1,PHF21A,CHCHD1,SLC3
BP1,CLASRP,MICAL1,UBE2I,SLC39A13,YIF1A,EIF3M,XBP1,PTK2,AUNIP,GUK1,FBXW9,HSBP1L1,IFI27L1,M
BP1,CLASRP,MICAL1,UBE2I,SLC39A13,YIF1A,EIF3M,XBP1,PTK2,AUNIP,GUK1,FBXW9,HSBP1L1,IFI27L1,M

,TKT,DENND1A,EPN1,ZCCHC9,GRAMD1A,MKRN1,DHRS4,MLST8,HMBS,MFGE8,AZI2,GOSR2,SGSM3,LAM
7B,APEH,NUTF2,PPOX,LSM12,CASC4,WNK1,C19ORF54,STRADA,ZFAND2B,CYB561,SHARPIN,STAU1,UPP1
7B,APEH,NUTF2,PPOX,LSM12,CASC4,WNK1,C19ORF54,STRADA,ZFAND2B,CYB561,SHARPIN,STAU1,UPP1

.,AUP1,SAP30BP,YIPF3,FOXRED1,RABL5,TRNT1,KXD1,ITGB1BP1,MED15,SH3YL1,IP6K2,CLASRP,C1ORF19
.,AUP1,SAP30BP,YIPF3,FOXRED1,RABL5,TRNT1,KXD1,ITGB1BP1,MED15,SH3YL1,IP6K2,CLASRP,C1ORF19

9,CD99L2,COASY,SRSF1,AUP1,SAP30BP,YIPF3,FOXRED1,TRNT1,KXD1,ITGB1BP1,MED15,CLASRP,UGGT2,
9,CD99L2,COASY,SRSF1,AUP1,SAP30BP,YIPF3,FOXRED1,TRNT1,KXD1,ITGB1BP1,MED15,CLASRP,UGGT2,
3,NUDT5,RBPMS,SNRPN,SPAG7,WDR62,IL1RAP,STYXL1,RFX5,POLD4,PLEKHA5,RANBP1,UBE2L6,VRK3,ZC
1,PHF19,LPCAT3,CD44,TM9SF1,MAGED2,SON,TET2,UQCRQ,HMGA2,CDC37,RPS16,CPSF3L,OBSL1,TTC21
1,PHF19,LPCAT3,CD44,TM9SF1,MAGED2,SON,TET2,UQCRQ,HMGA2,CDC37,RPS16,CPSF3L,OBSL1,TTC21
.36,MOB1A,NFAM1,CDK11A,LAMP2,DUT,SYNGR2,TMEM134,LRRFIP1,RAB2A,ZNF524,ELP6,TPM2,GPBP1
PG,CRIP1,PPIL2,PSMD4,LIG1,PSMA7,TMEM126B,XAF1,APEH,HYI,RPL21,TXNL4A,CAMLG,TCF4,UCK2,TMP
PG,CRIP1,PPIL2,PSMD4,LIG1,PSMA7,TMEM126B,XAF1,APEH,HYI,RPL21,TXNL4A,CAMLG,TCF4,UCK2,TMP
B,TMEM126B,APEH,HYI,CAPN10,RBM33,LYPLA2,PODXL,ATPAF2,LSM12,C17ORF62,PRMT1,CASC4,WNK1
B,TMEM126B,APEH,HYI,CAPN10,RBM33,LYPLA2,PODXL,ATPAF2,LSM12,C17ORF62,PRMT1,CASC4,WNK1
SME2,PHTF2,PABPC1,NAT9,BCAM,BLVRB,RABEP2,ZNF20,SLC33A1,RALGPS1,PDCD5,EPS15L1,POLR3H,AB
SME2,PHTF2,PABPC1,NAT9,BCAM,BLVRB,RABEP2,ZNF20,SLC33A1,RALGPS1,PDCD5,EPS15L1,POLR3H,AB
MT1,EBPL,TESC,IGSF8,NUDT5,RBPMS,NNMT,SNRPN,SPAG7,WDR62,IL1RAP,STYXL1,RFX5,POLD4,PLEKHA
MT1,EBPL,TESC,IGSF8,NUDT5,RBPMS,NNMT,SNRPN,SPAG7,WDR62,IL1RAP,STYXL1,RFX5,POLD4,PLEKHA

PL28,VIM,LIG1,CDC14B,APEH,RPL21,CAPN10,RBM33,LYPLA2,EPB41L4B,UCK2,LSM12,CASC4,STOML2,C1
PL28,VIM,LIG1,CDC14B,APEH,RPL21,CAPN10,RBM33,LYPLA2,EPB41L4B,UCK2,LSM12,CASC4,STOML2,C1
,NNMT,SNRPN,WDR62,STYXL1,RFX5,PLEKHA5,RANBP1,UBE2L6,VRK3,LZTFL1,PPM1K,TUBA1C,DST,ANTX
,NNMT,SNRPN,WDR62,STYXL1,RFX5,PLEKHA5,RANBP1,UBE2L6,VRK3,LZTFL1,PPM1K,TUBA1C,DST,ANTX
,CENPK,RPS6,GGPS1,RTEL1,SRSF1,AUP1,SAP30BP,YIPF3,FOXRED1,RABL5,KXD1,MED15,SH3YL1,IP6K2,CL
,CENPK,RPS6,GGPS1,RTEL1,SRSF1,AUP1,SAP30BP,YIPF3,FOXRED1,RABL5,KXD1,MED15,SH3YL1,IP6K2,CL
TXR2,STIP1,LRRC1,GRAMD1C,VCAN,PCBP2,LPCAT3,CD44,TM9SF1,SON,TET2,UQCRQ,SNF8,GRK6,HMGA
TXR2,STIP1,LRRC1,GRAMD1C,VCAN,PCBP2,LPCAT3,CD44,TM9SF1,SON,TET2,UQCRQ,SNF8,GRK6,HMGA
,YIF1A,PRMT2,EIF3M,TBX6,XBP1,PTK2,AUNIP,C14ORF2,GUK1,CENPN,PCYT2,FBXW9,HSBP1L1,AGPAT4,D
,YIF1A,PRMT2,EIF3M,TBX6,XBP1,PTK2,AUNIP,C14ORF2,GUK1,CENPN,PCYT2,FBXW9,HSBP1L1,AGPAT4,D
,YIF1A,PRMT2,EIF3M,TBX6,XBP1,PTK2,AUNIP,C14ORF2,GUK1,CENPN,PCYT2,FBXW9,HSBP1L1,AGPAT4,D
,YIF1A,PRMT2,EIF3M,TBX6,XBP1,PTK2,AUNIP,C14ORF2,GUK1,CENPN,PCYT2,FBXW9,HSBP1L1,AGPAT4,D
TIMM17B,XAF1,APEH,HYI,CAPN10,LYPLA2,NUTF2,PODXL,TCF4,ATPAF2,C17ORF62,PRMT1,CASC4,ZNF7,T
TIMM17B,XAF1,APEH,HYI,CAPN10,LYPLA2,NUTF2,PODXL,TCF4,ATPAF2,C17ORF62,PRMT1,CASC4,ZNF7,T
ANBP1,UBE2L6,VRK3,LZTFL1,PPM1K,TUBA1C,DST,ANTXR2,STIP1,LRRC1,GRAMD1C,VCAN,PCBP2,LPCAT3
X6,TMEM205,XBP1,PTK2,AUNIP,C14ORF2,GUK1,PCYT2,USE1,AGPAT4,MAF,CCDC25,DEF8,PPP1CC,RFX3,
X6,TMEM205,XBP1,PTK2,AUNIP,C14ORF2,GUK1,PCYT2,USE1,AGPAT4,MAF,CCDC25,DEF8,PPP1CC,RFX3,
,GRAMD1C,VCAN,PCBP2,LPCAT3,CD44,TM9SF1,SON,TET2,UQCRQ,SNF8,GRK6,HMGA2,CDC37,CPSF3L,T
NK1,ZNF7,TXNDC17,ZFAND2B,HNRNPC,CYB561,SHARPIN,UPP1,ZNF250,RPL36,MOB1A,LAMP2,CCNL2,HS
P1L1,IFI27L1,PPA1,ENDOV,CAMTA1,PPIL2,RPL28,CLDND1,LIG1,CDC14B,TIMM17B,APEH,HYI,TXNL4A,CAP
P1L1,IFI27L1,PPA1,ENDOV,CAMTA1,PPIL2,RPL28,CLDND1,LIG1,CDC14B,TIMM17B,APEH,HYI,TXNL4A,CAP
CK2,DBNDD2,NISCH,TMED4,RPS6,CD99L2,GGPS1,RTEL1,AUP1,YIPF3,FOXRED1,RABL5,TRNT1,PARD6G,S
CK2,DBNDD2,NISCH,TMED4,RPS6,CD99L2,GGPS1,RTEL1,AUP1,YIPF3,FOXRED1,RABL5,TRNT1,PARD6G,S
HNRNPUL1,TMEM199,UBE2D3,C1ORF63,TKT,DENND1A,EPN1,ITPA,ZCCHC9,GRAMD1A,MKRN1,SMUG1,

F8,HMGA2,CDC37,CPSF3L,TTC21B,PTPN13,UBR4,PSMA1,PHF21A,SLC39A11,FUZ,PSMA3,MUC1,METTL1

F8,HMGA2,CDC37,CPSF3L,TTC21B,PTPN13,UBR4,PSMA1,PHF21A,SLC39A11,FUZ,PSMA3,MUC1,METTL1

SF1B,EEF1B2,PPP1CC,EDF1,RFX3,DGKE,ENDOV,CAMTA1,CAPG,CRIP1,PPIL2,RPL28,VIM,LIG1,PSMA7,TM

SF1B,EEF1B2,PPP1CC,EDF1,RFX3,DGKE,ENDOV,CAMTA1,CAPG,CRIP1,PPIL2,RPL28,VIM,LIG1,PSMA7,TM

ST,ANTXR2,STIP1,LRRC1,GRAMD1C,VCAN,PCBP2,LPCAT3,CD44,TM9SF1,SON,TET2,SNF8,GRK6,HMGA2,

ST,ANTXR2,STIP1,LRRC1,GRAMD1C,VCAN,PCBP2,LPCAT3,CD44,TM9SF1,SON,TET2,SNF8,GRK6,HMGA2,

,FOXRED1,RABL5,TRNT1,KXD1,MED15,SH3YL1,CLASRP,JPH2,UGGT2,MICAL1,UBE2I,SLC39A13,PRMT2,T

,FOXRED1,RABL5,TRNT1,KXD1,MED15,SH3YL1,CLASRP,JPH2,UGGT2,MICAL1,UBE2I,SLC39A13,PRMT2,T

NL2,HSP90AB1,SYNGR2,CEP70,TMEM134,LRRFIP1,FAM219B,RAB2A,ZNF524,TPM2,GPBP1,ASPH,CHPF,

GA2,CDC37,OBSL1,TTC21B,UBR4,CHCHD1,SLC39A11,FUZ,RGS3,FAM3A,PSMA3,MUC1,METTL17,ECSIT,A

GA2,CDC37,OBSL1,TTC21B,UBR4,CHCHD1,SLC39A11,FUZ,RGS3,FAM3A,PSMA3,MUC1,METTL17,ECSIT,A

S3,FAM3A,PSMA3,MUC1,METTL17,ECSIT,ATG12,PPRC1,ZBTB44,NIP7,DUSP22,SLC25A10,TMX2,SPOCK2,

S3,FAM3A,PSMA3,MUC1,METTL17,ECSIT,ATG12,PPRC1,ZBTB44,NIP7,DUSP22,SLC25A10,TMX2,SPOCK2,

XRED1,TRNT1,KXD1,ITGB1BP1,SH3YL1,C1ORF198,MICAL1,UBE2I,YIF1A,PRMT2,EIF3M,TBX6,PTK2,CELF1

2,LPCAT3,CD44,TM9SF1,SON,TET2,SNF8,GRK6,HMGA2,CDC37,CPSF3L,TSG101,TTC21B,PTPN13,UBR4,P

,FAM104B,NCBP2,USP15,BCL7B,ECT2,IAH1,ZNF276,CEP63,SZT2,RQCD1,FDPS,RBM7,PKIG,ZNF345,ATP6

GLN2,SIRT5,RFXANK,SPINT2,REEP5,SMARCA4,MPC2,TYMS,DALRD3,BRF1,OARD1,FKBP5,MCFD2,ADAM1

GLN2,SIRT5,RFXANK,SPINT2,REEP5,SMARCA4,MPC2,TYMS,DALRD3,BRF1,OARD1,FKBP5,MCFD2,ADAM1

1BP1,SH3YL1,CLASRP,C1ORF198,UGGT2,MICAL1,UBE2I,SLC39A13,YIF1A,PRMT2,EIF3M,JUP,CDC42,TME

1BP1,SH3YL1,CLASRP,C1ORF198,UGGT2,MICAL1,UBE2I,SLC39A13,YIF1A,PRMT2,EIF3M,JUP,CDC42,TME

,LRRC1,GRAMD1C,VCAN,PCBP2,LPCAT3,CD44,TM9SF1,SON,TET2,SNF8,GRK6,HMGA2,CDC37,CPSF3L,TTC

NPLL,ZFAND4,HNRNPA1,FKBP3,ZNF506,SPINT2,REEP5,ASH2L,FTH1,PFDN5,TCTN1,MTHFS,MPC2,TXNRD2

TRMT112,PNPO,RPTOR,ZNF625,LYNX1,TWSG1,MAPK13,VPS28,ZSWIM7,HSPB1,MVD,SAT1,EMC4,IL32,SE

TRMT112,PNPO,RPTOR,ZNF625,LYNX1,TWSG1,MAPK13,VPS28,ZSWIM7,HSPB1,MVD,SAT1,EMC4,IL32,SE

UBE2L6,VRK3,ZC3H14,FIS1,SLC15A3,PPM1K,TUBA1C,IRF9,DST,ANTXR2,STIP1,LRRC1,PHF19,GRAMD1C,V

UBE2L6,VRK3,ZC3H14,FIS1,SLC15A3,PPM1K,TUBA1C,IRF9,DST,ANTXR2,STIP1,LRRC1,PHF19,GRAMD1C,V

22,SPOCK2,CENPK,RPL29,GGPS1,RTEL1,SRSF1,AUP1,FOXRED1,ITGB1BP1,SH3YL1,IP6K2,CLASRP,JPH2,U

22,SPOCK2,CENPK,RPL29,GGPS1,RTEL1,SRSF1,AUP1,FOXRED1,ITGB1BP1,SH3YL1,IP6K2,CLASRP,JPH2,U

D4,CLDND1,VIM,LIG1,PSMA7,TMEM126B,RPL21,TXNL4A,TCF4,TMPO,ATPAF2,LSM12,C17ORF62,PRMT1

D4,CLDND1,VIM,LIG1,PSMA7,TMEM126B,RPL21,TXNL4A,TCF4,TMPO,ATPAF2,LSM12,C17ORF62,PRMT1

P1,SAP30BP,FOXRED1,TRNT1,KXD1,ITGB1BP1,MED15,UGGT2,MICAL1,SLC39A13,YIF1A,PRMT2,EIF3M,

P1,SAP30BP,FOXRED1,TRNT1,KXD1,ITGB1BP1,MED15,UGGT2,MICAL1,SLC39A13,YIF1A,PRMT2,EIF3M,

1A,CHCHD1,FUZ,RGS3,MUC1,METTL17,ECSIT,PPRC1,RPS25,NLRP1,NIP7,TTC9C,COX4I1,DUSP22,SLC25A

1A,CHCHD1,FUZ,RGS3,MUC1,METTL17,ECSIT,PPRC1,RPS25,NLRP1,NIP7,TTC9C,COX4I1,DUSP22,SLC25A

AL1,HS2ST1,UBE2I,PRMT2,EIF3M,MAGT1,JUP,TBX6,TMEM205,PTK2,C14ORF2,USE1,AGPAT4,DEF8,PPP

AL1,HS2ST1,UBE2I,PRMT2,EIF3M,MAGT1,JUP,TBX6,TMEM205,PTK2,C14ORF2,USE1,AGPAT4,DEF8,PPP

LL5,LDHA,PSMA6,ACOT8,AP3B2,RPS9,SLC35G1,RCE1,ZNF410,RUVBL2,PCMT1,EBPL,TESC,IGSF8,NUDT5,

LL5,LDHA,PSMA6,ACOT8,AP3B2,RPS9,SLC35G1,RCE1,ZNF410,RUVBL2,PCMT1,EBPL,TESC,IGSF8,NUDT5,

PODXL,EPB41L4B,UCK2,ATPAF2,C17ORF62,PRMT1,CASC4,C19ORF54,ZNF7,STRADA,ZFAND2B,AP1S3,HN

9A13,PRMT2,CDC42,TBX6,TMEM205,PTK2,C14ORF2,GUK1,CENPN,ATF7,FBXW9,USE1,HSBP1L1,AGPAT

OV,CAMTA1,CAPG,CRIP1,PPIL2,VIM,LIG1,PSMA7,CDC14B,TMEM126B,CAPN10,NUTF2,PODXL,CAMLG,E

GA2,RPS16,CPSF3L,TSG101,OBSL1,PTPN13,UBR4,PSMA1,PHF21A,CHCHD1,FUZ,RGS3,FAM3A,MUC1,ME

GA2,RPS16,CPSF3L,TSG101,OBSL1,PTPN13,UBR4,PSMA1,PHF21A,CHCHD1,FUZ,RGS3,FAM3A,MUC1,ME

2,CPSF3L,OBSL1,TTC21B,PTPN13,UBR4,YLPM1,PHF21A,CHCHD1,SLC39A11,FAM3A,MUC1,ECSIT,EPB41L

A2,CPSF3L,OBSL1,TTC21B,PTPN13,UBR4,YLPM1,PHF21A,CHCHD1,SLC39A11,FAM3A,MUC1,ECSIT,EPB41L
D14,PCYT2,HSBP1L1,IFI27L1,ASF1B,CCDC25,EEF1B2,PPP1CC,PPA1,TNFRSF14,ENDOV,CAMTA1,PPIL2,PSM
RK6,HMGA2,CDC37,RPS16,CPSF3L,TSG101,TTC21B,UBR4,PSMA1,YLPM1,PHF21A,CHCHD1,FUZ,RGS3,FAM
RK6,HMGA2,CDC37,RPS16,CPSF3L,TSG101,TTC21B,UBR4,PSMA1,YLPM1,PHF21A,CHCHD1,FUZ,RGS3,FAM
PS28,DCTD,ZNF814,VWA2,EMC4,MCTS1,DDOST,LSR,DOCK10,PSME2,RRNAD1,DBNL,RAD1,TOMM40,SERI
PS28,DCTD,ZNF814,VWA2,EMC4,MCTS1,DDOST,LSR,DOCK10,PSME2,RRNAD1,DBNL,RAD1,TOMM40,SERI
1,ZNF223,FAM126A,RC3H2,OGFOD2,EIF3G,TSEN34,MPV17,ZNF512,RNF181,CHTOP,RPL35,SLC52A2,TRP
1,ZNF223,FAM126A,RC3H2,OGFOD2,EIF3G,TSEN34,MPV17,ZNF512,RNF181,CHTOP,RPL35,SLC52A2,TRP

L2,CLDND1,VIM,TIMM17B,APEH,CAPN10,LYPLA2,CAMLG,UCK2,ATPAF2,C17ORF62,PRMT1,CASC4,EXOSC
17,FKBP3,EGLN2,SIRT5,KCTD7,B3GAT3,ZNF506,SPINT2,PAFAH1B2,PTK7,ARHGEF1,DRG2,BRF1,ZFPL1,FAM
17,FKBP3,EGLN2,SIRT5,KCTD7,B3GAT3,ZNF506,SPINT2,PAFAH1B2,PTK7,ARHGEF1,DRG2,BRF1,ZFPL1,FAM
GA1,PSME2,PABPC1,DBNL,NAT9,BCAM,BLVRB,TOMM40,RABEP2,RALGPS1,ZNF687,PDCD5,NXN,EPS15L1
K,ASH2L,PFDN5,SMARCA4,COA3,ZNF343,SNRPF,PTK7,COPS4,OARD1,MFSD10,MRPS18C,MCFD2,RIC8A,S
13A,ARHGEF39,GSTK1,TTLL5,LDHA,PSMA6,ACOT8,AP3B2,RPS9,SLC35G1,RCE1,MRPL33,ZNF410,RUVBL2,
,CLDND1,TIMM17B,XAF1,CAMLG,EPB41L4B,TMPO,ATPAF2,LSM12,C17ORF62,PRMT1,EXOSC9,STOML2,C

LEKHA5,RANBP1,LZTFL1,FIS1,SLC15A3,PPM1K,TUBA1C,IRF9,DST,ANTXR2,STIP1,LRRC1,PHF19,VCAN,PCBP
LEKHA5,RANBP1,LZTFL1,FIS1,SLC15A3,PPM1K,TUBA1C,IRF9,DST,ANTXR2,STIP1,LRRC1,PHF19,VCAN,PCBP
OSPH10,STAU2,KRAS,GFOD1,METTL3,SSR3,RPL8,RBBP6,TJP1,RPL35,ANAPC13,NXF1,NT5E,CFLAR,RPL34,

A,MUC1,ECSIT,EPB41L5,ATG12,PPRC1,ZBTB44,NIP7,COX4I1,PLCB2,SLC25A10,DBNDD2,NISCH,TMED4,RPS
OV,PSMD4,RPL28,CLDND1,LIG1,CDC14B,TIMM17B,APEH,RPL21,CAPN10,RBM33,LYPLA2,EPB41L4B,UCK2
OV,PSMD4,RPL28,CLDND1,LIG1,CDC14B,TIMM17B,APEH,RPL21,CAPN10,RBM33,LYPLA2,EPB41L4B,UCK2
F48,ZFAND2B,HNRNPC,SLC9B2,HYPK,ZNF250,NFAM1,CDK11A,INO80E,MAPKAPK5,DGUOK,HSP90AB1,SY
F48,ZFAND2B,HNRNPC,SLC9B2,HYPK,ZNF250,NFAM1,CDK11A,INO80E,MAPKAPK5,DGUOK,HSP90AB1,SY
RF198,MICAL1,UBE2I,SLC39A13,YIF1A,PRMT2,CDC42,TBX6,TMEM205,PTK2,GUK1,CENPN,FBXW9,MAF,A
RF198,MICAL1,UBE2I,SLC39A13,YIF1A,PRMT2,CDC42,TBX6,TMEM205,PTK2,GUK1,CENPN,FBXW9,MAF,A
D17A,STXBP2,EEF1D,NAA10,LLGL2,CHEK1,COX17,SUMO3,ZFAND4,FKBP3,EGLN2,KCTD7,B3GAT3,RFXANK
2,MICAL1,HS2ST1,YIF1A,EIF3M,JUP,CDC42,TBX6,TMEM205,XBP1,PTK2,CELF1,AUNIP,C14ORF2,CENPN,C
2,MICAL1,HS2ST1,YIF1A,EIF3M,JUP,CDC42,TBX6,TMEM205,XBP1,PTK2,CELF1,AUNIP,C14ORF2,CENPN,C
F1,RFX3,PPA1,ENDOV,CAMTA1,PPIL2,RPL21,TXNL4A,PODXL,EPB41L4B,UCK2,PRMT1,EXOSC9,TXNDC17,A
F1,RFX3,PPA1,ENDOV,CAMTA1,PPIL2,RPL21,TXNL4A,PODXL,EPB41L4B,UCK2,PRMT1,EXOSC9,TXNDC17,A
S,NNMT,SPAG7,WDR62,IL1RAP,STYXL1,RFX5,POLD4,PLEKHA5,RANBP1,UBE2L6,VRK3,ZC3H14,SLC15A3,F
S,NNMT,SPAG7,WDR62,IL1RAP,STYXL1,RFX5,POLD4,PLEKHA5,RANBP1,UBE2L6,VRK3,ZC3H14,SLC15A3,F
2A,ZNF524,TPM2,GPBP1,CHPF,NIT2,LYNX1,ZSWIM7,GMPPB,MVD,LMBR1,RANGRF,CHCHD7,PSME2,PHTF

4,SLC9B2,SUGT1,COPZ1,CCNL2,MAPKAPK5,CEP70,LRRFIP1,FAM219B,RAB2A,ZNF524,ELP6,TPM2,GPBP1,
PCYT2,HSBP1L1,IFI27L1,ASF1B,DEF8,EEF1B2,PPP1CC,EDF1,RFX3,ENDOV,CAMTA1,CAPG,PPIL2,VIM,LIG1,
PCYT2,HSBP1L1,IFI27L1,ASF1B,DEF8,EEF1B2,PPP1CC,EDF1,RFX3,ENDOV,CAMTA1,CAPG,PPIL2,VIM,LIG1,
LEC1,SAT1,ERCC5,NAP1L1,DDOST,RANGRF,CHCHD7,MEIS2,PHTF2,PABPC1,NAT9,ZNF836,RBP1,RAD1,H2
2AFV,SERPINB6,TADA3,WAC,SLC33A1,ZNF706,ZNF687,EPS15L1,RAD23A,ILF3,PKP4,AGAP3,EEF1D,HSPH1
SP22,SLC25A10,TMX2,DBNDD2,SDF4,NISCH,TMED4,CD99L2,GGPS1,SRSF1,YIPF3,FOXRED1,TRNT1,PARD
SP22,SLC25A10,TMX2,DBNDD2,SDF4,NISCH,TMED4,CD99L2,GGPS1,SRSF1,YIPF3,FOXRED1,TRNT1,PARD

10,MRPL43,C9ORF64,IQCK,SETD4,MED8,NARF,RNMT,ABCC5,MRPL48,ZNF616,GANC,MBD1,CREM,CHFR,

10,MRPL43,C9ORF64,IQCK,SETD4,MED8,NARF,RNMT,ABCC5,MRPL48,ZNF616,GANC,MBD1,CREM,CHFR,
PHF21A,SLC39A11,FUZ,FAM3A,METTL17,EPB41L5,ATG12,PPRC1,RPS25,NLRP1,ZBTB44,TTC9C,COX4I1,D
PHF21A,SLC39A11,FUZ,FAM3A,METTL17,EPB41L5,ATG12,PPRC1,RPS25,NLRP1,ZBTB44,TTC9C,COX4I1,D
2,TADA3,WAC,ZNF706,PDCD5,NXN,KLK10,EPS15L1,RAD23A,ABHD17A,NT5C,ZCCHC4,MDH2,STXBP2,OA

IPF3,FOXRED1,TRNT1,ITGB1BP1,SH3YL1,IP6K2,SLC39A13,YIF1A,JUP,CDC42,TBX6,TMEM205,XBP1,PTK2,
IPF3,FOXRED1,TRNT1,ITGB1BP1,SH3YL1,IP6K2,SLC39A13,YIF1A,JUP,CDC42,TBX6,TMEM205,XBP1,PTK2,
9,USE1,EDF1,RFX3,DGKE,TNFRSF14,ENDOV,CAMTA1,CAPG,PPIL2,PSMD4,PSMA7,CDC14B,TIMM17B,TX

GADL1,RFX3,ENDOV,CAMTA1,PPIL2,PSMD4,VIM,LIG1,PSMA7,TMEM126B,APEH,TXNL4A,NUTF2,UCK2,T
GADL1,RFX3,ENDOV,CAMTA1,PPIL2,PSMD4,VIM,LIG1,PSMA7,TMEM126B,APEH,TXNL4A,NUTF2,UCK2,T
TCP1,CD74,TRPV3,PIGT,UBE2N,MFSD11,LFNG,SNX5,NOP2,FAM32A,ZSCAN20,RPAIN,PCID2,TGOLN2,NFY
TCP1,CD74,TRPV3,PIGT,UBE2N,MFSD11,LFNG,SNX5,NOP2,FAM32A,ZSCAN20,RPAIN,PCID2,TGOLN2,NFY
BP1,PTK2,C14ORF2,GUK1,CARD14,PCYT2,FBXW9,IFI27L1,MAF,ASF1B,CCDC25,RFX3,PPA1,TNFRSF14,END
B1,LYNX1,MPDU1,TWSG1,VPS28,ZSWIM7,ZNF814,ERLEC1,HHLA3,MVD,ERCC5,EMC4,MCTS1,SYNGR1,D
RRFIP1,RAB2A,ZNF524,TPM2,GPBP1,SIAH1,GGCX,ZEB2,ASPH,TRMT112,PNPO,STRIP2,ZNF625,NIT2,LYNX
CDC37,RPS16,CPSF3L,TSG101,OBSL1,TTC21B,YLPM1,PHF21A,CHCHD1,SLC39A11,FAM3A,MUC1,METTL1
CDC37,RPS16,CPSF3L,TSG101,OBSL1,TTC21B,YLPM1,PHF21A,CHCHD1,SLC39A11,FAM3A,MUC1,METTL1
5A3,PPM1K,TUBA1C,DST,ANTXR2,STIP1,LRRC1,PHF19,GRAMD1C,VCAN,LPCAT3,CD44,TM9SF1,MAGED2
5A3,PPM1K,TUBA1C,DST,ANTXR2,STIP1,LRRC1,PHF19,GRAMD1C,VCAN,LPCAT3,CD44,TM9SF1,MAGED2
K6,HMGA2,CDC37,RPS16,CPSF3L,TSG101,OBSL1,TTC21B,UBR4,PSMA1,PHF21A,RGS3,EPB41L5,ATG12,P
K6,HMGA2,CDC37,RPS16,CPSF3L,TSG101,OBSL1,TTC21B,UBR4,PSMA1,PHF21A,RGS3,EPB41L5,ATG12,P
D3,BRF1,OARD1,SLC25A23,FAM102A,MFSD10,PFKFB2,RIC8A,NFKBIA,GNB5,COMMD4,LFNG,ZNHIT3,XYL
MX2,SPOCK2,RDH11,DBNDD2,TMED4,CENPK,RPS6,RPL29,GGPS1,RTEL1,COASY,AUP1,YIPF3,FOXRED1,TR
2A,ELP6,RMDN1,GPBP1,TRMT112,PNPO,CHPF,MPDU1,TWSG1,UBXN1,ZNF814,GMPPB,MVD,SAT1,ERC
CAN,TM9SF1,SON,TET2,UQCRQ,SNF8,GRK6,HMGA2,CDC37,RPS16,CPSF3L,OBSL1,PTPN13,UBR4,YLPM1,
CAN,TM9SF1,SON,TET2,UQCRQ,SNF8,GRK6,HMGA2,CDC37,RPS16,CPSF3L,OBSL1,PTPN13,UBR4,YLPM1,
TUBGCP2,SSH3,RPL27,UBE2F,ZNF585A,C12ORF10,RNF4,MAGOH,ZRANB2,TXNDC12,SEC14L1,FANCI,SEC2
TUBGCP2,SSH3,RPL27,UBE2F,ZNF585A,C12ORF10,RNF4,MAGOH,ZRANB2,TXNDC12,SEC14L1,FANCI,SEC2

,PIGL,C19ORF48,PPAN,ACAD10,SHARPIN,SLC9B2,CARS,CDK11A,MDH1,MAPKAPK5,HSP90AB1,CEP70,TM
,PIGL,C19ORF48,PPAN,ACAD10,SHARPIN,SLC9B2,CARS,CDK11A,MDH1,MAPKAPK5,HSP90AB1,CEP70,TM
66,MARK2,TMEM223,ESRRA,GINS4,SEC24C,PPP1R7,SERINC5,MSRB1,TVP23B,LIN7B,TECPR2,MIB2,WDR
MA1,YIF1A,PRMT2,TBX6,GUK1,CENPN,ATF7,PCYT2,USE1,IFI27L1,ASF1B,CCDC25,EEF1B2,PPP1CC,RFX3,PP
MA1,YIF1A,PRMT2,TBX6,GUK1,CENPN,ATF7,PCYT2,USE1,IFI27L1,ASF1B,CCDC25,EEF1B2,PPP1CC,RFX3,PP
7B,HYI,NUTF2,PODXL,CAMLG,TCF4,WNK1,ZNF7,ARL6IP4,ZFAND2B,CYB561,SHARPIN,STAU1,SUGT1,UPP1
7B,HYI,NUTF2,PODXL,CAMLG,TCF4,WNK1,ZNF7,ARL6IP4,ZFAND2B,CYB561,SHARPIN,STAU1,SUGT1,UPP1
A10,TMX2,SPOCK2,DBNDD2,SDF4,NISCH,TMED4,CD99L2,GGPS1,COASY,SRSF1,AUP1,FOXRED1,TRNT1,K

NDOV,CAMTA1,PPIL2,PSMD4,VIM,LIG1,PSMA7,APEH,TXNL4A,NUTF2,TCF4,UCK2,TMPO,ATPAF2,LSM12,
NDOV,CAMTA1,PPIL2,PSMD4,VIM,LIG1,PSMA7,APEH,TXNL4A,NUTF2,TCF4,UCK2,TMPO,ATPAF2,LSM12,
P22,SPOCK2,THEM4,DBNDD2,SDF4,NISCH,TMED4,CENPK,RPS6,GGPS1,COASY,AUP1,SAP30BP,FOXRED1,
P22,SPOCK2,THEM4,DBNDD2,SDF4,NISCH,TMED4,CENPK,RPS6,GGPS1,COASY,AUP1,SAP30BP,FOXRED1,
,PSMA7,TIMM17B,NUTF2,PODXL,EPB41L4B,TCF4,PPOX,TMPO,ATPAF2,CASC4,ZNF7,PIGL,AP1S3,ACAD1
,PSMA7,TIMM17B,NUTF2,PODXL,EPB41L4B,TCF4,PPOX,TMPO,ATPAF2,CASC4,ZNF7,PIGL,AP1S3,ACAD1
T1,JUP,TBX6,AUNIP,GUK1,CENPN,CARD14,ATF7,FBXW9,ASF1B,EEF1B2,PPP1CC,GADL1,RFX3,DGKE,ENDO

T1,JUP,TBX6,AUNIP,GUK1,CENPN,CARD14,ATF7,FBXW9,ASF1B,EEF1B2,PPP1CC,GADL1,RFX3,DGKE,ENDO

,RPL28,VIM,LIG1,PSMA7,APEH,HYI,TXNL4A,LYPLA2,PODXL,CAMLG,TCF4,PPOX,TMPO,PRMT1,EXOSC9,W
,RPL28,VIM,LIG1,PSMA7,APEH,HYI,TXNL4A,LYPLA2,PODXL,CAMLG,TCF4,PPOX,TMPO,PRMT1,EXOSC9,W
1BP1,PARD6G,SH3YL1,IP6K2,UGGT2,MICAL1,HS2ST1,UBE2I,EIF3M,TBX6,AUNIP,C14ORF2,GUK1,FBXW9,
1BP1,PARD6G,SH3YL1,IP6K2,UGGT2,MICAL1,HS2ST1,UBE2I,EIF3M,TBX6,AUNIP,C14ORF2,GUK1,FBXW9,
,PCYT2,HSBP1L1,IFI27L1,ASF1B,DEF8,EEF1B2,PPP1CC,RFX3,DGKE,ENDOV,CAMTA1,CRIP1,PPIL2,RPL28,VI
P,JPH2,MICAL1,HS2ST1,UBE2I,PRMT2,TBX6,TMEM205,XBP1,PTK2,AUNIP,C14ORF2,GUK1,CENPN,PCYT2
M3A,PSMA3,METTL17,ATG12,PPRC1,RPS25,NLRP1,ZBTB44,NIP7,TTC9C,PLCB2,DUSP22,SLC25A10,TMX2,
M3A,PSMA3,METTL17,ATG12,PPRC1,RPS25,NLRP1,ZBTB44,NIP7,TTC9C,PLCB2,DUSP22,SLC25A10,TMX2,
B2,HYPK,UPP1,ZNF250,NFAM1,CDK11A,INO80E,MAPKAPK5,DGUOK,HSP90AB1,SYNGR2,TMEM134,FAM
B2,HYPK,UPP1,ZNF250,NFAM1,CDK11A,INO80E,MAPKAPK5,DGUOK,HSP90AB1,SYNGR2,TMEM134,FAM

,RPL21,TXNL4A,CAPN10,LYPLA2,CAMLG,TMPO,LSM12,C17ORF62,PRMT1,CASC4,ZNF7,TXNDC17,C19OR
,ZNF836,H2AFV,BLVRB,PPP3CB,RABEP2,TADA3,ZNF20,RALGPS1,ZNF687,PDCD5,GTF2IRD1,NXN,EPS15L
A3,WAC,ZNF687,POT1,ABHD17A,ILF3,AGAP3,EEF1D,HSPH1,RRP8,CARHSP1,CNST,LLGL2,SH3GLB1,CHEK
RD6G,SH3YL1,JPH2,UGGT2,MICAL1,UBE2I,YIF1A,MAGT1,JUP,CDC42,TBX6,TMEM205,XBP1,AUNIP,CENP
VRK3,LZTFL1,PPM1K,TUBA1C,DST,ANTXR2,STIP1,LRRC1,GRAMD1C,VCAN,PCBP2,LPCAT3,CD44,TM9SF1,
VRK3,LZTFL1,PPM1K,TUBA1C,DST,ANTXR2,STIP1,LRRC1,GRAMD1C,VCAN,PCBP2,LPCAT3,CD44,TM9SF1,

AGED2,SON,TET2,UQCRQ,SNF8,HMGA2,CDC37,RPS16,CPSF3L,OBSL1,TTC21B,PTPN13,UBR4,PSMA1,YLP
AGED2,SON,TET2,UQCRQ,SNF8,HMGA2,CDC37,RPS16,CPSF3L,OBSL1,TTC21B,PTPN13,UBR4,PSMA1,YLP
P30BP,KXD1,ITGB1BP1,MED15,PARD6G,IP6K2,UGGT2,MICAL1,UBE2I,SLC39A13,PRMT2,TBX6,PTK2,C14
3,CHCHD7,HMGA1,PSME2,PHTF2,PABPC1,DBNL,NAT9,BCAM,BLVRB,PPP3CB,RABEP2,TADA3,ZNF20,WA
1A,UBA52,VAPA,DHDDS,CEP63,SDHA,ZNF483,ABLIM2,ACHE,PEX16,SUFU,ZNF33A,MRPS33,EXOSC2,HNR
1A,UBA52,VAPA,DHDDS,CEP63,SDHA,ZNF483,ABLIM2,ACHE,PEX16,SUFU,ZNF33A,MRPS33,EXOSC2,HNR
,HMGA1,PABPC1,DBNL,NAT9,BCAM,BLVRB,PPP3CB,RABEP2,TADA3,ZNF20,WAC,RALGPS1,ZNF706,ZNF6

,CAMTA1,CAPG,PSMD4,RPL28,VIM,LIG1,PSMA7,HYI,TXNL4A,CAPN10,NUTF2,EPB41L4B,LSM12,PRMT1,V
PAT4,PPP1CC,EDF1,RFX3,PPA1,DGKE,ENDOV,CAMTA1,CAPG,CLDND1,VIM,LIG1,PSMA7,TMEM126B,XAF
PAT4,PPP1CC,EDF1,RFX3,PPA1,DGKE,ENDOV,CAMTA1,CAPG,CLDND1,VIM,LIG1,PSMA7,TMEM126B,XAF
NBP1,UBE2L6,LZTFL1,ZC3H14,FIS1,SLC15A3,TUBA1C,ANTXR2,STIP1,LRRC1,PHF19,GRAMD1C,VCAN,LPCA
NBP1,UBE2L6,LZTFL1,ZC3H14,FIS1,SLC15A3,TUBA1C,ANTXR2,STIP1,LRRC1,PHF19,GRAMD1C,VCAN,LPCA
HGDH,CC2D1B,ARPC5,GEMIN2,LYPLAL1,NDUFA11,THOC3,SH3GL1,BTF3,NDUFA3,C1ORF63,IDH3G,DNA
HGDH,CC2D1B,ARPC5,GEMIN2,LYPLAL1,NDUFA11,THOC3,SH3GL1,BTF3,NDUFA3,C1ORF63,IDH3G,DNA

B2,PPP1CC,EDF1,PPA1,DGKE,ENDOV,CAMTA1,RPL28,CLDND1,PSMA7,TMEM126B,RPL21,TXNL4A,LYPLA
B2,PPP1CC,EDF1,PPA1,DGKE,ENDOV,CAMTA1,RPL28,CLDND1,PSMA7,TMEM126B,RPL21,TXNL4A,LYPLA
T2,SLC9B2,SUGT1,UPP1,ZNF250,RPL36,RBM3,MDH1,ZNF197,COPZ1,CCNL2,MAPKAPK5,SYNGR2,TMEM1
T2,SLC9B2,SUGT1,UPP1,ZNF250,RPL36,RBM3,MDH1,ZNF197,COPZ1,CCNL2,MAPKAPK5,SYNGR2,TMEM1
,RABEP2,ZNF20,RALGPS1,PDCD5,EPS15L1,NOMO2,RAD23A,HSD17B10,ABHD17A,AGAP3,C11ORF48,CAR
,RABEP2,ZNF20,RALGPS1,PDCD5,EPS15L1,NOMO2,RAD23A,HSD17B10,ABHD17A,AGAP3,C11ORF48,CAR

ED1,RABL5,TRNT1,ITGB1BP1,MED15,PARD6G,IP6K2,CLASRP,JPH2,C1ORF198,UGGT2,EIF3M,MAGT1,CDC

ED1,RABL5,TRNT1,ITGB1BP1,MED15,PARD6G,IP6K2,CLASRP,JPH2,C1ORF198,UGGT2,EIF3M,MAGT1,CDC

A3,MUC1,ATG12,NLRP1,ZBTB44,DUSP22,SLC25A10,TMX2,NISCH,RPL29,GGPS1,RTEL1,COASY,SRSF1,AUP1

A3,MUC1,ATG12,NLRP1,ZBTB44,DUSP22,SLC25A10,TMX2,NISCH,RPL29,GGPS1,RTEL1,COASY,SRSF1,AUP1

5,OXLD1,ZNF223,METTL5,SLC35E1,OGFOD2,EIF3G,TSEN34,ARIH2,PICK1,FBXL18,TJP1,SLC52A2,NT5E,APC

ORF57,PEX26,BTN3A2,LY6E,ZNF223,STAU2,METTL3,ARIH2,PICK1,OCIAD1,TPT1,POLR2M,NME1,ANAPC13

PPAN,HNRNPC,ZNF644,SLC9B2,SUGT1,HYPK,UPP1,ZNF250,MDH1,ZNF197,INO80E,DGUOK,LRRFIP1,FAM

0,CDK11A,RBM3,COPZ1,INO80E,CCNL2,MAPKAPK5,CEP70,LRRFIP1,FAM219B,RAB2A,TPM2,GPBP1,GPAL

0,CDK11A,RBM3,COPZ1,INO80E,CCNL2,MAPKAPK5,CEP70,LRRFIP1,FAM219B,RAB2A,TPM2,GPBP1,GPAL

MA1,YLPM1,CHCHD1,FAM3A,PSMA3,EPB41L5,ATG12,PPRC1,RPS25,NLRP1,NIP7,TTC9C,COX4I1,TMX2,SP

MA1,YLPM1,CHCHD1,FAM3A,PSMA3,EPB41L5,ATG12,PPRC1,RPS25,NLRP1,NIP7,TTC9C,COX4I1,TMX2,SP

RPS15,ERLEC1,LRRC27,NAP1L1,SYNGR1,LSR,MTA3,MEIS2,PHTF2,PABPC1,NAT9,BCAM,ZNF836,RBP1,RAD

D5,GTF2IRD1,NXN,EPS15L1,POLR3H,ABHD17A,AGAP3,STXBP2,EEF1D,NAA10,LLGL2,CHEK1,COX17,ZFAND

D5,GTF2IRD1,NXN,EPS15L1,POLR3H,ABHD17A,AGAP3,STXBP2,EEF1D,NAA10,LLGL2,CHEK1,COX17,ZFAND

CAD10,CYB561,SHARPIN,STAU1,SLC9B2,SUGT1,UPP1,CARS,CDK11A,RBM3,MDH1,ZNF197,CCNL2,SYNGR

HNRNPC,CYB561,CBFA2T2,SLC9B2,UPP1,RPL36,RBM3,MDH1,ZNF197,COPZ1,HSP90AB1,SYNGR2,RAB2A,

HNRNPC,CYB561,CBFA2T2,SLC9B2,UPP1,RPL36,RBM3,MDH1,ZNF197,COPZ1,HSP90AB1,SYNGR2,RAB2A,

,HMGA1,PSME2,PABPC1,NAT9,BCAM,ZNF836,BLVRB,PPP3CB,RABEP2,TADA3,RALGPS1,ZNF706,ZNF687,

CAPG,CRIP1,RPL28,VIM,LIG1,CDC14B,TIMM17B,XAF1,APEH,HYI,CAPN10,LYPLA2,PODXL,ATPAF2,C17ORF

CAPG,CRIP1,RPL28,VIM,LIG1,CDC14B,TIMM17B,XAF1,APEH,HYI,CAPN10,LYPLA2,PODXL,ATPAF2,C17ORF

,PPRC1,RPS25,NLRP1,ZBTB44,NIP7,TTC9C,COX4I1,PLCB2,SLC25A10,SPOCK2,DBNDD2,TMED4,CENPK,RPS

,PPRC1,RPS25,NLRP1,ZBTB44,NIP7,TTC9C,COX4I1,PLCB2,SLC25A10,SPOCK2,DBNDD2,TMED4,CENPK,RPS

T1,KXD1,SH3YL1,IP6K2,CLASRP,C1ORF198,HS2ST1,YIF1A,PRMT2,MAGT1,AUNIP,GUK1,USE1,IFI27L1,CCDC

T1,KXD1,SH3YL1,IP6K2,CLASRP,C1ORF198,HS2ST1,YIF1A,PRMT2,MAGT1,AUNIP,GUK1,USE1,IFI27L1,CCDC

12,RPS25,NLRP1,ZBTB44,NIP7,TTC9C,COX4I1,DUSP22,SLC25A10,THEM4,RDH11,DBNDD2,NISCH,TMED4,

12,RPS25,NLRP1,ZBTB44,NIP7,TTC9C,COX4I1,DUSP22,SLC25A10,THEM4,RDH11,DBNDD2,NISCH,TMED4,

XNL4A,NUTF2,PPOX,TMPO,ATPAF2,LSM12,PRMT1,CASC4,STOML2,WNK1,PPAN,HNRNPC,ZNF644,STAU1

XNL4A,NUTF2,PPOX,TMPO,ATPAF2,LSM12,PRMT1,CASC4,STOML2,WNK1,PPAN,HNRNPC,ZNF644,STAU1

A2,CPSF3L,TSG101,OBSL1,TTC21B,PTPN13,UBR4,PSMA1,YLPM1,CHCHD1,SLC39A11,FUZ,FAM3A,PSMA3

A2,CPSF3L,TSG101,OBSL1,TTC21B,PTPN13,UBR4,PSMA1,YLPM1,CHCHD1,SLC39A11,FUZ,FAM3A,PSMA3

CKLF,AKT1S1,INO80C,GTF3A,ARRB2,DIP2A,MLST8,HMGN1,CDK2AP2,MYL12A,MAZ,C2ORF15,TBCA,DDX1

CKLF,AKT1S1,INO80C,GTF3A,ARRB2,DIP2A,MLST8,HMGN1,CDK2AP2,MYL12A,MAZ,C2ORF15,TBCA,DDX1

644,ZNF250,MOB1A,RBM3,ZNF197,INO80E,MAPKAPK5,HSP90AB1,SYNGR2,FAM219B,SIAH1,ZEB2,TRMT

32A,COBL,PIGH,C12ORF57,NUBP2,EIF4A1,STAU2,RPLP0,C6ORF132,KLHL7,METTL3,ARIH2,RPL8,TCF25,SL

ATP5J,TPM2,GPBP1,TRMT112,STRIP2,GABPB1,TWSG1,UBXN1,GMPPB,MVD,LMBR1,NAP1L1,PRKRIP1,RA

PRC1,RPS25,ZBTB44,COX4I1,PLCB2,SLC25A10,TMX2,DBNDD2,NISCH,TMED4,CD99L2,GGPS1,RTEL1,COA

PRC1,RPS25,ZBTB44,COX4I1,PLCB2,SLC25A10,TMX2,DBNDD2,NISCH,TMED4,CD99L2,GGPS1,RTEL1,COA

SHARPIN,STAU1,ZNF250,RPL36,MOB1A,LAMP2,CCNL2,DUT,SYNGR2,TMEM134,LRRFIP1,RAB2A,ZNF524,

1,ENDOV,PPIL2,TIMM17B,APEH,RPL21,TXNL4A,TMPO,LSM12,PRMT1,EXOSC9,TXNDC17,ARL6IP4,C19OR

1,ENDOV,PPIL2,TIMM17B,APEH,RPL21,TXNL4A,TMPO,LSM12,PRMT1,EXOSC9,TXNDC17,ARL6IP4,C19OR

,COASY,SRSF1,FOXRED1,TRNT1,ITGB1BP1,MED15,PARD6G,SH3YL1,IP6K2,C1ORF198,HS2ST1,UBE2I,SLC3

,COASY,SRSF1,FOXRED1,TRNT1,ITGB1BP1,MED15,PARD6G,SH3YL1,IP6K2,C1ORF198,HS2ST1,UBE2I,SLC3

7,SUMO3,FKBP3,EGLN2,KCTD7,B3GAT3,RFXANK,SPINT2,FTH1,TYMS,PTK7,TMEM39B,BRF1,ZFPL1,OARD1

PPA1,DGKE,TNFRSF14,VAMP8,CAMTA1,RPL28,PSMA7,TIMM17B,TMEM126B,RPL21,TXNL4A,CAPN10,PC

3SL1,UBR4,PSMA1,PHF21A,CHCHD1,SLC39A11,FUZ,RGS3,PSMA3,ECSIT,PPRC1,RPS25,NLRP1,ZBTB44,TTC
3SL1,UBR4,PSMA1,PHF21A,CHCHD1,SLC39A11,FUZ,RGS3,PSMA3,ECSIT,PPRC1,RPS25,NLRP1,ZBTB44,TTC
,PPP3CB,RABEP2,RALGPS1,ZNF706,PDCD5,GTF2IRD1,NXN,EPS15L1,POLR3H,NT5C,ILF3,AGAP3,STXBP2,E

PP1CC,GADL1,RFX3,DGKE,ENDOV,PPIL2,RPL28,LIG1,TXNL4A,CAPN10,LYPLA2,NUTF2,PODXL,PPOX,ATPAI
3,NSUN5,DAZAP2,OARD1,MFSD10,PFKFB2,RIC8A,SLC30A5,GNB5,TRPV3,NHP2,UBE2N,FAM32A,COX20,T
3,NSUN5,DAZAP2,OARD1,MFSD10,PFKFB2,RIC8A,SLC30A5,GNB5,TRPV3,NHP2,UBE2N,FAM32A,COX20,T
'6,ZNF248,TPT1,SMAGP,TSPAN31,TMEM9,NXF1,SLC39A6,ZNF593,GLYCTK,TRIM46,RBM42,CLASP2,CREB
S15L1,HSD17B10,ABHD17A,DDX41,ILF3,STXBP2,EEF1D,NAA10,C11ORF48,MED6,LLGL2,CD151,FKBP3,EG
S15L1,HSD17B10,ABHD17A,DDX41,ILF3,STXBP2,EEF1D,NAA10,C11ORF48,MED6,LLGL2,CD151,FKBP3,EG
2,CARS,ZNF250,CDK11A,MDH1,ZNF197,COPZ1,CCNL2,MAPKAPK5,DUT,CEP70,LRRFIP1,RAB2A,ZNF524,R
2,CARS,ZNF250,CDK11A,MDH1,ZNF197,COPZ1,CCNL2,MAPKAPK5,DUT,CEP70,LRRFIP1,RAB2A,ZNF524,R
2L6,VRK3,LZTFL1,ZC3H14,SLC15A3,PPM1K,TUBA1C,IRF9,ANTXR2,STIP1,LRRC1,PHF19,GRAMD1C,LPCAT3,
2L6,VRK3,LZTFL1,ZC3H14,SLC15A3,PPM1K,TUBA1C,IRF9,ANTXR2,STIP1,LRRC1,PHF19,GRAMD1C,LPCAT3,
X2,NISCH,RPL29,GGPS1,RTEL1,SRSF1,AUP1,YIPF3,FOXRED1,TRNT1,KXD1,ITGB1BP1,MED15,SH3YL1,IP6K
FB2,NFKBIA,GNB5,CBR1,LFNG,XYLT2,ZXDC,TNKS,SMIM4,NMNAT3,KPNA5,MPHOSPH10,MYL9,SLC35E1,G
MP2,ZNF197,INO80E,CCNL2,MAPKAPK5,DGUOK,DUT,TMEM134,LRRFIP1,FAM219B,RMDN1,GPBP1,GPA
17,COX5A,HNRNPLL,SUMO3,EGLN2,INTS4,RFXANK,SNRPF,TXNRD2,DAZAP2,RIC8A,LGALS3,TCP1,SCAP,PH
HGDH,ARPC5,MYL6,TLCD1,ST7,RPS3A,MOSPD3,TCP11L1,RHEB,MAGOHB,IDH3G,CREB3L2,RGS12,UCHL5,
3A1C,IRF9,DST,ANTXR2,STIP1,LRRC1,PHF19,GRAMD1C,VCAN,LPCAT3,TM9SF1,TET2,UQCRQ,SNF8,HMGA
3A1C,IRF9,DST,ANTXR2,STIP1,LRRC1,PHF19,GRAMD1C,VCAN,LPCAT3,TM9SF1,TET2,UQCRQ,SNF8,HMGA
T1,JUP,CDC42,TBX6,TMEM205,PTK2,AUNIP,CENPN,ATF7,PCYT2,USE1,MAF,ASF1B,DEF8,EEF1B2,PPP1CC
1134,FAM219B,RAB2A,RMDN1,GPBP1,GPALPP1,TRMT112,CHPF,MPDU1,ZSWIM7,DCTD,ERLEC1,GMPPB,
1134,FAM219B,RAB2A,RMDN1,GPBP1,GPALPP1,TRMT112,CHPF,MPDU1,ZSWIM7,DCTD,ERLEC1,GMPPB,
ROAP,VIM,LIG1,PSMA7,RPL21,TXNL4A,CAPN10,RBM33,NUTF2,PODXL,EPB41L4B,PPOX,ATPAF2,CASC4,ZN
ROAP,VIM,LIG1,PSMA7,RPL21,TXNL4A,CAPN10,RBM33,NUTF2,PODXL,EPB41L4B,PPOX,ATPAF2,CASC4,ZN
'B1,IL32,RANGRF,MTA3,HMGA1,DBNL,NAT9,RAD1,TOMM40,PPP3CB,RABEP2,ATP6V1F,WAC,PDCD5,EPS
'B1,IL32,RANGRF,MTA3,HMGA1,DBNL,NAT9,RAD1,TOMM40,PPP3CB,RABEP2,ATP6V1F,WAC,PDCD5,EPS
,ASPH,CHPF,NIT2,MAPK13,ZSWIM7,UBXN1,ERLEC1,GMPPB,SAT1,ERCC5,EMC4,IL32,SEMA3C,PRKRIP1,R
S15L1,POLR3H,ABHD17A,ILF3,AGAP3,STXBP2,EEF1D,NAA10,LLGL2,CHEK1,COX17,ZFAND4,FKBP3,NDUF
IB1,SYNGR1,RANGRF,CHCHD7,HMGA1,DBNL,RBP1,SERPINB6,WAC,RALGPS1,PDCD5,GTF2IRD1,EPS15L1,
IB1,SYNGR1,RANGRF,CHCHD7,HMGA1,DBNL,RBP1,SERPINB6,WAC,RALGPS1,PDCD5,GTF2IRD1,EPS15L1,
L1,RANGRF,CHCHD7,HMGA1,PSME2,PABPC1,NAT9,BCAM,ZNF836,BLVRB,PPP3CB,SERPINB6,RABEP2,TA
,UCK2,ATPAF2,CASC4,STOML2,C19ORF54,STAU1,CARS,RPL36,RBM3,INO80E,HSP90AB1,ZNF524,TPM2,S
,UCK2,ATPAF2,CASC4,STOML2,C19ORF54,STAU1,CARS,RPL36,RBM3,INO80E,HSP90AB1,ZNF524,TPM2,S
UZ,PSMA3,MUC1,METTL17,ECSIT,EPB41L5,ATG12,PPRC1,RPS25,ZBTB44,NIP7,TTC9C,COX4I1,DUSP22,SL
UZ,PSMA3,MUC1,METTL17,ECSIT,EPB41L5,ATG12,PPRC1,RPS25,ZBTB44,NIP7,TTC9C,COX4I1,DUSP22,SL
T2,UQCRQ,SNF8,GRK6,HMGA2,CDC37,RPS16,CPSF3L,TSG101,OBSL1,PTPN13,UBR4,YLPM1,PHF21A,CHC
T2,UQCRQ,SNF8,GRK6,HMGA2,CDC37,RPS16,CPSF3L,TSG101,OBSL1,PTPN13,UBR4,YLPM1,PHF21A,CHC
33A,FDPS,DYNC1I2,AP2S1,ZNF843,HAUS1,ZNF345,LAMTOR4,NDUFB10,ZNF586,ERAL1,BCAP29,DNAJB12
33A,FDPS,DYNC1I2,AP2S1,ZNF843,HAUS1,ZNF345,LAMTOR4,NDUFB10,ZNF586,ERAL1,BCAP29,DNAJB12
,TRMT112,CHPF,RPS20,NIT2,RPS15,DCTD,UBXN1,ERLEC1,HSPB1,SAT1,ERCC5,EMC4,LSR,CHCHD7,MEIS2,
RPV3,THADA,MROH1,SNX5,NUP133,TMEM147,ELP4,TSC22D3,ZXDC,COBL,ELOF1,PEX26,ARL16,HAX1,SLC
'1,UBE2L6,VRK3,LZTFL1,ZC3H14,FIS1,SLC15A3,PPM1K,TUBA1C,DST,LRRC1,PHF19,VCAN,LPCAT3,CD44,TM
'1,UBE2L6,VRK3,LZTFL1,ZC3H14,FIS1,SLC15A3,PPM1K,TUBA1C,DST,LRRC1,PHF19,VCAN,LPCAT3,CD44,TM

DX41,USP45,PKP4,ZCCHC4,STXBP2,CHCHD5,NAA10,SEC31A,ATP10D,EGFR,CASZ1,SUMO3,PHF20L1,LGM

DX41,USP45,PKP4,ZCCHC4,STXBP2,CHCHD5,NAA10,SEC31A,ATP10D,EGFR,CASZ1,SUMO3,PHF20L1,LGM

GCX,CHPF,STRIP2,RPS20,LYNX1,TWSG1,MAPK13,ZSWIM7,RPS15,VWA2,LRRC27,MVD,SAT1,LMBR1,NAP1

,ACHE,PEX16,SUFU,ZNF33A,MRPS33,EXOSC2,POLH,LAMTOR4,CCDC171,C12ORF73,MRPL43,SETD4,KIAA

,ACHE,PEX16,SUFU,ZNF33A,MRPS33,EXOSC2,POLH,LAMTOR4,CCDC171,C12ORF73,MRPL43,SETD4,KIAA

IT,ATG12,PPRC1,RPS25,ZBTB44,COX4I1,DUSP22,SLC25A10,SPOCK2,THEM4,DBNDD2,SDF4,NISCH,TMED

IT,ATG12,PPRC1,RPS25,ZBTB44,COX4I1,DUSP22,SLC25A10,SPOCK2,THEM4,DBNDD2,SDF4,NISCH,TMED

JFA3,C1ORF63,C5ORF24,SLC37A4,IDH3G,DNAJA3,CKLF,AKT1S1,INO80C,GTF3A,ARRB2,DIP2A,MLST8,CD

JFA3,C1ORF63,C5ORF24,SLC37A4,IDH3G,DNAJA3,CKLF,AKT1S1,INO80C,GTF3A,ARRB2,DIP2A,MLST8,CD

525,RPS20,GABPB1,VPS28,ZNF814,EMC4,PRKRIP1,DDOST,LSR,DOCK10,PSME2,PABPC1,RRNAD1,RBP1,TC

525,RPS20,GABPB1,VPS28,ZNF814,EMC4,PRKRIP1,DDOST,LSR,DOCK10,PSME2,PABPC1,RRNAD1,RBP1,TC

PALPP1,GGCX,CHPF,GABPB1,MPDU1,MAPK13,ZSWIM7,RPS15,ZNF814,GMPPB,MVD,IL32,NAP1L1,SERPI

PALPP1,GGCX,CHPF,GABPB1,MPDU1,MAPK13,ZSWIM7,RPS15,ZNF814,GMPPB,MVD,IL32,NAP1L1,SERPI

P4,C19ORF48,ZFAND2B,HNRNPC,HYPK,UPP1,ZNF250,NFAM1,CDK11A,RBM3,INO80E,MAPKAPK5,DGUOI

P4,C19ORF48,ZFAND2B,HNRNPC,HYPK,UPP1,ZNF250,NFAM1,CDK11A,RBM3,INO80E,MAPKAPK5,DGUOI

3L4,TJP1,TPT1,NTMT1,ANAPC13,TRPM6,SMAGP,TSPAN31,PPP2R5C,TMC7,GCDH,RALY,FAM69B,RER1,D

3L4,TJP1,TPT1,NTMT1,ANAPC13,TRPM6,SMAGP,TSPAN31,PPP2R5C,TMC7,GCDH,RALY,FAM69B,RER1,D

7L1,ASF1B,CCDC25,EEF1B2,PPP1CC,PDLIM5,RFX3,PPA1,DGKE,ENDOV,CAMTA1,RPL28,PSMA7,APEH,RPL

1,PHF19,GRAMD1C,VCAN,TM9SF1,MAGED2,SON,TET2,SNF8,HMGA2,CPSF3L,UBR4,PSMA1,CHCHD1,SLC

1,PHF19,GRAMD1C,VCAN,TM9SF1,MAGED2,SON,TET2,SNF8,HMGA2,CPSF3L,UBR4,PSMA1,CHCHD1,SLC

A3,MUC1,METTL17,ECSIT,EPB41L5,RPS25,ZBTB44,NIP7,TTC9C,SLC25A10,TMX2,SPOCK2,THEM4,RDH11,

3,PRKCSH,RPS4X,AK2,CEP63,ZNF783,LRRC23,PEX16,FBXO34,SLC39A14,ACSS1,CCT4,EXOSC8,TCFL5,UBE2

IT2,JUP,TMEM205,PTK2,AUNIP,C14ORF2,GUK1,CENPN,PCYT2,USE1,AGPAT4,MAF,ASF1B,CCDC25,DEF8,

APG,CLDND1,VIM,LIG1,PSMA7,TIMM17B,APEH,HYI,CAPN10,LYPLA2,NUTF2,PODXL,EPB41L4B,ATPAF2,C

APG,CLDND1,VIM,LIG1,PSMA7,TIMM17B,APEH,HYI,CAPN10,LYPLA2,NUTF2,PODXL,EPB41L4B,ATPAF2,C

TMEM134,LRRFIP1,RAB2A,ZNF524,ELP6,TPM2,GPBP1,SIAH1,ZEB2,TRMT112,CHPF,STRIP2,LYNX1,MAPK

TMEM134,LRRFIP1,RAB2A,ZNF524,ELP6,TPM2,GPBP1,SIAH1,ZEB2,TRMT112,CHPF,STRIP2,LYNX1,MAPK

,CASC4,ZNF7,ARL6IP4,PIGL,CYB561,STAU1,SLC9B2,SUGT1,UPP1,CARS,NFAM1,CDK11A,RBM3,MDH1,ZN

,APOLD1,RBM42,CNTRL,NUP62,SCN11A,OLFM1,IRF3,SCRN2,NDUFA11,AXIN1,TUBB3,MOSPD3,C5ORF24,

,APOLD1,RBM42,CNTRL,NUP62,SCN11A,OLFM1,IRF3,SCRN2,NDUFA11,AXIN1,TUBB3,MOSPD3,C5ORF24,

PRMT1,C19ORF54,TXNDC17,PIGL,ZFAND2B,AP1S3,CYB561,STAU1,UPP1,RPL36,RBM3,INO80E,SYNGR2,C

EDF1,RFX3,PPA1,DGKE,TNFRSF14,ENDOV,CAMTA1,CAPG,CLDND1,VIM,LIG1,PSMA7,TMEM126B,XAF1,A

EDF1,RFX3,PPA1,DGKE,TNFRSF14,ENDOV,CAMTA1,CAPG,CLDND1,VIM,LIG1,PSMA7,TMEM126B,XAF1,A

MM17B,APEH,HYI,CAPN10,NUTF2,PODXL,CAMLG,TCF4,PPOX,ATPAF2,C17ORF62,PRMT1,WNK1,C19ORF

IF250,MDH1,CCNL2,LRRFIP1,ZNF524,GPBP1,SIAH1,ASPH,TRMT112,RPTOR,STRIP2,ZNF625,RPS20,GABPB

IF250,MDH1,CCNL2,LRRFIP1,ZNF524,GPBP1,SIAH1,ASPH,TRMT112,RPTOR,STRIP2,ZNF625,RPS20,GABPB

EEF1D,NAA10,MED6,LLGL2,SSH2,FKBP3,EGLN2,SIRT5,KCTD7,ZNF506,SPINT2,FTH1,SMARCA4,PTK7,TME

EEF1D,NAA10,MED6,LLGL2,SSH2,FKBP3,EGLN2,SIRT5,KCTD7,ZNF506,SPINT2,FTH1,SMARCA4,PTK7,TME

ASC4,STOML2,WNK1,PPAN,HNRNPC,STAU1,UPP1,ZNF250,RBM3,MDH1,INO80E,CCNL2,CEP70,RAB2A,Z

ASC4,STOML2,WNK1,PPAN,HNRNPC,STAU1,UPP1,ZNF250,RBM3,MDH1,INO80E,CCNL2,CEP70,RAB2A,Z

,FAM32A,ZXDC,COBL,PRRC2B,ELOF1,C12ORF57,MPHOSPH10,NUBP2,EIF4A1,C6ORF132,ZMYM6,OGFOD

RPM4,GNB5,CBR1,COMMD4,CCDC40,UBE2N,MFSD11,ZNHIT3,PDCD6,MFF,ZNF81,FAM32A,COX20,TME

X4I1,PLCB2,DUSP22,SPOCK2,THEM4,DBNDD2,SDF4,TMED4,CENPK,CD99L2,COASY,SRSF1,AUP1,FOXRED

X4I1,PLCB2,DUSP22,SPOCK2,THEM4,DBNDD2,SDF4,TMED4,CENPK,CD99L2,COASY,SRSF1,AUP1,FOXRED

CH,CD99L2,GGPS1,COASY,SRSF1,AUP1,RABL5,KXD1,PARD6G,CLASRP,JPH2,C1ORF198,UGGT2,MICAL1,H

CH,CD99L2,GGPS1,COASY,SRSF1,AUP1,RABL5,KXD1,PARD6G,CLASRP,JPH2,C1ORF198,UGGT2,MICAL1,H

H2,C1ORF198,MICAL1,HS2ST1,SLC39A13,YIF1A,PRMT2,JUP,TBX6,TMEM205,PTK2,C14ORF2,GUK1,PCYT2

H2,C1ORF198,MICAL1,HS2ST1,SLC39A13,YIF1A,PRMT2,JUP,TBX6,TMEM205,PTK2,C14ORF2,GUK1,PCYT2

X,ATPAF2,C17ORF62,PRMT1,CASC4,WNK1,C19ORF54,ZNF7,PPAN,ZFAND2B,AP1S3,HNRNPC,CYB561,ZN

X,ATPAF2,C17ORF62,PRMT1,CASC4,WNK1,C19ORF54,ZNF7,PPAN,ZFAND2B,AP1S3,HNRNPC,CYB561,ZN

23A,HSD17B10,GGT5,SLC16A1,ILF3,AGAP3,EEF1D,CARHSP1,ATP10D,CAMK2D,SUMO3,PHF20L1,EGLN2,

23A,HSD17B10,GGT5,SLC16A1,ILF3,AGAP3,EEF1D,CARHSP1,ATP10D,CAMK2D,SUMO3,PHF20L1,EGLN2,

2ORF57,DCTN5,ERGIC1,EIF4A1,TMEM108,TSEN34,MPV17,PICK1,ECE1,ZNF512,TPT1,UBA1,ANAPC13,TM

2ORF57,DCTN5,ERGIC1,EIF4A1,TMEM108,TSEN34,MPV17,PICK1,ECE1,ZNF512,TPT1,UBA1,ANAPC13,TM

0C,AIFM2,SNRPA1,HMGN1,MFGE8,MYL12A,FAM104B,PLD1,FIG4,BTF3L4,C6ORF141,AK2,MLH1,PCMTD1

T1,UBE2I,SLC39A13,MAGT1,JUP,TBX6,TMEM205,PTK2,AUNIP,C14ORF2,PCYT2,FBXW9,USE1,HSBP1L1,AG

T1,UBE2I,SLC39A13,MAGT1,JUP,TBX6,TMEM205,PTK2,AUNIP,C14ORF2,PCYT2,FBXW9,USE1,HSBP1L1,AG

A1,DGKE,VAMP8,CAMTA1,CAPG,CRIP1,LIG1,PSMA7,CDC14B,TIMM17B,APEH,HYI,CAPN10,NUTF2,PODX

,VIM,LIG1,PSMA7,TMEM126B,XAF1,APEH,HYI,LYPLA2,NUTF2,PODXL,TCF4,UCK2,PPOX,TMPO,ATPAF2,C1

,VIM,LIG1,PSMA7,TMEM126B,XAF1,APEH,HYI,LYPLA2,NUTF2,PODXL,TCF4,UCK2,PPOX,TMPO,ATPAF2,C1

,FAM3A,MUC1,METTL17,ECSIT,EPB41L5,ATG12,RPS25,ZBTB44,NIP7,SLC25A10,TMX2,SPOCK2,THEM4,R

,FAM3A,MUC1,METTL17,ECSIT,EPB41L5,ATG12,RPS25,ZBTB44,NIP7,SLC25A10,TMX2,SPOCK2,THEM4,R

NFX1,DUSP18,HDAC10,MRPL51,ISOC2,FRMD8,MT1F,MRPL48,ANKRD10,ACOT9,MTHFD2L,RAP1B,ZNF61

NFX1,DUSP18,HDAC10,MRPL51,ISOC2,FRMD8,MT1F,MRPL48,ANKRD10,ACOT9,MTHFD2L,RAP1B,ZNF61

OSC9,TXNDC17,ARL6IP4,C19ORF48,ZFAND2B,HYPK,ZNF250,NFAM1,CDK11A,RBM3,INO80E,MAPKAPK5,

OSC9,TXNDC17,ARL6IP4,C19ORF48,ZFAND2B,HYPK,ZNF250,NFAM1,CDK11A,RBM3,INO80E,MAPKAPK5,

3,AGAP3,MDH2,STXBP2,EEF1D,HSPH1,RRP8,CARHSP1,MT1E,CNST,SUMO3,EGLN2,SIRT5,B3GAT3,RFXAN

FKBP3,NDUFV2,B3GAT3,GTF2IRD2,COPS4,ZFPL1,MFSD10,MRPS18C,GMPR2,RIC8A,SLC35B1,GNB5,THAD

FKBP3,NDUFV2,B3GAT3,GTF2IRD2,COPS4,ZFPL1,MFSD10,MRPS18C,GMPR2,RIC8A,SLC35B1,GNB5,THAD

PODXL,EPB41L4B,TCF4,UCK2,TMPO,ATPAF2,LSM12,C17ORF62,PRMT1,STOML2,C19ORF54,TXNDC17,AR

A1,RALGPS1,GTF2IRD1,NXN,EPS15L1,NUDT1,POLR3H,B2M,ABHD17A,ILF3,PKP4,EEF1D,NAA10,MED6,LL

A1,RALGPS1,GTF2IRD1,NXN,EPS15L1,NUDT1,POLR3H,B2M,ABHD17A,ILF3,PKP4,EEF1D,NAA10,MED6,LL

K2,PPOX,ATPAF2,C17ORF62,PRMT1,WNK1,STRADA,AP1S3,HNRNPC,CYB561,SHARPIN,UPP1,ZNF250,RPL

K2,PPOX,ATPAF2,C17ORF62,PRMT1,WNK1,STRADA,AP1S3,HNRNPC,CYB561,SHARPIN,UPP1,ZNF250,RPL

1,NIT2,LYNX1,ZSWIM7,RPS15,DCTD,ZNF814,ERLEC1,GMPPB,HSPB1,MVD,ERCC5,NAP1L1,DDOST,C12OR

A,RBM3,COPZ1,INO80E,CCNL2,DGUOK,HSP90AB1,FAM219B,RAB2A,ZNF524,ATP5J,TPM2,GPBP1,GPALP

A,RBM3,COPZ1,INO80E,CCNL2,DGUOK,HSP90AB1,FAM219B,RAB2A,ZNF524,ATP5J,TPM2,GPBP1,GPALP

,RBM3,INO80E,MAPKAPK5,DGUOK,HSP90AB1,CEP70,TMEM134,FAM219B,RAB2A,RMDN1,GPBP1,GPAL
,RBM3,INO80E,MAPKAPK5,DGUOK,HSP90AB1,CEP70,TMEM134,FAM219B,RAB2A,RMDN1,GPBP1,GPAL
RDX,IDH3G,DNAJA3,AKT1S1,NSFL1C,FARS2,DHRS4,ZBTB8OS,CARM1,PRKCSH,NFE2L1,BTF3L4,UBA52,KIF1
RDX,IDH3G,DNAJA3,AKT1S1,NSFL1C,FARS2,DHRS4,ZBTB8OS,CARM1,PRKCSH,NFE2L1,BTF3L4,UBA52,KIF1

L6IP5,RBBP6,NREP,CHTOP,BAX,TSPAN31,TMEM9,FAM192A,ADRA1A,ZNF593,GLYCTK,TCEAL3,RBM42,CL
RPS18C,GMPR2,PFKFB2,THADA,MROH1,LFNG,RPS11,SNX5,MRPS17,TMEM147,ELP4,PCID2,TGOLN2,POL
RPS18C,GMPR2,PFKFB2,THADA,MROH1,LFNG,RPS11,SNX5,MRPS17,TMEM147,ELP4,PCID2,TGOLN2,POL
ID4,CCDC40,GPR56,RPAIN,COX20,POLR1C,MDP1,STX10,USP4,C12ORF57,ARL16,EIF4A1,STAU2,FAM126A
C25A10,TMX2,SPOCK2,THEM4,DBNDD2,SDF4,NISCH,TMED4,RPS6,RPL29,RTEL1,AUP1,YIPF3,FOXRED1,TR
C25A10,TMX2,SPOCK2,THEM4,DBNDD2,SDF4,NISCH,TMED4,RPS6,RPL29,RTEL1,AUP1,YIPF3,FOXRED1,TR
T8,AP3B2,RPS9,SLC35G1,RCE1,MRPL33,ZNF410,RUVBL2,PCMT1,EBPL,TESC,IGSF8,NUDT5,RBPMS,NNMT,
T8,AP3B2,RPS9,SLC35G1,RCE1,MRPL33,ZNF410,RUVBL2,PCMT1,EBPL,TESC,IGSF8,NUDT5,RBPMS,NNMT,
SYP,CLASP2,RER1,NEK11,NKIRAS2,NUP62,ZNF675,GTF2H1,CLTA,BRD8,RBM4,THOC3,HNRNPUL1,SH3GL1
LC30A5,TRPV3,CBR1,PHC3,COMMD4,SNX5,ZNF81,TMEM147,SMIM4,DCTN5,NMNAT3,ARL16,BTN3A2,EI
LC30A5,TRPV3,CBR1,PHC3,COMMD4,SNX5,ZNF81,TMEM147,SMIM4,DCTN5,NMNAT3,ARL16,BTN3A2,EI

M147,ELP4,ACOT2,USP40,TGOLN2,PEX2,RPL37A,PIGH,DCTN5,NDUFS3,L3MBTL2,OXLD1,C6ORF132,ARIH
LA,AKT1S1,FARS2,B3GNTL1,REPS2,KAT8,KIF13B,CARM1,TTLL3,SYPL1,MDK,RNPS1,FIG4,IAH1,VAPA,AIG1,
OV,CAMTA1,RPL28,VIM,TIMM17B,HYI,RPL21,TXNL4A,NUTF2,PODXL,CAMLG,ATPAF2,LSM12,PRMT1,W
OV,CAMTA1,RPL28,VIM,TIMM17B,HYI,RPL21,TXNL4A,NUTF2,PODXL,CAMLG,ATPAF2,LSM12,PRMT1,W
,SLC37A4,AAMDC,RHEB,DCAF5,GRAMD1A,HMOX2,DIP2A,PRTFDC1,MLST8,HMGN1,MFGE8,GMFG,SGSM
,SLC37A4,AAMDC,RHEB,DCAF5,GRAMD1A,HMOX2,DIP2A,PRTFDC1,MLST8,HMGN1,MFGE8,GMFG,SGSM
ML2,C19ORF54,TXNDC17,ARL6IP4,C19ORF48,PPAN,ZFAND2B,HNRNPC,SHARPIN,ZNF644,SLC9B2,HYPK,
ML2,C19ORF54,TXNDC17,ARL6IP4,C19ORF48,PPAN,ZFAND2B,HNRNPC,SHARPIN,ZNF644,SLC9B2,HYPK,

H,HYI,CAPN10,NUTF2,PODXL,CAMLG,EPB41L4B,TCF4,PPOX,ATPAF2,C17ORF62,PRMT1,CASC4,STOML2,W
H,HYI,CAPN10,NUTF2,PODXL,CAMLG,EPB41L4B,TCF4,PPOX,ATPAF2,C17ORF62,PRMT1,CASC4,STOML2,W
AB2A,RMDN1,GPBP1,GPALPP1,PNPO,CHPF,MAPK13,ZSWIM7,ERLEC1,GMPPB,MVD,SAT1,ERCC5,EMC4,IL
AB2A,RMDN1,GPBP1,GPALPP1,PNPO,CHPF,MAPK13,ZSWIM7,ERLEC1,GMPPB,MVD,SAT1,ERCC5,EMC4,IL

PHTF2,DBNL,BCAM,RBP1,BLVRB,SERPINB6,SLC33A1,RALGPS1,ZNF706,ZNF687,PDCD5,GTF2IRD1,ANAPC
PHTF2,DBNL,BCAM,RBP1,BLVRB,SERPINB6,SLC33A1,RALGPS1,ZNF706,ZNF687,PDCD5,GTF2IRD1,ANAPC
15A3,PPM1K,TUBA1C,DST,LRRC1,PHF19,VCAN,LPCAT3,CD44,TM9SF1,MAGED2,SON,UQCRQ,GRK6,HMG
DUSP22,SLC25A10,DBNDD2,SDF4,TMED4,RPS6,CD99L2,GGPS1,RTEL1,COASY,AUP1,FOXRED1,RABL5,TRN
DUSP22,SLC25A10,DBNDD2,SDF4,TMED4,RPS6,CD99L2,GGPS1,RTEL1,COASY,AUP1,FOXRED1,RABL5,TRN
B3,MOSPD3,C5ORF24,RPS3,MAGOHB,CREB3L2,AIFM2,HMOX2,SPHK2,RBM6,HMGN1,NDUFB2,CDK2AP2
K1,MED6,SH3GLB1,SSH2,HNRNPLL,EGLN2,B3GAT3,RFXANK,ZNF506,REEP5,FTH1,PFDN5,TCTN1,TMEM10
K1,MED6,SH3GLB1,SSH2,HNRNPLL,EGLN2,B3GAT3,RFXANK,ZNF506,REEP5,FTH1,PFDN5,TCTN1,TMEM10
3,TMEM39B,PPIH,DALRD3,NSUN5,DAZAP2,ZFPL1,MFSD10,GMPR2,RIC8A,TCP1,CD74,PIGT,NHP2,UBE2N,
3,TMEM39B,PPIH,DALRD3,NSUN5,DAZAP2,ZFPL1,MFSD10,GMPR2,RIC8A,TCP1,CD74,PIGT,NHP2,UBE2N,
RUVBL2,PCMT1,TESC,IGSF8,NUDT5,RBPMS,SNRPN,SPAG7,WDR62,IL1RAP,STYXL1,RFX5,PLEKHA5,RANBP
RUVBL2,PCMT1,TESC,IGSF8,NUDT5,RBPMS,SNRPN,SPAG7,WDR62,IL1RAP,STYXL1,RFX5,PLEKHA5,RANBP

NDUFA7,CYB5D2,RABEPK,FDPS,EXOSC2,RBM7,PSMC5,AP2S1,CDK2AP1,CAMKMT,EXOSC8,DDX17,SNX13,
KPNA5,MPHOSPH10,MYL9,METTL5,C6ORF132,FAM126A,RC3H2,OGFOD2,CIZ1,EIF3G,FBXL18,TCF25,RNF
KPNA5,MPHOSPH10,MYL9,METTL5,C6ORF132,FAM126A,RC3H2,OGFOD2,CIZ1,EIF3G,FBXL18,TCF25,RNF
NF524,ATP5J,GPBP1,SIAH1,TRMT112,STRIP2,RPS20,GABPB1,TWSG1,MAPK13,ZSWIM7,UBXN1,ERLEC1,L
1,RC3H2,KLHL7,SSR3,MRPL55,PICK1,ECE1,OCIAD1,RBBP6,NREP,CHTOP,BAX,TSPAN31,TMEM9,NT5E,AD
1,RC3H2,KLHL7,SSR3,MRPL55,PICK1,ECE1,OCIAD1,RBBP6,NREP,CHTOP,BAX,TSPAN31,TMEM9,NT5E,AD
H2,KLHL7,SSR3,MRPL55,PICK1,ECE1,OCIAD1,RBBP6,NREP,CHTOP,BAX,TSPAN31,TMEM9,NT5E,ADRA1A,Z

P62,NADSYN1,ZNF511,IRF3,RNF130,D2HGDH,CC2D1B,ARPC5,TLCD1,NDUFA11,AXIN1,MOSPD3,SLC37A

ZNF524,ATP5J,GPBP1,GPALPP1,SIAH1,PNPO,RPTOR,ZNF625,MPDU1,VPS28,ZSWIM7,RPS15,UBXN1,ZNF
ZNF524,ATP5J,GPBP1,GPALPP1,SIAH1,PNPO,RPTOR,ZNF625,MPDU1,VPS28,ZSWIM7,RPS15,UBXN1,ZNF
PTOR,CHPF,STRIP2,GABPB1,LYNX1,MAPK13,RPS15,ERLEC1,LRRC27,LMBR1,NAP1L1,PRKRIP1,DDOST,SYN
PTOR,CHPF,STRIP2,GABPB1,LYNX1,MAPK13,RPS15,ERLEC1,LRRC27,LMBR1,NAP1L1,PRKRIP1,DDOST,SYN
NO80E,CCNL2,CEP70,RAB2A,ATP5J,TPM2,GPBP1,GPALPP1,SIAH1,CHPF,STRIP2,RPS20,GABPB1,LYNX1,M
NO80E,CCNL2,CEP70,RAB2A,ATP5J,TPM2,GPBP1,GPALPP1,SIAH1,CHPF,STRIP2,RPS20,GABPB1,LYNX1,M
KRD10,ACTG1,EHMT1,RAP1B,IFT81,UBE2A,PKN2,CREM,TUBGCP2,SWI5,MAP4,CORO1B,OAZ1,SEMA5A,F
B1BP1,PARD6G,SH3YL1,CLASRP,C1ORF198,UGGT2,MICAL1,JUP,CDC42,TMEM205,XBP1,PTK2,CELF1,AU
B1BP1,PARD6G,SH3YL1,CLASRP,C1ORF198,UGGT2,MICAL1,JUP,CDC42,TMEM205,XBP1,PTK2,CELF1,AU

5,MED15,PARD6G,SH3YL1,C1ORF198,UGGT2,MICAL1,UBE2I,SLC39A13,YIF1A,PRMT2,JUP,TBX6,TMEM2
5,MED15,PARD6G,SH3YL1,C1ORF198,UGGT2,MICAL1,UBE2I,SLC39A13,YIF1A,PRMT2,JUP,TBX6,TMEM2

34,RPL8,CREB3L4,TCF25,RNF181,TJP1,TPT1,SMAGP,SLC39A6,APOLD1,RALY,FAM69B,PSMD7,NADSYN1,
,ASF1B,EEF1B2,PPP1CC,EDF1,DGKE,ENDOV,CAPG,CRIP1,PPIL2,TROAP,CLDND1,APEH,HYI,TXNL4A,CAPN1
,ASF1B,EEF1B2,PPP1CC,EDF1,DGKE,ENDOV,CAPG,CRIP1,PPIL2,TROAP,CLDND1,APEH,HYI,TXNL4A,CAPN1
LASRP,JPH2,MICAL1,HS2ST1,UBE2I,SLC39A13,JUP,TBX6,TMEM205,XBP1,PTK2,C14ORF2,ATF7,PCYT2,FB
LASRP,JPH2,MICAL1,HS2ST1,UBE2I,SLC39A13,JUP,TBX6,TMEM205,XBP1,PTK2,C14ORF2,ATF7,PCYT2,FB
1,SUGT1,CDK11A,CCNL2,HSP90AB1,DUT,RAB2A,ELP6,TPM2,SIAH1,TSFM,ASPH,PNPO,GABPB1,NIT2,MP
NRP1,IFITM2,RHEB,MAGOHB,IDH3G,CREB3L2,UCHL5,GRAMD1A,AKT1S1,SPHK2,HMGN1,NDUFB2,NDUF

I,SLC39A13,YIF1A,PRMT2,JUP,TBX6,TMEM205,XBP1,PTK2,C14ORF2,GUK1,CENPN,PCYT2,HSBP1L1,MAF,
I,SLC39A13,YIF1A,PRMT2,JUP,TBX6,TMEM205,XBP1,PTK2,C14ORF2,GUK1,CENPN,PCYT2,HSBP1L1,MAF,
SMA7,TIMM17B,TMEM126B,RPL21,CAPN10,EPB41L4B,TCF4,PPOX,ATPAF2,LSM12,PRMT1,WNK1,C19O
SMA7,TIMM17B,TMEM126B,RPL21,CAPN10,EPB41L4B,TCF4,PPOX,ATPAF2,LSM12,PRMT1,WNK1,C19O
RF2,GUK1,IFI27L1,PDLIM5,RFX3,PPA1,DGKE,TNFRSF14,ENDOV,CAMTA1,CRIP1,PPIL2,RPL28,LIG1,PSMA
RF2,GUK1,IFI27L1,PDLIM5,RFX3,PPA1,DGKE,TNFRSF14,ENDOV,CAMTA1,CRIP1,PPIL2,RPL28,LIG1,PSMA
N5,DAZAP2,ZFPL1,MFSD10,GMPR2,RIC8A,TCP1,PIGT,NHP2,UBE2N,MFSD11,GPR56,NOP2,FAM32A,MA
N5,DAZAP2,ZFPL1,MFSD10,GMPR2,RIC8A,TCP1,PIGT,NHP2,UBE2N,MFSD11,GPR56,NOP2,FAM32A,MA
N,EPS15L1,ANAPC11,NOMO2,NT5C,USP45,SLC16A1,ILF3,AGAP3,STXBP2,EEF1D,CHEK1,SSH2,COX17,CA
N,EPS15L1,ANAPC11,NOMO2,NT5C,USP45,SLC16A1,ILF3,AGAP3,STXBP2,EEF1D,CHEK1,SSH2,COX17,CA
RAD23A,DDX41,SLC16A1,PKP4,PDE4DIP,MDH2,EEF1D,HSPH1,NAA10,RRP8,OAS3,SEC31A,MT1E,RPL18,A
RAD23A,DDX41,SLC16A1,PKP4,PDE4DIP,MDH2,EEF1D,HSPH1,NAA10,RRP8,OAS3,SEC31A,MT1E,RPL18,A

MD4,RPL28,LIG1,PSMA7,CDC14B,TIMM17B,HYI,CAPN10,UCK2,PRMT1,C19ORF54,PIGL,STRADA,PPAN,A
DK2AP2,NDUFA10,AZI2,MYL12A,U2AF1L4,ATL2,DDX10,PLD1,TMEM41A,EMC3,TAF9,RNPS1,DTD2,NMR

DK2AP2,NDUFA10,AZI2,MYL12A,U2AF1L4,ATL2,DDX10,PLD1,TMEM41A,EMC3,TAF9,RNPS1,DTD2,NMRA

OSPH10,EIF4A1,RC3H2,KLHL7,SSR3,PICK1,UBE2D4,ECE1,ZNF512,OCIAD1,ARL6IP5,SLMAP,RBBP6,NREP,CH

OSPH10,EIF4A1,RC3H2,KLHL7,SSR3,PICK1,UBE2D4,ECE1,ZNF512,OCIAD1,ARL6IP5,SLMAP,RBBP6,NREP,CH

5B,APEH,RPL21,CAPN10,PODXL,CAMLG,EPB41L4B,UCK2,PRMT1,WNK1,ZNF7,TXNDC17,ACAD10,CYB561,

YPK,UPP1,CARS,ZNF197,COPZ1,INO80E,HSP90AB1,DUT,ATP5J,TPM2,GPBP1,SIAH1,ASPH,PNPO,RPTOR,N

YPK,UPP1,CARS,ZNF197,COPZ1,INO80E,HSP90AB1,DUT,ATP5J,TPM2,GPBP1,SIAH1,ASPH,PNPO,RPTOR,N

17B,TMEM126B,RPL21,CAPN10,EPB41L4B,TCF4,PPOX,ATPAF2,LSM12,PRMT1,WNK1,C19ORF54,ARL6IP4

SLC9B2,UPP1,RPL36,NFAM1,RBM3,ZNF197,COPZ1,HSP90AB1,SYNGR2,RAB2A,ATP5J,ELP6,TPM2,GPALPI

SLC9B2,UPP1,RPL36,NFAM1,RBM3,ZNF197,COPZ1,HSP90AB1,SYNGR2,RAB2A,ATP5J,ELP6,TPM2,GPALPI

SLC9B2,UPP1,RPL36,NFAM1,RBM3,ZNF197,COPZ1,HSP90AB1,SYNGR2,RAB2A,ATP5J,ELP6,TPM2,GPALPI

ID1,FAM3A,MUC1,METTL17,EPB41L5,ATG12,RPS25,NIP7,SLC25A10,SPOCK2,RDH11,DBNDD2,SDF4,TME

ID1,FAM3A,MUC1,METTL17,EPB41L5,ATG12,RPS25,NIP7,SLC25A10,SPOCK2,RDH11,DBNDD2,SDF4,TME

F3A,DHRS4,NDUFB2,MFGE8,MYL12A,ZBTB8OS,SLC37A3,AKR1A1,PRKCSH,GPS2,FHL2,VAPA,CAMKK1,DH

F3A,DHRS4,NDUFB2,MFGE8,MYL12A,ZBTB8OS,SLC37A3,AKR1A1,PRKCSH,GPS2,FHL2,VAPA,CAMKK1,DH

SHARPIN,CBFA2T2,ZNF250,MOB1A,CDK11A,ZNF197,INO80E,DGUOK,RAB2A,TPM2,TSFM,TRMT112,PNP

SHARPIN,CBFA2T2,ZNF250,MOB1A,CDK11A,ZNF197,INO80E,DGUOK,RAB2A,TPM2,TSFM,TRMT112,PNP

S100A4,TXNRD2,PPIH,ARHGEF1,NSUN5,FKBP5,MFSD10,TCP1,SCAP,SLC30A5,TRPV3,THADA,CBR1,PHC3,N

ST,ANTXR2,STIP1,LRRC1,PHF19,GRAMD1C,VCAN,LPCAT3,CD44,TM9SF1,MAGED2,UQCRQ,SNF8,GRK6,CD

ST,ANTXR2,STIP1,LRRC1,PHF19,GRAMD1C,VCAN,LPCAT3,CD44,TM9SF1,MAGED2,UQCRQ,SNF8,GRK6,CD

,RPL28,PSMA7,TIMM17B,XAF1,RPL21,TXNL4A,LYPLA2,NUTF2,CAMLG,UCK2,ATPAF2,LSM12,PRMT1,WN

,RPL28,PSMA7,TIMM17B,XAF1,RPL21,TXNL4A,LYPLA2,NUTF2,CAMLG,UCK2,ATPAF2,LSM12,PRMT1,WN

,IP6K2,CLASRP,JPH2,UBE2I,KCNMA1,YIF1A,PRMT2,EIF3M,JUP,CDC42,XBP1,AUNIP,CENPN,ATF7,FBXW9,I

,IP6K2,CLASRP,JPH2,UBE2I,KCNMA1,YIF1A,PRMT2,EIF3M,JUP,CDC42,XBP1,AUNIP,CENPN,ATF7,FBXW9,I

BB3,MOSPD3,C5ORF24,RPS3,MAGOHB,CREB3L2,SMUG1,AIFM2,HMOX2,SPHK2,RBM6,HMGN1,NDUFB2,

MDH1,INO80E,MAPKAPK5,DUT,TMEM134,FAM219B,ELP6,GPBP1,TRMT112,CHPF,RPS20,NIT2,MPDU1,Z

,MTA3,MEIS2,PABPC1,PTPRB,NAT9,RAD1,RBMX,RABEP2,ZNF687,KLK10,NUDT1,B2M,USP45,AGAP3,PDE

,MTA3,MEIS2,PABPC1,PTPRB,NAT9,RAD1,RBMX,RABEP2,ZNF687,KLK10,NUDT1,B2M,USP45,AGAP3,PDE

MAPK13,UBXN1,HHLA3,LRRC27,SAT1,LMBR1,SERPINB1,LSR,MTA3,CHCHD7,MEIS2,HMGA1,PSME2,PHTF

RPL28,TIMM17B,TMEM126B,RPL21,TXNL4A,CAPN10,EPB41L4B,ATPAF2,LSM12,PRMT1,C19ORF54,ARL6I

PZ1,INO80E,CCNL2,DGUOK,HSP90AB1,DUT,TMEM134,FAM219B,RAB2A,ZNF524,ATP5J,RMDN1,GPBP1,S

PZ1,INO80E,CCNL2,DGUOK,HSP90AB1,DUT,TMEM134,FAM219B,RAB2A,ZNF524,ATP5J,RMDN1,GPBP1,S

2,NDUFA10,AZI2,MYL12A,U2AF1L4,ATL2,DDX10,PLD1,TMEM41A,EMC3,TAF9,RNPS1,DTD2,NMRAL1,ZN

ED4,RPS6,CD99L2,GGPS1,RTEL1,COASY,AUP1,FOXRED1,RABL5,PARD6G,SH3YL1,IP6K2,CLASRP,JPH2,UG

MAGT1,JUP,CDC42,TMEM205,PTK2,AUNIP,C14ORF2,GUK1,CENPN,CARD14,FBXW9,USE1,AGPAT4,IFI27L

MAGT1,JUP,CDC42,TMEM205,PTK2,AUNIP,C14ORF2,GUK1,CENPN,CARD14,FBXW9,USE1,AGPAT4,IFI27L

OT1,POT1,POLR3H,RAD23A,PKP4,EEF1D,HSPH1,OAS3,SEC31A,RPL18,ATP10D,SH3GLB1,PHF20L1,HNRNPA

OT1,POT1,POLR3H,RAD23A,PKP4,EEF1D,HSPH1,OAS3,SEC31A,RPL18,ATP10D,SH3GLB1,PHF20L1,HNRNPA

D1,NIPAL3,BAX,CDK4,NME1,ANAPC13,EEF2K,FAM192A,TCEAL3,GALT,VDAC1,ACAA1,GTF2H1,TAF1D,D2

D1,NIPAL3,BAX,CDK4,NME1,ANAPC13,EEF2K,FAM192A,TCEAL3,GALT,VDAC1,ACAA1,GTF2H1,TAF1D,D2

D1,NIPAL3,BAX,CDK4,NME1,ANAPC13,EEF2K,FAM192A,TCEAL3,GALT,VDAC1,ACAA1,GTF2H1,TAF1D,D2

2,PPP1CC,GADL1,PPA1,ENDOV,CAPG,CRIP1,PPIL2,TROAP,RPL28,CLDND1,LIG1,TIMM17B,RBM33,LYPLA
,TLCD1,GEMIN2,TUBB3,PSMD10,MOSPD3,C5ORF24,RPS3,MAGOHB,CREB3L2,PCTP,AIFM2,HMOX2,SPH
,TLCD1,GEMIN2,TUBB3,PSMD10,MOSPD3,C5ORF24,RPS3,MAGOHB,CREB3L2,PCTP,AIFM2,HMOX2,SPH

,PSMA1,FUZ,FAM3A,METTL17,ECSIT,ATG12,NLRP1,ZBTB44,NIP7,TTC9C,DUSP22,TMX2,SPOCK2,THEM4,
,PSMA1,FUZ,FAM3A,METTL17,ECSIT,ATG12,NLRP1,ZBTB44,NIP7,TTC9C,DUSP22,TMX2,SPOCK2,THEM4,
SPOCK2,THEM4,DBNDD2,SDF4,NISCH,TMED4,RPL29,CD99L2,GGPS1,AUP1,FOXRED1,TRNT1,KXD1,ITGB1
SPOCK2,THEM4,DBNDD2,SDF4,NISCH,TMED4,RPL29,CD99L2,GGPS1,AUP1,FOXRED1,TRNT1,KXD1,ITGB1
DDOST,RANGRF,LSR,CHCHD7,MEIS2,RBP1,RBMX,TOMM40,PPP3CB,RABEP2,ZNF20,RALGPS1,PDCD5,PO

D2B,HNRNPC,ZNF644,CBFA2T2,SLC9B2,HYPK,MOB1A,CDK11A,RBM3,MDH1,COPZ1,INO80E,CCNL2,DGU
D2B,HNRNPC,ZNF644,CBFA2T2,SLC9B2,HYPK,MOB1A,CDK11A,RBM3,MDH1,COPZ1,INO80E,CCNL2,DGU
M102A,PFKFB2,RIC8A,CCDC40,MFSD11,LFNG,XYLT2,PCID2,ZXDC,COBL,STX10,ERGIC1,NMNAT3,MYL9,C6
4,CBFA2T2,HYPK,MOB1A,CDK11A,RBM3,COPZ1,INO80E,CCNL2,DGUOK,HSP90AB1,FAM219B,RAB2A,ZN
4,CBFA2T2,HYPK,MOB1A,CDK11A,RBM3,COPZ1,INO80E,CCNL2,DGUOK,HSP90AB1,FAM219B,RAB2A,ZN
D3,ZNF396,MRPS24,MAGOHB,CREB3L2,VPS25,GRAMD1A,HMOX2,SPHK2,HMGN1,NDUFB2,NDUFA10,RP
D3,ZNF396,MRPS24,MAGOHB,CREB3L2,VPS25,GRAMD1A,HMOX2,SPHK2,HMGN1,NDUFB2,NDUFA10,RP
RK3,LZTFL1,ZC3H14,FIS1,SLC15A3,PPM1K,IRF9,ANTXR2,STIP1,LRRC1,PHF19,VCAN,LPCAT3,CD44,TM9SF
8,PPAN,HNRNPC,CYB561,SHARPIN,CBFA2T2,SLC9B2,SUGT1,CARS,NFAM1,INO80E,MAPKAPK5,DGUOK,H
8,PPAN,HNRNPC,CYB561,SHARPIN,CBFA2T2,SLC9B2,SUGT1,CARS,NFAM1,INO80E,MAPKAPK5,DGUOK,H
GCP2,NCK2,ETHE1,GSTO2,ARHGEF7,LRRC37A,TMEM229B,FDFT1,MAGOH,STAMBP,SEC14L1,CSRP2BP,U
TG12,PPRC1,RPS25,ZBTB44,NIP7,TTC9C,COX4I1,PLCB2,DUSP22,SLC25A10,THEM4,DBNDD2,SDF4,NISCH,
TG12,PPRC1,RPS25,ZBTB44,NIP7,TTC9C,COX4I1,PLCB2,DUSP22,SLC25A10,THEM4,DBNDD2,SDF4,NISCH,
,CDK2AP2,SGSM3,C2ORF15,FAM104B,ATL2,SNX3,TMEM41A,MDK,NFE2L1,SZT2,ZNF609,RAB26,NDUFA
,EIF2A,OLFM1,PAM16,FCHO2,TRAPPC2,TAF1D,RNF130,NAA38,TLCD1,NDUFA11,HNRNPUL1,RPS3A,TCP
,EIF2A,OLFM1,PAM16,FCHO2,TRAPPC2,TAF1D,RNF130,NAA38,TLCD1,NDUFA11,HNRNPUL1,RPS3A,TCP
UBGCP2,SWI5,OAZ1,WDR6,OPA3,LTA4H,MIEN1,LRRC37A,DBN1,MARK2,ESRRA,GINS4,UBA6,TMED1,DP

FA10,AZI2,MYL12A,U2AF1L4,ATL2,DDX10,PLD1,TMEM41A,EMC3,TAF9,RNPS1,DTD2,NMRAL1,ZNF483,L
FA10,AZI2,MYL12A,U2AF1L4,ATL2,DDX10,PLD1,TMEM41A,EMC3,TAF9,RNPS1,DTD2,NMRAL1,ZNF483,L

93,GLYCTK,RBM42,CLASP2,PCNP,CREBZF,EIF2A,RPL24,NADSYN1,ZNF511,GTF2H1,RNF130,MYL6,NAA38,
IP4,C19ORF48,CYB561,CBFA2T2,SLC9B2,UPP1,RPL36,NFAM1,RBM3,LAMP2,ZNF197,COPZ1,HSP90AB1,
IP4,C19ORF48,CYB561,CBFA2T2,SLC9B2,UPP1,RPL36,NFAM1,RBM3,LAMP2,ZNF197,COPZ1,HSP90AB1,
IP4,C19ORF48,CYB561,CBFA2T2,SLC9B2,UPP1,RPL36,NFAM1,RBM3,LAMP2,ZNF197,COPZ1,HSP90AB1,

L4,KIF13B,ATL2,DDX10,EMC3,RNPS1,AK2,AIG1,MLH1,DTD2,DHDDS,XPNPEP1,ZNF483,ZNF783,ACHE,FBX
HF20L1,HNRNPA1,CD151,FKBP3,NDUFV2,B3GAT3,GTF2IRD2,COPS4,TMEM39B,ZFPL1,TPM4,MFSD10,MF
HF20L1,HNRNPA1,CD151,FKBP3,NDUFV2,B3GAT3,GTF2IRD2,COPS4,TMEM39B,ZFPL1,TPM4,MFSD10,MF
,ALS2,IK,EXOSC8,DDX17,SLC31A1,RCAN1,C12ORF73,NSMF,ATP9B,IMMP1L,RAP1B,IFT81,COQ2,TUBGCP

MAF,CCDC25,DEF8,EEF1B2,PPP1CC,EDF1,RFX3,PPA1,DGKE,ENDOV,CAMTA1,CRIP1,PPIL2,RPL28,CLDND1

MAF,CCDC25,DEF8,EEF1B2,PPP1CC,EDF1,RFX3,PPA1,DGKE,ENDOV,CAMTA1,CRIP1,PPIL2,RPL28,CLDND1

1,POLR3H,RAD23A,GGT5,NT5C,SLC16A1,ILF3,AGAP3,MDH2,STXBP2,EEF1D,HSPH1,NAA10,RRP8,CARHSP

1,POLR3H,RAD23A,GGT5,NT5C,SLC16A1,ILF3,AGAP3,MDH2,STXBP2,EEF1D,HSPH1,NAA10,RRP8,CARHSP

K2,JPH2,C1ORF198,MICAL1,HS2ST1,UBE2I,SLC39A13,YIF1A,PRMT2,EIF3M,TBX6,TMEM205,XBP1,PTK2,C1

HLA3,MVD,ERCC5,IL32,LMBR1,MCTS1,NAP1L1,PRKRIP1,LSR,HMGA1,PHTF2,PABPC1,PTPRB,DBNL,BCAM,

RPC5,SIN3A,ST7,NDUFA11,MOSPD3,TCP11L1,SLC37A4,RHEB,MAGOHB,CREB3L2,EML3,VPS25,INO80C,D

RPC5,SIN3A,ST7,NDUFA11,MOSPD3,TCP11L1,SLC37A4,RHEB,MAGOHB,CREB3L2,EML3,VPS25,INO80C,D

P90AB1,CEP70,TMEM134,LRRFIP1,ZNF524,ATP5J,GPBP1,SIAH1,GGCX,ASPH,MPDU1,GMPPB,HSPB1,HHL

P90AB1,CEP70,TMEM134,LRRFIP1,ZNF524,ATP5J,GPBP1,SIAH1,GGCX,ASPH,MPDU1,GMPPB,HSPB1,HHL

S3,HAX1,SLC35E1,RC3H2,EIF3G,RPL8,ECE1,OCIAD1,RBBP6,ZNF248,NREP,CHTOP,TPT1,SMAGP,TSPAN31,

S3,HAX1,SLC35E1,RC3H2,EIF3G,RPL8,ECE1,OCIAD1,RBBP6,ZNF248,NREP,CHTOP,TPT1,SMAGP,TSPAN31,

RP1,ZBTB44,NIP7,PLCB2,SLC25A10,TMX2,SPOCK2,RDH11,DBNDD2,TMED4,RPS6,GGPS1,COASY,AUP1,YI

RP1,ZBTB44,NIP7,PLCB2,SLC25A10,TMX2,SPOCK2,RDH11,DBNDD2,TMED4,RPS6,GGPS1,COASY,AUP1,YI

1,TTC21B,PTPN13,UBR4,PSMA1,YLPM1,SLC39A11,FUZ,FAM3A,PSMA3,MUC1,METTL17,ECSIT,EPB41L5,A

1,TTC21B,PTPN13,UBR4,PSMA1,YLPM1,SLC39A11,FUZ,FAM3A,PSMA3,MUC1,METTL17,ECSIT,EPB41L5,A

RF54,C19ORF48,ZFAND2B,CBFA2T2,ZNF250,RPL36,CDK11A,INO80E,CCNL2,MAPKAPK5,DGUOK,HSP90AB

RF54,C19ORF48,ZFAND2B,CBFA2T2,ZNF250,RPL36,CDK11A,INO80E,CCNL2,MAPKAPK5,DGUOK,HSP90AB

NDC17,ARL6IP4,STRADA,AP1S3,CYB561,ZNF644,HYPK,CARS,ZNF250,NFAM1,CDK11A,MDH1,SYNGR2,LRR

NDC17,ARL6IP4,STRADA,AP1S3,CYB561,ZNF644,HYPK,CARS,ZNF250,NFAM1,CDK11A,MDH1,SYNGR2,LRR

NKIRAS2,ACAA1,ZNF675,PSMD7,GTF2H1,BFAR,THOC3,MGST2,C5ORF24,SLC37A4,ZNF396,RPS3,DNAJC2,

NKIRAS2,ACAA1,ZNF675,PSMD7,GTF2H1,BFAR,THOC3,MGST2,C5ORF24,SLC37A4,ZNF396,RPS3,DNAJC2,

2,UQCRQ,SNF8,GRK6,HMGA2,CDC37,RPS16,CPSF3L,TSG101,OBSL1,PTPN13,UBR4,YLPM1,PHF21A,CHCH

2,UQCRQ,SNF8,GRK6,HMGA2,CDC37,RPS16,CPSF3L,TSG101,OBSL1,PTPN13,UBR4,YLPM1,PHF21A,CHCH

TSFM,ASPH,TRMT112,RPTOR,ZNF625,GABPB1,ZSWIM7,DCTD,UBXN1,HHLA3,LRRC27,MVD,DDOST,SYNG

,MOB1A,CDK11A,RBM3,COPZ1,INO80E,CCNL2,DGUOK,HSP90AB1,FAM219B,RAB2A,ZNF524,ATP5J,GPBP

,MOB1A,CDK11A,RBM3,COPZ1,INO80E,CCNL2,DGUOK,HSP90AB1,FAM219B,RAB2A,ZNF524,ATP5J,GPBP

PSMA7,CDC14B,TIMM17B,TMEM126B,APEH,TXNL4A,EPB41L4B,UCK2,TMPO,PRMT1,EXOSC9,TXNDC17,

PSMA7,CDC14B,TIMM17B,TMEM126B,APEH,TXNL4A,EPB41L4B,UCK2,TMPO,PRMT1,EXOSC9,TXNDC17,

RC1,PHF19,GRAMD1C,TM9SF1,MAGED2,SON,TET2,GRK6,HMGA2,CDC37,CPSF3L,OBSL1,TTC21B,PTPN13,

RC1,PHF19,GRAMD1C,TM9SF1,MAGED2,SON,TET2,GRK6,HMGA2,CDC37,CPSF3L,OBSL1,TTC21B,PTPN13,

PPA1,DGKE,VAMP8,CRIP1,RPL28,PSMA7,TIMM17B,XAF1,RPL21,TXNL4A,LYPLA2,NUTF2,CAMLG,ATPAF2,

PPA1,DGKE,VAMP8,CRIP1,RPL28,PSMA7,TIMM17B,XAF1,RPL21,TXNL4A,LYPLA2,NUTF2,CAMLG,ATPAF2,

UP,TMEM205,PTK2,AUNIP,GUK1,ATF7,IFI27L1,MAF,PDLIM5,RFX3,PPA1,DGKE,CRIP1,PPIL2,TROAP,CLDN

UP,TMEM205,PTK2,AUNIP,GUK1,ATF7,IFI27L1,MAF,PDLIM5,RFX3,PPA1,DGKE,CRIP1,PPIL2,TROAP,CLDN

PBP1,SIAH1,GGCX,ZEB2,PNPO,RPS20,GABPB1,LYNX1,TWSG1,RPS15,ZNF814,ERLEC1,GMPPB,HSPB1,HHL

PBP1,SIAH1,GGCX,ZEB2,PNPO,RPS20,GABPB1,LYNX1,TWSG1,RPS15,ZNF814,ERLEC1,GMPPB,HSPB1,HHL

OSC9,WNK1,ZNF7,ARL6IP4,STRADA,PPAN,CYB561,SHARPIN,ZNF644,STAU1,SLC9B2,HYPK,RPL36,MOB1A,

M12,PRMT1,EXOSC9,TXNDC17,ARL6IP4,C19ORF48,ZFAND2B,HYPK,CARS,ZNF250,NFAM1,CDK11A,RBM3

M12,PRMT1,EXOSC9,TXNDC17,ARL6IP4,C19ORF48,ZFAND2B,HYPK,CARS,ZNF250,NFAM1,CDK11A,RBM3

PS15,UBXN1,ZNF814,GMPPB,VWA2,IL32,LMBR1,NAP1L1,SERPINB1,DDOST,C12ORF76,RANGRF,PABPC1,

PS15,UBXN1,ZNF814,GMPPB,VWA2,IL32,LMBR1,NAP1L1,SERPINB1,DDOST,C12ORF76,RANGRF,PABPC1,

O,RPTOR,NIT2,LYNX1,MPDU1,TWSG1,RPS15,UBXN1,ZNF814,GMPPB,VWA2,ERCC5,IL32,LMBR1,NAP1L1,

O,RPTOR,NIT2,LYNX1,MPDU1,TWSG1,RPS15,UBXN1,ZNF814,GMPPB,VWA2,ERCC5,IL32,LMBR1,NAP1L1,
2,CASC4,TXNDC17,PIGL,C19ORF48,STRADA,ZFAND2B,ACAD10,HNRNPC,CYB561,SHARPIN,SUGT1,CARS,Z
2,CASC4,TXNDC17,PIGL,C19ORF48,STRADA,ZFAND2B,ACAD10,HNRNPC,CYB561,SHARPIN,SUGT1,CARS,Z
L2,AKT1S1,SPHK2,ACTR10,HMGN1,NDUFB2,CDK2AP2,NDUFA10,RPS27L,AZI2,GOSR2,SPG21,KAT8,U2AF1
FAM192A,SLC39A6,MARS,TRIM46,LAMTOR5,GALT,VDAC1,ACAA1,NADSYN1,DCAKD,GTF2H1,D2HGDH,C
FAM192A,SLC39A6,MARS,TRIM46,LAMTOR5,GALT,VDAC1,ACAA1,NADSYN1,DCAKD,GTF2H1,D2HGDH,C
FAM192A,SLC39A6,MARS,TRIM46,LAMTOR5,GALT,VDAC1,ACAA1,NADSYN1,DCAKD,GTF2H1,D2HGDH,C
H14,FIS1,SLC15A3,TUBA1C,DST,ANTXR2,STIP1,LRRC1,PHF19,GRAMD1C,VCAN,PCBP2,TM9SF1,MAGED2,
H14,FIS1,SLC15A3,TUBA1C,DST,ANTXR2,STIP1,LRRC1,PHF19,GRAMD1C,VCAN,PCBP2,TM9SF1,MAGED2,

23A,DDX41,SLC16A1,PKP4,AGAP3,PDE4DIP,MDH2,EEF1D,HSPH1,NAA10,OAS3,SEC31A,MT1E,RPL18,ATP
23A,DDX41,SLC16A1,PKP4,AGAP3,PDE4DIP,MDH2,EEF1D,HSPH1,NAA10,OAS3,SEC31A,MT1E,RPL18,ATP
A1,DGKE,ENDOV,CRIP1,PPIL2,RPL28,PSMA7,CDC14B,CAPN10,LYPLA2,NUTF2,UCK2,ATPAF2,WNK1,ZNF7
RADA,PPAN,ZFAND2B,AP1S3,HNRNPC,CYB561,SHARPIN,ZNF644,STAU1,ZNF250,MOB1A,RBM3,LAMP2,Z
RADA,PPAN,ZFAND2B,AP1S3,HNRNPC,CYB561,SHARPIN,ZNF644,STAU1,ZNF250,MOB1A,RBM3,LAMP2,Z
GTF2H1,ST7,NDUFA11,AXIN1,SH3GL1,TKT,DNAJC2,RHEB,HPS1,RGS12,PCTP,AKT1S1,GTF3A,DHRS4,NDUF
GTF2H1,ST7,NDUFA11,AXIN1,SH3GL1,TKT,DNAJC2,RHEB,HPS1,RGS12,PCTP,AKT1S1,GTF3A,DHRS4,NDUF
L29,GGPS1,COASY,SRSF1,SAP30BP,YIPF3,FOXRED1,TRNT1,CLASRP,JPH2,MICAL1,SLC39A13,PRMT2,MAG
L29,GGPS1,COASY,SRSF1,SAP30BP,YIPF3,FOXRED1,TRNT1,CLASRP,JPH2,MICAL1,SLC39A13,PRMT2,MAG
HDDS,XPNPEP1,ZNF483,ZNF783,FBXO34,CYB5D2,ALS2,IK,EXOSC8,DDX17,SLC31A1,RCAN1,C12ORF73,NS
,SMARCC2,SLC30A5,TRPV3,CBR1,UBE2N,LFNG,TTC12,ZNHIT3,RPS11,SNX5,RAPGEF3,MRPS17,TMEM147
,SMARCC2,SLC30A5,TRPV3,CBR1,UBE2N,LFNG,TTC12,ZNHIT3,RPS11,SNX5,RAPGEF3,MRPS17,TMEM147
OR4,ATP6V0B,METTL23,C12ORF73,CGREF1,MCM7,PARP6,KIAA0513,RNMT,ACY1,AP4M1,SHMT1,COQ4,
OR4,ATP6V0B,METTL23,C12ORF73,CGREF1,MCM7,PARP6,KIAA0513,RNMT,ACY1,AP4M1,SHMT1,COQ4,
D3,GRAPL,PRRC2B,USP4,PEX26,NMNAT3,ARL16,ZNF223,OGFOD2,MRPL55,CREB3L4,ASH1L,CHTOP,ERRF
D3,GRAPL,PRRC2B,USP4,PEX26,NMNAT3,ARL16,ZNF223,OGFOD2,MRPL55,CREB3L4,ASH1L,CHTOP,ERRF
SLC9B2,SUGT1,CARS,NFAM1,CDK11A,RBM3,MDH1,ZNF197,CCNL2,SYNGR2,LRRFIP1,ZNF524,ATP5J,TPM
LG,ATPAF2,LSM12,PRMT1,EXOSC9,STOML2,WNK1,C19ORF54,PIGL,C19ORF48,ZFAND2B,ACAD10,HNRNP
LG,ATPAF2,LSM12,PRMT1,EXOSC9,STOML2,WNK1,C19ORF54,PIGL,C19ORF48,ZFAND2B,ACAD10,HNRNP

T,RPL31,H2AFY,TOMM7,WWP2,NPRL3,CEP152,SMOX,ATRX,GAPDH,AKAP8L,DNAJC10,TRIP10,HEATR1

,RNPS1,AK2,PCMTD1,MRPS16,ARL2,ZNF276,RPS23,NDUFA7,SELENBP1,SLC39A14,HNRNPM,DNAJC21,IK

LY,MARS,RBM42,RPL34,EIF2A,RPL24,PSPC1,SIN3A,RBM4,THOC3,HNRNPUL1,TP53,RPS3A,BTF3,EIF3C,RP

R3H,RAD23A,HSD17B10,DDX41,ILF3,ZCCHC4,MDH2,EEF1D,RRP8,OAS3,MED6,CARHSP1,RPL18,EGFR,CA

44,MAGED2,SON,TET2,SNF8,GRK6,HMGA2,CDC37,RPS16,CPSF3L,TSG101,OBSL1,TTC21B,PTPN13,UBR4

TSFM,ZEB2,PNPO,RPTOR,ZNF625,RPS20,GABPB1,MAPK13,RPS15,ZNF814,GMPPB,HSPB1,MVD,ERCC5,M

24,GPBP1,TSFM,ZEB2,PNPO,RPTOR,ZNF625,RPS20,GABPB1,MAPK13,RPS15,ZNF814,GMPPB,HSPB1,MV

A,RPS17,SEC13,EIF6,LMNA,AP4M1,IMMP1L,IFT81,BCAP29,MED1,RPL27,DYNC2LI1,TIMM9,OAZ1,RPS14

PINA1,PML,UBE2D3,RPS3,SYTL4,DENND1A,RDX,MAGOHB,CREB3L2,VPS25,SNRPD1,SPHK2,DHRS4,IFI27,

3A,HDAC8,PML,UBE2D3,RPS3,SYTL4,RDX,MAGOHB,DNAJA3,VPS25,NSFL1C,DHRS4,IFI27,TRAK1,GOSR2,

3A,HDAC8,PML,UBE2D3,RPS3,SYTL4,RDX,MAGOHB,DNAJA3,VPS25,NSFL1C,DHRS4,IFI27,TRAK1,GOSR2,N
CAPN2,MLH1,SUMO1,COMMD1,RPS23,RPL30,PSMD8,RQCD1,RAB26,SUFU,NUB1,OS9,GUSB,EXOSC2,RBM

DCTN5,PEX26,ERGIC1,KPNA5,RPLP0,HAX1,TMEM108,FAM126A,CIZ1,SSR3,MPV17,SSB,ARIH2,RPL8,PICK1

NUP62,ACAA1,RPL24,PSMD7,PAM16,NEDD4L,RNF130,BFAR,CLTA,LYPLAL1,AXIN1,TP53,PSMD10,RPS3A,F

1,PAK1,CSRP2BP,LUC7L3,PRPF31,NDUFA8,MSRB1,H2AFY,TRIM9,SRSF5,PSMD11,EIF3E,ATRX,ADD2,CRYA
TP53,PSMD10,SERPINA1,SH3GL1,PML,TMEM199,NDUFA3,EIF3C,COX19,RPS3,MRPS24,RDX,SNRPD1,DHR
MTOR5,GALT,PCNP,VDAC1,RPL34,ACAA1,RPL24,PSMD7,PAM16,NEDD4L,RNF130,BFAR,CLTA,AXIN1,TP53,

,BAX,RPL35,UBA1,CDK4,RNF8,PPP2R5C,NT5E,FAM192A,ADRA1A,GCDH,GLYCTK,LAMTOR5,GALT,PCNP,V
172,KDELR3,CLTA,THOC3,TP53,AP1G2,RPS3A,BTF3,PML,UBE2D3,RPS3,SYTL4,DENND1A,MAGOHB,VPS25
P62,ACAA1,RPL24,PAM16,IRF3,IFT172,KDELR3,CLTA,THOC3,TP53,AP1G2,RPS3A,BTF3,PML,UBE2D3,RPS3
PF,GBA2,PPIH,NSUN5,BRF1,TRPM4,MCFD2,LGALS3,TCP1,CD74,PARK7,CCDC40,NHP2,ZNHIT3,PDCD6,MF

ARS2,SPHK2,TRAK1,PRTFDC1,HMBS,RPS27L,RPS7,NCBP2,PLD1,AKR1A1,PIWIL4,NFE2L1,UBA52,RPS4X,AK

172,KDELR3,CLTA,THOC3,TP53,AP1G2,RPS3A,BTF3,PML,UBE2D3,RPS3,SYTL4,DENND1A,MAGOHB,VPS25
,IRF3,IFT172,KDELR3,CLTA,THOC3,TP53,AP1G2,RPS3A,BTF3,PML,UBE2D3,RPS3,SYTL4,DENND1A,MAGOI
LP0,HAX1,FAM126A,CIZ1,SSR3,MPV17,SSB,ARIH2,RPL8,PICK1,SLMAP,TJP1,UGT8,CHTOP,BAX,RPL35,DLG

GUOK,HSP90AB1,DUT,LRRFIP1,ZNF524,ATP5J,ELP6,GPBP1,TSFM,ZEB2,TRMT112,RPTOR,ZNF625,RPS20,G
TBCA,ECT2,TAF9,ARPC1A,UBA52,BMP1,SUMO1,ARL2,PRKCE,NDUFA7,GRHPR,NDUFS2,PSMC5,GTF3C5,AF
NF644,CBFA2T2,UPP1,CARS,ZNF250,RPL36,NFAM1,CDK11A,RBM3,ZNF197,INO80E,CCNL2,MAPKAPK5,D
IM16,ANAPC11,HSD17B10,SLC16A1,PDE4DIP,NAA10,RRP8,SEC31A,LLGL2,SH3GLB1,CHEK1,SSH2,COX17,

PK5,DGUOK,HSP90AB1,DUT,LRRFIP1,ZNF524,ATP5J,ELP6,GPBP1,TSFM,ZEB2,TRMT112,PNPO,RPTOR,ZNF

RNF149,RELA,OAZ2,APOC1,PDE1A,NUDT9,APOBEC3F,ATP13A2,UBA6,MVB12B,ADAMTS9,HGSNAT,GSK3

ACOT2,PCID2,SGCD,PEX2,TNKS,RPL37A,STX10,USP4,PEX26,BTN3A2,KPNA5,RPLP0,HAX1,FAM126A,DGAT
IAH1,TSFM,GGCX,ZEB2,ASPH,TRMT112,RPTOR,RPS20,MPDU1,TWSG1,MAPK13,VPS28,RPS15,UBXN1,ER
F3,HAUS4,COX20,COA1,NEK2,COBL,TNKS,USP4,NDUFS3,NUBP2,RPLP0,HAX1,DGAT1,EIF3G,LIMA1,PICK1
,CDC14B,TIMM17B,TMEM126B,APEH,TXNL4A,CAPN10,LYPLA2,PODXL,EPB41L4B,TCF4,ATPAF2,PRMT1,
AT1,MCTS1,MTA3,DCN,MEIS2,HMGA1,PSME2,PABPC1,ZNF836,H2AFV,RBMX,PPP3CB,BRIP1,TADA3,ZNF
K5,DGUOK,HSP90AB1,DUT,LRRFIP1,ZNF524,ATP5J,ELP6,GPBP1,TSFM,ZEB2,TRMT112,PNPO,RPTOR,ZNF

5A10,SPOCK2,THEM4,RDH11,DBNDD2,RPS6,RPL29,GGPS1,RTEL1,COASY,SRSF1,AUP1,SAP30BP,TRNT1,I

NF814,HSPB1,ERCC5,PRKRIP1,MTA3,DCN,MEIS2,HMGA1,PSME2,PABPC1,RRNAD1,ZNF836,RAD1,H2AFV
O80E,CCNL2,MAPKAPK5,DGUOK,HSP90AB1,DUT,LRRFIP1,ZNF524,ATP5J,ELP6,GPBP1,TSFM,ZEB2,TRMT

PL35,SLC52A2,SRSF7,NXF1,ZNF593,RPL34,NUP62,ACAA1,RPL24,PAM16,IRF3,IFT172,KDELR3,CLTA,SLC35
TA3,DCN,MEIS2,HMGA1,PSME2,PABPC1,ZNF836,RAD1,H2AFV,RBMX,PPP3CB,BRIP1,TADA3,ZNF20,WAC,

CTS1,NAP1L1,DDOST,MTA3,DCN,MEIS2,HMGA1,PSME2,PABPC1,ZNF836,RAD1,H2AFV,RBMX,PPP3CB,BF
10,LYPLA2,CAMLG,UCK2,PPOX,PRMT1,STOML2,WNK1,PIGL,STRADA,ZFAND2B,SHARPIN,UPP1,CARS,RPL

DK11A,RBM3,MDH1,ZNF197,CCNL2,MAPKAPK5,DGUOK,HSP90AB1,DUT,LRRFIP1,ZNF524,ATP5J,GPBP1,
2,PODXL,EPB41L4B,TCF4,ATPAF2,PRMT1,EXOSC9,STOML2,PPAN,HNRNPC,SHARPIN,CBFA2T2,STAU1,CD

CNL2,MAPKAPK5,DGUOK,HSP90AB1,DUT,LRRFIP1,ZNF524,ATP5J,GPBP1,TSFM,ZEB2,TRMT112,PNPO,RF

AMP2,INO80E,CCNL2,MAPKAPK5,DGUOK,HSP90AB1,RAB2A,SIAH1,TSFM,GGCX,ZEB2,ASPH,TRMT112,RP

F20,WAC,SLC33A1,ZNF706,GTF2IRD1,TRIM16,POLR3H,HSD17B10,DDX41,PSME1,ILF3,ZCCHC4,SGK1,RRF

DK11A,RBM3,MDH1,ZNF197,CCNL2,MAPKAPK5,DGUOK,HSP90AB1,DUT,LRRFIP1,ZNF524,ATP5J,GPBP1,

G12,PPRC1,RPS25,NLRP1,COX4I1,PLCB2,DUSP22,SLC25A10,SPOCK2,THEM4,RDH11,DBNDD2,NISCH,RPS
AFY,GSK3A,TOMM7,TRIM9,SURF4,NPRL3,CTNNB1,ZFYVE20,EXOSC10,SDCCAG8,ATRX,AKAP8L,ADD2,AN

3,RPS3,DNAJC2,CREB3L2,UCHL5,AKT1S1,SMUG1,INO80C,COMT,AIFM2,MLST8,HMGN1,RPS27L,GOSR2,SI

X3,PPA1,TNFRSF14,ENDOV,CAMTA1,PPIL2,PSMD4,RPL28,VIM,LIG1,PSMA7,CDC14B,APEH,RPL21,LYPLA2

3,HMGA2,RPS16,CPSF3L,TSG101,OBSL1,TTC21B,PTPN13,UBR4,PSMA1,YLPM1,PHF21A,CHCHD1,SLC39A1
PL21,TXNL4A,CAPN10,LYPLA2,NUTF2,EPB41L4B,UCK2,ATPAF2,PRMT1,EXOSC9,WNK1,TXNDC17,STRADA,
,PCBP2,LPCAT3,CD44,TM9SF1,MAGED2,SON,TET2,UQCRQ,SNF8,HMGA2,CDC37,RPS16,CPSF3L,TSG101,C
19,GRAMD1C,VCAN,PCBP2,LPCAT3,CD44,TM9SF1,MAGED2,SON,TET2,UQCRQ,SNF8,HMGA2,RPS16,CPSF
D44,TM9SF1,MAGED2,UQCRQ,SNF8,CDC37,RPS16,CPSF3L,TSG101,OBSL1,TTC21B,PTPN13,UBR4,PSMA1,
EKHA5,RANBP1,UBE2L6,VRK3,LZTFL1,ZC3H14,FIS1,SLC15A3,PPM1K,TUBA1C,IRF9,DST,ANTXR2,STIP1,LR
3,PPM1K,TUBA1C,IRF9,DST,ANTXR2,STIP1,LRRC1,PHF19,GRAMD1C,VCAN,PCBP2,LPCAT3,CD44,TM9SF1,

,PPIL2,PSMD4,VIM,LIG1,PSMA7,CDC14B,XAF1,APEH,TXNL4A,NUTF2,PODXL,TCF4,PPOX,ATPAF2,PRMT1,
,PPIL2,PSMD4,VIM,LIG1,PSMA7,CDC14B,XAF1,APEH,TXNL4A,NUTF2,PODXL,TCF4,PPOX,ATPAF2,PRMT1,
,PPIL2,PSMD4,VIM,LIG1,PSMA7,CDC14B,XAF1,APEH,TXNL4A,NUTF2,PODXL,TCF4,PPOX,ATPAF2,PRMT1,
BP1,PTK2,CELF1,AUNIP,CENPN,ATF7,FBXW9,HSBP1L1,MAF,ASF1B,PPP1CC,EDF1,RFX3,DGKE,ENDOV,CA

GRF,MTA3,DOCK10,HMGA1,PSME2,RBP1,RAD1,RBMX,BLVRB,PPP3CB,BRIP1,TADA3,WAC,ZNF687,GTF2I

0E,CCNL2,MAPKAPK5,HSP90AB1,DUT,TSFM,ZEB2,TRMT112,PNPO,RPTOR,RPS20,GABPB1,RPS15,ERCC5
OPZ1,INO80E,HSP90AB1,ATP5J,ELP6,TPM2,SIAH1,TPM3,TRMT112,RPTOR,RPS20,VPS28,ZSWIM7,RPS15,
OI,IMMP1L,MRPL48,AK9,COQ4,ERAL1,MTHFD2L,MRPL27,PRKCZ,SLC25A35,DNAJB12,COQ2,MRPS7,NDU
OI,IMMP1L,MRPL48,AK9,COQ4,ERAL1,MTHFD2L,MRPL27,PRKCZ,SLC25A35,DNAJB12,COQ2,MRPS7,NDU

S20,GABPB1,NIT2,RPS15,HSPB1,ERCC5,MCTS1,SERPINB1,PRKRIP1,MTA3,MEIS2,HMGA1,PABPC1,DBNL,
S20,GABPB1,NIT2,RPS15,HSPB1,ERCC5,MCTS1,SERPINB1,PRKRIP1,MTA3,MEIS2,HMGA1,PABPC1,DBNL,
NDUFS2,EXOSC2,PSMC5,HNRNPM,NDUFC1,TBC1D5,EXOSC8,UBE2L3,ACTB,RUVBL1,ZNFX1,HDAC10,NDU

MRPS16,ARL2,PRKCE,SDHA,NDUFA7,NDUFS2,MRPS33,ACSS1,MCUR1,NGDN,ME3,NDUFC1,QTRT1,DUSP

A23,MFSD10,MRPS18C,MCFD2,LGALS3,SLC35B1,SCAP,CD74,SLC30A5,THADA,PIGT,LFNG,PDCD6,XYLT2,M

ST,PRLR,EGFR,SH3GLB1,CAMK2D,HINT1,HNRNPA1,LGMN,B3GAT3,FTH1,PAFAH1B2,S100A4,COPS4,TPM

OLD4,PLEKHA5,RANBP1,UBE2L6,VRK3,LZTFL1,ZC3H14,FIS1,SLC15A3,PPM1K,TUBA1C,ANTXR2,STIP1,LRR

M1K,TUBA1C,ANTXR2,STIP1,LRRC1,PHF19,GRAMD1C,VCAN,LPCAT3,CD44,TM9SF1,MAGED2,SON,TET2,U

ORF198,UGGT2,MICAL1,UBE2I,SLC39A13,EIF3M,CDC42,TBX6,TMEM205,PTK2,C14ORF2,GUK1,CARD14,

,NUDT5,RBPMS,NNMT,SNRPN,SPAG7,WDR62,IL1RAP,STYXL1,RFX5,POLD4,PLEKHA5,RANBP1,UBE2L6,VR

,NUDT5,RBPMS,NNMT,SNRPN,SPAG7,WDR62,IL1RAP,STYXL1,RFX5,POLD4,PLEKHA5,RANBP1,UBE2L6,VR

1,POLD4,PLEKHA5,RANBP1,UBE2L6,VRK3,LZTFL1,ZC3H14,FIS1,SLC15A3,PPM1K,TUBA1C,IRF9,ANTXR2,S

1,POLD4,PLEKHA5,RANBP1,UBE2L6,VRK3,LZTFL1,ZC3H14,FIS1,SLC15A3,PPM1K,TUBA1C,IRF9,ANTXR2,S

H14,FIS1,SLC15A3,PPM1K,TUBA1C,DST,ANTXR2,STIP1,LRRC1,PHF19,VCAN,LPCAT3,CD44,TM9SF1,MAGE

37,RPS16,CPSF3L,TSG101,OBSL1,UBR4,YLPM1,PHF21A,FUZ,FAM3A,METTL17,EPB41L5,ATG12,RPS25,NIP

37,RPS16,CPSF3L,TSG101,OBSL1,UBR4,YLPM1,PHF21A,FUZ,FAM3A,METTL17,EPB41L5,ATG12,RPS25,NIP

P1,UBE2L6,VRK3,LZTFL1,ZC3H14,FIS1,SLC15A3,PPM1K,TUBA1C,IRF9,ANTXR2,STIP1,LRRC1,PHF19,GRAM

CLASRP,C1ORF198,UGGT2,MICAL1,UBE2I,YIF1A,EIF3M,CDC42,TBX6,TMEM205,CARD14,PCYT2,FBXW9,U

CLASRP,C1ORF198,UGGT2,MICAL1,UBE2I,YIF1A,EIF3M,CDC42,TBX6,TMEM205,CARD14,PCYT2,FBXW9,U

STOML2,C19ORF54,ZNF7,TXNDC17,ARL6IP4,PIGL,C19ORF48,PPAN,ZFAND2B,ACAD10,HNRNPC,SHARPIN

P7,COX4I1,DUSP22,SLC25A10,THEM4,CENPK,RPS6,RPL29,RTEL1,COASY,SRSF1,AUP1,SAP30BP,YIPF3,FO

P7,COX4I1,DUSP22,SLC25A10,THEM4,CENPK,RPS6,RPL29,RTEL1,COASY,SRSF1,AUP1,SAP30BP,YIPF3,FO

5A3,PPM1K,TUBA1C,DST,STIP1,LRRC1,PHF19,GRAMD1C,VCAN,PCBP2,LPCAT3,CD44,TM9SF1,MAGED2,S

5A3,PPM1K,TUBA1C,DST,STIP1,LRRC1,PHF19,GRAMD1C,VCAN,PCBP2,LPCAT3,CD44,TM9SF1,MAGED2,S
,RPS6,RPL29,RTEL1,COASY,AUP1,SAP30BP,YIPF3,FOXRED1,RABL5,TRNT1,KXD1,ITGB1BP1,MED15,PARD6
,RPS6,RPL29,RTEL1,COASY,AUP1,SAP30BP,YIPF3,FOXRED1,RABL5,TRNT1,KXD1,ITGB1BP1,MED15,PARD6
L1,UBE2I,SLC39A13,PRMT2,EIF3M,CDC42,TBX6,CARD14,PCYT2,FBXW9,USE1,HSBP1L1,IFI27L1,MAF,ASF
L1,UBE2I,SLC39A13,PRMT2,EIF3M,CDC42,TBX6,CARD14,PCYT2,FBXW9,USE1,HSBP1L1,IFI27L1,MAF,ASF
OX4I1,DUSP22,SLC25A10,THEM4,CENPK,RPS6,RPL29,RTEL1,COASY,SRSF1,AUP1,SAP30BP,YIPF3,FOXRED
OX4I1,DUSP22,SLC25A10,THEM4,CENPK,RPS6,RPL29,RTEL1,COASY,SRSF1,AUP1,SAP30BP,YIPF3,FOXRED
MS,NNMT,SNRPN,SPAG7,WDR62,IL1RAP,STYXL1,RFX5,POLD4,PLEKHA5,RANBP1,UBE2L6,VRK3,LZTFL1,Z
MS,NNMT,SNRPN,SPAG7,WDR62,IL1RAP,STYXL1,RFX5,POLD4,PLEKHA5,RANBP1,UBE2L6,VRK3,LZTFL1,Z
X4I1,DUSP22,SLC25A10,TMX2,THEM4,DBNDD2,CENPK,RPS6,RPL29,RTEL1,COASY,SRSF1,AUP1,SAP30BP,
X4I1,DUSP22,SLC25A10,TMX2,THEM4,DBNDD2,CENPK,RPS6,RPL29,RTEL1,COASY,SRSF1,AUP1,SAP30BP,
CRQ,SNF8,GRK6,HMGA2,CDC37,RPS16,CPSF3L,TSG101,OBSL1,UBR4,YLPM1,PHF21A,FUZ,FAM3A,PSMA3
CRQ,SNF8,GRK6,HMGA2,CDC37,RPS16,CPSF3L,TSG101,OBSL1,UBR4,YLPM1,PHF21A,FUZ,FAM3A,PSMA3
,LIG1,PSMA7,CDC14B,TMEM126B,APEH,RPL21,CAPN10,LYPLA2,LSM12,C17ORF62,PRMT1,STOML2,ZNF
,LIG1,PSMA7,CDC14B,TMEM126B,APEH,RPL21,CAPN10,LYPLA2,LSM12,C17ORF62,PRMT1,STOML2,ZNF
CDC37,RPS16,CPSF3L,TSG101,OBSL1,UBR4,YLPM1,PHF21A,FUZ,FAM3A,PSMA3,MUC1,METTL17,EPB41L
CDC37,RPS16,CPSF3L,TSG101,OBSL1,UBR4,YLPM1,PHF21A,FUZ,FAM3A,PSMA3,MUC1,METTL17,EPB41L
DC37,RPS16,CPSF3L,TSG101,OBSL1,UBR4,YLPM1,PHF21A,FUZ,FAM3A,PSMA3,METTL17,EPB41L5,ATG1
DC37,RPS16,CPSF3L,TSG101,OBSL1,UBR4,YLPM1,PHF21A,FUZ,FAM3A,PSMA3,METTL17,EPB41L5,ATG1
PM1K,TUBA1C,DST,ANTXR2,STIP1,LRRC1,PHF19,GRAMD1C,VCAN,LPCAT3,CD44,TM9SF1,SON,TET2,UQ
TSG101,UBR4,YLPM1,CHCHD1,FUZ,FAM3A,METTL17,EPB41L5,ATG12,RPS25,NIP7,COX4I1,DUSP22,SLC2
TSG101,UBR4,YLPM1,CHCHD1,FUZ,FAM3A,METTL17,EPB41L5,ATG12,RPS25,NIP7,COX4I1,DUSP22,SLC2
NDOV,CAMTA1,PPIL2,RPL28,LIG1,PSMA7,APEH,CAPN10,LYPLA2,NUTF2,ATPAF2,C17ORF62,PRMT1,STO
NTXR2,STIP1,LRRC1,PHF19,GRAMD1C,VCAN,LPCAT3,CD44,TM9SF1,SON,TET2,UQCRQ,SNF8,GRK6,HMGA
NTXR2,STIP1,LRRC1,PHF19,GRAMD1C,VCAN,LPCAT3,CD44,TM9SF1,SON,TET2,UQCRQ,SNF8,GRK6,HMGA
11A,RBM3,INO80E,CCNL2,MAPKAPK5,DGUOK,LRRFIP1,RAB2A,ZNF524,ATP5J,RMDN1,TRMT112,CHPF,
3,LZTFL1,ZC3H14,FIS1,SLC15A3,PPM1K,TUBA1C,ANTXR2,STIP1,LRRC1,PHF19,GRAMD1C,VCAN,LPCAT3,
MS,NNMT,SNRPN,SPAG7,WDR62,IL1RAP,STYXL1,RFX5,POLD4,PLEKHA5,RANBP1,UBE2L6,VRK3,LZTFL1,Z
MS,NNMT,SNRPN,SPAG7,WDR62,IL1RAP,STYXL1,RFX5,POLD4,PLEKHA5,RANBP1,UBE2L6,VRK3,LZTFL1,Z
,TRNT1,KXD1,ITGB1BP1,MED15,SH3YL1,IP6K2,CLASRP,C1ORF198,UGGT2,MICAL1,UBE2I,SLC39A13,PR
,TRNT1,KXD1,ITGB1BP1,MED15,SH3YL1,IP6K2,CLASRP,C1ORF198,UGGT2,MICAL1,UBE2I,SLC39A13,PR
,THEM4,CENPK,RPS6,RPL29,RTEL1,COASY,SRSF1,AUP1,SAP30BP,YIPF3,FOXRED1,RABL5,TRNT1,KXD1,IT
,THEM4,CENPK,RPS6,RPL29,RTEL1,COASY,SRSF1,AUP1,SAP30BP,YIPF3,FOXRED1,RABL5,TRNT1,KXD1,IT
ANTXR2,STIP1,LRRC1,VCAN,PCBP2,LPCAT3,CD44,TM9SF1,MAGED2,SON,UQCRQ,SNF8,GRK6,HMGA2,CD
ANTXR2,STIP1,LRRC1,VCAN,PCBP2,LPCAT3,CD44,TM9SF1,MAGED2,SON,UQCRQ,SNF8,GRK6,HMGA2,CD
ABL5,TRNT1,KXD1,ITGB1BP1,MED15,SH3YL1,IP6K2,CLASRP,C1ORF198,UGGT2,MICAL1,UBE2I,SLC39A13,
ABL5,TRNT1,KXD1,ITGB1BP1,MED15,SH3YL1,IP6K2,CLASRP,C1ORF198,UGGT2,MICAL1,UBE2I,SLC39A13,
GRK6,HMGA2,CDC37,RPS16,CPSF3L,TSG101,OBSL1,UBR4,YLPM1,PHF21A,CHCHD1,FUZ,FAM3A,METTL17
GRK6,HMGA2,CDC37,RPS16,CPSF3L,TSG101,OBSL1,UBR4,YLPM1,PHF21A,CHCHD1,FUZ,FAM3A,METTL17
X6,GUK1,CARD14,PCYT2,FBXW9,USE1,MAF,ASF1B,CCDC25,EEF1B2,RFX3,TNFRSF14,ENDOV,CAMTA1,C
X6,GUK1,CARD14,PCYT2,FBXW9,USE1,MAF,ASF1B,CCDC25,EEF1B2,RFX3,TNFRSF14,ENDOV,CAMTA1,C
15,SH3YL1,IP6K2,CLASRP,C1ORF198,UGGT2,MICAL1,UBE2I,SLC39A13,PRMT2,EIF3M,CDC42,TBX6,CARD
15,SH3YL1,IP6K2,CLASRP,C1ORF198,UGGT2,MICAL1,UBE2I,SLC39A13,PRMT2,EIF3M,CDC42,TBX6,CARD
FL1,ZC3H14,FIS1,SLC15A3,PPM1K,TUBA1C,IRF9,DST,ANTXR2,STIP1,LRRC1,PHF19,GRAMD1C,VCAN,LPCA
PAN,ZFAND2B,ACAD10,SHARPIN,SLC9B2,SUGT1,HYPK,CARS,ZNF250,CDK11A,RBM3,ZNF197,COPZ1,INO
K,LRRFIP1,ZNF524,ATP5J,RMDN1,TRMT112,CHPF,RPS20,MPDU1,DCTD,UBXN1,SAT1,ERCC5,EMC4,SERPI
BL5,TRNT1,KXD1,ITGB1BP1,MED15,SH3YL1,IP6K2,CLASRP,C1ORF198,UGGT2,MICAL1,UBE2I,SLC39A13,P

BL5,TRNT1,KXD1,ITGB1BP1,MED15,SH3YL1,IP6K2,CLASRP,C1ORF198,UGGT2,MICAL1,UBE2I,SLC39A13,P
TMX2,THEM4,DBNDD2,RPS6,RPL29,RTEL1,COASY,AUP1,SAP30BP,YIPF3,FOXRED1,RABL5,TRNT1,KXD1,IT
TMX2,THEM4,DBNDD2,RPS6,RPL29,RTEL1,COASY,AUP1,SAP30BP,YIPF3,FOXRED1,RABL5,TRNT1,KXD1,IT
GB1BP1,MED15,SH3YL1,IP6K2,CLASRP,C1ORF198,UGGT2,MICAL1,UBE2I,SLC39A13,PRMT2,EIF3M,CDC42
GB1BP1,MED15,SH3YL1,IP6K2,CLASRP,C1ORF198,UGGT2,MICAL1,UBE2I,SLC39A13,PRMT2,EIF3M,CDC42
),USE1,IFI27L1,ASF1B,CCDC25,EEF1B2,PDLIM5,RFX3,ENDOV,PPIL2,RPL28,CLDND1,LIG1,CDC14B,TIMM17
),USE1,IFI27L1,ASF1B,CCDC25,EEF1B2,PDLIM5,RFX3,ENDOV,PPIL2,RPL28,CLDND1,LIG1,CDC14B,TIMM17
CDC25,EEF1B2,RFX3,TNFRSF14,ENDOV,PPIL2,RPL28,LIG1,CDC14B,TIMM17B,APEH,RPL21,CAPN10,LYPLA
CDC25,EEF1B2,RFX3,TNFRSF14,ENDOV,PPIL2,RPL28,LIG1,CDC14B,TIMM17B,APEH,RPL21,CAPN10,LYPLA
XRED1,RABL5,TRNT1,KXD1,ITGB1BP1,MED15,SH3YL1,IP6K2,CLASRP,C1ORF198,UGGT2,MICAL1,UBE2I,SL
XRED1,RABL5,TRNT1,KXD1,ITGB1BP1,MED15,SH3YL1,IP6K2,CLASRP,C1ORF198,UGGT2,MICAL1,UBE2I,SL
GA2,CDC37,RPS16,CPSF3L,TSG101,TTC21B,UBR4,YLPM1,PHF21A,CHCHD1,FUZ,FAM3A,PSMA3,METTL17
GA2,CDC37,RPS16,CPSF3L,TSG101,TTC21B,UBR4,YLPM1,PHF21A,CHCHD1,FUZ,FAM3A,PSMA3,METTL17
F3,FOXRED1,RABL5,TRNT1,KXD1,ITGB1BP1,MED15,SH3YL1,IP6K2,CLASRP,C1ORF198,UGGT2,MICAL1,UB
F3,FOXRED1,RABL5,TRNT1,KXD1,ITGB1BP1,MED15,SH3YL1,IP6K2,CLASRP,C1ORF198,UGGT2,MICAL1,UB
BP2,EEF1D,HSPH1,CHCHD5,SEC31A,CNST,COX17,COX5A,HIGD1A,SIRT5,INTS4,ZNF506,PFDN5,PAFAH1B2,
5,NIP7,TTC9C,COX4I1,DUSP22,SLC25A10,TMX2,THEM4,DBNDD2,CENPK,RPS6,RPL29,RTEL1,COASY,SRSF1
5,NIP7,TTC9C,COX4I1,DUSP22,SLC25A10,TMX2,THEM4,DBNDD2,CENPK,RPS6,RPL29,RTEL1,COASY,SRSF1
,EIF3M,CDC42,TBX6,TMEM205,CARD14,PCYT2,FBXW9,USE1,HSBP1L1,MAF,ASF1B,CCDC25,EEF1B2,RFX3
,EIF3M,CDC42,TBX6,TMEM205,CARD14,PCYT2,FBXW9,USE1,HSBP1L1,MAF,ASF1B,CCDC25,EEF1B2,RFX3
AUP1,SAP30BP,YIPF3,FOXRED1,RABL5,TRNT1,KXD1,ITGB1BP1,MED15,IP6K2,CLASRP,C1ORF198,UGGT2,M
AUP1,SAP30BP,YIPF3,FOXRED1,RABL5,TRNT1,KXD1,ITGB1BP1,MED15,IP6K2,CLASRP,C1ORF198,UGGT2,M
P1,SAP30BP,YIPF3,FOXRED1,RABL5,TRNT1,KXD1,ITGB1BP1,MED15,SH3YL1,IP6K2,CLASRP,C1ORF198,UGG
P1,SAP30BP,YIPF3,FOXRED1,RABL5,TRNT1,KXD1,ITGB1BP1,MED15,SH3YL1,IP6K2,CLASRP,C1ORF198,UGG
3GAT3,ZNF506,SPINT2,MTHFS,PAFAH1B2,ZNF343,GBA2,TMEM39B,PPIH,DALRD3,NSUN5,MFSD10,MCFD
TRNT1,KXD1,ITGB1BP1,MED15,IP6K2,CLASRP,C1ORF198,UGGT2,MICAL1,UBE2I,SLC39A13,YIF1A,PRMT2,
TRNT1,KXD1,ITGB1BP1,MED15,IP6K2,CLASRP,C1ORF198,UGGT2,MICAL1,UBE2I,SLC39A13,YIF1A,PRMT2,
C1,ZBTB44,TTC9C,PLCB2,DUSP22,SLC25A10,TMX2,THEM4,DBNDD2,NISCH,TMED4,CD99L2,GGPS1,AUP1
C1,ZBTB44,TTC9C,PLCB2,DUSP22,SLC25A10,TMX2,THEM4,DBNDD2,NISCH,TMED4,CD99L2,GGPS1,AUP1
,TADA3,ZNF20,ATP6V1F,WAC,TRIM16,ANAPC11,ABHD17A,NT5C,MDH2,STXBP2,EEF1D,CHCHD5,MED6,S
B2,ZNF343,GBA2,PPIH,DALRD3,NSUN5,DRG2,ZFPL1,MFSD10,GMPR2,MCFD2,RIC8A,TCP1,THADA,CCDC4
PSF3L,TSG101,TTC21B,UBR4,YLPM1,PHF21A,CHCHD1,FUZ,FAM3A,PSMA3,METTL17,EPB41L5,ATG12,RPS
PSF3L,TSG101,TTC21B,UBR4,YLPM1,PHF21A,CHCHD1,FUZ,FAM3A,PSMA3,METTL17,EPB41L5,ATG12,RPS
M9SF1,SON,TET2,UQCRQ,SNF8,GRK6,HMGA2,CDC37,RPS16,CPSF3L,TSG101,OBSL1,TTC21B,PTPN13,PSMA
M9SF1,SON,TET2,UQCRQ,SNF8,GRK6,HMGA2,CDC37,RPS16,CPSF3L,TSG101,OBSL1,TTC21B,PTPN13,PSMA
PN,SPAG7,WDR62,IL1RAP,STYXL1,RFX5,POLD4,PLEKHA5,RANBP1,UBE2L6,VRK3,LZTFL1,ZC3H14,FIS1,SLC1
PN,SPAG7,WDR62,IL1RAP,STYXL1,RFX5,POLD4,PLEKHA5,RANBP1,UBE2L6,VRK3,LZTFL1,ZC3H14,FIS1,SLC1
F20,WAC,SLC33A1,KLK10,ANAPC11,ABHD17A,NT5C,MDH2,STXBP2,EEF1D,CHCHD5,NAA10,SEC31A,CNST,
QCRQ,SNF8,GRK6,HMGA2,CDC37,RPS16,CPSF3L,TSG101,PTPN13,UBR4,YLPM1,PHF21A,CHCHD1,FUZ,FAM
QCRQ,SNF8,GRK6,HMGA2,CDC37,RPS16,CPSF3L,TSG101,PTPN13,UBR4,YLPM1,PHF21A,CHCHD1,FUZ,FAM
,TUBA1C,ANTXR2,STIP1,LRRC1,PHF19,VCAN,LPCAT3,CD44,TM9SF1,MAGED2,SON,TET2,UQCRQ,SNF8,GR
,TUBA1C,ANTXR2,STIP1,LRRC1,PHF19,VCAN,LPCAT3,CD44,TM9SF1,MAGED2,SON,TET2,UQCRQ,SNF8,GR
CHPF,RPS20,MPDU1,DCTD,UBXN1,SAT1,ERCC5,EMC4,SERPINB1,DDOST,LSR,PSME2,PABPC1,RRNAD1,DB
EEF1B2,RFX3,ENDOV,PPIL2,RPL28,CLDND1,LIG1,CDC14B,TIMM17B,TMEM126B,APEH,RPL21,CAPN10,LYP
EEF1B2,RFX3,ENDOV,PPIL2,RPL28,CLDND1,LIG1,CDC14B,TIMM17B,TMEM126B,APEH,RPL21,CAPN10,LYP
F81,NOP2,FAM32A,ZSCAN20,COX20,TGOLN2,PEX2,POLR1C,TNKS,PIGH,ELOF1,STX10,C12ORF57,DCTN5,F
),ANAPC11,RAD23A,ABHD17A,NT5C,MDH2,STXBP2,EEF1D,CHCHD5,NAA10,SEC31A,CNST,PRLR,COX17,CA

F48,PPAN,AP1S3,ACAD10,CYB561,SHARPIN,SUGT1,HYPK,CARS,MOB1A,CDK11A,ZNF197,INO80E,CCNL2,
,RTEL1,COASY,SRSF1,AUP1,SAP30BP,YIPF3,FOXRED1,RABL5,TRNT1,KXD1,ITGB1BP1,MED15,SH3YL1,IP6
,RTEL1,COASY,SRSF1,AUP1,SAP30BP,YIPF3,FOXRED1,RABL5,TRNT1,KXD1,ITGB1BP1,MED15,SH3YL1,IP6
C9C,COX4I1,DUSP22,SLC25A10,TMX2,THEM4,DBNDD2,CENPK,RPS6,RPL29,RTEL1,COASY,SRSF1,AUP1,SA
C9C,COX4I1,DUSP22,SLC25A10,TMX2,THEM4,DBNDD2,CENPK,RPS6,RPL29,RTEL1,COASY,SRSF1,AUP1,SA
,C19ORF48,PPAN,ACAD10,HNRNPC,SUGT1,HYPK,CARS,CDK11A,RBM3,ZNF197,COPZ1,INO80E,CCNL2,MA
SF3L,TSG101,PTPN13,UBR4,PSMA1,YLPM1,PHF21A,CHCHD1,FUZ,RGS3,FAM3A,PSMA3,MUC1,METTL17,
SF3L,TSG101,PTPN13,UBR4,PSMA1,YLPM1,PHF21A,CHCHD1,FUZ,RGS3,FAM3A,PSMA3,MUC1,METTL17,
2,STXBP2,EEF1D,CHCHD5,SEC31A,RPL18,CNST,COX17,COX5A,HIGD1A,B3GAT3,ZNF506,PFDN5,PAFAH1B
3,LZTFL1,ZC3H14,FIS1,SLC15A3,PPM1K,TUBA1C,STIP1,LRRC1,PHF19,GRAMD1C,VCAN,LPCAT3,CD44,TM
3,LZTFL1,ZC3H14,FIS1,SLC15A3,PPM1K,TUBA1C,STIP1,LRRC1,PHF19,GRAMD1C,VCAN,LPCAT3,CD44,TM
,GMPR2,MCFD2,RIC8A,TCP1,THADA,CCDC40,PIGT,UBE2N,LFNG,ZNHIT3,PDCD6,FAM32A,ZSCAN20,COX2
B41L5,ATG12,PPRC1,ZBTB44,TTC9C,PLCB2,DUSP22,SLC25A10,TMX2,THEM4,RDH11,DBNDD2,NISCH,TM
B41L5,ATG12,PPRC1,ZBTB44,TTC9C,PLCB2,DUSP22,SLC25A10,TMX2,THEM4,RDH11,DBNDD2,NISCH,TM
,HNRNPLL,PHF20L1,HIGD1A,POLR2C,B3GAT3,RFXANK,ZNF506,SPINT2,PFDN5,PAFAH1B2,ZNF343,TXNRD
,HNRNPLL,PHF20L1,HIGD1A,POLR2C,B3GAT3,RFXANK,ZNF506,SPINT2,PFDN5,PAFAH1B2,ZNF343,TXNRD
PIN,SLC9B2,SUGT1,HYPK,CARS,ZNF250,CDK11A,RBM3,ZNF197,COPZ1,INO80E,CCNL2,DUT,LRRFIP1,ATP5
,PSMA3,METTL17,EPB41L5,ATG12,PPRC1,RPS25,ZBTB44,NIP7,COX4I1,PLCB2,DUSP22,SLC25A10,TMX2,
NAPC11,RAD23A,ABHD17A,NT5C,MDH2,STXBP2,EEF1D,CHCHD5,SEC31A,CARHSP1,COX17,HIGD1A,FKBP
1,PSMD7,NADSYN1,ZNF511,TRAPPC2,PIGO,CLTA,SLC35A2,THOC3,AXIN1,MOSPD3,TMEM199,EPN1,HPS1
,STOML2,C19ORF54,ZNF7,TXNDC17,ARL6IP4,PIGL,C19ORF48,PPAN,ACAD10,HNRNPC,SUGT1,HYPK,CAR
4,PIGL,PPAN,AP1S3,CYB561,SHARPIN,SUGT1,HYPK,CARS,ZNF250,RPL36,CDK11A,ZNF197,COPZ1,INO80
4,PIGL,PPAN,AP1S3,CYB561,SHARPIN,SUGT1,HYPK,CARS,ZNF250,RPL36,CDK11A,ZNF197,COPZ1,INO80
FDN5,PAFAH1B2,TXNRD2,GBA2,DALRD3,NSUN5,OARD1,MFSD10,MRPS18C,GMPR2,MCFD2,ZSCAN9,RIC
FDN5,PAFAH1B2,TXNRD2,GBA2,DALRD3,NSUN5,OARD1,MFSD10,MRPS18C,GMPR2,MCFD2,ZSCAN9,RIC
4,FBXW9,USE1,IFI27L1,CCDC25,EEF1B2,RFX3,TNFRSF14,ENDOV,CAMTA1,PPIL2,RPL28,CLDND1,LIG1,CD
4,FBXW9,USE1,IFI27L1,CCDC25,EEF1B2,RFX3,TNFRSF14,ENDOV,CAMTA1,PPIL2,RPL28,CLDND1,LIG1,CD
,MAGED2,SON,SNF8,GRK6,HMGA2,CDC37,RPS16,CPSF3L,TSG101,OBSL1,TTC21B,UBR4,PSMA1,YLPM1,C
,MAGED2,SON,SNF8,GRK6,HMGA2,CDC37,RPS16,CPSF3L,TSG101,OBSL1,TTC21B,UBR4,PSMA1,YLPM1,C
2,STXBP2,EEF1D,CHCHD5,SEC31A,CNST,COX17,COX5A,HIGD1A,SIRT5,INTS4,B3GAT3,ZNF506,PFDN5,C11
OPZ1,INO80E,CCNL2,LRRFIP1,ATP5J,RMDN1,TRMT112,CHPF,STRIP2,RPS20,GABPB1,NIT2,DCTD,UBXN1,E
1C,DST,ANTXR2,STIP1,LRRC1,PHF19,VCAN,LPCAT3,CD44,TM9SF1,MAGED2,SON,TET2,UQCRQ,GRK6,HM
1C,DST,ANTXR2,STIP1,LRRC1,PHF19,VCAN,LPCAT3,CD44,TM9SF1,MAGED2,SON,TET2,UQCRQ,GRK6,HM
41,NT5C,MDH2,STXBP2,EEF1D,CHCHD5,NAA10,MED6,SEC31A,CARHSP1,CNST,SSH2,COX17,HNRNPLL,HI
7,PDCD5,ANAPC11,RAD23A,ABHD17A,NT5C,MDH2,EEF1D,HSPH1,NAA10,MED6,SEC31A,CARHSP1,CNST
7,PDCD5,ANAPC11,RAD23A,ABHD17A,NT5C,MDH2,EEF1D,HSPH1,NAA10,MED6,SEC31A,CARHSP1,CNST
PCYT2,FBXW9,USE1,MAF,ASF1B,EEF1B2,RFX3,DGKE,ENDOV,CAMTA1,CAPG,PPIL2,PSMD4,LIG1,PSMA7,
PCYT2,FBXW9,USE1,MAF,ASF1B,EEF1B2,RFX3,DGKE,ENDOV,CAMTA1,CAPG,PPIL2,PSMD4,LIG1,PSMA7,
ZNF687,PDCD5,ANAPC11,RAD23A,ABHD17A,NT5C,MDH2,EEF1D,NAA10,MED6,SEC31A,CARHSP1,SSH2,C
ZNF687,PDCD5,ANAPC11,RAD23A,ABHD17A,NT5C,MDH2,EEF1D,NAA10,MED6,SEC31A,CARHSP1,SSH2,C
ARPIN,CBFA2T2,STAU1,SUGT1,HYPK,CARS,ZNF250,RPL36,CDK11A,ZNF197,COPZ1,INO80E,DGUOK,HSP9
ARPIN,CBFA2T2,STAU1,SUGT1,HYPK,CARS,ZNF250,RPL36,CDK11A,ZNF197,COPZ1,INO80E,DGUOK,HSP9
IL2,TROAP,LIG1,CDC14B,APEH,CAPN10,RBM33,LYPLA2,LSM12,C17ORF62,PRMT1,STOML2,C19ORF54,Z
IL2,TROAP,LIG1,CDC14B,APEH,CAPN10,RBM33,LYPLA2,LSM12,C17ORF62,PRMT1,STOML2,C19ORF54,Z
1,LRRC1,PHF19,VCAN,LPCAT3,CD44,TM9SF1,MAGED2,SON,TET2,UQCRQ,GRK6,HMGA2,CDC37,RPS16,C
1,LRRC1,PHF19,VCAN,LPCAT3,CD44,TM9SF1,MAGED2,SON,TET2,UQCRQ,GRK6,HMGA2,CDC37,RPS16,C
,DBNL,ZNF836,RAD1,TOMM40,RABEP2,TADA3,ZNF20,ATP6V1F,WAC,PDCD5,TRIM16,ANAPC11,POLR3H,

,DGUOK,LRRFIP1,ATP5J,RMDN1,TRMT112,CHPF,ZNF625,RPS20,GABPB1,DCTD,UBXN1,ERLEC1,HSPB1,SA

NF250,RPL36,CDK11A,ZNF197,INO80E,DGUOK,HSP90AB1,LRRFIP1,RAB2A,ZNF524,ATP5J,ELP6,TRMT112

NF250,RPL36,CDK11A,ZNF197,INO80E,DGUOK,HSP90AB1,LRRFIP1,RAB2A,ZNF524,ATP5J,ELP6,TRMT112

DH1,ZNF197,COPZ1,INO80E,CCNL2,DGUOK,CEP70,LRRFIP1,ATP5J,RMDN1,TRMT112,CHPF,STRIP2,RPS2

RPIN,STAU1,SUGT1,HYPK,CARS,ZNF250,CDK11A,RBM3,MDH1,ZNF197,COPZ1,INO80E,HSP90AB1,SYNG

RPIN,STAU1,SUGT1,HYPK,CARS,ZNF250,CDK11A,RBM3,MDH1,ZNF197,COPZ1,INO80E,HSP90AB1,SYNG

343,GBA2,PPIH,DALRD3,NSUN5,DRG2,OARD1,MFSD10,MCFD2,RIC8A,TCP1,SLC30A5,THADA,PIGT,UBE2

BA1C,STIP1,LRRC1,PHF19,GRAMD1C,VCAN,LPCAT3,CD44,TM9SF1,SON,UQCRQ,SNF8,GRK6,HMGA2,CDC

,ZNF7,TXNDC17,PPAN,ACAD10,CYB561,SHARPIN,SLC9B2,SUGT1,HYPK,CARS,ZNF250,CDK11A,RBM3,MD

RANBP1,VRK3,LZTFL1,ZC3H14,FIS1,SLC15A3,PPM1K,TUBA1C,DST,STIP1,PHF19,GRAMD1C,VCAN,LPCAT3,

POLR2C,B3GAT3,RFXANK,SPINT2,PFDN5,PAFAH1B2,TXNRD2,DALRD3,NSUN5,ZFPL1,OARD1,MFSD10,M

POLR2C,B3GAT3,RFXANK,SPINT2,PFDN5,PAFAH1B2,TXNRD2,DALRD3,NSUN5,ZFPL1,OARD1,MFSD10,M

1,UBR4,YLPM1,PHF21A,CHCHD1,FUZ,FAM3A,MUC1,METTL17,EPB41L5,ATG12,RPS25,NIP7,COX4I1,DUS

1,UBR4,YLPM1,PHF21A,CHCHD1,FUZ,FAM3A,MUC1,METTL17,EPB41L5,ATG12,RPS25,NIP7,COX4I1,DUS

1,UBE2I,SLC39A13,EIF3M,JUP,CDC42,TBX6,TMEM205,PTK2,CARD14,PCYT2,FBXW9,USE1,ASF1B,CCDC25

1,UBE2I,SLC39A13,EIF3M,JUP,CDC42,TBX6,TMEM205,PTK2,CARD14,PCYT2,FBXW9,USE1,ASF1B,CCDC25

1,UBE2I,SLC39A13,EIF3M,JUP,CDC42,TBX6,TMEM205,PTK2,CARD14,PCYT2,FBXW9,USE1,ASF1B,CCDC25

1,UBE2I,SLC39A13,EIF3M,JUP,CDC42,TBX6,TMEM205,PTK2,CARD14,PCYT2,FBXW9,USE1,ASF1B,CCDC25

ARHSP1,COX17,HNRNPLL,HIGD1A,POLR2C,B3GAT3,RFXANK,ZNF506,SPINT2,FTH1,PFDN5,PAFAH1B2,TX

ARHSP1,COX17,HNRNPLL,HIGD1A,POLR2C,B3GAT3,RFXANK,ZNF506,SPINT2,FTH1,PFDN5,PAFAH1B2,TX

ML2,ZNF7,TXNDC17,PPAN,ACAD10,CYB561,SHARPIN,SUGT1,HYPK,CARS,ZNF250,CDK11A,ZNF197,INO8

ML2,ZNF7,TXNDC17,PPAN,ACAD10,CYB561,SHARPIN,SUGT1,HYPK,CARS,ZNF250,CDK11A,ZNF197,INO8

L2,COX17,COX5A,HINT1,HIGD1A,POLR2C,RPL15,SIRT5,INTS4,SPINT2,PFDN5,PAFAH1B2,PPIH,DALRD3,N

P2,TADA3,PDCD5,KLK10,TRIM16,ANAPC11,RAD23A,ABHD17A,MDH2,EEF1D,HSPH1,CHCHD5,NAA10,SE

P2,TADA3,PDCD5,KLK10,TRIM16,ANAPC11,RAD23A,ABHD17A,MDH2,EEF1D,HSPH1,CHCHD5,NAA10,SE

1L5,ATG12,RPS25,ZBTB44,NIP7,TTC9C,COX4I1,DUSP22,SLC25A10,TMX2,THEM4,DBNDD2,RPS6,RPL29,R

1L5,ATG12,RPS25,ZBTB44,NIP7,TTC9C,COX4I1,DUSP22,SLC25A10,TMX2,THEM4,DBNDD2,RPS6,RPL29,R

LRD3,NSUN5,ZFPL1,OARD1,MFSD10,GMPR2,ZSCAN9,RIC8A,TCP1,THADA,UBE2N,LFNG,PDCD6,XYLT2,FA

LRD3,NSUN5,ZFPL1,OARD1,MFSD10,GMPR2,ZSCAN9,RIC8A,TCP1,THADA,UBE2N,LFNG,PDCD6,XYLT2,FA

,MED15,SH3YL1,IP6K2,CLASRP,C1ORF198,UGGT2,MICAL1,UBE2I,PRMT2,EIF3M,CDC42,TBX6,TMEM205,

,MED15,SH3YL1,IP6K2,CLASRP,C1ORF198,UGGT2,MICAL1,UBE2I,PRMT2,EIF3M,CDC42,TBX6,TMEM205,

101,PTPN13,UBR4,PSMA1,YLPM1,PHF21A,CHCHD1,FUZ,RGS3,FAM3A,PSMA3,ECSIT,ATG12,RPS25,ZBTB

101,PTPN13,UBR4,PSMA1,YLPM1,PHF21A,CHCHD1,FUZ,RGS3,FAM3A,PSMA3,ECSIT,ATG12,RPS25,ZBTB

EP2,TADA3,SLC33A1,ZNF687,PDCD5,ANAPC11,RAD23A,ABHD17A,MDH2,STXBP2,EEF1D,NAA10,SEC31A

EP2,TADA3,SLC33A1,ZNF687,PDCD5,ANAPC11,RAD23A,ABHD17A,MDH2,STXBP2,EEF1D,NAA10,SEC31A

RNT1,KXD1,ITGB1BP1,MED15,IP6K2,CLASRP,C1ORF198,UGGT2,MICAL1,UBE2I,SLC39A13,YIF1A,EIF3M,C

RNT1,KXD1,ITGB1BP1,MED15,IP6K2,CLASRP,C1ORF198,UGGT2,MICAL1,UBE2I,SLC39A13,YIF1A,EIF3M,C

FL1,ZC3H14,FIS1,SLC15A3,PPM1K,TUBA1C,ANTXR2,STIP1,LRRC1,PHF19,GRAMD1C,LPCAT3,CD44,TM9SF

,TSG101,OBSL1,UBR4,PSMA1,YLPM1,PHF21A,CHCHD1,SLC39A11,FUZ,FAM3A,PSMA3,MUC1,METTL17,E

,TSG101,OBSL1,UBR4,PSMA1,YLPM1,PHF21A,CHCHD1,SLC39A11,FUZ,FAM3A,PSMA3,MUC1,METTL17,E

44,PLCB2,DUSP22,TMX2,THEM4,DBNDD2,NISCH,TMED4,CD99L2,GGPS1,AUP1,YIPF3,FOXRED1,TRNT1,K

44,PLCB2,DUSP22,TMX2,THEM4,DBNDD2,NISCH,TMED4,CD99L2,GGPS1,AUP1,YIPF3,FOXRED1,TRNT1,K

,DRG2,OARD1,MFSD10,GMPR2,MCFD2,RIC8A,TCP1,PARK7,THADA,PIGT,UBE2N,LFNG,ZNHIT3,PDCD6,FA

ZBTB44,PLCB2,DUSP22,TMX2,THEM4,DBNDD2,NISCH,TMED4,CD99L2,GGPS1,SRSF1,AUP1,YIPF3,FOXRED

ZBTB44,PLCB2,DUSP22,TMX2,THEM4,DBNDD2,NISCH,TMED4,CD99L2,GGPS1,SRSF1,AUP1,YIPF3,FOXRED

EC31A,CARHSP1,COX17,HNRNPLL,PHF20L1,POLR2C,B3GAT3,PFDN5,MTHFS,PAFAH1B2,ZNF343,TXNRD2,

EC31A,CARHSP1,COX17,HNRNPLL,PHF20L1,POLR2C,B3GAT3,PFDN5,MTHFS,PAFAH1B2,ZNF343,TXNRD2,

OX17,PHF20L1,HIGD1A,POLR2C,B3GAT3,SPINT2,PFDN5,MTHFS,PAFAH1B2,ZNF343,TXNRD2,GBA2,NSUN

OX17,PHF20L1,HIGD1A,POLR2C,B3GAT3,SPINT2,PFDN5,MTHFS,PAFAH1B2,ZNF343,TXNRD2,GBA2,NSUN

ET2,UQCRQ,GRK6,HMGA2,CDC37,RPS16,CPSF3L,TSG101,OBSL1,TTC21B,UBR4,PSMA1,YLPM1,PHF21A,C

ET2,UQCRQ,GRK6,HMGA2,CDC37,RPS16,CPSF3L,TSG101,OBSL1,TTC21B,UBR4,PSMA1,YLPM1,PHF21A,C

LR1C,PIGH,C12ORF57,DCTN5,PEX26,NDUFS3,OXLD1,METTL5,SLC35E1,OGFOD2,METTL3,TSEN34,SSR3,AF

,CAPG,PPIL2,PSMD4,RPL28,LIG1,PSMA7,CDC14B,TMEM126B,APEH,TXNL4A,LYPLA2,NUTF2,TCF4,TMPO,

,CAPG,PPIL2,PSMD4,RPL28,LIG1,PSMA7,CDC14B,TMEM126B,APEH,TXNL4A,LYPLA2,NUTF2,TCF4,TMPO,

A5,RANBP1,UBE2L6,VRK3,LZTFL1,ZC3H14,FIS1,SLC15A3,PPM1K,TUBA1C,IRF9,ANTXR2,STIP1,LRRC1,PHF1

A5,RANBP1,UBE2L6,VRK3,LZTFL1,ZC3H14,FIS1,SLC15A3,PPM1K,TUBA1C,IRF9,ANTXR2,STIP1,LRRC1,PHF1

D14,PCYT2,FBXW9,USE1,HSBP1L1,ASF1B,CCDC25,EEF1B2,EDF1,RFX3,PPA1,TNFRSF14,ENDOV,VAMP8,CA

D14,PCYT2,FBXW9,USE1,HSBP1L1,ASF1B,CCDC25,EEF1B2,EDF1,RFX3,PPA1,TNFRSF14,ENDOV,VAMP8,CA

AP,STYXL1,RFX5,POLD4,PLEKHA5,RANBP1,UBE2L6,VRK3,LZTFL1,ZC3H14,FIS1,SLC15A3,PPM1K,TUBA1C,

AP,STYXL1,RFX5,POLD4,PLEKHA5,RANBP1,UBE2L6,VRK3,LZTFL1,ZC3H14,FIS1,SLC15A3,PPM1K,TUBA1C,

A,SIRT5,INTS4,B3GAT3,RFXANK,ZNF506,SPINT2,PFDN5,MTHFS,PAFAH1B2,ZNF343,GBA2,PPIH,ARHGEF1,

25A10,TMX2,THEM4,DBNDD2,SDF4,NISCH,TMED4,RPL29,CD99L2,GGPS1,AUP1,SAP30BP,FOXRED1,RABL

25A10,TMX2,THEM4,DBNDD2,SDF4,NISCH,TMED4,RPL29,CD99L2,GGPS1,AUP1,SAP30BP,FOXRED1,RABL

APC11,NT5C,MDH2,STXBP2,EEF1D,CHCHD5,SEC31A,CARHSP1,CNST,COX17,HIGD1A,FKBP3,RPL15,INTS4,

,UCK2,C17ORF62,STOML2,ZNF7,TXNDC17,PPAN,ZFAND2B,ACAD10,HNRNPC,CYB561,STAU1,SUGT1,HYP

,UCK2,C17ORF62,STOML2,ZNF7,TXNDC17,PPAN,ZFAND2B,ACAD10,HNRNPC,CYB561,STAU1,SUGT1,HYP

IDOV,VAMP8,CAMTA1,CAPG,PPIL2,TROAP,RPL28,LIG1,CDC14B,TIMM17B,APEH,RPL21,CAPN10,RBM33,

IDOV,VAMP8,CAMTA1,CAPG,PPIL2,TROAP,RPL28,LIG1,CDC14B,TIMM17B,APEH,RPL21,CAPN10,RBM33,

DK11A,RBM3,ZNF197,COPZ1,INO80E,CCNL2,HSP90AB1,TMEM134,LRRFIP1,ATP5J,ELP6,TRMT112,CHPF,

L1,CLASRP,C1ORF198,UGGT2,MICAL1,SLC39A13,EIF3M,CDC42,TMEM205,PTK2,AUNIP,C14ORF2,GUK1,C

L1,CLASRP,C1ORF198,UGGT2,MICAL1,SLC39A13,EIF3M,CDC42,TMEM205,PTK2,AUNIP,C14ORF2,GUK1,C

1,MFSD10,MRPS18C,GMPR2,ZSCAN9,RIC8A,TCP1,SMARCC2,THADA,UBE2N,LFNG,PDCD6,XYLT2,ZNF81,F

1,MFSD10,MRPS18C,GMPR2,ZSCAN9,RIC8A,TCP1,SMARCC2,THADA,UBE2N,LFNG,PDCD6,XYLT2,ZNF81,F

2N,NFATC4,PDCD6,FAM32A,ZSCAN20,COX20,COA1,TGOLN2,PEX2,TNKS,C12ORF57,DCTN5,PEX26,NDUF

32A,TMEM147,COA1,USP40,PCID2,PEX2,POLR1C,TNKS,C12ORF57,DCTN5,PEX26,ZNF223,OGFOD2,METT

32A,TMEM147,COA1,USP40,PCID2,PEX2,POLR1C,TNKS,C12ORF57,DCTN5,PEX26,ZNF223,OGFOD2,METT

THADA,PIGT,UBE2N,ZNF81,FAM32A,TMEM147,COA1,USP40,PCID2,ZXDC,PEX2,TNKS,RPL37A,STX10,C12

THADA,PIGT,UBE2N,ZNF81,FAM32A,TMEM147,COA1,USP40,PCID2,ZXDC,PEX2,TNKS,RPL37A,STX10,C12

RD1,MFSD10,MRPS18C,GMPR2,ZSCAN9,RIC8A,TCP1,SMARCC2,THADA,UBE2N,LFNG,PDCD6,ZNF81,FAM

RD1,MFSD10,MRPS18C,GMPR2,ZSCAN9,RIC8A,TCP1,SMARCC2,THADA,UBE2N,LFNG,PDCD6,ZNF81,FAM

,FBXL18,RNF181,TPT1,NME1,SLC52A2,TSPAN31,RALY,ZNF593,GLYCTK,CFLAR,TRIM46,CNTRL,RER1,EIF2A

1,SLC52A2,TSPAN31,RALY,GLYCTK,TRIM46,VDAC1,CNTRL,RER1,NEK11,NKIRAS2,ACAA1,TRAPPC2,PIGO,C

,ANAPC11,RAD23A,NT5C,MDH2,STXBP2,EEF1D,CHCHD5,SEC31A,CARHSP1,CNST,COX17,HIGD1A,FKBP3,

IT,EPB41L5,ATG12,PPRC1,ZBTB44,TTC9C,DUSP22,SLC25A10,DBNDD2,SDF4,NISCH,TMED4,RPL29,CD99L2

IT,EPB41L5,ATG12,PPRC1,ZBTB44,TTC9C,DUSP22,SLC25A10,DBNDD2,SDF4,NISCH,TMED4,RPL29,CD99L2

HPS1,ITPA,ZCCHC9,VPS25,SNRPD1,MKRN1,SMUG1,CHURC1,SNRPA1,AZI2,GOSR2,MYL12A,TATDN1,TBC1

AMD1C,LPCAT3,CD44,TM9SF1,SON,TET2,UQCRQ,SNF8,GRK6,HMGA2,CDC37,RPS16,CPSF3L,TSG101,OBS

5,ANAPC11,RAD23A,ABHD17A,NT5C,ZCCHC4,MDH2,STXBP2,EEF1D,NAA10,SEC31A,CARHSP1,COX17,HN

5,ANAPC11,RAD23A,ABHD17A,NT5C,ZCCHC4,MDH2,STXBP2,EEF1D,NAA10,SEC31A,CARHSP1,COX17,HN

7,AUP1,SAP30BP,FOXRED1,TRNT1,KXD1,ITGB1BP1,PARD6G,SH3YL1,IP6K2,CLASRP,JPH2,C1ORF198,UGGT

7,AUP1,SAP30BP,FOXRED1,TRNT1,KXD1,ITGB1BP1,PARD6G,SH3YL1,IP6K2,CLASRP,JPH2,C1ORF198,UGGT

3PMS,NNMT,SNRPN,SPAG7,WDR62,IL1RAP,STYXL1,RFX5,POLD4,PLEKHA5,RANBP1,UBE2L6,VRK3,LZTFL1,

3PMS,NNMT,SNRPN,SPAG7,WDR62,IL1RAP,STYXL1,RFX5,POLD4,PLEKHA5,RANBP1,UBE2L6,VRK3,LZTFL1

RPS25,ZBTB44,NIP7,TTC9C,COX4I1,DUSP22,SLC25A10,TMX2,THEM4,DBNDD2,NISCH,TMED4,CENPK,RPS

L1RAP,STYXL1,RFX5,POLD4,PLEKHA5,RANBP1,UBE2L6,VRK3,LZTFL1,ZC3H14,FIS1,SLC15A3,PPM1K,TUBA1
L1RAP,STYXL1,RFX5,POLD4,PLEKHA5,RANBP1,UBE2L6,VRK3,LZTFL1,ZC3H14,FIS1,SLC15A3,PPM1K,TUBA1
ZNF81,FAM32A,COA1,ACOT2,TGOLN2,PEX2,TNKS,PIGH,C12ORF57,DCTN5,PEX26,NDUFS3,OXLD1,EIF4A1
ZNF81,FAM32A,COA1,ACOT2,TGOLN2,PEX2,TNKS,PIGH,C12ORF57,DCTN5,PEX26,NDUFS3,OXLD1,EIF4A1
3,TRAPPC2,TAF1D,PIGO,CLTA,AXIN1,HNRNPUL1,TP53,TMEM199,UBE2D3,COX19,EPN1,HPS1,ITPA,ZCCH
L30,ZNF483,SZT2,PSMD8,PSMC5,LRP10,SLC31A1,ACTB,LAMTOR4,WDR45,C9ORF64,IQCK,SETD4,MED8,F
EH,TXNL4A,LYPLA2,NUTF2,TMPO,ATPAF2,C17ORF62,PRMT1,STOML2,WNK1,ZNF7,ARL6IP4,STRADA,PPA
EH,TXNL4A,LYPLA2,NUTF2,TMPO,ATPAF2,C17ORF62,PRMT1,STOML2,WNK1,ZNF7,ARL6IP4,STRADA,PPA
TYXL1,RFX5,POLD4,PLEKHA5,RANBP1,UBE2L6,VRK3,LZTFL1,ZC3H14,FIS1,SLC15A3,PPM1K,TUBA1C,ANTX
TYXL1,RFX5,POLD4,PLEKHA5,RANBP1,UBE2L6,VRK3,LZTFL1,ZC3H14,FIS1,SLC15A3,PPM1K,TUBA1C,ANTX

4,SLC15A3,PPM1K,TUBA1C,IRF9,ANTXR2,STIP1,LRRC1,PHF19,GRAMD1C,VCAN,LPCAT3,CD44,TM9SF1,SO
4,SLC15A3,PPM1K,TUBA1C,IRF9,ANTXR2,STIP1,LRRC1,PHF19,GRAMD1C,VCAN,LPCAT3,CD44,TM9SF1,SO
341L5,ATG12,PPRC1,RPS25,ZBTB44,PLCB2,DUSP22,TMX2,THEM4,DBNDD2,NISCH,TMED4,RPL29,CD99L2
341L5,ATG12,PPRC1,RPS25,ZBTB44,PLCB2,DUSP22,TMX2,THEM4,DBNDD2,NISCH,TMED4,RPL29,CD99L2
AA1,PSMD7,IRF3,TRAPPC2,GTF2H1,KDELR3,CLTA,SLC35A2,AXIN1,TP53,MOSPD3,TMEM199,C1ORF63,EP
K1,PCYT2,USE1,PPP1CC,RFX3,PPA1,DGKE,TNFRSF14,CAMTA1,CRIP1,PPIL2,CLDND1,LIG1,PSMA7,CDC14B,
K1,PCYT2,USE1,PPP1CC,RFX3,PPA1,DGKE,TNFRSF14,CAMTA1,CRIP1,PPIL2,CLDND1,LIG1,PSMA7,CDC14B,
,PSMA1,YLPM1,PHF21A,RGS3,FAM3A,METTL17,ECSIT,EPB41L5,ATG12,RPS25,ZBTB44,NIP7,TTC9C,COX4
RCC5,EMC4,DDOST,LSR,CHCHD7,MEIS2,PSME2,PABPC1,ZNF836,RBP1,TOMM40,SERPINB6,RABEP2,TAD
RCC5,EMC4,DDOST,LSR,CHCHD7,MEIS2,PSME2,PABPC1,ZNF836,RBP1,TOMM40,SERPINB6,RABEP2,TAD
D,ZNF687,TRIM16,ANAPC11,RAD23A,ABHD17A,NT5C,MDH2,EEF1D,HSPH1,CHCHD5,NAA10,MED6,SEC31
D,ZNF687,TRIM16,ANAPC11,RAD23A,ABHD17A,NT5C,MDH2,EEF1D,HSPH1,CHCHD5,NAA10,MED6,SEC31
TN5,PEX26,ZNF223,OGFOD2,EIF3G,SSR3,ARIH2,PICK1,FBXL18,NME1,SLC52A2,TSPAN31,MTIF2,FAM192A
TN5,PEX26,ZNF223,OGFOD2,EIF3G,SSR3,ARIH2,PICK1,FBXL18,NME1,SLC52A2,TSPAN31,MTIF2,FAM192A
CTK,TRIM46,RBM42,CNTRL,RER1,EIF2A,TRAPPC2,GTF2H1,PIGO,AXIN1,TP53,TMEM199,IFITM2,SLC25A24
ANTXR2,STIP1,LRRC1,VCAN,LPCAT3,CD44,TM9SF1,MAGED2,SON,TET2,SNF8,GRK6,HMGA2,CDC37,RPS16
ANTXR2,STIP1,LRRC1,VCAN,LPCAT3,CD44,TM9SF1,MAGED2,SON,TET2,SNF8,GRK6,HMGA2,CDC37,RPS16
ARS,ZNF250,RPL36,CDK11A,ZNF197,INO80E,DGUOK,HSP90AB1,RAB2A,ZNF524,ATP5J,TSFM,TRMT112,C
ARS,ZNF250,RPL36,CDK11A,ZNF197,INO80E,DGUOK,HSP90AB1,RAB2A,ZNF524,ATP5J,TSFM,TRMT112,C
30E,CCNL2,DGUOK,LRRFIP1,RAB2A,ATP5J,RMDN1,TRMT112,ZNF625,RPS20,MPDU1,RPS15,UBXN1,HSPB
R4,PSMA1,YLPM1,SLC39A11,FUZ,FAM3A,MUC1,METTL17,EPB41L5,PPRC1,NLRP1,ZBTB44,TTC9C,COX4I1,
R4,PSMA1,YLPM1,SLC39A11,FUZ,FAM3A,MUC1,METTL17,EPB41L5,PPRC1,NLRP1,ZBTB44,TTC9C,COX4I1,
DA,PIGT,UBE2N,MFSD11,LFNG,ZNHIT3,PDCD6,NOP2,FAM32A,COX20,PEX2,POLR1C,ELOF1,C12ORF57,DC
D6,XYLT2,FAM32A,COA1,USP40,TGOLN2,PEX2,POLR1C,C12ORF57,PEX26,OXLD1,EIF4A1,ZNF223,METTL5
P22,SLC25A10,SPOCK2,RDH11,NISCH,TMED4,CENPK,RPL29,CD99L2,GGPS1,RTEL1,SRSF1,AUP1,SAP30BP,
P22,SLC25A10,SPOCK2,RDH11,NISCH,TMED4,CENPK,RPL29,CD99L2,GGPS1,RTEL1,SRSF1,AUP1,SAP30BP,
5A10,THEM4,RDH11,DBNDD2,CD99L2,RTEL1,SRSF1,AUP1,SAP30BP,YIPF3,RABL5,ITGB1BP1,MED15,PARD
5A10,THEM4,RDH11,DBNDD2,CD99L2,RTEL1,SRSF1,AUP1,SAP30BP,YIPF3,RABL5,ITGB1BP1,MED15,PARD
SF1,AUP1,SAP30BP,YIPF3,FOXRED1,TRNT1,KXD1,ITGB1BP1,MED15,SH3YL1,C1ORF198,UGGT2,MICAL1,U
,PARD6G,SH3YL1,IP6K2,CLASRP,JPH2,C1ORF198,MICAL1,HS2ST1,UBE2I,PRMT2,EIF3M,CDC42,TBX6,TME
,PARD6G,SH3YL1,IP6K2,CLASRP,JPH2,C1ORF198,MICAL1,HS2ST1,UBE2I,PRMT2,EIF3M,CDC42,TBX6,TME
2,TBX6,TMEM205,PTK2,AUNIP,C14ORF2,CARD14,PCYT2,FBXW9,USE1,AGPAT4,MAF,ASF1B,CCDC25,EEF1
2,TBX6,TMEM205,PTK2,AUNIP,C14ORF2,CARD14,PCYT2,FBXW9,USE1,AGPAT4,MAF,ASF1B,CCDC25,EEF1
H2,C1ORF198,MICAL1,HS2ST1,UBE2I,SLC39A13,YIF1A,PRMT2,JUP,CDC42,TBX6,XBP1,PTK2,AUNIP,C14O
D8,EIF6,RNMT,PUF60,DHX30,AP4M1,MRPL27,GANC,DDX56,CHFR,SSH3,MED17,DHRS4L2,C12ORF10,RPS
2,LPCAT3,TM9SF1,MAGED2,SON,UQCRQ,SNF8,GRK6,HMGA2,CDC37,RPS16,CPSF3L,TSG101,OBSL1,TTC21

2,LPCAT3,TM9SF1,MAGED2,SON,UQCRQ,SNF8,GRK6,HMGA2,CDC37,RPS16,CPSF3L,TSG101,OBSL1,TTC21
PFDN5,PAFAH1B2,TXNRD2,GBA2,DALRD3,NSUN5,OARD1,MFSD10,MRPS18C,MCFD2,RIC8A,PARK7,SMAF
PFDN5,PAFAH1B2,TXNRD2,GBA2,DALRD3,NSUN5,OARD1,MFSD10,MRPS18C,MCFD2,RIC8A,PARK7,SMAF
TGB1BP1,MED15,PARD6G,SH3YL1,IP6K2,CLASRP,JPH2,C1ORF198,UGGT2,MICAL1,HS2ST1,UBE2I,SLC39A
TGB1BP1,MED15,PARD6G,SH3YL1,IP6K2,CLASRP,JPH2,C1ORF198,UGGT2,MICAL1,HS2ST1,UBE2I,SLC39A
UBA1C,IRF9,ANTXR2,STIP1,LRRC1,PHF19,GRAMD1C,VCAN,LPCAT3,CD44,TM9SF1,MAGED2,SON,TET2,GF
LRD3,NSUN5,DRG2,MFSD10,GMPR2,MCFD2,RIC8A,THADA,UBE2N,LFNG,ZNHIT3,NOP2,FAM32A,COX20,
IDUFS3,EIF4A1,METTL5,SLC35E1,OGFOD2,METTL3,TSEN34,SSR3,ARIH2,RPL8,PICK1,TPT1,SLC52A2,NTMT
51,GPS2,BTF3L4,ZNF581,SUMO1,ZNF483,SZT2,PSMD8,CYB5D2,PSMC5,ALS2,ACSS1,GTF3C5,POLH,LRP10,
25A10,TMX2,SPOCK2,DBNDD2,SDF4,NISCH,CENPK,RPS6,RPL29,CD99L2,GGPS1,SRSF1,AUP1,FOXRED1,RA
25A10,TMX2,SPOCK2,DBNDD2,SDF4,NISCH,CENPK,RPS6,RPL29,CD99L2,GGPS1,SRSF1,AUP1,FOXRED1,RA
L5,PPRC1,RPS25,ZBTB44,NIP7,TTC9C,COX4I1,PLCB2,SLC25A10,SPOCK2,DBNDD2,SDF4,NISCH,TMED4,CEN
L5,PPRC1,RPS25,ZBTB44,NIP7,TTC9C,COX4I1,PLCB2,SLC25A10,SPOCK2,DBNDD2,SDF4,NISCH,TMED4,CEN
LRRC1,PHF19,GRAMD1C,VCAN,PCBP2,LPCAT3,CD44,TM9SF1,SON,TET2,UQCRQ,SNF8,GRK6,HMGA2,CDC
,YIF1A,PRMT2,EIF3M,TBX6,TMEM205,XBP1,PTK2,AUNIP,C14ORF2,GUK1,PCYT2,USE1,AGPAT4,PPP1CC,E
,YIF1A,PRMT2,EIF3M,TBX6,TMEM205,XBP1,PTK2,AUNIP,C14ORF2,GUK1,PCYT2,USE1,AGPAT4,PPP1CC,E
BPMS,SPAG7,WDR62,IL1RAP,STYXL1,RFX5,POLD4,PLEKHA5,RANBP1,UBE2L6,VRK3,LZTFL1,ZC3H14,FIS1,S
BPMS,SPAG7,WDR62,IL1RAP,STYXL1,RFX5,POLD4,PLEKHA5,RANBP1,UBE2L6,VRK3,LZTFL1,ZC3H14,FIS1,S
I3A,PSMA3,MUC1,ECSIT,EPB41L5,ATG12,PPRC1,RPS25,ZBTB44,NIP7,TTC9C,COX4I1,DUSP22,SLC25A10,T
HF21A,SLC39A11,FUZ,FAM3A,METTL17,ECSIT,EPB41L5,ATG12,PPRC1,RPS25,NIP7,TTC9C,COX4I1,DUSP22
A,EIF3M,CDC42,TBX6,PTK2,GUK1,CARD14,PCYT2,FBXW9,CCDC25,EEF1B2,EDF1,RFX3,TNFRSF14,ENDOV,
A,EIF3M,CDC42,TBX6,PTK2,GUK1,CARD14,PCYT2,FBXW9,CCDC25,EEF1B2,EDF1,RFX3,TNFRSF14,ENDOV,
QL5,ACAA1,SKA2,TRAPPC2,TAF1D,PIGO,SLC35A2,AXIN1,HNRNPUL1,TP53,TMEM199,C1ORF63,SNAPC2,E
,ZNF511,IRF3,GTF2H1,PIGO,CLTA,ARPC5,SLC35A2,AXIN1,RPS3A,MOSPD3,TMEM199,C1ORF63,EPN1,HPS
,FBXW9,USE1,MAF,CCDC25,DEF8,EEF1B2,PPP1CC,EDF1,RFX3,DGKE,CAMTA1,CRIP1,PPIL2,PSMD4,RPL28,
CD99L2,GGPS1,RTEL1,COASY,SRSF1,AUP1,SAP30BP,YIPF3,FOXRED1,RABL5,TRNT1,KXD1,ITGB1BP1,MED1
CD99L2,GGPS1,RTEL1,COASY,SRSF1,AUP1,SAP30BP,YIPF3,FOXRED1,RABL5,TRNT1,KXD1,ITGB1BP1,MED1

T4,DEF8,PPP1CC,EDF1,RFX3,PPA1,DGKE,CAMTA1,CAPG,CRIP1,CLDND1,VIM,LIG1,PSMA7,CDC14B,TIMM1
T4,DEF8,PPP1CC,EDF1,RFX3,PPA1,DGKE,CAMTA1,CAPG,CRIP1,CLDND1,VIM,LIG1,PSMA7,CDC14B,TIMM1
T4,DEF8,PPP1CC,EDF1,RFX3,PPA1,DGKE,CAMTA1,CAPG,CRIP1,CLDND1,VIM,LIG1,PSMA7,CDC14B,TIMM1
T4,DEF8,PPP1CC,EDF1,RFX3,PPA1,DGKE,CAMTA1,CAPG,CRIP1,CLDND1,VIM,LIG1,PSMA7,CDC14B,TIMM1
16,CPSF3L,TSG101,OBSL1,PTPN13,UBR4,PSMA1,YLPM1,CHCHD1,FUZ,RGS3,FAM3A,PSMA3,ECSIT,PPRC1,
16,CPSF3L,TSG101,OBSL1,PTPN13,UBR4,PSMA1,YLPM1,CHCHD1,FUZ,RGS3,FAM3A,PSMA3,ECSIT,PPRC1,
2,CDC37,CPSF3L,TSG101,OBSL1,TTC21B,UBR4,YLPM1,PHF21A,FAM3A,PSMA3,METTL17,ECSIT,EPB41L5,A
RFX5,POLD4,PLEKHA5,RANBP1,UBE2L6,VRK3,LZTFL1,ZC3H14,FIS1,SLC15A3,PPM1K,TUBA1C,ANTXR2,STIF
RFX5,POLD4,PLEKHA5,RANBP1,UBE2L6,VRK3,LZTFL1,ZC3H14,FIS1,SLC15A3,PPM1K,TUBA1C,ANTXR2,STIF
HF19,GRAMD1C,VCAN,LPCAT3,CD44,TM9SF1,MAGED2,SON,TET2,UQCRQ,GRK6,HMGA2,CDC37,RPS16,C
SLC25A10,TMX2,RDH11,DBNDD2,SDF4,TMED4,CENPK,RPS6,RPL29,CD99L2,GGPS1,RTEL1,COASY,SRSF1,A
VCAN,LPCAT3,CD44,TM9SF1,SON,TET2,UQCRQ,SNF8,GRK6,HMGA2,CDC37,RPS16,CPSF3L,TSG101,PTPN
VCAN,LPCAT3,CD44,TM9SF1,SON,TET2,UQCRQ,SNF8,GRK6,HMGA2,CDC37,RPS16,CPSF3L,TSG101,PTPN
POCK2,DBNDD2,TMED4,RPS6,RPL29,GGPS1,RTEL1,COASY,SRSF1,AUP1,SAP30BP,YIPF3,FOXRED1,TRNT1,
POCK2,DBNDD2,TMED4,RPS6,RPL29,GGPS1,RTEL1,COASY,SRSF1,AUP1,SAP30BP,YIPF3,FOXRED1,TRNT1,
AMTA1,CRIP1,CLDND1,VIM,LIG1,TIMM17B,TMEM126B,APEH,HYI,CAPN10,NUTF2,PODXL,ATPAF2,LSM1
AMTA1,CRIP1,CLDND1,VIM,LIG1,TIMM17B,TMEM126B,APEH,HYI,CAPN10,NUTF2,PODXL,ATPAF2,LSM1
R4,YLPM1,PHF21A,CHCHD1,SLC39A11,FAM3A,PSMA3,MUC1,METTL17,ECSIT,ATG12,PPRC1,RPS25,ZBTB
,PPA1,DGKE,VAMP8,CAMTA1,CRIP1,PPIL2,RPL28,VIM,LIG1,TIMM17B,TMEM126B,APEH,HYI,CAPN10,LY

,PPA1,DGKE,VAMP8,CAMTA1,CRIP1,PPIL2,RPL28,VIM,LIG1,TIMM17B,TMEM126B,APEH,HYI,CAPN10,LY

1ORF63,HPS1,ITPA,DNAJA3,VPS25,NSFL1C,SMUG1,CHURC1,SNRPA1,B3GNTL1,AZI2,TATDN1,ZBTB8OS,TE

TPN13,UBR4,YLPM1,PHF21A,CHCHD1,FUZ,PSMA3,MUC1,ECSIT,EPB41L5,ATG12,PPRC1,RPS25,ZBTB44,N

F8,NUDT5,RBPMS,NNMT,SNRPN,SPAG7,WDR62,IL1RAP,STYXL1,RFX5,POLD4,PLEKHA5,RANBP1,UBE2L6,

F8,NUDT5,RBPMS,NNMT,SNRPN,SPAG7,WDR62,IL1RAP,STYXL1,RFX5,POLD4,PLEKHA5,RANBP1,UBE2L6,

25,ZBTB44,DUSP22,TMX2,SPOCK2,THEM4,RDH11,DBNDD2,NISCH,CENPK,GGPS1,SRSF1,AUP1,SAP30BP,

25,ZBTB44,DUSP22,TMX2,SPOCK2,THEM4,RDH11,DBNDD2,NISCH,CENPK,GGPS1,SRSF1,AUP1,SAP30BP,

7,LAMTOR5,RBM42,CLASP2,PCNP,RER1,NEK11,EIF2A,ACAA1,RPL24,PSMD7,SKA2,ZNF511,ZNF283,TRAP

CAL1,HS2ST1,UBE2I,YIF1A,PRMT2,EIF3M,TBX6,XBP1,PTK2,AUNIP,C14ORF2,GUK1,PCYT2,FBXW9,USE1,H

CAL1,HS2ST1,UBE2I,YIF1A,PRMT2,EIF3M,TBX6,XBP1,PTK2,AUNIP,C14ORF2,GUK1,PCYT2,FBXW9,USE1,H

TL17,ECSIT,EPB41L5,ATG12,PPRC1,NLRP1,ZBTB44,NIP7,TTC9C,COX4I1,PLCB2,DUSP22,SLC25A10,RDH11,

TL17,ECSIT,EPB41L5,ATG12,PPRC1,NLRP1,ZBTB44,NIP7,TTC9C,COX4I1,PLCB2,DUSP22,SLC25A10,RDH11,

ANBP1,UBE2L6,VRK3,LZTFL1,ZC3H14,FIS1,SLC15A3,PPM1K,TUBA1C,IRF9,ANTXR2,STIP1,LRRC1,PHF19,GR

ANBP1,UBE2L6,VRK3,LZTFL1,ZC3H14,FIS1,SLC15A3,PPM1K,TUBA1C,IRF9,ANTXR2,STIP1,LRRC1,PHF19,GR

22,SLC25A10,TMX2,SPOCK2,THEM4,DBNDD2,SDF4,NISCH,TMED4,RPS6,CD99L2,GGPS1,RTEL1,SRSF1,AU

42,TBX6,TMEM205,XBP1,PTK2,C14ORF2,GUK1,ATF7,PCYT2,FBXW9,USE1,HSBP1L1,IFI27L1,MAF,ASF1B,

42,TBX6,TMEM205,XBP1,PTK2,C14ORF2,GUK1,ATF7,PCYT2,FBXW9,USE1,HSBP1L1,IFI27L1,MAF,ASF1B,

42,TBX6,TMEM205,XBP1,PTK2,C14ORF2,GUK1,ATF7,PCYT2,FBXW9,USE1,HSBP1L1,IFI27L1,MAF,ASF1B,

42,TBX6,TMEM205,XBP1,PTK2,C14ORF2,GUK1,ATF7,PCYT2,FBXW9,USE1,HSBP1L1,IFI27L1,MAF,ASF1B,

42,TBX6,TMEM205,XBP1,PTK2,C14ORF2,GUK1,ATF7,PCYT2,FBXW9,USE1,HSBP1L1,IFI27L1,MAF,ASF1B,

42,TBX6,TMEM205,XBP1,PTK2,C14ORF2,GUK1,ATF7,PCYT2,FBXW9,USE1,HSBP1L1,IFI27L1,MAF,ASF1B,

1,DGKE,VAMP8,CAMTA1,CAPG,CRIP1,CLDND1,LIG1,TIMM17B,XAF1,APEH,HYI,CAPN10,LYPLA2,NUTF2,P

1,DGKE,VAMP8,CAMTA1,CAPG,CRIP1,CLDND1,LIG1,TIMM17B,XAF1,APEH,HYI,CAPN10,LYPLA2,NUTF2,P

IF1A,EIF3M,CDC42,TMEM205,PTK2,CARD14,PCYT2,FBXW9,USE1,IFI27L1,ASF1B,CCDC25,EEF1B2,EDF1,R

IF1A,EIF3M,CDC42,TMEM205,PTK2,CARD14,PCYT2,FBXW9,USE1,IFI27L1,ASF1B,CCDC25,EEF1B2,EDF1,R

P1S3,CYB561,CBFA2T2,SUGT1,HYPK,CARS,ZNF250,RPL36,CDK11A,ZNF197,COPZ1,INO80E,DGUOK,HSP9

P1S3,CYB561,CBFA2T2,SUGT1,HYPK,CARS,ZNF250,RPL36,CDK11A,ZNF197,COPZ1,INO80E,DGUOK,HSP9

G,SH3YL1,IP6K2,CLASRP,JPH2,C1ORF198,MICAL1,HS2ST1,UBE2I,SLC39A13,YIF1A,PRMT2,EIF3M,CDC42,T

G,SH3YL1,IP6K2,CLASRP,JPH2,C1ORF198,MICAL1,HS2ST1,UBE2I,SLC39A13,YIF1A,PRMT2,EIF3M,CDC42,T

CHC9,VPS25,SNRPD1,UCHL5,COMT,CHURC1,HMBS,AZI2,GOSR2,TATDN1,LAMTOR1,KAT8,U2AF1L4,BCL7

CHC9,VPS25,SNRPD1,UCHL5,COMT,CHURC1,HMBS,AZI2,GOSR2,TATDN1,LAMTOR1,KAT8,U2AF1L4,BCL7

HS2ST1,UBE2I,SLC39A13,YIF1A,PRMT2,EIF3M,JUP,TBX6,TMEM205,XBP1,PTK2,AUNIP,C14ORF2,GUK1,C

HS2ST1,UBE2I,SLC39A13,YIF1A,PRMT2,EIF3M,JUP,TBX6,TMEM205,XBP1,PTK2,AUNIP,C14ORF2,GUK1,C

LN2,PEX2,TNKS,RPL37A,STX10,C12ORF57,DCTN5,PEX26,NDUFS3,OXLD1,EIF4A1,ZNF223,ARIH2,PICK1,F

LN2,PEX2,TNKS,RPL37A,STX10,C12ORF57,DCTN5,PEX26,NDUFS3,OXLD1,EIF4A1,ZNF223,ARIH2,PICK1,F

A,PSMA3,MUC1,ECSIT,EPB41L5,ATG12,PPRC1,RPS25,ZBTB44,NIP7,COX4I1,PLCB2,SLC25A10,SPOCK2,DB

A,PSMA3,MUC1,ECSIT,EPB41L5,ATG12,PPRC1,RPS25,ZBTB44,NIP7,COX4I1,PLCB2,SLC25A10,SPOCK2,DB

MM40,RABEP2,TADA3,ATP6V1F,WAC,RALGPS1,KLK10,ANAPC11,POLR3H,RAD23A,HSD17B10,DDX41,NT

1,RTEL1,SRSF1,AUP1,YIPF3,FOXRED1,TRNT1,KXD1,ITGB1BP1,MED15,SH3YL1,IP6K2,CLASRP,JPH2,MICAL1

1,RTEL1,SRSF1,AUP1,YIPF3,FOXRED1,TRNT1,KXD1,ITGB1BP1,MED15,SH3YL1,IP6K2,CLASRP,JPH2,MICAL1

,AGPAT4,DEF8,PPP1CC,RFX3,PPA1,DGKE,CAMTA1,CAPG,CLDND1,VIM,LIG1,TIMM17B,XAF1,APEH,HYI,C

,AGPAT4,DEF8,PPP1CC,RFX3,PPA1,DGKE,CAMTA1,CAPG,CLDND1,VIM,LIG1,TIMM17B,XAF1,APEH,HYI,C

UNIP,CENPN,ATF7,PCYT2,FBXW9,HSBP1L1,IFI27L1,ASF1B,DEF8,EEF1B2,PPP1CC,EDF1,RFX3,PPA1,DGKE,

TA1,CRIP1,PPIL2,PSMD4,VIM,LIG1,PSMA7,TMEM126B,XAF1,APEH,HYI,TXNL4A,NUTF2,TCF4,UCK2,TMP

TA1,CRIP1,PPIL2,PSMD4,VIM,LIG1,PSMA7,TMEM126B,XAF1,APEH,HYI,TXNL4A,NUTF2,TCF4,UCK2,TMP

RK6,HMGA2,CDC37,CPSF3L,TSG101,OBSL1,TTC21B,PTPN13,UBR4,PSMA1,PHF21A,CHCHD1,SLC39A11,F

RK6,HMGA2,CDC37,CPSF3L,TSG101,OBSL1,TTC21B,PTPN13,UBR4,PSMA1,PHF21A,CHCHD1,SLC39A11,F

,TBC1D22A,KAT8,U2AF1L4,EMC3,PRKCSH,ZNF276,COMMD1,ZNF483,SZT2,PSMD8,ZNF606,BSCL2,DNAJC
,TBC1D22A,KAT8,U2AF1L4,EMC3,PRKCSH,ZNF276,COMMD1,ZNF483,SZT2,PSMD8,ZNF606,BSCL2,DNAJC
ET2,UQCRQ,SNF8,GRK6,HMGA2,CDC37,RPS16,CPSF3L,TSG101,OBSL1,TTC21B,PTPN13,PSMA1,PHF21A,C
ET2,UQCRQ,SNF8,GRK6,HMGA2,CDC37,RPS16,CPSF3L,TSG101,OBSL1,TTC21B,PTPN13,PSMA1,PHF21A,C
TPN13,UBR4,YLPM1,PHF21A,SLC39A11,FAM3A,PSMA3,MUC1,METTL17,ECSIT,EPB41L5,ATG12,PPRC1,R
TPN13,UBR4,YLPM1,PHF21A,SLC39A11,FAM3A,PSMA3,MUC1,METTL17,ECSIT,EPB41L5,ATG12,PPRC1,R
F8,GRK6,CDC37,RPS16,CPSF3L,TSG101,OBSL1,TTC21B,PTPN13,UBR4,PSMA1,YLPM1,CHCHD1,FUZ,FAM3
F8,GRK6,CDC37,RPS16,CPSF3L,TSG101,OBSL1,TTC21B,PTPN13,UBR4,PSMA1,YLPM1,CHCHD1,FUZ,FAM3
AN31,MTIF2,FAM192A,GLYCTK,CFLAR,TRIM46,C11ORF57,CREBZF,EIF2A,NKIRAS2,ACAA1,PSMD7,GTF2H1
AN31,MTIF2,FAM192A,GLYCTK,CFLAR,TRIM46,C11ORF57,CREBZF,EIF2A,NKIRAS2,ACAA1,PSMD7,GTF2H1
CENPK,GGPS1,RTEL1,SRSF1,AUP1,YIPF3,FOXRED1,RABL5,TRNT1,ITGB1BP1,PARD6G,SH3YL1,CLASRP,JPH2
CENPK,GGPS1,RTEL1,SRSF1,AUP1,YIPF3,FOXRED1,RABL5,TRNT1,ITGB1BP1,PARD6G,SH3YL1,CLASRP,JPH2
RFX3,PPA1,DGKE,VAMP8,CAMTA1,CRIP1,CLDND1,LIG1,PSMA7,TIMM17B,XAF1,APEH,HYI,CAPN10,NUTF
RFX3,PPA1,DGKE,VAMP8,CAMTA1,CRIP1,CLDND1,LIG1,PSMA7,TIMM17B,XAF1,APEH,HYI,CAPN10,NUTF
HARPIN,UPP1,ZNF250,RPL36,LAMP2,CCNL2,SYNGR2,CEP70,TMEM134,LRRFIP1,ZNF524,RMDN1,TPM2,G
HARPIN,UPP1,ZNF250,RPL36,LAMP2,CCNL2,SYNGR2,CEP70,TMEM134,LRRFIP1,ZNF524,RMDN1,TPM2,G
DCD5,NXN,TRIM16,ANAPC11,RAD23A,ABHD17A,ILF3,MDH2,STXBP2,EEF1D,HSPH1,CHCHD5,NAA10,SGK1
DCD5,NXN,TRIM16,ANAPC11,RAD23A,ABHD17A,ILF3,MDH2,STXBP2,EEF1D,HSPH1,CHCHD5,NAA10,SGK1
TA1,CAPG,CRIP1,CLDND1,VIM,LIG1,TIMM17B,TMEM126B,XAF1,APEH,HYI,CAPN10,NUTF2,PODXL,ATPAF
TA1,CAPG,CRIP1,CLDND1,VIM,LIG1,TIMM17B,TMEM126B,XAF1,APEH,HYI,CAPN10,NUTF2,PODXL,ATPAF
F62,PRMT1,CASC4,EXOSC9,STOML2,WNK1,C19ORF54,ZNF7,ARL6IP4,STRADA,HNRNPC,ZNF644,STAU1,S
F62,PRMT1,CASC4,EXOSC9,STOML2,WNK1,C19ORF54,ZNF7,ARL6IP4,STRADA,HNRNPC,ZNF644,STAU1,S
32,DUSP22,SLC25A10,THEM4,DBNDD2,NISCH,TMED4,CD99L2,GGPS1,COASY,AUP1,FOXRED1,TRNT1,ME
32,DUSP22,SLC25A10,THEM4,DBNDD2,NISCH,TMED4,CD99L2,GGPS1,COASY,AUP1,FOXRED1,TRNT1,ME
MD8,ZNF606,BSCL2,DNAJC24,ALS2,GTF3C5,AP2S1,ACTB,CCDC171,METTL23,SEC13,MRPL51,ZNF586,ME
MD8,ZNF606,BSCL2,DNAJC24,ALS2,GTF3C5,AP2S1,ACTB,CCDC171,METTL23,SEC13,MRPL51,ZNF586,ME
F197,INO80E,CCNL2,SYNGR2,LRRFIP1,FAM219B,RAB2A,ZNF524,ATP5J,ELP6,TSFM,GGCX,ZEB2,TPM3,CHP
N,MFSD11,PDCD6,GPR56,ENY2,NOP2,FAM32A,PEX2,POLR1C,TNKS,ELOF1,C12ORF57,DCTN5,PEX26,NDU
PC,CYB561,SHARPIN,UPP1,ZNF250,RPL36,MOB1A,LAMP2,CCNL2,HSP90AB1,SYNGR2,TMEM134,LRRFIP1
BE2N,LFNG,ZNF81,NOP2,FAM32A,RPAIN,COX20,PCID2,PEX2,PIGH,C12ORF57,DCTN5,NDUFS3,EIF4A1,ZN
BE2N,LFNG,ZNF81,NOP2,FAM32A,RPAIN,COX20,PCID2,PEX2,PIGH,C12ORF57,DCTN5,NDUFS3,EIF4A1,ZN
T,ATG12,PPRC1,ZBTB44,NIP7,TTC9C,COX4I1,DUSP22,SLC25A10,TMX2,SPOCK2,THEM4,DBNDD2,SDF4,NI
T,ATG12,PPRC1,ZBTB44,NIP7,TTC9C,COX4I1,DUSP22,SLC25A10,TMX2,SPOCK2,THEM4,DBNDD2,SDF4,NI
CAN,LPCAT3,CD44,TM9SF1,MAGED2,SON,TET2,UQCRQ,GRK6,HMGA2,CDC37,RPS16,CPSF3L,OBSL1,TTC2
CAN,LPCAT3,CD44,TM9SF1,MAGED2,SON,TET2,UQCRQ,GRK6,HMGA2,CDC37,RPS16,CPSF3L,OBSL1,TTC2
,COX4I1,DUSP22,SPOCK2,SDF4,TMED4,CENPK,RPS6,CD99L2,GGPS1,RTEL1,SRSF1,AUP1,FOXRED1,TRNT1
,COX4I1,DUSP22,SPOCK2,SDF4,TMED4,CENPK,RPS6,CD99L2,GGPS1,RTEL1,SRSF1,AUP1,FOXRED1,TRNT1
EF1B2,EDF1,RFX3,PPA1,ENDOV,VAMP8,PPIL2,RPL28,CLDND1,LIG1,PSMA7,CDC14B,TIMM17B,XAF1,APE
EF1B2,EDF1,RFX3,PPA1,ENDOV,VAMP8,PPIL2,RPL28,CLDND1,LIG1,PSMA7,CDC14B,TIMM17B,XAF1,APE
PL36,NFAM1,CCNL2,SYNGR2,TMEM134,LRRFIP1,FAM219B,RAB2A,ZNF524,TPM2,GPBP1,ASPH,STRIP2,N
PL36,NFAM1,CCNL2,SYNGR2,TMEM134,LRRFIP1,FAM219B,RAB2A,ZNF524,TPM2,GPBP1,ASPH,STRIP2,N
A52,ZNF276,RPL30,ZNF483,SZT2,PSMD8,ZNF606,BSCL2,DNAJC24,ALS2,GTF3C5,AP2S1,ACTB,METTL23,SE
A52,ZNF276,RPL30,ZNF483,SZT2,PSMD8,ZNF606,BSCL2,DNAJC24,ALS2,GTF3C5,AP2S1,ACTB,METTL23,SE
SF1,SON,TET2,SNF8,GRK6,HMGA2,CDC37,CPSF3L,TSG101,TTC21B,PTPN13,UBR4,PSMA1,YLPM1,CHCHD1
SF1,SON,TET2,SNF8,GRK6,HMGA2,CDC37,CPSF3L,TSG101,TTC21B,PTPN13,UBR4,PSMA1,YLPM1,CHCHD1
ON5,MTHFS,PAFAH1B2,COA3,TMEM39B,PPIH,DALRD3,NSUN5,DAZAP2,ZFPL1,OARD1,MFSD10,GMPR2,Z
ON5,MTHFS,PAFAH1B2,COA3,TMEM39B,PPIH,DALRD3,NSUN5,DAZAP2,ZFPL1,OARD1,MFSD10,GMPR2,Z

ADA3,ZNF20,RALGPS1,ZNF706,ZNF687,GTF2IRD1,KLK10,EPS15L1,POLR3H,RAD23A,ABHD17A,ILF3,AGAP3
ADA3,ZNF20,RALGPS1,ZNF706,ZNF687,GTF2IRD1,KLK10,EPS15L1,POLR3H,RAD23A,ABHD17A,ILF3,AGAP3
RRFIP1,RAB2A,ZNF524,TPM2,GPBP1,SIAH1,ASPH,CHPF,STRIP2,NIT2,LYNX1,GMPPB,MVD,IL32,SEMA3C,LI
RRFIP1,RAB2A,ZNF524,TPM2,GPBP1,SIAH1,ASPH,CHPF,STRIP2,NIT2,LYNX1,GMPPB,MVD,IL32,SEMA3C,LI
,SLC35E1,OGFOD2,METTL3,EIF3G,MPV17,ARIH2,PICK1,FBXL18,RNF181,KCNC4,TPT1,CDK4,TSPAN31,MTI
UGT1,HYPK,CARS,ZNF250,MOB1A,CDK11A,MDH1,ZNF197,INO80E,DGUOK,HSP90AB1,RAB2A,ATP5J,GPA
UGT1,HYPK,CARS,ZNF250,MOB1A,CDK11A,MDH1,ZNF197,INO80E,DGUOK,HSP90AB1,RAB2A,ATP5J,GPA
VRB,TOMM40,RABEP2,TADA3,ZNF20,RALGPS1,ZNF706,ZNF687,NXN,EPS15L1,RAD23A,ABHD17A,DDX41
VRB,TOMM40,RABEP2,TADA3,ZNF20,RALGPS1,ZNF706,ZNF687,NXN,EPS15L1,RAD23A,ABHD17A,DDX41
S25,NLRP1,ZBTB44,NIP7,TTC9C,COX4I1,DUSP22,SLC25A10,THEM4,RDH11,DBNDD2,SDF4,NISCH,TMED4,
5,ZBTB44,NIP7,COX4I1,DUSP22,SLC25A10,TMX2,SPOCK2,THEM4,DBNDD2,SDF4,NISCH,TMED4,RPS6,CD9
5,ZBTB44,NIP7,COX4I1,DUSP22,SLC25A10,TMX2,SPOCK2,THEM4,DBNDD2,SDF4,NISCH,TMED4,RPS6,CD9
10,MEIS2,PSME2,PABPC1,RRNAD1,RBP1,TOMM40,TADA3,ZNF687,PDCD5,ANAPC11,RAD23A,ABHD17A,
10,MEIS2,PSME2,PABPC1,RRNAD1,RBP1,TOMM40,TADA3,ZNF687,PDCD5,ANAPC11,RAD23A,ABHD17A,
198,MICAL1,HS2ST1,UBE2I,YIF1A,PRMT2,EIF3M,MAGT1,JUP,CDC42,TBX6,TMEM205,XBP1,AUNIP,GUK1,
198,MICAL1,HS2ST1,UBE2I,YIF1A,PRMT2,EIF3M,MAGT1,JUP,CDC42,TBX6,TMEM205,XBP1,AUNIP,GUK1,
C21B,UBR4,PSMA1,YLPM1,PHF21A,SLC39A11,FUZ,RGS3,FAM3A,PSMA3,METTL17,ECSIT,ATG12,PPRC1,R
C21B,UBR4,PSMA1,YLPM1,PHF21A,SLC39A11,FUZ,RGS3,FAM3A,PSMA3,METTL17,ECSIT,ATG12,PPRC1,R
D4,CENPK,RPS6,RPL29,CD99L2,RTEL1,SRSF1,AUP1,YIPF3,FOXRED1,TRNT1,KXD1,ITGB1BP1,PARD6G,SH3
L,C17ORF62,PRMT1,STOML2,C19ORF54,ZNF7,TXNDC17,PPAN,ACAD10,HNRNPC,CYB561,SHARPIN,CBFA2
L,C17ORF62,PRMT1,STOML2,C19ORF54,ZNF7,TXNDC17,PPAN,ACAD10,HNRNPC,CYB561,SHARPIN,CBFA2
R2,RAB2A,ZNF524,ATP5J,TPM2,GPALPP1,TSFM,TRMT112,PNPO,CHPF,RPS20,VPS28,DCTD,ZNF814,ERLEC
R2,RAB2A,ZNF524,ATP5J,TPM2,GPALPP1,TSFM,TRMT112,PNPO,CHPF,RPS20,VPS28,DCTD,ZNF814,ERLEC
TSG101,OBSL1,PTPN13,UBR4,PSMA1,YLPM1,PHF21A,FAM3A,ECSIT,EPB41L5,PPRC1,NLRP1,ZBTB44,NIP7,
1,UPP1,ZNF250,RPL36,MOB1A,LAMP2,CCNL2,HSP90AB1,DUT,SYNGR2,TMEM134,LRRFIP1,FAM219B,RAE
5,ATG12,PPRC1,RPS25,TTC9C,COX4I1,PLCB2,TMX2,THEM4,RDH11,TMED4,CENPK,RTEL1,COASY,AUP1,SA

5,MOB1A,NFAM1,LAMP2,CCNL2,HSP90AB1,DUT,SYNGR2,CEP70,TMEM134,LRRFIP1,RAB2A,ZNF524,ELP6
5,MOB1A,NFAM1,LAMP2,CCNL2,HSP90AB1,DUT,SYNGR2,CEP70,TMEM134,LRRFIP1,RAB2A,ZNF524,ELP6
UBA1C,IRF9,DST,ANTXR2,STIP1,LRRC1,PHF19,GRAMD1C,VCAN,TM9SF1,MAGED2,SON,TET2,UQCRQ,SNF8
UBA1C,IRF9,DST,ANTXR2,STIP1,LRRC1,PHF19,GRAMD1C,VCAN,TM9SF1,MAGED2,SON,TET2,UQCRQ,SNF8
FIS1,SLC15A3,PPM1K,TUBA1C,IRF9,ANTXR2,STIP1,LRRC1,PHF19,GRAMD1C,VCAN,LPCAT3,CD44,TM9SF1
FIS1,SLC15A3,PPM1K,TUBA1C,IRF9,ANTXR2,STIP1,LRRC1,PHF19,GRAMD1C,VCAN,LPCAT3,CD44,TM9SF1
F9,ANTXR2,STIP1,LRRC1,PHF19,GRAMD1C,VCAN,PCBP2,LPCAT3,CD44,TM9SF1,MAGED2,SON,TET2,UQC
2,GPBP1,SIAH1,GGCX,ZEB2,STRIP2,ZNF625,RPS20,LYNX1,TWSG1,RPS15,ERLEC1,GMPPB,SAT1,ERCC5,MCT
GGT2,MICAL1,UBE2I,SLC39A13,PRMT2,CDC42,TBX6,TMEM205,XBP1,PTK2,C14ORF2,GUK1,CENPN,ATF7,
1,UPP1,ZNF250,RPL36,MOB1A,LAMP2,CCNL2,HSP90AB1,DUT,SYNGR2,TMEM134,LRRFIP1,RAB2A,ZNF524
P2,RALGPS1,PDCD5,NXN,EPS15L1,NUDT1,ANAPC11,POLR3H,RAD23A,ABHD17A,NT5C,STXBP2,EEF1D,NA
AMTA1,CAPG,CLDND1,VIM,LIG1,CDC14B,TIMM17B,TMEM126B,XAF1,APEH,HYI,TXNL4A,CAPN10,LYPLA2
AMTA1,CAPG,CLDND1,VIM,LIG1,CDC14B,TIMM17B,TMEM126B,XAF1,APEH,HYI,TXNL4A,CAPN10,LYPLA2
AN31,PPP2R5C,FAM192A,RALY,ZNF593,GLYCTK,TRIM46,RBM42,RER1,NEK11,RPL34,NKIRAS2,ACAA1,PA
,PLEKHA5,RANBP1,UBE2L6,VRK3,LZTFL1,ZC3H14,FIS1,SLC15A3,PPM1K,TUBA1C,IRF9,DST,ANTXR2,STIP1,
,PLEKHA5,RANBP1,UBE2L6,VRK3,LZTFL1,ZC3H14,FIS1,SLC15A3,PPM1K,TUBA1C,IRF9,DST,ANTXR2,STIP1,
9A11,FUZ,FAM3A,MUC1,METTL17,ECSIT,EPB41L5,ATG12,PPRC1,ZBTB44,PLCB2,DUSP22,TMX2,RDH11,D
9A11,FUZ,FAM3A,MUC1,METTL17,ECSIT,EPB41L5,ATG12,PPRC1,ZBTB44,PLCB2,DUSP22,TMX2,RDH11,D
AF,PPP1CC,EDF1,RFX3,DGKE,ENDOV,CRIP1,PSMD4,LIG1,TIMM17B,HYI,CAPN10,CAMLG,EPB41L4B,TCF4,U
AF,PPP1CC,EDF1,RFX3,DGKE,ENDOV,CRIP1,PSMD4,LIG1,TIMM17B,HYI,CAPN10,CAMLG,EPB41L4B,TCF4,U

ITOR1,FAM104B,EMC3,GPS2,UBA52,AIG1,CAPN2,SDHA,SZT2,FBXO34,EXOSC2,MCUR1,AP2S1,ME3,SLC31

,ZNF250,RPL36,RBM3,LAMP2,INO80E,DGUOK,HSP90AB1,SYNGR2,LRRFIP1,FAM219B,RAB2A,ZNF524,AS

,ZNF250,RPL36,RBM3,LAMP2,INO80E,DGUOK,HSP90AB1,SYNGR2,LRRFIP1,FAM219B,RAB2A,ZNF524,AS

3,UGGT2,MICAL1,HS2ST1,KCNMA1,SLC39A13,YIF1A,EIF3M,CDC42,TBX6,PTK2,CELF1,CARD14,ATF7,FBXW

3,UGGT2,MICAL1,HS2ST1,KCNMA1,SLC39A13,YIF1A,EIF3M,CDC42,TBX6,PTK2,CELF1,CARD14,ATF7,FBXW

MICAL1,YIF1A,PRMT2,EIF3M,MAGT1,CDC42,TBX6,PTK2,AUNIP,CENPN,CARD14,PCYT2,USE1,ASF1B,CCDC

MICAL1,YIF1A,PRMT2,EIF3M,MAGT1,CDC42,TBX6,PTK2,AUNIP,CENPN,CARD14,PCYT2,USE1,ASF1B,CCDC

3H14,FIS1,TUBA1C,IRF9,DST,ANTXR2,STIP1,LRRC1,PHF19,GRAMD1C,PCBP2,LPCAT3,CD44,TM9SF1,MAG

.B,PTPN13,UBR4,PSMA1,YLPM1,PHF21A,CHCHD1,FUZ,FAM3A,PSMA3,METTL17,PPRC1,RPS25,NIP7,COX

.B,PTPN13,UBR4,PSMA1,YLPM1,PHF21A,CHCHD1,FUZ,FAM3A,PSMA3,METTL17,PPRC1,RPS25,NIP7,COX

,SIAH1,ASPH,TRMT112,NIT2,LYNX1,TWSG1,MAPK13,ZSWIM7,HSPB1,LRRC27,MVD,SAT1,SEMA3C,LMBR

O,LSM12,C17ORF62,PRMT1,CASC4,EXOSC9,WNK1,C19ORF54,ZNF7,ARL6IP4,STRADA,HNRNPC,CYB561,Z

O,LSM12,C17ORF62,PRMT1,CASC4,EXOSC9,WNK1,C19ORF54,ZNF7,ARL6IP4,STRADA,HNRNPC,CYB561,Z

,ZNF7,ARL6IP4,PIGL,ZFAND2B,CYB561,SHARPIN,UPP1,RPL36,MOB1A,NFAM1,RBM3,LAMP2,SYNGR2,CE

,ZNF7,ARL6IP4,PIGL,ZFAND2B,CYB561,SHARPIN,UPP1,RPL36,MOB1A,NFAM1,RBM3,LAMP2,SYNGR2,CE

HD17A,NT5C,ILF3,EEF1D,NAA10,RRP8,MED6,LLGL2,CHEK1,COX17,CAMK2D,SUMO3,ZFAND4,FKBP3,EGL

HD17A,NT5C,ILF3,EEF1D,NAA10,RRP8,MED6,LLGL2,CHEK1,COX17,CAMK2D,SUMO3,ZFAND4,FKBP3,EGL

5,RANBP1,UBE2L6,VRK3,LZTFL1,ZC3H14,FIS1,PPM1K,TUBA1C,IRF9,DST,ANTXR2,LRRC1,PHF19,GRAMD1C

5,RANBP1,UBE2L6,VRK3,LZTFL1,ZC3H14,FIS1,PPM1K,TUBA1C,IRF9,DST,ANTXR2,LRRC1,PHF19,GRAMD1C

9ORF54,ZNF7,PIGL,PPAN,HNRNPC,CYB561,SHARPIN,SLC9B2,CARS,MOB1A,CDK11A,RBM3,MDH1,LAMP2

9ORF54,ZNF7,PIGL,PPAN,HNRNPC,CYB561,SHARPIN,SLC9B2,CARS,MOB1A,CDK11A,RBM3,MDH1,LAMP2

R2,STIP1,LRRC1,GRAMD1C,VCAN,PCBP2,LPCAT3,CD44,TM9SF1,SON,TET2,UQCRQ,SNF8,GRK6,HMGA2,C

R2,STIP1,LRRC1,GRAMD1C,VCAN,PCBP2,LPCAT3,CD44,TM9SF1,SON,TET2,UQCRQ,SNF8,GRK6,HMGA2,C

LASRP,C1ORF198,UGGT2,MICAL1,HS2ST1,YIF1A,PRMT2,MAGT1,CDC42,PTK2,CELF1,GUK1,CENPN,ATF7,P

LASRP,C1ORF198,UGGT2,MICAL1,HS2ST1,YIF1A,PRMT2,MAGT1,CDC42,PTK2,CELF1,GUK1,CENPN,ATF7,P

2,CDC37,CPSF3L,TTC21B,PTPN13,UBR4,PSMA1,PHF21A,SLC39A11,FUZ,PSMA3,MUC1,METTL17,EPB41L5

2,CDC37,CPSF3L,TTC21B,PTPN13,UBR4,PSMA1,PHF21A,SLC39A11,FUZ,PSMA3,MUC1,METTL17,EPB41L5

EF8,PPP1CC,EDF1,GADL1,RFX3,PPA1,DGKE,TNFRSF14,VAMP8,CLDND1,VIM,LIG1,PSMA7,CDC14B,TIMM

EF8,PPP1CC,EDF1,GADL1,RFX3,PPA1,DGKE,TNFRSF14,VAMP8,CLDND1,VIM,LIG1,PSMA7,CDC14B,TIMM

EF8,PPP1CC,EDF1,GADL1,RFX3,PPA1,DGKE,TNFRSF14,VAMP8,CLDND1,VIM,LIG1,PSMA7,CDC14B,TIMM

EF8,PPP1CC,EDF1,GADL1,RFX3,PPA1,DGKE,TNFRSF14,VAMP8,CLDND1,VIM,LIG1,PSMA7,CDC14B,TIMM

XNDC17,PPAN,ZFAND2B,AP1S3,CYB561,CARS,ZNF250,RPL36,NFAM1,MDH1,LAMP2,COPZ1,CCNL2,HSP9

XNDC17,PPAN,ZFAND2B,AP1S3,CYB561,CARS,ZNF250,RPL36,NFAM1,MDH1,LAMP2,COPZ1,CCNL2,HSP9

,CD44,TM9SF1,SON,TET2,UQCRQ,SNF8,GRK6,HMGA2,CDC37,RPS16,CPSF3L,TSG101,TTC21B,PTPN13,UB

PPA1,DGKE,CAMTA1,CAPG,CRIP1,CLDND1,VIM,LIG1,TIMM17B,XAF1,APEH,HYI,CAPN10,NUTF2,PODXL,C

PPA1,DGKE,CAMTA1,CAPG,CRIP1,CLDND1,VIM,LIG1,TIMM17B,XAF1,APEH,HYI,CAPN10,NUTF2,PODXL,C

TC21B,PTPN13,UBR4,PSMA1,PHF21A,SLC39A11,FUZ,PSMA3,MUC1,METTL17,EPB41L5,ATG12,NLRP1,ZB

SP90AB1,SYNGR2,TMEM134,LRRFIP1,FAM219B,RAB2A,ZNF524,ATP5J,TPM2,SIAH1,GGCX,ZEB2,ASPH,TR

N10,LYPLA2,ATPAF2,C17ORF62,CASC4,STOML2,C19ORF54,ZNF7,C19ORF48,PPAN,HNRNPC,STAU1,SUG

N10,LYPLA2,ATPAF2,C17ORF62,CASC4,STOML2,C19ORF54,ZNF7,C19ORF48,PPAN,HNRNPC,STAU1,SUG

H3YL1,IP6K2,CLASRP,C1ORF198,UGGT2,MICAL1,HS2ST1,UBE2I,SLC39A13,PRMT2,JUP,TBX6,TMEM205,P

H3YL1,IP6K2,CLASRP,C1ORF198,UGGT2,MICAL1,HS2ST1,UBE2I,SLC39A13,PRMT2,JUP,TBX6,TMEM205,P

DHRS4,MLST8,HMBS,MFGE8,AZI2,GOSR2,SGSM3,RPS7,LAMTOR1,TTC39C,EMC3,GPS2,UBA52,AIG1,CAP

7,ATG12,NLRP1,ZBTB44,NIP7,TTC9C,DUSP22,SLC25A10,THEM4,RDH11,DBNDD2,NISCH,CENPK,GGPS1,C
7,ATG12,NLRP1,ZBTB44,NIP7,TTC9C,DUSP22,SLC25A10,THEM4,RDH11,DBNDD2,NISCH,CENPK,GGPS1,C
EM126B,TXNL4A,PODXL,CAMLG,EPB41L4B,UCK2,ATPAF2,LSM12,C17ORF62,PRMT1,WNK1,C19ORF54,Z
EM126B,TXNL4A,PODXL,CAMLG,EPB41L4B,UCK2,ATPAF2,LSM12,C17ORF62,PRMT1,WNK1,C19ORF54,Z
CDC37,CPSF3L,TTC21B,PTPN13,UBR4,PSMA1,PHF21A,SLC39A11,FUZ,PSMA3,MUC1,METTL17,EPB41L5,
CDC37,CPSF3L,TTC21B,PTPN13,UBR4,PSMA1,PHF21A,SLC39A11,FUZ,PSMA3,MUC1,METTL17,EPB41L5,
BX6,TMEM205,C14ORF2,GUK1,CENPN,FBXW9,USE1,HSBP1L1,AGPAT4,ASF1B,CCDC25,EEF1B2,PPP1CC,E
BX6,TMEM205,C14ORF2,GUK1,CENPN,FBXW9,USE1,HSBP1L1,AGPAT4,ASF1B,CCDC25,EEF1B2,PPP1CC,E
STRIP2,NIT2,LYNX1,ZSWIM7,GMPPB,LRRC27,MVD,LMBR1,NAP1L1,PRKRIP1,RANGRF,CHCHD7,HMGA1,P
TG12,PPRC1,ZBTB44,COX4I1,DUSP22,SLC25A10,TMX2,SPOCK2,THEM4,DBNDD2,SDF4,NISCH,TMED4,CE
TG12,PPRC1,ZBTB44,COX4I1,DUSP22,SLC25A10,TMX2,SPOCK2,THEM4,DBNDD2,SDF4,NISCH,TMED4,CE

,THEM4,SDF4,NISCH,TMED4,RPS6,RPL29,RTEL1,COASY,SRSF1,AUP1,YIPF3,FOXRED1,TRNT1,KXD1,ITGB1
,THEM4,SDF4,NISCH,TMED4,RPS6,RPL29,RTEL1,COASY,SRSF1,AUP1,YIPF3,FOXRED1,TRNT1,KXD1,ITGB1
,C14ORF2,ATF7,PCYT2,USE1,HSBP1L1,AGPAT4,IFI27L1,CCDC25,DEF8,EEF1B2,PPP1CC,EDF1,RFX3,PPA1,
SMA1,SLC39A11,FUZ,PSMA3,MUC1,METTL17,ATG12,NLRP1,ZBTB44,NIP7,DUSP22,SLC25A10,TMX2,RDH
V0B,C9ORF64,MED8,RNMT,MRPL48,MRPL27,C1QTNF6,DDX56,SSH3,RPL27,NCK2,ZNF655,TIMM9,MED1
9,GNB5,CBR1,PHC3,MROH1,EIF2B3,MFSD11,MOK,ZNF81,FAM32A,TMEM14B,ZXDC,SMIM4,RPL37A,ELO
9,GNB5,CBR1,PHC3,MROH1,EIF2B3,MFSD11,MOK,ZNF81,FAM32A,TMEM14B,ZXDC,SMIM4,RPL37A,ELO
M205,XBP1,AUNIP,C14ORF2,GUK1,CENPN,ATF7,PCYT2,FBXW9,HSBP1L1,IFI27L1,MAF,CCDC25,DEF8,EDF
M205,XBP1,AUNIP,C14ORF2,GUK1,CENPN,ATF7,PCYT2,FBXW9,HSBP1L1,IFI27L1,MAF,CCDC25,DEF8,EDF
21B,PTPN13,UBR4,PSMA1,PHF21A,SLC39A11,FUZ,PSMA3,MUC1,METTL17,EPB41L5,ATG12,NLRP1,ZBTB
,TYMS,GTF2IRD2,COPS4,DALRD3,BRF1,OARD1,SLC25A23,TPM4,MFSD10,RIC8A,SCAP,SMARCC2,SLC30A
MA3C,SYNGR1,C12ORF76,RANGRF,PABPC1,RRNAD1,DBNL,RBP1,TOMM40,SERPINB6,RABEP2,ATP6V1F,
MA3C,SYNGR1,C12ORF76,RANGRF,PABPC1,RRNAD1,DBNL,RBP1,TOMM40,SERPINB6,RABEP2,ATP6V1F,
CAN,LPCAT3,CD44,TM9SF1,SON,TET2,UQCRQ,SNF8,GRK6,HMGA2,RPS16,CPSF3L,TSG101,OBSL1,TTC21
CAN,LPCAT3,CD44,TM9SF1,SON,TET2,UQCRQ,SNF8,GRK6,HMGA2,RPS16,CPSF3L,TSG101,OBSL1,TTC21
GT2,HS2ST1,PRMT2,EIF3M,MAGT1,JUP,CDC42,TBX6,TMEM205,AUNIP,GUK1,CENPN,CARD14,ATF7,FBX
GT2,HS2ST1,PRMT2,EIF3M,MAGT1,JUP,CDC42,TBX6,TMEM205,AUNIP,GUK1,CENPN,CARD14,ATF7,FBX
,CASC4,EXOSC9,STOML2,WNK1,ZNF7,ARL6IP4,STRADA,AP1S3,HNRNPC,CYB561,ZNF644,CBFA2T2,STAU
,CASC4,EXOSC9,STOML2,WNK1,ZNF7,ARL6IP4,STRADA,AP1S3,HNRNPC,CYB561,ZNF644,CBFA2T2,STAU
AGT1,TBX6,TMEM205,XBP1,AUNIP,C14ORF2,GUK1,CENPN,CARD14,PCYT2,FBXW9,HSBP1L1,AGPAT4,IF
AGT1,TBX6,TMEM205,XBP1,AUNIP,C14ORF2,GUK1,CENPN,CARD14,PCYT2,FBXW9,HSBP1L1,AGPAT4,IF
10,SPOCK2,THEM4,RDH11,SDF4,NISCH,TMED4,CENPK,RPS6,RPL29,CD99L2,COASY,SRSF1,SAP30BP,YIPF3
10,SPOCK2,THEM4,RDH11,SDF4,NISCH,TMED4,CENPK,RPS6,RPL29,CD99L2,COASY,SRSF1,SAP30BP,YIPF3
1CC,EDF1,RFX3,PPA1,DGKE,CAMTA1,CAPG,CRIP1,PSMD4,VIM,LIG1,PSMA7,CDC14B,TIMM17B,TMEM12
1CC,EDF1,RFX3,PPA1,DGKE,CAMTA1,CAPG,CRIP1,PSMD4,VIM,LIG1,PSMA7,CDC14B,TIMM17B,TMEM12
BPMS,NNMT,SNRPN,SPAG7,WDR62,IL1RAP,STYXL1,RFX5,POLD4,PLEKHA5,RANBP1,UBE2L6,VRK3,LZTFL
BPMS,NNMT,SNRPN,SPAG7,WDR62,IL1RAP,STYXL1,RFX5,POLD4,PLEKHA5,RANBP1,UBE2L6,VRK3,LZTFL
IRNPC,CYB561,SHARPIN,STAU1,UPP1,ZNF250,RPL36,MOB1A,LAMP2,INO80E,CCNL2,HSP90AB1,SYNGR2,
4,MAF,ASF1B,CCDC25,PPP1CC,RFX3,DGKE,ENDOV,CAMTA1,CRIP1,PPIL2,TROAP,CLDND1,VIM,LIG1,PSM
B41L4B,TCF4,ATPAF2,WNK1,C19ORF54,ZNF7,PIGL,C19ORF48,STRADA,ZFAND2B,AP1S3,ACAD10,ZNF64
TL17,ECSIT,ATG12,PPRC1,RPS25,NLRP1,ZBTB44,NIP7,TTC9C,COX4I1,DUSP22,SLC25A10,TMX2,SPOCK2,
TL17,ECSIT,ATG12,PPRC1,RPS25,NLRP1,ZBTB44,NIP7,TTC9C,COX4I1,DUSP22,SLC25A10,TMX2,SPOCK2,
.5,ATG12,PPRC1,RPS25,NLRP1,ZBTB44,NIP7,TTC9C,COX4I1,PLCB2,DUSP22,SLC25A10,TMX2,DBNDD2,SD

.5,ATG12,PPRC1,RPS25,NLRP1,ZBTB44,NIP7,TTC9C,COX4I1,PLCB2,DUSP22,SLC25A10,TMX2,DBNDD2,SD
MD4,RPL28,LIG1,CDC14B,TIMM17B,TMEM126B,XAF1,CAPN10,TCF4,UCK2,TMPO,C17ORF62,WNK1,C19O
M3A,PSMA3,MUC1,METTL17,EPB41L5,RPS25,NIP7,TTC9C,COX4I1,SLC25A10,TMX2,THEM4,NISCH,TMED
M3A,PSMA3,MUC1,METTL17,EPB41L5,RPS25,NIP7,TTC9C,COX4I1,SLC25A10,TMX2,THEM4,NISCH,TMED
PINB6,ZNF20,SLC33A1,ZNF687,ANAPC11,NT5C,PDE4DIP,MDH2,NAA10,CARHSP1,PRLR,COX17,HNRNPLL,
PINB6,ZNF20,SLC33A1,ZNF687,ANAPC11,NT5C,PDE4DIP,MDH2,NAA10,CARHSP1,PRLR,COX17,HNRNPLL,
PM6,SMAGP,TSPAN31,APOLD1,ZNF593,GLYCTK,CFLAR,TRIM46,PCNP,RER1,RPL34,EIF2A,NKIRAS2,ACAA1
PM6,SMAGP,TSPAN31,APOLD1,ZNF593,GLYCTK,CFLAR,TRIM46,PCNP,RER1,RPL34,EIF2A,NKIRAS2,ACAA1

C9,C19ORF54,ARL6IP4,CYB561,SHARPIN,STAU1,UPP1,RPL36,MOB1A,RBM3,MDH1,COPZ1,SYNGR2,TMEN
M102A,MCFD2,PFKFB2,RIC8A,NFKBIA,CBR1,PIGT,EIF2B3,MFSD11,LFNG,XYLT2,MT2A,TMEM14B,PCID2,Z
M102A,MCFD2,PFKFB2,RIC8A,NFKBIA,CBR1,PIGT,EIF2B3,MFSD11,LFNG,XYLT2,MT2A,TMEM14B,PCID2,Z
,POLR3H,ABHD17A,STXBP2,EEF1D,NAA10,C11ORF48,MED6,LLGL2,COX17,CAMK2D,SUMO3,LGMN,CD15
LC30A5,THADA,CBR1,PHC3,COMMD4,ENY2,FAM32A,RPAIN,POLR1C,SMIM4,BTN3A2,ZMYM6,DGAT1,M
PCMT1,EBPL,TESC,IGSF8,NUDT5,RBPMS,NNMT,SNRPN,SPAG7,WDR62,IL1RAP,STYXL1,RFX5,POLD4,PLEK
C19ORF54,ZNF7,TXNDC17,PIGL,C19ORF48,STRADA,PPAN,ZFAND2B,AP1S3,ACAD10,CYB561,SHARPIN,ZN

P2,LPCAT3,TM9SF1,MAGED2,SON,TET2,UQCRQ,SNF8,GRK6,HMGA2,CDC37,RPS16,CPSF3L,TSG101,OBSL1
P2,LPCAT3,TM9SF1,MAGED2,SON,TET2,UQCRQ,SNF8,GRK6,HMGA2,CDC37,RPS16,CPSF3L,TSG101,OBSL1
EIF2A,PSPC1,TRAPPC2,SIN3A,BRD8,PLAC8,ST8SIA1,NDUFA11,TP53,AP1G2,BTF3,UBE2D3,AAMDC,DNAJC

56,RPL29,CD99L2,GGPS1,RTEL1,COASY,SRSF1,AUP1,YIPF3,FOXRED1,RABL5,TRNT1,KXD1,ITGB1BP1,MED1
2,LSM12,CASC4,STOML2,C19ORF54,ZNF7,PIGL,PPAN,HNRNPC,CYB561,SHARPIN,CBFA2T2,CARS,MOB1A,
2,LSM12,CASC4,STOML2,C19ORF54,ZNF7,PIGL,PPAN,HNRNPC,CYB561,SHARPIN,CBFA2T2,CARS,MOB1A,
YNGR2,TMEM134,RAB2A,RMDN1,GPBP1,GPALPP1,GGCX,TPM3,TRMT112,PNPO,CHPF,MPDU1,MAPK13,
YNGR2,TMEM134,RAB2A,RMDN1,GPBP1,GPALPP1,GGCX,TPM3,TRMT112,PNPO,CHPF,MPDU1,MAPK13,
ASF1B,CCDC25,EEF1B2,EDF1,GADL1,DGKE,TNFRSF14,ENDOV,CAMTA1,CAPG,CRIP1,PPIL2,TROAP,RPL28,L
ASF1B,CCDC25,EEF1B2,EDF1,GADL1,DGKE,TNFRSF14,ENDOV,CAMTA1,CAPG,CRIP1,PPIL2,TROAP,RPL28,L
K,SPINT2,PAFAH1B2,TYMS,PTK7,GTF2IRD2,BRF1,FAM102A,PFKFB2,RIC8A,NFKBIA,GNB5,COMMD4,CCDC
ARD14,ATF7,FBXW9,USE1,HSBP1L1,ASF1B,DEF8,EEF1B2,PPP1CC,EDF1,RFX3,PPA1,ENDOV,CAMTA1,CRIP
ARD14,ATF7,FBXW9,USE1,HSBP1L1,ASF1B,DEF8,EEF1B2,PPP1CC,EDF1,RFX3,PPA1,ENDOV,CAMTA1,CRIP
ARL6IP4,C19ORF48,ZFAND2B,ACAD10,SHARPIN,HYPK,ZNF250,NFAM1,CDK11A,RBM3,INO80E,MAPKAPK
ARL6IP4,C19ORF48,ZFAND2B,ACAD10,SHARPIN,HYPK,ZNF250,NFAM1,CDK11A,RBM3,INO80E,MAPKAPK
PPM1K,IRF9,ANTXR2,STIP1,LRRC1,PHF19,GRAMD1C,VCAN,LPCAT3,CD44,TM9SF1,MAGED2,SON,UQCRQ,
PPM1K,IRF9,ANTXR2,STIP1,LRRC1,PHF19,GRAMD1C,VCAN,LPCAT3,CD44,TM9SF1,MAGED2,SON,UQCRQ,
F2,PABPC1,NAT9,BCAM,ZNF836,BLVRB,TOMM40,RABEP2,TADA3,RALGPS1,PDCD5,EPS15L1,POLR3H,NO

GPALPP1,SIAH1,ZEB2,STRIP2,ZNF625,RPS20,LYNX1,MPDU1,TWSG1,VPS28,RPS15,ERLEC1,HHLA3,SAT1,L
PSMA7,TMEM126B,TXNL4A,LYPLA2,NUTF2,PODXL,CAMLG,EPB41L4B,TCF4,UCK2,TMPO,ATPAF2,LSM12,
PSMA7,TMEM126B,TXNL4A,LYPLA2,NUTF2,PODXL,CAMLG,EPB41L4B,TCF4,UCK2,TMPO,ATPAF2,LSM12,
AFV,RBMX,SERPINB6,RABEP2,TADA3,WAC,ZNF706,ZNF687,POT1,RAD23A,DDX41,ILF3,AGAP3,HSPH1,C1
.,NAA10,C11ORF48,OAS3,CNST,LLGL2,PRLR,SH3GLB1,CHEK1,COX17,HNRNPLL,SUMO3,HINT1,HNRNPA1,
6G,SH3YL1,IP6K2,CLASRP,JPH2,C1ORF198,UGGT2,MICAL1,HS2ST1,UBE2I,SLC39A13,PRMT2,JUP,CDC42,T
6G,SH3YL1,IP6K2,CLASRP,JPH2,C1ORF198,UGGT2,MICAL1,HS2ST1,UBE2I,SLC39A13,PRMT2,JUP,CDC42,T

ZNF655,DCTN6,OPA3,C12ORF10,SNRNP200,RPS14,RNF4,MIEN1,COX7C,CDKL3,C12ORF66,FDFT1,ZRANB

ZNF655,DCTN6,OPA3,C12ORF10,SNRNP200,RPS14,RNF4,MIEN1,COX7C,CDKL3,C12ORF66,FDFT1,ZRANB

USP22,SLC25A10,TMX2,SPOCK2,THEM4,RDH11,DBNDD2,NISCH,TMED4,CD99L2,GGPS1,RTEL1,COASY,SF

USP22,SLC25A10,TMX2,SPOCK2,THEM4,RDH11,DBNDD2,NISCH,TMED4,CD99L2,GGPS1,RTEL1,COASY,SF

AS3,SEC31A,CNST,SH3GLB1,SSH2,FKBP3,SIRT5,B3GAT3,SPINT2,ASH2L,PLAC9,PPIH,DALRD3,NSUN5,DAZA

C14ORF2,GUK1,CARD14,PCYT2,FBXW9,AGPAT4,IFI27L1,MAF,ASF1B,CCDC25,EEF1B2,RFX3,PPA1,TNFRSF

C14ORF2,GUK1,CARD14,PCYT2,FBXW9,AGPAT4,IFI27L1,MAF,ASF1B,CCDC25,EEF1B2,RFX3,PPA1,TNFRSF

NL4A,CAPN10,PODXL,EPB41L4B,C17ORF62,WNK1,ZNF7,TXNDC17,PPAN,ZFAND2B,AP1S3,ACAD10,STAU

MPO,LSM12,C17ORF62,PRMT1,CASC4,EXOSC9,STOML2,WNK1,ZNF7,ARL6IP4,STRADA,PPAN,HNRNPC,SH

MPO,LSM12,C17ORF62,PRMT1,CASC4,EXOSC9,STOML2,WNK1,ZNF7,ARL6IP4,STRADA,PPAN,HNRNPC,SH

C,PEX2,TNKS,RPL37A,STX10,USP4,C12ORF57,DCTN5,NDUFS3,ARL16,OXLD1,ZNF223,C6ORF132,SSR3,MF

C,PEX2,TNKS,RPL37A,STX10,USP4,C12ORF57,DCTN5,NDUFS3,ARL16,OXLD1,ZNF223,C6ORF132,SSR3,MF

OV,CAMTA1,CRIP1,PSMD4,TROAP,RPL28,LIG1,PSMA7,CDC14B,TIMM17B,TMEM126B,XAF1,RPL21,CAPN

OCK10,DCN,DBNL,BLVRB,RABEP2,TADA3,ATP6V1F,WAC,SLC33A1,RALGPS1,ZNF687,POT1,HSD17B10,NT

1,TWSG1,MAPK13,GMPPB,HSPB1,LRRC27,MVD,LMBR1,MCTS1,NAP1L1,SERPINB1,RANGRF,MTA3,CHCH

7,EPB41L5,ATG12,PPRC1,RPS25,ZBTB44,TTC9C,PLCB2,DUSP22,THEM4,DBNDD2,SDF4,NISCH,TMED4,CE

7,EPB41L5,ATG12,PPRC1,RPS25,ZBTB44,TTC9C,PLCB2,DUSP22,THEM4,DBNDD2,SDF4,NISCH,TMED4,CE

,SON,UQCRQ,GRK6,HMGA2,CDC37,RPS16,TSG101,OBSL1,PTPN13,UBR4,PSMA1,PHF21A,CHCHD1,SLC39

,SON,UQCRQ,GRK6,HMGA2,CDC37,RPS16,TSG101,OBSL1,PTPN13,UBR4,PSMA1,PHF21A,CHCHD1,SLC39

PRC1,RPS25,ZBTB44,TTC9C,COX4I1,PLCB2,DUSP22,SLC25A10,TMX2,SPOCK2,THEM4,SDF4,NISCH,RPS6,C

PRC1,RPS25,ZBTB44,TTC9C,COX4I1,PLCB2,DUSP22,SLC25A10,TMX2,SPOCK2,THEM4,SDF4,NISCH,RPS6,C

T2,COX20,ZXDC,COBL,TNKS,SMIM4,ERGIC1,NMNAT3,KPNA5,MPHOSPH10,MYL9,SLC35E1,OGFOD2,TSEN

NT1,KXD1,ITGB1BP1,MED15,SH3YL1,CLASRP,MICAL1,HS2ST1,UBE2I,KCNMA1,PRMT2,MAGT1,CDC42,TB

C5,EMC4,IL32,LMBR1,PRKRIP1,SYNGR1,C12ORF76,RANGRF,MEIS2,RRNAD1,NAT9,H2AFV,ZNF20,SLC33A1

PHF21A,CHCHD1,FUZ,RGS3,MUC1,METTL17,ECSIT,ATG12,PPRC1,RPS25,NLRP1,ZBTB44,NIP7,TTC9C,COX

PHF21A,CHCHD1,FUZ,RGS3,MUC1,METTL17,ECSIT,ATG12,PPRC1,RPS25,NLRP1,ZBTB44,NIP7,TTC9C,COX

EM134,TPM2,GGCX,CHPF,GMPPB,MVD,SAT1,MCTS1,PRKRIP1,C12ORF76,RANGRF,MTA3,RRNAD1,H2AF

EM134,TPM2,GGCX,CHPF,GMPPB,MVD,SAT1,MCTS1,PRKRIP1,C12ORF76,RANGRF,MTA3,RRNAD1,H2AF

A1,DGKE,ENDOV,CAMTA1,CAPG,CRIP1,PPIL2,RPL28,CLDND1,VIM,LIG1,PSMA7,TMEM126B,XAF1,APEH,F

A1,DGKE,ENDOV,CAMTA1,CAPG,CRIP1,PPIL2,RPL28,CLDND1,VIM,LIG1,PSMA7,TMEM126B,XAF1,APEH,F

L,CARS,CCNL2,DUT,CEP70,TMEM134,LRRFIP1,FAM219B,ATP5J,GPBP1,TSFM,TRMT112,MAPK13,RPS15,D

L,CARS,CCNL2,DUT,CEP70,TMEM134,LRRFIP1,FAM219B,ATP5J,GPBP1,TSFM,TRMT112,MAPK13,RPS15,D

XD1,ITGB1BP1,MED15,PARD6G,SH3YL1,JPH2,C1ORF198,UGGT2,MICAL1,HS2ST1,UBE2I,KCNMA1,SLC39

RMT1,CASC4,STOML2,WNK1,C19ORF48,PPAN,HNRNPC,ZNF644,STAU1,ZNF250,RBM3,MDH1,LAMP2,ZN

RMT1,CASC4,STOML2,WNK1,C19ORF48,PPAN,HNRNPC,ZNF644,STAU1,ZNF250,RBM3,MDH1,LAMP2,ZN

TRNT1,KXD1,ITGB1BP1,MED15,SH3YL1,CLASRP,UGGT2,UBE2I,KCNMA1,SLC39A13,PRMT2,EIF3M,MAGT1

TRNT1,KXD1,ITGB1BP1,MED15,SH3YL1,CLASRP,UGGT2,UBE2I,KCNMA1,SLC39A13,PRMT2,EIF3M,MAGT1

),CYB561,SHARPIN,STAU1,SLC9B2,SUGT1,UPP1,CARS,CDK11A,RBM3,MDH1,ZNF197,CCNL2,SYNGR2,CEP

),CYB561,SHARPIN,STAU1,SLC9B2,SUGT1,UPP1,CARS,CDK11A,RBM3,MDH1,ZNF197,CCNL2,SYNGR2,CEP

V,CAMTA1,CRIP1,PPIL2,PSMD4,RPL28,CLDND1,VIM,CDC14B,APEH,HYI,CAPN10,EPB41L4B,PPOX,TMPO,

V,CAMTA1,CRIP1,PPIL2,PSMD4,RPL28,CLDND1,VIM,CDC14B,APEH,HYI,CAPN10,EPB41L4B,PPOX,TMPO,

NK1,TXNDC17,ARL6IP4,PIGL,STRADA,ZFAND2B,AP1S3,ACAD10,CYB561,SHARPIN,ZNF644,SUGT1,UPP1,
NK1,TXNDC17,ARL6IP4,PIGL,STRADA,ZFAND2B,AP1S3,ACAD10,CYB561,SHARPIN,ZNF644,SUGT1,UPP1,
USE1,HSBP1L1,IFI27L1,MAF,CCDC25,RFX3,ENDOV,CAPG,CRIP1,PPIL2,RPL28,CLDND1,CDC14B,TIMM17B,
USE1,HSBP1L1,IFI27L1,MAF,CCDC25,RFX3,ENDOV,CAPG,CRIP1,PPIL2,RPL28,CLDND1,CDC14B,TIMM17B,
M,LIG1,CDC14B,TIMM17B,TMEM126B,TXNL4A,LYPLA2,NUTF2,PODXL,CAMLG,EPB41L4B,TCF4,UCK2,TM
,FBXW9,USE1,AGPAT4,CCDC25,EEF1B2,PPP1CC,EDF1,RFX3,PPA1,DGKE,ENDOV,CAMTA1,CRIP1,CLDND1,
,THEM4,SDF4,NISCH,TMED4,CENPK,RPS6,RPL29,CD99L2,GGPS1,COASY,SRSF1,AUP1,SAP30BP,YIPF3,FOX
,THEM4,SDF4,NISCH,TMED4,CENPK,RPS6,RPL29,CD99L2,GGPS1,COASY,SRSF1,AUP1,SAP30BP,YIPF3,FOX
219B,RAB2A,RMDN1,GPBP1,GPALPP1,TPM3,TRMT112,PNPO,CHPF,MPDU1,ZSWIM7,UBXN1,ERLEC1,GM
219B,RAB2A,RMDN1,GPBP1,GPALPP1,TPM3,TRMT112,PNPO,CHPF,MPDU1,ZSWIM7,UBXN1,ERLEC1,GM

F48,STRADA,AP1S3,HNRNPC,STAU1,SUGT1,ZNF250,RPL36,NFAM1,INO80E,HSP90AB1,SYNGR2,LRRFIP1,
1,RAD23A,ABHD17A,ILF3,EEF1D,NAA10,CNST,LLGL2,EGFR,CHEK1,COX17,CAMK2D,SUMO3,CD151,FKBP3
1,COX17,HNRNPLL,SUMO3,HNRNPA1,HIGD1A,RPL15,INTS4,ZNF506,MTHFS,SMARCA4,COA3,SNRPF,TYM
,FBXW9,USE1,HSBP1L1,IFI27L1,MAF,CCDC25,EEF1B2,EDF1,RFX3,PPA1,DGKE,TNFRSF14,ENDOV,CAMTA
SON,TET2,SNF8,HMGA2,CDC37,RPS16,CPSF3L,TTC21B,PTPN13,UBR4,PSMA1,SLC39A11,FUZ,PSMA3,MU
SON,TET2,SNF8,HMGA2,CDC37,RPS16,CPSF3L,TTC21B,PTPN13,UBR4,PSMA1,SLC39A11,FUZ,PSMA3,MU

1,PHF21A,CHCHD1,SLC39A11,FAM3A,MUC1,METTL17,ECSIT,EPB41L5,PPRC1,RPS25,NLRP1,ZBTB44,NI
1,PHF21A,CHCHD1,SLC39A11,FAM3A,MUC1,METTL17,ECSIT,EPB41L5,PPRC1,RPS25,NLRP1,ZBTB44,NI
RF2,GUK1,CENPN,CARD14,ATF7,PCYT2,USE1,MAF,EEF1B2,PPP1CC,EDF1,RFX3,PPA1,DGKE,CAMTA1,CA
C,RALGPS1,ZNF687,PDCD5,GTF2IRD1,NXN,EPS15L1,POLR3H,RAD23A,ABHD17A,ILF3,PKP4,AGAP3,EEF1
NPM,NIPA1,POLH,EXOSC8,ACTB,C12ORF73,MRPL43,SETD4,KIAA0513,ZSCAN30,EHMT1,CLIP4,CDK10,M
NPM,NIPA1,POLH,EXOSC8,ACTB,C12ORF73,MRPL43,SETD4,KIAA0513,ZSCAN30,EHMT1,CLIP4,CDK10,M
87,PDCD5,GTF2IRD1,NXN,EPS15L1,POLR3H,NOMO2,ABHD17A,ILF3,AGAP3,STXBP2,EEF1D,NAA10,LLGL2

VNK1,ZNF7,PIGL,C19ORF48,STRADA,PPAN,AP1S3,HNRNPC,CYB561,SHARPIN,SLC9B2,HYPK,CDK11A,MD
,APEH,HYI,TXNL4A,CAPN10,LYPLA2,NUTF2,PODXL,EPB41L4B,TCF4,UCK2,PPOX,TMPO,ATPAF2,C17ORF6
,APEH,HYI,TXNL4A,CAPN10,LYPLA2,NUTF2,PODXL,EPB41L4B,TCF4,UCK2,PPOX,TMPO,ATPAF2,C17ORF6
T3,CD44,MAGED2,TET2,UQCRQ,SNF8,HMGA2,CDC37,CPSF3L,OBSL1,PTPN13,UBR4,PSMA1,YLPM1,PHF2
T3,CD44,MAGED2,TET2,UQCRQ,SNF8,HMGA2,CDC37,CPSF3L,OBSL1,PTPN13,UBR4,PSMA1,YLPM1,PHF2
A3,CKLF,AKT1S1,INO80C,GTF3A,ARRB2,DIP2A,MLST8,HMGN1,CDK2AP2,B3GNTL1,MYL12A,SLK,MAZ,C2
A3,CKLF,AKT1S1,INO80C,GTF3A,ARRB2,DIP2A,MLST8,HMGN1,CDK2AP2,B3GNTL1,MYL12A,SLK,MAZ,C2

2,NUTF2,PODXL,CAMLG,EPB41L4B,UCK2,PPOX,TMPO,ATPAF2,LSM12,C17ORF62,PRMT1,STOML2,ZNF7,
2,NUTF2,PODXL,CAMLG,EPB41L4B,UCK2,PPOX,TMPO,ATPAF2,LSM12,C17ORF62,PRMT1,STOML2,ZNF7,
34,RAB2A,ZNF524,ELP6,RMDN1,TPM2,GPALPP1,SIAH1,ASPH,TRMT112,PNPO,NIT2,TWSG1,ZSWIM7,RP
34,RAB2A,ZNF524,ELP6,RMDN1,TPM2,GPALPP1,SIAH1,ASPH,TRMT112,PNPO,NIT2,TWSG1,ZSWIM7,RP
HSP1,LLGL2,SH3GLB1,CAMK2D,HNRNPLL,HINT1,HIGD1A,POLR2C,EGLN2,KCTD7,B3GAT3,ASH2L,MTHFS,
HSP1,LLGL2,SH3GLB1,CAMK2D,HNRNPLL,HINT1,HIGD1A,POLR2C,EGLN2,KCTD7,B3GAT3,ASH2L,MTHFS,

42,TBX6,TMEM205,AUNIP,CENPN,CARD14,FBXW9,USE1,IFI27L1,ASF1B,CCDC25,EEF1B2,PDLIM5,RFX3,PI
42,TBX6,TMEM205,AUNIP,CENPN,CARD14,FBXW9,USE1,IFI27L1,ASF1B,CCDC25,EEF1B2,PDLIM5,RFX3,PI
L,YIPF3,FOXRED1,TRNT1,KXD1,ITGB1BP1,MED15,SH3YL1,IP6K2,CLASRP,JPH2,HS2ST1,UBE2I,SLC39A13,YI
L,YIPF3,FOXRED1,TRNT1,KXD1,ITGB1BP1,MED15,SH3YL1,IP6K2,CLASRP,JPH2,HS2ST1,UBE2I,SLC39A13,YI
OLD1,RALY,ZNF593,GLYCTK,PCNP,VDAC1,RER1,DOK4,EIF2A,OLFM1,ZNF511,IRF3,PIGO,D2HGDH,KDELR3,
3,TSPAN31,MTIF2,MARS,GLYCTK,TCEAL3,ACAA1,ZNF283,TRAPPC2,DYNLT3,GTF2H1,TAF1D,PIGO,BFAR,RE
219B,RAB2A,TSFM,GGCX,ASPH,PNPO,CHPF,ZNF625,NIT2,ZSWIM7,DCTD,UBXN1,HSPB1,VWA2,MVD,SAT
LPP1,SIAH1,ZEB2,ASPH,CHPF,STRIP2,RPS20,GABPB1,LYNX1,MPDU1,ZSWIM7,RPS15,ERLEC1,GMPPB,SAT1
LPP1,SIAH1,ZEB2,ASPH,CHPF,STRIP2,RPS20,GABPB1,LYNX1,MPDU1,ZSWIM7,RPS15,ERLEC1,GMPPB,SAT1
OCK2,THEM4,RDH11,NISCH,TMED4,CENPK,RPS6,RPL29,CD99L2,GGPS1,RTEL1,COASY,SRSF1,TRNT1,KXD1
OCK2,THEM4,RDH11,NISCH,TMED4,CENPK,RPS6,RPL29,CD99L2,GGPS1,RTEL1,COASY,SRSF1,TRNT1,KXD1
D1,H2AFV,SERPINB6,WAC,ZNF706,ANAPC11,DDX41,ILF3,PKP4,AGAP3,EEF1D,HSPH1,C11ORF48,OAS3,CA
D4,FKBP3,NDUFV2,EGLN2,KCTD7,B3GAT3,SPINT2,TYMS,PTK7,GTF2IRD2,BRF1,FAM102A,TRPM4,PFKFB2,
D4,FKBP3,NDUFV2,EGLN2,KCTD7,B3GAT3,SPINT2,TYMS,PTK7,GTF2IRD2,BRF1,FAM102A,TRPM4,PFKFB2,
R2,CEP70,LRRFIP1,RAB2A,ZNF524,ATP5J,TPM2,GPALPP1,SIAH1,GGCX,TRMT112,ZNF625,GABPB1,LYNX1,
ATP5J,ELP6,TPM2,SIAH1,ASPH,PNPO,RPTOR,ZNF625,NIT2,LYNX1,MPDU1,RPS15,UBXN1,ZNF814,GMPPB
ATP5J,ELP6,TPM2,SIAH1,ASPH,PNPO,RPTOR,ZNF625,NIT2,LYNX1,MPDU1,RPS15,UBXN1,ZNF814,GMPPB
,PDCD5,GTF2IRD1,NXN,EPS15L1,POLR3H,RAD23A,ABHD17A,NT5C,AGAP3,STXBP2,EEF1D,NAA10,RRP8,LI

62,PRMT1,CASC4,ZNF7,ARL6IP4,PIGL,ZFAND2B,AP1S3,HNRNPC,CYB561,SHARPIN,UPP1,RPL36,MOB1A,N
62,PRMT1,CASC4,ZNF7,ARL6IP4,PIGL,ZFAND2B,AP1S3,HNRNPC,CYB561,SHARPIN,UPP1,RPL36,MOB1A,N
56,RPL29,CD99L2,RTEL1,COASY,SRSF1,AUP1,FOXRED1,ITGB1BP1,MED15,SH3YL1,JPH2,MICAL1,UBE2I,KC
56,RPL29,CD99L2,RTEL1,COASY,SRSF1,AUP1,FOXRED1,ITGB1BP1,MED15,SH3YL1,JPH2,MICAL1,UBE2I,KC
C25,EEF1B2,PPP1CC,EDF1,RFX3,PPA1,ENDOV,VAMP8,CAMTA1,CRIP1,PPIL2,RPL28,CLDND1,PSMA7,CDC1
C25,EEF1B2,PPP1CC,EDF1,RFX3,PPA1,ENDOV,VAMP8,CAMTA1,CRIP1,PPIL2,RPL28,CLDND1,PSMA7,CDC1
RPS6,RPL29,COASY,SRSF1,AUP1,SAP30BP,YIPF3,FOXRED1,RABL5,TRNT1,KXD1,ITGB1BP1,MED15,PARD6
RPS6,RPL29,COASY,SRSF1,AUP1,SAP30BP,YIPF3,FOXRED1,RABL5,TRNT1,KXD1,ITGB1BP1,MED15,PARD6
L,UPP1,ZNF250,CDK11A,RBM3,MDH1,INO80E,CCNL2,CEP70,RAB2A,ZNF524,ATP5J,TPM2,GPBP1,GPALPP
L,UPP1,ZNF250,CDK11A,RBM3,MDH1,INO80E,CCNL2,CEP70,RAB2A,ZNF524,ATP5J,TPM2,GPBP1,GPALPP
,MUC1,METTL17,ATG12,RPS25,NLRP1,ZBTB44,NIP7,TTC9C,COX4I1,DUSP22,SLC25A10,TMX2,THEM4,RD
,MUC1,METTL17,ATG12,RPS25,NLRP1,ZBTB44,NIP7,TTC9C,COX4I1,DUSP22,SLC25A10,TMX2,THEM4,RD
10,POLR2E,RNPS1,IAH1,PNRC1,ARPC1A,UBA52,VAPA,DHDDS,CEP63,SDHA,ZNF483,ABLIM2,ACHE,PEX16,
10,POLR2E,RNPS1,IAH1,PNRC1,ARPC1A,UBA52,VAPA,DHDDS,CEP63,SDHA,ZNF483,ABLIM2,ACHE,PEX16,
T112,ZNF625,GABPB1,NIT2,LYNX1,ZSWIM7,ZNF814,ERLEC1,HSPB1,MVD,ERCC5,LMBR1,SERPINB1,LSR,CH
MAP,TPT1,POLR2M,CDK4,SLC52A2,ERRFI1,ANAPC13,PPP2R5C,RALY,CFLAR,CLASP2,CNTRL,EIF2A,ZNF354
NGRF,LSR,CHCHD7,MEIS2,PABPC1,RRNAD1,NAT9,ZNF836,H2AFV,SERPINB6,TADA3,WAC,SLC33A1,ZNF6
SY,SRSF1,AUP1,SAP30BP,FOXRED1,TRNT1,KXD1,ITGB1BP1,MED15,PARD6G,SH3YL1,JPH2,C1ORF198,UG
SY,SRSF1,AUP1,SAP30BP,FOXRED1,TRNT1,KXD1,ITGB1BP1,MED15,PARD6G,SH3YL1,JPH2,C1ORF198,UG
ATP5J,TPM2,SIAH1,ZEB2,ASPH,TRMT112,CHPF,STRIP2,LYNX1,TWSG1,MAPK13,DCTD,GMPPB,LRRC27,LM
F48,ZFAND2B,HYPK,ZNF250,NFAM1,CDK11A,RBM3,INO80E,MAPKAPK5,DGUOK,HSP90AB1,SYNGR2,CEP
F48,ZFAND2B,HYPK,ZNF250,NFAM1,CDK11A,RBM3,INO80E,MAPKAPK5,DGUOK,HSP90AB1,SYNGR2,CEP
39A13,YIF1A,JUP,CDC42,TMEM205,XBP1,C14ORF2,GUK1,CARD14,PCYT2,FBXW9,HSBP1L1,IFI27L1,MAF,A
39A13,YIF1A,JUP,CDC42,TMEM205,XBP1,C14ORF2,GUK1,CARD14,PCYT2,FBXW9,HSBP1L1,IFI27L1,MAF,A
1,FAM102A,MFSD10,PFKFB2,RIC8A,NFKBIA,GNB5,MROH1,MFSD11,LFNG,XYLT2,PCID2,ZXDC,COBL,TNKS

DXL,EPB41L4B,ATPAF2,PRMT1,C19ORF54,ARL6IP4,C19ORF48,CYB561,CBFA2T2,SLC9B2,SUGT1,UPP1,ZN

9C,COX4I1,PLCB2,DUSP22,SLC25A10,RDH11,NISCH,TMED4,GGPS1,RTEL1,COASY,SRSF1,AUP1,SAP30BP,
9C,COX4I1,PLCB2,DUSP22,SLC25A10,RDH11,NISCH,TMED4,GGPS1,RTEL1,COASY,SRSF1,AUP1,SAP30BP,
EF1D,NAA10,RRP8,LLGL2,COX17,ZFAND4,FKBP3,NDUFV2,EGLN2,KCTD7,B3GAT3,RFXANK,SPINT2,REEP5,

F2,C17ORF62,PRMT1,STOML2,ZNF7,TXNDC17,ARL6IP4,C19ORF48,ZFAND2B,AP1S3,CYB561,SHARPIN,ZN
TNKS,MDP1,PIGH,ELOF1,KPNA5,MPHOSPH10,MYL9,METTL5,C6ORF132,SLC35E1,FAM126A,RC3H2,OGFC
TNKS,MDP1,PIGH,ELOF1,KPNA5,MPHOSPH10,MYL9,METTL5,C6ORF132,SLC35E1,FAM126A,RC3H2,OGFC
ZF,GLI4,NKIRAS2,ZNF511,GTF2H1,MYL6,BRD8,NAA38,RPS3A,MOSPD3,IFITM2,RHEB,MAGOHB,IDH3G,CF
LN2,SIRT5,KCTD7,RFXANK,ZNF506,SPINT2,SMARCA4,PTK7,TMEM39B,DALRD3,BRF1,ZFPL1,OARD1,SLC2
LN2,SIRT5,KCTD7,RFXANK,ZNF506,SPINT2,SMARCA4,PTK7,TMEM39B,DALRD3,BRF1,ZFPL1,OARD1,SLC2
MDN1,GPBP1,GPALPP1,SIAH1,GGCX,ZEB2,ASPH,TPM3,TRMT112,PNPO,STRIP2,ZNF625,RPS20,GABPB1,
MDN1,GPBP1,GPALPP1,SIAH1,GGCX,ZEB2,ASPH,TPM3,TRMT112,PNPO,STRIP2,ZNF625,RPS20,GABPB1,
,CD44,TM9SF1,MAGED2,TET2,SNF8,GRK6,CDC37,RPS16,TSG101,OBSL1,PTPN13,UBR4,YLPM1,PHF21A,SL
,CD44,TM9SF1,MAGED2,TET2,SNF8,GRK6,CDC37,RPS16,TSG101,OBSL1,PTPN13,UBR4,YLPM1,PHF21A,SL
2,CLASRP,SLC39A13,YIF1A,EIF3M,CDC42,TBX6,TMEM205,XBP1,PTK2,C14ORF2,GUK1,CENPN,CARD14,PC
FOD1,TSEN34,MRPL55,UBE2D4,ECE1,TCF25,TJP1,NXF1,SLC39A6,RALY,RBM42,FAM69B,RER1,GLI4,NUP6
LPP1,TSFM,ASPH,RPTOR,CHPF,STRIP2,ZNF625,RPS15,UBXN1,ERLEC1,LRRC27,MVD,ERCC5,MCTS1,DDOS
HF6,PARK7,GNB5,CBR1,COMMD4,CCDC40,MROH1,MFSD11,LFNG,SNX5,ENY2,ZSCAN20,TMEM147,TMEN
AKT1S1,AIFM2,HMOX2,SPHK2,HMGN1,NDUFB2,CDK2AP2,NDUFA10,AZI2,SPG21,KAT8,U2AF1L4,DDX10,
2,CDC37,CPSF3L,TSG101,OBSL1,PTPN13,UBR4,PHF21A,CHCHD1,SLC39A11,RGS3,FAM3A,MUC1,METTL1
2,CDC37,CPSF3L,TSG101,OBSL1,PTPN13,UBR4,PHF21A,CHCHD1,SLC39A11,RGS3,FAM3A,MUC1,METTL1
,RFX3,PPA1,DGKE,ENDOV,VAMP8,CRIP1,PPIL2,PSMD4,RPL28,LIG1,PSMA7,RPL21,TXNL4A,CAPN10,UCK2
,HHLA3,MVD,ERCC5,EMC4,IL32,LMBR1,MCTS1,DDOST,SYNGR1,RANGRF,PABPC1,RRNAD1,NAT9,H2AFV,
,HHLA3,MVD,ERCC5,EMC4,IL32,LMBR1,MCTS1,DDOST,SYNGR1,RANGRF,PABPC1,RRNAD1,NAT9,H2AFV,
NF7,ARL6IP4,PIGL,AP1S3,CYB561,SHARPIN,STAU1,SLC9B2,SUGT1,UPP1,CARS,NFAM1,CDK11A,RBM3,MD
NF7,ARL6IP4,PIGL,AP1S3,CYB561,SHARPIN,STAU1,SLC9B2,SUGT1,UPP1,CARS,NFAM1,CDK11A,RBM3,MD
15L1,ANAPC11,RAD23A,B2M,ABHD17A,USP45,ILF3,ZCCHC4,STXBP2,CHCHD5,MED6,CNST,COX5A,HNRN
15L1,ANAPC11,RAD23A,B2M,ABHD17A,USP45,ILF3,ZCCHC4,STXBP2,CHCHD5,MED6,CNST,COX5A,HNRN
ANGRF,CHCHD7,PABPC1,RRNAD1,RAD1,BLVRB,PPP3CB,RABEP2,SLC33A1,ZNF706,ZNF687,NOMO2,GGT5
2,EGLN2,KCTD7,B3GAT3,RFXANK,SPINT2,REEP5,PAFAH1B2,TYMS,PTK7,BRF1,SLC25A23,TPM4,FAM102
ANAPC11,NOMO2,RAD23A,HSD17B10,ABHD17A,DDX41,SLC16A1,AGAP3,STXBP2,EEF1D,C11ORF48,LLGL
ANAPC11,NOMO2,RAD23A,HSD17B10,ABHD17A,DDX41,SLC16A1,AGAP3,STXBP2,EEF1D,C11ORF48,LLGL
DA3,RALGPS1,ZNF706,ZNF687,PDCD5,GTF2IRD1,NXN,KLK10,EPS15L1,ANAPC11,ABHD17A,USP45,ILF3,E
IAH1,TRMT112,PNPO,RPTOR,RPS15,ERLEC1,MVD,EMC4,IL32,LMBR1,SERPINB1,CHCHD7,HMGA1,PSME
IAH1,TRMT112,PNPO,RPTOR,RPS15,ERLEC1,MVD,EMC4,IL32,LMBR1,SERPINB1,CHCHD7,HMGA1,PSME
C25A10,TMX2,SPOCK2,DBNDD2,NISCH,TMED4,CENPK,RPS6,RPL29,RTEL1,COASY,SRSF1,AUP1,SAP30BP,
C25A10,TMX2,SPOCK2,DBNDD2,NISCH,TMED4,CENPK,RPS6,RPL29,RTEL1,COASY,SRSF1,AUP1,SAP30BP,
HD1,RGS3,MUC1,METTL17,ECSIT,ATG12,PPRC1,RPS25,NLRP1,ZBTB44,NIP7,TTC9C,COX4I1,SLC25A10,SP
HD1,RGS3,MUC1,METTL17,ECSIT,ATG12,PPRC1,RPS25,NLRP1,ZBTB44,NIP7,TTC9C,COX4I1,SLC25A10,SP
,MPHOSPH9,SNAPC3,DYNC2LI1,SHPK,LSM14A,LTA4H,COX7C,LIN37,PHF12,ATP13A2,TXNDC12,FANCI,PR
,MPHOSPH9,SNAPC3,DYNC2LI1,SHPK,LSM14A,LTA4H,COX7C,LIN37,PHF12,ATP13A2,TXNDC12,FANCI,PR
,PSME2,PHTF2,RRNAD1,NAT9,RAD1,RBMX,BRIP1,SERPINB6,TADA3,ZNF20,ATP6V1F,WAC,PDCD5,KLK10,
35E1,DGAT1,TSEN34,MRPL55,ARIH2,RPL8,UBE2D4,ECE1,OCIAD1,TCF25,RBBP6,ZNF248,CHTOP,TPT1,UB
M9SF1,MAGED2,SON,UQCRQ,GRK6,HMGA2,CDC37,RPS16,CPSF3L,TSG101,OBSL1,TTC21B,PTPN13,UBR4
M9SF1,MAGED2,SON,UQCRQ,GRK6,HMGA2,CDC37,RPS16,CPSF3L,TSG101,OBSL1,TTC21B,PTPN13,UBR4

N,FKBP3,REEP5,FTH1,SMARCA4,PAFAH1B2,COA3,TYMS,TMEM39B,DALRD3,NSUN5,BRF1,OARD1,SLC25
N,FKBP3,REEP5,FTH1,SMARCA4,PAFAH1B2,COA3,TYMS,TMEM39B,DALRD3,NSUN5,BRF1,OARD1,SLC25

L1,MEIS2,HMGA1,PHTF2,DBNL,RBP1,H2AFV,RBMX,SERPINB6,RABEP2,TADA3,ATP6V1F,WAC,SLC33A1,Z
0513,NARF,EHMT1,CNTLN,MPHOSPH9,DPY30,MBD1,NDUFS7,DAD1,C15ORF40,TIMM9,UQCRB,DCTN6,
0513,NARF,EHMT1,CNTLN,MPHOSPH9,DPY30,MBD1,NDUFS7,DAD1,C15ORF40,TIMM9,UQCRB,DCTN6,
4,RPL29,RTEL1,SRSF1,SAP30BP,YIPF3,FOXRED1,RABL5,TRNT1,KXD1,ITGB1BP1,MED15,PARD6G,SH3YL1,I
4,RPL29,RTEL1,SRSF1,SAP30BP,YIPF3,FOXRED1,RABL5,TRNT1,KXD1,ITGB1BP1,MED15,PARD6G,SH3YL1,I
K2AP2,MYL12A,SLK,MAZ,C2ORF15,TBCA,DDX10,POLR2E,BCL7B,THOP1,ARPC1A,VAPA,MLH1,DHDDS,CEP
K2AP2,MYL12A,SLK,MAZ,C2ORF15,TBCA,DDX10,POLR2E,BCL7B,THOP1,ARPC1A,VAPA,MLH1,DHDDS,CEP
OMM40,SERPINB6,TADA3,SLC33A1,ZNF687,ANAPC11,PDE4DIP,MDH2,CHCHD5,NAA10,C11ORF48,OAS3,
OMM40,SERPINB6,TADA3,SLC33A1,ZNF687,ANAPC11,PDE4DIP,MDH2,CHCHD5,NAA10,C11ORF48,OAS3,

NB1,PRKRIP1,DDOST,SYNGR1,MTA3,DOCK10,PABPC1,BLVRB,SERPINB6,TADA3,ZNF706,ZNF687,PDCD5,
NB1,PRKRIP1,DDOST,SYNGR1,MTA3,DOCK10,PABPC1,BLVRB,SERPINB6,TADA3,ZNF706,ZNF687,PDCD5,
K,HSP90AB1,CEP70,TMEM134,FAM219B,RAB2A,RMDN1,GPBP1,GPALPP1,TRMT112,PNPO,CHPF,MAPK1
K,HSP90AB1,CEP70,TMEM134,FAM219B,RAB2A,RMDN1,GPBP1,GPALPP1,TRMT112,PNPO,CHPF,MAPK1
OK4,PSPC1,PAM16,SKA2,NEDD4L,FCHO2,IRF3,TRAPPC2,SIN3A,THOC3,TUBB3,MOSPD3,C1ORF63,RHEB,
OK4,PSPC1,PAM16,SKA2,NEDD4L,FCHO2,IRF3,TRAPPC2,SIN3A,THOC3,TUBB3,MOSPD3,C1ORF63,RHEB,
21,CAPN10,LYPLA2,NUTF2,PODXL,CAMLG,EPB41L4B,UCK2,PRMT1,CASC4,STOML2,WNK1,C19ORF54,ZN
39A11,FUZ,FAM3A,PSMA3,ECSIT,EPB41L5,ATG12,RPS25,NLRP1,ZBTB44,NIP7,TTC9C,COX4I1,DUSP22,SL
39A11,FUZ,FAM3A,PSMA3,ECSIT,EPB41L5,ATG12,RPS25,NLRP1,ZBTB44,NIP7,TTC9C,COX4I1,DUSP22,SL
NISCH,CENPK,RPL29,GGPS1,RTEL1,COASY,SRSF1,AUP1,SAP30BP,FOXRED1,TRNT1,KXD1,ITGB1BP1,MED1

PPP1CC,RFX3,PPA1,DGKE,TNFRSF14,CAMTA1,CRIP1,PPIL2,TROAP,VIM,LIG1,CDC14B,TIMM17B,XAF1,APE

17ORF62,PRMT1,CASC4,WNK1,ZNF7,ZFAND2B,AP1S3,CYB561,SHARPIN,ZNF644,STAU1,SLC9B2,HYPK,UP
17ORF62,PRMT1,CASC4,WNK1,ZNF7,ZFAND2B,AP1S3,CYB561,SHARPIN,ZNF644,STAU1,SLC9B2,HYPK,UP

13,RPS15,DCTD,GMPPB,LRRC27,LMBR1,NAP1L1,SERPINB1,SYNGR1,RANGRF,CHCHD7,MEIS2,HMGA1,PA
13,RPS15,DCTD,GMPPB,LRRC27,LMBR1,NAP1L1,SERPINB1,SYNGR1,RANGRF,CHCHD7,MEIS2,HMGA1,PA

F197,COPZ1,INO80E,CEP70,LRRFIP1,ZNF524,ATP5J,TPM2,GPALPP1,TSFM,TPM3,RPTOR,STRIP2,ZNF625,
,SLC37A4,SNAPC2,DNAJA3,ZCCHC9,SNRPD1,NSFL1C,TUSC3,ACTR10,TRAK1,CDK2AP2,B3GNTL1,SGSM3,Z
,SLC37A4,SNAPC2,DNAJA3,ZCCHC9,SNRPD1,NSFL1C,TUSC3,ACTR10,TRAK1,CDK2AP2,B3GNTL1,SGSM3,Z

EP70,TMEM134,LRRFIP1,FAM219B,ELP6,TPM2,GPALPP1,TSFM,RPTOR,CHPF,ZNF625,NIT2,MPDU1,TWS
PEH,HYI,TXNL4A,CAPN10,LYPLA2,PODXL,TCF4,TMPO,ATPAF2,C17ORF62,PRMT1,CASC4,ZNF7,TXNDC17,
PEH,HYI,TXNL4A,CAPN10,LYPLA2,PODXL,TCF4,TMPO,ATPAF2,C17ORF62,PRMT1,CASC4,ZNF7,TXNDC17,
54,ZNF7,TXNDC17,STRADA,ZFAND2B,AP1S3,HNRNPC,CYB561,STAU1,ZNF250,RPL36,HSP90AB1,DUT,SYN
31,LYNX1,MPDU1,RPS15,ERLEC1,HHLA3,IL32,NAP1L1,PRKRIP1,MTA3,MEIS2,RRNAD1,NAT9,ZNF836,RBP1
31,LYNX1,MPDU1,RPS15,ERLEC1,HHLA3,IL32,NAP1L1,PRKRIP1,MTA3,MEIS2,RRNAD1,NAT9,ZNF836,RBP1

M39B,BRF1,ZFPL1,OARD1,SLC25A23,FAM102A,GMPR2,MCFD2,PFKFB2,RIC8A,NFKBIA,TRPV3,PIGT,MROI
M39B,BRF1,ZFPL1,OARD1,SLC25A23,FAM102A,GMPR2,MCFD2,PFKFB2,RIC8A,NFKBIA,TRPV3,PIGT,MROI
NF524,ATP5J,TPM2,GPBP1,GPALPP1,SIAH1,CHPF,STRIP2,RPS20,GABPB1,LYNX1,MPDU1,TWSG1,ZSWIM7
NF524,ATP5J,TPM2,GPBP1,GPALPP1,SIAH1,CHPF,STRIP2,RPS20,GABPB1,LYNX1,MPDU1,TWSG1,ZSWIM7
2,TSEN34,ARIH2,KCNC4,ZNF248,TPT1,POLR2M,ANAPC13,SRSF7,GCDH,GLYCTK,CFLAR,SYP,CLASP2,CNTR

147,NEK2,PCID2,PEX2,SMIM4,ARL16,ZNF223,MYL9,HAX1,TMEM108,OGFOD2,EIF3G,TSEN34,MPV17,CI

1,KXD1,ITGB1BP1,MED15,PARD6G,SH3YL1,JPH2,MICAL1,HS2ST1,MAGT1,JUP,TBX6,TMEM205,XBP1,AUI
1,KXD1,ITGB1BP1,MED15,PARD6G,SH3YL1,JPH2,MICAL1,HS2ST1,MAGT1,JUP,TBX6,TMEM205,XBP1,AUI
S2ST1,UBE2I,KCNMA1,SLC39A13,YIF1A,PRMT2,TBX6,TMEM205,PTK2,AUNIP,PCYT2,USE1,HSBP1L1,AGP
S2ST1,UBE2I,KCNMA1,SLC39A13,YIF1A,PRMT2,TBX6,TMEM205,PTK2,AUNIP,PCYT2,USE1,HSBP1L1,AGP
,HSBP1L1,AGPAT4,MAF,ASF1B,EEF1B2,PPP1CC,EDF1,GADL1,RFX3,DGKE,ENDOV,CAMTA1,CAPG,CRIP1,R
,HSBP1L1,AGPAT4,MAF,ASF1B,EEF1B2,PPP1CC,EDF1,GADL1,RFX3,DGKE,ENDOV,CAMTA1,CAPG,CRIP1,R
F644,STAU1,MOB1A,NFAM1,MDH1,LAMP2,DGUOK,HSP90AB1,DUT,SYNGR2,CEP70,TMEM134,LRRFIP1,Z
F644,STAU1,MOB1A,NFAM1,MDH1,LAMP2,DGUOK,HSP90AB1,DUT,SYNGR2,CEP70,TMEM134,LRRFIP1,Z
SIRT5,KCTD7,RFXANK,MTHFS,TMEM101,PPIH,FKBP5,MFSD10,MRPS18C,SLC30A5,GNB5,TRPV3,CBR1,PH
SIRT5,KCTD7,RFXANK,MTHFS,TMEM101,PPIH,FKBP5,MFSD10,MRPS18C,SLC30A5,GNB5,TRPV3,CBR1,PH

C7,NT5E,FAM192A,SLC39A6,ZNF593,TCEAL3,NEK11,CREBZF,EIF2A,NKIRAS2,NUP62,ACAA1,PSPC1,PSMD
C7,NT5E,FAM192A,SLC39A6,ZNF593,TCEAL3,NEK11,CREBZF,EIF2A,NKIRAS2,NUP62,ACAA1,PSPC1,PSMD

,VTI1B,PRKCE,ABLIM2,STAG2,RQCD1,ZNF816,MCUR1,NGDN,KCNH2,POLH,CCT4,NDUFC1,ANXA2,SLC31

PAT4,ASF1B,CCDC25,DEF8,PPP1CC,EDF1,RFX3,PPA1,DGKE,ENDOV,CAMTA1,CAPG,CRIP1,VIM,LIG1,PSM
PAT4,ASF1B,CCDC25,DEF8,PPP1CC,EDF1,RFX3,PPA1,DGKE,ENDOV,CAMTA1,CAPG,CRIP1,VIM,LIG1,PSM
L,CAMLG,EPB41L4B,TCF4,PPOX,ATPAF2,C17ORF62,PRMT1,CASC4,WNK1,ZNF7,TXNDC17,STRADA,PPAN,
17ORF62,PRMT1,EXOSC9,STRADA,PPAN,ZFAND2B,AP1S3,HNRNPC,CYB561,SHARPIN,SUGT1,UPP1,ZNF25
17ORF62,PRMT1,EXOSC9,STRADA,PPAN,ZFAND2B,AP1S3,HNRNPC,CYB561,SHARPIN,SUGT1,UPP1,ZNF25

DH11,DBNDD2,NISCH,TMED4,CENPK,RPL29,CD99L2,RTEL1,AUP1,SAP30BP,FOXRED1,RABL5,TRNT1,KXD
DH11,DBNDD2,NISCH,TMED4,CENPK,RPL29,CD99L2,RTEL1,AUP1,SAP30BP,FOXRED1,RABL5,TRNT1,KXD
6,IFT81,COQ2,UBE2A,PKN2,CREM,SWI5,RNASEK,CORO1B,OAZ1,PHF20,REEP4,SHPK,DIDO1,B9D1,APOC
6,IFT81,COQ2,UBE2A,PKN2,CREM,SWI5,RNASEK,CORO1B,OAZ1,PHF20,REEP4,SHPK,DIDO1,B9D1,APOC
DGUOK,HSP90AB1,SYNGR2,CEP70,TMEM134,RAB2A,RMDN1,GPBP1,GPALPP1,TRMT112,CHPF,MPDU1,
DGUOK,HSP90AB1,SYNGR2,CEP70,TMEM134,RAB2A,RMDN1,GPBP1,GPALPP1,TRMT112,CHPF,MPDU1,
K,PFDN5,S100A4,TXNRD2,PPIH,ARHGEF1,NSUN5,FKBP5,TRPM4,MFSD10,SLC30A5,TRPV3,THADA,CBR1,
A,MROH1,RPS11,SNX5,NUP133,MT2A,MRPS17,TMEM147,ELP4,PCID2,NFYC,ZXDC,COBL,TNKS,ELOF1,AR
A,MROH1,RPS11,SNX5,NUP133,MT2A,MRPS17,TMEM147,ELP4,PCID2,NFYC,ZXDC,COBL,TNKS,ELOF1,AR
L6IP4,STRADA,ZFAND2B,CYB561,ZNF644,STAU1,CARS,RPL36,DUT,SYNGR2,LRRFIP1,RAB2A,TPM2,SIAH1,
GL2,CHEK1,HNRNPA1,FKBP3,NDUFV2,EGLN2,SIRT5,KCTD7,RFXANK,ZNF506,SPINT2,PAFAH1B2,PTK7,DR
GL2,CHEK1,HNRNPA1,FKBP3,NDUFV2,EGLN2,SIRT5,KCTD7,RFXANK,ZNF506,SPINT2,PAFAH1B2,PTK7,DR
36,MOB1A,CCNL2,HSP90AB1,DUT,SYNGR2,TMEM134,LRRFIP1,ZNF524,ATP5J,TPM2,SIAH1,GGCX,ZEB2,T
36,MOB1A,CCNL2,HSP90AB1,DUT,SYNGR2,TMEM134,LRRFIP1,ZNF524,ATP5J,TPM2,SIAH1,GGCX,ZEB2,T
F76,RANGRF,CHCHD7,PHTF2,RBP1,H2AFV,RBMX,BLVRB,TOMM40,RABEP2,WAC,KLK10,EPS15L1,POT1,TI
P1,SIAH1,TSFM,RPTOR,ZNF625,MPDU1,VPS28,ZSWIM7,RPS15,ZNF814,C12ORF76,HMGA1,NAT9,RAD1,
P1,SIAH1,TSFM,RPTOR,ZNF625,MPDU1,VPS28,ZSWIM7,RPS15,ZNF814,C12ORF76,HMGA1,NAT9,RAD1,

PP1,TRMT112,PNPO,CHPF,MAPK13,ZSWIM7,ERLEC1,GMPPB,MVD,SAT1,ERCC5,EMC4,IL32,LMBR1,SYNG

PP1,TRMT112,PNPO,CHPF,MAPK13,ZSWIM7,ERLEC1,GMPPB,MVD,SAT1,ERCC5,EMC4,IL32,LMBR1,SYNG

3A,VAPA,PSMD8,ZNF606,SUMF1,NDUFA7,CYB5D2,OS9,HNRNPM,NGDN,CAMKMT,EXOSC8,DDX17,SNX1

3A,VAPA,PSMD8,ZNF606,SUMF1,NDUFA7,CYB5D2,OS9,HNRNPM,NGDN,CAMKMT,EXOSC8,DDX17,SNX1

ASP2,VDAC1,CREBZF,EIF2A,RPL24,NADSYN1,ZNF511,GTF2H1,TAF1D,RNF130,D2HGDH,ARPC5,MYL6,SIN

R1C,USP4,LY6E,EIF4A1,SLC35E1,TMEM108,MPV17,SSB,RPL8,UBE2D4,ASH1L,TCF25,SLMAP,RBBP6,NIPA

R1C,USP4,LY6E,EIF4A1,SLC35E1,TMEM108,MPV17,SSB,RPL8,UBE2D4,ASH1L,TCF25,SLMAP,RBBP6,NIPA

,OGFOD2,KLHL7,MRPL55,ARIH2,RPL8,TCF25,ARL6IP5,SLMAP,TJP1,CDK4,ERRFI1,RNF8,CFLAR,TRIM46,C

NT1,ITGB1BP1,MED15,SH3YL1,C1ORF198,UGGT2,MICAL1,HS2ST1,KCNMA1,SLC39A13,YIF1A,PRMT2,EI

NT1,ITGB1BP1,MED15,SH3YL1,C1ORF198,UGGT2,MICAL1,HS2ST1,KCNMA1,SLC39A13,YIF1A,PRMT2,EI

SNRPN,SPAG7,WDR62,IL1RAP,STYXL1,RFX5,POLD4,PLEKHA5,RANBP1,UBE2L6,VRK3,LZTFL1,ZC3H14,FIS1

SNRPN,SPAG7,WDR62,IL1RAP,STYXL1,RFX5,POLD4,PLEKHA5,RANBP1,UBE2L6,VRK3,LZTFL1,ZC3H14,FIS1

,TMEM199,TCP11L1,NDUFA3,EIF3C,MGST2,AAMDC,RPS3,IDH3G,ZMAT1,CKLF,AKT1S1,INO80C,COMT,F

F4A1,C6ORF132,DNAJC12,TMEM108,ZMYM6,METTL3,SSR3,MPV17,MRPL55,SSB,ARIH2,CREB3L4,PICK1,

F4A1,C6ORF132,DNAJC12,TMEM108,ZMYM6,METTL3,SSR3,MPV17,MRPL55,SSB,ARIH2,CREB3L4,PICK1,

2,RPL8,CREB3L4,ECE1,ARL6IP5,ZNF248,UBA1,NME1,SMAGP,PPP2R5C,TMC7,MTIF2,GLYCTK,VDAC1,RER

CAMKK1,ZNF276,RQCD1,MCUR1,APRT,ANXA2,TCFL5,PPP1R13B,HDAC10,LARP1B,METTL23,TMEM53,PH

NK1,TXNDC17,ARL6IP4,ZFAND2B,ACAD10,SHARPIN,ZNF644,SUGT1,UPP1,ZNF250,NFAM1,CDK11A,LAM

NK1,TXNDC17,ARL6IP4,ZFAND2B,ACAD10,SHARPIN,ZNF644,SUGT1,UPP1,ZNF250,NFAM1,CDK11A,LAM

13,RPS7,APAF1,KIF13B,EPHA10,MAZ,CCDC24,NFE2L1,RNPS1,IAH1,BTF3L4,UBA52,AK2,CSF1,AIG1,LRRC5

13,RPS7,APAF1,KIF13B,EPHA10,MAZ,CCDC24,NFE2L1,RNPS1,IAH1,BTF3L4,UBA52,AK2,CSF1,AIG1,LRRC5

CARS,ZNF250,RPL36,CDK11A,RBM3,MDH1,CCNL2,MAPKAPK5,DGUOK,SYNGR2,CEP70,RAB2A,ATP5J,GPB

CARS,ZNF250,RPL36,CDK11A,RBM3,MDH1,CCNL2,MAPKAPK5,DGUOK,SYNGR2,CEP70,RAB2A,ATP5J,GPB

NK1,C19ORF54,ZNF7,PPAN,ZFAND2B,AP1S3,HNRNPC,CYB561,ZNF644,STAU1,RPL36,MOB1A,NFAM1,M

NK1,C19ORF54,ZNF7,PPAN,ZFAND2B,AP1S3,HNRNPC,CYB561,ZNF644,STAU1,RPL36,MOB1A,NFAM1,M

32,LMBR1,SYNGR1,RANGRF,CHCHD7,RRNAD1,RBP1,RAD1,H2AFV,SLC33A1,RALGPS1,ZNF687,NUDT1,PC

32,LMBR1,SYNGR1,RANGRF,CHCHD7,RRNAD1,RBP1,RAD1,H2AFV,SLC33A1,RALGPS1,ZNF687,NUDT1,PC

11,NT5C,ILF3,PKP4,AGAP3,STXBP2,HSPH1,C11ORF48,SGK1,SEC31A,EGFR,CASZ1,CHEK1,SUMO3,ZFAND4

11,NT5C,ILF3,PKP4,AGAP3,STXBP2,HSPH1,C11ORF48,SGK1,SEC31A,EGFR,CASZ1,CHEK1,SUMO3,ZFAND4

A2,CDC37,RPS16,CPSF3L,TSG101,OBSL1,TTC21B,PTPN13,UBR4,PSMA1,YLPM1,PHF21A,CHCHD1,SLC39A

NT1,PARD6G,SH3YL1,IP6K2,CLASRP,JPH2,UGGT2,MICAL1,UBE2I,SLC39A13,YIF1A,MAGT1,JUP,TMEM205,

NT1,PARD6G,SH3YL1,IP6K2,CLASRP,JPH2,UGGT2,MICAL1,UBE2I,SLC39A13,YIF1A,MAGT1,JUP,TMEM205,

,NDUFA10,AZI2,MYL12A,U2AF1L4,ATL2,DDX10,PLD1,TMEM41A,EMC3,TAF9,RNPS1,DTD2,NMRAL1,ZNF

1,C11ORF49,PTK7,TRPM4,TCP1,SCAP,MROH1,LFNG,TTC12,SNX5,MT2A,TMEM147,ELP4,ACOT2,USP40,T

1,C11ORF49,PTK7,TRPM4,TCP1,SCAP,MROH1,LFNG,TTC12,SNX5,MT2A,TMEM147,ELP4,ACOT2,USP40,T

MFSD11,GPR56,FAM32A,TMEM147,NFYC,PEX2,DCTN5,PEX26,NDUFS3,OXLD1,MPHOSPH10,ZNF223,RPL

MFSD11,GPR56,FAM32A,TMEM147,NFYC,PEX2,DCTN5,PEX26,NDUFS3,OXLD1,MPHOSPH10,ZNF223,RPL

1,UBE2L6,VRK3,LZTFL1,ZC3H14,FIS1,SLC15A3,PPM1K,TUBA1C,IRF9,DST,ANTXR2,LRRC1,PHF19,GRAMD

1,UBE2L6,VRK3,LZTFL1,ZC3H14,FIS1,SLC15A3,PPM1K,TUBA1C,IRF9,DST,ANTXR2,LRRC1,PHF19,GRAMD

UBE2L3,PABPN1,RPS15A,QTRT1,CCDC74B,RPS17,MRPL51,ZNF586,RBM43,MRPL48,COQ4,SLC25A35,DP
181,ADD1,NIPAL3,CHTOP,BAX,CDK4,SLC52A2,ANAPC13,SLC39A6,APOLD1,RBM42,CNTRL,NUP62,SCN11
181,ADD1,NIPAL3,CHTOP,BAX,CDK4,SLC52A2,ANAPC13,SLC39A6,APOLD1,RBM42,CNTRL,NUP62,SCN11
RRC27,MVD,LMBR1,PRKRIP1,SYNGR1,RANGRF,LSR,MTA3,CHCHD7,MEIS2,PABPC1,RRNAD1,NAT9,BCAM
RA1A,ZNF593,GLYCTK,RBM42,CLASP2,PCNP,NEK11,CREBZF,EIF2A,SCN11A,ACAA1,RPL24,NADSYN1,ZNF5
RA1A,ZNF593,GLYCTK,RBM42,CLASP2,PCNP,NEK11,CREBZF,EIF2A,SCN11A,ACAA1,RPL24,NADSYN1,ZNF5
NF593,GLYCTK,RBM42,CLASP2,NEK11,CREBZF,EIF2A,SCN11A,ACAA1,RPL24,NADSYN1,ZNF511,GTF2H1,T

,SNAPC2,DENND1A,RHEB,NQO2,AKT1S1,NSFL1C,TUSC3,DIP2A,MFGE8,B3GNTL1,REPS2,SPG21,KAT8,KIF

814,HHLA3,SAT1,ERCC5,IL32,NAP1L1,DDOST,C12ORF76,HMGA1,RAD1,RBMX,PPP3CB,BRIP1,RABEP2,ATI
814,HHLA3,SAT1,ERCC5,IL32,NAP1L1,DDOST,C12ORF76,HMGA1,RAD1,RBMX,PPP3CB,BRIP1,RABEP2,ATI
GR1,MTA3,CHCHD7,MEIS2,HMGA1,PSME2,PHTF2,DBNL,BCAM,PPP3CB,RABEP2,TADA3,ZNF20,WAC,RAL
GR1,MTA3,CHCHD7,MEIS2,HMGA1,PSME2,PHTF2,DBNL,BCAM,PPP3CB,RABEP2,TADA3,ZNF20,WAC,RAL
PDU1,TWSG1,ZSWIM7,RPS15,ERLEC1,GMPPB,HHLA3,ERCC5,LMBR1,MCTS1,NAP1L1,SERPINB1,DDOST,S
PDU1,TWSG1,ZSWIM7,RPS15,ERLEC1,GMPPB,HHLA3,ERCC5,LMBR1,MCTS1,NAP1L1,SERPINB1,DDOST,S
HF20,RELA,UNC119,PAK1,ESRRA,TMED1,SEC24C,SERINC5,C16ORF46,MSRB1,KIAA1958,RPE,TVP23B,LIN
IIP,GUK1,CENPN,CARD14,ATF7,USE1,HSBP1L1,IFI27L1,CCDC25,DEF8,EEF1B2,PPP1CC,PDLIM5,EDF1,RFX
IIP,GUK1,CENPN,CARD14,ATF7,USE1,HSBP1L1,IFI27L1,CCDC25,DEF8,EEF1B2,PPP1CC,PDLIM5,EDF1,RFX

)5,AUNIP,GUK1,CENPN,PCYT2,HSBP1L1,AGPAT4,MAF,ASF1B,CCDC25,DEF8,EEF1B2,PPP1CC,GADL1,RFX3
)5,AUNIP,GUK1,CENPN,PCYT2,HSBP1L1,AGPAT4,MAF,ASF1B,CCDC25,DEF8,EEF1B2,PPP1CC,GADL1,RFX3

ZNF511,ARPC5,SLC35A2,TLCD1,ALKBH6,NDUFA11,AXIN1,TUBB3,MOSPD3,BTF3,AAMDC,SNAPC2,SYTL4,
.0,PODXL,CAMLG,EPB41L4B,TMPO,STOML2,WNK1,C19ORF54,TXNDC17,PIGL,C19ORF48,ACAD10,CYB56
.0,PODXL,CAMLG,EPB41L4B,TMPO,STOML2,WNK1,C19ORF54,TXNDC17,PIGL,C19ORF48,ACAD10,CYB56
XW9,USE1,HSBP1L1,AGPAT4,MAF,ASF1B,DEF8,PPP1CC,EDF1,RFX3,PPA1,DGKE,ENDOV,CAMTA1,VIM,LIG
XW9,USE1,HSBP1L1,AGPAT4,MAF,ASF1B,DEF8,PPP1CC,EDF1,RFX3,PPA1,DGKE,ENDOV,CAMTA1,VIM,LIG
DU1,TWSG1,ZSWIM7,RPS15,DCTD,UBXN1,GMPPB,VWA2,LRRC27,MVD,LMBR1,SYNGR1,C12ORF76,MEIS
A10,AZI2,GOSR2,KAT8,U2AF1L4,ATL2,DDX10,EMC3,RNPS1,AIG1,ZNF483,GRN,ACHE,FBXO34,CYB5D2,RB

ASF1B,EEF1B2,PPP1CC,DGKE,CAPG,CRIP1,PPIL2,TROAP,CLDND1,LIG1,PSMA7,CDC14B,APEH,RPL21,TXNL
ASF1B,EEF1B2,PPP1CC,DGKE,CAPG,CRIP1,PPIL2,TROAP,CLDND1,LIG1,PSMA7,CDC14B,APEH,RPL21,TXNL
RF54,ARL6IP4,C19ORF48,HNRNPC,CYB561,CBFA2T2,SLC9B2,UPP1,ZNF250,RPL36,NFAM1,RBM3,MDH1,Z
RF54,ARL6IP4,C19ORF48,HNRNPC,CYB561,CBFA2T2,SLC9B2,UPP1,ZNF250,RPL36,NFAM1,RBM3,MDH1,Z
7,CDC14B,TIMM17B,APEH,TXNL4A,LYPLA2,PODXL,CAMLG,EPB41L4B,TCF4,UCK2,PPOX,TMPO,ATPAF2,C1
7,CDC14B,TIMM17B,APEH,TXNL4A,LYPLA2,PODXL,CAMLG,EPB41L4B,TCF4,UCK2,PPOX,TMPO,ATPAF2,C1
P3K8,TMEM147,PEX2,TNKS,DCTN5,NDUFS3,ARL16,OXLD1,MPHOSPH10,RPLP0,METTL5,C6ORF132,FAM1
P3K8,TMEM147,PEX2,TNKS,DCTN5,NDUFS3,ARL16,OXLD1,MPHOSPH10,RPLP0,METTL5,C6ORF132,FAM1
MK2D,HINT1,HNRNPA1,HIGD1A,NDUFV2,SIRT5,SPINT2,MTHFS,SMARCA4,PAFAH1B2,TYMS,RBFOX2,DAL
MK2D,HINT1,HNRNPA1,HIGD1A,NDUFV2,SIRT5,SPINT2,MTHFS,SMARCA4,PAFAH1B2,TYMS,RBFOX2,DAL
TP10D,SH3GLB1,CHEK1,COX5A,SUMO3,PHF20L1,HNRNPA1,CD151,FKBP3,NDUFV2,B3GAT3,REEP5,TXN
TP10D,SH3GLB1,CHEK1,COX5A,SUMO3,PHF20L1,HNRNPA1,CD151,FKBP3,NDUFV2,B3GAT3,REEP5,TXN

P1S3,ACAD10,STAU1,SUGT1,ZNF250,CDK11A,MDH1,LAMP2,MAPKAPK5,SYNGR2,CEP70,TMEM134,LRR
L1,ZNF483,LRRC23,SUMF1,FBXO34,CYB5D2,DNAJC24,ALS2,PKIG,EXOSC8,DDX17,SLC31A1,CCDC171,RP

AL1,ZNF483,LRRC23,SUMF1,FBXO34,CYB5D2,DNAJC24,ALS2,PKIG,EXOSC8,DDX17,SLC31A1,CCDC171,RPS

HTOP,BAX,ANAPC13,TSPAN31,TMEM9,FAM192A,ADRA1A,ZNF593,GLYCTK,TCEAL3,RBM42,CLASP2,VDAC

HTOP,BAX,ANAPC13,TSPAN31,TMEM9,FAM192A,ADRA1A,ZNF593,GLYCTK,TCEAL3,RBM42,CLASP2,VDAC

,SHARPIN,ZNF644,STAU1,SUGT1,UPP1,CARS,RPL36,CDK11A,MDH1,CCNL2,HSP90AB1,SYNGR2,TMEM134

IIT2,MPDU1,TWSG1,MAPK13,ZSWIM7,ZNF814,GMPPB,HSPB1,MVD,ERCC5,IL32,SEMA3C,LMBR1,NAP1L1

IIT2,MPDU1,TWSG1,MAPK13,ZSWIM7,ZNF814,GMPPB,HSPB1,MVD,ERCC5,IL32,SEMA3C,LMBR1,NAP1L1

,C19ORF48,HNRNPC,CYB561,CBFA2T2,SLC9B2,UPP1,ZNF250,RPL36,NFAM1,RBM3,ZNF197,COPZ1,CCNL2

P1,SIAH1,ASPH,PNPO,RPTOR,NIT2,LYNX1,MPDU1,TWSG1,RPS15,UBXN1,ZNF814,GMPPB,VWA2,ERCC5,IL

P1,SIAH1,ASPH,PNPO,RPTOR,NIT2,LYNX1,MPDU1,TWSG1,RPS15,UBXN1,ZNF814,GMPPB,VWA2,ERCC5,IL

P1,SIAH1,ASPH,PNPO,RPTOR,NIT2,LYNX1,MPDU1,TWSG1,RPS15,UBXN1,ZNF814,GMPPB,VWA2,ERCC5,IL

D4,CENPK,GGPS1,COASY,SRSF1,SAP30BP,RABL5,TRNT1,KXD1,ITGB1BP1,MED15,SH3YL1,JPH2,C1ORF198

D4,CENPK,GGPS1,COASY,SRSF1,SAP30BP,RABL5,TRNT1,KXD1,ITGB1BP1,MED15,SH3YL1,JPH2,C1ORF198

DDS,ARL2,CEP63,RPS23,ZNF783,IFRD2,ACHE,SUMF1,NDUFA7,CCNG2,RABEPK,GRHPR,SLC39A14,HNRNP

DDS,ARL2,CEP63,RPS23,ZNF783,IFRD2,ACHE,SUMF1,NDUFA7,CCNG2,RABEPK,GRHPR,SLC39A14,HNRNP

D,CHPF,LYNX1,RPS15,UBXN1,ZNF814,ERCC5,EMC4,LMBR1,MCTS1,PRKRIP1,DDOST,LSR,MTA3,DOCK10,P

D,CHPF,LYNX1,RPS15,UBXN1,ZNF814,ERCC5,EMC4,LMBR1,MCTS1,PRKRIP1,DDOST,LSR,MTA3,DOCK10,P

MFSD11,LFNG,SNX5,NOP2,RPAIN,TMEM147,SMIM4,PRRC2B,ARL16,BTN3A2,MPHOSPH10,C6ORF132,ZM

C37,RPS16,CPSF3L,TSG101,OBSL1,TTC21B,PTPN13,YLPM1,PHF21A,CHCHD1,SLC39A11,FAM3A,PSMA3,N

C37,RPS16,CPSF3L,TSG101,OBSL1,TTC21B,PTPN13,YLPM1,PHF21A,CHCHD1,SLC39A11,FAM3A,PSMA3,N

K1,C19ORF54,PIGL,C19ORF48,ZFAND2B,HNRNPC,ZNF644,CBFA2T2,HYPK,UPP1,MOB1A,CDK11A,RBM3,N

K1,C19ORF54,PIGL,C19ORF48,ZFAND2B,HNRNPC,ZNF644,CBFA2T2,HYPK,UPP1,MOB1A,CDK11A,RBM3,N

FI27L1,MAF,ASF1B,DEF8,EEF1B2,PPP1CC,RFX3,PPA1,TNFRSF14,VAMP8,CAMTA1,CAPG,PSMD4,LIG1,PSM

FI27L1,MAF,ASF1B,DEF8,EEF1B2,PPP1CC,RFX3,PPA1,TNFRSF14,VAMP8,CAMTA1,CAPG,PSMD4,LIG1,PSM

,CDK2AP2,NDUFA10,AZI2,MYL12A,U2AF1L4,ATL2,DDX10,PLD1,TMEM41A,EMC3,TAF9,RNPS1,DTD2,NM

SWIM7,DCTD,HSPB1,LRRC27,MVD,SERPINB1,DDOST,SYNGR1,MEIS2,HMGA1,PABPC1,RRNAD1,PPP3CB,S

4DIP,EEF1D,HSPH1,CHCHD5,C11ORF48,CHEK1,CAMK2D,COX5A,HNRNPA1,FKBP3,RPL15,SIRT5,KCTD7,ZN

4DIP,EEF1D,HSPH1,CHCHD5,C11ORF48,CHEK1,CAMK2D,COX5A,HNRNPA1,FKBP3,RPL15,SIRT5,KCTD7,ZN

2,PABPC1,BCAM,PPP3CB,TADA3,WAC,RALGPS1,ZNF687,PDCD5,GTF2IRD1,NXN,EPS15L1,RAD23A,HSD17

IP4,C19ORF48,PPAN,CYB561,CBFA2T2,SLC9B2,UPP1,ZNF250,RPL36,NFAM1,RBM3,MDH1,ZNF197,COPZ1

IAH1,ASPH,PNPO,ZNF625,MPDU1,VPS28,ZSWIM7,UBXN1,ZNF814,HHLA3,SAT1,ERCC5,IL32,NAP1L1,DD

IAH1,ASPH,PNPO,ZNF625,MPDU1,VPS28,ZSWIM7,UBXN1,ZNF814,HHLA3,SAT1,ERCC5,IL32,NAP1L1,DD

F483,LRRC23,SUMF1,FBXO34,CYB5D2,DNAJC24,ALS2,PKIG,EXOSC8,DDX17,SLC31A1,RPS17,RCAN1,NSM

T2,MICAL1,UBE2I,SLC39A13,YIF1A,MAGT1,JUP,PTK2,C14ORF2,ATF7,PCYT2,USE1,AGPAT4,IFI27L1,MAF,

L1,ASF1B,CCDC25,EEF1B2,EDF1,PPA1,DGKE,CAMTA1,PPIL2,RPL28,LIG1,CDC14B,TIMM17B,TMEM126B,R

L1,ASF1B,CCDC25,EEF1B2,EDF1,PPA1,DGKE,CAMTA1,PPIL2,RPL28,LIG1,CDC14B,TIMM17B,TMEM126B,R

A1,FKBP3,NDUFV2,B3GAT3,REEP5,TCTN1,GTF2IRD2,COPS4,TMEM39B,TPM4,MFSD10,MRPS18C,RIC8A,S

A1,FKBP3,NDUFV2,B3GAT3,REEP5,TCTN1,GTF2IRD2,COPS4,TMEM39B,TPM4,MFSD10,MRPS18C,RIC8A,S

HGDH,CC2D1B,ARPC5,ST7,SH3GL1,DNAJC2,MRPS24,RHEB,CREB3L2,RGS12,DNAJA3,CKLF,AKT1S1,COMT,

HGDH,CC2D1B,ARPC5,ST7,SH3GL1,DNAJC2,MRPS24,RHEB,CREB3L2,RGS12,DNAJA3,CKLF,AKT1S1,COMT,

HGDH,CC2D1B,ARPC5,ST7,SH3GL1,DNAJC2,MRPS24,RHEB,CREB3L2,RGS12,DNAJA3,CKLF,AKT1S1,COMT,

2,NUTF2,PODXL,LSM12,C17ORF62,STOML2,C19ORF54,ARL6IP4,STRADA,ACAD10,HNRNPC,CYB561,SHAR
K2,RBM6,HMGN1,NDUFB2,CDK2AP2,NDUFA10,AZI2,MYL12A,U2AF1L4,ATL2,DDX10,PLD1,TMEM41A,EM
K2,RBM6,HMGN1,NDUFB2,CDK2AP2,NDUFA10,AZI2,MYL12A,U2AF1L4,ATL2,DDX10,PLD1,TMEM41A,EM

RDH11,DBNDD2,SDF4,NISCH,TMED4,CENPK,RPS6,RPL29,CD99L2,GGPS1,RTEL1,SRSF1,AUP1,SAP30BP,YI
RDH11,DBNDD2,SDF4,NISCH,TMED4,CENPK,RPS6,RPL29,CD99L2,GGPS1,RTEL1,SRSF1,AUP1,SAP30BP,YI
1BP1,PARD6G,SH3YL1,IP6K2,CLASRP,JPH2,C1ORF198,MICAL1,HS2ST1,UBE2I,SLC39A13,YIF1A,PRMT2,MA
1BP1,PARD6G,SH3YL1,IP6K2,CLASRP,JPH2,C1ORF198,MICAL1,HS2ST1,UBE2I,SLC39A13,YIF1A,PRMT2,MA
T1,POLR3H,ILF3,AGAP3,NAA10,RRP8,SEC31A,EGFR,SH3GLB1,SSH2,HNRNPLL,SUMO3,ZFAND4,HNRNPA1,

OK,HSP90AB1,DUT,FAM219B,RAB2A,ZNF524,ATP5J,RMDN1,GPBP1,GPALPP1,SIAH1,PNPO,RPTOR,ZNF62
OK,HSP90AB1,DUT,FAM219B,RAB2A,ZNF524,ATP5J,RMDN1,GPBP1,GPALPP1,SIAH1,PNPO,RPTOR,ZNF62
ORF132,GFOD1,TSEN34,LIMA1,ECE1,TCF25,KCNC4,TJP1,CHTOP,TPT1,SMAGP,NXF1,SYP,FAM69B,GLI4,N
F524,ATP5J,GPBP1,GPALPP1,SIAH1,PNPO,RPTOR,ZNF625,MPDU1,VPS28,ZSWIM7,RPS15,UBXN1,ZNF81
F524,ATP5J,GPBP1,GPALPP1,SIAH1,PNPO,RPTOR,ZNF625,MPDU1,VPS28,ZSWIM7,RPS15,UBXN1,ZNF81
S27L,SPG21,KIF13B,DDX10,PLD1,TMEM41A,EMC3,DTD2,DHDDS,NMRAL1,PRKCE,ZNF483,LRRC23,ZNF60
S27L,SPG21,KIF13B,DDX10,PLD1,TMEM41A,EMC3,DTD2,DHDDS,NMRAL1,PRKCE,ZNF483,LRRC23,ZNF60
1,MAGED2,SON,UQCRQ,SNF8,GRK6,HMGA2,CDC37,RPS16,CPSF3L,TSG101,OBSL1,TTC21B,PTPN13,YLPM
SP90AB1,CEP70,RAB2A,ZNF524,GGCX,TRMT112,RPTOR,RPS20,GABPB1,NIT2,TWSG1,VPS28,ZSWIM7,RPS
SP90AB1,CEP70,RAB2A,ZNF524,GGCX,TRMT112,RPTOR,RPS20,GABPB1,NIT2,TWSG1,VPS28,ZSWIM7,RPS

,TMED4,RTEL1,COASY,SRSF1,YIPF3,FOXRED1,KXD1,ITGB1BP1,MED15,PARD6G,SH3YL1,IP6K2,CLASRP,JPH
,TMED4,RTEL1,COASY,SRSF1,YIPF3,FOXRED1,KXD1,ITGB1BP1,MED15,PARD6G,SH3YL1,IP6K2,CLASRP,JPH
7,DYNC1I2,NIPA1,ATP11B,PABPN1,NDUFB10,ATPAF1,ISOC2,WDR20,C9ORF64,IQCK,MED8,RNMT,DHX30
11L1,UBE2D3,C5ORF24,SNAPC2,IDH3G,CREB3L2,CKLF,AKT1S1,SPHK2,CHURC1,HMGN1,NDUFA10,AZI2,S
11L1,UBE2D3,C5ORF24,SNAPC2,IDH3G,CREB3L2,CKLF,AKT1S1,SPHK2,CHURC1,HMGN1,NDUFA10,AZI2,S
P9,PTGR1,C8ORF82,MSRB1,SURF4,GBGT1,CDK5RAP3,LIN7B,PTMS,EXOSC10,TECPR2,GCA,SLC41A3

RRC23,SUMF1,FBXO34,CYB5D2,DNAJC24,ALS2,IK,PKIG,EXOSC8,DDX17,SLC31A1,RPS17,RCAN1,NSMF,RN
RRC23,SUMF1,FBXO34,CYB5D2,DNAJC24,ALS2,IK,PKIG,EXOSC8,DDX17,SLC31A1,RPS17,RCAN1,NSMF,RN

TLCD1,RPS3A,MOSPD3,TCP11L1,MAGOHB,IDH3G,CREB3L2,AKT1S1,AIFM2,HMOX2,SPHK2,ACTR10,HMG
SYNGR2,RAB2A,ATP5J,ELP6,TPM2,GPALPP1,SIAH1,ASPH,PNPO,RPTOR,NIT2,LYNX1,MPDU1,TWSG1,RPS1
SYNGR2,RAB2A,ATP5J,ELP6,TPM2,GPALPP1,SIAH1,ASPH,PNPO,RPTOR,NIT2,LYNX1,MPDU1,TWSG1,RPS1
SYNGR2,RAB2A,ATP5J,ELP6,TPM2,GPALPP1,SIAH1,ASPH,PNPO,RPTOR,NIT2,LYNX1,MPDU1,TWSG1,RPS1

O34,CYB5D2,ALS2,IK,EXOSC8,DDX17,SLC31A1,RCAN1,C12ORF73,MPND,NSMF,ATP9B,IMMP1L,RAP1B,IF
RPS18C,GMPR2,THADA,COMMD4,MROH1,RPS11,MFF,SNX5,NUP133,TMEM147,ELP4,NFYC,ZXDC,COBL,T
RPS18C,GMPR2,THADA,COMMD4,MROH1,RPS11,MFF,SNX5,NUP133,TMEM147,ELP4,NFYC,ZXDC,COBL,T
2,SWI5,SNRNP40,DCTN6,WDR6,OPA3,SNRNP200,ARHGEF7,AHI1,LIN37,C12ORF66,MAGOH,PAK1,GINS4,

,LIG1,PSMA7,CDC14B,XAF1,APEH,HYI,RPL21,TXNL4A,CAPN10,LYPLA2,PODXL,CAMLG,EPB41L4B,UCK2,P

,LIG1,PSMA7,CDC14B,XAF1,APEH,HYI,RPL21,TXNL4A,CAPN10,LYPLA2,PODXL,CAMLG,EPB41L4B,UCK2,P

1,MT1E,CNST,LLGL2,SUMO3,EGLN2,SIRT5,B3GAT3,RFXANK,PFDN5,S100A4,TXNRD2,PPIH,NSUN5,FKBP5,

1,MT1E,CNST,LLGL2,SUMO3,EGLN2,SIRT5,B3GAT3,RFXANK,PFDN5,S100A4,TXNRD2,PPIH,NSUN5,FKBP5,

L4ORF2,CENPN,PCYT2,FBXW9,USE1,HSBP1L1,MAF,ASF1B,DEF8,PPP1CC,EDF1,GADL1,RFX3,PPA1,DGKE,T

ZNF836,RBMX,BLVRB,TOMM40,ATP6V1F,WAC,NXN,POT1,TRIM16,RAD23A,HSD17B10,GGT5,DDX41,US

HRS4,DIP2A,NAA15,MLST8,NDUFB2,HMBS,NDUFA10,MFGE8,MYL12A,TBC1D22A,CARM1,C1QC,CCDC24

HRS4,DIP2A,NAA15,MLST8,NDUFB2,HMBS,NDUFA10,MFGE8,MYL12A,TBC1D22A,CARM1,C1QC,CCDC24

A3,LRRC27,ERCC5,LMBR1,SERPINB1,MTA3,CHCHD7,PABPC1,RRNAD1,NAT9,RAD1,PPP3CB,SERPINB6,TA

A3,LRRC27,ERCC5,LMBR1,SERPINB1,MTA3,CHCHD7,PABPC1,RRNAD1,NAT9,RAD1,PPP3CB,SERPINB6,TA

,TMEM9,NXF1,NT5E,SLC39A6,ADRA1A,ZNF593,GLYCTK,FAM131A,TRIM46,RBM42,CLASP2,PCNP,CREBZF

,TMEM9,NXF1,NT5E,SLC39A6,ADRA1A,ZNF593,GLYCTK,FAM131A,TRIM46,RBM42,CLASP2,PCNP,CREBZF

PF3,TRNT1,KXD1,ITGB1BP1,MED15,PARD6G,SH3YL1,IP6K2,CLASRP,JPH2,C1ORF198,MICAL1,KCNMA1,E

PF3,TRNT1,KXD1,ITGB1BP1,MED15,PARD6G,SH3YL1,IP6K2,CLASRP,JPH2,C1ORF198,MICAL1,KCNMA1,E

TG12,PPRC1,RPS25,ZBTB44,TTC9C,COX4I1,DUSP22,TMX2,SPOCK2,THEM4,NISCH,CENPK,GGPS1,COASY,

TG12,PPRC1,RPS25,ZBTB44,TTC9C,COX4I1,DUSP22,TMX2,SPOCK2,THEM4,NISCH,CENPK,GGPS1,COASY,

1,TMEM134,FAM219B,RAB2A,GPBP1,GPALPP1,SIAH1,TRMT112,PNPO,CHPF,NIT2,ZSWIM7,RPS15,DCTD

1,TMEM134,FAM219B,RAB2A,GPBP1,GPALPP1,SIAH1,TRMT112,PNPO,CHPF,NIT2,ZSWIM7,RPS15,DCTD

FIP1,FAM219B,ZNF524,ATP5J,SIAH1,ASPH,TRMT112,RPTOR,STRIP2,RPS20,GABPB1,MPDU1,UBXN1,ERL

FIP1,FAM219B,ZNF524,ATP5J,SIAH1,ASPH,TRMT112,RPTOR,STRIP2,RPS20,GABPB1,MPDU1,UBXN1,ERL

DENND1A,EPN1,CREB3L2,EML3,ZMAT1,VPS25,SNRPD1,SLC6A9,TRAK1,SNRPA1,HMBS,RPS27L,MFGE8,Z

DENND1A,EPN1,CREB3L2,EML3,ZMAT1,VPS25,SNRPD1,SLC6A9,TRAK1,SNRPA1,HMBS,RPS27L,MFGE8,Z

D1,RGS3,FAM3A,ECSIT,ATG12,PPRC1,RPS25,NLRP1,ZBTB44,NIP7,TTC9C,COX4I1,PLCB2,SLC25A10,SPOC

D1,RGS3,FAM3A,ECSIT,ATG12,PPRC1,RPS25,NLRP1,ZBTB44,NIP7,TTC9C,COX4I1,PLCB2,SLC25A10,SPOC

R1,DCN,PHTF2,PABPC1,PTPRB,RABEP2,ATP6V1F,WAC,SLC33A1,PDCD5,TRIM16,POLR3H,RAD23A,ABHD

1,GPALPP1,SIAH1,PNPO,RPTOR,ZNF625,MPDU1,VPS28,ZSWIM7,RPS15,UBXN1,ZNF814,HHLA3,SAT1,ER

1,GPALPP1,SIAH1,PNPO,RPTOR,ZNF625,MPDU1,VPS28,ZSWIM7,RPS15,UBXN1,ZNF814,HHLA3,SAT1,ER

ARL6IP4,ZFAND2B,HYPK,UPP1,ZNF250,NFAM1,CDK11A,RBM3,INO80E,MAPKAPK5,DGUOK,HSP90AB1,C

ARL6IP4,ZFAND2B,HYPK,UPP1,ZNF250,NFAM1,CDK11A,RBM3,INO80E,MAPKAPK5,DGUOK,HSP90AB1,C

UBR4,PSMA1,PHF21A,CHCHD1,SLC39A11,FAM3A,PSMA3,MUC1,ECSIT,EPB41L5,PPRC1,RPS25,NLRP1,ZB

UBR4,PSMA1,PHF21A,CHCHD1,SLC39A11,FAM3A,PSMA3,MUC1,ECSIT,EPB41L5,PPRC1,RPS25,NLRP1,ZB

LSM12,PRMT1,EXOSC9,STOML2,WNK1,C19ORF54,ARL6IP4,PIGL,C19ORF48,STRADA,ZFAND2B,AP1S3,H

LSM12,PRMT1,EXOSC9,STOML2,WNK1,C19ORF54,ARL6IP4,PIGL,C19ORF48,STRADA,ZFAND2B,AP1S3,H

D1,VIM,PSMA7,CDC14B,TMEM126B,RPL21,TXNL4A,CAPN10,LYPLA2,CAMLG,TCF4,UCK2,TMPO,ATPAF2,

D1,VIM,PSMA7,CDC14B,TMEM126B,RPL21,TXNL4A,CAPN10,LYPLA2,CAMLG,TCF4,UCK2,TMPO,ATPAF2,

A3,MVD,LMBR1,PRKRIP1,RANGRF,LSR,MTA3,MEIS2,HMGA1,RRNAD1,ZNF836,RBP1,ATP6V1F,SLC33A1,Z

A3,MVD,LMBR1,PRKRIP1,RANGRF,LSR,MTA3,MEIS2,HMGA1,RRNAD1,ZNF836,RBP1,ATP6V1F,SLC33A1,Z

MDH1,LAMP2,CCNL2,MAPKAPK5,SYNGR2,LRRFIP1,RAB2A,ZNF524,GPBP1,GPALPP1,SIAH1,TSFM,ZEB2,A

,INO80E,MAPKAPK5,DGUOK,HSP90AB1,SYNGR2,CEP70,TMEM134,RAB2A,RMDN1,GPBP1,GPALPP1,TP

,INO80E,MAPKAPK5,DGUOK,HSP90AB1,SYNGR2,CEP70,TMEM134,RAB2A,RMDN1,GPBP1,GPALPP1,TP

RRNAD1,RAD1,BLVRB,ZNF20,SLC33A1,ZNF706,PDCD5,GTF2IRD1,NUDT1,ANAPC11,POLR3H,RAD23A,GG

RRNAD1,RAD1,BLVRB,ZNF20,SLC33A1,ZNF706,PDCD5,GTF2IRD1,NUDT1,ANAPC11,POLR3H,RAD23A,GG

,SERPINB1,DDOST,C12ORF76,RANGRF,PABPC1,RRNAD1,RAD1,BLVRB,ZNF20,SLC33A1,ZNF706,PDCD5,GT

,SERPINB1,DDOST,C12ORF76,RANGRF,PABPC1,RRNAD1,RAD1,BLVRB,ZNF20,SLC33A1,ZNF706,PDCD5,GT
NF250,RBM3,ZNF197,MAPKAPK5,RAB2A,ZNF524,ATP5J,RMDN1,GPBP1,GPALPP1,GGCX,PNPO,STRIP2,R
NF250,RBM3,ZNF197,MAPKAPK5,RAB2A,ZNF524,ATP5J,RMDN1,GPBP1,GPALPP1,GGCX,PNPO,STRIP2,R

C2D1B,ARPC5,ST7,NDUFA11,MGST2,SYTL4,DNAJC2,MRPS24,DNAJA3,ZMAT1,AKT1S1,INO80C,COMT,GT
C2D1B,ARPC5,ST7,NDUFA11,MGST2,SYTL4,DNAJC2,MRPS24,DNAJA3,ZMAT1,AKT1S1,INO80C,COMT,GT
C2D1B,ARPC5,ST7,NDUFA11,MGST2,SYTL4,DNAJC2,MRPS24,DNAJA3,ZMAT1,AKT1S1,INO80C,COMT,GT
SON,GRK6,HMGA2,CDC37,RPS16,CPSF3L,TSG101,OBSL1,PTPN13,UBR4,YLPM1,PHF21A,CHCHD1,FAM3A
SON,GRK6,HMGA2,CDC37,RPS16,CPSF3L,TSG101,OBSL1,PTPN13,UBR4,YLPM1,PHF21A,CHCHD1,FAM3A

10D,SH3GLB1,CHEK1,COX5A,SUMO3,PHF20L1,HNRNPA1,CD151,FKBP3,NDUFV2,EGLN2,B3GAT3,REEP5,
10D,SH3GLB1,CHEK1,COX5A,SUMO3,PHF20L1,HNRNPA1,CD151,FKBP3,NDUFV2,EGLN2,B3GAT3,REEP5,
,STRADA,PPAN,ZFAND2B,AP1S3,ACAD10,SHARPIN,STAU1,ZNF250,RPL36,MOB1A,CDK11A,MDH1,LAMP
NF197,CCNL2,DGUOK,DUT,SYNGR2,TMEM134,LRRFIP1,RAB2A,ZNF524,GPBP1,GGCX,ZEB2,PNPO,RPTOR,
NF197,CCNL2,DGUOK,DUT,SYNGR2,TMEM134,LRRFIP1,RAB2A,ZNF524,GPBP1,GGCX,ZEB2,PNPO,RPTOR,
B2,HMBS,MFGE8,MYL12A,FAM104B,SLC37A3,PRKCSH,IAH1,BTF3L4,UBA52,FHL2,VAPA,CAMKK1,ARL2,R
B2,HMBS,MFGE8,MYL12A,FAM104B,SLC37A3,PRKCSH,IAH1,BTF3L4,UBA52,FHL2,VAPA,CAMKK1,ARL2,R
T1,CDC42,AUNIP,GUK1,CARD14,FBXW9,USE1,HSBP1L1,IFI27L1,ASF1B,EEF1B2,PPP1CC,GADL1,RFX3,DGK
T1,CDC42,AUNIP,GUK1,CARD14,FBXW9,USE1,HSBP1L1,IFI27L1,ASF1B,EEF1B2,PPP1CC,GADL1,RFX3,DGK
MF,ATP9B,IMMP1L,RAP1B,IFT81,COQ2,TUBGCP2,SWI5,SNRNP40,DCTN6,WDR6,OPA3,SNRNP200,ARHG
,TGOLN2,PEX2,STX10,PEX26,NDUFS3,RNASE1,OXLD1,EIF4A1,DGAT1,OGFOD2,GFOD1,ARIH2,RPL8
,TGOLN2,PEX2,STX10,PEX26,NDUFS3,RNASE1,OXLD1,EIF4A1,DGAT1,OGFOD2,GFOD1,ARIH2,RPL8
ACOT9,MRPL27,MFSD1,DNAJB12,FCGRT,PKN2,RNASEK,SSH3,CORO1B,PHF20,WDR6,MRPL36,SYNJ2,MIE
ACOT9,MRPL27,MFSD1,DNAJB12,FCGRT,PKN2,RNASEK,SSH3,CORO1B,PHF20,WDR6,MRPL36,SYNJ2,MIE
I1,RNF8,PPP2R5C,TMEM9,FAM192A,SLC39A6,ADRA1A,APOLD1,FAM131A,RPL34,CREBZF,SCN11A,SKA2,
I1,RNF8,PPP2R5C,TMEM9,FAM192A,SLC39A6,ADRA1A,APOLD1,FAM131A,RPL34,CREBZF,SCN11A,SKA2,
2,GPALPP1,SIAH1,TSFM,GGCX,ASPH,TPM3,RPTOR,STRIP2,ZNF625,GABPB1,LYNX1,MPDU1,MAPK13,VPS
C,CYB561,ZNF644,CBFA2T2,HYPK,ZNF250,MOB1A,CDK11A,RBM3,COPZ1,INO80E,CCNL2,DGUOK,HSP90
C,CYB561,ZNF644,CBFA2T2,HYPK,ZNF250,MOB1A,CDK11A,RBM3,COPZ1,INO80E,CCNL2,DGUOK,HSP90

,HAUS1,DDX17,ACTB,PABPN1,RPS15A,QTRT1,CCDC171,RCAN1,NDUFB10,CLSPN,MCM7,C9ORF64,EIF6,

| |
|---|
| 'S3,DNAJC2,MRPS24,RDX,MAGOHB,ZCCHC9,SNRPD1,UCHL5,MKRN1,GTF3A,FARS2,RBM6,SNRPA1,NAA1 |
| 5Z1,HNRNPLL,HNRNPA1,POLR2C,FKBP3,RPL15,RFXANK,ZNF506,ASH2L,SMARCA4,S100A4,ZNF343,SNRPF |
| ,PSMA1,YLPM1,PHF21A,CHCHD1,FUZ,RGS3,FAM3A,PSMA3,MUC1,METTL17,ECSIT,EPB41L5,ATG12,PPR( |
| CTS1,NAP1L1,PRKRIP1,MTA3,DCN,MEIS2,HMGA1,PHTF2,PABPC1,ZNF836,RAD1,H2AFV,RBMX,BRIP1,ZNI |
| D,ERCC5,MCTS1,NAP1L1,PRKRIP1,MTA3,DCN,MEIS2,HMGA1,PHTF2,PABPC1,ZNF836,RAD1,H2AFV,RBM) |
| ,AP4S1,CLIP1,OAZ2,MAGOH,GLE1,PAK1,ATP13A2,TMED1,SEC24C,RPL31,GSK3A,TOMM7,EDEM2,ITGB2,S |
| ACTR10,TRAK1,GOSR2,SGSM3,TBC1D22A,RPS7,LAMTOR1,U2AF1L4,KIF13B,NCBP2,SNX3,ECT2,RNPS1,UB |
| MYL12A,SGSM3,TBC1D22A,RPS7,LAMTOR1,U2AF1L4,KIF13B,CARM1,NCBP2,SNX3,ECT2,RNPS1,UBA52,KI |

MYL12A,SGSM3,TBC1D22A,RPS7,LAMTOR1,U2AF1L4,KIF13B,CARM1,NCBP2,SNX3,ECT2,RNPS1,UBA52,KI

L,SLMAP,UGT8,CHTOP,POLR2M,BAX,RPL35,DLG4,SRSF7,NXF1,ADRA1A,ZNF593,TRIM46,LAMTOR5,CLASP

PML,TMEM199,UBE2D3,MGST2,RPS3,RDX,MAGOHB,ITPA,VPS25,UCHL5,NSFL1C,SMUG1,COMT,HMOX2,

RS4,MLST8,NDUFB3,NDUFB2,NDUFA10,RPS27L,GMFG,REPS2,GOSR2,PSMG2,ATL2,PSMG4,TBCA,ECT2,TA
,PSMD10,RPS3A,PML,TMEM199,UBE2D3,MGST2,RPS3,RHEB,RDX,MAGOHB,ITPA,VPS25,UCHL5,GRAMD1

VDAC1,RPL34,NUP62,ACAA1,RPL24,PSMD7,PAM16,NEDD4L,RNF130,BFAR,CLTA,LYPLAL1,AXIN1,TP53,PSI
,SPHK2,DHRS4,IFI27,ARRB2,TRAK1,GOSR2,SGSM3,TBC1D22A,RPS7,U2AF1L4,KIF13B,NCBP2,SNX3,MDK,E
,SYTL4,DENND1A,RDX,MAGOHB,VPS25,SPHK2,DHRS4,IFI27,ARRB2,WDPCP,TRAK1,GOSR2,SGSM3,TBC1D
F,NOP2,RAPGEF3,HAUS4,COX20,COA1,NEK2,COBL,TNKS,USP4,NDUFS3,MPHOSPH10,NUBP2,RPLP0,HAX

2,VAPA,CAPN2,DHDDS,MRPS16,NMRAL1,RPS23,RPL30,RQCD1,ACHE,NDUFA7,DNAJC24,MRPS33,ACSS1,

,SPHK2,DHRS4,IFI27,ARRB2,TRAK1,GOSR2,SGSM3,TBC1D22A,RPS7,U2AF1L4,KIF13B,NCBP2,SNX3,MDK,E
HB,VPS25,SPHK2,DHRS4,IFI27,ARRB2,TRAK1,GOSR2,SGSM3,TBC1D22A,RPS7,U2AF1L4,KIF13B,NCBP2,SN
4,SRSF7,NXF1,ZNF593,TRIM46,LAMTOR5,CLASP2,RER1,RPL34,NUP62,ACAA1,RPL24,PAM16,NEDD4L,FCI

GABPB1,TWSG1,ZSWIM7,RPS15,DCTD,ZNF814,GMPPB,HSPB1,MVD,ERCC5,PRKRIP1,MTA3,DCN,MEIS2,HI
P2S1,NDUFC1,ANXA2,RUVBL1,SEC13,NDUFB10,ATPAF1,SFSWAP,MCM7,EIF6,DHX30,SHMT1,ERAL1,RAP1
GUOK,HSP90AB1,DUT,LRRFIP1,ZNF524,ATP5J,ELP6,GPBP1,TSFM,ZEB2,TRMT112,PNPO,RPTOR,ZNF625,F
,CAMK2D,HNRNPA1,HIGD1A,NDUFV2,SIRT5,ASH2L,TCTN1,SMARCA4,COA3,PTK7,GBA2,COPS4,TPM4,MC

F625,RPS20,GABPB1,TWSG1,ZSWIM7,RPS15,DCTD,ZNF814,GMPPB,HSPB1,MVD,ERCC5,PRKRIP1,MTA3,D

T1,CIZ1,SSR3,MPV17,SSB,ARIH2,RPL8,LIMA1,PICK1,SLMAP,TJP1,UGT8,CHTOP,BAX,RPL35,DLG4,SRSF7,N
LEC1,HSPB1,MVD,MCTS1,SERPINB1,PRKRIP1,DDOST,MTA3,DCN,PSME2,PABPC1,PTPRB,DBNL,NAT9,PPP
,ADD1,KCNC4,TJP1,UGT8,BAX,DLG4,TRPM6,EEF2K,CFLAR,CLASP2,CNTRL,EIF2A,NUP62,RPL24,FCHO2,IF
EXOSC9,STOML2,PPAN,HNRNPC,SHARPIN,CBFA2T2,STAU1,CDK11A,LAMP2,INO80E,MAPKAPK5,DGUOK,
20,WAC,SLC33A1,ZNF706,GTF2IRD1,NUDT1,POT1,TRIM16,POLR3H,GGT5,DDX41,NT5C,PSME1,ILF3,ZCC
625,RPS20,GABPB1,TWSG1,ZSWIM7,RPS15,DCTD,ZNF814,GMPPB,HSPB1,MVD,ERCC5,PRKRIP1,MTA3,D

TGB1BP1,MED15,SH3YL1,IP6K2,CLASRP,UGGT2,MICAL1,HS2ST1,UBE2I,PRMT2,EIF3M,MAGT1,JUP,CDC4

,RBMX,PPP3CB,BRIP1,TADA3,ZNF20,WAC,SLC33A1,ZNF706,GTF2IRD1,NUDT1,POT1,TRIM16,POLR3H,RA
112,PNPO,RPTOR,ZNF625,RPS20,GABPB1,TWSG1,ZSWIM7,RPS15,DCTD,ZNF814,GMPPB,HSPB1,MVD,ER

A2,THOC3,TP53,AP1G2,RPS3A,BTF3,PML,SLC25A24,UBE2D3,RPS3,SYTL4,DENND1A,MAGOHB,VPS25,SP
,SLC33A1,ZNF706,GTF2IRD1,POT1,TRIM16,POLR3H,ABHD17A,GGT5,DDX41,PSME1,ILF3,ZCCHC4,EEF1D,

IP1,TADA3,ZNF20,WAC,SLC33A1,ZNF706,GTF2IRD1,POT1,TRIM16,POLR3H,ABHD17A,GGT5,DDX41,PSM
36,CDK11A,RBM3,LAMP2,INO80E,CCNL2,MAPKAPK5,DGUOK,HSP90AB1,DUT,RAB2A,ATP5J,SIAH1,TSFM,

TSFM,ZEB2,TRMT112,PNPO,RPTOR,CHPF,ZNF625,RPS20,GABPB1,MPDU1,TWSG1,RPS15,DCTD,ZNF814,
K11A,LAMP2,INO80E,MAPKAPK5,DGUOK,HSP90AB1,SYNGR2,CEP70,RAB2A,ATP5J,TPM2,SIAH1,ZEB2,AS

TOR,CHPF,ZNF625,RPS20,GABPB1,MPDU1,TWSG1,RPS15,DCTD,ZNF814,GMPPB,HSPB1,MVD,SAT1,MCT

TOR,CHPF,RPS20,MPDU1,TWSG1,MAPK13,VPS28,RPS15,UBXN1,ERLEC1,HSPB1,MVD,MCTS1,SERPINB1,

8,OAS3,MED6,CARHSP1,RPL18,EGFR,CASZ1,CHEK1,CAMK2D,HNRNPLL,HINT1,PHF20L1,HNRNPA1,HIGD

TSFM,ZEB2,TRMT112,PNPO,RPTOR,CHPF,ZNF625,RPS20,GABPB1,MPDU1,TWSG1,RPS15,DCTD,ZNF814,

,RPL29,GGPS1,RTEL1,COASY,SRSF1,AUP1,SAP30BP,FOXRED1,TRNT1,ITGB1BP1,MED15,SH3YL1,IP6K2,C

LK,APAF1,CARM1,POLR2E,ECT2,NFE2L1,TAF9,GPS2,UBA52,MLH1,SUMO1,PRKCE,COMMD1,CEP63,SZT2,

,CAMLG,TCF4,TMPO,PRMT1,EXOSC9,STOML2,WNK1,ZNF7,PIGL,STRADA,ZFAND2B,HNRNPC,SHARPIN,ZN

1,RGS3,PSMA3,MUC1,METTL17,ECSIT,EPB41L5,ATG12,PPRC1,RPS25,NLRP1,ZBTB44,NIP7,COX4I1,DUSP

AP1S3,HNRNPC,SHARPIN,STAU1,SUGT1,UPP1,CARS,RPL36,MOB1A,MDH1,COPZ1,MAPKAPK5,DGUOK,H

OBSL1,TTC21B,PTPN13,UBR4,PSMA1,YLPM1,PHF21A,CHCHD1,SLC39A11,FUZ,RGS3,PSMA3,MUC1,METT

F3L,TSG101,OBSL1,TTC21B,PTPN13,UBR4,PSMA1,YLPM1,PHF21A,CHCHD1,SLC39A11,FUZ,RGS3,PSMA3,

YLPM1,CHCHD1,SLC39A11,FUZ,RGS3,PSMA3,MUC1,METTL17,ECSIT,EPB41L5,ATG12,RPS25,NLRP1,NIP7

RC1,PHF19,GRAMD1C,VCAN,PCBP2,LPCAT3,CD44,TM9SF1,MAGED2,SON,TET2,UQCRQ,SNF8,HMGA2,CD

MAGED2,SON,TET2,UQCRQ,SNF8,HMGA2,CDC37,RPS16,CPSF3L,TSG101,OBSL1,TTC21B,PTPN13,UBR4,

EXOSC9,STOML2,ARL6IP4,STRADA,PPAN,ACAD10,HNRNPC,HYPK,UPP1,RPL36,MOB1A,RBM3,LAMP2,IN

EXOSC9,STOML2,ARL6IP4,STRADA,PPAN,ACAD10,HNRNPC,HYPK,UPP1,RPL36,MOB1A,RBM3,LAMP2,IN

EXOSC9,STOML2,ARL6IP4,STRADA,PPAN,ACAD10,HNRNPC,HYPK,UPP1,RPL36,MOB1A,RBM3,LAMP2,IN

MTA1,CAPG,PPIL2,PSMD4,VIM,LIG1,PSMA7,CDC14B,XAF1,APEH,TXNL4A,NUTF2,PODXL,TCF4,TMPO,ATP

RD1,POT1,TRIM16,ANAPC11,POLR3H,RAD23A,USP45,PSME1,ILF3,EEF1D,HSPH1,SGK1,RRP8,OAS3,MED6

PRKRIP1,RANGRF,MTA3,DOCK10,MEIS2,HMGA1,PSME2,RBP1,RAD1,H2AFV,RBMX,BLVRB,PPP3CB,BRIP

UBXN1,HSPB1,ERCC5,EMC4,MCTS1,PRKRIP1,DDOST,LSR,MTA3,DCN,HMGA1,PSME2,PABPC1,PTPRB,NAT

FS7,IFI6,DCTN1,TIMM9,BIN1,UQCRB,PHF20,MRPL36,CLIP1,COX7C,NDUFS4,SLC25A45,CPNE1,NUDT9,GL

FS7,IFI6,DCTN1,TIMM9,BIN1,UQCRB,PHF20,MRPL36,CLIP1,COX7C,NDUFS4,SLC25A45,CPNE1,NUDT9,GL

RBP1,RAD1,H2AFV,RBMX,PPP3CB,RABEP2,TADA3,POT1,ANAPC11,POLR3H,RAD23A,SLC16A1,ILF3,PKP4,

RBP1,RAD1,H2AFV,RBMX,PPP3CB,RABEP2,TADA3,POT1,ANAPC11,POLR3H,RAD23A,SLC16A1,ILF3,PKP4,

JFB10,MPND,PPIE,RNMT,SMARCE1,DPY30,UBE2A,MED1,NDUFS7,SNRNP40,DYNC2LI1,DAD1,DCTN1,UQ

18,PPP1R13B,NDUFB10,ATPAF1,MRPL51,MRPL43,DHX30,IMMP1L,MRPL48,COQ4,ERAL1,ACOT9,MTHFD

MFF,SNX5,NUP133,COX20,MRPS17,TMEM147,TMEM14B,COA1,TGOLN2,PEX2,TNKS,PIGH,STX10,DCTN5,

4,FKBP5,MFSD10,MCFD2,LGALS3,TCP1,SYTL1,SCAP,CD74,PARK7,SLC30A5,CBR1,PIGT,UBE2N,LFNG,RPS1

C1,PHF19,GRAMD1C,VCAN,LPCAT3,CD44,TM9SF1,SON,TET2,UQCRQ,SNF8,GRK6,HMGA2,CDC37,RPS16,

QCRQ,SNF8,GRK6,HMGA2,CDC37,RPS16,CPSF3L,TSG101,OBSL1,TTC21B,PTPN13,UBR4,PSMA1,YLPM1,P

PCYT2,FBXW9,USE1,IFI27L1,ASF1B,CCDC25,EEF1B2,RFX3,DGKE,ENDOV,CAMTA1,PPIL2,PSMD4,RPL28,LI

K3,LZTFL1,ZC3H14,FIS1,SLC15A3,PPM1K,TUBA1C,ANTXR2,STIP1,LRRC1,PHF19,GRAMD1C,VCAN,LPCAT3

K3,LZTFL1,ZC3H14,FIS1,SLC15A3,PPM1K,TUBA1C,ANTXR2,STIP1,LRRC1,PHF19,GRAMD1C,VCAN,LPCAT3

TIP1,LRRC1,PHF19,GRAMD1C,VCAN,LPCAT3,CD44,TM9SF1,MAGED2,SON,TET2,UQCRQ,SNF8,GRK6,HM

TIP1,LRRC1,PHF19,GRAMD1C,VCAN,LPCAT3,CD44,TM9SF1,MAGED2,SON,TET2,UQCRQ,SNF8,GRK6,HM

D2,SON,TET2,UQCRQ,GRK6,HMGA2,CDC37,RPS16,CPSF3L,TSG101,OBSL1,TTC21B,PTPN13,UBR4,PSMA1

7,TTC9C,COX4I1,DUSP22,SLC25A10,TMX2,THEM4,CENPK,RPS6,RPL29,RTEL1,COASY,SRSF1,AUP1,SAP30

7,TTC9C,COX4I1,DUSP22,SLC25A10,TMX2,THEM4,CENPK,RPS6,RPL29,RTEL1,COASY,SRSF1,AUP1,SAP30

D1C,LPCAT3,CD44,TM9SF1,SON,TET2,UQCRQ,SNF8,GRK6,HMGA2,CDC37,RPS16,CPSF3L,TSG101,OBSL1,

SE1,HSBP1L1,IFI27L1,MAF,ASF1B,CCDC25,EEF1B2,RFX3,TNFRSF14,ENDOV,CAMTA1,PPIL2,RPL28,CLDND

SE1,HSBP1L1,IFI27L1,MAF,ASF1B,CCDC25,EEF1B2,RFX3,TNFRSF14,ENDOV,CAMTA1,PPIL2,RPL28,CLDND

,SLC9B2,SUGT1,HYPK,CARS,ZNF250,CDK11A,RBM3,ZNF197,COPZ1,INO80E,CCNL2,DGUOK,SYNGR2,LRR

RED1,RABL5,TRNT1,KXD1,ITGB1BP1,MED15,SH3YL1,IP6K2,CLASRP,C1ORF198,UGGT2,MICAL1,UBE2I,SL

RED1,RABL5,TRNT1,KXD1,ITGB1BP1,MED15,SH3YL1,IP6K2,CLASRP,C1ORF198,UGGT2,MICAL1,UBE2I,SL

ON,TET2,UQCRQ,SNF8,GRK6,CDC37,RPS16,CPSF3L,TSG101,OBSL1,TTC21B,PTPN13,UBR4,PSMA1,YLPM1

ON,TET2,UQCRQ,SNF8,GRK6,CDC37,RPS16,CPSF3L,TSG101,OBSL1,TTC21B,PTPN13,UBR4,PSMA1,YLPM1,

G,SH3YL1,IP6K2,CLASRP,C1ORF198,UGGT2,MICAL1,UBE2I,SLC39A13,PRMT2,EIF3M,CDC42,TBX6,CARD1

G,SH3YL1,IP6K2,CLASRP,C1ORF198,UGGT2,MICAL1,UBE2I,SLC39A13,PRMT2,EIF3M,CDC42,TBX6,CARD1

1B,CCDC25,EEF1B2,RFX3,TNFRSF14,ENDOV,CAMTA1,PPIL2,RPL28,CLDND1,LIG1,CDC14B,TIMM17B,TME

1B,CCDC25,EEF1B2,RFX3,TNFRSF14,ENDOV,CAMTA1,PPIL2,RPL28,CLDND1,LIG1,CDC14B,TIMM17B,TME

1,RABL5,TRNT1,KXD1,ITGB1BP1,MED15,PARD6G,SH3YL1,IP6K2,CLASRP,C1ORF198,UGGT2,MICAL1,UBE

1,RABL5,TRNT1,KXD1,ITGB1BP1,MED15,PARD6G,SH3YL1,IP6K2,CLASRP,C1ORF198,UGGT2,MICAL1,UBE

3H14,FIS1,SLC15A3,PPM1K,TUBA1C,DST,ANTXR2,STIP1,LRRC1,PHF19,VCAN,LPCAT3,CD44,TM9SF1,MA

3H14,FIS1,SLC15A3,PPM1K,TUBA1C,DST,ANTXR2,STIP1,LRRC1,PHF19,VCAN,LPCAT3,CD44,TM9SF1,MA

YIPF3,FOXRED1,RABL5,TRNT1,KXD1,ITGB1BP1,MED15,SH3YL1,IP6K2,CLASRP,C1ORF198,UGGT2,MICAL1

YIPF3,FOXRED1,RABL5,TRNT1,KXD1,ITGB1BP1,MED15,SH3YL1,IP6K2,CLASRP,C1ORF198,UGGT2,MICAL1

,METTL17,EPB41L5,ATG12,RPS25,NIP7,TTC9C,COX4I1,DUSP22,SLC25A10,TMX2,SPOCK2,THEM4,CENPK,

,METTL17,EPB41L5,ATG12,RPS25,NIP7,TTC9C,COX4I1,DUSP22,SLC25A10,TMX2,SPOCK2,THEM4,CENPK,

7,TXNDC17,ARL6IP4,C19ORF48,PPAN,ACAD10,CYB561,SHARPIN,SUGT1,HYPK,CARS,ZNF250,CDK11A,RB

7,TXNDC17,ARL6IP4,C19ORF48,PPAN,ACAD10,CYB561,SHARPIN,SUGT1,HYPK,CARS,ZNF250,CDK11A,RB

5,ATG12,RPS25,NIP7,TTC9C,COX4I1,DUSP22,SLC25A10,THEM4,CENPK,RPS6,RPL29,RTEL1,COASY,SRSF1,

5,ATG12,RPS25,NIP7,TTC9C,COX4I1,DUSP22,SLC25A10,THEM4,CENPK,RPS6,RPL29,RTEL1,COASY,SRSF1,

2,RPS25,NIP7,TTC9C,COX4I1,DUSP22,SLC25A10,TMX2,THEM4,DBNDD2,CENPK,RPS6,RPL29,RTEL1,COAS

2,RPS25,NIP7,TTC9C,COX4I1,DUSP22,SLC25A10,TMX2,THEM4,DBNDD2,CENPK,RPS6,RPL29,RTEL1,COAS

RQ,SNF8,GRK6,HMGA2,CDC37,RPS16,CPSF3L,TSG101,OBSL1,TTC21B,PTPN13,UBR4,YLPM1,PHF21A,CH

5A10,THEM4,CENPK,RPS6,RPL29,RTEL1,COASY,SRSF1,AUP1,SAP30BP,YIPF3,FOXRED1,RABL5,TRNT1,KXD

5A10,THEM4,CENPK,RPS6,RPL29,RTEL1,COASY,SRSF1,AUP1,SAP30BP,YIPF3,FOXRED1,RABL5,TRNT1,KXD

L2,C19ORF54,ZNF7,TXNDC17,ARL6IP4,C19ORF48,PPAN,ZFAND2B,AP1S3,ACAD10,HNRNPC,CYB561,SH

2,CDC37,RPS16,CPSF3L,TSG101,OBSL1,TTC21B,PTPN13,UBR4,PSMA1,YLPM1,PHF21A,CHCHD1,SLC39A1

2,CDC37,RPS16,CPSF3L,TSG101,OBSL1,TTC21B,PTPN13,UBR4,PSMA1,YLPM1,PHF21A,CHCHD1,SLC39A1

PS20,NIT2,MPDU1,RPS15,UBXN1,ERLEC1,HSPB1,SAT1,ERCC5,EMC4,LMBR1,MCTS1,SERPINB1,DDOST,L

D44,TM9SF1,SON,TET2,UQCRQ,SNF8,GRK6,HMGA2,CDC37,RPS16,CPSF3L,TSG101,OBSL1,TTC21B,PTPN

3H14,SLC15A3,PPM1K,TUBA1C,DST,ANTXR2,STIP1,LRRC1,PHF19,GRAMD1C,VCAN,LPCAT3,CD44,TM9S

3H14,SLC15A3,PPM1K,TUBA1C,DST,ANTXR2,STIP1,LRRC1,PHF19,GRAMD1C,VCAN,LPCAT3,CD44,TM9S

T2,EIF3M,CDC42,TBX6,TMEM205,CARD14,PCYT2,FBXW9,USE1,HSBP1L1,MAF,ASF1B,CCDC25,EEF1B2,R

T2,EIF3M,CDC42,TBX6,TMEM205,CARD14,PCYT2,FBXW9,USE1,HSBP1L1,MAF,ASF1B,CCDC25,EEF1B2,R

GB1BP1,MED15,SH3YL1,IP6K2,CLASRP,C1ORF198,UGGT2,MICAL1,UBE2I,SLC39A13,PRMT2,EIF3M,CDC4

GB1BP1,MED15,SH3YL1,IP6K2,CLASRP,C1ORF198,UGGT2,MICAL1,UBE2I,SLC39A13,PRMT2,EIF3M,CDC4

C37,RPS16,CPSF3L,TSG101,OBSL1,UBR4,YLPM1,PHF21A,CHCHD1,FUZ,FAM3A,PSMA3,METTL17,ECSIT,E

C37,RPS16,CPSF3L,TSG101,OBSL1,UBR4,YLPM1,PHF21A,CHCHD1,FUZ,FAM3A,PSMA3,METTL17,ECSIT,E

,PRMT2,EIF3M,CDC42,TBX6,TMEM205,PTK2,CARD14,PCYT2,FBXW9,USE1,HSBP1L1,IFI27L1,MAF,ASF1B,

,PRMT2,EIF3M,CDC42,TBX6,TMEM205,PTK2,CARD14,PCYT2,FBXW9,USE1,HSBP1L1,IFI27L1,MAF,ASF1B,

,EPB41L5,ATG12,RPS25,NIP7,TTC9C,COX4I1,DUSP22,SLC25A10,TMX2,THEM4,DBNDD2,CENPK,RPS6,RP

,EPB41L5,ATG12,RPS25,NIP7,TTC9C,COX4I1,DUSP22,SLC25A10,TMX2,THEM4,DBNDD2,CENPK,RPS6,RP

PG,PPIL2,RPL28,LIG1,PSMA7,CDC14B,TIMM17B,APEH,RPL21,CAPN10,LYPLA2,NUTF2,LSM12,C17ORF62

PG,PPIL2,RPL28,LIG1,PSMA7,CDC14B,TIMM17B,APEH,RPL21,CAPN10,LYPLA2,NUTF2,LSM12,C17ORF62

14,PCYT2,FBXW9,USE1,HSBP1L1,MAF,ASF1B,CCDC25,EEF1B2,RFX3,TNFRSF14,ENDOV,CAMTA1,CAPG,P

14,PCYT2,FBXW9,USE1,HSBP1L1,MAF,ASF1B,CCDC25,EEF1B2,RFX3,TNFRSF14,ENDOV,CAMTA1,CAPG,P

T3,TM9SF1,MAGED2,SON,TET2,UQCRQ,SNF8,GRK6,HMGA2,CDC37,RPS16,CPSF3L,TSG101,OBSL1,TTC21

0E,CCNL2,DGUOK,LRRFIP1,ZNF524,ATP5J,RMDN1,TRMT112,CHPF,STRIP2,RPS20,GABPB1,NIT2,MPDU1,

NB1,DDOST,LSR,PSME2,PABPC1,RRNAD1,DBNL,NAT9,RAD1,TOMM40,RABEP2,TADA3,ZNF20,ATP6V1F,

RMT2,EIF3M,CDC42,TBX6,CARD14,PCYT2,FBXW9,USE1,MAF,ASF1B,CCDC25,EEF1B2,PDLIM5,RFX3,END

RMT2,EIF3M,CDC42,TBX6,CARD14,PCYT2,FBXW9,USE1,MAF,ASF1B,CCDC25,EEF1B2,PDLIM5,RFX3,END
GB1BP1,MED15,PARD6G,SH3YL1,IP6K2,CLASRP,C1ORF198,UGGT2,MICAL1,UBE2I,SLC39A13,YIF1A,PRM
GB1BP1,MED15,PARD6G,SH3YL1,IP6K2,CLASRP,C1ORF198,UGGT2,MICAL1,UBE2I,SLC39A13,YIF1A,PRM
,TBX6,PTK2,CARD14,PCYT2,FBXW9,USE1,HSBP1L1,MAF,ASF1B,CCDC25,EEF1B2,RFX3,TNFRSF14,ENDOV,
,TBX6,PTK2,CARD14,PCYT2,FBXW9,USE1,HSBP1L1,MAF,ASF1B,CCDC25,EEF1B2,RFX3,TNFRSF14,ENDOV,
B,APEH,RPL21,CAPN10,LYPLA2,NUTF2,EPB41L4B,UCK2,LSM12,C17ORF62,PRMT1,CASC4,STOML2,ZNF7,
B,APEH,RPL21,CAPN10,LYPLA2,NUTF2,EPB41L4B,UCK2,LSM12,C17ORF62,PRMT1,CASC4,STOML2,ZNF7,
2,C17ORF62,PRMT1,STOML2,C19ORF54,ZNF7,TXNDC17,ARL6IP4,C19ORF48,PPAN,ZFAND2B,HNRNPC,C
2,C17ORF62,PRMT1,STOML2,C19ORF54,ZNF7,TXNDC17,ARL6IP4,C19ORF48,PPAN,ZFAND2B,HNRNPC,C
C39A13,YIF1A,PRMT2,EIF3M,CDC42,TBX6,PTK2,C14ORF2,CARD14,PCYT2,FBXW9,USE1,HSBP1L1,MAF,A
C39A13,YIF1A,PRMT2,EIF3M,CDC42,TBX6,PTK2,C14ORF2,CARD14,PCYT2,FBXW9,USE1,HSBP1L1,MAF,A
,EPB41L5,ATG12,RPS25,ZBTB44,NIP7,TTC9C,COX4I1,DUSP22,SLC25A10,TMX2,SPOCK2,THEM4,DBNDD2,
,EPB41L5,ATG12,RPS25,ZBTB44,NIP7,TTC9C,COX4I1,DUSP22,SLC25A10,TMX2,SPOCK2,THEM4,DBNDD2,
E2I,SLC39A13,PRMT2,EIF3M,CDC42,TBX6,PTK2,CARD14,PCYT2,FBXW9,USE1,HSBP1L1,IFI27L1,MAF,ASF1
E2I,SLC39A13,PRMT2,EIF3M,CDC42,TBX6,PTK2,CARD14,PCYT2,FBXW9,USE1,HSBP1L1,IFI27L1,MAF,ASF1
ZNF343,GBA2,PPIH,DALRD3,NSUN5,ZFPL1,MFSD10,MCFD2,RIC8A,TCP1,PIGT,UBE2N,MFSD11,LFNG,ZNH
,AUP1,SAP30BP,YIPF3,FOXRED1,RABL5,TRNT1,KXD1,ITGB1BP1,MED15,SH3YL1,IP6K2,CLASRP,C1ORF19
,AUP1,SAP30BP,YIPF3,FOXRED1,RABL5,TRNT1,KXD1,ITGB1BP1,MED15,SH3YL1,IP6K2,CLASRP,C1ORF19
,DGKE,ENDOV,CAMTA1,CAPG,PPIL2,PSMD4,LIG1,PSMA7,CDC14B,TMEM126B,APEH,RPL21,TXNL4A,CAP
,DGKE,ENDOV,CAMTA1,CAPG,PPIL2,PSMD4,LIG1,PSMA7,CDC14B,TMEM126B,APEH,RPL21,TXNL4A,CAP
MICAL1,UBE2I,SLC39A13,YIF1A,EIF3M,CDC42,TBX6,TMEM205,CARD14,PCYT2,FBXW9,USE1,MAF,ASF1B,
MICAL1,UBE2I,SLC39A13,YIF1A,EIF3M,CDC42,TBX6,TMEM205,CARD14,PCYT2,FBXW9,USE1,MAF,ASF1B,
GT2,MICAL1,UBE2I,SLC39A13,YIF1A,PRMT2,EIF3M,CDC42,TBX6,TMEM205,PTK2,CARD14,PCYT2,FBXW9,
GT2,MICAL1,UBE2I,SLC39A13,YIF1A,PRMT2,EIF3M,CDC42,TBX6,TMEM205,PTK2,CARD14,PCYT2,FBXW9,
D2,RIC8A,TCP1,PARK7,THADA,PIGT,UBE2N,LFNG,ZNHIT3,XYLT2,ZNF81,FAM32A,ZSCAN20,COX20,COA1,T
,EIF3M,CDC42,TBX6,TMEM205,CARD14,PCYT2,FBXW9,USE1,HSBP1L1,MAF,ASF1B,CCDC25,EEF1B2,RFX3
,EIF3M,CDC42,TBX6,TMEM205,CARD14,PCYT2,FBXW9,USE1,HSBP1L1,MAF,ASF1B,CCDC25,EEF1B2,RFX3
,YIPF3,FOXRED1,TRNT1,KXD1,ITGB1BP1,PARD6G,SH3YL1,IP6K2,CLASRP,JPH2,C1ORF198,MICAL1,HS2ST
,YIPF3,FOXRED1,TRNT1,KXD1,ITGB1BP1,PARD6G,SH3YL1,IP6K2,CLASRP,JPH2,C1ORF198,MICAL1,HS2ST
EC31A,CARHSP1,CNST,COX17,COX5A,PHF20L1,HIGD1A,FKBP3,RPL15,INTS4,B3GAT3,ZNF506,PFDN5,C11
0,PIGT,UBE2N,LFNG,ZNHIT3,PDCD6,FAM32A,ZSCAN20,COX20,PEX2,POLR1C,PIGH,STX10,C12ORF57,DCT
25,ZBTB44,NIP7,TTC9C,COX4I1,DUSP22,SLC25A10,TMX2,SPOCK2,THEM4,DBNDD2,TMED4,RPS6,RPL29,
25,ZBTB44,NIP7,TTC9C,COX4I1,DUSP22,SLC25A10,TMX2,SPOCK2,THEM4,DBNDD2,TMED4,RPS6,RPL29,
A1,YLPM1,PHF21A,RGS3,FAM3A,PSMA3,METTL17,ECSIT,EPB41L5,ATG12,RPS25,ZBTB44,NIP7,TTC9C,CO
A1,YLPM1,PHF21A,RGS3,FAM3A,PSMA3,METTL17,ECSIT,EPB41L5,ATG12,RPS25,ZBTB44,NIP7,TTC9C,CO
L5A3,PPM1K,TUBA1C,IRF9,ANTXR2,STIP1,LRRC1,PHF19,GRAMD1C,VCAN,LPCAT3,CD44,TM9SF1,SON,TE
L5A3,PPM1K,TUBA1C,IRF9,ANTXR2,STIP1,LRRC1,PHF19,GRAMD1C,VCAN,LPCAT3,CD44,TM9SF1,SON,TE
,COX17,COX5A,HIGD1A,INTS4,B3GAT3,ZNF506,PFDN5,PAFAH1B2,ZNF343,GBA2,PPIH,DALRD3,NSUN5,D
M3A,PSMA3,METTL17,ECSIT,EPB41L5,ATG12,RPS25,ZBTB44,NIP7,TTC9C,COX4I1,DUSP22,SLC25A10,TM
M3A,PSMA3,METTL17,ECSIT,EPB41L5,ATG12,RPS25,ZBTB44,NIP7,TTC9C,COX4I1,DUSP22,SLC25A10,TM
K6,HMGA2,CDC37,RPS16,CPSF3L,TSG101,OBSL1,TTC21B,PTPN13,UBR4,YLPM1,PHF21A,CHCHD1,SLC39
K6,HMGA2,CDC37,RPS16,CPSF3L,TSG101,OBSL1,TTC21B,PTPN13,UBR4,YLPM1,PHF21A,CHCHD1,SLC39
NL,NAT9,RAD1,TOMM40,RABEP2,TADA3,ZNF20,ATP6V1F,WAC,KLK10,TRIM16,ANAPC11,RAD23A,ABHD
LA2,LSM12,C17ORF62,PRMT1,CASC4,STOML2,C19ORF54,ZNF7,TXNDC17,ARL6IP4,PIGL,C19ORF48,PPA
LA2,LSM12,C17ORF62,PRMT1,CASC4,STOML2,C19ORF54,ZNF7,TXNDC17,ARL6IP4,PIGL,C19ORF48,PPA
EX26,NDUFS3,ARL16,OXLD1,STAU2,METTL5,SLC35E1,OGFOD2,METTL3,TSEN34,SSR3,ARIH2,RPL8,PICK1
AMK2D,HNRNPLL,HIGD1A,POLR2C,SIRT5,INTS4,B3GAT3,SPINT2,PFDN5,MTHFS,SMARCA4,C11ORF49,PA

MAPKAPK5,DGUOK,LRRFIP1,ZNF524,ATP5J,RMDN1,GGCX,TRMT112,CHPF,RPS20,NIT2,MPDU1,RPS15,D
K2,CLASRP,C1ORF198,UGGT2,MICAL1,UBE2I,SLC39A13,PRMT2,EIF3M,CDC42,TBX6,C14ORF2,GUK1,CAR
K2,CLASRP,C1ORF198,UGGT2,MICAL1,UBE2I,SLC39A13,PRMT2,EIF3M,CDC42,TBX6,C14ORF2,GUK1,CAR
AP30BP,YIPF3,FOXRED1,RABL5,TRNT1,KXD1,ITGB1BP1,MED15,SH3YL1,IP6K2,CLASRP,C1ORF198,UGGT2,
AP30BP,YIPF3,FOXRED1,RABL5,TRNT1,KXD1,ITGB1BP1,MED15,SH3YL1,IP6K2,CLASRP,C1ORF198,UGGT2,
APKAPK5,DGUOK,HSP90AB1,DUT,LRRFIP1,ZNF524,ATP5J,ELP6,RMDN1,TRMT112,CHPF,RPS20,MPDU1,D
,EPB41L5,ATG12,RPS25,NIP7,TTC9C,COX4I1,DUSP22,SLC25A10,TMX2,SPOCK2,THEM4,DBNDD2,RPS6,RP
,EPB41L5,ATG12,RPS25,NIP7,TTC9C,COX4I1,DUSP22,SLC25A10,TMX2,SPOCK2,THEM4,DBNDD2,RPS6,RP
2,ZNF343,GBA2,PPIH,DALRD3,NSUN5,DRG2,ZFPL1,MFSD10,GMPR2,MCFD2,RIC8A,TCP1,SLC30A5,THAD
9SF1,SON,TET2,UQCRQ,SNF8,GRK6,HMGA2,CDC37,RPS16,CPSF3L,TSG101,OBSL1,TTC21B,PTPN13,UBR4,
9SF1,SON,TET2,UQCRQ,SNF8,GRK6,HMGA2,CDC37,RPS16,CPSF3L,TSG101,OBSL1,TTC21B,PTPN13,UBR4,
0,PEX2,POLR1C,PIGH,STX10,C12ORF57,DCTN5,PEX26,NDUFS3,OXLD1,EIF4A1,METTL5,SLC35E1,OGFOD
IED4,CD99L2,GGPS1,AUP1,YIPF3,FOXRED1,TRNT1,KXD1,ITGB1BP1,PARD6G,SH3YL1,IP6K2,CLASRP,JPH2,
IED4,CD99L2,GGPS1,AUP1,YIPF3,FOXRED1,TRNT1,KXD1,ITGB1BP1,PARD6G,SH3YL1,IP6K2,CLASRP,JPH2,
2,GBA2,DALRD3,NSUN5,OARD1,MFSD10,MRPS18C,GMPR2,MCFD2,ZSCAN9,RIC8A,TCP1,PARK7,SMARC
2,GBA2,DALRD3,NSUN5,OARD1,MFSD10,MRPS18C,GMPR2,MCFD2,ZSCAN9,RIC8A,TCP1,PARK7,SMARC
J,RMDN1,TRMT112,CHPF,STRIP2,RPS20,GABPB1,NIT2,DCTD,UBXN1,ERLEC1,HSPB1,SAT1,ERCC5,EMC4,
SPOCK2,THEM4,DBNDD2,TMED4,CENPK,RPS6,RPL29,CD99L2,GGPS1,RTEL1,COASY,SRSF1,AUP1,YIPF3,FC
3,RPL15,SIRT5,INTS4,B3GAT3,RFXANK,ZNF506,SPINT2,PFDN5,C11ORF49,PAFAH1B2,ZNF343,GBA2,PPIH,
L,ITPA,ZCCHC9,VPS25,SNRPD1,MKRN1,SMUG1,HMOX2,CHURC1,SNRPA1,CDK2AP2,AZI2,GOSR2,MYL12A
S,CDK11A,RBM3,ZNF197,COPZ1,INO80E,CCNL2,MAPKAPK5,DGUOK,HSP90AB1,DUT,LRRFIP1,ZNF524,AT
E,CCNL2,MAPKAPK5,DGUOK,HSP90AB1,LRRFIP1,RAB2A,ATP5J,TSFM,TRMT112,CHPF,RPS20,GABPB1,DC
E,CCNL2,MAPKAPK5,DGUOK,HSP90AB1,LRRFIP1,RAB2A,ATP5J,TSFM,TRMT112,CHPF,RPS20,GABPB1,DC
8A,TCP1,PARK7,SMARCC2,UBE2N,LFNG,PDCD6,ZNF81,FAM32A,TMEM147,COA1,USP40,PCID2,TSC22D3
8A,TCP1,PARK7,SMARCC2,UBE2N,LFNG,PDCD6,ZNF81,FAM32A,TMEM147,COA1,USP40,PCID2,TSC22D3
C14B,TIMM17B,APEH,CAPN10,LYPLA2,EPB41L4B,UCK2,C17ORF62,PRMT1,STOML2,C19ORF54,ZNF7,TXN
C14B,TIMM17B,APEH,CAPN10,LYPLA2,EPB41L4B,UCK2,C17ORF62,PRMT1,STOML2,C19ORF54,ZNF7,TXN
HCHD1,SLC39A11,FUZ,FAM3A,PSMA3,MUC1,METTL17,ECSIT,EPB41L5,ATG12,PPRC1,RPS25,ZBTB44,NIP
HCHD1,SLC39A11,FUZ,FAM3A,PSMA3,MUC1,METTL17,ECSIT,EPB41L5,ATG12,PPRC1,RPS25,ZBTB44,NIP
ORF49,PAFAH1B2,ZNF343,GBA2,PPIH,DALRD3,NSUN5,DRG2,ZFPL1,MFSD10,MCFD2,RIC8A,TCP1,THADA
RLEC1,HSPB1,SAT1,ERCC5,MCTS1,NAP1L1,SERPINB1,DDOST,LSR,PSME2,PHTF2,PABPC1,RRNAD1,DBNL,
IGA2,CDC37,RPS16,CPSF3L,TSG101,OBSL1,TTC21B,PTPN13,UBR4,PSMA1,YLPM1,PHF21A,CHCHD1,SLC39
IGA2,CDC37,RPS16,CPSF3L,TSG101,OBSL1,TTC21B,PTPN13,UBR4,PSMA1,YLPM1,PHF21A,CHCHD1,SLC39
GD1A,INTS4,ZNF506,SPINT2,PFDN5,MTHFS,SMARCA4,C11ORF49,PAFAH1B2,ZNF343,PTK7,GBA2,PPIH,D
,CHEK1,COX17,HNRNPLL,PHF20L1,HIGD1A,POLR2C,B3GAT3,RFXANK,SPINT2,FTH1,PFDN5,PAFAH1B2,TX
,CHEK1,COX17,HNRNPLL,PHF20L1,HIGD1A,POLR2C,B3GAT3,RFXANK,SPINT2,FTH1,PFDN5,PAFAH1B2,TX
APEH,TXNL4A,CAPN10,LYPLA2,ATPAF2,C17ORF62,PRMT1,STOML2,ZNF7,TXNDC17,ARL6IP4,C19ORF48,S
APEH,TXNL4A,CAPN10,LYPLA2,ATPAF2,C17ORF62,PRMT1,STOML2,ZNF7,TXNDC17,ARL6IP4,C19ORF48,S
COX17,HNRNPLL,PHF20L1,HIGD1A,POLR2C,B3GAT3,RFXANK,ZNF506,SPINT2,FTH1,PFDN5,PAFAH1B2,TX
COX17,HNRNPLL,PHF20L1,HIGD1A,POLR2C,B3GAT3,RFXANK,ZNF506,SPINT2,FTH1,PFDN5,PAFAH1B2,TX
0AB1,RAB2A,ATP5J,ELP6,TSFM,TRMT112,PNPO,CHPF,RPS20,VPS28,DCTD,UBXN1,ZNF814,HSPB1,VWA2,
0AB1,RAB2A,ATP5J,ELP6,TSFM,TRMT112,PNPO,CHPF,RPS20,VPS28,DCTD,UBXN1,ZNF814,HSPB1,VWA2,
NF7,TXNDC17,PPAN,ACAD10,CYB561,SHARPIN,SUGT1,HYPK,CARS,ZNF250,CDK11A,ZNF197,INO80E,CCN
NF7,TXNDC17,PPAN,ACAD10,CYB561,SHARPIN,SUGT1,HYPK,CARS,ZNF250,CDK11A,ZNF197,INO80E,CCN
PSF3L,TSG101,OBSL1,TTC21B,PTPN13,UBR4,PSMA1,YLPM1,PHF21A,CHCHD1,SLC39A11,FUZ,FAM3A,MU
PSF3L,TSG101,OBSL1,TTC21B,PTPN13,UBR4,PSMA1,YLPM1,PHF21A,CHCHD1,SLC39A11,FUZ,FAM3A,MU
,RAD23A,NT5C,MDH2,STXBP2,EEF1D,CHCHD5,MED6,SEC31A,CARHSP1,RPL18,CNST,COX17,PHF20L1,HI

T1,ERCC5,SERPINB1,PRKRIP1,DDOST,LSR,PSME2,PABPC1,RRNAD1,DBNL,RAD1,TOMM40,RABEP2,TADA
,CHPF,MPDU1,RPS15,DCTD,UBXN1,ZNF814,HSPB1,SAT1,ERCC5,EMC4,MCTS1,DDOST,LSR,MEIS2,RRNAD
,CHPF,MPDU1,RPS15,DCTD,UBXN1,ZNF814,HSPB1,SAT1,ERCC5,EMC4,MCTS1,DDOST,LSR,MEIS2,RRNAD
,GABPB1,NIT2,LYNX1,ERLEC1,HSPB1,SAT1,ERCC5,MCTS1,NAP1L1,SERPINB1,DDOST,LSR,MTA3,PSME2,P
2,RAB2A,ATP5J,RMDN1,TRMT112,CHPF,RPS20,GABPB1,ZNF814,ERLEC1,HSPB1,MVD,ERCC5,EMC4,DDO
2,RAB2A,ATP5J,RMDN1,TRMT112,CHPF,RPS20,GABPB1,ZNF814,ERLEC1,HSPB1,MVD,ERCC5,EMC4,DDO
N,LFNG,ZNHIT3,FAM32A,ZSCAN20,COX20,COA1,PEX2,POLR1C,STX10,C12ORF57,DCTN5,PEX26,NDUFS3,
37,RPS16,CPSF3L,TSG101,OBSL1,TTC21B,PTPN13,UBR4,PSMA1,YLPM1,PHF21A,CHCHD1,FUZ,RGS3,FAM
H1,INO80E,CCNL2,MAPKAPK5,DGUOK,DUT,SYNGR2,LRRFIP1,RAB2A,ATP5J,ELP6,RMDN1,ASPH,TRMT112
CD44,TM9SF1,MAGED2,SON,TET2,UQCRQ,SNF8,GRK6,HMGA2,CDC37,RPS16,CPSF3L,TSG101,OBSL1,TT
PS18C,GMPR2,ZSCAN9,RIC8A,TCP1,SMARCC2,THADA,UBE2N,MFSD11,LFNG,PDCD6,GPR56,ENY2,ZNF81
PS18C,GMPR2,ZSCAN9,RIC8A,TCP1,SMARCC2,THADA,UBE2N,MFSD11,LFNG,PDCD6,GPR56,ENY2,ZNF81
P22,SLC25A10,TMX2,SPOCK2,THEM4,TMED4,CENPK,RPS6,RPL29,GGPS1,RTEL1,COASY,AUP1,SAP30BP,Y
P22,SLC25A10,TMX2,SPOCK2,THEM4,TMED4,CENPK,RPS6,RPL29,GGPS1,RTEL1,COASY,AUP1,SAP30BP,Y
,EEF1B2,PPP1CC,RFX3,DGKE,ENDOV,VAMP8,CAPG,PPIL2,PSMD4,TROAP,RPL28,LIG1,PSMA7,CDC14B,TI
,EEF1B2,PPP1CC,RFX3,DGKE,ENDOV,VAMP8,CAPG,PPIL2,PSMD4,TROAP,RPL28,LIG1,PSMA7,CDC14B,TI
,EEF1B2,PPP1CC,RFX3,DGKE,ENDOV,VAMP8,CAPG,PPIL2,PSMD4,TROAP,RPL28,LIG1,PSMA7,CDC14B,TI
,EEF1B2,PPP1CC,RFX3,DGKE,ENDOV,VAMP8,CAPG,PPIL2,PSMD4,TROAP,RPL28,LIG1,PSMA7,CDC14B,TI
RD2,GBA2,DALRD3,NSUN5,OARD1,MFSD10,MRPS18C,GMPR2,MCFD2,ZSCAN9,RIC8A,TCP1,PARK7,SM
RD2,GBA2,DALRD3,NSUN5,OARD1,MFSD10,MRPS18C,GMPR2,MCFD2,ZSCAN9,RIC8A,TCP1,PARK7,SM
E,CCNL2,MAPKAPK5,DGUOK,HSP90AB1,LRRFIP1,RAB2A,ZNF524,ATP5J,ELP6,TRMT112,CHPF,RPS20,DCT
E,CCNL2,MAPKAPK5,DGUOK,HSP90AB1,LRRFIP1,RAB2A,ZNF524,ATP5J,ELP6,TRMT112,CHPF,RPS20,DCT
SUN5,ZFPL1,OARD1,MFSD10,GMPR2,MCFD2,RIC8A,TCP1,PARK7,THADA,COMMD4,PIGT,UBE2N,NFATC4,
31A,CARHSP1,CNST,COX17,HIGD1A,POLR2C,RPL15,SPINT2,PFDN5,PAFAH1B2,COA3,DALRD3,NSUN5,ZF
31A,CARHSP1,CNST,COX17,HIGD1A,POLR2C,RPL15,SPINT2,PFDN5,PAFAH1B2,COA3,DALRD3,NSUN5,ZF
TEL1,COASY,SRSF1,AUP1,SAP30BP,YIPF3,FOXRED1,RABL5,TRNT1,KXD1,ITGB1BP1,MED15,SH3YL1,IP6K2,
TEL1,COASY,SRSF1,AUP1,SAP30BP,YIPF3,FOXRED1,RABL5,TRNT1,KXD1,ITGB1BP1,MED15,SH3YL1,IP6K2,
M32A,TMEM147,COA1,USP40,PCID2,PEX2,POLR1C,TNKS,RPL37A,C12ORF57,DCTN5,PEX26,NDUFS3,OXL
M32A,TMEM147,COA1,USP40,PCID2,PEX2,POLR1C,TNKS,RPL37A,C12ORF57,DCTN5,PEX26,NDUFS3,OXL
PTK2,CARD14,PCYT2,FBXW9,USE1,HSBP1L1,IFI27L1,MAF,ASF1B,CCDC25,EEF1B2,RFX3,PPA1,ENDOV,VA
PTK2,CARD14,PCYT2,FBXW9,USE1,HSBP1L1,IFI27L1,MAF,ASF1B,CCDC25,EEF1B2,RFX3,PPA1,ENDOV,VA
4,NIP7,TTC9C,COX4I1,DUSP22,SLC25A10,TMX2,SPOCK2,THEM4,DBNDD2,RPS6,RPL29,GGPS1,RTEL1,CO
4,NIP7,TTC9C,COX4I1,DUSP22,SLC25A10,TMX2,SPOCK2,THEM4,DBNDD2,RPS6,RPL29,GGPS1,RTEL1,CO
,CARHSP1,COX17,HNRNPLL,HIGD1A,POLR2C,SIRT5,B3GAT3,RFXANK,SPINT2,PFDN5,MTHFS,PAFAH1B2,T
,CARHSP1,COX17,HNRNPLL,HIGD1A,POLR2C,SIRT5,B3GAT3,RFXANK,SPINT2,PFDN5,MTHFS,PAFAH1B2,T
DC42,TBX6,CARD14,PCYT2,FBXW9,USE1,MAF,ASF1B,CCDC25,EEF1B2,EDF1,RFX3,TNFRSF14,ENDOV,VA
DC42,TBX6,CARD14,PCYT2,FBXW9,USE1,MAF,ASF1B,CCDC25,EEF1B2,EDF1,RFX3,TNFRSF14,ENDOV,VA
1,MAGED2,SON,TET2,UQCRQ,SNF8,GRK6,HMGA2,CDC37,RPS16,CPSF3L,TSG101,OBSL1,TTC21B,PTPN13
CSIT,PPRC1,RPS25,NIP7,TTC9C,COX4I1,SLC25A10,TMX2,SPOCK2,THEM4,DBNDD2,TMED4,RPS6,RPL29,G
CSIT,PPRC1,RPS25,NIP7,TTC9C,COX4I1,SLC25A10,TMX2,SPOCK2,THEM4,DBNDD2,TMED4,RPS6,RPL29,G
XD1,ITGB1BP1,PARD6G,SH3YL1,IP6K2,CLASRP,JPH2,C1ORF198,MICAL1,HS2ST1,UBE2I,KCNMA1,SLC39A
XD1,ITGB1BP1,PARD6G,SH3YL1,IP6K2,CLASRP,JPH2,C1ORF198,MICAL1,HS2ST1,UBE2I,KCNMA1,SLC39A
M32A,ZSCAN20,COX20,COA1,PEX2,POLR1C,STX10,C12ORF57,DCTN5,PEX26,NDUFS3,OXLD1,EIF4A1,ST
1,TRNT1,KXD1,ITGB1BP1,PARD6G,SH3YL1,IP6K2,CLASRP,JPH2,C1ORF198,MICAL1,HS2ST1,UBE2I,KCNM
1,TRNT1,KXD1,ITGB1BP1,PARD6G,SH3YL1,IP6K2,CLASRP,JPH2,C1ORF198,MICAL1,HS2ST1,UBE2I,KCNM
GBA2,NSUN5,MFSD10,MRPS18C,MCFD2,ZSCAN9,RIC8A,PARK7,MROH1,UBE2N,LFNG,ZNHIT3,PDCD6,ZN
GBA2,NSUN5,MFSD10,MRPS18C,MCFD2,ZSCAN9,RIC8A,PARK7,MROH1,UBE2N,LFNG,ZNHIT3,PDCD6,ZN

5,OARD1,MFSD10,GMPR2,ZSCAN9,RIC8A,TCP1,PARK7,UBE2N,LFNG,PDCD6,ZNF81,FAM32A,TMEM147,C
5,OARD1,MFSD10,GMPR2,ZSCAN9,RIC8A,TCP1,PARK7,UBE2N,LFNG,PDCD6,ZNF81,FAM32A,TMEM147,C
HCHD1,SLC39A11,FUZ,FAM3A,MUC1,METTL17,ECSIT,EPB41L5,ATG12,PPRC1,RPS25,NLRP1,ZBTB44,NIP7
HCHD1,SLC39A11,FUZ,FAM3A,MUC1,METTL17,ECSIT,EPB41L5,ATG12,PPRC1,RPS25,NLRP1,ZBTB44,NIP7
RIH2,RPL8,PICK1,FBXL18,RNF181,TPT1,NME1,SLC52A2,NTMT1,TSPAN31,MTIF2,RALY,ZNF593,GLYCTK,TR
ATPAF2,C17ORF62,PRMT1,STOML2,WNK1,ZNF7,ARL6IP4,STRADA,PPAN,AP1S3,ACAD10,CYB561,SHARPI
ATPAF2,C17ORF62,PRMT1,STOML2,WNK1,ZNF7,ARL6IP4,STRADA,PPAN,AP1S3,ACAD10,CYB561,SHARPI
19,GRAMD1C,VCAN,LPCAT3,CD44,TM9SF1,MAGED2,SON,TET2,UQCRQ,GRK6,HMGA2,CDC37,CPSF3L,TSG
19,GRAMD1C,VCAN,LPCAT3,CD44,TM9SF1,MAGED2,SON,TET2,UQCRQ,GRK6,HMGA2,CDC37,CPSF3L,TSG
MTA1,CAPG,PPIL2,TROAP,LIG1,CDC14B,TIMM17B,TMEM126B,APEH,LYPLA2,NUTF2,CAMLG,LSM12,C17
MTA1,CAPG,PPIL2,TROAP,LIG1,CDC14B,TIMM17B,TMEM126B,APEH,LYPLA2,NUTF2,CAMLG,LSM12,C17
IRF9,DST,ANTXR2,STIP1,LRRC1,PHF19,GRAMD1C,VCAN,PCBP2,LPCAT3,CD44,TM9SF1,MAGED2,SON,TET
IRF9,DST,ANTXR2,STIP1,LRRC1,PHF19,GRAMD1C,VCAN,PCBP2,LPCAT3,CD44,TM9SF1,MAGED2,SON,TET
DALRD3,NSUN5,DRG2,ZFPL1,OARD1,MFSD10,MCFD2,RIC8A,TCP1,THADA,PIGT,UBE2N,LFNG,ZNHIT3,FAM
5,TRNT1,KXD1,ITGB1BP1,PARD6G,SH3YL1,IP6K2,CLASRP,JPH2,C1ORF198,UGGT2,MICAL1,HS2ST1,UBE2I
5,TRNT1,KXD1,ITGB1BP1,PARD6G,SH3YL1,IP6K2,CLASRP,JPH2,C1ORF198,UGGT2,MICAL1,HS2ST1,UBE2I
B3GAT3,RFXANK,ZNF506,PFDN5,C11ORF49,PAFAH1B2,ZNF343,GBA2,PPIH,ARHGEF1,DALRD3,NSUN5,DR
K,CARS,ZNF250,RBM3,MDH1,ZNF197,COPZ1,INO80E,DGUOK,HSP90AB1,SYNGR2,RAB2A,ZNF524,ATP5J,F
K,CARS,ZNF250,RBM3,MDH1,ZNF197,COPZ1,INO80E,DGUOK,HSP90AB1,SYNGR2,RAB2A,ZNF524,ATP5J,F
LYPLA2,LSM12,C17ORF62,PRMT1,STOML2,ZNF7,TXNDC17,ARL6IP4,PIGL,C19ORF48,PPAN,CYB561,SHAR
LYPLA2,LSM12,C17ORF62,PRMT1,STOML2,ZNF7,TXNDC17,ARL6IP4,PIGL,C19ORF48,PPAN,CYB561,SHAR
ZNF625,RPS20,GABPB1,DCTD,UBXN1,EMC4,PRKRIP1,DDOST,LSR,CHCHD7,PSME2,PABPC1,RRNAD1,DBNL
CARD14,FBXW9,USE1,MAF,ASF1B,CCDC25,EEF1B2,PPP1CC,RFX3,DGKE,ENDOV,CAMTA1,CAPG,PPIL2,PSM
CARD14,FBXW9,USE1,MAF,ASF1B,CCDC25,EEF1B2,PPP1CC,RFX3,DGKE,ENDOV,CAMTA1,CAPG,PPIL2,PSM
FAM32A,TMEM147,COA1,USP40,PCID2,PEX2,POLR1C,TNKS,STX10,C12ORF57,DCTN5,PEX26,ZNF223,OGF
FAM32A,TMEM147,COA1,USP40,PCID2,PEX2,POLR1C,TNKS,STX10,C12ORF57,DCTN5,PEX26,ZNF223,OGF
S3,ARL16,OXLD1,EIF3G,ARIH2,PICK1,FBXL18,ZNF512,RNF181,CHTOP,NTMT1,TSPAN31,GCDH,RALY,ZNF5
L3,EIF3G,SSR3,ARIH2,PICK1,FBXL18,NME1,SLC52A2,NTMT1,TSPAN31,FAM192A,ZNF593,GLYCTK,CFLAR,
L3,EIF3G,SSR3,ARIH2,PICK1,FBXL18,NME1,SLC52A2,NTMT1,TSPAN31,FAM192A,ZNF593,GLYCTK,CFLAR,
ORF57,DCTN5,PEX26,NDUFS3,ARL16,OXLD1,ZNF223,OGFOD2,METTL3,EIF3G,ARIH2,PICK1,FBXL18,ZNF5
ORF57,DCTN5,PEX26,NDUFS3,ARL16,OXLD1,ZNF223,OGFOD2,METTL3,EIF3G,ARIH2,PICK1,FBXL18,ZNF5
M32A,TMEM147,COA1,USP40,PCID2,PEX2,POLR1C,TNKS,STX10,C12ORF57,DCTN5,PEX26,OXLD1,ZNF223,
M32A,TMEM147,COA1,USP40,PCID2,PEX2,POLR1C,TNKS,STX10,C12ORF57,DCTN5,PEX26,OXLD1,ZNF223,
A,ACAA1,TRAPPC2,TAF1D,PIGO,KDELR3,AXIN1,HNRNPUL1,TP53,TMEM199,UBE2D3,C1ORF63,TKT,EPN1,I
CLTA,AXIN1,TMEM199,C1ORF63,TKT,EPN1,HPS1,ITPA,EML3,ZCCHC9,VPS25,SNRPD1,UCHL5,CHURC1,SNF
RPL15,SIRT5,INTS4,B3GAT3,RFXANK,ZNF506,PFDN5,C11ORF49,PAFAH1B2,ZNF343,GBA2,PPIH,ARHGEF1,
2,GGPS1,RTEL1,AUP1,YIPF3,FOXRED1,TRNT1,KXD1,PARD6G,SH3YL1,IP6K2,CLASRP,JPH2,C1ORF198,MICA
2,GGPS1,RTEL1,AUP1,YIPF3,FOXRED1,TRNT1,KXD1,PARD6G,SH3YL1,IP6K2,CLASRP,JPH2,C1ORF198,MICA
D22A,LAMTOR1,NCBP2,TTLL3,BCL7B,IAH1,ZNF581,ZNF276,RPL30,SZT2,PSMC5,IK,LRP10,SLC31A1,ACTB,
L1,TTC21B,PTPN13,UBR4,PSMA1,YLPM1,PHF21A,CHCHD1,SLC39A11,FAM3A,PSMA3,MUC1,ECSIT,EPB41
RNPLL,HINT1,CD151,HIGD1A,POLR2C,B3GAT3,RFXANK,SPINT2,FTH1,PFDN5,MTHFS,PAFAH1B2,TXNRD2,
RNPLL,HINT1,CD151,HIGD1A,POLR2C,B3GAT3,RFXANK,SPINT2,FTH1,PFDN5,MTHFS,PAFAH1B2,TXNRD2,
T2,MICAL1,HS2ST1,UBE2I,SLC39A13,YIF1A,PRMT2,EIF3M,MAGT1,JUP,TBX6,TMEM205,XBP1,PTK2,AUNIP
T2,MICAL1,HS2ST1,UBE2I,SLC39A13,YIF1A,PRMT2,EIF3M,MAGT1,JUP,TBX6,TMEM205,XBP1,PTK2,AUNIP
,ZC3H14,FIS1,SLC15A3,PPM1K,TUBA1C,DST,ANTXR2,STIP1,PHF19,GRAMD1C,VCAN,LPCAT3,CD44,TM9SF
,ZC3H14,FIS1,SLC15A3,PPM1K,TUBA1C,DST,ANTXR2,STIP1,PHF19,GRAMD1C,VCAN,LPCAT3,CD44,TM9SF
6,CD99L2,RTEL1,COASY,AUP1,FOXRED1,RABL5,TRNT1,KXD1,ITGB1BP1,PARD6G,SH3YL1,CLASRP,C1ORF1

1C,IRF9,DST,ANTXR2,STIP1,LRRC1,PHF19,GRAMD1C,VCAN,PCBP2,LPCAT3,CD44,TM9SF1,MAGED2,SON,T
1C,IRF9,DST,ANTXR2,STIP1,LRRC1,PHF19,GRAMD1C,VCAN,PCBP2,LPCAT3,CD44,TM9SF1,MAGED2,SON,T
1,ZNF223,METTL5,SLC35E1,OGFOD2,METTL3,SSR3,MPV17,ARIH2,PICK1,FBXL18,ZNF512,POLR2M,SLC52
1,ZNF223,METTL5,SLC35E1,OGFOD2,METTL3,SSR3,MPV17,ARIH2,PICK1,FBXL18,ZNF512,POLR2M,SLC52
C9,VPS25,SNRPD1,MKRN1,SMUG1,CHURC1,SNRPA1,AZI2,GOSR2,MYL12A,TBC1D22A,RPS7,LAMTOR1,U
RNMT,AP4M1,MRPL27,ZNF616,GANC,DDX56,CHFR,SSH3,ZNF655,C15ORF40,MED17,C12ORF10,RPS14,R
N,AP1S3,ACAD10,CYB561,SHARPIN,SUGT1,HYPK,CARS,ZNF250,RPL36,CDK11A,LAMP2,ZNF197,INO80E,C
N,AP1S3,ACAD10,CYB561,SHARPIN,SUGT1,HYPK,CARS,ZNF250,RPL36,CDK11A,LAMP2,ZNF197,INO80E,C
R2,STIP1,LRRC1,PHF19,GRAMD1C,VCAN,LPCAT3,CD44,TM9SF1,MAGED2,SON,TET2,UQCRQ,SNF8,GRK6,
R2,STIP1,LRRC1,PHF19,GRAMD1C,VCAN,LPCAT3,CD44,TM9SF1,MAGED2,SON,TET2,UQCRQ,SNF8,GRK6,

N,TET2,UQCRQ,SNF8,GRK6,HMGA2,RPS16,CPSF3L,TSG101,OBSL1,TTC21B,PTPN13,UBR4,PSMA1,YLPM1
N,TET2,UQCRQ,SNF8,GRK6,HMGA2,RPS16,CPSF3L,TSG101,OBSL1,TTC21B,PTPN13,UBR4,PSMA1,YLPM1
,GGPS1,SRSF1,AUP1,SAP30BP,YIPF3,FOXRED1,TRNT1,KXD1,PARD6G,SH3YL1,IP6K2,CLASRP,JPH2,C1ORF
,GGPS1,SRSF1,AUP1,SAP30BP,YIPF3,FOXRED1,TRNT1,KXD1,PARD6G,SH3YL1,IP6K2,CLASRP,JPH2,C1ORF
N1,HPS1,ITPA,DNAJA3,VPS25,SNRPD1,DHRS4,IFI27,SMIM12,AZI2,TATDN1,ZBTB8OS,TBC1D22A,U2AF1L
,TIMM17B,XAF1,APEH,HYI,CAPN10,NUTF2,TCF4,ATPAF2,LSM12,C17ORF62,PRMT1,CASC4,WNK1,C19OR
,TIMM17B,XAF1,APEH,HYI,CAPN10,NUTF2,TCF4,ATPAF2,LSM12,C17ORF62,PRMT1,CASC4,WNK1,C19OR
I1,DUSP22,SLC25A10,SPOCK2,THEM4,DBNDD2,SDF4,NISCH,TMED4,CENPK,RPL29,CD99L2,SRSF1,AUP1,S
A3,ZNF687,PDCD5,KLK10,TRIM16,ANAPC11,POLR3H,RAD23A,ABHD17A,PDE4DIP,MDH2,STXBP2,EEF1D,
A3,ZNF687,PDCD5,KLK10,TRIM16,ANAPC11,POLR3H,RAD23A,ABHD17A,PDE4DIP,MDH2,STXBP2,EEF1D,
LA,CARHSP1,CNST,COX17,HNRNPLL,PHF20L1,HIGD1A,POLR2C,B3GAT3,RFXANK,ZNF506,SPINT2,FTH1,PF
LA,CARHSP1,CNST,COX17,HNRNPLL,PHF20L1,HIGD1A,POLR2C,B3GAT3,RFXANK,ZNF506,SPINT2,FTH1,PF
A,ZNF593,GLYCTK,CFLAR,TRIM46,C11ORF57,RBM42,CREBZF,EIF2A,NKIRAS2,SCN11A,RECQL5,ACAA1,ZNF
A,ZNF593,GLYCTK,CFLAR,TRIM46,C11ORF57,RBM42,CREBZF,EIF2A,NKIRAS2,SCN11A,RECQL5,ACAA1,ZNF
,MGST2,EPN1,HPS1,ITPA,ZCCHC9,VPS25,SNRPD1,GTF3A,CHURC1,AZI2,GOSR2,MYL12A,TATDN1,TBC1D2
,CPSF3L,TSG101,OBSL1,TTC21B,PTPN13,UBR4,PHF21A,CHCHD1,FUZ,RGS3,FAM3A,PSMA3,MUC1,METT
,CPSF3L,TSG101,OBSL1,TTC21B,PTPN13,UBR4,PHF21A,CHCHD1,FUZ,RGS3,FAM3A,PSMA3,MUC1,METT
HPF,MPDU1,VPS28,UBXN1,ZNF814,HSPB1,SAT1,ERCC5,EMC4,MCTS1,PRKRIP1,DDOST,LSR,DOCK10,MEI
HPF,MPDU1,VPS28,UBXN1,ZNF814,HSPB1,SAT1,ERCC5,EMC4,MCTS1,PRKRIP1,DDOST,LSR,DOCK10,MEI
1,VWA2,ERCC5,LMBR1,NAP1L1,SERPINB1,DDOST,LSR,MTA3,MEIS2,PABPC1,RRNAD1,DBNL,NAT9,ZNF83
,DUSP22,SPOCK2,THEM4,RDH11,DBNDD2,NISCH,TMED4,CENPK,GGPS1,RTEL1,SRSF1,AUP1,SAP30BP,YIP
,DUSP22,SPOCK2,THEM4,RDH11,DBNDD2,NISCH,TMED4,CENPK,GGPS1,RTEL1,SRSF1,AUP1,SAP30BP,YIP
TN5,PEX26,NDUFS3,OXLD1,EIF4A1,ZNF223,METTL5,SLC35E1,OGFOD2,METTL3,TSEN34,SSR3,ARIH2,PIC
,C6ORF132,SLC35E1,OGFOD2,METTL3,TSEN34,SSR3,MPV17,ARIH2,PICK1,TCF25,RNF181,TPT1,UBA1,SL
YIPF3,FOXRED1,TRNT1,KXD1,ITGB1BP1,MED15,SH3YL1,IP6K2,CLASRP,JPH2,UGGT2,HS2ST1,UBE2I,YIF1A
YIPF3,FOXRED1,TRNT1,KXD1,ITGB1BP1,MED15,SH3YL1,IP6K2,CLASRP,JPH2,UGGT2,HS2ST1,UBE2I,YIF1A
6G,CLASRP,C1ORF198,UGGT2,MICAL1,HS2ST1,UBE2I,SLC39A13,YIF1A,PRMT2,CDC42,TBX6,TMEM205,P
6G,CLASRP,C1ORF198,UGGT2,MICAL1,HS2ST1,UBE2I,SLC39A13,YIF1A,PRMT2,CDC42,TBX6,TMEM205,P
JBE2I,SLC39A13,YIF1A,PRMT2,EIF3M,JUP,CDC42,XBP1,PTK2,GUK1,CENPN,CARD14,PCYT2,FBXW9,USE1,
M205,XBP1,PTK2,AUNIP,C14ORF2,GUK1,PCYT2,FBXW9,USE1,AGPAT4,MAF,CCDC25,DEF8,PPP1CC,EDF1
M205,XBP1,PTK2,AUNIP,C14ORF2,GUK1,PCYT2,FBXW9,USE1,AGPAT4,MAF,CCDC25,DEF8,PPP1CC,EDF1
1B2,PPP1CC,EDF1,RFX3,PPA1,DGKE,TNFRSF14,ENDOV,VAMP8,CAMTA1,CAPG,PPIL2,PSMD4,TROAP,LIG1
1B2,PPP1CC,EDF1,RFX3,PPA1,DGKE,TNFRSF14,ENDOV,VAMP8,CAMTA1,CAPG,PPIL2,PSMD4,TROAP,LIG1
RF2,GUK1,ATF7,PCYT2,FBXW9,USE1,AGPAT4,ASF1B,DEF8,EEF1B2,PPP1CC,EDF1,RFX3,PPA1,DGKE,ENDO
S14,RNF4,MAGOH,ZRANB2,TXNDC12,SEC14L1,FANCI,DPP9,SEC24C,C8ORF82,ZNF850,C16ORF46,TVP23B
1B,PTPN13,UBR4,PSMA1,YLPM1,PHF21A,CHCHD1,FUZ,FAM3A,PSMA3,MUC1,ECSIT,EPB41L5,ATG12,PPR

1B,PTPN13,UBR4,PSMA1,YLPM1,PHF21A,CHCHD1,FUZ,FAM3A,PSMA3,MUC1,ECSIT,EPB41L5,ATG12,PPR

RCC2,UBE2N,SNX5,ZNF81,FAM32A,COX20,TMEM147,COA1,USP40,PCID2,TSC22D3,PEX2,TNKS,PIGH,ELO

RCC2,UBE2N,SNX5,ZNF81,FAM32A,COX20,TMEM147,COA1,USP40,PCID2,TSC22D3,PEX2,TNKS,PIGH,ELO

13,PRMT2,EIF3M,JUP,TBX6,TMEM205,XBP1,PTK2,AUNIP,C14ORF2,GUK1,ATF7,PCYT2,USE1,AGPAT4,PP

13,PRMT2,EIF3M,JUP,TBX6,TMEM205,XBP1,PTK2,AUNIP,C14ORF2,GUK1,ATF7,PCYT2,USE1,AGPAT4,PP

RK6,HMGA2,CDC37,RPS16,CPSF3L,TSG101,OBSL1,TTC21B,PTPN13,UBR4,YLPM1,PHF21A,CHCHD1,SLC39

TMEM147,PEX2,POLR1C,PIGH,C12ORF57,DCTN5,PEX26,NDUFS3,METTL5,SLC35E1,DGAT1,OGFOD2,MET

1,ANAPC13,TSPAN31,MTIF2,RALY,GLYCTK,TRIM46,C11ORF57,RER1,CREBZF,NKIRAS2,RECQL5,ACAA1,RP

LAMTOR4,RPS17,WDR45,C9ORF64,IQCK,SETD4,MED8,EIF6,RNMT,DHX30,AP4M1,COQ4,ACTG1,MRPL27

BL5,TRNT1,KXD1,MED15,SH3YL1,JPH2,C1ORF198,UGGT2,MICAL1,UBE2I,SLC39A13,YIF1A,JUP,TBX6,XBP

BL5,TRNT1,KXD1,MED15,SH3YL1,JPH2,C1ORF198,UGGT2,MICAL1,UBE2I,SLC39A13,YIF1A,JUP,TBX6,XBP

NPK,RPS6,RPL29,CD99L2,GGPS1,RTEL1,COASY,SRSF1,AUP1,SAP30BP,YIPF3,FOXRED1,RABL5,TRNT1,KXD1

NPK,RPS6,RPL29,CD99L2,GGPS1,RTEL1,COASY,SRSF1,AUP1,SAP30BP,YIPF3,FOXRED1,RABL5,TRNT1,KXD1

37,RPS16,CPSF3L,TSG101,OBSL1,TTC21B,PTPN13,UBR4,PSMA1,PHF21A,SLC39A11,FUZ,FAM3A,MUC1,M

DF1,RFX3,PPA1,DGKE,VAMP8,CAMTA1,CAPG,CRIP1,CLDND1,VIM,LIG1,TIMM17B,XAF1,APEH,HYI,CAPN1

DF1,RFX3,PPA1,DGKE,VAMP8,CAMTA1,CAPG,CRIP1,CLDND1,VIM,LIG1,TIMM17B,XAF1,APEH,HYI,CAPN1

SLC15A3,PPM1K,TUBA1C,IRF9,DST,ANTXR2,STIP1,LRRC1,PHF19,GRAMD1C,VCAN,PCBP2,LPCAT3,CD44,T

SLC15A3,PPM1K,TUBA1C,IRF9,DST,ANTXR2,STIP1,LRRC1,PHF19,GRAMD1C,VCAN,PCBP2,LPCAT3,CD44,T

MX2,SPOCK2,THEM4,DBNDD2,TMED4,RPL29,CD99L2,GGPS1,RTEL1,COASY,SRSF1,AUP1,SAP30BP,RABL

2,SLC25A10,SPOCK2,THEM4,RDH11,DBNDD2,TMED4,CENPK,CD99L2,GGPS1,RTEL1,SRSF1,AUP1,SAP30B

,VAMP8,CAPG,PPIL2,TROAP,RPL28,LIG1,CDC14B,TIMM17B,APEH,RPL21,CAPN10,RBM33,LYPLA2,PODXL,

,VAMP8,CAPG,PPIL2,TROAP,RPL28,LIG1,CDC14B,TIMM17B,APEH,RPL21,CAPN10,RBM33,LYPLA2,PODXL,

1,ITPA,ZCCHC9,VPS25,SNRPD1,DCAF5,MKRN1,HMOX2,CHURC1,SNRPA1,CDK2AP2,AZI2,GOSR2,MYL12A

,VIM,TIMM17B,RPL21,TXNL4A,CAPN10,LYPLA2,CAMLG,EPB41L4B,UCK2,TMPO,ATPAF2,LSM12,C17ORF6

15,SH3YL1,IP6K2,CLASRP,JPH2,C1ORF198,UGGT2,MICAL1,UBE2I,SLC39A13,EIF3M,JUP,CDC42,XBP1,PTK2

15,SH3YL1,IP6K2,CLASRP,JPH2,C1ORF198,UGGT2,MICAL1,UBE2I,SLC39A13,EIF3M,JUP,CDC42,XBP1,PTK2

17B,XAF1,APEH,HYI,CAPN10,LYPLA2,NUTF2,PODXL,TCF4,ATPAF2,C17ORF62,PRMT1,CASC4,C19ORF54,Z

17B,XAF1,APEH,HYI,CAPN10,LYPLA2,NUTF2,PODXL,TCF4,ATPAF2,C17ORF62,PRMT1,CASC4,C19ORF54,Z

17B,XAF1,APEH,HYI,CAPN10,LYPLA2,NUTF2,PODXL,TCF4,ATPAF2,C17ORF62,PRMT1,CASC4,C19ORF54,Z

17B,XAF1,APEH,HYI,CAPN10,LYPLA2,NUTF2,PODXL,TCF4,ATPAF2,C17ORF62,PRMT1,CASC4,C19ORF54,Z

RPS25,NIP7,COX4I1,DUSP22,TMX2,SPOCK2,RDH11,DBNDD2,SDF4,NISCH,TMED4,CENPK,RPS6,RPL29,CD

RPS25,NIP7,COX4I1,DUSP22,TMX2,SPOCK2,RDH11,DBNDD2,SDF4,NISCH,TMED4,CENPK,RPS6,RPL29,CD

ATG12,RPS25,ZBTB44,NIP7,TTC9C,COX4I1,PLCB2,DUSP22,SLC25A10,SPOCK2,THEM4,DBNDD2,NISCH,TM

1,LRRC1,PHF19,GRAMD1C,VCAN,LPCAT3,CD44,TM9SF1,MAGED2,SON,TET2,UQCRQ,SNF8,GRK6,HMGA

1,LRRC1,PHF19,GRAMD1C,VCAN,LPCAT3,CD44,TM9SF1,MAGED2,SON,TET2,UQCRQ,SNF8,GRK6,HMGA

PSF3L,TSG101,OBSL1,TTC21B,PTPN13,UBR4,YLPM1,PHF21A,CHCHD1,SLC39A11,FUZ,FAM3A,MUC1,MET

UP1,YIPF3,FOXRED1,RABL5,TRNT1,KXD1,ITGB1BP1,MED15,SH3YL1,CLASRP,MICAL1,HS2ST1,UBE2I,YIF1

13,UBR4,YLPM1,PHF21A,CHCHD1,SLC39A11,FUZ,FAM3A,PSMA3,MUC1,METTL17,ECSIT,EPB41L5,ATG12

13,UBR4,YLPM1,PHF21A,CHCHD1,SLC39A11,FUZ,FAM3A,PSMA3,MUC1,METTL17,ECSIT,EPB41L5,ATG12

KXD1,ITGB1BP1,MED15,SH3YL1,IP6K2,CLASRP,JPH2,MICAL1,HS2ST1,UBE2I,KCNMA1,SLC39A13,YIF1A,P

KXD1,ITGB1BP1,MED15,SH3YL1,IP6K2,CLASRP,JPH2,MICAL1,HS2ST1,UBE2I,KCNMA1,SLC39A13,YIF1A,P

2,C17ORF62,PRMT1,CASC4,WNK1,ZNF7,ARL6IP4,PPAN,ZFAND2B,CYB561,SHARPIN,ZNF644,UPP1,ZNF25

2,C17ORF62,PRMT1,CASC4,WNK1,ZNF7,ARL6IP4,PPAN,ZFAND2B,CYB561,SHARPIN,ZNF644,UPP1,ZNF25

44,NIP7,PLCB2,SLC25A10,TMX2,SPOCK2,RDH11,DBNDD2,SDF4,TMED4,CENPK,RPS6,RPL29,CD99L2,GGP

LA2,NUTF2,PODXL,ATPAF2,LSM12,C17ORF62,PRMT1,CASC4,ARL6IP4,PIGL,C19ORF48,PPAN,ZFAND2B,C

PLA2,NUTF2,PODXL,ATPAF2,LSM12,C17ORF62,PRMT1,CASC4,ARL6IP4,PIGL,C19ORF48,PPAN,ZFAND2B,C
3C1D22A,APAF1,KAT8,NCBP2,TTLL3,RNPS1,ZNF581,SSC5D,ZNF483,SZT2,LRRC23,NDUFA7,CYB5D2,DYNC1
IP7,TTC9C,COX4I1,PLCB2,SLC25A10,THEM4,DBNDD2,SDF4,NISCH,TMED4,CENPK,RPS6,RPL29,CD99L2,GG
VRK3,LZTFL1,ZC3H14,FIS1,SLC15A3,PPM1K,TUBA1C,IRF9,DST,ANTXR2,STIP1,LRRC1,PHF19,GRAMD1C,VC
VRK3,LZTFL1,ZC3H14,FIS1,SLC15A3,PPM1K,TUBA1C,IRF9,DST,ANTXR2,STIP1,LRRC1,PHF19,GRAMD1C,VC
YIPF3,RABL5,ITGB1BP1,MED15,PARD6G,SH3YL1,CLASRP,JPH2,C1ORF198,UGGT2,MICAL1,UBE2I,KCNMA
YIPF3,RABL5,ITGB1BP1,MED15,PARD6G,SH3YL1,CLASRP,JPH2,C1ORF198,UGGT2,MICAL1,UBE2I,KCNMA
PC2,GTF2H1,PIGO,D2HGDH,KDELR3,CLTA,SLC35A2,SNX29,AXIN1,TMEM199,SLC25A24,C1ORF63,EIF3C,H
SBP1L1,AGPAT4,DEF8,PPP1CC,EDF1,RFX3,PPA1,DGKE,CAMTA1,CAPG,CRIP1,CLDND1,VIM,LIG1,TIMM17B
SBP1L1,AGPAT4,DEF8,PPP1CC,EDF1,RFX3,PPA1,DGKE,CAMTA1,CAPG,CRIP1,CLDND1,VIM,LIG1,TIMM17B
,DBNDD2,TMED4,CENPK,RPS6,CD99L2,GGPS1,RTEL1,COASY,SRSF1,RABL5,TRNT1,KXD1,ITGB1BP1,MED1
,DBNDD2,TMED4,CENPK,RPS6,CD99L2,GGPS1,RTEL1,COASY,SRSF1,RABL5,TRNT1,KXD1,ITGB1BP1,MED1
AMD1C,VCAN,LPCAT3,CD44,TM9SF1,MAGED2,SON,TET2,UQCRQ,SNF8,GRK6,HMGA2,CDC37,RPS16,CPS
AMD1C,VCAN,LPCAT3,CD44,TM9SF1,MAGED2,SON,TET2,UQCRQ,SNF8,GRK6,HMGA2,CDC37,RPS16,CPS
P1,SAP30BP,YIPF3,TRNT1,KXD1,ITGB1BP1,PARD6G,SH3YL1,IP6K2,CLASRP,UGGT2,MICAL1,UBE2I,YIF1A,
CDC25,DEF8,EEF1B2,PPP1CC,PDLIM5,EDF1,RFX3,DGKE,ENDOV,CAMTA1,CRIP1,PPIL2,RPL28,CLDND1,LI
CDC25,DEF8,EEF1B2,PPP1CC,PDLIM5,EDF1,RFX3,DGKE,ENDOV,CAMTA1,CRIP1,PPIL2,RPL28,CLDND1,LI
CDC25,DEF8,EEF1B2,PPP1CC,PDLIM5,EDF1,RFX3,DGKE,ENDOV,CAMTA1,CRIP1,PPIL2,RPL28,CLDND1,LI
CDC25,DEF8,EEF1B2,PPP1CC,PDLIM5,EDF1,RFX3,DGKE,ENDOV,CAMTA1,CRIP1,PPIL2,RPL28,CLDND1,LI
CDC25,DEF8,EEF1B2,PPP1CC,PDLIM5,EDF1,RFX3,DGKE,ENDOV,CAMTA1,CRIP1,PPIL2,RPL28,CLDND1,LI
ODXL,TCF4,ATPAF2,C17ORF62,PRMT1,CASC4,ZNF7,PPAN,ZFAND2B,AP1S3,CYB561,SHARPIN,STAU1,SU
ODXL,TCF4,ATPAF2,C17ORF62,PRMT1,CASC4,ZNF7,PPAN,ZFAND2B,AP1S3,CYB561,SHARPIN,STAU1,SU
FX3,PPA1,DGKE,ENDOV,VAMP8,CAMTA1,PPIL2,RPL28,CLDND1,LIG1,CDC14B,TIMM17B,APEH,RPL21,TX
FX3,PPA1,DGKE,ENDOV,VAMP8,CAMTA1,PPIL2,RPL28,CLDND1,LIG1,CDC14B,TIMM17B,APEH,RPL21,TX
DAB1,SYNGR2,RAB2A,ATP5J,TSFM,TRMT112,CHPF,ZNF625,GABPB1,DCTD,UBXN1,ZNF814,HSPB1,VWA2,
DAB1,SYNGR2,RAB2A,ATP5J,TSFM,TRMT112,CHPF,ZNF625,GABPB1,DCTD,UBXN1,ZNF814,HSPB1,VWA2,
BX6,TMEM205,XBP1,PTK2,AUNIP,C14ORF2,GUK1,CENPN,PCYT2,USE1,IFI27L1,PPP1CC,EDF1,RFX3,PPA1,
BX6,TMEM205,XBP1,PTK2,AUNIP,C14ORF2,GUK1,CENPN,PCYT2,USE1,IFI27L1,PPP1CC,EDF1,RFX3,PPA1,

ENPN,PCYT2,USE1,HSBP1L1,EEF1B2,PPP1CC,EDF1,RFX3,PPA1,DGKE,CAMTA1,CAPG,CRIP1,CLDND1,LIG1,
ENPN,PCYT2,USE1,HSBP1L1,EEF1B2,PPP1CC,EDF1,RFX3,PPA1,DGKE,CAMTA1,CAPG,CRIP1,CLDND1,LIG1,
3XL18,ZNF512,RNF181,CHTOP,POLR2M,FAM192A,SLC39A6,ZNF593,GLYCTK,CFLAR,PCNP,VDAC1,CNTRL,
3XL18,ZNF512,RNF181,CHTOP,POLR2M,FAM192A,SLC39A6,ZNF593,GLYCTK,CFLAR,PCNP,VDAC1,CNTRL,
NDD2,NISCH,TMED4,CENPK,RPS6,RPL29,CD99L2,GGPS1,RTEL1,COASY,SRSF1,AUP1,YIPF3,FOXRED1,RAB
NDD2,NISCH,TMED4,CENPK,RPS6,RPL29,CD99L2,GGPS1,RTEL1,COASY,SRSF1,AUP1,YIPF3,FOXRED1,RAB
SC,MDH2,STXBP2,EEF1D,HSPH1,SEC31A,ATP10D,CNST,SSH2,COX17,CAMK2D,COX5A,HNRNPLL,HINT1,HI
L,HS2ST1,YIF1A,PRMT2,EIF3M,CDC42,TBX6,TMEM205,XBP1,AUNIP,GUK1,CENPN,ATF7,FBXW9,MAF,ASF
L,HS2ST1,YIF1A,PRMT2,EIF3M,CDC42,TBX6,TMEM205,XBP1,AUNIP,GUK1,CENPN,ATF7,FBXW9,MAF,ASF
APN10,NUTF2,PODXL,EPB41L4B,ATPAF2,C17ORF62,PRMT1,CASC4,WNK1,ZNF7,C19ORF48,STRADA,PPAN
APN10,NUTF2,PODXL,EPB41L4B,ATPAF2,C17ORF62,PRMT1,CASC4,WNK1,ZNF7,C19ORF48,STRADA,PPAN
ENDOV,CAMTA1,CRIP1,PPIL2,PSMD4,RPL28,VIM,LIG1,PSMA7,TIMM17B,TMEM126B,APEH,TXNL4A,NUT
O,ATPAF2,LSM12,C17ORF62,PRMT1,CASC4,EXOSC9,STOML2,WNK1,C19ORF54,ZNF7,ARL6IP4,STRADA,P
O,ATPAF2,LSM12,C17ORF62,PRMT1,CASC4,EXOSC9,STOML2,WNK1,C19ORF54,ZNF7,ARL6IP4,STRADA,P
UZ,RGS3,FAM3A,METTL17,ECSIT,EPB41L5,PPRC1,RPS25,NLRP1,NIP7,TTC9C,COX4I1,SLC25A10,SPOCK2,T
UZ,RGS3,FAM3A,METTL17,ECSIT,EPB41L5,PPRC1,RPS25,NLRP1,NIP7,TTC9C,COX4I1,SLC25A10,SPOCK2,T

C24,ALS2,GTF3C5,AP2S1,IK,ACTB,CCDC171,METTL23,SEC13,MRPL51,ZNF586,MED8,RNMT,DHX30
C24,ALS2,GTF3C5,AP2S1,IK,ACTB,CCDC171,METTL23,SEC13,MRPL51,ZNF586,MED8,RNMT,DHX30
HCHD1,SLC39A11,FUZ,FAM3A,MUC1,ECSIT,EPB41L5,ATG12,PPRC1,RPS25,ZBTB44,NIP7,TTC9C,COX4I1,P
HCHD1,SLC39A11,FUZ,FAM3A,MUC1,ECSIT,EPB41L5,ATG12,PPRC1,RPS25,ZBTB44,NIP7,TTC9C,COX4I1,P
PS25,ZBTB44,TTC9C,COX4I1,PLCB2,DUSP22,SLC25A10,TMX2,SPOCK2,DBNDD2,SDF4,NISCH,TMED4,CENF
PS25,ZBTB44,TTC9C,COX4I1,PLCB2,DUSP22,SLC25A10,TMX2,SPOCK2,DBNDD2,SDF4,NISCH,TMED4,CENF
A,PSMA3,MUC1,ECSIT,EPB41L5,ATG12,PPRC1,RPS25,NLRP1,ZBTB44,NIP7,TTC9C,COX4I1,PLCB2,SLC25A1
A,PSMA3,MUC1,ECSIT,EPB41L5,ATG12,PPRC1,RPS25,NLRP1,ZBTB44,NIP7,TTC9C,COX4I1,PLCB2,SLC25A1
L,BFAR,SNX29,TP53,SH3GL1,BTF3,TMEM199,SLC25A24,UBE2D3,ZNF396,SNAPC2,EPN1,ITPA,ZCCHC9,VPS
L,BFAR,SNX29,TP53,SH3GL1,BTF3,TMEM199,SLC25A24,UBE2D3,ZNF396,SNAPC2,EPN1,ITPA,ZCCHC9,VPS
2,UGGT2,HS2ST1,YIF1A,PRMT2,EIF3M,CDC42,TBX6,TMEM205,XBP1,PTK2,CELF1,AUNIP,C14ORF2,GUK1,C
2,UGGT2,HS2ST1,YIF1A,PRMT2,EIF3M,CDC42,TBX6,TMEM205,XBP1,PTK2,CELF1,AUNIP,C14ORF2,GUK1,C
2,PODXL,TCF4,ATPAF2,C17ORF62,PRMT1,CASC4,C19ORF54,ZNF7,TXNDC17,STRADA,PPAN,ZFAND2B,AP1
2,PODXL,TCF4,ATPAF2,C17ORF62,PRMT1,CASC4,C19ORF54,ZNF7,TXNDC17,STRADA,PPAN,ZFAND2B,AP1
PBP1,GPALPP1,SIAH1,ASPH,CHPF,STRIP2,NIT2,LYNX1,TWSG1,VPS28,ZSWIM7,HSPB1,LRRC27,MVD,LMBR
PBP1,GPALPP1,SIAH1,ASPH,CHPF,STRIP2,NIT2,LYNX1,TWSG1,VPS28,ZSWIM7,HSPB1,LRRC27,MVD,LMBR
,SEC31A,CARHSP1,RPL18,ATP10D,LLGL2,EGFR,SSH2,COX17,HIGD1A,POLR2C,NDUFV2,RPL15,INTS4,B3GA
,SEC31A,CARHSP1,RPL18,ATP10D,LLGL2,EGFR,SSH2,COX17,HIGD1A,POLR2C,NDUFV2,RPL15,INTS4,B3GA
F2,C17ORF62,PRMT1,CASC4,ZNF7,PPAN,ZFAND2B,AP1S3,HNRNPC,CYB561,SHARPIN,SUGT1,UPP1,ZNF25
F2,C17ORF62,PRMT1,CASC4,ZNF7,PPAN,ZFAND2B,AP1S3,HNRNPC,CYB561,SHARPIN,SUGT1,UPP1,ZNF25
LC9B2,ZNF250,MOB1A,CDK11A,RBM3,MDH1,ZNF197,CCNL2,TMEM134,LRRFIP1,ZNF524,ELP6,RMDN1,S
LC9B2,ZNF250,MOB1A,CDK11A,RBM3,MDH1,ZNF197,CCNL2,TMEM134,LRRFIP1,ZNF524,ELP6,RMDN1,S
D15,PARD6G,SH3YL1,IP6K2,CLASRP,JPH2,C1ORF198,MICAL1,HS2ST1,UBE2I,SLC39A13,YIF1A,PRMT2,EIF3
D15,PARD6G,SH3YL1,IP6K2,CLASRP,JPH2,C1ORF198,MICAL1,HS2ST1,UBE2I,SLC39A13,YIF1A,PRMT2,EIF3

PF,STRIP2,RPS20,NIT2,MPDU1,RPS15,ERLEC1,LRRC27,ERCC5,LMBR1,NAP1L1,DDOST,MEIS2,PHTF2,RRNA
UFS3,OXLD1,ZNF223,OGFOD2,METTL3,ARIH2,CREB3L4,PICK1,FBXL18,RNF181,CDK4,NME1,TSPAN31,MTI
,FAM219B,RAB2A,ZNF524,ATP5J,TPM2,SIAH1,GGCX,ZEB2,ASPH,TRMT112,PNPO,CHPF,STRIP2,NIT2,LYNX
F223,STAU2,METTL5,SLC35E1,OGFOD2,TSEN34,RPL8,FBXL18,RNF181,TJP1,TPT1,BAX,UBA1,SLC52A2,NT
F223,STAU2,METTL5,SLC35E1,OGFOD2,TSEN34,RPL8,FBXL18,RNF181,TJP1,TPT1,BAX,UBA1,SLC52A2,NT
ISCH,TMED4,CENPK,RPS6,RPL29,CD99L2,RTEL1,SRSF1,AUP1,YIPF3,FOXRED1,RABL5,TRNT1,KXD1,ITGB1B
ISCH,TMED4,CENPK,RPS6,RPL29,CD99L2,RTEL1,SRSF1,AUP1,YIPF3,FOXRED1,RABL5,TRNT1,KXD1,ITGB1B
1B,PTPN13,UBR4,YLPM1,PHF21A,CHCHD1,SLC39A11,FUZ,RGS3,FAM3A,MUC1,METTL17,EPB41L5,ATG12
1B,PTPN13,UBR4,YLPM1,PHF21A,CHCHD1,SLC39A11,FUZ,RGS3,FAM3A,MUC1,METTL17,EPB41L5,ATG12
,ITGB1BP1,SH3YL1,IP6K2,CLASRP,JPH2,MICAL1,HS2ST1,YIF1A,PRMT2,EIF3M,MAGT1,JUP,CDC42,TBX6,TI
,ITGB1BP1,SH3YL1,IP6K2,CLASRP,JPH2,MICAL1,HS2ST1,YIF1A,PRMT2,EIF3M,MAGT1,JUP,CDC42,TBX6,TI
H,RPL21,RBM33,LYPLA2,NUTF2,CAMLG,C17ORF62,PRMT1,CASC4,STOML2,C19ORF54,ZNF7,TXNDC17,A
H,RPL21,RBM33,LYPLA2,NUTF2,CAMLG,C17ORF62,PRMT1,CASC4,STOML2,C19ORF54,ZNF7,TXNDC17,A
IT2,TWSG1,GMPPB,MVD,IL32,SEMA3C,LMBR1,SERPINB1,MTA3,CHCHD7,HMGA1,PABPC1,NAT9,BCAM,Z
IT2,TWSG1,GMPPB,MVD,IL32,SEMA3C,LMBR1,SERPINB1,MTA3,CHCHD7,HMGA1,PABPC1,NAT9,BCAM,Z

,SLC39A11,FUZ,RGS3,PSMA3,MUC1,METTL17,ECSIT,EPB41L5,ATG12,NLRP1,ZBTB44,NIP7,DUSP22,SLC2
,SLC39A11,FUZ,RGS3,PSMA3,MUC1,METTL17,ECSIT,EPB41L5,ATG12,NLRP1,ZBTB44,NIP7,DUSP22,SLC2
SCAN9,RIC8A,TCP1,SMARCC2,SLC30A5,TRPV3,PIGT,UBE2N,LFNG,PDCD6,FAM32A,ZSCAN20,COX20,NFYC
SCAN9,RIC8A,TCP1,SMARCC2,SLC30A5,TRPV3,PIGT,UBE2N,LFNG,PDCD6,FAM32A,ZSCAN20,COX20,NFYC

3,EEF1D,CHCHD5,MED6,CASZ1,COX5A,PHF20L1,HNRNPA1,POLR2C,FKBP3,EGLN2,RPL15,SIRT5,KCTD7,B3
3,EEF1D,CHCHD5,MED6,CASZ1,COX5A,PHF20L1,HNRNPA1,POLR2C,FKBP3,EGLN2,RPL15,SIRT5,KCTD7,B3
MBR1,NAP1L1,SERPINB1,CHCHD7,HMGA1,PSME2,PHTF2,PABPC1,NAT9,BCAM,BLVRB,TOMM40,RABEP2
MBR1,NAP1L1,SERPINB1,CHCHD7,HMGA1,PSME2,PHTF2,PABPC1,NAT9,BCAM,BLVRB,TOMM40,RABEP2
IF2,FAM192A,RALY,ZNF593,GLYCTK,FAM131A,CFLAR,DOK4,EIF2A,RECQL5,PSMD7,ZNF283,GTF2H1,PIGO
ALPP1,TSFM,TPM3,TRMT112,PNPO,RPTOR,CHPF,RPS20,VPS28,DCTD,ZNF814,HSPB1,ERCC5,EMC4,MCTS1
ALPP1,TSFM,TPM3,TRMT112,PNPO,RPTOR,CHPF,RPS20,VPS28,DCTD,ZNF814,HSPB1,ERCC5,EMC4,MCTS1
,ILF3,AGAP3,EEF1D,CHCHD5,MED6,CASZ1,COX5A,SUMO3,PHF20L1,HNRNPA1,POLR2C,FKBP3,EGLN2,RP
,ILF3,AGAP3,EEF1D,CHCHD5,MED6,CASZ1,COX5A,SUMO3,PHF20L1,HNRNPA1,POLR2C,FKBP3,EGLN2,RP
,CENPK,RPS6,RPL29,CD99L2,GGPS1,RTEL1,COASY,SRSF1,AUP1,FOXRED1,RABL5,TRNT1,ITGB1BP1,MED15
99L2,GGPS1,RTEL1,SRSF1,AUP1,SAP30BP,YIPF3,TRNT1,KXD1,ITGB1BP1,PARD6G,SH3YL1,IP6K2,CLASRP,U
99L2,GGPS1,RTEL1,SRSF1,AUP1,SAP30BP,YIPF3,TRNT1,KXD1,ITGB1BP1,PARD6G,SH3YL1,IP6K2,CLASRP,U
,MDH2,EEF1D,CHCHD5,NAA10,MED6,SEC31A,CARHSP1,COX17,HNRNPLL,HINT1,HIGD1A,POLR2C,B3GAT
,MDH2,EEF1D,CHCHD5,NAA10,MED6,SEC31A,CARHSP1,COX17,HNRNPLL,HINT1,HIGD1A,POLR2C,B3GAT
,CENPN,FBXW9,USE1,HSBP1L1,IFI27L1,ASF1B,CCDC25,DEF8,EEF1B2,EDF1,GADL1,RFX3,DGKE,CAMTA1,PI
,CENPN,FBXW9,USE1,HSBP1L1,IFI27L1,ASF1B,CCDC25,DEF8,EEF1B2,EDF1,GADL1,RFX3,DGKE,CAMTA1,PI
PS25,ZBTB44,NIP7,TTC9C,COX4I1,DUSP22,SLC25A10,TMX2,SPOCK2,THEM4,RDH11,DBNDD2,SDF4,TMEI
PS25,ZBTB44,NIP7,TTC9C,COX4I1,DUSP22,SLC25A10,TMX2,SPOCK2,THEM4,RDH11,DBNDD2,SDF4,TMEI
YL1,IP6K2,CLASRP,JPH2,C1ORF198,MICAL1,HS2ST1,UBE2I,KCNMA1,SLC39A13,YIF1A,PRMT2,EIF3M,TBX6
2T2,STAU1,SUGT1,HYPK,CARS,ZNF250,MOB1A,CDK11A,ZNF197,COPZ1,INO80E,MAPKAPK5,DGUOK,HSP9
2T2,STAU1,SUGT1,HYPK,CARS,ZNF250,MOB1A,CDK11A,ZNF197,COPZ1,INO80E,MAPKAPK5,DGUOK,HSP9
C1,VWA2,ERCC5,EMC4,MCTS1,PRKRIP1,DDOST,LSR,DOCK10,HMGA1,PSME2,RRNAD1,DBNL,ZNF836,TON
C1,VWA2,ERCC5,EMC4,MCTS1,PRKRIP1,DDOST,LSR,DOCK10,HMGA1,PSME2,RRNAD1,DBNL,ZNF836,TON
,TTC9C,COX4I1,PLCB2,DUSP22,SLC25A10,TMX2,SPOCK2,THEM4,RDH11,DBNDD2,SDF4,NISCH,CENPK,RP
32A,ZNF524,ATP5J,TPM2,SIAH1,GGCX,ZEB2,ASPH,TRMT112,PNPO,CHPF,STRIP2,ZNF625,NIT2,LYNX1,TW
AP30BP,YIPF3,FOXRED1,RABL5,TRNT1,MED15,SH3YL1,IP6K2,CLASRP,C1ORF198,UGGT2,MICAL1,SLC39A1

5,TPM2,GPBP1,SIAH1,GGCX,ZEB2,ASPH,TRMT112,CHPF,STRIP2,NIT2,LYNX1,MAPK13,RPS15,DCTD,LRRC2
5,TPM2,GPBP1,SIAH1,GGCX,ZEB2,ASPH,TRMT112,CHPF,STRIP2,NIT2,LYNX1,MAPK13,RPS15,DCTD,LRRC2
8,GRK6,HMGA2,CDC37,RPS16,CPSF3L,OBSL1,TTC21B,PTPN13,UBR4,PSMA1,YLPM1,PHF21A,CHCHD1,FU
8,GRK6,HMGA2,CDC37,RPS16,CPSF3L,OBSL1,TTC21B,PTPN13,UBR4,PSMA1,YLPM1,PHF21A,CHCHD1,FU
,SON,TET2,UQCRQ,SNF8,GRK6,HMGA2,CDC37,RPS16,CPSF3L,TSG101,PTPN13,UBR4,PSMA1,YLPM1,PHF
,SON,TET2,UQCRQ,SNF8,GRK6,HMGA2,CDC37,RPS16,CPSF3L,TSG101,PTPN13,UBR4,PSMA1,YLPM1,PHF
CRQ,SNF8,GRK6,HMGA2,CDC37,RPS16,CPSF3L,TSG101,OBSL1,TTC21B,PTPN13,UBR4,PSMA1,PHF21A,CH
TS1,NAP1L1,PRKRIP1,DDOST,CHCHD7,MEIS2,PHTF2,PABPC1,DBNL,ZNF836,RBP1,H2AFV,RBMX,SERPINB
,PCYT2,FBXW9,HSBP1L1,IFI27L1,ASF1B,DEF8,EEF1B2,PPP1CC,EDF1,RFX3,DGKE,ENDOV,CAMTA1,CAPG,C
4,ATP5J,TPM2,SIAH1,GGCX,ZEB2,ASPH,TRMT112,PNPO,CHPF,STRIP2,NIT2,LYNX1,TWSG1,MAPK13,ZSWI
AA10,RRP8,ATP10D,LLGL2,CHEK1,COX17,SUMO3,CD151,FKBP3,EGLN2,KCTD7,B3GAT3,RFXANK,SPINT2,F
2,NUTF2,PODXL,EPB41L4B,TCF4,UCK2,TMPO,ATPAF2,C17ORF62,PRMT1,CASC4,WNK1,ZNF7,STRADA,ZFA
2,NUTF2,PODXL,EPB41L4B,TCF4,UCK2,TMPO,ATPAF2,C17ORF62,PRMT1,CASC4,WNK1,ZNF7,STRADA,ZFA
M16,GTF2H1,BFAR,D2HGDH,CLTA,NAA38,SLC35A2,SCRN2,PLAC8,ST8SIA1,AXIN1,RPS3A,BTF3,TMEM199
LRRC1,PHF19,LPCAT3,CD44,TM9SF1,MAGED2,SON,TET2,UQCRQ,SNF8,GRK6,HMGA2,CDC37,RPS16,CPSI
LRRC1,PHF19,LPCAT3,CD44,TM9SF1,MAGED2,SON,TET2,UQCRQ,SNF8,GRK6,HMGA2,CDC37,RPS16,CPSI
BNDD2,SDF4,NISCH,TMED4,CENPK,RPL29,CD99L2,GGPS1,COASY,SRSF1,AUP1,FOXRED1,TRNT1,PARD6G
BNDD2,SDF4,NISCH,TMED4,CENPK,RPL29,CD99L2,GGPS1,COASY,SRSF1,AUP1,FOXRED1,TRNT1,PARD6G
JCK2,C17ORF62,WNK1,C19ORF54,ZNF7,ARL6IP4,PIGL,C19ORF48,STRADA,PPAN,CYB561,SLC9B2,ZNF250
JCK2,C17ORF62,WNK1,C19ORF54,ZNF7,ARL6IP4,PIGL,C19ORF48,STRADA,PPAN,CYB561,SLC9B2,ZNF250

LA1,MRPL51,SFSWAP,MED8,RNMT,PUF60,ANKRD10,SLC25A35,DPAGT1,DDX56,MBD1,ZNF655,DAD1,SN
PH,NIT2,TWSG1,MAPK13,VPS28,HSPB1,IL32,SEMA3C,SERPINB1,RANGRF,CHCHD7,PHTF2,PABPC1,DBNL,
PH,NIT2,TWSG1,MAPK13,VPS28,HSPB1,IL32,SEMA3C,SERPINB1,RANGRF,CHCHD7,PHTF2,PABPC1,DBNL,

/9,USE1,EEF1B2,EDF1,GADL1,RFX3,PPA1,ENDOV,CAPG,PPIL2,RPL28,CLDND1,VIM,LIG1,CDC14B,TMEM12
/9,USE1,EEF1B2,EDF1,GADL1,RFX3,PPA1,ENDOV,CAPG,PPIL2,RPL28,CLDND1,VIM,LIG1,CDC14B,TMEM12

25,DEF8,EEF1B2,PPP1CC,EDF1,RFX3,PPA1,DGKE,TNFRSF14,CAMTA1,PPIL2,PSMD4,RPL28,CLDND1,TIMM
25,DEF8,EEF1B2,PPP1CC,EDF1,RFX3,PPA1,DGKE,TNFRSF14,CAMTA1,PPIL2,PSMD4,RPL28,CLDND1,TIMM
ED2,SON,UQCRQ,SNF8,GRK6,HMGA2,CDC37,RPS16,CPSF3L,TSG101,TTC21B,PTPN13,UBR4,PSMA1,YLPM
4I1,DUSP22,TMX2,SPOCK2,RDH11,DBNDD2,CENPK,RPL29,GGPS1,RTEL1,COASY,SRSF1,AUP1,YIPF3,FOXR
4I1,DUSP22,TMX2,SPOCK2,RDH11,DBNDD2,CENPK,RPL29,GGPS1,RTEL1,COASY,SRSF1,AUP1,YIPF3,FOXR
1,SYNGR1,C12ORF76,RANGRF,CHCHD7,PABPC1,RRNAD1,NAT9,BCAM,ZNF836,BLVRB,TOMM40,RABEP2,
NF644,STAU1,SUGT1,HYPK,ZNF250,CDK11A,MDH1,INO80E,CCNL2,TMEM134,LRRFIP1,ZNF524,ATP5J,RN
NF644,STAU1,SUGT1,HYPK,ZNF250,CDK11A,MDH1,INO80E,CCNL2,TMEM134,LRRFIP1,ZNF524,ATP5J,RN
P70,TMEM134,LRRFIP1,RAB2A,RMDN1,TPM2,GPBP1,GPALPP1,SIAH1,ZEB2,ASPH,CHPF,STRIP2,TWSG1,D
P70,TMEM134,LRRFIP1,RAB2A,RMDN1,TPM2,GPBP1,GPALPP1,SIAH1,ZEB2,ASPH,CHPF,STRIP2,TWSG1,D
N2,SIRT5,KCTD7,B3GAT3,ZNF506,SPINT2,PAFAH1B2,S100A4,PTK7,ARHGEF1,DRG2,BRF1,ZFPL1,FAM102
N2,SIRT5,KCTD7,B3GAT3,ZNF506,SPINT2,PAFAH1B2,S100A4,PTK7,ARHGEF1,DRG2,BRF1,ZFPL1,FAM102
C,VCAN,LPCAT3,CD44,TM9SF1,MAGED2,SON,TET2,UQCRQ,SNF8,GRK6,HMGA2,CDC37,RPS16,CPSF3L,TS
C,VCAN,LPCAT3,CD44,TM9SF1,MAGED2,SON,TET2,UQCRQ,SNF8,GRK6,HMGA2,CDC37,RPS16,CPSF3L,TS

2,ZNF197,COPZ1,INO80E,CCNL2,MAPKAPK5,HSP90AB1,DUT,TMEM134,LRRFIP1,ZNF524,RMDN1,ASPH,T
2,ZNF197,COPZ1,INO80E,CCNL2,MAPKAPK5,HSP90AB1,DUT,TMEM134,LRRFIP1,ZNF524,RMDN1,ASPH,T
DC37,RPS16,CPSF3L,TSG101,TTC21B,PTPN13,UBR4,PSMA1,YLPM1,CHCHD1,SLC39A11,FUZ,PSMA3,MUC
DC37,RPS16,CPSF3L,TSG101,TTC21B,PTPN13,UBR4,PSMA1,YLPM1,CHCHD1,SLC39A11,FUZ,PSMA3,MUC
CYT2,USE1,AGPAT4,IFI27L1,ASF1B,CCDC25,EEF1B2,PPP1CC,EDF1,RFX3,DGKE,TNFRSF14,VAMP8,CAMTA
CYT2,USE1,AGPAT4,IFI27L1,ASF1B,CCDC25,EEF1B2,PPP1CC,EDF1,RFX3,DGKE,TNFRSF14,VAMP8,CAMTA
,ATG12,NLRP1,ZBTB44,NIP7,TTC9C,DUSP22,SLC25A10,TMX2,THEM4,RDH11,DBNDD2,NISCH,CENPK,RPL
,ATG12,NLRP1,ZBTB44,NIP7,TTC9C,DUSP22,SLC25A10,TMX2,THEM4,RDH11,DBNDD2,NISCH,CENPK,RPL
17B,TMEM126B,APEH,RPL21,TXNL4A,CAPN10,LYPLA2,NUTF2,PODXL,CAMLG,UCK2,PPOX,ATPAF2,PRMT
17B,TMEM126B,APEH,RPL21,TXNL4A,CAPN10,LYPLA2,NUTF2,PODXL,CAMLG,UCK2,PPOX,ATPAF2,PRMT
17B,TMEM126B,APEH,RPL21,TXNL4A,CAPN10,LYPLA2,NUTF2,PODXL,CAMLG,UCK2,PPOX,ATPAF2,PRMT
17B,TMEM126B,APEH,RPL21,TXNL4A,CAPN10,LYPLA2,NUTF2,PODXL,CAMLG,UCK2,PPOX,ATPAF2,PRMT
0AB1,DUT,SYNGR2,CEP70,TMEM134,LRRFIP1,RAB2A,ZNF524,ATP5J,TPM2,GPBP1,SIAH1,GGCX,ZEB2,AS
0AB1,DUT,SYNGR2,CEP70,TMEM134,LRRFIP1,RAB2A,ZNF524,ATP5J,TPM2,GPBP1,SIAH1,GGCX,ZEB2,AS
R4,PSMA1,YLPM1,CHCHD1,SLC39A11,FUZ,PSMA3,MUC1,METTL17,EPB41L5,NLRP1,ZBTB44,NIP7,TTC9C
AMLG,EPB41L4B,TCF4,PPOX,ATPAF2,C17ORF62,PRMT1,CASC4,WNK1,ZNF7,PPAN,ZFAND2B,AP1S3,HNR
AMLG,EPB41L4B,TCF4,PPOX,ATPAF2,C17ORF62,PRMT1,CASC4,WNK1,ZNF7,PPAN,ZFAND2B,AP1S3,HNR
TB44,NIP7,TTC9C,DUSP22,SLC25A10,TMX2,THEM4,RDH11,DBNDD2,NISCH,CENPK,RPL29,CD99L2,GGPS1
MT112,PNPO,STRIP2,NIT2,LYNX1,TWSG1,MAPK13,ZSWIM7,DCTD,GMPPB,HSPB1,LRRC27,MVD,LMBR1,N
T1,HYPK,RPL36,MOB1A,NFAM1,CDK11A,ZNF197,COPZ1,INO80E,CCNL2,DGUOK,DUT,ATP5J,ELP6,RMDN1
T1,HYPK,RPL36,MOB1A,NFAM1,CDK11A,ZNF197,COPZ1,INO80E,CCNL2,DGUOK,DUT,ATP5J,ELP6,RMDN1
TK2,AUNIP,C14ORF2,CENPN,ATF7,PCYT2,FBXW9,USE1,HSBP1L1,AGPAT4,MAF,ASF1B,PPP1CC,EDF1,GAD
TK2,AUNIP,C14ORF2,CENPN,ATF7,PCYT2,FBXW9,USE1,HSBP1L1,AGPAT4,MAF,ASF1B,PPP1CC,EDF1,GAD
N2,SDHA,SZT2,FBXO34,EXOSC2,RBM7,MCUR1,AP2S1,ME3,LRP10,PKIG,SLC31A1,MRPL51,SFSWAP,MED8

OASY,SRSF1,AUP1,SAP30BP,YIPF3,FOXRED1,TRNT1,KXD1,ITGB1BP1,MED15,SH3YL1,IP6K2,CLASRP,JPH2,

OASY,SRSF1,AUP1,SAP30BP,YIPF3,FOXRED1,TRNT1,KXD1,ITGB1BP1,MED15,SH3YL1,IP6K2,CLASRP,JPH2,

NF7,TXNDC17,STRADA,AP1S3,ZNF644,CARS,ZNF250,RPL36,NFAM1,MDH1,INO80E,CCNL2,MAPKAPK5,HS

NF7,TXNDC17,STRADA,AP1S3,ZNF644,CARS,ZNF250,RPL36,NFAM1,MDH1,INO80E,CCNL2,MAPKAPK5,HS

ATG12,NLRP1,ZBTB44,NIP7,TTC9C,DUSP22,SLC25A10,TMX2,THEM4,RDH11,DBNDD2,NISCH,CENPK,RPL2

ATG12,NLRP1,ZBTB44,NIP7,TTC9C,DUSP22,SLC25A10,TMX2,THEM4,RDH11,DBNDD2,NISCH,CENPK,RPL2

DF1,RFX3,DGKE,TNFRSF14,ENDOV,VAMP8,CAMTA1,PPIL2,TROAP,RPL28,CLDND1,VIM,LIG1,PSMA7,TME

DF1,RFX3,DGKE,TNFRSF14,ENDOV,VAMP8,CAMTA1,PPIL2,TROAP,RPL28,CLDND1,VIM,LIG1,PSMA7,TME

SME2,PABPC1,RRNAD1,DBNL,NAT9,BCAM,BLVRB,PPP3CB,RABEP2,TADA3,ZNF20,ATP6V1F,RALGPS1,ZNF

NPK,RPS6,RPL29,GGPS1,RTEL1,COASY,SRSF1,AUP1,YIPF3,FOXRED1,TRNT1,KXD1,ITGB1BP1,MED15,SH3Y

NPK,RPS6,RPL29,GGPS1,RTEL1,COASY,SRSF1,AUP1,YIPF3,FOXRED1,TRNT1,KXD1,ITGB1BP1,MED15,SH3Y

BP1,SH3YL1,IP6K2,CLASRP,C1ORF198,UGGT2,MICAL1,HS2ST1,UBE2I,KCNMA1,SLC39A13,YIF1A,PRMT2,E

BP1,SH3YL1,IP6K2,CLASRP,C1ORF198,UGGT2,MICAL1,HS2ST1,UBE2I,KCNMA1,SLC39A13,YIF1A,PRMT2,E

DGKE,TNFRSF14,VAMP8,CAMTA1,PSMD4,RPL28,PSMA7,TIMM17B,TMEM126B,HYI,RPL21,TXNL4A,CAPN

11,DBNDD2,NISCH,CENPK,RPS6,RPL29,GGPS1,RTEL1,COASY,SRSF1,AUP1,SAP30BP,YIPF3,FOXRED1,TRNT

7,SHPK,SLC25A45,CDKL3,MARK2,SEC14L1,CSRP2BP,SEC24C,ZNF850,NDUFA8,ZNF195,RPE,EDEM2,TVP23

F1,DCTN5,ARL16,LY6E,TMEM108,KRAS,GFOD1,KLHL7,SSB,RPL8,LIMA1,UBE2D4,ECE1,ASH1L,TCF25,ZNF2

F1,DCTN5,ARL16,LY6E,TMEM108,KRAS,GFOD1,KLHL7,SSB,RPL8,LIMA1,UBE2D4,ECE1,ASH1L,TCF25,ZNF2

F1,GADL1,PPA1,DGKE,TNFRSF14,ENDOV,CAMTA1,PPIL2,PSMD4,CLDND1,VIM,LIG1,PSMA7,CDC14B,TME

F1,GADL1,PPA1,DGKE,TNFRSF14,ENDOV,CAMTA1,PPIL2,PSMD4,CLDND1,VIM,LIG1,PSMA7,CDC14B,TME

44,TTC9C,DUSP22,SLC25A10,TMX2,RDH11,DBNDD2,NISCH,CENPK,RPL29,GGPS1,RTEL1,COASY,SRSF1,A

5,THADA,PHC3,CCDC40,PIGT,EIF2B3,NFATC4,MFSD11,ZNHIT3,RPS11,MOK,SNX5,NUP133,RAPGEF3,HAU

,RALGPS1,ZNF687,PDCD5,NXN,KLK10,POLR3H,RAD23A,HSD17B10,ABHD17A,GGT5,DDX41,SLC16A1,PKP

,RALGPS1,ZNF687,PDCD5,NXN,KLK10,POLR3H,RAD23A,HSD17B10,ABHD17A,GGT5,DDX41,SLC16A1,PKP

,PTPN13,UBR4,PSMA1,YLPM1,PHF21A,CHCHD1,SLC39A11,FUZ,FAM3A,MUC1,ECSIT,EPB41L5,ATG12,PP

,PTPN13,UBR4,PSMA1,YLPM1,PHF21A,CHCHD1,SLC39A11,FUZ,FAM3A,MUC1,ECSIT,EPB41L5,ATG12,PP

W9,ASF1B,CCDC25,EEF1B2,PPP1CC,EDF1,GADL1,RFX3,ENDOV,CAMTA1,PPIL2,PSMD4,CLDND1,VIM,LIG1

W9,ASF1B,CCDC25,EEF1B2,PPP1CC,EDF1,GADL1,RFX3,ENDOV,CAMTA1,PPIL2,PSMD4,CLDND1,VIM,LIG1

1,SUGT1,HYPK,UPP1,MOB1A,CDK11A,RBM3,MDH1,LAMP2,COPZ1,CCNL2,MAPKAPK5,LRRFIP1,FAM219B

1,SUGT1,HYPK,UPP1,MOB1A,CDK11A,RBM3,MDH1,LAMP2,COPZ1,CCNL2,MAPKAPK5,LRRFIP1,FAM219B

I27L1,MAF,CCDC25,PPP1CC,RFX3,PPA1,ENDOV,CAMTA1,PPIL2,PSMA7,TIMM17B,APEH,RPL21,TXNL4A,R

I27L1,MAF,CCDC25,PPP1CC,RFX3,PPA1,ENDOV,CAMTA1,PPIL2,PSMA7,TIMM17B,APEH,RPL21,TXNL4A,R

,FOXRED1,TRNT1,KXD1,ITGB1BP1,MED15,SH3YL1,IP6K2,CLASRP,UGGT2,MICAL1,SLC39A13,YIF1A,PRMT

,FOXRED1,TRNT1,KXD1,ITGB1BP1,MED15,SH3YL1,IP6K2,CLASRP,UGGT2,MICAL1,SLC39A13,YIF1A,PRMT

6B,APEH,HYI,CAPN10,LYPLA2,NUTF2,PODXL,TCF4,ATPAF2,C17ORF62,PRMT1,CASC4,WNK1,C19ORF54,Z

6B,APEH,HYI,CAPN10,LYPLA2,NUTF2,PODXL,TCF4,ATPAF2,C17ORF62,PRMT1,CASC4,WNK1,C19ORF54,Z

1,ZC3H14,FIS1,SLC15A3,PPM1K,TUBA1C,IRF9,ANTXR2,STIP1,LRRC1,PHF19,GRAMD1C,VCAN,PCBP2,LPCA

1,ZC3H14,FIS1,SLC15A3,PPM1K,TUBA1C,IRF9,ANTXR2,STIP1,LRRC1,PHF19,GRAMD1C,VCAN,PCBP2,LPCA

CEP70,TMEM134,LRRFIP1,RAB2A,ZNF524,ATP5J,TPM2,GPBP1,SIAH1,GGCX,ZEB2,ASPH,CHPF,LYNX1,TW

7,TMEM126B,APEH,RBM33,LYPLA2,NUTF2,UCK2,TMPO,ATPAF2,C17ORF62,PRMT1,ZNF7,TXNDC17,ARL

4,STAU1,SLC9B2,HYPK,ZNF250,MOB1A,CDK11A,MDH1,COPZ1,CCNL2,SYNGR2,TMEM134,LRRFIP1,RAB2

HEM4,RDH11,DBNDD2,SDF4,NISCH,TMED4,RPS6,GGPS1,RTEL1,SRSF1,AUP1,SAP30BP,RABL5,TRNT1,KX

HEM4,RDH11,DBNDD2,SDF4,NISCH,TMED4,RPS6,GGPS1,RTEL1,SRSF1,AUP1,SAP30BP,RABL5,TRNT1,KX

F4,NISCH,TMED4,CENPK,RPS6,RPL29,CD99L2,GGPS1,COASY,SRSF1,AUP1,YIPF3,FOXRED1,TRNT1,PARD6

F4,NISCH,TMED4,CENPK,RPS6,RPL29,CD99L2,GGPS1,COASY,SRSF1,AUP1,YIPF3,FOXRED1,TRNT1,PARD6

RF54,ZNF7,TXNDC17,PIGL,STRADA,ZFAND2B,HNRNPC,ZNF644,HYPK,UPP1,ZNF250,MOB1A,RBM3,ZNF1

4,CENPK,RPS6,RPL29,RTEL1,COASY,SRSF1,AUP1,SAP30BP,YIPF3,TRNT1,KXD1,ITGB1BP1,MED15,SH3YL1,

4,CENPK,RPS6,RPL29,RTEL1,COASY,SRSF1,AUP1,SAP30BP,YIPF3,TRNT1,KXD1,ITGB1BP1,MED15,SH3YL1,

HINT1,POLR2C,INTS4,B3GAT3,RFXANK,SPINT2,FTH1,PFDN5,MTHFS,PAFAH1B2,TXNRD2,GBA2,PPIH,DAL

HINT1,POLR2C,INTS4,B3GAT3,RFXANK,SPINT2,FTH1,PFDN5,MTHFS,PAFAH1B2,TXNRD2,GBA2,PPIH,DAL

,FCHO2,GTF2H1,BFAR,SLC35A2,PML,TMEM199,SLC25A24,OXR1,UBE2D3,C1ORF63,ZNF396,EPN1,HPS1,

,FCHO2,GTF2H1,BFAR,SLC35A2,PML,TMEM199,SLC25A24,OXR1,UBE2D3,C1ORF63,ZNF396,EPN1,HPS1,

M134,LRRFIP1,RAB2A,GPALPP1,SIAH1,TSFM,ASPH,TRMT112,CHPF,STRIP2,ZNF625,RPS20,LYNX1,TWSG1,

XDC,COBL,TNKS,SMIM4,ERGIC1,MYL9,C6ORF132,SLC35E1,TSEN34,RPL8,LIMA1,UBE2D4,ECE1,TCF25,KC

XDC,COBL,TNKS,SMIM4,ERGIC1,MYL9,C6ORF132,SLC35E1,TSEN34,RPL8,LIMA1,UBE2D4,ECE1,TCF25,KC

1,FKBP3,EGLN2,SIRT5,KCTD7,B3GAT3,ZNF506,SPINT2,FTH1,PAFAH1B2,TYMS,PTK7,TMEM39B,DALRD3,

ETTL3,SSR3,PICK1,RNF181,KCNC4,RPL35,ANAPC13,TMC7,SRSF7,MTIF2,NXF1,SLC39A6,RALY,MARS,GLYC

HA5,RANBP1,UBE2L6,VRK3,LZTFL1,ZC3H14,FIS1,SLC15A3,PPM1K,TUBA1C,IRF9,DST,ANTXR2,STIP1,LRRC

F644,SLC9B2,RPL36,MOB1A,CDK11A,MDH1,INO80E,HSP90AB1,DUT,SYNGR2,TMEM134,LRRFIP1,FAM21

,PTPN13,UBR4,PSMA1,YLPM1,PHF21A,SLC39A11,FUZ,PSMA3,MUC1,EPB41L5,RPS25,NLRP1,ZBTB44,NI

,PTPN13,UBR4,PSMA1,YLPM1,PHF21A,SLC39A11,FUZ,PSMA3,MUC1,EPB41L5,RPS25,NLRP1,ZBTB44,NI

2,RHEB,IDH3G,EML3,CKLF,MKRN1,INO80C,FARS2,EMC9,CD37,ARRB2,RBM6,SNRPA1,HMGN1,CDK2AP2,

15,PARD6G,SH3YL1,JPH2,UGGT2,MICAL1,UBE2I,YIF1A,EIF3M,MAGT1,JUP,CDC42,TBX6,TMEM205,PTK2,

CDK11A,RBM3,MDH1,LAMP2,ZNF197,COPZ1,INO80E,CCNL2,MAPKAPK5,DGUOK,HSP90AB1,DUT,TMEM

CDK11A,RBM3,MDH1,LAMP2,ZNF197,COPZ1,INO80E,CCNL2,MAPKAPK5,DGUOK,HSP90AB1,DUT,TMEM

ZSWIM7,ERLEC1,GMPPB,HHLA3,MVD,SAT1,ERCC5,EMC4,IL32,NAP1L1,PRKRIP1,DDOST,SYNGR1,RANGR

ZSWIM7,ERLEC1,GMPPB,HHLA3,MVD,SAT1,ERCC5,EMC4,IL32,NAP1L1,PRKRIP1,DDOST,SYNGR1,RANGR

IG1,TMEM126B,XAF1,APEH,RPL21,EPB41L4B,UCK2,PPOX,TMPO,ATPAF2,PRMT1,EXOSC9,STOML2,ZNF7,

IG1,TMEM126B,XAF1,APEH,RPL21,EPB41L4B,UCK2,PPOX,TMPO,ATPAF2,PRMT1,EXOSC9,STOML2,ZNF7,

40,MROH1,NFATC4,MFSD11,LFNG,XYLT2,TMEM14B,PCID2,ZXDC,COBL,SMIM4,ERGIC1,NMNAT3,MPHOS

1,PSMD4,RPL28,CLDND1,LIG1,TMEM126B,HYI,RPL21,TXNL4A,CAMLG,TCF4,UCK2,TMPO,LSM12,C17OR

1,PSMD4,RPL28,CLDND1,LIG1,TMEM126B,HYI,RPL21,TXNL4A,CAMLG,TCF4,UCK2,TMPO,LSM12,C17OR

5,DGUOK,HSP90AB1,SYNGR2,CEP70,TMEM134,RAB2A,RMDN1,GPBP1,GPALPP1,GGCX,ASPH,CHPF,MPD

5,DGUOK,HSP90AB1,SYNGR2,CEP70,TMEM134,RAB2A,RMDN1,GPBP1,GPALPP1,GGCX,ASPH,CHPF,MPD

,GRK6,HMGA2,CDC37,RPS16,CPSF3L,TSG101,OBSL1,PTPN13,UBR4,YLPM1,PHF21A,CHCHD1,SLC39A11,F

,GRK6,HMGA2,CDC37,RPS16,CPSF3L,TSG101,OBSL1,PTPN13,UBR4,YLPM1,PHF21A,CHCHD1,SLC39A11,F

MO2,ABHD17A,NT5C,USP45,ILF3,STXBP2,EEF1D,NAA10,RRP8,MED6,LLGL2,CHEK1,COX17,COX5A,SUMO

MBR1,MCTS1,NAP1L1,DDOST,SYNGR1,CHCHD7,MEIS2,PHTF2,DBNL,NAT9,ZNF836,RBP1,H2AFV,RBMX,S

PRMT1,CASC4,C19ORF54,ZNF7,TXNDC17,PIGL,STRADA,AP1S3,ACAD10,ZNF644,HYPK,UPP1,CARS,ZNF25

PRMT1,CASC4,C19ORF54,ZNF7,TXNDC17,PIGL,STRADA,AP1S3,ACAD10,ZNF644,HYPK,UPP1,CARS,ZNF25

1ORF48,OAS3,MED6,CARHSP1,CNST,LLGL2,PRLR,CHEK1,COX17,HNRNPLL,SUMO3,HNRNPA1,HIGD1A,E

HIGD1A,RPL15,INTS4,ASH2L,SMARCA4,SNRPF,TYMS,ZFPL1,OARD1,TPM4,TRPM4,MRPS18C,RIC8A,LGAL

BX6,TMEM205,PTK2,AUNIP,C14ORF2,GUK1,CENPN,PCYT2,FBXW9,USE1,AGPAT4,MAF,DEF8,PPP1CC,ED

BX6,TMEM205,PTK2,AUNIP,C14ORF2,GUK1,CENPN,PCYT2,FBXW9,USE1,AGPAT4,MAF,DEF8,PPP1CC,ED

RSF1,AUP1,YIPF3,FOXRED1,RABL5,KXD1,ITGB1BP1,PARD6G,SH3YL1,JPH2,MICAL1,HS2ST1,UBE2I,YIF1A,EI
RSF1,AUP1,YIPF3,FOXRED1,RABL5,KXD1,ITGB1BP1,PARD6G,SH3YL1,JPH2,MICAL1,HS2ST1,UBE2I,YIF1A,EI
P2,ZFPL1,MFSD10,ZSCAN9,RIC8A,LGALS3,TCP1,SLC35B1,SMARCC2,NFKBIA,PHC3,PIGT,UBE2N,MFSD11,L

14,ENDOV,CAMTA1,CRIP1,PSMD4,TROAP,RPL28,LIG1,PSMA7,CDC14B,TIMM17B,TMEM126B,XAF1,RPL2
14,ENDOV,CAMTA1,CRIP1,PSMD4,TROAP,RPL28,LIG1,PSMA7,CDC14B,TIMM17B,TMEM126B,XAF1,RPL2
1,UPP1,CARS,NFAM1,RBM3,MDH1,ZNF197,CCNL2,DUT,CEP70,TMEM134,LRRFIP1,RAB2A,ZNF524,ELP6,(

HARPIN,ZNF644,STAU1,SLC9B2,UPP1,ZNF250,RPL36,RBM3,MDH1,INO80E,CCNL2,MAPKAPK5,CEP70,LRR
HARPIN,ZNF644,STAU1,SLC9B2,UPP1,ZNF250,RPL36,RBM3,MDH1,INO80E,CCNL2,MAPKAPK5,CEP70,LRR
PV17,ARIH2,PICK1,ZNF512,RNF181,NIPAL3,CHTOP,RPL35,TSPAN31,PPP2R5C,SLC39A6,ZNF593,GLYCTK,R
PV17,ARIH2,PICK1,ZNF512,RNF181,NIPAL3,CHTOP,RPL35,TSPAN31,PPP2R5C,SLC39A6,ZNF593,GLYCTK,R
N10,LYPLA2,TMPO,C17ORF62,WNK1,ZNF7,TXNDC17,ARL6IP4,ZFAND2B,AP1S3,HNRNPC,ZNF644,SLC9B2,
5C,AGAP3,MDH2,HSPH1,OAS3,SEC31A,CNST,SSH2,COX17,HNRNPLL,ZFAND4,LGMN,FKBP3,NDUFV2,INTS
D7,HMGA1,PSME2,PABPC1,DBNL,NAT9,BCAM,ZNF836,BLVRB,PPP3CB,RABEP2,TADA3,ZNF20,WAC,RAL(
NPK,CD99L2,GGPS1,COASY,SRSF1,AUP1,FOXRED1,RABL5,TRNT1,KXD1,ITGB1BP1,PARD6G,SH3YL1,IP6K2
NPK,CD99L2,GGPS1,COASY,SRSF1,AUP1,FOXRED1,RABL5,TRNT1,KXD1,ITGB1BP1,PARD6G,SH3YL1,IP6K2
A11,FUZ,FAM3A,PSMA3,MUC1,METTL17,ATG12,PPRC1,RPS25,NLRP1,ZBTB44,TTC9C,COX4I1,DUSP22,TN
A11,FUZ,FAM3A,PSMA3,MUC1,METTL17,ATG12,PPRC1,RPS25,NLRP1,ZBTB44,TTC9C,COX4I1,DUSP22,TN
CD99L2,GGPS1,AUP1,FOXRED1,RABL5,TRNT1,KXD1,PARD6G,SH3YL1,UGGT2,MICAL1,HS2ST1,UBE2I,SLC3
CD99L2,GGPS1,AUP1,FOXRED1,RABL5,TRNT1,KXD1,PARD6G,SH3YL1,UGGT2,MICAL1,HS2ST1,UBE2I,SLC3
N34,RPL8,CREB3L4,ECE1,TCF25,RNF181,TJP1,SLC52A2,NXF1,SLC39A6,APOLD1,RALY,RBM42,FAM69B,REF
BX6,TMEM205,XBP1,PTK2,C14ORF2,GUK1,PCYT2,FBXW9,USE1,HSBP1L1,MAF,ASF1B,PPP1CC,RFX3,PPA1,
L,ZNF687,GTF2IRD1,KLK10,NUDT1,POT1,TRIM16,POLR3H,RAD23A,PSME1,SLC16A1,ILF3,PKP4,MDH2,STX
4I1,DUSP22,SLC25A10,SPOCK2,RDH11,SDF4,NISCH,TMED4,CENPK,RPS6,RPL29,CD99L2,GGPS1,COASY,SI
4I1,DUSP22,SLC25A10,SPOCK2,RDH11,SDF4,NISCH,TMED4,CENPK,RPS6,RPL29,CD99L2,GGPS1,COASY,SI

FV,RBMX,PPP3CB,RABEP2,WAC,RALGPS1,GTF2IRD1,NXN,TRIM16,RAD23A,HSD17B10,ABHD17A,PKP4,C1
FV,RBMX,PPP3CB,RABEP2,WAC,RALGPS1,GTF2IRD1,NXN,TRIM16,RAD23A,HSD17B10,ABHD17A,PKP4,C1

RPL21,CAPN10,NUTF2,CAMLG,EPB41L4B,TCF4,UCK2,PPOX,ATPAF2,C17ORF62,PRMT1,WNK1,C19ORF54,
RPL21,CAPN10,NUTF2,CAMLG,EPB41L4B,TCF4,UCK2,PPOX,ATPAF2,C17ORF62,PRMT1,WNK1,C19ORF54,
CTD,GMPPB,HHLA3,SAT1,LMBR1,SYNGR1,RANGRF,MTA3,CHCHD7,HMGA1,PHTF2,PABPC1,RRNAD1,NA
CTD,GMPPB,HHLA3,SAT1,LMBR1,SYNGR1,RANGRF,MTA3,CHCHD7,HMGA1,PHTF2,PABPC1,RRNAD1,NA
A13,YIF1A,PRMT2,JUP,CDC42,TBX6,XBP1,PTK2,C14ORF2,CENPN,ATF7,PCYT2,FBXW9,USE1,HSBP1L1,AGP

NF197,INO80E,CCNL2,CEP70,FAM219B,RAB2A,ZNF524,ATP5J,ELP6,TPM2,GPBP1,GPALPP1,SIAH1,GGCX,C
NF197,INO80E,CCNL2,CEP70,FAM219B,RAB2A,ZNF524,ATP5J,ELP6,TPM2,GPBP1,GPALPP1,SIAH1,GGCX,C
L,XBP1,PTK2,C14ORF2,GUK1,CENPN,CARD14,ATF7,PCYT2,FBXW9,USE1,AGPAT4,IFI27L1,MAF,CCDC25,DE
L,XBP1,PTK2,C14ORF2,GUK1,CENPN,CARD14,ATF7,PCYT2,FBXW9,USE1,AGPAT4,IFI27L1,MAF,CCDC25,DE
70,LRRFIP1,RAB2A,ZNF524,ATP5J,TPM2,GPALPP1,SIAH1,GGCX,TRMT112,STRIP2,ZNF625,GABPB1,LYNX1
70,LRRFIP1,RAB2A,ZNF524,ATP5J,TPM2,GPALPP1,SIAH1,GGCX,TRMT112,STRIP2,ZNF625,GABPB1,LYNX1
ATPAF2,C17ORF62,PRMT1,CASC4,EXOSC9,STOML2,WNK1,ZNF7,TXNDC17,ARL6IP4,PIGL,PPAN,AP1S3,HN

ATPAF2,C17ORF62,PRMT1,CASC4,EXOSC9,STOML2,WNK1,ZNF7,TXNDC17,ARL6IP4,PIGL,PPAN,AP1S3,HN

CARS,ZNF250,RPL36,NFAM1,RBM3,MDH1,LAMP2,ZNF197,INO80E,CCNL2,MAPKAPK5,HSP90AB1,CEP70,T
CARS,ZNF250,RPL36,NFAM1,RBM3,MDH1,LAMP2,ZNF197,INO80E,CCNL2,MAPKAPK5,HSP90AB1,CEP70,T
,TMEM126B,CAPN10,NUTF2,EPB41L4B,UCK2,PPOX,EXOSC9,STOML2,C19ORF54,ZNF7,STRADA,AP1S3,HN
,TMEM126B,CAPN10,NUTF2,EPB41L4B,UCK2,PPOX,EXOSC9,STOML2,C19ORF54,ZNF7,STRADA,AP1S3,HN
IPO,ATPAF2,LSM12,C17ORF62,PRMT1,CASC4,C19ORF54,ZNF7,ARL6IP4,PIGL,STRADA,AP1S3,ACAD10,ZN
,VIM,LIG1,PSMA7,TIMM17B,XAF1,APEH,HYI,TXNL4A,CAPN10,NUTF2,PODXL,EPB41L4B,TCF4,UCK2,ATPA
RED1,RABL5,TRNT1,KXD1,ITGB1BP1,MED15,SH3YL1,C1ORF198,UGGT2,MICAL1,HS2ST1,UBE2I,KCNMA1,
RED1,RABL5,TRNT1,KXD1,ITGB1BP1,MED15,SH3YL1,C1ORF198,UGGT2,MICAL1,HS2ST1,UBE2I,KCNMA1
MPPB,HHLA3,MVD,SAT1,ERCC5,EMC4,IL32,PRKRIP1,DDOST,SYNGR1,RANGRF,MEIS2,PABPC1,RRNAD1,NA
MPPB,HHLA3,MVD,SAT1,ERCC5,EMC4,IL32,PRKRIP1,DDOST,SYNGR1,RANGRF,MEIS2,PABPC1,RRNAD1,NA

ZNF524,ATP5J,TPM2,GPBP1,SIAH1,CHPF,STRIP2,GABPB1,LYNX1,ZSWIM7,ZNF814,ERLEC1,LRRC27,MVD,I
,EGLN2,SIRT5,KCTD7,B3GAT3,ZNF506,SPINT2,C11ORF49,PAFAH1B2,PTK7,GTF2IRD2,BRF1,ZFPL1,FAM10
MS,ZFPL1,OARD1,TPM4,MRPS18C,LGALS3,TCP1,PHF6,GNB5,COMMD4,NFATC4,MFSD11,SNX5,FAM32A,R
1,CAPG,PSMD4,RPL28,VIM,LIG1,PSMA7,TMEM126B,APEH,HYI,TXNL4A,CAPN10,NUTF2,EPB41L4B,PPOX,
C1,METTL17,ECSIT,NLRP1,ZBTB44,NIP7,TTC9C,COX4I1,DUSP22,SLC25A10,SPOCK2,THEM4,RDH11,DBND
C1,METTL17,ECSIT,NLRP1,ZBTB44,NIP7,TTC9C,COX4I1,DUSP22,SLC25A10,SPOCK2,THEM4,RDH11,DBND

P7,TTC9C,COX4I1,DUSP22,SLC25A10,TMX2,THEM4,DBNDD2,SDF4,NISCH,RPS6,RPL29,RTEL1,COASY,AUP
P7,TTC9C,COX4I1,DUSP22,SLC25A10,TMX2,THEM4,DBNDD2,SDF4,NISCH,RPS6,RPL29,RTEL1,COASY,AUP
PG,CRIP1,PPIL2,RPL28,CLDND1,LIG1,PSMA7,CDC14B,APEH,HYI,TXNL4A,CAPN10,PODXL,CAMLG,EPB41L4
),NAA10,LLGL2,CHEK1,COX17,ZFAND4,FKBP3,NDUFV2,EGLN2,KCTD7,B3GAT3,SPINT2,FTH1,SMARCA4,PA
PHOSPH9,DPY30,MBD1,NDUFS7,TPM1,DAD1,C15ORF40,DCTN1,UQCRB,DCTN6,RNF149,HBS1L,B9D1,OA
PHOSPH9,DPY30,MBD1,NDUFS7,TPM1,DAD1,C15ORF40,DCTN1,UQCRB,DCTN6,RNF149,HBS1L,B9D1,OA
,CHEK1,COX17,SUMO3,ZFAND4,FKBP3,NDUFV2,EGLN2,KCTD7,B3GAT3,SPINT2,PAFAH1B2,PTK7,GTF2IR

H1,LAMP2,ZNF197,INO80E,CCNL2,MAPKAPK5,HSP90AB1,DUT,CEP70,ZNF524,ELP6,GPBP1,GGCX,ZEB2,TR
2,PRMT1,WNK1,STRADA,ZFAND2B,AP1S3,HNRNPC,CYB561,SHARPIN,STAU1,UPP1,ZNF250,RPL36,MOB1
2,PRMT1,WNK1,STRADA,ZFAND2B,AP1S3,HNRNPC,CYB561,SHARPIN,STAU1,UPP1,ZNF250,RPL36,MOB1
1A,CHCHD1,SLC39A11,FUZ,FAM3A,MUC1,METTL17,EPB41L5,ATG12,PPRC1,ZBTB44,COX4I1,PLCB2,DUSP
1A,CHCHD1,SLC39A11,FUZ,FAM3A,MUC1,METTL17,EPB41L5,ATG12,PPRC1,ZBTB44,COX4I1,PLCB2,DUSP
ORF15,TBCA,DDX10,POLR2E,BCL7B,IAH1,PNRC1,ARPC1A,VAPA,SSC5D,DHDDS,CEP63,RPS23,SDHA,ZNF4
ORF15,TBCA,DDX10,POLR2E,BCL7B,IAH1,PNRC1,ARPC1A,VAPA,SSC5D,DHDDS,CEP63,RPS23,SDHA,ZNF4

TXNDC17,ARL6IP4,C19ORF48,STRADA,ZFAND2B,AP1S3,HNRNPC,CYB561,STAU1,SLC9B2,SUGT1,UPP1,ZN
TXNDC17,ARL6IP4,C19ORF48,STRADA,ZFAND2B,AP1S3,HNRNPC,CYB561,STAU1,SLC9B2,SUGT1,UPP1,ZN
S15,DCTD,UBXN1,ZNF814,HSPB1,VWA2,LRRC27,EMC4,IL32,RANGRF,HMGA1,PABPC1,RAD1,RBMX,BLVR
S15,DCTD,UBXN1,ZNF814,HSPB1,VWA2,LRRC27,EMC4,IL32,RANGRF,HMGA1,PABPC1,RAD1,RBMX,BLVR
SNRPF,GTF2IRD2,DRG2,BRF1,OARD1,TRPM4,MFSD10,PHF6,PARK7,ADAM19,SMARCC2,NFKBIA,SLC30A5
SNRPF,GTF2IRD2,DRG2,BRF1,OARD1,TRPM4,MFSD10,PHF6,PARK7,ADAM19,SMARCC2,NFKBIA,SLC30A5

PA1,DGKE,ENDOV,CAPG,PPIL2,RPL28,CLDND1,LIG1,TIMM17B,TMEM126B,XAF1,APEH,HYI,RPL21,LYPLA2

PA1,DGKE,ENDOV,CAPG,PPIL2,RPL28,CLDND1,LIG1,TIMM17B,TMEM126B,XAF1,APEH,HYI,RPL21,LYPLA2

IF1A,EIF3M,CDC42,TBX6,TMEM205,XBP1,PTK2,C14ORF2,GUK1,CENPN,CARD14,PCYT2,IFI27L1,MAF,ASF1

IF1A,EIF3M,CDC42,TBX6,TMEM205,XBP1,PTK2,C14ORF2,GUK1,CENPN,CARD14,PCYT2,IFI27L1,MAF,ASF1

,TLCD1,PLAC8,THOC3,AXIN1,MOSPD3,IFITM2,SLC25A24,EIF3C,AAMDC,EPN1,HPS1,SNRPD1,DCAF5,GRAN

3M4,PLAC8,HNRNPUL1,TP53,TMEM199,C1ORF63,SNAPC2,EPN1,MRPS24,CREB3L2,ZCCHC9,MKRN1,SMU

1,ERCC5,LMBR1,NAP1L1,SERPINB1,RANGRF,MTA3,CHCHD7,MEIS2,HMGA1,PHTF2,PABPC1,NAT9,TOMM

L,ERCC5,LMBR1,MCTS1,NAP1L1,DDOST,SYNGR1,RANGRF,CHCHD7,MEIS2,PHTF2,NAT9,ZNF836,RAD1,H2A

L,ERCC5,LMBR1,MCTS1,NAP1L1,DDOST,SYNGR1,RANGRF,CHCHD7,MEIS2,PHTF2,NAT9,ZNF836,RAD1,H2A

1,ITGB1BP1,PARD6G,SH3YL1,C1ORF198,UGGT2,MICAL1,HS2ST1,YIF1A,PRMT2,MAGT1,TMEM205,XBP1,

1,ITGB1BP1,PARD6G,SH3YL1,C1ORF198,UGGT2,MICAL1,HS2ST1,YIF1A,PRMT2,MAGT1,TMEM205,XBP1,

RHSP1,LLGL2,CHEK1,COX17,CAMK2D,HNRNPLL,SUMO3,HIGD1A,RPL15,INTS4,B3GAT3,SPINT2,SMARCA4

RIC8A,NFKBIA,CCDC40,MFSD11,LFNG,XYLT2,TMEM14B,PCID2,ZXDC,COBL,PRRC2B,STX10,ERGIC1,NMNA

RIC8A,NFKBIA,CCDC40,MFSD11,LFNG,XYLT2,TMEM14B,PCID2,ZXDC,COBL,PRRC2B,STX10,ERGIC1,NMNA

MPDU1,MAPK13,ZSWIM7,RPS15,ZNF814,ERLEC1,GMPPB,HSPB1,VWA2,MVD,LMBR1,NAP1L1,PRKRIP1,R

3,VWA2,ERCC5,IL32,LMBR1,NAP1L1,SERPINB1,DDOST,C12ORF76,RANGRF,LSR,PABPC1,RRNAD1,RAD1,BL

3,VWA2,ERCC5,IL32,LMBR1,NAP1L1,SERPINB1,DDOST,C12ORF76,RANGRF,LSR,PABPC1,RRNAD1,RAD1,BL

LGL2,CHEK1,COX17,SUMO3,ZFAND4,FKBP3,NDUFV2,EGLN2,SIRT5,KCTD7,B3GAT3,RFXANK,SPINT2,PAFA

NFAM1,LAMP2,HSP90AB1,DUT,SYNGR2,CEP70,TMEM134,LRRFIP1,ZNF524,TPM2,GPBP1,GPALPP1,SIAH1

NFAM1,LAMP2,HSP90AB1,DUT,SYNGR2,CEP70,TMEM134,LRRFIP1,ZNF524,TPM2,GPBP1,GPALPP1,SIAH1

NMA1,YIF1A,MAGT1,JUP,CDC42,TBX6,TMEM205,XBP1,PTK2,CELF1,AUNIP,CENPN,CARD14,ATF7,PCYT2,I

NMA1,YIF1A,MAGT1,JUP,CDC42,TBX6,TMEM205,XBP1,PTK2,CELF1,AUNIP,CENPN,CARD14,ATF7,PCYT2,I

L4B,XAF1,APEH,RPL21,TXNL4A,CAPN10,LYPLA2,NUTF2,CAMLG,TMPO,ATPAF2,LSM12,PRMT1,WNK1,C19

L4B,XAF1,APEH,RPL21,TXNL4A,CAPN10,LYPLA2,NUTF2,CAMLG,TMPO,ATPAF2,LSM12,PRMT1,WNK1,C19

G,SH3YL1,IP6K2,CLASRP,MICAL1,UBE2I,YIF1A,EIF3M,CDC42,TMEM205,XBP1,PTK2,CENPN,CARD14,PCYT

G,SH3YL1,IP6K2,CLASRP,MICAL1,UBE2I,YIF1A,EIF3M,CDC42,TMEM205,XBP1,PTK2,CENPN,CARD14,PCYT

P1,SIAH1,CHPF,STRIP2,RPS20,GABPB1,LYNX1,MPDU1,TWSG1,ZSWIM7,RPS15,ERLEC1,GMPPB,HHLA3,SAT

P1,SIAH1,CHPF,STRIP2,RPS20,GABPB1,LYNX1,MPDU1,TWSG1,ZSWIM7,RPS15,ERLEC1,GMPPB,HHLA3,SAT

H11,NISCH,TMED4,CENPK,RPS6,COASY,SRSF1,AUP1,SAP30BP,YIPF3,FOXRED1,RABL5,TRNT1,ITGB1BP1,N

H11,NISCH,TMED4,CENPK,RPS6,COASY,SRSF1,AUP1,SAP30BP,YIPF3,FOXRED1,RABL5,TRNT1,ITGB1BP1,N

SUFU,ZNF33A,MRPS33,EXOSC2,HNRNPM,NIPA1,POLH,EXOSC8,ACTB,CCDC171,C12ORF73,MRPL43,SETD

SUFU,ZNF33A,MRPS33,EXOSC2,HNRNPM,NIPA1,POLH,EXOSC8,ACTB,CCDC171,C12ORF73,MRPL43,SETD

ICHD7,PSME2,RRNAD1,TOMM40,SERPINB6,TADA3,RALGPS1,PDCD5,GTF2IRD1,NXN,ANAPC11,B2M,DDX

4A,NUP62,IRF3,SIN3A,RBM4,TLCD1,NDUFA11,HNRNPUL1,TP53,RPS3A,BTF3,AAMDC,RHEB,RDX,MAGOHB

87,NXN,KLK10,POT1,ANAPC11,PSME1,SLC16A1,ILF3,PKP4,ZCCHC4,MED6,LLGL2,EGFR,CHEK1,CAMK2D,C

GT2,MICAL1,HS2ST1,UBE2I,KCNMA1,YIF1A,PRMT2,JUP,CDC42,TBX6,XBP1,PTK2,C14ORF2,CENPN,CARD1

GT2,MICAL1,HS2ST1,UBE2I,KCNMA1,YIF1A,PRMT2,JUP,CDC42,TBX6,XBP1,PTK2,C14ORF2,CENPN,CARD1

MBR1,NAP1L1,SERPINB1,SYNGR1,RANGRF,LSR,MTA3,CHCHD7,MEIS2,HMGA1,PSME2,PABPC1,RRNAD1,N

70,TMEM134,FAM219B,RAB2A,RMDN1,GPBP1,GPALPP1,TRMT112,PNPO,CHPF,MPDU1,MAPK13,ZSWIM

70,TMEM134,FAM219B,RAB2A,RMDN1,GPBP1,GPALPP1,TRMT112,PNPO,CHPF,MPDU1,MAPK13,ZSWIM

ASF1B,EEF1B2,PPP1CC,EDF1,RFX3,DGKE,ENDOV,CAMTA1,CAPG,CRIP1,PPIL2,PSMD4,RPL28,PSMA7,TMEM

ASF1B,EEF1B2,PPP1CC,EDF1,RFX3,DGKE,ENDOV,CAMTA1,CAPG,CRIP1,PPIL2,PSMD4,RPL28,PSMA7,TMEM

,ERGIC1,NMNAT3,MPHOSPH10,MYL9,OGFOD2,TSEN34,RPL8,CREB3L4,LIMA1,UBE2D4,TCF25,TJP1,CHTO

NF250,RPL36,NFAM1,RBM3,MDH1,ZNF197,COPZ1,INO80E,CCNL2,MAPKAPK5,HSP90AB1,DUT,SYNGR2,R

FOXRED1,RABL5,TRNT1,KXD1,ITGB1BP1,MED15,SH3YL1,CLASRP,JPH2,UGGT2,MICAL1,HS2ST1,UBE2I,KCI
FOXRED1,RABL5,TRNT1,KXD1,ITGB1BP1,MED15,SH3YL1,CLASRP,JPH2,UGGT2,MICAL1,HS2ST1,UBE2I,KCI
,PAFAH1B2,TYMS,PTK7,DRG2,BRF1,FAM102A,PFKFB2,RIC8A,NFKBIA,GNB5,CBR1,NHP2,MFSD11,LFNG,PI

F644,SLC9B2,UPP1,ZNF250,COPZ1,CCNL2,MAPKAPK5,DGUOK,HSP90AB1,SYNGR2,LRRFIP1,RAB2A,ATP5J
D2,GFOD1,CIZ1,EIF3G,FBXL18,TCF25,RNF181,ADD1,NIPAL3,CHTOP,BAX,CDK4,SLC52A2,ANAPC13,SLC39
D2,GFOD1,CIZ1,EIF3G,FBXL18,TCF25,RNF181,ADD1,NIPAL3,CHTOP,BAX,CDK4,SLC52A2,ANAPC13,SLC39
REB3L2,UCHL5,AKT1S1,NSFL1C,SPHK2,ACTR10,HMGN1,NDUFB2,CDK2AP2,NDUFA10,RPS27L,AZI2,SPG21
5A23,FAM102A,GMPR2,MCFD2,PFKFB2,RIC8A,NFKBIA,SLC30A5,TRPV3,PIGT,EIF2B3,UBE2N,LFNG,ZNHIT3
5A23,FAM102A,GMPR2,MCFD2,PFKFB2,RIC8A,NFKBIA,SLC30A5,TRPV3,PIGT,EIF2B3,UBE2N,LFNG,ZNHIT3
YNX1,MPDU1,TWSG1,VPS28,ZSWIM7,RPS15,DCTD,ERLEC1,HSPB1,MVD,SEMA3C,MCTS1,NAP1L1,PRKRI
YNX1,MPDU1,TWSG1,VPS28,ZSWIM7,RPS15,DCTD,ERLEC1,HSPB1,MVD,SEMA3C,MCTS1,NAP1L1,PRKRI
C39A11,FUZ,RGS3,FAM3A,ECSIT,EPB41L5,ATG12,RPS25,NLRP1,ZBTB44,NIP7,TTC9C,COX4I1,PLCB2,DUSI
C39A11,FUZ,RGS3,FAM3A,ECSIT,EPB41L5,ATG12,RPS25,NLRP1,ZBTB44,NIP7,TTC9C,COX4I1,PLCB2,DUSI
CYT2,IFI27L1,MAF,ASF1B,RFX3,PPA1,DGKE,TNFRSF14,CAMTA1,CRIP1,PPIL2,PSMD4,RPL28,LIG1,PSMA7,T
52,NADSYN1,ZNF511,ZNF283,IRF3,D2HGDH,CC2D1B,TLCD1,SNX29,NDUFA11,AXIN1,MOSPD3,SLC37A4,SI
T,RANGRF,LSR,MTA3,CHCHD7,PSME2,PABPC1,RRNAD1,PTPRB,DBNL,NAT9,RBP1,RAD1,H2AFV,RABEP2,T
M14B,NEK2,ACOT2,COBL,PIGH,ELOF1,C12ORF57,NMNAT3,NUBP2,STAU2,RPLP0,OGFOD2,KLHL7,METTL3
SLC37A3,EMC3,TAF9,RNPS1,IAH1,DTD2,CAMKK1,DHDDS,ARL2,NMRAL1,ZNF483,ACHE,PEX16,FBXO34,N
7,EPB41L5,ATG12,RPS25,NLRP1,ZBTB44,NIP7,TTC9C,COX4I1,PLCB2,DUSP22,TMX2,SPOCK2,THEM4,DBN
7,EPB41L5,ATG12,RPS25,NLRP1,ZBTB44,NIP7,TTC9C,COX4I1,PLCB2,DUSP22,TMX2,SPOCK2,THEM4,DBN
,ATPAF2,PRMT1,PIGL,STRADA,PPAN,ZFAND2B,AP1S3,ACAD10,STAU1,SUGT1,CARS,ZNF250,RPL36,CDK1
TOMM40,ZNF20,SLC33A1,ZNF687,POT1,RAD23A,B2M,ABHD17A,DDX41,SLC16A1,PKP4,MDH2,EEF1D,HS
TOMM40,ZNF20,SLC33A1,ZNF687,POT1,RAD23A,B2M,ABHD17A,DDX41,SLC16A1,PKP4,MDH2,EEF1D,HS
H1,ZNF197,COPZ1,INO80E,CEP70,LRRFIP1,ZNF524,ATP5J,TPM2,GPALPP1,SIAH1,TSFM,TPM3,PNPO,RPT
H1,ZNF197,COPZ1,INO80E,CEP70,LRRFIP1,ZNF524,ATP5J,TPM2,GPALPP1,SIAH1,TSFM,TPM3,PNPO,RPT
IPLL,HNRNPA1,CD151,FKBP3,NDUFV2,RPL15,INTS4,ZNF506,REEP5,ASH2L,FTH1,MTHFS,MPC2,TXNRD2,G
IPLL,HNRNPA1,CD151,FKBP3,NDUFV2,RPL15,INTS4,ZNF506,REEP5,ASH2L,FTH1,MTHFS,MPC2,TXNRD2,G
,SLC16A1,MDH2,STXBP2,EEF1D,NAA10,C11ORF48,SEC31A,CARHSP1,RPL18,EGFR,SH3GLB1,COX5A,SUM
A,TRPM4,PFKFB2,RIC8A,NFKBIA,GNB5,COMMD4,CCDC40,MROH1,MFSD11,LFNG,XYLT2,PCID2,ZXDC,COE
2,SH3GLB1,COX17,CAMK2D,COX5A,SUMO3,HINT1,HIGD1A,EGLN2,RPL15,B3GAT3,ASH2L,MTHFS,SNRPF
2,SH3GLB1,COX17,CAMK2D,COX5A,SUMO3,HINT1,HIGD1A,EGLN2,RPL15,B3GAT3,ASH2L,MTHFS,SNRPF
EF1D,NAA10,RRP8,LLGL2,EGFR,CHEK1,COX17,SUMO3,FKBP3,EGLN2,KCTD7,B3GAT3,SPINT2,PAFAH1B2,T
2,RRNAD1,DBNL,BCAM,RBMX,TOMM40,PPP3CB,ZNF20,WAC,SLC33A1,ZNF687,GTF2IRD1,NXN,HSD17B1
2,RRNAD1,DBNL,BCAM,RBMX,TOMM40,PPP3CB,ZNF20,WAC,SLC33A1,ZNF687,GTF2IRD1,NXN,HSD17B1
YIPF3,FOXRED1,RABL5,TRNT1,KXD1,ITGB1BP1,MED15,IP6K2,C1ORF198,UGGT2,MICAL1,UBE2I,TBX6,XBI
YIPF3,FOXRED1,RABL5,TRNT1,KXD1,ITGB1BP1,MED15,IP6K2,C1ORF198,UGGT2,MICAL1,UBE2I,TBX6,XBI
OCK2,THEM4,RDH11,SDF4,NISCH,TMED4,CENPK,RPS6,RPL29,CD99L2,GGPS1,COASY,SRSF1,AUP1,FOXRE
OCK2,THEM4,RDH11,SDF4,NISCH,TMED4,CENPK,RPS6,RPL29,CD99L2,GGPS1,COASY,SRSF1,AUP1,FOXRE
PF31,PPP1R7,GSK3A,ZNF599,SRSF5,ABCB8,PSMD11,SDCCAG8,N4BP2L2,AAK1,ACP6,MIB2,TSTA3,RANGA
PF31,PPP1R7,GSK3A,ZNF599,SRSF5,ABCB8,PSMD11,SDCCAG8,N4BP2L2,AAK1,ACP6,MIB2,TSTA3,RANGA
POT1,ANAPC11,POLR3H,DDX41,NT5C,AGAP3,ZCCHC4,MDH2,STXBP2,EEF1D,CHCHD5,NAA10,C11ORF48,
A1,SMAGP,TSPAN31,SRSF7,TMEM9,SLC39A6,ZNF593,GLYCTK,RBM42,CLASP2,CREBZF,GLI4,NADSYN1,ZN
,PSMA1,YLPM1,PHF21A,CHCHD1,SLC39A11,FUZ,FAM3A,PSMA3,MUC1,EPB41L5,ATG12,RPS25,NLRP1,ZE
,PSMA1,YLPM1,PHF21A,CHCHD1,SLC39A11,FUZ,FAM3A,PSMA3,MUC1,EPB41L5,ATG12,RPS25,NLRP1,ZE

A23,FAM102A,MFSD10,LGALS3,NFKBIA,GNB5,TRPV3,CBR1,COMMD4,NHP2,MROH1,UBE2N,MFSD11,NC

A23,FAM102A,MFSD10,LGALS3,NFKBIA,GNB5,TRPV3,CBR1,COMMD4,NHP2,MROH1,UBE2N,MFSD11,NC

NF706,EPS15L1,RAD23A,B2M,ILF3,PKP4,AGAP3,EEF1D,HSPH1,C11ORF48,RRP8,CARHSP1,RPL18,ATP10D

MRPL36,RNF149,HBS1L,B9D1,HNRNPL,ENTPD6,ZNF652,SEC24C,NDUFA8,TOMM7,NPRL3,C17ORF97,PAP

MRPL36,RNF149,HBS1L,B9D1,HNRNPL,ENTPD6,ZNF652,SEC24C,NDUFA8,TOMM7,NPRL3,C17ORF97,PAP

P6K2,CLASRP,JPH2,UGGT2,MICAL1,HS2ST1,UBE2I,SLC39A13,YIF1A,PRMT2,MAGT1,JUP,CDC42,XBP1,PTK

P6K2,CLASRP,JPH2,UGGT2,MICAL1,HS2ST1,UBE2I,SLC39A13,YIF1A,PRMT2,MAGT1,JUP,CDC42,XBP1,PTK

63,SDHA,ZNF483,ABLIM2,PEX16,SUFU,ZNF33A,EXOSC2,PSMC5,HNRNPM,NIPA1,APRT,EXOSC8,TCFL5,AC

63,SDHA,ZNF483,ABLIM2,PEX16,SUFU,ZNF33A,EXOSC2,PSMC5,HNRNPM,NIPA1,APRT,EXOSC8,TCFL5,AC

MED6,CARHSP1,PRLR,COX17,HINT1,POLR2C,B3GAT3,FTH1,PFDN5,MTHFS,PAFAH1B2,GBA2,PPIH,NSUN

MED6,CARHSP1,PRLR,COX17,HINT1,POLR2C,B3GAT3,FTH1,PFDN5,MTHFS,PAFAH1B2,GBA2,PPIH,NSUN

NXN,KLK10,POT1,TRIM16,ANAPC11,RAD23A,DDX41,PKP4,AGAP3,STXBP2,HSPH1,NAA10,RRP8,SEC31A,C

NXN,KLK10,POT1,TRIM16,ANAPC11,RAD23A,DDX41,PKP4,AGAP3,STXBP2,HSPH1,NAA10,RRP8,SEC31A,C

3,ZSWIM7,ERLEC1,GMPPB,MVD,SAT1,ERCC5,EMC4,IL32,LMBR1,NAP1L1,SYNGR1,RANGRF,CHCHD7,PAB

3,ZSWIM7,ERLEC1,GMPPB,MVD,SAT1,ERCC5,EMC4,IL32,LMBR1,NAP1L1,SYNGR1,RANGRF,CHCHD7,PAB

VPS25,UCHL5,TUSC3,SMUG1,DIP2A,NDUFA10,RPS7,U2AF1L4,KIF13B,CARM1,FAM104B,NCBP2,SYPL1,G

VPS25,UCHL5,TUSC3,SMUG1,DIP2A,NDUFA10,RPS7,U2AF1L4,KIF13B,CARM1,FAM104B,NCBP2,SYPL1,G

F7,TXNDC17,ARL6IP4,PIGL,C19ORF48,PPAN,ZFAND2B,SHARPIN,SLC9B2,SUGT1,CARS,MOB1A,RBM3,LAM

C25A10,TMX2,SPOCK2,RDH11,SDF4,NISCH,TMED4,CENPK,RPS6,CD99L2,GGPS1,RTEL1,SRSF1,AUP1,SAP3

C25A10,TMX2,SPOCK2,RDH11,SDF4,NISCH,TMED4,CENPK,RPS6,CD99L2,GGPS1,RTEL1,SRSF1,AUP1,SAP3

L5,SH3YL1,IP6K2,CLASRP,C1ORF198,UGGT2,HS2ST1,UBE2I,MAGT1,JUP,CDC42,TBX6,TMEM205,XBP1,AU

EH,HYI,LYPLA2,NUTF2,PODXL,CAMLG,EPB41L4B,TCF4,PPOX,ATPAF2,C17ORF62,PRMT1,CASC4,WNK1,C1

P1,MOB1A,CDK11A,CCNL2,MAPKAPK5,HSP90AB1,DUT,SYNGR2,CEP70,TMEM134,LRRFIP1,ZNF524,ELP6

P1,MOB1A,CDK11A,CCNL2,MAPKAPK5,HSP90AB1,DUT,SYNGR2,CEP70,TMEM134,LRRFIP1,ZNF524,ELP6

BPC1,BCAM,H2AFV,BLVRB,PPP3CB,RABEP2,TADA3,RALGPS1,ZNF687,PDCD5,GTF2IRD1,NXN,KLK10,EPS1

BPC1,BCAM,H2AFV,BLVRB,PPP3CB,RABEP2,TADA3,RALGPS1,ZNF687,PDCD5,GTF2IRD1,NXN,KLK10,EPS1

GABPB1,VPS28,ZSWIM7,RPS15,DCTD,ZNF814,ERLEC1,HSPB1,VWA2,LRRC27,MVD,ERCC5,NAP1L1,PRKRIP

BTB8OS,SLK,APAF1,KAT8,U2AF1L4,CARM1,CCDC24,PSMG4,TBCA,SNX3,BCL7B,PNRC1,UBA52,CAMKK1,X

BTB8OS,SLK,APAF1,KAT8,U2AF1L4,CARM1,CCDC24,PSMG4,TBCA,SNX3,BCL7B,PNRC1,UBA52,CAMKK1,X

G1,MAPK13,UBXN1,ZNF814,GMPPB,HSPB1,MVD,ERCC5,PRKRIP1,SYNGR1,C12ORF76,RANGRF,LSR,PABP

C19ORF48,STRADA,ZFAND2B,AP1S3,HNRNPC,CYB561,SHARPIN,STAU1,UPP1,ZNF250,MOB1A,ZNF197,CC

C19ORF48,STRADA,ZFAND2B,AP1S3,HNRNPC,CYB561,SHARPIN,STAU1,UPP1,ZNF250,MOB1A,ZNF197,CC

IGR2,TMEM134,LRRFIP1,RAB2A,ZNF524,ATP5J,TPM2,SIAH1,ZEB2,ASPH,TRMT112,STRIP2,ZNF625,NIT2,L

L,RAD1,H2AFV,BLVRB,TOMM40,SERPINB6,RABEP2,TADA3,WAC,ZNF706,ZNF687,PDCD5,DDX41,NT5C,ILF

L,RAD1,H2AFV,BLVRB,TOMM40,SERPINB6,RABEP2,TADA3,WAC,ZNF706,ZNF687,PDCD5,DDX41,NT5C,ILF

H1,EIF2B3,LFNG,ZNHIT3,XYLT2,RAPGEF3,TMEM14B,ZXDC,TNKS,SMIM4,ERGIC1,KPNA5,MYL9,SLC35E1,TS

H1,EIF2B3,LFNG,ZNHIT3,XYLT2,RAPGEF3,TMEM14B,ZXDC,TNKS,SMIM4,ERGIC1,KPNA5,MYL9,SLC35E1,TS

7,RPS15,ERLEC1,GMPPB,HHLA3,SAT1,ERCC5,LMBR1,MCTS1,NAP1L1,SERPINB1,DDOST,SYNGR1,RANGRF,

7,RPS15,ERLEC1,GMPPB,HHLA3,SAT1,ERCC5,LMBR1,MCTS1,NAP1L1,SERPINB1,DDOST,SYNGR1,RANGRF,

L,RER1,CREBZF,DCAKD,IRF3,TAF1D,RNF130,CLTA,NAA38,RBM4,SCRN2,LYPLAL1,ST8SIA1,TP53,PSMD10,C

REB3L4,ECE1,ASH1L,SLMAP,NIPAL3,TPT1,BAX,CDK4,SLC52A2,TSPAN31,NXF1,FAM192A,CFLAR,TRIM46,L

NIP,C14ORF2,GUK1,ATF7,PCYT2,USE1,AGPAT4,EDF1,GADL1,RFX3,PPA1,DGKE,TNFRSF14,VAMP8,CAMTA

NIP,C14ORF2,GUK1,ATF7,PCYT2,USE1,AGPAT4,EDF1,GADL1,RFX3,PPA1,DGKE,TNFRSF14,VAMP8,CAMTA

AT4,MAF,CCDC25,PPP1CC,RFX3,PPA1,DGKE,TNFRSF14,VAMP8,CAMTA1,CAPG,CRIP1,VIM,LIG1,CDC14B,

AT4,MAF,CCDC25,PPP1CC,RFX3,PPA1,DGKE,TNFRSF14,VAMP8,CAMTA1,CAPG,CRIP1,VIM,LIG1,CDC14B,

PL28,LIG1,PSMA7,TIMM17B,XAF1,APEH,HYI,TXNL4A,CAPN10,LYPLA2,NUTF2,CAMLG,EPB41L4B,PPOX,TM

PL28,LIG1,PSMA7,TIMM17B,XAF1,APEH,HYI,TXNL4A,CAPN10,LYPLA2,NUTF2,CAMLG,EPB41L4B,PPOX,TM

ZNF524,TPM2,GPBP1,SIAH1,GGCX,ZEB2,ASPH,TRMT112,PNPO,RPTOR,CHPF,STRIP2,ZNF625,RPS20,MAPI

ZNF524,TPM2,GPBP1,SIAH1,GGCX,ZEB2,ASPH,TRMT112,PNPO,RPTOR,CHPF,STRIP2,ZNF625,RPS20,MAPI

C3,COMMD4,MFF,SNX5,COX20,TMEM147,COA1,SMIM4,NMNAT3,BTN3A2,OXLD1,MPHOSPH10,C6ORF1

C3,COMMD4,MFF,SNX5,COX20,TMEM147,COA1,SMIM4,NMNAT3,BTN3A2,OXLD1,MPHOSPH10,C6ORF1

7,SKA2,GTF2H1,TAF1D,CLTA,NAA38,ZNF69,NRP1,AXIN1,AP1G2,MOSPD3,TMEM199,DNAJC2,MRPS24,EN

7,SKA2,GTF2H1,TAF1D,CLTA,NAA38,ZNF69,NRP1,AXIN1,AP1G2,MOSPD3,TMEM199,DNAJC2,MRPS24,EN

A1,CCDC74B,ZNFX1,DUSP18,PLEKHA8,CLSPN,ISOC2,MCM7,PPIE,RCN1,IQCK,PARP6,AP4M1,ATP9B,AK9,C

A7,TIMM17B,APEH,HYI,TXNL4A,CAPN10,LYPLA2,NUTF2,PODXL,CAMLG,EPB41L4B,TCF4,PPOX,TMPO,ATI

A7,TIMM17B,APEH,HYI,TXNL4A,CAPN10,LYPLA2,NUTF2,PODXL,CAMLG,EPB41L4B,TCF4,PPOX,TMPO,ATI

ZFAND2B,AP1S3,HNRNPC,CYB561,SHARPIN,STAU1,RPL36,MOB1A,LAMP2,CCNL2,HSP90AB1,DUT,SYNGR

0,RPL36,MOB1A,CCNL2,HSP90AB1,DUT,SYNGR2,TMEM134,LRRFIP1,ZNF524,ATP5J,TPM2,GPALPP1,SIAH

0,RPL36,MOB1A,CCNL2,HSP90AB1,DUT,SYNGR2,TMEM134,LRRFIP1,ZNF524,ATP5J,TPM2,GPALPP1,SIAH

1,ITGB1BP1,SH3YL1,IP6K2,CLASRP,JPH2,UGGT2,MICAL1,HS2ST1,UBE2I,KCNMA1,SLC39A13,YIF1A,PRMT

1,ITGB1BP1,SH3YL1,IP6K2,CLASRP,JPH2,UGGT2,MICAL1,HS2ST1,UBE2I,KCNMA1,SLC39A13,YIF1A,PRMT

MAPK13,ZSWIM7,DCTD,ERLEC1,GMPPB,HHLA3,VWA2,MVD,SAT1,ERCC5,EMC4,IL32,LMBR1,MCTS1,DDC

MAPK13,ZSWIM7,DCTD,ERLEC1,GMPPB,HHLA3,VWA2,MVD,SAT1,ERCC5,EMC4,IL32,LMBR1,MCTS1,DDC

PHC3,MFSD11,LFNG,SNX5,RPAIN,COX20,TMEM147,GRAPL,SMIM4,PRRC2B,DCTN5,ARL16,BTN3A2,MPH

L16,EIF4A1,SLC35E1,RC3H2,KLHL7,MRPL55,ARIH2,LIMA1,PICK1,UBE2D4,ECE1,OCIAD1,TCF25,RBBP6,ZN

L16,EIF4A1,SLC35E1,RC3H2,KLHL7,MRPL55,ARIH2,LIMA1,PICK1,UBE2D4,ECE1,OCIAD1,TCF25,RBBP6,ZN

GGCX,ASPH,TRMT112,CHPF,LYNX1,MPDU1,TWSG1,MAPK13,ZSWIM7,RPS15,DCTD,HSPB1,LRRC27,SAT1,

G2,BRF1,ZFPL1,FAM102A,MCFD2,PFKFB2,RIC8A,PHF6,SMARCC2,NFKBIA,COMMD4,PIGT,EIF2B3,NFATC4,

G2,BRF1,ZFPL1,FAM102A,MCFD2,PFKFB2,RIC8A,PHF6,SMARCC2,NFKBIA,COMMD4,PIGT,EIF2B3,NFATC4,

TRMT112,CHPF,STRIP2,GABPB1,MPDU1,TWSG1,MAPK13,ZSWIM7,RPS15,DCTD,UBXN1,LRRC27,SEMA3C,

TRMT112,CHPF,STRIP2,GABPB1,MPDU1,TWSG1,MAPK13,ZSWIM7,RPS15,DCTD,UBXN1,LRRC27,SEMA3C,

RIM16,RAD23A,B2M,ILF3,AGAP3,MDH2,SEC31A,CNST,PRLR,CHEK1,COX17,COX5A,HNRNPLL,RPL15,B3GA

RBMX,PPP3CB,RABEP2,ATP6V1F,WAC,PDCD5,EPS15L1,ANAPC11,NOMO2,RAD23A,HSD17B10,NT5C,USP

RBMX,PPP3CB,RABEP2,ATP6V1F,WAC,PDCD5,EPS15L1,ANAPC11,NOMO2,RAD23A,HSD17B10,NT5C,USP

GR1,RANGRF,CHCHD7,RRNAD1,RBP1,RAD1,H2AFV,RALGPS1,ZNF687,NUDT1,POT1,ANAPC11,POLR3H,RA
GR1,RANGRF,CHCHD7,RRNAD1,RBP1,RAD1,H2AFV,RALGPS1,ZNF687,NUDT1,POT1,ANAPC11,POLR3H,RA
L3,UBE2L3,PABPN1,C12ORF73,ATPAF1,ZNF586,ISOC2,PPIE,XYLB,IMMP1L,COQ4,ANKRD10,GANC,DPAGT1
L3,UBE2L3,PABPN1,C12ORF73,ATPAF1,ZNF586,ISOC2,PPIE,XYLB,IMMP1L,COQ4,ANKRD10,GANC,DPAGT1

3A,NAA38,TLCD1,GEMIN2,TUBB3,MOSPD3,C5ORF24,RPS3,MAGOHB,CREB3L2,VPS25,AIFM2,HMOX2,SP
L3,TMC7,CFLAR,C11ORF57,RBM42,VDAC1,CNTRL,RER1,CREBZF,GLI4,NUP62,OLFM1,PSMD7,PAM16,IRF3
L3,TMC7,CFLAR,C11ORF57,RBM42,VDAC1,CNTRL,RER1,CREBZF,GLI4,NUP62,OLFM1,PSMD7,PAM16,IRF3
NTRL,FAM69B,RER1,CREBZF,RPL24,PSMD7,NADSYN1,SKA2,NEDD4L,RNF130,PIGO,BFAR,D2HGDH,KDELR3
F3M,MAGT1,JUP,TBX6,XBP1,PTK2,CELF1,C14ORF2,GUK1,ATF7,PCYT2,FBXW9,USE1,AGPAT4,IFI27L1,MAF
F3M,MAGT1,JUP,TBX6,XBP1,PTK2,CELF1,C14ORF2,GUK1,ATF7,PCYT2,FBXW9,USE1,AGPAT4,IFI27L1,MAF
,SLC15A3,PPM1K,TUBA1C,IRF9,DST,ANTXR2,STIP1,LRRC1,PHF19,GRAMD1C,VCAN,PCBP2,LPCAT3,CD44,
,SLC15A3,PPM1K,TUBA1C,IRF9,DST,ANTXR2,STIP1,LRRC1,PHF19,GRAMD1C,VCAN,PCBP2,LPCAT3,CD44,
ARS2,EMC9,NDUFA10,REPS2,ZBTB8OS,KAT8,TBCA,SYPL1,USP15,BCL7B,PIWIL4,PRKCSH,CAPN2,ACVR1B,
ZNF512,OCIAD1,TCF25,RNF181,SLMAP,RBBP6,ADD1,BAX,CDK4,NME1,NTMT1,ANAPC13,EEF2K,FAM192
ZNF512,OCIAD1,TCF25,RNF181,SLMAP,RBBP6,ADD1,BAX,CDK4,NME1,NTMT1,ANAPC13,EEF2K,FAM192

1,CREBZF,NKIRAS2,ACAA1,OLFM1,ZNF675,FCHO2,ZNF511,IRF3,TRAPPC2,GTF2H1,KDELR3,CLTA,MYL6,PL
PT1,PUF60,MT1F,ATP9B,AK9,ANKRD10,ACTG1,EHMT1,RAP1B,IFT81,UBE2A,PKN2,CREM,CHFR,TUBGCP2
P2,INO80E,CCNL2,HSP90AB1,CEP70,TMEM134,LRRFIP1,ELP6,GPBP1,GPALPP1,SIAH1,CHPF,GABPB1,MPD
P2,INO80E,CCNL2,HSP90AB1,CEP70,TMEM134,LRRFIP1,ELP6,GPBP1,GPALPP1,SIAH1,CHPF,GABPB1,MPD
,NMRAL1,CEP63,ABLIM2,NSRP1,BSCL2,FBXO34,NUB1,ZNF816,KCNH2,LRP10,APRT,HAUS1,EXOSC8,SLC3
,NMRAL1,CEP63,ABLIM2,NSRP1,BSCL2,FBXO34,NUB1,ZNF816,KCNH2,LRP10,APRT,HAUS1,EXOSC8,SLC3
P1,ZEB2,ASPH,PNPO,CHPF,MPDU1,MAPK13,ZSWIM7,RPS15,UBXN1,ZNF814,ERLEC1,HHLA3,MVD,ERCC5
P1,ZEB2,ASPH,PNPO,CHPF,MPDU1,MAPK13,ZSWIM7,RPS15,UBXN1,ZNF814,ERLEC1,HHLA3,MVD,ERCC5

DH1,LAMP2,HSP90AB1,DUT,SYNGR2,CEP70,TMEM134,LRRFIP1,RAB2A,ZNF524,TPM2,GPBP1,SIAH1,GG
DH1,LAMP2,HSP90AB1,DUT,SYNGR2,CEP70,TMEM134,LRRFIP1,RAB2A,ZNF524,TPM2,GPBP1,SIAH1,GG
DT1,ANAPC11,POLR3H,RAD23A,DDX41,SLC16A1,PKP4,AGAP3,PDE4DIP,MDH2,EEF1D,HSPH1,NAA10,RRP
DT1,ANAPC11,POLR3H,RAD23A,DDX41,SLC16A1,PKP4,AGAP3,PDE4DIP,MDH2,EEF1D,HSPH1,NAA10,RRP

,NDUFV2,RPL15,INTS4,KCTD7,ZNF506,REEP5,ASH2L,PFDN5,SMARCA4,COA3,TYMS,GTF2IRD2,PPIH,DALR
,NDUFV2,RPL15,INTS4,KCTD7,ZNF506,REEP5,ASH2L,PFDN5,SMARCA4,COA3,TYMS,GTF2IRD2,PPIH,DALR
A11,FUZ,FAM3A,PSMA3,MUC1,EPB41L5,ATG12,RPS25,NLRP1,ZBTB44,NIP7,TTC9C,COX4I1,DUSP22,SLC25
,PTK2,C14ORF2,ATF7,PCYT2,USE1,AGPAT4,IFI27L1,MAF,DEF8,EEF1B2,PPP1CC,EDF1,PPA1,DGKE,TNFRSF1
,PTK2,C14ORF2,ATF7,PCYT2,USE1,AGPAT4,IFI27L1,MAF,DEF8,EEF1B2,PPP1CC,EDF1,PPA1,DGKE,TNFRSF1
483,LRRC23,SUMF1,FBXO34,CYB5D2,DNAJC24,ZNF816,ALS2,IK,PKIG,EXOSC8,DDX17,SLC31A1,CCDC171,
GOLN2,PEX2,RPL37A,PIGH,DCTN5,NDUFS3,L3MBTL2,OXLD1,EIF4A1,C6ORF132,DGAT1,ARIH2,RPL8,CREB
GOLN2,PEX2,RPL37A,PIGH,DCTN5,NDUFS3,L3MBTL2,OXLD1,EIF4A1,C6ORF132,DGAT1,ARIH2,RPL8,CREB
P0,METTL5,C6ORF132,ZMYM6,SSR3,MPV17,ARIH2,PICK1,ZNF512,TCF25,RNF181,NIPAL3,CHTOP,NME1,
P0,METTL5,C6ORF132,ZMYM6,SSR3,MPV17,ARIH2,PICK1,ZNF512,TCF25,RNF181,NIPAL3,CHTOP,NME1,
1C,LPCAT3,CD44,TM9SF1,MAGED2,SON,TET2,UQCRQ,SNF8,GRK6,HMGA2,RPS16,OBSL1,TTC21B,UBR4,P
1C,LPCAT3,CD44,TM9SF1,MAGED2,SON,TET2,UQCRQ,SNF8,GRK6,HMGA2,RPS16,OBSL1,TTC21B,UBR4,P

AGT1,MRPS7,MED1,NDUFS7,CHFR,HYKK,SHPK,DIDO1,ZNF585A,RSBN1L,LSM14A,AP4S1,COX7C,AHI1,GA
A,OLFM1,NEDD4L,IRF3,SCRN2,NDUFA11,AXIN1,TUBB3,MOSPD3,UBE2D3,C5ORF24,SLC37A4,SNAPC2,DN
A,OLFM1,NEDD4L,IRF3,SCRN2,NDUFA11,AXIN1,TUBB3,MOSPD3,UBE2D3,C5ORF24,SLC37A4,SNAPC2,DN
I,ZNF836,RBP1,H2AFV,SERPINB6,TADA3,WAC,ZNF706,ZNF687,ANAPC11,RAD23A,PSME1,ILF3,PKP4,EEF1
511,GTF2H1,TAF1D,RNF130,D2HGDH,MYL6,SIN3A,NAA38,TLCD1,GEMIN2,TUBB3,RPS3A,MOSPD3,C5ORF
511,GTF2H1,TAF1D,RNF130,D2HGDH,MYL6,SIN3A,NAA38,TLCD1,GEMIN2,TUBB3,RPS3A,MOSPD3,C5ORF
TAF1D,RNF130,D2HGDH,MYL6,SIN3A,NAA38,TLCD1,GEMIN2,TUBB3,RPS3A,MOSPD3,C5ORF24,SLC37A4,

13B,CARM1,MAZ,CCDC24,TTLL3,MDK,FIG4,ACVR1B,CAMKK1,ZNF276,XPNPEP1,ABLIM2,CCNG2,NUB1,ZN

P6V1F,WAC,PDCD5,EPS15L1,ANAPC11,NOMO2,RAD23A,HSD17B10,NT5C,USP45,ILF3,ZCCHC4,MDH2,STX
P6V1F,WAC,PDCD5,EPS15L1,ANAPC11,NOMO2,RAD23A,HSD17B10,NT5C,USP45,ILF3,ZCCHC4,MDH2,STX
LGPS1,ZNF706,ZNF687,GTF2IRD1,NXN,KLK10,EPS15L1,ANAPC11,POLR3H,ABHD17A,DDX41,SLC16A1,ILF3
LGPS1,ZNF706,ZNF687,GTF2IRD1,NXN,KLK10,EPS15L1,ANAPC11,POLR3H,ABHD17A,DDX41,SLC16A1,ILF3
YNGR1,RANGRF,CHCHD7,MEIS2,PHTF2,PABPC1,DBNL,NAT9,RBP1,H2AFV,TOMM40,SERPINB6,TADA3,ZN
YNGR1,RANGRF,CHCHD7,MEIS2,PHTF2,PABPC1,DBNL,NAT9,RBP1,H2AFV,TOMM40,SERPINB6,TADA3,ZN

3,PPA1,DGKE,TNFRSF14,ENDOV,VAMP8,CAMTA1,CAPG,PPIL2,PSMD4,RPL28,CLDND1,PSMA7,CDC14B,TI
3,PPA1,DGKE,TNFRSF14,ENDOV,VAMP8,CAMTA1,CAPG,PPIL2,PSMD4,RPL28,CLDND1,PSMA7,CDC14B,TI

3,DGKE,TNFRSF14,ENDOV,CAMTA1,CAPG,CRIP1,CLDND1,LIG1,PSMA7,APEH,HYI,TXNL4A,CAPN10,LYPLA2
3,DGKE,TNFRSF14,ENDOV,CAMTA1,CAPG,CRIP1,CLDND1,LIG1,PSMA7,APEH,HYI,TXNL4A,CAPN10,LYPLA2

DENND1A,MRPS24,RHEB,NQO2,AKT1S1,TUSC3,HMGN1,MFGE8,B3GNTL1,REPS2,KAT8,KIF13B,CARM1,M
1,STAU1,HYPK,CARS,ZNF250,CDK11A,MDH1,COPZ1,CCNL2,DGUOK,HSP90AB1,DUT,SYNGR2,GPBP1,GPA
1,STAU1,HYPK,CARS,ZNF250,CDK11A,MDH1,COPZ1,CCNL2,DGUOK,HSP90AB1,DUT,SYNGR2,GPBP1,GPA
51,PSMA7,APEH,HYI,CAPN10,LYPLA2,NUTF2,PODXL,CAMLG,EPB41L4B,TCF4,UCK2,TMPO,ATPAF2,CASC4,
51,PSMA7,APEH,HYI,CAPN10,LYPLA2,NUTF2,PODXL,CAMLG,EPB41L4B,TCF4,UCK2,TMPO,ATPAF2,CASC4,
2,HMGA1,PABPC1,RAD1,H2AFV,BLVRB,TOMM40,PPP3CB,SERPINB6,ATP6V1F,ZNF706,PDCD5,GTF2IRD1
M7,ALS2,EXOSC8,DDX17,SLC31A1,ZNFX1,RCAN1,C12ORF73,NSMF,ATP9B,IMMP1L,RAP1B,PRKCZ,COQ2,

L4A,CAPN10,LYPLA2,NUTF2,CAMLG,EPB41L4B,UCK2,TMPO,ATPAF2,PRMT1,CASC4,STOML2,WNK1,ZNF7,
L4A,CAPN10,LYPLA2,NUTF2,CAMLG,EPB41L4B,UCK2,TMPO,ATPAF2,PRMT1,CASC4,STOML2,WNK1,ZNF7,
NF197,COPZ1,CCNL2,MAPKAPK5,HSP90AB1,SYNGR2,TMEM134,RAB2A,ZNF524,ATP5J,ELP6,RMDN1,TPN
NF197,COPZ1,CCNL2,MAPKAPK5,HSP90AB1,SYNGR2,TMEM134,RAB2A,ZNF524,ATP5J,ELP6,RMDN1,TPN
17ORF62,PRMT1,CASC4,EXOSC9,STOML2,C19ORF54,TXNDC17,PPAN,ZFAND2B,HNRNPC,CYB561,SUGT1,I
17ORF62,PRMT1,CASC4,EXOSC9,STOML2,C19ORF54,TXNDC17,PPAN,ZFAND2B,HNRNPC,CYB561,SUGT1,I
L26A,SSR3,ARIH2,PICK1,ZNF512,TCF25,RNF181,NIPAL3,CHTOP,TSPAN31,PPP2R5C,SRSF7,MTIF2,ZNF593,
L26A,SSR3,ARIH2,PICK1,ZNF512,TCF25,RNF181,NIPAL3,CHTOP,TSPAN31,PPP2R5C,SRSF7,MTIF2,ZNF593,
RD3,DAZAP2,DRG2,BRF1,SLC25A23,MFSD10,MCFD2,RIC8A,SCAP,PHF6,PARK7,SMARCC2,CCDC40,PIGT,U
RD3,DAZAP2,DRG2,BRF1,SLC25A23,MFSD10,MCFD2,RIC8A,SCAP,PHF6,PARK7,SMARCC2,CCDC40,PIGT,U
RD2,GTF2IRD2,COPS4,TMEM39B,ZFPL1,TPM4,MFSD10,MRPS18C,GMPR2,SLC30A5,THADA,COMMD4,MF
RD2,GTF2IRD2,COPS4,TMEM39B,ZFPL1,TPM4,MFSD10,MRPS18C,GMPR2,SLC30A5,THADA,COMMD4,MF

FIP1,ZNF524,ELP6,SIAH1,GGCX,ZNF625,NIT2,MPDU1,VPS28,ZSWIM7,DCTD,UBXN1,ERLEC1,GMPPB,HSPB
517,RCAN1,NSMF,RNMT,ATP9B,COQ4,RAP1B,CLIP4,IFT81,COQ2,CREM,TUBGCP2,SWI5,RNASEK,SNRNP4

517,RCAN1,NSMF,RNMT,ATP9B,COQ4,RAP1B,CLIP4,IFT81,COQ2,CREM,TUBGCP2,SWI5,RNASEK,SNRNP4

C1,CREBZF,EIF2A,RPL24,NADSYN1,ZNF511,GTF2H1,TAF1D,RNF130,D2HGDH,ARPC5,MYL6,SIN3A,NAA38,

C1,CREBZF,EIF2A,RPL24,NADSYN1,ZNF511,GTF2H1,TAF1D,RNF130,D2HGDH,ARPC5,MYL6,SIN3A,NAA38,

4,LRRFIP1,TPM2,GPBP1,SIAH1,TRMT112,PNPO,STRIP2,ZNF625,LYNX1,MPDU1,TWSG1,MAPK13,RPS15,D

1,SERPINB1,DDOST,DCN,HMGA1,PABPC1,RRNAD1,DBNL,ZNF836,RAD1,H2AFV,BLVRB,SERPINB6,RABEP2,

1,SERPINB1,DDOST,DCN,HMGA1,PABPC1,RRNAD1,DBNL,ZNF836,RAD1,H2AFV,BLVRB,SERPINB6,RABEP2,

2,MAPKAPK5,HSP90AB1,SYNGR2,TMEM134,RAB2A,ZNF524,ATP5J,ELP6,RMDN1,TPM2,GPALPP1,SIAH1,A

L32,LMBR1,NAP1L1,SERPINB1,DDOST,C12ORF76,RANGRF,PHTF2,PABPC1,RRNAD1,DBNL,ZNF836,RAD1,B

L32,LMBR1,NAP1L1,SERPINB1,DDOST,C12ORF76,RANGRF,PHTF2,PABPC1,RRNAD1,DBNL,ZNF836,RAD1,B

L32,LMBR1,NAP1L1,SERPINB1,DDOST,C12ORF76,RANGRF,PHTF2,PABPC1,RRNAD1,DBNL,ZNF836,RAD1,B

3,MICAL1,HS2ST1,UBE2I,KCNMA1,YIF1A,EIF3M,JUP,CDC42,TMEM205,PTK2,CELF1,AUNIP,C14ORF2,CENP

3,MICAL1,HS2ST1,UBE2I,KCNMA1,YIF1A,EIF3M,JUP,CDC42,TMEM205,PTK2,CELF1,AUNIP,C14ORF2,CENP

M,ACSS1,MCUR1,ME3,QTRT1,LAMTOR4,ATP6V0B,METTL23,C12ORF73,CGREF1,MCM7,SETD4,PARP6,KIA

M,ACSS1,MCUR1,ME3,QTRT1,LAMTOR4,ATP6V0B,METTL23,C12ORF73,CGREF1,MCM7,SETD4,PARP6,KIA

SME2,PABPC1,PTPRB,RBP1,TOMM40,BRIP1,SERPINB6,RABEP2,TADA3,SLC33A1,RALGPS1,PDCD5,NXN,N

SME2,PABPC1,PTPRB,RBP1,TOMM40,BRIP1,SERPINB6,RABEP2,TADA3,SLC33A1,RALGPS1,PDCD5,NXN,N

IYM6,OGFOD2,GFOD1,SSR3,MRPL55,SSB,ARIH2,CREB3L4,TCF25,NIPAL3,NREP,BAX,CDK4,NME1,TMC7,EE

METTL17,ECSIT,EPB41L5,ATG12,RPS25,ZBTB44,TTC9C,COX4I1,DUSP22,SLC25A10,TMX2,SPOCK2,THEM4,

METTL17,ECSIT,EPB41L5,ATG12,RPS25,ZBTB44,TTC9C,COX4I1,DUSP22,SLC25A10,TMX2,SPOCK2,THEM4,

MDH1,ZNF197,INO80E,CCNL2,DGUOK,HSP90AB1,FAM219B,RAB2A,ZNF524,ATP5J,ELP6,GPBP1,SIAH1,ZE

MDH1,ZNF197,INO80E,CCNL2,DGUOK,HSP90AB1,FAM219B,RAB2A,ZNF524,ATP5J,ELP6,GPBP1,SIAH1,ZE

MA7,TMEM126B,HYI,RPL21,LYPLA2,CAMLG,TCF4,TMPO,ATPAF2,LSM12,C17ORF62,CASC4,EXOSC9,STOM

MA7,TMEM126B,HYI,RPL21,LYPLA2,CAMLG,TCF4,TMPO,ATPAF2,LSM12,C17ORF62,CASC4,EXOSC9,STOM

RAL1,ZNF483,LRRC23,SUMF1,FBXO34,CYB5D2,DNAJC24,ZNF816,ALS2,IK,PKIG,EXOSC8,DDX17,SLC31A1,

SERPINB6,WAC,SLC33A1,ZNF706,PDCD5,GTF2IRD1,NXN,NUDT1,POT1,ANAPC11,NOMO2,RAD23A,ABHD1

NF506,FTH1,MTHFS,C11ORF49,PAFAH1B2,ZNF343,MPC2,TYMS,GTF2IRD2,GBA2,DALRD3,NSUN5,DAZAP2

NF506,FTH1,MTHFS,C11ORF49,PAFAH1B2,ZNF343,MPC2,TYMS,GTF2IRD2,GBA2,DALRD3,NSUN5,DAZAP2

YB10,B2M,GGT5,NT5C,PKP4,MDH2,RRP8,MED6,MT1E,EGFR,CASZ1,SSH2,EGLN2,KCTD7,RFXANK,SMARCA

L,CCNL2,MAPKAPK5,HSP90AB1,SYNGR2,RAB2A,ATP5J,ELP6,TPM2,GPALPP1,SIAH1,ASPH,TRMT112,PNPO

DST,C12ORF76,DCN,HMGA1,PTPRB,ZNF836,RAD1,RBMX,PPP3CB,RABEP2,TADA3,ATP6V1F,WAC,RALGPS

DST,C12ORF76,DCN,HMGA1,PTPRB,ZNF836,RAD1,RBMX,PPP3CB,RABEP2,TADA3,ATP6V1F,WAC,RALGPS

F,RNMT,ATP9B,COQ4,RAP1B,CLIP4,IFT81,COQ2,CREM,TUBGCP2,SWI5,RNASEK,SNRNP40,UQCRB,DCTN6,

DEF8,EEF1B2,PPP1CC,EDF1,PPA1,DGKE,TNFRSF14,ENDOV,CAMTA1,CRIP1,VIM,PSMA7,CDC14B,TIMM17

PL21,CAPN10,RBM33,LYPLA2,UCK2,C17ORF62,PRMT1,CASC4,STOML2,TXNDC17,PIGL,ZFAND2B,HNRNP

PL21,CAPN10,RBM33,LYPLA2,UCK2,C17ORF62,PRMT1,CASC4,STOML2,TXNDC17,PIGL,ZFAND2B,HNRNP

LC35B1,SLC30A5,THADA,COMMD4,MROH1,RPS11,MFF,SNX5,NUP133,MT2A,TMEM147,ELP4,NFYC,ZXD

LC35B1,SLC30A5,THADA,COMMD4,MROH1,RPS11,MFF,SNX5,NUP133,MT2A,TMEM147,ELP4,NFYC,ZXD

,GTF3A,EMC9,DHRS4,SMIM12,NAA15,NDUFB2,B3GNTL1,MYL12A,ZBTB8OS,SPG21,TBC1D22A,LAMTOR1

,GTF3A,EMC9,DHRS4,SMIM12,NAA15,NDUFB2,B3GNTL1,MYL12A,ZBTB8OS,SPG21,TBC1D22A,LAMTOR1

,GTF3A,EMC9,DHRS4,SMIM12,NAA15,NDUFB2,B3GNTL1,MYL12A,ZBTB8OS,SPG21,TBC1D22A,LAMTOR1

PIN,ZNF644,STAU1,SUGT1,HYPK,CARS,RPL36,CDK11A,ZNF197,DGUOK,HSP90AB1,RAB2A,ZNF524,ATP5J,

C3,TAF9,RNPS1,DTD2,NMRAL1,ZNF483,NSRP1,LRRC23,SUMF1,FBXO34,CYB5D2,DNAJC24,ZNF816,ALS2,

C3,TAF9,RNPS1,DTD2,NMRAL1,ZNF483,NSRP1,LRRC23,SUMF1,FBXO34,CYB5D2,DNAJC24,ZNF816,ALS2,

PF3,FOXRED1,TRNT1,KXD1,ITGB1BP1,MED15,PARD6G,SH3YL1,IP6K2,CLASRP,JPH2,C1ORF198,MICAL1,YI

PF3,FOXRED1,TRNT1,KXD1,ITGB1BP1,MED15,PARD6G,SH3YL1,IP6K2,CLASRP,JPH2,C1ORF198,MICAL1,YI

AGT1,TBX6,TMEM205,XBP1,PTK2,C14ORF2,ATF7,PCYT2,USE1,AGPAT4,IFI27L1,ASF1B,DEF8,PPP1CC,EDF1

AGT1,TBX6,TMEM205,XBP1,PTK2,C14ORF2,ATF7,PCYT2,USE1,AGPAT4,IFI27L1,ASF1B,DEF8,PPP1CC,EDF1

,LGMN,FKBP3,INTS4,B3GAT3,RFXANK,ASH2L,SMARCA4,PAFAH1B2,TYMS,COPS4,ARHGEF1,TPM4,MFSD1

5,MPDU1,VPS28,ZSWIM7,RPS15,UBXN1,ZNF814,HHLA3,LRRC27,SAT1,ERCC5,IL32,LMBR1,NAP1L1,DDOS

5,MPDU1,VPS28,ZSWIM7,RPS15,UBXN1,ZNF814,HHLA3,LRRC27,SAT1,ERCC5,IL32,LMBR1,NAP1L1,DDOS

UP62,SCN11A,PSMD7,NADSYN1,ZNF511,D2HGDH,KDELR3,CC2D1B,ARPC5,TLCD1,ST8SIA1,SNX29,AXIN1,

4,HHLA3,SAT1,ERCC5,IL32,NAP1L1,DDOST,C12ORF76,DOCK10,HMGA1,RAD1,RBMX,PPP3CB,BRIP1,RABE

4,HHLA3,SAT1,ERCC5,IL32,NAP1L1,DDOST,C12ORF76,DOCK10,HMGA1,RAD1,RBMX,PPP3CB,BRIP1,RABE

6,FBXO34,CYB5D2,DNAJC24,ALS2,IK,PKIG,ANXA2,DDX17,SNX13,SLC31A1,ZNFX1,RCAN1,NSMF,NARF,PU

6,FBXO34,CYB5D2,DNAJC24,ALS2,IK,PKIG,ANXA2,DDX17,SNX13,SLC31A1,ZNFX1,RCAN1,NSMF,NARF,PU

1,PHF21A,CHCHD1,SLC39A11,FUZ,FAM3A,PSMA3,MUC1,METTL17,EPB41L5,ATG12,PPRC1,RPS25,NLRP

S15,GMPPB,HHLA3,VWA2,MVD,EMC4,LMBR1,MCTS1,NAP1L1,PRKRIP1,DDOST,LSR,DCN,PSME2,PHTF2,P

S15,GMPPB,HHLA3,VWA2,MVD,EMC4,LMBR1,MCTS1,NAP1L1,PRKRIP1,DDOST,LSR,DCN,PSME2,PHTF2,P

H2,C1ORF198,UGGT2,MICAL1,HS2ST1,UBE2I,SLC39A13,YIF1A,JUP,TBX6,TMEM205,XBP1,PTK2,AUNIP,C1

H2,C1ORF198,UGGT2,MICAL1,HS2ST1,UBE2I,SLC39A13,YIF1A,JUP,TBX6,TMEM205,XBP1,PTK2,AUNIP,C1

,XYLB,IMMP1L,ERAL1,ARMCX4,BCAP29,CNTLN,DDX56,PPARD,TPM1,DAD1,BIN1,SEMA5A,MRPL36,REEP

PG21,RPS7,U2AF1L4,CARM1,EPHA10,MAZ,C2ORF15,TTC39C,TBCA,DDX10,PLD1,POLR2E,BCL7B,GRK4,G

PG21,RPS7,U2AF1L4,CARM1,EPHA10,MAZ,C2ORF15,TTC39C,TBCA,DDX10,PLD1,POLR2E,BCL7B,GRK4,G

IMT,ATP9B,COQ4,RAP1B,CLIP4,IFT81,COQ2,CREM,TUBGCP2,SWI5,RNASEK,SNRNP40,UQCRB,DCTN6,PH

IMT,ATP9B,COQ4,RAP1B,CLIP4,IFT81,COQ2,CREM,TUBGCP2,SWI5,RNASEK,SNRNP40,UQCRB,DCTN6,PH

N1,NDUFB2,CDK2AP2,NDUFA10,RPS27L,AZI2,GOSR2,SPG21,KAT8,U2AF1L4,ATL2,DDX10,PLD1,EMC3,TA

5,UBXN1,ZNF814,GMPPB,VWA2,ERCC5,IL32,LMBR1,NAP1L1,SERPINB1,DDOST,C12ORF76,RANGRF,PAB

5,UBXN1,ZNF814,GMPPB,VWA2,ERCC5,IL32,LMBR1,NAP1L1,SERPINB1,DDOST,C12ORF76,RANGRF,PAB

5,UBXN1,ZNF814,GMPPB,VWA2,ERCC5,IL32,LMBR1,NAP1L1,SERPINB1,DDOST,C12ORF76,RANGRF,PAB

FT81,COQ2,NDUFS7,TUBGCP2,SWI5,SNRNP40,DCTN6,WDR6,OPA3,SNRNP200,ARHGEF7,AHI1,LIN37,C12

TNKS,ELOF1,L3MBTL2,ARL16,EIF4A1,SLC35E1,RC3H2,DGAT1,KLHL7,MRPL55,ARIH2,LIMA1,PICK1,UBE2D

TNKS,ELOF1,L3MBTL2,ARL16,EIF4A1,SLC35E1,RC3H2,DGAT1,KLHL7,MRPL55,ARIH2,LIMA1,PICK1,UBE2D

POX,TMPO,ATPAF2,C17ORF62,WNK1,ZNF7,ARL6IP4,AP1S3,ACAD10,HNRNPC,CYB561,SHARPIN,ZNF644,S

POX,TMPO,ATPAF2,C17ORF62,WNK1,ZNF7,ARL6IP4,AP1S3,ACAD10,HNRNPC,CYB561,SHARPIN,ZNF644,S

,MFSD10,MCFD2,SLC30A5,TRPV3,THADA,CBR1,PHC3,EIF2B3,MFSD11,LFNG,XYLT2,SNX5,NOP2,RPAIN,TM

,MFSD10,MCFD2,SLC30A5,TRPV3,THADA,CBR1,PHC3,EIF2B3,MFSD11,LFNG,XYLT2,SNX5,NOP2,RPAIN,TM

NFRSF14,ENDOV,CAMTA1,CRIP1,TROAP,LIG1,PSMA7,APEH,HYI,CAPN10,LYPLA2,NUTF2,PODXL,EPB41L4

P45,PSME1,SLC16A1,ILF3,PKP4,AGAP3,HSPH1,NAA10,C11ORF48,MED6,MT1E,ATP10D,LLGL2,SH3GLB1,C

,SYPL1,SLC37A3,EMC3,THOP1,GRK4,NAAA,C6ORF141,CSF1,VAPA,AIG1,ACVR1B,PCMTD1,LRRC59,CAMK

,SYPL1,SLC37A3,EMC3,THOP1,GRK4,NAAA,C6ORF141,CSF1,VAPA,AIG1,ACVR1B,PCMTD1,LRRC59,CAMK

DA3,ZNF20,SLC33A1,RALGPS1,ZNF706,NXN,ANAPC11,ILF3,MDH2,HSPH1,NAA10,C11ORF48,CARHSP1,LL

DA3,ZNF20,SLC33A1,RALGPS1,ZNF706,NXN,ANAPC11,ILF3,MDH2,HSPH1,NAA10,C11ORF48,CARHSP1,LL

,RECQL5,RPL24,ZNF675,ZNF511,DYNLT3,GTF2H1,BFAR,D2HGDH,KDELR3,ARPC5,MYL6,TLCD1,ST7,MOSP

,RECQL5,RPL24,ZNF675,ZNF511,DYNLT3,GTF2H1,BFAR,D2HGDH,KDELR3,ARPC5,MYL6,TLCD1,ST7,MOSP

F3M,MAGT1,JUP,CDC42,TMEM205,XBP1,PTK2,C14ORF2,GUK1,CARD14,ATF7,PCYT2,FBXW9,HSBP1L1,A

F3M,MAGT1,JUP,CDC42,TMEM205,XBP1,PTK2,C14ORF2,GUK1,CARD14,ATF7,PCYT2,FBXW9,HSBP1L1,A

SRSF1,SAP30BP,FOXRED1,RABL5,TRNT1,KXD1,MED15,PARD6G,SH3YL1,JPH2,UGGT2,MICAL1,HS2ST1,UB

SRSF1,SAP30BP,FOXRED1,RABL5,TRNT1,KXD1,MED15,PARD6G,SH3YL1,JPH2,UGGT2,MICAL1,HS2ST1,UB

,UBXN1,ZNF814,ERLEC1,GMPPB,MVD,IL32,NAP1L1,SYNGR1,C12ORF76,RANGRF,PABPC1,RRNAD1,PTPRE

,UBXN1,ZNF814,ERLEC1,GMPPB,MVD,IL32,NAP1L1,SYNGR1,C12ORF76,RANGRF,PABPC1,RRNAD1,PTPRE

EC1,NAP1L1,SYNGR1,LSR,MTA3,MEIS2,PABPC1,RRNAD1,NAT9,BCAM,ZNF836,H2AFV,BLVRB,PPP3CB,WA

EC1,NAP1L1,SYNGR1,LSR,MTA3,MEIS2,PABPC1,RRNAD1,NAT9,BCAM,ZNF836,H2AFV,BLVRB,PPP3CB,WA

BTB8OS,TBC1D22A,C1QC,FAM104B,TTC39C,BCL7B,TMEM41A,ZNF581,ACSM3,AK2,SSC5D,DHDDS,XPNPE

BTB8OS,TBC1D22A,C1QC,FAM104B,TTC39C,BCL7B,TMEM41A,ZNF581,ACSM3,AK2,SSC5D,DHDDS,XPNPE

K2,THEM4,RDH11,SDF4,NISCH,TMED4,CENPK,RPS6,RPL29,CD99L2,GGPS1,RTEL1,COASY,SRSF1,AUP1,FO

K2,THEM4,RDH11,SDF4,NISCH,TMED4,CENPK,RPS6,RPL29,CD99L2,GGPS1,RTEL1,COASY,SRSF1,AUP1,FO

17A,NT5C,PKP4,PDE4DIP,STXBP2,HSPH1,SGK1,SEC31A,CARHSP1,COX17,HNRNPLL,HINT1,HIGD1A,FKBP3,

CC5,IL32,NAP1L1,DDOST,C12ORF76,DOCK10,HMGA1,DBNL,ZNF836,RAD1,RBMX,PPP3CB,BRIP1,RABEP2,

CC5,IL32,NAP1L1,DDOST,C12ORF76,DOCK10,HMGA1,DBNL,ZNF836,RAD1,RBMX,PPP3CB,BRIP1,RABEP2,

EP70,TMEM134,RAB2A,RMDN1,GPBP1,GPALPP1,TRMT112,PNPO,CHPF,NIT2,MPDU1,TWSG1,MAPK13,Z

EP70,TMEM134,RAB2A,RMDN1,GPBP1,GPALPP1,TRMT112,PNPO,CHPF,NIT2,MPDU1,TWSG1,MAPK13,Z

TB44,TTC9C,COX4I1,DUSP22,SLC25A10,TMX2,SPOCK2,SDF4,NISCH,TMED4,CENPK,CD99L2,GGPS1,RTEL1

TB44,TTC9C,COX4I1,DUSP22,SLC25A10,TMX2,SPOCK2,SDF4,NISCH,TMED4,CENPK,CD99L2,GGPS1,RTEL1

NRNPC,CYB561,SHARPIN,ZNF644,CBFA2T2,SUGT1,HYPK,MOB1A,CDK11A,RBM3,MDH1,COPZ1,INO80E,C

NRNPC,CYB561,SHARPIN,ZNF644,CBFA2T2,SUGT1,HYPK,MOB1A,CDK11A,RBM3,MDH1,COPZ1,INO80E,C

C17ORF62,EXOSC9,STOML2,WNK1,C19ORF54,ARL6IP4,PPAN,ZFAND2B,AP1S3,HNRNPC,CYB561,ZNF644,

C17ORF62,EXOSC9,STOML2,WNK1,C19ORF54,ARL6IP4,PPAN,ZFAND2B,AP1S3,HNRNPC,CYB561,ZNF644,

NF687,PDCD5,NXN,EPS15L1,ANAPC11,NOMO2,B2M,DDX41,SLC16A1,ILF3,AGAP3,MDH2,EEF1D,CHCHD

NF687,PDCD5,NXN,EPS15L1,ANAPC11,NOMO2,B2M,DDX41,SLC16A1,ILF3,AGAP3,MDH2,EEF1D,CHCHD

SPH,STRIP2,GABPB1,TWSG1,ZSWIM7,RPS15,DCTD,ERLEC1,HHLA3,LRRC27,MVD,LMBR1,PRKRIP1,SYNGR

M3,TRMT112,PNPO,CHPF,MAPK13,ZSWIM7,ERLEC1,GMPPB,HHLA3,MVD,SAT1,ERCC5,EMC4,IL32,LMBR1,

M3,TRMT112,PNPO,CHPF,MAPK13,ZSWIM7,ERLEC1,GMPPB,HHLA3,MVD,SAT1,ERCC5,EMC4,IL32,LMBR1,

T5,DDX41,NT5C,SLC16A1,ILF3,AGAP3,MDH2,STXBP2,EEF1D,HSPH1,NAA10,RRP8,CARHSP1,MT1E,CNST,L

T5,DDX41,NT5C,SLC16A1,ILF3,AGAP3,MDH2,STXBP2,EEF1D,HSPH1,NAA10,RRP8,CARHSP1,MT1E,CNST,L

F2IRD1,NUDT1,ANAPC11,POLR3H,RAD23A,GGT5,NT5C,PSME1,SLC16A1,ILF3,AGAP3,MDH2,STXBP2,EEF

TF2IRD1,NUDT1,ANAPC11,POLR3H,RAD23A,GGT5,NT5C,PSME1,SLC16A1,ILF3,AGAP3,MDH2,STXBP2,EEF
PS20,GABPB1,TWSG1,VPS28,ZSWIM7,ZNF814,ERLEC1,HHLA3,LRRC27,MVD,SAT1,ERCC5,EMC4,IL32,SEM
PS20,GABPB1,TWSG1,VPS28,ZSWIM7,ZNF814,ERLEC1,HHLA3,LRRC27,MVD,SAT1,ERCC5,EMC4,IL32,SEM

F3A,DHRS4,ARRB2,SMIM12,NDUFB2,HMBS,B3GNTL1,SGSM3,SPG21,TBC1D22A,LAMTOR1,USP15,EMC3,
F3A,DHRS4,ARRB2,SMIM12,NDUFB2,HMBS,B3GNTL1,SGSM3,SPG21,TBC1D22A,LAMTOR1,USP15,EMC3,
F3A,DHRS4,ARRB2,SMIM12,NDUFB2,HMBS,B3GNTL1,SGSM3,SPG21,TBC1D22A,LAMTOR1,USP15,EMC3,
,MUC1,EPB41L5,ATG12,PPRC1,RPS25,NLRP1,ZBTB44,NIP7,TTC9C,PLCB2,DUSP22,SLC25A10,SPOCK2,DBI
,MUC1,EPB41L5,ATG12,PPRC1,RPS25,NLRP1,ZBTB44,NIP7,TTC9C,PLCB2,DUSP22,SLC25A10,SPOCK2,DBI

GTF2IRD2,COPS4,TMEM39B,ZFPL1,TPM4,MFSD10,MRPS18C,GMPR2,SLC30A5,THADA,COMMD4,MROH1
GTF2IRD2,COPS4,TMEM39B,ZFPL1,TPM4,MFSD10,MRPS18C,GMPR2,SLC30A5,THADA,COMMD4,MROH1
2,INO80E,CCNL2,MAPKAPK5,CEP70,TMEM134,LRRFIP1,RAB2A,ZNF524,ELP6,SIAH1,GGCX,ASPH,TPM3,ST
,CHPF,STRIP2,RPS20,GABPB1,NIT2,MAPK13,VPS28,ZSWIM7,RPS15,DCTD,UBXN1,ERLEC1,GMPPB,LRRC2
,CHPF,STRIP2,RPS20,GABPB1,NIT2,MAPK13,VPS28,ZSWIM7,RPS15,DCTD,UBXN1,ERLEC1,GMPPB,LRRC2
PS23,RPL30,ZNF783,IFRD2,ACHE,SUMF1,NDUFA7,CCNG2,SELENBP1,HNRNPM,ACSS1,MCUR1,ME3,LAMT
PS23,RPL30,ZNF783,IFRD2,ACHE,SUMF1,NDUFA7,CCNG2,SELENBP1,HNRNPM,ACSS1,MCUR1,ME3,LAMT
E,TNFRSF14,ENDOV,VAMP8,CAMTA1,CAPG,PPIL2,PSMD4,TROAP,RPL28,CLDND1,LIG1,TIMM17B,TMEM
E,TNFRSF14,ENDOV,VAMP8,CAMTA1,CAPG,PPIL2,PSMD4,TROAP,RPL28,CLDND1,LIG1,TIMM17B,TMEM

N1,ABHD14B,C12ORF66,NUDT9,UNC119,ZRANB2,CSRP2BP,MVB12B,ANKHD1,SSBP4,TRIM9,WWP2,LIN
N1,ABHD14B,C12ORF66,NUDT9,UNC119,ZRANB2,CSRP2BP,MVB12B,ANKHD1,SSBP4,TRIM9,WWP2,LIN
FCHO2,IRF3,RNF130,KDELR3,PLAC8,HNRNPUL1,TP53,AP1G2,MOSPD3,SERPINA1,TMEM199,ZNF396,AA
FCHO2,IRF3,RNF130,KDELR3,PLAC8,HNRNPUL1,TP53,AP1G2,MOSPD3,SERPINA1,TMEM199,ZNF396,AA
28,ZSWIM7,RPS15,ZNF814,ERLEC1,GMPPB,VWA2,MVD,ERCC5,NAP1L1,RANGRF,LSR,MEIS2,PSME2,PABP
AB1,FAM219B,RAB2A,ZNF524,ATP5J,ELP6,GPBP1,GPALPP1,SIAH1,PNPO,RPTOR,ZNF625,MPDU1,MAPK1
AB1,FAM219B,RAB2A,ZNF524,ATP5J,ELP6,GPBP1,GPALPP1,SIAH1,PNPO,RPTOR,ZNF625,MPDU1,MAPK1

NARF,DHX30,MFSD1,DUSP11,PON2,DNAJB12,DDX56,TUBGCP2,PPARD,RPL27,MAP4,OAZ1,WDR6,DIDO1,

2L,MRPL27,SLC25A35,FLAD1,PON2,COQ2,MRPS7,NDUFS7,IFI

P2,FAM32A,PCID2,ZXDC,PIGH,NMNAT3,KPNA5,MPHOSPH1
P2,FAM32A,PCID2,ZXDC,PIGH,NMNAT3,KPNA5,MPHOSPH1

| Gene | Chr_Num | Exon_Num | Splice | Novel_Spli | Strand_Ch | Exon_Start | Exon_Stop | Mutation_ | Mutation_ |
|---|---|---|---|---|---|---|---|---|---|
| SDR39U1 | 14 | 3 | AP | 0 | - | 24909989 | 24910132 | 24910070 | 24910073 |
| RPL15 | 3 | 1 | AD | 0 | + | 23962968 | 23965759 | 23960714 | 23960715 |
| RPL15 | 3 | 3 | AD | 0 | + | 23960687 | 23962302 | 23960714 | 23960715 |
| SDHA | 5 | 2 | ES | 0 | + | 251107 | 251158 | 228334 | 228335 |
| SDHA | 5 | 5 | ES | 0 | + | 251107 | 251218 | 228334 | 228335 |
| SDHA | 5 | 14 | ES | 0 | + | 256449 | 257189 | 228334 | 228335 |
| SDHA | 5 | 13 | ES | 0 | + | 254508 | 254621 | 228334 | 228335 |
| SNAPC3 | 9 | 9 | AT | 0 | + | 15459717 | 15461627 | 15447239 | 15447240 |
| ACAA1 | 3 | 3 | ES | 0 | - | 38175443 | 38175500 | 38175489 | 38175489 |
| ACAA1 | 3 | 4 | ES | 0 | - | 38175364 | 38175500 | 38175489 | 38175489 |
| CDC37 | 19 | 2 | ES | 0 | - | 10514054 | 10514271 | 10505909 | 10505909 |
| CPSF3L | 1 | 2 | ES | 0 | - | 1256376 | 1256476 | 1255909 | 1255909 |
| CPSF3L | 1 | 5 | ES | 0 | - | 1255836 | 1255909 | 1255909 | 1255909 |
| CPSF3L | 1 | 4 | ES | 0 | - | 1255836 | 1255909 | 1255909 | 1255909 |
| CPSF3L | 1 | 3 | ES | 0 | - | 1257288 | 1257364 | 1255909 | 1255909 |
| DDOST | 1 | 2 | ES | 0 | - | 20987374 | 20987484 | 20987419 | 20987419 |
| DHPS | 19 | 1 | AP | 0 | - | 12790975 | 12791139 | 12790297 | 12790297 |
| DHPS | 19 | 4 | ES | 0 | - | 12790437 | 12790533 | 12790297 | 12790297 |
| ITSN1 | 21 | 32 | AT | 0 | + | 35237455 | 35237638 | 35191585 | 35191585 |
| MGRN1 | 16 | 7 | AA | 0 | + | 4715100 | 4715152 | 4702743 | 4702743 |
| MGRN1 | 16 | 17 | AA | 0 | + | 4738797 | 4741347 | 4702743 | 4702743 |
| NGDN | 14 | 10 | RI | 0 | + | 23946712 | 23946769 | 23945486 | 23945486 |
| NPRL3 | 16 | 2 | ES | 0 | - | 188149 | 188428 | 169169 | 169169 |
| NPRL3 | 16 | 1 | ES | 0 | - | 188641 | 188672 | 169169 | 169169 |
| NPRL3 | 16 | 3 | ES | 0 | - | 169125 | 169254 | 169169 | 169169 |
| NPRL3 | 16 | 5 | ES | 0 | - | 169125 | 169254 | 169169 | 169169 |
| NPRL3 | 16 | 4 | ES | 0 | - | 169125 | 169254 | 169169 | 169169 |
| OSGEP | 14 | 4 | RI | 0 | - | 20916265 | 20916330 | 20916117 | 20916117 |
| PPIL2 | 22 | 22 | RI | 0 | + | 22049683 | 22052201 | 22049265 | 22049265 |
| SMARCA4 | 19 | 32 | ES | 0 | + | 11169464 | 11169565 | 11134266 | 11134266 |
| ZNF141 | 4 | 4 | AT | 0 | + | 376884 | 377227 | 367133 | 367133 |
| BMP1 | 8 | 5 | AA | 0 | + | 22069783 | 22069837 | 22069241 | 22069241 |
| CNTN4 | 3 | 4 | AP | 0 | + | 3085276 | 3085388 | 3084848 | 3084848 |
| ELOF1 | 19 | 1 | AP | 0 | - | 11666160 | 11666281 | 11664626 | 11664626 |
| SH3GL1 | 19 | 4 | ES | 0 | - | 4365570 | 4365622 | 4364084 | 4364084 |
| SH3GL1 | 19 | 5 | ES | 0 | - | 4364085 | 4364117 | 4364084 | 4364084 |
| IL32 | 16 | 5 | RI | 0 | + | 3118181 | 3118240 | 3117423 | 3117423 |
| IL32 | 16 | 1 | RI | 0 | + | 3118119 | 3118240 | 3117423 | 3117423 |
| IL32 | 16 | 2 | RI | 0 | + | 3117378 | 3118011 | 3117423 | 3117423 |
| ACTB | 7 | 3 | ES | 1 | - | 5568792 | 5569031 | 5568999 | 5568999 |
| ACTB | 7 | 4 | ES | 0 | - | 5567912 | 5569031 | 5568999 | 5568999 |
| BMP1 | 8 | 5 | AA | 0 | + | 22068326 | 22068500 | 22059428 | 22059428 |
| BMP1 | 8 | 5 | AA | 0 | + | 22068326 | 22068500 | 22067082 | 22067082 |
| BMP1 | 8 | 17 | AA | 0 | + | 22064367 | 22064494 | 22059428 | 22059428 |
| BMP1 | 8 | 18 | AA | 0 | + | 22064816 | 22065029 | 22059428 | 22059428 |
| BMP1 | 8 | 18 | AA | 0 | + | 22066958 | 22067208 | 22067082 | 22067082 |

| | | | | | | | | | |
|---|---|---|---|---|---|---|---|---|---|
| **C20orf96** | 20 | 3 | ES | 0 | - | 259813 | 259971 | 259966 | 259966 |
| **CAMTA1** | 1 | 3 | ES | 1 | + | 7796404 | 7796493 | 7737765 | 7737765 |
| **CAMTA1** | 1 | 5 | AT | 0 | + | 7797315 | 7797507 | 7737765 | 7737765 |
| **CAMTA1** | 1 | 5 | AT | 0 | + | 7797315 | 7797507 | 7796575 | 7796575 |
| **CAMTA1** | 1 | 3 | ES | 1 | + | 7815698 | 7815728 | 7796575 | 7796575 |
| **CARM1** | 19 | 16 | ES | 0 | + | 11032291 | 11033448 | 11024705 | 11024705 |
| **CARM1** | 19 | 4 | ES | 0 | + | 11031510 | 11031622 | 11024705 | 11024705 |
| **CD44** | 11 | 3 | ES | 0 | + | 35236399 | 35236461 | 35227738 | 35227738 |
| **CD44** | 11 | 10 | ES | 0 | + | 35250676 | 35251213 | 35227738 | 35227738 |
| **CD44** | 11 | 6 | ES | 0 | + | 35243201 | 35243279 | 35227738 | 35227738 |
| **CD44** | 11 | 9 | ES | 1 | + | 35232793 | 35233725 | 35227738 | 35227738 |
| **CD44** | 11 | 8 | ES | 1 | + | 35243201 | 35243279 | 35227738 | 35227738 |
| **CD44** | 11 | 11 | ES | 1 | + | 35227659 | 35227790 | 35227738 | 35227738 |
| **CD44** | 11 | 14 | ES | 0 | + | 35232793 | 35232996 | 35227738 | 35227738 |
| **CDC37** | 19 | 2 | ES | 0 | - | 10514054 | 10514271 | 10506754 | 10506754 |
| **CDCA3** | 12 | 5 | AT | 0 | - | 6958487 | 6958593 | 6958344 | 6958344 |
| **CNTLN** | 9 | 7 | AT | 0 | + | 17298188 | 17302048 | 17298326 | 17298326 |
| **DRG2** | 17 | 6 | RI | 0 | + | 18010468 | 18010676 | 18003681 | 18003681 |
| **DRG2** | 17 | 8 | AA | 0 | + | 18005235 | 18005311 | 18003681 | 18003681 |
| **DRG2** | 17 | 3 | AD | 0 | + | 18003883 | 18003973 | 18003681 | 18003681 |
| **DRG2** | 17 | 5 | AD | 0 | + | 18002947 | 18003749 | 18003681 | 18003681 |
| **DRG2** | 17 | 12 | AD | 0 | + | 18009747 | 18009800 | 18003681 | 18003681 |
| **EIF3C** | 16 | 2 | AP | 0 | + | 28746253 | 28746340 | 28734551 | 28734551 |
| **FAM195A** | 16 | 3 | ES | 0 | + | 697783 | 697884 | 697444 | 697444 |
| **FAM195A** | 16 | 4 | ES | 0 | + | 698120 | 698474 | 697444 | 697444 |
| **FDFT1** | 8 | 2 | AP | 0 | + | 11689027 | 11689179 | 11687884 | 11687884 |
| **GBA2** | 9 | 15 | RI | 0 | - | 35737745 | 35737936 | 35737268 | 35737268 |
| **GBA2** | 9 | 14 | RI | 0 | - | 35738229 | 35738371 | 35737268 | 35737268 |
| **GIPC1** | 19 | 2 | ES | 0 | - | 14602468 | 14602555 | 14593528 | 14593528 |
| **GIPC1** | 19 | 3 | ES | 0 | - | 14602468 | 14602555 | 14593528 | 14593528 |
| **GIPC1** | 19 | 4 | ES | 0 | - | 14593668 | 14593818 | 14593528 | 14593528 |
| **GTF2H1** | 11 | 1 | AP | 0 | + | 18373075 | 18373516 | 18359773 | 18359773 |
| **GTF3A** | 13 | 1 | AD | 0 | + | 28008747 | 28010546 | 28009581 | 28009581 |
| **HMHA1** | 19 | 15 | AP | 0 | + | 1080891 | 1081063 | 1080691 | 1080691 |
| **HMHA1** | 19 | 15 | AP | 0 | + | 1080891 | 1081063 | 1074171 | 1074171 |
| **HNRNPC** | 14 | 2 | AD | 0 | - | 21736782 | 21736827 | 21702191 | 21702191 |
| **HNRNPC** | 14 | 1 | AP | 0 | - | 21737457 | 21737585 | 21702191 | 21702191 |
| **HNRNPC** | 14 | 3 | ES | 0 | - | 21702112 | 21702388 | 21702191 | 21702191 |
| **HNRNPC** | 14 | 4 | ES | 0 | - | 21702112 | 21702388 | 21702191 | 21702191 |
| **HNRNPC** | 14 | 5 | ES | 0 | - | 21702112 | 21702388 | 21702191 | 21702191 |
| **HNRNPM** | 19 | 11 | ES | 0 | + | 8551908 | 8551959 | 8551214 | 8551214 |
| **HNRNPM** | 19 | 3 | ES | 0 | + | 8550487 | 8551289 | 8551214 | 8551214 |
| **IAH1** | 2 | 7 | AT | 0 | + | 9634831 | 9634994 | 9621566 | 9621566 |
| **IQCK** | 16 | 9 | AT | 0 | + | 19868777 | 19868907 | 19838353 | 19838353 |
| **ITSN1** | 21 | 32 | AT | 0 | + | 35237455 | 35237638 | 35202044 | 35202044 |
| **KIAA1328** | 18 | 12 | AT | 0 | + | 34811623 | 34812141 | 34802046 | 34802046 |
| **LRP10** | 14 | 7 | RI | 0 | + | 23346149 | 23346262 | 23345402 | 23345402 |

| | | | | | | | | | |
|---|---|---|---|---|---|---|---|---|---|
| **LSM14A** | 19 | 3 | ES | 0 | + | 34718274 | 34720187 | 34710699 | 34710699 |
| **LSM14A** | 19 | 11 | ES | 0 | + | 34718270 | 34720420 | 34710699 | 34710699 |
| **LSM14A** | 19 | 5 | ES | 0 | + | 34710611 | 34710782 | 34710699 | 34710699 |
| **MAP3K8** | 10 | 9 | AT | 0 | + | 30749635 | 30750761 | 30747153 | 30747153 |
| **MEIS2** | 15 | 2 | AP | 0 | - | 37390168 | 37390400 | 37390266 | 37390266 |
| **MGRN1** | 16 | 17 | AA | 0 | + | 4738797 | 4741347 | 4732890 | 4732890 |
| **MLST8** | 16 | 4 | AA | 0 | + | 2256498 | 2257111 | 2256400 | 2256400 |
| **MLST8** | 16 | 5 | AA | 0 | + | 2257036 | 2257111 | 2256400 | 2256400 |
| **MLST8** | 16 | 3 | AA | 0 | + | 2256359 | 2256410 | 2256400 | 2256400 |
| **MLST8** | 16 | 8 | AA | 0 | + | 2258451 | 2258614 | 2256400 | 2256400 |
| **MRPL51** | 12 | 2 | AP | 0 | - | 6602028 | 6602138 | 6601500 | 6601500 |
| **MYL9** | 20 | 3 | ES | 0 | + | 35177480 | 35178226 | 35177499 | 35177499 |
| **NPRL3** | 16 | 2 | ES | 0 | - | 188149 | 188428 | 136767 | 136767 |
| **NPRL3** | 16 | 2 | ES | 0 | - | 188149 | 188428 | 150396 | 150396 |
| **NPRL3** | 16 | 5 | ES | 0 | - | 162621 | 162774 | 136767 | 136767 |
| **NPRL3** | 16 | 5 | ES | 0 | - | 162621 | 162774 | 150396 | 150396 |
| **NPRL3** | 16 | 4 | ES | 0 | - | 160523 | 160604 | 136767 | 136767 |
| **NPRL3** | 16 | 4 | ES | 0 | - | 160523 | 160604 | 150396 | 150396 |
| **NPRL3** | 16 | 1 | ES | 0 | - | 188641 | 188672 | 136767 | 136767 |
| **NPRL3** | 16 | 1 | ES | 0 | - | 188641 | 188672 | 150396 | 150396 |
| **NPRL3** | 16 | 3 | ES | 0 | - | 169125 | 169254 | 136767 | 136767 |
| **NPRL3** | 16 | 3 | ES | 0 | - | 169125 | 169254 | 150396 | 150396 |
| **NPRL3** | 16 | 7 | ES | 0 | - | 160523 | 160604 | 136767 | 136767 |
| **NPRL3** | 16 | 7 | ES | 0 | - | 160523 | 160604 | 150396 | 150396 |
| **NRP1** | 10 | 5 | ES | 0 | - | 33545244 | 33545399 | 33515104 | 33515104 |
| **PPIL2** | 22 | 22 | RI | 0 | + | 22049683 | 22052201 | 22039067 | 22039067 |
| **PRKCSH** | 19 | 12 | AD | 0 | + | 11558508 | 11558604 | 11557939 | 11557939 |
| **PRKCSH** | 19 | 2 | AD | 0 | + | 11559729 | 11559803 | 11557939 | 11557939 |
| **PRLR** | 5 | 10 | AT | 0 | - | 35066051 | 35066204 | 35065452 | 35065452 |
| **QDPR** | 4 | 2 | ES | 0 | - | 17492285 | 17492368 | 17488811 | 17488811 |
| **QDPR** | 4 | 3 | ES | 0 | - | 17506002 | 17506098 | 17488811 | 17488811 |
| **QDPR** | 4 | 6 | ES | 0 | - | 17493855 | 17493963 | 17488811 | 17488811 |
| **QDPR** | 4 | 7 | ES | 0 | - | 17488813 | 17488859 | 17488811 | 17488811 |
| **RBBP6** | 16 | 18 | AT | 0 | + | 24582197 | 24584181 | 24581479 | 24581479 |
| **RCAN1** | 21 | 5 | AP | 0 | - | 35888786 | 35890554 | 35890400 | 35890400 |
| **RFXANK** | 19 | 5 | ES | 0 | + | 19309466 | 19309532 | 19308954 | 19308954 |
| **RIC8A** | 11 | 2 | AA | 0 | + | 209407 | 209989 | 209565 | 209565 |
| **RIC8A** | 11 | 6 | AA | 0 | + | 212416 | 212511 | 209565 | 209565 |
| **RIC8A** | 11 | 3 | AA | 0 | + | 209407 | 210000 | 209565 | 209565 |
| **RRP8** | 11 | 3 | RI | 0 | - | 6622156 | 6622832 | 6622651 | 6622651 |
| **SERPINB1** | 6 | 3 | ES | 0 | - | 2838817 | 2838920 | 2838132 | 2838132 |
| **SMARCA4** | 19 | 32 | ES | 0 | + | 11169464 | 11169565 | 11107017 | 11107017 |
| **SMARCA4** | 19 | 32 | ES | 0 | + | 11169464 | 11169565 | 11152114 | 11152114 |
| **SMARCA4** | 19 | 32 | ES | 0 | + | 11170429 | 11170561 | 11170476 | 11170476 |
| **SMARCA4** | 19 | 32 | ES | 0 | + | 11170429 | 11170561 | 11170509 | 11170509 |
| **SSBP4** | 19 | 1 | AP | 0 | + | 18539858 | 18541354 | 18538218 | 18538218 |
| **STXBP2** | 19 | 1 | AP | 0 | + | 7709400 | 7709638 | 7707366 | 7707366 |

| | | | | | | | | | |
|---|---|---|---|---|---|---|---|---|---|
| **STXBP2** | 19 | 5 | AA | 0 | + | 7707315 | 7707422 | 7707366 | 7707366 |
| **TARDBP** | 1 | 3 | ES | 0 | + | 11085167 | 11085265 | 11082189 | 11082189 |
| **TARDBP** | 1 | 4 | ES | 0 | + | 11085608 | 11085766 | 11082189 | 11082189 |
| **TBC1D5** | 3 | 2 | AP | 0 | - | 17688453 | 17688548 | 17208310 | 17208310 |
| **TCP11L1** | 11 | 11 | AT | 0 | + | 33094020 | 33094390 | 33094069 | 33094069 |
| **THOP1** | 19 | 1 | AP | 0 | + | 2806978 | 2807050 | 2790603 | 2790603 |
| **TMED1** | 19 | 1 | AP | 0 | - | 10946685 | 10947274 | 10945733 | 10945733 |
| **TMEM147** | 19 | 5 | RI | 0 | + | 36037847 | 36037931 | 36037431 | 36037431 |
| **TMEM147** | 19 | 1 | RI | 0 | + | 36036558 | 36038142 | 36037431 | 36037431 |
| **TNKS** | 8 | 1 | AP | 0 | + | 9627642 | 9627772 | 9605704 | 9605704 |
| **TNKS** | 8 | 1 | AP | 0 | + | 9627642 | 9627772 | 9623249 | 9623249 |
| **TWSG1** | 18 | 7 | AT | 0 | + | 9399344 | 9400815 | 9359999 | 9359999 |
| **TYMS** | 18 | 2 | ES | 0 | + | 671380 | 671451 | 670816 | 670816 |
| **TYMS** | 18 | 3 | ES | 0 | + | 670692 | 670867 | 670816 | 670816 |
| **TYMS** | 18 | 4 | ES | 0 | + | 671380 | 671451 | 670816 | 670816 |
| **VIM** | 10 | 7 | RI | 0 | + | 17278293 | 17278378 | 17276772 | 17276772 |
| **VIM** | 10 | 7 | RI | 0 | + | 17278293 | 17278378 | 17277377 | 17277377 |
| **WDR83** | 19 | 6 | AA | 0 | + | 12786337 | 12786642 | 12786384 | 12786384 |
| **ZCCHC4** | 4 | 13 | AT | 0 | + | 25370660 | 25371164 | 25366651 | 25366651 |
| **ZNF345** | 19 | 4 | AT | 0 | + | 37383720 | 37383827 | 37368905 | 37368905 |
| **ZSCAN20** | 1 | 8 | AT | 0 | + | 33959818 | 33961199 | 33957115 | 33957115 |
| **LYPLA2** | 1 | 8 | RI | 0 | + | 24120714 | 24121171 | 24121164 | 24121164 |
| **ACTB** | 7 | 3 | ES | 1 | - | 5569166 | 5569294 | 5569258 | 5569260 |
| **ACACA** | 17 | 1 | AP | 0 | - | 35550315 | 35550399 | 35454098 | 35454098 |
| **ACACA** | 17 | 1 | AP | 0 | - | 35550315 | 35550399 | 35446008 | 35446008 |
| **ACACA** | 17 | 1 | AP | 0 | - | 35550315 | 35550399 | 35486405 | 35486405 |
| **ACACA** | 17 | 1 | AP | 0 | - | 35550315 | 35550399 | 35549147 | 35549147 |
| **ACSM3** | 16 | 15 | AT | 0 | + | 20808208 | 20808478 | 20787326 | 20787326 |
| **ACSM3** | 16 | 15 | AT | 0 | + | 20808208 | 20808478 | 20781387 | 20781387 |
| **ACSS1** | 20 | 12 | ES | 0 | - | 24993237 | 24993300 | 24993297 | 24993297 |
| **AK2** | 1 | 2 | ES | 0 | - | 33490043 | 33490168 | 33487216 | 33487216 |
| **AK2** | 1 | 3 | ES | 0 | - | 33497145 | 33497262 | 33487216 | 33487216 |
| **AK2** | 1 | 5 | ES | 0 | - | 33487194 | 33487304 | 33487216 | 33487216 |
| **AKAP8L** | 19 | 4 | RI | 0 | - | 15514469 | 15514526 | 15511996 | 15511996 |
| **APP** | 21 | 2 | AP | 0 | - | 27484296 | 27484463 | 27394204 | 27394204 |
| **APP** | 21 | 2 | AP | 0 | - | 27484296 | 27484463 | 27354787 | 27354787 |
| **APP** | 21 | 4 | ES | 0 | - | 27423316 | 27423509 | 27394204 | 27394204 |
| **APP** | 21 | 4 | ES | 0 | - | 27423316 | 27423509 | 27354787 | 27354787 |
| **APP** | 21 | 9 | ES | 0 | - | 27354657 | 27354790 | 27354787 | 27354787 |
| **ARRB2** | 17 | 15 | AA | 0 | + | 4624241 | 4624791 | 4619793 | 4619793 |
| **ARRB2** | 17 | 15 | AA | 0 | + | 4624241 | 4624791 | 4621608 | 4621608 |
| **ARRB2** | 17 | 15 | AA | 0 | + | 4624241 | 4624791 | 4621244 | 4621244 |
| **ASF1B** | 19 | 4 | AT | 0 | - | 14231647 | 14231769 | 14231410 | 14231410 |
| **ASH2L** | 8 | 1 | AP | 0 | + | 37993186 | 37993284 | 37964684 | 37964684 |
| **ASH2L** | 8 | 1 | AP | 0 | + | 37993186 | 37993284 | 37974234 | 37974234 |
| **ASH2L** | 8 | 15 | ES | 0 | + | 37996334 | 37996393 | 37964684 | 37964684 |
| **ASH2L** | 8 | 15 | ES | 0 | + | 37996334 | 37996393 | 37974234 | 37974234 |

| | | | | | | | | | |
|---|---|---|---|---|---|---|---|---|---|
| **ASH2L** | 8 | 4 | ES | 0 | + | 37967897 | 37967985 | 37964684 | 37964684 |
| **ASH2L** | 8 | 4 | ES | 0 | + | 38001257 | 38001594 | 37974234 | 37974234 |
| **BFAR** | 16 | 3 | ES | 0 | + | 14742245 | 14742449 | 14738367 | 14738367 |
| **BFAR** | 16 | 4 | ES | 0 | + | 14743661 | 14743830 | 14738367 | 14738367 |
| **BFAR** | 16 | 3 | ES | 0 | + | 14755749 | 14755922 | 14749057 | 14749057 |
| **BFAR** | 16 | 4 | ES | 0 | + | 14758726 | 14758925 | 14749057 | 14749057 |
| **BFAR** | 16 | 5 | ES | 0 | + | 14758726 | 14758928 | 14738367 | 14738367 |
| **BFAR** | 16 | 5 | ES | 0 | + | 14758726 | 14758928 | 14749057 | 14749057 |
| **BMP1** | 8 | 5 | AA | 0 | + | 22049562 | 22049664 | 22037910 | 22037910 |
| **BMP1** | 8 | 15 | AA | 0 | + | 22054753 | 22054933 | 22037910 | 22037910 |
| **BMP1** | 8 | 15 | AA | 0 | + | 22054753 | 22054933 | 22053064 | 22053064 |
| **BMP1** | 8 | 15 | AA | 0 | + | 22054753 | 22054933 | 22054897 | 22054897 |
| **BMP1** | 8 | 5 | AA | 0 | + | 22068326 | 22068500 | 22064900 | 22064900 |
| **BMP1** | 8 | 5 | AA | 0 | + | 22068326 | 22068500 | 22053064 | 22053064 |
| **BMP1** | 8 | 5 | AA | 0 | + | 22068326 | 22068500 | 22064400 | 22064400 |
| **BMP1** | 8 | 5 | AA | 0 | + | 22068326 | 22068500 | 22054897 | 22054897 |
| **BMP1** | 8 | 17 | AA | 0 | + | 22064367 | 22064494 | 22037910 | 22037910 |
| **BMP1** | 8 | 17 | AA | 0 | + | 22064367 | 22064494 | 22053064 | 22053064 |
| **BMP1** | 8 | 17 | AA | 0 | + | 22064367 | 22064494 | 22064400 | 22064400 |
| **BMP1** | 8 | 17 | AA | 0 | + | 22064367 | 22064494 | 22054897 | 22054897 |
| **BMP1** | 8 | 18 | AA | 0 | + | 22064816 | 22065029 | 22037910 | 22037910 |
| **BMP1** | 8 | 18 | AA | 0 | + | 22064816 | 22065029 | 22064900 | 22064900 |
| **BMP1** | 8 | 18 | AA | 0 | + | 22064816 | 22065029 | 22053064 | 22053064 |
| **BMP1** | 8 | 18 | AA | 0 | + | 22064816 | 22065029 | 22064400 | 22064400 |
| **BMP1** | 8 | 18 | AA | 0 | + | 22064816 | 22065029 | 22054897 | 22054897 |
| **BMP1** | 8 | 17 | AA | 0 | + | 22064816 | 22065029 | 22064900 | 22064900 |
| **BTN3A2** | 6 | 12 | AT | 0 | + | 26376025 | 26378539 | 26370581 | 26370581 |
| **C17orf97** | 17 | 2 | AT | 0 | + | 270463 | 270881 | 263004 | 263004 |
| **CAMTA1** | 1 | 3 | ES | 1 | + | 7796404 | 7796493 | 7724340 | 7724340 |
| **CAMTA1** | 1 | 3 | ES | 1 | + | 7796404 | 7796493 | 7724600 | 7724600 |
| **CAMTA1** | 1 | 3 | ES | 1 | + | 7796404 | 7796493 | 7700507 | 7700507 |
| **CAMTA1** | 1 | 5 | AT | 0 | + | 7797315 | 7797507 | 7724340 | 7724340 |
| **CAMTA1** | 1 | 5 | AT | 0 | + | 7797315 | 7797507 | 7724600 | 7724600 |
| **CAMTA1** | 1 | 5 | AT | 0 | + | 7797315 | 7797507 | 7700507 | 7700507 |
| **CAMTA1** | 1 | 3 | ES | 1 | + | 7815698 | 7815728 | 7811265 | 7811265 |
| **CAMTA1** | 1 | 5 | AT | 0 | + | 7812519 | 7812593 | 7811265 | 7811265 |
| **CARM1** | 19 | 16 | ES | 0 | + | 11032291 | 11033448 | 11019791 | 11019791 |
| **CARM1** | 19 | 16 | ES | 0 | + | 11032291 | 11033448 | 11022882 | 11022882 |
| **CARM1** | 19 | 4 | ES | 0 | + | 11019779 | 11019883 | 11019791 | 11019791 |
| **CARM1** | 19 | 4 | ES | 0 | + | 11031510 | 11031622 | 11022882 | 11022882 |
| **CASZ1** | 1 | 21 | AT | 0 | - | 10696661 | 10700116 | 10699493 | 10699493 |
| **CCDC25** | 8 | 5 | ES | 0 | - | 27610029 | 27610104 | 27605688 | 27605688 |
| **CCDC25** | 8 | 2 | ES | 0 | - | 27622721 | 27622768 | 27605688 | 27605688 |
| **CCDC25** | 8 | 3 | ES | 0 | - | 27621803 | 27622050 | 27605688 | 27605688 |
| **CCDC25** | 8 | 7 | ES | 0 | - | 27605377 | 27605796 | 27605688 | 27605688 |
| **CCDC25** | 8 | 8 | ES | 0 | - | 27605270 | 27605796 | 27605688 | 27605688 |
| **CD44** | 11 | 3 | ES | 0 | + | 35236399 | 35236461 | 35218302 | 35218302 |

| | | | | | | | | | |
|---|---|---|---|---|---|---|---|---|---|
| **CD44** | 11 | 3 | ES | 0 | + | 35236399 | 35236461 | 35218359 | 35218359 |
| **CD44** | 11 | 10 | ES | 0 | + | 35250676 | 35251213 | 35218302 | 35218302 |
| **CD44** | 11 | 10 | ES | 0 | + | 35250676 | 35251213 | 35250775 | 35250775 |
| **CD44** | 11 | 10 | ES | 0 | + | 35250676 | 35251213 | 35218359 | 35218359 |
| **CD44** | 11 | 6 | ES | 0 | + | 35243201 | 35243279 | 35218302 | 35218302 |
| **CD44** | 11 | 6 | ES | 0 | + | 35243201 | 35243279 | 35218359 | 35218359 |
| **CD44** | 11 | 9 | ES | 1 | + | 35232793 | 35233725 | 35218302 | 35218302 |
| **CD44** | 11 | 9 | ES | 1 | + | 35232793 | 35233725 | 35218359 | 35218359 |
| **CD44** | 11 | 8 | ES | 1 | + | 35223218 | 35223334 | 35218302 | 35218302 |
| **CD44** | 11 | 8 | ES | 1 | + | 35223218 | 35223334 | 35218359 | 35218359 |
| **CD44** | 11 | 11 | ES | 1 | + | 35227659 | 35227790 | 35218302 | 35218302 |
| **CD44** | 11 | 11 | ES | 1 | + | 35227659 | 35227790 | 35218359 | 35218359 |
| **CD44** | 11 | 9 | ES | 1 | + | 35250676 | 35253948 | 35250775 | 35250775 |
| **CD44** | 11 | 8 | ES | 1 | + | 35250676 | 35250880 | 35250775 | 35250775 |
| **CD44** | 11 | 14 | ES | 0 | + | 35232793 | 35232996 | 35218302 | 35218302 |
| **CD44** | 11 | 14 | ES | 0 | + | 35232793 | 35232996 | 35218359 | 35218359 |
| **CD59** | 11 | 1 | AP | 0 | - | 33757928 | 33758025 | 33731808 | 33731808 |
| **CD59** | 11 | 1 | AP | 0 | - | 33757928 | 33758025 | 33739011 | 33739011 |
| **CLN3** | 16 | 9 | ES | 0 | - | 28493947 | 28493993 | 28493652 | 28493652 |
| **CLN3** | 16 | 4 | ES | 0 | - | 28499912 | 28499983 | 28493652 | 28493652 |
| **CLN3** | 16 | 7 | ES | 0 | - | 28493798 | 28493866 | 28493652 | 28493652 |
| **CLN3** | 16 | 10 | ES | 0 | - | 28493648 | 28493703 | 28493652 | 28493652 |
| **CLN3** | 16 | 8 | ES | 0 | - | 28497899 | 28497971 | 28493652 | 28493652 |
| **CLN3** | 16 | 11 | ES | 0 | - | 28493648 | 28493703 | 28493652 | 28493652 |
| **CLN3** | 16 | 5 | ES | 0 | - | 28495327 | 28495439 | 28493652 | 28493652 |
| **CLN3** | 16 | 6 | ES | 0 | - | 28497668 | 28497811 | 28493652 | 28493652 |
| **CLN3** | 16 | 14 | ES | 0 | - | 28493426 | 28493703 | 28493652 | 28493652 |
| **CNTN4** | 3 | 4 | AP | 0 | + | 2968982 | 2969315 | 2908487 | 2908487 |
| **CNTN4** | 3 | 4 | AP | 0 | + | 2968982 | 2969315 | 2928774 | 2928774 |
| **CNTN4** | 3 | 4 | AP | 0 | + | 3085276 | 3085388 | 3067840 | 3067840 |
| **CNTN4** | 3 | 4 | AP | 0 | + | 3085276 | 3085388 | 3078896 | 3078896 |
| **CPSF3L** | 1 | 5 | ES | 0 | - | 1254676 | 1254904 | 1254756 | 1254756 |
| **CPSF3L** | 1 | 2 | ES | 0 | - | 1256376 | 1256476 | 1254756 | 1254756 |
| **CPSF3L** | 1 | 4 | ES | 0 | - | 1255836 | 1255909 | 1254756 | 1254756 |
| **CPSF3L** | 1 | 6 | ES | 0 | - | 1254783 | 1254904 | 1254756 | 1254756 |
| **CPSF3L** | 1 | 3 | ES | 0 | - | 1257288 | 1257364 | 1254756 | 1254756 |
| **CSRP2BP** | 20 | 1 | AP | 0 | + | 18166339 | 18168678 | 18162409 | 18162409 |
| **CSRP2BP** | 20 | 1 | AP | 0 | + | 18166339 | 18168678 | 18143582 | 18143582 |
| **DHDDS** | 1 | 4 | AA | 0 | + | 26769222 | 26769364 | 26769270 | 26769270 |
| **DHRS4** | 14 | 6 | ES | 0 | + | 24436420 | 24436448 | 24429144 | 24429144 |
| **DHRS4** | 14 | 6 | ES | 0 | + | 24436420 | 24436448 | 24429127 | 24429127 |
| **DHRS4** | 14 | 4 | ES | 0 | + | 24434971 | 24435041 | 24429144 | 24429144 |
| **DHRS4** | 14 | 4 | ES | 0 | + | 24434971 | 24435041 | 24429127 | 24429127 |
| **DHRS4** | 14 | 3 | ES | 0 | + | 24429111 | 24429212 | 24429144 | 24429144 |
| **DHRS4** | 14 | 3 | ES | 0 | + | 24429111 | 24429212 | 24429127 | 24429127 |
| **DHRS4** | 14 | 5 | ES | 0 | + | 24436420 | 24436475 | 24429144 | 24429144 |
| **DHRS4** | 14 | 5 | ES | 0 | + | 24436420 | 24436475 | 24429127 | 24429127 |

| | | | | | | | | | |
|---|---|---|---|---|---|---|---|---|---|
| **DHRS4L2** | 14 | 4 | ES | 0 | + | 24473546 | 24473601 | 24470603 | 24470603 |
| **DHRS4L2** | 14 | 7 | ES | 0 | + | 24473546 | 24473601 | 24470603 | 24470603 |
| **DHRS4L2** | 14 | 5 | ES | 0 | + | 24473546 | 24473601 | 24470603 | 24470603 |
| **DHRS4L2** | 14 | 6 | ES | 0 | + | 24473546 | 24473566 | 24470603 | 24470603 |
| **DHRS4L2** | 14 | 8 | ES | 0 | + | 24473546 | 24473601 | 24470603 | 24470603 |
| **DRG2** | 17 | 6 | RI | 0 | + | 18010468 | 18010676 | 18002358 | 18002358 |
| **DRG2** | 17 | 6 | RI | 0 | + | 18010468 | 18010676 | 18007126 | 18007126 |
| **DRG2** | 17 | 8 | AA | 0 | + | 18005235 | 18005311 | 18002358 | 18002358 |
| **DRG2** | 17 | 5 | AD | 0 | + | 18002947 | 18003037 | 18002358 | 18002358 |
| **DRG2** | 17 | 3 | AD | 0 | + | 18003883 | 18003973 | 18002358 | 18002358 |
| **DRG2** | 17 | 8 | AA | 0 | + | 18009747 | 18009800 | 18007126 | 18007126 |
| **DRG2** | 17 | 12 | AD | 0 | + | 18009747 | 18009800 | 18002358 | 18002358 |
| **DRG2** | 17 | 12 | AD | 0 | + | 18009747 | 18009800 | 18007126 | 18007126 |
| **DRG2** | 17 | 5 | AD | 0 | + | 18009747 | 18009800 | 18007126 | 18007126 |
| **DRG2** | 17 | 3 | AD | 0 | + | 18007115 | 18007203 | 18007126 | 18007126 |
| **DUSP22** | 6 | 7 | RI | 0 | + | 348769 | 348841 | 345913 | 345913 |
| **EIF3C** | 16 | 2 | AP | 0 | + | 28746253 | 28746340 | 28734529 | 28734529 |
| **ELP4** | 11 | 10 | ES | 0 | + | 31671663 | 31671769 | 31541617 | 31541617 |
| **ELP4** | 11 | 11 | ES | 0 | + | 31736179 | 31736442 | 31541617 | 31541617 |
| **ENTPD6** | 20 | 3 | AA | 0 | + | 25206135 | 25206369 | 25205943 | 25205943 |
| **ENTPD6** | 20 | 3 | AA | 0 | + | 25206135 | 25206369 | 25205907 | 25205907 |
| **ENTPD6** | 20 | 3 | AA | 0 | + | 25206135 | 25206369 | 25203540 | 25203540 |
| **ENTPD6** | 20 | 14 | AA | 0 | + | 25206135 | 25207288 | 25205943 | 25205943 |
| **ENTPD6** | 20 | 14 | AA | 0 | + | 25206135 | 25207288 | 25205907 | 25205907 |
| **ENTPD6** | 20 | 14 | AA | 0 | + | 25206135 | 25207288 | 25203540 | 25203540 |
| **ERAL1** | 17 | 3 | AD | 0 | + | 27185594 | 27185842 | 27185169 | 27185169 |
| **EXOSC8** | 13 | 9 | AA | 0 | + | 37582866 | 37582972 | 37582916 | 37582916 |
| **FARS2** | 6 | 2 | AP | 0 | + | 5771524 | 5771709 | 5771561 | 5771561 |
| **FDFT1** | 8 | 2 | AP | 0 | + | 11683588 | 11683724 | 11660380 | 11660380 |
| **FDFT1** | 8 | 2 | AP | 0 | + | 11683588 | 11683724 | 11667339 | 11667339 |
| **GGT5** | 22 | 5 | AA | 0 | - | 24628019 | 24628176 | 24622171 | 24622171 |
| **GGT5** | 22 | 5 | AA | 0 | - | 24628019 | 24628176 | 24622114 | 24622114 |
| **GPR125** | 4 | 21 | AT | 0 | - | 22390711 | 22390806 | 22390718 | 22390718 |
| **GPR125** | 4 | 21 | AT | 0 | - | 22390711 | 22390806 | 22390256 | 22390256 |
| **GPR125** | 4 | 21 | AT | 0 | - | 22390711 | 22390806 | 22390383 | 22390383 |
| **GTF2H1** | 11 | 1 | AP | 0 | + | 18387195 | 18388590 | 18387396 | 18387396 |
| **GTF3A** | 13 | 1 | AD | 0 | + | 28003827 | 28004103 | 28001293 | 28001293 |
| **HMHA1** | 19 | 15 | AP | 0 | + | 1080891 | 1081063 | 1068512 | 1068512 |
| **HNRNPC** | 14 | 8 | ES | 0 | - | 21679969 | 21680082 | 21680019 | 21680019 |
| **HNRNPC** | 14 | 3 | ES | 0 | - | 21698478 | 21698525 | 21680019 | 21680019 |
| **HNRNPC** | 14 | 2 | AD | 0 | - | 21736782 | 21736827 | 21680019 | 21680019 |
| **HNRNPC** | 14 | 1 | AP | 0 | - | 21737457 | 21737585 | 21680019 | 21680019 |
| **HNRNPC** | 14 | 4 | ES | 0 | - | 21699117 | 21699231 | 21680019 | 21680019 |
| **HNRNPC** | 14 | 6 | ES | 0 | - | 21698478 | 21698525 | 21680019 | 21680019 |
| **HNRNPC** | 14 | 7 | ES | 0 | - | 21679969 | 21680082 | 21680019 | 21680019 |
| **HNRNPC** | 14 | 5 | ES | 0 | - | 21698478 | 21698525 | 21680019 | 21680019 |
| **HNRNPM** | 19 | 11 | ES | 0 | + | 8550487 | 8550858 | 8548057 | 8548057 |

| | | | | | | | | | |
|---|---|---|---|---|---|---|---|---|---|
| **HNRNPM** | 19 | 11 | ES | 0 | + | 8550487 | 8550858 | 8528515 | 8528515 |
| **HNRNPM** | 19 | 11 | ES | 0 | + | 8551908 | 8551959 | 8550883 | 8550883 |
| **HNRNPM** | 19 | 11 | ES | 0 | + | 8551908 | 8551959 | 8550903 | 8550903 |
| **HNRNPM** | 19 | 3 | ES | 0 | + | 8532419 | 8532468 | 8528515 | 8528515 |
| **HNRNPM** | 19 | 3 | ES | 0 | + | 8550487 | 8551289 | 8548057 | 8548057 |
| **HNRNPM** | 19 | 3 | ES | 0 | + | 8550487 | 8551289 | 8550883 | 8550883 |
| **HNRNPM** | 19 | 3 | ES | 0 | + | 8550487 | 8551289 | 8550903 | 8550903 |
| **HSPH1** | 13 | 4 | ES | 0 | - | 31729651 | 31729791 | 31724241 | 31724241 |
| **HSPH1** | 13 | 5 | ES | 0 | - | 31726989 | 31727088 | 31724241 | 31724241 |
| **HSPH1** | 13 | 7 | ES | 0 | - | 31725084 | 31725328 | 31724241 | 31724241 |
| **HSPH1** | 13 | 6 | ES | 0 | - | 31725746 | 31725879 | 31724241 | 31724241 |
| **ILF3** | 19 | 14 | AA | 0 | + | 10793813 | 10793938 | 10787961 | 10787961 |
| **ILF3** | 19 | 14 | AA | 0 | + | 10793813 | 10793938 | 10781680 | 10781680 |
| **ILF3** | 19 | 14 | AA | 0 | + | 10794593 | 10797165 | 10794144 | 10794144 |
| **ILF3** | 19 | 17 | AA | 0 | + | 10794593 | 10794646 | 10794144 | 10794144 |
| **ILF3** | 19 | 17 | AA | 0 | + | 10794593 | 10794646 | 10787961 | 10787961 |
| **ILF3** | 19 | 17 | AA | 0 | + | 10794593 | 10794646 | 10781680 | 10781680 |
| **INO80C** | 18 | 3 | ES | 0 | - | 33060417 | 33060527 | 33060428 | 33060428 |
| **INO80C** | 18 | 4 | ES | 0 | - | 33060417 | 33060527 | 33060428 | 33060428 |
| **INO80E** | 16 | 7 | ES | 0 | + | 30013757 | 30014334 | 30007925 | 30007925 |
| **INO80E** | 16 | 6 | AA | 0 | + | 30012533 | 30014334 | 30007925 | 30007925 |
| **INO80E** | 16 | 9 | ES | 0 | + | 30014971 | 30014991 | 30007925 | 30007925 |
| **INO80E** | 16 | 8 | ES | 0 | + | 30014757 | 30014847 | 30007925 | 30007925 |
| **INO80E** | 16 | 10 | ES | 0 | + | 30016542 | 30017113 | 30007925 | 30007925 |
| **ITGB1BP1** | 2 | 3 | ES | 0 | - | 9554307 | 9554385 | 9547660 | 9547660 |
| **ITGB1BP1** | 2 | 5 | ES | 0 | - | 9552398 | 9552534 | 9547660 | 9547660 |
| **ITSN1** | 21 | 32 | AT | 0 | + | 35237455 | 35237638 | 35237530 | 35237530 |
| **ITSN1** | 21 | 32 | AT | 0 | + | 35237455 | 35237638 | 35094941 | 35094941 |
| **ITSN1** | 21 | 32 | AT | 0 | + | 35237455 | 35237638 | 35208915 | 35208915 |
| **ITSN1** | 21 | 32 | AT | 0 | + | 35254547 | 35254759 | 35247768 | 35247768 |
| **ITSN1** | 21 | 32 | AT | 0 | + | 35254547 | 35254759 | 35254750 | 35254750 |
| **KIAA1328** | 18 | 12 | AT | 0 | + | 34811623 | 34812141 | 34802090 | 34802090 |
| **LPCAT3** | 12 | 10 | RI | 0 | - | 7086296 | 7086424 | 7086376 | 7086376 |
| **LRP10** | 14 | 7 | RI | 0 | + | 23346149 | 23350918 | 23346384 | 23346384 |
| **LSM14A** | 19 | 3 | ES | 0 | + | 34687539 | 34687668 | 34685499 | 34685499 |
| **LSM14A** | 19 | 3 | ES | 0 | + | 34687539 | 34687668 | 34663663 | 34663663 |
| **LSM14A** | 19 | 3 | ES | 0 | + | 34718274 | 34720187 | 34710328 | 34710328 |
| **LSM14A** | 19 | 11 | ES | 0 | + | 34718270 | 34720420 | 34685499 | 34685499 |
| **LSM14A** | 19 | 11 | ES | 0 | + | 34718270 | 34720420 | 34663663 | 34663663 |
| **LSM14A** | 19 | 11 | ES | 0 | + | 34718270 | 34720420 | 34710328 | 34710328 |
| **LSM14A** | 19 | 5 | ES | 0 | + | 34706501 | 34706566 | 34685499 | 34685499 |
| **LSM14A** | 19 | 5 | ES | 0 | + | 34706501 | 34706566 | 34663663 | 34663663 |
| **LSM14A** | 19 | 5 | ES | 0 | + | 34710611 | 34710782 | 34710328 | 34710328 |
| **LSR** | 19 | 5 | ES | 0 | + | 35757262 | 35757435 | 35753550 | 35753550 |
| **LSR** | 19 | 2 | ES | 0 | + | 35757582 | 35757870 | 35753550 | 35753550 |
| **LSR** | 19 | 4 | ES | 0 | + | 35757262 | 35757330 | 35753550 | 35753550 |
| **LSR** | 19 | 3 | ES | 0 | + | 35758012 | 35758607 | 35753550 | 35753550 |

| | | | | | | | | | |
|---|---|---|---|---|---|---|---|---|---|
| **LSR** | 19 | 8 | ES | 0 | + | 35758012 | 35758637 | 35753550 | 35753550 |
| **LSR** | 19 | 1 | ES | 0 | + | 35757399 | 35757435 | 35753550 | 35753550 |
| **LSR** | 19 | 7 | ES | 0 | + | 35757742 | 35757870 | 35753550 | 35753550 |
| **MAGOHB** | 12 | 2 | ES | 0 | - | 10763221 | 10763279 | 10762505 | 10762505 |
| **MAGOHB** | 12 | 4 | ES | 0 | - | 10762430 | 10762540 | 10762505 | 10762505 |
| **MAPK13** | 6 | 5 | ES | 0 | + | 36108765 | 36109042 | 36106684 | 36106684 |
| **MAPK13** | 6 | 9 | ES | 0 | + | 36107071 | 36107827 | 36106684 | 36106684 |
| **MAPK13** | 6 | 8 | ES | 0 | + | 36106656 | 36106832 | 36106684 | 36106684 |
| **MAZ** | 16 | 2 | AP | 0 | + | 29819551 | 29819614 | 29819148 | 29819148 |
| **MED15** | 22 | 18 | RI | 0 | + | 20940855 | 20941918 | 20937634 | 20937634 |
| **MED15** | 22 | 18 | RI | 0 | + | 20940855 | 20941918 | 20939239 | 20939239 |
| **MED15** | 22 | 18 | RI | 0 | + | 20940855 | 20941918 | 20921060 | 20921060 |
| **MEIS2** | 15 | 2 | AP | 0 | - | 37387765 | 37387815 | 37385817 | 37385817 |
| **METTL3** | 14 | 3 | RI | 0 | - | 21971316 | 21971720 | 21971467 | 21971467 |
| **MGRN1** | 16 | 17 | AA | 0 | + | 4738797 | 4741347 | 4733904 | 4733904 |
| **MIB2** | 1 | 10 | AA | 0 | + | 1562676 | 1562824 | 1560722 | 1560722 |
| **MIB2** | 1 | 5 | AA | 0 | + | 1560666 | 1560808 | 1560722 | 1560722 |
| **MLH1** | 3 | 20 | AT | 0 | + | 37091977 | 37092330 | 37053334 | 37053334 |
| **MOSPD2** | X | 15 | AA | 0 | + | 14937794 | 14939458 | 14929376 | 14929376 |
| **MYL9** | 20 | 3 | ES | 0 | + | 35177480 | 35178226 | 35177521 | 35177521 |
| **NIPA1** | 15 | 2 | AP | 0 | - | 23062022 | 23062319 | 23049053 | 23049053 |
| **NOP2** | 12 | 4 | ES | 0 | - | 6675695 | 6675783 | 6670153 | 6670153 |
| **NOP2** | 12 | 4 | ES | 0 | - | 6675695 | 6675783 | 6672578 | 6672578 |
| **NOP2** | 12 | 3 | ES | 0 | - | 6675943 | 6675988 | 6670153 | 6670153 |
| **NOP2** | 12 | 3 | ES | 0 | - | 6675943 | 6675988 | 6672578 | 6672578 |
| **NOP2** | 12 | 5 | ES | 0 | - | 6670097 | 6670236 | 6670153 | 6670153 |
| **NOP2** | 12 | 5 | ES | 0 | - | 6675267 | 6675502 | 6672578 | 6672578 |
| **OSGEP** | 14 | 4 | RI | 0 | - | 20916265 | 20916330 | 20916132 | 20916132 |
| **PCNA** | 20 | 2 | AP | 0 | - | 5100224 | 5100560 | 5100411 | 5100411 |
| **PDCD5** | 19 | 5 | RI | 0 | + | 33077764 | 33077835 | 33076792 | 33076792 |
| **PDCD5** | 19 | 1 | AP | 0 | + | 33074864 | 33078331 | 33076792 | 33076792 |
| **PHF12** | 17 | 12 | RI | 0 | - | 27234608 | 27234690 | 27233281 | 27233281 |
| **PLEKHA8** | 7 | 14 | AT | 0 | + | 30118206 | 30124338 | 30094411 | 30094411 |
| **PLEKHA8** | 7 | 14 | AT | 0 | + | 30118206 | 30124338 | 30100541 | 30100541 |
| **PPARD** | 6 | 9 | AT | 0 | + | 35393609 | 35395970 | 35393666 | 35393666 |
| **PRKCSH** | 19 | 2 | AD | 0 | + | 11561401 | 11561780 | 11560084 | 11560084 |
| **PRLR** | 5 | 10 | AT | 0 | - | 35066051 | 35066204 | 35065420 | 35065420 |
| **PSMD11** | 17 | 13 | RI | 0 | + | 30807507 | 30810337 | 30796059 | 30796059 |
| **QDPR** | 4 | 2 | ES | 0 | - | 17510894 | 17510986 | 17510920 | 17510920 |
| **QDPR** | 4 | 3 | ES | 0 | - | 17510894 | 17510986 | 17510920 | 17510920 |
| **RABEP2** | 16 | 7 | ES | 0 | - | 28922209 | 28922307 | 28920040 | 28920040 |
| **RABEP2** | 16 | 3 | ES | 0 | - | 28931107 | 28931264 | 28920040 | 28920040 |
| **RABEP2** | 16 | 2 | ES | 0 | - | 28931107 | 28931264 | 28920040 | 28920040 |
| **RAD23A** | 19 | 7 | AA | 0 | + | 13060089 | 13060222 | 13059574 | 13059574 |
| **RAD23A** | 19 | 4 | AA | 0 | + | 13059895 | 13059973 | 13059574 | 13059574 |
| **RBBP6** | 16 | 18 | AT | 0 | + | 24582197 | 24584181 | 24581622 | 24581622 |
| **RBBP6** | 16 | 18 | AT | 0 | + | 24582197 | 24584181 | 24580170 | 24580170 |

| | | | | | | | | | |
|---|---|---|---|---|---|---|---|---|---|
| RBM42 | 19 | 5 | ES | 0 | + | 36128344 | 36128586 | 36128403 | 36128403 |
| RBM42 | 19 | 8 | ES | 0 | + | 36128344 | 36128586 | 36128403 | 36128403 |
| RBM42 | 19 | 9 | ES | 0 | + | 36128344 | 36128587 | 36128403 | 36128403 |
| RHBDF1 | 16 | 18 | AT | 0 | - | 108059 | 108758 | 108512 | 108512 |
| RHBDF1 | 16 | 18 | AT | 0 | - | 108059 | 108758 | 108725 | 108725 |
| RNF8 | 6 | 7 | ES | 1 | + | 37348926 | 37349130 | 37348968 | 37348968 |
| RNF8 | 6 | 7 | ES | 1 | + | 37348926 | 37349130 | 37336474 | 37336474 |
| RNH1 | 11 | 4 | AA | 0 | - | 500535 | 500654 | 494985 | 494985 |
| RNH1 | 11 | 5 | AA | 0 | - | 499015 | 499185 | 494985 | 494985 |
| RNH1 | 11 | 1 | AP | 0 | - | 504427 | 504552 | 494985 | 494985 |
| RNH1 | 11 | 6 | AA | 0 | - | 498763 | 498933 | 494985 | 494985 |
| RRP8 | 11 | 3 | RI | 0 | - | 6622156 | 6622832 | 6622389 | 6622389 |
| RRP8 | 11 | 3 | RI | 0 | - | 6622156 | 6622832 | 6621780 | 6621780 |
| RRP8 | 11 | 3 | RI | 0 | - | 6622156 | 6622832 | 6622562 | 6622562 |
| RRP8 | 11 | 5 | RI | 0 | - | 6621173 | 6621812 | 6621780 | 6621780 |
| SDHA | 5 | 2 | ES | 0 | + | 251107 | 251158 | 235284 | 235284 |
| SDHA | 5 | 5 | ES | 0 | + | 251107 | 251218 | 235284 | 235284 |
| SDHA | 5 | 14 | ES | 0 | + | 256449 | 257189 | 235284 | 235284 |
| SDHA | 5 | 14 | ES | 0 | + | 256449 | 257189 | 251561 | 251561 |
| SDHA | 5 | 5 | ES | 0 | + | 256449 | 256794 | 251561 | 251561 |
| SDHA | 5 | 13 | ES | 0 | + | 254508 | 254621 | 235284 | 235284 |
| SDHA | 5 | 13 | ES | 0 | + | 254508 | 254621 | 251561 | 251561 |
| SDHA | 5 | 2 | ES | 0 | + | 256545 | 256794 | 251561 | 251561 |
| SH2B1 | 16 | 3 | AD | 0 | + | 28878652 | 28878753 | 28877932 | 28877932 |
| SH2B1 | 16 | 3 | AD | 0 | + | 28878652 | 28878753 | 28877903 | 28877903 |
| SH2B1 | 16 | 3 | AD | 0 | + | 28884742 | 28885523 | 28883991 | 28883991 |
| SH2B1 | 16 | 9 | AD | 0 | + | 28884491 | 28884590 | 28883991 | 28883991 |
| SH2B1 | 16 | 9 | AD | 0 | + | 28884491 | 28884590 | 28877932 | 28877932 |
| SH2B1 | 16 | 9 | AD | 0 | + | 28884491 | 28884590 | 28877903 | 28877903 |
| SHMT1 | 17 | 4 | ES | 0 | - | 18256986 | 18257131 | 18232655 | 18232655 |
| SHMT1 | 17 | 9 | ES | 0 | - | 18236480 | 18236602 | 18232655 | 18232655 |
| SHMT1 | 17 | 8 | ES | 0 | - | 18238873 | 18238989 | 18232655 | 18232655 |
| SMARCA4 | 19 | 32 | ES | 0 | + | 11169464 | 11169565 | 11143994 | 11143994 |
| SMARCA4 | 19 | 32 | ES | 0 | + | 11169464 | 11169565 | 11105645 | 11105645 |
| SNAPC3 | 9 | 9 | AT | 0 | + | 15459717 | 15461627 | 15422996 | 15422996 |
| SSBP4 | 19 | 1 | AP | 0 | + | 18542449 | 18542516 | 18541673 | 18541673 |
| SYTL1 | 1 | 6 | AA | 0 | + | 27677749 | 27677762 | 27677325 | 27677325 |
| SYTL1 | 1 | 6 | AA | 0 | + | 27677749 | 27677762 | 27677441 | 27677441 |
| TBC1D5 | 3 | 2 | AP | 0 | - | 17688453 | 17688548 | 17418045 | 17418045 |
| THOP1 | 19 | 1 | AP | 0 | + | 2806978 | 2807050 | 2794884 | 2794884 |
| TMEM159 | 16 | 3 | ES | 0 | + | 21181758 | 21181926 | 21181905 | 21181905 |
| TMEM159 | 16 | 5 | ES | 0 | + | 21190796 | 21191935 | 21181905 | 21181905 |
| TMEM159 | 16 | 6 | ES | 0 | + | 21190796 | 21191859 | 21181905 | 21181905 |
| TMEM66 | 8 | 2 | ES | 0 | - | 29927158 | 29927575 | 29923589 | 29923589 |
| TMEM66 | 8 | 5 | ES | 0 | - | 29923504 | 29923655 | 29923589 | 29923589 |
| TMEM66 | 8 | 6 | ES | 0 | - | 29923504 | 29923655 | 29923589 | 29923589 |
| TMEM66 | 8 | 1 | ES | 0 | - | 29940378 | 29940580 | 29923589 | 29923589 |

| | | | | | | | | | | |
|---|---|---|---|---|---|---|---|---|---|---|
| **TMEM98** | 17 | 8 | AT | 0 | + | | 31267804 | 31269785 | 31266550 | 31266550 |
| **TMEM98** | 17 | 8 | AT | 0 | + | | 31267804 | 31269785 | 31263420 | 31263420 |
| **TNKS** | 8 | 1 | AP | 0 | + | | 9627642 | 9627772 | 9627692 | 9627692 |
| **TNKS** | 8 | 1 | AP | 0 | + | | 9627642 | 9627772 | 9538260 | 9538260 |
| **TNKS** | 8 | 1 | AP | 0 | + | | 9627642 | 9627772 | 9623794 | 9623794 |
| **TXNDC17** | 17 | 1 | AD | 0 | + | | 6544628 | 6544818 | 6544417 | 6544417 |
| **TXNDC17** | 17 | 2 | AD | 0 | + | | 6545073 | 6545154 | 6544417 | 6544417 |
| **TYMS** | 18 | 3 | ES | 0 | + | | 672860 | 673135 | 672975 | 672975 |
| **USE1** | 19 | 5 | RI | 0 | + | | 17330022 | 17330196 | 17326260 | 17326260 |
| **USE1** | 19 | 7 | AT | 0 | + | | 17330440 | 17330627 | 17326260 | 17326260 |
| **WAC** | 10 | 2 | AP | 0 | + | | 28878665 | 28878780 | 28878735 | 28878735 |
| **WAC** | 10 | 2 | AP | 0 | + | | 28899628 | 28899750 | 28897207 | 28897207 |
| **WDR62** | 19 | 25 | RI | 0 | + | | 36592566 | 36592676 | 36572414 | 36572414 |
| **WDR62** | 19 | 25 | RI | 0 | + | | 36592566 | 36592676 | 36590347 | 36590347 |
| **WDR83** | 19 | 7 | AA | 0 | + | | 12784016 | 12784130 | 12784068 | 12784068 |
| **WDR83** | 19 | 6 | AA | 0 | + | | 12786337 | 12786642 | 12784068 | 12784068 |
| **ZCCHC4** | 4 | 13 | AT | 0 | + | | 25370660 | 25371164 | 25370758 | 25370758 |
| **ZNF141** | 4 | 4 | AT | 0 | + | | 376884 | 377227 | 367319 | 367319 |
| **ZNF141** | 4 | 4 | AT | 0 | + | | 376884 | 377227 | 367212 | 367212 |
| **ZNF141** | 4 | 4 | AT | 0 | + | | 376884 | 377227 | 367647 | 367647 |
| **ZNF345** | 19 | 4 | AT | 0 | + | | 37383720 | 37383827 | 37368703 | 37368703 |
| **ZNF345** | 19 | 4 | AT | 0 | + | | 37383720 | 37383827 | 37368427 | 37368427 |
| **ZNF345** | 19 | 4 | AT | 0 | + | | 37383720 | 37383827 | 37368396 | 37368396 |
| **ZNF345** | 19 | 4 | AT | 0 | + | | 37383720 | 37383827 | 37368910 | 37368910 |
| **ZNF506** | 19 | 3 | ES | 0 | - | | 19916840 | 19916935 | 19905821 | 19905821 |
| **ZNF850** | 19 | 5 | AT | 0 | - | | 37234380 | 37241706 | 37239046 | 37239046 |
| **ZSCAN20** | 1 | 8 | AT | 0 | + | | 33959818 | 33961199 | 33960721 | 33960721 |
| **ZSCAN20** | 1 | 8 | AT | 0 | + | | 33959818 | 33961199 | 33960823 | 33960823 |
| **CGREF1** | 2 | 7 | AT | 0 | - | | 27324152 | 27324261 | 27321998 | 27321998 |
| **CGREF1** | 2 | 7 | AT | 0 | - | | 27324152 | 27324261 | 27321993 | 27321993 |
| **B9D1** | 17 | 8 | AT | 0 | - | | 19247103 | 19247170 | 19246523 | 19246523 |
| **BMP1** | 8 | 5 | AA | 0 | + | | 22069783 | 22069837 | 22069260 | 22069260 |
| **C1orf86** | 1 | 4 | ES | 1 | - | | 2134875 | 2135069 | 2121108 | 2121108 |
| **C1orf86** | 1 | 6 | ES | 0 | - | | 2120990 | 2121220 | 2121108 | 2121108 |
| **CDC37** | 19 | 2 | ES | 0 | - | | 10514054 | 10514271 | 10502151 | 10502151 |
| **CIRBP** | 19 | 7 | RI | 0 | + | | 1274306 | 1274479 | 1272618 | 1272618 |
| **CIRBP** | 19 | 8 | AA | 0 | + | | 1274648 | 1274806 | 1272618 | 1272618 |
| **CIRBP** | 19 | 9 | RI | 0 | + | | 1274648 | 1274772 | 1272618 | 1272618 |
| **CIRBP** | 19 | 5 | AP | 0 | + | | 1271980 | 1274439 | 1272618 | 1272618 |
| **DHRS4L2** | 14 | 4 | ES | 0 | + | | 24475095 | 24475612 | 24475125 | 24475125 |
| **DHRS4L2** | 14 | 8 | ES | 0 | + | | 24475095 | 24475610 | 24475125 | 24475125 |
| **DHRS4L2** | 14 | 5 | ES | 0 | + | | 24475100 | 24475610 | 24475125 | 24475125 |
| **DHRS4L2** | 14 | 6 | ES | 0 | + | | 24475095 | 24475610 | 24475125 | 24475125 |
| **DHRS4L2** | 14 | 7 | ES | 0 | + | | 24475077 | 24475610 | 24475125 | 24475125 |
| **ELP4** | 11 | 10 | ES | 0 | + | | 31804941 | 31805329 | 31805078 | 31805078 |
| **ELP4** | 11 | 11 | ES | 0 | + | | 31804941 | 31805319 | 31805078 | 31805078 |
| **ERRFI1** | 1 | 3 | AT | 0 | - | | 8075368 | 8075460 | 8073211 | 8073211 |

| | | | | | | | | | |
|---|---|---|---|---|---|---|---|---|---|
| **GGT5** | 22 | 10 | AA | 0 | - | 24620964 | 24621074 | 24615921 | 24615921 |
| **GGT5** | 22 | 5 | AA | 0 | - | 24628019 | 24628176 | 24615921 | 24615921 |
| **ILK** | 11 | 1 | RI | 0 | + | 6625041 | 6632102 | 6631921 | 6631921 |
| **MAP3K8** | 10 | 9 | AT | 0 | + | 30749635 | 30750761 | 30749820 | 30749820 |
| **PPARD** | 6 | 9 | AT | 0 | + | 35393609 | 35395970 | 35393974 | 35393974 |
| **QDPR** | 4 | 2 | ES | 0 | - | 17492285 | 17492368 | 17488658 | 17488658 |
| **QDPR** | 4 | 3 | ES | 0 | - | 17506002 | 17506098 | 17488658 | 17488658 |
| **QDPR** | 4 | 6 | ES | 0 | - | 17493855 | 17493963 | 17488658 | 17488658 |
| **QDPR** | 4 | 7 | ES | 0 | - | 17488813 | 17488859 | 17488658 | 17488658 |
| **RNMT** | 18 | 12 | AA | 0 | + | 13762038 | 13764556 | 13760059 | 13760059 |
| **SLC39A6** | 18 | 10 | AT | 0 | - | 33688495 | 33689708 | 33689468 | 33689468 |
| **TADA3** | 3 | 2 | AP | 0 | - | 9833900 | 9833913 | 9821956 | 9821956 |
| **TLCD1** | 17 | 2 | AP | 0 | - | 27053409 | 27053824 | 27051417 | 27051417 |
| **TP53I3** | 2 | 1 | RI | 0 | - | 24308527 | 24308855 | 24300349 | 24300349 |
| **ZNF599** | 19 | 4 | AT | 0 | - | 35258170 | 35258316 | 35249888 | 35249888 |
| **BFAR** | 16 | 3 | ES | 0 | + | 14742245 | 14742449 | 14738159 | 14738159 |
| **BFAR** | 16 | 4 | ES | 0 | + | 14743661 | 14743830 | 14738159 | 14738159 |
| **BFAR** | 16 | 5 | ES | 0 | + | 14758726 | 14758928 | 14738159 | 14738159 |
| **LRP10** | 14 | 7 | RI | 0 | + | 23346149 | 23346262 | 23341345 | 23341345 |
| **MAPK13** | 6 | 5 | ES | 0 | + | 36103530 | 36103638 | 36098345 | 36098345 |
| **MAPK13** | 6 | 8 | ES | 0 | + | 36104433 | 36104547 | 36098345 | 36098345 |
| **MAPK13** | 6 | 9 | ES | 0 | + | 36107071 | 36107827 | 36098345 | 36098345 |
| **RPAIN** | 17 | 5 | ES | 0 | + | 5335862 | 5336340 | 5323023 | 5323023 |
| **RPAIN** | 17 | 4 | ES | 0 | + | 5329556 | 5329619 | 5323023 | 5323023 |
| **RPAIN** | 17 | 6 | ES | 0 | + | 5331391 | 5332985 | 5323023 | 5323023 |
| **WAC** | 10 | 2 | AP | 0 | + | 28822927 | 28822963 | 28822471 | 28822472 |
| **BMP1** | 8 | 5 | AA | 0 | + | 22049562 | 22049664 | 22035193 | 22035193 |
| **BMP1** | 8 | 15 | AA | 0 | + | 22054753 | 22054933 | 22035193 | 22035193 |
| **BMP1** | 8 | 5 | AA | 0 | + | 22068326 | 22068500 | 22058842 | 22058842 |
| **BMP1** | 8 | 17 | AA | 0 | + | 22064367 | 22064494 | 22035193 | 22035193 |
| **BMP1** | 8 | 17 | AA | 0 | + | 22064367 | 22064494 | 22058842 | 22058842 |
| **BMP1** | 8 | 18 | AA | 0 | + | 22064816 | 22065029 | 22035193 | 22035193 |
| **BMP1** | 8 | 18 | AA | 0 | + | 22064816 | 22065029 | 22058842 | 22058842 |
| **DHRS4L2** | 14 | 4 | ES | 0 | + | 24475095 | 24475612 | 24473710 | 24473710 |
| **DHRS4L2** | 14 | 8 | ES | 0 | + | 24475095 | 24475610 | 24473710 | 24473710 |
| **DHRS4L2** | 14 | 6 | ES | 0 | + | 24473546 | 24473725 | 24473710 | 24473710 |
| **DHRS4L2** | 14 | 5 | ES | 0 | + | 24475100 | 24475610 | 24473710 | 24473710 |
| **DHRS4L2** | 14 | 7 | ES | 0 | + | 24475077 | 24475610 | 24473710 | 24473710 |
| **ERRFI1** | 1 | 3 | AT | 0 | - | 8075368 | 8075460 | 8074543 | 8074543 |
| **GMPR2** | 14 | 10 | RI | 0 | + | 24707452 | 24708448 | 24702249 | 24702249 |
| **GMPR2** | 14 | 2 | RI | 0 | + | 24702425 | 24702546 | 24702249 | 24702249 |
| **LYPLA2** | 1 | 8 | RI | 0 | + | 24120714 | 24121171 | 24121145 | 24121145 |
| **NRP1** | 10 | 5 | ES | 0 | - | 33545244 | 33545399 | 33495125 | 33495125 |
| **RBBP6** | 16 | 18 | AT | 0 | + | 24582197 | 24584181 | 24560342 | 24560342 |
| **CIRBP** | 19 | 7 | RI | 0 | + | 1274306 | 1274479 | 1270152 | 1270152 |
| **CIRBP** | 19 | 5 | AP | 0 | + | 1271980 | 1272050 | 1270152 | 1270152 |
| **CIRBP** | 19 | 8 | AA | 0 | + | 1274648 | 1274806 | 1270152 | 1270152 |

| | | | | | | | | | |
|---|---|---|---|---|---|---|---|---|---|
| **CIRBP** | 19 | 9 | RI | 0 | + | 1274648 | 1274772 | 1270152 | 1270152 |
| **TNKS** | 8 | 1 | AP | 0 | + | 9627642 | 9627772 | 9599253 | 9599253 |

| Variant_Cl | Variant_Ty | CONTEXT | BIOTYPE | DOMAINS | IMPACT |
|---|---|---|---|---|---|
| Frame_Shi | DEL | AGAAAAA( | protein_co | hmmpanth | HIGH |
| Frame_Shi | INS | GGGCTCTG | protein_co | hmmpanth | HIGH |
| Frame_Shi | INS | GGGCTCTG | protein_co | hmmpanth | HIGH |
| Frame_Shi | INS | GGAGTATT | protein_co | hmmpanth | HIGH |
| Frame_Shi | INS | GGAGTATT | protein_co | hmmpanth | HIGH |
| Frame_Shi | INS | GGAGTATT | protein_co | hmmpanth | HIGH |
| Frame_Shi | INS | GGAGTATT | protein_co | hmmpanth | HIGH |
| Frame_Shi | INS | ATCAGCAA | protein_co | hmmpanth | HIGH |
| Nonsense_ | SNP | AGGCTGCA | protein_co | Superfamil | HIGH |
| Nonsense_ | SNP | AGGCTGCA | protein_co | Superfamil | HIGH |
| Nonsense_ | SNP | CTCCTCCA( | protein_co | hmmpanth | HIGH |
| Nonsense_ | SNP | GCGTCGCT | protein_co | hmmpanth | HIGH |
| Nonsense_ | SNP | GCGTCGCT | protein_co | hmmpanth | HIGH |
| Nonsense_ | SNP | GCGTCGCT | protein_co | hmmpanth | HIGH |
| Nonsense_ | SNP | GCGTCGCT | protein_co | hmmpanth | HIGH |
| Nonsense_ | SNP | GAATTCCC | protein_co | hmmpanth | HIGH |
| Nonsense_ | SNP | GGACTCTG | protein_co | hmmpanth | HIGH |
| Nonsense_ | SNP | GGACTCTG | protein_co | hmmpanth | HIGH |
| Nonsense_ | SNP | TAAAGCGA | protein_co | hmmpanth | HIGH |
| Nonsense_ | SNP | GCCTGGA( | protein_co | hmmpanth | HIGH |
| Nonsense_ | SNP | GCCTGGA( | protein_co | hmmpanth | HIGH |
| Nonsense_ | SNP | TAGAACGA | protein_co | Low_comp | HIGH |
| Nonsense_ | SNP | AAATCGCA | protein_co | hmmpanth | HIGH |
| Nonsense_ | SNP | AAATCGCA | protein_co | hmmpanth | HIGH |
| Nonsense_ | SNP | AAATCGCA | protein_co | hmmpanth | HIGH |
| Nonsense_ | SNP | AAATCGCA | protein_co | hmmpanth | HIGH |
| Nonsense_ | SNP | AAATCGCA | protein_co | hmmpanth | HIGH |
| Nonsense_ | SNP | GGCTCGCT | protein_co | Superfamil | HIGH |
| Nonsense_ | SNP | CCCCGGA( | protein_co | . | HIGH |
| Nonsense_ | SNP | TGCTCCGA | protein_co | Superfamil | HIGH |
| Nonsense_ | SNP | ATAAGCGA | protein_co | PROSITE_p | HIGH |
| Nonstop_M | SNP | AAGTGAC( | protein_co | . | HIGH |
| Splice_Site | SNP | GCCACGTA | protein_co | . | HIGH |
| Splice_Site | SNP | CAGATCTG | protein_co | . | HIGH |
| Splice_Site | SNP | CAGCACCT | protein_co | . | HIGH |
| Splice_Site | SNP | CAGCACCT | protein_co | . | HIGH |
| Missense_ | SNP | TAATGCTC | protein_co | . | LOW |
| Missense_ | SNP | TAATGCTC | protein_co | . | LOW |
| Missense_ | SNP | TAATGCTC | protein_co | . | LOW |
| Silent | SNP | ACATAGGA | protein_co | hmmpanth | LOW |
| Silent | SNP | ACATAGGA | protein_co | hmmpanth | LOW |
| Silent | SNP | AAGCACGA | protein_co | PROSITE_p | LOW |
| Silent | SNP | GGCTACG( | protein_co | PROSITE_p | LOW |
| Silent | SNP | AAGCACGA | protein_co | PROSITE_p | LOW |
| Silent | SNP | AAGCACGA | protein_co | PROSITE_p | LOW |
| Silent | SNP | GGCTACG( | protein_co | PROSITE_p | LOW |

| | | | | | |
|---|---|---|---|---|---|
| Silent | SNP | CTGAGCGA | protein_co | . | LOW |
| Silent | SNP | CCTTCCTC( | protein_co | hmmpanth | LOW |
| Silent | SNP | CCTTCCTC( | protein_co | hmmpanth | LOW |
| Silent | SNP | CCACCCTA | protein_co | PROSITE_p | LOW |
| Silent | SNP | CCACCCTA | protein_co | PROSITE_p | LOW |
| Silent | SNP | CTCCACGC | protein_co | PROSITE_p | LOW |
| Silent | SNP | CTCCACGC | protein_co | PROSITE_p | LOW |
| Silent | SNP | TTCTTCAA( | protein_co | hmmpanth | LOW |
| Silent | SNP | TTCTTCAA( | protein_co | hmmpanth | LOW |
| Silent | SNP | TTCTTCAA( | protein_co | hmmpanth | LOW |
| Silent | SNP | TTCTTCAA( | protein_co | hmmpanth | LOW |
| Silent | SNP | TTCTTCAA( | protein_co | hmmpanth | LOW |
| Silent | SNP | TTCTTCAA( | protein_co | hmmpanth | LOW |
| Silent | SNP | TTCTTCAA( | protein_co | hmmpanth | LOW |
| Silent | SNP | TTGCCGCC | protein_co | Coiled-coil | LOW |
| Silent | SNP | ACTTAGG( | protein_co | . | LOW |
| Silent | SNP | CACACAG( | protein_co | hmmpanth | LOW |
| Silent | SNP | GGTCTCTG | protein_co | PROSITE_p | LOW |
| Silent | SNP | GGTCTCTG | protein_co | PROSITE_p | LOW |
| Silent | SNP | GGTCTCTG | protein_co | PROSITE_p | LOW |
| Silent | SNP | GGTCTCTG | protein_co | PROSITE_p | LOW |
| Silent | SNP | GGTCTCTG | protein_co | PROSITE_p | LOW |
| Silent | SNP | AAGAAGCA | protein_co | Low_comp | LOW |
| Silent | SNP | AATCGTGT | protein_co | hmmpanth | LOW |
| Silent | SNP | AATCGTGT | protein_co | hmmpanth | LOW |
| Silent | SNP | CTTTCGAG | protein_co | Superfami | LOW |
| Silent | SNP | TTGTGCTG | protein_co | PIRSF_dom | LOW |
| Silent | SNP | TTGTGCTG | protein_co | PIRSF_dom | LOW |
| Silent | SNP | GCCTCGG( | protein_co | hmmpanth | LOW |
| Silent | SNP | GCCTCGG( | protein_co | hmmpanth | LOW |
| Silent | SNP | GCCTCGG( | protein_co | hmmpanth | LOW |
| Silent | SNP | AGTTCTTC | protein_co | hmmpanth | LOW |
| Silent | SNP | AAATCTCG | protein_co | Low_comp | LOW |
| Silent | SNP | TTTAAGAA | protein_co | hmmpanth | LOW |
| Silent | SNP | CACTGCTG | protein_co | hmmpanth | LOW |
| Silent | SNP | TGAACGAA | protein_co | Superfami | LOW |
| Silent | SNP | TGAACGAA | protein_co | Superfami | LOW |
| Silent | SNP | TGAACGAA | protein_co | Superfami | LOW |
| Silent | SNP | TGAACGAA | protein_co | Superfami | LOW |
| Silent | SNP | TGAACGAA | protein_co | Superfami | LOW |
| Silent | SNP | AGCTTCGC | protein_co | Low_comp | LOW |
| Silent | SNP | AGCTTCGC | protein_co | Low_comp | LOW |
| Silent | SNP | CAGTGCAT | protein_co | hmmpanth | LOW |
| Silent | SNP | GTTCGCTG | protein_co | . | LOW |
| Silent | SNP | GAGAGCT( | protein_co | PROSITE_p | LOW |
| Silent | SNP | CAGCGCTA | protein_co | . | LOW |
| Silent | SNP | GAGACGT( | protein_co | PROSITE_p | LOW |

| | | | | | |
|---|---|---|---|---|---|
| Silent | SNP | AATGCCGA | protein_co | hmmpanth | LOW |
| Silent | SNP | AATGCCGA | protein_co | hmmpanth | LOW |
| Silent | SNP | AATGCCGA | protein_co | hmmpanth | LOW |
| Silent | SNP | CCCTCCTA | protein_co | PROSITE_p | LOW |
| Silent | SNP | TAGTGCTG | protein_co | hmmpanth | LOW |
| Silent | SNP | GTGGACG | protein_co | hmmpanth | LOW |
| Silent | SNP | ATGATTGC | protein_co | PROSITE_p | LOW |
| Silent | SNP | ATGATTGC | protein_co | PROSITE_p | LOW |
| Silent | SNP | ATGATTGC | protein_co | PROSITE_p | LOW |
| Silent | SNP | ATGATTGC | protein_co | PROSITE_p | LOW |
| Silent | SNP | AGGTTGTG | protein_co | hmmpanth | LOW |
| Silent | SNP | GACCACCT | protein_co | PROSITE_p | LOW |
| Silent | SNP | AGCACGCT | protein_co | hmmpanth | LOW |
| Silent | SNP | TCGATGGC | protein_co | hmmpanth | LOW |
| Silent | SNP | AGCACGCT | protein_co | hmmpanth | LOW |
| Silent | SNP | TCGATGGC | protein_co | hmmpanth | LOW |
| Silent | SNP | AGCACGCT | protein_co | hmmpanth | LOW |
| Silent | SNP | TCGATGGC | protein_co | hmmpanth | LOW |
| Silent | SNP | AGCACGCT | protein_co | hmmpanth | LOW |
| Silent | SNP | TCGATGGC | protein_co | hmmpanth | LOW |
| Silent | SNP | AGCACGCT | protein_co | hmmpanth | LOW |
| Silent | SNP | TCGATGGC | protein_co | hmmpanth | LOW |
| Silent | SNP | AGCACGCT | protein_co | hmmpanth | LOW |
| Silent | SNP | TCGATGGC | protein_co | hmmpanth | LOW |
| Silent | SNP | TCCCCGTT | protein_co | PROSITE_p | LOW |
| Silent | SNP | GACCCGTC | protein_co | hmmpanth | LOW |
| Silent | SNP | GACCGCGT | protein_co | hmmpanth | LOW |
| Silent | SNP | GACCGCGT | protein_co | hmmpanth | LOW |
| Silent | SNP | CCGGGCTT | protein_co | . | LOW |
| Silent | SNP | ATTAGGCT | protein_co | hmmpanth | LOW |
| Silent | SNP | ATTAGGCT | protein_co | hmmpanth | LOW |
| Silent | SNP | ATTAGGCT | protein_co | hmmpanth | LOW |
| Silent | SNP | ATTAGGCT | protein_co | hmmpanth | LOW |
| Silent | SNP | GGCGTAGA | protein_co | hmmpanth | LOW |
| Silent | SNP | ATCGGCGT | protein_co | hmmpanth | LOW |
| Silent | SNP | GAGAGCGC | protein_co | Superfamil | LOW |
| Silent | SNP | GCTGACAT | protein_co | hmmpanth | LOW |
| Silent | SNP | GCTGACAT | protein_co | hmmpanth | LOW |
| Silent | SNP | GCTGACAT | protein_co | hmmpanth | LOW |
| Silent | SNP | GTGGGGG | protein_co | hmmpanth | LOW |
| Silent | SNP | CCTTTGAC | protein_co | Superfamil | LOW |
| Silent | SNP | GCTGCCCA | protein_co | hmmpanth | LOW |
| Silent | SNP | AAGGACGA | protein_co | hmmpanth | LOW |
| Silent | SNP | GTGCGGCA | protein_co | Superfamil | LOW |
| Silent | SNP | GAAGGCGA | protein_co | hmmpanth | LOW |
| Silent | SNP | GCCCAGAC | protein_co | Prints_dom | LOW |
| Silent | SNP | CTGGACGA | protein_co | Low_comp | LOW |

| | | | | | |
|---|---|---|---|---|---|
| Silent | SNP | CTGGACGA | protein_co | Low_comp | LOW |
| Silent | SNP | GCGCAGTC | protein_co | PROSITE_p | LOW |
| Silent | SNP | GCGCAGTC | protein_co | PROSITE_p | LOW |
| Silent | SNP | CCCTGGCC | protein_co | Low_comp | LOW |
| Silent | SNP | CATCAGAA | protein_co | Pfam_dom | LOW |
| Silent | SNP | AAGGCGCT | protein_co | hmmpanth | LOW |
| Silent | SNP | TCGGAGAT | protein_co | PROSITE_p | LOW |
| Silent | SNP | AGATGCTG | protein_co | Transmem | LOW |
| Silent | SNP | AGATGCTG | protein_co | Transmem | LOW |
| Silent | SNP | CAGACGCC | protein_co | PROSITE_p | LOW |
| Silent | SNP | CACCGACA | protein_co | PROSITE_p | LOW |
| Silent | SNP | GAAGGCAA | protein_co | hmmpanth | LOW |
| Silent | SNP | AACATCGC | protein_co | HAMAP:M | LOW |
| Silent | SNP | AACATCGC | protein_co | HAMAP:M | LOW |
| Silent | SNP | AACATCGC | protein_co | HAMAP:M | LOW |
| Silent | SNP | CGGAGACA | protein_co | Coiled-coil | LOW |
| Silent | SNP | GAAGGCGA | protein_co | hmmpanth | LOW |
| Silent | SNP | CAGTCGCT | protein_co | hmmpanth | LOW |
| Silent | SNP | TGGATCCA | protein_co | PROSITE_p | LOW |
| Silent | SNP | CTAATCCA | protein_co | PROSITE_p | LOW |
| Silent | SNP | ACAGATGC | protein_co | hmmpanth | LOW |
| Splice_Reg | SNP | TTAATCCC | protein_co | . | LOW |
| In_Frame_ | DEL | CGTTGTCG | protein_co | hmmpanth | MODERATE |
| Missense_ | SNP | CAGGATAT | protein_co | hmmpanth | MODERATE |
| Missense_ | SNP | CATAAGCC | protein_co | hmmpanth | MODERATE |
| Missense_ | SNP | ACTGTGCA | protein_co | PROSITE_p | MODERATE |
| Missense_ | SNP | ATTAGCAC | protein_co | hmmpanth | MODERATE |
| Missense_ | SNP | TGCCCAGC | protein_co | Superfamil | MODERATE |
| Missense_ | SNP | TGCTACGT | protein_co | . | MODERATE |
| Missense_ | SNP | GTGGTCGC | protein_co | hmmpanth | MODERATE |
| Missense_ | SNP | CAGTCCGA | protein_co | HAMAP:M | MODERATE |
| Missense_ | SNP | CAGTCCGA | protein_co | HAMAP:M | MODERATE |
| Missense_ | SNP | CAGTCCGA | protein_co | HAMAP:M | MODERATE |
| Missense_ | SNP | GGTCCGCC | protein_co | hmmpanth | MODERATE |
| Missense_ | SNP | GGTGGCAA | protein_co | Low_comp | MODERATE |
| Missense_ | SNP | TTGTAGGA | protein_co | hmmpanth | MODERATE |
| Missense_ | SNP | GGTGGCAA | protein_co | Low_comp | MODERATE |
| Missense_ | SNP | TTGTAGGA | protein_co | hmmpanth | MODERATE |
| Missense_ | SNP | TTGTAGGA | protein_co | hmmpanth | MODERATE |
| Missense_ | SNP | TCATCGCC | protein_co | Superfamil | MODERATE |
| Missense_ | SNP | CAACTCCA | protein_co | Low_comp | MODERATE |
| Missense_ | SNP | CCCAGCCT | protein_co | Superfamil | MODERATE |
| Missense_ | SNP | TGTCCATG | protein_co | hmmpanth | MODERATE |
| Missense_ | SNP | TACCTCGT | protein_co | hmmpanth | MODERATE |
| Missense_ | SNP | GTGGGAAT | protein_co | hmmpanth | MODERATE |
| Missense_ | SNP | TACCTCGT | protein_co | hmmpanth | MODERATE |
| Missense_ | SNP | GTGGGAAT | protein_co | hmmpanth | MODERATE |

| | | | | | |
|---|---|---|---|---|---|
| Missense_ | SNP | TACCTCGT | protein_co | hmmpanth | MODERATE |
| Missense_ | SNP | GTGGGAAT | protein_co | hmmpanth | MODERATE |
| Missense_ | SNP | CTGCCGTC | protein_co | PROSITE_p | MODERATE |
| Missense_ | SNP | CTGCCGTC | protein_co | PROSITE_p | MODERATE |
| Missense_ | SNP | TCTCTGGG | protein_co | hmmpanth | MODERATE |
| Missense_ | SNP | TCTCTGGG | protein_co | hmmpanth | MODERATE |
| Missense_ | SNP | CTGCCGTC | protein_co | PROSITE_p | MODERATE |
| Missense_ | SNP | TCTCTGGG | protein_co | hmmpanth | MODERATE |
| Missense_ | SNP | GCACAGG( | protein_co | PROSITE_p | MODERATE |
| Missense_ | SNP | GCACAGG( | protein_co | PROSITE_p | MODERATE |
| Missense_ | SNP | ACCCCGGC | protein_co | PROSITE_p | MODERATE |
| Missense_ | SNP | ACACCGTG | protein_co | PROSITE_p | MODERATE |
| Missense_ | SNP | AGGCCCCC | protein_co | PROSITE_p | MODERATE |
| Missense_ | SNP | ACCCCGGC | protein_co | PROSITE_p | MODERATE |
| Missense_ | SNP | CAGTGGTA | protein_co | PROSITE_p | MODERATE |
| Missense_ | SNP | ACACCGTG | protein_co | PROSITE_p | MODERATE |
| Missense_ | SNP | GCACAGG( | protein_co | PROSITE_p | MODERATE |
| Missense_ | SNP | ACCCCGGC | protein_co | PROSITE_p | MODERATE |
| Missense_ | SNP | CAGTGGTA | protein_co | PROSITE_p | MODERATE |
| Missense_ | SNP | ACACCGTG | protein_co | PROSITE_p | MODERATE |
| Missense_ | SNP | GCACAGG( | protein_co | PROSITE_p | MODERATE |
| Missense_ | SNP | AGGCCCCC | protein_co | PROSITE_p | MODERATE |
| Missense_ | SNP | ACCCCGGC | protein_co | PROSITE_p | MODERATE |
| Missense_ | SNP | CAGTGGTA | protein_co | PROSITE_p | MODERATE |
| Missense_ | SNP | ACACCGTG | protein_co | PROSITE_p | MODERATE |
| Missense_ | SNP | AGGCCCCC | protein_co | PROSITE_p | MODERATE |
| Missense_ | SNP | GTGAAGG( | protein_co | hmmpanth | MODERATE |
| Missense_ | SNP | TAAGTCCT | protein_co | . | MODERATE |
| Missense_ | SNP | TCCCAGCA | protein_co | Low_comp | MODERATE |
| Missense_ | SNP | AGACGCG( | protein_co | hmmpanth | MODERATE |
| Missense_ | SNP | ATCGAGGA | protein_co | PROSITE_p | MODERATE |
| Missense_ | SNP | TCCCAGCA | protein_co | Low_comp | MODERATE |
| Missense_ | SNP | AGACGCG( | protein_co | hmmpanth | MODERATE |
| Missense_ | SNP | ATCGAGGA | protein_co | PROSITE_p | MODERATE |
| Missense_ | SNP | ACGCACTT | protein_co | hmmpanth | MODERATE |
| Missense_ | SNP | ACGCACTT | protein_co | hmmpanth | MODERATE |
| Missense_ | SNP | GCTACCTG | protein_co | PROSITE_p | MODERATE |
| Missense_ | SNP | CTGTGGCT | protein_co | PROSITE_p | MODERATE |
| Missense_ | SNP | GCTACCTG | protein_co | PROSITE_p | MODERATE |
| Missense_ | SNP | CTGTGGCT | protein_co | PROSITE_p | MODERATE |
| Missense_ | SNP | CTTCTCGT( | protein_co | hmmpanth | MODERATE |
| Missense_ | SNP | TTCACGAT | protein_co | Coiled-coil | MODERATE |
| Missense_ | SNP | TTCACGAT | protein_co | Coiled-coil | MODERATE |
| Missense_ | SNP | TTCACGAT | protein_co | Coiled-coil | MODERATE |
| Missense_ | SNP | TTCACGAT | protein_co | Coiled-coil | MODERATE |
| Missense_ | SNP | TTCACGAT | protein_co | Coiled-coil | MODERATE |
| Missense_ | SNP | GATGAGCA | protein_co | Low_comp | MODERATE |

| | | | | | |
|---|---|---|---|---|---|
| Missense_ | SNP | TTGGTTTT | protein_co | hmmpanth | MODERATE |
| Missense_ | SNP | GATGAGC | protein_co | Low_comp | MODERATE |
| Missense_ | SNP | CAGGAAA | protein_co | hmmpanth | MODERATE |
| Missense_ | SNP | TTGGTTTT | protein_co | hmmpanth | MODERATE |
| Missense_ | SNP | GATGAGC | protein_co | Low_comp | MODERATE |
| Missense_ | SNP | TTGGTTTT | protein_co | hmmpanth | MODERATE |
| Missense_ | SNP | GATGAGC | protein_co | Low_comp | MODERATE |
| Missense_ | SNP | TTGGTTTT | protein_co | hmmpanth | MODERATE |
| Missense_ | SNP | GATGAGC | protein_co | Low_comp | MODERATE |
| Missense_ | SNP | TTGGTTTT | protein_co | hmmpanth | MODERATE |
| Missense_ | SNP | GATGAGC | protein_co | Low_comp | MODERATE |
| Missense_ | SNP | TTGGTTTT | protein_co | hmmpanth | MODERATE |
| Missense_ | SNP | CAGGAAA | protein_co | hmmpanth | MODERATE |
| Missense_ | SNP | CAGGAAA | protein_co | hmmpanth | MODERATE |
| Missense_ | SNP | GATGAGC | protein_co | Low_comp | MODERATE |
| Missense_ | SNP | TTGGTTTT | protein_co | hmmpanth | MODERATE |
| Missense_ | SNP | ACGTTAG | protein_co | hmmpanth | MODERATE |
| Missense_ | SNP | GCAGGCT | protein_co | Cleavage_ | MODERATE |
| Missense_ | SNP | CCAGCGG | protein_co | Superfami | MODERATE |
| Missense_ | SNP | CCAGCGG | protein_co | Superfami | MODERATE |
| Missense_ | SNP | CCAGCGG | protein_co | Superfami | MODERATE |
| Missense_ | SNP | CCAGCGG | protein_co | Superfami | MODERATE |
| Missense_ | SNP | CCAGCGG | protein_co | Superfami | MODERATE |
| Missense_ | SNP | CCAGCGG | protein_co | Superfami | MODERATE |
| Missense_ | SNP | CCAGCGG | protein_co | Superfami | MODERATE |
| Missense_ | SNP | CCAGCGG | protein_co | Superfami | MODERATE |
| Missense_ | SNP | CCAGCGG | protein_co | Superfami | MODERATE |
| Missense_ | SNP | TAATCGC | protein_co | Superfami | MODERATE |
| Missense_ | SNP | AGCAAGG | protein_co | Superfami | MODERATE |
| Missense_ | SNP | AGAGAGT | protein_co | Superfami | MODERATE |
| Missense_ | SNP | CACAGCG | protein_co | Superfami | MODERATE |
| Missense_ | SNP | GGCCTCG | protein_co | hmmpanth | MODERATE |
| Missense_ | SNP | GGCCTCG | protein_co | hmmpanth | MODERATE |
| Missense_ | SNP | GGCCTCG | protein_co | hmmpanth | MODERATE |
| Missense_ | SNP | GGCCTCG | protein_co | hmmpanth | MODERATE |
| Missense_ | SNP | GGCCTCG | protein_co | hmmpanth | MODERATE |
| Missense_ | SNP | GTATCGCT | protein_co | hmmpanth | MODERATE |
| Missense_ | SNP | TGACCGAT | protein_co | hmmpanth | MODERATE |
| Missense_ | SNP | TCTACGCA | protein_co | Superfami | MODERATE |
| Missense_ | SNP | TCCTAGTC | protein_co | hmmpanth | MODERATE |
| Missense_ | SNP | TCATGGA | protein_co | hmmpanth | MODERATE |
| Missense_ | SNP | TCCTAGTC | protein_co | hmmpanth | MODERATE |
| Missense_ | SNP | TCATGGA | protein_co | hmmpanth | MODERATE |
| Missense_ | SNP | TCCTAGTC | protein_co | hmmpanth | MODERATE |
| Missense_ | SNP | TCATGGA | protein_co | hmmpanth | MODERATE |
| Missense_ | SNP | TCCTAGTC | protein_co | hmmpanth | MODERATE |
| Missense_ | SNP | TCATGGA | protein_co | hmmpanth | MODERATE |

| | | | | | |
|---|---|---|---|---|---|
| Missense_ | SNP | CAGTCCTT | protein_co | Superfamil | MODERATE |
| Missense_ | SNP | CAGTCCTT | protein_co | Superfamil | MODERATE |
| Missense_ | SNP | CAGTCCTT | protein_co | Superfamil | MODERATE |
| Missense_ | SNP | CAGTCCTT | protein_co | Superfamil | MODERATE |
| Missense_ | SNP | CAGTCCTT | protein_co | Superfamil | MODERATE |
| Missense_ | SNP | TCCTGGAC | protein_co | PROSITE_p | MODERATE |
| Missense_ | SNP | CATGAAG( | protein_co | PROSITE_p | MODERATE |
| Missense_ | SNP | TCCTGGAC | protein_co | PROSITE_p | MODERATE |
| Missense_ | SNP | TCCTGGAC | protein_co | PROSITE_p | MODERATE |
| Missense_ | SNP | TCCTGGAC | protein_co | PROSITE_p | MODERATE |
| Missense_ | SNP | CATGAAG( | protein_co | PROSITE_p | MODERATE |
| Missense_ | SNP | TCCTGGAC | protein_co | PROSITE_p | MODERATE |
| Missense_ | SNP | CATGAAG( | protein_co | PROSITE_p | MODERATE |
| Missense_ | SNP | CATGAAG( | protein_co | PROSITE_p | MODERATE |
| Missense_ | SNP | CATGAAG( | protein_co | PROSITE_p | MODERATE |
| Missense_ | SNP | TGAGAGCT | protein_co | PROSITE_p | MODERATE |
| Missense_ | SNP | GAAACGG( | protein_co | Low_comp | MODERATE |
| Missense_ | SNP | TAGCCGTT | protein_co | Low_comp | MODERATE |
| Missense_ | SNP | TAGCCGTT | protein_co | Low_comp | MODERATE |
| Missense_ | SNP | GAGCAAA( | protein_co | hmmpanth | MODERATE |
| Missense_ | SNP | CGTCAGCC | protein_co | hmmpanth | MODERATE |
| Missense_ | SNP | CAGGACG( | protein_co | hmmpanth | MODERATE |
| Missense_ | SNP | GAGCAAA( | protein_co | hmmpanth | MODERATE |
| Missense_ | SNP | CGTCAGCC | protein_co | hmmpanth | MODERATE |
| Missense_ | SNP | CAGGACG( | protein_co | hmmpanth | MODERATE |
| Missense_ | SNP | GCATCACC | protein_co | HAMAP:M | MODERATE |
| Missense_ | SNP | AACAGGA/ | protein_co | hmmpanth | MODERATE |
| Missense_ | SNP | CGTACCGC | protein_co | Superfamil | MODERATE |
| Missense_ | SNP | GAGTTCTA | protein_co | hmmpanth | MODERATE |
| Missense_ | SNP | GCCAGGT( | protein_co | Superfamil | MODERATE |
| Missense_ | SNP | CCCCCGGC | protein_co | Superfamil | MODERATE |
| Missense_ | SNP | CGTGCCTG | protein_co | Superfamil | MODERATE |
| Missense_ | SNP | GGGTGCGT | protein_co | Pfam_dom | MODERATE |
| Missense_ | SNP | AAGCCCCA | protein_co | Transmem | MODERATE |
| Missense_ | SNP | ATGATTTA | protein_co | Pfam_dom | MODERATE |
| Missense_ | SNP | CACGGCGT | protein_co | hmmpanth | MODERATE |
| Missense_ | SNP | GAGCCGCC | protein_co | Superfamil | MODERATE |
| Missense_ | SNP | TCAAGCG( | protein_co | . | MODERATE |
| Missense_ | SNP | GGAGAGA | protein_co | PIRSF_dom | MODERATE |
| Missense_ | SNP | GGAGAGA | protein_co | PIRSF_dom | MODERATE |
| Missense_ | SNP | GGAGAGA | protein_co | PIRSF_dom | MODERATE |
| Missense_ | SNP | GGAGAGA | protein_co | PIRSF_dom | MODERATE |
| Missense_ | SNP | GGAGAGA | protein_co | PIRSF_dom | MODERATE |
| Missense_ | SNP | GGAGAGA | protein_co | PIRSF_dom | MODERATE |
| Missense_ | SNP | GGAGAGA | protein_co | PIRSF_dom | MODERATE |
| Missense_ | SNP | GGAGAGA | protein_co | PIRSF_dom | MODERATE |
| Missense_ | SNP | TAATGCAC | protein_co | hmmpanth | MODERATE |

| | | | | | |
|---|---|---|---|---|---|
| Missense_ | SNP | AAAAGCTC | protein_co | PROSITE_p | MODERATE |
| Missense_ | SNP | CGAGCGCA | protein_co | hmmpanth | MODERATE |
| Missense_ | SNP | TGGAGCGC | protein_co | hmmpanth | MODERATE |
| Missense_ | SNP | AAAAGCTC | protein_co | PROSITE_p | MODERATE |
| Missense_ | SNP | TAATGCAC | protein_co | hmmpanth | MODERATE |
| Missense_ | SNP | CGAGCGCA | protein_co | hmmpanth | MODERATE |
| Missense_ | SNP | TGGAGCGC | protein_co | hmmpanth | MODERATE |
| Missense_ | SNP | TTGAGATG | protein_co | hmmpanth | MODERATE |
| Missense_ | SNP | TTGAGATG | protein_co | hmmpanth | MODERATE |
| Missense_ | SNP | TTGAGATG | protein_co | hmmpanth | MODERATE |
| Missense_ | SNP | TTGAGATG | protein_co | hmmpanth | MODERATE |
| Missense_ | SNP | AGAGAAGA | protein_co | SMART_do | MODERATE |
| Missense_ | SNP | ATGACCGC | protein_co | PROSITE_p | MODERATE |
| Missense_ | SNP | ACACCCCT | protein_co | hmmpanth | MODERATE |
| Missense_ | SNP | ACACCCCT | protein_co | hmmpanth | MODERATE |
| Missense_ | SNP | AGAGAAGA | protein_co | SMART_do | MODERATE |
| Missense_ | SNP | ATGACCGC | protein_co | PROSITE_p | MODERATE |
| Missense_ | SNP | GTTGGGAT | protein_co | hmmpanth | MODERATE |
| Missense_ | SNP | GTTGGGAT | protein_co | hmmpanth | MODERATE |
| Missense_ | SNP | AGGTGTCC | protein_co | Coiled-coil | MODERATE |
| Missense_ | SNP | AGGTGTCC | protein_co | Coiled-coil | MODERATE |
| Missense_ | SNP | AGGTGTCC | protein_co | Coiled-coil | MODERATE |
| Missense_ | SNP | AGGTGTCC | protein_co | Coiled-coil | MODERATE |
| Missense_ | SNP | AGGTGTCC | protein_co | Coiled-coil | MODERATE |
| Missense_ | SNP | TTCCCGCC | protein_co | Low_comp | MODERATE |
| Missense_ | SNP | TTCCCGCC | protein_co | Low_comp | MODERATE |
| Missense_ | SNP | GCACAGCT | protein_co | PROSITE_p | MODERATE |
| Missense_ | SNP | ACCTGTTT | protein_co | PROSITE_p | MODERATE |
| Missense_ | SNP | CCACAGAC | protein_co | PROSITE_p | MODERATE |
| Missense_ | SNP | AAGGAGAA | protein_co | Coiled-coil | MODERATE |
| Missense_ | SNP | AAAATGTA | protein_co | PROSITE_p | MODERATE |
| Missense_ | SNP | TTTCCGAC | protein_co | . | MODERATE |
| Missense_ | SNP | GAATAGTA | protein_co | Transmem | MODERATE |
| Missense_ | SNP | CACAGGTC | protein_co | . | MODERATE |
| Missense_ | SNP | GTGAGCCA | protein_co | hmmpanth | MODERATE |
| Missense_ | SNP | CCTTGCCA | protein_co | hmmpanth | MODERATE |
| Missense_ | SNP | GAGGGCG | protein_co | Low_comp | MODERATE |
| Missense_ | SNP | GTGAGCCA | protein_co | hmmpanth | MODERATE |
| Missense_ | SNP | CCTTGCCA | protein_co | hmmpanth | MODERATE |
| Missense_ | SNP | GAGGGCG | protein_co | Low_comp | MODERATE |
| Missense_ | SNP | GTGAGCCA | protein_co | hmmpanth | MODERATE |
| Missense_ | SNP | CCTTGCCA | protein_co | hmmpanth | MODERATE |
| Missense_ | SNP | GAGGGCG | protein_co | Low_comp | MODERATE |
| Missense_ | SNP | GCTACGTC | protein_co | Low_comp | MODERATE |
| Missense_ | SNP | GCTACGTC | protein_co | Low_comp | MODERATE |
| Missense_ | SNP | GCTACGTC | protein_co | Low_comp | MODERATE |
| Missense_ | SNP | GCTACGTC | protein_co | Low_comp | MODERATE |

| | | | | | |
|---|---|---|---|---|---|
| Missense_ | SNP | GCTACGTC | protein_co | Low_comp | MODERATE |
| Missense_ | SNP | GCTACGTC | protein_co | Low_comp | MODERATE |
| Missense_ | SNP | GCTACGTC | protein_co | Low_comp | MODERATE |
| Missense_ | SNP | ATTCTCTT( | protein_co | hmmpanth | MODERATE |
| Missense_ | SNP | ATTCTCTT( | protein_co | hmmpanth | MODERATE |
| Missense_ | SNP | CTGGAGCT | protein_co | PROSITE_p | MODERATE |
| Missense_ | SNP | CTGGAGCT | protein_co | PROSITE_p | MODERATE |
| Missense_ | SNP | CTGGAGCT | protein_co | PROSITE_p | MODERATE |
| Missense_ | SNP | TCTCCCGG | protein_co | Superfamil | MODERATE |
| Missense_ | SNP | ACCTGAGT | protein_co | Low_comp | MODERATE |
| Missense_ | SNP | TGTCTTCA | protein_co | hmmpanth | MODERATE |
| Missense_ | SNP | CGCAAGCT | protein_co | hmmpanth | MODERATE |
| Missense_ | SNP | AAGTTCTT | protein_co | hmmpanth | MODERATE |
| Missense_ | SNP | AACTGGCA | protein_co | Low_comp | MODERATE |
| Missense_ | SNP | TGGCCGA( | protein_co | hmmpanth | MODERATE |
| Missense_ | SNP | AGCACGG( | protein_co | hmmpanth | MODERATE |
| Missense_ | SNP | AGCACGG( | protein_co | hmmpanth | MODERATE |
| Missense_ | SNP | AGGCATTA | protein_co | hmmpanth | MODERATE |
| Missense_ | SNP | TATAGCTA | protein_co | hmmpanth | MODERATE |
| Missense_ | SNP | CCACCATG | protein_co | PROSITE_p | MODERATE |
| Missense_ | SNP | GGAGTCGA | protein_co | hmmpanth | MODERATE |
| Missense_ | SNP | CAGCCGAT | protein_co | PROSITE_p | MODERATE |
| Missense_ | SNP | ATTCAGAA | protein_co | hmmpanth | MODERATE |
| Missense_ | SNP | CAGCCGAT | protein_co | PROSITE_p | MODERATE |
| Missense_ | SNP | ATTCAGAA | protein_co | hmmpanth | MODERATE |
| Missense_ | SNP | CAGCCGAT | protein_co | PROSITE_p | MODERATE |
| Missense_ | SNP | ATTCAGAA | protein_co | hmmpanth | MODERATE |
| Missense_ | SNP | CTCTACCA | protein_co | Superfamil | MODERATE |
| Missense_ | SNP | CTTGAGGA | protein_co | HAMAP:M | MODERATE |
| Missense_ | SNP | GCAAGATA | protein_co | hmmpanth | MODERATE |
| Missense_ | SNP | GCAAGATA | protein_co | hmmpanth | MODERATE |
| Missense_ | SNP | CTGCAGCA | protein_co | hmmpanth | MODERATE |
| Missense_ | SNP | CAAGTTGC | protein_co | hmmpanth | MODERATE |
| Missense_ | SNP | TGGATCTT | protein_co | hmmpanth | MODERATE |
| Missense_ | SNP | CCTGCGTG | protein_co | hmmpanth | MODERATE |
| Missense_ | SNP | AGGTGCG( | protein_co | hmmpanth | MODERATE |
| Missense_ | SNP | CCTTGGCA | protein_co | . | MODERATE |
| Missense_ | SNP | CAAAGCTC | protein_co | hmmpanth | MODERATE |
| Missense_ | SNP | CGAGTCTG | protein_co | hmmpanth | MODERATE |
| Missense_ | SNP | CGAGTCTG | protein_co | hmmpanth | MODERATE |
| Missense_ | SNP | CAACCGCT | protein_co | Coiled-coil | MODERATE |
| Missense_ | SNP | CAACCGCT | protein_co | Coiled-coil | MODERATE |
| Missense_ | SNP | CAACCGCT | protein_co | Coiled-coil | MODERATE |
| Missense_ | SNP | TGGCCGC( | protein_co | PROSITE_p | MODERATE |
| Missense_ | SNP | TGGCCGC( | protein_co | PROSITE_p | MODERATE |
| Missense_ | SNP | AAGTGCG( | protein_co | hmmpanth | MODERATE |
| Missense_ | SNP | ATTCAGCC | protein_co | Low_comp | MODERATE |

| | | | | | |
|---|---|---|---|---|---|
| Missense_ | SNP | GGAATCTG | protein_co | hmmpanth | MODERATE |
| Missense_ | SNP | GGAATCTG | protein_co | hmmpanth | MODERATE |
| Missense_ | SNP | GGAATCTG | protein_co | hmmpanth | MODERATE |
| Missense_ | SNP | TTTCCGGT | protein_co | hmmpanth | MODERATE |
| Missense_ | SNP | GCAGGCCA | protein_co | Transmem | MODERATE |
| Missense_ | SNP | ACTGTATC | protein_co | Superfamil | MODERATE |
| Missense_ | SNP | GAGAACTA | protein_co | PIRSF_dom | MODERATE |
| Missense_ | SNP | GCAGGCTG | protein_co | hmmpanth | MODERATE |
| Missense_ | SNP | GCAGGCTG | protein_co | hmmpanth | MODERATE |
| Missense_ | SNP | GCAGGCTG | protein_co | hmmpanth | MODERATE |
| Missense_ | SNP | GCAGGCTG | protein_co | hmmpanth | MODERATE |
| Missense_ | SNP | GCGAAGAT | protein_co | hmmpanth | MODERATE |
| Missense_ | SNP | CAAAGCGG | protein_co | hmmpanth | MODERATE |
| Missense_ | SNP | CCAGCCGG | protein_co | hmmpanth | MODERATE |
| Missense_ | SNP | CAAAGCGG | protein_co | hmmpanth | MODERATE |
| Missense_ | SNP | ATCACGTC | protein_co | hmmpanth | MODERATE |
| Missense_ | SNP | ATCACGTC | protein_co | hmmpanth | MODERATE |
| Missense_ | SNP | ATCACGTC | protein_co | hmmpanth | MODERATE |
| Missense_ | SNP | GGGCGCGG | protein_co | hmmpanth | MODERATE |
| Missense_ | SNP | GGGCGCGG | protein_co | hmmpanth | MODERATE |
| Missense_ | SNP | ATCACGTC | protein_co | hmmpanth | MODERATE |
| Missense_ | SNP | GGGCGCGG | protein_co | hmmpanth | MODERATE |
| Missense_ | SNP | GGGCGCGG | protein_co | hmmpanth | MODERATE |
| Missense_ | SNP | AGTGGCGG | protein_co | hmmpanth | MODERATE |
| Missense_ | SNP | TGTCCGAG | protein_co | Low_comp | MODERATE |
| Missense_ | SNP | TGTCCTTG | protein_co | PROSITE_p | MODERATE |
| Missense_ | SNP | TGTCCTTG | protein_co | PROSITE_p | MODERATE |
| Missense_ | SNP | AGTGGCGG | protein_co | hmmpanth | MODERATE |
| Missense_ | SNP | TGTCCGAG | protein_co | Low_comp | MODERATE |
| Missense_ | SNP | CAGTGCTG | protein_co | HAMAP:M | MODERATE |
| Missense_ | SNP | CAGTGCTG | protein_co | HAMAP:M | MODERATE |
| Missense_ | SNP | CAGTGCTG | protein_co | HAMAP:M | MODERATE |
| Missense_ | SNP | CCACCGCA | protein_co | Superfamil | MODERATE |
| Missense_ | SNP | ACGAGCGG | protein_co | SMART_do | MODERATE |
| Missense_ | SNP | TACGCGCG | protein_co | hmmpanth | MODERATE |
| Missense_ | SNP | CAGCCCCG | protein_co | Pfam_dom | MODERATE |
| Missense_ | SNP | GCTGAGCC | protein_co | PROSITE_p | MODERATE |
| Missense_ | SNP | AGCCCCGG | protein_co | hmmpanth | MODERATE |
| Missense_ | SNP | ATGTAGAA | protein_co | PROSITE_p | MODERATE |
| Missense_ | SNP | AGGACGTG | protein_co | hmmpanth | MODERATE |
| Missense_ | SNP | CTCTGCTG | protein_co | Low_comp | MODERATE |
| Missense_ | SNP | CTCTGCTG | protein_co | Low_comp | MODERATE |
| Missense_ | SNP | CTCTGCTG | protein_co | Low_comp | MODERATE |
| Missense_ | SNP | CTATTCCA | protein_co | Pfam_dom | MODERATE |
| Missense_ | SNP | CTATTCCA | protein_co | Pfam_dom | MODERATE |
| Missense_ | SNP | CTATTCCA | protein_co | Pfam_dom | MODERATE |
| Missense_ | SNP | CTATTCCA | protein_co | Pfam_dom | MODERATE |

| | | | | | |
|---|---|---|---|---|---|
| Missense_ | SNP | TGGACGCA | protein_co | hmmpanth | MODERATE |
| Missense_ | SNP | TTCAGCCA | protein_co | hmmpanth | MODERATE |
| Missense_ | SNP | CGCCTCCA | protein_co | PROSITE_p | MODERATE |
| Missense_ | SNP | TAATGGCT | protein_co | PROSITE_p | MODERATE |
| Missense_ | SNP | TGAGCGAC | protein_co | PROSITE_p | MODERATE |
| Missense_ | SNP | TATGAGGA | protein_co | hmmpanth | MODERATE |
| Missense_ | SNP | TATGAGGA | protein_co | hmmpanth | MODERATE |
| Missense_ | SNP | AACTATTA | protein_co | HAMAP:M | MODERATE |
| Missense_ | SNP | TATCCCGC | protein_co | Coiled-coil | MODERATE |
| Missense_ | SNP | TATCCCGC | protein_co | Coiled-coil | MODERATE |
| Missense_ | SNP | TTGTCGAA | protein_co | PROSITE_p | MODERATE |
| Missense_ | SNP | CTCCAACA | protein_co | Low_comp | MODERATE |
| Missense_ | SNP | CATTCGCT | protein_co | PROSITE_p | MODERATE |
| Missense_ | SNP | CAAGCGCA | protein_co | hmmpanth | MODERATE |
| Missense_ | SNP | GCGAGCGT | protein_co | PROSITE_p | MODERATE |
| Missense_ | SNP | GCGAGCGT | protein_co | PROSITE_p | MODERATE |
| Missense_ | SNP | GAGGGAA | protein_co | hmmpanth | MODERATE |
| Missense_ | SNP | GAGAGCGC | protein_co | PROSITE_p | MODERATE |
| Missense_ | SNP | TAAACATA | protein_co | PROSITE_p | MODERATE |
| Missense_ | SNP | TCATACTT | protein_co | PROSITE_p | MODERATE |
| Missense_ | SNP | CTTTAGAA | protein_co | PROSITE_p | MODERATE |
| Missense_ | SNP | TGAATGCA | protein_co | PROSITE_p | MODERATE |
| Missense_ | SNP | ATCGGCGC | protein_co | PROSITE_p | MODERATE |
| Missense_ | SNP | CCATACTG | protein_co | PROSITE_p | MODERATE |
| Missense_ | SNP | AGGCTTTG | protein_co | PROSITE_p | MODERATE |
| Missense_ | SNP | GTGTGTAC | protein_co | PROSITE_p | MODERATE |
| Missense_ | SNP | CCACACTG | protein_co | PROSITE_p | MODERATE |
| Missense_ | SNP | CTACAAAT | protein_co | PROSITE_p | MODERATE |
| 3'Flank | SNP | GGCAACGC | protein_co | . | MODIFIER |
| 3'Flank | SNP | ATGGGGG | protein_co | . | MODIFIER |
| 3'UTR | SNP | CTATTCTG | protein_co | . | MODIFIER |
| 3'UTR | SNP | AGGGGCG | protein_co | . | MODIFIER |
| 3'UTR | SNP | AGCGTGTC | protein_co | . | MODIFIER |
| 3'UTR | SNP | AGCGTGTC | protein_co | . | MODIFIER |
| 3'UTR | SNP | CGGGCGA | protein_co | . | MODIFIER |
| 3'UTR | SNP | CCTCGTTA | protein_co | . | MODIFIER |
| 3'UTR | SNP | CCTCGTTA | protein_co | . | MODIFIER |
| 3'UTR | SNP | CCTCGTTA | protein_co | . | MODIFIER |
| 3'UTR | SNP | CCTCGTTA | protein_co | . | MODIFIER |
| 3'UTR | SNP | GGCATCGT | protein_co | . | MODIFIER |
| 3'UTR | SNP | GGCATCGT | protein_co | . | MODIFIER |
| 3'UTR | SNP | GGCATCGT | protein_co | . | MODIFIER |
| 3'UTR | SNP | GGCATCGT | protein_co | . | MODIFIER |
| 3'UTR | SNP | GGCATCGT | protein_co | . | MODIFIER |
| 3'UTR | SNP | GGATTCCT | protein_co | . | MODIFIER |
| 3'UTR | SNP | GGATTCCT | protein_co | . | MODIFIER |
| 3'UTR | SNP | TCAAACTG | protein_co | . | MODIFIER |

| | | | | | |
|---|---|---|---|---|---|
| 3'UTR | SNP | CAGCTCTG | protein_co | . | MODIFIER |
| 3'UTR | SNP | CAGCTCTG | protein_co | . | MODIFIER |
| 3'UTR | SNP | GGCCTCTG | protein_co | . | MODIFIER |
| 3'UTR | SNP | TGGTTCTG | protein_co | . | MODIFIER |
| 3'UTR | SNP | CCCACGAT | protein_co | . | MODIFIER |
| 3'UTR | SNP | ACCACAAA | protein_co | . | MODIFIER |
| 3'UTR | SNP | ACCACAAA | protein_co | . | MODIFIER |
| 3'UTR | SNP | ACCACAAA | protein_co | . | MODIFIER |
| 3'UTR | SNP | ACCACAAA | protein_co | . | MODIFIER |
| 3'UTR | SNP | TATTTCTCT | protein_co | . | MODIFIER |
| 3'UTR | SNP | TAAACGCT | protein_co | . | MODIFIER |
| 3'UTR | SNP | ACAGGCCA | protein_co | . | MODIFIER |
| 3'UTR | SNP | CTCAGAAG | protein_co | . | MODIFIER |
| 3'UTR | SNP | CTCTGGAG | protein_co | . | MODIFIER |
| 3'UTR | SNP | CACCACTA | protein_co | . | MODIFIER |
| 5'UTR | SNP | TTCTACGT | protein_co | . | MODIFIER |
| 5'UTR | SNP | TTCTACGT | protein_co | . | MODIFIER |
| 5'UTR | SNP | TTCTACGT | protein_co | . | MODIFIER |
| 5'UTR | SNP | CCGCCGCC | protein_co | . | MODIFIER |
| 5'UTR | SNP | GCCGAGAT | protein_co | . | MODIFIER |
| 5'UTR | SNP | GCCGAGAT | protein_co | . | MODIFIER |
| 5'UTR | SNP | GCCGAGAT | protein_co | . | MODIFIER |
| 5'UTR | SNP | GCACAGCC | protein_co | . | MODIFIER |
| 5'UTR | SNP | GCACAGCC | protein_co | . | MODIFIER |
| 5'UTR | SNP | GCACAGCC | protein_co | . | MODIFIER |
| 5'UTR | INS | CGCTCTCC | protein_co | . | MODIFIER |
| Intron | SNP | TCTTCCTGA | protein_co | . | MODIFIER |
| Intron | SNP | TCTTCCTGA | protein_co | . | MODIFIER |
| Intron | SNP | CTGGCCGC | protein_co | . | MODIFIER |
| Intron | SNP | TCTTCCTGA | protein_co | . | MODIFIER |
| Intron | SNP | CTGGCCGC | protein_co | . | MODIFIER |
| Intron | SNP | TCTTCCTGA | protein_co | . | MODIFIER |
| Intron | SNP | CTGGCCGC | protein_co | . | MODIFIER |
| Intron | SNP | AGGCTCTC | protein_co | . | MODIFIER |
| Intron | SNP | AGGCTCTC | protein_co | . | MODIFIER |
| Intron | SNP | AGGCTCTC | protein_co | . | MODIFIER |
| Intron | SNP | AGGCTCTC | protein_co | . | MODIFIER |
| Intron | SNP | AGGCTCTC | protein_co | . | MODIFIER |
| Intron | SNP | TGCTTCCA | protein_co | . | MODIFIER |
| Intron | SNP | AGTACCTG | protein_co | . | MODIFIER |
| Intron | SNP | AGTACCTG | protein_co | . | MODIFIER |
| Intron | SNP | TGCCTCTC | protein_co | . | MODIFIER |
| Intron | SNP | GATTCGCA | protein_co | . | MODIFIER |
| Intron | SNP | CATTGCTT | protein_co | . | MODIFIER |
| RNA | SNP | CGATTCCC | protein_co | . | MODIFIER |
| RNA | SNP | CGATTCCC | protein_co | . | MODIFIER |
| RNA | SNP | CGATTCCC | protein_co | . | MODIFIER |

| | | | | | |
|---|---|---|---|---|---|
| RNA | SNP | CGATTCCC | protein_co | . | MODIFIER |
| RNA | SNP | TGTGGCTC | protein_co | . | MODIFIER |

| Genes | Name.Gene | UniProt.ID | Uniprot.Title | PDB.ID | HGNC.ID | Species |
|---|---|---|---|---|---|---|
| **ACAA1** | 3-ketoacyl-CoA thiolase, | P09110 | THIK_HUMAN | 2IIK | HGNC:82 | Humans |
| **ACHE** | Acetylcholinesterase | P22303 | ACES_HUMAN | 1B41; 1F8U; 1PU\ | HGNC:108 | Humans |
| **ACHE** | Acetylcholinesterase | P22303 | ACES_HUMAN | 1B41; 1F8U; 1PU\ | HGNC:108 | Humans |
| **ACHE** | Acetylcholinesterase | P22303 | ACES_HUMAN | 1B41; 1F8U; 1PU\ | HGNC:108 | Humans |
| **ACHE** | Acetylcholinesterase | P22303 | ACES_HUMAN | 1B41; 1F8U; 1PU\ | HGNC:108 | Humans |
| **ACHE** | Acetylcholinesterase | P22303 | ACES_HUMAN | 1B41; 1F8U; 1PU\ | HGNC:108 | Humans |
| **ACHE** | Acetylcholinesterase | P22303 | ACES_HUMAN | 1B41; 1F8U; 1PU\ | HGNC:108 | Humans |
| **ACHE** | Acetylcholinesterase | P22303 | ACES_HUMAN | 1B41; 1F8U; 1PU\ | HGNC:108 | Humans |
| **ACHE** | Acetylcholinesterase | P22303 | ACES_HUMAN | 1B41; 1F8U; 1PU\ | HGNC:108 | Humans |
| **ACHE** | Acetylcholinesterase | P22303 | ACES_HUMAN | 1B41; 1F8U; 1PU\ | HGNC:108 | Humans |
| **ACHE** | Acetylcholinesterase | P22303 | ACES_HUMAN | 1B41; 1F8U; 1PU\ | HGNC:108 | Humans |
| **ACHE** | Acetylcholinesterase | P22303 | ACES_HUMAN | 1B41; 1F8U; 1PU\ | HGNC:108 | Humans |
| **ACHE** | Acetylcholinesterase | P22303 | ACES_HUMAN | 1B41; 1F8U; 1PU\ | HGNC:108 | Humans |
| **ACHE** | Acetylcholinesterase | P22303 | ACES_HUMAN | 1B41; 1F8U; 1PU\ | HGNC:108 | Humans |
| **ACHE** | Acetylcholinesterase | P22303 | ACES_HUMAN | 1B41; 1F8U; 1PU\ | HGNC:108 | Humans |
| **ACHE** | Acetylcholinesterase | P22303 | ACES_HUMAN | 1B41; 1F8U; 1PU\ | HGNC:108 | Humans |
| **ADRA1A** | Alpha-1A adrenergic rece | P35348 | ADA1A_HUMAN | | HGNC:277 | Humans |
| **ADRA1A** | Alpha-1A adrenergic rece | P35348 | ADA1A_HUMAN | | HGNC:277 | Humans |
| **ADRA1A** | Alpha-1A adrenergic rece | P35348 | ADA1A_HUMAN | | HGNC:277 | Humans |
| **ADRA1A** | Alpha-1A adrenergic rece | P35348 | ADA1A_HUMAN | | HGNC:277 | Humans |
| **ADRA1A** | Alpha-1A adrenergic rece | P35348 | ADA1A_HUMAN | | HGNC:277 | Humans |
| **ADRA1A** | Alpha-1A adrenergic rece | P35348 | ADA1A_HUMAN | | HGNC:277 | Humans |
| **ADRA1A** | Alpha-1A adrenergic rece | P35348 | ADA1A_HUMAN | | HGNC:277 | Humans |
| **ADRA1A** | Alpha-1A adrenergic rece | P35348 | ADA1A_HUMAN | | HGNC:277 | Humans |
| **ADRA1A** | Alpha-1A adrenergic rece | P35348 | ADA1A_HUMAN | | HGNC:277 | Humans |
| **ADRA1A** | Alpha-1A adrenergic rece | P35348 | ADA1A_HUMAN | | HGNC:277 | Humans |
| **ADRA1A** | Alpha-1A adrenergic rece | P35348 | ADA1A_HUMAN | | HGNC:277 | Humans |
| **ADRA1A** | Alpha-1A adrenergic rece | P35348 | ADA1A_HUMAN | | HGNC:277 | Humans |
| **ADRA1A** | Alpha-1A adrenergic rece | P35348 | ADA1A_HUMAN | | HGNC:277 | Humans |
| **ADRA1A** | Alpha-1A adrenergic rece | P35348 | ADA1A_HUMAN | | HGNC:277 | Humans |
| **ADRA1A** | Alpha-1A adrenergic rece | P35348 | ADA1A_HUMAN | | HGNC:277 | Humans |
| **ADRA1A** | Alpha-1A adrenergic rece | P35348 | ADA1A_HUMAN | | HGNC:277 | Humans |
| **ADRA1A** | Alpha-1A adrenergic rece | P35348 | ADA1A_HUMAN | | HGNC:277 | Humans |
| **ADRA1A** | Alpha-1A adrenergic rece | P35348 | ADA1A_HUMAN | | HGNC:277 | Humans |
| **ADRA1A** | Alpha-1A adrenergic rece | P35348 | ADA1A_HUMAN | | HGNC:277 | Humans |
| **ADRA1A** | Alpha-1A adrenergic rece | P35348 | ADA1A_HUMAN | | HGNC:277 | Humans |
| **ADRA1A** | Alpha-1A adrenergic rece | P35348 | ADA1A_HUMAN | | HGNC:277 | Humans |
| **ADRA1A** | Alpha-1A adrenergic rece | P35348 | ADA1A_HUMAN | | HGNC:277 | Humans |
| **ADRA1A** | Alpha-1A adrenergic rece | P35348 | ADA1A_HUMAN | | HGNC:277 | Humans |
| **ADRA1A** | Alpha-1A adrenergic rece | P35348 | ADA1A_HUMAN | | HGNC:277 | Humans |
| **ADRA1A** | Alpha-1A adrenergic rece | P35348 | ADA1A_HUMAN | | HGNC:277 | Humans |
| **ADRA1A** | Alpha-1A adrenergic rece | P35348 | ADA1A_HUMAN | | HGNC:277 | Humans |
| **ADRA1A** | Alpha-1A adrenergic rece | P35348 | ADA1A_HUMAN | | HGNC:277 | Humans |
| **ADRA1A** | Alpha-1A adrenergic rece | P35348 | ADA1A_HUMAN | | HGNC:277 | Humans |
| **ADRA1A** | Alpha-1A adrenergic rece | P35348 | ADA1A_HUMAN | | HGNC:277 | Humans |
| **ADRA1A** | Alpha-1A adrenergic rece | P35348 | ADA1A_HUMAN | | HGNC:277 | Humans |

| | | | | | | |
|---|---|---|---|---|---|---|
| **ADRA1A** | Alpha-1A adrenergic rece | P35348 | ADA1A_HUMAN | | HGNC:277 | Humans |
| **ADRA1A** | Alpha-1A adrenergic rece | P35348 | ADA1A_HUMAN | | HGNC:277 | Humans |
| **ADRA1A** | Alpha-1A adrenergic rece | P35348 | ADA1A_HUMAN | | HGNC:277 | Humans |
| **ADRA1A** | Alpha-1A adrenergic rece | P35348 | ADA1A_HUMAN | | HGNC:277 | Humans |
| **ADRA1A** | Alpha-1A adrenergic rece | P35348 | ADA1A_HUMAN | | HGNC:277 | Humans |
| **ADRA1A** | Alpha-1A adrenergic rece | P35348 | ADA1A_HUMAN | | HGNC:277 | Humans |
| **ADRA1A** | Alpha-1A adrenergic rece | P35348 | ADA1A_HUMAN | | HGNC:277 | Humans |
| **ADRA1A** | Alpha-1A adrenergic rece | P35348 | ADA1A_HUMAN | | HGNC:277 | Humans |
| **ADRA1A** | Alpha-1A adrenergic rece | P35348 | ADA1A_HUMAN | | HGNC:277 | Humans |
| **ADRA1A** | Alpha-1A adrenergic rece | P35348 | ADA1A_HUMAN | | HGNC:277 | Humans |
| **ADRA1A** | Alpha-1A adrenergic rece | P35348 | ADA1A_HUMAN | | HGNC:277 | Humans |
| **ADRA1A** | Alpha-1A adrenergic rece | P35348 | ADA1A_HUMAN | | HGNC:277 | Humans |
| **ADRA1A** | Alpha-1A adrenergic rece | P35348 | ADA1A_HUMAN | | HGNC:277 | Humans |
| **ADRA1A** | Alpha-1A adrenergic rece | P35348 | ADA1A_HUMAN | | HGNC:277 | Humans |
| **ADRA1A** | Alpha-1A adrenergic rece | P35348 | ADA1A_HUMAN | | HGNC:277 | Humans |
| **ADRA1A** | Alpha-1A adrenergic rece | P35348 | ADA1A_HUMAN | | HGNC:277 | Humans |
| **ADRA1A** | Alpha-1A adrenergic rece | P35348 | ADA1A_HUMAN | | HGNC:277 | Humans |
| **ADRA1A** | Alpha-1A adrenergic rece | P35348 | ADA1A_HUMAN | | HGNC:277 | Humans |
| **ADRA1A** | Alpha-1A adrenergic rece | P35348 | ADA1A_HUMAN | | HGNC:277 | Humans |
| **ADRA1A** | Alpha-1A adrenergic rece | P35348 | ADA1A_HUMAN | | HGNC:277 | Humans |
| **ADRA1A** | Alpha-1A adrenergic rece | P35348 | ADA1A_HUMAN | | HGNC:277 | Humans |
| **ADRA1A** | Alpha-1A adrenergic rece | P35348 | ADA1A_HUMAN | | HGNC:277 | Humans |
| **ADRA1A** | Alpha-1A adrenergic rece | P35348 | ADA1A_HUMAN | | HGNC:277 | Humans |
| **ADRA1A** | Alpha-1A adrenergic rece | P35348 | ADA1A_HUMAN | | HGNC:277 | Humans |
| **ANXA2** | Annexin A2 | P07355 | ANXA2_HUMAN | 1W7B; 1XJL; 2HYU | HGNC:537 | Humans |
| **ANXA5** | Annexin A5 | P08758 | ANXA5_HUMAN | 1ANW; 1ANX; 1AV | HGNC:543 | Humans |
| **APP** | Amyloid beta A4 protein | P05067 | A4_HUMAN | 1AAP; 1AMB; 1AN | HGNC:620 | Humans |
| **APP** | Amyloid beta A4 protein | P05067 | A4_HUMAN | 1AAP; 1AMB; 1AN | HGNC:620 | Humans |
| **APP** | Amyloid beta A4 protein | P05067 | A4_HUMAN | 1AAP; 1AMB; 1AN | HGNC:620 | Humans |
| **ARL2** | ADP-ribosylation factor-li | P36404 | ARL2_HUMAN | 3DOE; 3DOF | HGNC:693 | Humans |
| **BLVRB** | Flavin reductase (NADPH | P30043 | BLVRB_HUMAN | 1HDO; 1HE2; 1HE | HGNC:1063 | Humans |
| **CAMLG** | Calcium signal-modulatin | P49069 | CAMLG_HUMAN | | HGNC:1471 | Humans |
| **CD33** | Myeloid cell surface antig | P20138 | CD33_HUMAN | | HGNC:1659 | Humans |
| **CD44** | CD44 antigen | P16070 | CD44_HUMAN | 1POZ; 1UUH; 2I83 | HGNC:1681 | Humans |
| **CDK4** | Cyclin-dependent kinase | P11802 | CDK4_HUMAN | 1LD2; 2W96; 2W9 | HGNC:1773 | Humans |
| **CDK4** | Cyclin-dependent kinase | P11802 | CDK4_HUMAN | 1LD2; 2W96; 2W9 | HGNC:1773 | Humans |
| **CDK4** | Cyclin-dependent kinase | P11802 | CDK4_HUMAN | 1LD2; 2W96; 2W9 | HGNC:1773 | Humans |
| **COMT** | Catechol O-methyltransfe | P21964 | COMT_HUMAN | 3A7E; 3BWM; 3BV | HGNC:2228 | Humans |
| **COMT** | Catechol O-methyltransfe | P21964 | COMT_HUMAN | 3A7E; 3BWM; 3BV | HGNC:2228 | Humans |
| **COMT** | Catechol O-methyltransfe | P21964 | COMT_HUMAN | 3A7E; 3BWM; 3BV | HGNC:2228 | Humans |
| **CSF2RA** | Granulocyte-macrophage | P15509 | CSF2R_HUMAN | 4NKQ; 4RS1 | HGNC:2435 | Humans |
| **CSF2RA** | Granulocyte-macrophage | P15509 | CSF2R_HUMAN | 4NKQ; 4RS1 | HGNC:2435 | Humans |
| **EGFR** | Epidermal growth factor | P00533 | EGFR_HUMAN | 1DNQ; 1DNR; 1IV | HGNC:3236 | Humans |
| **EGFR** | Epidermal growth factor | P00533 | EGFR_HUMAN | 1DNQ; 1DNR; 1IV | HGNC:3236 | Humans |
| **EGFR** | Epidermal growth factor | P00533 | EGFR_HUMAN | 1DNQ; 1DNR; 1IV | HGNC:3236 | Humans |
| **EGFR** | Epidermal growth factor | P00533 | EGFR_HUMAN | 1DNQ; 1DNR; 1IV | HGNC:3236 | Humans |
| **EGFR** | Epidermal growth factor | P00533 | EGFR_HUMAN | 1DNQ; 1DNR; 1IV | HGNC:3236 | Humans |

| | | | | | | |
|---|---|---|---|---|---|---|
| **EGFR** | Epidermal growth factor | P00533 | EGFR_HUMAN | 1DNQ; 1DNR; 1IV | HGNC:3236 | Humans |
| **EGFR** | Epidermal growth factor | P00533 | EGFR_HUMAN | 1DNQ; 1DNR; 1IV | HGNC:3236 | Humans |
| **EGFR** | Epidermal growth factor | P00533 | EGFR_HUMAN | 1DNQ; 1DNR; 1IV | HGNC:3236 | Humans |
| **EGFR** | Epidermal growth factor | P00533 | EGFR_HUMAN | 1DNQ; 1DNR; 1IV | HGNC:3236 | Humans |
| **EGFR** | Epidermal growth factor | P00533 | EGFR_HUMAN | 1DNQ; 1DNR; 1IV | HGNC:3236 | Humans |
| **EGFR** | Epidermal growth factor | P00533 | EGFR_HUMAN | 1DNQ; 1DNR; 1IV | HGNC:3236 | Humans |
| **EGFR** | Epidermal growth factor | P00533 | EGFR_HUMAN | 1DNQ; 1DNR; 1IV | HGNC:3236 | Humans |
| **EGFR** | Epidermal growth factor | P00533 | EGFR_HUMAN | 1DNQ; 1DNR; 1IV | HGNC:3236 | Humans |
| **EGFR** | Epidermal growth factor | P00533 | EGFR_HUMAN | 1DNQ; 1DNR; 1IV | HGNC:3236 | Humans |
| **FDPS** | Farnesyl pyrophosphate s | P14324 | FPPS_HUMAN | 1YQ7; 1YV5; 1ZW | HGNC:3631 | Humans |
| **FDPS** | Farnesyl pyrophosphate s | P14324 | FPPS_HUMAN | 1YQ7; 1YV5; 1ZW | HGNC:3631 | Humans |
| **FDPS** | Farnesyl pyrophosphate s | P14324 | FPPS_HUMAN | 1YQ7; 1YV5; 1ZW | HGNC:3631 | Humans |
| **FDPS** | Farnesyl pyrophosphate s | P14324 | FPPS_HUMAN | 1YQ7; 1YV5; 1ZW | HGNC:3631 | Humans |
| **FDPS** | Farnesyl pyrophosphate s | P14324 | FPPS_HUMAN | 1YQ7; 1YV5; 1ZW | HGNC:3631 | Humans |
| **FDPS** | Farnesyl pyrophosphate s | P14324 | FPPS_HUMAN | 1YQ7; 1YV5; 1ZW | HGNC:3631 | Humans |
| **FTH1** | Ferritin heavy chain | P02794 | FRIH_HUMAN | 1FHA; 2CEI; 2CHI; | HGNC:3976 | Humans |
| **FTH1** | Ferritin heavy chain | P02794 | FRIH_HUMAN | 1FHA; 2CEI; 2CHI; | HGNC:3976 | Humans |
| **GANC** | Neutral alpha-glucosidase | Q8TET4 | GANC_HUMAN | | HGNC:4139 | Humans |
| **GAPDH** | Glyceraldehyde-3-phosph | P04406 | G3P_HUMAN | 1U8F; 1ZNQ; 2FEH | HGNC:4141 | Humans |
| **GGCX** | Vitamin K-dependent gan | P38435 | VKGC_HUMAN | | HGNC:4247 | Humans |
| **GGCX** | Vitamin K-dependent gan | P38435 | VKGC_HUMAN | | HGNC:4247 | Humans |
| **GGCX** | Vitamin K-dependent gan | P38435 | VKGC_HUMAN | | HGNC:4247 | Humans |
| **GGPS1** | Geranylgeranyl pyrophos | O95749 | GGPPS_HUMAN | 2Q80 | HGNC:4249 | Humans |
| **GGPS1** | Geranylgeranyl pyrophos | O95749 | GGPPS_HUMAN | 2Q80 | HGNC:4249 | Humans |
| **HDAC10** | Histone deacetylase 10 | Q969S8 | HDA10_HUMAN | | HGNC:1812 | Humans |
| **HDAC10** | Histone deacetylase 10 | Q969S8 | HDA10_HUMAN | | HGNC:1812 | Humans |
| **HDAC8** | Histone deacetylase 8 | Q9BY41 | HDAC8_HUMAN | 1T64; 1T67; 1T69 | HGNC:1331 | Humans |
| **HDAC8** | Histone deacetylase 8 | Q9BY41 | HDAC8_HUMAN | 1T64; 1T67; 1T69 | HGNC:1331 | Humans |
| **HDAC9** | Histone deacetylase 9 | Q9UKV0 | HDAC9_HUMAN | | HGNC:1406 | Humans |
| **HDAC9** | Histone deacetylase 9 | Q9UKV0 | HDAC9_HUMAN | | HGNC:1406 | Humans |
| **HDAC9** | Histone deacetylase 9 | Q9UKV0 | HDAC9_HUMAN | | HGNC:1406 | Humans |
| **HSD17B10** | 3-hydroxyacyl-CoA dehyd | Q99714 | HCD2_HUMAN | 1F67; 1SO8; 1U7T | HGNC:4800 | Humans |
| **IFNAR1** | Interferon alpha/beta rec | P17181 | INAR1_HUMAN | 3S98; 3SE3; 3SE4; | HGNC:5432 | Humans |
| **IFNAR1** | Interferon alpha/beta rec | P17181 | INAR1_HUMAN | 3S98; 3SE3; 3SE4; | HGNC:5432 | Humans |
| **IFNAR1** | Interferon alpha/beta rec | P17181 | INAR1_HUMAN | 3S98; 3SE3; 3SE4; | HGNC:5432 | Humans |
| **IFNAR1** | Interferon alpha/beta rec | P17181 | INAR1_HUMAN | 3S98; 3SE3; 3SE4; | HGNC:5432 | Humans |
| **IFNAR1** | Interferon alpha/beta rec | P17181 | INAR1_HUMAN | 3S98; 3SE3; 3SE4; | HGNC:5432 | Humans |
| **IFNAR1** | Interferon alpha/beta rec | P17181 | INAR1_HUMAN | 3S98; 3SE3; 3SE4; | HGNC:5432 | Humans |
| **IFNAR1** | Interferon alpha/beta rec | P17181 | INAR1_HUMAN | 3S98; 3SE3; 3SE4; | HGNC:5432 | Humans |
| **IFNAR1** | Interferon alpha/beta rec | P17181 | INAR1_HUMAN | 3S98; 3SE3; 3SE4; | HGNC:5432 | Humans |
| **IFNAR1** | Interferon alpha/beta rec | P17181 | INAR1_HUMAN | 3S98; 3SE3; 3SE4; | HGNC:5432 | Humans |
| **IFNAR1** | Interferon alpha/beta rec | P17181 | INAR1_HUMAN | 3S98; 3SE3; 3SE4; | HGNC:5432 | Humans |
| **IFNAR2** | Interferon alpha/beta rec | P48551 | INAR2_HUMAN | 1N6U; 1N6V; 2HY | HGNC:5433 | Humans |
| **IFNAR2** | Interferon alpha/beta rec | P48551 | INAR2_HUMAN | 1N6U; 1N6V; 2HY | HGNC:5433 | Humans |
| **IFNAR2** | Interferon alpha/beta rec | P48551 | INAR2_HUMAN | 1N6U; 1N6V; 2HY | HGNC:5433 | Humans |
| **IFNAR2** | Interferon alpha/beta rec | P48551 | INAR2_HUMAN | 1N6U; 1N6V; 2HY | HGNC:5433 | Humans |
| **IFNAR2** | Interferon alpha/beta rec | P48551 | INAR2_HUMAN | 1N6U; 1N6V; 2HY | HGNC:5433 | Humans |

| | | | | | | |
|---|---|---|---|---|---|---|
| **IFNAR2** | Interferon alpha/beta re | P48551 | INAR2_HUMAN | 1N6U; 1N6V; 2HY | HGNC:5433 | Humans |
| **IFNAR2** | Interferon alpha/beta re | P48551 | INAR2_HUMAN | 1N6U; 1N6V; 2HY | HGNC:5433 | Humans |
| **IFNAR2** | Interferon alpha/beta re | P48551 | INAR2_HUMAN | 1N6U; 1N6V; 2HY | HGNC:5433 | Humans |
| **IFNAR2** | Interferon alpha/beta re | P48551 | INAR2_HUMAN | 1N6U; 1N6V; 2HY | HGNC:5433 | Humans |
| **IFNAR2** | Interferon alpha/beta re | P48551 | INAR2_HUMAN | 1N6U; 1N6V; 2HY | HGNC:5433 | Humans |
| **KCNH2** | Potassium voltage-gated | Q12809 | KCNH2_HUMAN | 1BYW; 1UJL; 2L0V | HGNC:6251 | Humans |
| **KCNH2** | Potassium voltage-gated | Q12809 | KCNH2_HUMAN | 1BYW; 1UJL; 2L0V | HGNC:6251 | Humans |
| **KCNH2** | Potassium voltage-gated | Q12809 | KCNH2_HUMAN | 1BYW; 1UJL; 2L0V | HGNC:6251 | Humans |
| **KCNH2** | Potassium voltage-gated | Q12809 | KCNH2_HUMAN | 1BYW; 1UJL; 2L0V | HGNC:6251 | Humans |
| **KCNH2** | Potassium voltage-gated | Q12809 | KCNH2_HUMAN | 1BYW; 1UJL; 2L0V | HGNC:6251 | Humans |
| **KCNH2** | Potassium voltage-gated | Q12809 | KCNH2_HUMAN | 1BYW; 1UJL; 2L0V | HGNC:6251 | Humans |
| **KCNH2** | Potassium voltage-gated | Q12809 | KCNH2_HUMAN | 1BYW; 1UJL; 2L0V | HGNC:6251 | Humans |
| **KCNH2** | Potassium voltage-gated | Q12809 | KCNH2_HUMAN | 1BYW; 1UJL; 2L0V | HGNC:6251 | Humans |
| **KCNH2** | Potassium voltage-gated | Q12809 | KCNH2_HUMAN | 1BYW; 1UJL; 2L0V | HGNC:6251 | Humans |
| **KCNMA1** | Calcium-activated potass | Q12791 | KCMA1_HUMAN | 2K44; 3MT5; 3NA | HGNC:6284 | Humans |
| **KCNMA1** | Calcium-activated potass | Q12791 | KCMA1_HUMAN | 2K44; 3MT5; 3NA | HGNC:6284 | Humans |
| **KCNMA1** | Calcium-activated potass | Q12791 | KCMA1_HUMAN | 2K44; 3MT5; 3NA | HGNC:6284 | Humans |
| **KCNMA1** | Calcium-activated potass | Q12791 | KCMA1_HUMAN | 2K44; 3MT5; 3NA | HGNC:6284 | Humans |
| **KCNMA1** | Calcium-activated potass | Q12791 | KCMA1_HUMAN | 2K44; 3MT5; 3NA | HGNC:6284 | Humans |
| **KRAS** | GTPase KRas | P01116 | RASK_HUMAN | 1D8D; 1D8E; 1KZ( | HGNC:6407 | Humans |
| **KRAS** | GTPase KRas | P01116 | RASK_HUMAN | 1D8D; 1D8E; 1KZ( | HGNC:6407 | Humans |
| **LDHA** | L-lactate dehydrogenase | P00338 | LDHA_HUMAN | 1I10; 4AJP; 4JNK; | HGNC:6535 | Humans |
| **MAP4** | Microtubule-associated p | P27816 | MAP4_HUMAN | | HGNC:6862 | Humans |
| **MAP4** | Microtubule-associated p | P27816 | MAP4_HUMAN | | HGNC:6862 | Humans |
| **MDH2** | Malate dehydrogenase, r | P40926 | MDHM_HUMAN | 2DFD | HGNC:6971 | Humans |
| **NISCH** | Nischarin | Q9Y2I1 | NISCH_HUMAN | 3P0C | HGNC:1800 | Humans |
| **NISCH** | Nischarin | Q9Y2I1 | NISCH_HUMAN | 3P0C | HGNC:1800 | Humans |
| **NNMT** | Nicotinamide N-methyltr | P40261 | NNMT_HUMAN | 2IIP; 3ROD | HGNC:7861 | Humans |
| **PDE1A** | Calcium/calmodulin-depe | P54750 | PDE1A_HUMAN | 1LXQ | HGNC:8774 | Humans |
| **PDXK** | Pyridoxal kinase | O00764 | PDXK_HUMAN | 2AJP; 2F7K; 2YXT; | HGNC:8819 | Humans |
| **PDXK** | Pyridoxal kinase | O00764 | PDXK_HUMAN | 2AJP; 2F7K; 2YXT; | HGNC:8819 | Humans |
| **PPARD** | Peroxisome proliferator-a | Q03181 | PPARD_HUMAN | 1GWX; 1Y0S; 2AW | HGNC:9235 | Humans |
| **PPARD** | Peroxisome proliferator-a | Q03181 | PPARD_HUMAN | 1GWX; 1Y0S; 2AW | HGNC:9235 | Humans |
| **PPARD** | Peroxisome proliferator-a | Q03181 | PPARD_HUMAN | 1GWX; 1Y0S; 2AW | HGNC:9235 | Humans |
| **PPARD** | Peroxisome proliferator-a | Q03181 | PPARD_HUMAN | 1GWX; 1Y0S; 2AW | HGNC:9235 | Humans |
| **PRKCE** | Protein kinase C epsilon t | Q02156 | KPCE_HUMAN | 2WH0 | HGNC:9401 | Humans |
| **PRKCZ** | Protein kinase C zeta typ | Q05513 | KPCZ_HUMAN | | HGNC:9412 | Humans |
| **PRLR** | Prolactin receptor | P16471 | PRLR_HUMAN | 1BP3; 2LFG; 3D48 | HGNC:9446 | Humans |
| **PRLR** | Prolactin receptor | P16471 | PRLR_HUMAN | 1BP3; 2LFG; 3D48 | HGNC:9446 | Humans |
| **RELA** | Transcription factor p65 | Q04206 | TF65_HUMAN | 1NFI; 2LSP; 2O61; | HGNC:9955 | Humans |
| **SCN11A** | Sodium channel protein t | Q9UI33 | SCNBA_HUMAN | | HGNC:1058 | Humans |
| **SCN11A** | Sodium channel protein t | Q9UI33 | SCNBA_HUMAN | | HGNC:1058 | Humans |
| **SCN11A** | Sodium channel protein t | Q9UI33 | SCNBA_HUMAN | | HGNC:1058 | Humans |
| **SCN11A** | Sodium channel protein t | Q9UI33 | SCNBA_HUMAN | | HGNC:1058 | Humans |
| **SCN11A** | Sodium channel protein t | Q9UI33 | SCNBA_HUMAN | | HGNC:1058 | Humans |
| **SCN11A** | Sodium channel protein t | Q9UI33 | SCNBA_HUMAN | | HGNC:1058 | Humans |
| **SCN11A** | Sodium channel protein t | Q9UI33 | SCNBA_HUMAN | | HGNC:1058 | Humans |

| | | | | | | |
|---|---|---|---|---|---|---|
| **SCN11A** | Sodium channel protein t | Q9UI33 | SCNBA_HUMAN | | HGNC:1058 | Humans |
| **SCN11A** | Sodium channel protein t | Q9UI33 | SCNBA_HUMAN | | HGNC:1058 | Humans |
| **SCN11A** | Sodium channel protein t | Q9UI33 | SCNBA_HUMAN | | HGNC:1058 | Humans |
| **SCN11A** | Sodium channel protein t | Q9UI33 | SCNBA_HUMAN | | HGNC:1058 | Humans |
| **SCN8A** | Sodium channel protein t | Q9UQD0 | SCN8A_HUMAN | | HGNC:1059 | Humans |
| **SCN8A** | Sodium channel protein t | Q9UQD0 | SCN8A_HUMAN | | HGNC:1059 | Humans |
| **SCN8A** | Sodium channel protein t | Q9UQD0 | SCN8A_HUMAN | | HGNC:1059 | Humans |
| **SCN8A** | Sodium channel protein t | Q9UQD0 | SCN8A_HUMAN | | HGNC:1059 | Humans |
| **SCN8A** | Sodium channel protein t | Q9UQD0 | SCN8A_HUMAN | | HGNC:1059 | Humans |
| **SCN8A** | Sodium channel protein t | Q9UQD0 | SCN8A_HUMAN | | HGNC:1059 | Humans |
| **SCN8A** | Sodium channel protein t | Q9UQD0 | SCN8A_HUMAN | | HGNC:1059 | Humans |
| **SCN8A** | Sodium channel protein t | Q9UQD0 | SCN8A_HUMAN | | HGNC:1059 | Humans |
| **SCN8A** | Sodium channel protein t | Q9UQD0 | SCN8A_HUMAN | | HGNC:1059 | Humans |
| **SCN8A** | Sodium channel protein t | Q9UQD0 | SCN8A_HUMAN | | HGNC:1059 | Humans |
| **SCN8A** | Sodium channel protein t | Q9UQD0 | SCN8A_HUMAN | | HGNC:1059 | Humans |
| **SDHA** | Succinate dehydrogenase | P31040 | SDHA_HUMAN | | HGNC:1068 | Humans |
| **SHMT1** | Serine hydroxymethyltra | P34896 | GLYC_HUMAN | 1BJ4 | HGNC:1085 | Humans |
| **SIRT5** | NAD-dependent protein | Q9NXA8 | SIR5_HUMAN | 2B4Y; 2NYR; 3RIG | HGNC:1493 | Humans |
| **SLC52A2** | Solute carrier family 52, r | Q9HAB3 | S52A2_HUMAN | | HGNC:3022 | Humans |
| **SMOX** | Spermine oxidase | Q9NWM0 | SMOX_HUMAN | | HGNC:1586 | Humans |
| **SMS** | Spermine synthase | P52788 | SPSY_HUMAN | 3C6K; 3C6M | HGNC:1112 | Humans |
| **TRPV3** | Transient receptor poten | Q8NET8 | TRPV3_HUMAN | | HGNC:1808 | Humans |
| **TRPV3** | Transient receptor poten | Q8NET8 | TRPV3_HUMAN | | HGNC:1808 | Humans |
| **TRPV3** | Transient receptor poten | Q8NET8 | TRPV3_HUMAN | | HGNC:1808 | Humans |
| **TUBB3** | Tubulin beta-3 chain | Q13509 | TBB3_HUMAN | | HGNC:2077 | Humans |
| **TYMS** | Thymidylate synthase | P04818 | TYSY_HUMAN | 1HVY; 1HW3; 1HV | HGNC:1244 | Humans |
| **TYMS** | Thymidylate synthase | P04818 | TYSY_HUMAN | 1HVY; 1HW3; 1HV | HGNC:1244 | Humans |
| **TYMS** | Thymidylate synthase | P04818 | TYSY_HUMAN | 1HVY; 1HW3; 1HV | HGNC:1244 | Humans |
| **TYMS** | Thymidylate synthase | P04818 | TYSY_HUMAN | 1HVY; 1HW3; 1HV | HGNC:1244 | Humans |
| **TYMS** | Thymidylate synthase | P04818 | TYSY_HUMAN | 1HVY; 1HW3; 1HV | HGNC:1244 | Humans |
| **TYMS** | Thymidylate synthase | P04818 | TYSY_HUMAN | 1HVY; 1HW3; 1HV | HGNC:1244 | Humans |
| **TYMS** | Thymidylate synthase | P04818 | TYSY_HUMAN | 1HVY; 1HW3; 1HV | HGNC:1244 | Humans |
| **TYMS** | Thymidylate synthase | P04818 | TYSY_HUMAN | 1HVY; 1HW3; 1HV | HGNC:1244 | Humans |
| **TYMS** | Thymidylate synthase | P04818 | TYSY_HUMAN | 1HVY; 1HW3; 1HV | HGNC:1244 | Humans |
| **TYMS** | Thymidylate synthase | P04818 | TYSY_HUMAN | 1HVY; 1HW3; 1HV | HGNC:1244 | Humans |
| **VKORC1** | Vitamin K epoxide reduct | Q9BQB6 | VKOR1_HUMAN | | HGNC:2366 | Humans |
| **VKORC1** | Vitamin K epoxide reduct | Q9BQB6 | VKOR1_HUMAN | | HGNC:2366 | Humans |
| **VKORC1** | Vitamin K epoxide reduct | Q9BQB6 | VKOR1_HUMAN | | HGNC:2366 | Humans |
| **VKORC1** | Vitamin K epoxide reduct | Q9BQB6 | VKOR1_HUMAN | | HGNC:2366 | Humans |
| **VKORC1** | Vitamin K epoxide reduct | Q9BQB6 | VKOR1_HUMAN | | HGNC:2366 | Humans |
| **VKORC1** | Vitamin K epoxide reduct | Q9BQB6 | VKOR1_HUMAN | | HGNC:2366 | Humans |

| Name.Drug | CAS.Numb | Drug.Type | PubChem.S | Drug.IDs |
|---|---|---|---|---|
| Trimetazidi | 5011-34-7 | SmallMole | 3.1E+08 | DB09069 |
| Edrophoniu | 312-48-1 | SmallMole | 46507530 | DB01010 |
| Demecariu | 16505-84-3 | SmallMole | 46507824 | DB00944 |
| Rivastigmir | 123441-03 | SmallMole | 46507452 | DB00989 |
| Tacrine | 321-64-2 | SmallMole | 46505487 | DB00382 |
| Physostigm | 57-47-6 | SmallMole | 46506998 | DB00981 |
| Neostigmir | 59-99-4 | SmallMole | 46509161 | DB01400 |
| Pralidoxim | 6735-59-7 | SmallMole | 46509152 | DB00733 |
| Pyridostign | 155-97-5 | SmallMole | 46506129 | DB00545 |
| Minaprine | 25905-77-5 | SmallMole | 46505380 | DB00805 |
| Donepezil | 120014-06 | SmallMole | 46504803 | DB00843 |
| Dimetacrin | 4757-55-5 | SmallMole | 3.1E+08 | DB08996 |
| Gallamine t | 65-29-2 | SmallMole | 46508651 | DB00483 |
| Ambenoniu | 7648-98-8 | SmallMole | 46508143 | DB01122 |
| Isoflurophа | 55-91-4 | SmallMole | 46504499 | DB00677 |
| Galantamir | 357-70-0 | SmallMole | 46505659 | DB00674 |
| Droperidol | 548-73-2 | SmallMole | 46505291 | DB00450 |
| Alfuzosin | 81403-80-7 | SmallMole | 46508512 | DB00346 |
| Nicergoline | 27848-84-6 | SmallMole | 46508741 | DB00699 |
| Metaramin | 54-49-9 | SmallMole | 46505593 | DB00610 |
| Ergotamine | 113-15-5 | SmallMole | 46507632 | DB00696 |
| Carvedilol | 72956-09-3 | SmallMole | 46505146 | DB01136 |
| Tetryzoline | 84-22-0 | SmallMole | 3.1E+08 | DB06764 |
| Epinephrin | 51-43-4 | SmallMole | 46509097 | DB00668 |
| Isomethep | 503-01-5 | SmallMole | 99443258 | DB06706 |
| Phendimet | 634-03-7 | SmallMole | 46506122 | DB01579 |
| Propiverine | 60569-19-9 | SmallMole | 3.48E+08 | DB12278 |
| Prazosin | 19216-56-9 | SmallMole | 46508594 | DB00457 |
| Benzphetar | 156-08-1 | SmallMole | 46506102 | DB00865 |
| Silodosin | 160970-54 | SmallMole | 1.75E+08 | DB06207 |
| Dapiprazol | 72822-12-9 | SmallMole | 46508859 | DB00298 |
| Thioridazin | 50-52-2 | SmallMole | 46509070 | DB00679 |
| Ergometrin | 60-79-7 | SmallMole | 46506922 | DB01253 |
| Doxazosin | 74191-85-8 | SmallMole | 46506825 | DB00590 |
| Chlorprom | 50-53-3 | SmallMole | 46508395 | DB00477 |
| Lorpiprazol | 108785-69 | SmallMole | 3.1E+08 | DB09195 |
| Oxymetazo | 1491-59-4 | SmallMole | 46505622 | DB00935 |
| Tamsulosin | 106133-20 | SmallMole | 46507763 | DB00706 |
| Dronedaro | 141626-36 | SmallMole | 1.75E+08 | DB04855 |
| Arotinolol | 68377-92-4 | SmallMole | 3.1E+08 | DB09204 |
| Racepineph | 329-65-7 | SmallMole | 3.48E+08 | DB11124 |
| Ergoloid m | 8067-24-1 | SmallMole | 46505963 | DB01049 |
| Dihydroerg | 25447-65-8 | SmallMole | NA | DB11273 |
| Phenoxybe | 59-96-1 | SmallMole | 46507191 | DB00925 |
| Norepinepl | 51-41-2 | SmallMole | 46506201 | DB00368 |
| Mephenter | 100-92-5 | SmallMole | 46505918 | DB01365 |

| | | | | |
|---|---|---|---|---|
| Xylometazo | 526-36-3 | SmallMolec | 99443248 | DB06694 |
| Cirazoline | 59939-16-1 | SmallMolec | 3.1E+08 | DB09202 |
| Tolazoline | 59-98-3 | SmallMolec | 46505641 | DB00797 |
| DL-Methyle | 1201-56-5 | SmallMolec | NA | DB11278 |
| Moxisylyte | 54-32-0 | SmallMolec | 3.1E+08 | DB09205 |
| Phenylephr | 59-42-7 | SmallMolec | 46506961 | DB00388 |
| Tiapride | 51012-32-9 | SmallMolec | 3.48E+08 | DB13025 |
| Trifluopera | 117-89-5 | SmallMolec | 46507961 | DB00831 |
| Phentolam | 50-60-2 | SmallMolec | 46506535 | DB00692 |
| 4-Methoxy | 64-13-1 | SmallMolec | 46507471 | DB01472 |
| Midodrine | 42794-76-3 | SmallMolec | 46507373 | DB00211 |
| Dihydroerg | 17479-19-5 | SmallMolec | NA | DB13345 |
| Thioproper | 316-81-4 | SmallMolec | 46507111 | DB01622 |
| Dosulepin | 113-53-1 | SmallMolec | 3.1E+08 | DB09167 |
| Droxidopa | 23651-95-8 | SmallMolec | 99443239 | DB06262 |
| Labetalol | 36894-69-6 | SmallMolec | 46505511 | DB00598 |
| Methoxam | 390-28-3 | SmallMolec | 46506264 | DB00723 |
| Flupentixol | 2709-56-0 | SmallMolec | 46507816 | DB00875 |
| Naphazolin | 835-31-4 | SmallMolec | 99443263 | DB06711 |
| Dipivefrin | 52365-63-6 | SmallMolec | 46504716 | DB00449 |
| Dobutamin | 34368-04-2 | SmallMolec | 46505241 | DB00841 |
| Apraclonid | 66711-21-5 | SmallMolec | 46505614 | DB00964 |
| Pseudoeph | 90-82-4 | SmallMolec | 46508190 | DB00852 |
| Terazosin | 63590-64-7 | SmallMolec | 46509129 | DB01162 |
| Fluocinolor | 67-73-2 | SmallMolec | 46506244 | DB00591 |
| Fluocinolor | 67-73-2 | SmallMolec | 46506244 | DB00591 |
| Florbetabe | 902143-01- | SmallMolec | 3.1E+08 | DB09148 |
| Florbetapir | 956103-76- | SmallMolec | 3.1E+08 | DB09149 |
| Flutemetar | 765922-62- | SmallMolec | 3.1E+08 | DB09151 |
| Tagraxofus | 2055491-0 | BiotechDru | NA | DB14731 |
| Riboflavin | 83-88-5 | SmallMolec | 46507342 | DB00140 |
| Cyclosporir | 59865-13-3 | SmallMolec | 46508198 | DB00091 |
| Gemtuzum | 220578-59- | BiotechDru | 46505767 | DB00056 |
| Hyaluronic | 9004-61-9 | SmallMolec | 3.1E+08 | DB08818 |
| Ribociclib | 1211441-9 | SmallMolec | 3.48E+08 | DB11730 |
| Abemacicli | 1231929-9 | SmallMolec | 3.48E+08 | DB12001 |
| Palbociclib | 571190-30- | SmallMolec | 3.1E+08 | DB09073 |
| Entacapone | 130929-57- | SmallMolec | 46508734 | DB00494 |
| Tolcapone | 134308-13- | SmallMolec | 46504932 | DB00323 |
| Opicapone | 923287-50- | SmallMolec | NA | DB11632 |
| Foreskin keratinocyte ( | | BiotechDru | NA | DB10772 |
| Sargramost | 123774-72- | BiotechDru | 46507000 | DB00020 |
| Cetuximab | 205923-56- | BiotechDru | 46507042 | DB00002 |
| Gefitinib | 184475-35- | SmallMolec | 46508649 | DB00317 |
| Neratinib | 698387-09- | SmallMolec | 3.48E+08 | DB11828 |
| Osimertinib | 1421373-6 | SmallMolec | 3.1E+08 | DB09330 |
| Dacomitini | 1110813-3 | SmallMolec | 3.48E+08 | DB11963 |

| | | | | |
|---|---|---|---|---|
| Lapatinib | 231277-92- | SmallMole | 46507141 | DB01259 |
| Necitumum | 906805-06- | BiotechDru | 3.48E+08 | DB09559 |
| Olmutinib | 1353550-1 | SmallMole | 3.48E+08 | DB13164 |
| Panitumum | 339177-26- | BiotechDru | 46505063 | DB01269 |
| Foreskin keratinocyte ( | | BiotechDru | NA | DB10772 |
| Erlotinib | 183321-74- | SmallMole | 46508021 | DB00530 |
| Afatinib | 850140-72- | SmallMole | 1.75E+08 | DB08916 |
| Brigatinib | 1197953-5 | SmallMole | 3.48E+08 | DB12267 |
| Zalutumum | 667901-13- | BiotechDru | 3.48E+08 | DB12202 |
| Ibandronat | 114084-78- | SmallMole | 46508134 | DB00710 |
| Zoledronic | 118072-93- | SmallMole | 46507310 | DB00399 |
| Risedronic | 105462-24- | SmallMole | 46507526 | DB00884 |
| Alendronic | 66376-36-1 | SmallMole | 46507199 | DB00630 |
| Minodroni | 180064-38- | SmallMole | NA | DB06548 |
| Pamidroni | 40391-99-9 | SmallMole | 46504823 | DB00282 |
| Ferric pyro | 1802359-9 | SmallMole | NA | DB13995 |
| Iron Dextra | 9004-66-4 | SmallMole | 46506136 | DB00893 |
| Miglitol | 72432-03-2 | SmallMole | 46504492 | DB00491 |
| Xanthinol | 2530-97-4 | SmallMole | 3.1E+08 | DB09092 |
| Anisindione | 117-37-3 | SmallMole | 46504660 | DB01125 |
| Phylloquin | 84-80-0 | SmallMole | 46504528 | DB01022 |
| Menadione | 58-27-5 | SmallMole | 46505447 | DB00170 |
| Zoledronic | 118072-93- | SmallMole | 46507310 | DB00399 |
| Minodroni | 180064-38- | SmallMole | NA | DB06548 |
| Panobinost | 404950-80- | SmallMole | 3.1E+08 | DB06603 |
| Belinostat | 866323-14- | SmallMole | 3.48E+08 | DB05015 |
| Belinostat | 866323-14- | SmallMole | 3.48E+08 | DB05015 |
| Panobinost | 404950-80- | SmallMole | 3.1E+08 | DB06603 |
| Valproic ac | 99-66-1 | SmallMole | 46505925 | DB00313 |
| Panobinost | 404950-80- | SmallMole | 3.1E+08 | DB06603 |
| Belinostat | 866323-14- | SmallMole | 3.48E+08 | DB05015 |
| Omega-3-carboxylic ac | | SmallMole | 3.48E+08 | DB09568 |
| Interferon | 76543-88-9 | BiotechDru | 46506712 | DB00034 |
| Interferon | 145258-61- | BiotechDru | 46504899 | DB00060 |
| Peginterfer | 198153-51- | BiotechDru | 46504860 | DB00008 |
| Interferon | 145155-23- | BiotechDru | 46504458 | DB00068 |
| Interferon | 74899-72-2 | BiotechDru | 46506227 | DB00011 |
| Peginterfer | 215647-85- | BiotechDru | 46506669 | DB00022 |
| Interferon | 98530-12-2 | BiotechDru | 46507075 | DB00105 |
| Interferon | 118390-30- | BiotechDru | 46505985 | DB00069 |
| Human inte | 74899-71-1 | BiotechDru | NA | DB14999 |
| Interferon alfa-n3 | | BiotechDru | 46508902 | DB00018 |
| Interferon | 76543-88-9 | BiotechDru | 46506712 | DB00034 |
| Interferon | 145258-61- | BiotechDru | 46504899 | DB00060 |
| Interferon | 98530-12-2 | BiotechDru | 46507075 | DB00105 |
| Interferon | 145155-23- | BiotechDru | 46504458 | DB00068 |
| Peginterfer | 198153-51- | BiotechDru | 46504860 | DB00008 |

| | | | | |
|---|---|---|---|---|
| Peginterfer | 215647-85- | BiotechDru | 46506669 | DB00022 |
| Interferon | 118390-30- | BiotechDru | 46505985 | DB00069 |
| Interferon | 74899-72-2 | BiotechDru | 46506227 | DB00011 |
| Interferon alfa-n3 | | BiotechDru | 46508902 | DB00018 |
| Nylidrin | 447-41-6 | SmallMole | 3.48E+08 | DB06152 |
| Ibutilide | 122647-31- | SmallMole | 46507904 | DB00308 |
| Dofetilide | 115256-11- | SmallMole | 46509127 | DB00204 |
| Amiodaron | 1951-25-3 | SmallMole | 46507387 | DB01118 |
| Propafenor | 54063-53-5 | SmallMole | 46504529 | DB01182 |
| Phenytoin | 57-41-0 | SmallMole | 46508847 | DB00252 |
| Pimozide | 2062-78-4 | SmallMole | 46507096 | DB01100 |
| Dronedaro | 141626-36- | SmallMole | 1.75E+08 | DB04855 |
| Sotalol | 3930-20-9 | SmallMole | 46505012 | DB00489 |
| Halofantrin | 69756-53-2 | SmallMole | 46506753 | DB01218 |
| Chlorzoxaz | 95-25-0 | SmallMole | 46507755 | DB00356 |
| Bendroflun | 73-48-3 | SmallMole | 46508672 | DB00436 |
| Trimebutin | 39133-31-8 | SmallMole | 3.1E+08 | DB09089 |
| Cromoglici | 16110-51-3 | SmallMole | 46504509 | DB01003 |
| Hydrochlor | 58-93-5 | SmallMole | 46504440 | DB00999 |
| MRTX849 | | SmallMole | NA | DB15568 |
| AMG-510 | 2252403-5 | SmallMole | NA | DB15569 |
| Stiripentol | 49763-96-4 | SmallMole | 3.1E+08 | DB09118 |
| Docetaxel | 114977-28- | SmallMole | 46506766 | DB01248 |
| Paclitaxel | 33069-62-4 | SmallMole | 46506910 | DB01229 |
| Xanthinol | 2530-97-4 | SmallMole | 3.1E+08 | DB09092 |
| Agmatine | 306-60-5 | SmallMole | 1.75E+08 | DB08838 |
| Moxonidin | 75438-57-2 | SmallMole | 3.1E+08 | DB09242 |
| Niacin | 59-67-6 | SmallMole | 46507508 | DB00627 |
| Trapidil | 15421-84-8 | SmallMole | 3.1E+08 | DB09283 |
| Pyridoxine | 65-23-6 | SmallMole | 46508560 | DB00165 |
| Pyridoxal | 66-72-8 | SmallMole | 46506282 | DB00147 |
| Treprostini | 81846-19-7 | SmallMole | 46504572 | DB00374 |
| Glycerin | 56-81-5 | SmallMole | 3.48E+08 | DB09462 |
| Bezafibrate | 41859-67-0 | SmallMole | 46509188 | DB01393 |
| Icosapent | 10417-94-4 | SmallMole | 46506267 | DB00159 |
| Tamoxifen | 10540-29-1 | SmallMole | 46505515 | DB00675 |
| Tamoxifen | 10540-29-1 | SmallMole | 46505515 | DB00675 |
| Fluoxymest | 76-43-7 | SmallMole | 46508867 | DB01185 |
| Somatotro | 12629-01-5 | BiotechDru | 46506010 | DB00052 |
| SC-236 | 170569-86- | SmallMole | NA | DB14059 |
| Pramocain | 140-65-8 | SmallMole | 3.1E+08 | DB09345 |
| Dichlorobe | 1777-82-8 | SmallMole | NA | DB13269 |
| Lamotrigin | 84057-84-1 | SmallMole | 46505408 | DB00555 |
| Amylocaine | 644-26-8 | SmallMole | 3.1E+08 | DB09088 |
| Topiramate | 97240-79-4 | SmallMole | 46508334 | DB00273 |
| Zonisamide | 68291-97-4 | SmallMole | 46505278 | DB00909 |
| Tetracaine | 94-24-6 | SmallMole | 3.1E+08 | DB09085 |

| | | | | |
|---|---|---|---|---|
| Brivaraceta | 357336-20- | SmallMole | 3.48E+08 | DB05541 |
| Propoxycai | 86-43-1 | SmallMole | 3.1E+08 | DB09342 |
| Cocaine | 50-36-2 | SmallMole | 46506326 | DB00907 |
| Carbamaze | 298-46-4 | SmallMole | 46507583 | DB00564 |
| Carbamaze | 298-46-4 | SmallMole | 46507583 | DB00564 |
| Brivaraceta | 357336-20- | SmallMole | 3.48E+08 | DB05541 |
| Cocaine | 50-36-2 | SmallMole | 46506326 | DB00907 |
| Propoxycai | 86-43-1 | SmallMole | 3.1E+08 | DB09342 |
| Dichlorobe | 1777-82-8 | SmallMole | NA | DB13269 |
| Amylocaine | 644-26-8 | SmallMole | 3.1E+08 | DB09088 |
| Tetracaine | 94-24-6 | SmallMole | 3.1E+08 | DB09085 |
| Topiramate | 97240-79-4 | SmallMole | 46508334 | DB00273 |
| Phenytoin | 57-41-0 | SmallMole | 46508847 | DB00252 |
| Lamotrigin | 84057-84-1 | SmallMole | 46505408 | DB00555 |
| Pramocain | 140-65-8 | SmallMole | 3.1E+08 | DB09345 |
| Ubidecaren | 303-98-0 | SmallMole | 3.1E+08 | DB09270 |
| Mimosine | 500-44-7 | SmallMole | 46505895 | DB01055 |
| Suramin | 145-63-1 | SmallMole | 46507013 | DB04786 |
| gamma-Hy | 591-81-1 | SmallMole | 46507548 | DB01440 |
| Spermine | 71-44-3 | SmallMole | 46507215 | DB00127 |
| Spermine | 71-44-3 | SmallMole | 46507215 | DB00127 |
| Camphor | 464-49-3 | SmallMole | 46508429 | DB01744 |
| (S)-campho | 464-48-2 | SmallMole | NA | DB11345 |
| Levomenth | 2216-51-5 | SmallMole | 46506513 | DB00825 |
| Ixabepilone | 219989-84- | SmallMole | 99443224 | DB04845 |
| Fluorourac | 51-21-8 | SmallMole | 46508557 | DB00544 |
| Methotrex | 59-05-2 | SmallMole | 46507678 | DB00563 |
| Pralatrexat | 146464-95- | SmallMole | 1.75E+08 | DB06813 |
| Pemetrexe | 137281-23- | SmallMole | 46505640 | DB00642 |
| Capecitabi | 154361-50- | SmallMole | 46508686 | DB01101 |
| Raltitrexed | 112887-68- | SmallMole | 46504880 | DB00293 |
| Floxuridine | 50-91-9 | SmallMole | 46508645 | DB00322 |
| Trifluridine | 70-00-8 | SmallMole | 46506192 | DB00432 |
| Tegafur-ura | 74578-38-4 | SmallMole | 3.1E+08 | DB09327 |
| Tegafur | 17902-23-7 | SmallMole | 3.1E+08 | DB09256 |
| Acenocoun | 152-72-7 | SmallMole | 46507631 | DB01418 |
| Dicoumaro | 66-76-2 | SmallMole | 46505536 | DB00266 |
| Warfarin | 81-81-2 | SmallMole | 46504711 | DB00682 |
| Phenproco | 435-97-2 | SmallMole | 46506423 | DB00946 |
| Phenindion | 83-12-5 | SmallMole | 46505018 | DB00498 |
| Menadione | 58-27-5 | SmallMole | 46505447 | DB00170 |
